# The proteome of agroinfiltrated *Nicotiana benthamiana* is shaped by extensive protein processing

**DOI:** 10.1101/2023.11.02.565301

**Authors:** Kaijie Zheng, Joy C. Lyu, Emma L. Thomas, Mariana Schuster, Nattapong Sanguankiattichai, Sabrina Ninck, Farnusch Kaschani, Markus Kaiser, Renier A. L. van der Hoorn

**Affiliations:** Key Laboratory of Soybean Molecular Design Breeding, Northeast Institute of Geography and Agroecology, Chinese Academy of Sciences, Changchun 130102, China; The Plant Chemetics Laboratory, Department of Biology, University of Oxford, Oxford OX1 3RD, UK; ZMB Chemical Biology, Faculty of Biology, University of Duisburg-Essen, Essen 45141, Germany; Leibniz Institute of Plant Biochemistry, Halle 06120, Germany

**Keywords:** protein processing, protein migration, molecular weight, agroinfiltration, *Nicotiana benthamiana*

## Abstract

Processing by proteases irreversibly regulates the fate of plant proteins and hampers the production of recombinant protein in plants, yet only few processing events have been described in agroinfiltrated *Nicotiana benthamiana*, which has emerged as a favorite transient protein expression platform in plant science and molecular pharming. Here, we used in-gel digests and mass spectrometry to monitor the migration and topography of 5,040 plant proteins of agroinfiltrated *N. benthamiana* within a protein gel. By plotting the peptides over the gel slices, we generated peptographs that reveal where which part of each protein was detected within the protein gel. These data uncovered that 60% of the detected proteins have proteoforms that migrate at lower than predicted molecular weights, implicating extensive proteolytic processing. For instance, this analysis confirms the proteolytic removal and degradation of autoinhibitory prodomains of most but not all proteases, and revealed differential processing within pectinemethylesterase and lipase families. This analysis also uncovered intricate processing of glycosidases and uncovered that ectodomain shedding might be common for a diverse range of receptor-like kinases. Transient expression of double-tagged candidate proteins confirmed various processing events *in vivo*. This extensive proteomic dataset can be investigated further and demonstrates that most plant proteins are proteolytically processed and implicates an extensive proteolytic machinery shaping the proteome of agroinfiltrated *N. benthamiana*.

## Introduction

Agroinfiltration of *Nicotiana benthamiana* is routinely used in plant science and has emerged as a powerful protein expression platform in molecular pharming. Yet, the yields or recombinant protein production are hampered by processing and degradation caused by endogenous plant proteases. The *N. benthamiana* genome encodes for >1,200 putative proteases (Jutras et al., 2020), but little is known on their endogenous substrates. Most processing events may inactivate proteins, but protein cleavages can also regulate proteins for instance by releasing active proteins from their precursors (Dissmeyer et al., 2018). Protein processing can also change the subcellular location of proteins or their ability to interact with other components inside and outside the cell. Understanding protein processing in plants is limited by the absence of global studies on protein processing that reveal endogenous protein substrates.

Several global studies on protein processing include the use of shotgun proteomics on isotope-labeled Arabidopsis cell cultures to monitor life times of plant proteins (Nelson et al., 2014); the use of Terminal Amine Isotope Labelling of Samples (TAILS), e.g. to monitor the role of PRT6 in the N-terminome of *Arabidopsis thaliana* (Zhang et al., 2018); and the use of COmbined FRactional DIagonal Chromatography (COFRADIC), e.g. to identify substrates of metacaspase-9, based on the detection of N-termini (Tsiatsiani et al., 2013). These, and similar powerful proteomic methods resulted in many candidate protease substrates that remain to be confirmed. Candidate substrates identified by TAILS, COFRADIC and shotgun proteomics are challenging to confirm because neo-termini can originate from proteins that are only incompletely processed or already degraded. In addition, neo-termini that appear near the termini of proteins are difficult to confirm by detecting shifts in molecular weight (MW).

Here, we take a different, complementary approach to identify candidate protease substrates by a shift in migration in protein gels. We use the PRotein TOpography and Migration Analysis Platform (PROTOMAP) to identify proteins that migrate in the protein gel at a different than predicted MW. PROTOMAP is a proteomic analysis approach that charts tryptic peptides detected by mass spectrometry (MS) from gel slices of a 1D-SDS-PAGE gel in two dimensions: the protein sequence (x-axis) and the gel slice at which the peptide was detected (y-scale) (Dix et al., 2008). By including MS data from all gel slices of an 1D-gel and including replicates, this PROTOMAP approach results in a global overview of where within the protein gel, which part of the protein is migrating. This analysis generates the same type of information as western blots, but then for all detectable proteins simultaneously. Thus, in contrast to other proteomic techniques, PROTOMAP will display changes in the migration of proteins within protein gels.

PROTOMAP was originally used to identify 261 processing events during apoptosis (Dix et al., 2008) and was used to demonstrate that phosphorylation sculpts the apoptotic proteome (Dix et al., 2012). PROTOMAP has also been used to profile protein processing during infection with the malaria parasite (Bowyer et al., 2011), but hasn’t been used in plant science because of the high demand on machine time. For instance, the generation of PROTOMAP data for a single proteome involves the analysis of ∼100 in-gel tryptic digests by mass spectrometry.

Here, we sought to generate a PROTOMAP dataset from agroinfiltrated *N. benthamiana*, taking advantage of its recently improved genome annotation (Ranawaka et al., 2023). Transient expression by agroinfiltration of *N. benthamiana* is frequently used in plant science to study e.g. subcellular localization and protein-protein interactions, and in molecular pharming to produce pharmaceutical proteins and metabolites (Eidenberger et al., 2023). However, the proteolytic machinery of *N. benthamiana* includes >1,000 putative proteases that hamper molecular pharming and research in plant science (Jutras et al., 2020) and therefore the characterization of protein processing in agroinfiltrated leaves will benefit both plant science and molecular pharming. In addition, by analyzing processing in agroinfiltrated leaves, we should be able to verify processing events upon transient expression of reporter-tagged precursor proteins.

## Results

To generate the PROTOMAP dataset, we separated the soluble proteomes of extracts from agroinfiltrated leaves in adjacent lanes on a 4-12% gradient Bis-Tris protein gel and cut each lane into 24 gel slices, covering a range of 10-180 kDa indicated by the flanking marker lanes (Supplemental **Figure S1**). We digested the proteins in these gel slices with trypsin and analysed the released peptides by LC-MS/MS (**Figure 1A**). We annotated the spectra to predicted tryptic peptides generated *in silico* from the recently released LAB3.60 annotation of the *N. benthamiana* genome (Ranawaka et al., 2023). The annotation to the LAB3.60 drastically increased the frequency of unique peptides when compared to our earlier NbDE database (Kourelis et al., 2019, Supplemental **Figure S2**). A total of 36,967 different tryptic peptides derived from 5,040 plant proteins were detected in these 96 gel slices, making this the largest proteomics dataset available for agroinfiltrated *N. benthamiana* leaves (Supplemental **File S1**). Almost 82% of the detected proteins were identified with more than one single peptide (**Figure 1B**). The 919 proteins that were identified by a single peptide are retained in this analysis because their detection within the gel is often consistent with the predicted MW and the migration of homologous proteins (Supplemental **Figure S3**).

**Figure 1.**
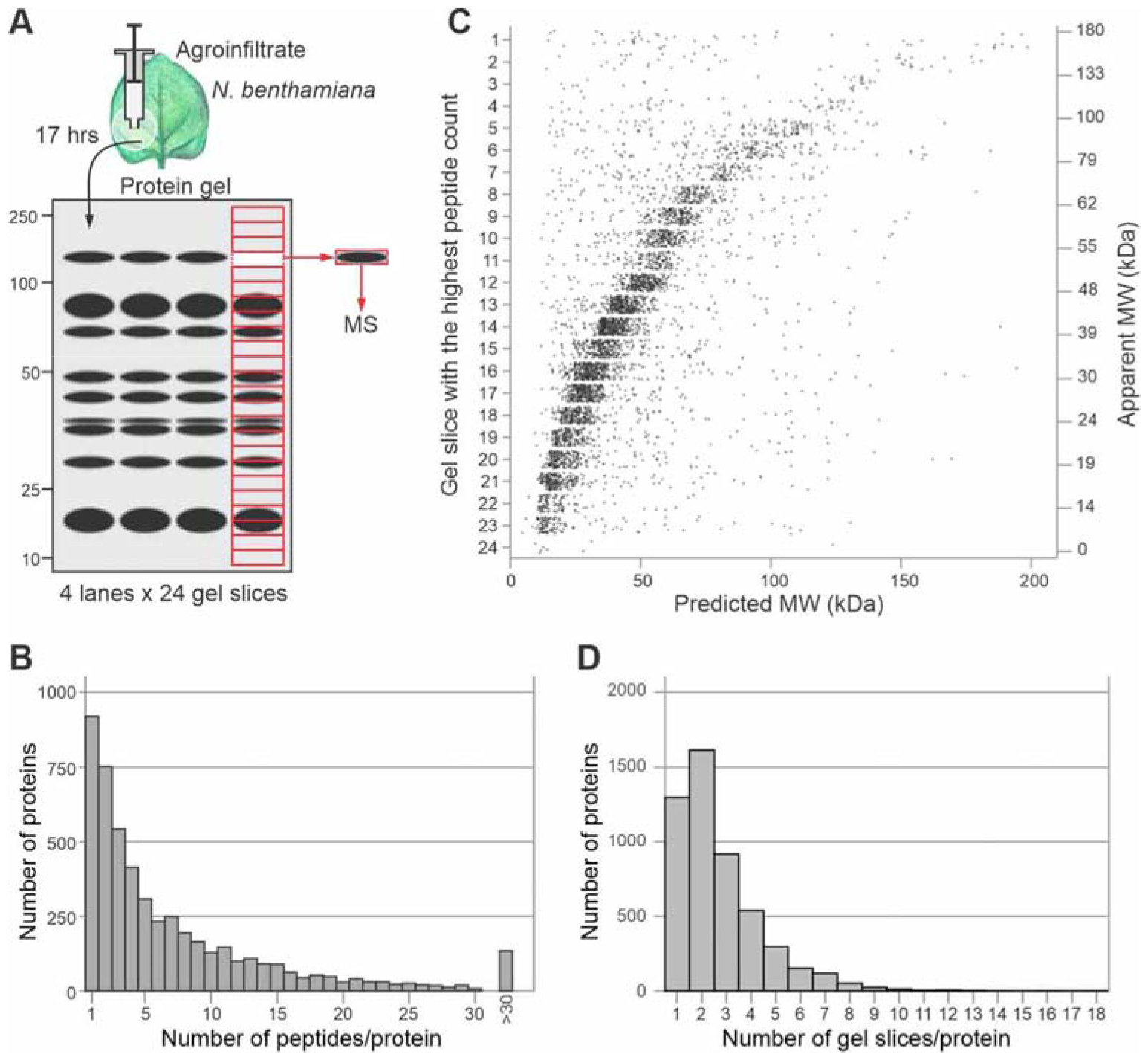
Detection of 5,040 plant proteins in agroinfiltrated *N. benthamiana* leaves. **(A)** Total extracts from agroinfiltrated *N. benthamiana* leaves were separated by SDS-PAGE and the lane was cut into 24 slices. Proteins in each gel slice were digested with trypsin and analysed by LC-MS/MS. **(B)** Number of peptides per protein. Spectra were mapped to the LAB3.60 annotation of the *N. benthamiana* proteome and the number of unique and non-unique peptides were plotted for each protein. **(C)** Correlation between predicted and apparent MW. For each protein, the apparent MW was plotted against the predicted MW. The predicted MW was calculated after subtracting the putative signal peptide. **(D)** Number of gel slices in which each protein was detected.

We next plotted the predicted MW on the x-axis against the gel slice with the highest number of peptides (y-axis) for each protein. Since we are interested in secreted proteins, we also calculated the predicted MW of the protein without its SignalP-predicted signal peptide (SP, Teufel et al., 2022). This graph shows a linear correlation between the predicted and apparent MW (r = 0.399, p = 2.2e^-16^) (**Figure 1C**). Interestingly, there are also 1,052 proteins from which most peptides were found at MW 10% below the predicted MW. Likewise, 1,059 proteins have the highest peptide count at MW 10% above the predicted MW. Another 59 proteins have their highest peptide count in slice 1 (top slice), which is either caused by protein insolubility or the protein being too large to migrate into the polyacrylamide gel.

Most proteins (74.3%) are not detected in a single slice (**Figure 1D**). For 79.5% of these 3,744 proteins (2,977 proteins), peptides from the same protein were detected in adjacent gel slices, which is presumably caused by diffuse protein signals within the protein gel. For 3,011 proteins, however, peptides were also detected at two or more gel slices distant from the gel slice with the main count, indicating that we are detecting multiple proteoforms for 59.7% of the 5,040 detected proteins.

### Building an upgraded PROTOMAP dataset

To map the distributions of the peptides over the detected protein (x-axis) and the gel slice (y-axis), we produced an R script that generates peptographs for each protein, similar to how PROTOMAP was built originally (Dix et al., 2008), but with several important upgrades (**Figure 2**). First, we highlight all unique peptides with red edges. Second, we depict in how many of the four replicates the peptide was detected in the thickness of the peptide box to signify robustly detected peptides. The peptides are also depicted translucent to show partially overlapping peptides. Third, we depict the average signal intensity of each peptide using the greyscale to highlight peptides detected at higher ion intensities. Fourth, we added horizontal lines to indicate the gel slice in which the protein is expected to accumulate, based on the predicted molecular weight for the protein without SignalP-predicted signal peptide and predicted *N*-glycans (red horizontal line) or with predicted *N*-glycans (blue horizontal line), based on the presence of sequons (NxS/T, +3 kDa each). *O*-glycosylation was not considered as it is less common and less predictable when compared to *N*-glycosylation. Fifth, instead of the average spectral counts (Dix et al., 2008), we plotted the total peptide count for all four replicates for each gel slice in the graph on the right to indicate gel slices containing the strongest protein signals. Finally, we highlight the position of putative *N*-glycosylation sites (NxS/T sequons) in the protein, together with InterPro-predicted PFAM domains (Mistry et al., 2021), SignalP-predicted signal peptides (Teufel et al., 2022), and TMHMM-predicted transmembrane domains (Krogh et al., 2001). And as with the original PROTOMAP, we provide the annotation of the putative protein function (top), an explanation of the PFAM domains, and the protein sequence (bottom). These peptographs have been generated for every detected protein in a PDF format that can be searched for accession numbers, PFAM domains, keywords and sequences (Supplemental **File S2**). The R script to produce these peptographs is provided as Supplemental **File S3**.

**Figure 2.**
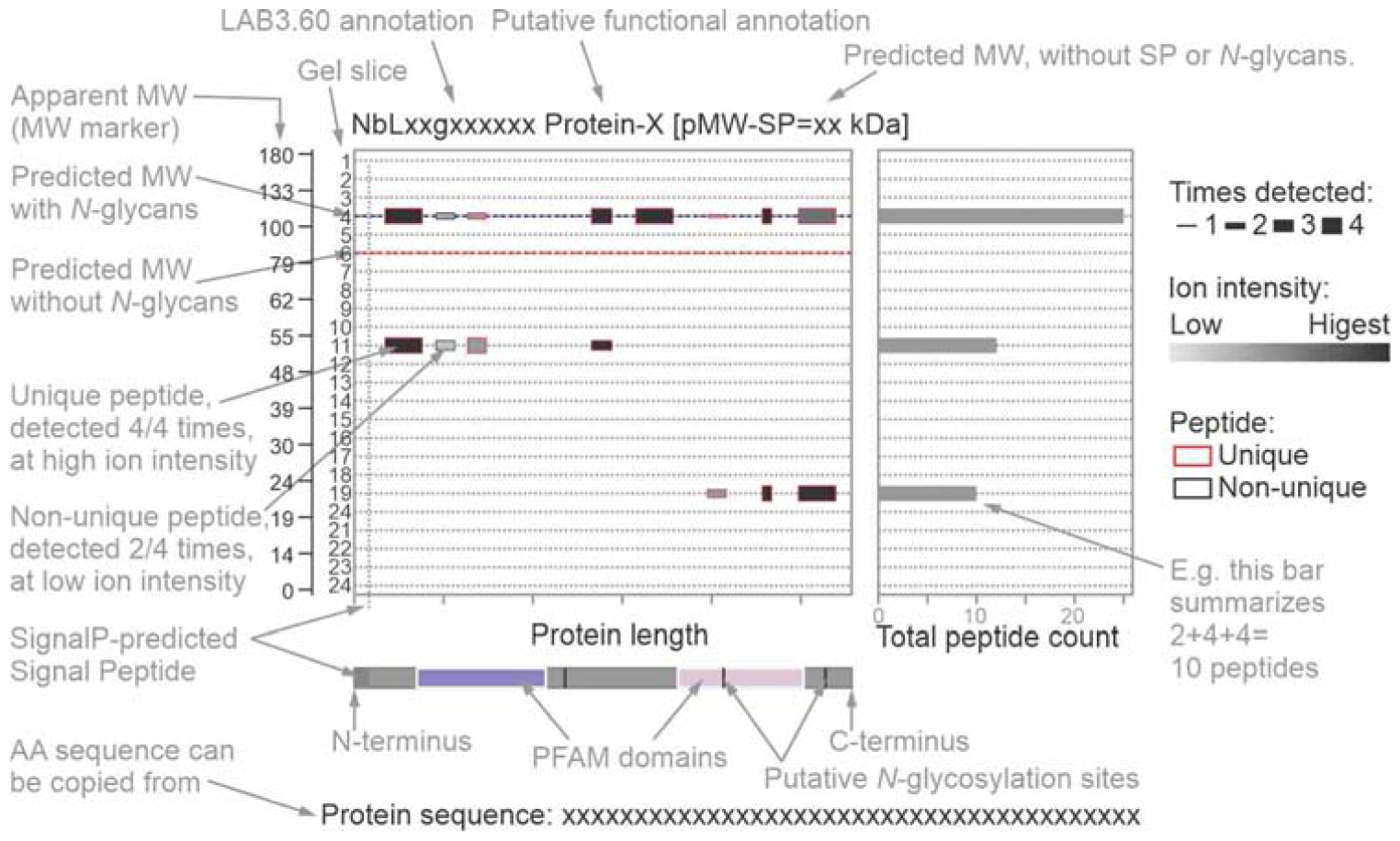
Peptographs display topology and migration for each protein. For each protein the detected peptides were mapped onto the protein (x-axis) for each gel slide (y-axis). The number of peptides detected in a gel slice is summarized in the bar graph on the left. The apparent MW marker estimated from the protein gel is added to the left. The relative ion intensities of the peptides are indicated in grey scale and the number of times of the 4 replicates they were detected is indicated by the height of the peptide box. Unique peptides have a red outline. The red dash line indicated the predicted MW without SignalP-predicted signal peptide or predicted *N*-glycans; the blue dash line indicates the MW without SP but with *N*-glycans, based on the presence of sequons (NxS/T). The header on the top summarizes the protein identifier, putative function, and predicted molecular weight without SP and without predicted *N*-glycans. The bar on the bottom represents the protein from N- to C-terminus, with the PFAM domains, putative SP and transmembrane (TM) domain, and the position of the putative *N*-glycosylation sites (NxS/T). The protein sequence is given at the bottom of the peptograph to be copied for further analysis.

### Secreted proteases accumulate as mature enzymes lacking autoinhibitory domains

To investigate if the PROTOMAP dataset displays known protein processing events, we mined the dataset for secreted proteases that carry an autoinhibitory prodomain that is removed upon activation, resulting in a mature protease with a lower MW. Using searches for the PFAM code of the catalytic domain for proteins detected with more than a single peptide, we identified 24 subtilisin-like proteases (SBTs, family S08, PF00082); 16 papain-like cysteine proteases (PLCPs, family C01, PF00112); 14 pepsin-like aspartic proteases (family A01, PF14541); and three vacuolar processing enzymes (VPEs, family C13, PF01650). Most of these proteases accumulate at a MW that is predicted for the mature proteases and not their proenzymes (**Figure 3A**), indicating that these enzymes do no longer carry their autoinhibitory prodomain. The position of the identified peptides within the protease precursors confirms this pattern: most peptides originate from the protease domains (**Figure 3B**). The absence of peptides from the prodomain in any gel slice indicates that these domains are degraded *in vivo*, presumably to avoid inhibition *in trans*.

**Figure 3.**
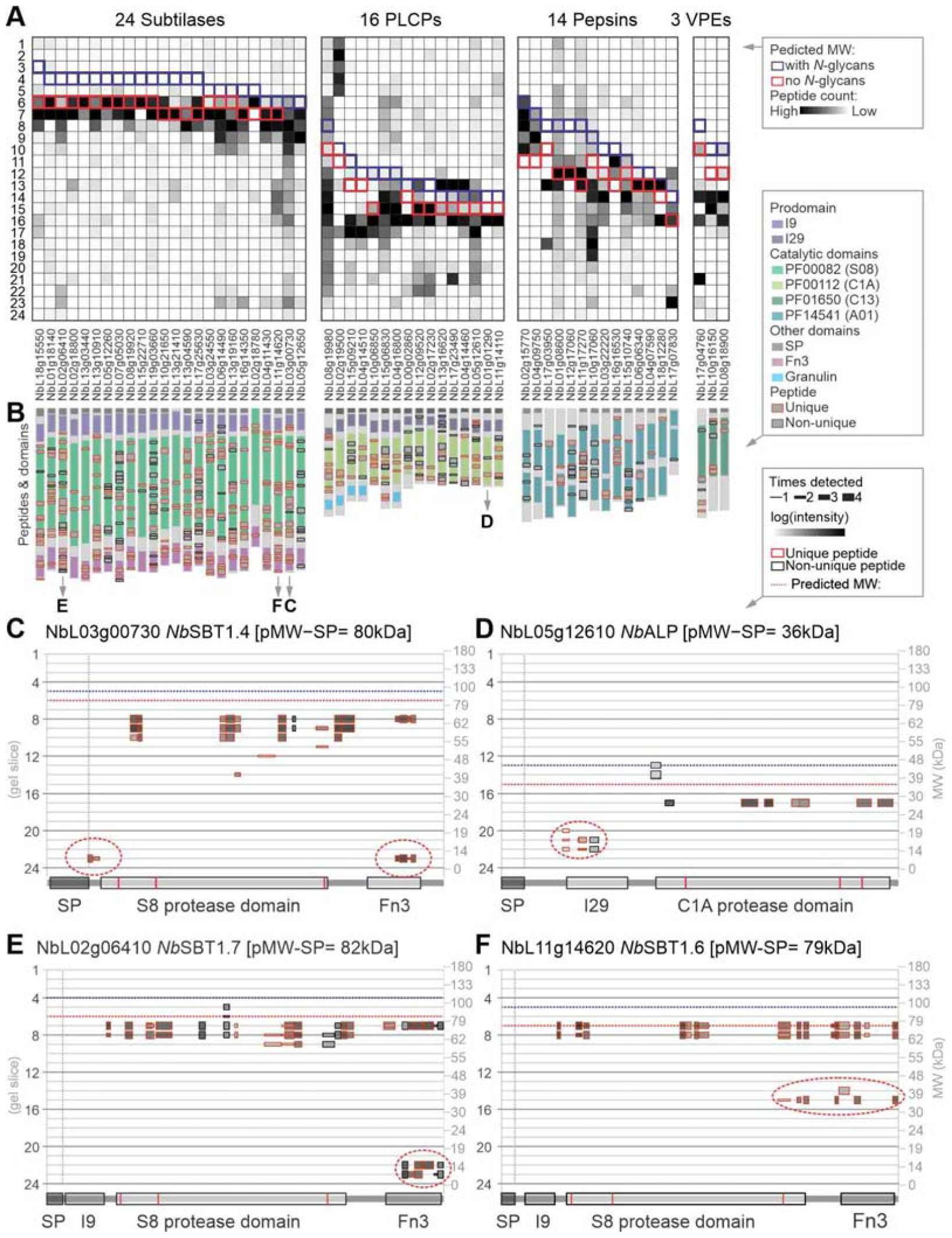
PROTOMAP displays migration and processing of mature proteases. **(A)** Peptographs of proteases with known autoinhibitory prodomains were extracted using PFAM families PF00082 (S08 proteases/subtilases); PF00112 (C01 proteases/papains); PF14541 (A01 Asp proteases/pepsins); and PF01650 (C13 legumains/VPEs). The peptides were counted per gel slice and presented as a heat map illustrating the distribution of the peptides. The outlines indicate the gel slice where the full-length pro-protease is predicted to migrate without SP but with (blue) or without (red) predicted *N*-glycans. Proteases were ranked on their predicted MW. *Nb*TPP2 (NbL03g11790) was omitted from this analysis because it is a distinct cytoplasmic subtilase that lacks an autoinhibitory prodomain, and NbL04g13700 was omitted because it was detected by a single peptide. **(B)** Most peptides of detected proteases are not from the autoinhibitory prodomains. The position of all detected peptides (grey) is shown on top of the protein architectures, with the outline indicating unique (red) and non-unique (black) peptides. **(C-F)** Example peptographs revealing proteoforms of two subtilases. Peptograph features are explained in Figure 2. Catalytic residues are highlighted red within the catalytic domains.

Notable exceptions are subtilases NbL03g00730 (*Nb*SBT1.4, **Figure 3C**), NbL17g25630, NbL06g14490 and NbL16g14350, for which peptides were detected of the prodomain, but these peptides were only detected at 15 kDa (Supplemental **File S2**), indicating that these prodomains have been removed by processing but that they persist. Likewise, peptides for the prodomain were also detected at 15 kDa for PLCP NbL05g12610, an aleurain-like protease (*Nb*ALP, **Figure 3D**), suggesting that also this prodomain persists after release. By contrast, peptides of the prodomain were also detected for PLCPs NbL12g09520, NbL13g16620 and NbL04g14460 but these peptides all migrate at the predicted MW of these proteins (Supplemental **File S2**), indicating that they are from precursor proteases. Another exception is the detection of peptides from the C-terminal inhibitory domain of VPE NbL17g4760, which were detected both at the predicted MW and at 20 kDa (Supplemental **File S2**), indicating that only part of this protease is processed but that the inhibitory domain can persist separately.

Interestingly, peptograph of subtilase NbL02g06410 (*Nb*SBT1.7, **Figure 3E**), also displays C-terminal peptides corresponding to the fibronectin type-III domain (PF17766) at 12 kDa, indicating that this domain is released from these subtilases. Released Fn3 domains were also detected for subtilases NbL16g14350, NbL06g14490, NbL03g00730 and NbL13g19160. By contrast, the peptograph of subtilase NbL11g14620 (*Nb*SBT1.6, **Figure 3F**) displays multiple C-terminal peptides at 35 kDa, indicating that this subtilase is processed more upstream in the protease domain, severing the catalytic Ser from the catalytic Asp and His residues (**Figure 3F**). It is unclear at this stage what the consequences of these cleavages are. One distinct subtilase not included in Figure 3 is tripeptidyl peptidase (*Nb*TPP2, NbL03g11790), which was identified at 150 kDa with >250 peptides covering its entirety (Supplemental **File S2**), consistent with the high abundance and high predicted MW (147 kDa) of this cytoplasmic subtilase.

### SCPL proteases/acyltransferases are differentially processed into functional heterodimers

The PROTOMAP dataset contains peptographs of 18 Serine Carboxypeptidase-Like (SCPL) proteins with more than a single peptide, carrying catalytic domain PF00450. SCPLs can be carboxypeptidases or acyltransferases involved in secondary metabolism (Mikowski and Strack, 2004). The peptograph of *Nb*SCPL20a (NbL09g13440, see phylogeny in Supplemental **Figure S4**) shows that this 51 kDa enzyme is cleaved into an N-terminal 37 kDa fragment in slice 15, and a C-terminal 19 kDa fragment in slice 20 (**Figure 4A**). Peptographs of 11 additional SCPLs also indicate that they accumulate as cleaved proteins (Supplemental **File S2**; NbL01g02560, NbL02g23560, NbL06g01240, NbL07g00400, NbL07g17710, NbL09g11980, NbL09g13440, NbL13g00950, NbL13g01230, NbL14g14490, NbL16g05780 and NbL18g14840). These data are consistent with processing of SCPLs into a large and small subunit that function together as a disulfide-linked heterodimer. Oat SCPL1, for instance, is cleaved into 29 and 19 kDa subunits that stay together as a disulfide-linked heterodimer that mediate a step in the biosynthesis of antimicrobial avenacins (Mugford et al., 2009). Likewise, brassinosteroid insensitive-1 suppressor BRS1 of Arabidopsis is processed into 34 and 19 kDa subunits, resulting in a heterodimer acting as secreted carboxypeptidase (Zhou and Li, 2005). It is unclear at this stage if SCPL processing is required for their activity and which endopeptidase cleaves these SCPLs.

**Figure 4.**
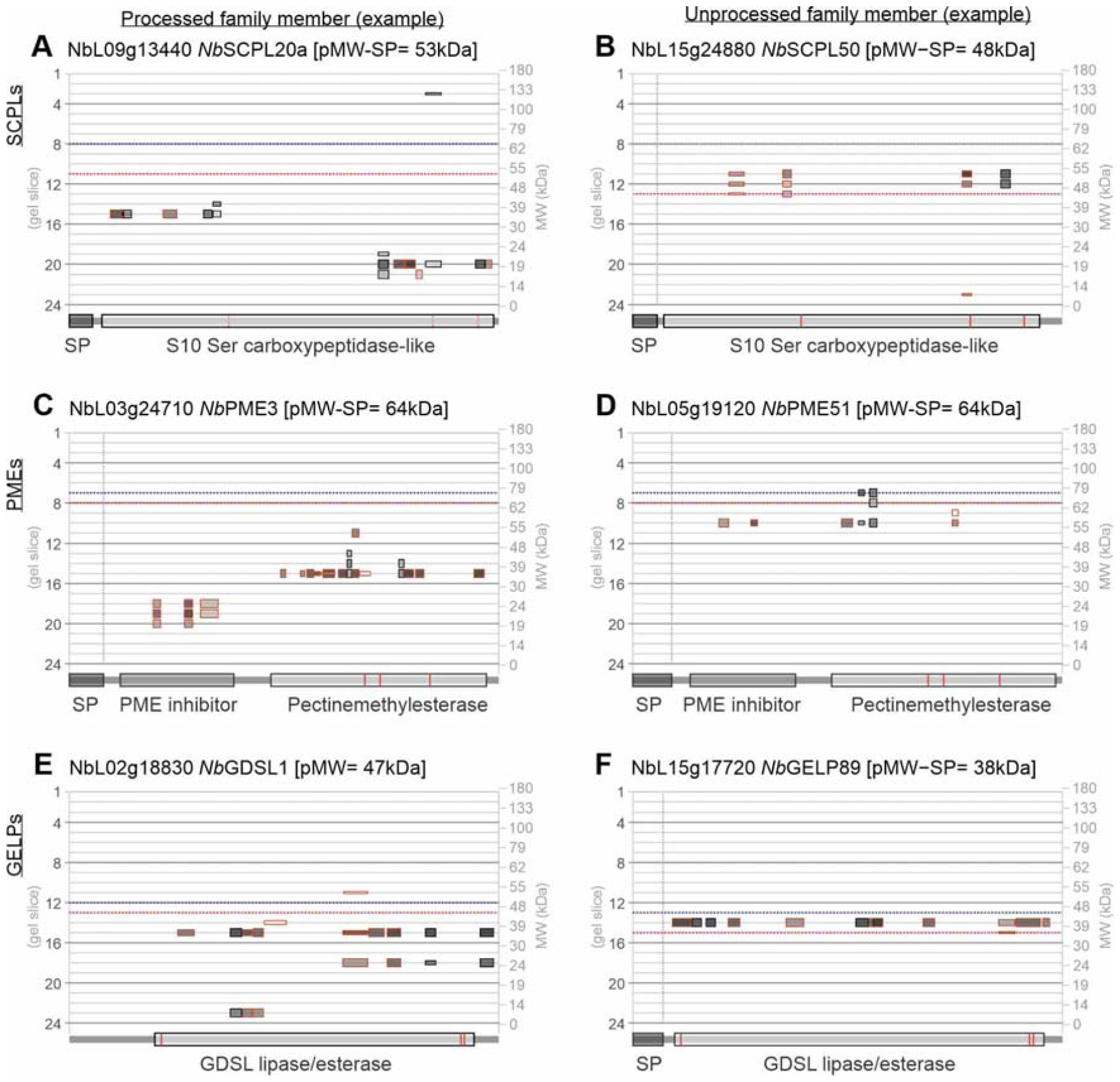
Differential processing of SCPLs, PMEs and GELPs. Examples of processed **(A, C, E)** and unprocessed **(B, D, F)** precursors of Ser carboxypeptidase-like proteins (SCPLs), Pectinemethylesterases (PMEs) and GDSL-type lipase/esterase-like proteins (GELPs). Catalytic residues are highlighted red within the catalytic domains. Peptograph features are explained in Figure 2. Catalytic residues are highlighted red within the catalytic domains.

By contrast, peptides from *Nb*SCPL50 (NbL15g24880, see phylogeny in Supplemental **Figure S4**) are mostly in gel slice 11 at 50 kDa, consistent with an uncleaved predicted MW of 48 kDa (**Figure 4B**). Peptographs of five additional SCPLs show peptides mostly at their predicted MW, indicating that these SCPLs are not processed (Supplemental **File S2**; NbL01g18230, NbL02g04500, NbL04g07300, NbL05g22050, NbL07g00690, NbL12g02780, NbL14g17630 and NbL15g24880). The differential processing of SCPLs is thought to be caused by the length and sequence of the linker peptide located between the two subunits in the SCPL sequence (Mugford et al., 2009). Shorter linker peptides are generally not processed. Indeed, alignment of the 18 detected SCPLs, ranked on the length of the linker region reveals that uncleaved SCPLs have a short linker region (4-15 residues), in contrast to cleaved SCPLs that have longer linker regions (32-65 residues, Supplemental **Figure S5**). The alignment also shows that these cleaved SCPLs have highly polymorphic linker sequences, although many carry dibasic motifs that might be targeted by subtilases that have furin-like activities (Supplemental **Figure S5**).

### PMEs are differentially processed to remove the inhibitor domain

The PROTOMAP dataset contains 12 type-I pectinemethylesterases (PMEs) with two or more peptides. Type-I PMEs consist of a signal peptide for secretion, followed by a 20 kDa inhibitory domain (PF04043) and the 35 kDa PME domain (PF01095). Eight of the detected PMEs migrate in protein gels as processed, mature enzymes at 35-45 kDa. *Nb*PME3 (NbL03g24710, see phylogeny in Supplemental **Figure S6**), for instance, shows peptides of the catalytic domain at 40 kDa and peptides of the inhibitory domain at 20 kDa (**Figure 4C**). By contrast to the eight processed PMEs, the other four PMEs are detected close to the predicted MW. For instance, peptides from both the inhibitory and catalytic domains of *Nb*PME51 (NbL05g19120), are detected at 60 kDa, close to the predicted MW of 62 kDa, confirming that this PME accumulates as an uncleaved precursor (**Figure 4D**).

The removal of the inhibitory domain is required to activate and secrete PMEs, which then regulate the direction of cell growth by targeted cell wall relaxation (Wolf et al., 2009;). A dibasic motif (e.g. RRLL) at the junction between the inhibitory and catalytic domains is required for cleavage (Wolf et al., 2009) and Arabidopsis SBT3.3 was found to be responsible for processing PME17 (Senechal et al., 2014). Interestingly, all eight processed PMEs detected by PROTOMAP contain either RRLL (4x), RKLL (2x), RVLL or RTLL at the putative cleavage region, whilst four unprocessed PMEs lack basic residues in this region with the exception of *Nb*PME40 (NbL01g19610), which carries RALL (Supplemental **Figure S7**).

### GELPs are differentially processed, separating its catalytic triad

The PROTOMAP dataset contains 33 GDSL esterases/lipases (GELPs, PF00657, see phylogeny in Supplemental **Figure S8**), with two or more peptides. GELPs are lipolytic enzymes that contain a 35 kDa catalytic domain that contain the catalytic Ser in an N-terminal GDSL motif (Shen et al., 2002). Peptographs of eight GELPs (e.g. NbL02g18830, *Nb*GDSL1, **Figure 4E**) show additional peptides at 25 kDa from the C-terminal half and peptides at 13 kDa from the N-terminal half of the catalytic domain. This indicates that these GELPs are processed into two subunits that are stable, possibly by remaining in a heterodimeric state. Interestingly, such processing would separate the catalytic triad, which consists of the catalytic Ser residue in the N-terminus, and the catalytic Asp and His residues in the C-terminus (**Figure 4E**). Although several plant GELPs have been biochemically characterized (e.g. Zhang et al., 2017; Tang et al., 2020), their processing *in planta* has not been reported before. By contrast, the other 25 detected GELPs only show peptides at the predicted MW from the entire catalytic domain (e.g. NbL15g17720, *Nb*GELP89, **Figure 4F**), indicating that they are not processed. The mechanisms and consequences underlying this differential processing remain to be identified.

### Glycosidases are often cleaved from their C-terminal domains

The PROTOMAP dataset contains peptographs of 96 glycosyl hydrolases (GHs, also called glycosidases) that mostly act in the endomembrane system and apoplast to modify the glycans in the cell wall, membrane surface and proteins (Leonard et al., 2009). Many GHs carry carbohydrate binding motifs (CBM) that enable substrate selection and localization. Interestingly, the peptographs of GHs often indicate that the CBM or other C-terminal domains is processed and can be detected at a low molecular weight, indicating that it is not degraded. β-galactosidase-1 (*Nb*BGAL1, NbL07g00850, family GH35) for instance, carries a galactose-binding lectin domain (PF02140) at the C-terminus, which is detected at 15, 28 and 40 kDa in the *Nb*BGAL1 peptograph (**Figure 5A**). Peptides from the catalytic domain (PF01301) of *Nb*BGAL1 migrate at the predicted MW (90 kDa) and at 48 kDa, consistent with activity-based labeling of this proteoform in our previous experiments (Chandrasekar et al., 2016; Buscaill et al., 2019). The four other detected BGALs of family GH35 (NbL03g01030, NbL02g14360, NbL13g19810 and NbL03g00970), all have similar peptographs, indicating that the processing of the C-terminal lectin domain is common for the GH35 family.

**Figure 5.**
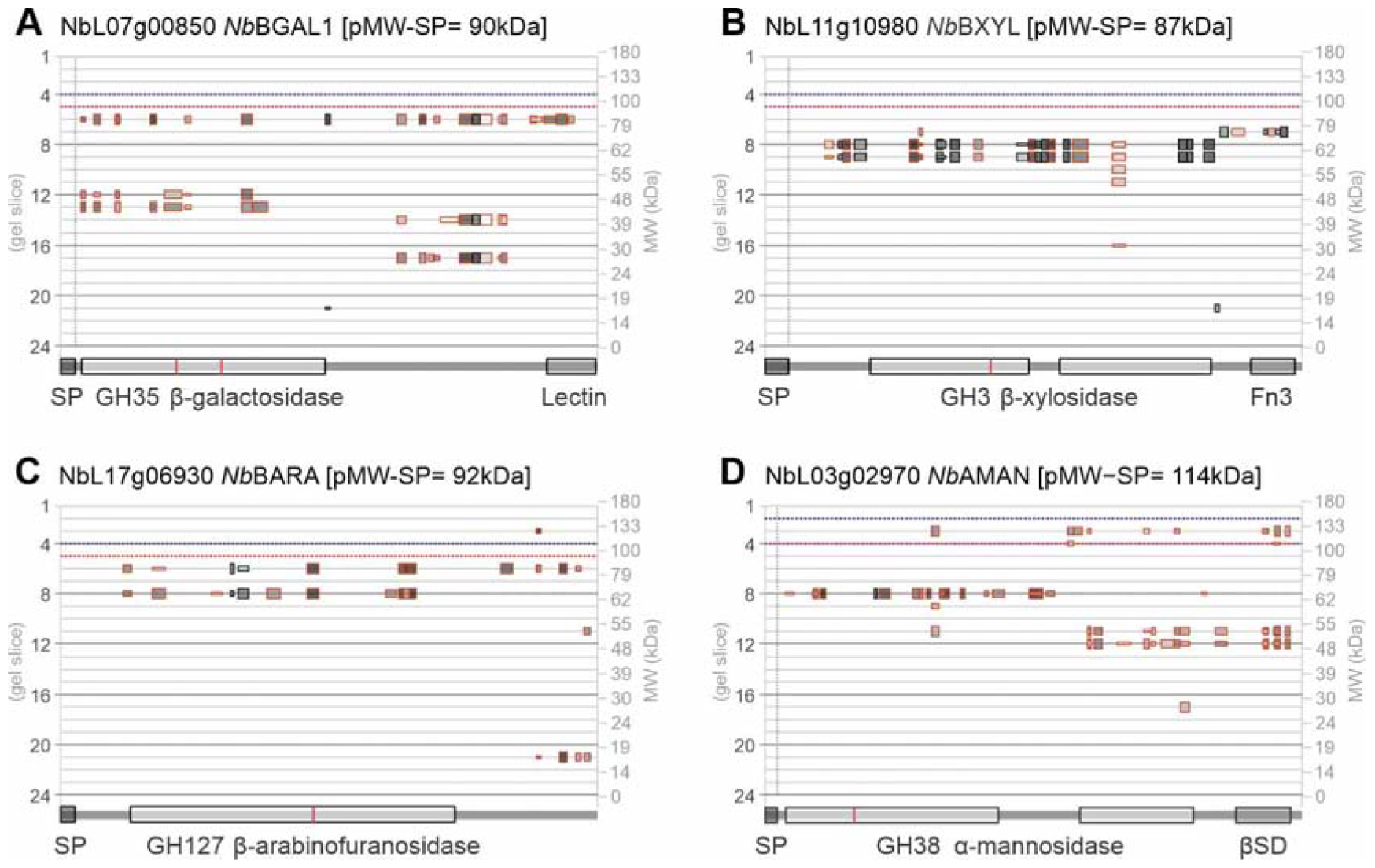
Fascinating processing of various glycosyl hydrolases (glycosidases). Shown peptographs are from: **(A)** β-galactosidase-1 *Nb*BGAL1; **(B)** putative β-xylosidase *Nb*BXYL; **(C)** putative β-arabinofuranosidase *Nb*BARA; and **(D)** putative α-mannosidase *Nb*AMAN. Peptograph features are explained in Figure 2. Catalytic residues are highlighted red within the catalytic domains.

Likewise, putative β-xylosidase (*Nb*BXYL, NbL11g10980, family GH3) carries a C-terminal fibronectin III-like domain (Fn3, PF14310) that can be detected at 75 kDa as part of the full-length protein (**Figure 5B**). In addition to the full-length protein, peptides of the catalytic domain are also detected at 60 kDa but this gel slice lacks peptides of the Fn3 domain, indicating that this proteoform is C-terminally truncated. Although no peptides of the Fn3 domain can be detected for the four other GH3-family β-xylosidase carrying an Fn3 domain (NbL11g20540, NbL04g10020, NbL15g17830, and NbL01g14480), they all show peptides from only the catalytic domain migrating at 60-65 kDa, below the predicted MW, indicating that the C-terminal Fn3 domain has been removed from all these GH3 β-xylosidases. The function of the Fn3 domain in GH3 β-xylosidase is unknown but it might be involved in anchoring the enzyme on large polymeric substrates and in thermostability (Pozzo et al., 2010). Interestingly, we detected the processing of a different Fn3 domain (PF17766) from the C-terminus of three subtilases (see subtilase section).

Likewise, putative β-arabinofuranosidase NbL17g06930 (*Nb*BARA, family GH127), is detected at 88 kDa, close to its predicted MW of 92 kDa, but the catalytic GH127 domain also migrates at 65 kDa. This shorter proteoform lacks peptides from the C-terminal region, whilst this C-terminal region is detected at 15 kDa (**Figure 5C**). This indicates that the C-terminal region is removed from this GH127 glycosidase. The C-terminal region has homology to two β-sandwich domains that are important for homodimerization of a bacterial GH127 enzyme (Ito et al., 2014), but it is unknown how this processing might influence GH127 function.

Several other glycosidases show interesting processing events. The peptograph of heparanase-like NbL13g02440 (*Nb*HEPA, family GH79) for instance, has peptides covering the most of the protein at 58 kDa, consistent with its predicted MW (56 kDa), but also peptides of the catalytic domain (PF03662) at 35 kDa and even a smaller fragment at 15 kDa (Supplemental **File S2**), indicating that this glycosidase is processed. In addition, the peptograph of putative α-mannosidase NbL03g02970 (*Nb*AMAN, family GH38) shows peptides across the protein at 130 kDa, close to its predicted MW (114 kDa), but much more and intense peptides of the N-terminal GH38 domain (PF01074) at 65 kDa and peptides of the C-terminal GH38 domain (PF07748) at 50 kDa (**Figure 5D**), indicating that most of this hydrolase is cleaved into two subunits. This peptograph is consistent with the observation that a homologous GH38 α-mannosidase from tomato fruits contains 70 and 47 kDa subunits (Hossain et al., 2009).

### Transient expression of double-tagged precursors confirms processing in vivo

To confirm the existence of proteoforms of secreted proteins in vivo, we transiently expressed several extracellular proteins with suspected processing by agroinfiltration with tags on both sides of the protein (**Figure 6A**). We added an N-terminal RFP and a C-terminal GFP protein to detect expression before protein extraction and to ensure that also processing near the N- and C-terminus can be detected by western blots. We also added an N-terminal FLAG tag and C-terminal His tag because we found that antibodies against these tags are more sensitive than anti-RFP/GFP antibodies. The double-tagged protein was fused behind the signal peptide of PR1a to ensure entry into the secretory pathway. These double tagged constructs were driven by the 2x35S promotor and contain an intron to exclude bacterial expression.

**Figure 6.**
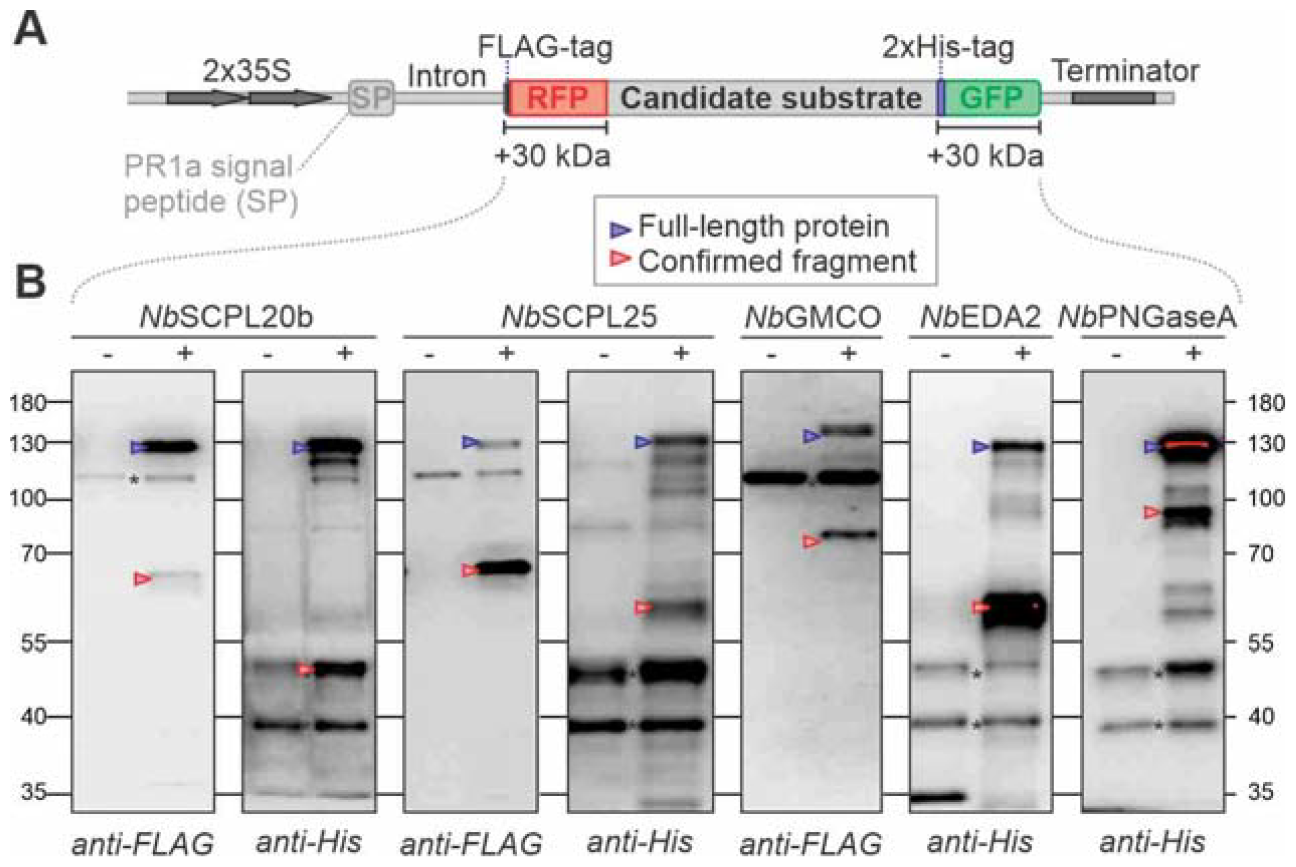
Several processing events are confirmed upon transient expression. **(A)** The coding region of candidate proteins without its own signal peptide were cloned in-between sequences encoding SP-FLAG-RFP and 2xHis-GFP. These coding sequences are located on the T-DNA of binary vectors and are driven by a 2x35S promoter and terminated by a 35S terminator and carry an intron to avoid protein expression by bacteria. **(B)** Western blots of double tagged proteins confirm several processing events *in vivo*. The double tagged candidate proteins were transiently expressed by agroinfiltration in *N. benthamiana* and extracted in gel loading buffer for western blot analysis with anti-FLAG and anti-His antibodies to detect the N- and C-termini of the fusion proteins, respectively. *, background signal, also seen in the empty vector control.

We first selected two SCPLs that are processed according to the PROTOMAP dataset. Peptides of *Nb*SCPL20b (NbL09g13440, see phylogeny in Supplemental **Figure S4**) were detected at 37 kDa (N-terminal part) and 20 kDa (C-terminal part) (Supplemental **Figure S9A**). Both fragments were confirmed with double tagged SCPL20b, with tagged fragments accumulating at 65 and 45 kDa, respectively (**Figure 6B**). Likewise, the peptograph of *Nb*SCPL25 (NbL02g23560) displayed peptides at 40, 30, and 25 kDa, indicating full-length, N- and C-terminal fragments, respectively (Supplemental **Figure S9B**). Western blot analyses of double-tagged SCPL25 confirmed the accumulation of both the tagged N- and C-terminal fragments, accumulating at 65 and 55 kDa, respectively (**Figure 6B**). These data are consistent with the processing of SCPLs as explained above.

The peptograph of glucose-methanol-choline oxidoreductase (*Nb*GMCO, NbL10g21430) shows peptides at 64 kDa, representing the predicted full-length protein lacking its signal peptide, but also at 50 kDa, with peptides from the N-terminal half, and at 28 kDa with peptides from the C-terminal half (Supplemental **Figure S9C**). This indicates that *Nb*GMCO is cleaved between the N-terminal FAD-binding domain and the substrate binding domain. To confirm this observation, we double-tagged GMCO and monitored its accumulation upon transient expression by agroinfiltration. Western blot analysis confirmed the accumulation of the N-terminal processing product, accumulating at 75 kDa (**Figure 6B**), but the C-terminal processing product could not be detected, possibly because of another processing event at the junction with the 2xHis-GFP reporter tag.

The peptograph of the putative serine protease *Nb*EDA2 (NbL06g09140, S28 family, PF05577) shows most peptides at 53 kDa, consistent with its predicted MW (52 kDa), but also shows N-terminal peptides at 28 kDa and C-terminal peptides at 25 kDa (Supplemental **Figure S9D**). Western blot analysis of transiently expressed double-tagged *Nb*EDA2 confirmed the accumulation of the His-GFP-tagged C-terminal fragment at 60 kDa (**Figure 6B**), whereas the N-terminal fragment was not detected. Interestingly, this cleavage would separate the catalytic triad since the catalytic Ser residues locates in the N-terminal half, and the catalytic Asp and His residues at the C-terminus. However, as with SCPLs, this cleavage might result in a functional heterodimer.

Finally, the peptograph of putative Golgi-localized peptide-N4-(N-acetyl-beta-glucosaminyl) asparagine amidase A (*Nb*PNGaseA, NbL17g03090), shows peptides of the majority of the protein at 58 kDa, which might be the full-length protein (62 kDa), and both C- and N-terminal peptides at 45 kDa, indicating that this protein is processed (Supplemental **Figure S9E**). Western blot analysis of transiently expressed double tagged *Nb*PNGaseA confirmed the accumulation of a His-GFP-tagged C-terminal fragment at 80 kDa, consistent with the peptograph (**Figure 6B**). Taken together, these western blot experiments confirmed that the cleavages observed in the peptographs occur *in vivo* and can be detected by transient expression of double-tagged precursors proteins.

### Detection of extracellular domains of RLKs indicates ectodomain shedding

Interestingly, the PROTOMAP dataset also contains peptides of receptor-like kinases (RLKs). This was unexpected because the sample preparation included the removal of insoluble material by centrifugation before boiling in lithium dodecyl sulfate. To investigate this further, we searched the data for RLKs using PFAM identifiers for the protein kinase domain (PF00069 and PF07714) preceded by a signal peptide and/or transmembrane domain. This search delivered 37 putative RLKs, each with at least two peptides, including likely orthologs of the brassinosteroid receptor BRI1/SR160 (Wang et al., 2018); chitin receptor CERK1 (Miya et al., 2007); RALF1 receptor FERONIA (Haruta et al., 2014); cell wall sensor HERK1 (Guo et al., 2009); BAK1-regulator BIR2 (Halter et al., 2014); and auxin co-receptor TMK1 (Friml et al., 2022).

Notably, most RLK-derived peptides were found in gel slices much below the gel slice that is predicted to contain full-length, *N*-glycosylated RLKs (**Figure 7A**), indicating that these RLKs are truncated. This truncation is consistently 50-70 kDa, irrespective of the type of RLK. Further inspection of the detected peptides revealed that nearly all RLK-derived peptides originate from ectodomains and only few peptides are from kinase domains, despite being a substantial part of the RLK protein (**Figure 7B**). Putative brassinosteroid receptor BRI1/SR160 (NbL15g00030) for instance, is detected by peptides exclusively from the ectodomain at 90 kDa, 100 kDa below its predicted MW when fully *N*-glycosylated (**Figure 7C**). Likewise, LRR-RLK NbL03g04950 is detected with only ectodomain peptides at 80 kDa, 77 kDa below its predicted MW when fully *N*-glycosylated (**Figure 7D**). Taken together, these data indicate that we detected a wide range of ectodomains that were cleaved from the membrane, releasing a soluble extracellular domain. The remaining TM-kinase domains probably remained in the membrane fraction that was pelleted during sample preparation. These data suggest that ectodomains of RLKs representing different RLK subfamilies are shed from the membrane.

**Figure 7.**
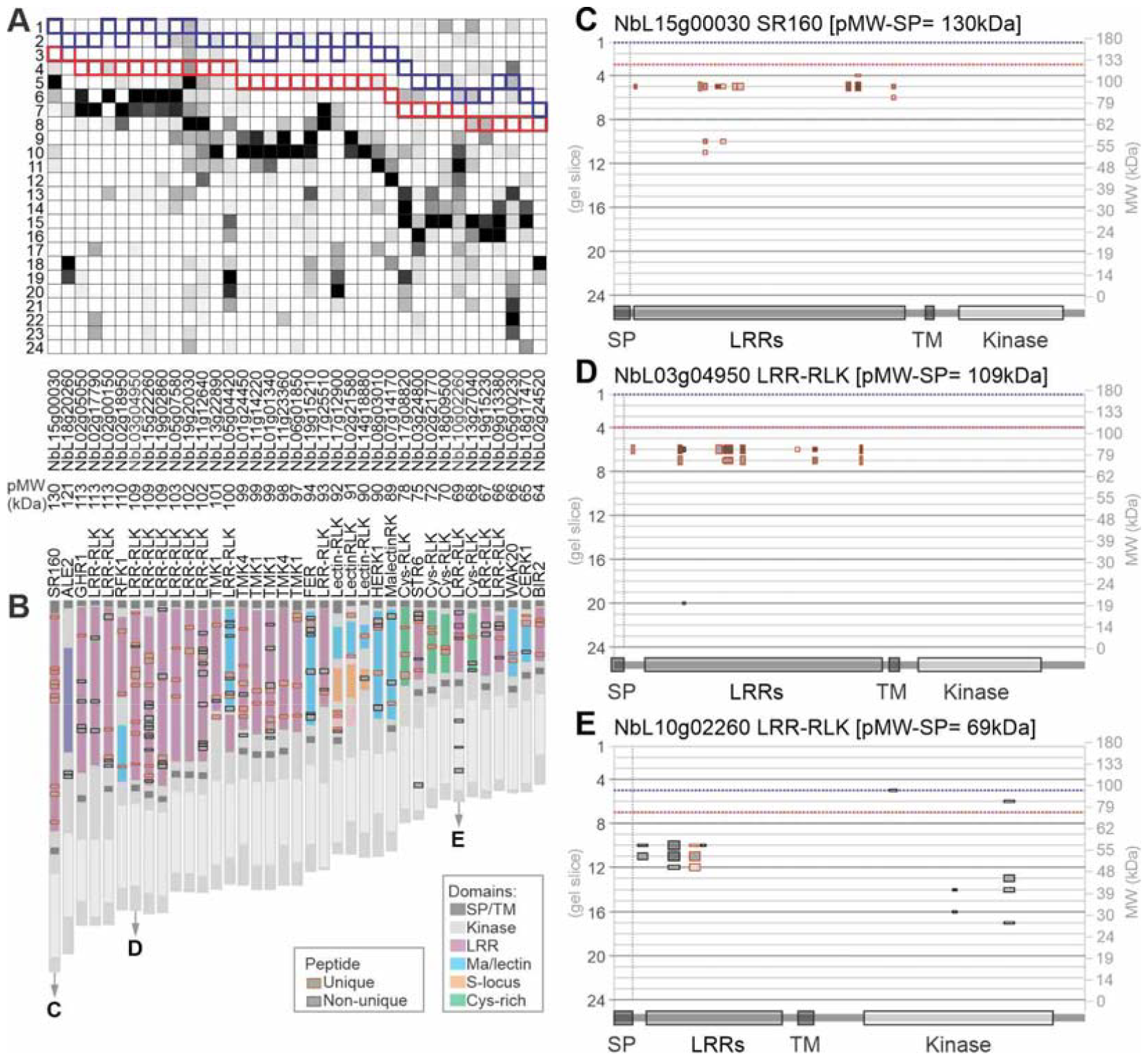
Detection of processed ectodomains suggest frequent receptor shedding in plants. **(A)** Peptides from RLKs migrate below their predicted MW. Peptides detected from RLKs are summarized for each gel slice, with highest coverage and ion intensity (black), medium ion intensity (dark grey) and low ion intensities (light grey). RLKs were extracted by selecting peptographs that contain protein kinase domains PF07714 or PF00069, preceded by a signal peptide (SP, not shown) and/or transmembrane domain (TM). RLKs identified by a single peptide were omitted. RLKs were ranked on their predicted MW. Outlines is the gel slice where the full-length RLK is predicted migrate without SP but including predicted *N*-glycans (blue) or without predicted *N*-glycans (blue). **(B)** Peptides of detected transmembrane receptors are almost exclusively from the ectodomains. The position of the detected peptides is shown as grey boxes on top of the domain architecture as unique (red outline) or non-unique (black outline). **(C-E)** Example peptographs of processed RLKs. Peptograph features are explained in Figure 2.

A notable exception is LRR-RLK NbL10g02260 (**Figure 7E**), which has a predicted MW of 90 kDa including *N*-glycans, but for which peptides derived from the kinase domain were detected at 40-44 kDa, corresponding to the predicted MW of the cytoplasmic domain. The peptograph also shows peptides of the ectodomain at 55 kDa, consistent with ectodomain shedding. These data suggest that this LRR-RLK is cleaved on both sides of the membrane, releasing both a soluble ectodomain, and a soluble kinase domain.

## Discussion

Our conceptually simple PROTOMAP experiment has uncovered 2,977 endogenous plant proteins that have multiple signals in protein gels. Whilst most are novel and implicate extensive protein processing in agroinfiltrated *N. benthamiana* leaves, these processing events are consistent with the literature. Nine processing events were confirmed by western blot analysis by transiently expressing double-tagged proteins. The PROTOMAP dataset is an extensive source of putative endogenous protease substrates to unravel the proteolytic machinery of this protein production platform in the future.

The PROTOMAP dataset is a rich source of proteoforms in agroinfiltrated leaves, most of which have not yet been reported before. This PROTOMAP dataset outperforms more modern approaches to detect protein processing such as COFRADIC and TAILS for several reasons. First, although TAILS and COFRADIC will reveal cleavage sites, they do not display the molecular weight of the precursor, meaning that the identified neo-termini can also originate from further degradation products. Second, it is unclear from TAILS and COFRADIC datasets if the detected processing event is a minority or majority processing event. PROTOMAP, by contrast, displays the relative abundance of the cleaved products and precursor, forcing the focus on major cleavage events that cause a shift of the protein in the gel. Third, PROTOMAP displays many more putative cleavage events. Our own TAILS analysis on the exact same proteomes only revealed 688 cleavage events in 1038 protein groups, and 31% of these proteins were not even detected in the PROTOMAP dataset (Thomas, 2020). Unfortunately, the RAW datasets of these TAILS samples were not maintained and could not be annotated to LAB3.60 and included in this study. This study nevertheless indicates that this PROTOMAP dataset contains highly valuable protein processing and abundance information that could not have been generated through modern approaches that are sometimes presumed to be better. Although the cleavage sites remain to be revealed, this dataset will focus further research on major processing events that can easily be detected by a shift in protein gels.

The PROTOMAP dataset is an extensive source of proteoforms that can be explored further for protein processing and abundance in agroinfiltrated *N. benthamiana* leaves. We mostly focused our further analysis on secreted proteins and have highlighted several phenomena in this dataset, which are briefly discussed below.

### Zymogen maturation

PROTOMAP analysis uncovered numerous processing events that are consistent with the literature. For instance, the prodomain of proteases was found to be removed in our dataset for papains, subtilases, legumains and pepsins. It is interestingly to note that we did not detect peptides of most of the prodomains, indicating that they are degraded as soon as they are released from the precursor protein. The differential stability of different parts of secreted proteins indicates mechanisms in extracellular protein homeostasis that are not yet understood.

### Differential processing within protein families

We also detected differential processing of SCPLs, PMEs and GELPs. Processing of these proteins is consistent with the literature on family members. In SCPLs, cleavage occurs in a polymorphic linker region and cleaved proteins continue to function as heterodimers (Mugford et al., 2009). The PMEs require processing at a dibasic RKxL/RRxL-like motif to remove the autoinhibitory prodomain (Wolf et al., 2009). Processing of GELPs separate their catalytic triad but the cleavage site is unknown, and it is unknown if this affects the function of these enzymes.

### Processing of glycosidases

We also discovered that glycosidases are processed. We detected the separation from the C-terminal domain in three different glycosidase families, and we highlighted the processing of glycosidases of the GH79 and GH38 families. It is unclear how these processing events affect glycosidase function, but the removal of CBDs is likely to affect the localization and specificity of these enzymes.

### RLK processing

Interestingly, the PROTOMAP dataset contains peptides of 37 RLKs suggesting receptor processing. Processing leads to the regulation of the activity of receptors via activation or deactivation but also to the generation of proteoforms with novel functions (Lichtenthaler et al., 2018). In most cases, peptides from ectodomains but not of the kinase domains were found. These data are consistent with ectodomain shedding that has been reported widely in mammals (Lichtenthaler et al., 2018). In plants, by contrast, only few ectodomain shedding events have been reported: chitin receptor CERK1 in Arabidopsis is cleaved in the second LysM motif of its ectodomain generating a N-terminal 33 kDa fragment. The *cerk1-4* mutant harbors a point mutation that impedes the generation of this N-terminal shedding product and these mutants are still able to perceive chitin but show enhanced cell death upon biotic stress (Petutschnig et al., 2014). Likewise, processing in the extracellular malectin-like domain of SYMRK from *Lotus japonicus* generates a truncated version of the receptor that still contains an extracellular LRR domain but is less stable and more prompt to form a complex with the Nod factor receptor 5 (NFR5) thereby activating symbiotic development (Antolin-Llovera et al., 2014).

We also found one LRR-RLK (NbL10g02260) that not only shows putative ectodomain shedding, but also a kinase migrating at 40-44 kDa, indicating that this kinase is released from the membrane. Research on Notch receptors in mammals has shown that ectodomain shedding can induce subsequent processing and release of a cytosolic proteoform (De Strooper et al., 1999). In plants, the proteolytic generation of cytosolic proteoforms of RLKs has been reported in four cases (Xa21 (Park & Ronald, 2012), BAK1 (Zhou et al., 2019), TMK1 (Gu et al., 2022) and FERONIA (Chen et al., 2023)). But in none of these cases, an ectodomain shedding event as reported preceding cytosolic processing How processing of the detected RLKs regulates signaling in plants is an exciting topic for future investigations.

### Double-tagged processing reporters

We used western blot analysis of transiently expressed double-tagged proteins to confirm several of the observed cleavage events. We noticed that some processing events were not detected by western blot of double-tagged precursors. There are several reasons for this. First, all proteins also accumulated as full-length proteins, even though these were often not detected in the peptographs. This is probably caused by the continuous overproduction of these proteins, causing either an overdose when compared to the processing enzymes, or an accumulation of the precursor before their processing. Second, not all processing events were detected with double tagged vectors. This could be caused by additional processing events closer to the N- and C-termini resulting in the removal of the reporter tags. The bulky reporter tags may also have caused misprocessing or mislocalization of the tagged protein. We nevertheless were able to confirm processing of several proteins, testifying that these cleavages occur *in vivo* and enabling us to use these proteins in the future as reporters to identify the proteases responsible for these cleavage events.

### Further use of PROTOMAP data

Our PROTOMAP dataset can be mined further by searching the peptographs in the PDF file (Supplemental **Figure S2**) with identifiers, PFAM domains, sequences, and functional names. A PDF editor can be used to quickly extract all peptographs with shared PFAM codes, identifiers or functional names. Extra care should be taken with the interpretation of peptographs showing very few peptides, but these peptographs are usually consistent with peptographs of other members of the same protein family. It should be stressed that the different proteoforms in peptographs may also result from alternative splicing and alternative initiation of transcription and translation. These cases can be identified using alternative gene models and mapping RNA seq reads on genome sequences. In addition, our PROTOMAP dataset lacks information of cleavage sites because it only contains tryptic peptides. Our searches for half-tryptic peptides reduced the confidence threshold, which drastically reduced the number of tryptic peptides and did not deliver many reliable cleavage events.

In conclusion, this PROTOMAP dataset revealed many fascinating novel proteoforms of plant proteins. We discovered that 59.7% of the proteins exist in multiple proteoforms that run at different apparent MWs in protein gels and are presumably the result of protein processing by proteases *in vivo*. This includes prodomain removal, the separation of catalytic triads and receptor shedding. Processing is often consistent within protein families but there are also interesting exceptions. These data indicate extensive protein processing pathways in agroinfiltrated *N. benthamiana* and identified substrates that will facilitate future studies on protein processing in *N. benthamiana*.

## Materials and methods

### Plant growth and agroinfiltration

*Nicotiana benthamiana* (LAB strain) plants were grown in a growth cabinet under long day cycle with 16 hours of light (80 μmol/m^2^s), 22°C and 8 hours dark at 20°C. This experiment was originally designed to be the control to detect the earliest protein processing events during the hypersensitive response (HR) triggered by co-expressing potato *Rx* with the coat protein of PVX, which induces tissue collapse at 21 hours post agroinfiltration (21 hpa), but the chosen timepoint (17 hpa) was too early to detect HR-related processing or even to detect Rx or CP by LC-MS/MS (Supplemental **File S1**). The control samples used for this experiment consisted of a 1:1 mixture of frozen leaf powders from leaves expressing *Rx* with leaves expressing *CP*.

*Agrobacterium tumefaciens* C58C1 containing binary vectors with either *Rx* from *Solanum tuberosum* or Coat Protein from Potato Virus X (Bendahmane et al., 1999) were grown overnight in Luria-Bertani medium supplemented with 100 μg/ml rifampicin and kanamycin at 28°C with 220 rpm overnight. Cultures were pelleted and washed with infiltration buffer (10 mM MES, 10 mM MgCl_2_, pH 5.7). Cells were resuspended in infiltration buffer to final OD_600_ 0.5 and acetosyringone added to 100 μM. The top two fully expanded leaves of 4-5 weeks old plants were agroinfiltrated with either Rx or CP alone (non-HR samples). Leaves were harvested at 17 hours post infiltration and ground with liquid nitrogen. Ground frozen tissue from non-HR induced leaves agroinfiltrated with *Rx* or *CP* alone was mixed in a 1:1 ratio.

### Protein extraction and gel electrophoresis

Extraction buffer (100 mM HEPES pH7.5, 5 mM DTT, MS SAFE Protease and Phosphatase Inhibitor (Sigma)) and leaf powder were mixed in a 1:1 ratio (i.e. 300 μg powder in 300 μl buffer). Extract was sonicated (Diagenode Biorupter) at full power for 10 minutes at 4°C. The samples were cleared by centrifugation at 11,000 x *g* for 20 mins at 4°C and the retained supernatant centrifuged for a further five minutes. Protein concentration was determined using Bradford Assay and the cleared lysate was mixed with lithium dodecyl sulfate loading buffer (ThermoFisher) and DTT to a final concentration of 200 mM and then boiled at 90°C for 10 min. 50 μg of sample was then loaded onto an SDS PAGE pre-cast 4-12% Bis Tris gel (ThermoFisher) and separated with MOPS buffer (ThermoFisher). Each sample lane was sliced into 24 2.5 mm sections guided by a background grid (Supplemental **Figure S1**). The migration of protein ladder (Multicolor Broad Range Protein Ladder, Thermo Scientific) was used to estimate MWs of proteins in each gel slice (Supplemental **Figure S1**).

Gel slices were washed twice in HPLC grade water and then in 100 mM ammonium bicarbonate. Proteins were reduced by incubation with 10 mM TCEP at 62°C for 30 min and then alkylated in 55 mM iodoacetamide for 30 min in the dark. Gel slices were dehydrated by four washes in 50% acetonitrile (ACN) and a final wash in 100% ACN. Remaining solution was removed in a vacuum centrifuge (Eppendorf). Proteins were then digested with addition of 400 ng of trypsin in 25 mM ammonium bicarbonate and incubation overnight at 37°C. All solutions were prepared in 100 mM ammonium bicarbonate. Peptide containing solution was eluted by successive addition of 5% formic acid in ACN washes. Peptides were dried in a vacuum centrifuge (Eppendorf) and desalted on home-made C18 stage tips as previously described (Rappsilber et al., 2007). Peptides were resuspended in 0.1% (v/v) formic acid for loading onto the LC-MS/MS.

### Proteomic analysis

Samples were analysed with an Orbitrap Elite (Thermo) coupled to an Easy-nLC 1000 liquid chromatography (LC) system (Thermo) (Michalski et al., 2012). The LC was run in a single column mode with an analytical column of a fused silica capillary (75 μm × 22 cm) with an integrated PicoFrit emitter (New Objective) packed in-house with Reprosil-Pur 120 C18-AQ 1.9 μm resin (Dr. Maisch). The analytical column was encased by a column oven (PRSO-V1; Sonation) at 45°C and attached to a nanospray flex ion source (ThermoFisher). All solvents were of UPLC grade (Sigma-Aldrich). Peptides were directly loaded onto the analytical column with a maximum flow rate that would not exceed the set pressure limit of 980 bar (usually around 0.6-1.0 μl/min). Peptides were separated on the running a 70 min gradient of solvent A and solvent B (start with 7% B; gradient 7% to 35% B for 60 min; gradient 35% to 80% B for 5 min and 80% B for 5 min) at a flow rate of 300 nl/min.

Xcalibur software (version 2.2 SP1.48) was used for data acquisition with the mass spectrometer in positive ion mode. Precursor ion scanning was performed in the Orbitrap analyser (FTMS; Fourier Transform Mass Spectrometry) in the scan range of m/z 300-1500 and at a resolution of 60000 with the internal lock mass option turned on (lock mass was 445.120025 m/z, polysiloxane) (Olsen et al., 2005). Product ion spectra were recorded in a data dependent fashion in the ion trap (ITMS) in a variable scan range and at a rapid scan rate (Wideband activation was turned on). The ionization potential (spray voltage) was set to 1.8 kV. Peptides were analysed using a repeating cycle consisting of a full precursor ion scan (3.0 × 10^6^ ions or 50 ms) followed by 12 product ion scans (1.0 ×10^4^ ions or 50 ms) where peptides are isolated based on their intensity in the full survey scan (threshold of 500 counts) for tandem mass spectrum (MS2) generation that permits peptide sequencing and identification. Collision induced dissociation (CID) energy was set to 35% for the generation of MS2 spectra. During MS2 data acquisition dynamic ion exclusion was set to 120 seconds with a maximum list of excluded ions consisting of 500 members and a repeat count of one. Ion injection time prediction, preview mode for the FTMS, monoisotopic precursor selection and charge state screening were enabled. Only charge states higher than 1 were considered for fragmentation.

Raw spectra were analysed with Andromeda search engine (Cox et al., 2011) in Max Quant v 1.5.5.30 (Cox and Mann, 2008). Primarily default parameters were selected including LFQ and match between runs. Spectra were searched against the LAB3.60 *N. benthamiana* proteome database (Ranawaka et al., 2023). Permitted modifications included static carbamidomethylation and cysteines, variable N-terminal acetylation and methionine oxidation and trypsin specific digestion. The instrument type in Andromeda searches was set to Orbitrap and the precursor mass tolerance was set to ±20 ppm (first search) and ±4.5 ppm (main search). The MS/MS match tolerance was set to ±0.5 Da. The peptide spectrum match FDR and the protein FDR were set to 0.01 (based on target-decoy approach). Minimum peptide length was 7 amino acids. For protein quantification unique and razor peptides were allowed. Modified peptides were allowed for quantification. The minimum score for modified peptides was 40. Label-free protein quantification was switched on, and unique and razor peptides were considered for quantification with a minimum ratio count of 2. Retention times were recalibrated based on the built-in nonlinear time-rescaling algorithm. MS/MS identifications were transferred between LC-MS/MS runs with the “match between runs” option in which the maximal match time window was set to 0.7 min and the alignment time window set to 20 min. The quantification is based on the “value at maximum” of the extracted ion current. At least two quantitation events were required for a quantifiable protein.

### PROTOMAP analysis

Peptide reads were analysed in R with customized scripts (Supplemental **File S3**). Contaminants and non-plant sequences were removed prior to any processing. To suppress noise signals of ions that were detected across many gel slices caused by high ionisation efficiencies in contrast to other peptides of the same proteins, we removed peptides from gel slices that were detected at <10% of the total ion intensity of that peptide across all gel slices. For each protein, peptides were plotted along the sequence across each gel slice on peptograph with ggplot2 package (Wickham, 2011). The number of peptides from each gel slice were plotted alongside as a reference of signal strength from the slice. Signal peptides, transmembrane domains and PFAM domain annotation were predicted by SignalP5.0 (Teufel et al., 2022), TMHMM (Krogh et al., 2001), and InterPro (Mistry et al., 2021), respectively, and mapped along the sequences. MWs of proteins were predicted using the Peptides package (Osorio et al., 2015). The sequons were predicted by scanning through the sequence for NxS/T, and an additional 3 kDa was added for each sequon to the predicted MW to calculate the *N*-glycosylated MW.

### Molecular cloning

To generate the double-tagged binary cloning vector, the *2x35S* promoter and *SP*(*PR1a*)-intron fusion, a fragment encoding FLAG-RFP, and a fragment encoding 2xHis-GFP fused to the *35S* terminator was amplified from template plasmids listed in Supplemental **Table S1** using primers listed in Supplemental **Table S2**. The PCR products were digested with restriction enzymes listed in Supplemental **Table S2** and cloned into *pJK001c* (Paulus et al., 2000) using EcoRI and PmeI restriction enzymes, resulting in *pKZ45*, a binary vector carrying an open reading frame (ORF) encoding SP-FLAG-RFP-2xHis-GFP (SP-FR-HG). Full length ORFs of each target gene without the region encoding the endogenous signal peptide were amplified by PCR from cDNA generated from mRNA isolated from wild-type *N. benthamiana* seedlings, using primers listed in Supplemental **Table S2**. These fragments were cloned into *pKZ45* restriction enzymes indicated in the primer names (Supplemental **Table S2**), resulting in binary vectors for expression of double tagged SCPL25 (pKZ47); EDA2 (pKZ49); GMC (pKZ54); PNGaseA (pKZ60) and SCPL20b (pKZ67), listed in Supplemental **Table S1**.

### Western blot analysis

Total protein was isolated from *N. benthamiana* leaves at 3-5□days after agroinfiltration. Three 8 mm diameter discs were taken per leaf and ground in liquid nitrogen. The frozen leaf powder was mixed with 200 μl 2×gel loading buffer containing 100 mM Tris-HCl pH6.8, 200 mM DTT, 4% SDS, 0.2% bromophenol blue and 20% glycerol. Samples were vortexed vigorously and immediately heated at 95°C for 8 min. Heated samples were briefly vortexed and centrifugated at 10,000 x *g* for 1 minute. 5 μl supernatant was loaded onto 10% SDS polyacrylamide gels and the proteins were transferred onto PVDF membranes using the Trans-Blot Turbo RTA Transfer Kit (1704275, Bio-rad). Membranes were blocked in PBS-T (tablets 524650-1EA, Sigma-Aldrich) containing 0.05% Tween-20 and 5% skim milk for 1 hour, and then with antibody (1:5000 anti-His-HRP (130-092-785, Miltenyi Biotec) or anti-FLAG-HRP (Ab49763, Abcam)) overnight at 4°C and washed with PBS-T three times and PBS one time. To detect the signals, the membrane was incubated with SuperSignal West Femto Maximum Sensitivity Substrate (34096, ThermoFisher) and chemiluminescent signals were detected with ImageQuant LAS 4000 (GE Healthcare).

### Phylogenetic analysis

Proteome sequences of *N. benthamiana* and *A. thaliana* were obtained from LAB3.60 (Ranawaka et al., 2023) and Araport11 (Cheng et al., 2017) annotations, respectively. Protein sequences from these proteomes were annotated for PFAM (Mistry et al., 2021) protein families and domains using InterProScan 5 (Jones et al., 2014). To select proteins to construct a phylogenetic tree for each family of interest, all proteins annotated with a PFAM ID of each family: serine carboxypeptidase-like (SCPL) (PF00450), pectin methylesterase (PME) (PF01095) and GDSL-like lipase/acylhydrolase (GELP) (PF00657), were selected. Next, to identify additional family members that might have been missed, this PFAM-identified protein set was used as a query to search against proteomes of both LAB3.60 an Araport11 using BLAST+ (Camacho et al., 2009), keeping hits with higher than 80% query coverage. The PFAM-identified and BLAST-identified proteins were combined to generate the final protein set for each family. Then, protein sequences were aligned using MAFFT 7 (Katoh and Standley, 2013) with the L-INS-i algorithm. The resulting multiple sequence alignment was used to construct a maximum likelihood phylogenetic tree using IQ-TREE 2 (Minh et al., 2020) with WAG amino acid substitution model, empirical amino acid frequencies (+F), allowance for invariant sites (+I) and Gamma distributed evolutionary rate heterogeneity among sites with four discrete categories (+G4). Branch support values were calculated using ultrafast bootstrap with 1000 replications. Phylogenetic trees were visualised using iTOL (Letunic and Bork, 2021). Trees were rooted at midpoint.

## Funding

This project was financially supported by the China Scholarship and the Northeast Institute of Geography and Agroecology, Chinese Academy of Sciences (KZ); Interdisciplinary Bioscience DTC DDT00060 (ET) and DDT00230 (JL); BBSRC 19RM3 project BB/015128/1 (NS); BBSRC 18RM1 project BB/S003193/1 ‘Pip1S’ (MS); ERG-2013-CoG project 616449 ‘GreenProteases’ (RH); and ERC-2020-AdG project 101019324 ‘ExtraImmune’ (RH).

## Acknowledgements

We like to thank Urszula Pyzio for excellent plant care, Sarah Rodgers and Caroline O’Brien for excellent technical support, and Pitter Huesgen and Fatih Demir for TAILS analysis.

## Author contributions

RH conceived and managed the project; ET produced the protomap samples; KZ mined the dataset to identify processing events and confirmed processing with double-tagged precursors; FK and MK generated proteomics data; ET and JL analysed MS data; NS supported PROTOMAP analysis and build phylogenetic trees; MS analyzed receptor processing; RH wrote the article with input from all co-authors.

## Data availability

The peptide file that was used as an input for the PROTOMAP is provided as Supplemental **File S1**. The code that was used to generate the PROTOMAP is in Supplemental **File S3** and available on GitHub (https://github.com/joyylyu/protomap.git). Uncropped images are provided in Supplemental **File S4**.

## Notes

### Competing Interest Statement

The authors have declared no competing interest.

### Summary of Updates

This manuscript has been adjusted to stress the importance of the used materials, the superiority over TAILS data is discussed, and the multimerisation sections have been removed.

