## Supplementary material for "The proteome of agroinfiltrated *Nicotiana benthamiana* is shaped by extensive protein processing": File S2

NbL19g24480.1 tubulin_alpha−3_chain [MW = 49.78 kDa]

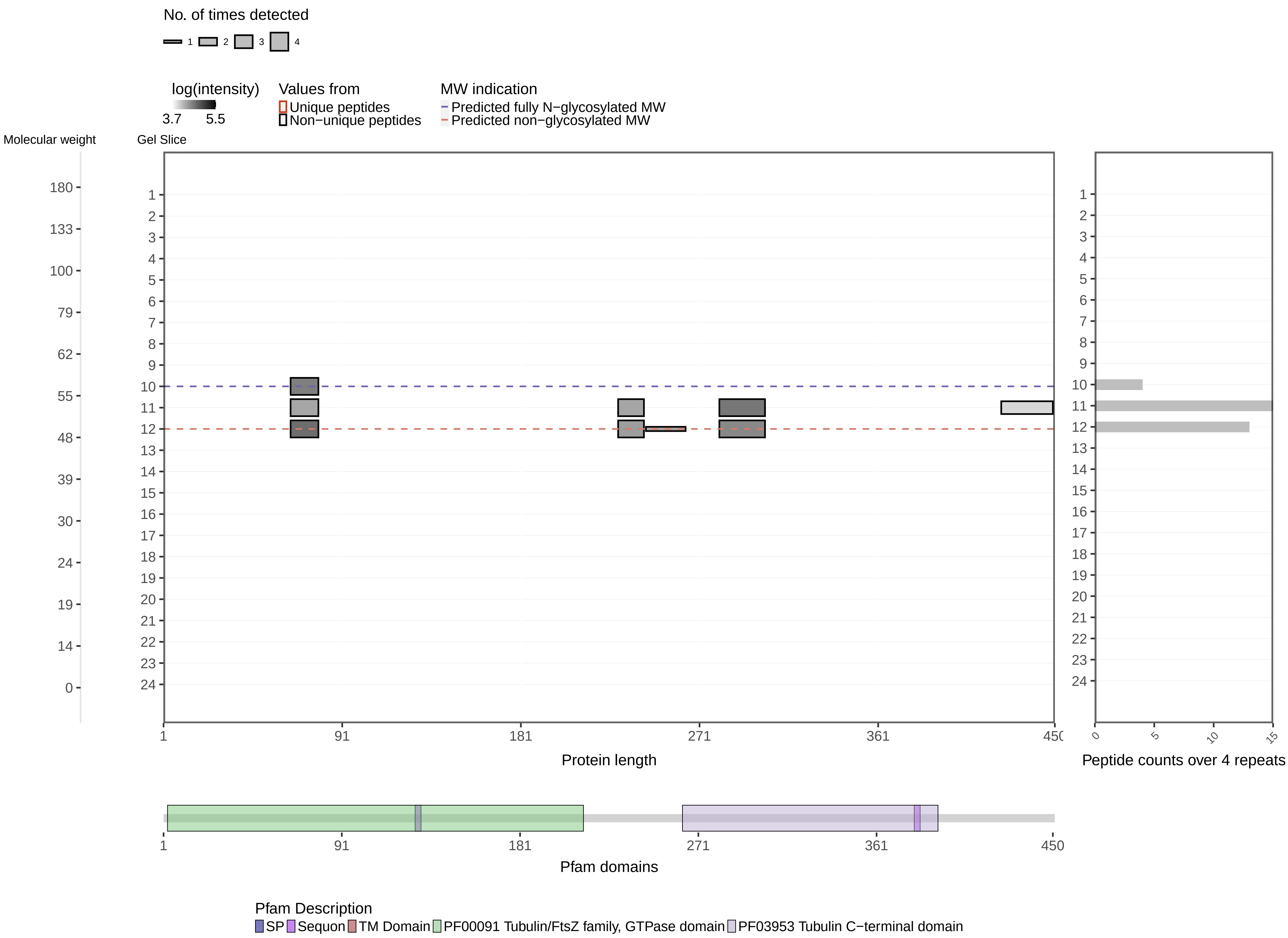


Protein sequence:

CRRSLDIERPTYTNLNRLISQIISSLTTSLRFDGAINVDITEFQTNLVPYPRIHFMLSSYAPVISAEKAFHEQLSVPEITNAVFEPSSMMAKCDPRHGKYMACCLMYRGDVVPKDVNAAVATIKTKRTVQFVDWCPTGFKCGINYQPPTVVPGGDLAKVQRAVCMISNNTAVAEVFSRIDHKFDLMYAKRAFVHWYVGEGMEEGEFSEAREDLAALEKDYEEVGAEGVDDEEDGDEY*

### NbL19g24460.1 importin−5 [MW = 123.81 kDa]

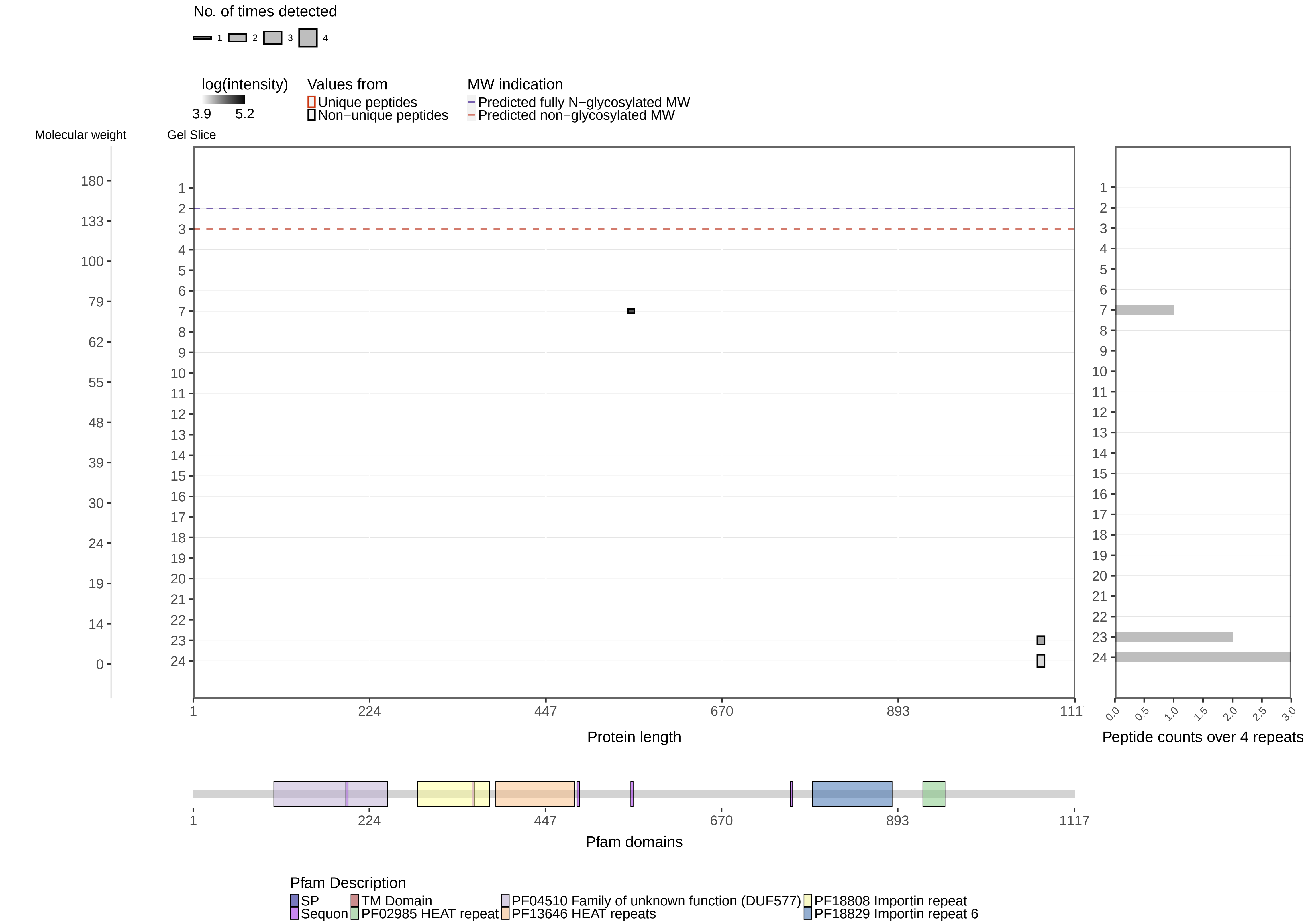


Protein sequence:

.KTILVNATDKSNRMLRAKAMECISLVGMAVGKDKFRDDAKQVMEVLMSLQGSQMETDDPTTSYMLQAWARLCKCLGQDFLPYMSVVMPPLLQSAQLKPDVTISSADSDNELDDSDDDSMETITLGDKRIGIKTSVLEEKATACNMLCCYADELKEGFYPWIDQVAPTLVPLLKFYFHEEVRKAAVSAMPELLRSAKLAVEKGIAQGRNESYVKQLSDYIVPALVQALHKEPDTEICASMLDALNECLQISGPLLDEGQVRSIVDEIKQVIMASSSRKRERAERAKAEDFDAEE

NbL19g24420.1 probable_acyl−activating_enzyme_18_peroxisomal [MW = 80.71 kDa]

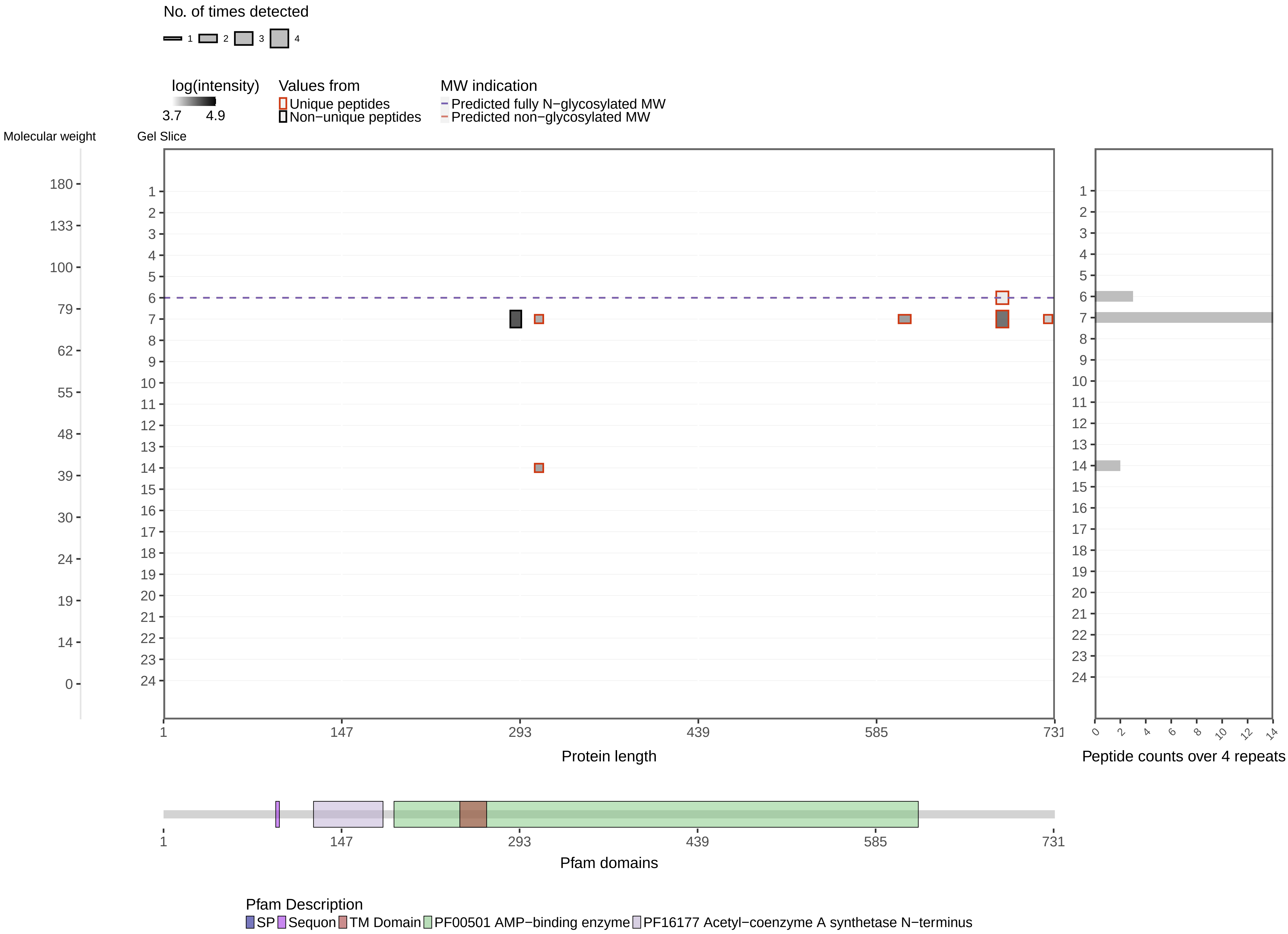


Protein sequence:

VHQSIDSVTNILFSSGTTGDPKAIPWTHLSPIRCGAESWANLDVRAGDVSCWPTNLGWVMGPILIYSCFLTGATLALYHGSPLDHGFGKFVQDAGVTILGTVPSLVKTWKSTGCMKGLDWTRIRMFASTGEASNVDDDLWLSSRVDYKPVIECCGGTELSSSYIMGNLLQPQAFGTFSSASMSTGFVILDEDGRPYPDDQACVGEVGLFPTYMGATDRLLNADHEKVYFKGMPMYKDMKLRRHGDILKRTVGGYLVVQGRADDTMNLGGIKTSSIEIERVCDRADVCILETV

### NbL19g24380.1 NADH_dehydrogenase_[ubiquinone]_1_alpha_subcomplex_subunit_6−like [MW = 18.5 kDa]

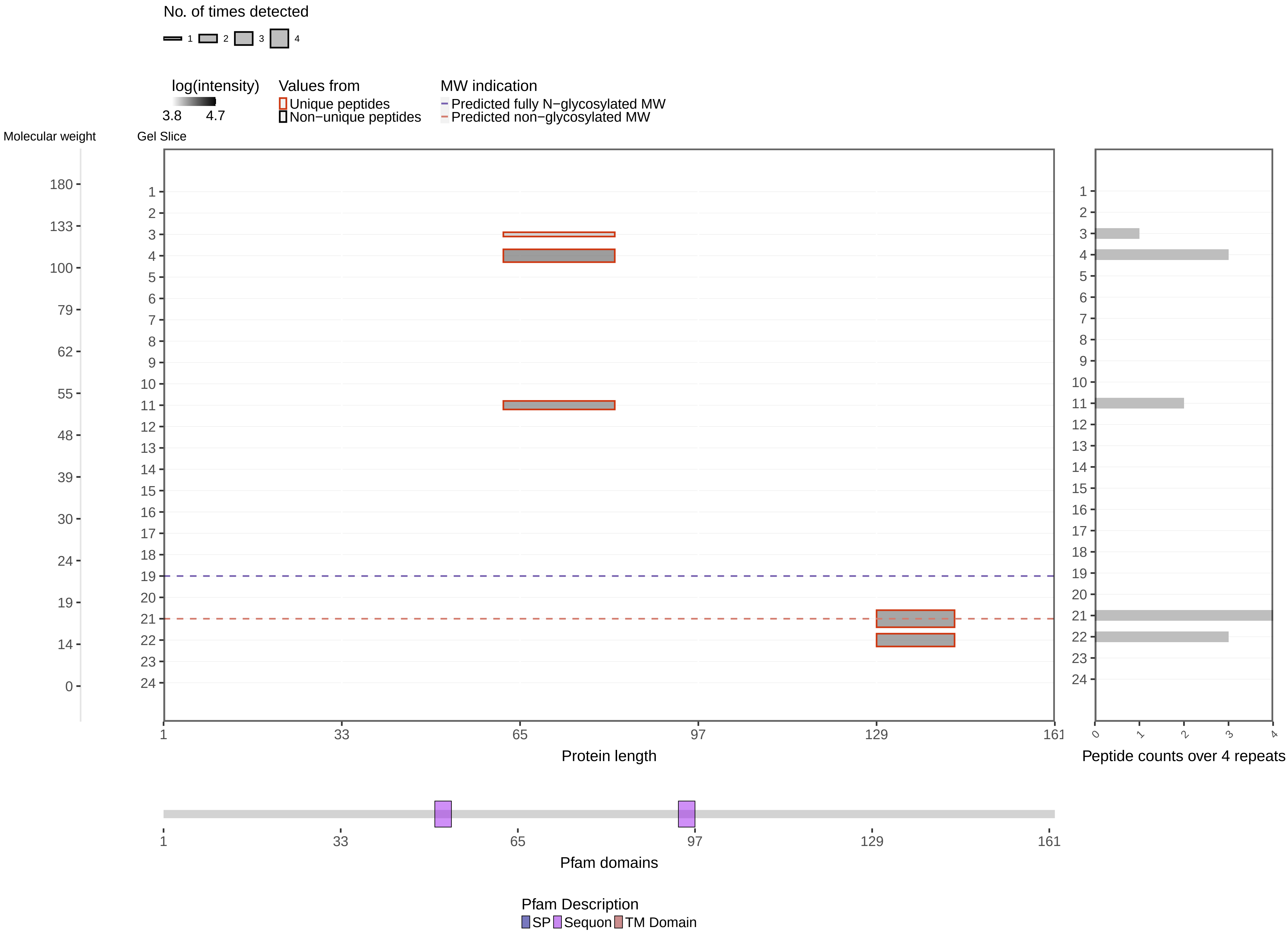


Protein sequence:

IYNLYDVVSPSQLRSAVAAEVRKNSNVTNPKVIDMLLFKGMEELKNFVDHSKQRHHIVGKYVVGNQGLVQDLEAKDQGSSNFLKNFYSSNYF*

### NbL19g24250.1 50S_ribosomal_protein_L6_chloroplastic [MW = 25.13 kDa]

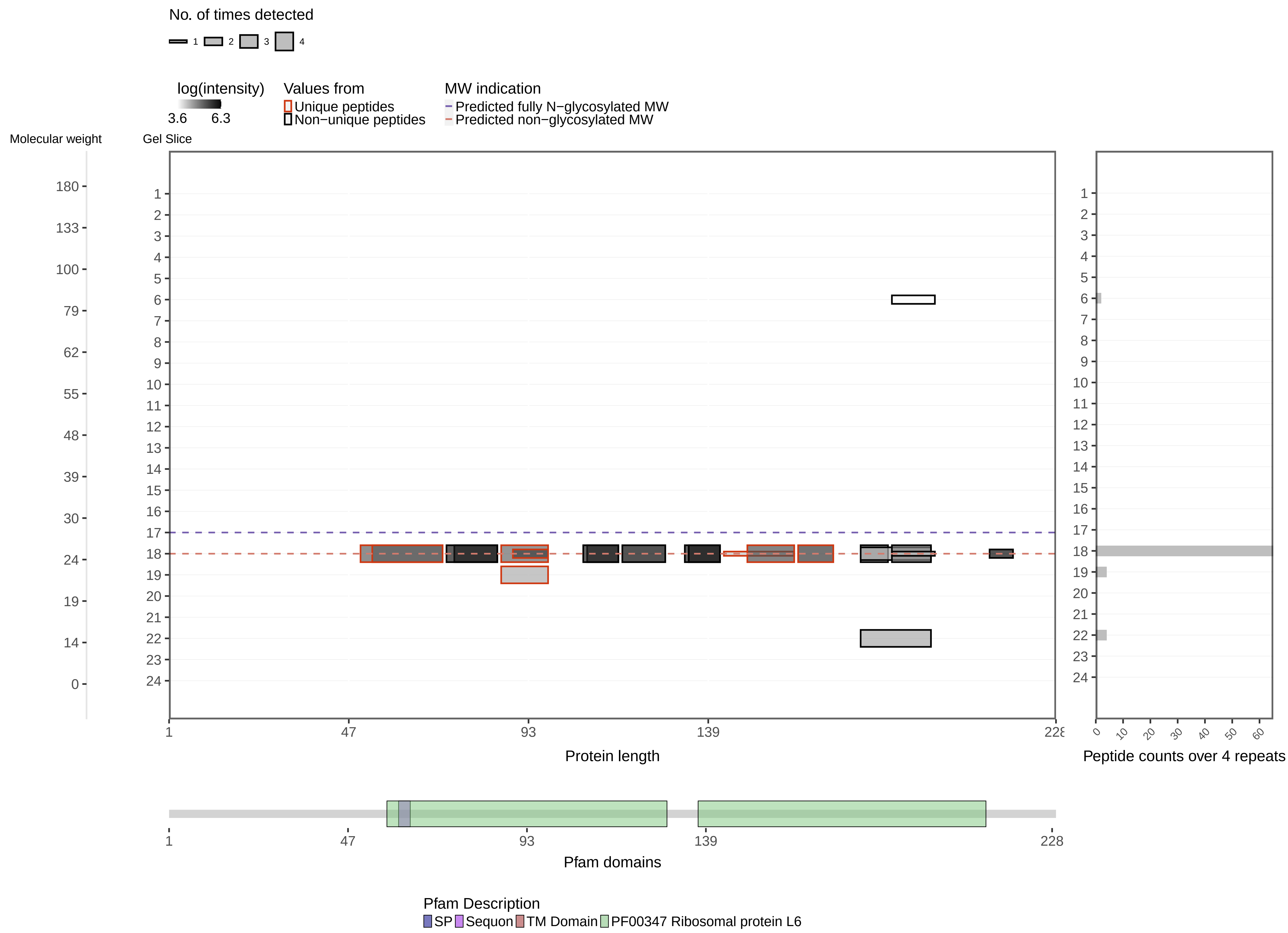


Protein sequence:

AVETRRANQMHGLFRTLTDNMVVGVSKGFEKKLQLVGVGYRATVEGKDIVLNLGFSHPVKMEIPDGLQVKVEENTRITVSGYDKSEIGQFAASIRKWRPPEPYKGKGVKYADEIVRRKEGKAGKKK*

### NbL19g24150.1 probable_nucleoside_diphosphate_kinase_5 [MW − SP = 17.87 kDa]

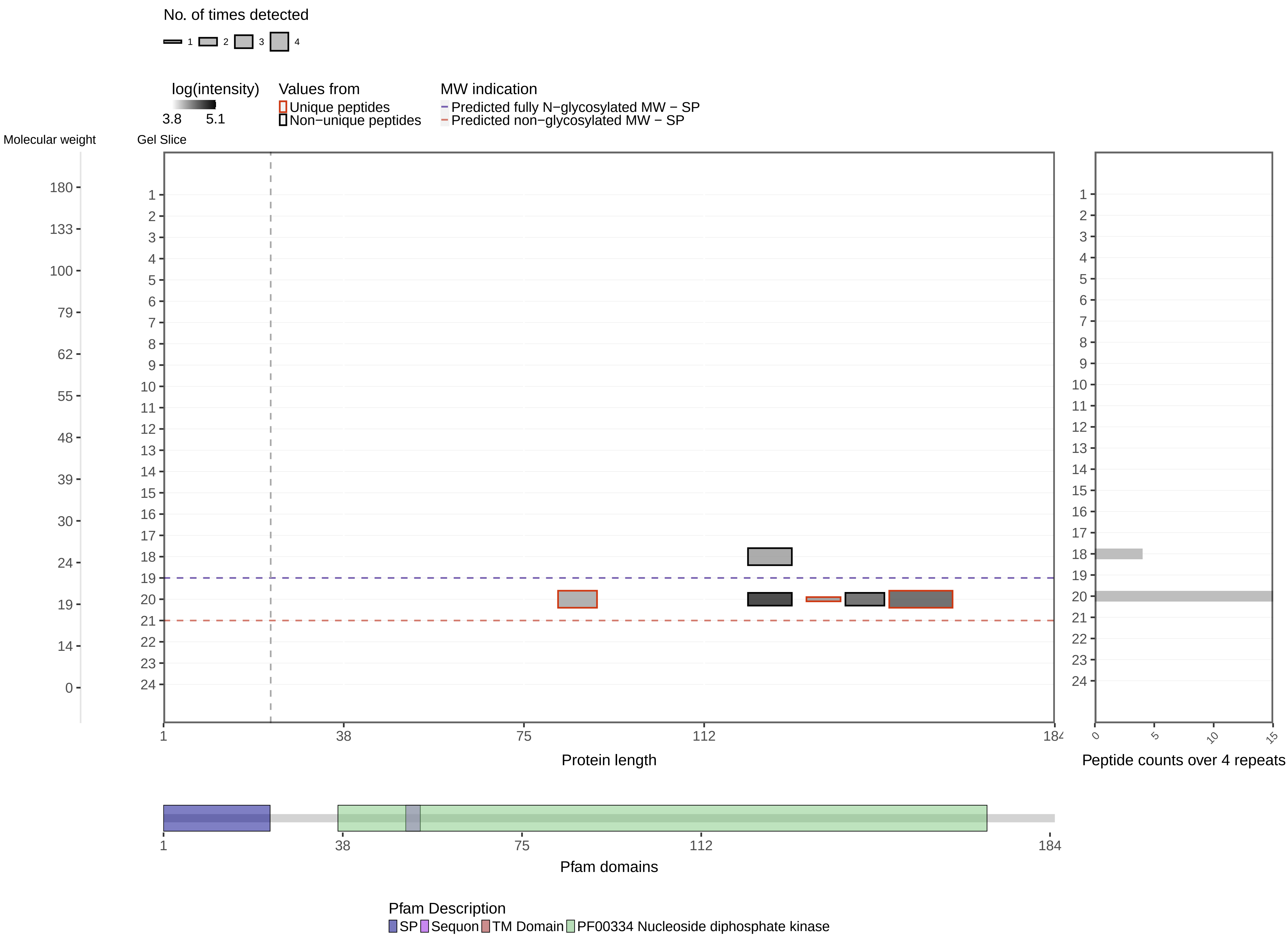


Protein sequence:

NFYAEHSSRSFFPSLVEYMASGPVLIMVLEKGNAIADWRALIGPTDPLKAKVTHPYSVRAICGLDLQKNCVHGSDSPQSAAREISFFFERTSSGHVSKHDEL*

### NbL19g24090.1 serine−−glyoxylate_aminotransferase−like [MW = 46.92 kDa]

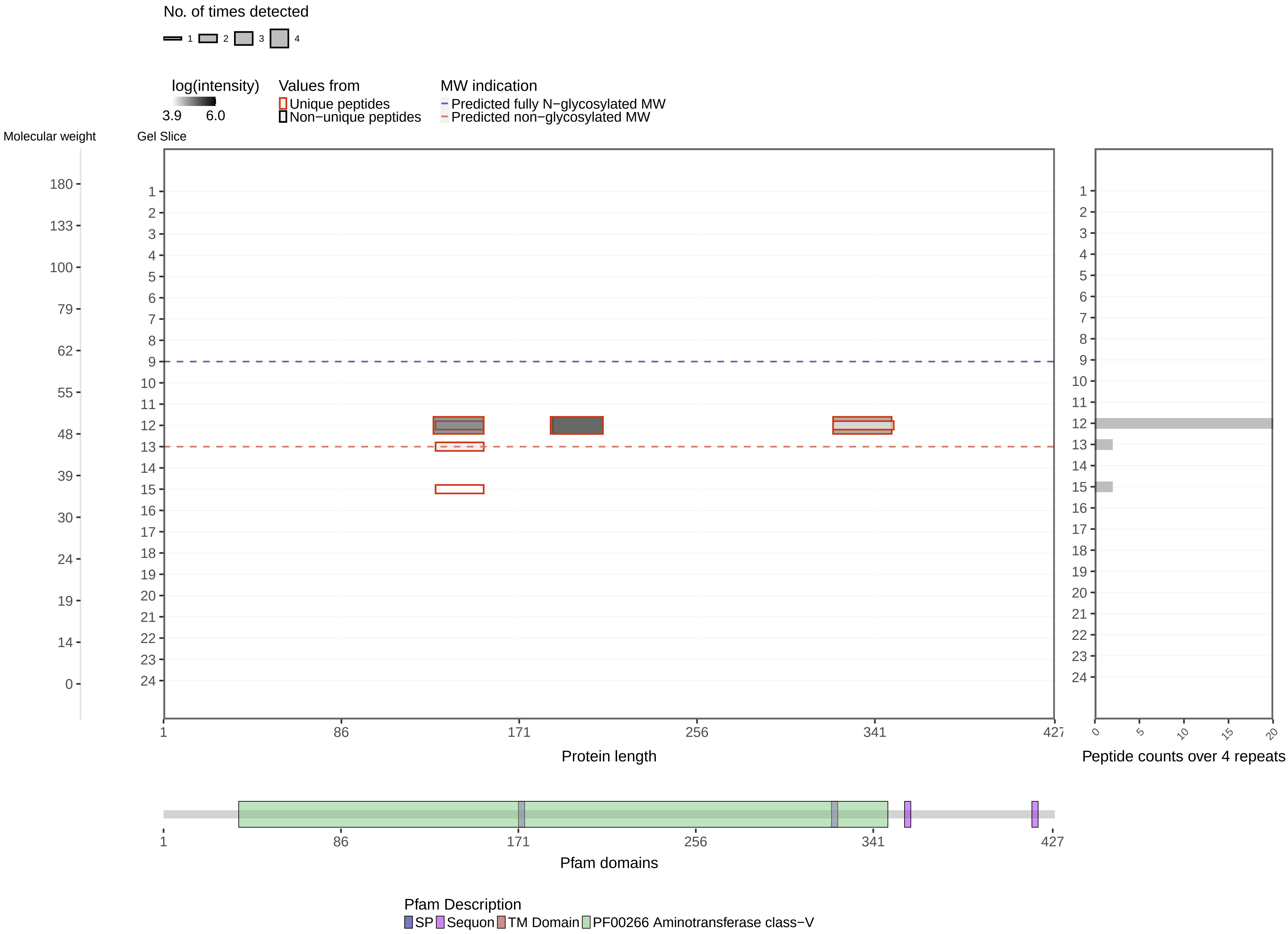


Protein sequence:

'SSICALDFRMDEWGVDVALTGSQKALSLPTGLGIVCASPKALEAAKTAKSVRVFFDWSDYLKFYKMGTYWPYTPSIQLLYGLRAALDLLFEEGLENVIARHARLGKATRLAVEAWGLKNCTQKEEWFSDTVTAVVVPPYIDSAEIVKRAWKRYNLSLGLGLNKVAGKVFRIGHLGNLNELQLLGCLAGVEMVLKDVGYPVKFGSGVAAASTFLQNSTPLIPSRI*

### NbL19g23780.1 DEAD−box_ATP−dependent_RNA_helicase_20 [MW = 55.72 kDa]

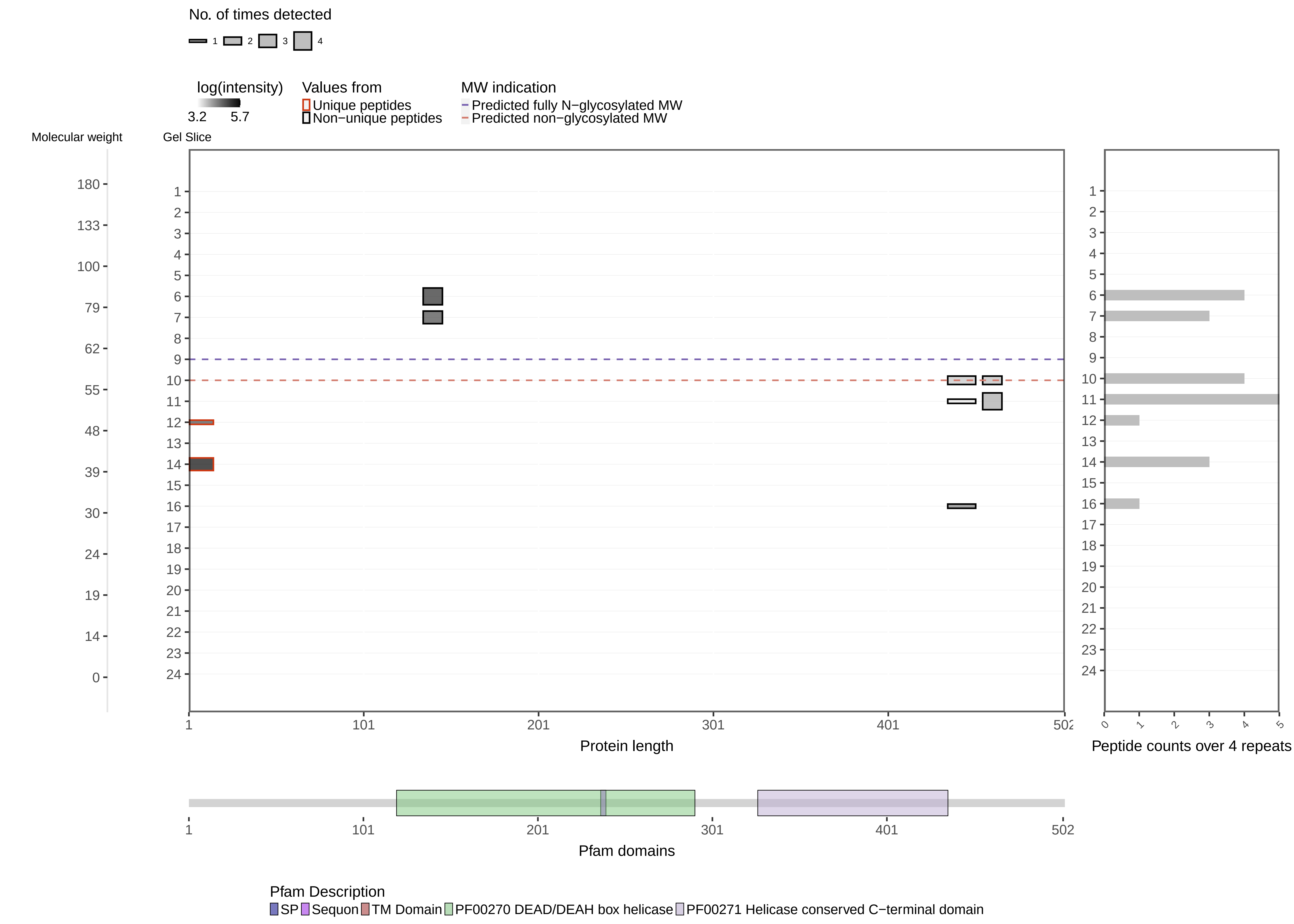


Protein sequence:

NLRRVTYLVLDEADRMLDMGFEPQIRKIVSQIRPDRQTLYWSATWPKEVEQLARQFLFNAYKVVIGSADLKANHAIRQHVEIVSESQKYNKLVKLLEDIMDGSRILIFMDTKKGCDQVTRQLRMDGWPALSIHGDKSQAERDWVLSEFKAGKSPIMTATDVAARGLDVKDVKFVINYDFPGSLEDYVHRIGRTGRAGASGTAYTYFTAANARFAKDLINILEEAGQKVSPELAKMGRGAPPPPGLGGFRDRGRGYGGNRQWS*

NbL19g23730.1 60S_ribosomal_protein_L28−1−like [MW = 15.8 kDa]

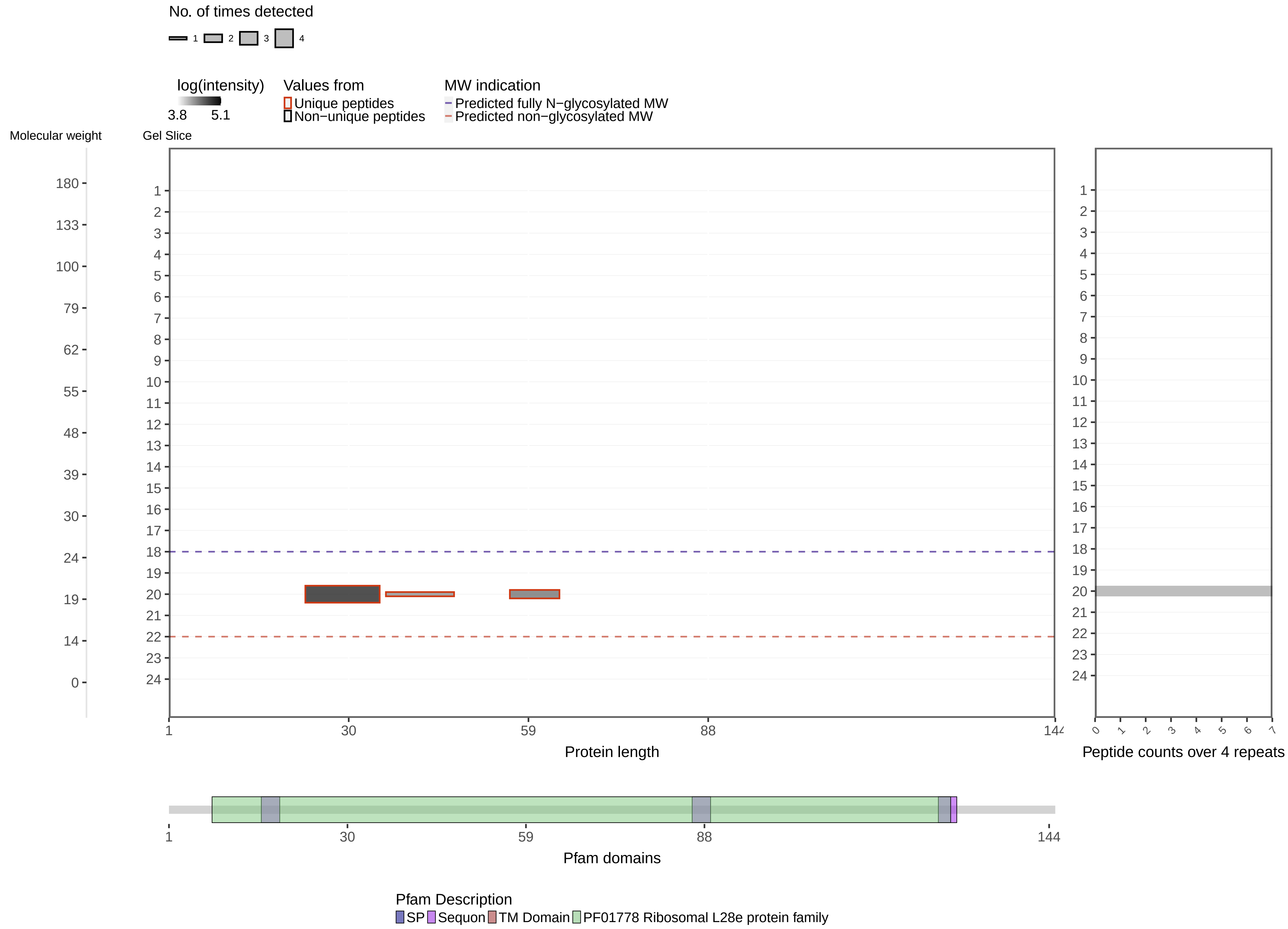


Protein sequence:

AGGKDQSVLLATTKTKKQNKPSSLLNKSAMKKEFRRMAKAVTNQVADNYYRPDLKKAALARLSAVNRSLKVAKSGVKKRNRQA*

NbL19g23700.1 probable_voltage−gated_potassium_channel_subunit_beta [MW = 36.94 kDa]

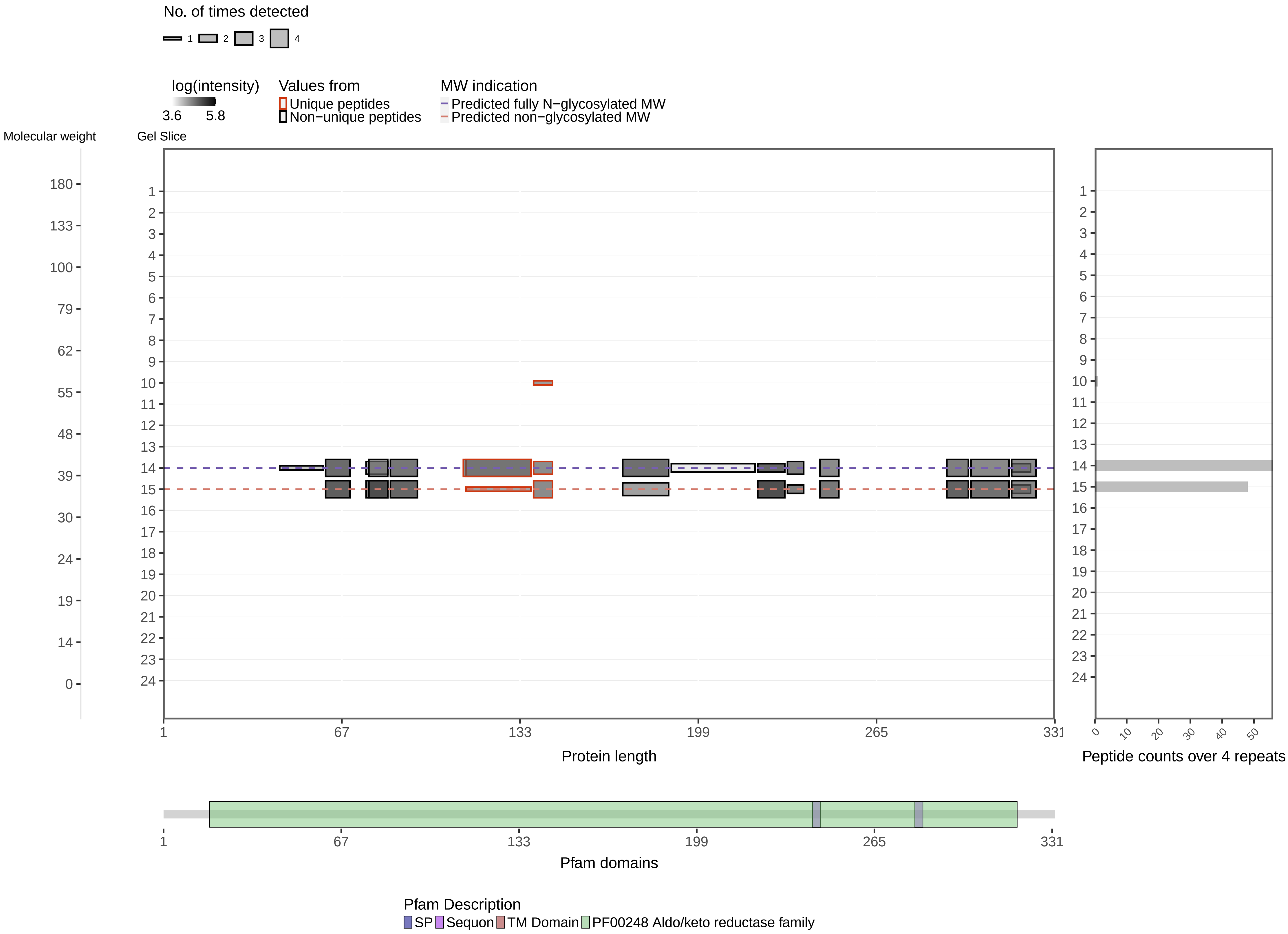


Protein sequence:

WGTSEWSAOOITEAWGVAORLDLVGPIVEOPEYNLLSRHKVESEYLPLYSNYGIGLTTWSPLASGVLTGKYNAGNIPADSRFALENYKNLANRSLVDDVLRKVDGLKPIAESLGVPLPOLAIAWCAANPNVSSVITGATKEYOIEENMKAINVIPMLTPAVMERIEAVVOSKPKRODSYR*

### NbL19g23450.1 actin−related_protein_2/3_complex_subunit_4 [MW = 19.84 kDa]

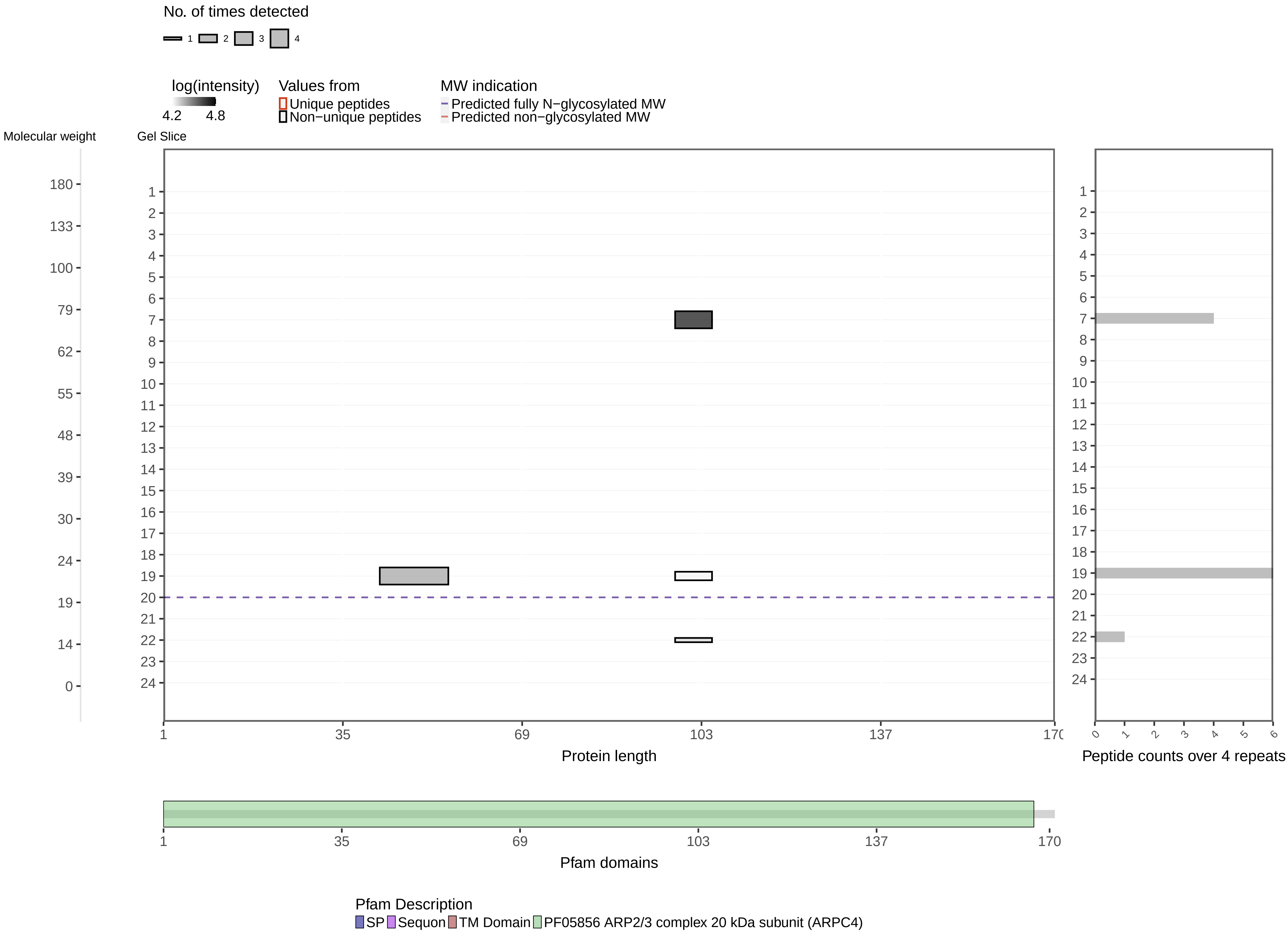


Protein sequence:

RISLKVKQSDELENIITKKFLRFLSMRAEAFQVLRRKPVQGYDISFLITNYHCEEMQKQKLIDFIVQFMEDIDKEISELKLSVNTRGRLVATEFLKQFI*

### NbL19g23400.1 serine_hydroxymethyltransferase_4 [MW = 20.68 kDa]

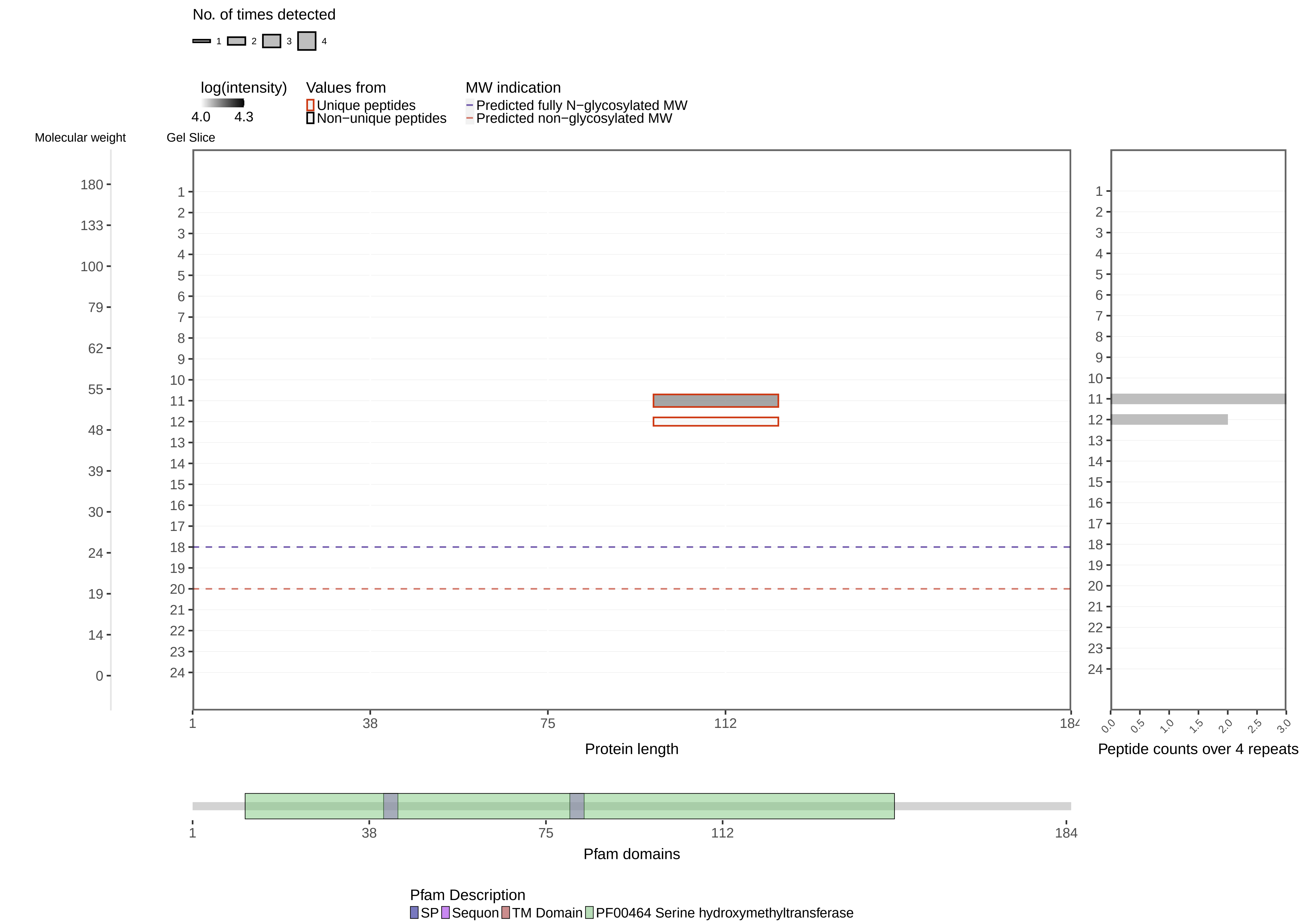


Protein sequence:

ENLTRSRALQAFHVDPTKWGVNVQPYSGSTANFAAYTAVLNPHDRIMGLDLPSGGHLTHGYYTSGGKKIQLLQFILRVYLISWIQKLGILIMIDWKRRHWILGLS*

### NbL19g23180.1 signal_peptide_peptidase−like_3 [MW − SP = 55.03 kDa]

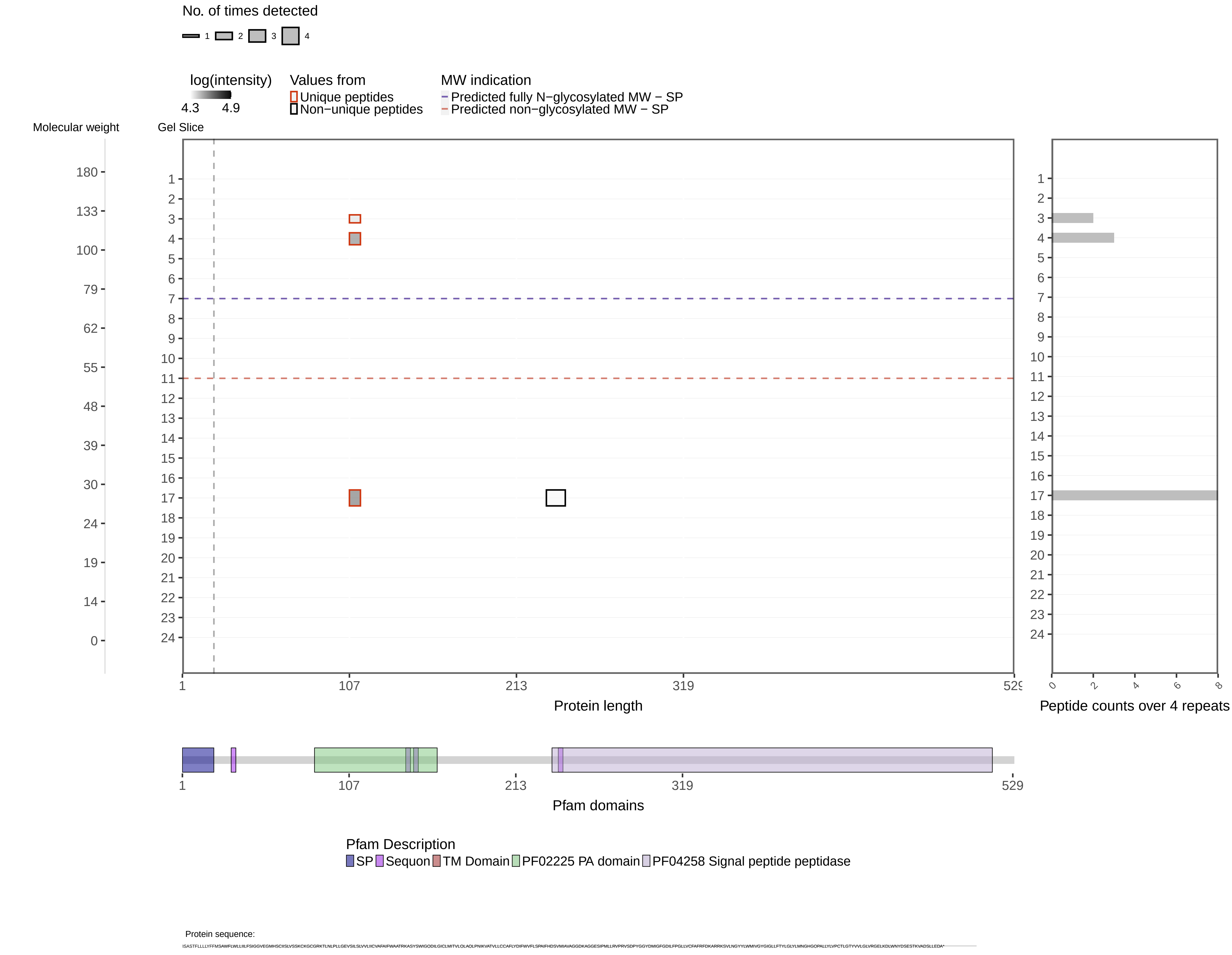

### NbL19g23020.1 5'−nucleotidase_domain−containing_protein_4 [MW = 71.44 kDa]

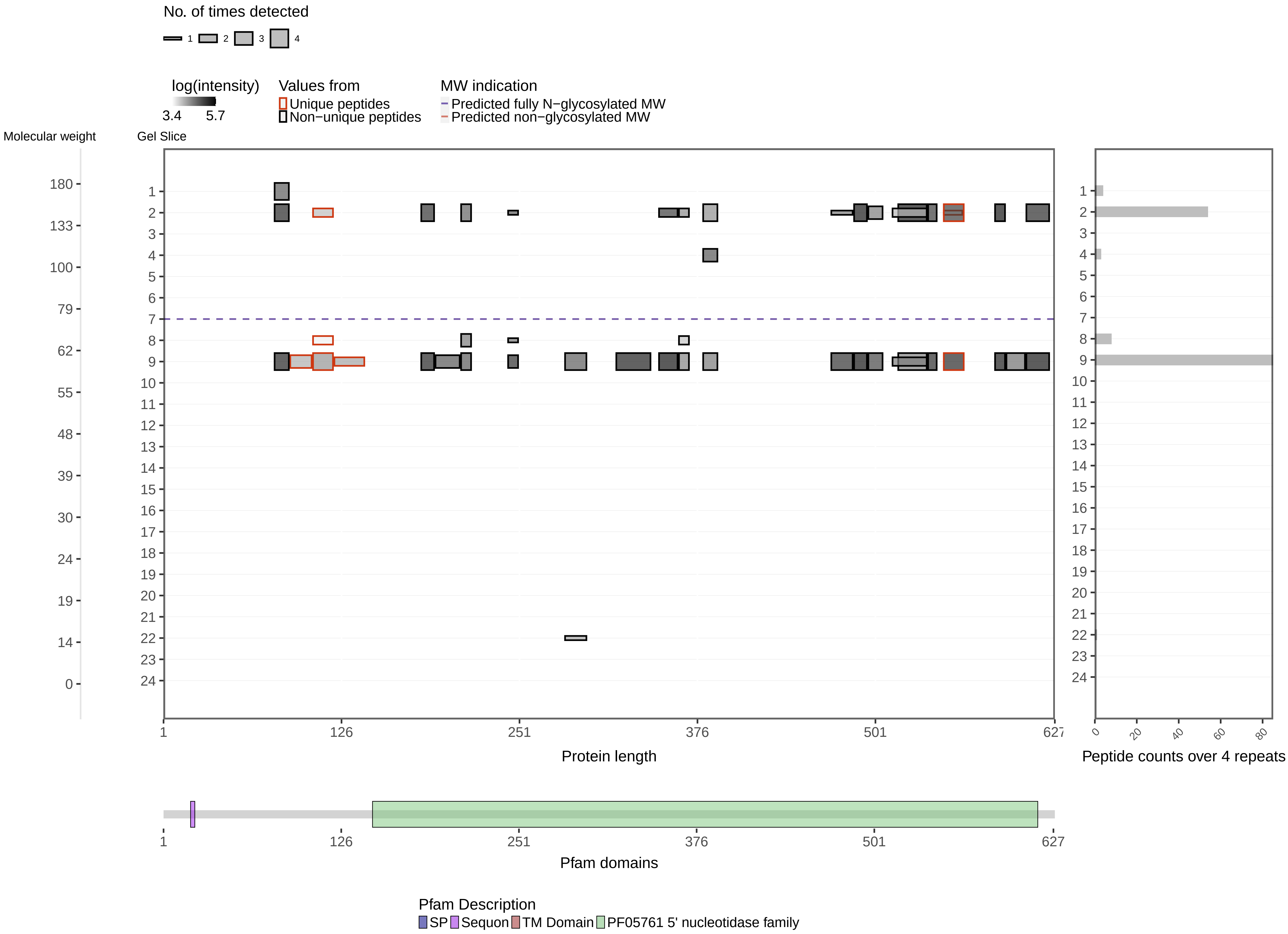


Protein sequence:

NbL19g22910.1 novel_plant_SNARE_13 [MW = 30.41 kDa]

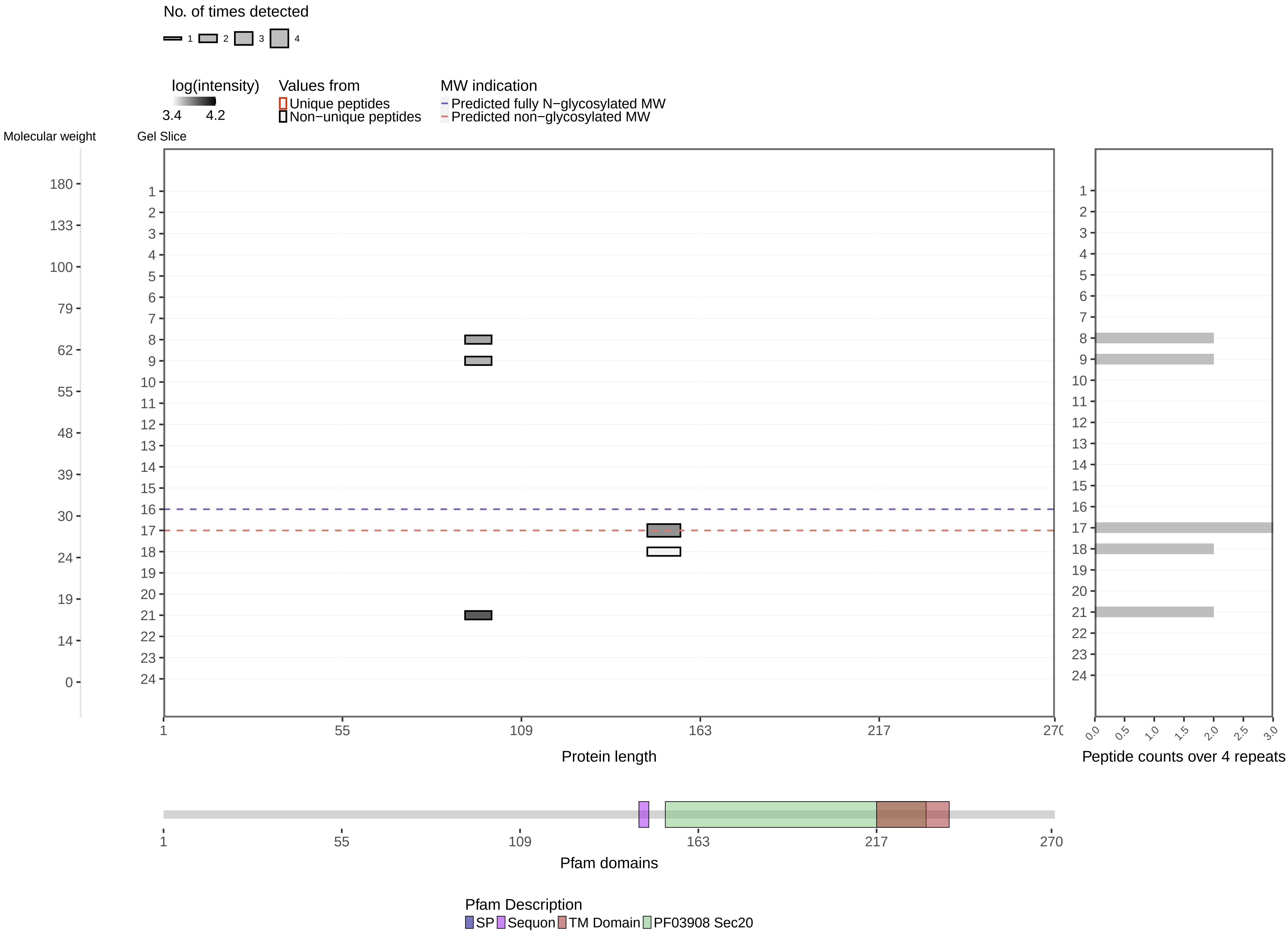


Protein sequence:
ASEPTAEENVQMASAMSNQELISAGNKTMDETDQAIERSKQVVHQTIEVGTQTATTLKGQTDQMGRVVNELDTIQFSIKKASQLVKEIGRQVATDKCIMLFLFLIVCGVVAIIVVKIVNPHNKDIRDIPGLAPPAPSRRLLYLKSGQDFM*

### NbL19g22860.1 S−adenosylmethionine_synthase_2 [MW = 43.04 kDa]

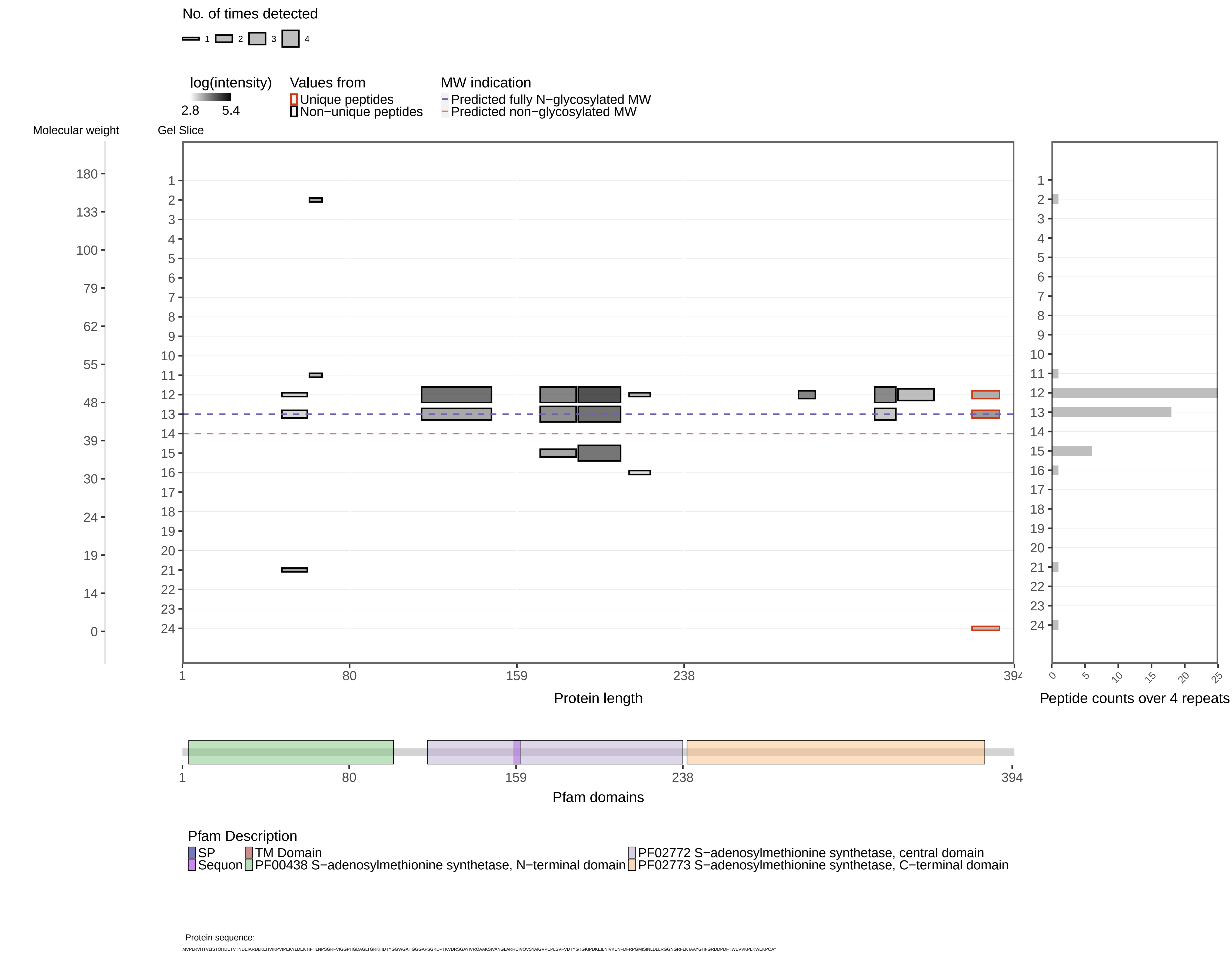

### NbL19g22800.1 ADP−ribosylation_factor_1−like [MW = 20.67 kDa]

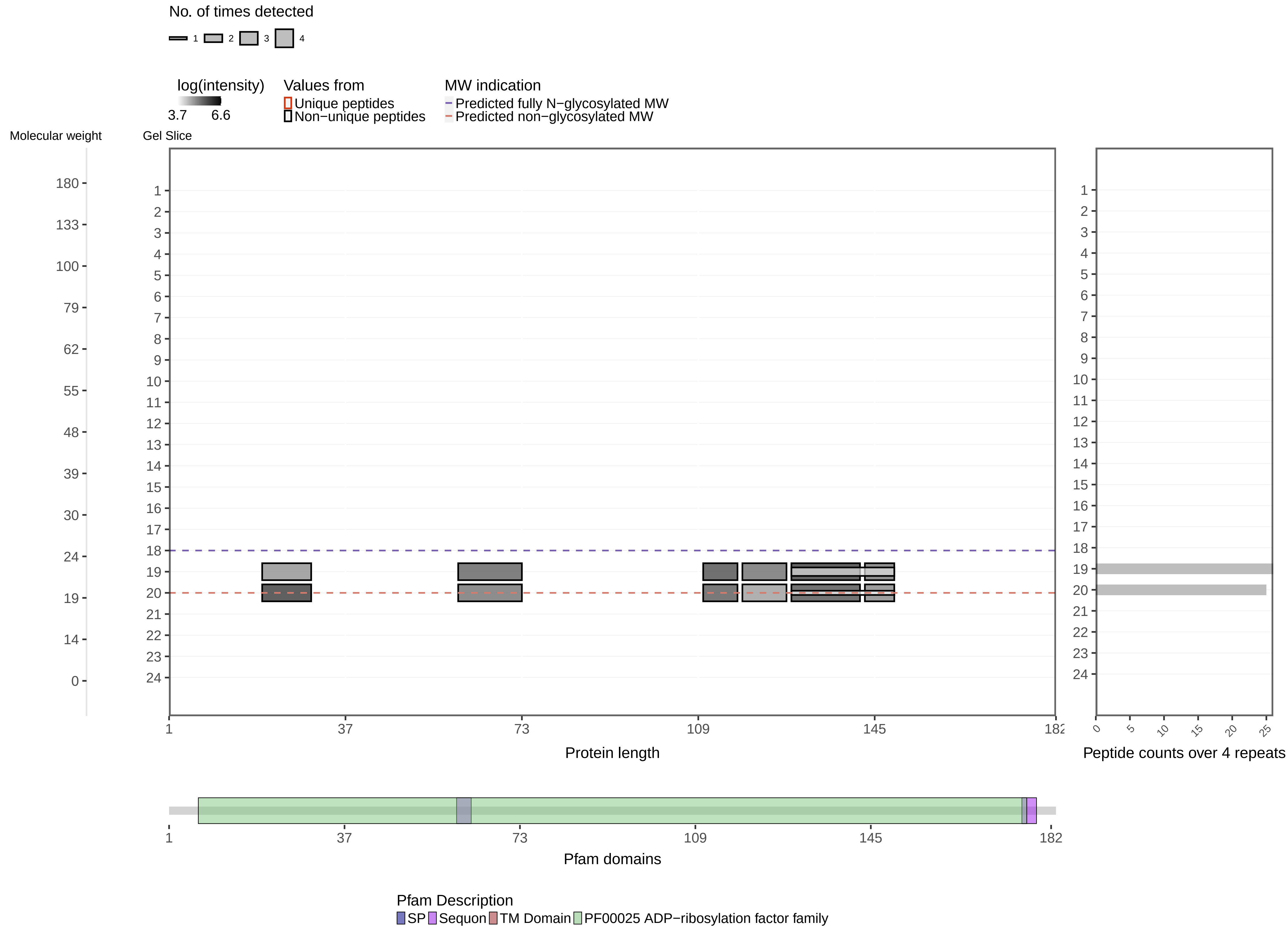


Protein sequence:

HYFQNTQGLIFVVDSNDRDRVVEARDELHRMLNEDELREAVLLVFANKQDLPNAMNAAEITDKLGLHSLRQRHWYIQSTCATSGEGLYEGLDWLSNNISNKA*

### NbL19g22780.1 dihydrolipoyl_dehydrogenase_mitochondrial [MW = 16.05 kDa]

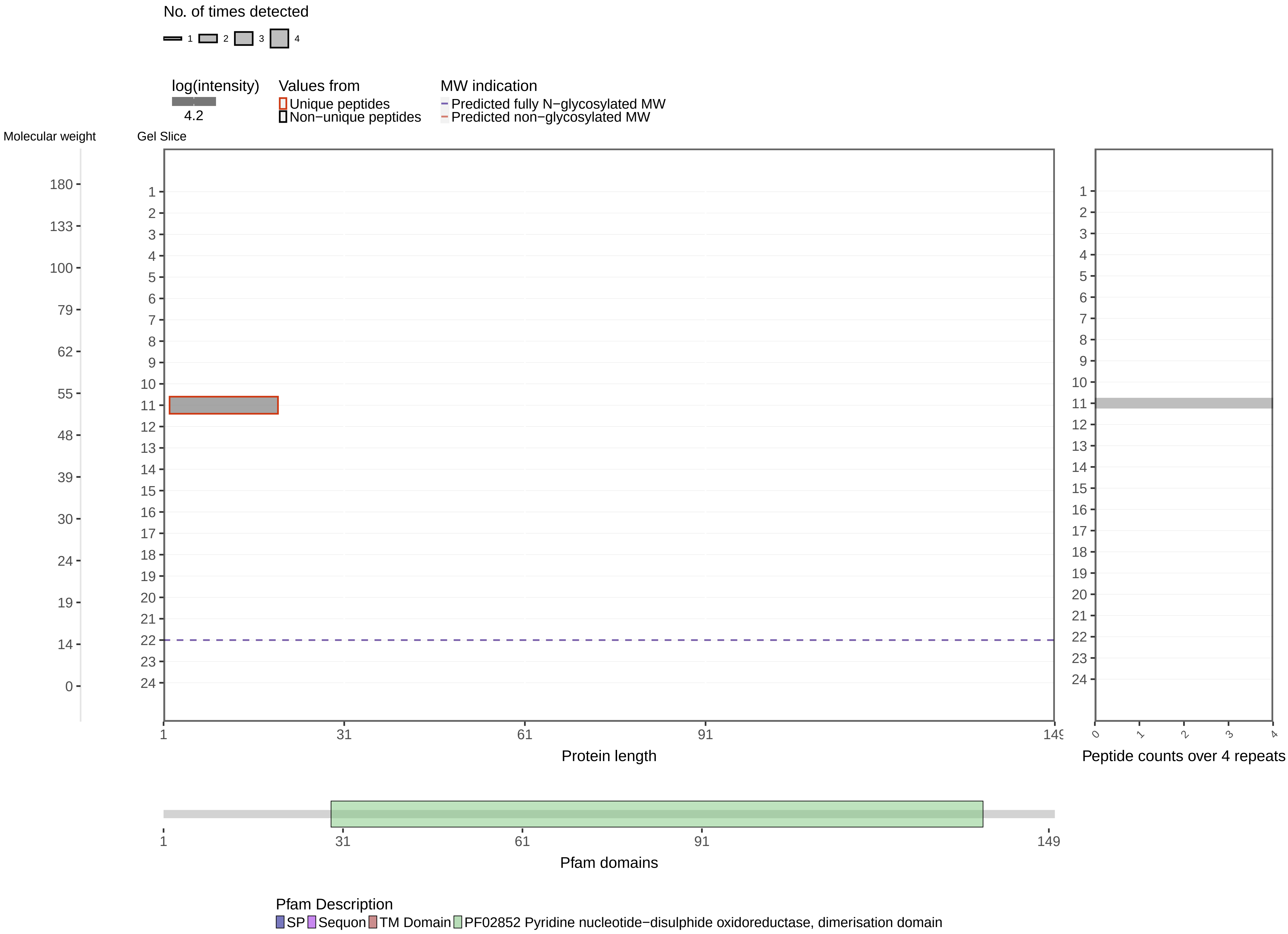


Protein sequence:

KFPFLANSRAKAIDDAEGLVKILAEKESDKILGVHIMSPNTGELIHEAVLALQYGASSEDIARTCHAHPTMSEALKEAAMATYDKPIHM*

NbL19g22710.1 cell_wall_/_vacuolar_inhibitor_of_fructosidase_1−like [MW − SP = 15.75 kDa]

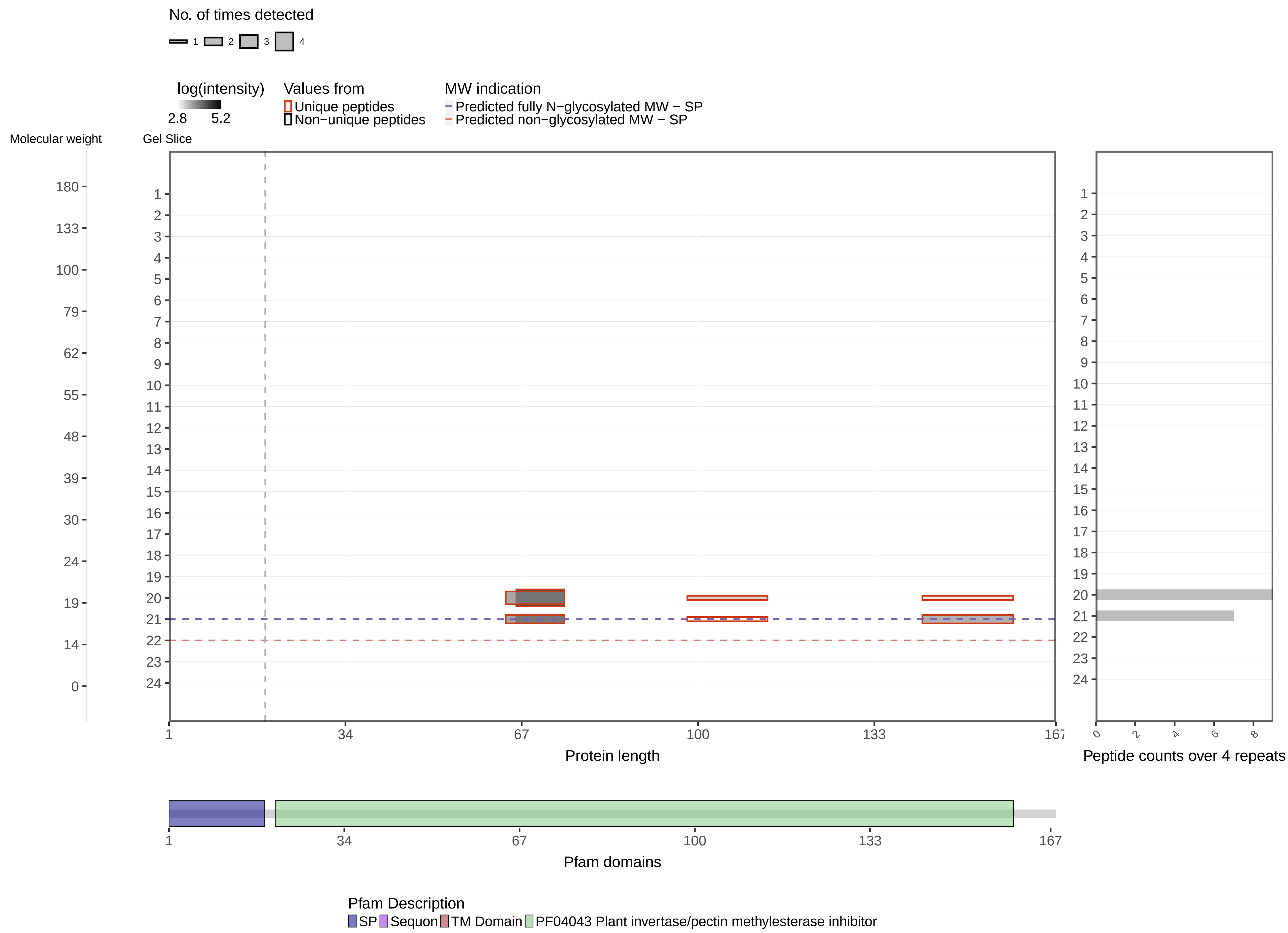


Protein sequence:

/TISKLRHSNPPAAWKGPLKNCAISYKVILTTSLPEAIEALTKGDPKFAEDGMVGSSGDAQECEEYFKGSKSPLSALNVAVHQLSDVGRAIVRNLL*

NbL19g22640.1 ATP−citrate_synthase_beta_chain_protein_2 [MW = 65.63 kDa]

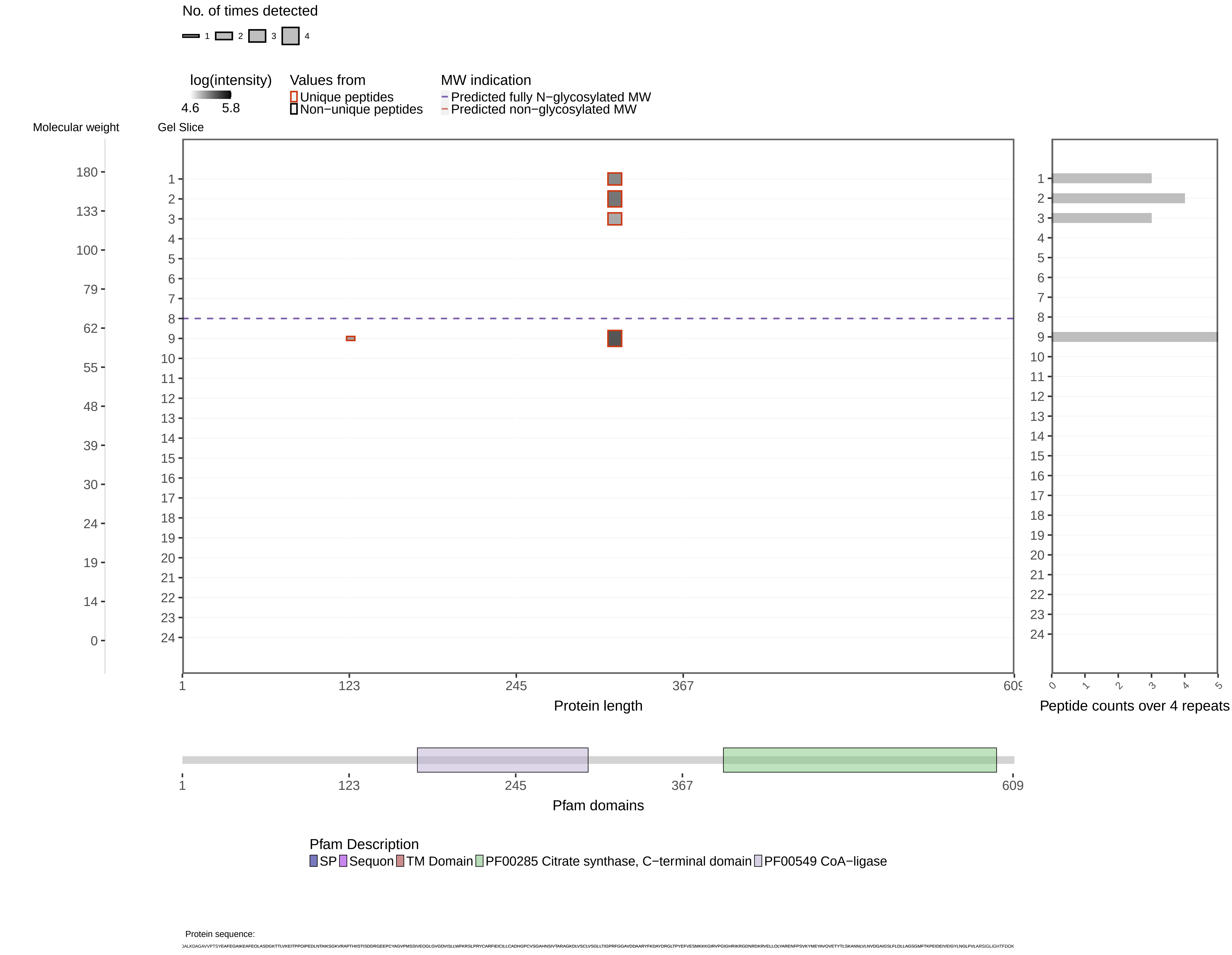

NbL19g22510.1 uncharacterized_LOC109241286 [MW = 35.45 kDa]

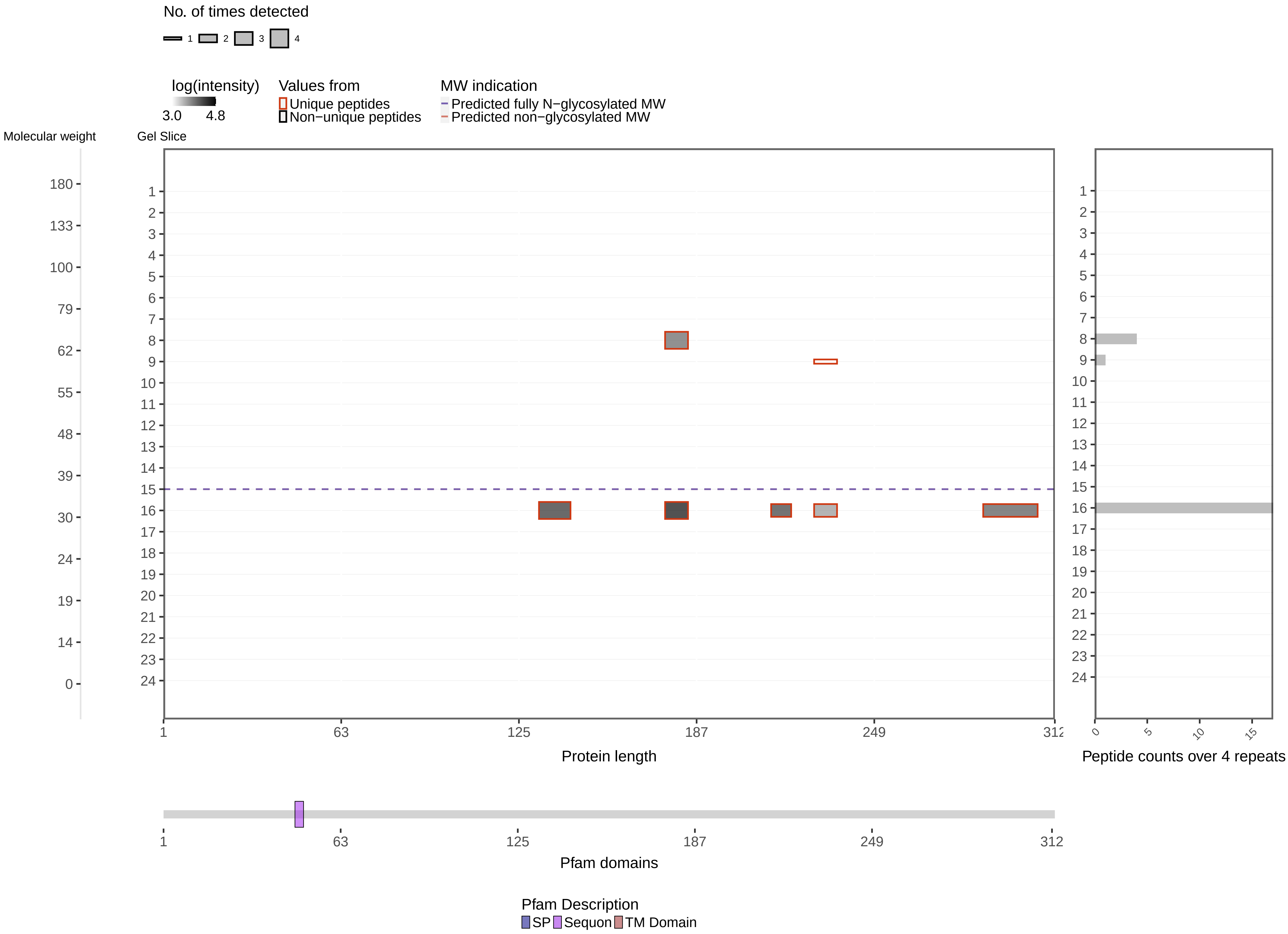


Protein sequence:

ERGIANPDTKEISSLSVEPCEGETLIVTLFEIQRSEIPAFIEREHEFRFLAVIPETLNGLFYINPAIICARYSDEEYLKTRCRGDKGIFFQLYGRYNIDKIWRDDILPCRVYLRHCVLAAQNLGDIAYDNFLDHTFLGDRRTTIREYLATTGSGIMEEEPPELLKYRYGG*

### NbL19g22470.1 glyoxysomal_fatty_acid_beta−oxidation_multifunctional_protein_MFP−a [MW = 78.86 kDa]

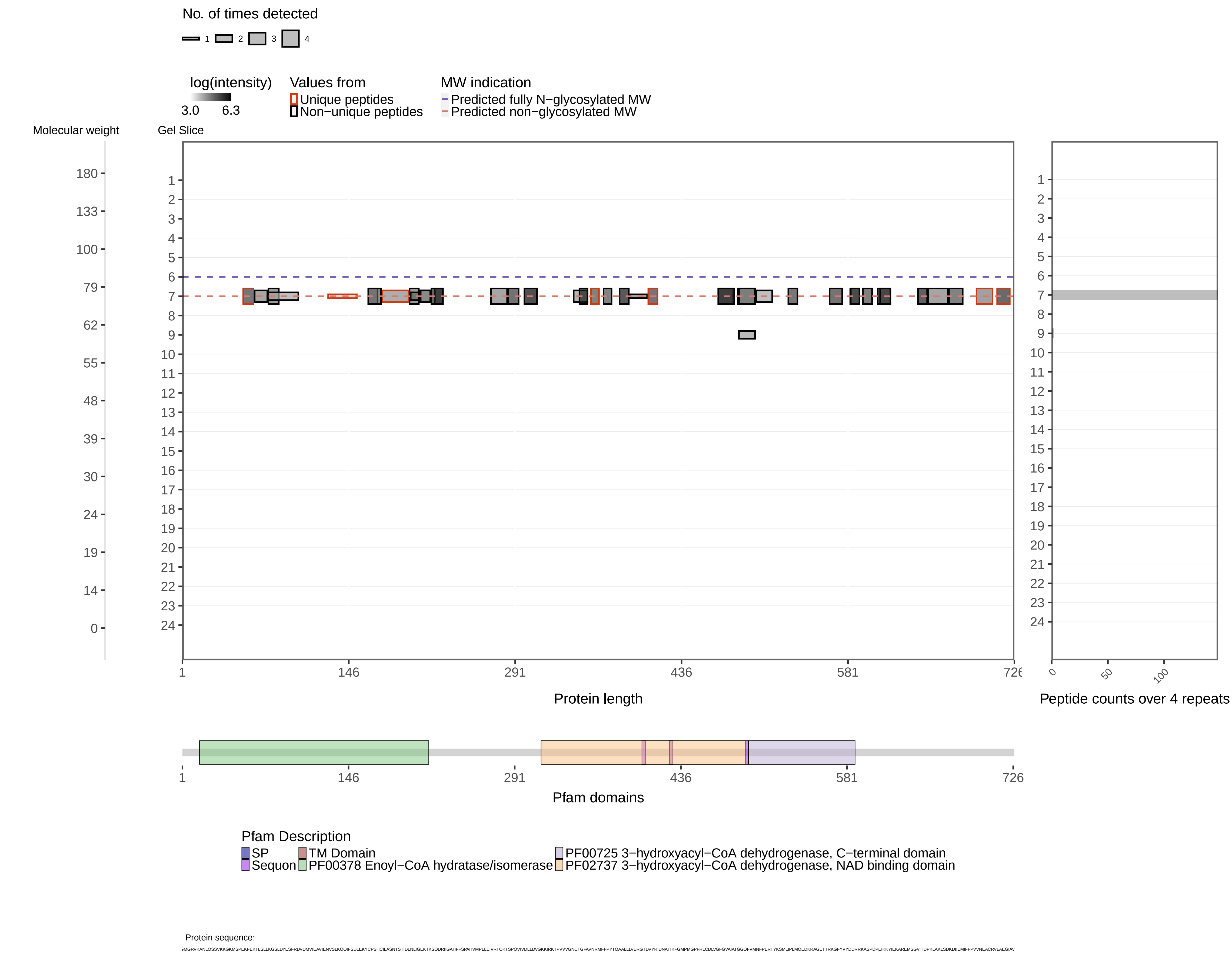

### NbL19g22450.1 eukaryotic_translation_initiation_factor_2_subunit_gamma−like [MW = 50.74 kDa]

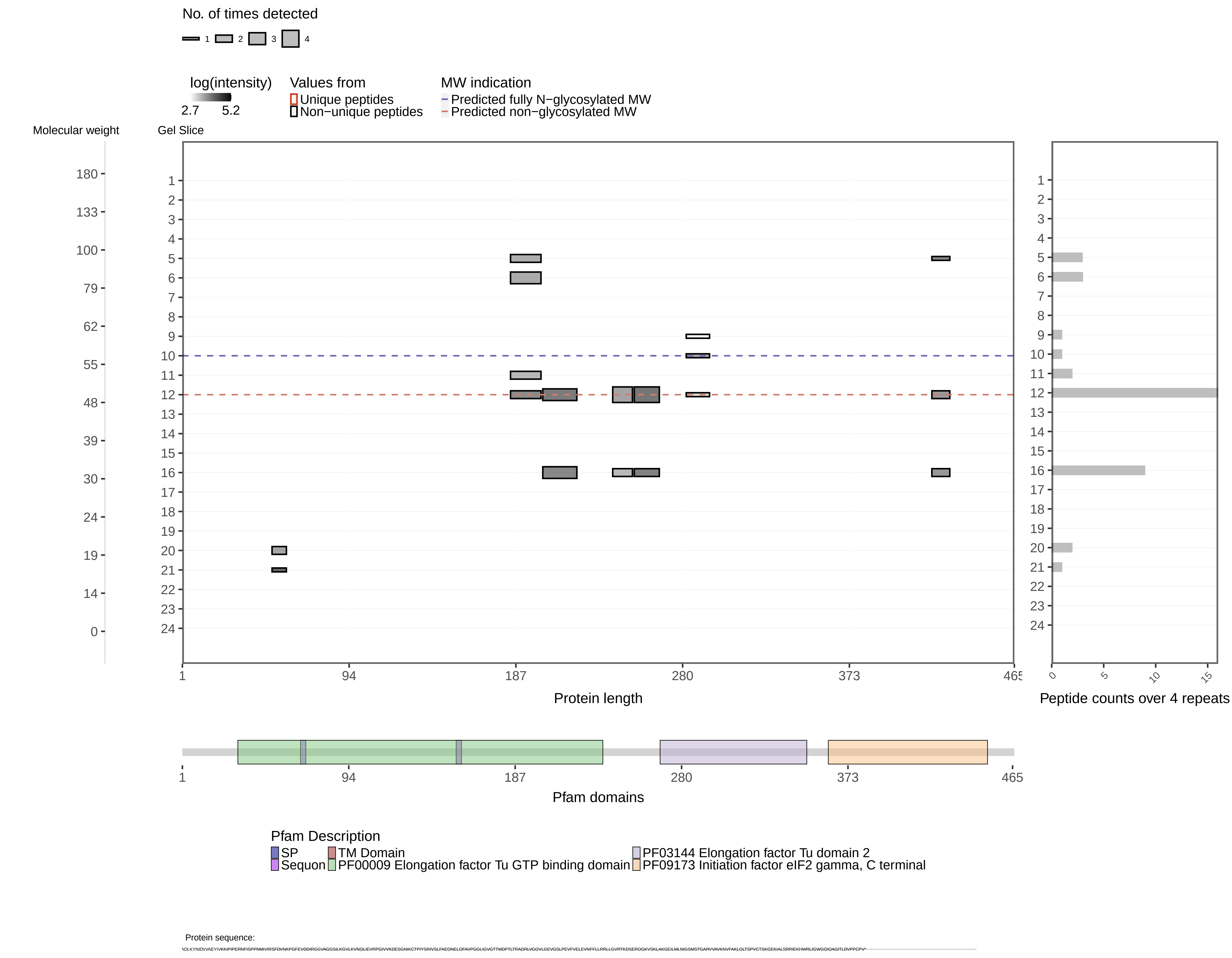

NbL19g22340.1 ATP_synthase_subunit_O_mitochondrial−like [MW = 25.84 kDa]

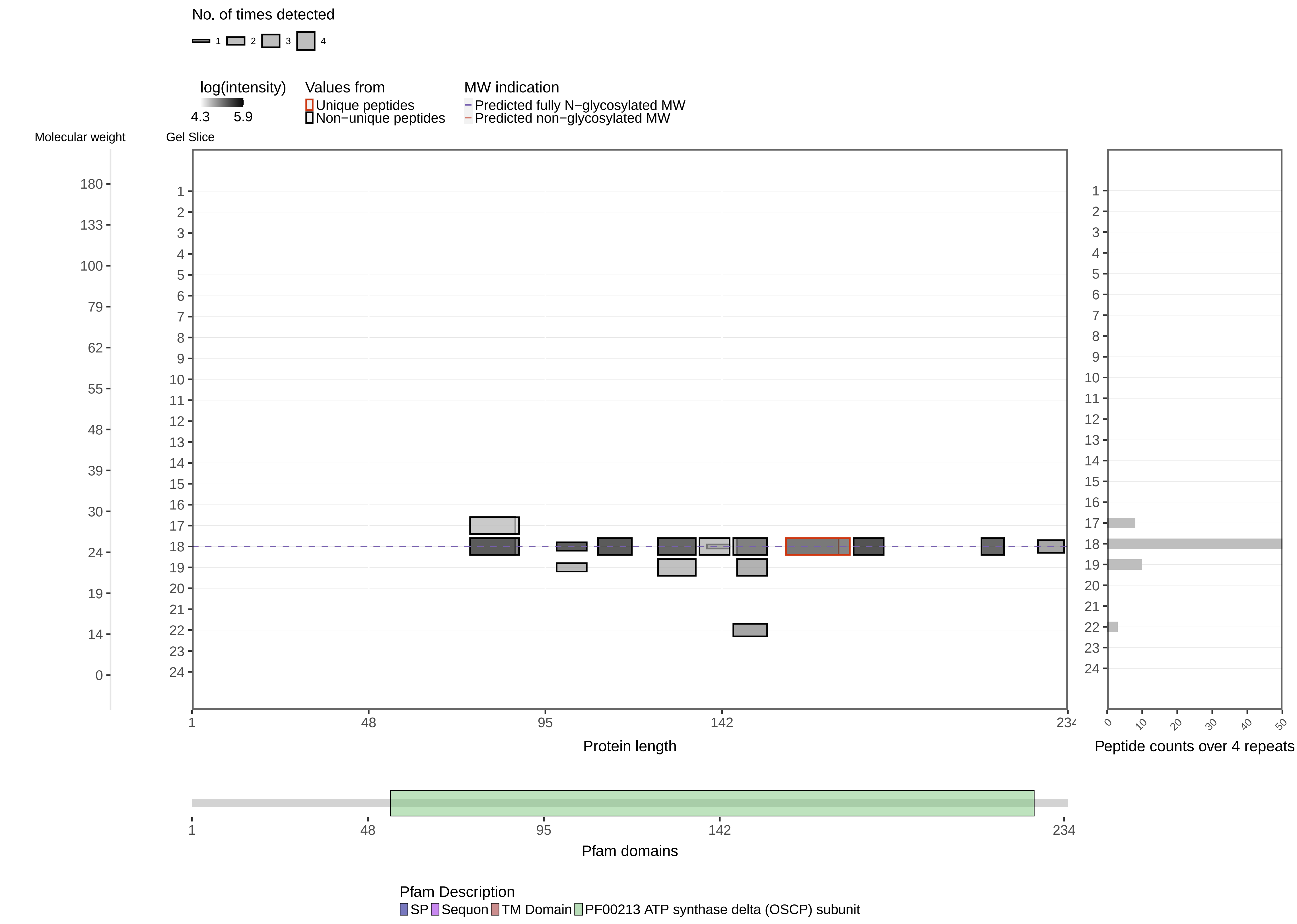


Protein sequence:
TDTRVKAISDICAQAKFGDLTRNFLVVLAENGRLKHIDAIAKRFAELTMAHKGELKAIVTTVIPLPPEEEKELKETLQDVMGKGKKVNIEQKIDPSILGGLVIEFGQKVFDMSIRTRARQMERLLKEPLNF*

NbL19g22330.1 heat_shock_factor−binding_protein_1−like [MW = 12.5 kDa]

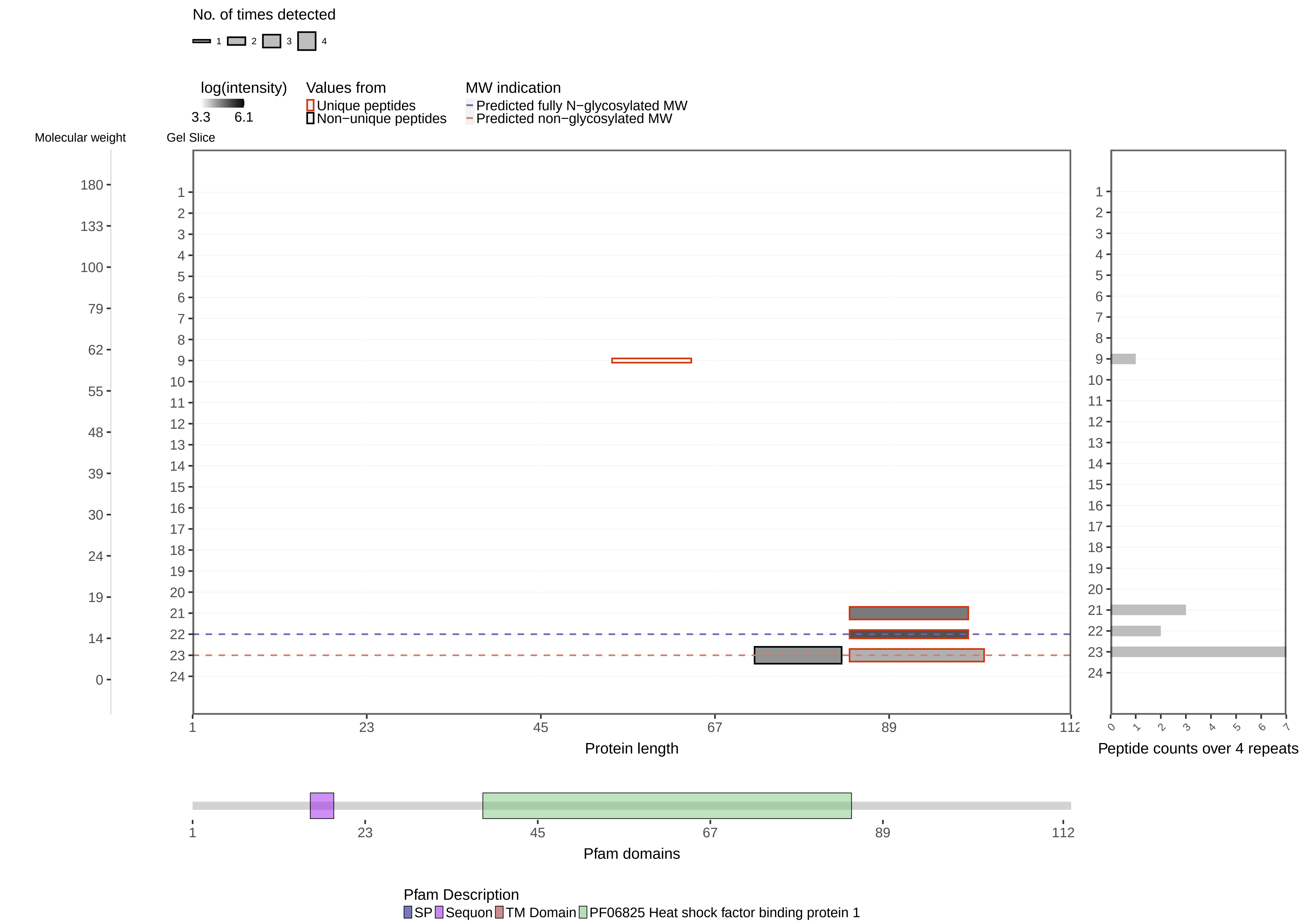


Protein sequence:

QNLLQQMQTRFQTMSESIILKIDEMGNRIDELEQSINDLRAEMGQDGAPSPSATLKPREEPKTSDDNA*

### NbL19g22230.1 lipoyl_synthase_chloroplastic [MW = 40.88 kDa]

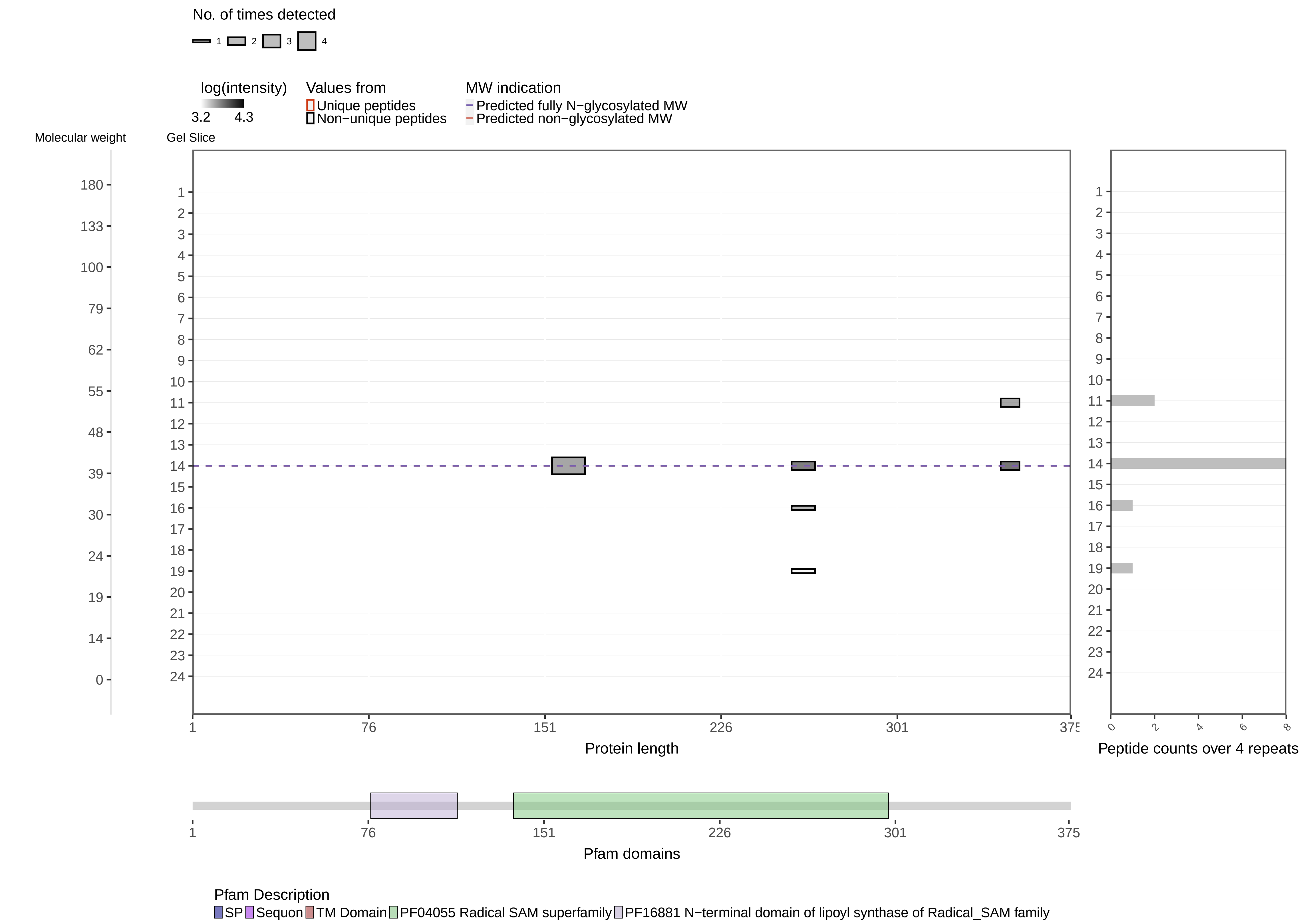


Protein sequence:

/GVDYIVLTSVDRDDLPDGGSGHFAETVKAMKLLKPDIMVECLTSDFRGDLSAVSTLVHSGLDVFAHNIETVKRLQRIVRDPRAGYEQSLSVLKHAKLDKEGMITKTSIMLGLGETDDELKEAMADLRAIDVDILTLGQYLQPTPLHLTVKEYVTPEKFAFWKEYGESIGFRYVASGPLVRSSYRAGELFVQTMVKERTRNA*

### NbL19g22140.1 signal_recognition_particle_9_kDa_protein [MW = 12.04 kDa]

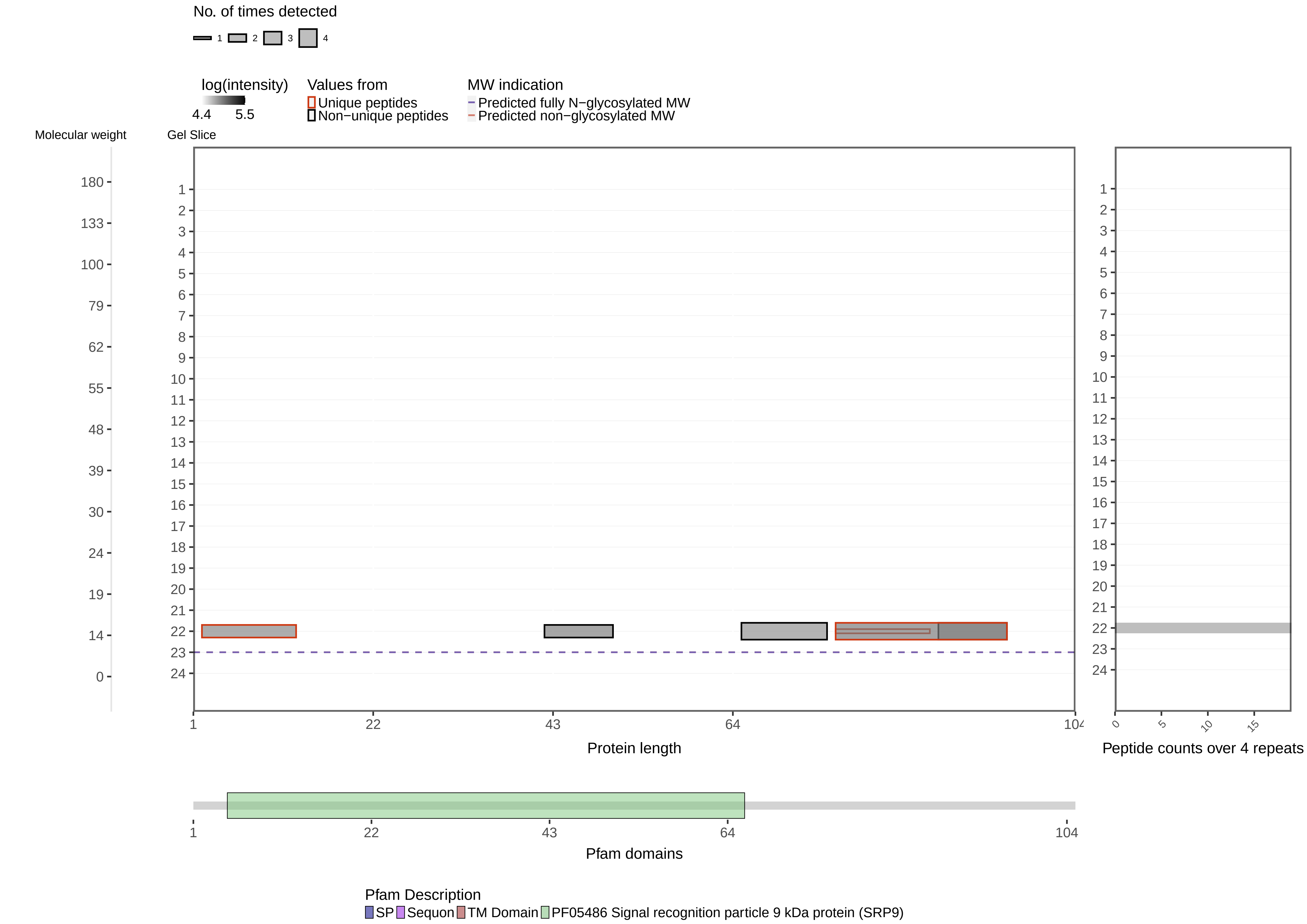


Protein sequence:

VLKVTDDTECIKFRTDQAQDAKKMEKLNNIFFTLMARGPDVDISEVGGKEQMEAQAPKKGRGRKQ*

NbL19g21970.1 single−stranded_DNA−binding_protein_mitochondrial−like [MW = 26.14 kDa]

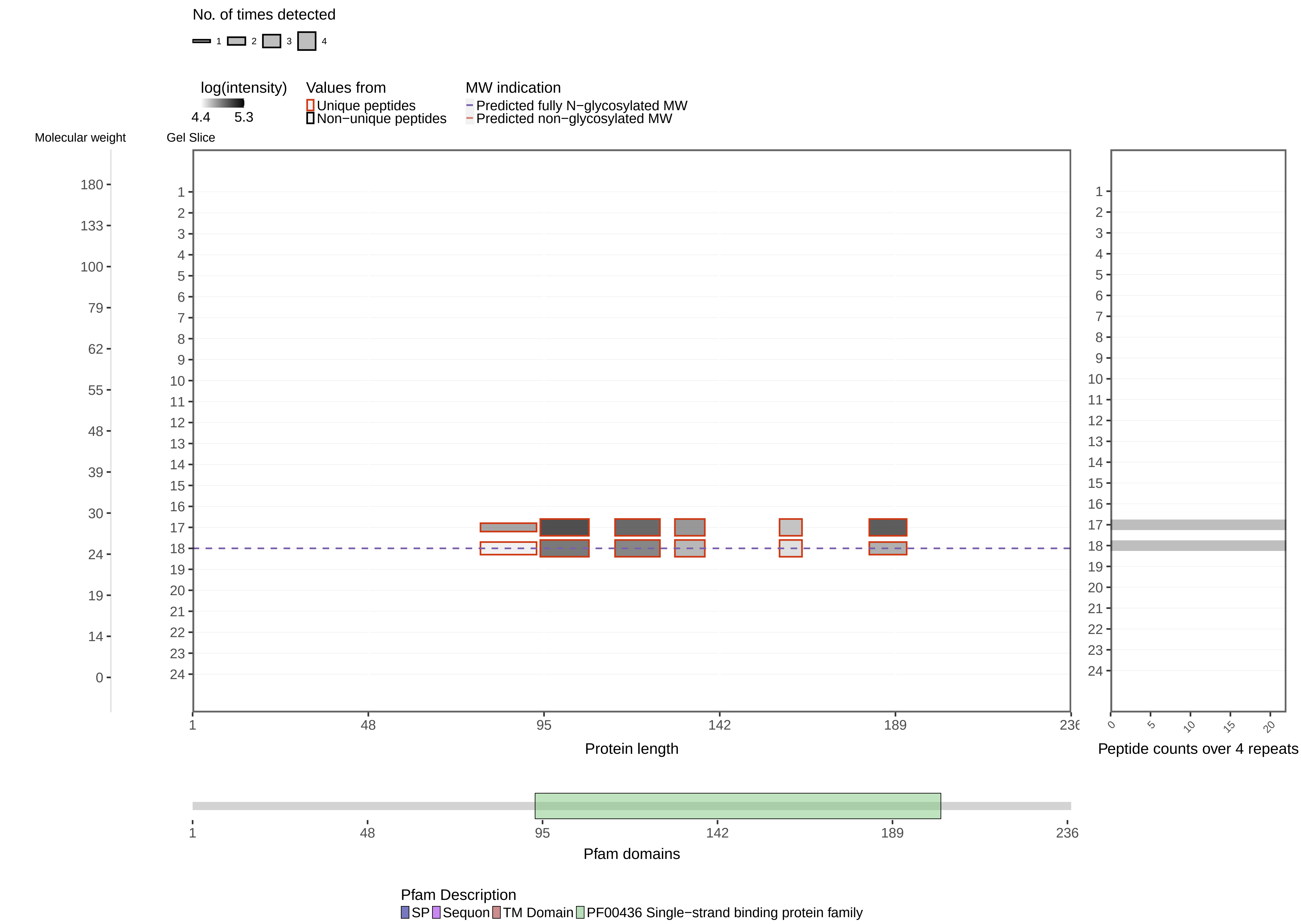


Protein sequence:

)KKLKSGRSVTLLSLGTGGIRNNRRPFDNEEPREYANRCAVQWHRVSVYPERLGEMAMKSIVPGSILYVEGNLESKIFTDPITGLVRRIREVLYEEMVVWFSLGMGMKPSSHHKQKLKGLVITESLRVSL*

### NbL19g21950.1 probable_acyl−activating_enzyme_17_peroxisomal [MW = 78.95 kDa]

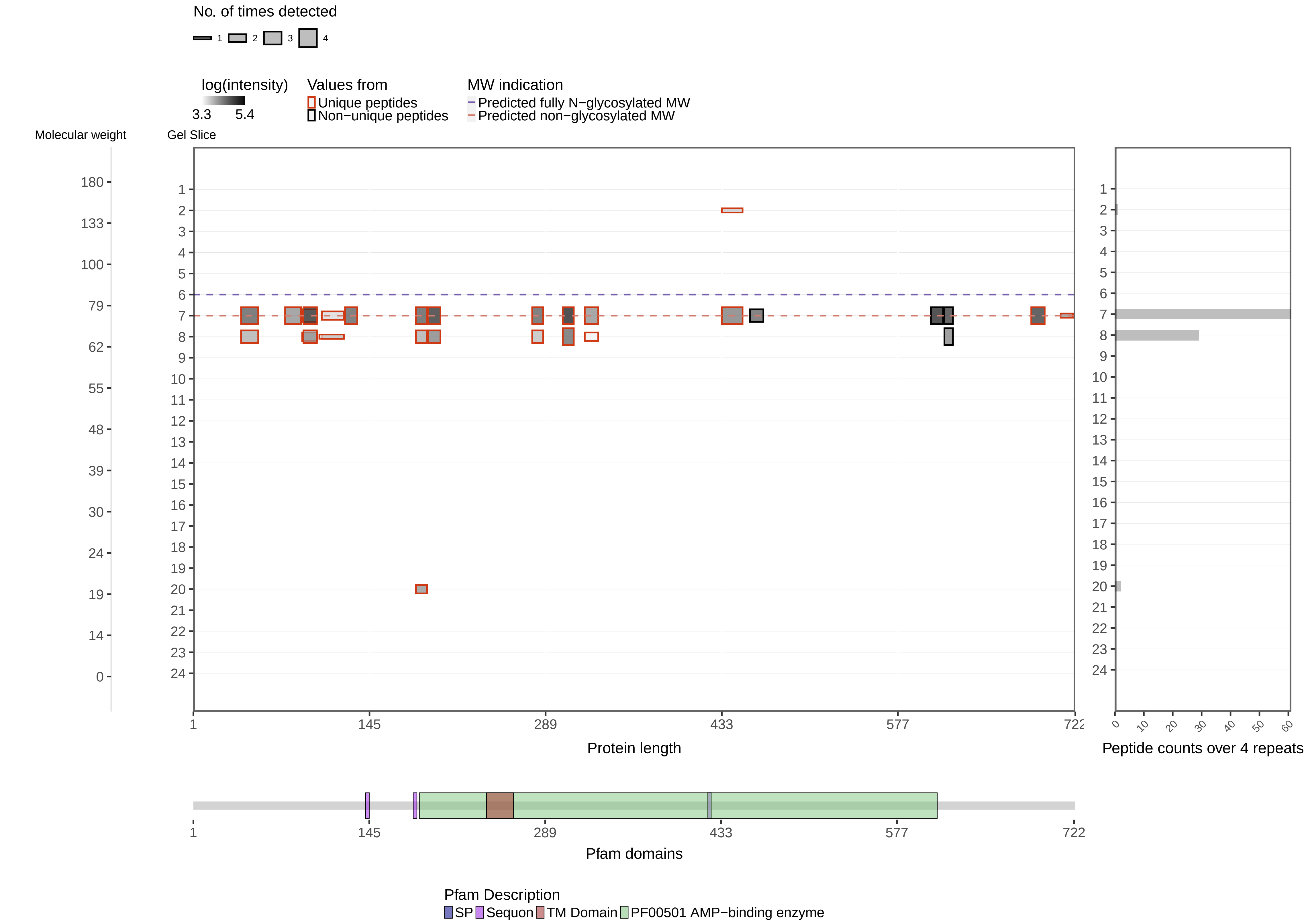


Protein sequence:
FTNILFSSGTTGEPKAIPWSLLSHFKAAADGWCHMDIKKGDVVAWPTNLGWMMGPWLVYASLLNGASIALYNGSPLGSGFAKFVQDAEVTMLGVIPSIVRTWKAKNSPDGFDWSTIRCFGSTGEASSVDEYLWLMGRAQYKPIIEYCGGTEIGGSFVSGSLLQPQSLAAFSTAVMGCSLHILGDDGLPIPSDVPGIGELALGPLMFGASSTLLNADHNEIYFKGMPVLNGKVLRRHGDVFERTSKGYYHAHGRADDTMNLGGIKVSSLEIERICNAADENILEAAAVGVPPA

NbL19g21920.1 H/ACA_ribonucleoprotein_complex_subunit_2−like_protein [MW = 17.07 kDa]

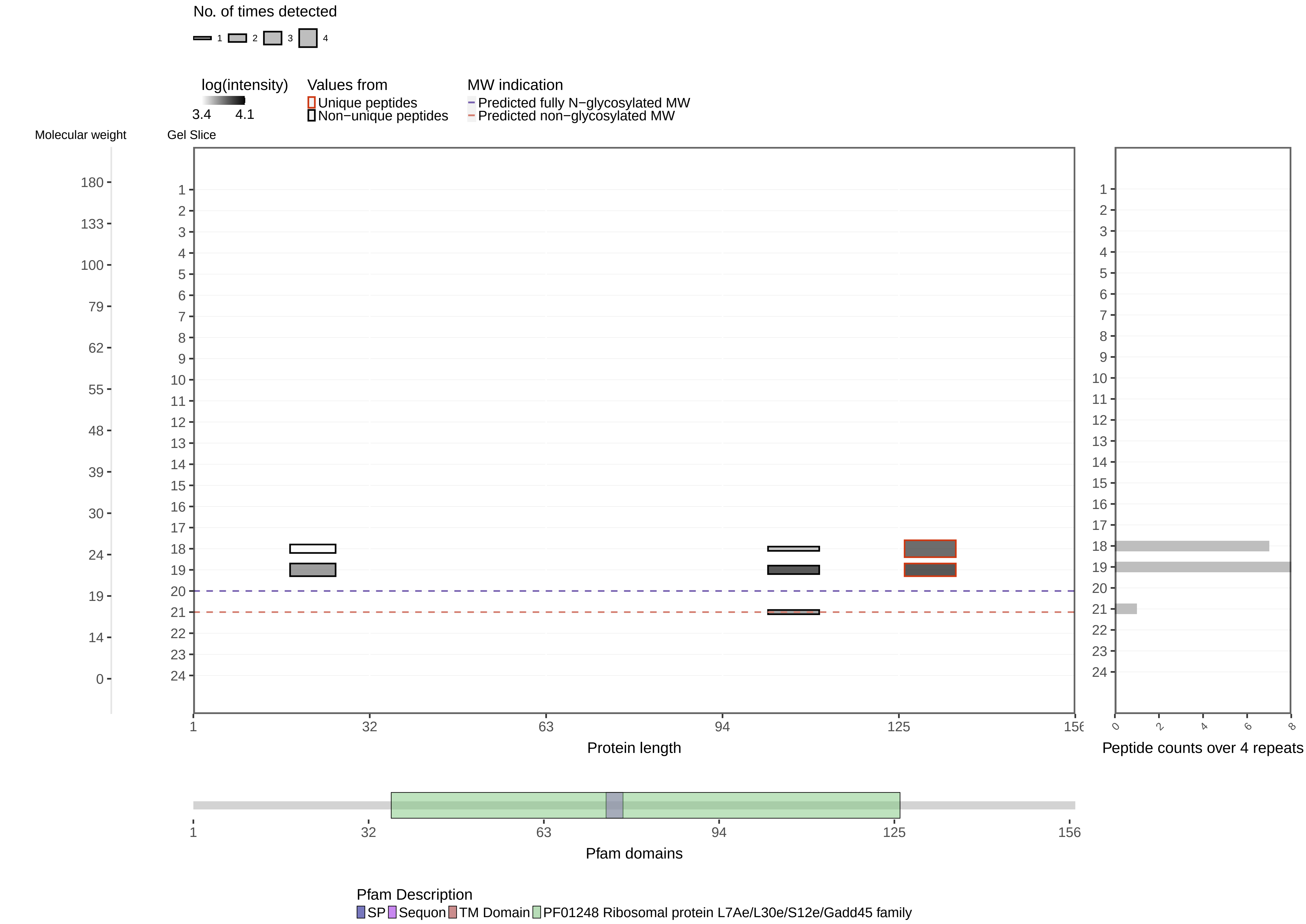


Protein sequence:

QKGVCVIAGNISPIDVITHVPILCEEAEIPYVYVSSKEDLANAGATKRPTCCVLVQTKPAKEELTQEDQEKLKADYDQVVSEIGEMASSMF*

### NbL19g21910.1 agmatine_deiminase−like [MW = 41.57 kDa]

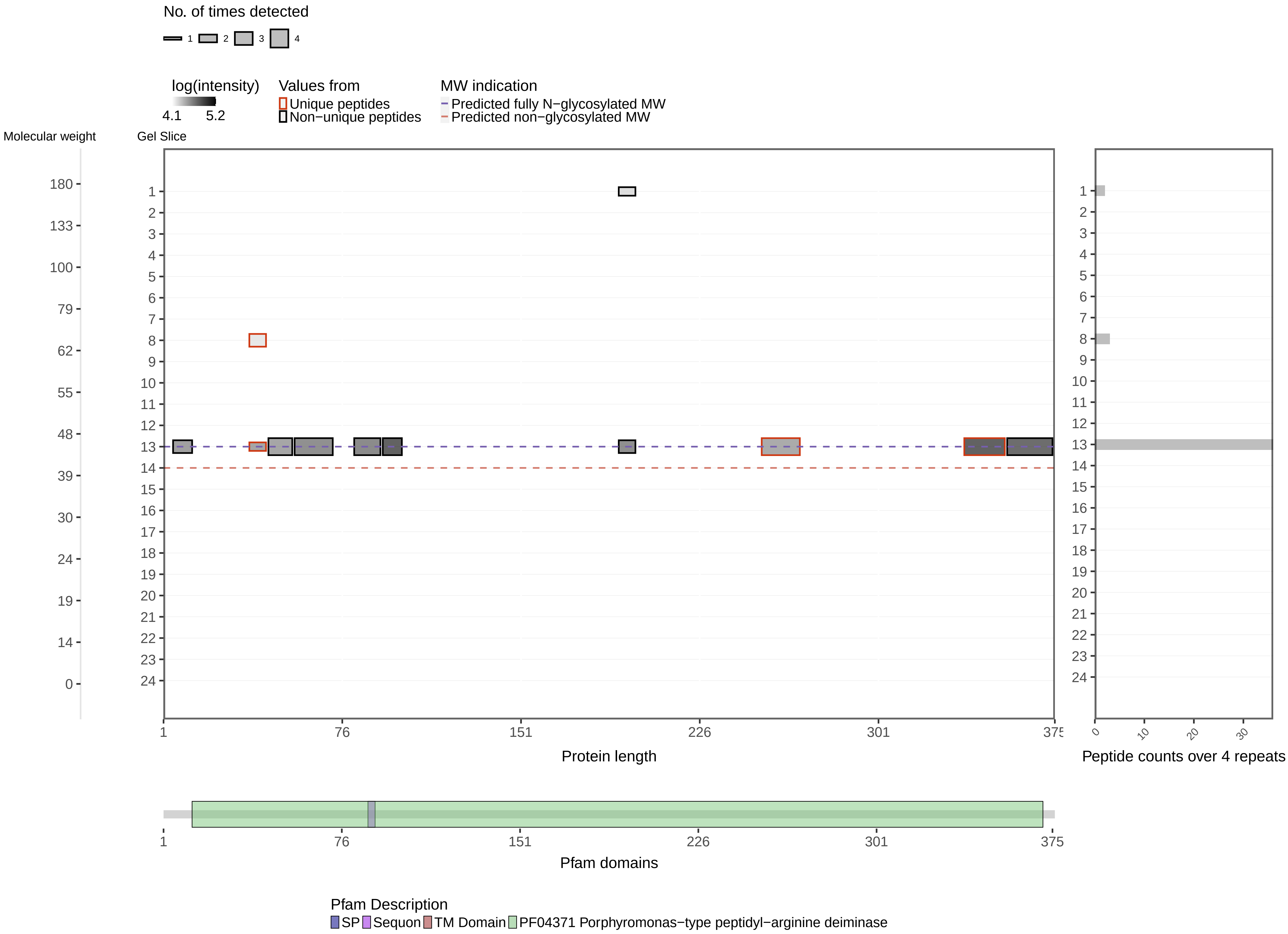


Protein sequence:

TCLTTEECLLNKNRNPHLSKEQIENELKAYLGVRKIIWLPRGLFGDDDTNGHIDNMCCFVKPGVVLLSWTDDHSDPQYERSIEALSILETTSDANGRKLQVIKLHVPGPLCMTDEEADGIFQDGDAKPRTSGTRLAASYVNFYIANGAIITPQFGDKKWDEEAIRVLSLAFPDHEVVGVEGAREIVLGGGNIHCITQQQPSGP*

### NbL19g21890.1 uncharacterized_LOC109206323 [MW = 35.25 kDa]

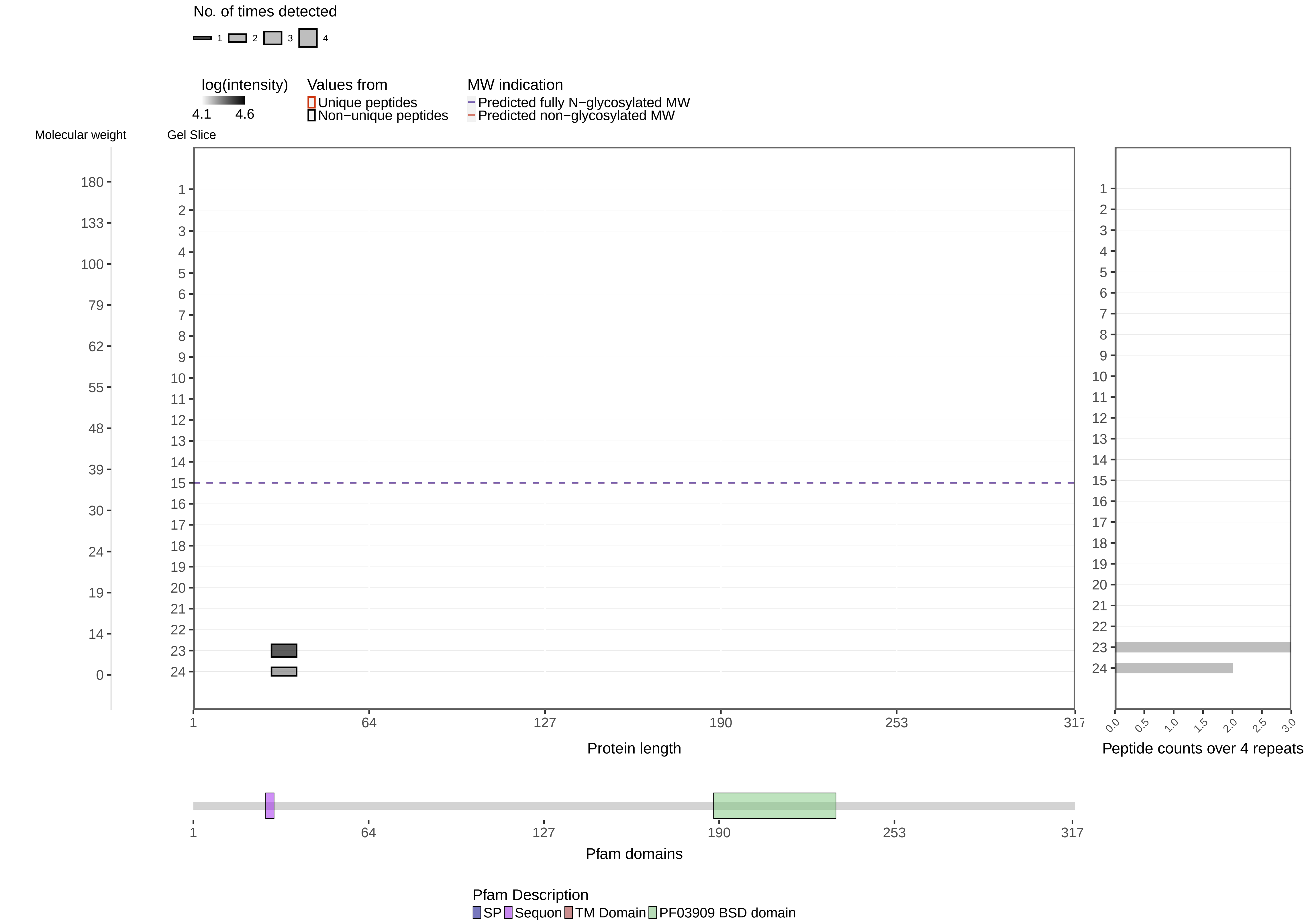


Protein sequence:

TDDLREFVKGITINTFQDFPLEDDSVISDIPTVSNVRQDLTEFQEKHAKLVLQSVKEISKLRYELCPRIMRERKFWRIYFILVNSHVAPYEKKYMEEAKIKSVEKAQVESEKQLSSAGTTSKPVVEAAPQTNKKVASSTSDQDLDVFLLGEDSDDGPDDGDDAFDDDFDKI*

NbL19g21790.1 psbP−like_protein_1_chloroplastic [MW = 28.07 kDa]

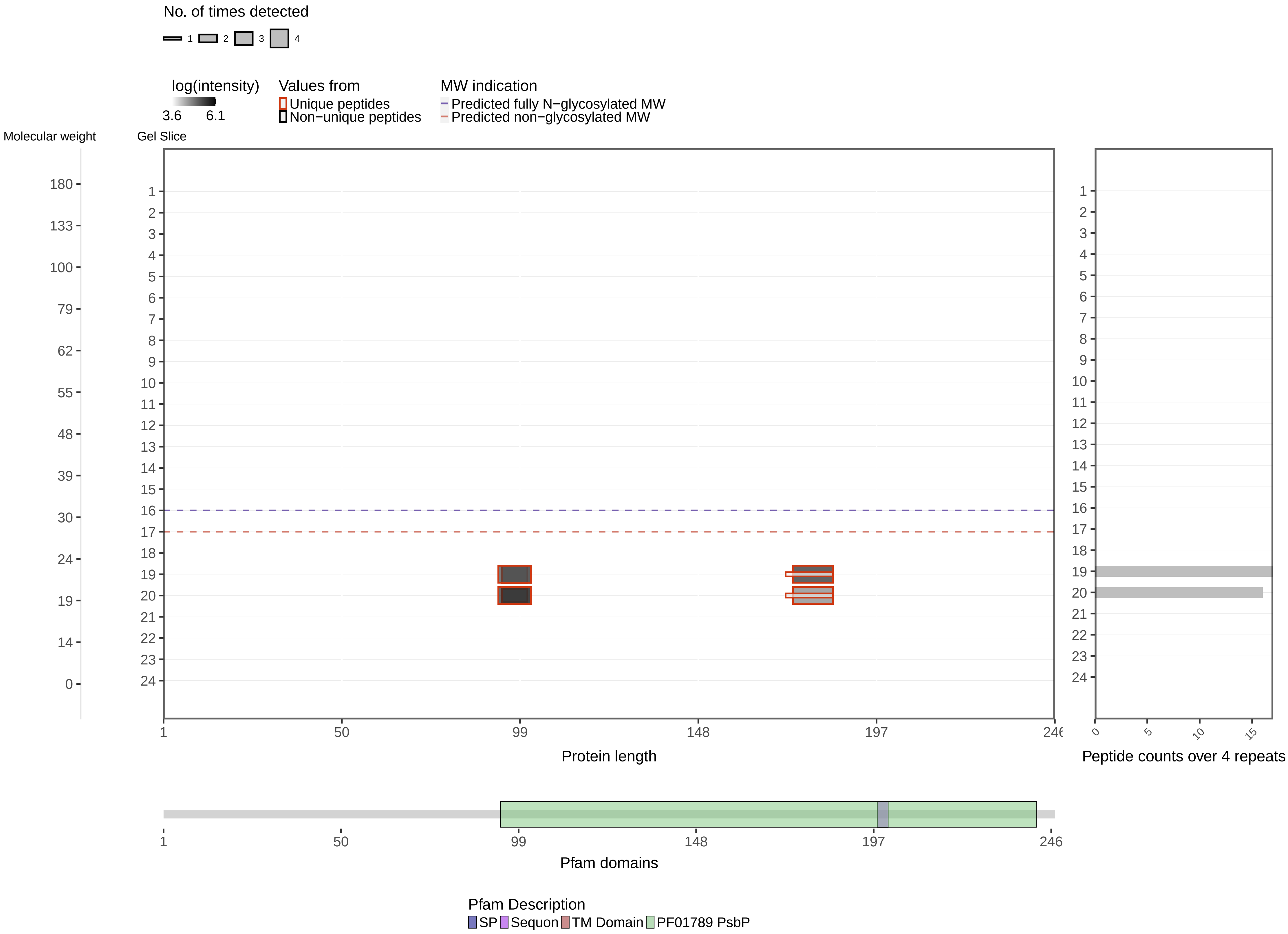


Protein sequence:

PFGWQEVVVEGQDKVFKDVIEPLESVSVNVIPTSKQDIRDFGSPQEVAETLIKKFLSSPSQKTKLIEASEHDIEGKAYYTFEFVAQAPNFTRHGLTAVCIGNGKFYTLTTGANERRWEKMKDRLHTVVDSFQIFNV*

NbL19g21560.1 2−alkenal_reductase_(NADP(+)−dependent)−like [MW = 37.8 kDa]

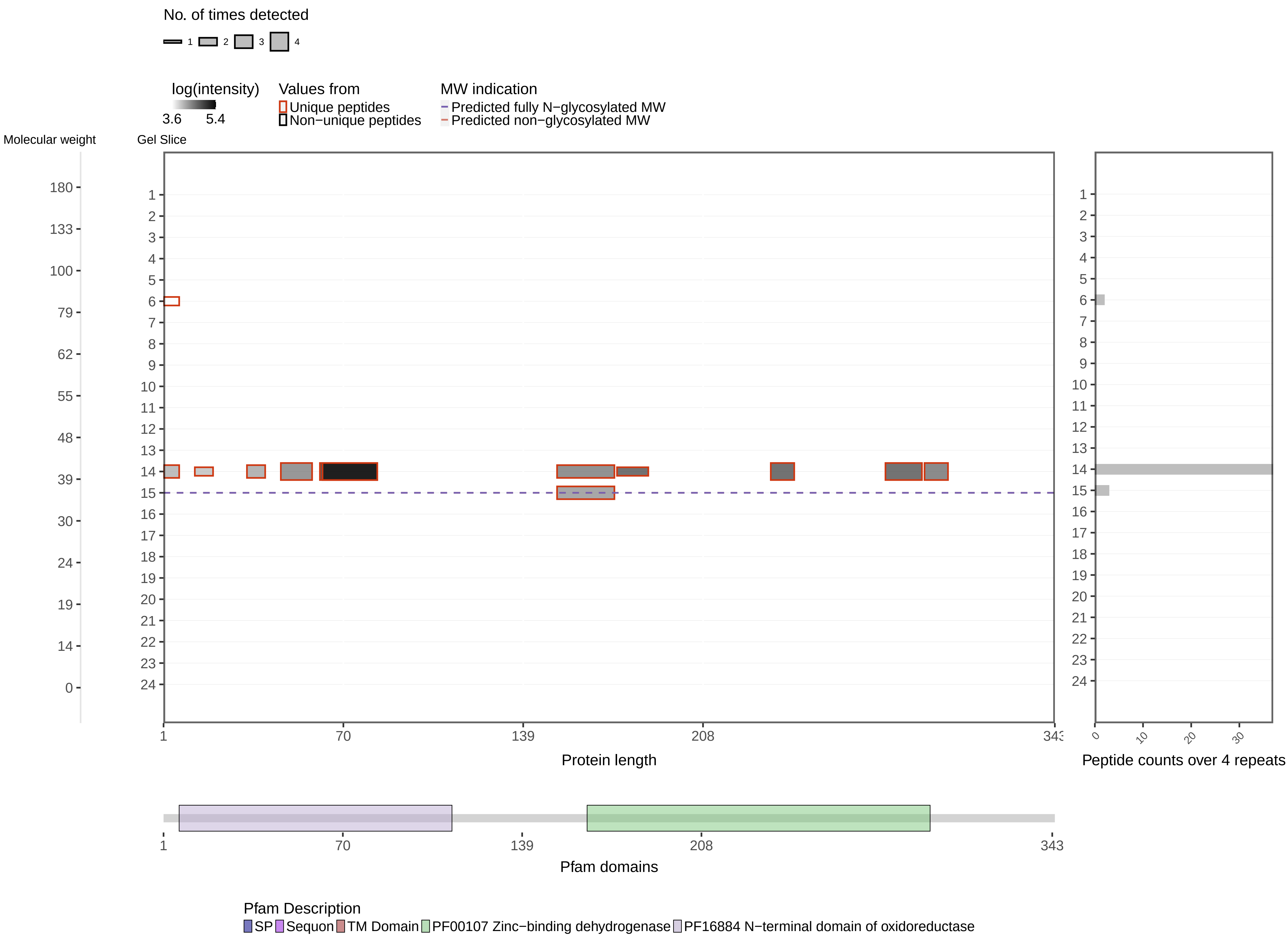


Protein sequence:
AASGAVGQLVGQFAKLLGCYVVGSAGTKEKVDLLKNKFGFDEAFNYKEEQDLSAALKRYFPDGIDIYFENVGGKMLDAVLLNMRVHGRIAVCGFISQYNLGQPEGVHNLFCLISKRVRMQGFLAADYFPLYQKFVEMVTPHIKEGKVTYVEDIAEGIESAPGALVGLFSGRNVGKQLVRVSRE*

NbL19g21520.1 UDP−glycosyltransferase_74E2−like [MW = 50.54 kDa]

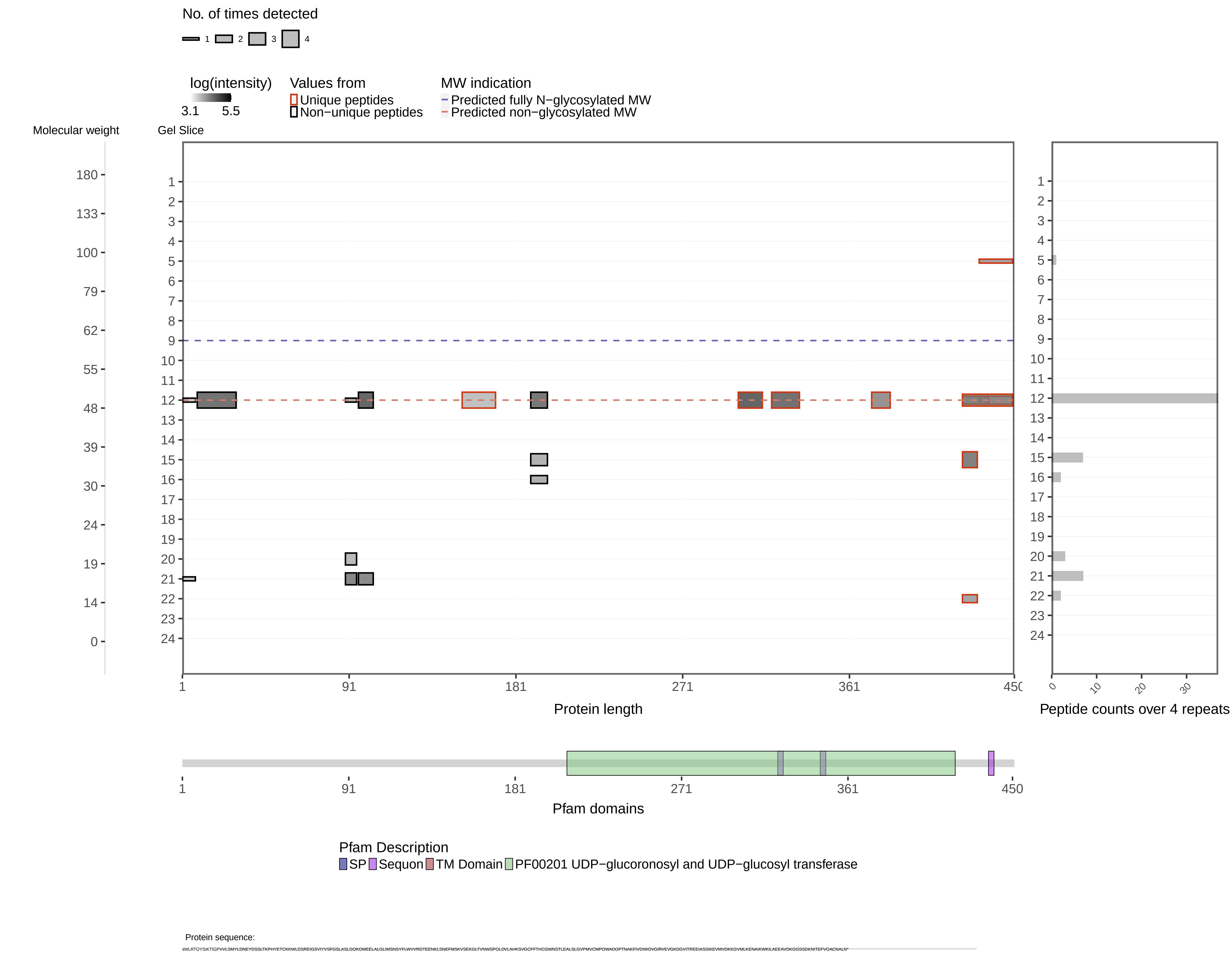

### NbL19g21430.1 pathogenesis−related_protein_STH−2−like [MW = 17.8 kDa]

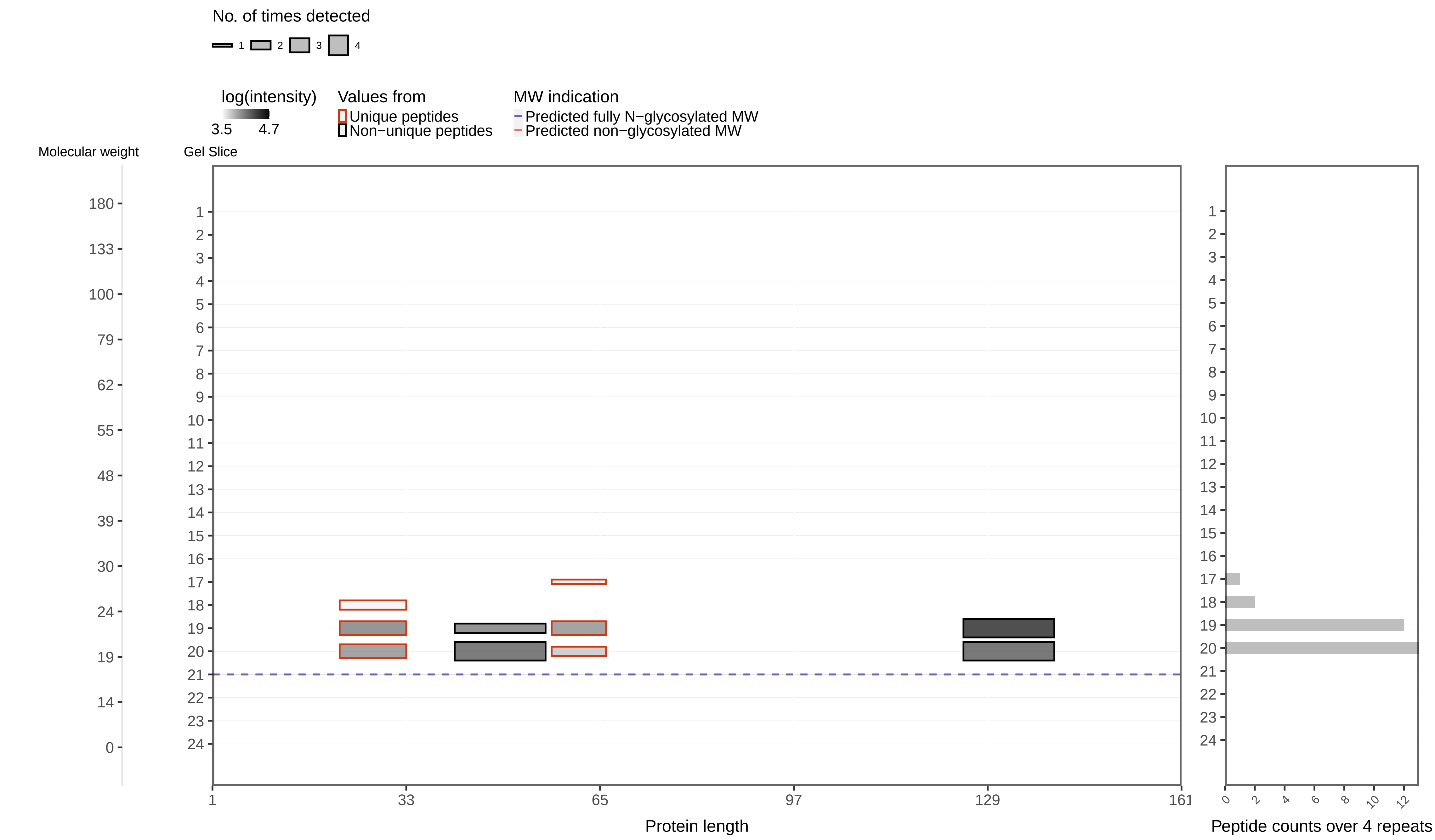


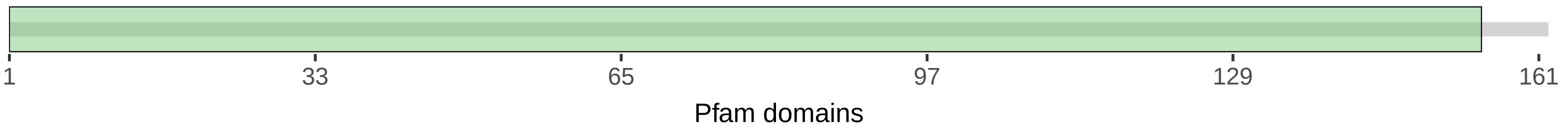


Pfam Description

SP Sequon TM Domain PF00407 Pathogenesis−related protein Bet v 1 family

Protein sequence:

LKHKIHVVDDKNMVIKYSLIEGEVLGENLEYISYVVSFEGSDNGGCICKTISNYHTKDDFVVKEEEIKEGEEKAVELFKAVEAYLLANPSSYA*

### NbL19g21380.1 carboxyl−terminal−processing_peptidase_2_chloroplastic−like [MW = 59.91 kDa]

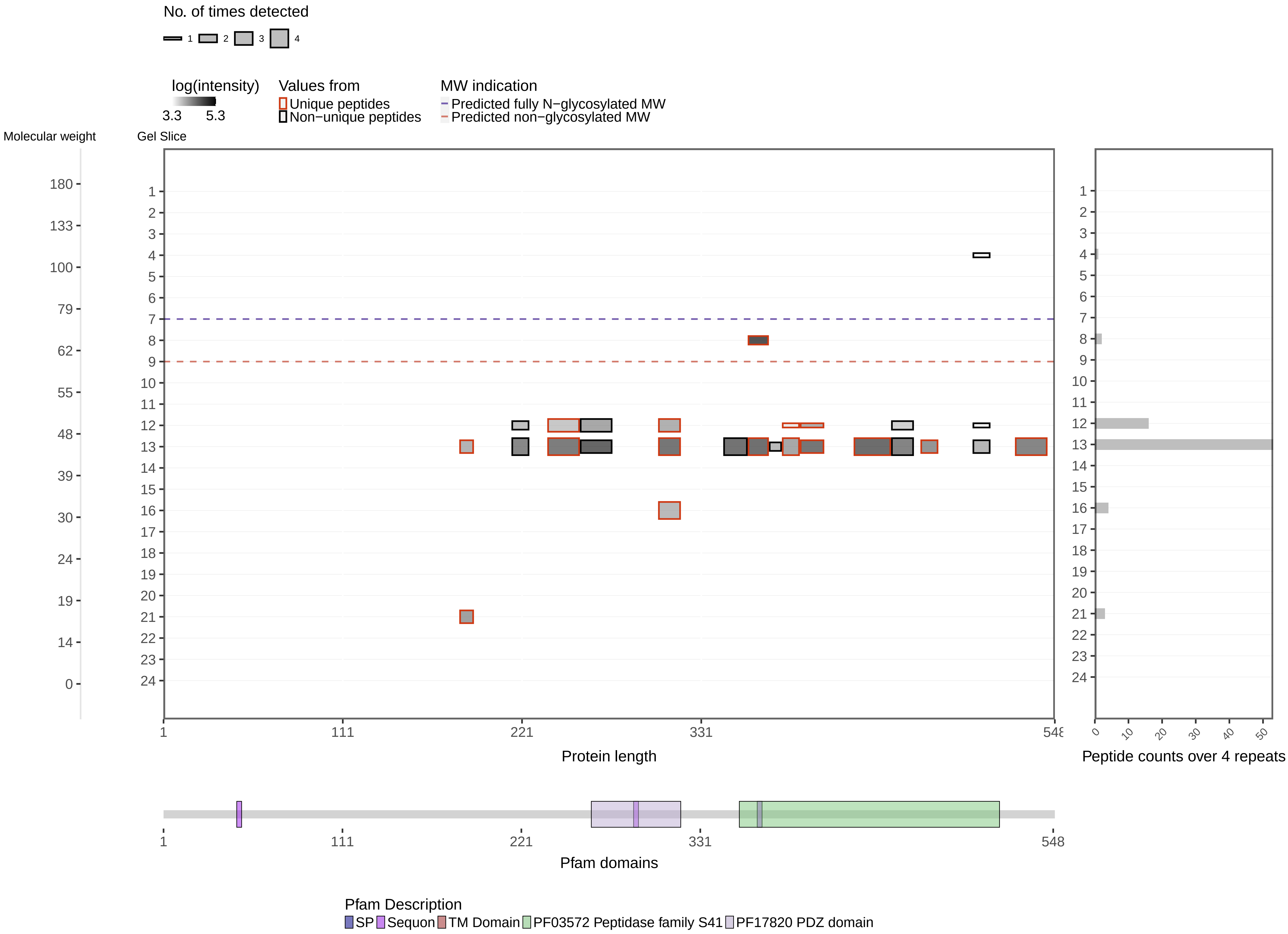


Protein sequence:

PGLVVVSASPGGPADRAGISSGDIILEIDNSSTENMGIYDAAERLQGPEGSGVELTVRRGSETRNLPLIREKVSLNPVKSRICKLPTGGDDAPQIGYIKLSTFNQNASGAVKEAIEALRKSNVNAFVLDLRDNSGGLFPEGVEIAKIWLNKGVIVYICDSRGVRDIYDTDGSNVVAASEPLAVLVNKGTASASEILAGALKDNKRALIFGEPTYGKGKIQSVFQLSDGSGLAVTVARYETPAHNDIDKVGVIPDHPLPASFPKDDESFCNCLQNPASACHINRVELF*

NbL19g21310.1 trans−resveratrol_di−O−methyltransferase−like [MW = 40.72 kDa]

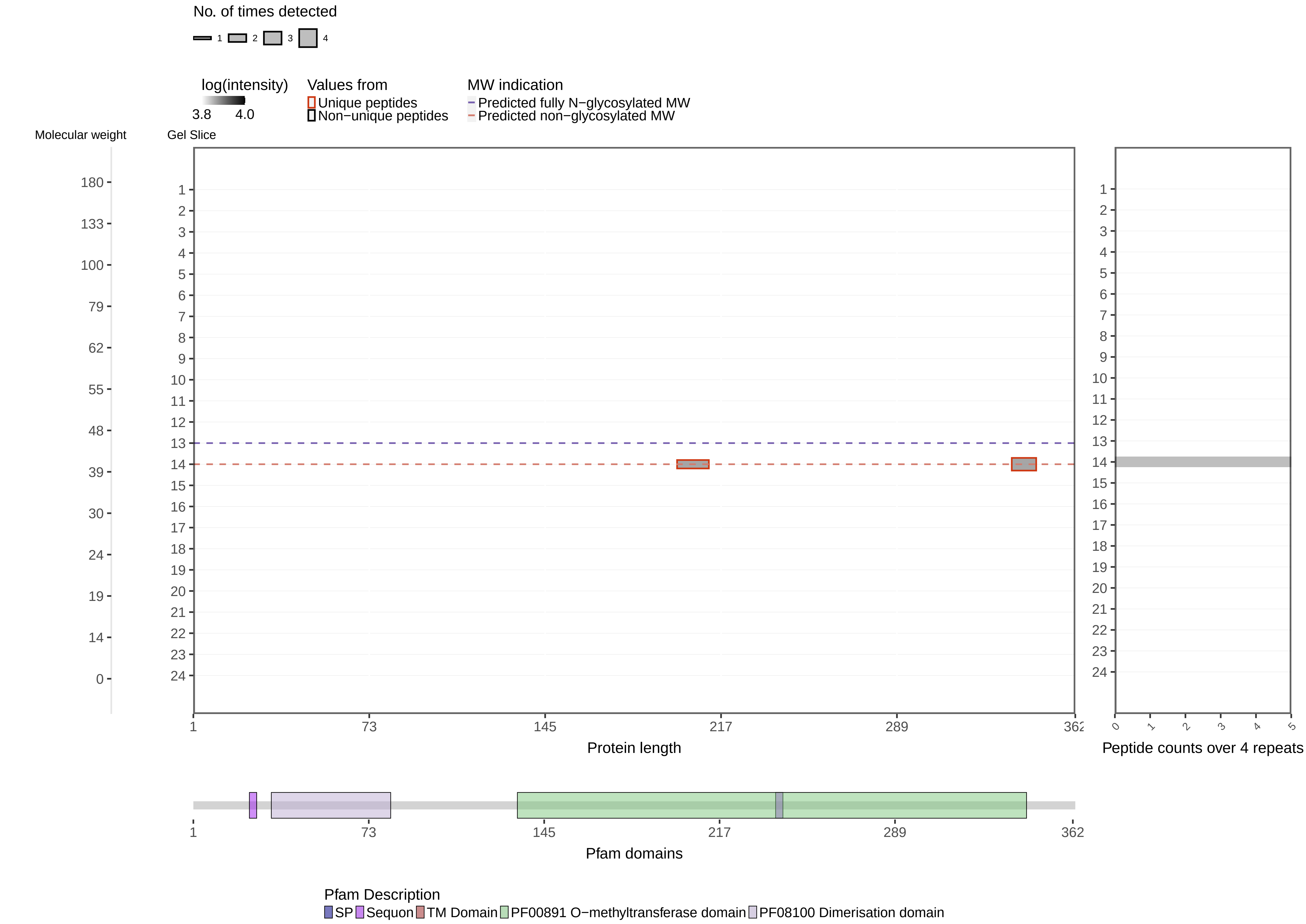


Protein sequence:

NCLFNEAMASDARLIISIMIENCKEVFEGLKSLVDVGGGTGTVAKAIADAFPKTNCIVFDLPHVIEDCEGRKNLSYVGGDMFKSIPSADAVLLKWILHDWNDEECIQILKKCKEAIPSKENGGKVLIIDMVLDKEKGDHKSFETQLFWDMLMMAFNSGKERTQQDWAKLFSTAGFSDYKIIPVLGLRSIIEVYP*

### NbL19g21260.1 transcription_factor_MYB1R1 [MW = 40.48 kDa]

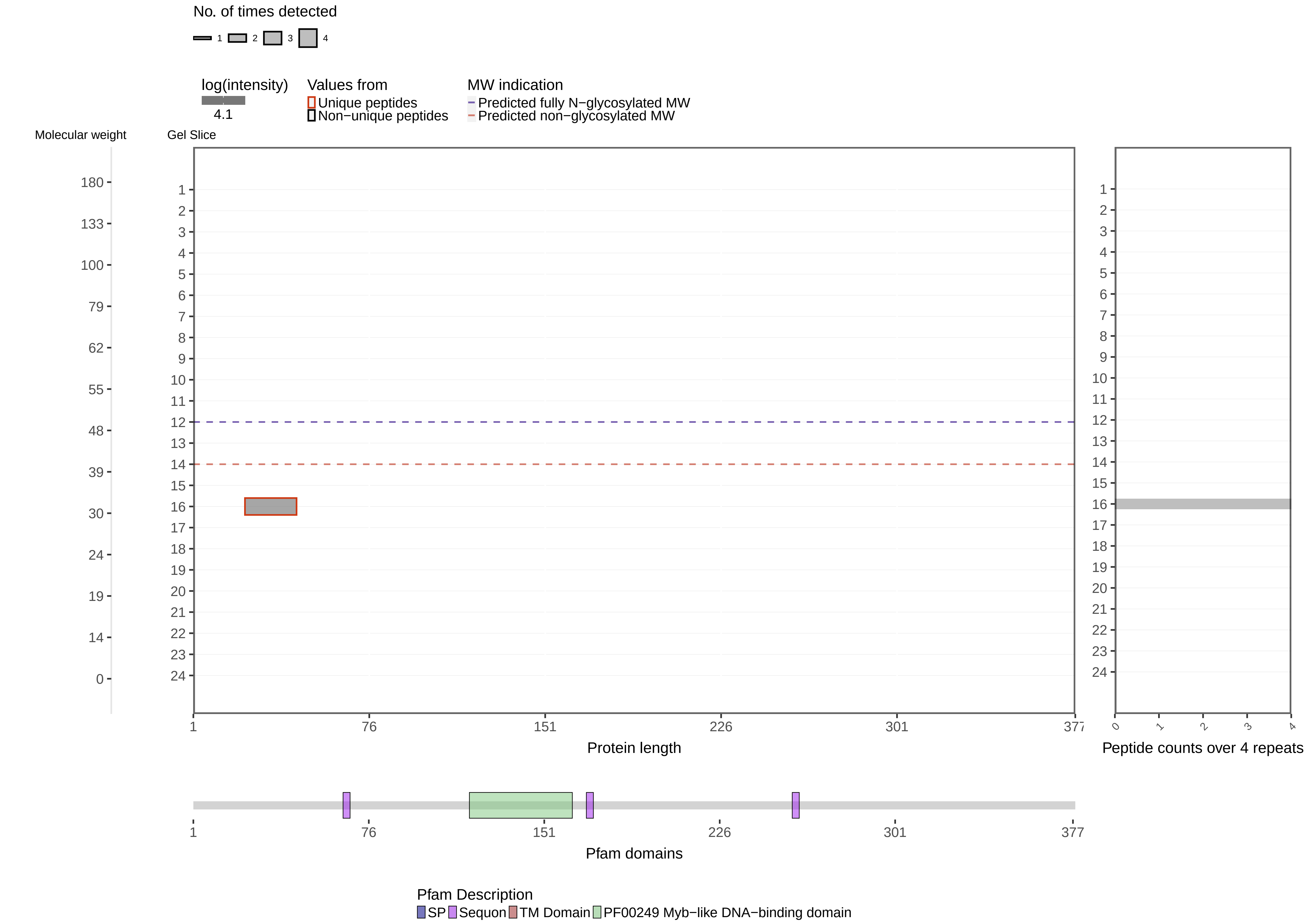


Protein sequence:

ATRRKRRSSLFDIVADTGTDASHPLSEEQFMLPPRAIESDKEHLAPSATKAIESDFADTLPSLDLSLKSDFESMETTASEPVEETKLNTTTDEIPSVYPAFFPAYIPVPYSFWPSNAVPVPEDRGSEPSHHQILKPIPTVPKEPVNVDELVGMSQLTLADAGSGHIEPSPLSLKLTAEPSRQSAFHASTPVKNREITKGETAPIQAV*

NbL19g21240.1 NADP−dependent_malic_enzyme_chloroplastic−like [MW = 70.51 kDa]

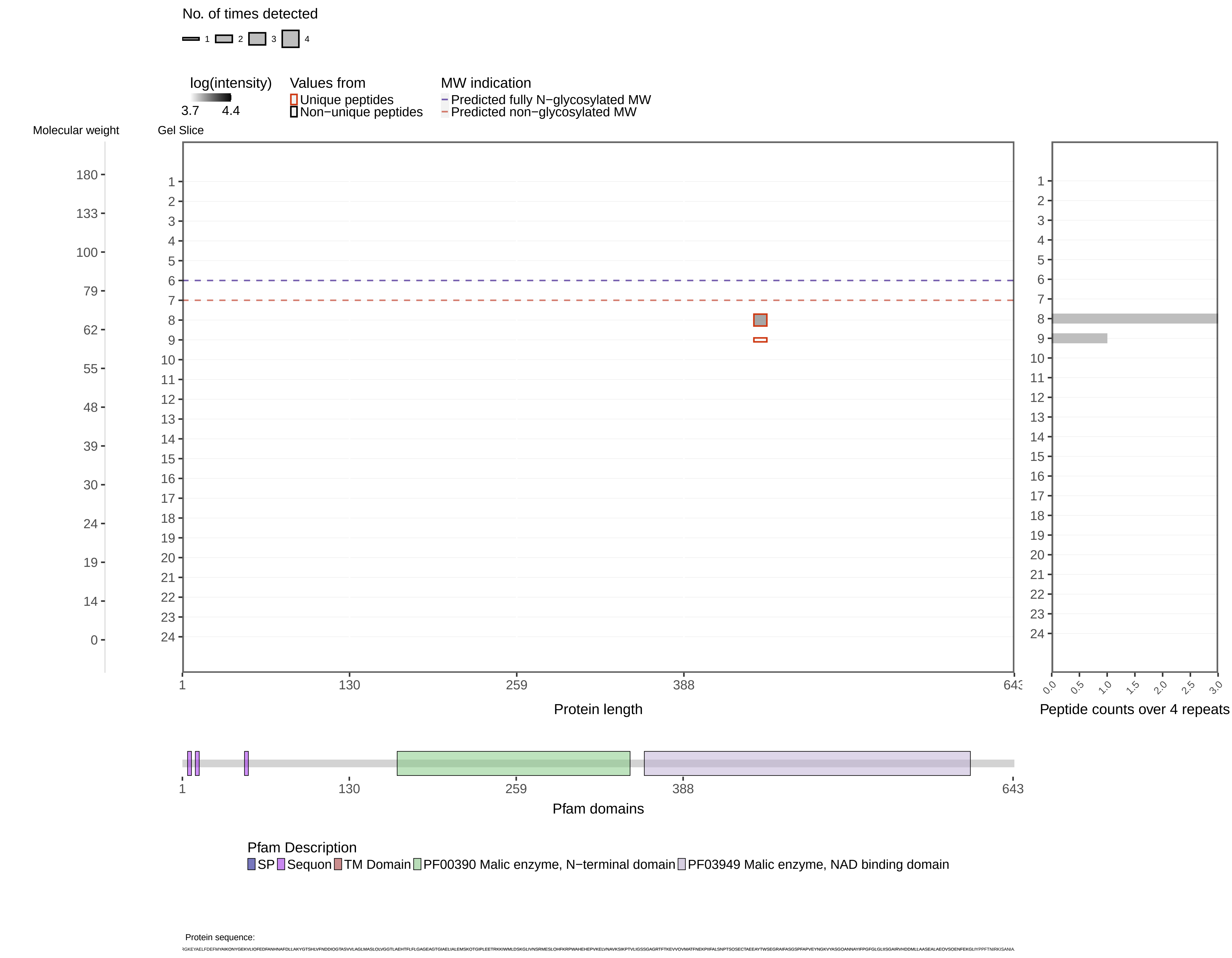

### NbL19g21140.1 aquaporin_TIP2−1−like [MW = 24.99 kDa]

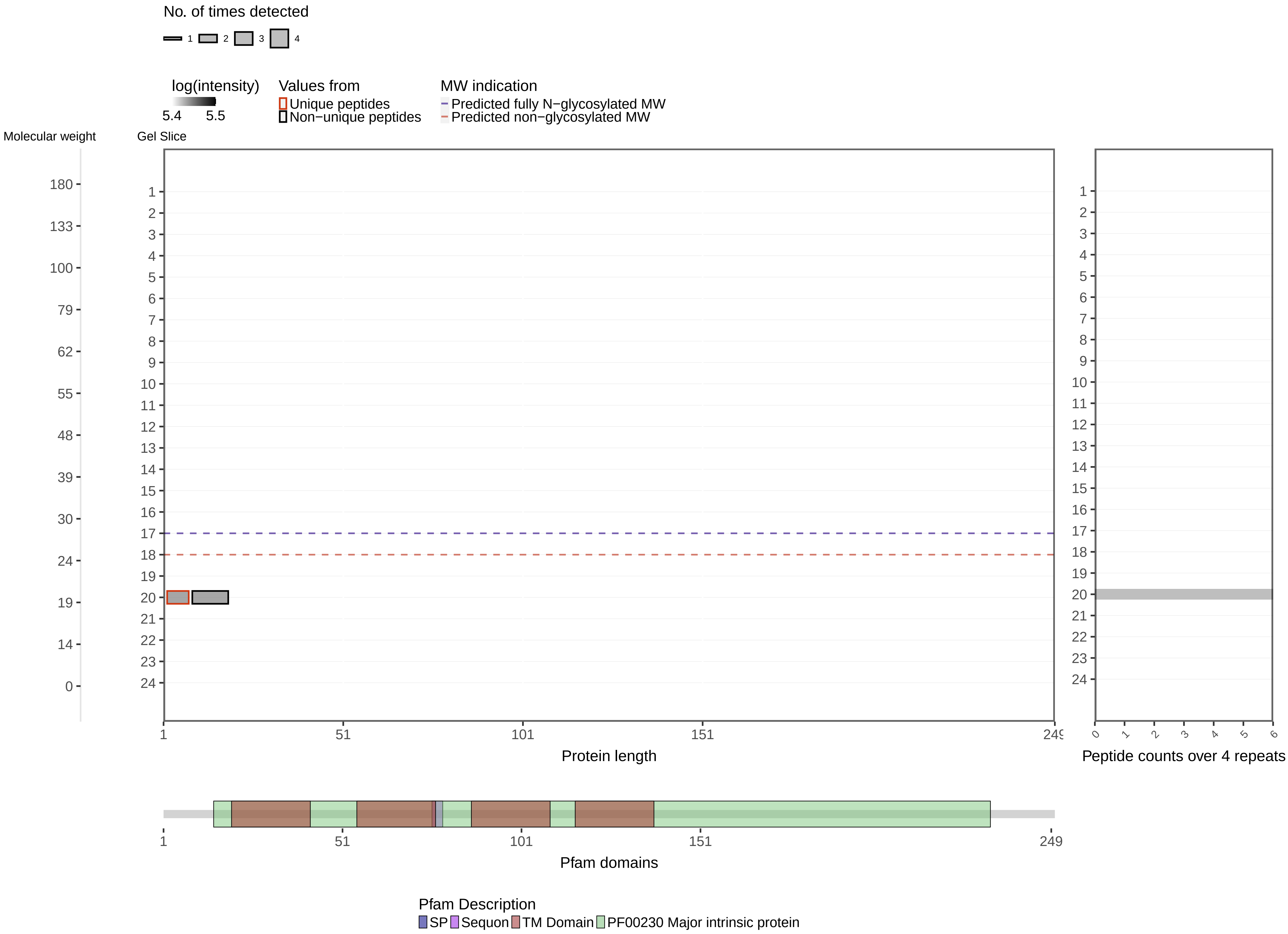


Protein sequence:

ATAASYLLKVVTGGLAVPIHSVAAGVGAAEGVVMEIIITFALVYTVYATAADPKKGSLGTIAPIAIGFIVGANILAAGPFSGGSMNPARSFGPAVASGNFAGHWIYWVGPLVGGGLAGLIYSNVFMNNDHAPLSTEF*

NbL19g21130.1 alpha−glucan_phosphorylase_H_isozyme [MW = 80.92 kDa]

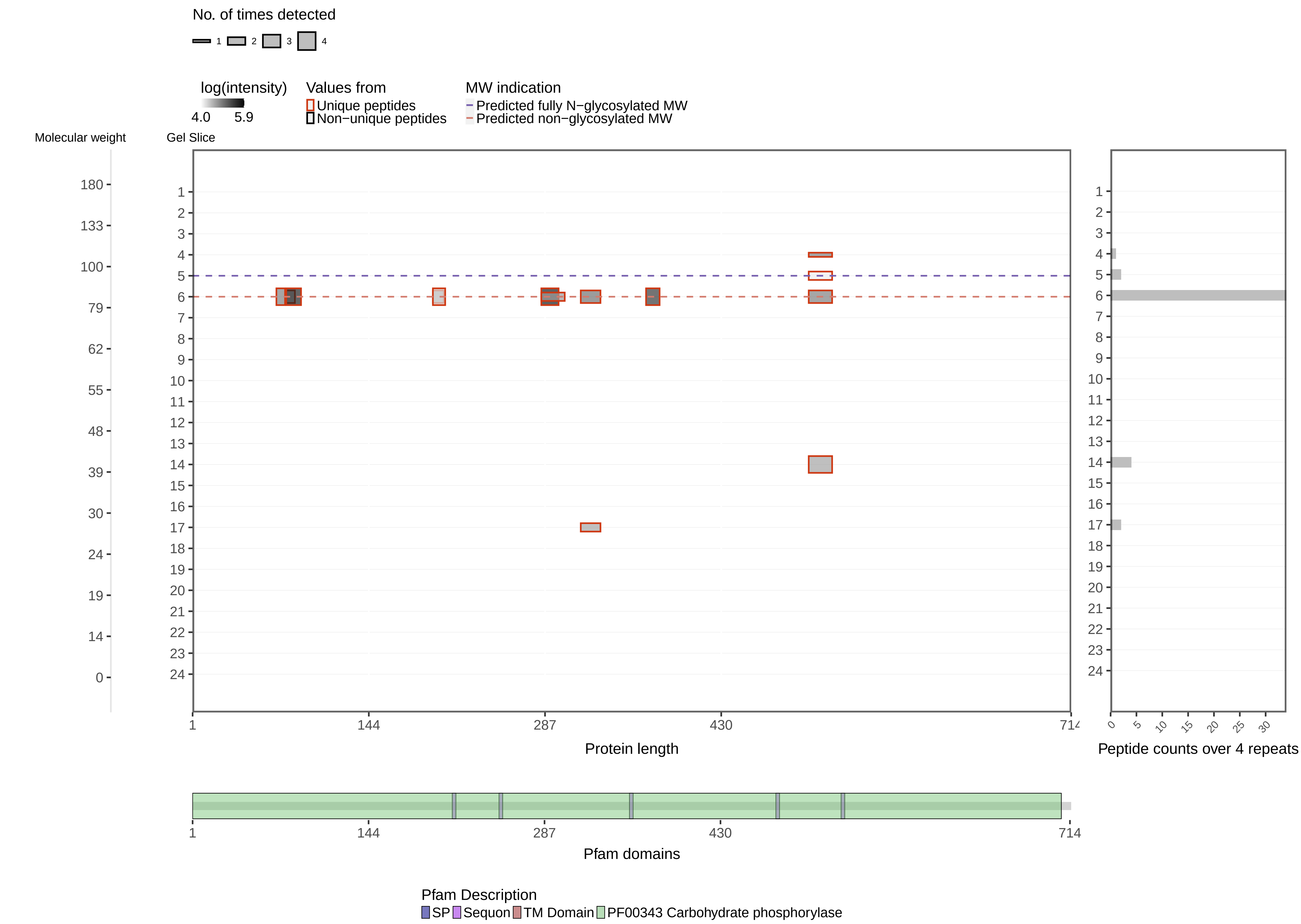


Protein sequence:

LFADYVSVWPTKFQNKTNGITPRRWIRFCSPELSNIITKWLKTDQWVTNLELLANLREFADNAELHAEWESAKMANKQRLAQYVLRVTGVSIDPDTLFDIQVKRIHEYKRQLLNVLGVVYRYKKLKEMSPQERNNTTPRTVMIGGKAFATYTNAKRIVKLVTDVGDIINSDPDINNYLKMVFVPNYNVSVAEMLIPGSEISQHISTAGMEASGTSNMKFALNGCLIIGTLDGANVEIREEIGEENFFLFGATADEVPRLRKEREDGLFKPDPRFEEAKQFIRSGAFGTYDYNPLLES

NbL19g21110.1 neutral_ceramidase−like [MW − SP = 81.65 kDa]

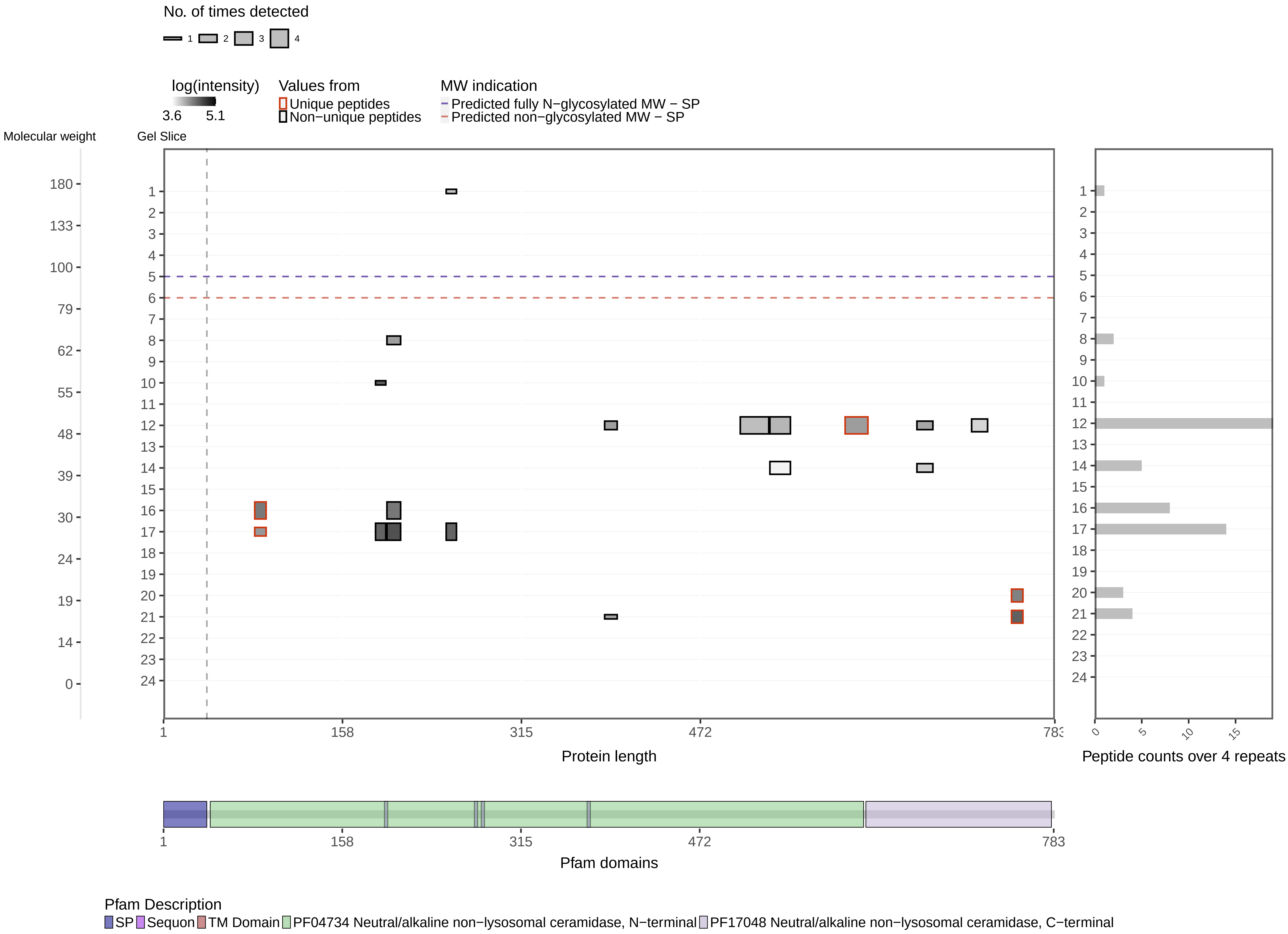


Protein sequence:

CGGKNELCYGRGPGYPDEFESTRIIGERQFKKAVELFNKATEQVKGKVDFRHTYVDFSNLEVTIPKEGAGIETAKTCPAAMGFAFAAGTTDGPGAFDFQQGDDKGNVFWRLVRNLLKKPGSEQIKCQHPKPILLDTGEMKEPYDWAPSILPVQILRIGQLVILSVPGEFSTMAGRRLRDAVKMVLTSGGTKEFDSNIHVVIAGLTNTYSQYITTFEEYQIQRYEGASTLYGPHTLSGYIQQFKTLAAALITGQTLQAGPQPPNLLGKQIGLLPPVVMDATPLGSKFGDLITDVPQ

NbL19g21070.1 50S_ribosomal_protein_L4_chloroplastic [MW = 30.86 kDa]

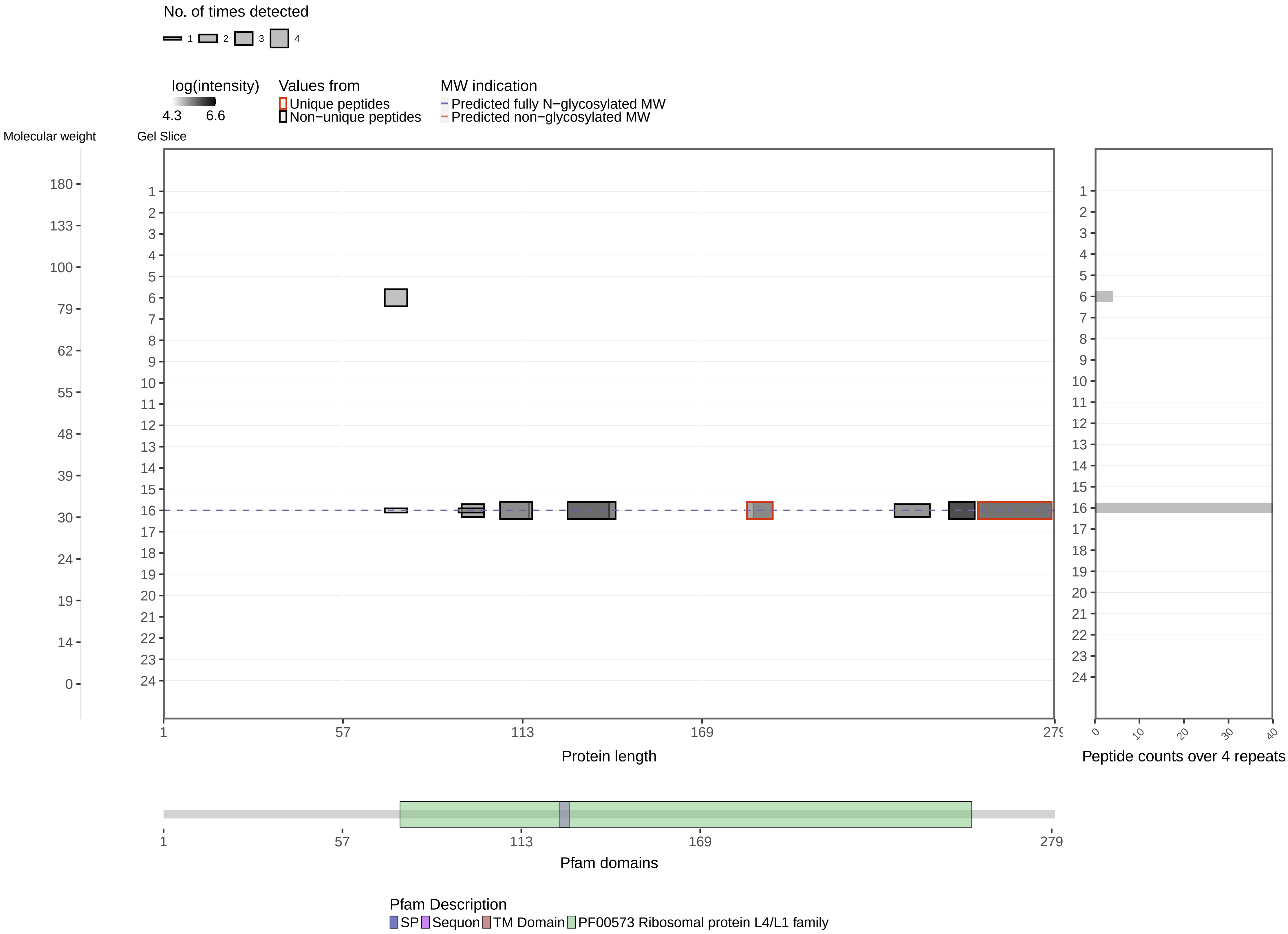


Protein sequence:

PLRPGGGVVFGPKPKDWSVKINKKEKRLAISTALSSASENTIVVEEFNDKFEKPKTKEFIDLMRRWGLDPKEKSLFLMTEVSDNVILSSRNIGTLKMLTPRTLNLFDILDSEKLVLTKSAVDFLNERYGDENEWEDEEKDDQEDNDGGGER*

NbL19g20980.1 probable_aquaporin_PIP−type_pTOM75 [MW = 30.83 kDa]

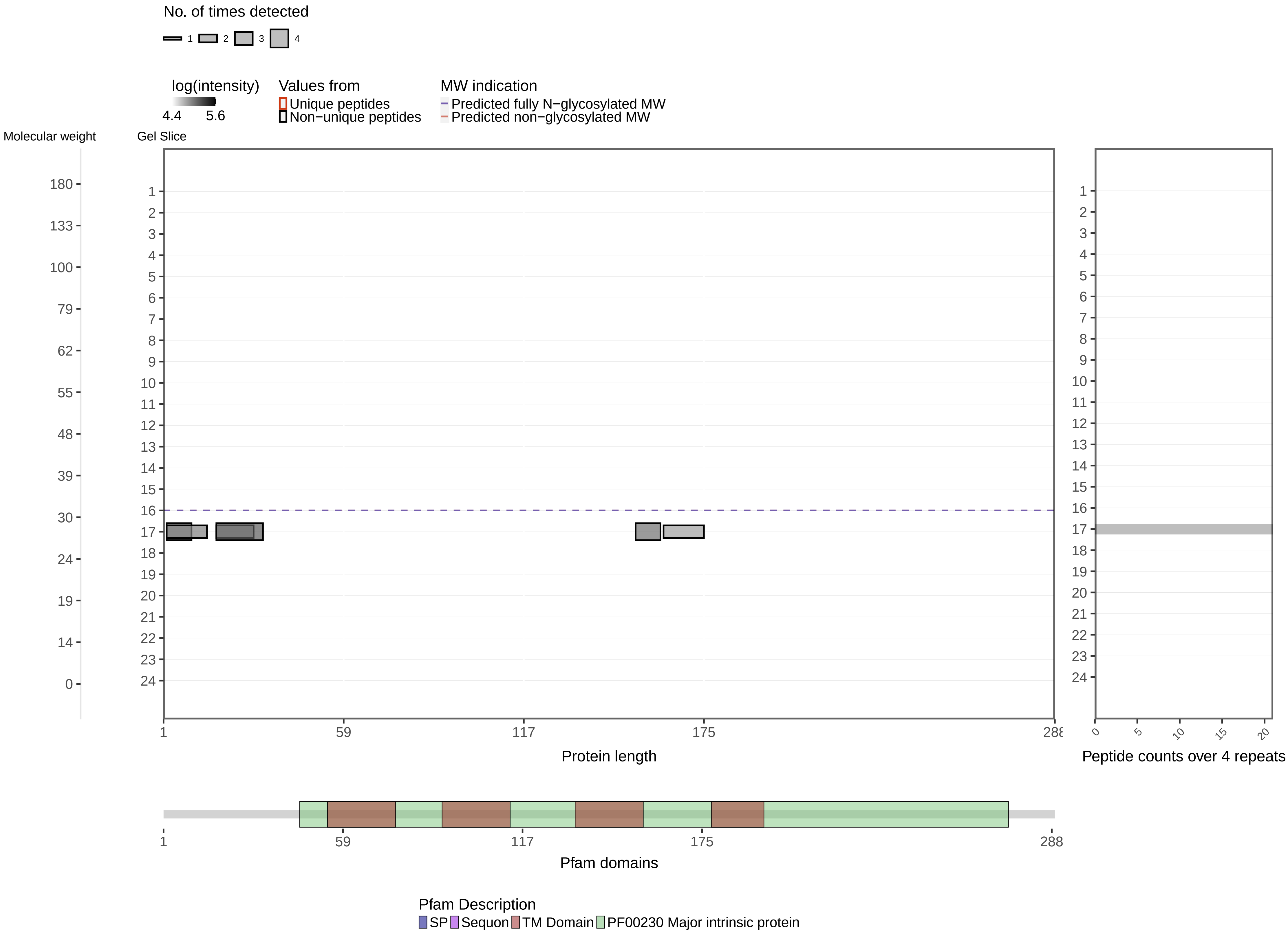


Protein sequence:

TRALFYMVMQCLGAICGAGVVKGFMVGPYQRLGGGANVVQPGYTKGDGLGAEIIGTFVLVYTVFSATDAKRNARDSHVPILAPLPIGFAVFLVHLATIPITGTGINPARSLGAAIIFNHDQAWDDHWIFWVGPFIGAALAAVYHQIIIRAIPFKTKS*

NbL19g20910.1 putative_cytochrome_c_oxidase_subunit_5b−like [MW = 18.76 kDa]

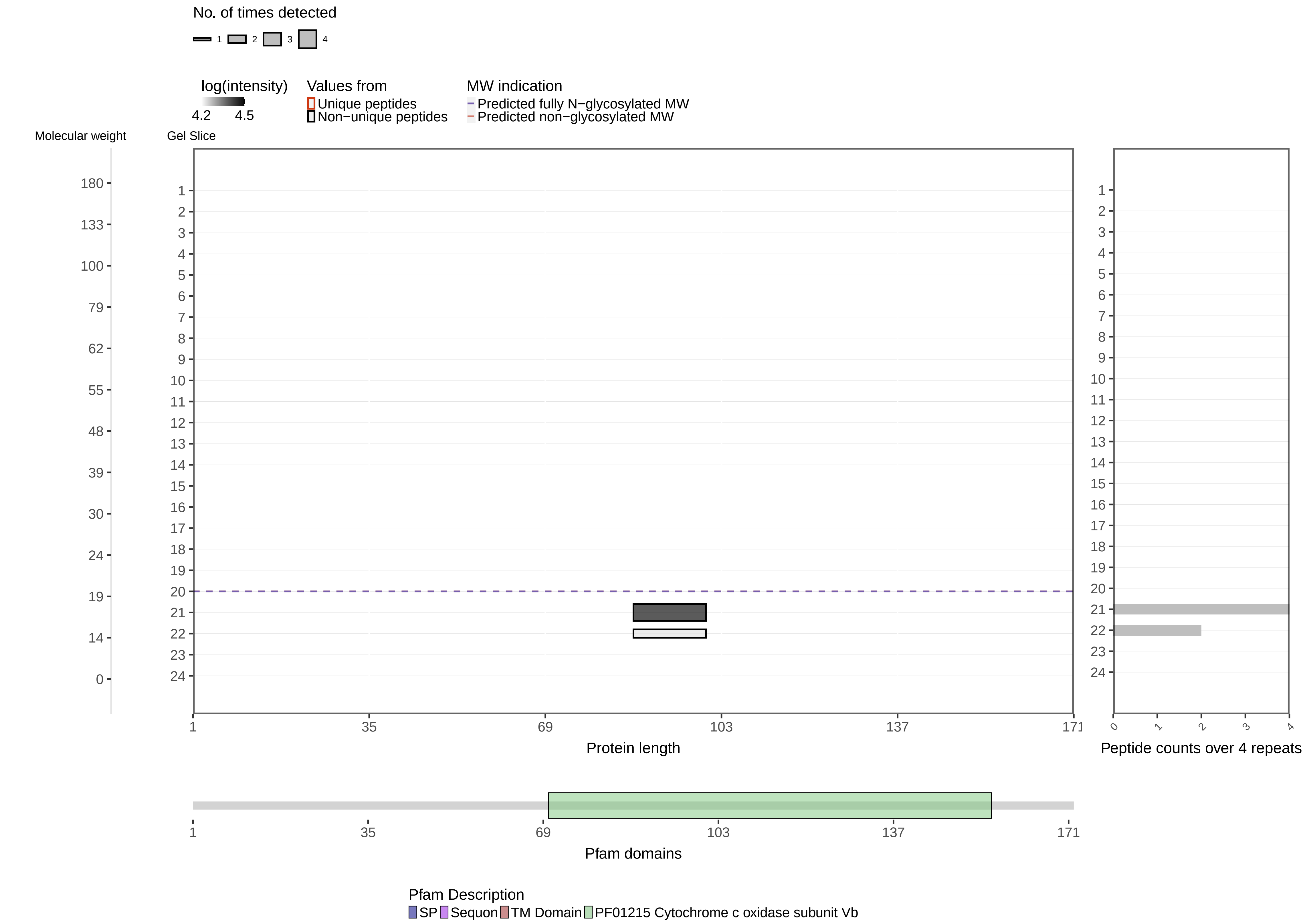


Protein sequence:

REELEAELQGRELLDINFPEGPFGTKEAPAVVKSYYDRRIVGCPGGEGEDEHDVVWFWLEKGKPHECPVCSQYFELEVVGPGGPPDGHGSDDEGHH*

### NbL19g20880.1 uncharacterized_LOC109242909 [MW = 37.21 kDa]

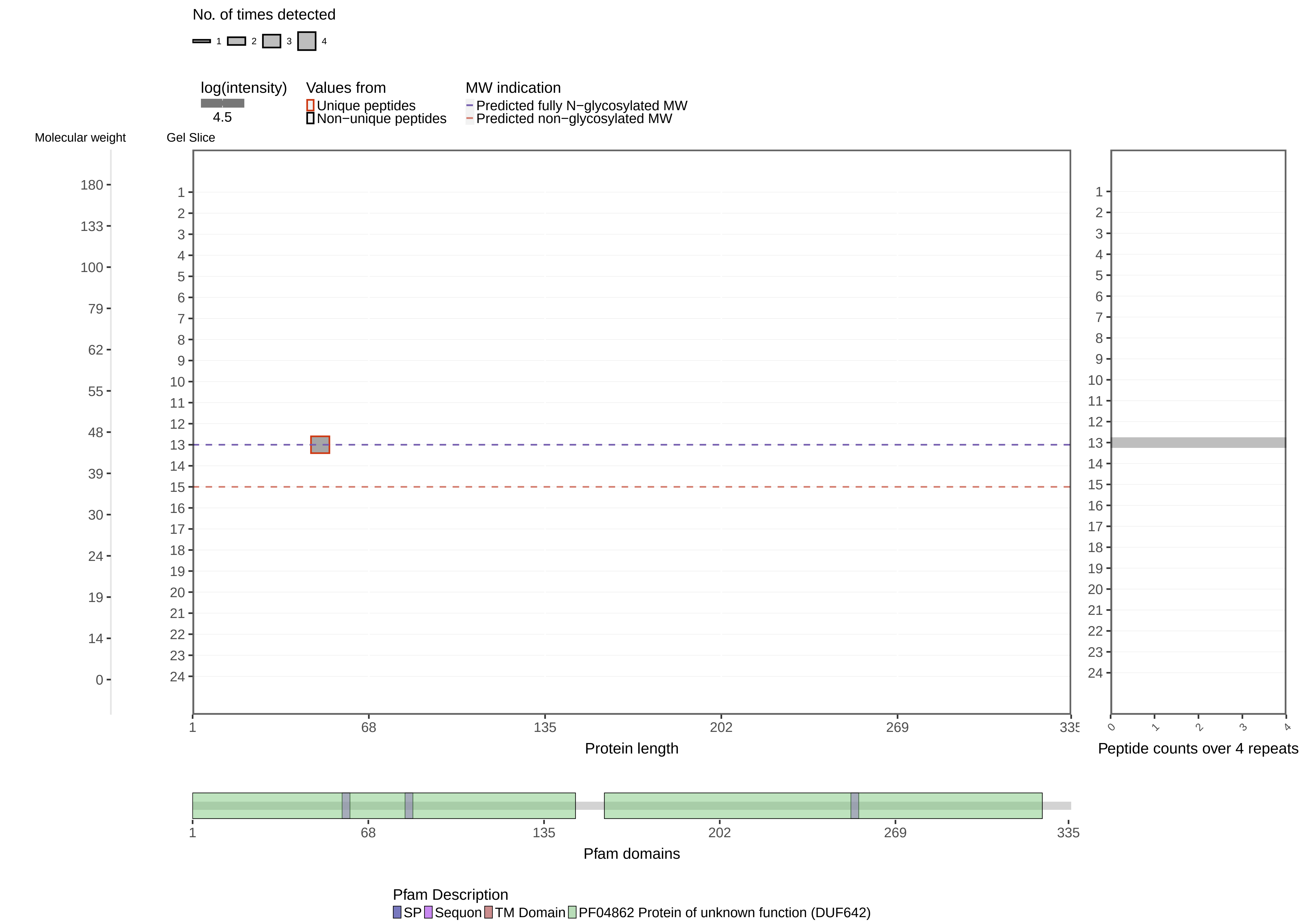


Protein sequence:

RPKGNMLKNGNFEEGPYIFPNTNWGILIPPNIEDDHCPLPGWIIESLKAVKYVDAEHFAVPEGKRAVELVAGRESAIAQEVITKAGKIYDLIFSFGDANNSCEGSMLIEAFAGKVALQVPYMSKGKGGSKRAKLRFTAISRRTRVRFLSTNYHMKSDNSGSLCGPVIDDVRLVSVRNPRIP*

NbL19g20830.1 coatomer_subunit_beta'−2−like [MW = 103.36 kDa]

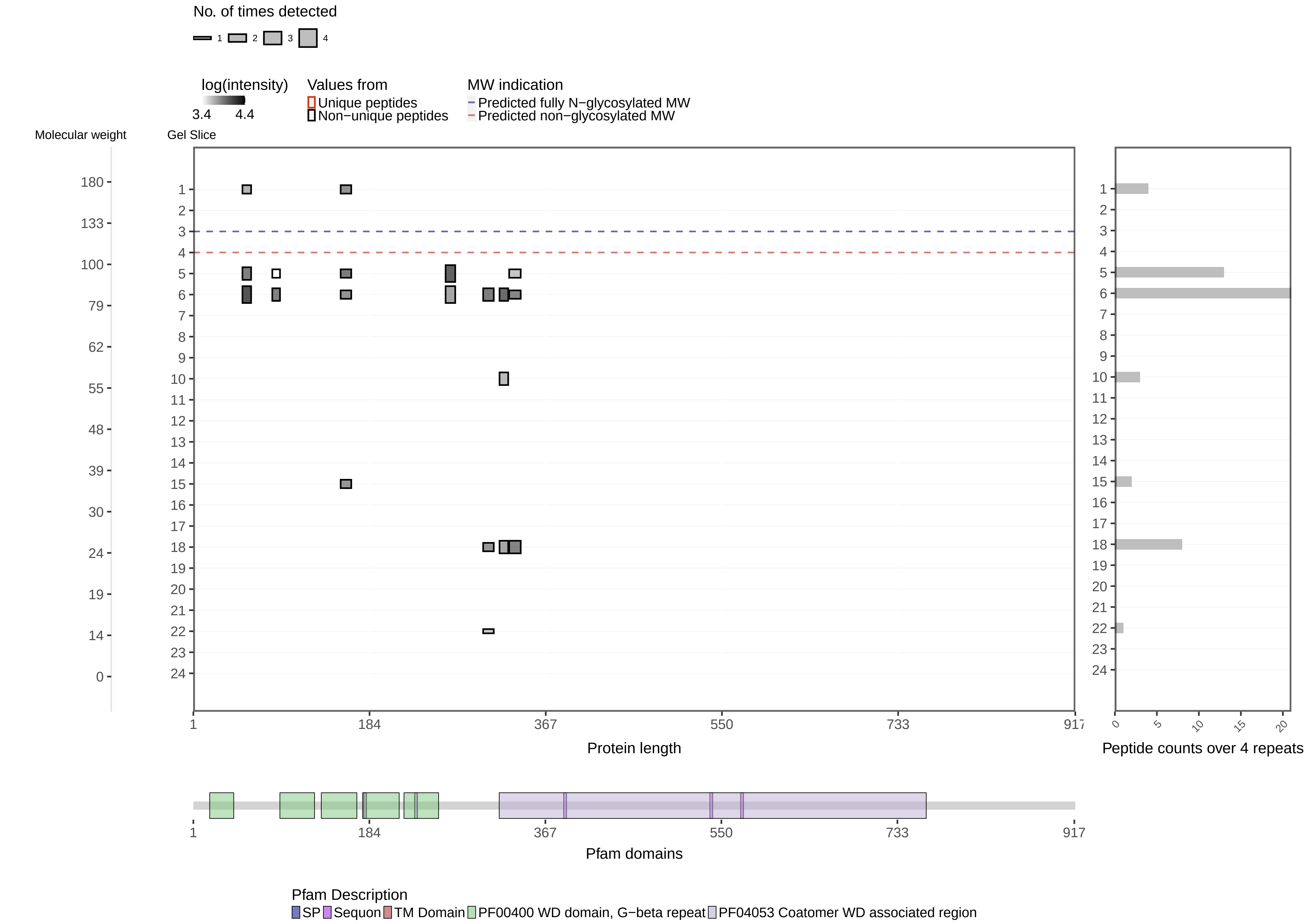


Protein sequence:

DFICFYDWADCRLIRRIDVNVKNLYWADSGDLVAIASDASFYILKYNRDVVSAHFDSGRSVDEOGVEEAFELLNEINERVRTGIWVGDCFIYNNSSWRLNYCVGGEVTTMFHLDRPMYLLGYLANOSRVFLIDKEFNVVGYTLLLGLIEYKTLVMRGDWDRANEVLPSIPKEHHNSVARFLESRGMIEEALEVATDLDYRFELAIOLGKLDIAKEIAVVAOSESKWKOLGDLAMSSGRLETAEECLKHANDLSGLLLLYSSLGDAEGITELASFAKEHGKNNVAFLCMFLLGKVDE

### NbL19g20750.1 heat_shock_70_kDa_protein_15−like [MW = 94.36 kDa]

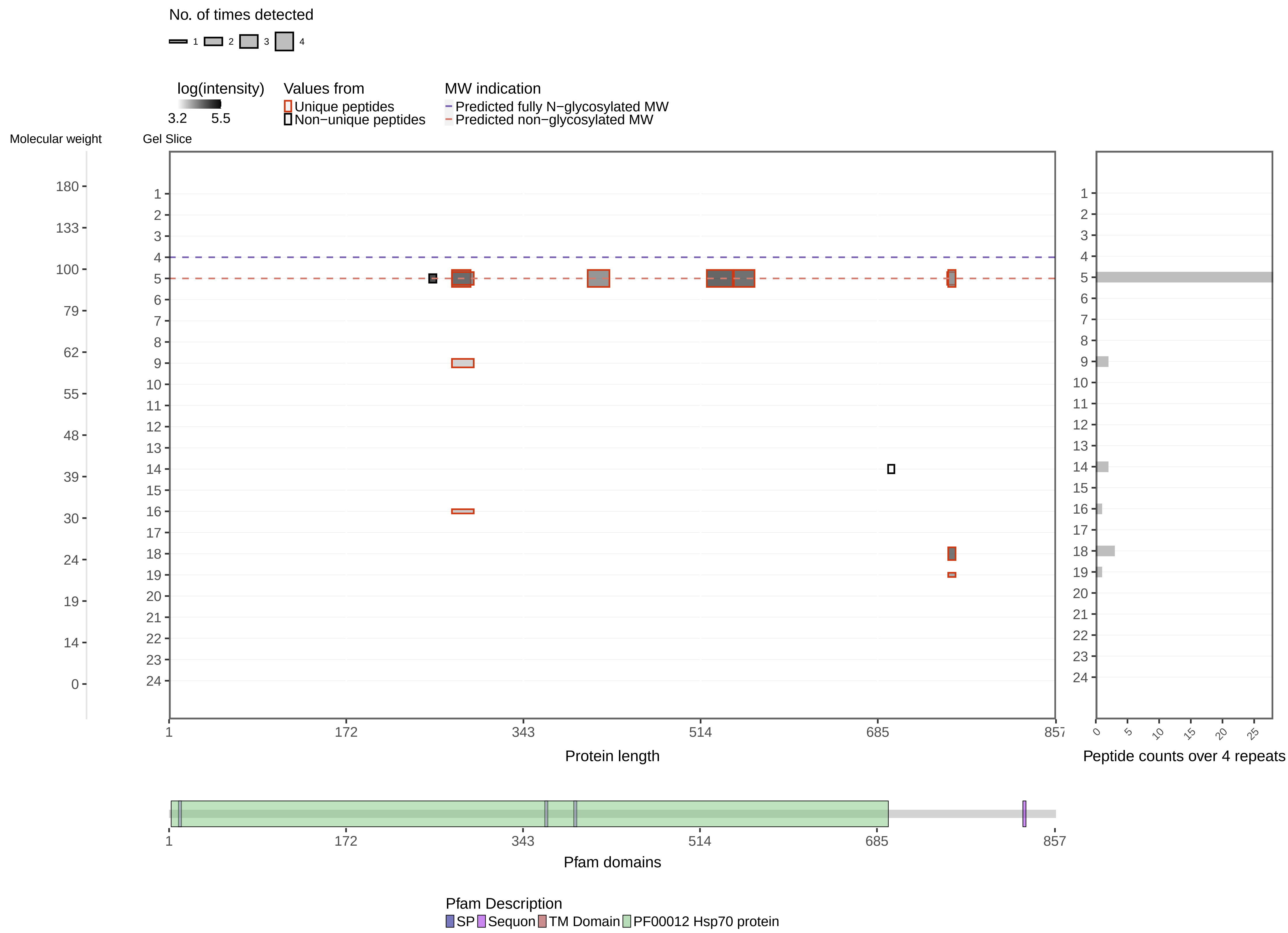


Protein sequence:

IVFPKGNPIPSVKALTFYRSGTFTIDVQYADVSELQAPPKISTYTIGPFQSAKGERAKLKVKARLNLHGVVSVDSATLLEEEEVEVPVVREAVKEPAKMEMDEASGDAAHSTTSESDVNIQDAKAAGDASGAENGVPESGDKPVKMETDAKVGTTKKKVKKTSVPLTEIVYGAMAAADVQKAVEKEFEMALQDRVMEETKDKKNAVEAYVYDMRNKLSDKYQEFVIDSEREQFVAKLQEVEDWLYEDGEDETKGVYISKLEELKKQGDPIEQRYKEYTERGSVVDQFVYCIN

### NbL19g20710.1 hydroxyphenylpyruvate_reductase−like [MW = 34.11 kDa]

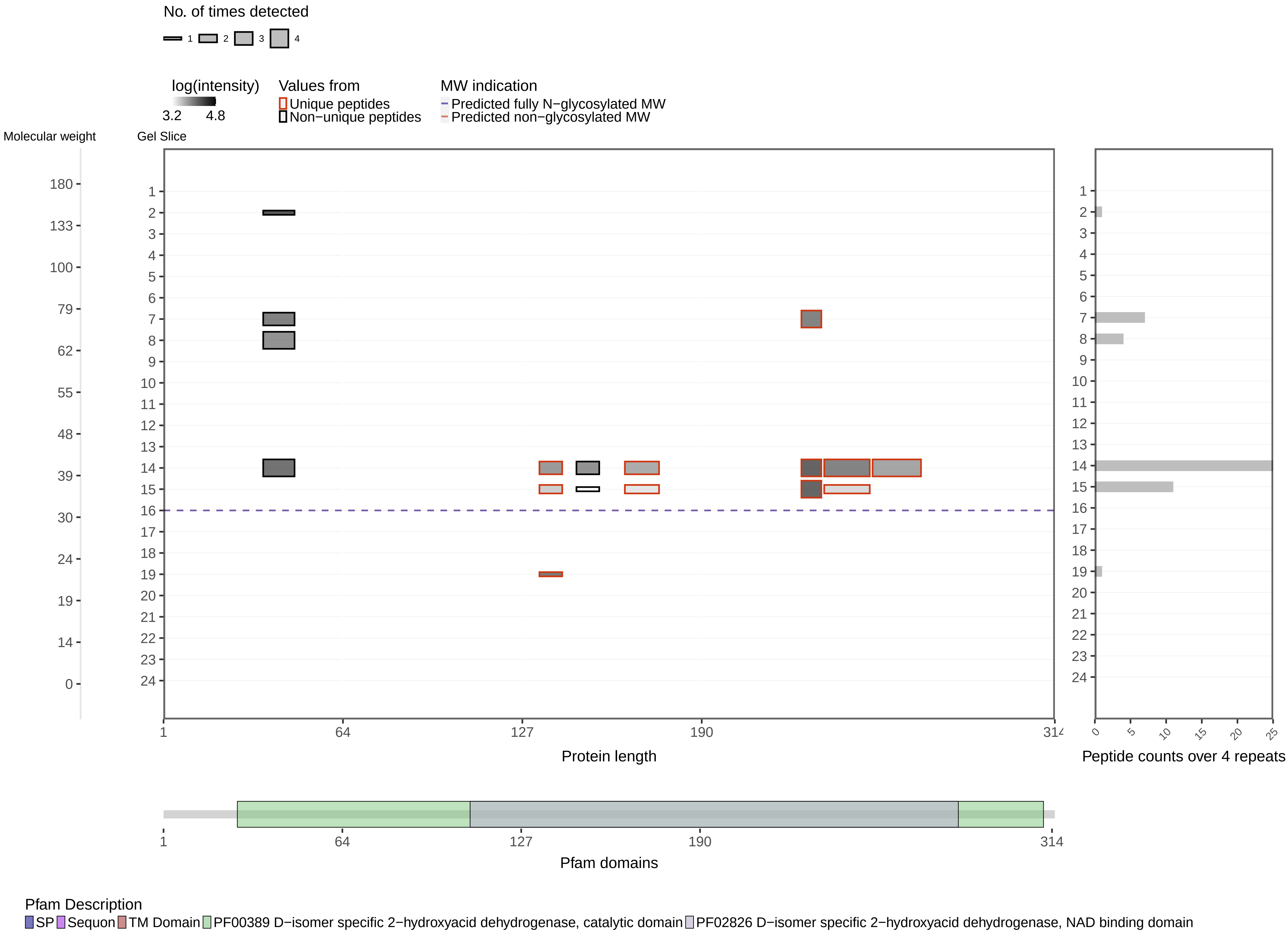


Protein sequence:

FSGKLVGIIGLGRIGSAIAKRAEAFGCPVSYHSRSKKPNINYKYYSNVVDLAENSEILIVACALTEETQHIVNRKVLDALGPKGILINIGRGAHVDESELVAAIIEGRIGGAGLDVYQTEPEVPERFFGLENVVLLSHAASDTVETCTSMADLVVANLEAHFLQKPLLTPVL*

### NbL19g20690.1 malonate−−CoA_ligase−like [MW = 67.75 kDa]

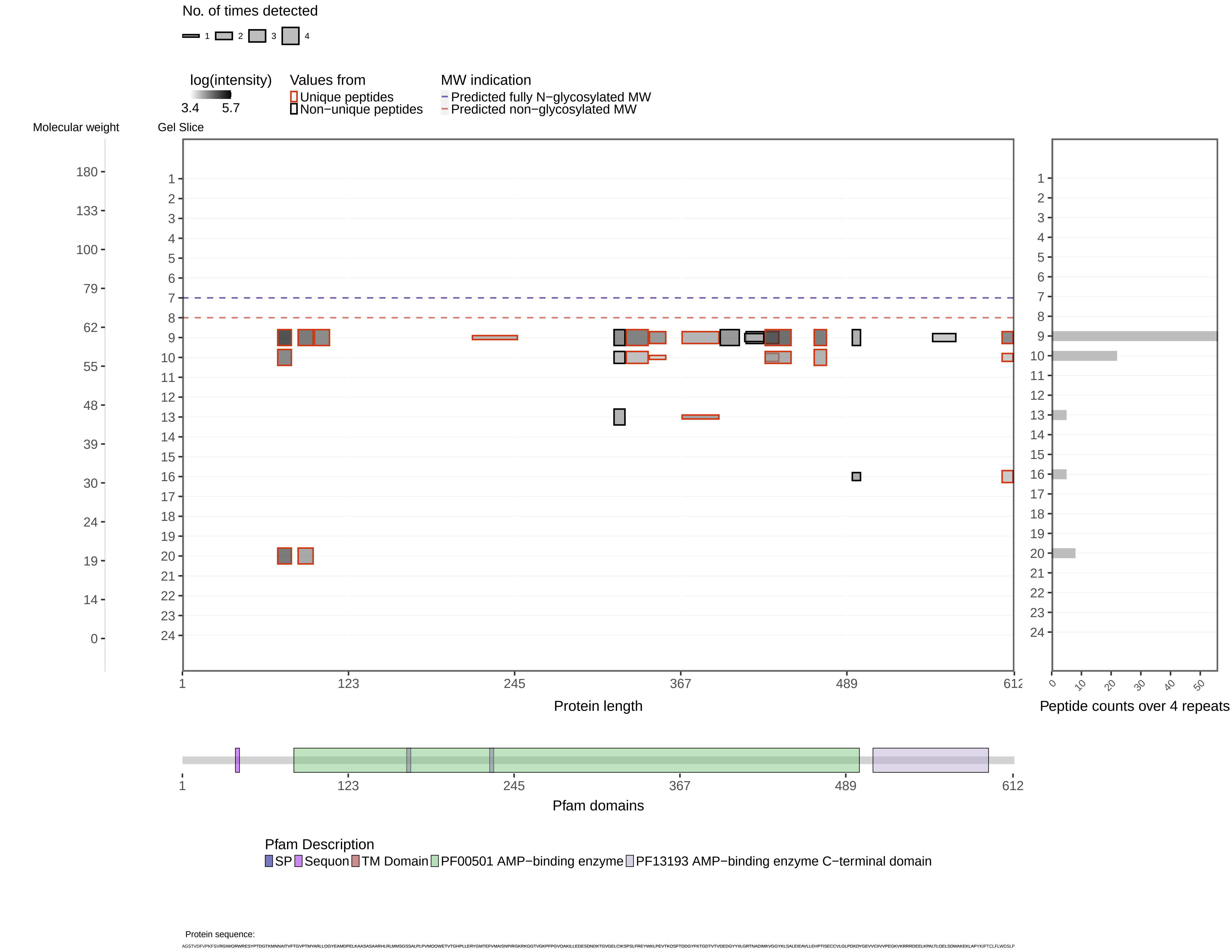

### NbL19g20450.1 cinnamoyl−CoA_reductase_1 [MW = 35.78 kDa]

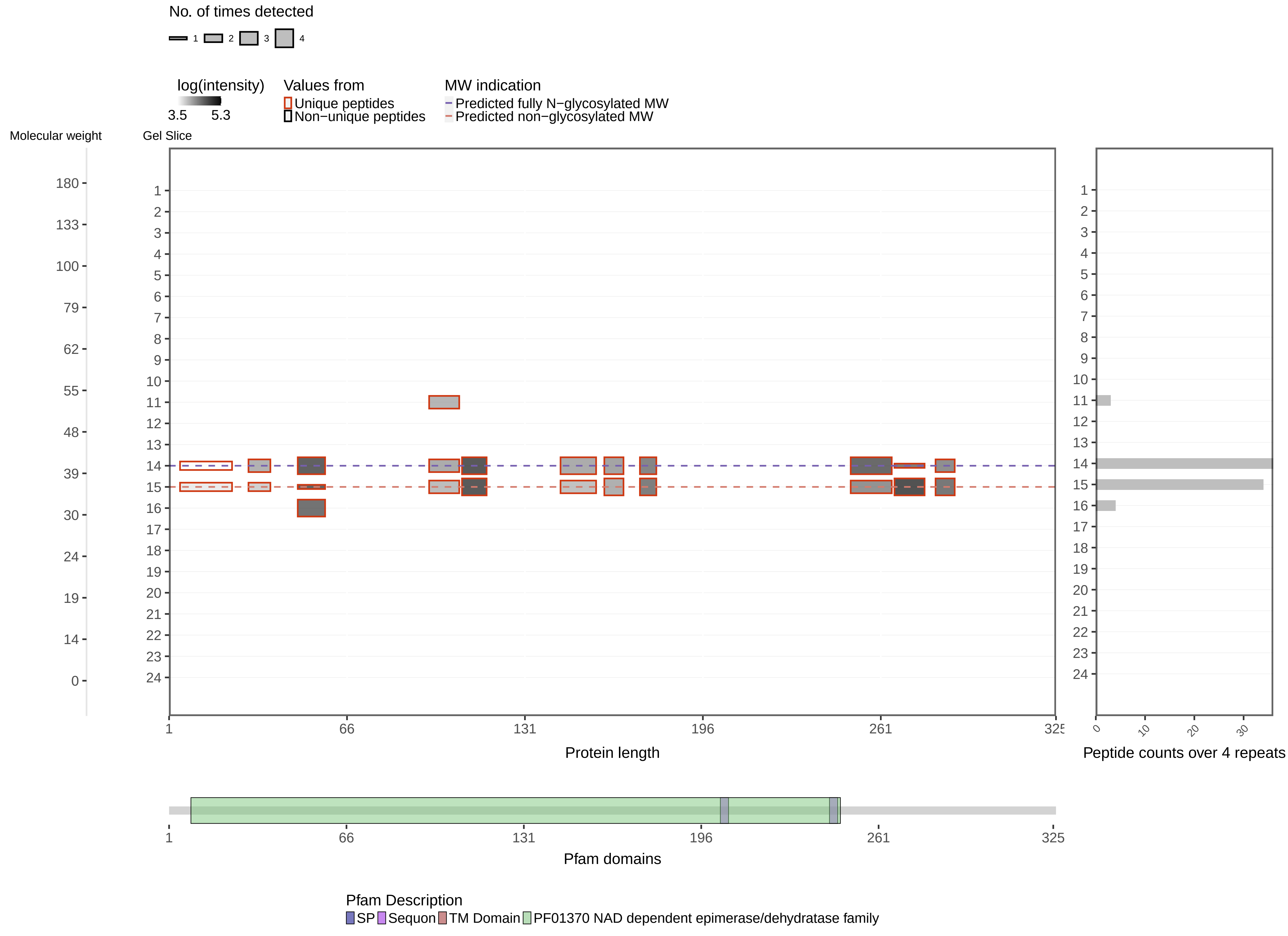


Protein sequence:

[illegible]TDIEYCKQKGVWYPLSKTLAEKASWEFAKEKDLDIVVVNPGTVMGPILPPTLNASMLMMLRLLQGCTTAYEDFFMGLVHVKDVALAHILVYENISAKGRHLCVEAITHYGDFAAKVAELYPEYNIPRLPKDTQPGLLRAKDGGKKLMDLGLQFIDMEQIIRDSVESLKMKGFMS*

NbL19g20340.1 peptidyl−prolyl_cis−trans_isomerase_CYP20−1−like [MW − SP = 19.67 kDa]

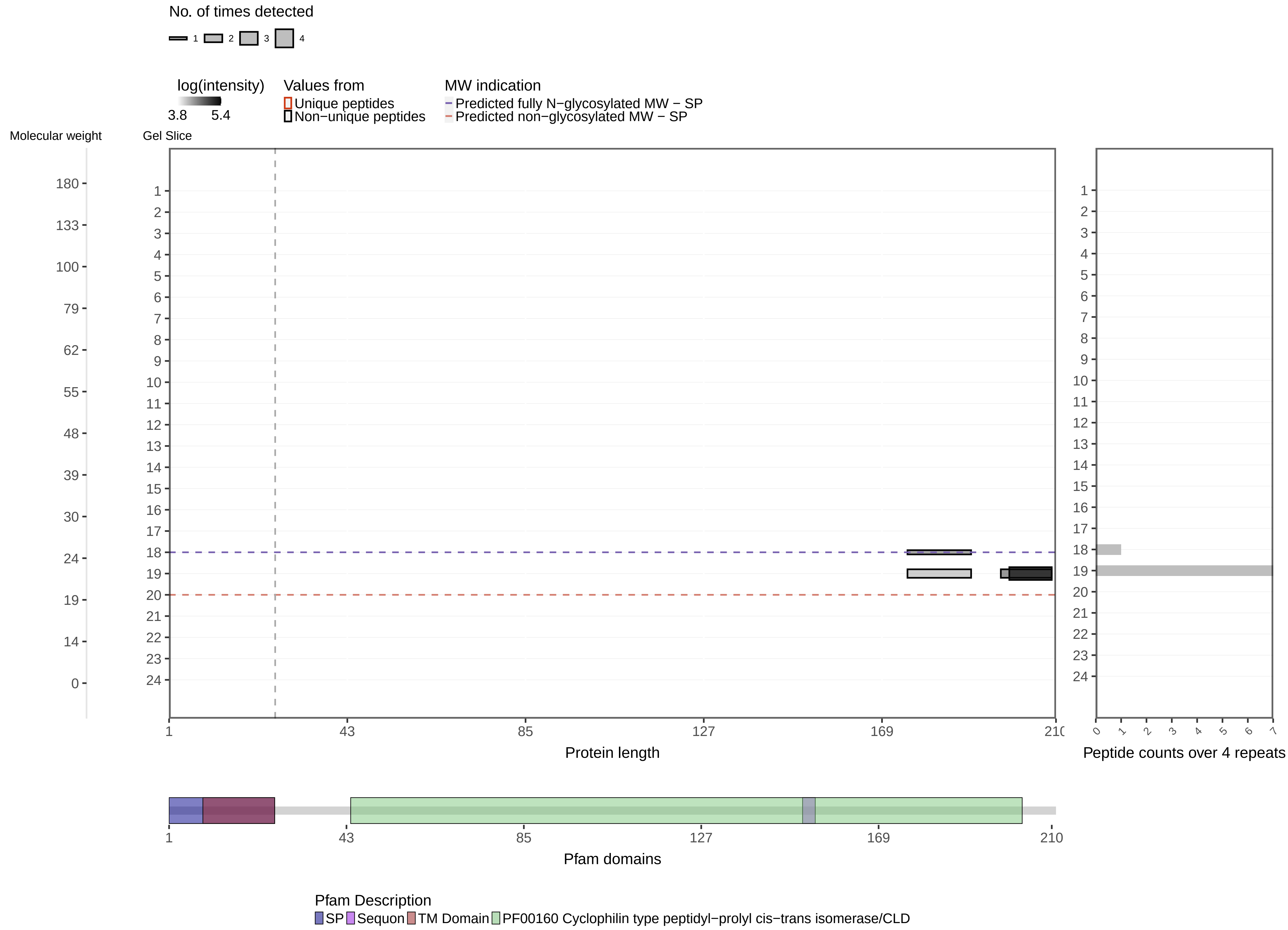


Protein sequence:

;STFHRIIPSFMLQGGDFTLGDGRGGESIYGEKFADENFKIKHTGPGLLSMANAGSDTNGSQFFITTVTTSWLDGRHVVFGKVLSGMDVVYKVEAEGRQSGTPKSKVVIAESGELPL*

### NbL19g20110.1 probable_glutathione_S−transferase_parA [MW = 28.58 kDa]

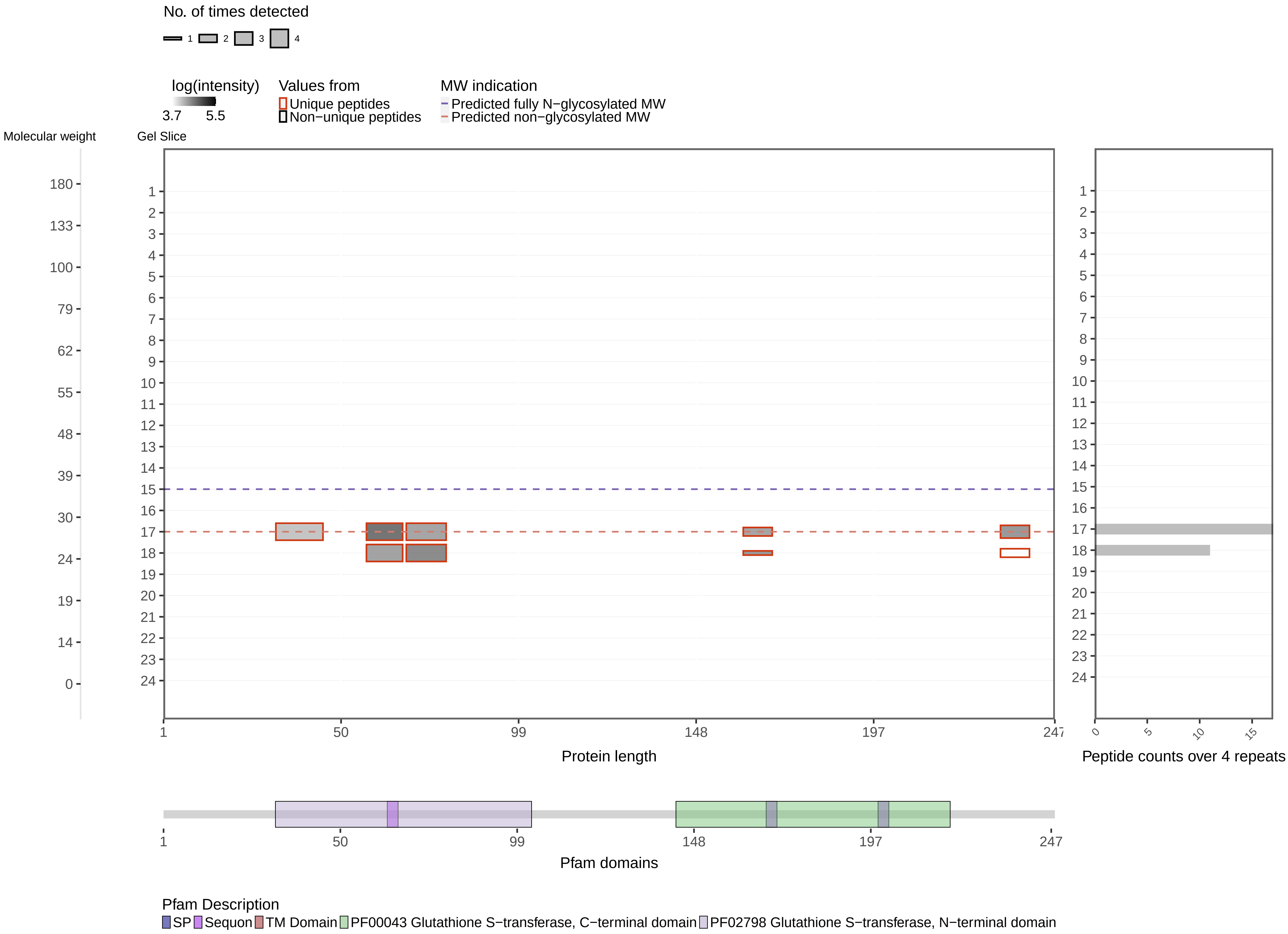


Protein sequence:

SDPYEKSRARFWADYIDKKIYSTGRRVWSGKGEDQEEAKKEFLEIFKTLEGELGNKTYFGGDNLGFVDVALVPFTSWFYSYETFANFSIEAECPKLVVWAKRCMENECVSKSLPHPHKIYDFVLELKHKLGLA*

NbL19g20030.1 probable_LRR_receptor−like_serine/threonine−protein_kinase_At1g06840 [MW − SP = 102.38 kDa]

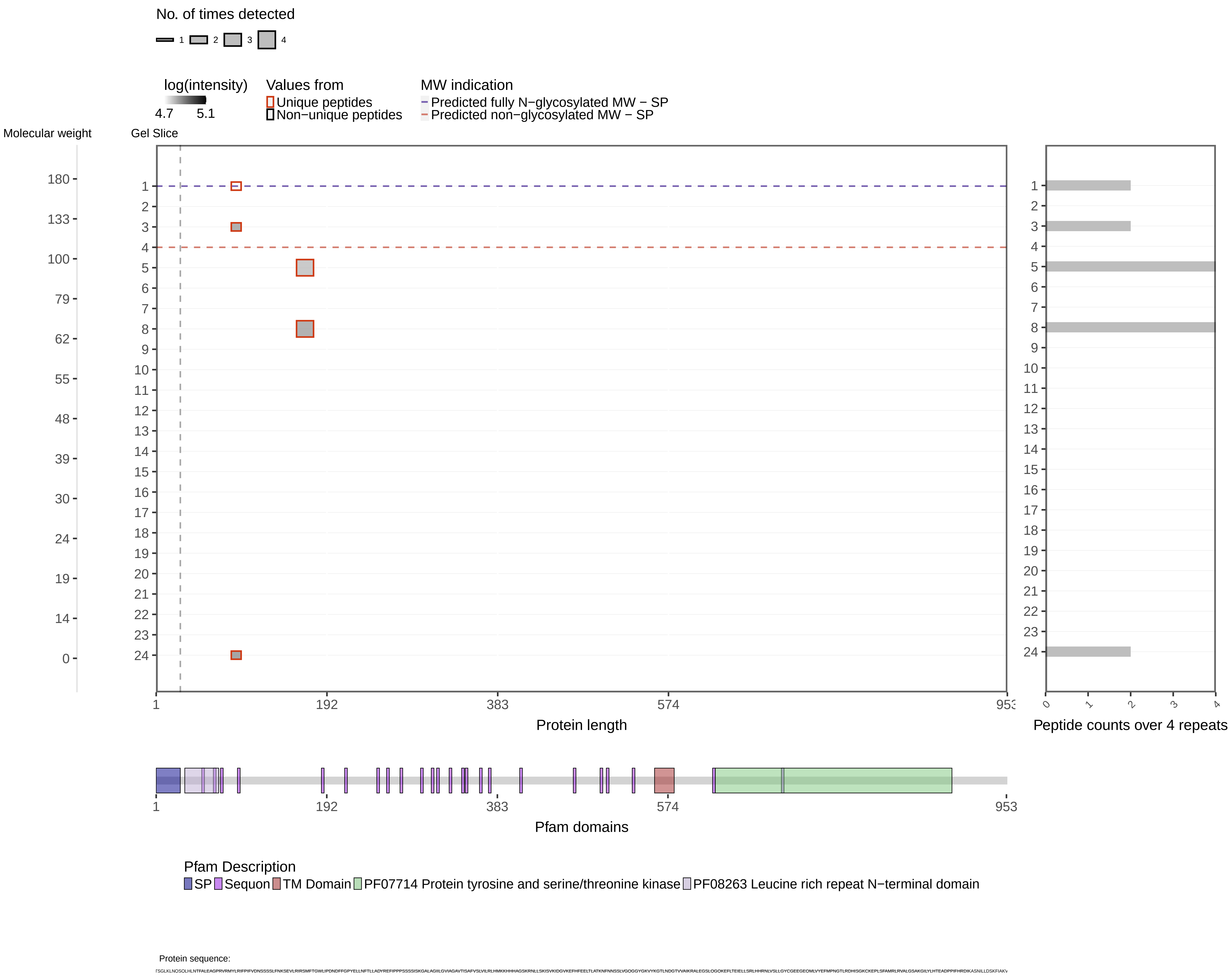

### NbL19g19930.1 bis(5'−adenosyl)−triphosphatase−like [MW = 18.11 kDa]

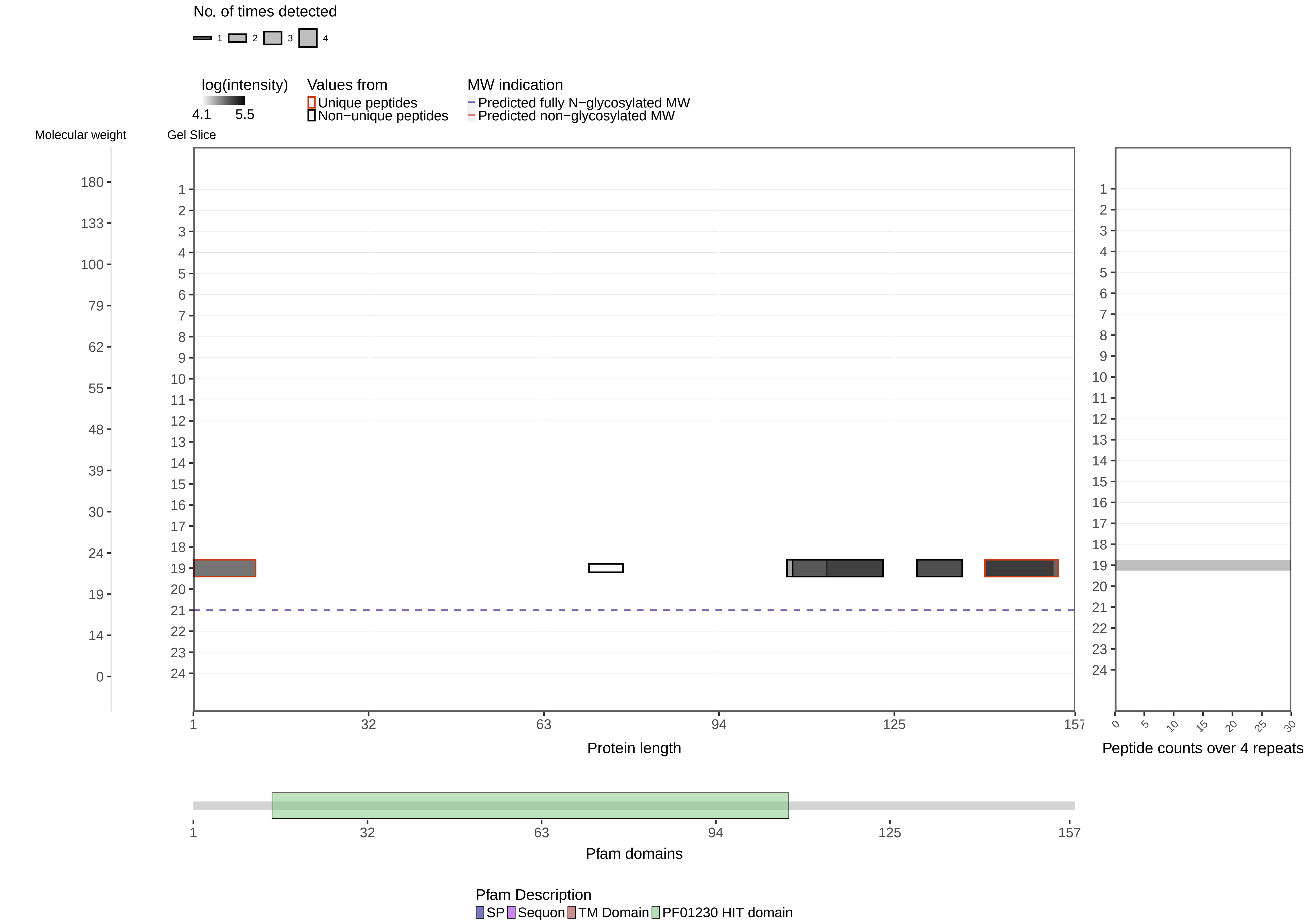


Protein sequence:

QKVGKQLESYHKASSLTFAIQDGPQAGQTVPHVHIHIIPRKSGDFEKNDEIYDALDVKEKEMKQSLDLDKERKDRSIEVMAEEADEYRKLF*

### NbL19g19920.1 50S_ribosomal_protein_L17_chloroplastic−like [MW = 22.38 kDa]

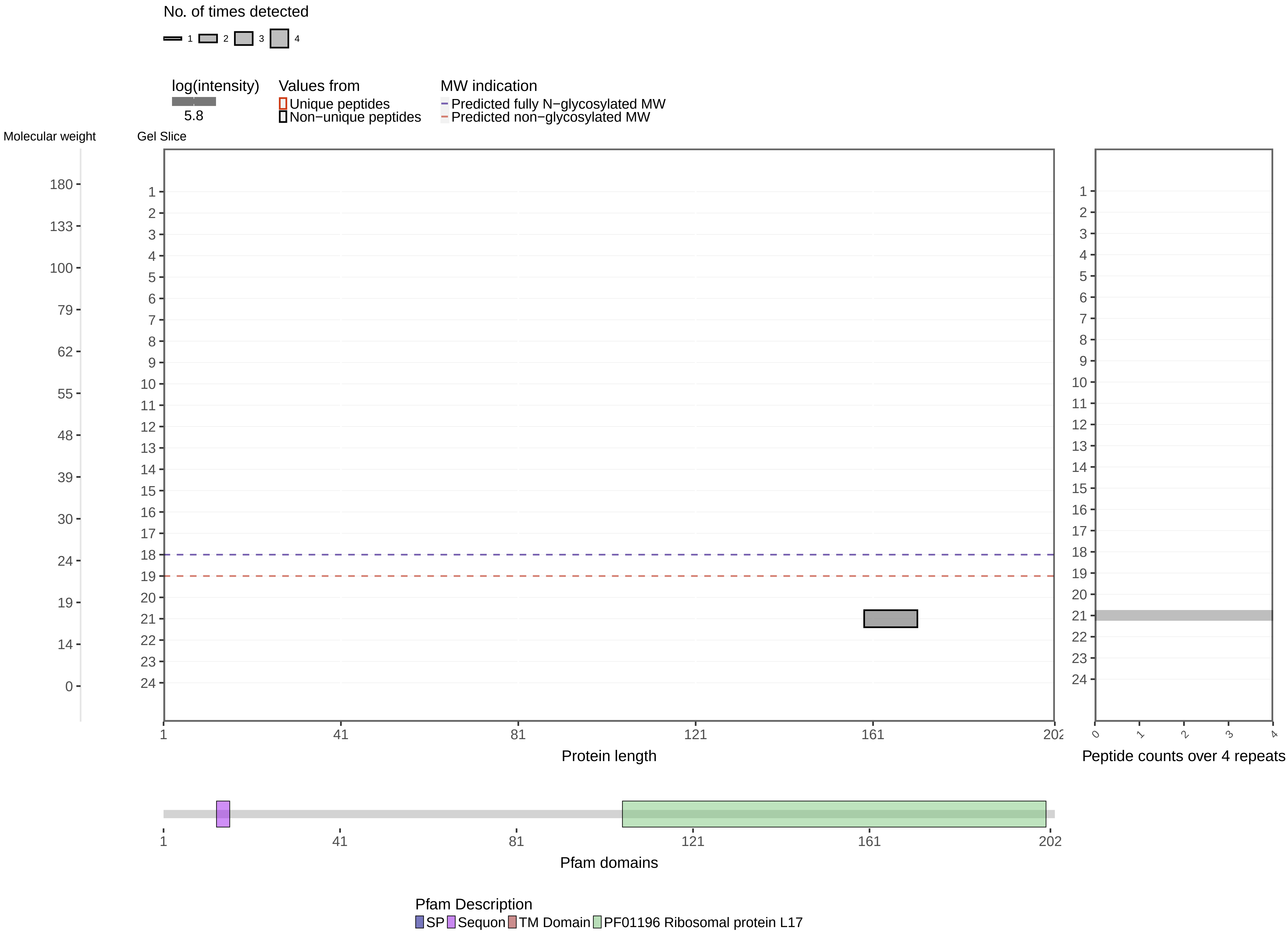


Protein sequence:

HGRKVPKLNRPPDQRRALLRGLTTQLLKHGRIKTTKARARAVRKYVDKMITMAKDGTLHKRRQALGFIYEKQIVHALFAEVPERYGERNGGYTRIIRTLPRRGDNAPMAYIELV*

### NbL19g19880.1 CBS_domain−containing_protein_CBSX3_mitochondrial [MW = 22.57 kDa]

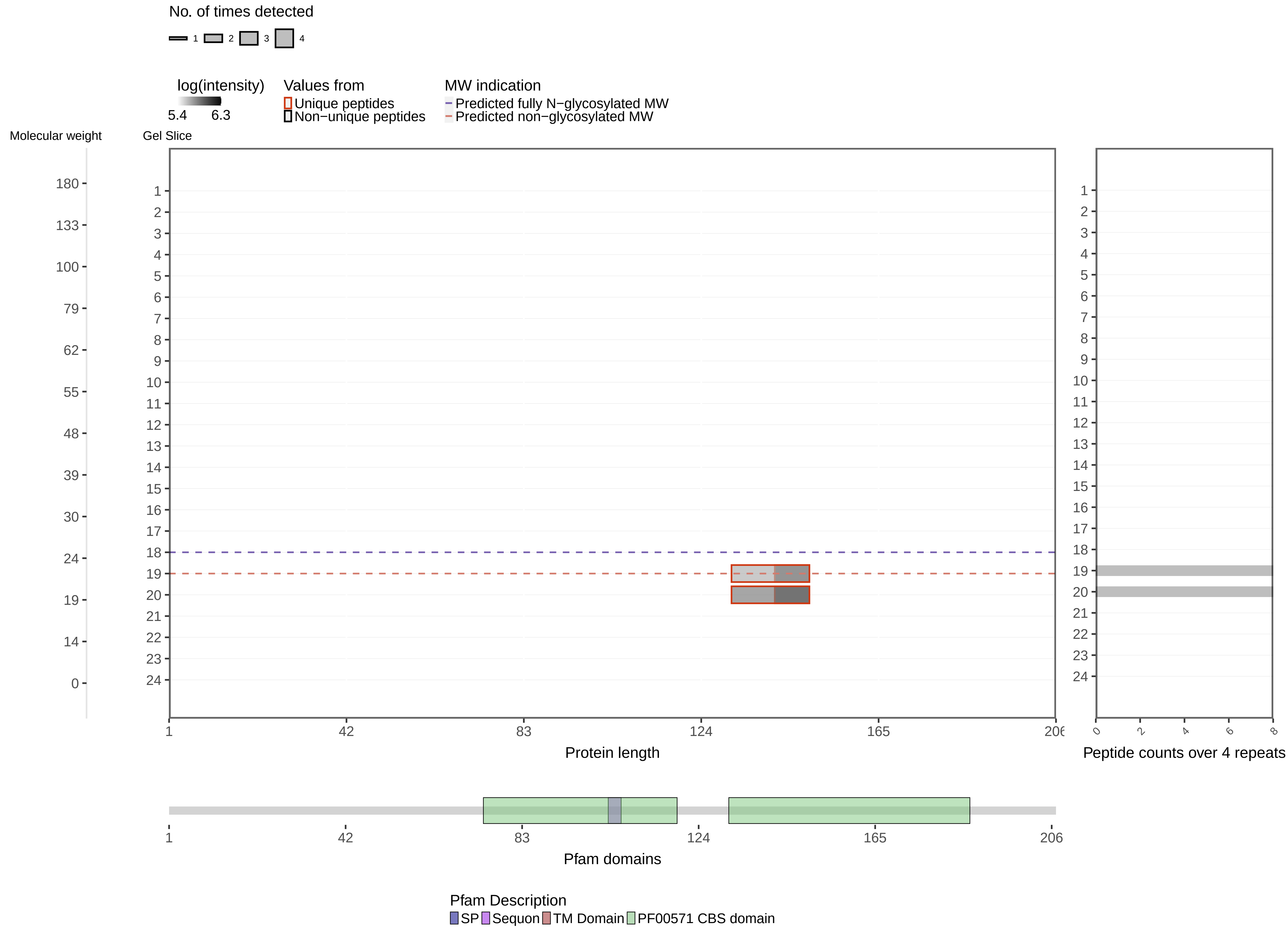


Protein sequence:

QHNVGALVVVKPGENKSIAGIITERDYLRKIIVQGRSSKSTKVGDIMTEENKLITVTPETKVLKAMQLMTDNRIRHIPVIDQTGMIGMVSIGDVVRAVVSEHREELNRLNAFIQGGY*

NbL19g19850.1 glyoxylate/succinic_semialdehyde_reductase_1 [MW = 30.79 kDa]

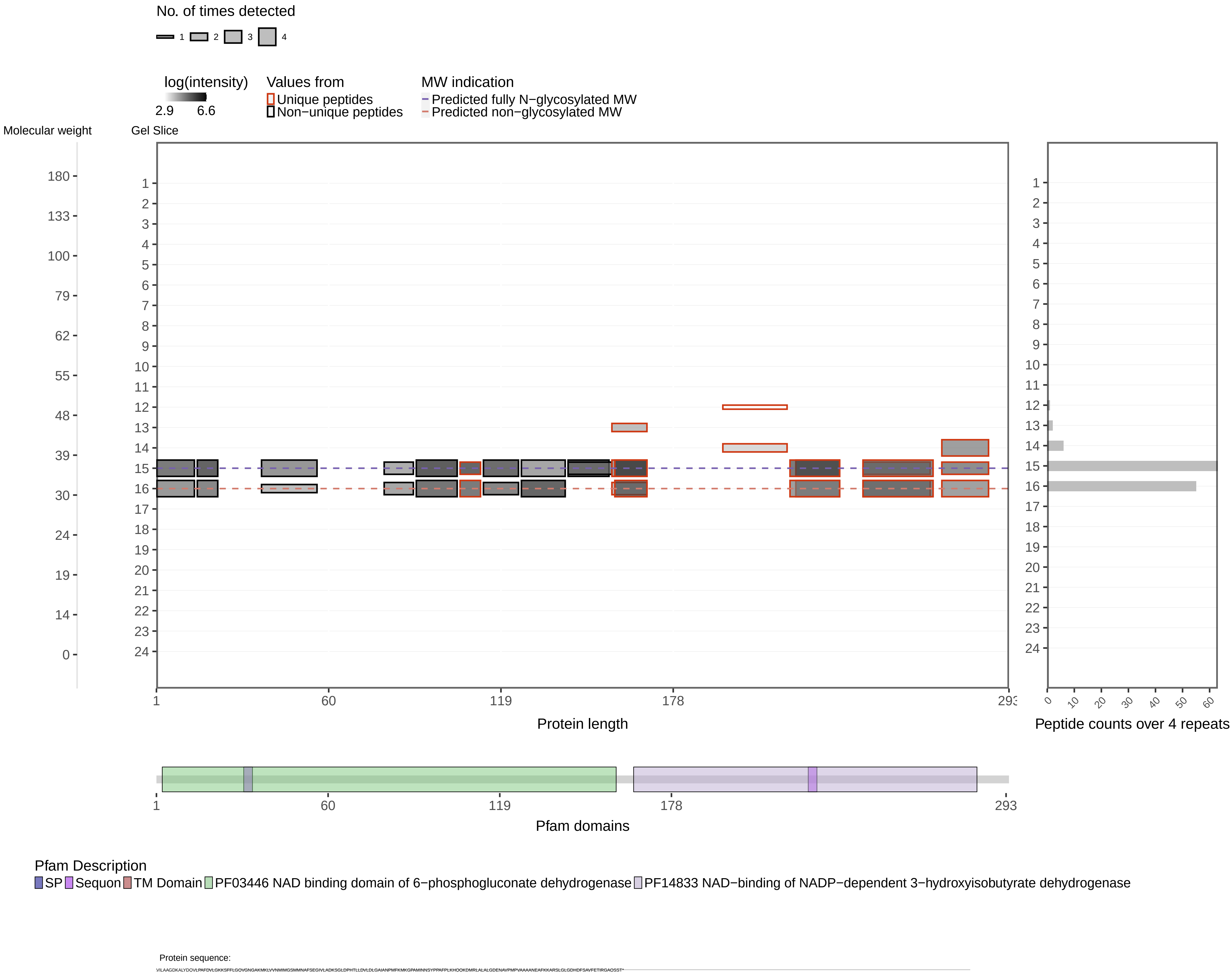

NbL19g19790.1 auxilin−related_protein_2−like [MW = 105.52 kDa]

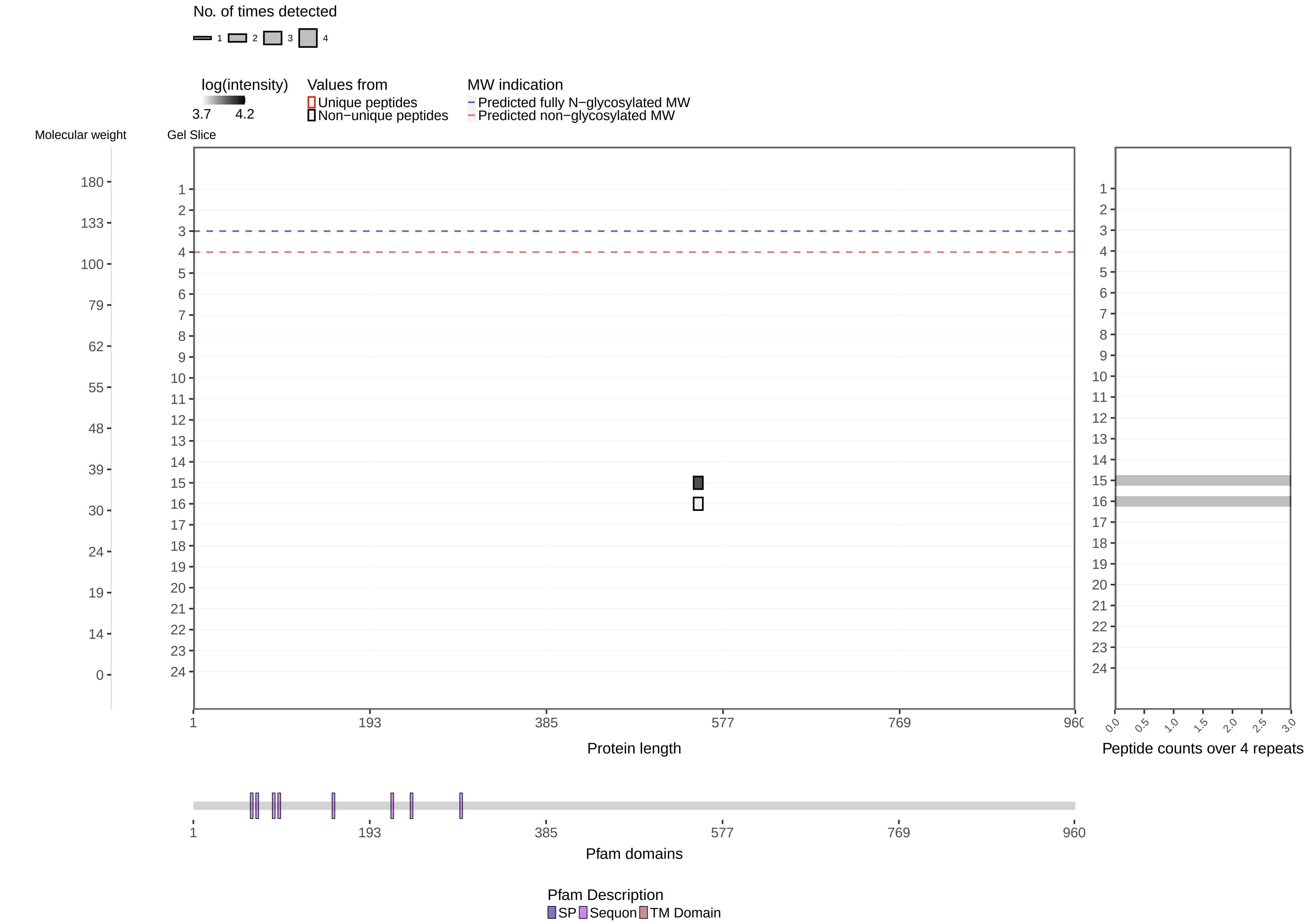


Protein sequence:

### NbL19g19750.1 60S_ribosomal_protein_L5−like [MW = 34.54 kDa]

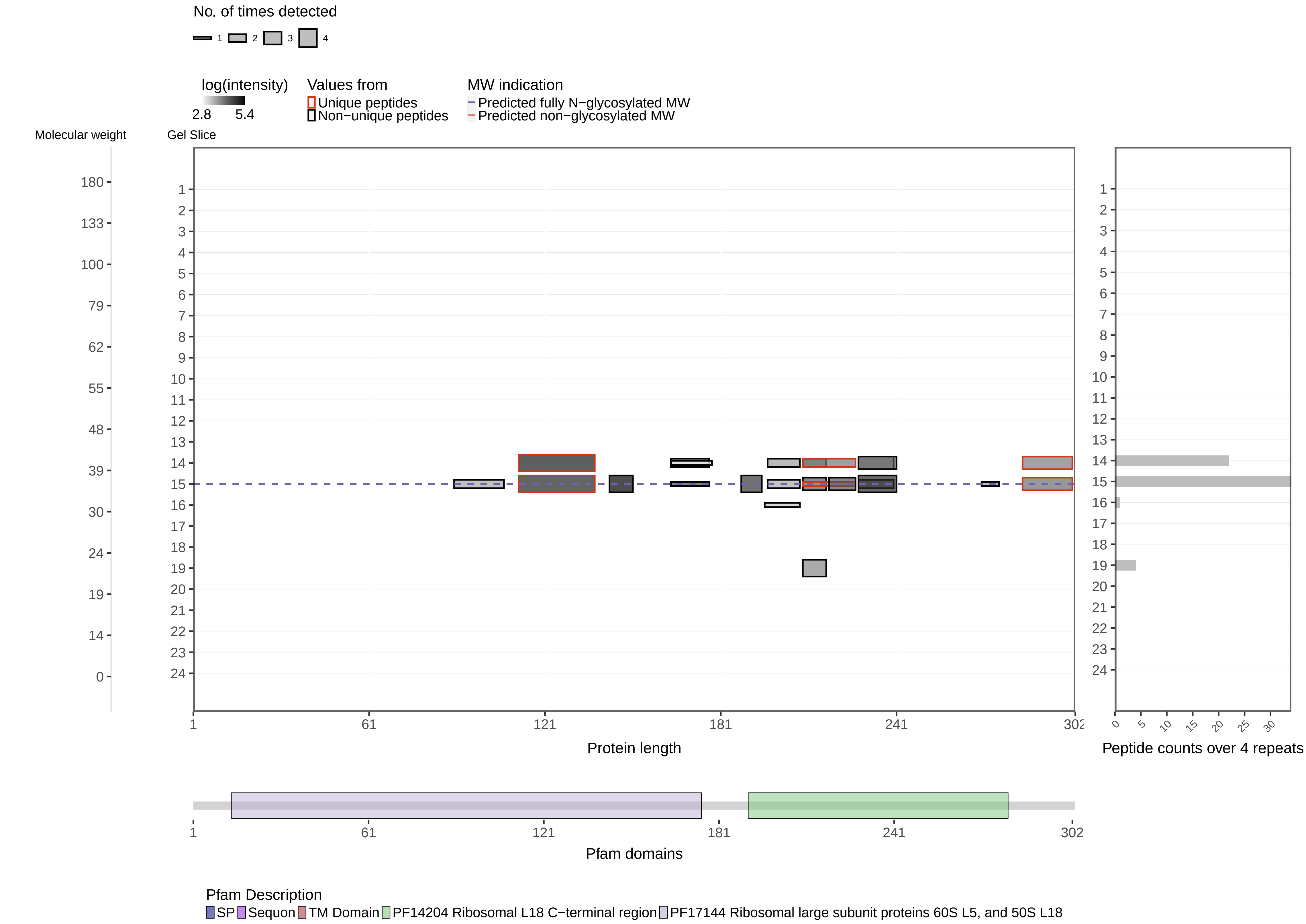


Protein sequence:

### NbL19g19720.1 ribonuclease_2−like [MW − SP = 26.42 kDa]

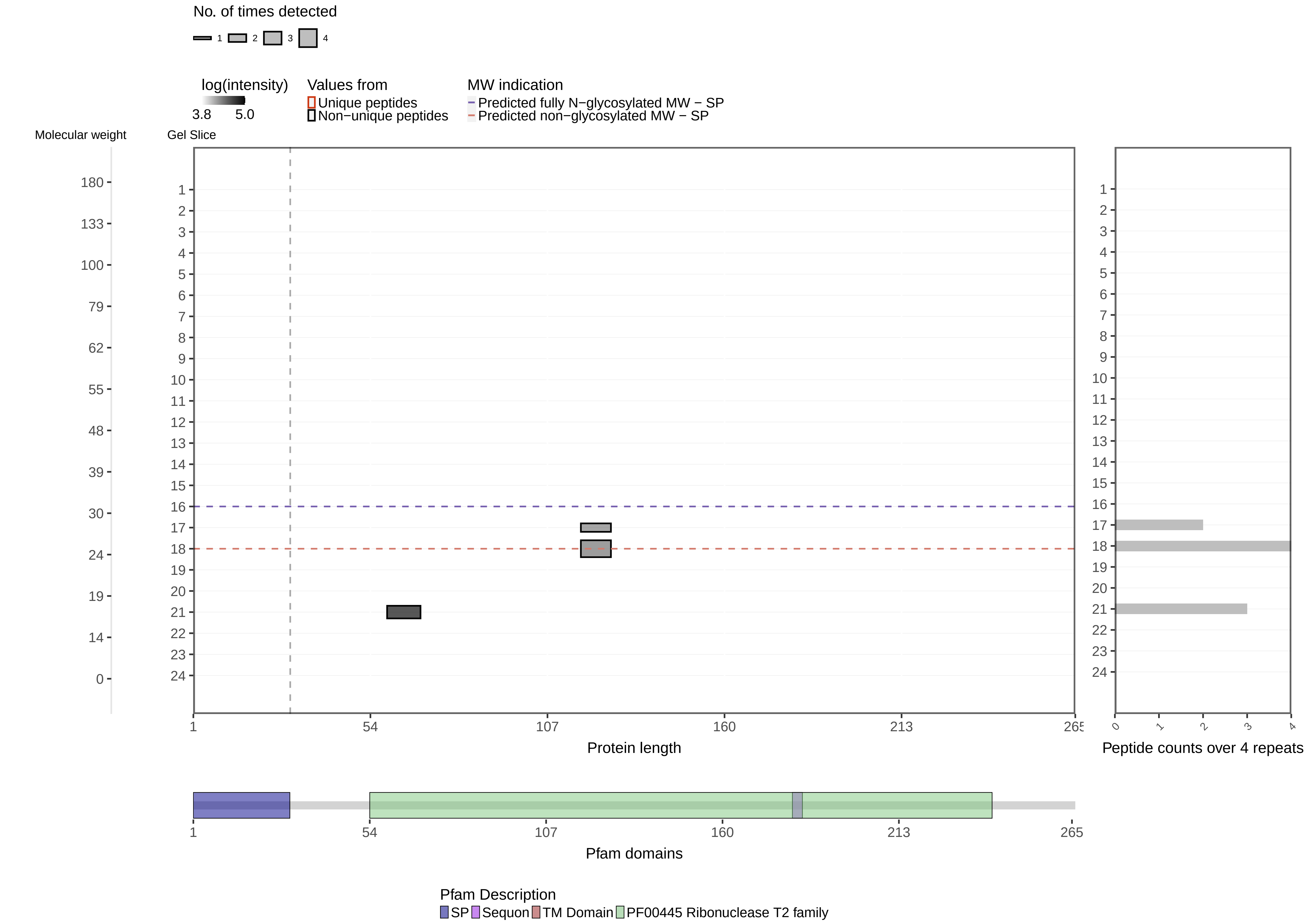


Protein sequence:

STLLEPLRKYWPSLSCGSPRSCHHRKGPFWAHEVEKHGTCAYPVVHDEYEFFLTTLNVYFKYNVTEVLFEAGYVPSDSEKYPLGGIISSIENAFHATPEVTCSGDALEELRICFYKNFEPRDCAHDTSALSRGSCPQYVSLPAHGS*

### NbL19g19710.1 ATP−dependent_RNA_helicase_DHX36 [MW = 116.31 kDa]

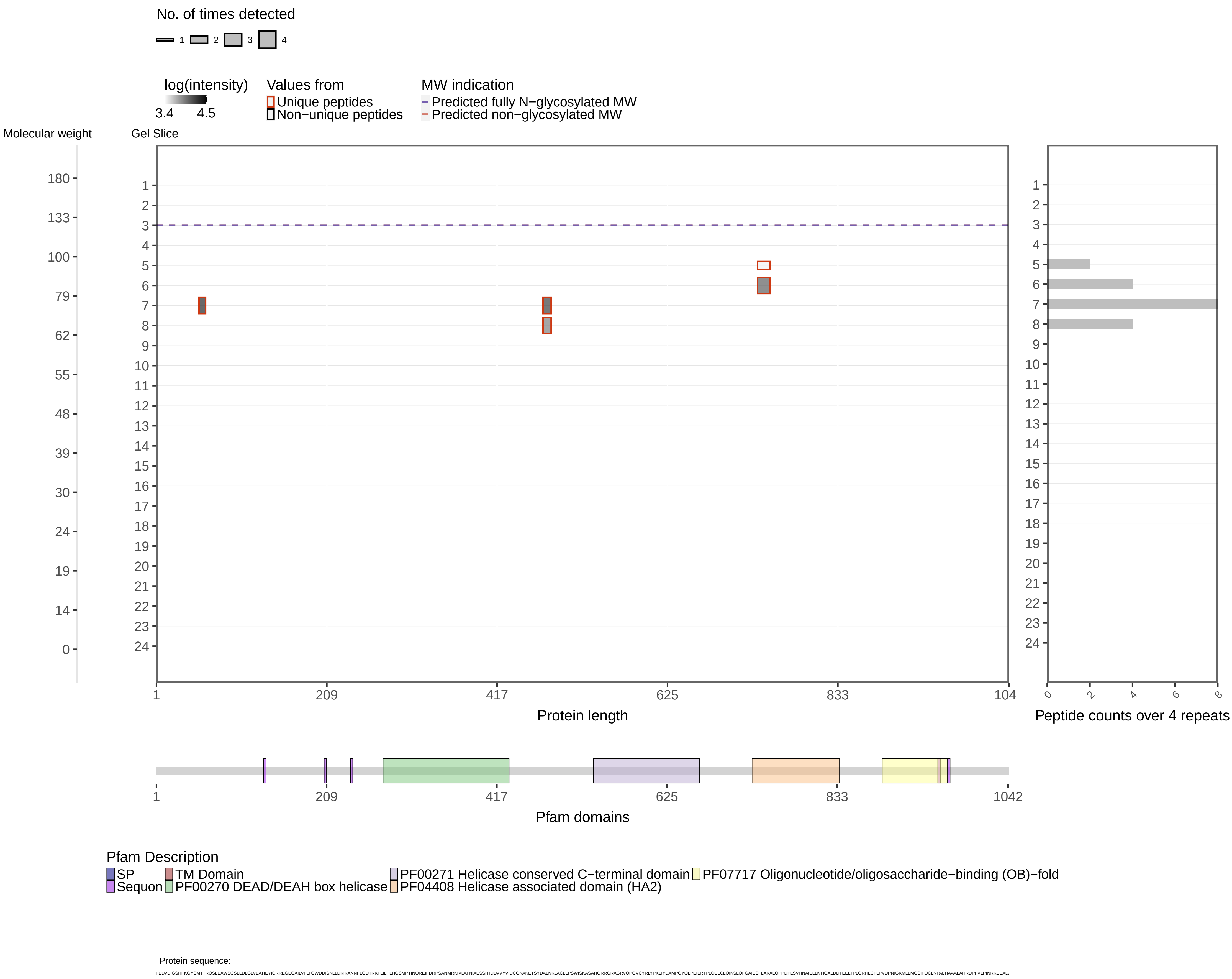

### NbL19g19470.1 leucine−−tRNA_ligase_cytoplasmic−like [MW = 131.56 kDa]

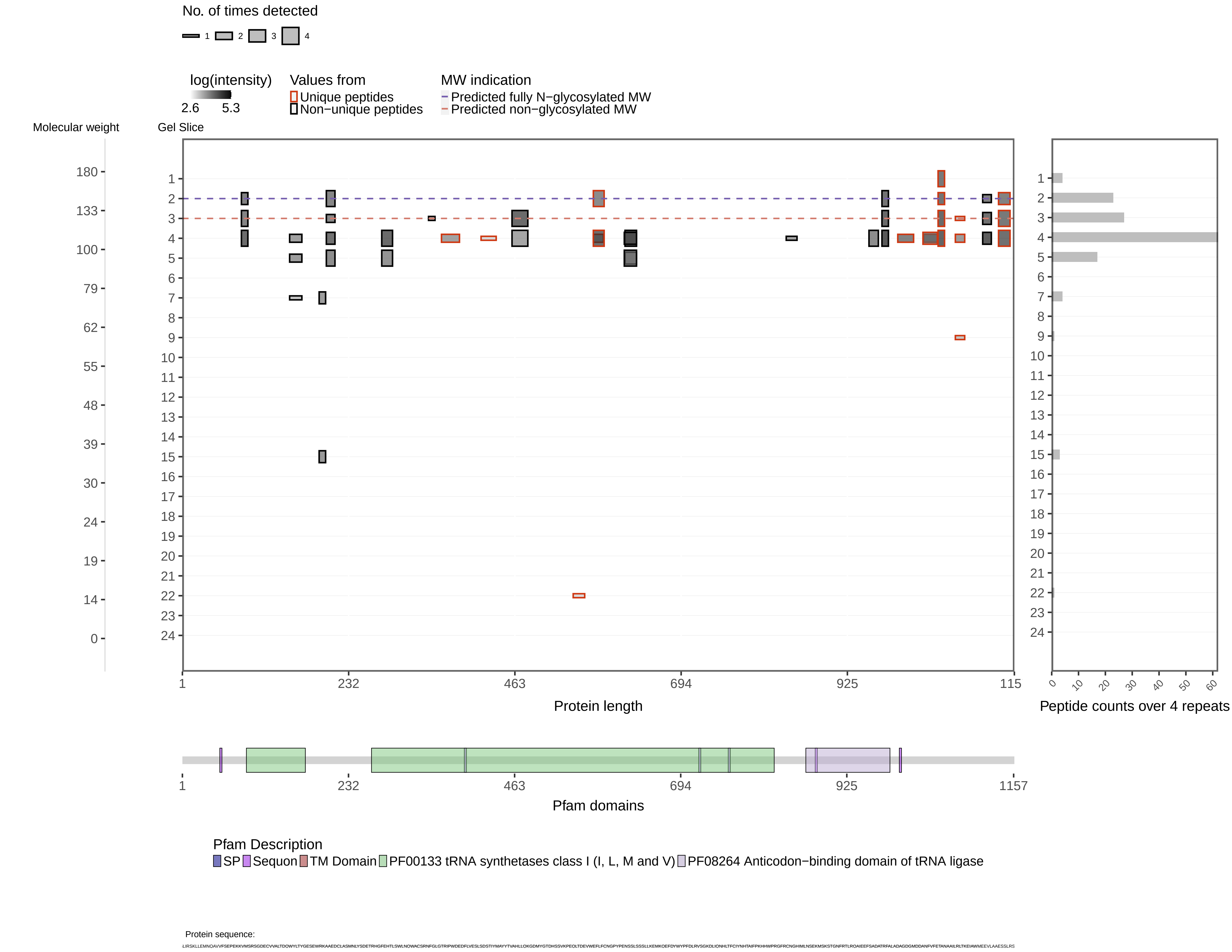

NbL19g19230.1 ras−related_protein_RABD1−like [MW = 29.38 kDa]

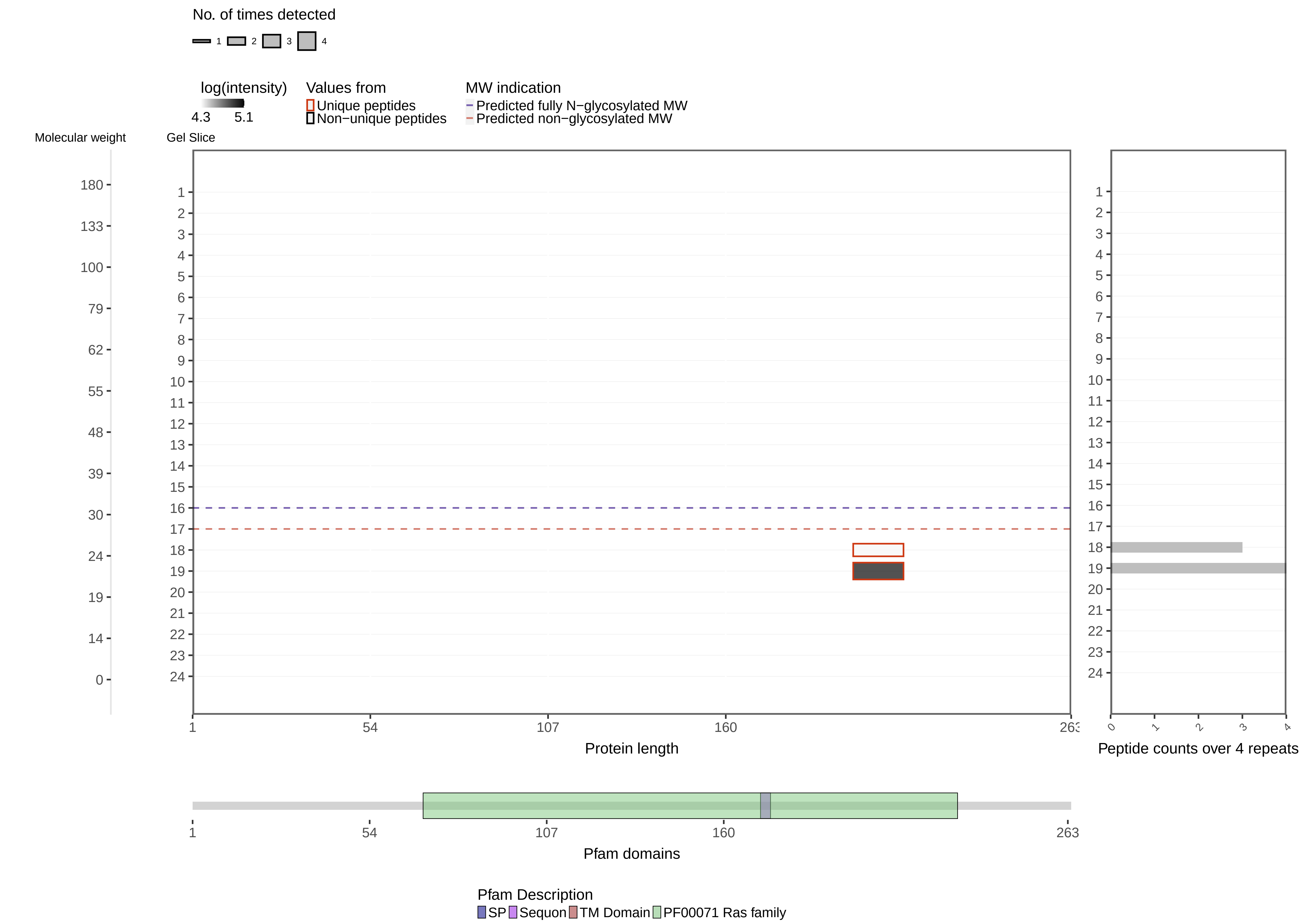


Protein sequence:

QIWDTAGQERFRTITSSYYRGAHGIIIVYDVTEKESFDNVKQWLSEIDRYANESVCKLLVGNKCDLVENKVVDTQTAKAFADELGIPFIETSAKDSINVEQAFLTMAGEIKKKMGSQPAGAKNSGGGVQIKGHPIEQKSNCCG*

### NbL19g19130.1 cleavage_and_polyadenylation_specificity_factor_subunit_3−I [MW = 77.37 kDa]

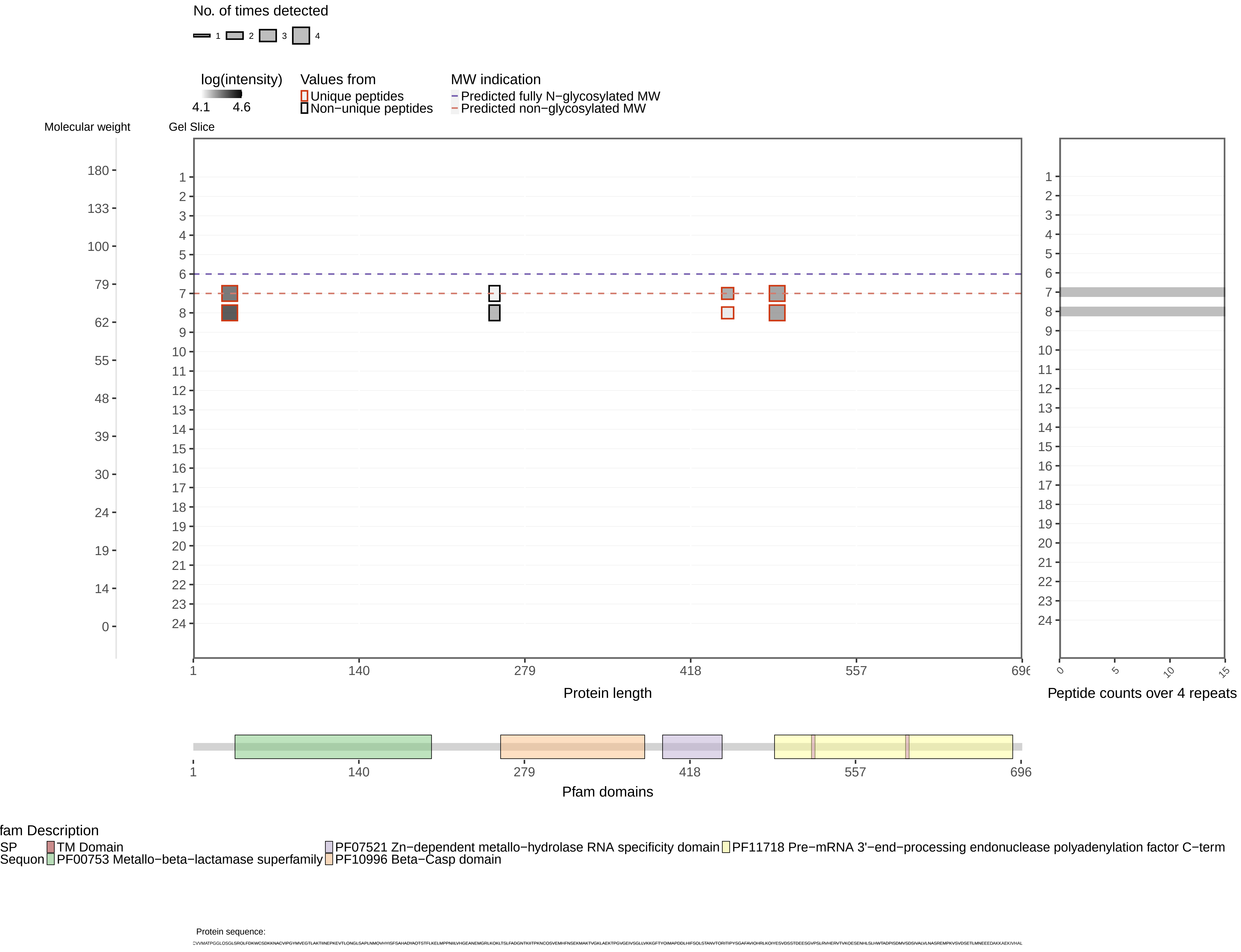

NbL19g19040.1 monodehydroascorbate_reductase [MW = 47.05 kDa]

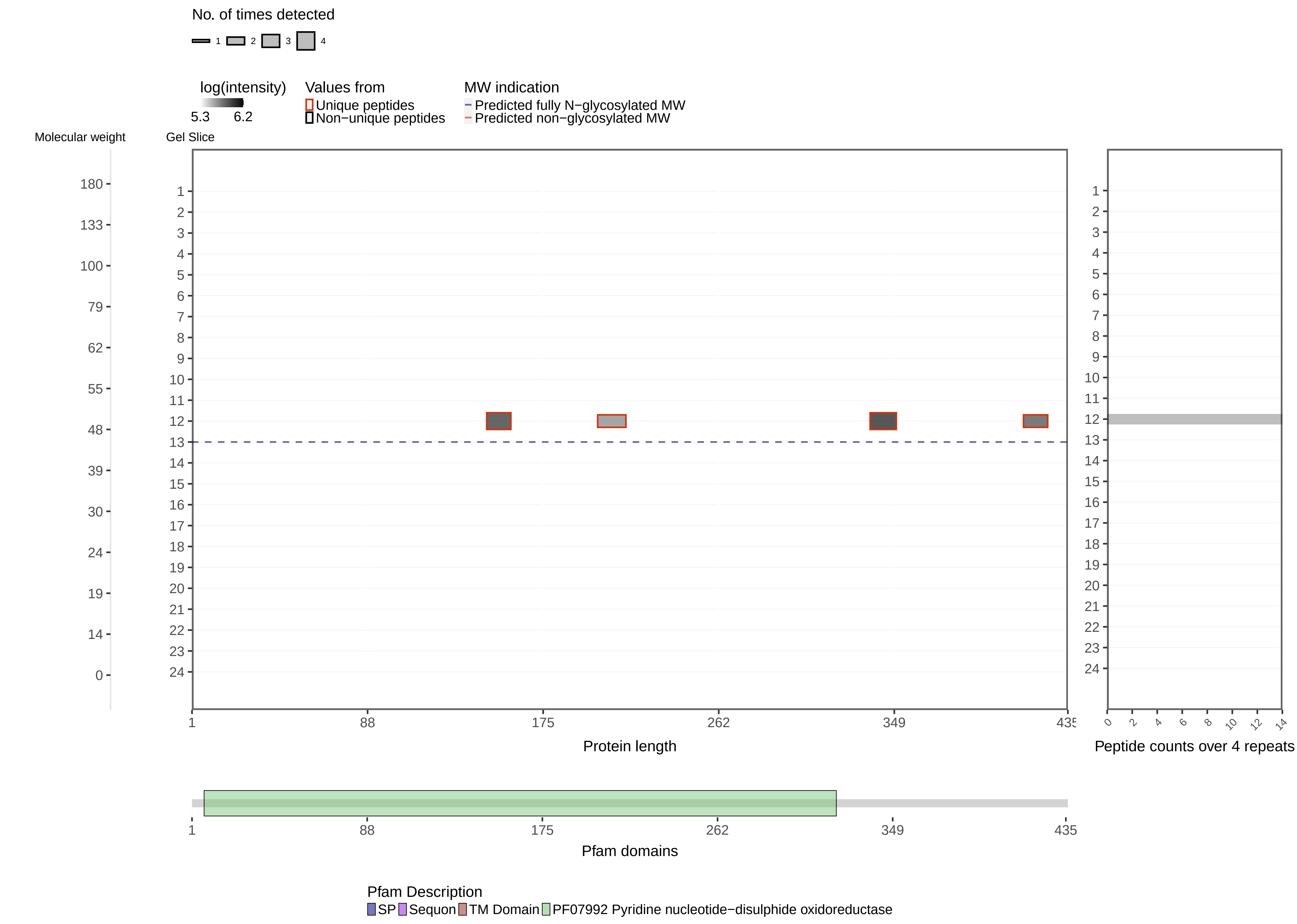


Protein sequence:

GIAAFYEGYYRNKGVNIIKGTVAVGFDTHPNGEVKEVKLKDGRVLEADIVVVGVGARPLTTLFKGQVAEEKGGIKTDAFFKTSVPDVYAVGDVATFPLKMYSDIRRVEHVDHSRKSAEHAVKAIFAREQGKSIDEYDYLPYFYSRAFDLSWQFYGDNVGETVLFGDADPNSATHKFGQYWIQDGKVVGVFLESGTPEENKAIAKVAKVQPLVSGLDQLAKEGLSFASKI*

NbL19g19020.1 cell_division_protein_FtsZ_homolog_2−2_chloroplastic−like [MW = 49.13 kDa]

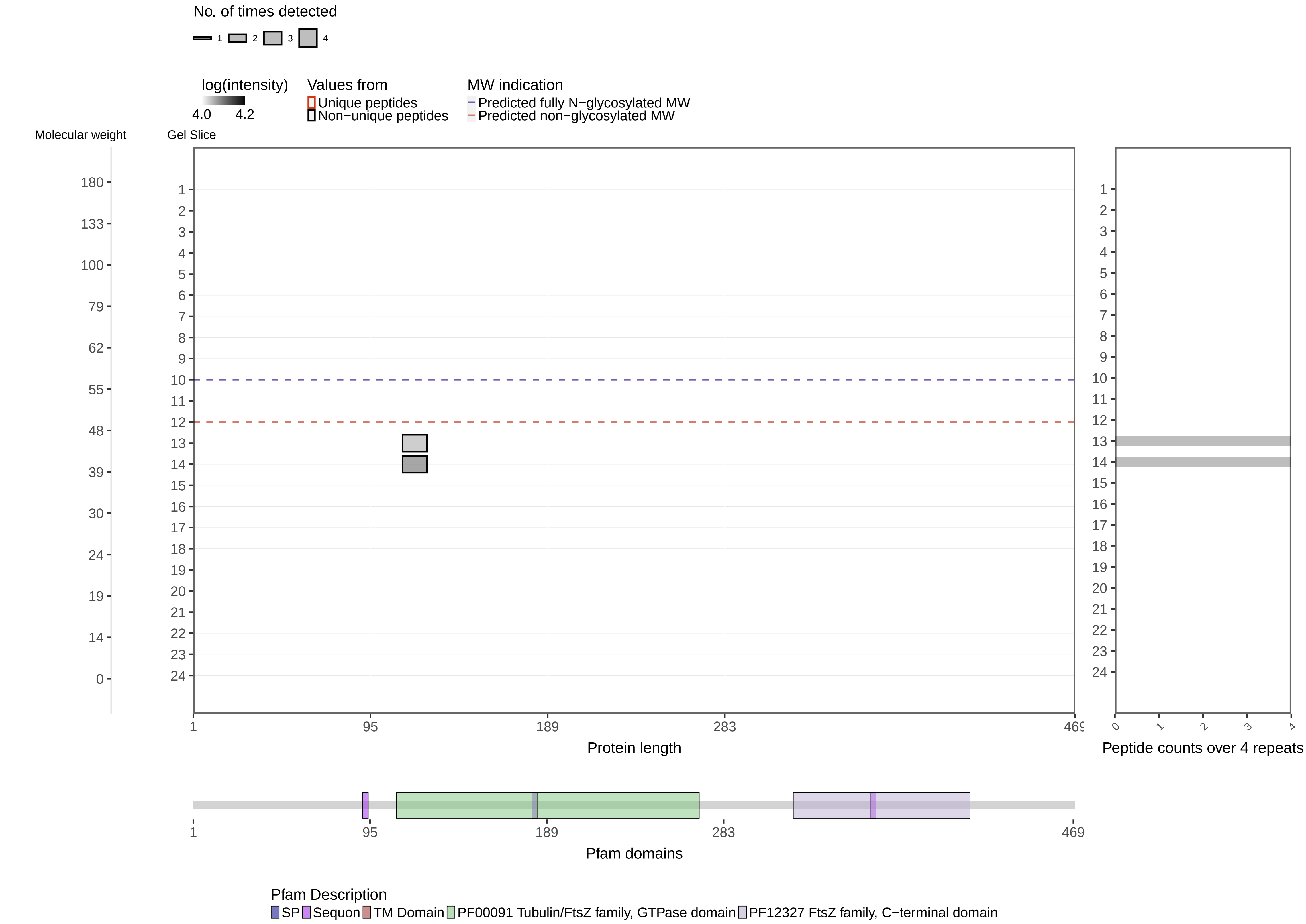


Protein sequence:

NbL19g18920.1 fatty_acyl−CoA_reductase_2−like [MW = 27.94 kDa]

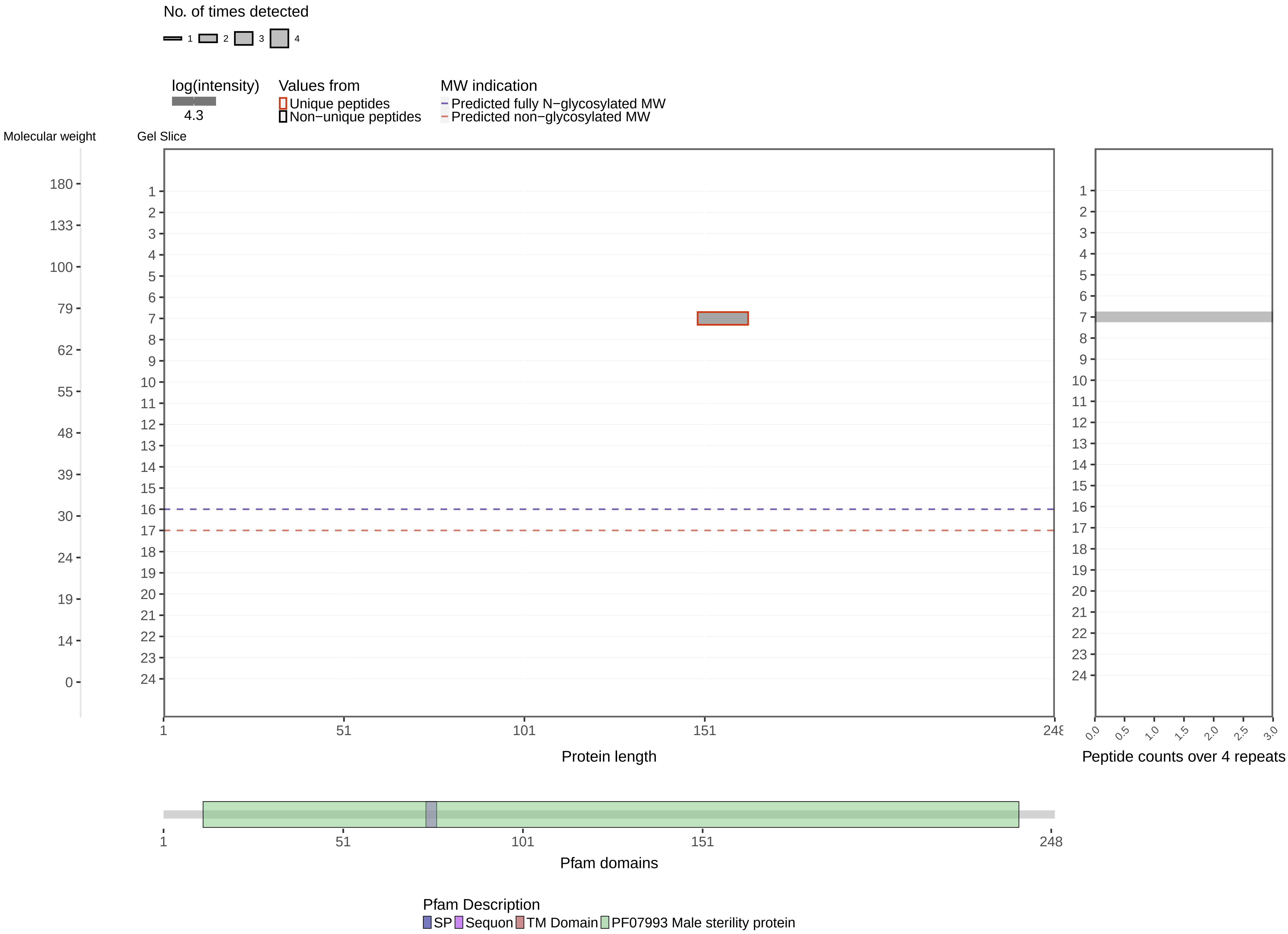


Protein sequence:

STAYANGEREGLILEKPFTMGESITKEKITSFSPFTKFPSLNAMSEMDLVSKLKNAITTNNGLEQIMKDTGIERAKLYGWQDTYSFTKAIGEMMIDSMREEIPILILRPSVVASSFREPFPGWIQGFRCRPSISQG*

NbL19g18850.1 probable_uridine_nucleosidase_1 [MW = 36.72 kDa]

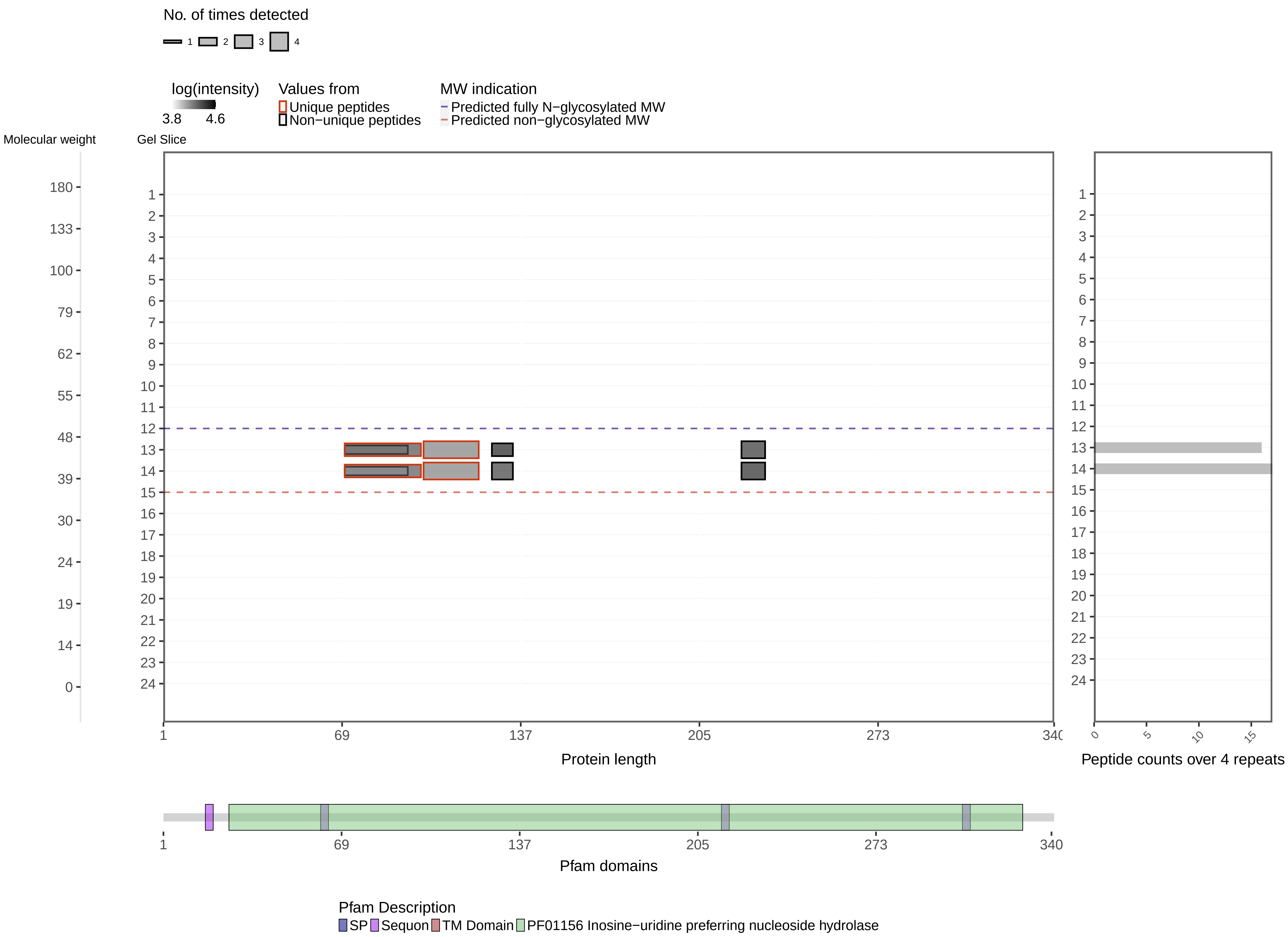


Protein sequence:

RDSTFASKVKRVVVLGGSFFALGNVNPAAEANIYGDPEAADIAFTSGANIDIVGINITTOVKLTDADLEELROSEGKHAKFLCNMCKFYRDWHVKSDGVYGIFLHDPVSFVALVRPELFTFKKGVVRVETOGLCIGHTLMDOGLKKWNTSNPWTDYSPVSVAWTVDVDEVLNYVKKMLMKP*

### NbL19g18760.1 glucosamine_6−phosphate_N−acetyltransferase [MW = 17.87 kDa]

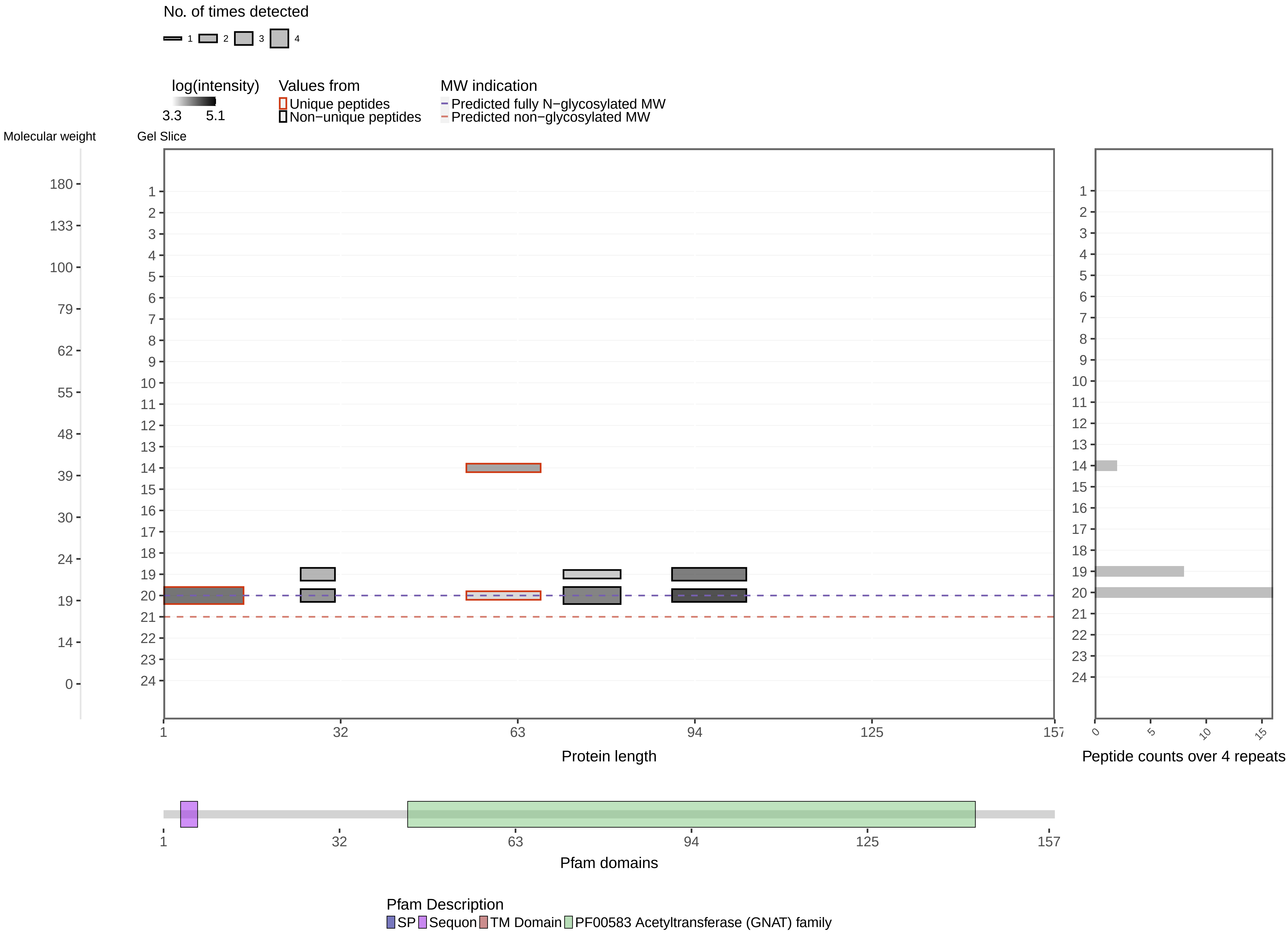


Protein sequence:

KSGKIVATGSVFIEKKFVRNCGKAGHIEDVVVDSSTRGMQLGKRIVEYLANHAYSMGCYKVILDCTEENRPFYEKCGFKKKEIQMVKYFV*

### NbL19g18720.1 uncharacterized_LOC109242020 [MW = 39.71 kDa]

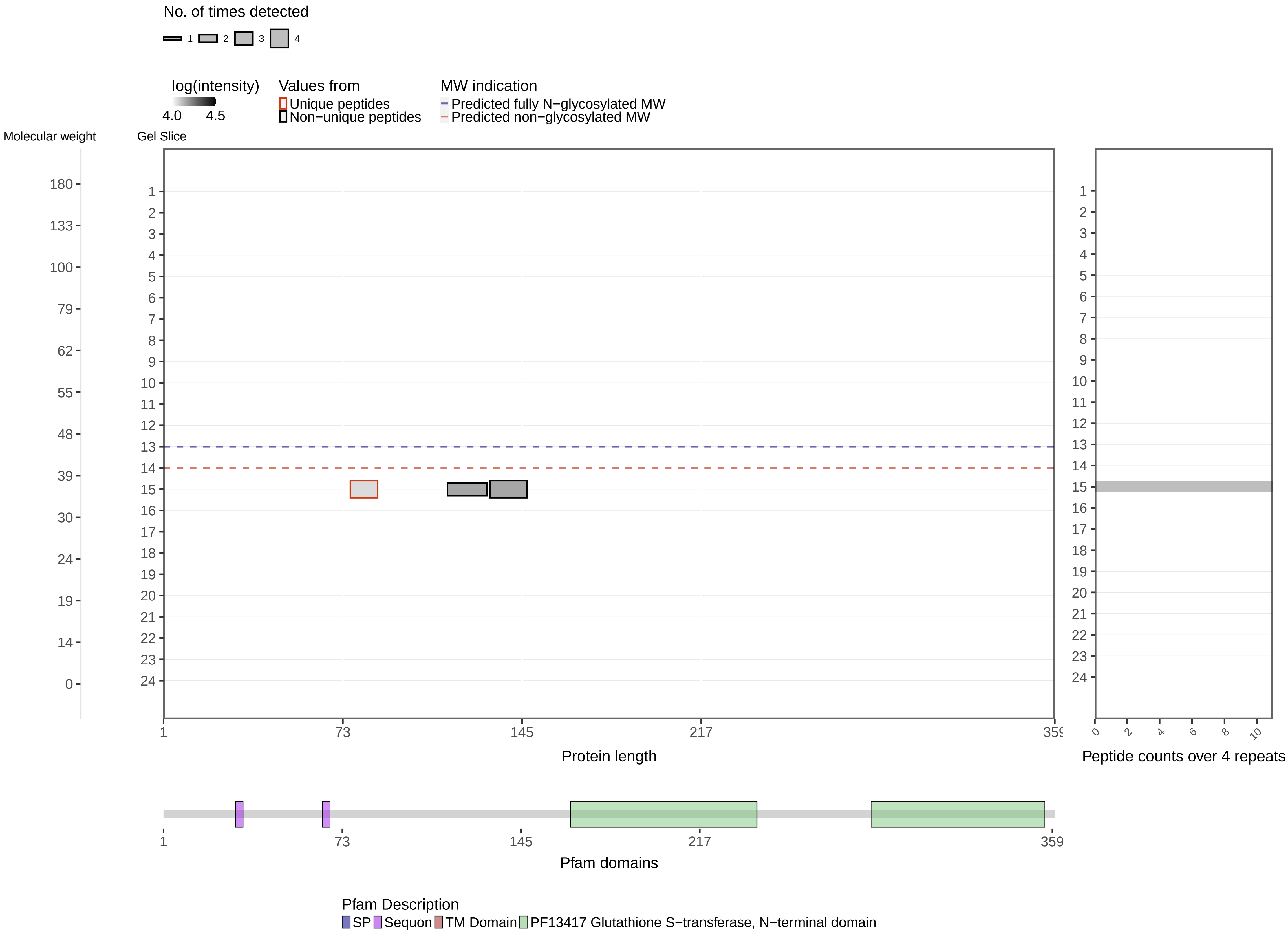


Protein sequence:

ESCPFCRKVREIVAVLDLDVLYYPCPRNGPNFRPKVGQMGGKQQFPYMVDPNTGVEMYESDEIIKYLVGKYGDGNVPLMLSLGLLTTLTAGFAMIGRMGKGSSYRPSKLPPKPLEIWAYEPSPFCKVVREVLVELELPHILHSCARGSPKRQVLYERVGHFQVPYLEDPNTGVQMFESADIVDYLQATYAL*

### NbL19g18710.1 alpha_carbonic_anhydrase_1_chloroplastic [MW − SP = 27.99 kDa]

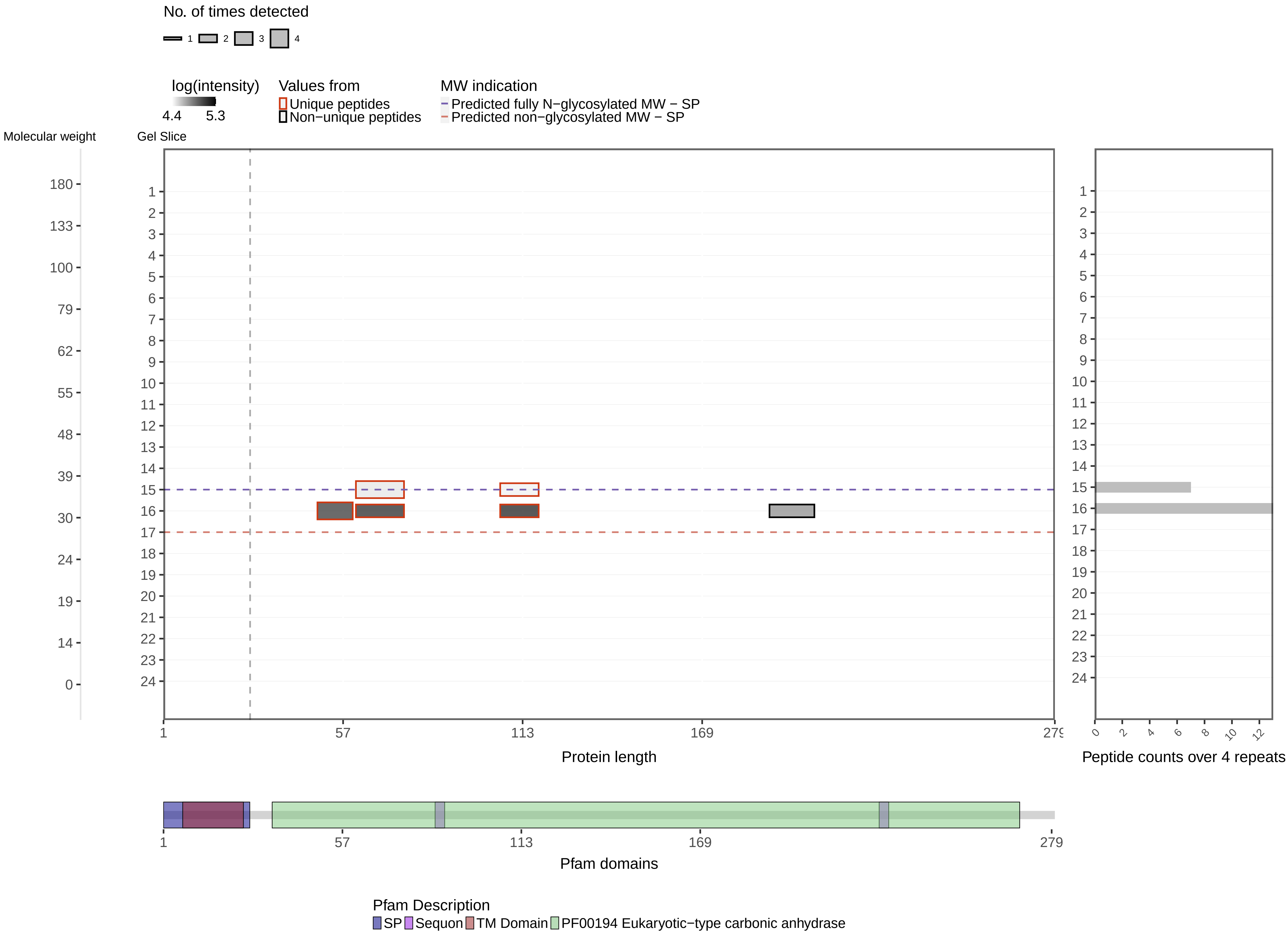


Protein sequence:

VHSPSEHRLDGTQYAAELHLVHFAEDNSIAVLATLFHIGHPDPLMTKLQNKLNELARDVASQKETQIQLGTIDTHEIRKQSHKYYSYTGSLTTPPCSENVSWHILGKVRSISREQVEALRAPLDSRCKRNARPVQPLNGRQIQMYEVQPNVN*

NbL19g18680.1 heterogeneous_nuclear_ribonucleoprotein_R [MW = 53.02 kDa]

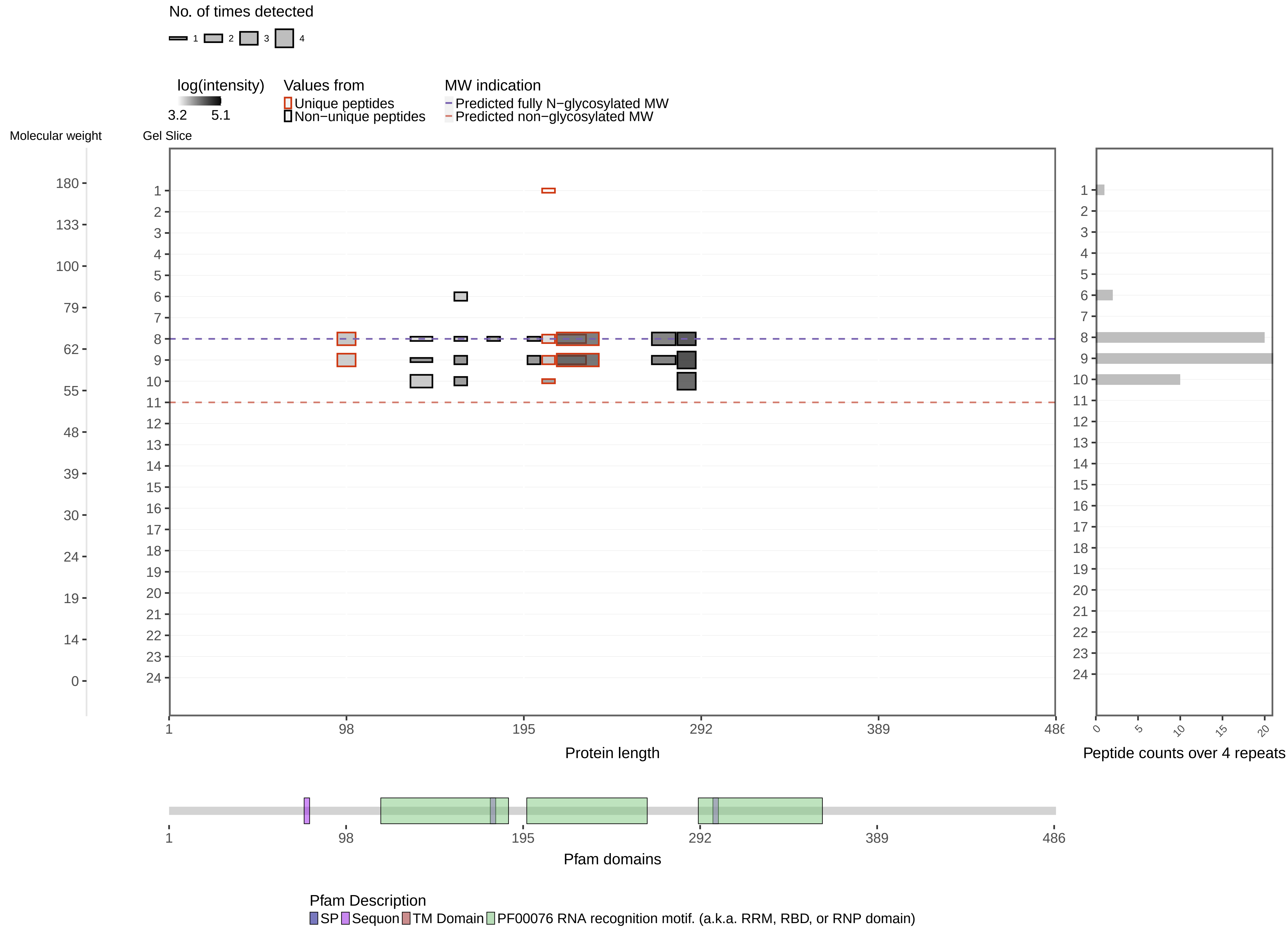


Protein sequence:

NSGRNRGFAFIEYYNNACAEYSRQKMSNSDFKLDDNAPTVSWADPKNAETAASAQVKAVYVKNLPKNITQDKIKELFEHHGKITKVVLPPAKPGQEQSRYGFVHFAERSSAMKALKNTERYEIDGKILECSLAKPQADQKSSGGSSSQKGPTLPTYPPRIGYGMVGAPYGALSAGYGGAGVGQPLVYGRGASPAGLAMMPMLLPDGRIGYVLQQPGVMHTPLPASRGGRSGGGGSSGGRRGDSGRGRSRYNPY*

NbL19g18640.1 signal_recognition_particle_54_kDa_protein_chloroplastic [MW = 61.91 kDa]

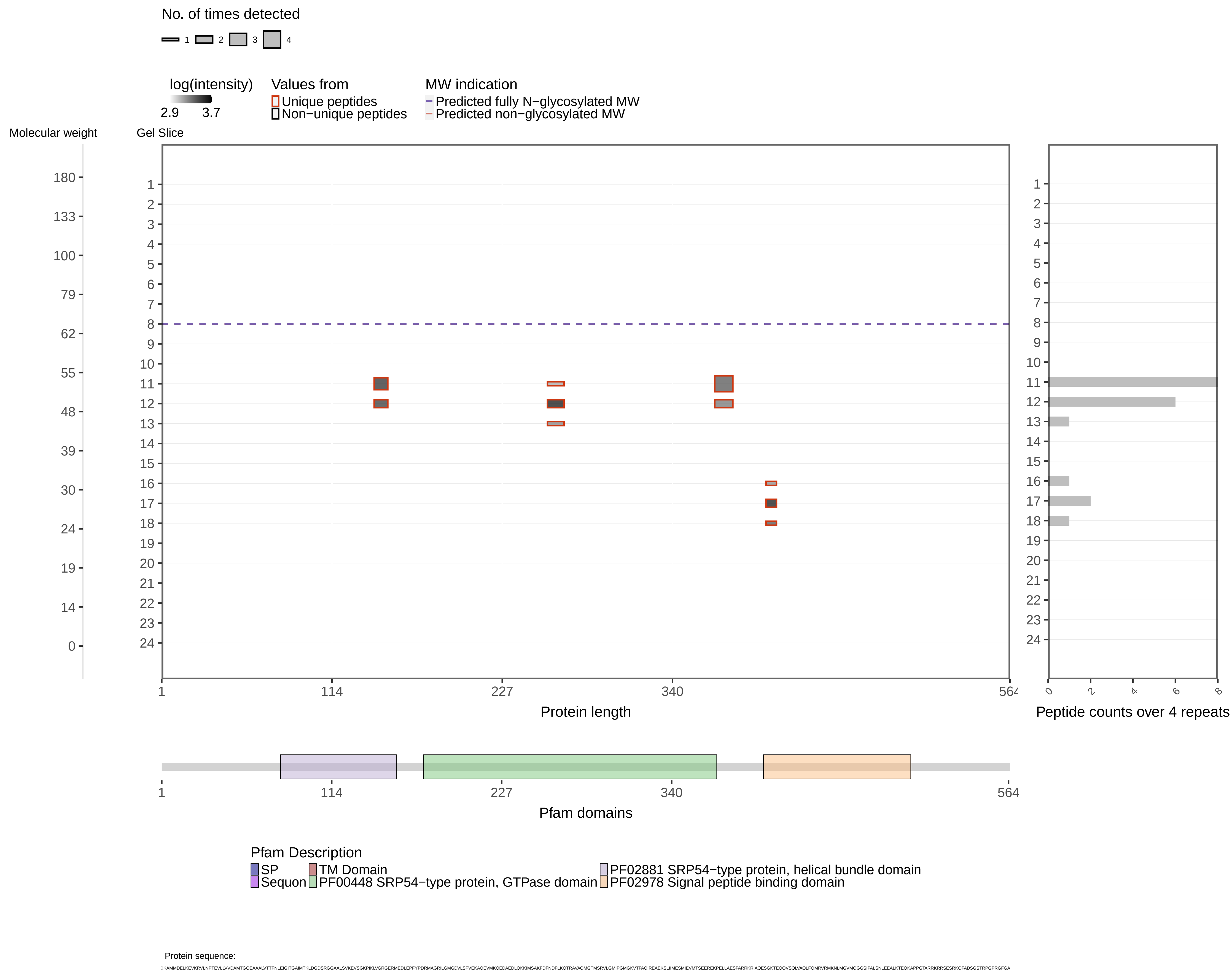

### NbL19g18570.1 1−aminocyclopropane−1−carboxylate_oxidase_homolog_1−like [MW = 40.94 kDa]

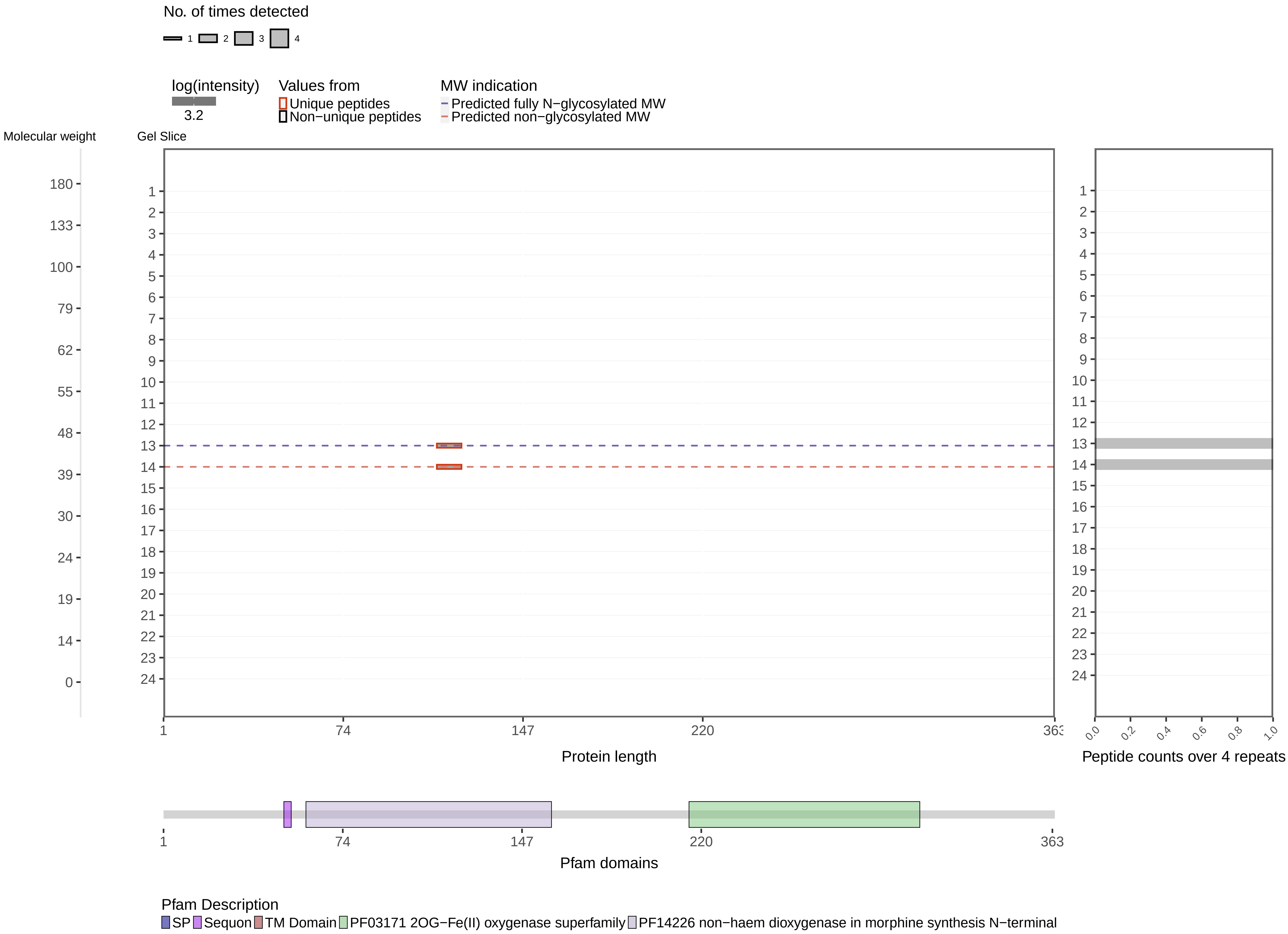


Protein sequence:

LPPICRDIIIKYSEEVKKLGNSLFELLAEGLGLNRNHLTDMDCDKGLSVVCHYYPACPEPELTLGASKHADDGFITLLLQDHIGGLQVLHQNHWVDVPPTPGALVVNIADLLQLISNDKFKSVEHRVLANHIGPRISIACFFSTFLLASSRLYEPIKELLSEDNPPKYRETTIHEYAAYFTAKGLDGTSALHHFRL*

NbL19g18490.1 uncharacterized_LOC109237744 [MW = 21.26 kDa]

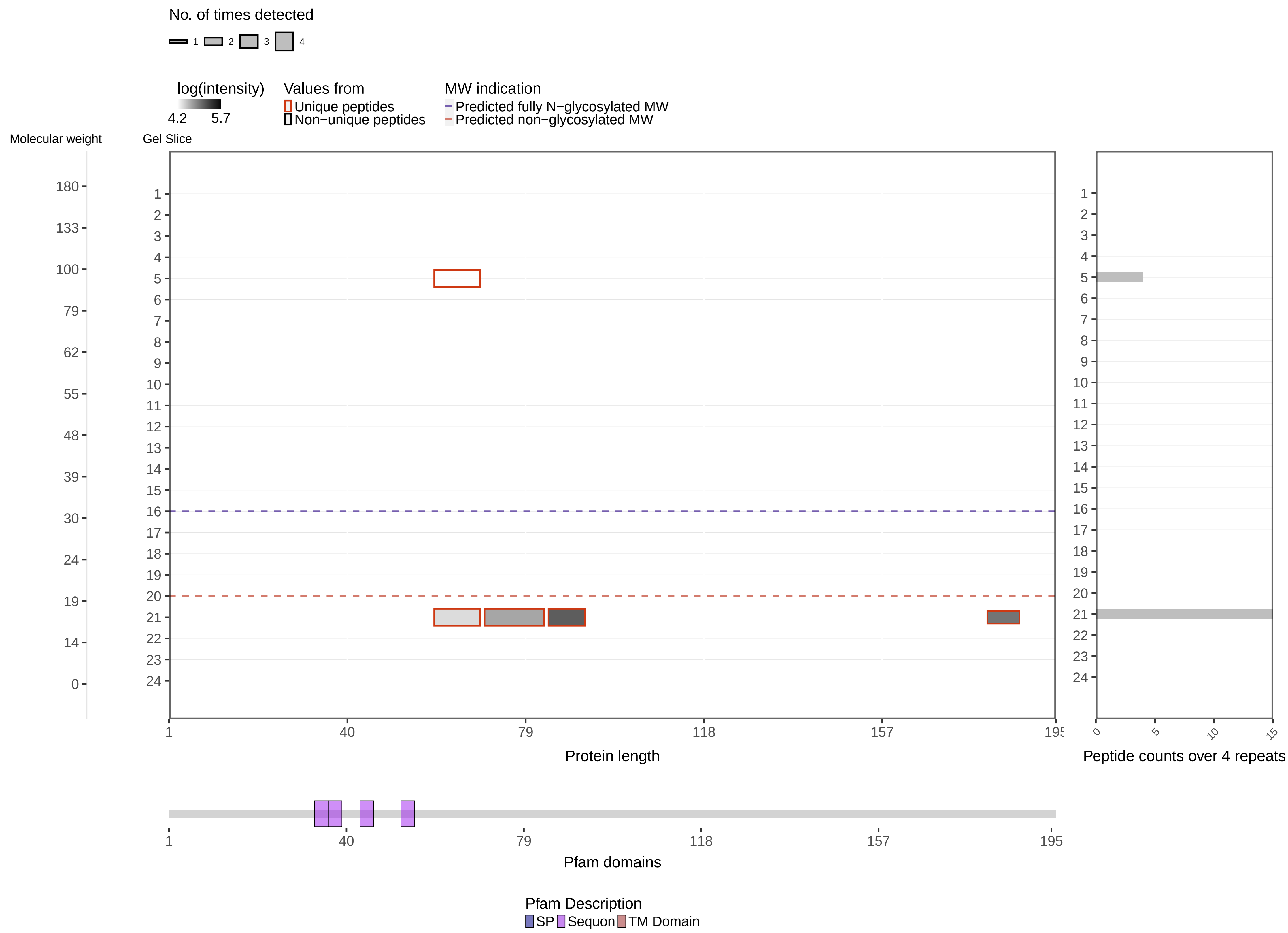


Protein sequence:

LPLDIDVARFETLLFQWANSLCQGAMLPLPVPLKVDKIPRGARLGFITVEDAQTEVLVYIDCMVFPATDNSPPIFRAIRNGPSRDKAPPGEPRIMRSLLGALQKSVEIARV*

### NbL19g18460.1 beta−fructofuranosidase_insoluble_isoenzyme_1−like [MW = 66.5 kDa]

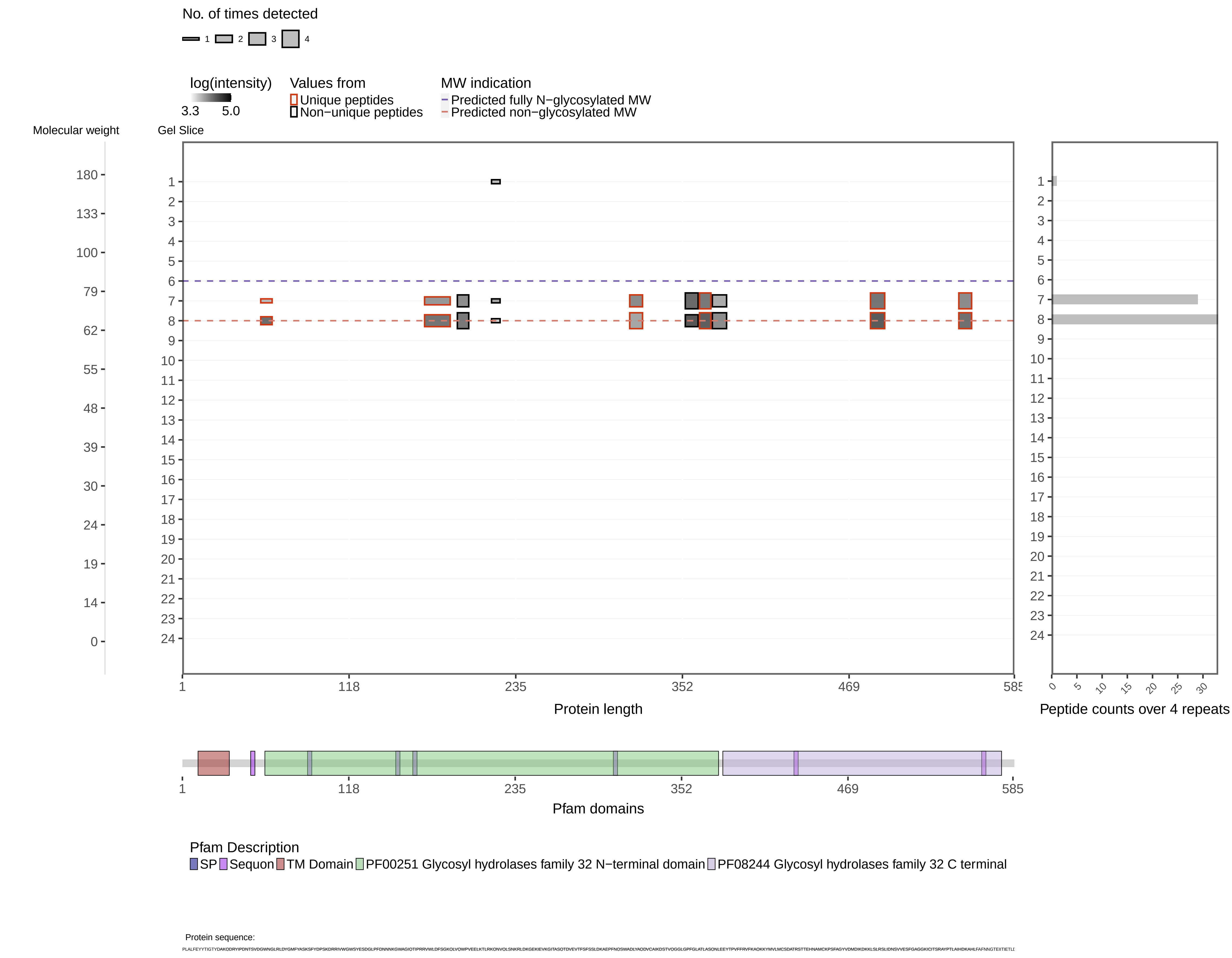

### NbL19g18360.1 peptidyl−prolyl_cis−trans_isomerase_CYP18−2 [MW = 18.16 kDa]

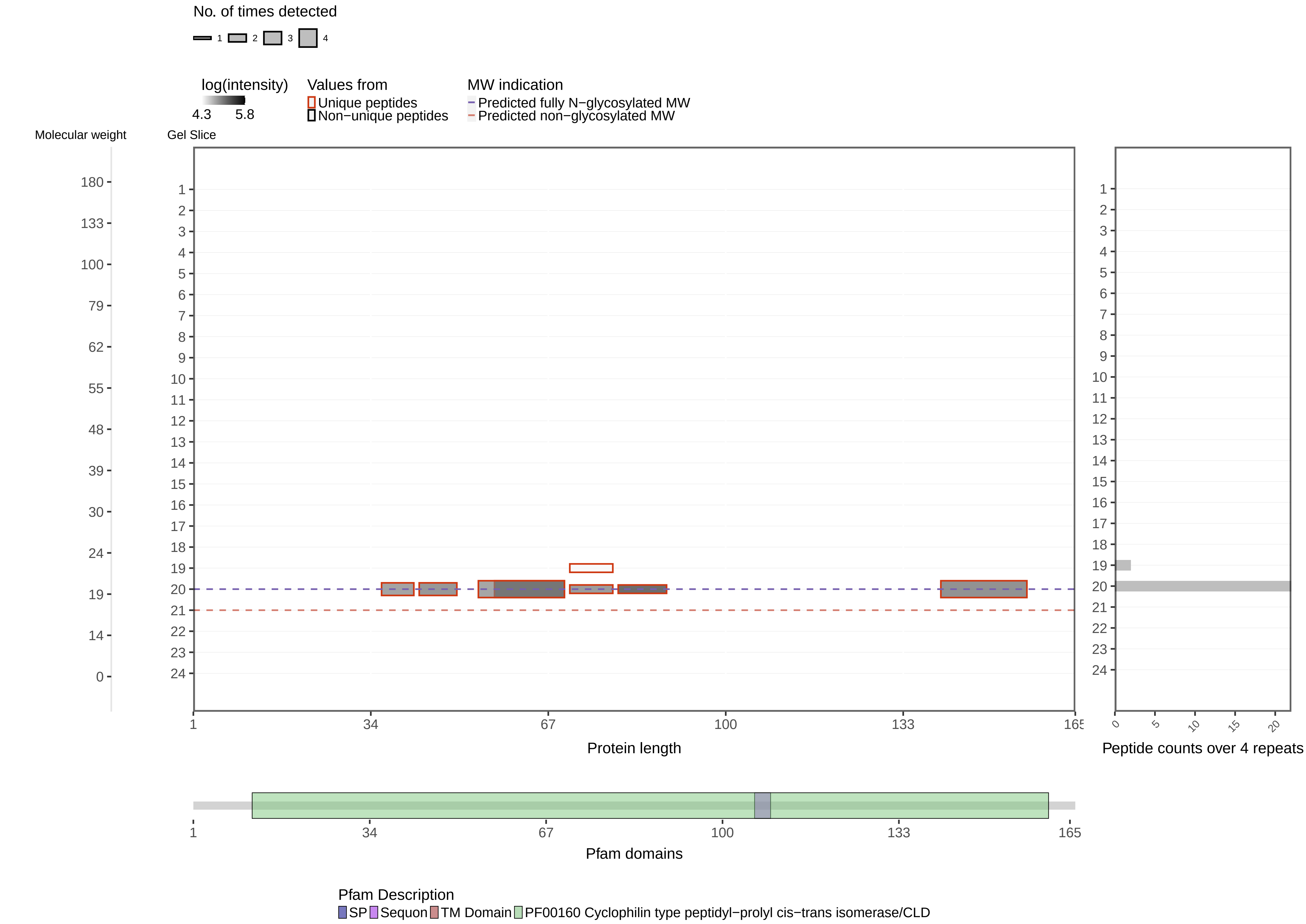


Protein sequence:

GRGGESIYGAKFDDEINQQLKHTGAGIISMANAGPNTNGSQFFITLAPAQSLDGKHTIFGRVCRGMEIIKRLGSVQTDNTDRPIHDVKILRTTVKD*

NbL19g18330.1 endoglucanase_8−like [MW = 55.82 kDa]

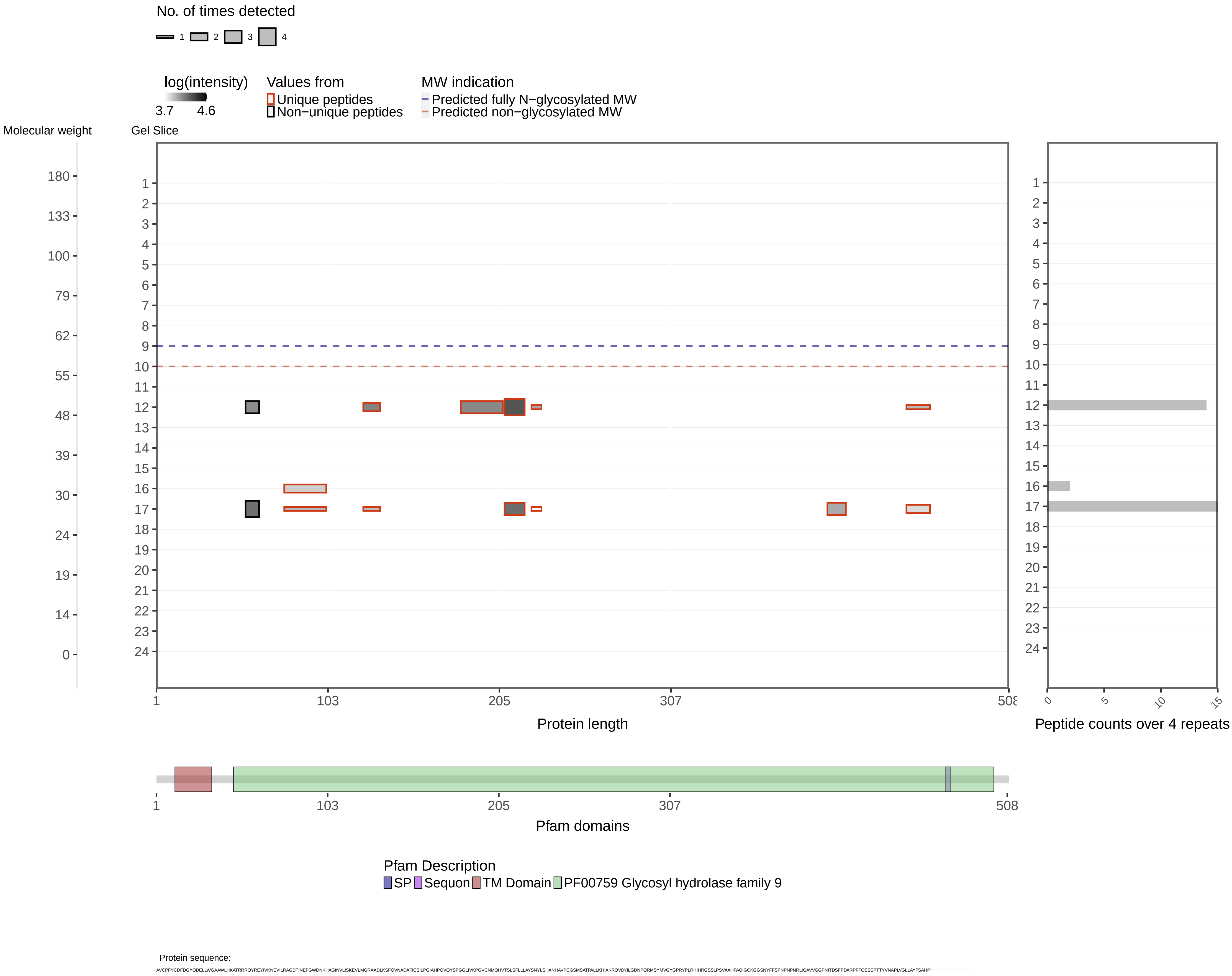

NbL19g18320.1 lysosomal_Pro−X_carboxypeptidase−like [MW − SP = 55.3 kDa]

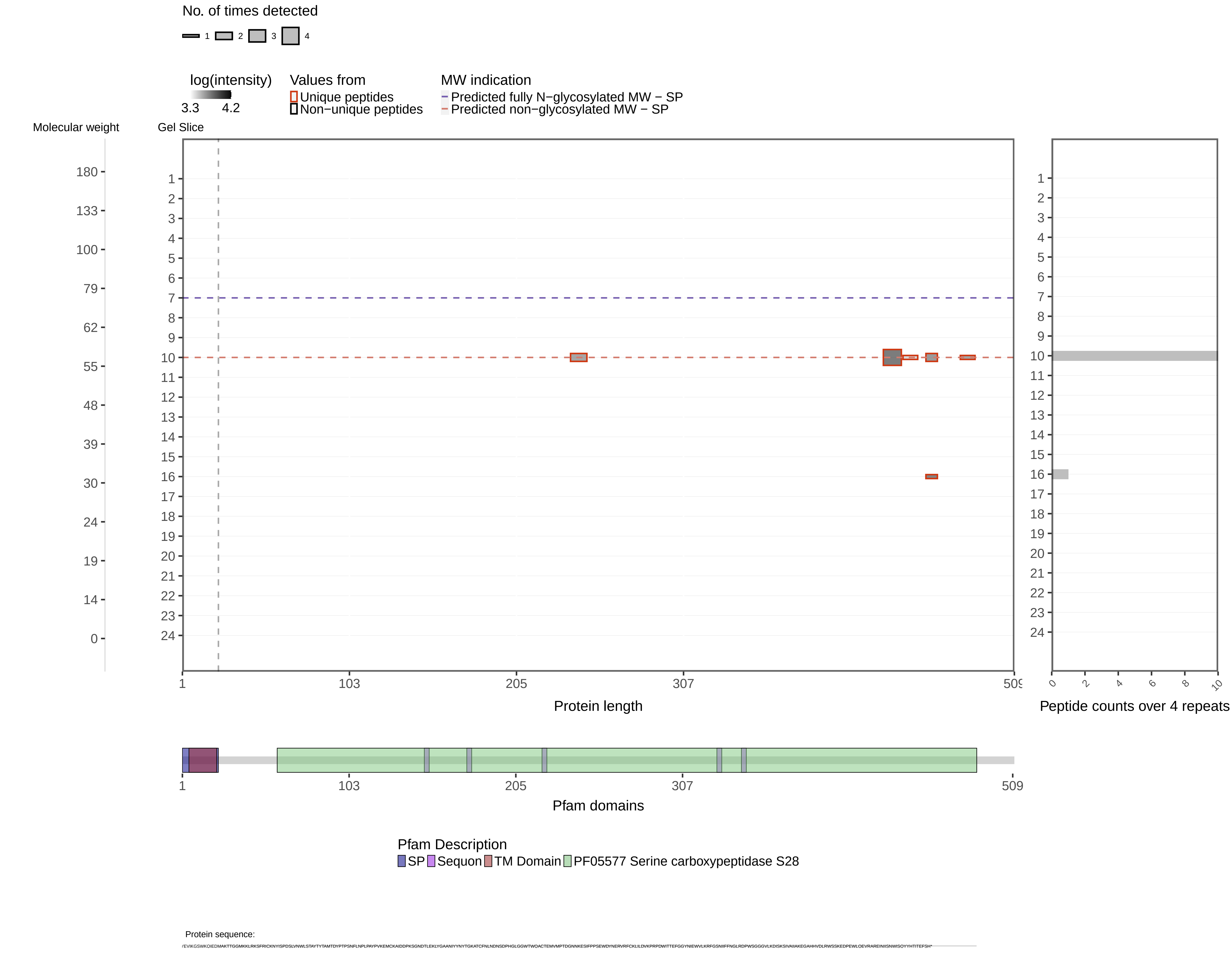

### NbL19g18260.1 glutathione_S−transferase_L3−like [MW = 36.16 kDa]

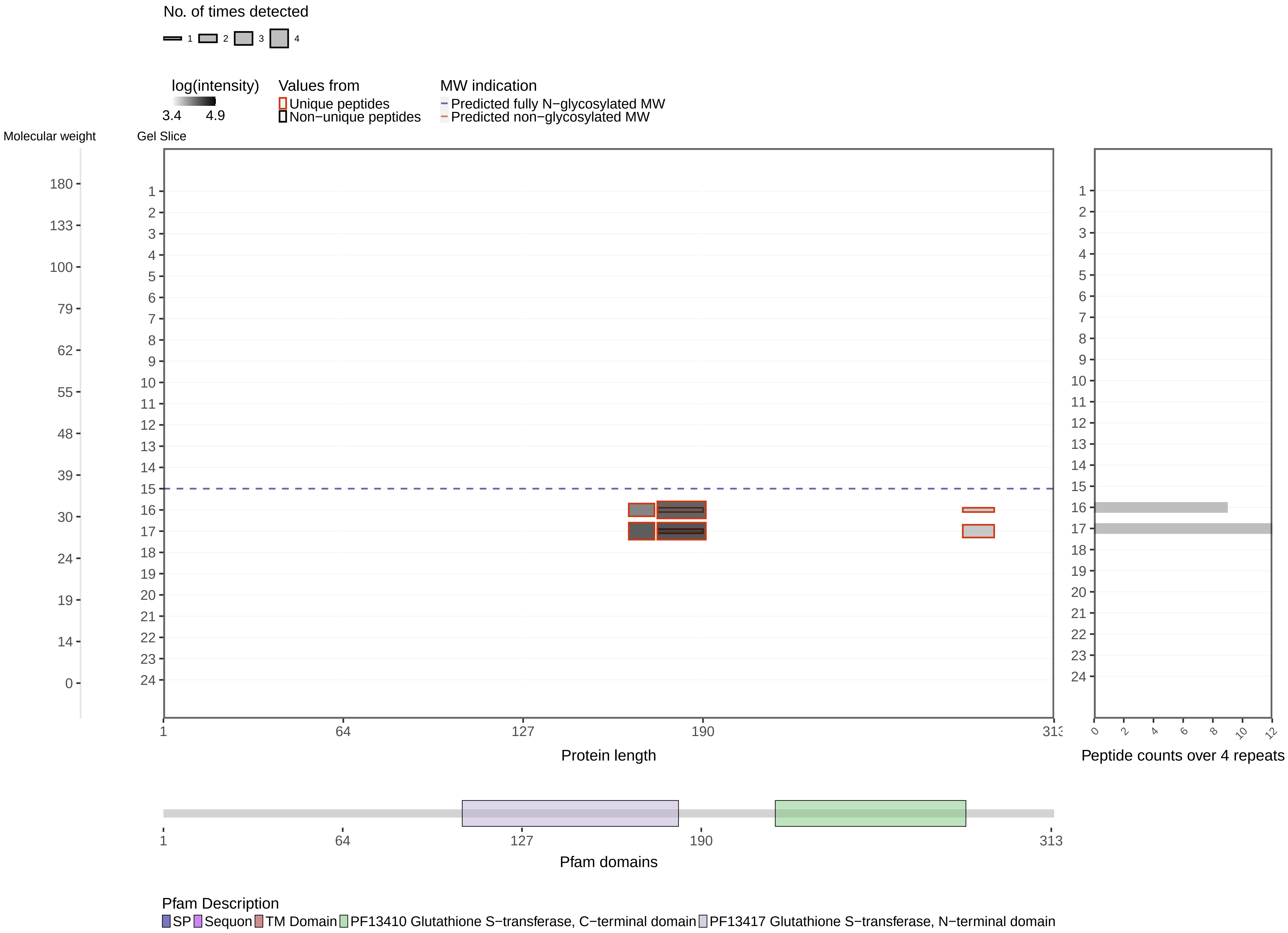


Protein sequence:

YKEKVYAENKVPSLEHNNKVIGESLDLMKYVDSNFEGPSLLPDDPKKQKFAEELIAYSDTFVKEVYGSFKKDAQILAGAQFDYLEKALEKFDDGPFFLGQFSQVDIAYAPFIERFQIFLQEVFNYDITSARPKLAKWIEEMSNIDGYKQTKVLDPKRLVEYYKNRFMA*

NbL19g18130.1 uncharacterized_LOC107831931 [MW = 26.85 kDa]

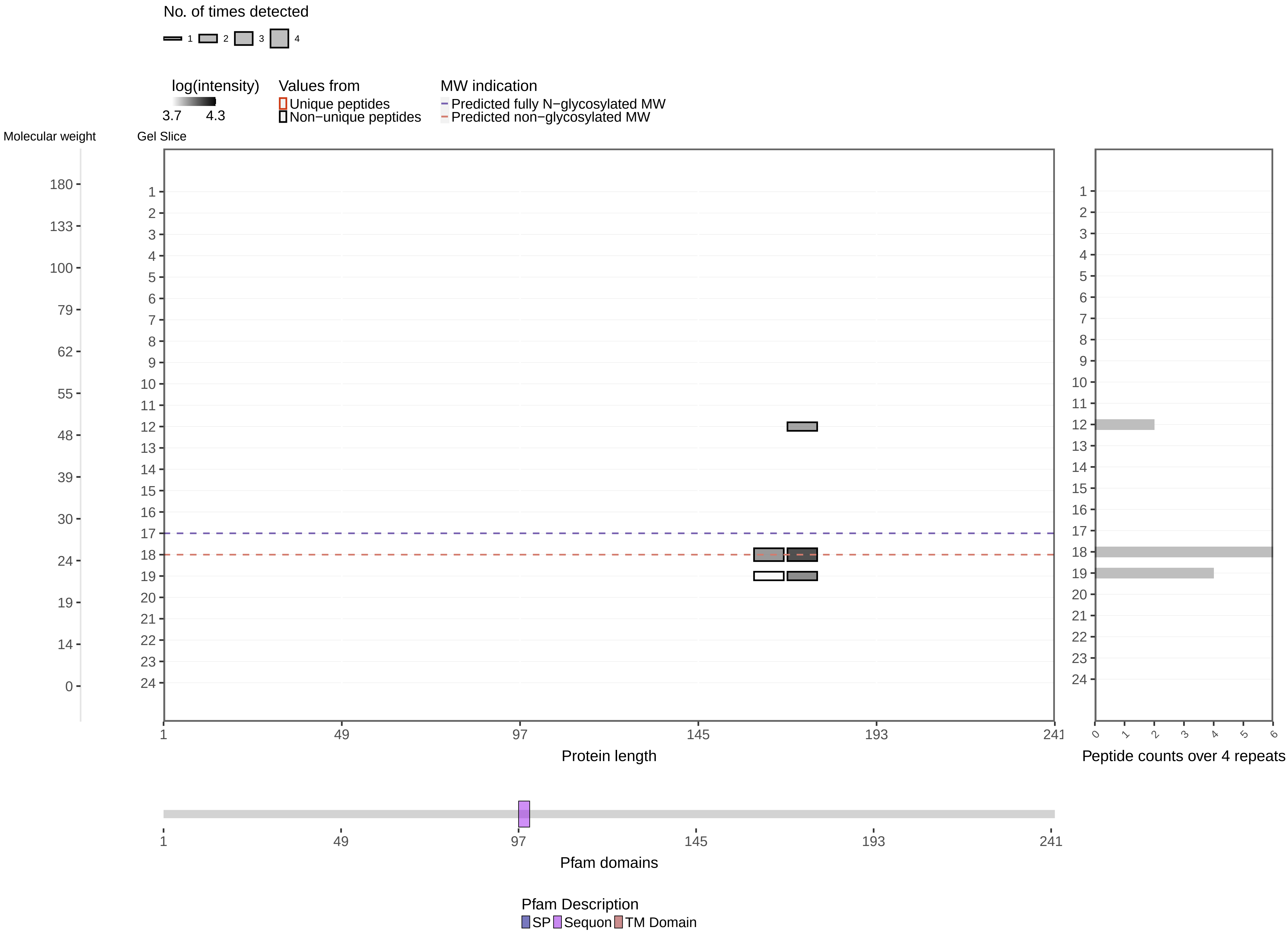


Protein sequence:

YNPNYAPVAAMIQFPTRNAYDASLRAVVRTSRLFRMERADRQQWDTLPNYDGKTILLQGIPRNALADDVERFLSGCQYDGSSIQIFARQQRGFDRPPARVALVQFPSQALAAHAFITKNRGFVLNNQIAVRLLQ*

### NbL19g18070.1 probable_pectate_lyase_4 [MW = 43.89 kDa]

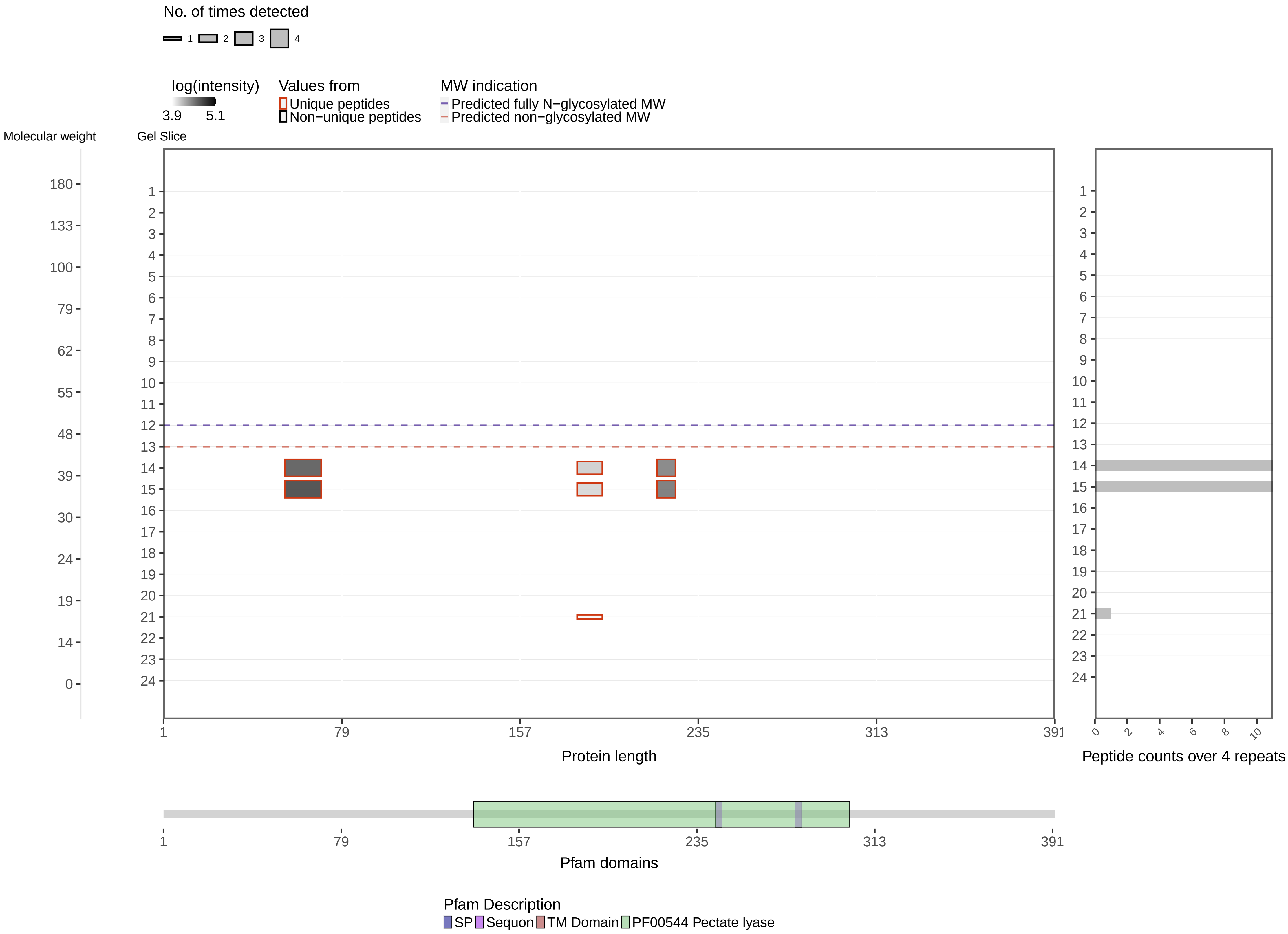


Protein sequence:

GSDVDAIQIKPKSRHIWIDRCSLSDYDDGLIDITRESTDITVSRCHFSKHDKTMLIGGDPSNCTDRCMRVTIHHCFFDGTRQRHPRVRFGKVHLYNNYTRNWGIYAVCASVESQIYSQCNIYEAGQKKVAFKYLTEKAADKEEGCTGSIKSEDDLFVCGTQAGLLPTCSESNIFNPCEFHQTWTVERPNDALKHYLQHCTGWQSIPRPN*

### NbL19g18010.1 protein_SLOW_GREEN_1_chloroplastic−like [MW = 40.2 kDa]

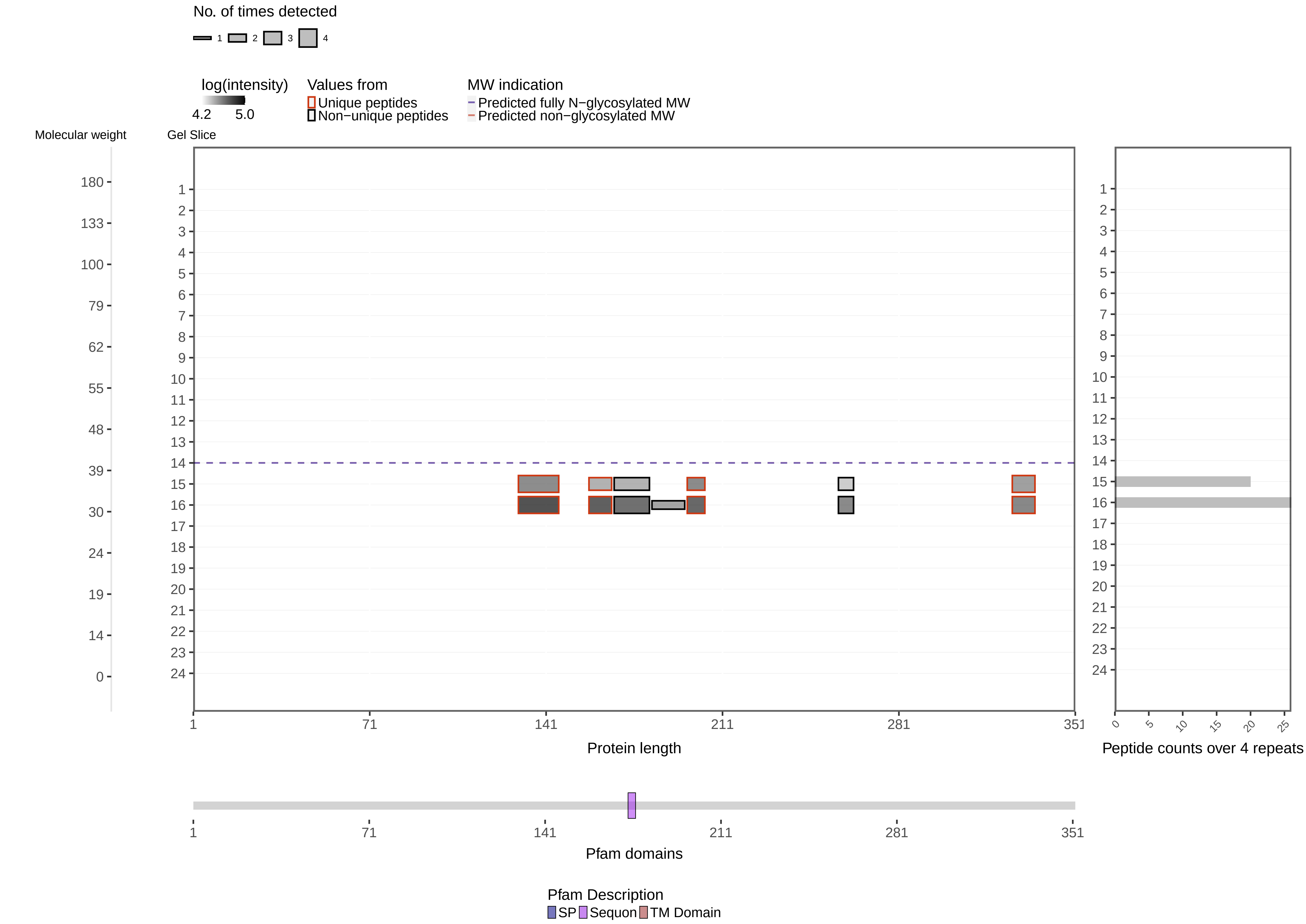


Protein sequence:

'IDKLIEVEPNESEWPLMKSHLYVNIGEVELAKVGFNEILKKDPFRVEAYHGLVMAASQDESIDDLKGIEKMIEEGMKLCKKENKKTDLRDFKLLLAQIRVIEGKYEDALKVYGELVKEEPRDFRPYLCQGIIYTLLRKSSEAEKCFEKYRRLVPQGHPYARYFDENMMATKVFAQKVENERAGSKS*

### NbL19g17980.1 60S_ribosomal_protein_L23a−like [MW = 17.32 kDa]

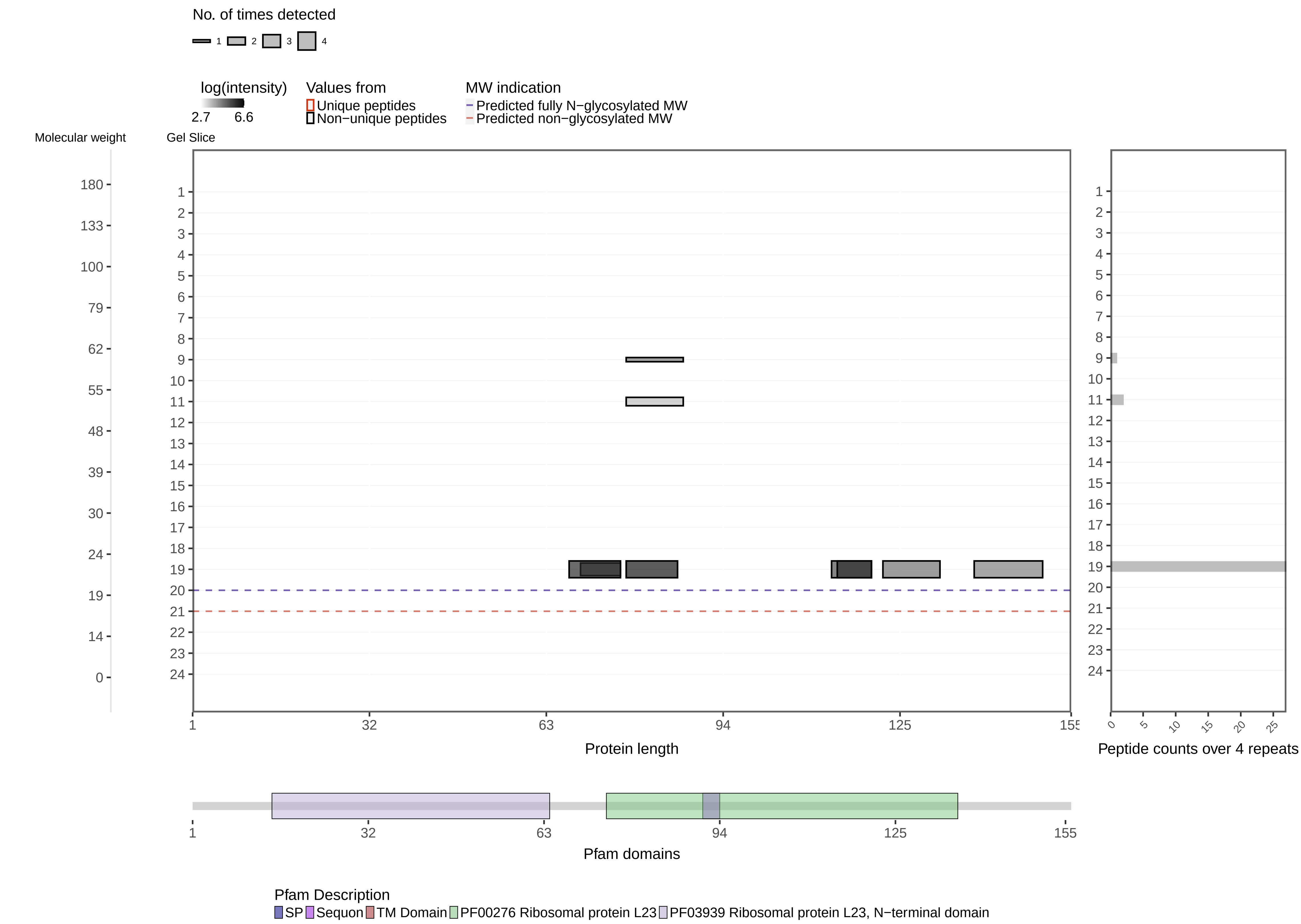


Protein sequence:

SAPGRNKLDQYGILKYPLTTESAMKKIEDNNTLVFIVDIKADKKKIKDAVKKMYDIQTKKVNTLIRPDGTKKAYVRLTPDYDALDVANKIGII*

### NbL19g17970.1 diaminopimelate_epimerase_chloroplastic−like [MW = 39.6 kDa]

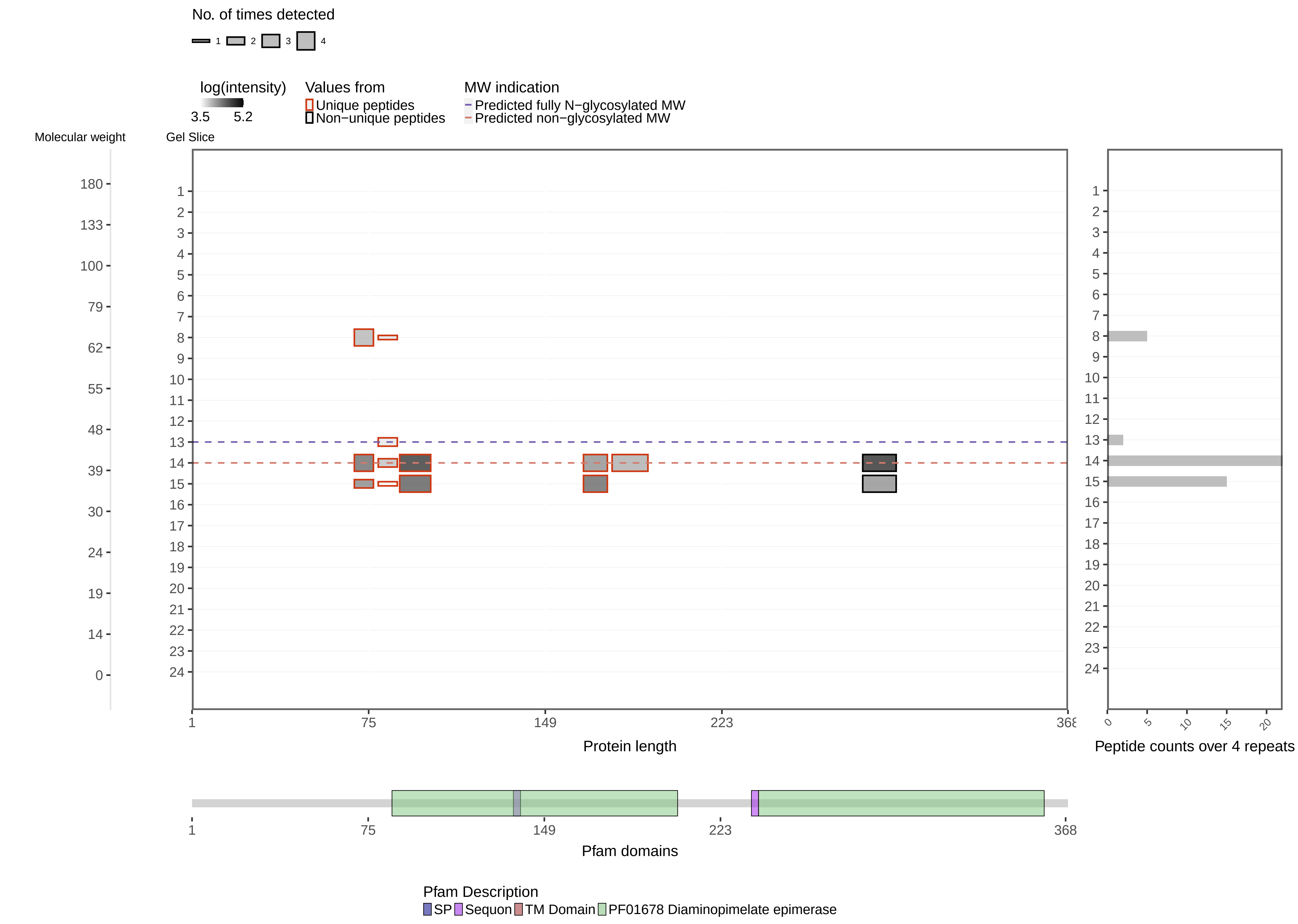


Protein sequence:

GKRSFTIHTGAGLIVPEIREDGKVRVDMGEPILKASDVPTKLPPNRDQSVVKARLDVDGSAWNVTCVSMGNPHCVTFGTEQSQDLQVDELNLADIGPKFEHHVMFPARTNTEFVQVFSPTHLKMRVWERGAGATLACGTGACAVVVAAVLEDRAARRCTVDLPGGPLDIEWSEKDNHIYMTGPAEVVFYGSVPL*

NbL19g17900.1 probable_23−bisphosphoglycerate−independent_phosphoglycerate_mutase [MW = 52.92 kDa]

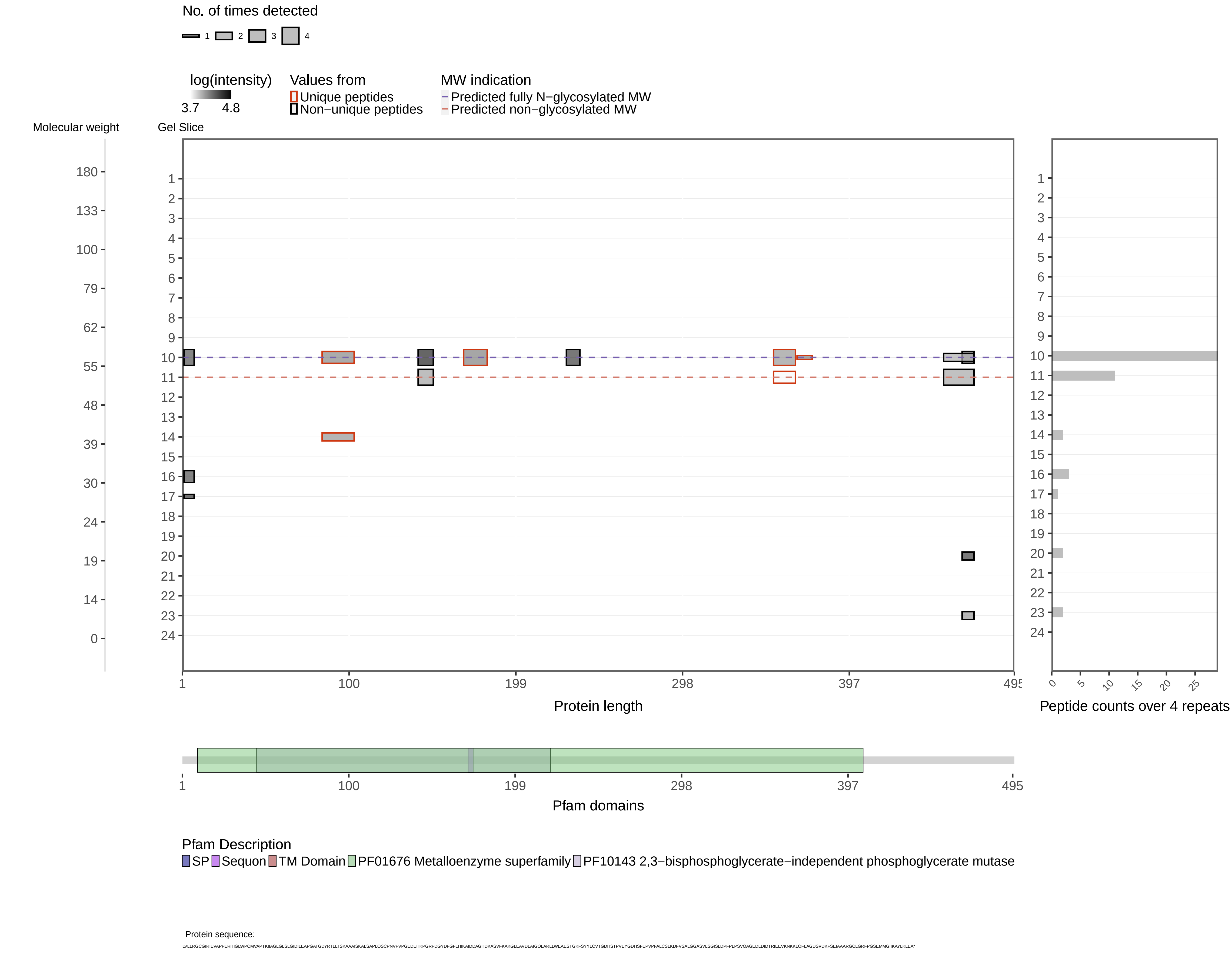

NbL19g17890.1 soluble_inorganic_pyrophosphatase_4 [MW = 24.46 kDa]

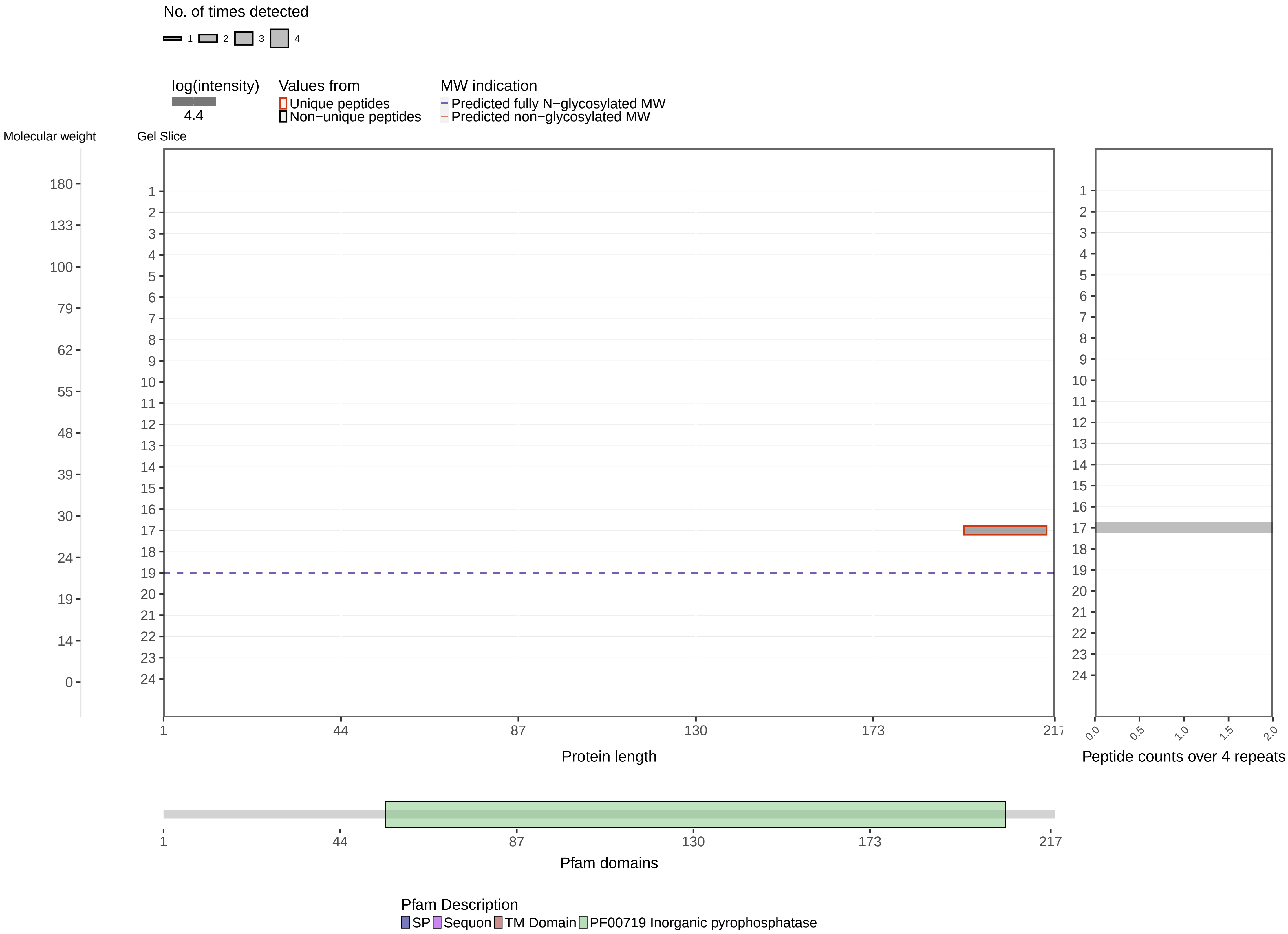


Protein sequence:

IPRTLCEDSDPLDVLVIMQEPVLPGCFLRAKAIGLMPMIDQGEKDDKIIAVCADDPEYKEYTDIKELPPHRLAEIRRFFEDYKKNENKEVAVNDFLPATKAFQAVQHSQDLYADYIVESLRR*

### NbL19g17870.1 histone_H2A−like [MW = 15.31 kDa]

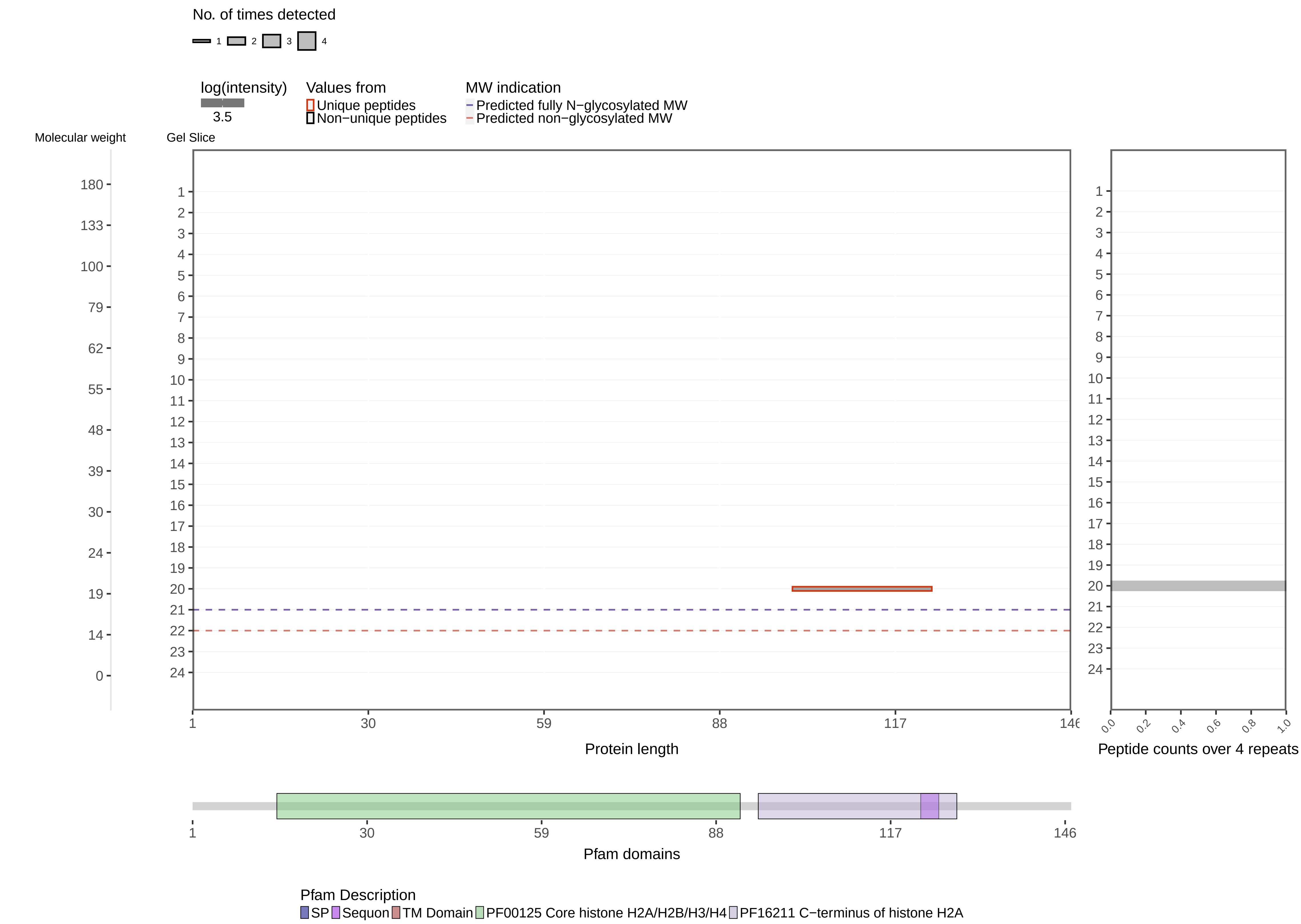


Protein sequence:

/LEYLAAEVLELAGNAARDNKKTRIIPRHVLLAVRNDEELGKLLAGVTIAHGGVLPNINPVLLPNKSDKAGKEPTKSPTKATKSPKKA*

### NbL19g17810.1 actin−depolymerizing_factor_1 [MW = 16 kDa]

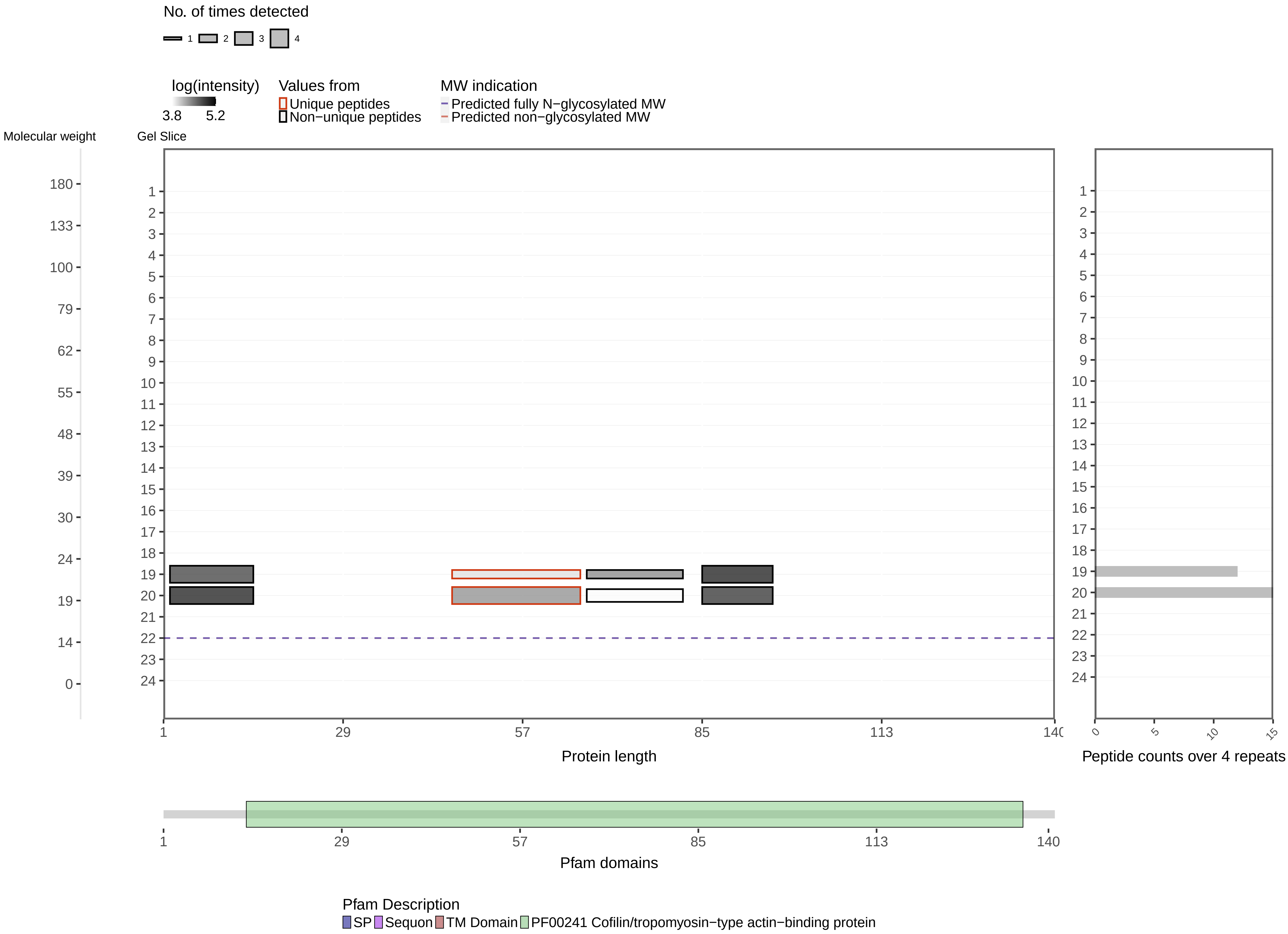


Protein sequence:

AASLPDNECRYAVYDFDFVTAENCQKSRIFFIAWSPDTSRVRSKMIYASSKDRFKRELDGIQVELQATDPTEMGLDVIKSRAN*

NbL19g17550.1 heat_shock_cognate_70_kDa_protein_2−like [MW = 71.13 kDa]

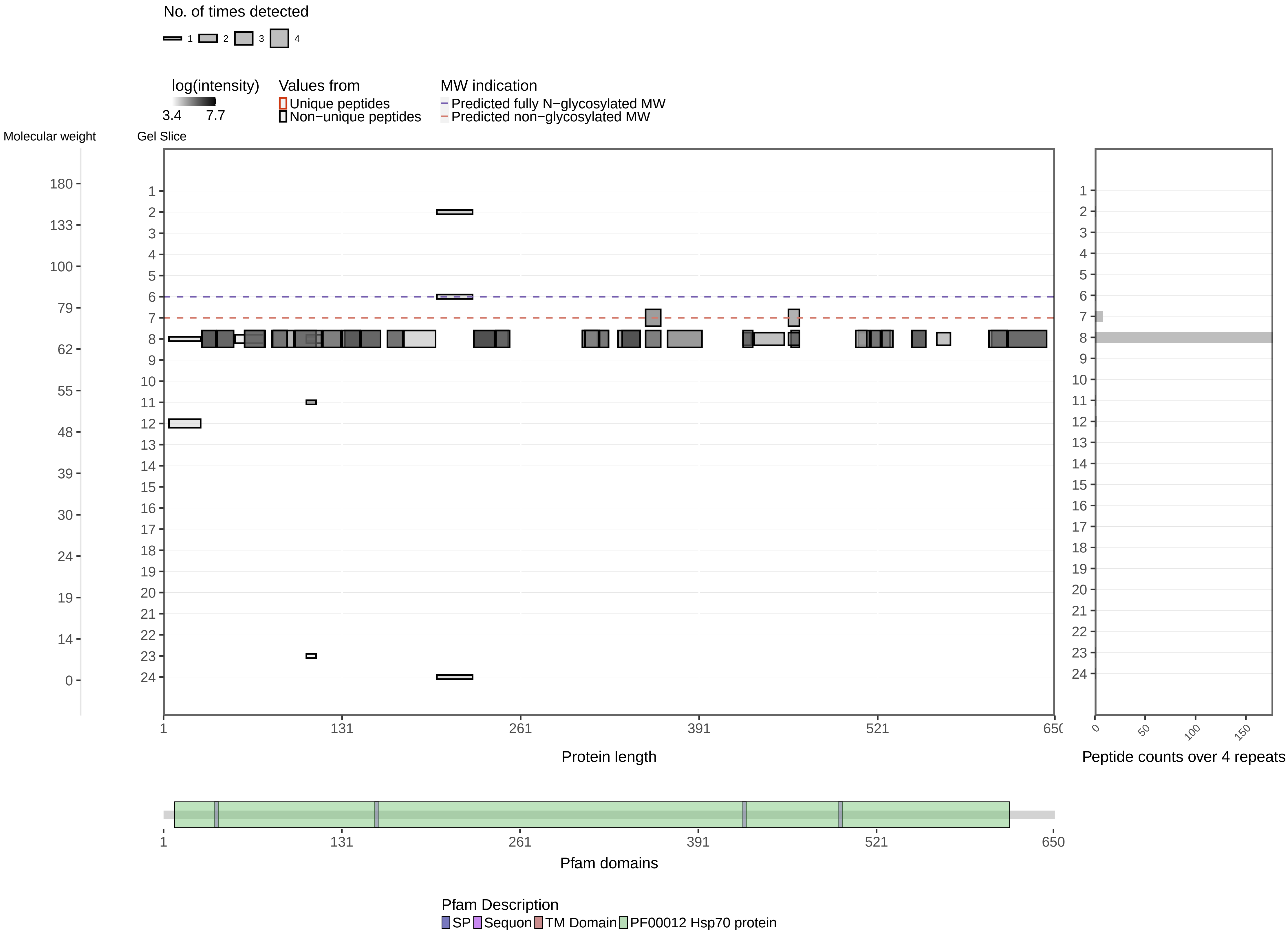


Protein sequence:

LFRKCMEPVEKCLRDAKMDKSTIHDVVLVGGSTRIPKVQQLLQDFFNGKELCKSINPDEAVAYGAAVQAAILSGEGNEKVQDLLLLDVTPLSLGLETAGGVMTVLIPRNTTIPTKKEQVFSTYSDNQPGVLIQVYEGERTRTRDNNLLGKFELSGIPPAPRGVPQITVCFDIDANGILNVSAEDKTTGQKNKITITNDKGRLSKEEIEKMVQEAEKYKSEDEEHKKKVEAKNALENYAYNMRNTVKDEKIASKLPEADKKKIEDAIESAIQWLDANQLAESDEFEDKMKELESICN

NbL19g17240.1 ribulose_bisphosphate_carboxylase/oxygenase_activase_1_chloroplastic [MW = 52.72 kDa]

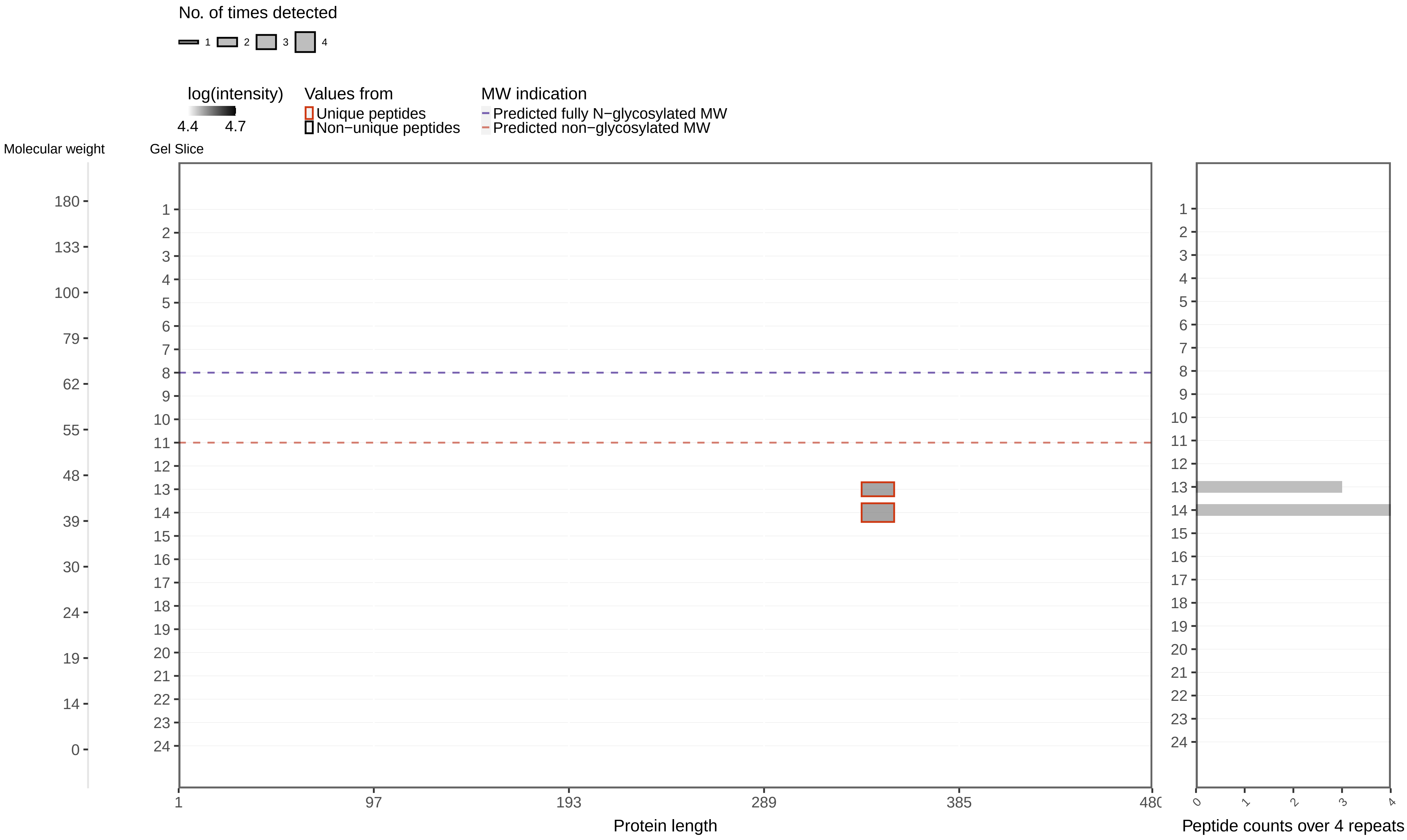


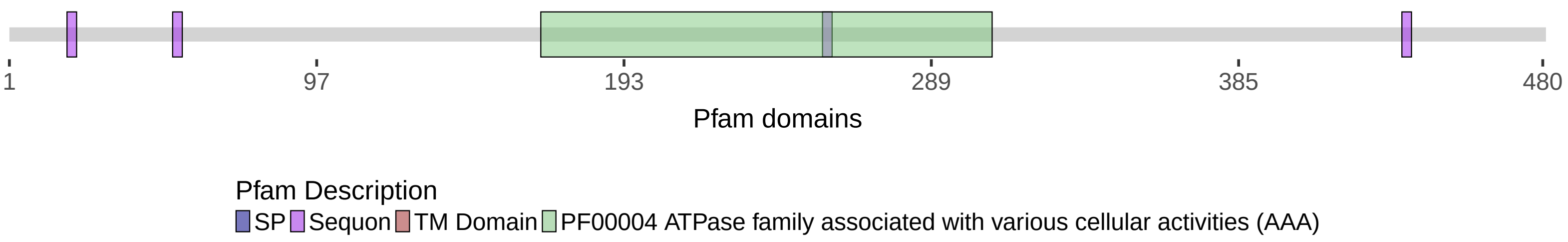


Protein sequence:

CLFINDLDAGAGRMGGTTQYTVNNQMVNATLMNIADNPTNVQLPGMYNKQENARVPIIVTGNDFSTLYAPLIRDGRMEKFYWAPTREDRIGVCTGIFRTDNVPAEDVVKIVDNFPGQSIDFFGALRARVYDDEVRKWVSGTGIEKIGDKLLNSFDGPPTFEQPKMTIEKLLEYGNMLVQEQENVKRVQLADKYLKEAALGDANADAINNGSFFTRKEAQHVDVPVGEGSTDSIATKLDSNSMAEDGSYLFKF*

### NbL19g17140.1 aldo−keto_reductase_family_4_member_C10−like [MW = 35.23 kDa]

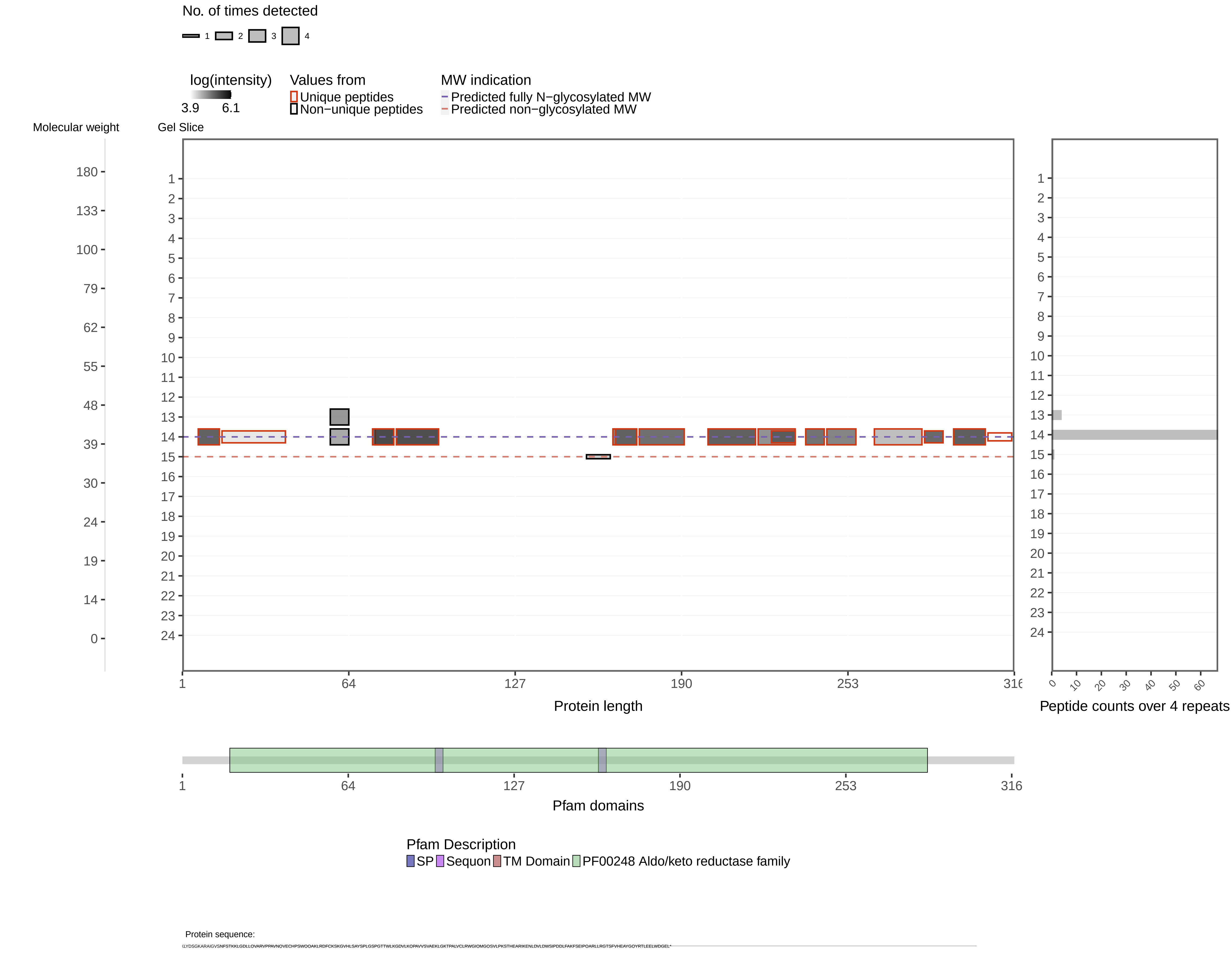

### NbL19g17100.1 uracil_phosphoribosyltransferase [MW = 36.74 kDa]

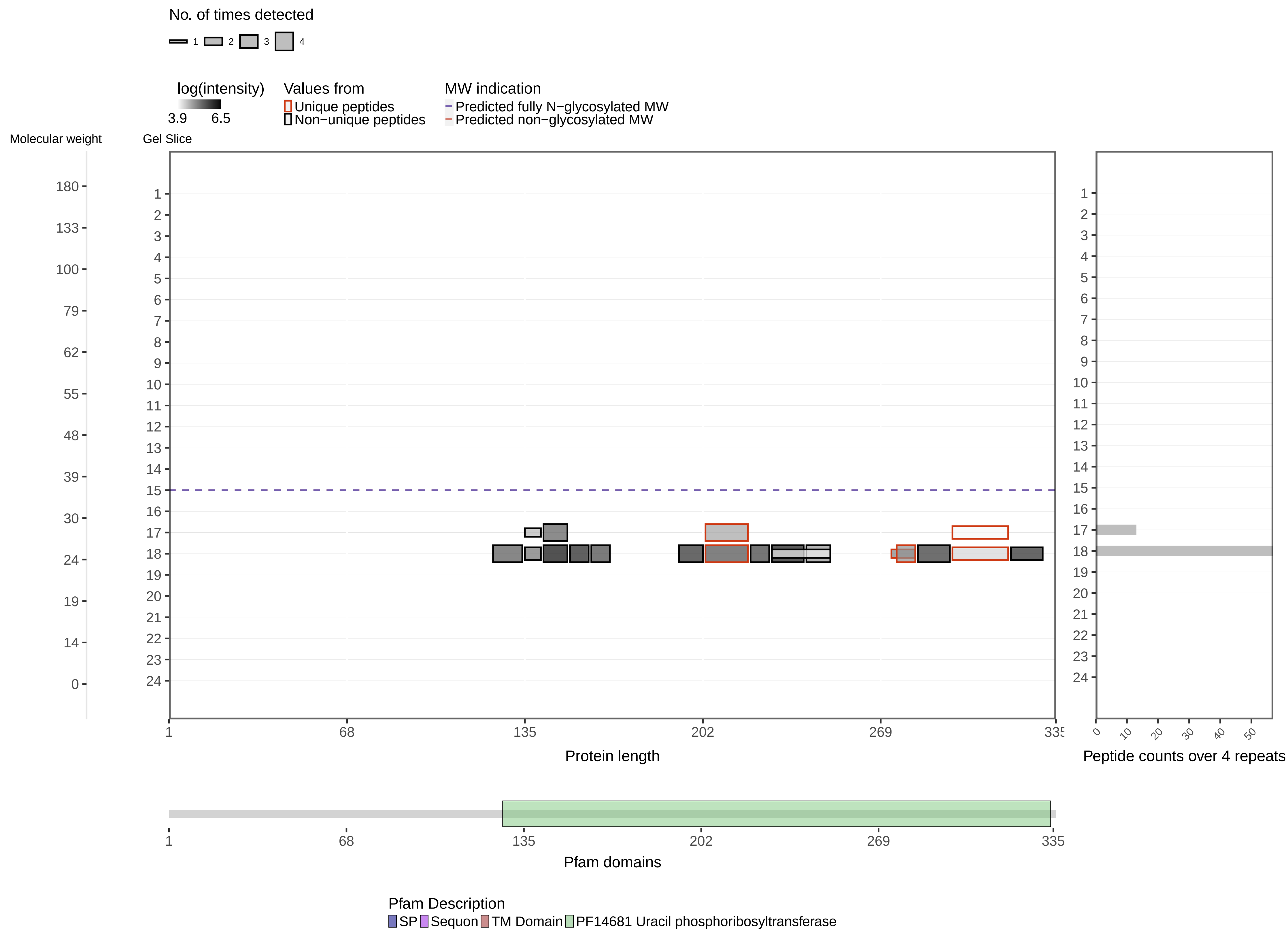


Protein sequence:

[illegible]MSELGRLLMYEASRDWLPIITGEIQSPMGVASVEFVDPREPVAIVPILRAGLALAEHASSILPATKTYHLGISRNEETLQPSVYLNKLPDKFPEGSRVFVVDPMLATGGTIVAAIDLIKERGVDNSQIKVICAVGAPPALQKLSENFPGLHVYAGILDPTVNDKGFIIPGLGDAGDRSFGT*

NbL19g17000.1 NHP2−like_protein_1 [MW = 13.91 kDa]

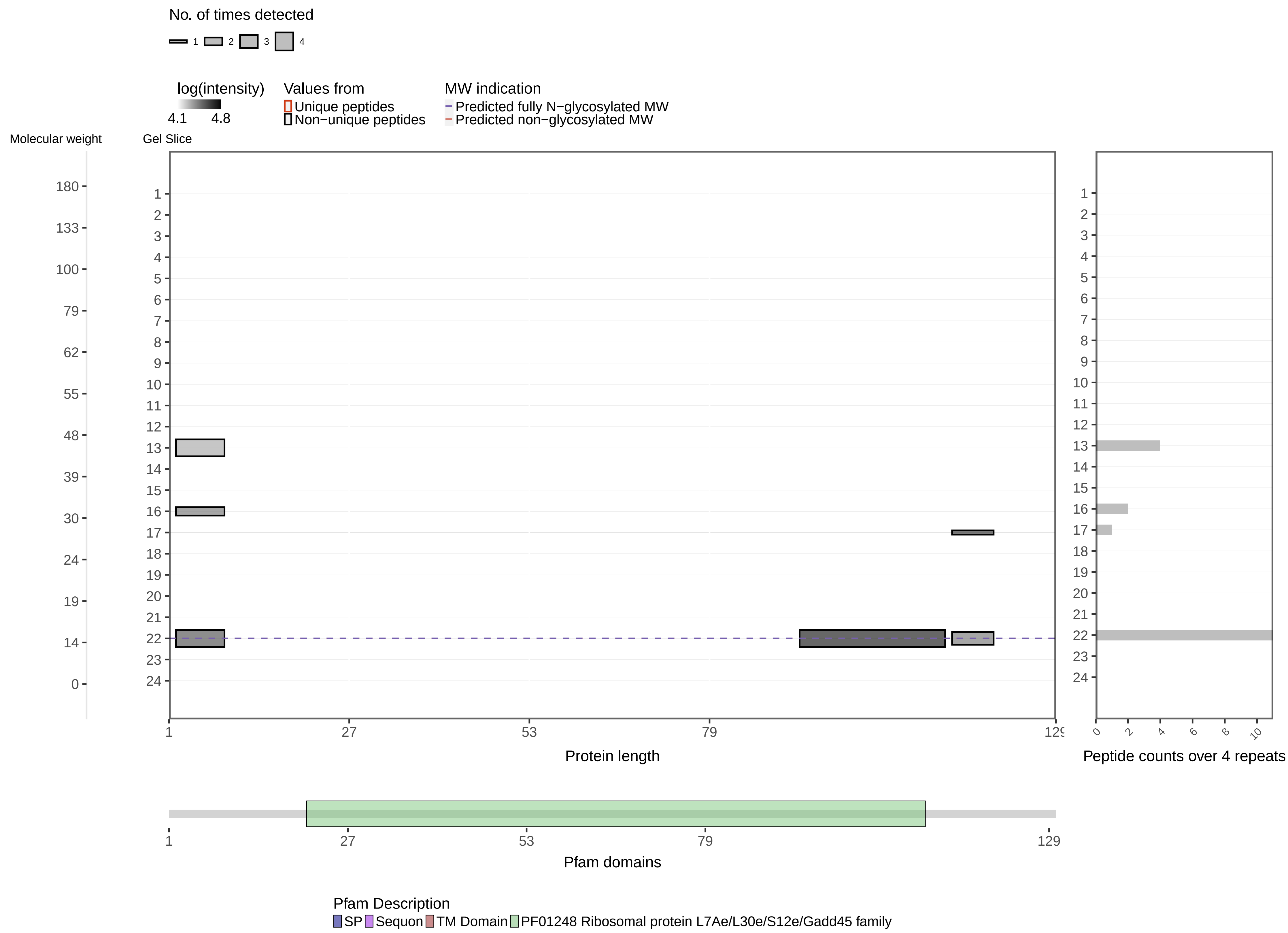


Protein sequence:

SEFVVMAADTEPLEILLHLPLLAEDKNVPYVFVPSKQALGRACGVTRPVIACSVTSNEGSQLKSQIQQLKDAIEKLLI*

### NbL19g16990.1 probable_proteasome_inhibitor [MW = 31.83 kDa]

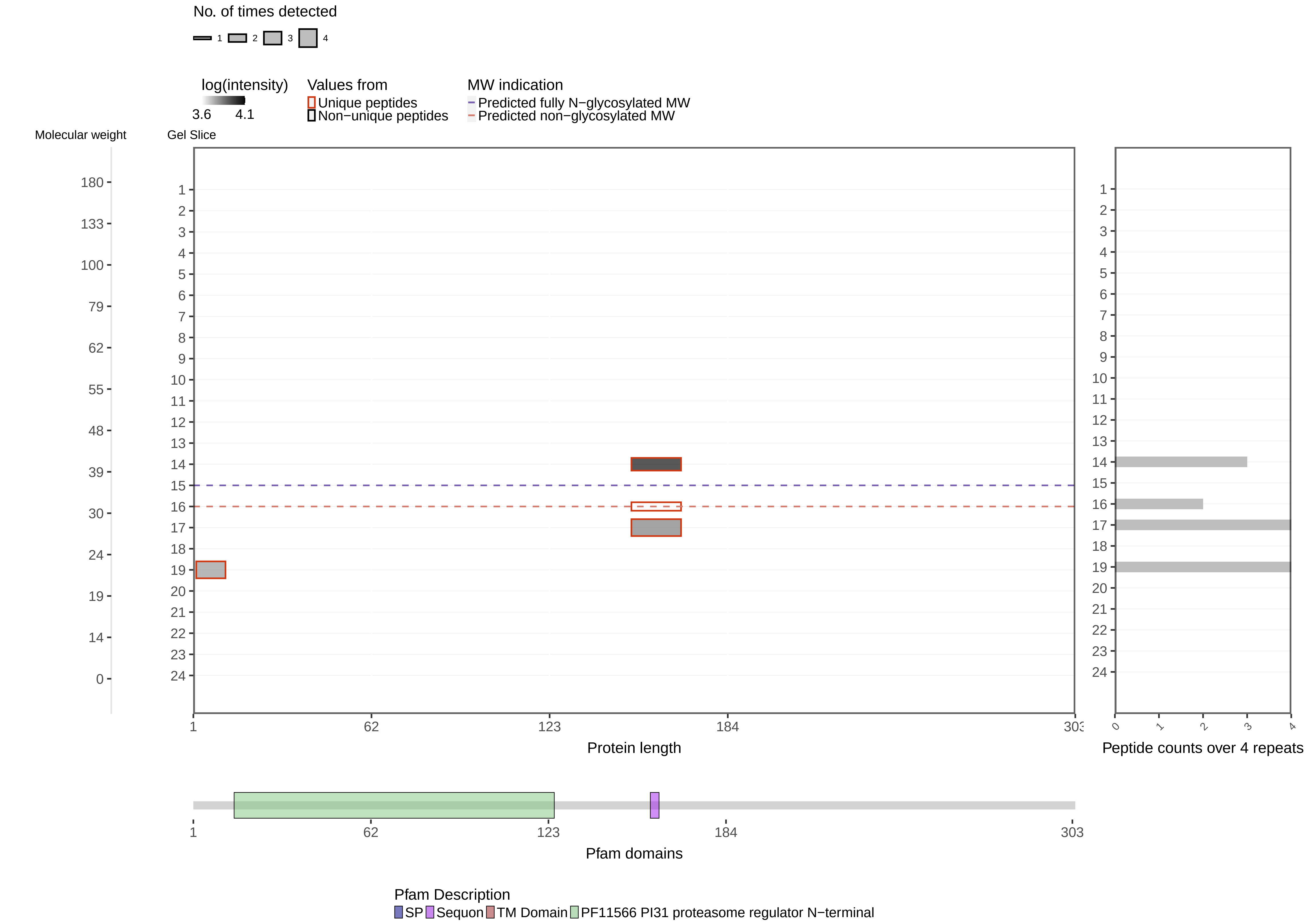


Protein sequence:

ELNKEVLSKYSGSSVANSSTQTSSSGKGGLGDEPSAERRSDYPYQPESSFPSGYVVPPIPGSGGGDLFPGPGAGVYPTRGGSGGGSMLVGPNDPRFFGGMGGDPRLPGGLPGVPPGARFDPYGPPDVPGFEPGRFIRDPRRPGGGRPHPDLPHFGGDSDFI*

### NbL19g16950.1 probable_glutathione_S−transferase [MW = 25.47 kDa]

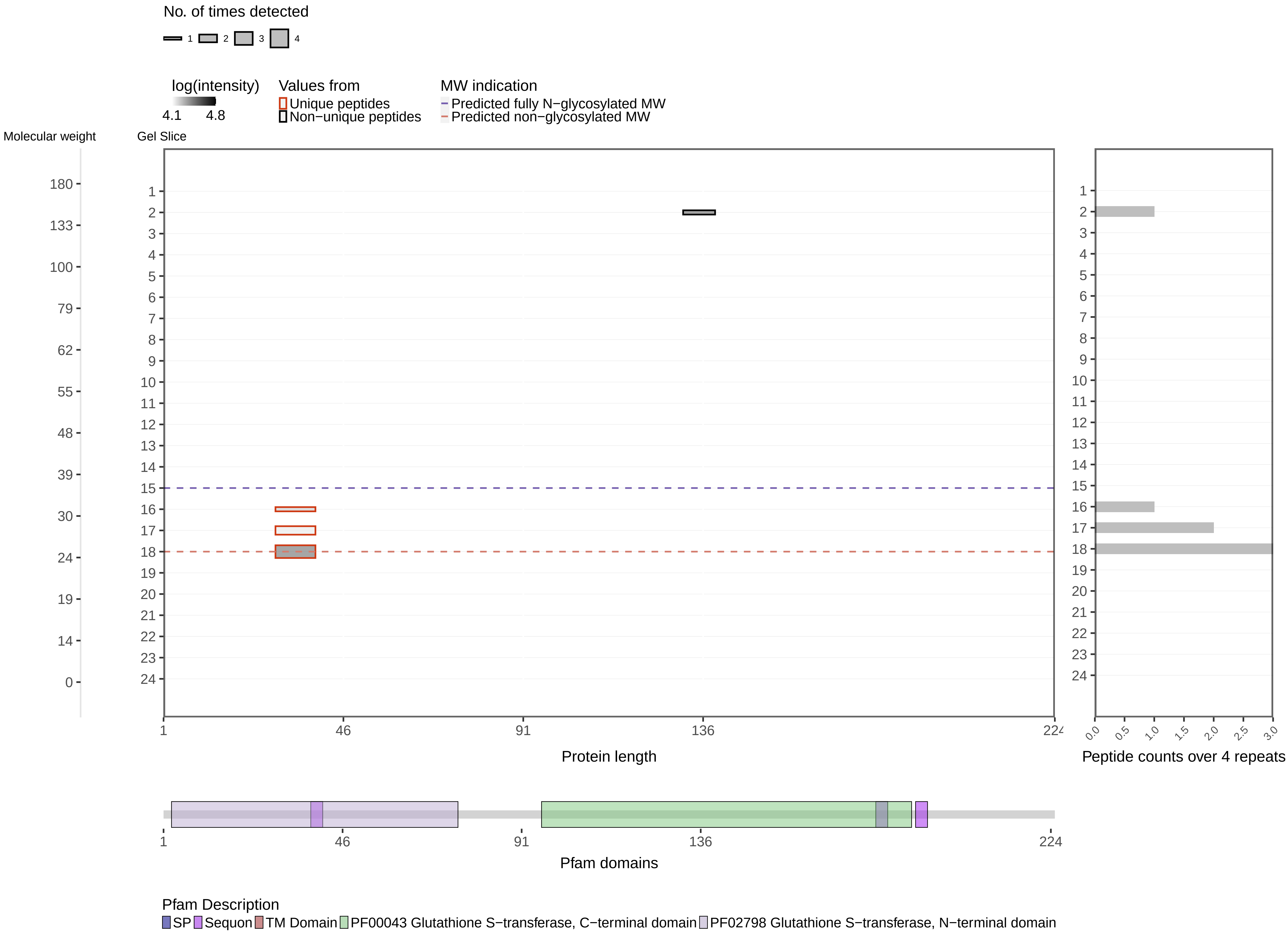


Protein sequence:

.DDKVTVIVNTFLSKGEEQEKAKEQVWEMLKVLDNELKDKKFLVGDKFGFADIAANLVAFWLGVFEEASGIVLMTSEKFPNFSKWRDEYTNCSQLKESLPPRDELLAFFRARSQAATTSASAPK*

NbL19g16910.1 probable_glutathione_S−transferase [MW = 25.04 kDa]

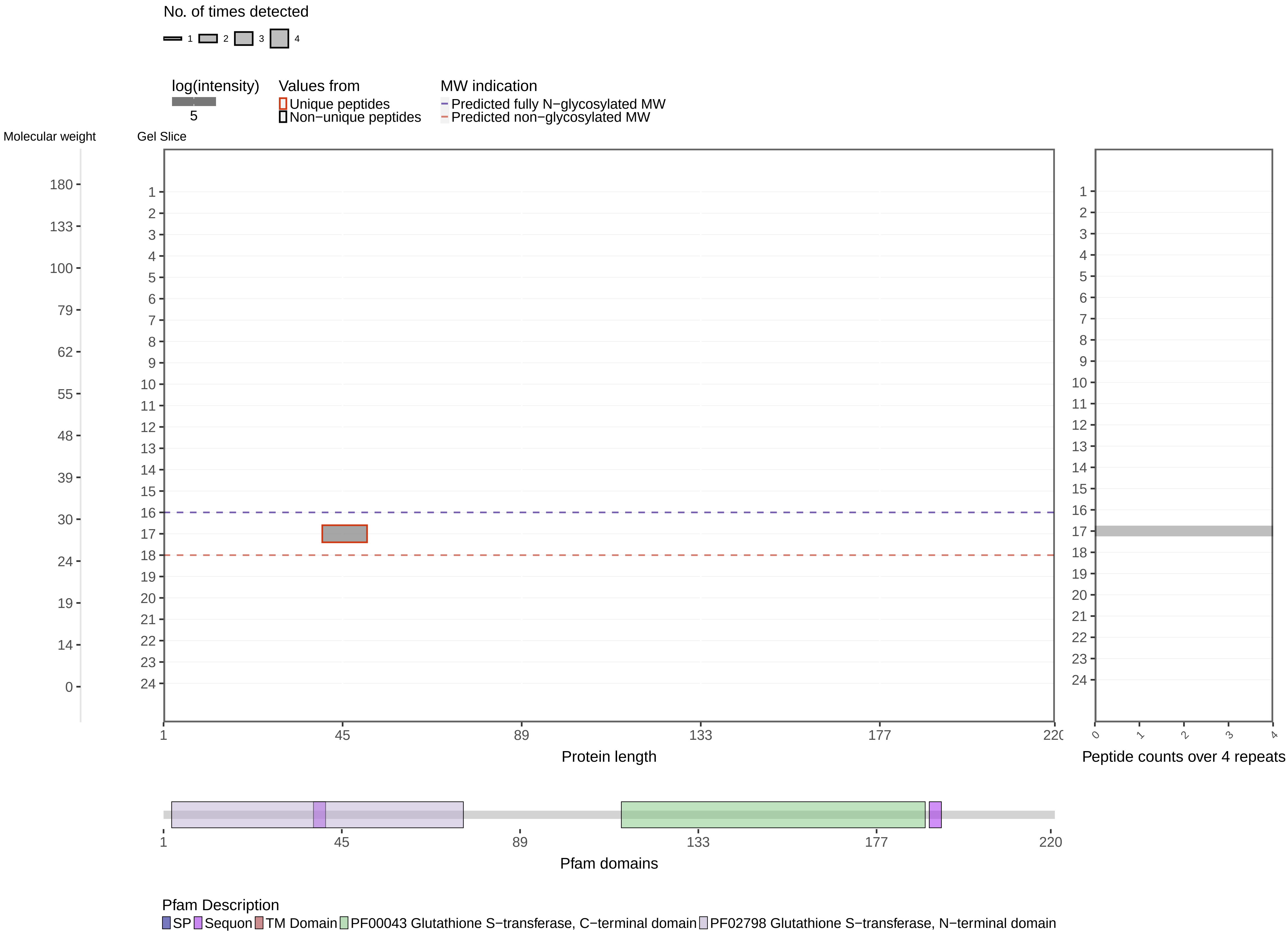


Protein sequence:

FEEKGASVGRSFFLKGEEQEKAKEQVCEMLKVLDNEVKDKKFFVGDKFGFADIAANAAALWLGVLEEASCVVLVTREKFPNFCAWRDEYINCSQNKEYLPPRDELLALFKARFQAAAAAP*

NbL19g16900.1 probable_glutathione_S−transferase [MW = 25.38 kDa]

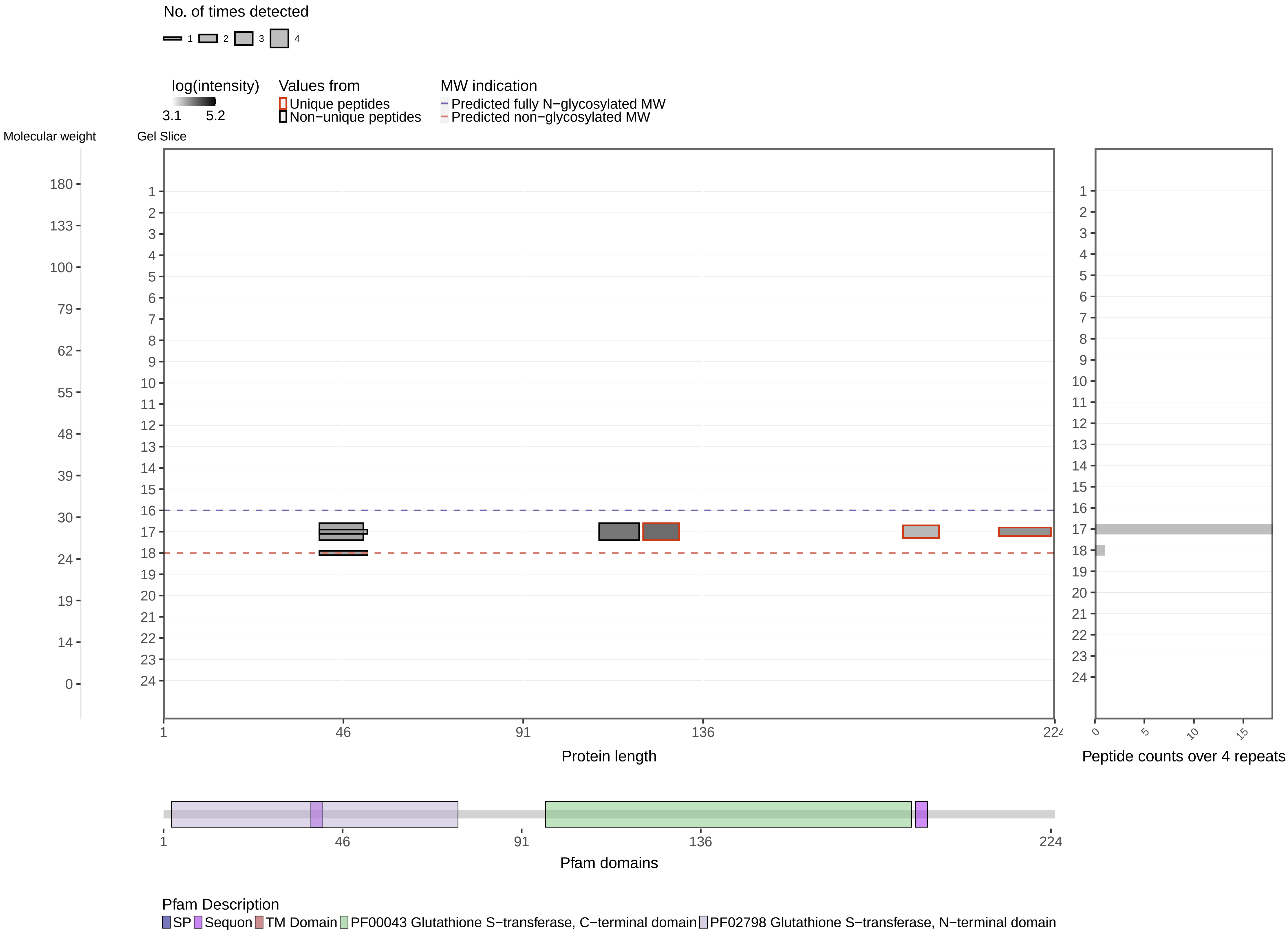


Protein sequence:

LEDKVRAVGKSFFLKGEEQEKSKEDAFEILKILDNELKDKKFFTGDKIGLADIAANFVGLWLGVFEEASGIILVTREKFPNFCAWRNEYINCSQNKEYLPPRDELVAHFKAHFQAAAAAAAAPK*

### NbL19g16890.1 probable_glutathione_S−transferase [MW = 24.92 kDa]

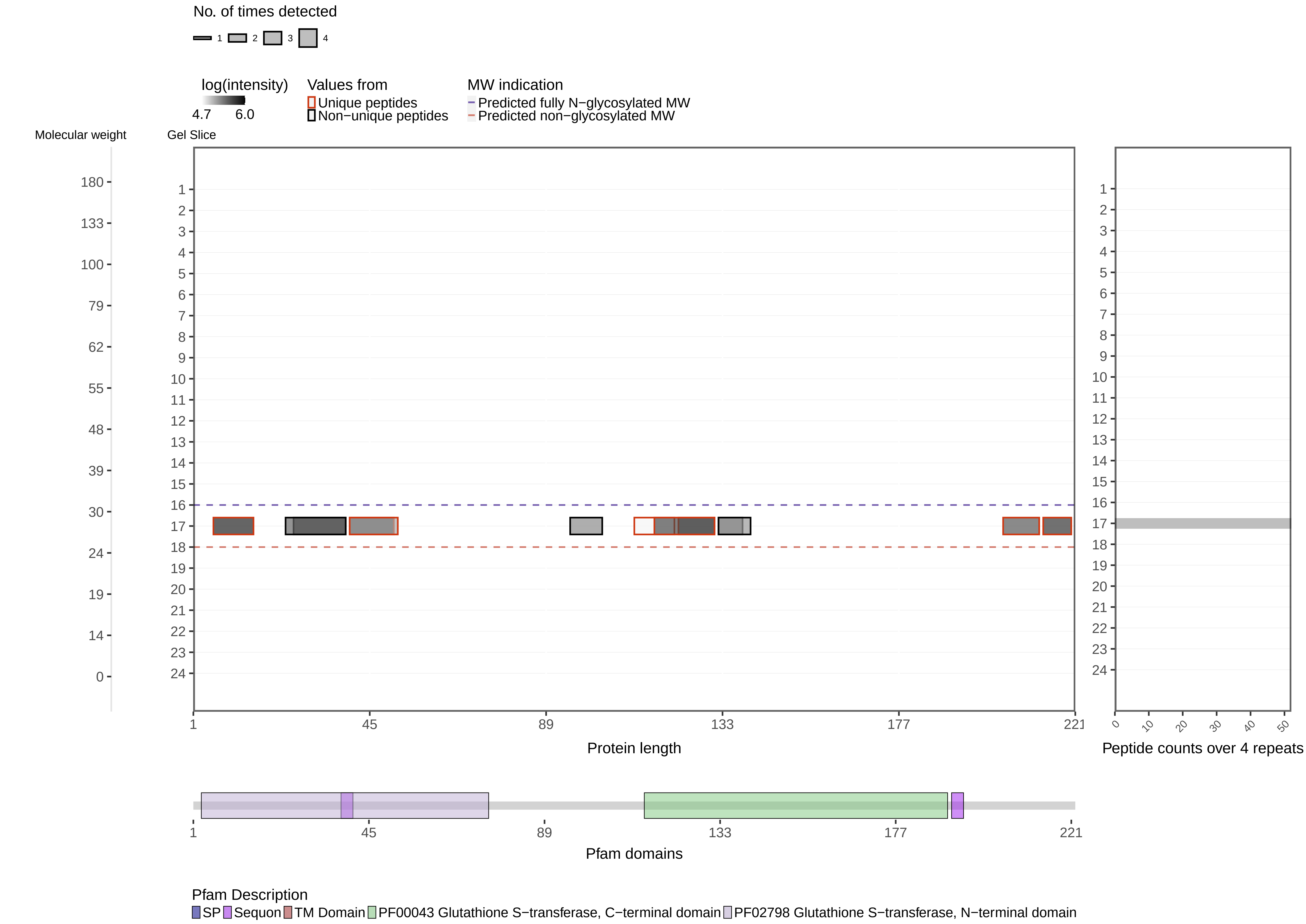


Protein sequence:

NFLDVKCITTIGKALFGKGEESDKAIEECGELLKILDNELKDKKFFVGDNFGYADIAANLMAFWLGILEEASGVILVTNEKFPNFCAWKDEYINCSHVKEHLPSRDALLSHFQPRFQTAAAPK*

NbL19g16860.1 universal_stress_protein_PHOS32−like [MW = 18.45 kDa]

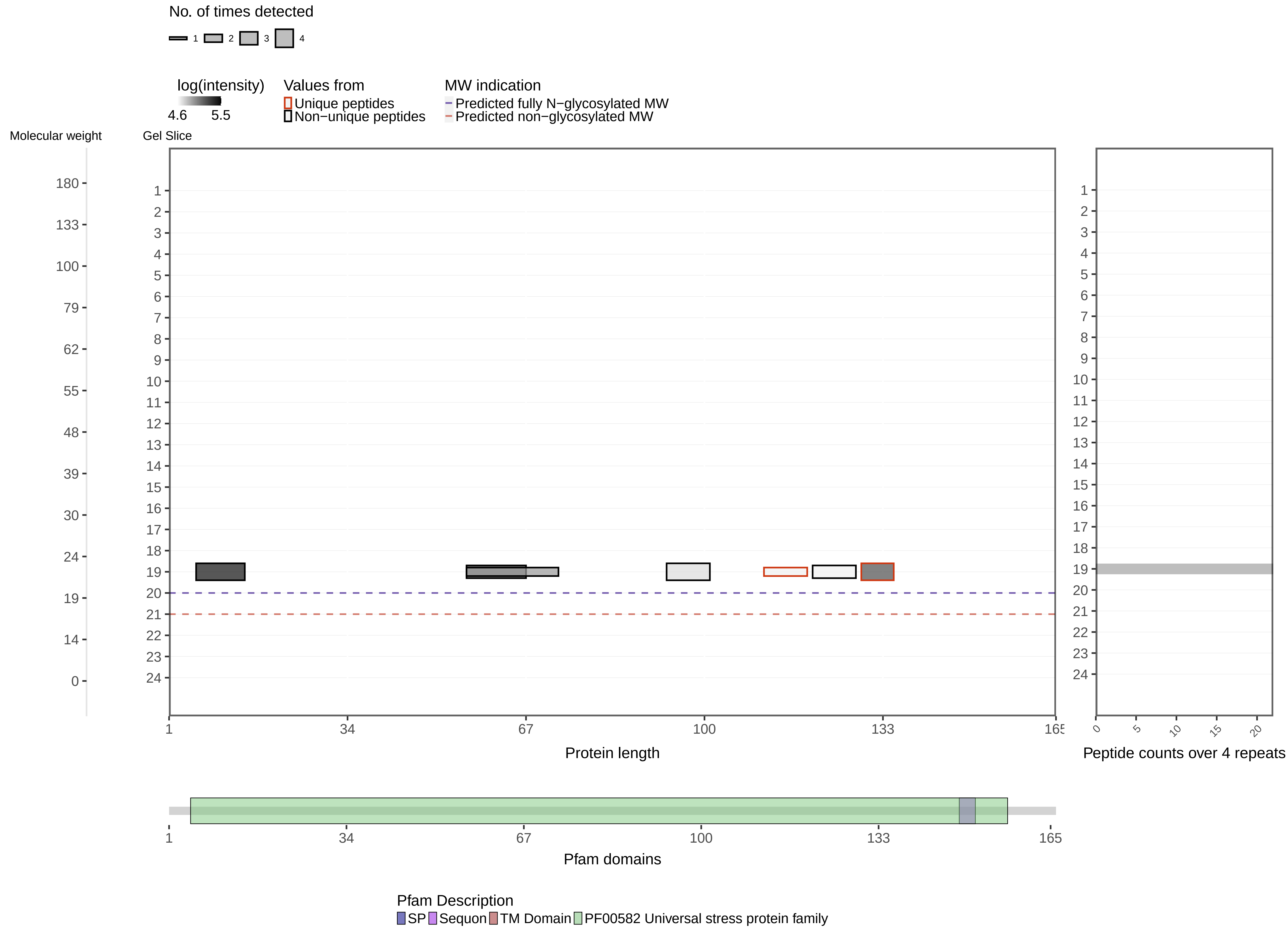


Protein sequence:

EVMKKYDVETDIDVLDLLDTATRQKEITVVTKLYWGDAREKLIDATEDLKLDSLVMGSRGLGTLQRIIMGSVTNYVLTNATCAVTVVKDPDFHKH*

NbL19g16750.1 fructose−16−bisphosphatase_chloroplastic−like [MW = 44.29 kDa]

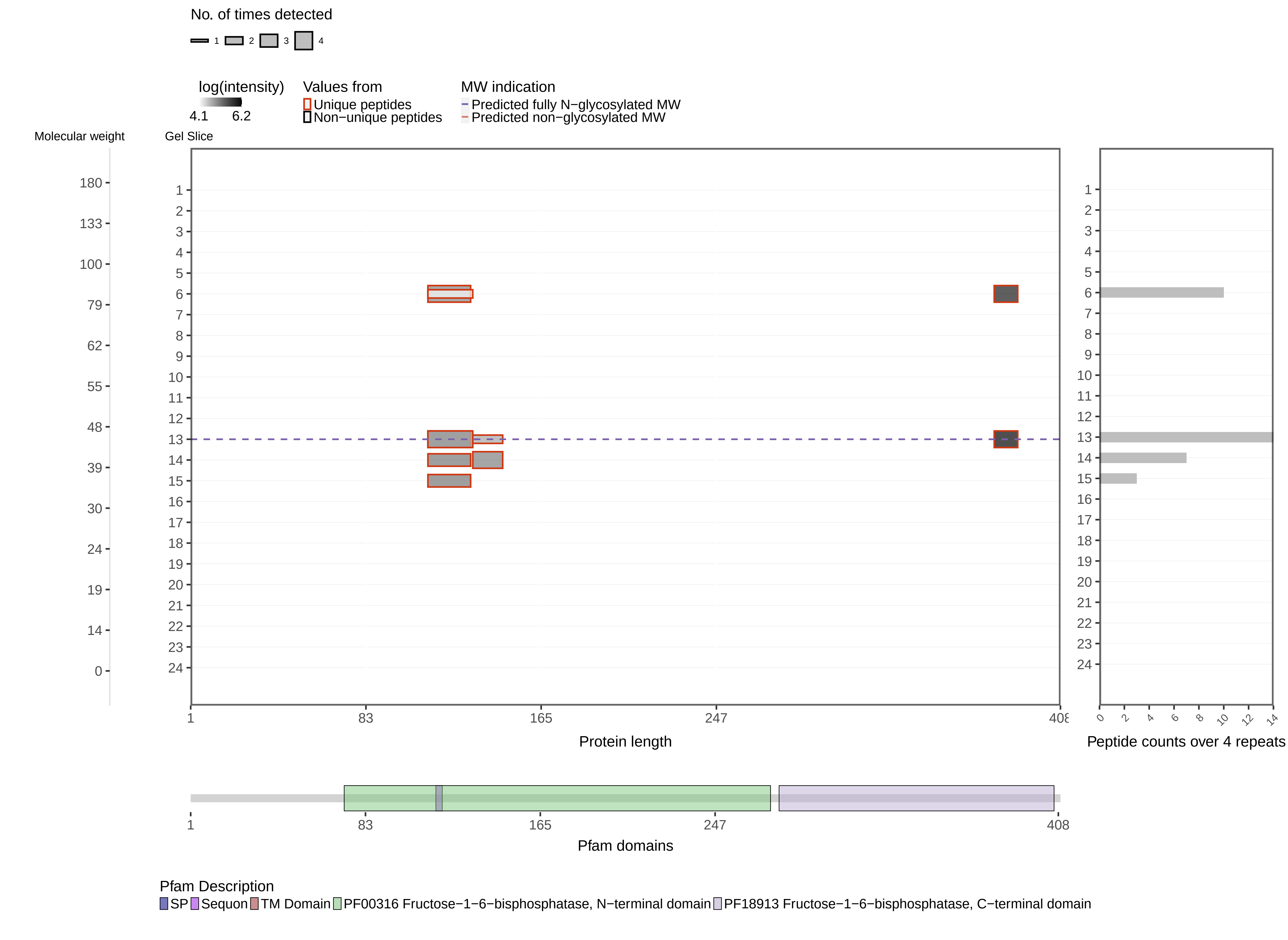


Protein sequence:

TGSIFGIYSPNDECLADHGDDSTLDNVEQRCIVNVCQPGSNLLAAGYCMYSSSVIFVVTLGNGVFAFNLDPMYGEFVMTQENIQIPKSGKIYSFNEGNYQLWDDKLKKYIDDLKDPGPSGKPYSARYIGSLVGDFHRTLLYGGIYGYPRDKKSKNGKLRLLYECAPMSFLVEQAGGKGSDGHQRILDIEPTEIHQRVPLYIGSTEEVEKLEKYLS*

NbL19g16550.1 proteasome_subunit_alpha_type−4−like [MW = 27.31 kDa]

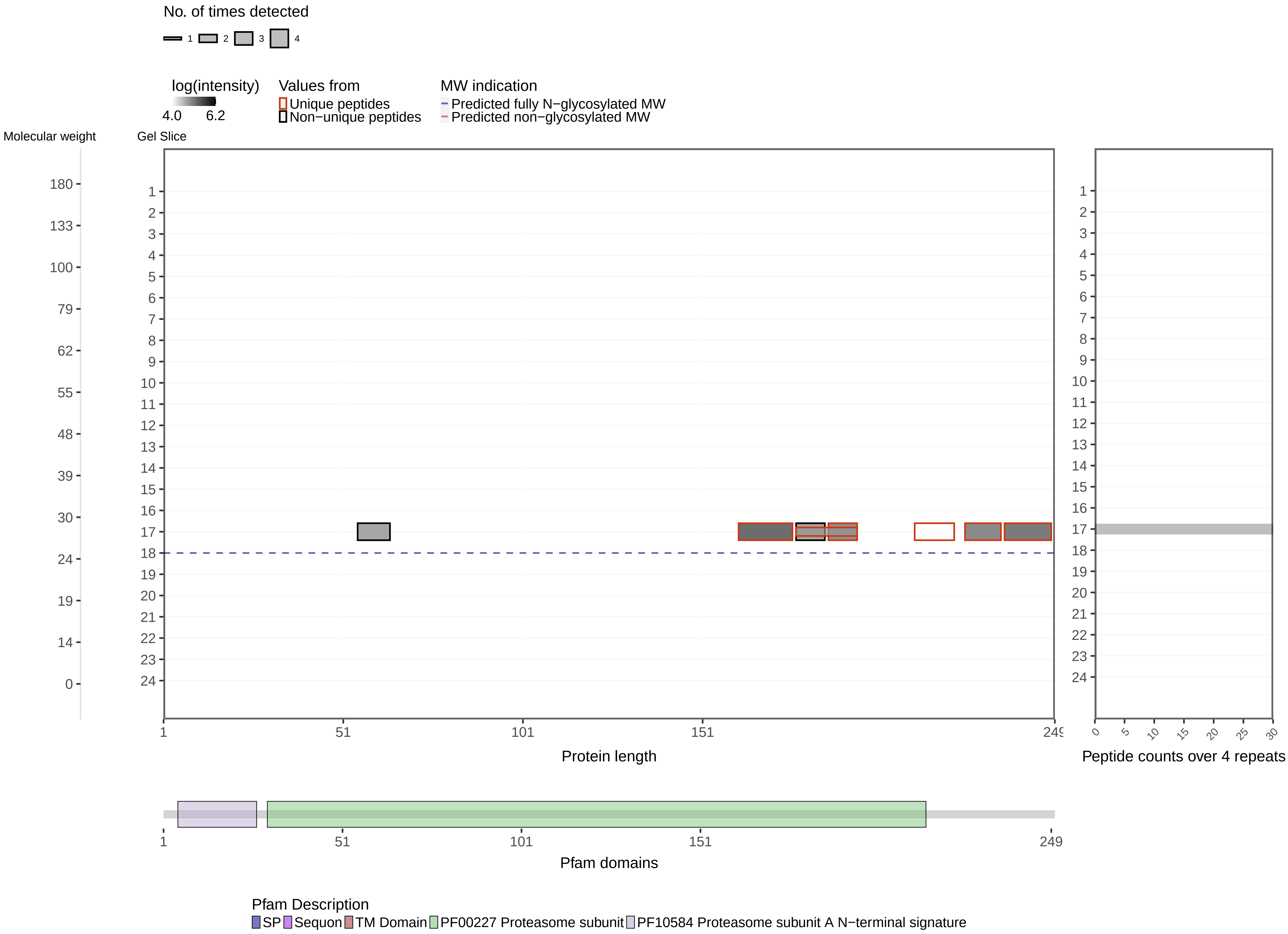


Protein sequence:

LCDTKQGYTQFGGLRPFGVSFLFAGWDKNYGFQLYMSDPSGNYGGWKAAATGANNQAAQSMLKQDYKDEITREEAVHLVLKVLSKTMDSTSLTSEKLELAEVFLVNGKVKYQVHSPESLNKLLVQYGLTQPAADA*

### NbL19g16410.1 uncharacterized_acetyltransferase_At3g50280 [MW = 53.21 kDa]

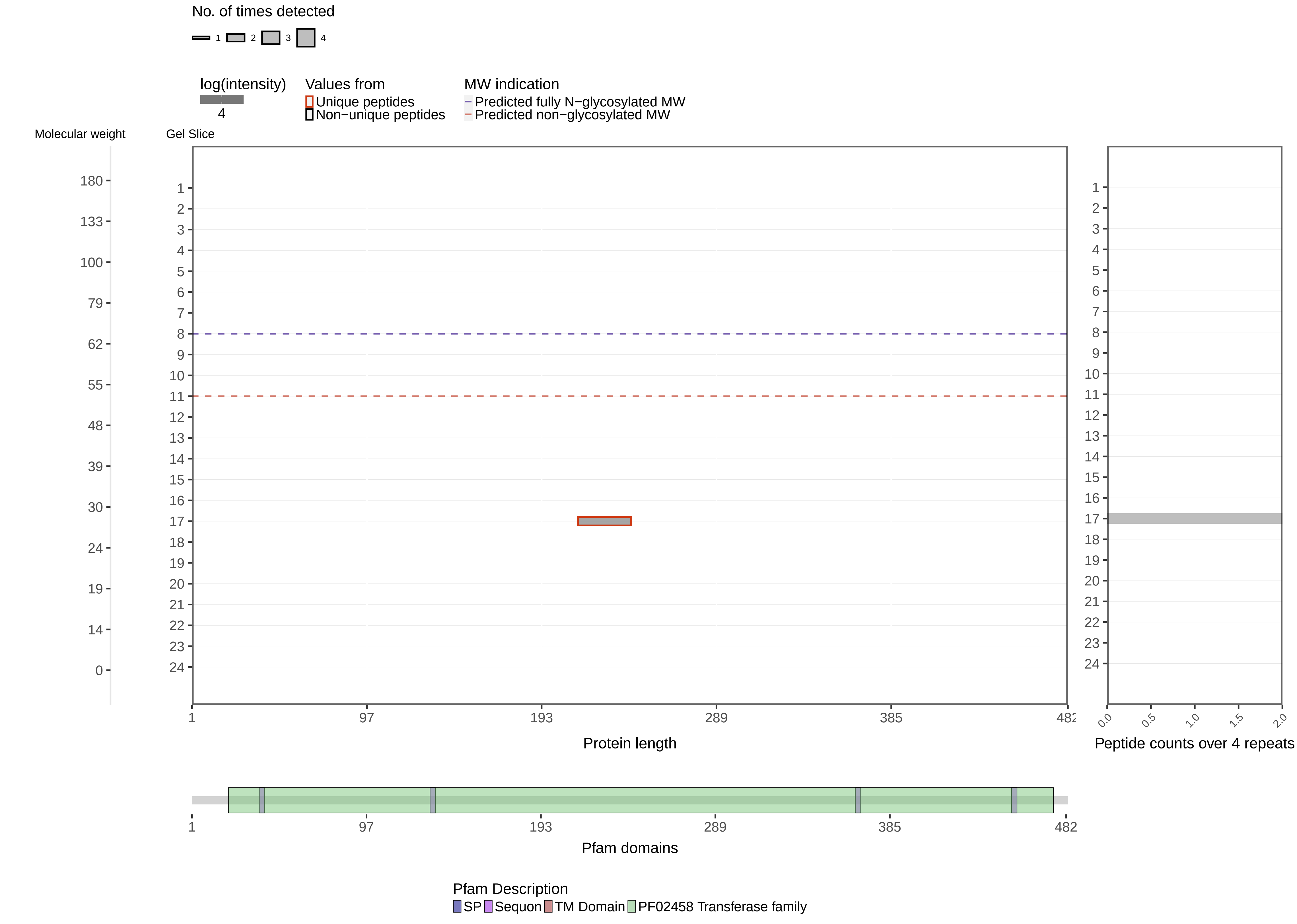


Protein sequence:

FSGDEPIREKIFHFSREAILKLKFRANNKMAVNGQNGKTNGKITPVNNEPALQNGAAVKYTASEISSFQSLSAQLWRSVTRARKLDGSKTTTFRMAVNCRHRLEPKMEPLYFGNAIQSIPTVASVNELLTNDLNWSANLLHQNVVAHDDATVRRGVKDWESNPRLFPLGNFDGAMITMGSSPRFPMYDNDFGWGRPVAVRSGMANKFDGKISAFPGREGNGTVDLEVVLAPETMAGLEEDMEFVQYVSY*

### NbL19g16340.1 uncharacterized_LOC104085387 [MW = 46.63 kDa]

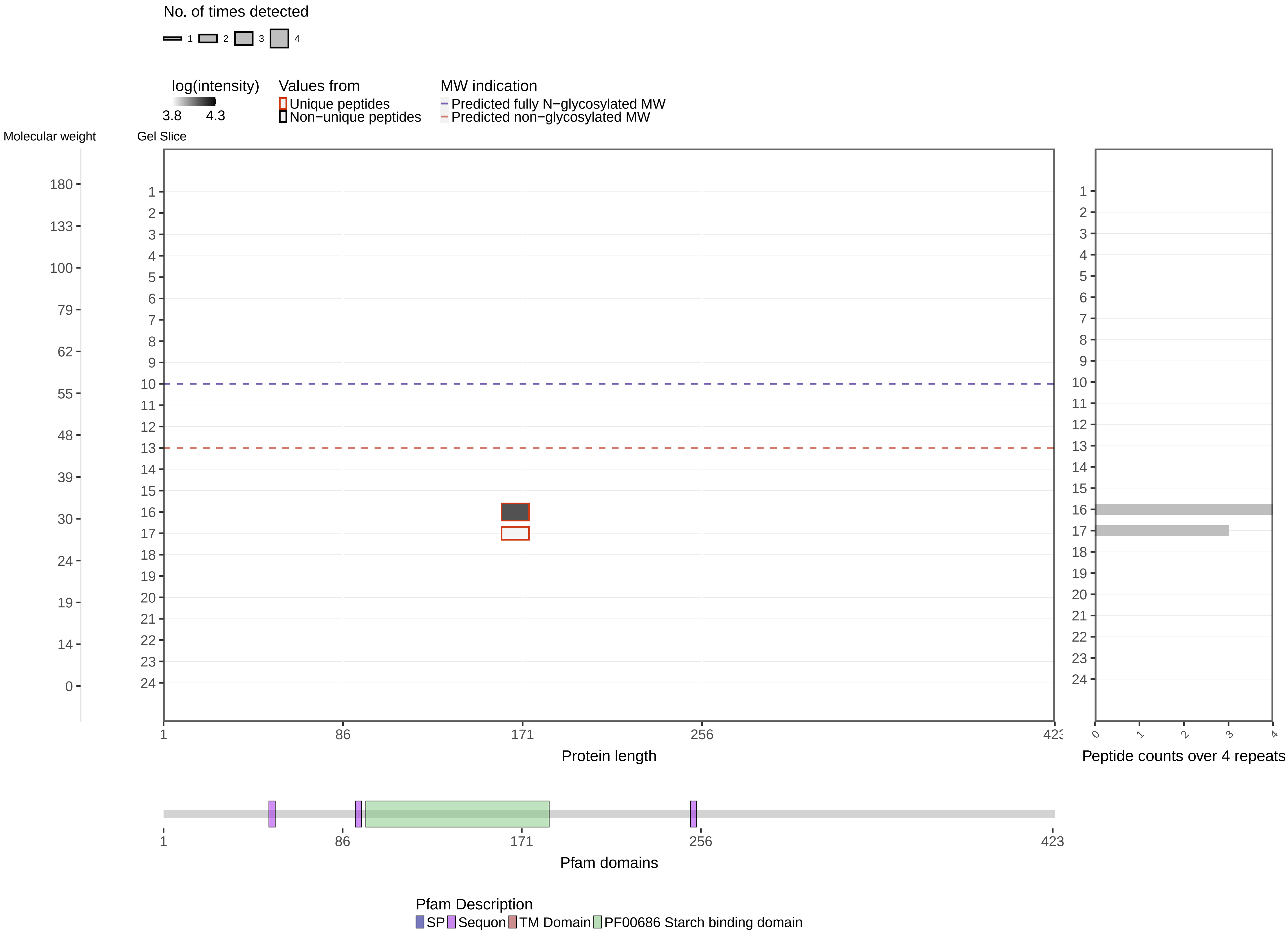


Protein sequence:

[illegible]TIVEEEPAAEQSKESAANPEILIAENLVPPTVVDLEDDVSEGKSNEVLAMVAENITEVNENTIIEAKAIGKNGMVFNDEVLSSKEESILVASETVPVLVPSLTQVLDMDKVIAETLLLDEVQMNKVVAEGSLGLKSKEFNVPELSLNEEVATDLTHPRETPEMKLNVKQEVRGNEDIEKSELVEGGDWPNGKPVEEIFDSDMQWGKRTLQKFFANLGFFKVELEG*

### NbL19g16270.1 chlorophyll_a−b_binding_protein_CP29.1_chloroplastic−like [MW = 31.08 kDa]

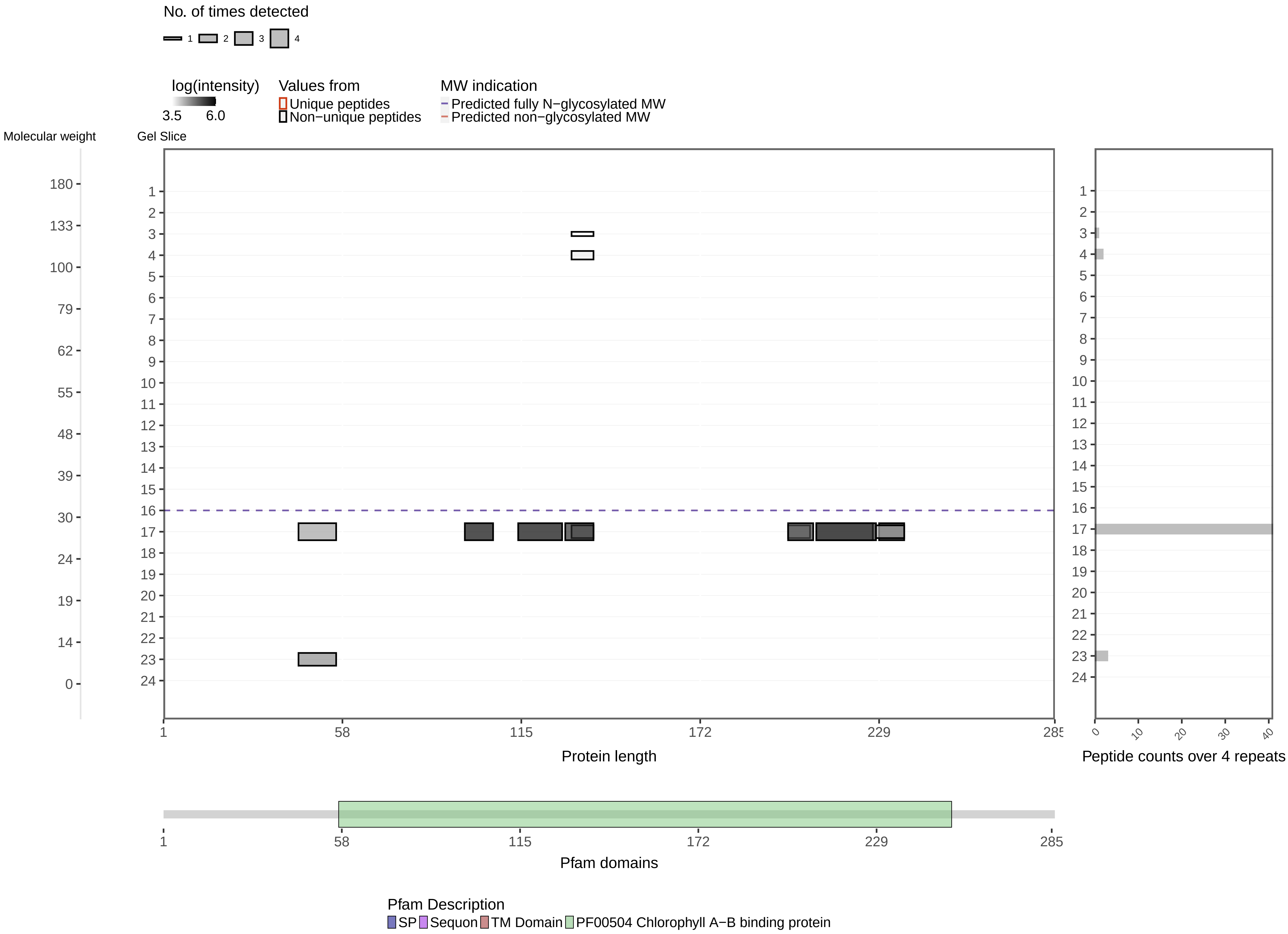


Protein sequence:

RFRECELIHGRWAMLATLGALTVEWLTGITWQDAGKVELVEGSSYLGQPLPFSITTLIWIEVLVIGYIEFQRNAELDPEKRLYPGGSFFDPLGLAADPEKKATLQLAEIKHARLAMVAFLGFAVQAAATGKGPLNNWATHLSDPLHTTIFDTFGFFS*

### NbL19g16130.1 dnaJ_homolog_subfamily_B_member_4−like [MW = 40.35 kDa]

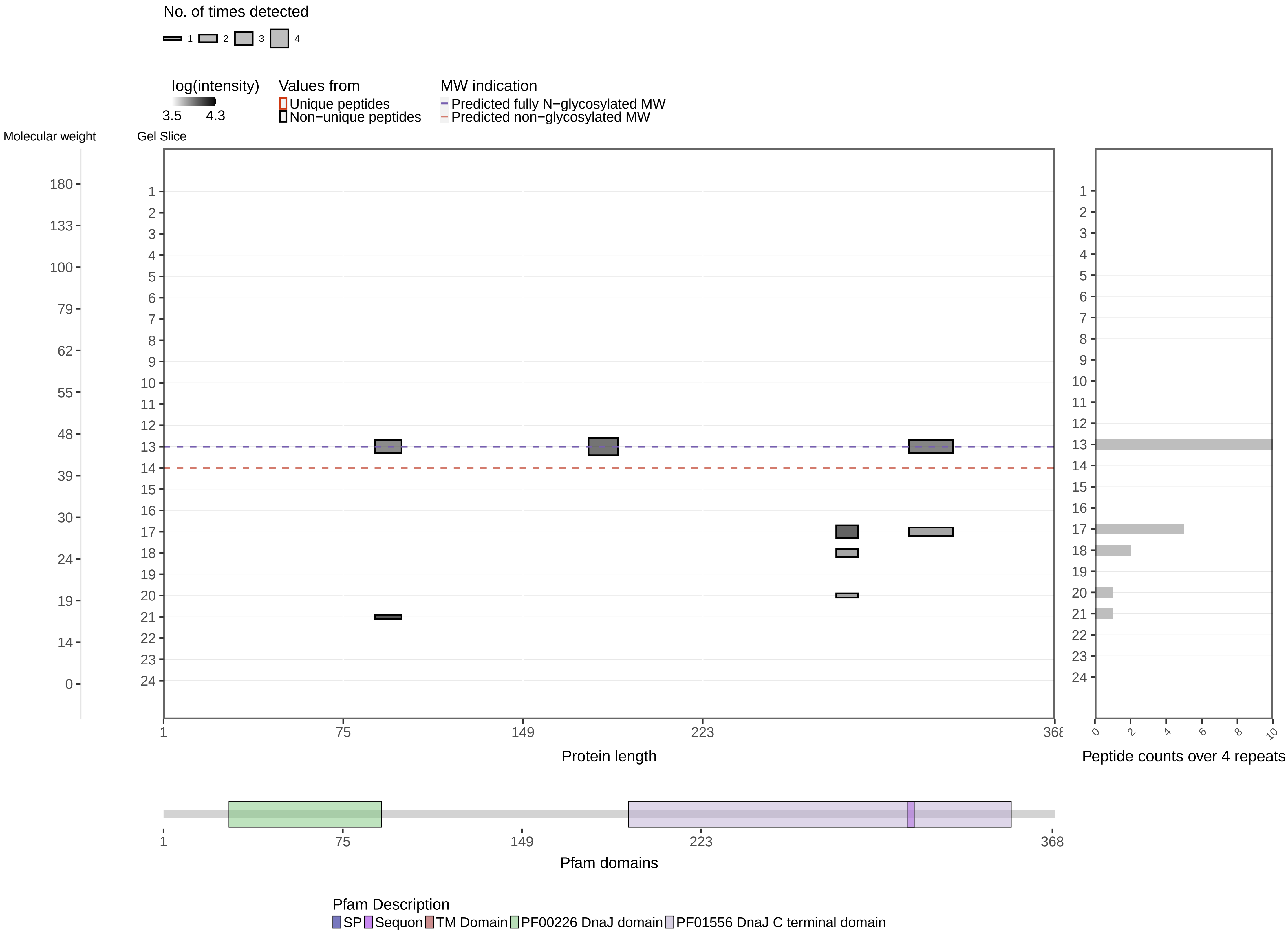


Protein sequence:

GMFGEDIFTSFRNAAGEGASSNVPRKAAPIERTLPCSLEDLYKGTTKKMKISREVSDATGRPSTTEEILTIDIKPGWKKGTKITFPEKGNEORGIIPSDLVFIIDEKPHSVFKRDGNDLVVTOKISLVEALTGYTAOITTLDGRNLTIPINSIISPNYEEVIKGEGMPIPKEPSKIGNLRVKFNIKFPSKLTSEOKAGIKRYLT*

NbL19g16020.1 bifunctional_monothiol_glutaredoxin−S16_chloroplastic [MW = 35.84 kDa]

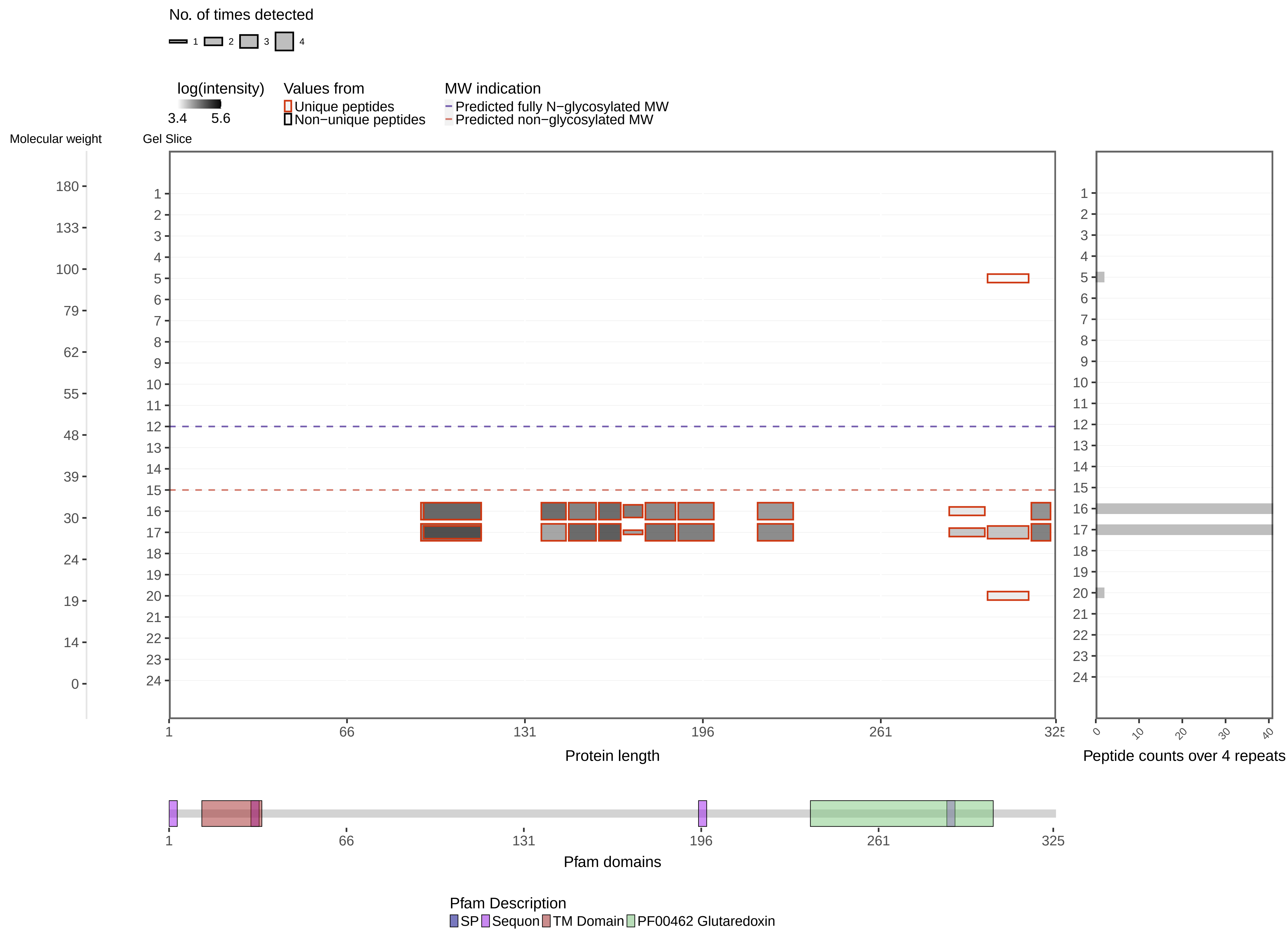


Protein sequence:

CSSVKVGVVDEPNRIALTESWKSWMEEHIAATGKVPPGNEPGNSTWVRQPPKKKTDLRLTPGRHVQLTVPLEDLIDRLVKENKVVAFIKGSRSAPQCGFSQRVVAMLESEGVDYESVDVLDEEYNYGLRETLKNYSNWPTFPQIFVKGELVGGCDILTSMYEKGELATLFKS*

NbL19g15680.1 ubiquitin−activating_enzyme_E1_1−like [MW = 121.21 kDa]

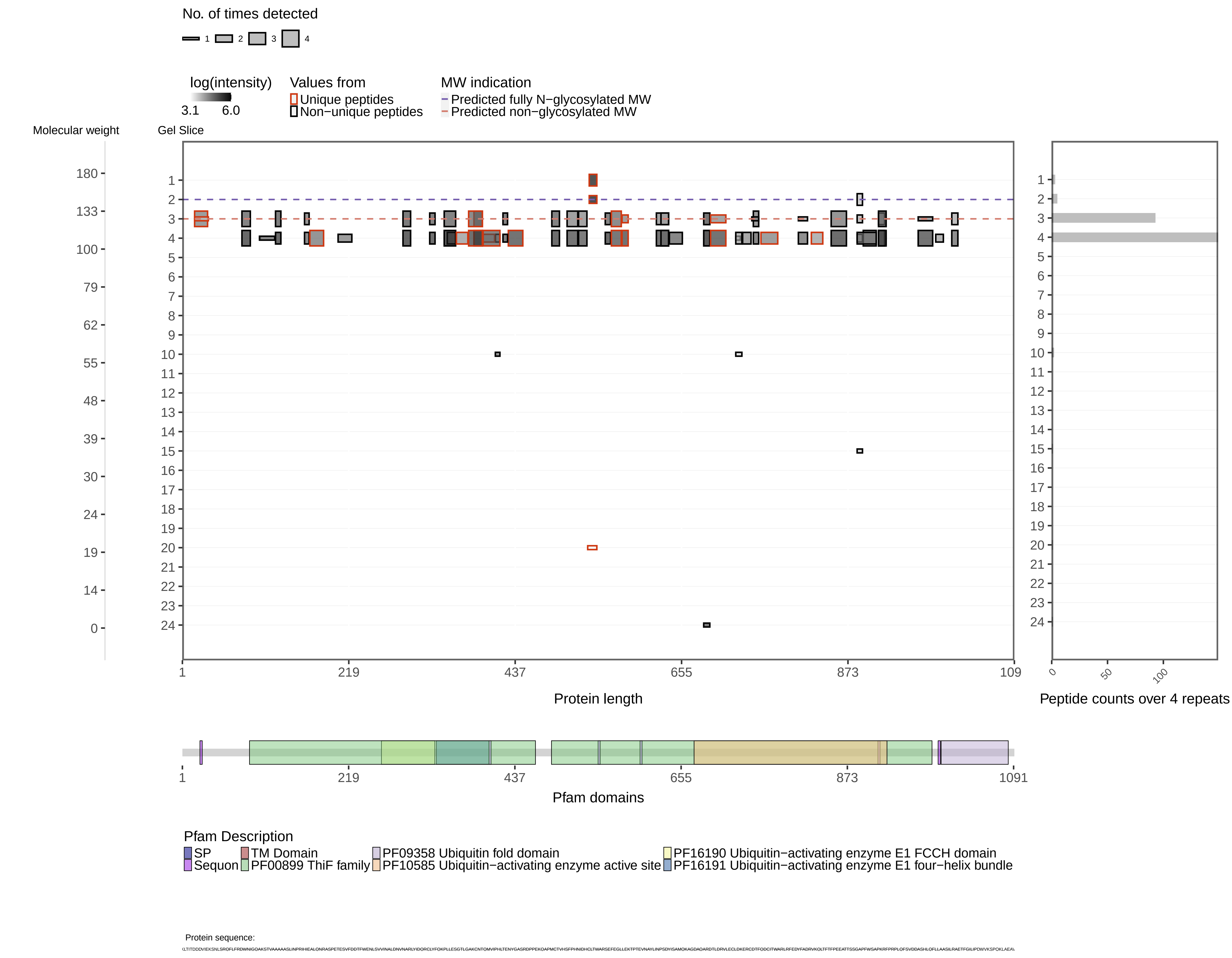

### NbL19g15670.1 uncharacterized_LOC107798940 [MW = 37.57 kDa]

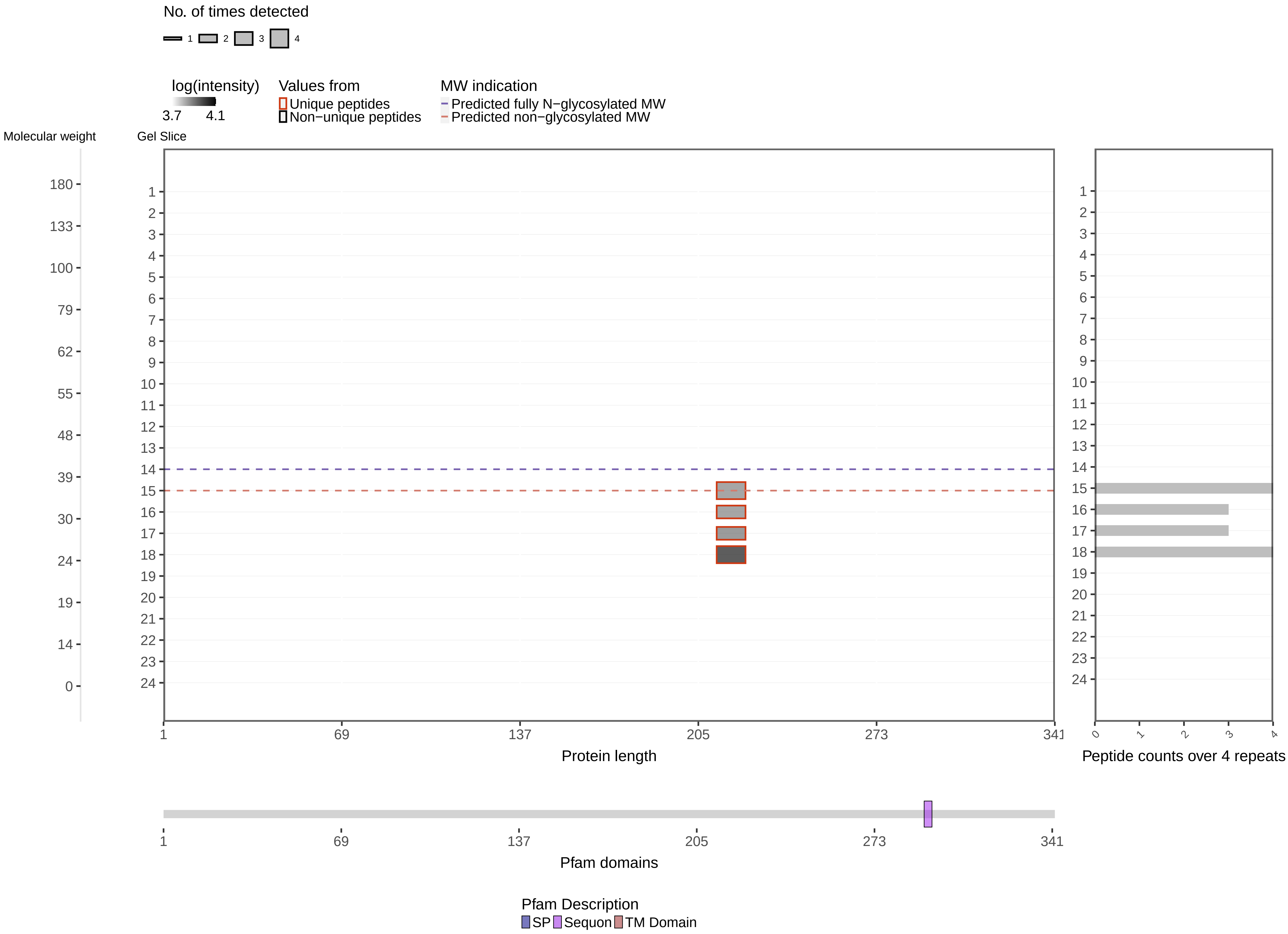


Protein sequence:

[illegible]SHSFTPQNAPYLLSLPPSLFLLLTTPFHEQLIHTHEYIAFQYRISTDSFDPKSLDIINIGPSFRSHYGSFSPISPFPLISCDLRGPNAMAEDEKVESLVSIKRGLKDQKELDLCAEGFEIGRLNKLMGSDASNYTAELEHLYDKMLVKLDSLARLVETSSAKVLEQENHNMKLRYKVAGLE*

NbL19g15230.1 probable_inactive_receptor_kinase_At5g58300 [MW − SP = 67.45 kDa]

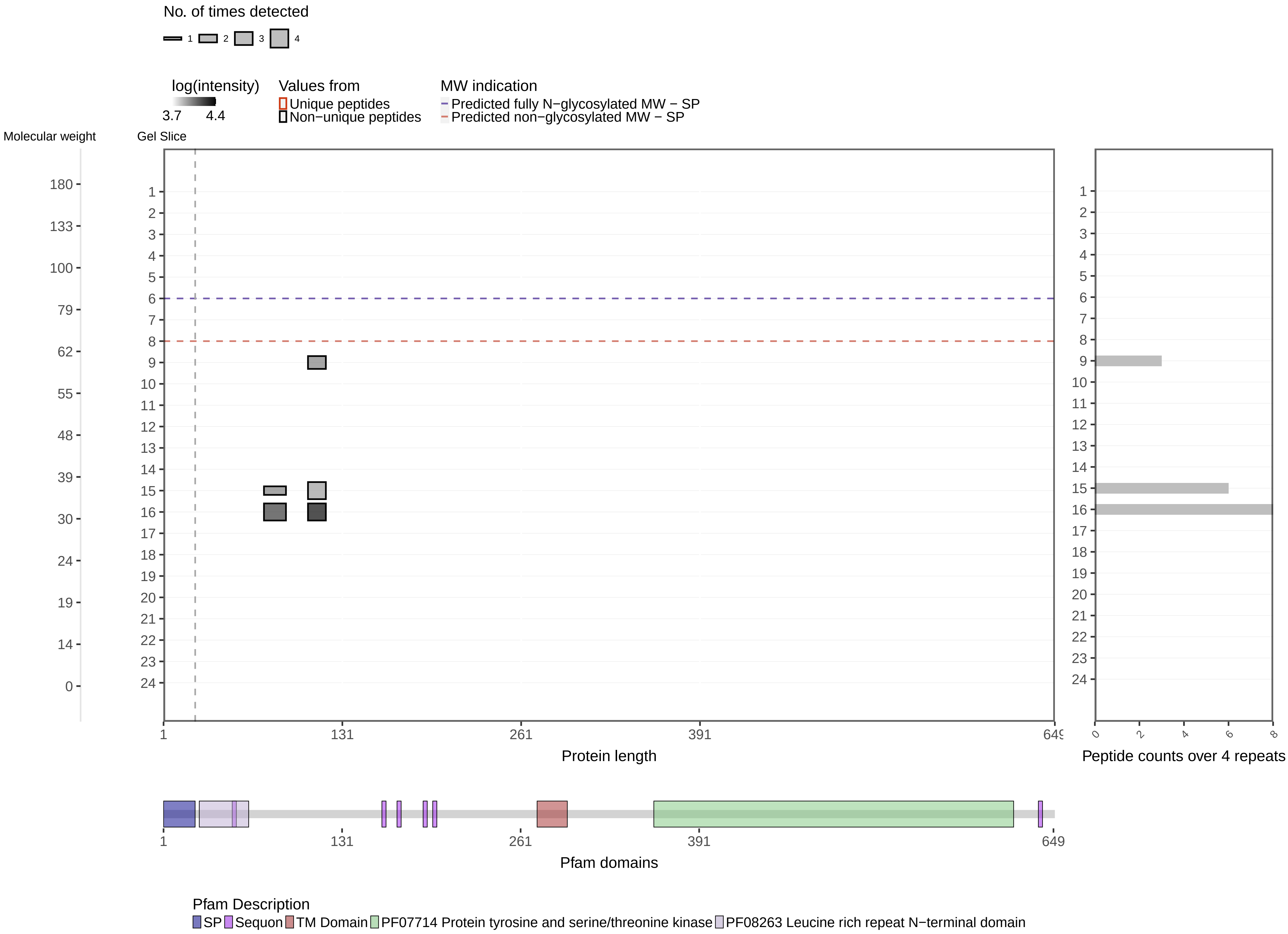


Protein sequence:

GSPKQTEDFSSGVQAAEKNKLVFFGGCSFNFDLEDLLRASAQVLGKGSYGTTYKAILEEGTTVVVKRLKEVVVGKREFEQQMEIIGTVAQHGNVVALRAYYFSKDEKLLIYDHVPAGSLSTRMHGNRELGRTLSWESRLRIAHGAASGIAHIHSVAGGKLIHGNIRSSNVLLTHDNSGCISDVGLTPLMGSPTIPSRSAGYRAPEVIETRKCTQKSDVYSFGVLLLELLTGKAPVQPPGHDEVVDLPRWVQSVVREEWTAEVFDAELIKFQNIEEEMVQMLQIAMACVAKVPET

### NbL19g15210.1 receptor−like_protein_kinase_FERONIA [MW − SP = 93.8 kDa]

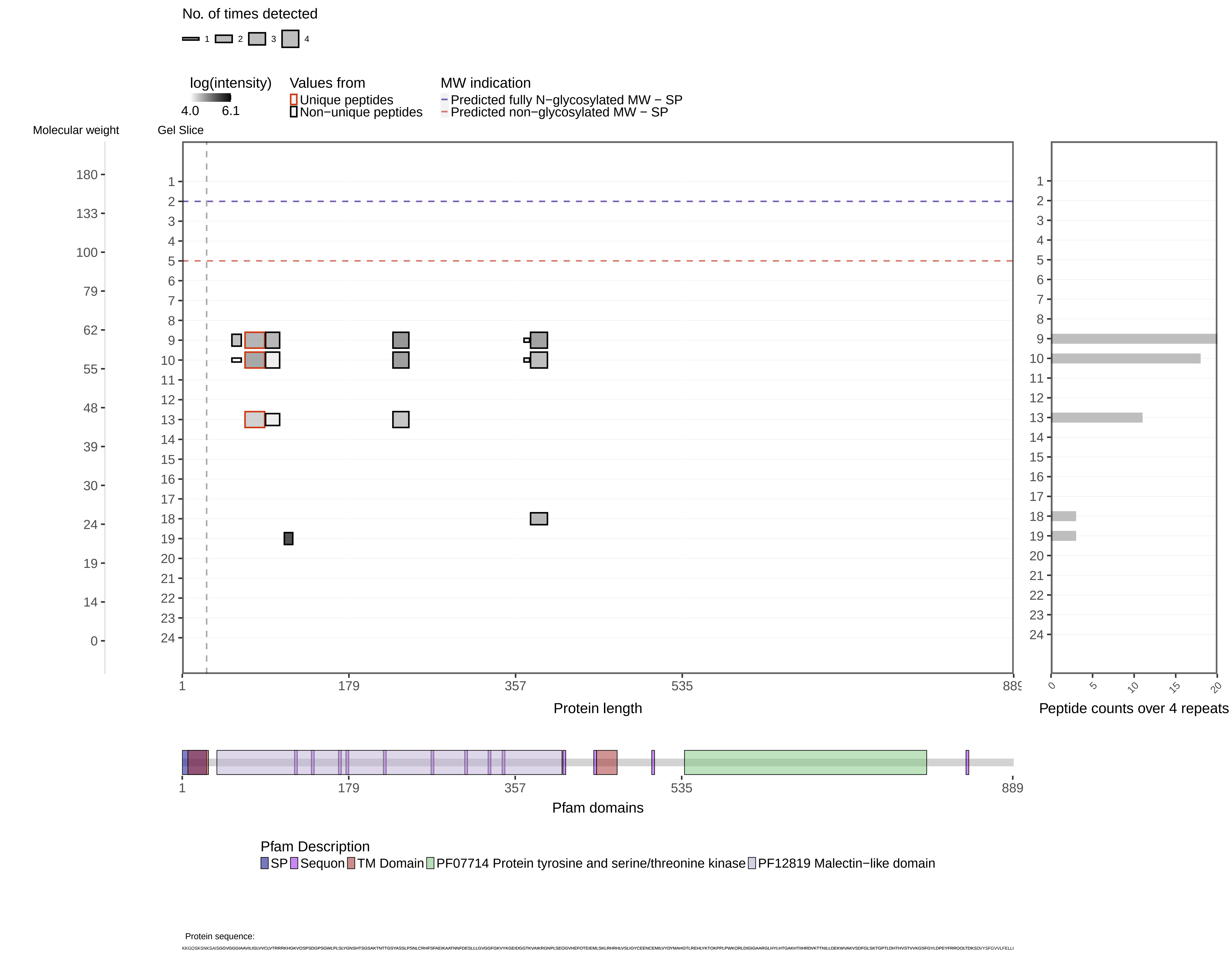

NbL19g15060.1 non−specific_lipid−transfer_protein_1−like [MW − SP = 9.78 kDa]

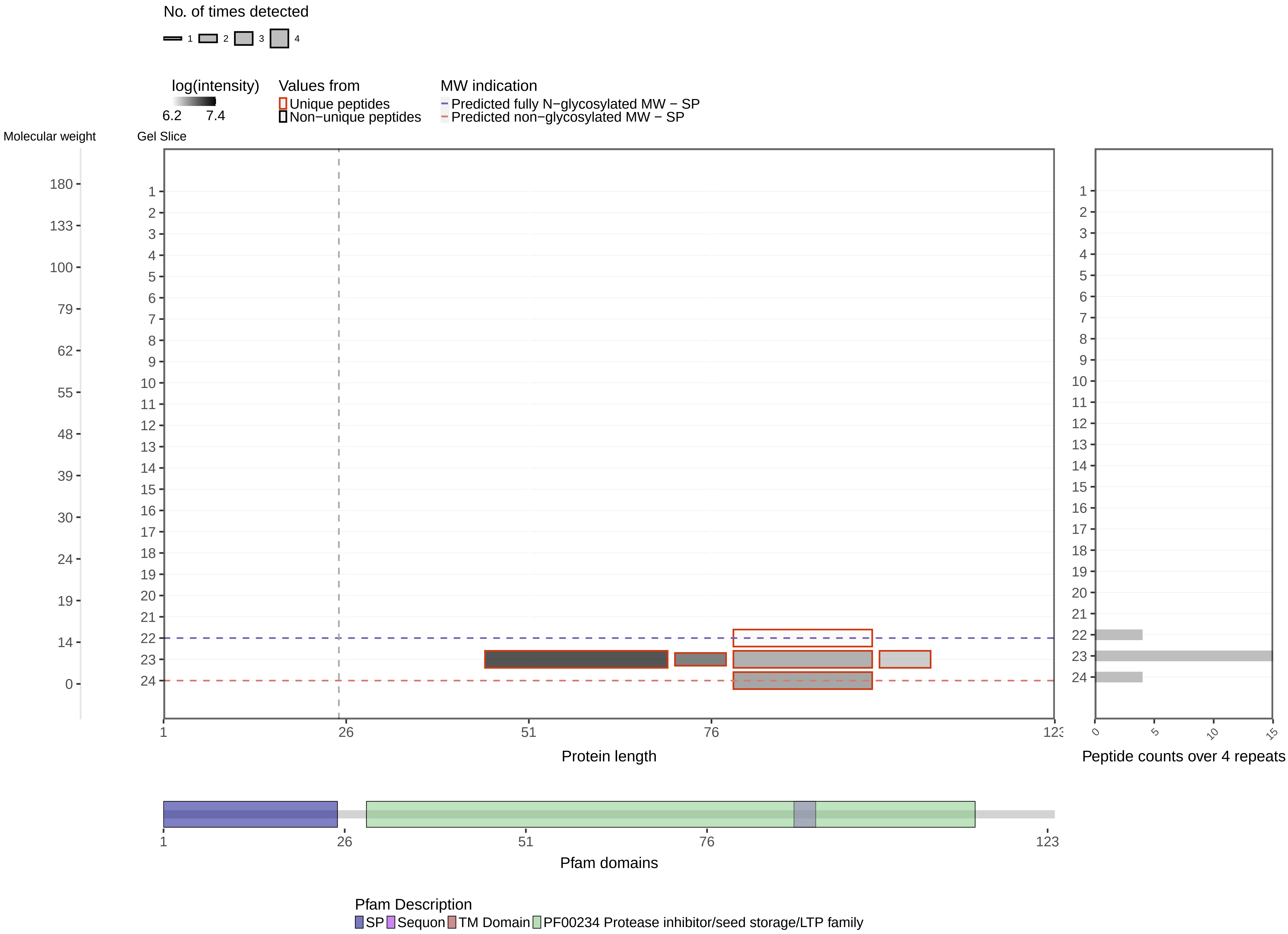


Protein sequence:

AIPAPCCSGINSLNNQATSTPDRQTACSCIKSAAAGISGINLSLAGSLPSKCGVNLPYKISPSIDCSTYVSNTHF*

NbL19g15050.1 non−specific_lipid−transfer_protein_1−like [MW − SP = 11.38 kDa]

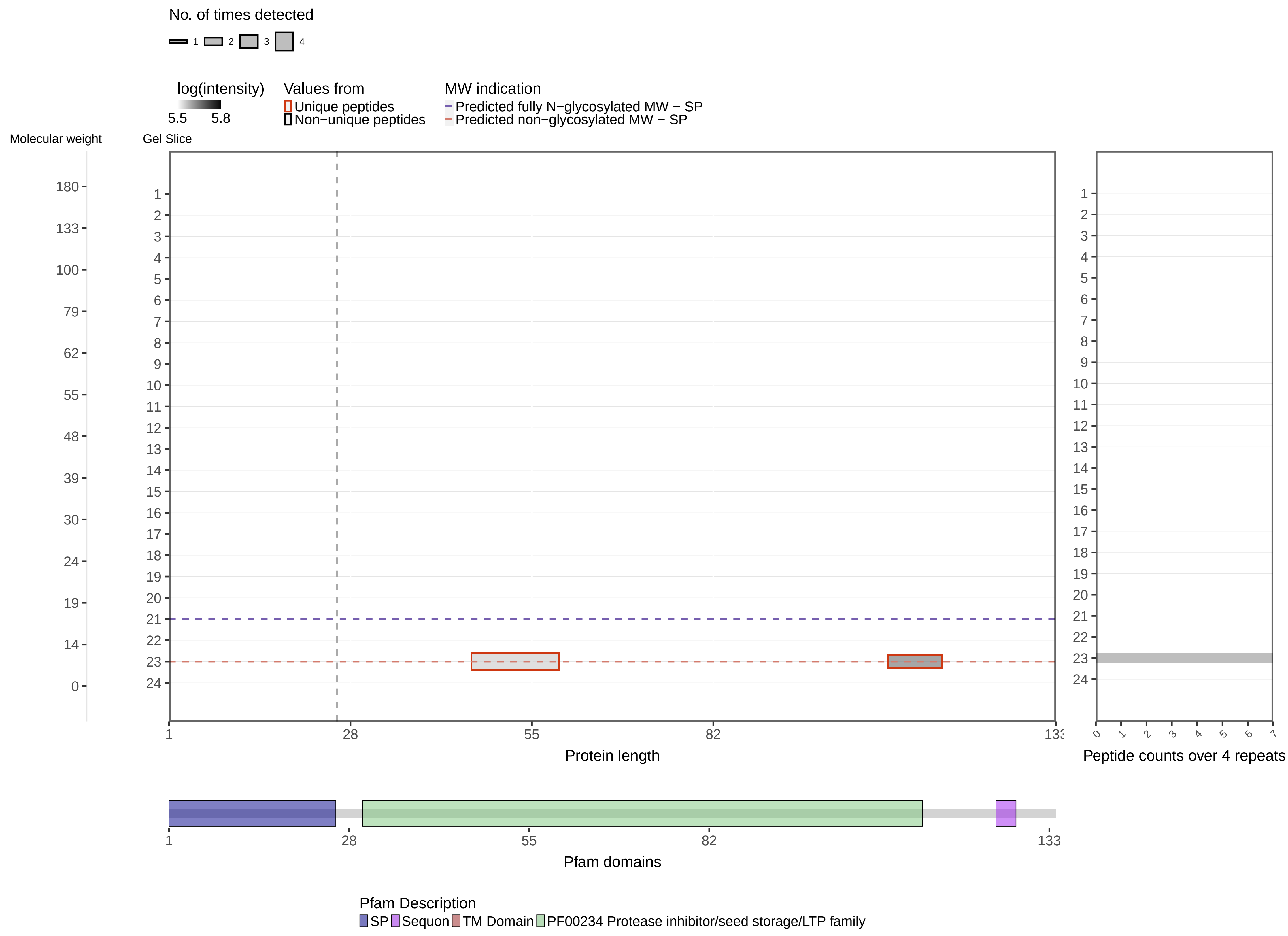


Protein sequence:

MCCNGIRSLFRAASTTQDRQTVCNCMKSAARVISGVNLNLAAALPSECGVNIPYKISPTIDCARYIYIYSFLNISQHFCA*

NbL19g15040.1 non−specific_lipid−transfer_protein_1−like [MW − SP = 14.85 kDa]

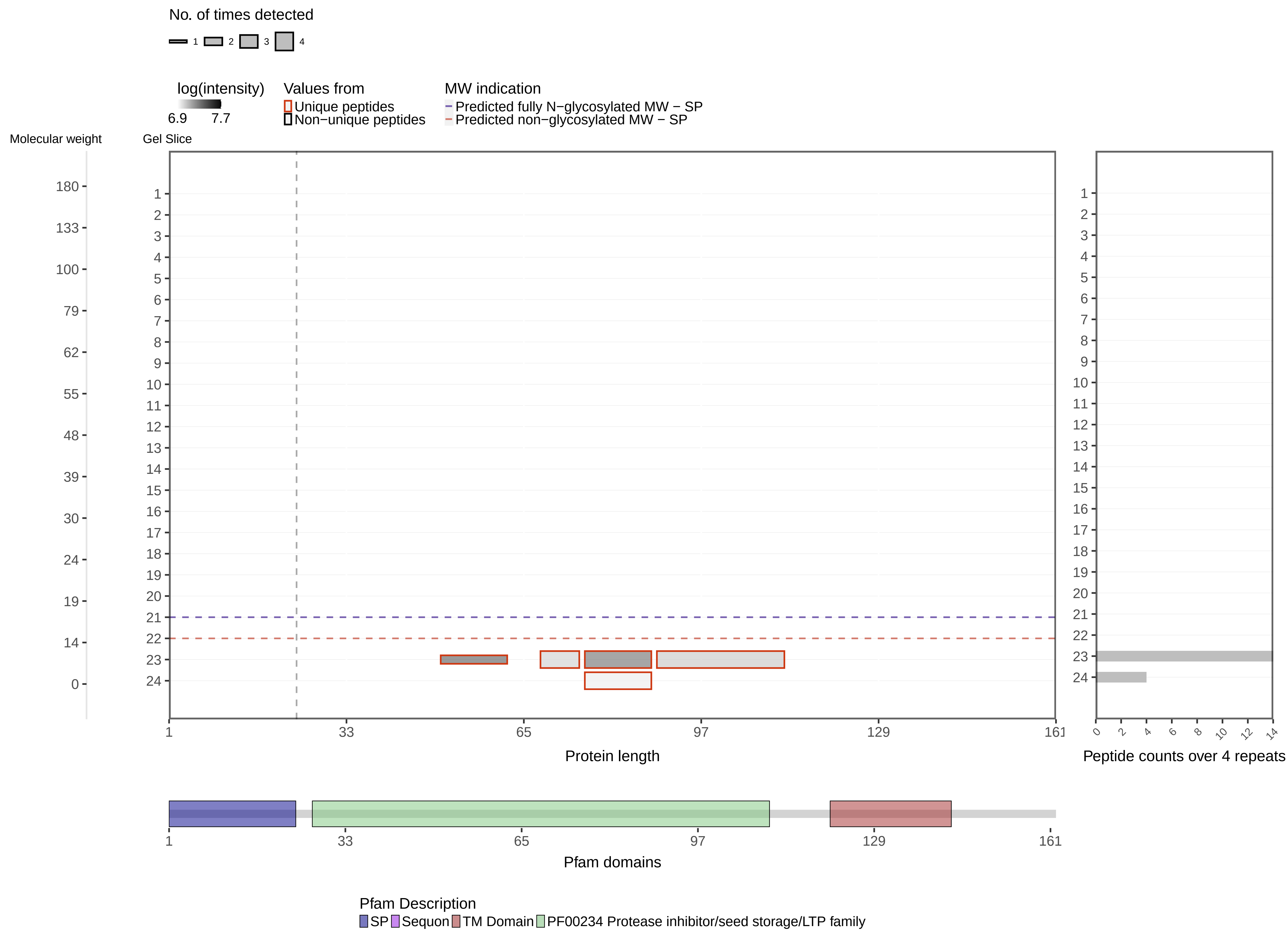


Protein sequence:

TQDRQTACTCLKSAAAAISGIDLGKAAGLPSTCGVNIPYAISPSTDCSKYLTFFLSHLILILYSICFIFSYPLNFLFRLFKQITRRIHYLFFSSRYK*

NbL19g14840.1 phosphoglucomutase_cytoplasmic−like [MW = 63.45 kDa]

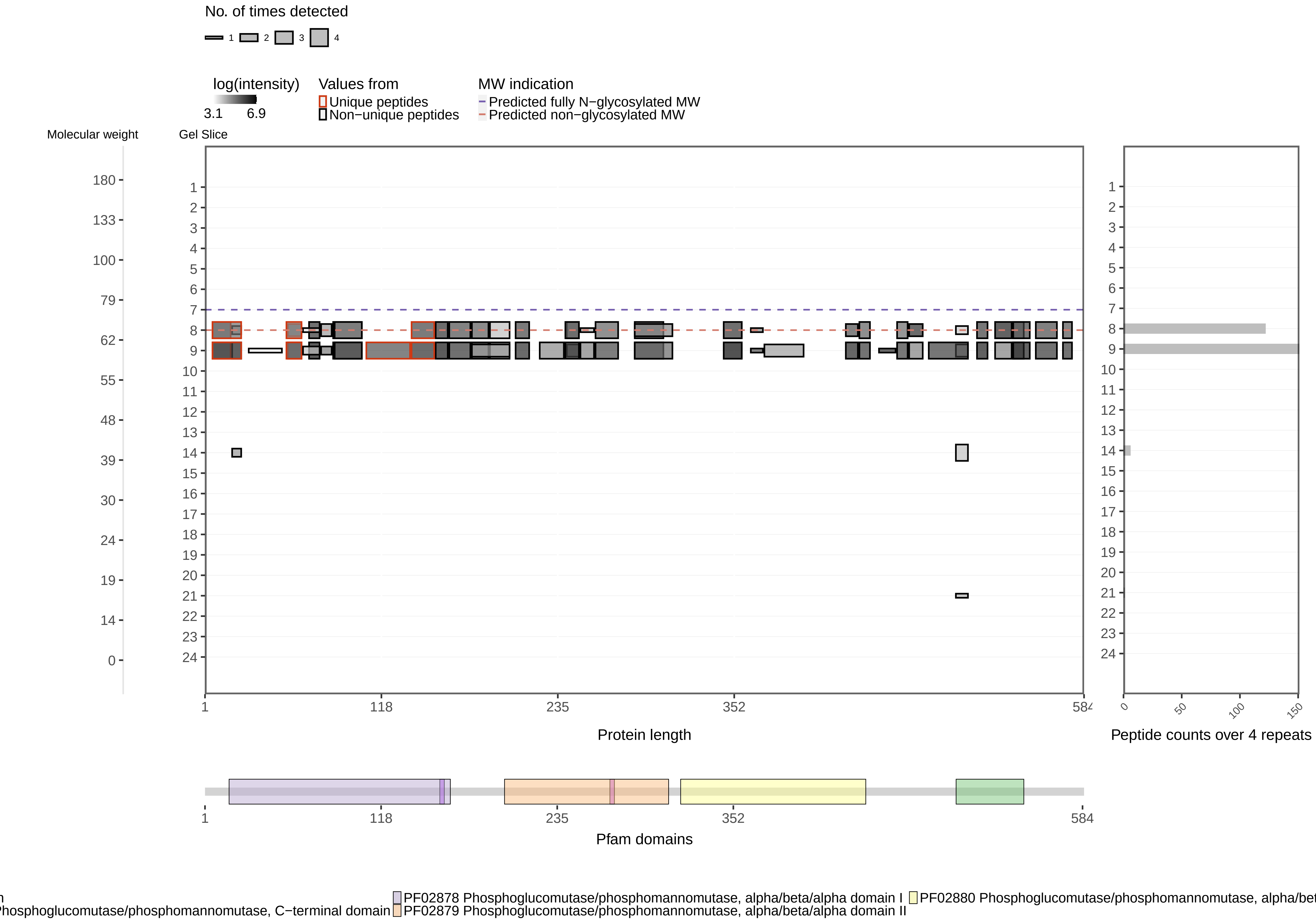


Protein sequence:

ARMGLSKSHSEPNPPEFGAAADGDADRNMILGKRFFVTPSDSVAIIAANAVEAIPYFSGGLKGVARSMPTSAALDVVAKSLNLKFYEVPTGWKFFGNLMDAGMCSICGEESFGTGSDHIREKDGIWAVLAWLSILADKNKDNLGGEKLVSVEDIVRQHWATYGRHYYTRYDYENVDAAGAKELMAHLVKLQSSLDEVNKIIKGIHSDVSNVVHGDEFEYKDPVDSSISKHQGIRYLFEDGSRLVFRLSGTGSEGATIRVYIEQYEKDPSKIGRDSQEALAPLVEVAIKLSKMQE

NbL19g14660.1 heat_shock_70_kDa_protein_16 [MW = 84.35 kDa]

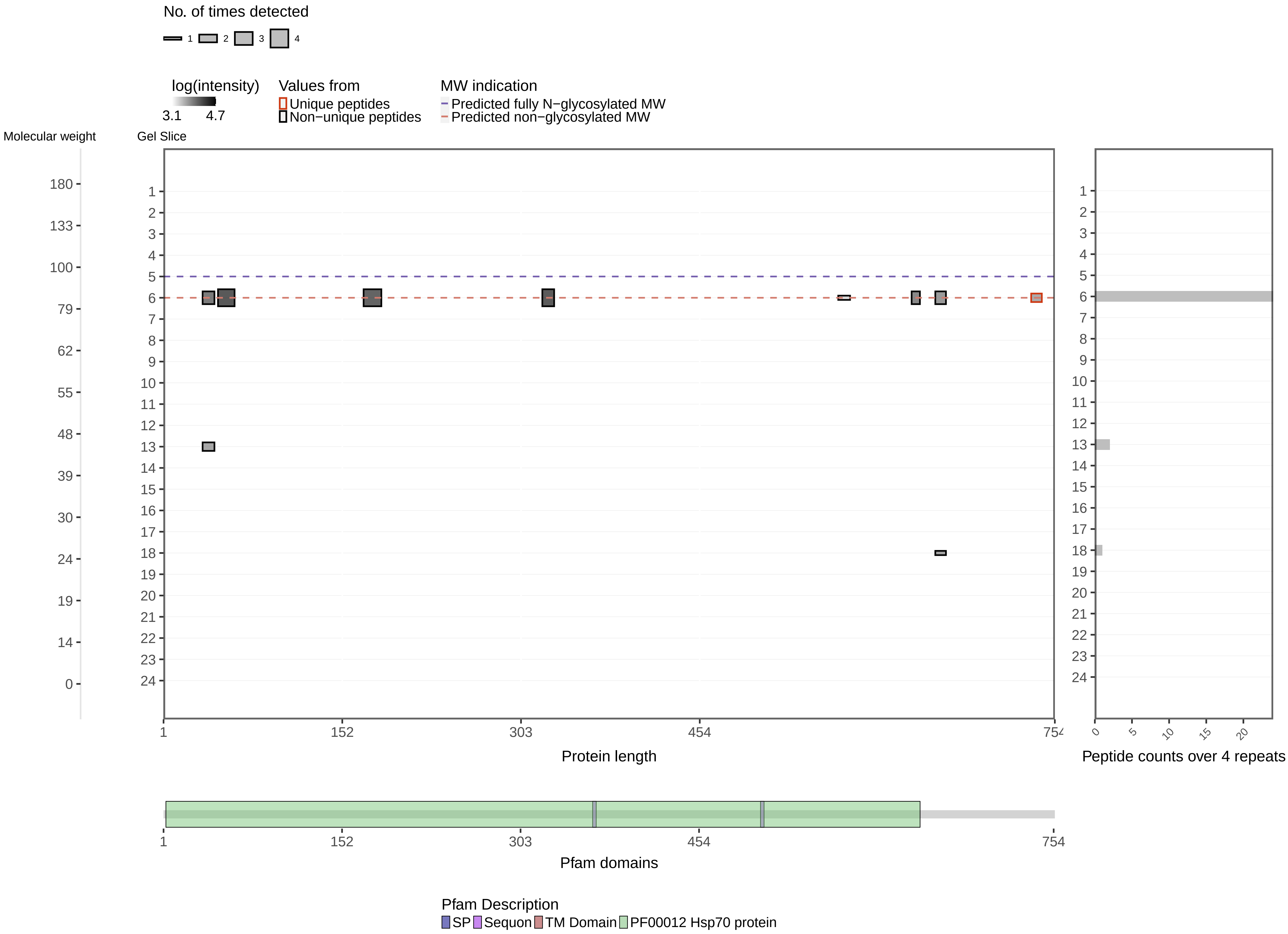


Protein sequence:

/ARGCALQCAMLSPIFRVREYEVQDSFPFSIGFASDEGPVCTLSNGILFPRGHSFPSMKVLTLQRSTSFYLEAFYTNQNELPPGVSDKISKSTIGPFQVPHSEKAKIKVKIQLNLHGIVTVESAWLIKDQTSHSTSENNTDTHAEDMERDDTRKSMAVKRQDIPVSESVDGGMTLAELSQAQEKECHLAEQDIKVERTKDKKNTLEAYVYETRNKLLNTYRSFATDSEREGISCNLQQTEEWLYEDGDDESEHVYAEKLDDLKKMVDPVENRYKEEETRAQATRNLLNSIVEYI

NbL19g14650.1 UPF0051_protein_ABCI8_chloroplastic−like [MW = 63.93 kDa]

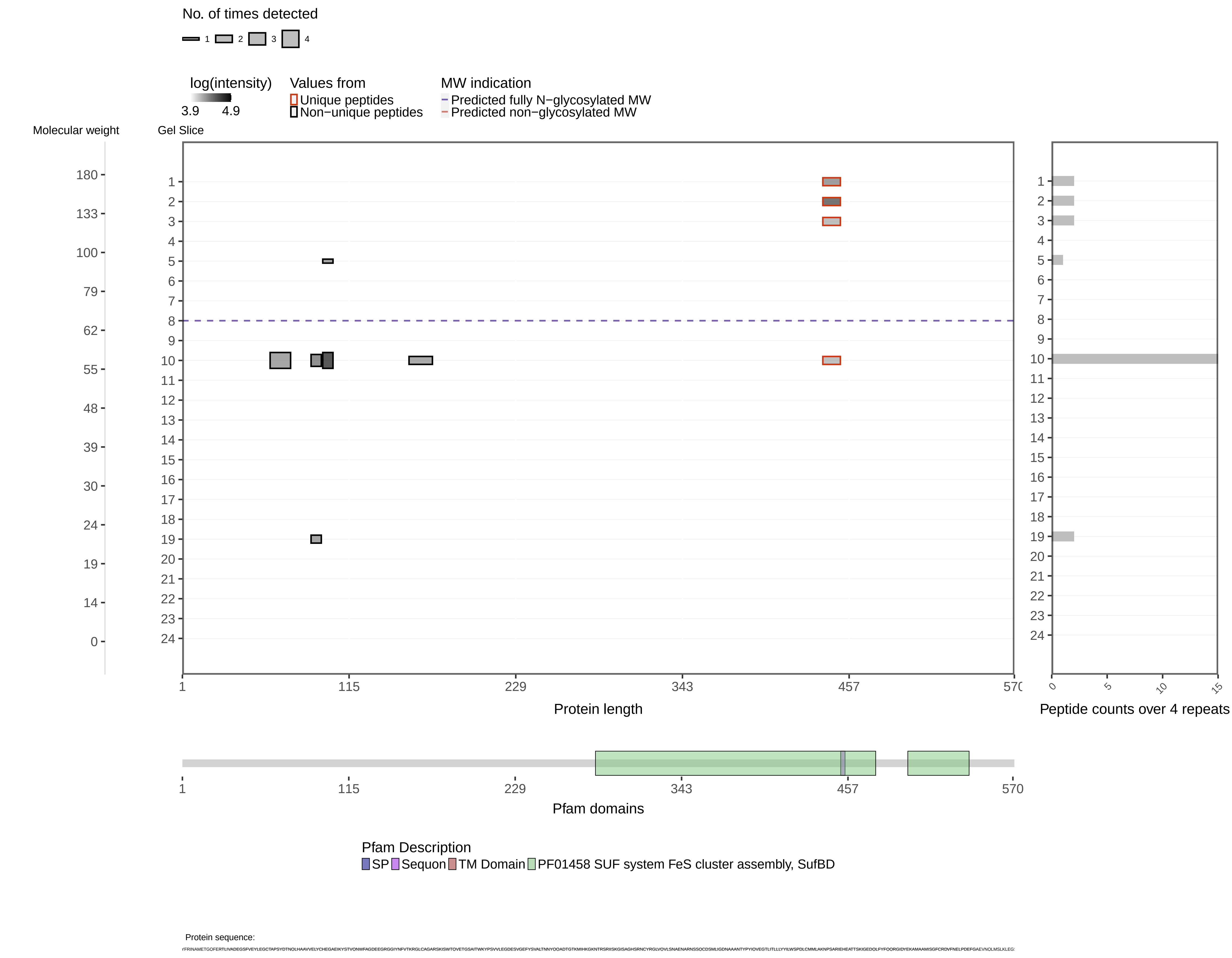

NbL19g14480.1 zinc_finger_CCCH_domain−containing_protein_14−like [MW = 30.73 kDa]

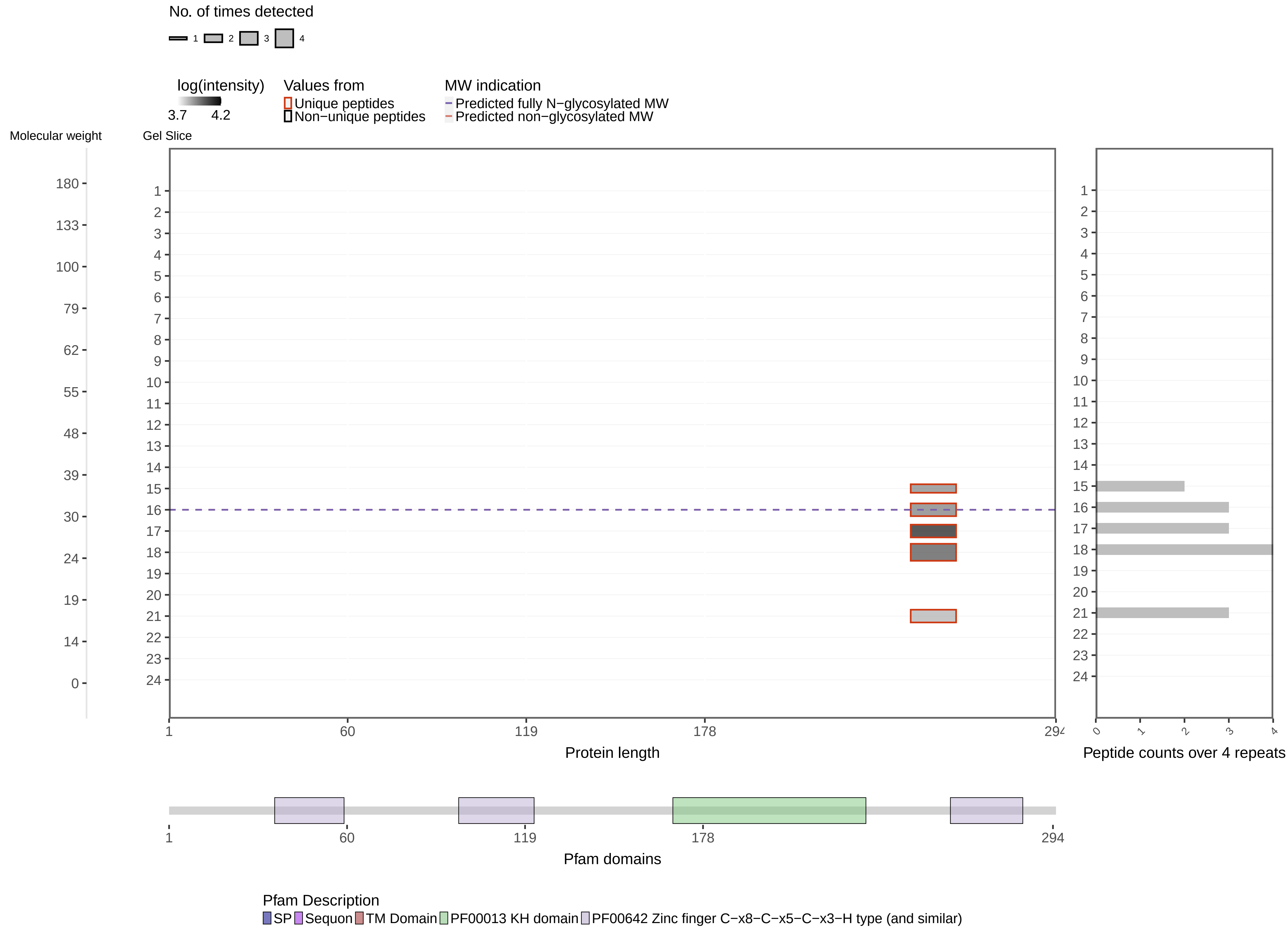


Protein sequence:

QEEPRAMGVGPVPGRFGGRIEPAVSGPAASFGTSATAKISVDASLAGAIIGKGGVHSKQICRQTGAKLAIRDHETDKNLRNIELEGTFEQISQASAMVRELINSLGSVGCPGRTPVVPGGAAPPGSNFKTKLCENFAKGSCTFGDRCHFAHGAAELRKTGV*

### NbL19g14410.1 60S_ribosomal_protein_L21−2 [MW = 21.71 kDa]

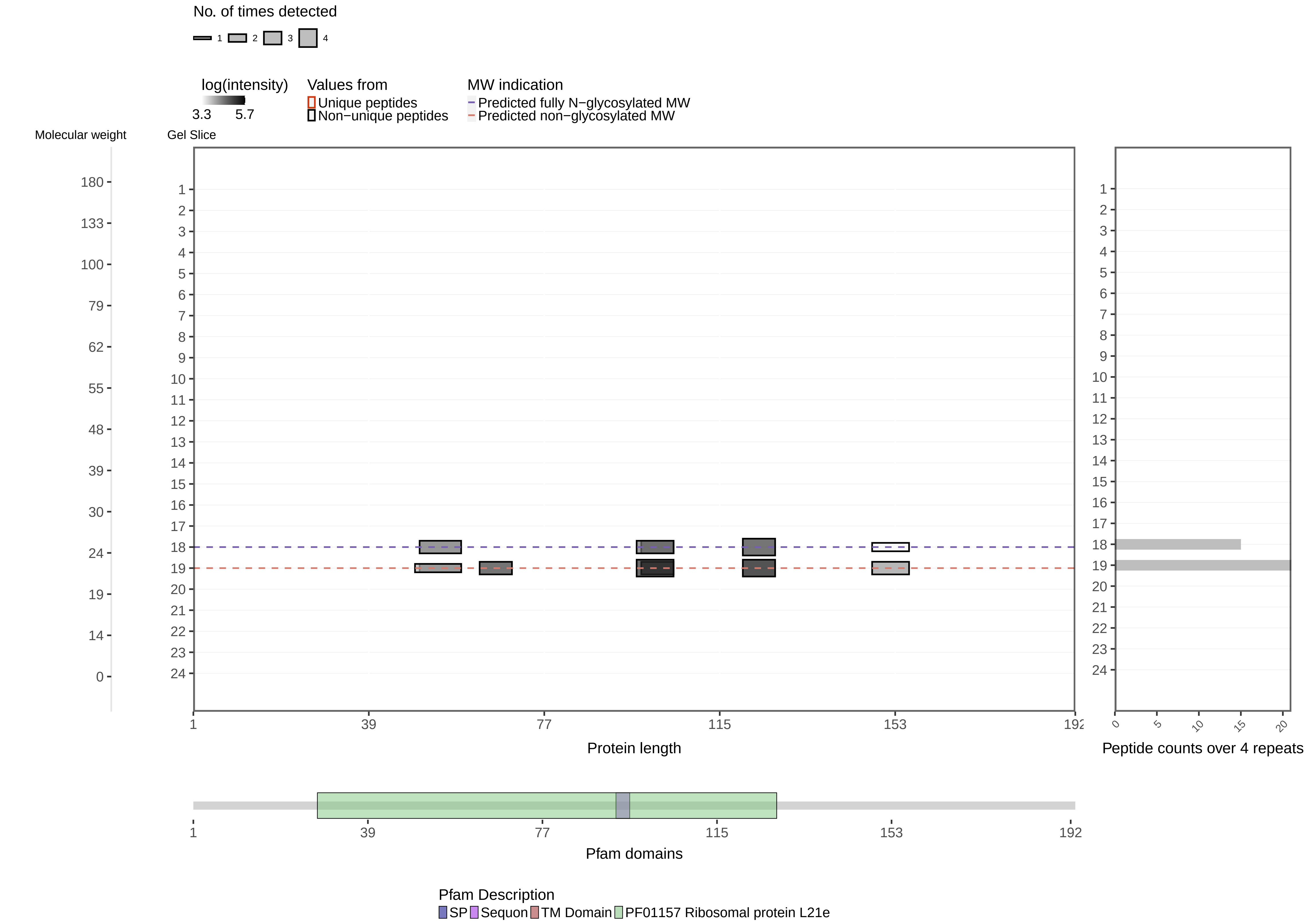


Protein sequence:

FYHGRTGRIWNVTKRAVGVEVNKQVRNKILRKRIHVRIEHVQLSRCTEEVKERIKRNDQLKAEAKARGEVISTKRQPLGPNPGFMVEGATLETVTPIPYDVVNDLKGGY*

NbL19g14270.1 asparagine−−tRNA_ligase_cytoplasmic_1−like [MW = 64.92 kDa]

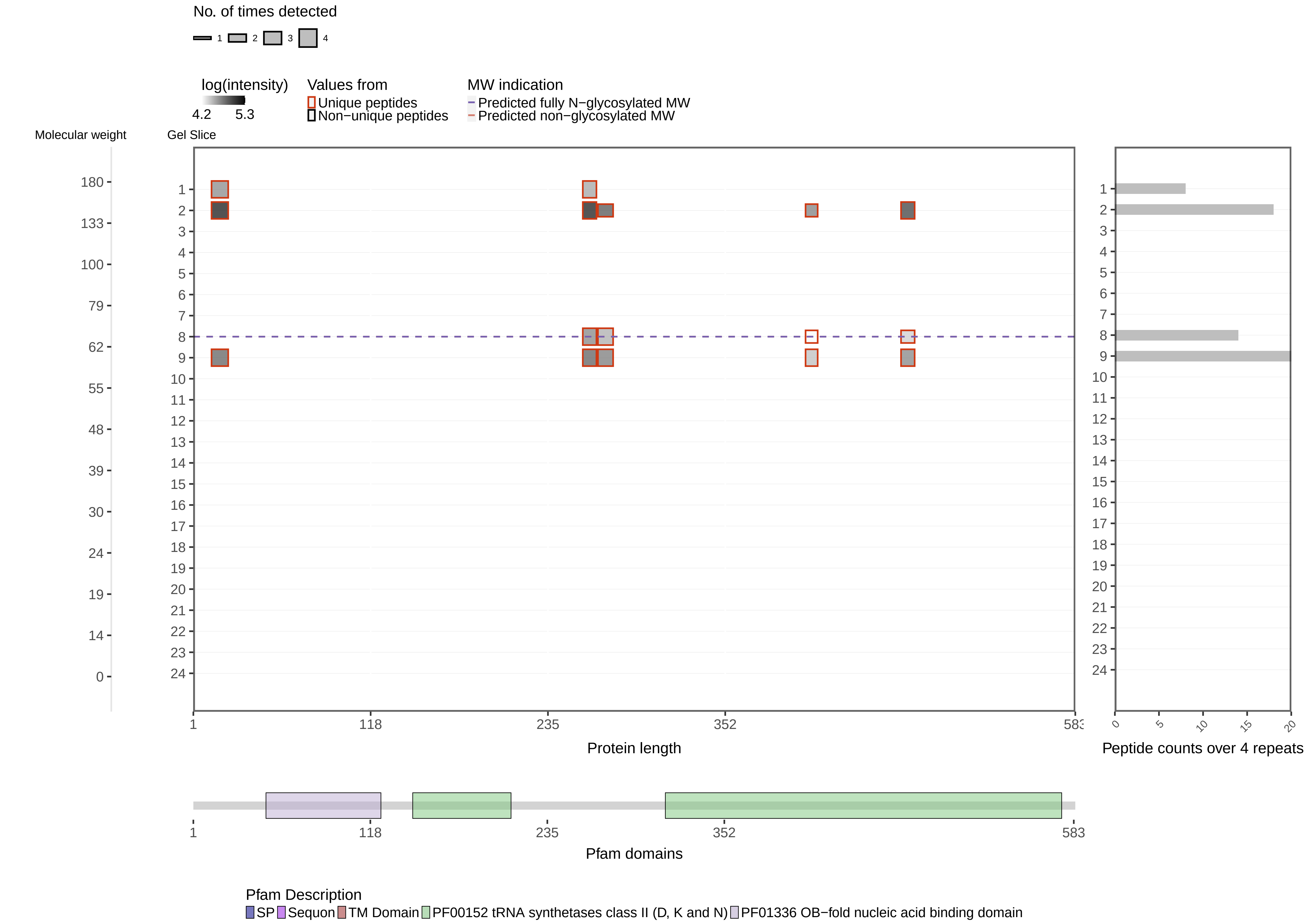


Protein sequence:

KENLLKLQERFKLGERCRLSAGIPKKDGKVDYSEDFFARQAFLTVSGQLQVETYACALSSVYTFGPTFRAEQSHTSRHLAEFCMVEPEIAFADLQDDMNCAEAYVKFLCQWLLDHCLDDMEFITKFIDKDALNRLRMVASSNCYRITYTEAIAILEVAAKVKKFENKVEWGIDLASEHERYLTEEHFKAPVIVFSYPKGIKAFYMKVNEDKKTVAAMDLLVPKVGELIGGSQREEDYEVLRTRILEMGLPLEPYEWYLDLRRYGTVKHSGFGLGFERMILFATGIENIRDVIPFPR

### NbL19g14080.1 bifunctional_protein_FolD_2−like [MW = 32.94 kDa]

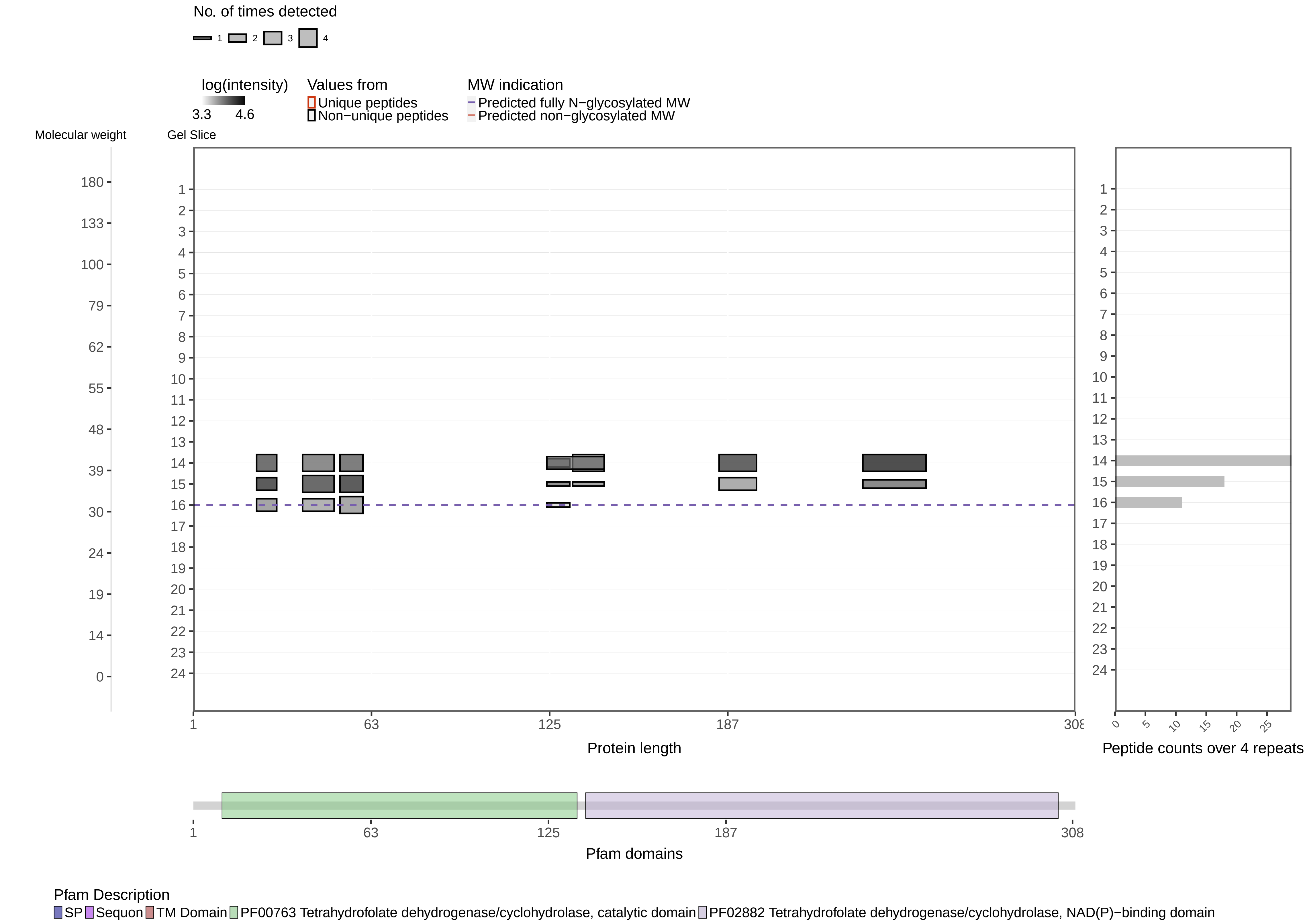


Protein sequence:

NIGKLAMKGREPLFLPCTPKGCIELLHRSGISIKGKNAVVVGRSNIVGLPVSLLLLKEDATVTIVHSRTKEPEKIVREADIIIAAAGQAMMIKGSWIKPGAAVIDVGTNAVDDPTRKSGYRLVGDVDFQEACEVAGCITPVPGGVGPMTVAMLLKNTLDGAKRVIEK*

NbL19g14030.1 nascent_polypeptide−associated_complex_subunit_alpha−like_protein [MW = 22.1 kDa]

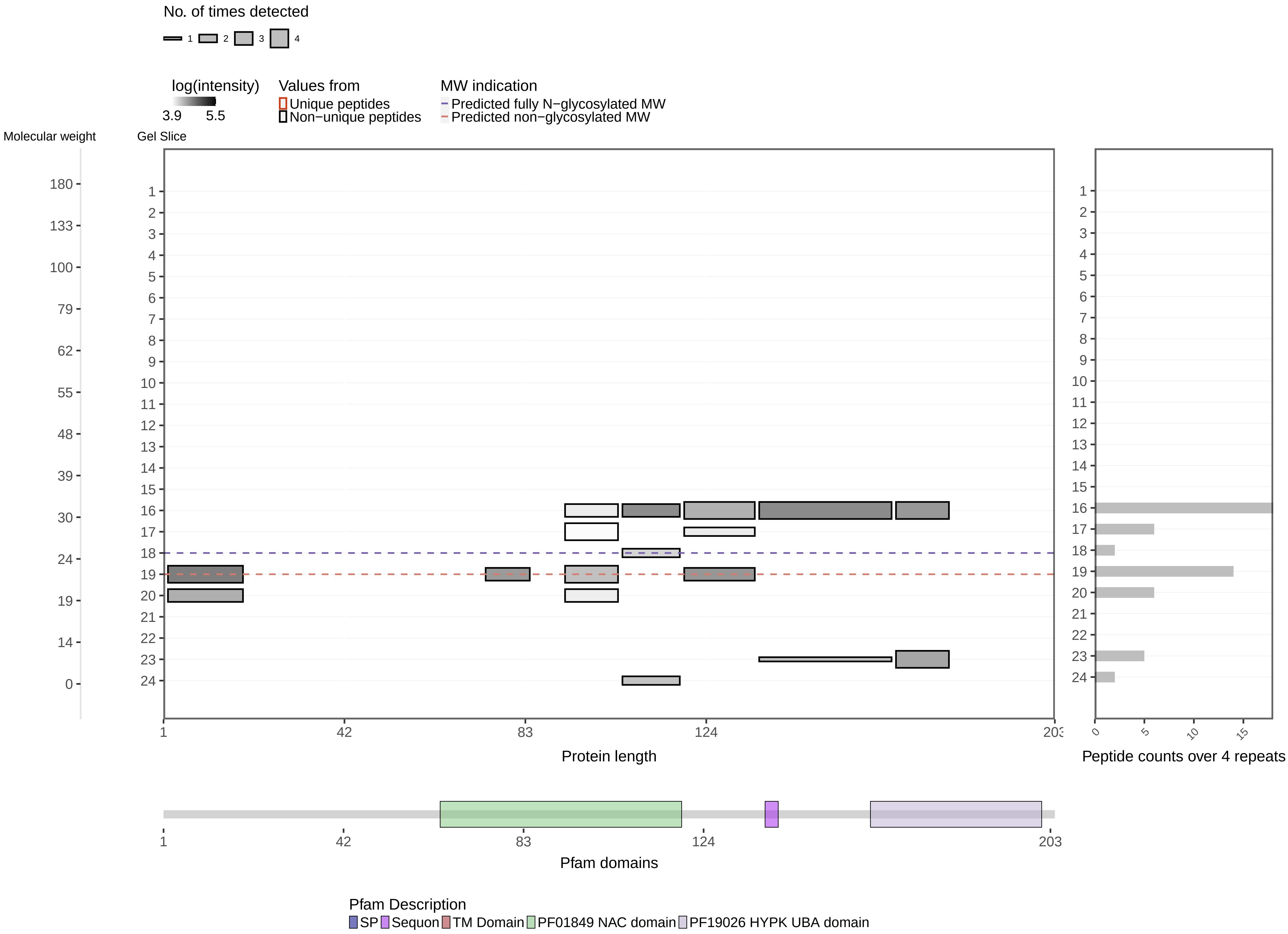


Protein sequence:
VKKSKNILFVISKPDVFKSPNSDTYVIFGEAKIEDLSSQLQTQAAEQFKAPNLSNVISKPEPSTVAQDDEDVDETGVEPKDIELVMTQAGVSRVKAVKALKAADGDIVSAIMELTN*

NbL19g13790.1 thioredoxin_reductase_NTRC−like [MW = 59.44 kDa]

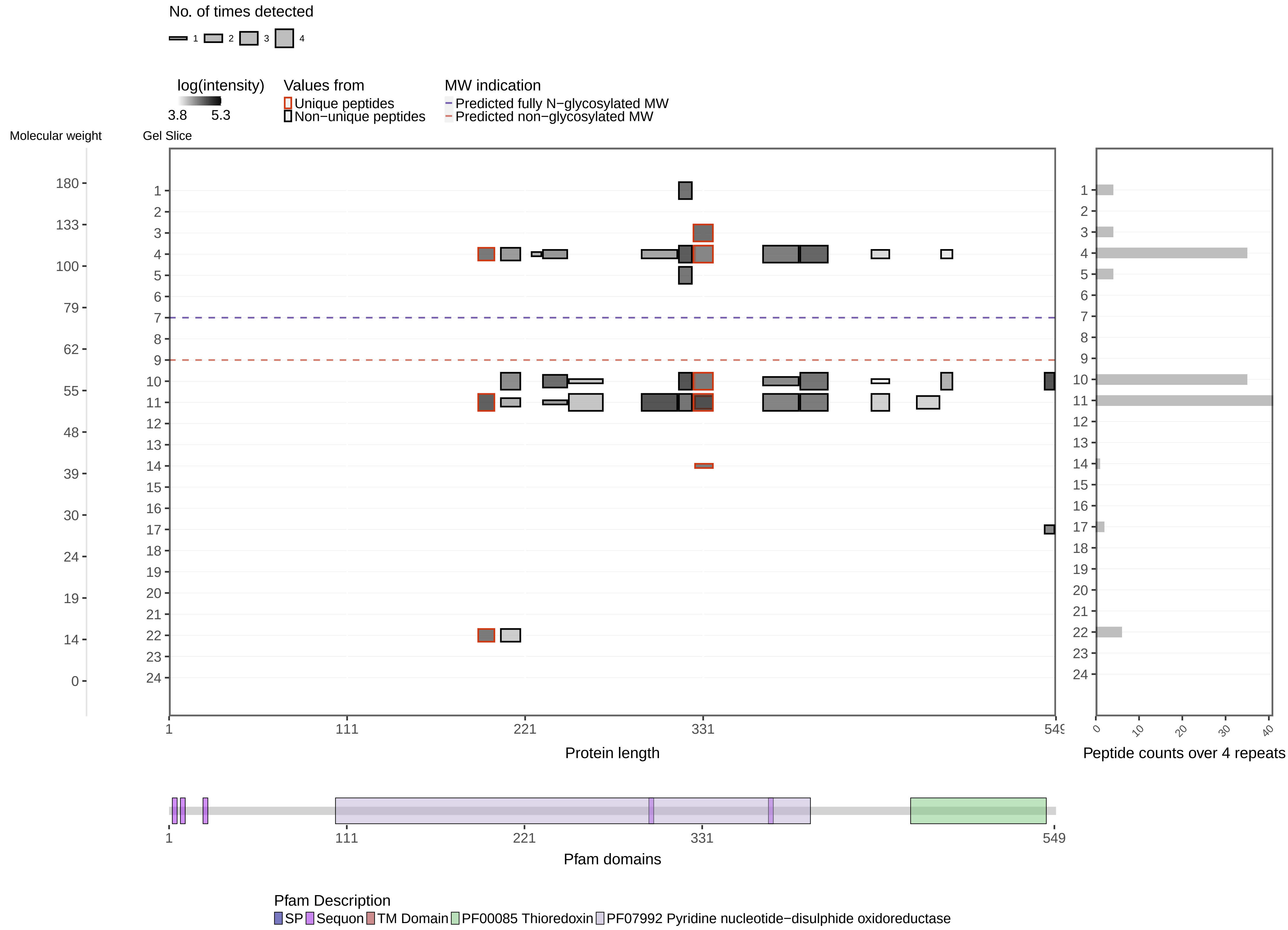


Protein sequence:

TEEAIYLTKYARHVHLLVRKDQLRASRAMQDRVFNNPNITVHFNTETVDVVSNPKGQMSGILIRKADTQEESVLEAKGLFYGIGHSPNSQLLEGQVELDSAGYILVKEGTANTSVEGVFAAGDVQDHEWRQAITAAGSGCVAALSVERYLTSKNLLIEFHQPHTEEVKKELTQKDVQEGFDITLTKHKGQYALRKLYHESPRIICVLYTSPTCGPCRTLKPILSKVIDEFDQHVHFVEIDITEDPEIAEAAGIMGTPCVQFFKNKEMLRTVSGVKMKREYRELIEANK*

NbL19g13710.1 probable_calcium−binding_protein_CML49 [MW = 35.43 kDa]

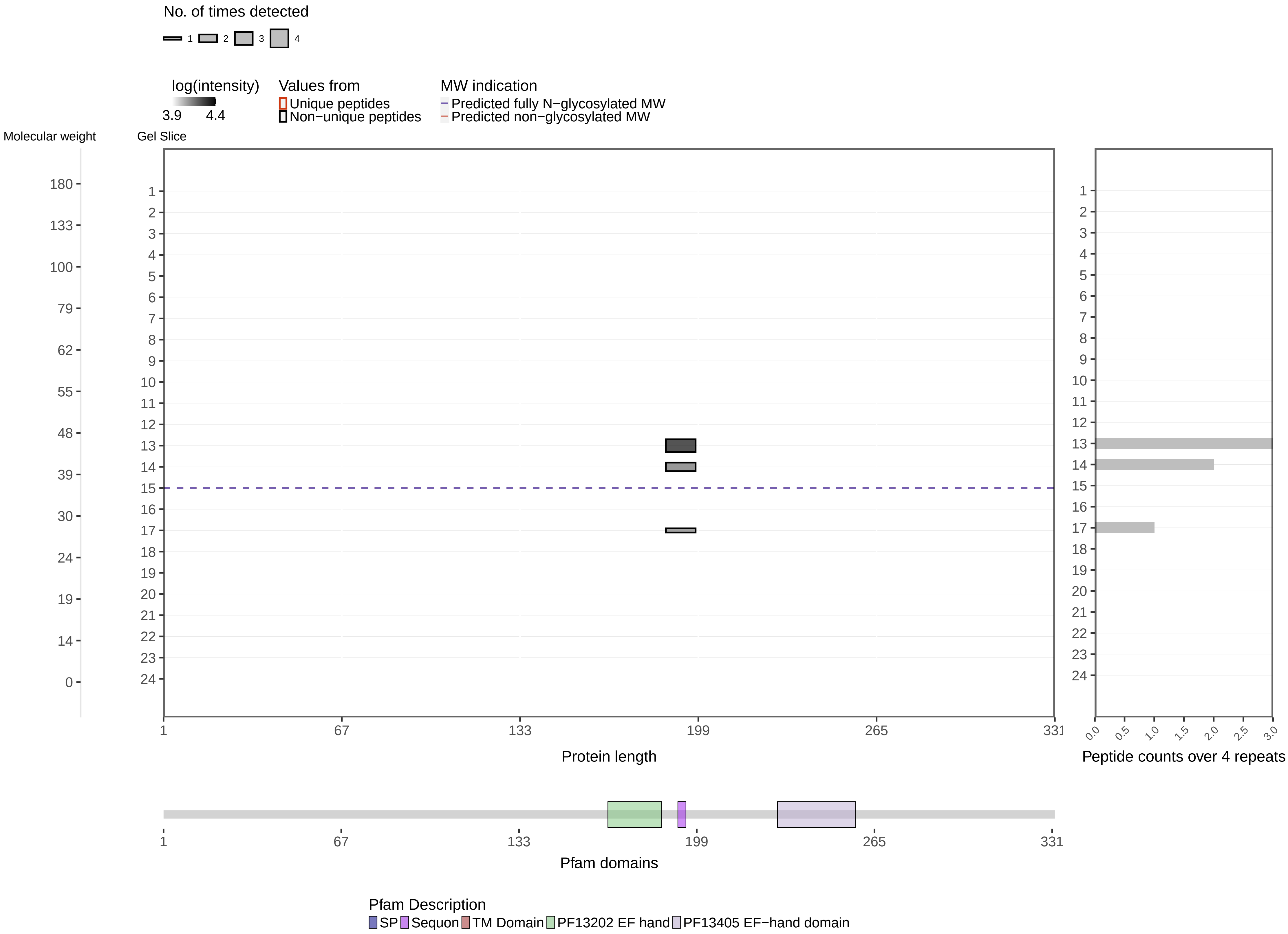


Protein sequence:

FASLVPSAFPPGTDPNVIACFQLADQDGSGLIDDKELQKALSSYNQSFSLRTVHLLMYLFTNTNTRKIGPKEFTSVFYSLQEWRGIFERFDRDRSGRIDSSELRDALLSLGYAVSPPILDLLVSKFDKTGGKNKAIEYDNFIECCLTVKGLTDKFKEKDASYSGSATFSYESFMLTVLPFLIA*

### NbL19g13540.1 extensin−like [MW = 29.83 kDa]

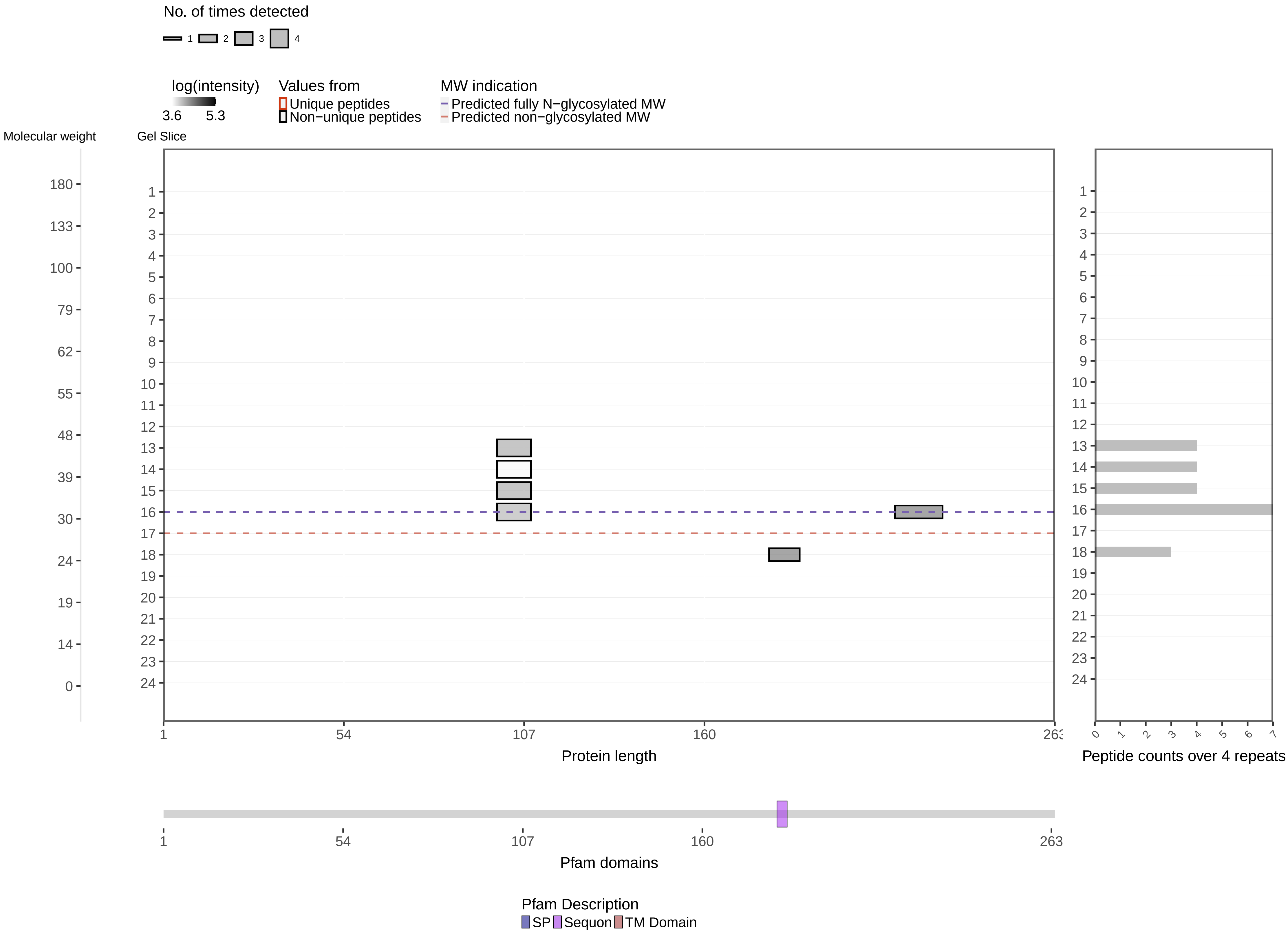


Protein sequence:

PPPPQPPSYSSVEHVSHESESESAHRHHDHHRFQPHVPSFFHHETSSHPELIDKPSFRVYTKADPNYSLTIRDGKVVLASSDPSDPFQHWYKAEKYSTKVKDEEGFPSFALVNKAAGLAMKHSVGASHLCSSPLTTLMFLMRQSCG*

### NbL19g13520.1 SNW/SKI−interacting_protein−like [MW = 67.2 kDa]

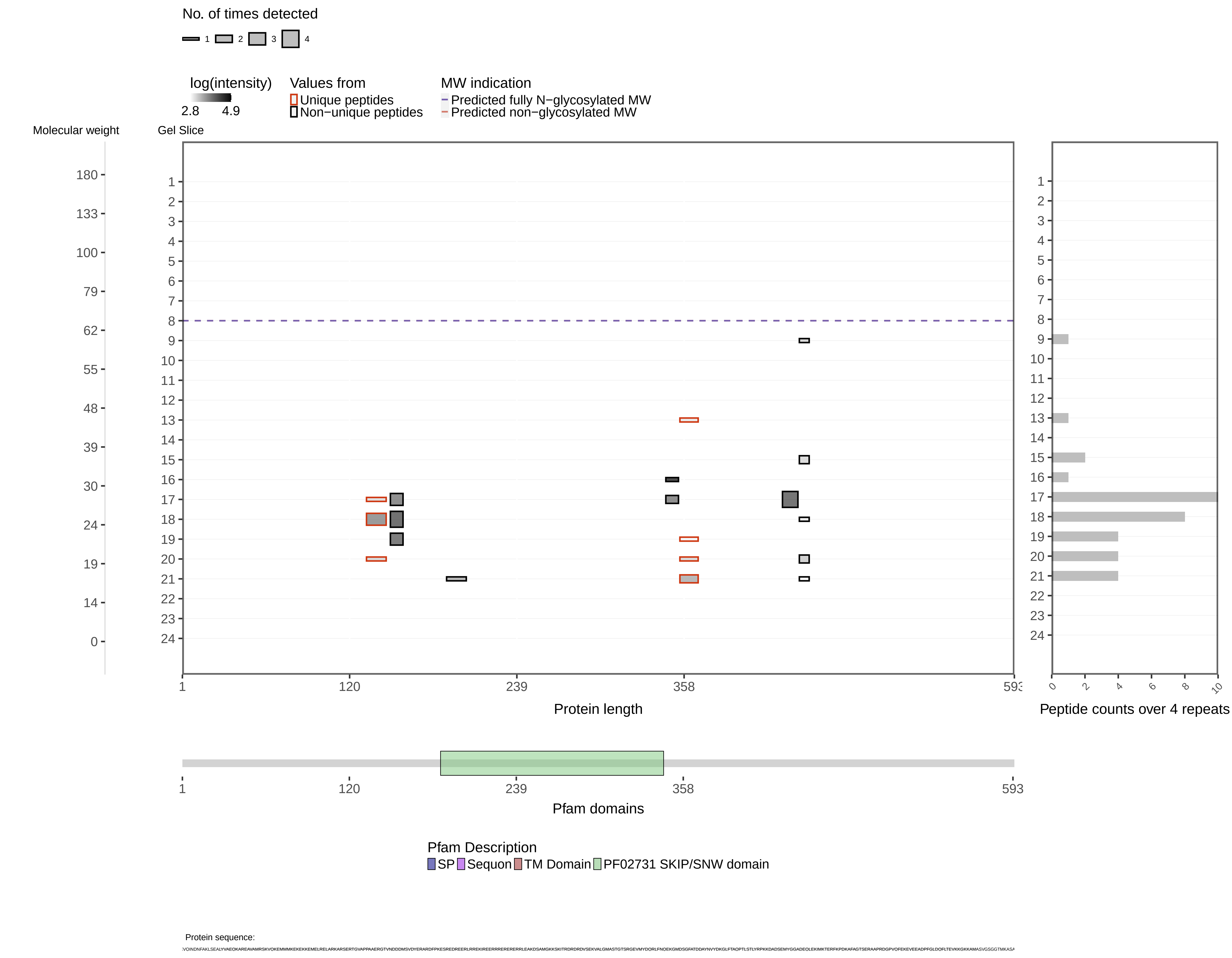

### NbL19g13500.1 cystathionine_beta−lyase_chloroplastic [MW = 51.35 kDa]

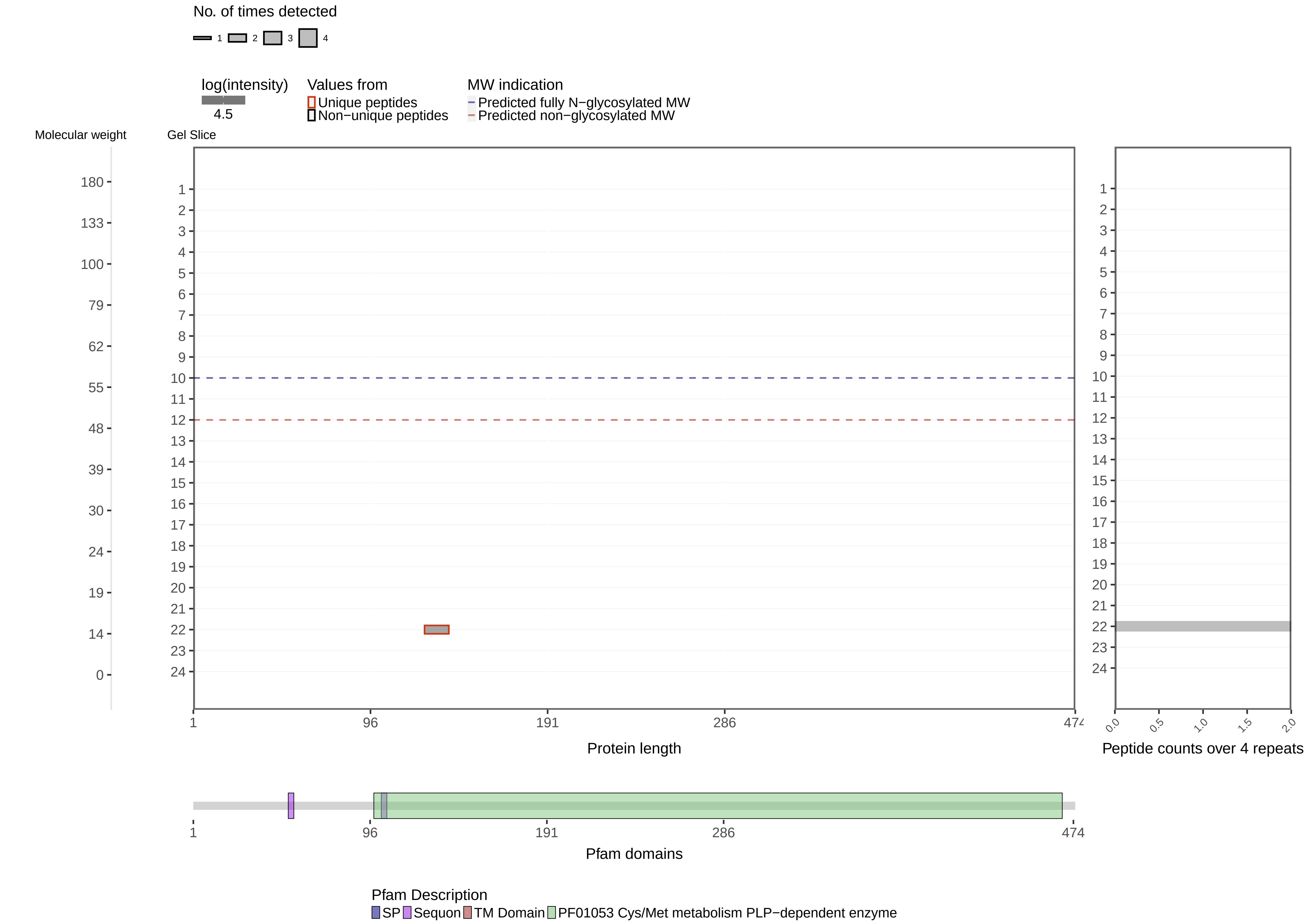


Protein sequence:

SAINPOTKLVWLESPTNPROOICDIRKIAKIAHAHGALVLVDNSIMSPVFSHPLELGADIVMHSATKFISGHSDLMAGVLAVRGESLAKEIYFLONSEGAGLAPFDCWLCLRGIKTMALRVEKOOENAOKIAEFLSSHPRVKKVNYAGLPDHPGRSLHFSOARGAGSVLSFLTGSLALSKHVAEATKYFSITVSFGSVKSLISLPCFMSHASIPVEVREARGLTEDLVRISVGIEDVNDLIDDLDYALRTGPA*

### NbL19g13430.1 protein_STRICTOSIDINE_SYNTHASE−LIKE_10−like [MW = 40.88 kDa]

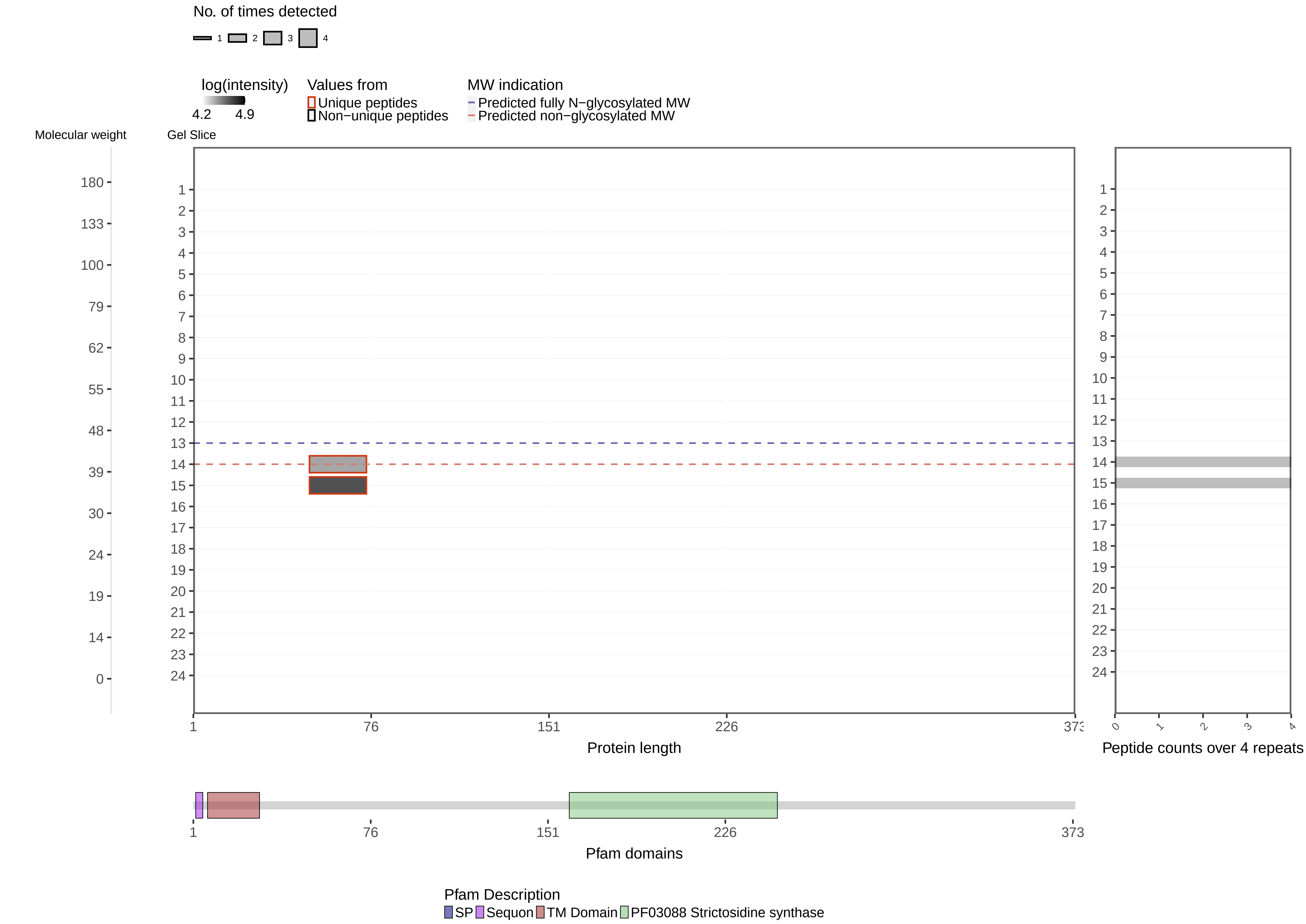


Protein sequence:

DTSTQFQRRQFIASVVIGDRTGRLMKYDKSTKAVTVLLRGLAFANGVALSKDKSFVLVAETSTGRIVKYWLKGPDAGKHATFADLPGYPDNIRRNSRGEFWVALHSKRSLLARLVTSNSWLGRTLLKLPFTFQQLHYLLVGGQPHATAIKLSEEGQVLEVLEDVESKTLKFLSEVEEKKGKLWIGSVLVPFVGVYELS*

### NbL19g13410.1 methylthioribose−1−phosphate_isomerase [MW = 39.6 kDa]

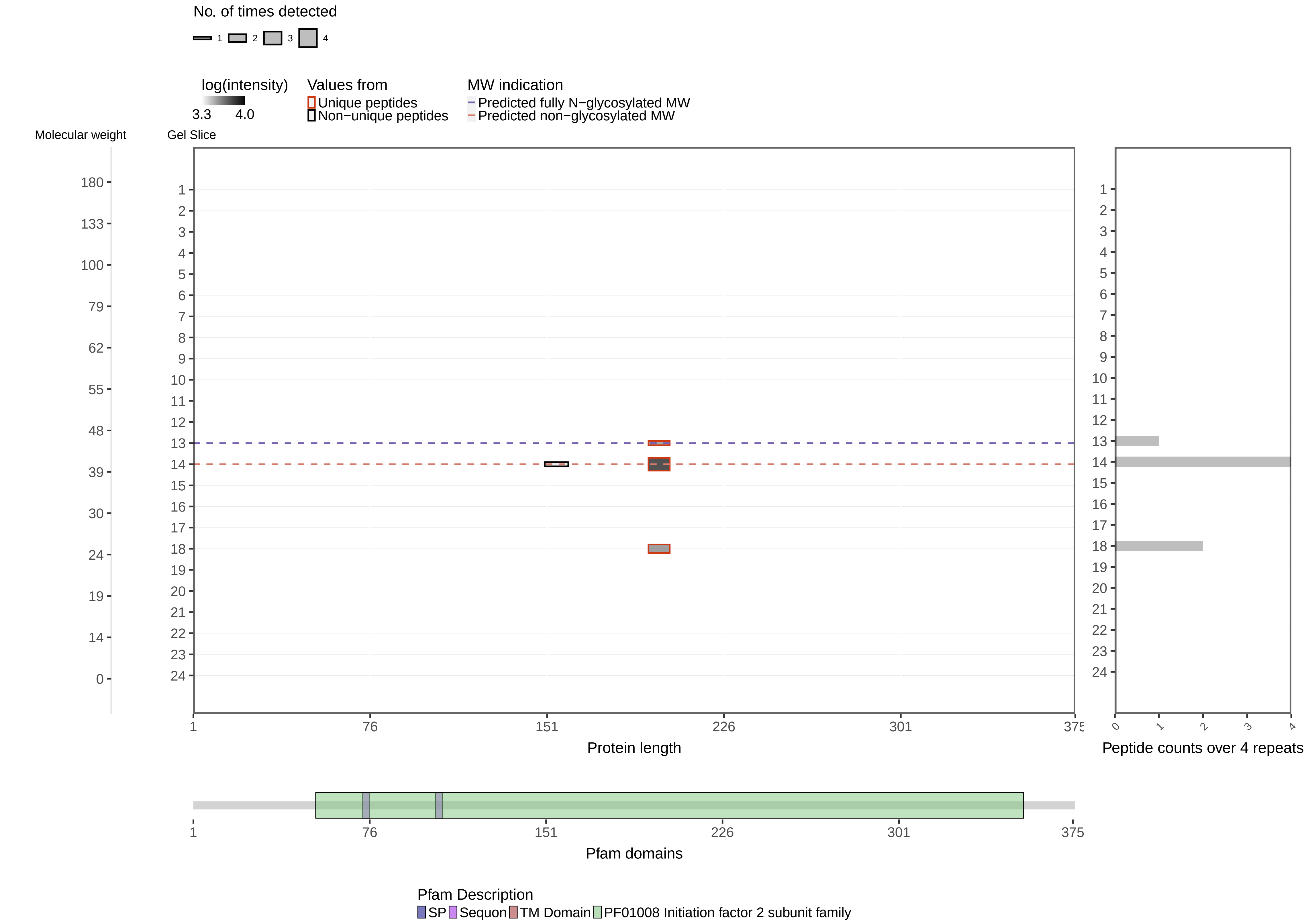


Protein sequence:

HCNTGSLATAGYGTALGVIRALYTDGVLERAYCTETRPFNOGSRLTAFELVHENIPATLIADSAAAALMNAGRVKAVIVGADRVAGNGDTANKIGTYNLAVSAMHHGIPFYVAAPLTSIDLSLSSGOEIVIEERSPKELLHTRGGLGEOVAASGISVWNPAFDVTPANLISGIITEKGVITKKGTDTFDIAEFVRKTSGNL*

### NbL19g13350.1 NAD−dependent_malic_enzyme_62_kDa_isoform_mitochondrial [MW = 70.05 kDa]

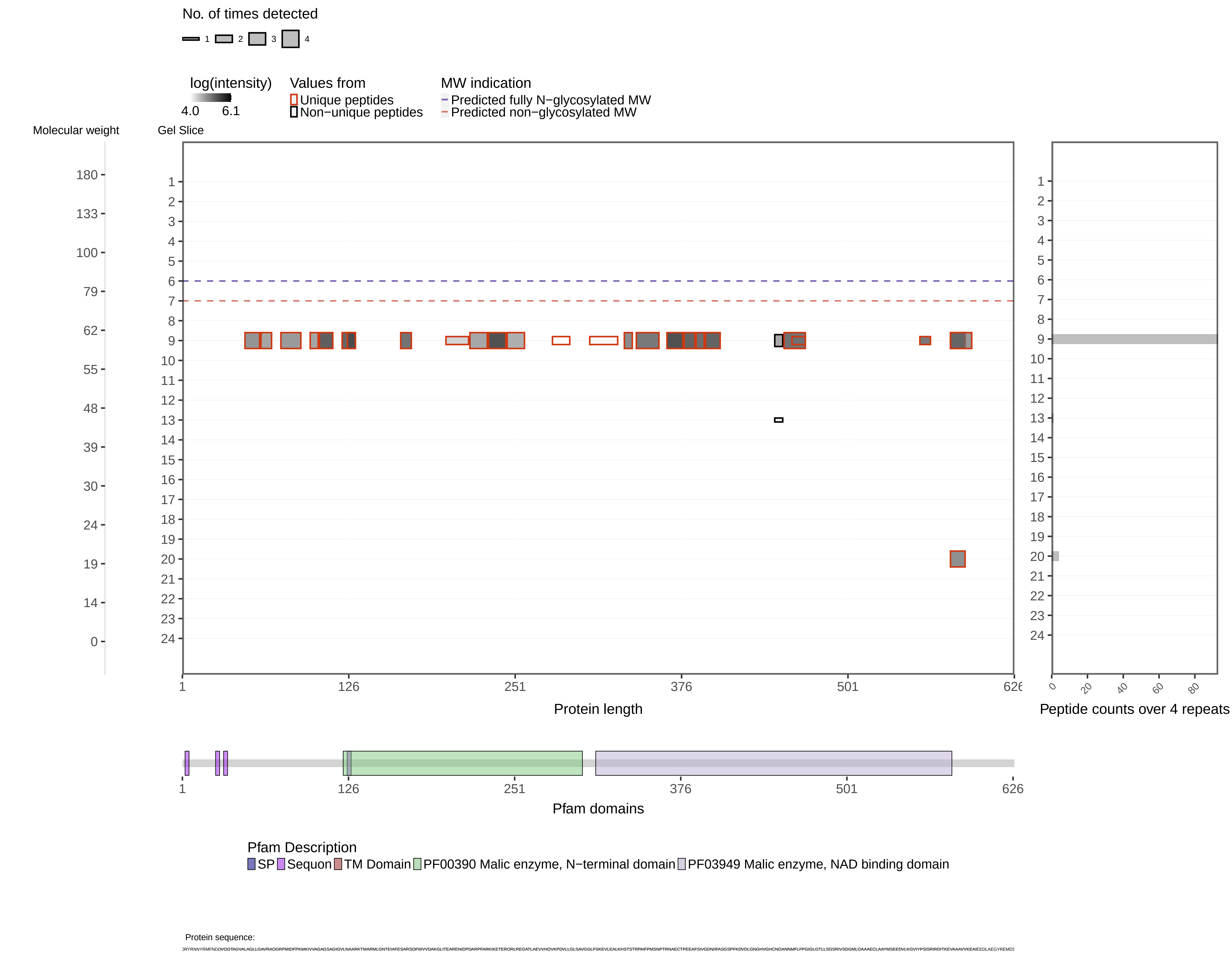

### NbL19g13340.1 haloacid_dehalogenase−like_hydrolase_domain−containing_protein_Sgpp [MW = 31.65 kDa]

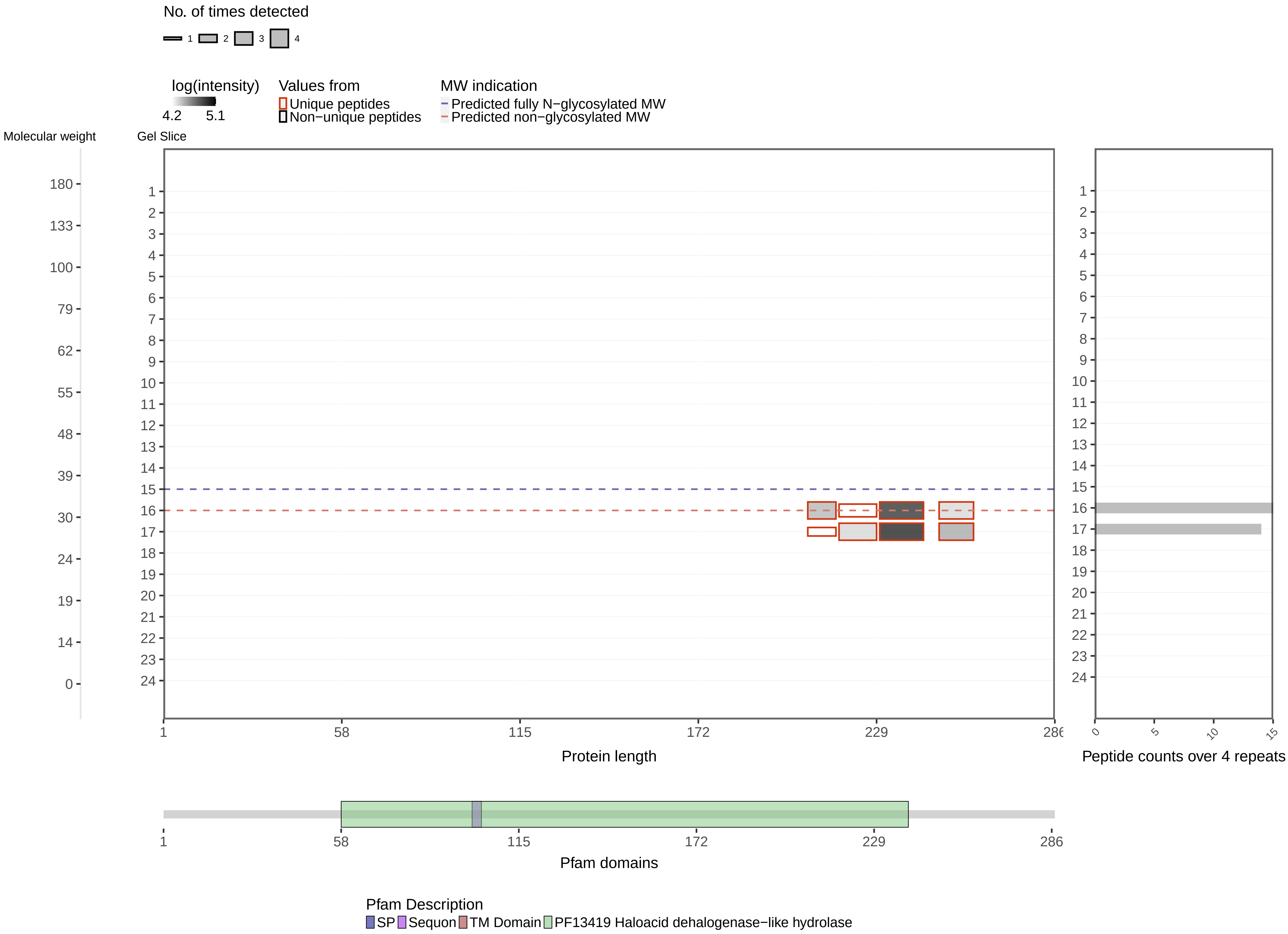


Protein sequence:

EDMFRKIASEQLKPLNGLEEVCKWIEDHCLKRAAVTNAPRANAELIISMLGLADFFELLVIGSECERAKPFPDPYLKALHELGVSPKHAFVFEDSISGIKAGVAAGMPVVGLGLRNPEKLLSEAGATFVIKDFKDSKLWTALEELETEVDSEKLKT*

NbL19g13270.1 eukaryotic_translation_initiation_factor_2A [MW = 57.89 kDa]

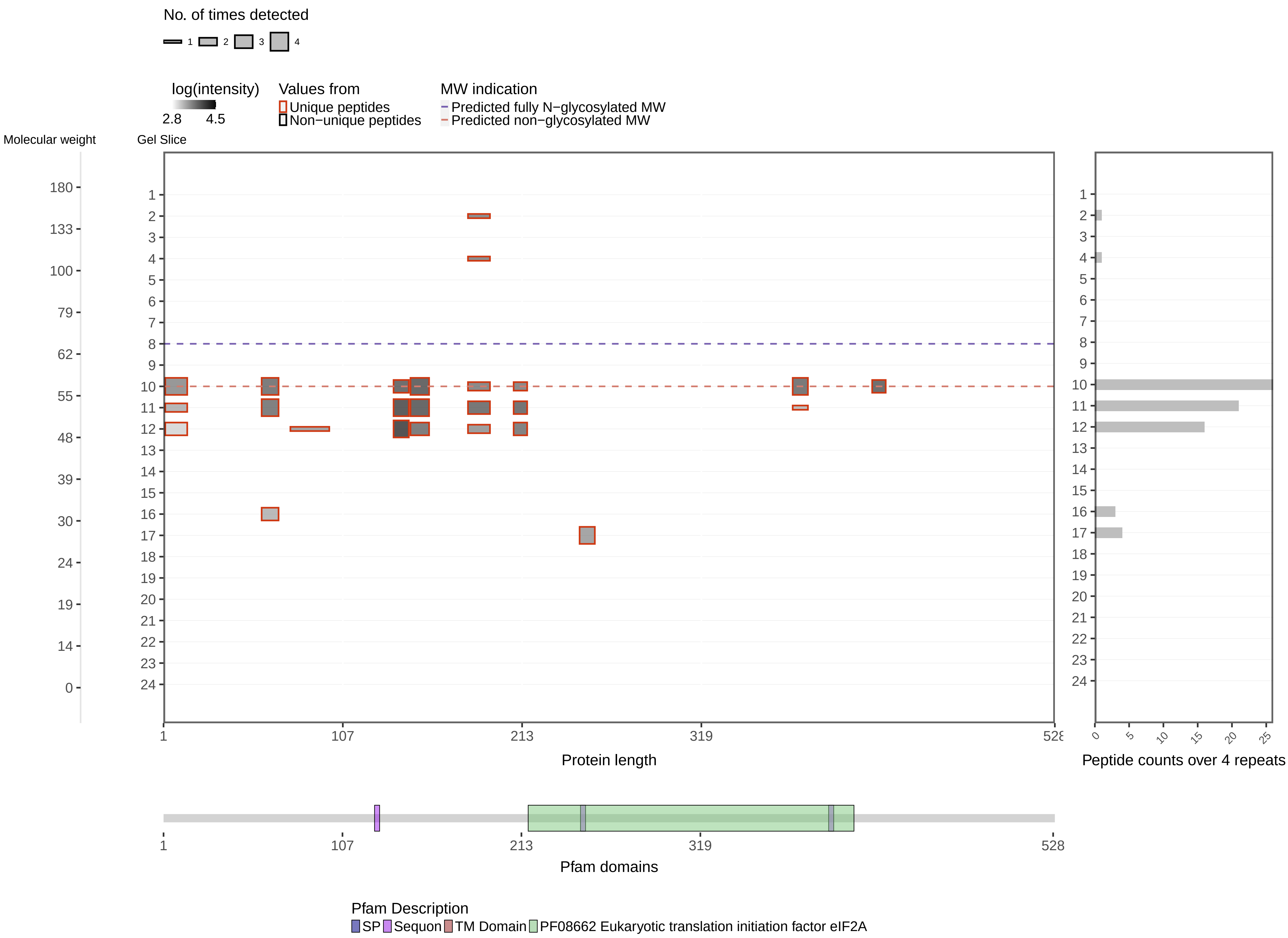


Protein sequence:

ESKLNYLTTDGTHEGLVPLRKDGPIHDVQWSYSGKEFAVVYGFMPAMATVFDKKCNPLLELGTGPYNTIRWNPLGNFICLAGFGNLPGDMAFWDYKEKKQLGTTRAELSVTSEWSPDGRYFMTATTAPRLQVDNGVKIFHHNGSLYFKKMFDRLYQVDWKPESVEKFGGIEDLVKAVGSVKMDEAKAPGKGSKATQAVPKATQAIPKATPANPPAAKPAAYRPPHAKVAAAVQEELFGGSSSGELSKNALKNKKKREKQRERKQTEGASGTGGS*

### NbL19g13190.1 2−isopropylmalate_synthase_A−like [MW = 71.24 kDa]

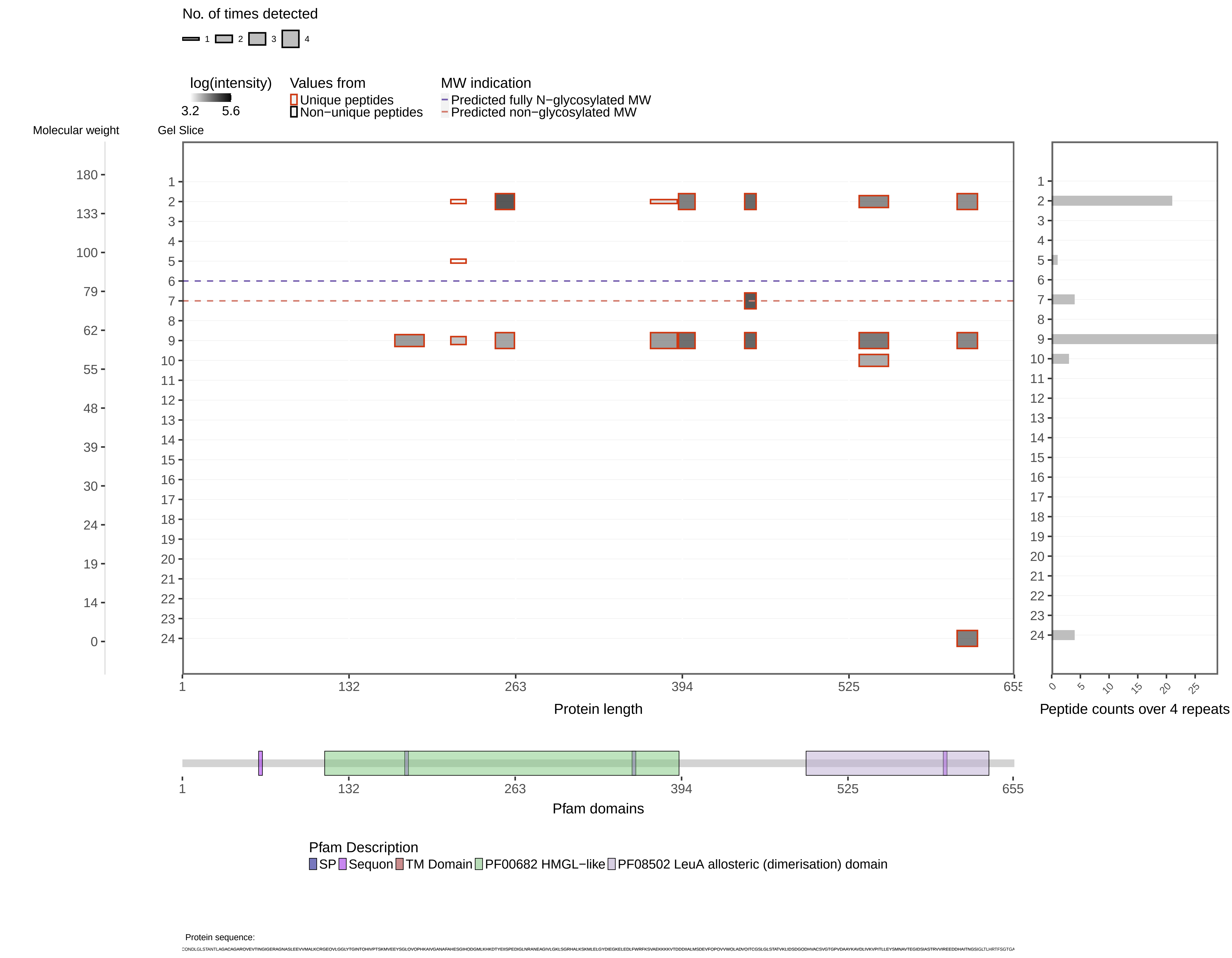

### NbL19g13030.1 beta−galactosidase [MW = 127.97 kDa]

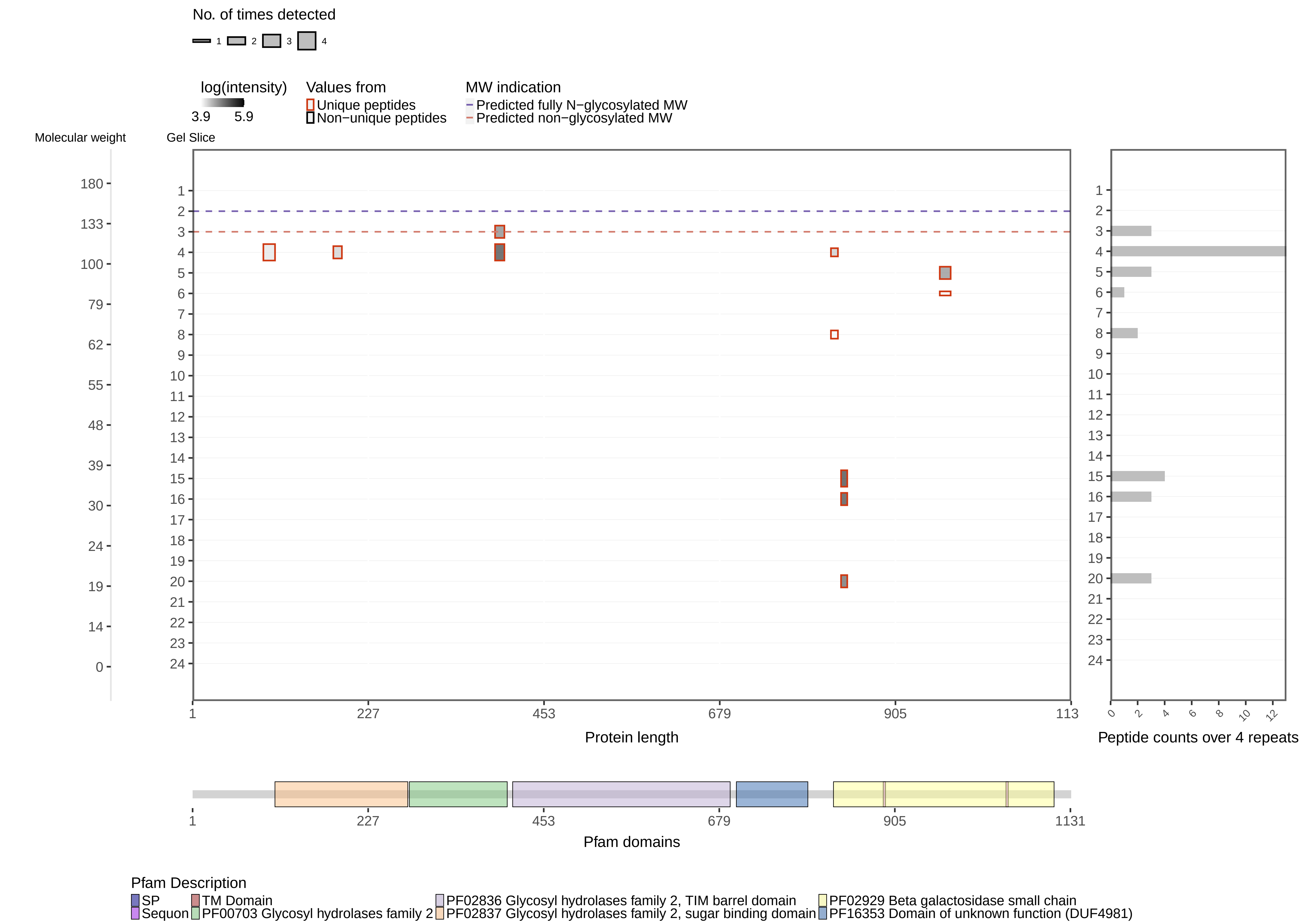


Protein sequence:

### NbL19g13020.1 60S_ribosomal_protein_L9−1−like [MW = 22.65 kDa]

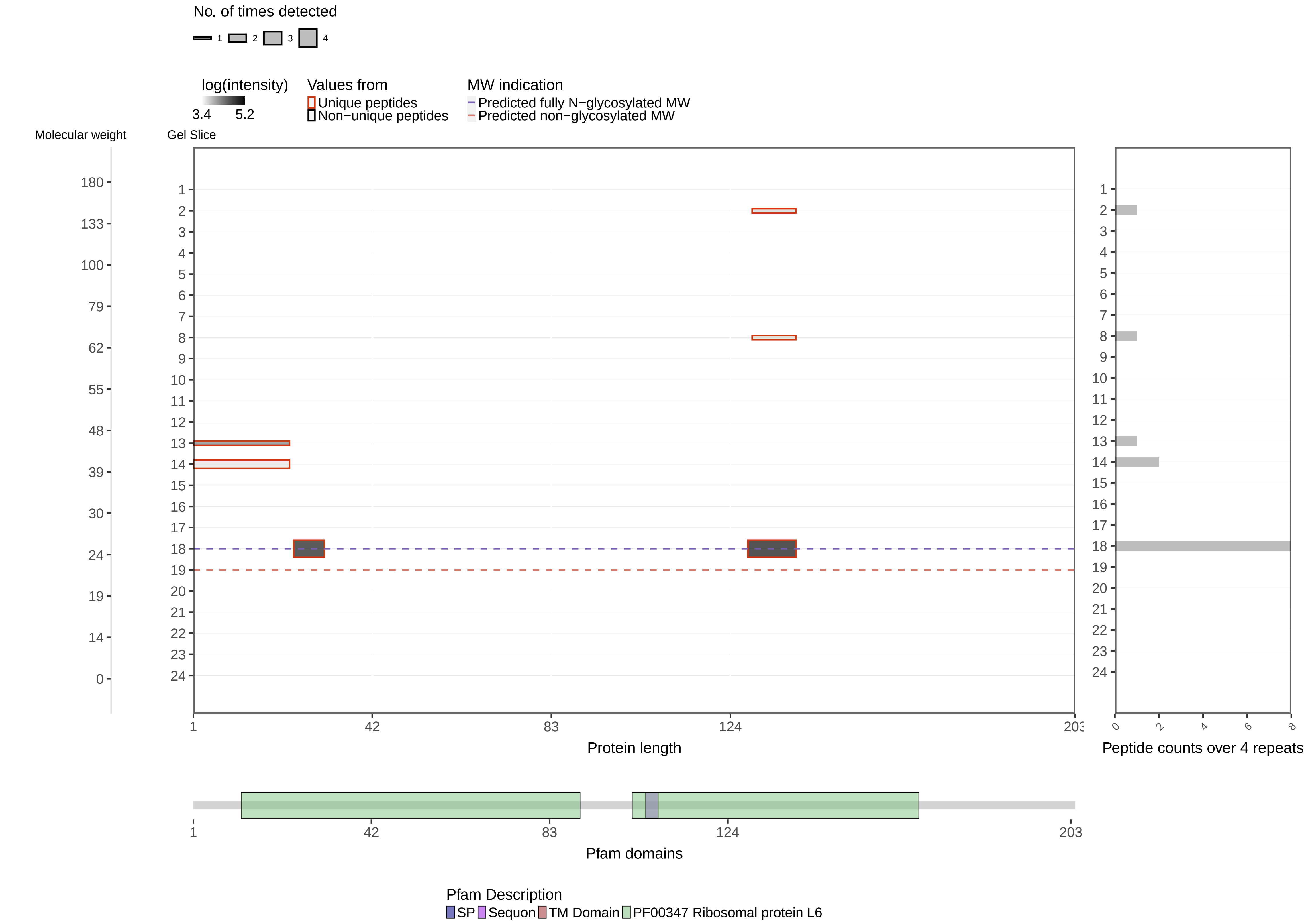


Protein sequence:

KGYRYKMRFVYAHFPINASITGGNKAIEIRNFLGEKKVRKVDMLEGVTVVRSEKVKDELVLDGNDIELVSRSAALINQVKLLFYSSGQCISALYSTFSSSALWISSPFFDRNAM*

### NbL19g12950.1 hydroxymethylglutaryl−CoA_synthase−like [MW = 51.11 kDa]

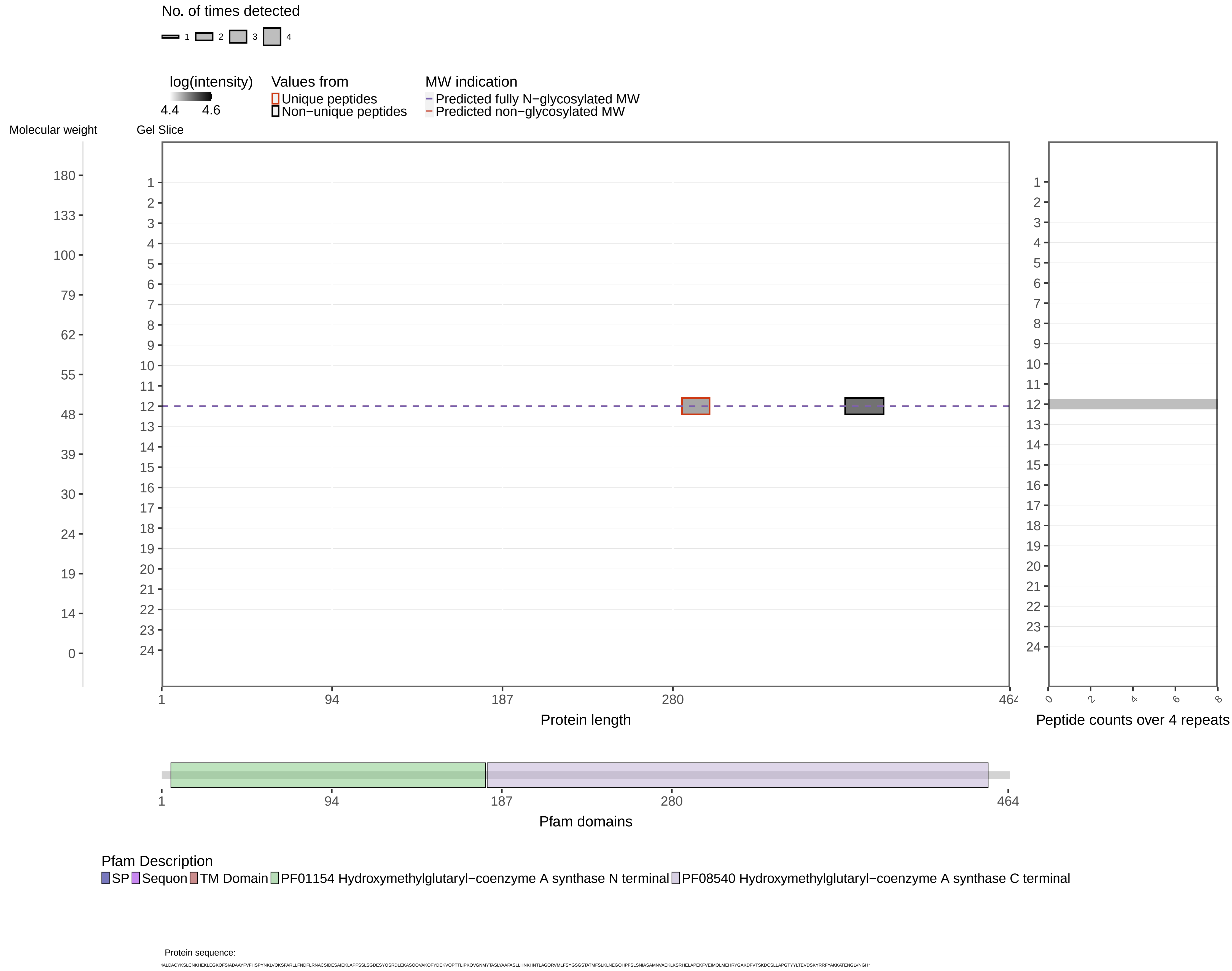

NbL19g12910.1 actin−104 [MW = 41.75 kDa]

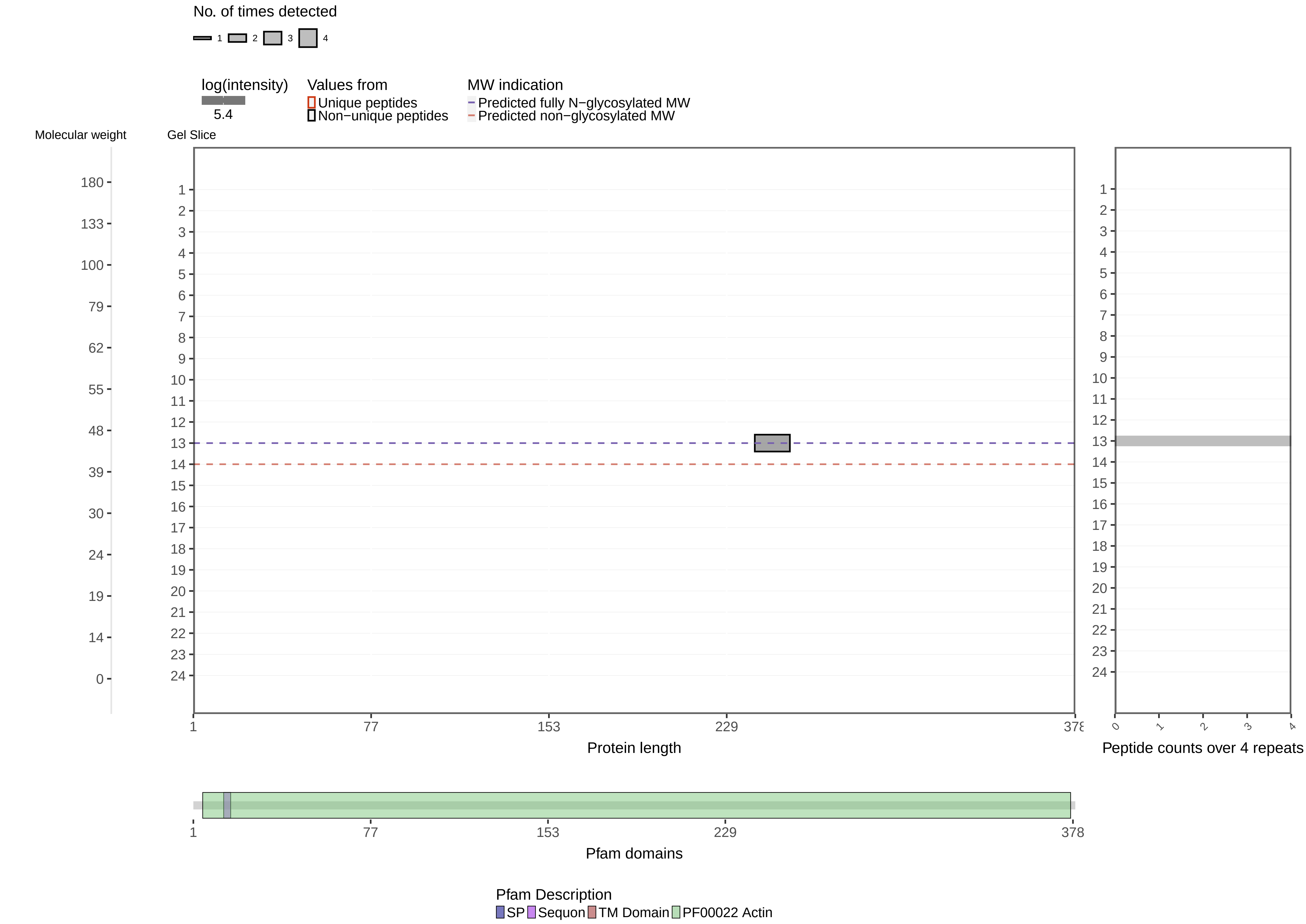


Protein sequence:

PHAILRLDLAGRDLTDYLMKILTERGYMFTTSAEREIVRDVKEKLSYVALDFEQELETAKSSSSVEKSYELPDGQIITIGAERFRCPEVLFQPSLIGMEAAGIHETTYNSIMKCDVDIRKDLYGNIVLSGGSTMFPGIADRMSKEITALAPSSMKIKVVAPPERKYSVWIGGSILASLSTFQQMWISKAEYDESGPAIVHRKCF*

NbL19g12890.1 ATP−dependent_Clp_protease_ATP−binding_subunit_ClpA_homolog_CD4B_chloroplastic [MW = 102.3 kDa]

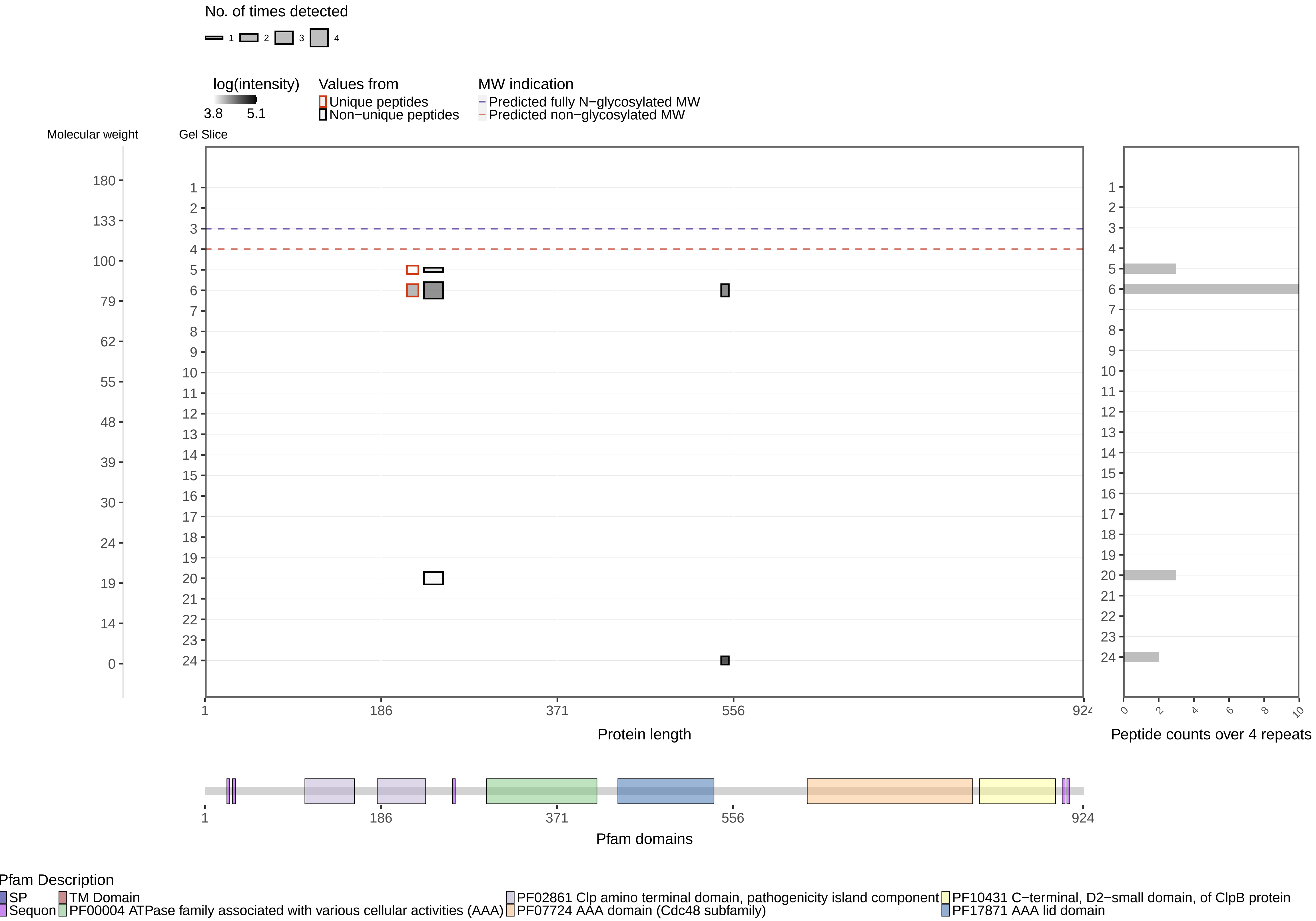


Protein sequence:

YEIHHKLRYTDEALEAAAQLSYQYISDRFLPDKAIDLIDEAGSRVRLRHAQLPEEAKELEKELRQITKEKNEAVRGQDFEKAGELRDREMDLKAQITALIDKNKEMNKAESEAGDTGPLVTEADIQHIVSSWTGIPVEKVSTDESDRLLKMEETLHTRIIGQDEAVKAISRAIRRARVGLKNPNRPIASFIFSGPTGVGKSELAKSLAAYYFGSEEAMIRLDMSEFMERHTVSKLIGSPPGYVGYTEGGQLTEAVRRRPYTVVLFDEIEKAHPDVFNMMLQILEDGRLTDSKGRTVD

NbL19g12830.1 guanine_nucleotide−binding_protein_subunit_beta−like_protein [MW = 35.91 kDa]

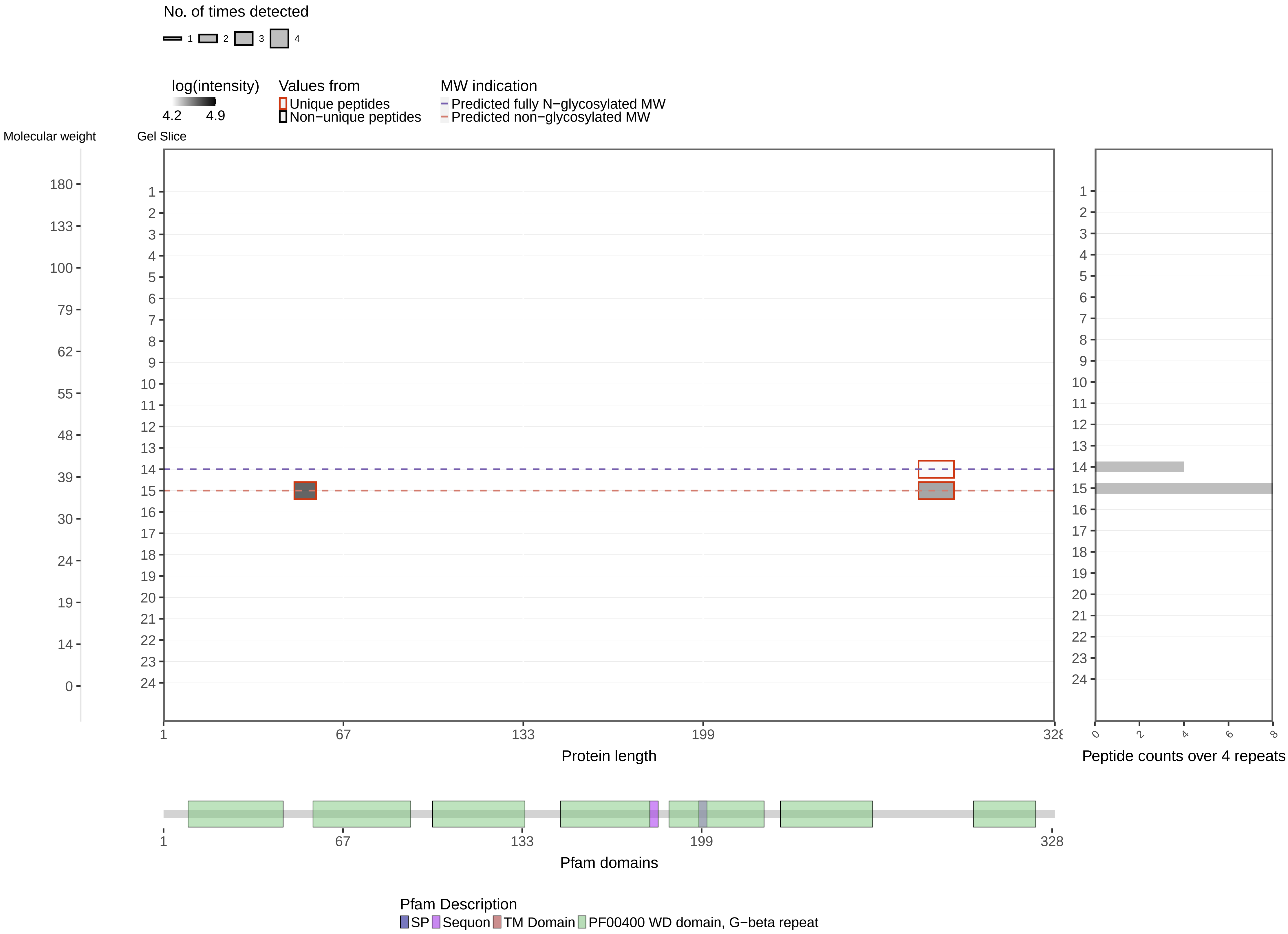


Protein sequence:

SDWVSCVRFSPNNIQPTIVSGSWDRTVKIWNLTNCKLRATLAGHTGYVNTTAVSPDGSLCASGGKDGVILLWDLAEGKKLYSLESGSIIHSLCFSPNRYWLCAATESSIKIWDLESKTIVDDLKVDLKQETEMSAEGTASGKNKVVIYCTSLGWSADGSTLFSGYTDGLIRVWGIGRY*

### NbL19g12590.1 actin−related_protein_4 [MW = 48.2 kDa]

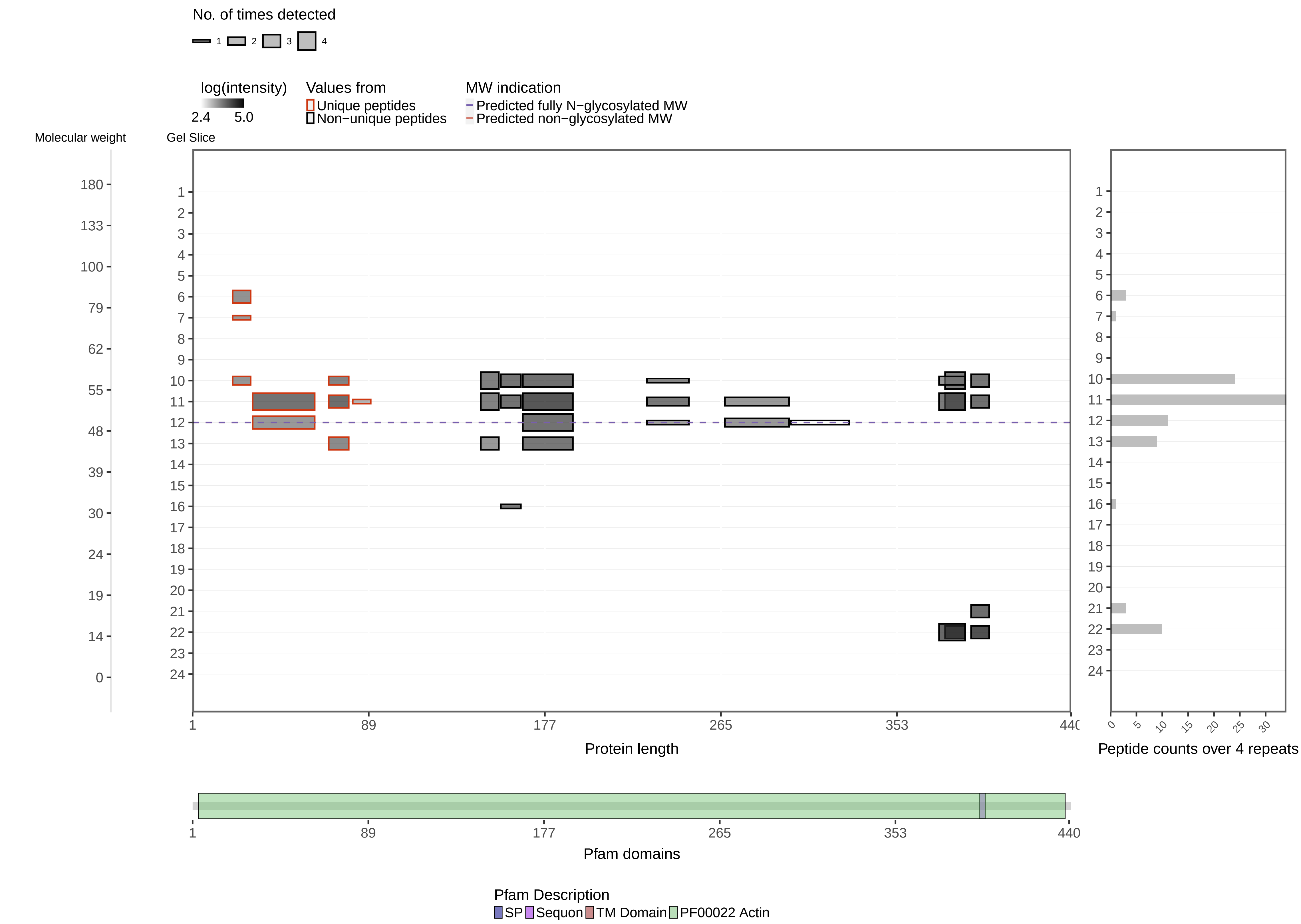


Protein sequence:

TDCLIKSLEQKGITIKPRYSFKRKEIRPGEFQTVDLDFPDTTESYKLYCQRAIASDIKECVSRAPDTPYDDSSYSNIPTTSYELPDGQTIEVGADRFKIPDIIFNPSLVQTIPGMESFAETAASLRGLPQMVIESINKCDVDIRRELFSSILCLIQIAGGTASMQQLKERLEKDLLEESPQAARVKVLASGNATERRFSVWIGGSILASLGSFQQMWFSKSEFEEHGASYVQRKCP*

### NbL19g12520.1 glucan_endo−13−beta−glucosidase_11−like [MW − SP = 42.58 kDa]

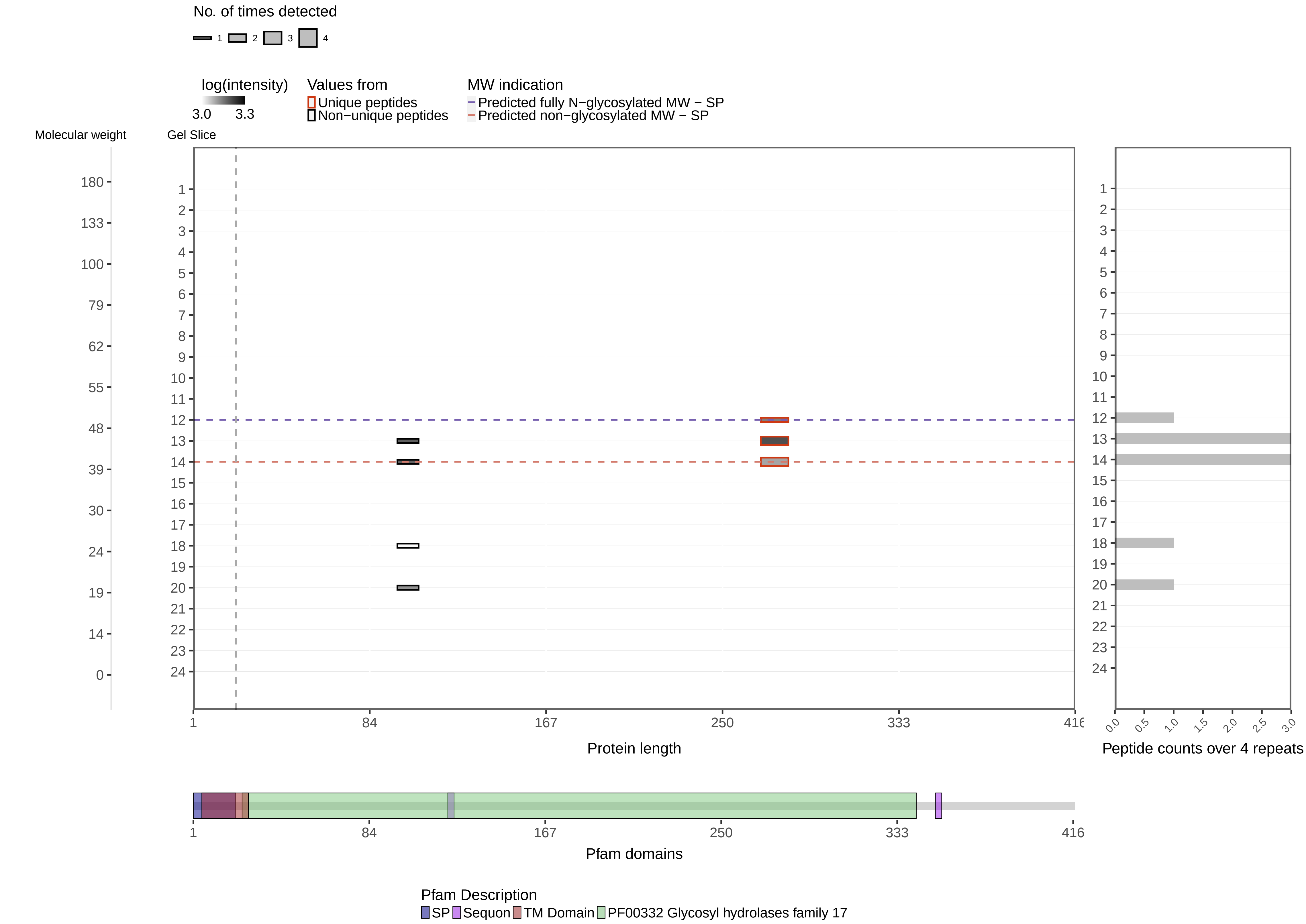


Protein sequence:

YPYFAYKANPKQVPLDYVLFQPNPGNVDPVTNLHYDNMLYAQIDAVHSALALIGYKNVCVQISETGWPSKGDGEEVGATPDNARKYNCNLIKLVSQNKGTPLRPNTNLNIYVFALFNENLKPGPTSERNYGLFKPDGSPSYPLGFAGINAVGTNSSSSRSPAAGTGSSTSTSSSTPSWNSPDGYLSITSDSGRIQFCWKSLVLRLHIAGLISLLFSQL*

### NbL19g12500.1 V−type_proton_ATPase_catalytic_subunit_A−like [MW = 68.67 kDa]

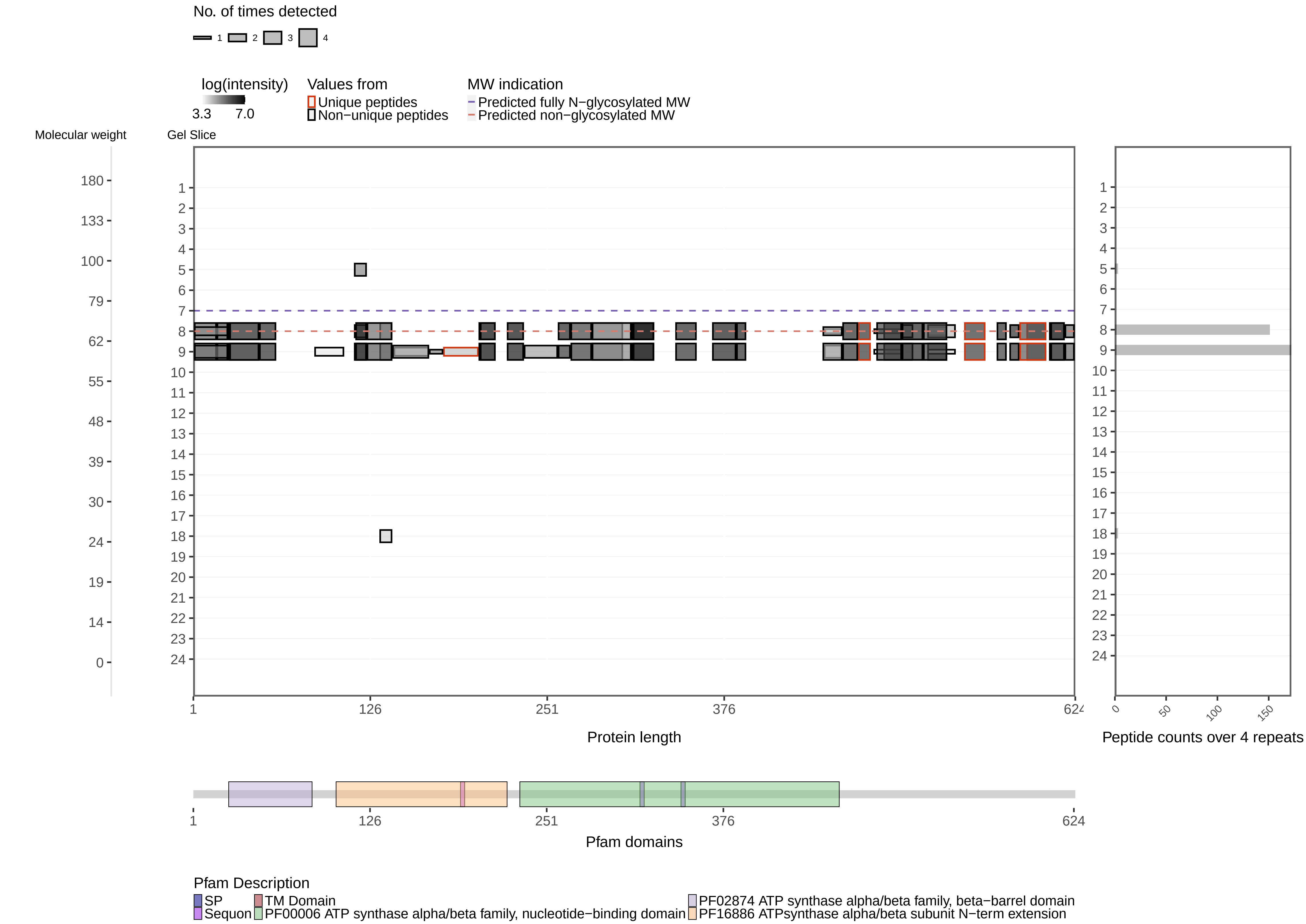


Protein sequence:

TLPDGREESVMKRTTLVANTSNMPVAAREASIYTGITIAEYFRDMGYNVSMMADSTSRWAEALREISGRLAEMPADSGYPAYLAARLASFYERAGKVKCLGGPERTGSVTIVGAVSPPGGDFSDPVTSATLGIVQVFWGLDKKLAQRKHFPSVNWLISYSKYSGALESFYEKFDADFIDIRTKAREVLQREDDLNEIVQLVGKDALAETDKITLETAKLLREDYLAQNAFTPYDKFCPFYKSVWMLRNIIHFYNLANQAVERGAGMDGQKITYTLIKHRLGDLFYRLVSQKFEDI

### NbL19g12370.1 UDP−glucose_4−epimerase_GEPI48−like [MW = 38.37 kDa]

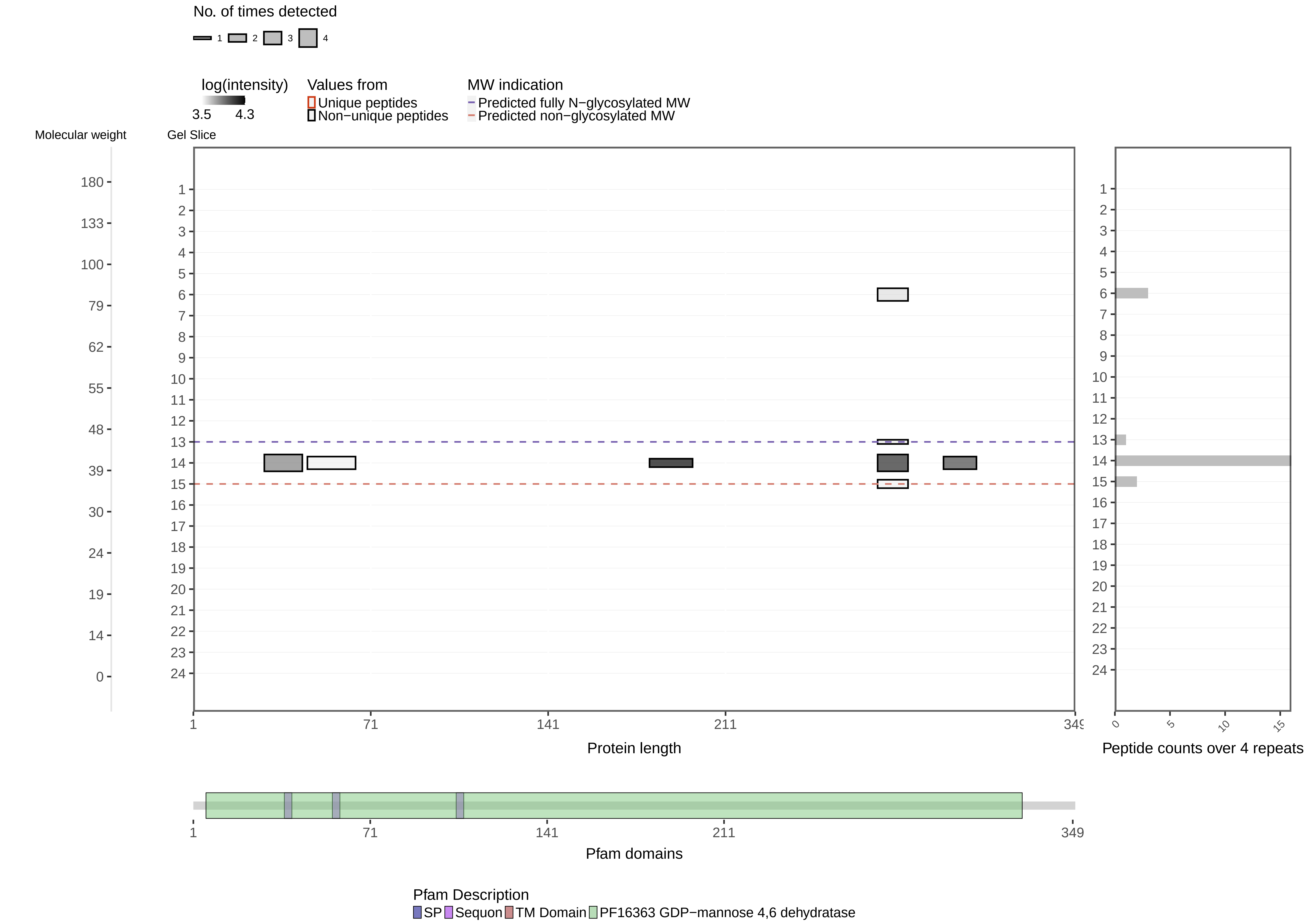


Protein sequence:

CRDVYHSDHEWKVILLRYFNPIGAHPSGYIGDDPRGIPNNLMPFVOOVAVGRRPALTVYGTDYPTKDGTAVRDYIHVVDLADGHIAALOKMFDPSTGCEVYNLGTGKGSSVLEVVSAFEKASGKKIPLVMSSRRPGDAEIVYAATEKAERELNWRAKYGIDEMCRDOWNWASKNPYGYEGSEESN*

### NbL19g12240.1 6−phosphogluconate_dehydrogenase_decarboxylating_2_chloroplastic [MW = 54.27 kDa]

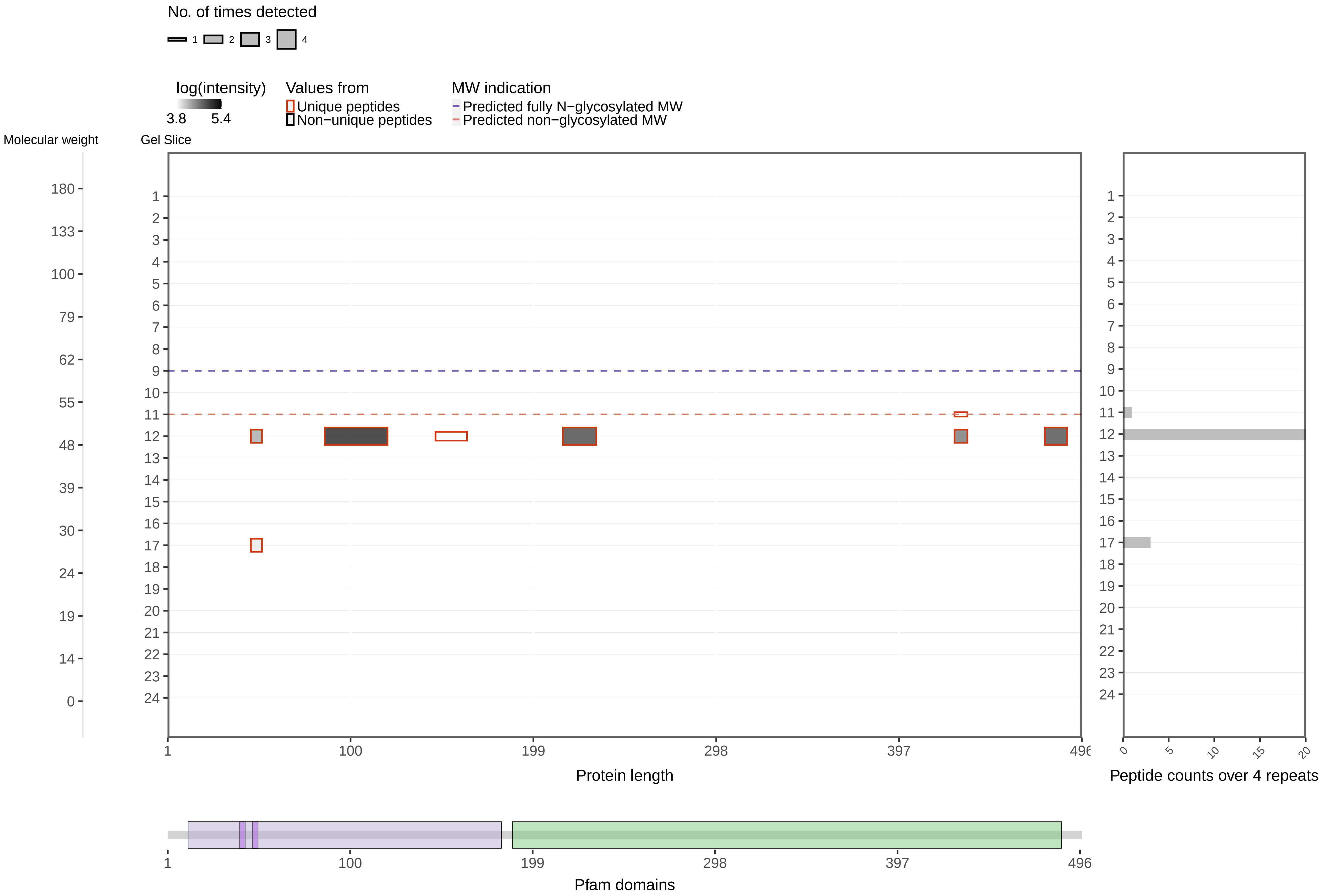


Pfam Description

SP Sequon TM Domain PF00393 6−phosphogluconate dehydrogenase, C−terminal domain PF03446 NAD binding domain of 6−phosphogluconate dehydrogenase

Protein sequence:

LESFLVEITADIFNVKDEKTGSGELVDKILDKTGMKGTGKWTVQQAAELSIAAPTIAASLDGRYMSGLKEEREDAARIFKKEGLKEEMNFDVNGASVDKKRLIDDVRQALYASKICSYAQGMNLLRAKSAEKGWNLNLGEMARIWKGGCIIRAVFLDRIKKAYQRNPGLANLMVDPEFAREMVQRQAAWRRVVGLAIQRGISVPGMSASLQYFDTYRRSRLPANLVQAQRDYFGAHTYERVDMPGAYHTEWSKLARKARV*

NbL19g12230.1 uncharacterized_LOC109239217 [MW = 39.96 kDa]

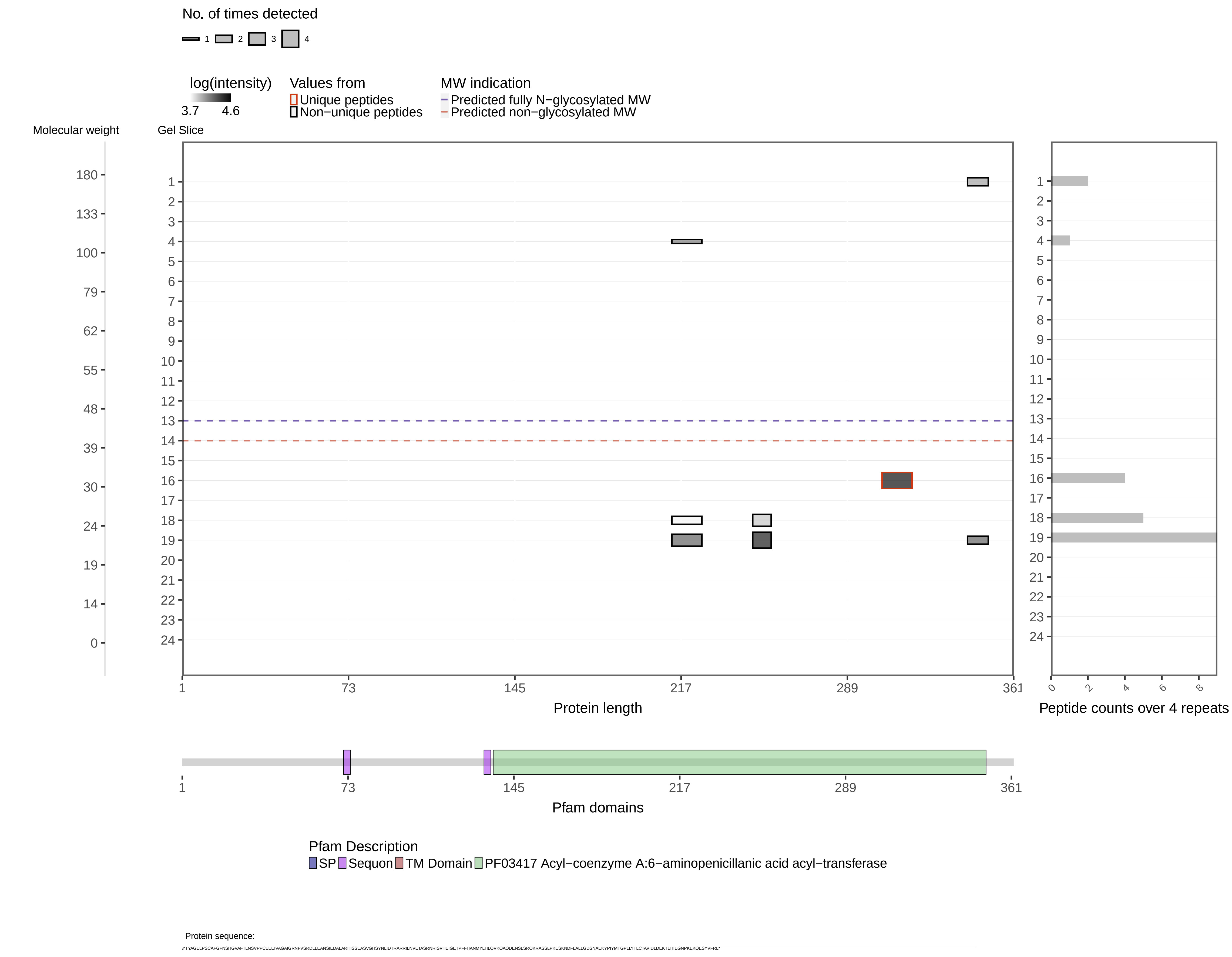

NbL19g12210.1 uncharacterized_LOC109238591 [MW = 47.18 kDa]

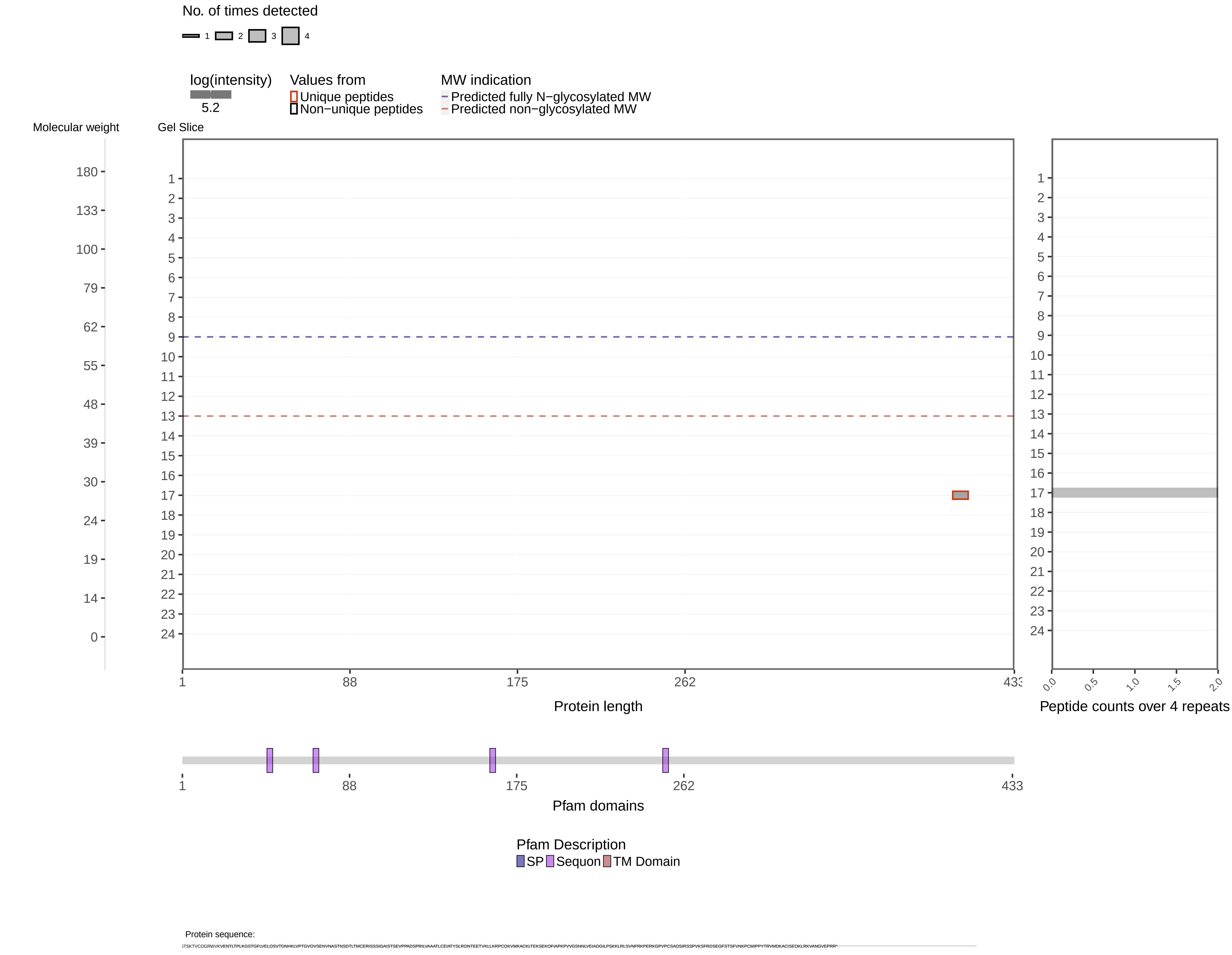

### NbL19g12170.1 glucan_endo−13−beta−glucosidase_1 [MW − SP = 53.34 kDa]

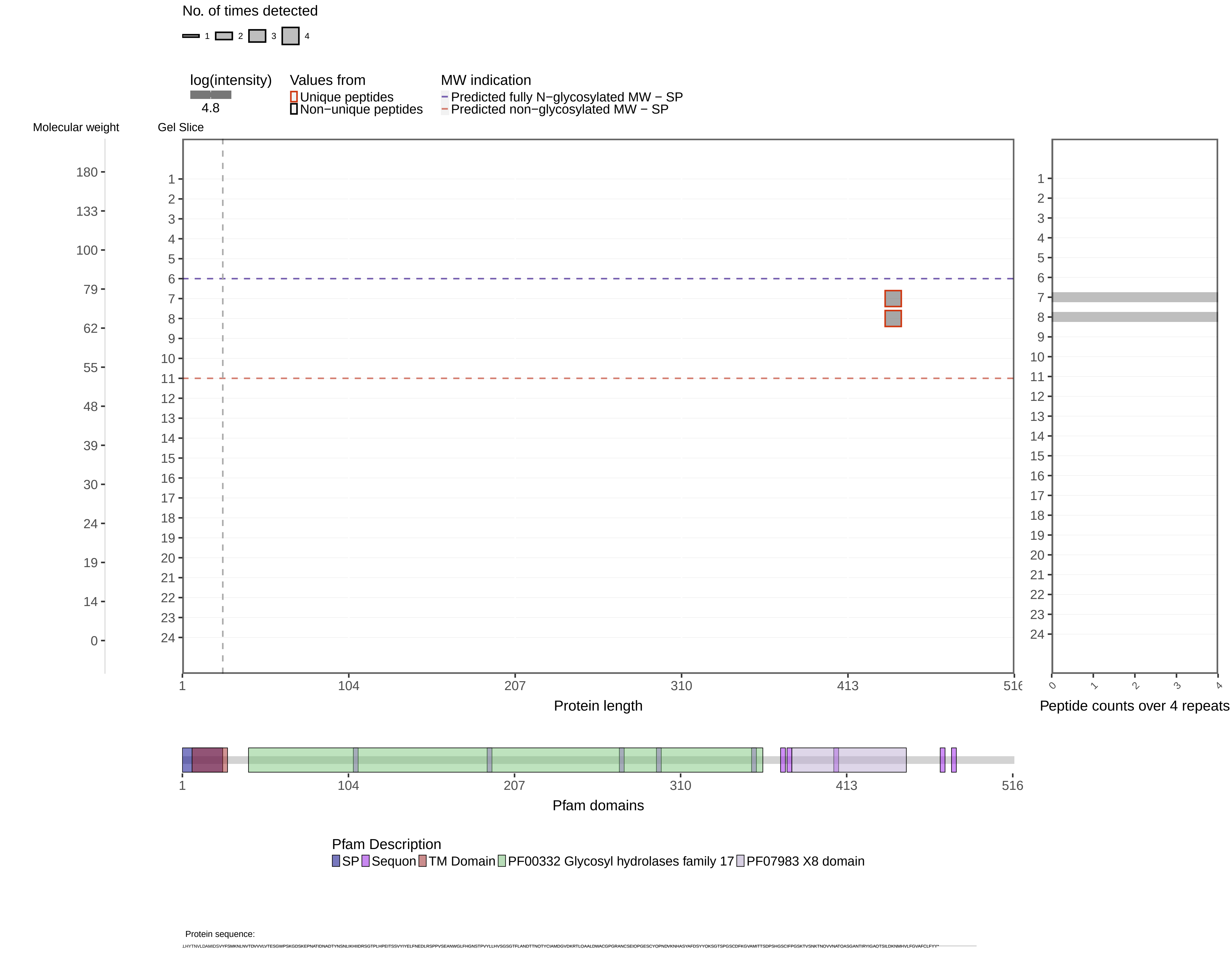

### NbL19g12120.1 aminomethyltransferase_mitochondrial [MW = 44.12 kDa]

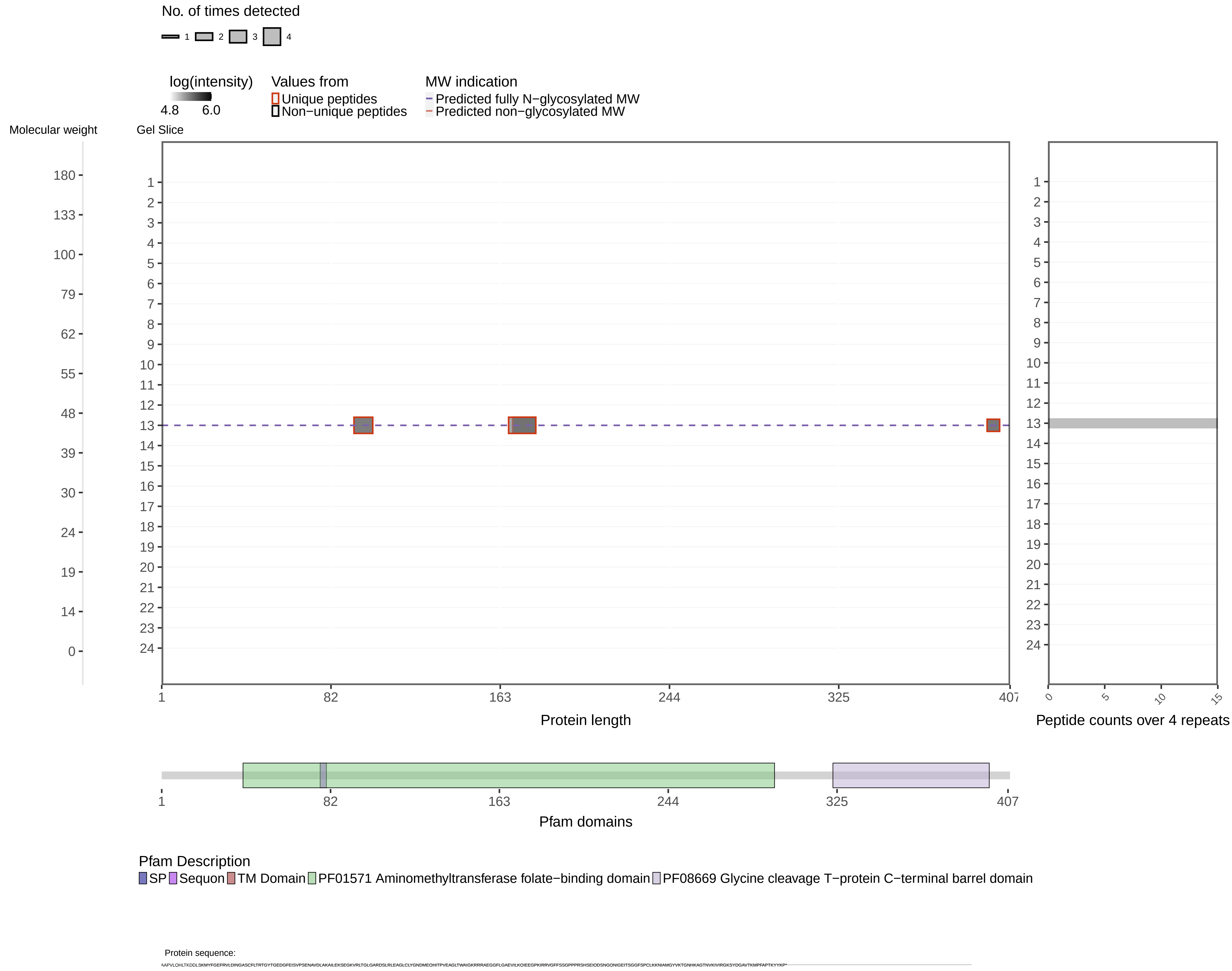

### NbL19g12110.1 deoxyhypusine_synthase−like [MW = 27.25 kDa]

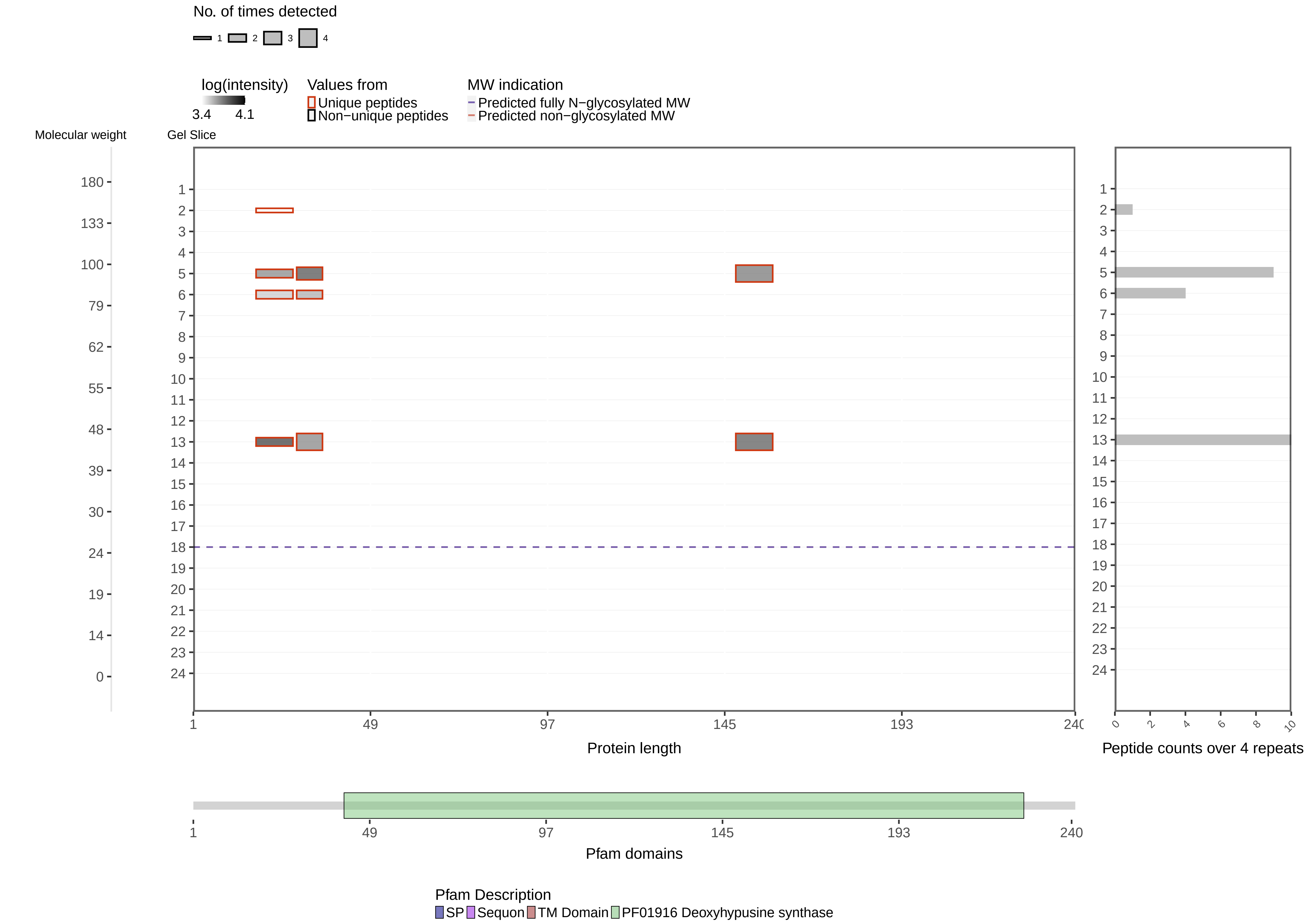


Protein sequence:

NLVSSGVRDTIRYLVOHRMVDVVVTTAGGIEEDLIKCLAPTYKGDFSLPGAVLRSKGLNRIGNLLVPNDNYCKFENWIIPIFDOMYEEOIKEKVLWTPSKVIARLGKEINDETSYLYWAYKVNFOSTSLSLLFDKI

NbL19g12040.1 proline_synthase_co−transcribed_bacterial_homolog_protein [MW = 30.44 kDa]

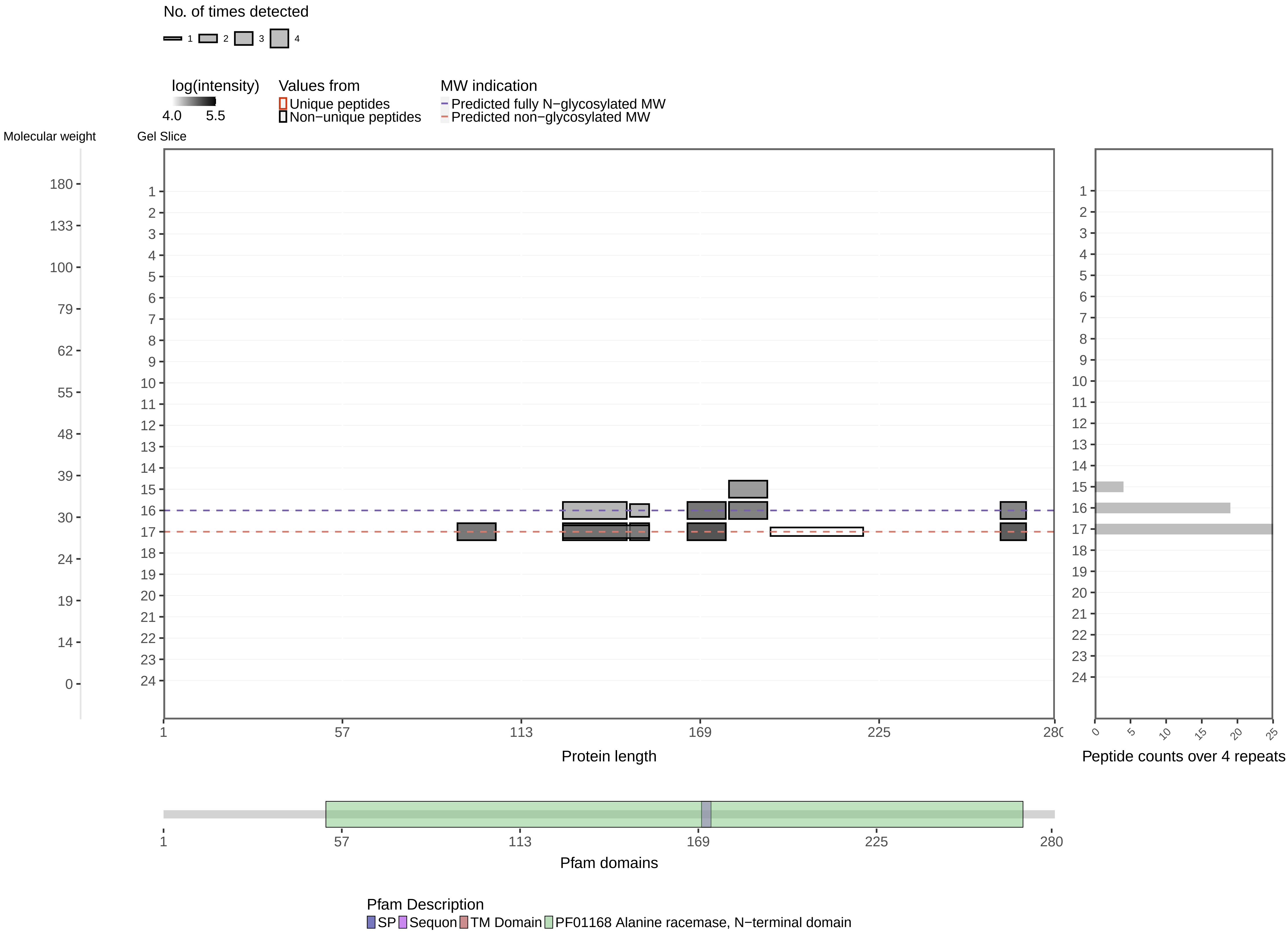


Protein sequence:

‹PLLTGVPNLAMVETVDDEKIANQLNRVAGNIGRKPLKVFIQVNTSGEETKSGVEPDGCLELVKHVTSNCPNLEFCGLMTIGMPDYTSTPENFKTLVRCRSEVCKALEIPEGQCELSMGMSGDFELAVEMGSTNVRVGSTIFGAREYPKKQTN*

### NbL19g11990.1 leucine−−tRNA_ligase_chloroplastic/mitochondrial−like [MW = 113.85 kDa]

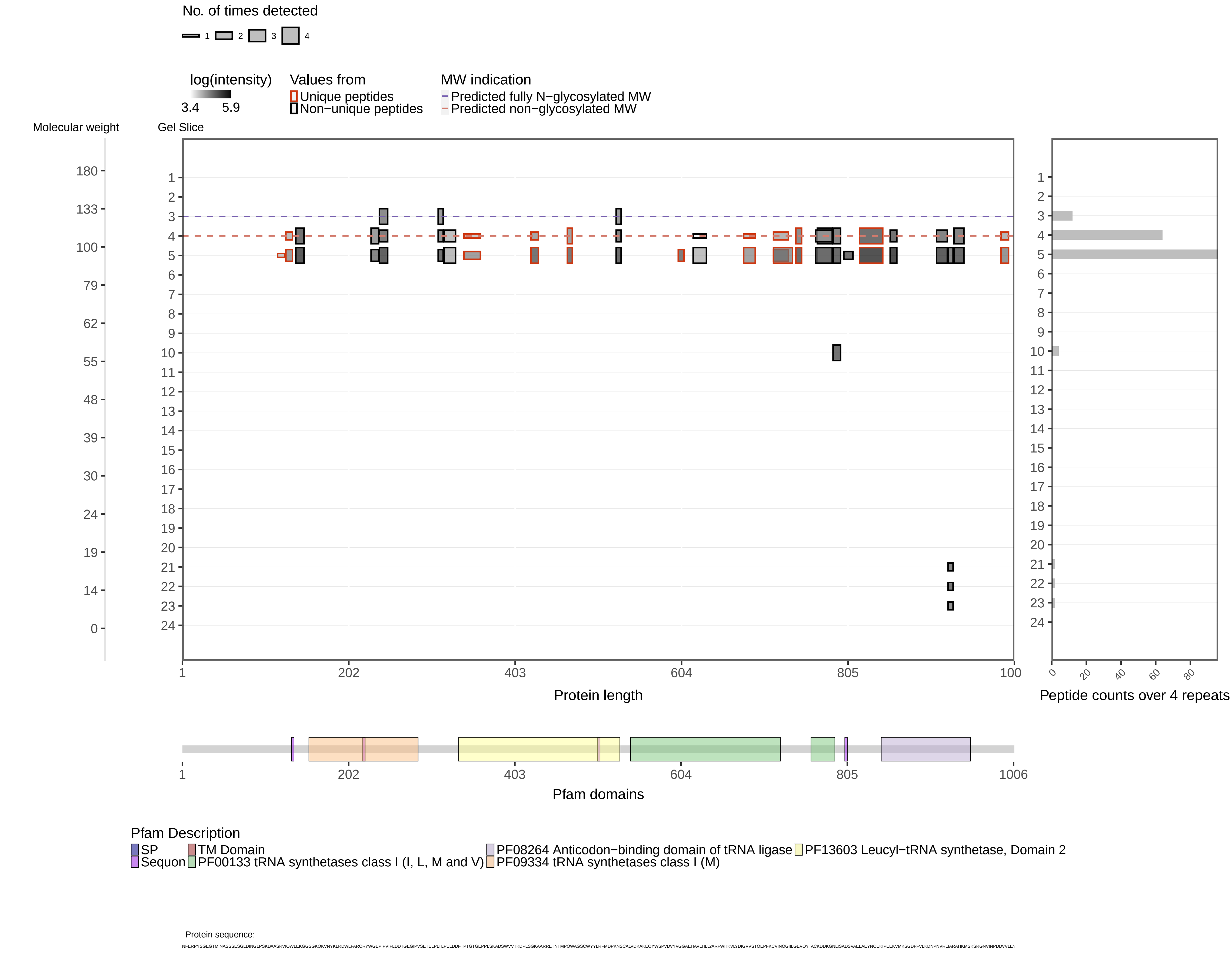

NbL19g11940.1 light−induced_protein_chloroplastic [MW = 34.93 kDa]

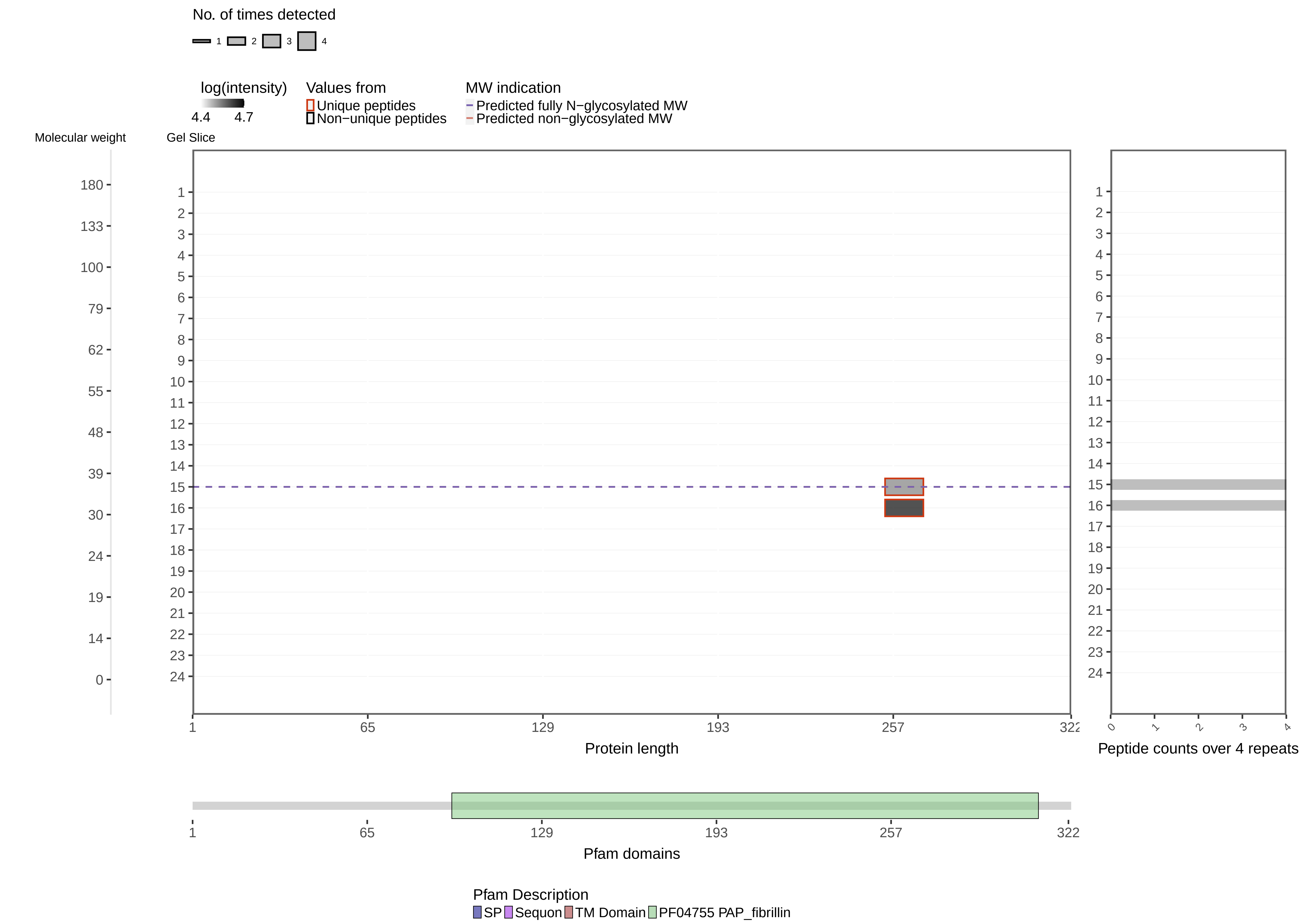


Protein sequence:

GKWILAYTSFSGLFPLLSRGTLPLVRVEEISQTIDSETFAVQNCVVFAGPLATTSITTNAKFEVRSPKRVQIKFDEGIIGTPQLTDSIELPENVEFLGQKIDLSPFKGLINSVQYTASSVAKSISSQPPIKIPISNSNAQSWLLTTYLDDELRISRGDGGSVFVLIKEGSPLLKP*

### NbL19g11930.1 pyrophosphate−−fructose_6−phosphate_1−phosphotransferase_subunit_beta [MW = 65.17 kDa]

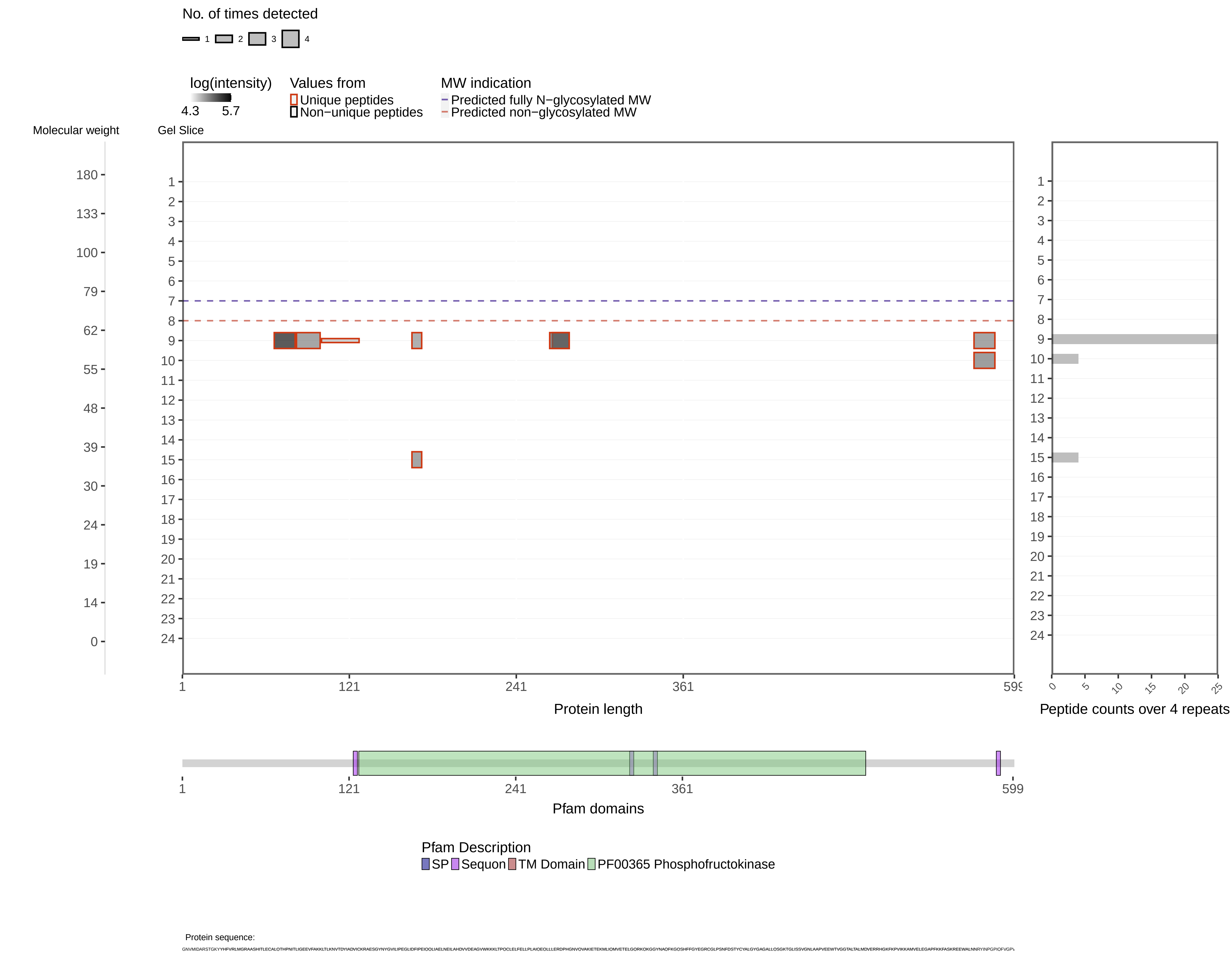

### NbL19g11880.1 UBA_and_UBX_domain−containing_protein_At4g15410−like [MW = 32.69 kDa]

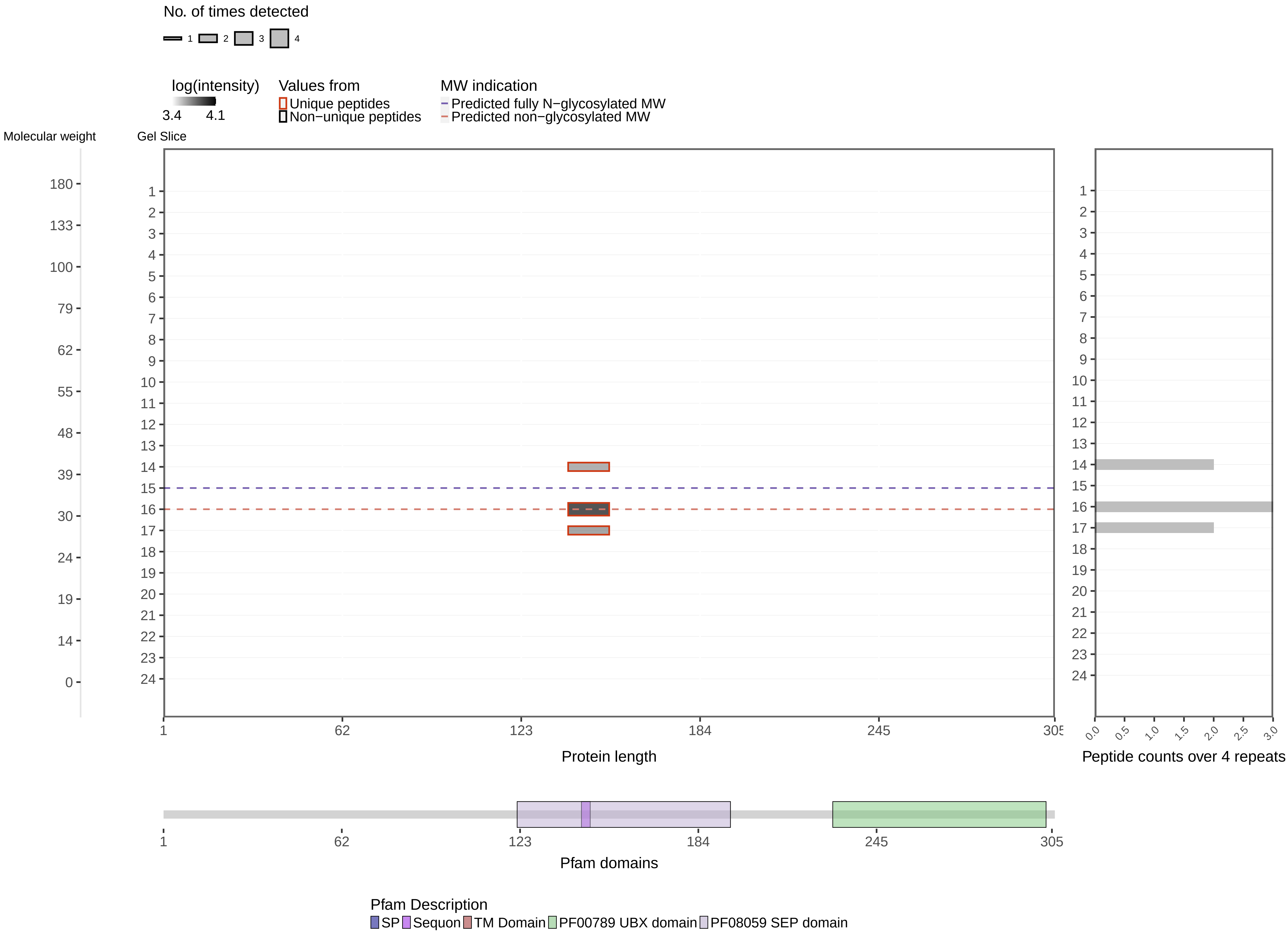


Protein sequence:

RLDDPENASFLESITKSECPRELAPADTRSOVHCNLIRRDENCPEPEKROVPFOGVGRTLGSSSTPATSEPSASVPTSTASAPTSILDDSLPSTSLOVRLADGTRMVAHFNFOOTVGDIRAFIDASRPSNTRAYLLOTVGFPPRILSDPSOTIEOAGLANSVVIOKF*

NbL19g11820.1 1−aminocyclopropane−1−carboxylate_oxidase−like [MW = 36.5 kDa]

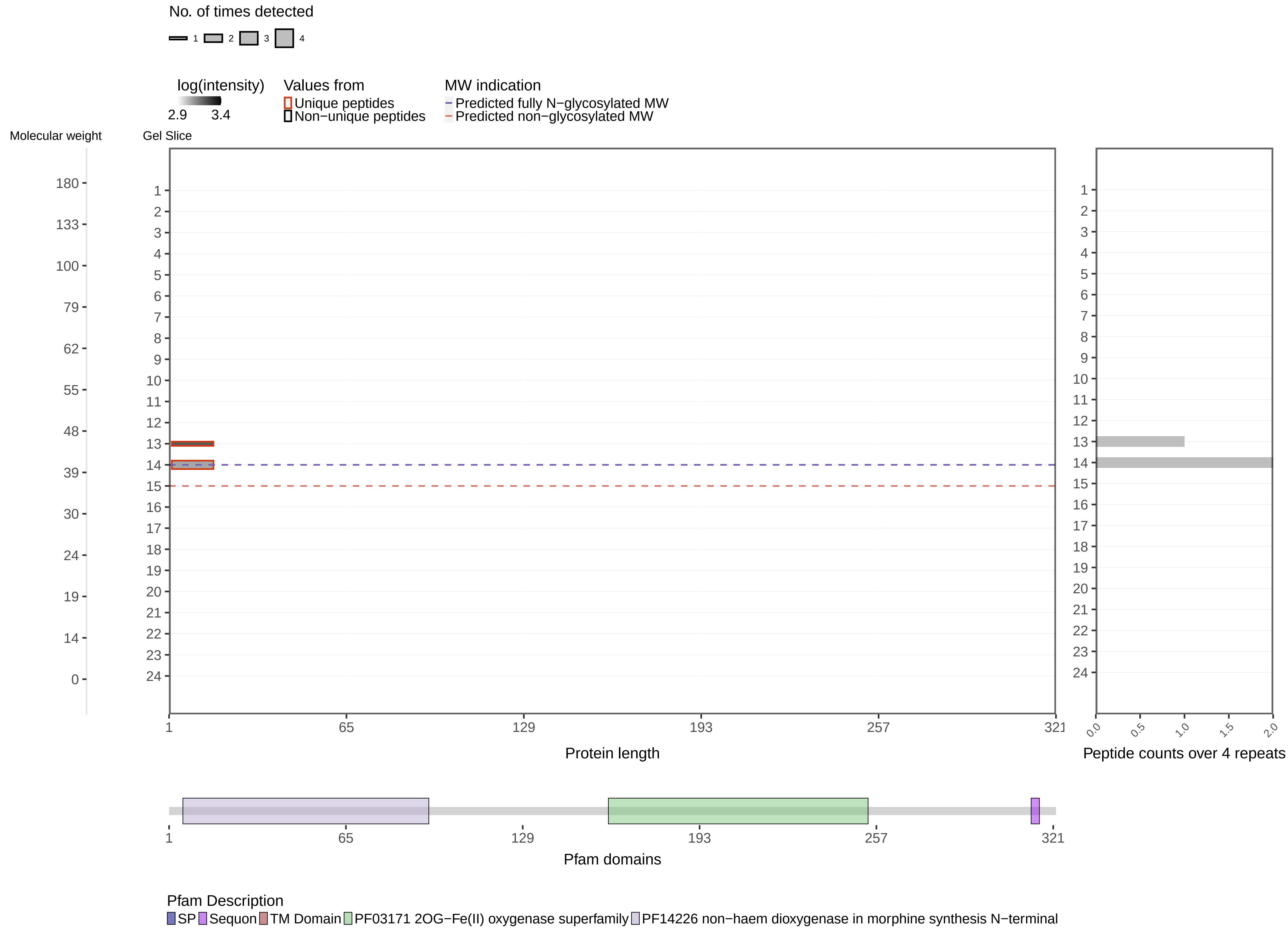


Protein sequence:

KVFYGLKGPTFGTKVSNYPPCPKPELIKGLRAHTDAGGLILLFQDDKVSGLQLLKDGNWIDVPPMKHSIVINLGDQLEVITNGRYKSIEHRVIAQQDGTRMSIASFYNPGSDAVIFPAPELVEKAEEENKLKYPKFVFEDYMKLYAGLKFQAKEPRFEAMKAVETTVNISPIETV*

### NbL19g11780.1 expansin−A13−like [MW − SP = 26.55 kDa]

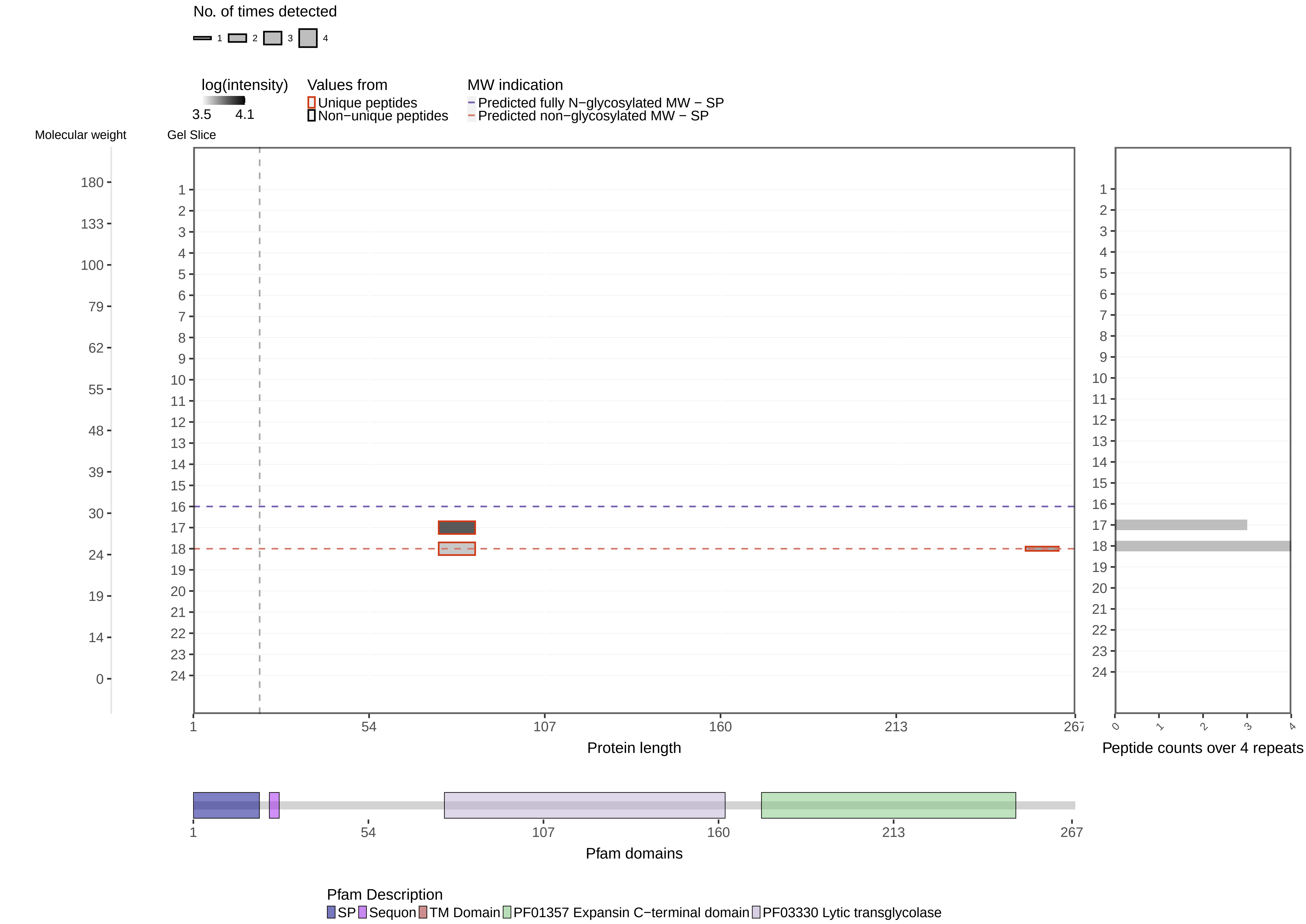


Protein sequence:

NYGFDVDGGGHCNPPNAHFVLPIEAFEKIAIWKASNMPIQYRRIKCRKEGGVRFTMGGAGIFLSVLVSNVAGAGDIVAAKIKGSRTGWLPMGRNWGQNWHINADLKNQPLSFEITSSDGVTLTSYNVAPKIWNFGQTFEGKQFGS*

### NbL19g11740.1 DEAD−box_ATP−dependent_RNA_helicase_21 [MW = 83.1 kDa]

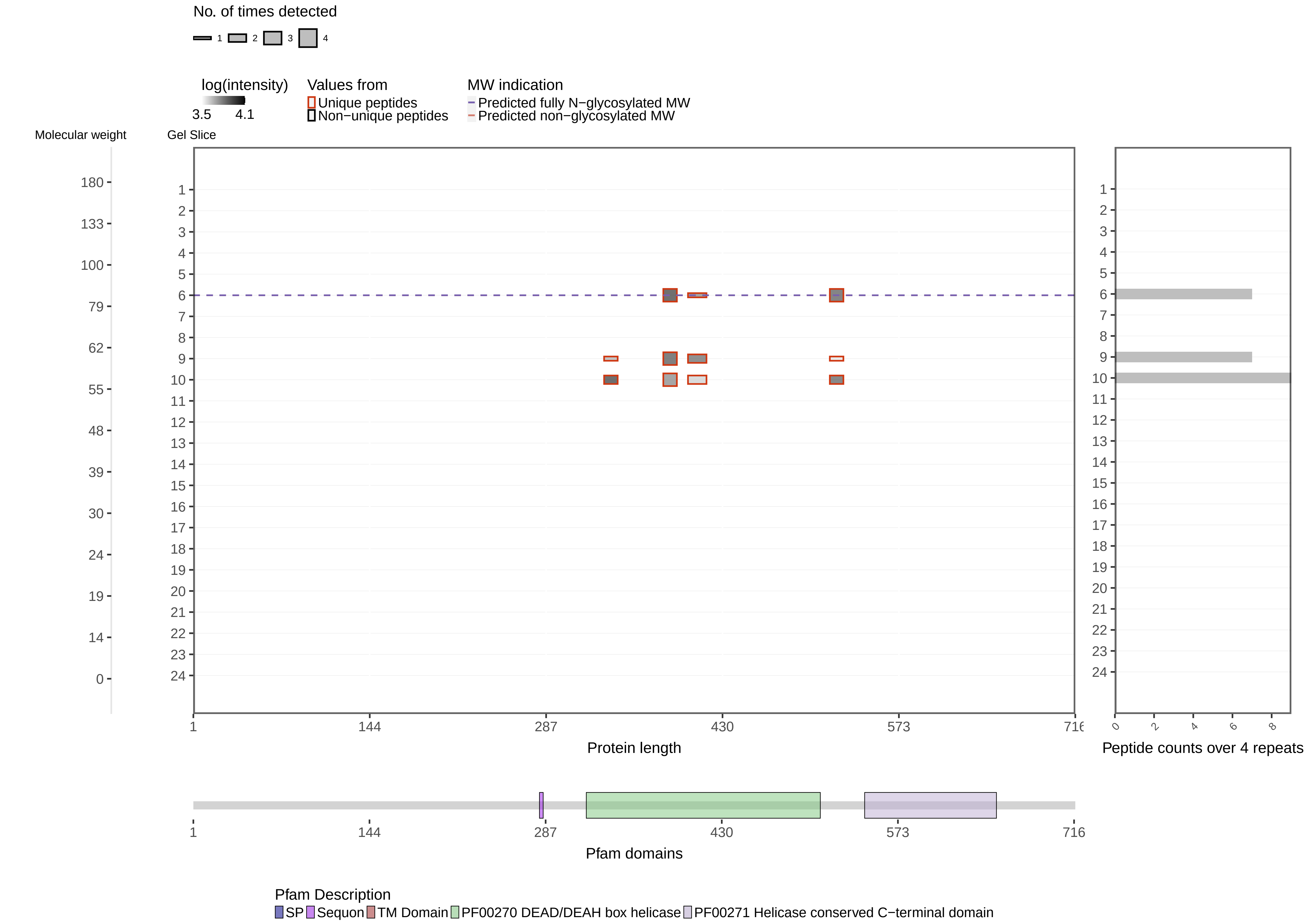


Protein sequence:

NbL19g11550.1 coatomer_subunit_beta−1 [MW = 98 kDa]

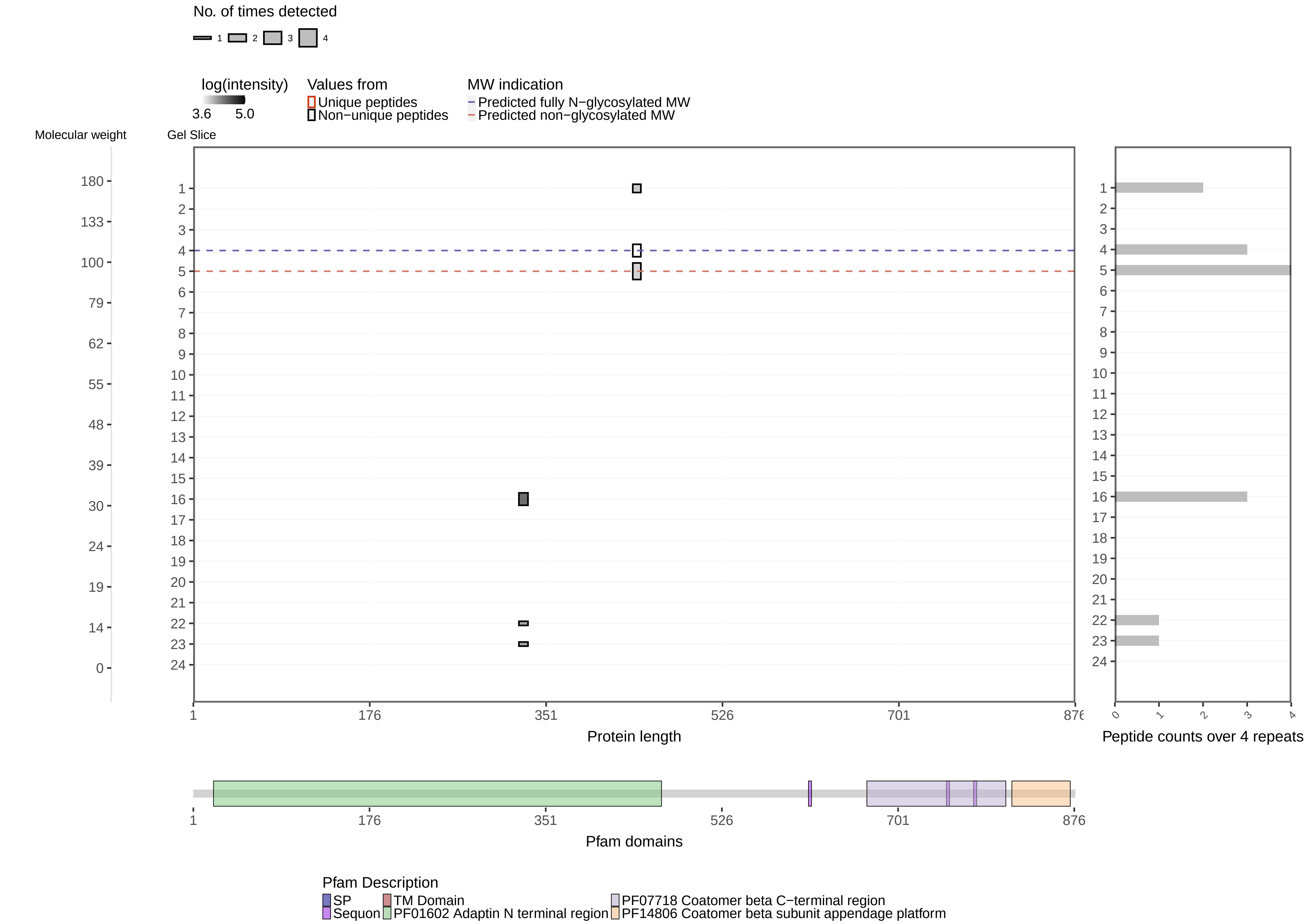


Protein sequence:

[illegible]

NbL19g11520.1 protein_YeeZ−like [MW = 39.32 kDa]

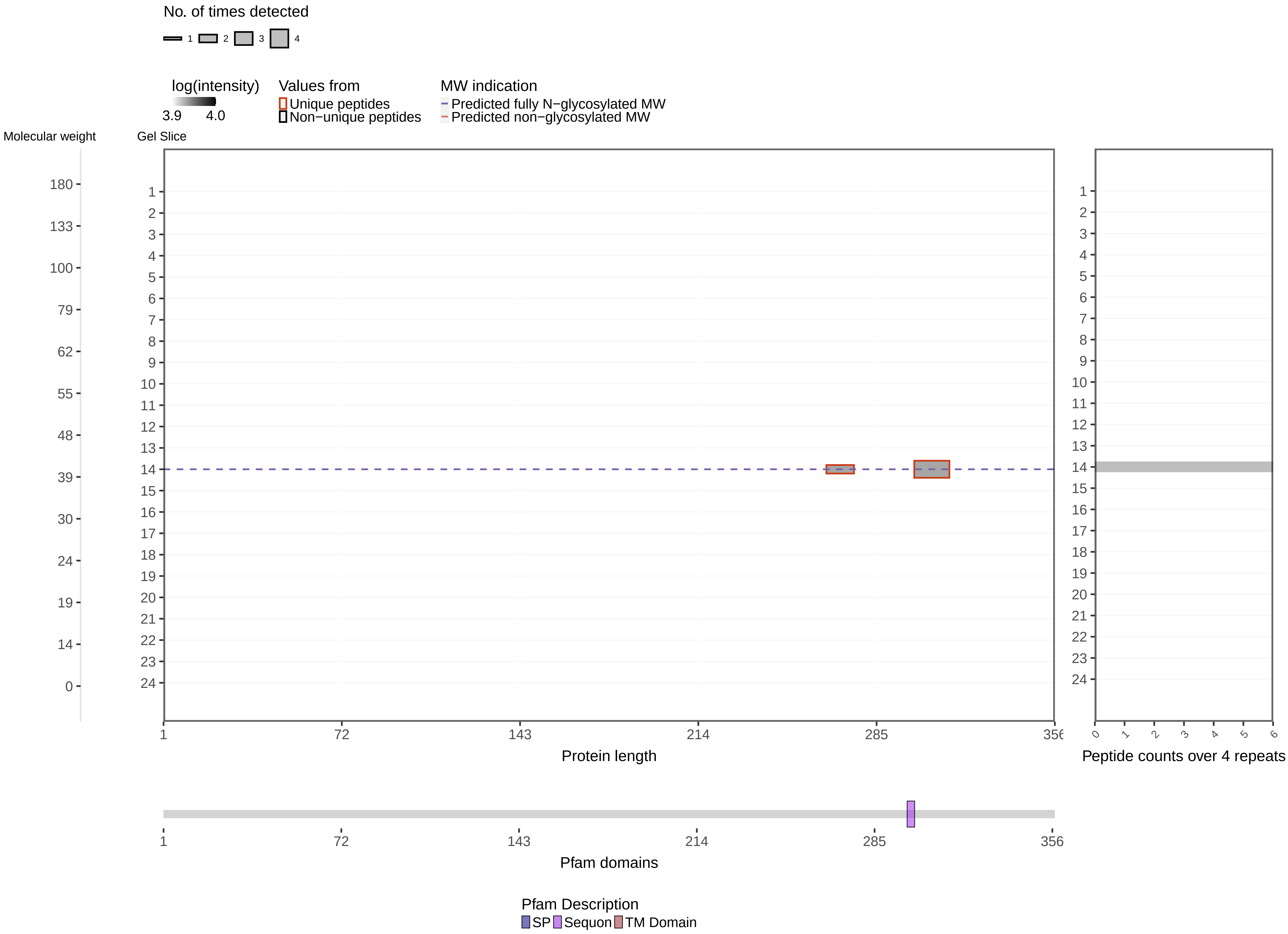


Protein sequence:

GGAWVDEKFPPRPITESAKARMAAEEGWLHLACDVGVTAQIFRLGGIYGPGRSAIDTILKQESLSKGQVMRFSRHYTSRIHVADICQTLKASIQRPSPGKIYNVVDDDPAPRELVFTFARNLVEKKWPGHFTSNKSSEDAASLNPQGASRGEKRVSNKRIKSELAVRLLYPSYESGLRSIIEHMENPFRQS*

### NbL19g11360.1 putative_endo−13(4)−beta−glucanase_2 [MW = 80.36 kDa]

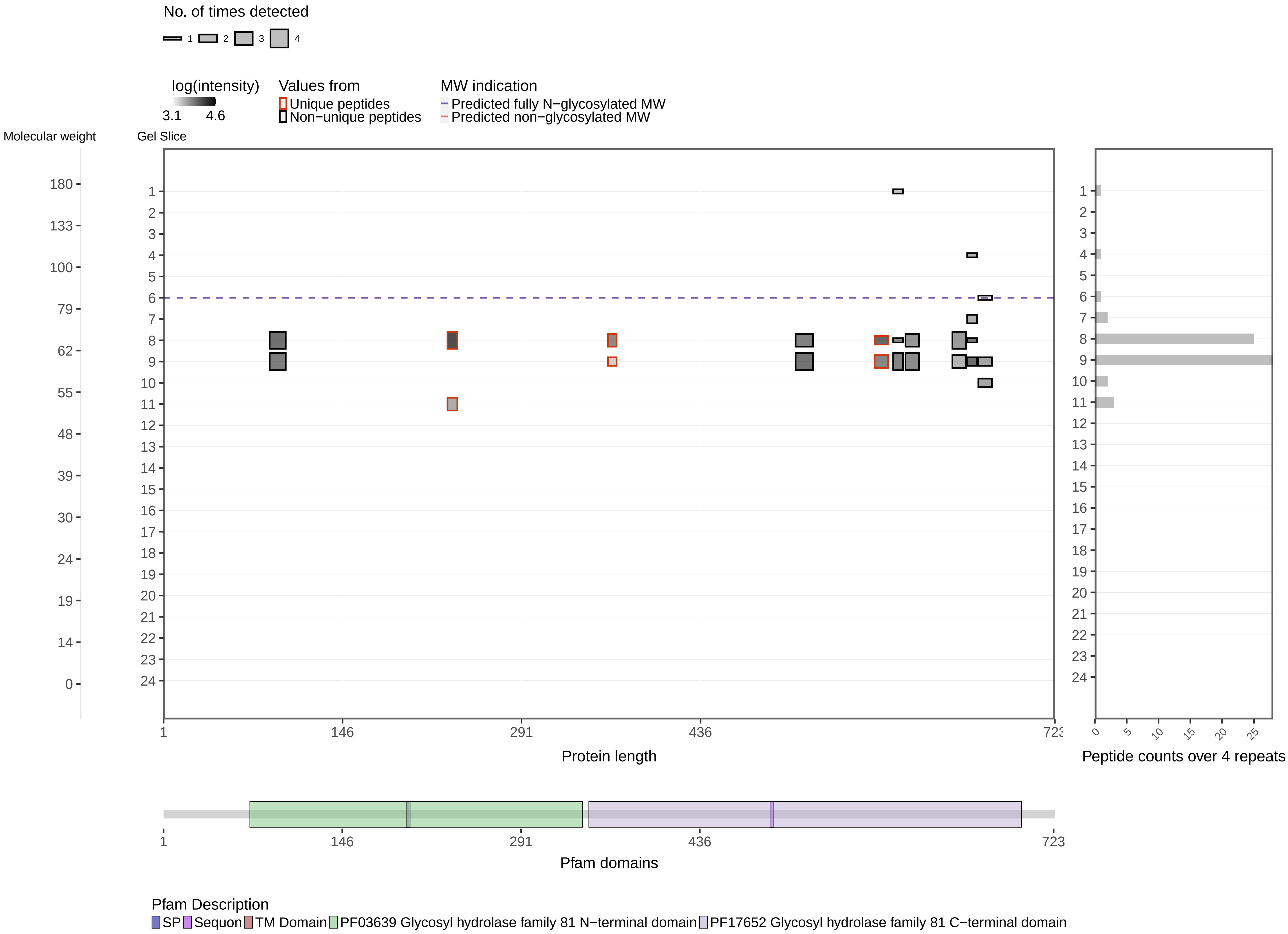


Protein sequence:

SEIIDALNKDVSLLDSKGISTTSSYFYGKLIARAARLALIAEEVCYLDVIPTIRKFLKDTVEPWLEGTFEANGFFYESKWGGIVTKQGALDSGADFGFGVYNDHHFHIGHFLYAIAVLAKIDPIWGRKYRPQAYALMADFMNLSRRANSKYARLRCFDLWKLHSWAGGLTEFADGRNQESTSEAVNAYYSAALMGLAYGDTNLVAIGSTLSAFEIHSAQTWWHVKEGSTLYGDEFTKDNRVVGVLWSNKRDSGLWFAPAEWRECRLGIQVLPILPITEVLFSDVQFVKQLVEWTL

NbL19g11280.1 60S_ribosomal_protein_L22−3−like [MW = 13.87 kDa]

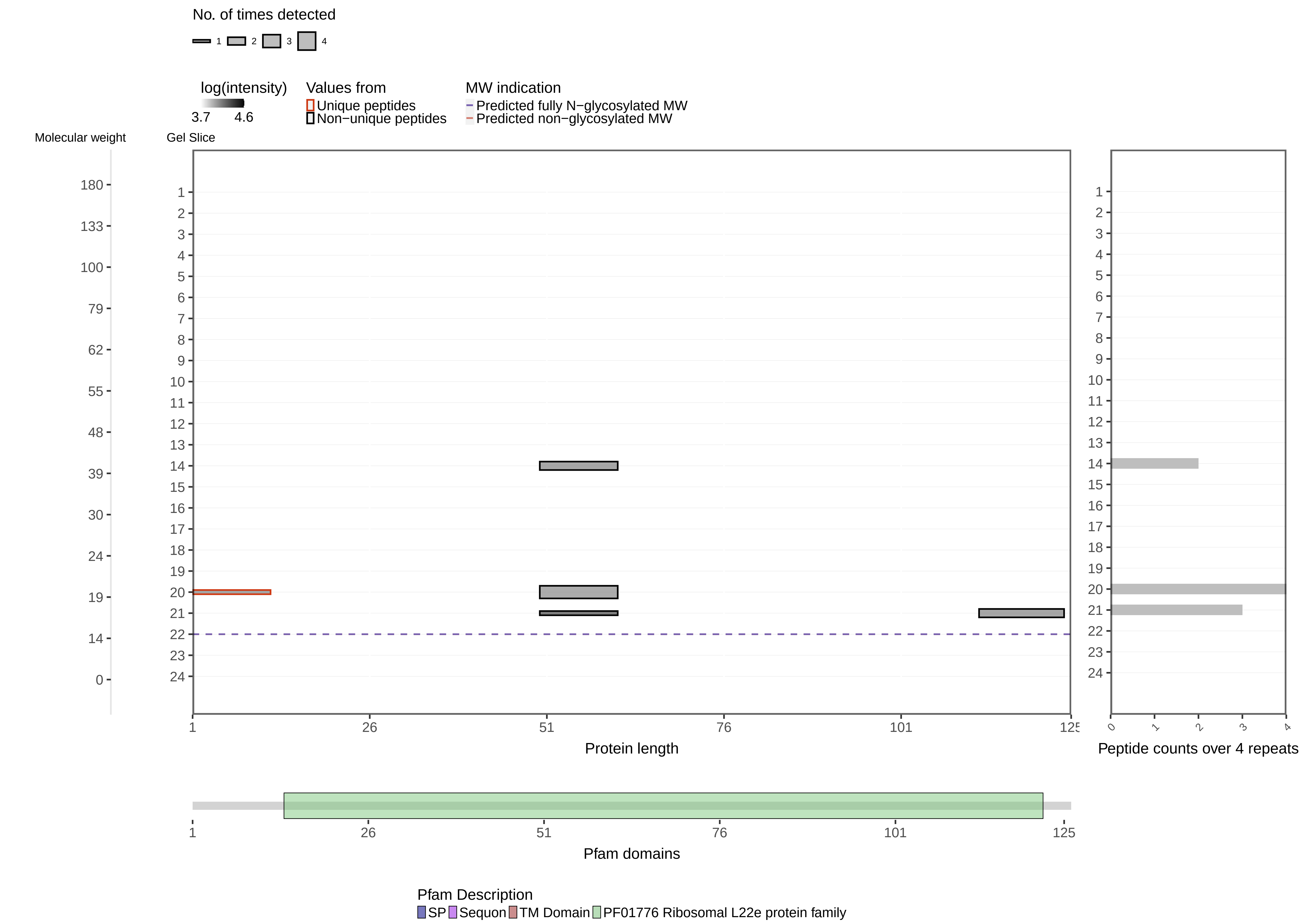


Protein sequence:

:AGALGDSVTVTRDKNTITVTSNSAFSKRYLKYLVKKYLKKNNVRDWLRVIAANKDKNLYELRYFNIAETEADEED*

NbL19g10960.1 uncharacterized_protein_YwbO [MW = 26.45 kDa]

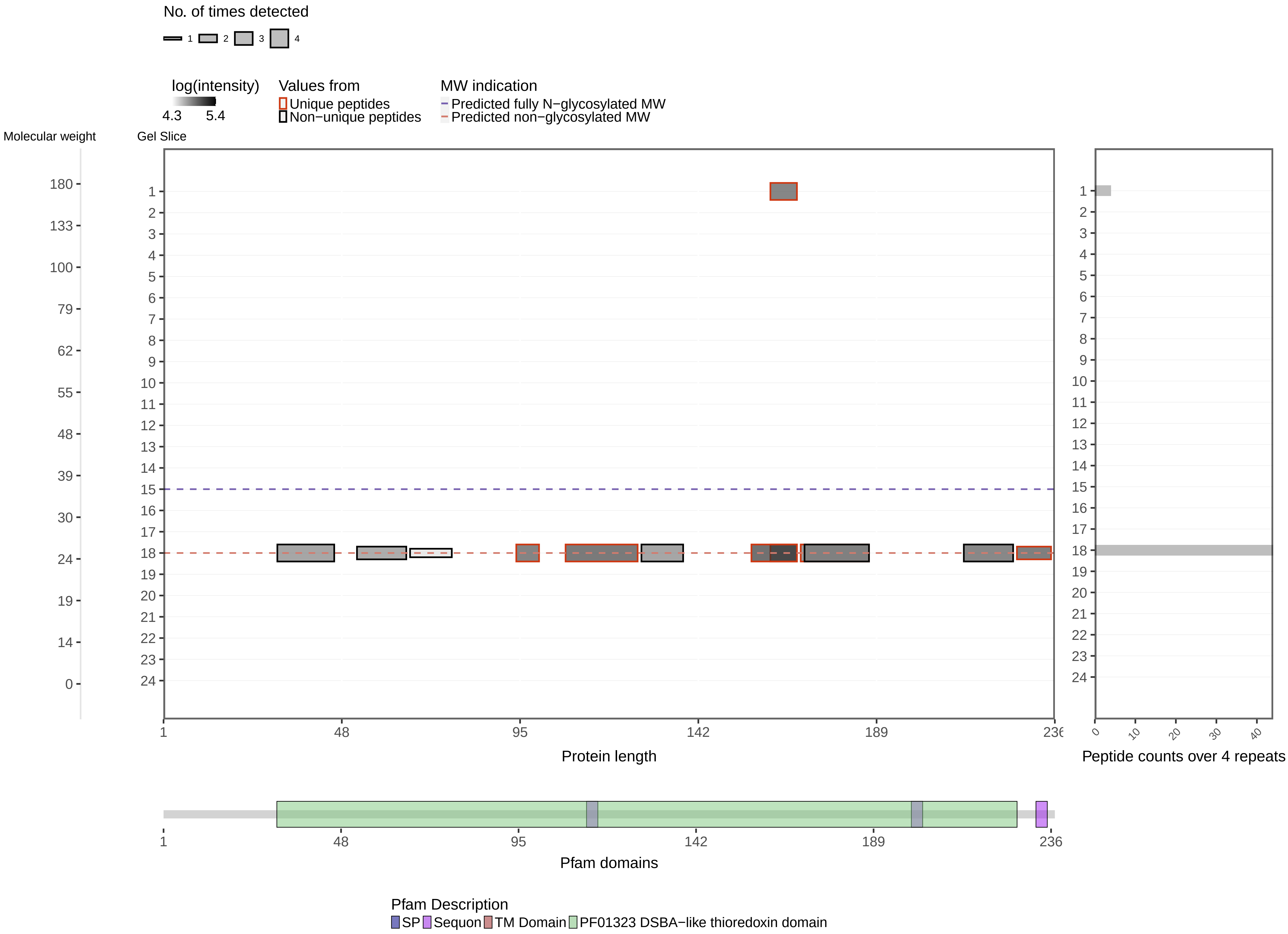


Protein sequence:

KGLGLEYNMSGVTGSSLDSHRLLYFAGQQGLDKQNNLAEELFLGYFTQGRYIGDREFLVEAARKVGVEGAADFLEDPNNGLKEVNEELQQYSSNISGVPHFVLNGKYQLSGGQPPESFIRAFQAASNITA*

### NbL19g10920.1 probable_tRNA_(guanine(26)−N(2))−dimethyltransferase_1 [MW = 65.92 kDa]

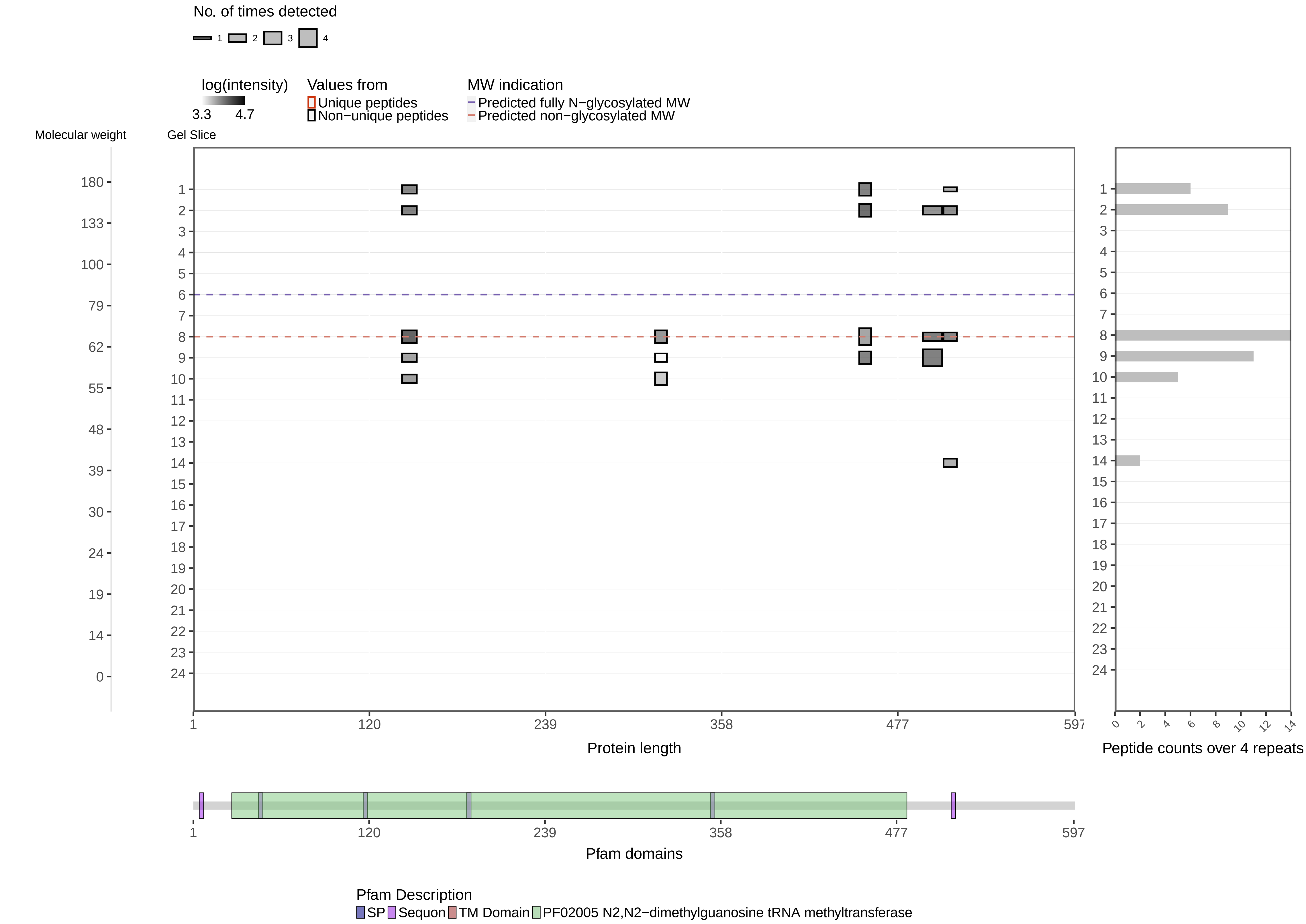


Protein sequence:

### NbL19g10780.1 uncharacterized_LOC109206144 [MW = 14.58 kDa]

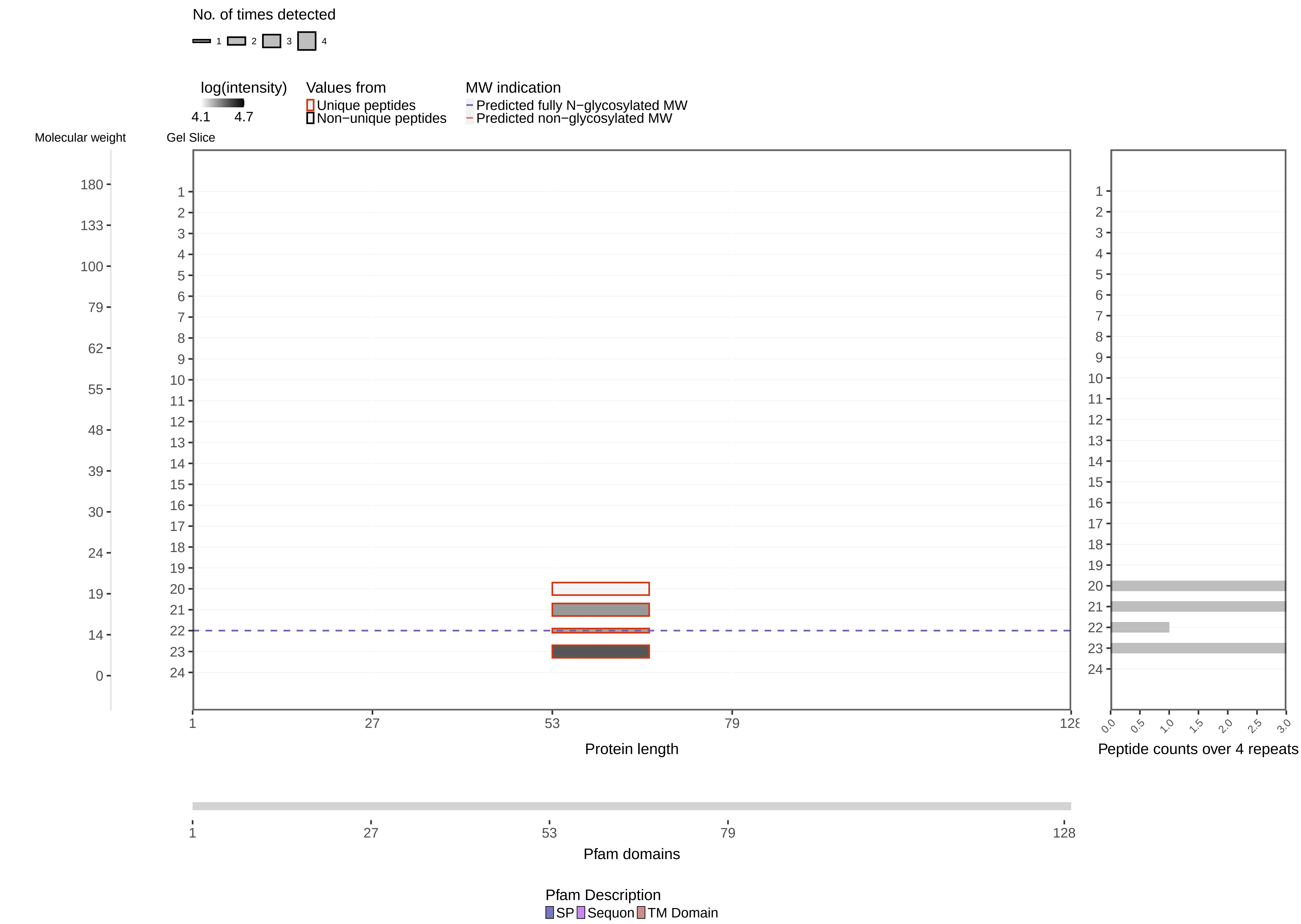


Protein sequence:

KAVVPTSDPETVFDIKYFPRDQRRNRPPIRRTFLNKADVEKMMMEKKTFDVNDFPQPYLTAKVEEDMDAIGGGYQK*

### NbL19g10690.1 glycine−rich_RNA−binding_protein_3_mitochondrial−like [MW = 31.67 kDa]

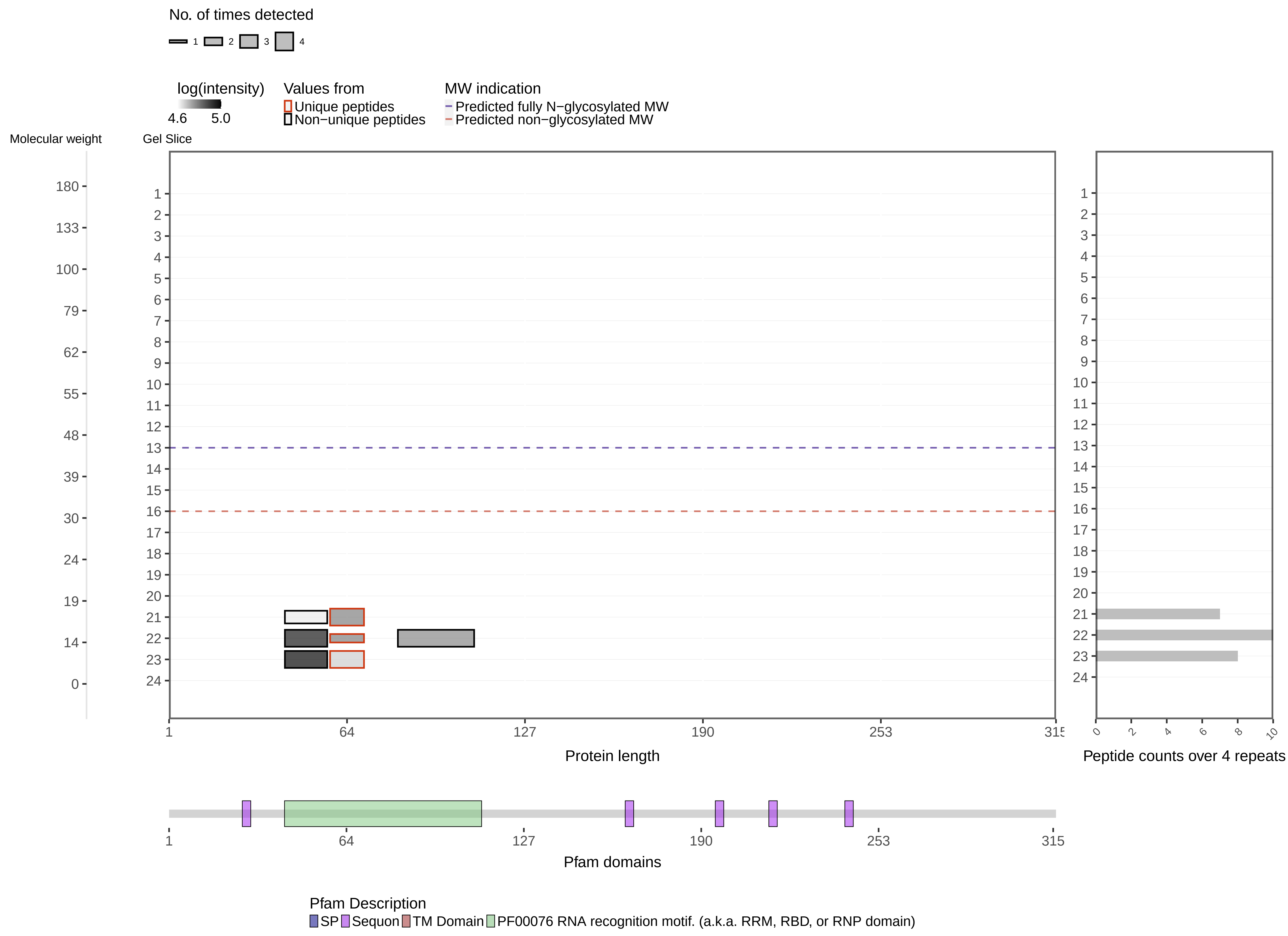


Protein sequence:

GYAASNYGGGGGYGNSSGFNYGGEGGNSAGAGGYPTNNYGGGFSAGNTSAGGYGSYGGGSNYGNDNSSPAEGGNYGGGVGYGSASYGNIPNSNYSGQGNSFSGGYDGGNSGNGFGESPGASFNGGQEQVSVDQGTQSVNEDIEQETSEGNYRDDEDEPKDFANTRG*

### NbL19g10590.1 UDP−glucose_6−dehydrogenase_3 [MW = 53.07 kDa]

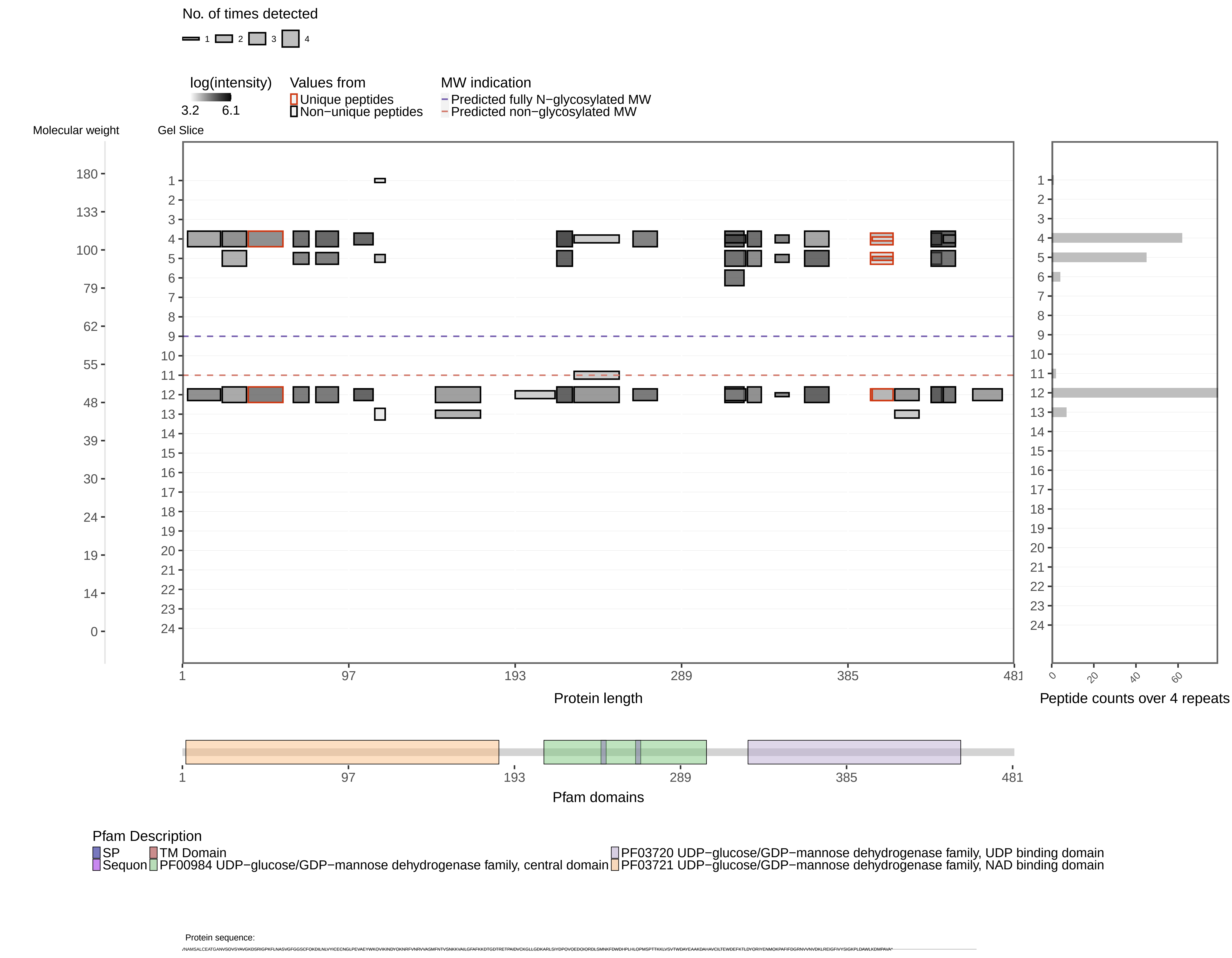

### NbL19g10540.1 chaperone_protein_ClpB3_chloroplastic−like [MW = 109.73 kDa]

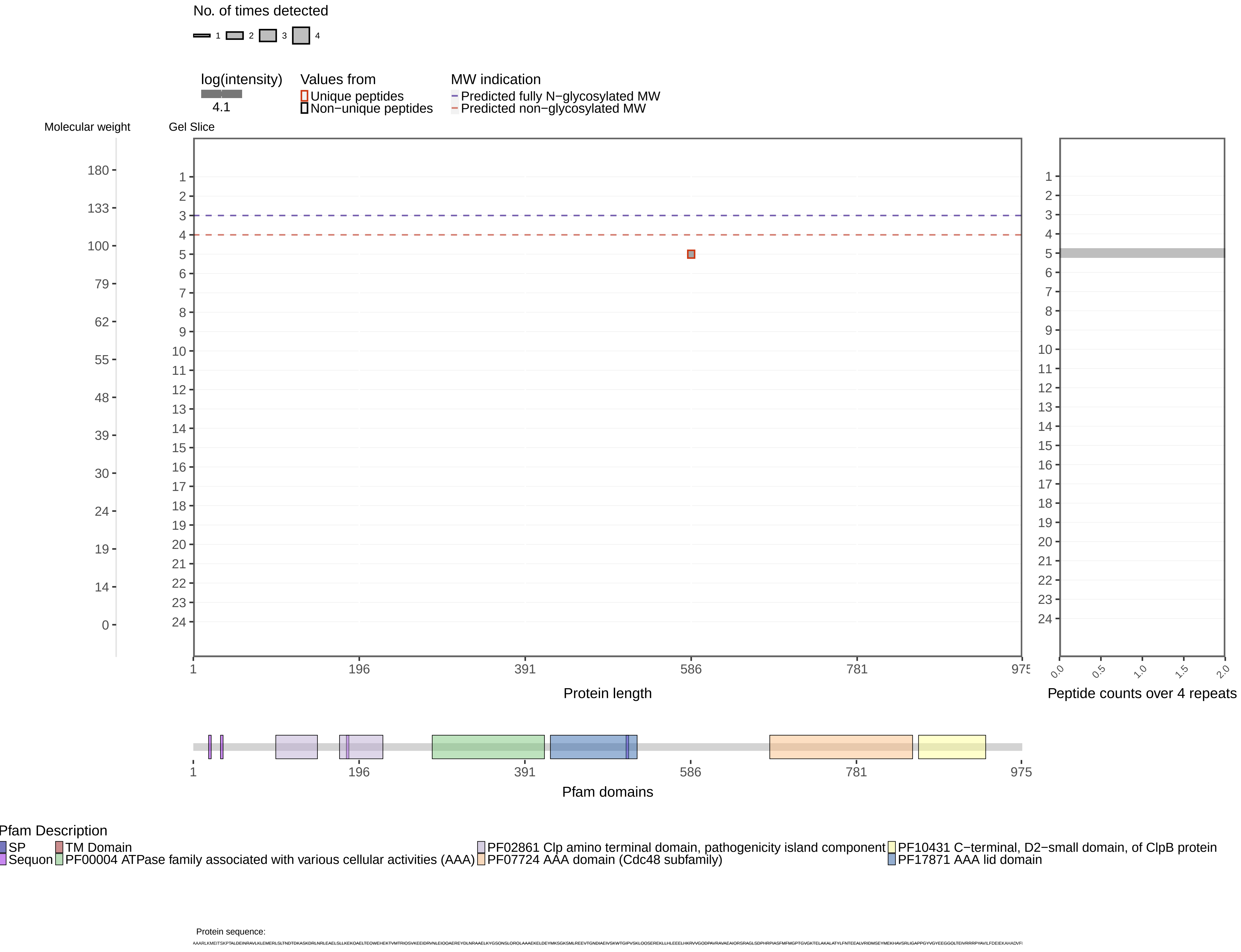

NbL19g10360.1 chorismate_mutase_1_chloroplastic−like [MW = 40.12 kDa]

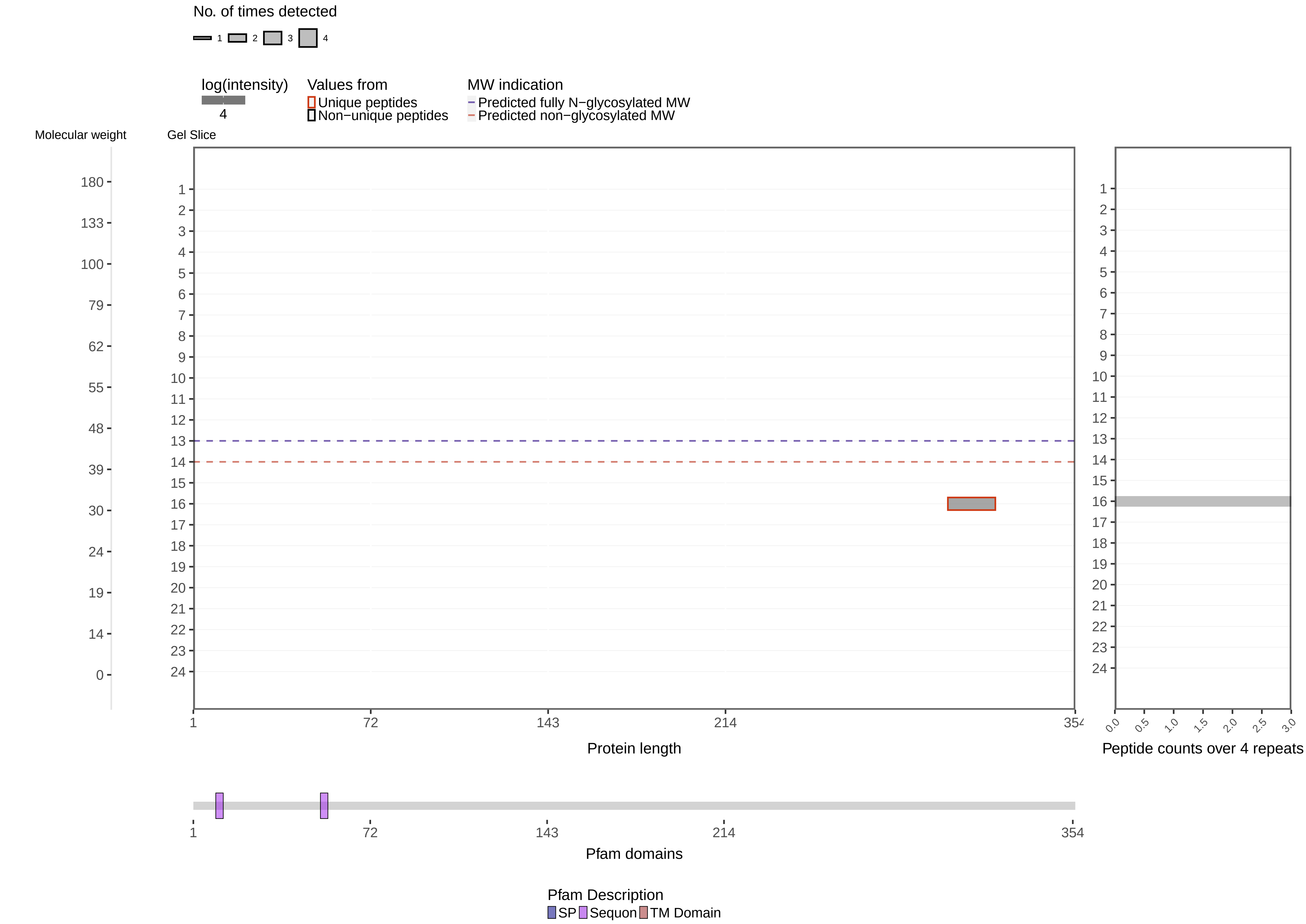


Protein sequence:

IAKVGRYKSPDEHPFFFPKELPEPMLPPMOFPKVLHSVADSININVKIWEMYFKNLLPRLVKEGDDGNFGSTAVCDTICLOALSKRIHYGKFVAEAKFRASPDVYKAAIRAODRNGLMDLLFYPTVEEAVTRRVEMKTRTYGOEININGPENGVDPVYKIKPSLVAELYGDWIMPLTKEVOVEYLLRRLD*

NbL19g10350.1 lamin−like_protein [MW − SP = 16.24 kDa]

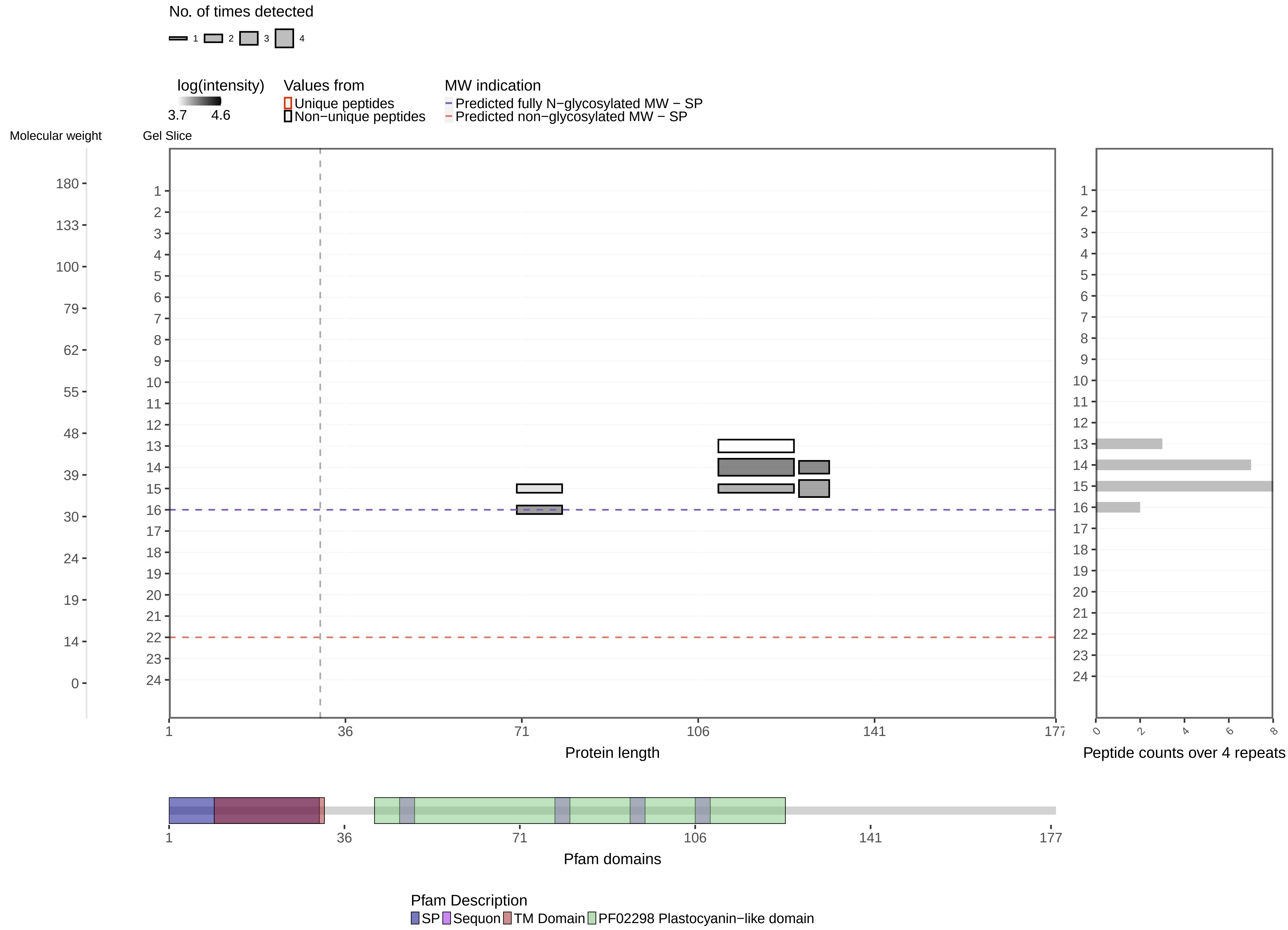


Protein sequence:

VLELNKTNYENCITDHPLHNYTTGAGRDVVPLNVTKTYYFASSGGLCYGGMKVAIHVEKAPPPPKAAPVRSGSTNVLTSFTGOILIPALFATSAVWNAFLRLW*

### NbL19g10340.1 uncharacterized_LOC107762835 [MW = 46.13 kDa]

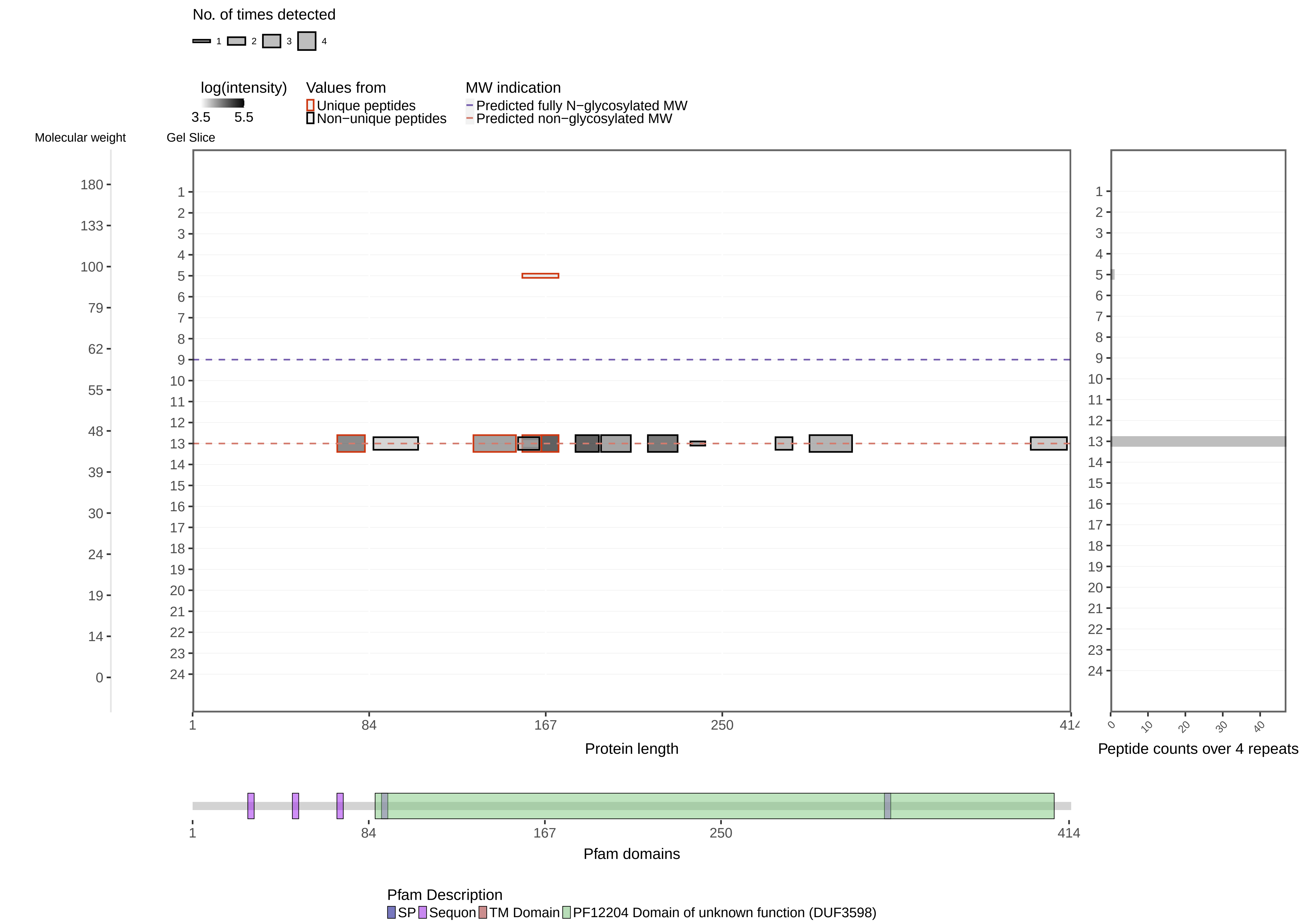


Protein sequence:

[illegible]GVLGEDDIGEESPRNLKLPSKKPSIVCENCLYSLEKDKRVRAFHIMDPKGILEMLLVFLEERGNKEAIPPSFDNFKEDSERIMPHLGTWKGHSLTKRTGVYGATITEAETTAVLEMNKDGQLVQDITSTSGASNVTTNVHWTGTISGNLVTFHGGFQLTLLPGGIYMGYPSDVANSVQESKSFHLELCWLESPGKRQRLIRTYDVEGLAVSSTYFIESKI*

NbL19g10230.1 uncharacterized_LOC104214001 [MW = 29.72 kDa]

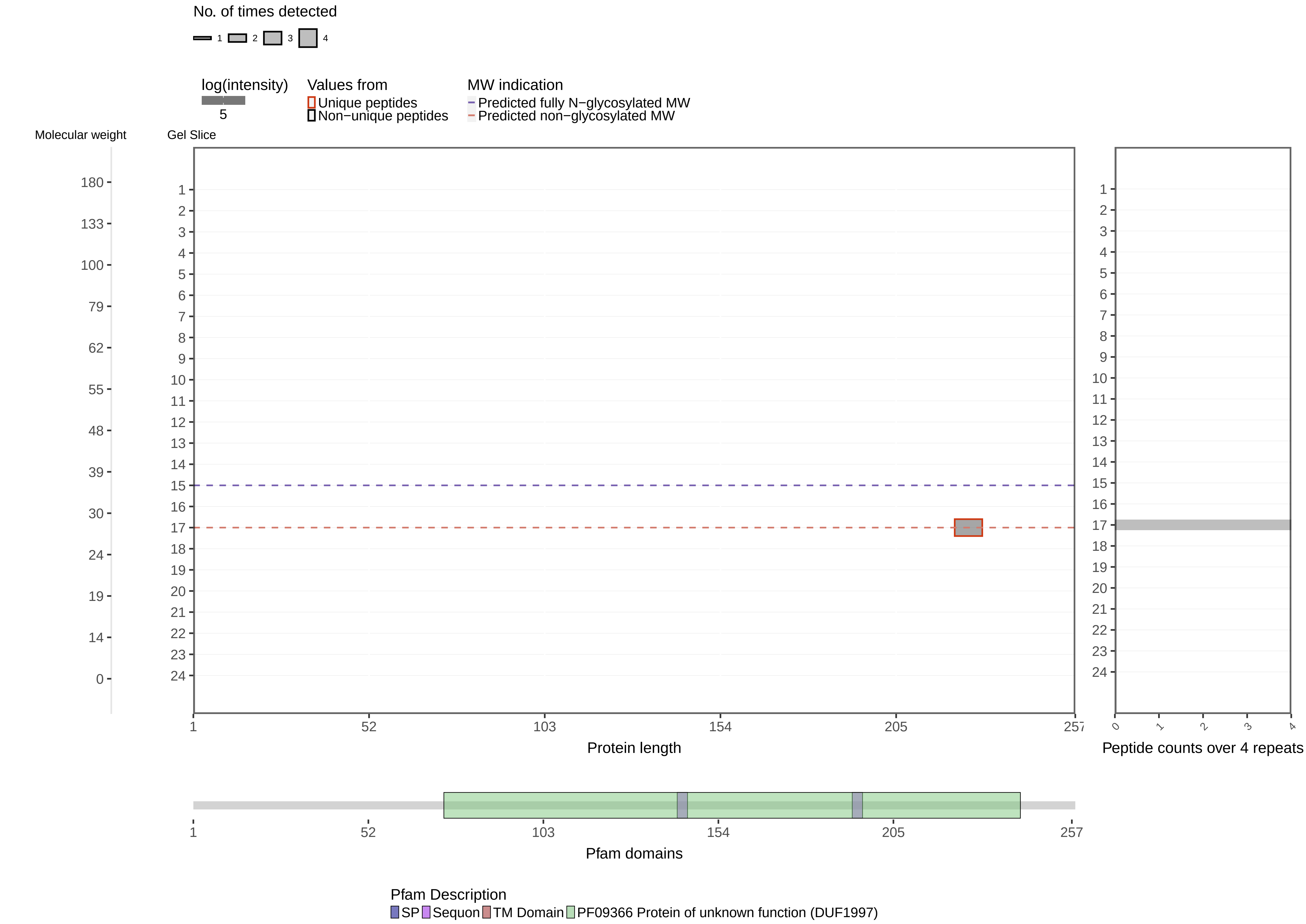


Protein sequence:

LTAWPVVNMRIRYKTKGKEYPSGVPTNTSTVLELKIIKWDLEGVTEGKNKPSEFRINMEGALYPDRRGTRSRIKGHFNMSLSFAPPPMLALVPPHVHQHVTQSVMKNIGESMQHKVRDNLLADYTKFKKENYKNIEGEDRK*

NbL19g10220.1 uncharacterized_protein_At5g39570 [MW = 27.42 kDa]

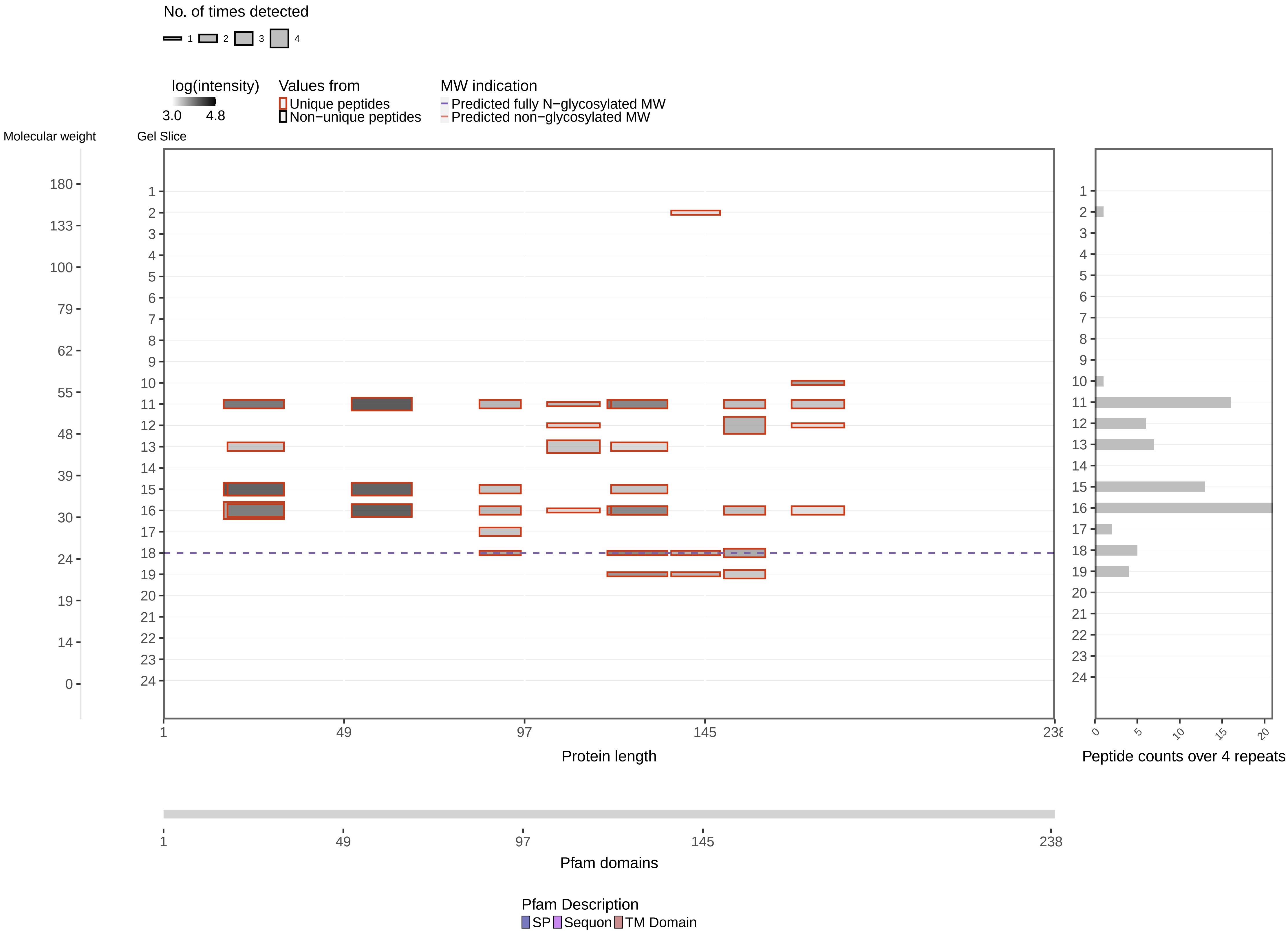


Protein sequence:
EPAPEYGSGYRRKSEYEEPSSEYGSGYERKTEPEEDGSGGYGRKPSYGQDEEGERRPSYGRSSYQTEEGEGYERPRYGRSEEEDYRRPSYERRGDDDDEGYGRKKYGDDDSDDDEEKKQHYKHHHRKHYDD*

NbL19g10170.1 cystathionine_gamma−synthase_1_chloroplastic−like [MW = 58.47 kDa]

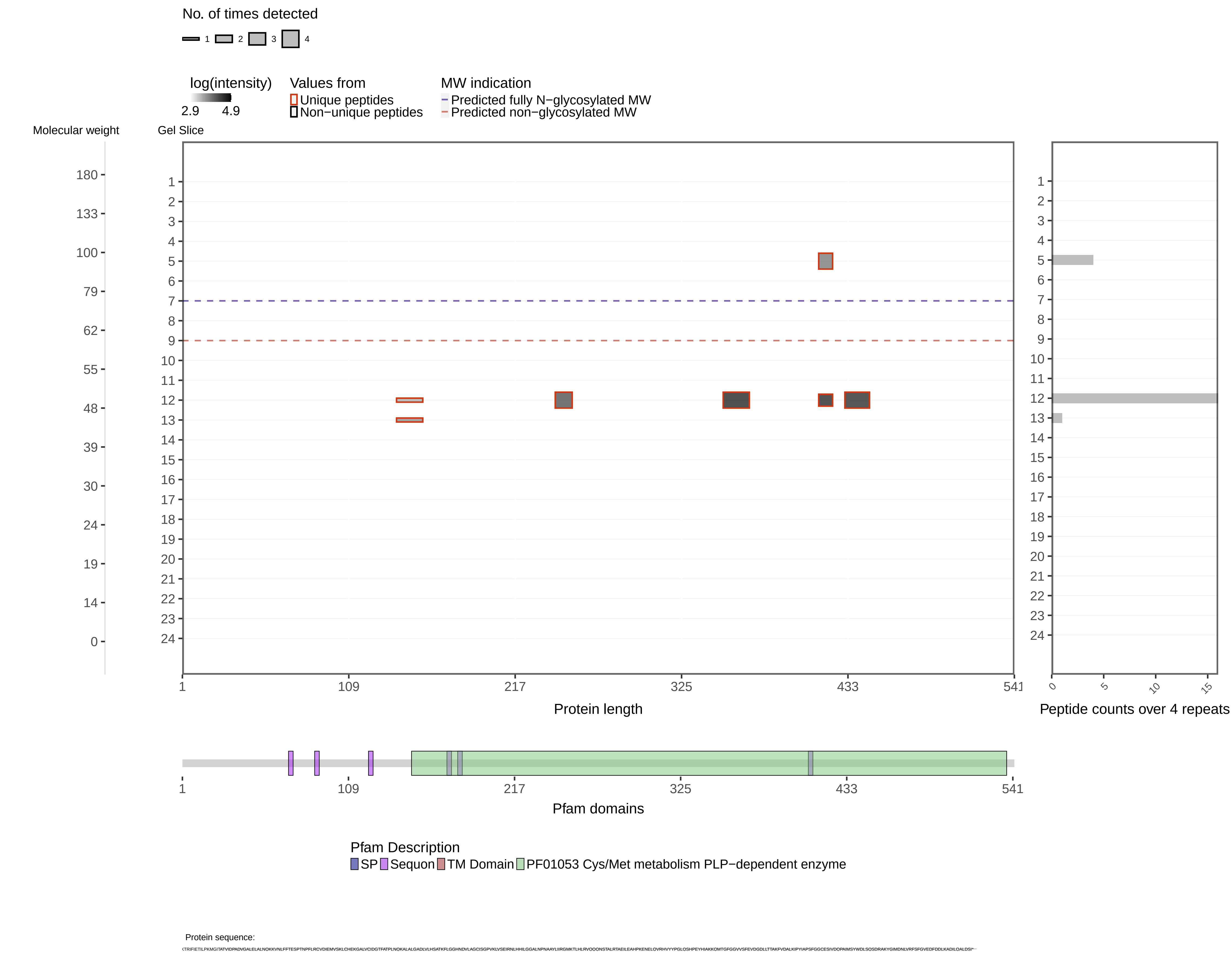

NbL19g10090.1 expansin−A10−like [MW − SP = 23.98 kDa]

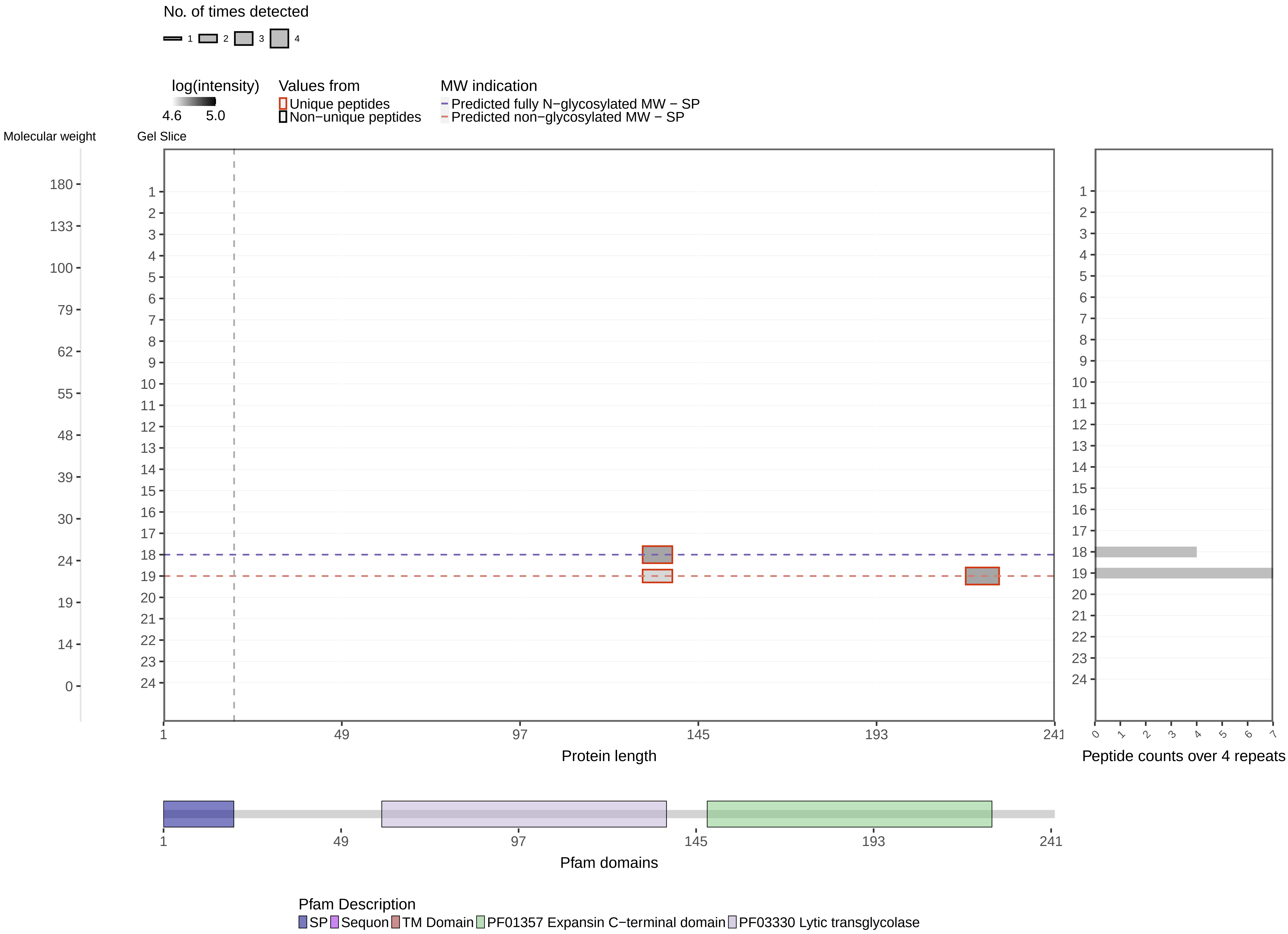


Protein sequence:

DPPNHHFDLSQPVYLRIAQYRSGIVPVAYRRVPCRRRGGIRFTINGHSYFNLVLVTNVGGAGDVRSLYIKGSRTQWQPMSRNWGQNWQNNAYLNGQSLSFKVTTSDGRTVVSYNAAPRSWSFGQTFTGGQFR*

NbL19g10060.1 granule−bound_starch_synthase_2_chloroplastic/amyloplastic [MW = 87.9 kDa]

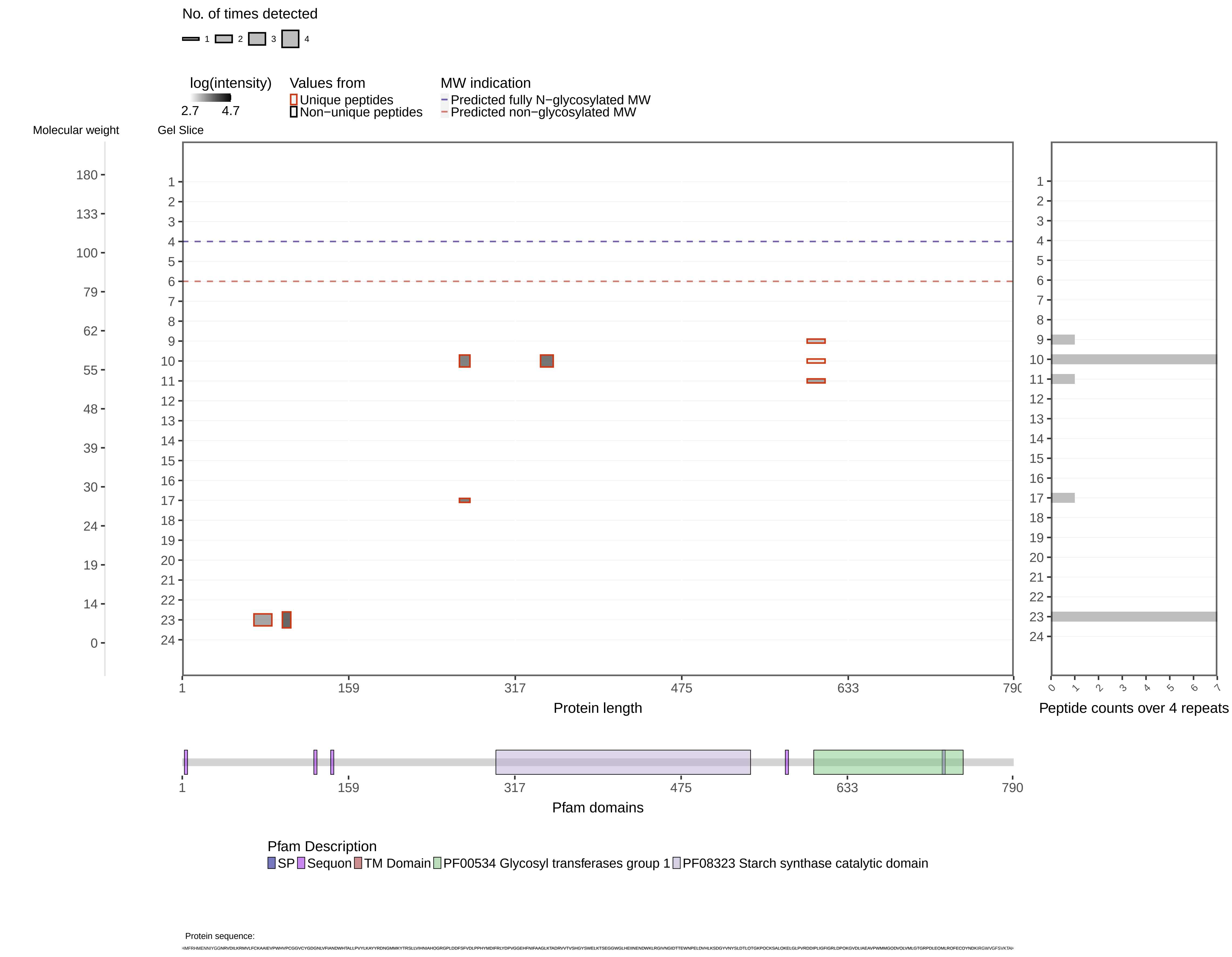

NbL19g09940.1 tubulin_alpha_chain−like [MW = 49.65 kDa]

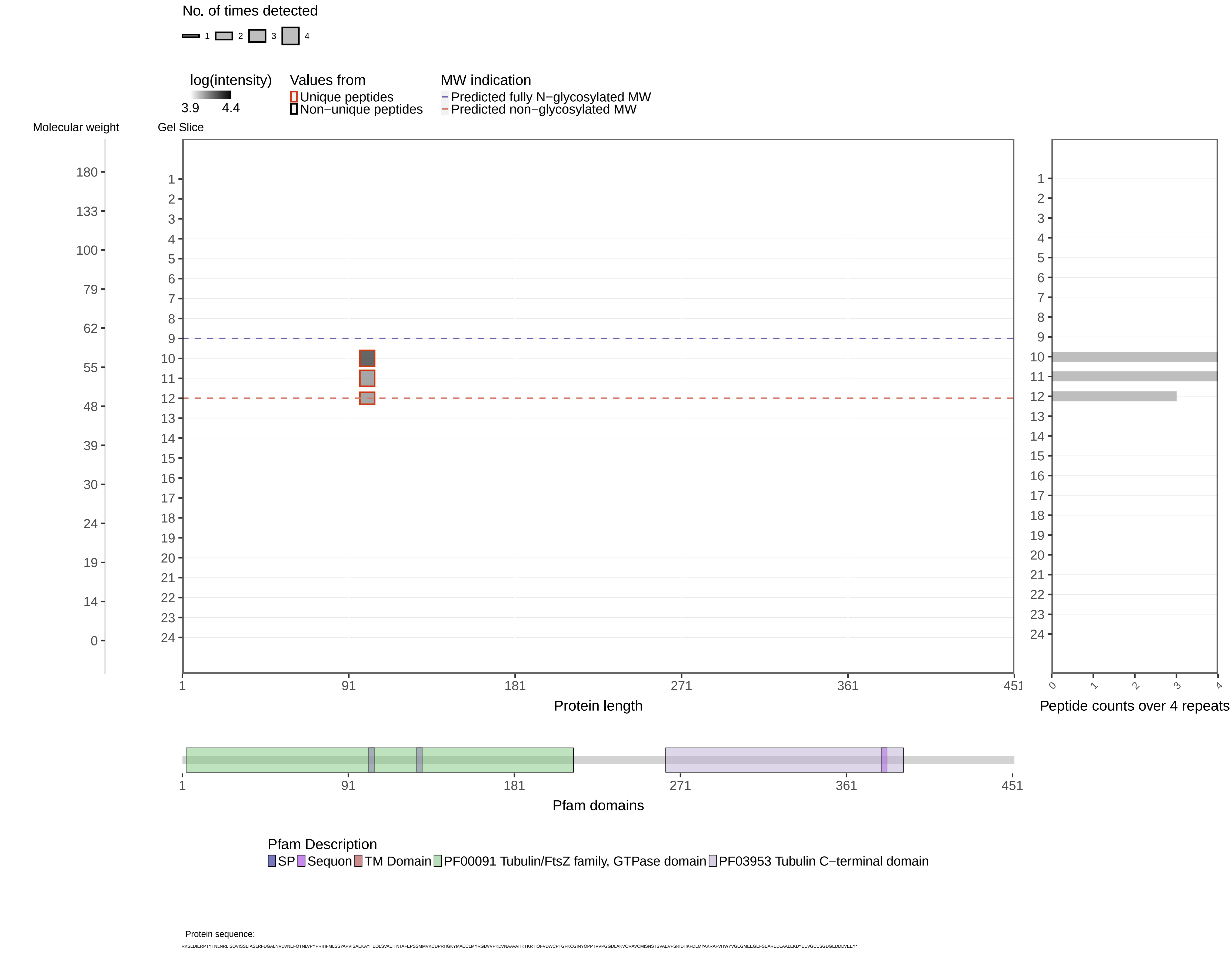

NbL19g09860.1 aldose_1−epimerase−like [MW − SP = 37.14 kDa]

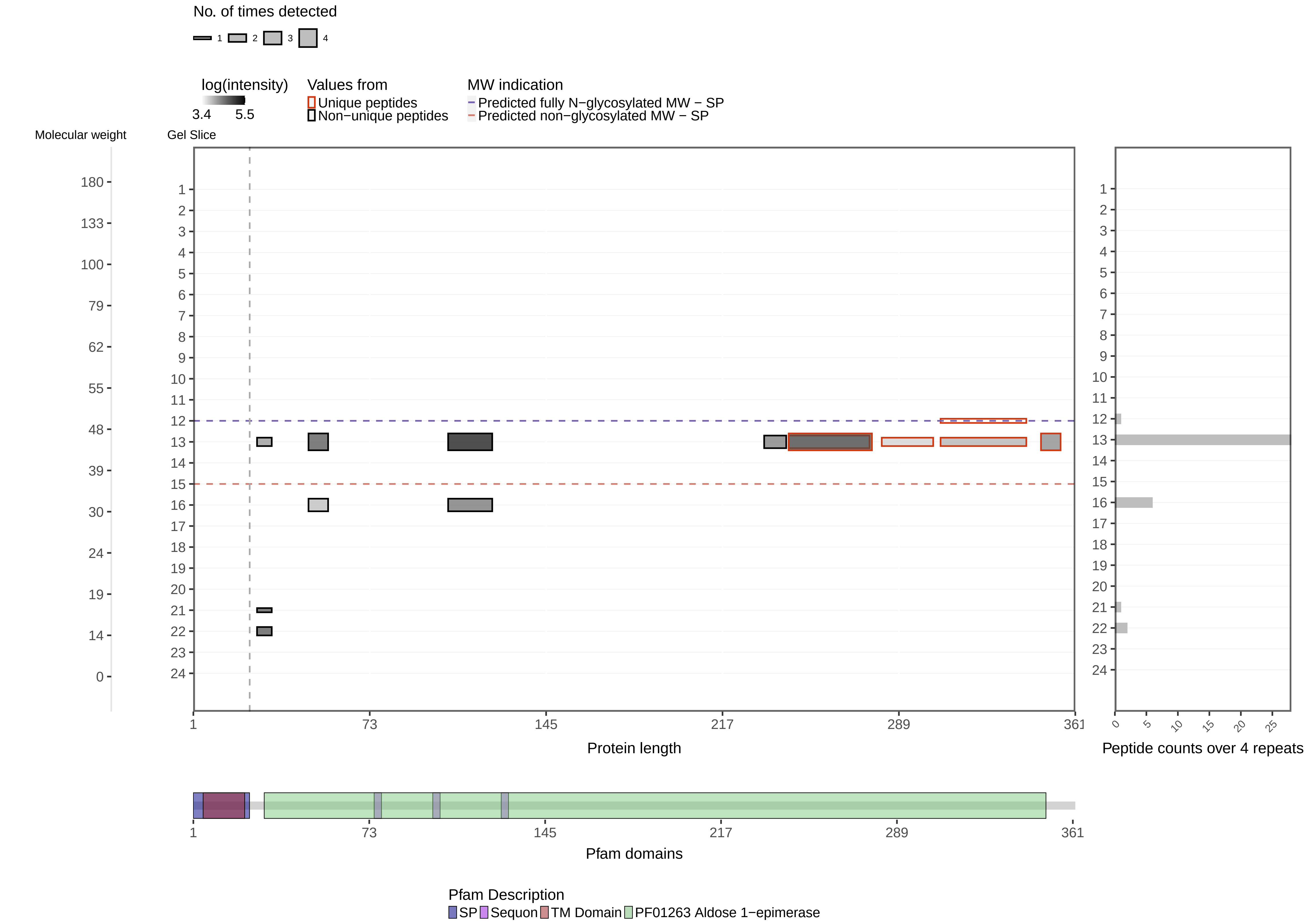


Protein sequence:

YKLSVVMKAKALNKATPVNLAQHTYWNLGGHSSGDILSNIVQIFASHITPVDEHFIPTGEISPVKNTPYDFLKPREVGSQINELQSGYDINYALDSTEEMKPVAIVYDKKSGRVMDLQASAPGVQFYTANSLIQKGKGGYVYQPHAALCFETQGFPDAVNHPNFPSTIVIPGKAYTHKMLYTFSIMKYWNSY*

### NbL19g09830.1 protein_IQ−DOMAIN_31−like [MW = 62.22 kDa]

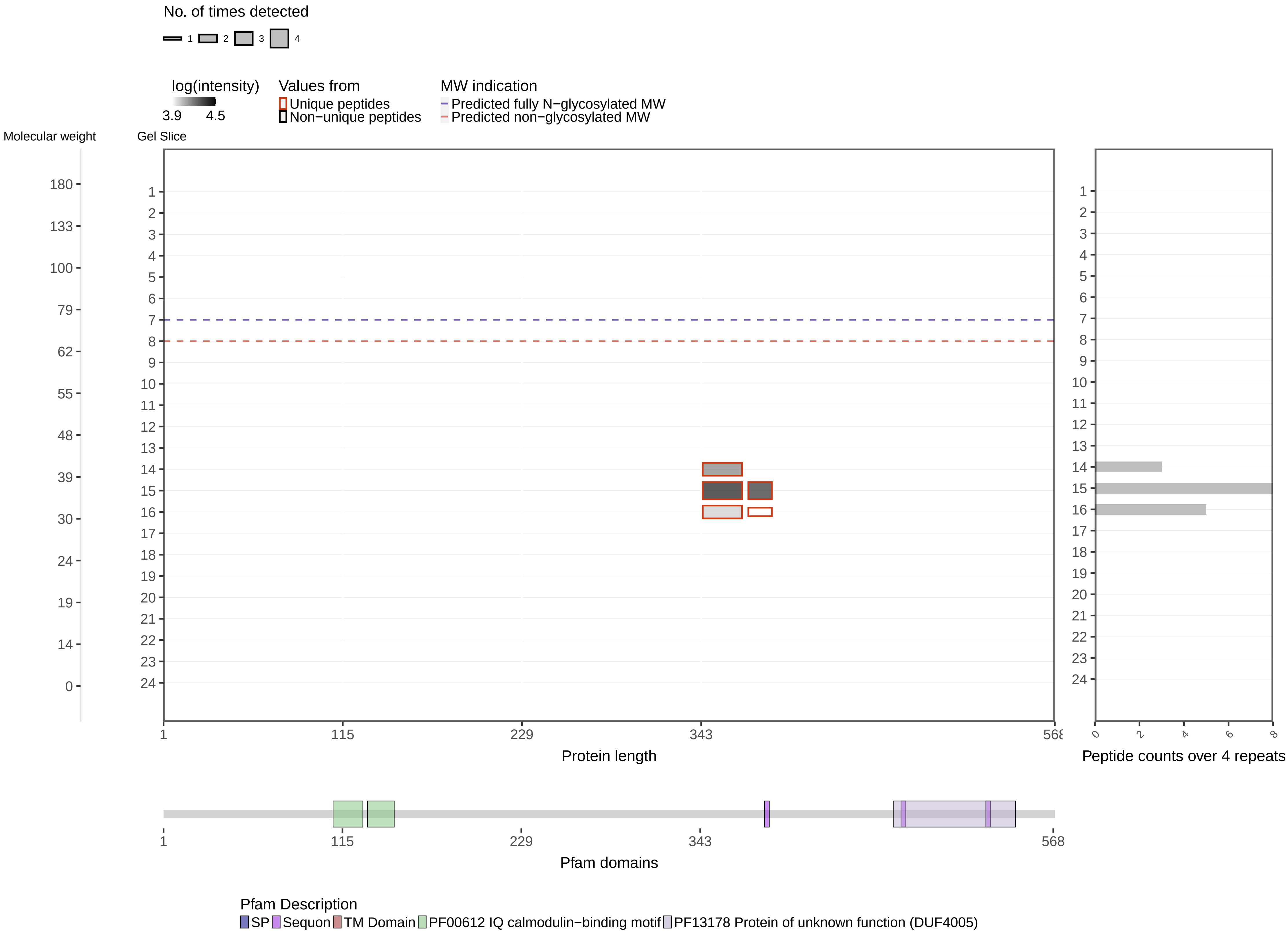


Protein sequence:

RSKRSVKKVLTSANGDAYAVSSSEPEKTKRYPRKFSHPHIEPVLDQPQNELEKVKRNLRKVSAALATPSERTETESEKVQETPNLAQAQAQAQAQAKTSKSSALDVEQIVVNSSEKTRDSAPEIEKLAVTEAPLPVAVDEPSNELLDHPTTEQQQLEDVGNTANSPVINEELSSMEDHTAKERTRRRKSLPPKQENSENISHNTPNVPSYMAATQSAKAKLKAQGSAKVSEDGAENGFVRRHSLPSSTNGKFNTSSPRVQRPVQANGKAGNKNRAMSSSKDAPSSEKVL

### NbL19g09720.1 thioredoxin_H2−like [MW = 15.57 kDa]

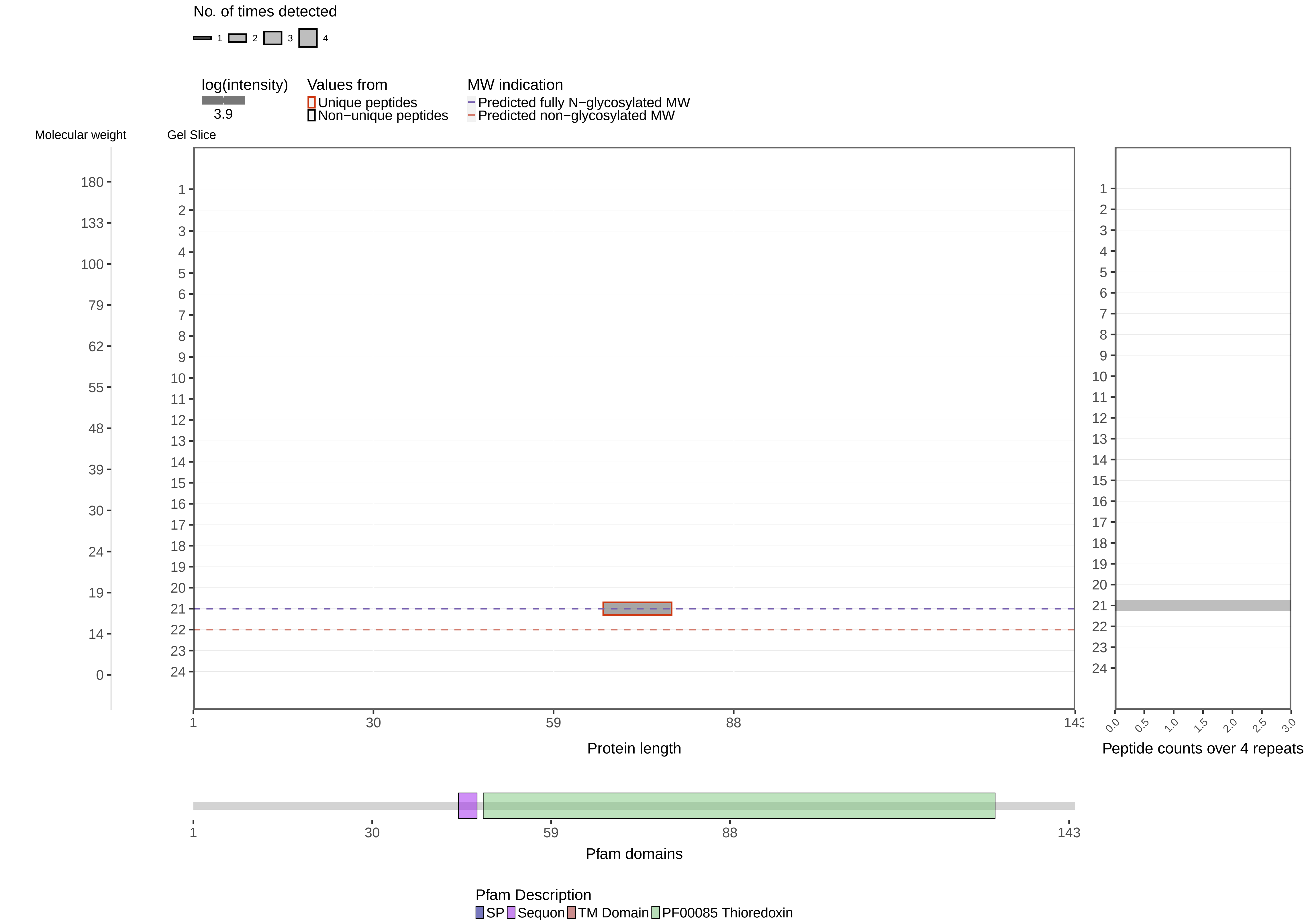


Protein sequence:

ATWCGPCKMMEPVINAMSAKYTDVDFVKIDVDELSDVAQEFGVQAMPTFLLLKQGKEVERVVGAKKDELEKKILKHREAPKFAA*

NbL19g09710.1 inositol_hexakisphosphate_and_diphosphoinositol−pentakisphosphate_kinase_2−like [MW = 124.18 kDa]

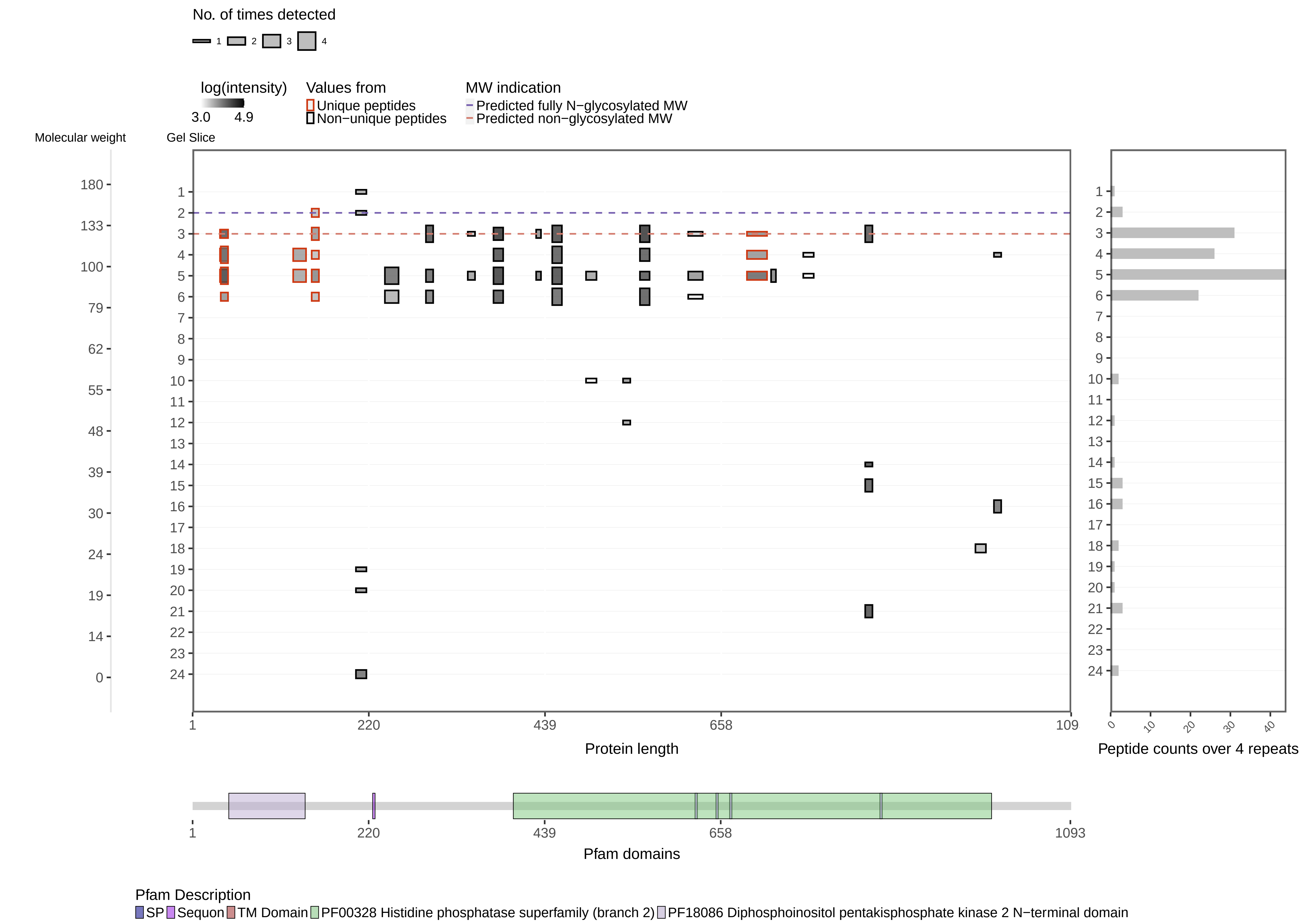


Protein sequence:

[illegible]

### NbL19g09510.1 signal_recognition_particle_43_kDa_protein_chloroplastic [MW = 41.73 kDa]

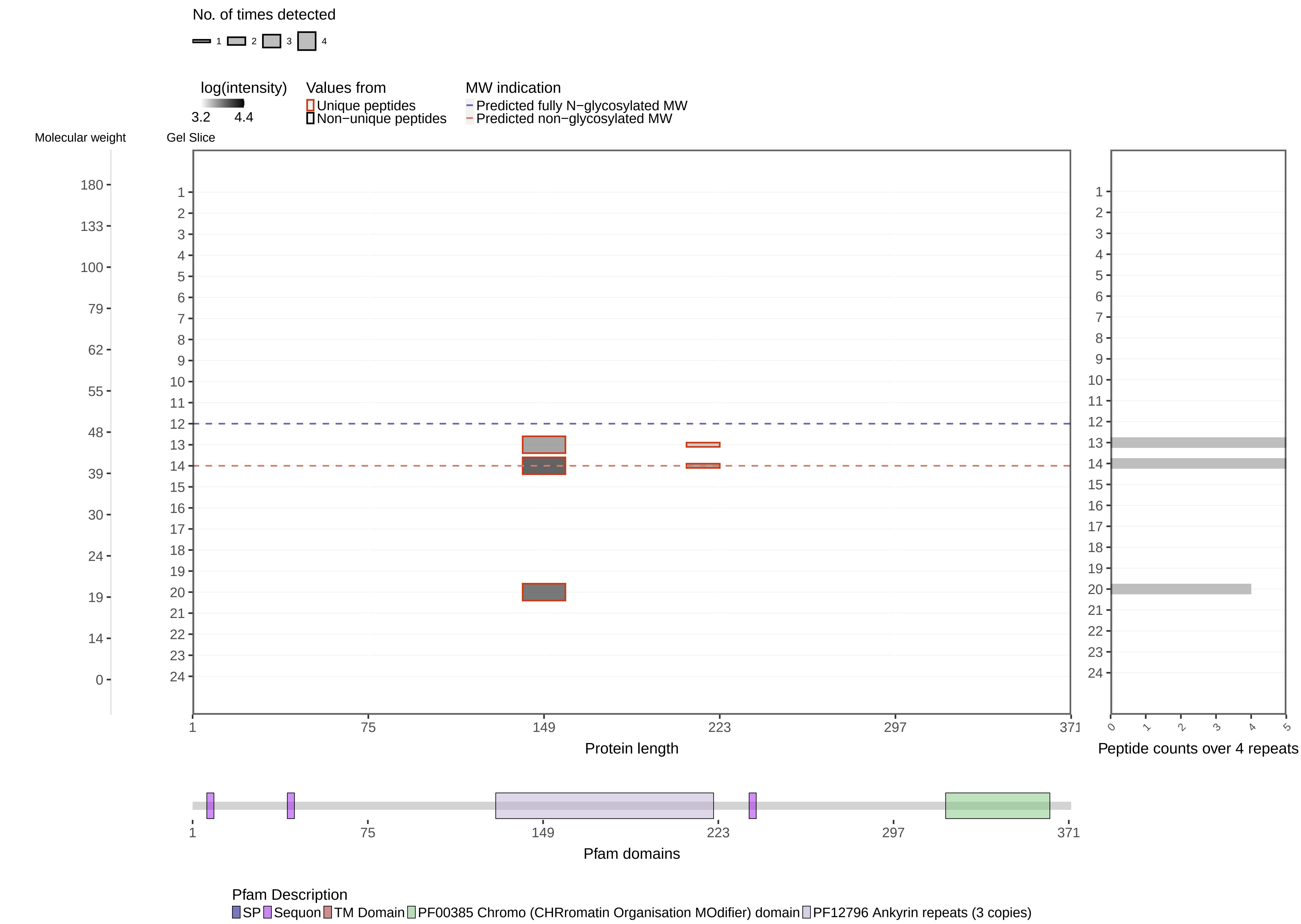


Protein sequence:

[illegible]VKLLAEAGADVDYRDRNGGLTALHMAAGYVKPDVAKLLIDLGADPEVQDYRGQTPLSLARMILNQTPKGNPMQFARRLGLENVVRILEDAIFEYVTVEEILEKRGKGENVEYLVKWKDGEDNEWVKAWVISEDLVRDFEAGLEYAVAECILEKREGEDGKGEYLVKWTDIEEATWEPEENVDPLLIEDFEKGQQNVVS*

NbL19g09400.1 protein_transport_protein_SEC13_homolog_B [MW = 32.52 kDa]

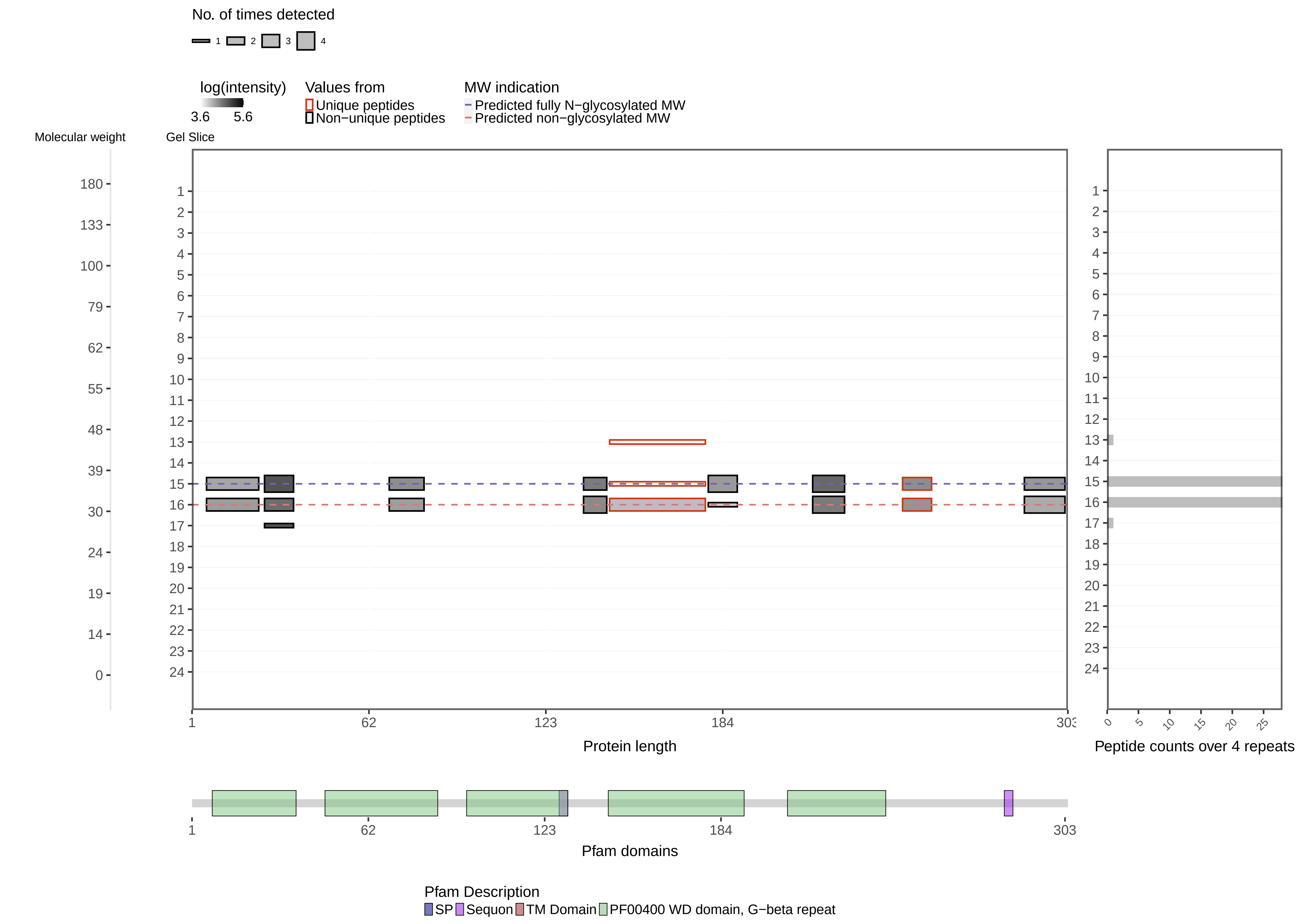


Protein sequence:

DTTRIDQAHPVGVTSVSWAPSMAPGALVGSGALEPVQKLASGGCDNTVKVWKLYNGIWKMDCFPALQMHTNWVRDVAWAPNLGLPKSTIASASEDGTVVIWTVAKEGDQWEGNVLKDFKTPVWRVSWSLTGNLLAVAAGDNNVTLWKEAVDGEWQQASTVDQ*

NbL19g09300.1 ferredoxin−thioredoxin_reductase_catalytic_chain_chloroplastic−like [MW = 16.61 kDa]

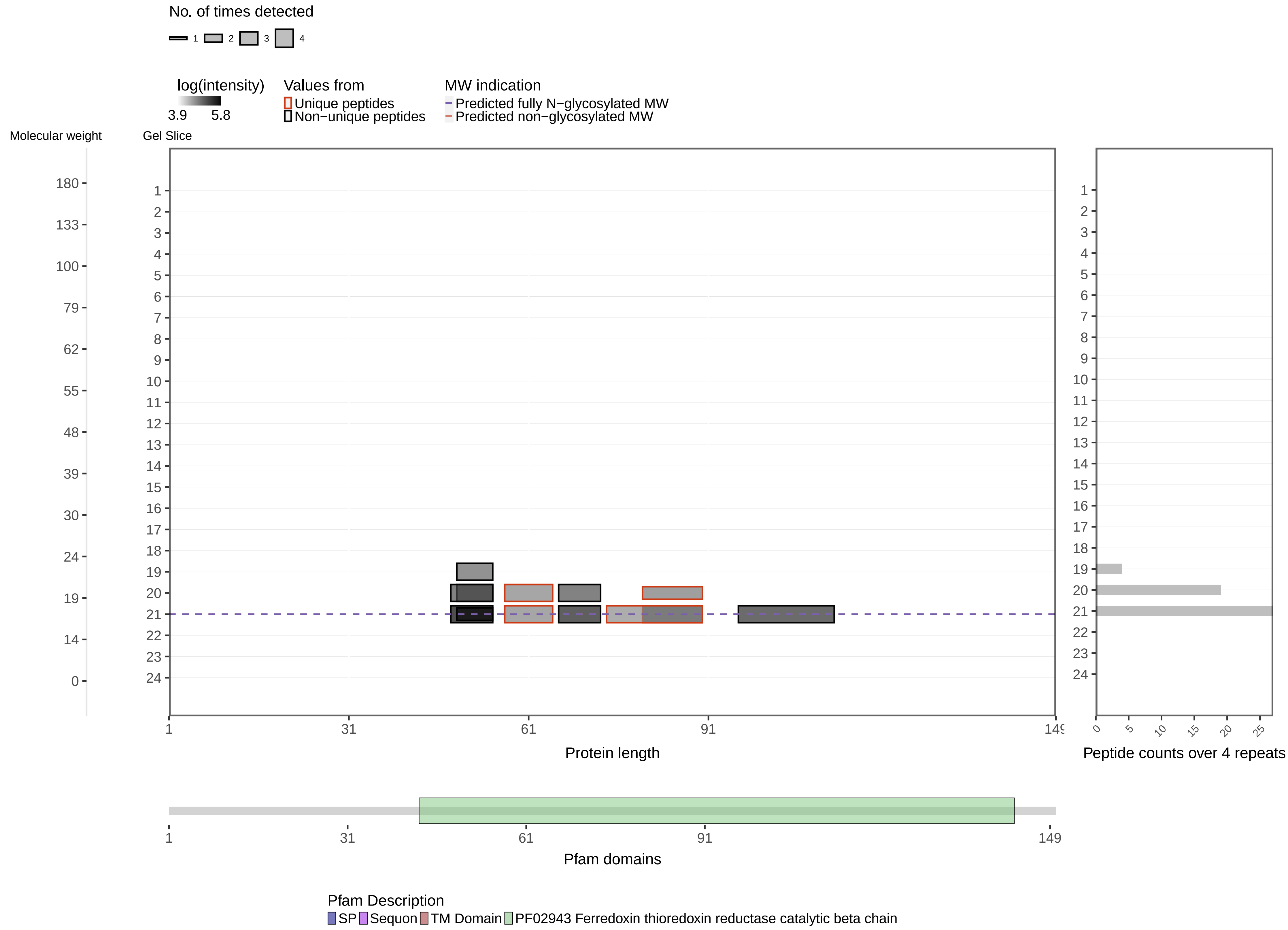


Protein sequence:

MDKGVTSVVIKGLAEHKDTLGAPLCPCRHYDDKAAEAQQGFWNCPCVPMRERKECHCMLFLTPDNDFAGEEQAISMEEIKETTANM*

### NbL19g09270.1 peroxidase_63−like [MW − SP = 34.17 kDa]

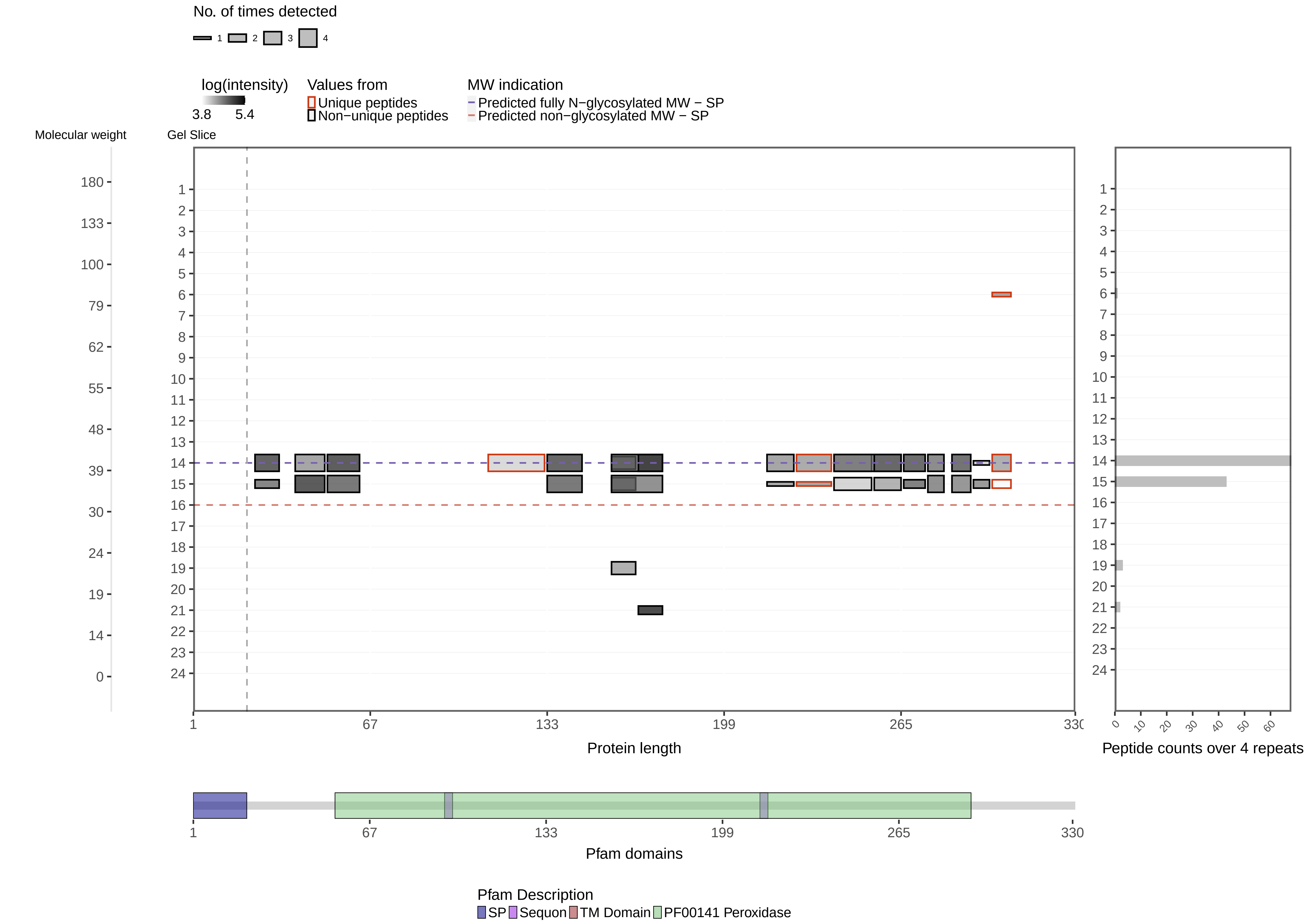


Protein sequence:

SKASLVEGNLPRPTMPMDOIIKIFESRGFSTOEMVALSGAHTIGFSHCKEFNSDLYNYNKTSOADPSYNPRFAOALENACSNSOKDPTLSVFNDIMTPNKFDNMYYONLPKGLGLLSSDRGLFSDPRTRIHVEEYVRDONAFFKAFAAAMOKLSEHGVKIGKHGEIRHRCDAFNN*

### NbL19g08990.1 formate_dehydrogenase_mitochondrial [MW = 42.46 kDa]

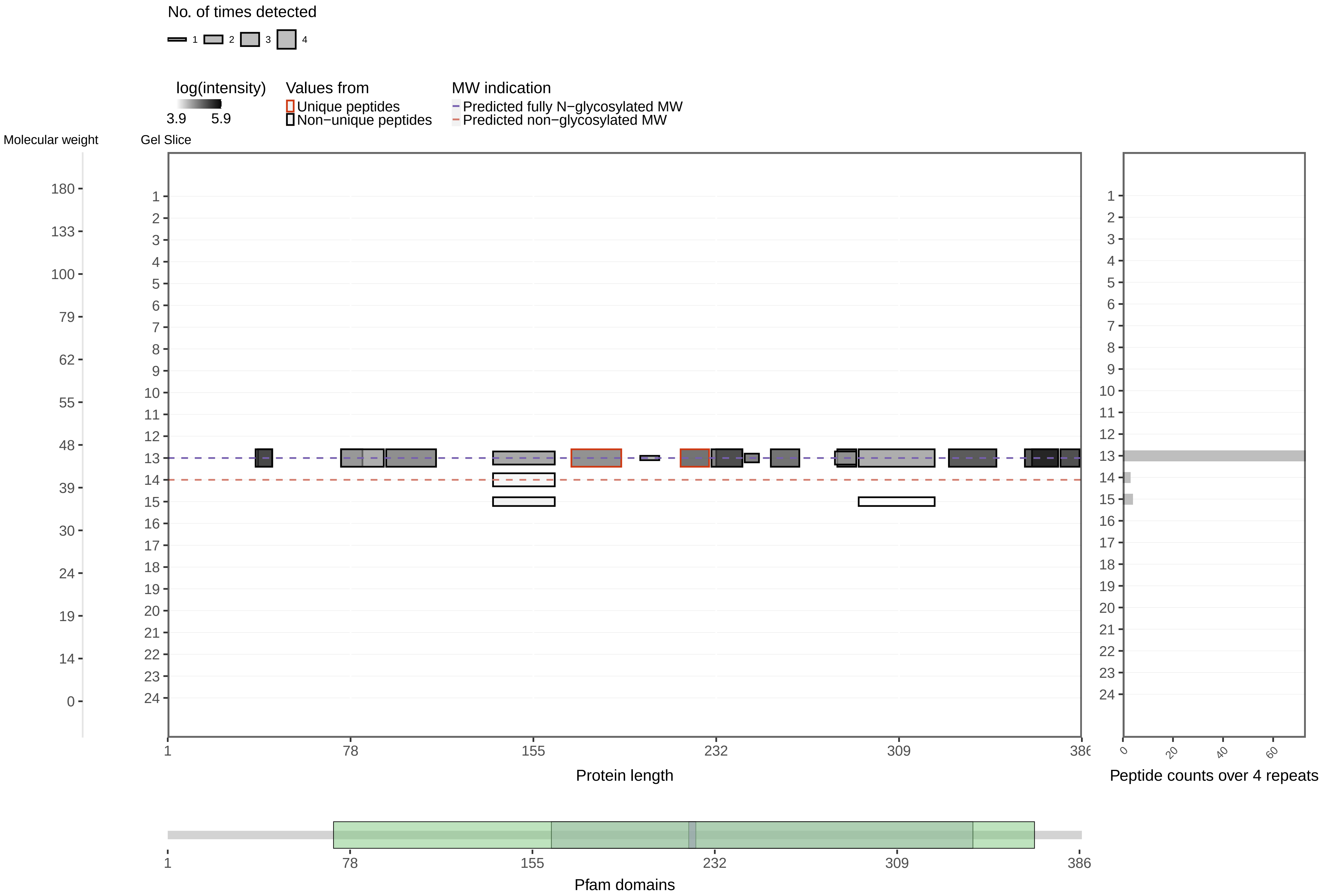


Pfam Description

SP | Sequon | TM Domain | PF00389 D−isomer specific 2−hydroxyacid dehydrogenase, catalytic domain | PF02826 D−isomer specific 2−hydroxyacid dehydrogenase, NAD binding domain

Protein sequence:

EWNVAGIAYRAYDLEGKTVGTVGAGRIGRLLLQRLKPFNCSLLYHDRVKMDSELENQIGAKFEEDLDKMLSKCDIVVVNTPLTEKTKGMFDKERIAKMKKGVLIVNNARGAIMDTQAVVDACNSGHIAGYSGDVWYPQPAPKDHPWRYMPNQAMTPHISGTTIDAQLRYAAGTKDMLDRYFKGEDFPAQNYIVKDGELAPQYR*

NbL19g08980.1 protein_farnesyltransferase_subunit_beta−like [MW = 49.86 kDa]

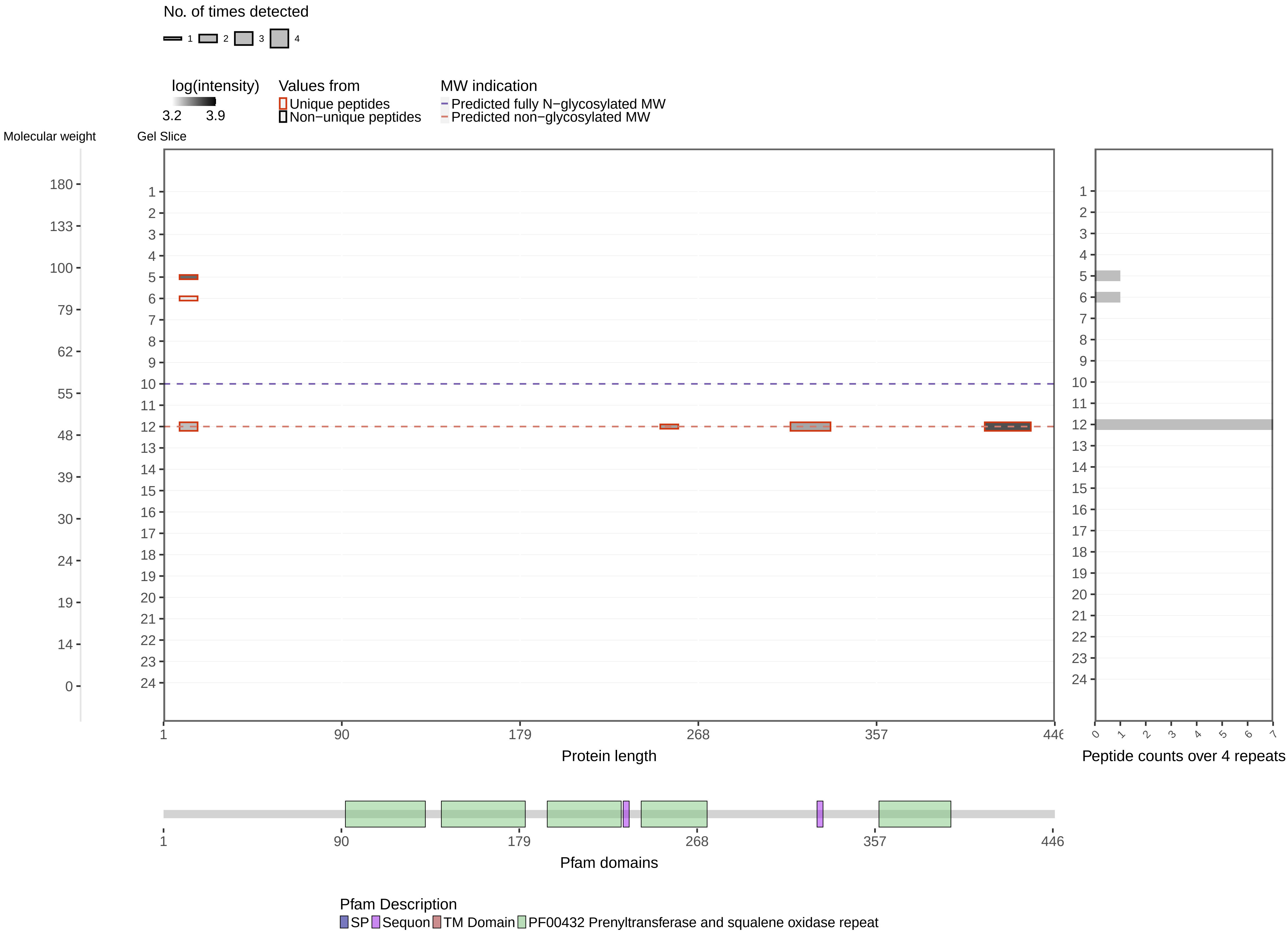


Protein sequence:

SEAHGGYTFCGLAAMILINETNRLDLPRLIDWMVFRQGVEGGFQGRTNKLVDGCYSFWQGAVAFLVQRLKSIVYEQLGLSNELSAESADESSESELSDEEKHLEGTSSHGQEGQENASDPTKIADTGYDFVNRRIAMQPVFDSFYLQQYVLLCSQIDGGFRDKPGKGRDHYHTCYCLSGLSIAQYSWTNEADAPPLPKDVFGPYSQNLLEQIHPLYNVVLDRYYEARRFFSCL*

### NbL19g08930.1 carbonic_anhydrase_chloroplastic [MW = 35.96 kDa]

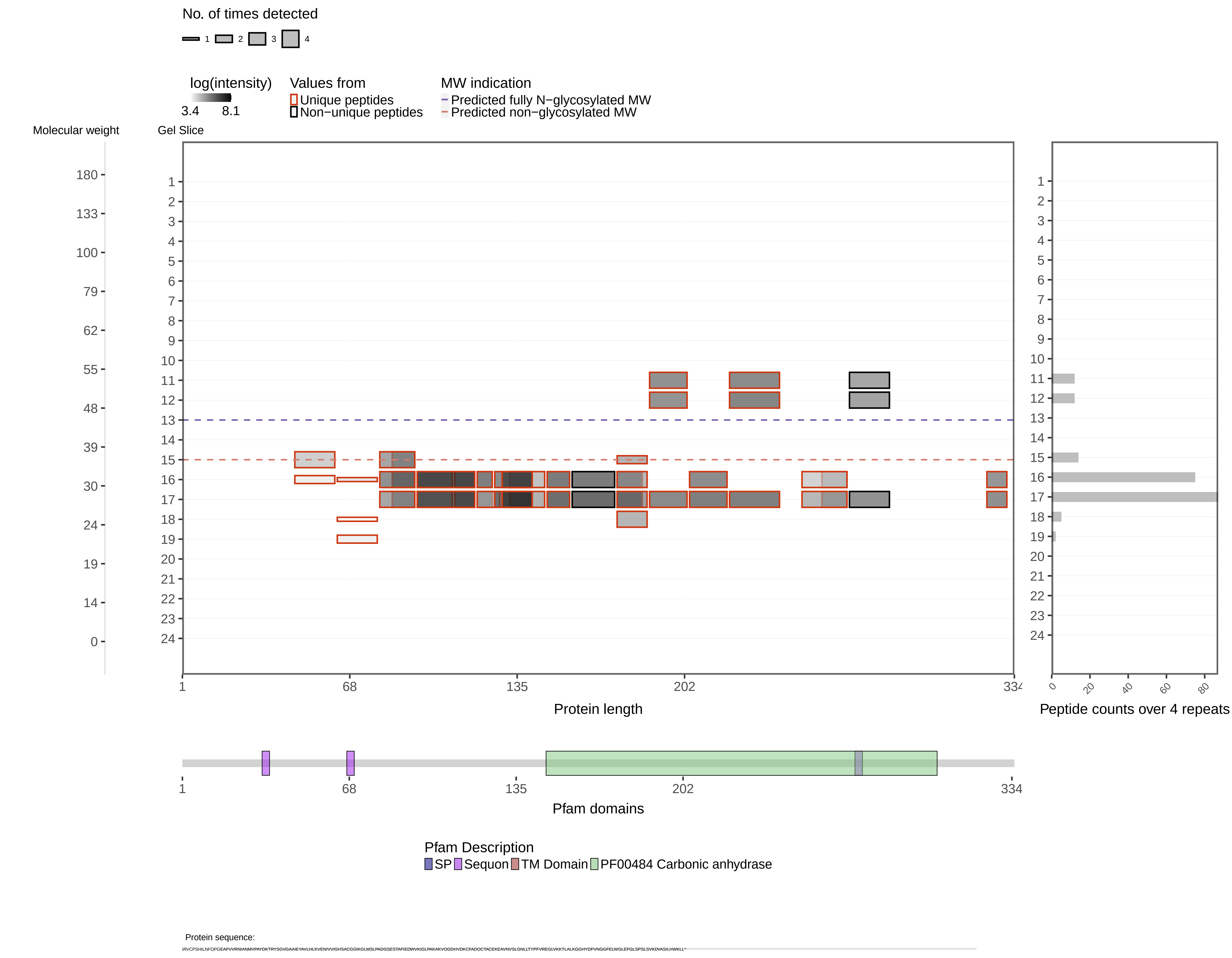

### NbL19g08840.1 50S_ribosomal_protein_L12_chloroplastic [MW = 22.18 kDa]

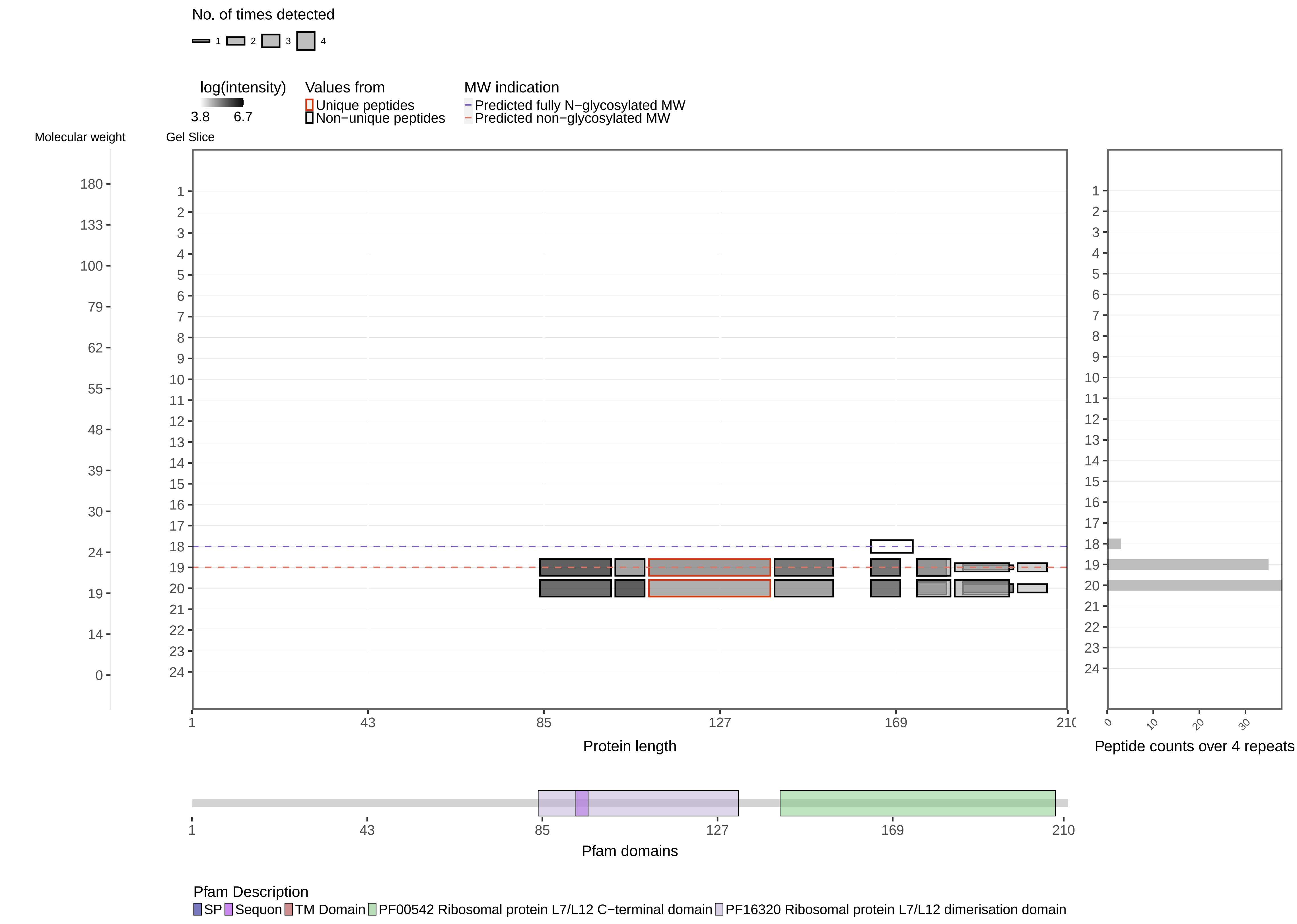


Protein sequence:

SNLTLADAQKLVEYLQDKLGVTAASFAPAAVVAAPGAAAEAPAVVEEKTEFDVVIDEVPSNARIATIKAVRALTNLALKEAKELIEGLPKKFKEGVSKDEAEDAKKQLEEAGAKVSIA*

NbL19g08740.1 glutaredoxin [MW = 11.29 kDa]

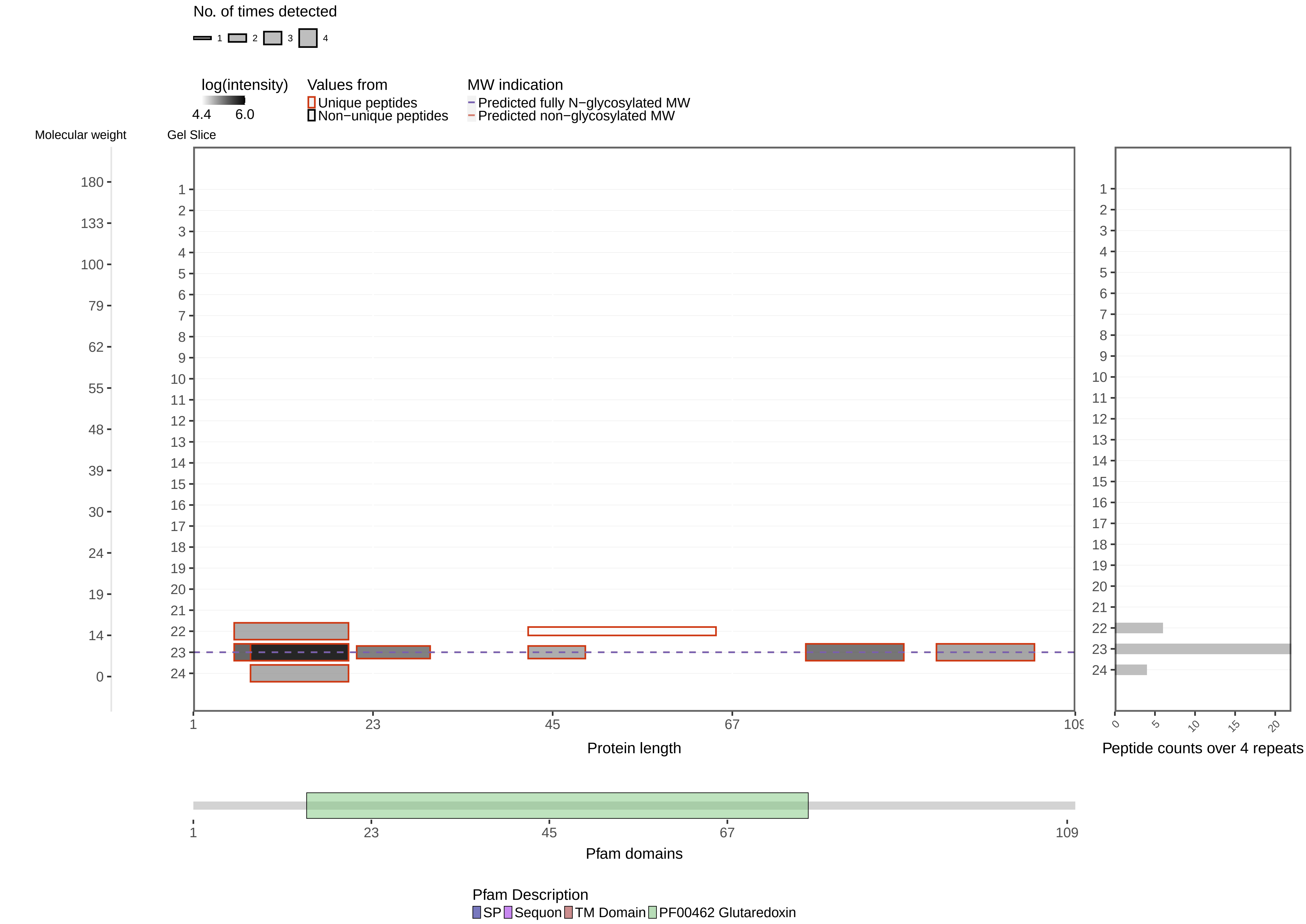


Protein sequence:

AVELDTEKDGSEIQAALAEWTGQRTVPNVFIGGKHIGGCDATTALHREGKLVPLLTEAGALAKSSSA*

NbL19g08650.1 sirohydrochlorin_ferrochelatase_chloroplastic [MW = 22.61 kDa]

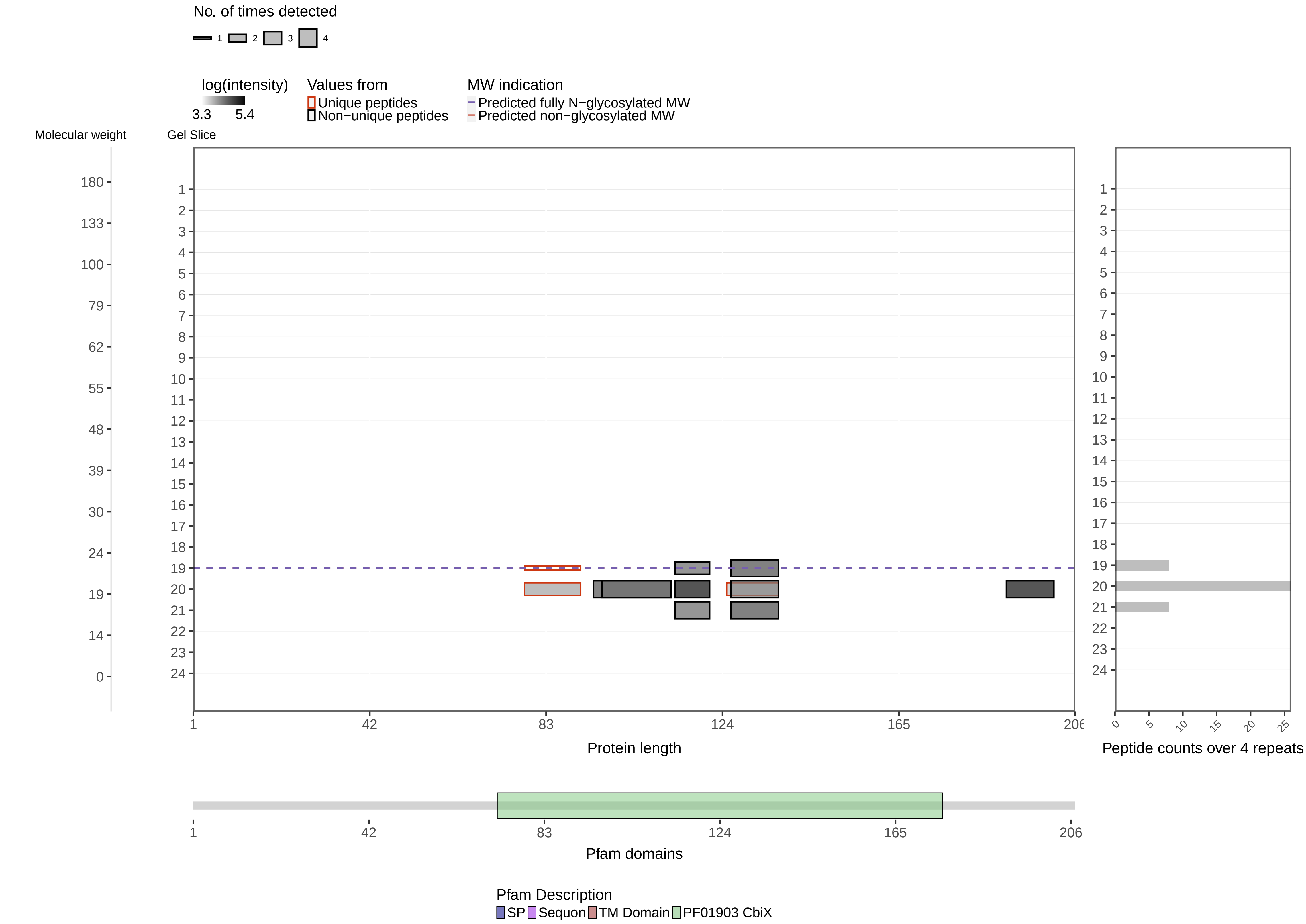


Protein sequence:

FKERTKYPIVEPAHMELAEPSIRDAFSLCVQRGARRVIVSPFFLSPGRHWNQDIPSLTAEAAKEHPGVSYVITAPLGLHELLVDVVNDRIKHCLNHIAGKADECSVCAGTGKCQLR*

### NbL19g08600.1 50S_ribosomal_protein_L5_chloroplastic [MW = 29.53 kDa]

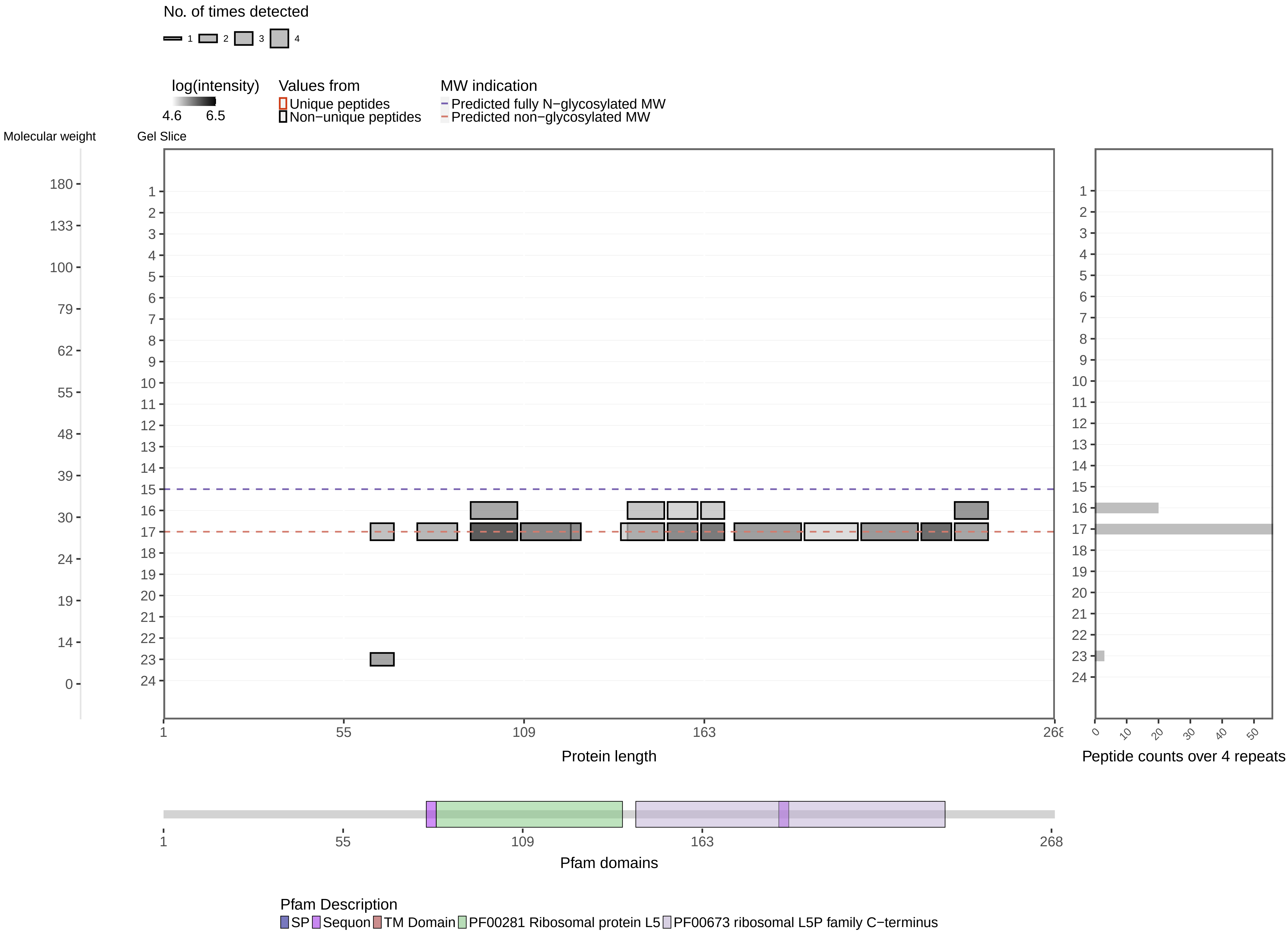


Protein sequence:

RPVKTRAKNAIATFKIREGQPLGIAVTLRGNVMFSFLDRLINLGLPRTRDFQGVNPNSFDGHGNYSIGFREQSVFPELSYDALGKPRGMDVCITTTAETDKEAHRLLALMGMPFREGAVGQTNVVRKKKLRGHHFDSKAKQRSRR*

### NbL19g08550.1 titin [MW = 418.08 kDa]

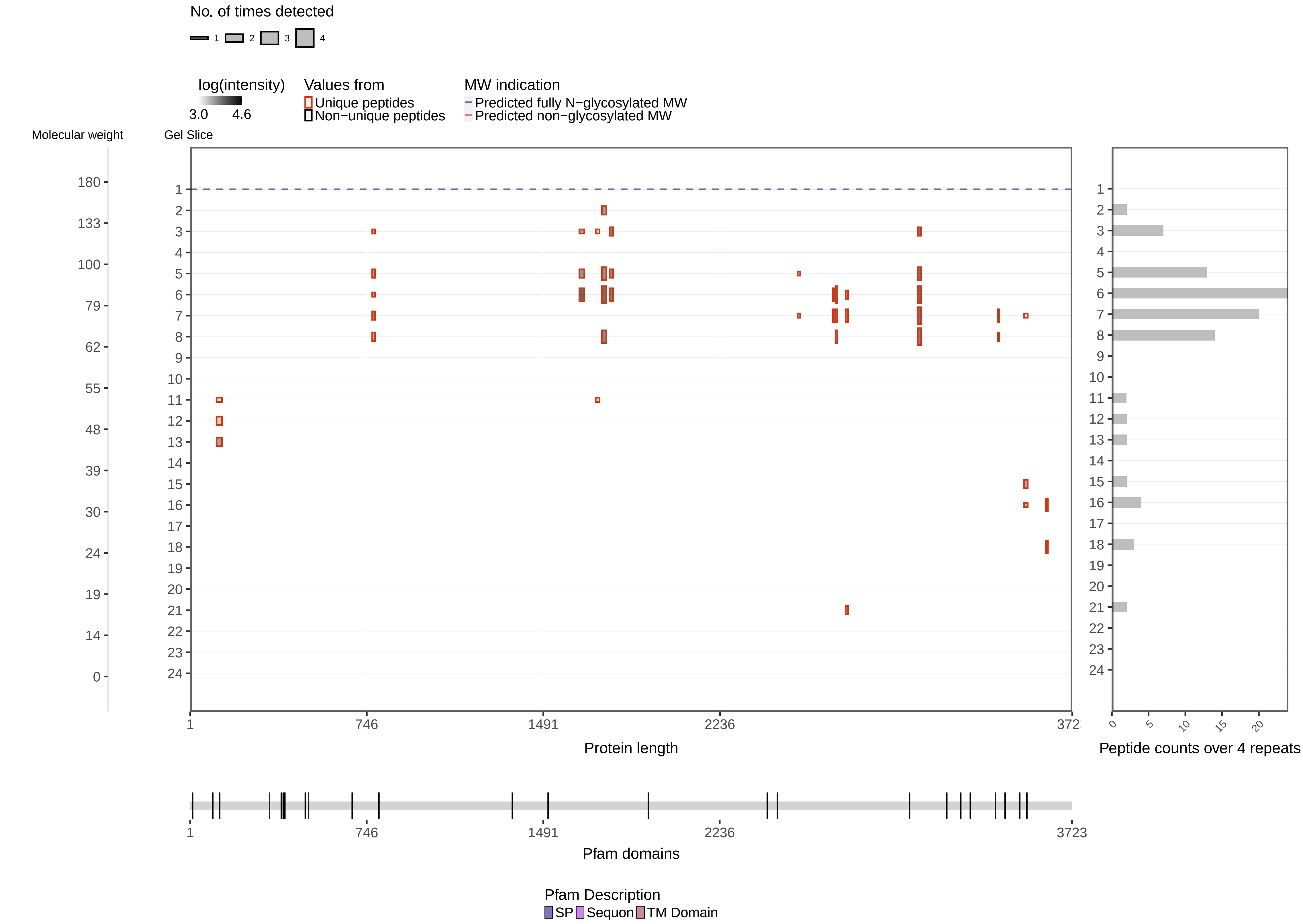


Protein sequence:

VGEKLEEPVVEQTPAGQAWEETPKELAFEEKPEEPAVEETPEELAVEKNPDKTAVGEKHEKVTLLEKNIPIIELNEKNDSSTPCNTNNDAADYLAVEEKPEEPAIEKKTEEQMAKEKLEEPNVEDKPEKSVIEEKSEKLATEETLEEQAGEETPEALAVEEKPDESAVEETVEELAIEQKPKETATEKEKPETVTLLEKNMPVVDTKDKINSEIPCNANNDVSDYPAVEAKPEELAVETKYEEPPAVILEKPAELAVEETPEKTVVEEKPENLAAREETLEEPALEEIPEEPTVEEKPI

### NbL19g08480.1 RNA−binding_protein_squid [MW = 39.48 kDa]

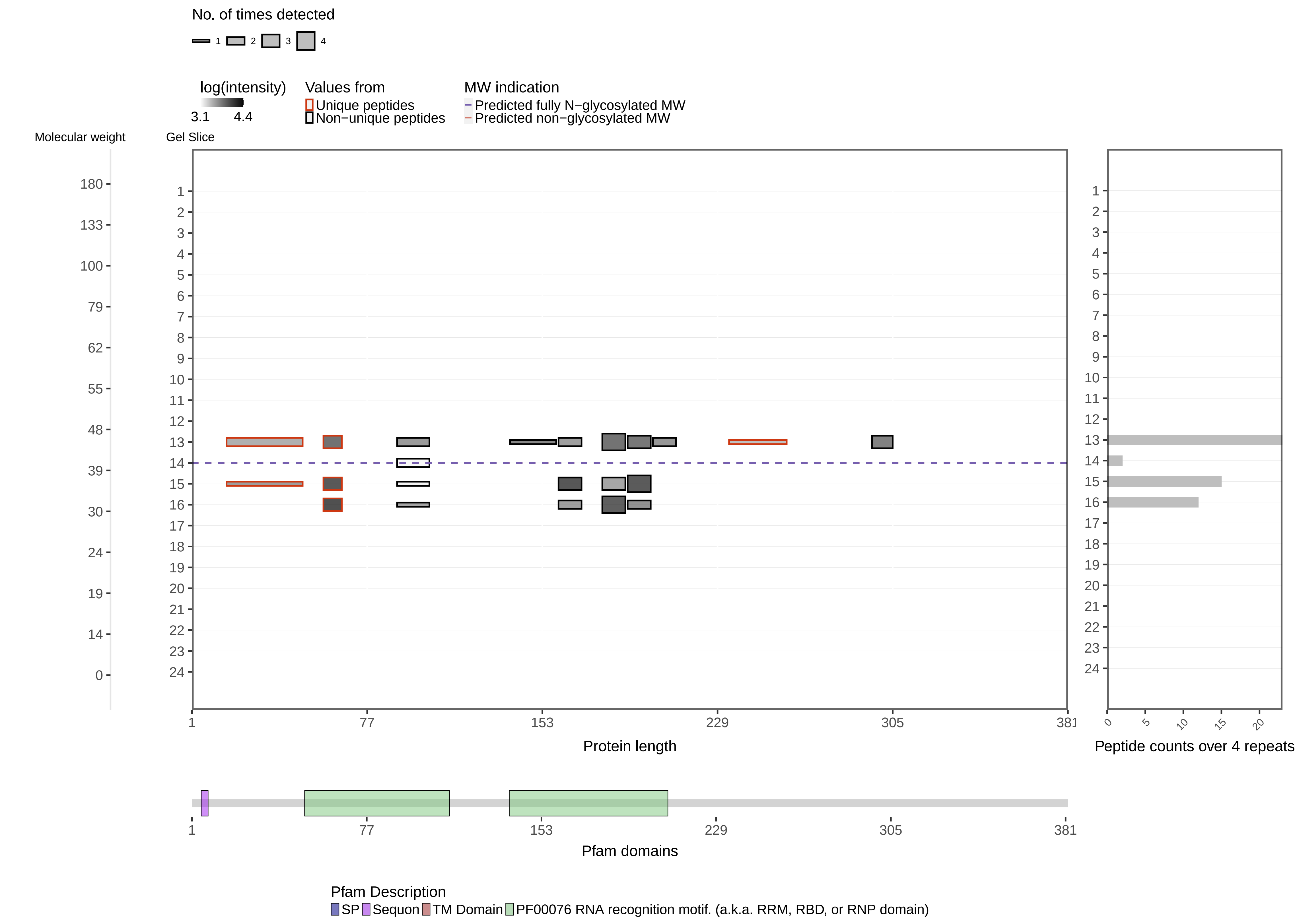


Protein sequence:

FDSEKSVEDILANGNRLDFAGTQVEIKKAEPKRPNAPLPATSKRQGHPRASFGGGFSDAYGGYGDSGFGAGGPYRSGGPYGGRSSAYGGYSGAGMGGYGAYGGSGAYGGFREEASFGYSSRYGGAFGRGYDVGGGYGGPSEGYGGYAAGSGGYGSGYGAGLGGGYGGGDGTGGSMYGSSRGGYGGAGSGRYHPYGR*

### NbL19g08390.1 glucuronokinase_1−like [MW = 39.74 kDa]

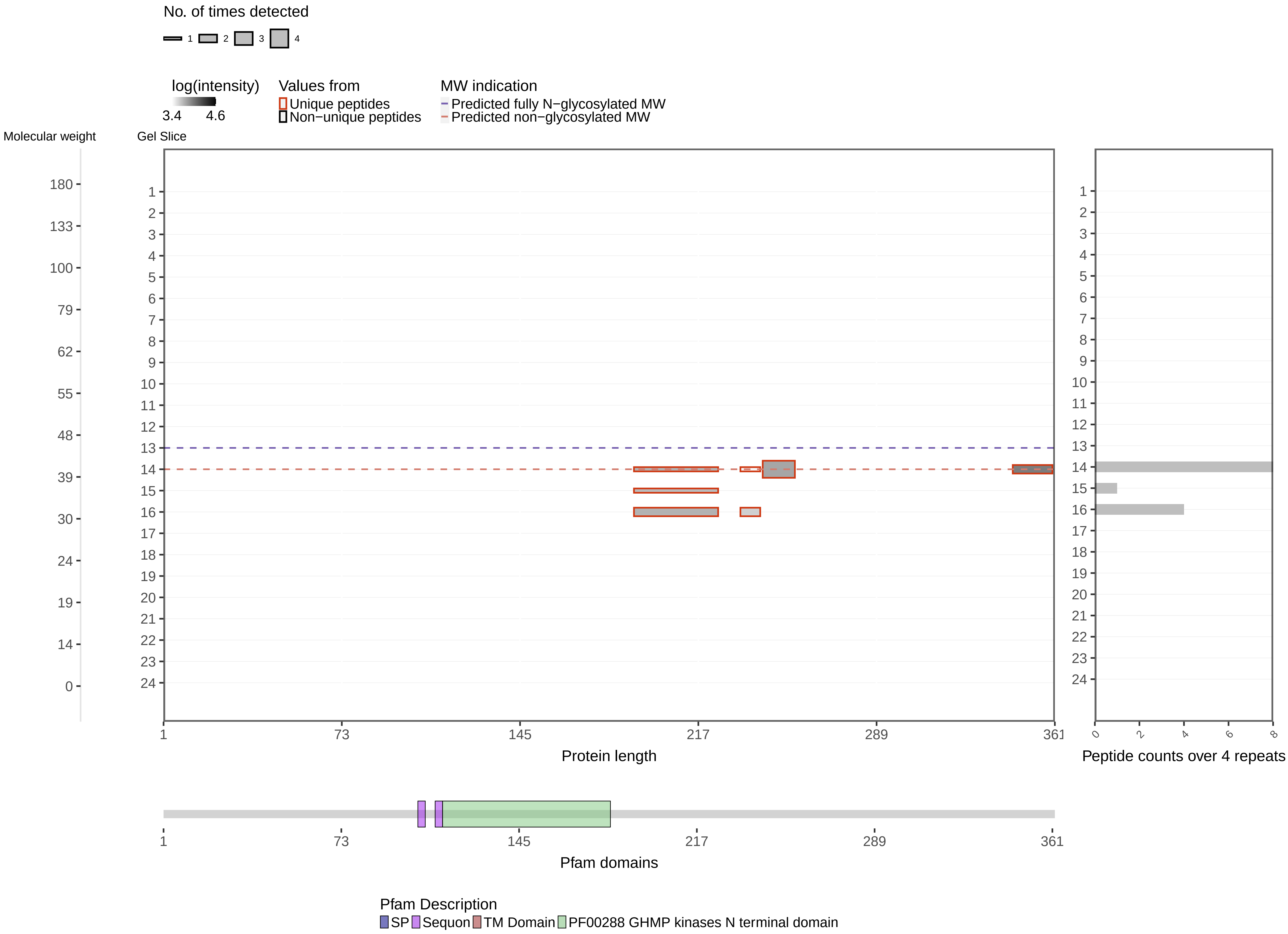


Protein sequence:

GLQDRVAQVYGGLVYMDFGKNHMDELGHGIYTPVDVDLLPPLYLIYAENPSDSGKVHSTVRQRWLDGDEFIRSTMEEVANIAAEGRKALLEKDYLKLAALMNHNFDLRRLMFGDDALGAMNIEMVEIARRIGAASKFTGSGGAVVVFCPDGPSQVKQLEDACHSAGFTFQPIKVMPSFLSEVDLQTPGSH*

### NbL19g08250.1 ras−related_protein_RABB1b [MW = 23.34 kDa]

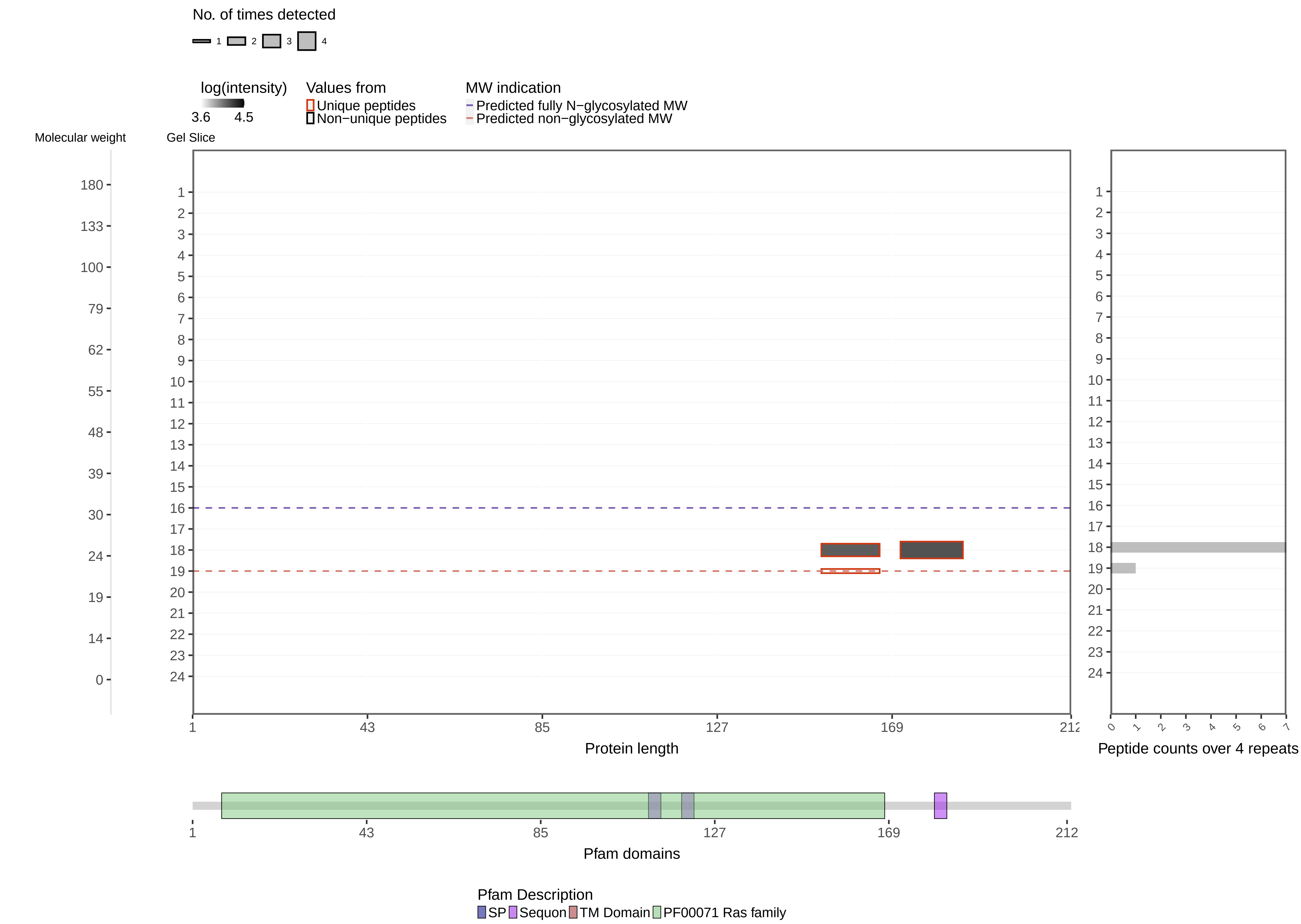


Protein sequence:

NHLASWLEDARQHANPNMTIMLVGNKSDLSHRRAVSKEEGEQFAKENGLLFLEASARTAQNVEEAFIQTAAKILQKIQEGVFDVSNESSGIKVGYGRTQGPAGPRDGAVAQRGGCCG*

### NbL19g08220.1 uncharacterized_LOC104120577 [MW = 33.63 kDa]

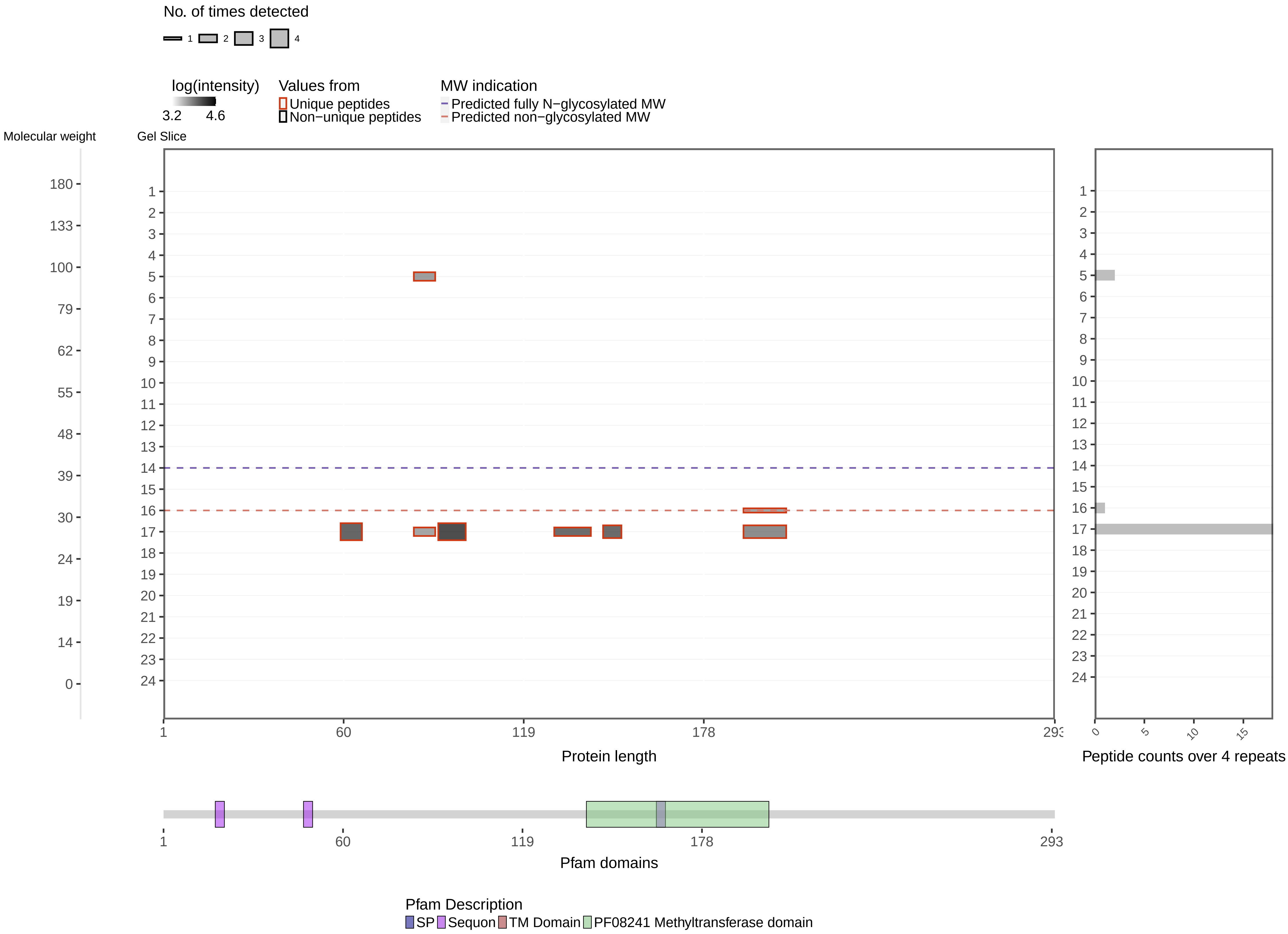


Protein sequence:

NAQELAKNPRLDYFIVKDLNQDQKLEFENCSFDAVLCTVSVQYLQQPEKVFAEVFRVLKPGGVFIVSFSNRLFYEKVISAWRDSTGYGRVQLVVQYFQCVEGFTQPEVIRKLPTGKDGVAQQNKLPSNWIMQLTGLLSASSDPFYAVVAYRNFKPVHE*

NbL19g08070.1 glucose−6−phosphate_1−dehydrogenase_cytoplasmic_isoform [MW = 26.69 kDa]

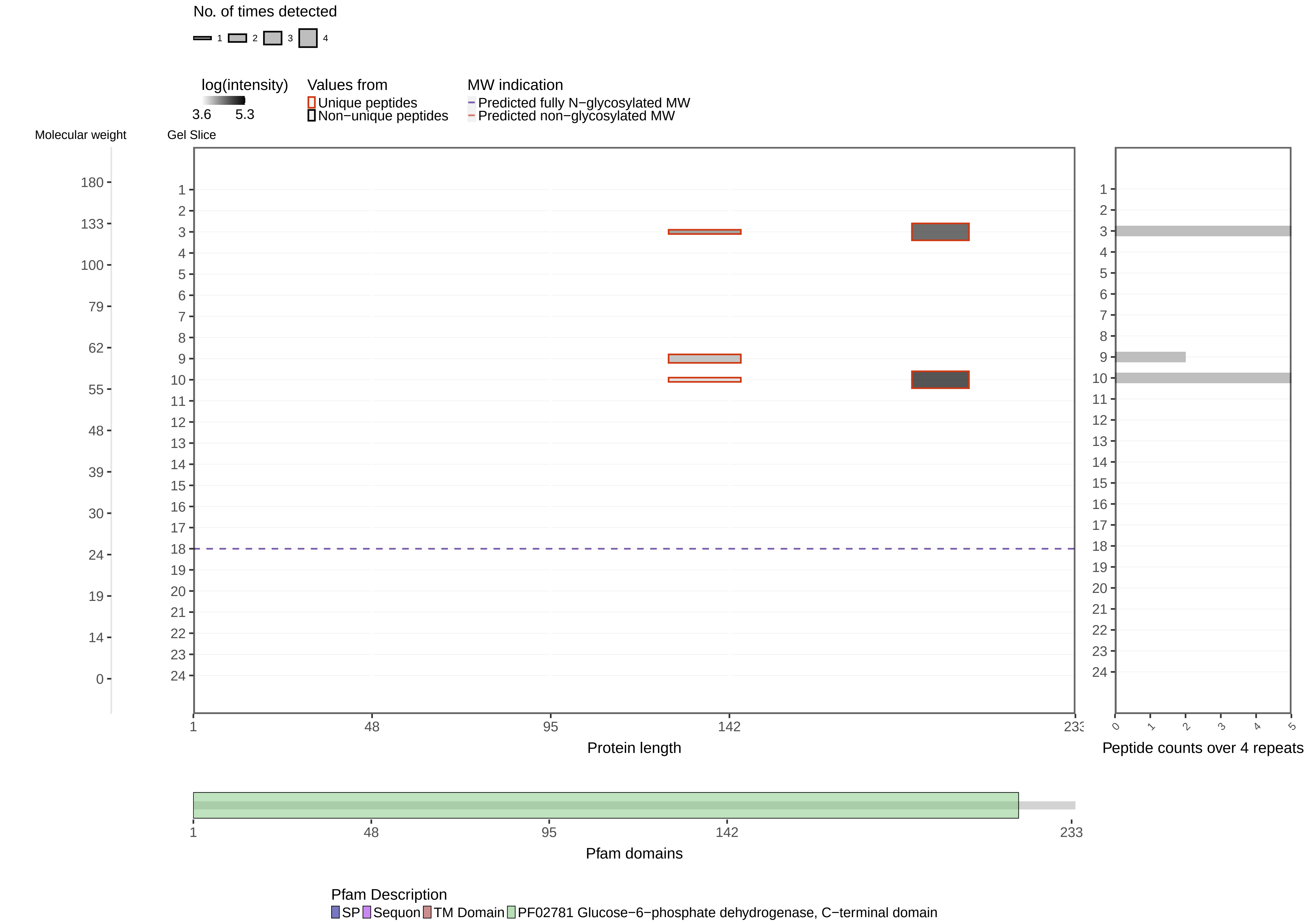


Protein sequence:

GRNEFVIRLQPSEAMYMKLTVKKPGLEMSTVQSELDLSYGQRYQGVVIPEAYERLILDTIRGDQQHFVRRDELKAAWEIFTPLLHRIDDGEVKPIPYKPGSRGPAEADELLQNVGYVQTHGYIWIPPTL*

NbL19g07750.1 nuclear_transport_factor_2−like [MW = 24.87 kDa]

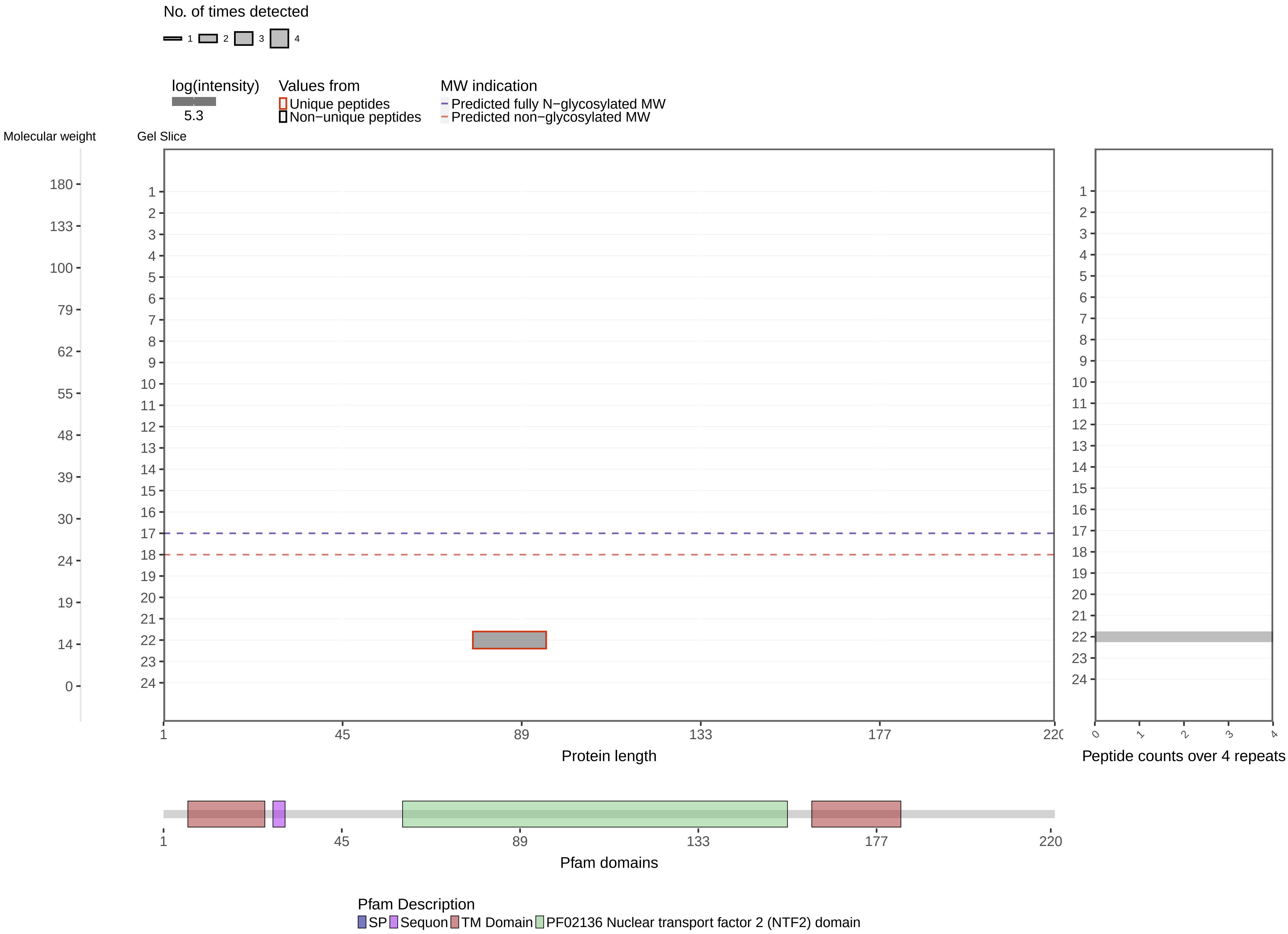


Protein sequence:

QGAQHIVAKLTSLPFQQCKHQINTVDCQPSGPIGGMLVFVSGNLQLPGEEHVLKFSQVFSPLLRALLLLFHVLFIIIGDLLMYFAISKSIIFNRLSKENGWNFDFLFRTPVDFPFRWGRMFKV*

NbL19g07710.1 uncharacterized_protein_YciO−like [MW = 33.36 kDa]

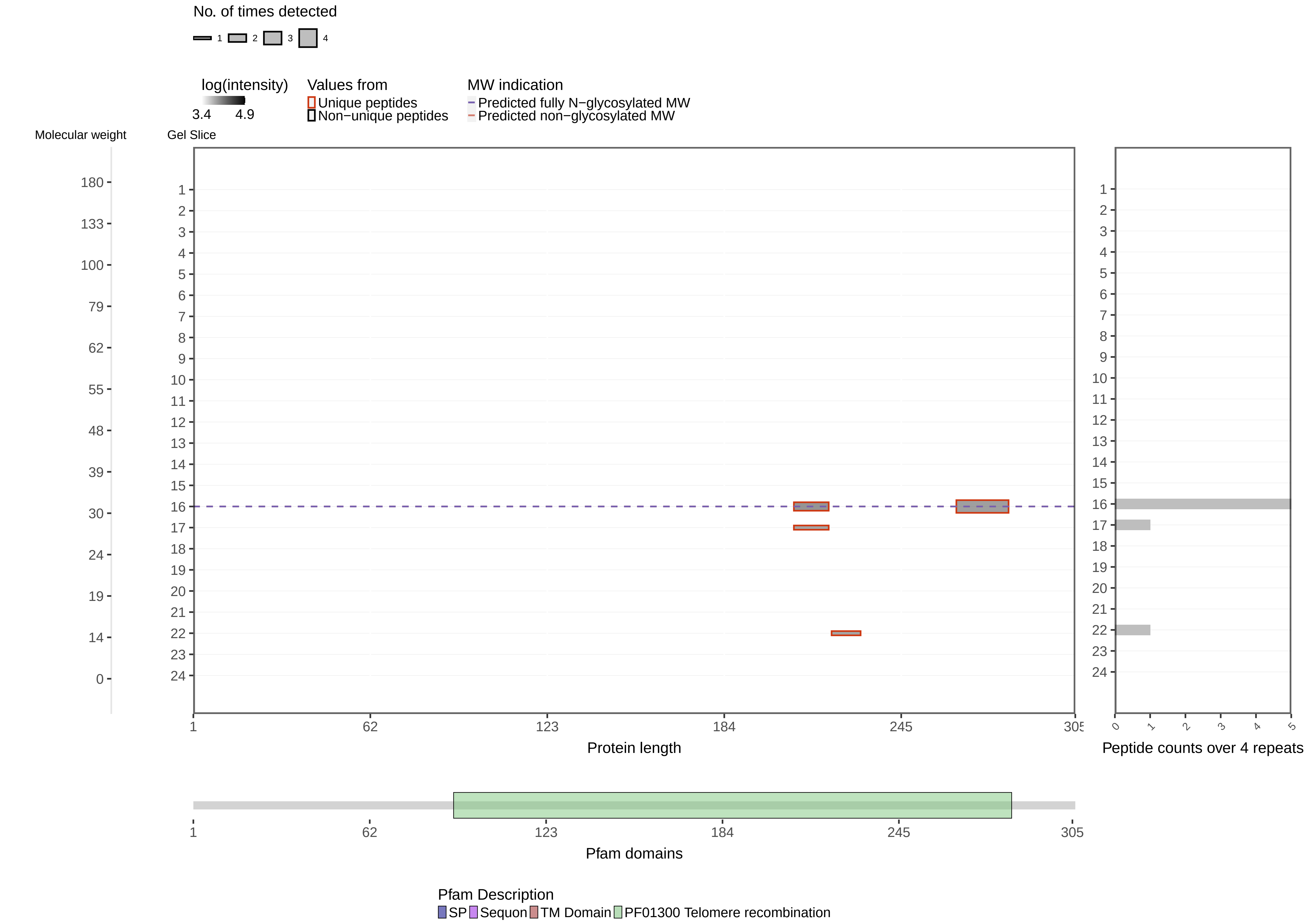


Protein sequence:

SFRDIDTYTTGFSSGNGQGLTDIFRAVKHCLPGPYTFILTASKQLPKQCTRYGTAASRYISRKNVGVRIPDDPVCQAILEKMDGPLISTSVKSPEENEWILDPVIIADVYGPEGLDFVVDAGLRVADPSTVVDMTASSPHLIRQGKGPKQPWMIAEDDDSARDGN*

### NbL19g07620.1 basic_blue_protein [MW = 20.95 kDa]

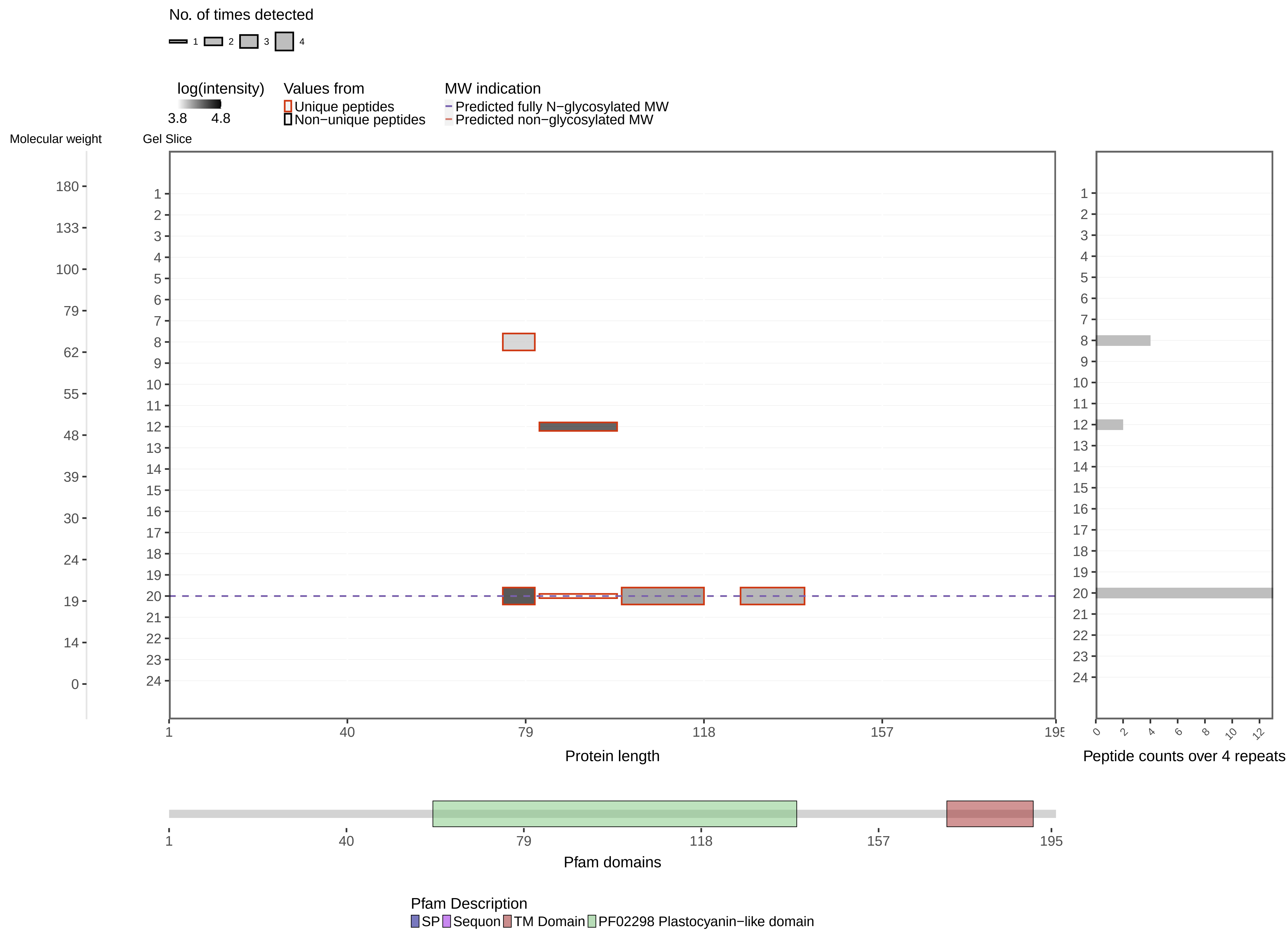


Protein sequence:

PGLHSVVEVEGESAYKSCDTSSSVNSMSAGNDVVKLNKPGTRYFACGTAGHCDQGMKLKITTVTGNAPSNQAATSSNPSSSSVADSRRFSTAFFTFVAAIFTIQMALVFQL*

NbL19g07360.1 mitochondrial_phosphate_carrier_protein_3_mitochondrial−like [MW = 39.22 kDa]

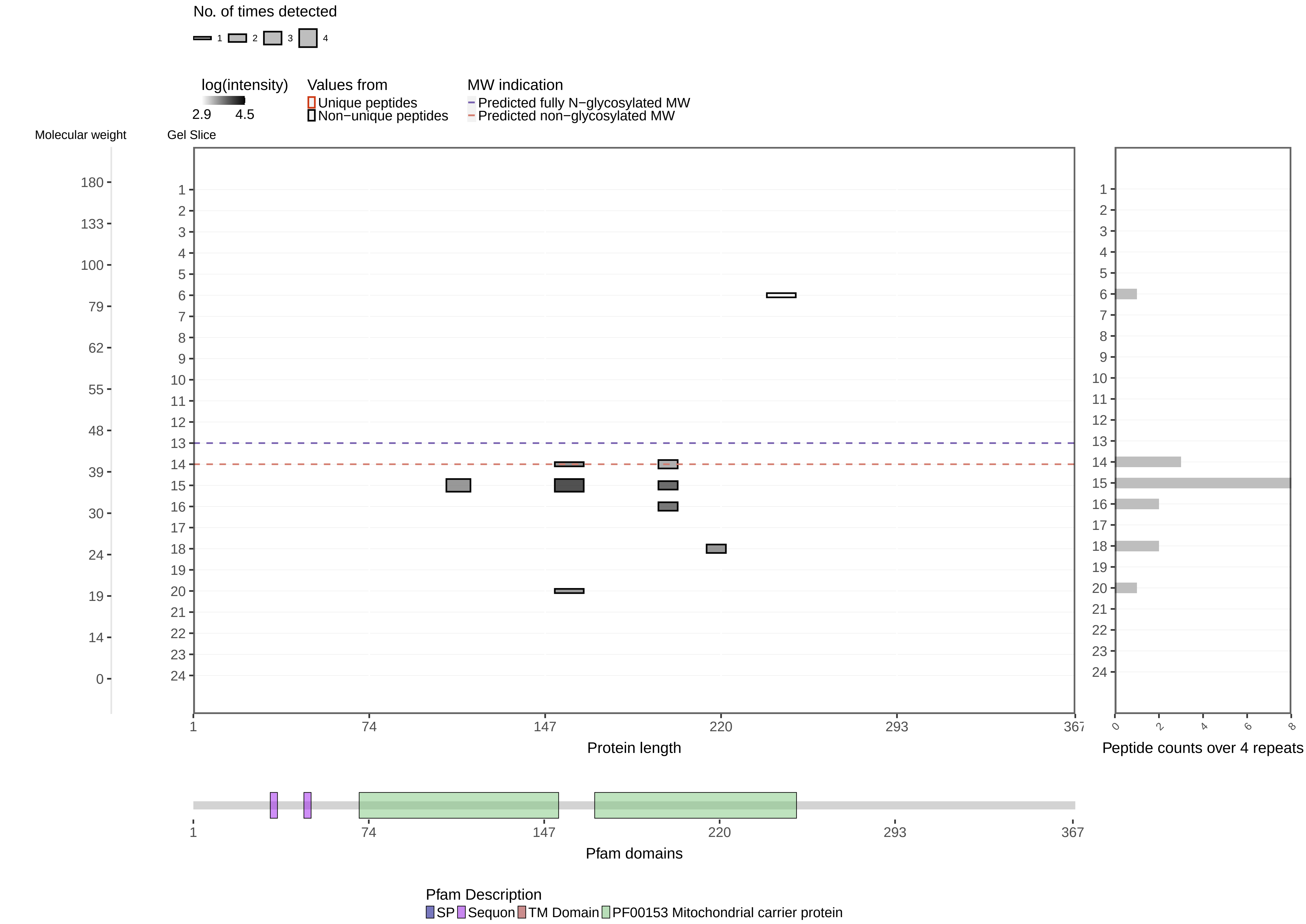


Protein sequence:

AGSASAEVIADIALCPFEAVKVRVQTQPGFARGLSDGLPKFVRSEGTMGLYKGLVPLWGRQIPYTMMKFASFETIVEMIYKHAIPKPKNECSKTMQLGISFAGGYIAGVFCAIVSHPADNLVSFLNNAKGATVGDAVKKMGVVGLFTRGLPLRIVMIGTLTGAQWGIYDAFKVFVGLPTTGGVTPAAPAASEVAKV*

NbL19g07210.1 acetyl−coenzyme_A_synthetase_chloroplastic/glyoxysomal [MW = 86.21 kDa]

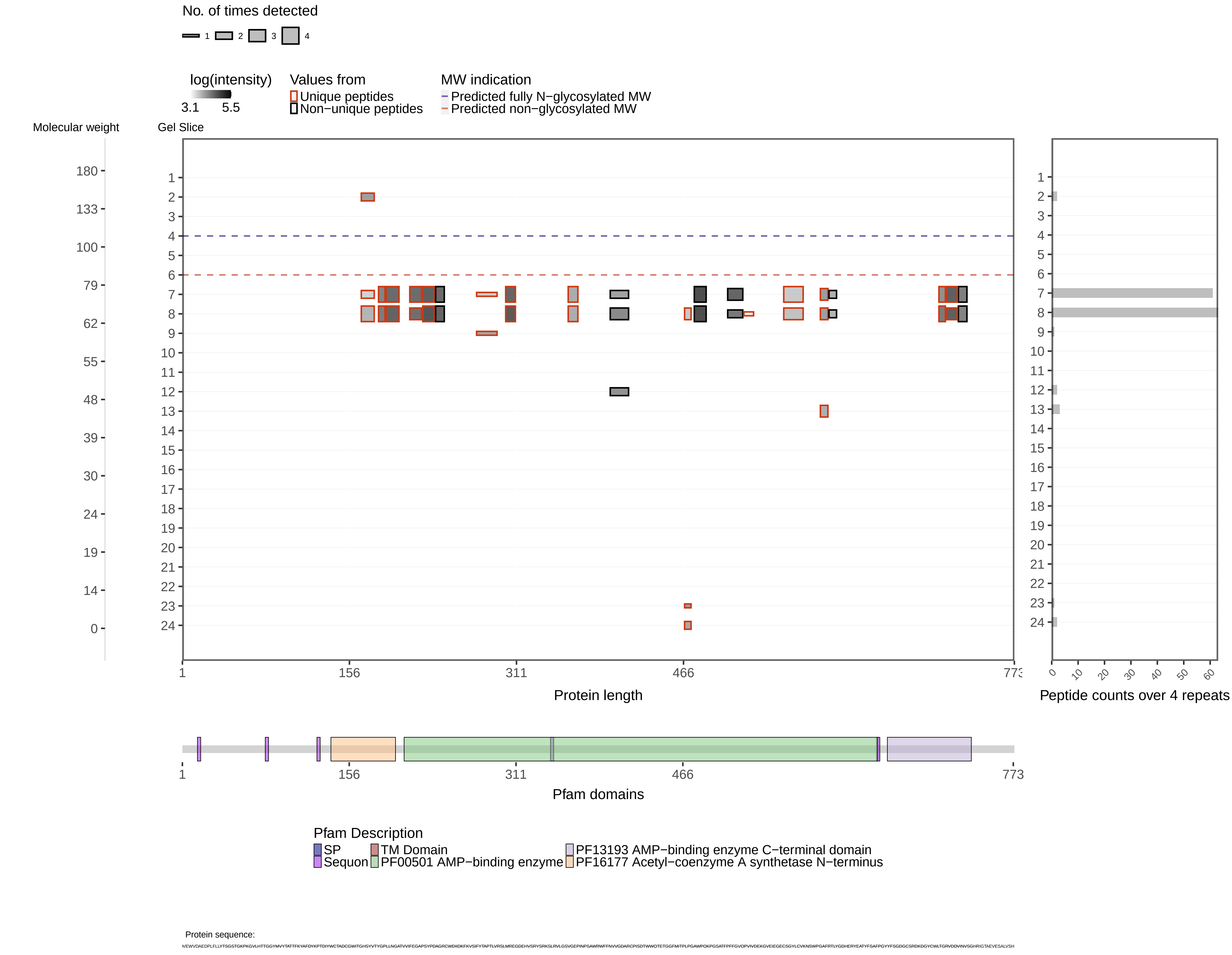

### NbL19g07200.1 beta−glucosidase_18−like [MW − SP = 57.03 kDa]

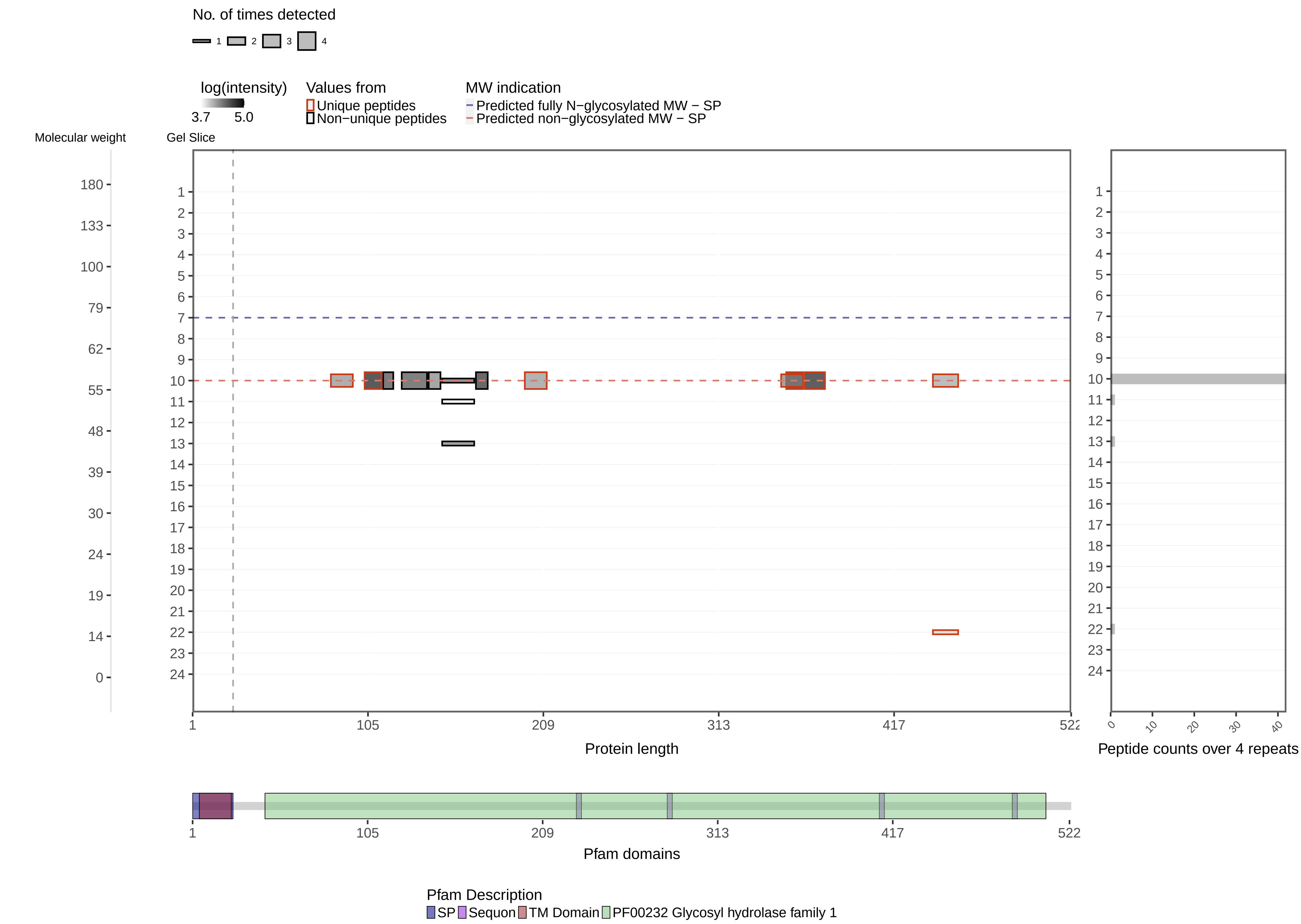


Protein sequence:

SHAAAVSIYRTRYQKRQGGMIGISLDIQWYEPYSNSSEDIAATQRARSFYVNWFLDPIILGRYPMEMVQILGSHLPDFSVNDLSKLSYGLDFIGINYYTAKYIKDCLYSACEHGKTWSEGSYFVTTEKDGVYIGQPTEVDWLFLYPQGMKKIVMYMKDRFNNTPMVITENGIAENENLNPSITDTLNDVHRVNYMHSYLNSLANAIREGADVRGYFVWSLLDNFEWLDGYRLRFGLHYVNFTNLERTPKFSATRYKELMYNFHIQLETHTAQK*

### NbL19g07190.1 phosphoribosylamine−−glycine_ligase−like [MW = 55.25 kDa]

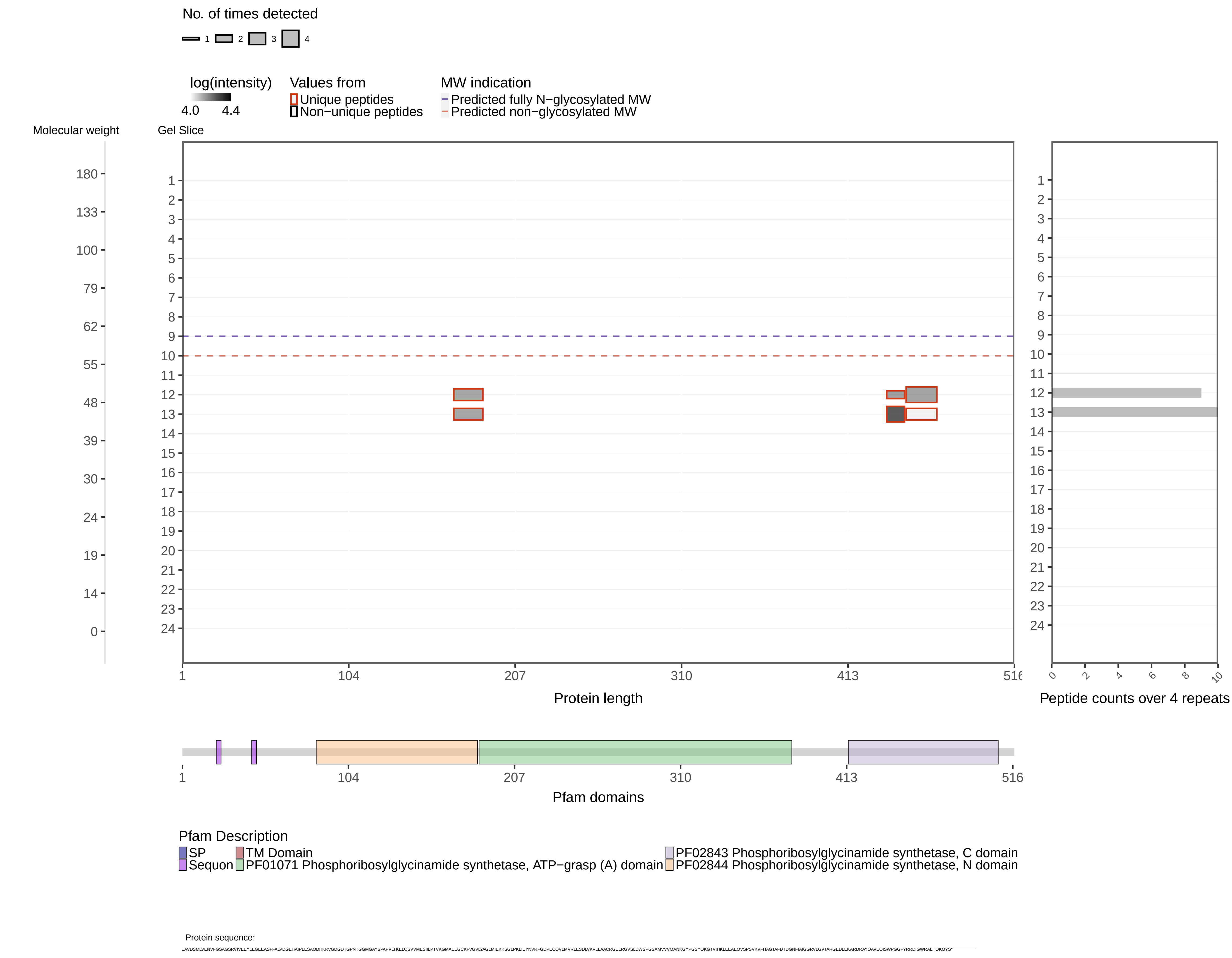

NbL19g07040.1 oxygen−evolving_enhancer_protein_3−2_chloroplastic−like [MW = 24.19 kDa]

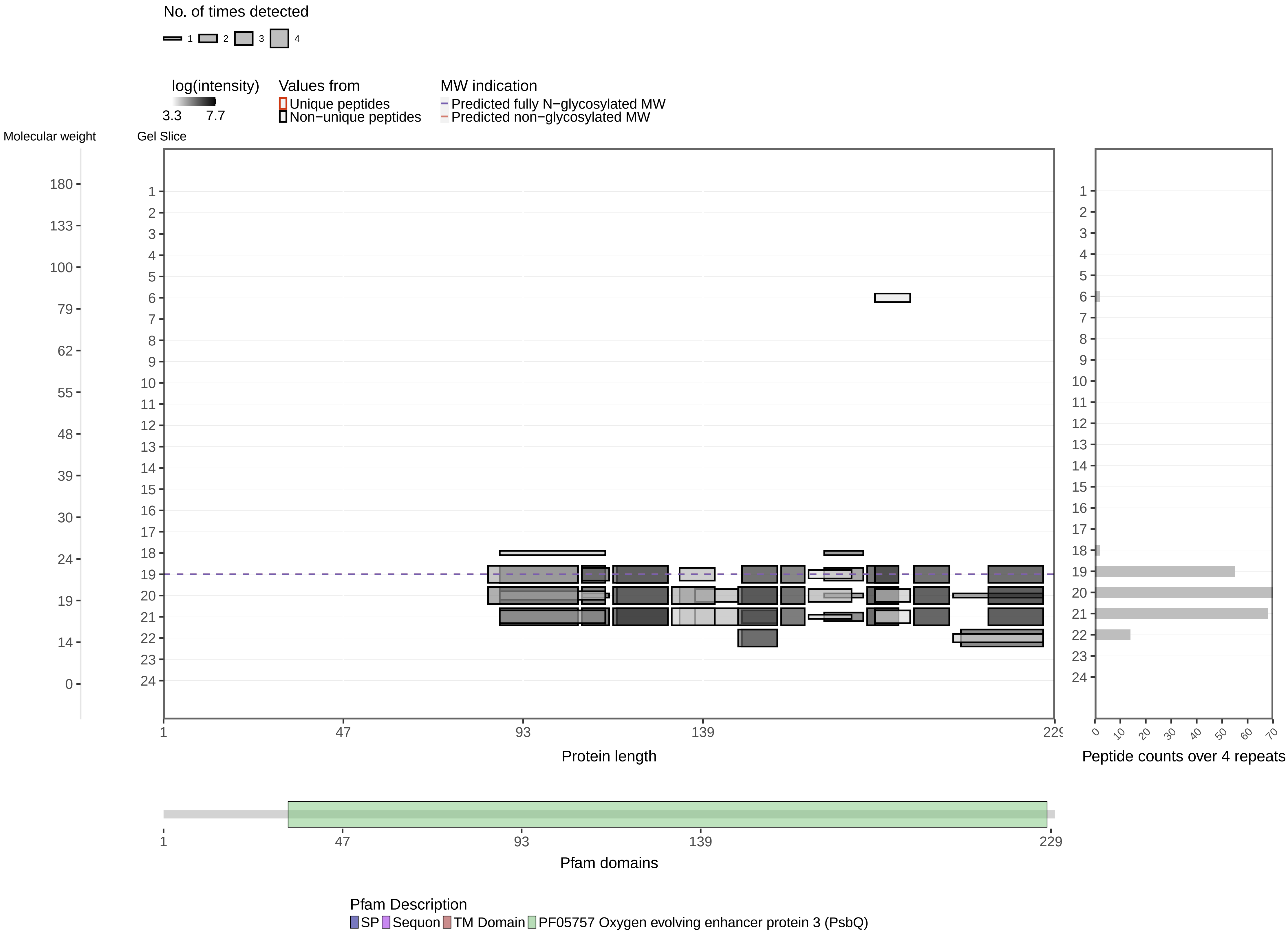


Protein sequence:

LNSDEARDFGLPLKKRFYLQPLTPAEAAQRVKDSAKEIVSVKNFIDKKAWPYVQNDLRLRAEYLRYDLKTVISAKPKEEKGKLQDLTGKLFKTISDLDHAAKTKNSPEAEKYYAETVSTLNDVLAKLG*

### NbL19g06980.1 (−)−alpha−terpineol_synthase−like [MW = 62.89 kDa]

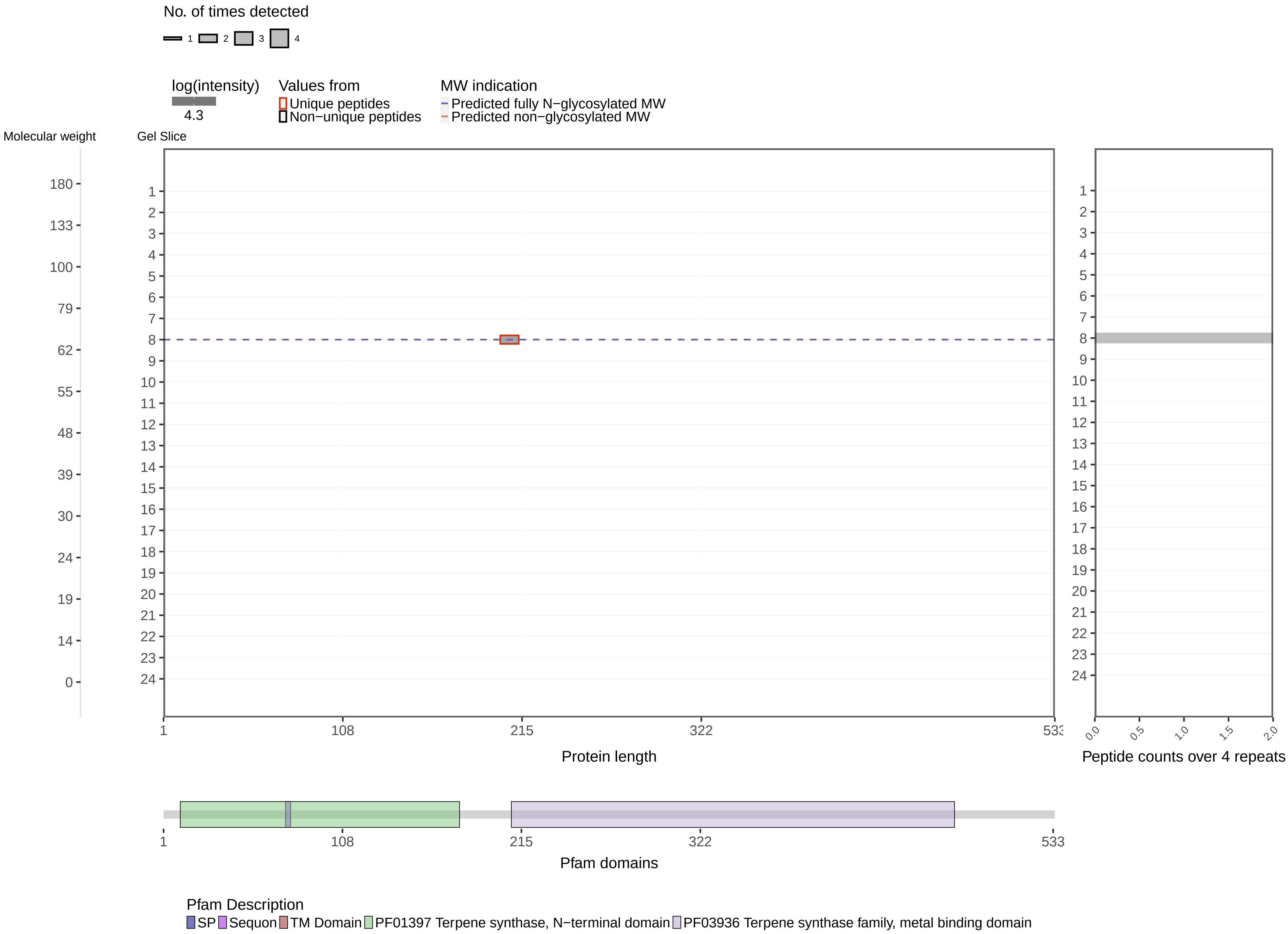


Protein sequence:

RLVEGFLWAVGFTPNPQFGYCRKFSTKLSVLLTTIDDIYDVYGTMDELELFTDIVDRWDINAIERLPDYMKICFLALFNSMNELAYDILKEQGFSIISHIRKQWANLCKAYLLEVKWYQSRYTPSLNEFLRNAWITNTGPVLIMHAYFCITNPIKEEELECLKNYPAIIESPSLILRLANDLATSPDEIKKGDYLKSIHCYMHESGRSEENARNYINKLIDQTWKKMNKDILRGSSFSKDFRTTAMNLARIAQCMYQHGDGFGIPDRETKDRIRSLFFEPIPLS*

### NbL19g06930.1 DAG_protein_chloroplastic−like [MW = 25.42 kDa]

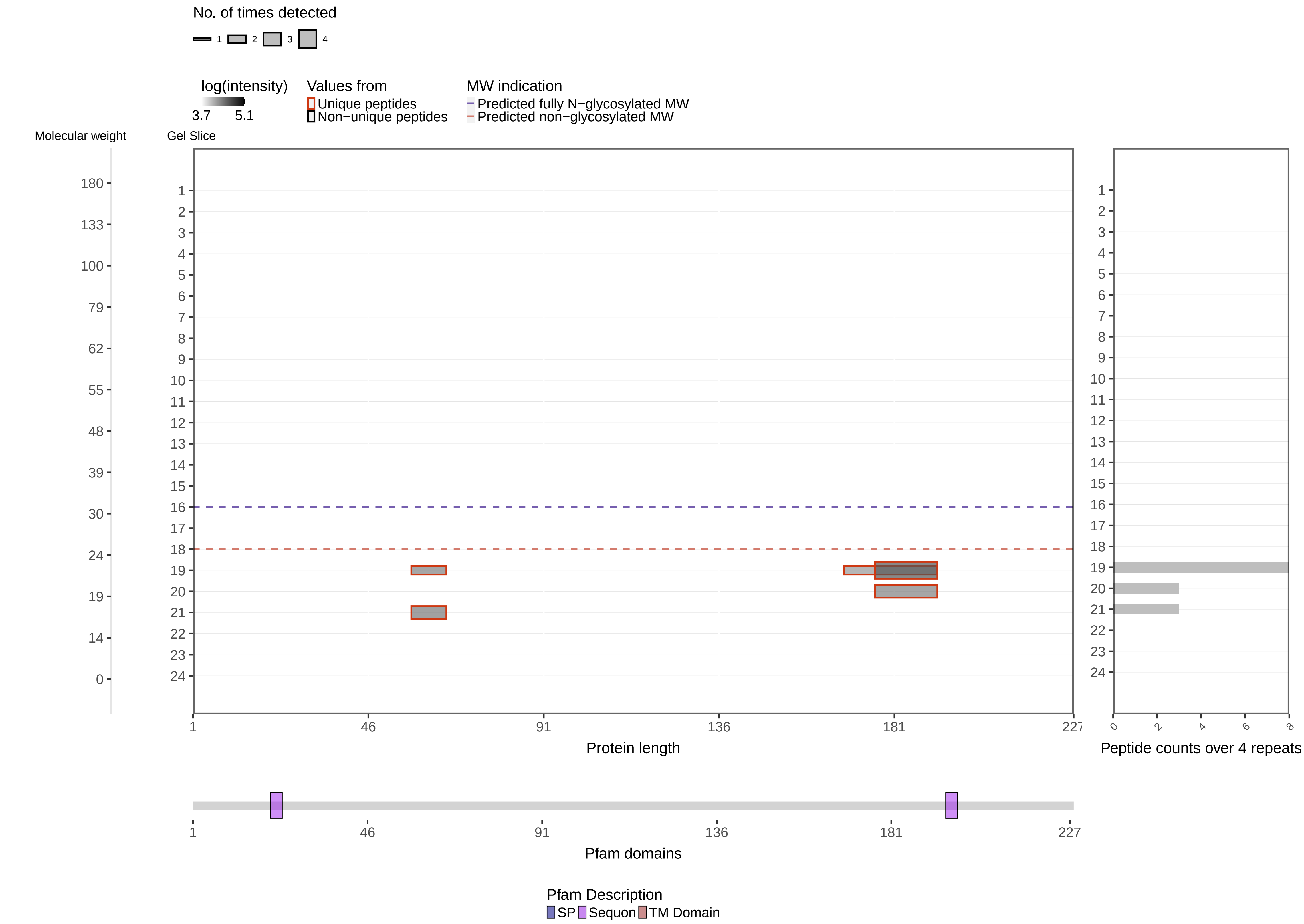


Protein sequence:

EQMIDTYLNTLATVLGSMEEAKKNMYAFSTTTYTGFQCTVSEETSEKFKGLPGVLWVLPDSYIDVKNKDYGGDKYINGEIIPCQYPTYQPKQNNRTRSKSKAYVRRRDGPPPERTKQEAAPESSS*

### NbL19g06910.1 probable_NAD(P)H_dehydrogenase_(quinone)_FQR1−like_1 [MW = 24.95 kDa]

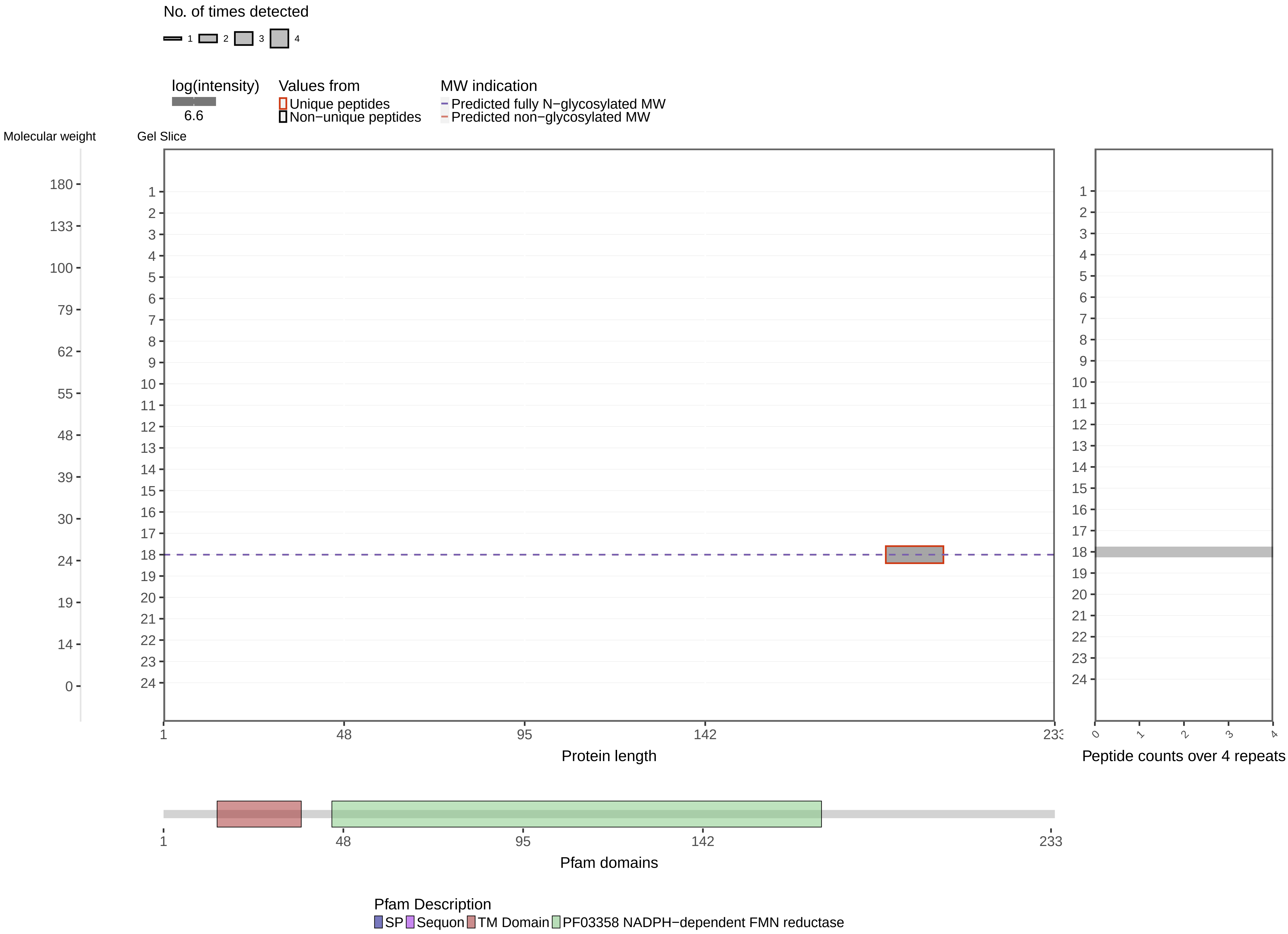


Protein sequence:

GFPTRFGMMAAQFKAFLDATGGLWRTQQLAGKPAGLFYSTGSQGGGQETTPLTAITQLVHHGMIFVPIGYTFGGGMFEMENIKGGSPYGSGTYAGDGSRQPTELEMEQAFHQGKYIATITKKLKGSA*

### NbL19g06780.1 suberization−associated_anionic_peroxidase−like [MW − SP = 35.98 kDa]

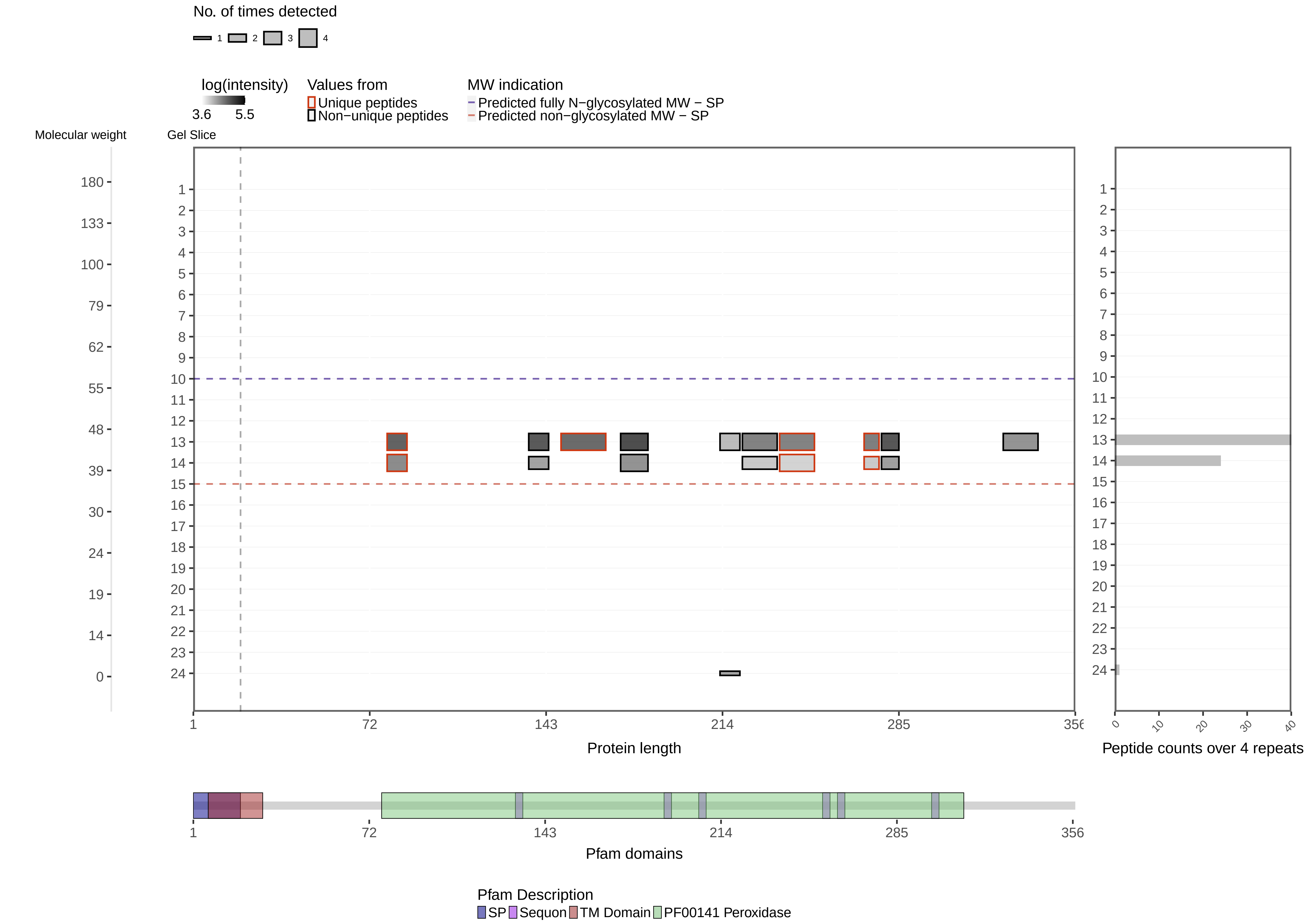


Protein sequence:

:DSVVKLGGOGYNVALGRRDARSANFTGALTOLPAPFDNLTVOLOKFNDKNFNAREMVALAGAHTVGFTRCAVACNSNFVNPAARLOCNCSVPONDTNLORLDRTPAVFDRVYFODLNRNOGILFSDOVLTGNTTTAAIVTTYSNNGAVFLGDFAAAMIKMGNLPPSPGAOLEIRDVCSRVNPSSVASM*

### NbL19g06760.1 suberization−associated_anionic_peroxidase−like [MW − SP = 36.4 kDa]

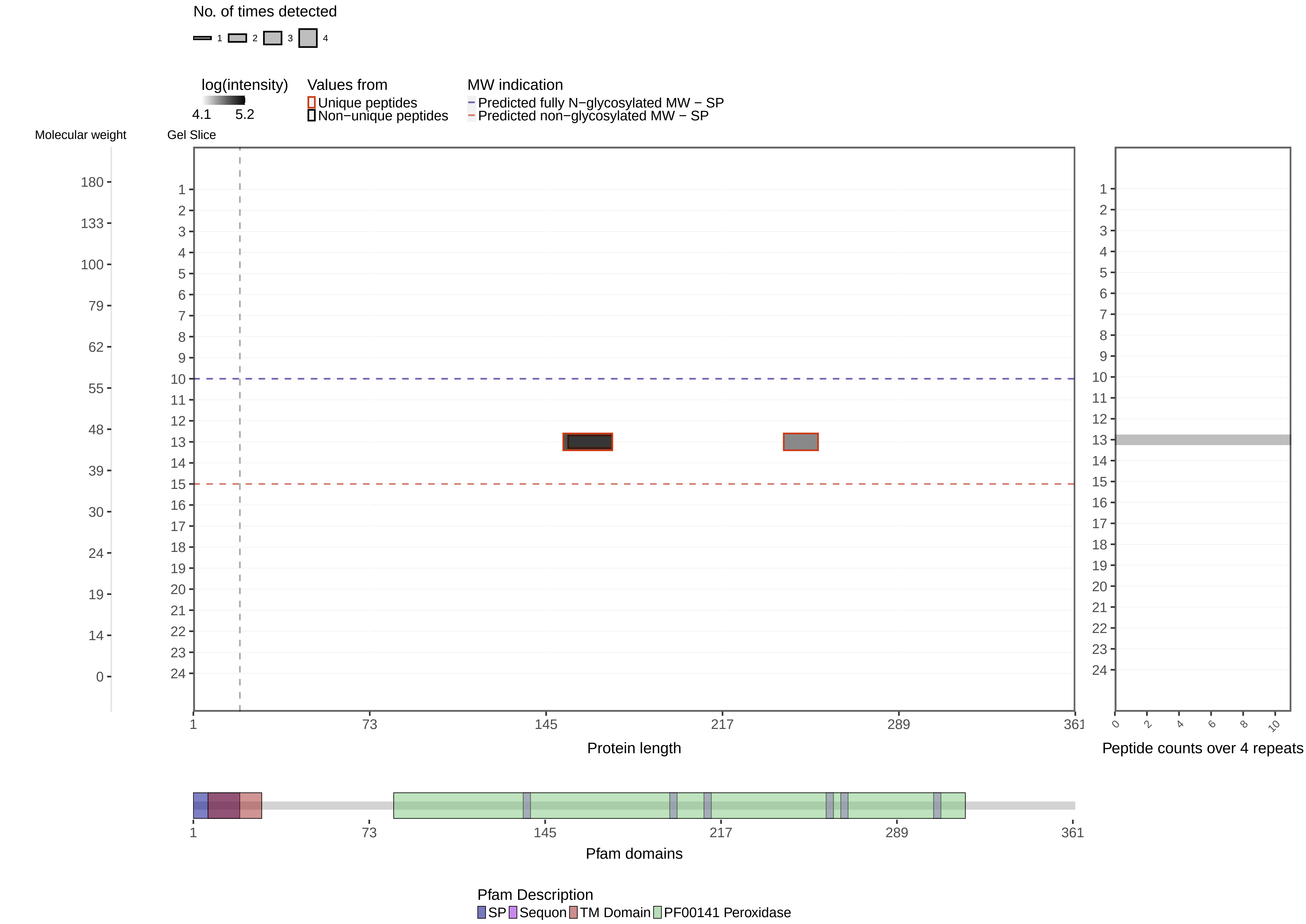


Protein sequence:

ARDSVVKLGGOGYNVALGRRDARAANFTGALTOLPAPFDNLTVOLRKFNDKNFNAREMVALAGAHTVGFTRCVTACNSNNVNPAARLOCNCSVTONDTNLOOLDTTPAVFDRVYFODLNRNOGILFSDOVLTGNTTTAAIVTTYSNNGAVFLGDFAAAMIKMGNLPPSPGAOLEIRDVCSRVNPSSVASM*

### NbL19g06700.1 uncharacterized_LOC104219407 [MW = 26.2 kDa]

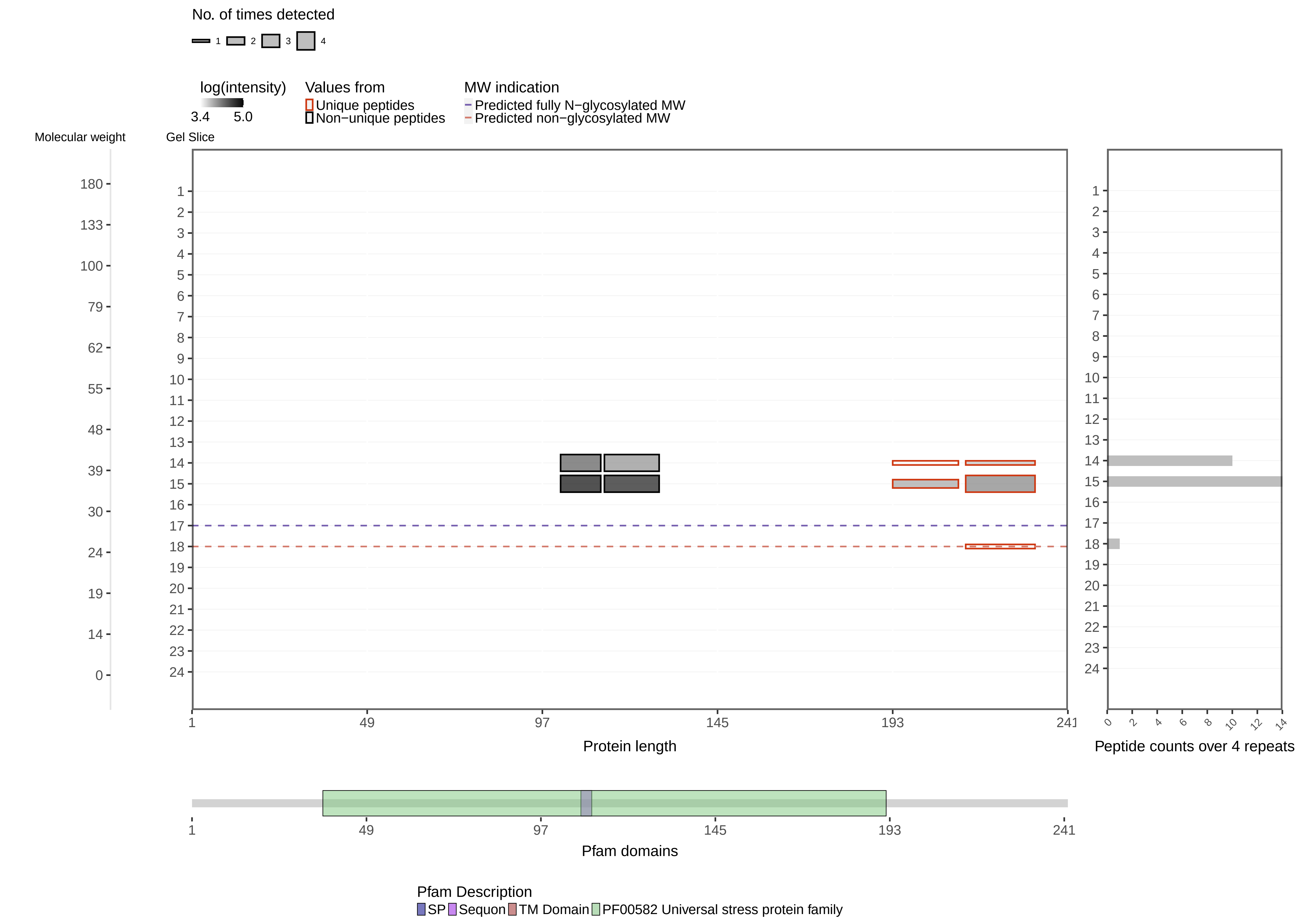


Protein sequence:

NFTTTKANDLAQPLVEGNIPFKIHIVKDHDMKERLCLEVERLGLSAVIMGSRGFGASRRSIKGGRLGSVSDYCVHHSVCPVVVVRYPDDKDGVAADDANEGSVKKETDLHPVPEDEPIYHDAEDKAADLEKAS*

### NbL19g06660.1 uncharacterized_LOC107758959 [MW = 44.9 kDa]

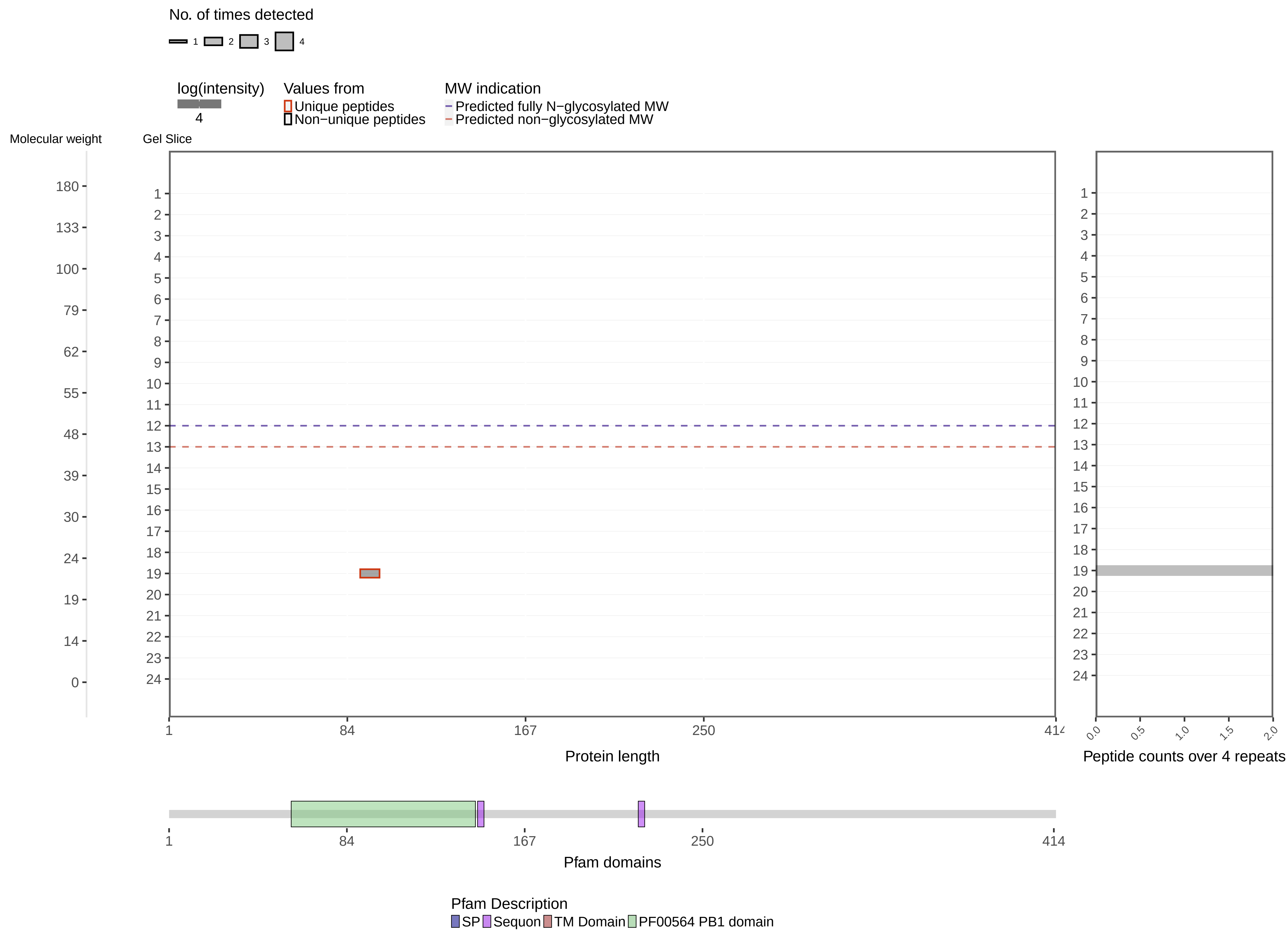


Protein sequence:

PDYLFGLDAADDPPRESRFGKIKNISDPGSPAPPIVSSPYCSVSSTMTSSAMTMIPDLPPVKTKPDKPNPVTESKETPVQRQQPSYPGSPLWHYPAPTVQTIPVYYVPDPVHQPGSIPVQSIPMRAPYVHPFPVPPNQLPVGYPPVSSIGQIYSGMRPVMGVDPNEFSAAARVVTEGTNQQMYYGFRNSGVVPGYGGLAGEEMQGPGNDGKAGRVSQ*

NbL19g06540.1 putative_PAP−specific_phosphatase_mitochondrial [MW = 44.21 kDa]

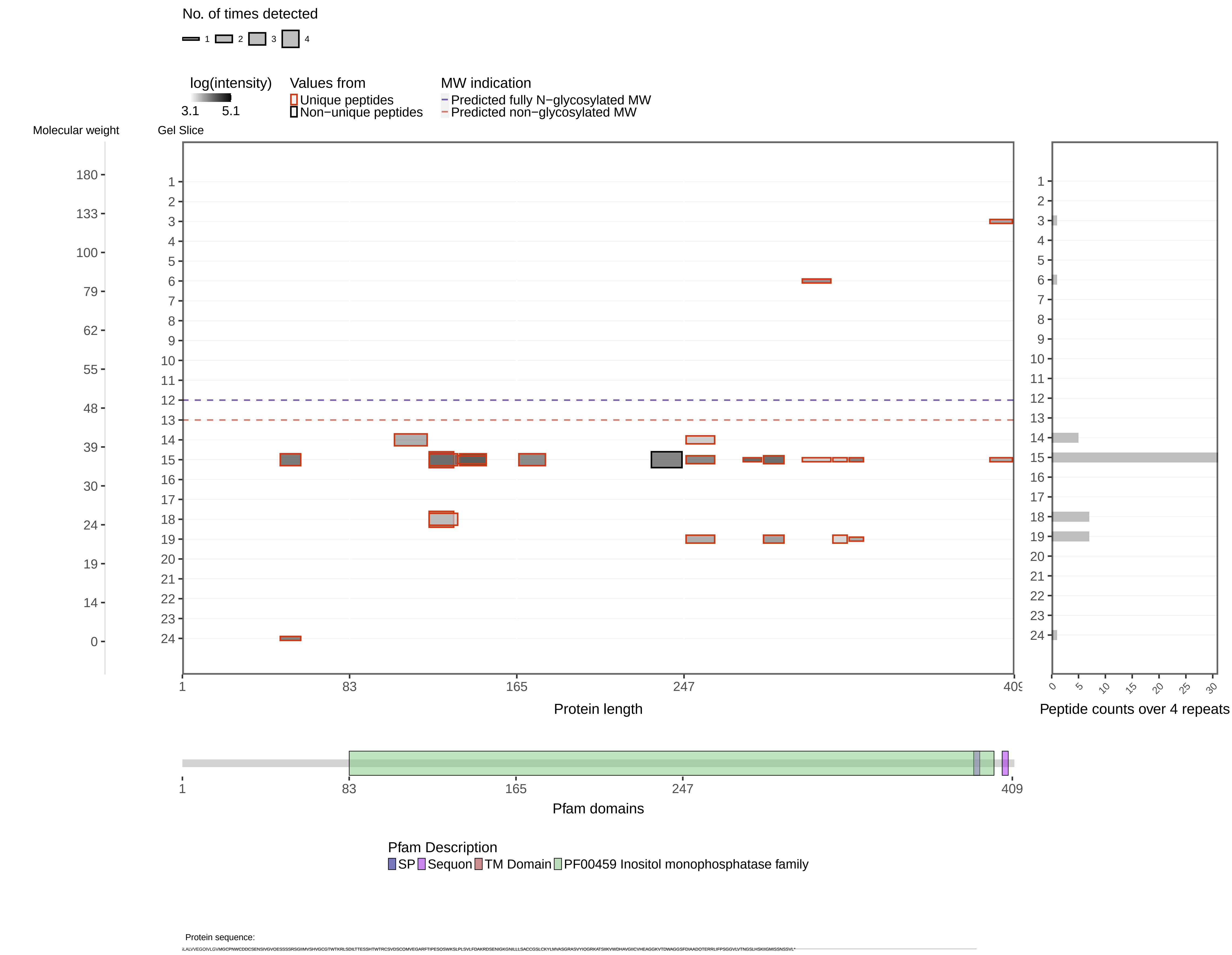

### NbL19g06530.1 vesicle−associated_protein_4−2 [MW = 30.44 kDa]

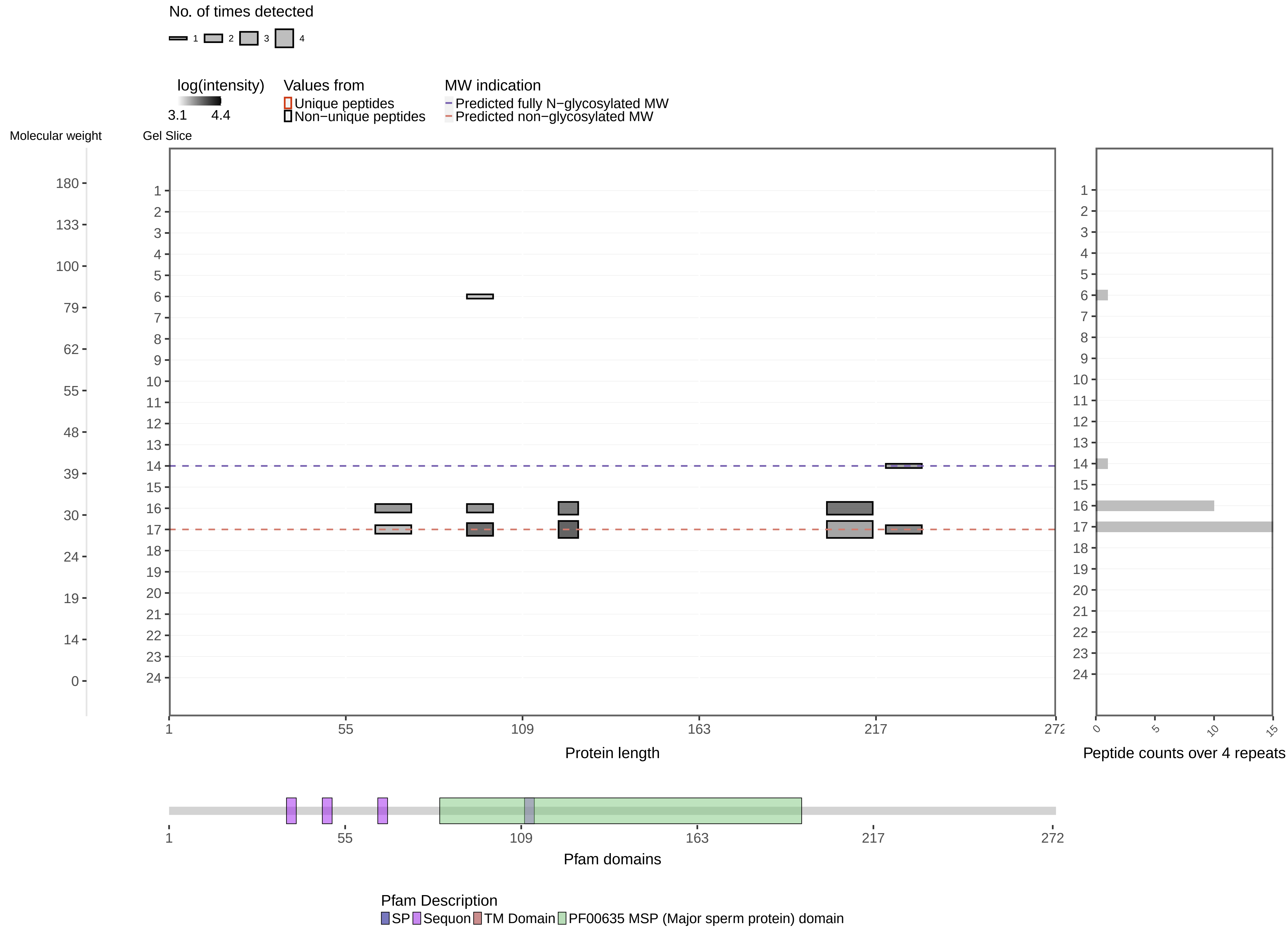


Protein sequence:

TAPKSCFMRPPGAILAPGESIIATVFKFVEHPENNEKPMDQKSKVKFKIMSLKVKGPMDYVPELFDEQKDQVAVEQILRVIFLDVERPSPAFEKLKRQLAEADAALEARKKPAEDTGPKIIGEGLVIDEWKERRERYLARQQVEGVDSV*

NbL19g06420.1 bifunctional_riboflavin_kinase/FMN_phosphatase−like [MW = 42.83 kDa]

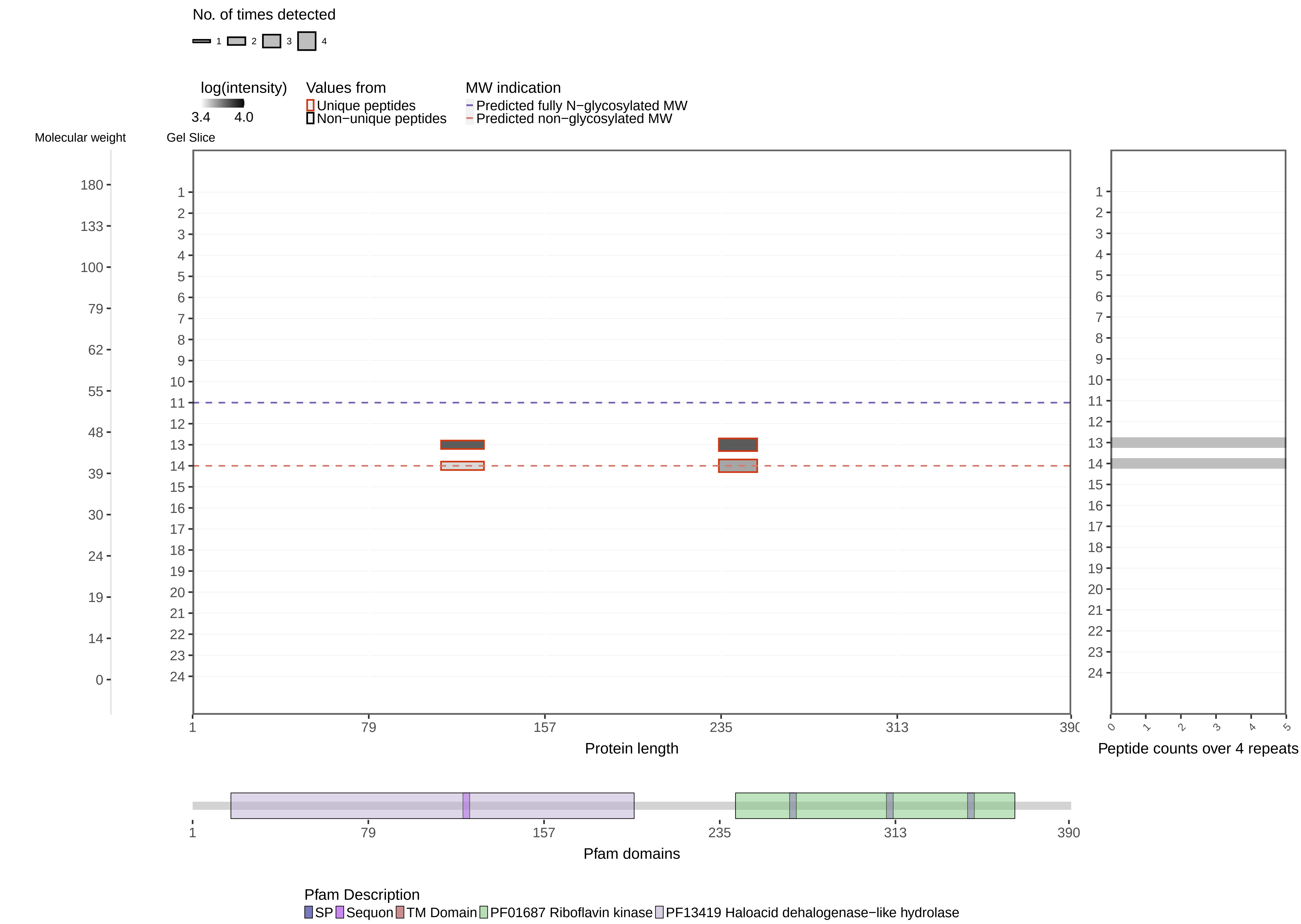


Protein sequence:

PGVTAGKAAGMAVVAVPSLAKQSHLYTSADEVINSLLDLQLEKWGLPAFQDRIEGTLPLEPWYIGGPIIKGFGRGSKVLGIPTANLSPEGYSAILSEHPAGVYFGWAGLSARGVYKMVMSIGWNPYFNNTEKAIEPWLLHDFNEDFYGEELHLVVVGYIRPEANFSSLEALIAKIHEDRKIAERALELPQYLKYKDDPYLKISLHQDN*

### NbL19g06320.1 calmodulin−binding_protein_60_B−like [MW = 69.31 kDa]

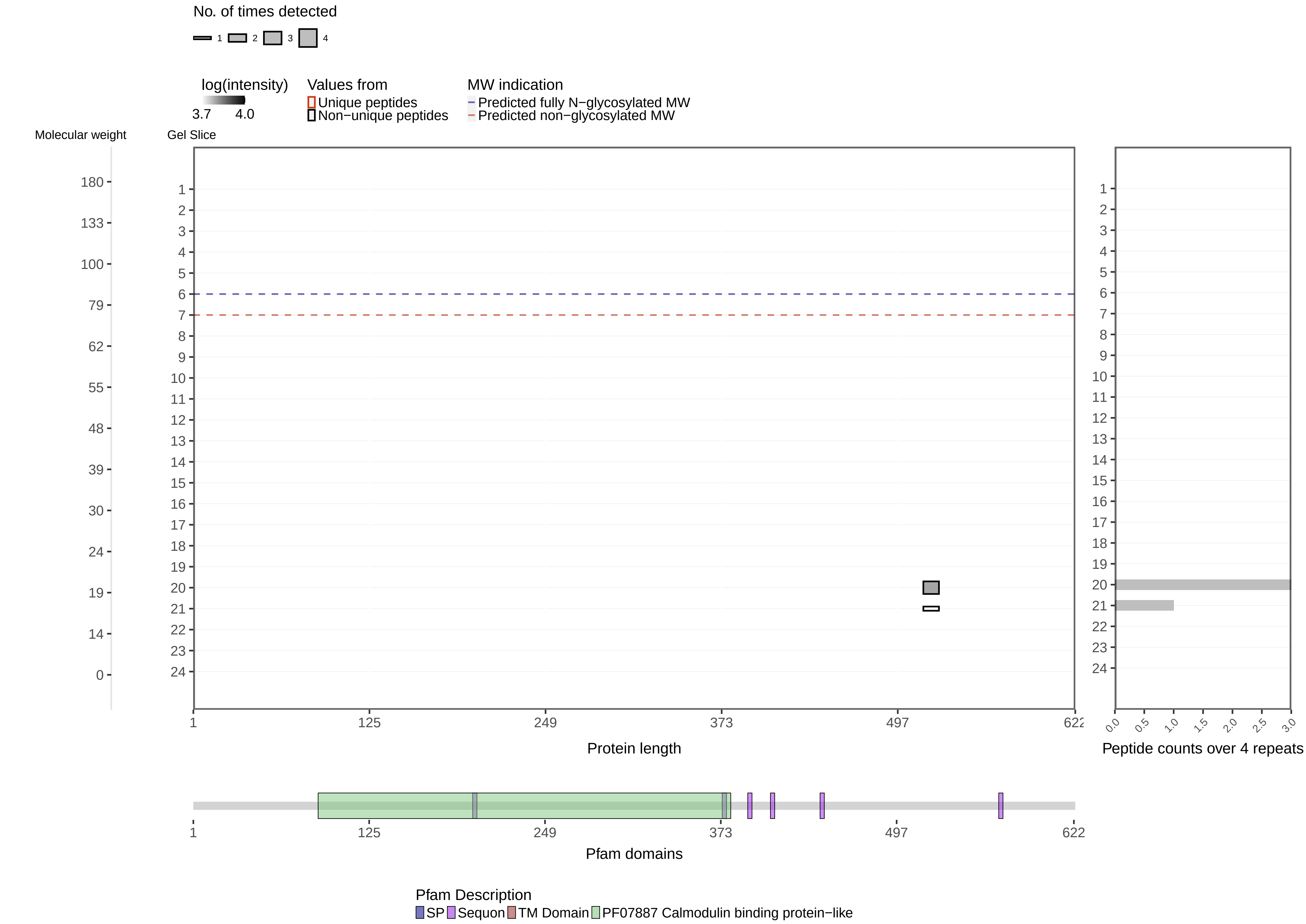


Protein sequence:

MWDALVEHAKTCVLGGKLYVYYPDDMRNVGVVFNNIYELCGLISGGQYHSVDSLSENQKVYVDTLVKKAYDNWTHVVEYDGKSLLSLGQNKTSVASQNVLTIGSQNHSTSFDQQINLPSLPASISSEQPAMNIGLNMGGYNESLSSRYTMQPQRMNLNGNMQLNGASFPPQNHLLGTSQQAQPHGSESMLALRPPQPSMSSYFAASSPNPYKGAEDFLTEEEIRMRSHEMLENEDMQHLLRIFSMGQGHASSSGEENYQYGSSYMPNMSSTFGFDEDRTRSSGK

### NbL19g06300.1 eukaryotic_translation_initiation_factor_3_subunit_J−A−like [MW = 25.18 kDa]

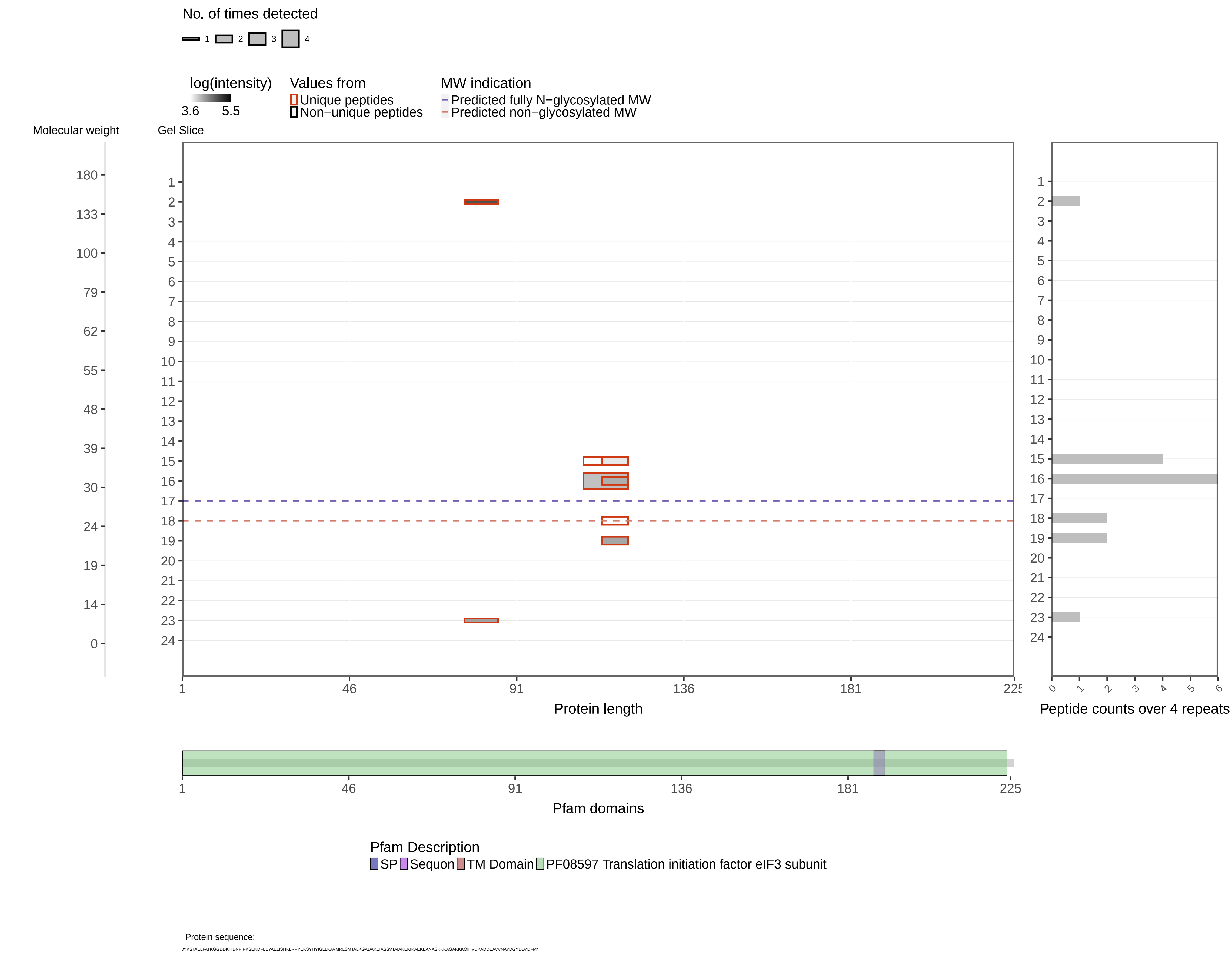

### NbL19g06240.1 glucan_endo−13−beta−glucosidase−like [MW − SP = 48.89 kDa]

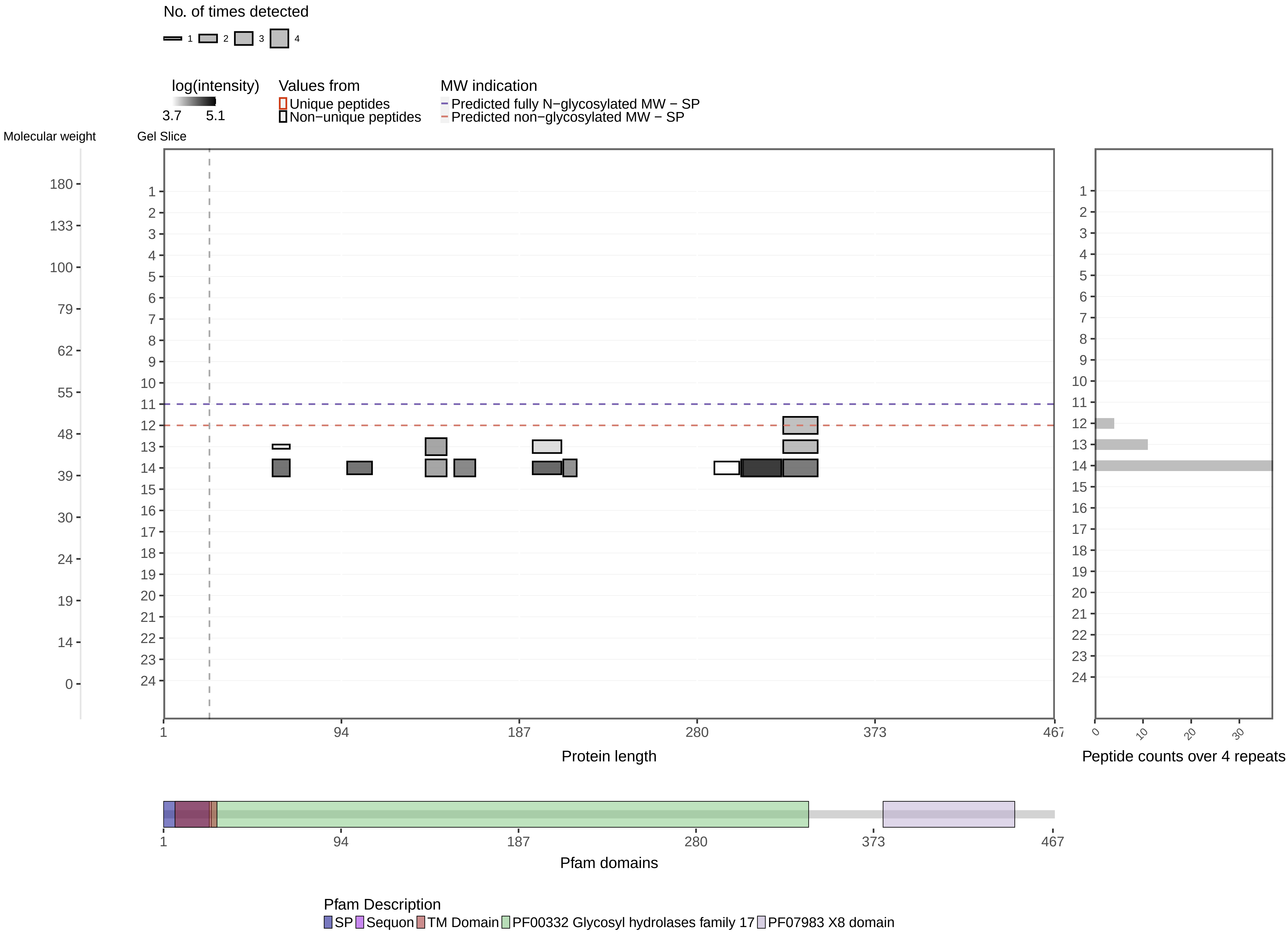


Protein sequence:

FSKRWYTNSFDMLLDAVYMSMVRLKYSDVEIVAAETGWPSQGESFEPQCTVENAASYNGGLLRKYNSGTGTPLMPHRKIETYIFGLFNENTKPGSTAERNFGLFRPDFTPVYNIGVLKGEQAQPTPALPAPRNGNGNKGQPTSPVRPAESKKFCMPKAEATDAQLQSNINYVCSQGVDCTPIQAGGPCFNPNTIRSHAAFVMNSYYQREGRNAFNCDFAGTGVVASADPSKSHSELKLFLC*

### NbL19g06220.1 endonuclease_4−like [MW − SP = 31.06 kDa]

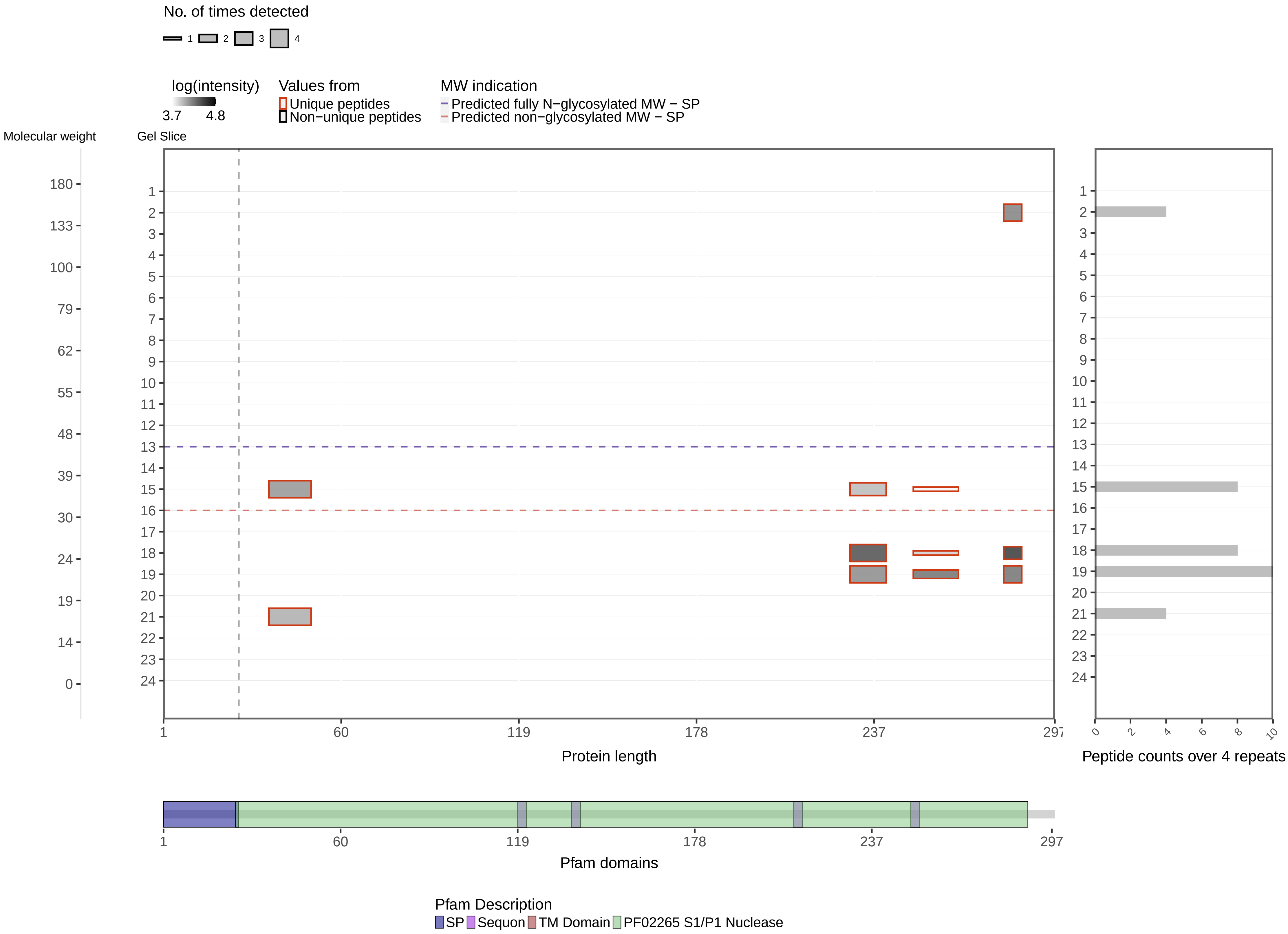


Protein sequence:

EKYNLTEALMFLSHFVGDVHQPLHVGFTGDVGGNTIVVRWYRRKTNLHHVWDTMIIESALKTYYKSDLTLMIQALLRNITDEWSDDVPSWENCKKMVCPDPYASESVRLACKFAYRNATPGSTLGDDYFLSRLPVVEKRLAQSGIRLAEVLNRIFSKQPFTAQ*

NbL19g05870.1 clathrin_light_chain_1−like [MW = 34.84 kDa]

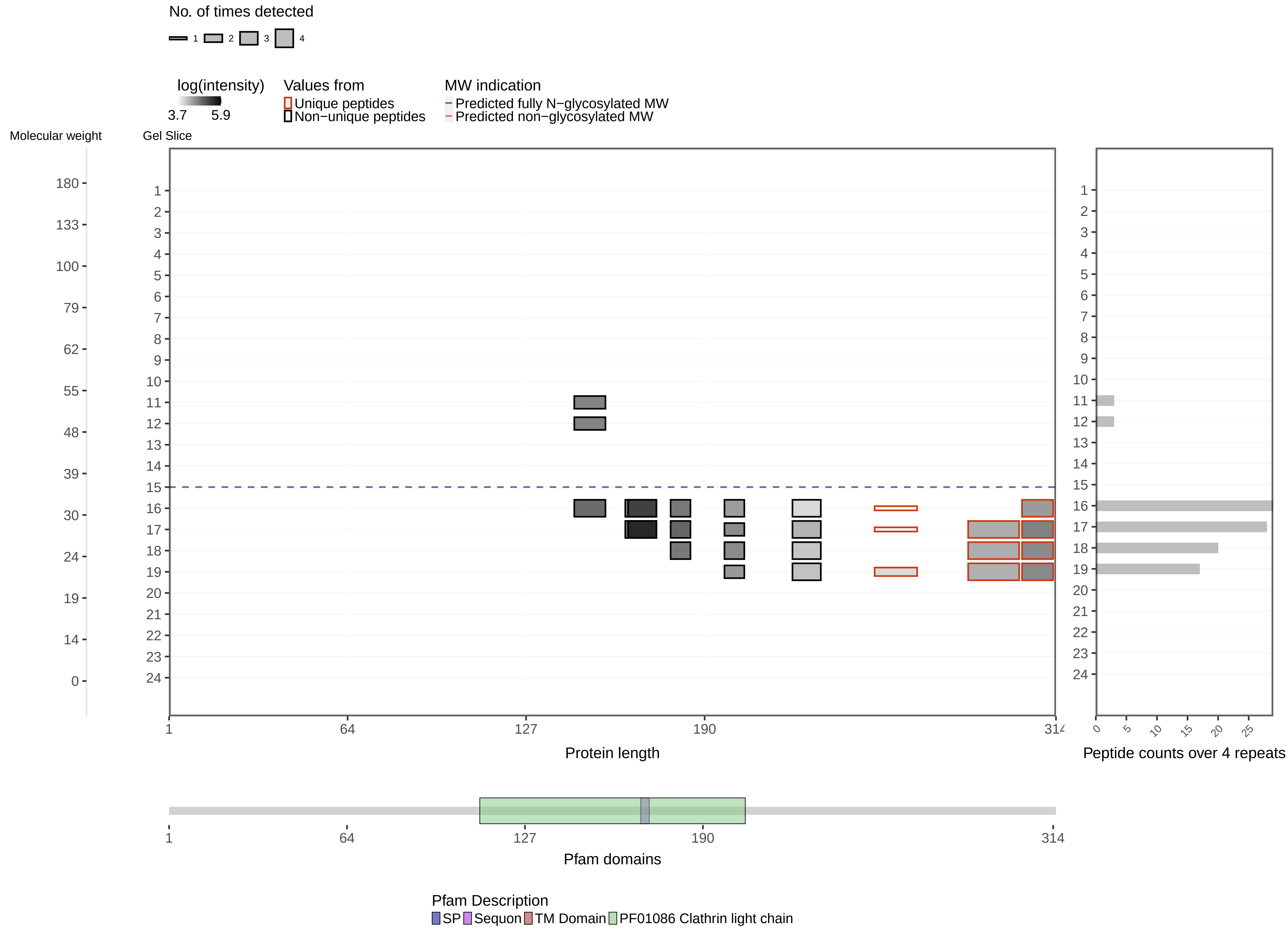


Protein sequence:

IRNQIIEEGEEYKKAFYEKRKLTVETNQSTNREKEKLYLSNQEKFHKEADKQYWKAIAELIPKEVATIEKKGRKKDQEKKPSVTVVQGPKPGKPTELSRMRHILLKLKHSPPFHMIPPPPPPAKDAKAGKDGKDAKVKDSKNAKDEAPNGHSDAPPTDAKVTQTSEEPAAAE*

NbL19g05830.1 30S_ribosomal_protein_S5_chloroplastic [MW = 36.85 kDa]

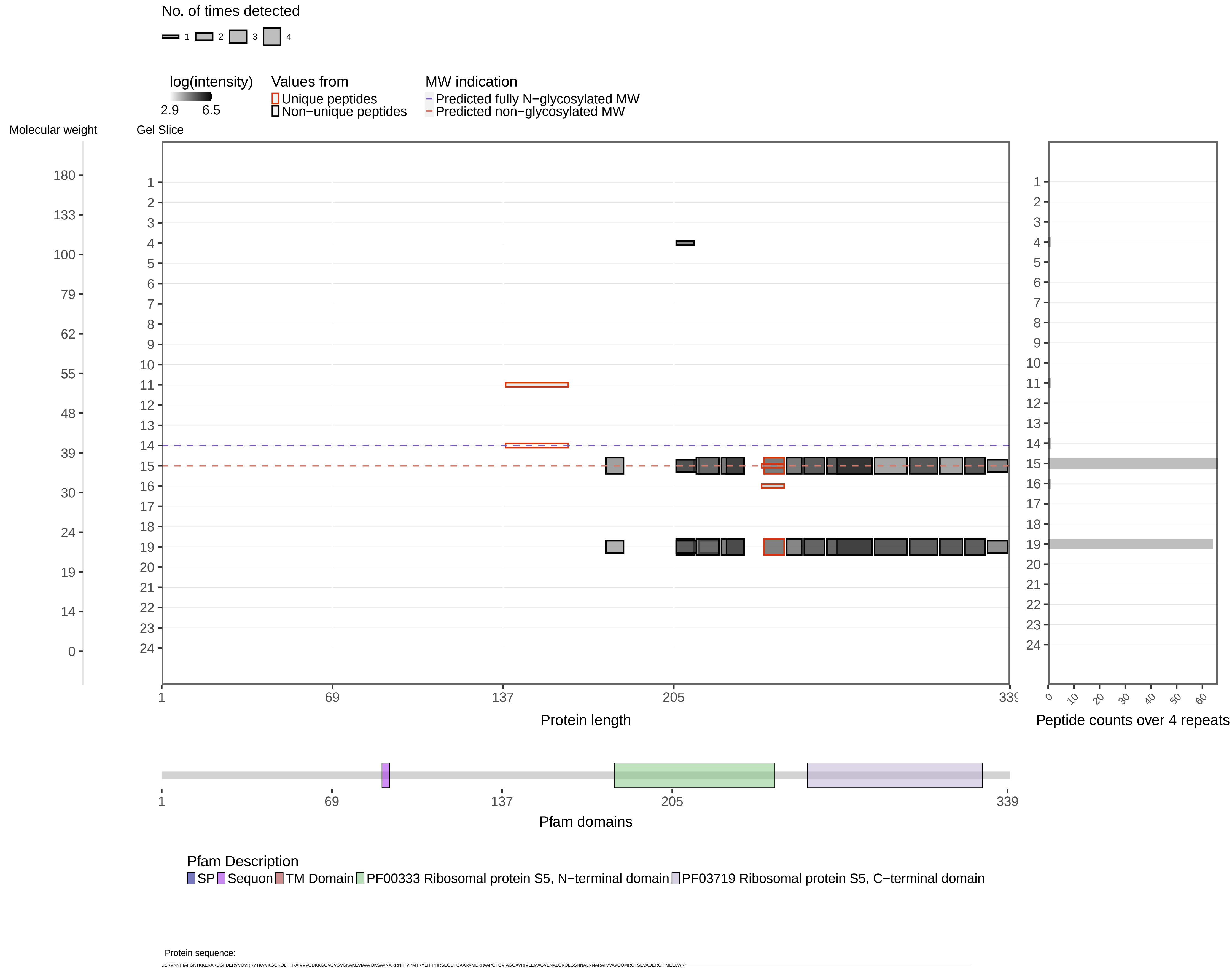

NbL19g05770.1 pistil−specific_extensin−like_protein [MW − SP = 25.44 kDa]

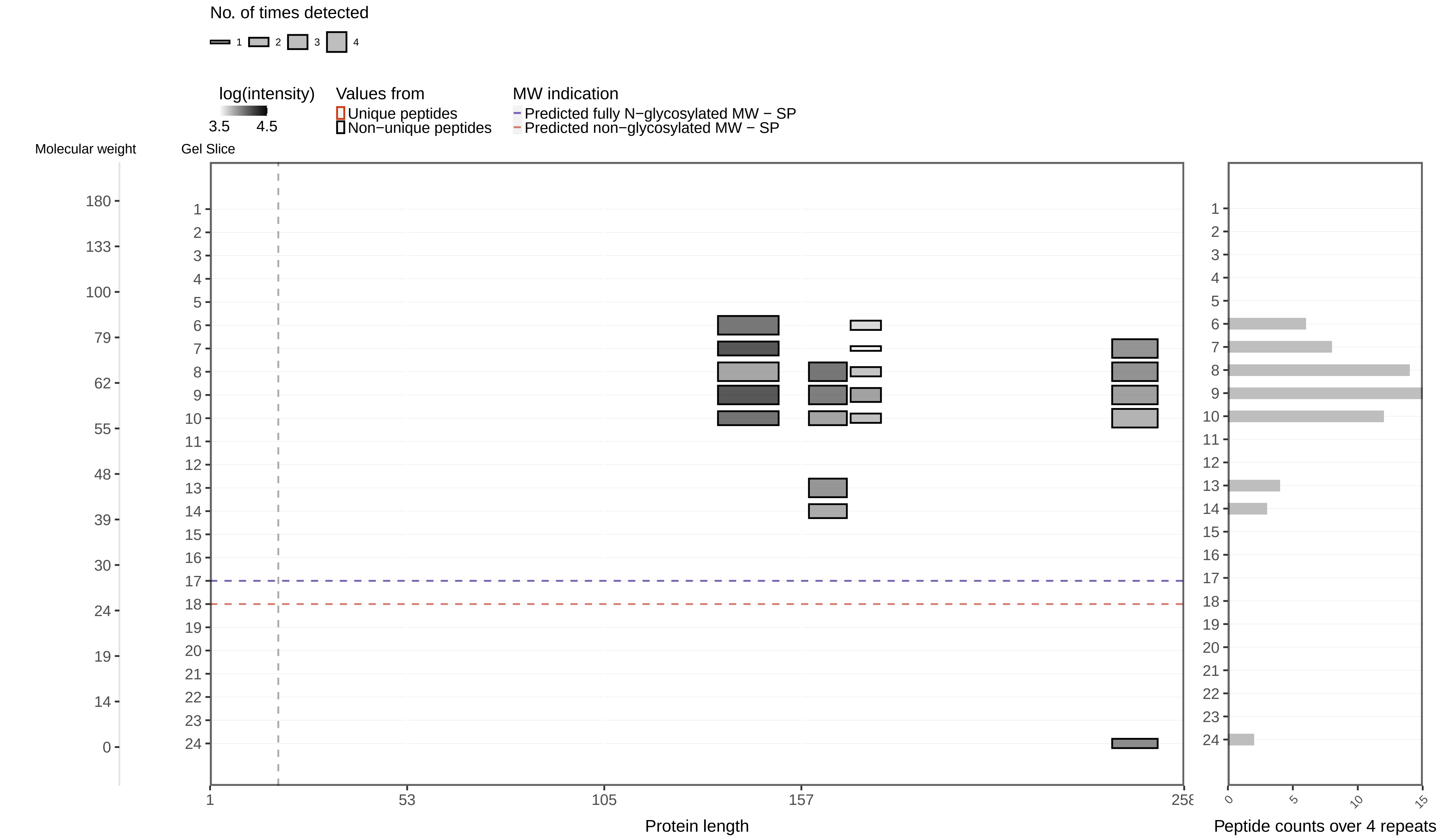


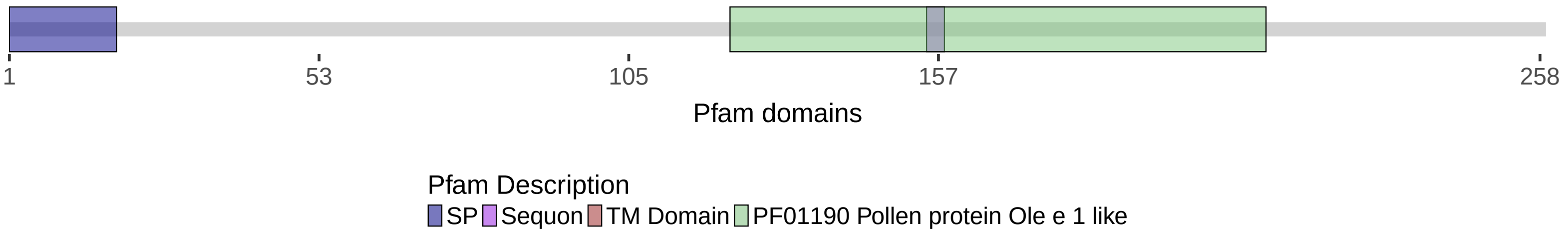


Protein sequence:

RNFVGVRGVVYCKSCKYRGIDTLLGASPIQGAVVKLACNNTKYHLTALGTTDKNGYFFIQPKWLTTAGYHKCKVFLAKSPKAECSVPTNFHNGQAGAMLVPAPLPPPSPMISKPVDSETGVKLFNVGPFAFEPSKNLPCKQ*

### NbL19g05700.1 eukaryotic_translation_initiation_factor_3_subunit_D−like [MW = 65.58 kDa]

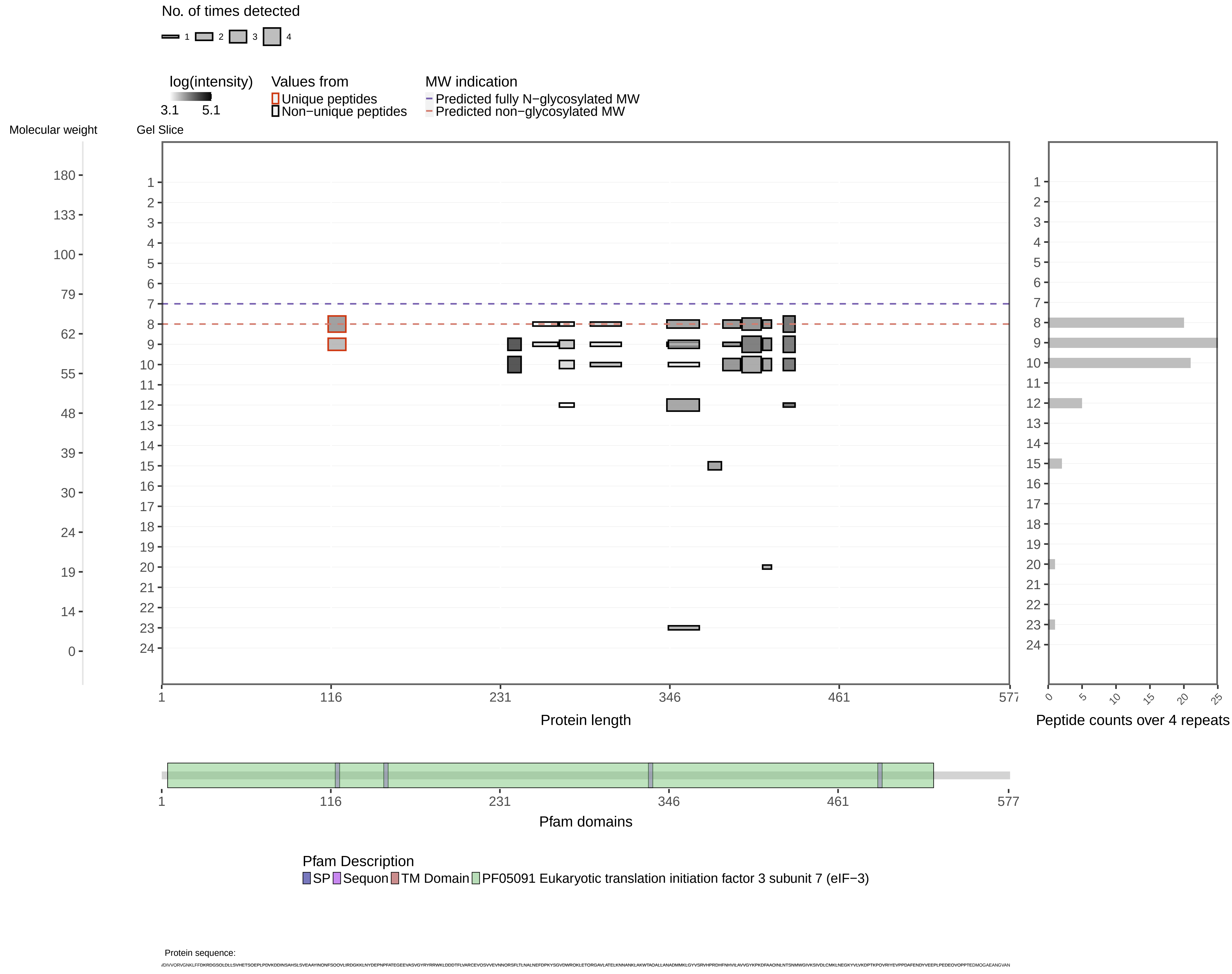

NbL19g05590.1 metal−independent_phosphoserine_phosphatase−like [MW = 34.03 kDa]

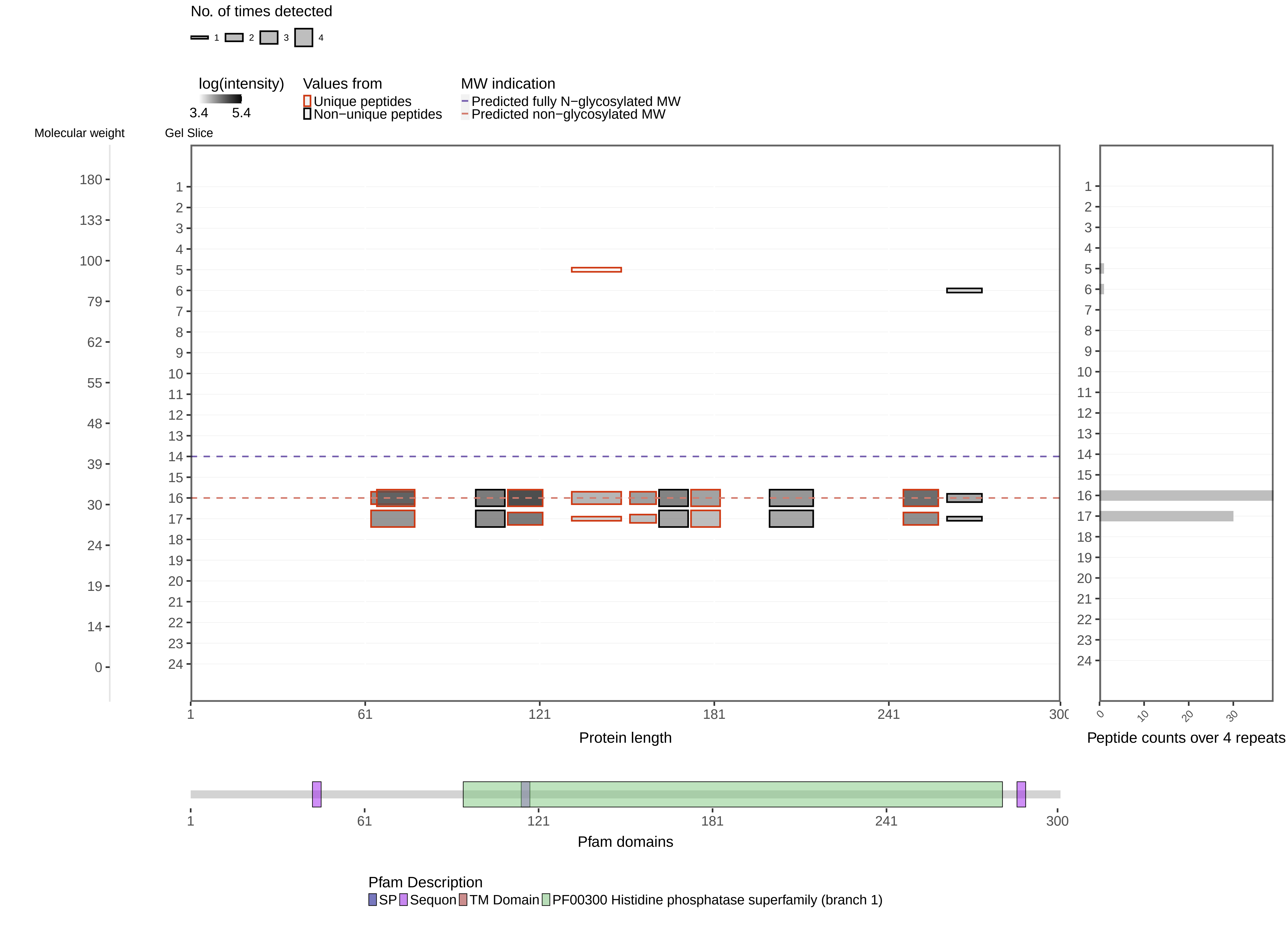


Protein sequence:
MHFDQCFSSPISRAKTTAEIMWQGKEEPLVFLDSLKEAHLYFLEGMKNEDAKRIYPKEYTTWREDPSNFCVDGVYPLQKLWGRAYEAWEEILLTPGENFLVVTHKSILRALICTALGLGPERFRSVDINNGGLCVFKFNKQGEAMLHSLNMTAHMYTDHIYHY*

### NbL19g05540.1 molybdopterin_synthase_sulfur_carrier_subunit [MW = 13.33 kDa]

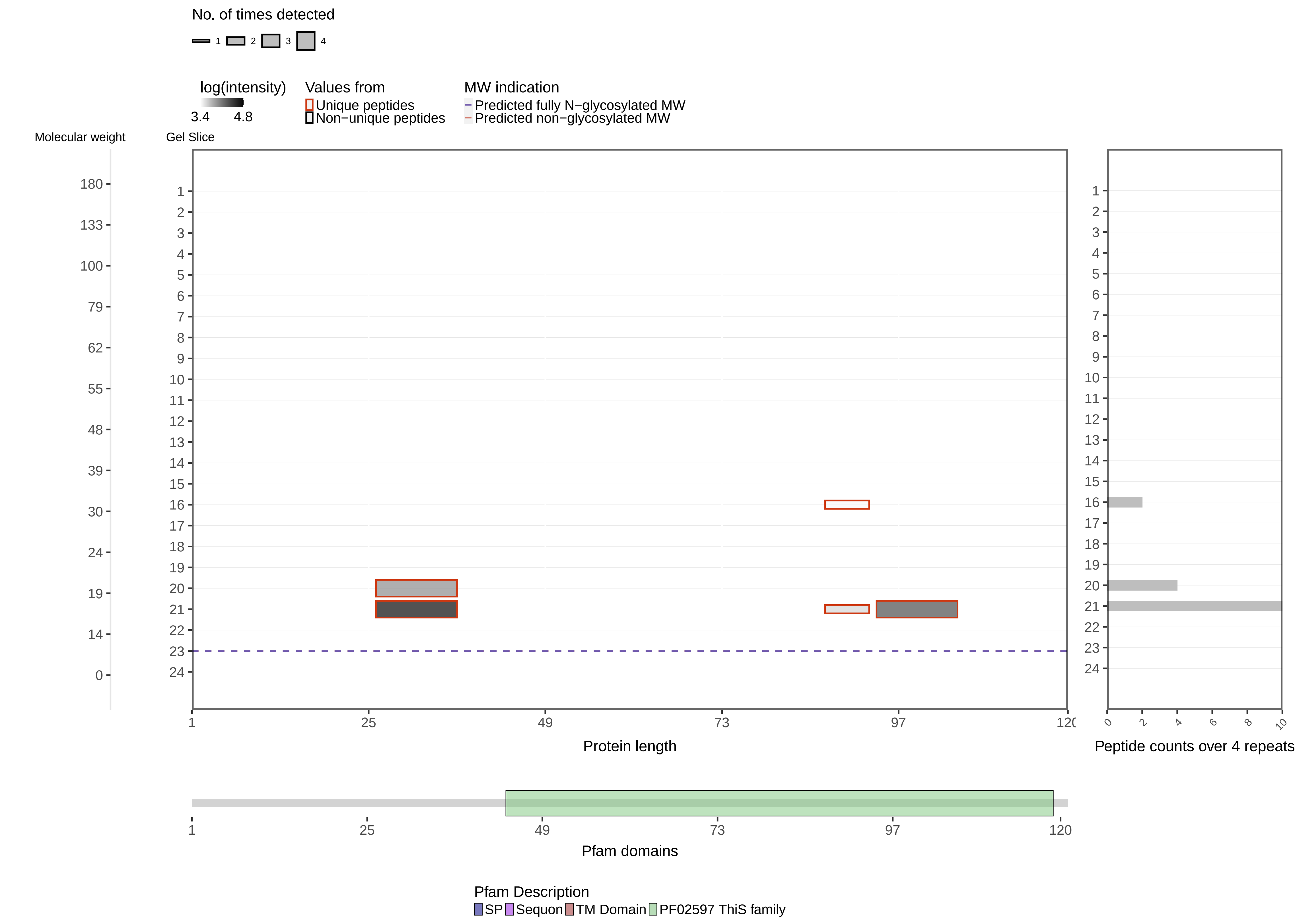


Protein sequence:

ARARDLTGMTEMPLEVSYGSTAGDCLNKVISRFPRLEEIRGCMLLARNEEYTPESTTVKDGDELAIIPPISGG*

NbL19g05470.1 14−dihydroxy−2−naphthoyl−CoA_thioesterase_1−like [MW = 17.1 kDa]

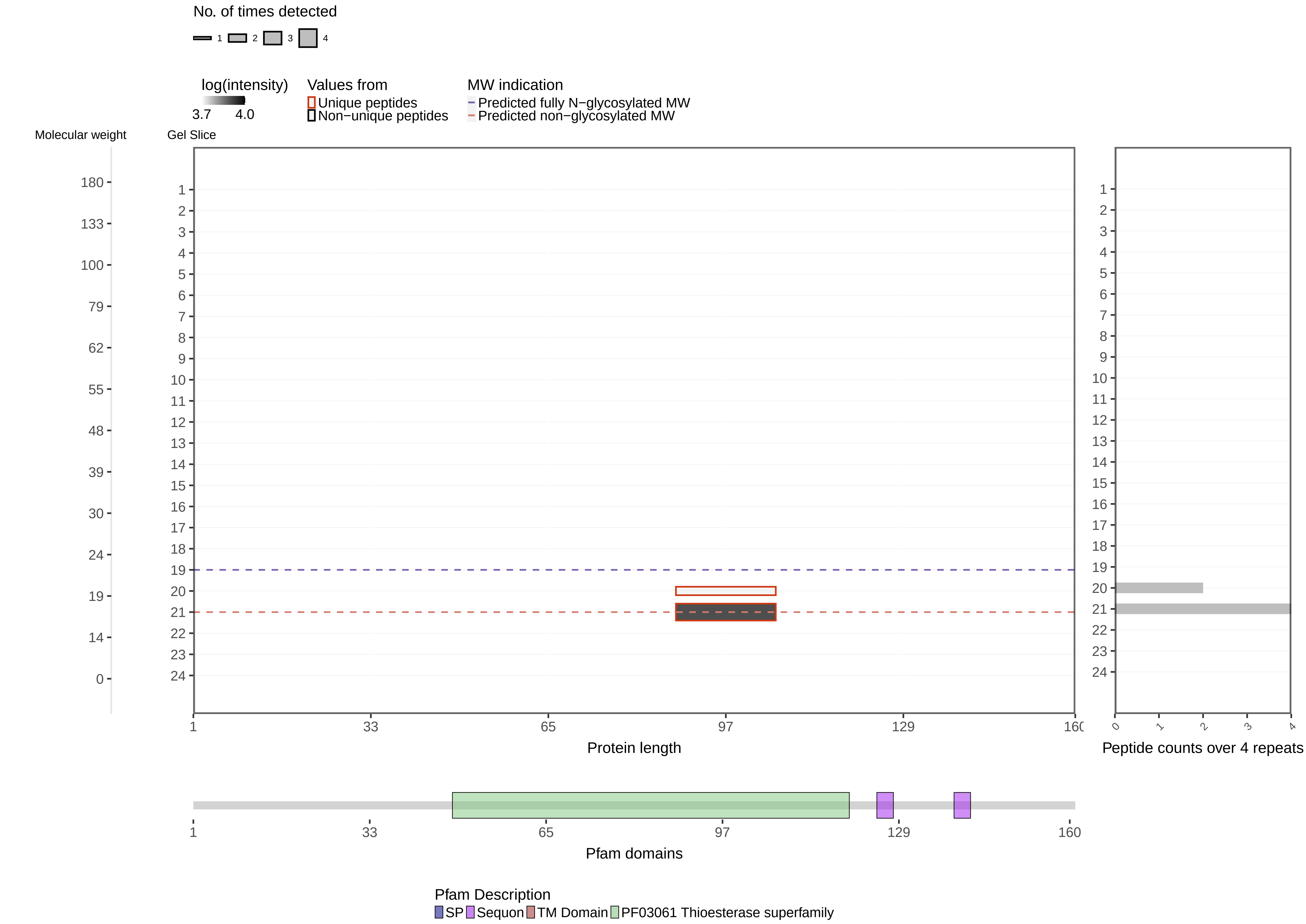


Protein sequence:

AHMASGFGRVAGVQLSIHHLKSAHIGDLVFAEAVPVNIGKTIHVWEVCIWKIDPNNEENKTLIASSRVTLKANMSVPENAKDAAVNLKKYAKL*

NbL19g05160.1 uncharacterized_protein_At4g28440−like [MW = 17.54 kDa]

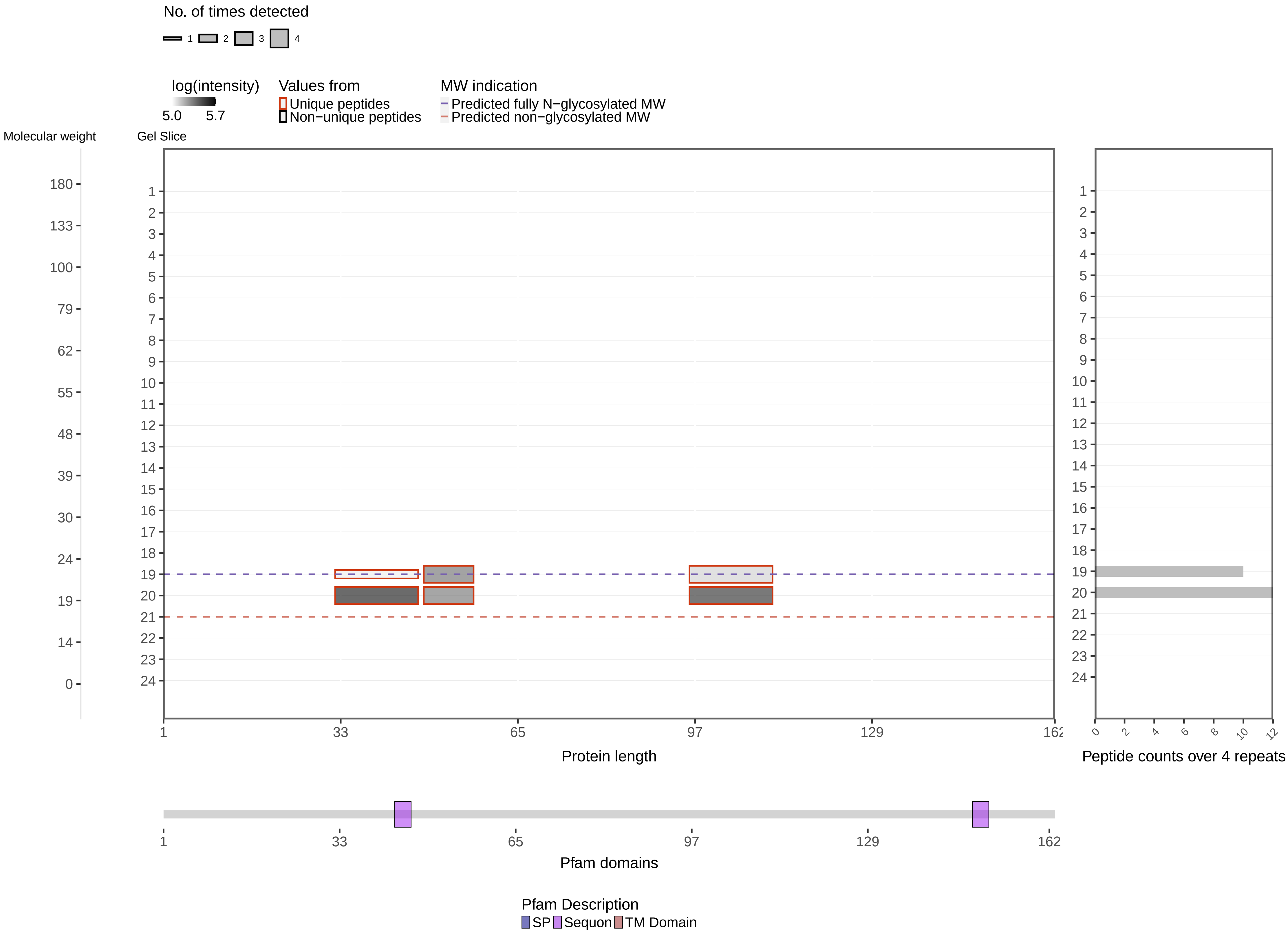


Protein sequence:

RVPARPQQNTRISECLVGDESGSILFTARNDQVDMMKPDATIILRNAKIDMFKGSMRLAVDKWGRVEVAAEPATFVVNEENNLSLVEYELVNVEE*

### NbL19g05100.1 glycogen_phosphorylase_1−like [MW = 92.51 kDa]

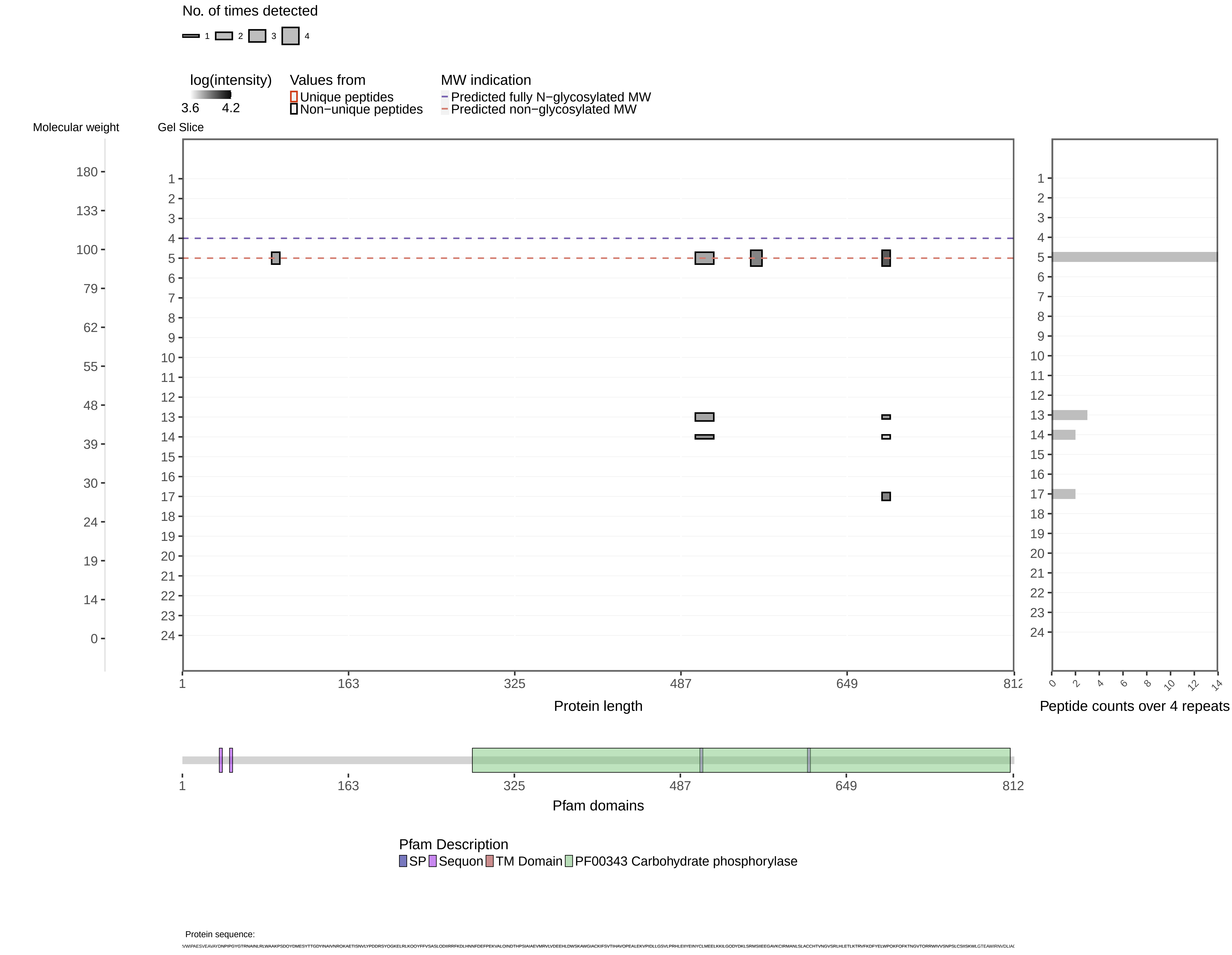

### NbL19g04820.1 GDSL_esterase/lipase_At5g45950 [MW − SP = 37.79 kDa]

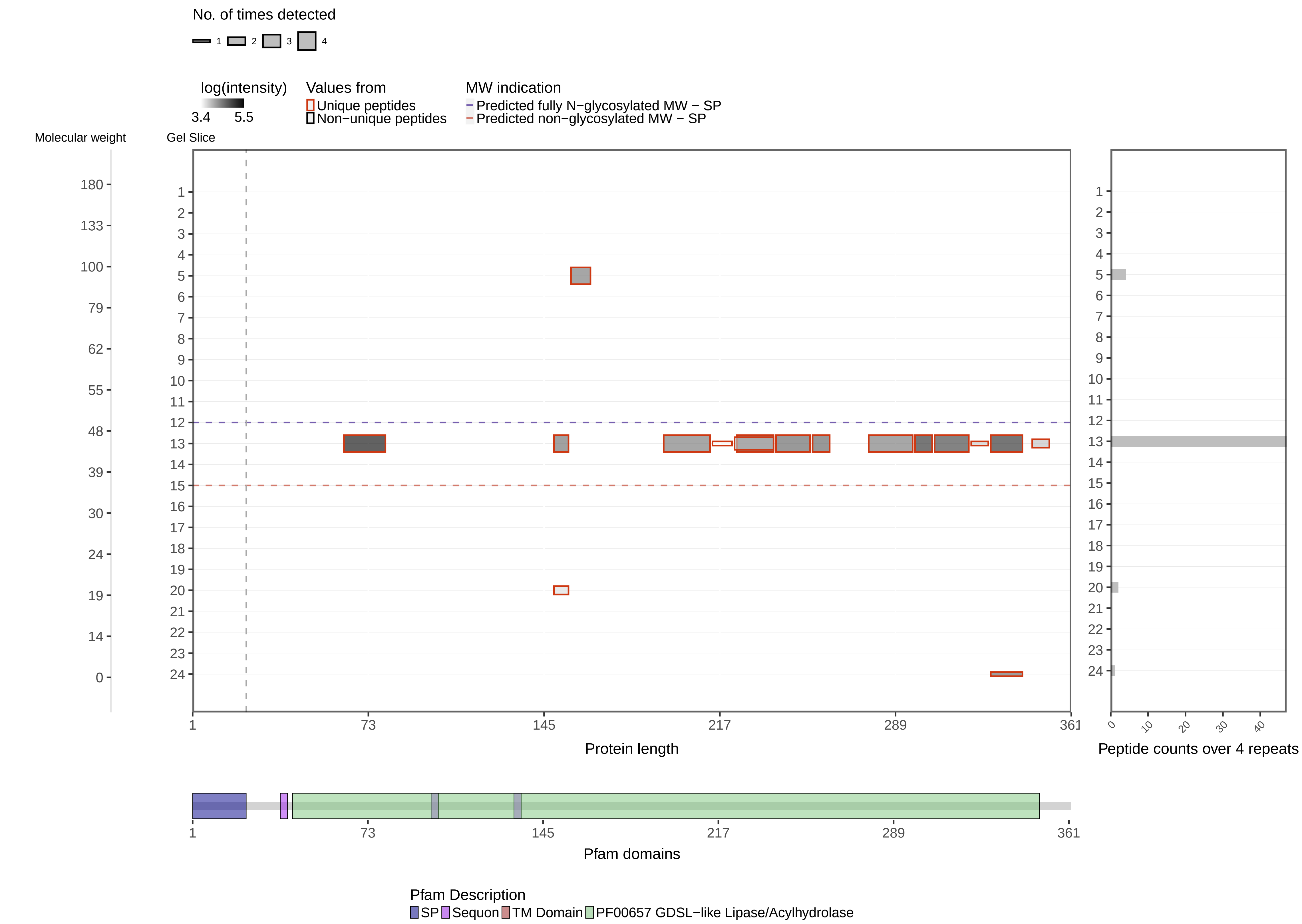


Protein sequence:

MKNALYILSMGTNDFLQNYFVEVERSKQYSVEQYQNFLIHSMFTHVKDIIHGVGARRLAVVGVPPLGCEPLIRTIRDDETKCDDDLNKVAFSFNLKLKRELQTLKRLFGIKASFIDIYSIIQEAVQNPHKFGFEETSKGCCGTGTTEYGETCKGLTTCADRTQFVFWDAVHPSEKMHKIMADEALKSINADLFS*

### NbL19g04710.1 GDSL_esterase/lipase_At5g45920 [MW = 26.32 kDa]

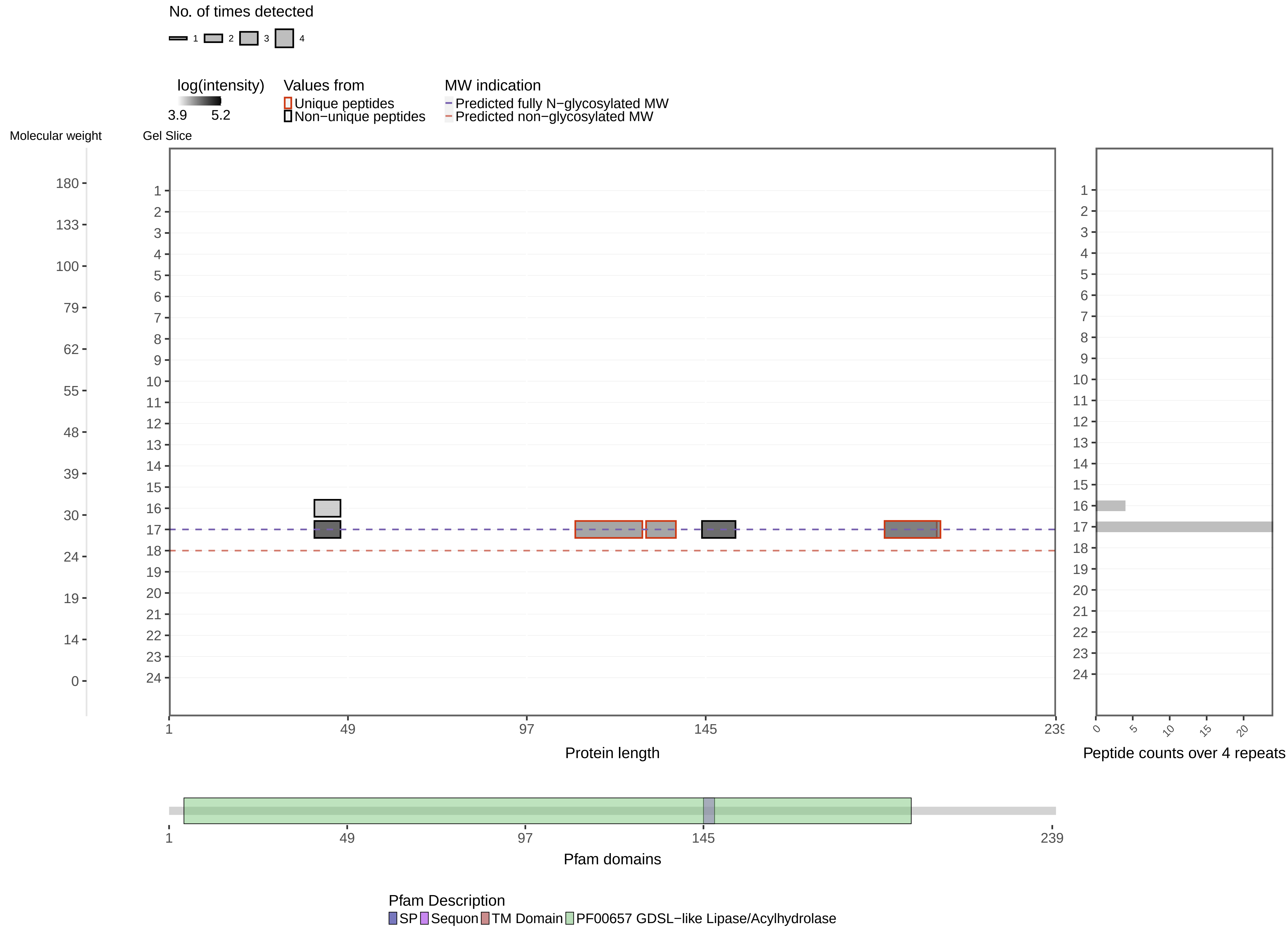


Protein sequence:

FLKGRWPTTQIVLISPPPIDEPARLLYPFIENKLGLPERTNETAGNYAKASLTVAAECGVLAVDLWTRMQQIPGWQTACLSDGLHLSKTGNEIVFEEVVEALKKKGLSVETLAVDLPVINEIDPNDPLKSFEGI*

### NbL19g04660.1 GDSL_esterase/lipase_At1g28600−like [MW = 43.02 kDa]

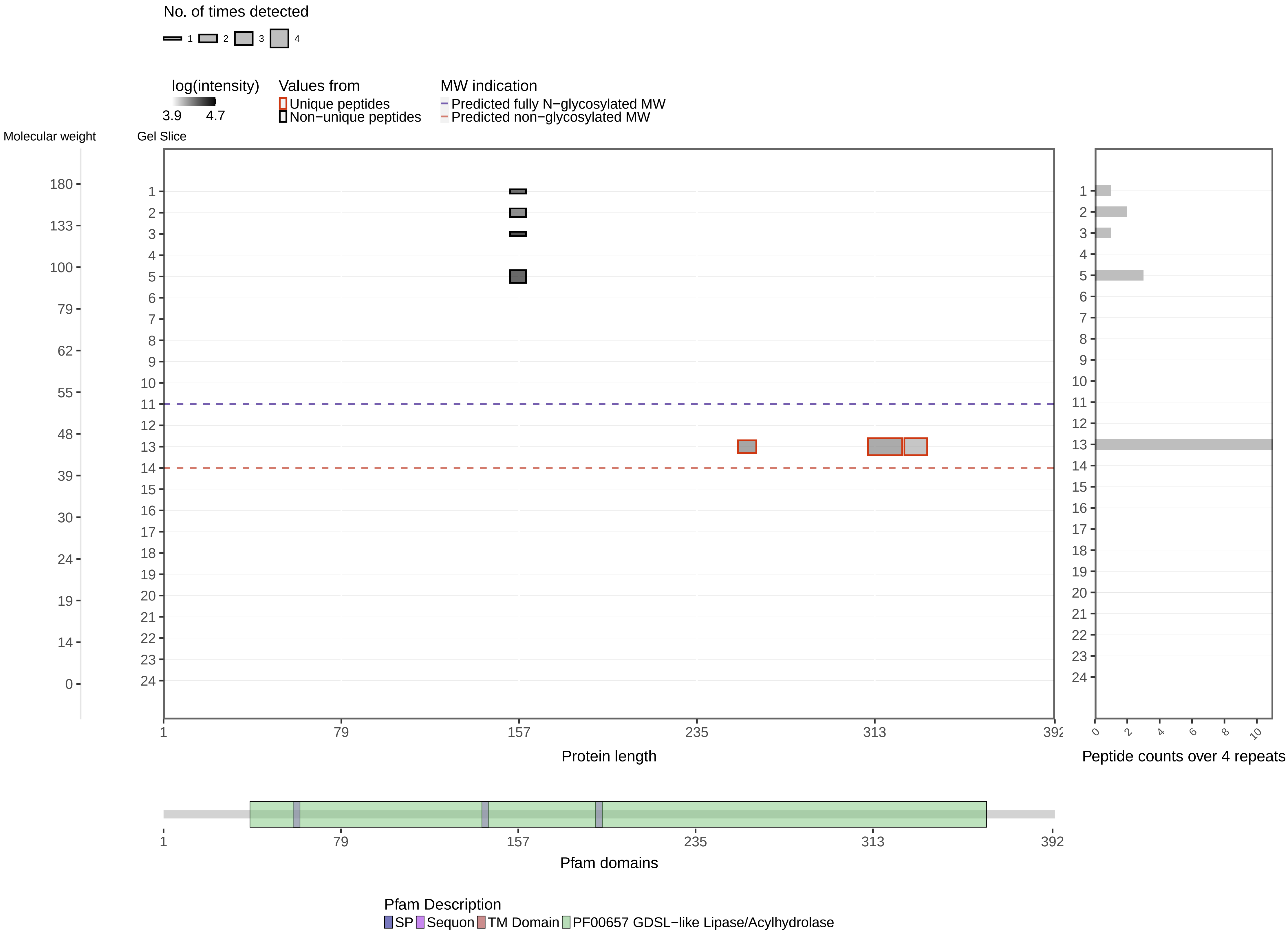


Protein sequence:

DFNYGFLGNSTKEEVESYVPAVINTISSAIQELIELGASTLVVPGDLPIGCSTAYLTKFMHSDKENYDPKTGCINWLNRFSQHYNELLQKELHLLRELYPSATIIYADYYNAAMQFYVHPMSHGFTKGALDACCGAGGPYNFKFSAVCGDPPARNICSDTSVYASWDGVHFTEAAYKLIARGLLEGTFTFPPLPKIFTPFNPIANHFDS*

NbL19g04650.1 phospholipase_A1−II_1−like [MW − SP = 44.01 kDa]

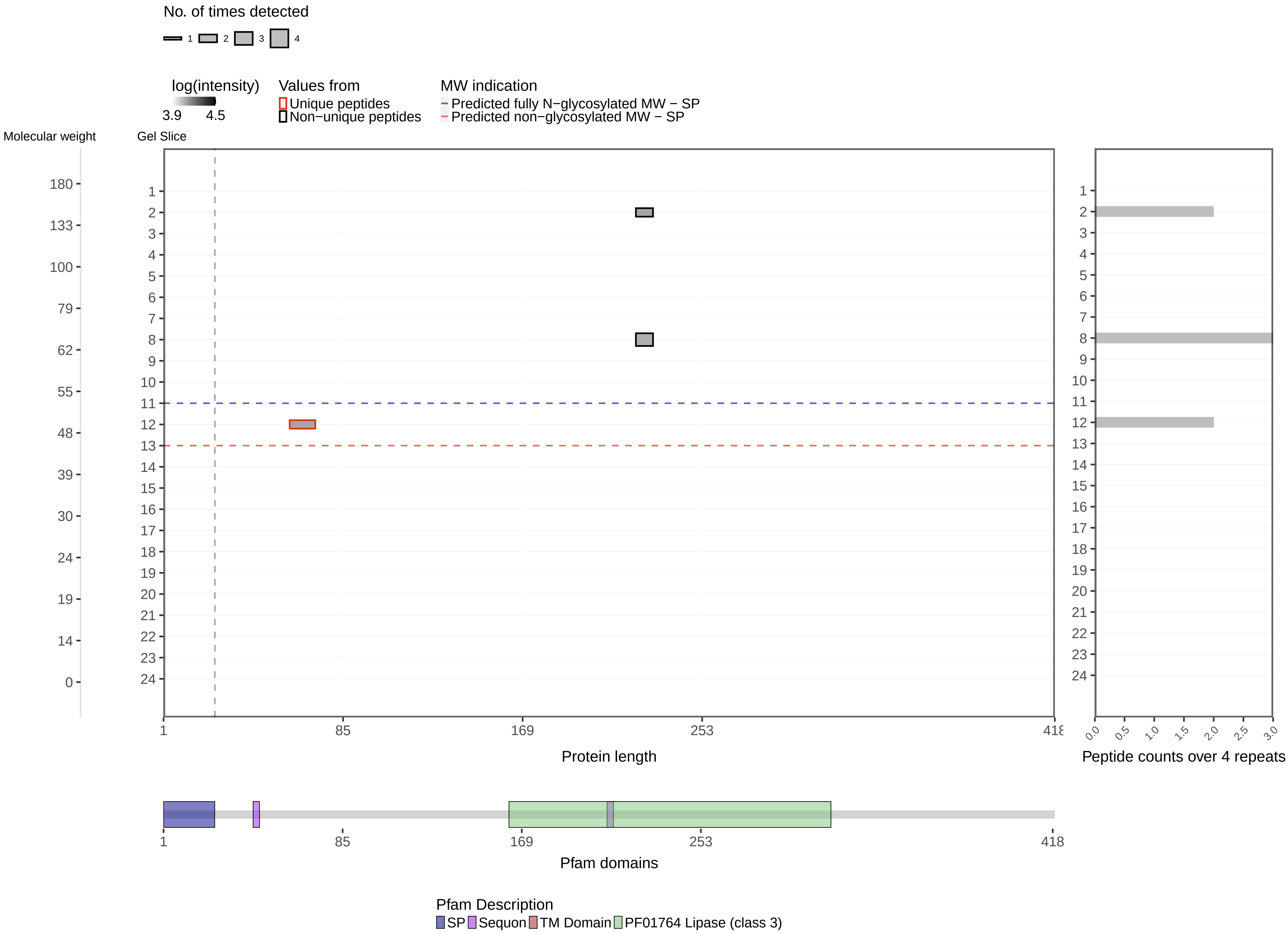


Protein sequence:

IHGWYSVYTSNDSRSPFSKASARDQVLGEVQRLVEQYKNEEITITVTGHSMGAAVATLNAVDIVSNGFNKGFPVTAFLFASPRVGDQNFKKTFSKLVNLRALRVRNAPDIVPNYPLLGYSDVGVQLDIDTTKSTYLKGPGDESSWHNMDCYMHGVAGNQGSKGGFKLEVQHDISVVNKYLDALKDEYGVPASWWVEKNKGMVQQQDGSWILMDHEDDDL*

### NbL19g04630.1 protein_trichome_birefringence−like_37 [MW − SP = 39.09 kDa]

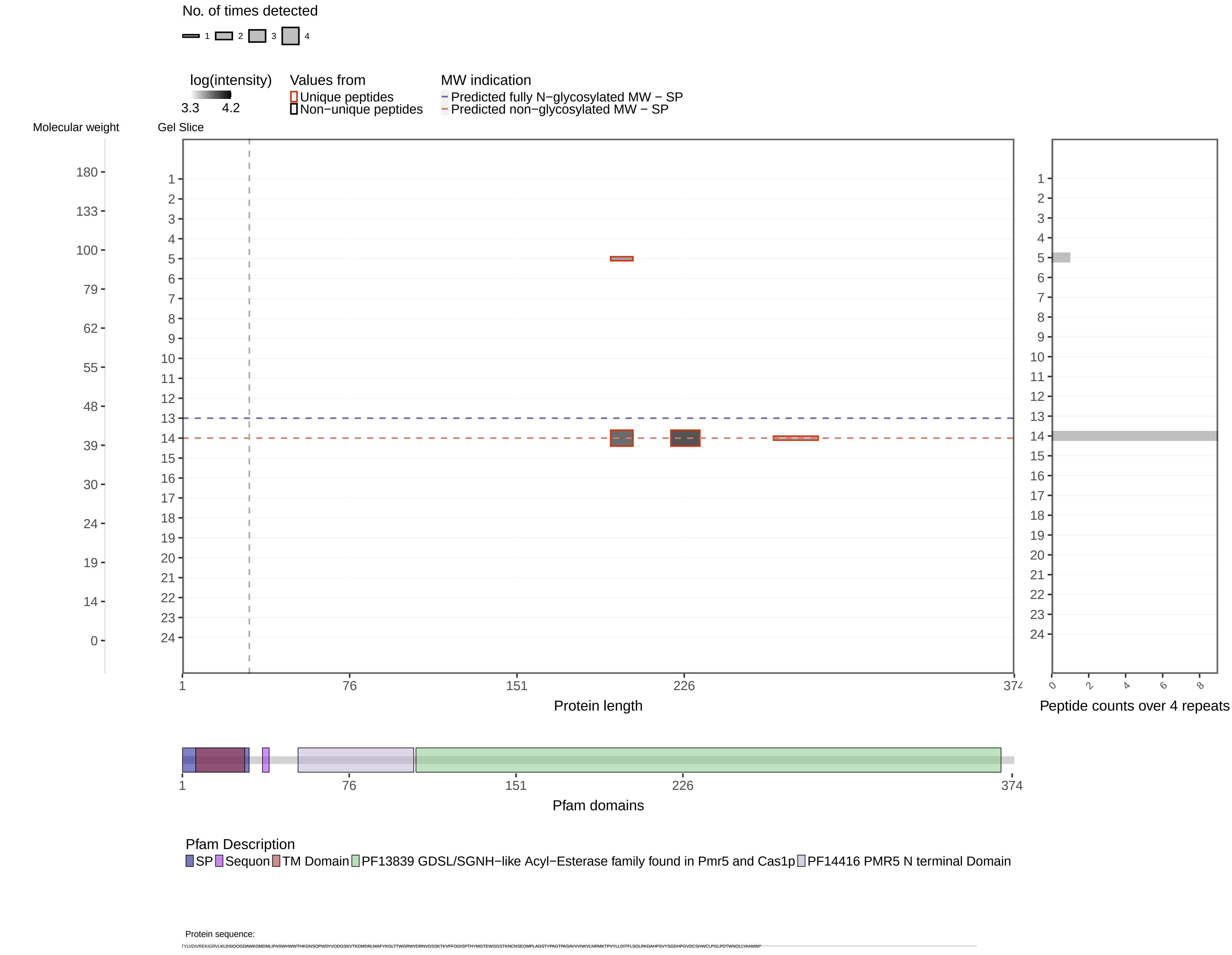

NbL19g04120.1 ras−related_protein_RABA1d−like [MW = 24.14 kDa]

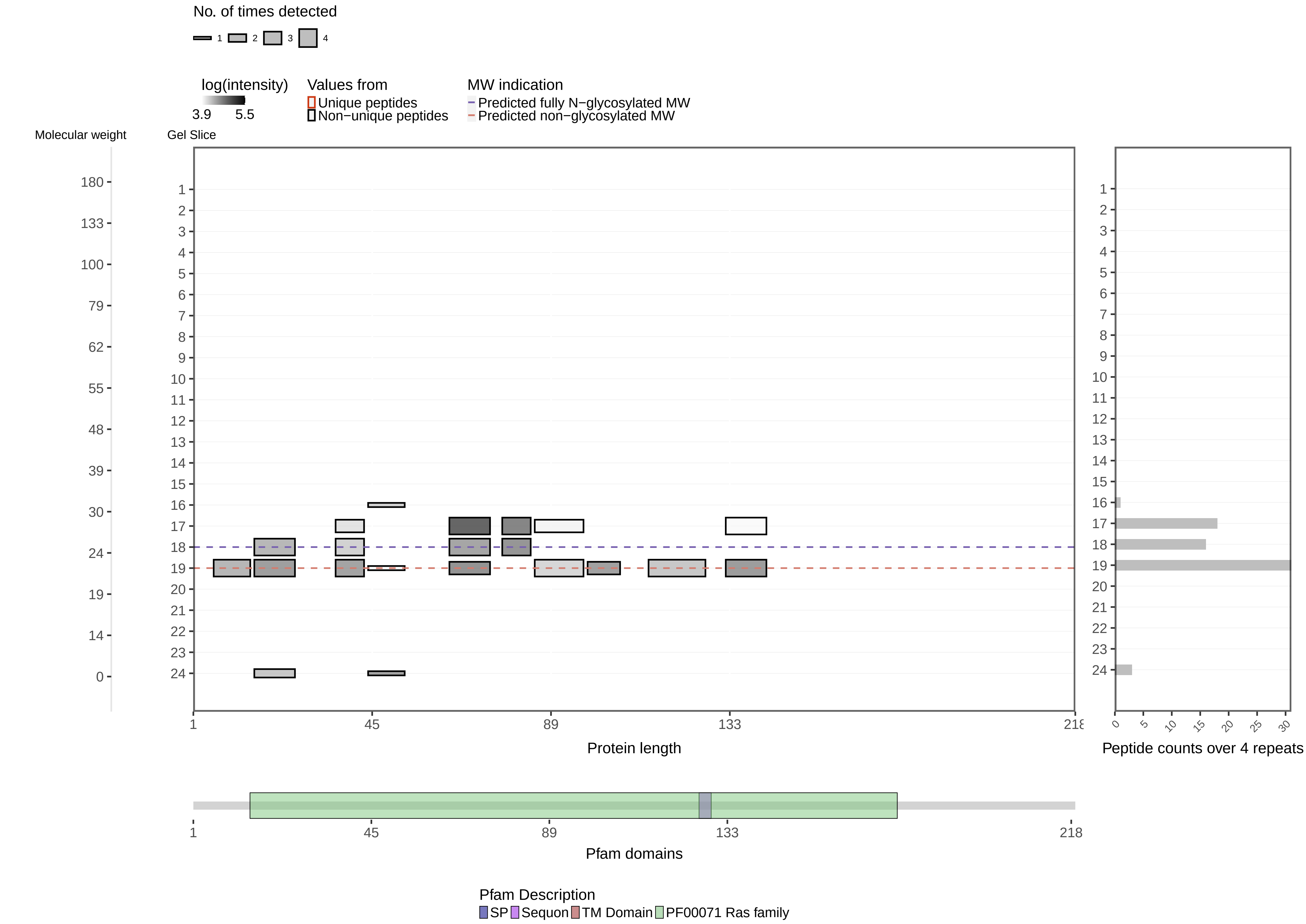


Protein sequence:

RHSTFENVERWLKELRDHTDPNIVVMLVGNKSDLRHLVAVSTEDGKSFAKKESLYFMETSALEATNVDNAFAEVLTEIYHVVSRKAMENGENGNANVPPQGEKIDIGKEVSDVKKTGCCST*

### NbL19g03830.1 histidinol_dehydrogenase_chloroplastic−like [MW = 55.4 kDa]

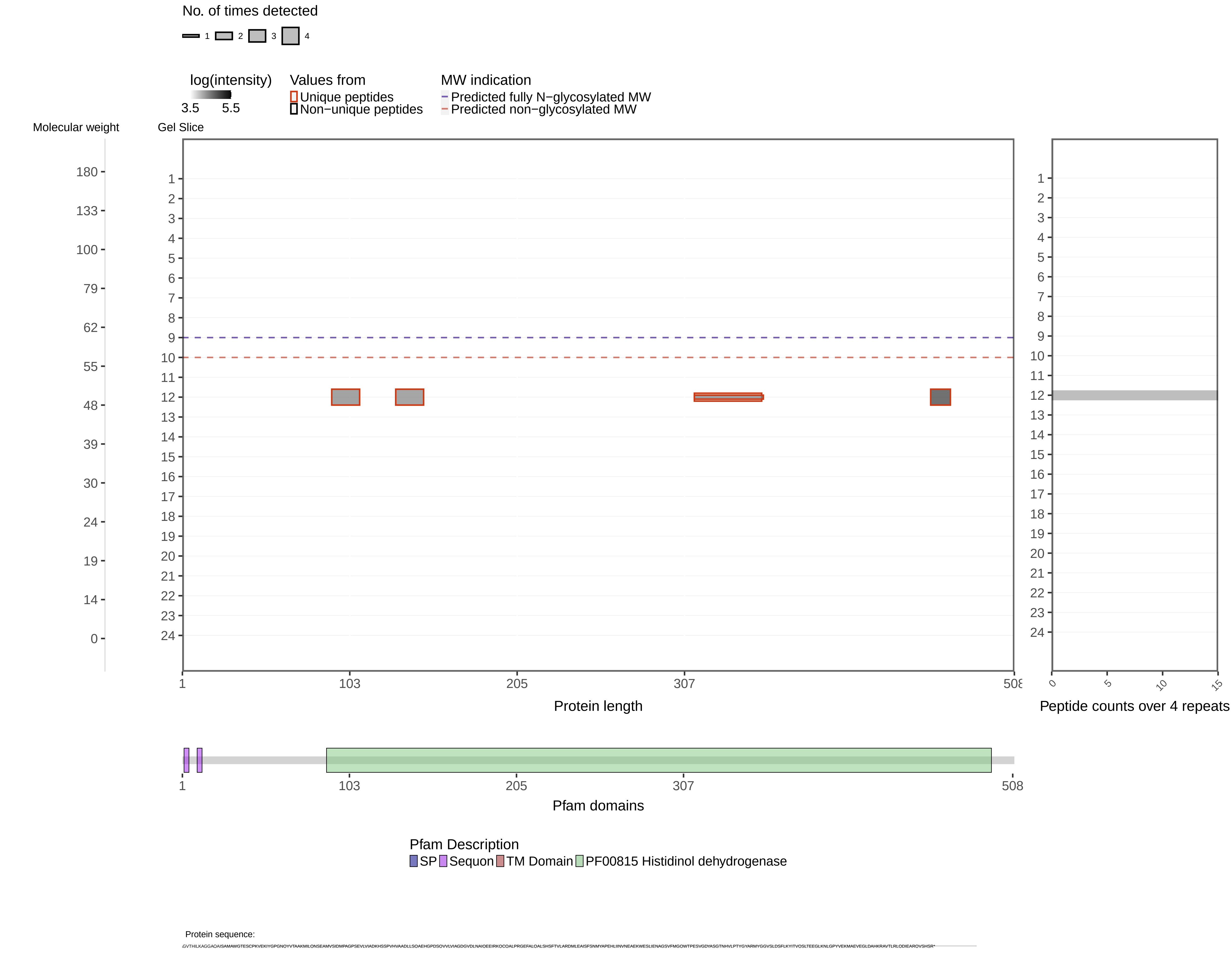

### NbL19g03780.1 MACPF_domain−containing_protein_CAD1 [MW = 63.38 kDa]

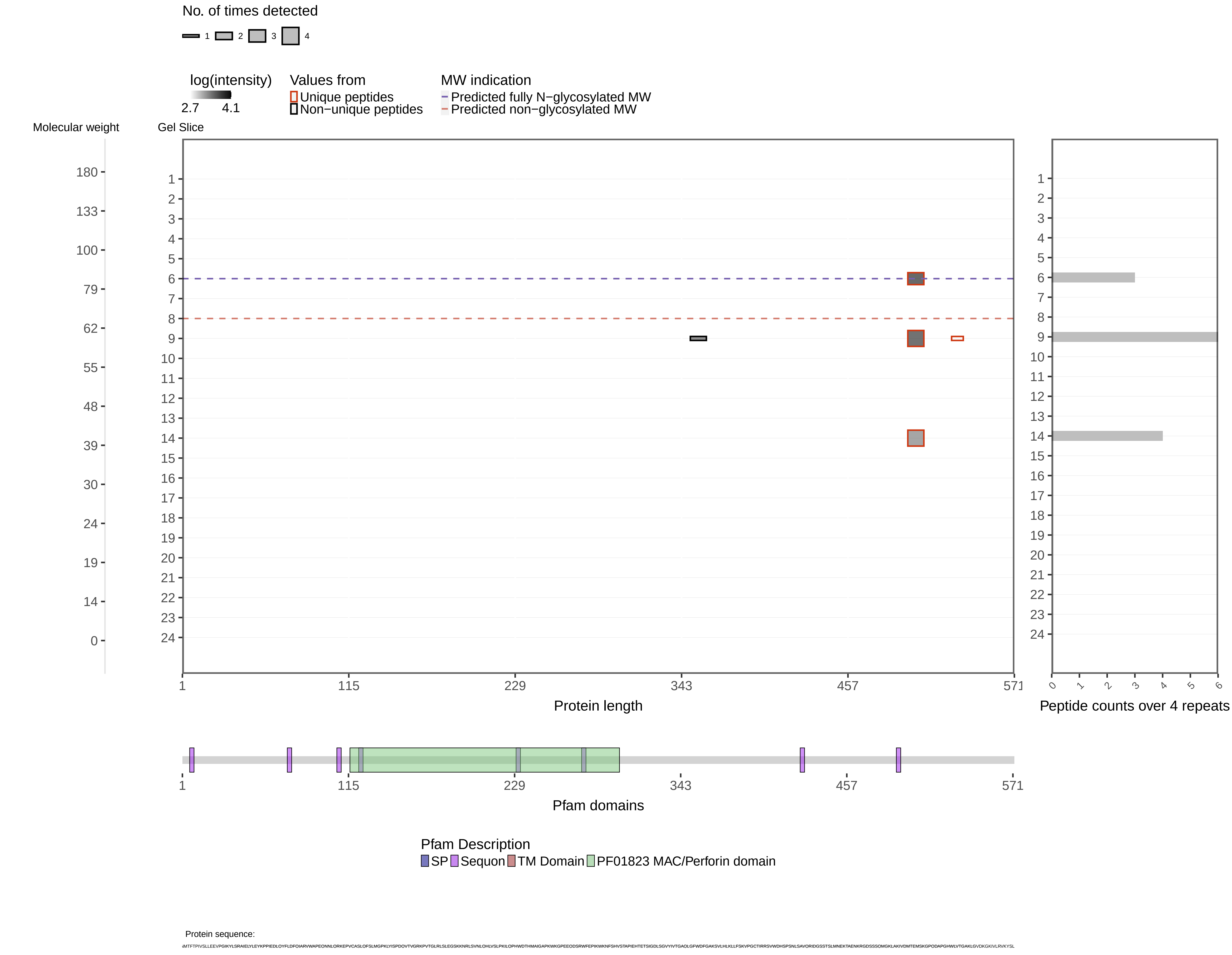

NbL19g03760.1 serine/threonine−protein_kinase_HT1−like [MW = 53.32 kDa]

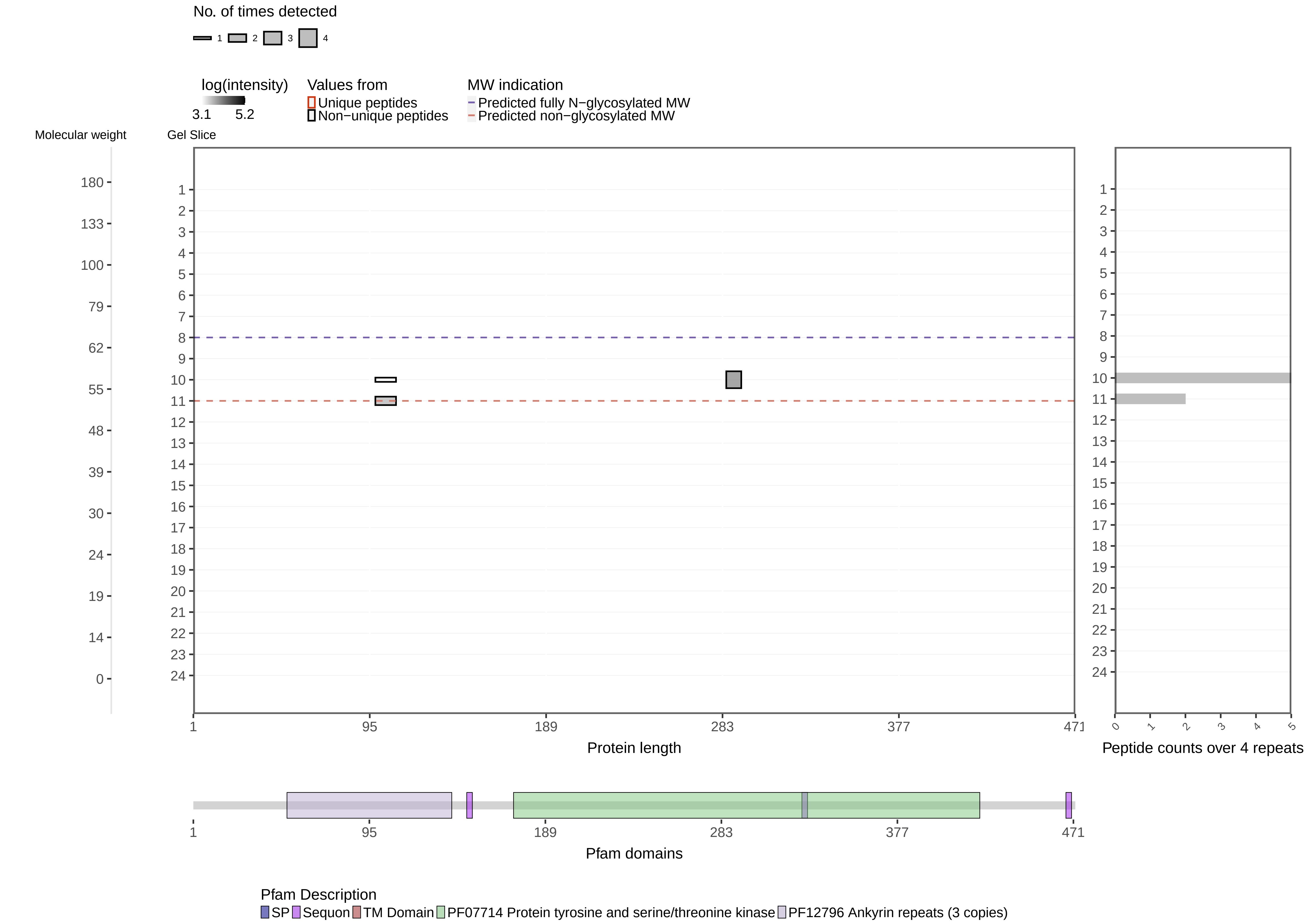


Protein sequence:

LGAVTQSSPMMIVTEYLPKGDLHAYLMKEGPLRPTKAIRFAMDIARGLNYLHEIRPEAIIHRDLEPSNILRDDTGHLKVADFGVSKLLKVTNRVKEDKPLTCDNTSCRYVAPEVFKNEEYDTKVDVFSFALILQEMIEGCPPFHAKIESEVAKCYAAKERPPFKAPGKFYAHGLRELIEECWNEKPALRPSFKQIIPRLESIYTKFGHKRRWKVRPMKCFQNFEAMWKKDHSSLSSRNGGSSRSNRSI*

NbL19g03730.1 uncharacterized_LOC104210613 [MW − SP = 37.21 kDa]

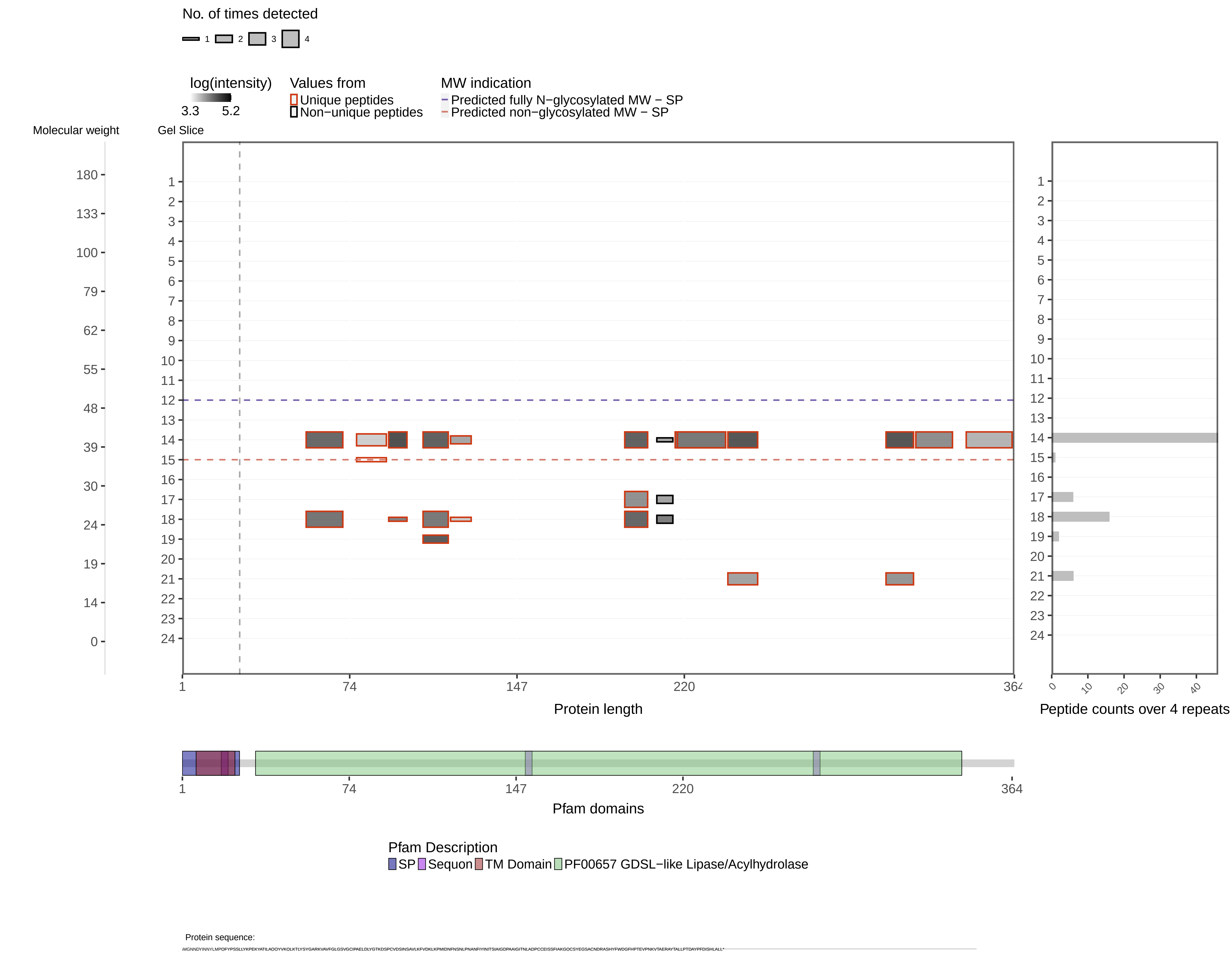

### NbL19g03700.1 GDSL_esterase/lipase_At1g29670 [MW − SP = 37.18 kDa]

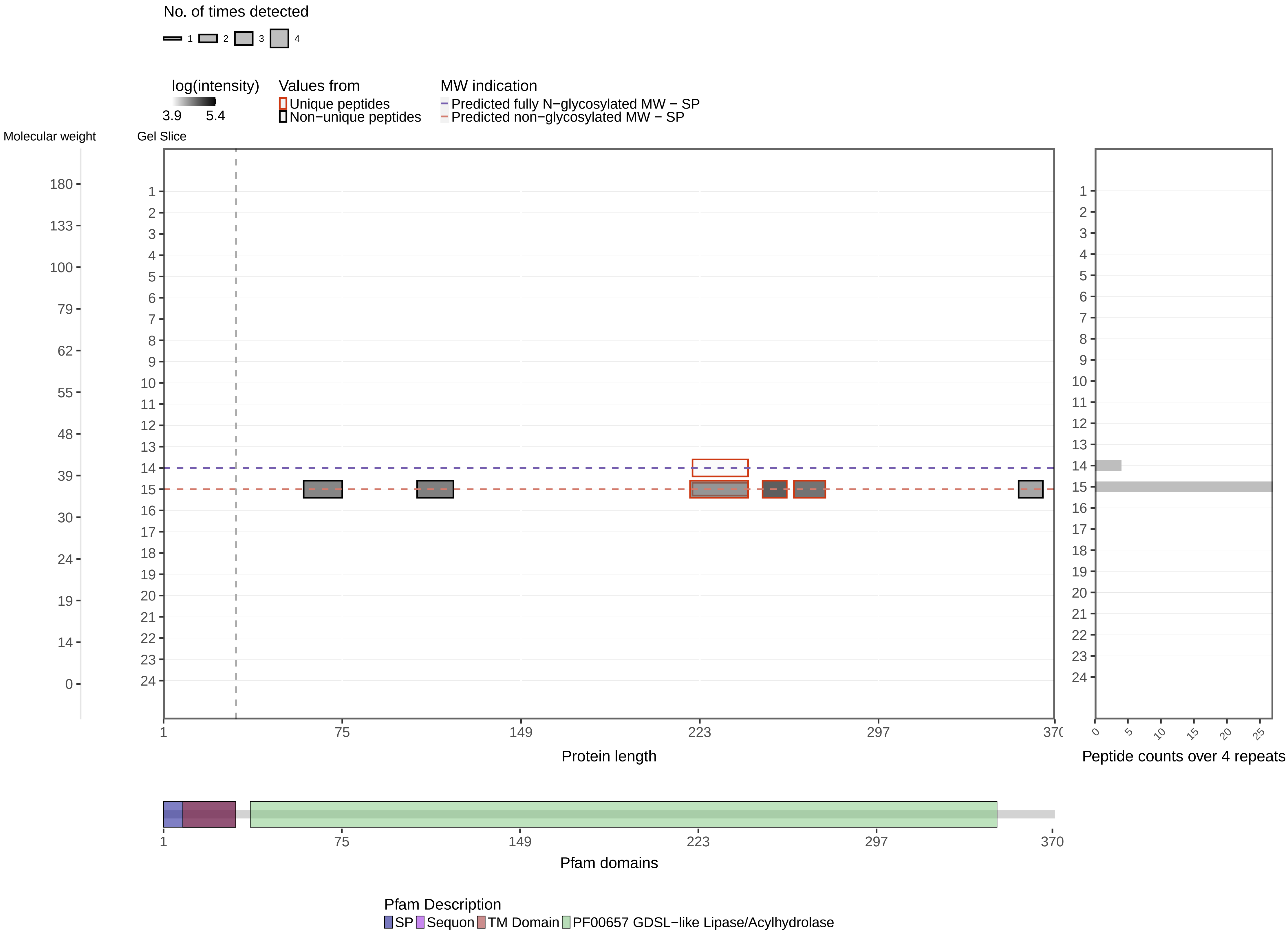


Protein sequence:

GSNDYLNNYFMPLYYPTSROFTPEOYASVLIOOYTOOLRILYNNGARKFALIGVGOIGCSPNALAONSPDGRTCVORINVANOLFNSKLKALVDNLNGNTPDAKFIYINAYGIFODLIDNPSAFGLSVTNAGCCGVGRNNGOITCLPFORPCRNRNOYLFWDAFHPTEVANIVVGRRSYRAOKSSDAYPFDISRLAOL*

NbL19g03690.1 GDSL_esterase/lipase_At5g45670−like [MW = 43.6 kDa]

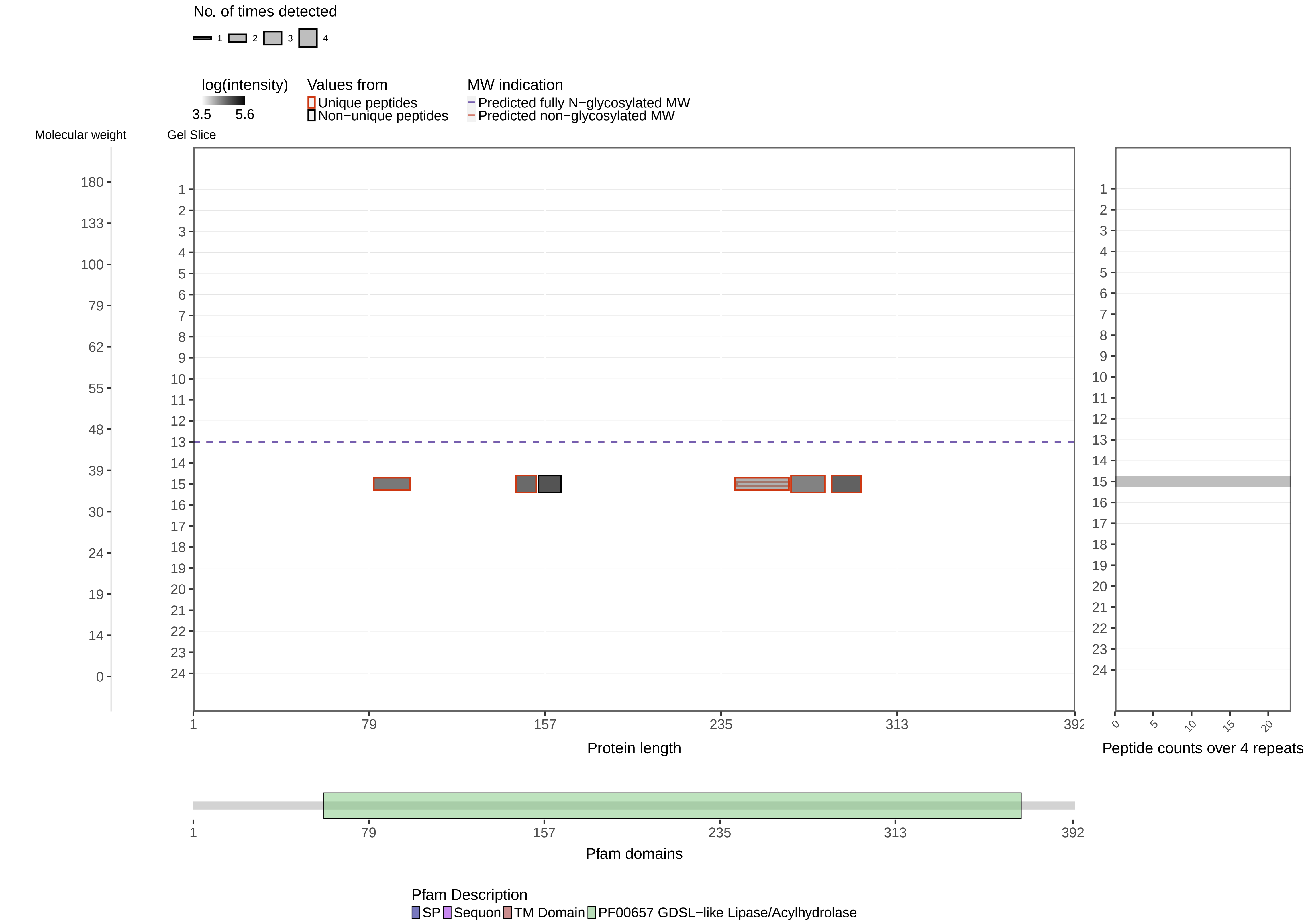


Protein sequence:

YLSKCIYSIGLGSNDYLNNYFMPLYYSTSRQFNPEQYADVLIQQYTQQLKTLYDYGARKFVLIGVGQIGCSPNSLAQNSPDGRTCVQNINVANQIFNNRLRGVVGDFNSNTPDAKFIYINAYDIFQDLIDNPSAFGFRVTNAGCCGVGRNNGQITCLPLQNPCPNRDEYLFWDAFHPGEAANVIVGRRSYRAERPSDSYPFDIQRLAQL*

NbL19g03660.1 subtilisin−like_protease_SBT5.6 [MW − SP = 80.79 kDa]

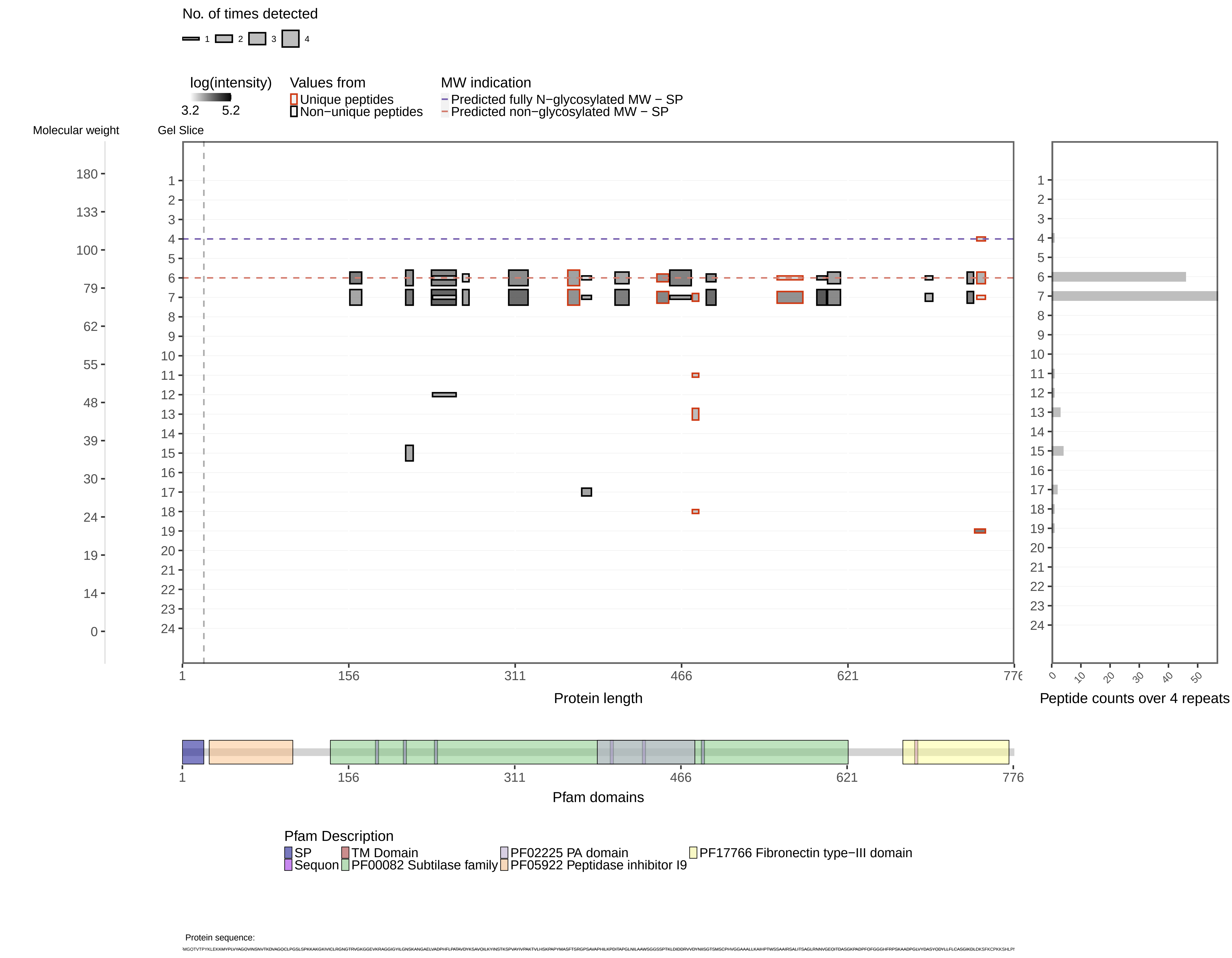

NbL19g03370.1 carboxyl−terminal−processing_peptidase_1_chloroplastic [MW = 50.46 kDa]

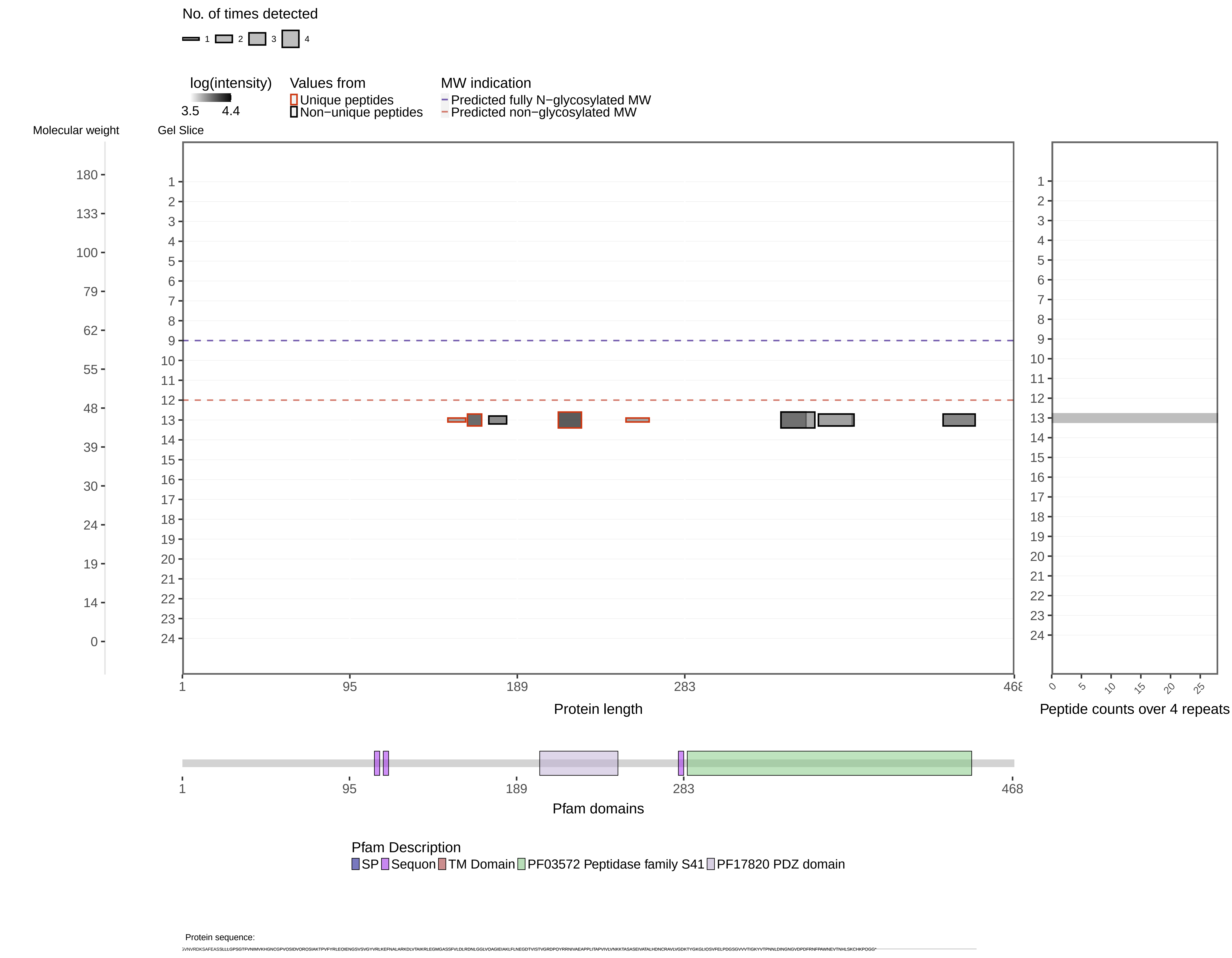

NbL19g03350.1 uncharacterized_LOC109236735 [MW = 89.52 kDa]

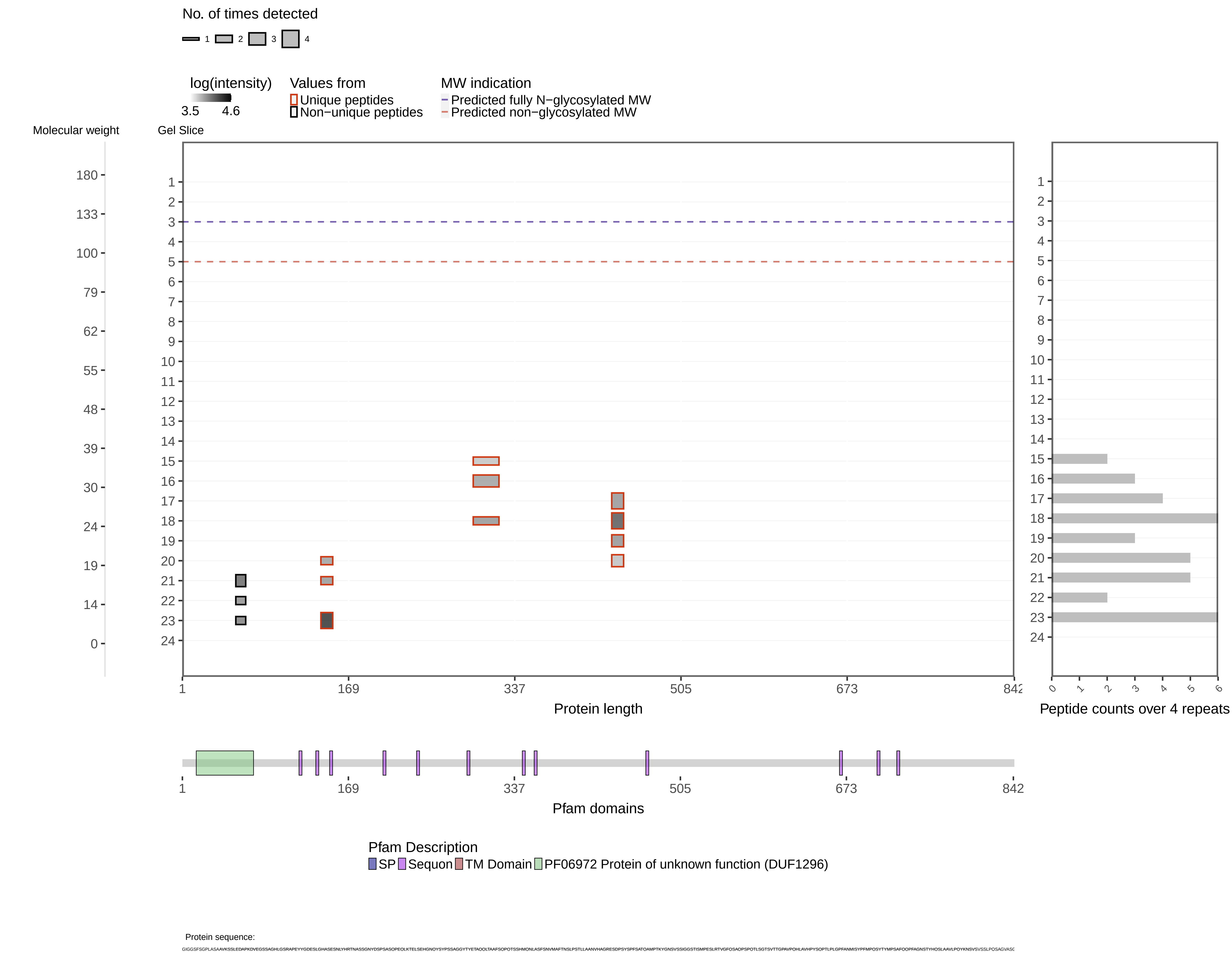

NbL19g03130.1 nuclear_pore_complex_protein_GP210 [MW − SP = 210.28 kDa]

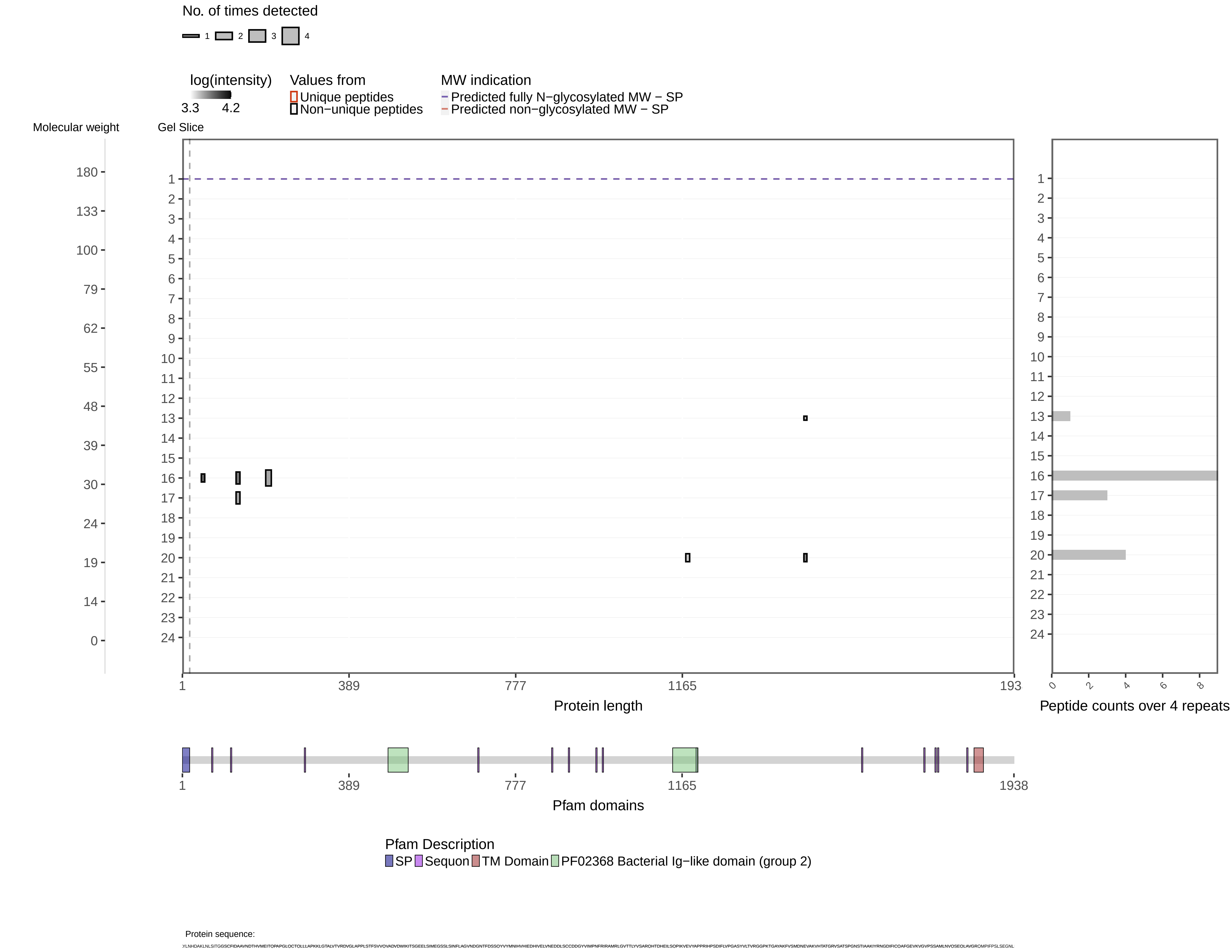

### NbL19g03120.1 uncharacterized_protein_ycf23−like [MW = 31.66 kDa]

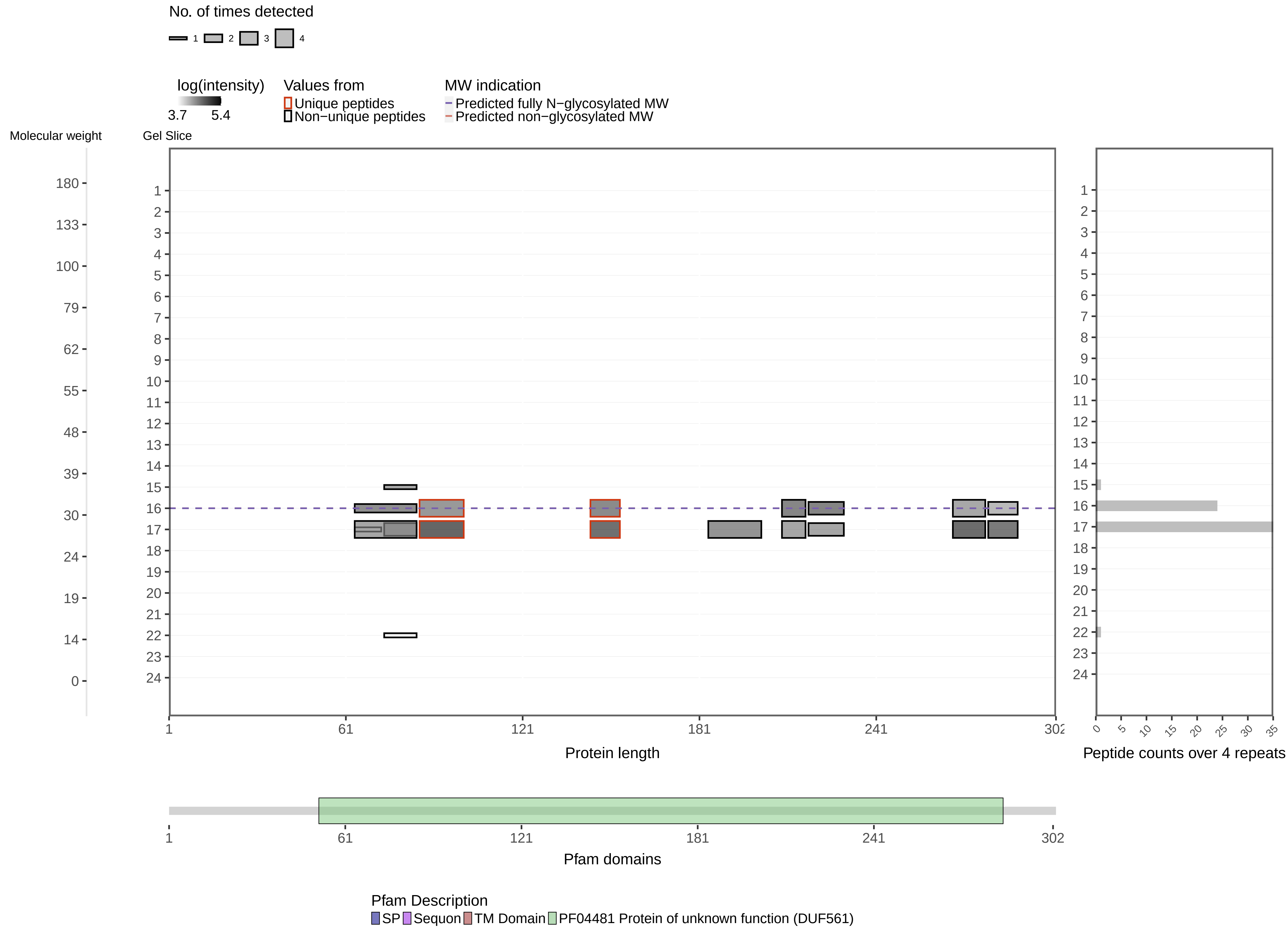


Protein sequence:

DSFYEKGLVFTPEQILKLTKETKRILPSVTLSVTVPHTLSLPDQIKLAELLEQEGVDIIQTEGGKCSTPSKAGVLGLIEKATPTLAAAYSISRAVNIPVMCSSGLSAVTAPMAITAGAAGVGVGSAINKLNDQVAMIAEVKSIAHSLGLSTKTQNLFEGSSLRL*

### NbL19g03110.1 3−oxoacyl−[acyl−carrier−protein]_synthase_I_chloroplastic [MW = 49.99 kDa]

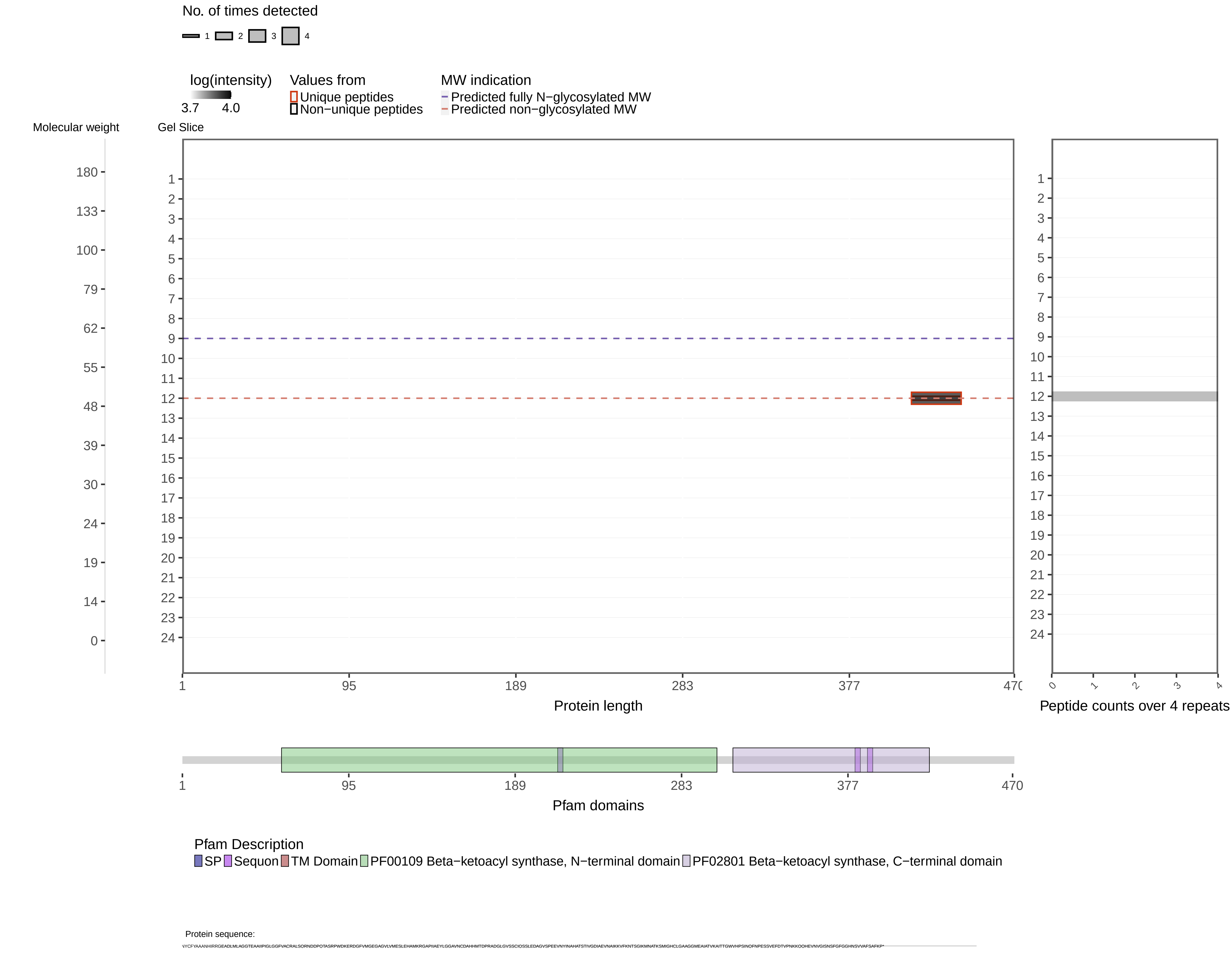

### NbL19g03100.1 DNA_replication_licensing_factor_MCM3_homolog_2 [MW = 94.57 kDa]

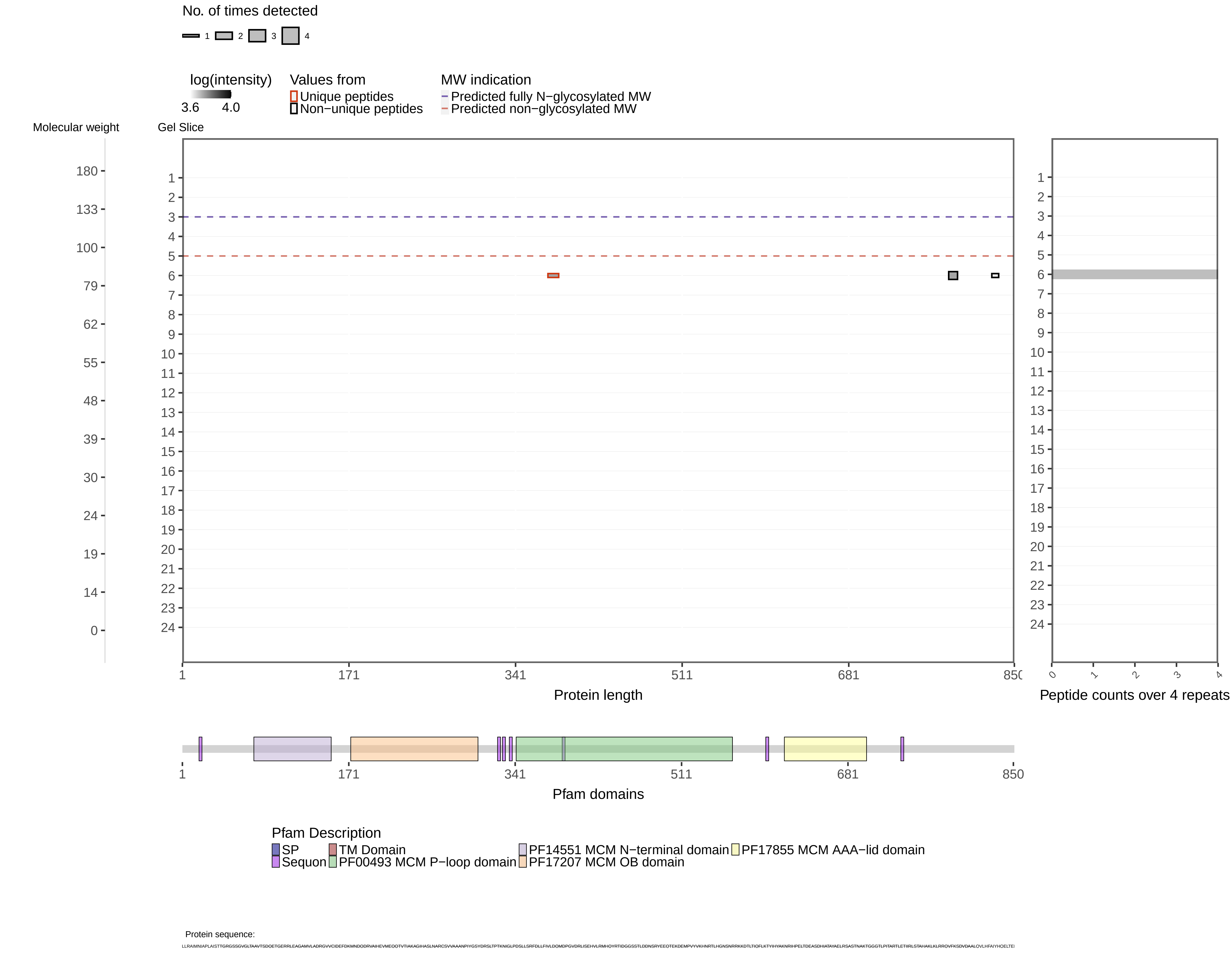

NbL19g03090.1 uncharacterized_LOC104243949 [MW = 50.11 kDa]

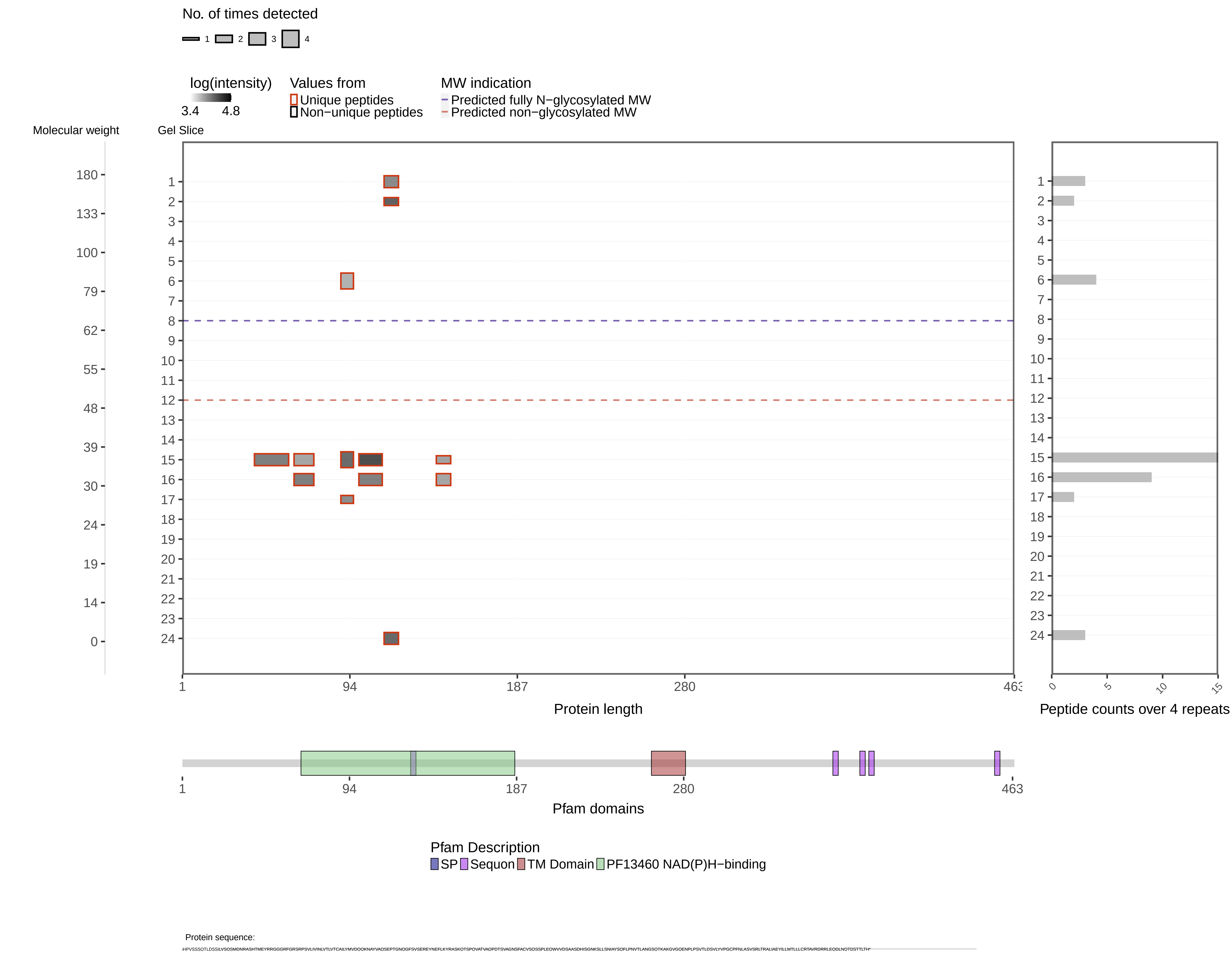

NbL19g03000.1 urease_accessory_protein_G−like [MW = 30.67 kDa]

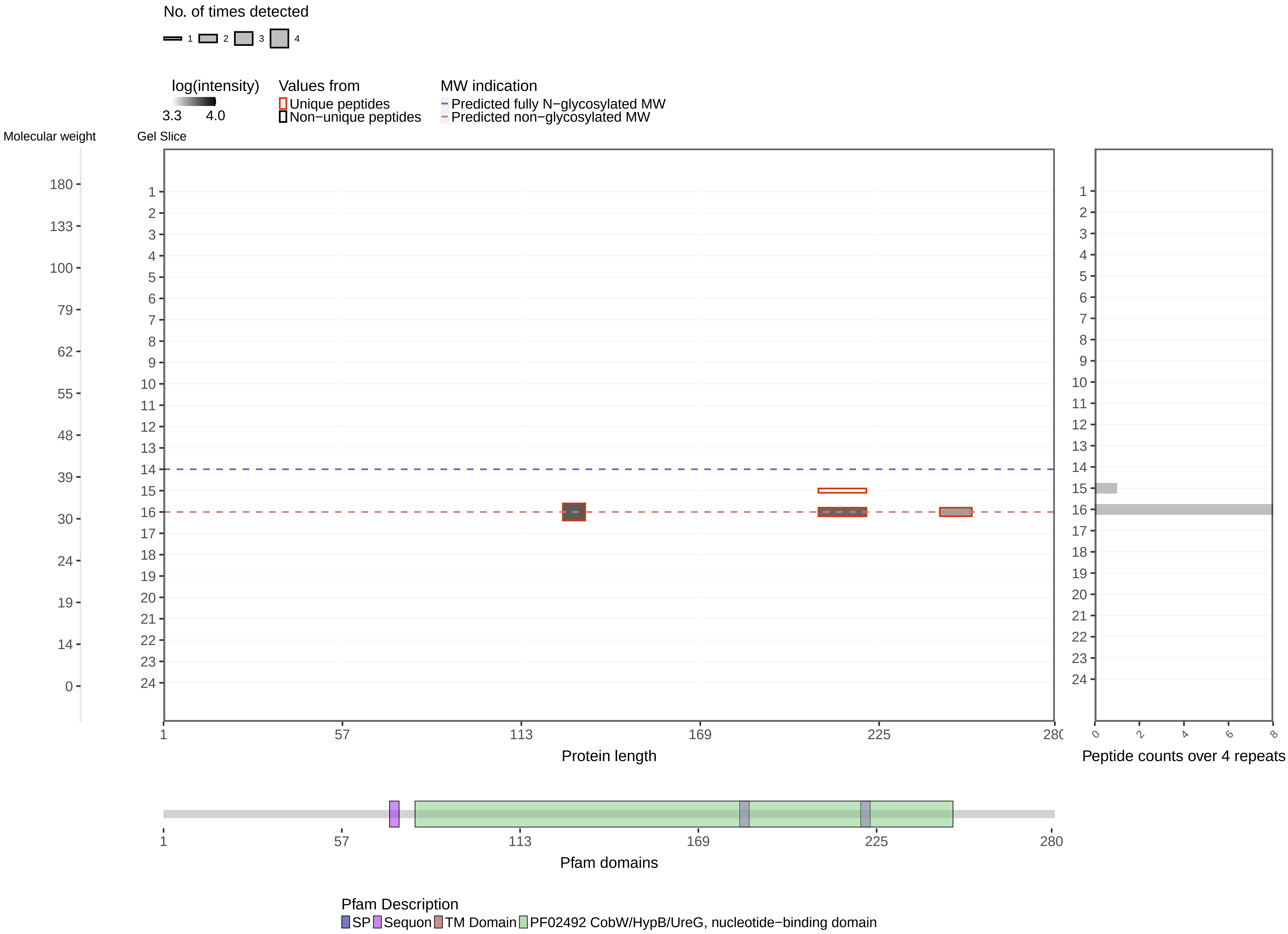


Protein sequence:

KHGALPEERIRAVETGGCPHAAIREDISINLGPLEELSNLYKADILLCESGGDNLAANFSRELADYIIYIIDVSGGDKIPRKGGPGITQADLLVINKTDLAAAVGADLSVMERDALRMRDGGPFVFAQVKHEVGVEDIVNHILQAWEVATGNKKH*

NbL19g02860.1 leucine−rich_repeat_receptor−like_serine/threonine−protein_kinase_BAM1 [MW − SP = 108.91 kDa]

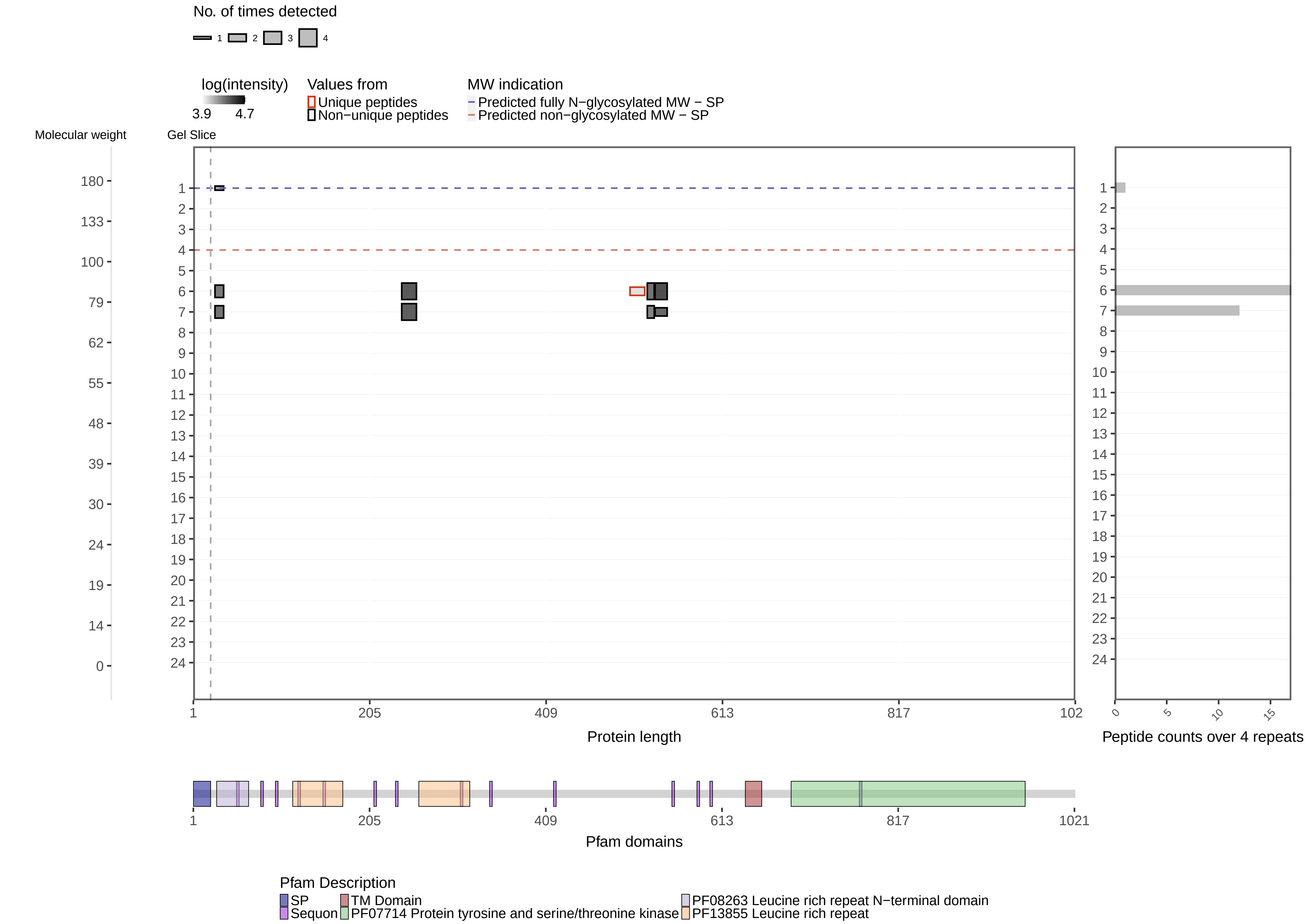


Protein sequence:

### NbL19g02790.1 COMPASS−like_H3K4_histone_methylase_component_WDR5A [MW = 35.45 kDa]

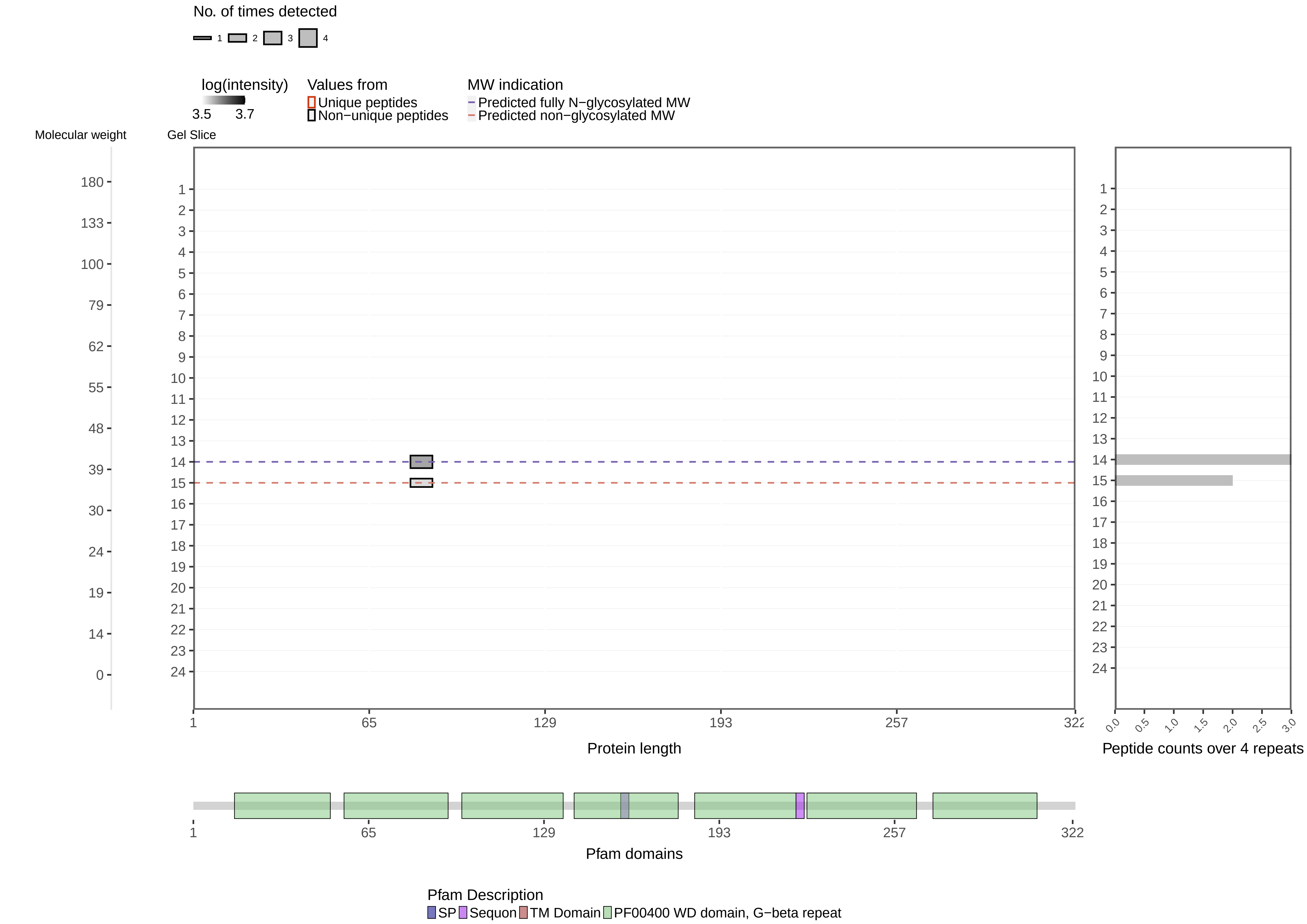


Protein sequence:

PAHSDPVTAVNFSRDGLLIVSSSYDGLCRIWDASTGHCMKTIIDDENPPVSFVKFSPNGKFILVGTLDNTLRLWNFSTGKVLKTYTGHANSKYCVSSTFSITDGKYIVSGSEDNCVYLWELQSRKIVQKLEGHTDTVISVACHPTQNMIASGSLGNDKTVKIWTQAQREDSEDMT*

NbL19g02730.1 probable_beta−D−xylosidase_6 [MW − SP = 83.95 kDa]

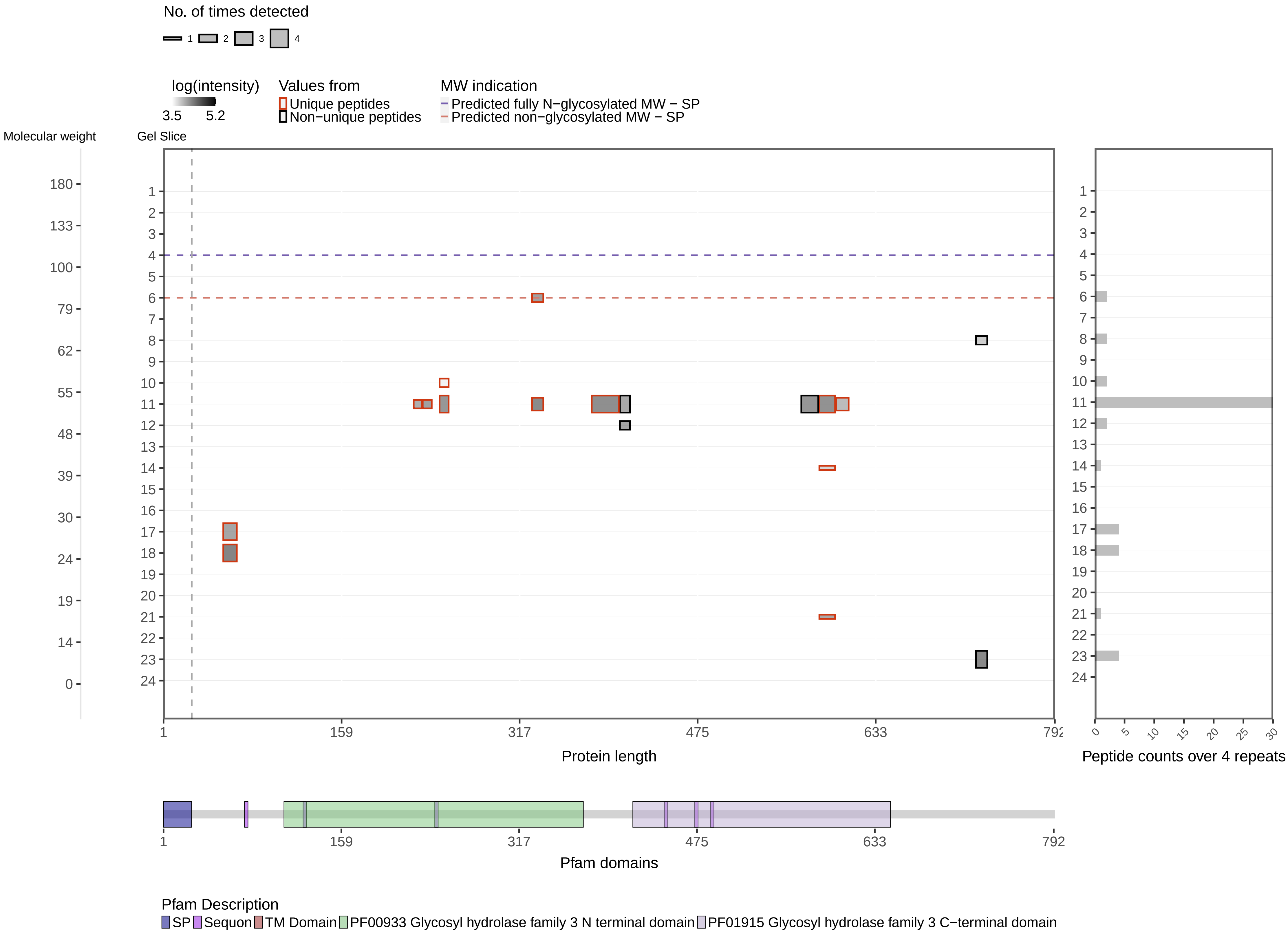


Protein sequence:

.GLFDGNPADGQFANFGAEDVCSSKHLNLALDAARQGIVLLKNDQKFLPLDKTSVSTLAIVGPMANVSSPGGTYSGVPCKLKSIREGFHRHINRTLYAAGCLDVGCNSTSGFQDAISIAKEADYVIVAAGLDLSEETEDHDRYSLLLPGQQANLVTTLAAASKKPIILVLTGGGPVDVSFAEKDPRIASILWVAYPGETGGKALSEIVFGYQNPGGKLPVTWYLESFTKVPMTNMNMRADPSNGYPGRSYRFYTGDVLYGFGHGLSYTSFSSQLLSAPSRLSLSLAKSNRKRSIL/

### NbL19g02580.1 probable_fructokinase−6_chloroplastic [MW = 42.05 kDa]

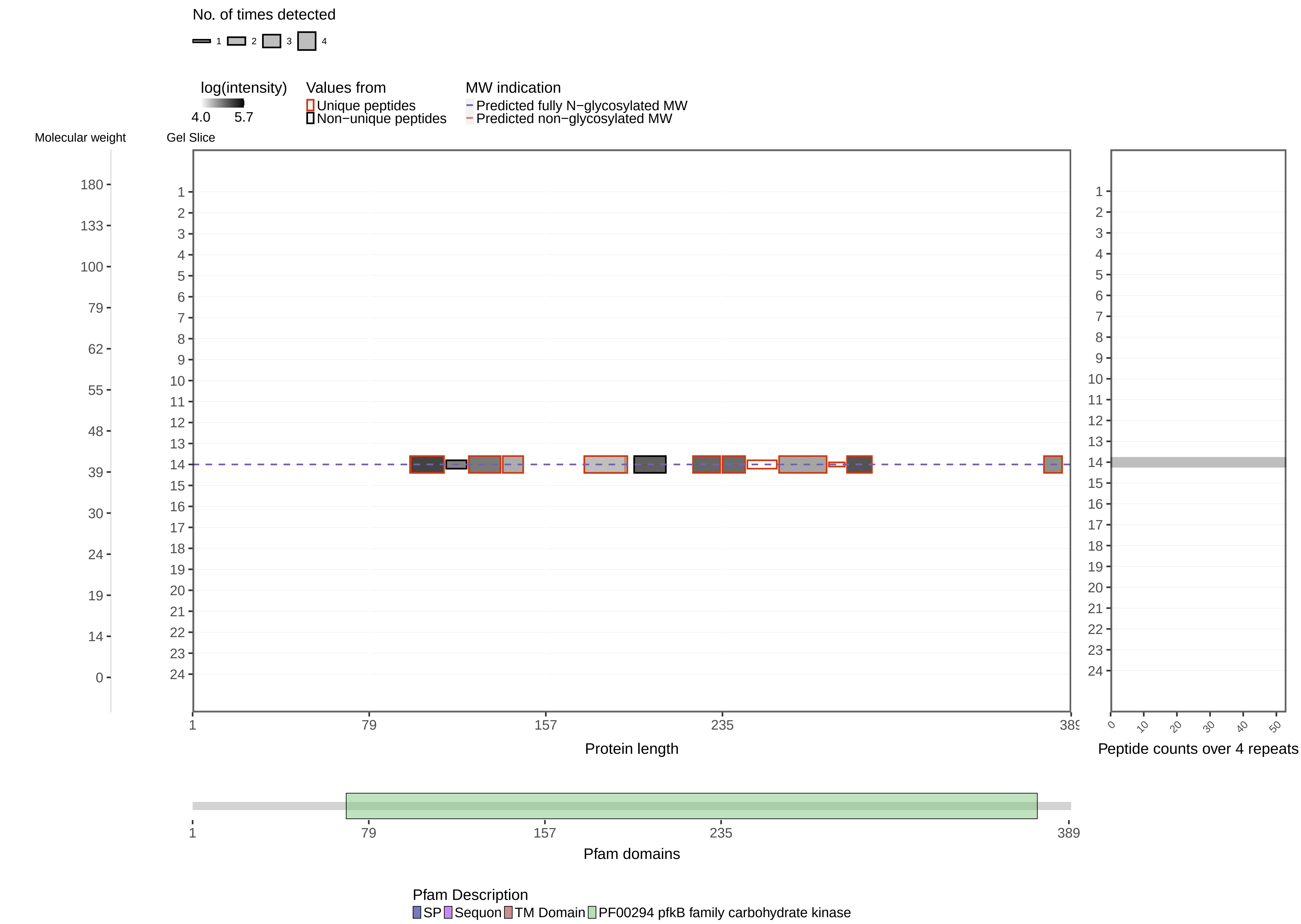


Protein sequence:

DMLLQEDELDLELIRKAKIFHYGSISLITEPCRSAHIAATRAAKDAGVILSYDPNLRLPLWPSAEIAREGILSIWETADIIKISEEEISFLTQGEDPYDDNVVRKLYHPNLKLLLVTEGPEGCRYYTKDFSGRVKGLKVDAVDTTGAGDAFVAGILSQLASDVSLLQDEDKLRDALCFANACGALTVMERGAIPALPTREVVLNSLLKSVS*

### NbL19g02480.1 protease_Do−like_7 [MW = 122.68 kDa]

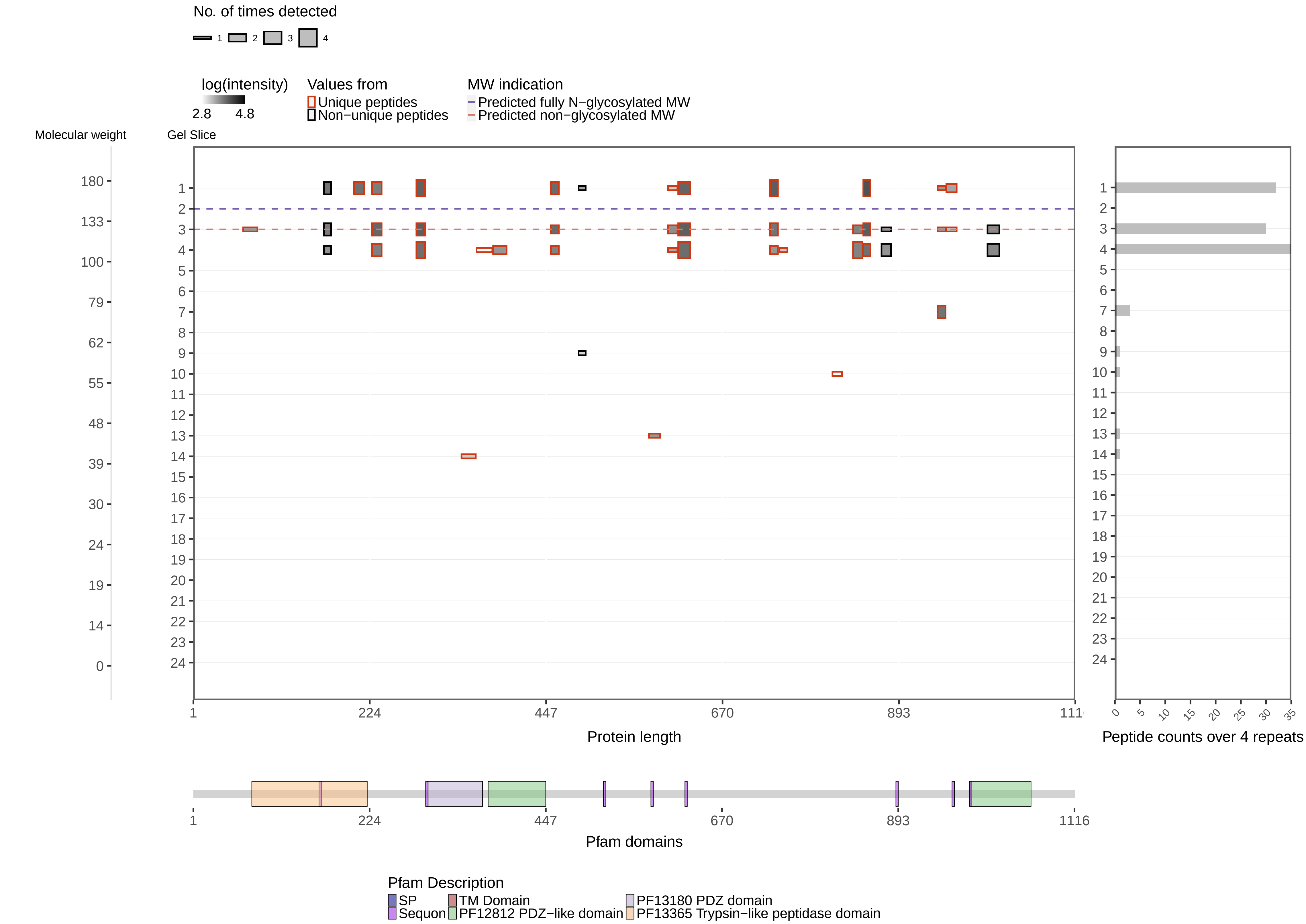


Protein sequence:
[illegible]

### NbL19g02450.1 pyridoxal_kinase [MW = 37.43 kDa]

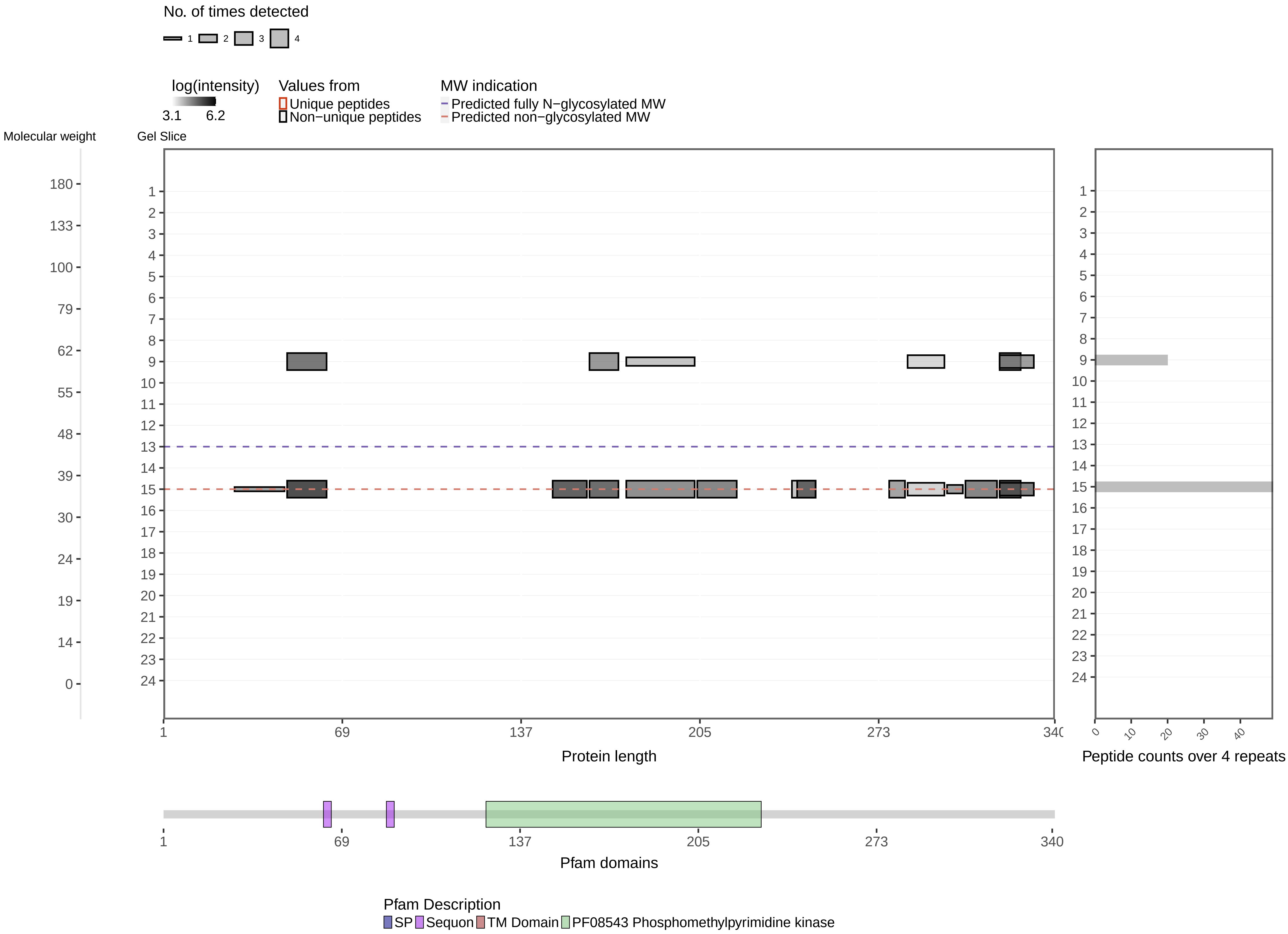


Protein sequence:

VMGDEGKLYVPPELVAVYREKVVPVASMLTPNOFEAEOLTGSSITSEKDGOEACNILHAAGPSKVVITSINIDGNLLLIGSHOKEKGOSPEOFKIVIPKIPAYFTGTGDLMTALLLGWSNKYPNDLDKAAELAVSSLOALLLRTLADYOKAGYDCOSSSLEIRLIOSODNIRNPEVKYRAKRYI*

NbL19g02400.1 methionyl−tRNA_formyltransferase−like [MW = 38.92 kDa]

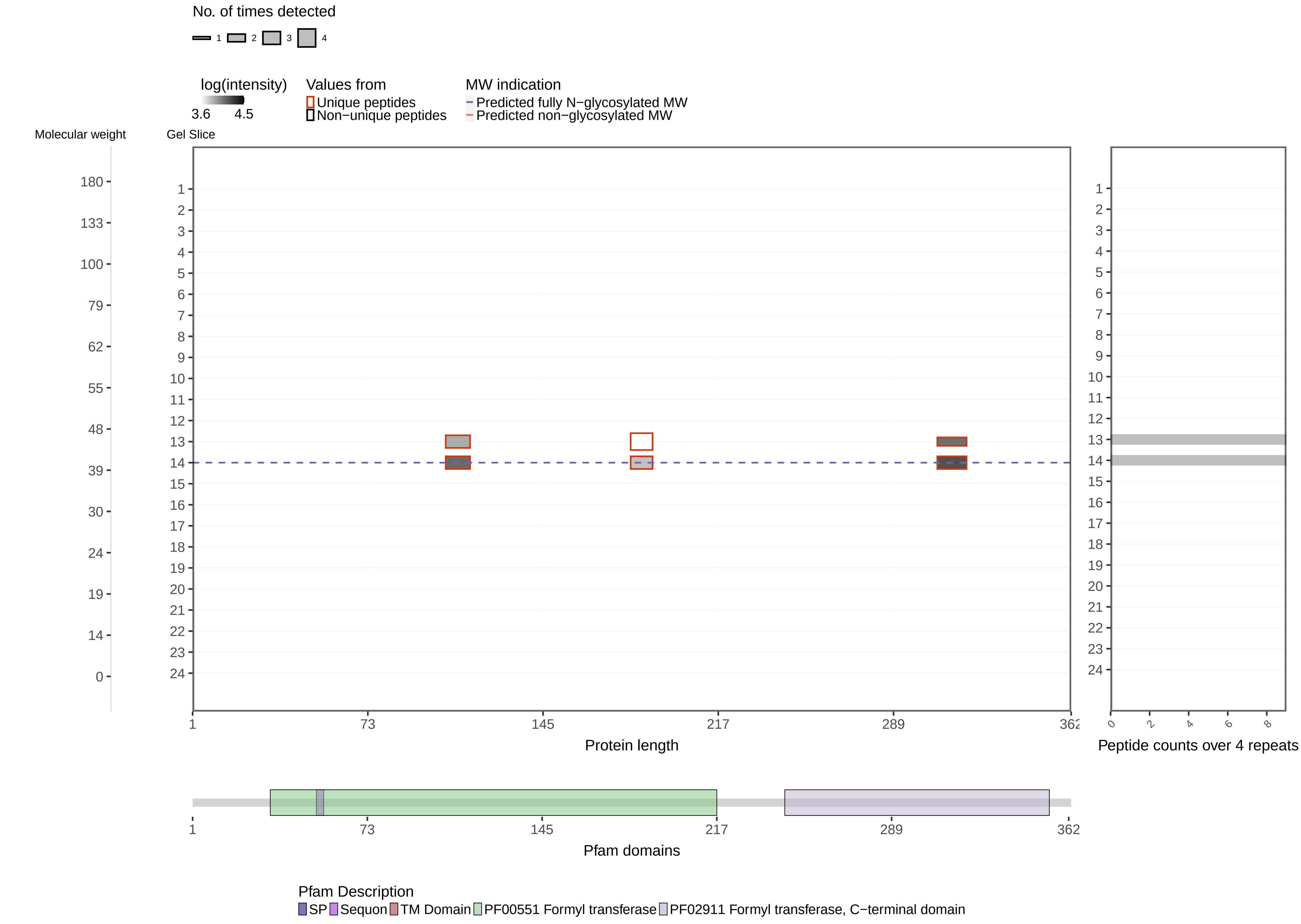


Protein sequence:

GVKETGVSLAYTVRKLDAGPVIARERVTVDDQIKAPDLLDLLFALGSKLLIRELPSIFDGSASGRAEEQDDSKATLAPKITPEESWLSFDEDAQVLHDKVRAFAGWPGTRSRLMVIDPSSGESSSIELKIITTRIYGGIRSQDIETNDVLFTKGSLVIPCGGGTALEVLELQLPGKRAVNAASFWNGLRGQILKR*

### NbL19g02380.1 ATP−dependent_Clp_protease_proteolytic_subunit_3_chloroplastic [MW = 20.67 kDa]

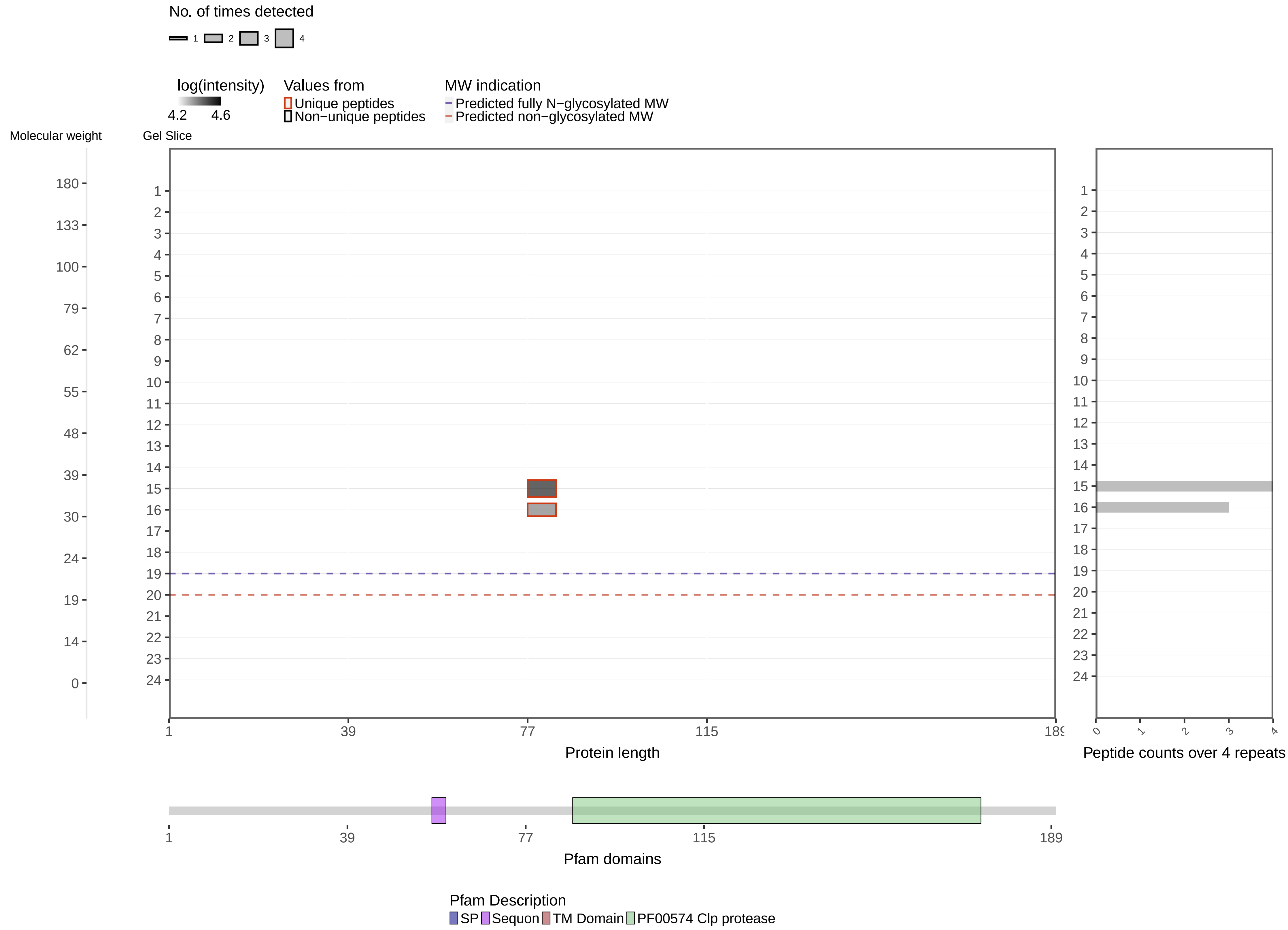


Protein sequence:

PRFEELDTTNMLLRQRIIFLGSQVDDMTADFVISQLLFLDAEDQKKDIRLIINSPGGSVTAGMGIYDAMKMCKADVSTICMGLAASMGAFSSHLAARERGTACRTQG*

NbL19g02300.1 ATP_synthase_subunit_beta_mitochondrial−like [MW = 59.82 kDa]

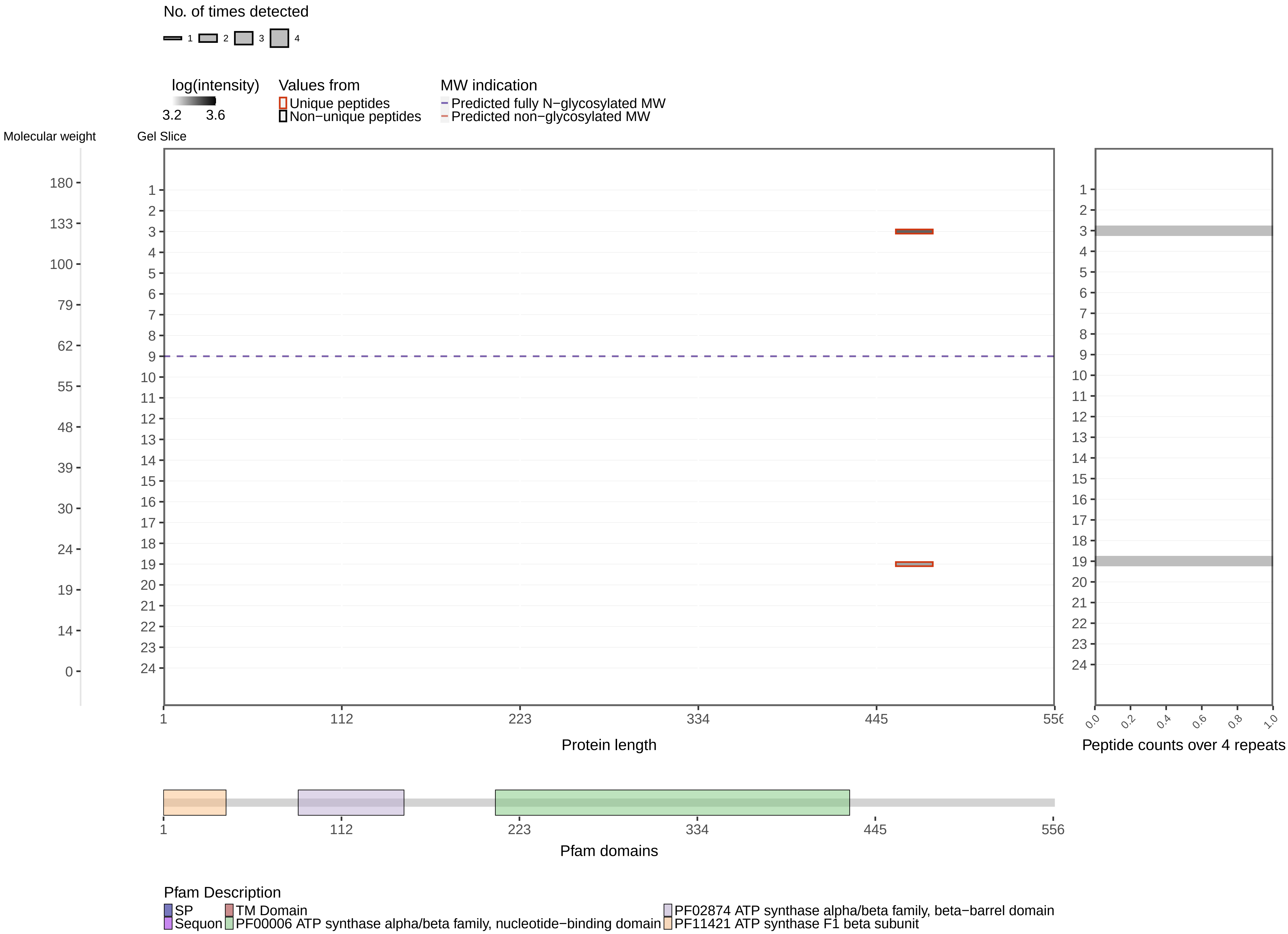


Protein sequence:

GNDLYREMIESGVIKLGDKQGESKCALVYGQMNEPPGARARVGLTGLTVAEHFRDAEGQDVLLFIDNIFRFTQANSEVSALLGRIPSAVGYQPTLATDLGGLQERITTTKKGSITSVQAIYVPADDLTDPAPATTFAHLDATTVLSRQISELGIYPAVDPLDSTSRMLSPHILGEDHYNTARGVQKVLQNYKNLQDIIAILGMDELSEDDKLTVARARKIQRFLSQPFHVAEVFTGAPGKYVDLKESIQSFQQGVLDGKYDDLSEQSFYMVGGIEEVIAKAEKIAKESATS*

### NbL19g02270.1 2−hydroxyacyl−CoA_lyase−like [MW = 61.41 kDa]

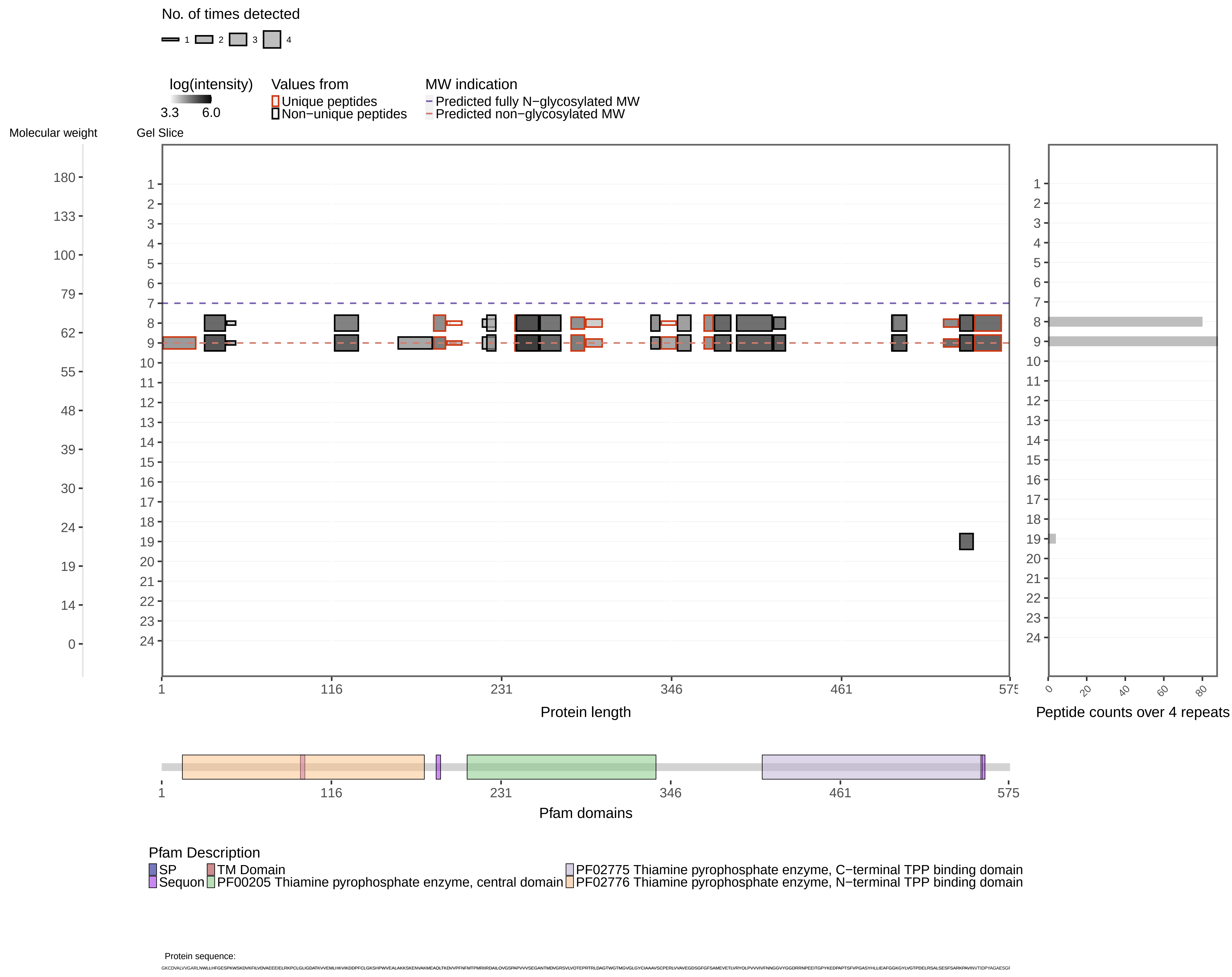

NbL19g01990.1 chloroplastic_group_IIB_intron_splicing_facilitator_CRS2_chloroplastic−like [MW = 26.86 kDa]

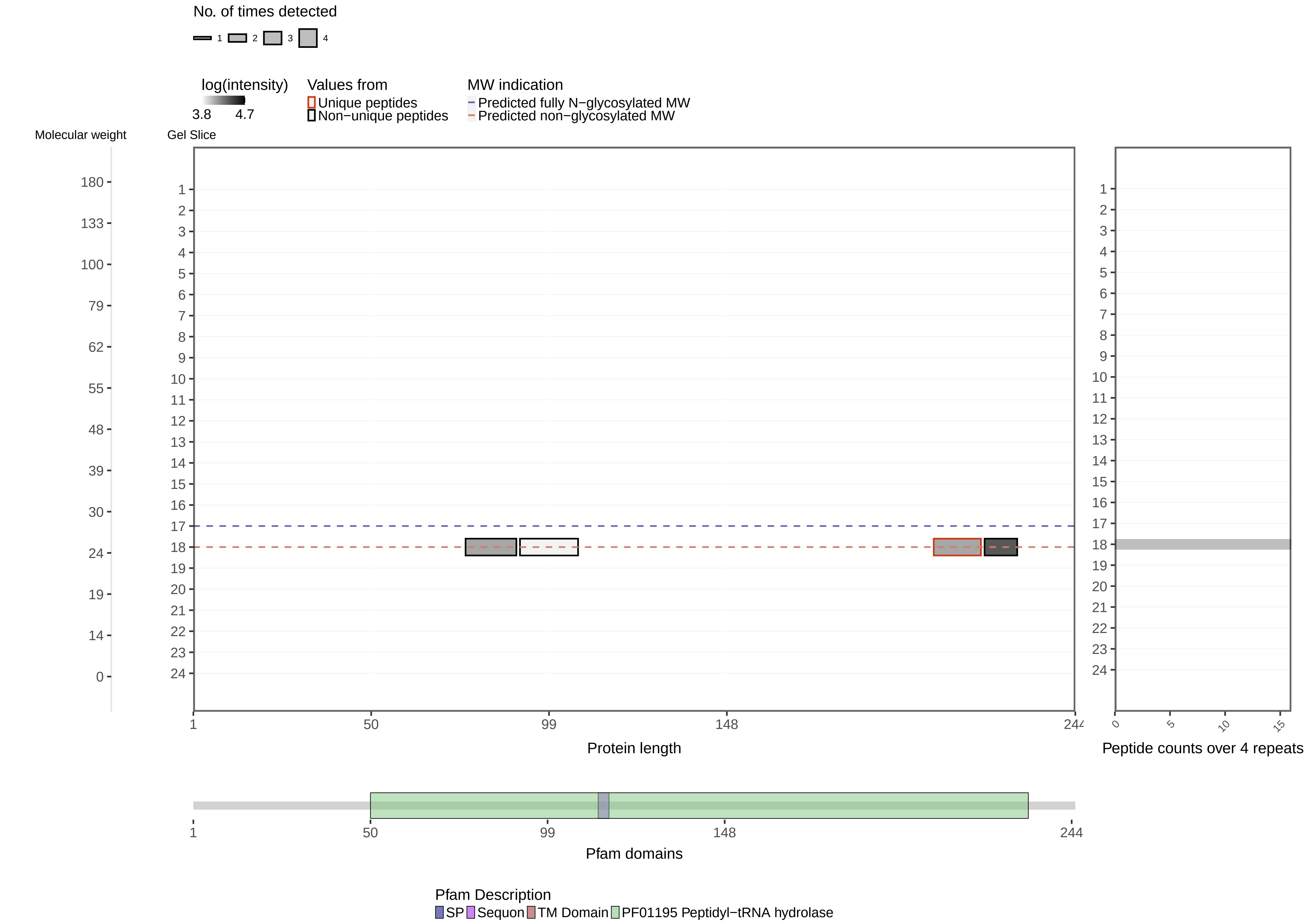


Protein sequence:

YMNFSGESVGPLAAYYQVPLRHILLVYDEMSLPNGVLRLQPKGGHGHHNGVKSVMEHLDGRREFPRFCIGIGNPPGTMDMKAYLLQKFSDIERKQVDAAFDQGVEAVRTVVLEGFGSRISRFNIGQKYKYHKV*

### NbL19g01960.1 ribulose_bisphosphate_carboxylase_small_chain_S41_chloroplastic−like [MW = 20.37 kDa]

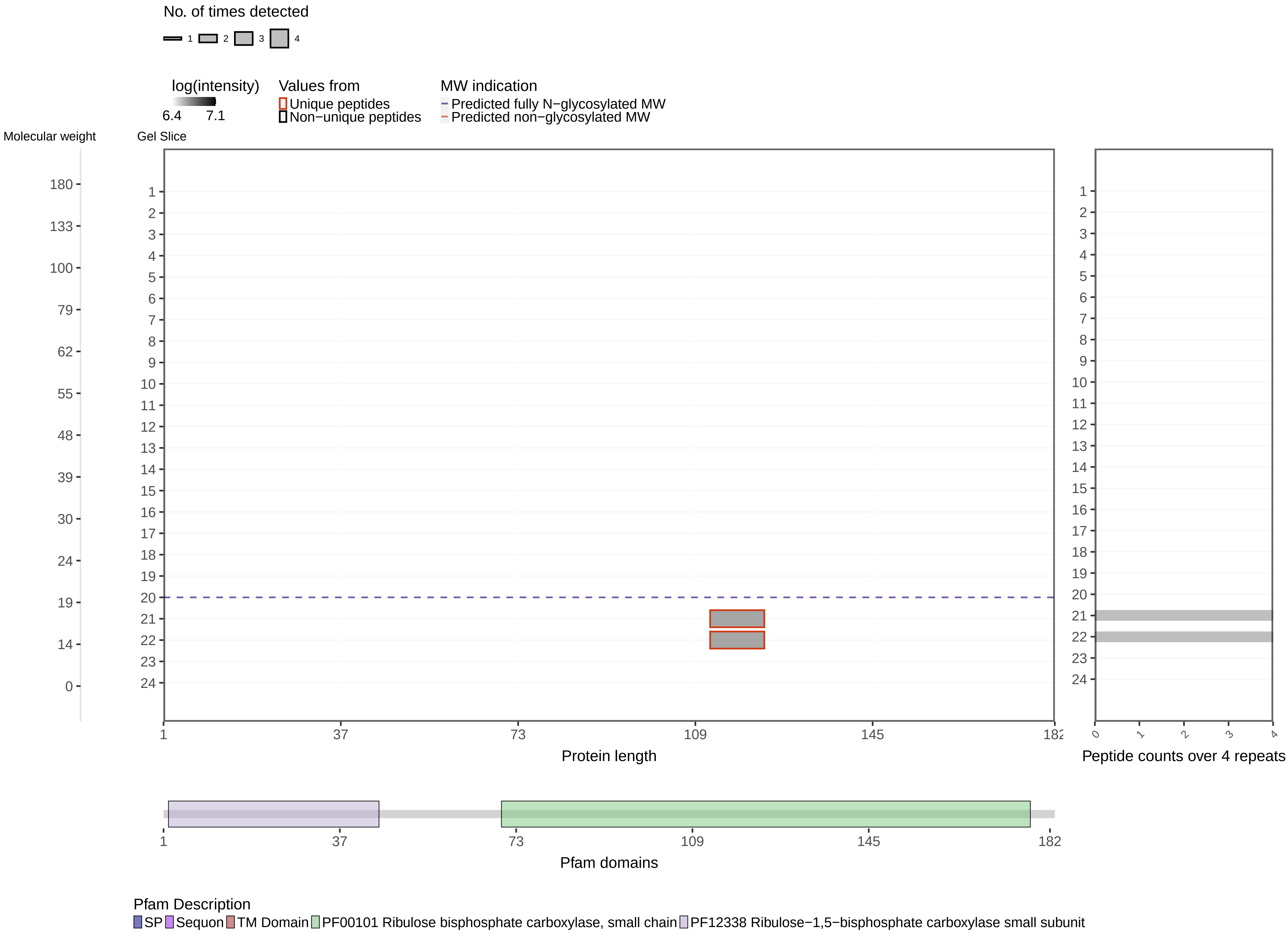


Protein sequence:

[illegible]LSVEQLLSEIEYLLKNGWVPCLEFETEHGFVYREHHHSPGYYDGRYWTMWKLPMFGCTDATQVLAEVEEAKKAYPQAWVRIIGFDNVRQVQCISFIAYKPEGY*

### NbL19g01770.1 T−complex_protein_1_subunit_zeta_1−like [MW = 58.97 kDa]

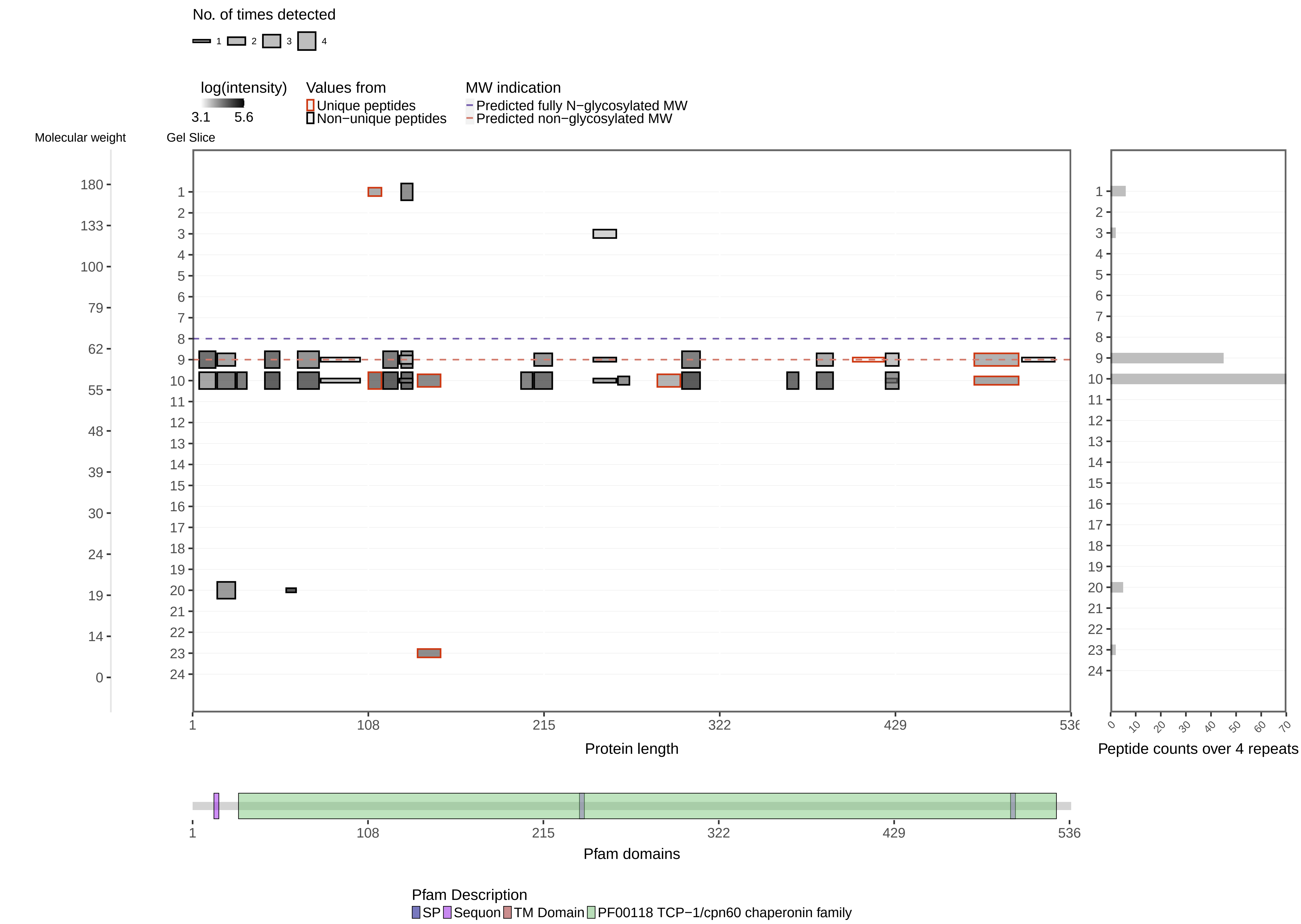


Protein sequence:

NAEQREAMVAAERRSVDERVQKIIDLKNKVCSGNENNFVVINQKGIDPPSLDILARAGIIALRRAKRRNMERLVLACGGDAVNSVDDLTPECLGWAGLVYEHVLGEEKYTFVENVKNPHSCTILIKGPNDHTIAQIKDAVRDGLRAVKNTIEDEAVVLGAGAFEVAARQYLINEVKKTVKGRAQLGVEAFANALLVIPKTLAENSGLDTQDVIIALTGEHDKGNIVGLNQHTGEPIDPQMEGIFDNYSVKRQIVNSGPVIASQLLLVDEVIRAGRNMRKPT*

NbL19g01710.1 allene_oxide_cyclase_4_chloroplastic−like [MW = 30.28 kDa]

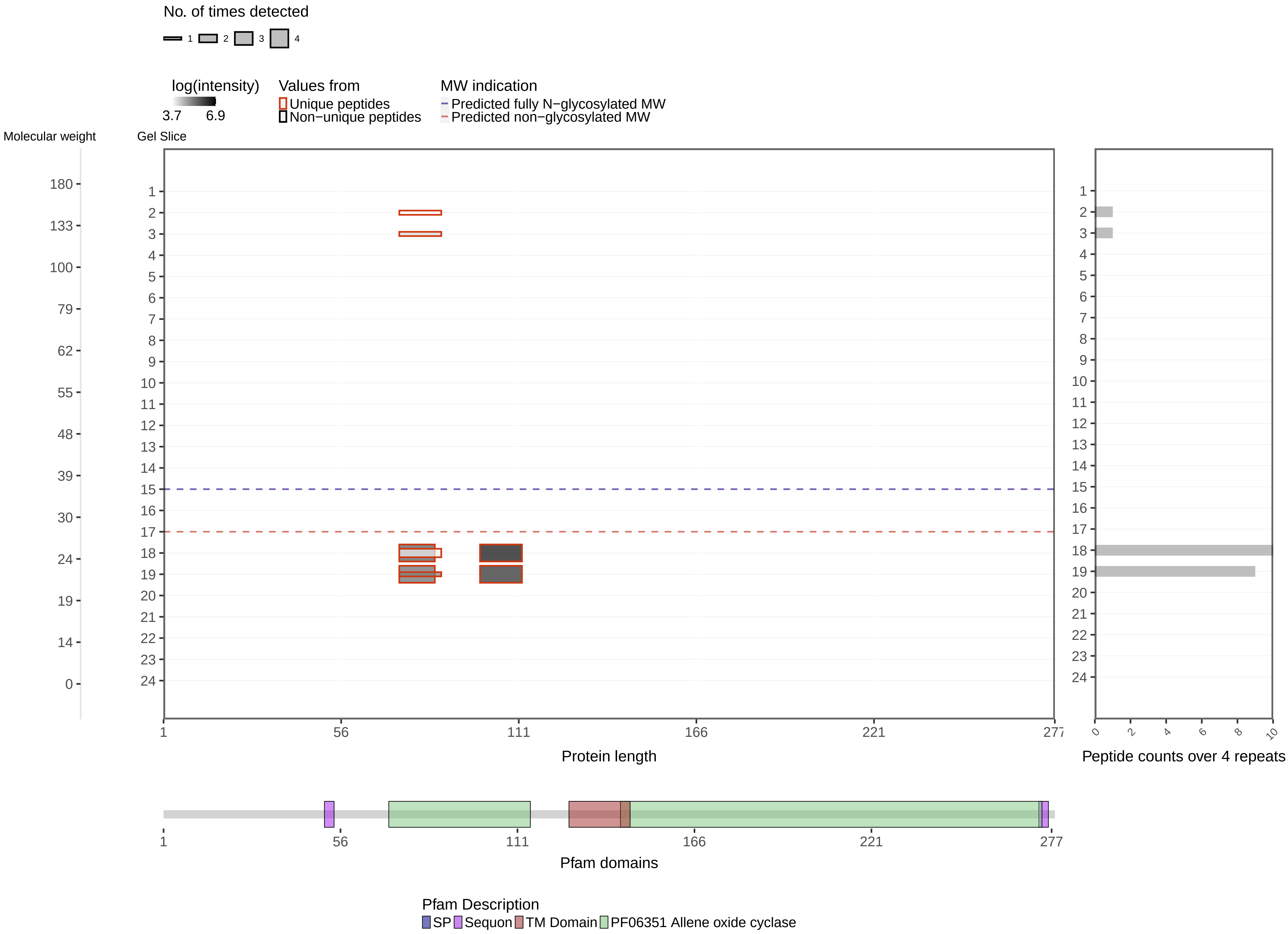


Protein sequence:

[illegible]VKKRIFSLYFILMLILYIIFQLYTGDLKKRIGITAGLCILIKHEEEKKGDRYEAIYSFYFGDYGQIAVQGSYLTYEDTYLAVTGGSGIFAGVSGQVKLQQIIFPFKLFYTFYLKGIPDLPSELLVTAVPPSPTVEPAPEAKACEAGATVKNFTN*

### NbL19g01520.1 60S_acidic_ribosomal_protein_P2−2−like [MW = 14.1 kDa]

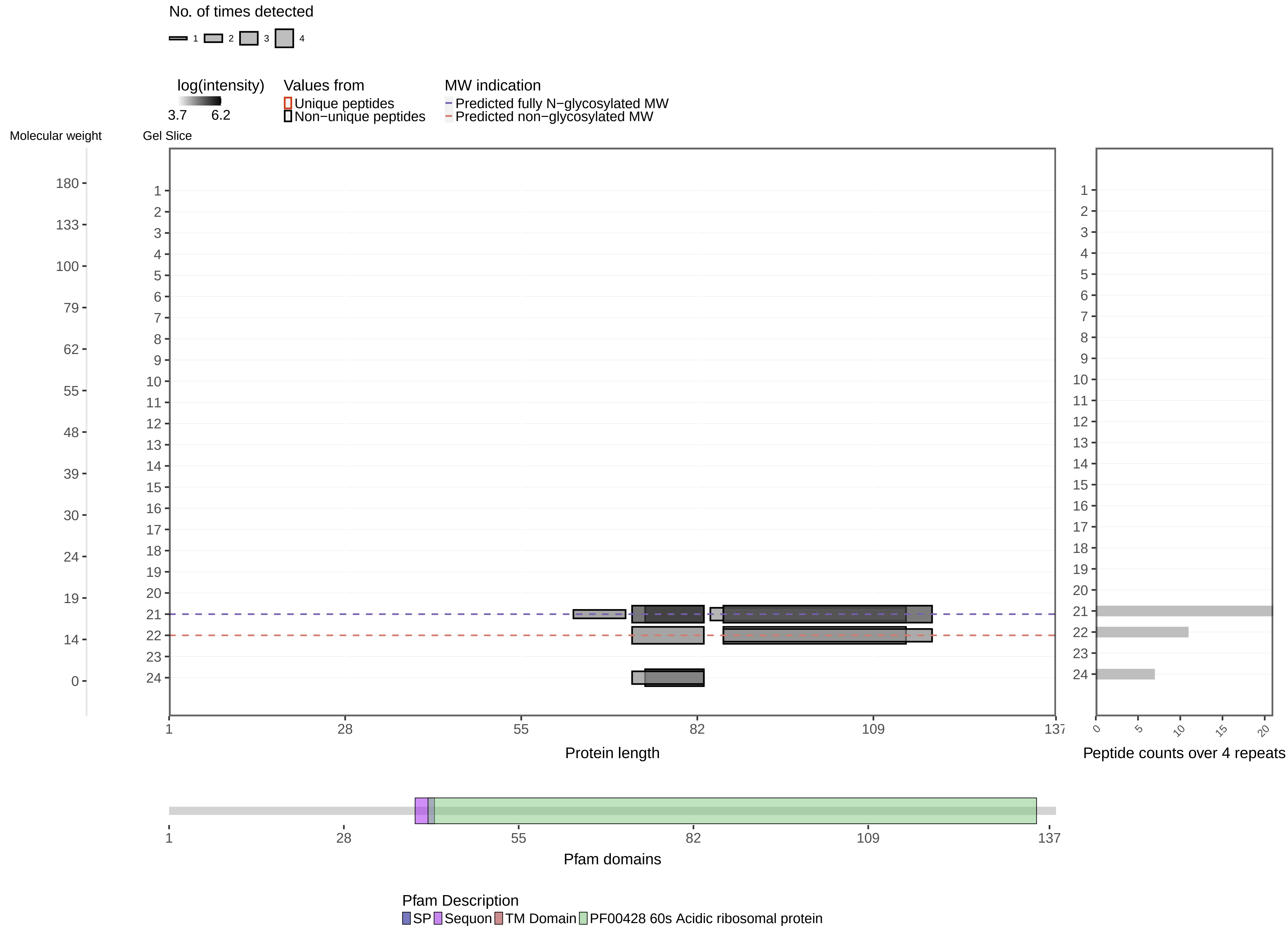


Protein sequence:

AEADDDRIELLLSQVKGKDITELIAAGREKLASVPAGGGGVAVAAPTGGAAAAAPAEEKKEEKKVEEKEESDDDMGFSLFD*

### NbL19g01430.1 succinate_dehydrogenase_[ubiquinone]_flavoprotein_subunit_1_mitochondrial [MW = 69.39 kDa]

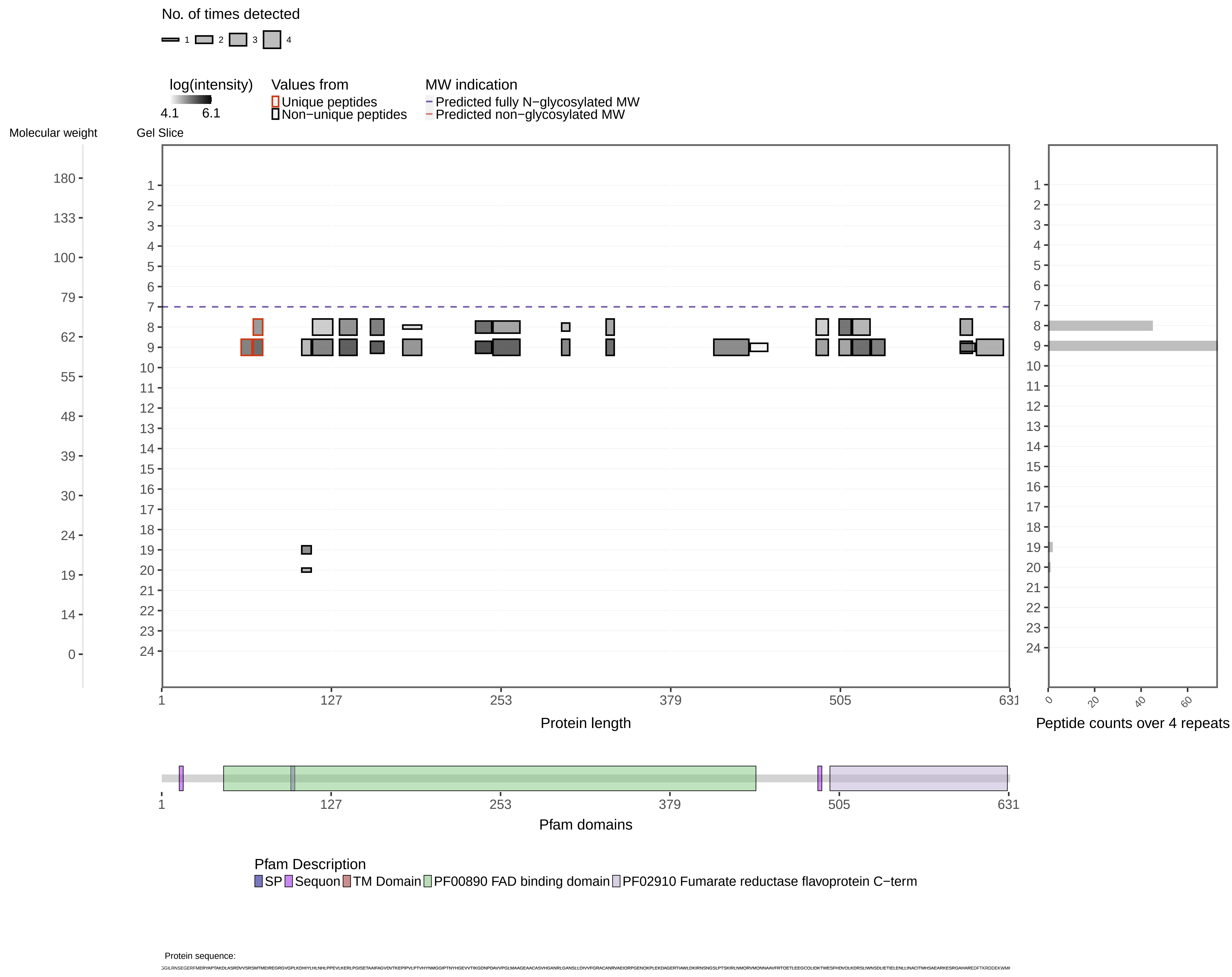

### NbL19g01370.1 maf−like_protein_DDB_G0281937 [MW = 29.27 kDa]

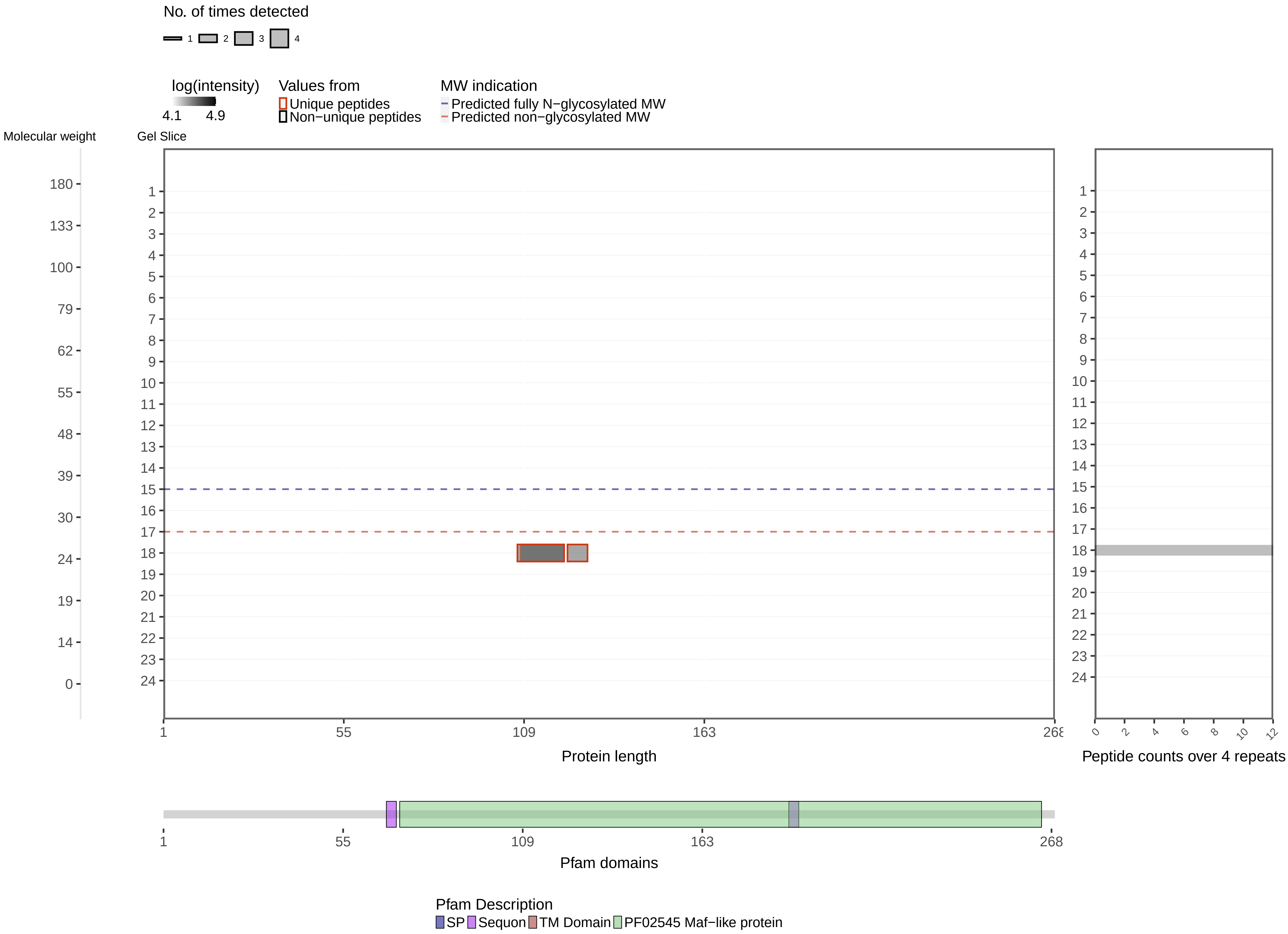


Protein sequence:

KADAIMSRLETTDHLEENTHPTLLITADTVAVYQGIIREKPSSKEEARQFIKSYSGGQATVVGSVVVTNLTKGIRKGAWERSEVYFHEIPDEVIDSLIEEGIILNVAGGLMLEHPLTLPFVDTVIGTADHVMGLPRSLTEKLIQEAL*

### NbL19g01220.1 laccase−11−like [MW − SP = 59.37 kDa]

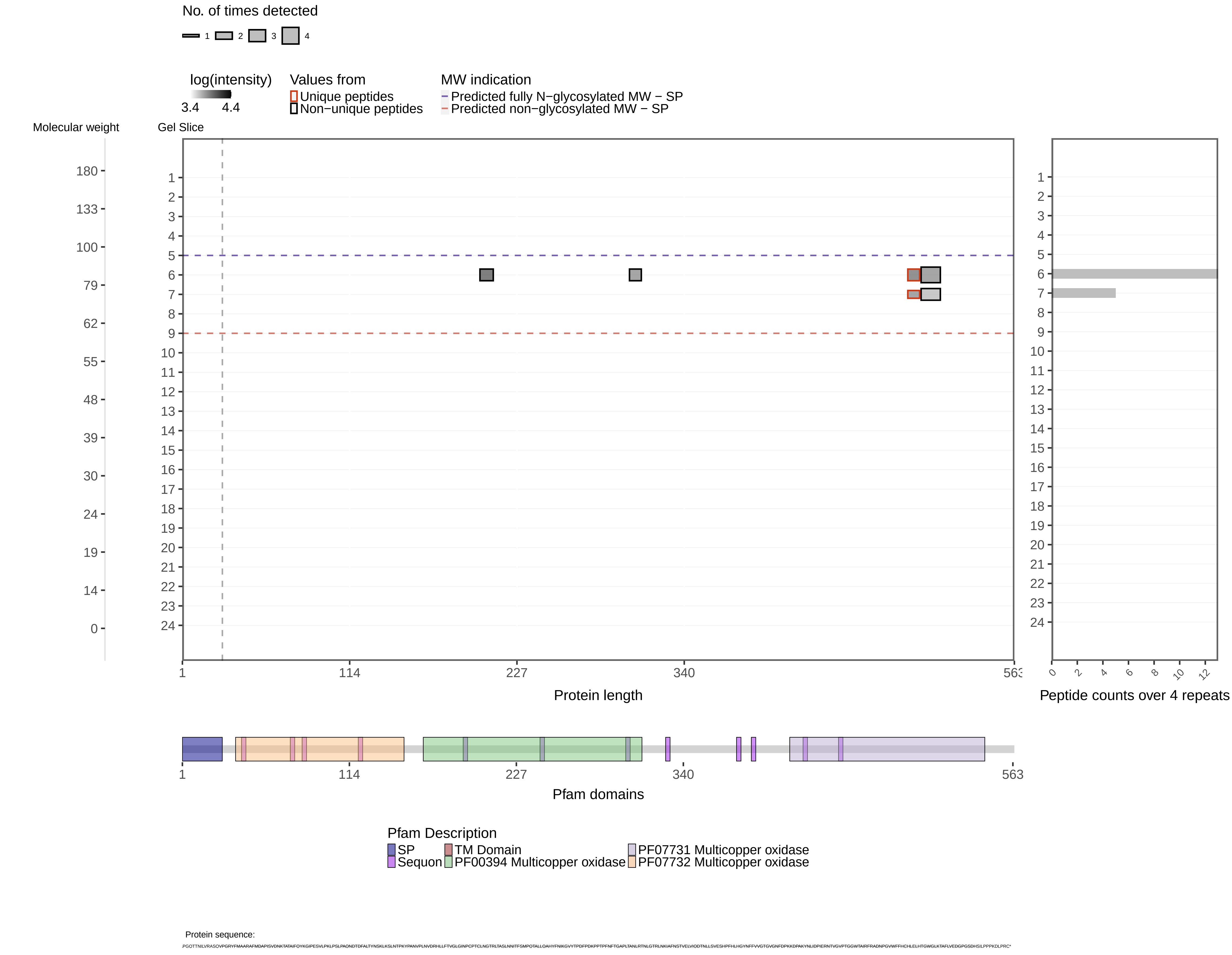

### NbL19g01150.1 soluble_inorganic_pyrophosphatase−like [MW = 24.07 kDa]

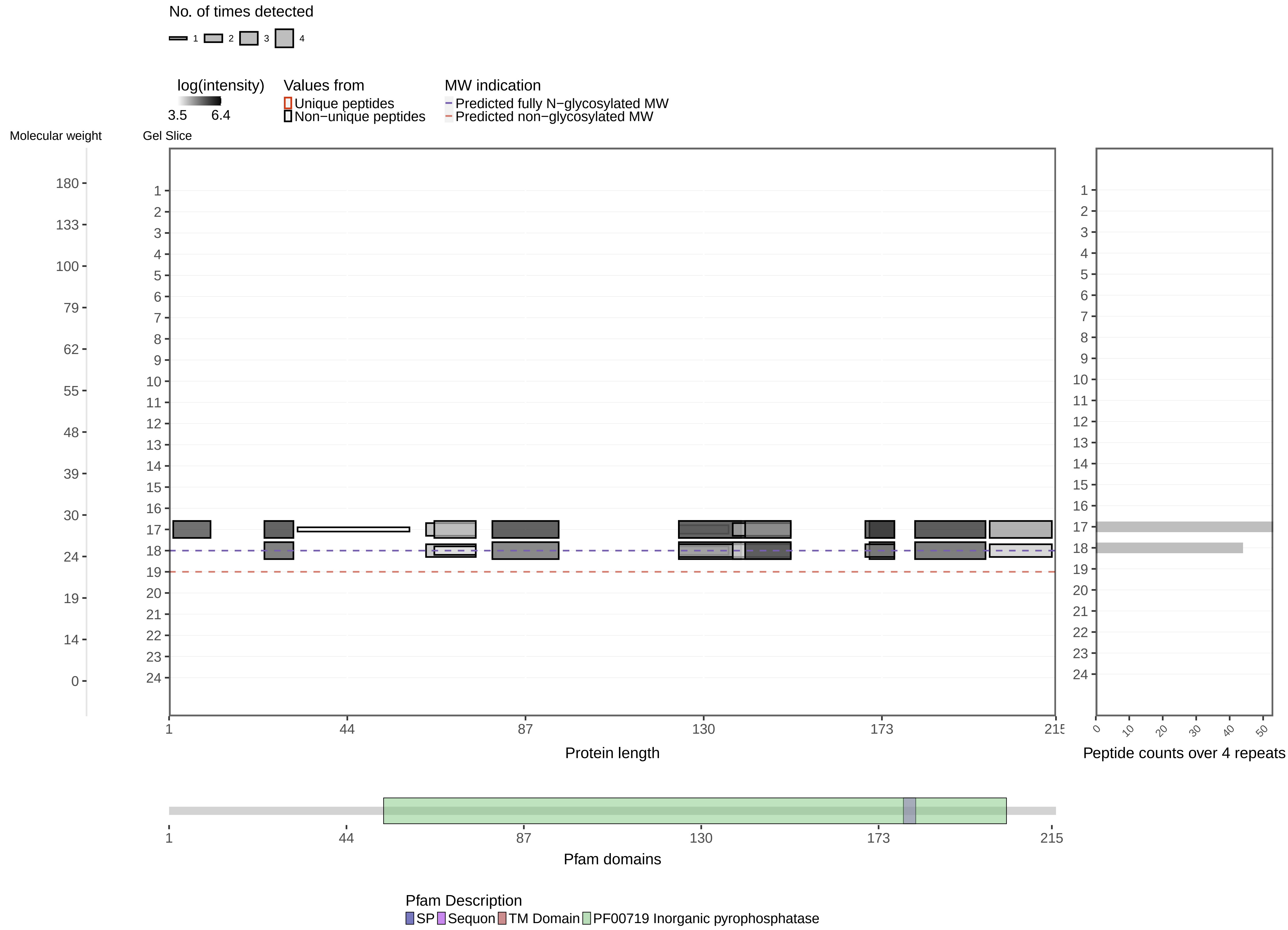


Protein sequence:

PRTICEDSDPMDVLVLMQEPVLPSTFLRARAIGLMPMIDQGEKDDKIIAVCADDPEFRHYSDIKELPPHRLAEIRRFFEDYKKNENKSVAVEDFLPAEAAVDAIKYSMDLYASYIVESLRK*

### NbL19g01070.1 uncharacterized_LOC109238053 [MW = 53.01 kDa]

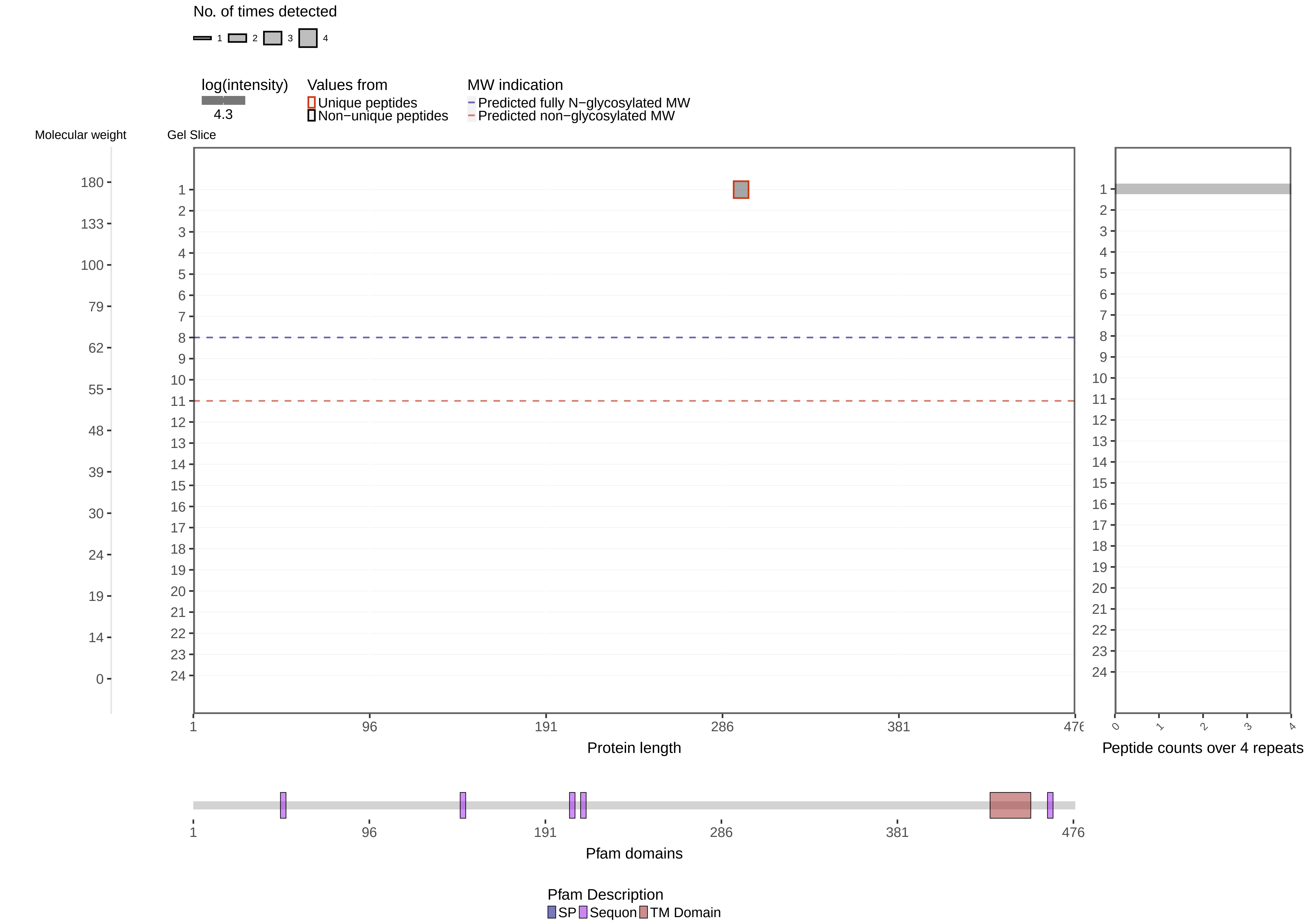


Protein sequence:

DDFFDPQESLSVKSNGESENNGGLQRSVSAMTPLGEFYDAWEELSSENGPQIPTHDFENELREIRLSLLMEIEKRKHVEEALSNMQTQWQRIREQLSLVGLNFPPCPVAGDGLDNEQLDDQVEDLFQQVHIVRFVSDSIGRGIAKAEVEIEMEAQIESKNIEMARLWDKLNYYEAVNREMSQRNQEAIETARRLRHIRKRRQKKWIWGSIAATITLGSAVLAWSYFSAERASSSSNQSHSLEGDSTSEL*

### NbL19g00880.1 peroxidase_72−like [MW − SP = 33.81 kDa]

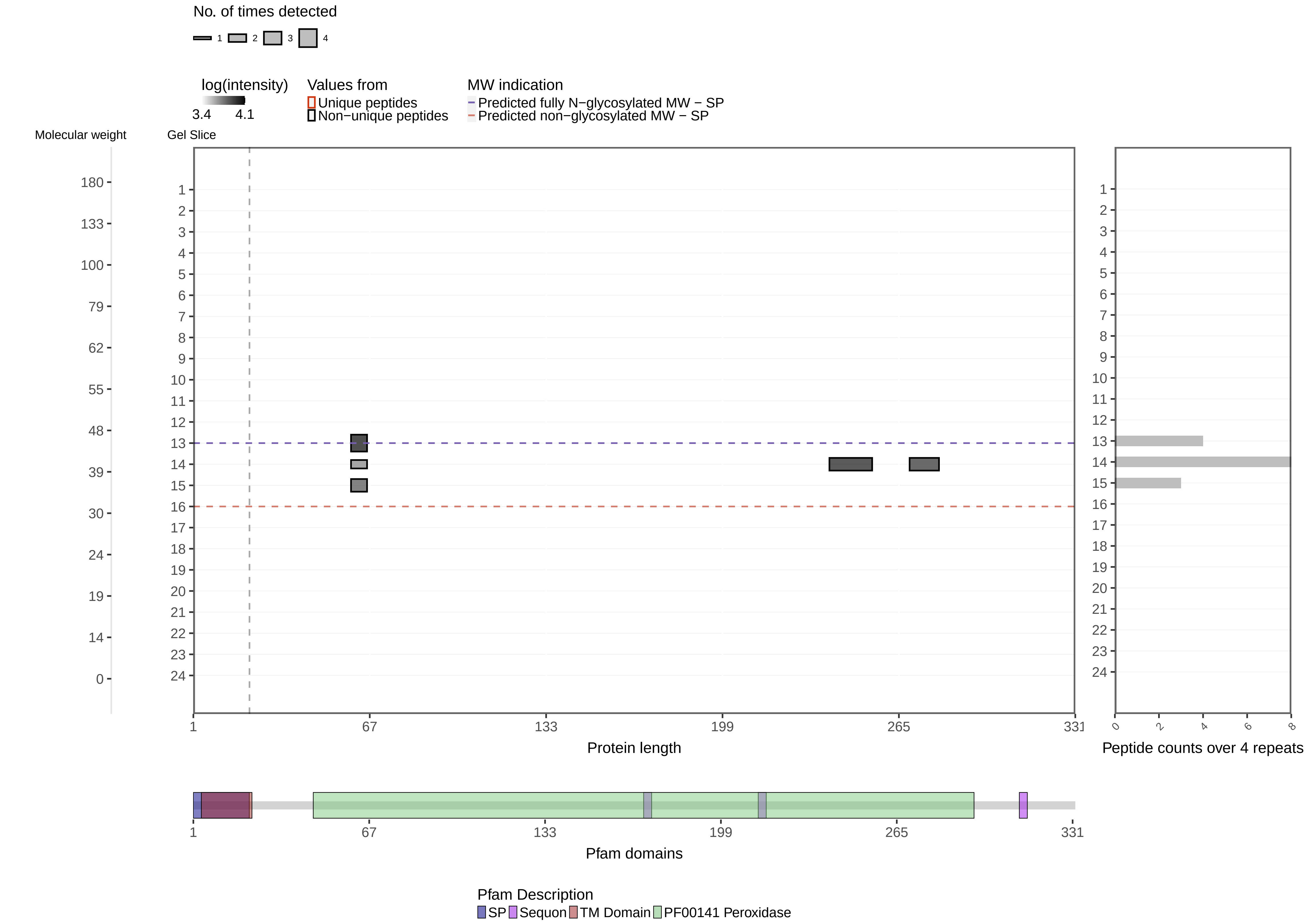


Protein sequence:

SRSASLSGSNNNIPAPNNTFDTILSKFKRQGLDLVDLVALSGSHTIGNSRCTSFRQRLYNQSGNNQPDSTLDQSYAAQLRNRCPKSGGDQNLFFLDFVSPTKFDNSYFKNLLASKGLLNSDQVLATKSQASLALVKQYAENNALFFEHFAKSMVKMGNISPLTGSSGEIRKNCRKMN*

### NbL19g00840.1 CDK5RAP1−like_protein [MW = 70.67 kDa]

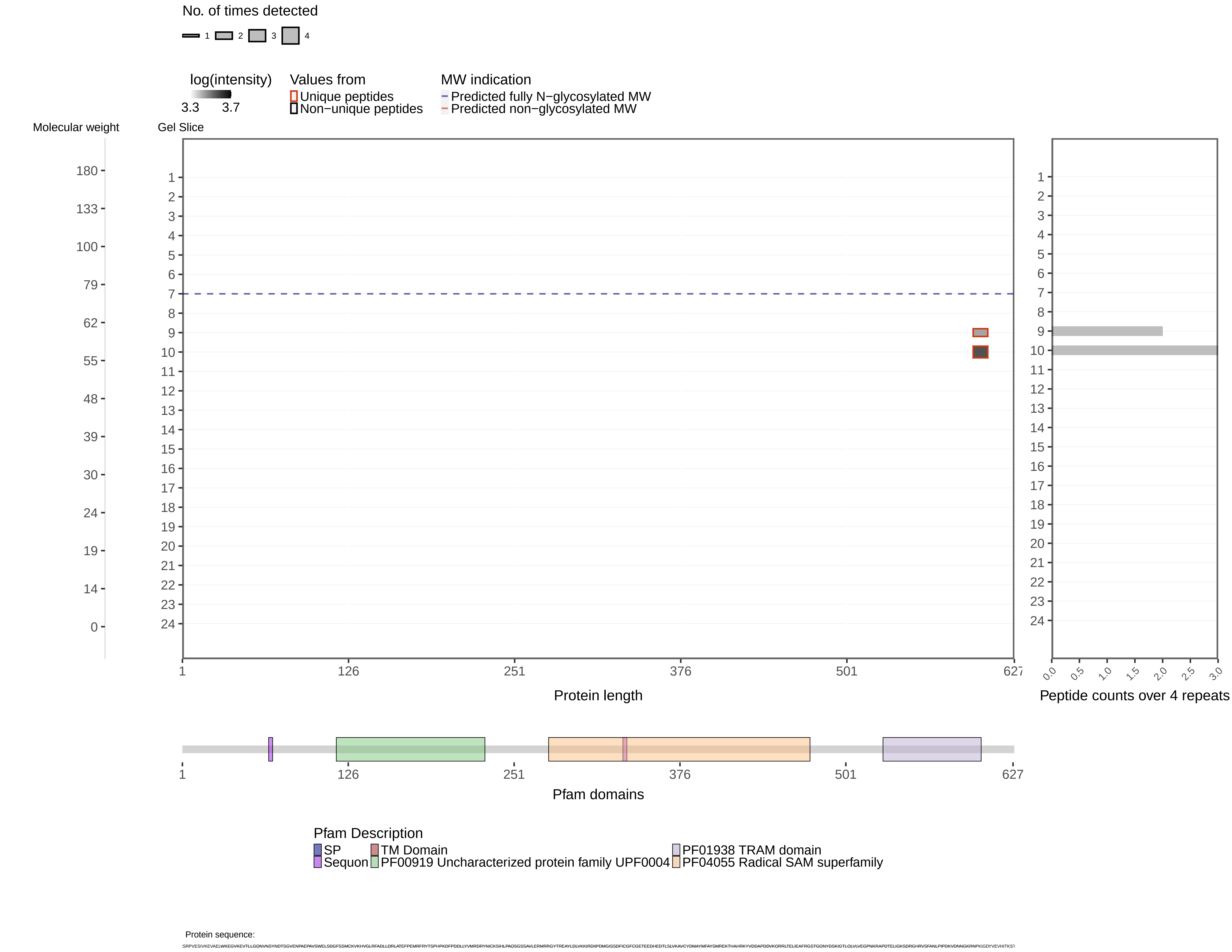

### NbL19g00810.1 thioredoxin_H−type_2 [MW = 12.99 kDa]

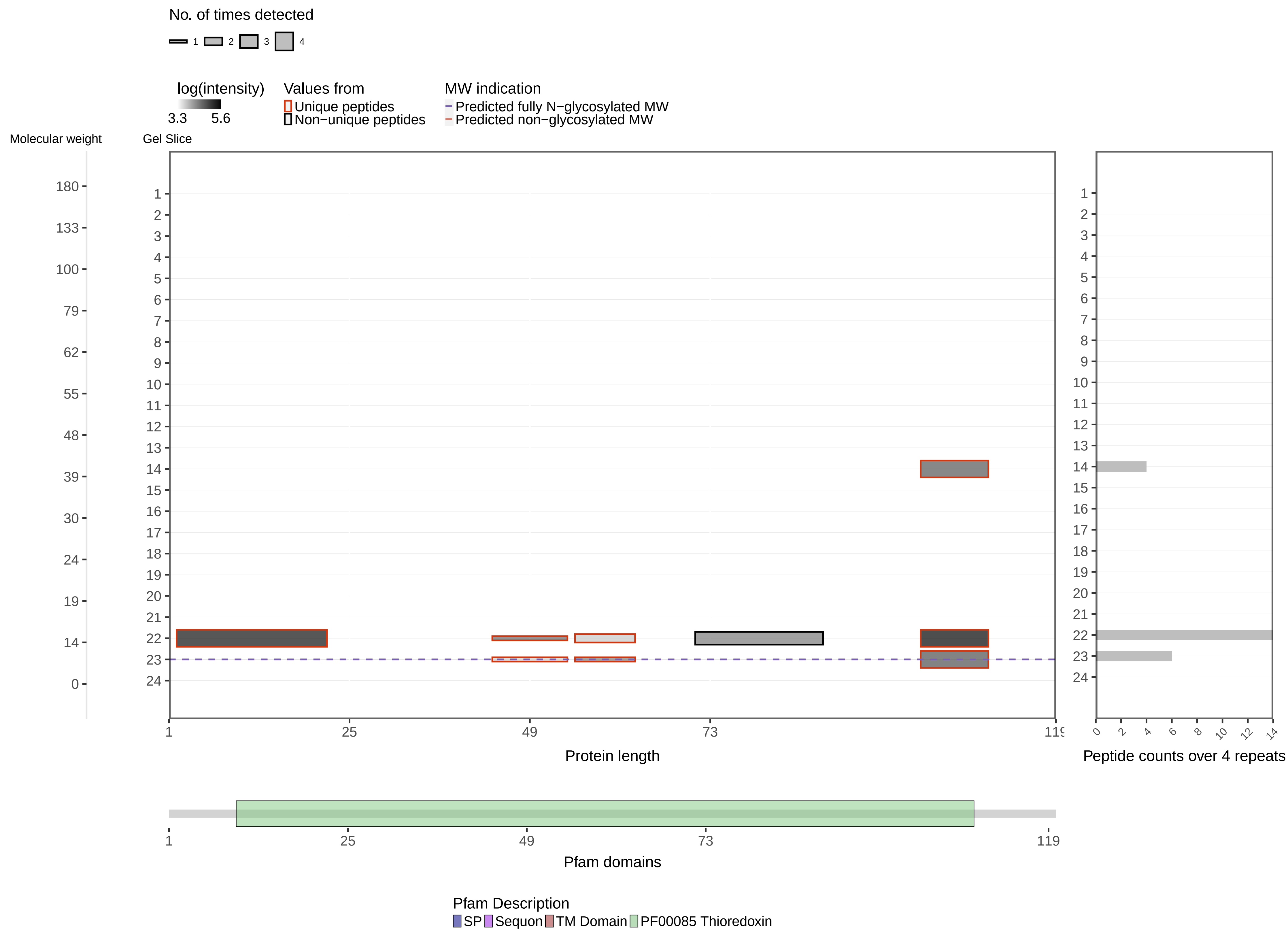


Protein sequence:

APFFAELAKKIPTVTFLKVDVDELKSVATDWAVEAMPTFMFLKEGKIVDKVVGAKKDELQQTIAKHISGTSTA*

### NbL19g00680.1 fructose−bisphosphate_aldolase_1_chloroplastic−like [MW = 42.75 kDa]

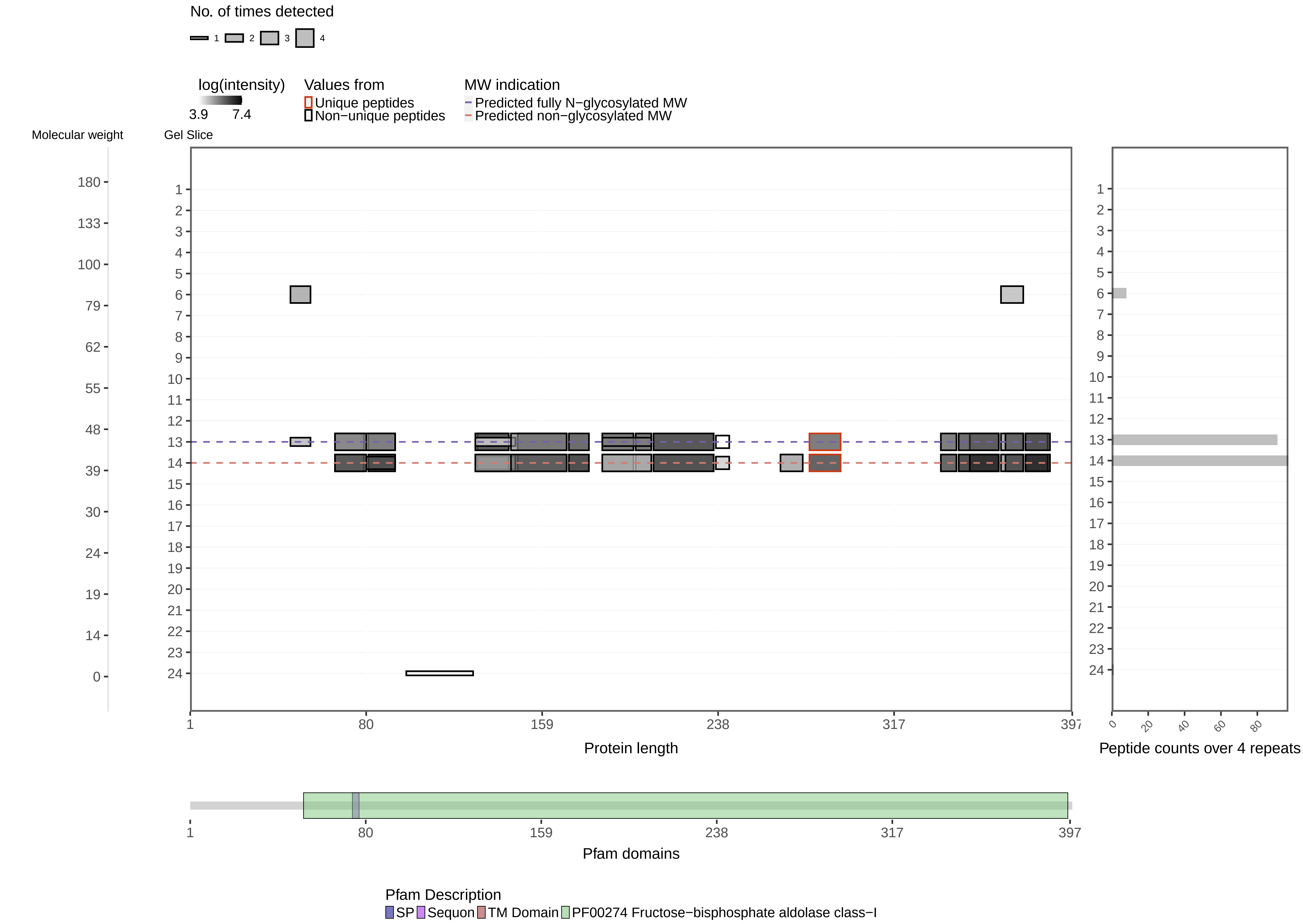


Protein sequence:

VVSIPNGPSALAVKEAAWGLARYAAISQDNGLVPIVEPEILLDGEHNIDRTFEVAKKVWAEVFFYLAENNVMFEGILLKPSMVTPGAECKERATPHQVADYTLNLLRQRIPPAVPGIMFLSGGQSEVEATLNLNAMNQGPNPWHVSFSYARALQNTCLKTWGGRPENVQAAQEALLVRAKANSLAQLGKYTGEGESEEAKKGMFVKGYVY*

### NbL19g00300.1 50S_ribosomal_protein_L3_chloroplastic [MW = 30.03 kDa]

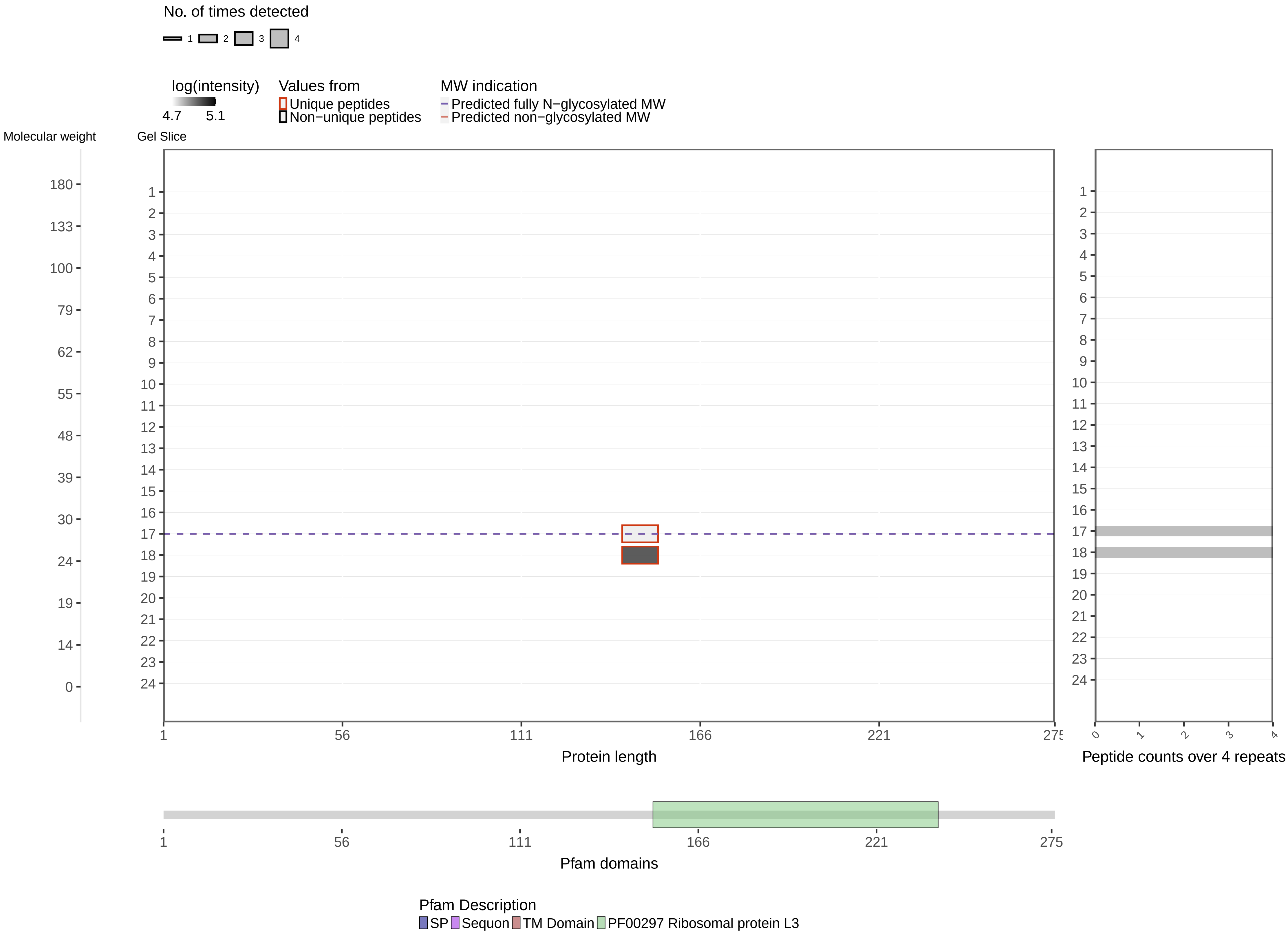


Protein sequence:

HLEKSGIIPLRHLQEFRLQCIDGFEVTQKLDFGELFKEGDLVDVSGTTIGKGFQGGIKRHNFKRGQMTHGSKSHRQLGSIGAGTTPGRVYKGKKMPGRMGGTKRKIRKLKIVKIDDQLNIIMIKGALPGKPGNLLRIAPAKIVGKNIPKN*

NbL19g00260.1 zinc_finger_CCCH_domain−containing_protein_44−like [MW = 30.62 kDa]

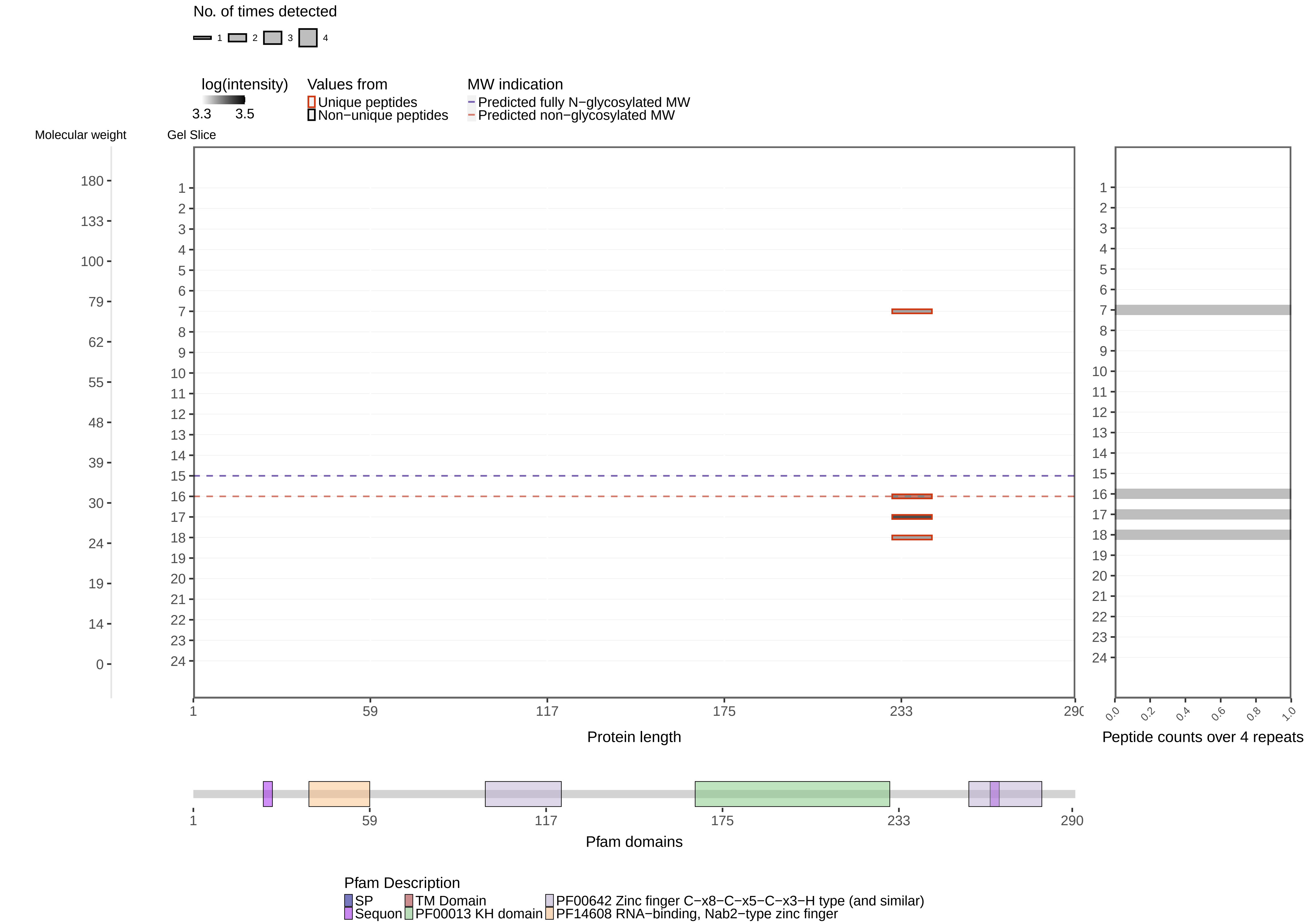


Protein sequence:

/APAYDDFQAAGPPSRFPGYHKPSQPSPAATNFGASATTTISIDAALAGTIIGRNGVNSKQICRLTGVKLSVKDHETDSNRRNIELQGTFDQINQASAMVREIISNVGTPTGGPKNTGPFARGPAGPFKTKMCSNFSKGSCTFGEKCHFAHGASELQKPLA*

NbL19g00190.1 protein_GrpE−like [MW = 42.46 kDa]

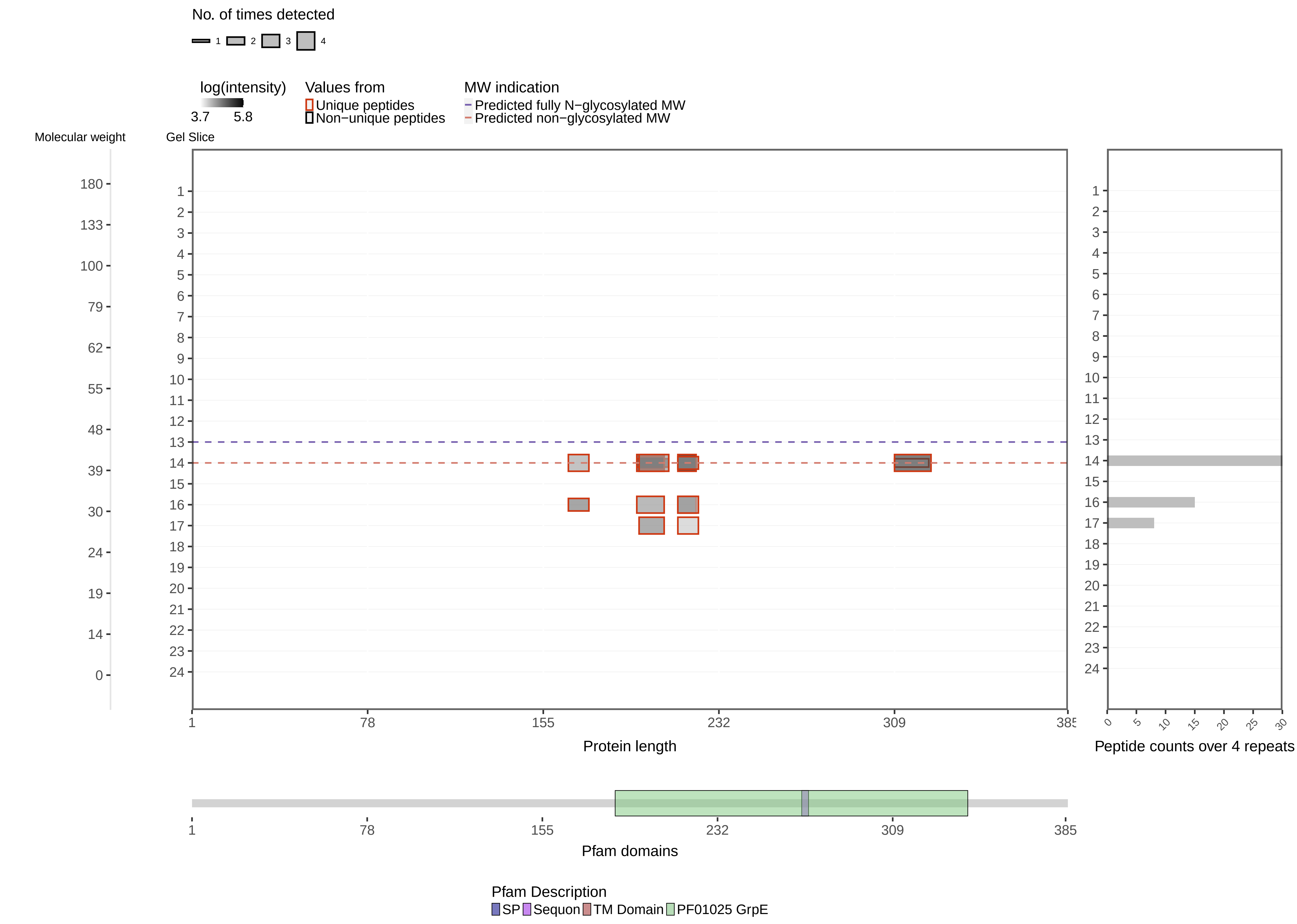


Protein sequence:

ISLKSIENEKIELQRKVASLTEELSSERDRVLRISADFDNFRKRTERERLSLVTNAQGEVVEKLLSVLDNFERAKMQIKVATEGEEKINNSYQSISKQFGEILGSLGVETVDTVGKPFDPLLHEAIMREDSEEFEEGVILEEYRKGFKLGDRLLRPSMVKVSAGPGPAKPETAEPKEEQHKVEEKSEEGTAETAGDEGTGEGGN*

### NbL19g00170.1 replication_protein_A_14_kDa_subunit_B−like [MW = 14.75 kDa]

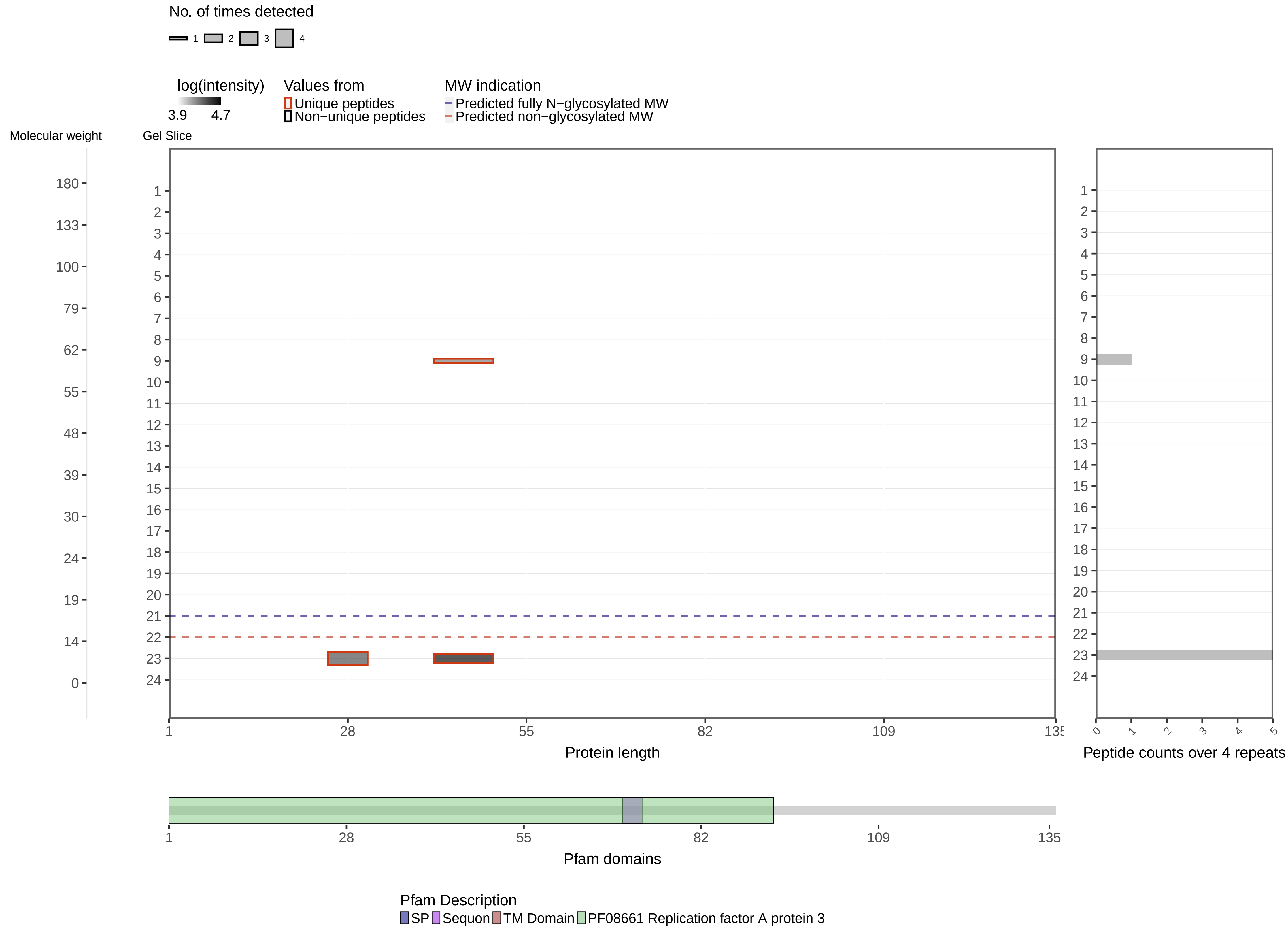


Protein sequence:

PGPLTTFVEVIGIADSNQSIRAEIWSNFGDTLGITLDHSMNNFDNVLSHDVCRWWNLYDCFLCLLCVGSALALKSFTTNDL*

NbL19g00150.1 BRASSINOSTEROID_INSENSITIVE_1−associated_receptor_kinase_1−like [MW − SP = 21.02 kDa]

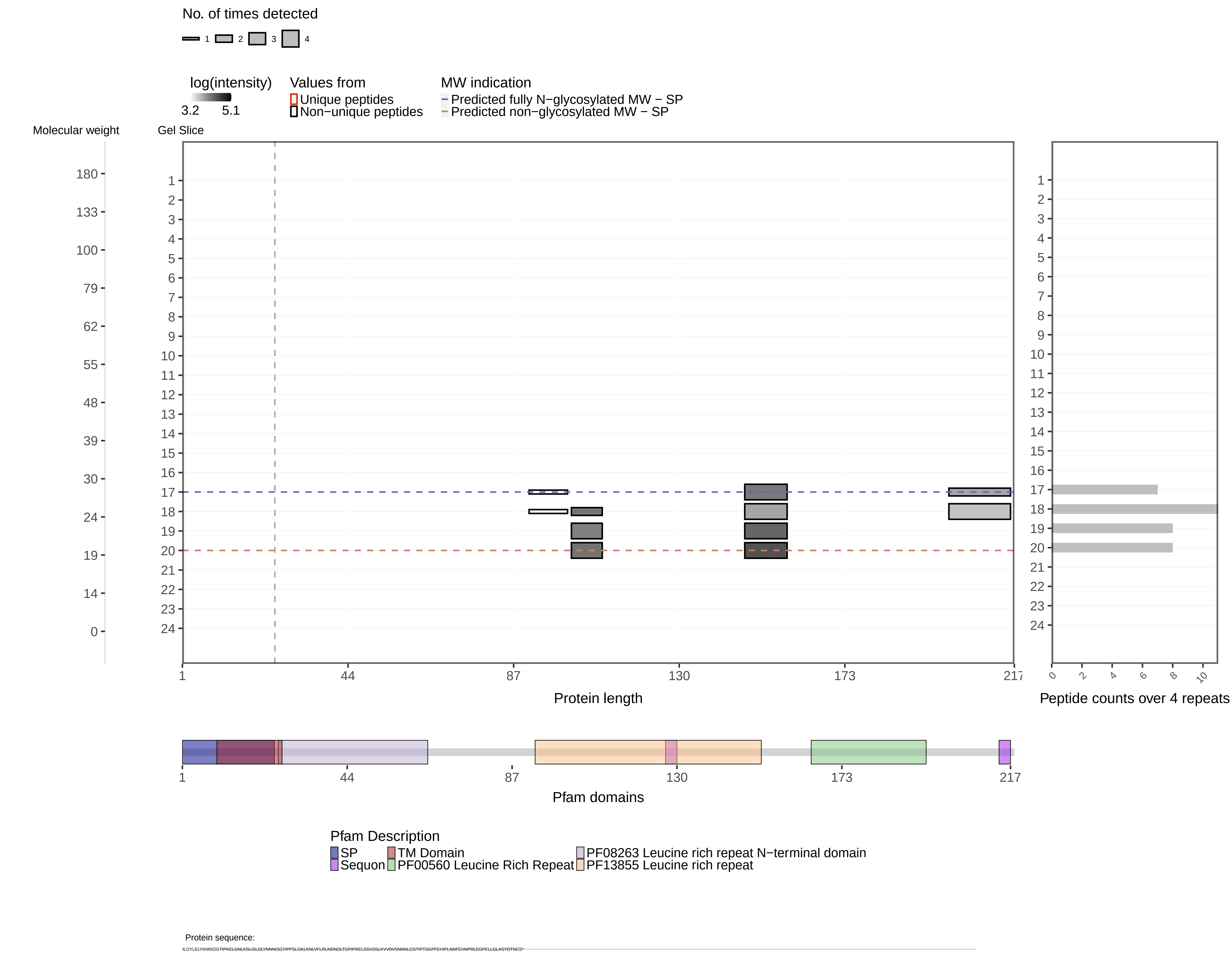

NbL19g00140.1 glycine−rich_RNA−binding_protein_2_mitochondrial−like [MW = 29.7 kDa]

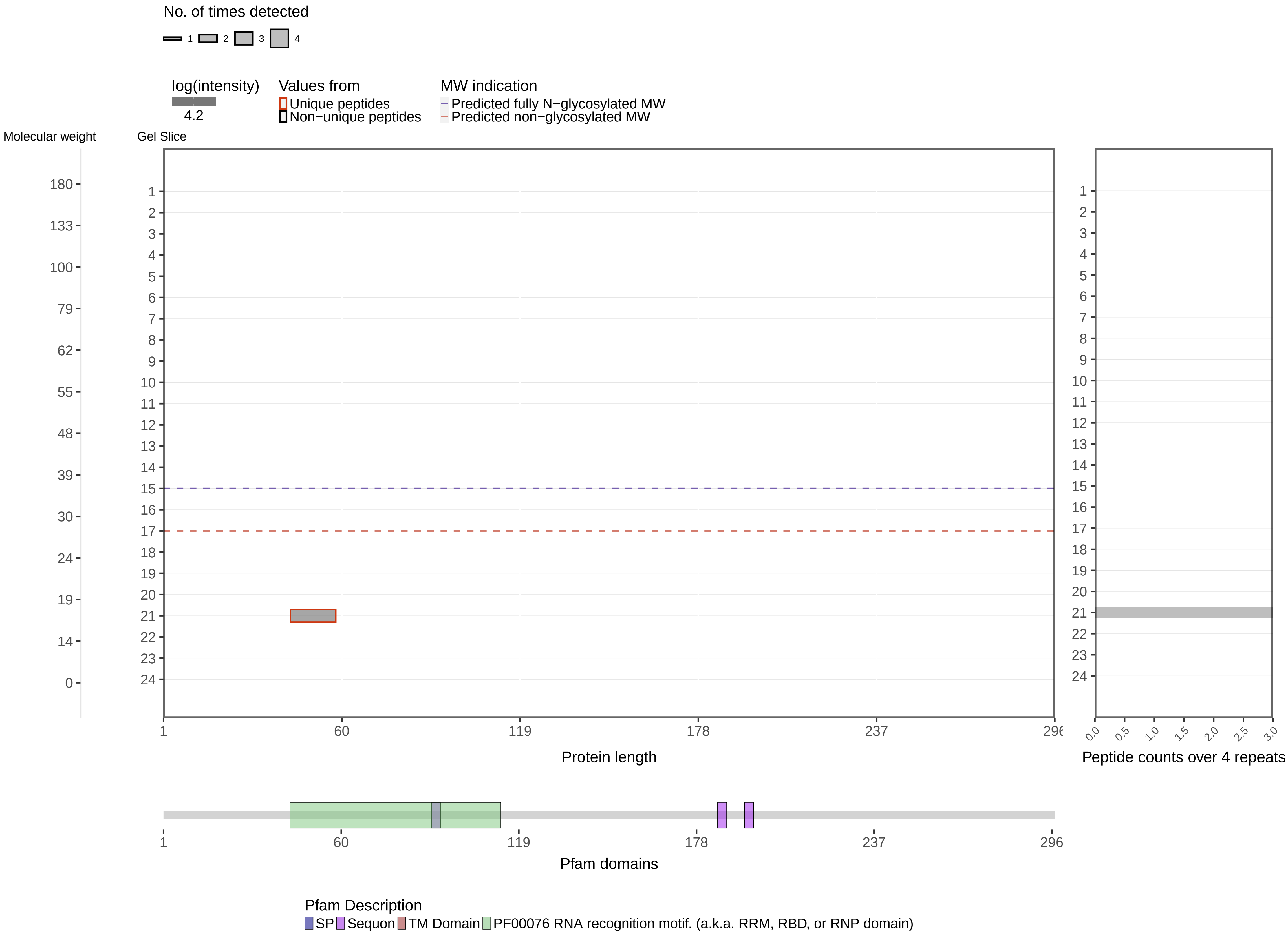


Protein sequence:

GGGFGAGGYGGGGNYGAGGYGGGGNYGAGGYGGGGNYGSGGGGDANVTGSYGANNYTSGESSFGFGTSGGNNFPSGDSYASENQNLDAAAAGSGDTYGTGEAASNSFTSSFGSNDQFGNVDTSNQAETGDVNSPVDAGEDYRDDNVESGDYANRRA*

NbL19g00110.1 50S_ribosomal_protein_5_chloroplastic−like [MW = 15.98 kDa]

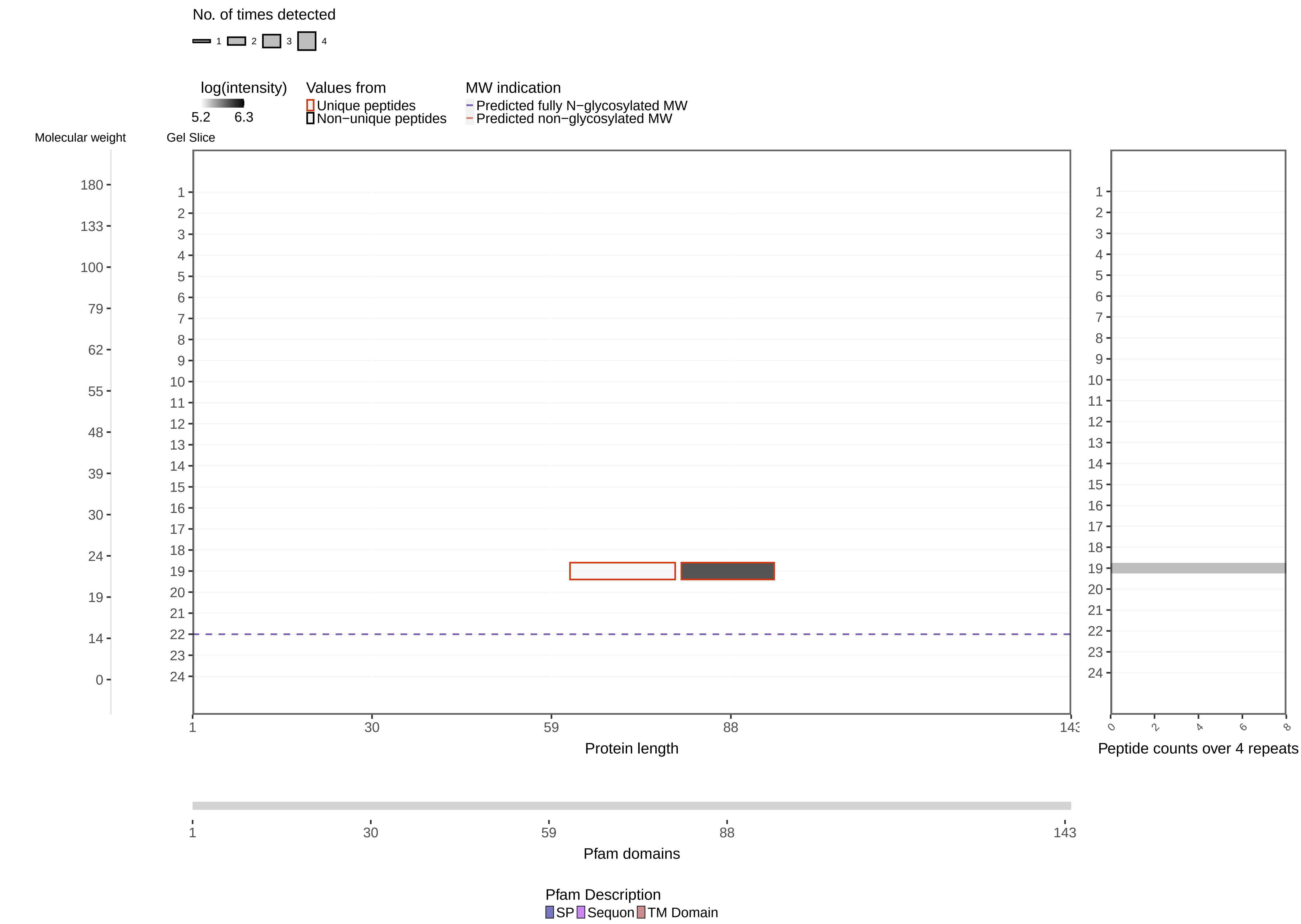


Protein sequence:

KAAADIDSVGSDNPEPAPKNNEESVPVENLPLESKLQQKLDQKMKMKLAKKIRLRRKRLVRKRHLRKKGRWPPSKMKKNKNV*

NbL19g00100.1 splicing_factor_3B_subunit_1 [MW = 141.25 kDa]

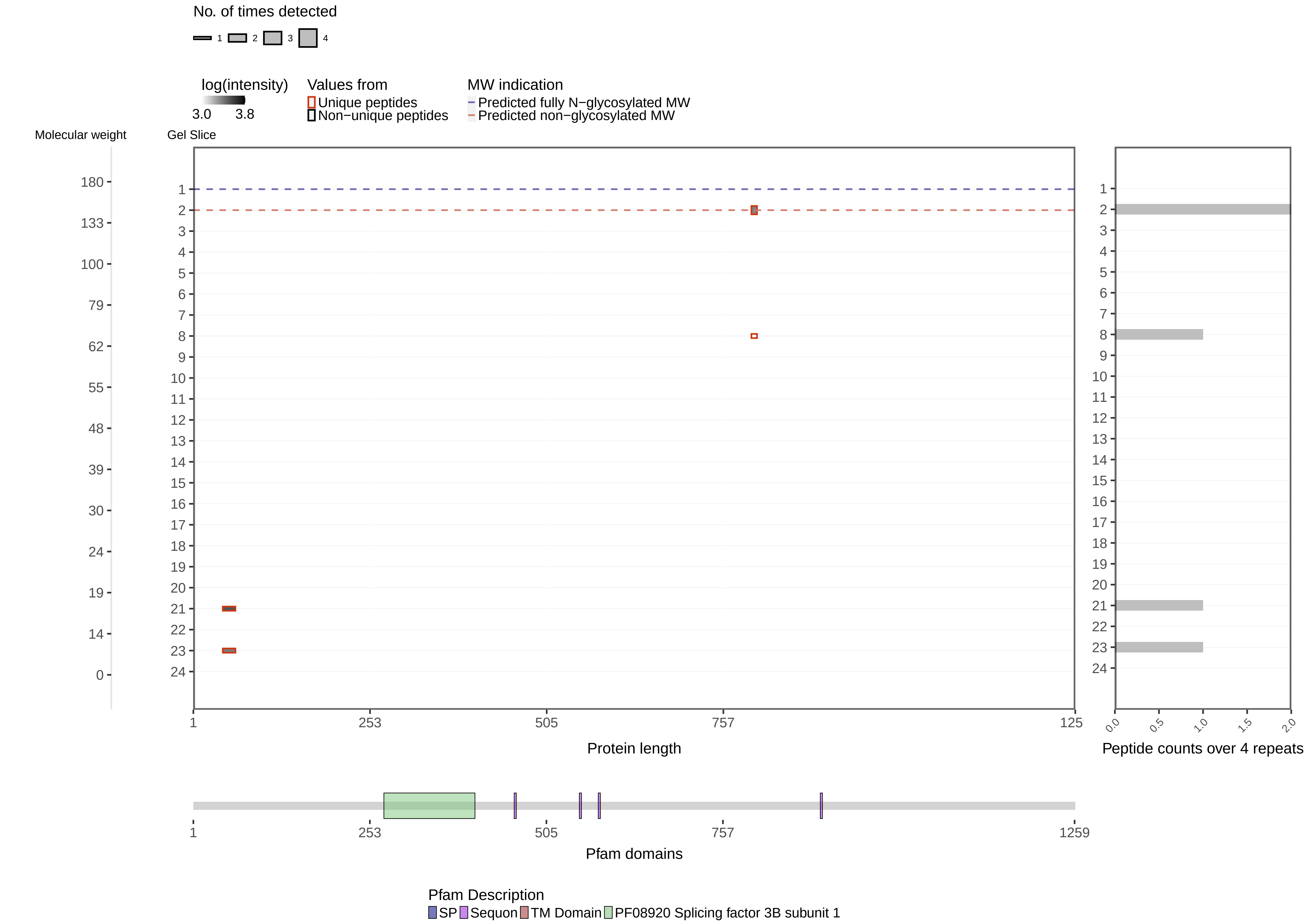


Protein sequence:

KSWQARHTGIKIVQQIAILIGCAVLPHLRSLVEIIEHGLNDENQKVRTITALSLAALAEAAAPYGIESFDSVLKPLWKGIRSHRGKVLAAFLKAIGFIIPLMDAVYASYYTKEVMVILIREFQSPDEEMKKIVLKVVKQCVSTEGVEPDYIRQDILPEFFRNFWVRRMALDRRNYKQLVETTVEIANKVGVADIVGRIVEDLKDESEPYRRMVMETIEKVVANLGASDIDARLEELLIDGILYAFQEQTSDDANVMLNGFGAVVNALGQRVKPYLPQICGTIKWRLNNKSAKVRQQAADLI

### NbL19g00040.1 phytanoyl−CoA_dioxygenase [MW = 32.09 kDa]

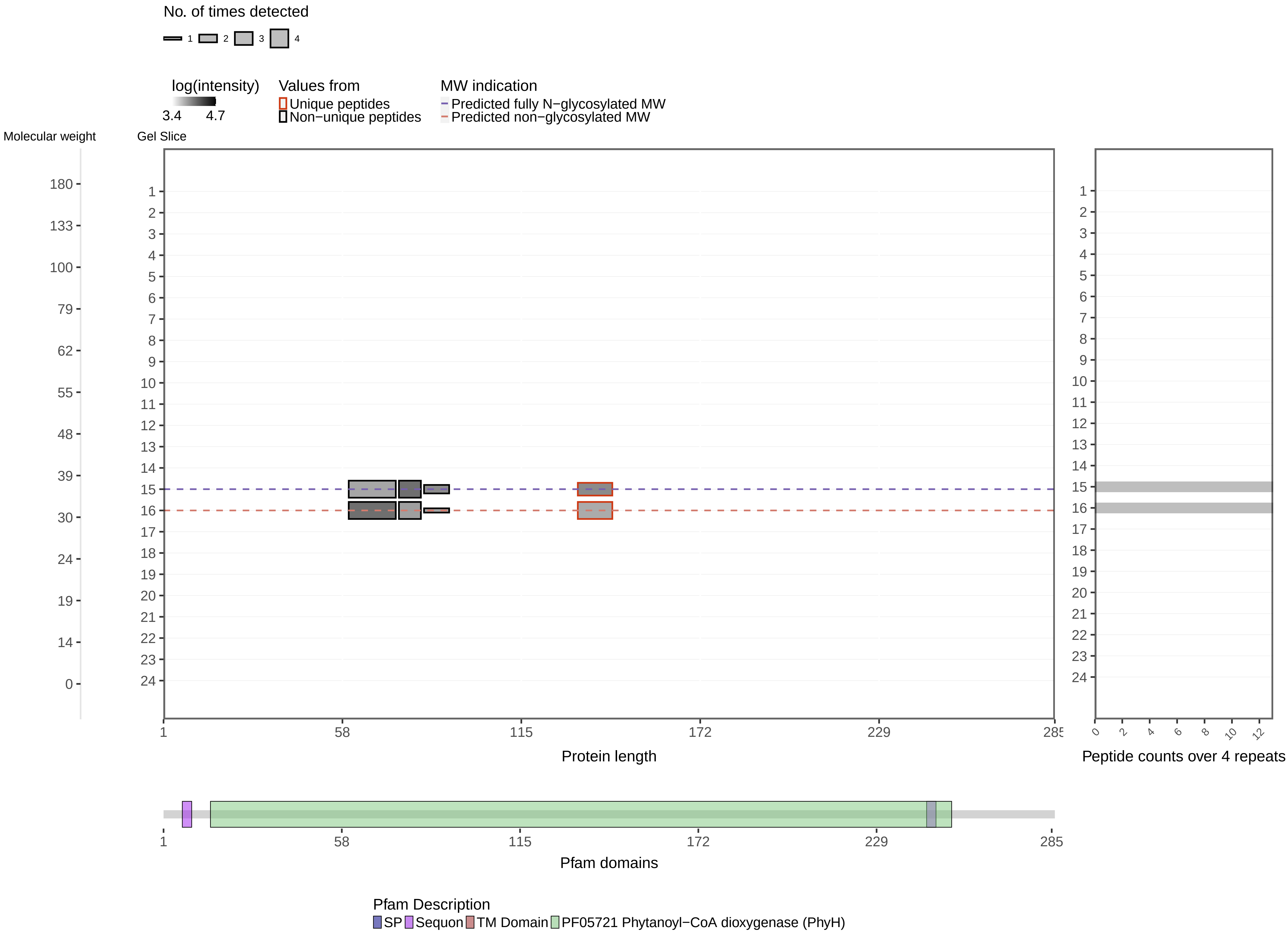


Protein sequence:

LGYRSPVVIQSMYIFKQPGIGGEVVPHQDNSFLYTEPTTCTGLWLALEDATIINGCLWAIPGSHKDGLVRRFLRDENGVHFDKPSPCYYQKDFIPLEVKAGSLVVIHGDLIHQSFENKSSKSRHAYSLHVVDTNGCKWAEDNWIRRNVDPVPLYSS*

### NbL18g20630.1 xaa−Pro_dipeptidase [MW = 55.79 kDa]

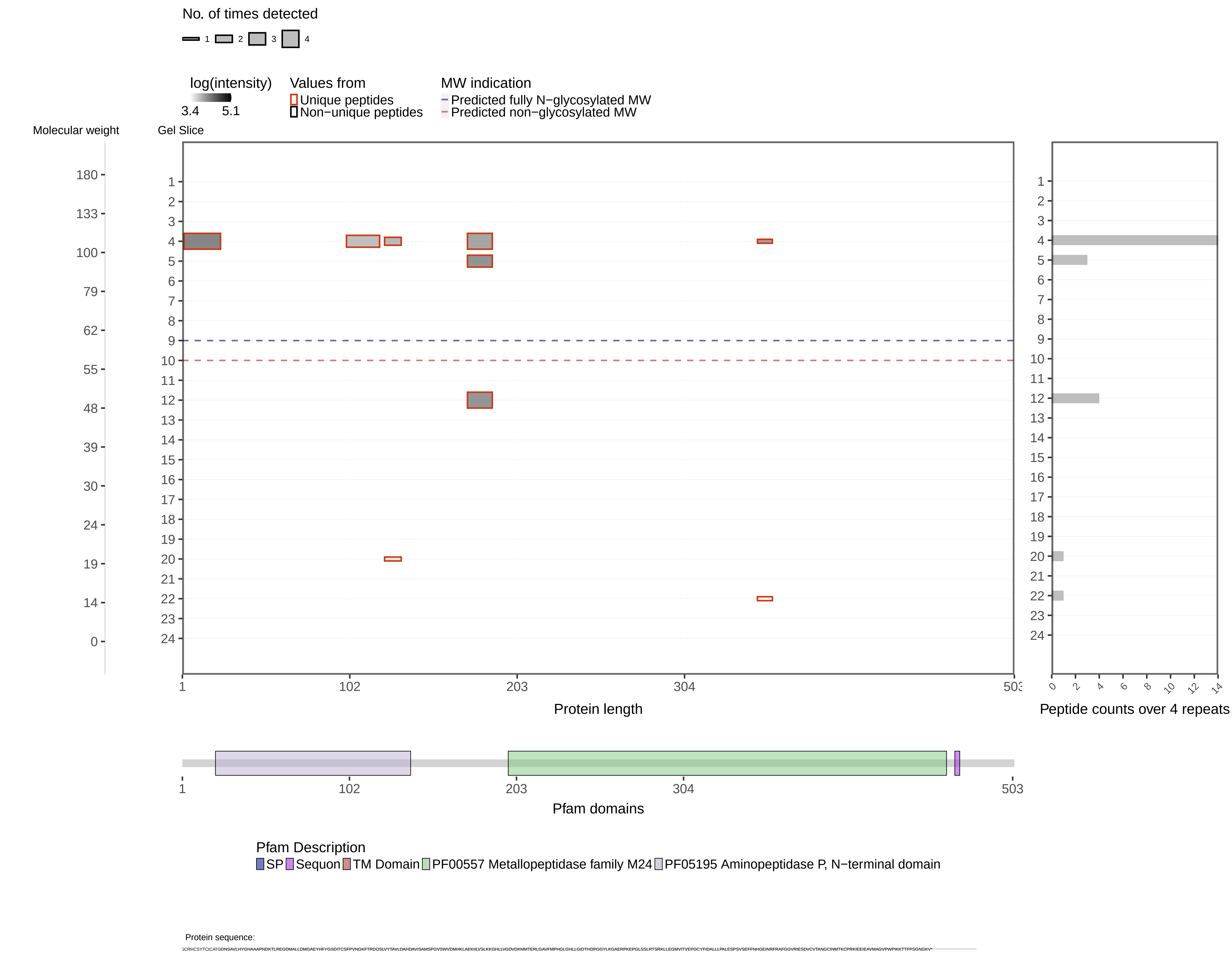

NbL18g20450.1 uncharacterized_LOC107790078 [MW = 34.94 kDa]

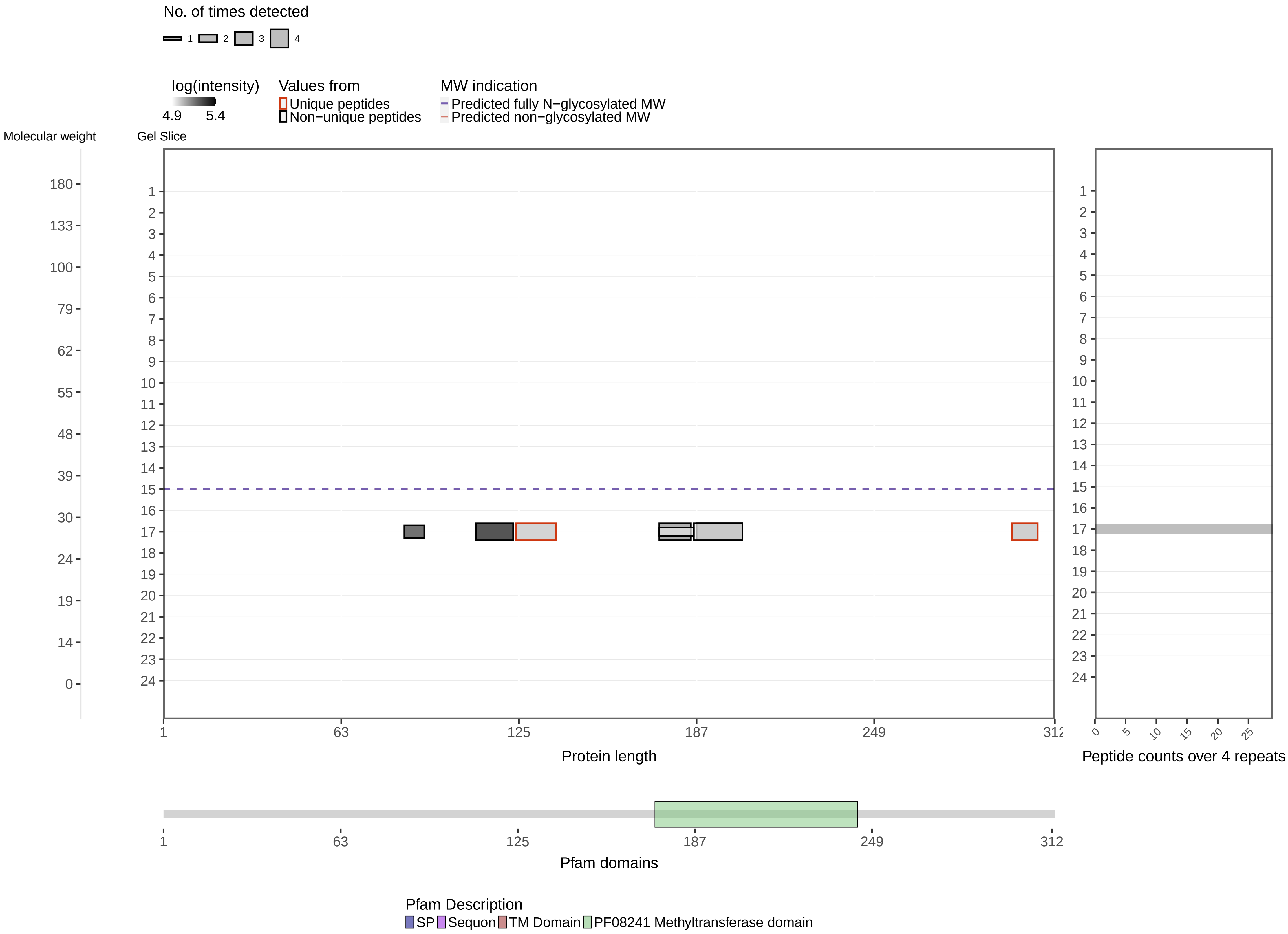


Protein sequence:

VLPPSNAPGVAILDMCSSWVSHYPAGYKQERIVGMGLNEEELKRNPVLTEYVVQDLNSNPKLPFGDNTFDVITNVVSVDYLTKPLDVFKEMSRILKHGGLAIMSFSNRCFFTKAISIWTSTGDADHVMIVGSYFHYAGGFEPPQAVDISPNPGRSDPMYIVYSRKIATA*

### NbL18g20430.1 cytidine_deaminase_1−like [MW = 35.49 kDa]

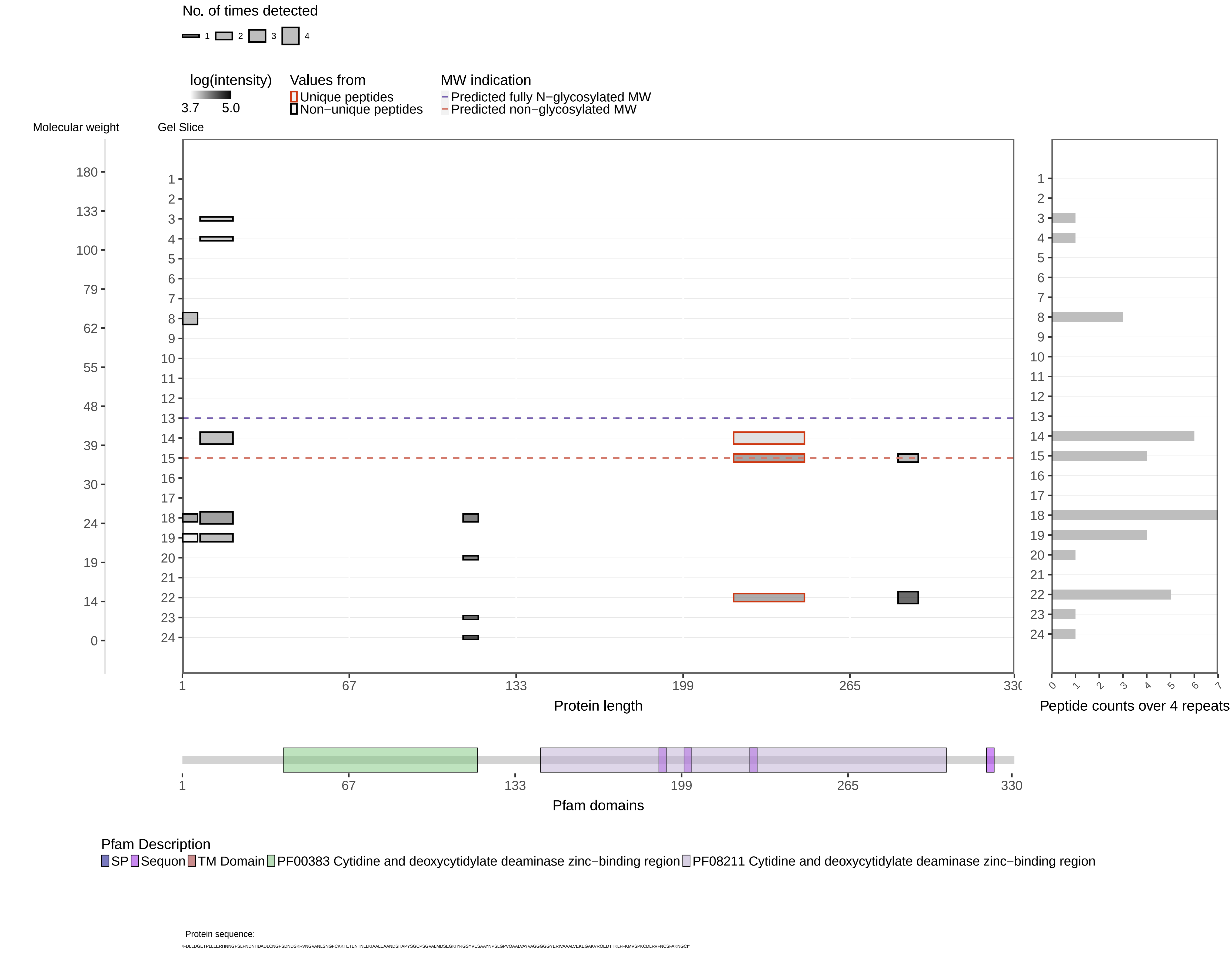

NbL18g20390.1 ectonucleotide_pyrophosphatase/phosphodiesterase_family_member_3−like [MW = 54.85 kDa]

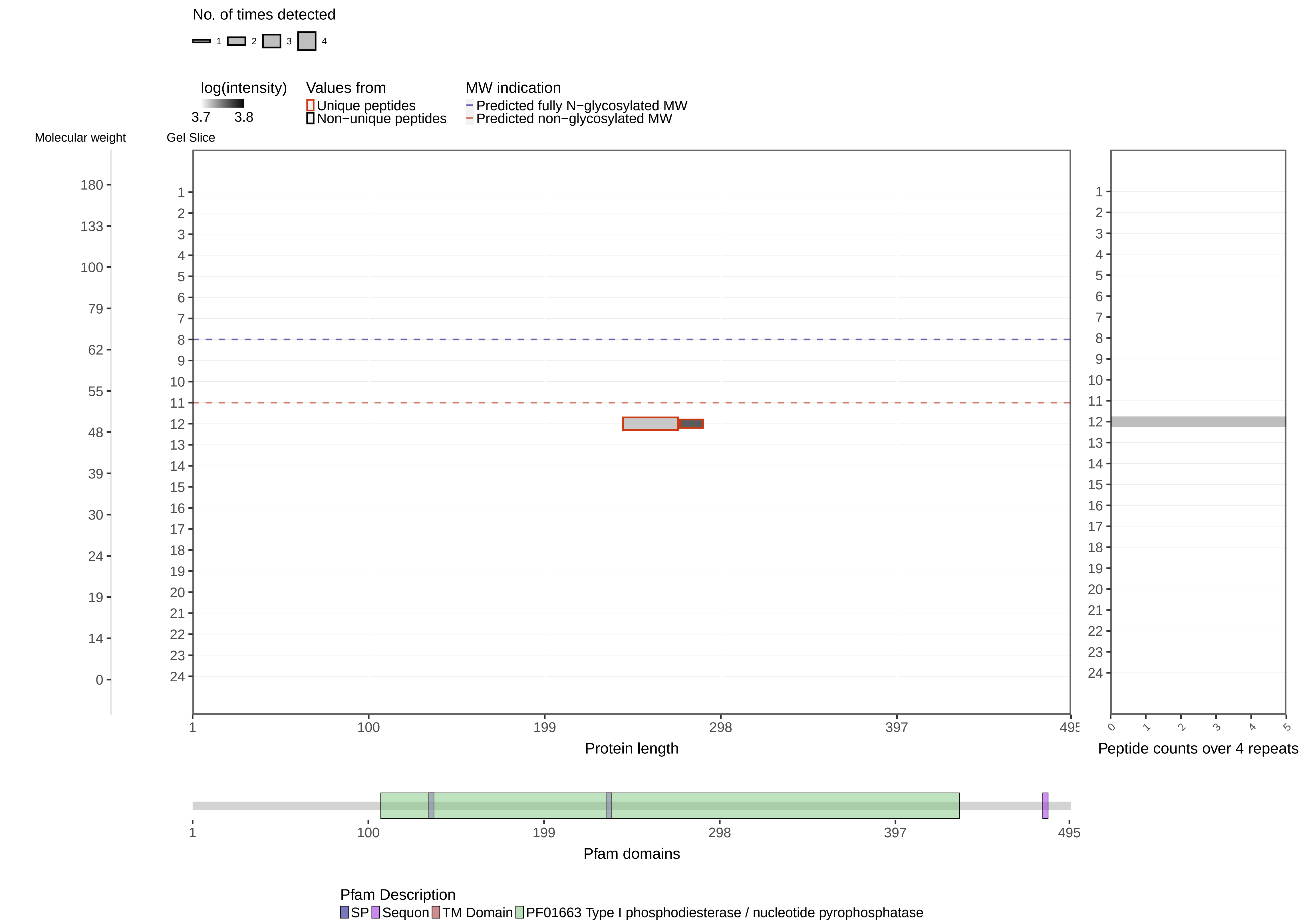


Protein sequence:

GSVPFEERVDTVLNSFDLPIDEIPSFMTLYFEDPDHQGHKVGPDDPQITEAVARVDAMIGRLIRGLEERGVFEDVNIIMVGDHGMVGTCDKKLIFLEDLAPWIEISKDWIQSYSPLLSIRPPASYSAKDVVTKMNEGLKSGKVQNGQYLKVYLKEELPARLHYFASDRIPPIIGLIDESFKVEQKSSKRKECGGSHGYDNAFFSMRTIFIAHGPRFARGSKVTSFENVQIYNLVTTILNIQGASNNGTTSFPKTILLPSH*

### NbL18g20290.1 WD−40_repeat−containing_protein_MSI4−like [MW = 57.46 kDa]

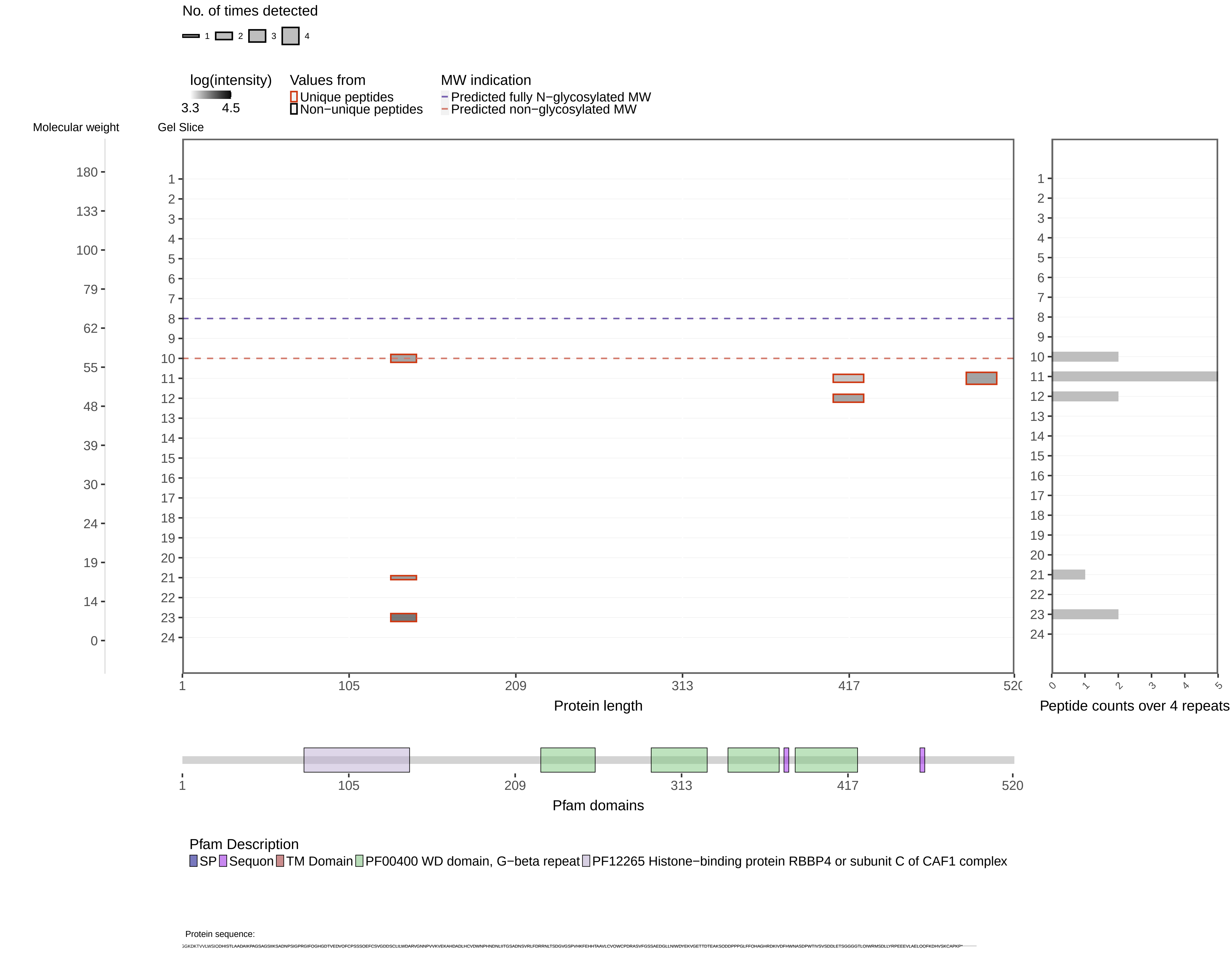

NbL18g20260.1 receptor−like_serine/threonine−protein_kinase_ALE2 [MW − SP = 120.55 kDa]

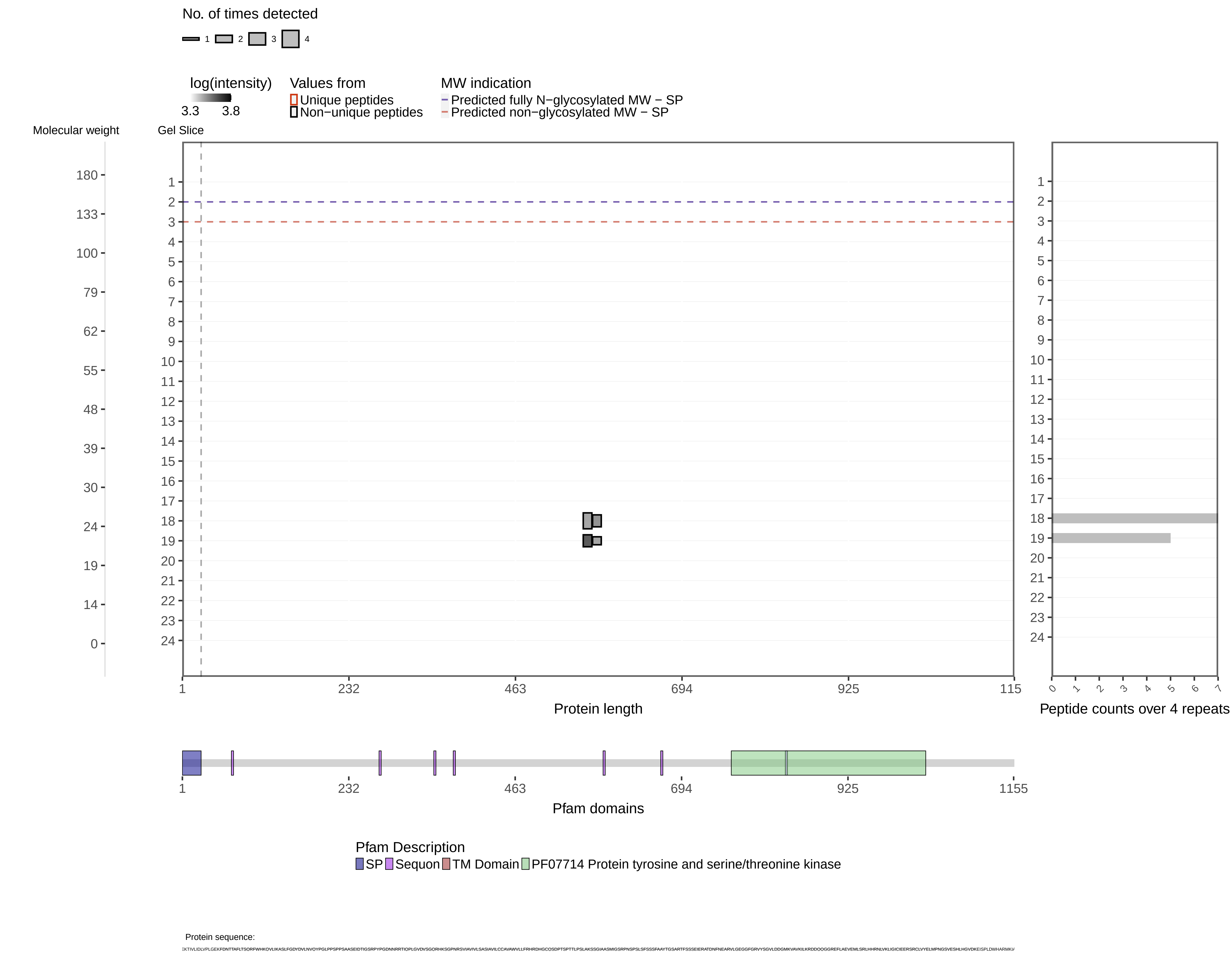

### NbL18g20240.1 translation_machinery−associated_protein_22−like [MW = 20.68 kDa]

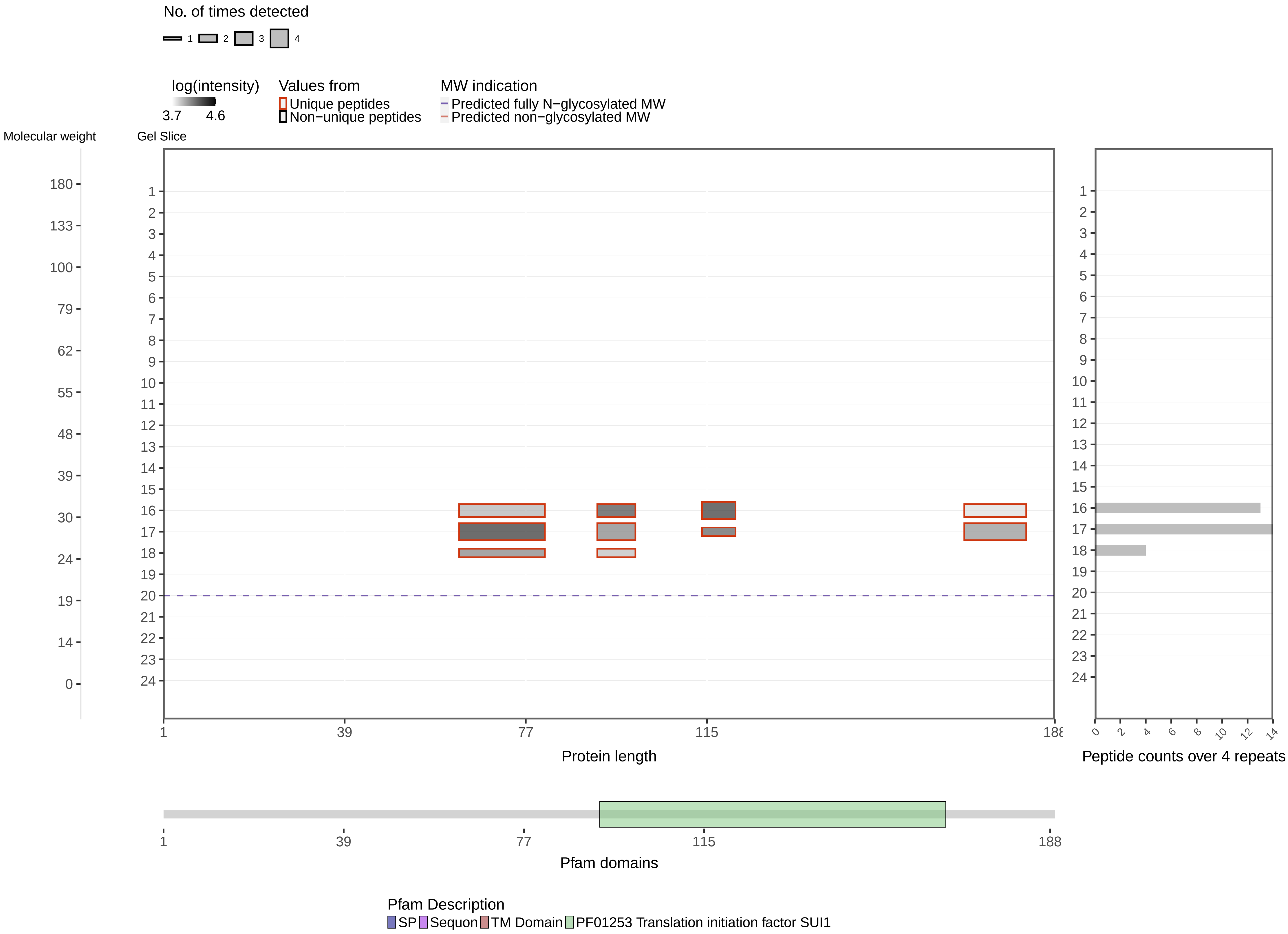


Protein sequence:

EEVKRLPGGKIKKKDKQEIIIEKVTRNKRKSITTIKGLELFGVKLSDASKKLGKKFATGASVVKGPTEKEQIDVQGDIFYDIVDFITETWRDVPESAIFFIEDGKKVPAA*

### NbL18g19830.1 WD_repeat−containing_protein_VIP3−like [MW = 34.79 kDa]

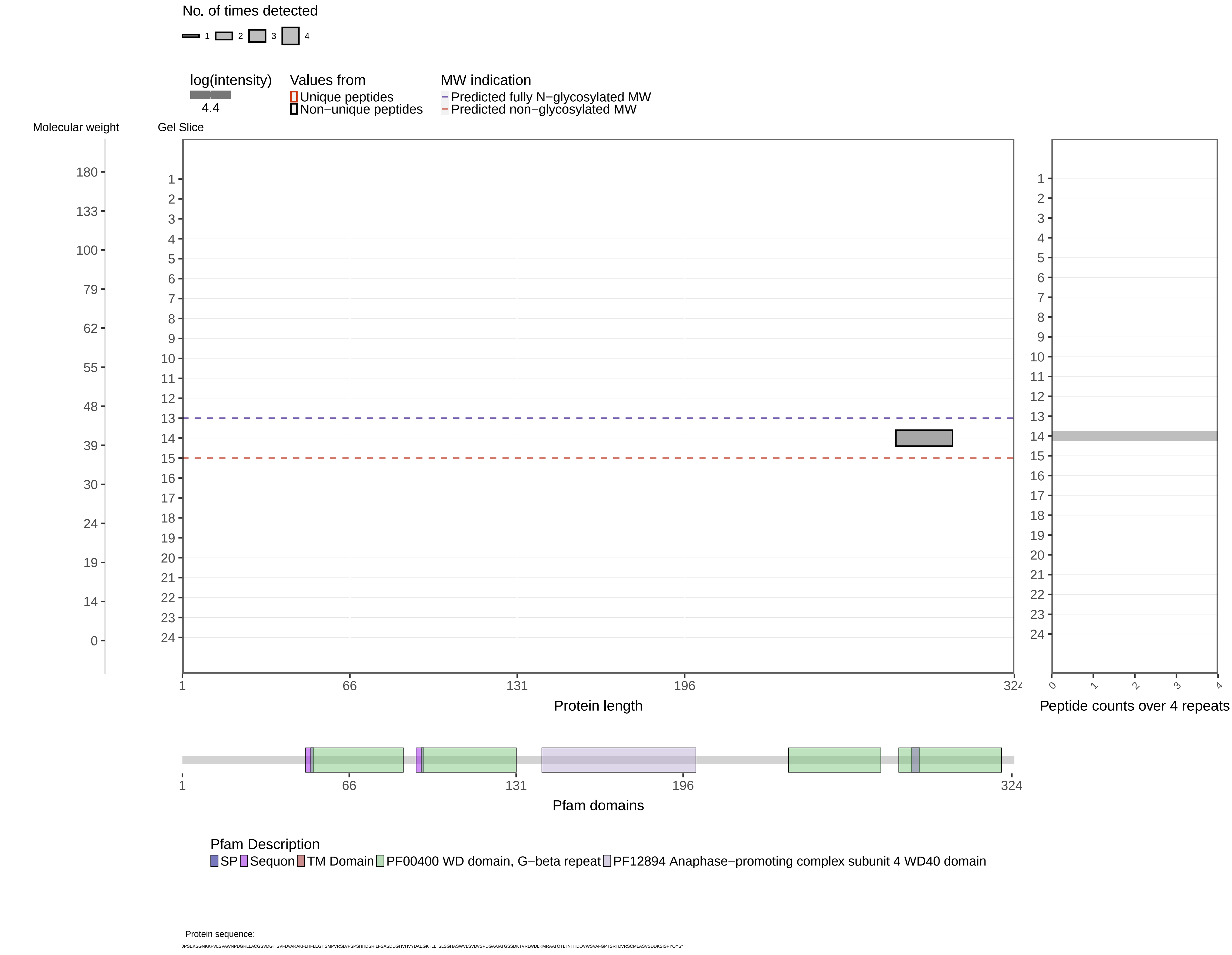

### NbL18g19750.1 PHD_finger_protein_ALFIN−LIKE_4−like [MW = 27.73 kDa]

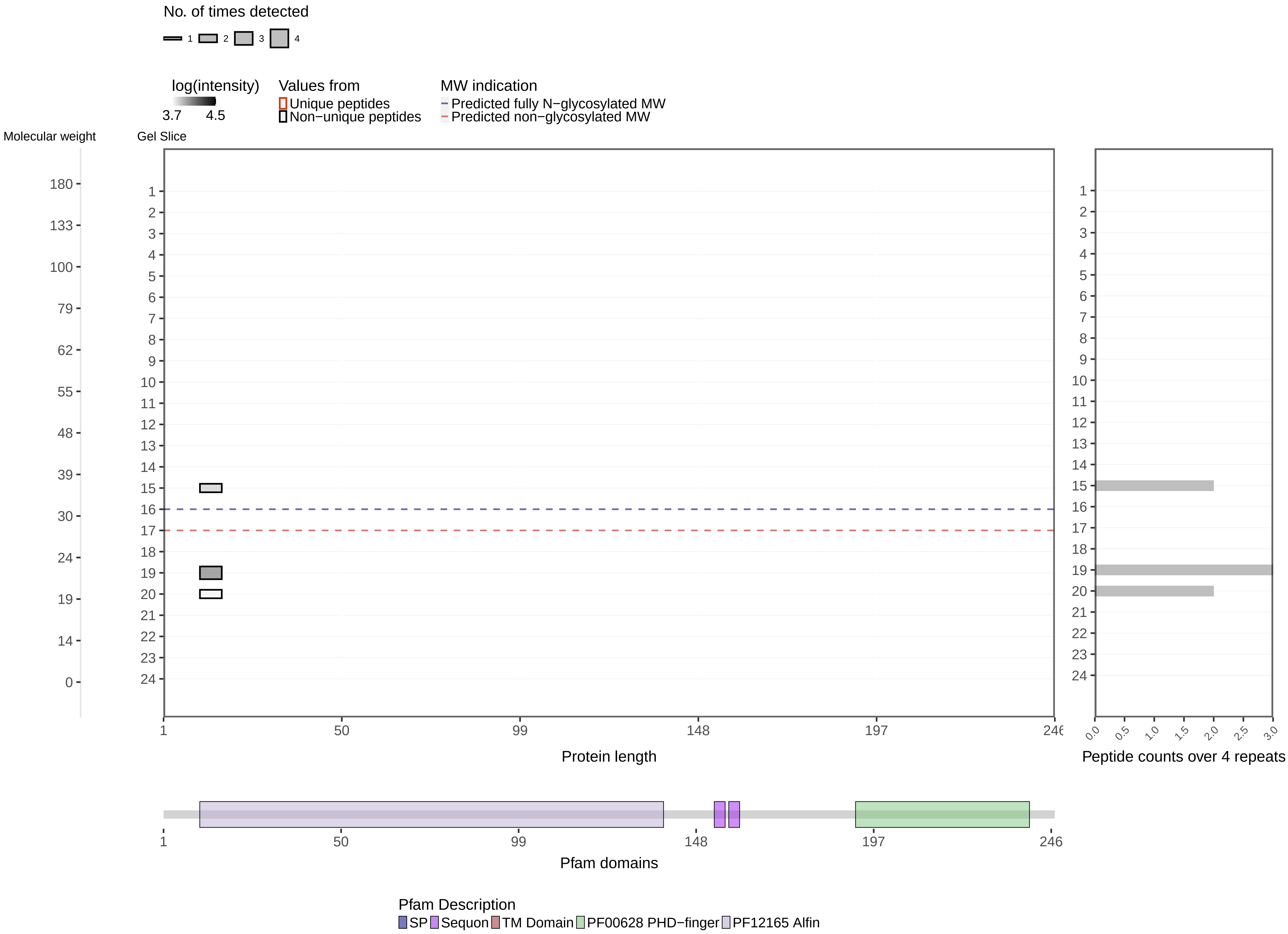


Protein sequence:

[illegible]ARFGFDKTDRKCLFNMINDLPTIYEVVTGVAKKQVKDRSTISNHSSNKSKSNSKAGKYSKAHVDEDDGLDEEEDEEHGDTLCGACGENYASDEFWICCDICERWFHGKCVKITPARAEHIKQYKCPSCSNKRSCP*

### NbL18g19710.1 40S_ribosomal_protein_S3−1−like [MW = 26.43 kDa]

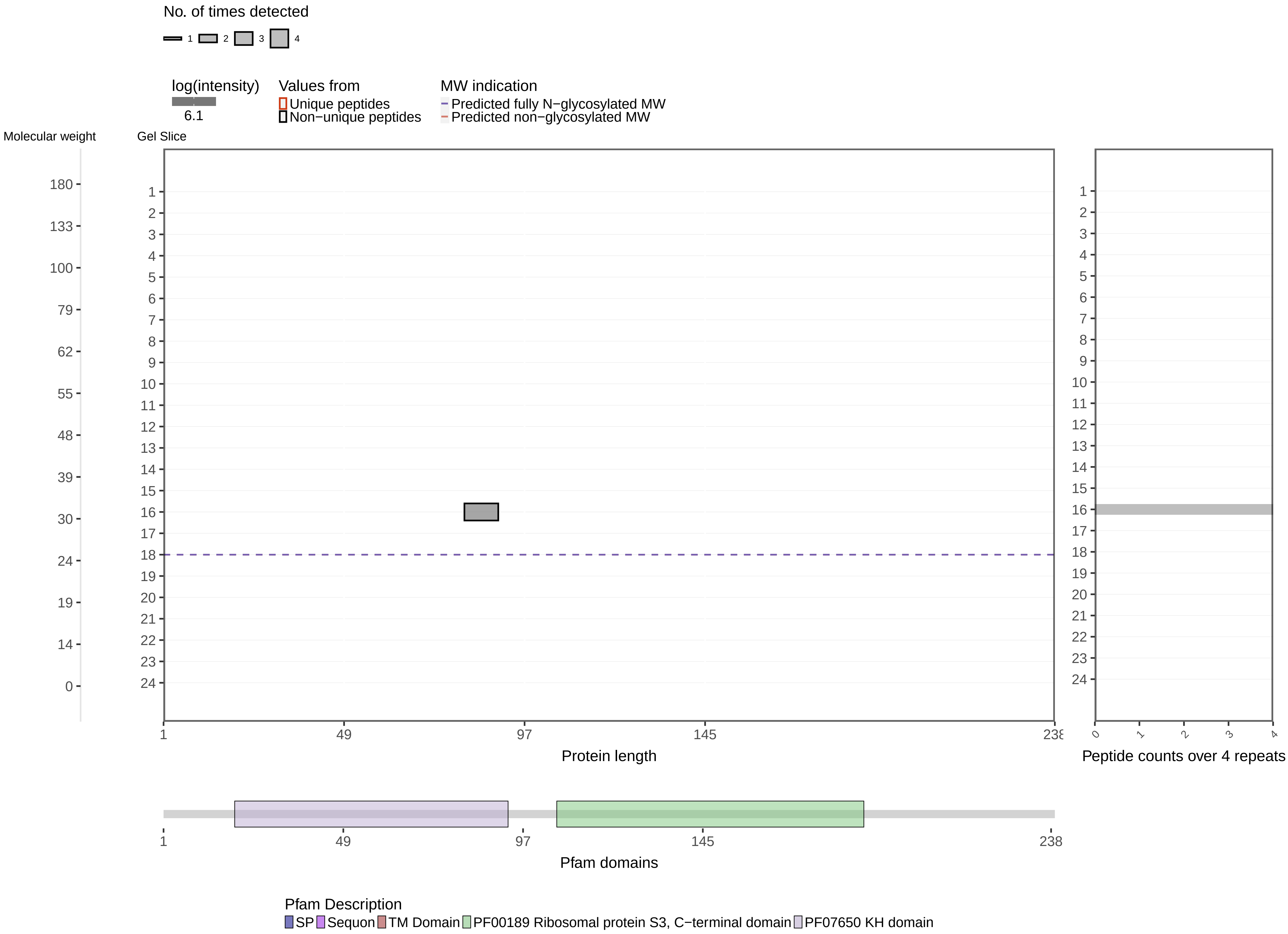


Protein sequence:

RYKLLGGLAVRRACYGVLRFIMESGAKGCEVIVSGKLRAQRAKSMKFKDGYMISSGQPVKEYIDSAVRHILMRQGVLGIKVKIMLDWDPKGKQGPTTPLPDLVTIHPPKEEEEYNIPPPAAPVDIEVSVPIS*

### NbL18g19670.1 stress−response_A/B_barrel_domain−containing_protein_UP3 [MW = 31.82 kDa]

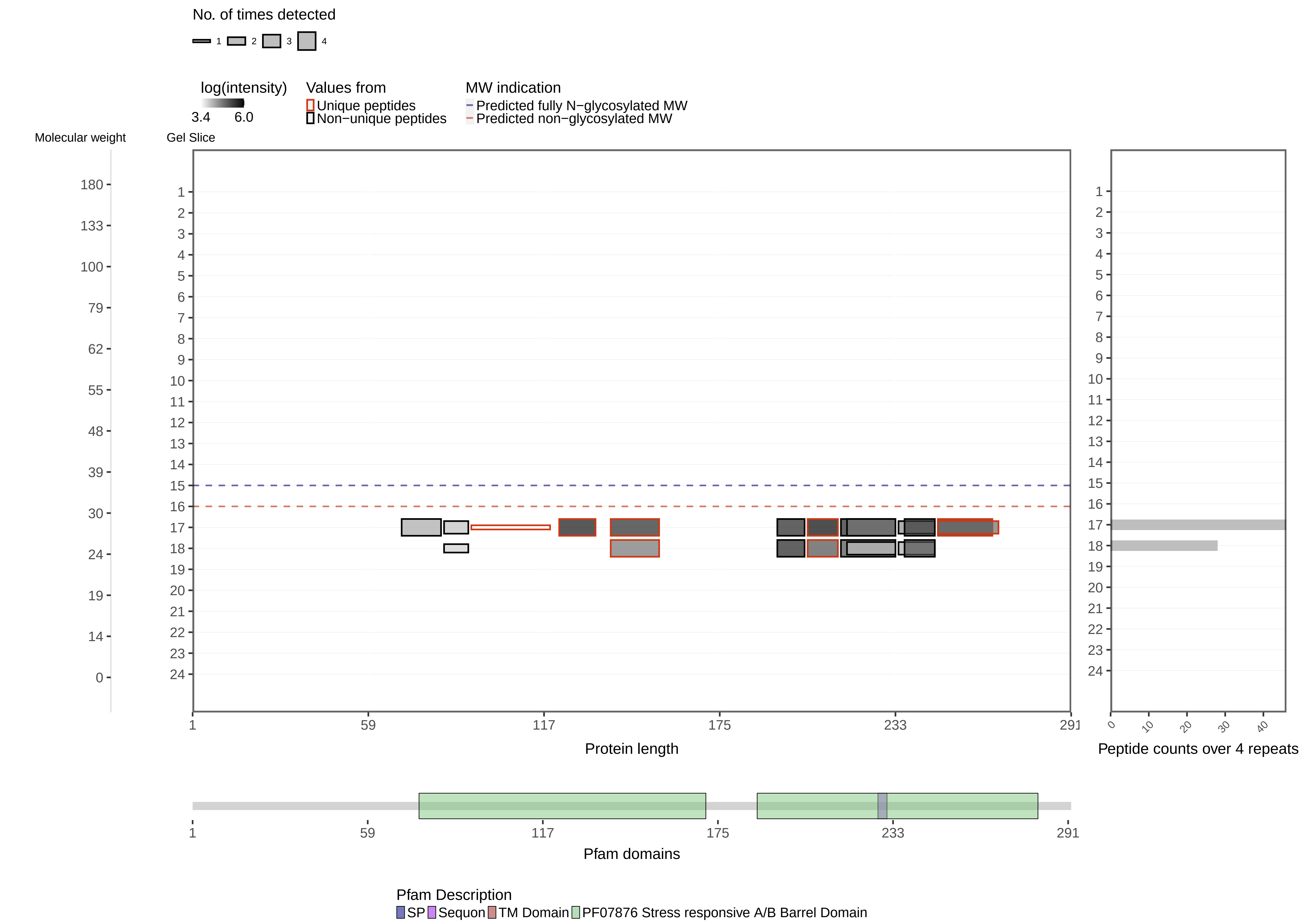


Protein sequence:

SRYKSKSDLADYSAHPDHVSVVKEYVLPVVDDIMAVDWIPTDFSGPIEVPPGSALRVTFLKLKENLGENEKSQILNAVGGIKEKFPGIEQLTVGENFSPGRAKGFSIASIAVFKGVNELEALEAQTELANEQKEKVRDFLDGVVVVDYAVPPTQSASL*

### NbL18g19380.1 protein_trichome_birefringence−like_14 [MW = 55.35 kDa]

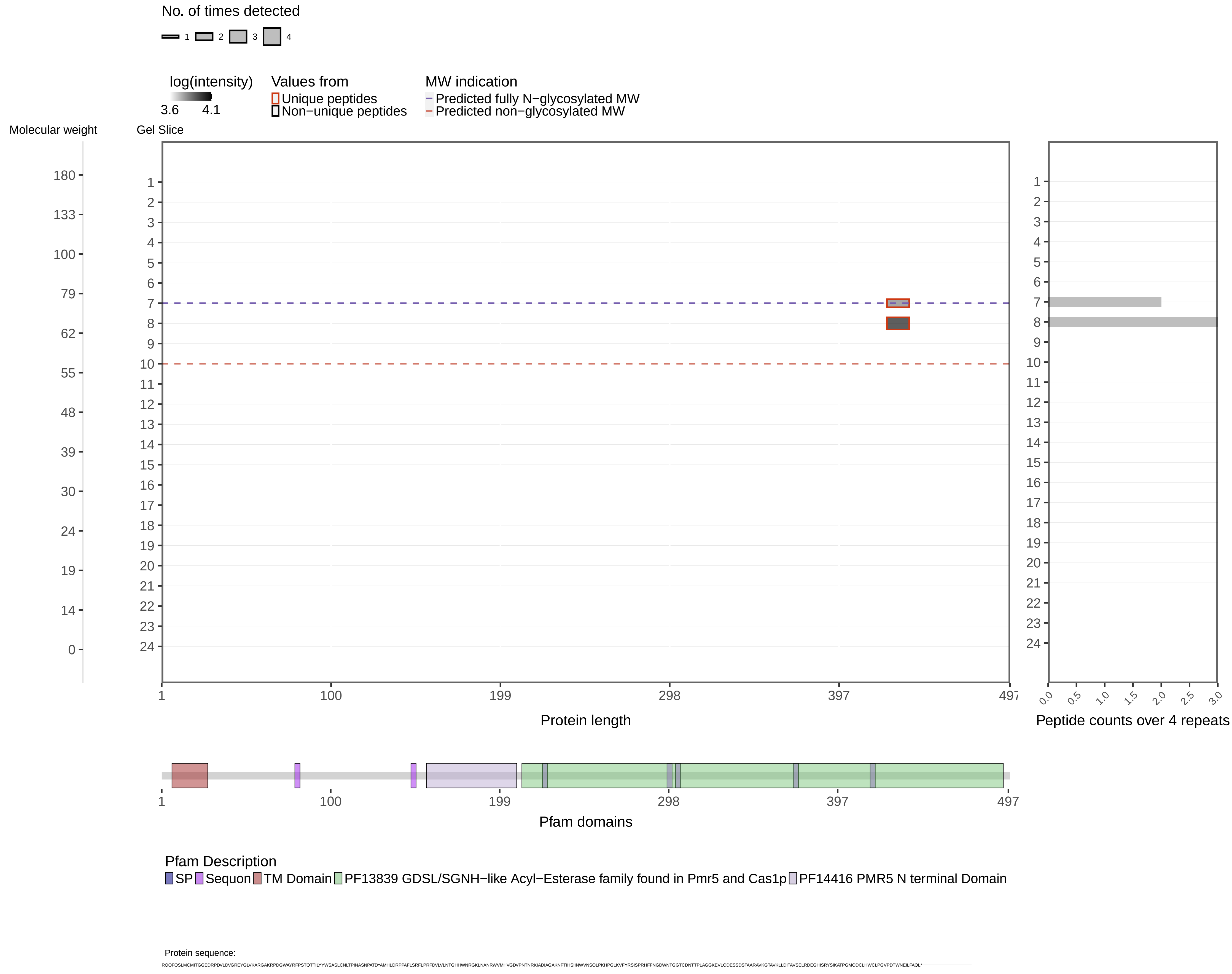

NbL18g19260.1 beta−galactosidase_15−like [MW − SP = 89.71 kDa]

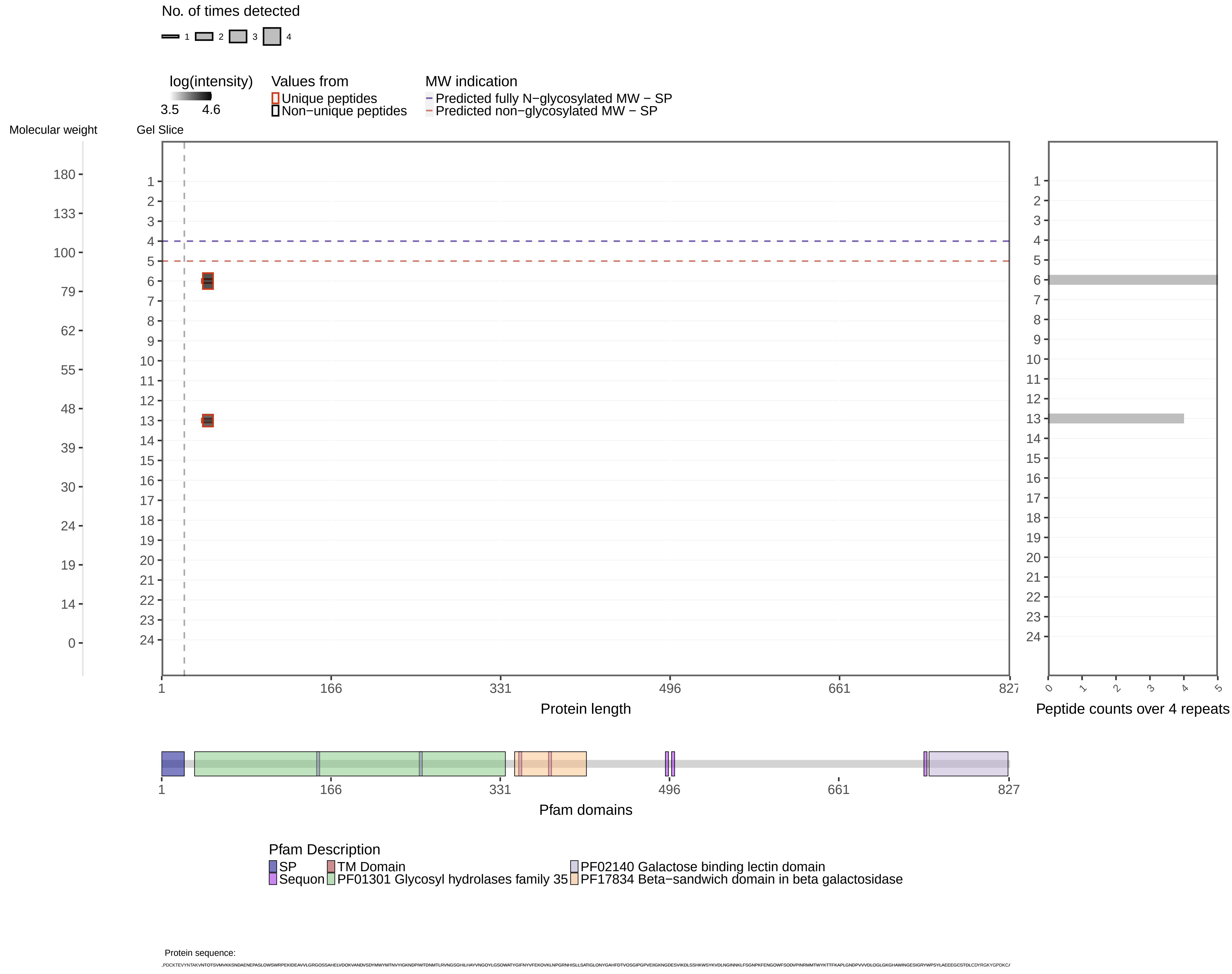

NbL18g19210.1 3−oxoacyl−[acyl−carrier−protein]_synthase_II_chloroplastic−like [MW = 53.04 kDa]

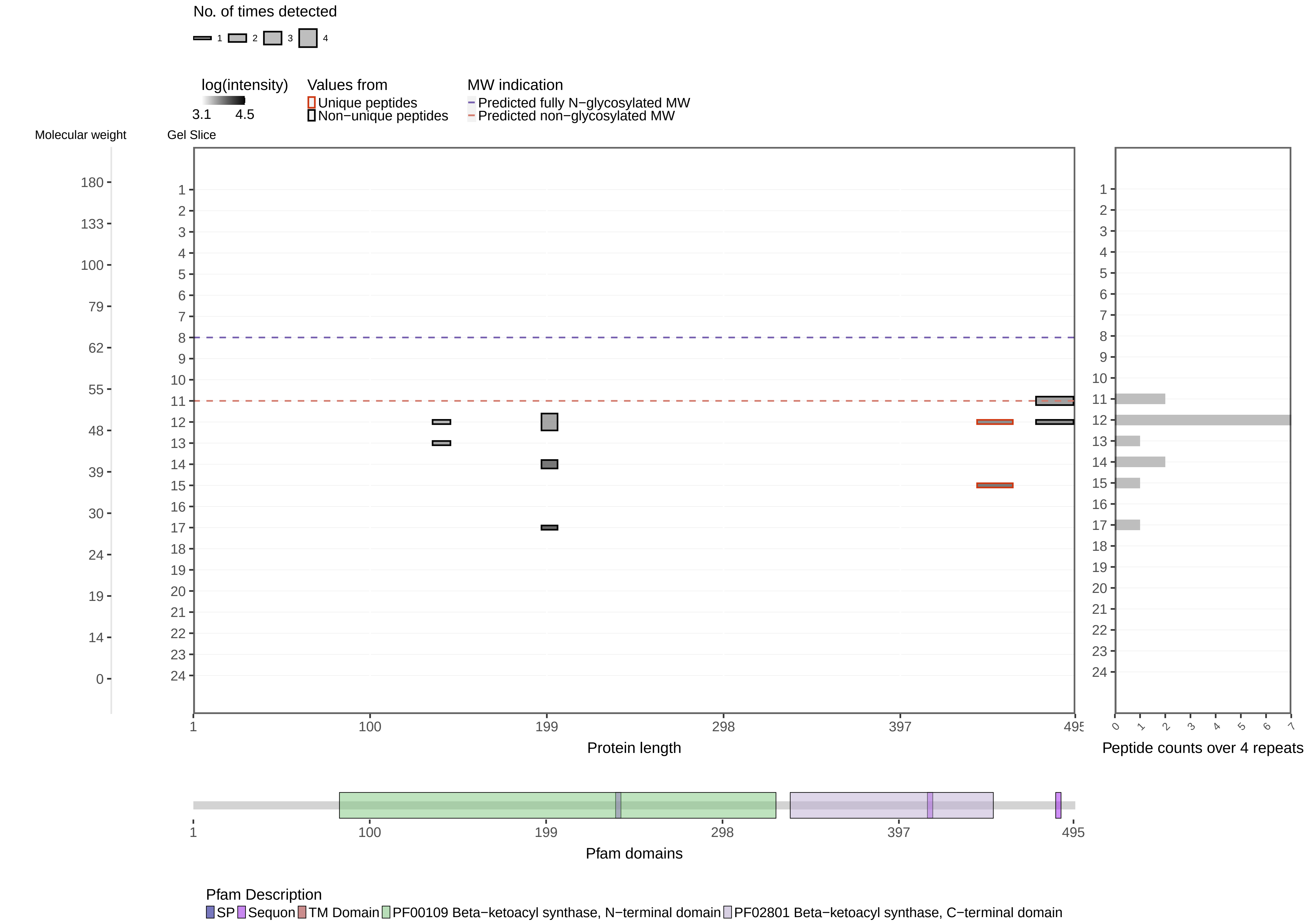


Protein sequence:

[illegible]LGWMGPNYSISTACATSNFCILNAANHIIRGEADLMLCGGSDAAIIPIGLGGFVACRALSQRNGDPARASRPWDSNRDGFVMGEGAGVLVLEELEHAKQRGATIYAEFLGGSFTCDAYHMTEPHPEGTGVILCIEKALAQSGVSKEDVNYINAHATSTLAGDIKEYQALLYCFGQNPELRVNSTKSMIGHLLGAAGAVEAVATIQAIRTGWVHPNINLENPDEGVDTTKLVGPIKERLDIKVALSNSFGFGGHNSSILFAPYK*

### NbL18g19190.1 21_kDa_protein−like [MW = 16.77 kDa]

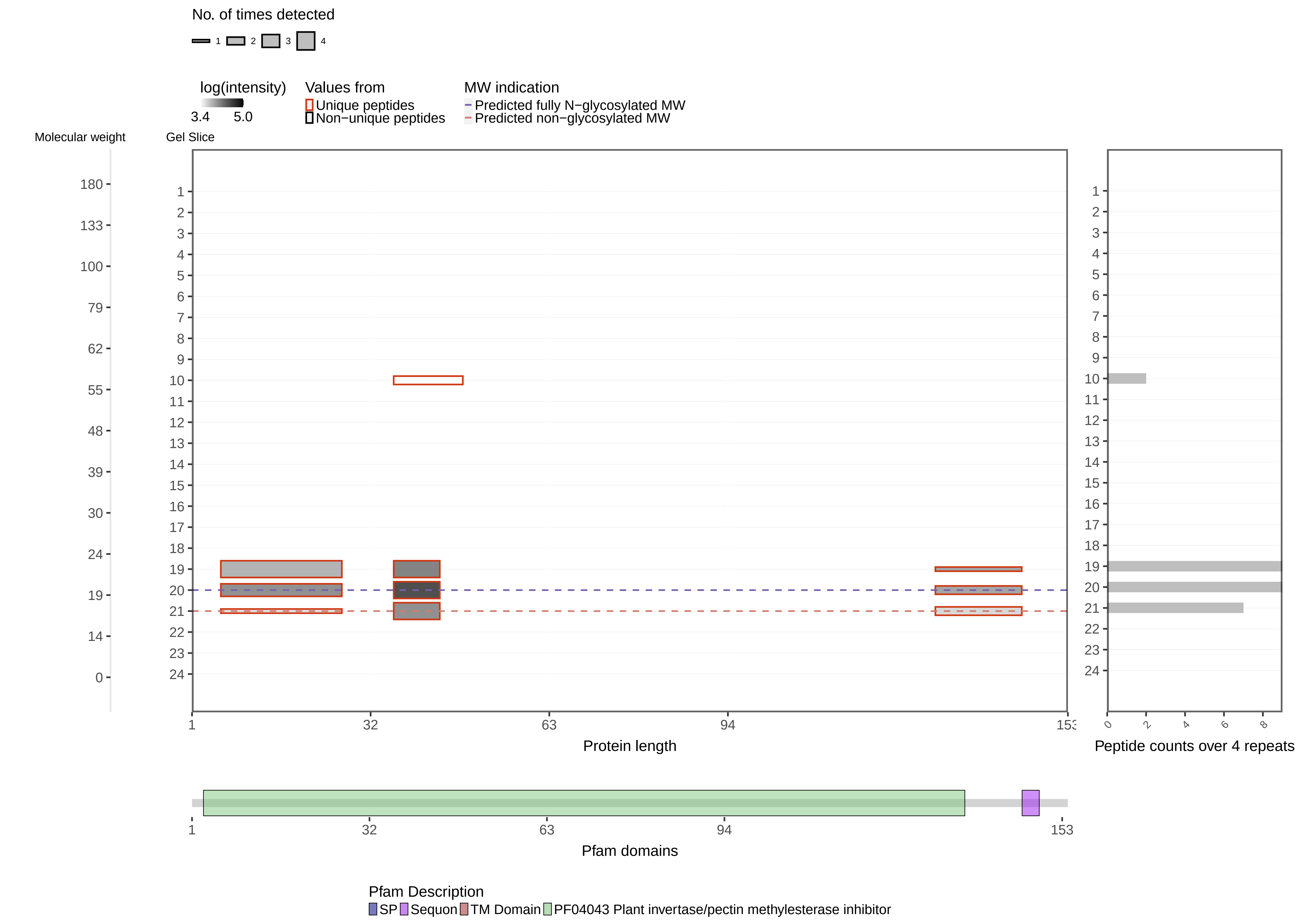


Protein sequence:

DSMDELSKTLSELKHLSKGNAFKWQMSNLETWVSAALTNEDTCIDGFKEIDGKLRSDVKRKITNVARVTSNALYLINQLDDSRNKTAAHP*

NbL18g19130.1 peroxiredoxin_Q_chloroplastic−like [MW = 23.59 kDa]

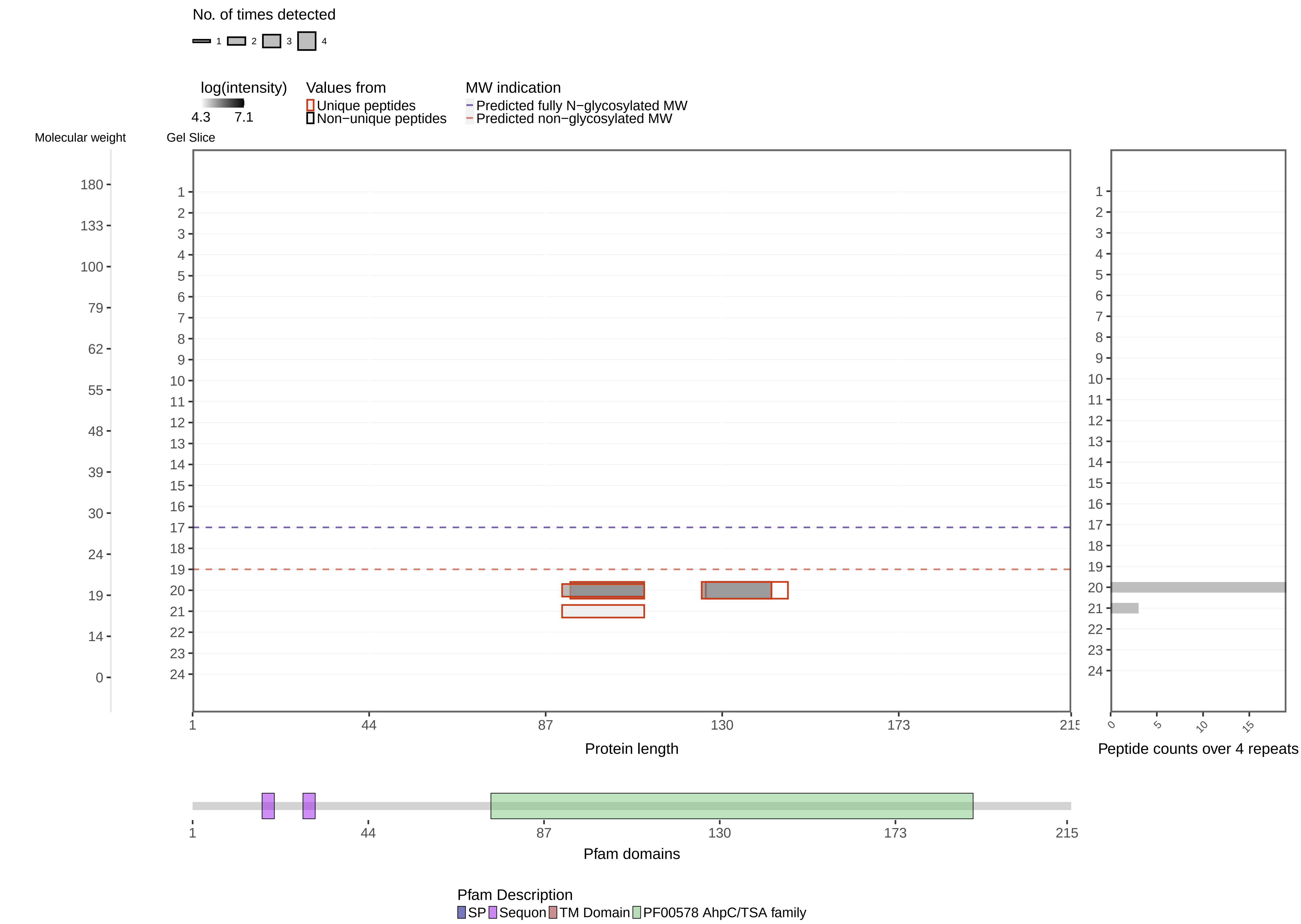


Protein sequence:

VVVYFYPADETPGCTKQACAFRDSYEKFKKAGAQVVGISGDDPASHKAFAKKYRLPFTLLCDEGNKVRKEWGIPGDLFGTLPGRQTYVLDKNGVVQLIYNNQFQPEKHIDETLKLLQSL*

### NbL18g19090.1 sucrose_synthase−like [MW = 91.19 kDa]

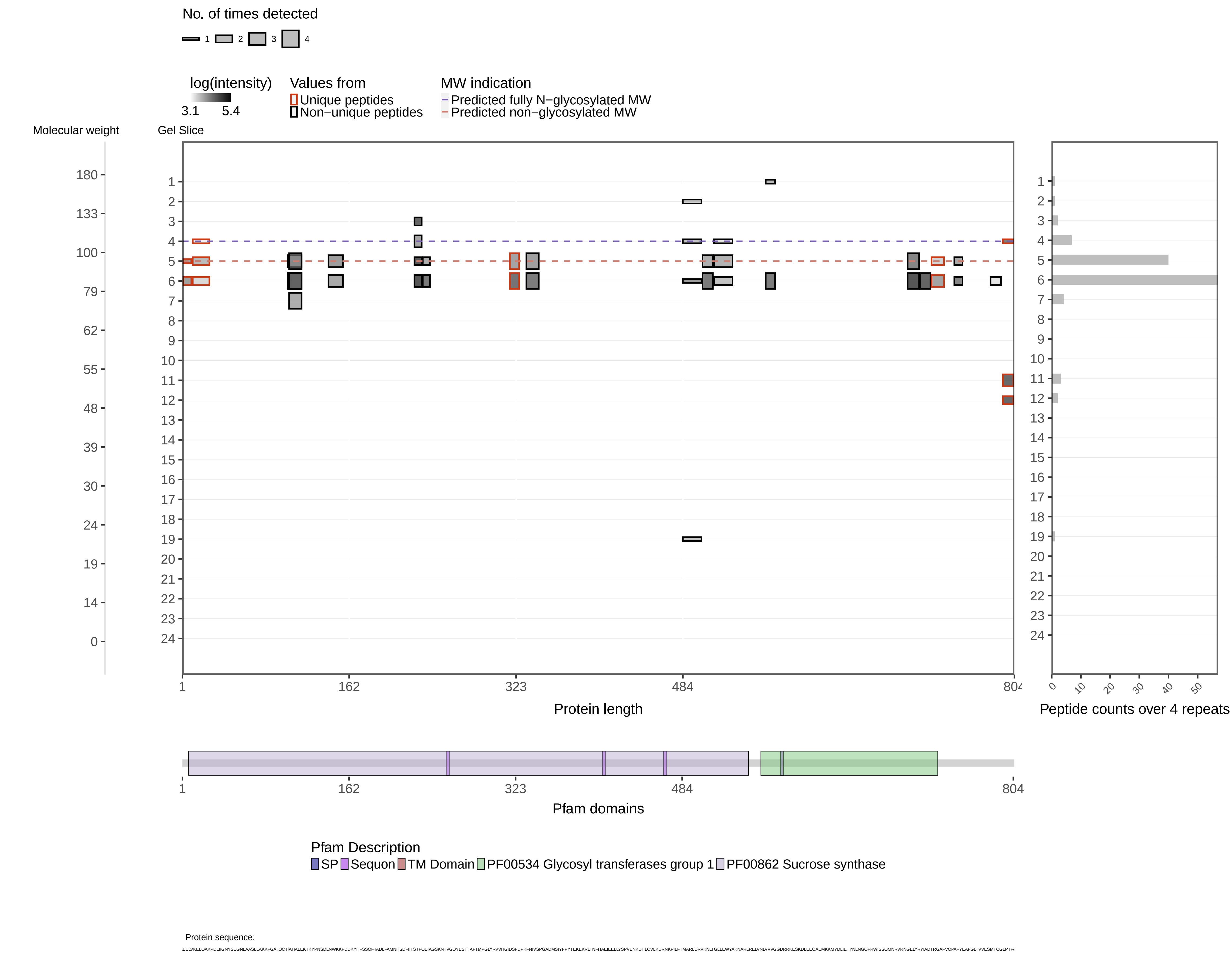

NbL18g19080.1 sucrose_synthase−like [MW = 92.49 kDa]

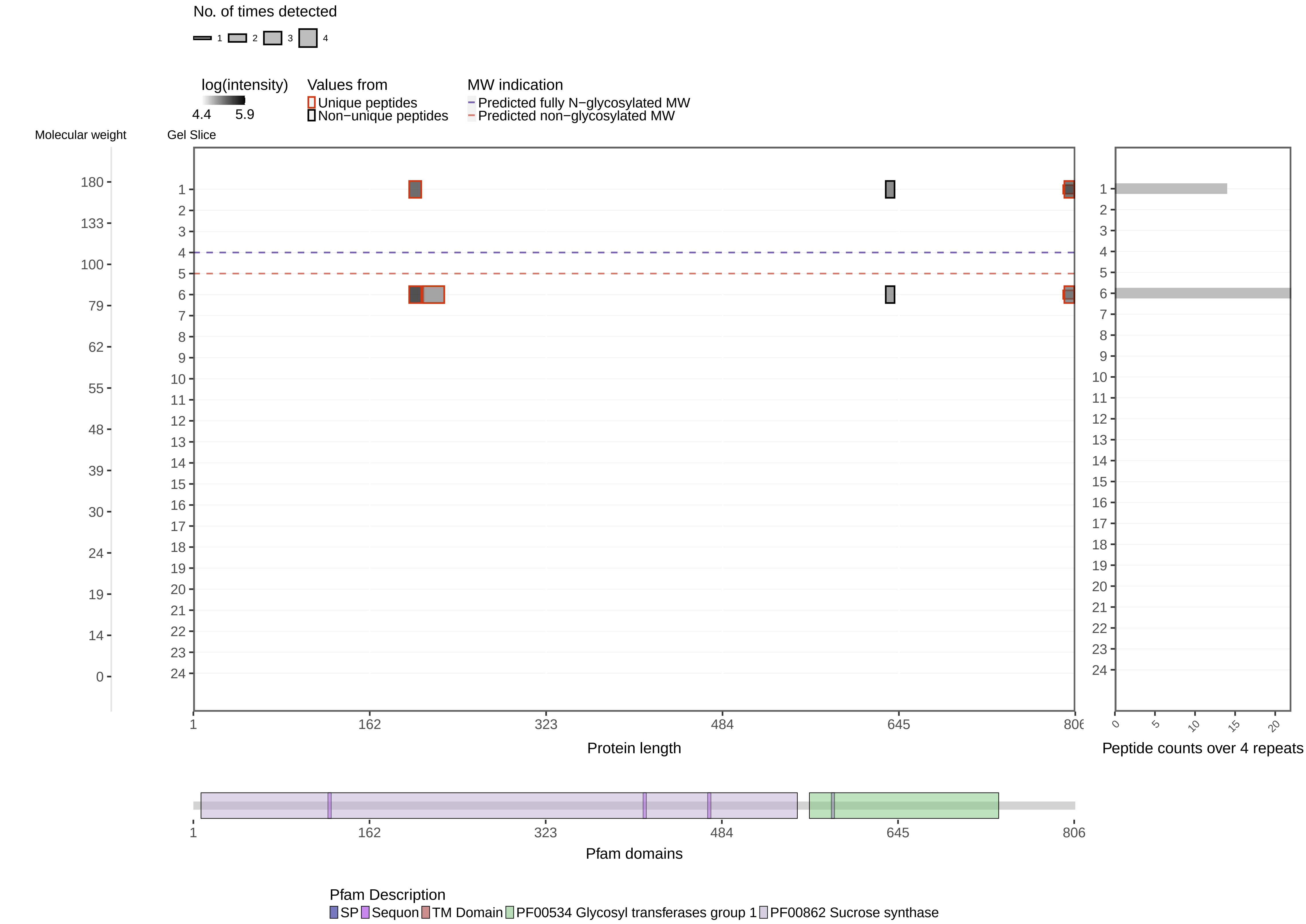


Protein sequence:

TFTEDVAKEIAAELQAKPDLIIGNYSEGNLAASLLAHKLGVTQCTIAHALEKTKYPDSDIYLKKFDEKYHFSAQFTADLIAMNHTDFIITSTFQEIAGSKDTVGQYESHMAFTMPGLYRVVHGIDVFDPKFNIVSPGADMNLYFPYFEKEKRLTAYHPEIEELLFSDVENDEHTCVLKDRNKPIIFTMARLDRVKNLTGLVELYAKNPRLRELVNLVVVGGDRRKESKDLEEQAEMKKMYELIKTHNLNGQFRWISSQMNRVRNGELYRYIADTRGAFVQPAFYEAFGLTVVEAMTCGI

NbL18g19060.1 eukaryotic_translation_initiation_factor_3_subunit_K [MW = 26.2 kDa]

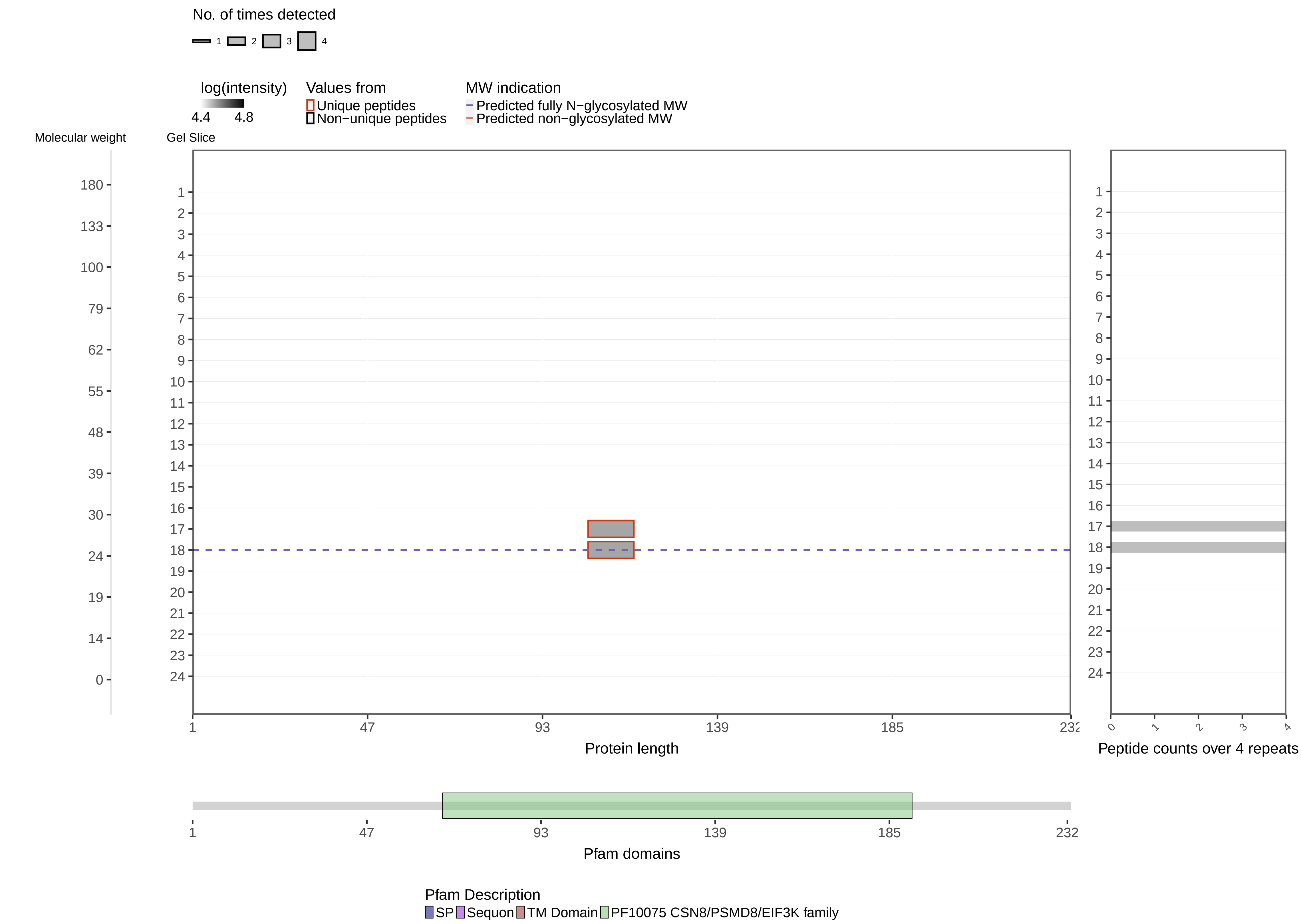


Protein sequence:

[illegible]FKTLIVLSHYLETARFRQFWDEAAKSRHIVEVVPGFEQAILAYAVHVLSITYQKVPRAIVAEAINIEGLSLDKFLEHQVSNNGWVIEKGQGKGQLITLPPTEFNHPELKKNTADSVPLEHVTRIFPILG*

### NbL18g19050.1 microtubule−associated_protein_TORTIFOLIA1−like [MW = 92.53 kDa]

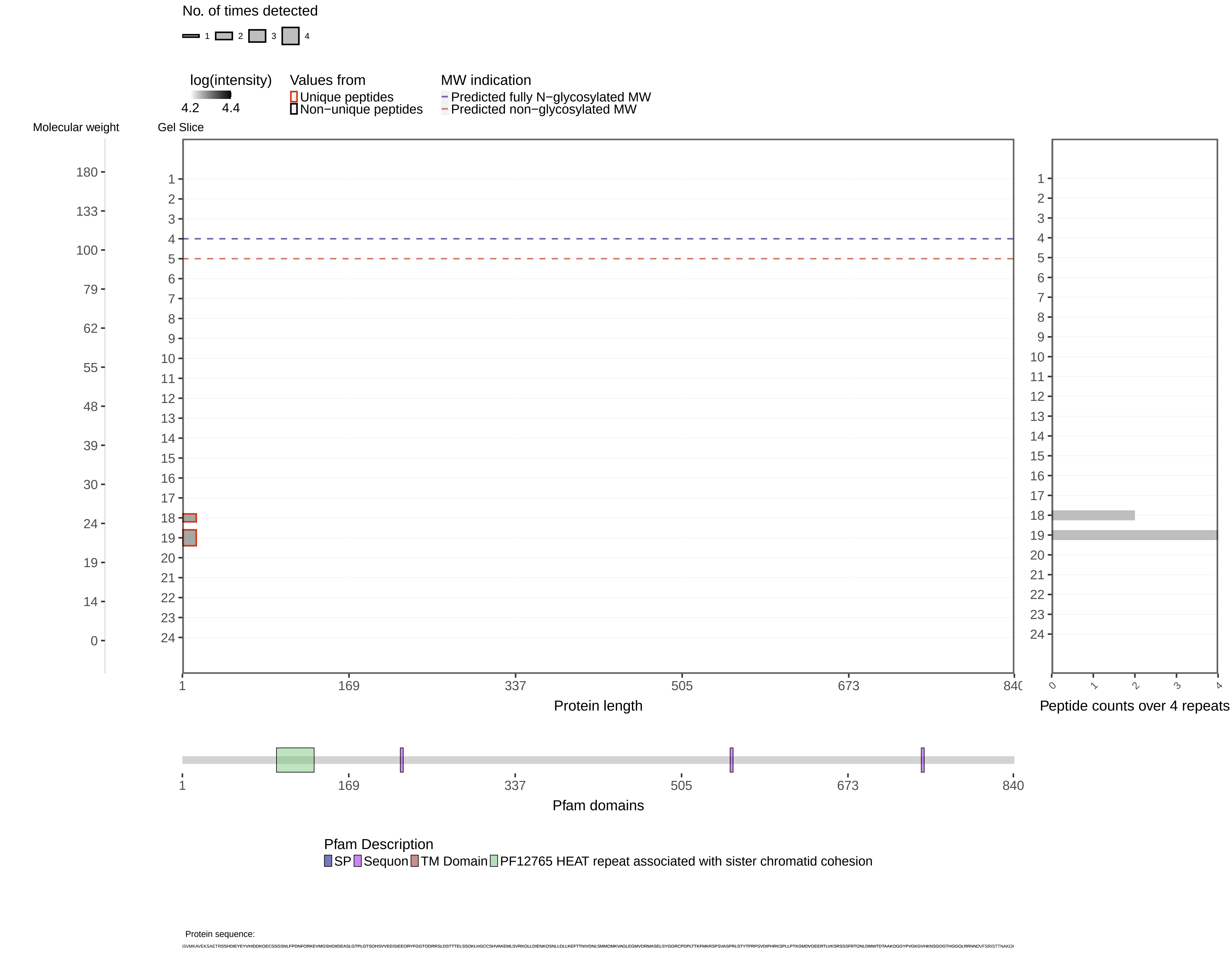

### NbL18g19030.1 prefoldin_subunit_1 [MW = 14.83 kDa]

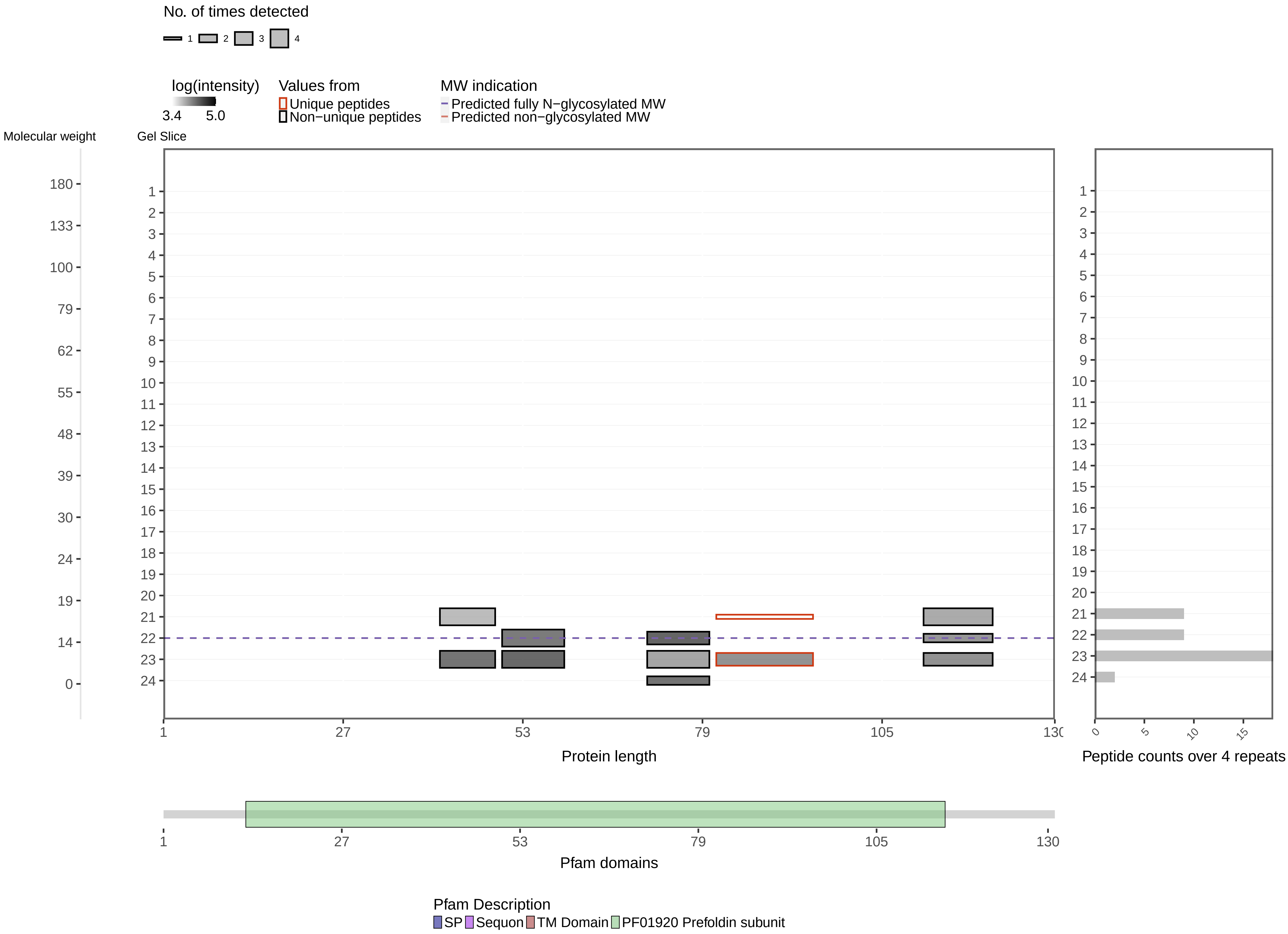


Protein sequence:

SDDTNTYKAIGMRFILEPKTVLMDEQEQKLKDSETAIASLQTSKEYLEKHLAEVENNLRELLQQDPGLARQIMTMSVA*

NbL18g18900.1 pyruvate_dehydrogenase_E1_component_subunit_alpha−3_chloroplastic−like [MW = 47.9 kDa]

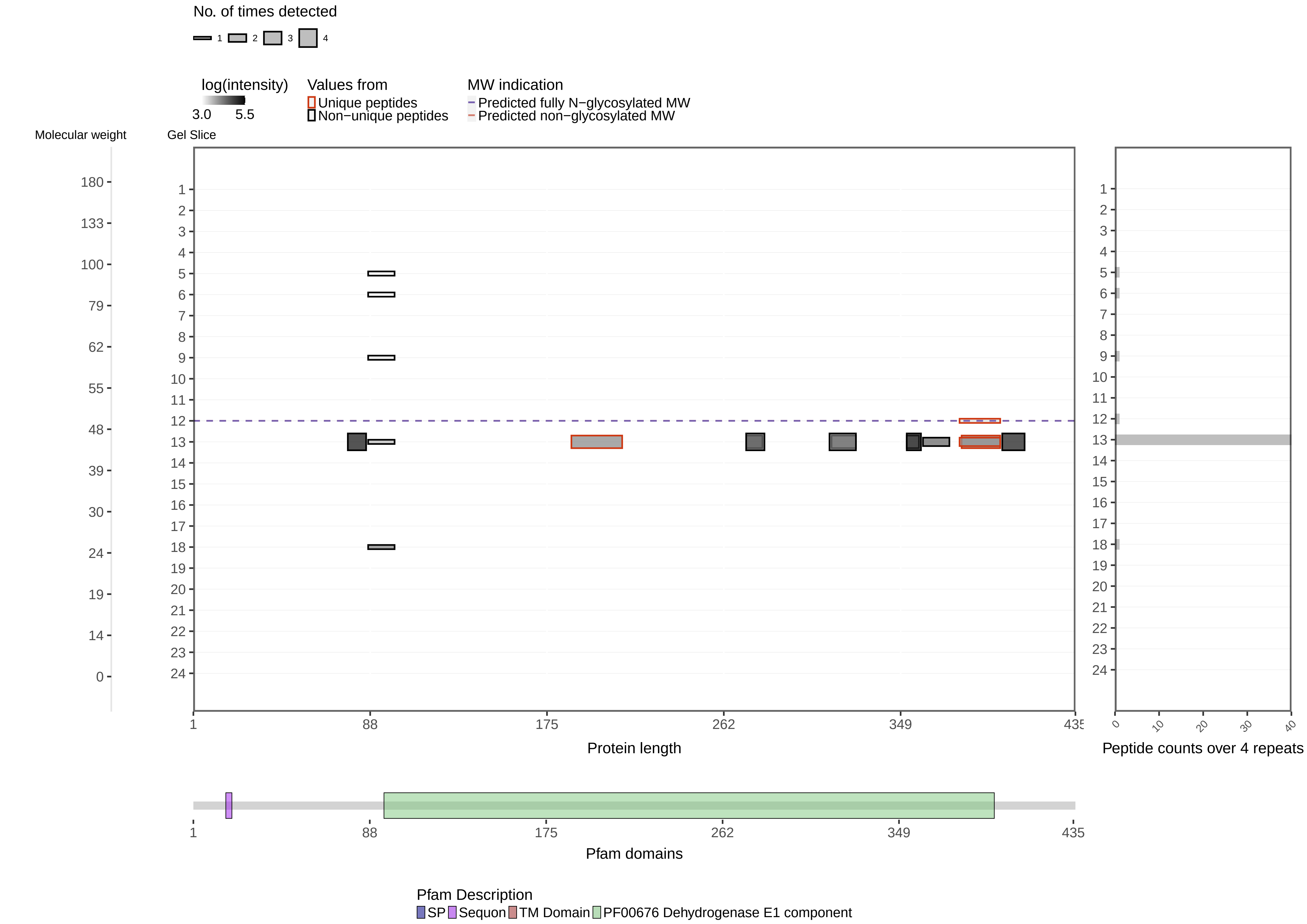


Protein sequence:

AGAAFTSKYRREVLKEADCDHVTLAFFGDGTCNNGQFYECLNMAALWKLPIIFVVENNLWAIGMSHLRSTSDPEIWKKGPAFGMPGVHVDGMDVLKVREVAKEAIGRARRGEGPTLVECETYRFRGHSLADPDELRDPAEKNHYATRDPITALKKYMFENNLVSEAELKAIDKKIDEVVEESVEFADASPVPARSQLLENVFADPKGFGIGPDGRYRCEDPKFTEGTAHV*

### NbL18g18770.1 gamma_aminobutyrate_transaminase_1_mitochondrial [MW = 41.62 kDa]

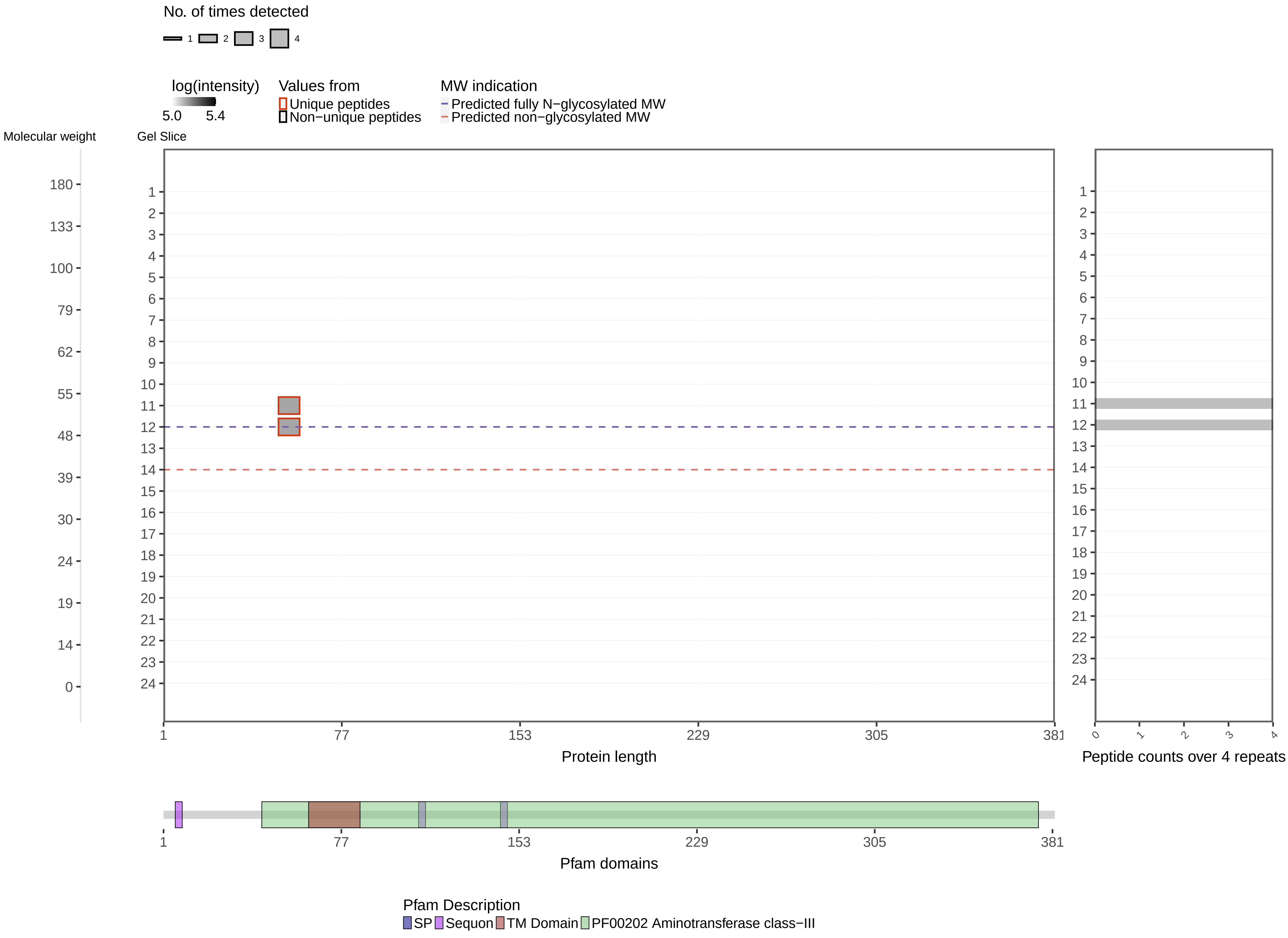


Protein sequence:

YHGSTLISASLTGTLTAPIIGAIICOMYRPGETEEEFSTRLAKNVEDLILKEGPETIAAFIAEPVMGAGGVIPPPATYFDKIOAVVKKYDILFIADEVICAFGRLGTMFGCDKYNIKPDLVSLAKALSSAYMPIGAVLVSPEVSDVIYSOSNKLGSFSHGFTLSGHPVSCAVALEALKIYKERNMVEKVNRISPKFOEGLKAFLTVP*

NbL18g18760.1 myosin_heavy_chain_cardiac_muscle_isoform−like [MW = 146.68 kDa]

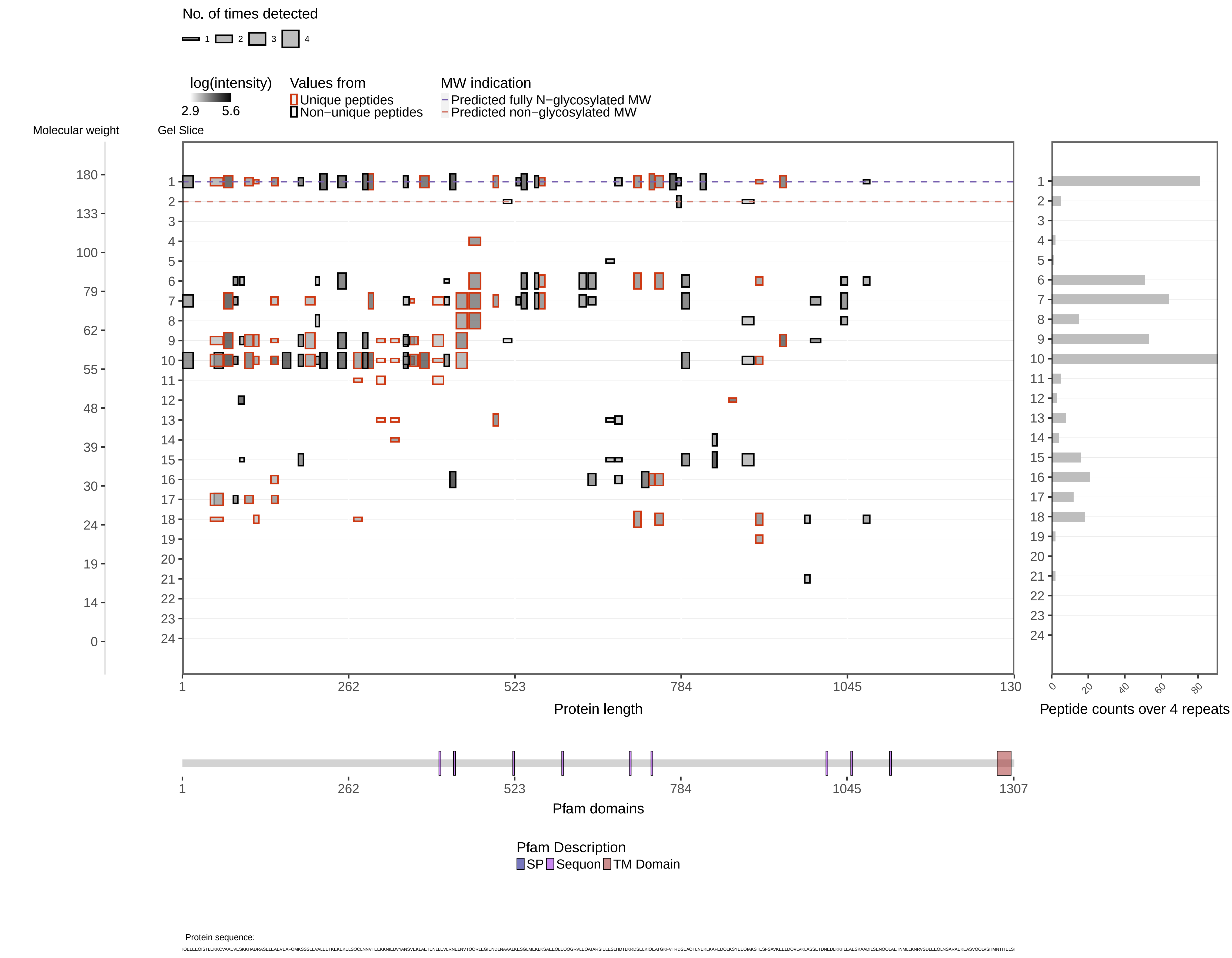

NbL18g18740.1 60S_ribosomal_protein_L27−3−like [MW = 15.68 kDa]

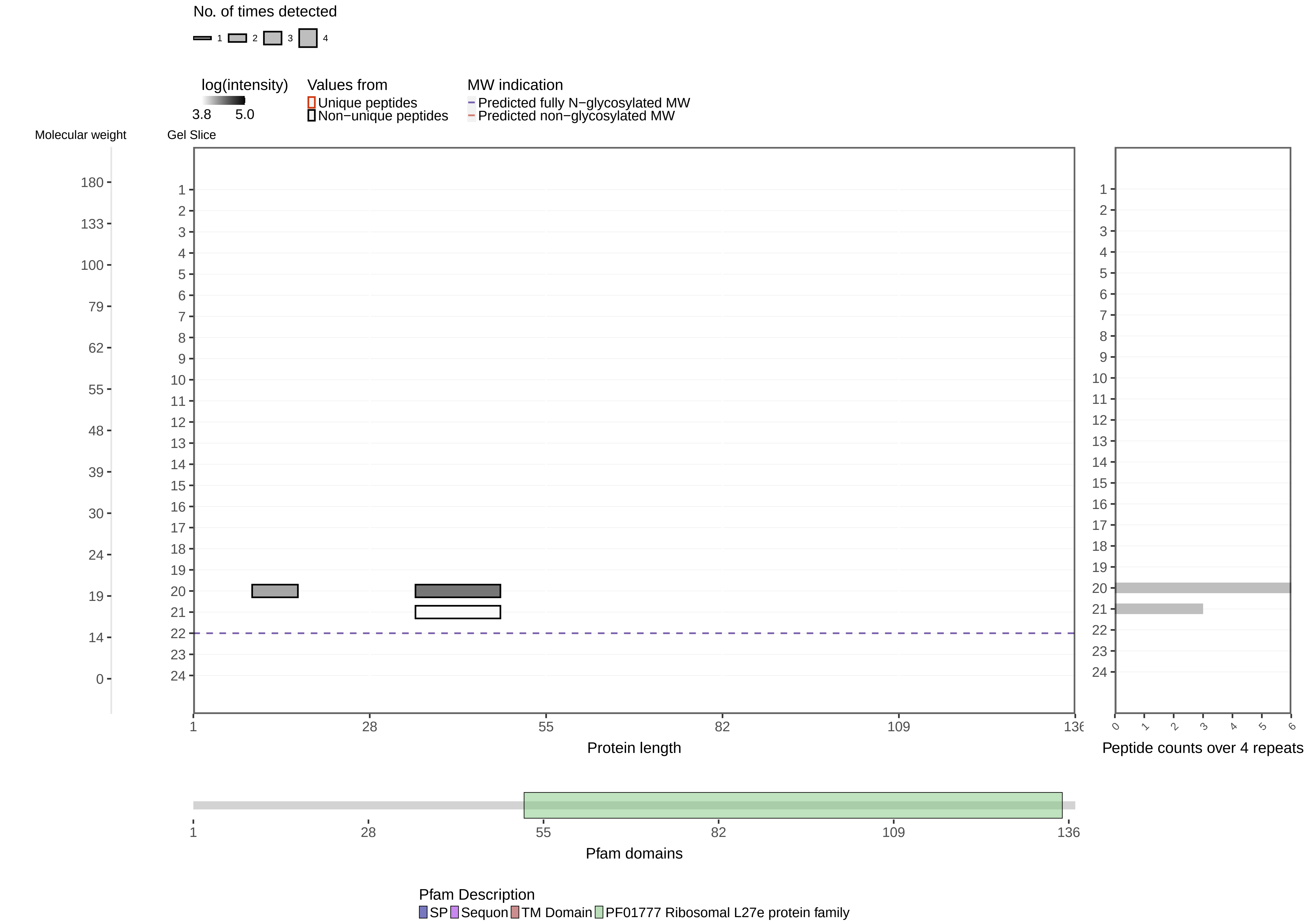


Protein sequence:

RKDSDKKQAKKSRVKTFIKLVNYNHIMPTRYTLDVDLKEAVTVDCLQSRDKKVTAAKEAKAKFEERFKTGKNRWFFSKLRF*

### NbL18g18640.1 heat_shock_70_kDa_protein_17 [MW − SP = 96.92 kDa]

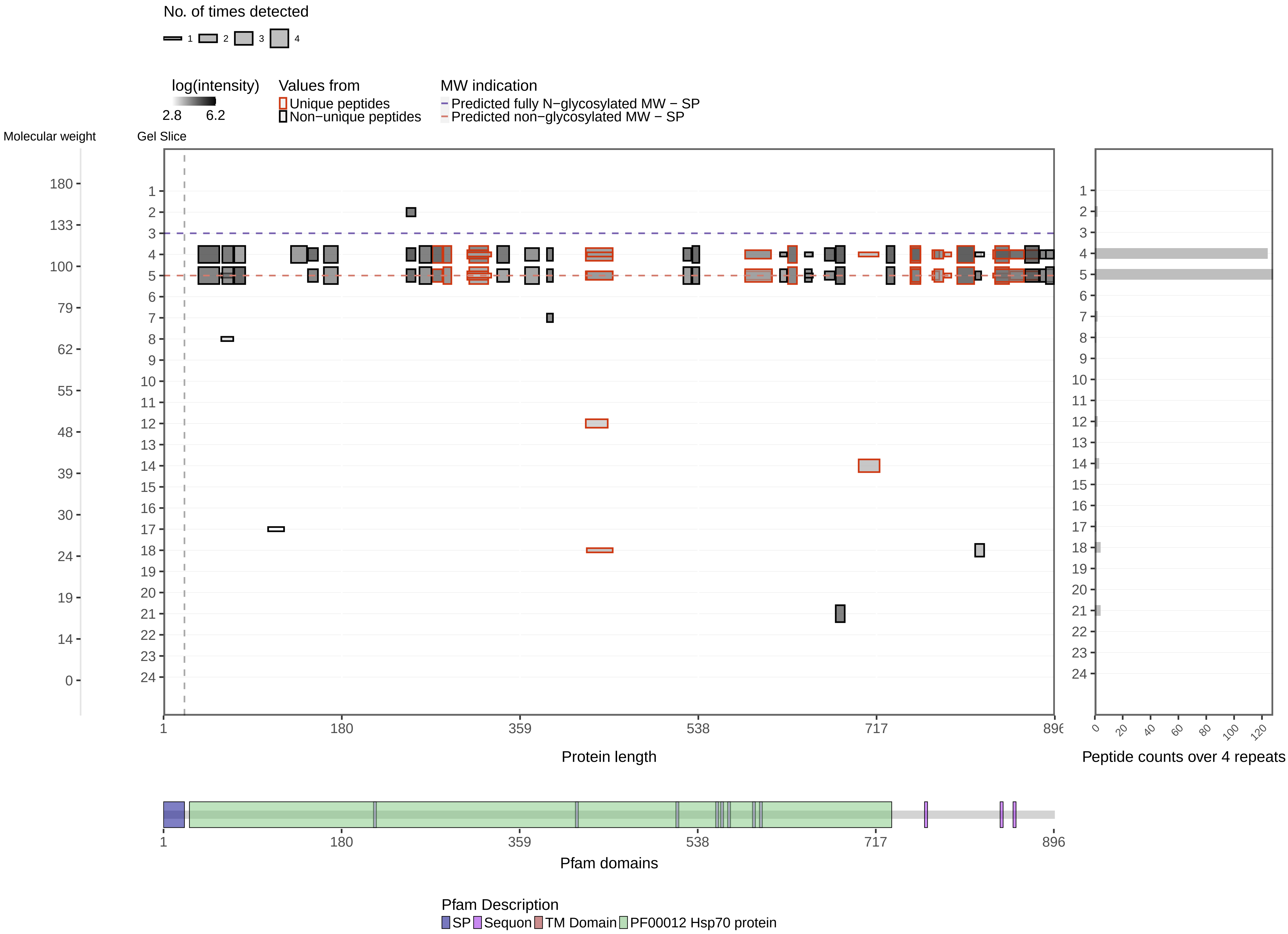


Protein sequence:

EVDGADLPKDESTKQLIIPRMKKLPSKMFRSIVHNKDFEVSLAYESDDFLPPGTPSLTFAQYAVSGLADTSEKYASLNLSAPIKANLHFSLSRSGIFSLDRADAVIEITEWVEAPRRNLTVDNSTSASTNTSTEGSPSNTEESSEKLNADDSNSNNSDPSANDSSTTSLSIEKKLKKRTFRVPLKIDEKTAGPGASLLKESLNEAKSKLEALDKKDEERRRTAELKNSLEGYIYDTRDKLESEDFAKISTSQERQSFIEKLDEVQEWLYTDGEDASATQFKKRLDELKAIGDPIFFRHK

### NbL18g18610.1 uncharacterized_oxidoreductase_At1g06690_chloroplastic [MW = 40.32 kDa]

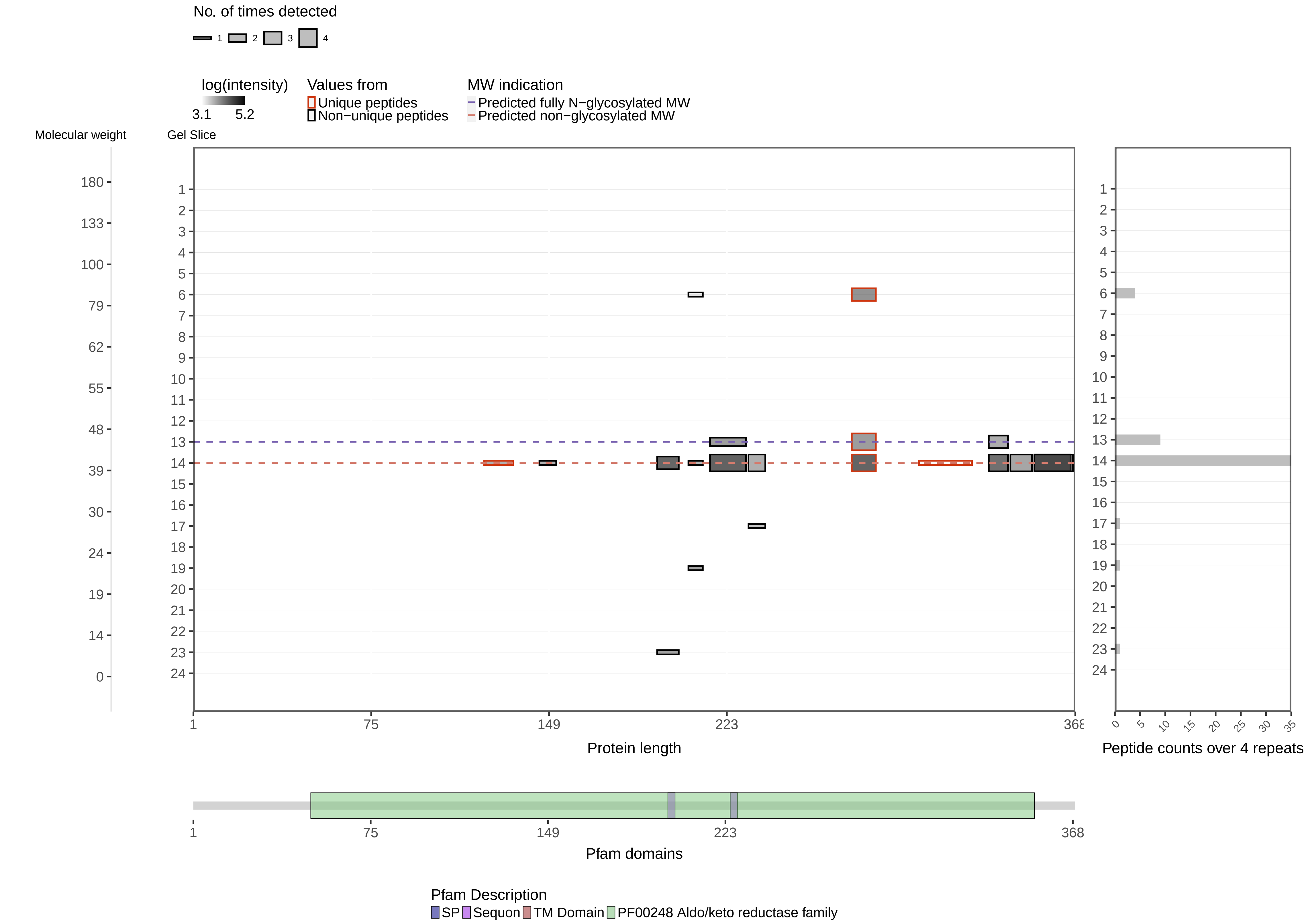


Protein sequence:

/PGIWGNEGYIDGLGDAVEQGLVKAVGVSNYSEKRLRAAYEQLKRRGIPLASNQVNYSLIYRFPEENGVKAACDELGVTLIAYSPIAQGALTGKYTPENPPTGPRGQIYTPEFLTKLQPIINRIKVIGESYSKTPTQVVLNWLIAQDNVVPIPGAKNADQAKEFAGALGWRLTQDEVDELRSLALDVKPVTGFPVEKL*

NbL18g18480.1 23−bisphosphoglycerate−independent_phosphoglycerate_mutase−like [MW = 61.06 kDa]

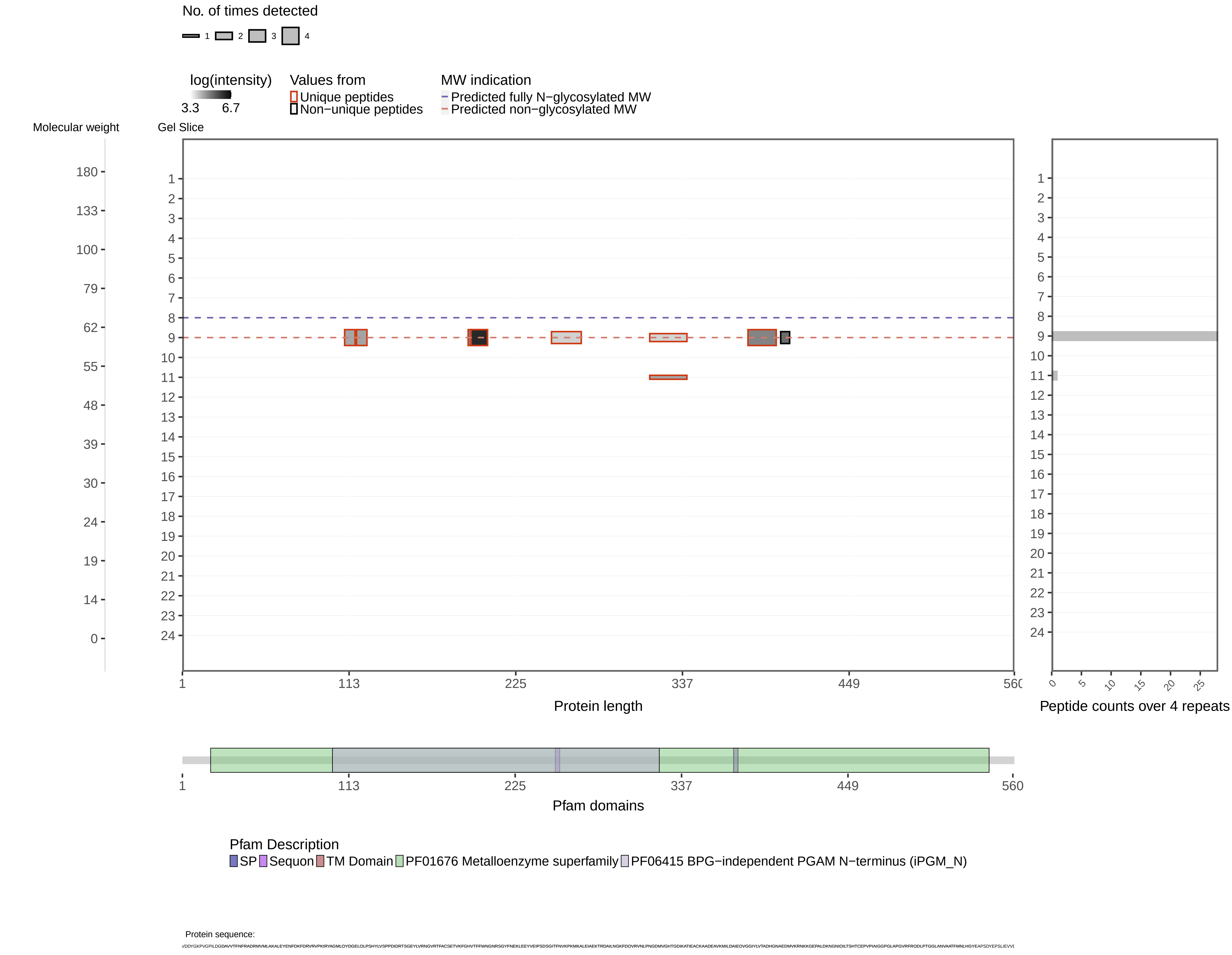

### NbL18g18460.1 enhancer_protein_2−1_chloroplastic−like [MW = 28.55 kDa]

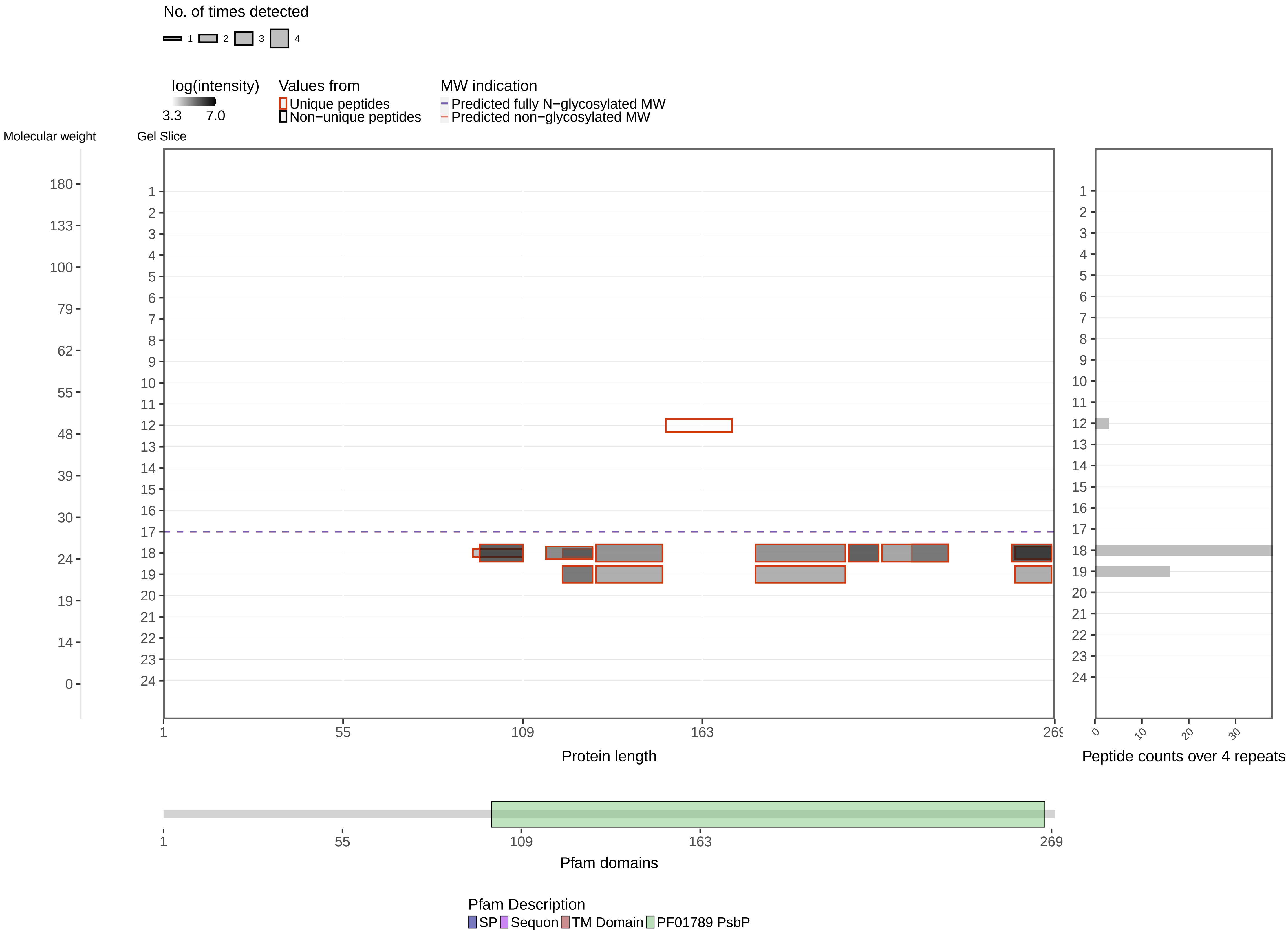


Protein sequence:

EVEFPGQVLRYEDNFDSTSNLIVTVTPTDKKSITGYGSPEEFLSQVDFLLGKQAYFGKTDSEGGFESGAVATANLLEASSTTVGGKEYYILSVLTRTADGDEGGKHQLISATVNDGKLYICKAQAGDKRWFKGARKFVENAATSFSVA*

NbL18g18270.1 uncharacterized_LOC107810187 [MW = 26.64 kDa]

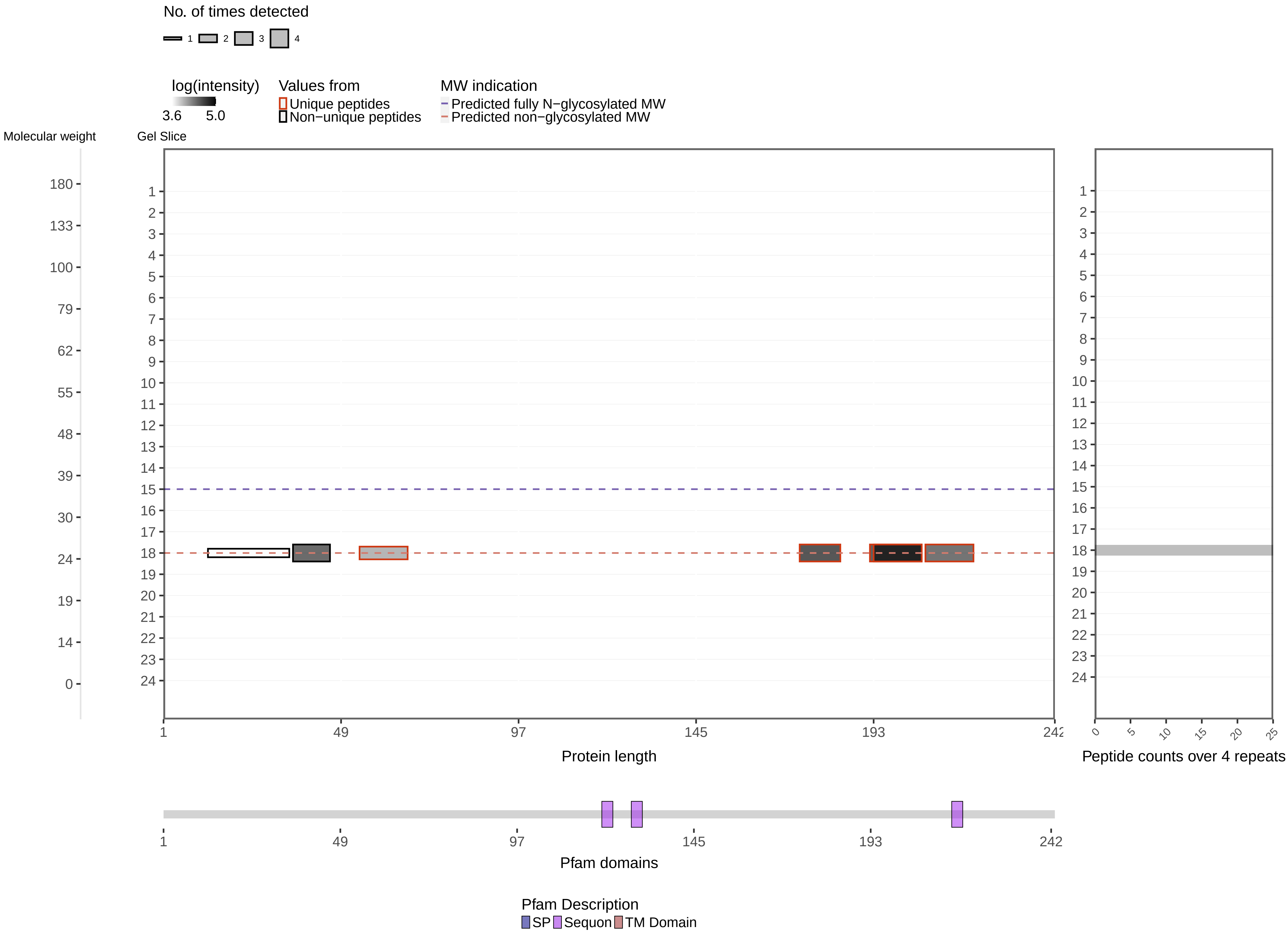


Protein sequence:
PFLVTDTKLKMENRTSVGVDNISVYGLQVGASSGLTKALVTNLQSSPPSLQIFHEQLACLLQLVRVLNLPTVVIIGKKGQKLHRKTSEDELEVIHEIGRHLASFSSLNFSAEKIAWDATKSSRETQEPWRALYG*

### NbL18g18210.1 uncharacterized_LOC104231634 [MW = 66.42 kDa]

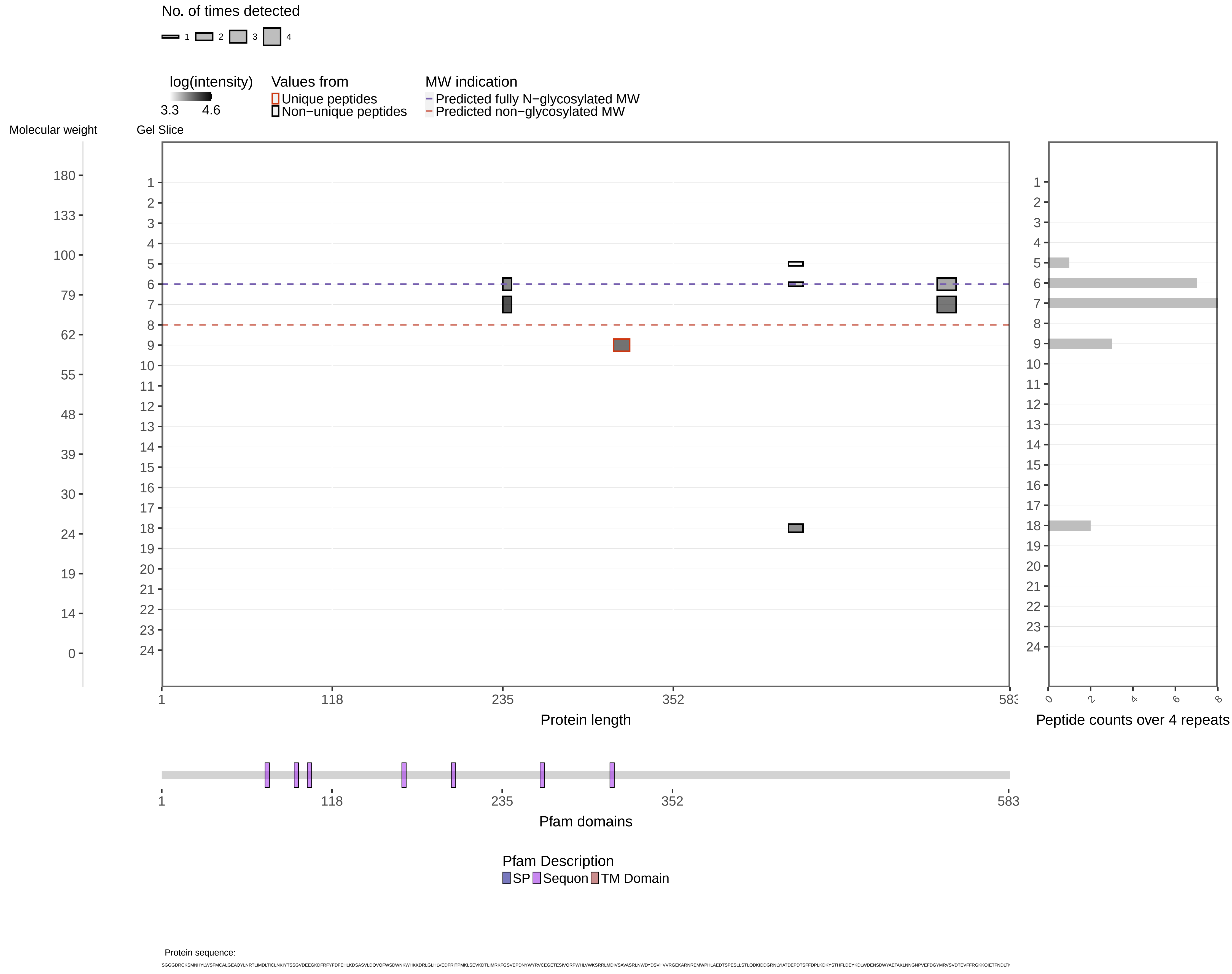

### NbL18g18180.1 heterogeneous_nuclear_ribonucleoprotein_1−like [MW = 35.55 kDa]

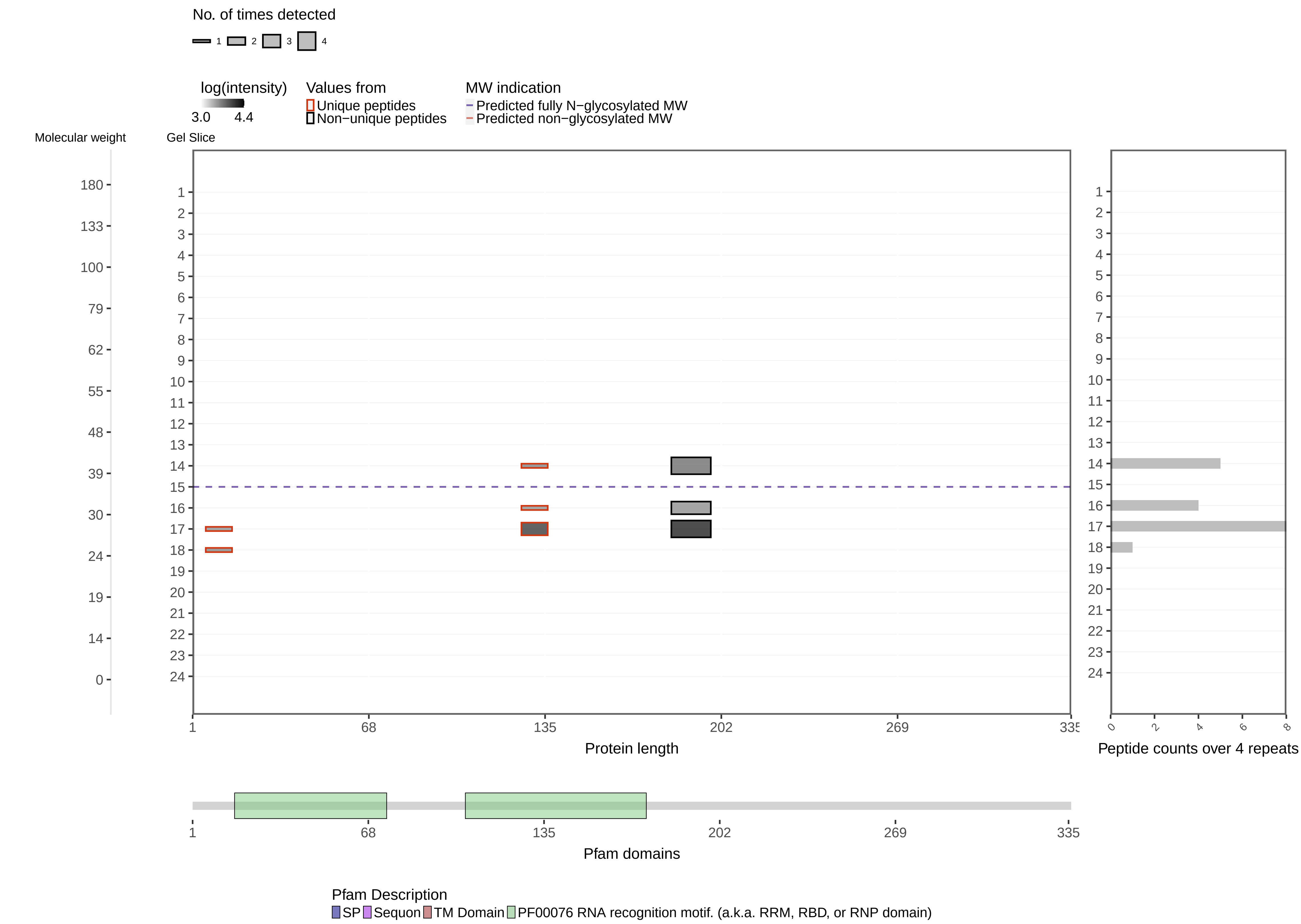


Protein sequence:

AEGNRIDMMGTQVEIKKAEPKKPSNPASAPAYGSDSRGRGYGDSYGGFGNSYSGFGSGGFGPASYRSFGARFGEYGYGGGEFGGRYGDFGGYRGDPSLGYSSRFGSYAAGFGGGYGGSGLGSYGRGGGYGGYGGVGPDASYESGPGAGYGGPGGLYGTRAGYSGSGRYHPYSR*

NbL18g18080.1 acetyl−CoA_acetyltransferase_cytosolic_1 [MW = 41.28 kDa]

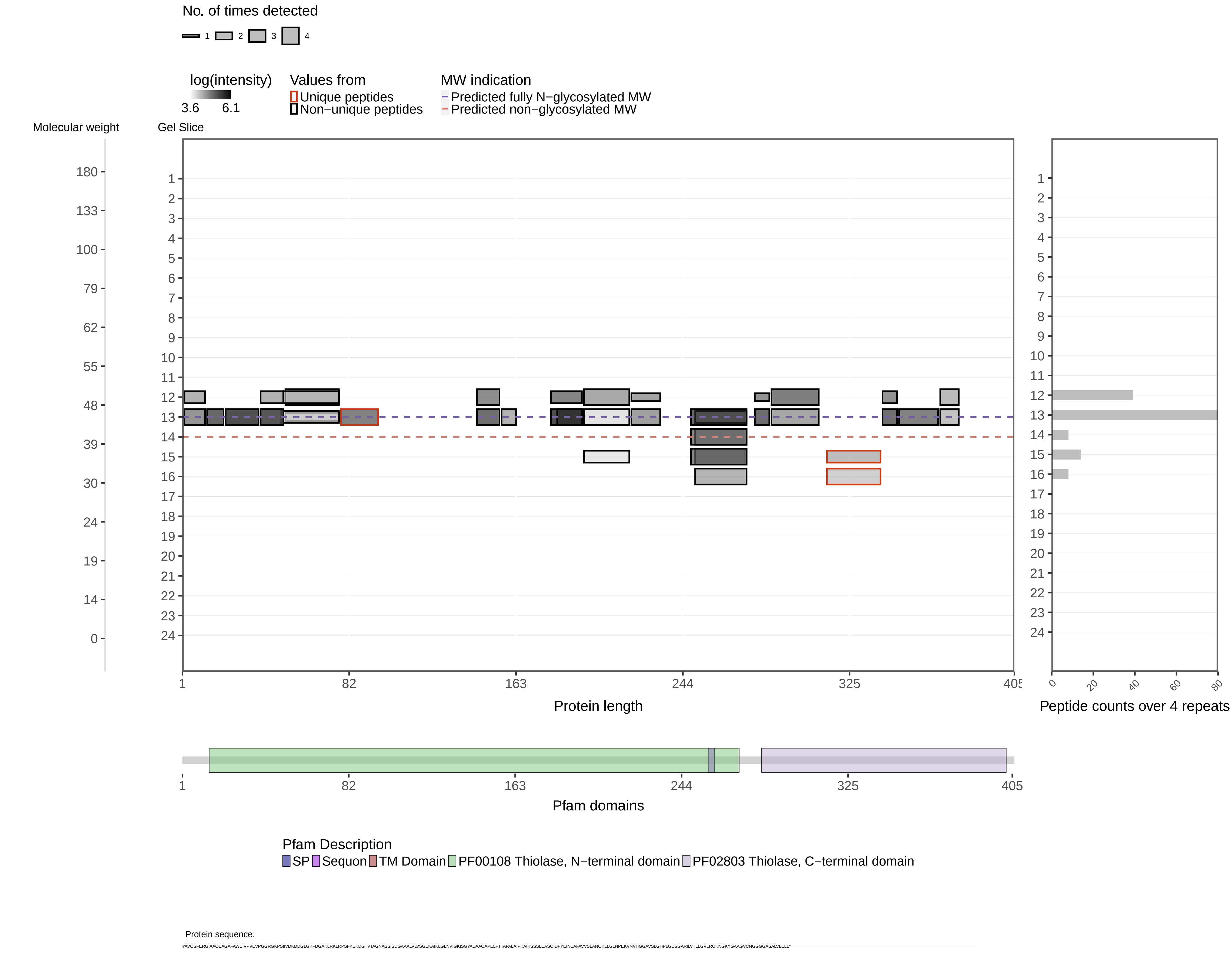

### NbL18g18030.1 fasciclin−like_arabinogalactan_protein_2 [MW − SP = 48.02 kDa]

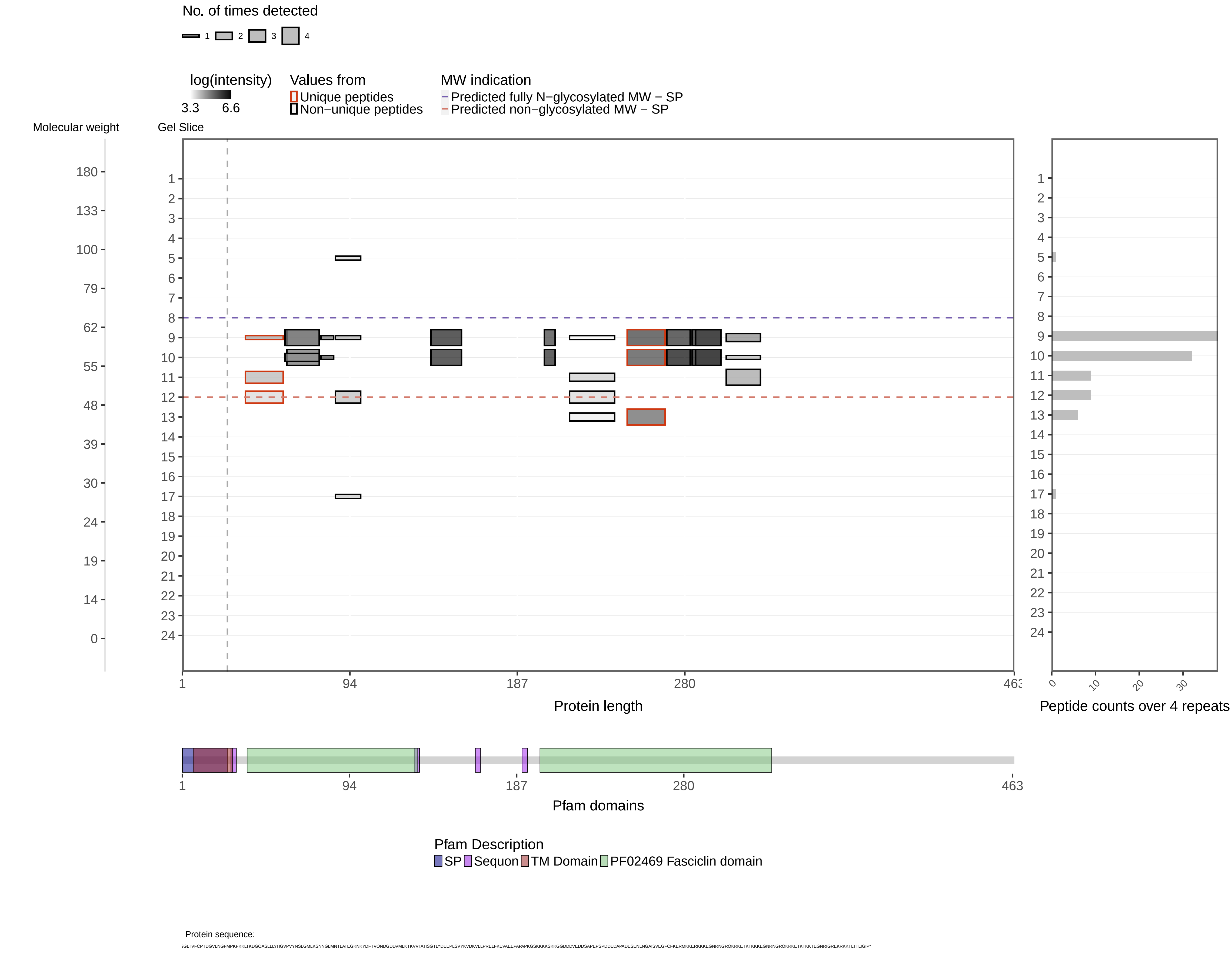

### NbL18g17790.1 levodione_reductase−like [MW = 29.73 kDa]

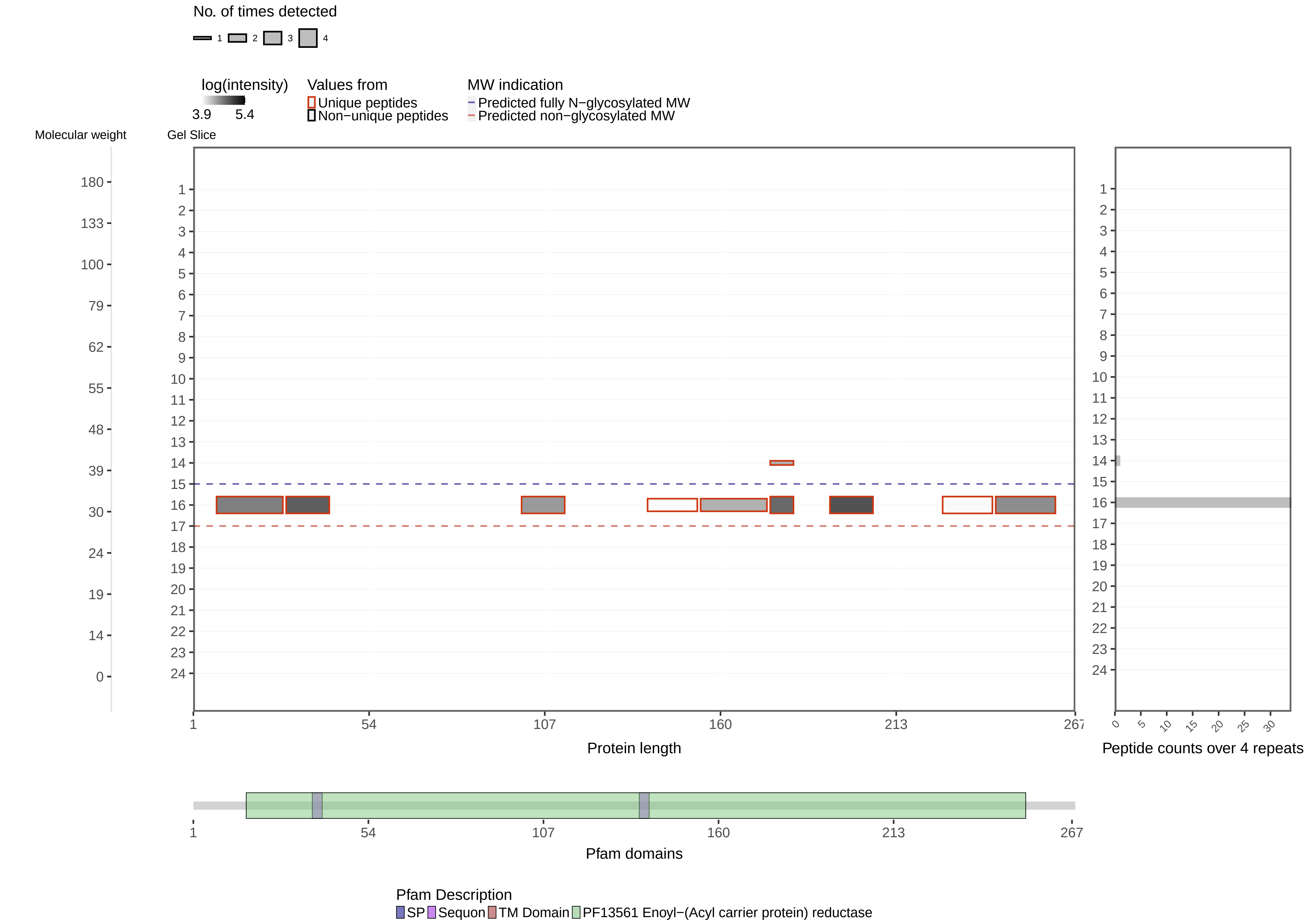


Protein sequence:

AGWYLLKCIGKRMRDNKSGGSIVFLTSIIGAERGIYQGAAAYGSCAAGMQQLVRLSALEMGKHQIRVNGISRGLHLEDEYPMSVGKERAEKLTKEAAPLNRWLDVRKDLASTVIYLISDDSRYMTGTSIFVDGAQSLVRPRMRSYM*

NbL18g17720.1 chlorophyll_a−b_binding_protein_37_chloroplastic−like [MW = 28.6 kDa]

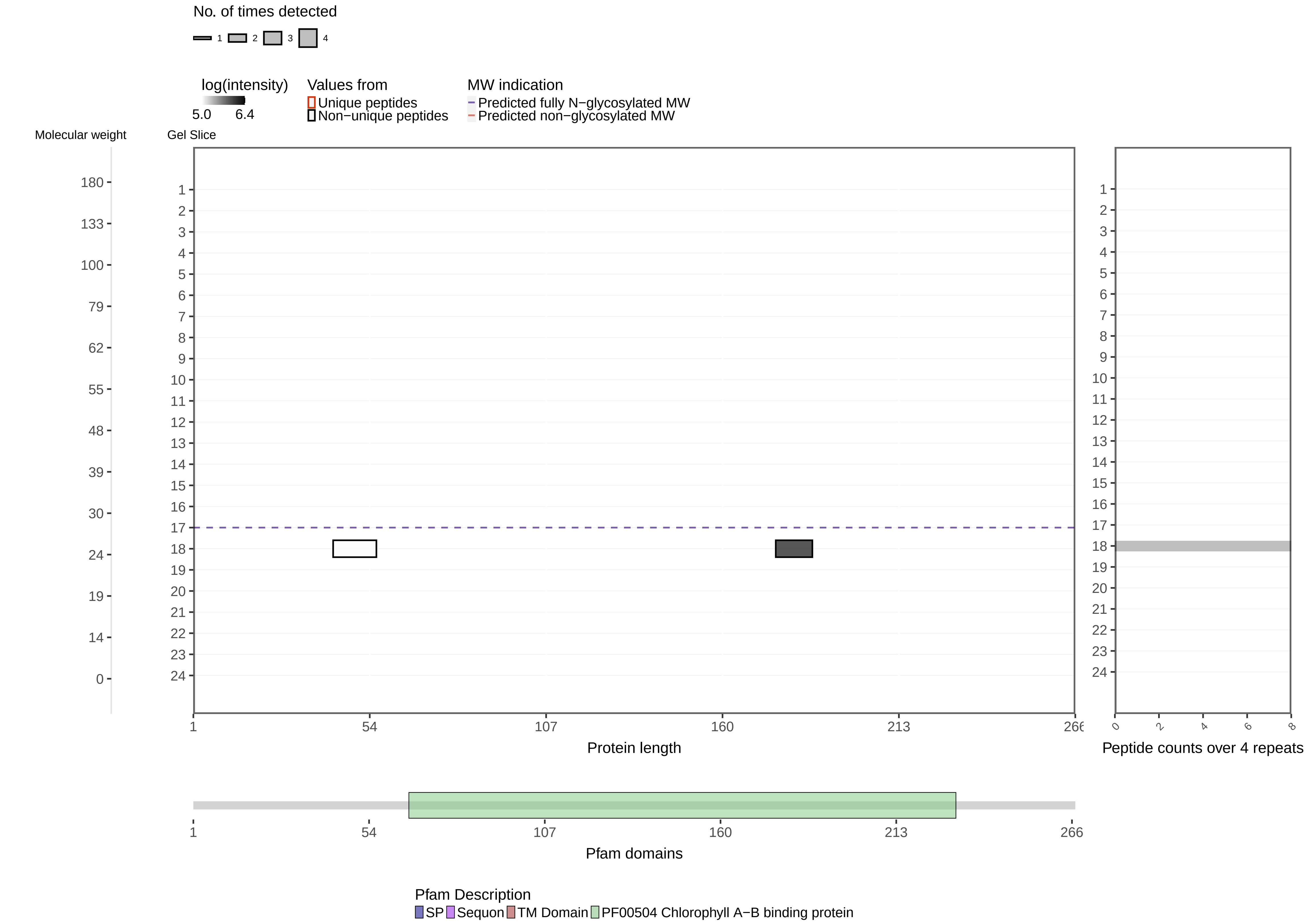


Protein sequence:

[illegible]KNGVKFGEAVWFKAGAQIFSEGGLDYLGNPNLVHAQSILAIWACQVVLVGLIEGYRVGGGPLGEGLDKIYPGGAFDPLGLADDPEAFAELKVKEIKNGRLAMFSMFGFFVQAIVTGKGPIENLFDHINDPVANNAWAFATNFVPGK*

NbL18g17540.1 cytochrome_b561_and_DOMON_domain−containing_protein_At5g47530−like [MW − SP = 41.28 kDa]

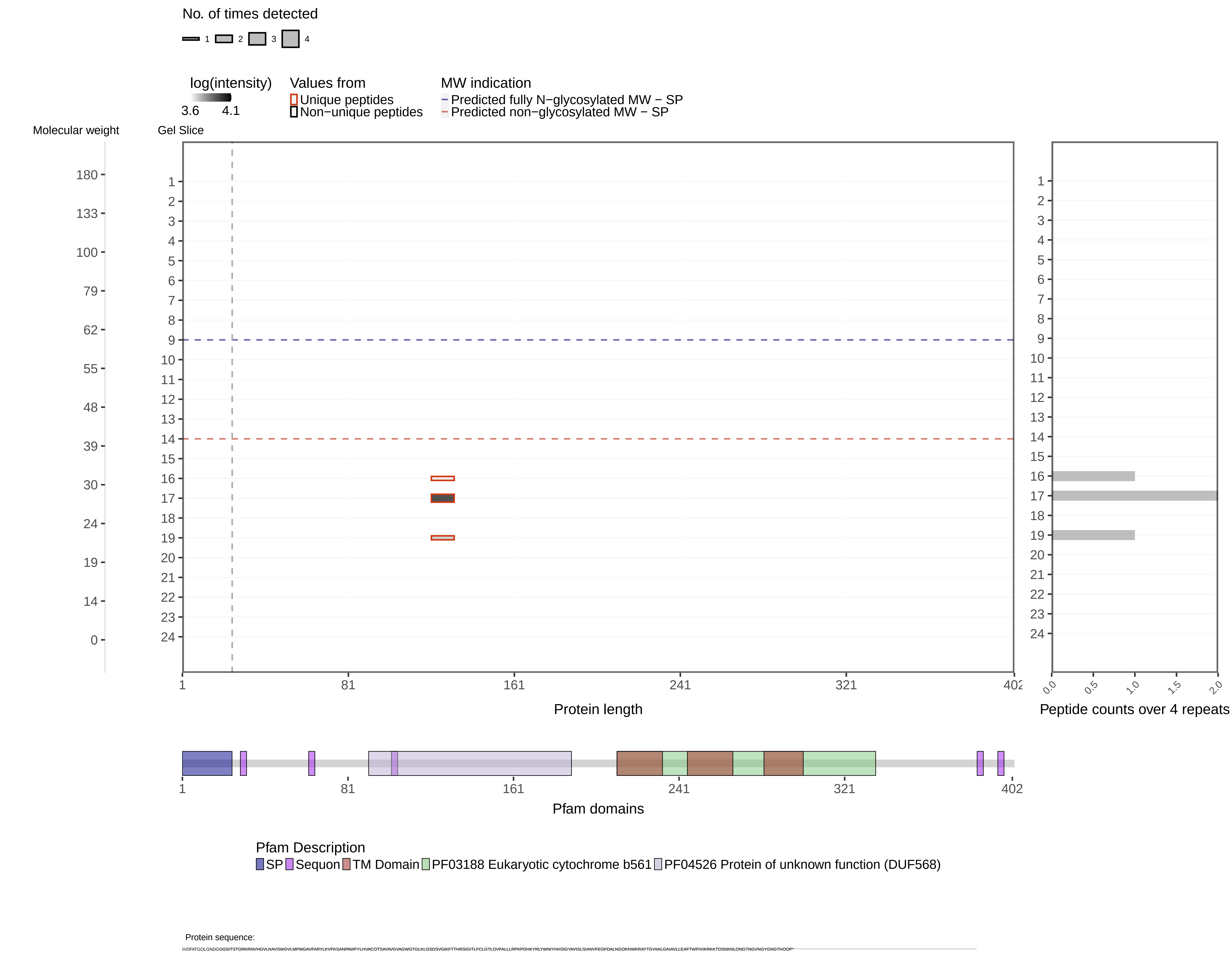

### NbL18g17530.1 threonine−−tRNA_ligase_chloroplastic/mitochondrial_2 [MW = 77.82 kDa]

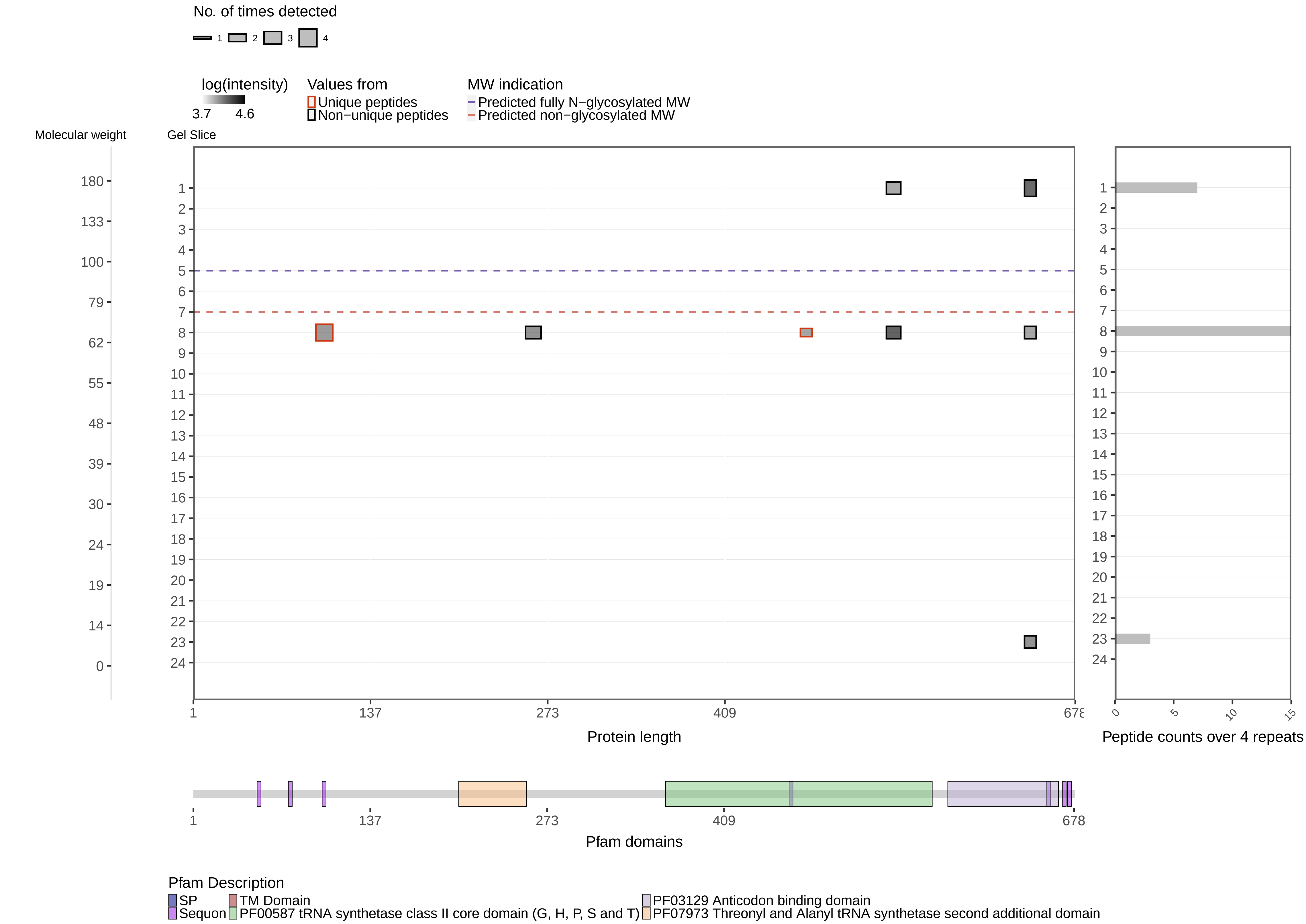


Protein sequence:

### NbL18g17520.1 fasciclin−like_arabinogalactan_protein_7 [MW − SP = 25.75 kDa]

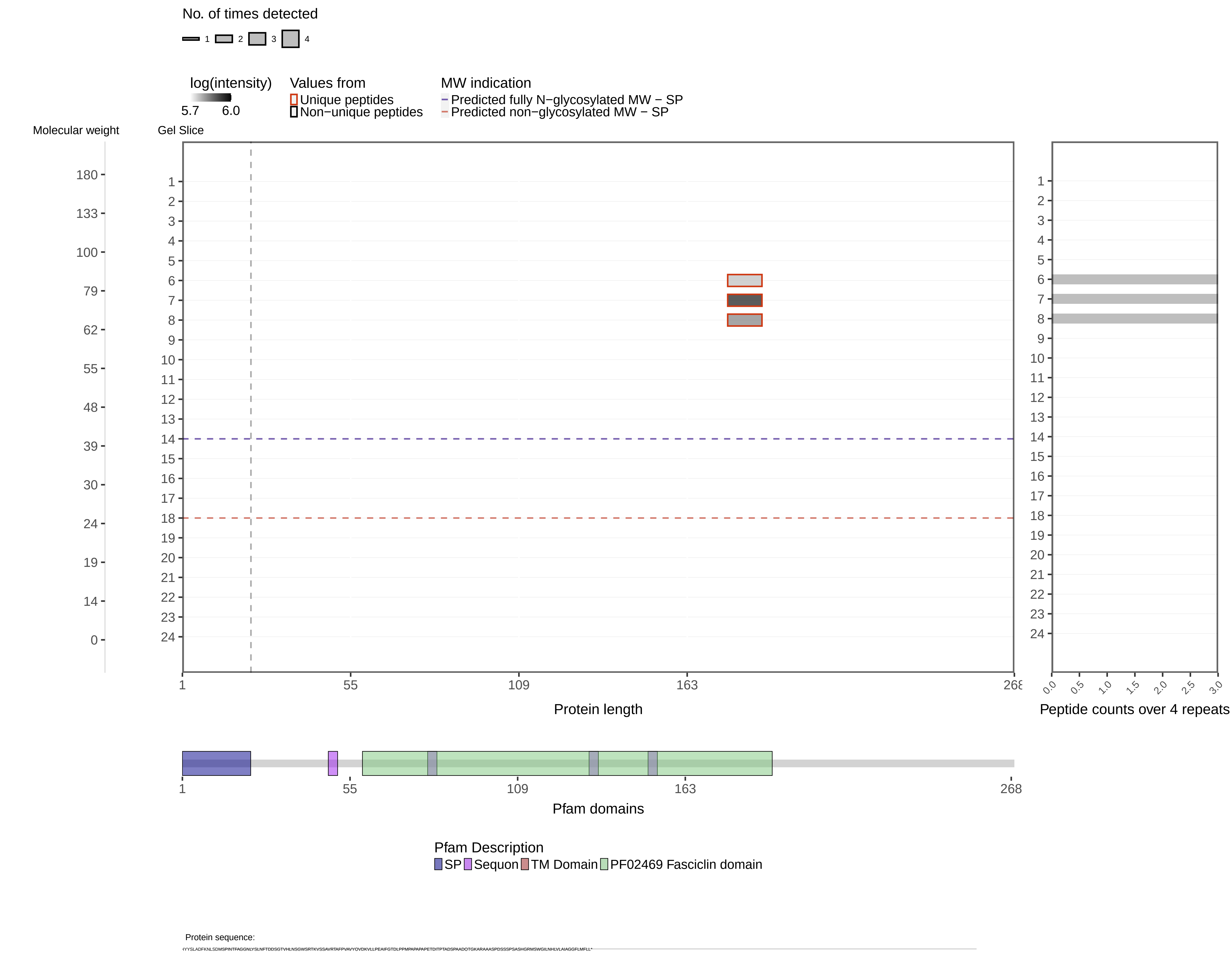

### NbL18g17470.1 chitin_elicitor_receptor_kinase_1−like [MW − SP = 65.34 kDa]

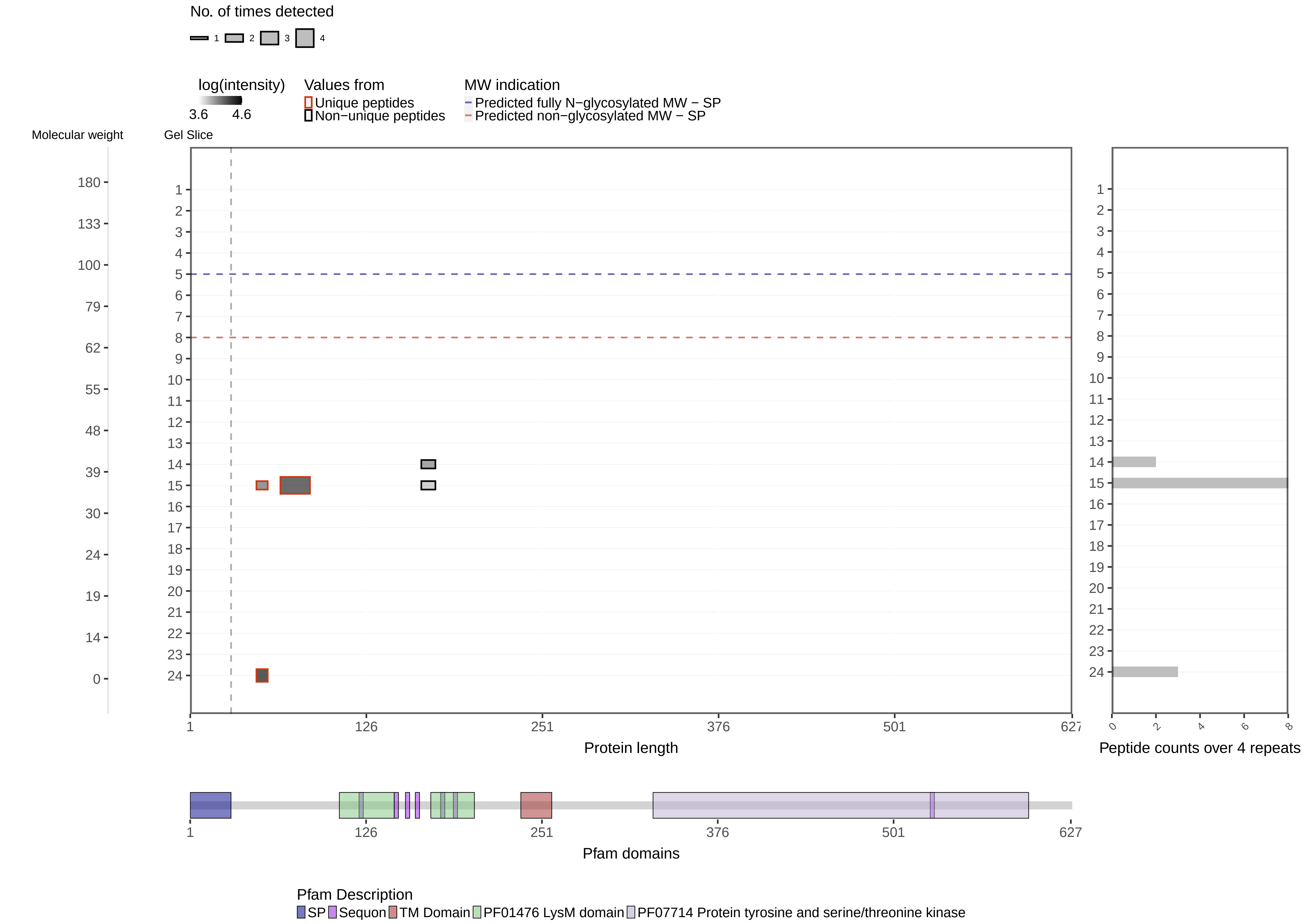


Protein sequence:

SPELSGITVDKSVEFTYEELATATNDFSIANKIGQGGFGAVYYAELRGEKAAIKKMDMEATREFLAELKVLTHVHHLNLVRLIGYCVEGSLFLVYEYVENGHIGQHLRGTGRDPLPWSTRVQIALDSARGLEYIHEHTVPVYIHRDIKTANILIDKNFHAKVADFGLTKLTEVGSSSLQTRLVGTFGYMPPEYAQYGDVSPKVDVYAFGVVLYELISAKEAIVKPNESVTESKGLVGLFEEVLNQPEPDEDLRKLVDPRLGNDYPLDSVRKMAQLAKACTHENPLIRPSMRSIVVALMT

### NbL18g17430.1 uncharacterized_LOC107779001 [MW = 16.98 kDa]

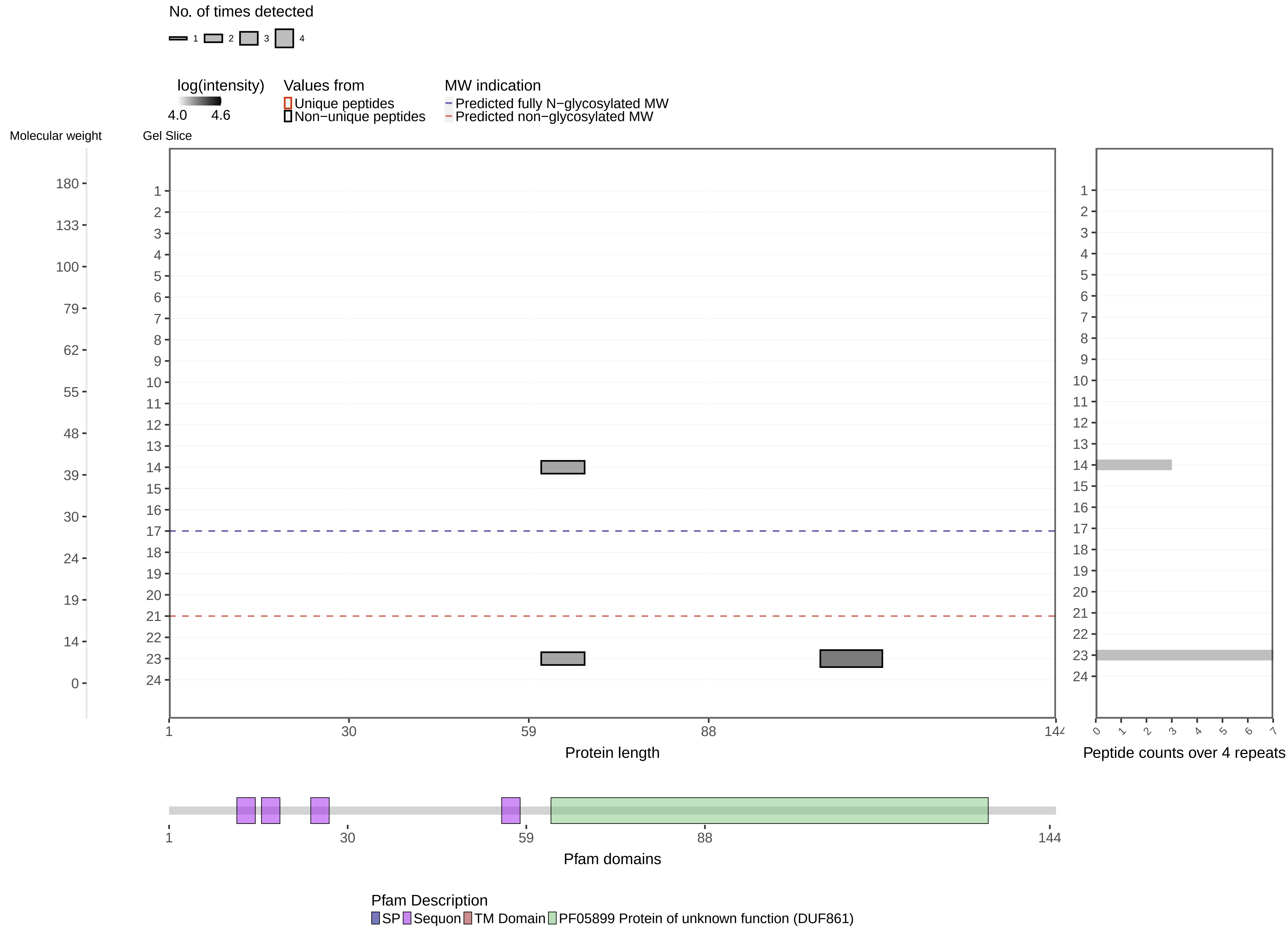


Protein sequence:

LMELGVPRWSMWKTGKCKLPWDWHVDQLVYIEEGEVRVVPEGSQRFMQFVAGDLVRYPKWFEADLYFNDFYQERYSFRAYGDS*

NbL18g17300.1 small_ubiquitin−related_modifier_2 [MW = 12.06 kDa]

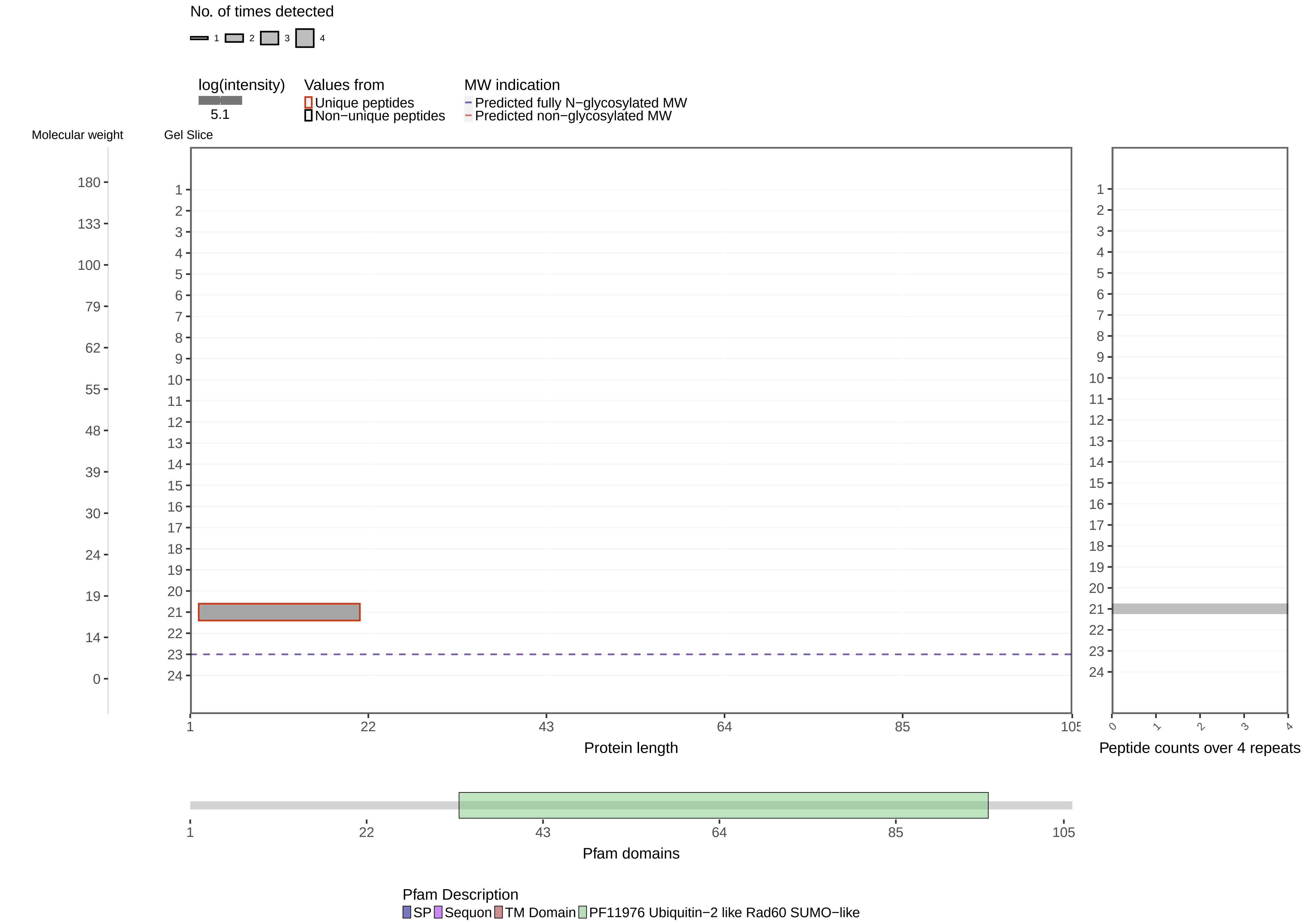


Protein sequence:

FRIKRSTQLKKLMNAYCDRQSVDFNSIAFLFDGRRLRAEQTPDELEMEDGDEIDAMLHQTGGSTV*

### NbL18g17280.1 nifU−like_protein_4_mitochondrial [MW = 30.8 kDa]

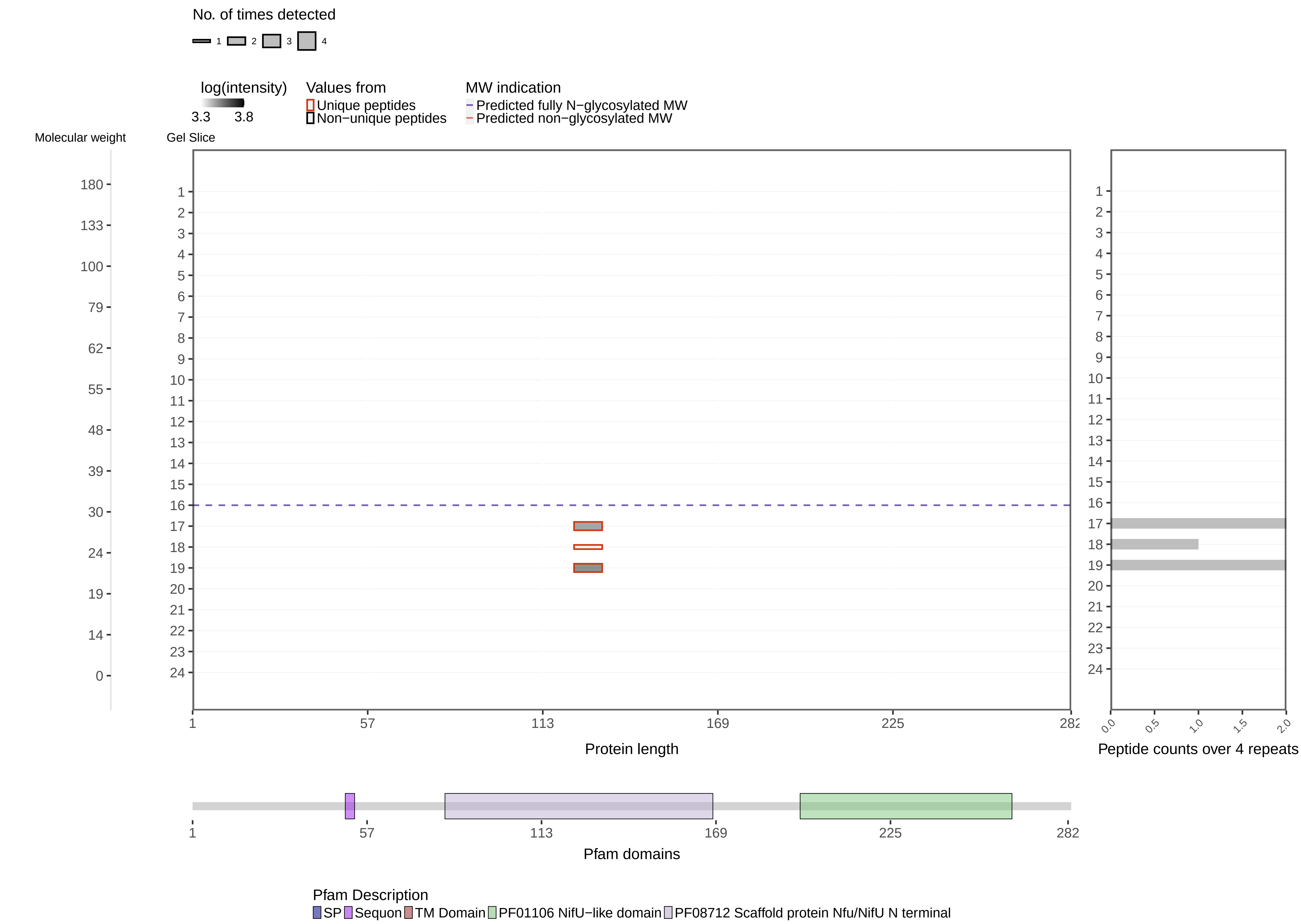


Protein sequence:

GISRVFFGSDFVTVTKSEDASWDFLKPEIFAAIMDFYSSGKPLFLDSSTAASMDTAIHEDDSETVAMIKELLETRIRPAVQDDGGDIVYRGFDLDRGIVKLKMQGACSGCPSSSVTLKSGIENMLMHYVPEVKGVEQEMDDDEEHGLTGVPYE*

### NbL18g17220.1 protein_disulfide_isomerase−like_2−3 [MW − SP = 44.7 kDa]

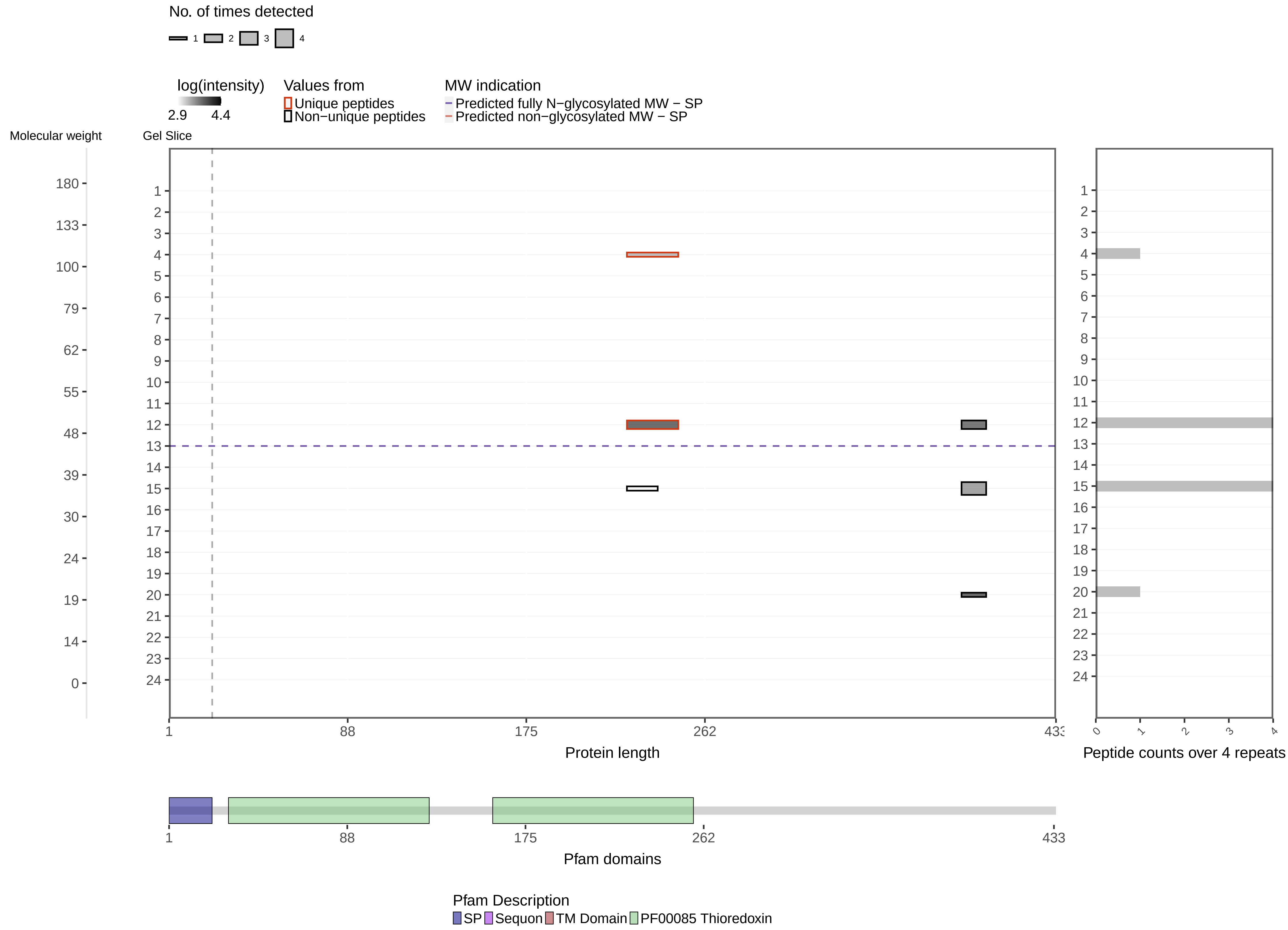


Protein sequence:

GKVKLGHVDCDAEKSLMSRYNVQGFPTILVFGADKESPVPYEGARTAAAIESFGIEQLETNVAPPEVIELTSPDVMEEKCNSAAICFVSFLPDILDSKAEGRNKYLEMLLSVAEKFKRNPYSYVWVAAGNQPDLEKHVGVGGYGYPAMVALNVKKGVYAPLKSAFQRQPIIDFVKEAGLGGKGNLPLAATPSIVKTEAWDGKDGEIIEEDEFSLEELMGDDTRNKDEL*

NbL18g17210.1 GDSL_esterase/lipase_At4g26790−like [MW − SP = 36.16 kDa]

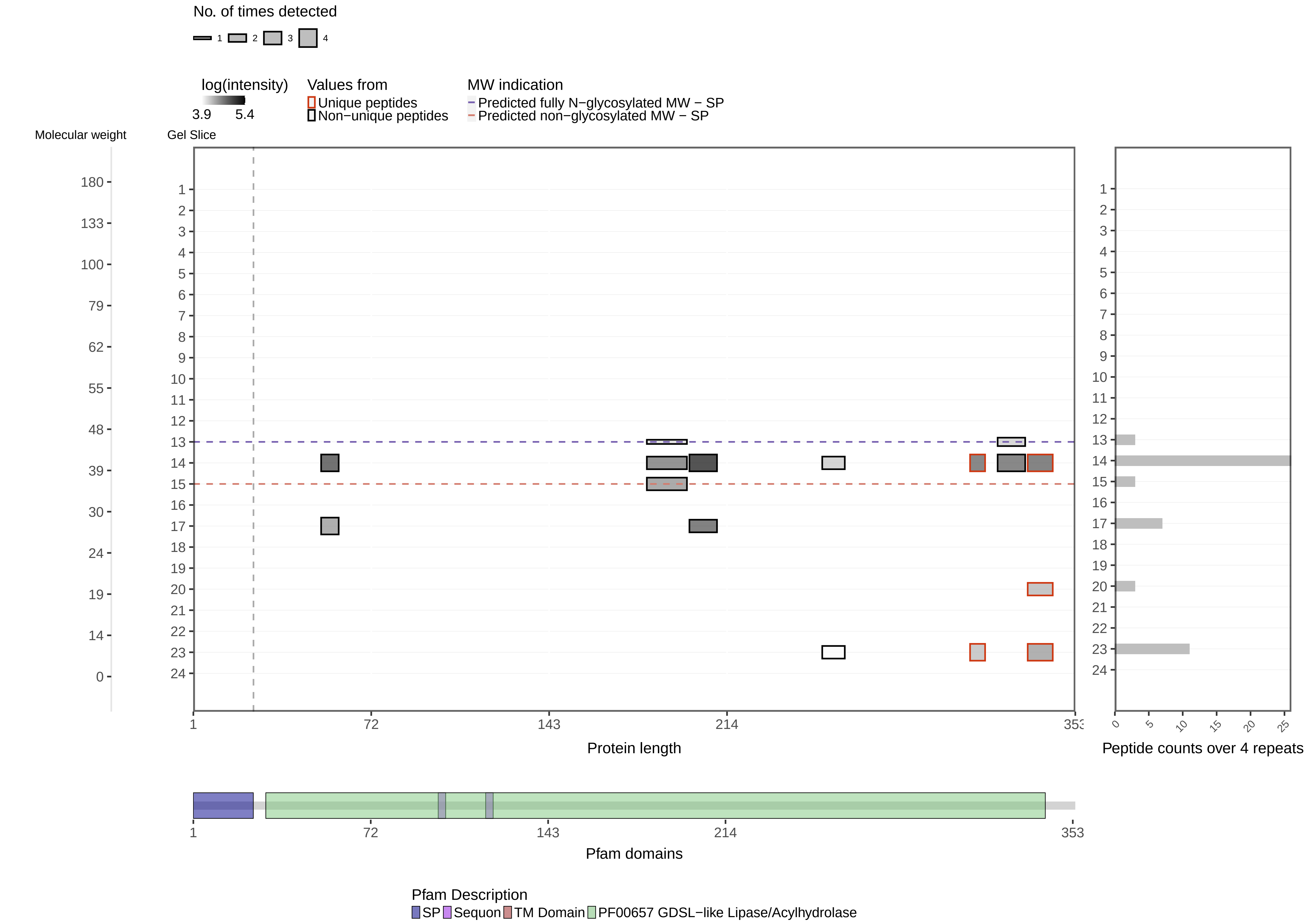


Protein sequence:

GTNDFLENYYTMPORRSOYTVDOYOTFLIGIAKNFVTNLYNLGARKISLGGLPPMGCMPLERTSNVANGNECLESYNVVAMNFNDKLSGLVIELNKELSGIOVVLSNPYNIMLOMIKNPSSYGFEVASLACCATGLFEMGYACDRYNLFTCKDANKYIFWDAFHPTEKTDGIIADHVVKTALSKFLA*

### NbL18g17030.1 1−aminocyclopropane−1−carboxylate_oxidase [MW = 35.58 kDa]

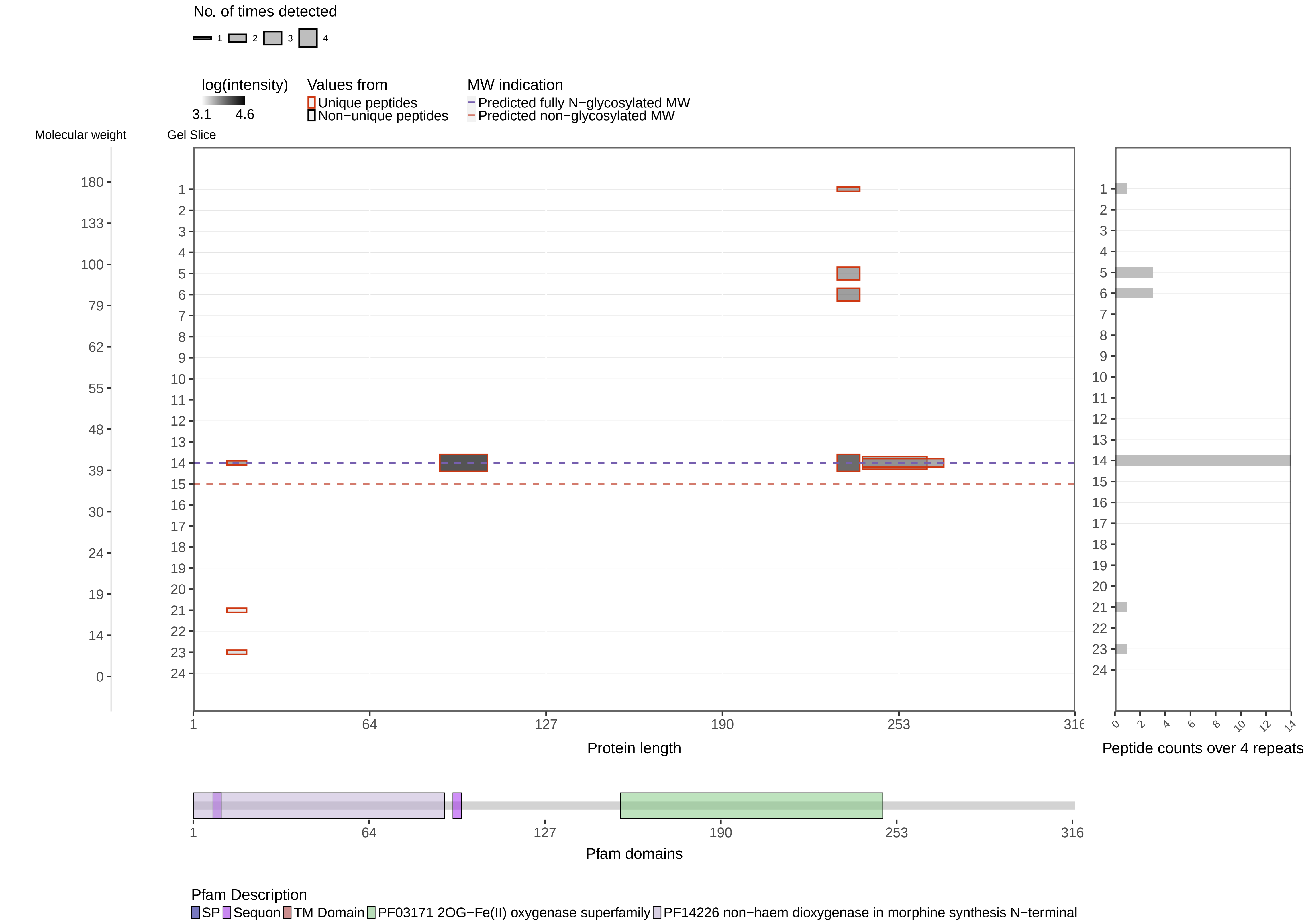


Protein sequence:

[illegible]TKGPNFGTKVSNYPPCPKPDLIKGLRAHTDAGGIILLFQDDKVSGLQLLKDGQWIDVPPMRHSIVVNLGDQLEVITNGKYKSVMHRVITQTDGTRMSLASFYNPGGDAVIFPAPALVEKDAEESKAVYPKFVFDDYMKLYAGLKFQAKEPRFEAMIKAMETGVKCDPIATA*

### NbL18g17000.1 uncharacterized_LOC107024435 [MW − SP = 10.94 kDa]

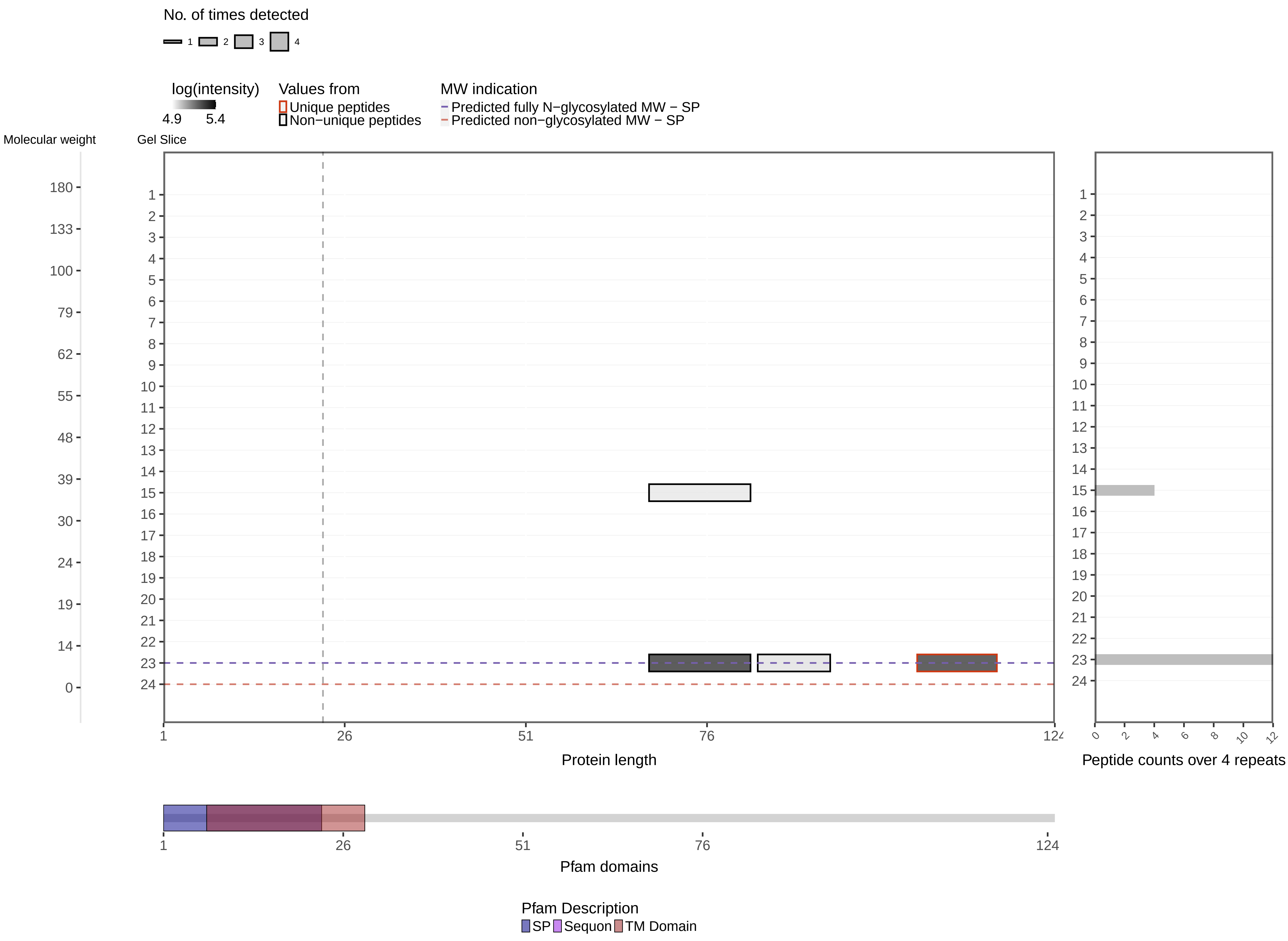


Protein sequence:

FPTEESPLVSVDLCCELVKSVTSVSESEQHYDLKECLCTAATINRIVVGRDRRVSKVIGSCTGETTTRYRCFSDI*

### NbL18g16990.1 benzyl_alcohol_O−benzoyltransferase−like [MW = 50.78 kDa]

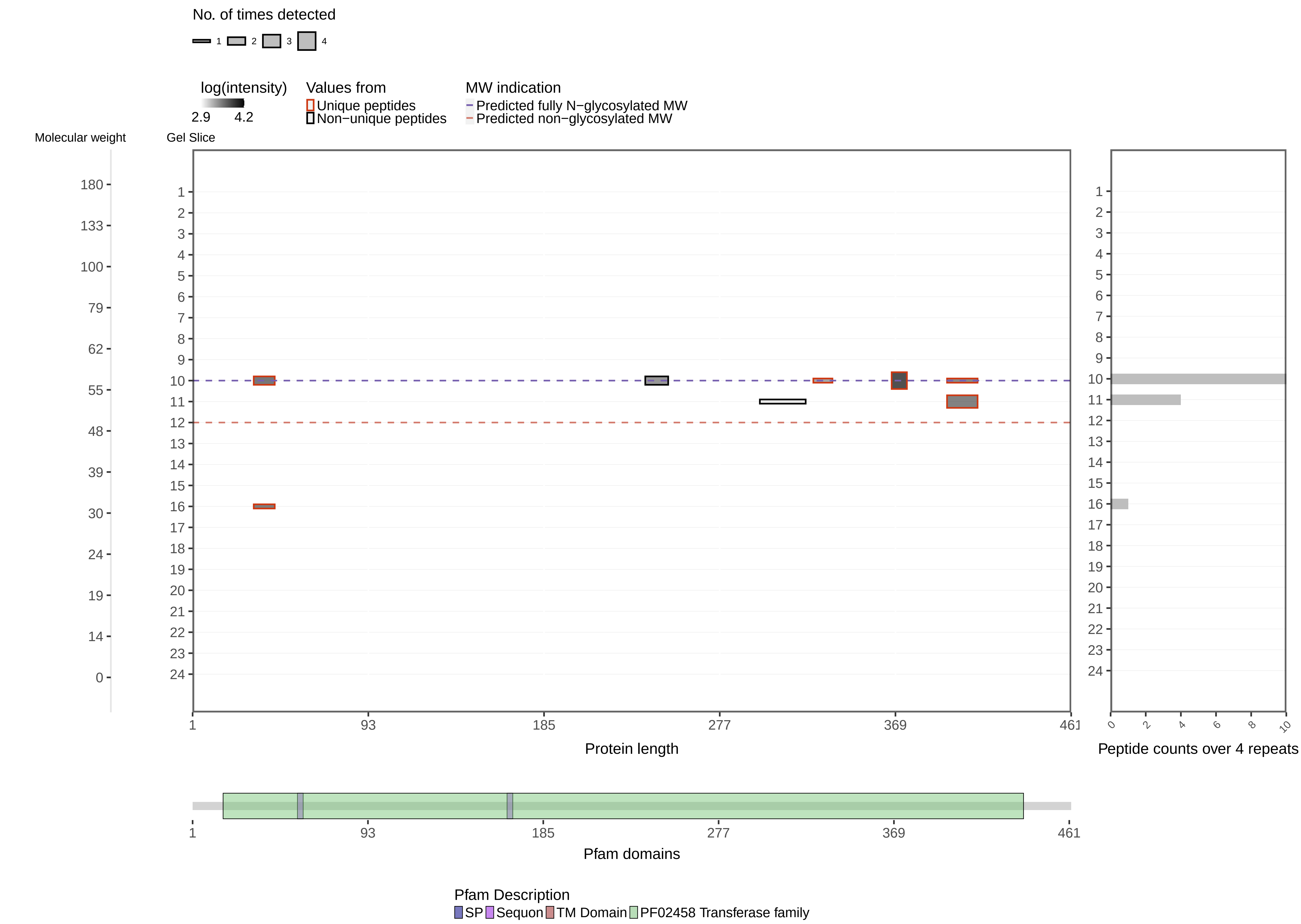


Protein sequence:

### NbL18g16950.1 probable_lipid−A−disaccharide_synthase_mitochondrial [MW = 51.85 kDa]

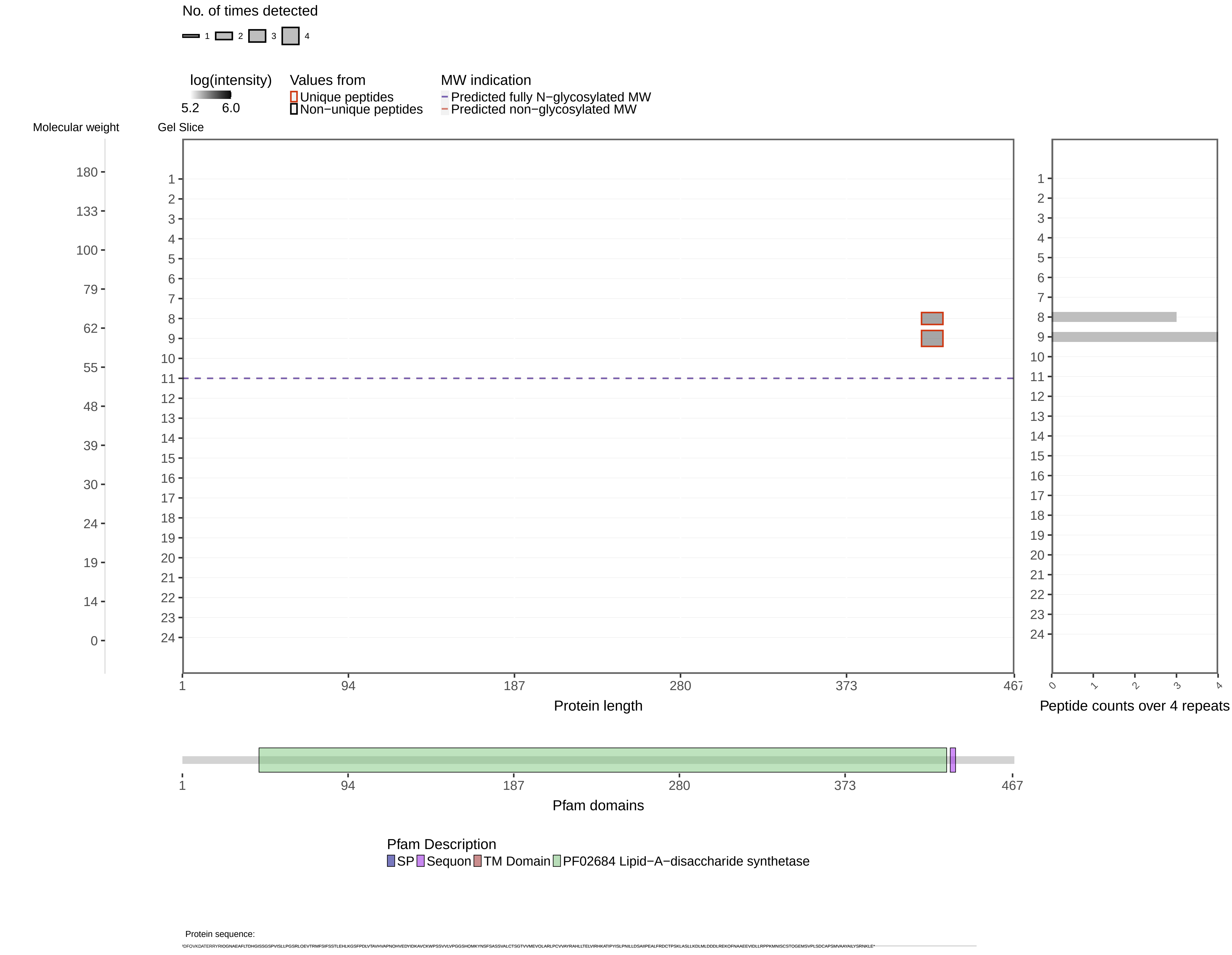

### NbL18g16710.1 universal_stress_protein_PHOS32−like [MW = 23.77 kDa]

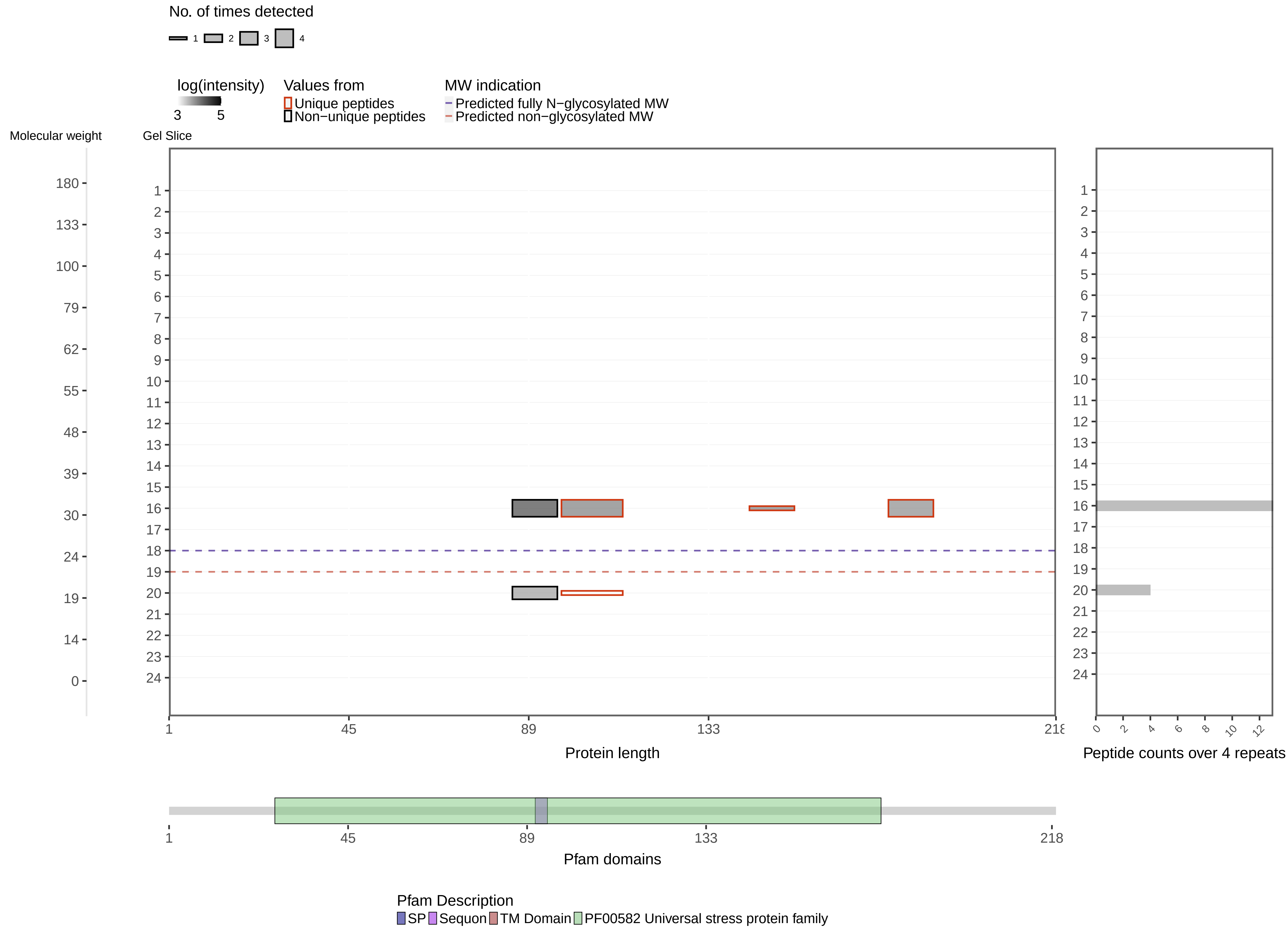


Protein sequence:

ATQLALPLVEANVPYKIHIVKDRDMKERLCLEVEGLGLSAMIMGSRGFGATSSGFISKGKLGSVSDYCVHHCICPVVVVRYPQDDGDGGVEKKAVVEAELPPVPEEEVEYHHDDDELTKGG*

### NbL18g16280.1 aconitate_hydratase_cytoplasmic [MW = 108.85 kDa]

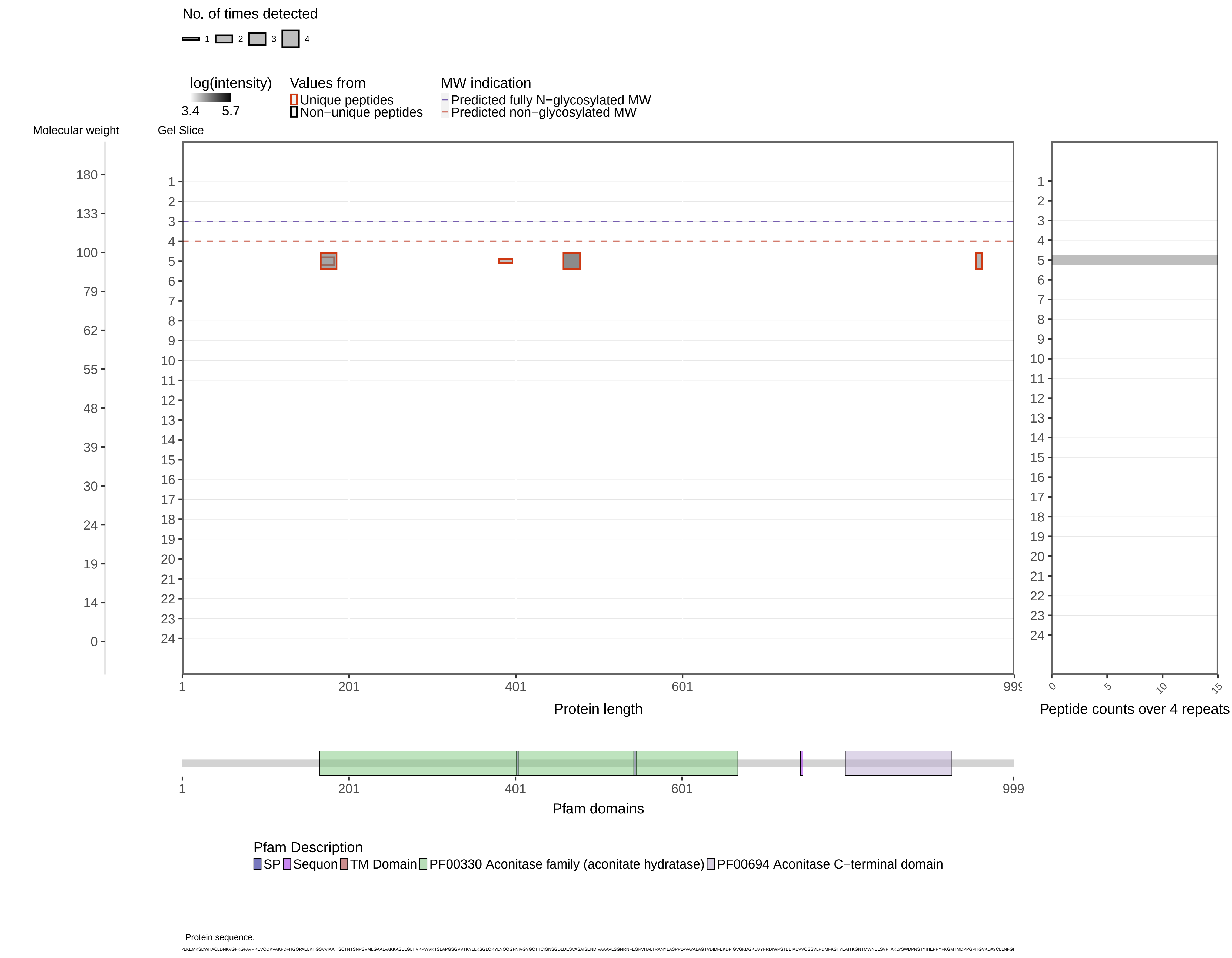

### NbL18g16090.1 cyprosin−like [MW − SP = 53.42 kDa]

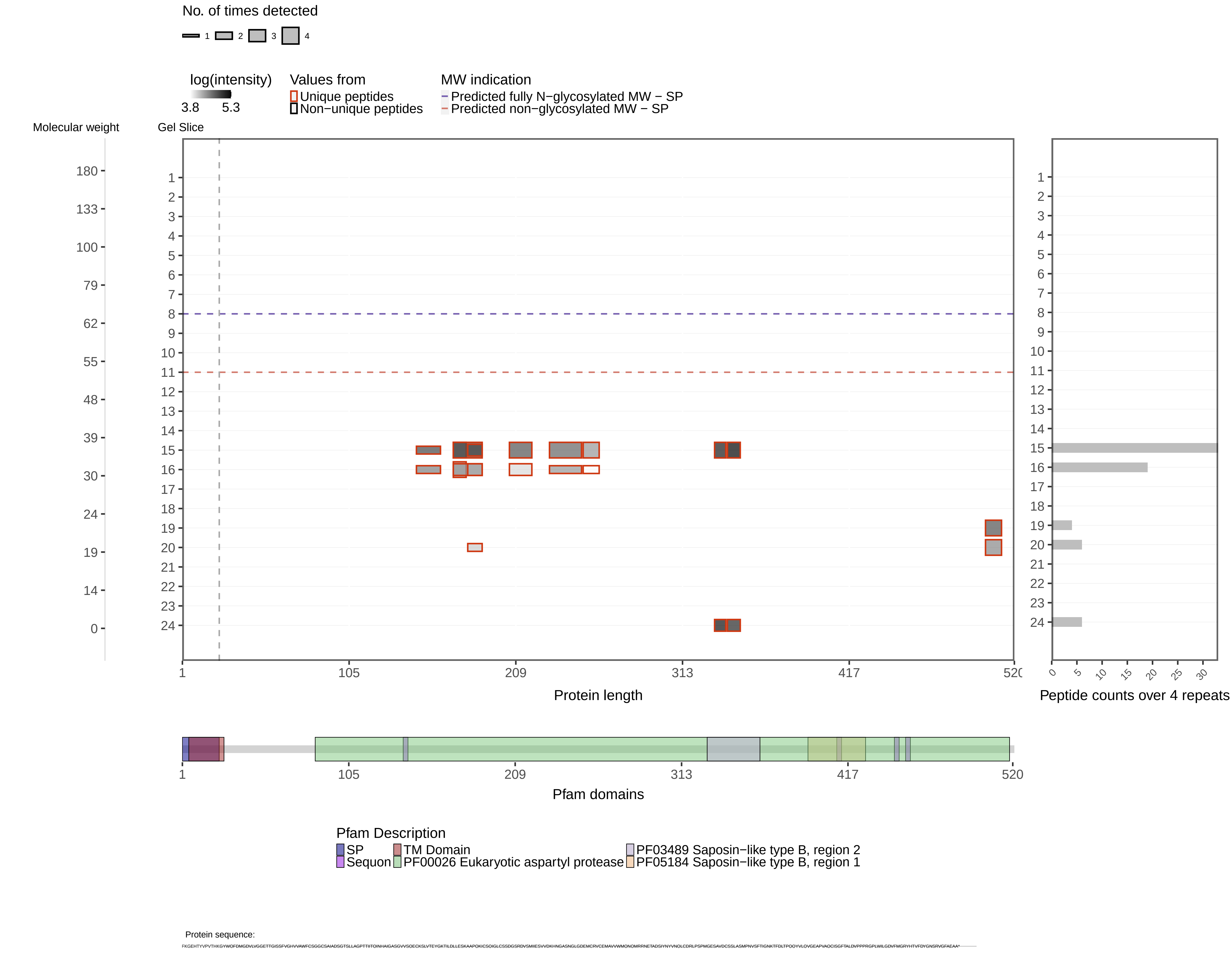

### NbL18g15780.1 lipoyl_synthase_mitochondrial [MW = 47.75 kDa]

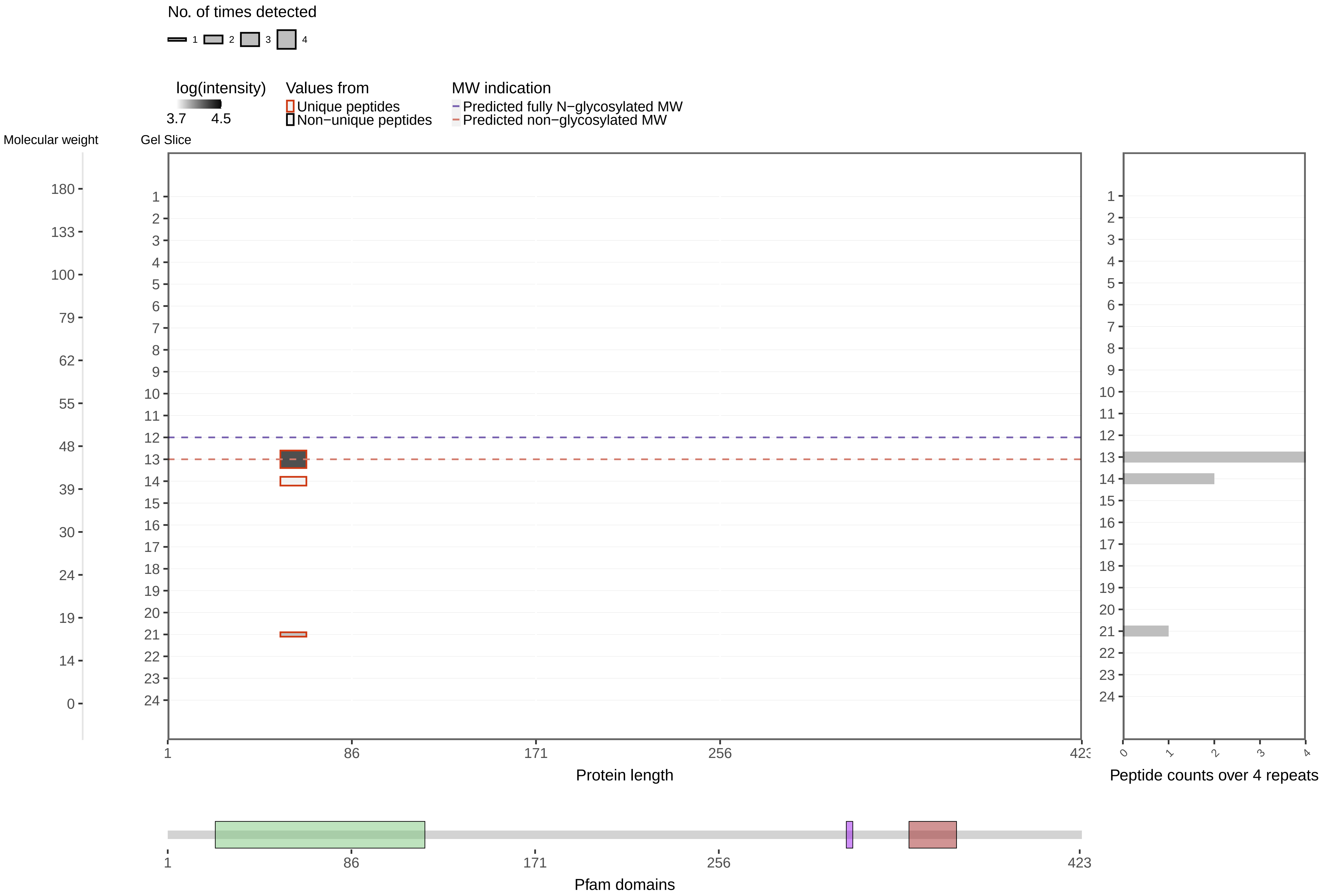


Pfam Description
SP Sequon TM Domain PF16881 N−terminal domain of lipoyl synthase of Radical_SAM family

Protein sequence:

### NbL18g15600.1 polyadenylate−binding_protein_2−like [MW = 71.84 kDa]

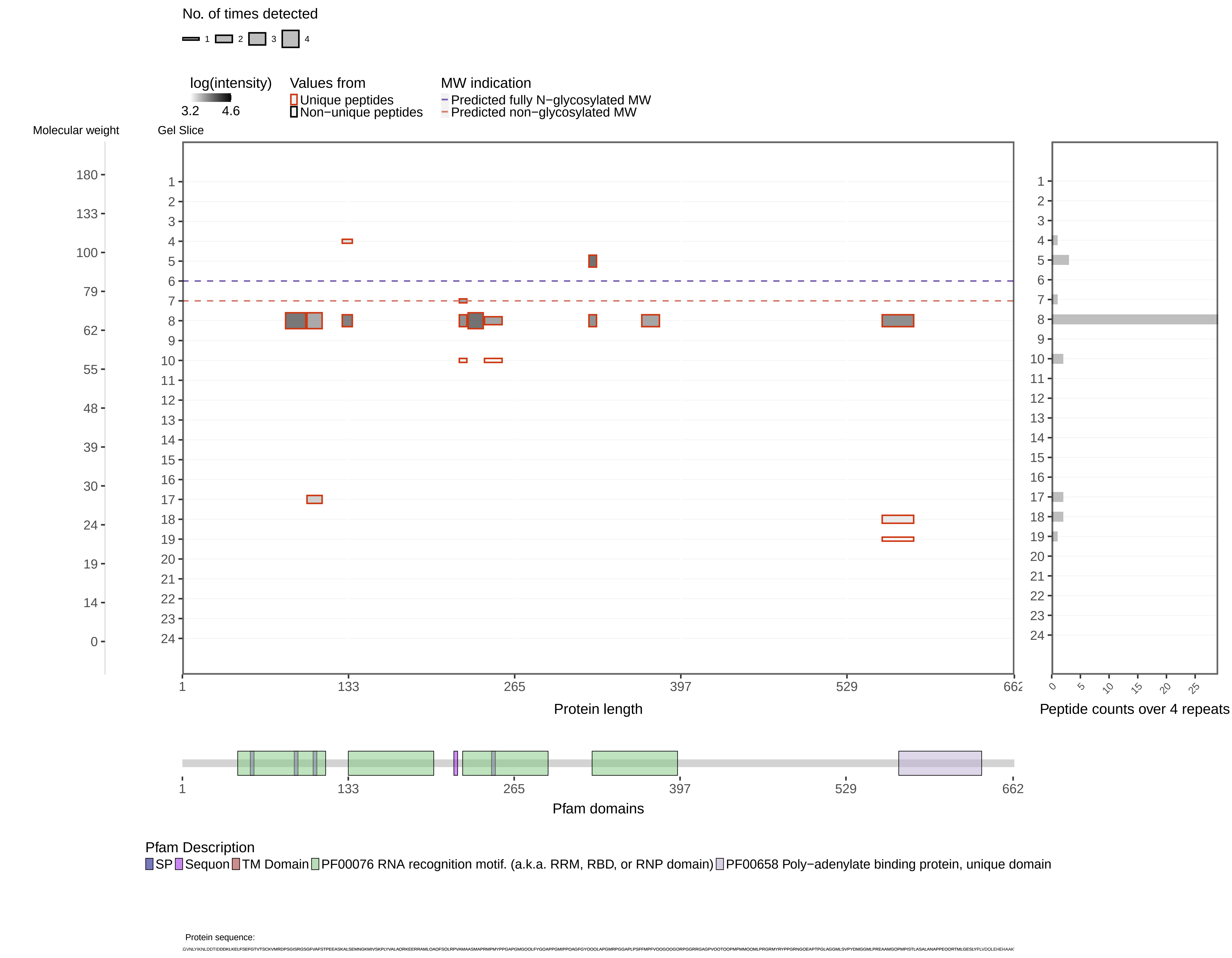

NbL18g15550.1 subtilisin−like_protease [MW = 87.81 kDa]

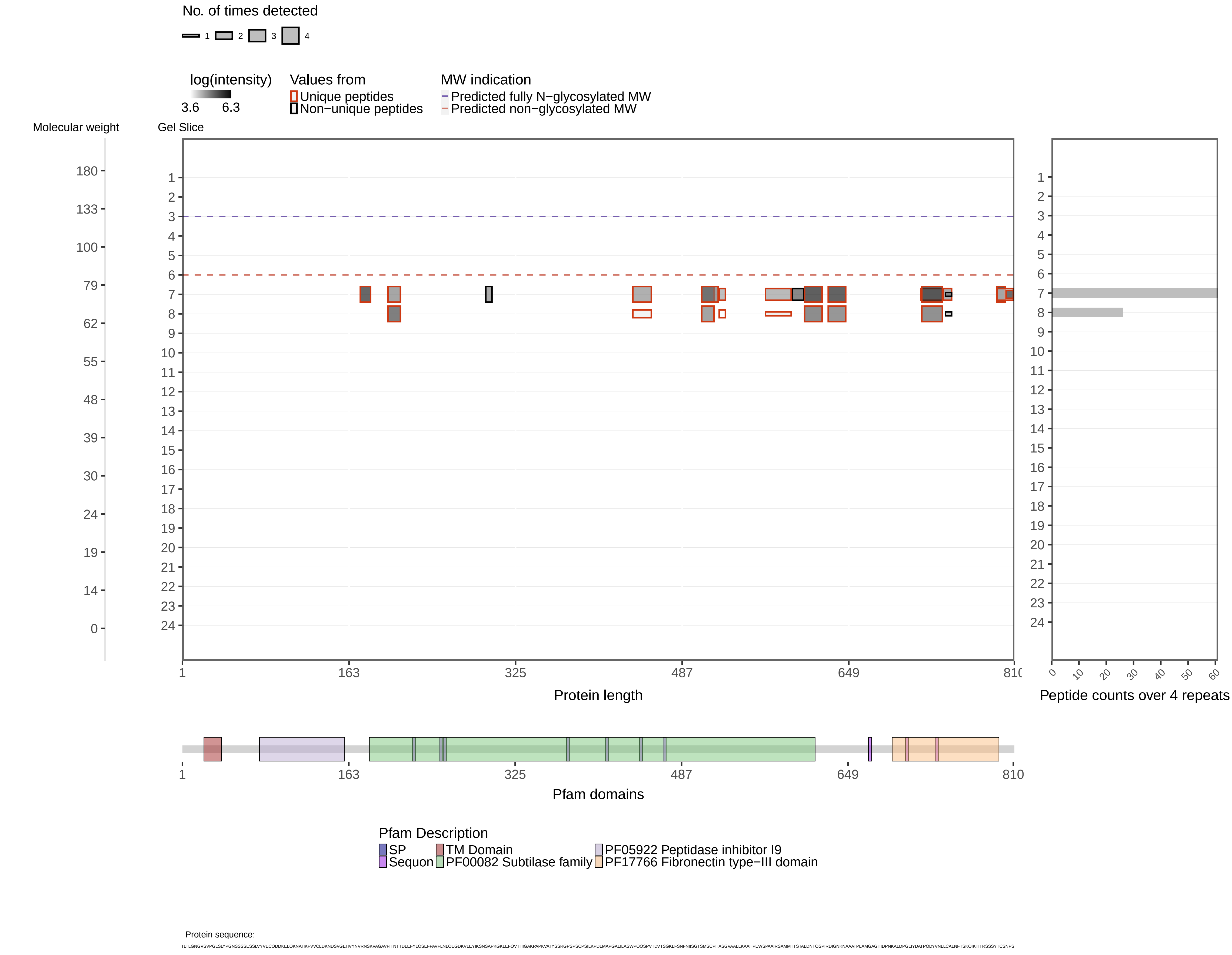

### NbL18g15390.1 snakin−1 [MW − SP = 7.07 kDa]

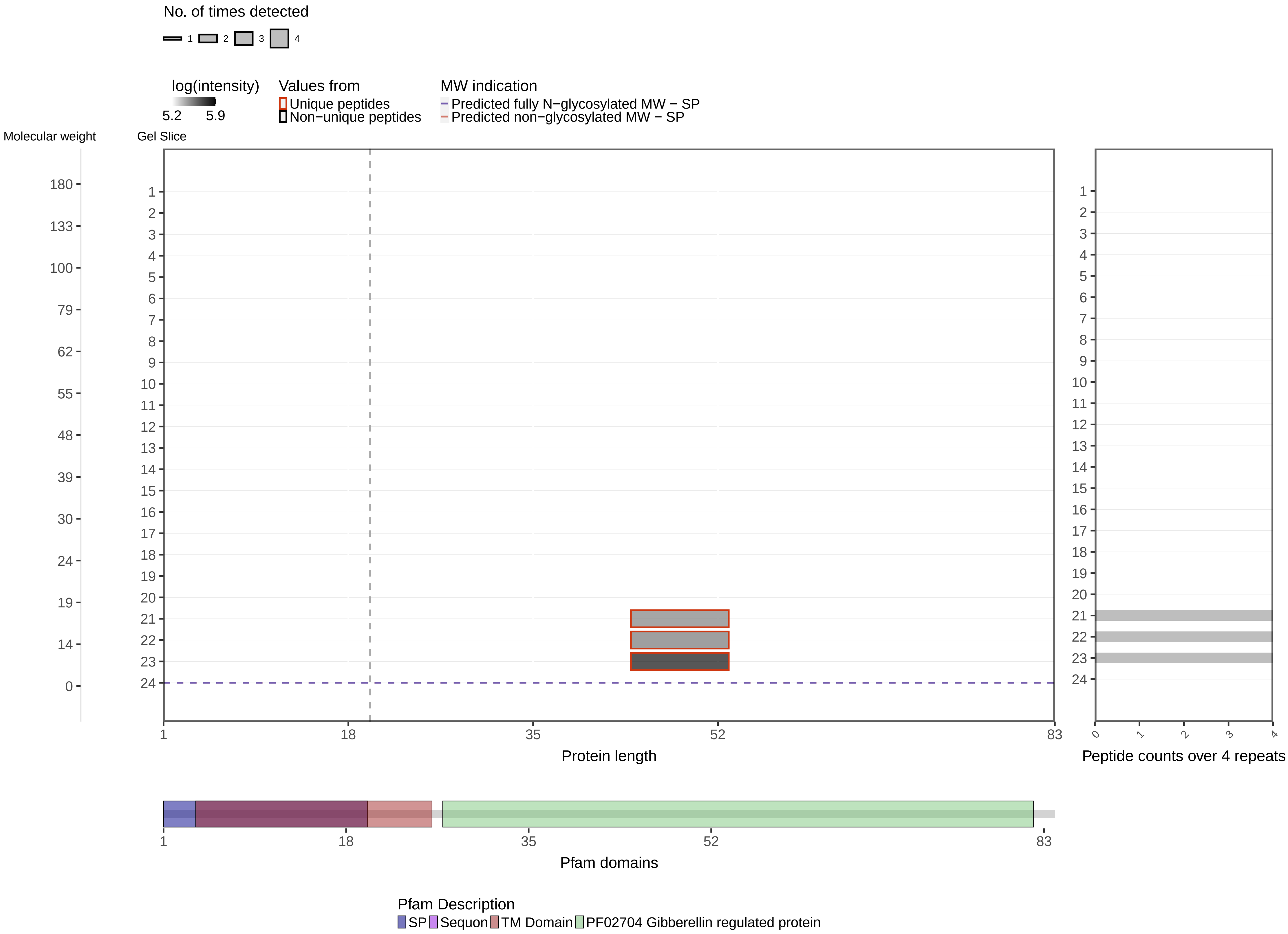


Protein sequence:

RCAKAGVMERCLKYCGICCEECKCVPCGTYGNNHECPRYRDKKNNKGNPKCP*

### NbL18g15140.1 adenine_phosphoribosyltransferase_1−like [MW = 30.08 kDa]

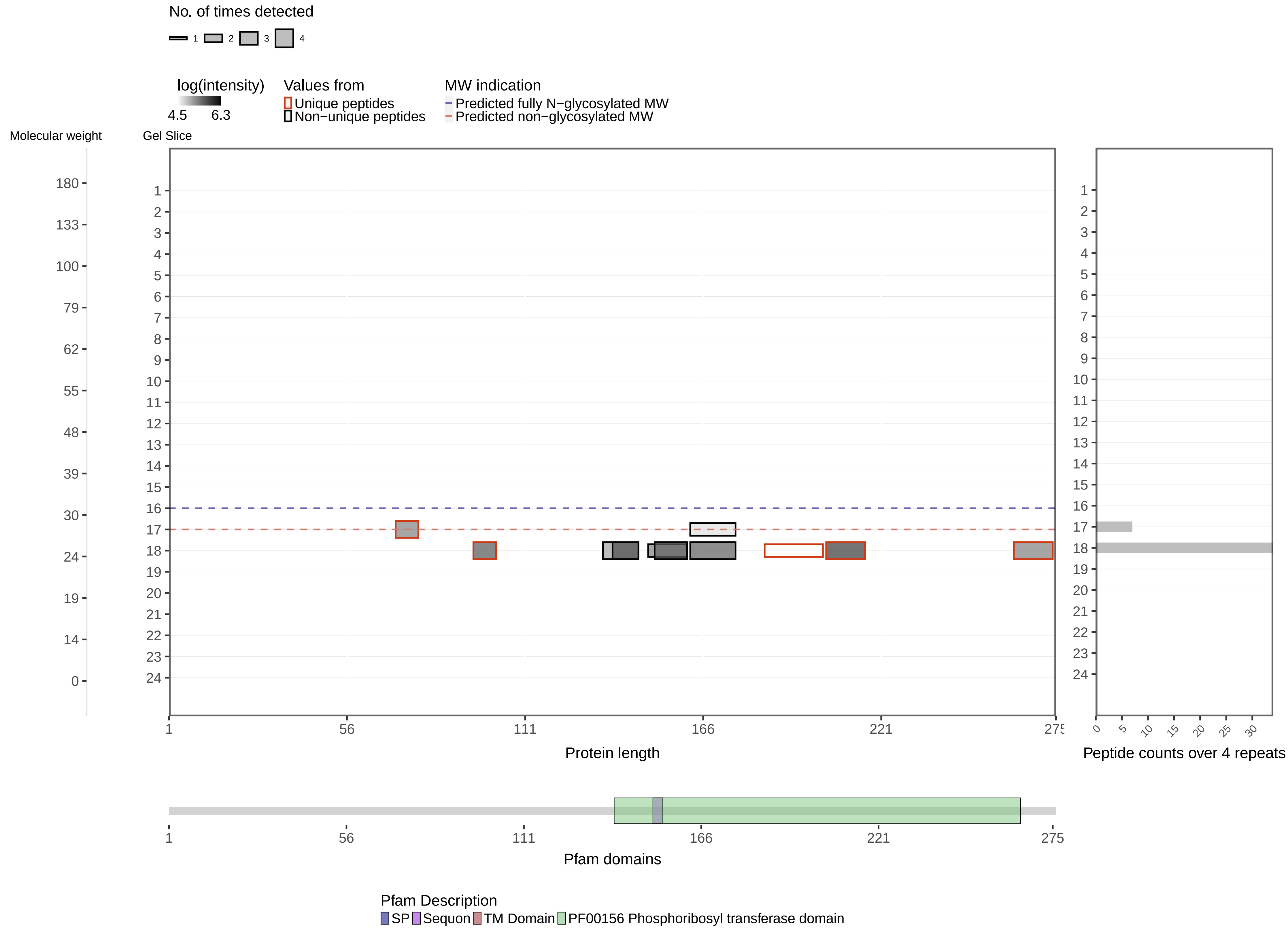


Protein sequence:

DITTLLLDTKAFKDTIDLFVERYKDKNISVVAGIEARGFIFGPPIALAIGAKFVPMRKPKKLPGEVISEEYSLEYGTDKIEMHVGAVQAGERALVVDDLIATGGTLSAAIRLLERVGVEVVECACLIELPELKGRDRLGDKPLFVLVSST*

NbL18g14970.1 LIM_domain−containing_protein_WLIM1−like [MW = 21.62 kDa]

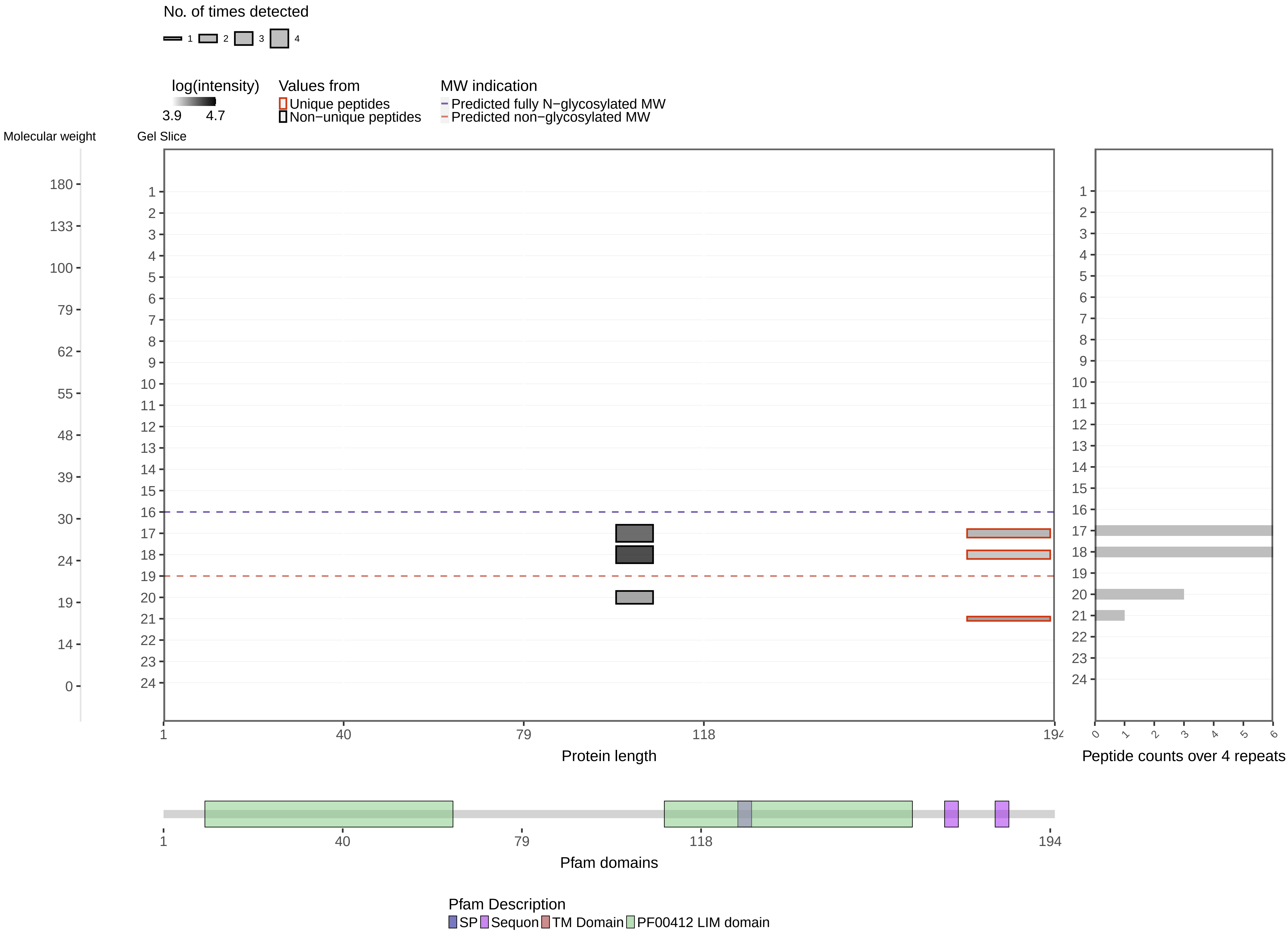


Protein sequence:

KPQKPIDSEKPQVAKVTSMFGGTREKCFGCKKTVYPTEKVSVNGTPYHKSCFKCSHGGCVISPSNYIAHEGRLYCKHHHIQLIKEKGNLSKLEGDHEQNSTTTTEVTAES*

NbL18g14840.1 serine_carboxypeptidase−like_7 [MW − SP = 49.86 kDa]

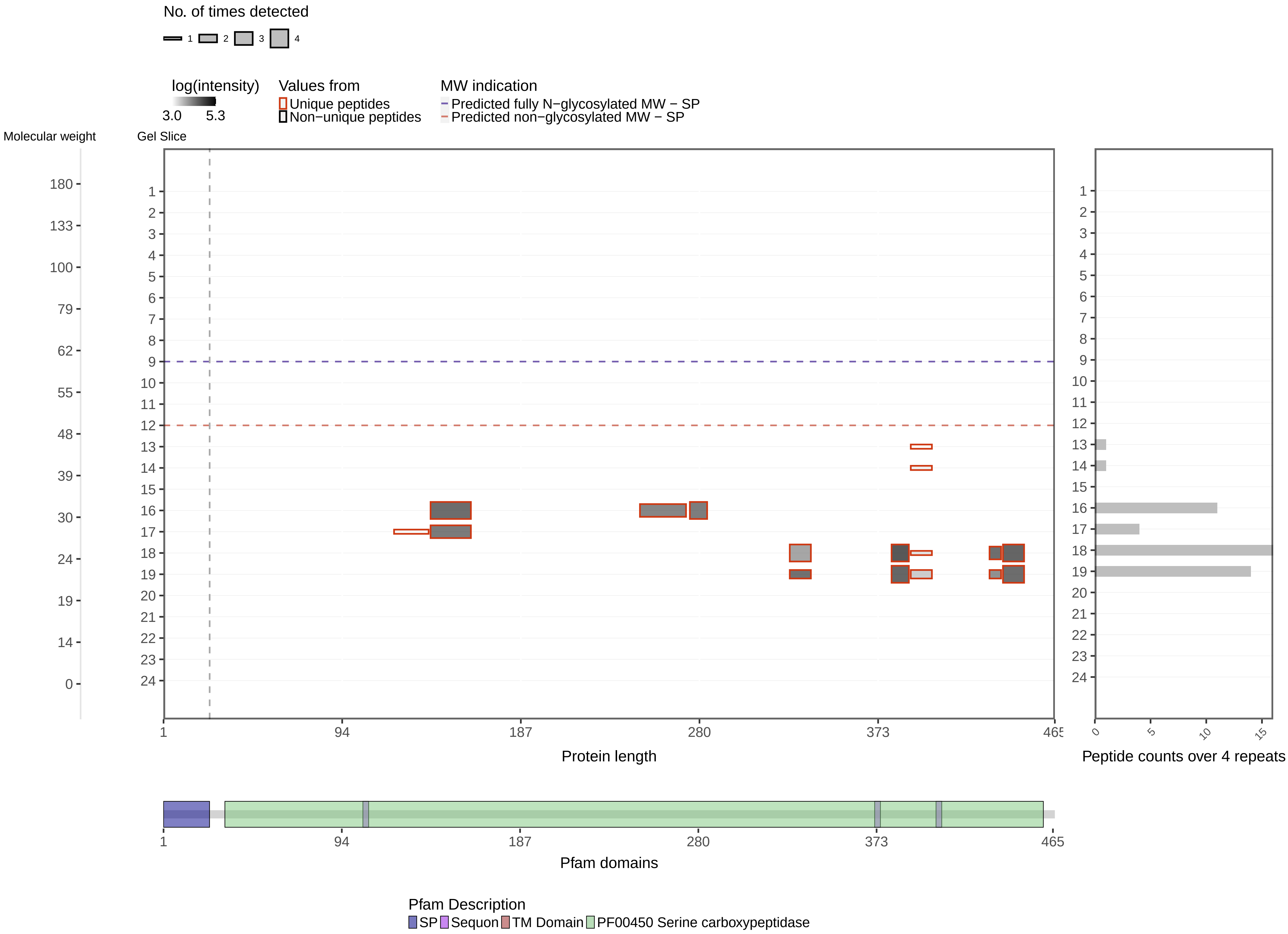


Protein sequence:

YEIPFAYGMGLISDELYESLKRNCKGEYQNTDPSNTQCLQDGHTFHELLKRINNPHILEPKCQFASPKPHLLFGQRRSLNVKFHQLNNPQQLPALKCRNDWYKHSTHWADDGQVREALHIRKGTIGKWVRCASLQYQTTIMSSIPYHANLSAKGYRSLIYSGDHDKVVTFLSTQAWIKSLNYSIVDDWRPWIVDNQVAGYTRSYSNRMTFATVKGAGHTAPEYKPRECLAMLKRWMSCKPL*

### NbL18g14580.1 branched−chain−amino−acid_aminotransferase_2_chloroplastic−like [MW = 42.81 kDa]

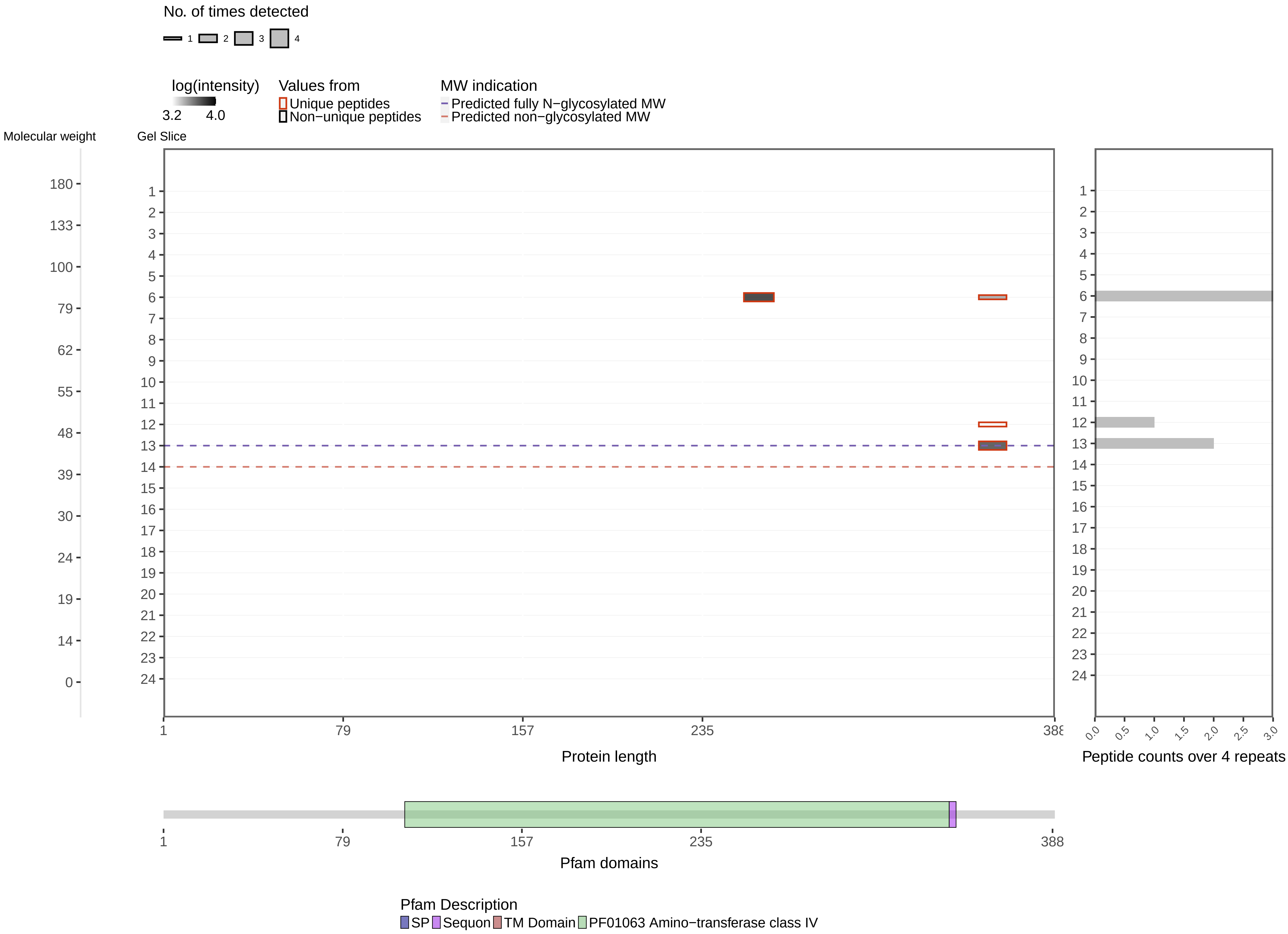


Protein sequence:

MGSGAVLGVAPAPEYTFLVYSCPVGNYFKDGTEALSLYVEEEFHRSALGGAGRIKSITIYAPVMKAMRNAKGKGFTDVLYLDSVNNKYIEEASSSNIFLVKGKVISTPKAKGTILEGVTKKSAIDIAHDLGYQVEERAIEAEELNTADEVFCTGTALGVAVVGNITYKGKKIEYKISSEMISQQLNSRLVGIQRGTIEDKRNWMVEVK*

### NbL18g14390.1 xylulose_kinase−like [MW = 61.44 kDa]

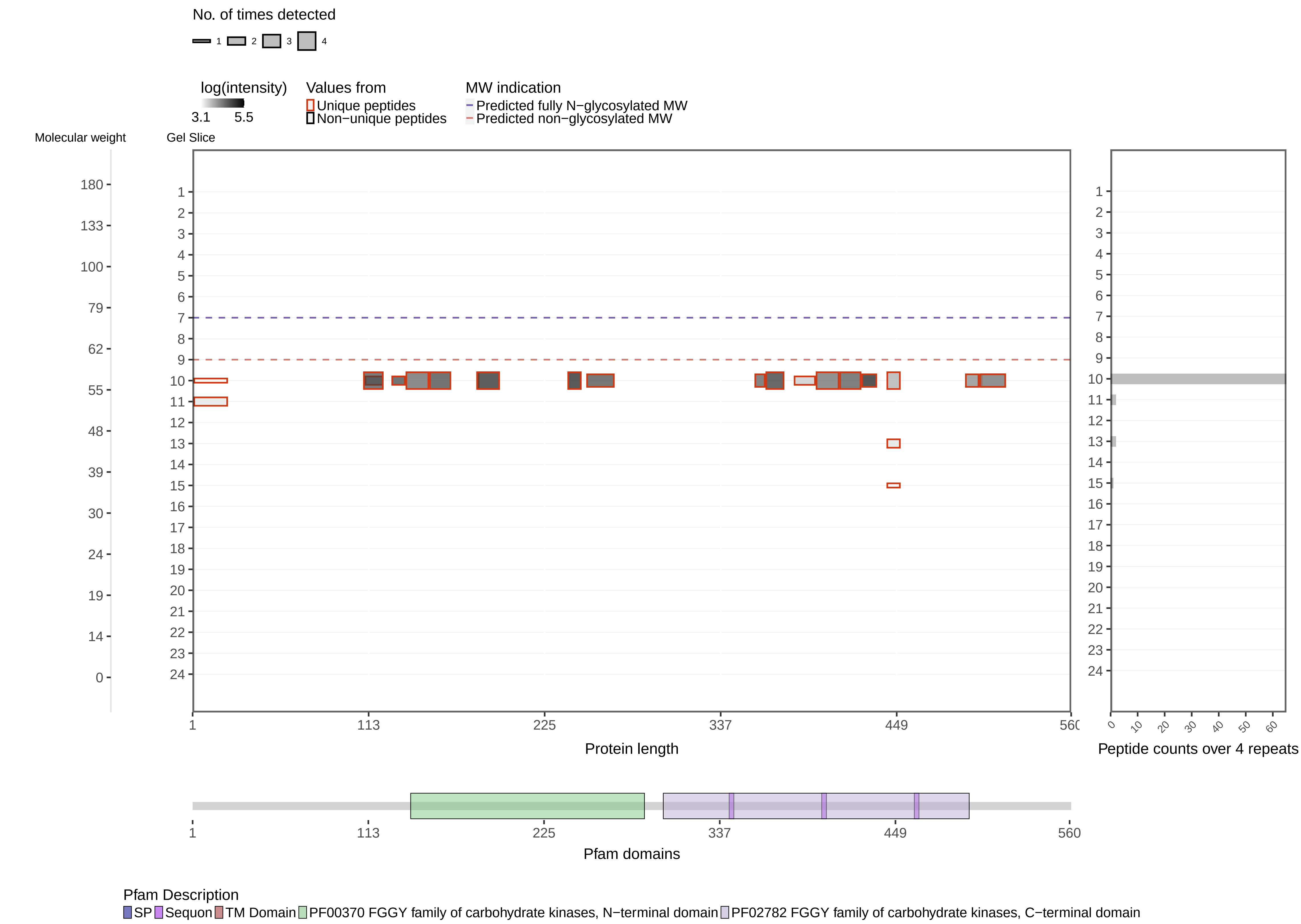


Protein sequence:

ERYHFSKDCLIIQWSGDNPNSLAGLTLNTPGDLAVSLGTSDTVFGITKDHNPSLEGHVFPNPVDTESYMVMLCYKNGSLAREDIRNLYAEKSWDVFNKYLQQTPPLNSGRLGFYYKDFEILPPLPVGFHRYILENFTGDSLDGIKEHEVQEFDPPSEVRAIVEGQFLSKRAHAERFGMPSPPKRIIATGGASANSSILSLIASIFGCDVYTVQQPDSASLGAALRAAHGWLCSKKGNFVPISSMYMDNMDKTSLNCKLAATAGDQKLVAKYAMLMKKRIEIENRLVEKLGRL*

### NbL18g14320.1 60S_ribosomal_protein_L14−1 [MW = 17.07 kDa]

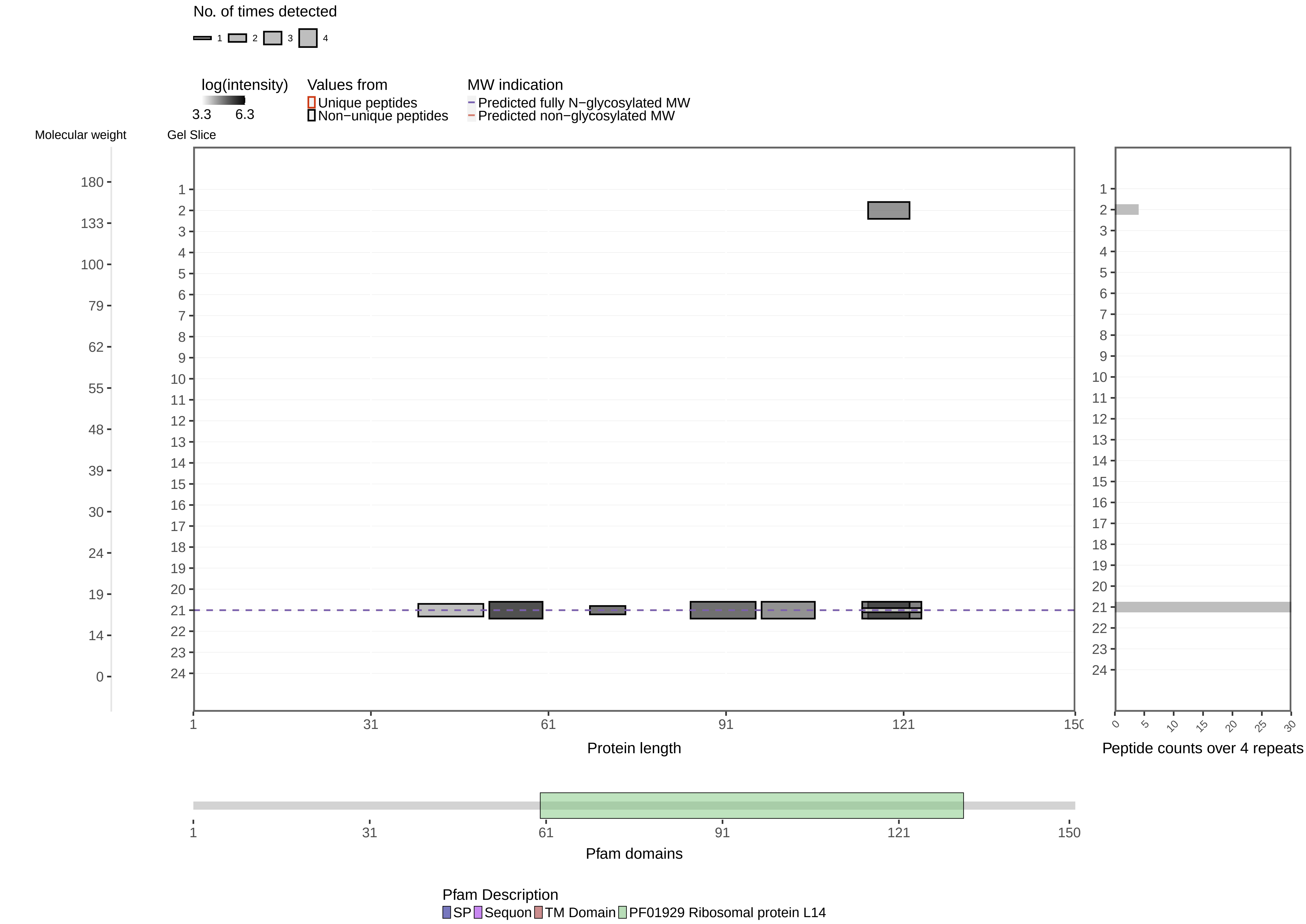


Protein sequence:

SQINFKRLSLTDIKIDIKRIPKKKTLVAAMDAADVKNKWESSSWGRKLIVQKRRASLNDFDRFKLMLAKIKRAGVVRQELAKLKKETVA*

### NbL18g14290.1 E3_ubiquitin−protein_ligase_UPL2−like [MW = 403.04 kDa]

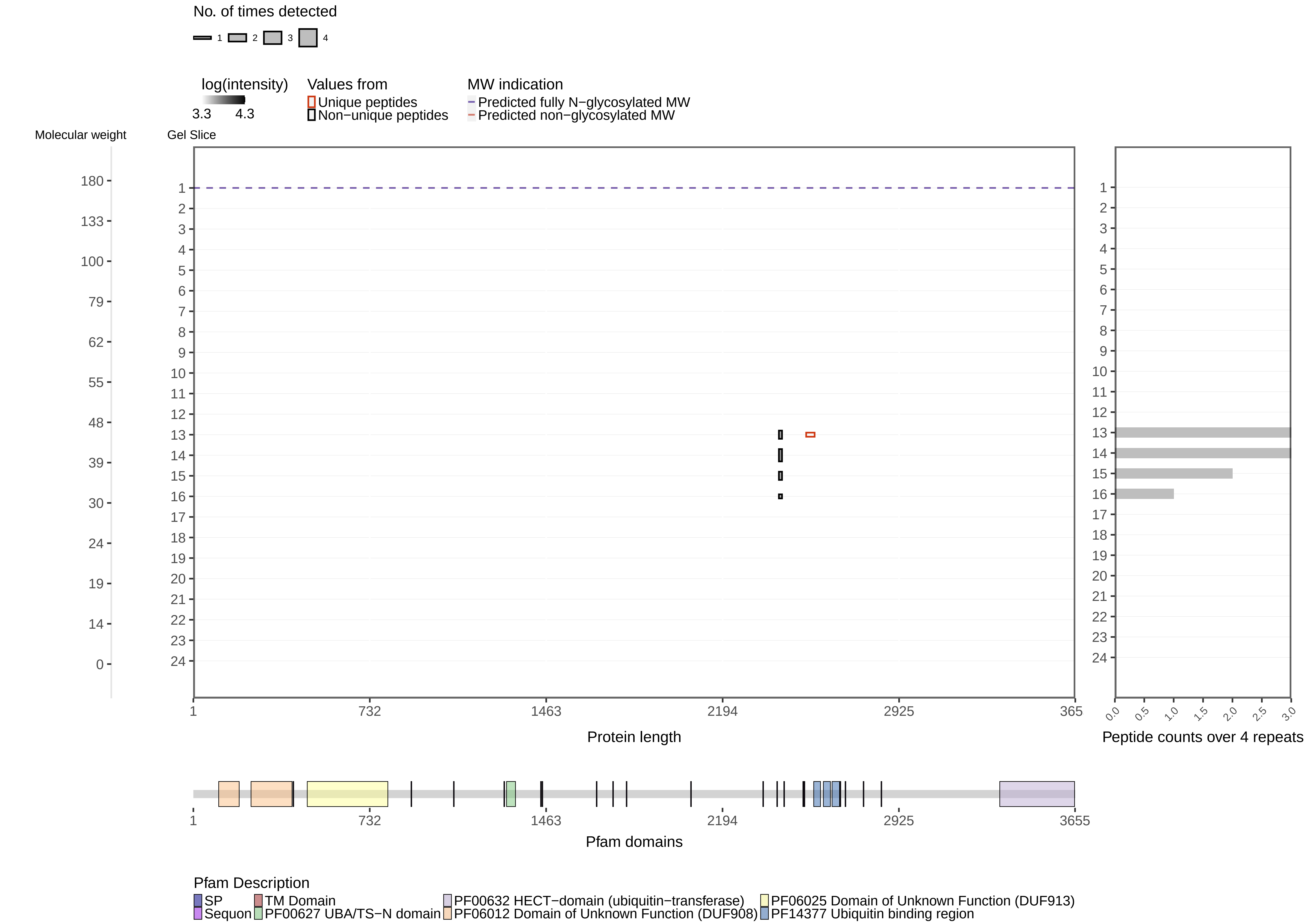


Protein sequence:

NbL18g14060.1 phosphoglycerate_mutase−like_protein_1 [MW = 31.4 kDa]

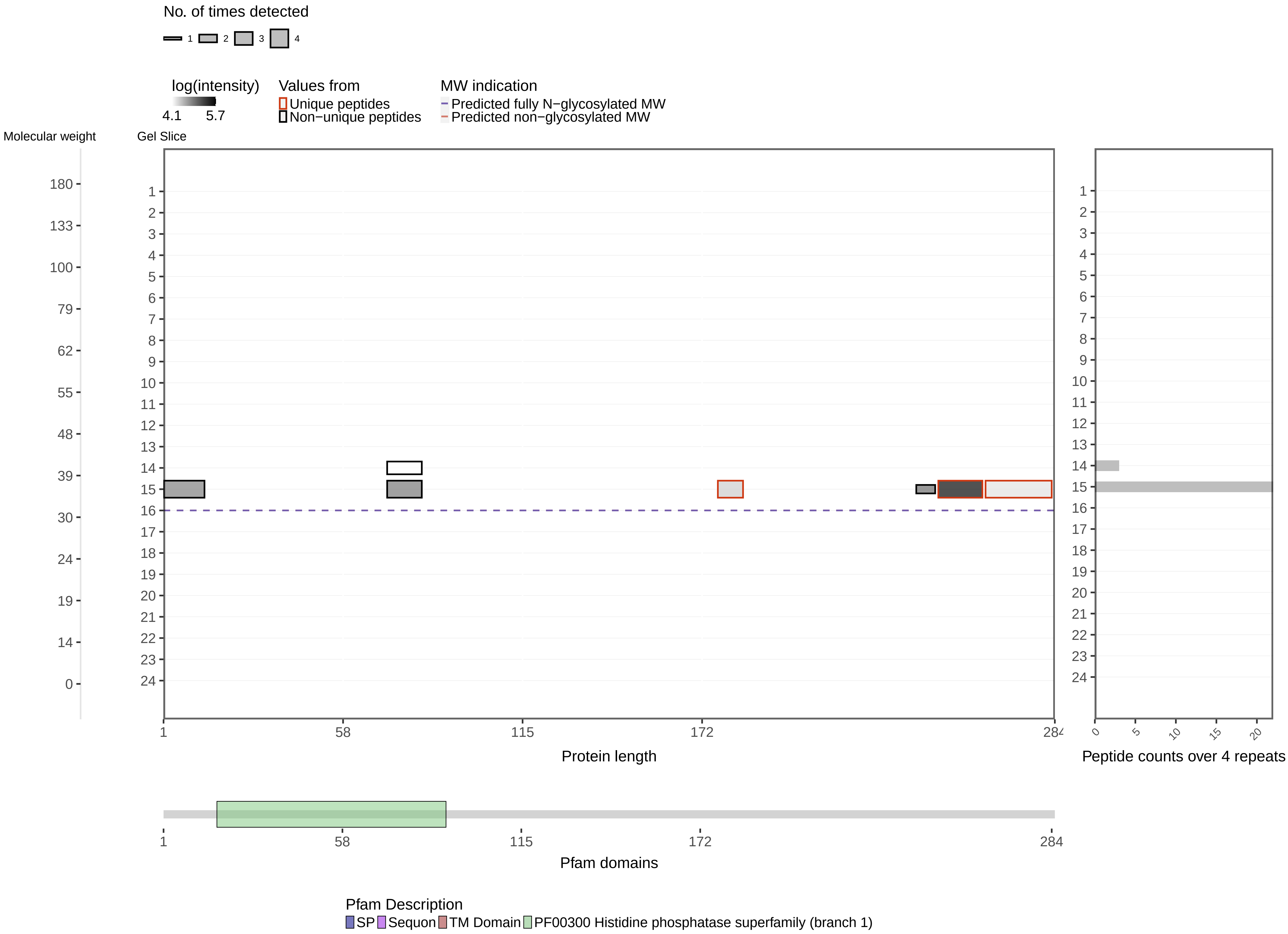


Protein sequence:
AVEGCREHLGVHWCDKRRSISEYKPLFPAIDFSLIECDDDVLWEPDIREPNEHLAARGMEFLRWLWTRKEKEIAVVTHSGFLIHTLSAFGNDCHPDVKNEICRPFNNCELRSMVIIDRSMIGSDSSTTNYPGKIPSGVDVPSDNTYINHPDGLSN*

### NbL18g13960.1 phosphoenolpyruvate_carboxykinase_[ATP]−like [MW = 82.61 kDa]

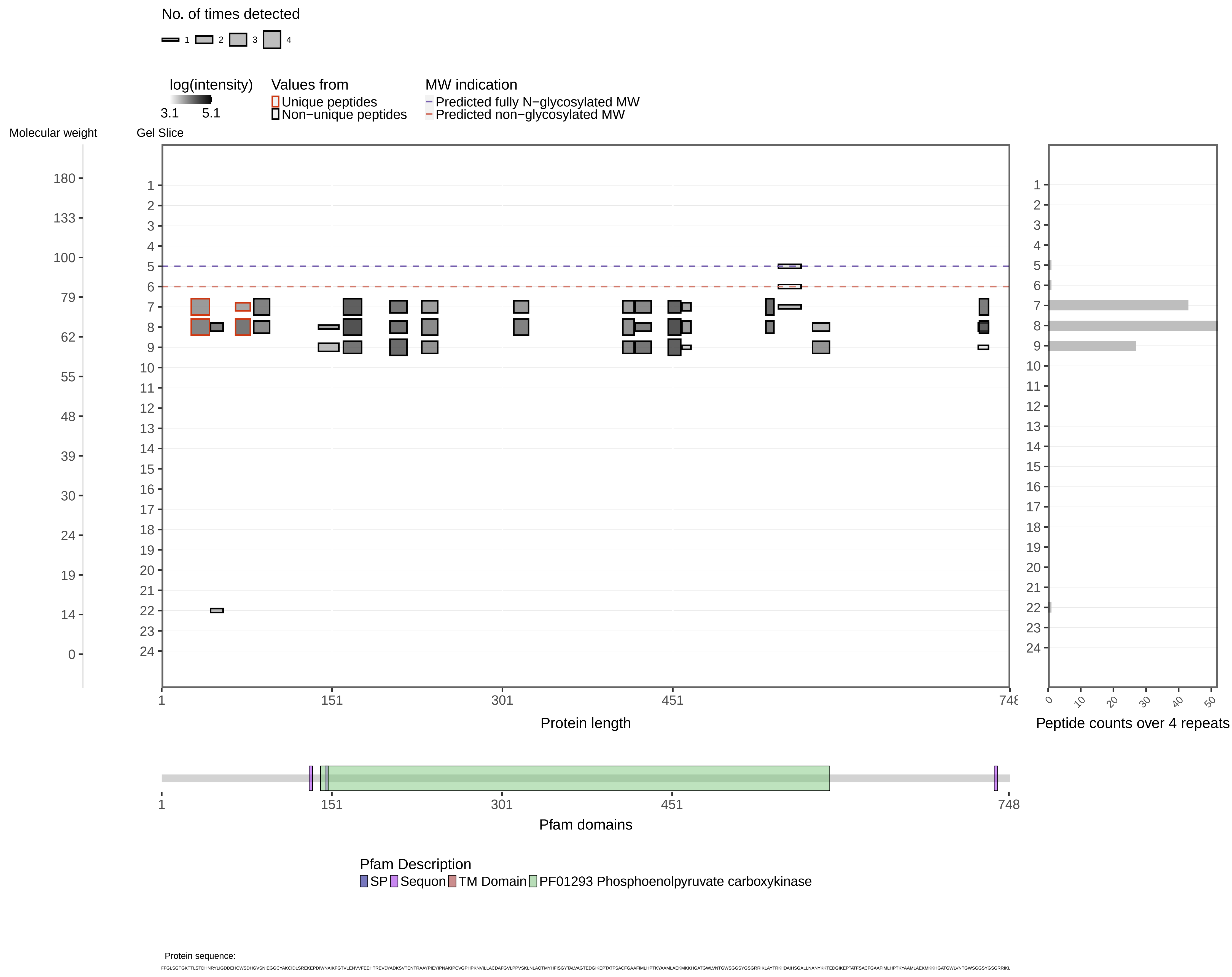

### NbL18g13950.1 ubiquitin−fold_modifier−conjugating_enzyme_1 [MW = 13.64 kDa]

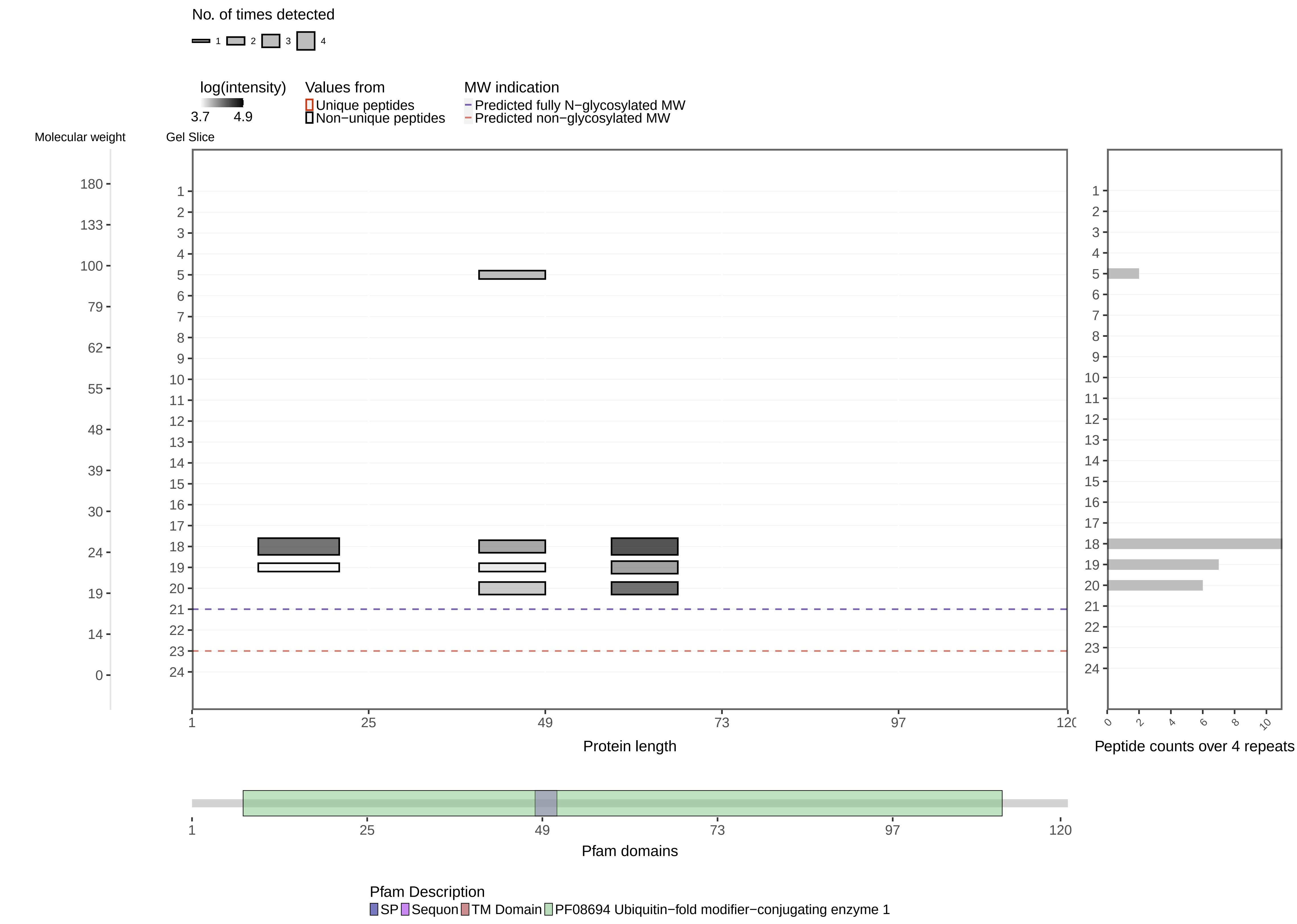


Protein sequence:

INKSKDNDWFRISAANPEGTRWKGKCWYVHNLLKYEFDLOFDIPVTYPATAPELELPELDGKTEKVTSFLCVF*

### NbL18g13800.1 geranylgeranyl_transferase_type−2_subunit_alpha_1 [MW = 81.33 kDa]

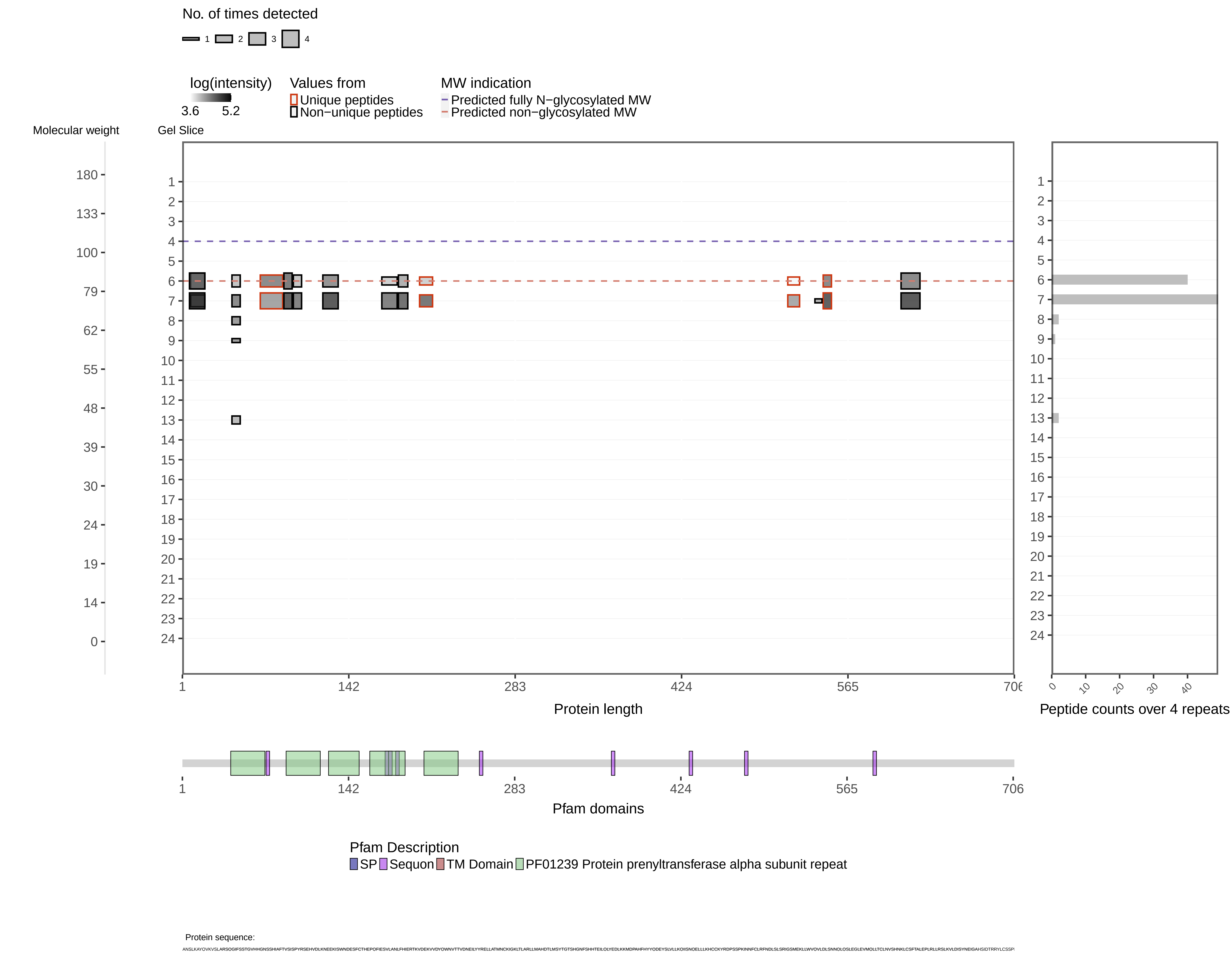

### NbL18g13760.1 NADPH−−cytochrome_P450_reductase [MW = 76.81 kDa]

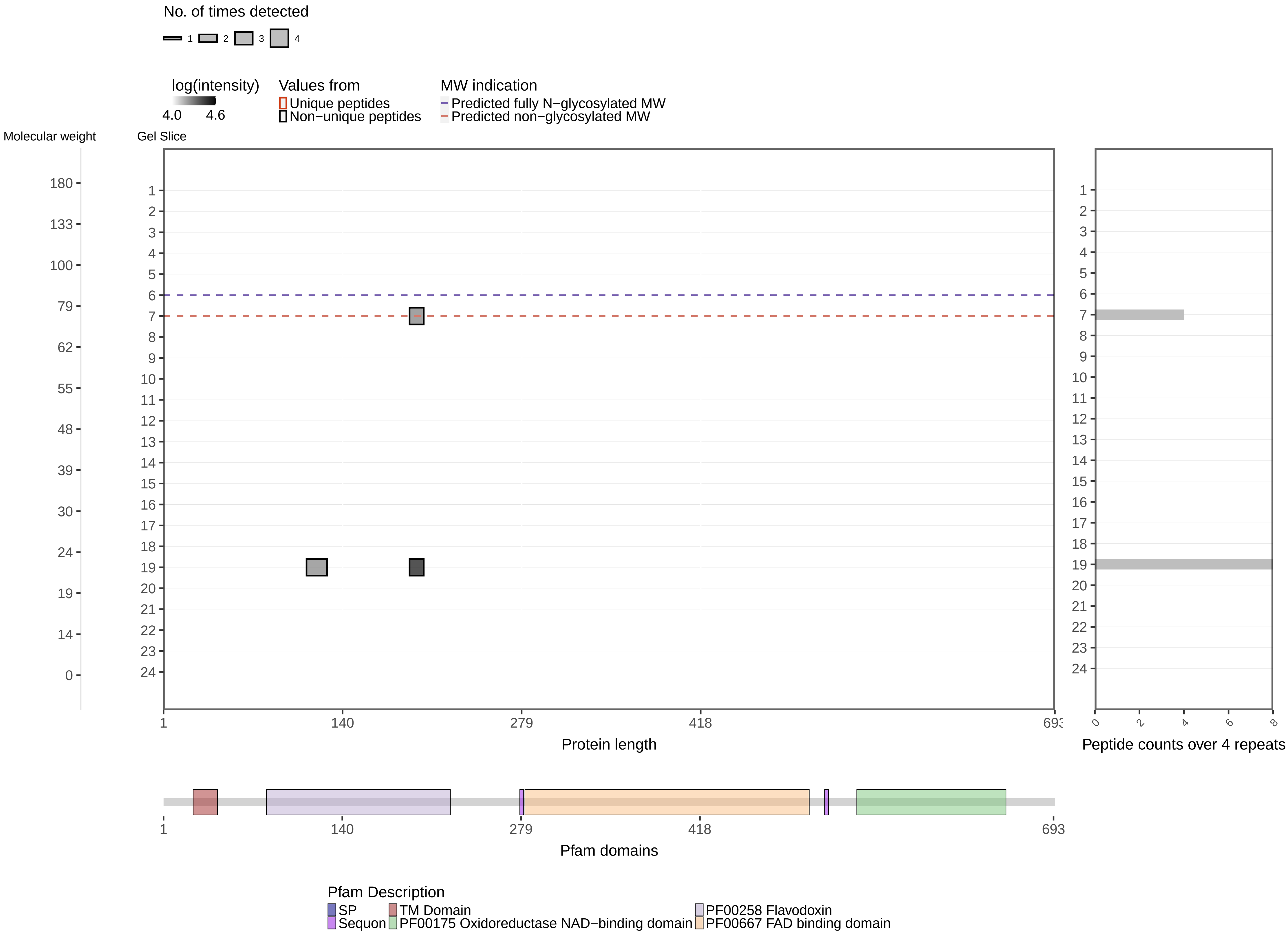


Protein sequence:

EIVEEAARLLGQSLDLIFSIHTDKEDGTALGGSLPPPFPGPCTLRAALARFADLLNPPRKATLVALAAHAAEPSEAEKLKFLASPQGKDEYSAWVVASQRSLLEVMAEFPSAKPPLGVFFAAVAPRLQPRYYSISSSPRFAPARIHVTCALVYGPTPTGRIHKGVCSTWMKNAVPSEKSQNCSSAPIFIRPSNFKLPVDPSVPIVMVGPGTGLAPFRGFLQERAALKEDGAQLGPAFLFFGCRNRRMDFIYEEELQSFVKQGVISELIIAFSREGPQKEYVQHKMLEKLMTYEQA

### NbL18g13540.1 glutamate_dehydrogenase_B [MW = 44.47 kDa]

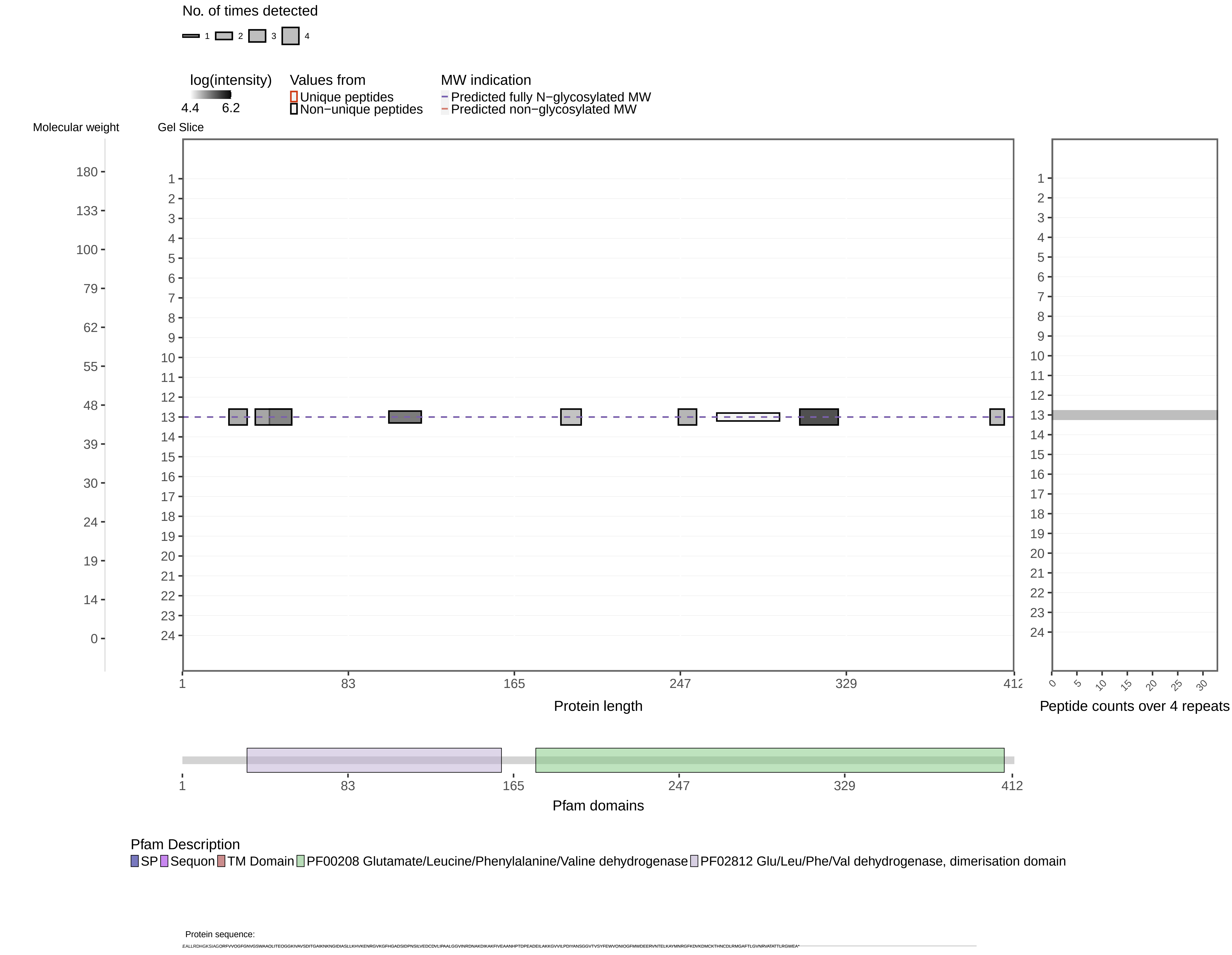

NbL18g13530.1 protein_ASPARTIC_PROTEASE_IN_GUARD_CELL_2 [MW − SP = 43.94 kDa]

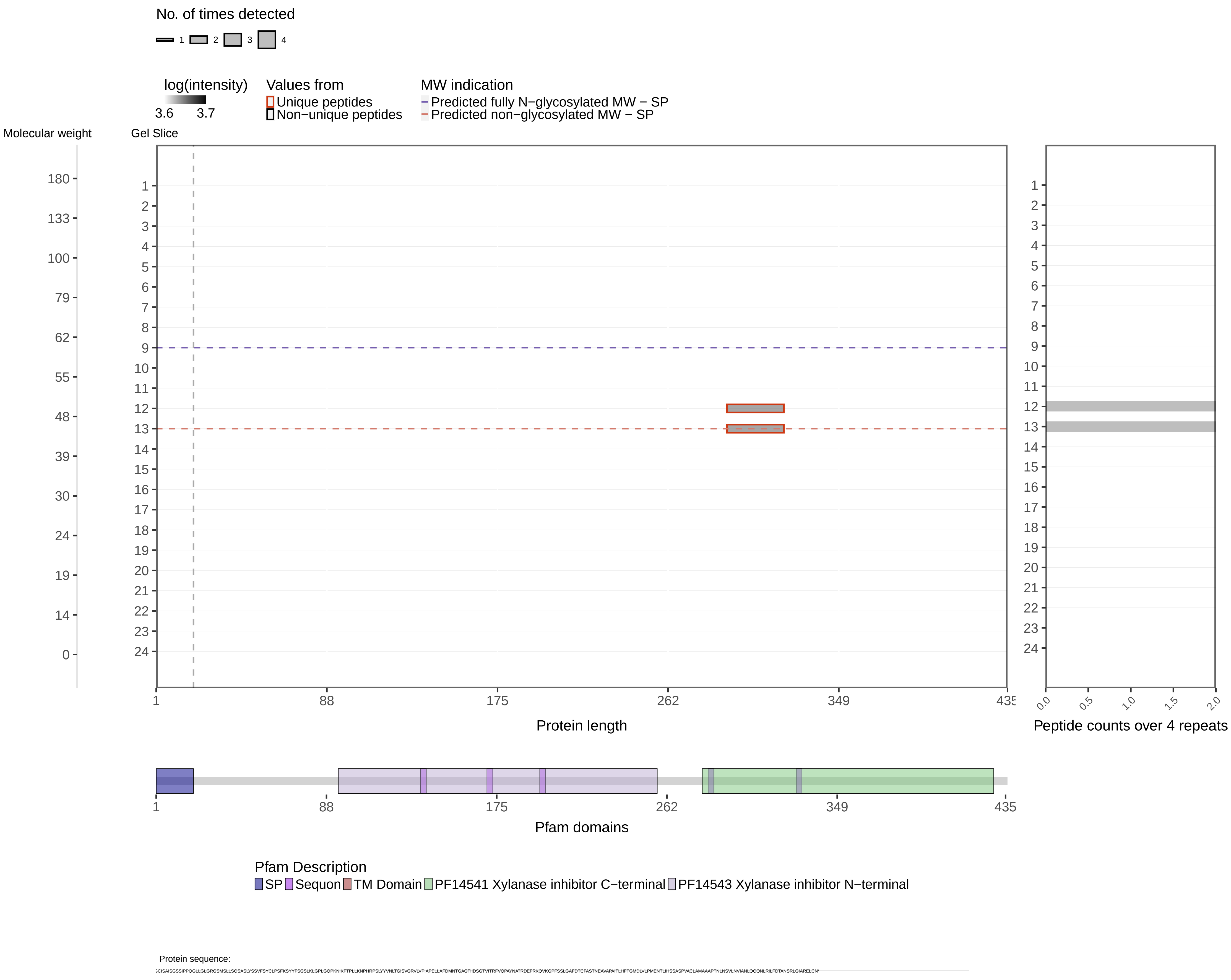

NbL18g13490.1 glucose−6−phosphate_isomerase_1_chloroplastic [MW = 68.04 kDa]

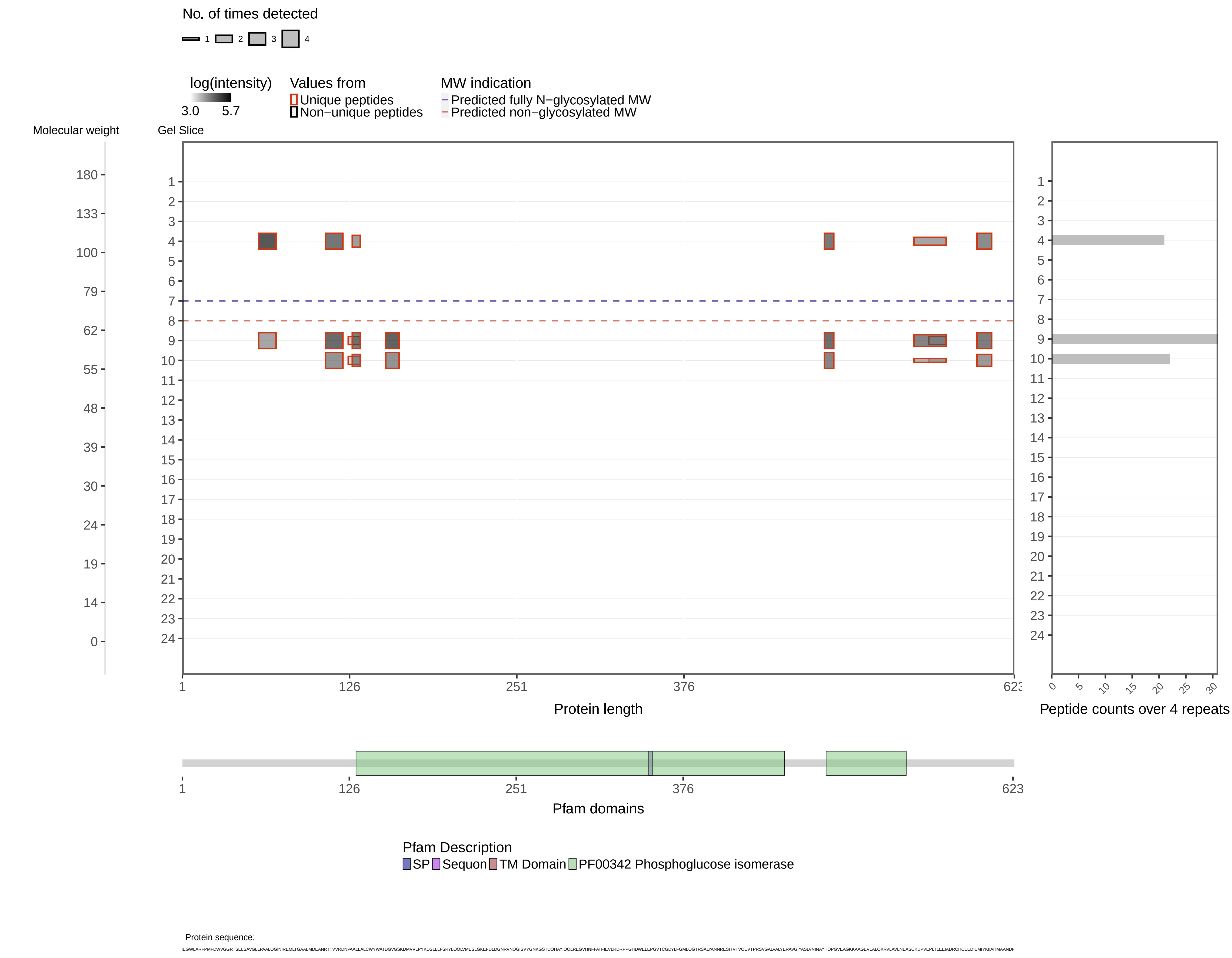

### NbL18g13430.1 14−3−3_protein_10 [MW = 32.9 kDa]

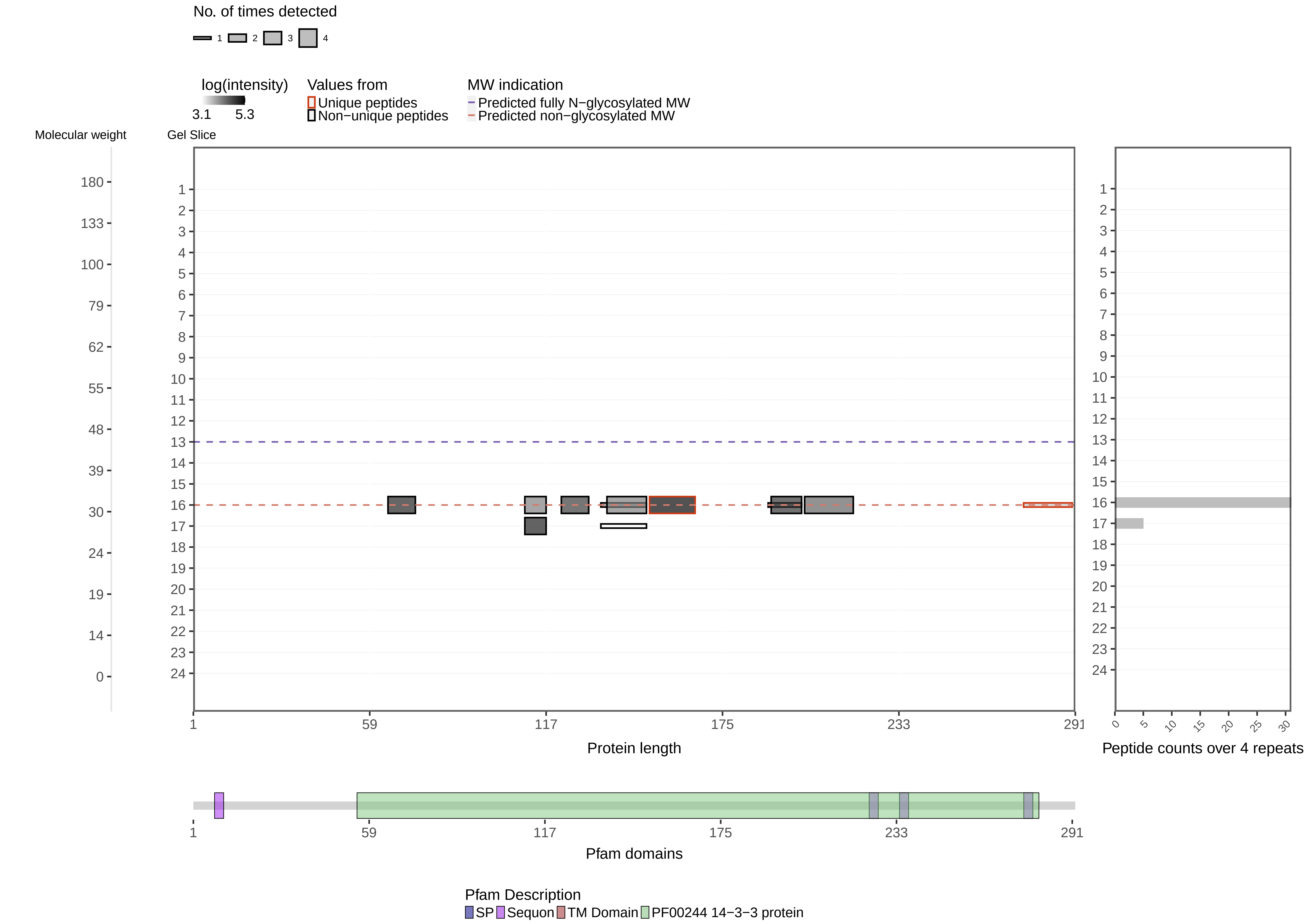


Protein sequence:

RGKVENELTEVCAGILKLLESNLEPSASTGESRVFYLKMKGDYYRYLAEFKVGDERKOAAEDTMNSYKAAOEIALADLPPTHPIRLGLALNFSVFYFEILNSSDKACSMAKOAFEEAIAELDTLGEESYKDSTLIMOLLRDNLTLWTSDAODOLDES*

### NbL18g13420.1 protein_CutA_chloroplastic [MW = 19.92 kDa]

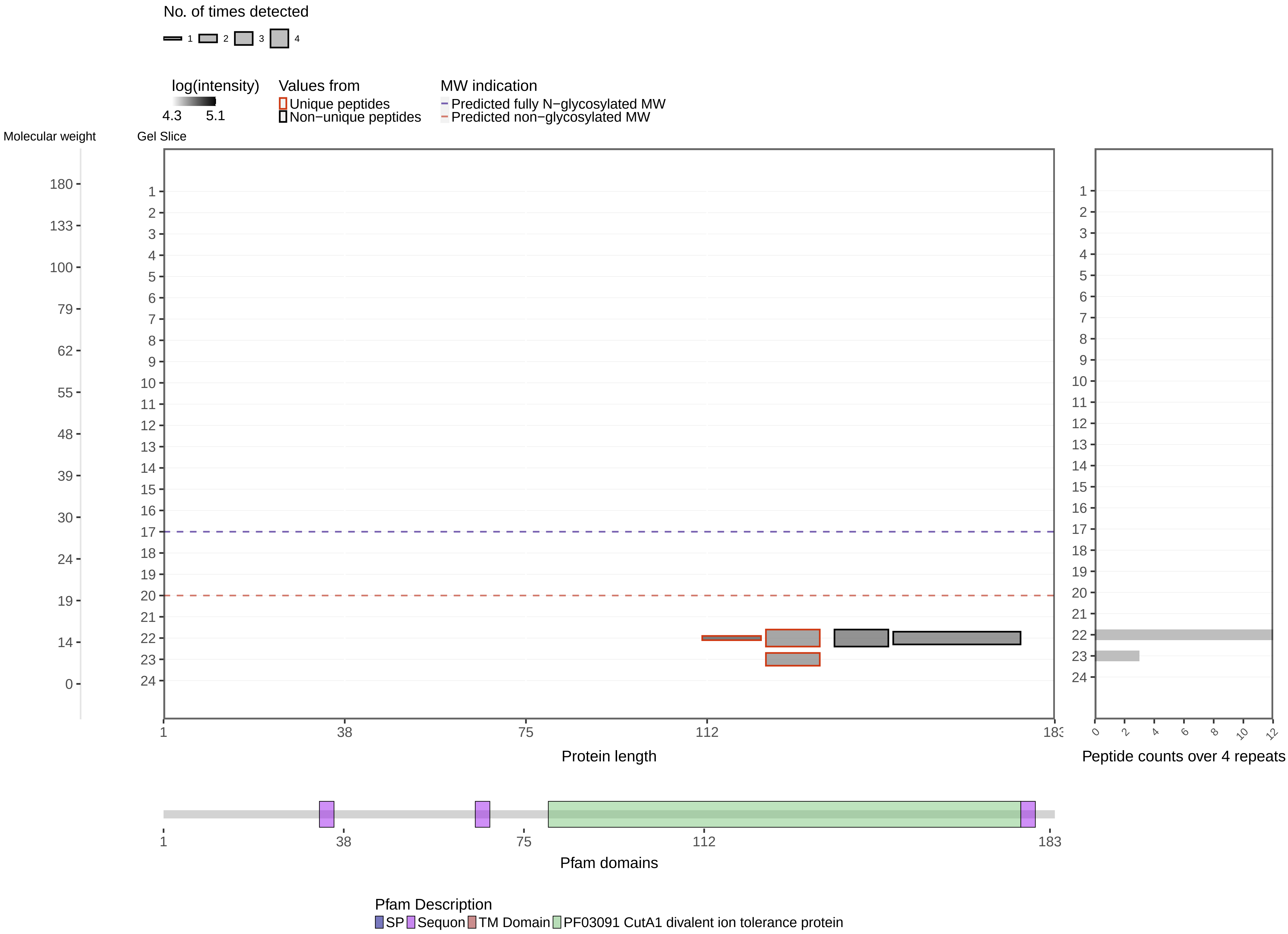


Protein sequence:

SIVVYVTVPNKELGKKLAGSIVKEKLASCVNRIPGVESVYEWEGKVQTDSEELLIIKTRESLLEALTEHVKANHEYDVPEVIAMPIVGGSPQYLEWLKNSTREK*

NbL18g13260.1 28_kDa_ribonucleoprotein_chloroplastic−like [MW = 32.77 kDa]

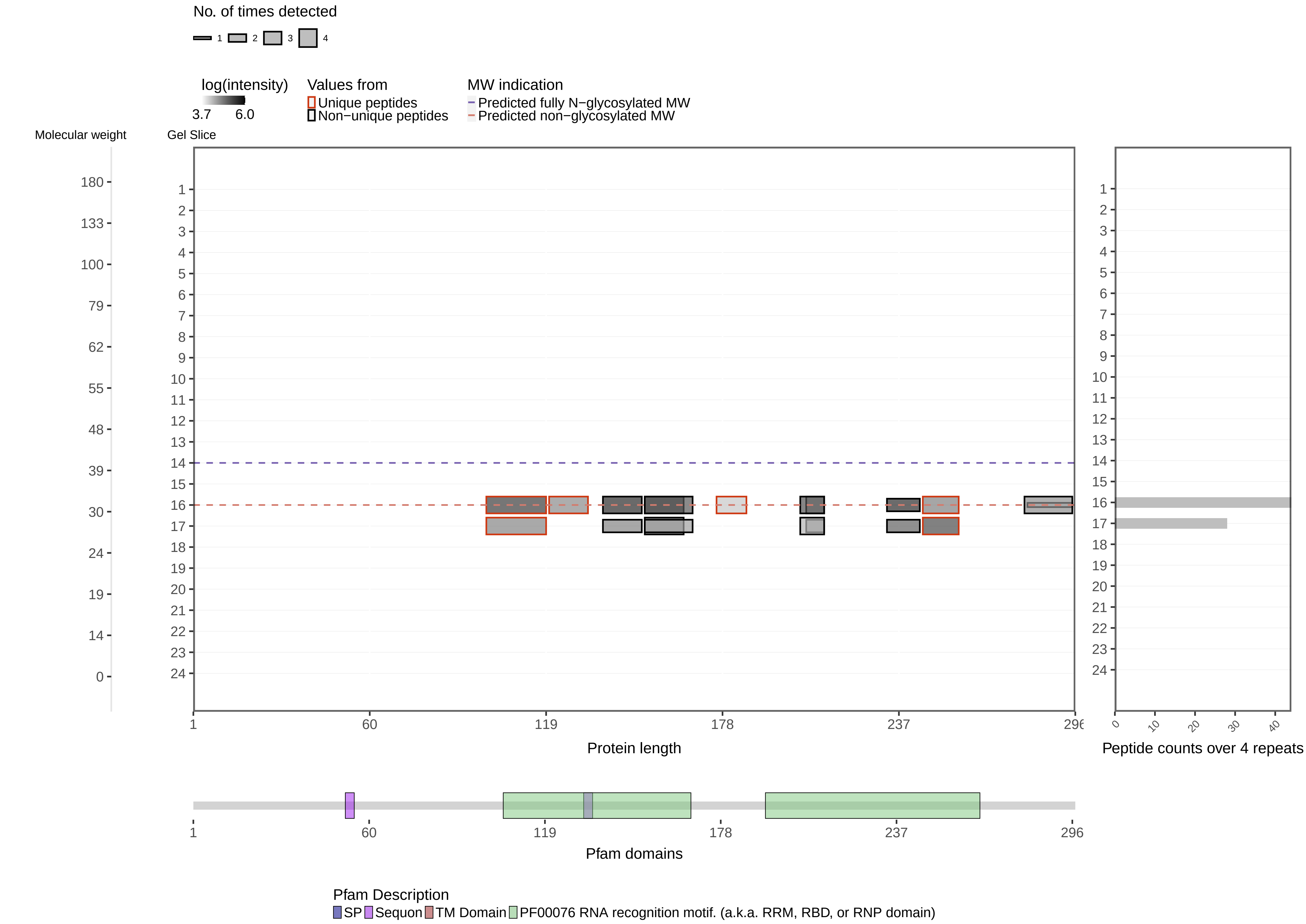


Protein sequence:

SRGLVFVTMGSHEEAKAVLENLEAYELEGRPLRLAWAKPKTVKPSSPLPSKPLPVHNVFVANLHFEARSKDLMEFFKANGADVVSAEVIFNSDNPRRSAGYGFVSFNTKVEADSALSSFEGKEFMGRPIRVAHSKRFLRGETKKILQSQEVSSELISVAE*

NbL18g13160.1 thylakoid_lumenal_29_kDa_protein_chloroplastic [MW = 37.52 kDa]

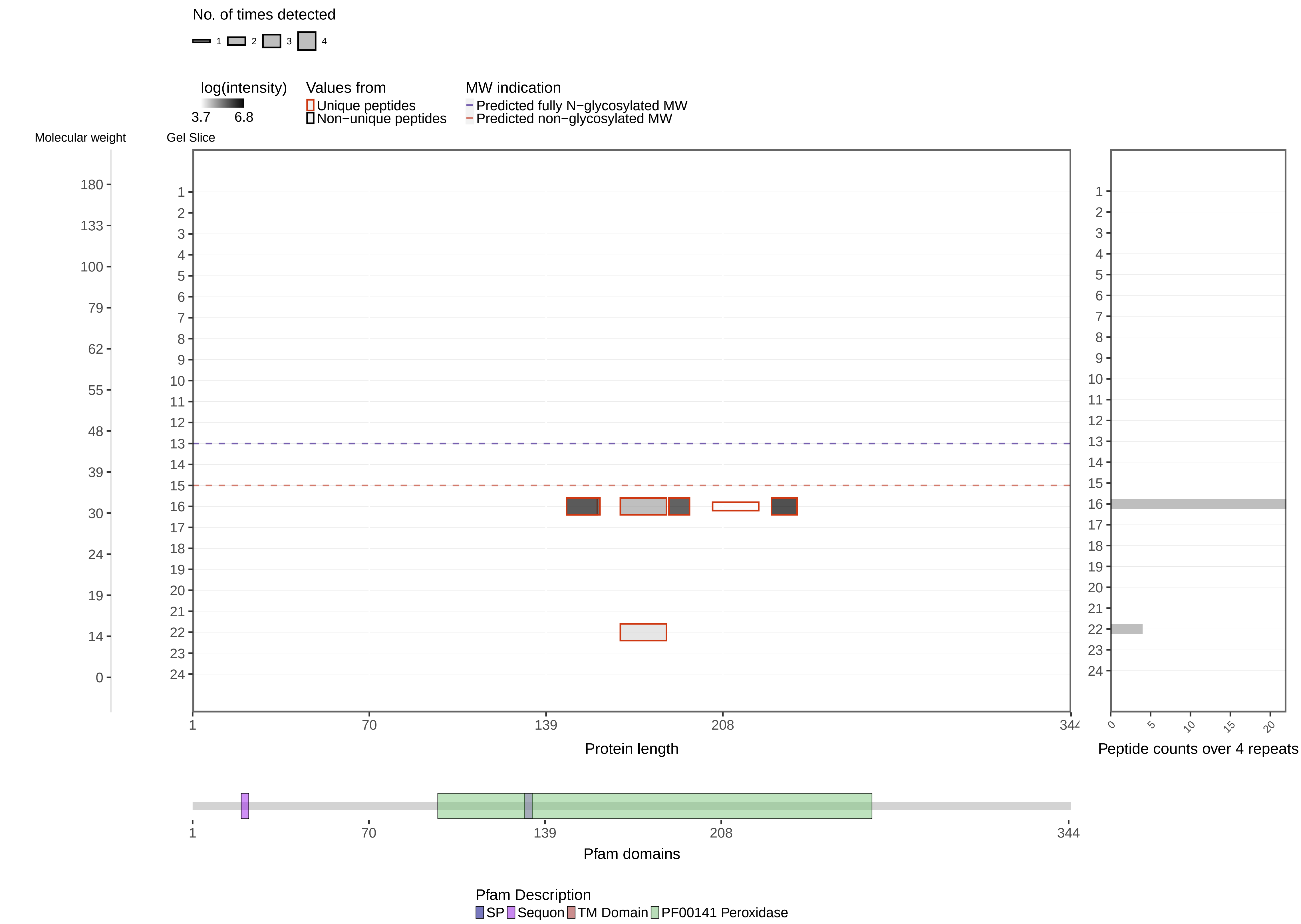


Protein sequence:

IDADSKGGPISYADLIQLAAQSALKSTFLASAIRKCGGNVEKGSLLYSAYGSNGQWGLFDRQFGRSDAQEPDPEGRVPQWDKASVQEMKDKFKAVGFGPRQLAVMSAFIGPDQDATEALLANDPEVIPWVQKYQRSRETVSRTDYEVDLITTLTKLSCLGQQINYEAYSYPVQKIDVTKLKL*

NbL18g13150.1 serine−−tRNA_ligase_chloroplastic/mitochondrial [MW = 59.96 kDa]

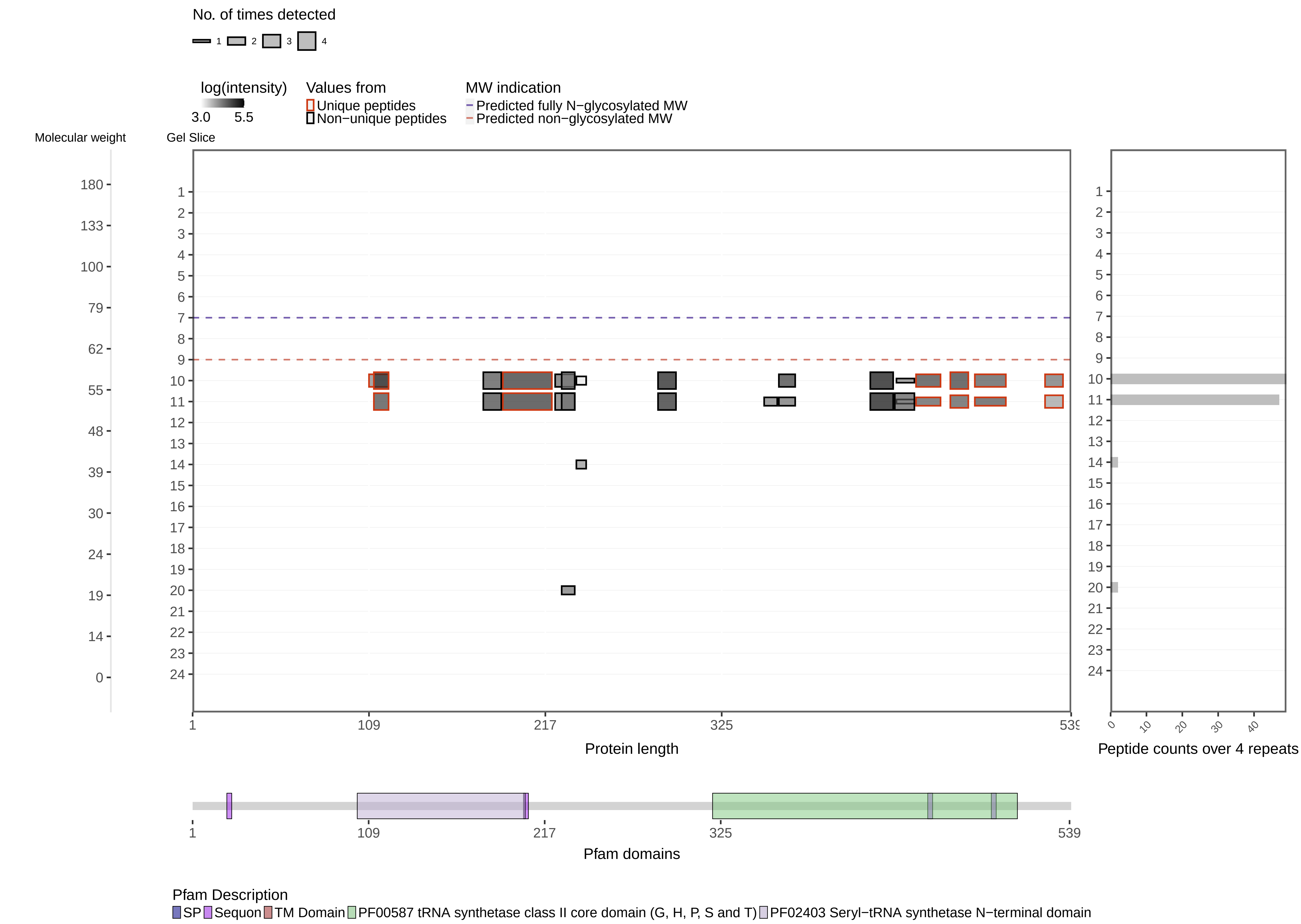


Protein sequence:

'SGSKFYYLKNEAVILEMALVNWAVSEAMKRGFTPLTTPEIVRSSVVEKCGFQPRGTNTQVYSIENSDQCLIGTAEIPVGGIHMDSILSESSLPLKYVAFSHCFRTEAGAAGAATRGLYRVHQFSKVEMFILCRPDESDSYHEELIEIEEDLFSSLGFHFKTLDMASQDLGAPAYRKFDVEAWMPGLGRYGEISSASNCTDYQSRRLGIRYRPEPSASSSPKKGKNTAPTQFVHTLNATACAVPRMIVCLLENFQQEDGSVVIPEPLRSFMGGLQIIKPKHIQN*

### NbL18g13120.1 cytochrome_c_oxidase_subunit_6b−1−like [MW = 19.83 kDa]

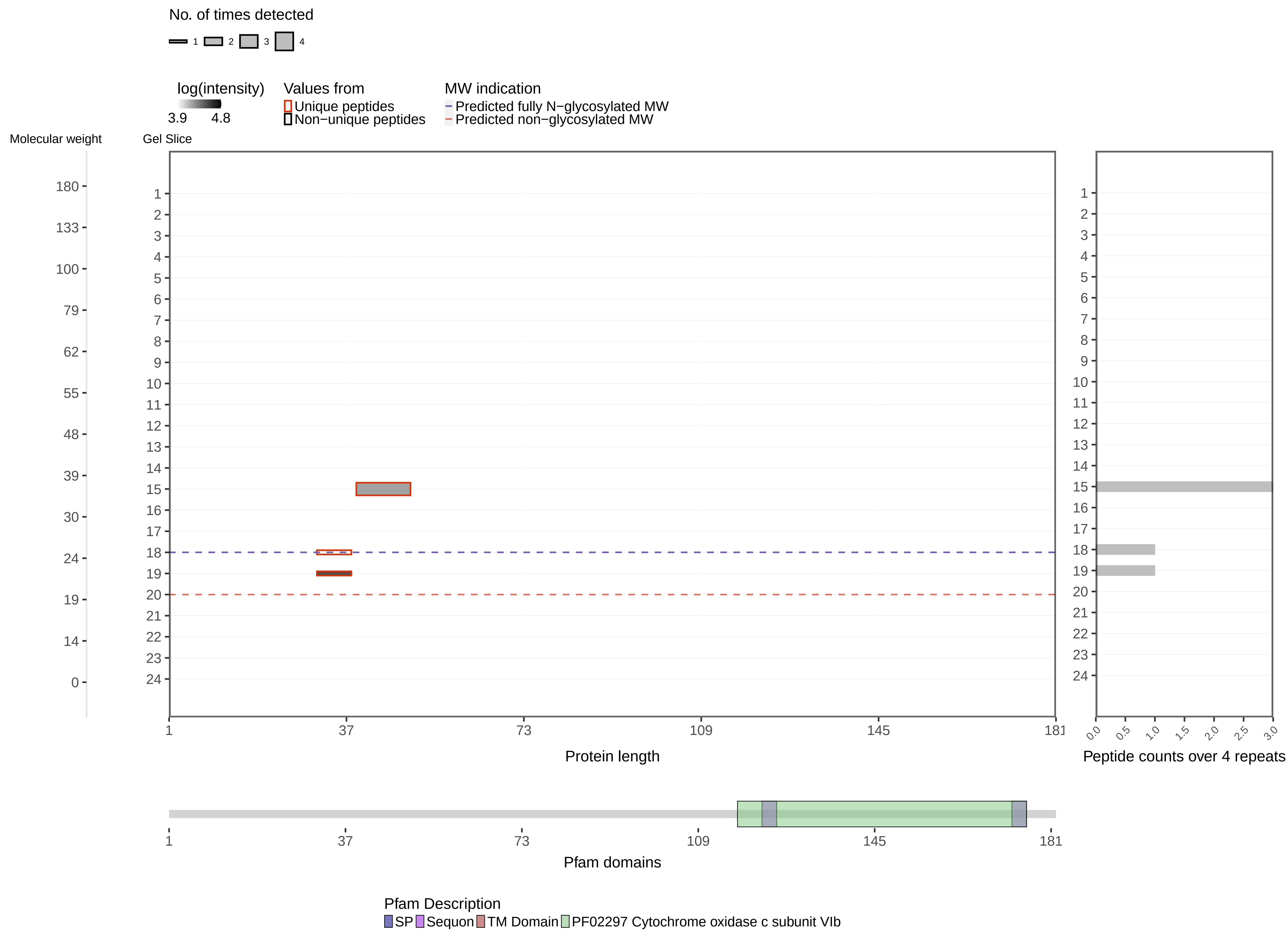


Protein sequence:

VEASSDESSESTEDETSGNEESAVETPEIKLETAPADFRFPTTNQTRHCFTRYVEYHRCVAAKGEGAPECDKFAKYYRSLCPGEWIDRWNEQRENGTFPGPL*

### NbL18g13010.1 probable_carboxylesterase_2 [MW = 36.95 kDa]

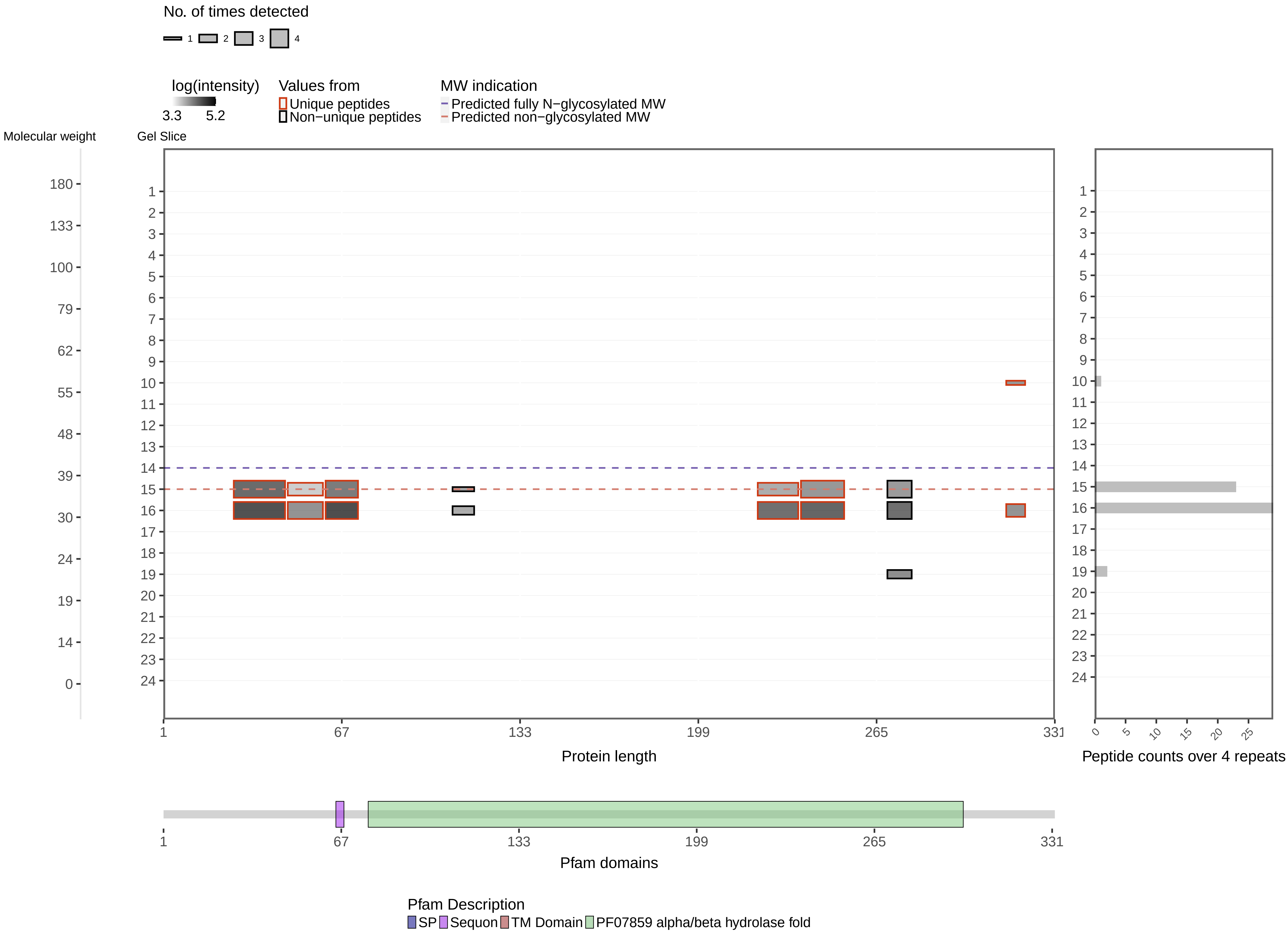


Protein sequence:

ANFGRVFLSGDSAGANIAHNLAMAAGSVEPELMVDIIGVALVHPYFWGSGSIGSEALYPDKKAHVDRLWAFVCPSNPDNDDPRINPMFEGAPSLTSLGCRRVLIIVAEKDILKDRGWAYYQALGRSGWMGVVVIQETEGEDHVFHLNNLQCQKSKDLIKNLAHFFNREMPHFLKIKH*

NbL18g12750.1 ornithine_carbamoyltransferase_chloroplastic−like [MW = 43.22 kDa]

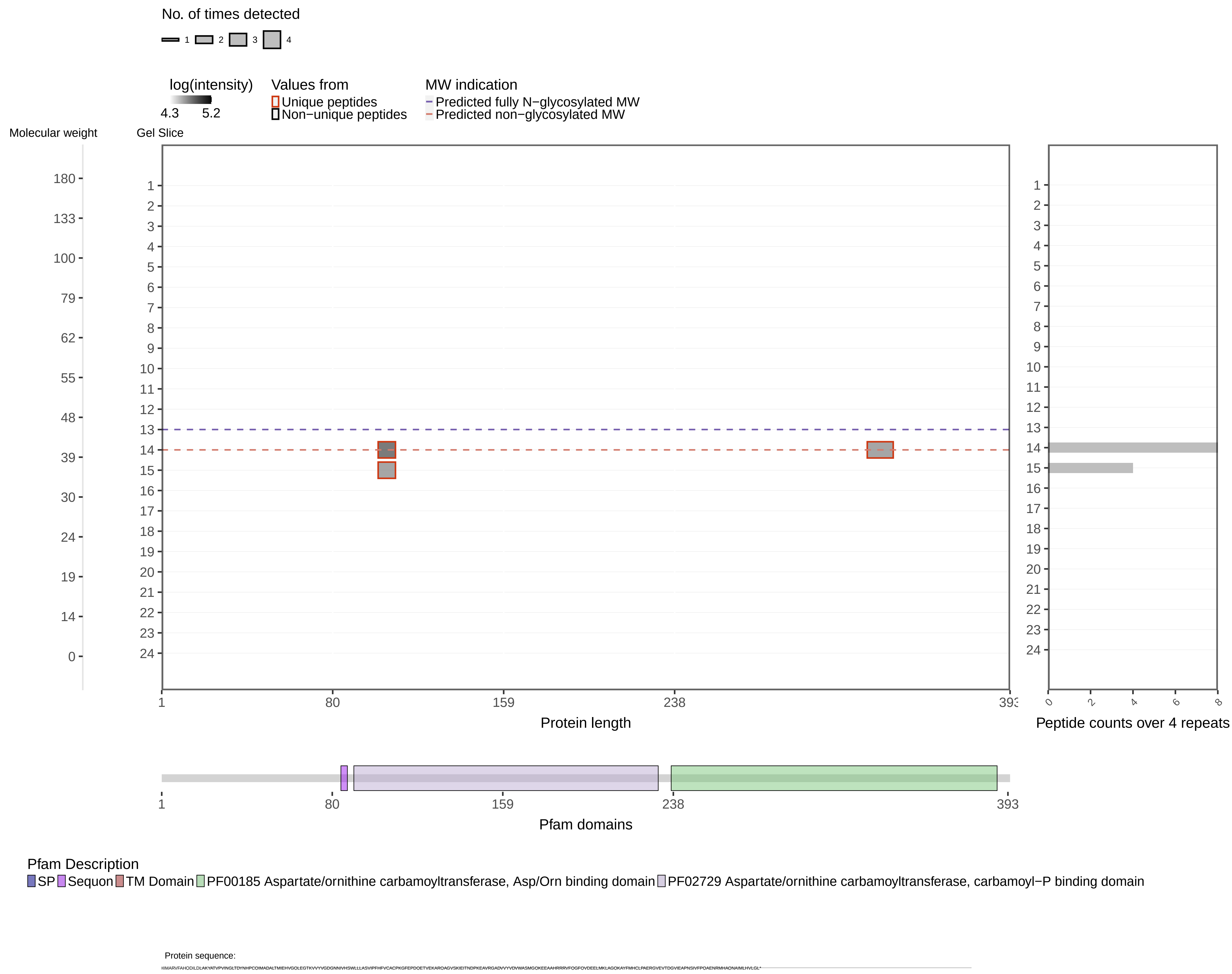

### NbL18g12390.1 alcohol_dehydrogenase−like_7 [MW = 41.88 kDa]

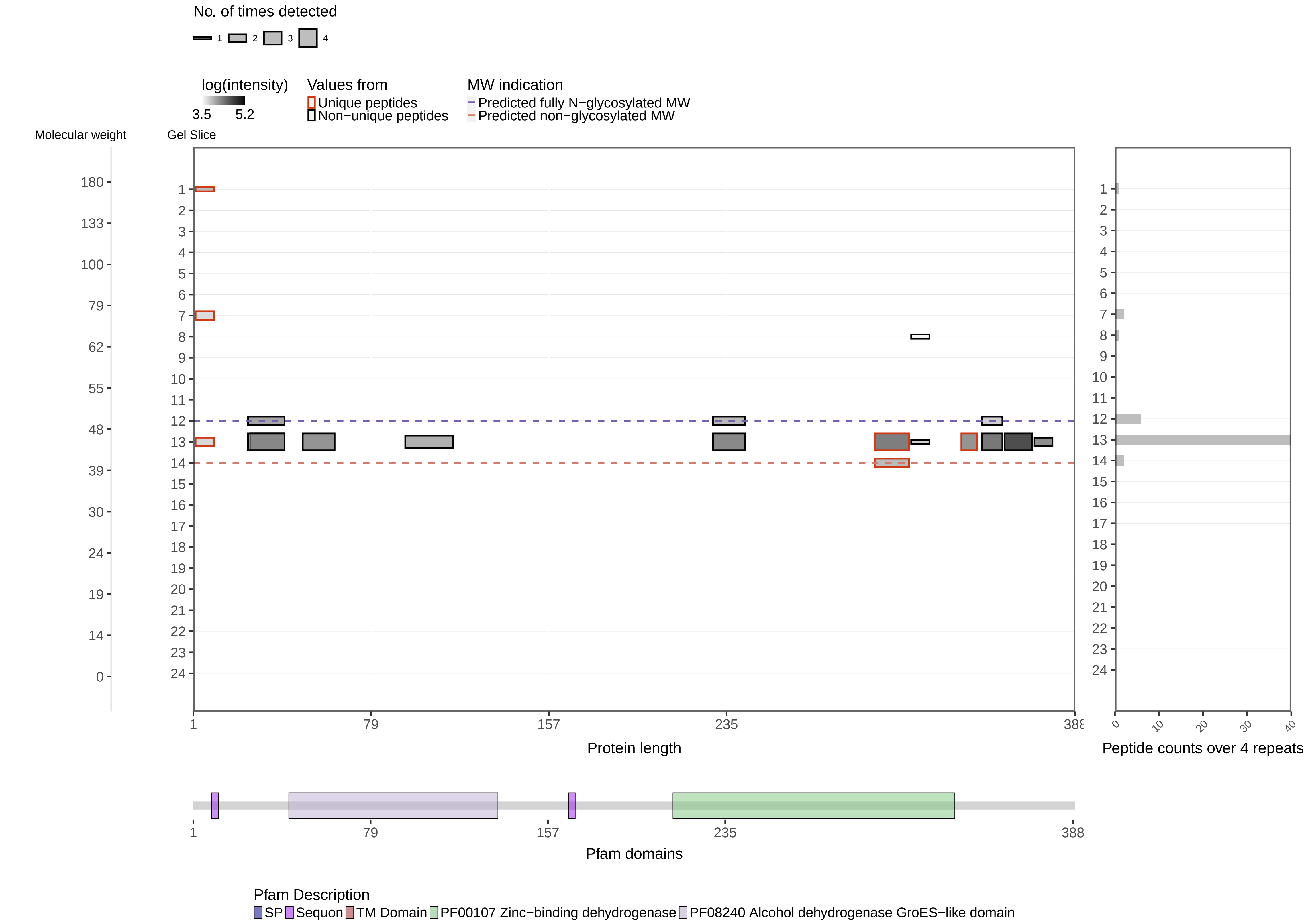


Protein sequence:

FSCGVSTGLGAAFKTANVEPGSTVVIFGLGAIGLAVAEGARICGATRIIGVDINSDKFEIGKOFGVTDFVNSNSFGDKPISOVINEMTNGGADYCFECVGLGTVVOEAYACCRKGWGKTIVLGVDKPGADLTFKSFDVLHSGKTLMGSLFGGLKPKSDIPFLVKRYLDKELOLDKFVTHEVSFODINKAFDLLAOGKSLRCVIWMD*

NbL18g12280.1 aspartic_proteinase_Asp1 [MW − SP = 42.69 kDa]

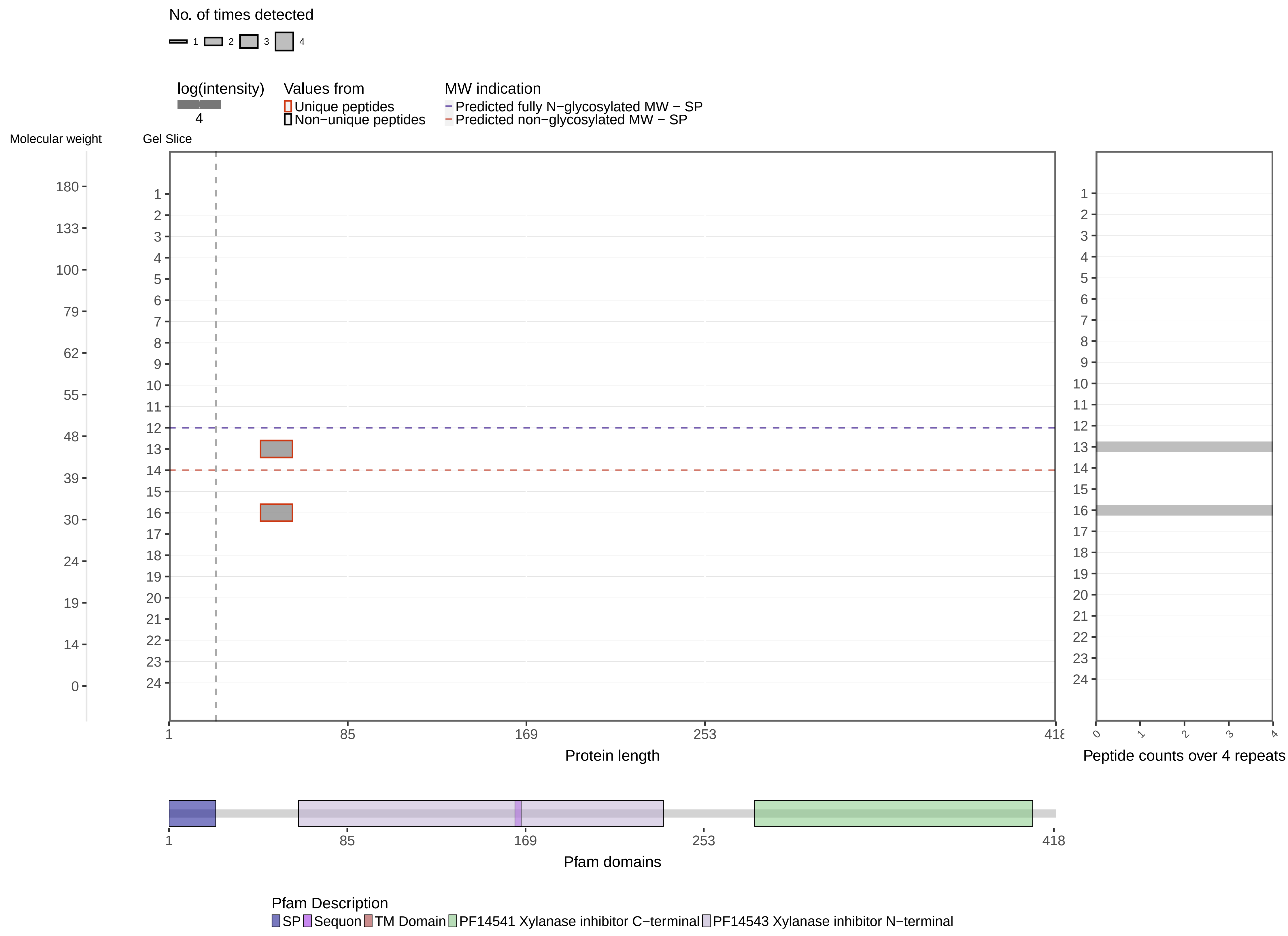


Protein sequence:

GILGLGNGKSSVLSQLSSLGLIRNVVGHCLSGQGGGFLFFGDDVLPSSGIVWTPIVRASSEKHYSLGPADLLFSGQATGIKGIPIIFDSGSTYSYFNSKAYKILVSSIKKDVNTKQLTDAVDDKSLPVCWKGSKSFKSIHDAKSFFKPLTLSFRKAKNVQLQLPPEAYLVRTEHGNICLGILDGTEVGLGNYNLIGDISMVDKMVIYDNEKQQIGWVPANCNKLP*

### NbL18g12270.1 catalase_isozyme_1 [MW = 56.85 kDa]

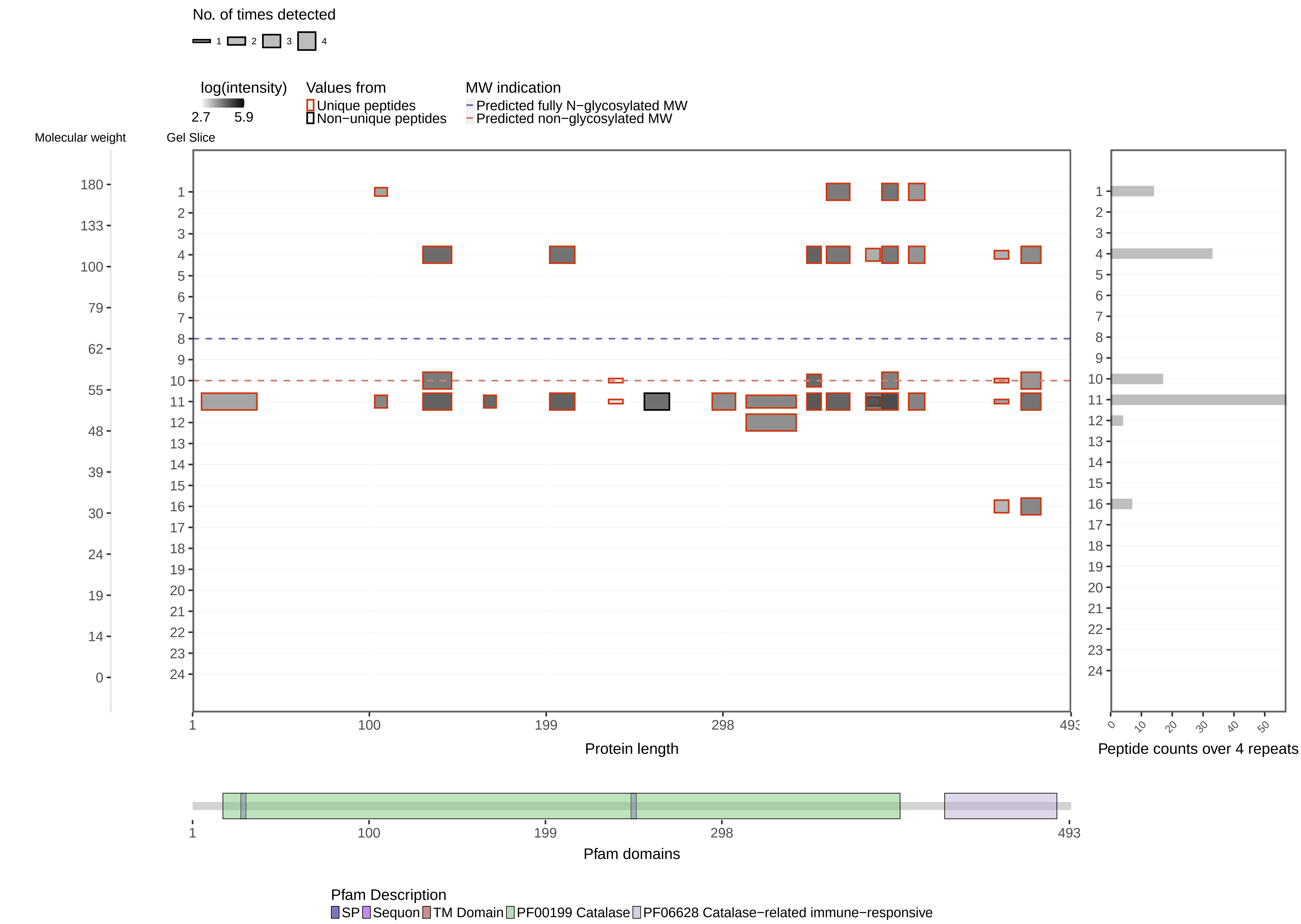


Protein sequence:

MSEEEAIRVGGTNHSHATKDLYDSIAAGNYPEWKLFIQIMDTEDVDKFDFDPLDVTKTWPEDILPLMPVGRLVLNRNIDNFFAENEQLAFNPGHIVPGLYYSEDKLLQTRIFAYADTQRHRIGPNYMQLPVNAPKCAHHNNHRDGAMNFMHRDEEVDYLPSRFDPCRHAEQYPIPSRVLTGRREMCVIEKENNFKQAGERYRSWEPDRQDRYVSKWVEHLSDPRVTYEIRSIWISYLSQADKSCGQKVASRLTLKPTM*

### NbL18g12180.1 psbP_domain−containing_protein_3_chloroplastic−like [MW = 28.34 kDa]

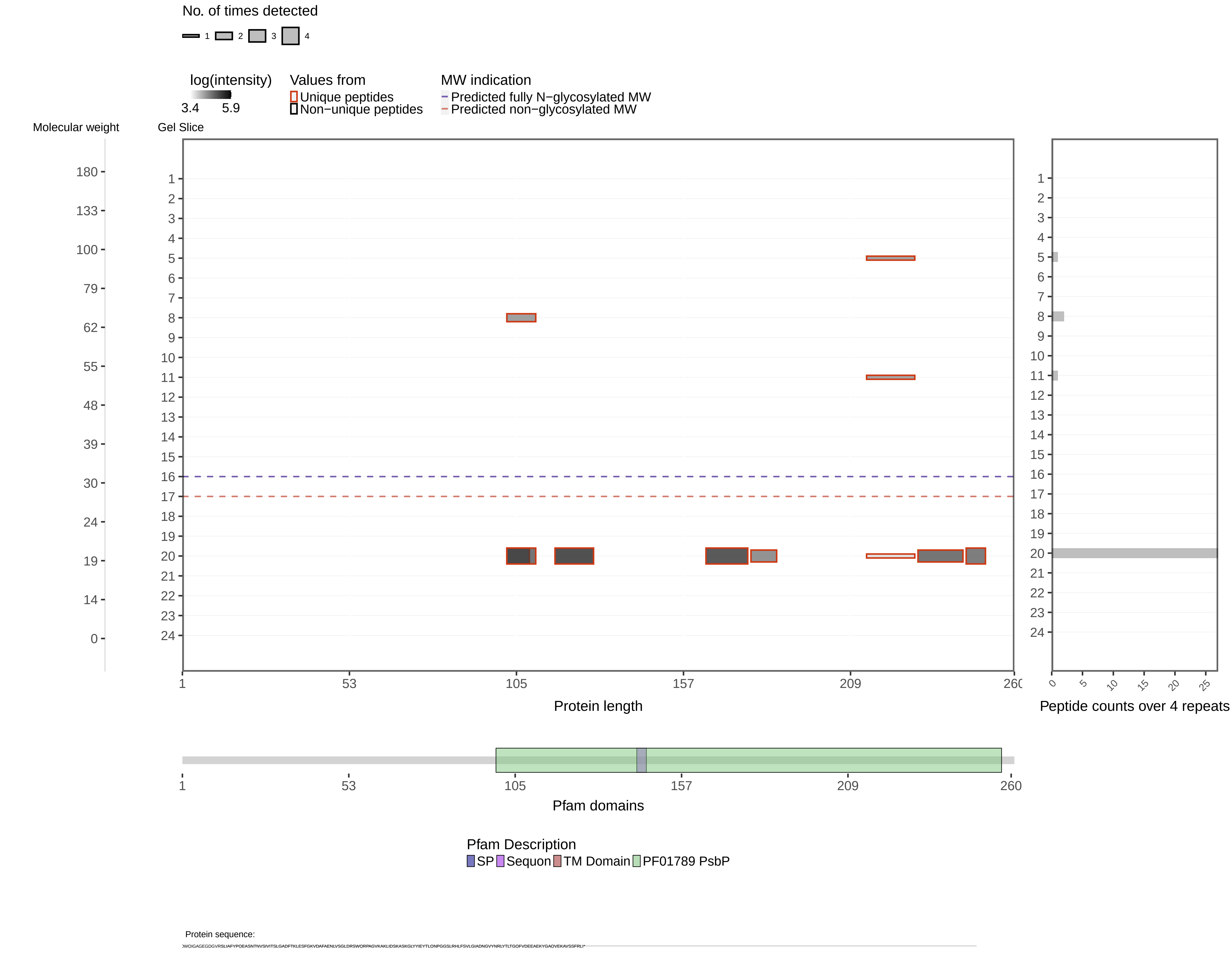

### NbL18g12170.1 pyrophosphate−−fructose_6−phosphate_1−phosphotransferase_subunit_alpha−like [MW = 67.18 kDa]

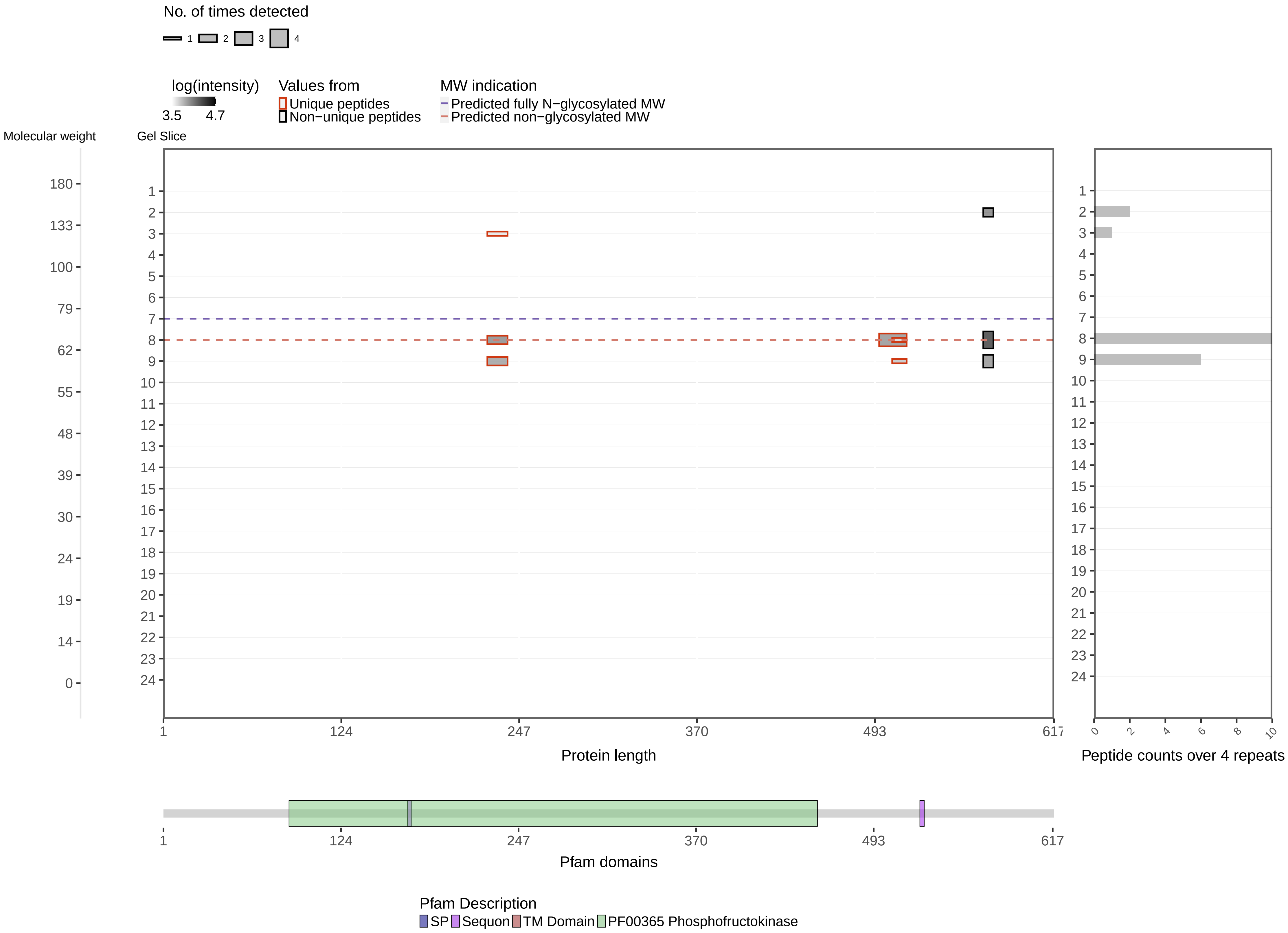


Protein sequence:

KLTLFDITNKICDAVQARAEQDKNHGVILLPEGLIESIPEVYALLQEIHGLLRQGVSADKISSQLSPWASALFEFLPPFIKKQLLLYPESDDSAQLSQIETEKLIAHLVETEMNKRLKEGTYKGKKFNAICHFFGYQARGSLPSKFDCDYAYVLGHICYHILAAGLNGYMATVTNLKNHVNKWRCAAAPITAMMTVKRYGRGPGNAAIGKPAVHPTTVDLKGKAYELLSQNATKFLLDDVYRNPGPLQFDGPGADAKAVTLCVEDQDYMGRIKKLQEYLDKVRTIVKPGCSQDVLKA

### NbL18g12160.1 UPF0047_protein_YjbQ [MW = 22.5 kDa]

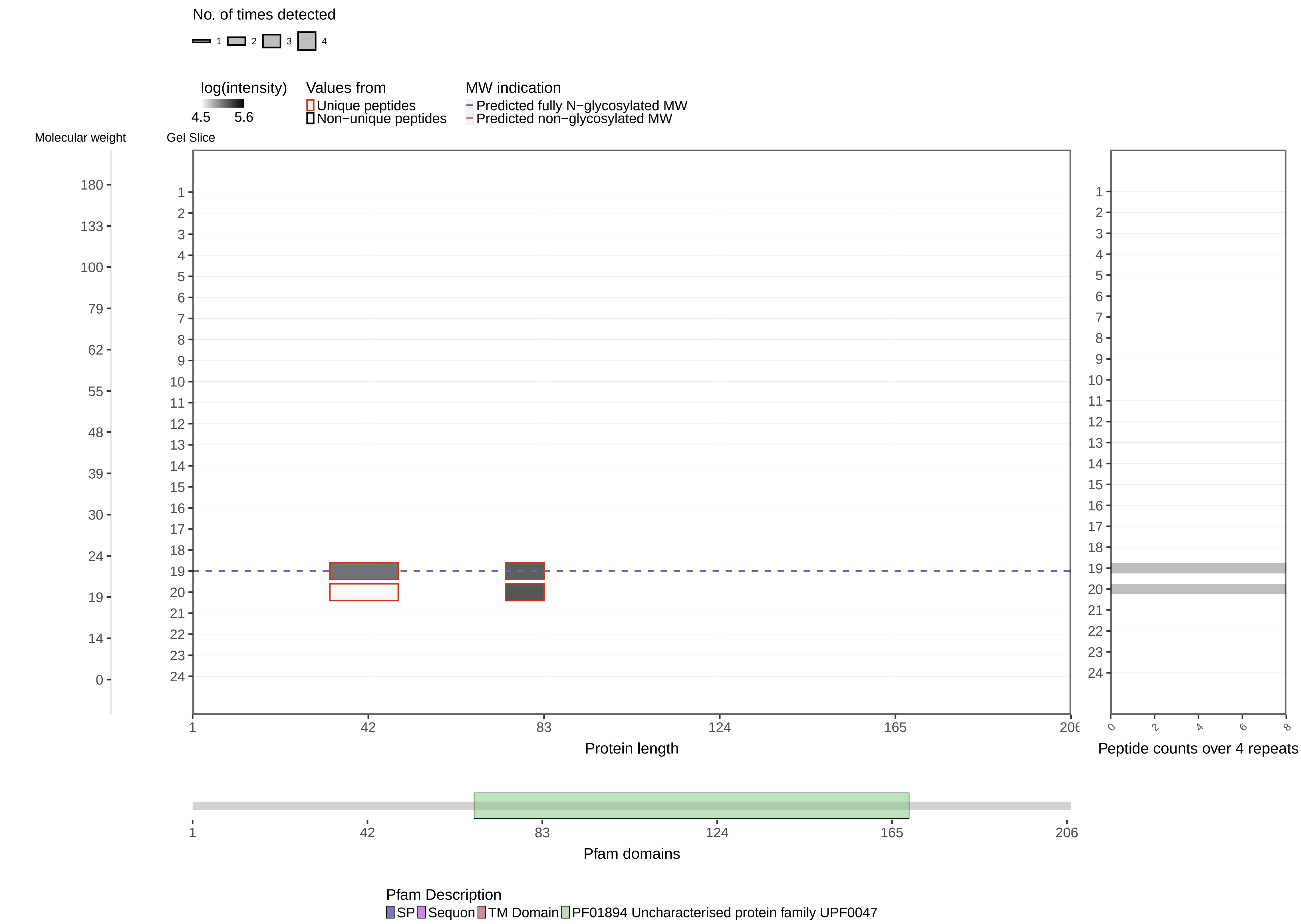


Protein sequence:

FLQHTSASLTINENYDSDVRDDTETFLNRVVPEGISAPWKHTLEGPDDMPAHIKSSMFGCTLTSEIPITDGKLNMGTWQYLPSSAEEEPWIQLGIWLCEHRDAATPRRIVITLNGM*

NbL18g12080.1 tryptophan−−tRNA_ligase_chloroplastic/mitochondrial [MW = 49.77 kDa]

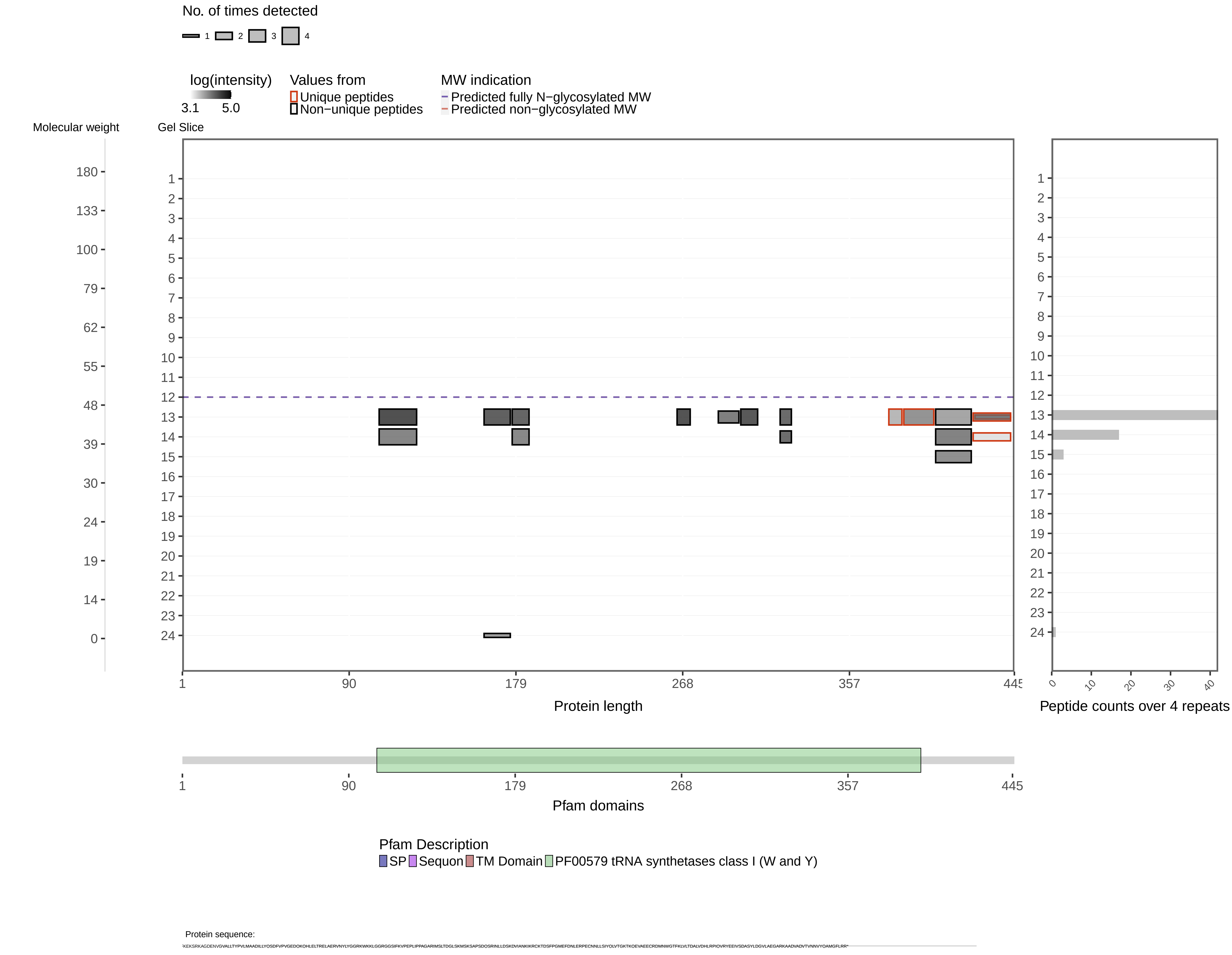

### NbL18g12010.1 serine_hydroxymethyltransferase_3_chloroplastic−like [MW = 31.26 kDa]

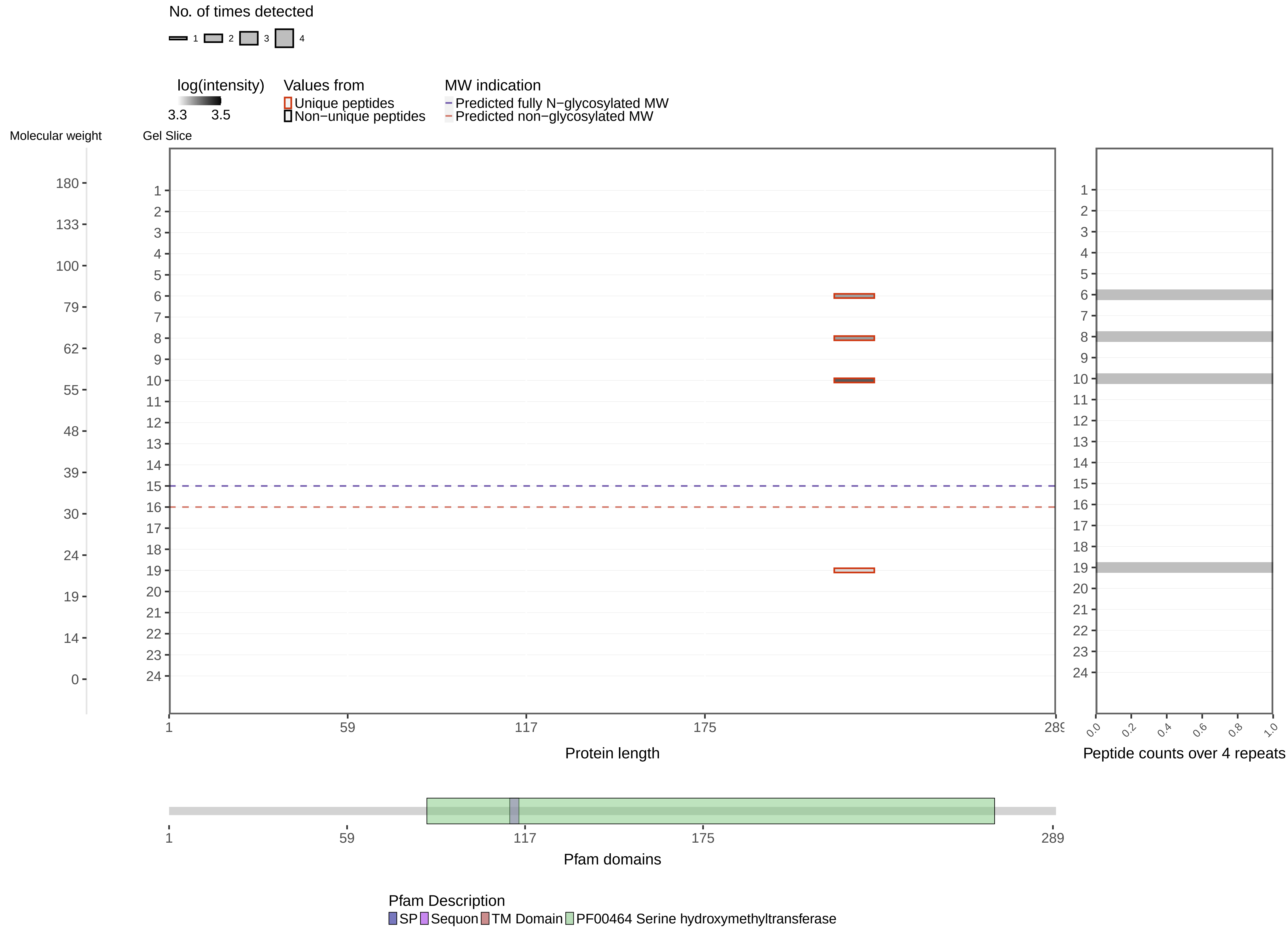


Protein sequence:

EGLPGKRYYGGNEYIDELEILCQERALAAFNLDGKQWGVNVQPLSGSPANFEVYTAILNPHDRIMGLDLPHGGHLSHGFMTPKRRVSGTSIYFESMPYRLDESSGLVDYDMLEKTASLFRPKLIIAGASAYPRDFDYPRSQAGLNCFIRSSAGQRGC*

### NbL18g11940.1 far_upstream_element−binding_protein_2 [MW = 67.71 kDa]

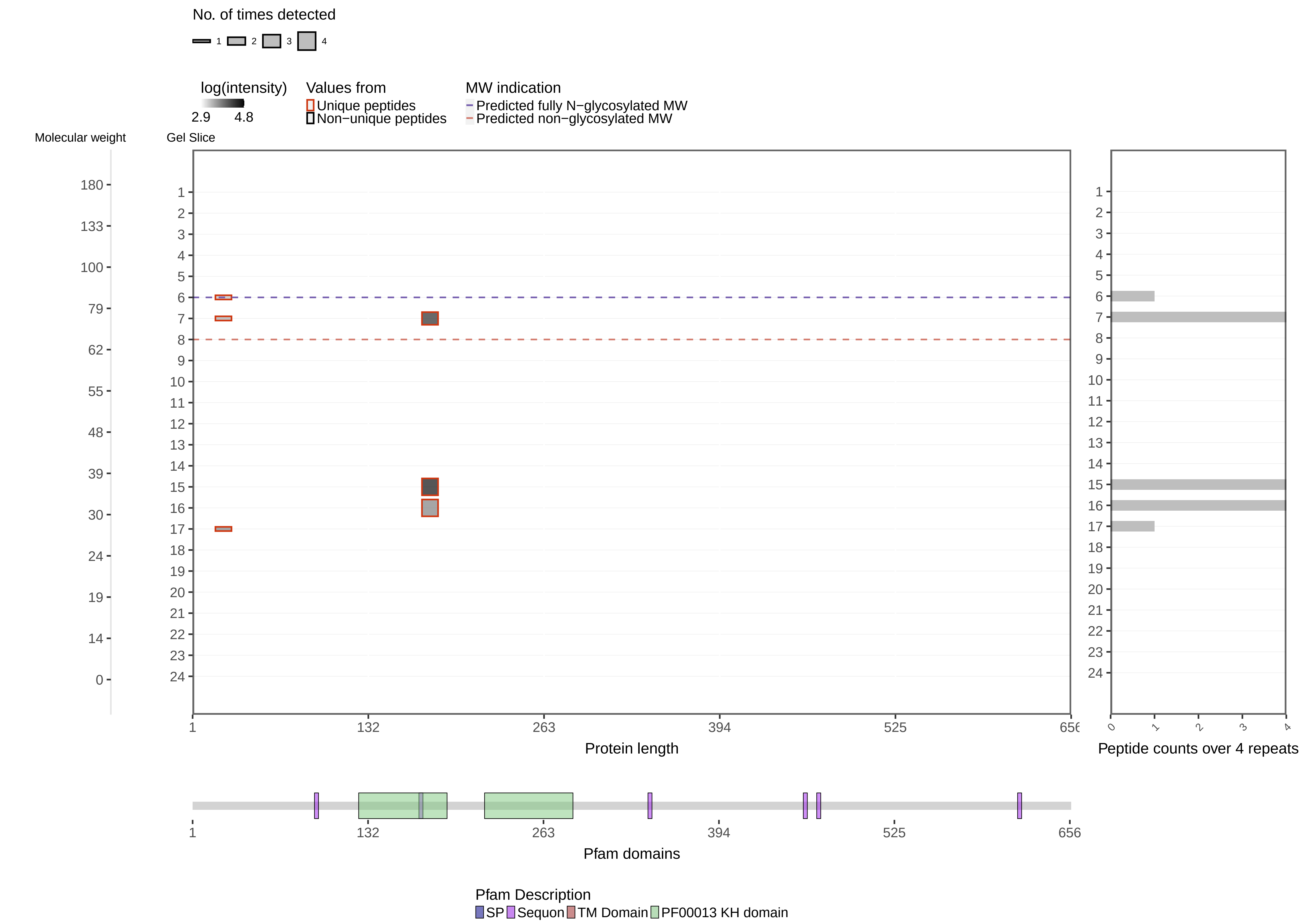


Protein sequence:

QPGYGYVQPGAYPGPSPQYNTSQQPYPGYPPQPASAGYASGWDQTASQNQQSQGGGYDYYNQQAPPQQQQTPGGTAASTDNSAYGYNQLSASGYSQGQAYSQDGYGGYHAPAPQSGYQGTGYEQQQGYSSAAGYGNASNPTSEGQNSSYGTQGDGSQAPPPVQSSATGQQGSYPSQGSTQAGYGMPPTSQGGYGTPPPAGYAASYGAPQTQKPSPSQPAYGQPQQSPSAQGGYAQPPPAQPGYPQQAAQSGYALPDSGTLRPPASGYGAPTAQPGY

NbL18g11750.1 60S_ribosomal_protein_L7−2 [MW = 30.82 kDa]

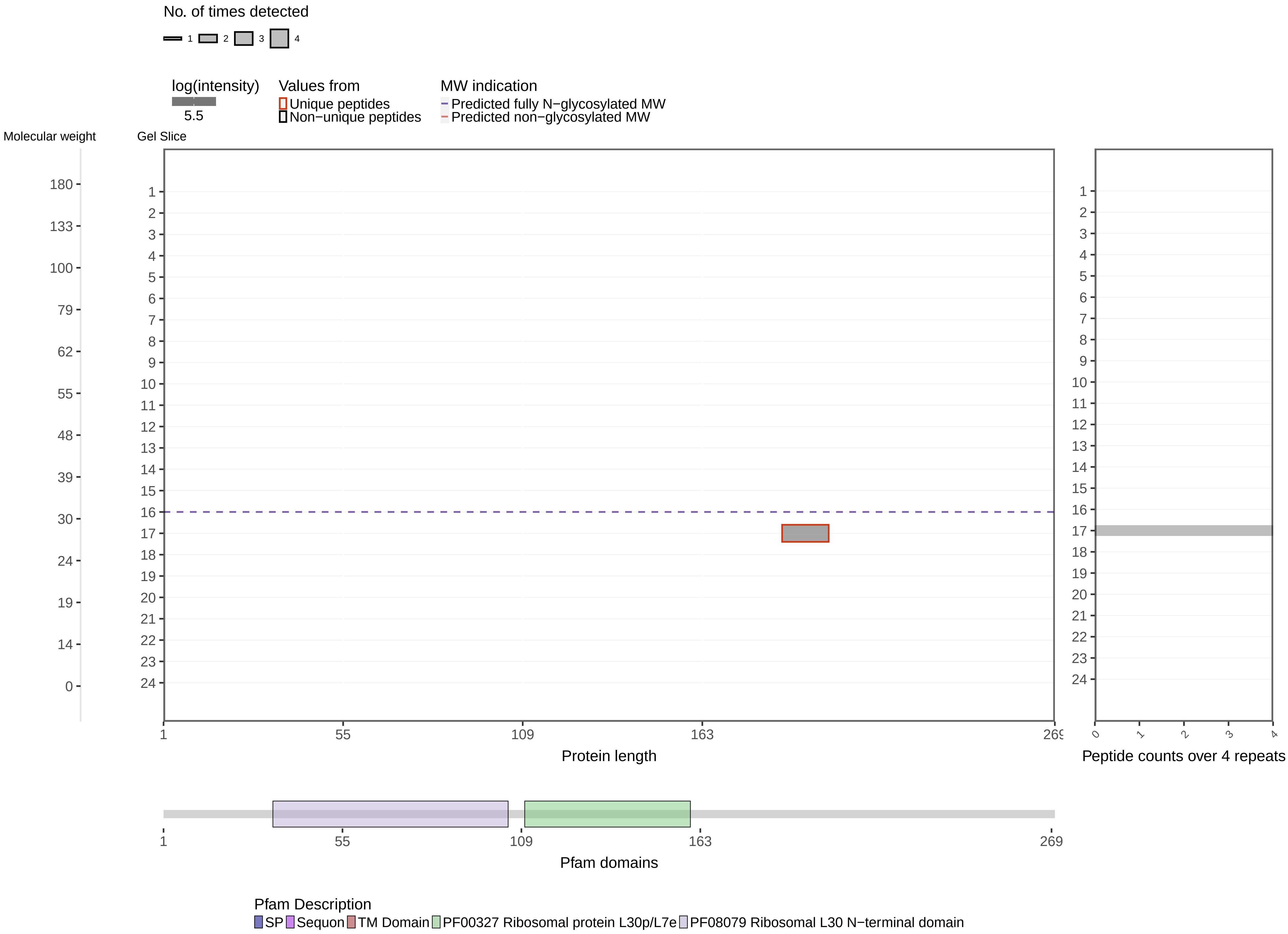


Protein sequence:

MAPQTKKILQLLRLRQIFNGVFLKVNKATVNMLHRVEPYVTYGYPNLKSVRELIYKRGYGKLDKQRIALTDNSIIEQALGKHGIICAEDLVHEIMTVGPHFKEANNFLWPFQLKAPLGGLKKKRNHYVEGGDAGNRENYINELIRRMN*

NbL18g11660.1 40S_ribosomal_protein_S6−like [MW = 28.32 kDa]

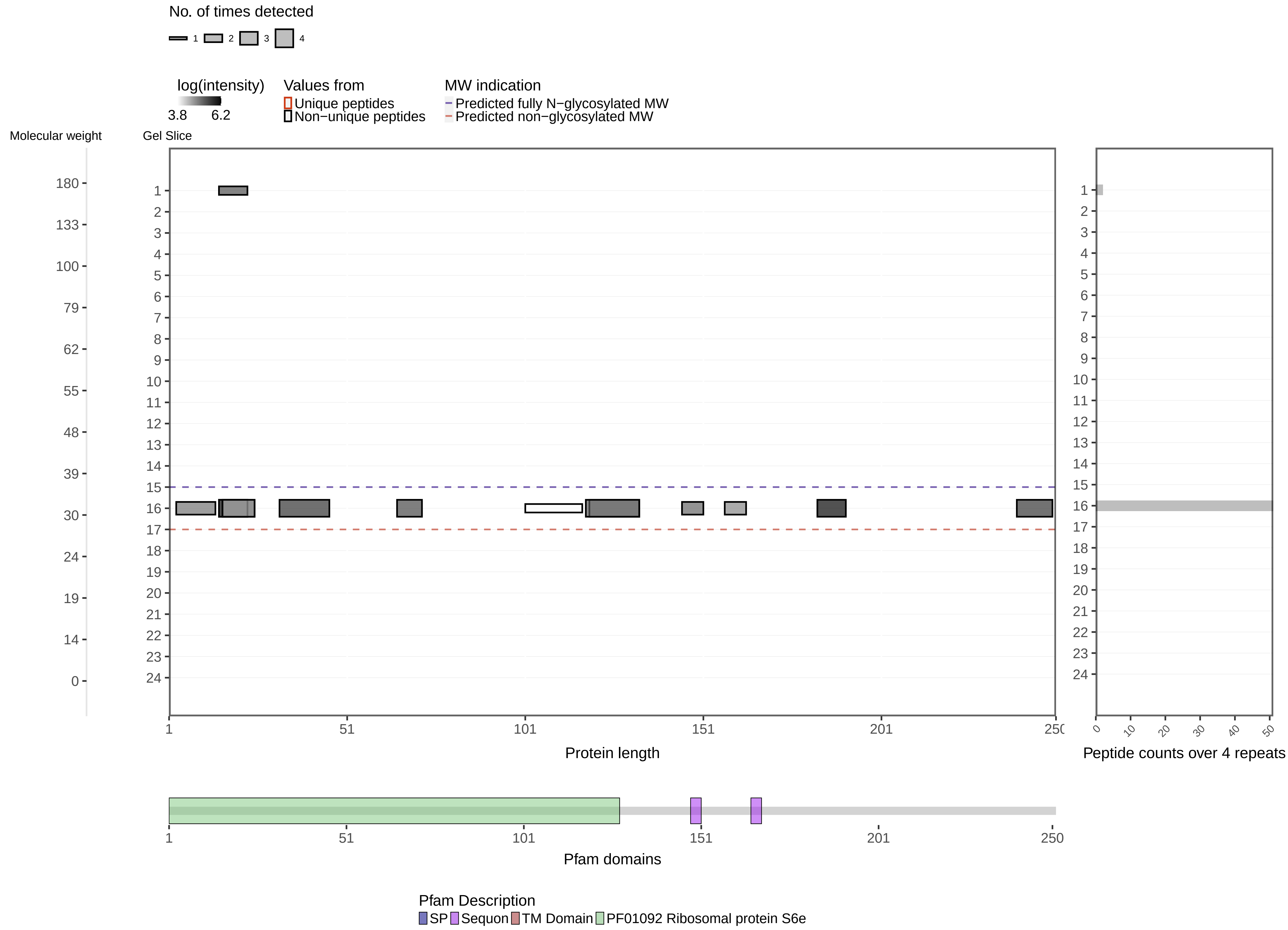


Protein sequence:

/LNLVIVKKGENDLPGLTDTEKPRMRGPKRASKIRKLFNLSKEDDVRKYVNTYRRNFTTKTGKKASKAPKIQRLVTPLTLQRKRARIADKKKRIAKAKSEAAEYQKLLATRLKEQRERRSESLAKKRSRLSAASKPSIAA*

NbL18g11560.1 probable_protein_phosphatase_2C_59 [MW = 31.49 kDa]

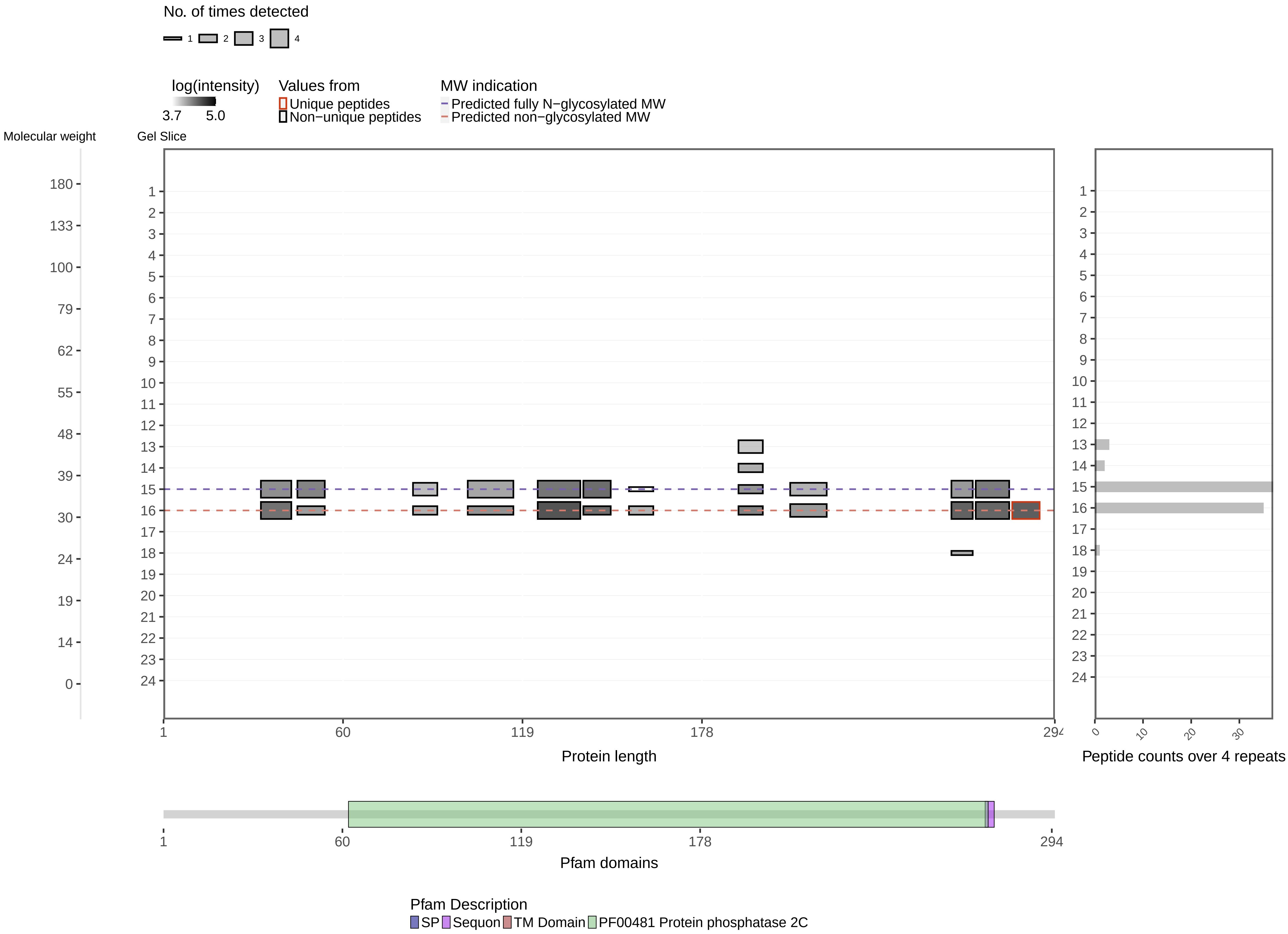


Protein sequence:

.VGDRLLVANVGDSRAVICRGGTAIAVSRDHKPDQTDERQRIEDAGGFVMWAGTWRVGGVLAVSRAFGDRLLKQYVVADPEIQEEKVDGSLEFLILASDGLWDVVTNEEAVSMVKPIQDPEEAAKRLMQEAYQRGSADNITIVVVRFLGNQEGSSRGVSA*

### NbL18g11540.1 apyrase_2−like [MW = 51.2 kDa]

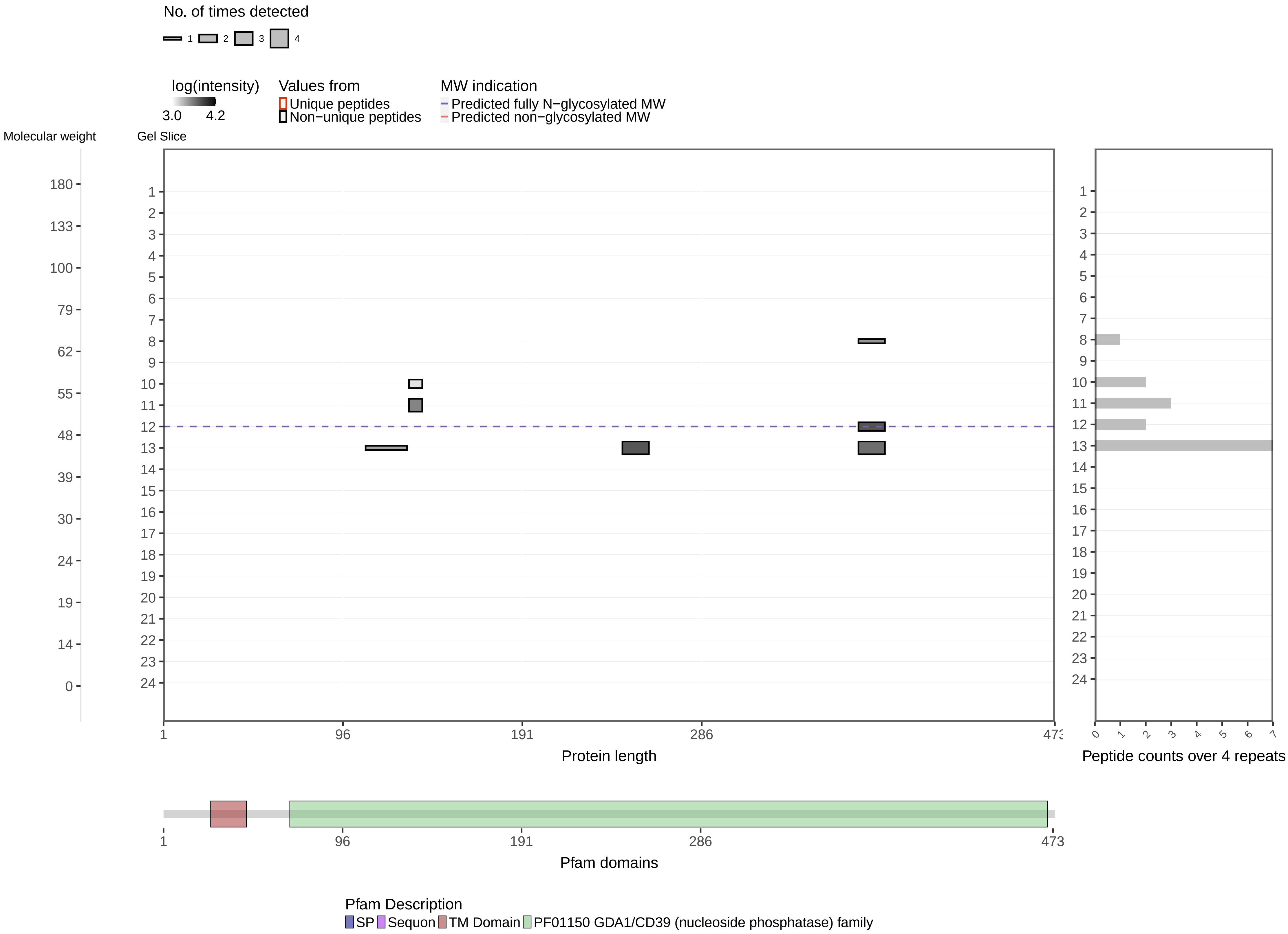


Protein sequence:

SVQMAYAIAESDAQKAPKVSDGEDTYVQEMYLKGTKYYLYVHSYLHYGLLAARAEILKATDESGSPCILGGYHGSYKYGGAVYPASAMSQGSSLSSCREVSLKALKVNEPTCTHMKCTFGGVWNGGGGDGQKNMFVASFFFDRAAEAGMINPSLAVAKVRPVDYESAAQRACETGLEGAKSAFPRVDPDNLPYLCMDLVYQYTLLVDGFGLDPQQEITLVKKVQYKNSLVEAAWPLGSAIEVASSLT*

NbL18g11510.1 67−dimethyl−8−ribityllumazine_synthase_chloroplastic−like [MW = 25.46 kDa]

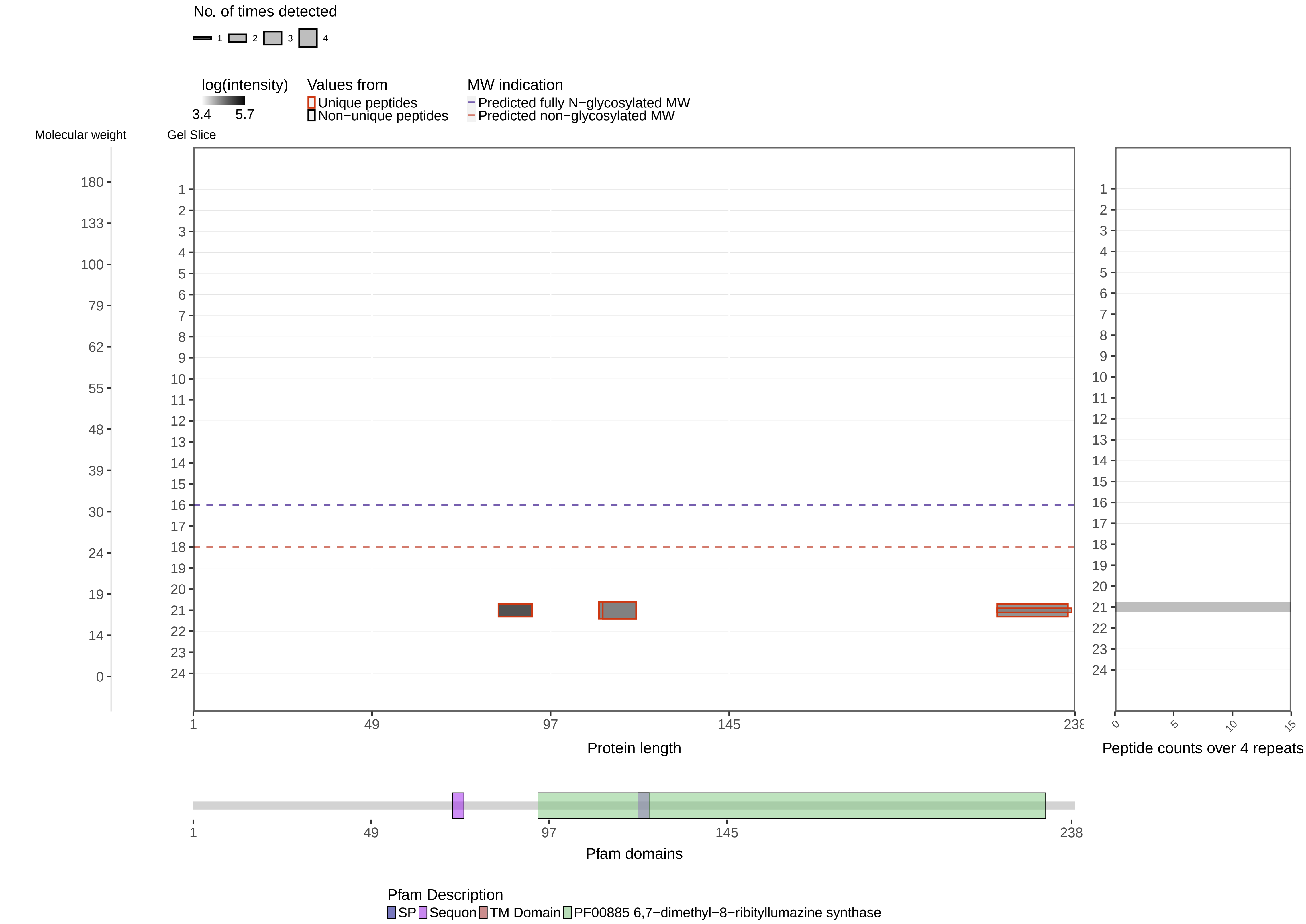


Protein sequence:

LITKKLLEGALETFKNYSVREEDIDVVWVPGCFEIGVVAQQLGKSQKYQAILCIGAVIRGDTSHYDAVVNAATSGVLSAGLNSGTPCIFGVLTCDTLEQAFNRVGGKAGNKGAETALTAIEMASLFEHHLKV*

NbL18g11320.1 uncharacterized_LOC109236326 [MW = 30.24 kDa]

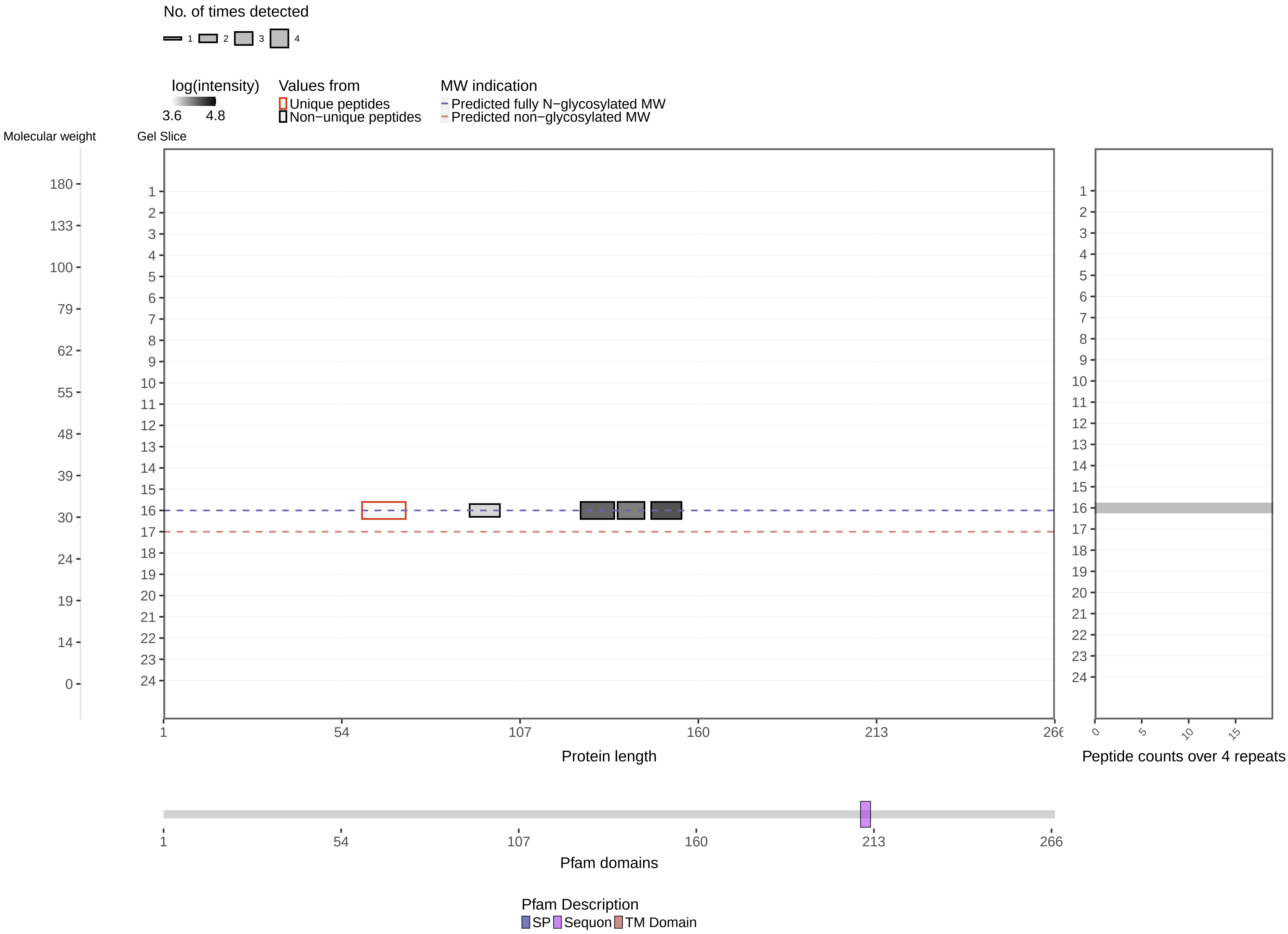


Protein sequence:

LYSKVFQGVGDFSLRENDYVFGNRKFGGNAQSITKGRWIHHTSFLWDYEMINMAYLKLPKRAPEYRQARDHSDFICRMKDYLSRQEFINRTISTLDSHFSVTSLELKSFDCPDDSKFMPSSRLLGKQELEESFESESGNVILQSM*

NbL18g11300.1 prolycopene_isomerase_1_chloroplastic [MW = 61.73 kDa]

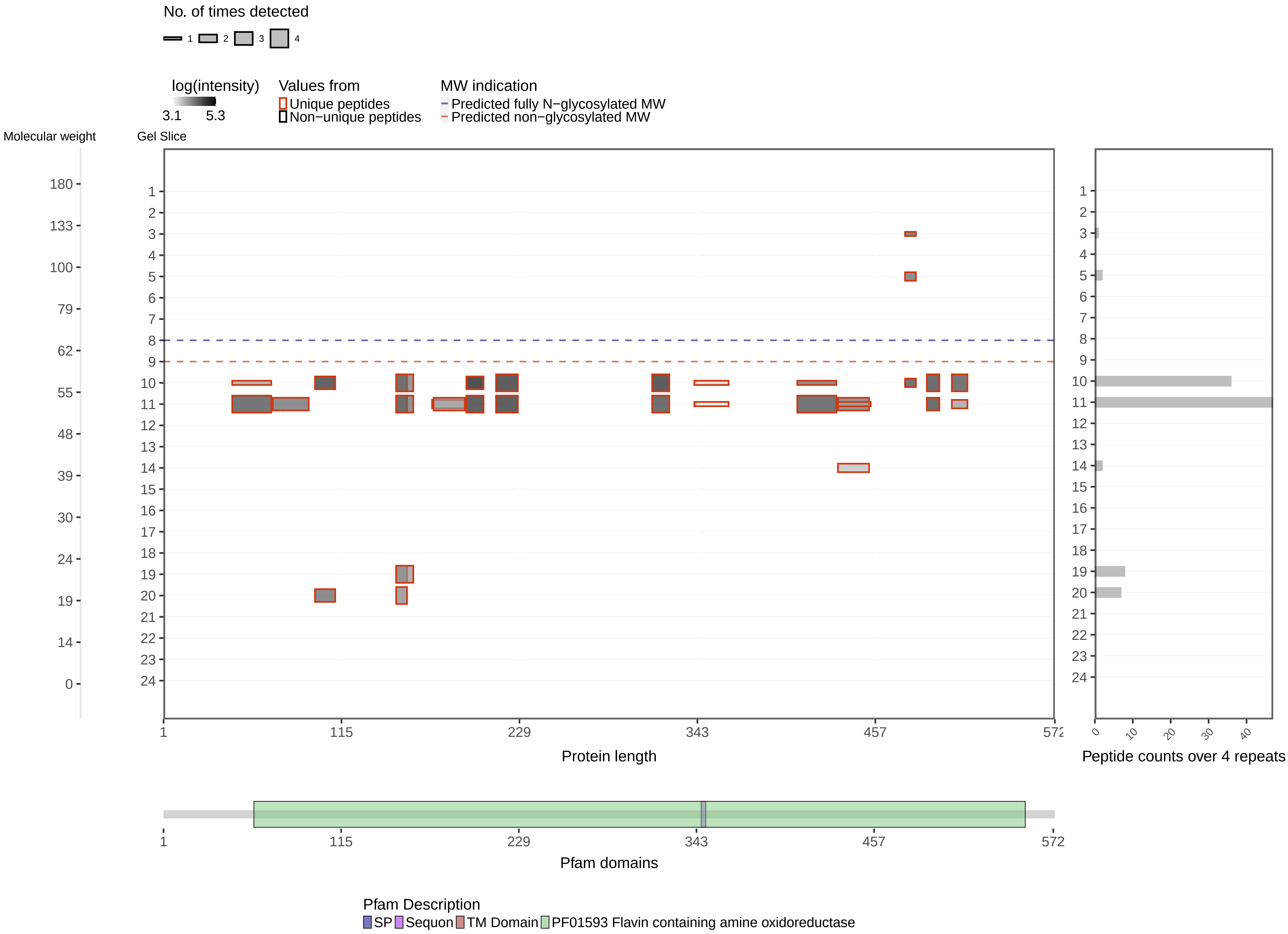


Protein sequence:

FSEWYKPGCTLEYPLQGSGAIVDALVRGLQKFGGRISLKSHVENIVVENGRAVGVKLRGGQFVRAKKAVVSNASMWDTLSLLPLEVVPKSYRDNIKSTPQCESFMHLHLGFDAEGIRDDLGIHHIIVNDWDRGVDADQNVVLISVPSVLSPNLAPPGKHILHAYTPGTEPFEIWEGLDRRSNEYKNLKAERSEVMWRAVEKALGPGFNREKCEVKLVGTPLTHKRFLRRNRGTYGPAILAGKGTFPGHSTPIPQLWCCGDSTFPGIGVPAVAASGSIVANSLVSVAEHSQLL[illegible]

NbL18g11220.1 glutathione_S−transferase_U9−like [MW = 25.5 kDa]

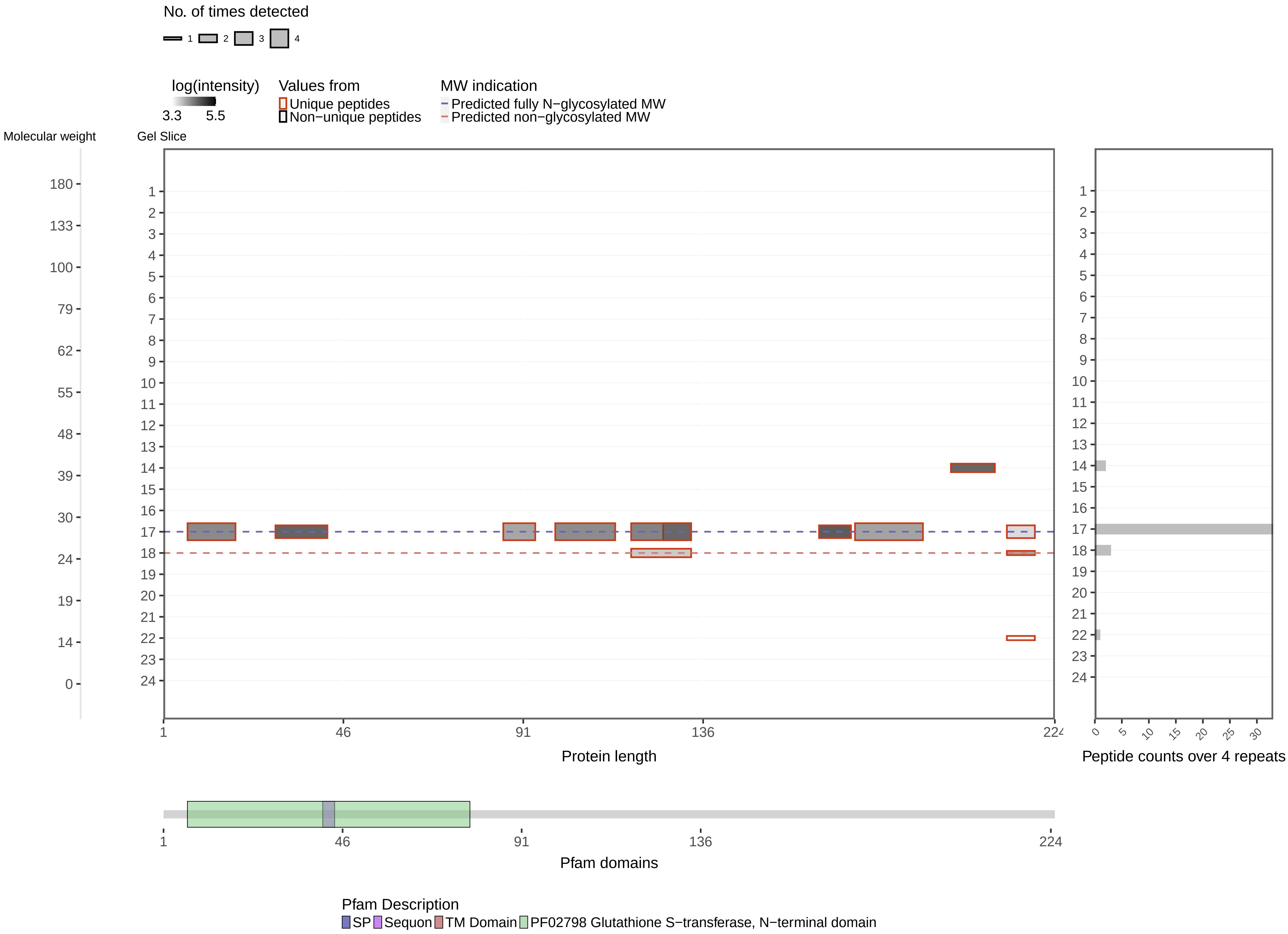


Protein sequence:

RFWANYIDQHLIPSTAKLWKSTGEEQEKALEEVLEKIRILEKGIPEIHGNEFGLLQILLVTTFGSHKAYEDVLGVKILVPENNPLLSSCVTSLKELPLVKEICPPYDMLLSKLKIYRENALKSTTH*

NbL18g11190.1 flavonol_synthase/flavanone_3−hydroxylase−like [MW = 36.18 kDa]

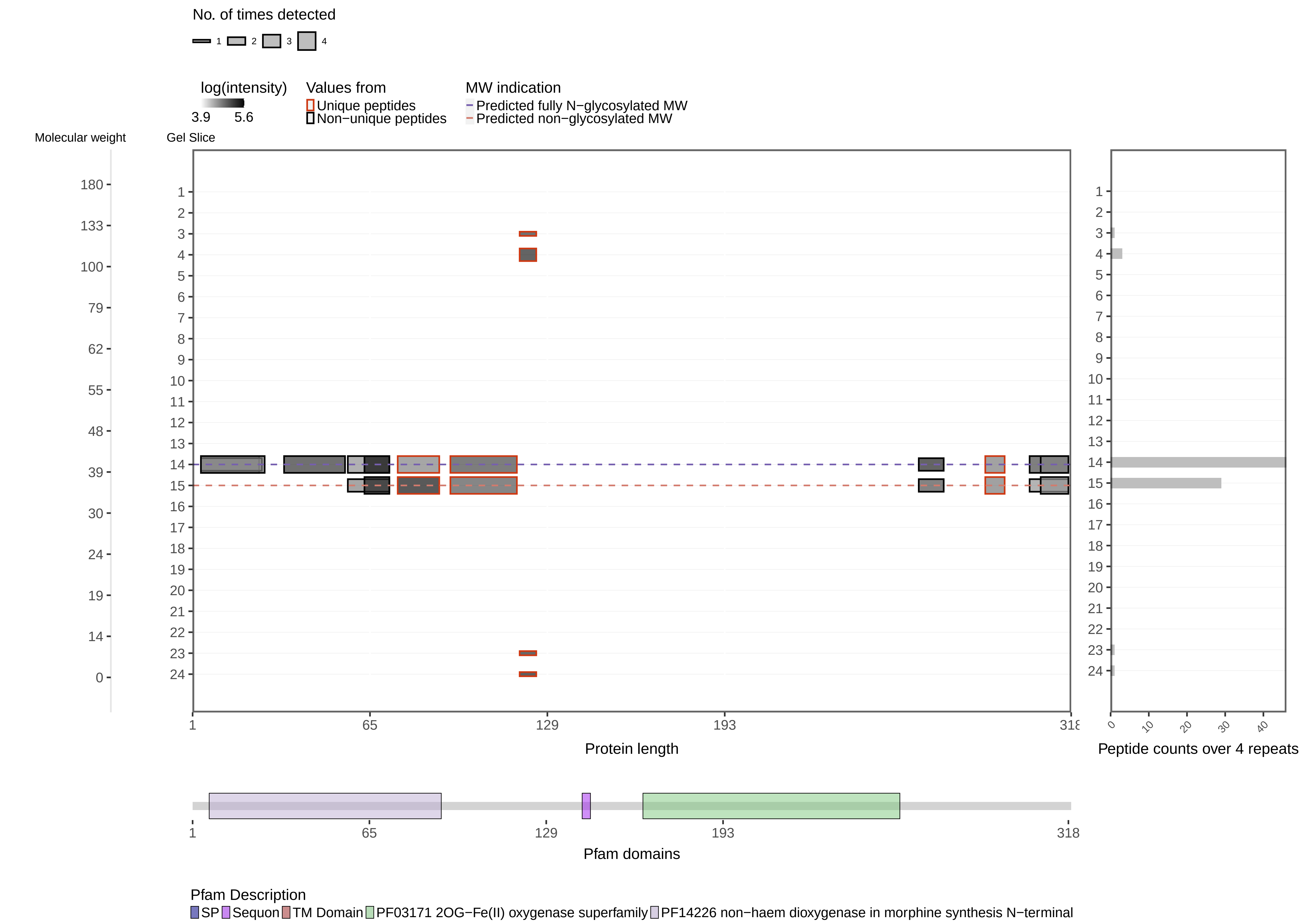


Protein sequence:

LPPNCLKDYNNDRCADLLLSCRYFPATEMEDNGASSHQDPVVLGLIFQDDAGGLEVLIDGEWIPVTPIEGALVFHICDVLQVLTNDKFVSPFHQVLRPKGKSRYVFVFFYNLNGEKWVEPLPQFSKEIGEPPKYRRFQYKDYFGAWVKEKLNPPVRHEDKTSLITFASATN*

### NbL18g11080.1 flavonol_synthase/flavanone_3−hydroxylase−like [MW = 36.12 kDa]

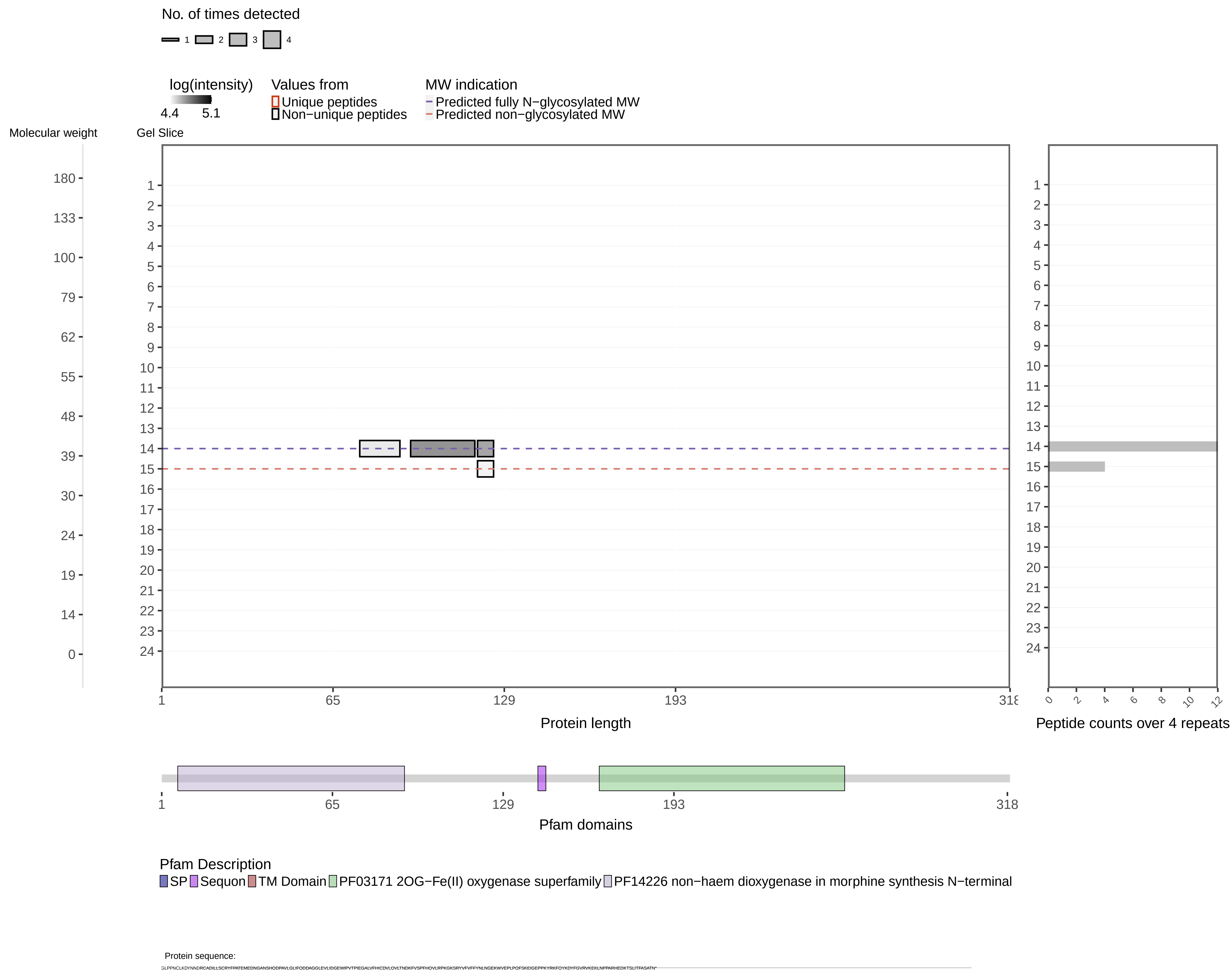

### NbL18g10900.1 uncharacterized_LOC107799956 [MW = 31.48 kDa]

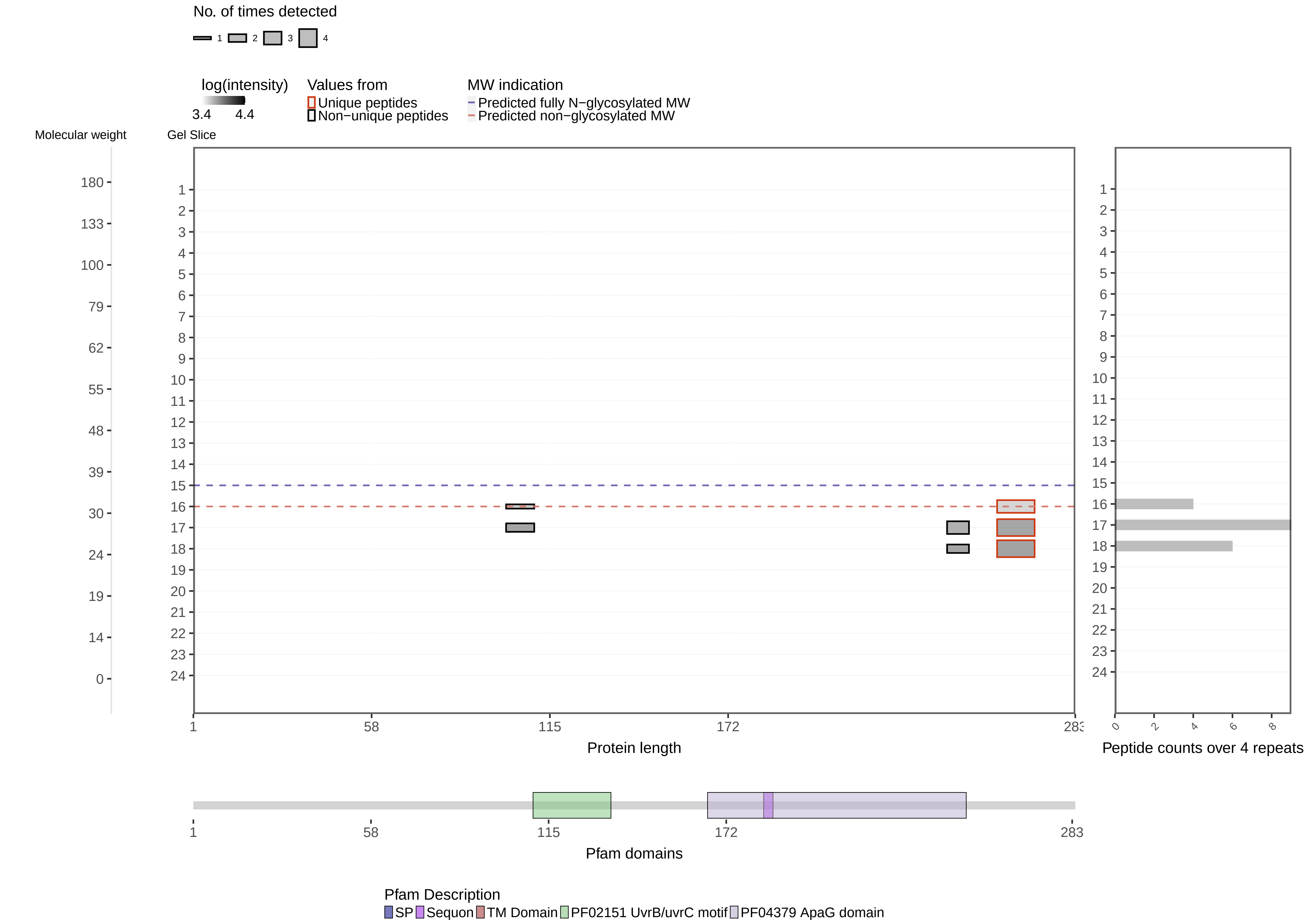


Protein sequence:

[illegible]ARYRDELKDVAPHSLLKCSSDATTLGIRVQVRSVYIEGRSHPSKGLYFFAYRIRITNNSDRPVQLLRRHWIITDANGKTENVWGIGVIGEQPVILPNTGFEYSSACPLGTPNGRMEGDFEMKHIDKVGSRTFNVAIAPFSLSKVGDGSDTFSDGS*

NbL18g10890.1 60S_ribosomal_protein_L44 [MW = 12 kDa]

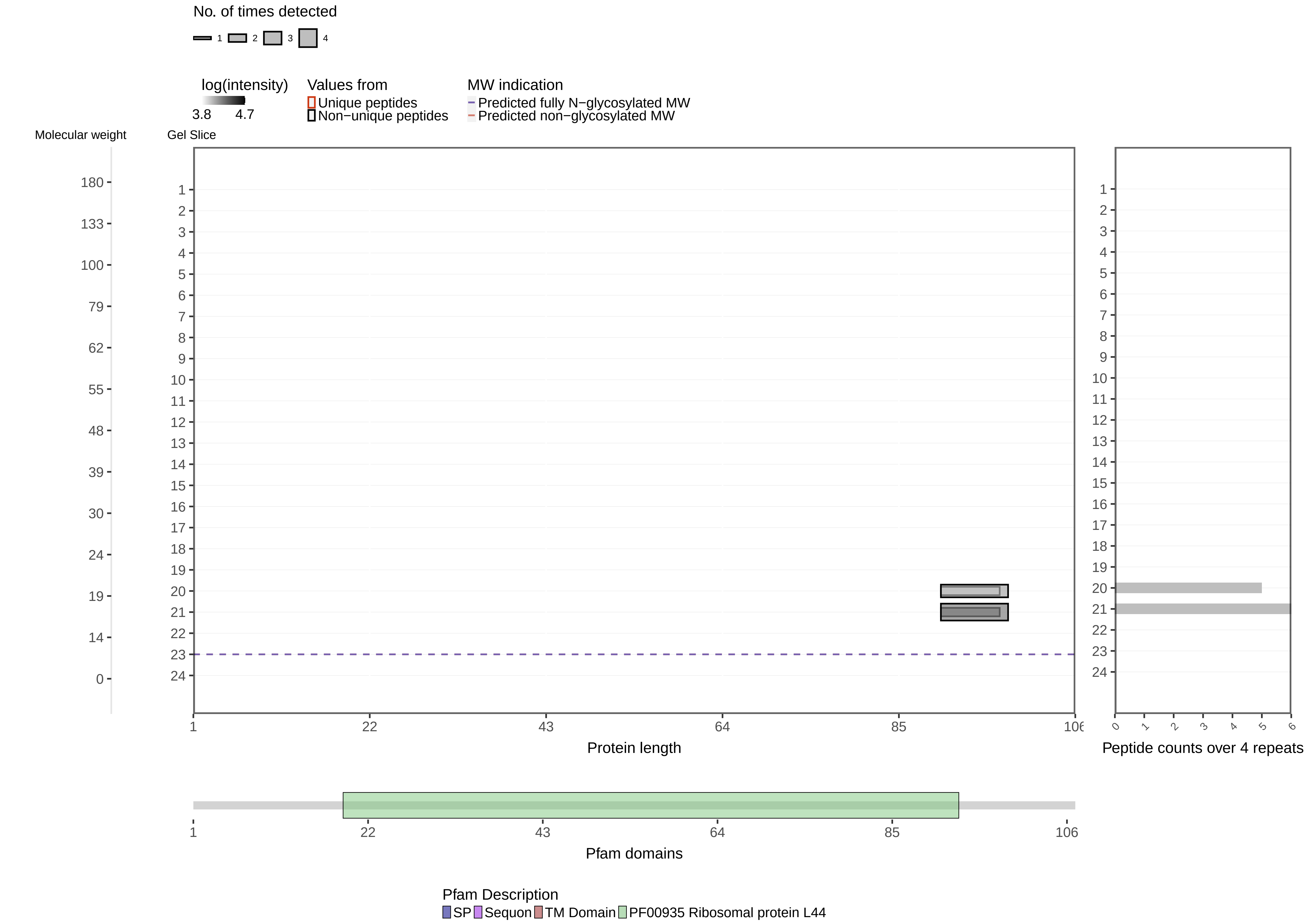


Protein sequence:

KRRYDRKQSGYGGQTKPVFHKKAKTTKKIVLRLQCQGCKHVSQHPIKRCKHFEIGGDKKGKGTSLF*

NbL18g10730.1 indole−3−glycerol_phosphate_synthase_chloroplastic−like [MW = 45.02 kDa]

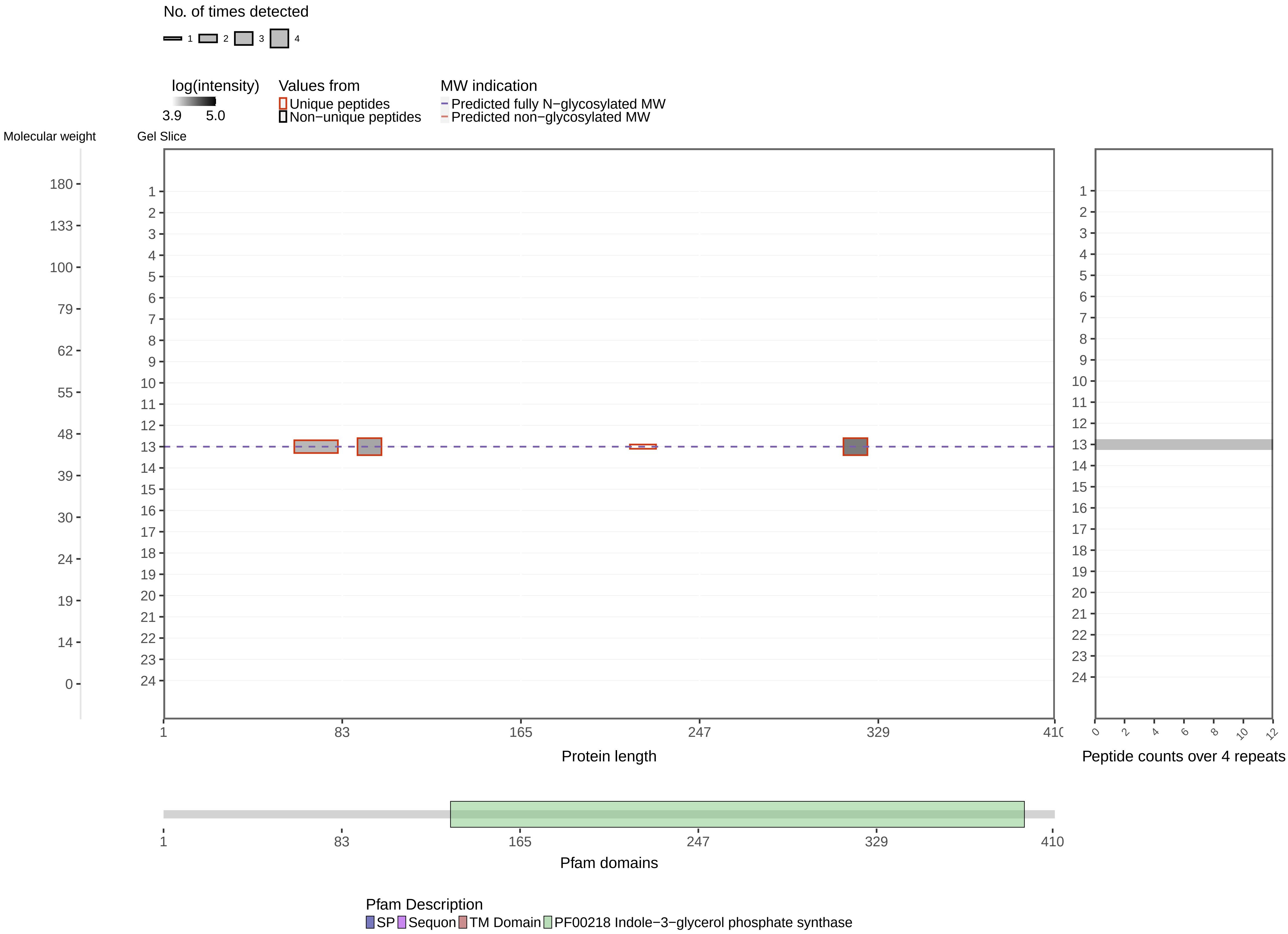


Protein sequence:

KASPSRGVLREDFNPVAIAKAYEKGGAACLSVLTDEKYFQGSFENLEAIRNAGVECPLLCKEFVVDAWQIYYARIKGADAVLLIAAVLPDLDIMYMTKICKLLGLTALVEVHDEREMDRVLRIDGIELIGINNRDLGTFKVDISNTKNLLEGERGKRIHEEGIIVVGESGLFTPADVAYVQEAGVKAVLVGESLVKQKDPAKGIRQLFGKDISCEGPSI*

### NbL18g10720.1 31_kDa_ribonucleoprotein_chloroplastic [MW = 34.51 kDa]

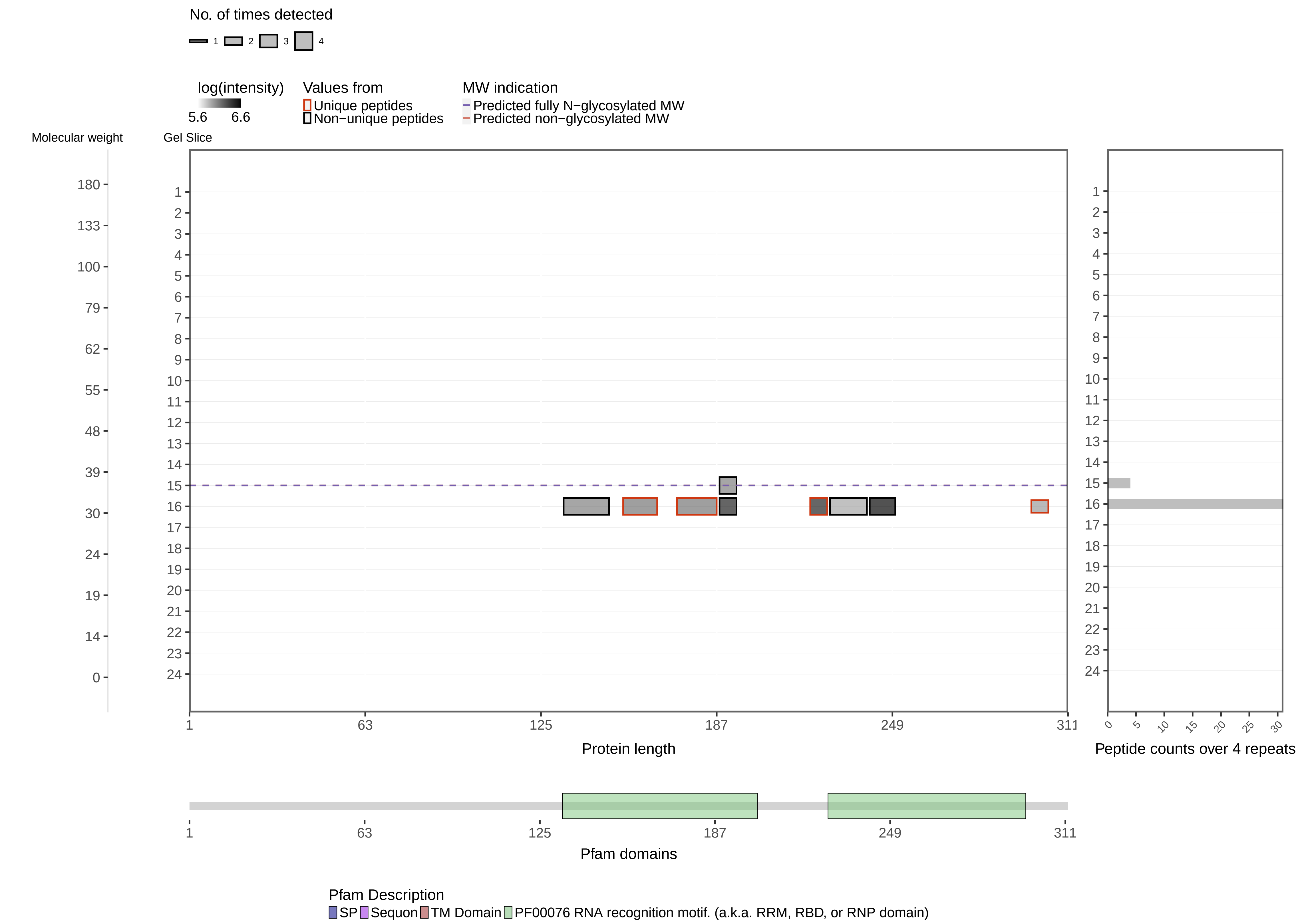


Protein sequence:

GLARLFEKAGVVEIAEVIYNRDTDQSRGFGFVTMSTVEEAEKAVEMYNRYDVNGRLLTVNKAARRGERPERPPRTFEQSYRIYVGNIPWGIDDARLEQLFSEHGKVVSARVVYDRETGRSRGFGFVTMASEAEMSDAIANLDGQSLDGRTIRVNVAEDRSRRNTF*

NbL18g10610.1 adenylate_kinase_4−like [MW = 26.74 kDa]

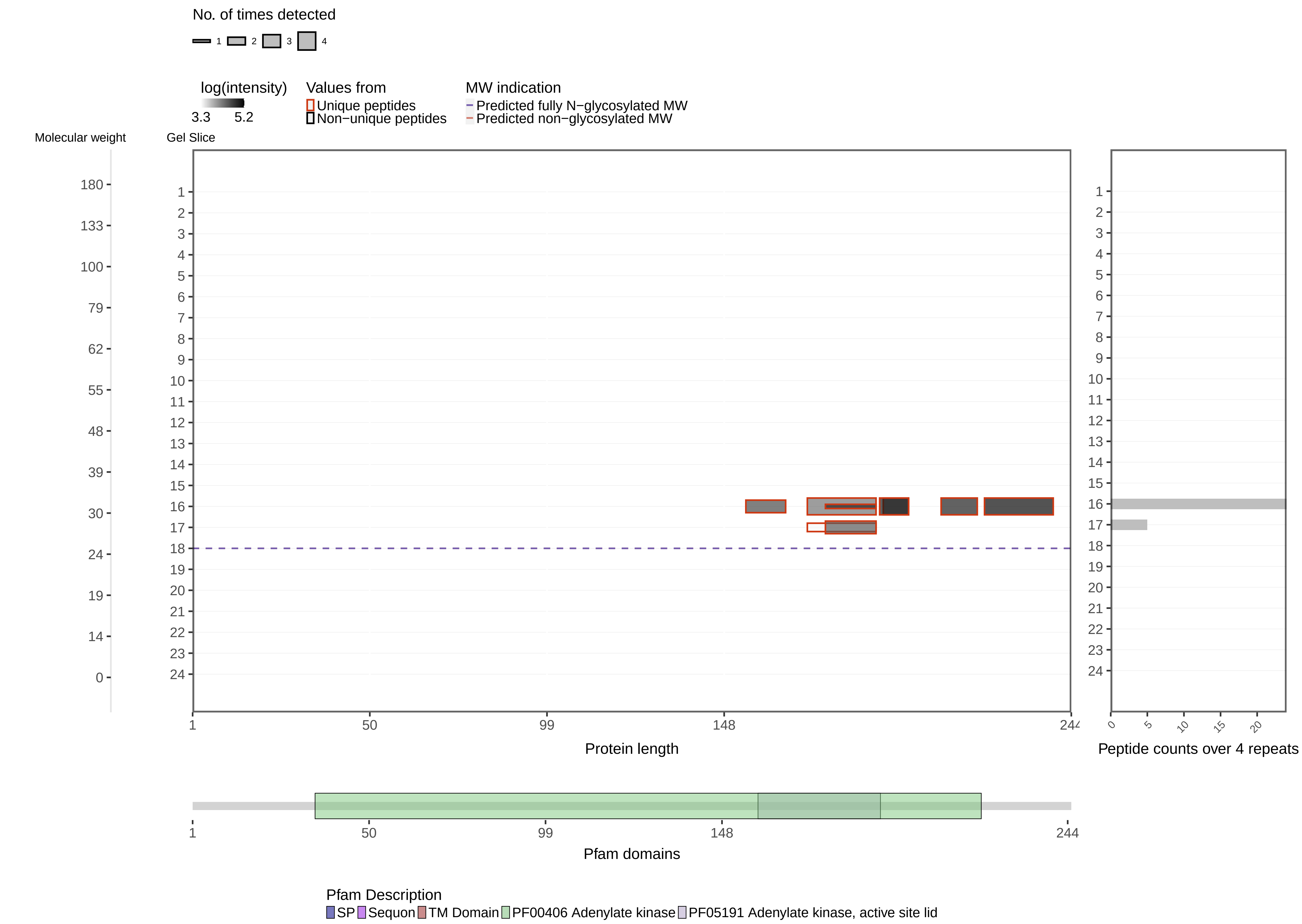


Protein sequence:

KGFILDGFPRTVVQAEKLDEMLQNRGTKVDKVLNFAIDDAILEERITGLWIHPASGRSYHTKFAPPKVPGIDDITGEPLIQRKDDTAAVFKSRLESFHRQTEPVIDYYVKKNMVVNLPAEKPPQTVTSEVKKALS*

### NbL18g10590.1 uncharacterized_LOC107817829 [MW = 10.95 kDa]

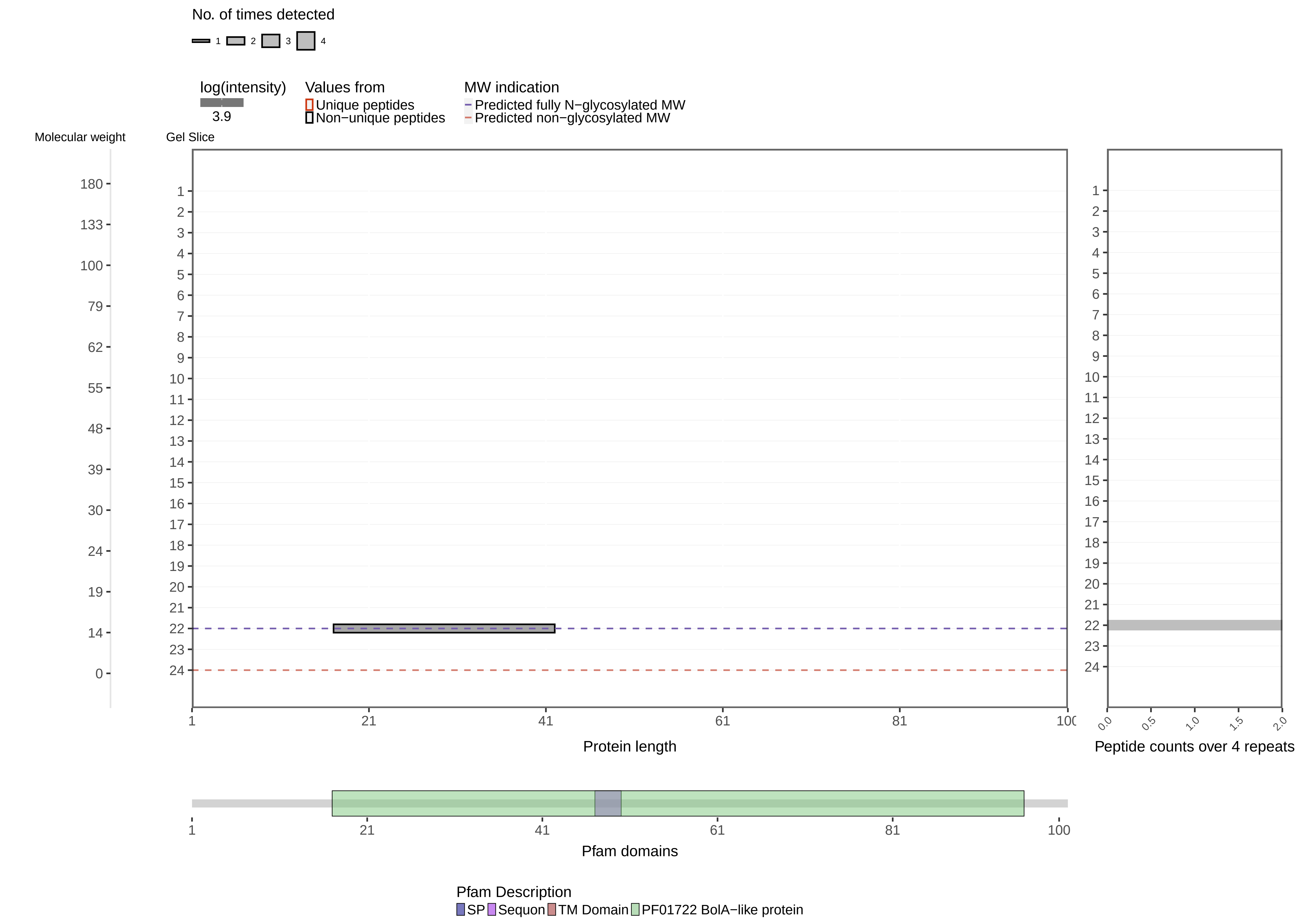


Protein sequence:

GHAAVKESGTNETHFNVKIVSSKFDGQNLVKRHRMVYELLSDELQSGLHALSIIAKTPKEAGM*

NbL18g10550.1 pyruvate_dehydrogenase_E1_component_subunit_beta−1_mitochondrial−like [MW = 40.55 kDa]

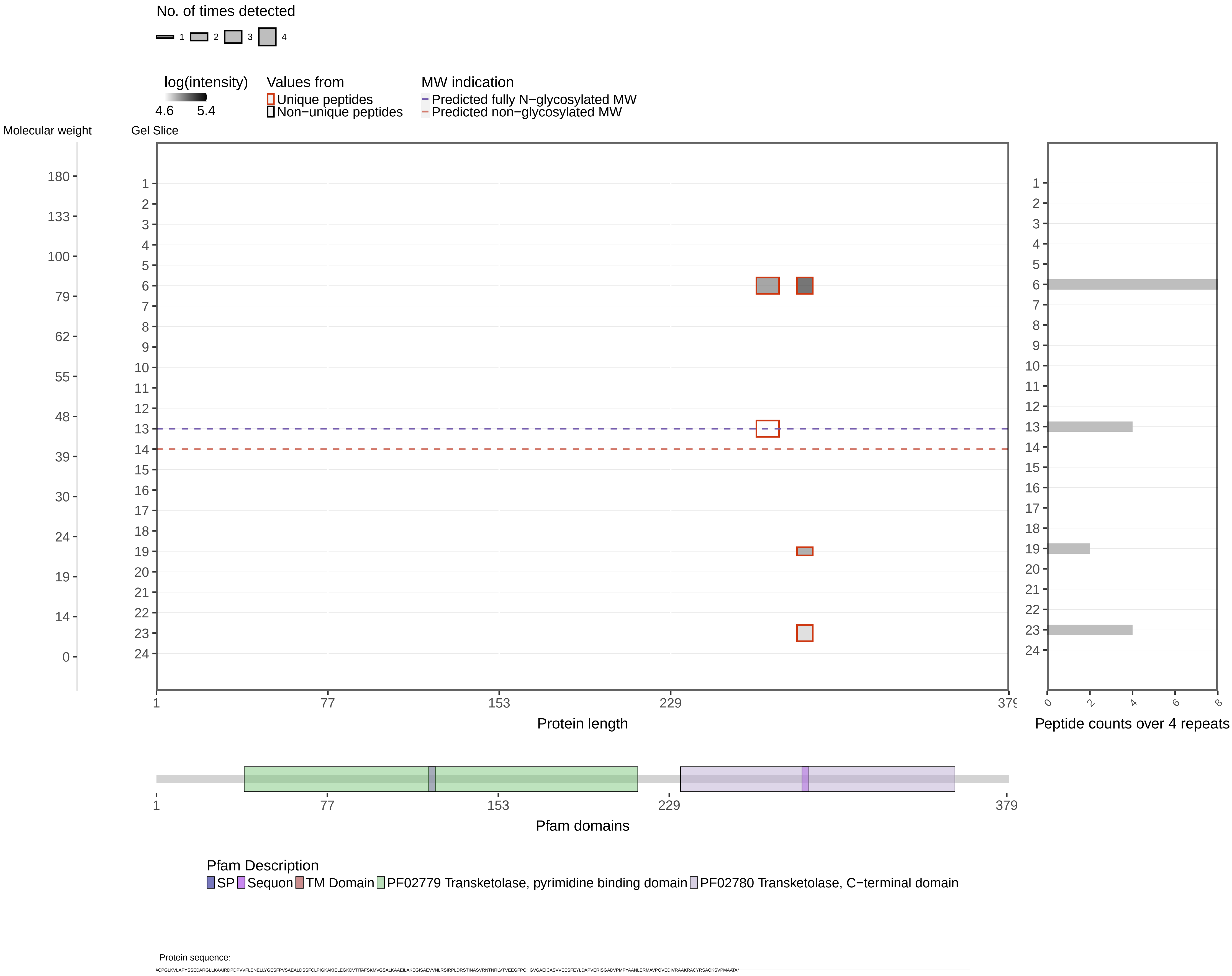

### NbL18g10400.1 protein_IQ−DOMAIN_14−like [MW = 54.31 kDa]

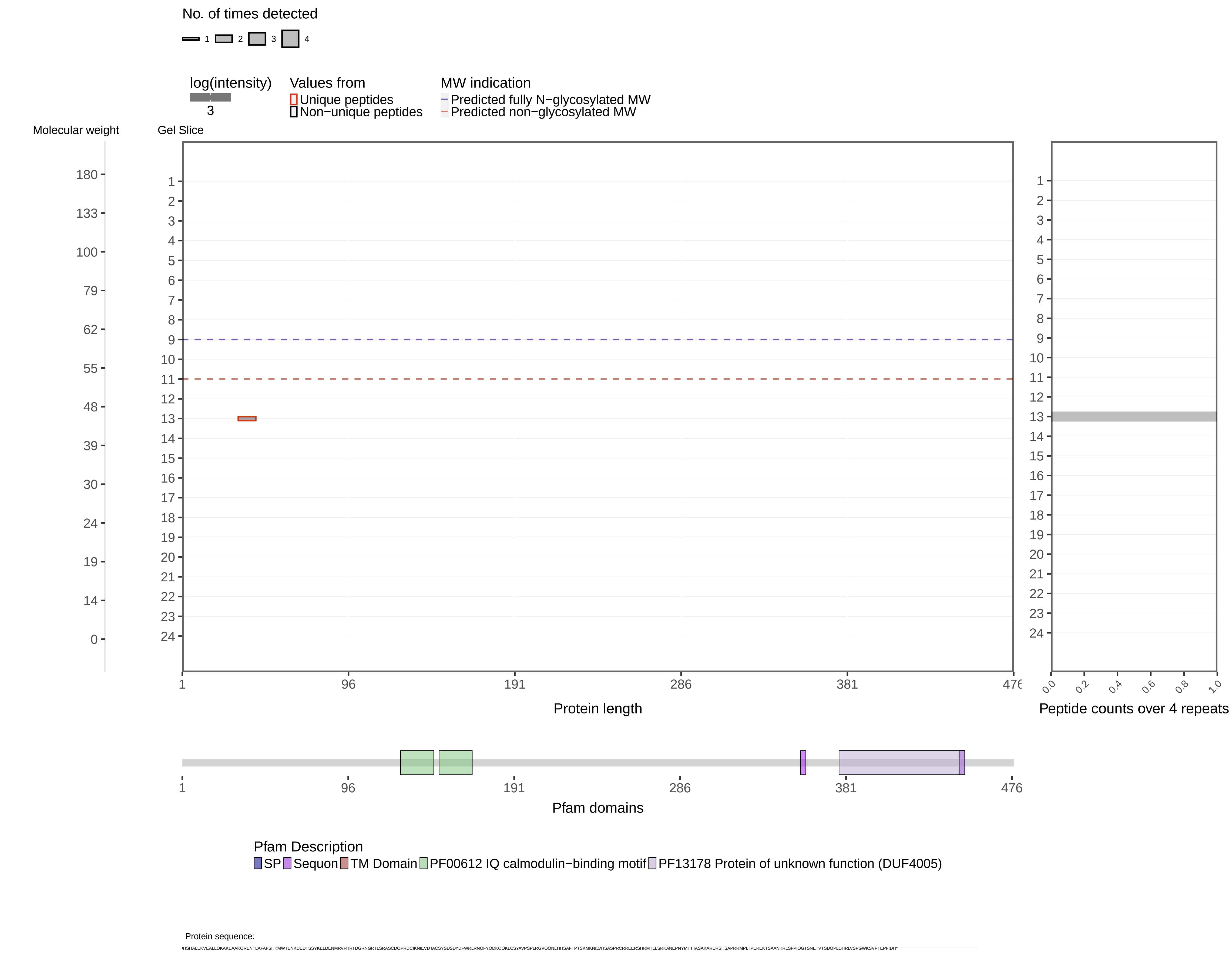

NbL18g10370.1 uncharacterized_LOC104217540 [MW − SP = 68.67 kDa]

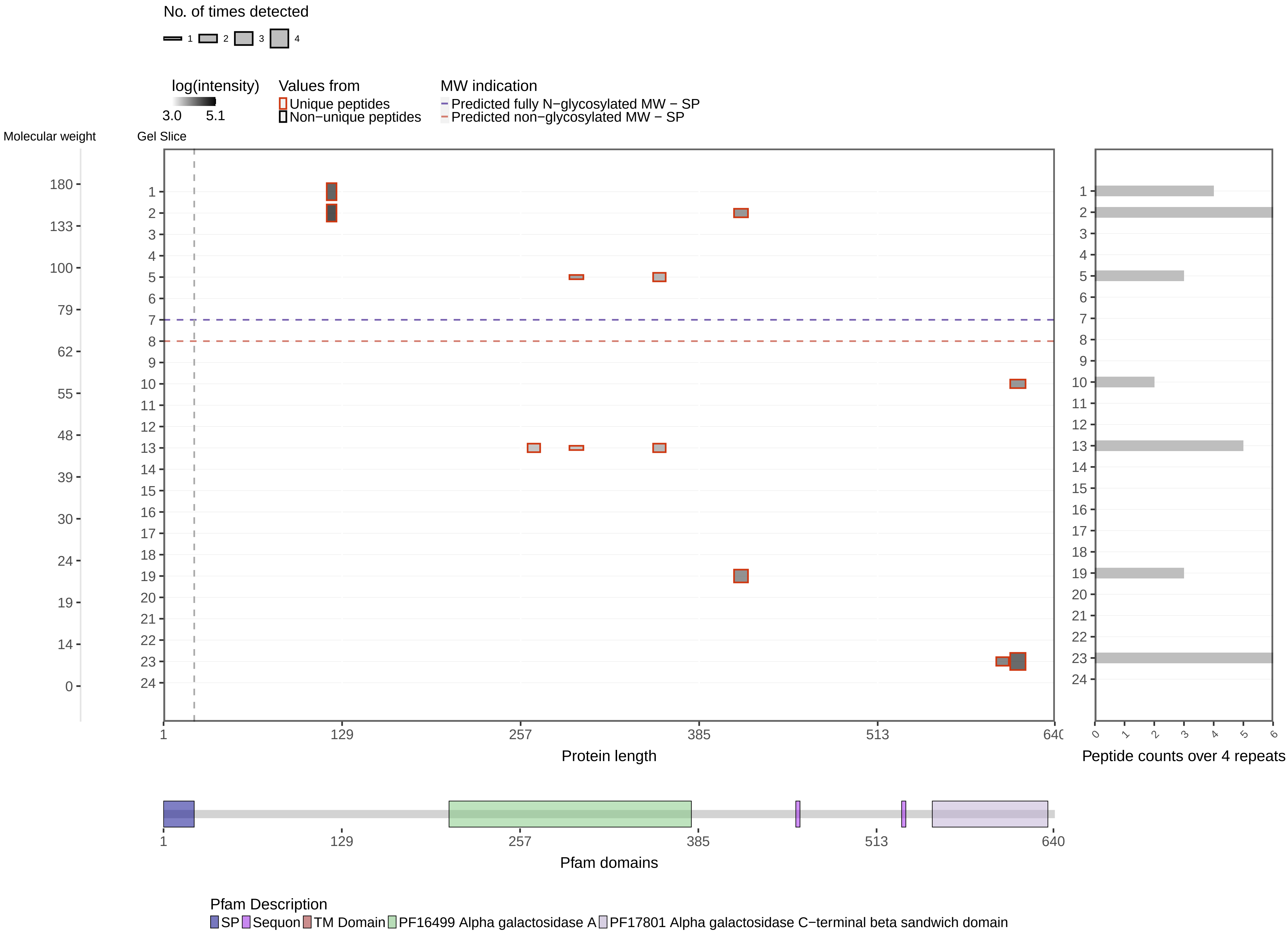


Protein sequence:

KSWPDLDMLPLGWLTDAGSNLGPHRNCRLNLDEQRTQVTLWSMVRSPLIFGGDMRKLDDTTFNLLTNPTLLEINWFSSNNMEFRYVTGSLSSPGKHGLTHNSKNEGKTSVSETRVLALRSCKDVKANGWSTKVLDNDVAQVCWEENSNNHTSPFCLYKRKAPSLSEGEMVYNHQSHGKLHLLVTERTELCFSASSNRKLTSKESSRGSFSRCRSHANQIWELNNNGTLKNSYSGLCASLDIVKASPGGVRSWLATGRRGEIYLAFFNLNNQVTDMSAKISDIAKAIPA[illegible]

### NbL18g10340.1 uncharacterized_LOC109235488 [MW = 13.05 kDa]

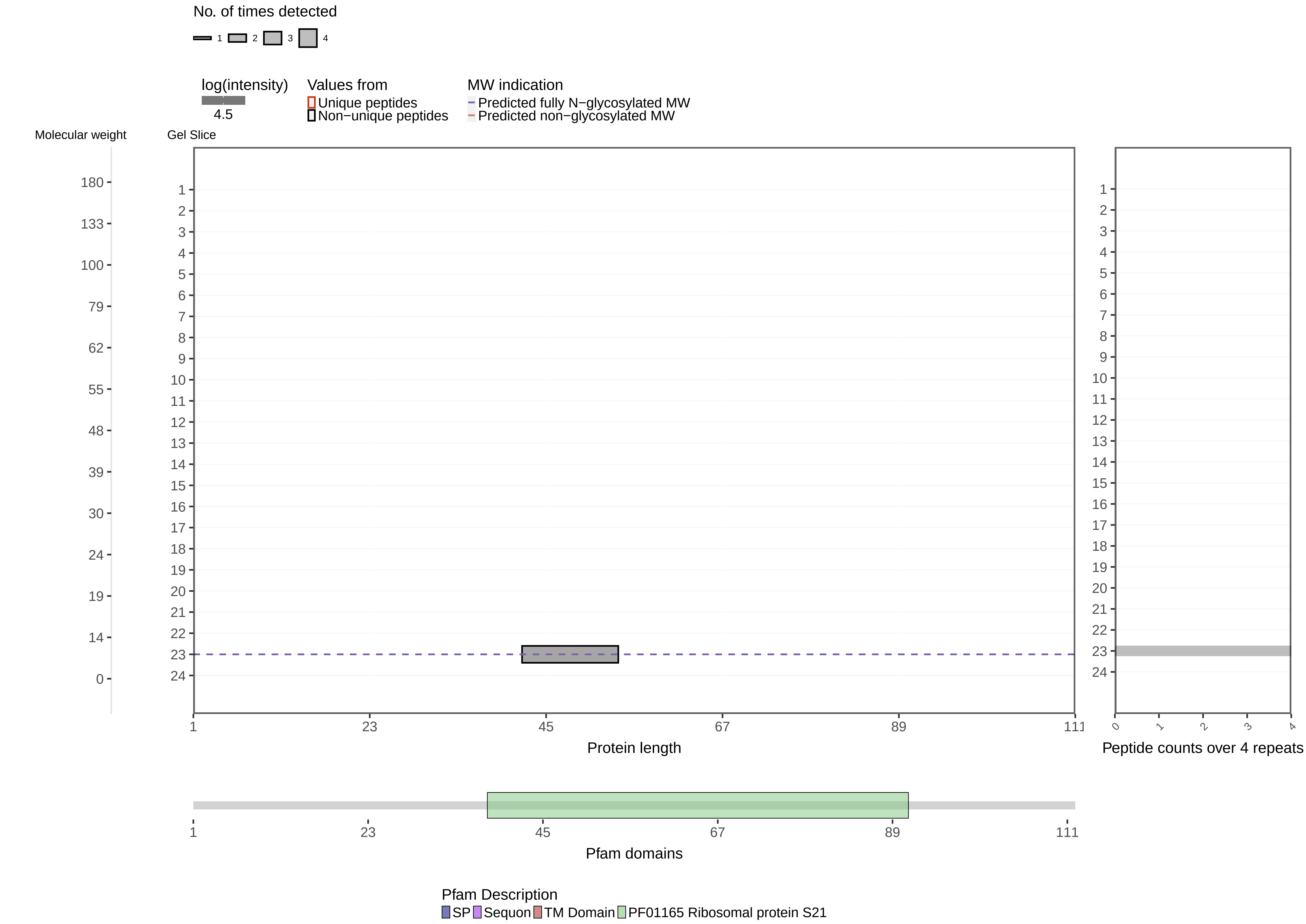


Protein sequence:

NNNLDQALFLMQRKMQSSGMERLIKREQLHHIKNSEKRVLARKLLERKLRSQDLARKLKSILIKKVRGL*

NbL18g10270.1 uncharacterized_LOC107778361 [MW − SP = 59.56 kDa]

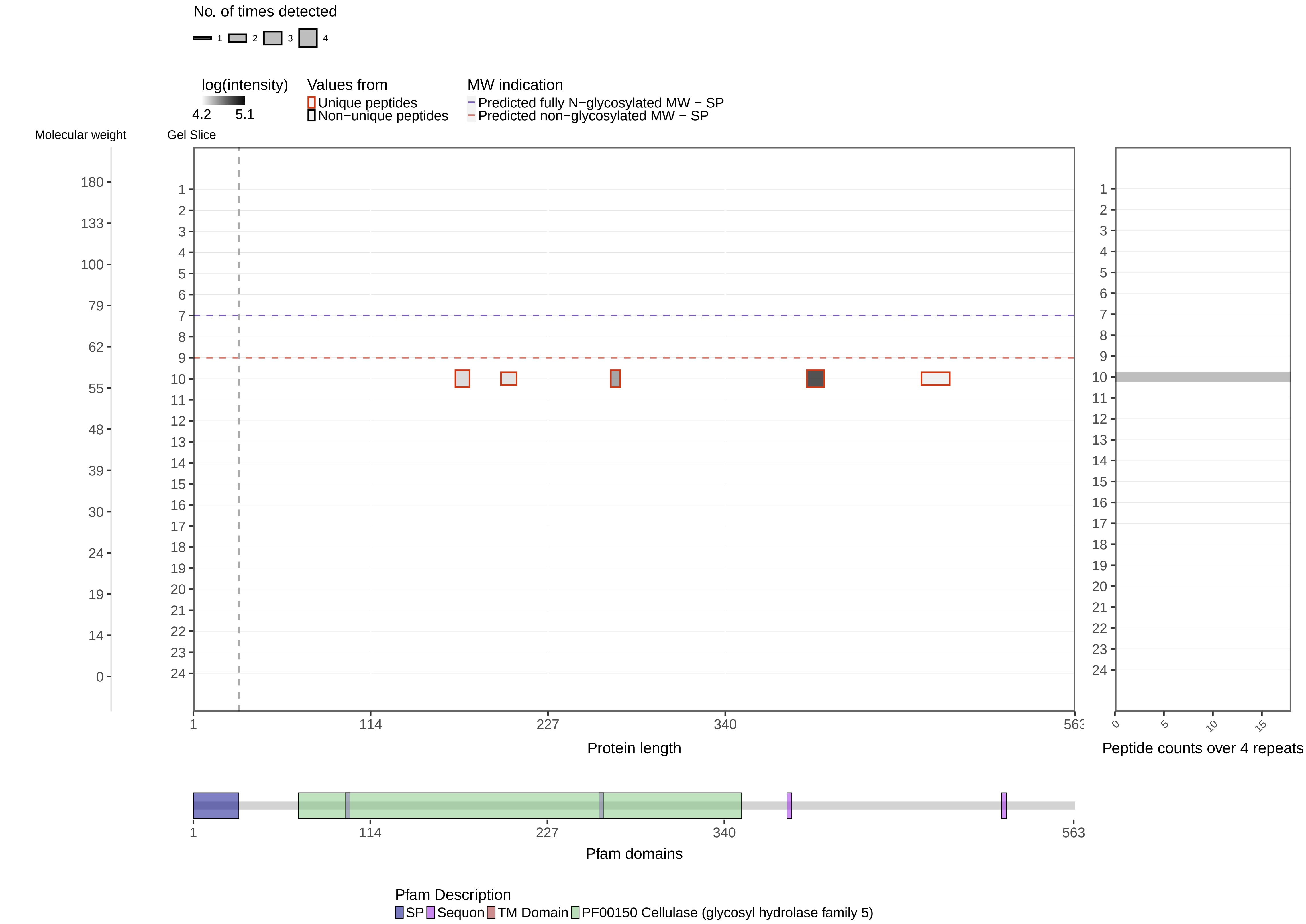


Protein sequence:

### NbL18g10240.1 tumor_protein [MW = 18.76 kDa]

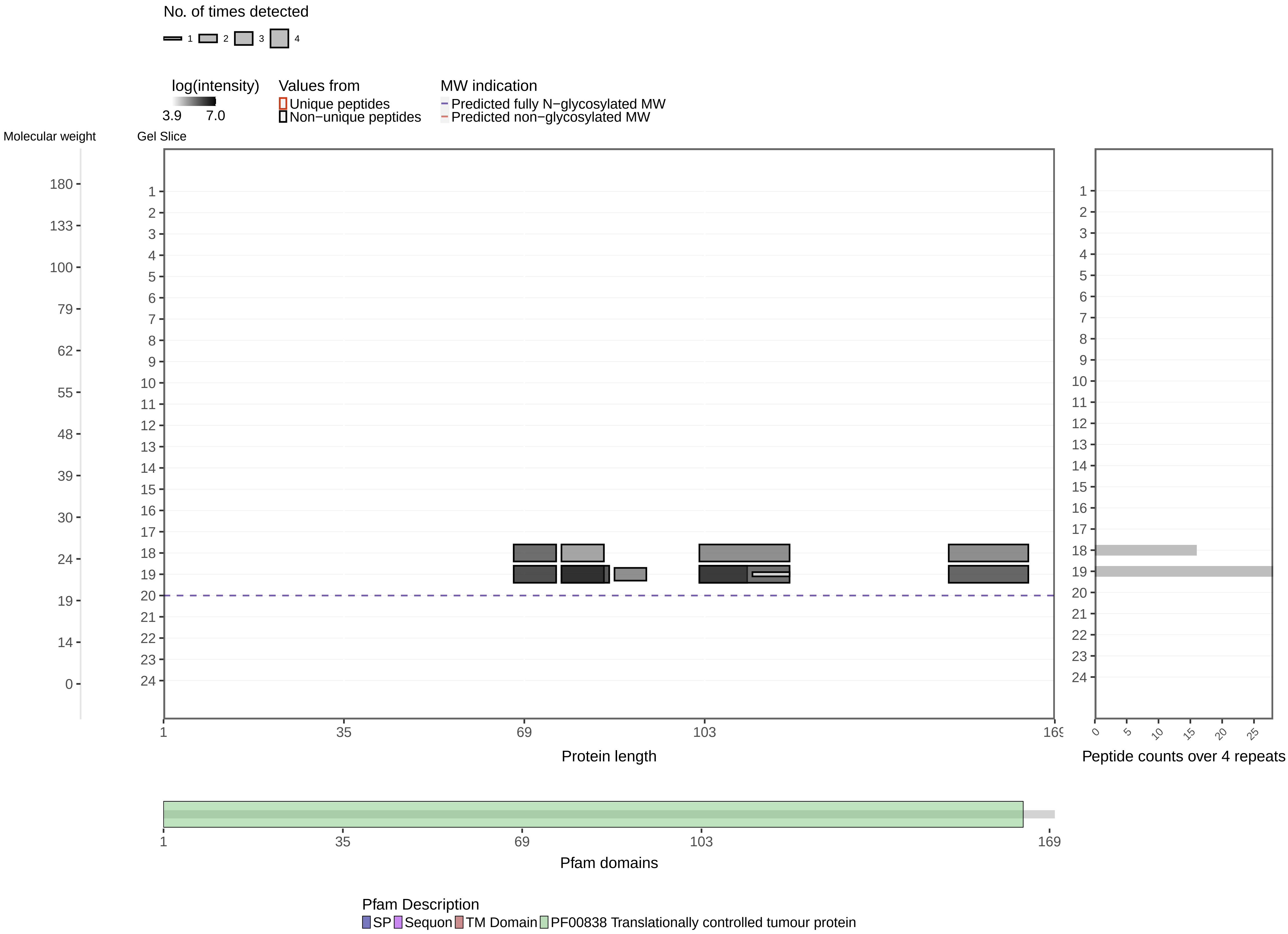


Protein sequence:

VDTFRLQEQPSFDKKQFVAYMKKYIKSLTPKLGAEQEEVFKNNIQGATKYLLSKLSDLQFFVGESMADDTGMVFAYYKDGATDPTFLYLAHGLKEVKC*

### NbL18g10190.1 putative_protease_Do−like_14 [MW = 46.37 kDa]

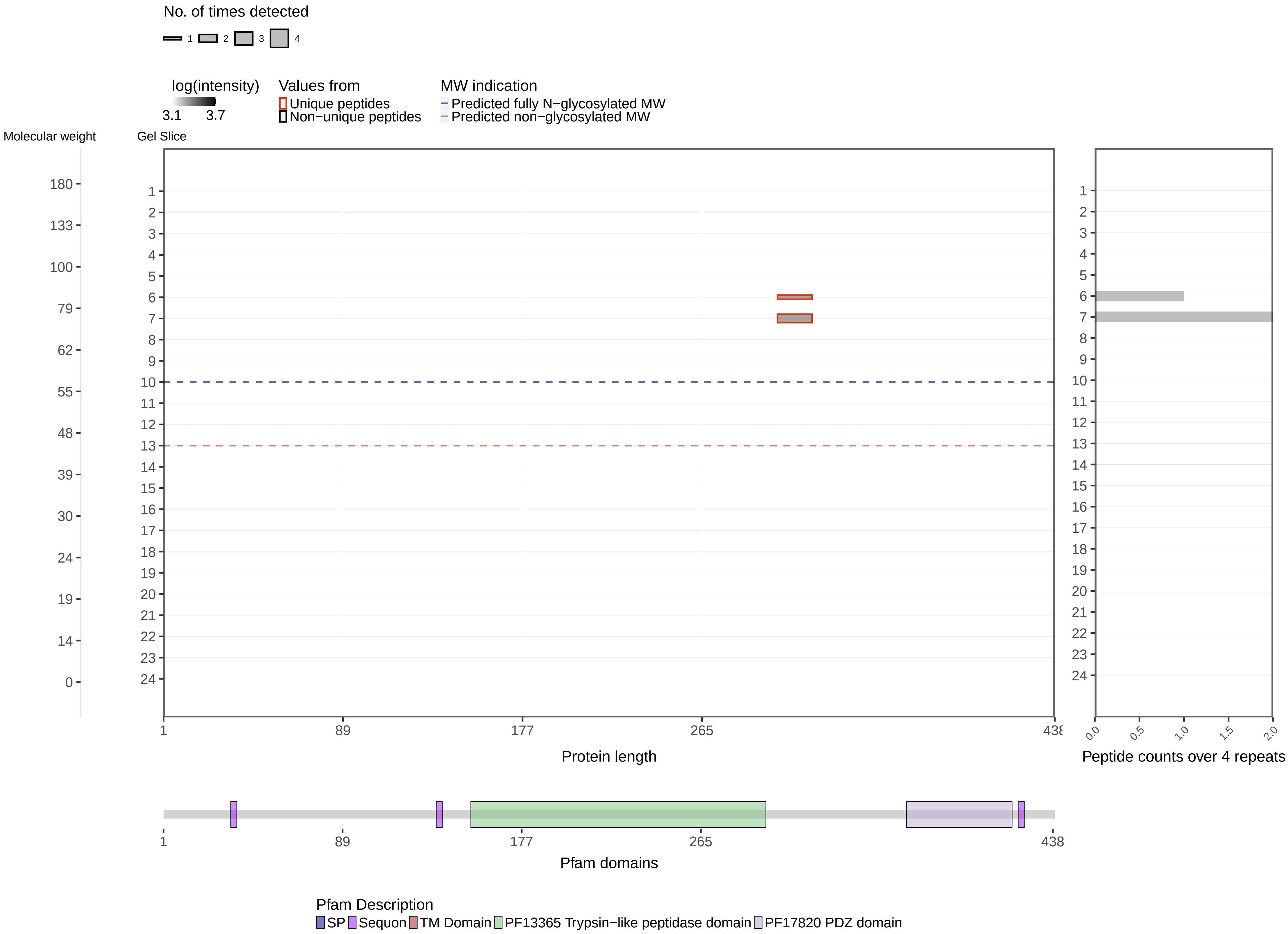


Protein sequence:

DIAIVKIKSKTPLPTAKLGTSNMLRPGDWVVAMGCPLSLQNTITAGIVSCVDRKSSDLGLGGMRREYLQTDCAINAGNSGGPLVNVDGEVIGVNIMKVLAADGLGFSVPIDSVTKIIDHFKKRGRVIRPWLGLKMLDLNDMVVAQLQERDPSFPKVNKGVLVSMVTPGSPADCAGFHPSDVVVEFDGKPVGSIKEIIDIMGENLGKQLTAVVKRANNATATLTVIPEEANPDM*

NbL18g10090.1 amidase_1 [MW = 45.14 kDa]

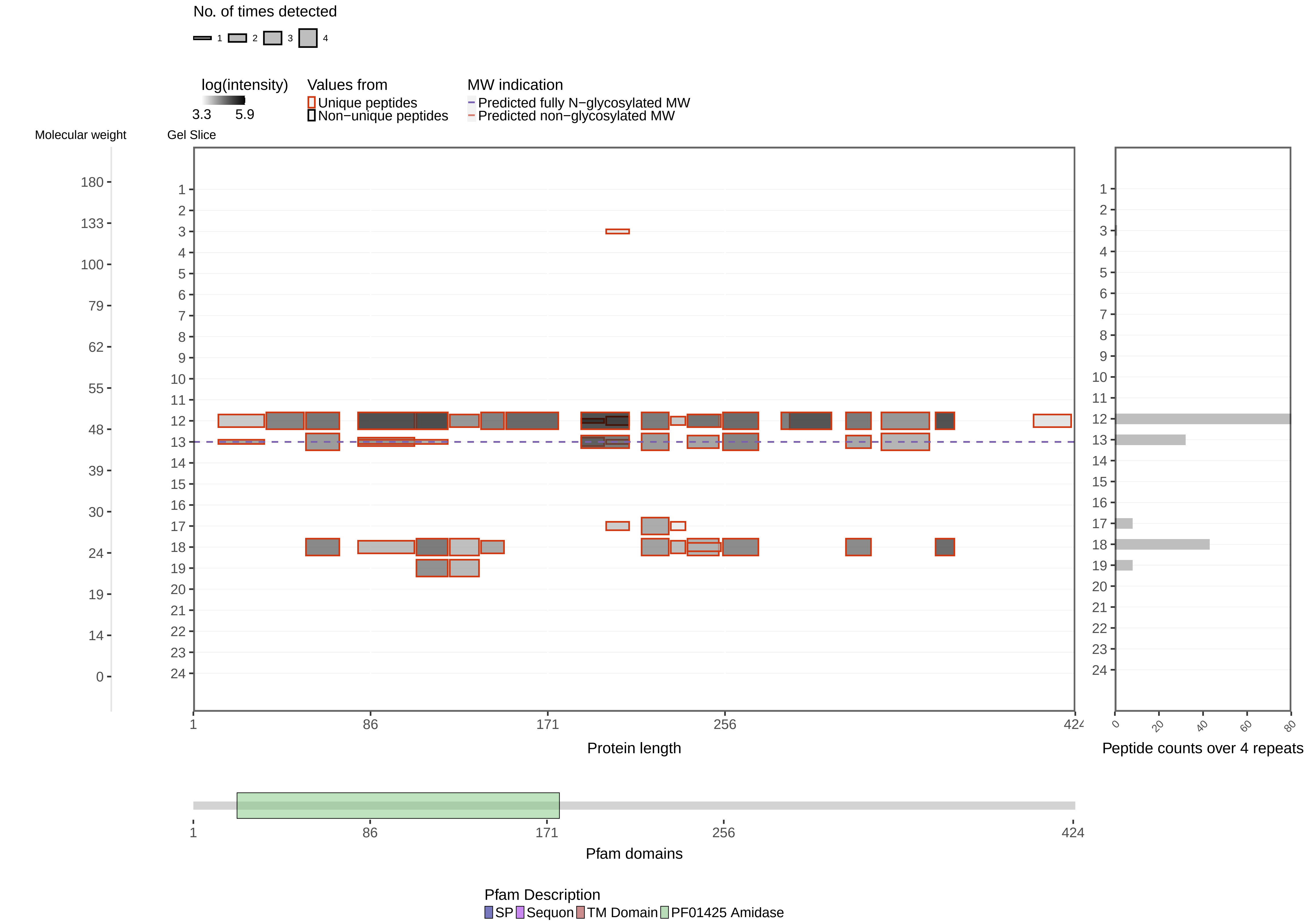


Protein sequence:

/RPTNIIIAEDCFKLLDSKSNQLIEVLVNSVEKLYGSHMIKYVSVGDYIEGNVPSLKKFMTPGTGNDESYIPSLLRLSAAMRLLQRYEFKENHGEWVSTVKPSLGPGIAERVWEALKATDENIDVCHSVKAELRAALTALLGDSGILAIPTVPGPPPKLRSETSALEGFRVKAFSLLSIAGVSGFCQVSIPLGMQDNLPISVSLLVKHGSDWLLLDAVEAIHNVLKI*

### NbL18g09960.1 eukaryotic_translation_initiation_factor_4B1−like [MW = 70 kDa]

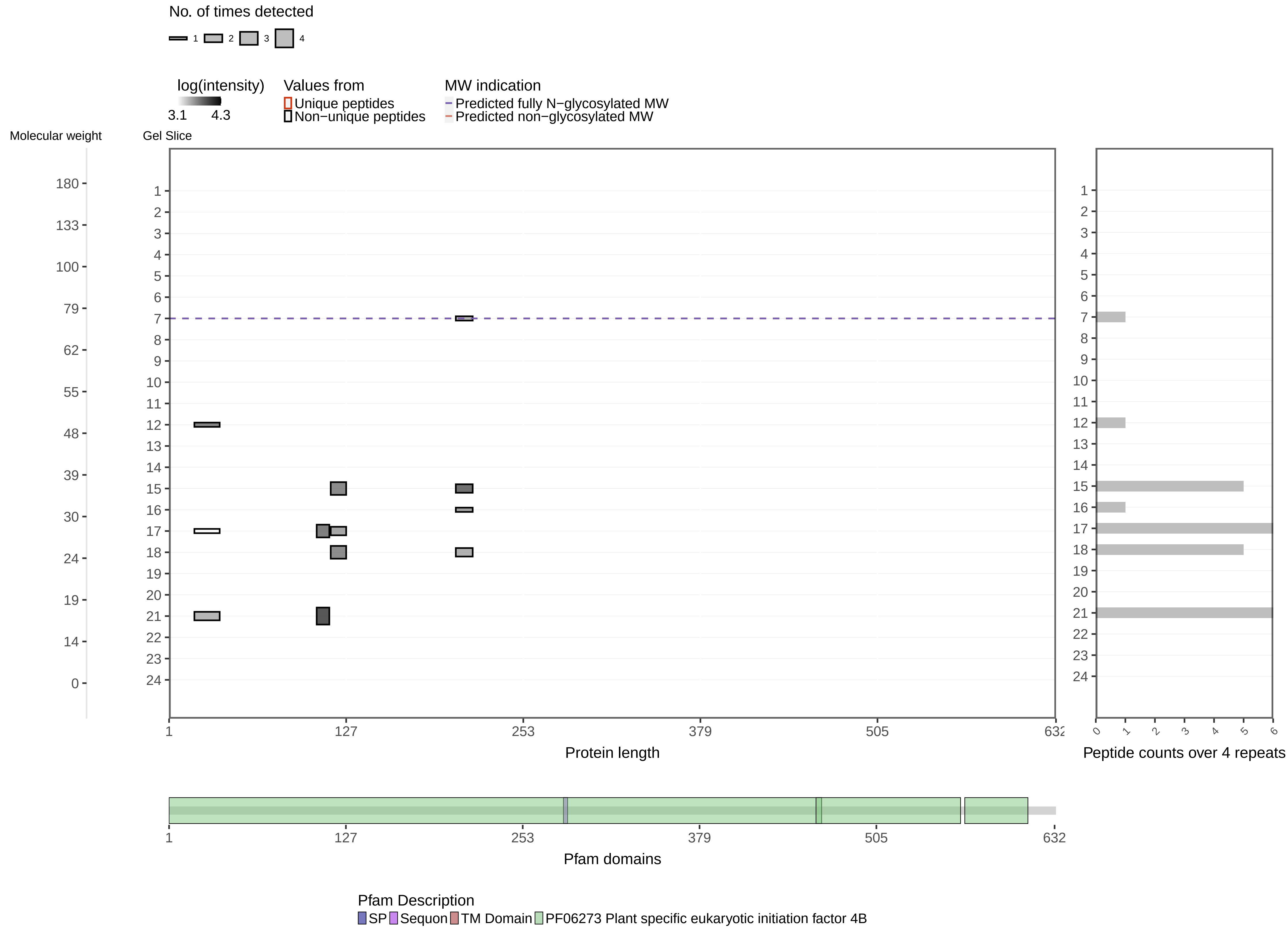


Protein sequence:

EEVLAEKGLDWKKLDMEIEVKKGSRPTSSQSSRPGSSHSAPSEGPAVLQGGVMEVKPKSKVNPFRDAKPREVLLEEKGLDWKKIDLELEHRRVDRPETEEEKNLKEEIEHLRKEAMQSSGHDQTNLQNVINPKDRDLELLIRDLDDKVRFGQKRPGSASGRERNLKEEIEQLRKEAMQSSGQDQTNLQNLINQKERELELLIRDFDVKVRFSQKHIQRPGSGSGRVTGLPERSLSQSASYEDRRAGEQIDRPRSRGGDAWARPTDPMERPRSRGGVDSWARPNEQV

### NbL18g09950.1 GDSL_esterase/lipase_At3g26430−like [MW − SP = 39.69 kDa]

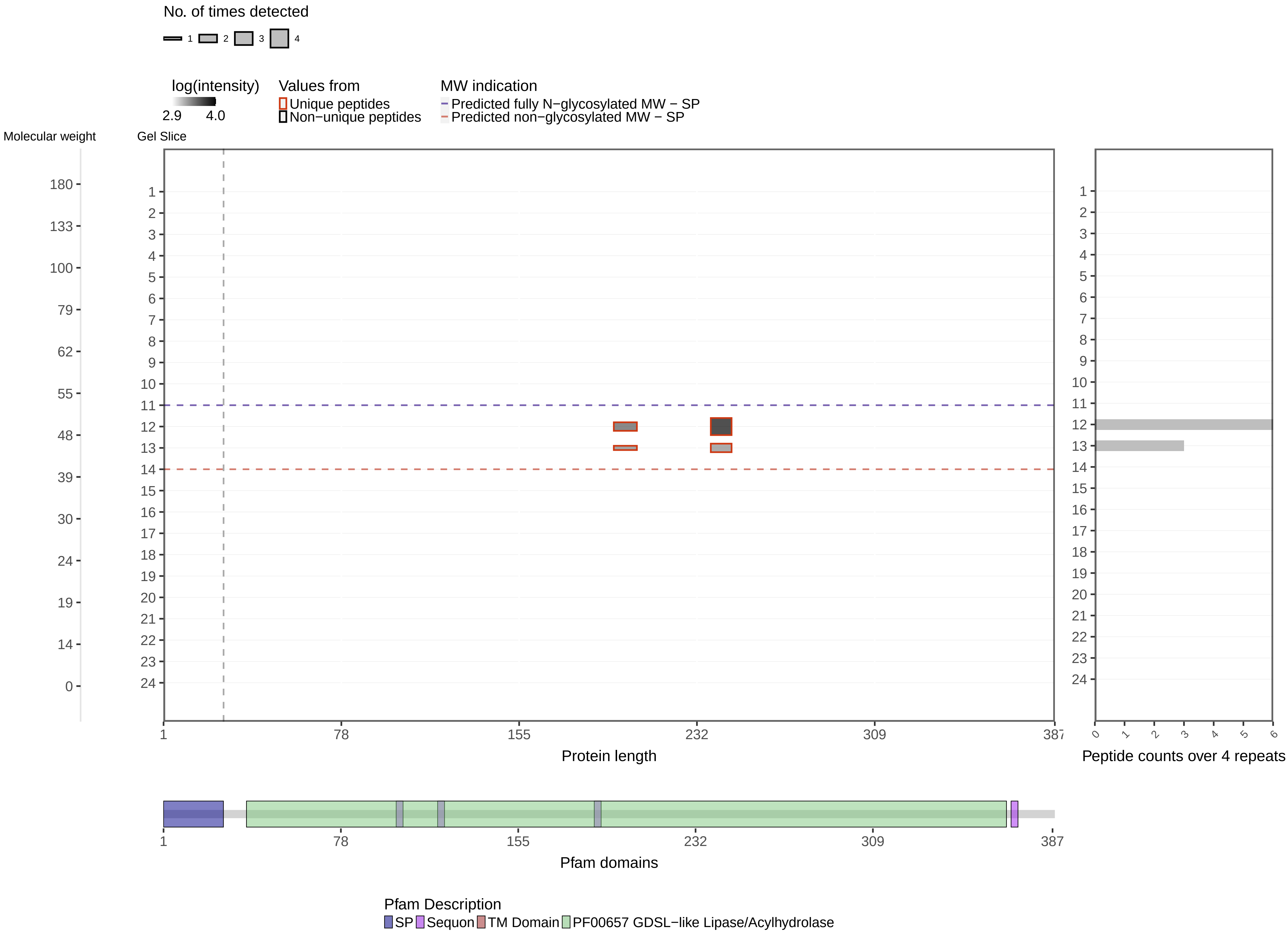


Protein sequence:

GYFLNMSTDQVRAYVPDLMDQFRTVIKDIYDEGGRYFWIHNTGPVGCLPYILDRLLITAAQIDKAGCASPFNKVAQYFNAKLKETVVQLRKDLPLAALTYVDVYSAKYELISHANKHGFEHPLRACCGHGGKYNYNMNHGCGSKIKVKGKEIMLGKSCKDPSVRINWDGVHYTEAANRWVFEKIVNGSYSDPSLPLNMACHKQ*

NbL18g09690.1 peroxisomal_primary_amine_oxidase−like [MW = 86.25 kDa]

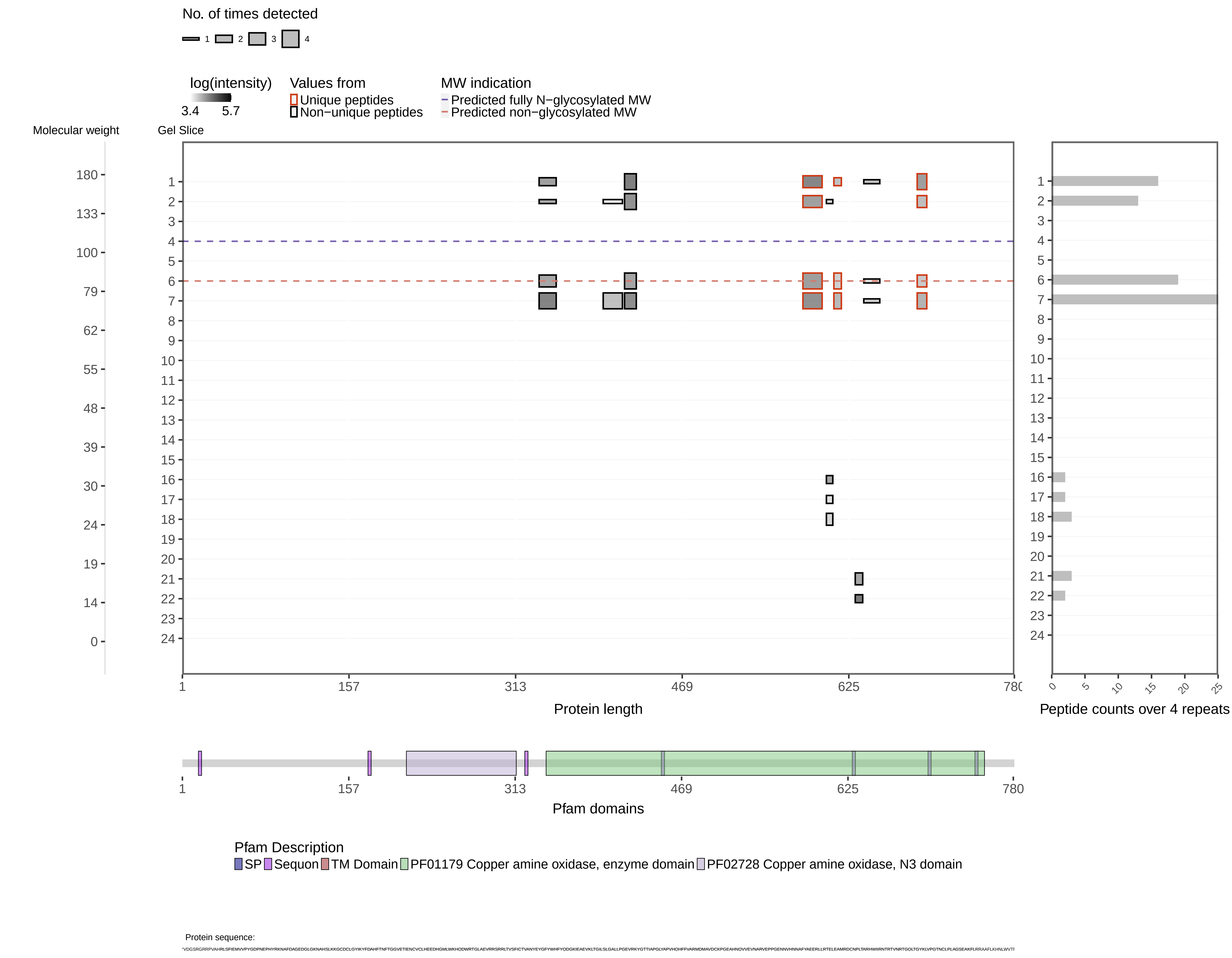

NbL18g09650.1 glutamate−−glyoxylate_aminotransferase_2 [MW = 53.48 kDa]

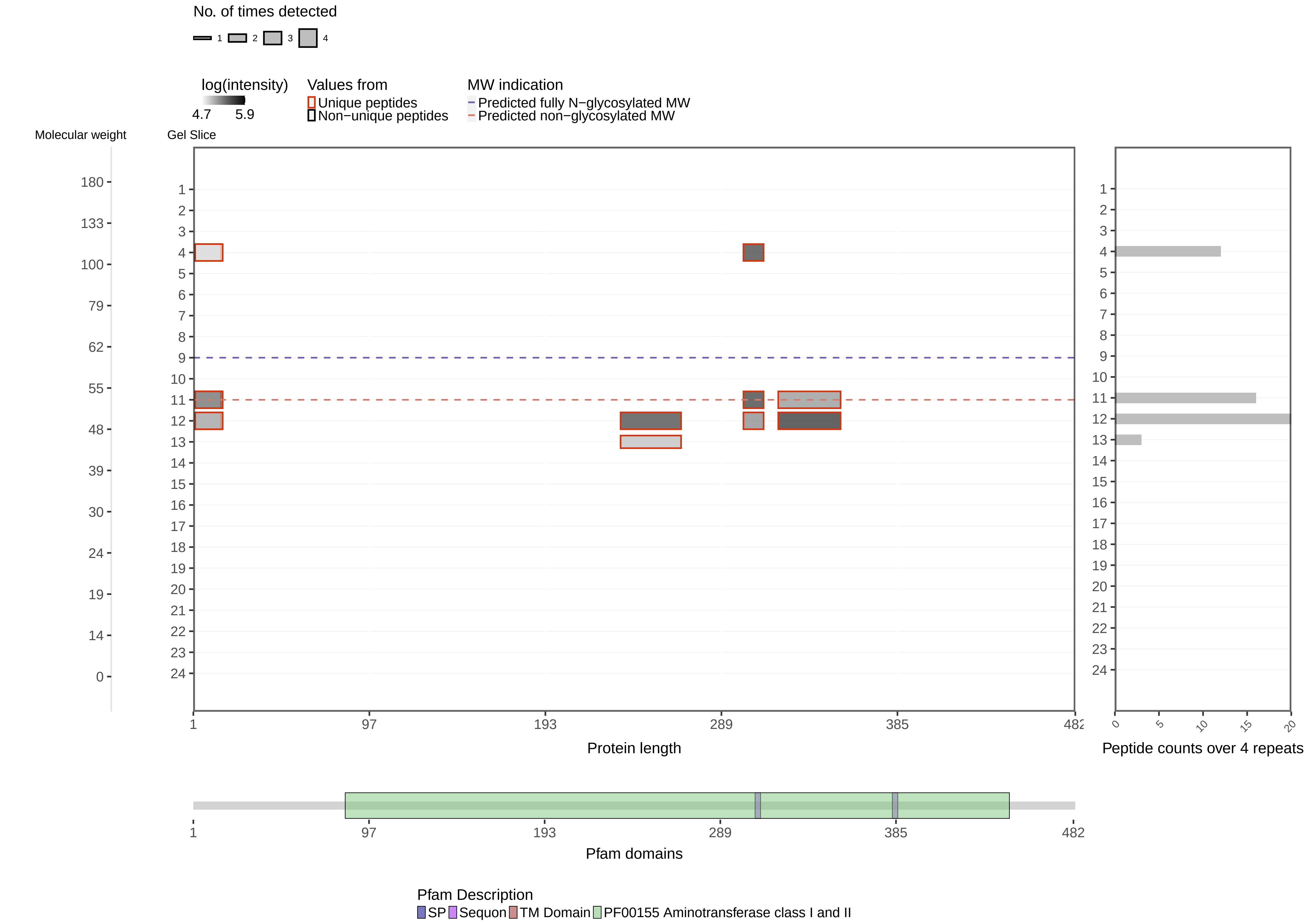


Protein sequence:

ANLKQIIQFCHQESLVLLGDEVYQQNIYQDERPFISARKVLLDMGPPISKELQLVSFHTVSKGYWGECGQRGGYFEMTNISPKSVEEIYKVASISLSPNVPGQIFMGLMVNPPKPGDISYDQYVRESKGILESLRKRAHMMTDGFNSCRNVVCNFTEGAMYSFPQIRLPPKAIEAAKKLGKVPDVFYCLRLLEATGISTVPGSGFGQKEGVFHLRTTILPAEEDMPAIMSSFKKFNDEFMEQYEDHRGYSRM*

### NbL18g09500.1 cysteine−rich_receptor−like_protein_kinase_2 [MW − SP = 69.77 kDa]

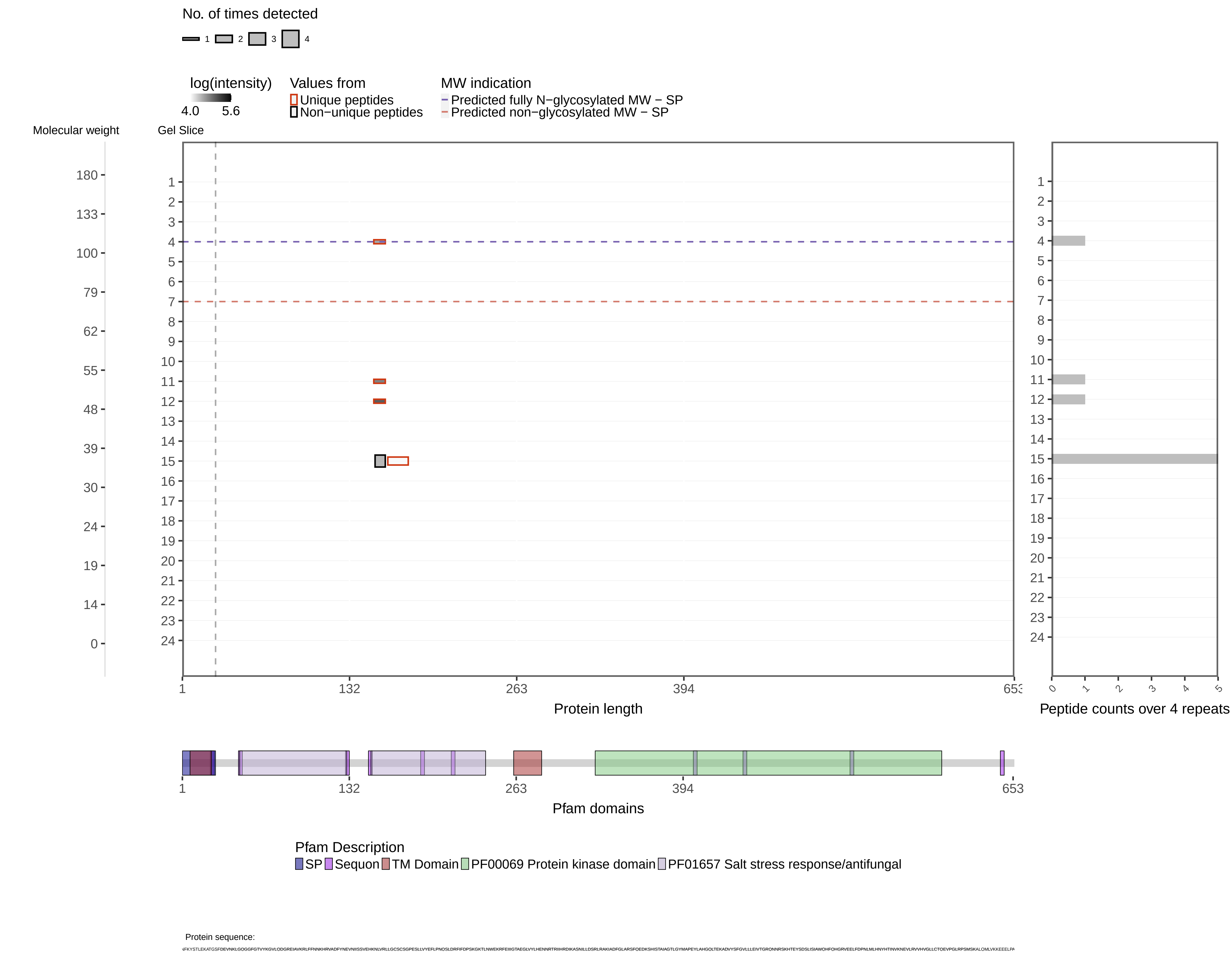

NbL18g09260.1 ATP−citrate_synthase_alpha_chain_protein_2 [MW = 46.5 kDa]

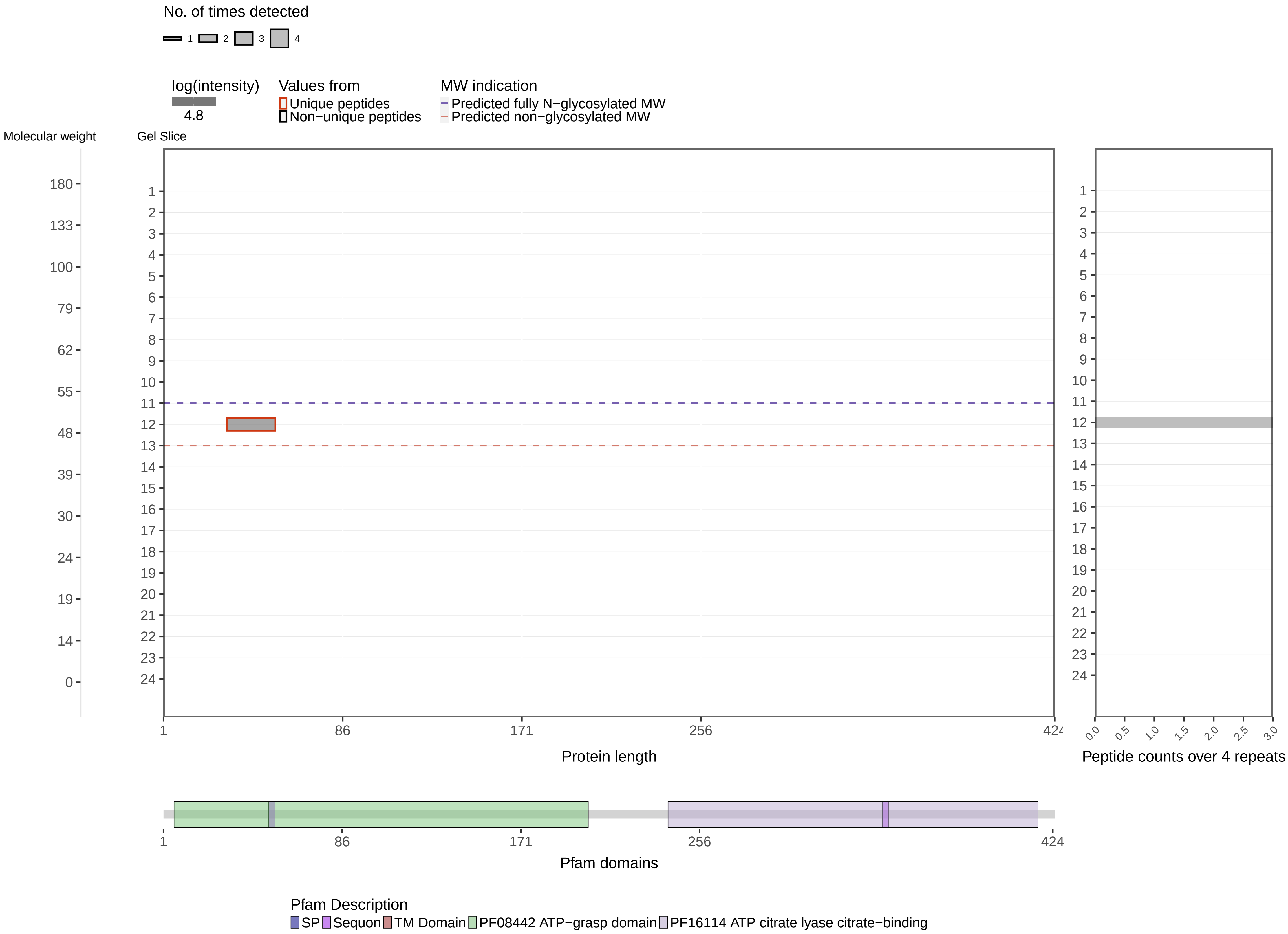


Protein sequence:

INPFALVEGKPYPLDMRGELDDTAAFKNFKKWGNIEFPLPFGRVMSATESFIHGLDEKTSASLKFTVLNPKGRIWTMVAGGGASVIYADTVGDLGYASELGNYAEYSGAPNEEEVLQYARVVIDCATANPDGRKRALVIGGGIANFTDVAATFSGIIRALKEKESKLKAARMQIFVRRGGPNYQKGLAKMRSLGEEIGIPIEVYGPEATMTGICKQAIECITAAA*

NbL18g09140.1 2−methylene−furan−3−one_reductase−like [MW = 41.11 kDa]

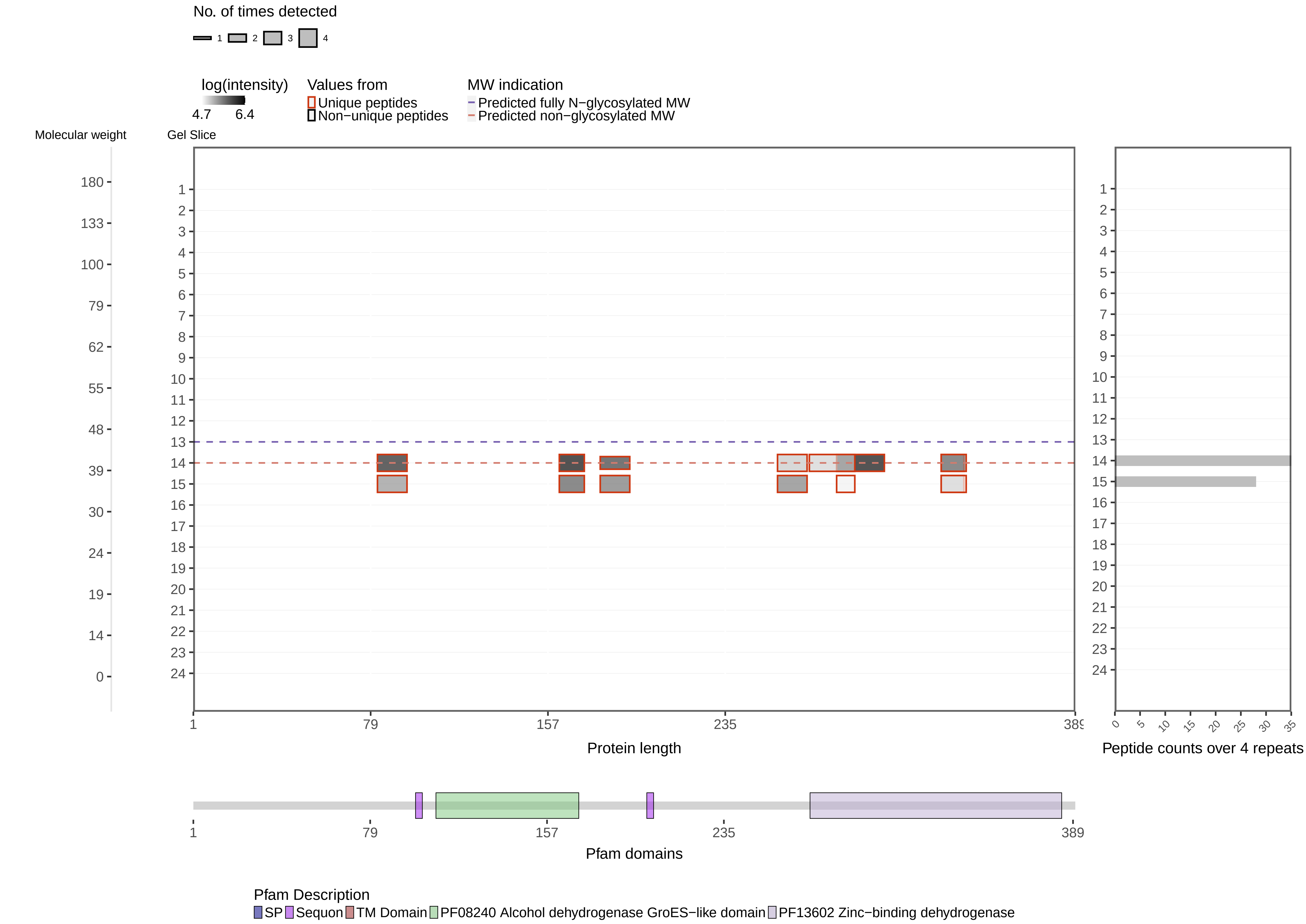


Protein sequence:

QFGSLAEFTAVEEKLVALKPKNLSFAEAASLPLAIETAYEGLEKAGFSAGKSILVLGGAGGVGSLVIQVAKHVFGASKVAATSSTGKLELLKSIGADLAIDYTKENFENLPEKFDVVYDTVGQGEKAVKAVKEGGSVVVLTGAVTPPGFRFVVTSNGEMLKKLKPYLESGKVKPVIDPKGPFPFDKVVEAFSYLETGRATGKVVIHPIP*

### NbL18g09130.1 carbonic_anhydrase_2−like [MW = 33.23 kDa]

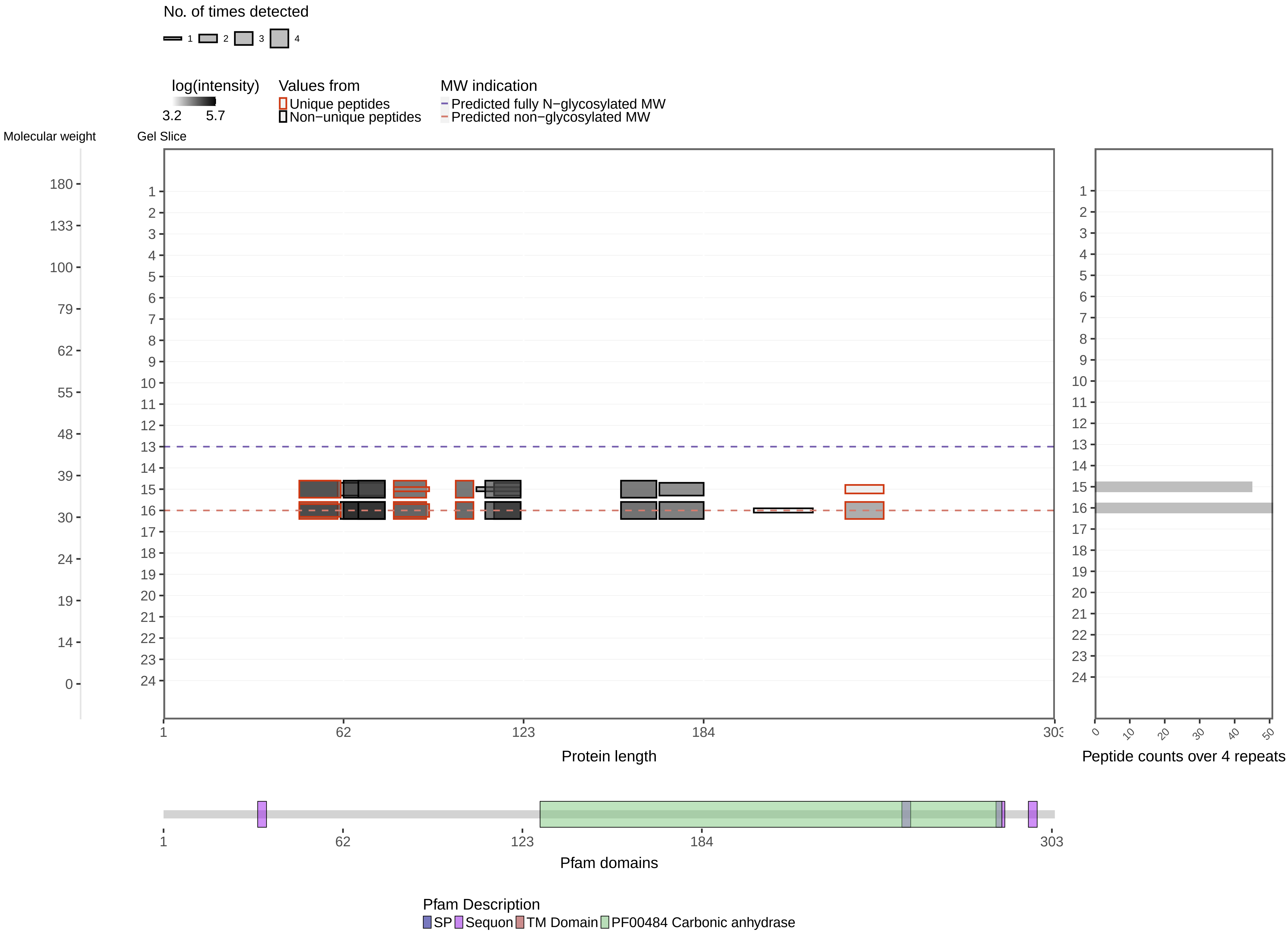


Protein sequence:

NbL18g09040.1 RNA−binding_protein_25 [MW = 106.19 kDa]

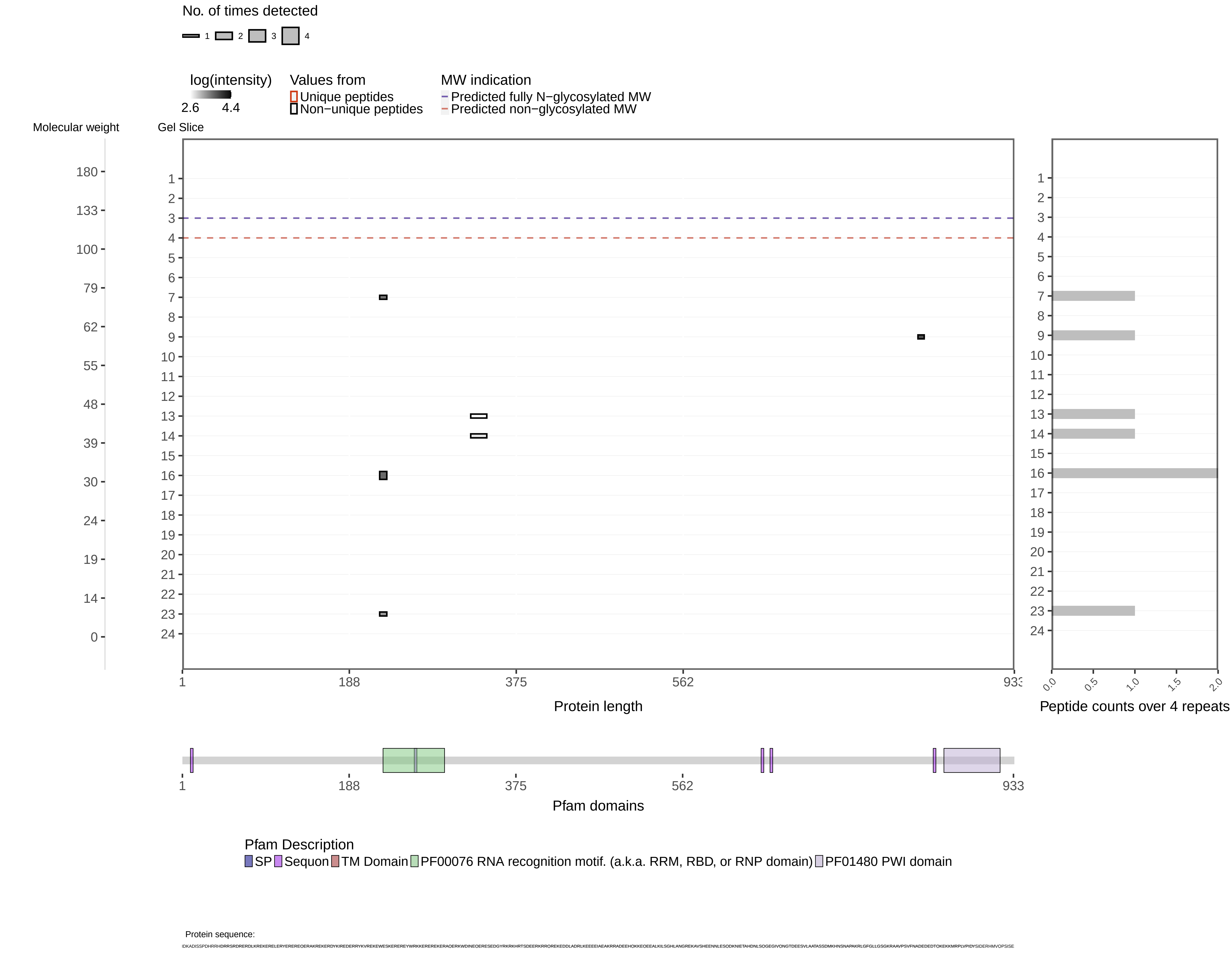

NbL18g08970.1 aldehyde_dehydrogenase_family_2_member_B7_mitochondrial−like [MW = 58.53 kDa]

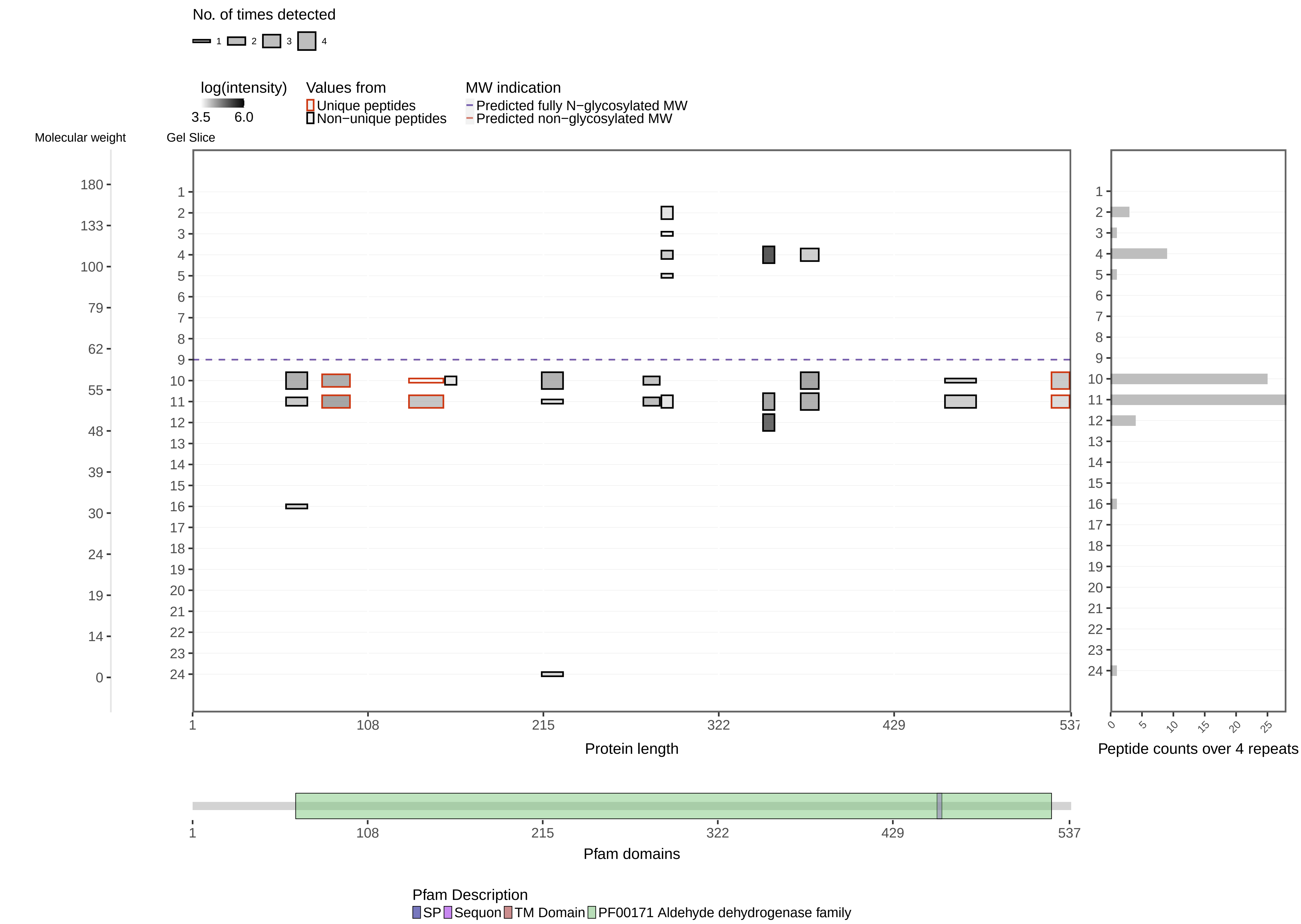


Protein sequence:

FGPTAGAPLCSHMDVDKLAFTGSTDTGKAILSLAAKSNLKPVTLELGGKSPFIVCEDADIDTAVEQAHFALFFNQGQCCCAGSRTYVHEKVYDEFLEKAKARALKRTVGDPFKSGTEQGPQIDSKQFDKIMNYIRCGIDSGATLETGGERLGERGYYIKPTVFSNVKDDMLIAQDEIFGPVQSILKFKDLDDVIRRANNSRYGLAAGVFTQNIDTANTLTRALRVGTVWVNCFDTFDATIPFGGYKMSGHGREKGEYSLKNYLQVKAVVTPLNNPAWL*

### NbL18g08950.1 synthase [MW = 38.14 kDa]

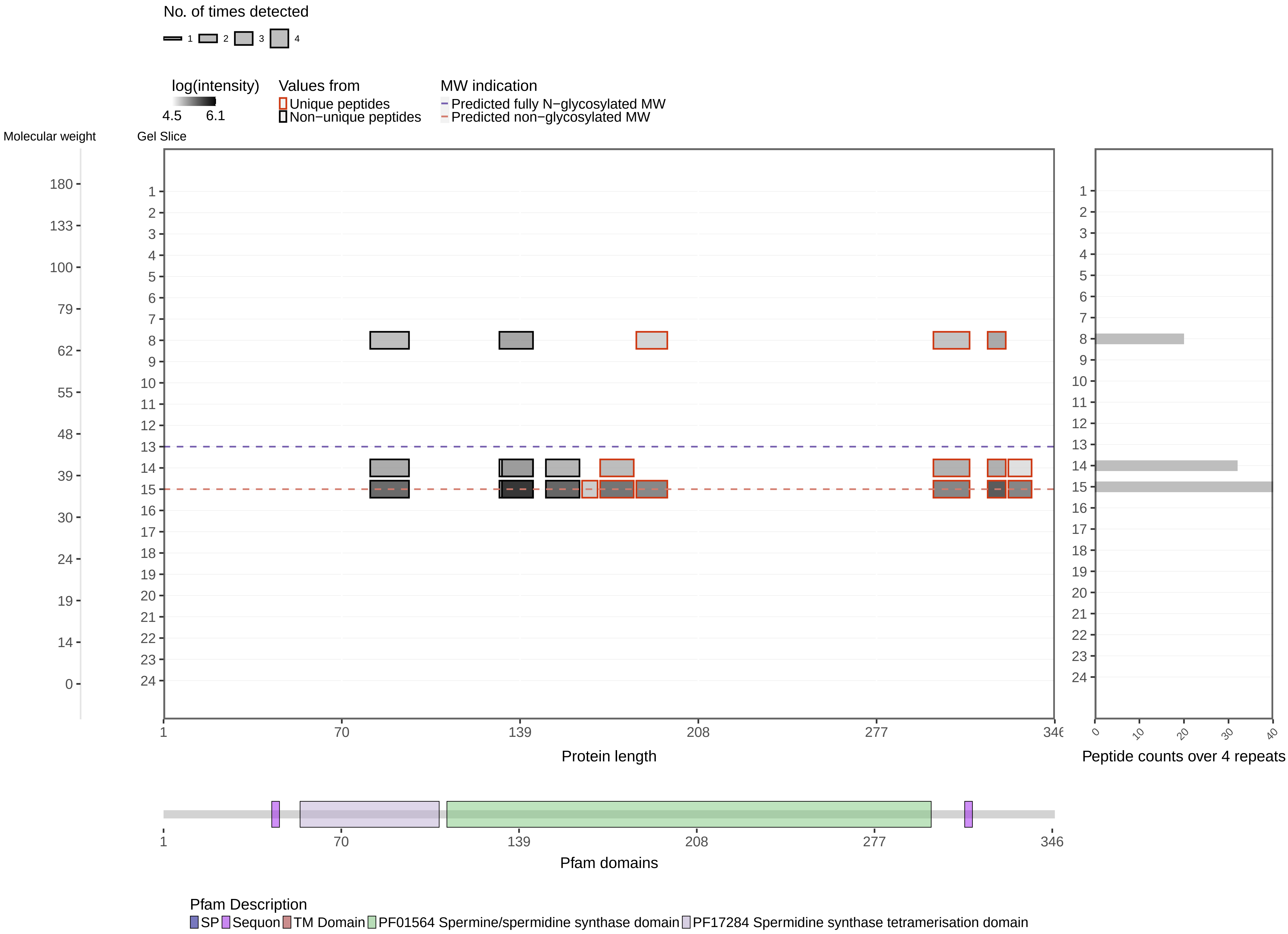


Protein sequence:

CEIDKMVVEVSKQFFPDVAVGYEDPRVNLHIGDGVAFLKNVAAGTYDAVIVDSSDPIGPAQELFEKPFFESIARALRPGGVVSTQAESIWLHMHIIEEIVANCRLIFKGSVNYAWTTVPTYPSGMIGFMLCSTEGPAVDFKNPINPIDEDDVSHNKTLGPLKFYNSELHKASFCLPSFAKRVIESKGK*

### NbL18g08850.1 AP−1_complex_subunit_gamma−2−like [MW = 96.03 kDa]

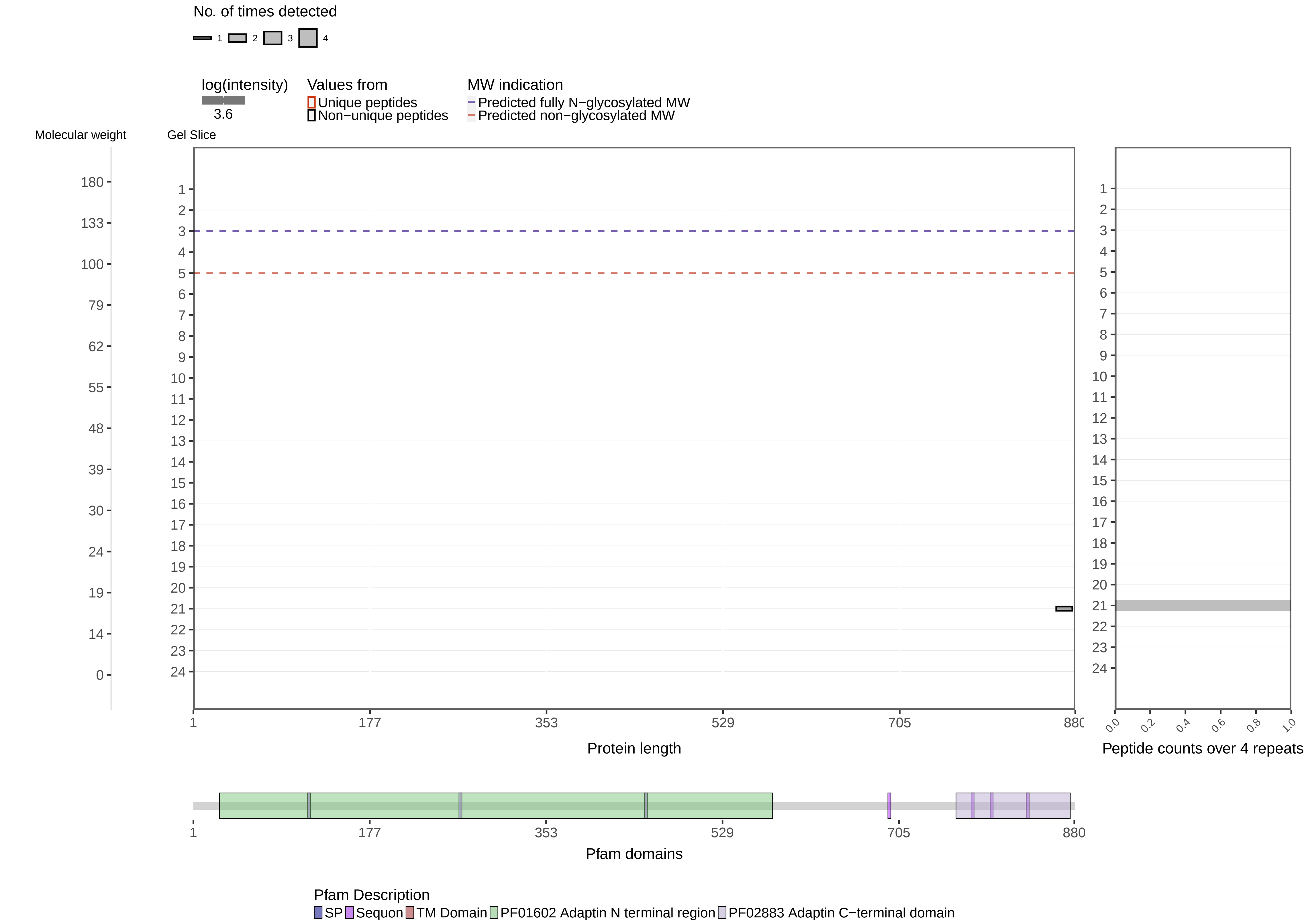


Protein sequence:

PEAGNYVKDEVWHSLIVVITNASNLHGYAVRSLYRAVQAAGEQETLIRVAVWCIGEYGDMLVNNAGRLDIEEPLTVTESDAVDVVETSIKSHSFDLTTQAMCLIALLKLSSRFPSCSQRINDIIVQYKGCFVLELQQRAIEFNSVIERHQNIRPSLVERMPVLDAATYSGKKAGSLPAAGSTSQGVSVNLPNGVAKPSATPLVDLLDLSSDDVAAPSSSGGDFLQDLLGVDLAPVSSQSGTNQAQKSGTDVLLDLLSIGTPPANSSPSTTQVSPSNVDIRSPVDLLDRLSSPSAPS\

NbL18g08830.1 threonine−−tRNA_ligase_mitochondrial [MW = 53.09 kDa]

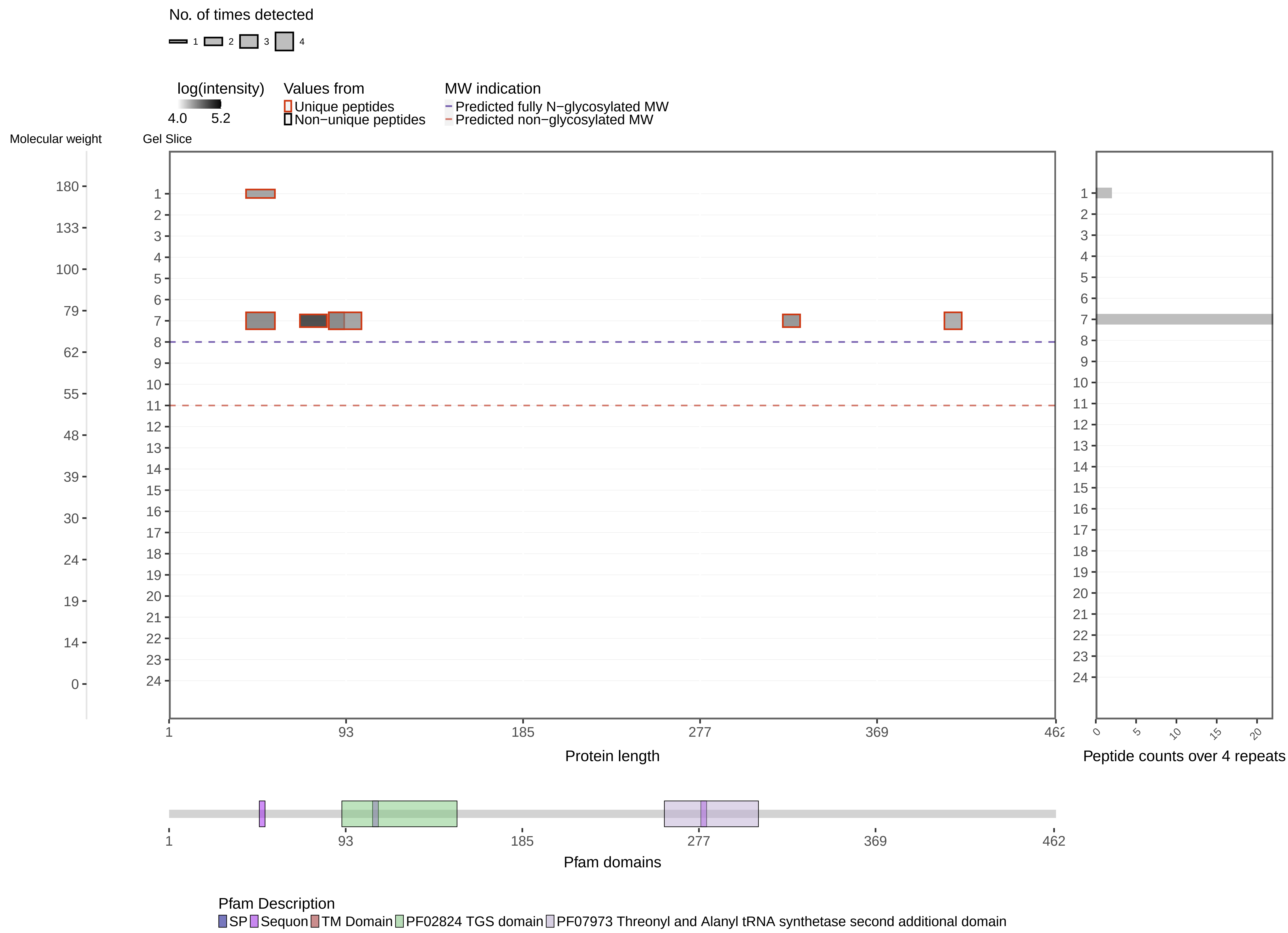


Protein sequence:

NbL18g08760.1 MLP−like_protein_423 [MW = 16.82 kDa]

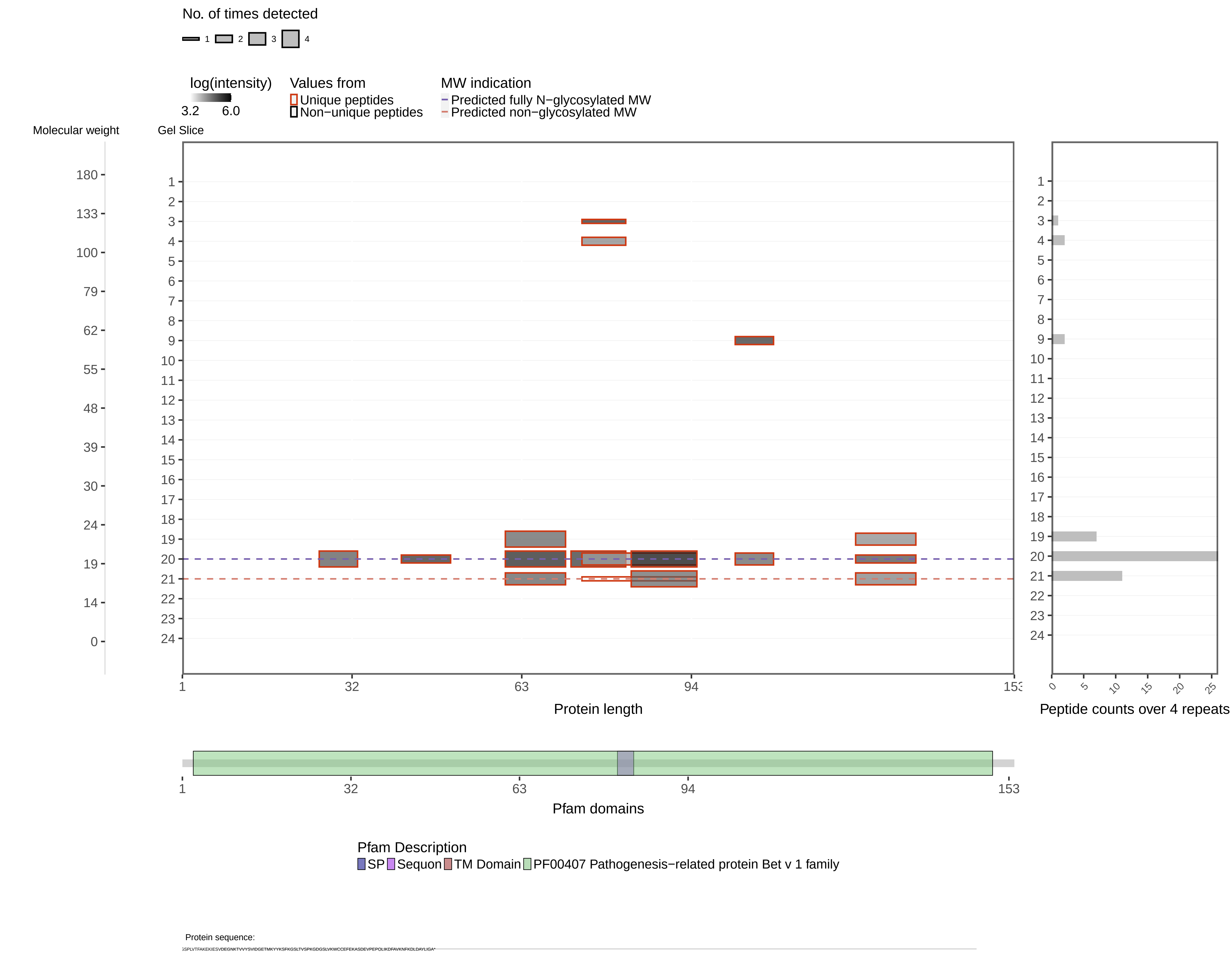

NbL18g08740.1 30S_ribosomal_protein_S13_chloroplastic [MW = 21.93 kDa]

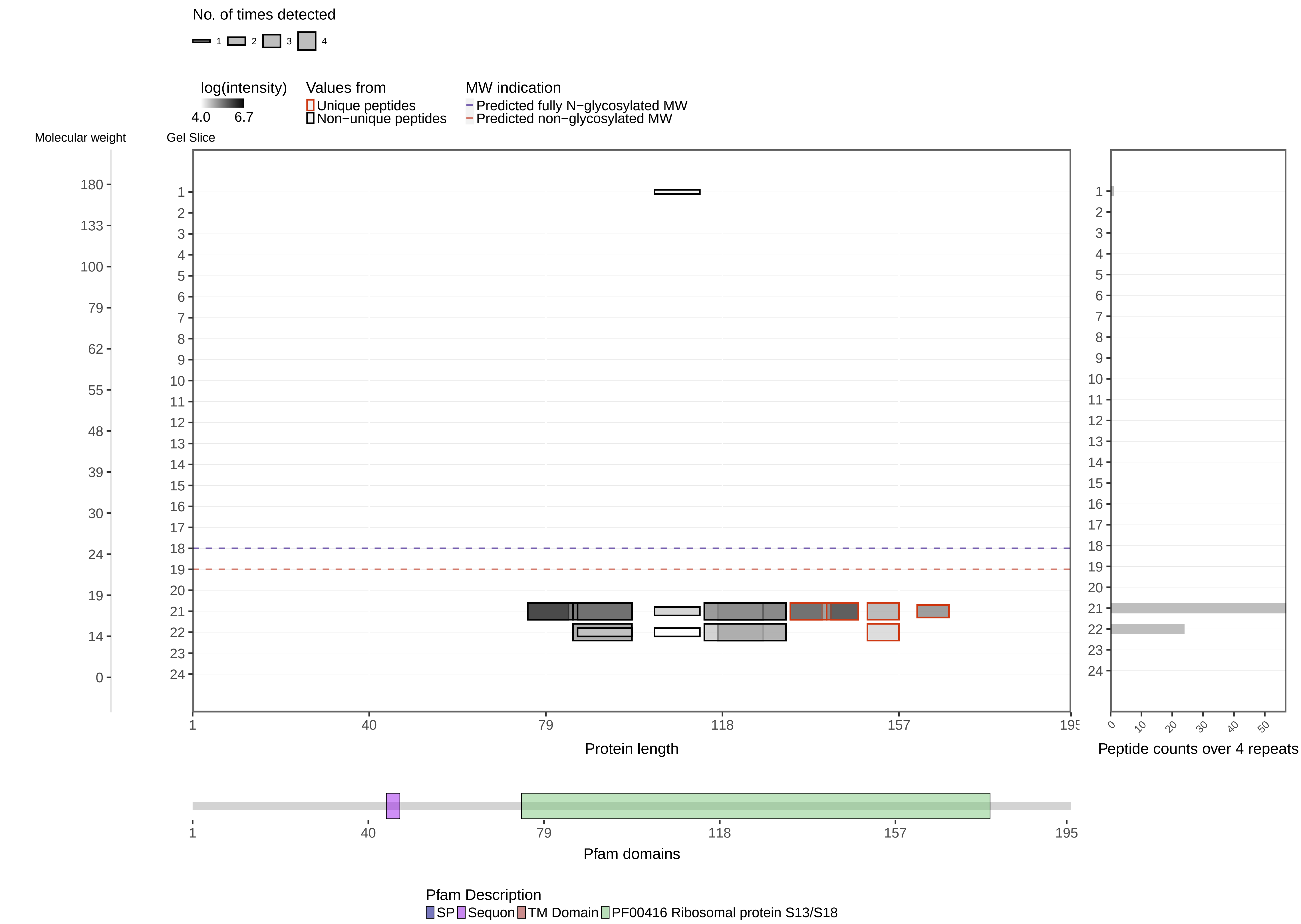


Protein sequence:

RVEFALQYIHGIGRTSARQILVDLQIENKIMKDFSEEELITLRDEVSKYMIEGDLRRFNALAIRRLKEIQCYRGVRHIQGLPCRGQRTKNNCRTLKGKKVAIAGKKKAPR*

### NbL18g08630.1 uncharacterized_LOC107812748 [MW = 40.41 kDa]

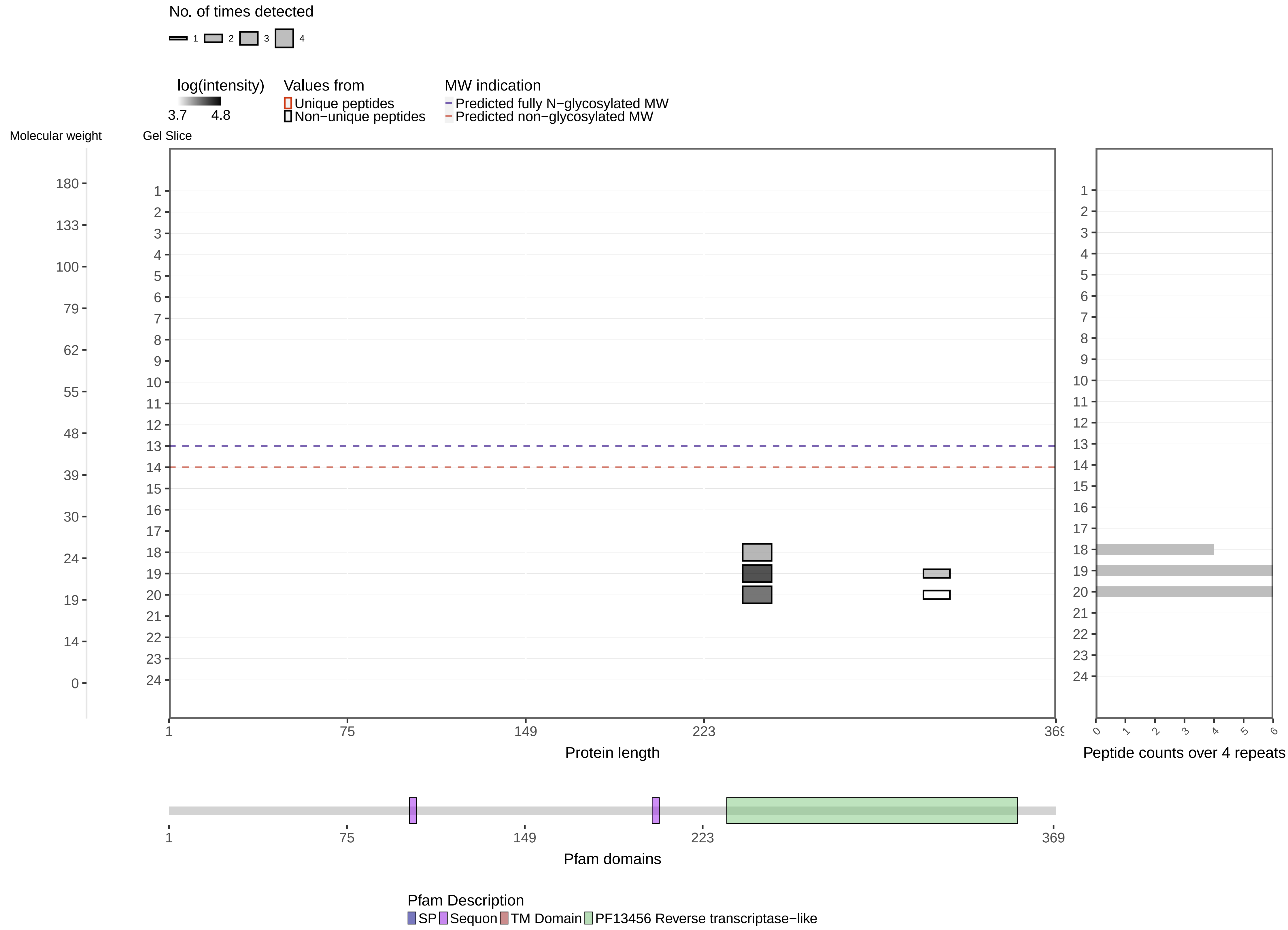


Protein sequence:

KSGTSDHLPKKRSQEAMWSEYADAVGSAVIPNDSIRKHVKIEHHKGDQVLALPSGQRSCTLEFDGASKGNPGQAGAGAVIRADDGSLTCRLREGLGVATSNHAEYRAFILGLKYALSKGFTNIRVQGDSKLVCMQIQGLWKVKNQNIATVFEQAKQLKERFLTFRIIHVLRESNSDADQQANLAVELAEGQIQEEIEK*

NbL18g08550.1 uncharacterized_protein_At4g28440−like [MW = 13.82 kDa]

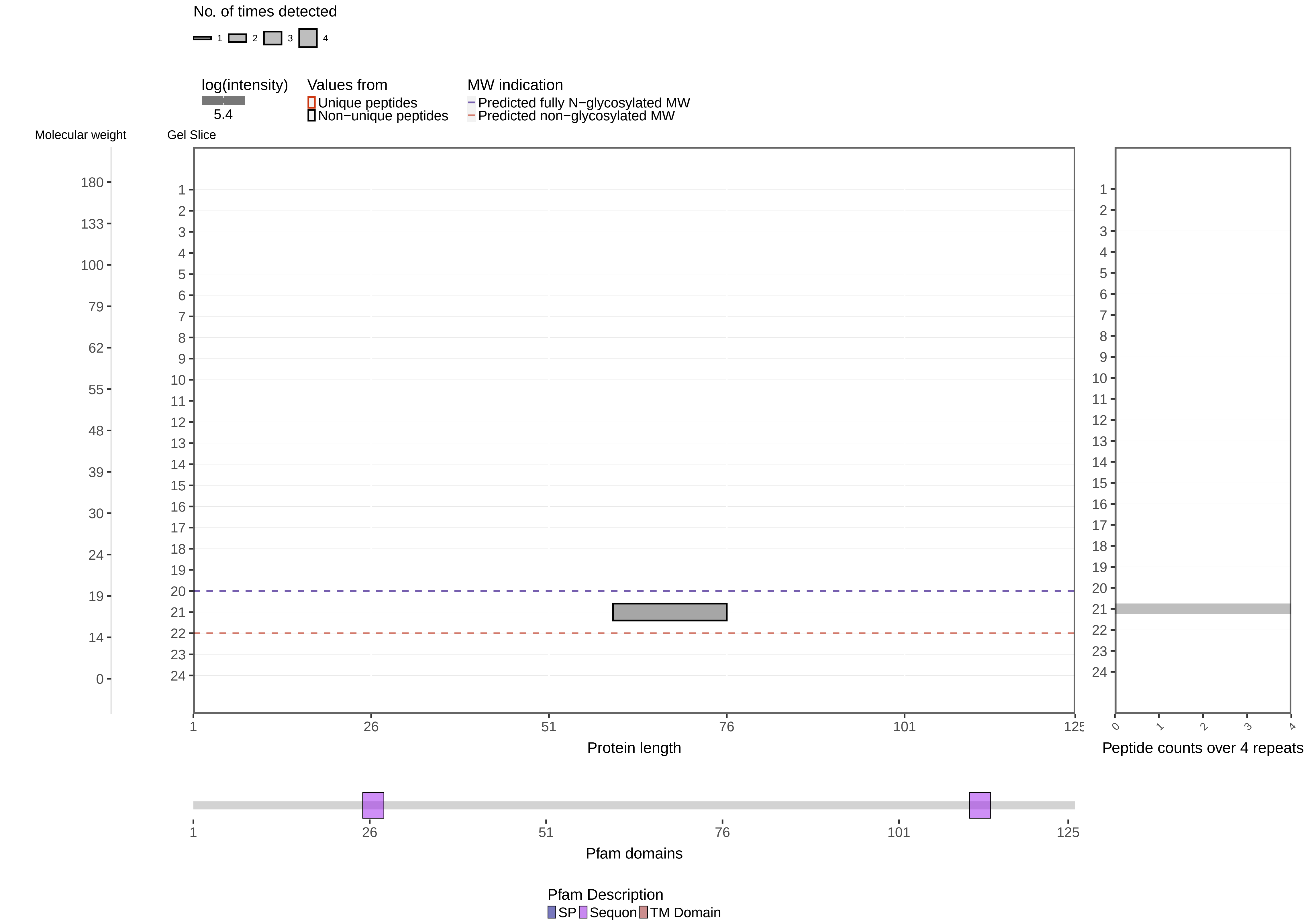


Protein sequence:

DETGIIIFTARNDQVDLIQEGSTLVLTKAKVDMFKGSMRLAVDRFGRIEVGQPASFSVDQDINMSLIEFERVDVVV*

NbL18g08290.1 pyruvate_dehydrogenase_E1_component_subunit_alpha_mitochondrial−like [MW = 45.9 kDa]

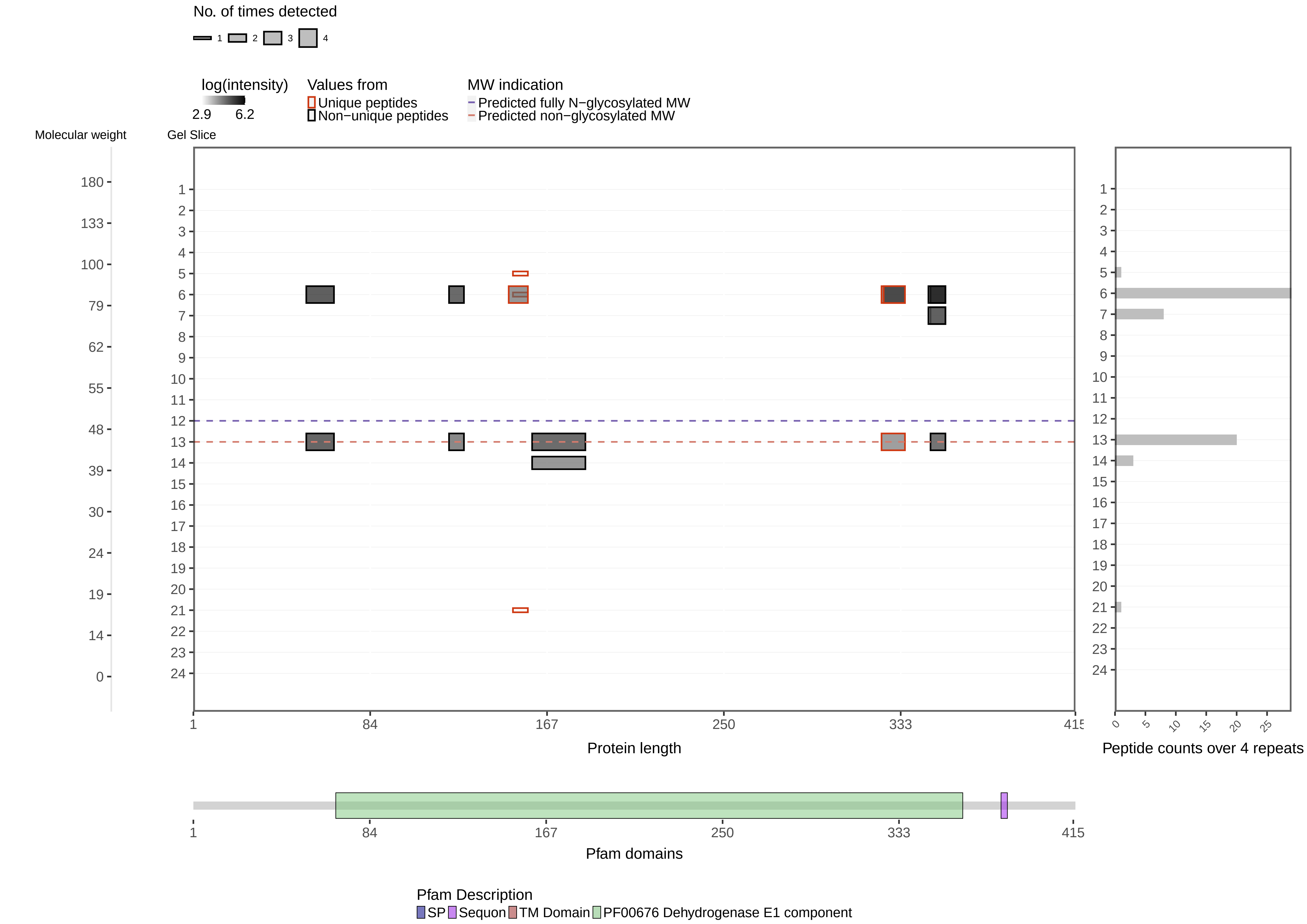


Protein sequence:

MYGDGAANQGQLFEALNMAALWDLPAILVCENNHYGMGTAEWRAAKSPAYYKRGDYVPGLKVDGMDALAVKQACKFAKEHALKNGPIILEMDTYRYHGHSMSDPGSTYRTRDEISGVRQERDPIERIRKLILAHDISTEKELKDIEKEKRKIVDEAIAKAKESPMPDPSELFTNVYVKGFGVEVCNCSNHQFSLLNYLRATVSSRNQVMIAQFSLEHII*

### NbL18g08060.1 HIPL1_protein−like [MW = 78.01 kDa]

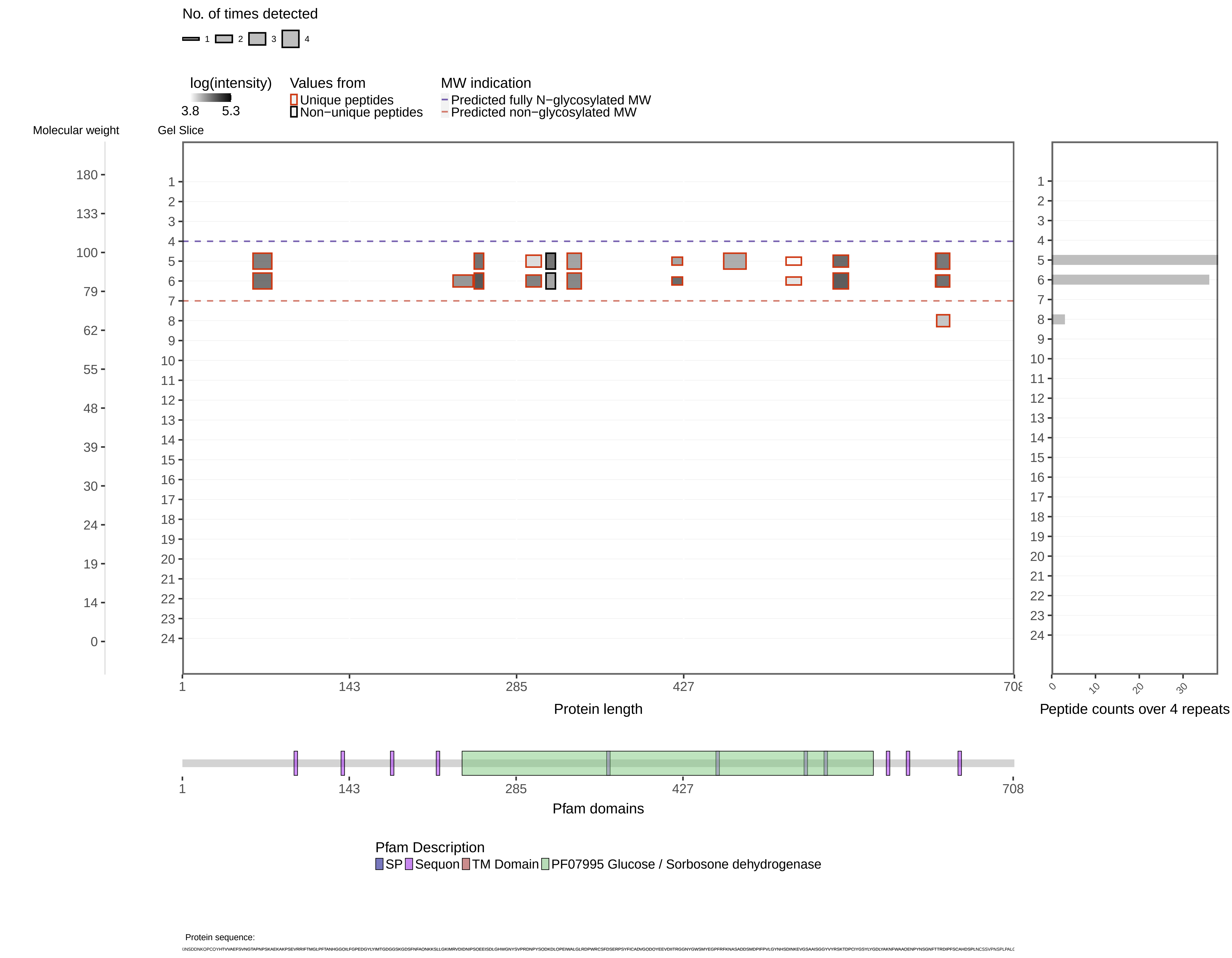

### NbL18g08050.1 glutathione_S−transferase_U17−like [MW = 25.33 kDa]

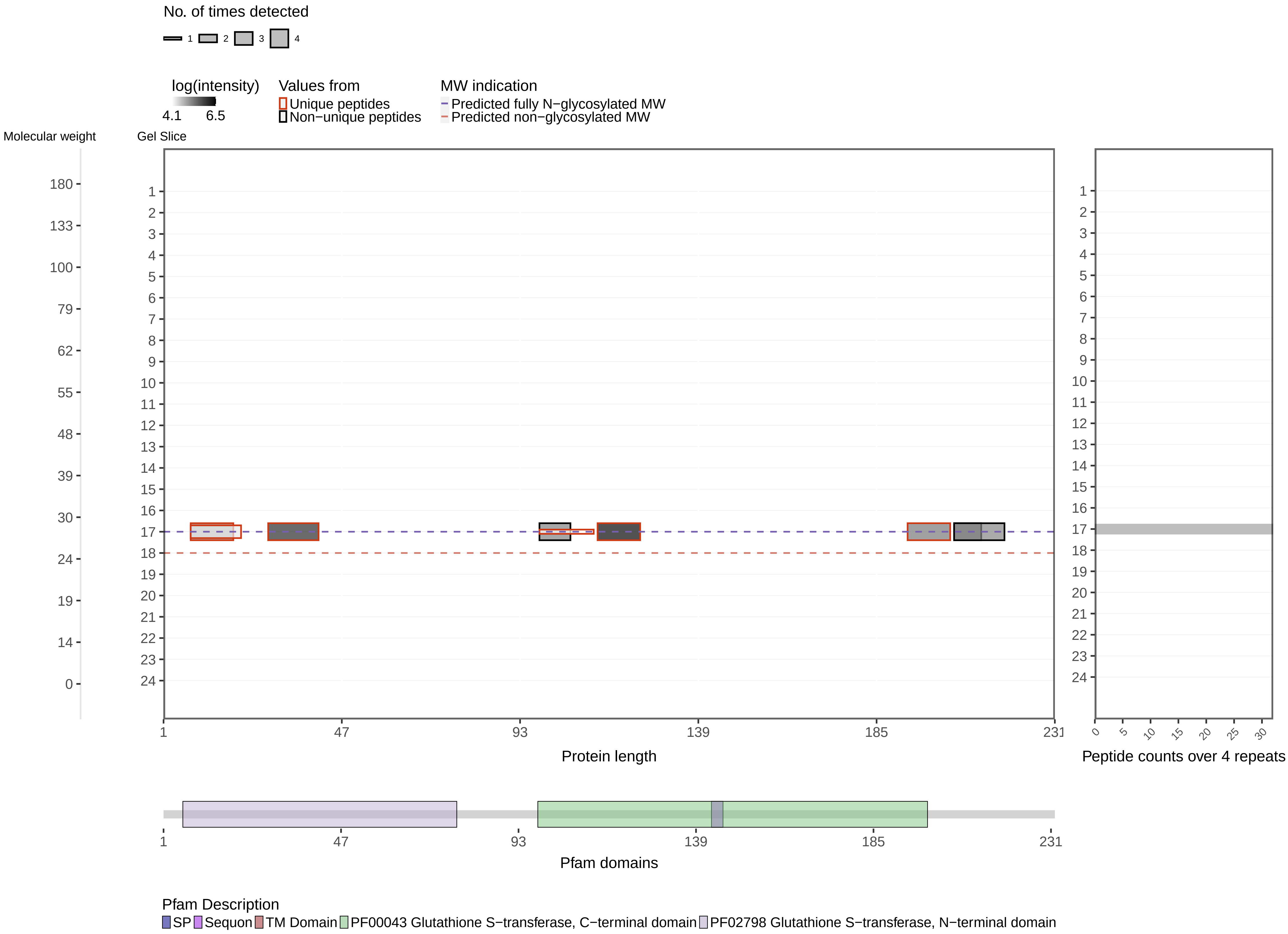


Protein sequence:

KWFPAMRGIAAAQGEDAQKAATEPVVEGLVLLEDAFKNCSEGNKFFGGDKIGYLDIALGCFLGWMRVTEKLNNVKLLDEVKTPGLCKWAEDFCADSAVKDVMPATDKLAEAAKVIMAKIRAQASS*

### NbL18g08000.1 tetratricopeptide_repeat_protein_38 [MW = 57.38 kDa]

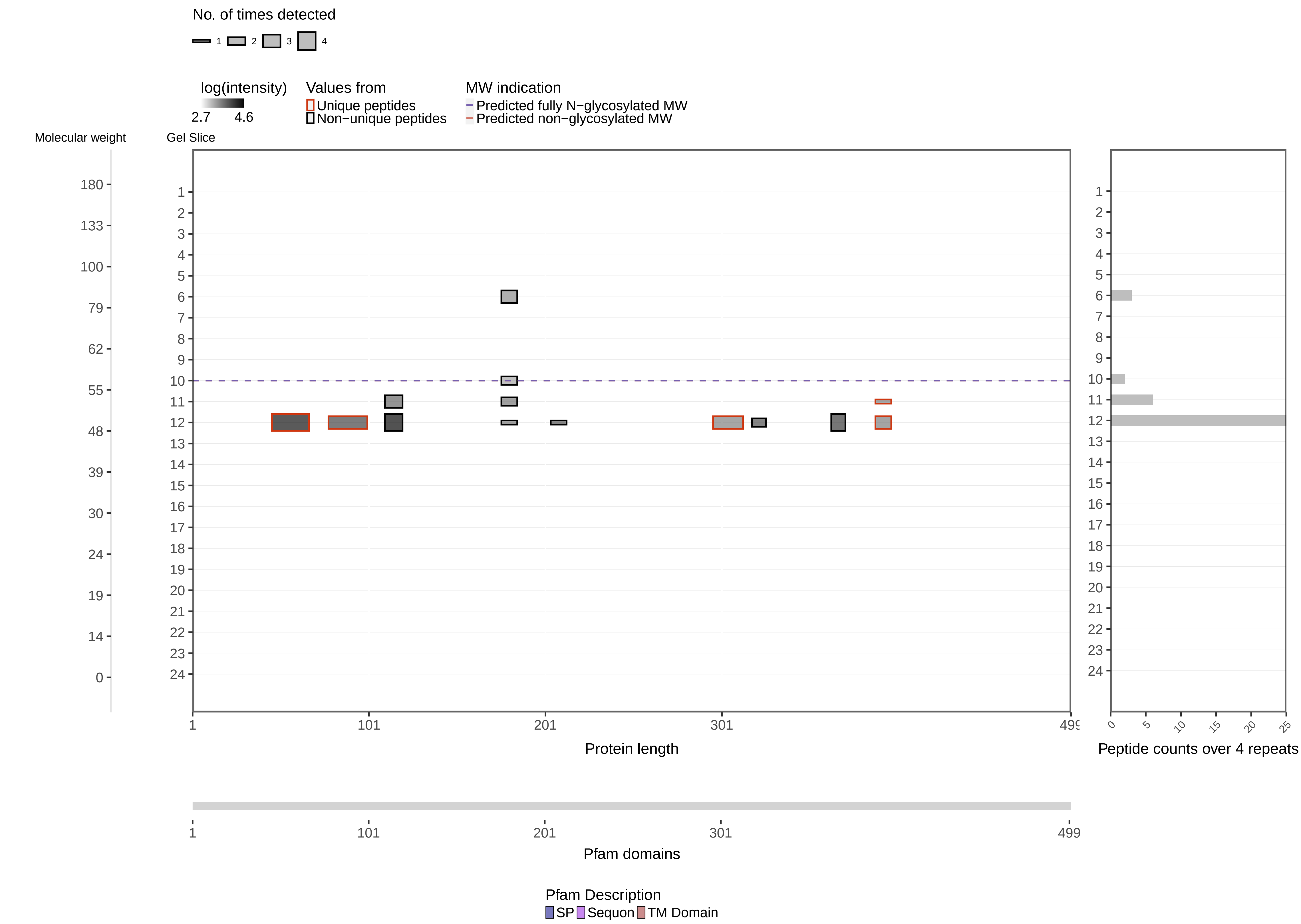


Protein sequence:

[illegible]EAVQFMEKCSRTWSSLSSFMYTHNWWHVALCYLEGHSQMEKVRDVYDQNIWKELERSDASPAEVYLNVVGLLLRVYVRGGINVFGDRLKILADCLTNKAFWYLEWHLDVLVVWALSFTGEVSKAEELLEGLKSRMSKMTKKKRQHMQRAMLLAEALFKYGKGENERALELLGEPFDAINYKIIGASDEQLDVFNEVWITMLLNSGQVTKAIQAIERQLKKREGTPFLWRLLEKSYSMSRKQEAIDAGKKAQALEAAYFD*

### NbL18g07960.1 uncharacterized_protein_At1g27050 [MW = 24.87 kDa]

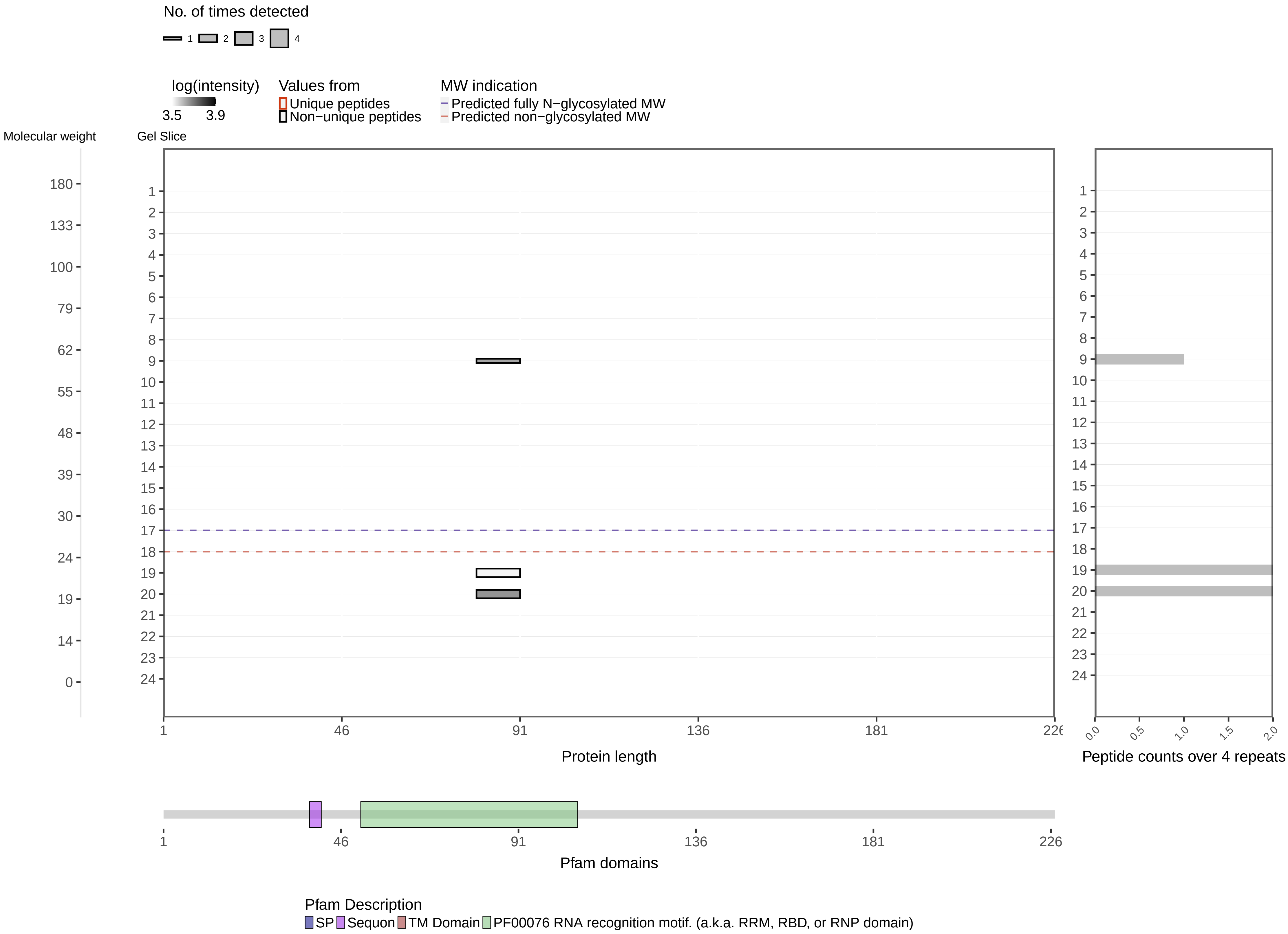


Protein sequence:

[illegible]NAAVGSSFPITLHSKPVQVIWATDHVPQWKEGVMRKDDLSTSTVASKLVRAEIPLSRHGRGNRLGSAIVNPKDEDNDNAGNHDRGHVSTRLGEPDNAGDNGRGHVSTRLVEPYKGREIVAYDDIL*

### NbL18g07870.1 uncharacterized_LOC104243986 [MW = 42.67 kDa]

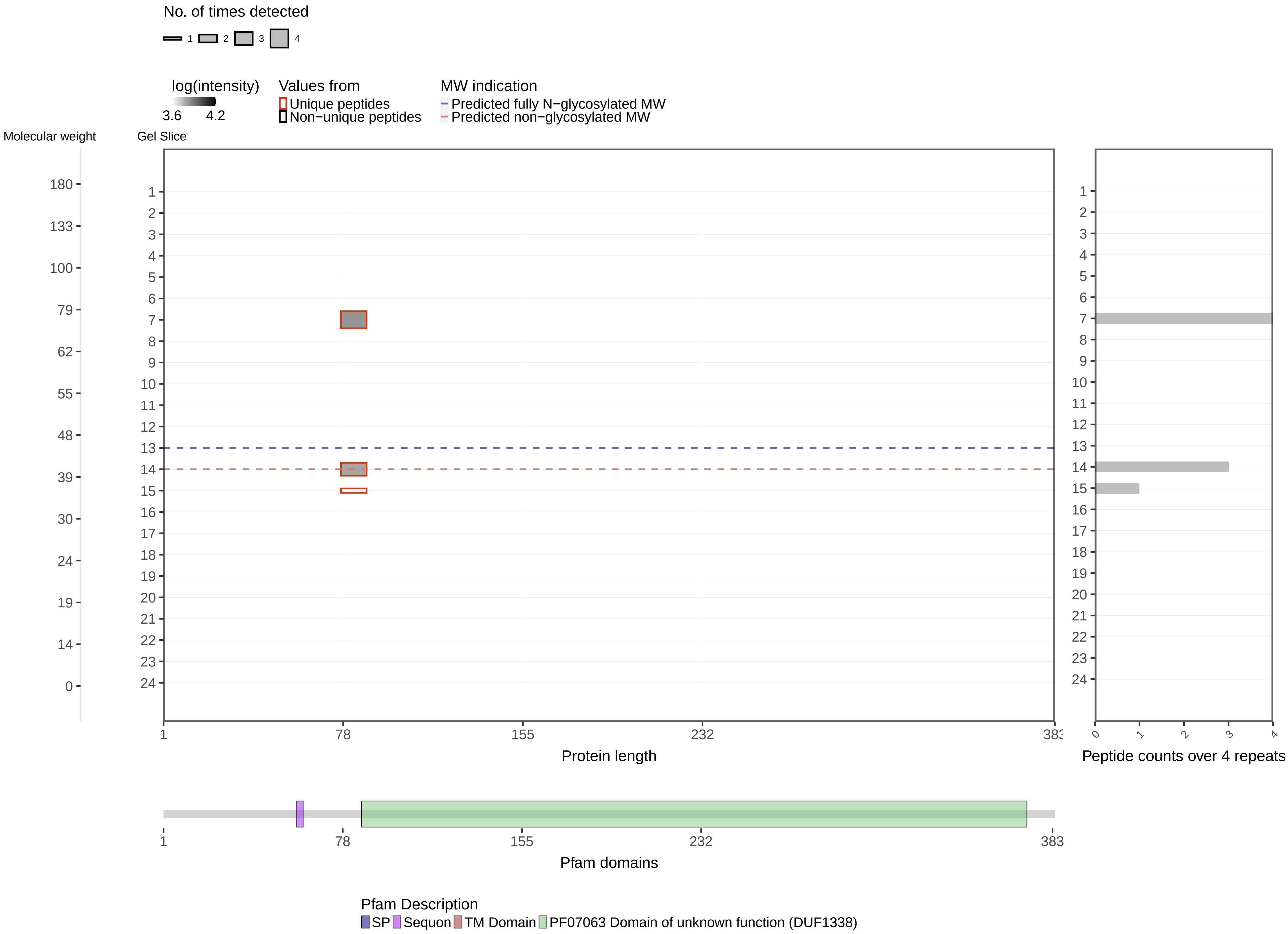


Protein sequence:

RGSGVNGPLPRIFISELLVGQLSPEAQDIVKKYTDISQCGKEYAALASAFGTLTWEKPFYSEFQQLARESEYAAWTLVNGYALNHVTISTHRLTSNLRSIGNLNQFIEKNGFNLNSEGGVLKVSPDGLLLQSSTVADSTSFEFSDGITEAVPFSYIEFAERLVLPQYKDLPPEKVEEFHRRDGFEVGNADKIFESTSKDQLTRRAA*

NbL18g07850.1 single−stranded_DNA−binding_protein_WHY1_chloroplastic−like [MW = 29.43 kDa]

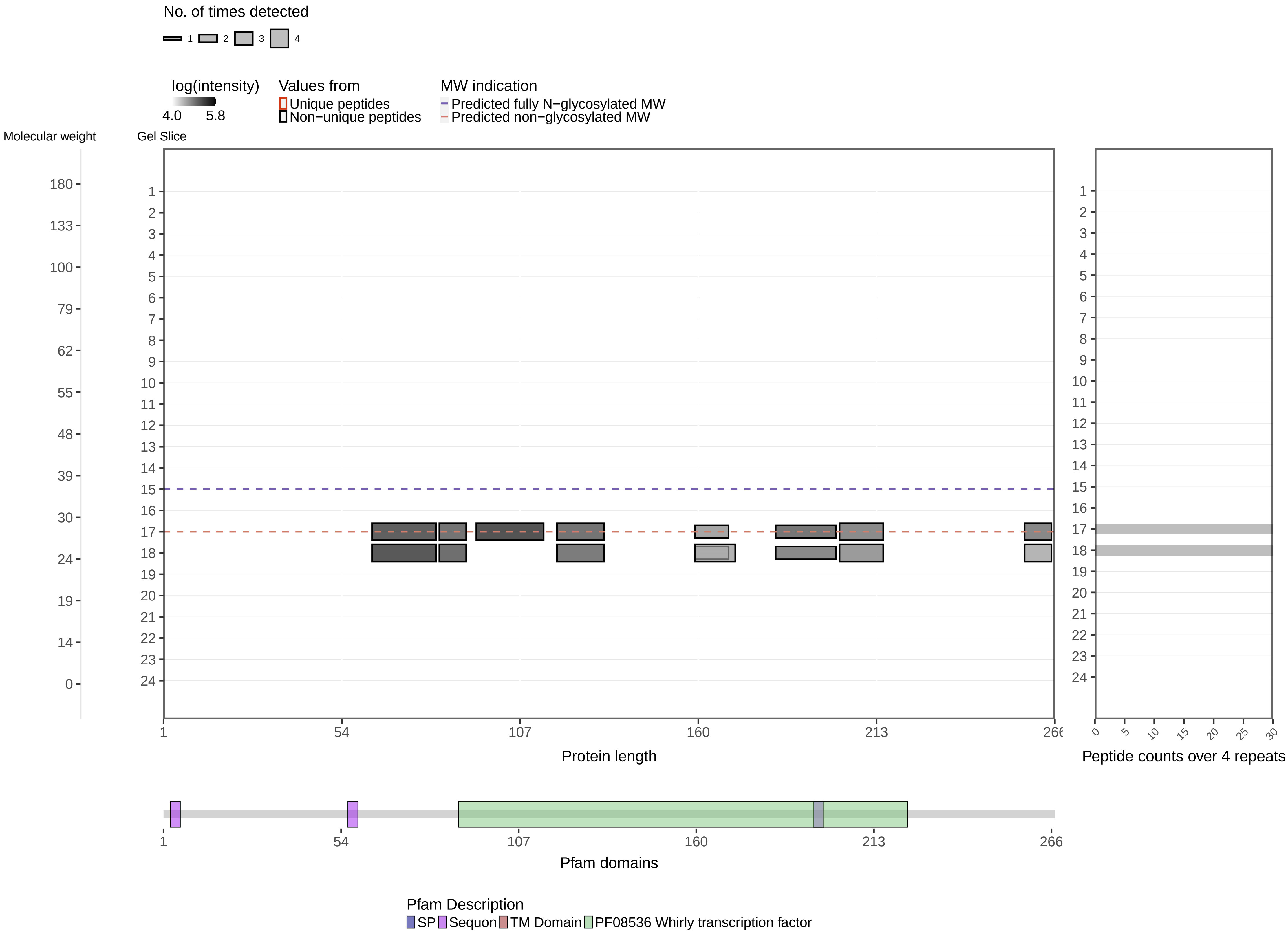


Protein sequence:

FVLLQFAPAAGVRQYDWGRKQVFSLSVTEMGSLISFGAKDSCEFFHDPNKGRSDEGKVRKVLKVEPLPDGSGHFFNLSVQNKLINLDENIYIPVTKAEFAVLISAFNFVVPYLLGWHTAVNSFKPEDASRSNNANPRSGADLEWSR*

NbL18g07810.1 fumarate_hydratase_1_mitochondrial [MW = 50.36 kDa]

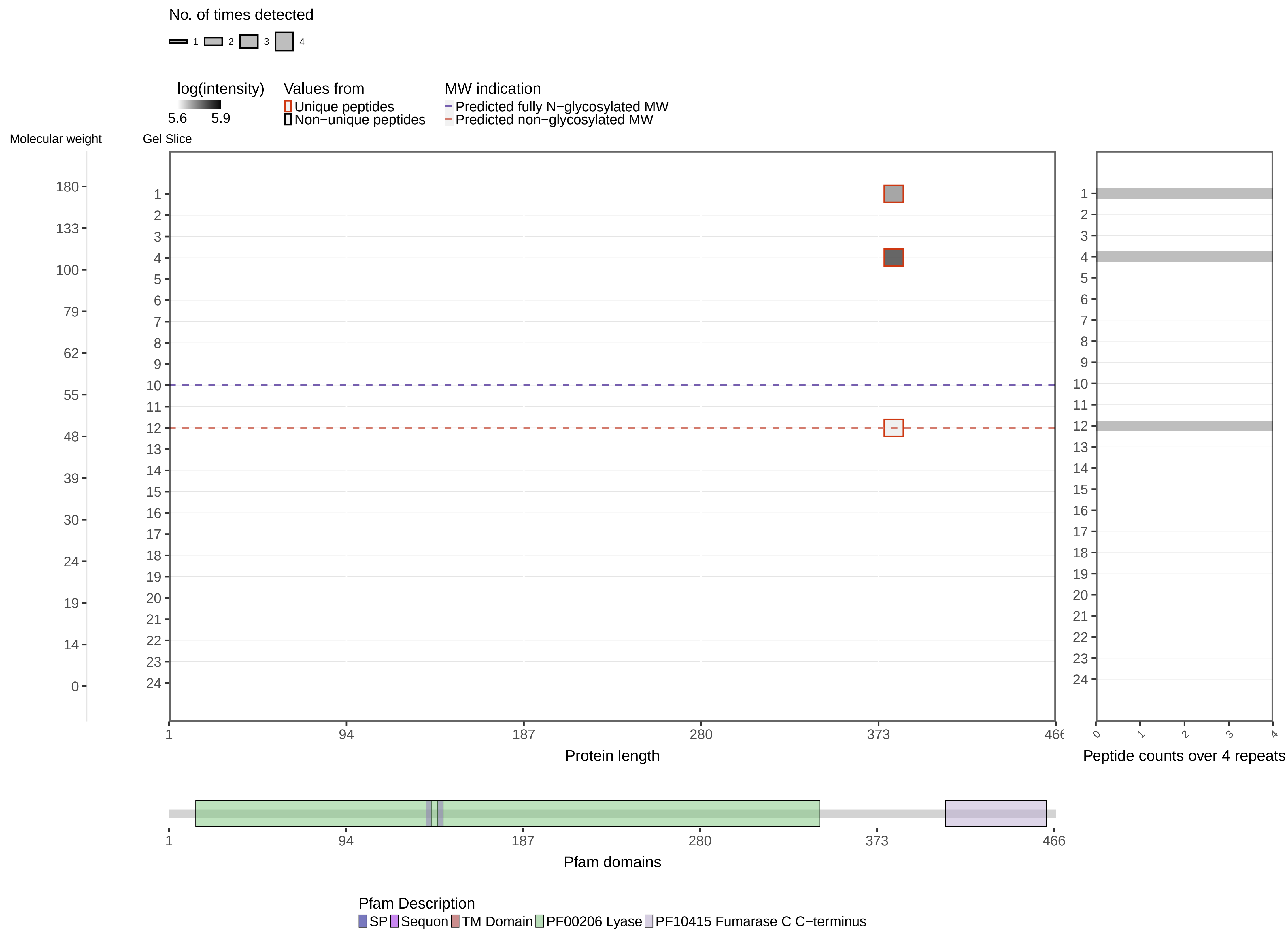


Protein sequence:

LPHMYQIAQGGTAVGTGLNTKKGFDIKIAAAVAEETNLPFVTAENKFEALAAHDAFVETSGAVNTLAASLMKVGNDIRFLGSGPRCGLGELILPENEPGSSIMPGKVNPTQCEALTMVCAQVMGNHVAVTIGGSNGHFELNVFKPMIANALLHSVRLLGDASASFEKNCVRGIQANRERIDKLLHESLMLVTCLNPKIGYDIAAAVAKKAHKEGTSLKDAALNLGVLKSEEFDQFVVPEKMIGPTD*

NbL18g07620.1 40S_ribosomal_protein_S9−2−like [MW = 23.01 kDa]

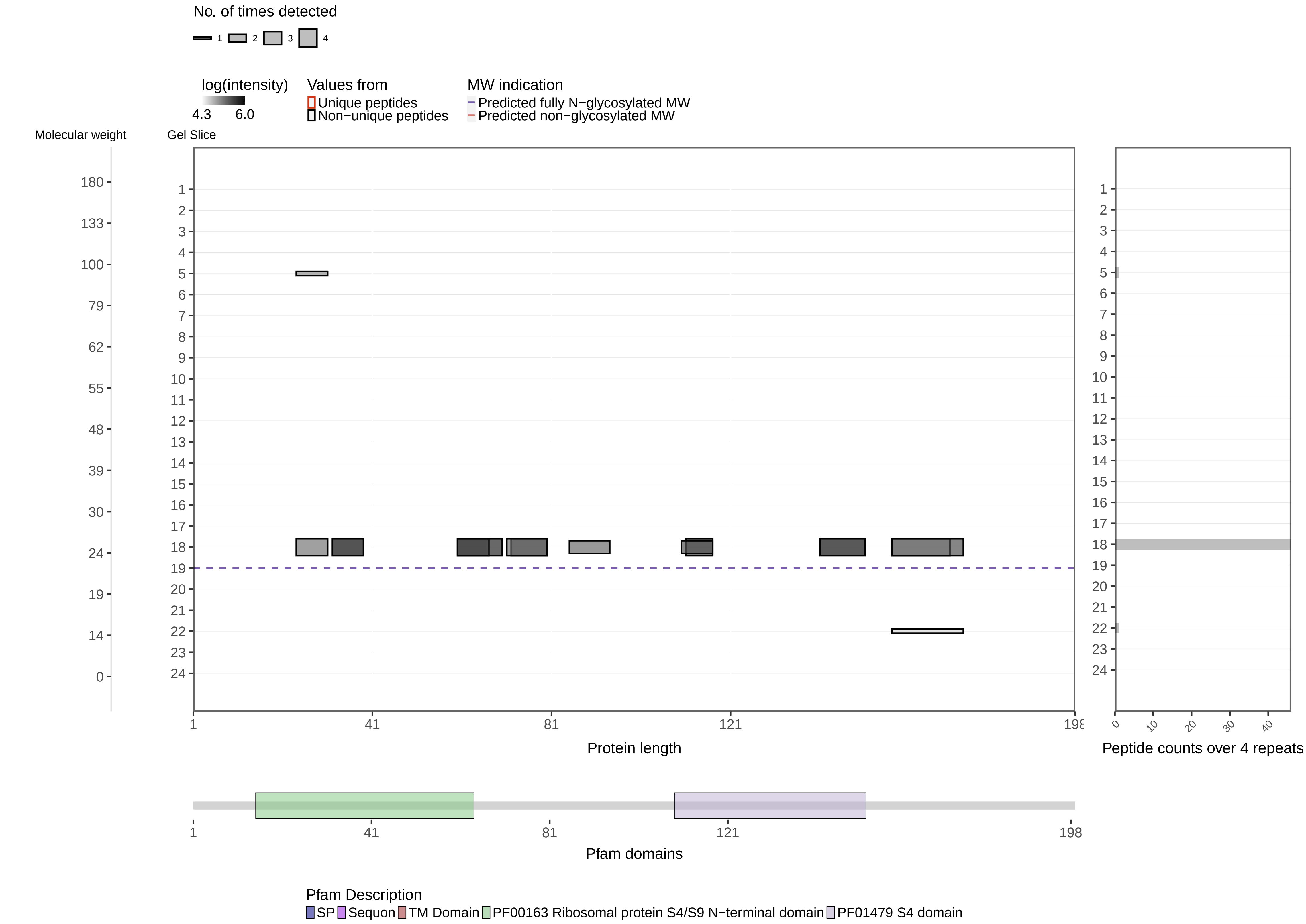


Protein sequence:

LLDESQNKLDYVLSLTVENFLERRLQTLVFKTGMAKSIHHARVLIRQRHIRVGRQVVNVPSFMVRVDSQKHIDFSLTSPFGGGRPGRVKRKNQKAAAKKASGGDGDEEDEE*

### NbL18g07420.1 60S_ribosomal_protein_L34 [MW = 13.77 kDa]

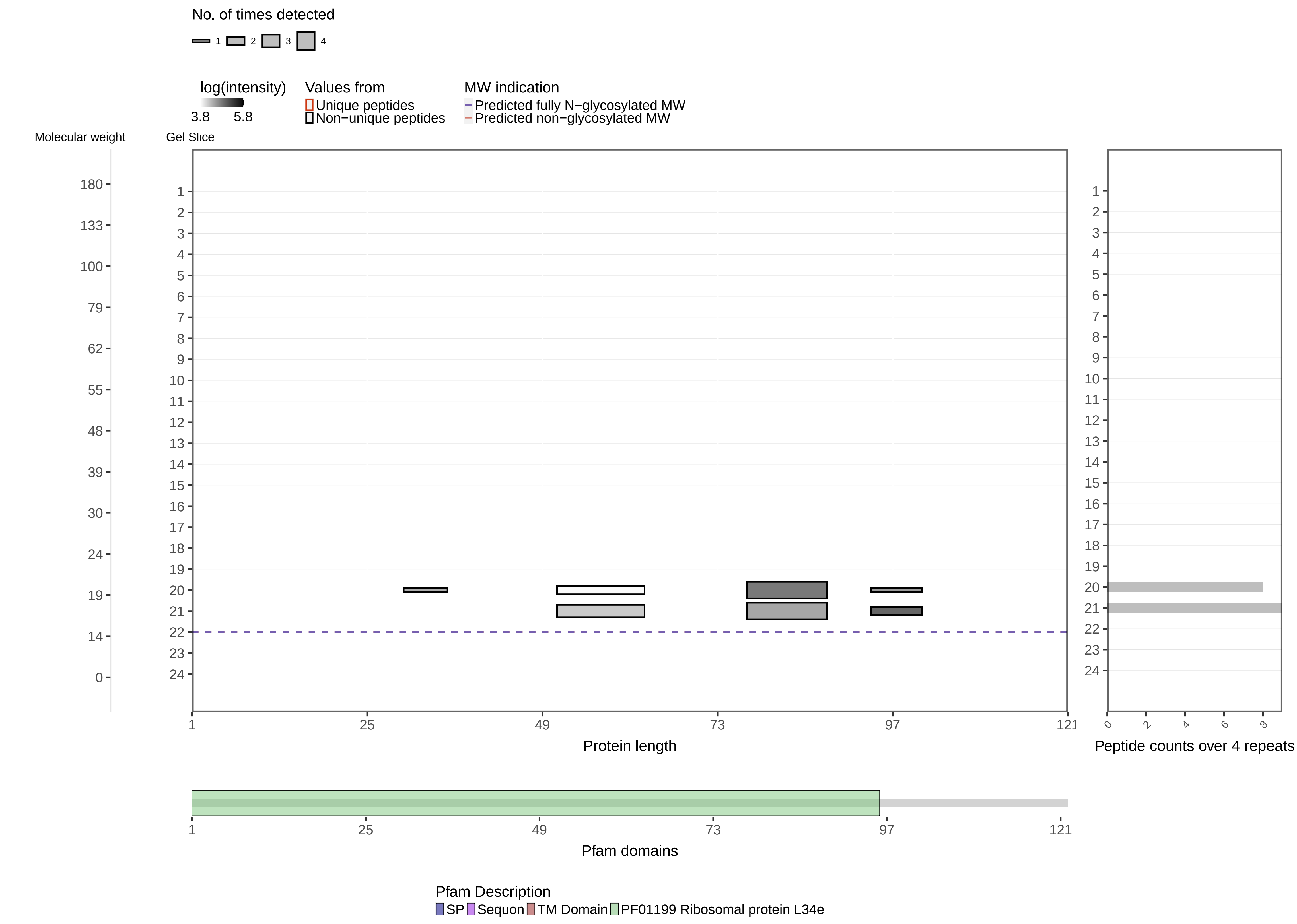


Protein sequence:

/TGKRIQGIPHLRPTEYKRSRLSRNRRTVNRAYGGVLSGSAVRERIIRAFLVEEQKIVKKVLKIQKAKEKLASKS*

NbL18g07140.1 intracellular_ribonuclease_LX−like [MW = 32.47 kDa]

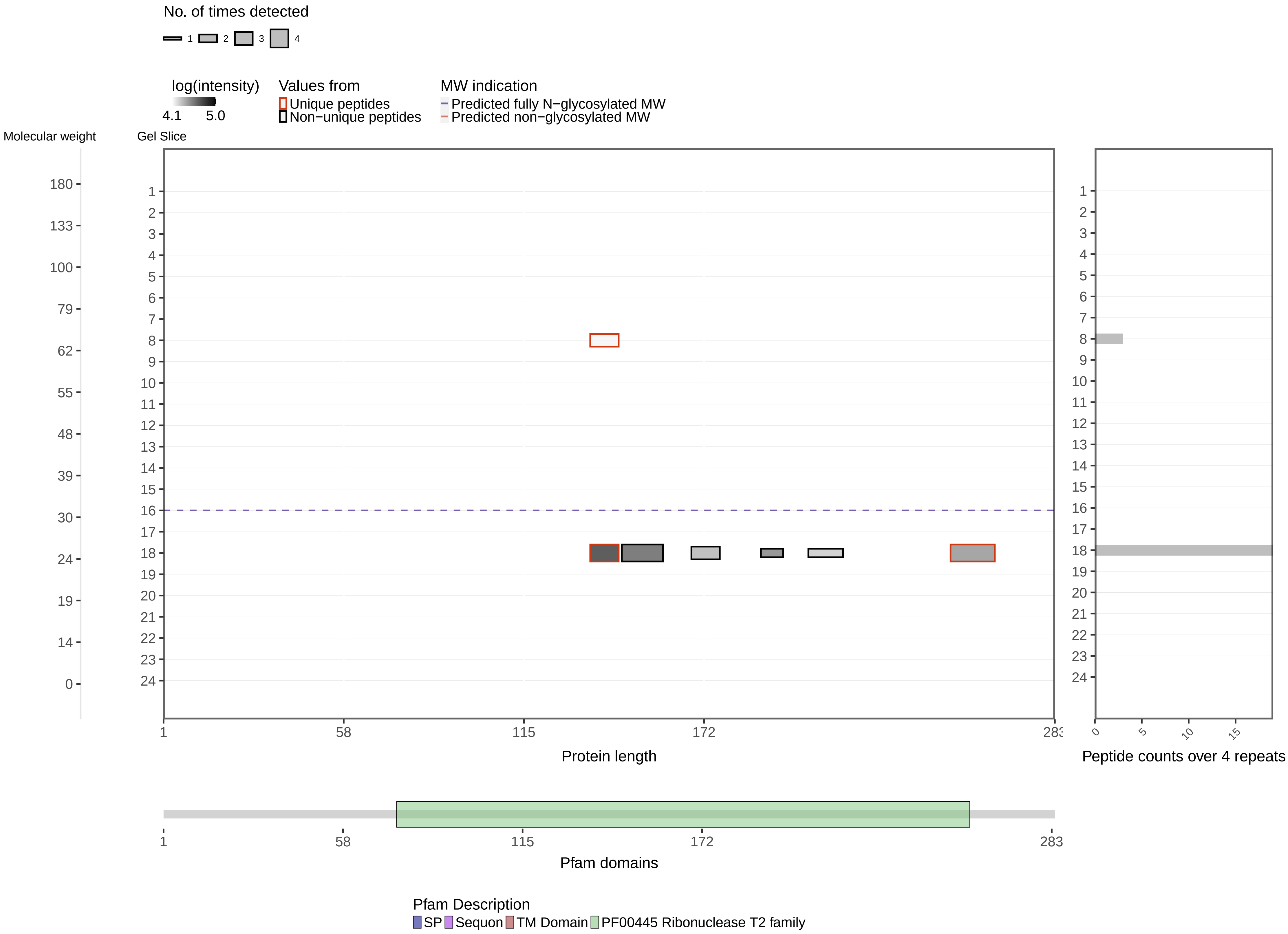


Protein sequence:

SLDESKISDLISTMEKNWPSLACPSSDGVRFWSHEWLKHGTCSALGERAYFQAALDFRKKSNLLENLKNAGITPRNGEHYSLESIKKAIEEGVGHSPYIECNVDTQGNHQIYQVYMCVDKTATDFIDCPVFPHGRGCGSKIEFPPFSSDHAEF*

NbL18g06980.1 ribulose−15_bisphosphate_carboxylase/oxygenase_large_subunit_N−methyltransferase_chloroplastic−like [MW = 55.99 kDa]

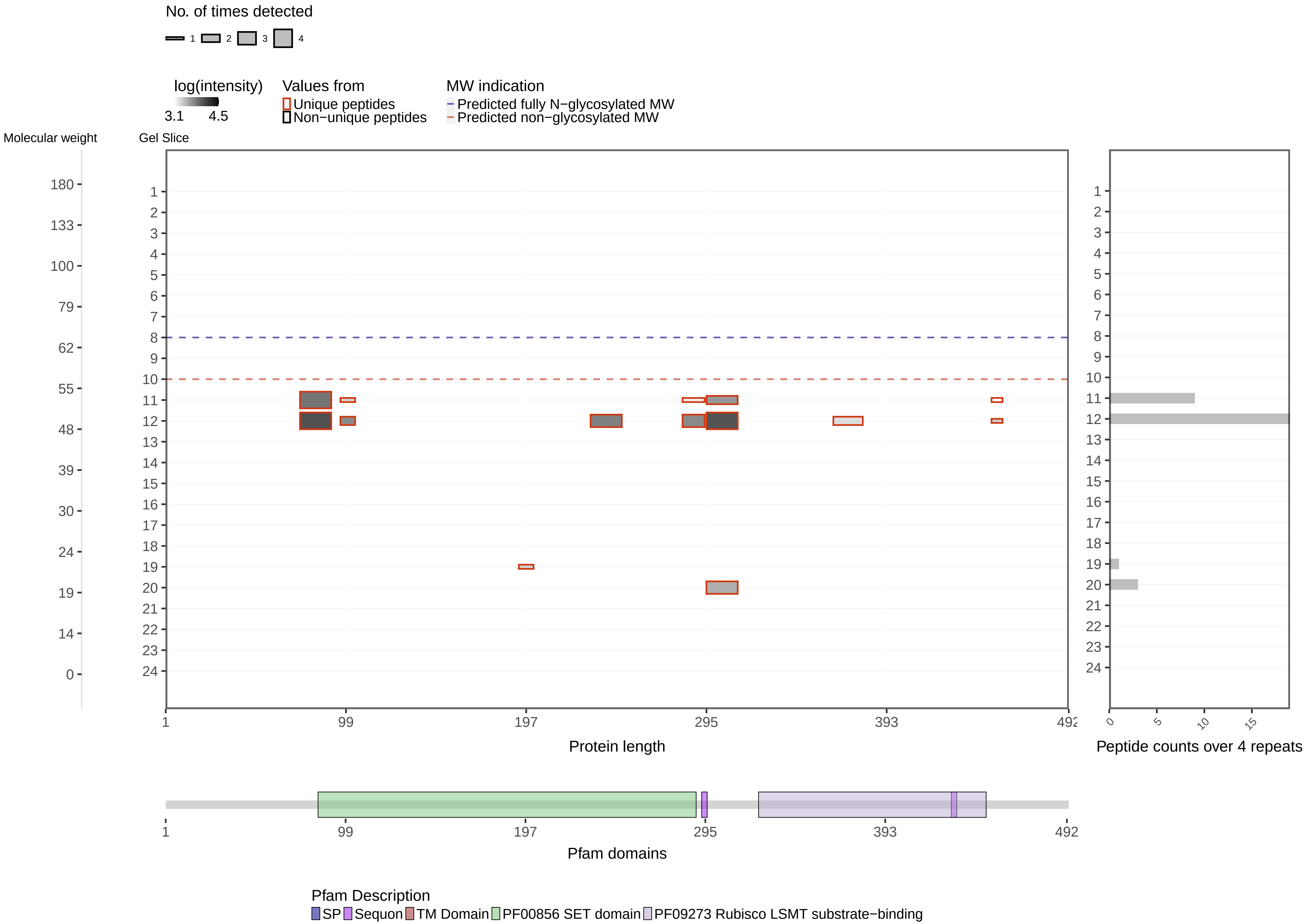


Protein sequence:

RNQNLILVPFADLTNHNARVTTEDHAHEVRGPAGLFSWDLLFSLRSPLKLKAGDQLFIQYDLNKSNADMALDYGFIEPSSARDAFTLTLEISESDEFYGDKLDIAETNGIGETAYFDIKIGQSLPPTMIPYLRLVALGGTDAFLLESIFRNSVWGHLGLPVSRANEELICKVVRDACKSALSGYHTTIEEDEKLMEQGNLSTRLQLAVGIRLGEKKVLKQIDNIFRERELELDELEYYGERRLKDLGLVGEQGDIIFWEPK*

### NbL18g06970.1 uncharacterized_LOC109214564 [MW = 19.96 kDa]

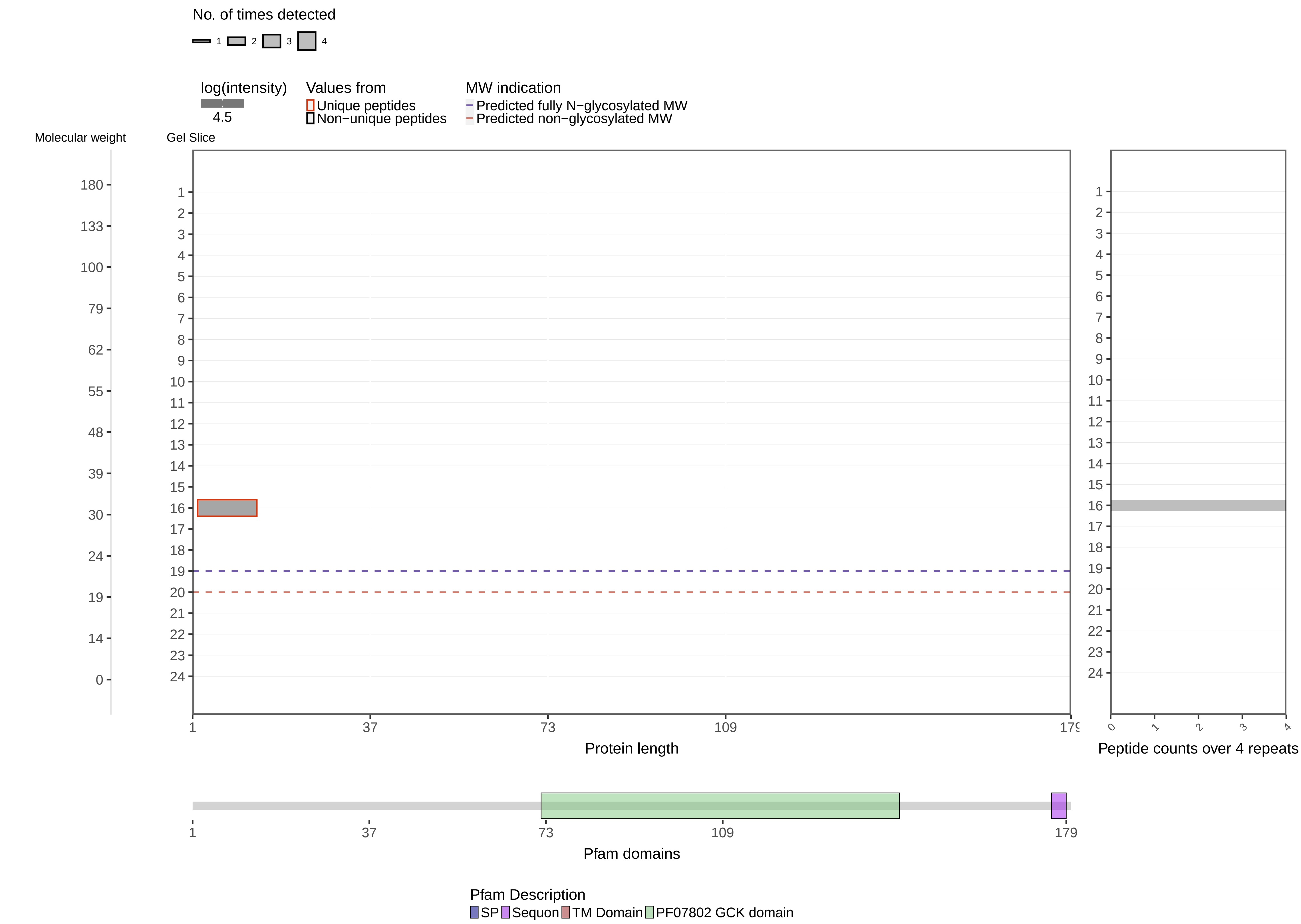


Protein sequence:

CLFMKGGGCKDEFIDWEKCIEEGEKNKEDIVEKCFEVTSALKKCMEAHSDYYAPIFQAEKAAEAELEKELDKEKEKEEQRNDADALGSVSVDPTLDGNSNSS*

### NbL18g06930.1 extracellular_ribonuclease_LE−like [MW − SP = 22.51 kDa]

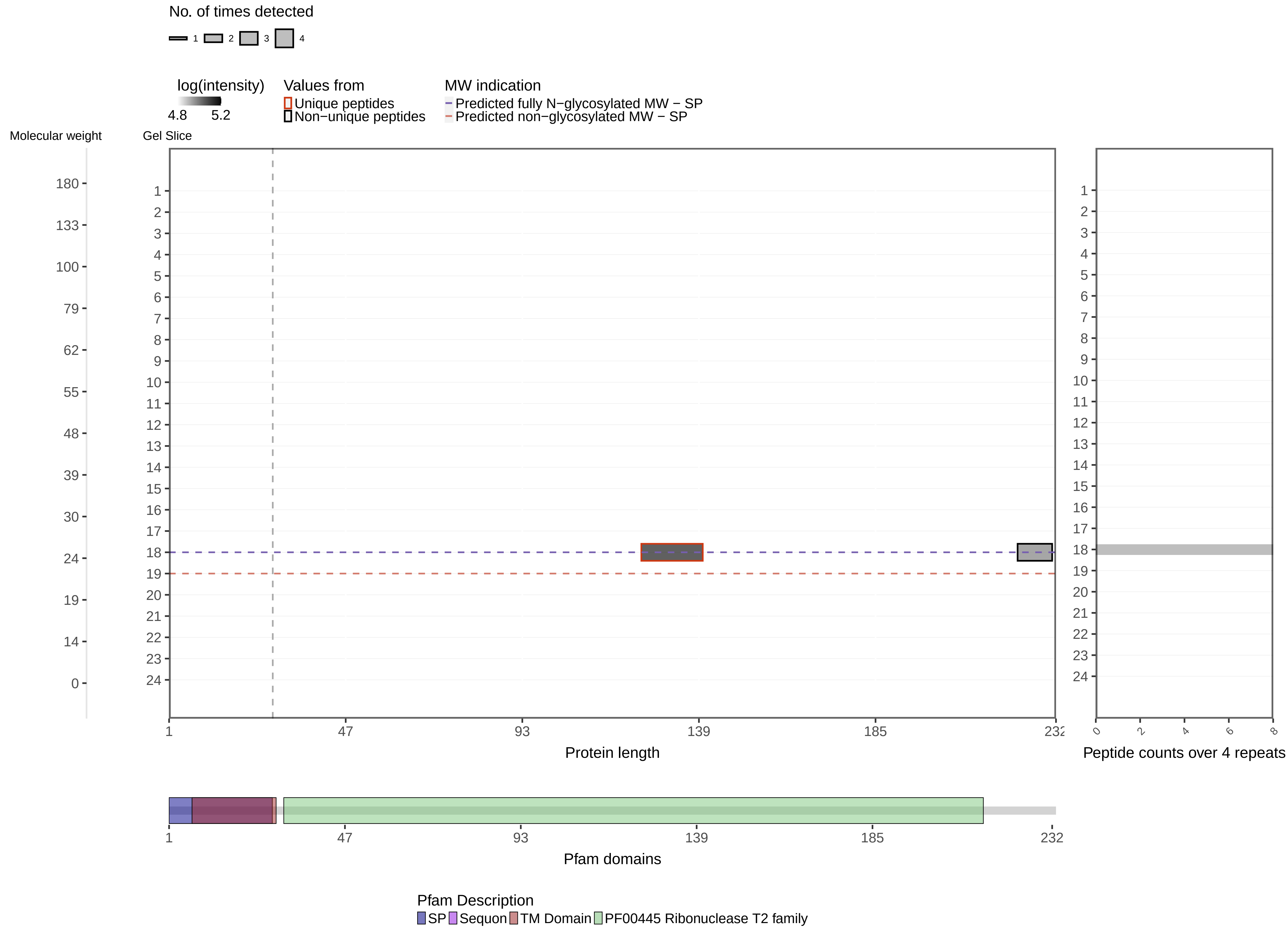


Protein sequence:

IWPTLACPSGTGSAFWSHEWQKHGTCAENVFDQHGYFMKALDLKNQINLLEILQGAGINPDGGFYSLNSVKNAIRSAIGYTPGIECNVDESGNSQLYQVYICVDGSGSNLIECPVFPRGKCGSSIEFPTF*

NbL18g06660.1 alpha−14_glucan_phosphorylase_L−2_isozyme_chloroplastic/amyloplastic [MW = 111.4 kDa]

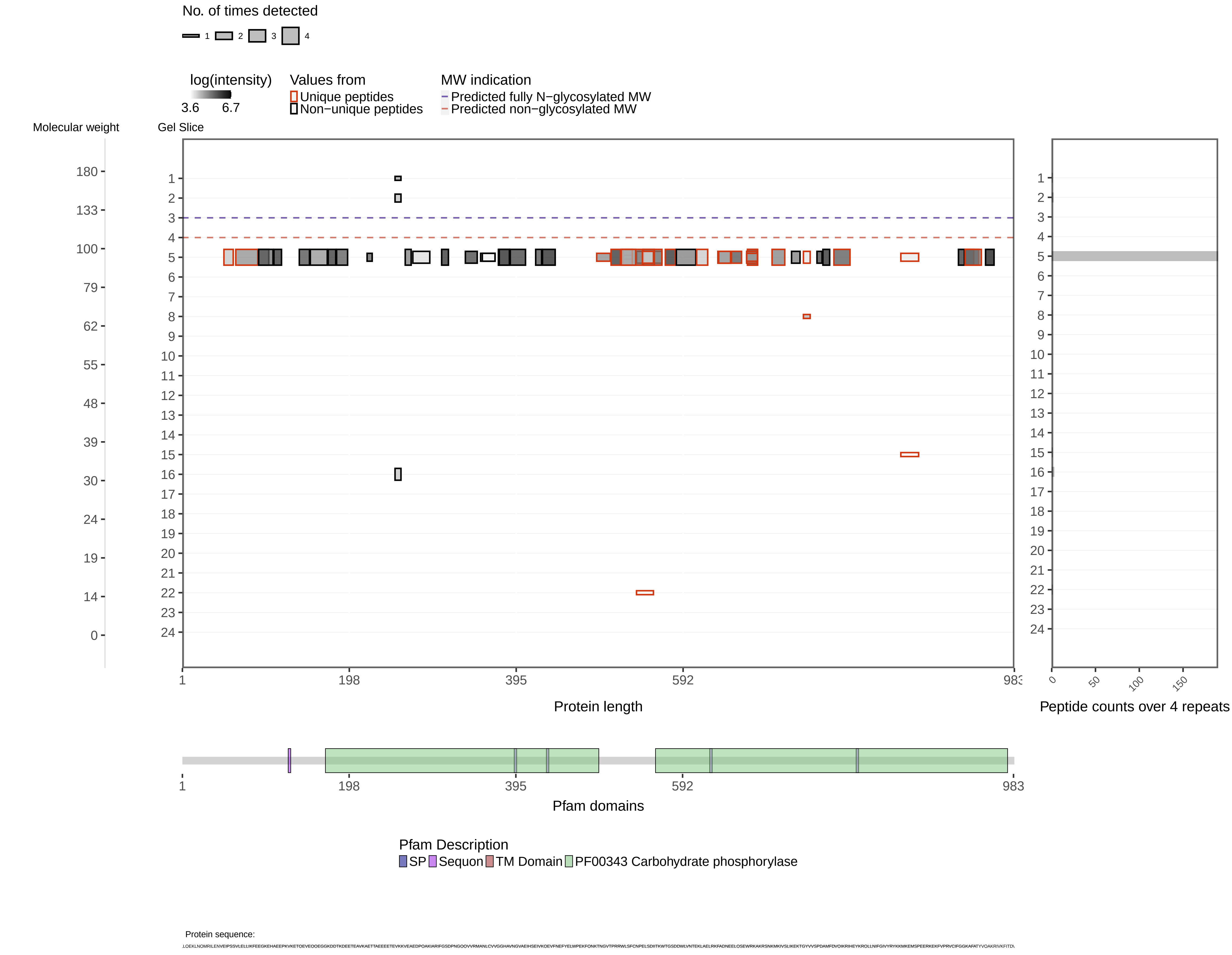

NbL18g06610.1 probable_pectinesterase/pectinesterase_inhibitor_41 [MW = 12.6 kDa]

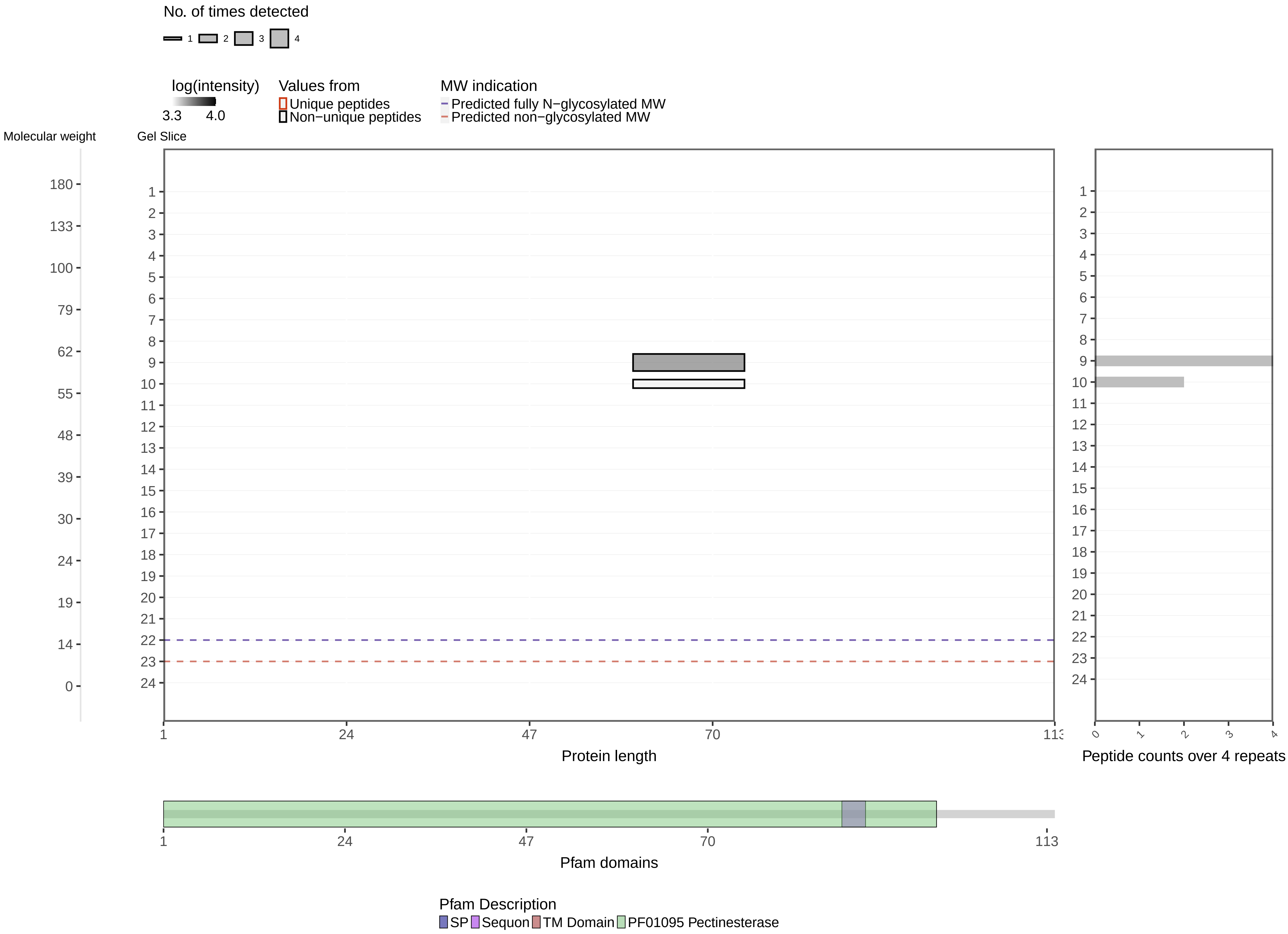


Protein sequence:

FIFGNAAAVFQNCNLYPRLPLPNQFNAITAQGRTDINQNTGISIHNCTITPSCGQRTVASTSNLLSRYEQI*

NbL18g06590.1 probable_mitochondrial−processing_peptidase_subunit_beta_mitochondrial [MW = 57.17 kDa]

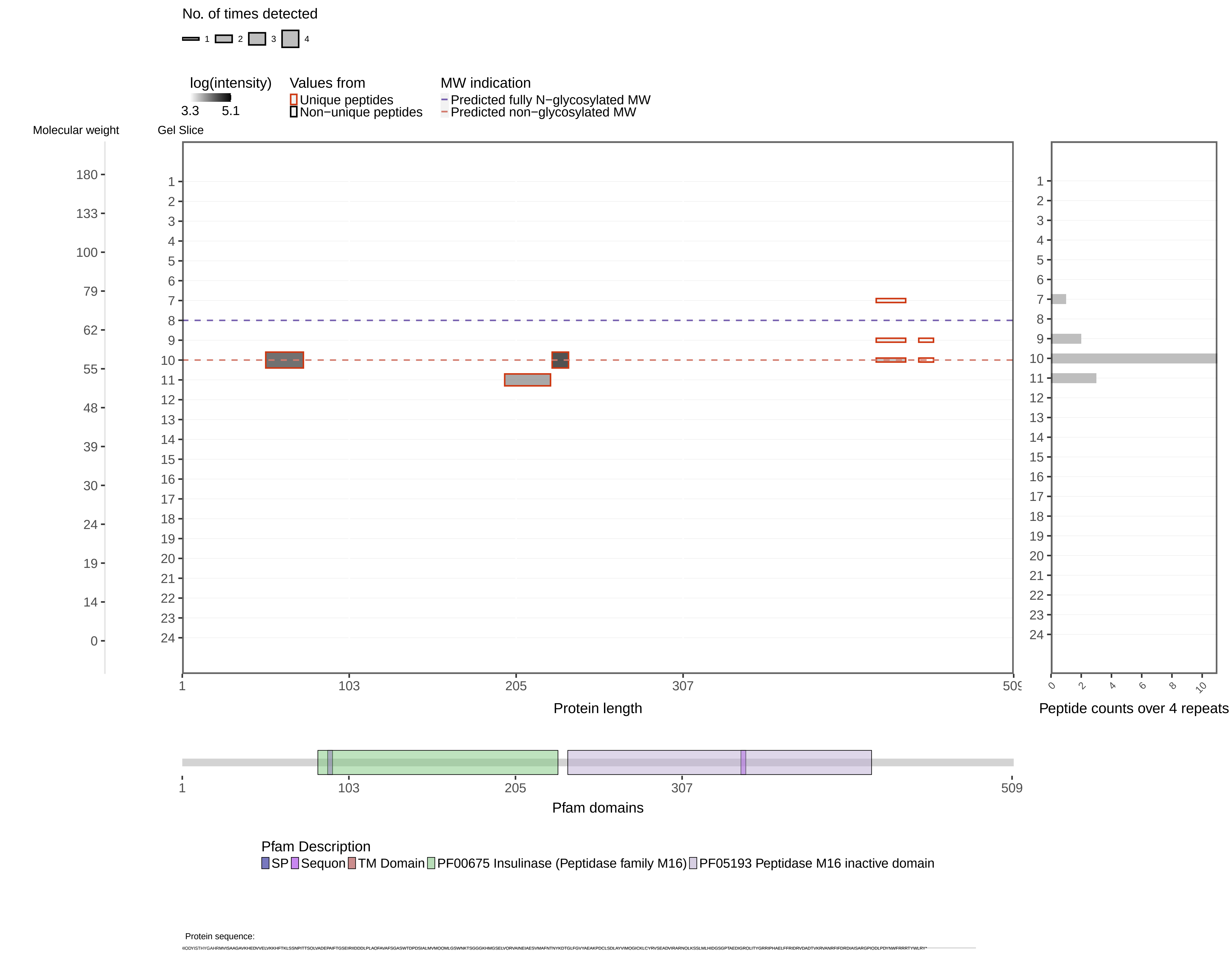

### NbL18g06520.1 pentatricopeptide_repeat−containing_protein_At1g26460_mitochondrial [MW = 69.29 kDa]

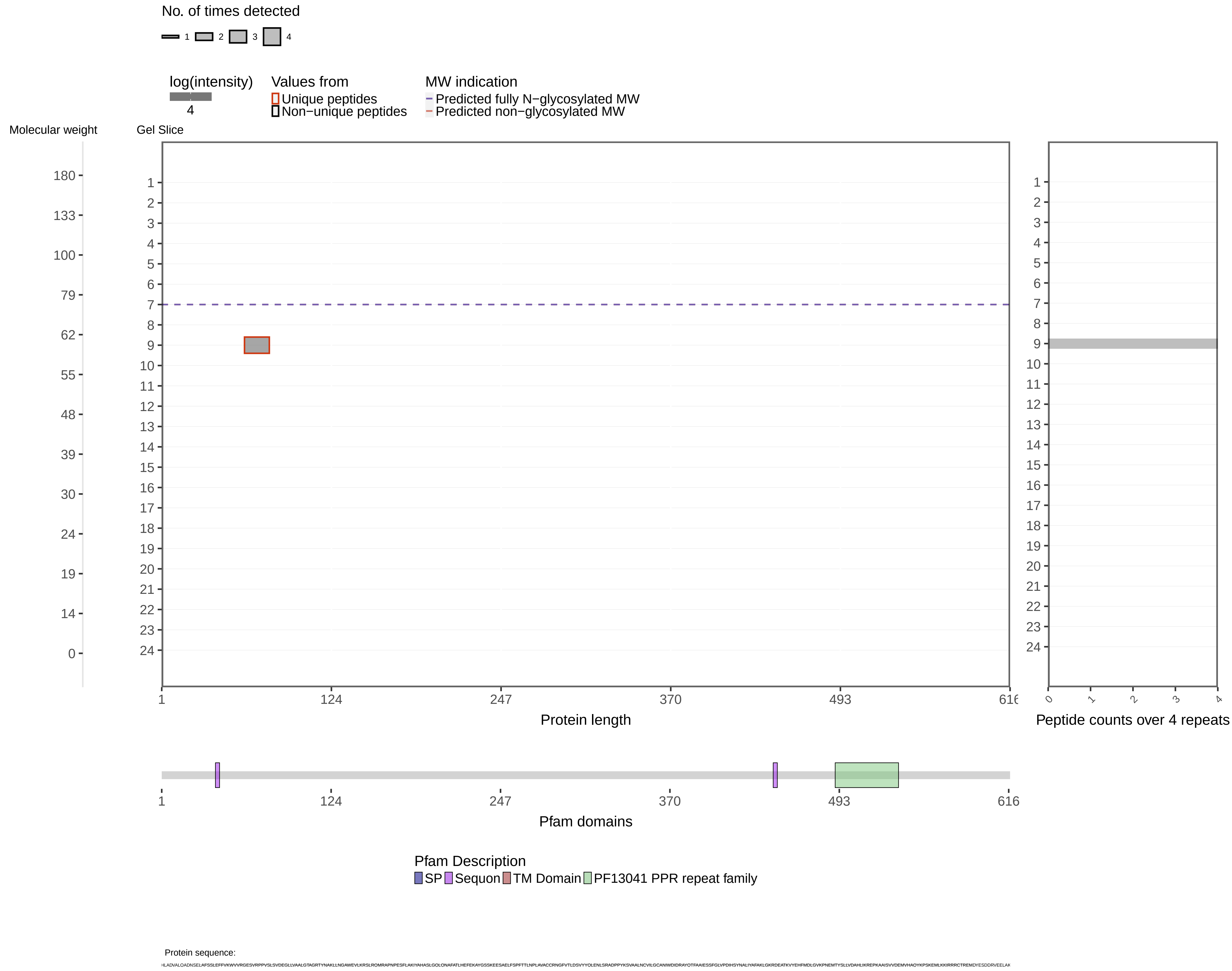

NbL18g06440.1 probable_inactive_purple_acid_phosphatase_1 [MW − SP = 66.47 kDa]

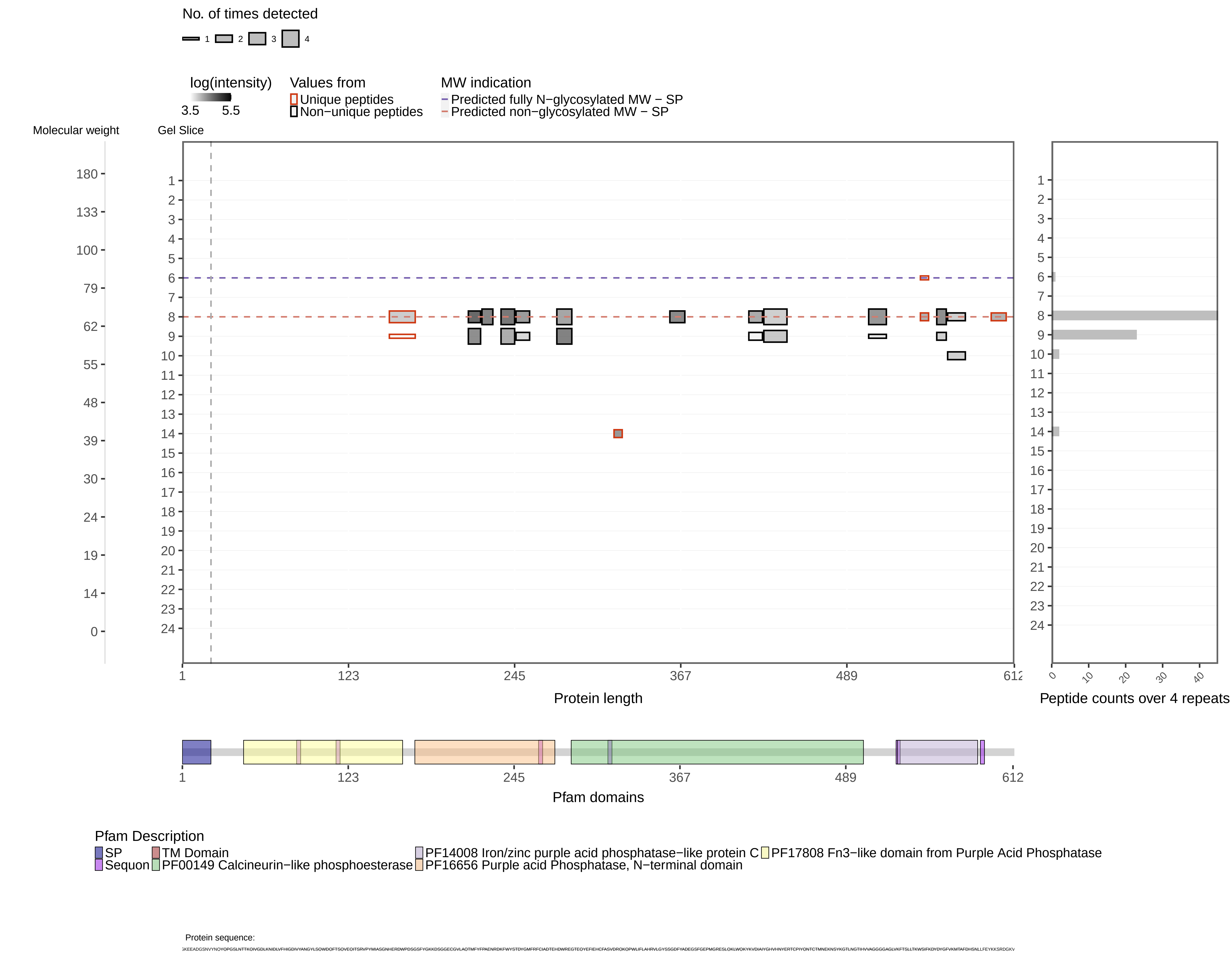

### NbL18g06260.1 probable_6−phosphogluconolactonase_1 [MW = 29.2 kDa]

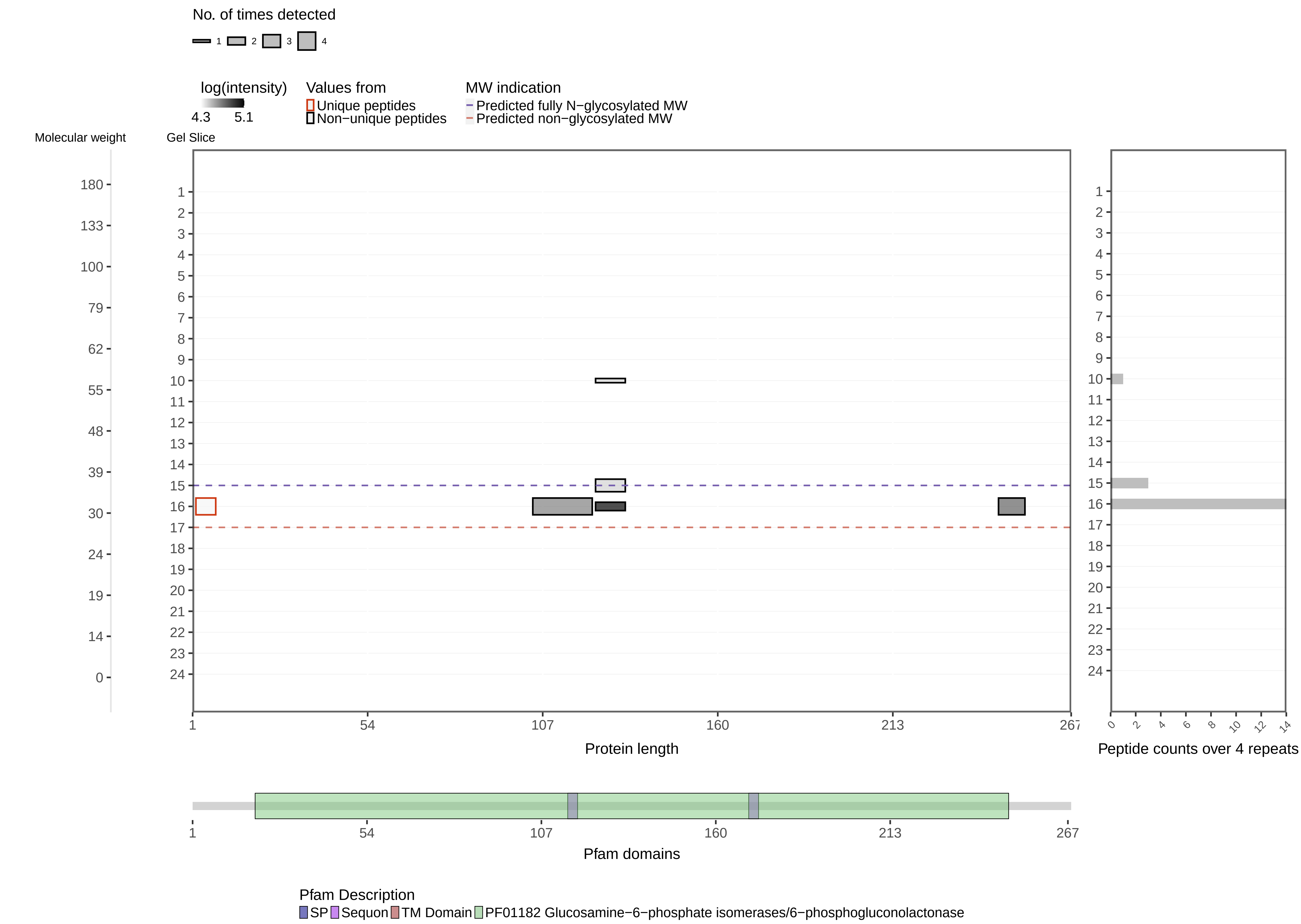


Protein sequence:

SAEKAAEDYEFVIRQLVRTRVISVSDISDCPKFDLILLGMGQDGHVASLFPNHSILDEKEEWVTFLTDSPKPPPERISFTLPVINSASNVAIVATGSSKAEAVHLAIDDDGPECPKSPAKMVQPSKGNLVWFLDKAAASKLDGAKFSE*

### NbL18g06250.1 uncharacterized_LOC104234369 [MW = 72.8 kDa]

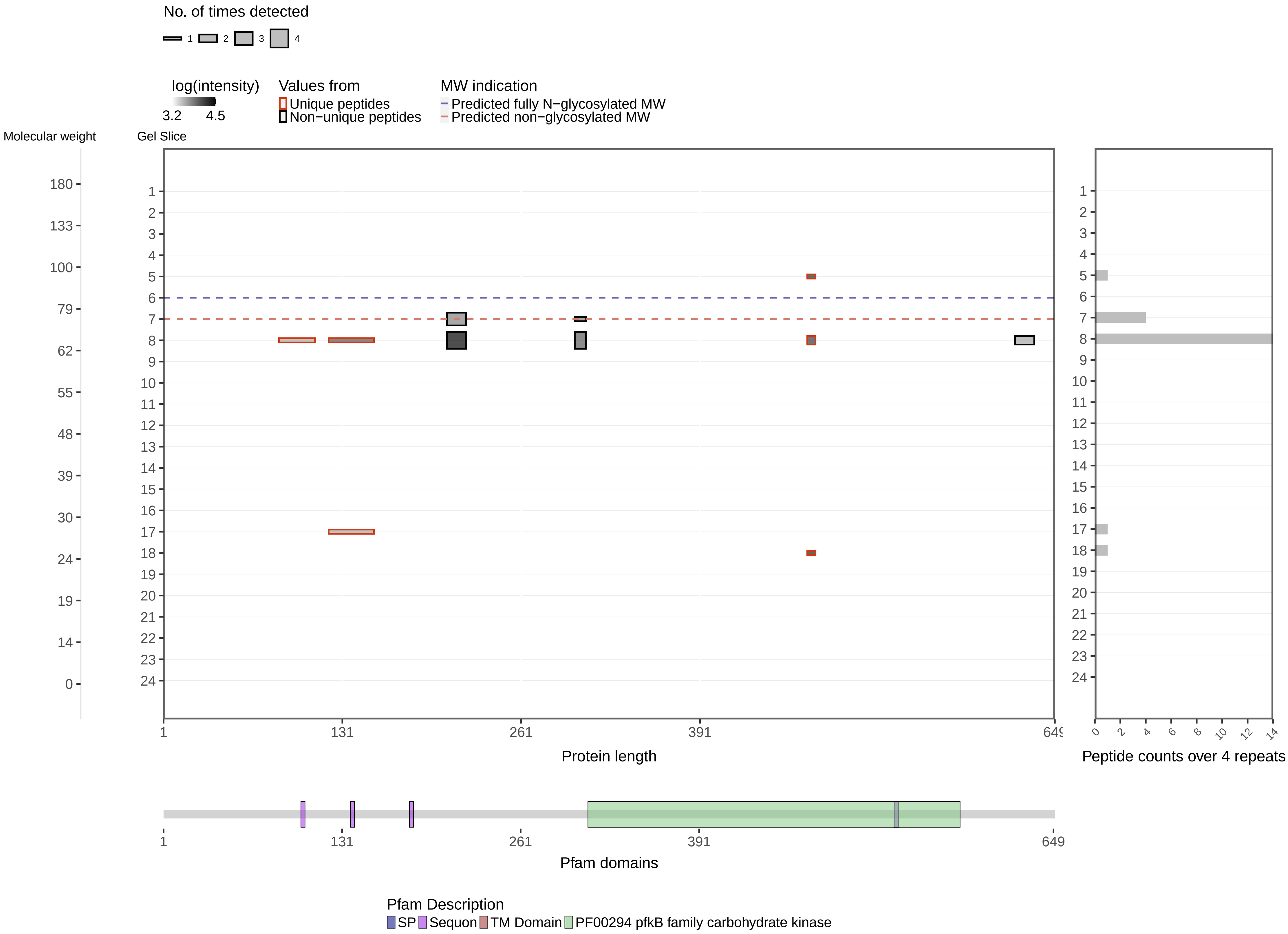


Protein sequence:

RMKDAIWDPEKFTRSPGGCSSNVAVALASLGGKVAFMGKLGDDDFGQSLVYFMNINKVQTRSVRLDSKKATAITHMKIGKRGGLRMTTTKPSAEDSLLKSEINIDVLKEAKMFYFNTFSLLDPNMRLTTLRATKISKKLGGVVFYDVNLPFPLWESGDKAKTFIQQARDLADIIEVTKQELEFLCEIKPSERFDTKDNDRSKFTHYPPEVIAPLWHENLKILFVTNGTSKIHYYTKEHNGAVLGLEDVPLTPYTSDMSASGDGIIAGIIRMLTVQPHLMTDKGYLERTLKYAISCGV

### NbL18g06200.1 alpha−14−glucan−protein_synthase_[UDP−forming]_1 [MW = 41.18 kDa]

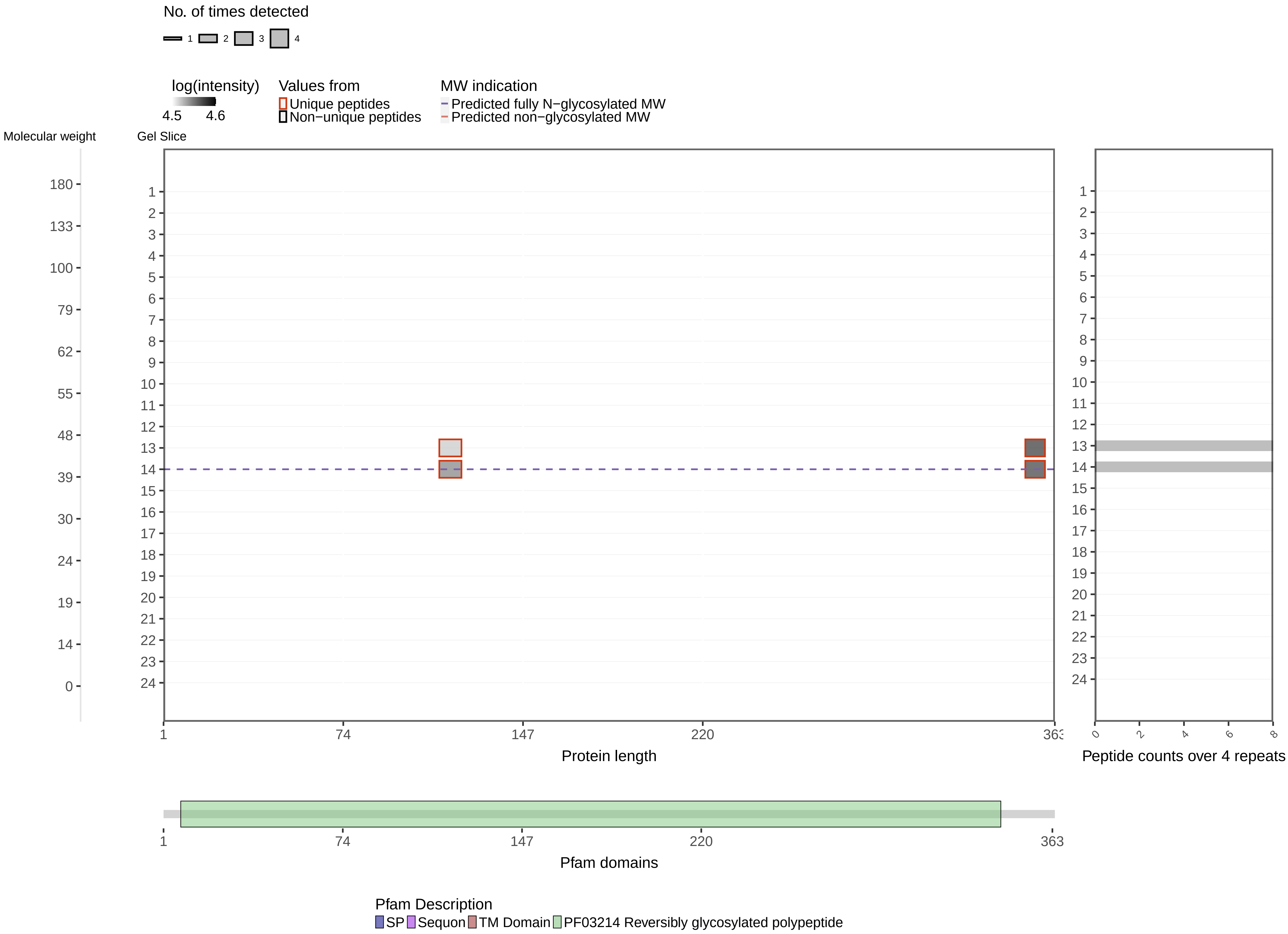


Protein sequence:

PDYDAPTQLVKPHERNTRYVDAVMTIPKGTLFPMCGMNLAFDRELIGPAMYFGLMGDGQPIGRYDDMWAGWCTKVICDHLGLGIKTGLPYIWHSKASNPFVNLKKEYNGIFWQEEIIPFFQAATFPKECKTVQQCYLELSKQVKDKLSKIDPYFTKLGDAMVTWIEAWDELNPTGENLAKLSIADGPAKAAK*

### NbL18g05980.1 chalcone−−flavonone_isomerase [MW = 26.41 kDa]

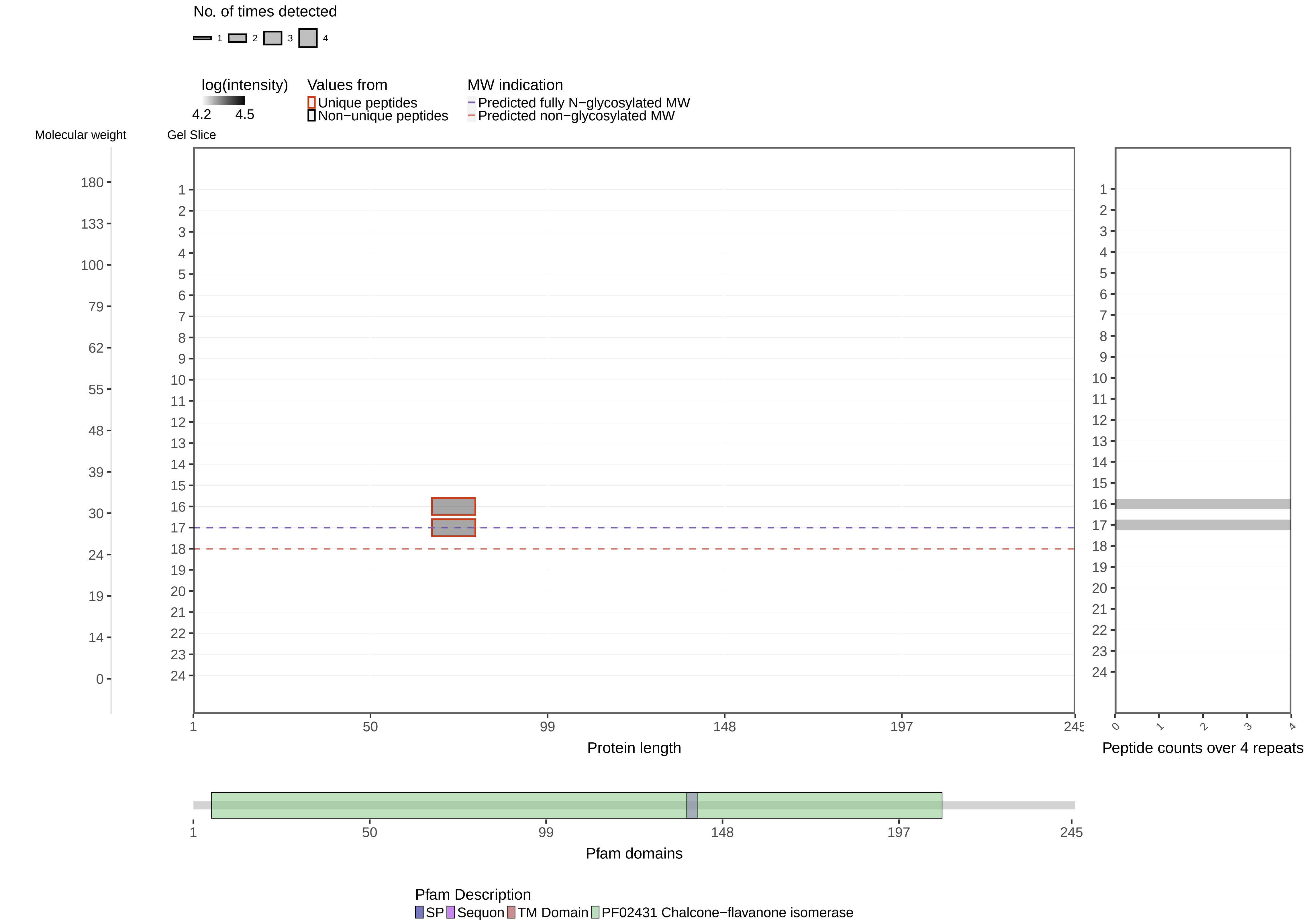


Protein sequence:

ENCVAHWKAIGTYTDAESQAIEEFLNIFQNETFPLGASILFTQSPVGALTISFIKDDTITGTGNAVIENKQLSEAVLESIIGKHGVSPAAKCSIAERVSSLLKKSYGEVPVCEKPGNGKSSDPVNEDKPTIAEIGV*

### NbL18g05950.1 uncharacterized_LOC104233992 [MW = 29.66 kDa]

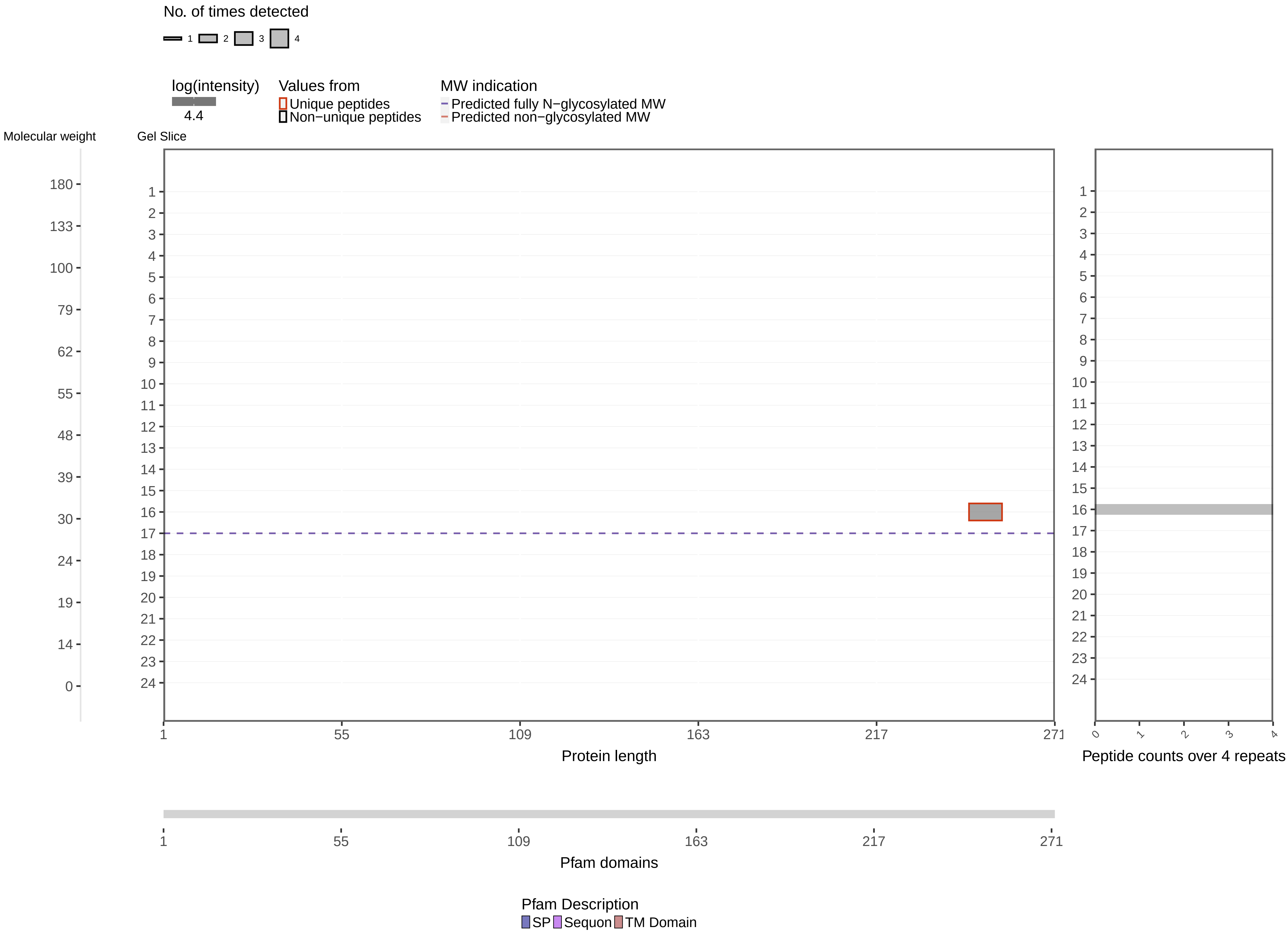


Protein sequence:

VANLLIGEQIIPEKVLILDSIRSNNFRGKLSPDEAFAFKLESSAERKGKADGHQDSPLVKCADYLPSGSVVDGLAAALLSRCQLKKIRGTLCVSWPEAGISVISLVKSLLLKDVLSGVNHKNVGDLEEEFLKLGRNKDFLLDSELYT*

### NbL18g05920.1 6−phosphogluconate_dehydrogenase_decarboxylating_3 [MW = 54.05 kDa]

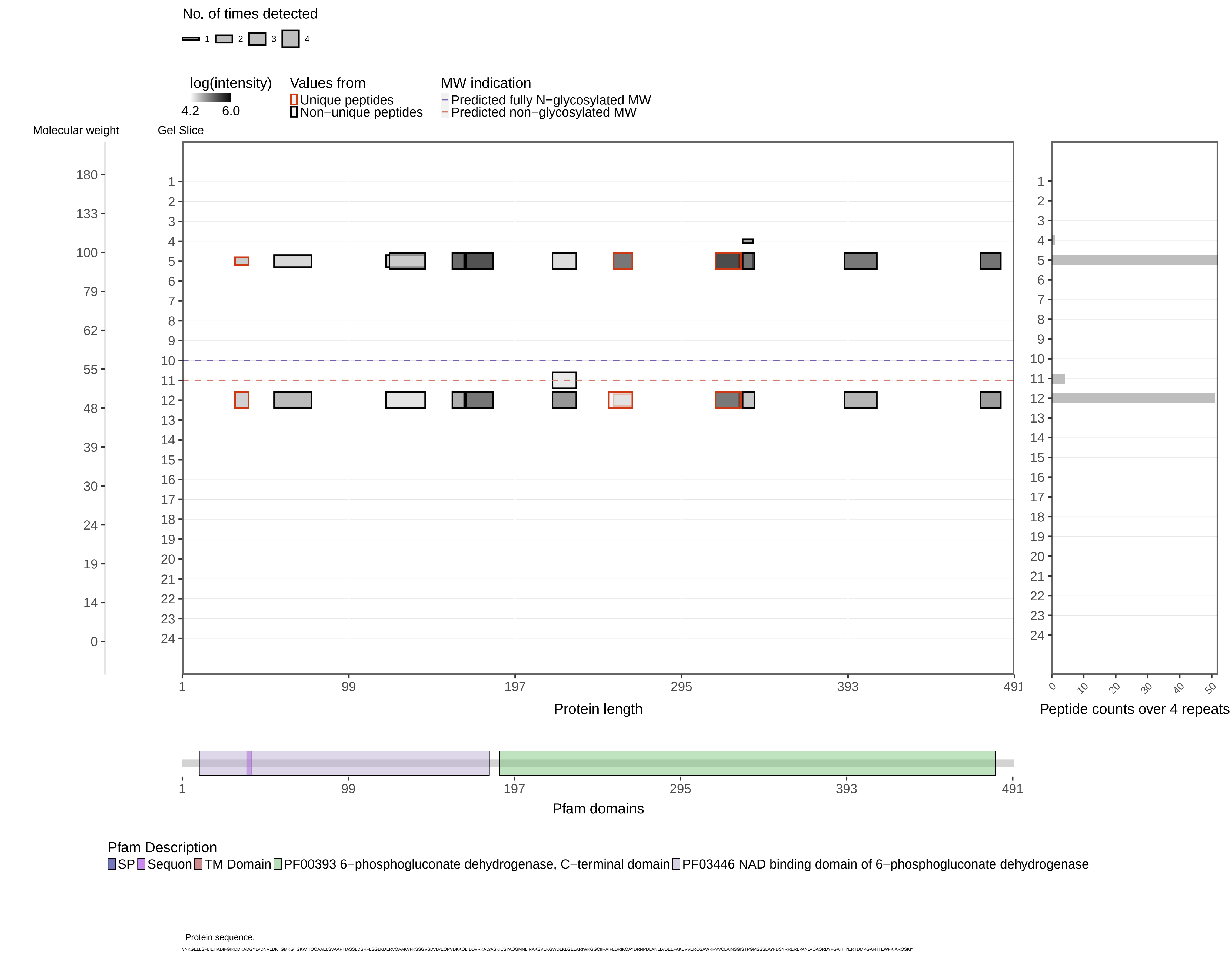

### NbL18g05770.1 heat_shock_protein_90−5_chloroplastic−like [MW = 51.78 kDa]

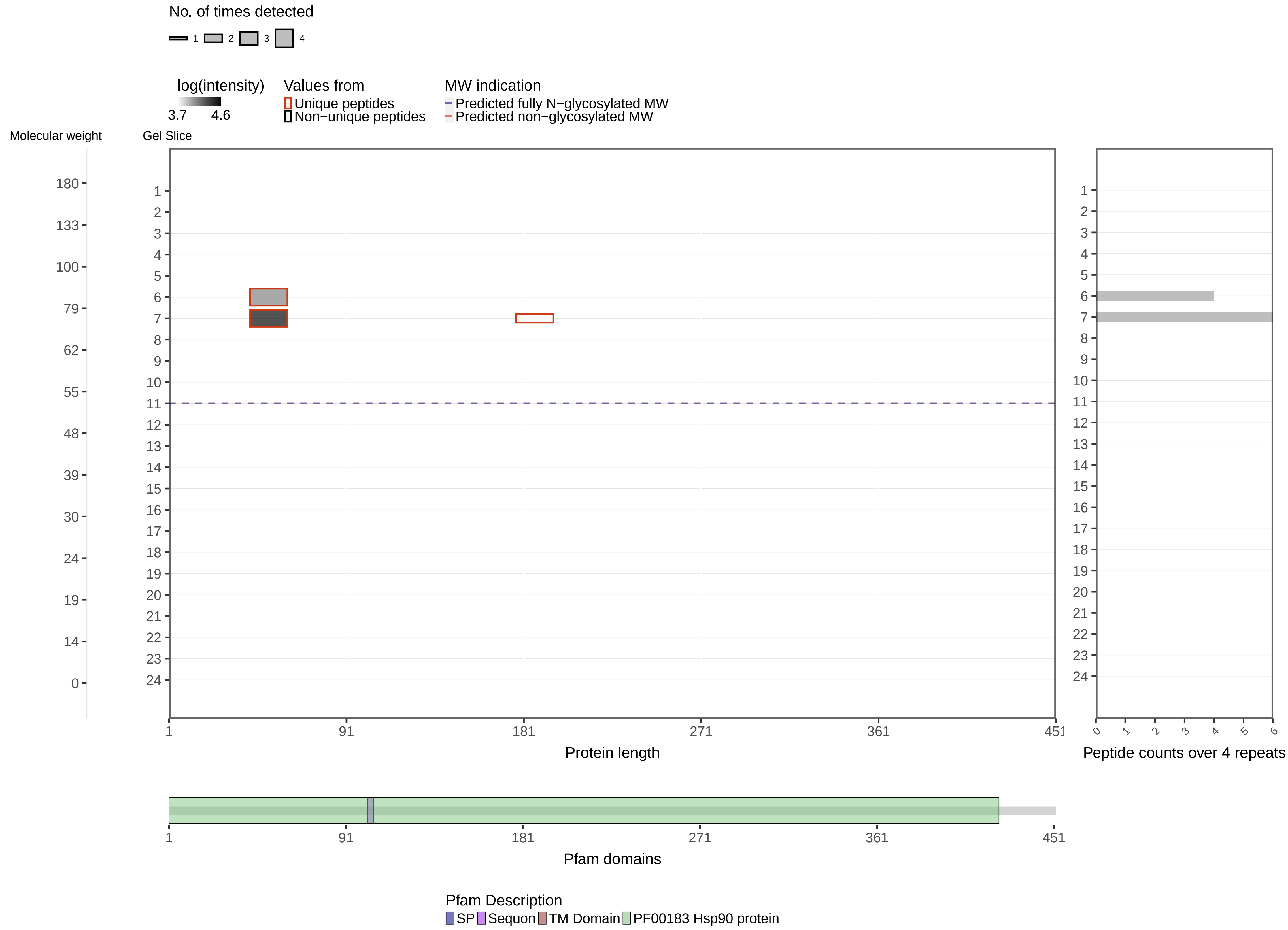


Protein sequence:

EKLVQKGIEVLYLVEPIDEVAIQNLQTYKEKKFVDISKEDLELDDDDEVKEREAKQEYNLLCDWIKQQLGDKVAKVQVSKRLSSSPCVLVSGKFGWSANMERLMRAQTLGDTSTLEFMRGRRILEVNPDHPIVKDLNAACKNAPDSSDAKRAVELLYDTALISSGFTPDSPAELGNKIYDMMAMALGGRWGRSEEDESETSVETSTESDAGSAEVSEVQVVEPSEVRTESNDYWE*

### NbL18g05620.1 phosphopantothenate−−cysteine_ligase_2−like [MW = 36.24 kDa]

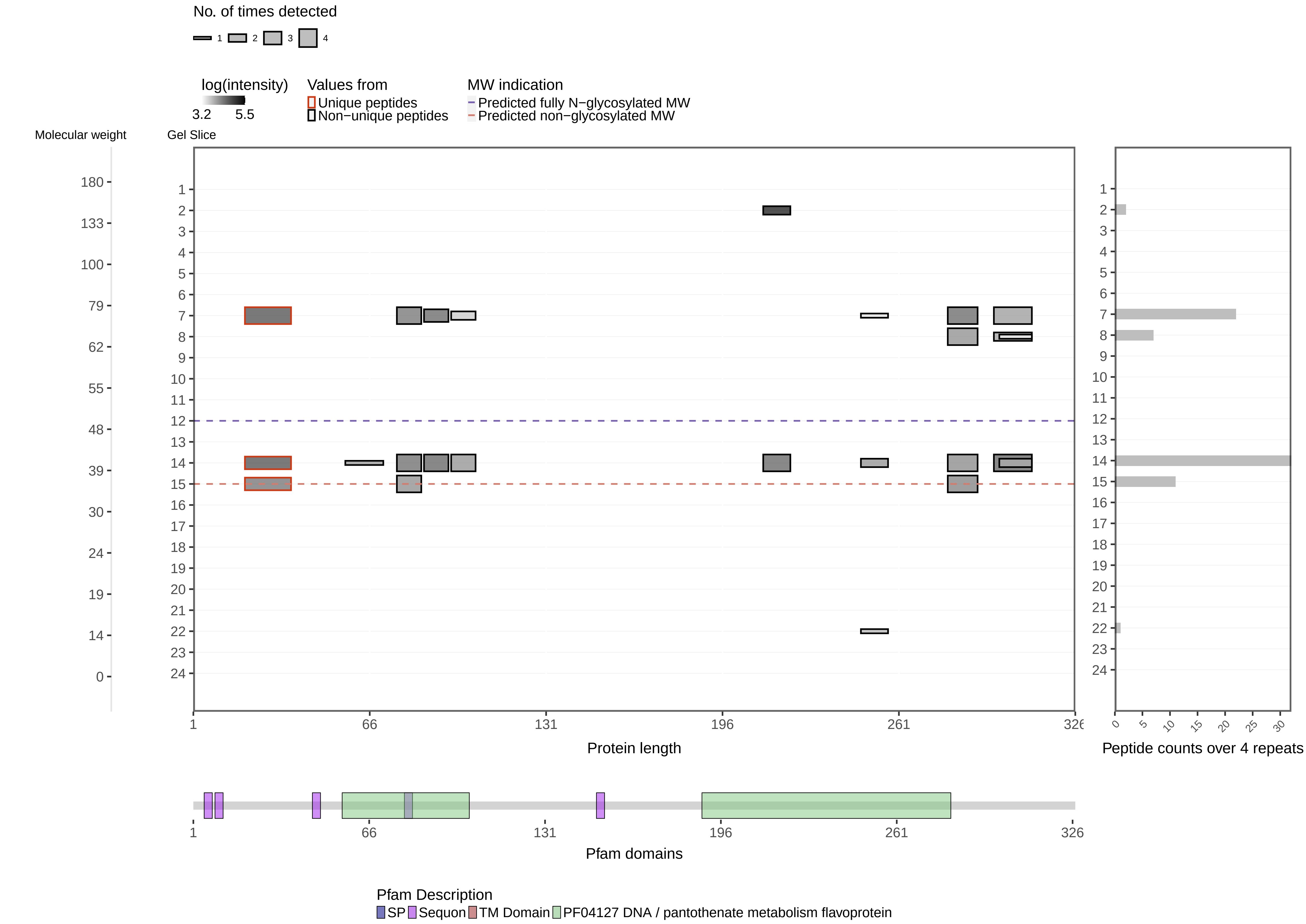


Protein sequence:

NRSAVNGGFLLKLPFTTIFEYLQILRLIAGSLRSRGPSGMFYLAAAVSDFYVPWESMTVHKIQSGSGPLDMRLAQVPKMLYVLRNEWAPVAFCISFKLETDTDILLGKADMALKKYKMHMVVANELSTRKEEVIVVTEQEKITVRRDKTQAGADVEDPLIKLVMDWHSAYIKHSDA*

NbL18g05470.1 4−coumarate−−CoA_ligase−like_9 [MW = 58.35 kDa]

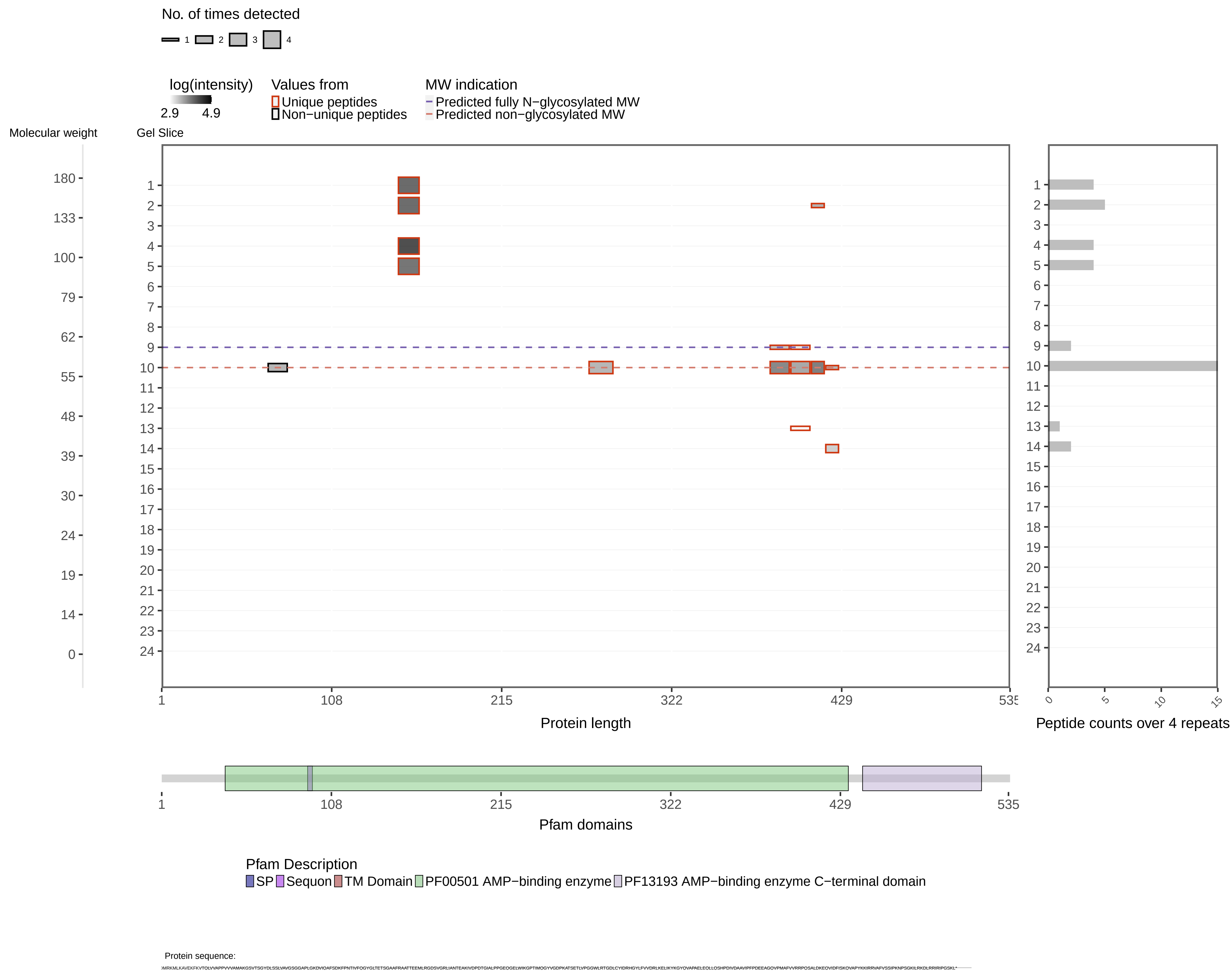

NbL18g05350.1 uncharacterized_LOC107789946 [MW = 14.02 kDa]

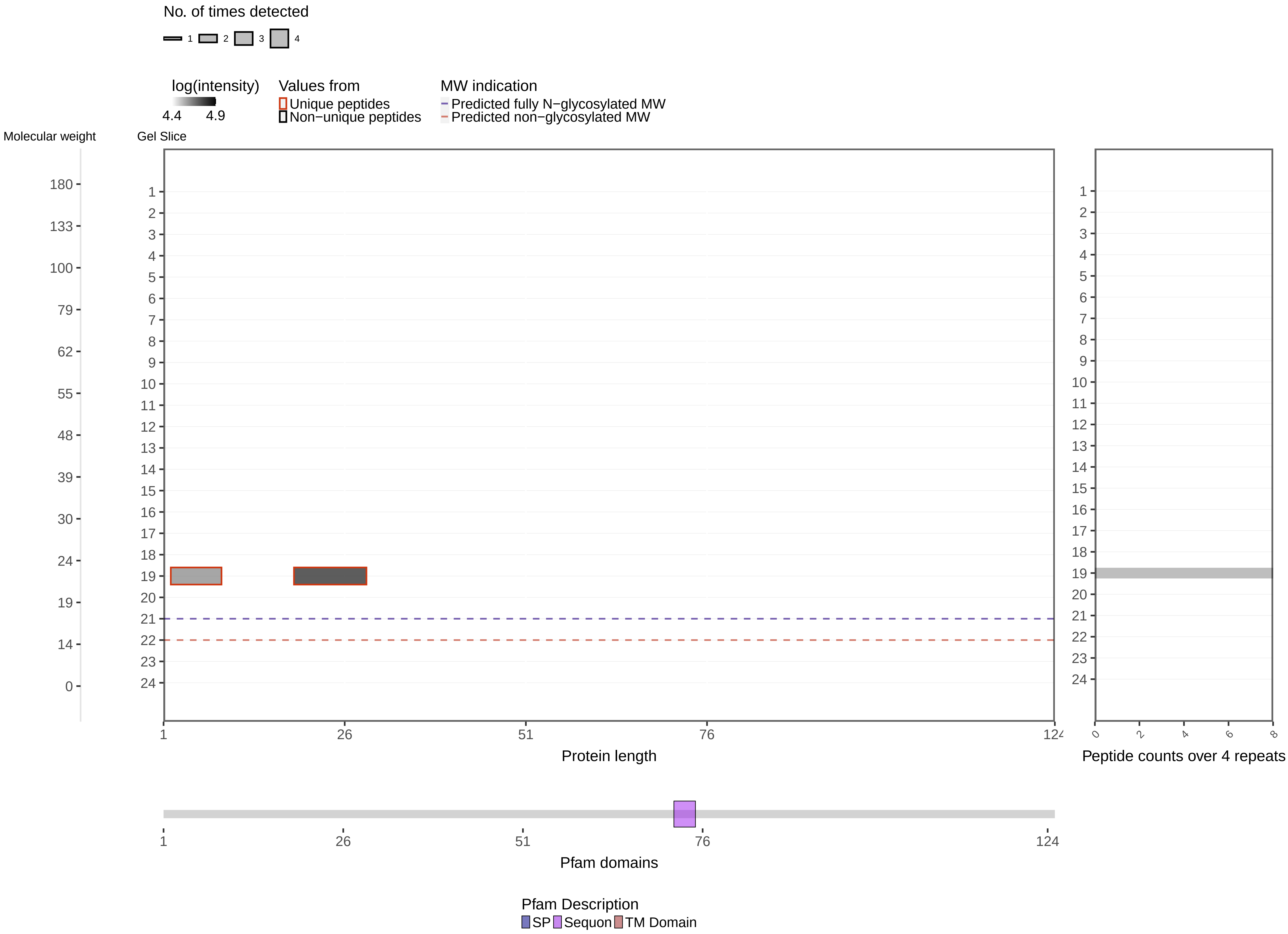


Protein sequence:

EEIFSEMGIDPRFGLACLGKINMTYESDRDLMIHFYEFVAKEEMACEEAELGPDGFAERLKMQQNLQSSNWRC*

### NbL18g05300.1 ATP_synthase_subunit_alpha_chloroplastic [MW = 60.15 kDa]

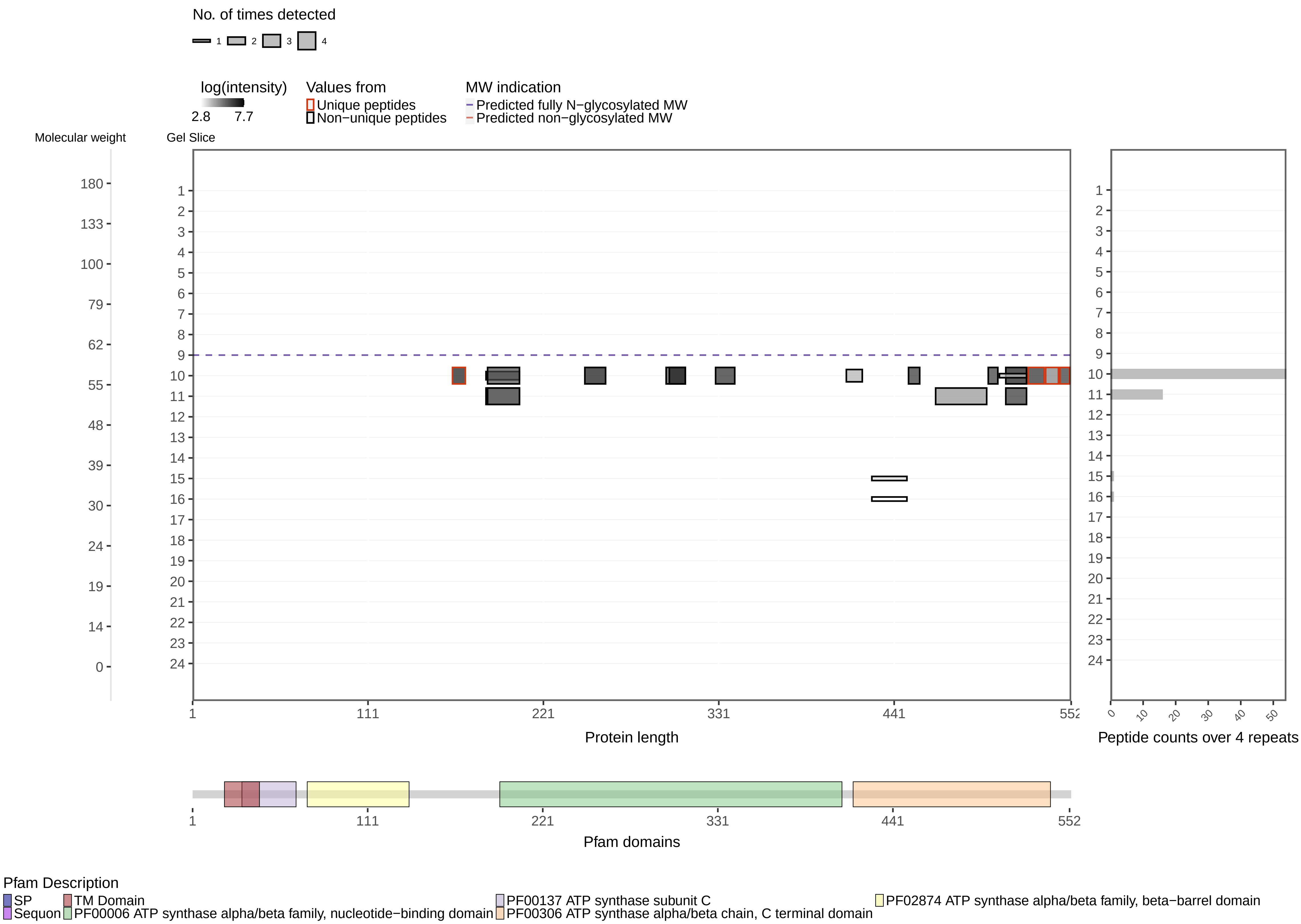


Protein sequence:

EYTIVVAETADSPATLQYLAPYTGAALAEYFMYRERHTLIIYDDLSKQAQAYRQMSLLLRRPPGREAYPGDVFYLHSRLLERAAKLSSSLGEGSMTALPIVETQSGDVSAYIPTNVISITDGQIFLSADLFNSGIRPAINVGISVSRVGSAAQIKAMKQVAGKLKLELAQFAELEAFAQFASDLDKATQNQLARGQRLRELLKQSQSAPLTVEEQIMTIYTGTNGYLDSLEVGQVRKFLVELRTYLKTNKPQFQEIISSTKTFTEEAEALLKEAIQEQMDRFILQEQA*

NbL18g05240.1 50S_ribosomal_protein_L27_chloroplastic [MW = 20.57 kDa]

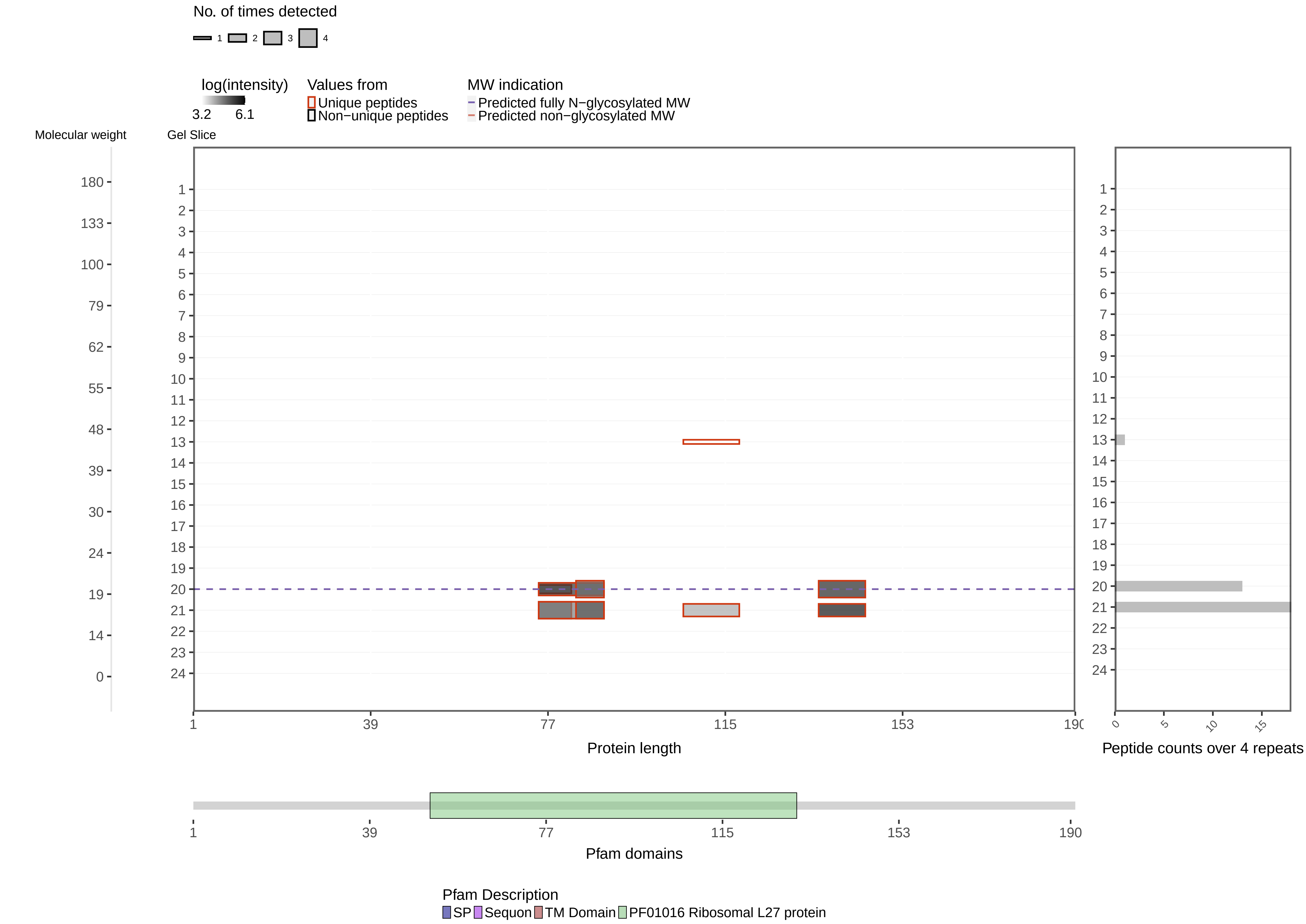


Protein sequence:

KPGSIIVRQRGTKFHPGKNVGLGKDHTIFSLIDGLVKFEKFGPDRKKISVYPREVQPENPNSYRNRKRESFRVQRERRKARREGVILQPQSILASAPESSEDTPSPSC*

NbL18g04880.1 ATP_sulfurylase_1_chloroplastic−like [MW = 52.15 kDa]

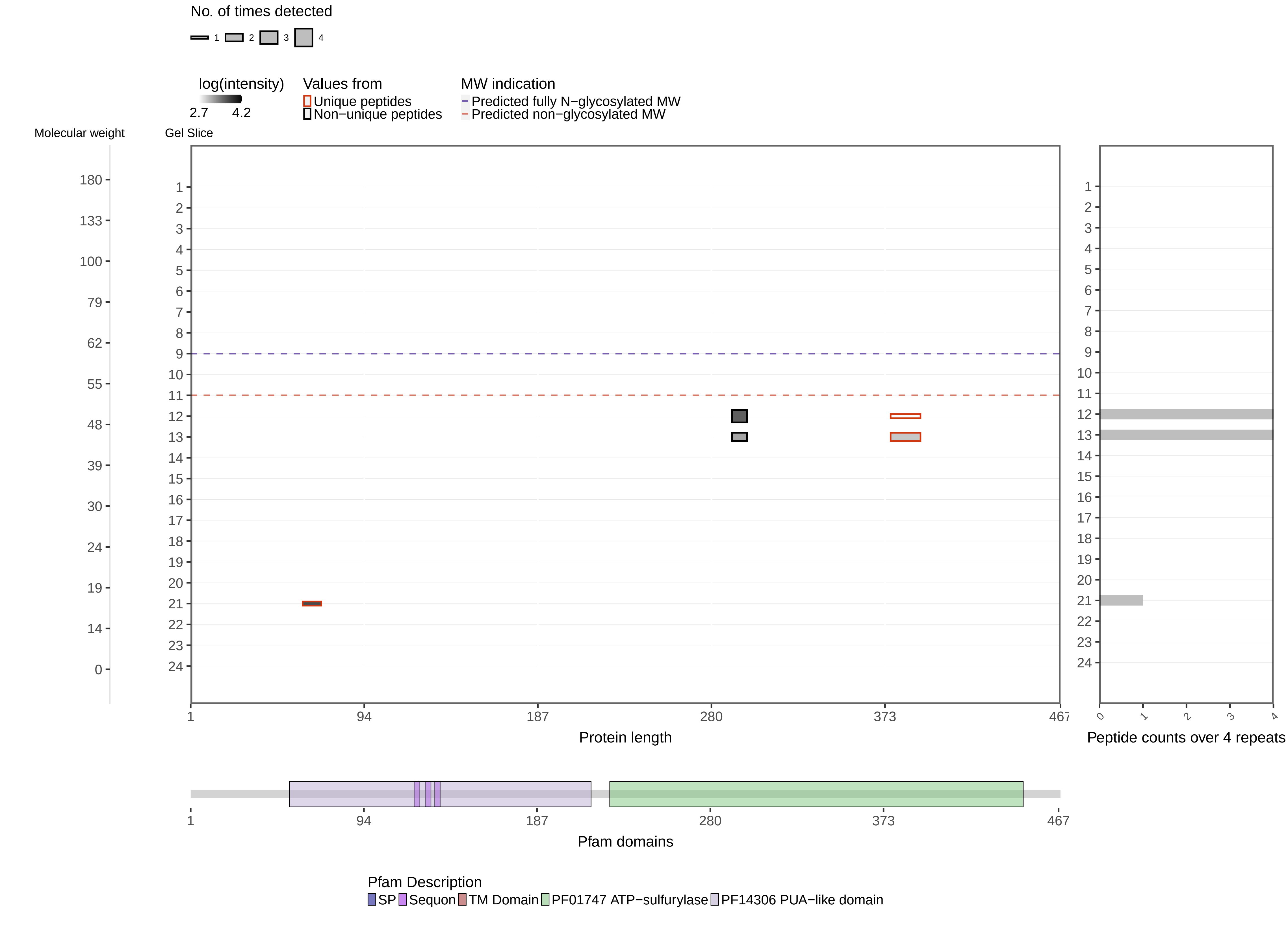


Protein sequence:

### NbL18g04860.1 2−methyl−6−phytyl−14−hydroquinone_methyltransferase_chloroplastic−like [MW = 38.07 kDa]

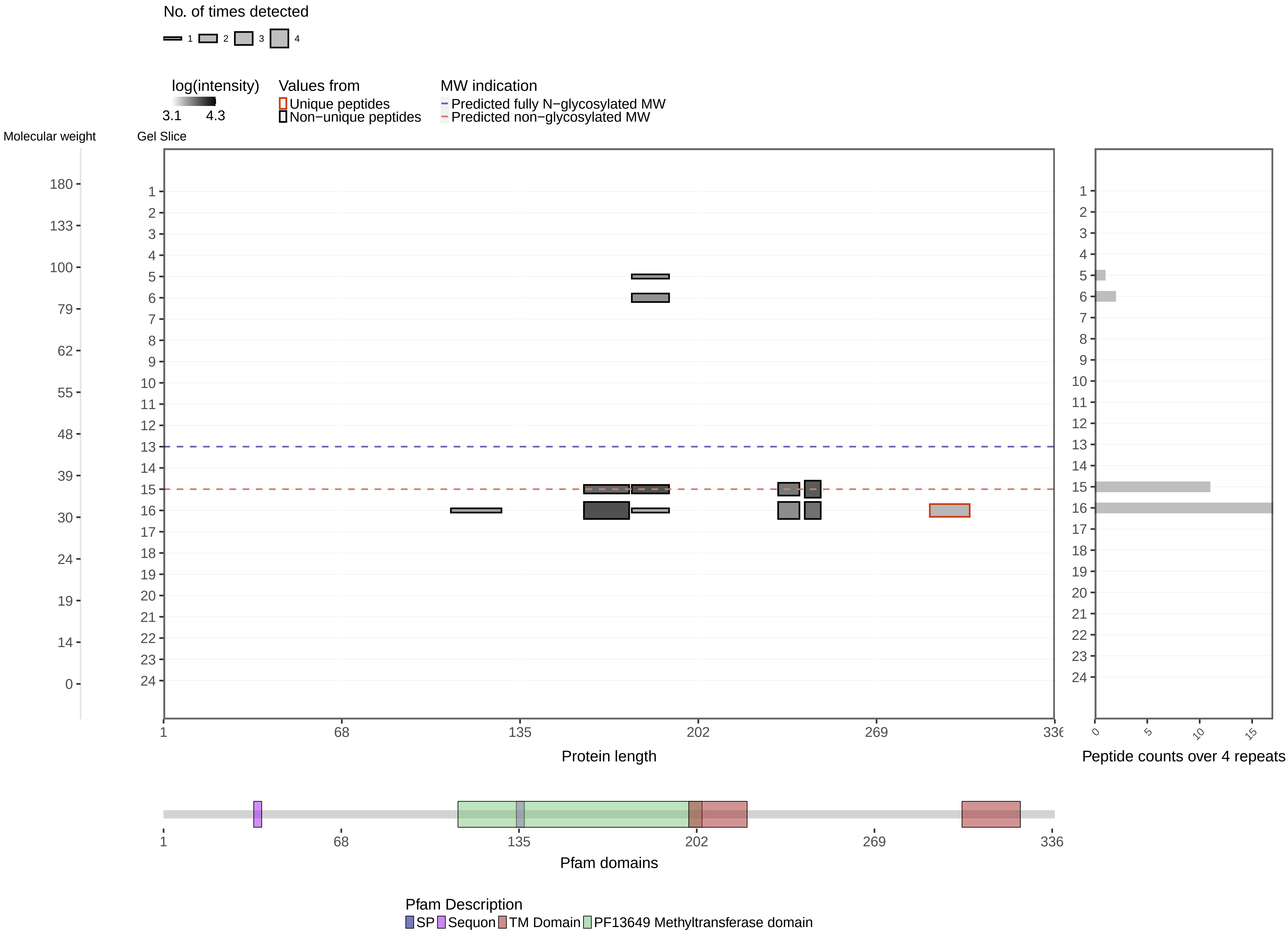


Protein sequence:

KECKILEGDAEDLPFPTDTFDRYVSAGSIEYWPDPQRGIKEAYRVLTIGGVACLIGPVYPSFWLSRFFADMWMLFPKEEEYIEWFKKAGFAQVKLKRIGPKWYRGVRRHGLIMGCSVTGVKPYFGESPLQLGPKVEDVSKPVNPFVFLVRFLLGITAATYYVLVPIYMWLKDQITPKGQPI*

### NbL18g04830.1 pyruvate_kinase_isozyme_A_chloroplastic−like [MW = 63.11 kDa]

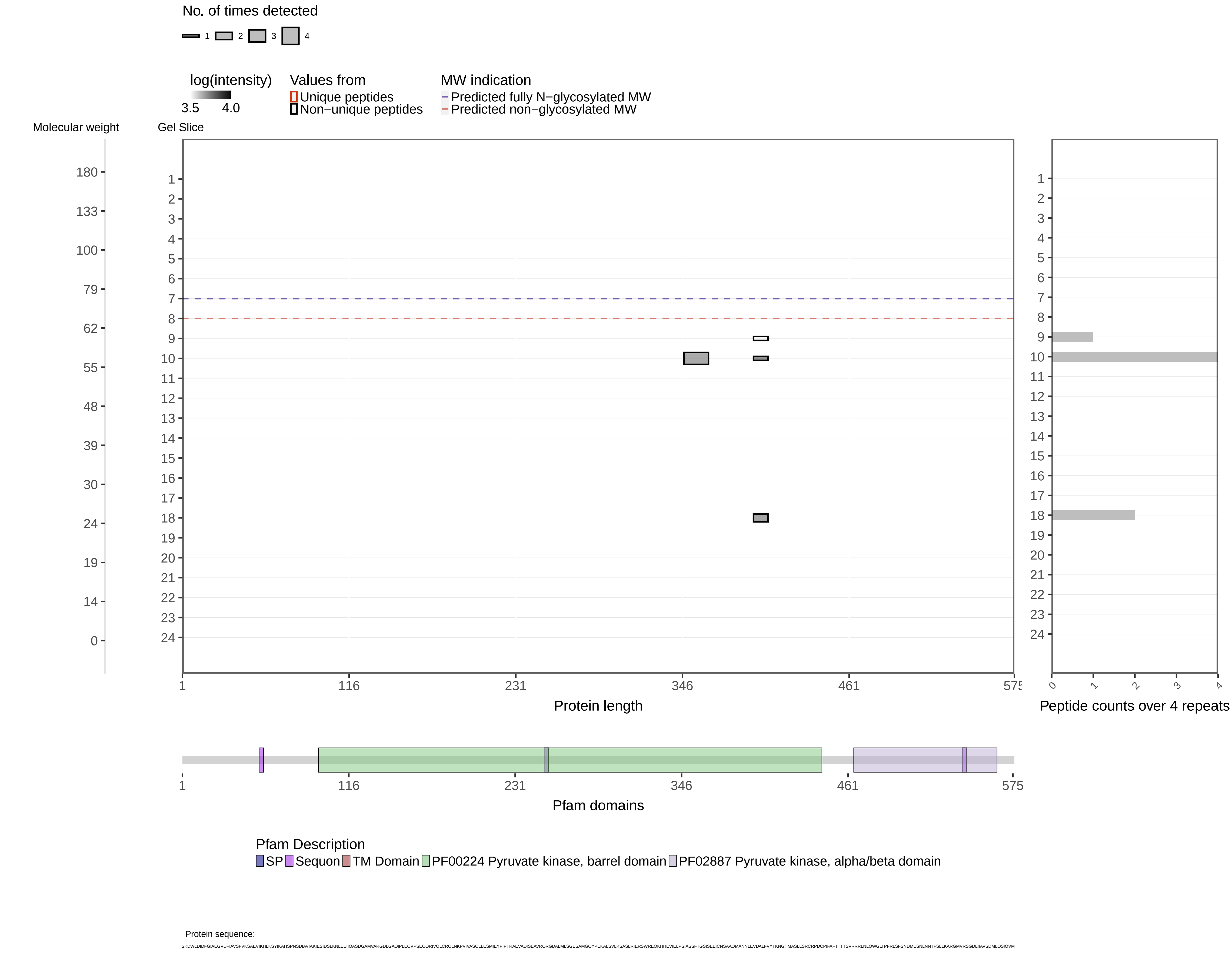

NbL18g04770.1 acylamino−acid−releasing_enzyme−like [MW = 66.34 kDa]

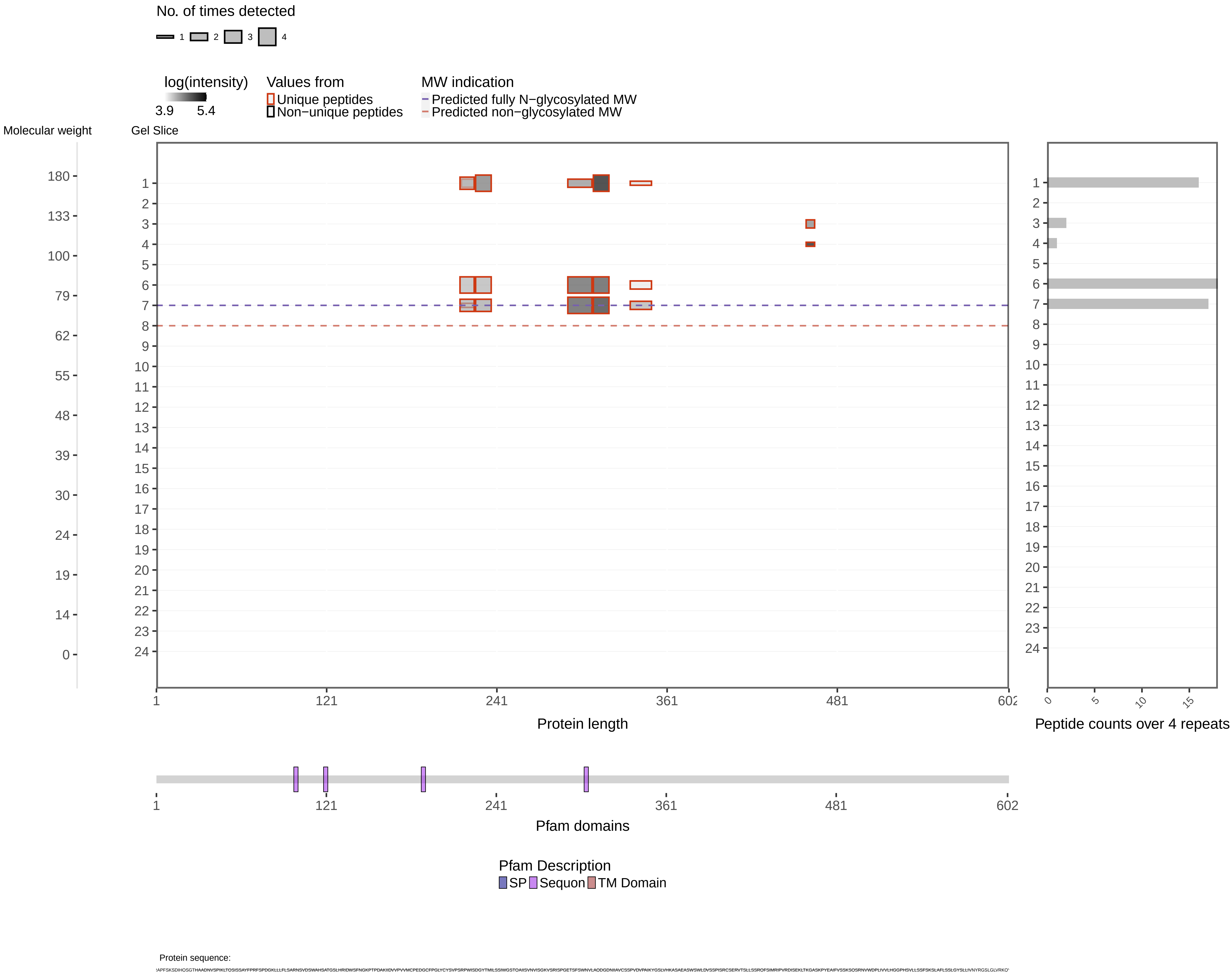

NbL18g04730.1 histone_H2AX−like [MW = 14.92 kDa]

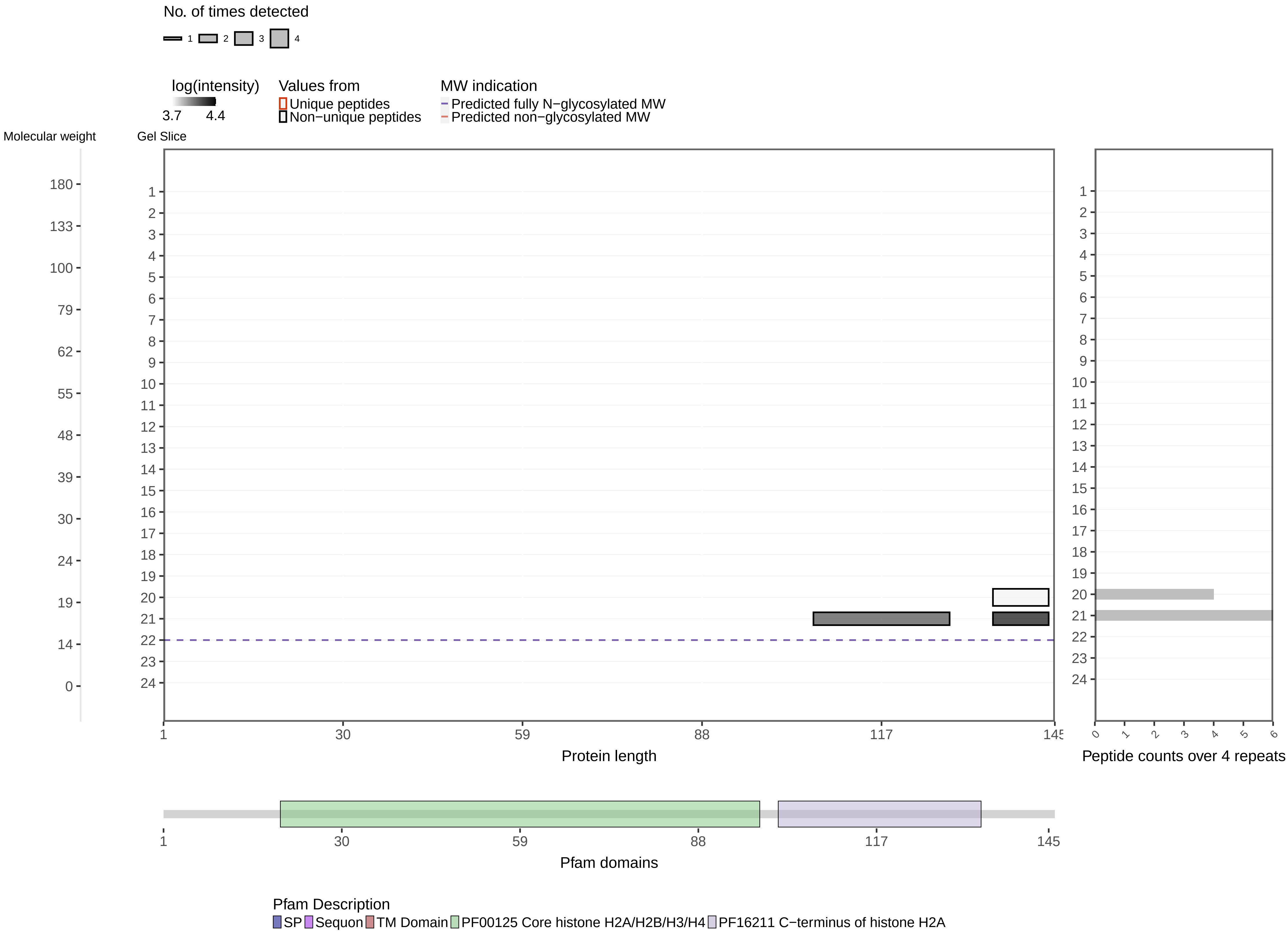


Protein sequence:

PVYLSAVLEYLAAEVLELAGNAARDNKKNRIVPRHIQLAVRNDEELSKLLGQVTIANGGVMPNIHQNLLPKKAGSGKGDIGSASQEF*

### NbL18g04640.1 ADP−ribosylation_factor−like_protein_5 [MW = 20.94 kDa]

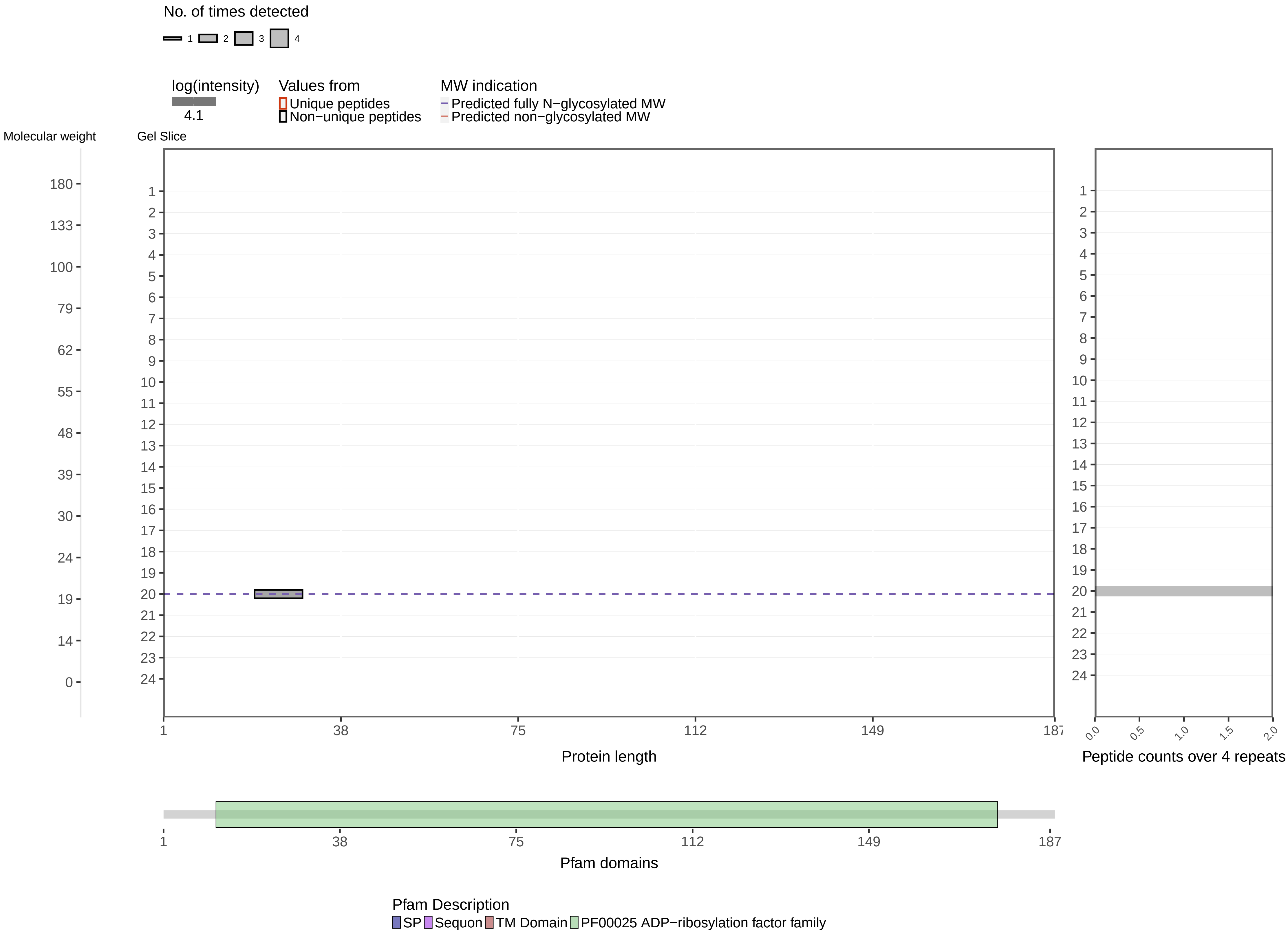


Protein sequence:

ATYYRGTHAVIAVIDSTDRARISIMKDELFRLLPHEDLQSAVLLVFANKQDLKDAMTPAEITDALSLHSIKNHDWHIQACSALTGDGLYDGLGWIAQRVPAMKIDGQK*

### NbL18g04600.1 acylamino−acid−releasing_enzyme−like [MW = 91.52 kDa]

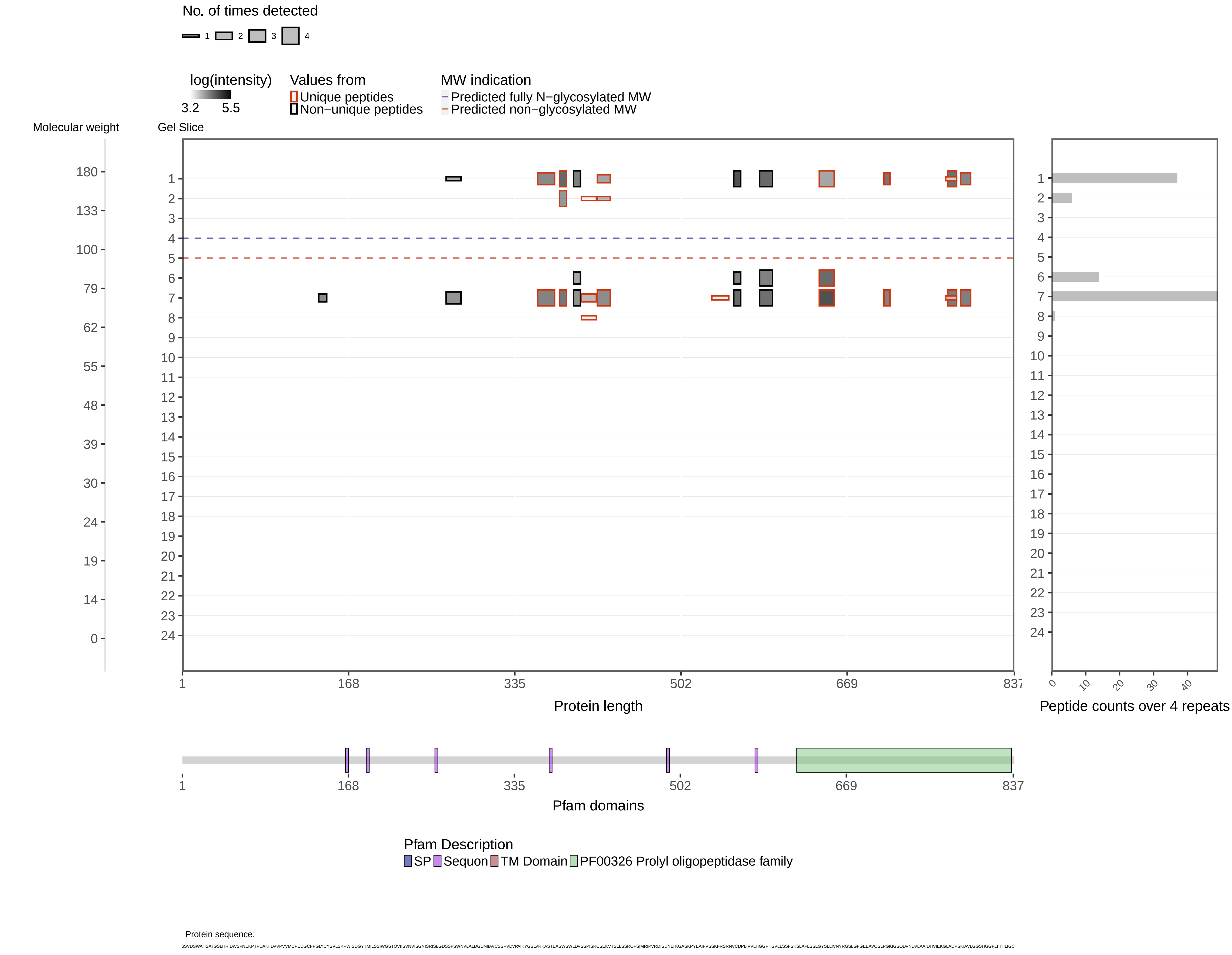

NbL18g03900.1 chlorophyll_a−b_binding_protein_50_chloroplastic−like [MW = 28.41 kDa]

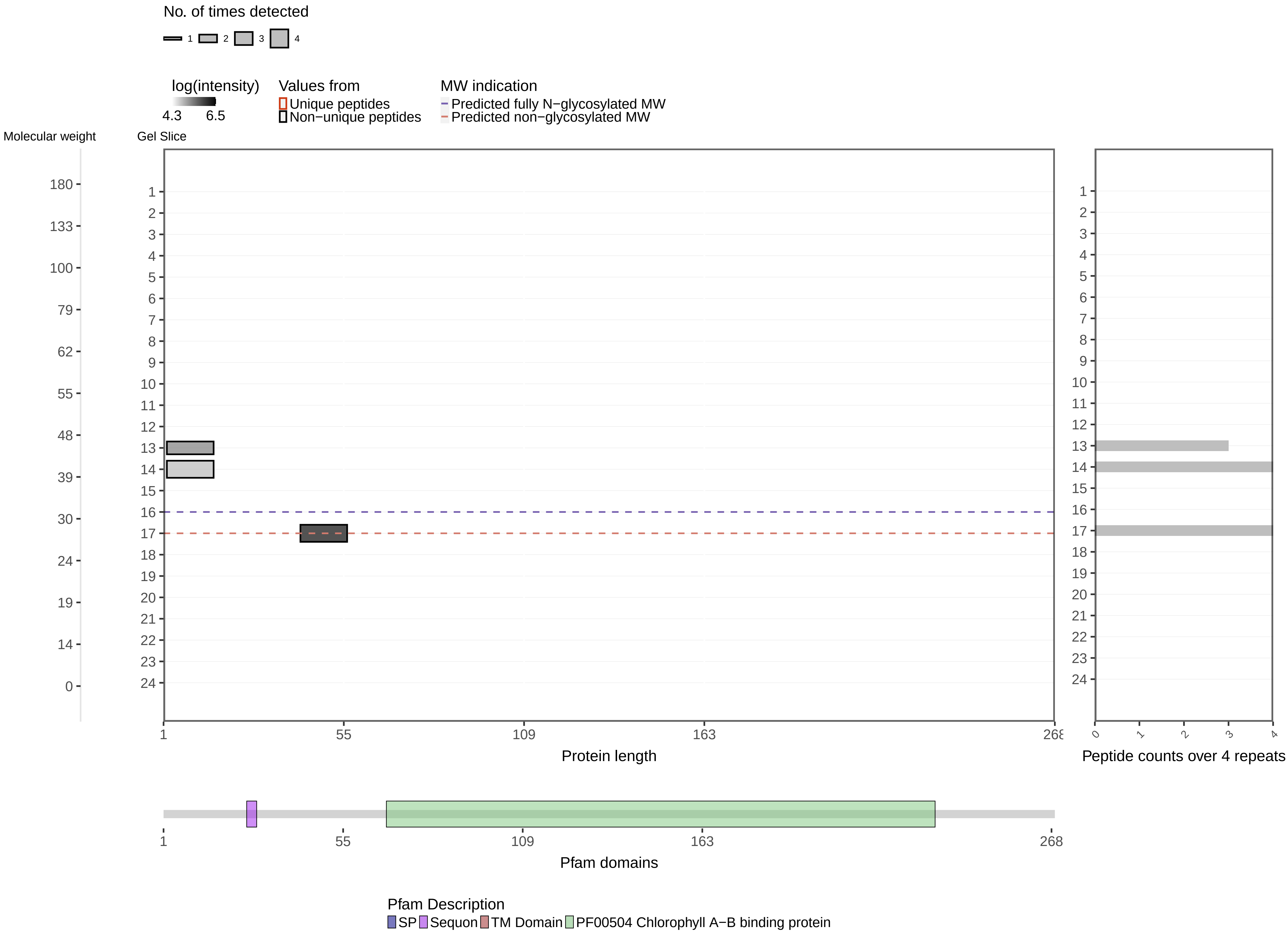


Protein sequence:

ARNGVKFGEAVWFKAGSQIFSEGGLDYLGNPSLVHAQSILAIWACQVVLMGAVEGYRVAGGPLGEVVDPLYPGGSFDPLGLADDPEAFAELKVKEIKNGRLAMFSMFGFFVQAIVTGKGPLENLADHLTDPVYYNAWAYATNFVPGK*

NbL18g03850.1 chlorophyll_a−b_binding_protein_40_chloroplastic [MW = 28.7 kDa]

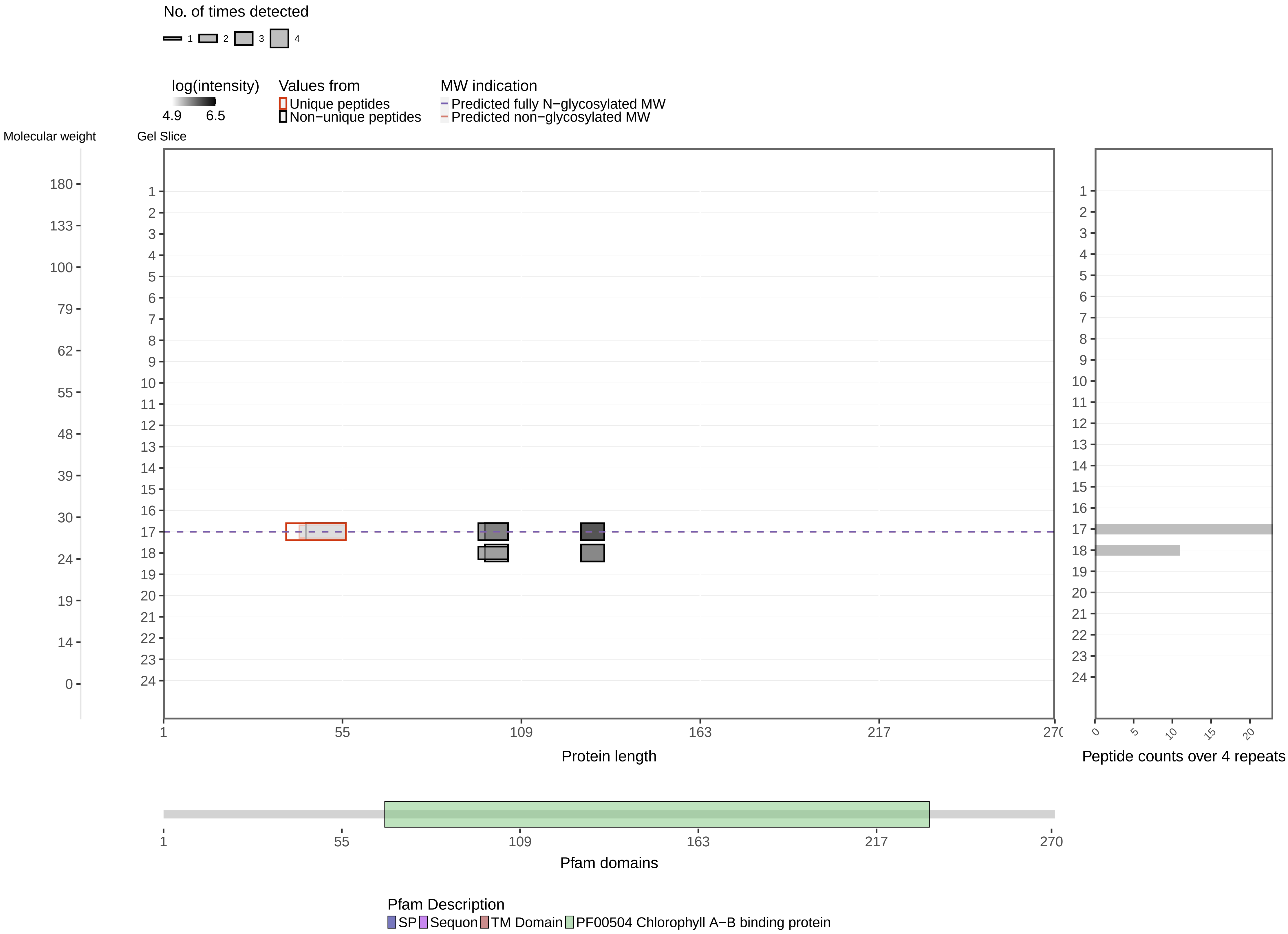


Protein sequence:

RNGVKFGEAVWFKAGSOIFSEGGLDYLGNPSLVHAOSILAIWACOVVLMGAVEGYRVAGGPLGEVVDPLYPGGSFDPLGLAEDPEAFAELKVKEIKNGRLAMFSMFGFFVOAIVTGKGPLENLADHLADPVNNNAWAYATYYLPLLEN*

### NbL18g03810.1 3−isopropylmalate_dehydratase_large_subunit_chloroplastic−like [MW = 55.21 kDa]

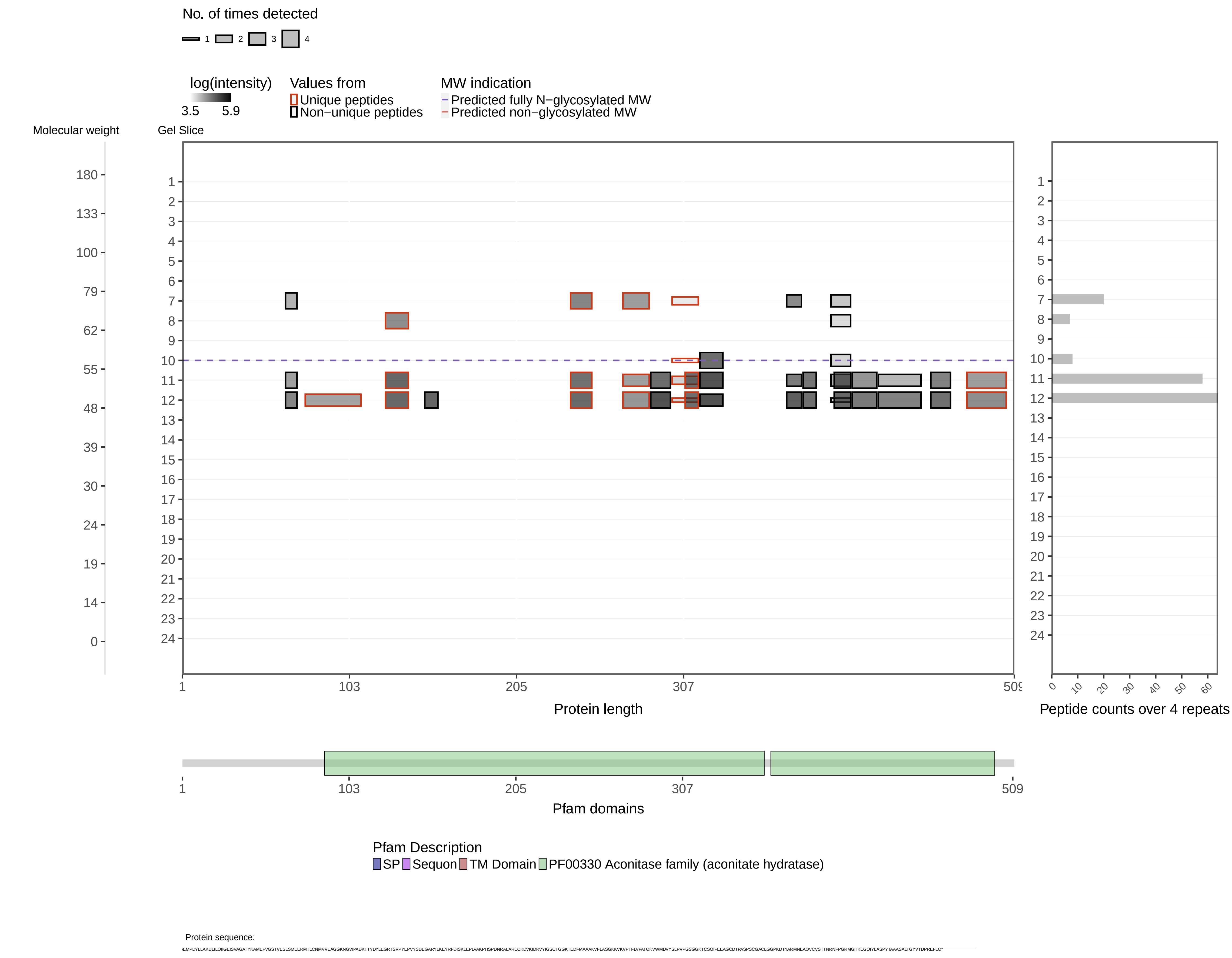

### NbL18g03510.1 pleiotropic_drug_resistance_protein_1 [MW = 140.5 kDa]

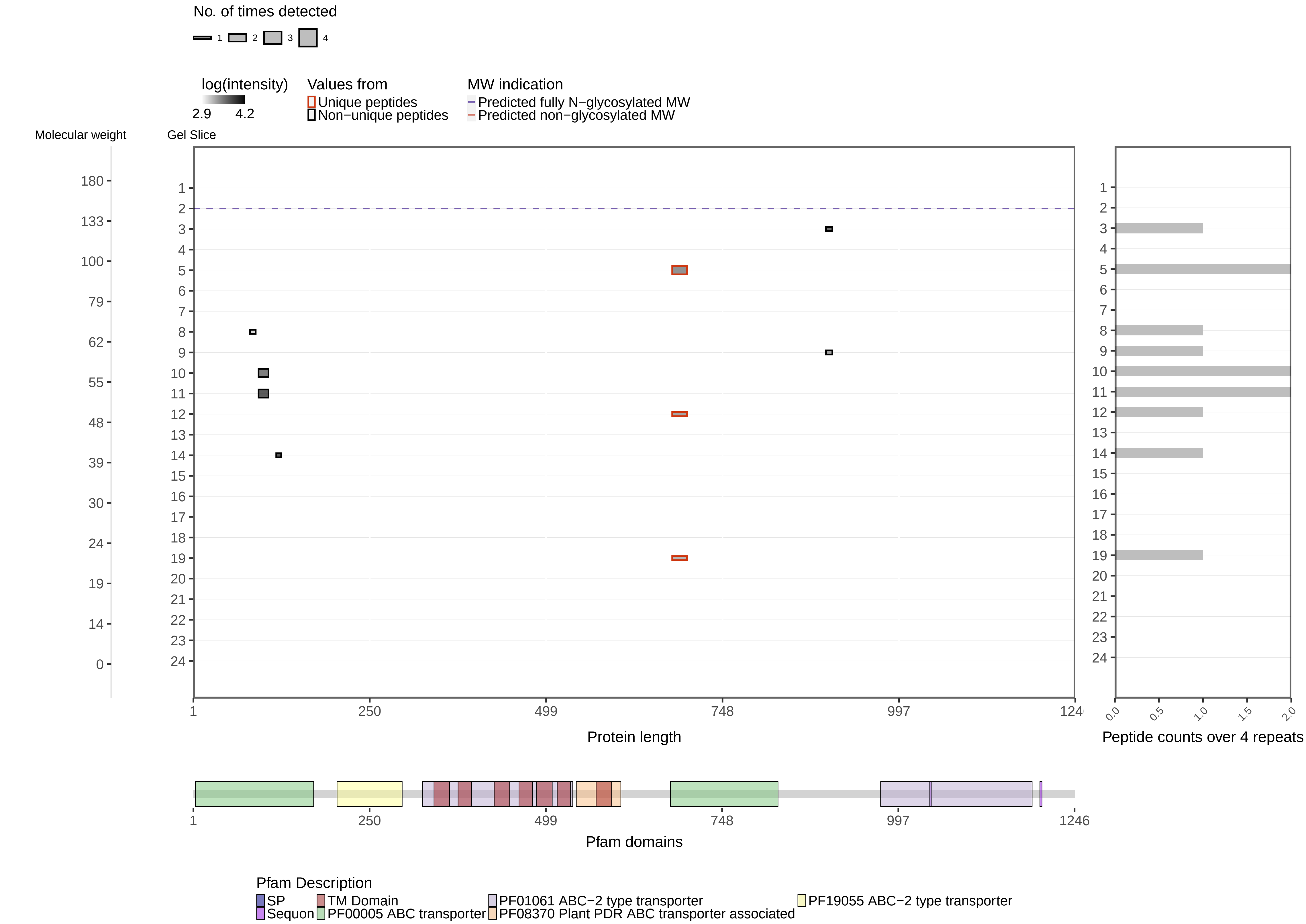


Protein sequence:

NENTESSPQITSTKEGDSVNENKKKGMVLPFDPHSITFDEVVYSVDMPPEMRESGTSENRLVLLKSVSGAFRPGVLTALMGVSGAGKTTLMDVLAGRKTGGYIDGSIKISGYPKKQETFARISGYCEQNDIHSPYVTVFESLVYSAWLRLPQDVDEQKRMMFVEEVMDLVELTSLRSALVGLPGVNGLSTEQRKRLTIAVELVANPSIIFMDEPTSGLDARAAAIVMRAVRNTVDTGRTVVCTIHQPSIDIFEAFDELFLMKRGGQEIYVGPLGRQSCHLIKYFESIPGVSKIVE[illegible]

### NbL18g03310.1 histone_H2B.3 [MW = 15.96 kDa]

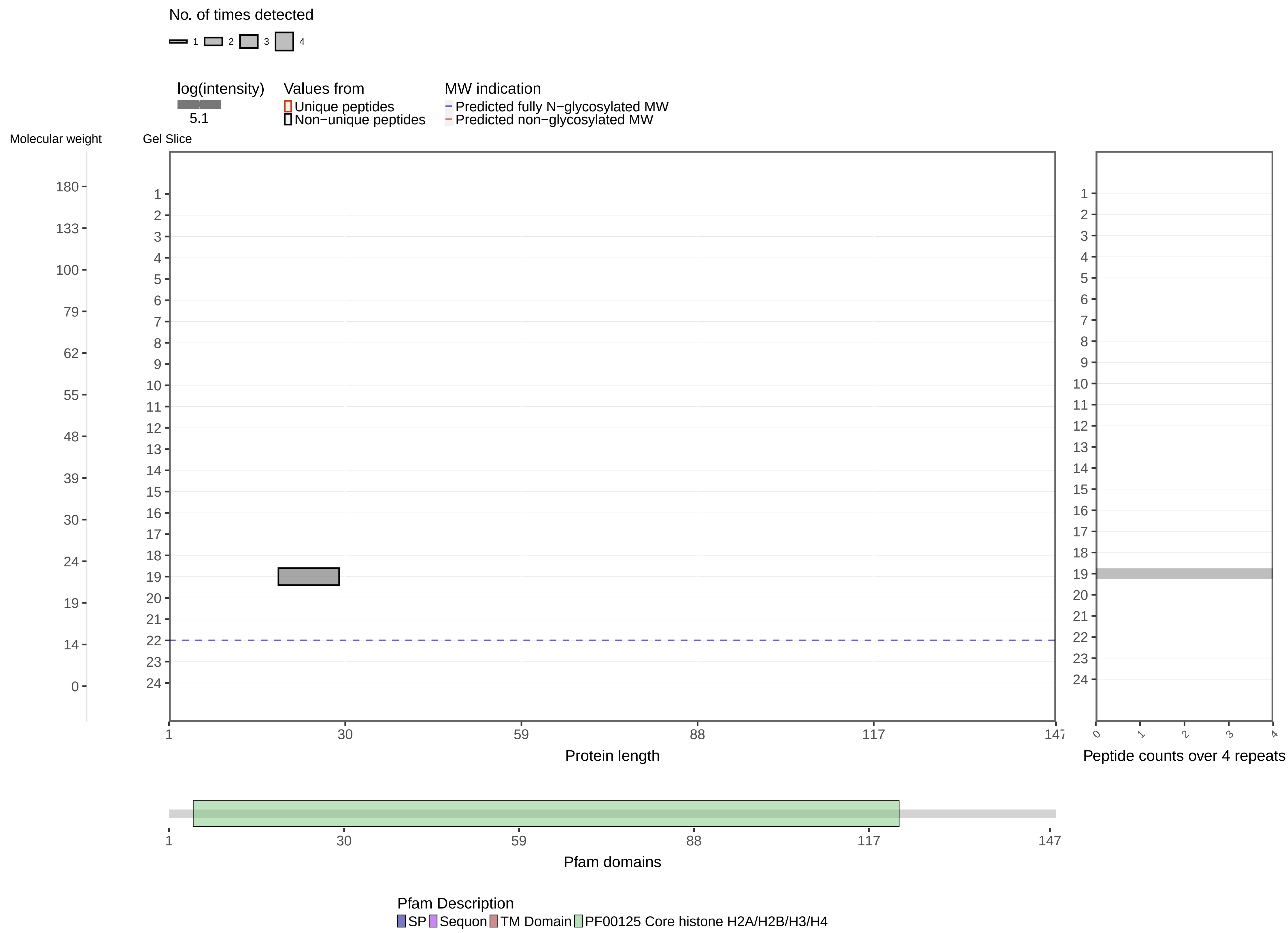


Protein sequence:

YKIYIFKVLKQVHPDIGISSKAMGIMNSFINDIFEKLAQESSRLARYNKKPTITSREIQTAVRLVLPGELAKHAVSEGTKAVTKFTSS*

NbL18g03130.1 bark_storage_protein_A−like [MW − SP = 34.47 kDa]

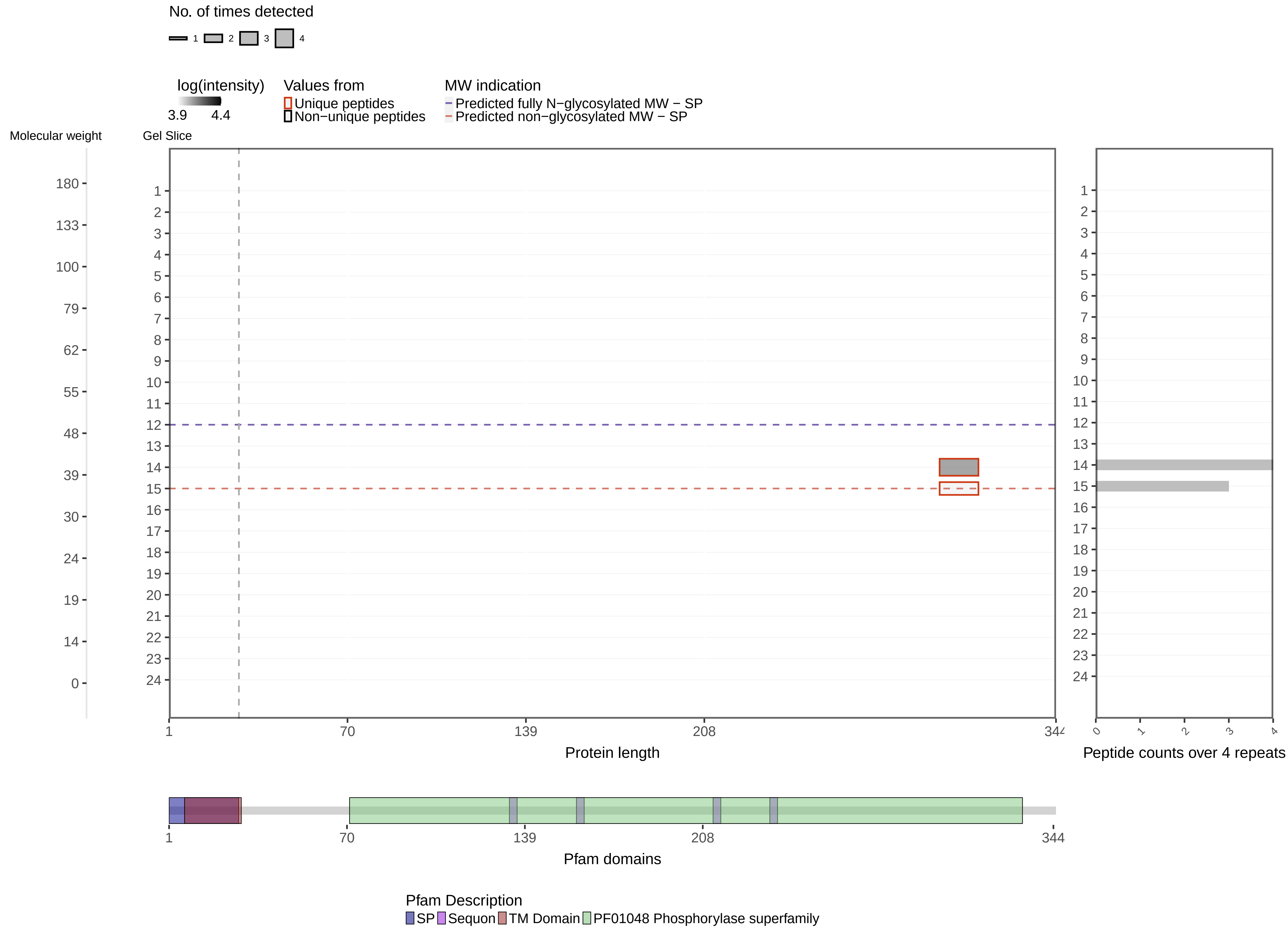


Protein sequence:

TLEPNDFAOFDFKNYNAPKGGDNKLGRVGYSTEOFYSTSGEINVPORPVWFNITKNWLNVASHLOGMKLEOCANSTMCLPOKPKLVVGLKGATSNFFIDNAAYTDFLFKTFGVTSLDMESSAVVMTCLSNGYPVIAIRGVSDLAGTOEGDNTIRLFGPLAALNTAKVVISFVKSLPGNYISKF*

NbL18g03090.1 pyruvate_kinase_1_cytosolic−like [MW = 57.62 kDa]

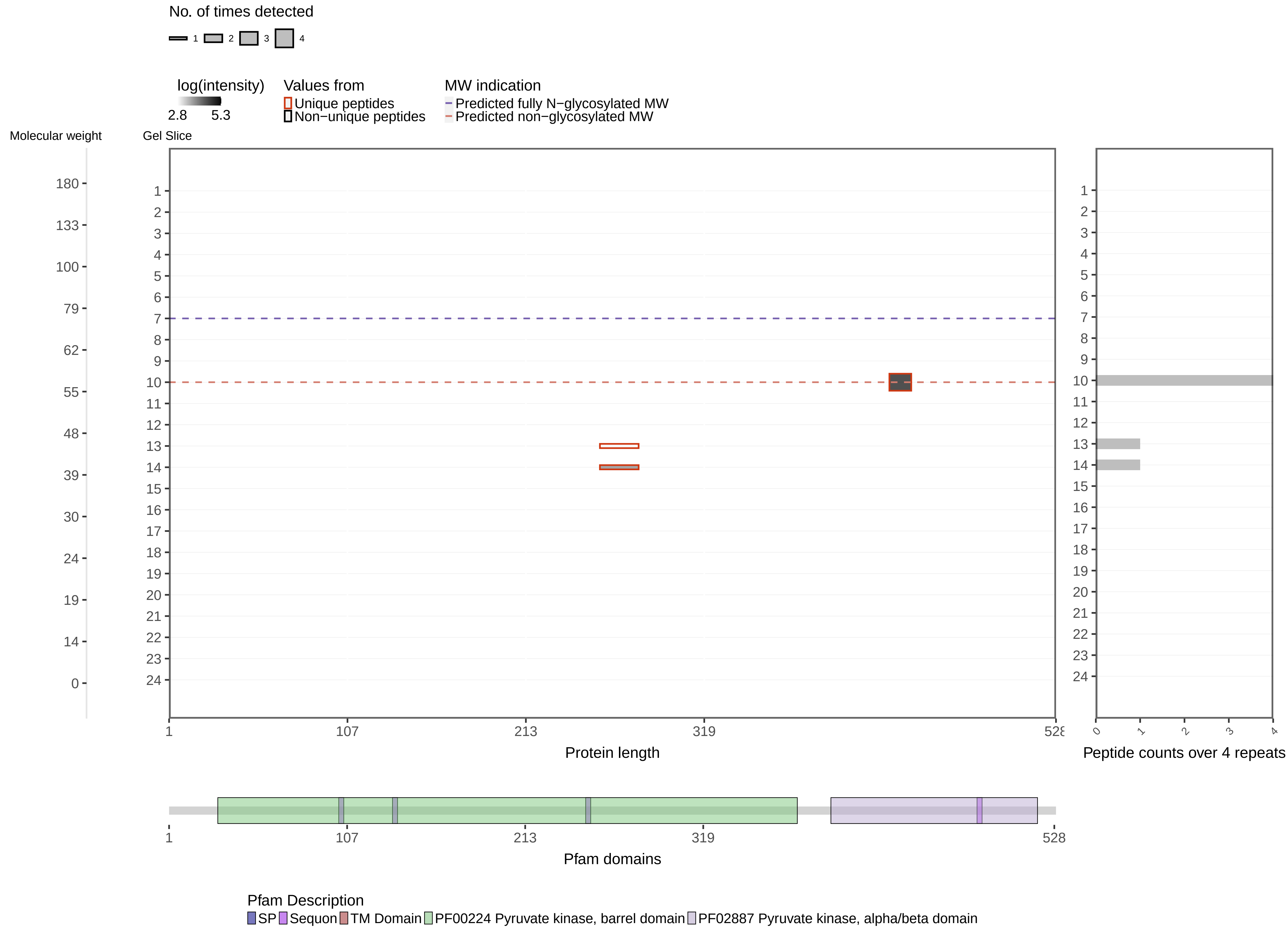


Protein sequence:

TQIFAKIENVEGMANFDEILEEADGIILSRGNLGIDLPPEKVFLFQKAAVHKCNMAGKPAVVTRVVDSMTDNLRPTRAEATDVANAVLDGSDAILLGAETLRGLYPVETISIVGKICAEAEKVYNQDVYFKRAVKYVGEPMTHLESIASSAVRAAIKVKASVIICFTSSGRAARLIAKYRPPMPVLSVVIPRLKTNQLRWTFSGAFQARQSLIVRGLFPILADPRHPAESTNATDESVLKVALDHGKTAGVIKPHDRVVVFQKVGDSSVVKIIELEN*

### NbL18g02860.1 neutral/alkaline_invertase_3_chloroplastic−like [MW = 73.53 kDa]

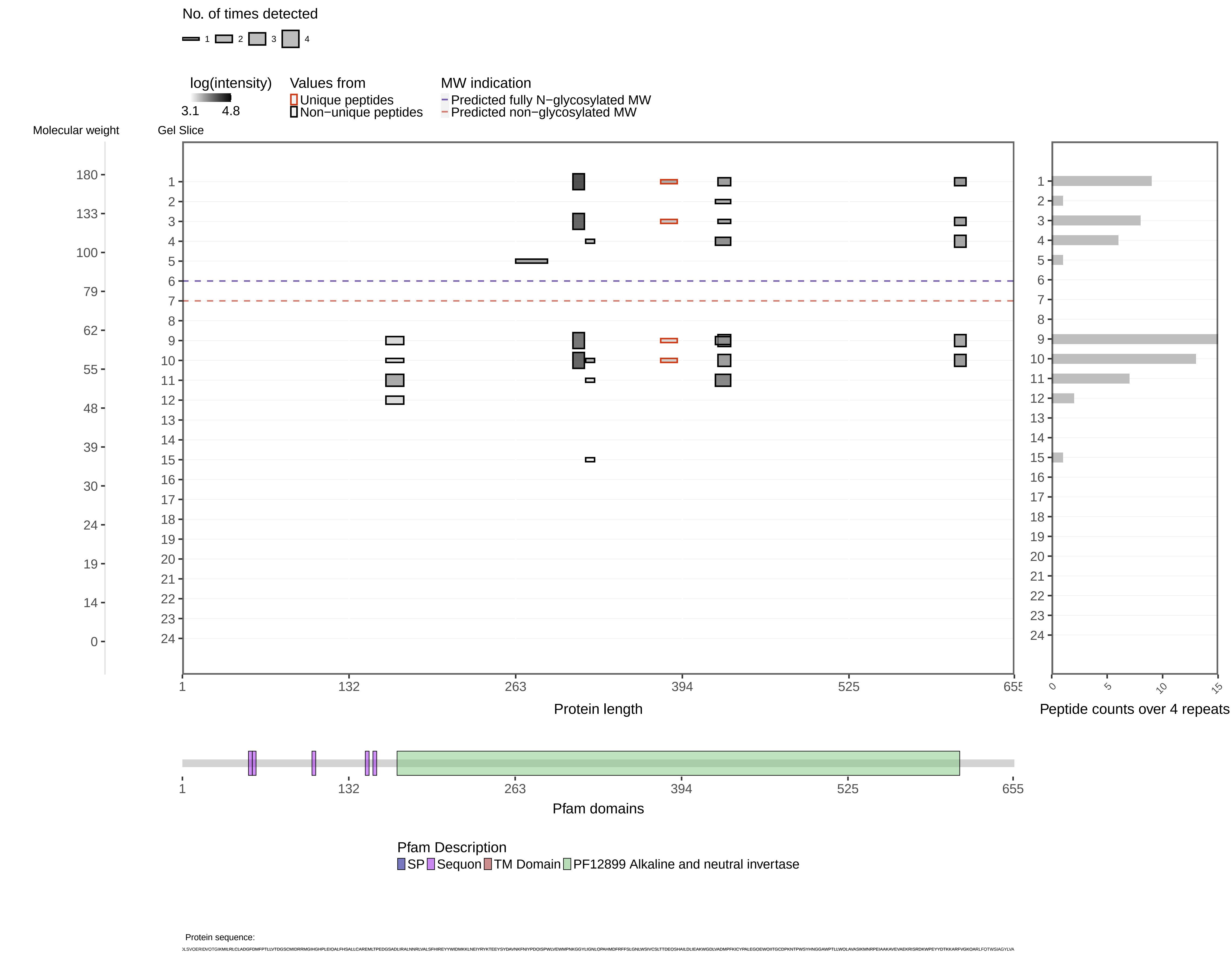

NbL18g02750.1 nifU−like_protein_4_mitochondrial [MW = 30.86 kDa]

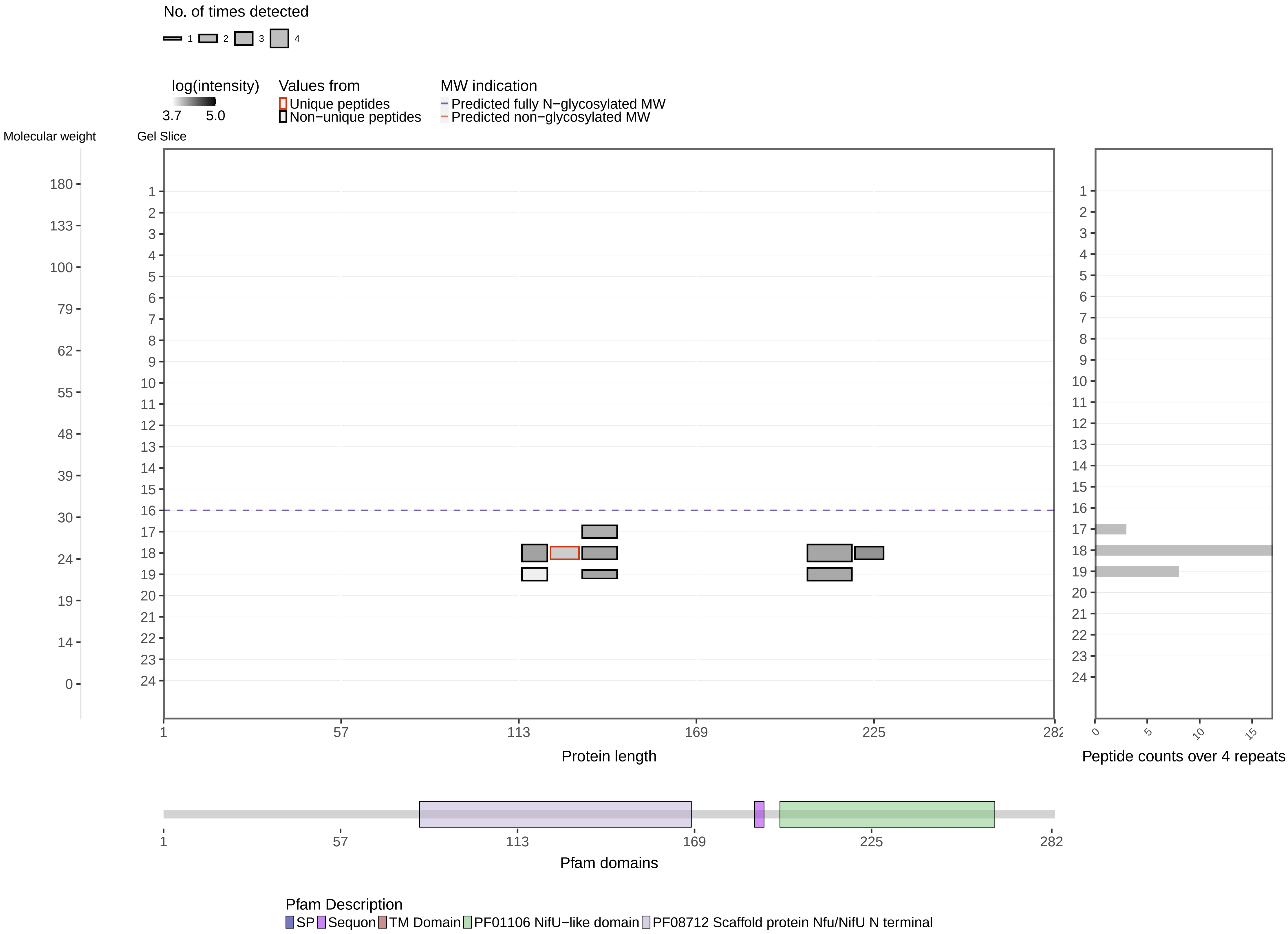


Protein sequence:

GITRVFFGSDFVTVTKSDDASWDFLKPEIFAAIMDFYSSGKPLFLDSNTAASMDTAIHENDSETVAMIKELLETRIRPAVQDDGGDIVYRGFDLDTGIVKLKMQGACSGCPSSSVTLKSGIENMLMHYVPEVKGVEQEMDEDEEPGLTGAPYE*

NbL18g02710.1 probable_aspartyl_aminopeptidase [MW = 59.77 kDa]

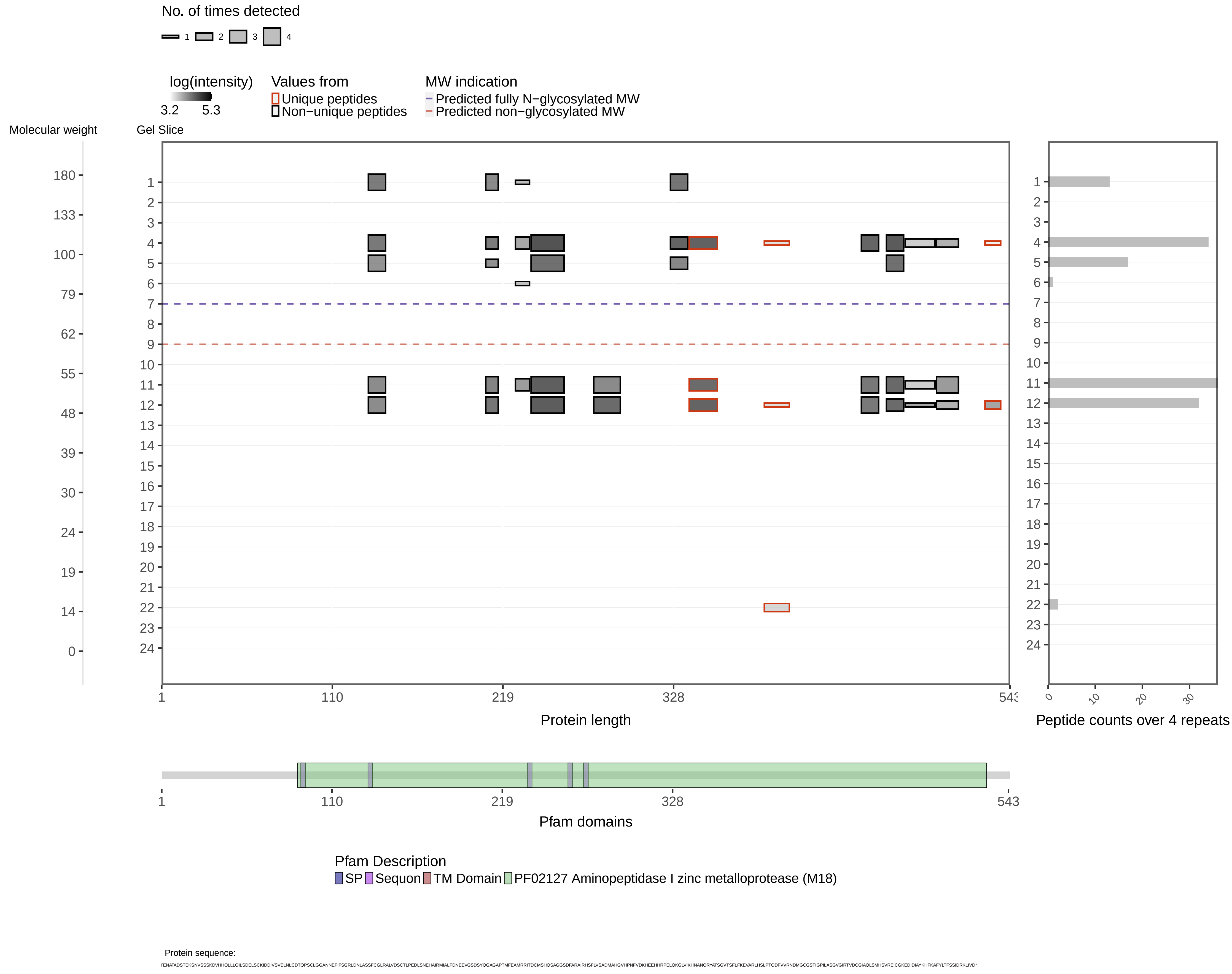

NbL18g02680.1 diphosphomevalonate_decarboxylase_MVD2−like [MW = 46.55 kDa]

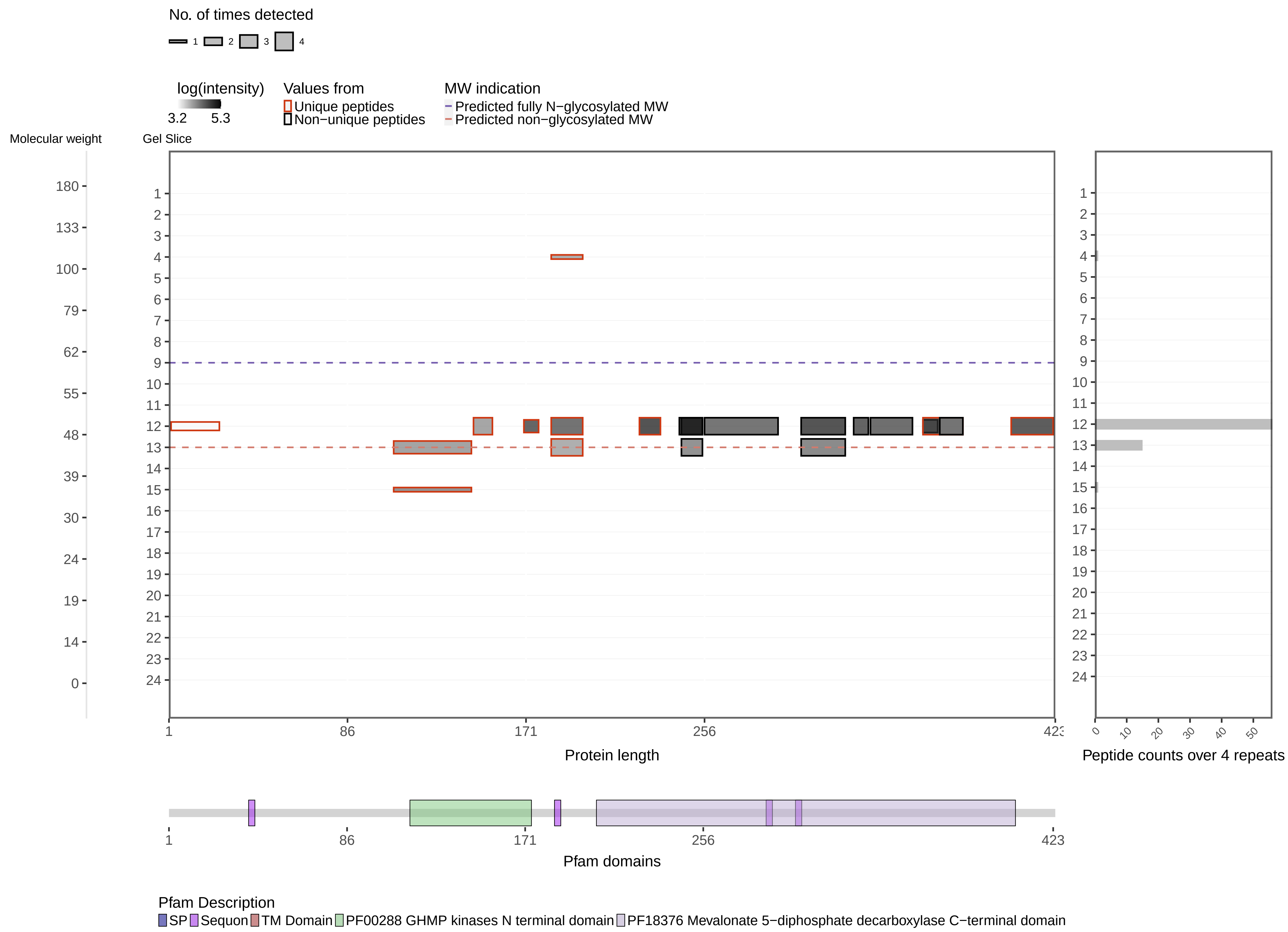


Protein sequence:

EKHWDELVIIIAVVSSRQKETSSTSGMRDTVETSALIERRAKEVVPKRIIQMEEAIQNRDFPTFAQLTCADSNQFHAVCMDTSPPIFYMNDTSHRVIGCVEKWNRSEGTPQVAYTFDAGPNAVLIARNRKAATLMLQRLLFHFPPNSDTDLNSYVIGDKSILKDAGIQDIKDVEALSPPPEIKDKVPAQKYKGEISYFICTRPGRGPVLLTDDSLALLNPETGLPK*

NbL18g02620.1 ACT_domain−containing_protein_ACR12−like [MW = 35.4 kDa]

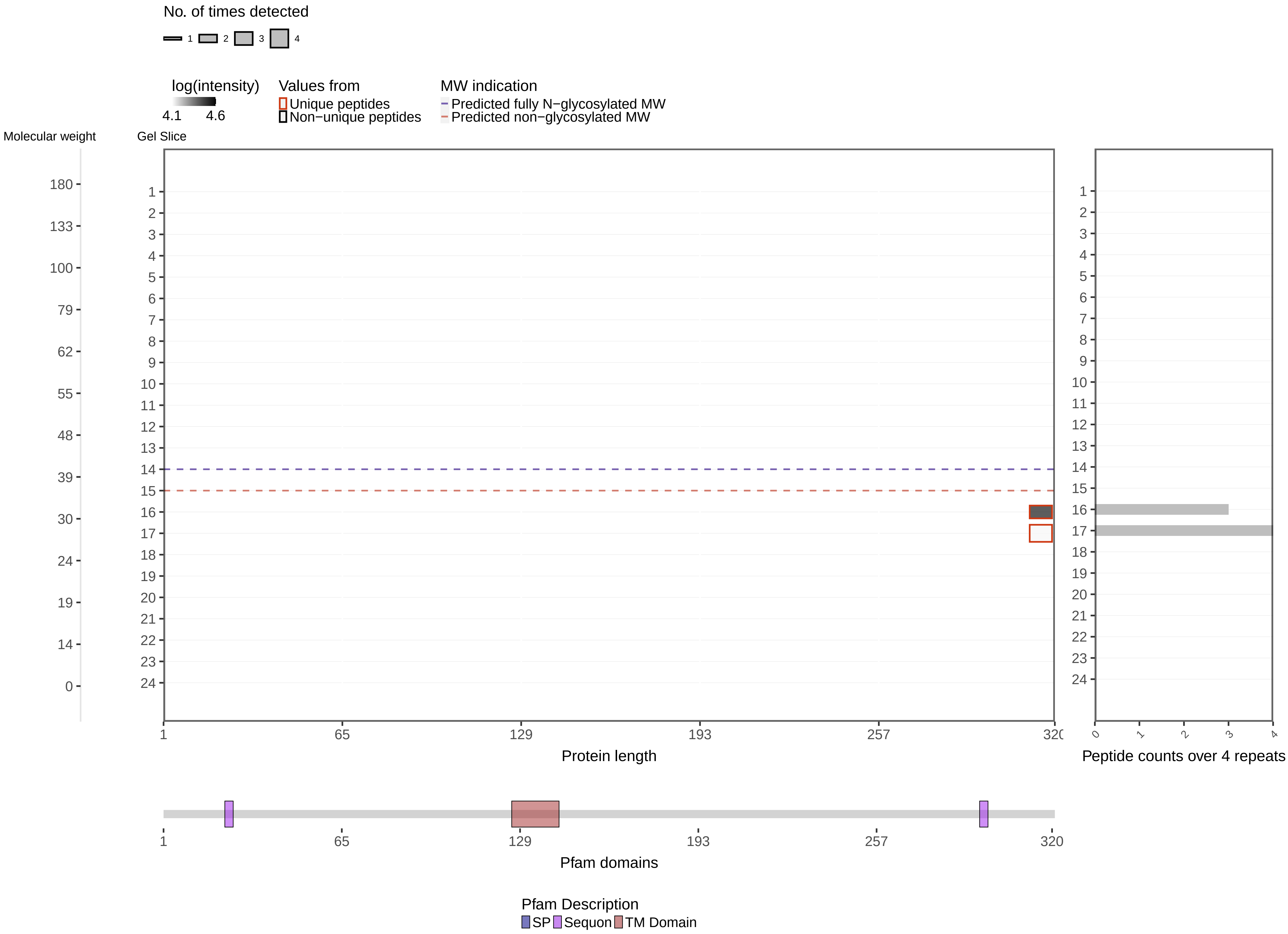


Protein sequence:

LKDLGLDVVKGIVTTEGSVKKTKFLITRLDTGRKVEDPDLLERIRLTIISNLIKYHPESSEQLAMGEAFGIKAPEKKPDVDIATHIHVKDDGSKRSILHVETADRSGLLVEVVKIMADINIDVESAEIDTEGLVAKDKFYVSYRGAALNSSLSQVLVNCLRYFLRRPETDEESY*

### NbL18g02590.1 ferredoxin_root_R−B2−like [MW = 16 kDa]

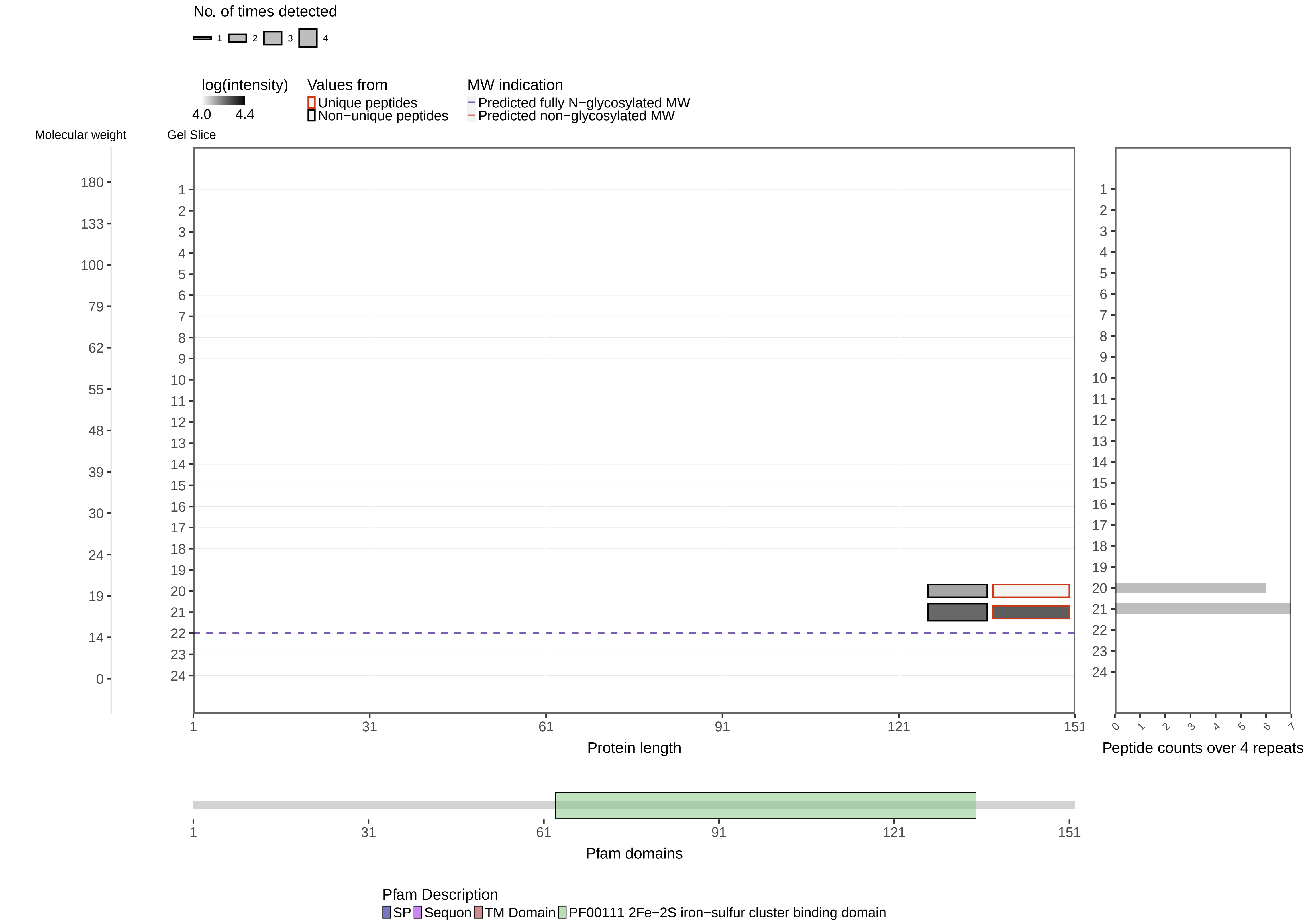


Protein sequence:

DGTENEIEVSDDQYILDAAEDAGLELPYSCRAGSCCTCAGQLVSGSVDQSEGAFLDDDQMEKGYLLTCISYPRSDCVIHTNKEEEVH*

NbL18g02520.1 5−epi−aristolochene_synthase−like [MW = 63.22 kDa]

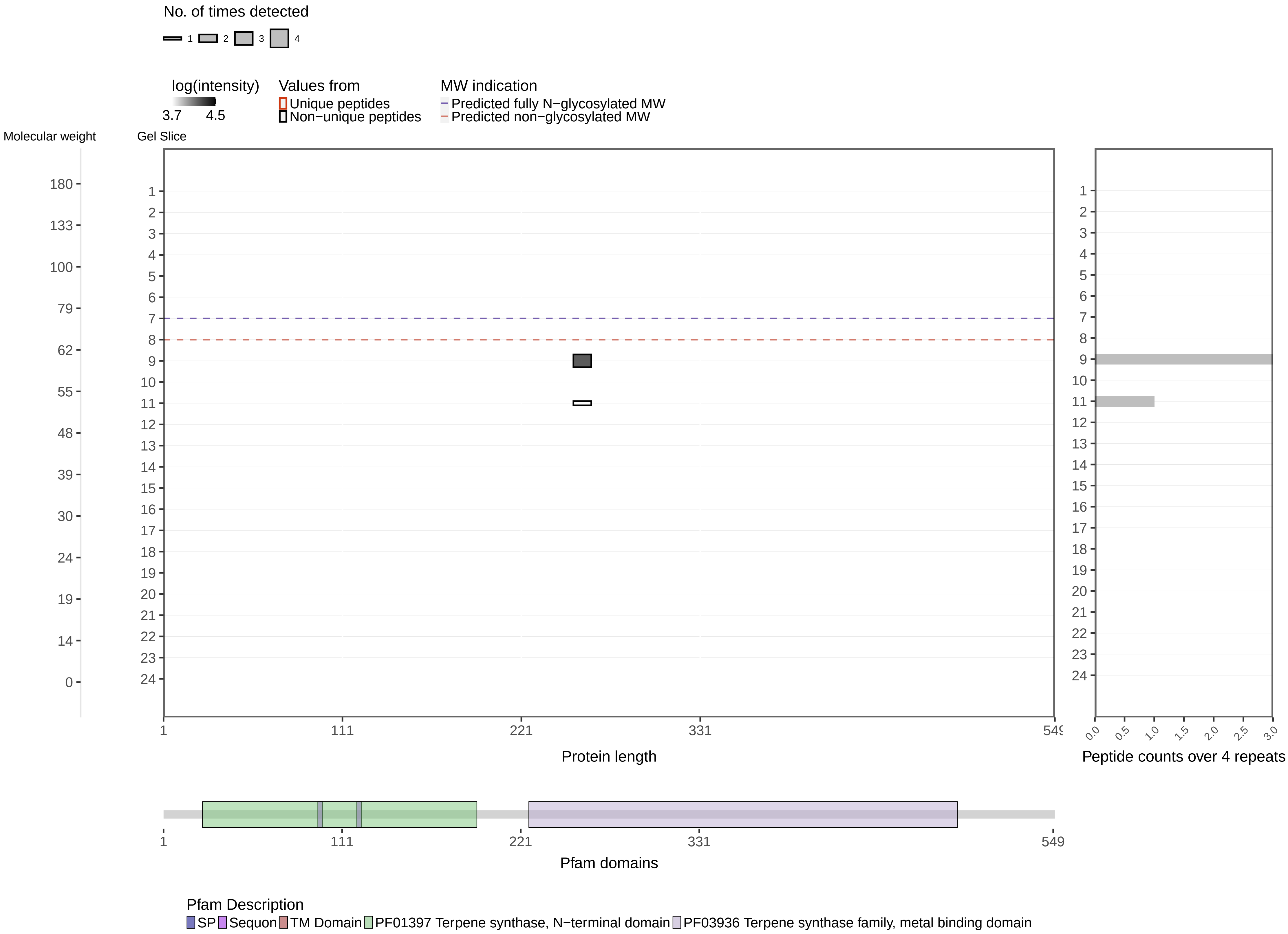


Protein sequence:

YARDRVVECYFWALGVYFEPQYSKARVMLVKTISMISIVDDTFDAYGTVKELEVYTNAIQRWDINEIDRLPDYMKLSYKAILDLYNDYEKELSSAGRSHLVCHAIERMKEVVRNYNVESTWFIEGYKPPVTEYLSNALATTTYYYLATTSYMGMKSVTERDFEWLSKNPKILEASVIICRVIDDTATYEVEKSRGQIATGIECCMRDYGISTEEAMTKFQEMAEAAWKDLNEGLLRPTPVSTEFLSRILNLARIVEVTYIHNLDGYTHPEKVLKPHIIALLVDSIEV*

### NbL18g02510.1 5−epi−aristolochene_synthase−like [MW = 63.24 kDa]

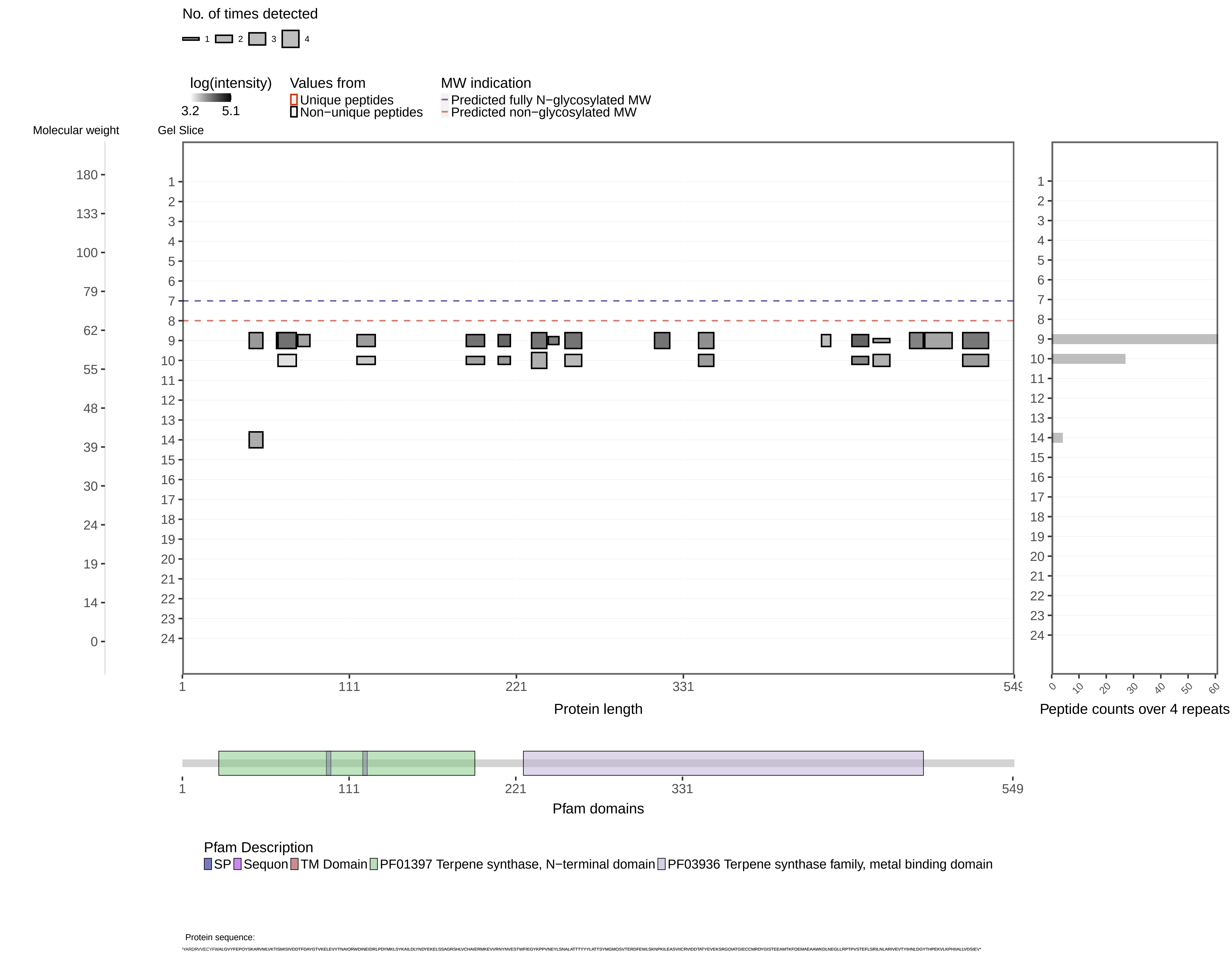

NbL18g02400.1 nudix_hydrolase_26_chloroplastic−like [MW = 26.02 kDa]

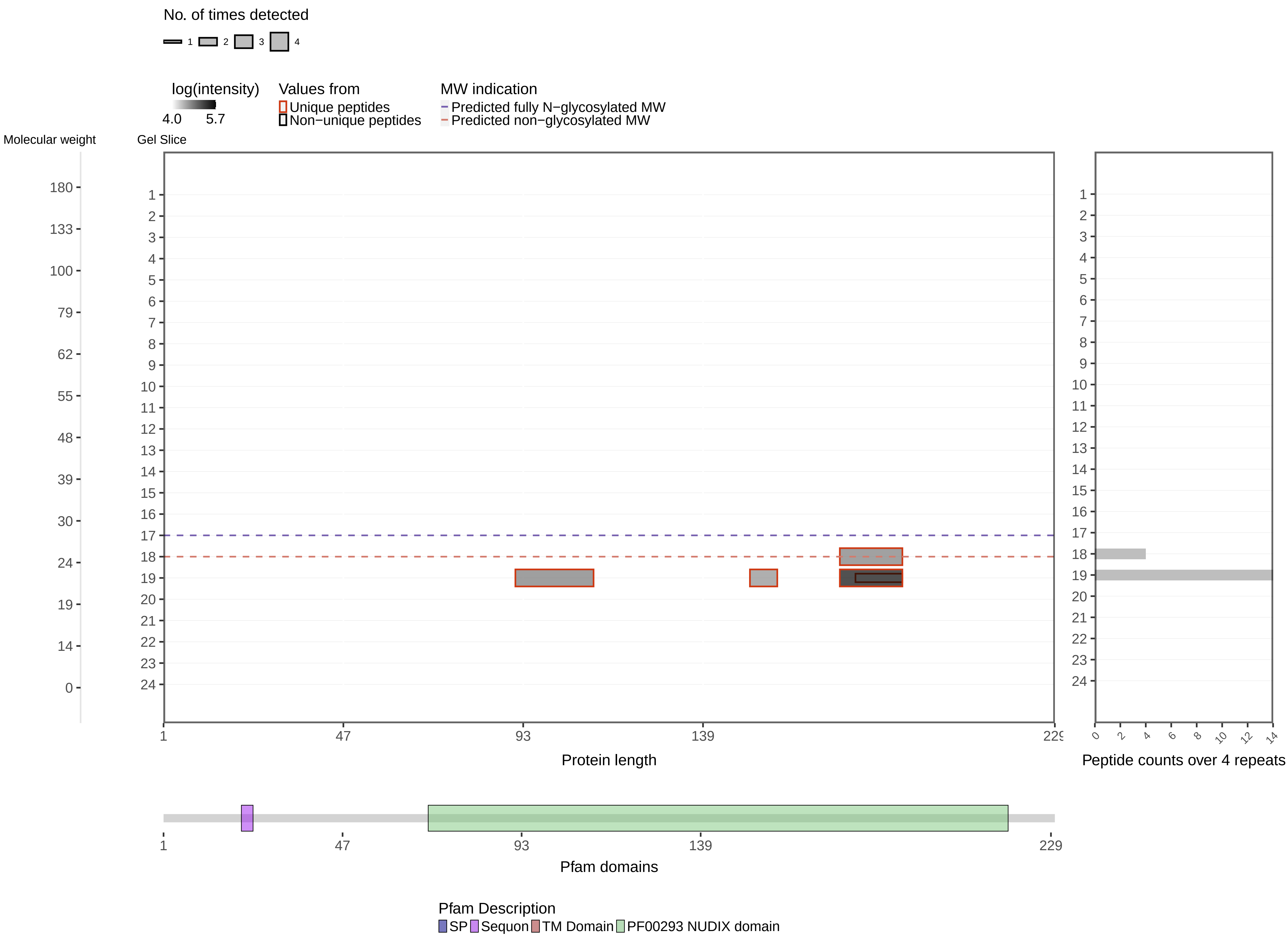


Protein sequence:
VDESEDPRNAAIRELREETGVTSAEIVAEAPHWLTYDFPPDVREKLRHQWGSDWKGQAQKCMENSRFLFKFTGKDEEINLLGDGTEKAEFGEWSWMSPEQVVEFAADFKKPVYKEVLSVFSQYFL*

### NbL18g02360.1 vacuolar_protein_sorting−associated_protein_60.1 [MW = 25.95 kDa]

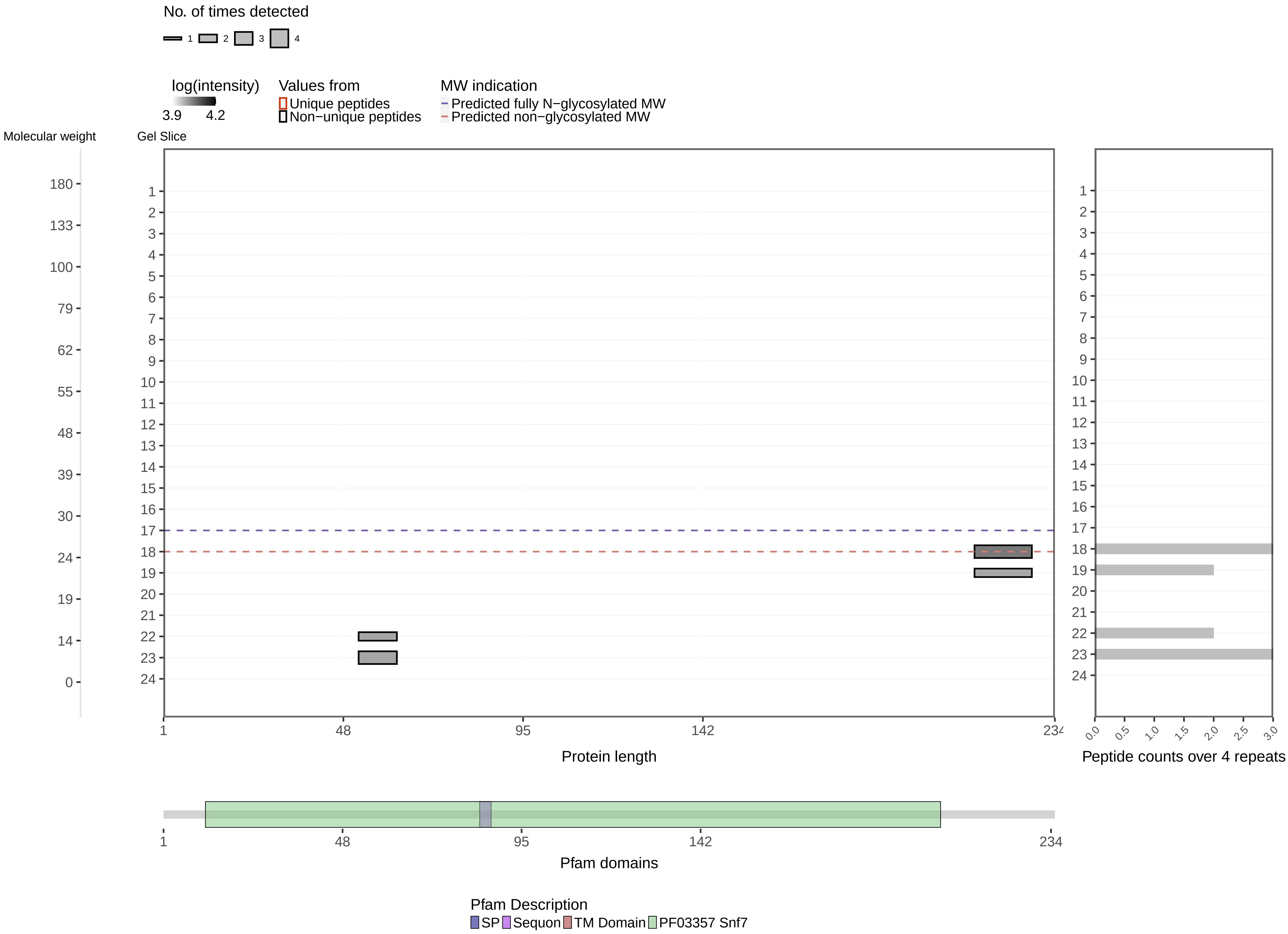


Protein sequence:

)TMSALKSANKELKGMMKTVKINDIDNLQDEMMDLMDVSNEIQESLGRSYSVPDDIDEEELMGELDALEADMDTEGEGVPSYLQPEPDLDAELSLPSAPTGHAPMPAGRANPQADDELGLPAVPRASIRG*

### NbL18g02330.1 dnaJ_protein_homolog [MW = 46.56 kDa]

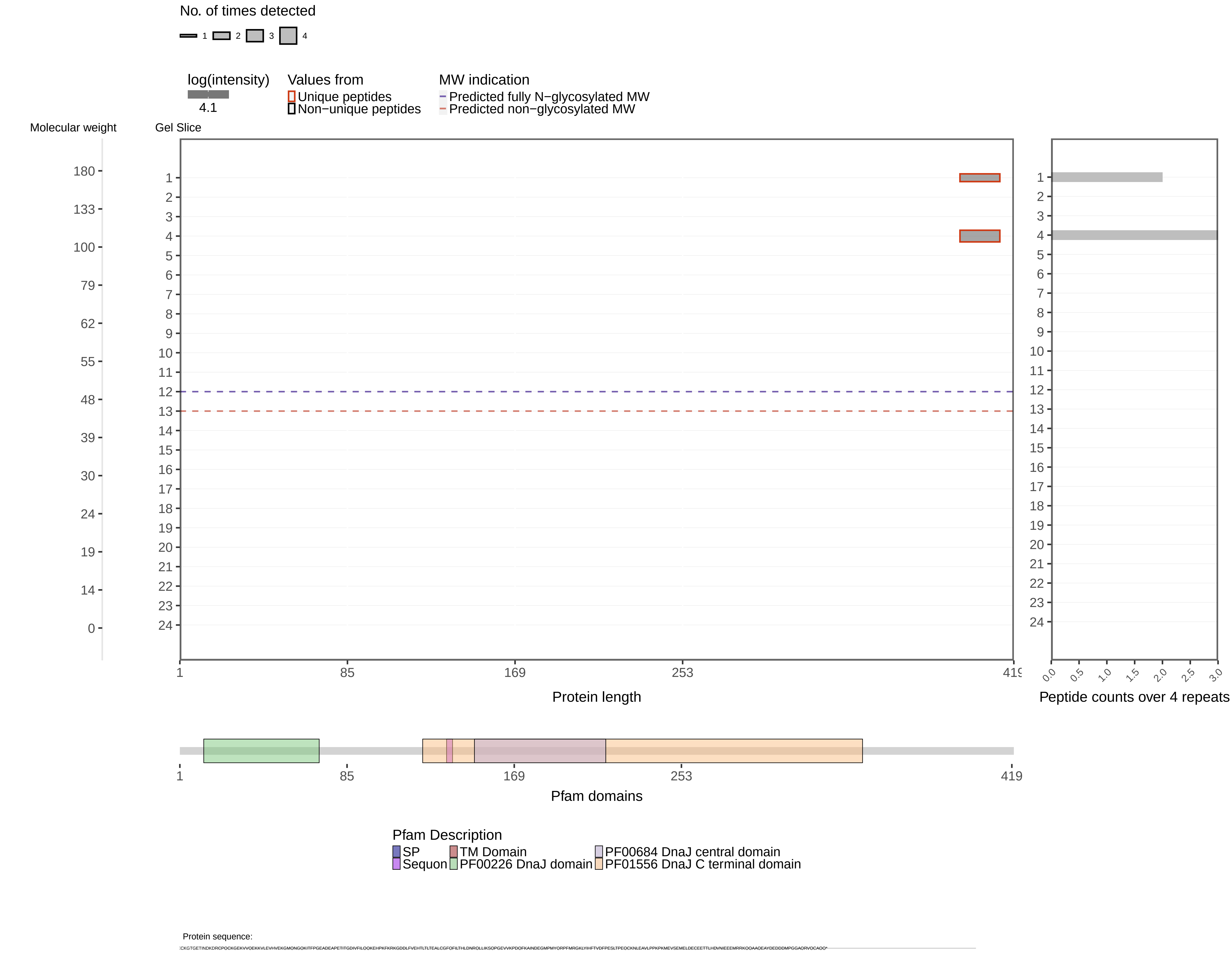

NbL18g02290.1 mitochondrial_acidic_protein_MAM33 [MW = 25.26 kDa]

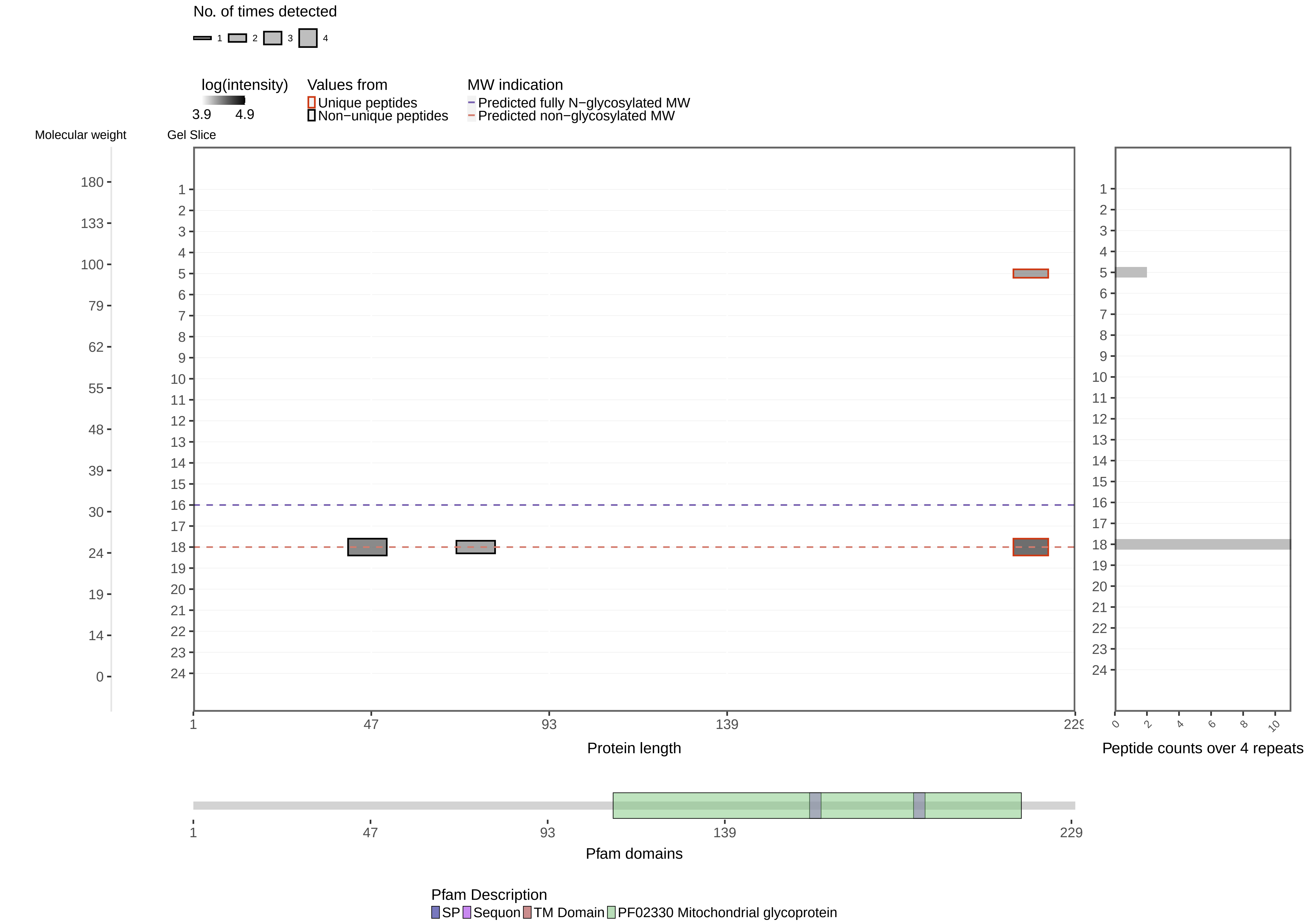


Protein sequence:

GGDEEEDGINQLFLHVDISKPGQKKSLHFLCGLYPDALGIHSVSLRPKAESTGFLAVPTNYSGPVFQDIDEAMRDALHRFIEERGINESLFPFLQAWLYVKDHRNLMRWFKTVGSLVNDIKQGSSHA*

NbL18g02240.1 ubiquitin_carboxyl−terminal_hydrolase_12−like [MW = 130.72 kDa]

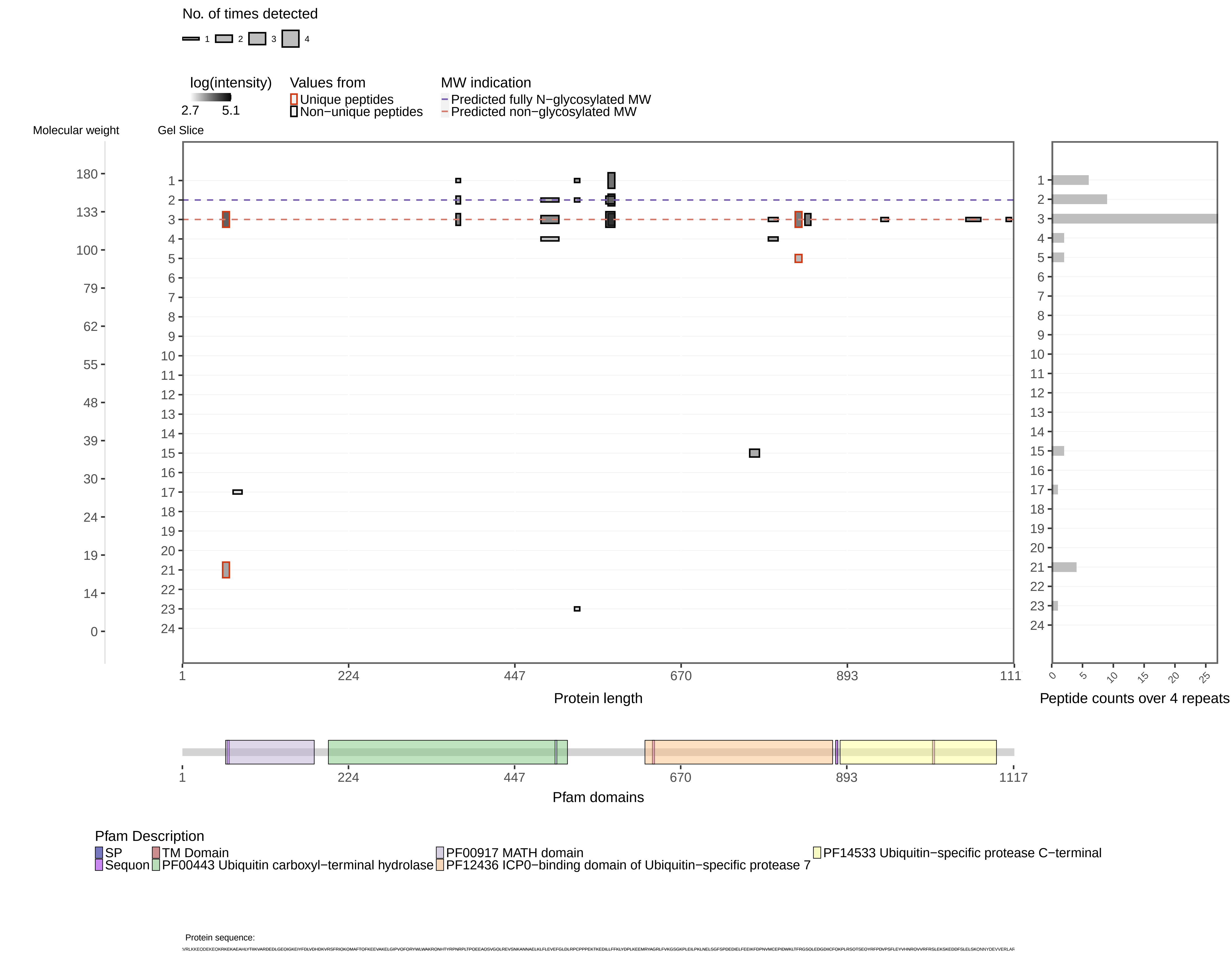

### NbL18g02180.1 GDSL_esterase/lipase_At5g33370−like [MW = 41.52 kDa]

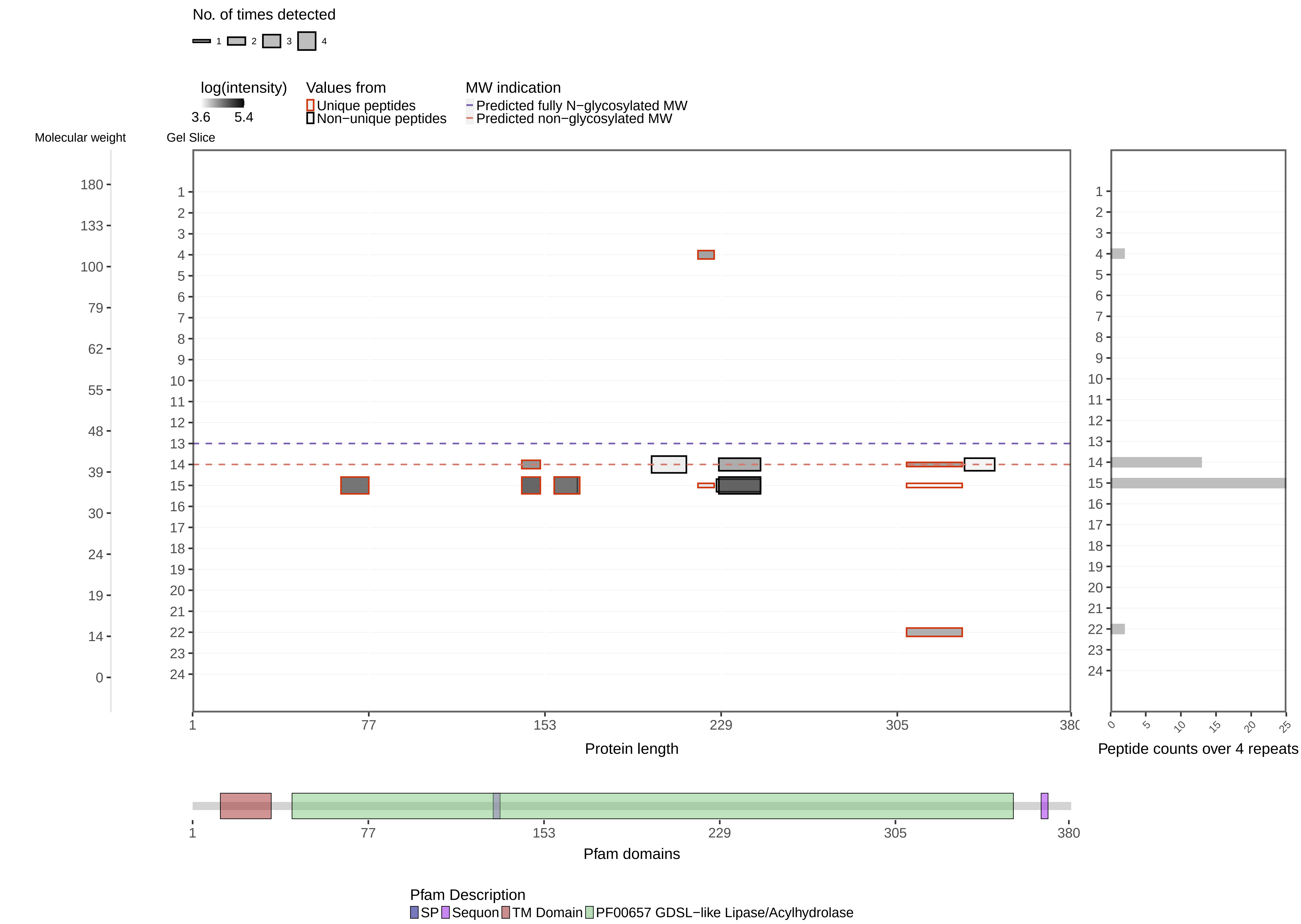


Protein sequence:

GNDFVNNYYLVPNSARSROYALPDYVPFLISEYRKILRRVYDLGGRRVLVTGTGPLGCVPAELAORSRNGECAVELOOAAALFNPOLTOMLODLNTELGSNIFIAANTOOMHIDFITNPOAYGFITSKVACCGOGPYNGLGLCTPLSNLCPNRDVFAFWDPFHPSERANKIIVOOIMTGSSDYMNPMNLSTIMAMDSMA*

### NbL18g02050.1 HMG−Y−related_protein_A−like [MW = 18.23 kDa]

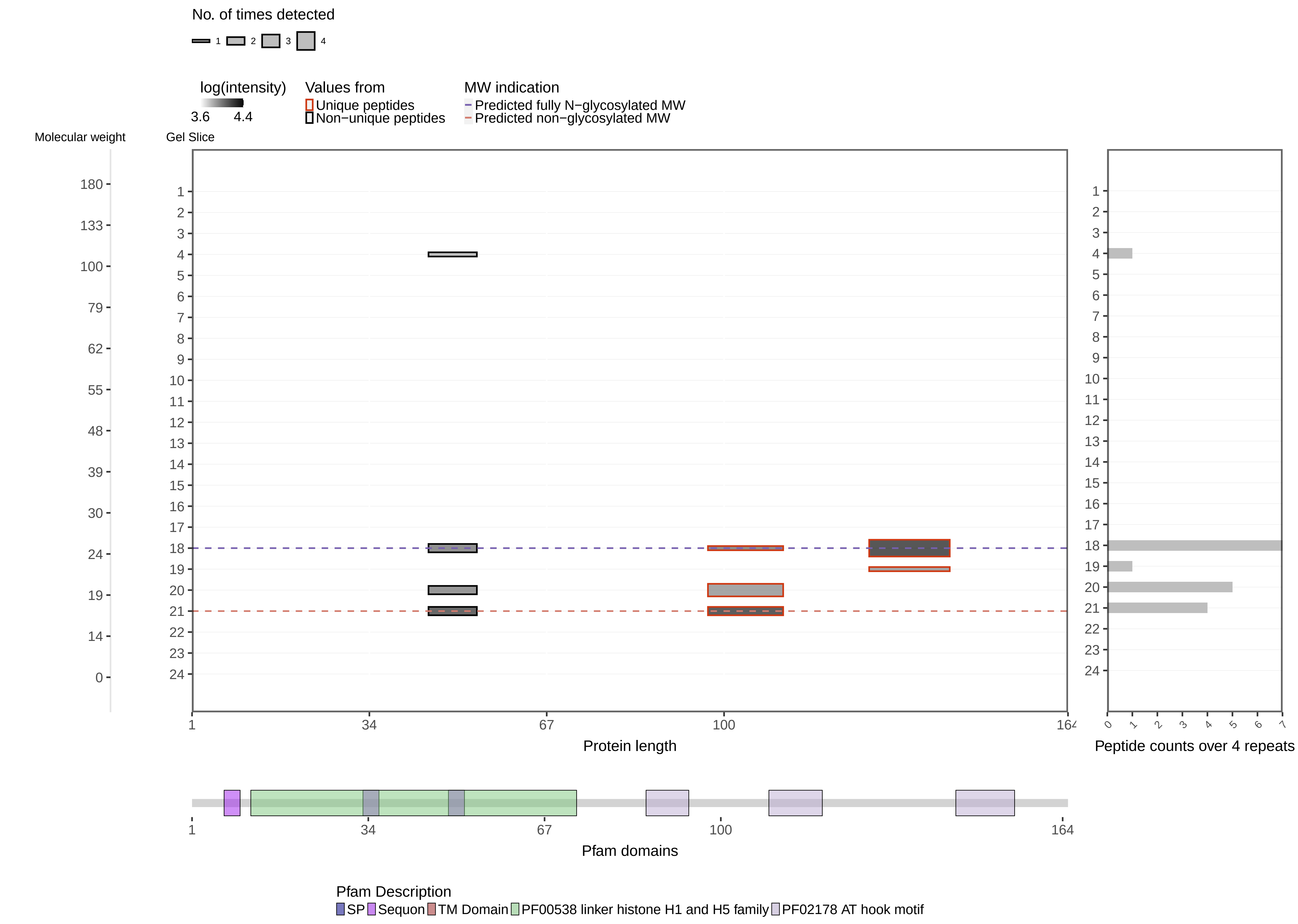


Protein sequence:

FLKNNYIKPGPNVPPKRGRGRPPKPKTTLPQDTEIAASKPRGRPKKDPNAPPTPKKLKPTIVPINSANPVSKTGRPRGRPRKVKPQSAQNGVE*

### NbL18g01810.1 probable_isoleucine−−tRNA_ligase_cytoplasmic [MW = 135.05 kDa]

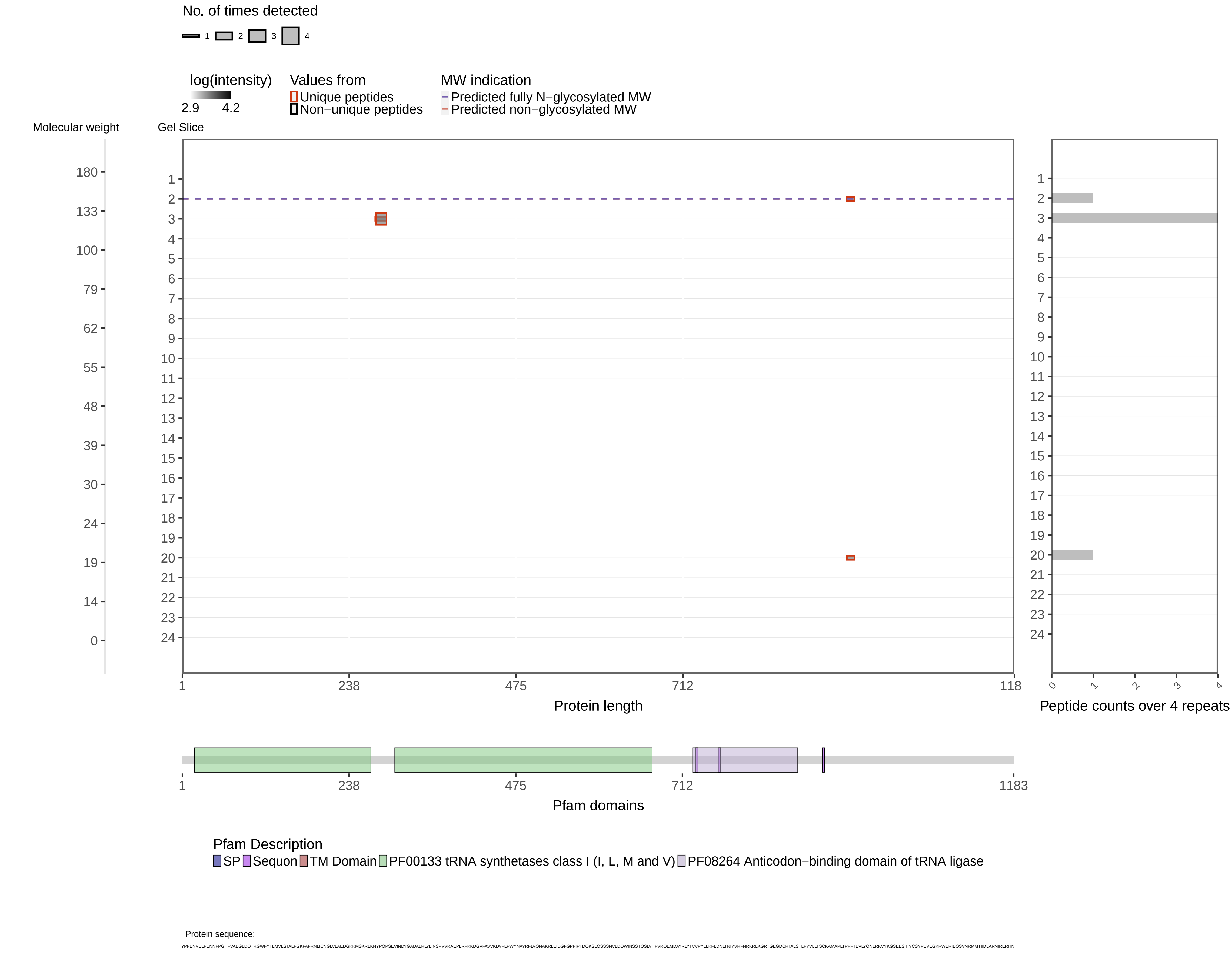

### NbL18g01780.1 60S_ribosomal_protein_L6−like [MW = 25.45 kDa]

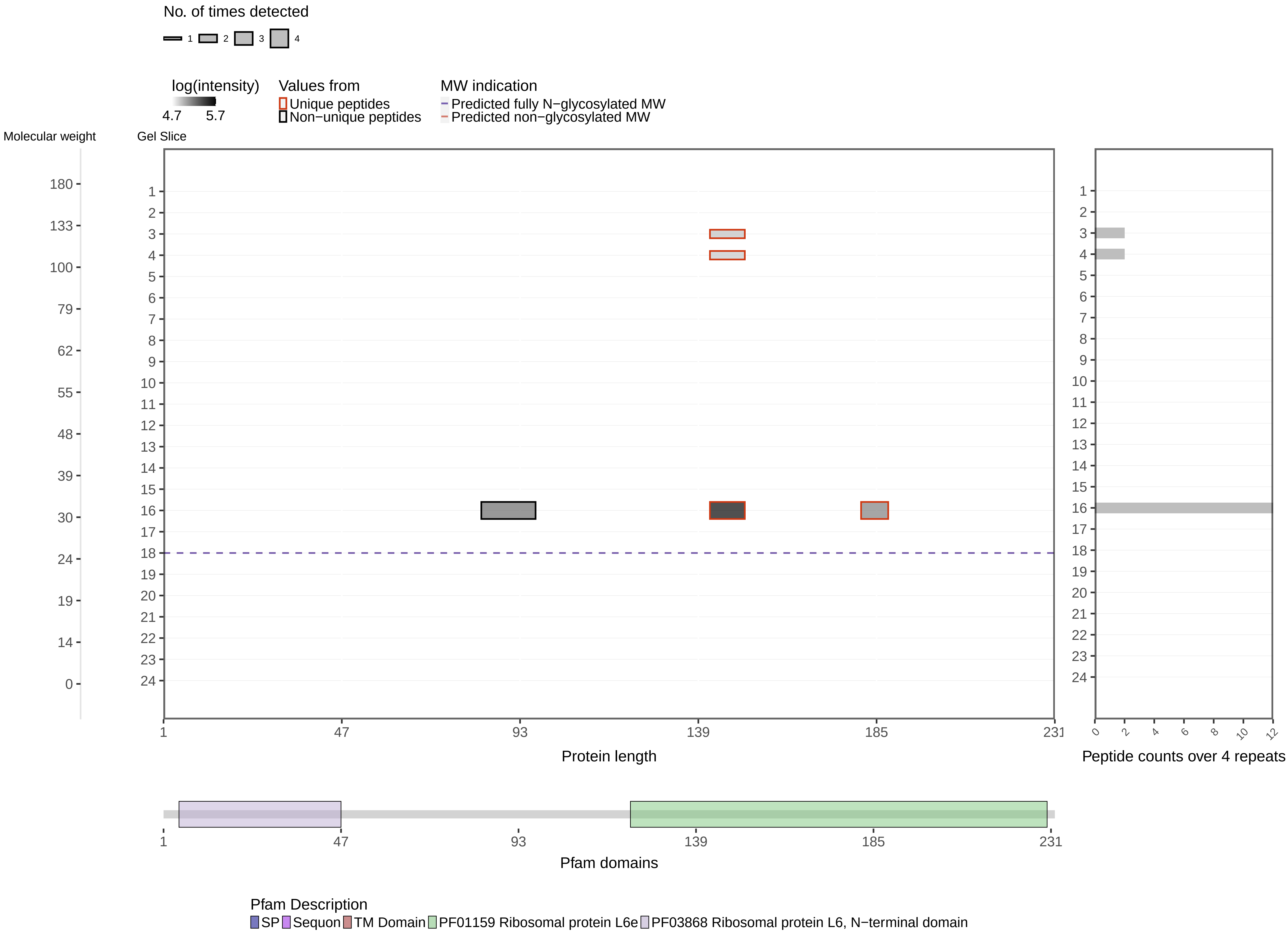


Protein sequence:

VFLKQLKSGLLLVSGPFKLNGVPLRRVNQAYVIGTSTKVDISGVSVEKFDDKYFAKQAEKKQKKGEGEFFEDKKEEKKNVLPQEKKDDQKAVDEALIKAVECVPELKAYLSARFSLKSGMKPHELVF*

NbL18g01770.1 phosphoribosylglycinamide_formyltransferase_chloroplastic [MW = 33.07 kDa]

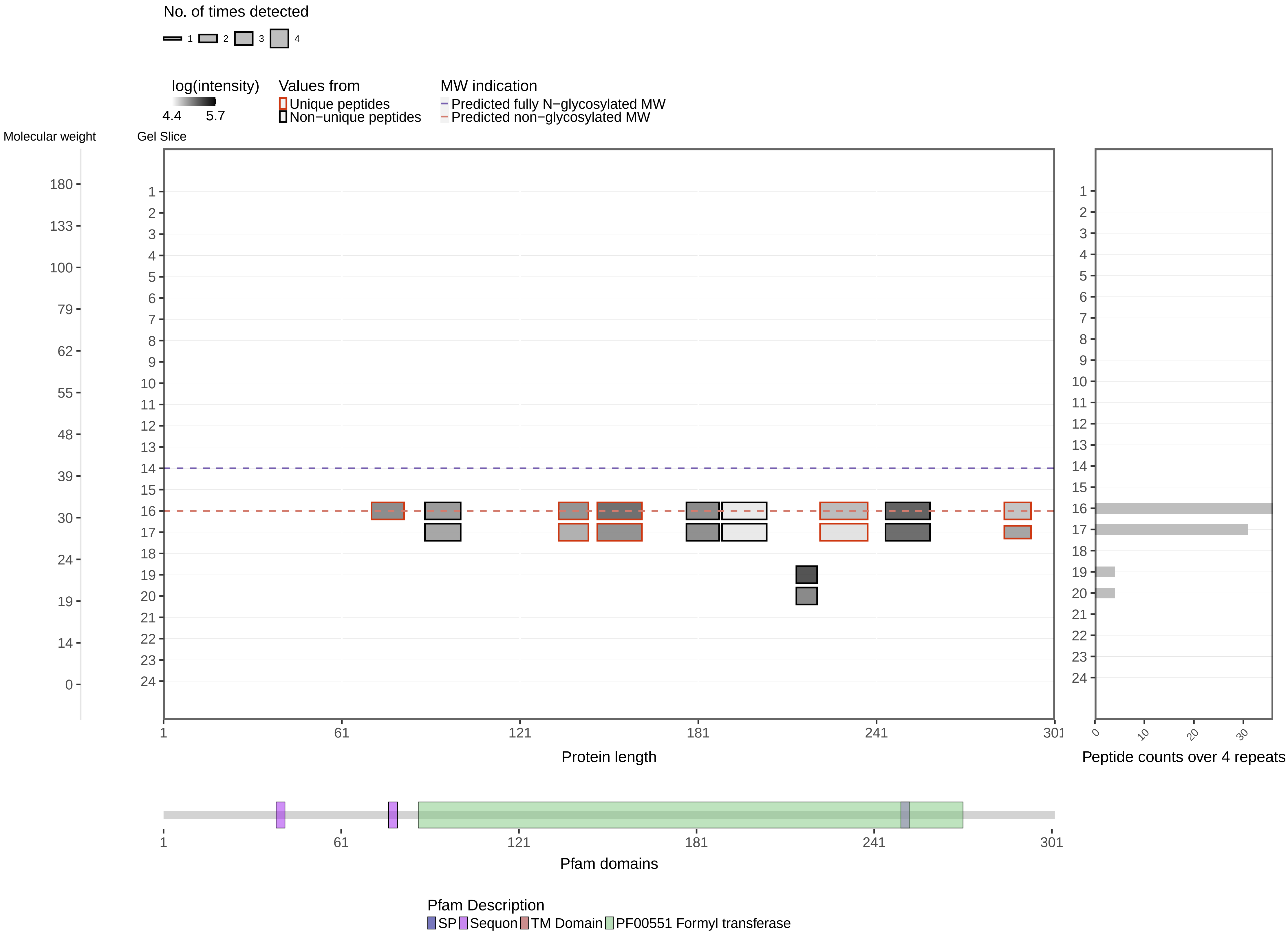


Protein sequence:
IPVILFPKAKGTSEGLSEEDLVGALRTYNIDFILLAGYLKLIPTELVQAFPRSIFNIHPSLLPAFGGKGYYGMKVHKAVIASGARYSGPTIHYVDEHYDTGRILAQRVVPVLANDTADTLAARVLQEEHKLYVEVAAALCEERIVWRQDDVPLIQSKEDPNQYK*

NbL18g01740.1 acyl−protein_thioesterase_1−like [MW = 28.01 kDa]

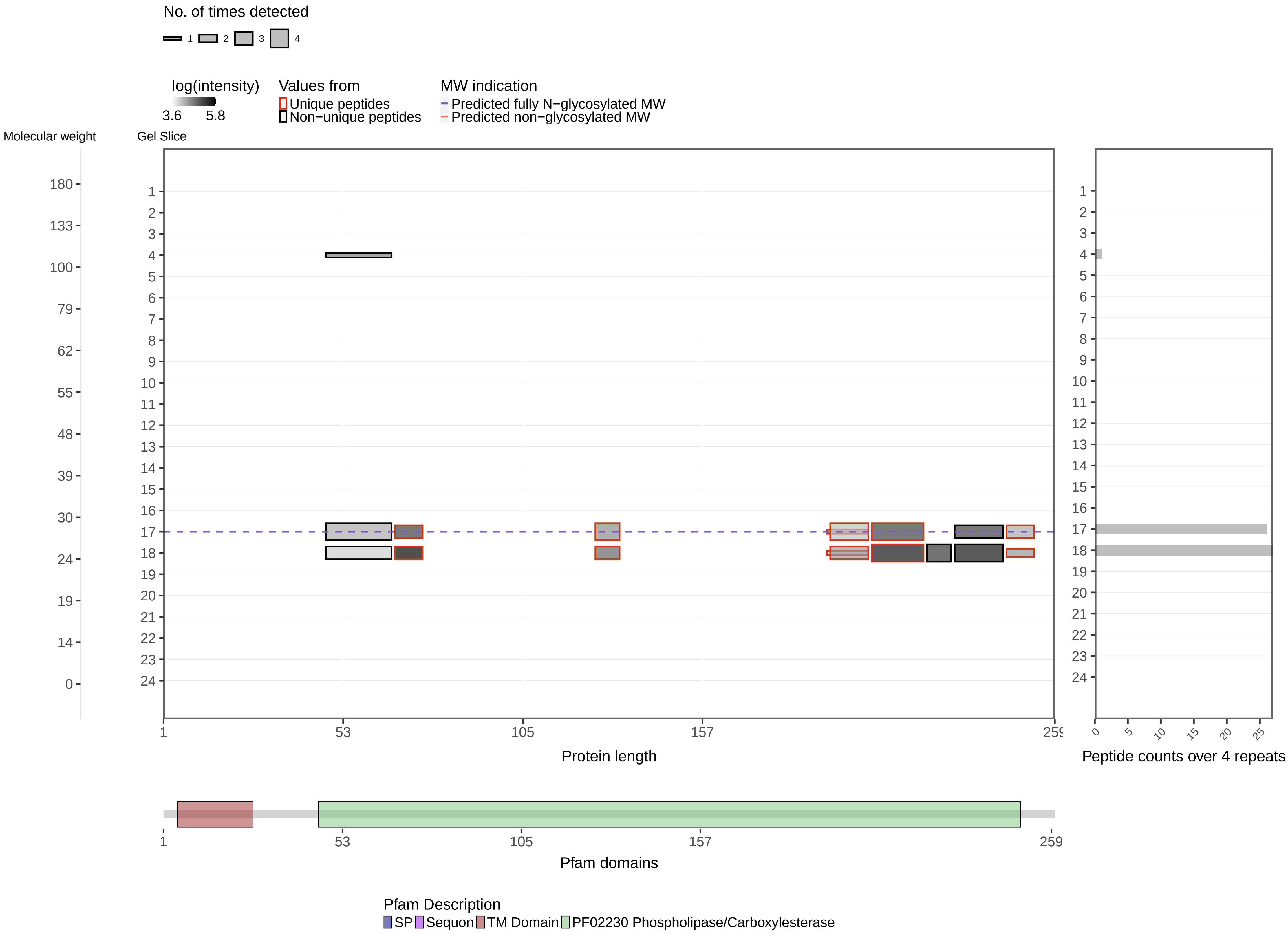


Protein sequence:

SGLLKAVKNIHAMIDKEIDAGTNPSNVFICGFSQGGALTLASVLLYPKTLGGGAVFSGWVPFNSAILEKQTPEAKKTPILWSHGIADRTVLFEAGQAGPPFLERAGMTCEFKAYPGLAHSINNEELKNLELWIKPRLQSNS*

NbL18g01690.1 tropinone_reductase−like_3 [MW = 27.18 kDa]

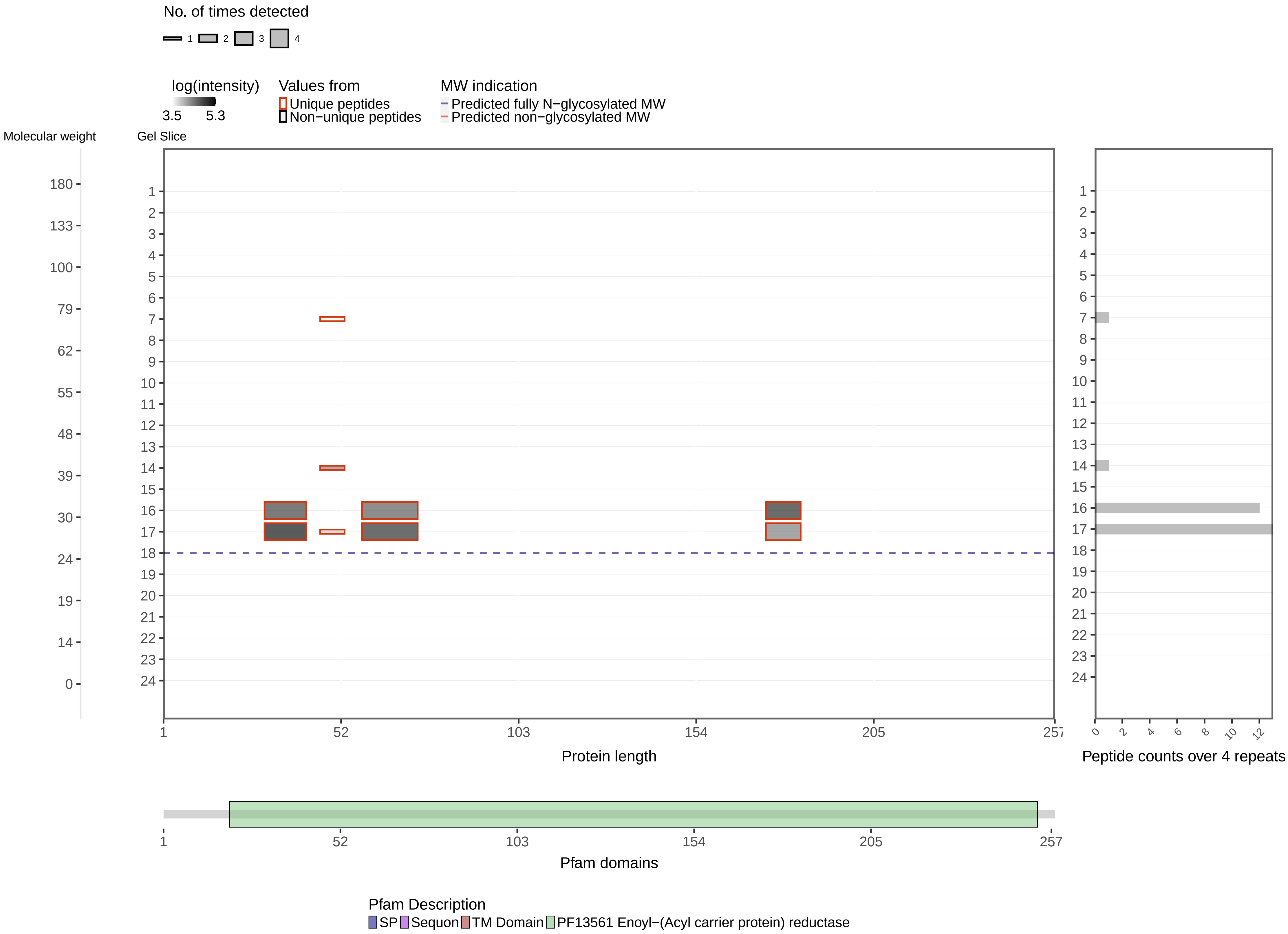


Protein sequence:

WDINVKAAILLLQHLQDAAPCLQKGSSVVIISSIAGYTPPAAMGMYGVTKTALLGLTKALAAEMAPNTRVNCIAPGFVPTHFAEFITGNANVRRELEAKTLLNRLGTTQDMAAATAYLASDDAAYVTGETLVVAGGMHARL*

### NbL18g01640.1 UTP−−glucose−1−phosphate_uridylyltransferase [MW = 54.7 kDa]

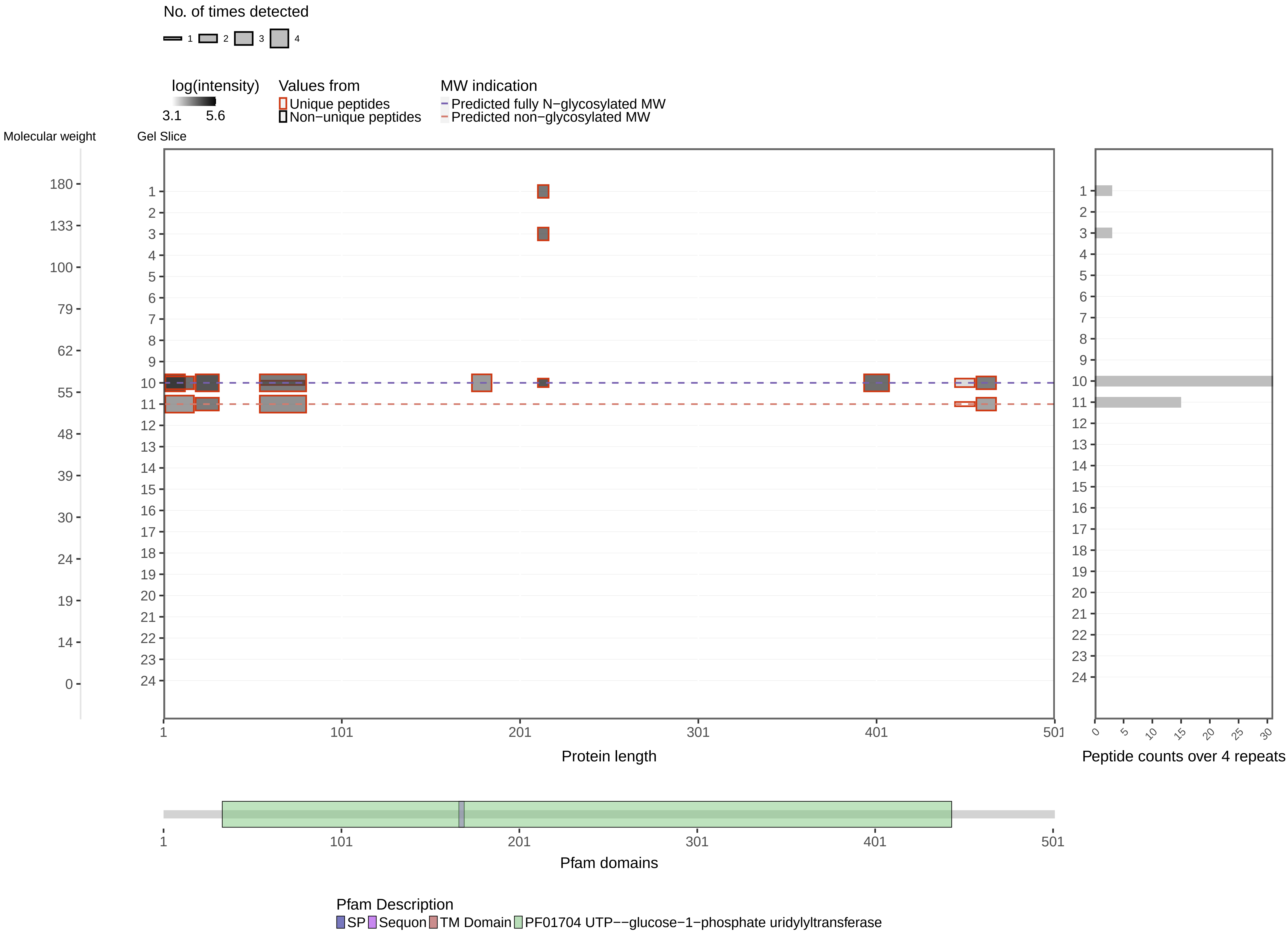


Protein sequence:

VDLKILNHLIQNKNEYCMEVTPKTLADVKGGTLISYEGKVQLLEIAQVPDEHVNEFKSIEKFKIFNTNNLWVNLNAINRLVEADALKMEIIPNPKEVDGVKVLQLETAAGAAIKFFDRAIGVNVPRSRFLPVKATSDLLLVQSDLYTLTDEGYVIRNPARSDPSNPSIELGPEFKKVANFLGRFKSIPSIIELDSLKVTGDVWFGSGITLKGIVIVAAISGVKLEIPDGSVIANKVIQFTLAPNGAARYRFTSFFDPVTSIFRISMD*

### NbL18g01600.1 isocitrate_dehydrogenase_[NADP]−like [MW = 57.88 kDa]

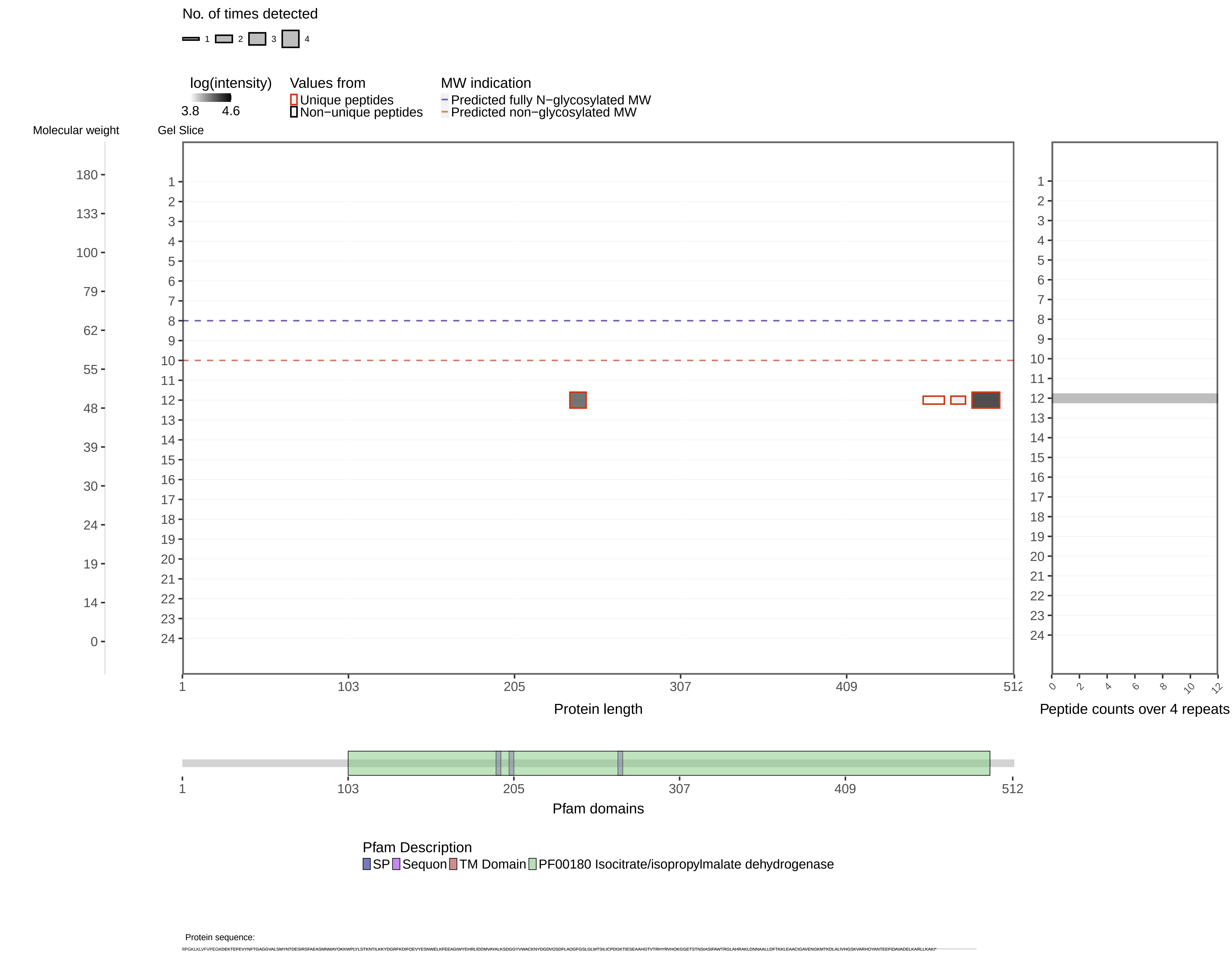

NbL18g01590.1 glutamate_decarboxylase [MW = 55.86 kDa]

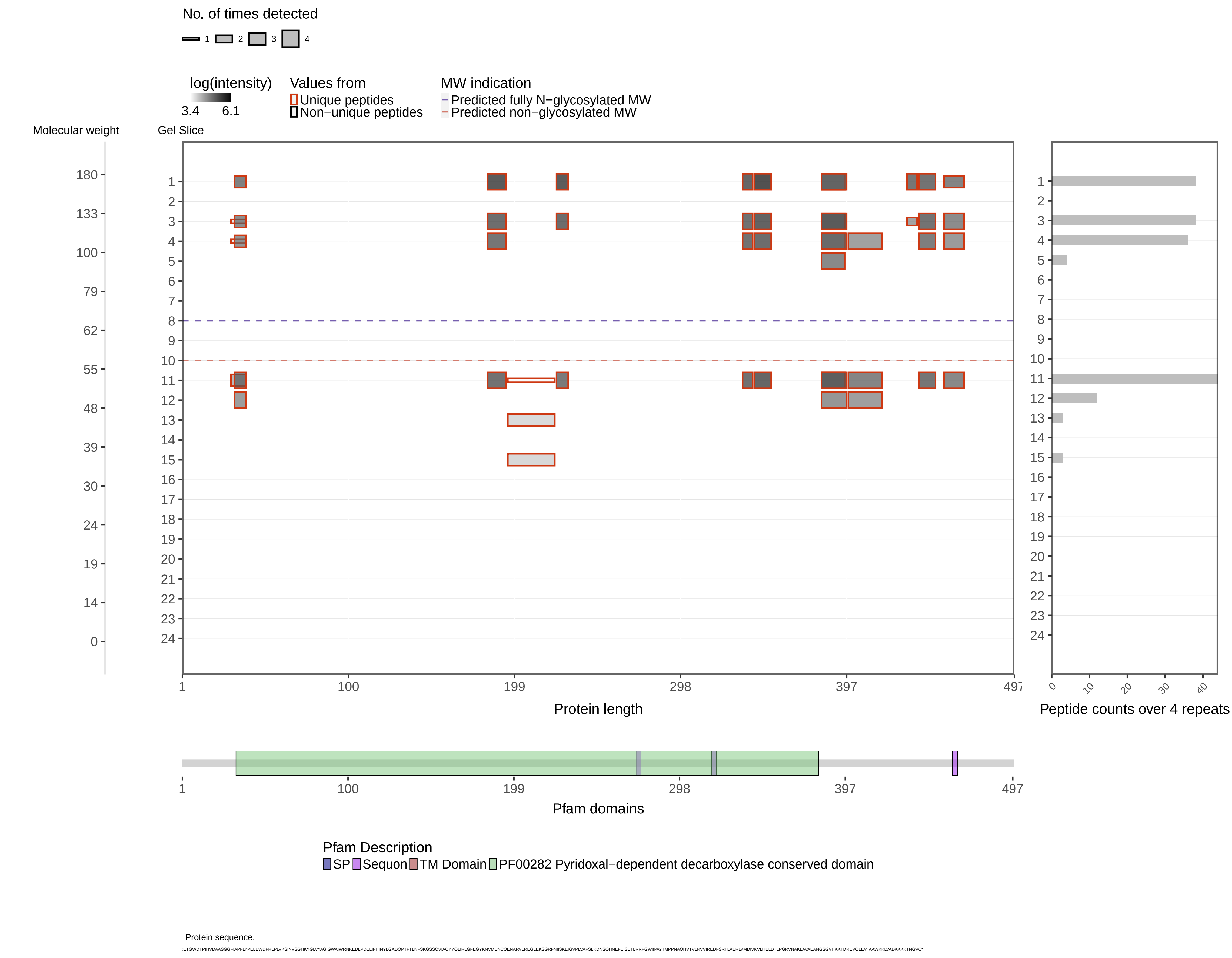

### NbL18g01390.1 30S_ribosomal_protein_3−1_chloroplastic−like [MW = 22.47 kDa]

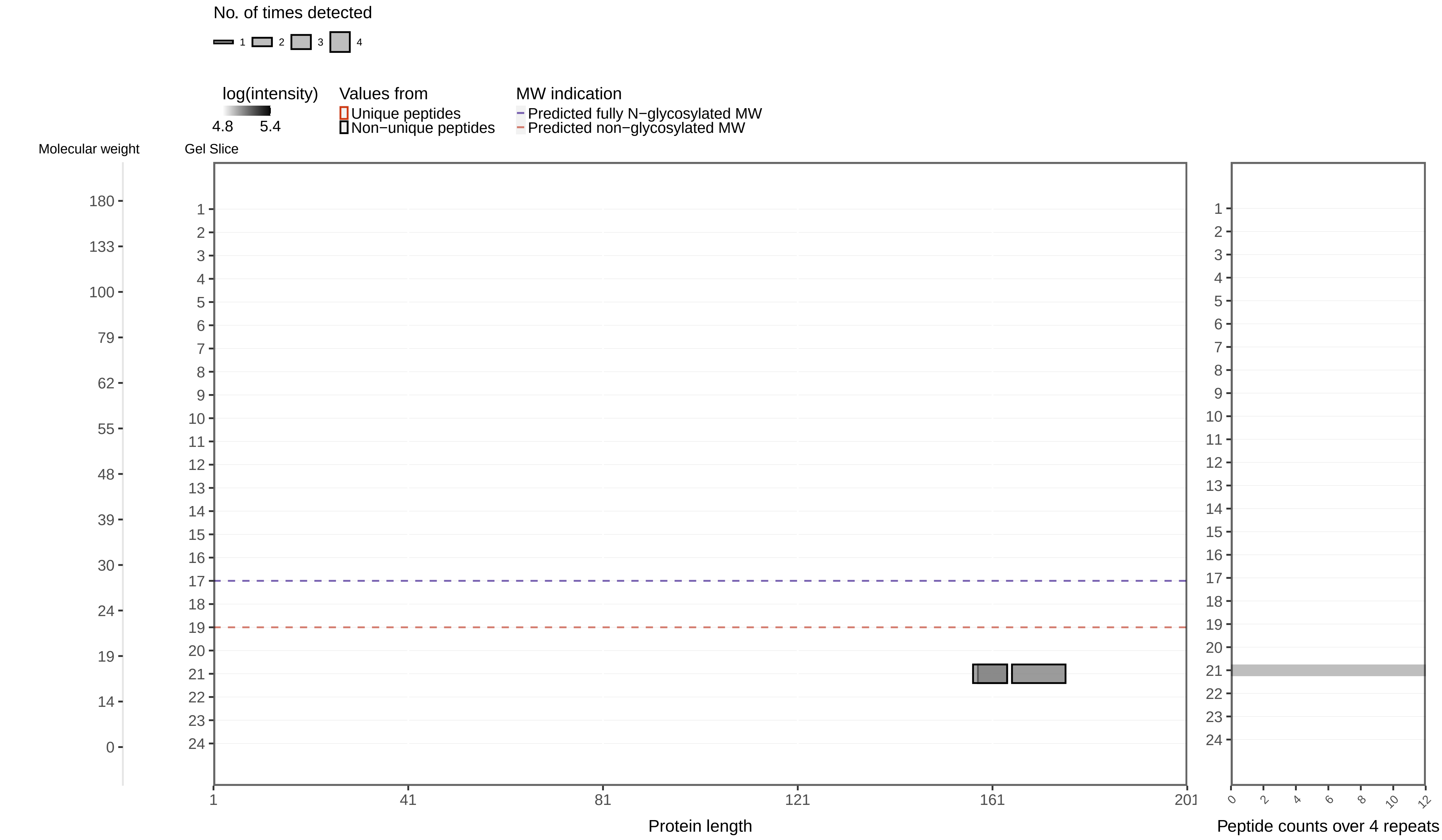


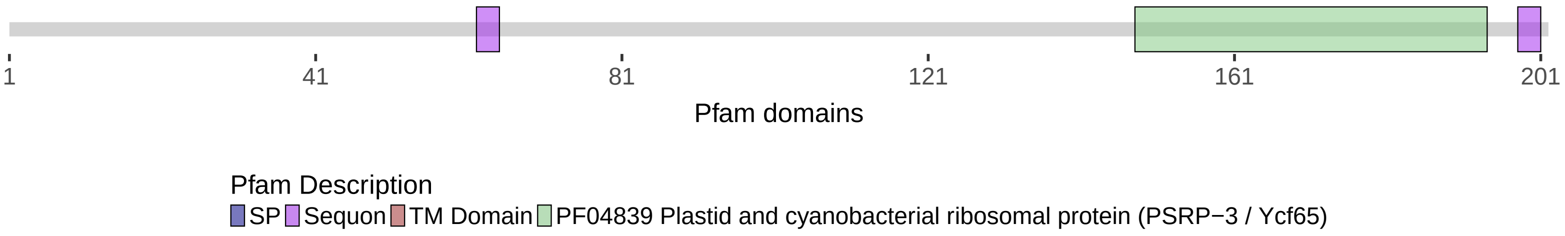


Protein sequence:

TETELVTTDTSDSVVQEIENTDSKKKVIEKPRLVLKFIWMEKNIGLGLDQVLPGHGSVPLSPYFFWPRKDAWEELKTTLESKPWISQKQMIILLNQATDIINLWQQSGGNLS*

NbL18g01300.1 alpha−xylosidase_1−like [MW − SP = 102.24 kDa]

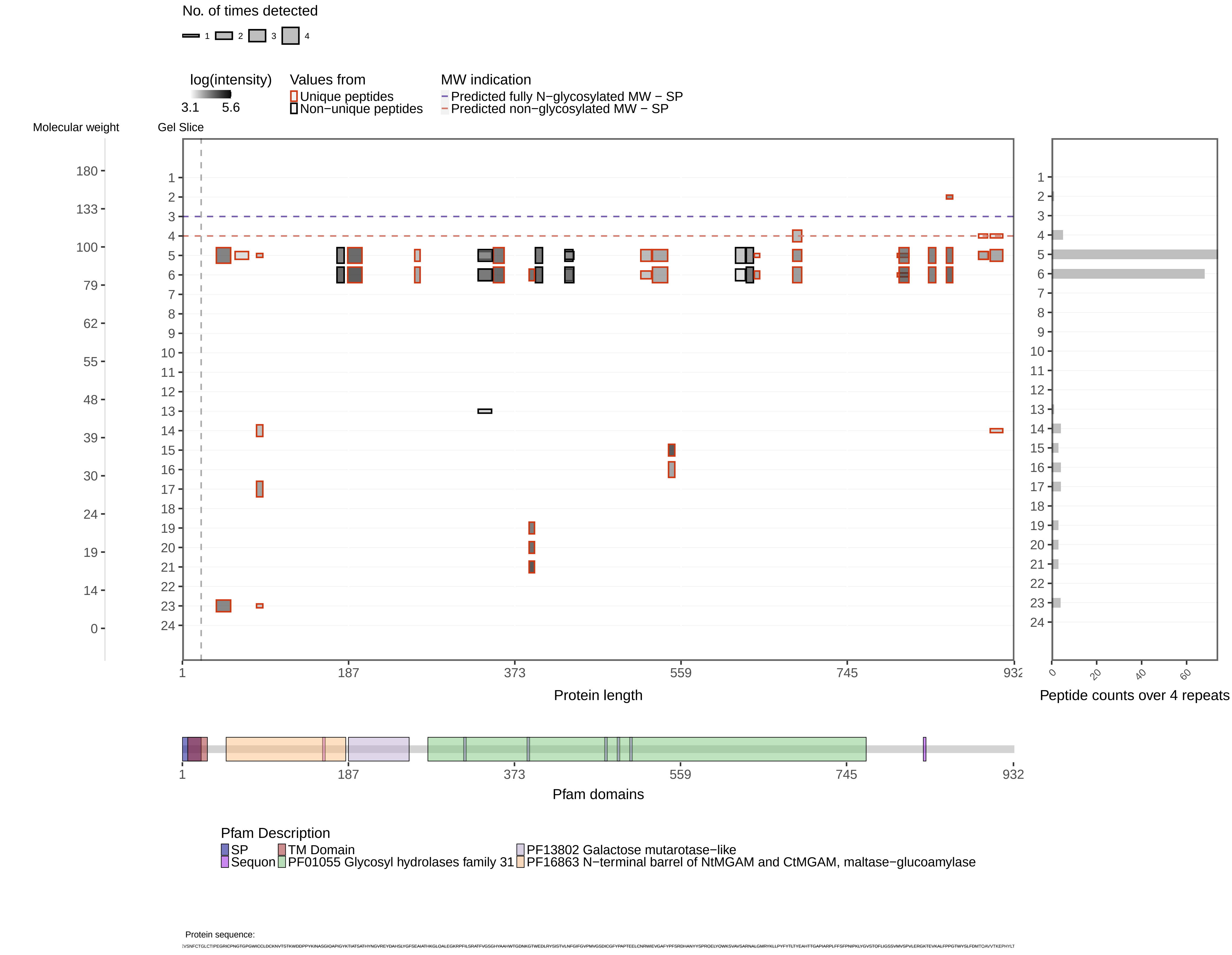

### NbL18g01100.1 glutamine−−tRNA_ligase [MW = 90.51 kDa]

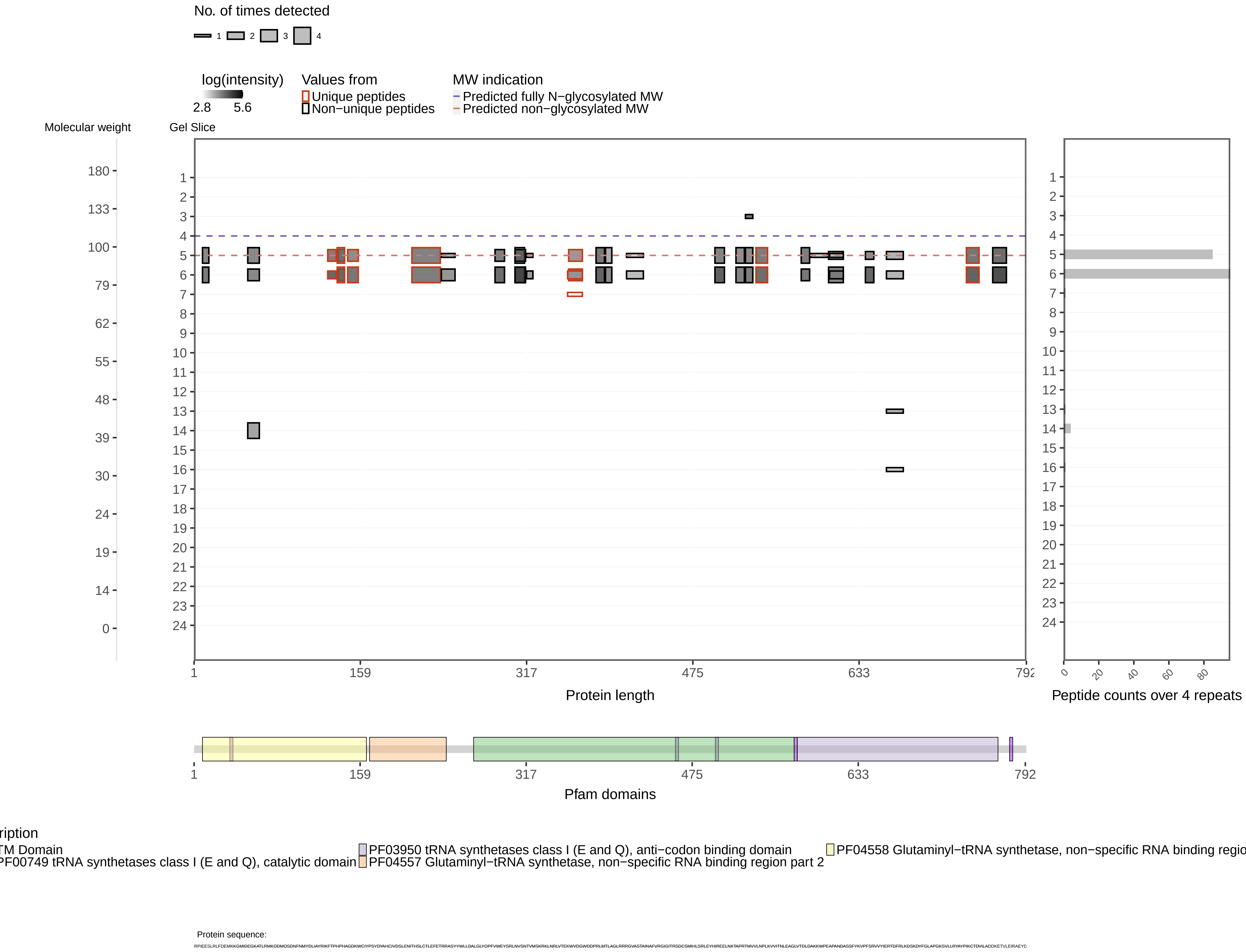

NbL18g01080.1 cysteine−−tRNA_ligase_2_cytoplasmic−like [MW = 63.8 kDa]

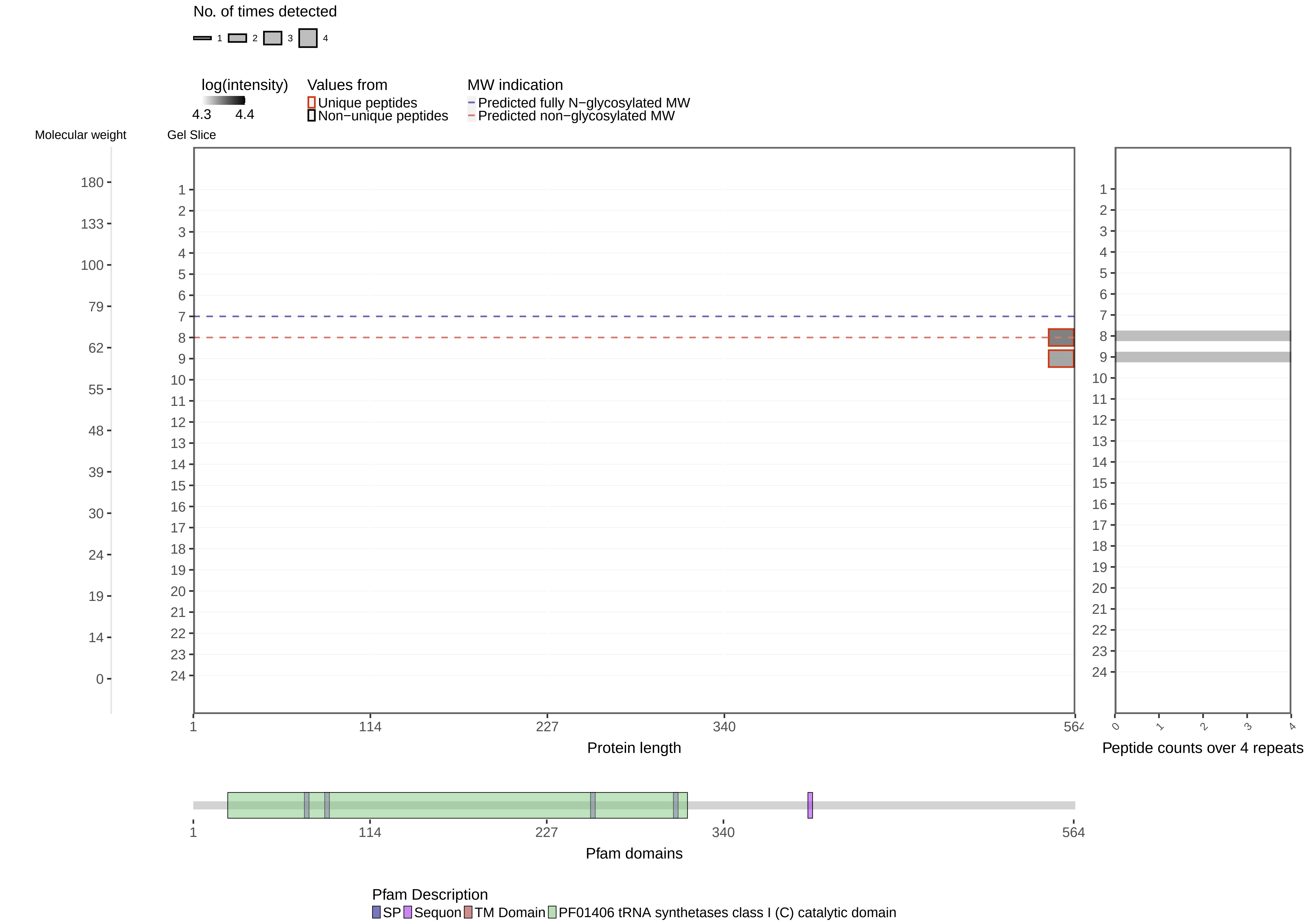


Protein sequence:

IDDEKMSKSLGNFFTIREVTRLYHPIALRYFLLGMHYRSPVNYSISOIEIASESVFYLYOTLHDCEEALSVFOOGTETETEVRGVRVSPOAOECIKKLRNELETKLSDDLHTPTILNAALOEALRLMNSSLNMLKKKOOKKOOLSAVLSLAELLKEVKAVLDVLGLLSSSTCAEVLOOFKERALKRAELTEEDILHAIEERTLARKNKEFARSDOIRSDLVAKGIALMDIGTETVWRPCVRPEPEOPAAPVOOEOSAAAVOKEKPVAPSOOKOSAAAPOKEOPAAPLOOEOPAVRI

### NbL18g00530.1 fruit_protein_pKIWI502−like [MW = 33.43 kDa]

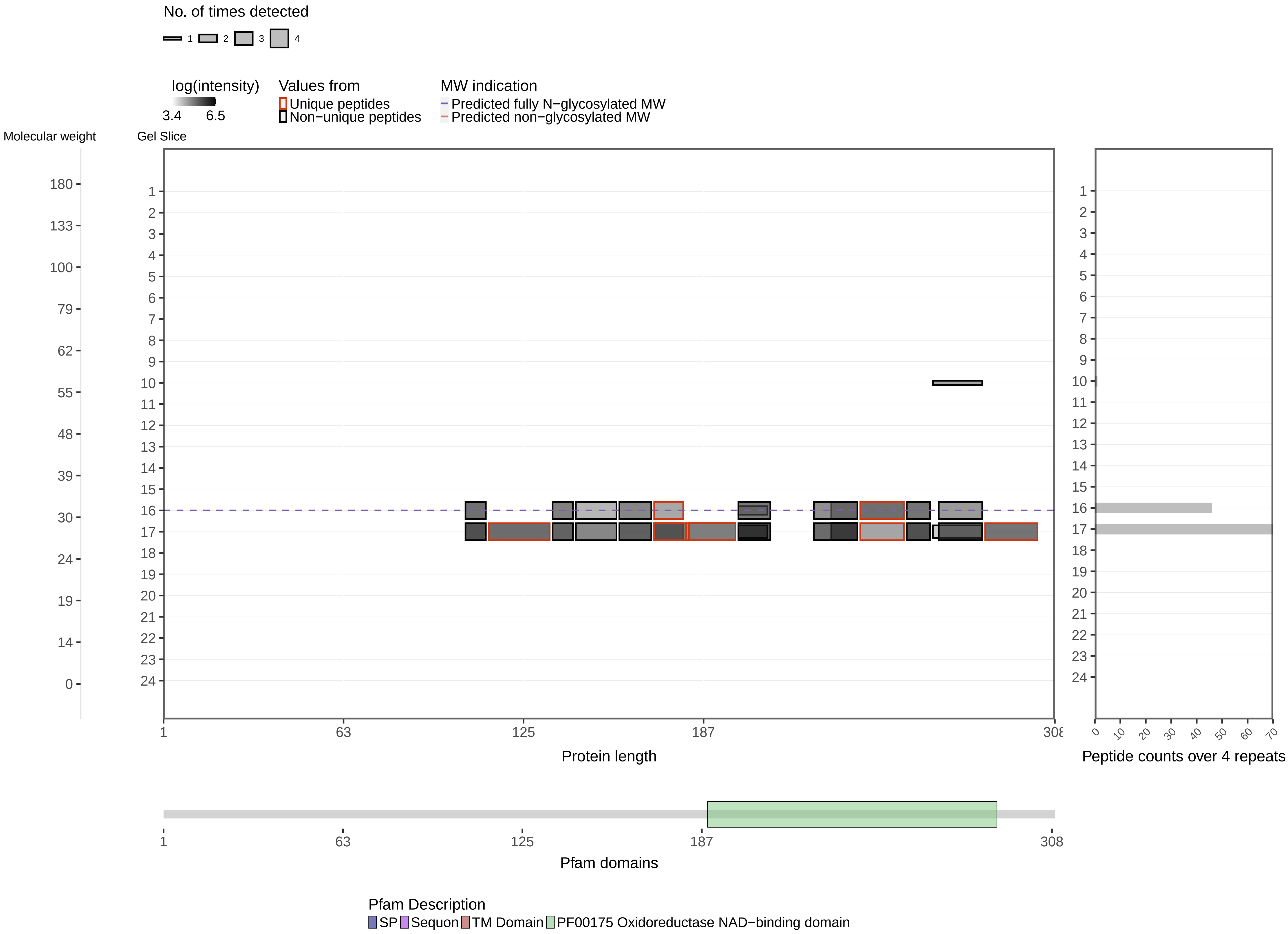


Protein sequence:

SISGSTAELLCGLQKGDVVELSQVMGKGFDLDQISPPKKYQTILIFATGSGISPIRSLIEAGFSADKRSDVRLYYGARNLKRMAYQDRFENWASSGVKVVPVLSQPDDAWTGERGFVQAAFARAKNIFSPQSTGAVLCGQKQMAEEVTALLLADGVSTEKILKNF*

NbL18g00520.1 ATP_synthase_subunit_beta_mitochondrial−like [MW = 66.27 kDa]

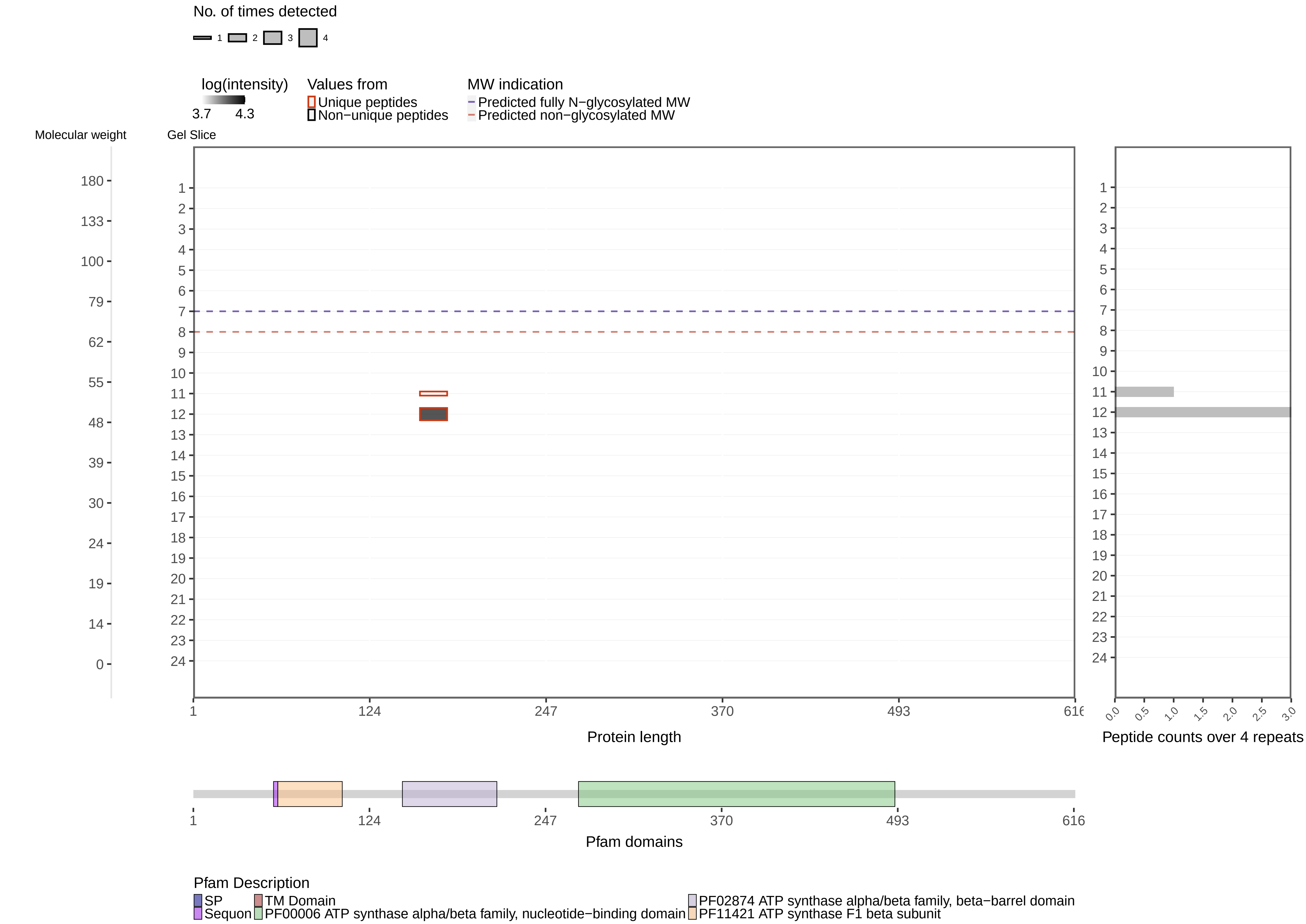


Protein sequence:

TVLIMELINNVAKAHGGFSVFAGVGERTREGNDLYREMIESGVIKLGDKQSESKCALVYGQMNEPPGARARVGLTGLTVAEHFRDAEGQDVLLFIDNIFRFTQANSEVSALLGRIPSAVGYQPTLATDLGGLQERITTTKKGSITSVQAIYVPADDLTDPAPATTFAHLDATTVLSRQISELGIYPAVDPLDSTSRMLSPHILGEDHYNTARGVQKVLQNYKNLQDIIAILGMDELSEDDKMTVARARKIQRFLSQPFHVAEVFTGAPGKYVDLKESINSFQGVLDGKYDDLSEQSFY

NbL18g00010.1 60S_ribosomal_protein_L31−like [MW = 13.91 kDa]

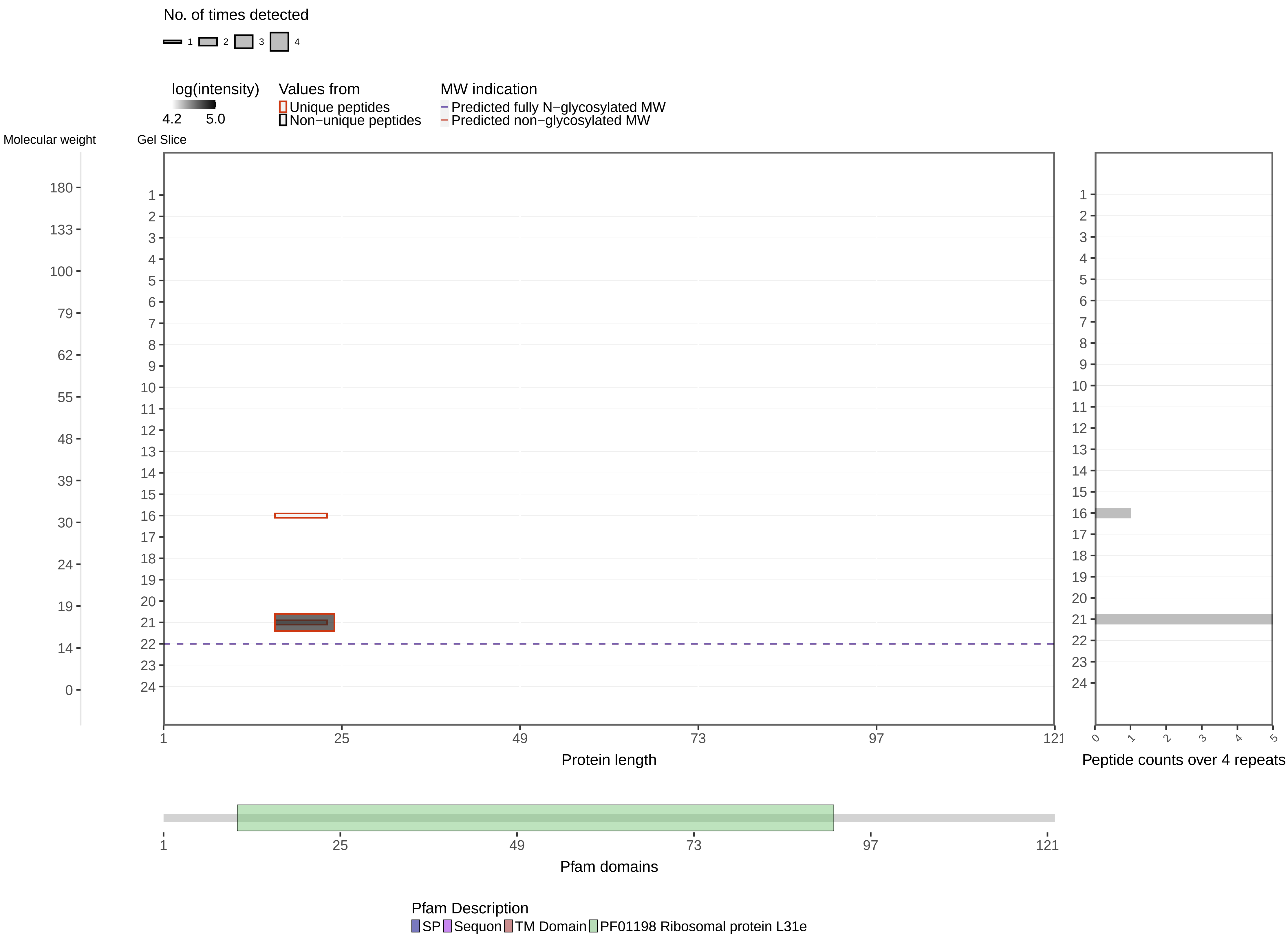


Protein sequence:

KAMGTKDVRVDVKLNKHIWSRGIRSVPRRVRVRIARKRNDDEDAKEELYSLVTVTEIPPEGLKGLGTKIIEDED*

### NbL17g28270.1 nuclear_pore_complex_protein_NUP98A [MW = 105.87 kDa]

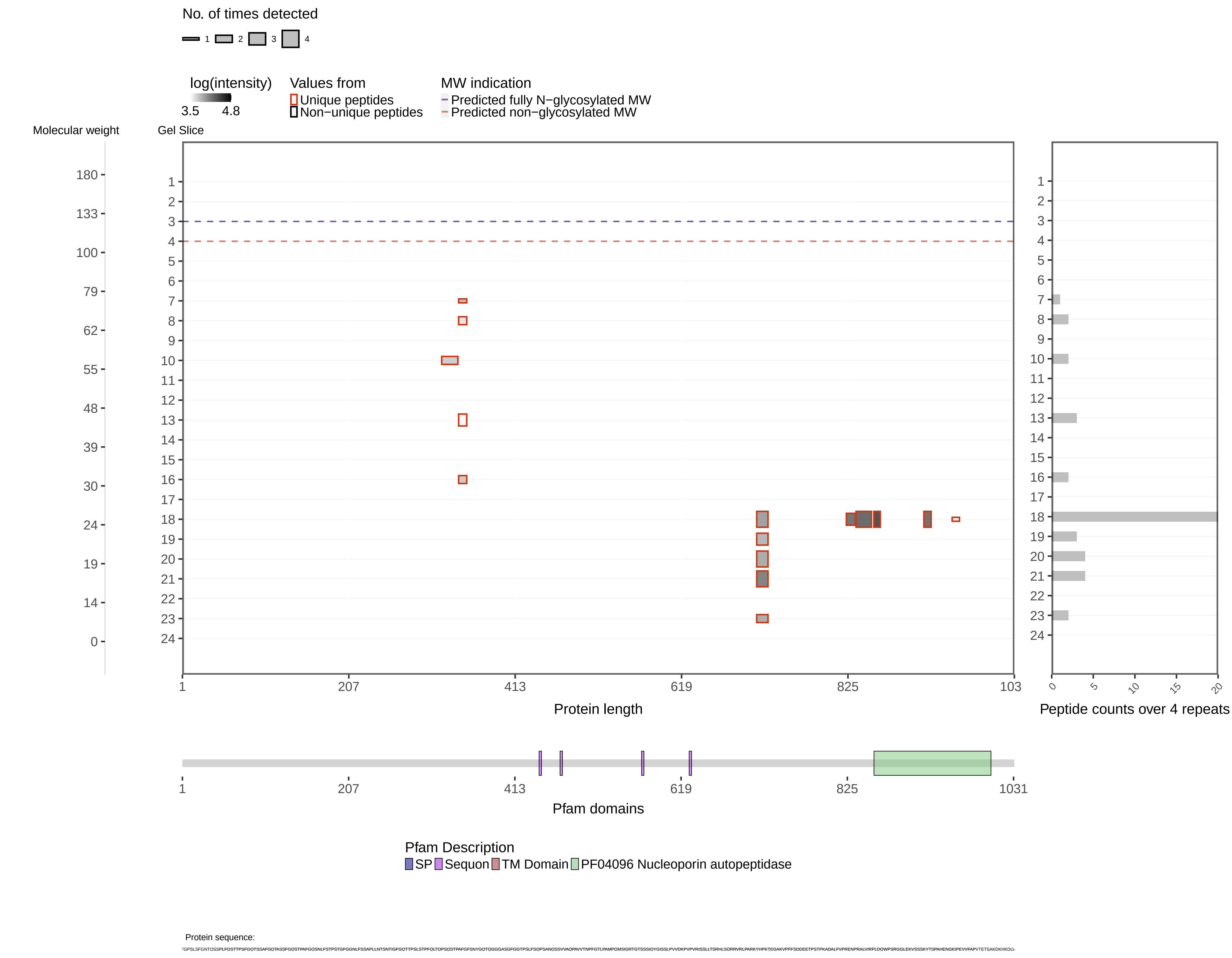

### NbL17g28220.1 soluble_inorganic_pyrophosphatase_PPA1−like [MW = 24.12 kDa]

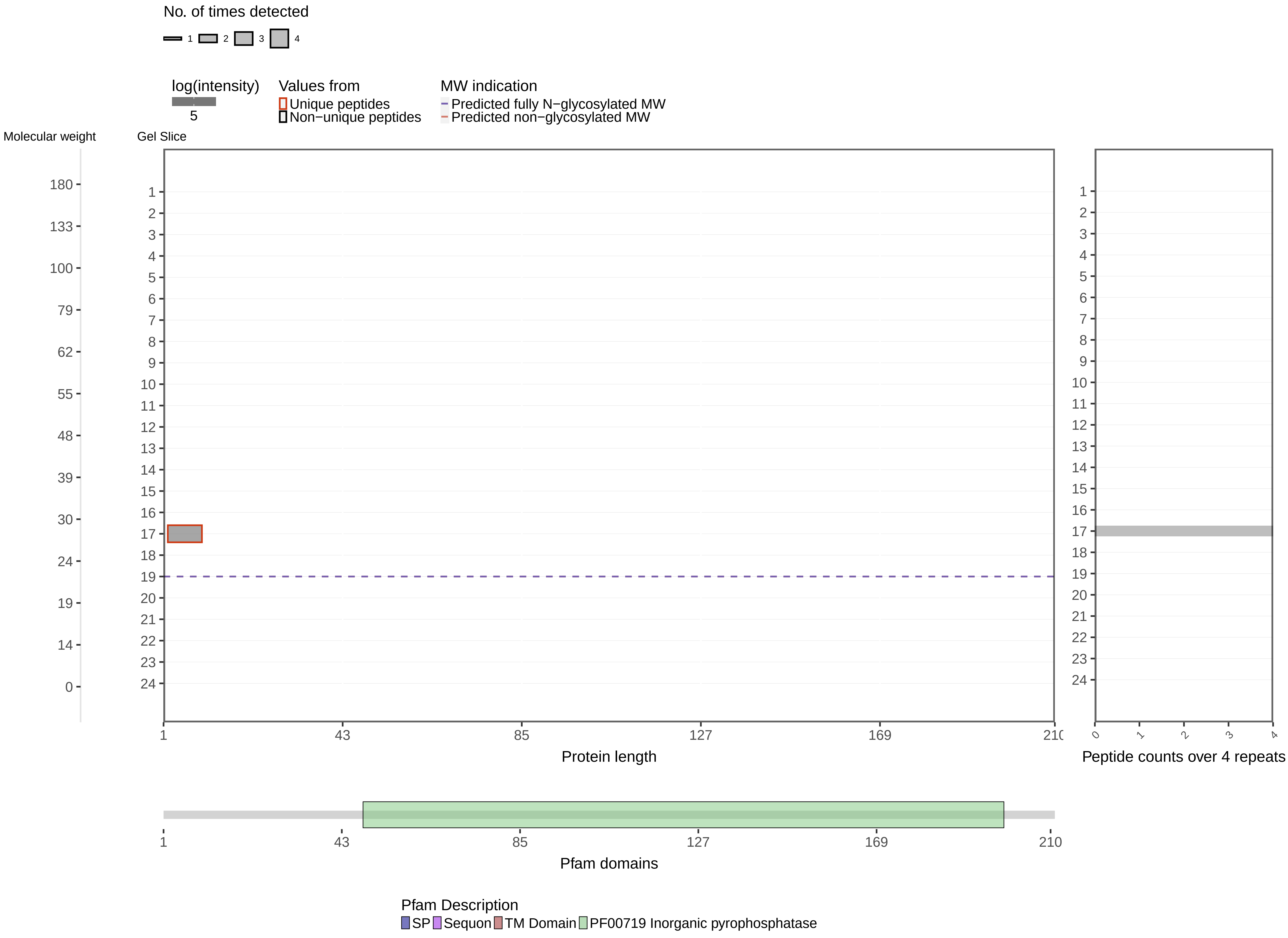


Protein sequence:

TLCEDNDPMDVLVLMQEPVLPGCFLRARAIGLMPMIDQGEKDDKIIAVCADDPEYRHYTDINQLAPHRLAEIRRFFEDYKKLENKEVAVNEFLPPNTAVDAIQYSMDLYAEYILQTLRK*

### NbL17g28120.1 putative_glucose−6−phosphate_1−epimerase [MW = 34.21 kDa]

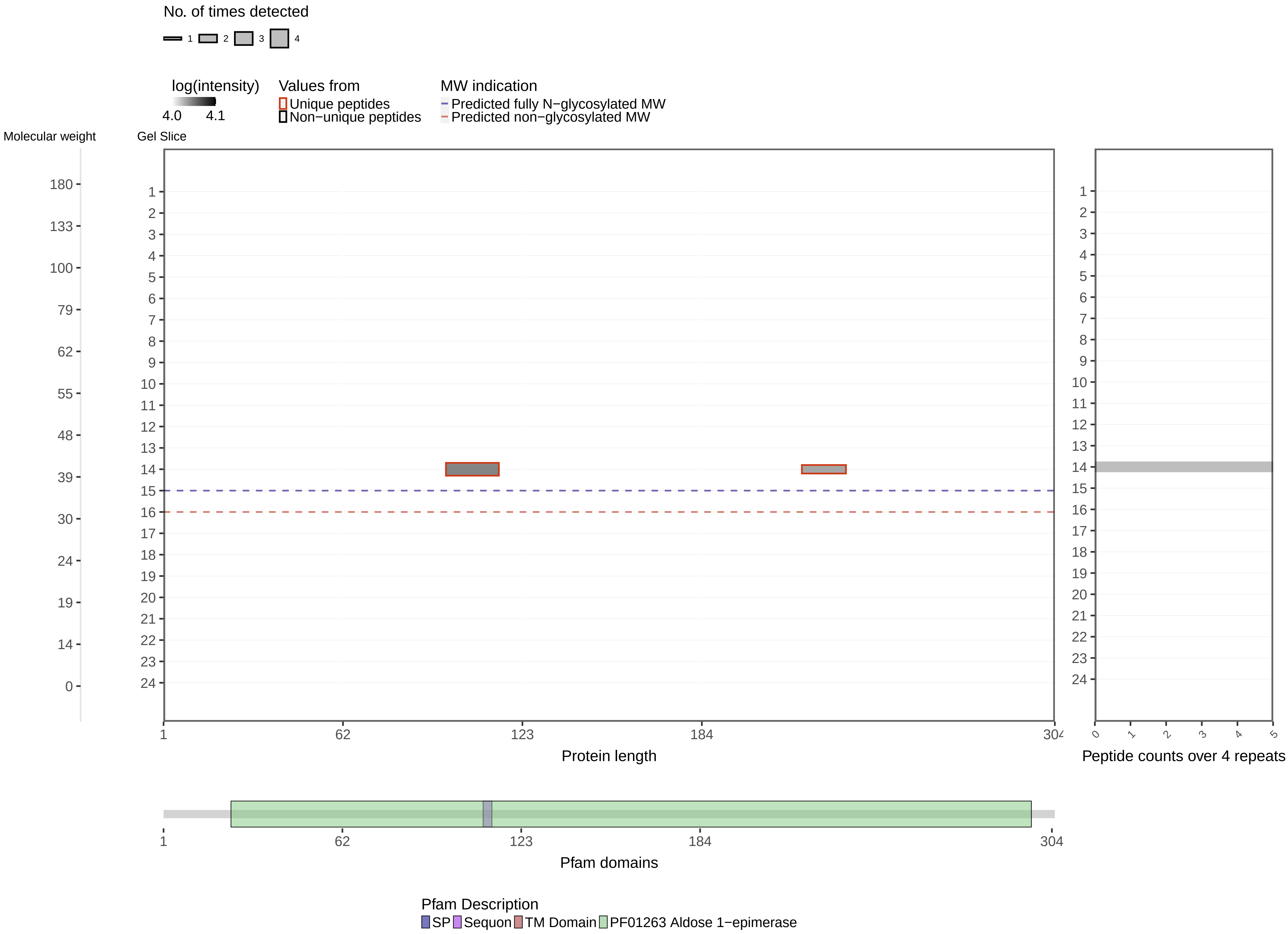


Protein sequence:

EFRLRVTLAVDGSLTLISRIRNVNCKPFSFSIAYHTYFALSDISEVRLEGLETLDYLDNLCNRERFTEQGDALTFETEVDRVYLTSSDVIAIFDHEKKRTFVIKREGLPDVVVWNPWEKKSKTIADFGDDEYRHMLCVDGAAIEKPITLKPGEEWTGRLELSVMPSS*

NbL17g28080.1 myosin−11−like [MW = 159.57 kDa]

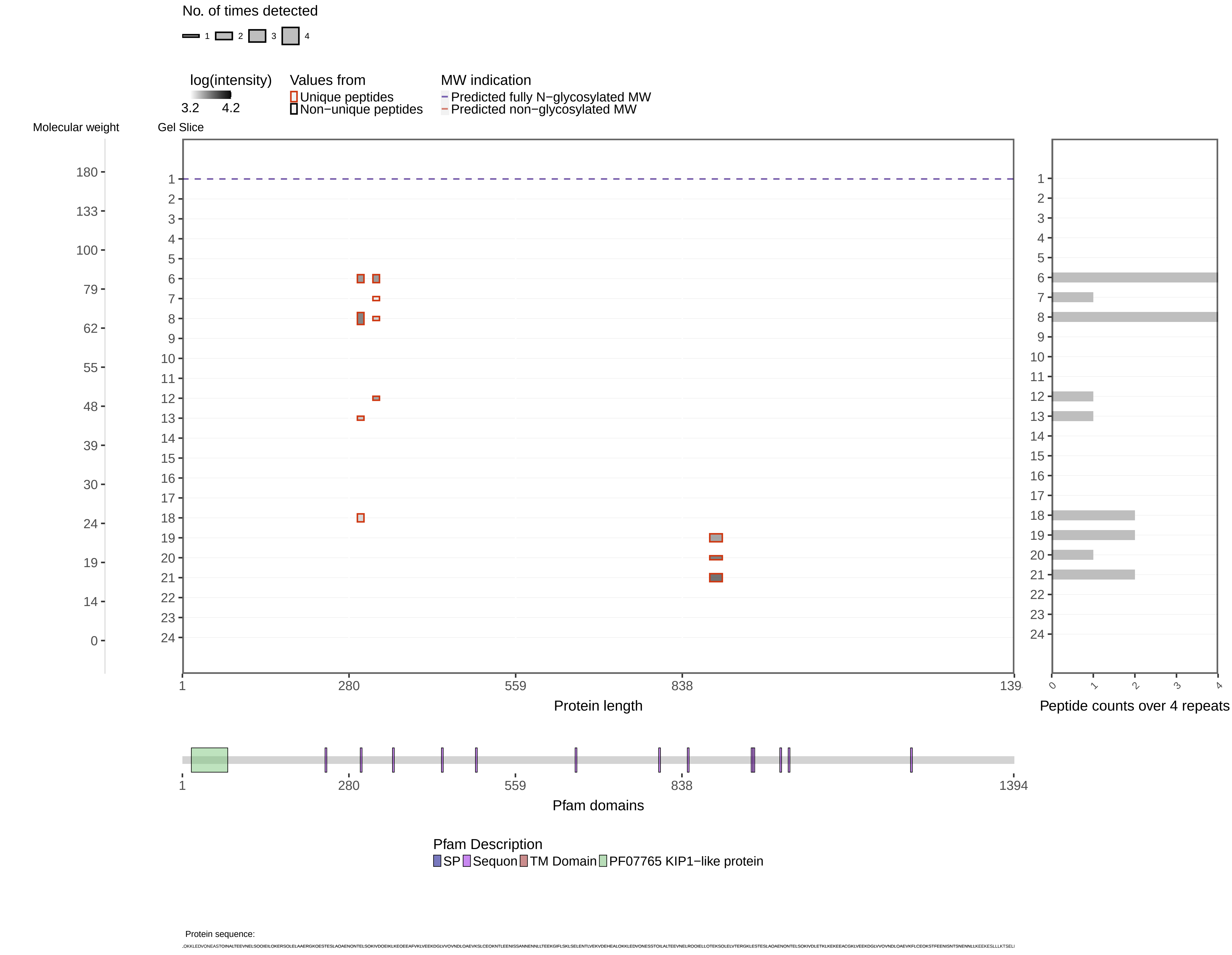

### NbL17g27940.1 probable_polygalacturonase [MW = 53.73 kDa]

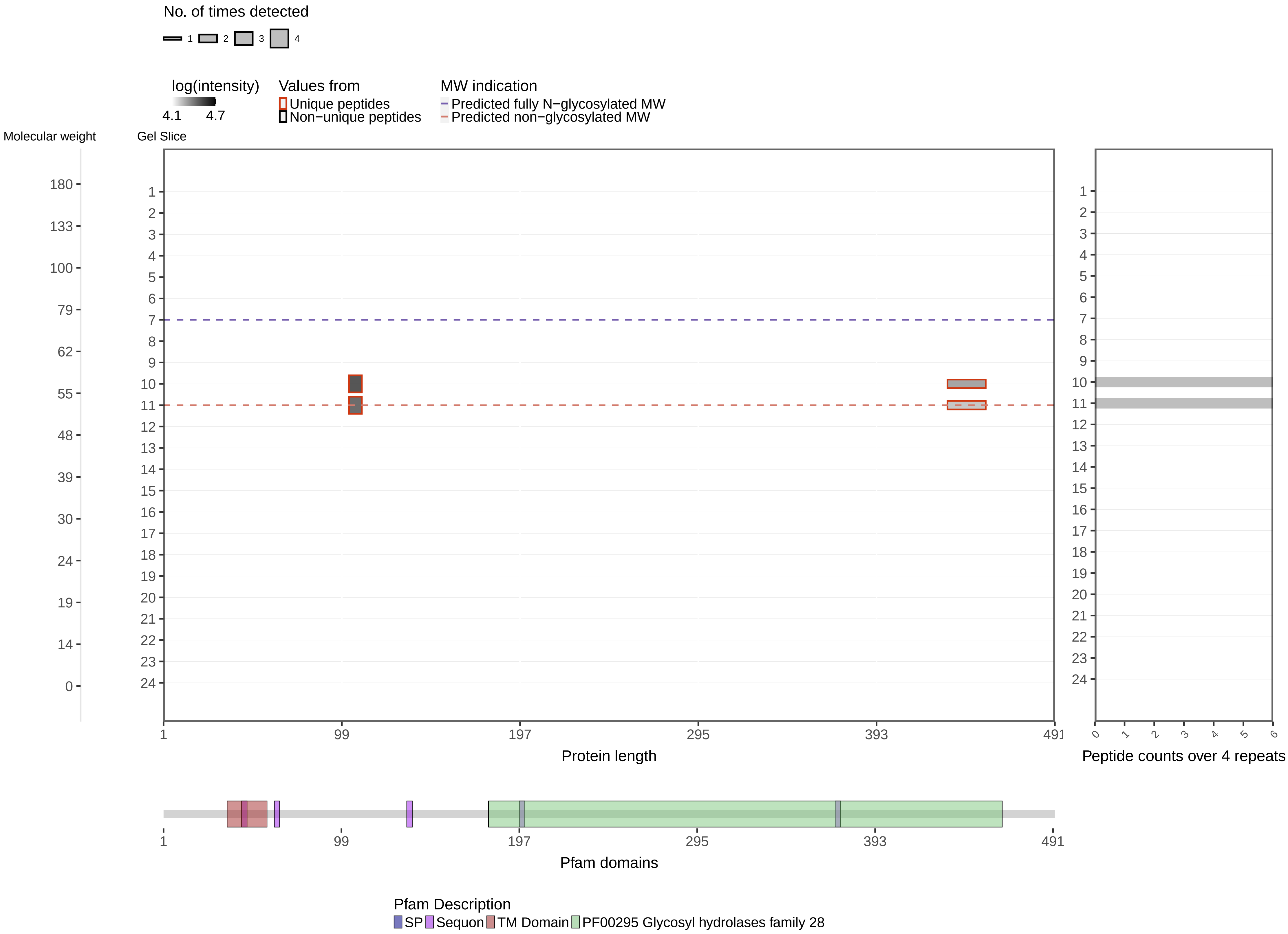


Protein sequence:

RGIIISNVIFKNSPFWNIHPVYCSNVIIRYVTILAPADSPNTDGIDPDSSSHVCIEDSYISVGDDLVAVKSGWDQYGIAYGRPSHGITIRRITGSSPFAGIAVGSETSGGIVDVLAEHINLFNMGVGIHIKTNIGRGGMIRNITVSNVYMENTRTGIKIAGDVGDHPDEKFNPNALPIVKGIKIKDVWGEKVLQAGLIRGLKNSPFSGICLSNINLHGNKPGPRNSPWKCSDVSGAAIQVSPWPCSELTSSQQTGACSTSF*

### NbL17g27850.1 long_chain_acyl−CoA_synthetase_4−like [MW = 25.45 kDa]

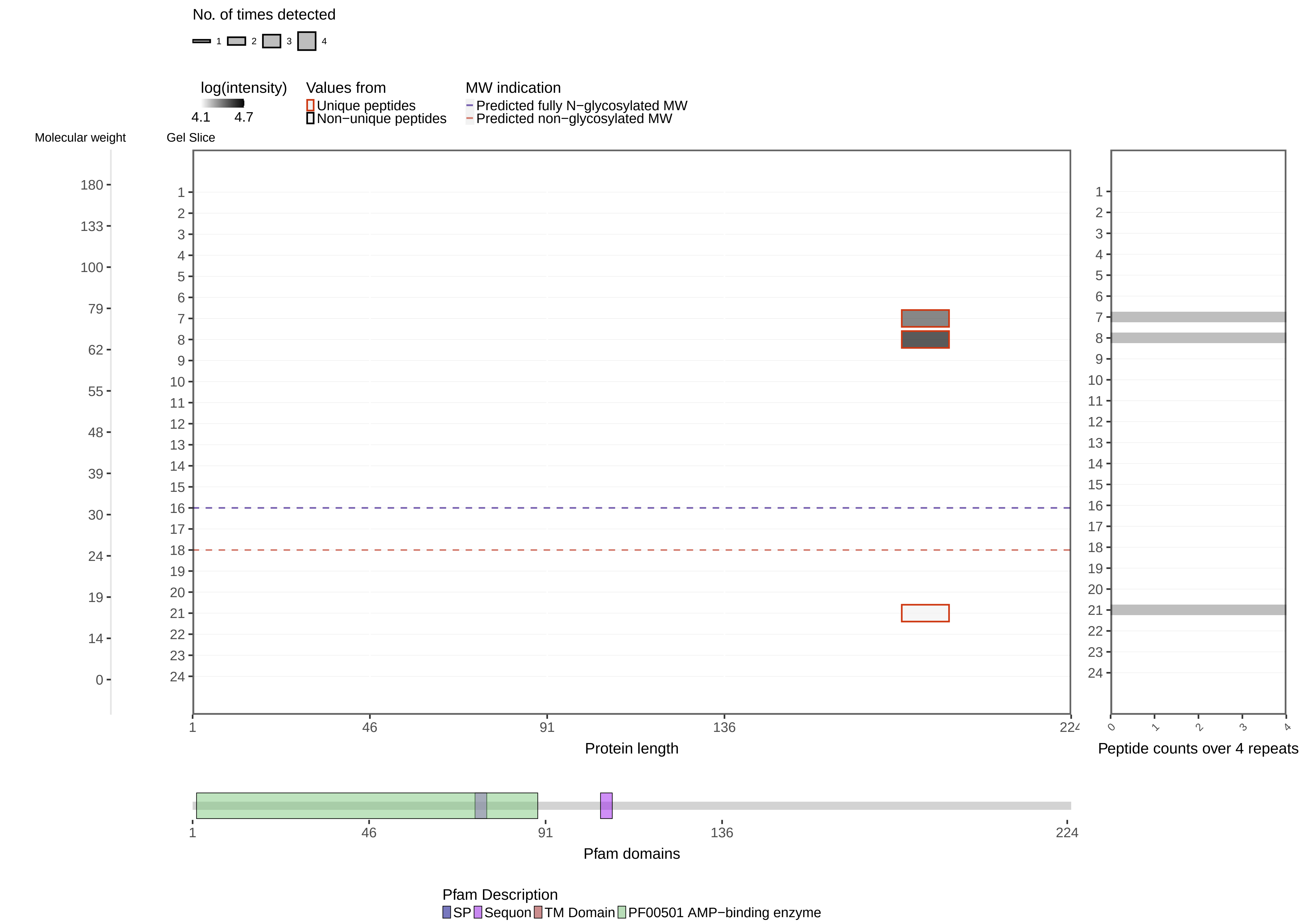


Protein sequence:

NLENIYGNSSLIDSIWIYGNSFESFLVAVLNPNQRAVEQWAQQNGISGDFDSLCENPKVKEYILGELTKAGKEKKLKGFEFIKAVHLDAQPFDMERDLLTPTFKKKRPQLLKYYKVISLLICWWML*

### NbL17g27770.1 nucleoside_diphosphate_kinase_3−like [MW = 25.75 kDa]

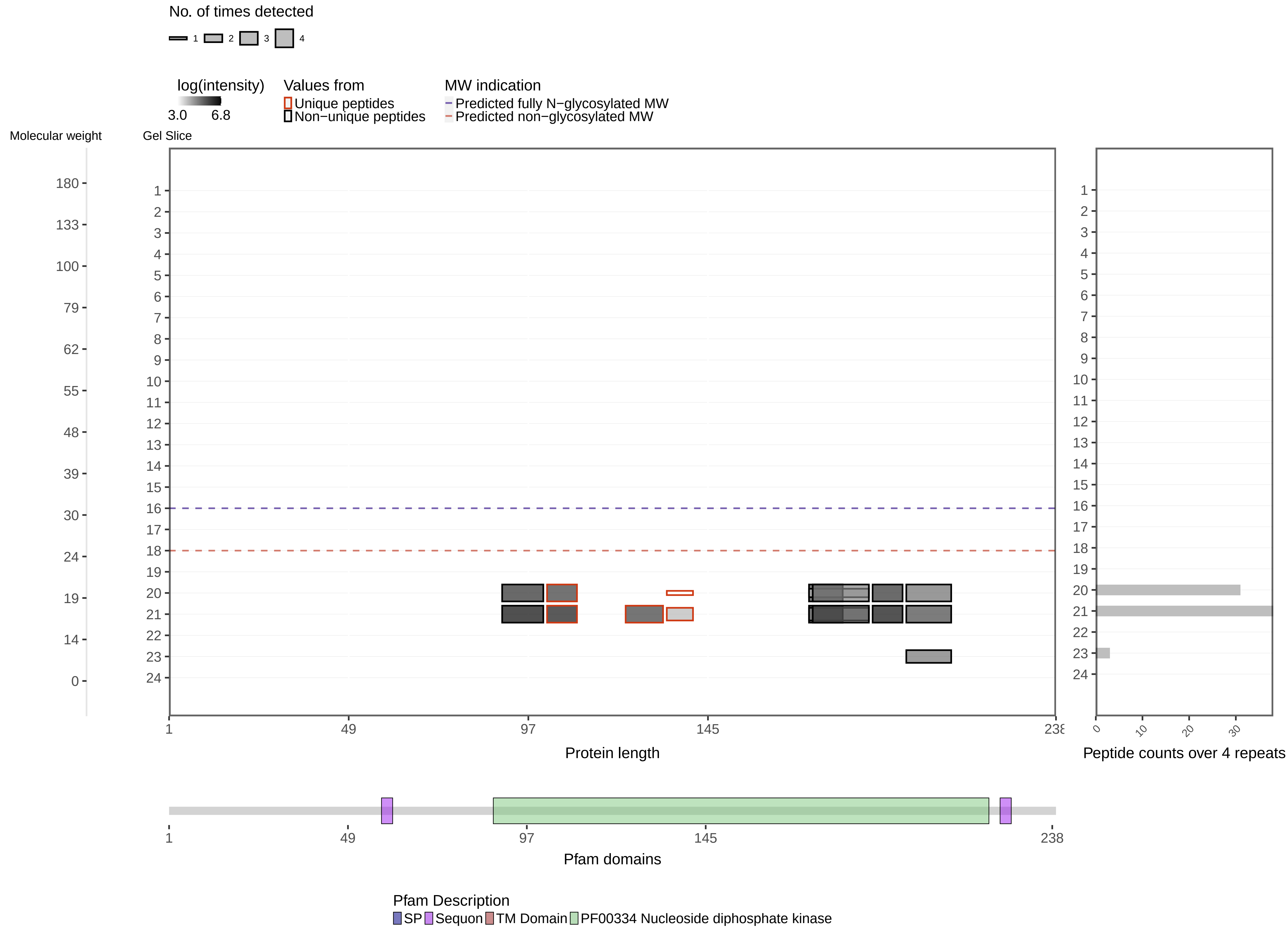


Protein sequence:

SEIIARFERKGFKLVAVKIVIPSKEFAOKHYHDLSERPFFNGLCDFLSSGPVLAMVWEGEGVIRYGRKLIGATDPOKSEPGTIRGDLAVVVGRNIIHGSDGPETAKDEINLWFKPEELVNYTSNSERWVYGDN*

NbL17g27710.1 uncharacterized_LOC109210112 [MW = 22.28 kDa]

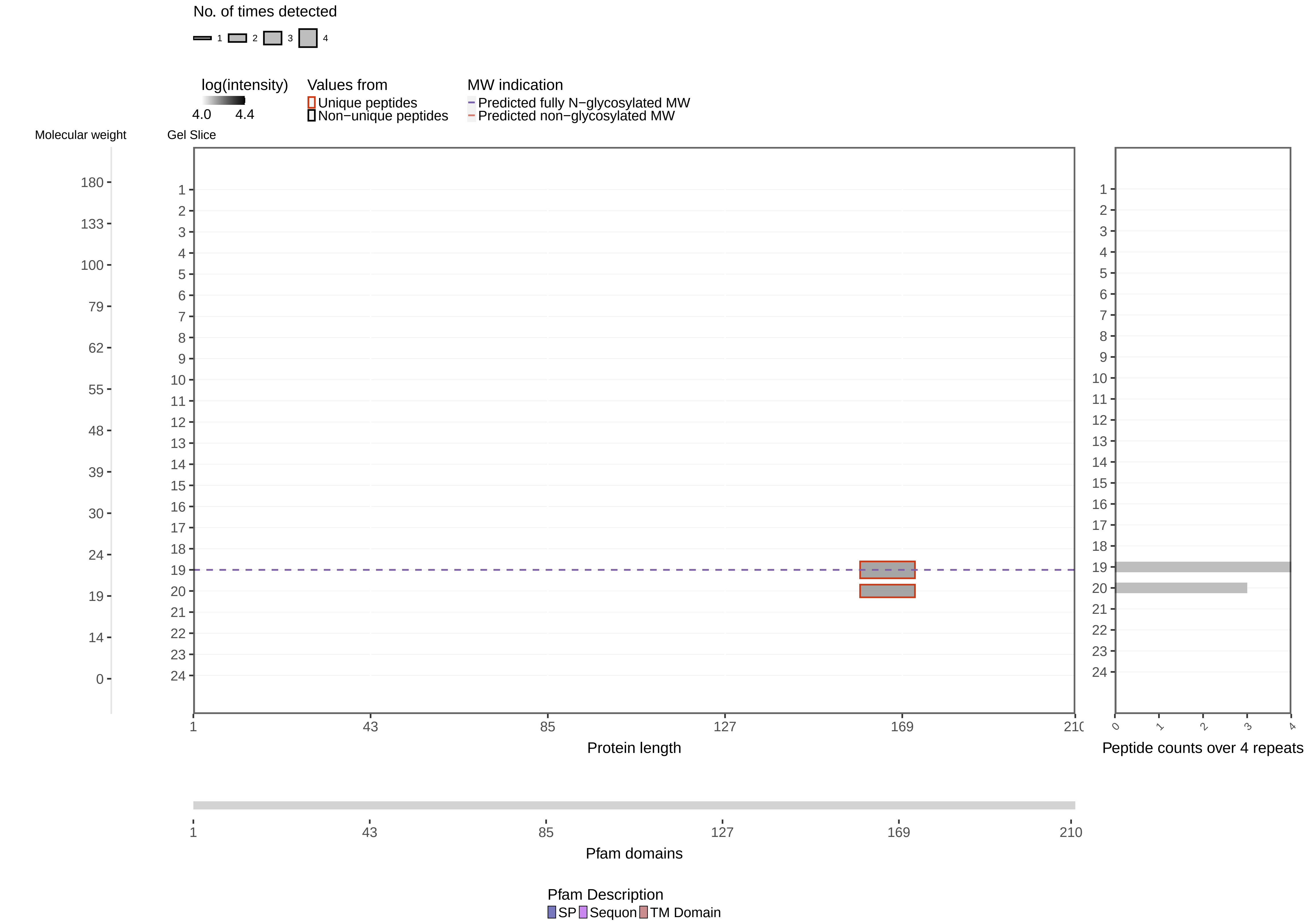


Protein sequence:

INQESTSVLPPGACQACGREEIERGCNGEGRIQGGIATVPGFGWWPIKAYRPCPAFVASGGRYRRFGQSMDEVVSGSGRGGRASSAGTDAEVQSSLFCRKPSRFGLNLNGIAPVRH*

### NbL17g27580.1 signal_recognition_particle_19_kDa_protein−like [MW = 15.29 kDa]

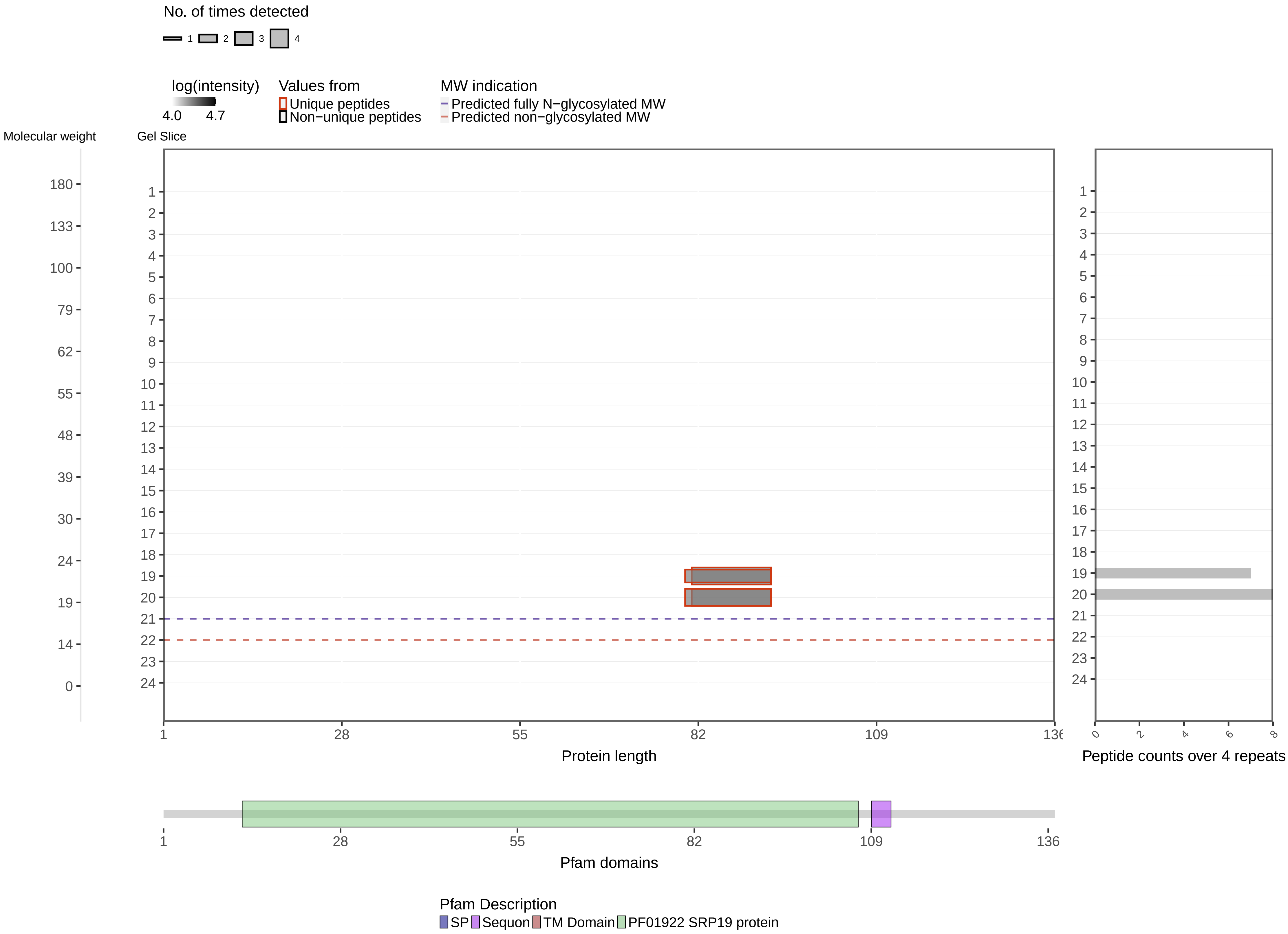


Protein sequence:

AIELDKAYPRDFMQRGRVRVLLKKEDGTTYNPAISSRKQLMIHVAELVRKHPNRTKKQEPASSSAAGSSKSGKGGKKKR*

### NbL17g27510.1 luminal−binding_protein [MW − SP = 70.53 kDa]

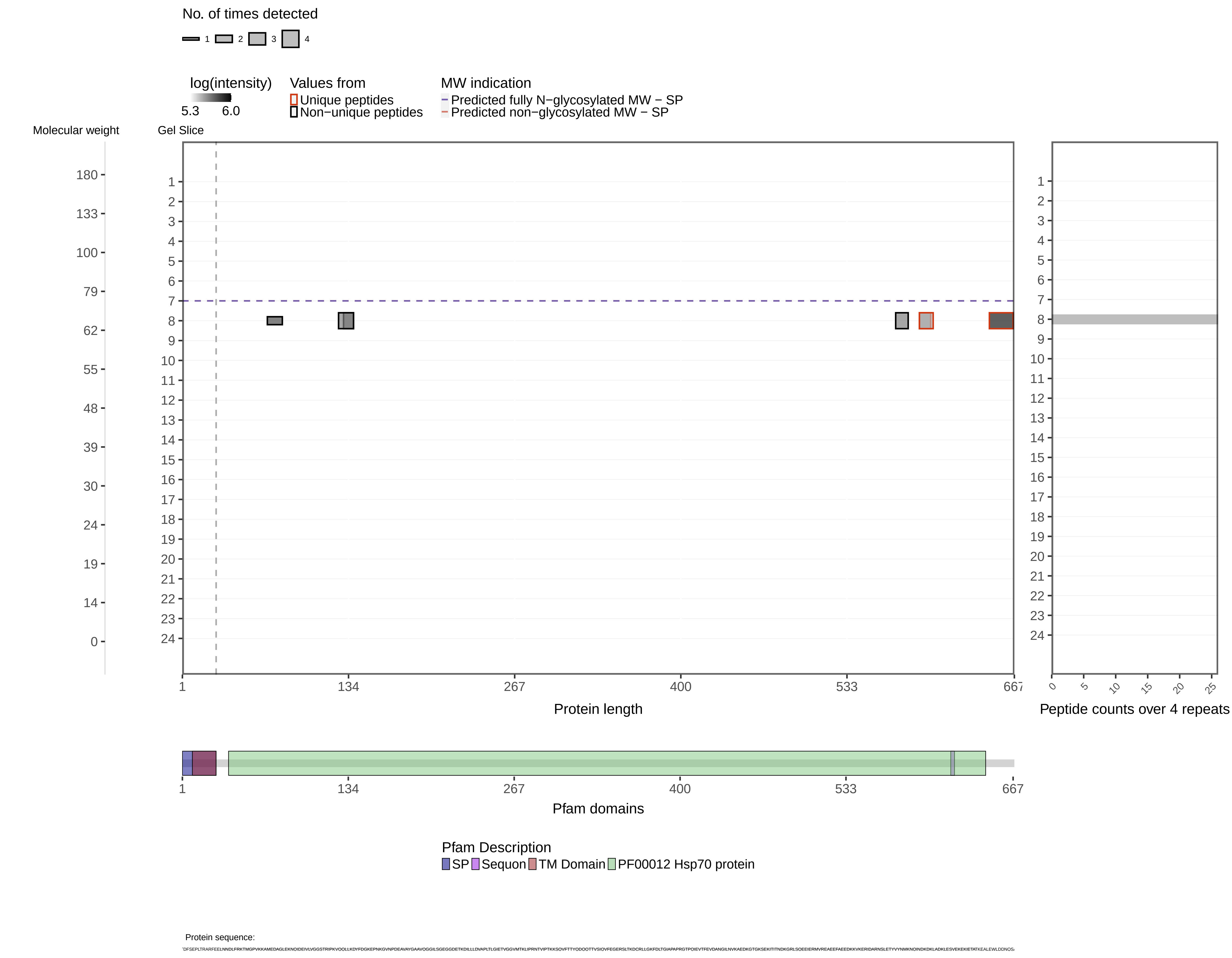

### NbL17g27460.1 probable_trehalase [MW − SP = 63.48 kDa]

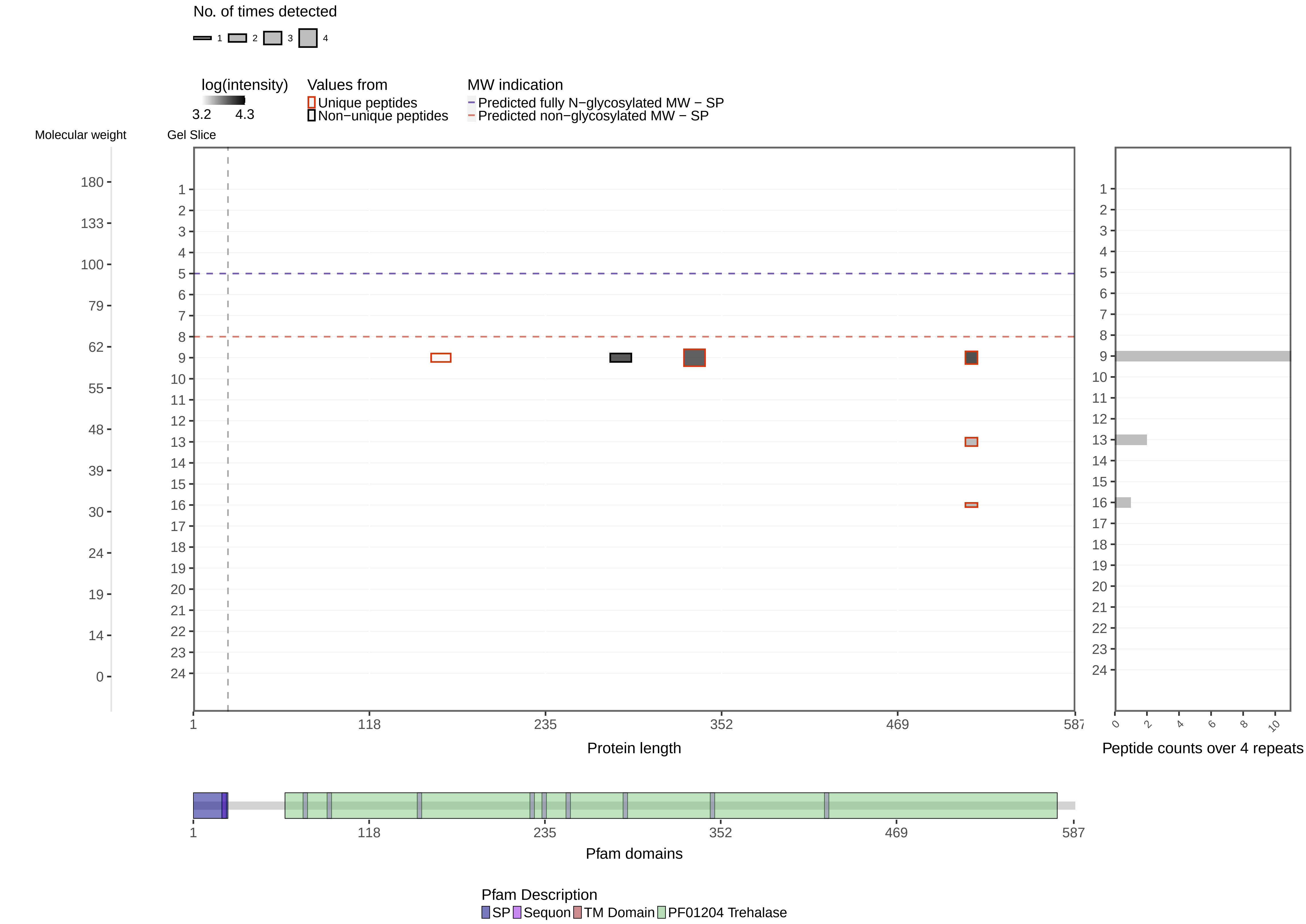


Protein sequence:

QGSNHSLSRYYAMWNKPRPESSTIDGETASKLPNICEKRQLYRDVASAAESGWDFSSRWMRNQSDLTTTSTTSIIPVDLNAFLLKMELDIAFLANLVGESSIIAHFTEASQNRQKAMNSILWNTEMGQWLDYWLDNSNTSEDIYKWEDLHQNKQSFASNFVPLWAELLNSDDITTQKVVQSLKSSGLLQPAGIATTLSNTGQQWDFPNGWPPLQHIIIEGLSRSRLEEAKALAEDIAVRWIRTNYVAYKKTGAMHEKYDVTKCGAYGGGGEYVPQTGFGWANGVILALLEE

NbL17g27220.1 uncharacterized_LOC107784855 [MW = 64.25 kDa]

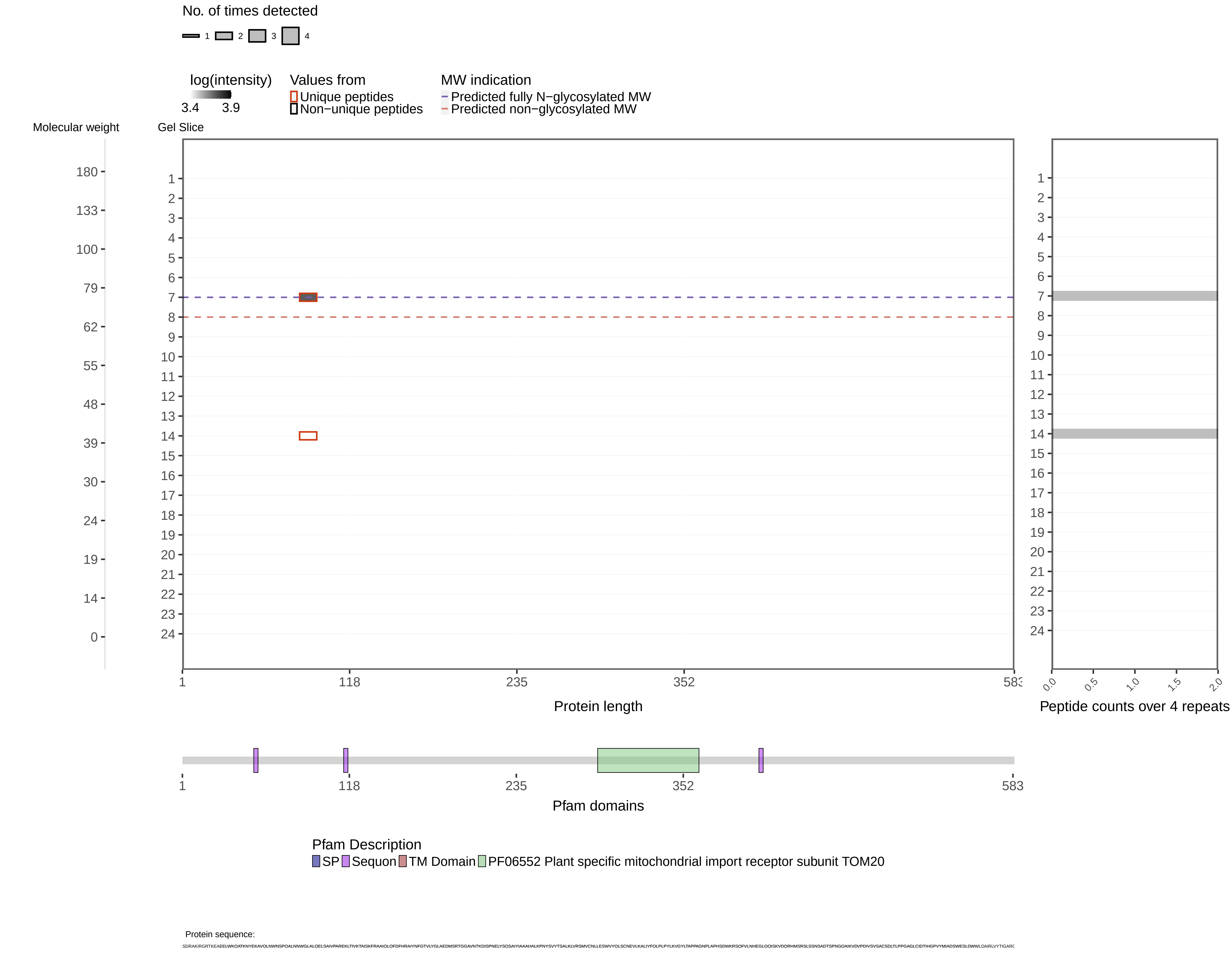

### NbL17g27030.1 granule−bound_starch_synthase_1_chloroplastic/amyloplastic−like [MW = 34.66 kDa]

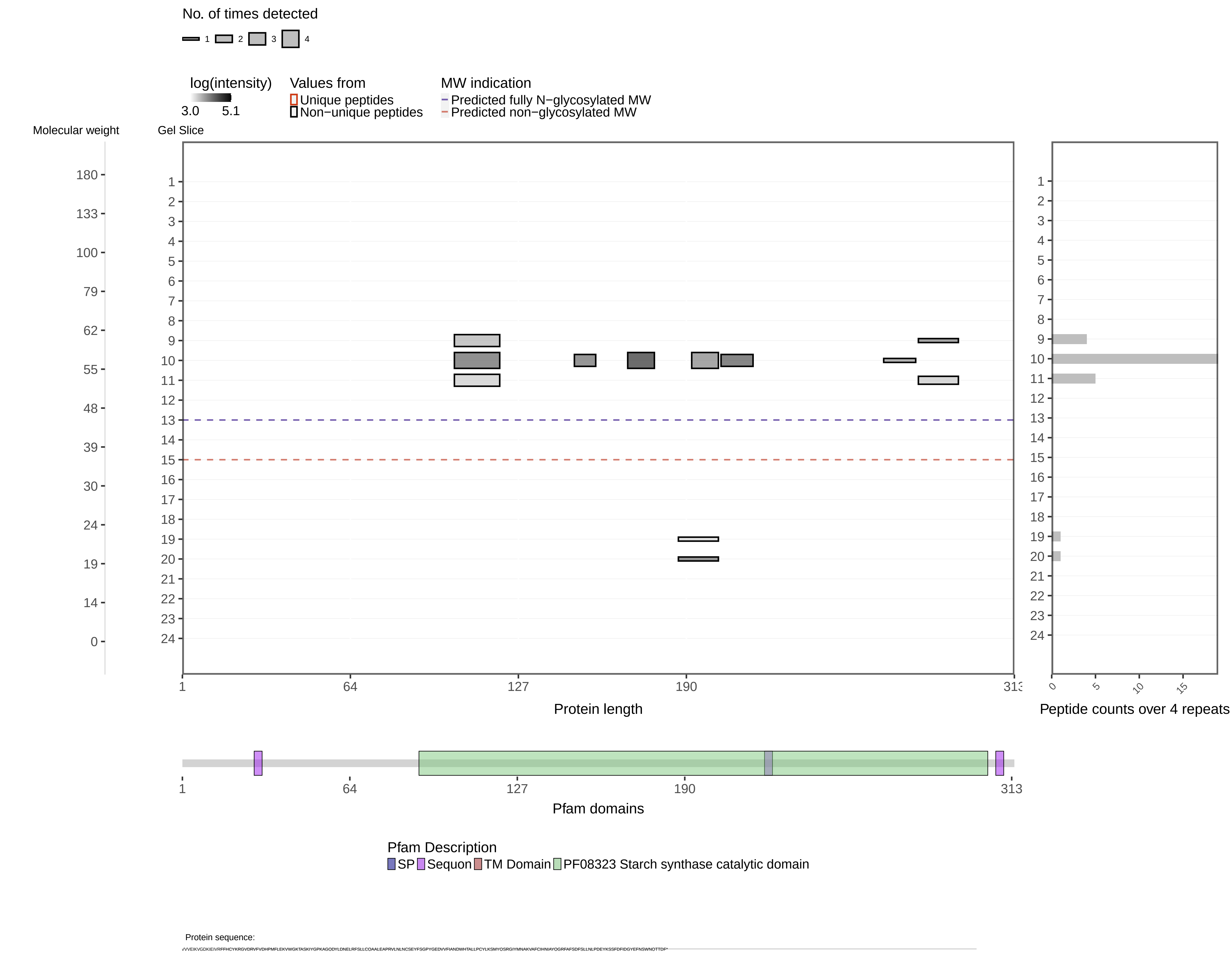

### NbL17g26920.1 reticulon−like_protein_B4 [MW = 30.05 kDa]

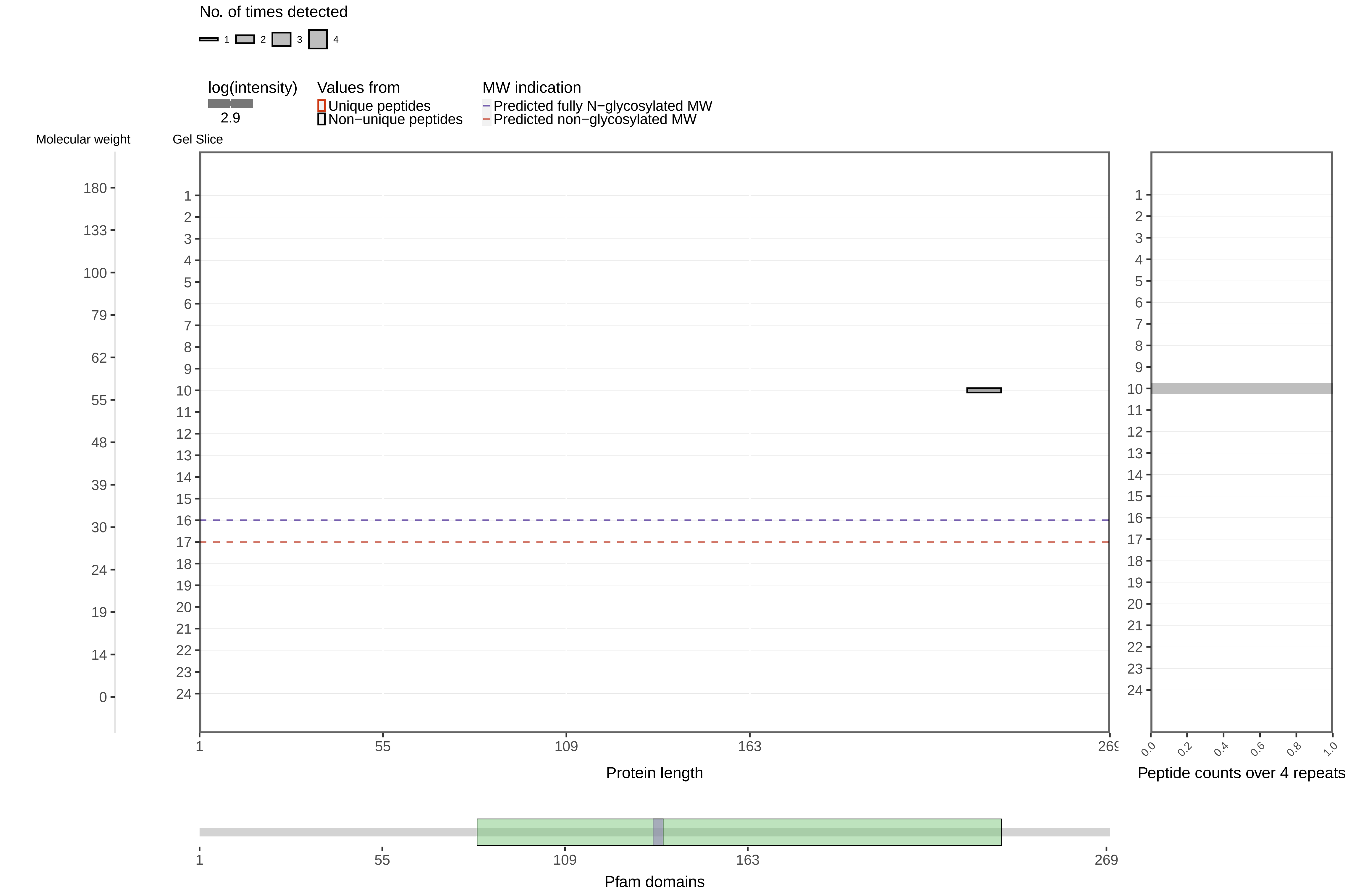


Protein sequence:

LCHGLIVALAGLFLWSNASAFIHKSAPHIPEVHIPEKPVLEFATGLRNEINRAFAVLREIAAGRDLKMFLMVIGGVWILSVLGSCCDFLSLLYIIVVVLHTVPVLYEKYEDOIDSFAEKAFHELKKOYGVFDAKVLSKIPRGPLRDKKKA*

NbL17g26860.1 endoglucanase_18−like [MW = 57.18 kDa]

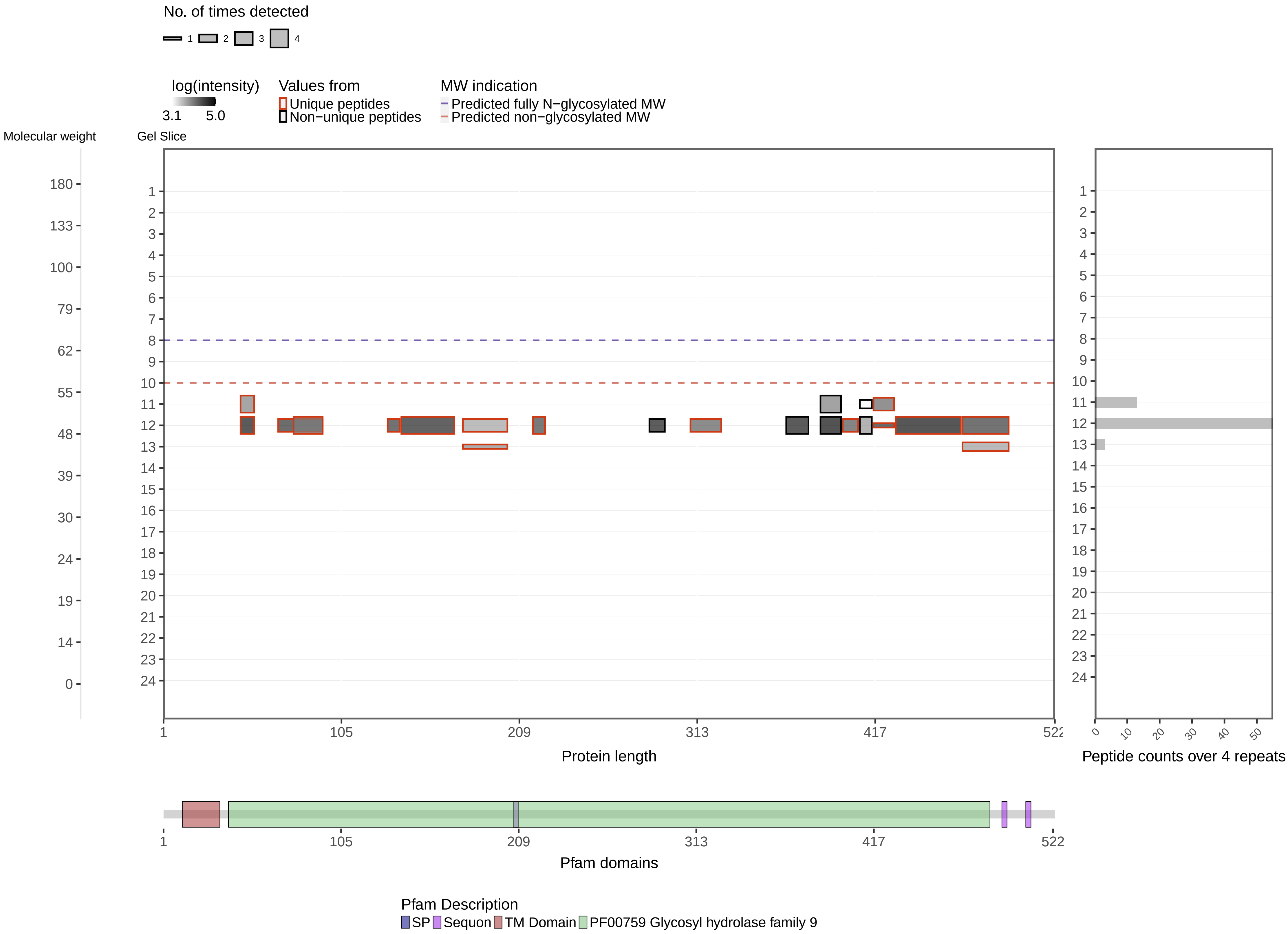


Protein sequence:

VLYKAGGGNNYLNYASSNQGWSQVATEFSWDNKFAGAQTLLAKEFLNGKSSLEKFKKDADSFICALMPGSSSIQIKTTPGGLLYTRDSSNLQYVTGATMVFSMYSKVLETAGKGGVTCGSVTFSTSKIKSFAKSQVDYILGNNPLKMSYMVGFGSKYPTQLHHRASSIPSIYNHPTRVGCNDGYSSWYNSNNPNPNIHVGAIVGGPNSGDQFADSRSDYSHSEPTTYMNAAFVGSVAALINQSKVEQSLQMPQLNKTTFYDRQSTKNILY*

### NbL17g26730.1 mitogen−activated_protein_kinase_homolog_MMK2−like [MW = 43.77 kDa]

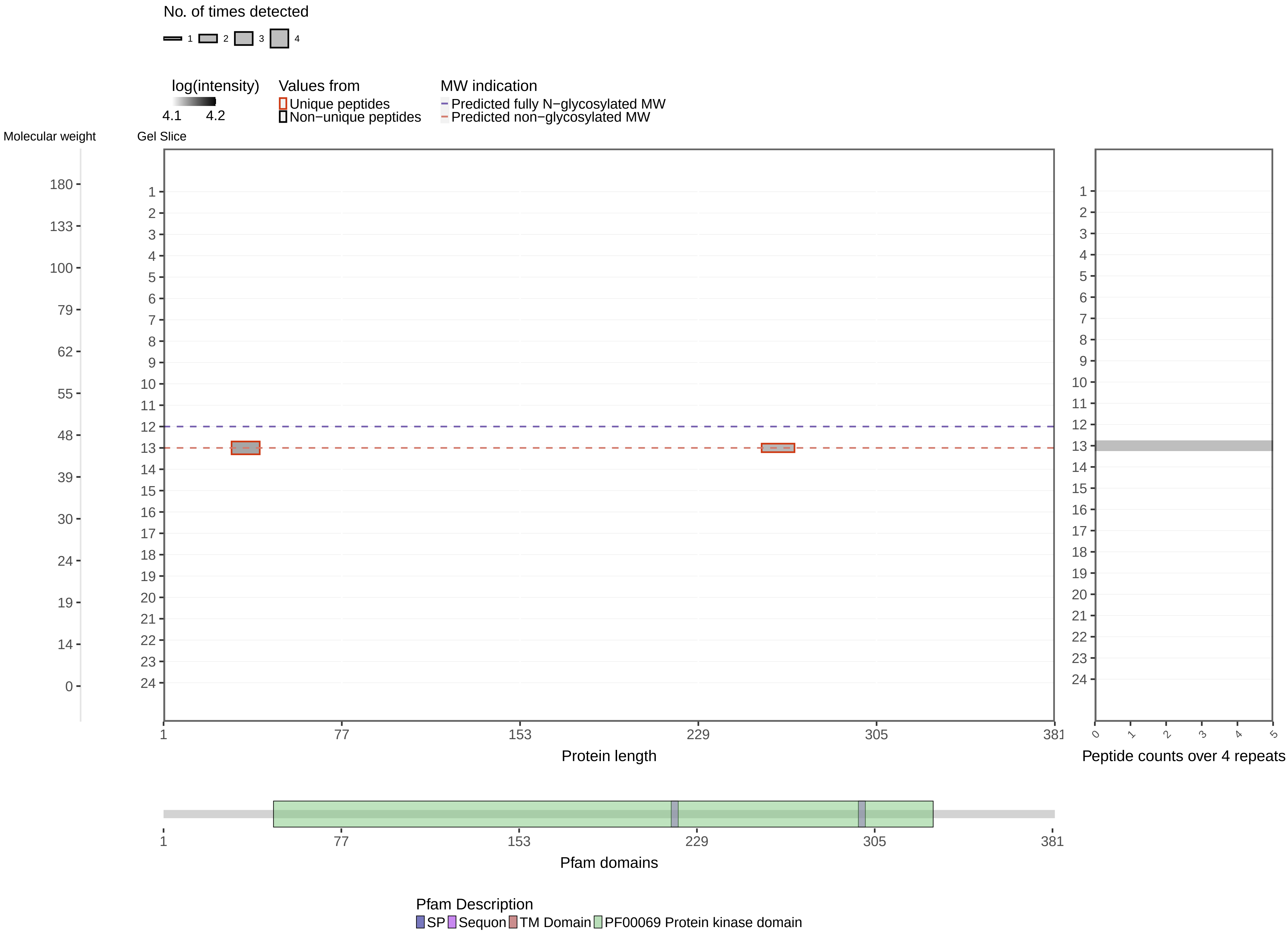


Protein sequence:

NANCDLKIGDFGLARTTSETDFMTEYVVTRWYRAPELLLNCSEYTAAIDVWSVGCILGEIMTREPLFPGKDYVHQLRLITELLGSPDDASLRFLRSDNARRYVRQLPQCPKQQFSARFPNMSTLAIDLLEKMLVFDPTRRITVDEALCHPFLSSLHDINDEPICPRPFSFDFEQPSITEENIKELIWKESVKFYPDFVELKI*

### NbL17g26610.1 beta−adaptin−like_protein_B [MW = 99.81 kDa]

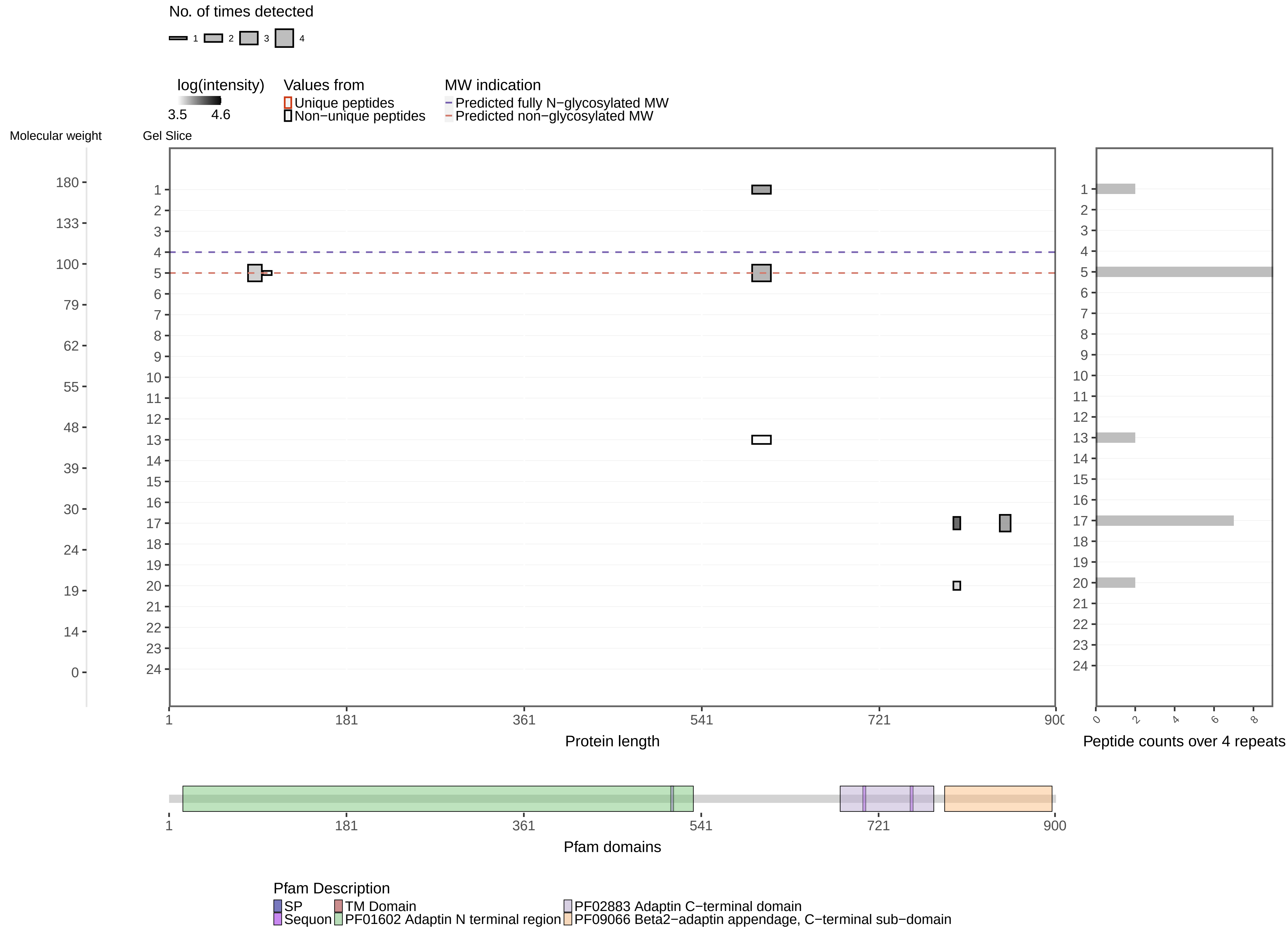


Protein sequence:

NbL17g26530.1 puromycin−sensitive_aminopeptidase [MW = 100.13 kDa]

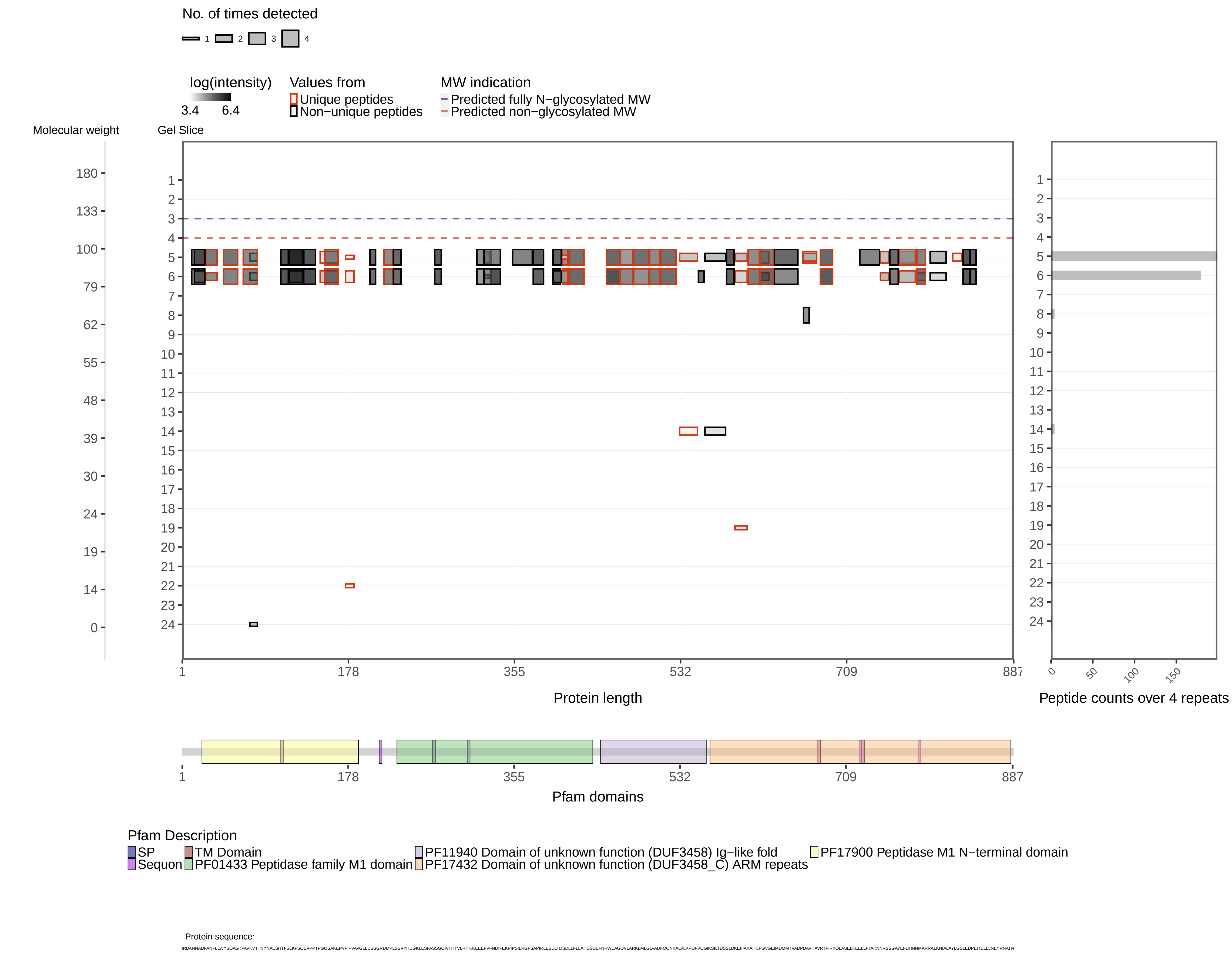

NbL17g26470.1 short−chain_dehydrogenase_TIC_32_chloroplastic−like [MW = 34.51 kDa]

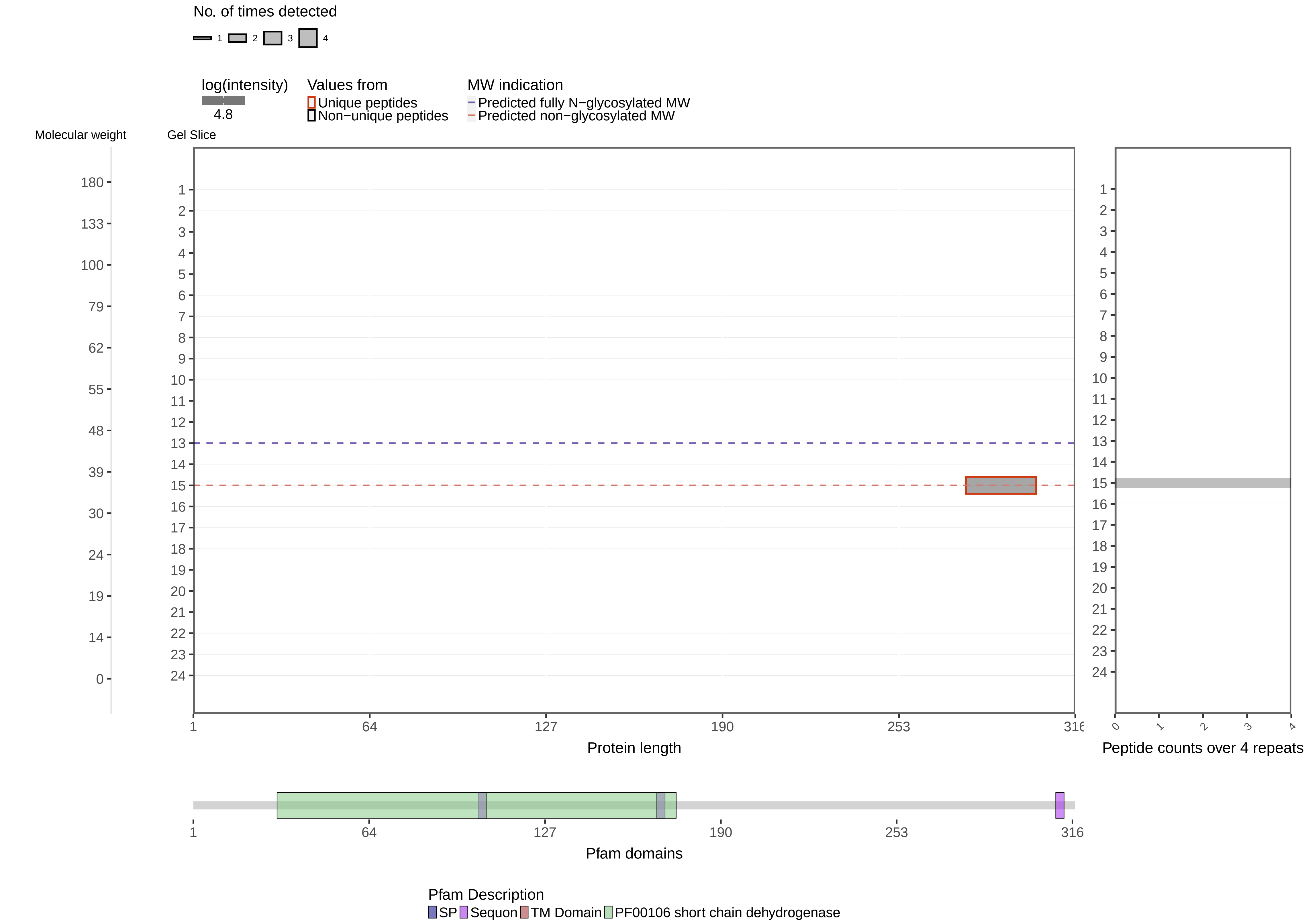


Protein sequence:
TNLLLENLKNTAQQSHKEGRIVNVSSMGHKFTYREGIRFDKINDKDSYTSWYAYGQSKLANILHANELARRFKDEGLEITANSLHPGAINTNLMRHQNLIEGIVNWIGKYFIKDIPQGAATTCYVALHPQVKGISGEYFSDSNVATPTSHATDTELAKKLWDFSLNLTKPQ*

NbL17g26460.1 aquaporin_PIP−type_pTOM75 [MW = 30.76 kDa]

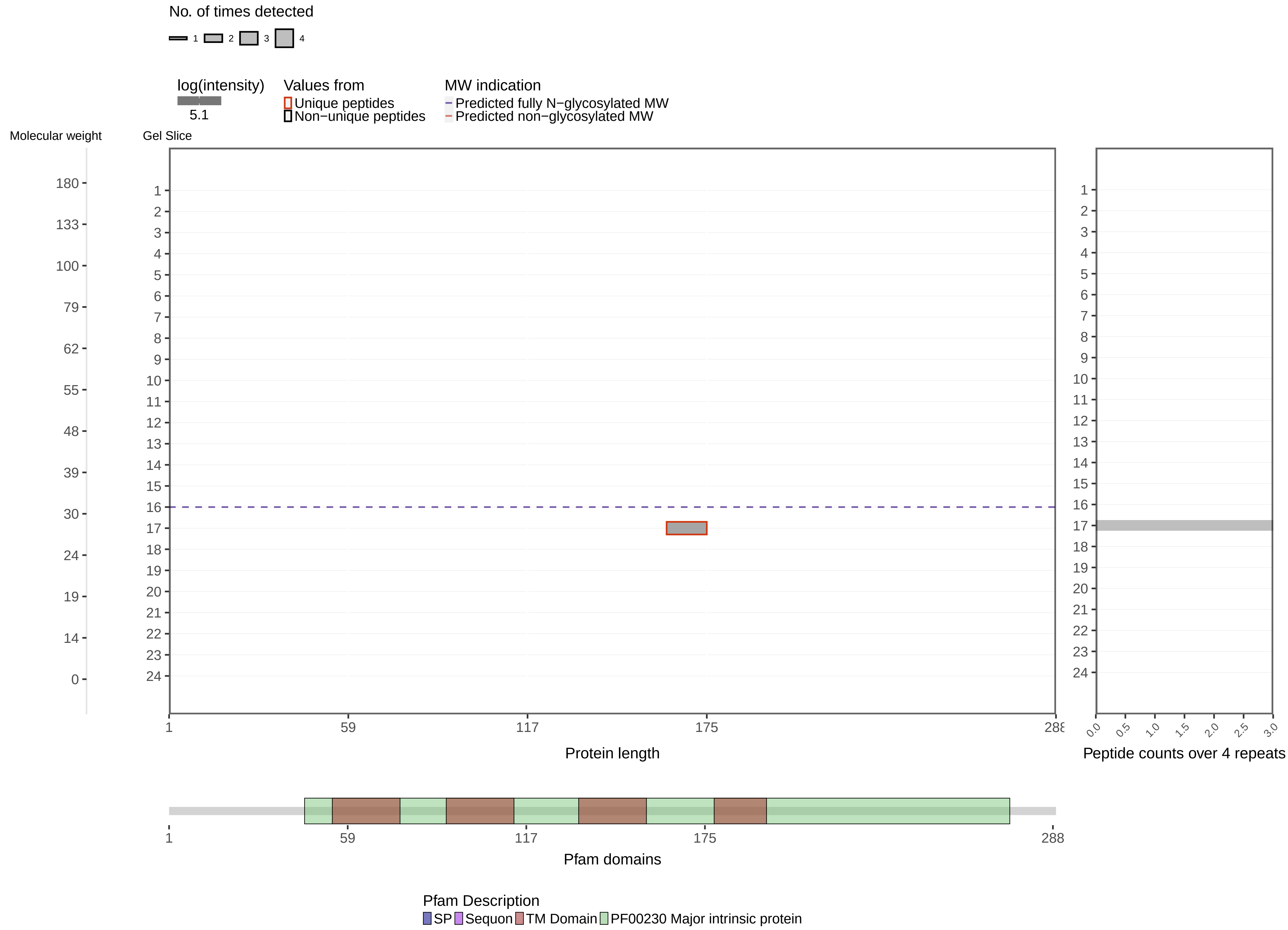


Protein sequence:

LTRAIFYIVMQCLGAICGAGVVKGFMVGPYQRLGGGANIVNPGYTKGDGLGAEIIGTFVLVYTVFSATDAKRNARDSHVPILAPLPIGFAVFLVHLATIPITGTGINPARSLGAAIIYNNDHAWDDHWIFWVGPFIGAALAAVYHQIIIRAIPFHKSS*

### NbL17g26400.1 GMP_synthase_[glutamine−hydrolyzing] [MW = 64.12 kDa]

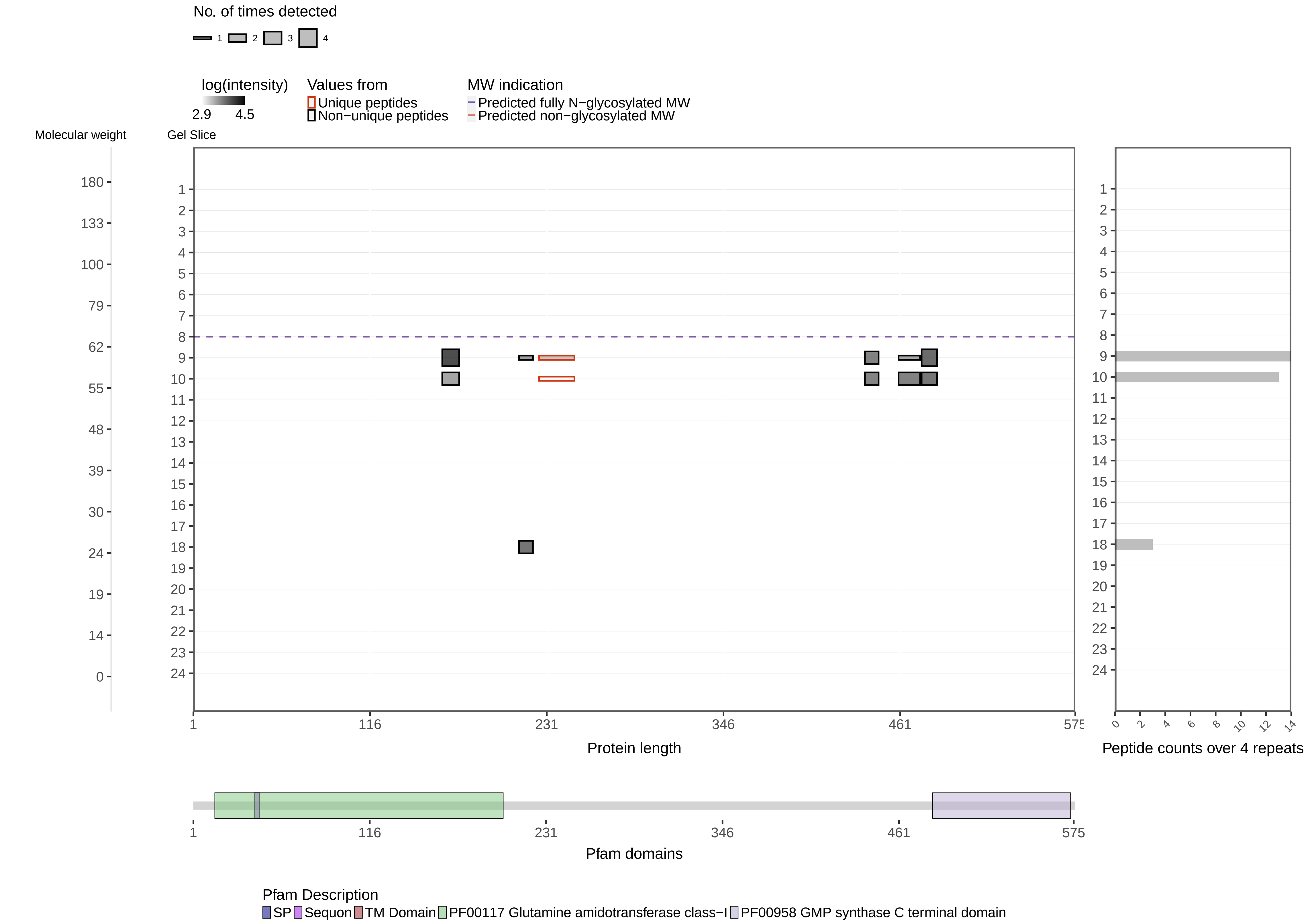


Protein sequence:

ERERVMDLFEKRLHLPVTCVDATEEFLSKLKGVTEPEMKRKIIGKEFINIFDLFSHDVEKKVGKKPTYLVQGTLYPDVIESCPPPGSGRTHSHTIKSHHNVGGLPKDMKLKLIEPLKLLFKDEVSFCLLKLLHASLHYFFLTLQIGLYVDLSCTPLISQVRELGKILDISEDFLKRHPFPGPGLAVRIPGDVTAGNSLDILRQVDEIFIQSIREAKIYDEIWQAFAVFLPVKTVGVQGDQRTHSHAVALRAVTSQDGMTADWYYFDFKFLDDVSRKICNSVRGVNRVLLDITSKPPSTIE

NbL17g26310.1 glutamate−−cysteine_ligase_chloroplastic [MW = 58.98 kDa]

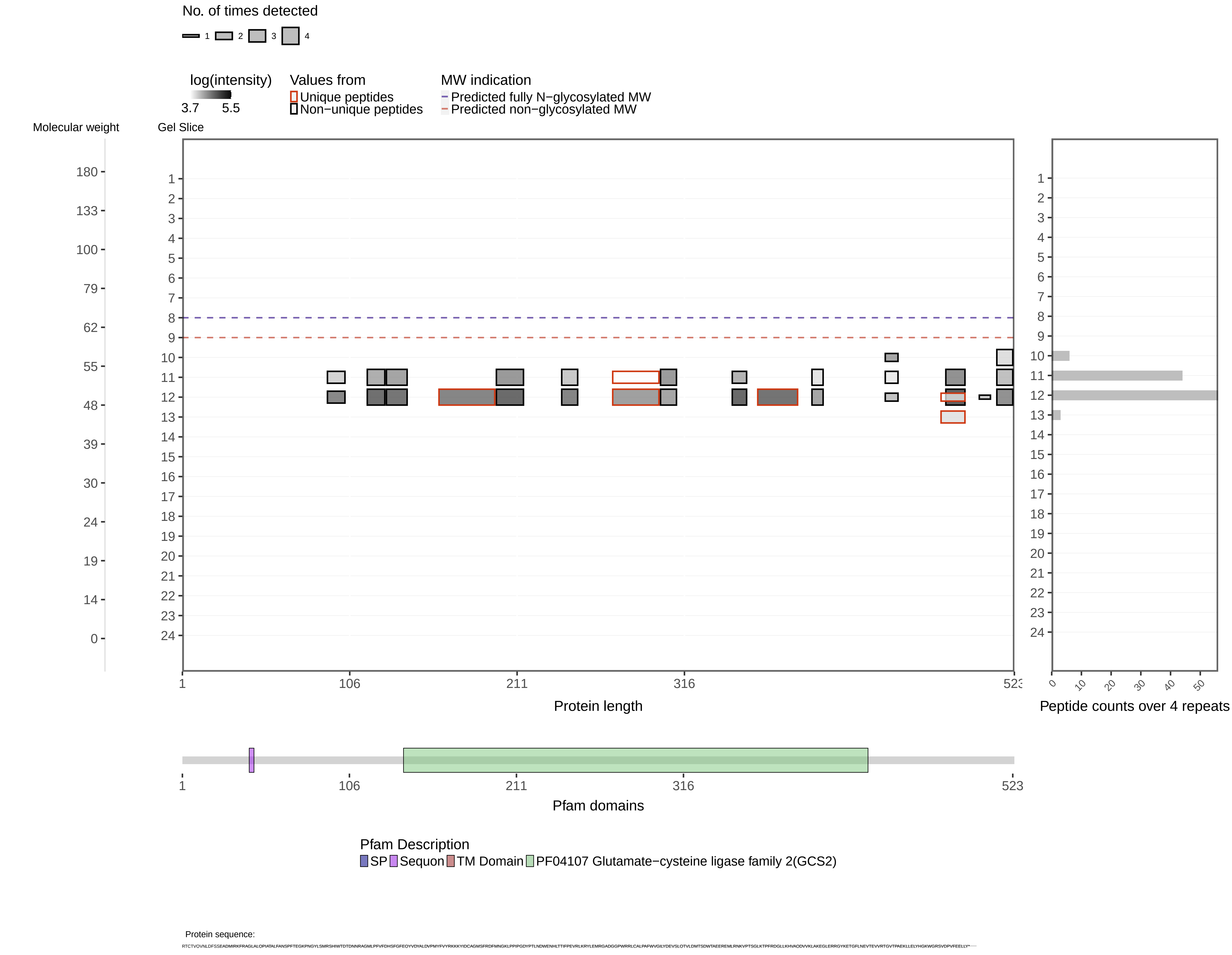

NbL17g26250.1 probable_phospholipid_hydroperoxide_glutathione_peroxidase [MW = 25.99 kDa]

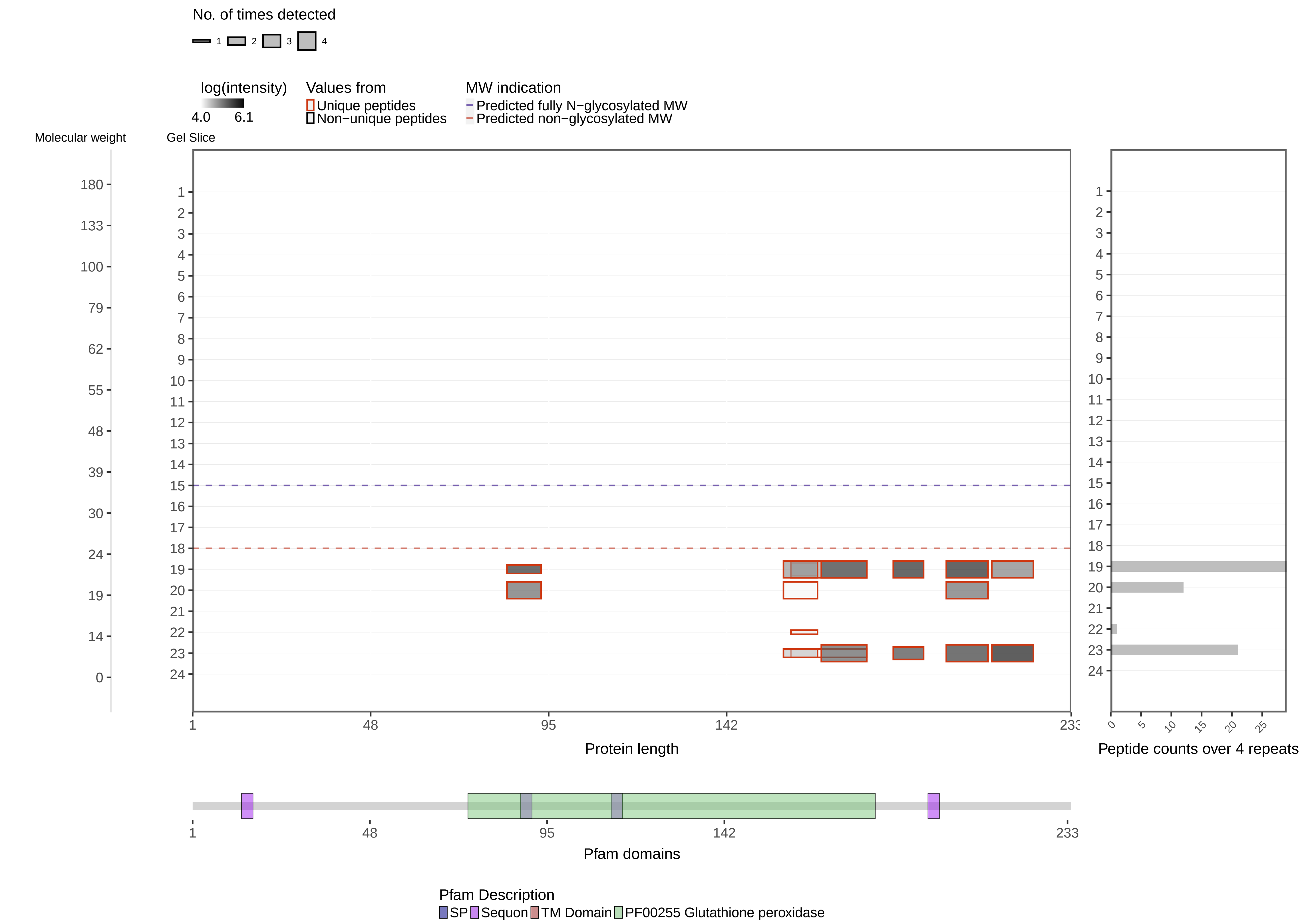


Protein sequence:

CGLTNSNYTDMTEIYKKYKDQGLEILAFPCNQFGGQEPGSIEEIQNMVCTRFKAEYPIFDKVDVNGDNAAPLYKFLKSSKGGFFGDSIKWNFSKFLVDKEGNVVDRYSPTTAPASMEKDIKKLLGVA*

### NbL17g26230.1 probable_phosphopantothenoylcysteine_decarboxylase [MW = 23.09 kDa]

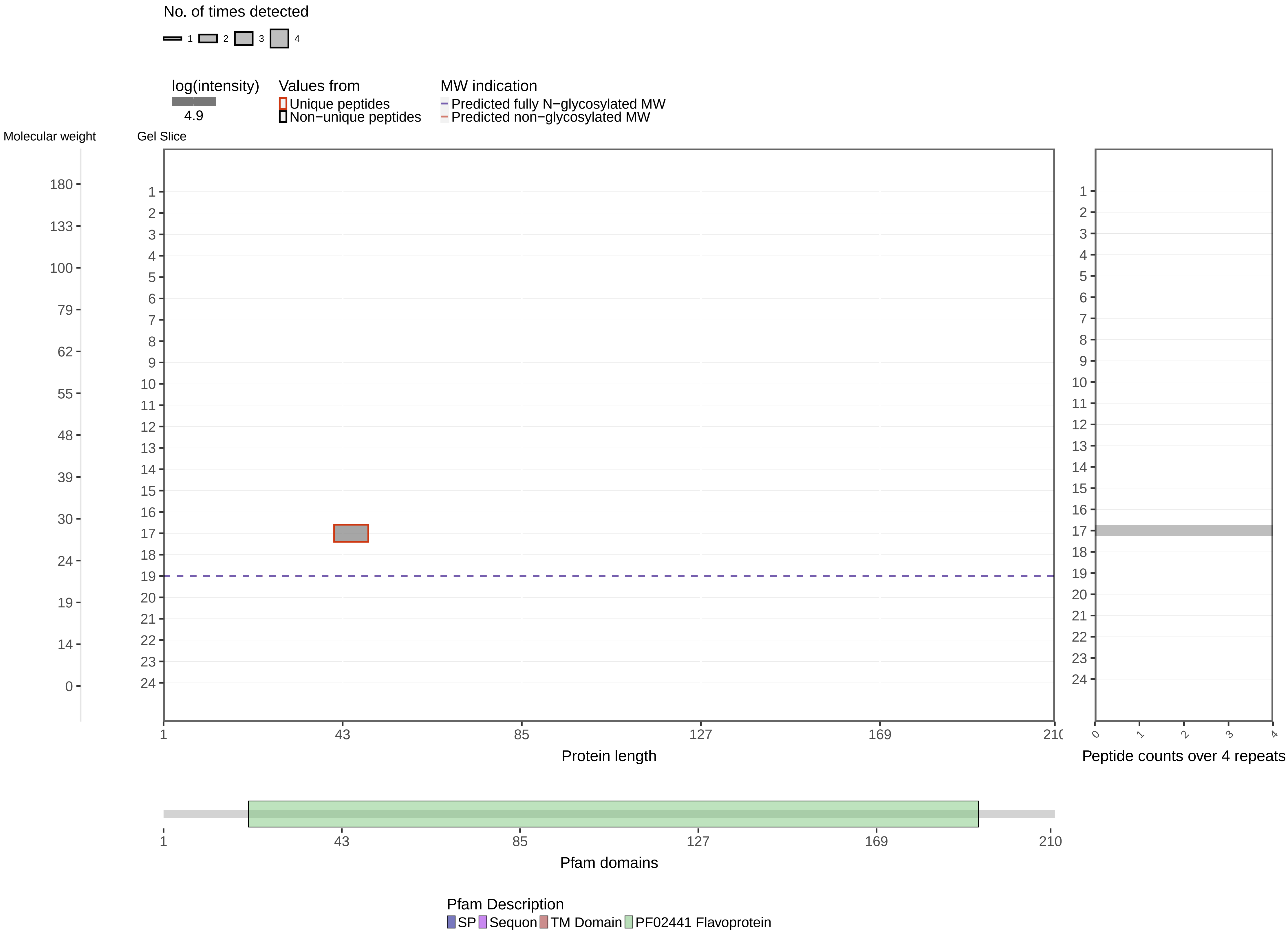


Protein sequence:

ELRRWADIMIIAPLSANTLGKIAGGLCDNLLTCIVRAWDYDKPLFVAPAMNTLMWNNPFTEKHLMVIDELGISLIPPVSKRLACGDYGNGAMAEPSLIYQAVRIYYDAQLRSGGSNVA*

### NbL17g26060.1 elicitor−responsive_protein_3 [MW = 16.07 kDa]

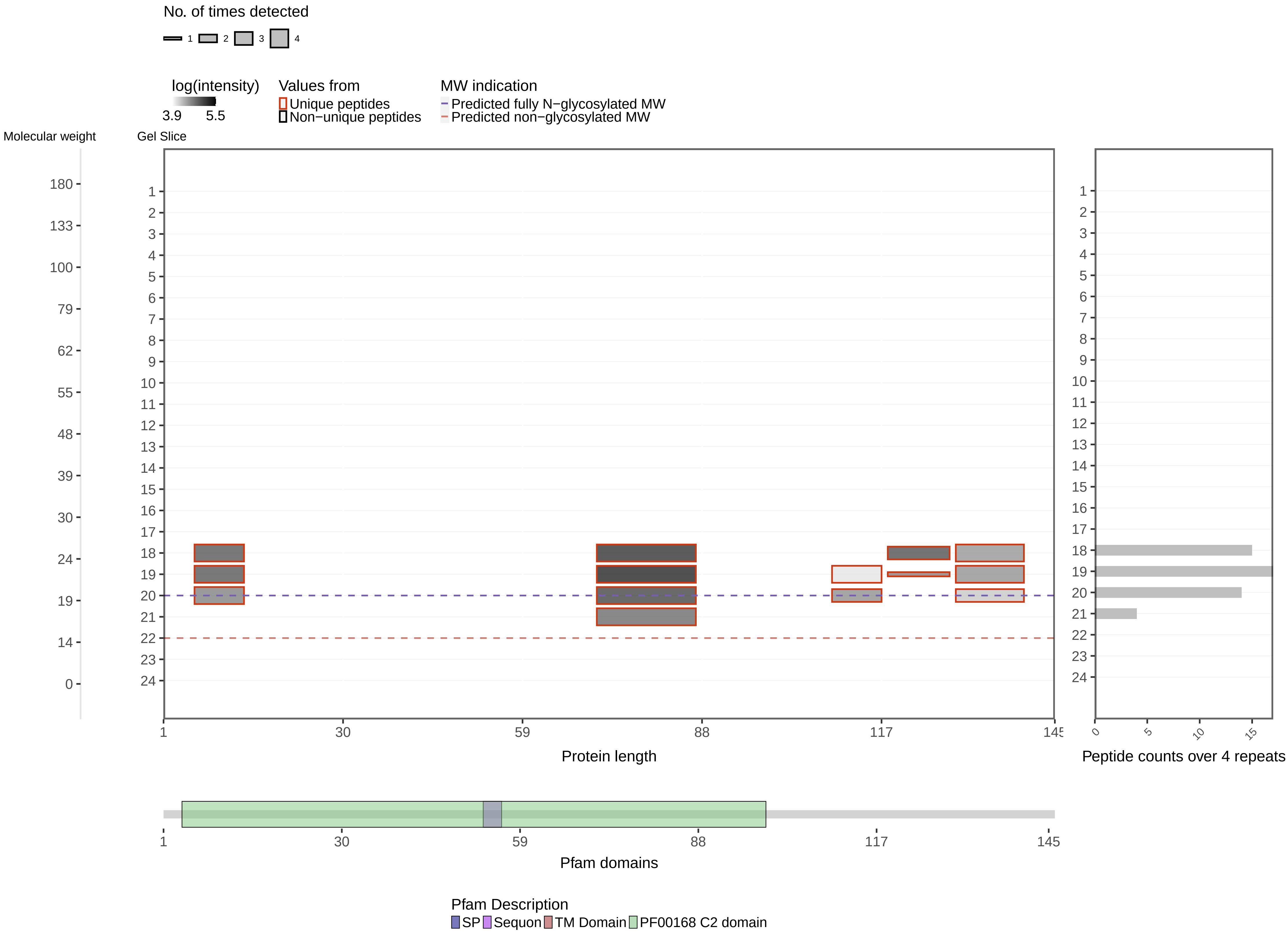


Protein sequence:

FSISEGVEELFLKIMDSDSVGEDDFVGEAKIPIEPVCSEGSIPTTAYNVVKNEEYCGEIRVSLTFTPQEERDYEQEESYGGWKESTY*

NbL17g26040.1 bifunctional_UDP−glucose_4−epimerase_and_UDP−xylose_4−epimerase_1 [MW = 39.3 kDa]

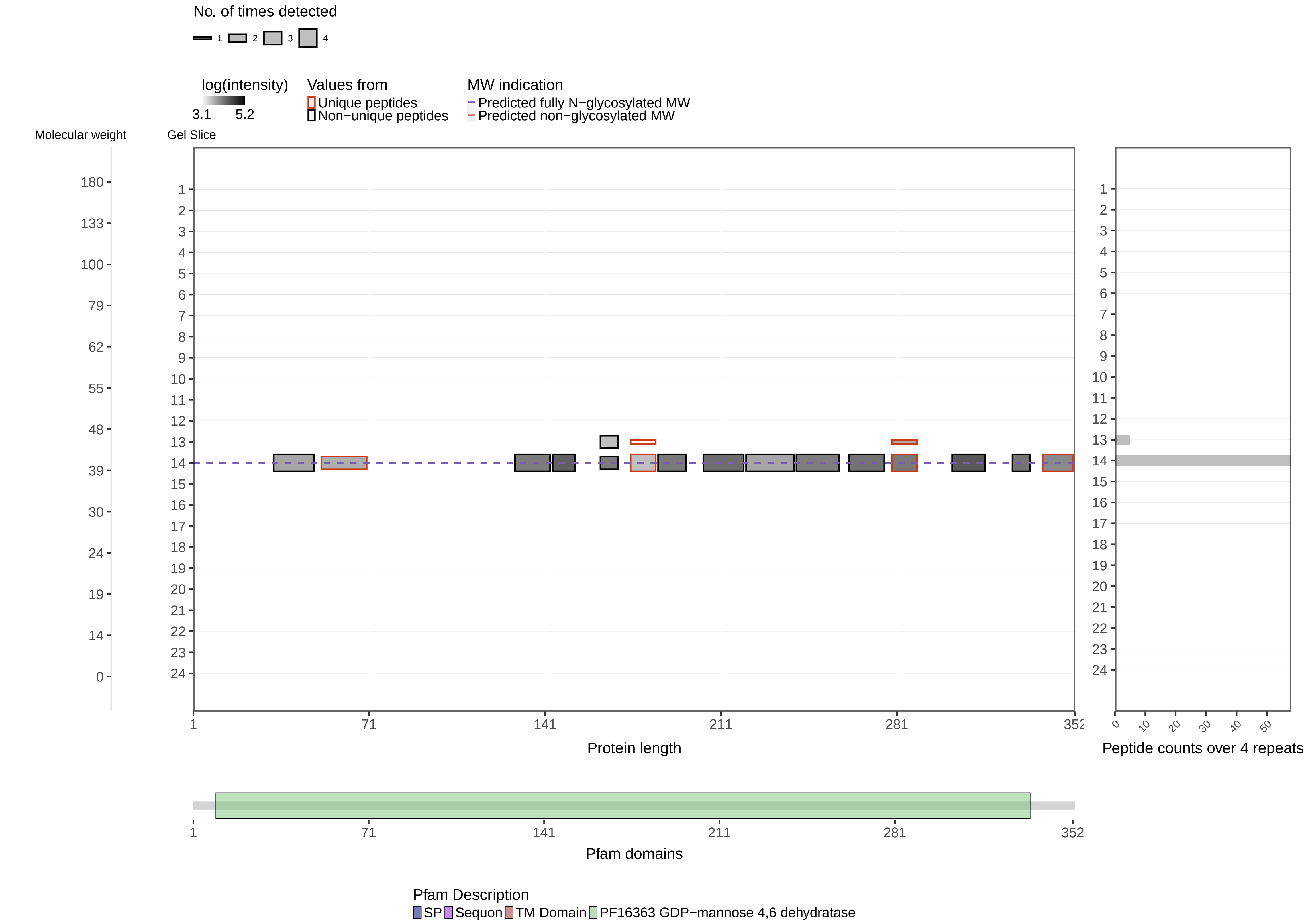


Protein sequence:

.EEIARDIQKADAEWSIILLRYFNPVGAHESGKLGEDPRGIPNNLMPYIQQVAVGRLPELNVYGTDYPTPDGTAIRDYIHVMDLADGHVVALQRLLKQNDLGCVAYNLGTGKGKSVLEMVAAFERASGKKIPLKMCPRRPGDATAVYASTEKAEKELGWKAKYGIDEMCRDQWKWASQNPWGYQSKP*

NbL17g25940.1 plant_UBX_domain−containing_protein_8−like [MW = 66.83 kDa]

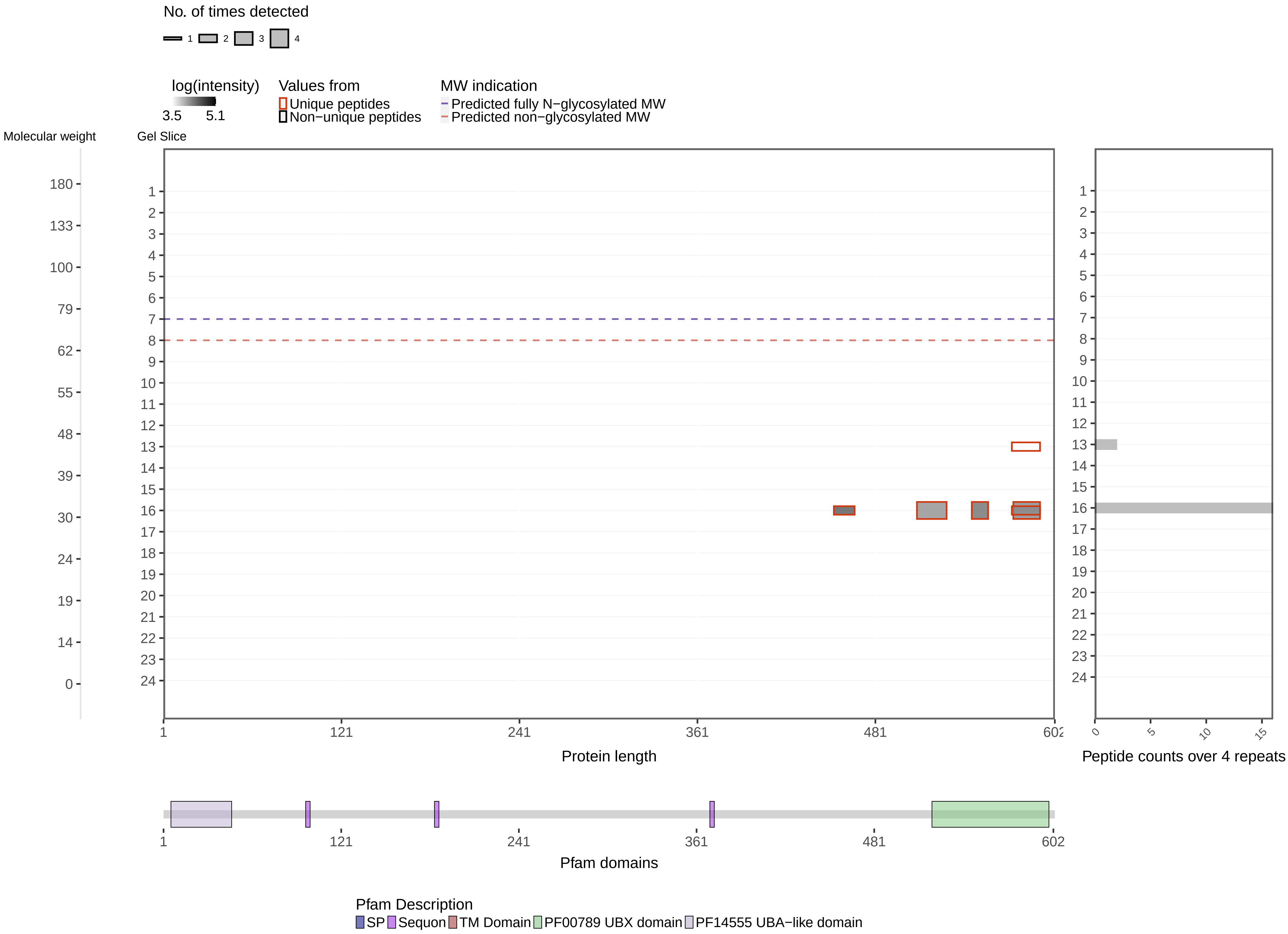


Protein sequence:

EGYKPAEEDERGKSITSNGRQFRSEVGSSSVPDEAEDLEEEPLVWRRRKPISSGSETAQDIEEHVGSPLSSPRQHDNLNRSPRNGADFPSDEWGGISSLEHDEAVMLEAALFGGIPEGSGYRLPNAPHQLMQNGVEGPMGPYQWHMPRPPSPSLVAQRLLREQQDDEYHTALQADREKELKAKEEAEAALEEKRQEEEESRRKAEEEKEMERQLAAKEASLPQEPTADNENAVNLVVRMPDGSRRGRRFLKSDRLQYLFNYIDVGRAVKPGTYRLVRPYPRRAFSD

NbL17g25820.1 bifunctional_dTDP−4−dehydrorhamnose_35−epimerase/dTDP−4−dehydrorhamnose_reductase [MW = 33.47 kDa]

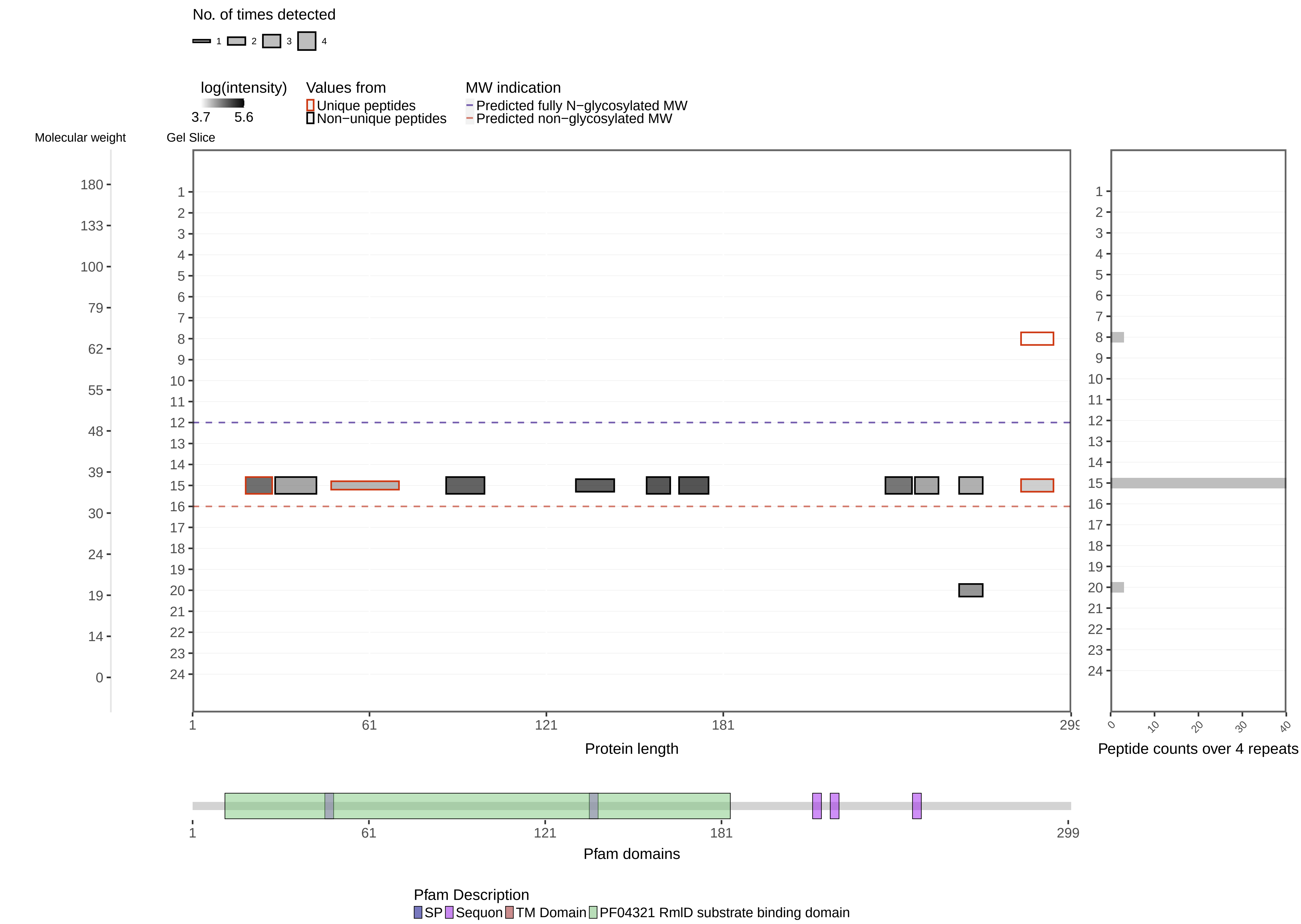


Protein sequence:

NFTGSFYSKTKAMVEELLKNYDNVCTLRVRMPISADLTNPRNFITKITRYDKVVDIPNSMTILDELLPISLEMAKRNLTGIWNFTNPGVVSHNEILEMYRDYVDPSFSWKNFTLEEQAKVIVAPRSNNELDASKLSKEFPEMKSIKESLIEYVFKPNRKTPVA*

### NbL17g25810.1 phospholipase_D_beta_1−like [MW = 95.71 kDa]

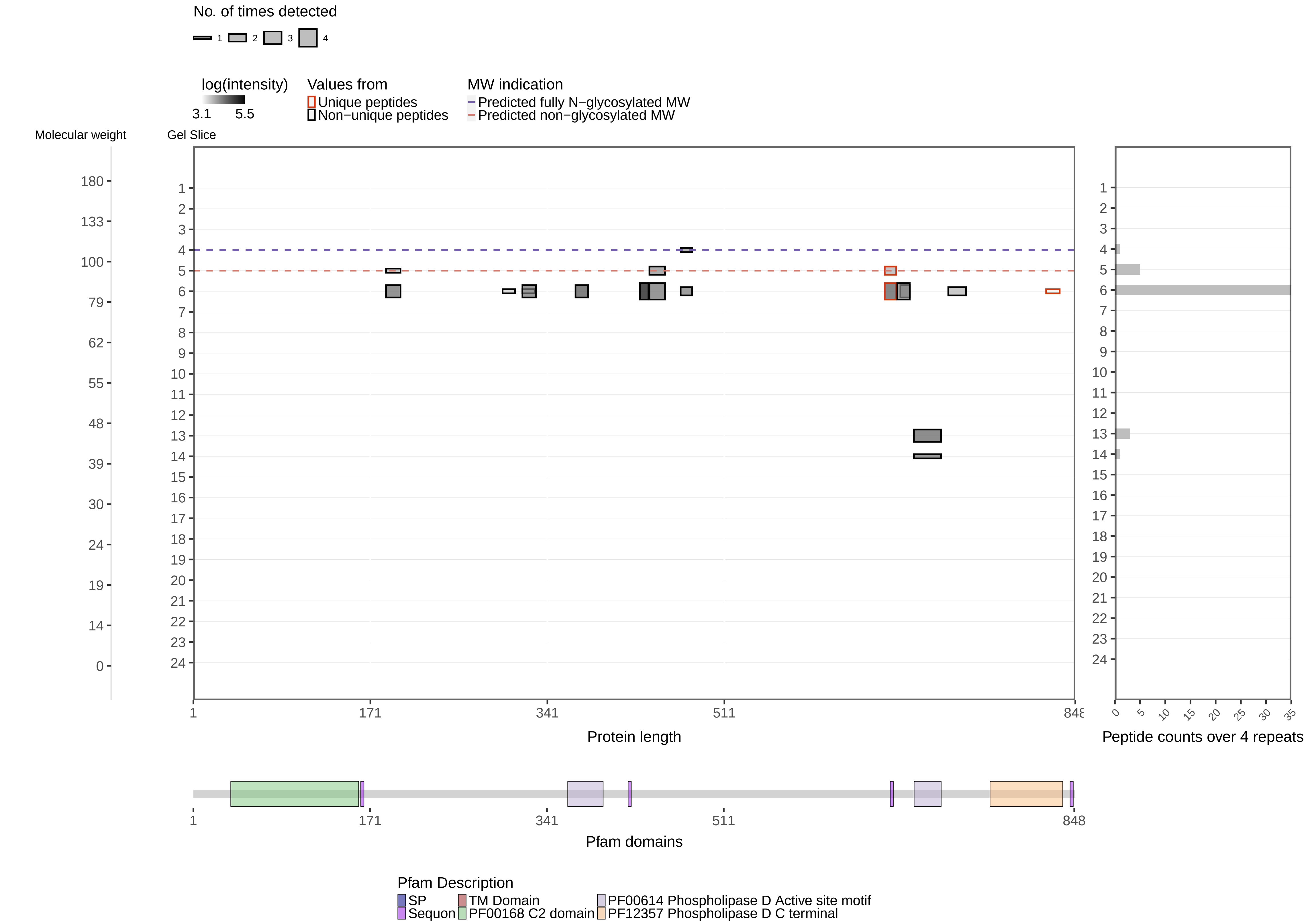


Protein sequence:

### NbL17g25740.1 uncharacterized_LOC109240160 [MW = 138.05 kDa]

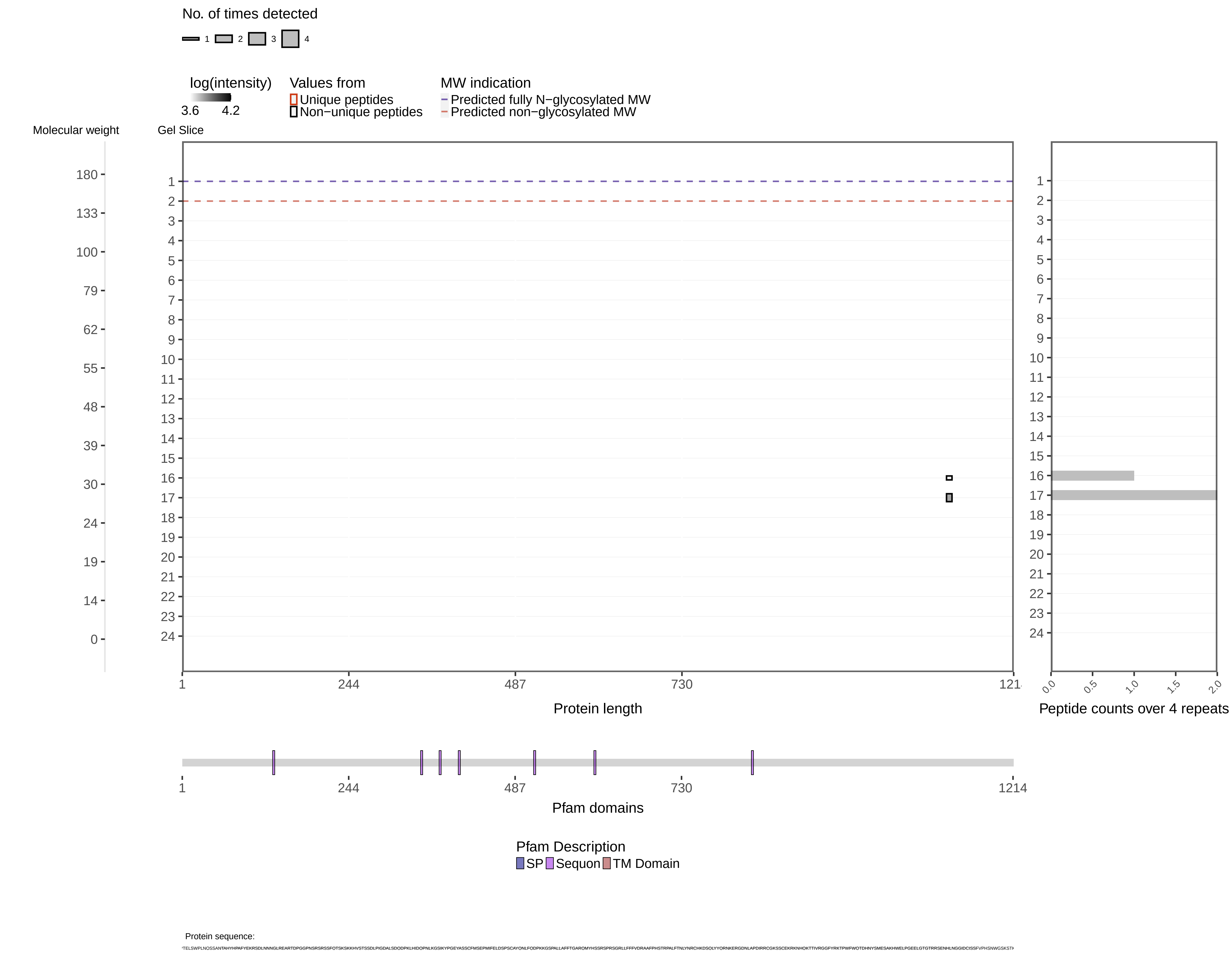

NbL17g25630.1 subtilisin−like_protease_SBT1.2 [MW − SP = 77.37 kDa]

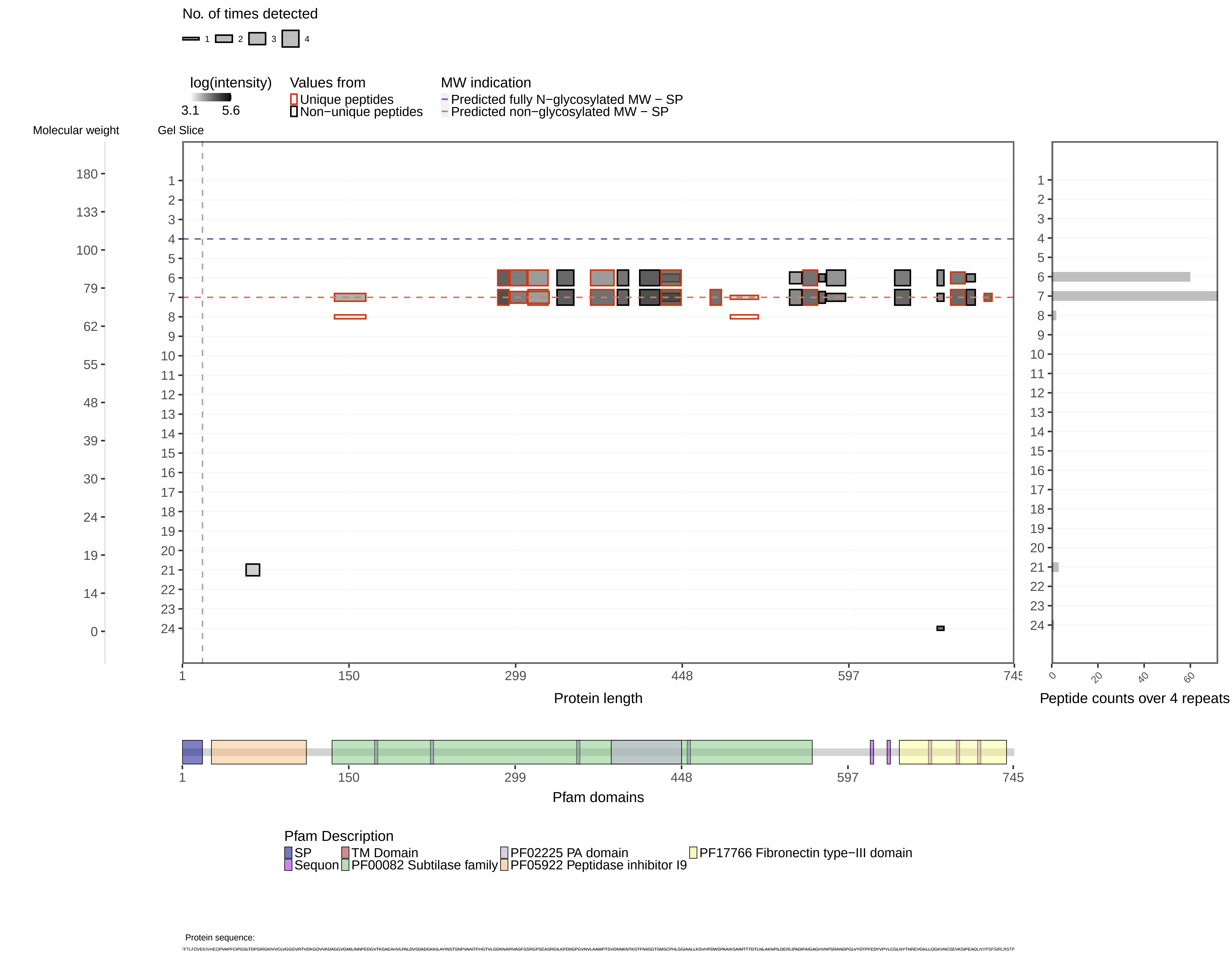

NbL17g25620.1 copper_chaperone_for_superoxide_dismutase_chloroplastic/cytosolic [MW = 34.1 kDa]

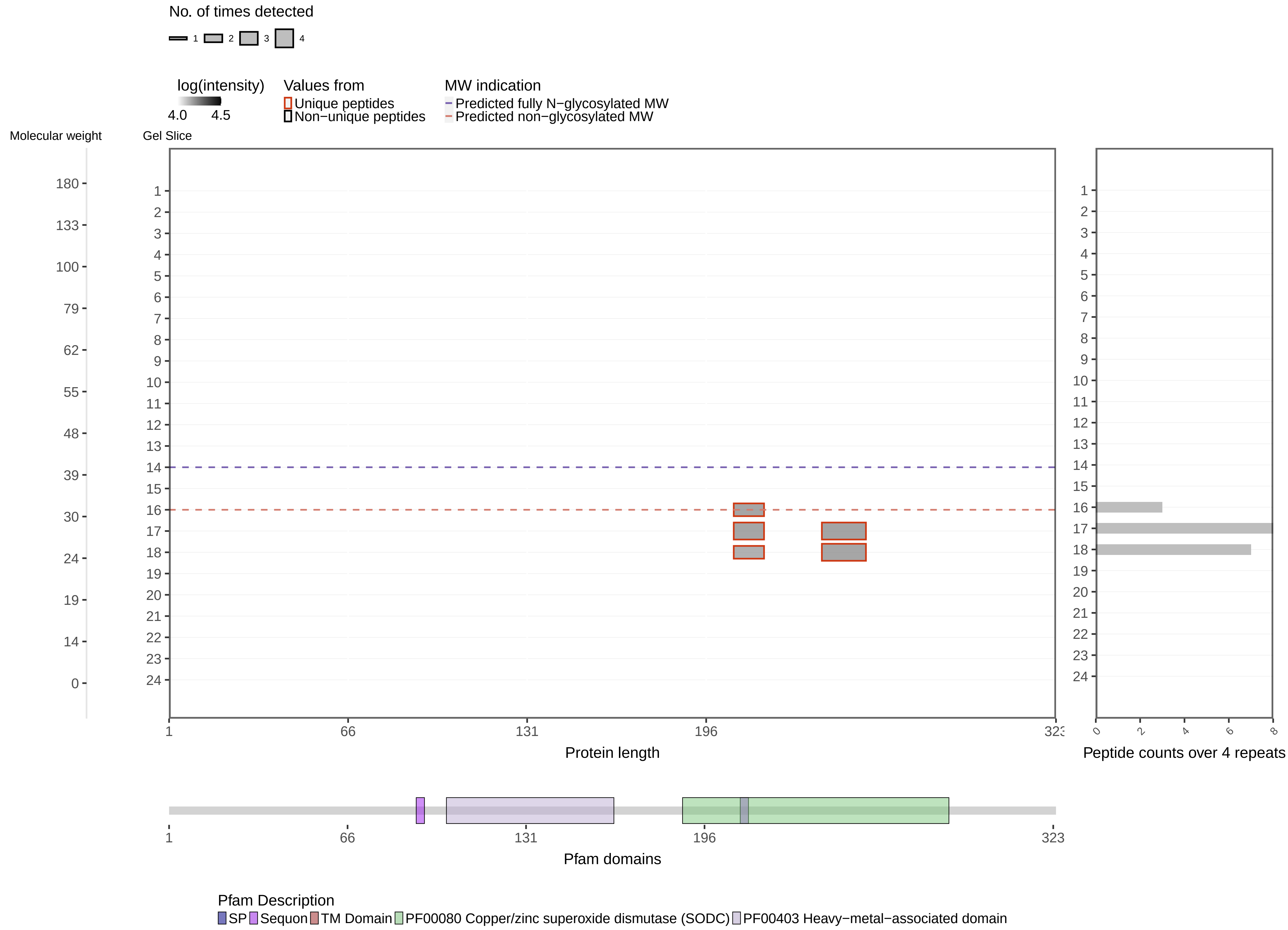


Protein sequence:

SSPVKILTEALEQTGRKARLIGQGVPDDFLISAAVAEFKGSDIFGVVRLAQVNIELTRIEANFSGLSPGKHAWSINEFGDLTRGAASTGKLYSPPVGDLGTLQVDGKGEAFYSGAKEKLRVADLIGRAIAVYSTEDKSDPGLKAAVIARSAGVGENYKKLCTCDGTTIWEATNKL*

### NbL17g25610.1 enoyl−CoA_delta_isomerase_1_peroxisomal [MW = 26.85 kDa]

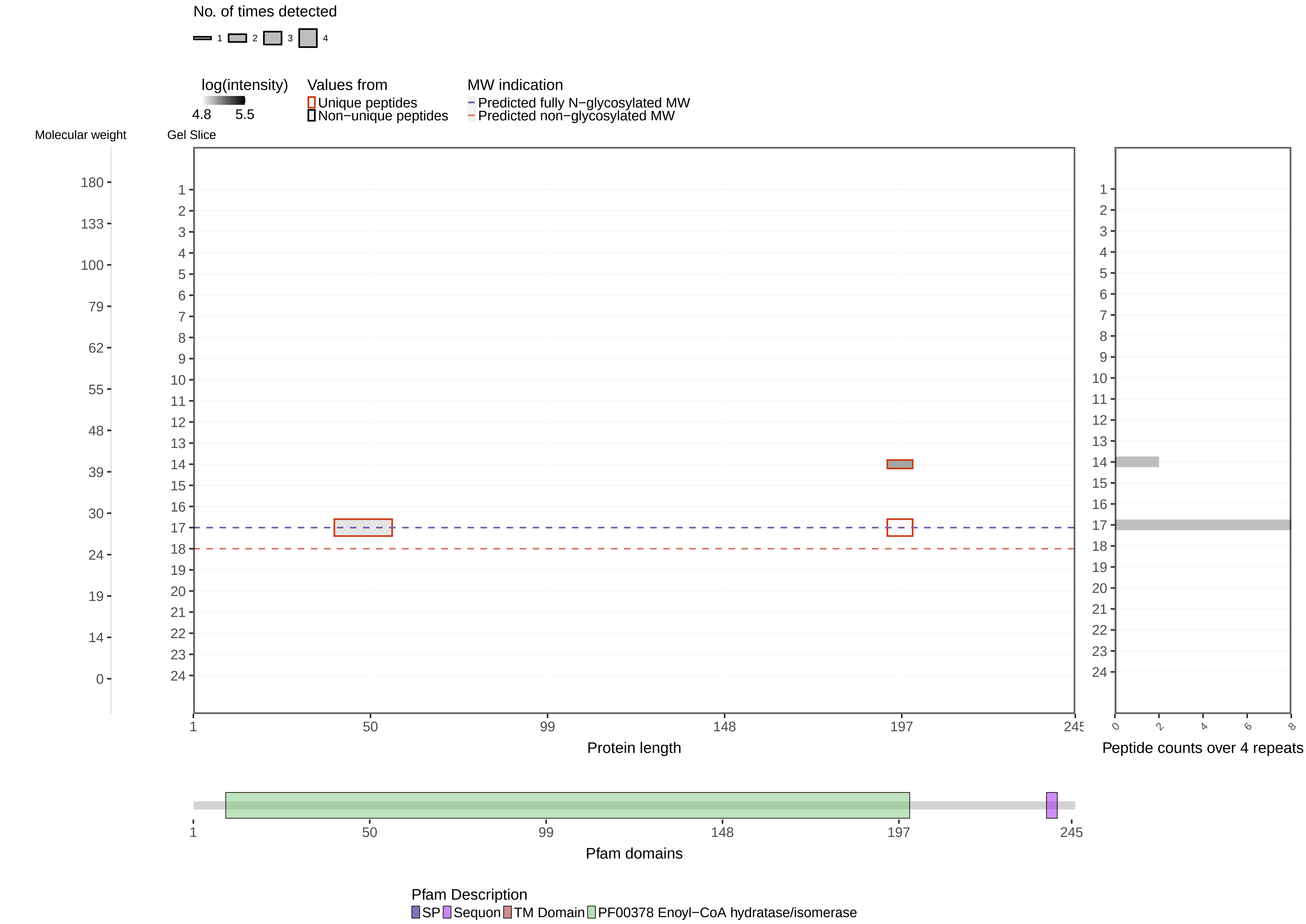


Protein sequence:

-SHDYVLMRKDQGFLCMSELDIGYKIPIWFSVLLKCKVGSPAVWREVVLKSAKMTAEMGVEMAIVDSAHSGAEATVEAAVRLGEELVSRNWDGNVYADCRKTLYVQLLNSLGSDETVGHYGEEEKPNKTLSRL*

NbL17g25510.1 probable_LRR_receptor−like_serine/threonine−protein_kinase_At1g12460 [MW − SP = 92.52 kDa]

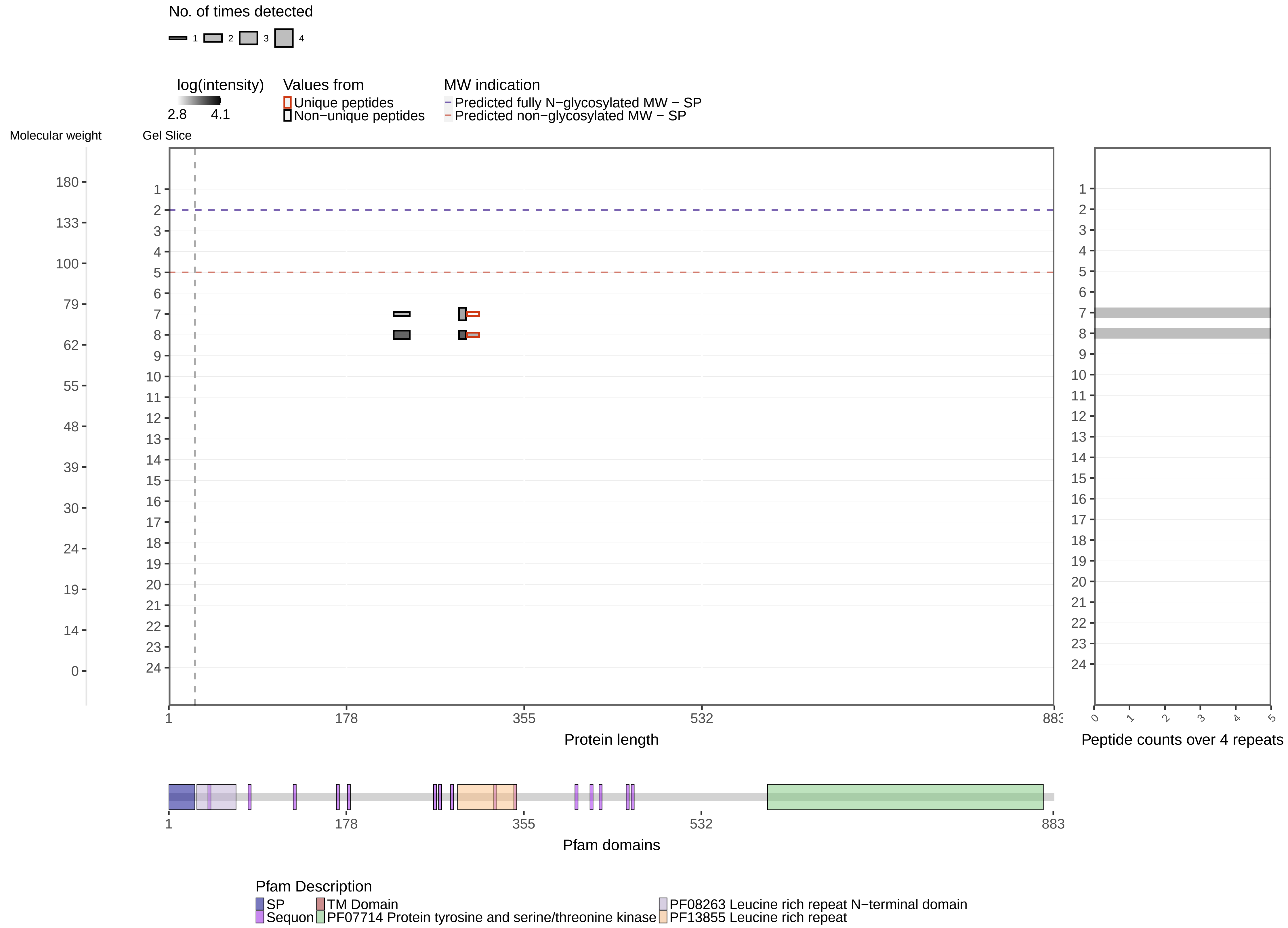


Protein sequence:

NLSNLHFLDLSQNLLSGSIPVALGNLHNLTHFNISYNLLSGAIPSIESIOOFGPSAFFHNSGLCGAPLDVSCYPSGIAPAKGKPRLSASAIVAIVAAAVILTGVCLITIINMKARRRRRRADETIIVESTPLASTESNVIIGKLVLFSKTLPSKYEDWEAGTKALLDKESLIGGGTIGSVYRTSFEGGVSLAVKKLETLGRIRNQDEFEQEIGRLGTLHHPNLVALQGYYWSSSMQLILSEFVPNGNLYNNLHGLSYPGTSTGVGNPELNWPRRFQIAVGTARALAYLHHDCKPPVLHLNV

### NbL17g25390.1 ATP−dependent_Clp_protease_ATP−binding_subunit_CLPT2_chloroplastic−like [MW = 29.13 kDa]

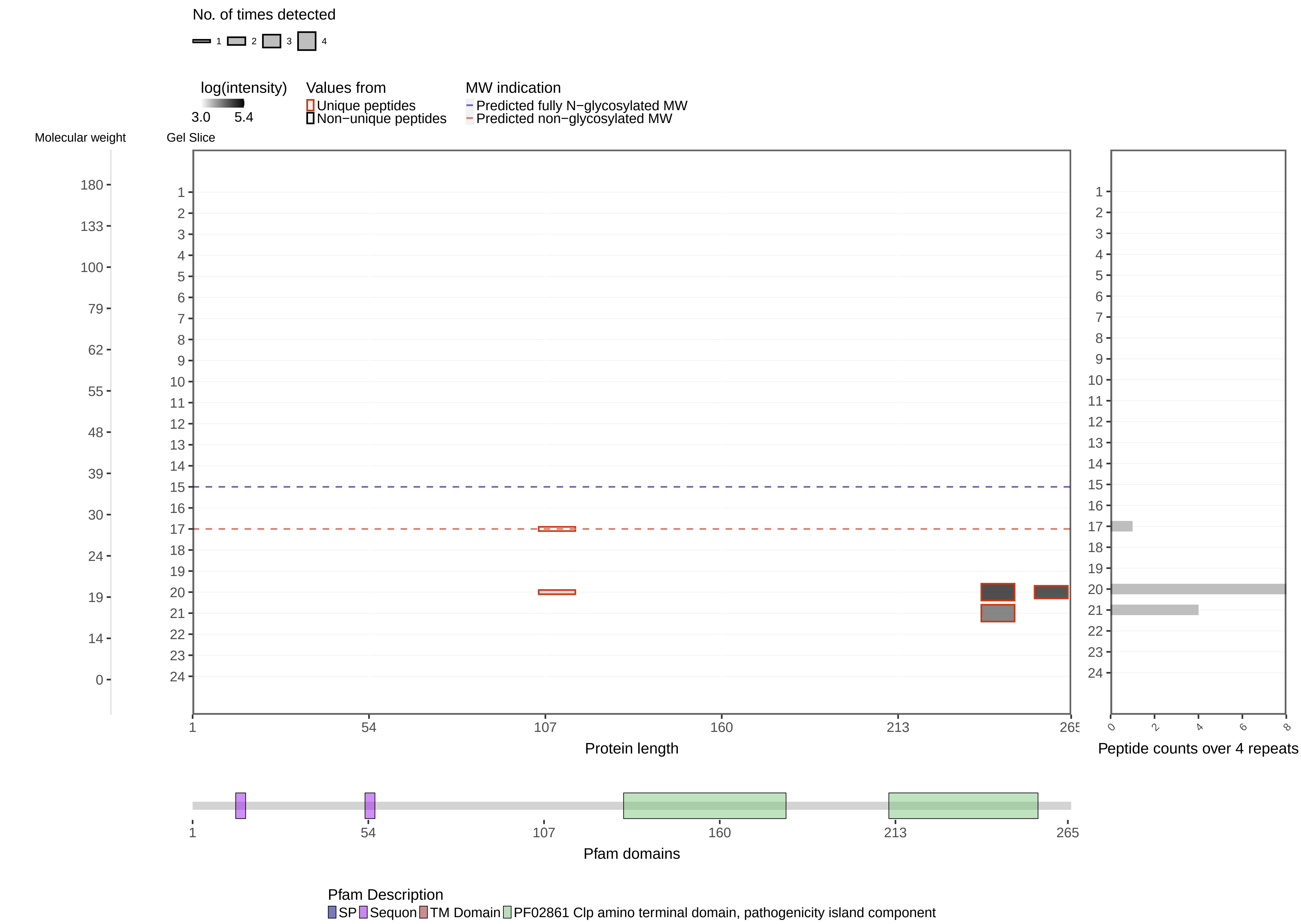


Protein sequence:

KAIKSFAMGELEARKLKYPTTGTEALLMGILIEGTNFASKYLRANSITLFKVREETIKVLGKADMWFFSPEHPPLTEDAQKALDWALDEKLKSSDNGEITTTHLLLGVWSQVGSPGYKILSALGFNDEKAQELKNVISEPGFVDD*

### NbL17g25360.1 ATP−dependent_Clp_protease_proteolytic_subunit−related_protein_2_chloroplastic−like [MW = 36 kDa]

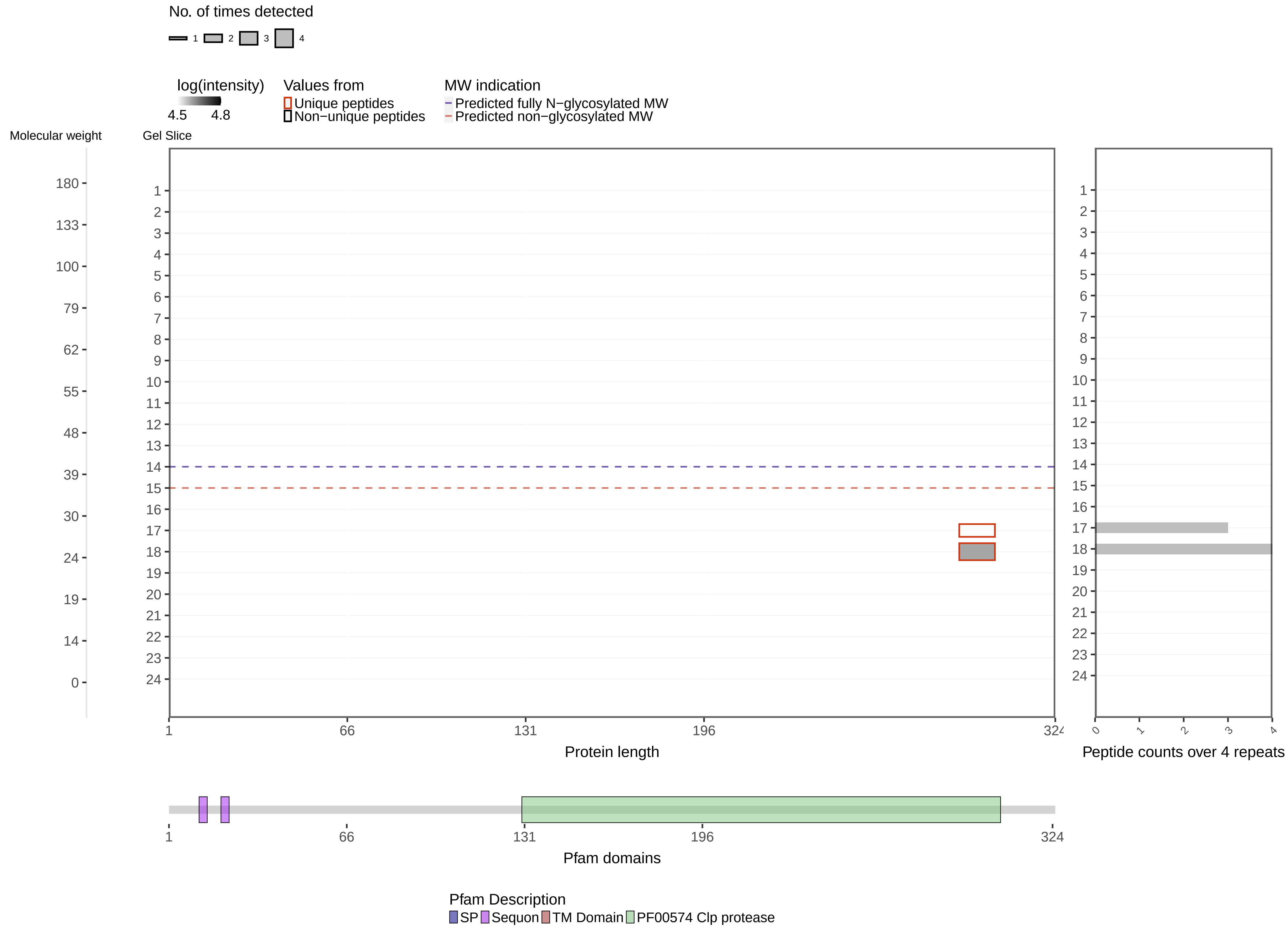


Protein sequence:

QHIDEEFSNQILATMLYLDSIDDSKKLYLYINGPGGDLTPSMAIYDTMQSLKSAVGTHCVGYAYNLAGFLLAAGEKGNRFAMPLSRIALQSPAGAARGQADDIRNEADELLRIRDYLFKELAEKTGQPVEKVHKDLSRMKRFNAQEALEYGLIDRIVRPPRIKADAPRKDTTAGLG*

NbL17g25240.1 peptidyl−tRNA_hydrolase_ICT1_mitochondrial [MW = 26.3 kDa]

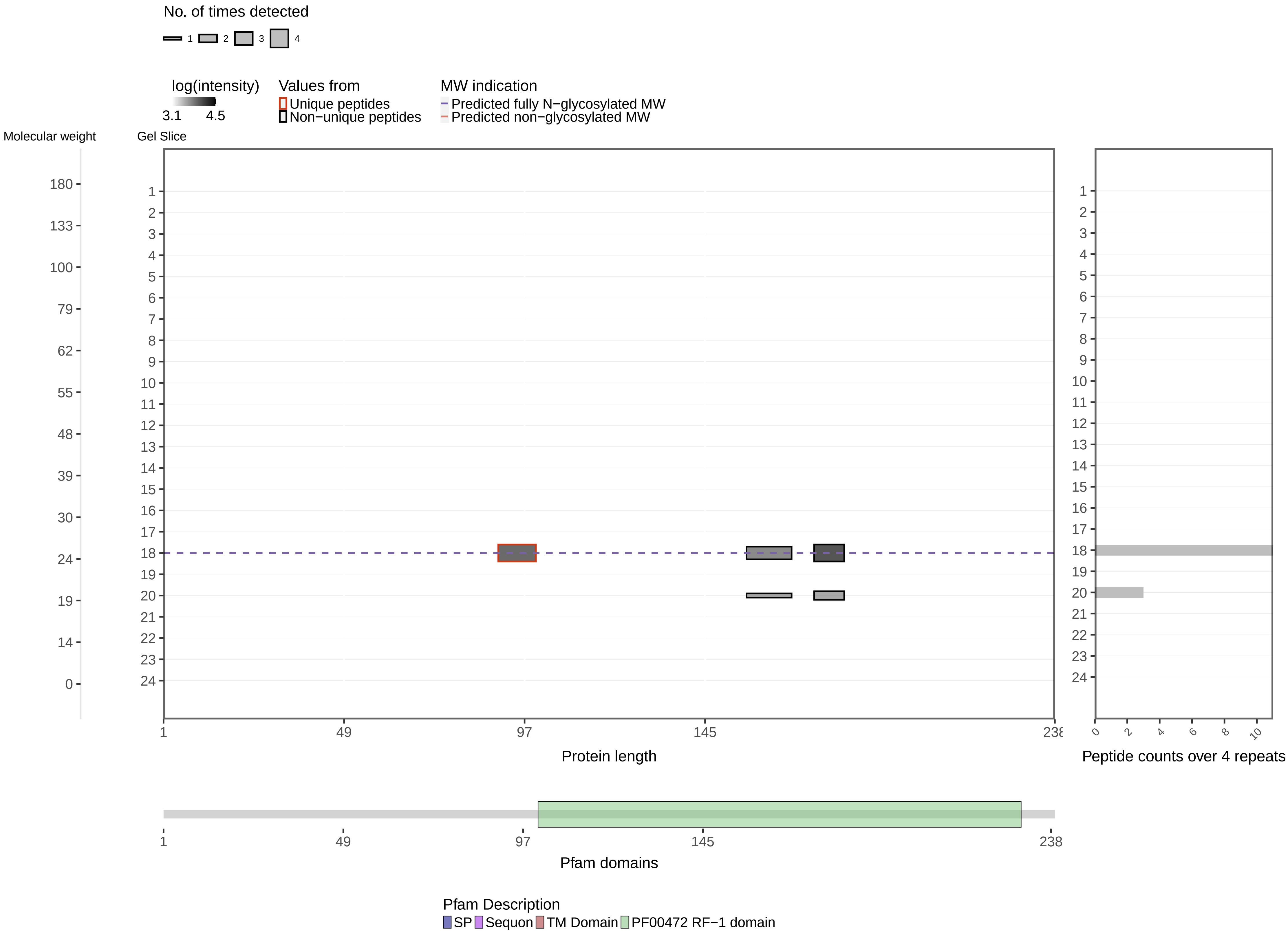


Protein sequence:

/NFARSGGPGGQNVNKVNTKVDMRFNVKNAYWLSDRVREKILQMEKNRINKDGELVISSTKTRTQKGNIEDALSKLQAIIDAASYVPPPPSEEQVKRIAKLAAIGERKRLDNKKAQSQKKAIRRSKDSWD*

### NbL17g25180.1 primary_amine_oxidase−like [MW = 83.54 kDa]

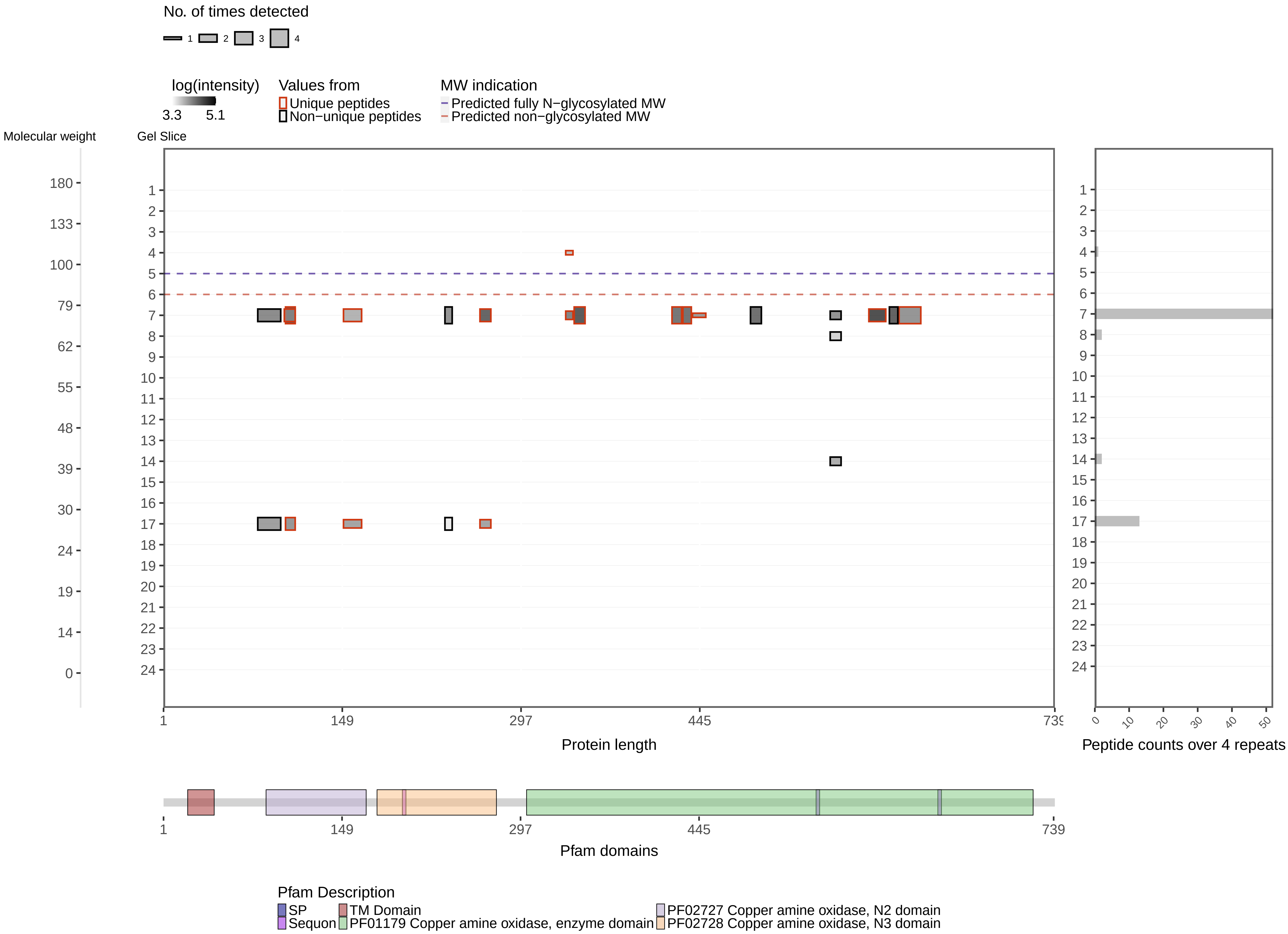


Protein sequence:

### NbL17g25150.1 FAM10_family_protein_At4g22670−like [MW = 44.7 kDa]

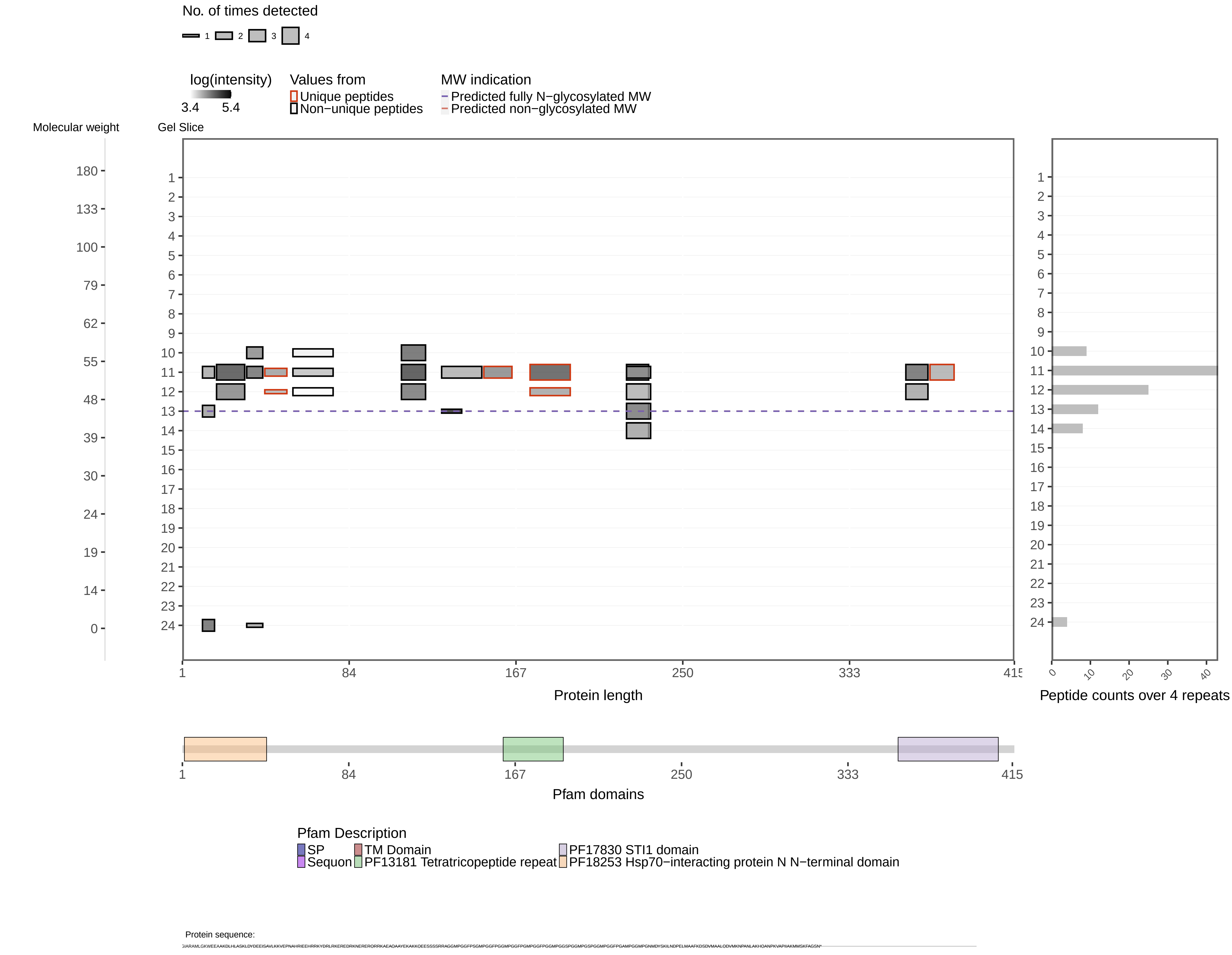

NbL17g25120.1 uncharacterized_LOC104104568 [MW = 28.48 kDa]

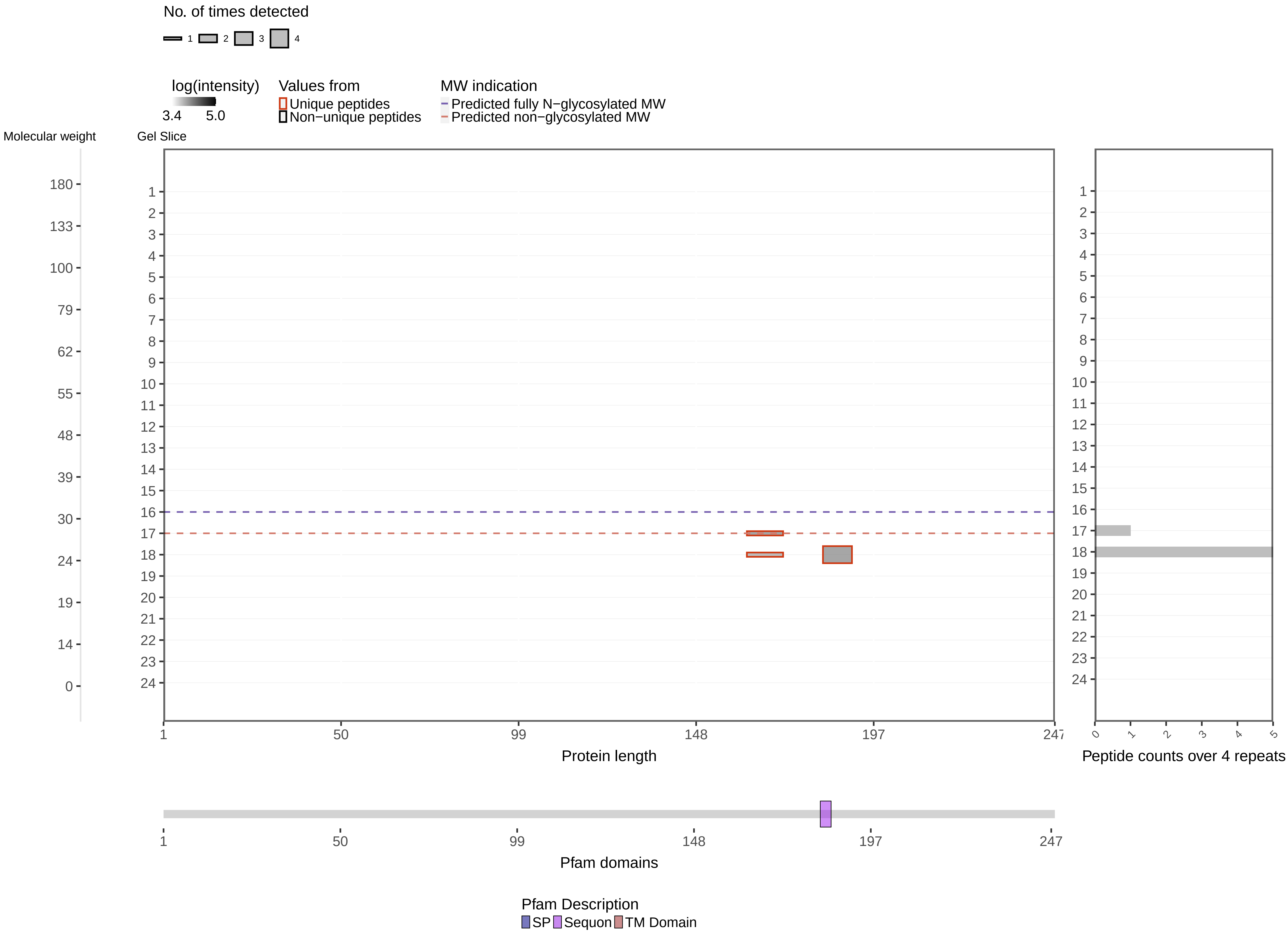


Protein sequence:

EAQEIEFEIARLRPRLNKEFFDHLQVEVGKLRFSMTKTQEVEDRLIELEALQKALQEATEAFDKMEADIVATKKNLTEILTSKDVKATLLDLVERNELNRPLLTLLDENIATAHSVNQKQAAEYMEKLRGLVLKYLTV*

### NbL17g24960.1 protein_IQ−DOMAIN_32 [MW = 97.86 kDa]

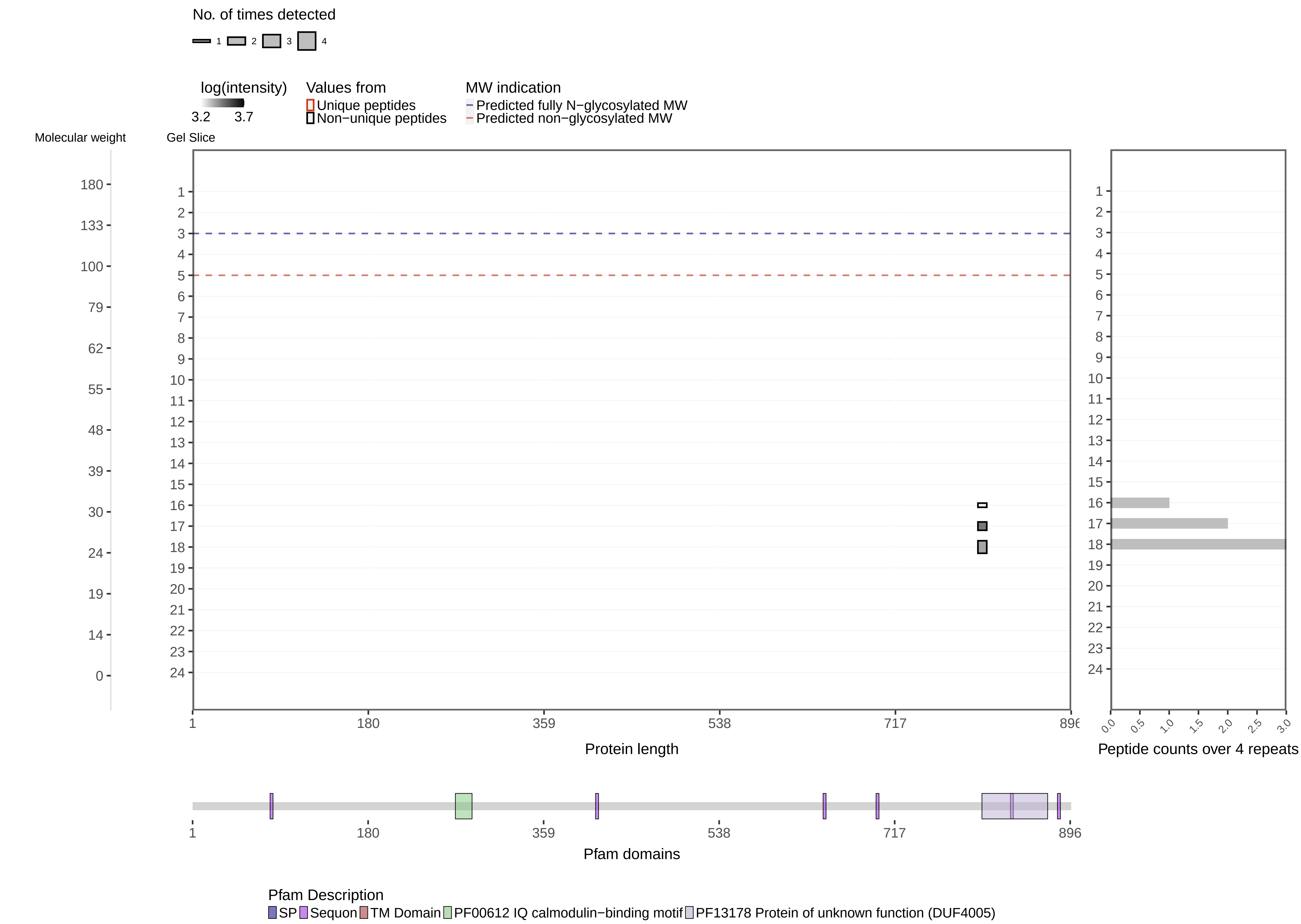


Protein sequence:

APLLESDEHLITYDADSLEVQADIPVPPQPQNVDETNSRDDNFCSIPTQYKESELLPESETNSFPAKTEADSDDTHSLTVEPPETESKQVVHGSRKASNPAFIAAQTKFEELTSAAKSTKISSLSNHKTEDESSEDTFSSITDQSFGARETAMSEASVPHSTRAQVGGSECGTELSISSTLDSPDRFEVGAREFEQELKPSYYQTGHNKSNGYPHIGDDNINDLLHSDNVQTGTEDPTDNANSELKDVMVSSDLSHEEKPENNSVDVQIEQEAKTDRLYKVSPDASPRSHVT

### NbL17g24900.1 5'−3'_exoribonuclease_3−like [MW = 122.11 kDa]

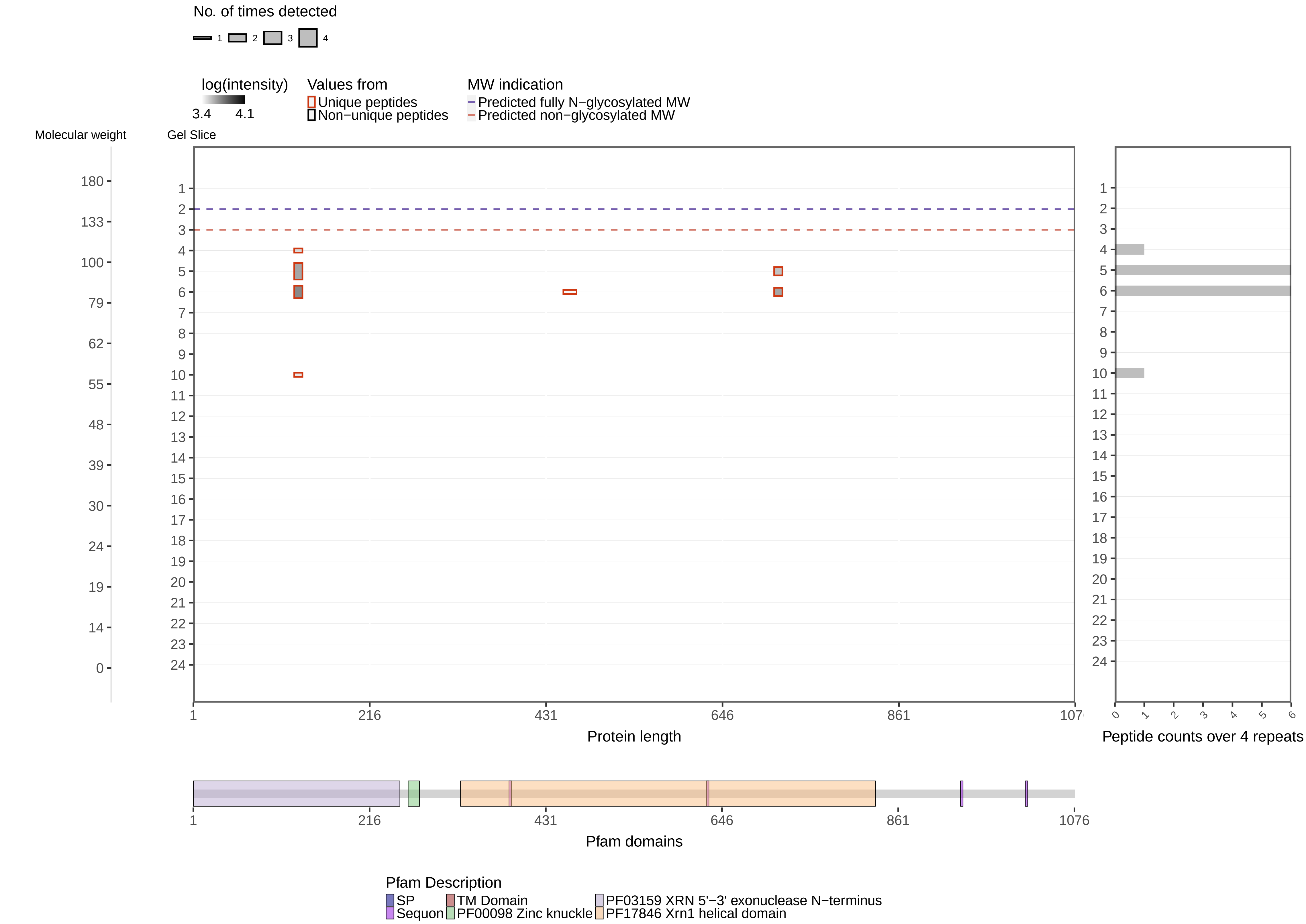


Protein sequence:

### NbL17g24880.1 endoglucanase_2 [MW = 57.17 kDa]

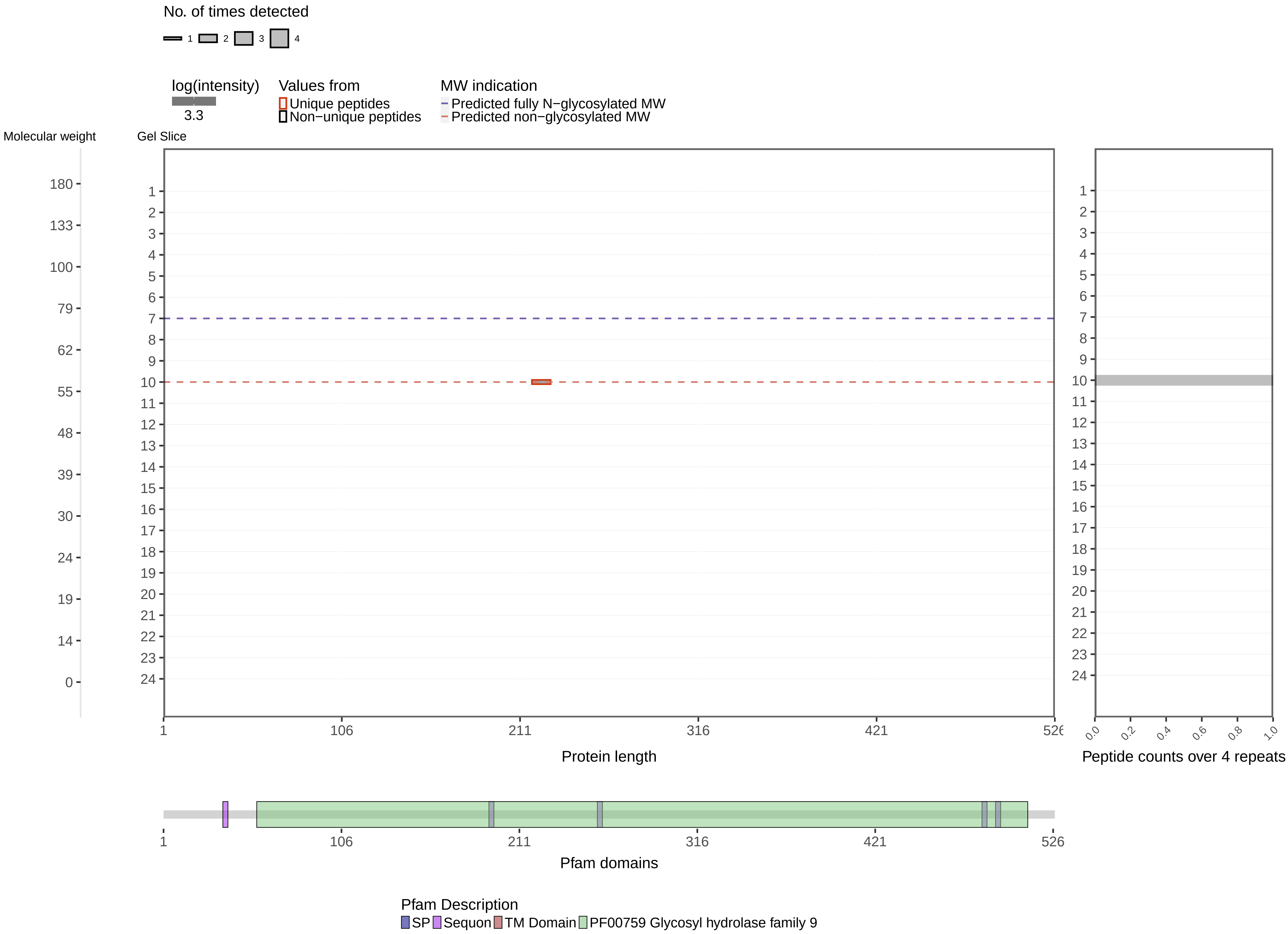


Protein sequence:

PEVATYYNSTGYGDELLWAASWLYHATGDQSYYDYATGKNADSFGNFGNPAWFSWDNKLAGTQVLLSRVSFFDSKGTSNSETLQEYRKTAEAVMCGLLPKSPTATSSRTDSGLIWISEWNALQHPVASAFLAVLYSDYMLTSRTAKITCNGDAYTPSDLRKFAISQADYVLGDNPSKMSYLIGYGDNYPQYVHHRGASIPKDADTGCKDGWKWLDSTEPNPNVATGALVGGPFLNETYIDSRNNSMQAEPTTYNSAVIVGLLSGLVTTSSVVQSFT*

### NbL17g24850.1 probable_diphthine_methyl_ester_synthase [MW = 30.19 kDa]

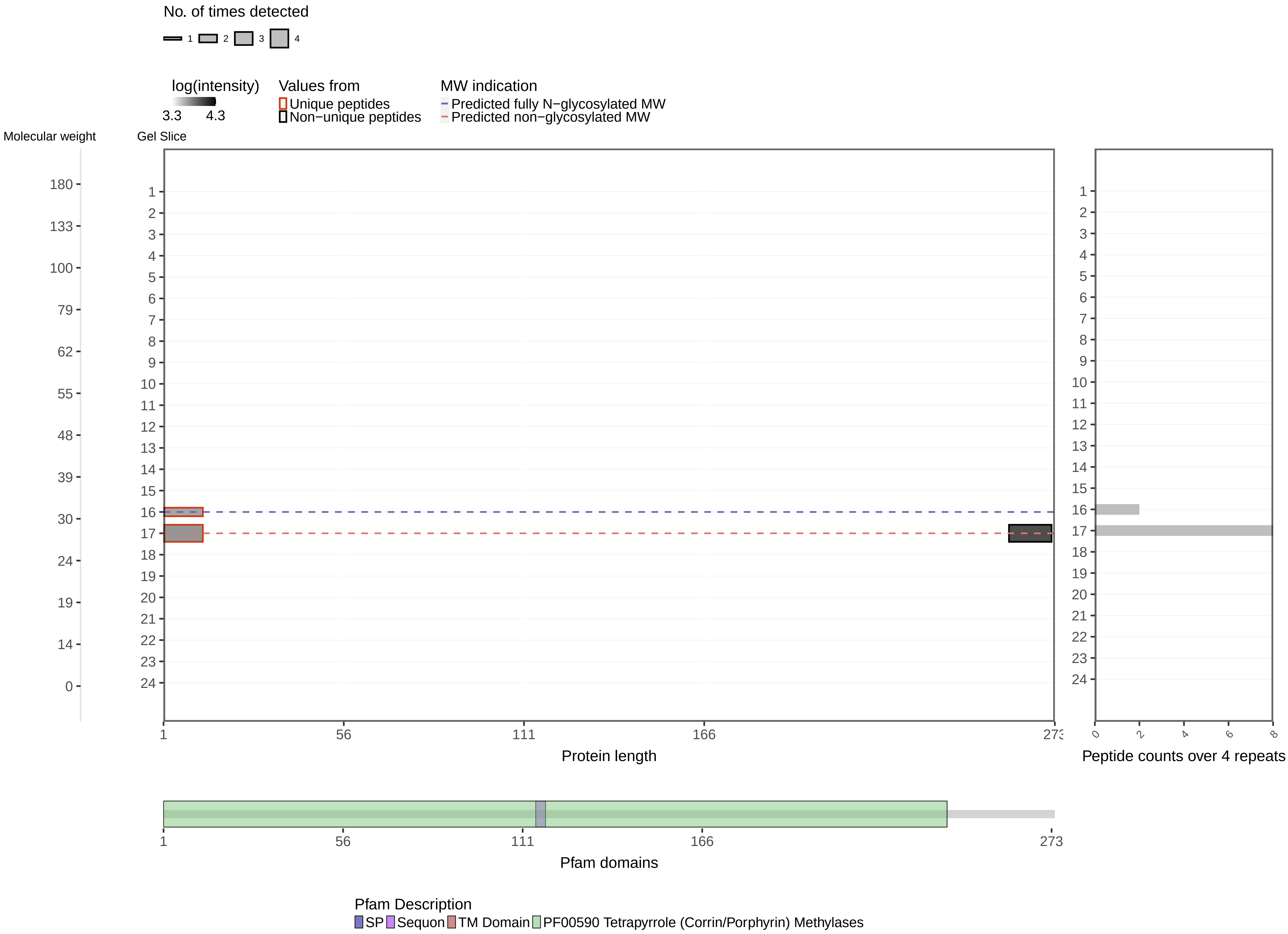


Protein sequence:

CGLQLYHYGETVSIPFFTETWRPDSFYEKIRENRKLGLHTLCLLDIRVKEPTIESLCRGKKQYEPARFMTVNTAIEQLQEVEEARGESVYGENTICVGFARLGSEDQKVVAGSMKQLLTVDFGAPLHCLVIVGKTHPVEEEMLEFYGL*

NbL17g24780.1 alkylated_DNA_repair_protein_alkB_homolog_8−like [MW = 38.52 kDa]

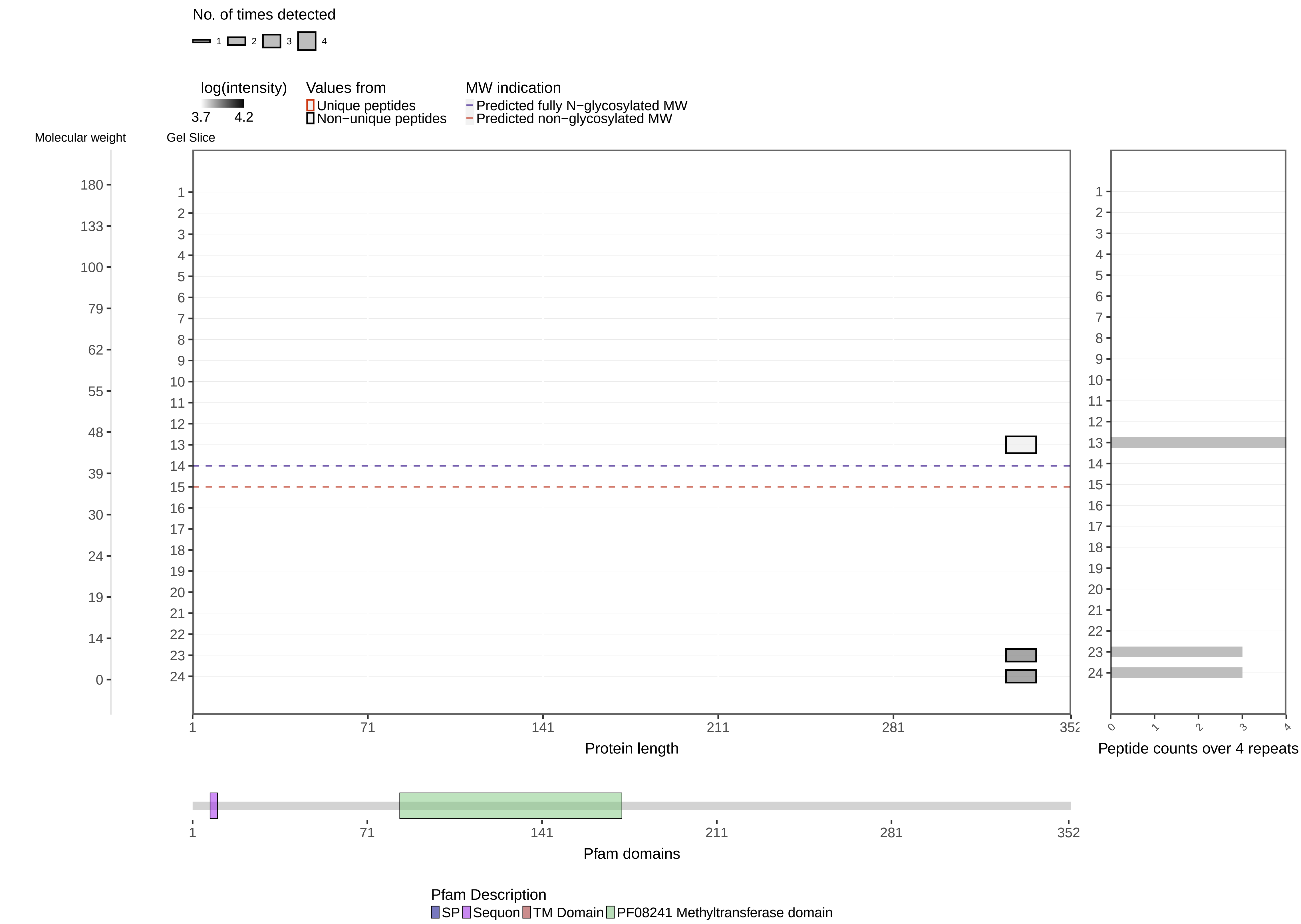


Protein sequence:

VVKKGGCILITVWAREQEDTSLVEKWTPLNQRYIAEWIGPGSPRVRTPSSPCTLESIPEAEENGAGEQLKDLHAKSSKVKSDEVMDSISQDKCRSLLSGSGKGYAEQQEFFVPWHLPYHRAEVSGASAVALASGLAKKDDKKGSVVYNRYYHVFSEGELERLVSGLDNAILVDRFYDKSNWCIILFE*

### NbL17g24570.1 GDSL_esterase/lipase_EXL3 [MW − SP = 36.69 kDa]

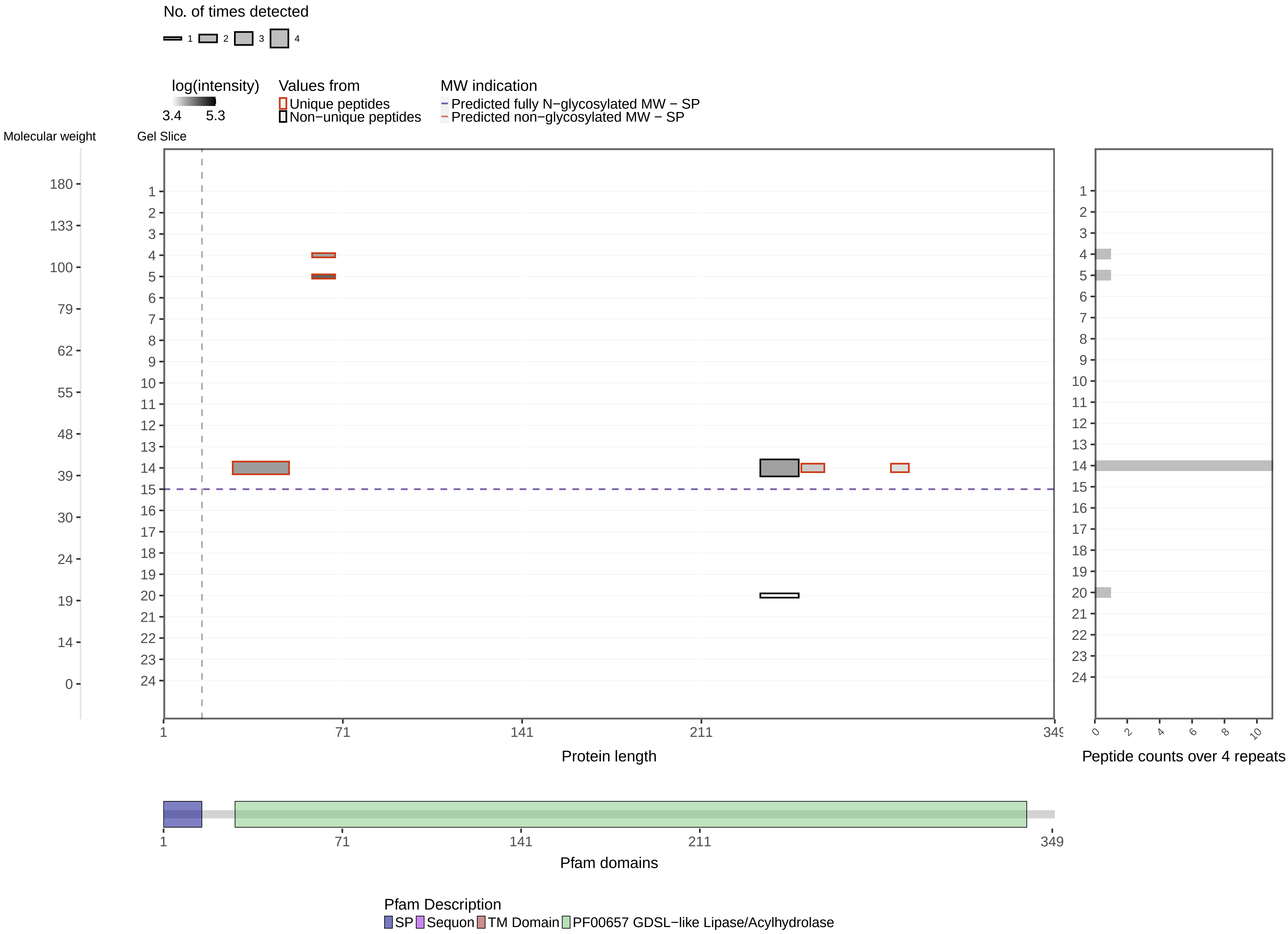


Protein sequence:

/VAGSDDIANTYFTAGIRLKQDINSYTNFMVDKVSEFLQELYKLGARKVGFFGIPPLGCLPSQRTLAGGFNRVCVEEYNQAAELANTKLSAELNSLSKYLPQSKLVFIDIYNPILDIIVNPQKYGLEEVEKGCCGTGKIEVTLLCNKFSGTCEDDTKYFFWDSYHPTEKGYRILVDQILKKYINNFL*

### NbL17g24520.1 uncharacterized_LOC109224900 [MW − SP = 13.94 kDa]

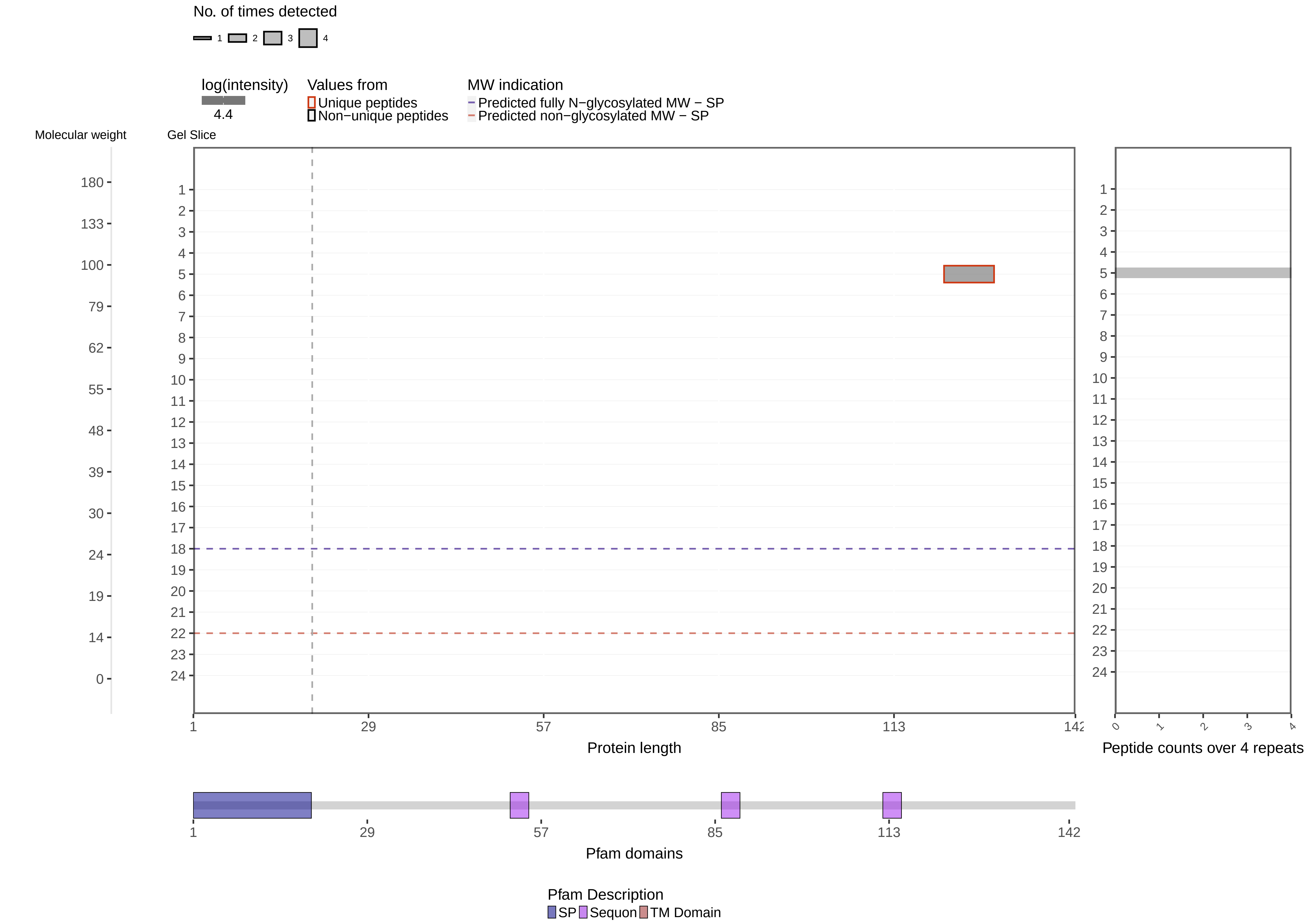


Protein sequence:

IVDNRRYNELDVSMEILNLKTELHSETENNTSCNTIMEKECDNLMGSETVISNENVSSSVRRKFDLICELDKFLIIQSNFMSRN*

### NbL17g24350.1 expansin−A11−like [MW − SP = 25.21 kDa]

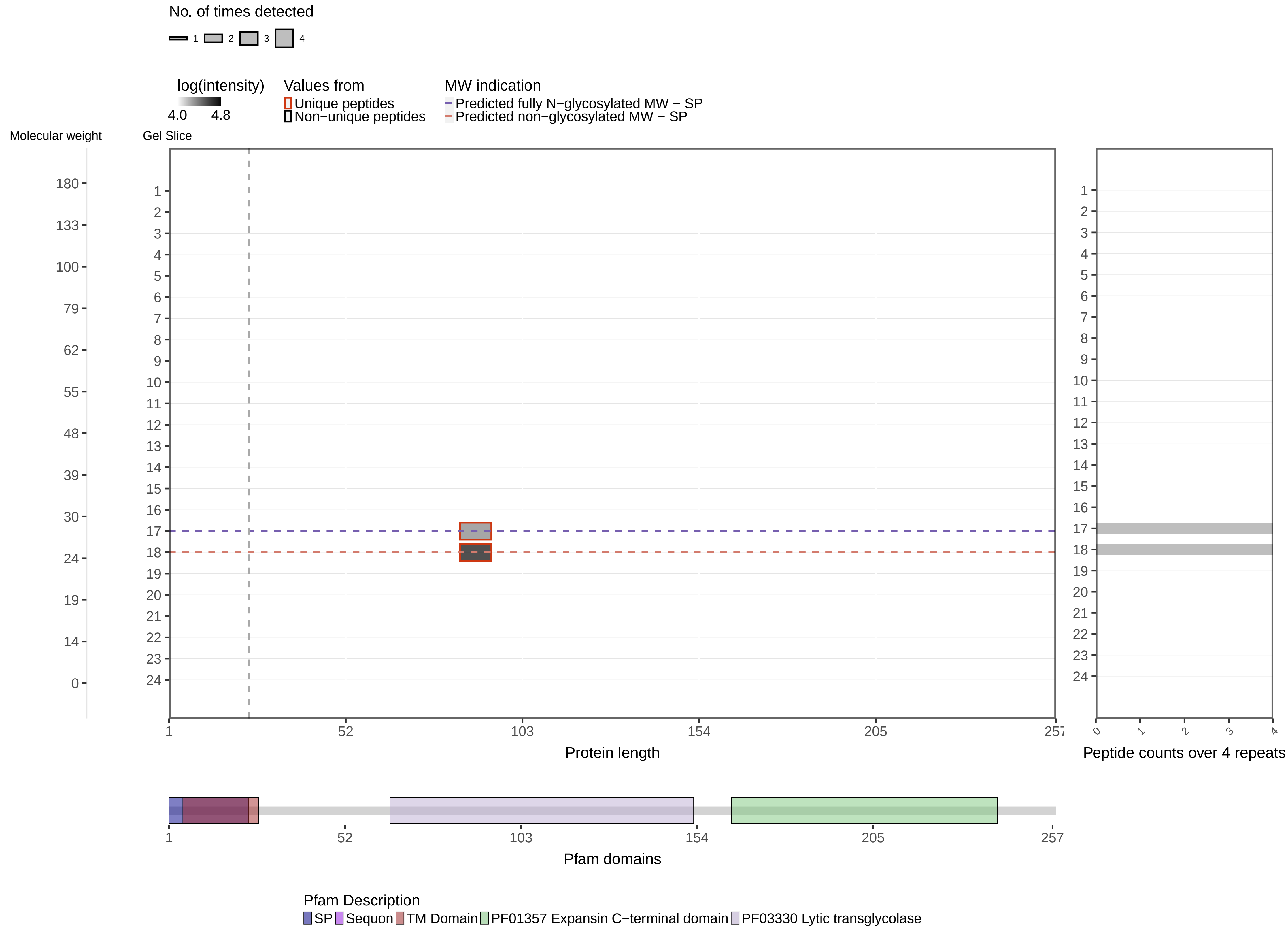


Protein sequence:

PSYNGGWCNPPRPHFDMAQPAWENIGIYRGGIVPVLYQRIPCVKKGGVRFTINGRNYFELLMISNVGGAGSAQSVQIKGSKTNWMTMSRNWGANWQSNSYLNGQPLSFKVTTTDGVTKTFLNAVPSNWHFGQTYSTSINF*

NbL17g24340.1 ribonuclease_P_protein_subunit_p25−like_protein [MW = 28.13 kDa]

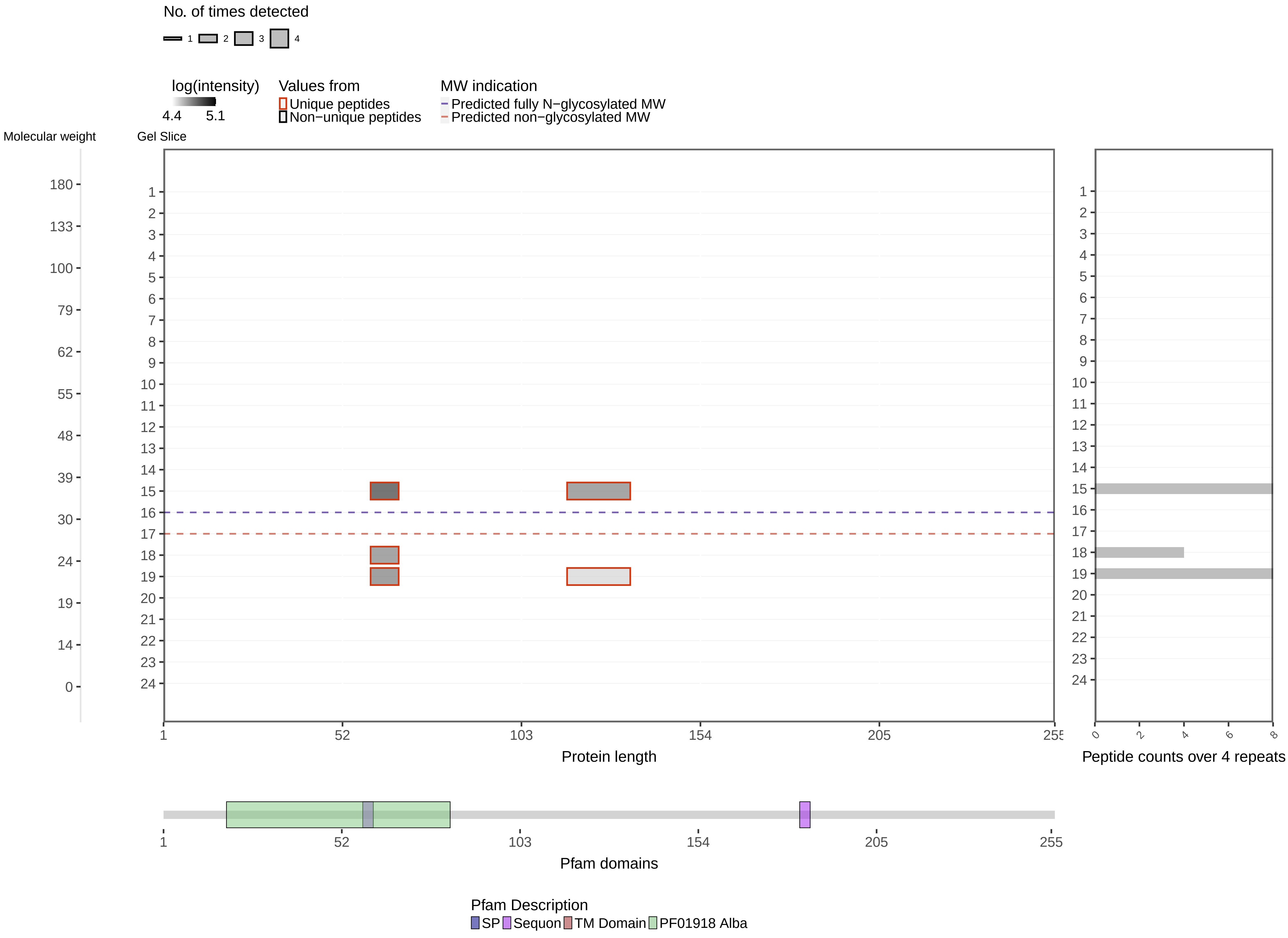


Protein sequence:

/GYQPPLPADQVKVLTDFDYDGEGSPSGGRRGRGGRGRGRFRGFAGNGFMSAEYDDGGYDRNRSYGRGRSRGRGRSFRGRGRGGYNGPQDTQQDAGFYNQEAPMQGRGRGRGRGTRGRGRGFRTTGPIHGGGA*

NbL17g24230.1 thioredoxin−like_protein_CDSP32_chloroplastic [MW = 33.5 kDa]

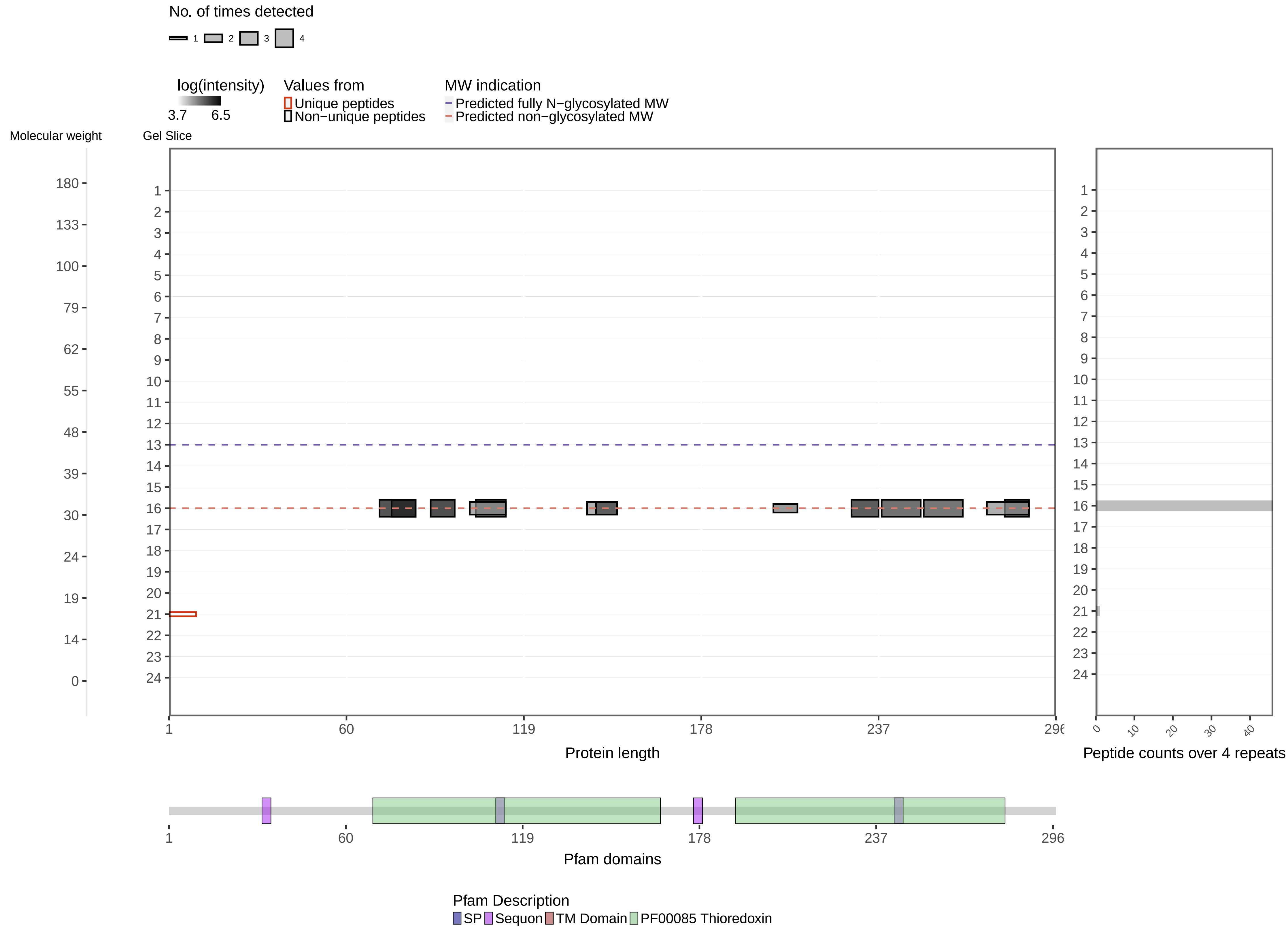


Protein sequence:

REKINKVPHFNFYKSMEKIHEEEGIGPDQLAGDVLYYGDNHSEVVQLHSREDVEKLIQDHKIDQKLIVLDVGLKHCGPCVKVYPTVIKLSRQMADTVVFARMNGDENDSCMQFLKDMDVVEVPTFLFIREGEICGRYVGSGKGELIGEILRYQGVRVTY*

### NbL17g24160.1 tRNA_pseudouridine_synthase_A_mitochondrial [MW = 64.29 kDa]

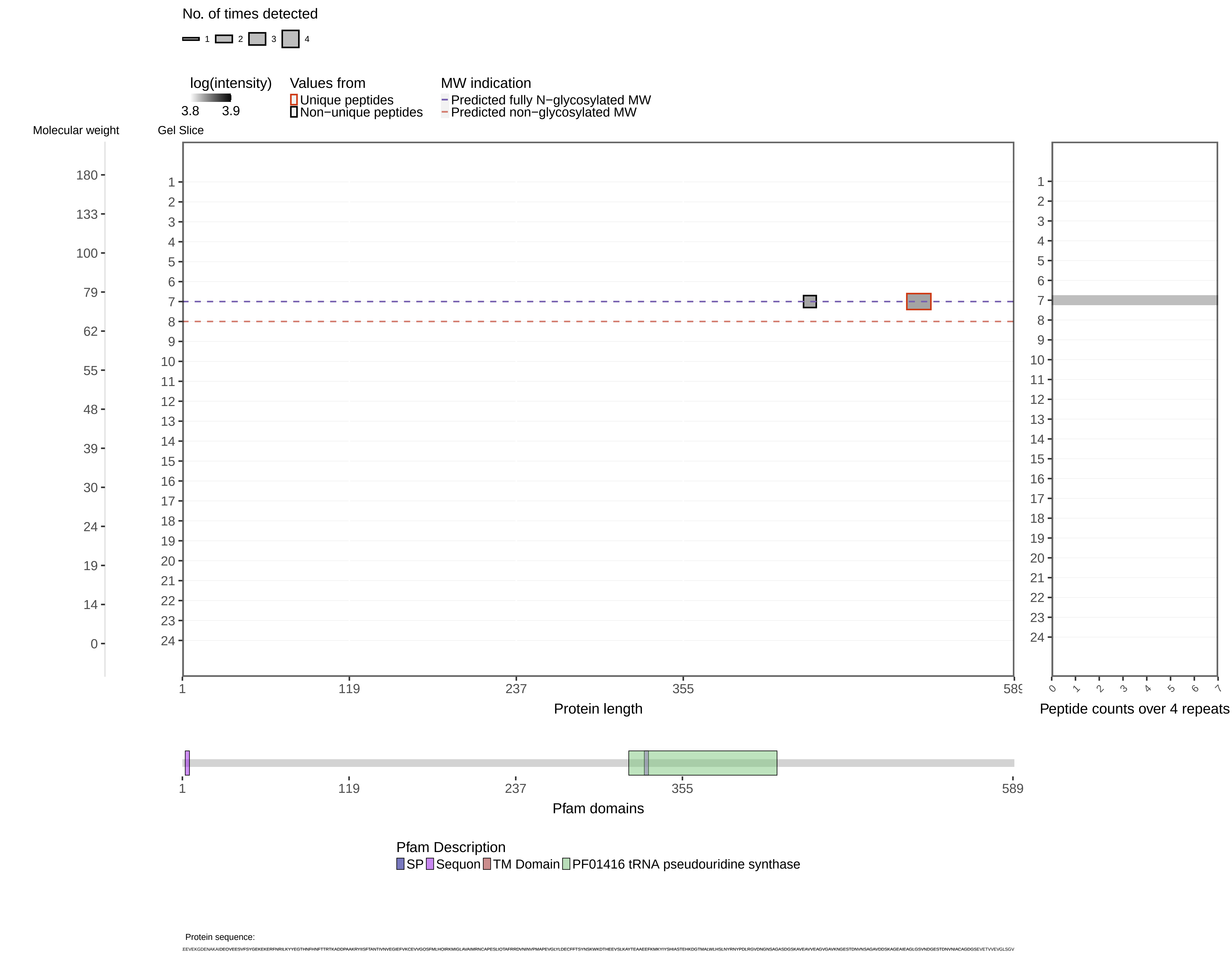

NbL17g24120.1 enoyl−CoA_hydratase_2_peroxisomal−like [MW = 34.08 kDa]

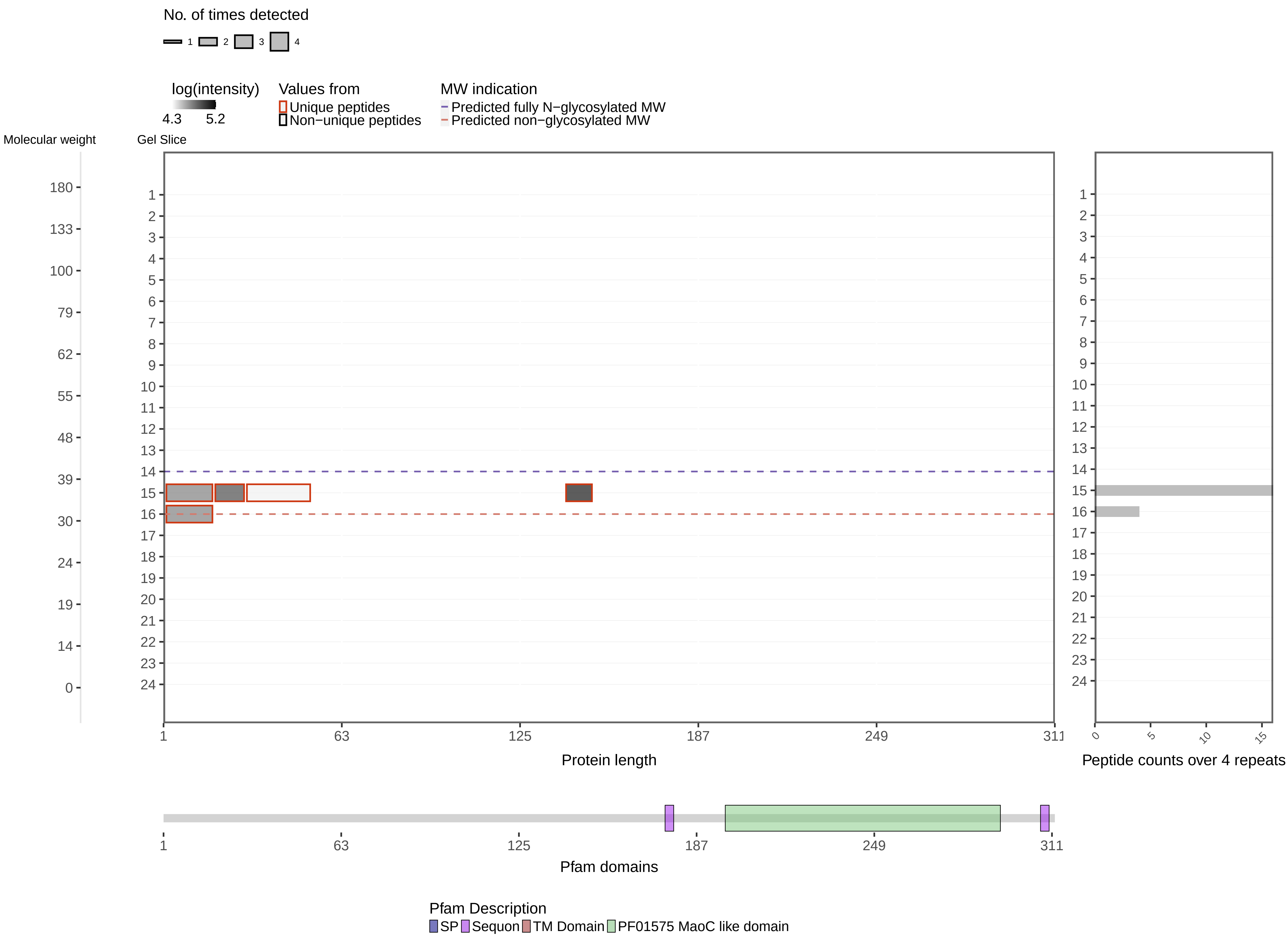


Protein sequence:

SVEPLCMNRMSIYLRGSGGFSKSSRPYSYSRNPSNQTAFPKIPKSQPFAVFEGCTQASQALLYRLSGDYNPLHSDPLVASIAGFSRPILHGLCTLGFAIRAIIGRICRGDHTLIKSISGRFLLHVYPGETLITEMWPEASRVIYQVKVKERNRAVLLGFVDLDRINSSL*

### NbL17g24100.1 L−ascorbate_oxidase_homolog [MW − SP = 57.77 kDa]

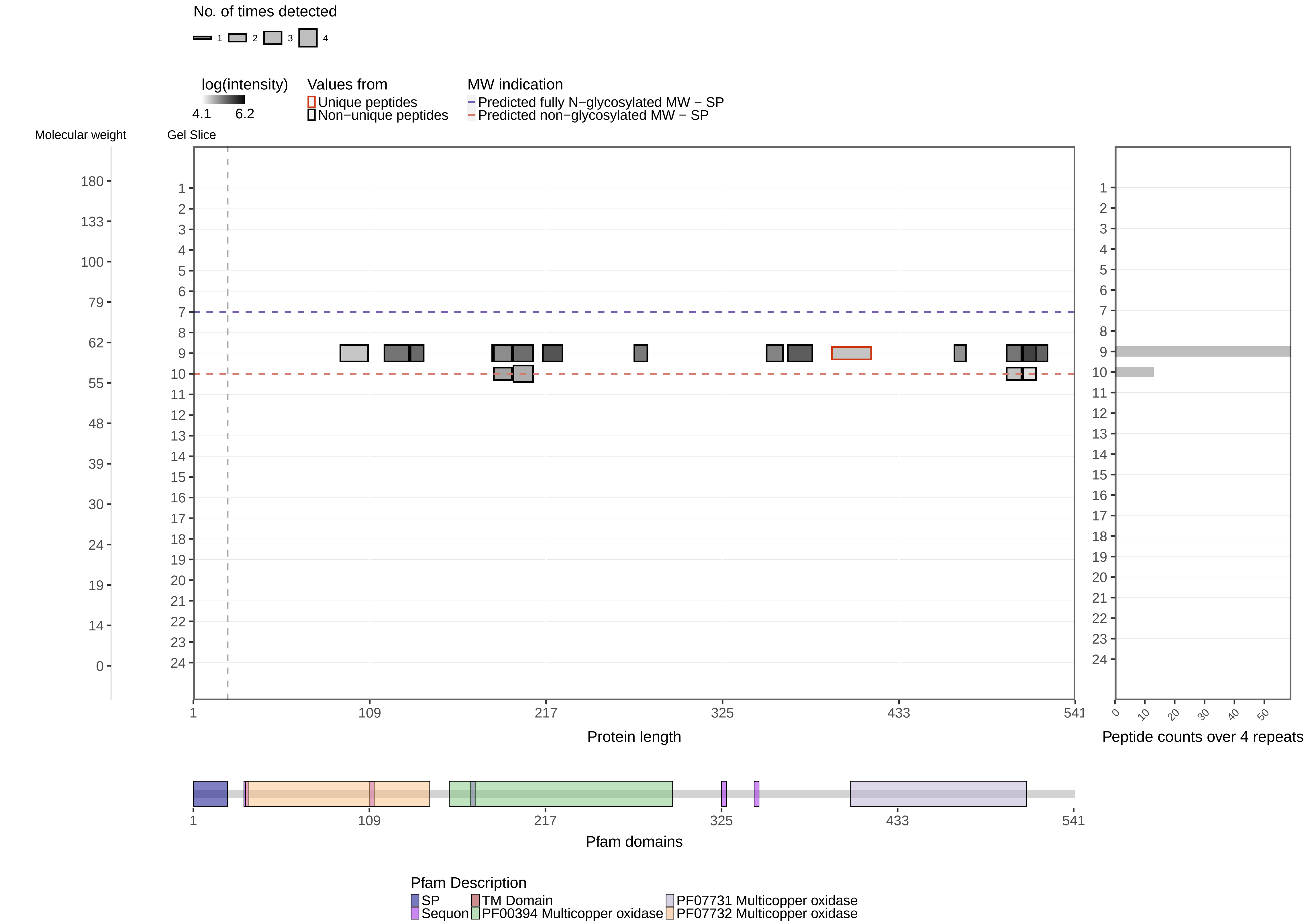


Protein sequence:

VLITADQEAKDHYIVVSSRFTSQVLTTTGVLHYSNSNTPVSGPPPGGPTIQIDWSLNQARSIRTNLSASGPRPNPQGSYHYGMINTTRTIRLASSAGQVNGKQRYAVNSVSFVPLDTPLKLLDYFKVGGFRVGSISDAPSGGGIFLDTSVLGADYRQFIEIVFENNEDIVQSWHLNGYSFWVVGMDGGQWTQASRSGYNLRDAVARCTTQVYPKSWTAIYIALDNVGMWNLRTEFWARQYLGQQLYMRVYTTSTSLRDEYPIPRNARLCGKVAGRHTRPL*

NbL17g24020.1 ATP−dependent_zinc_metalloprotease_FTSH_chloroplastic [MW = 76.02 kDa]

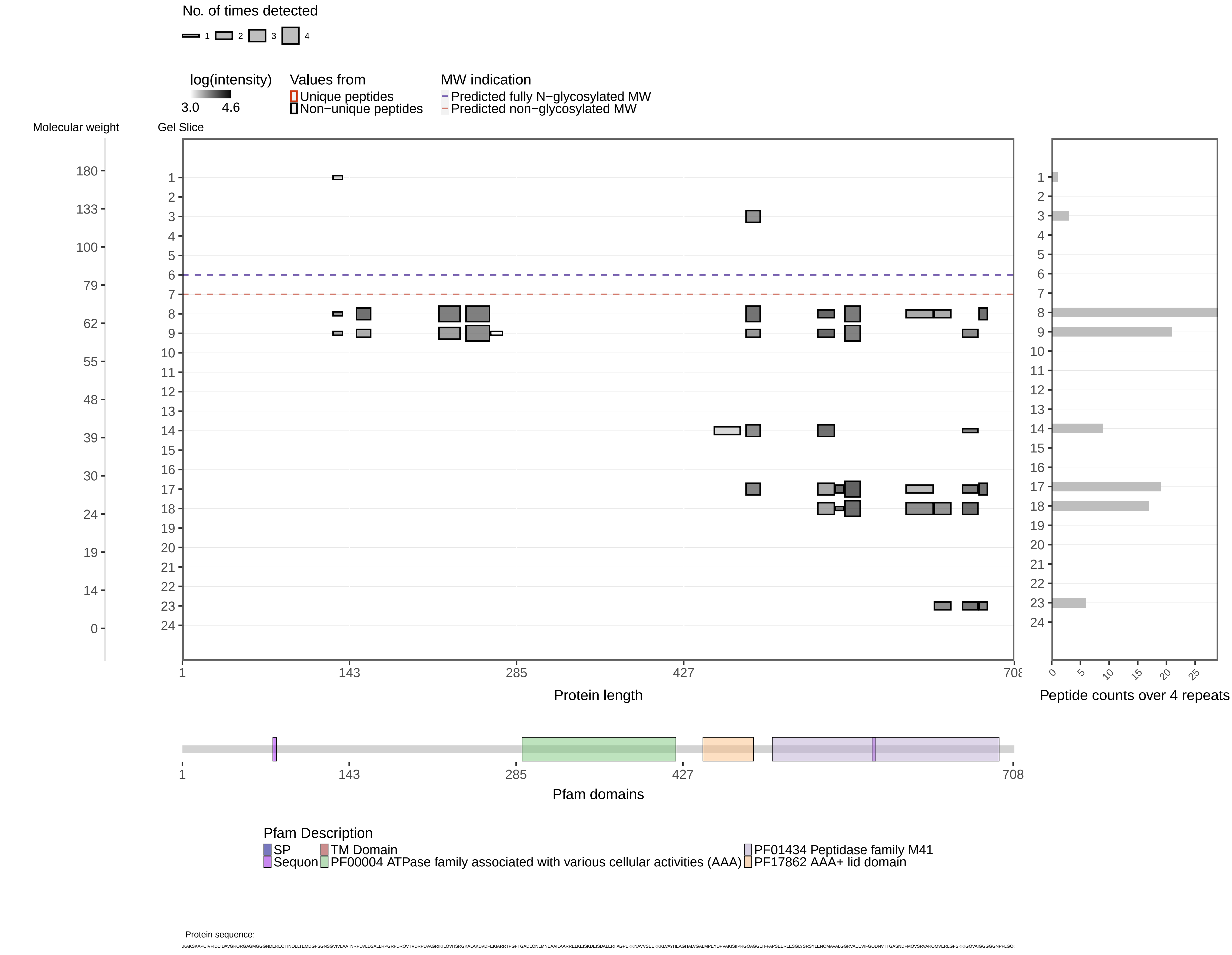

### NbL17g23970.1 uncharacterized_LOC107816208 [MW = 37.39 kDa]

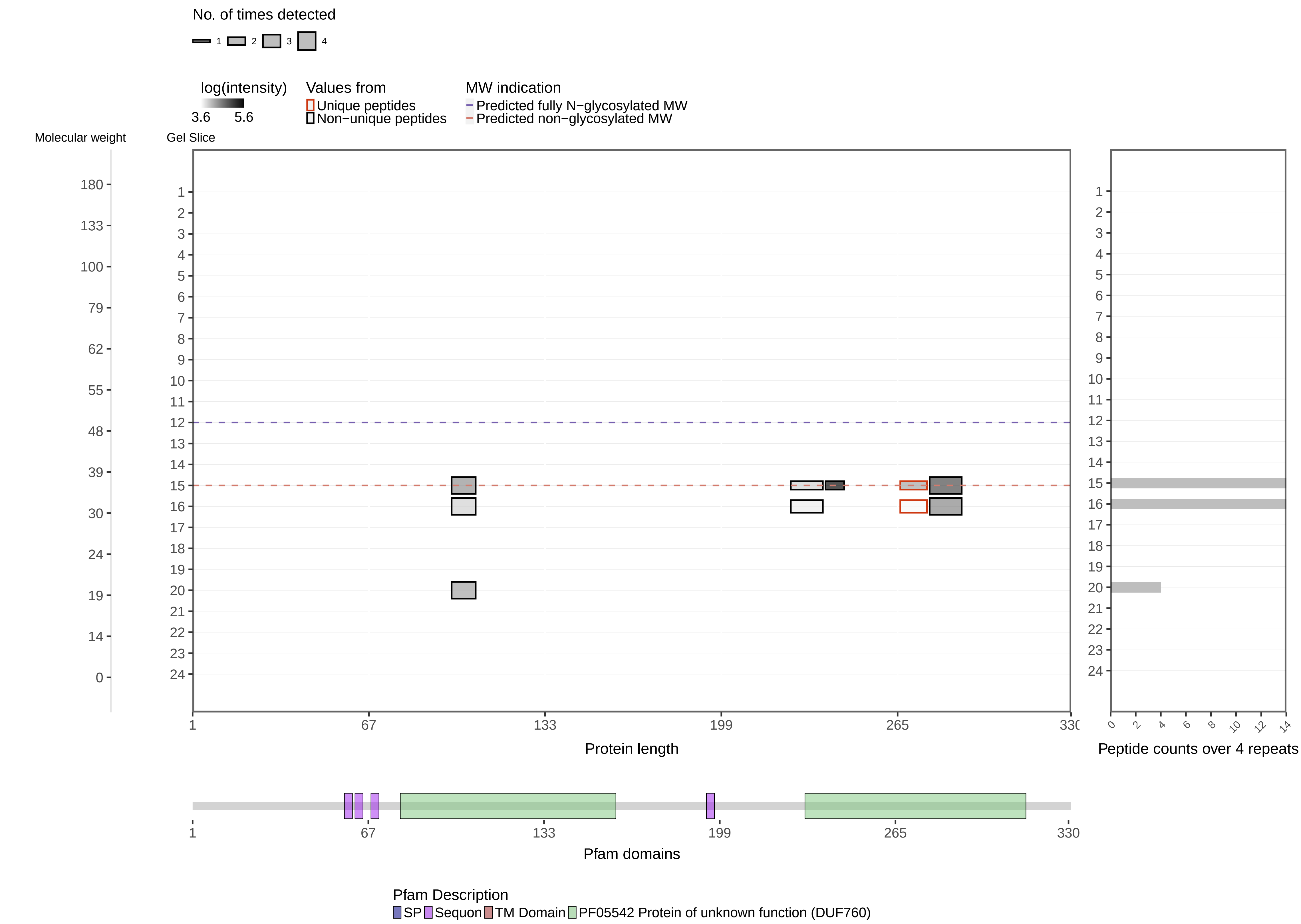


Protein sequence:

AQFRLELQQSLEQAALPEAQEEKDSPDYASGTQKKVTGEVIRWSNVTGPEKIDALKYIELLEAEVEELNRQMERKSGSGYNELLEYLKTLEPQNIKELTSTAGEDVVLAMNSFIKRLVAVSDPSQMKTSVTETSAPELAKLLYWLMVVGYSIRNIEVRFDMERVLGTPPKLAELPPGENI*

NbL17g23910.1 14−alpha−glucan−branching_enzyme−like [MW = 98.34 kDa]

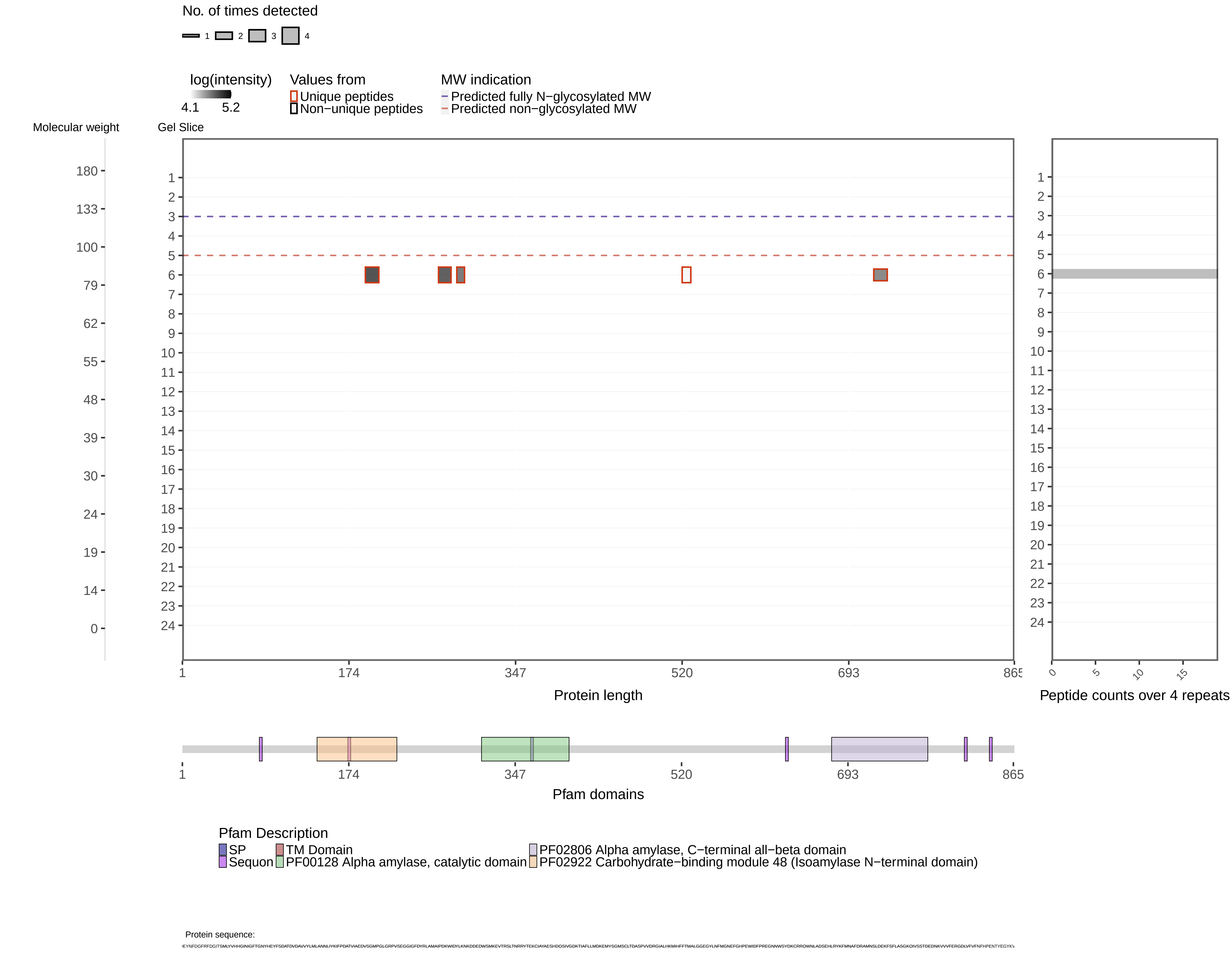

### NbL17g23850.1 WD_repeat−containing_protein_LWD1 [MW = 38.97 kDa]

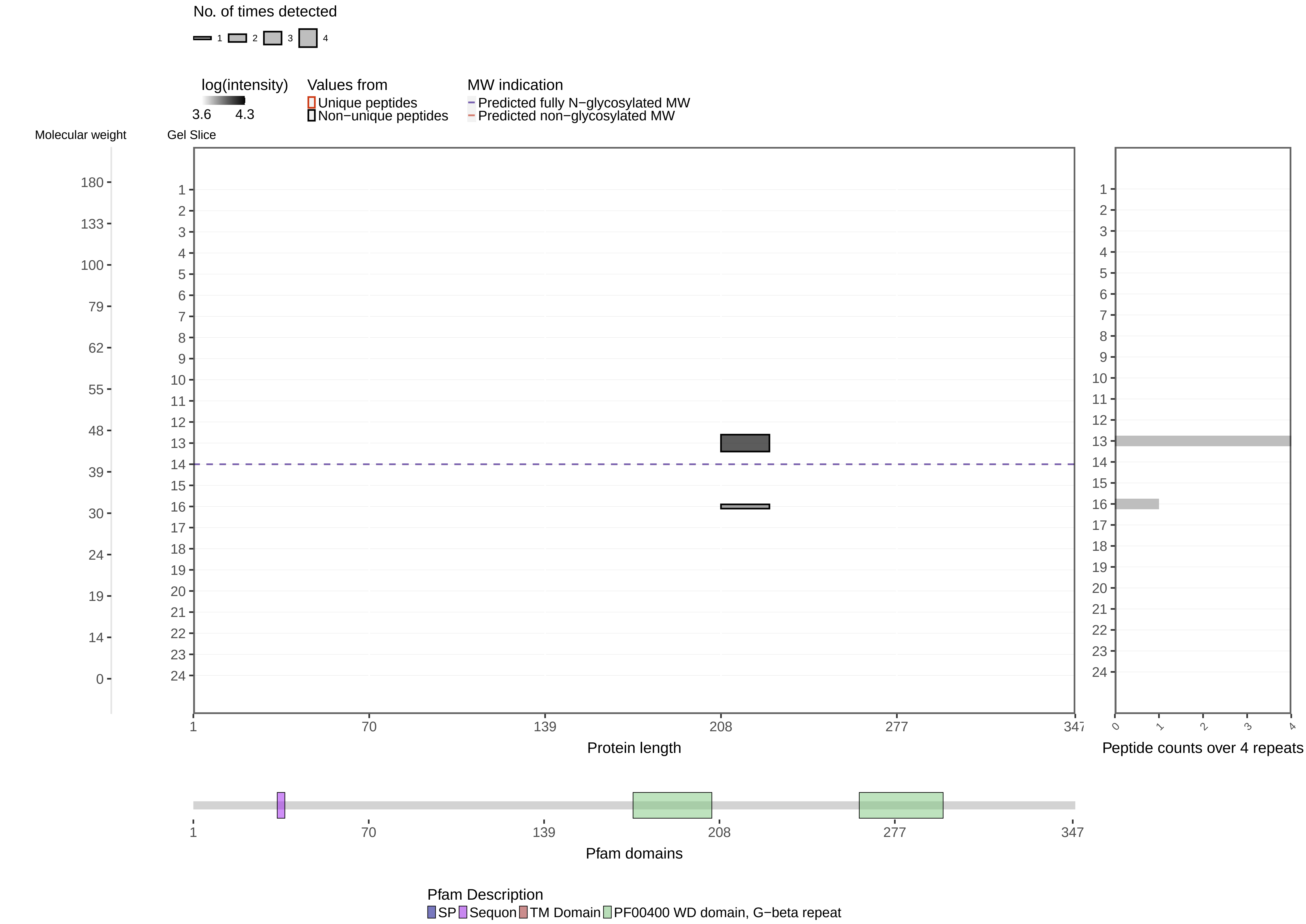


Protein sequence:

TCTIWDIERETVDTQLIAHDKEVYDIAWGGVGVFASVSADGSVRVFDLRDKEHSTIIYESSEPDTPLVRLGWNKQDPRYMATIIMDSAKVVVLDIRFPTLPVVELQRHQASVNAIAWAPHSSCHICTAGDDSQALIWDLSSMGQPIEGGLDPILAYTAGAEIEQLQWSSSQPDWVAIAFSNKLQILRV*

### NbL17g23740.1 epidermal_growth_factor_receptor_substrate_15−like_1 [MW = 107.52 kDa]

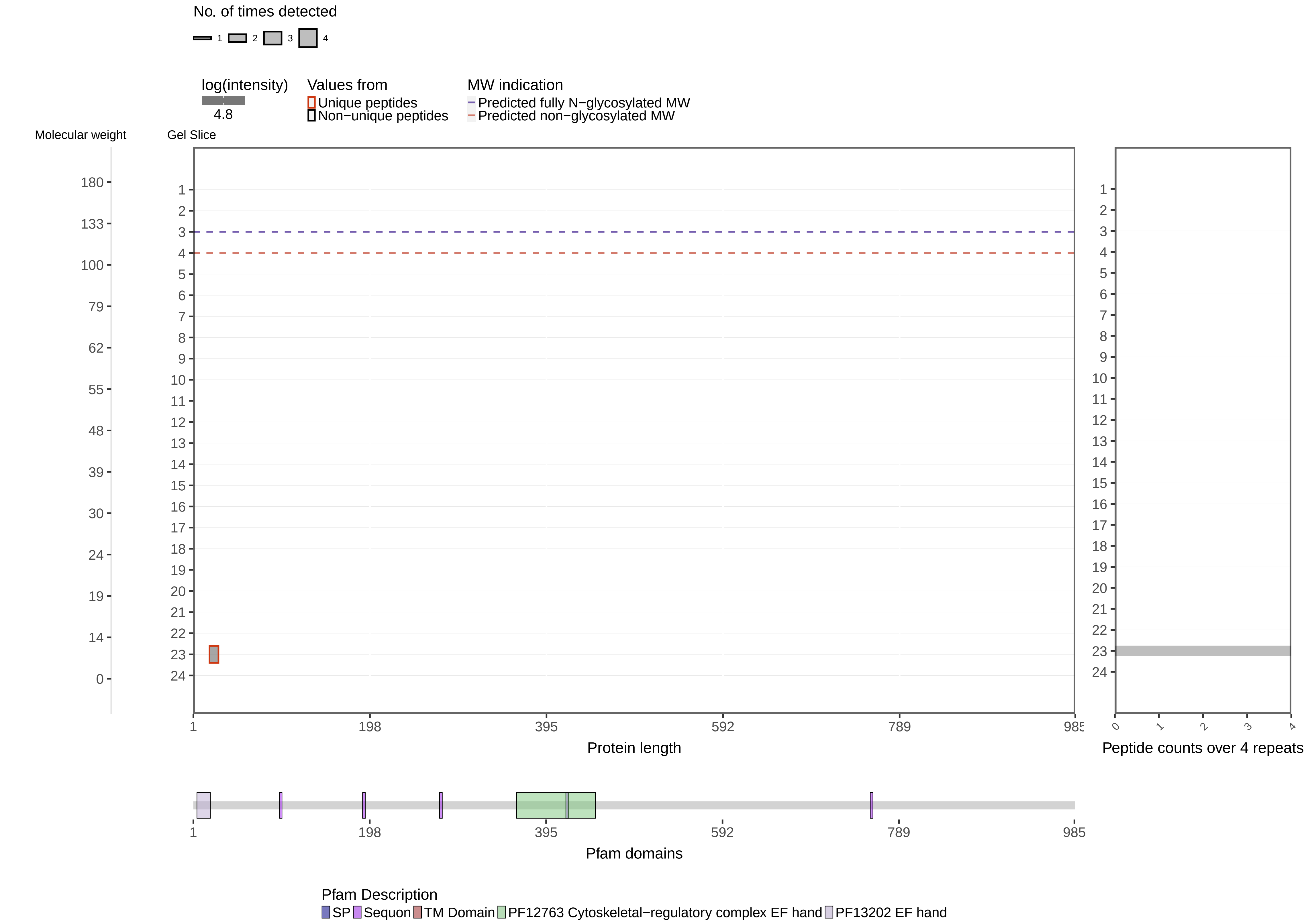


Protein sequence:

[illegible]

### NbL17g23690.1 uncharacterized_LOC109224789 [MW = 17.59 kDa]

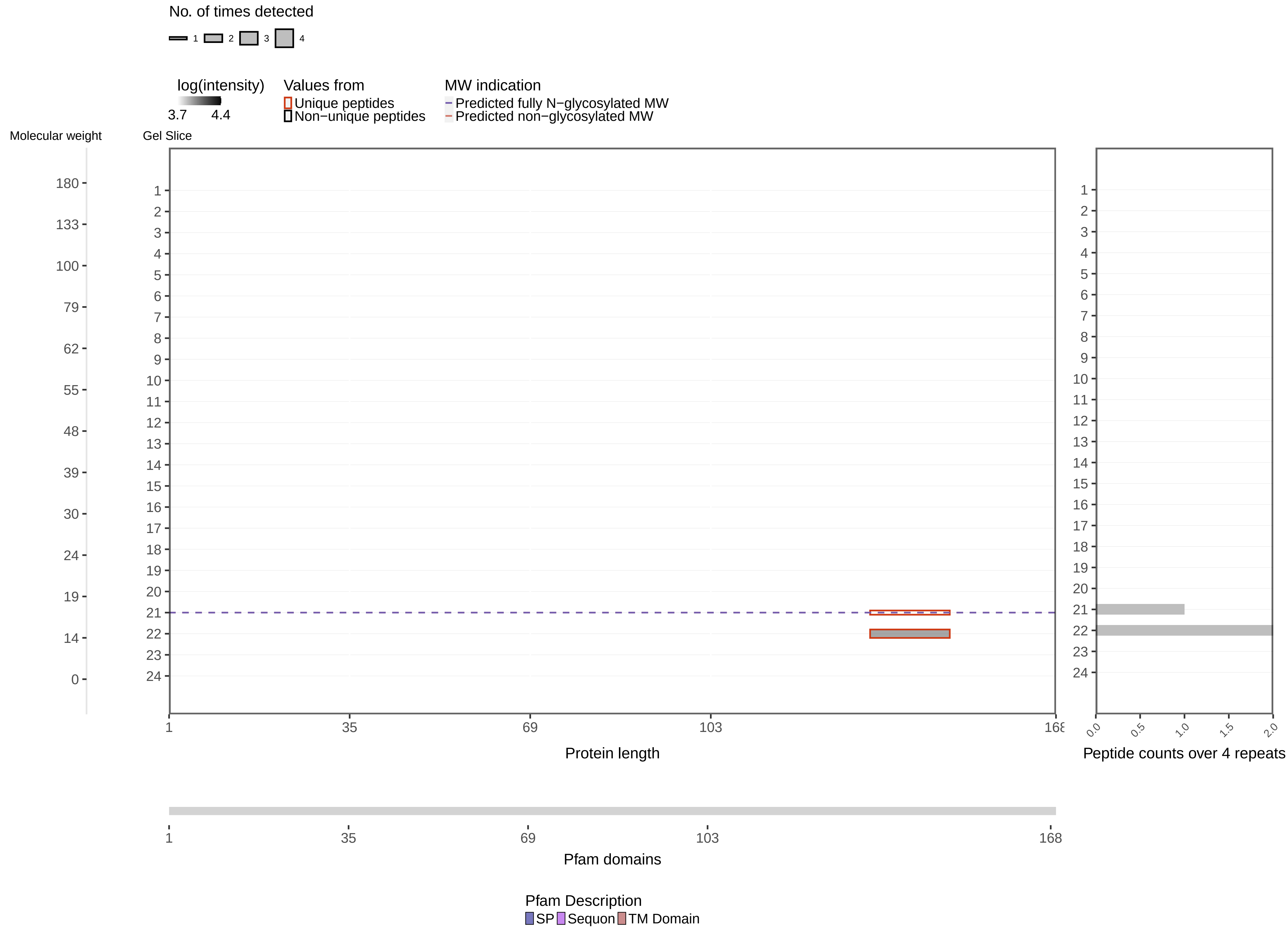


Protein sequence:

AGIFVGGFVLGGLIVGTLGCVFALQISKALAGADKKDLMRKLPKFIYDEDKALEKQRKKLAEKIDQLNSAIDDISTQLRSDDTPNGAAVNSDDVEAVV*

### NbL17g23640.1 protein_canopy_homolog_1−like [MW − SP = 18.78 kDa]

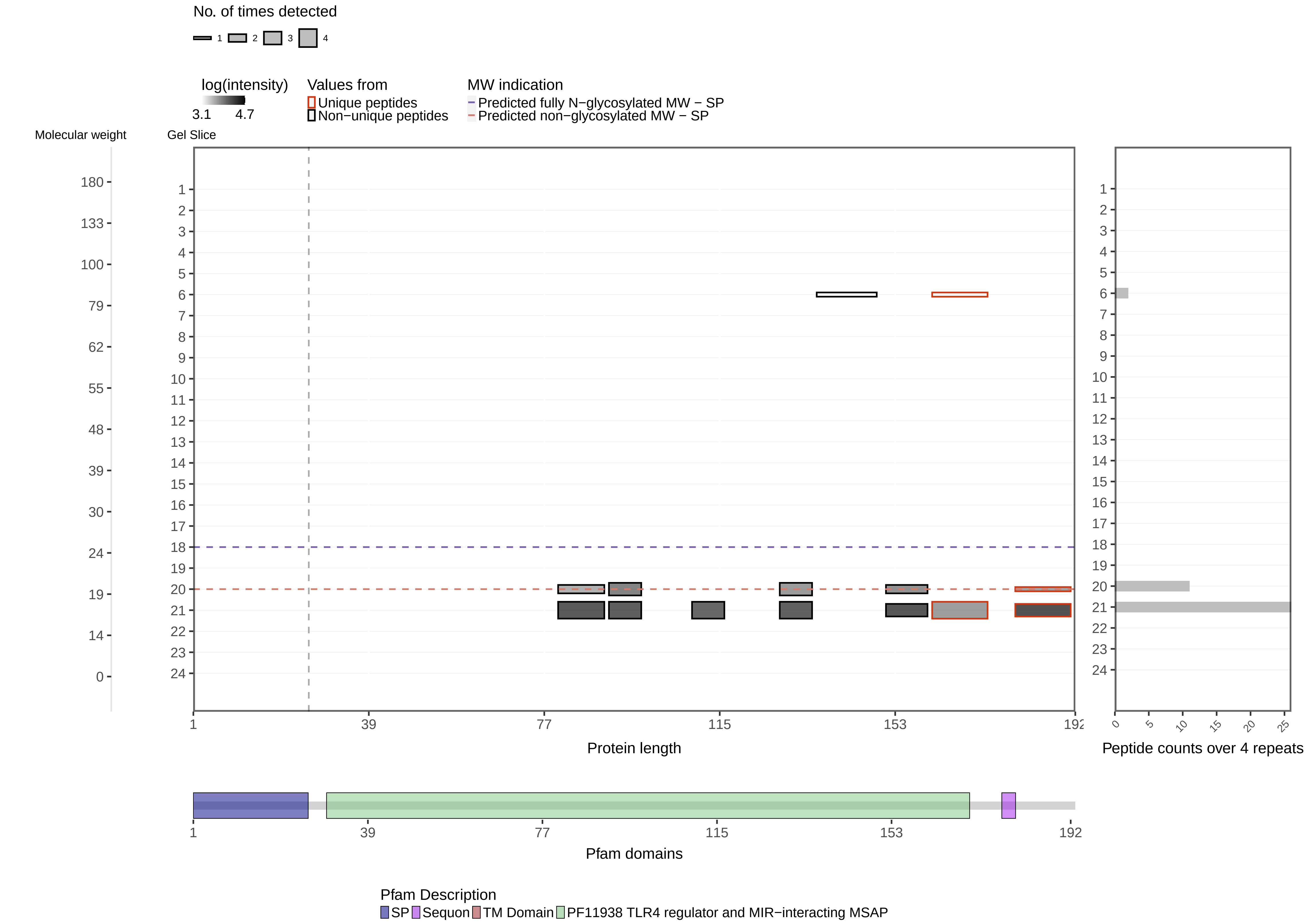


Protein sequence:

DLCEKMQDYTLEKVDSSTKTWIKVNNWDLLKTNKQEARAHSKAISSFCGRLLEQTEDDFTELMKKGSVQVGDVSKVLCEDLSNYCNGKSSSNKTVGDENEDADREL*

NbL17g23590.1 peptidyl−prolyl_cis−trans_isomerase_FKBP18_chloroplastic [MW = 24.73 kDa]

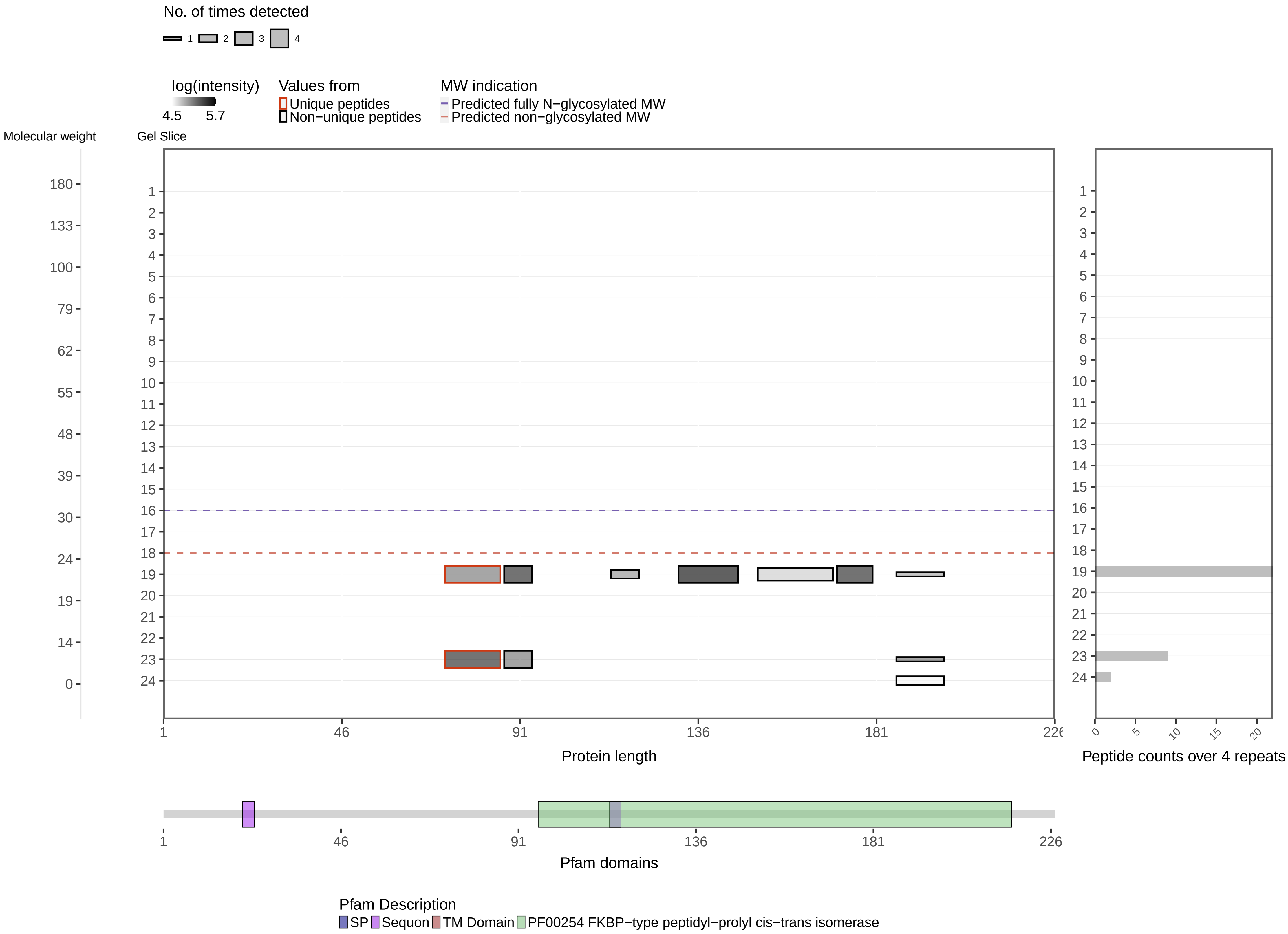


Protein sequence:

STVQVHFECLYRNITAVSSRESKLLAGNRIISQPYEFQVGAPPGKERKRDFVDNPNGLFSAQAAPKPPKAMYTITEGMKVGGKRTVIVPPEAGYGSRGMNEIPPGATFDLNIELLEVKKPEGKQ*

### NbL17g23490.1 xylem_cysteine_proteinase_1−like [MW − SP = 37.04 kDa]

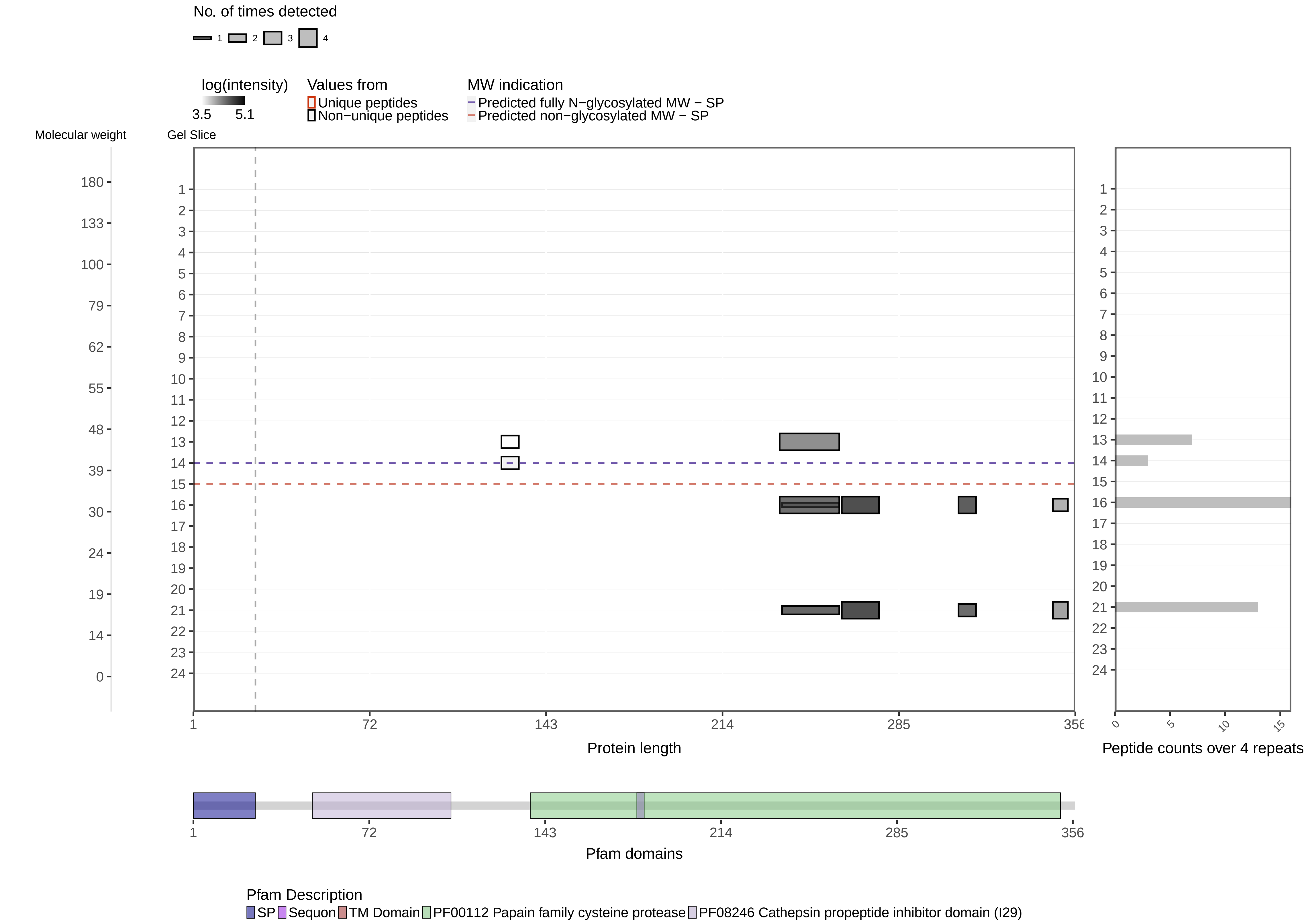


Protein sequence:

AAVEGINQIKTGNLTSLSEQELIDCDTTYNNGCNGGLMDYAFQFIISNGGLHKEDDYPYLMEEGTCNEKRDESEVVTIDGYHDVPANDEQSLLKALANQPLSVAIEASGRDFQFYSGGVFDGHCGTELDHGVAAVGYGTTKGADYIIVKNSWGPKWGEKGFIRMKRNTGKAEGKCGINKMVSFPVKKK*

### NbL17g23450.1 increased_DNA_methylation_3 [MW = 19.91 kDa]

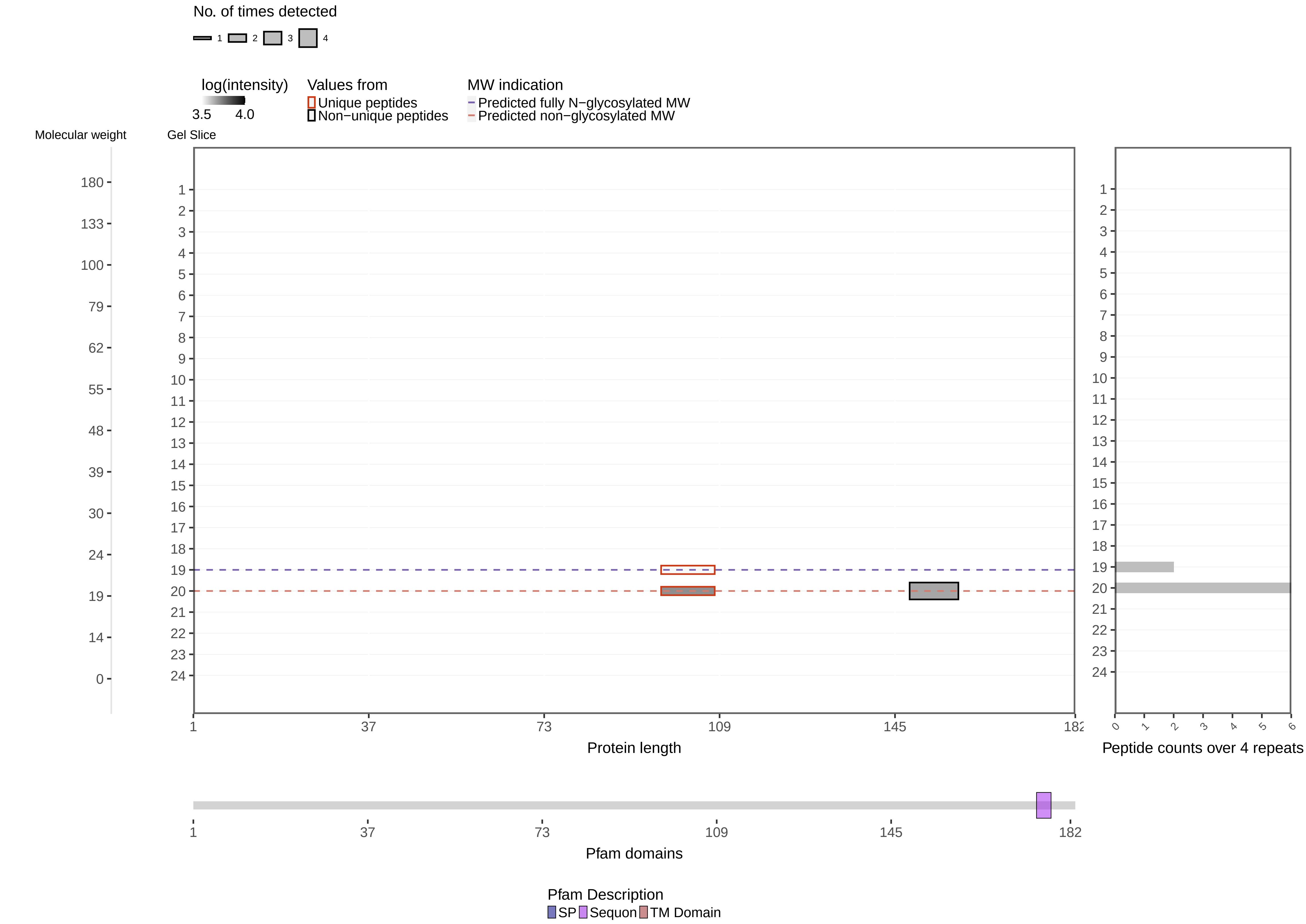


Protein sequence:

GVRNKCNIKCDIQREGRVRIEGVITESDVLKNSPKDYEMKVQQLSPPGPFTVSFNLPGPVDPRLCSRRFRSDGILEVIVLKYRIPIVSAEGLPENWYNGSFPAP*

### NbL17g23380.1 cinnamoyl−CoA_reductase_2−like [MW = 37.37 kDa]

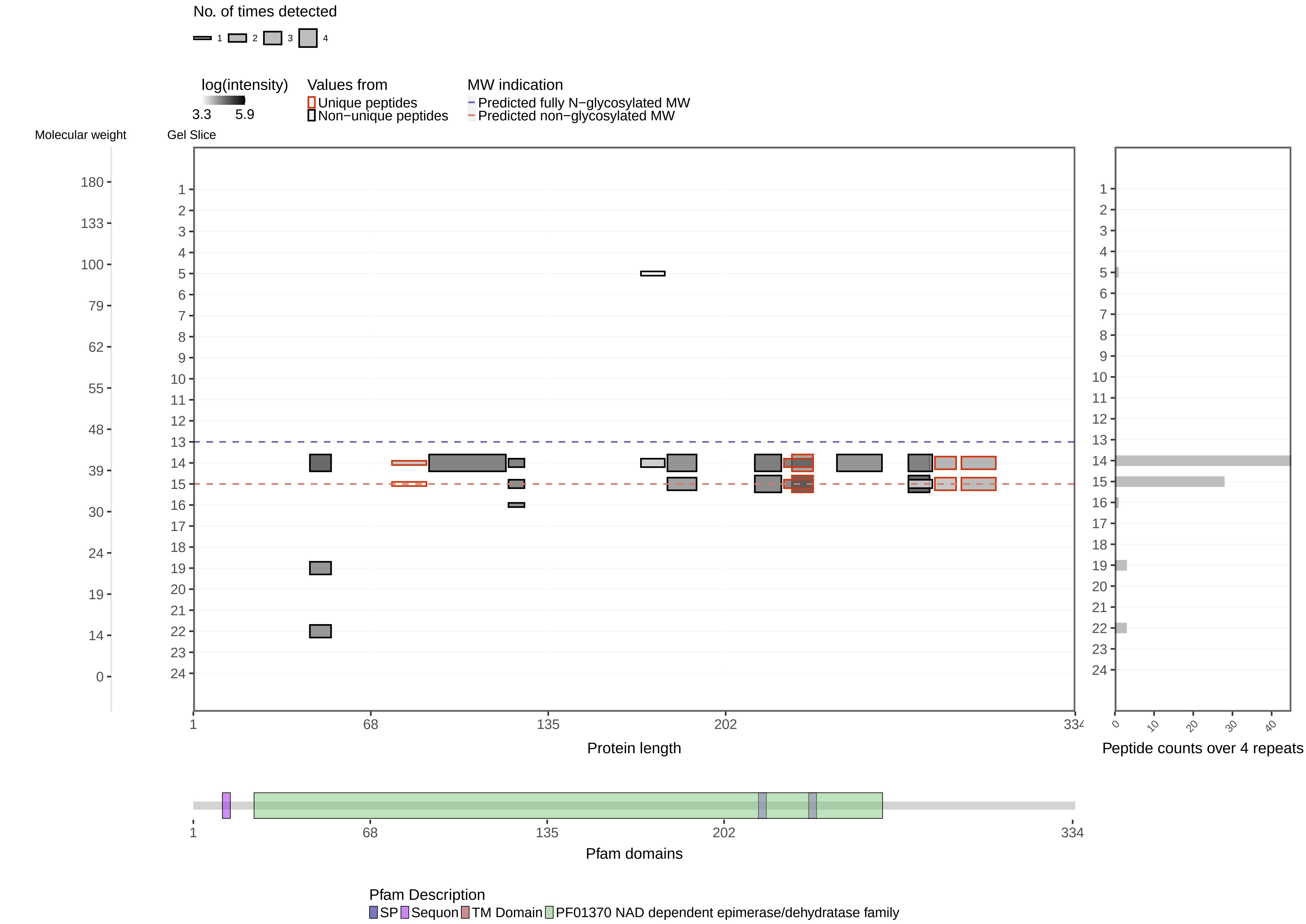


Protein sequence:

)VKDETCWSDSEYCKATNNWYSCSKTMAETEAWSYAKQSGLDLVTVLPTIVWGPMLQKTTNASSLILIKLLKEGYEELENKTRFIIDVRDLAEALVLVYEKPEAEGRHICMAHKAKSQDLVDMLKRYYPNYNYPKRFTDVREEGSYSTEKLQKLGWKCRPLEETLIDAVESYKQAGILD*

### NbL17g23280.1 pyrophosphate−−fructose_6−phosphate_1−phosphotransferase_subunit_alpha [MW = 67.36 kDa]

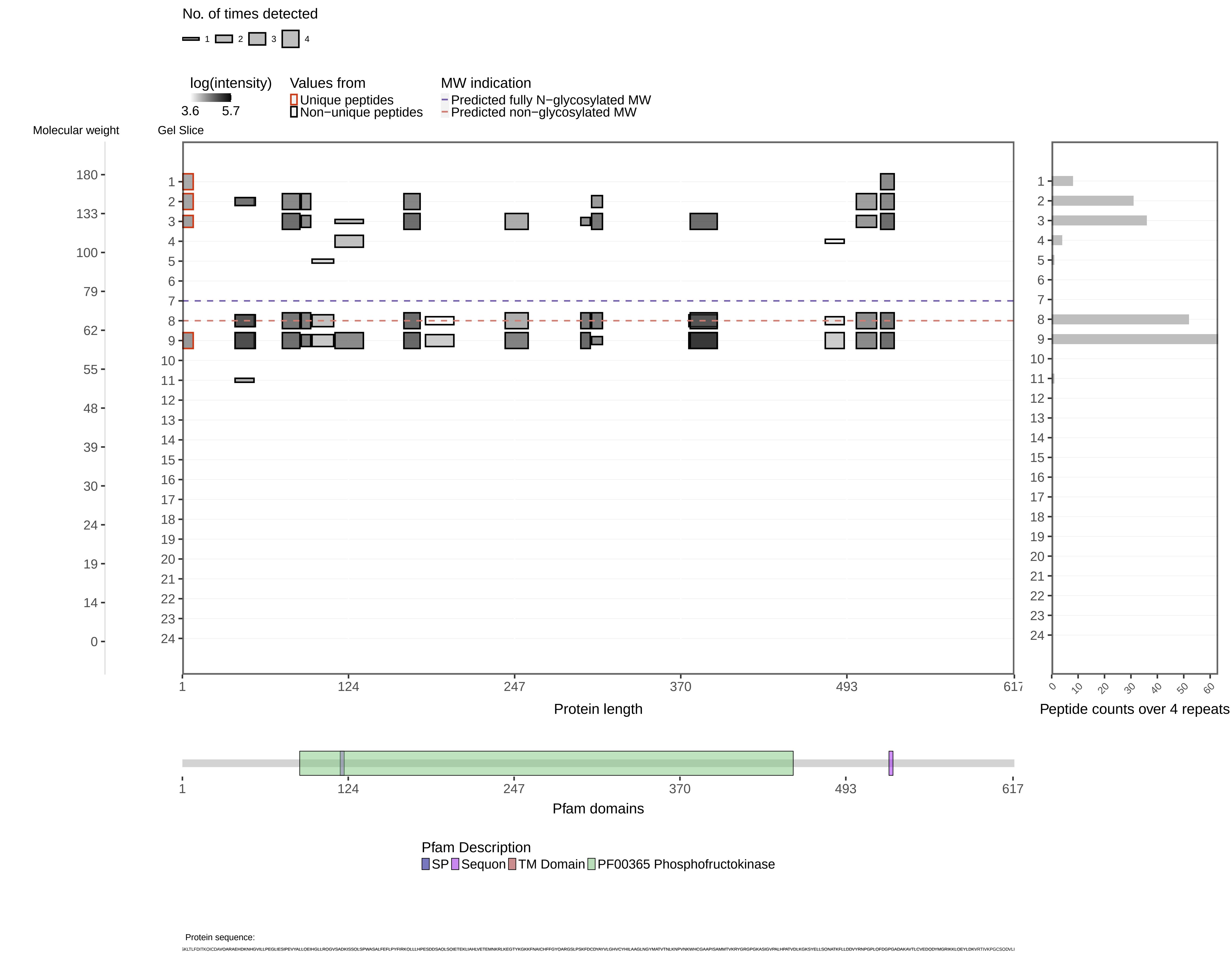

NbL17g23180.1 methylcrotonoyl−CoA_carboxylase_beta_chain_mitochondrial [MW = 61.59 kDa]

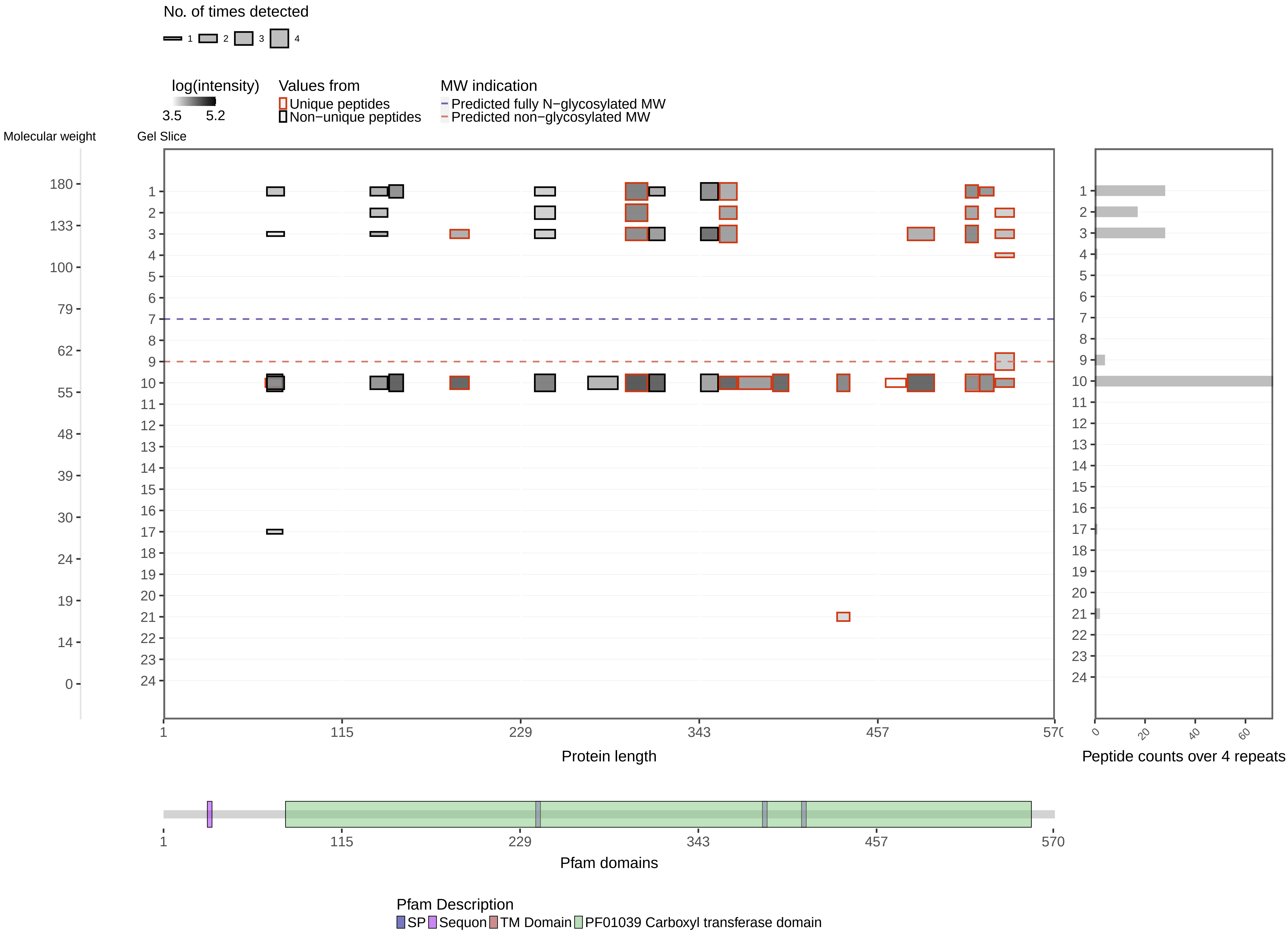


Protein sequence:
SGVSDYFAHDELHALAIGRNIVKNLHMAGGPEASGQTRADYKEPLYDVKELRTIAPTDLKQPFDVRSVIARIVDGSEFDEFKKLYGTTLVTGFGRICGQPVGILGNNGILFNESAQKGAHFIELCAQRNIPLIFLQNITGFMVGSKSEASGIAKAGAKMVMAVSCAKVPKITVVIGGSFGAGNYAMCGRAYSPNFMFFWPNARISVMGGPQAAGVLAQVEKATKKKKGIQWAKEEEEKFKAEVVEAYDREGSPYYATARLWDDGIIDPADTRKILSLCISASLNRGPETTKFGVFR

### NbL17g23170.1 thioredoxin_M3_chloroplastic−like [MW = 19.95 kDa]

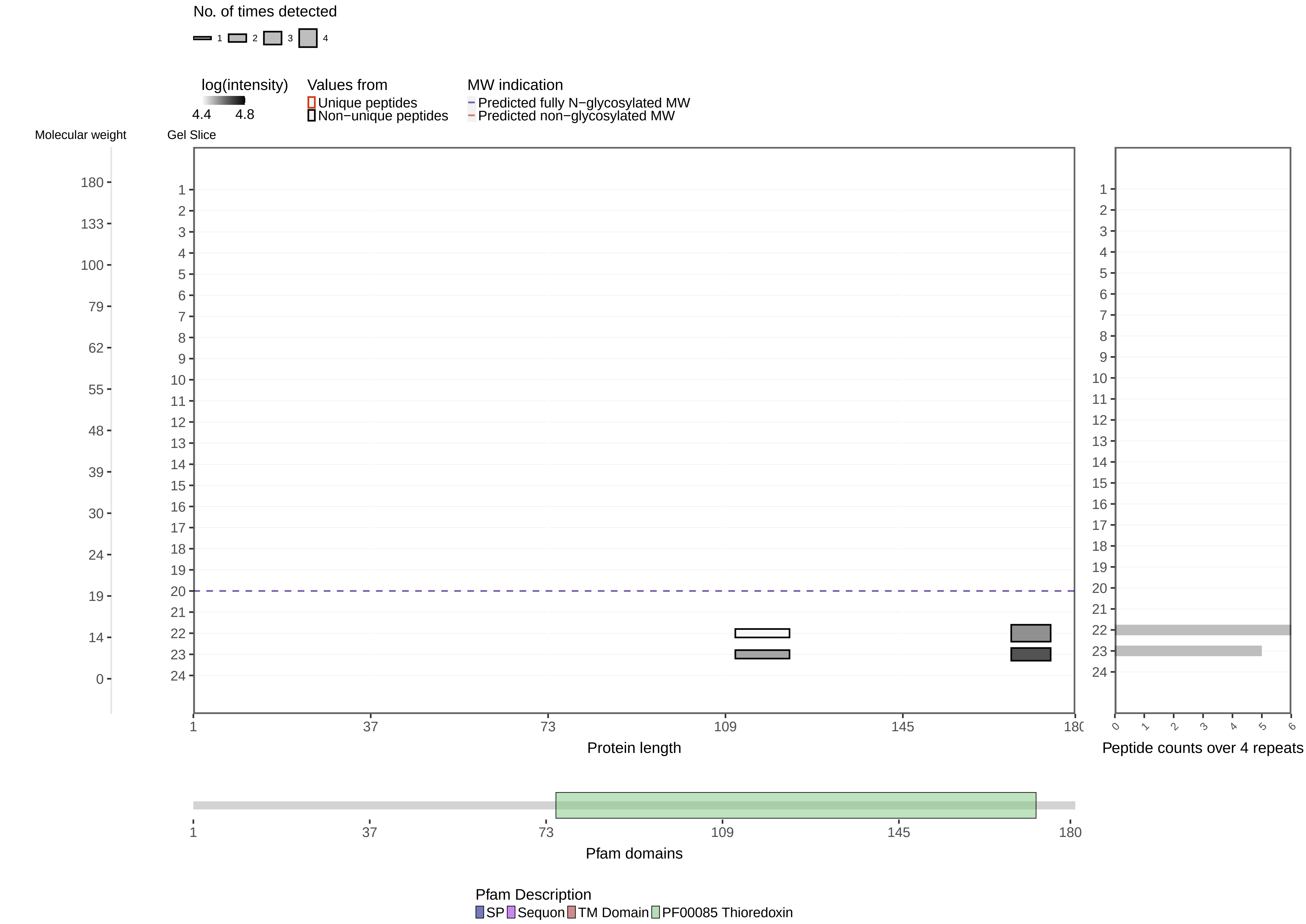


Protein sequence:

EVVTAKSWEKLILSSDTLVLVEFHATWCGPCQMVHRVIDEIAAEYSGRLKCYVLDADSESQIAEDYDIKAVPVVLLFKNGEKCDSVIGTMPKEFYVASIERLLAS*

### NbL17g23140.1 PI−PLC_X_domain−containing_protein_At5g67130−like [MW = 52.03 kDa]

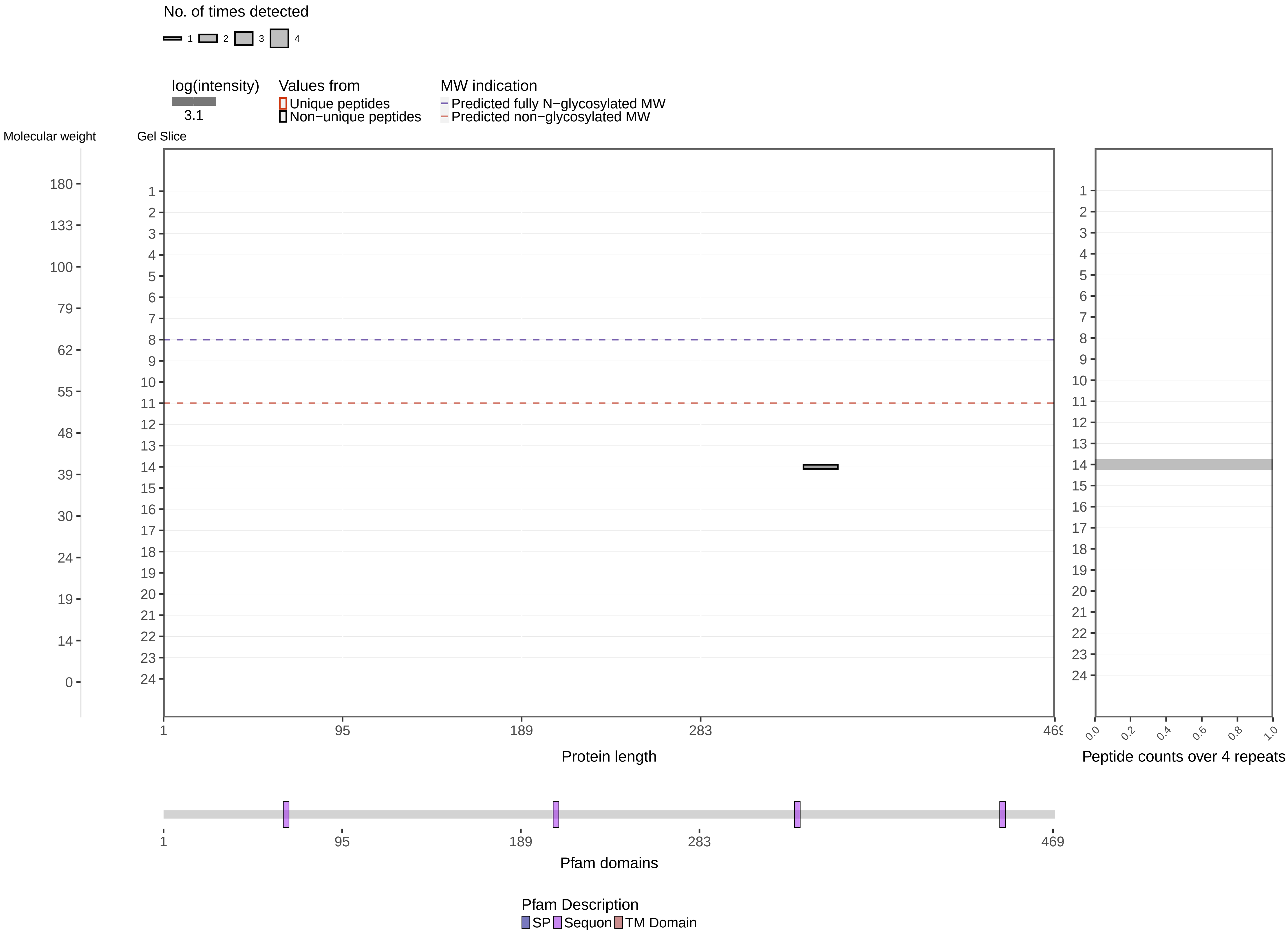


Protein sequence:

EANPSDIVTVIIEDYVRSPNGLTKLFDAAGLRKFWFPVSRMPKNGGEWPTVDDMVQRNQRLVVFTSKSAKEPSEGIAYEWRYVVENQYGNGGMKPGSCPNRAESSAMDNKTKSLVLMNYFPDAPDFAQACKHNSAPLINMMNTCHEAAGKRWPNFIAVDFYKRSDGGGAPEAVDVANGHLVCGCTNIASCKEHMTYGICELPEPRISPVTPAAAAKNETSFSNFNCQPKLQWLLGTTLMSFFLL*

NbL17g23090.1 caffeoyl−CoA_O−methyltransferase_6 [MW = 27.78 kDa]

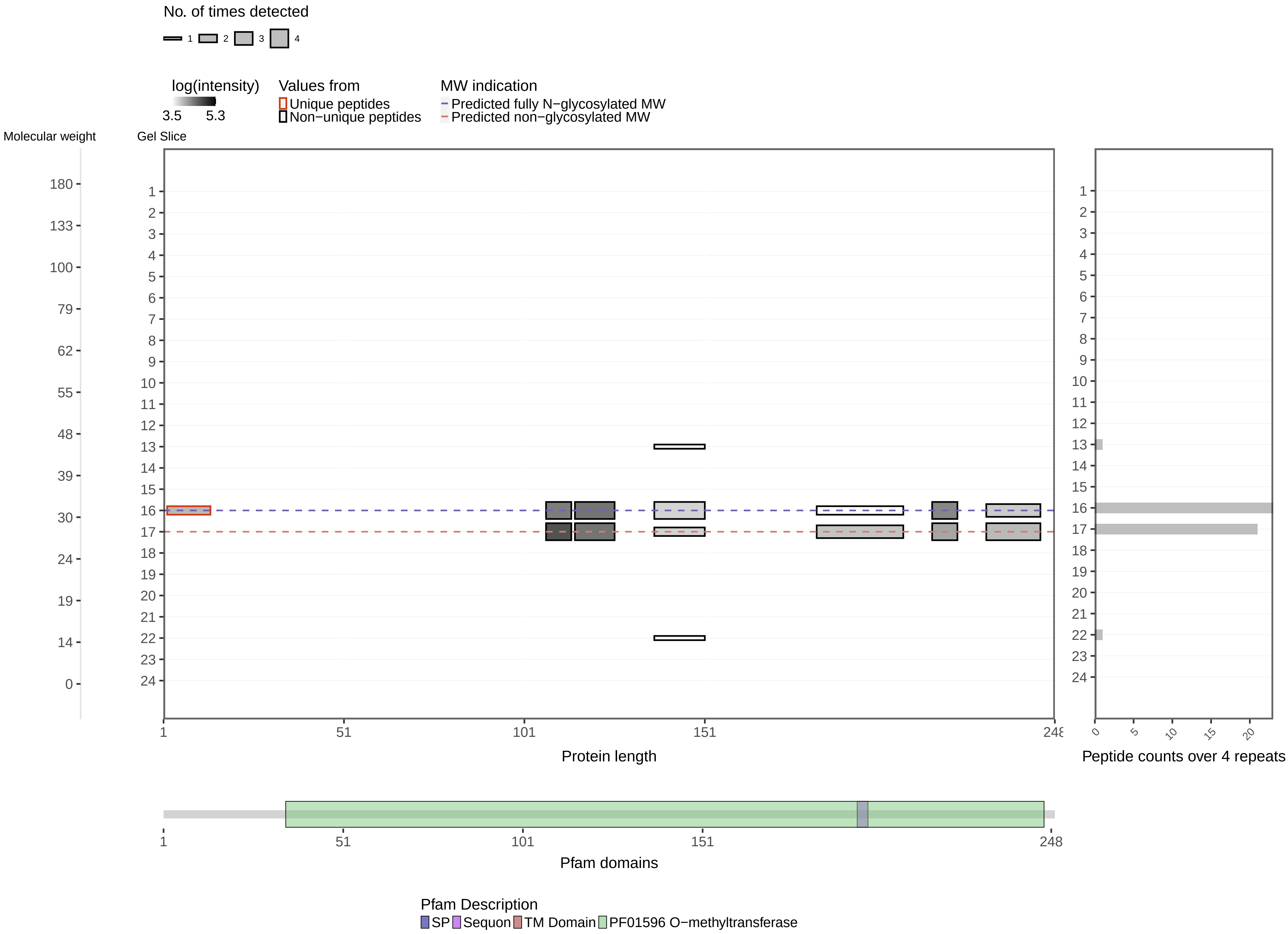


Protein sequence:

MDINRENYEIGLPVIQKAGVAHKIDFREGPALPVLDLMVEDKNNHGTYDFIFVDADKDNYINYHKRIIELVKVGGVIGYDNTLWNGSVAAPPDAPMRKYVRYYRDFVLELNKALAADPRIEICMLPVGDGITLCRRIN*

NbL17g23020.1 CBS_domain−containing_protein_CBSX1_chloroplastic−like [MW = 26.4 kDa]

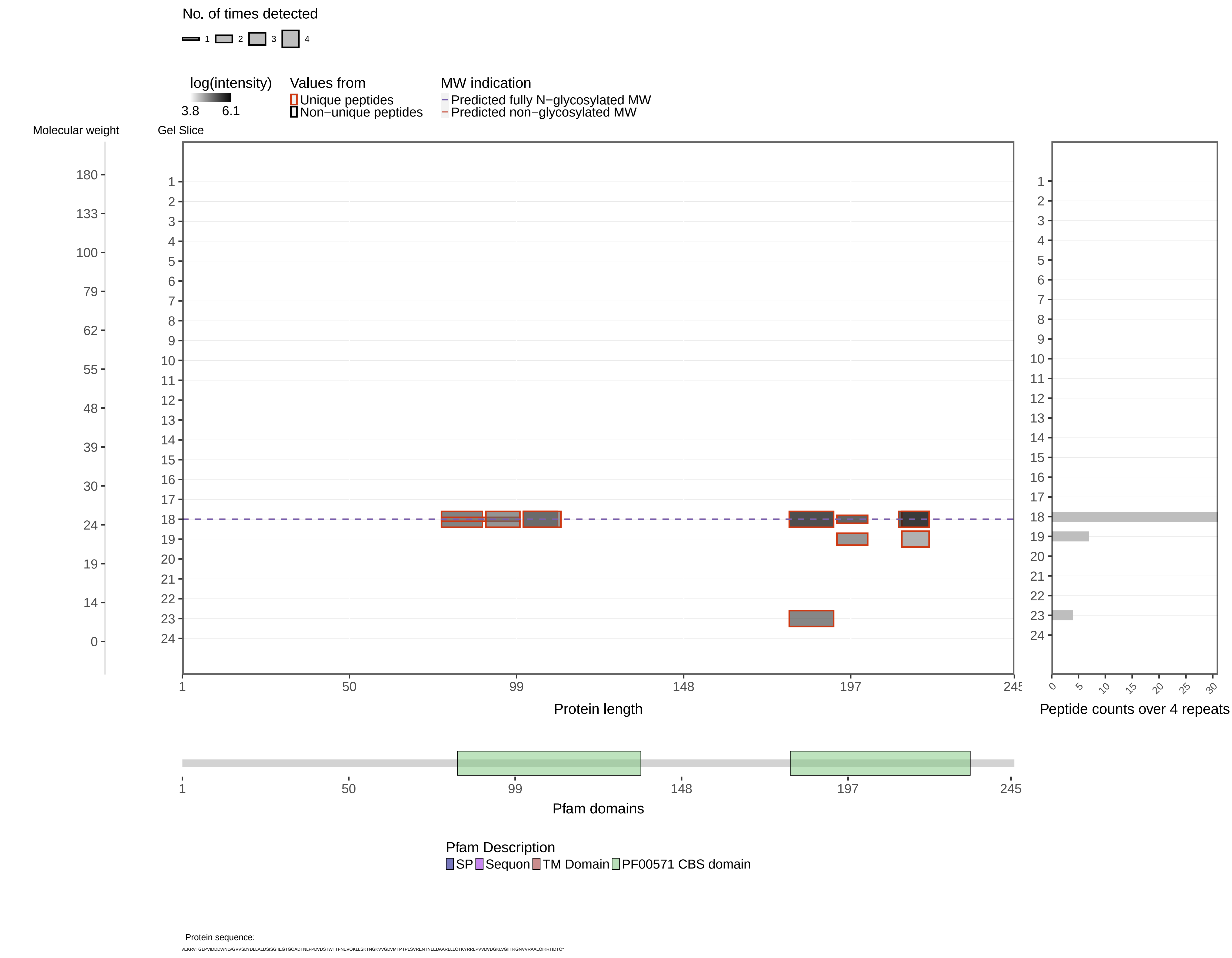

### NbL17g22990.1 scopoletin_glucosyltransferase [MW = 53.77 kDa]

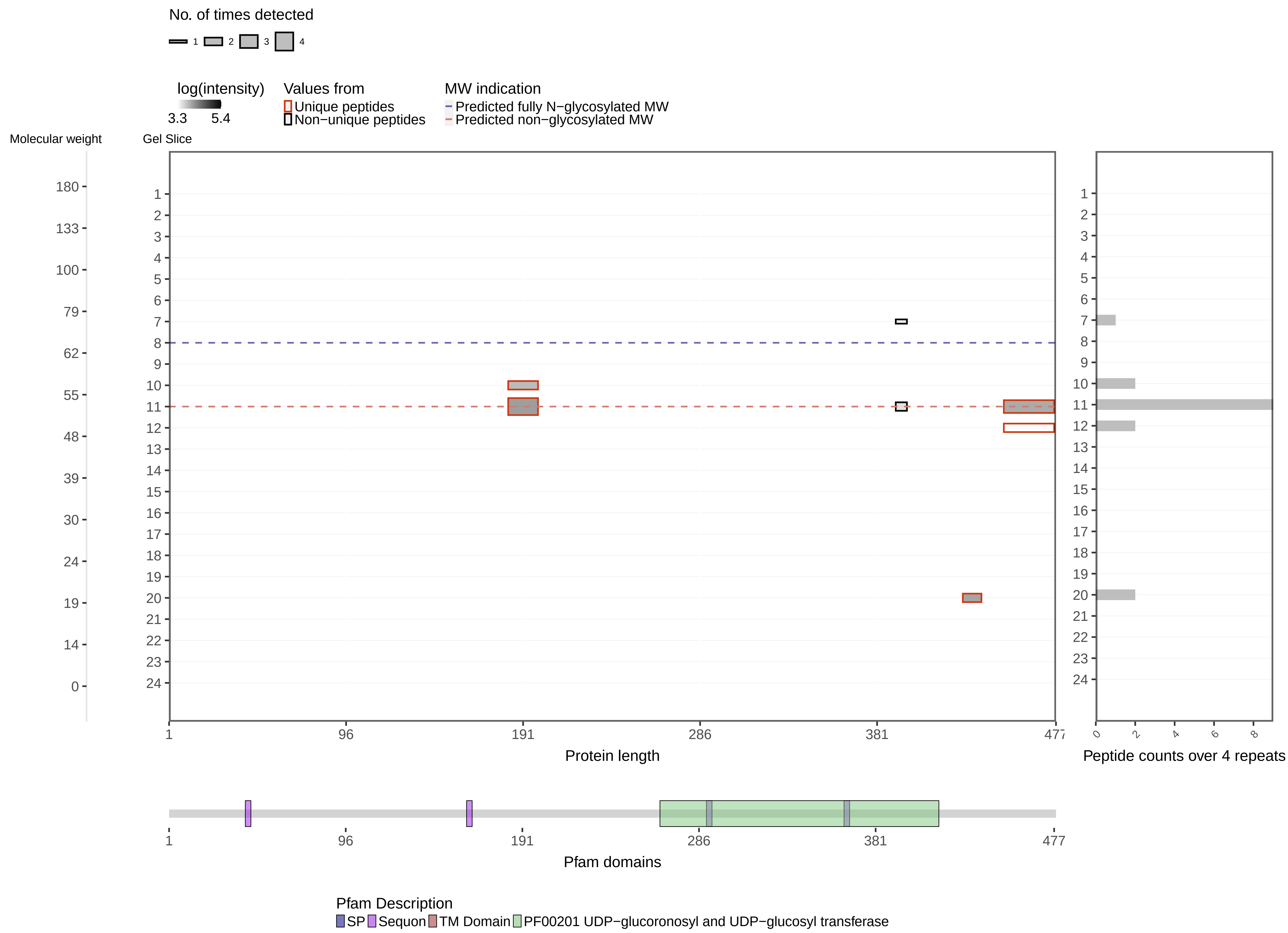


Protein sequence:

HYTKVLGRRAWAIGPLSMCNRDIEDKAERGKKSSIDKHECLKWLDSKKPSSVVYICFGSVANFTASQLHELAMGIEASGQEFIWVVRTELDNEDWLPEGFEERTREKGLIIRGWAPQVVILDHESVGAFVTHCGWNSTLEGVSGGVPMVTWPVFAEQFFNEKLVTEVLRTGADVGSIQWKRSASEGVKREAIAKAIQRVMVSEEAEGFRNRAKAYKEMARKAVEEGGSSYTGLTTLLEDISTYSSTGH*

NbL17g22970.1 phosphatidylinositol_345−trisphosphate_3−phosphatase_and_protein−tyrosine−phosphatase_PTEN2A [MW = 46.86 kDa]

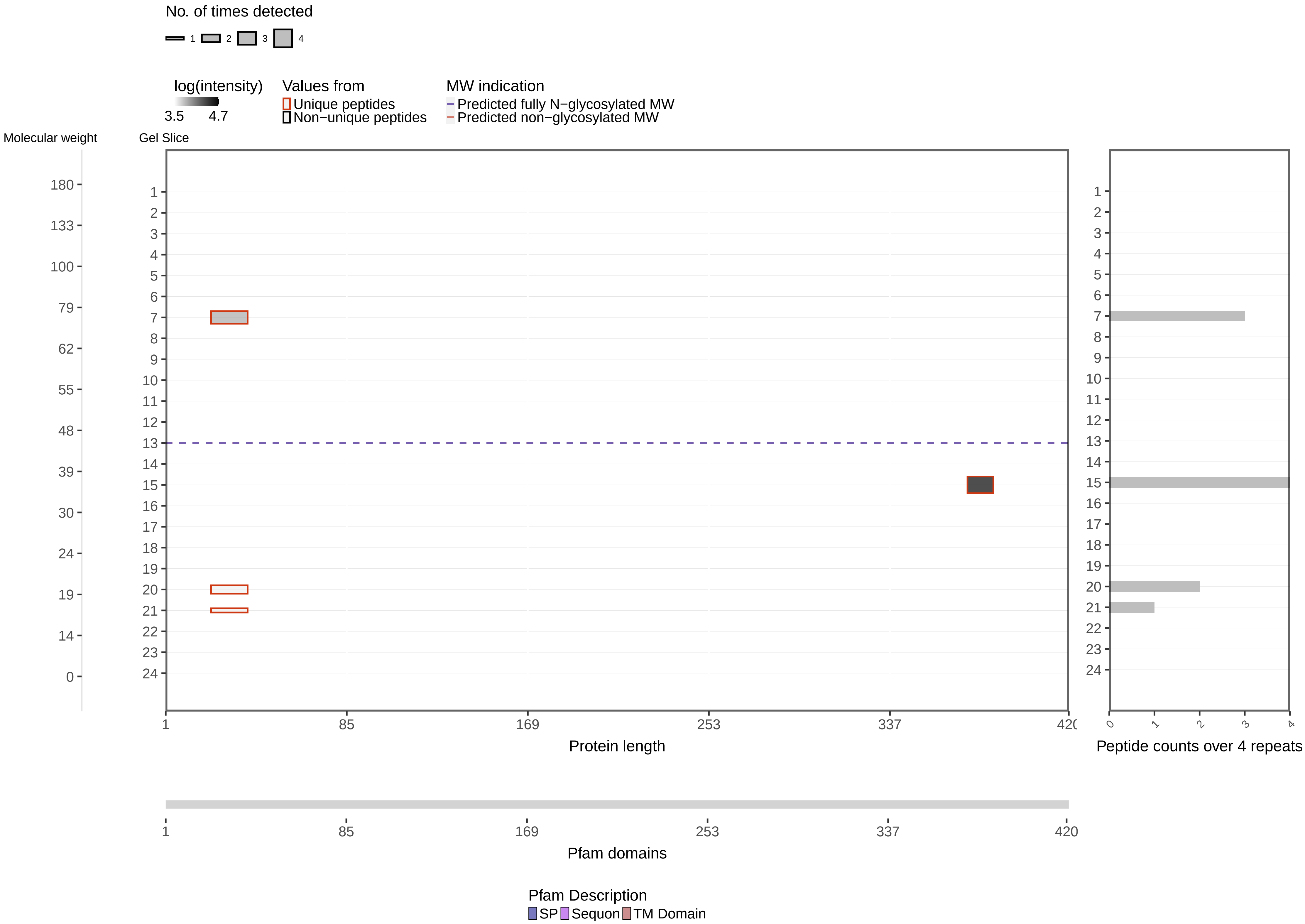


Protein sequence:

YKVYNLCSERLYDASLFEGKVACFPFDDHNCPPIQLIISFCQSAYSWLKEDIENVVVVHCKAGMARTGLMISSLLLYLKLSYMVFLQFFPTAEESMDYYNQKRCVDGKGLVLPSQIRYVKYFERILMYFNGENQPGRRCMLRGFRLHRCPYWIRPSITVSDHNGVLFSTKKHPRTKELSPEDFWFSAPKKGVMVFALPGEPGLAELAGDFKIHFHDGQGDFYW*

### NbL17g22960.1 elicitor−responsive_protein_1−like [MW = 26.28 kDa]

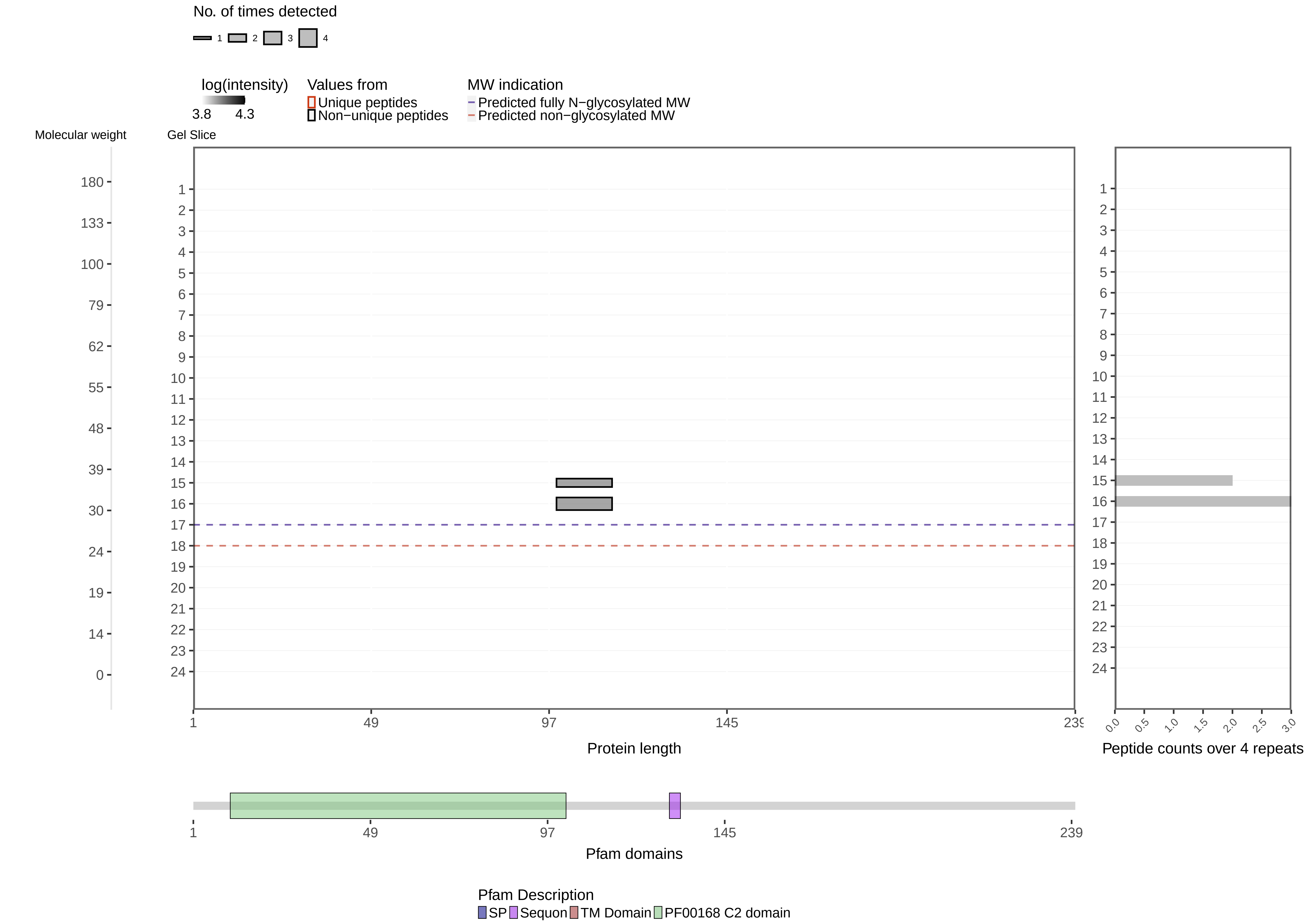


Protein sequence:

AWPIQTKTGRHAGEVRLIMHYANASNPATSYAPSAPPYAAPQPQAPMYSVPPPHLASYPPTAAYQTPSPYPAYPPHSAAYPPSPYPPPQTTAYPPAYPPPSAYPPVSYPPPSQDYSYPPGPYQGTYPPRPY*

### NbL17g22930.1 cyclase−like_protein_2 [MW − SP = 27.88 kDa]

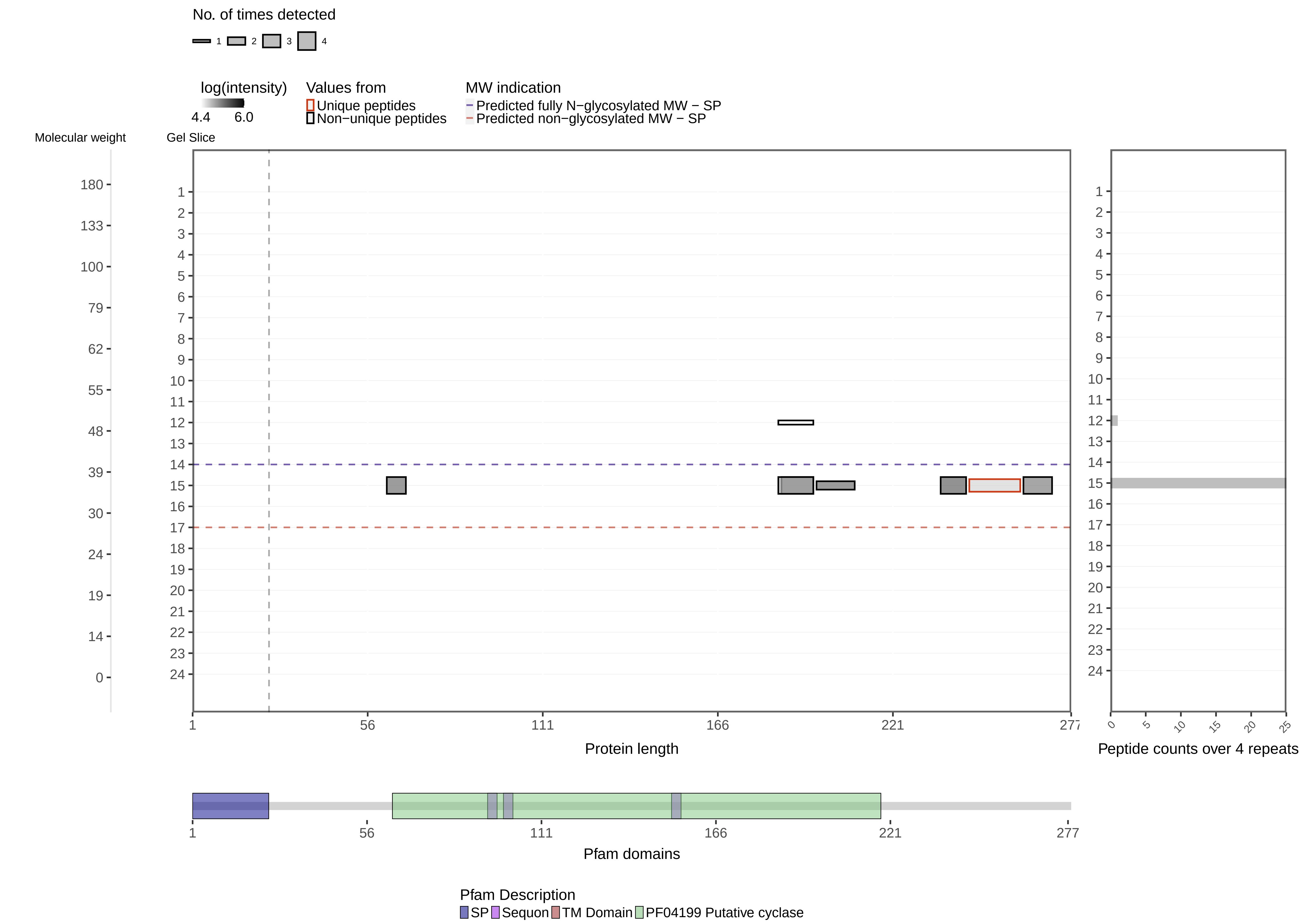


Protein sequence:

YFDAGFDIDTLDLEVLNGPALVVDVPRDKNLTADVMQSLHIPKGVKRVLFRTLNTDRRLMWKKAFDTSYVGFMKDGAQWLVDNTDIKLVGIDYLSVAAFDDLIPSHLVFLKSREIILVEGLKLDDIETGIYNVHCLPLRLLGAEGSPIRCILIK*

### NbL17g22910.1 stress_enhanced_protein_1_chloroplastic [MW = 14.74 kDa]

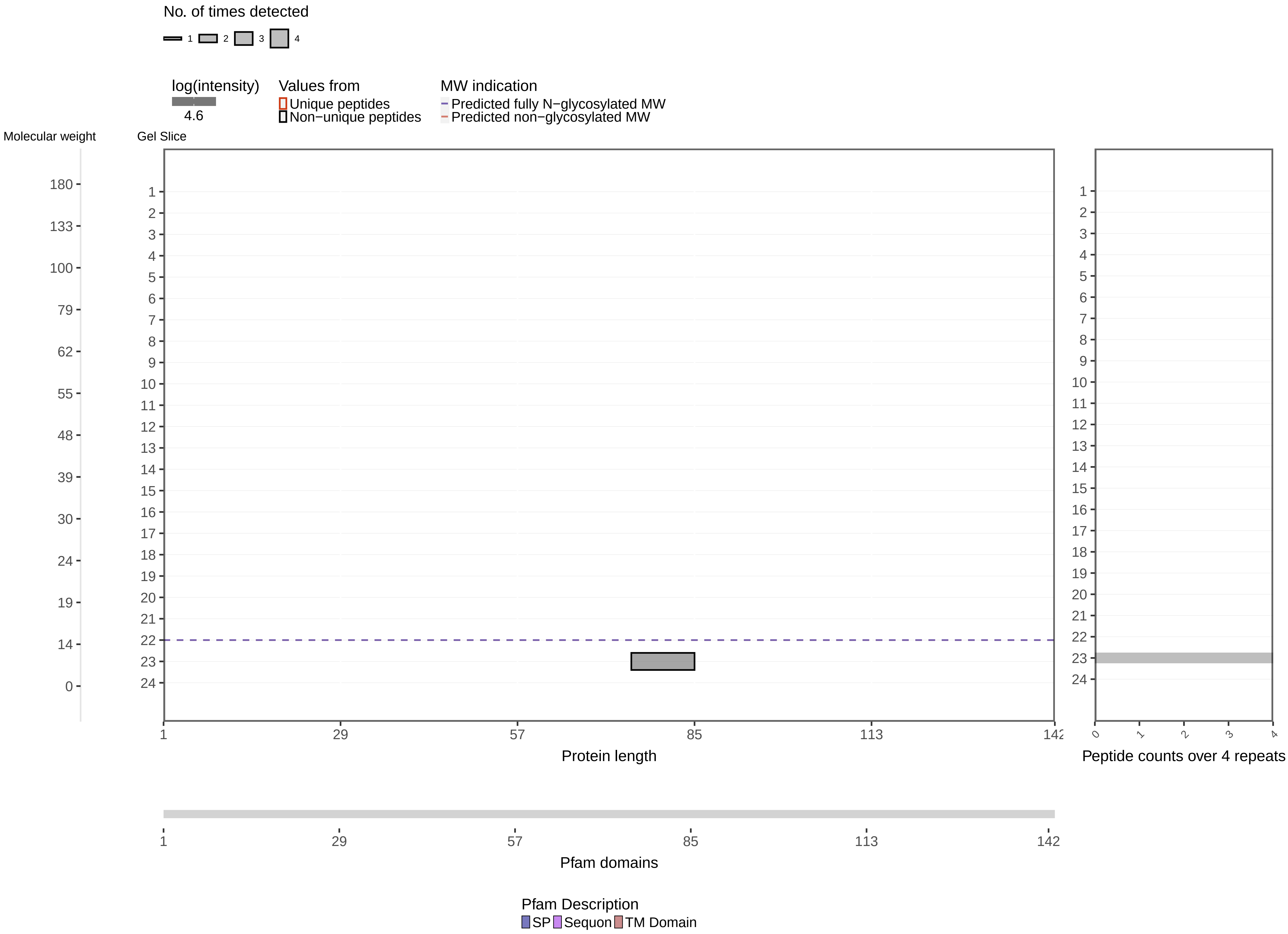


Protein sequence:

AIPSRATSVSIRCGQSTKEESNLDVWLGRSAMVGFAVAISVEIATGKGLLENFGVTGPLPTVALAVTALVGILTAVFIFQSASKN*

### NbL17g22820.1 methylthioribose_kinase [MW = 48.02 kDa]

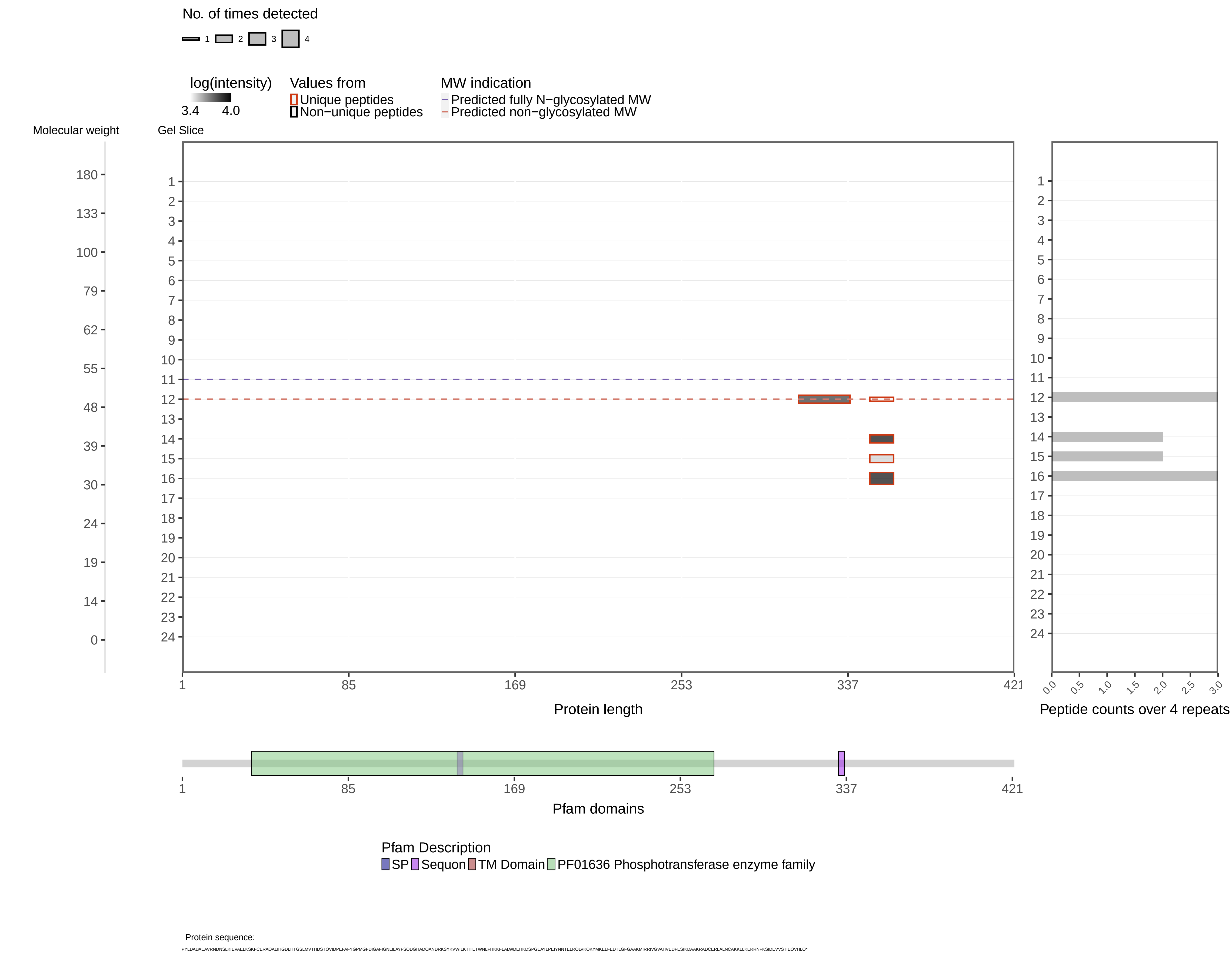

NbL17g22800.1 uncharacterized_LOC104232799 [MW − SP = 22.74 kDa]

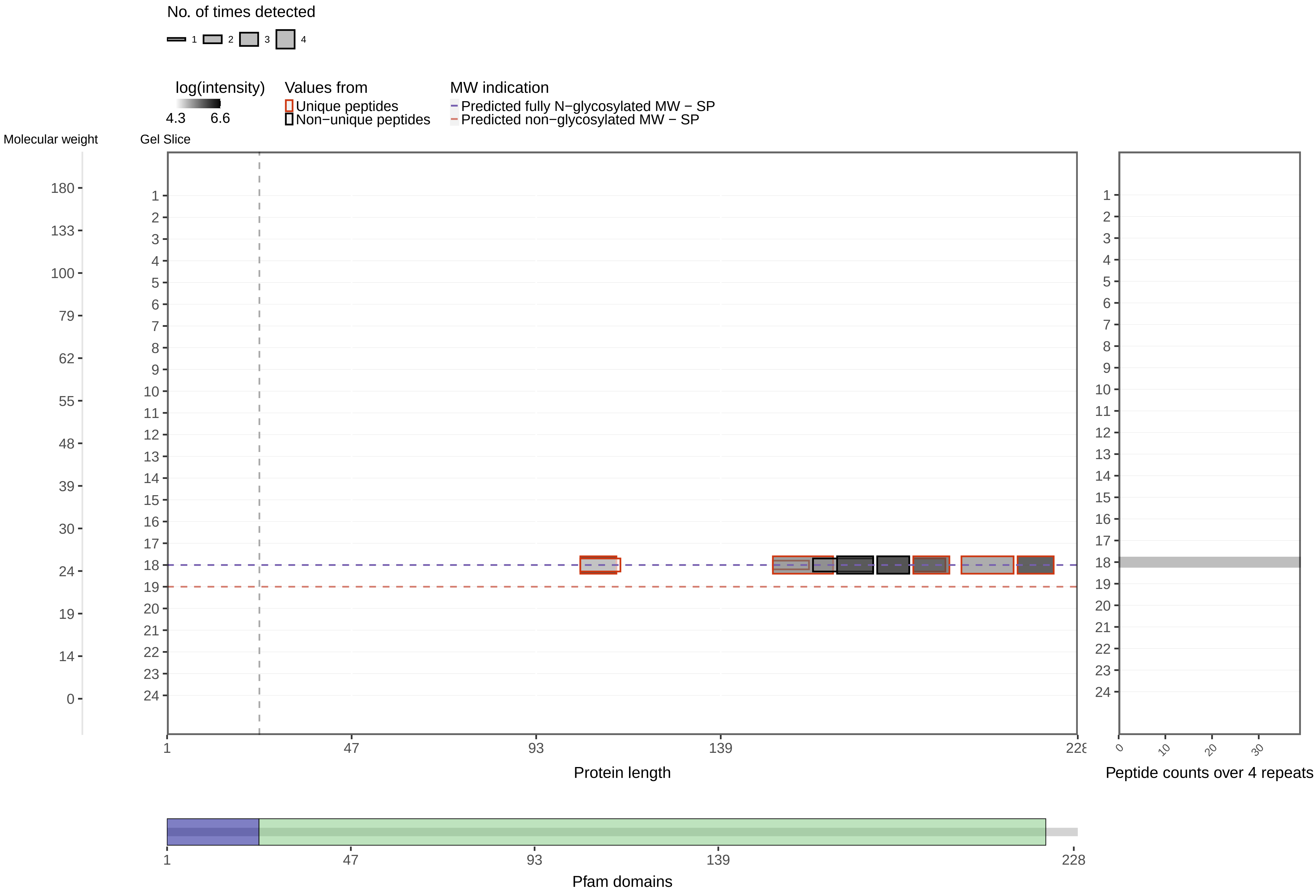


Protein sequence:
YIQGYSGDVRREITGVLYHESTHVWQWNGNGGAPGGLIEGIADYVRLKAGFGPSHWVKPGQGDRWDQGYDVTARFLDYCNSLRNGFVAQLNKKMRTGYSNQFFVDLLGKTVDQLWNDYKAKFRA*

NbL17g22790.1 uncharacterized_LOC101249214 [MW − SP = 23.22 kDa]

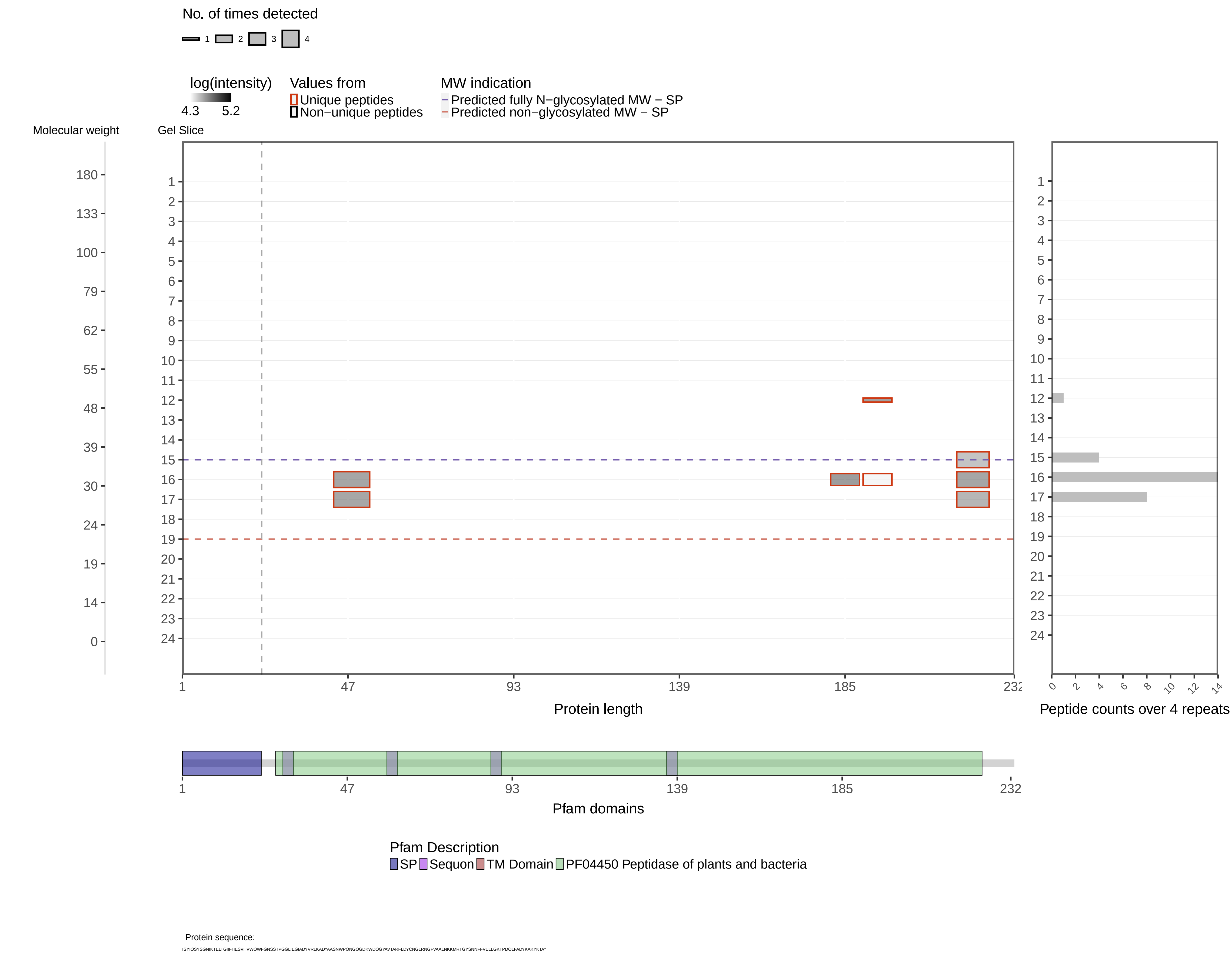

NbL17g22750.1 LYR_motif−containing_protein_At3g19508 [MW = 10.37 kDa]

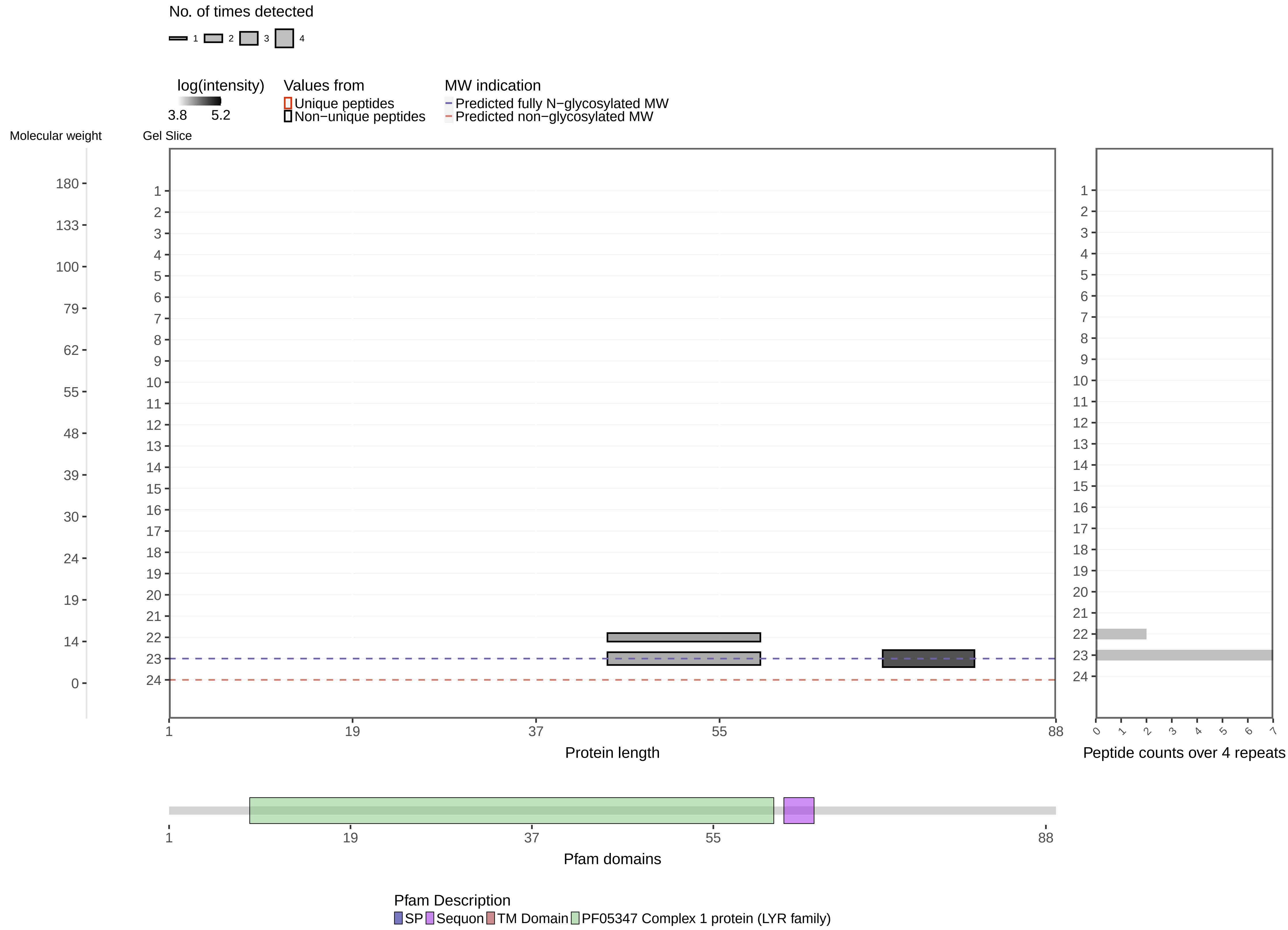


Protein sequence:

AKYARENFVNYREIDSNDPNALQELLQRTYNHSLWVLNKYSVDQSAADRLKDMCTA*

NbL17g22700.1 TIP41−like_protein [MW = 33.97 kDa]

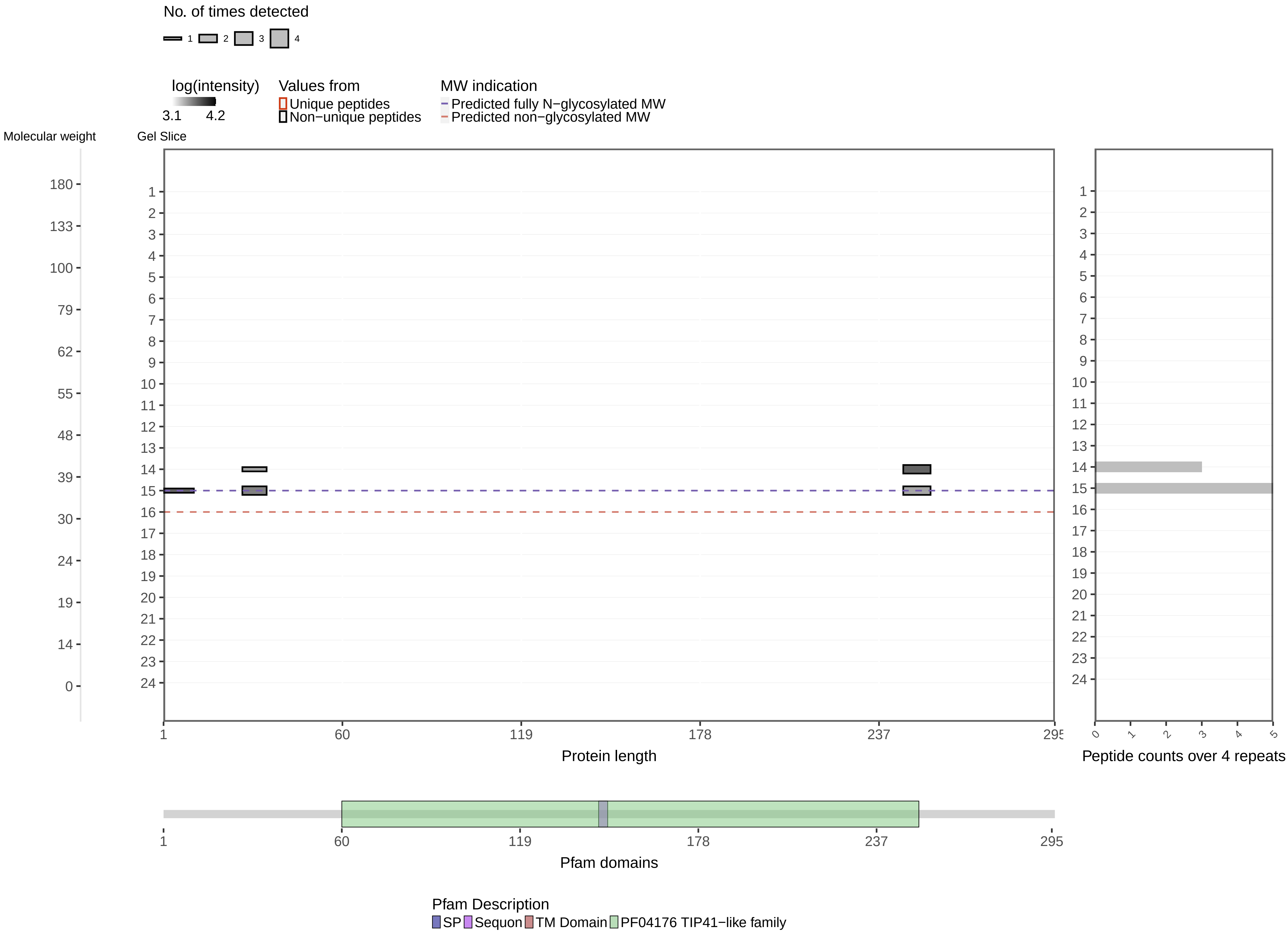


Protein sequence:

FDTNAERVRGLSNESSCSLQWEDCMEKIDLVALASKEPILFYDEIILYEDELADNGISLLTVKVRVMPSGWFLLLRFWLRVDGVLMRLRDTRLHCLFGEHMKSVILRESCWREATFQALSSKGFPSDCAAYSDPSTISERLPIMMQKTQKLSIEIPCKWCLE*

NbL17g22490.1 mitotic_checkpoint_protein_BUB3.1−like [MW = 37.76 kDa]

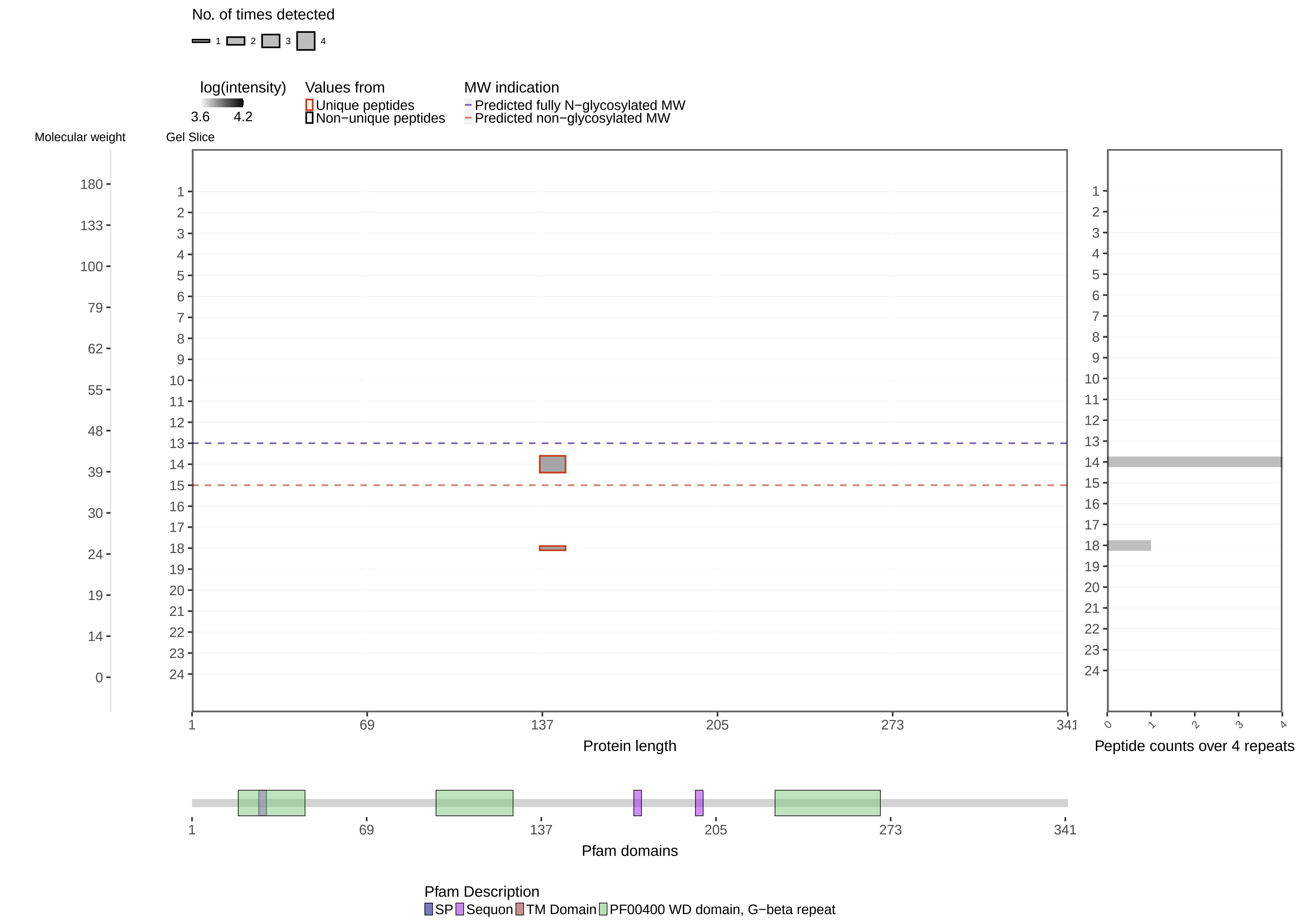


Protein sequence:

LVVATAGRHVNVYDLRNMSOPEORRESSLKYOTRCVRCYPNGTGYALSSVEGRVAMEFFDLSEAGOSKKYAFKCHRKTEAGRDIVYPVNAIAFHPIYGTFATGGCDGYVNVWDGNNKKRLYOYPKYPSSIAALSFSRDGRLLAVASSYTFEEGEKPHEPDAIIVRSVNEVEVKPKPKVLPNPTT*

NbL17g22140.1 aspartate−−tRNA_ligase_chloroplastic/mitochondrial [MW = 75.22 kDa]

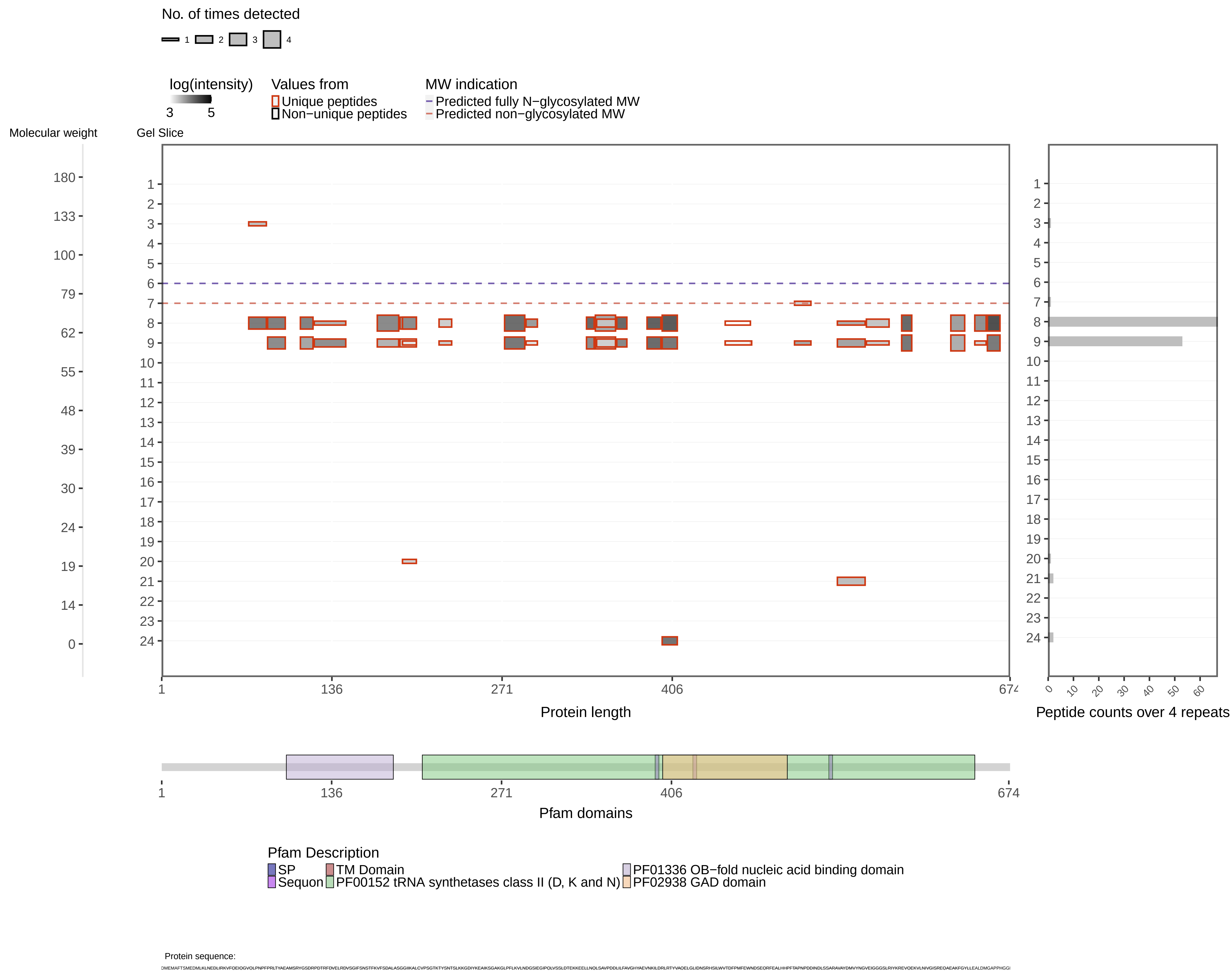

NbL17g21950.1 L−galactose_dehydrogenase [MW = 34.48 kDa]

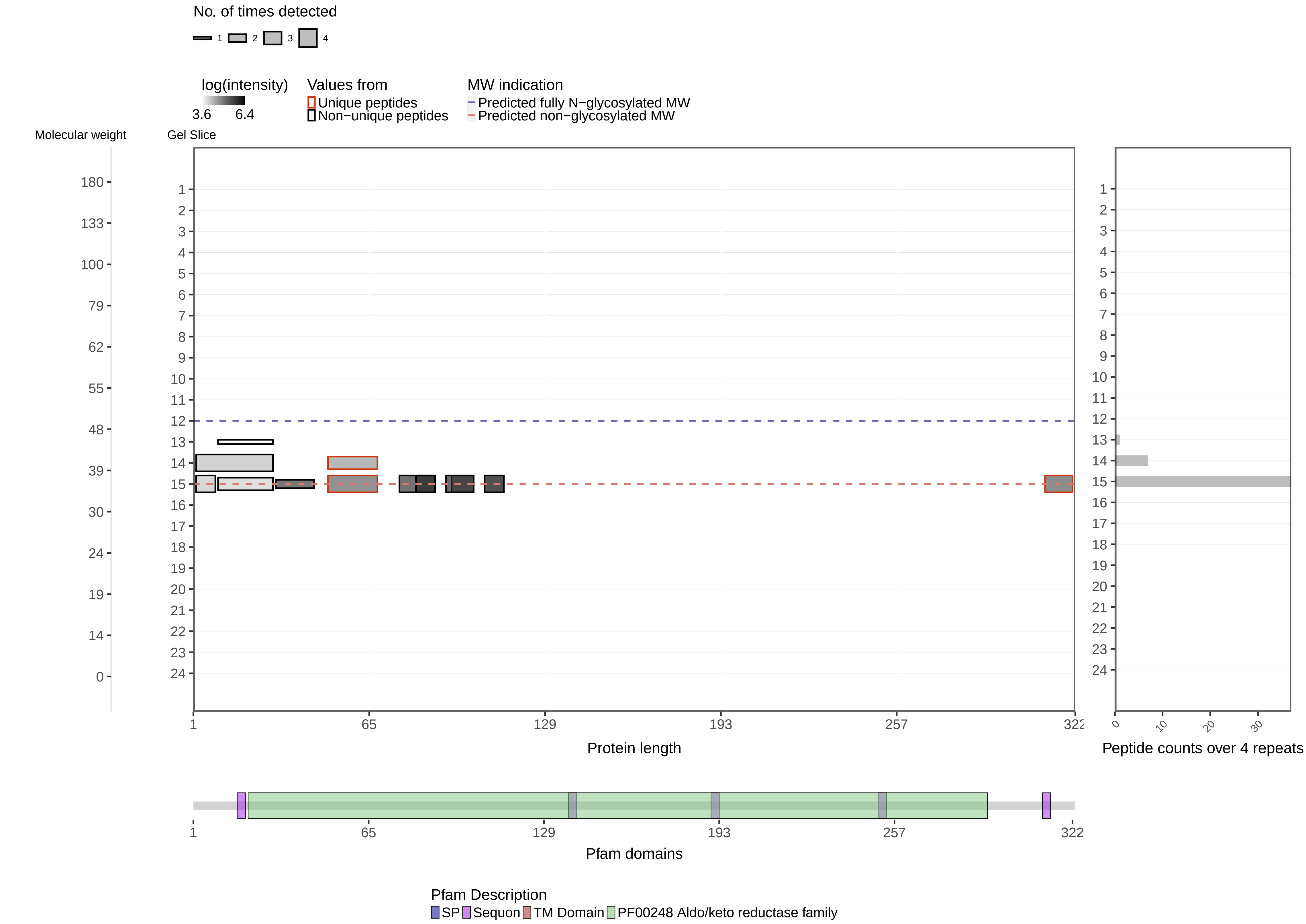


Protein sequence:

KLKQAGKIRFIGITGLPLGIFTYVLDRVPPGTVDVILSYCHYSINDSTLEDLLPYLKSKGVGVISASPLAMGLLTEAGAPEWHPASSELKAACRAAVDHCKGKGKNISKLALQYSLANTGISSVLVGMKSVKEVEENIAAALELATAGMDEETLSEITDILKPVKNQSWPSGIQQS*

NbL17g21930.1 soluble_inorganic_pyrophosphatase_6_chloroplastic−like [MW = 32.4 kDa]

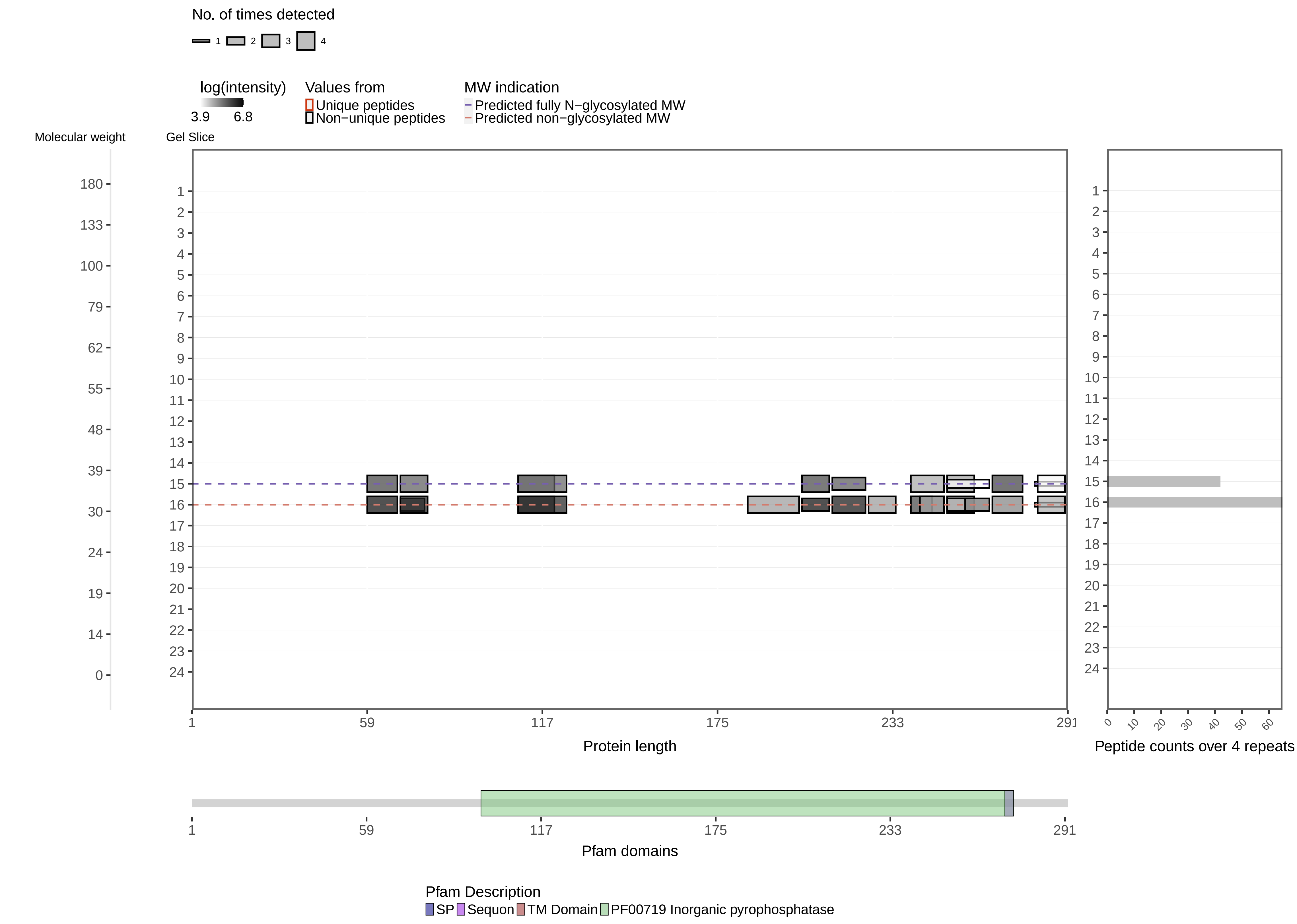


Protein sequence:

YPYNIHWNYGLLPQTWEDPSFANSEVEGAFGDNDPVDVVEIGDSRGKIGQVLKVKPLAALAMIDEGELDWKIVAISLDDPRASLVNDVDDVEKHFPGTLTAIRDWFRDYKIPDGKPANRFALGNKPANKDYALKVITETNESWAKLVKRSIPAGELSLV*

### NbL17g21910.1 peptide_methionine_sulfoxide_reductase_B5−like [MW = 19.39 kDa]

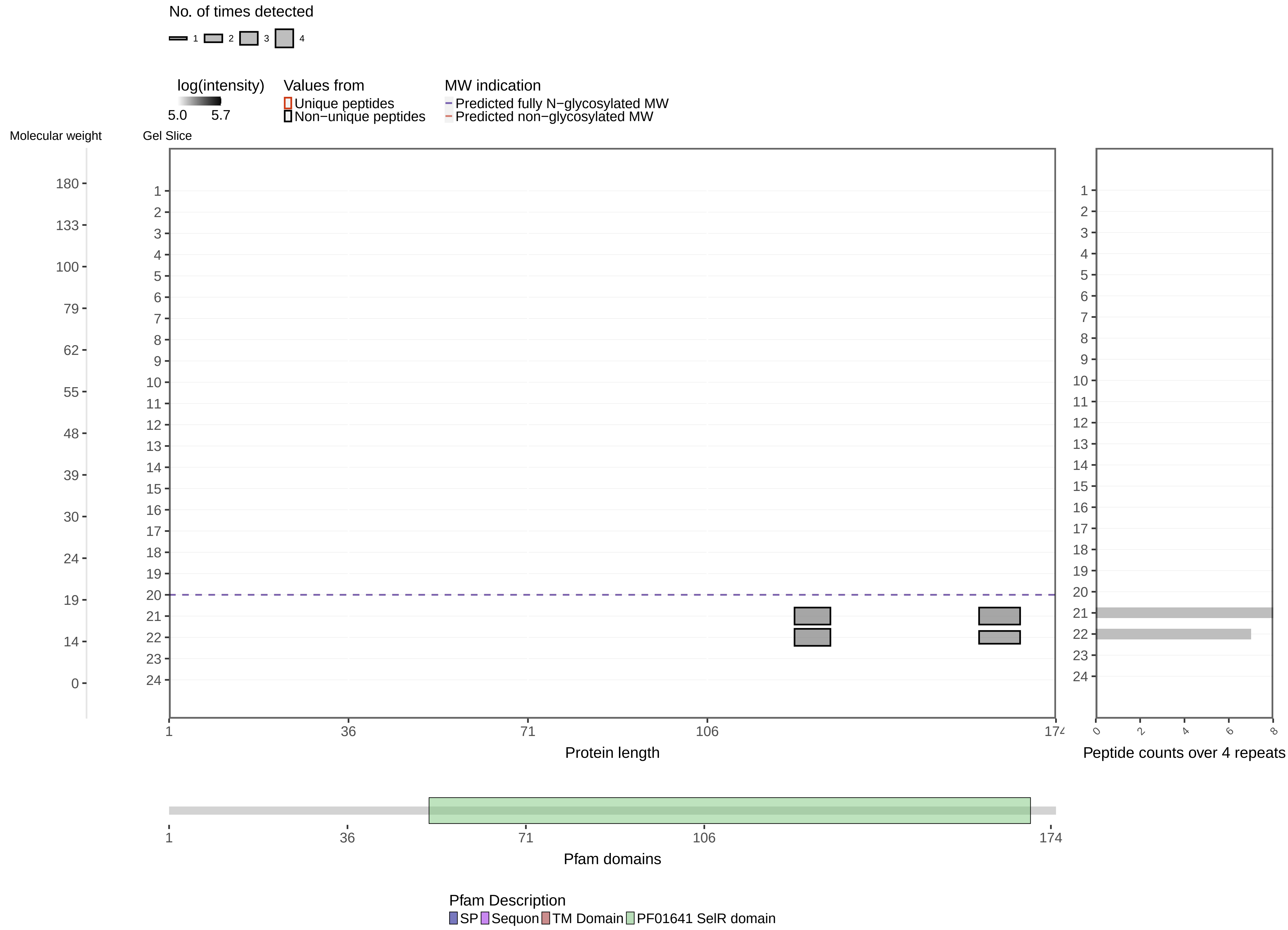


Protein sequence:

GEYNKFFGVGTYLCAGCGTPIYRSATKFNSPCGWPSFYEGLPGAINRNPDPDGVRMEITCAACGGHLGHVFKGEGFRTPTNERHCVNSISLKFKPPNF*

### NbL17g21840.1 uncharacterized_LOC107824885 [MW = 53.39 kDa]

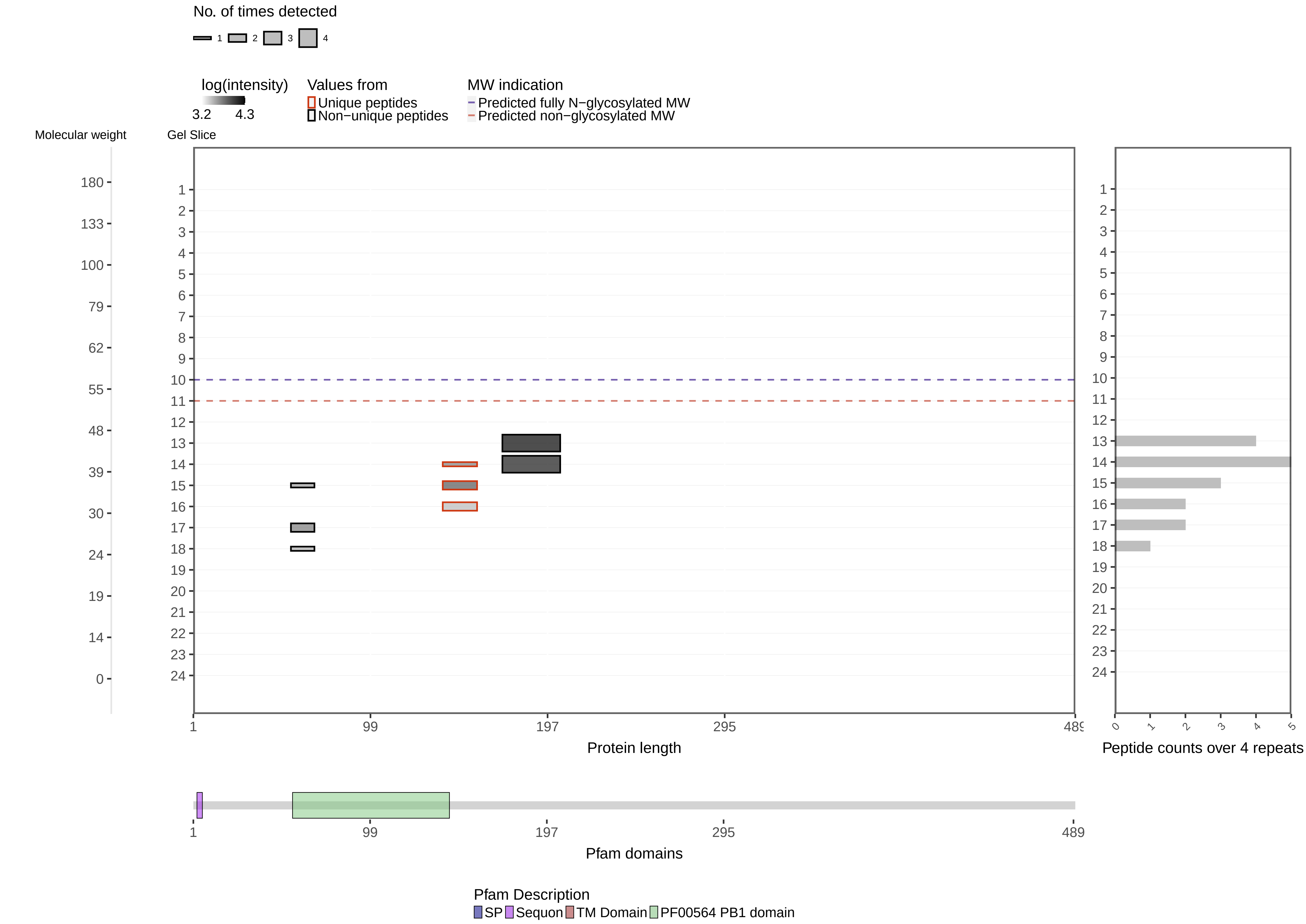


Protein sequence:

GTDPIQKHIQDLQRLHLEEQQNMYLRKSDDNLAAAFTGGHDYYVQKVPEKIAPAATVPGTVPAQVGYWQEKQVPGGVHPANSLGTEQPVYMIQAPAGAYHQHQHPQFVRPMAGPPGQGYYAVQRMPQQEPYRDHQPVYNVMPSVPQAVPTVAATLPSQGGPKVAGYSEGYGMVRPATTMGVGVADAGGYTQMVYDGGVGRQMYYPPQGGVMAPPPQQPQYQAMPPQAVVSSDMRTQEAAKVVPVKVNQCNS*

### NbL17g21730.1 psbP_domain−containing_protein_1_chloroplastic−like [MW = 39.6 kDa]

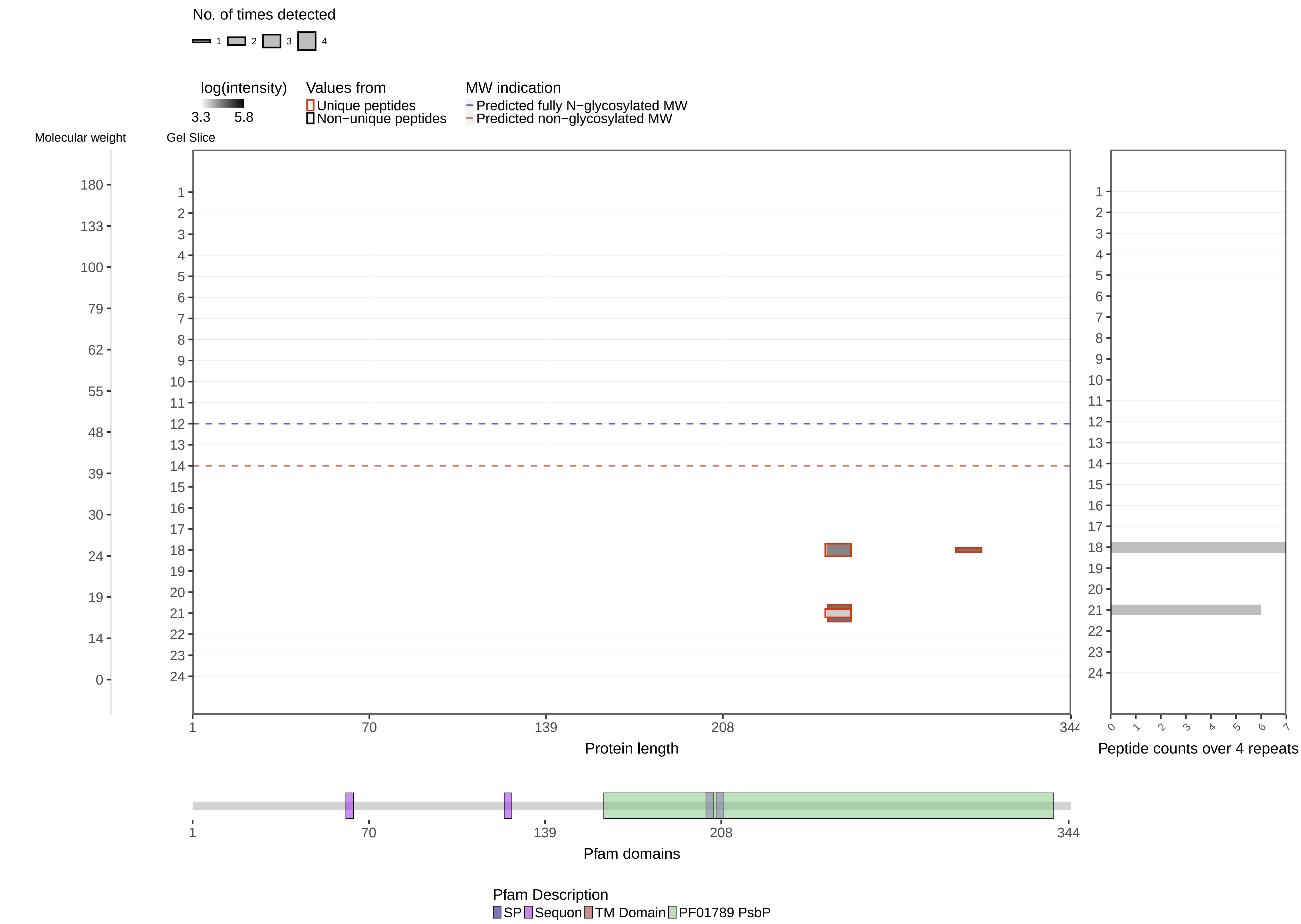


Protein sequence:

STEFREYIDTFDGYSFMYPRNWIQVRGAGADIFFRDPLVLDENLSMNISSPSSSAYKSVEDLGPPEKAGKGVLRQYLTEFMSTRLGVRRESSVLSTSSRVADDGKLYYEVEVNIKSYANNNELAVMPQDRVARLEWDRRYLSVLGVENNRLYELRIQTPEKTFADEESDIRAIMDSFRVNKIEV*

NbL17g21710.1 methylmalonate−semialdehyde_dehydrogenase_[acylating]_mitochondrial [MW = 56.75 kDa]

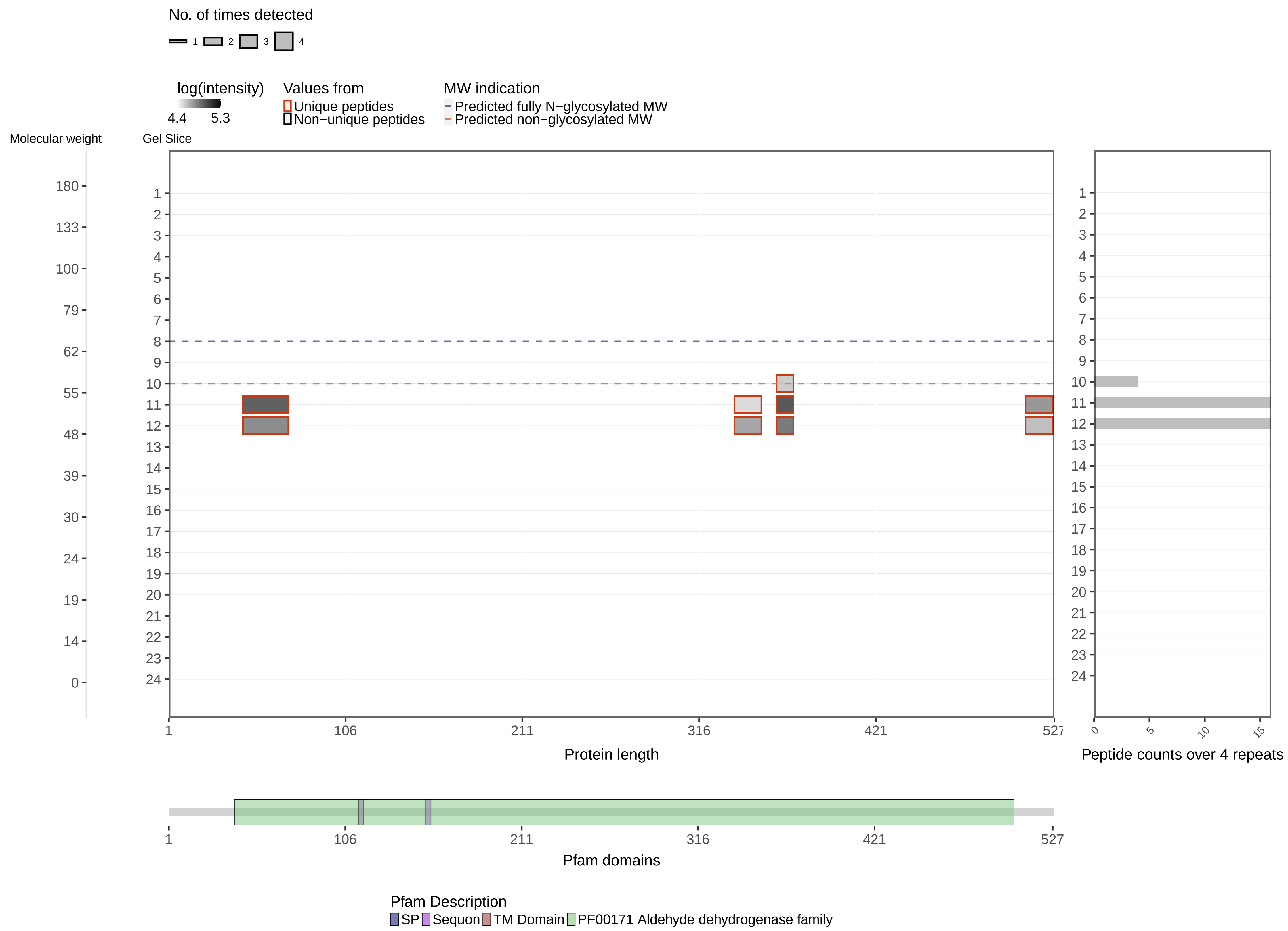


Protein sequence:

VGSNQAGMHIYARASAKGKRVQSNMGAKNHGVVMPDANIDSTVNALVAAGFGAAGQRCMALSTVVFVGDSKPWEEKLLERAKALKVSGGTELDADLGPVISKQAKERVCRLVQSGVDSGARLLLDGRDIVVPGYEKGNFVGPTILSDVTPDMECYKEEIFGPVLICMQANSLDEAINIVNKNKWGNGAAIFTTSGVAARKFQTEIEAGQVGINVPIPVPLPFFSFTGSKASFAGDLNFYGKAGVQFYTQIKTITQQWKDLPGGSGVSLAMPTSQK*

NbL17g21680.1 gamma−glutamyl_peptidase_5−like [MW = 27.3 kDa]

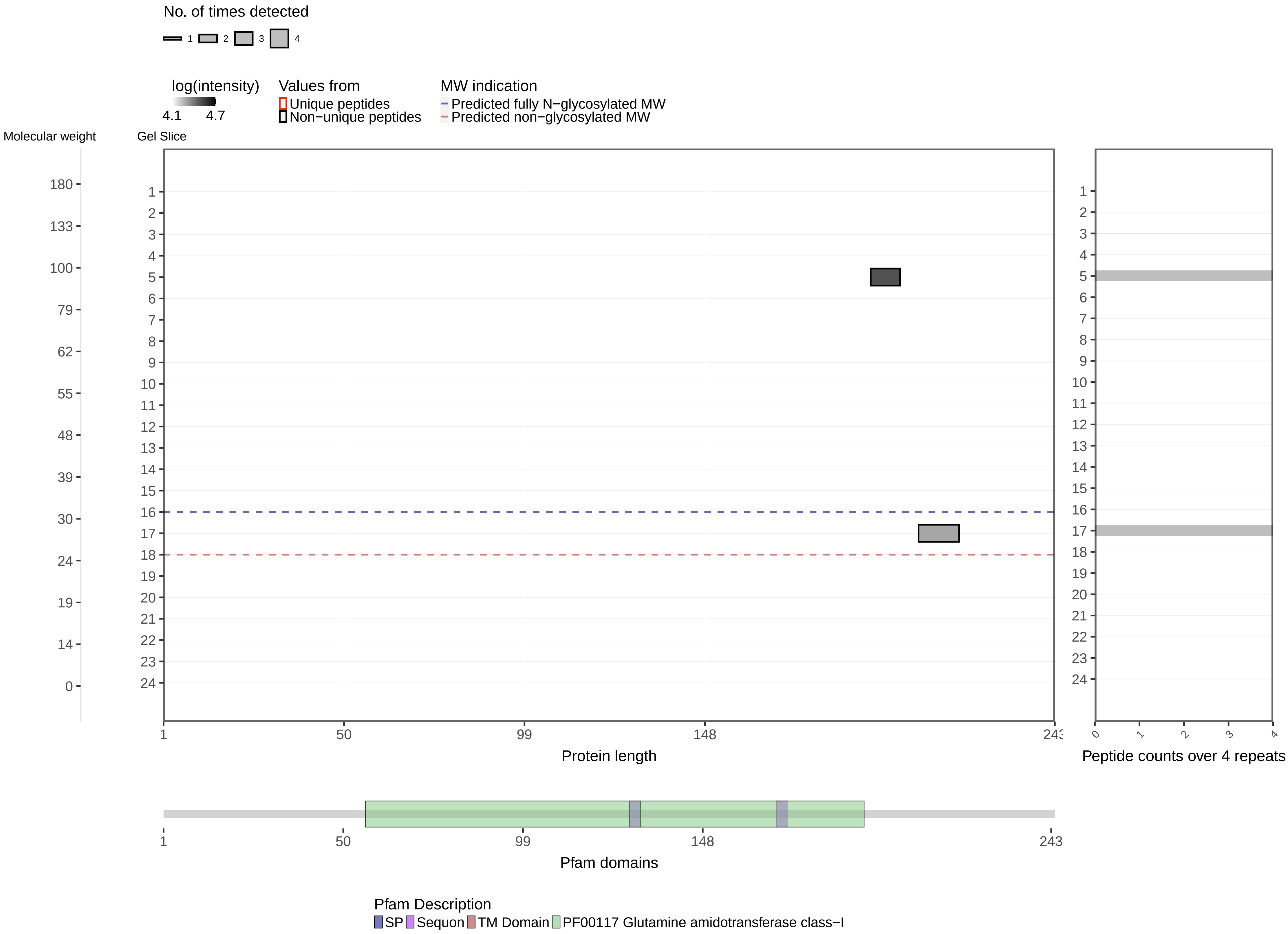


Protein sequence:
SLGGVIERATTGWDIGVTTVNLSTSKLFKTLNLPAFMQVIECHRDEIRELPPKAEVMAWSNKTGIEMFRYGNHIMGIQGHPEYTKDILLHLIDRLLQRNLIEESIADVAKAKVEGCEPDREAWKKLCVSFLKGRL*

NbL17g21630.1 magnesium−dependent_phosphatase_1 [MW = 21.52 kDa]

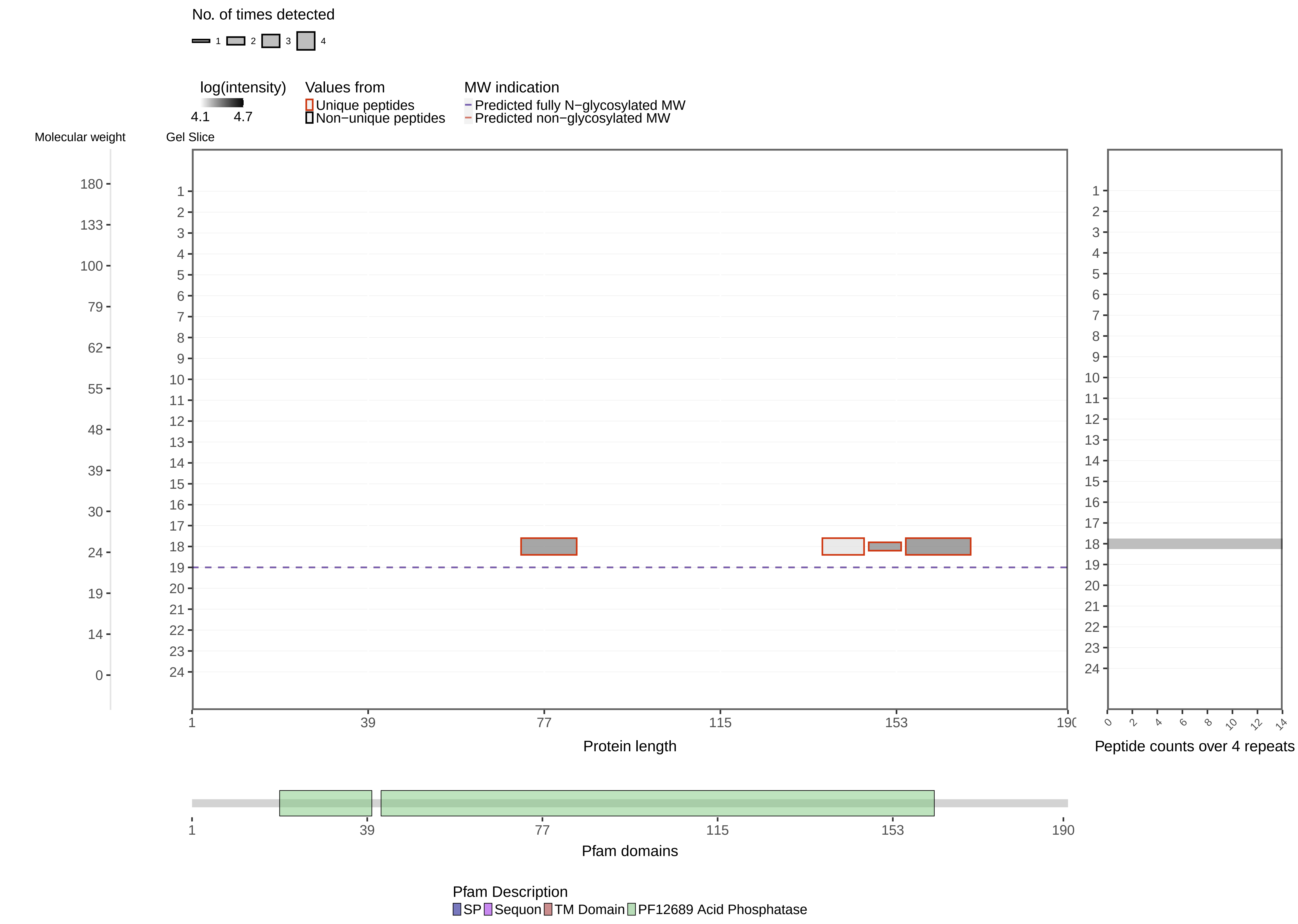


Protein sequence:

QKLEIKSMFVAQEIFSSWTHKTEHFQKINRKTAVPYSDMLFFDDENRNIQAVSKMGVTSILVGKGVNIGAFRQGLSEFVQNSASVEKNKERWRKFSKEPGSSKSDTQ*

NbL17g21460.1 3−oxoacyl−[acyl−carrier−protein]_synthase_mitochondrial [MW = 50.01 kDa]

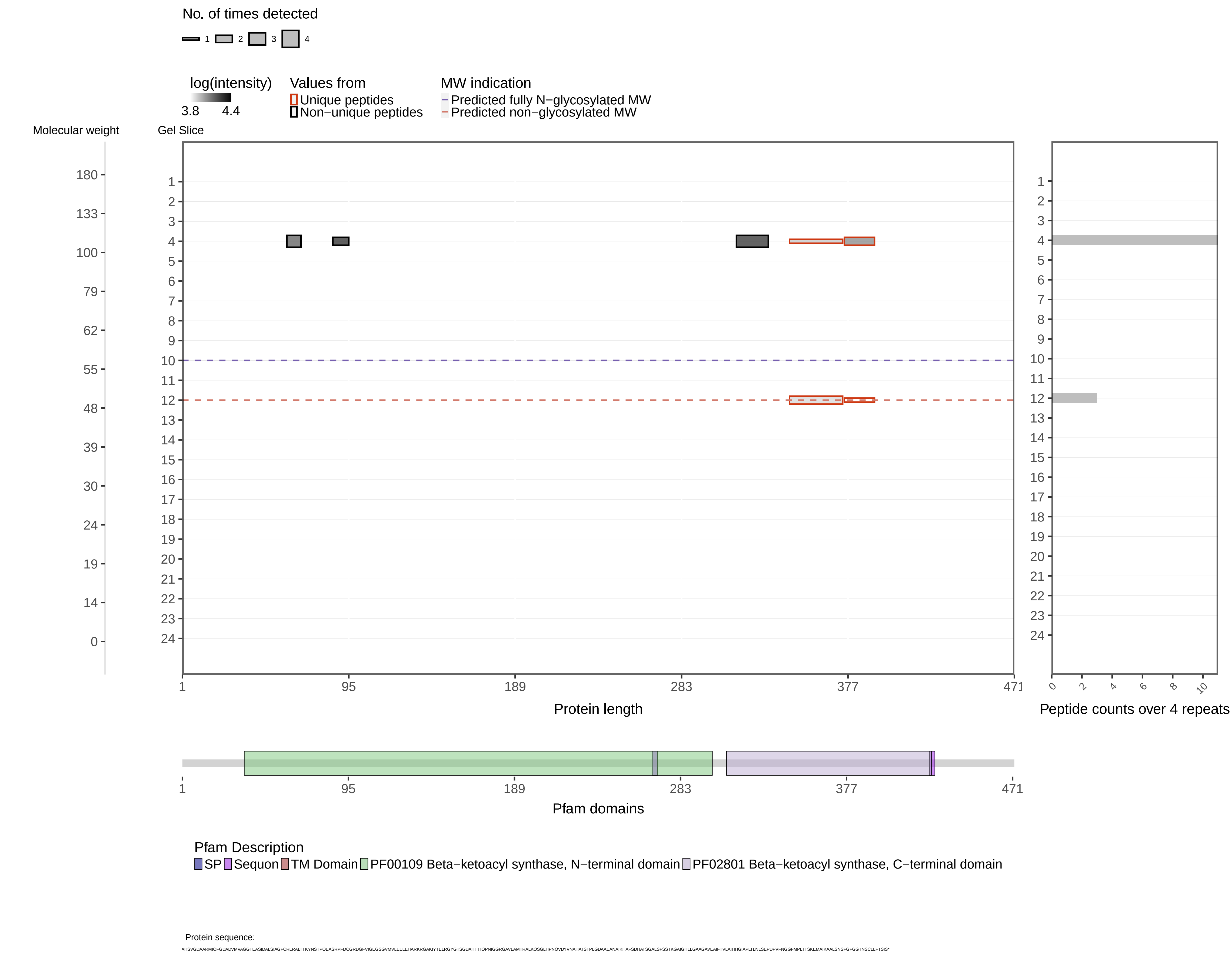

NbL17g21130.1 uncharacterized_LOC104219300 [MW = 71.17 kDa]

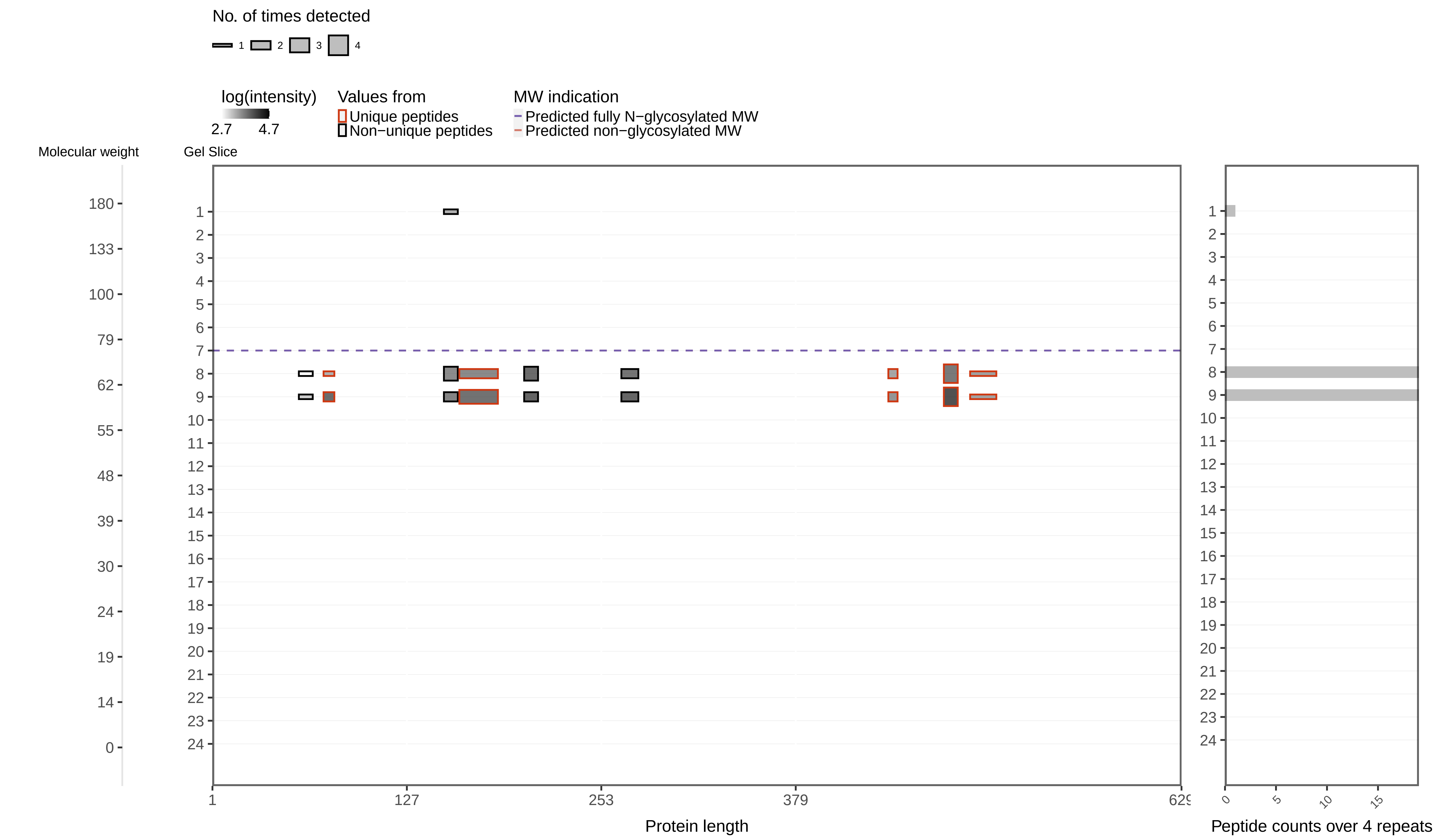


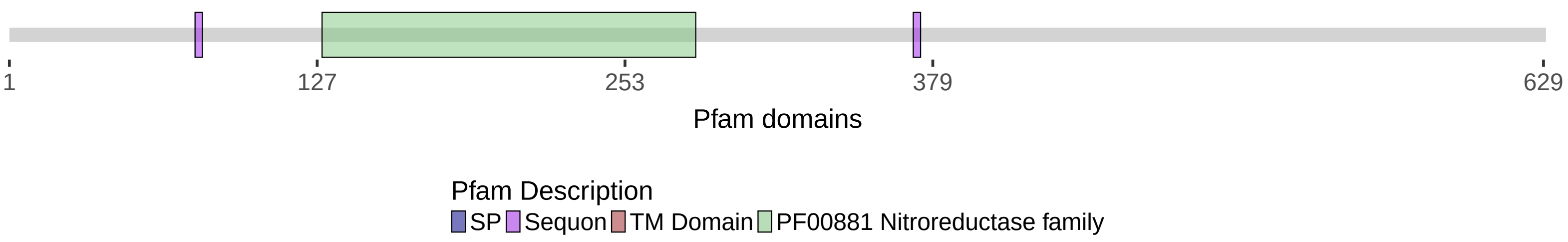


Protein sequence:

EHPDCVLLVFPSDVNEFEVDYKELSFGISELSGLDWNGKPNVLSKEHICWDIIYRTAEAAKKPLTMFNVSVVDPFQTSGTISESSYKDLSLRELVRKRRSAVDMDGFTEMGSDTFYQILLHCMPSGSRDGEKHVRQLALPFRSLAWDSEIHAALFVHRVVGLPSGLYFLVRNENHLDDLKKATRAEFKWVKPDDCPDDLPLYELARGDCRELSKRLSCHQDIASDGCFSLGMIAHFEPTLRNKGSWMYPRLFWETGVLGQVLYLEAHAVGISATGIGCFFDDPVHEVLGLKG

NbL17g21120.1 arginase_1_mitochondrial [MW = 37.13 kDa]

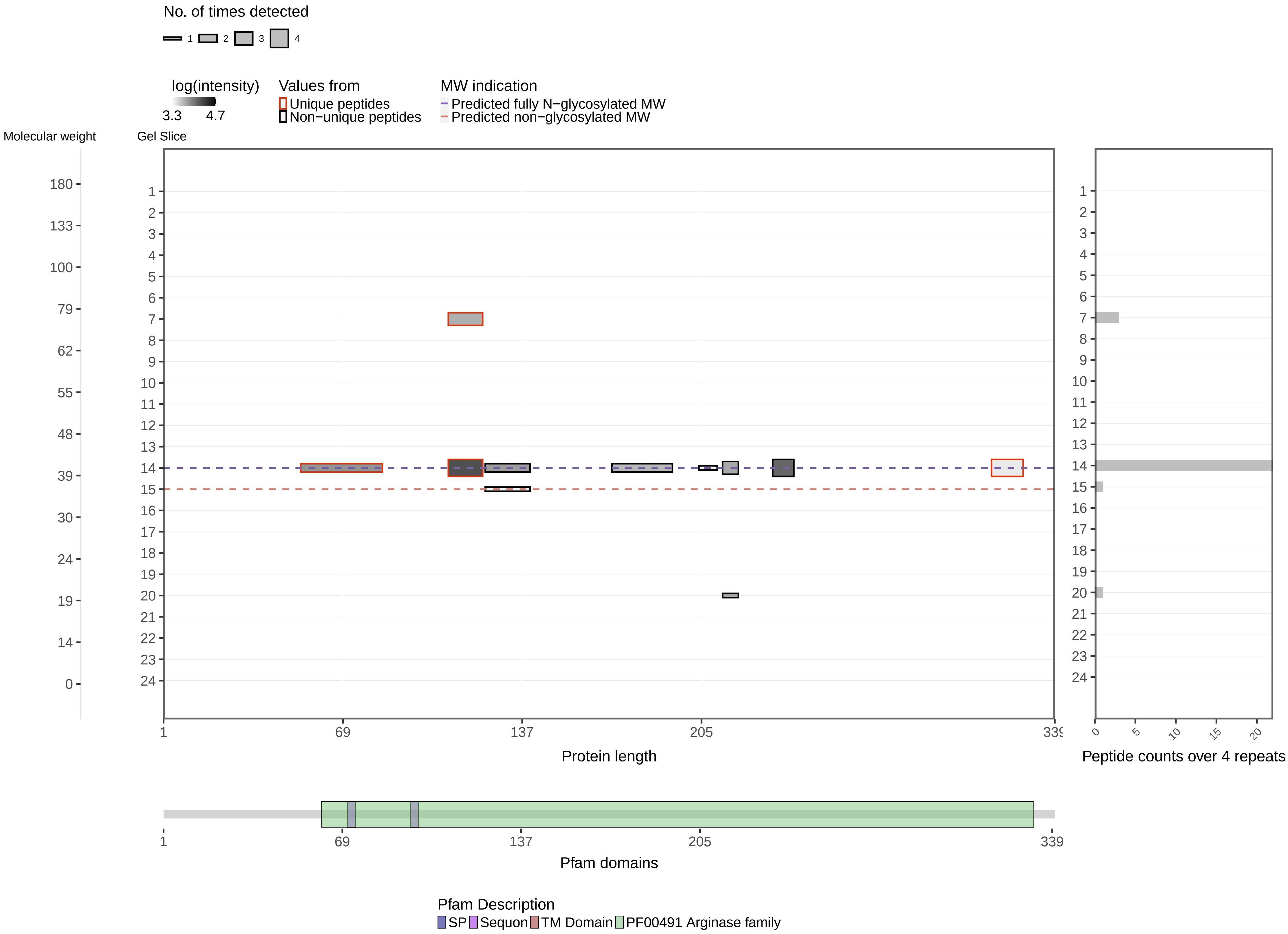


Protein sequence:

HSISYPVVRAVSEKLGGPIDILHLDAHPDIYHSFEGNKYSHASSFARIMEGGYVRRLLQVGIRSITKEGREQGKRFGVEQYEMRTFSRDRQFLENLKLGEGVKGVYISVDIDCMDPAFAPGVSHIEPGGLSFRDVLNILHNLQADVVGADVVEFNPQRDTVDGMTAMVAAKLVRELTAKISK*

NbL17g21090.1 3−phosphoshikimate_1−carboxyvinyltransferase_1_chloroplastic [MW = 56.45 kDa]

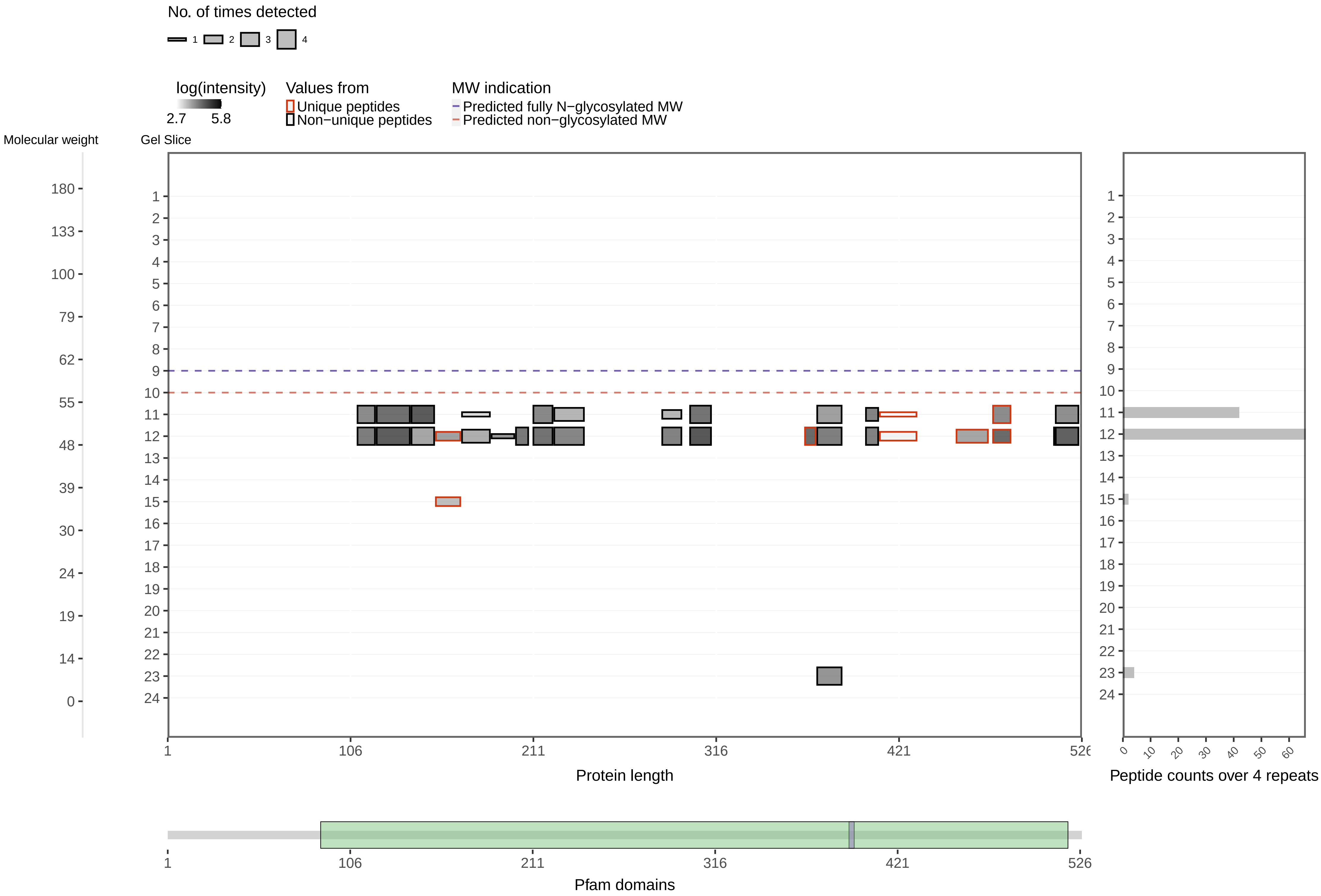


Pfam Description
SP Sequon TM Domain PF00275 EPSP synthase (3−phosphoshikimate 1−carboxyvinyltransferase)

Protein sequence:
KVKLSGSISSQYLTALLMAAPLALGDVEIEIIDKLISVPYVEMTLKLMERFGVSVEHTSSWDKFLVRGGQKYKSPGKAYVEGDASSASYFLAGAAVTGGTVTVEGCGTSSLQGDVKFAEVLEKMGAEVTWTENSVTVKGPPRNSSGMKHLRAVDVNMNKMPDVAMTLAVVALFADSPTAIRDVASWRVKETERMIAICTELRKLGATVVEGPDYCIITPPEKLKVAEIDTYDDHRMAMAFSLAACADVPVTIKDPGCTRKTFPNYFDVLQQYSKH*

### NbL17g21060.1 40S_ribosomal_protein_S13 [MW = 17.08 kDa]

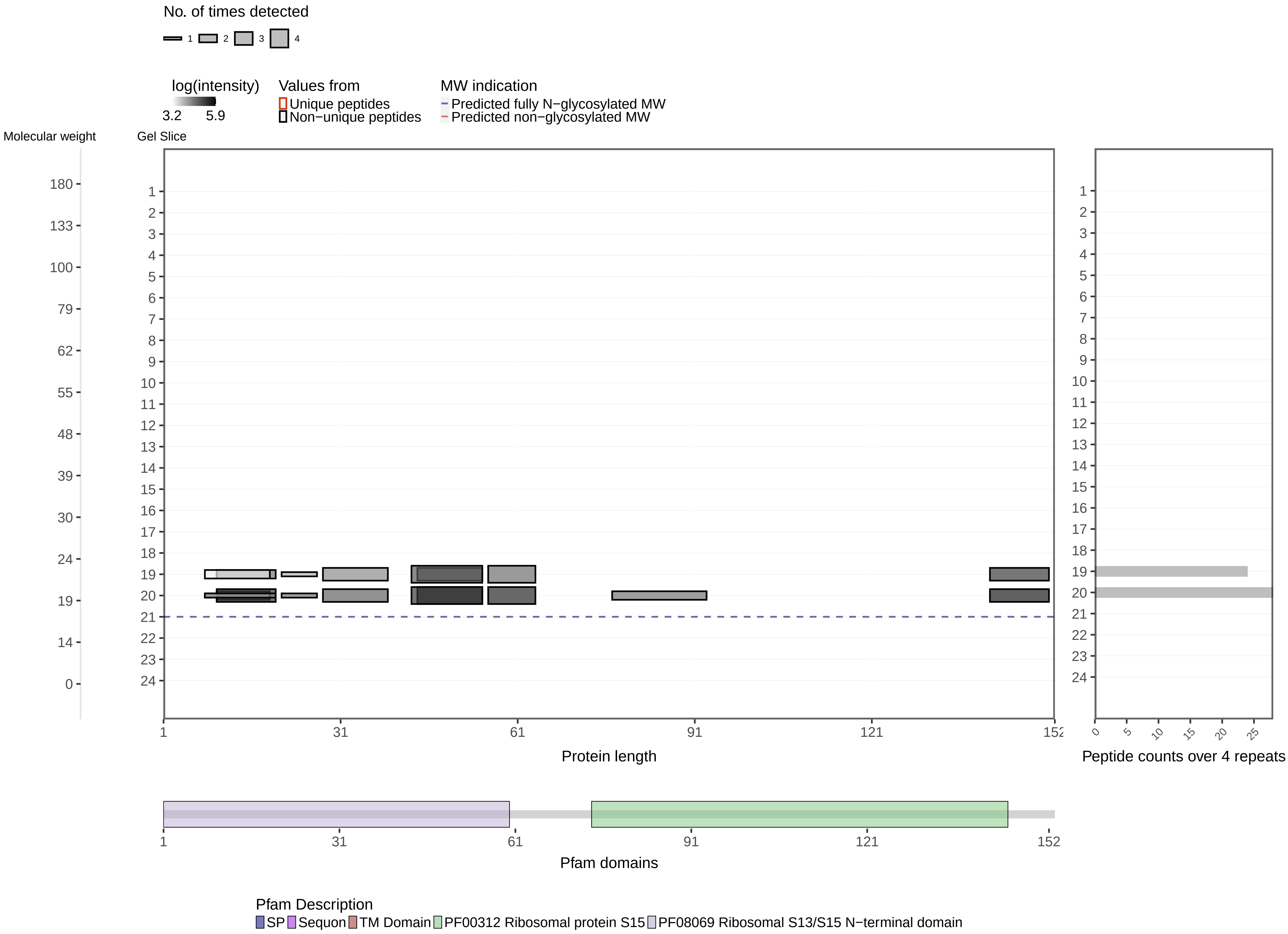


Protein sequence:

AQVKSITGSKILRILKAHGLAPEIPEDLYHLIKKAVAIRKHLERNRKDKDSKFRLILVESRIHRLARYYKKTKKLPPVWKYESTTASTLVA*

### NbL17g20970.1 transducin_beta−like_protein_2 [MW = 47.19 kDa]

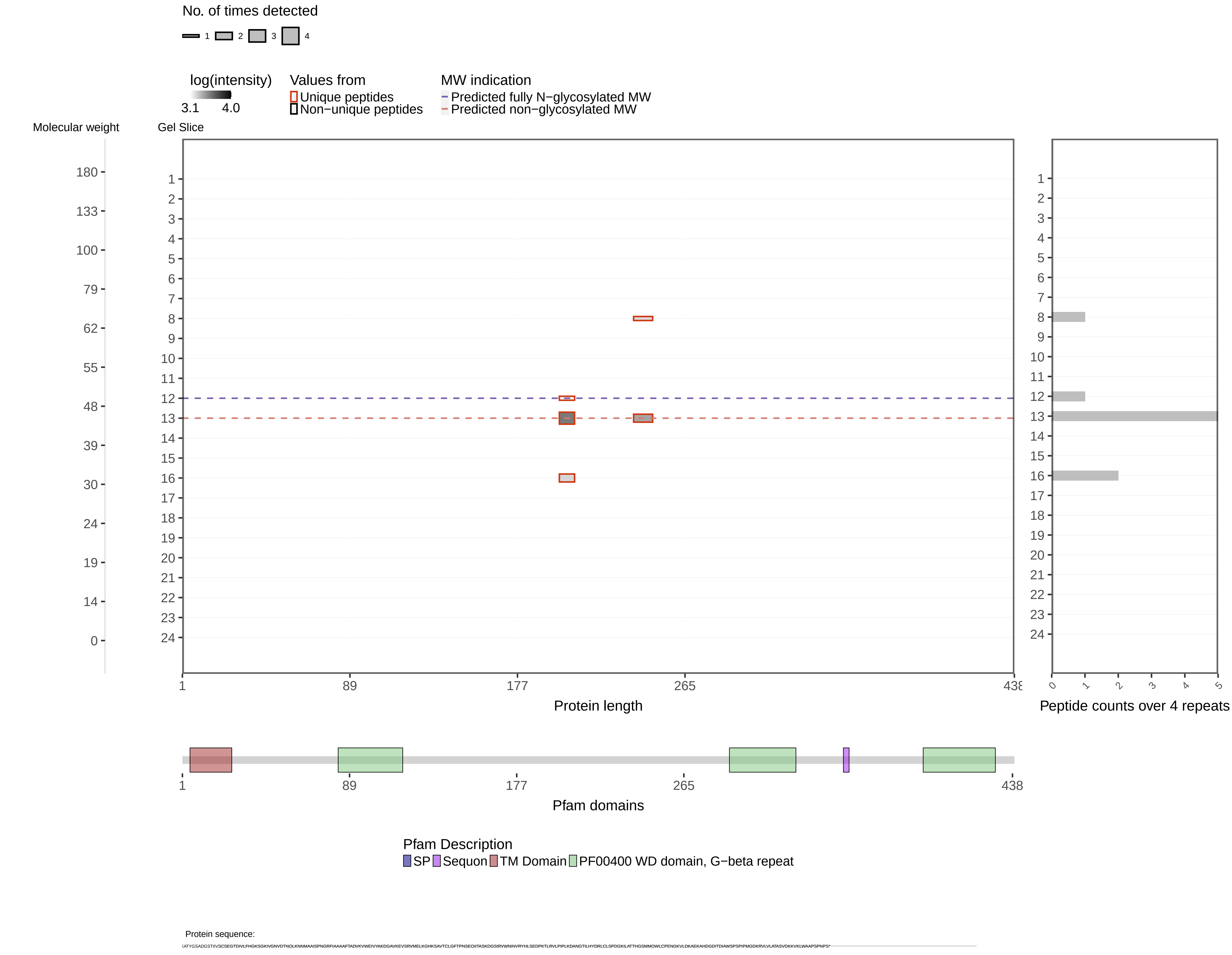

NbL17g20730.1 fasciclin−like_arabinogalactan_protein_8 [MW − SP = 41.53 kDa]

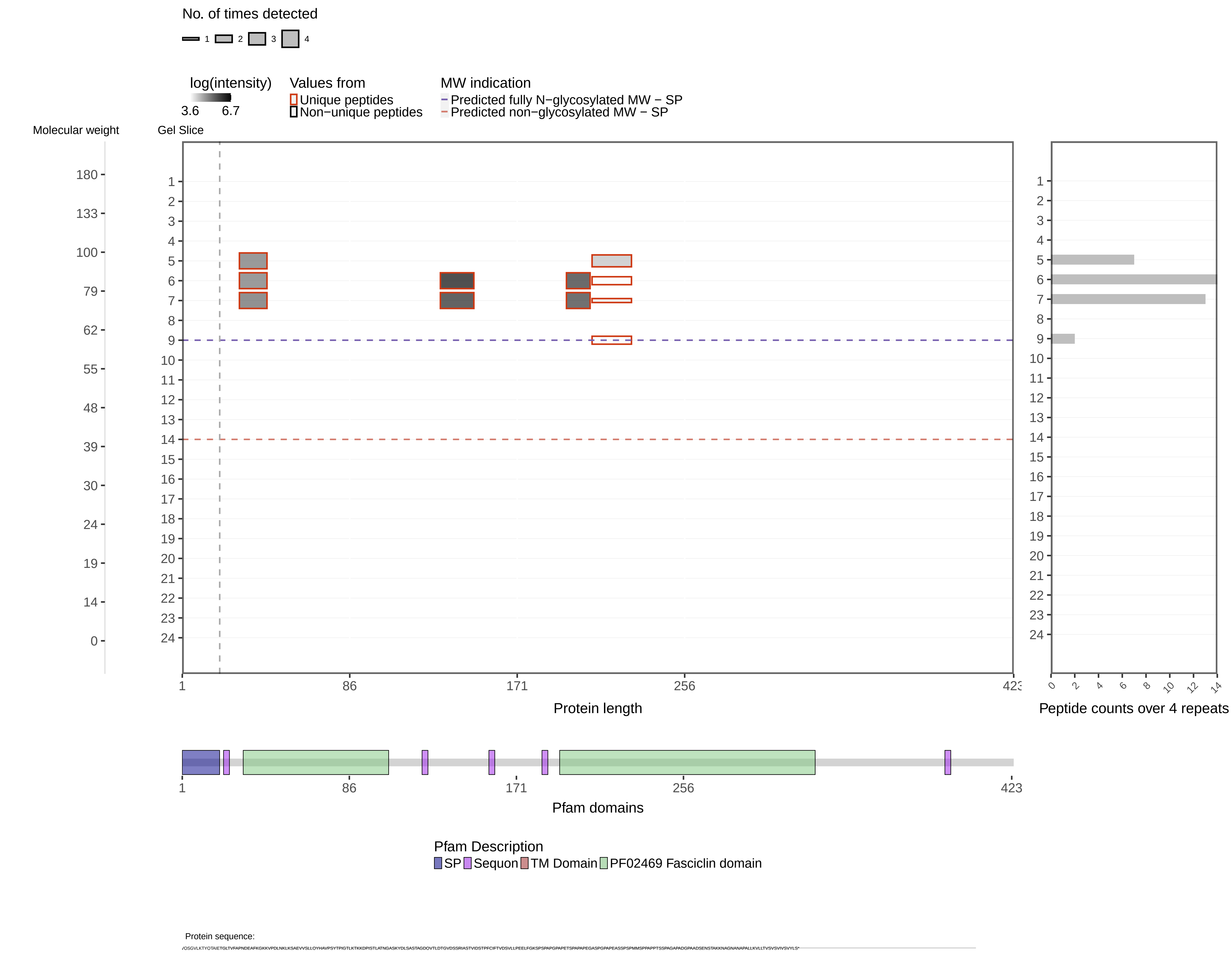

### NbL17g20710.1 uncharacterized_LOC109229227 [MW = 18.6 kDa]

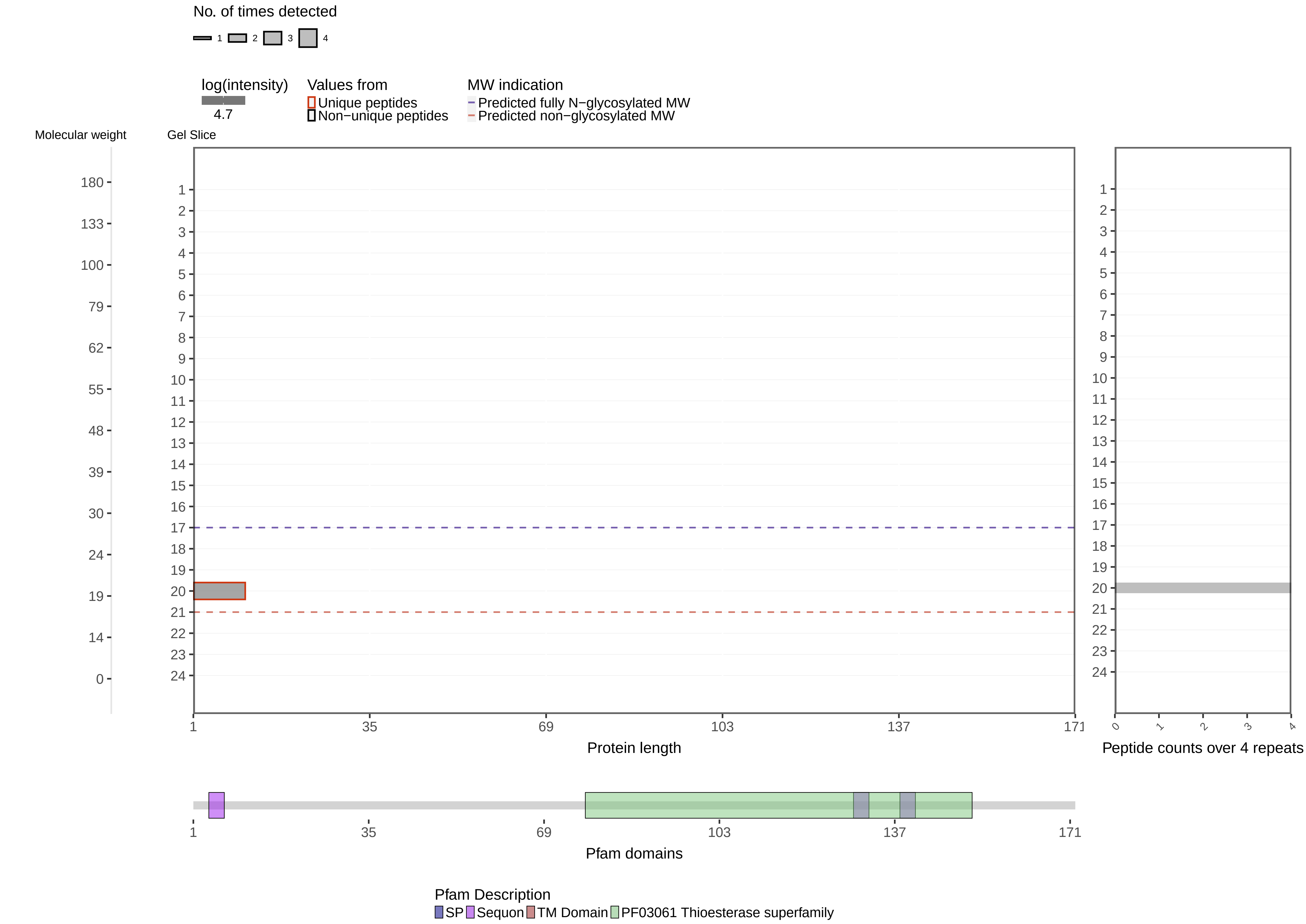


Protein sequence:

PIMNIYGSLHGGAVGDVTVRIATACARTIVGKDKKLFLGELSISYLSGAPRNAEVIVNATIIRSGRNLTVVAVDFRVKESEKLSYISHATFYHMPVASL*

### NbL17g20460.1 protein−ribulosamine_3−kinase_chloroplastic [MW = 37.49 kDa]

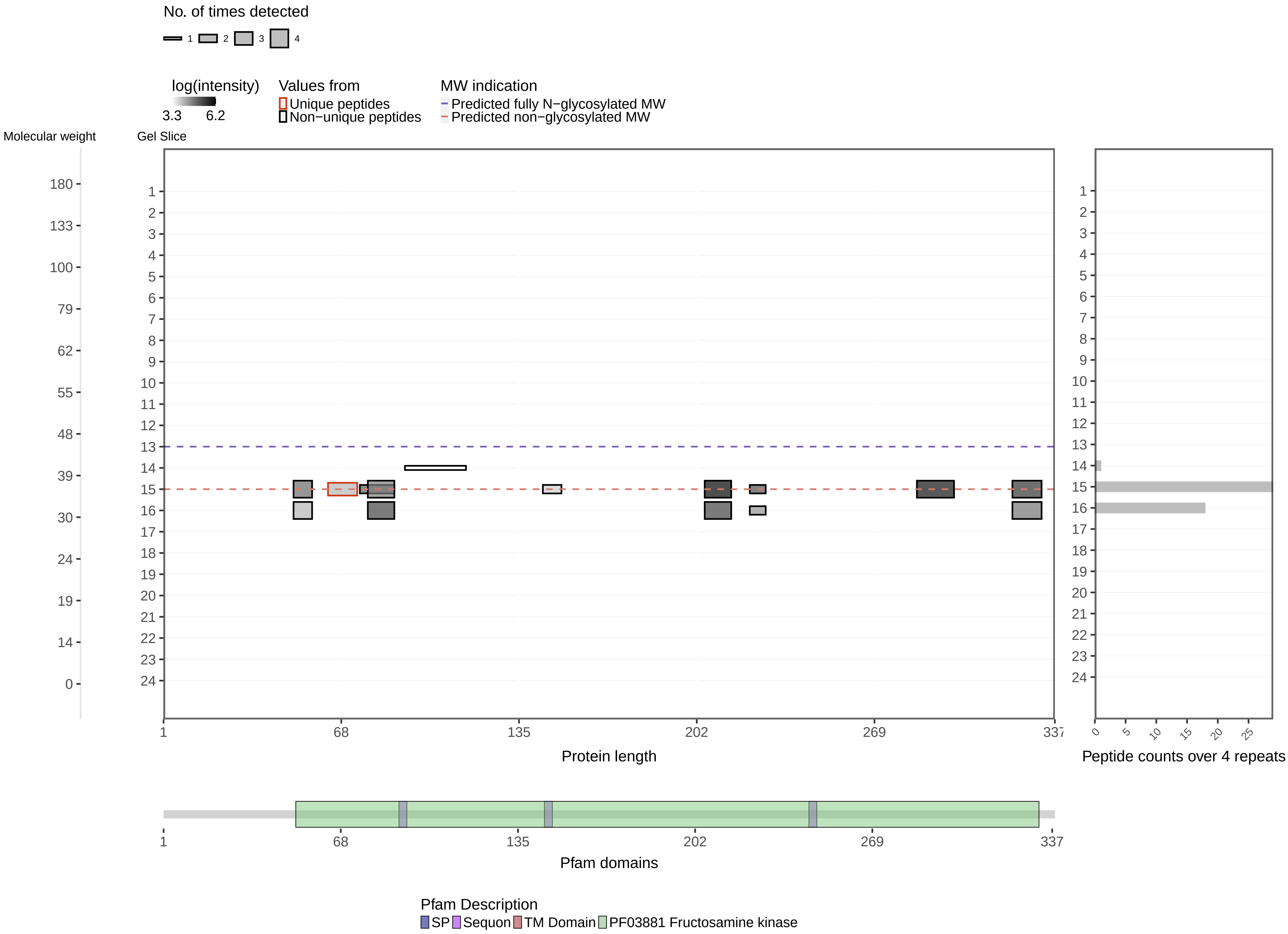


Protein sequence:

KATKSEKGFGFHVENTIGSTPQINTWTSDWIEFYAELRLGYQLKLAREQYGDSTIYERGQRLAKNLGPLFKNAVIEPCLLHGDLWSGNITYDQNGEPVILDPACYYGHNEAEFGMSWCAGFGGDFYKAYFQVMPEQPGFEQRRDLYMLYHYLNHYNLFGSGYRSSAMSIIDDYLRMLNV*

### NbL17g20370.1 ATPase_GET3−like [MW = 40.49 kDa]

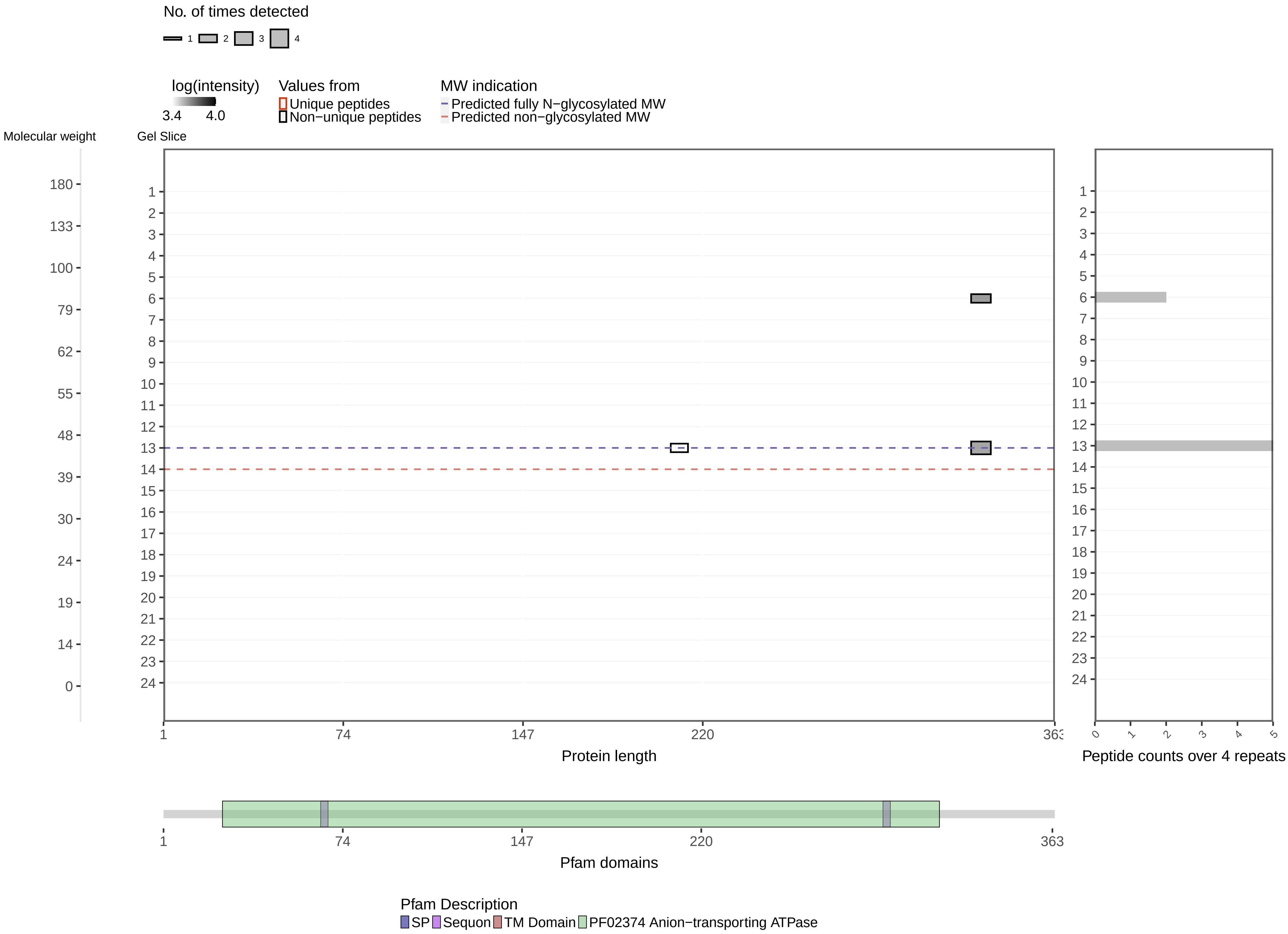


Protein sequence:

KVMSLKSRFGGVLSOMTRLLGVDEEFGEAAILGKLEGMKEIIEEVNKOFKDPDLTTFICVCIPEFLSLYETERLVOELAKFEIDTHNILINOVIFDEEVVESKLLKARMRMOOKYLDOFYMLYDDFNITKLPLLPOEVCGVEALKGFSHNFITPYKPFLARGSVEELENRVASLKEOLKDAEAELEKVRKGKOKV*

### NbL17g20360.1 ubiquitin−related_modifier_1_homolog_1−like [MW = 11.14 kDa]

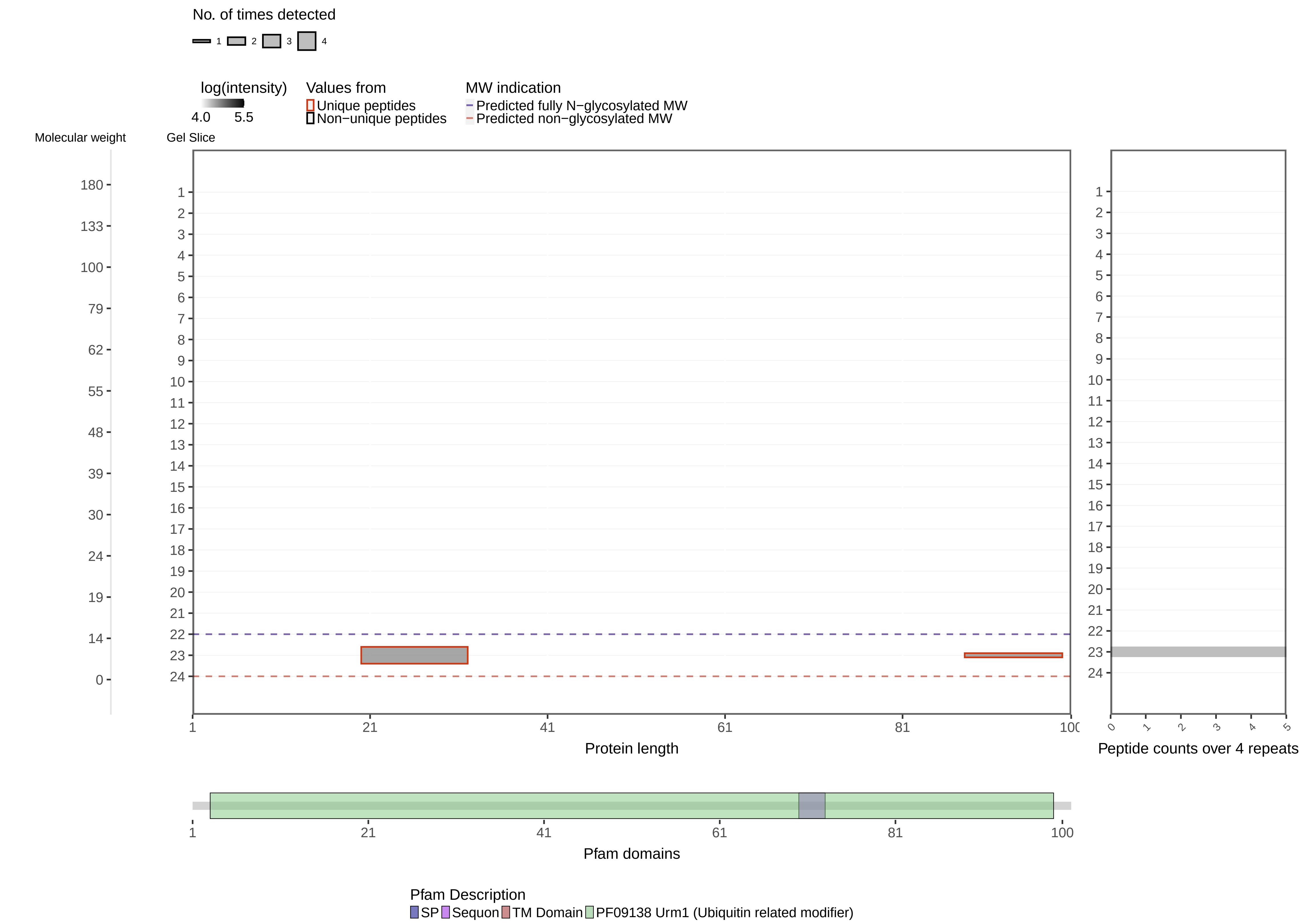


Protein sequence:

NLLTWVRTNLIKERPEMFMKGDSVRPGVLVLVNDTDWELSGQLETVLEEKDNVVFISTLHGG*

NbL17g20300.1 uncharacterized_LOC104238971 [MW = 66.74 kDa]

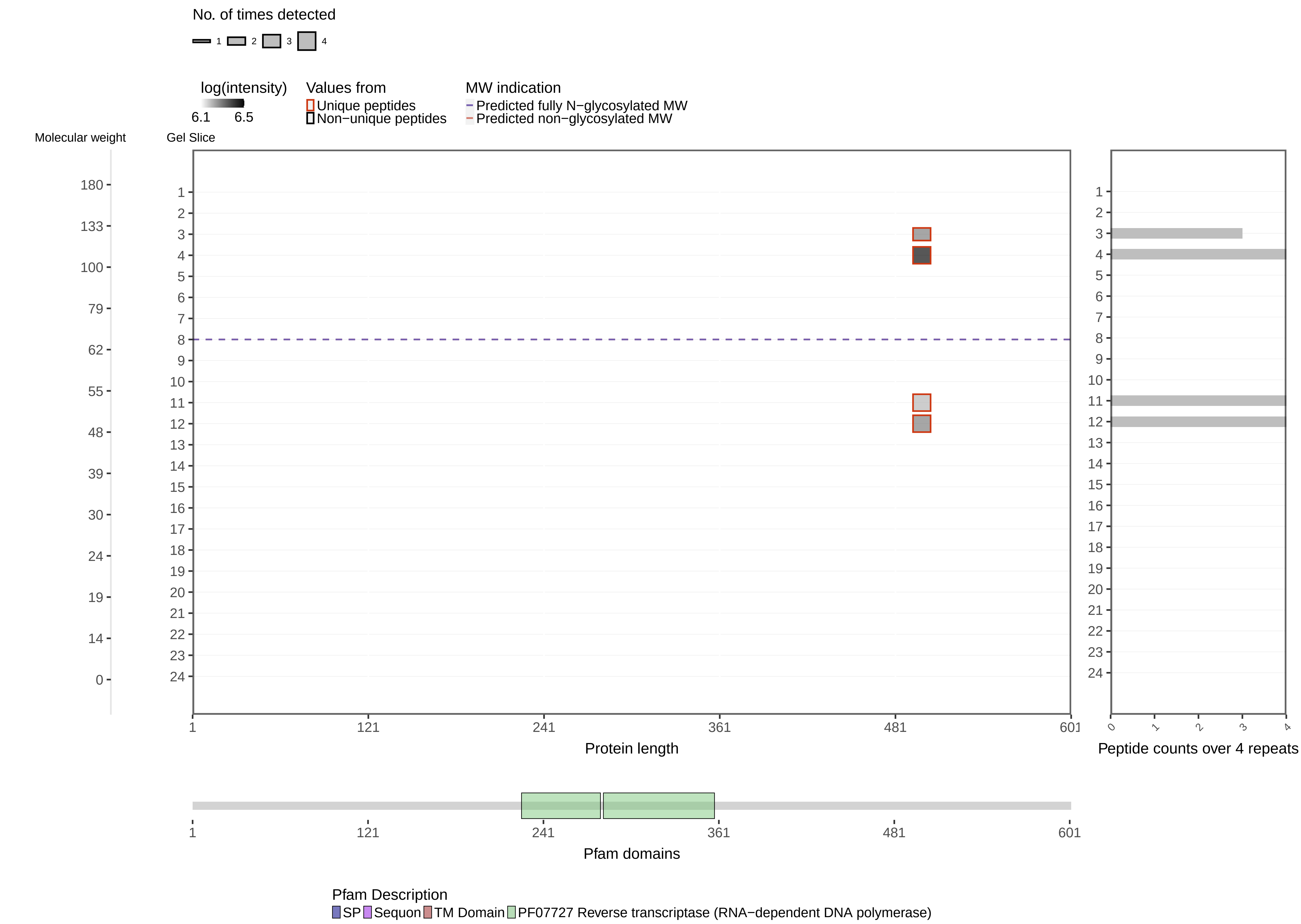


Protein sequence:

VDDIVITGNDQDGITKLKQHLFQHFQTKDLGTLKYFLGIEVAQSSSGIVISQRKYALDILEETGMTGCKPVDTPMDPNSKLLPGHGEPLNDPERYRRLVGKLNYLTATRPDISFPVSVVSQFMGSPCDSHWDAVVRILRYIKSTPGKGLLFEDRGYEQIVGYSDADWAGSPSDRRSTSGYCVLVGGNLVSSKSKKQNVVARSSAEAEYRAMAIATCELVWVKQLLKELKFGDISQMGLVCDNQAALHIASNPVFHERTKHIEIDCHFIREKILSGDIATKFVKSNDQLADIFTKSLT

### NbL17g20230.1 glyoxylate/hydroxypyruvate_reductase_HPR3−like [MW = 35.7 kDa]

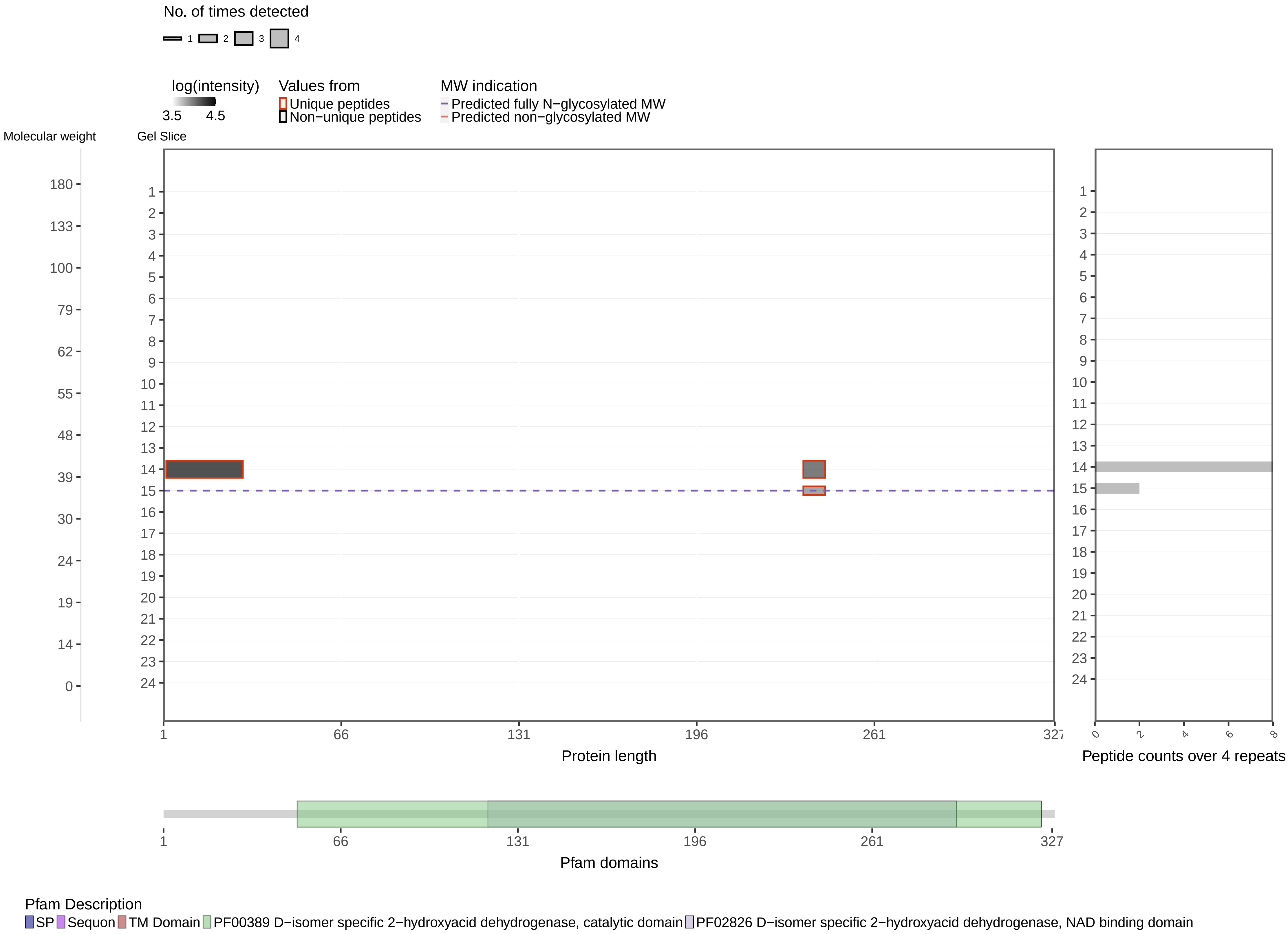


Protein sequence:

.CSKVGGRKVGIVGLGSIGLKVAKRLEAFGCIISYQSRKKKPVSYPFYPDVHELATKCDVLVICCALTEQTRHLINKEVLLALGKEGVIINIARGSVINEMELVQCLVQGEIAGVGLDVFENEPNVPEDLFSLDNVVLTRHVAACTEDSFRDLYELVSGNLEAFFLNKPLLSPVLDD*

### NbL17g20220.1 glyoxylate/hydroxypyruvate_reductase_HPR3−like [MW = 35.38 kDa]

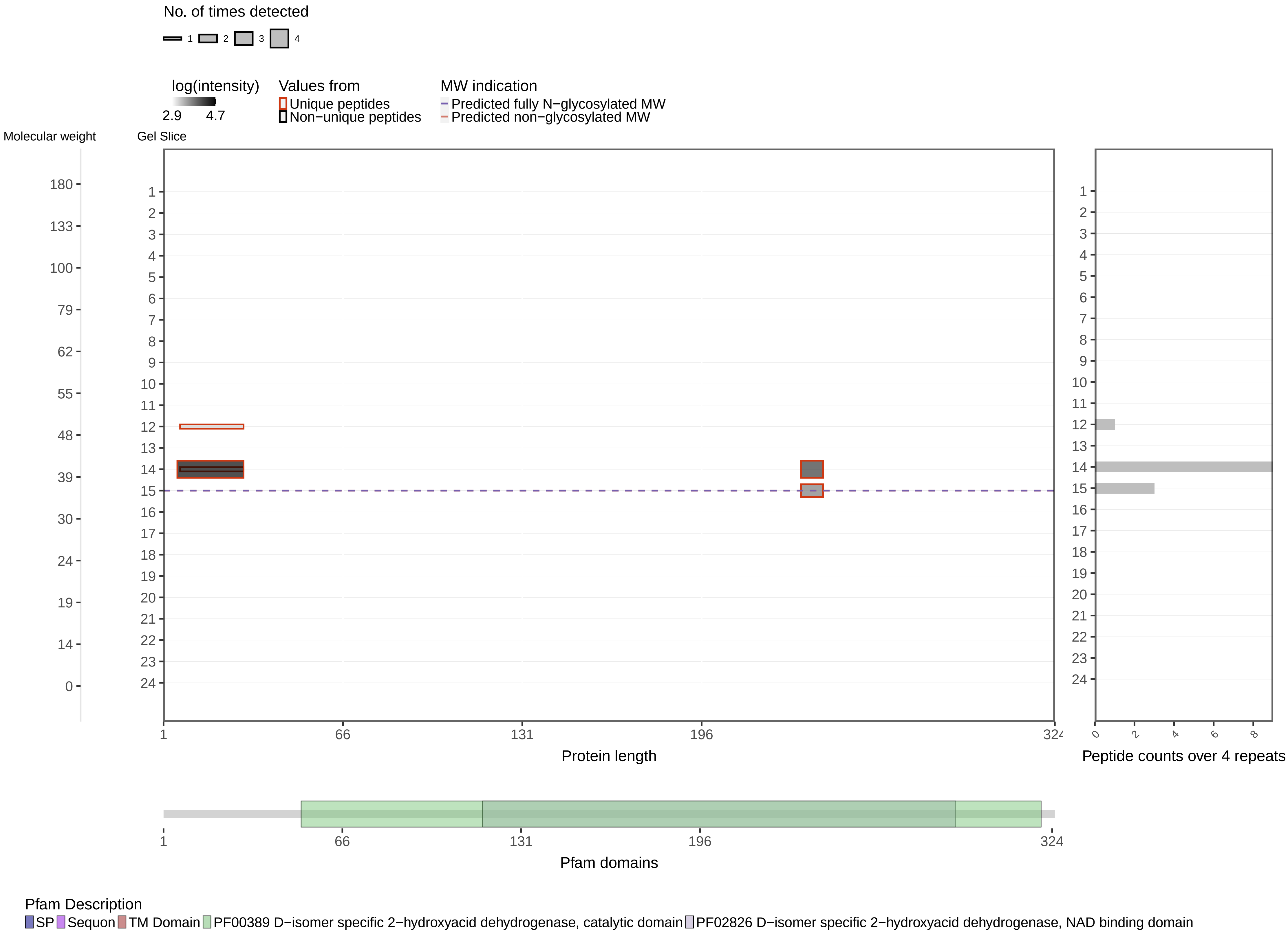


Protein sequence:

LASKVGGRKVGIVGLGSIGLEVAKRLEAFGCIISYQSRKKKPVVYPFYPDVHELATKCDVLVICCALTDQTHHLINKEVLLALGKEGIIINIARGAIINEMELVQCLVQGEIAGAGLDVFENEPNVPEELLSLDNVVLSQHVAFLTEDSFRDLYELMSGNLEAFFLNKPLLSPVFDD*

NbL17g20130.1 methionine_adenosyltransferase_2_subunit_beta−like [MW = 35.65 kDa]

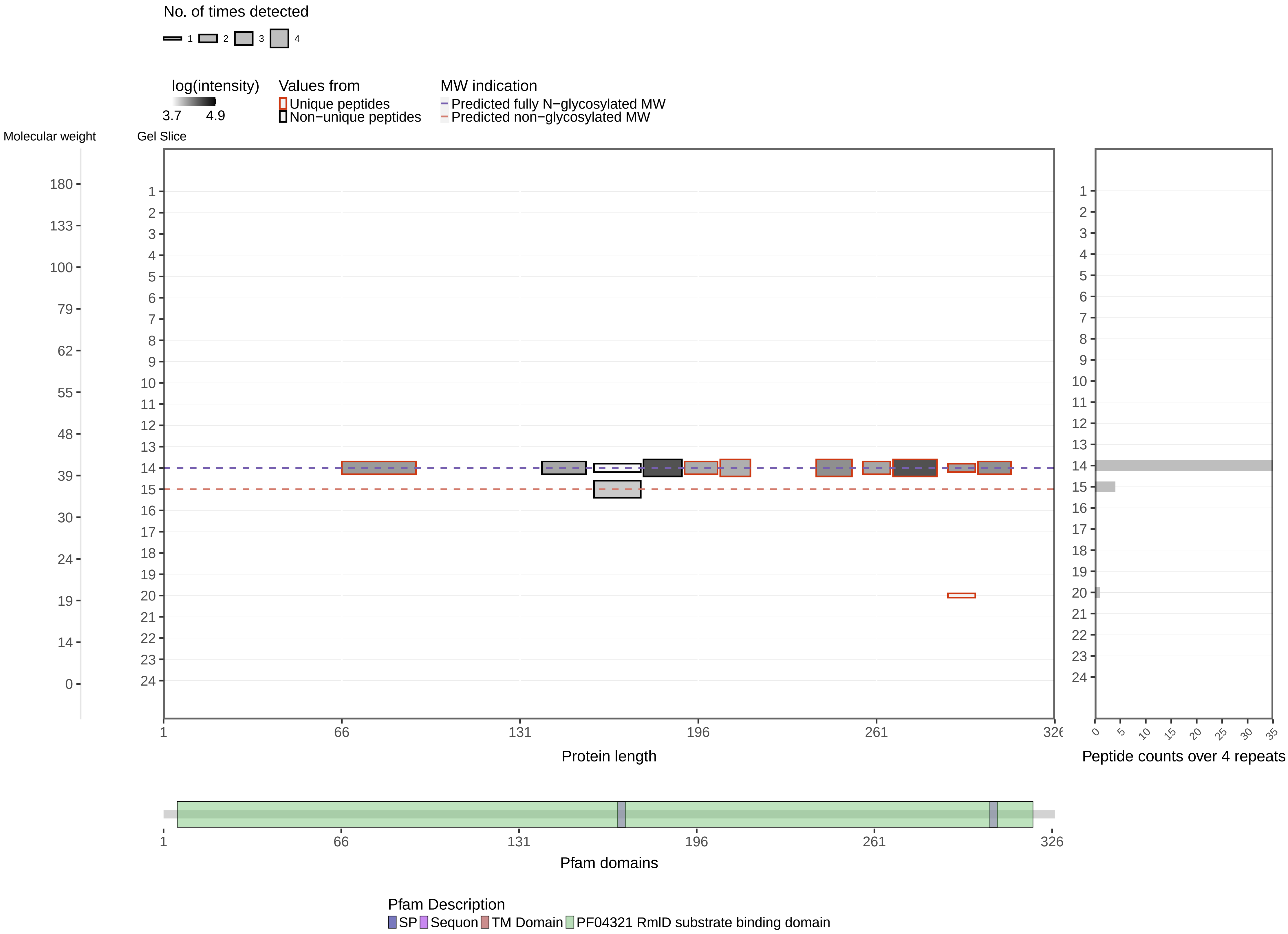


Protein sequence:

VNVYGRSKVAAEQFISANYSNFAILRSSIIYGPQTVSPVPKSLPVQWMDSVLAKGDAMEFFHDEFRCPIYVKDLVTIVWTLTNRWISGREPIQLLLNVGGPDRVSRVQMAEAVAHIRGYSLSLIKPVSSSTVNRGVKSPADISMDITRLIQTLNMSTTTFRDGVKLTLEAELASTS*

NbL17g20120.1 NAD−dependent_malic_enzyme_59_kDa_isoform_mitochondrial [MW = 66.34 kDa]

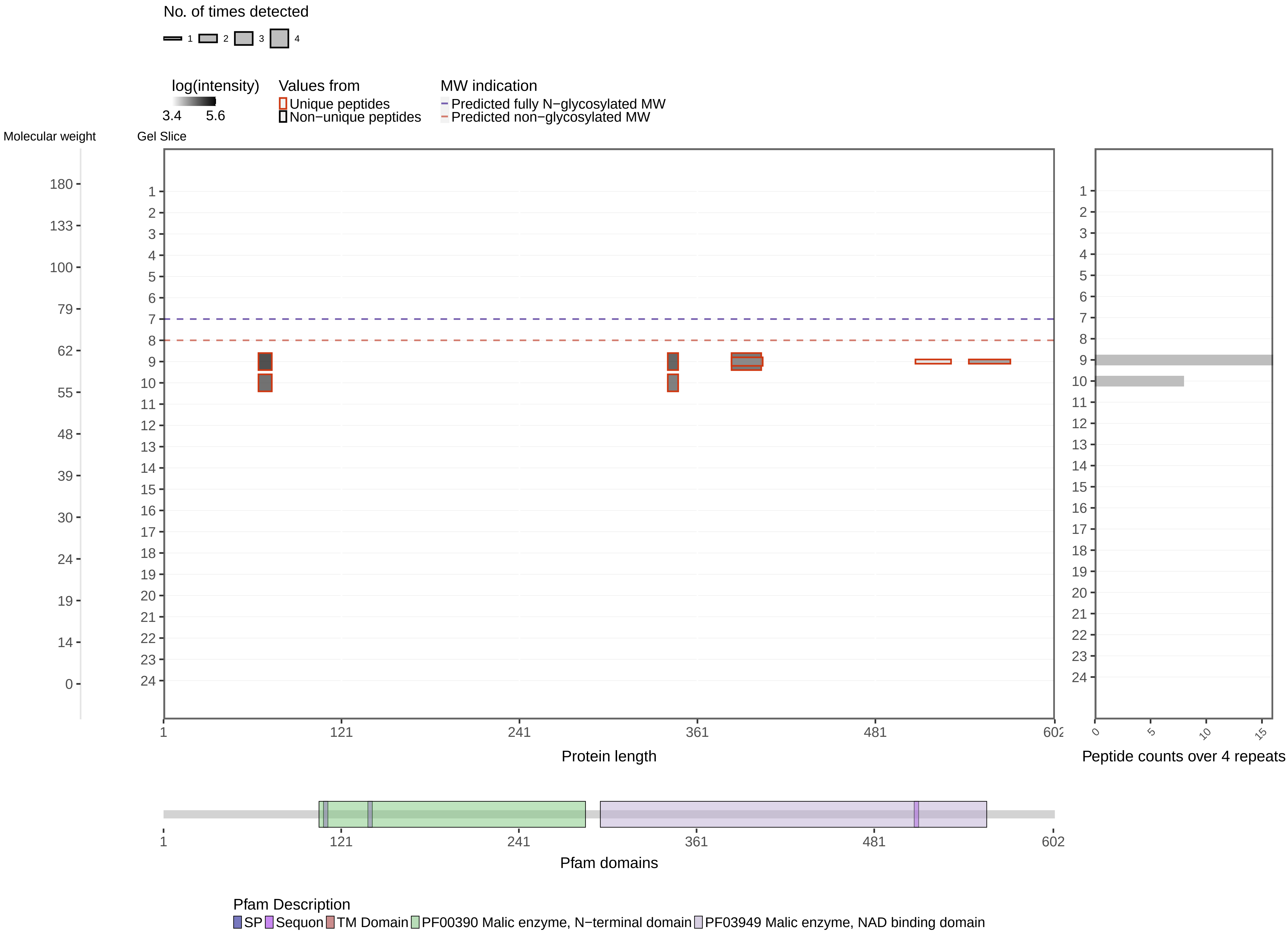


Protein sequence:

KFCMFNDDIQGTAGVALAGLLGTVRAQGRPLTDFANQKIVVVGAGSAGLGVLKMALQTVSRMAGPSADPHFFLLDKNGLITKHRKDIDPAALPFAKAQHEIEGLGLQEGAGLLEVVKKVKPHVLLGLSGVGGIFHEEVLRAMRESDSVRPAIFAMSNPTNNAECCPVDAFKLAGENIVFASGSPFENVDLGNGKIGHVNQANNMYLFPGIGLGALLSGARNISDTMLEAAAECLANYMSDDEIKRGILYPSINDIRDITAEVGAAVLQAAVAEDLAEGHGDVGARELQHMSKEE

### NbL17g20110.1 50S_ribosomal_protein_HLP_mitochondrial−like [MW = 18.03 kDa]

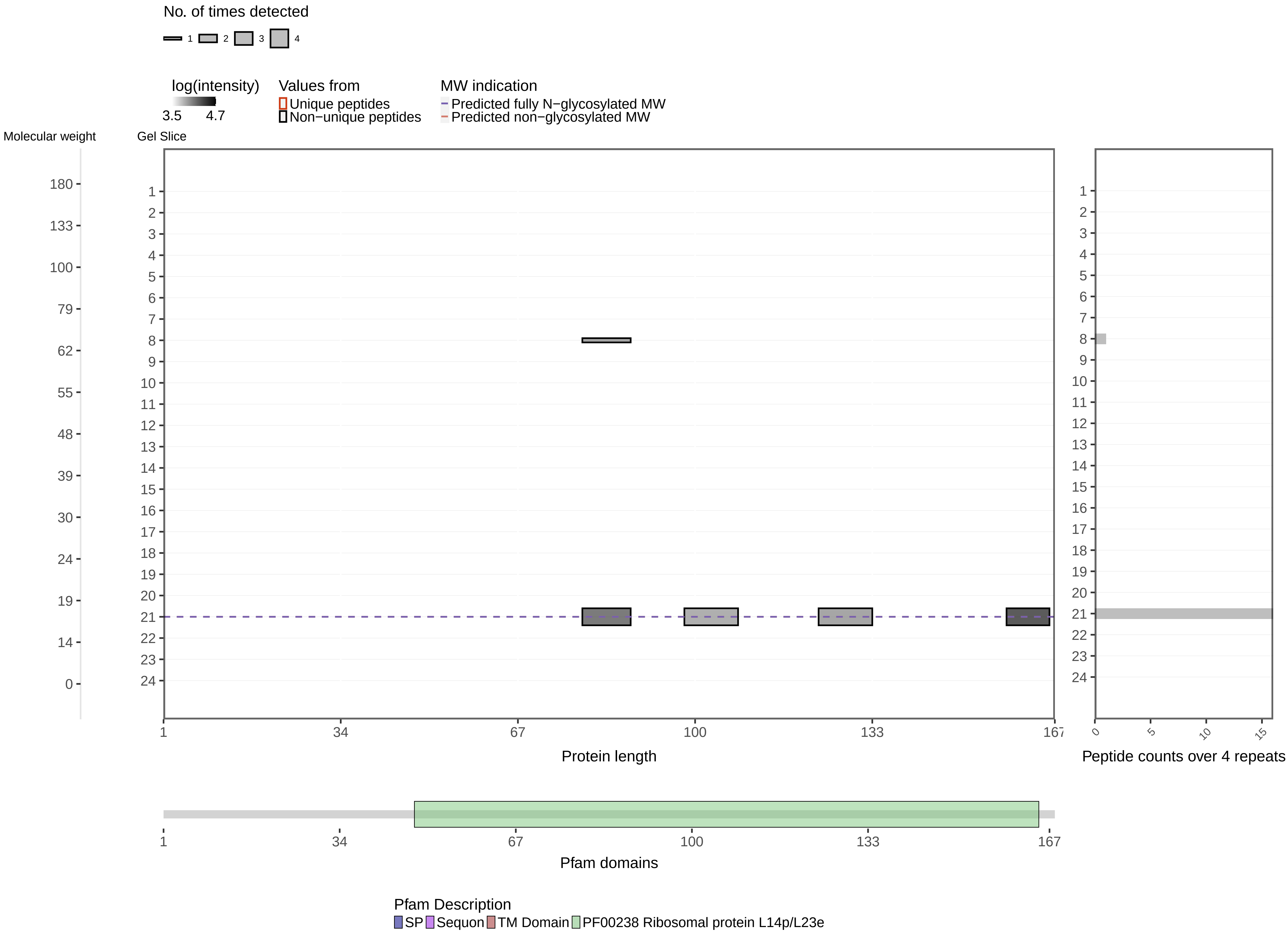


Protein sequence:

ALKGKKGARLGDTIVCSVKEAQPGGKVKKGDVHYGVVVRAAMPRGRCDGSEVKFDDNAVVLINKHGEPIGTRVFGPVPHELRKKKHVKILSLAEHIA*

NbL17g20090.1 (+)−neomenthol_dehydrogenase−like [MW = 33.42 kDa]

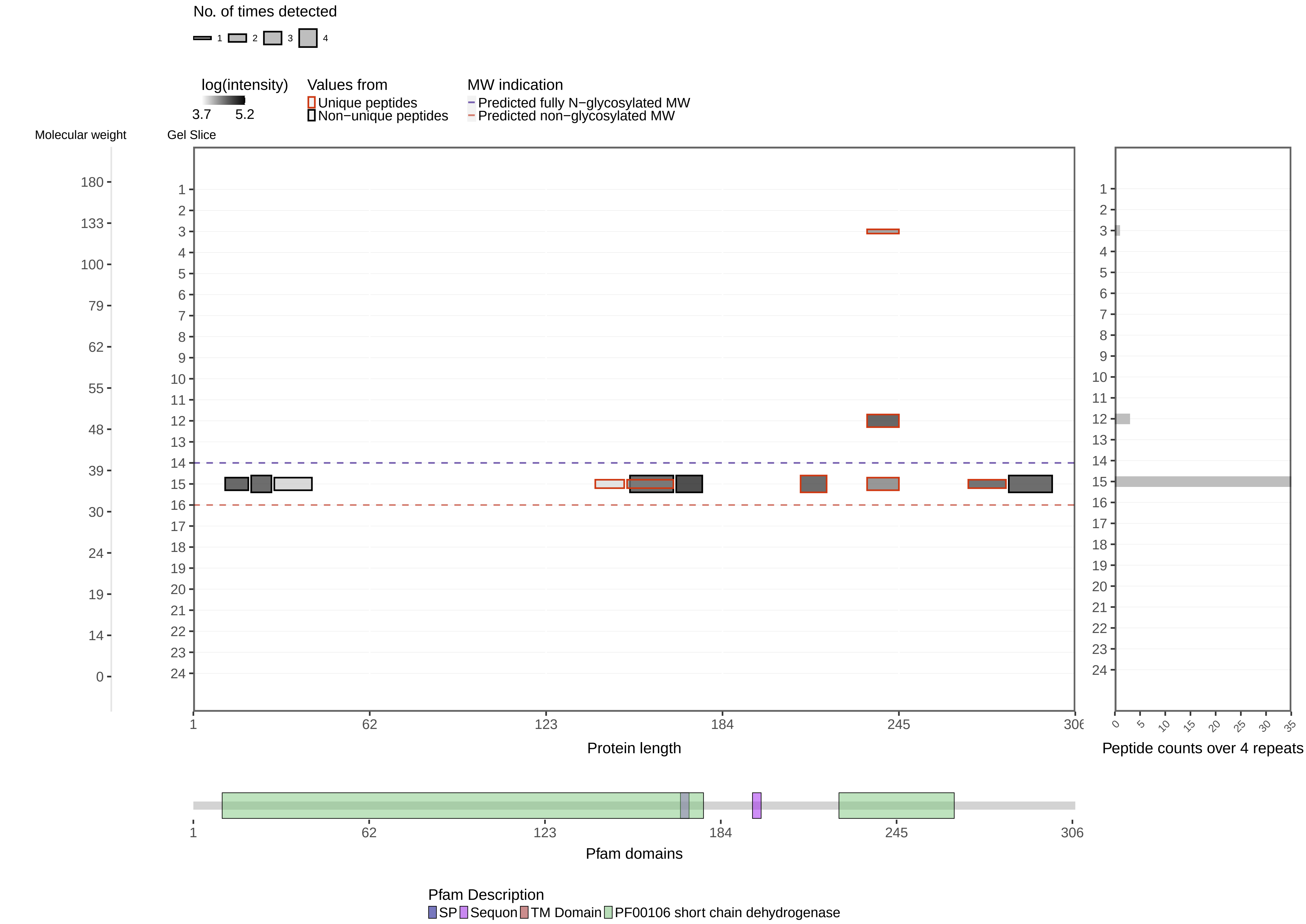


Protein sequence:

LETNYYGVKRMTEAFIPLLQLSNSPRIVNVSSSMGRLRNLKHEWAKGVLSDSENLSEEKIEEVIAQYLKDYKEDSLQAKGWPSFLSAYTLSKAAVNAYTSIMAKKHPSIQINCVCPGFVKTDMNYRRGLLSIEEGAESPVRLAMQPADGPSGLFFDRKEVSSFE*

### NbL17g20040.1 bifunctional_protein_FolD_4_chloroplastic−like [MW = 40.12 kDa]

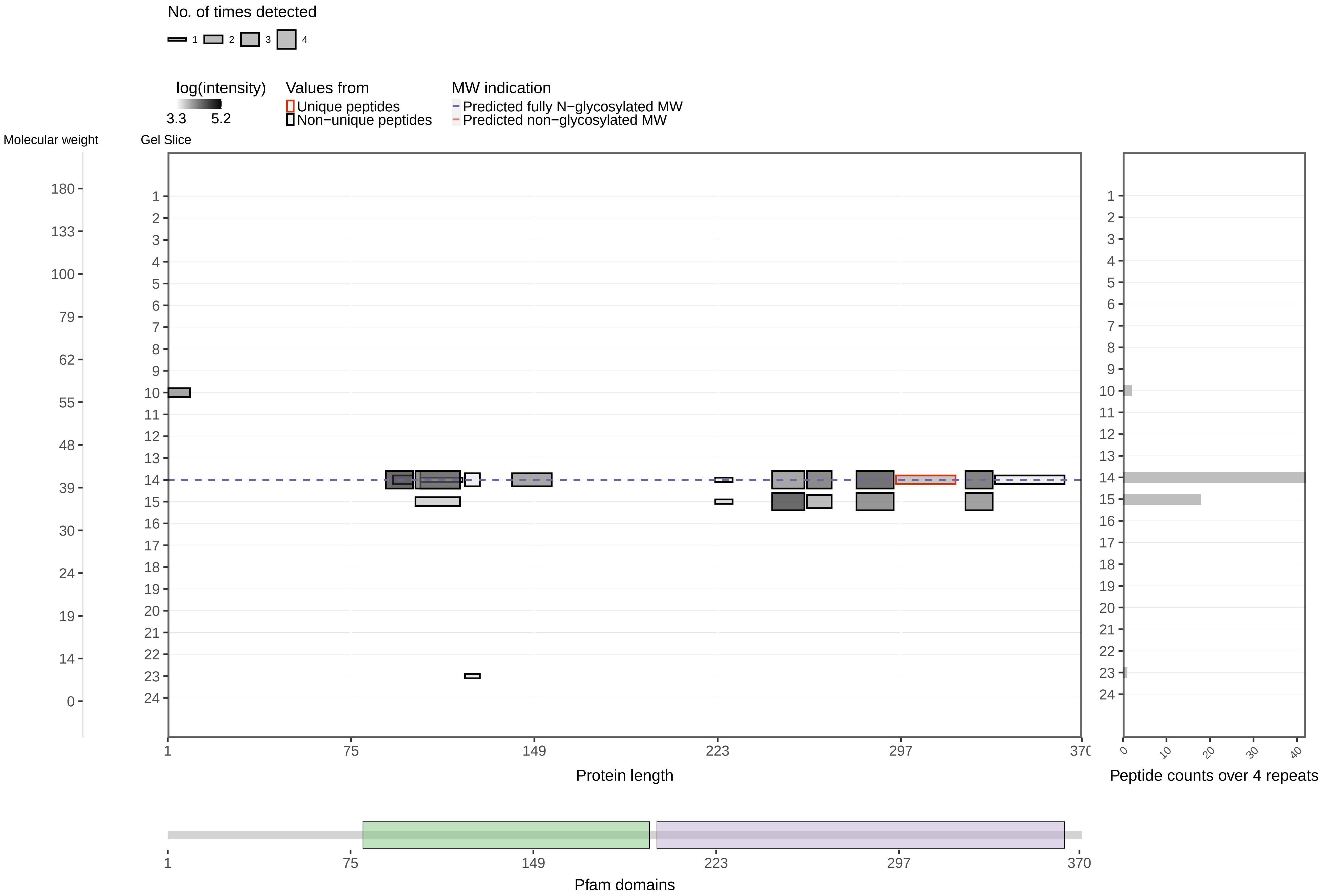


Pfam Description

SP Sequon TM Domain PF00763 Tetrahydrofolate dehydrogenase/cyclohydrolase, catalytic domain PF02882 Tetrahydrofolate dehydrogenase/cyclohydrolase, NAD(P)−binding domain

Protein sequence:

VQLPLPSHMNEQNILNAVCIEKDVDGFHPLNIGRLAMRGREPLFVPCTPKGCIELLHRYNVEIKGKRAVVVGRSNIVGTPAALLLQREDATVSIVHSRTKKPEEITRQADIIISAVGQPNMVRGSWIKPGAVIIDVGINPVEDATNTRGYRLVGDVCYEEASKVASAITPVPGGVGPMTIAMLLSNTLLSAKRIHNFN*

NbL17g19770.1 small_nuclear_ribonucleoprotein−associated_protein_B'−like [MW = 33.61 kDa]

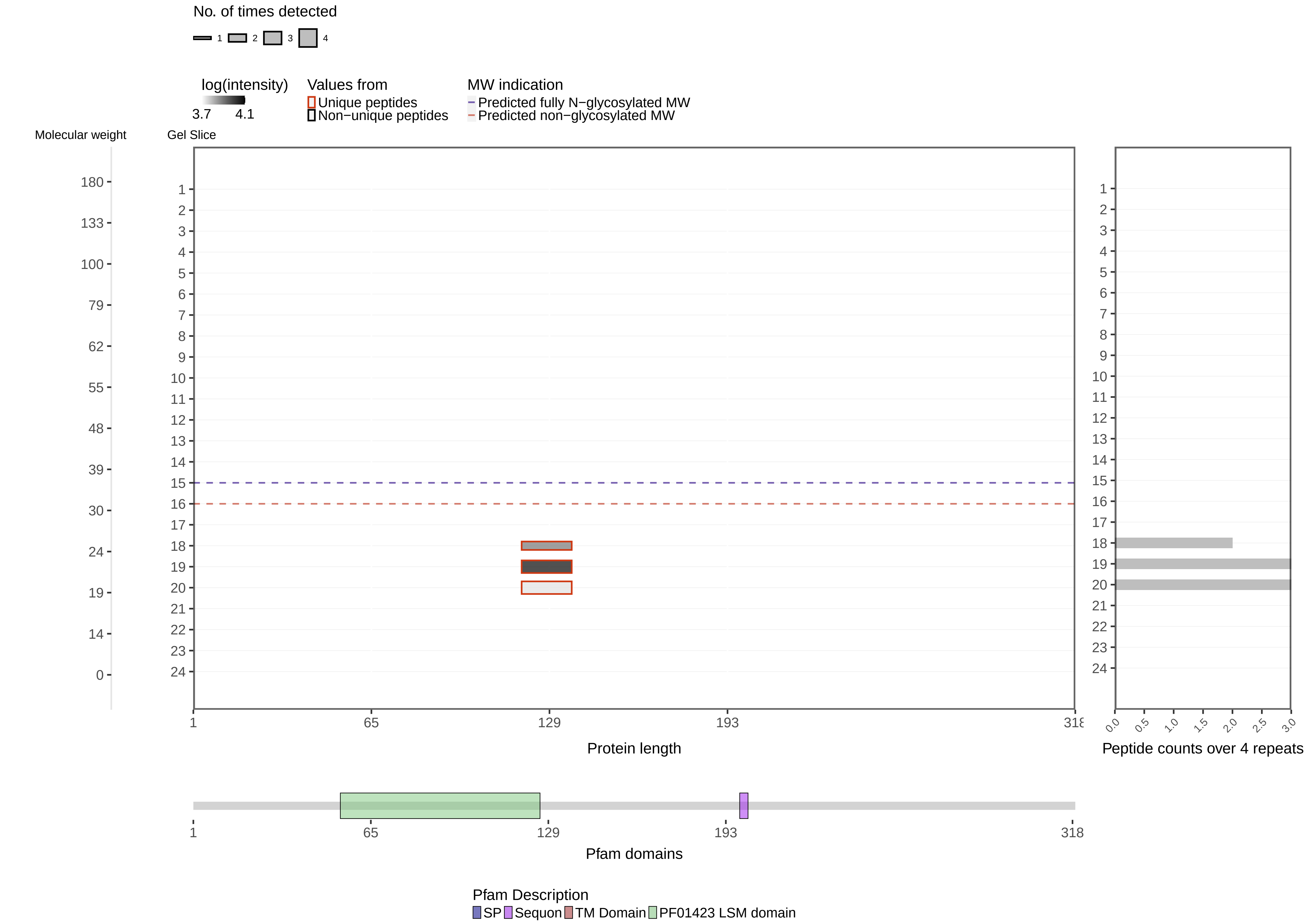


Protein sequence:

LGRAAGRGVPTGPLVHAOPGLAGPVRGVGGPAPGMMOPOISRPPMPNVSAPPMNYPPPVMRPPGGPPMOMOMORPGGPPPPMAPPPOFRPGGGPPOFPPOFGORPMVPPPPMMRGPPPPGAPPRPGMPGGPPGOPPRPGMPPPPSGGOVPVYGPPRSGMPPPPNOO*

### NbL17g19510.1 uncharacterized_LOC104224003 [MW = 30.32 kDa]

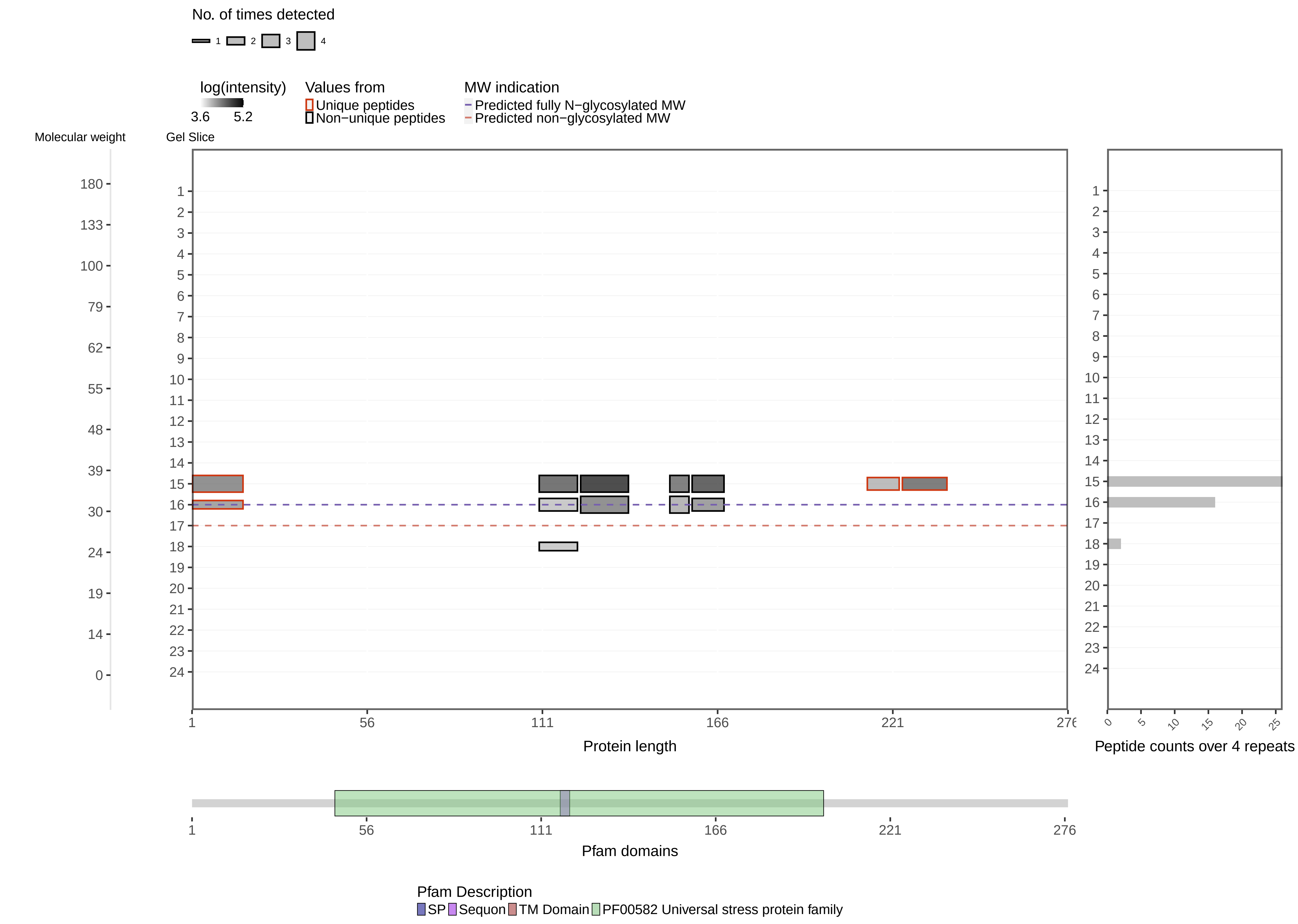


Protein sequence:

AQPLVDAQIPYKIHIVKDHDMKERLCLEVERLGLSAVIMGSRGFGATKRGNDGRLGSVSDYCVRHCACPVVVVRYPDEKDGGNAARKPVVSMASAAKENEEETAYHDASDDRQGKLRKNCCNCWFVDSYTNSLLVSFKSLHERGKWILC*

NbL17g19330.1 reductase−like [MW = 42.98 kDa]

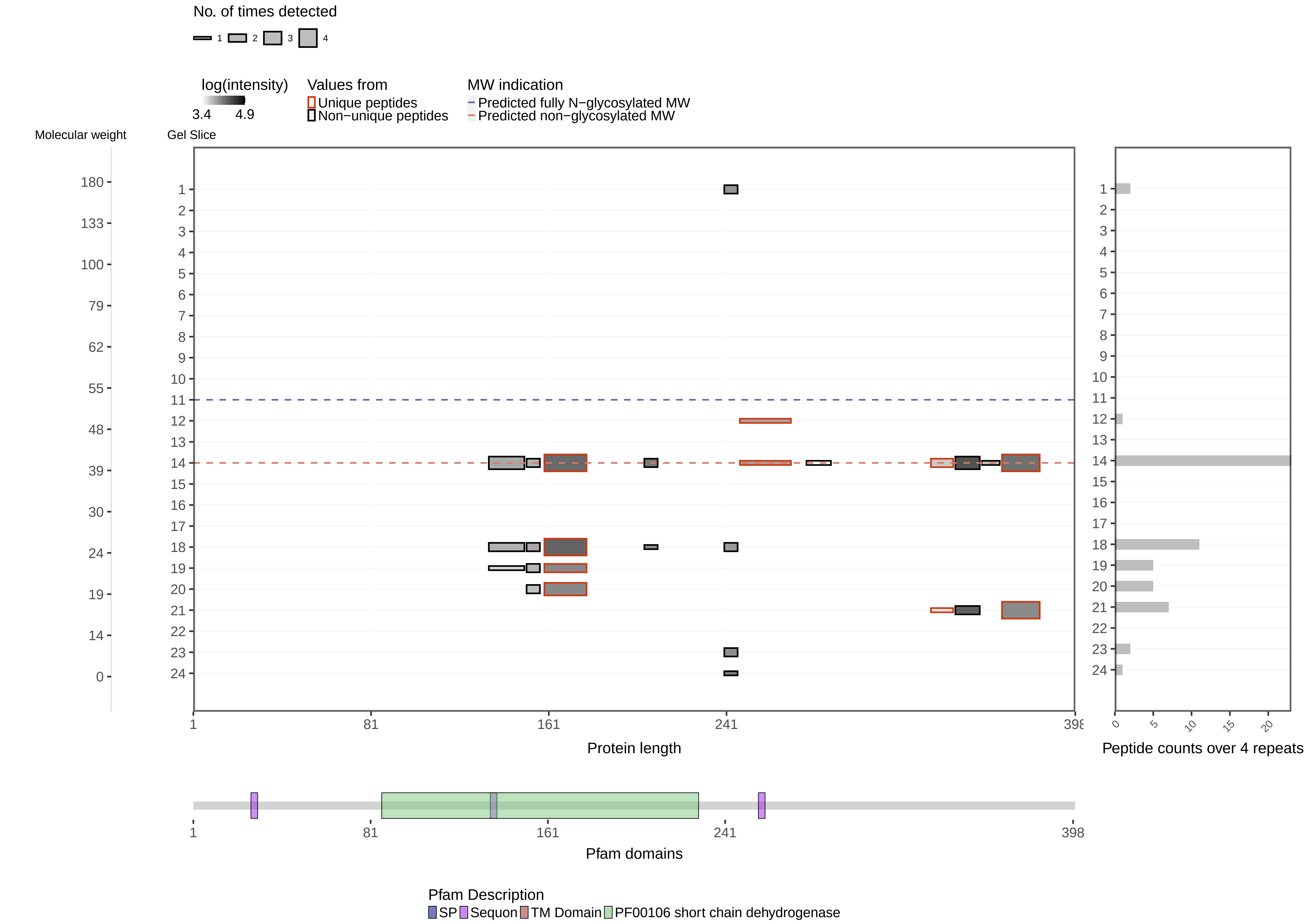


Protein sequence:

GFELSVGTNHLGHFLLSRLLLDDLKQSDYPSKRLIIVGSITGNTNTLAGNVPPKANLGDLRGMAGGLNGINSSAMIDGGEFDGAKAYKDSKVCNMLTMQEFHRRYHEETGITFASLYPGCIATTGLFREHIPLFRLLFPPFQKYITKGFVSETEAGKRLAQVVSDPSLTKSGVYWSWNKDSASFENQLSEEASDVEKARKVWEVSEKLVGLV*

### NbL17g19280.1 vesicle−fusing_ATPase [MW = 82.27 kDa]

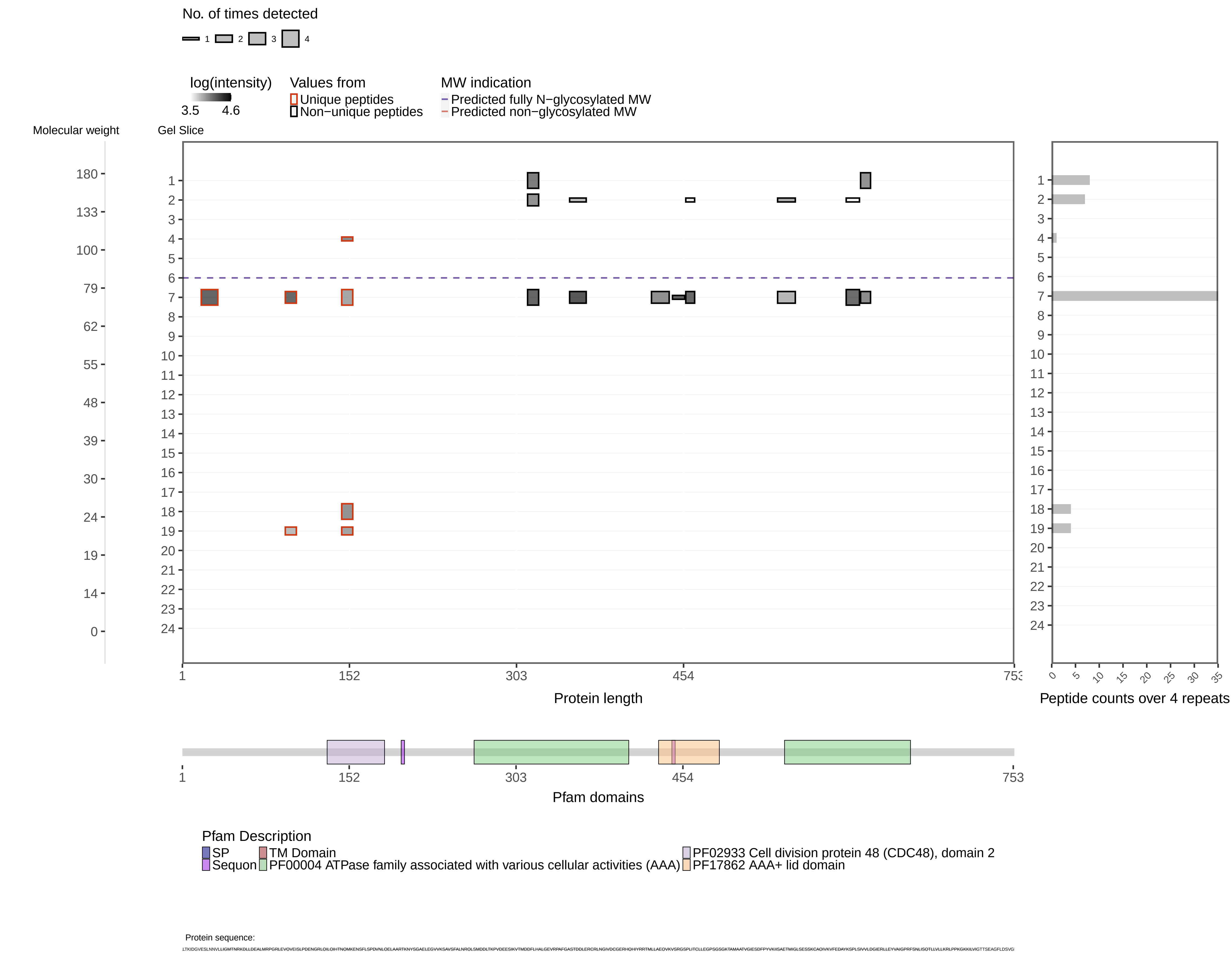

### NbL17g19230.1 probable_bifunctional_methylthioribulose−1−phosphate_dehydratase/enolase−phosphatase_E1_1 [MW = 58.16 kDa]

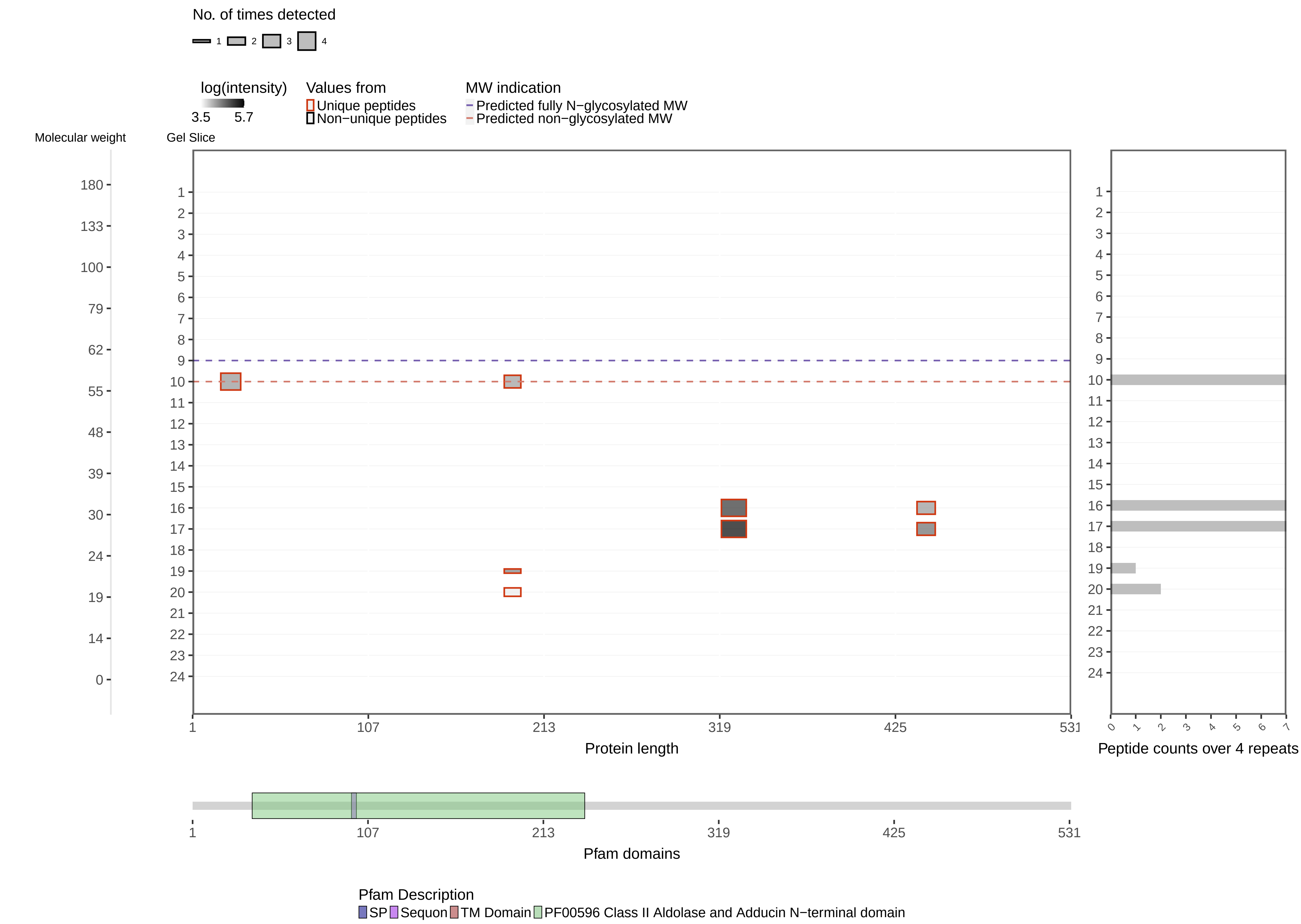


Protein sequence:

PTHGPIHNSNGALSITRNVKTSAKAGGLSLNGDNEPLRRCIVLDIEGTTTPISFVTDVLFPYAFDNVGRHLDATYDSAETQEDIKLLRAQVQEDLENGVVGAVPIPSDDAGKEEVIAALVTNVEAMIKADRKITALKELQGHIWQTGFQNNELEAVVFDDVPEALERWAALGLKVYIYSSGSRLAQRLIFGYTNYGDLRKYLCGFFDTTVGNKRETKSYTEITASLGVDNPSEILFVTDVYQEAIAAKAAGLEVIISVRPGNGPLPENHGFRTITSFSEI*

### NbL17g19220.1 early_nodulin−like_protein_1 [MW = 24.18 kDa]

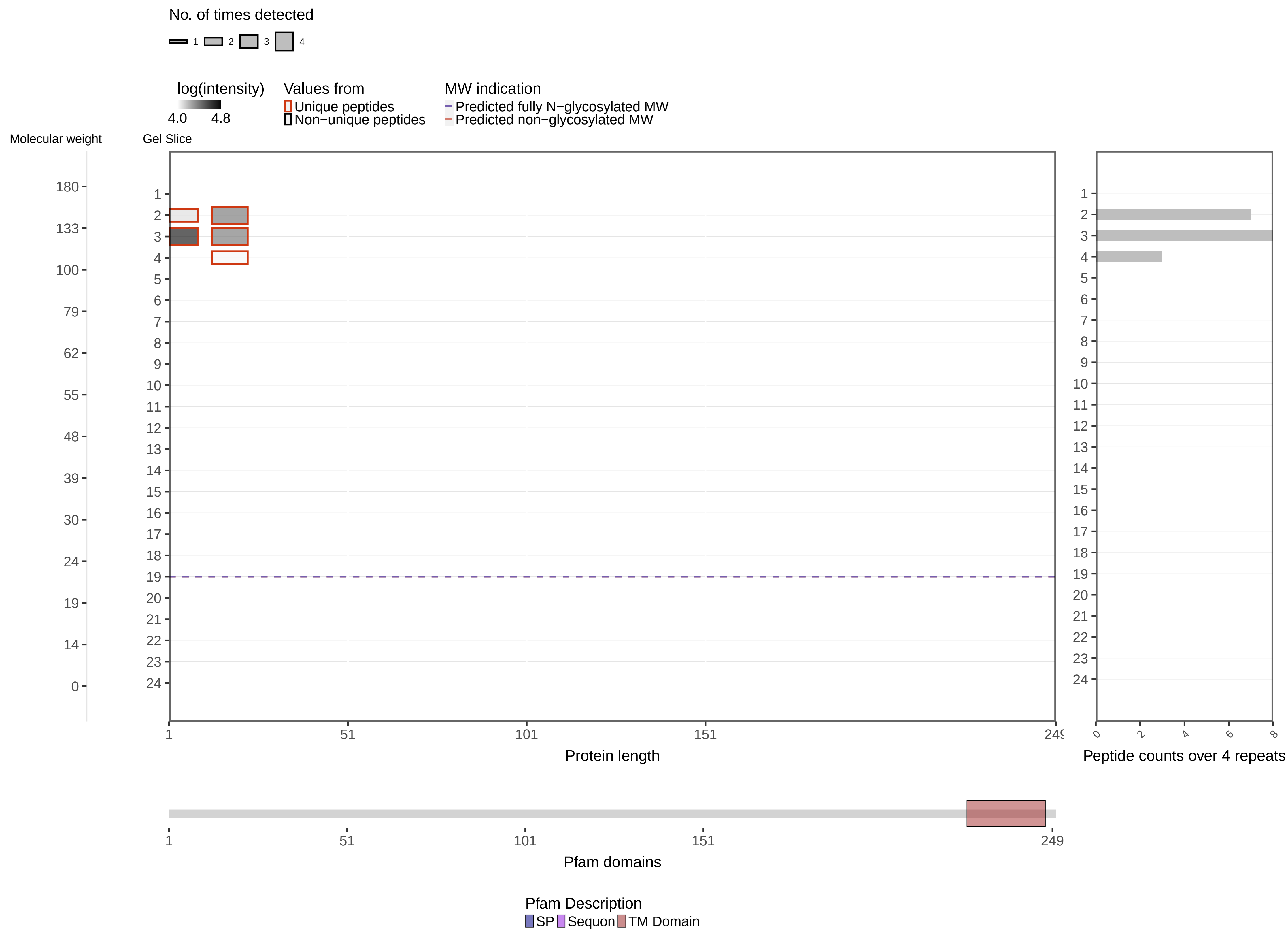


Protein sequence:

SPGNAPTPANSPATSPSTSHAPSINSPTSSPSPATSPAPARISPSGDITAPSPSSSGSSSSPTSSPPASGTPESPGKPTPGSIAPATGGGNSPPVDIQSPTGSSTNSKNSAIKAFTPSAVLMSLVLTIFLGGFVVFP*

NbL17g19210.1 uncharacterized_oxidoreductase_At4g09670−like [MW = 40.46 kDa]

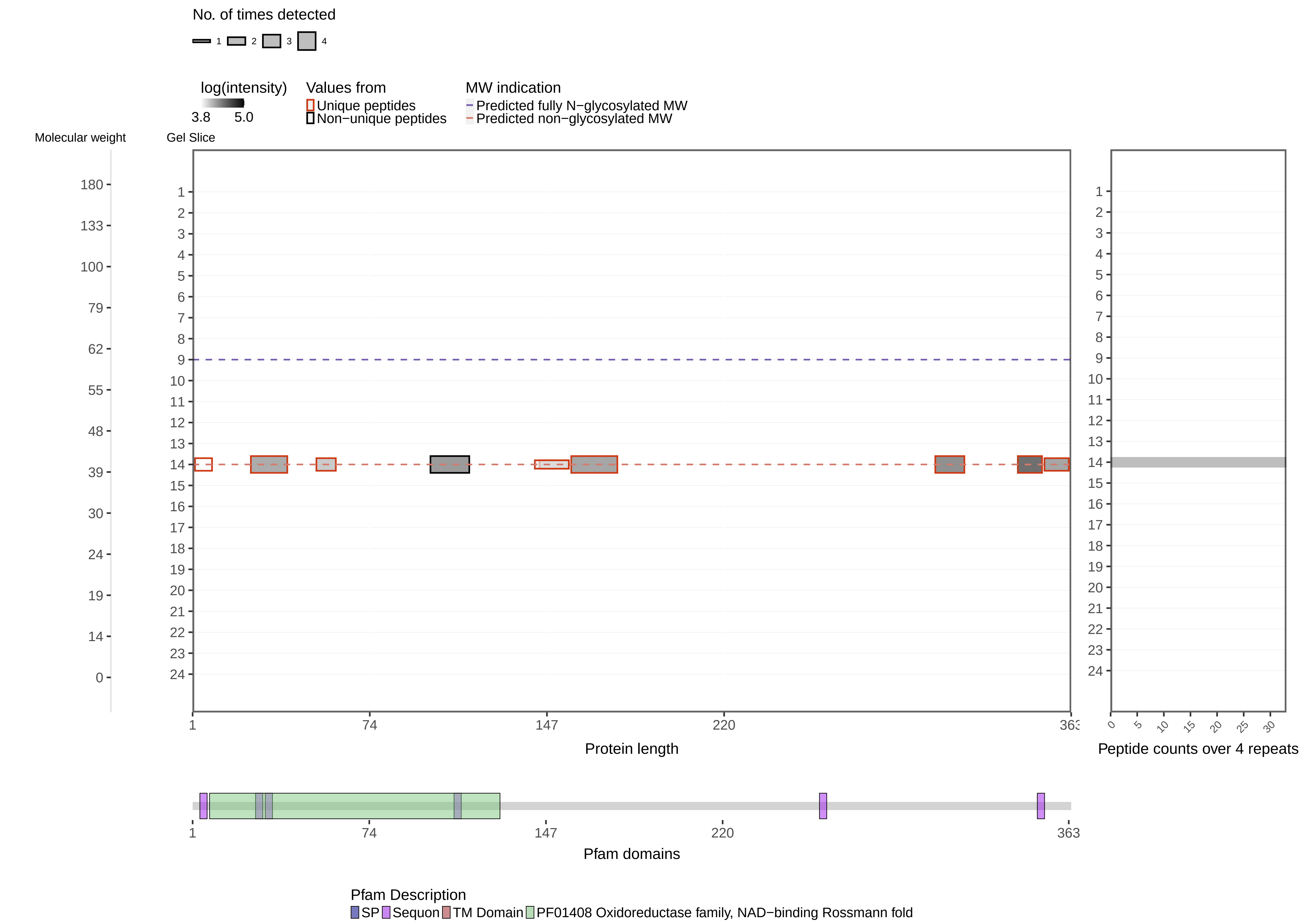


Protein sequence:

/FLENNIRVKPDLDALGALGDAGWYCIGAILWAMNQNLPTTVTALPTVARNSAGVILTCSASLHWEKEETVATFYCSFVSHETMDLIVYGSNGTFYLYDFIIPMDENSASFSFTSGAKFVDRHIGWNMKPRAVEVTSKFPQEASMIQEFSRLVKAIEVSRSKPESKWPHISRVTQLVLDAVNKSIDLGFQPVHI*

### NbL17g19090.1 phosphoribulokinase_chloroplastic−like [MW = 45.15 kDa]

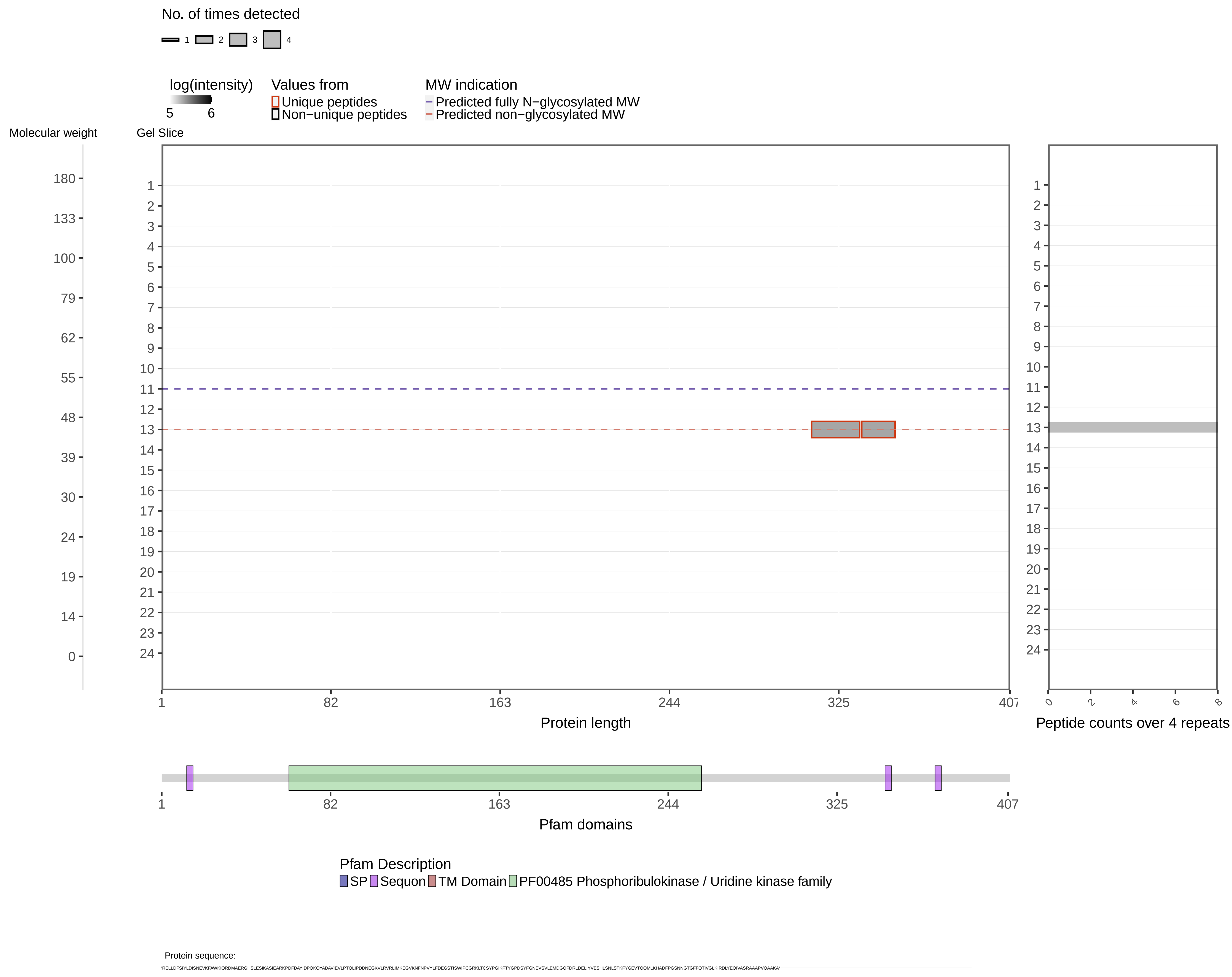

NbL17g18810.1 tocopherol_O−methyltransferase_chloroplastic [MW = 41.53 kDa]

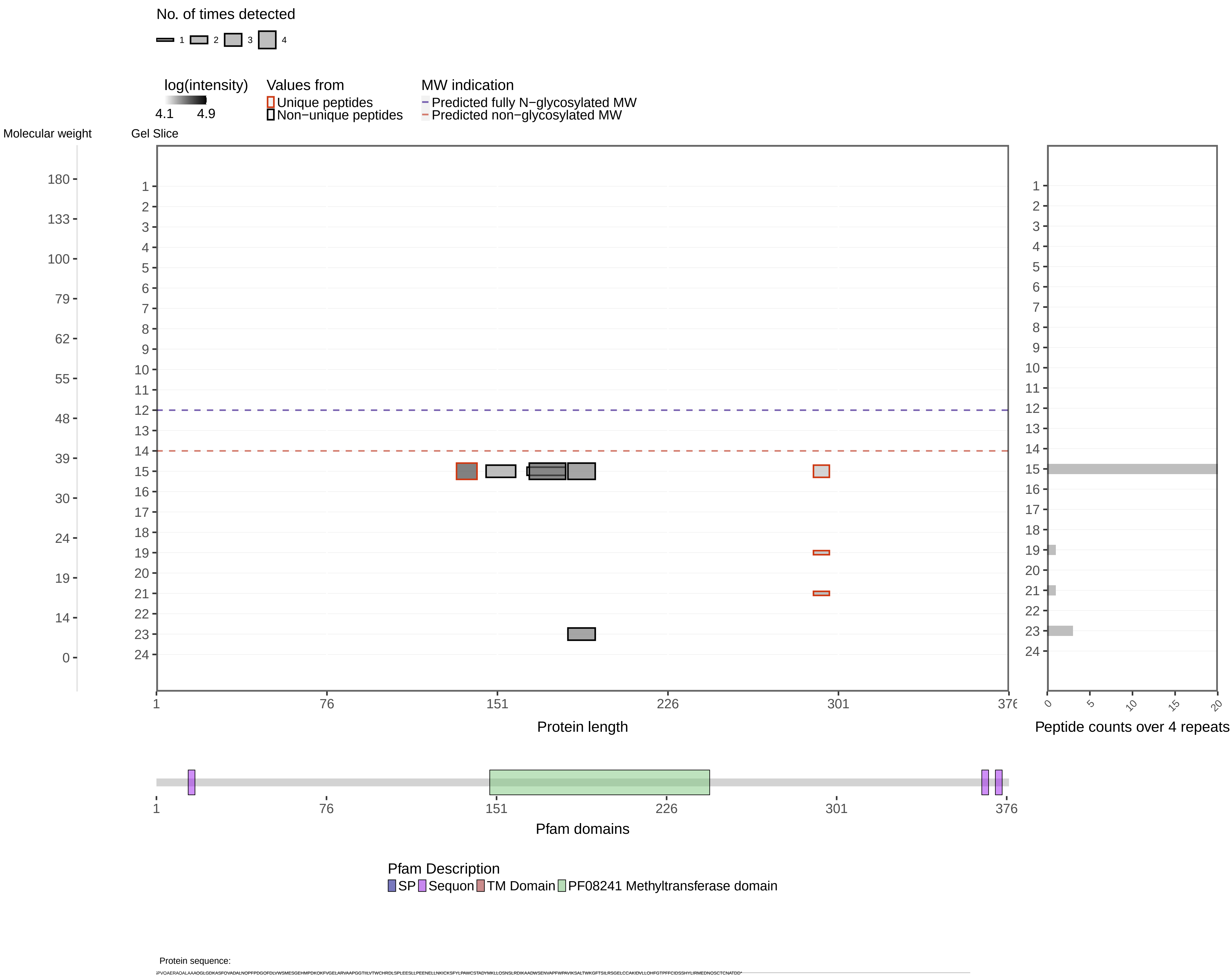

### NbL17g18710.1 probable_ADP−ribosylation_factor_GTPase−activating_protein_AGD8 [MW = 43.71 kDa]

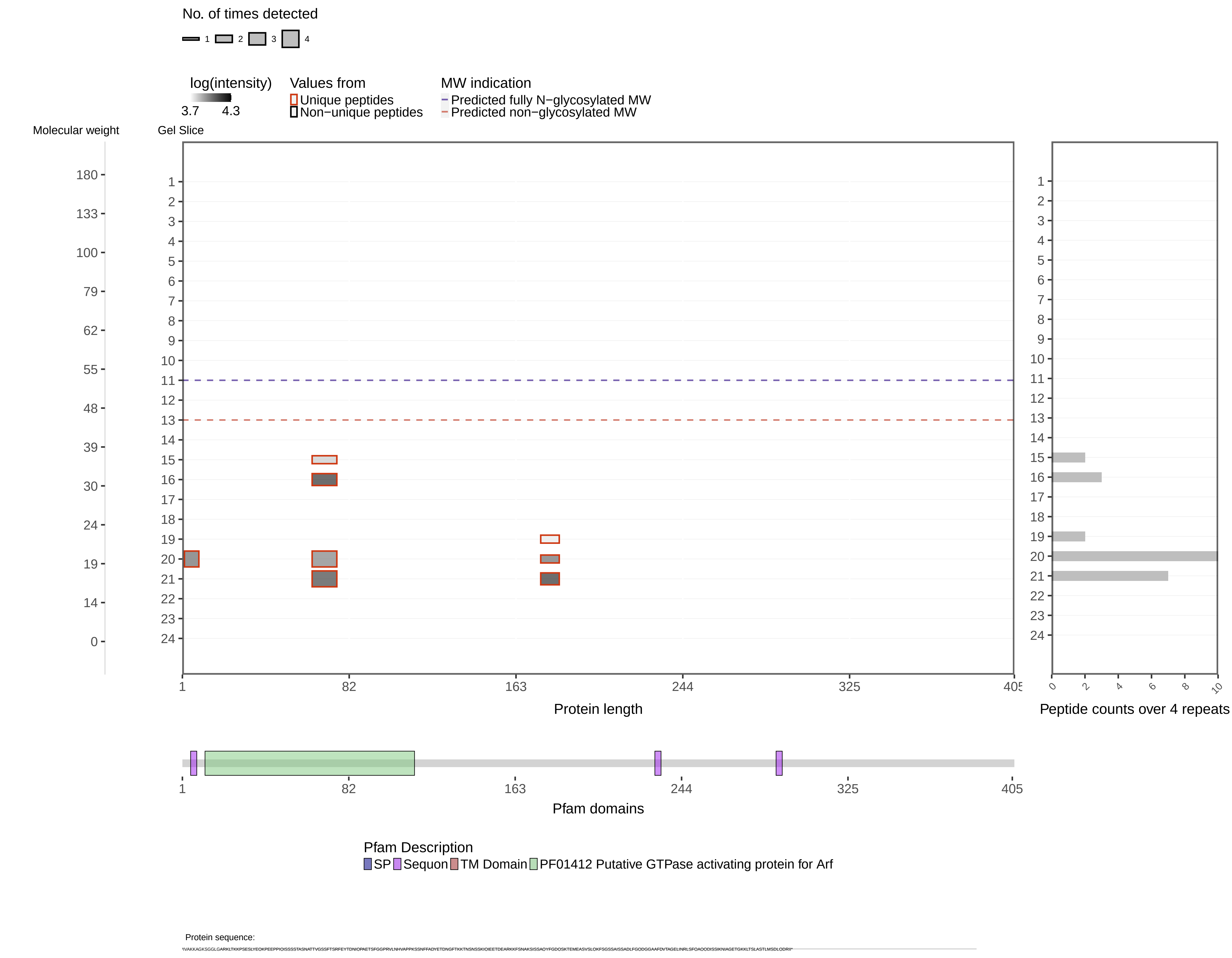

NbL17g18440.1 phosphoglucomutase_cytoplasmic [MW = 63.29 kDa]

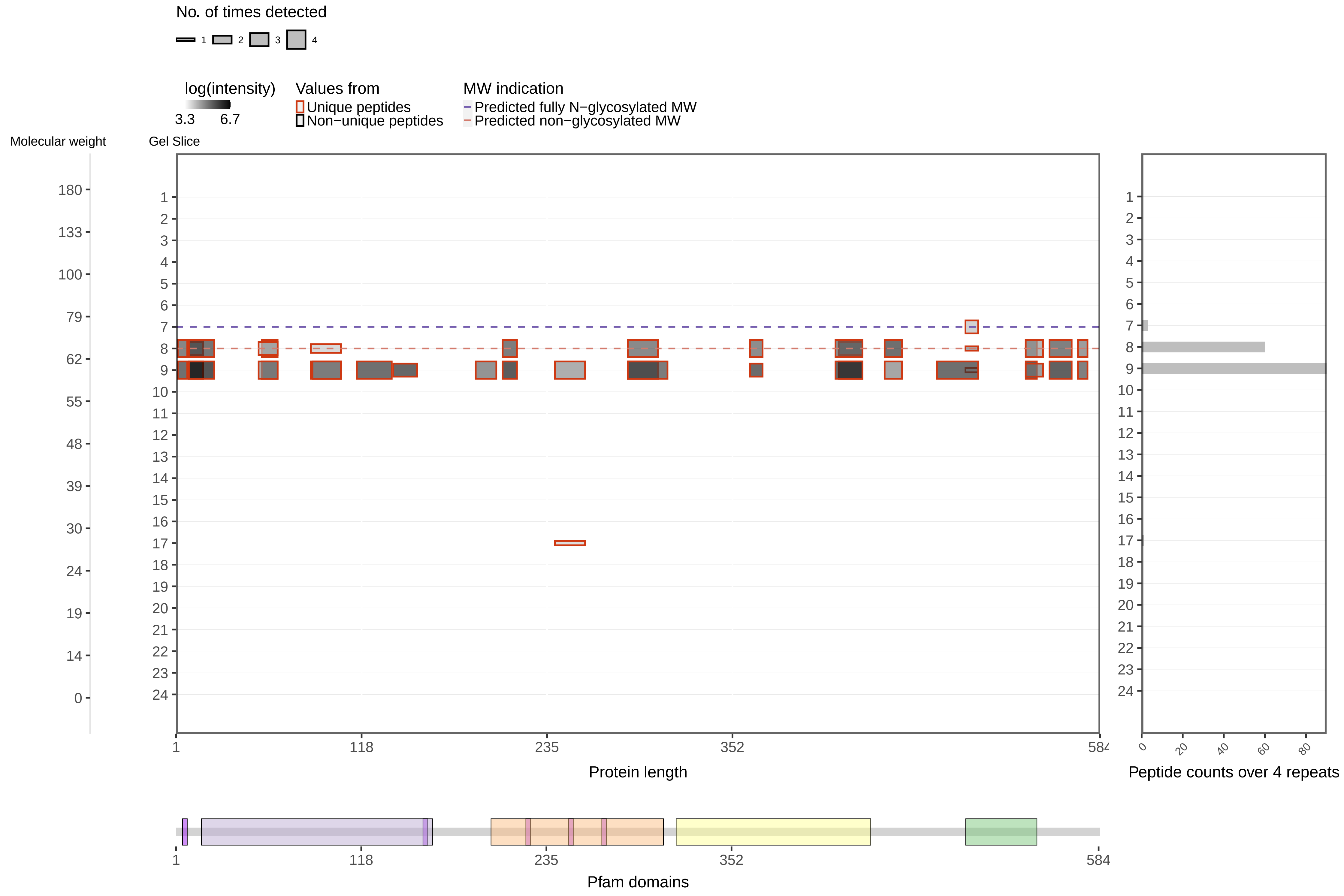


n
PF02878 Phosphoglucomutase/phosphomannomutase, alpha/beta/alpha domain I
PF02880 Phosphoglucomutase/phosphomannomutase, alpha/bet
hosphoglucomutase/phosphomannomutase, C−terminal domain
PF02879 Phosphoglucomutase/phosphomannomutase, alpha/beta/alpha domain II

Protein sequence:

NbL17g18180.1 villin−4−like [MW = 109.1 kDa]

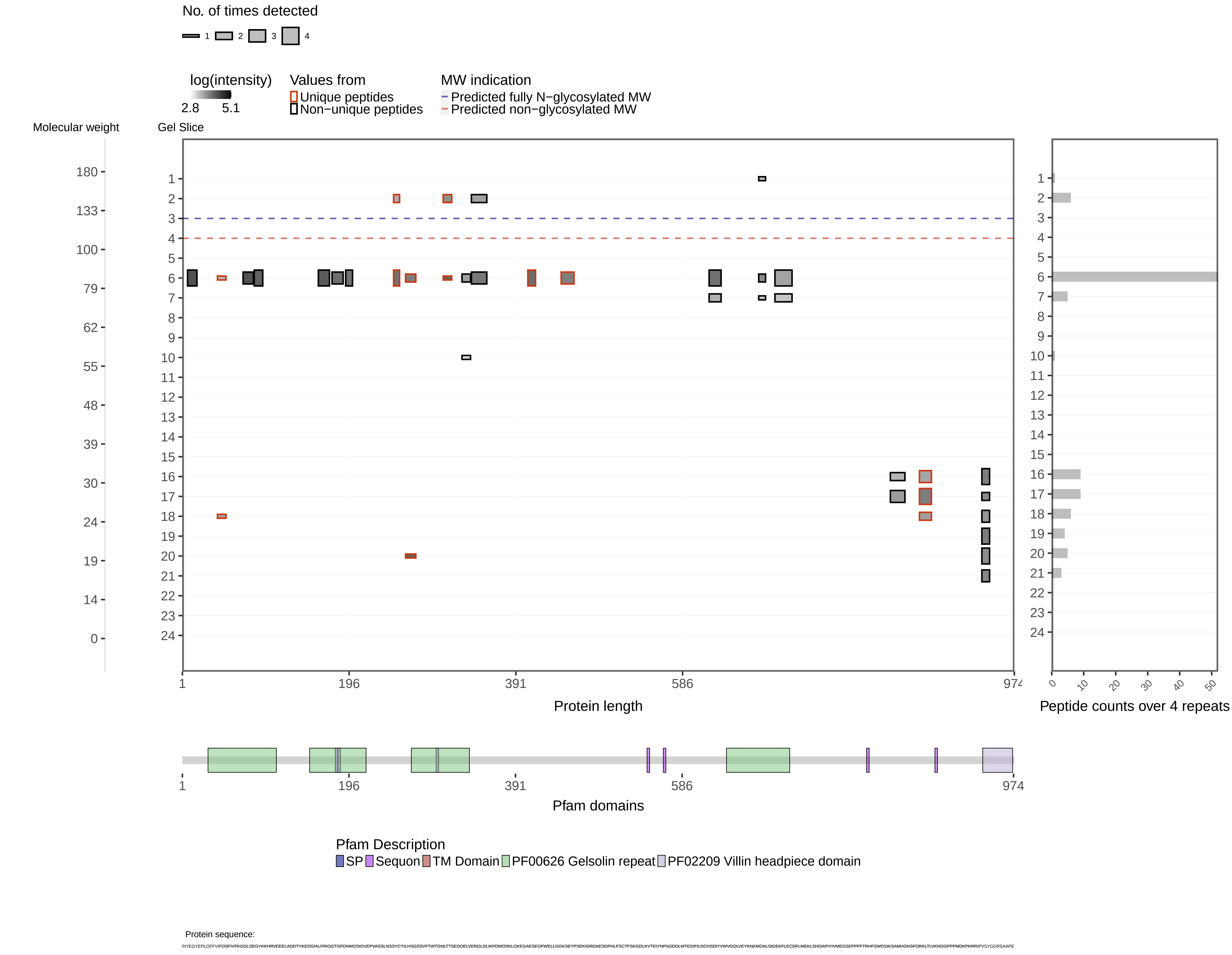

NbL17g18130.1 thioredoxin−like_protein_Clot [MW = 18.85 kDa]

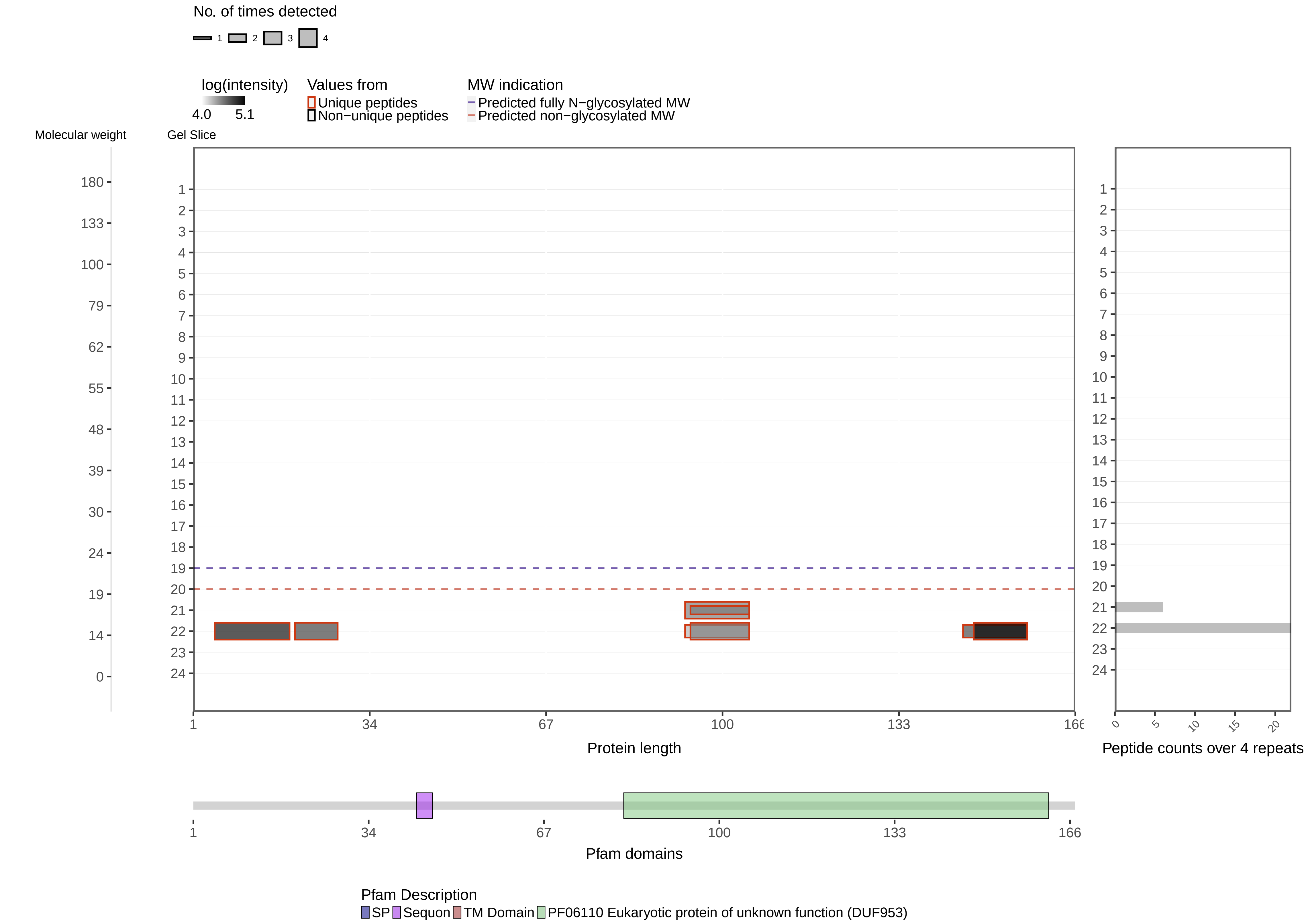


Protein sequence:

LHLLQASRKNTDCVRAEPVIYKKMEESAEDIALLRAYVGDRPTWRTPQHPWRVDPRFNLKGVPTLIRWENDATKGRLEDSEAHLAHKIDSLLIGN*

### NbL17g18120.1 30S_ribosomal_protein_2_chloroplastic−like [MW = 27.7 kDa]

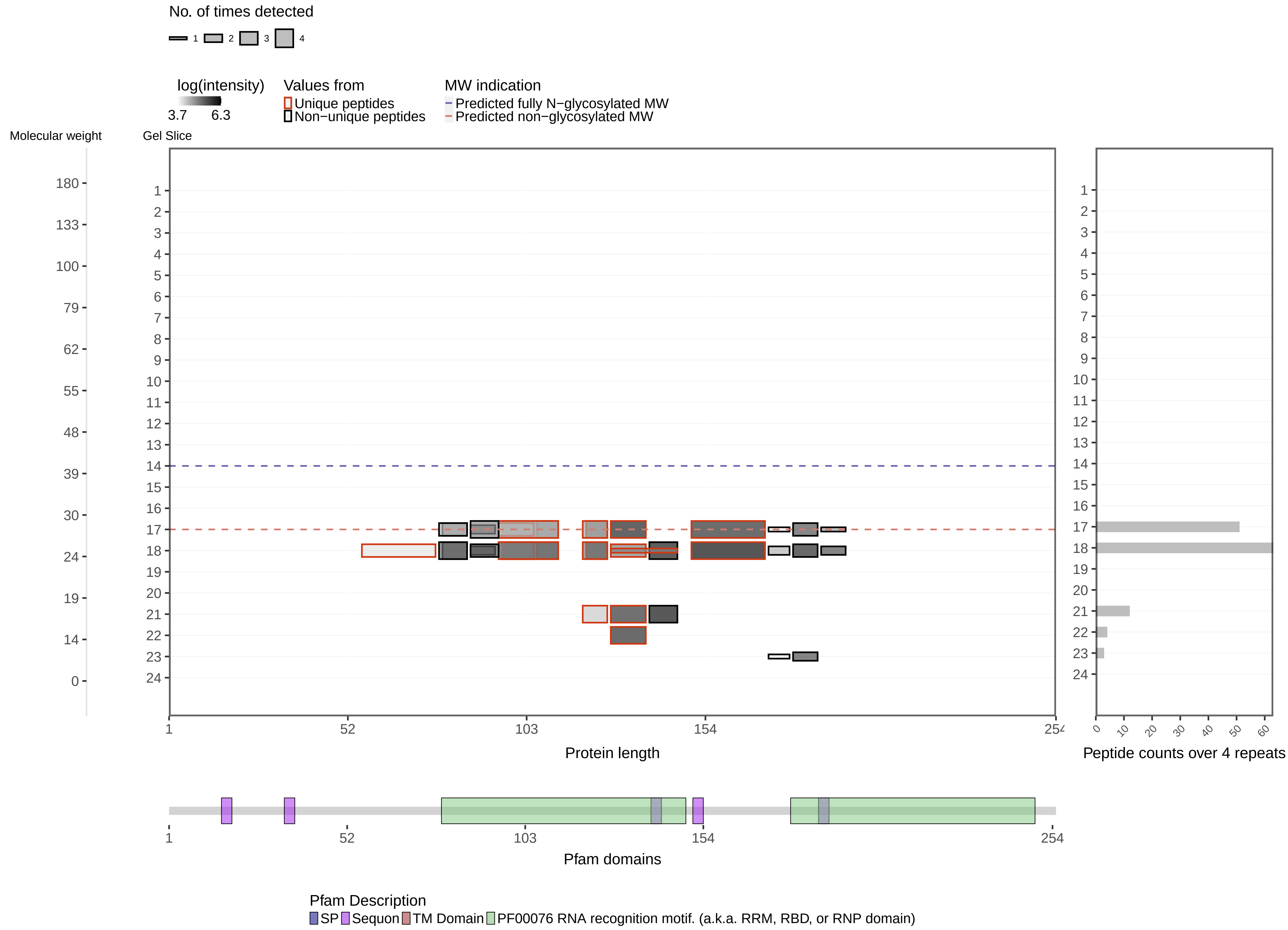


Protein sequence:

RSRRFGFVTMKTVEDASAAVEKLNGTEIGGREIKVNITEKPLSSVGSSLLQSDDSKFVDSPHKVYVGNLAKNVTSESLKNFFSEKGNVLSAKVSRVPGTSKSSGYGFVSFSSEEDVEAAISSLNNALLDGQKIRVNKA*

NbL17g18020.1 V−type_proton_ATPase_subunit_D−like [MW = 29.14 kDa]

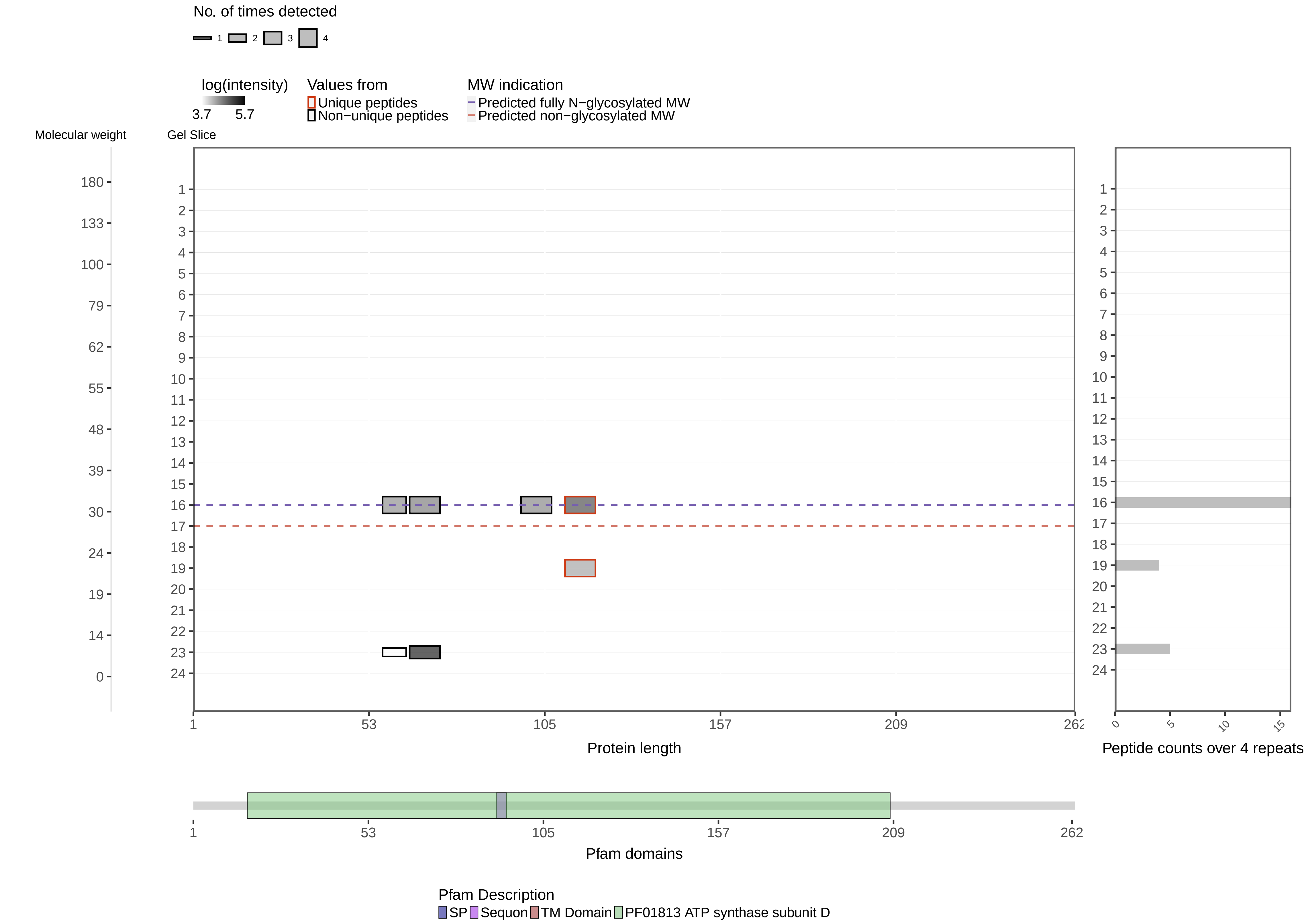


Protein sequence:

ETKNDLTGLARGGQQVQACRAAYVKSIELLVELASLQTSFLTLDEAIKTTNRRVNALENVVKPRLENTVLYIKGELDELEREDFFRLKKIQGYKKREVERQMLASKQYAEEKAAEEISLKRGISLGTAHNLLSHASQKDEDIIF*

NbL17g17800.1 ras−related_protein_RABH1b [MW = 23.11 kDa]

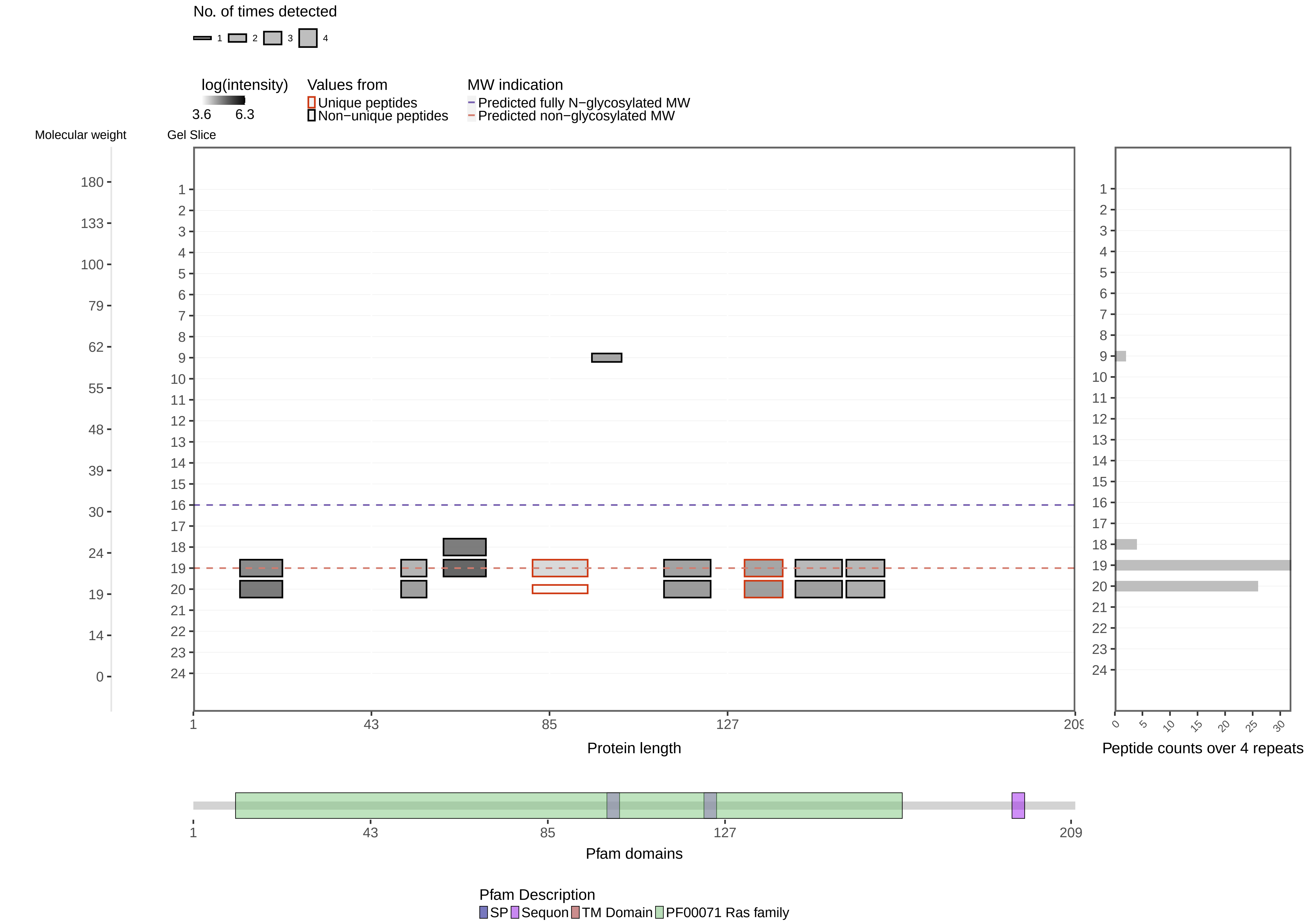


Protein sequence:

SRQSFLNTSKWIEEVRTERGSDVIIVLVGNKTDLVEKRQVSIEEAEAKARELNVMFIETSAKAGFNIKPLFRKIAAALPGMETLSSAKQEDMVDVNLKSSNANASQSQAQSGGCAC*

### NbL17g17730.1 heterogeneous_nuclear_ribonucleoprotein_R−like [MW = 88.06 kDa]

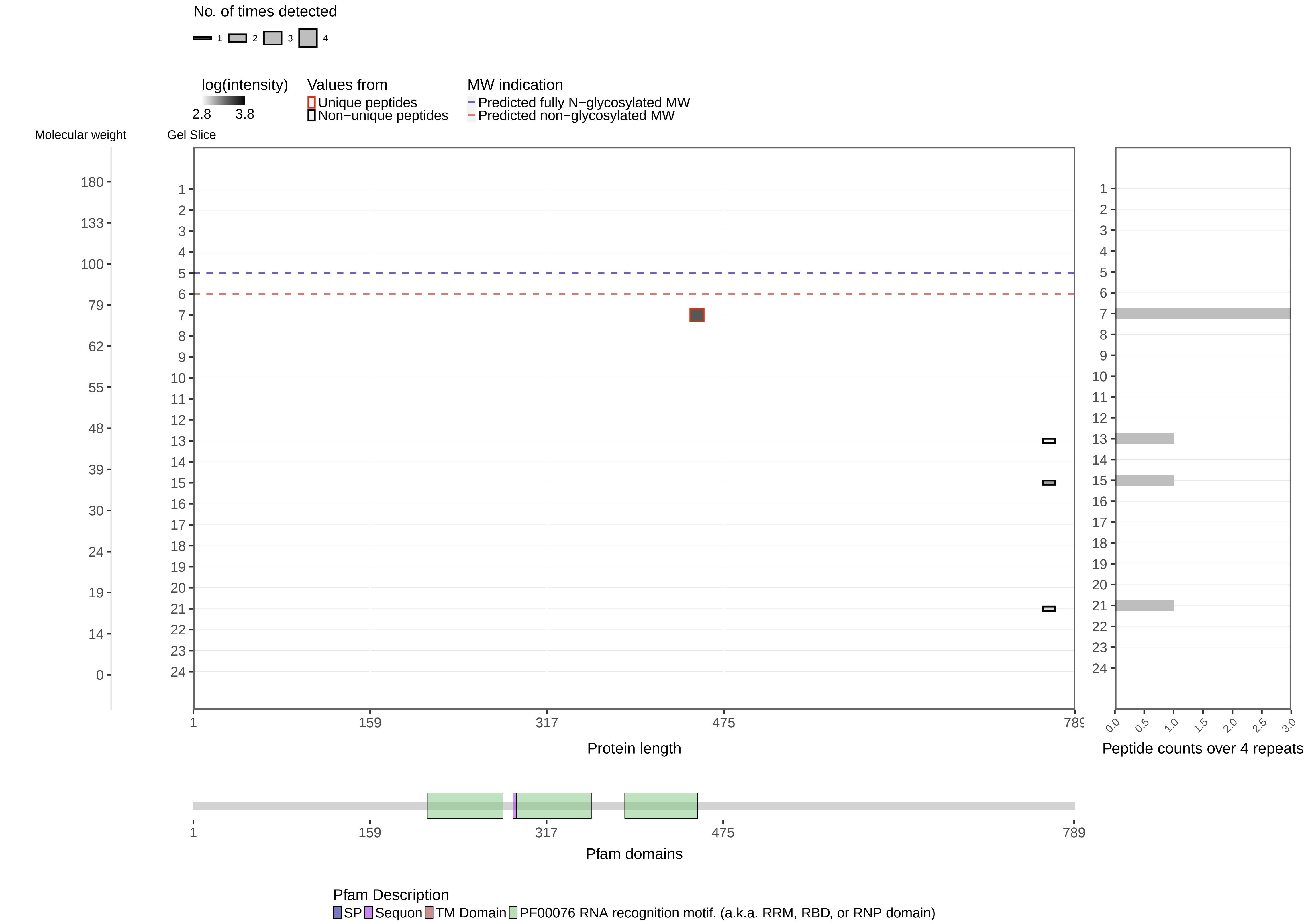


Protein sequence:

VFVDGLSASWDEDRVQELLKEYGKIEKVELARNMPSAKRKDFGFVTFDTHDAAVTCAKSINNEELGEGDNRVKVRARLSRPLQRGRGKYGGRGDLRPRHMPLRGLRAPWGRTVQHSRPIRGTRVSTRLPPVVGRGFKRPAMLRDRRAVMALPPRGRPMAPPPSSRSYERRPPVPSYAKSSLKREYGRREEIPPRSRALAEYPSRVHSGRRASYSDEYSSRDSGYPEMPRGTHSAGRRAYVDDGYGQRFERPPPSYRDGRDREYDSMSGSKRPYSAVDDVHPR

NbL17g17700.1 nucleoside_diphosphate_kinase_1−like [MW = 15.41 kDa]

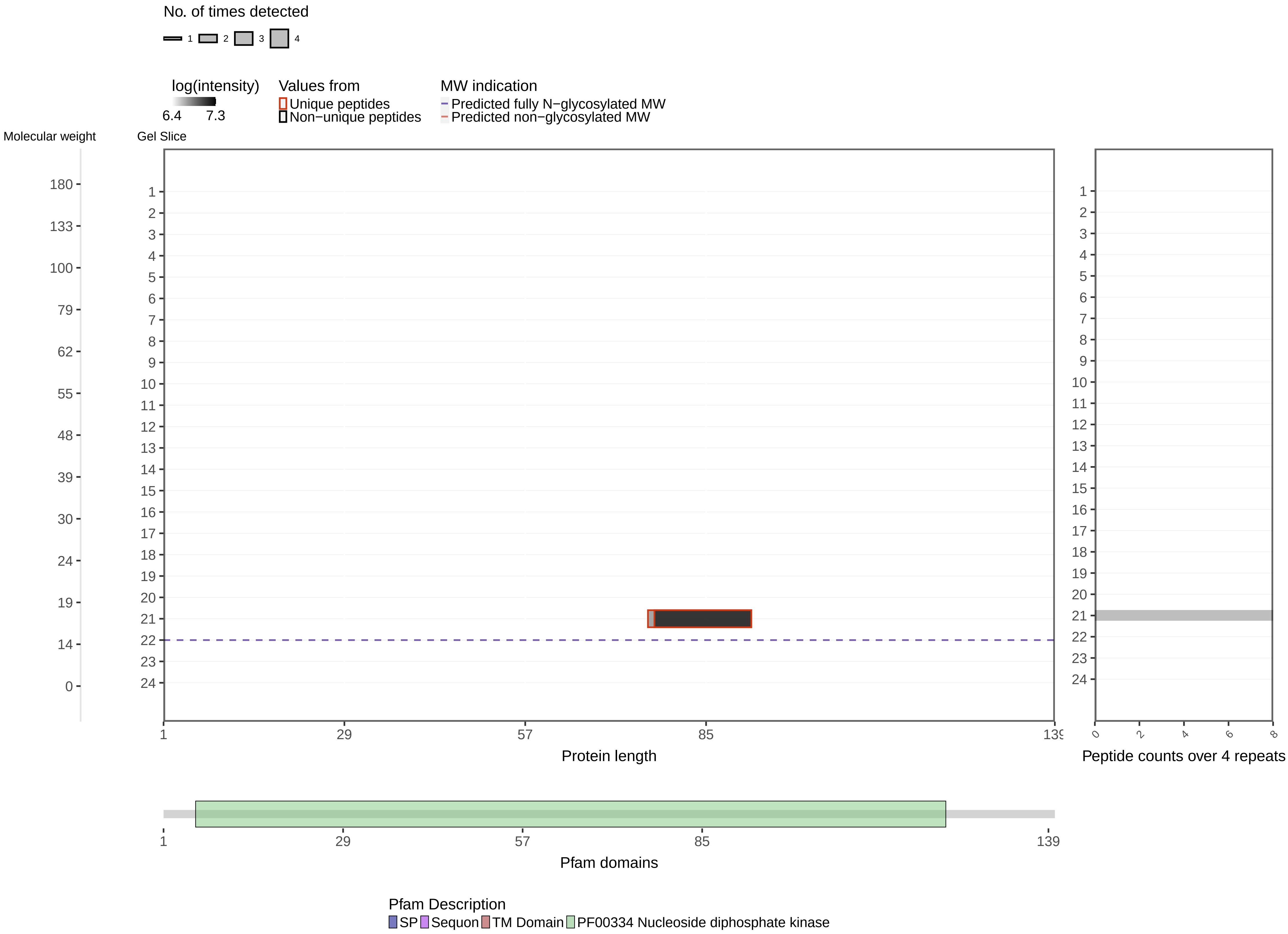


Protein sequence:

SGPVVAMIWEGKGVISTGRKLIGATNPLESAPGTIRGDYTIDIGRNVIHGSDSVDSAQKEKDLWFPEGINWWRPSQHSWIYE*

NbL17g17680.1 uncharacterized_LOC107820933 [MW = 56.8 kDa]

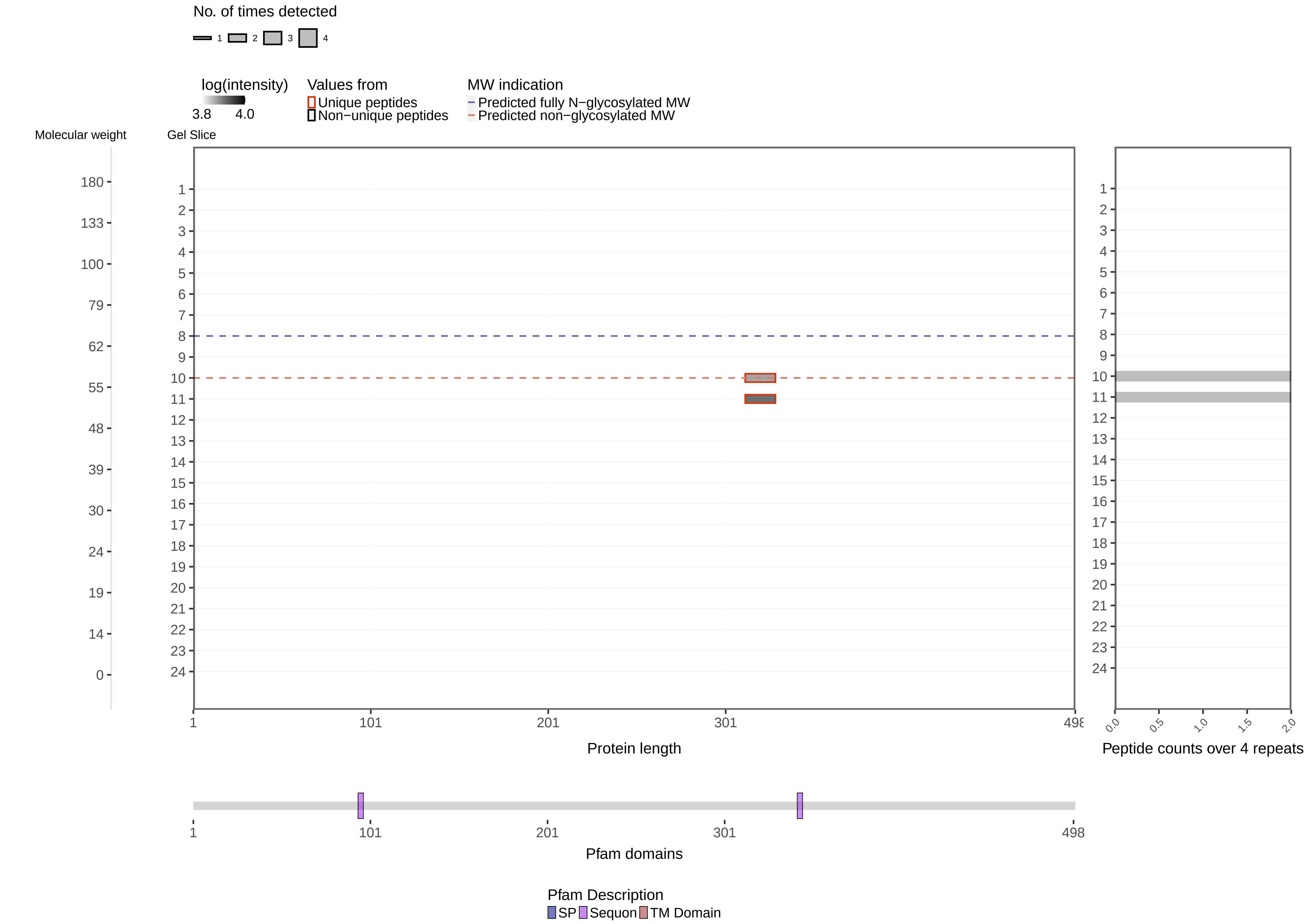


Protein sequence:

SRHQKLPKFESFDFEKSDIMEEDIMGMEPGLVFFEDGSYSRGPVDIPVGELDESKYYISPTFKFEQCLVKGCHKRLRIVHTIDFSNGGSDIQILRVAVYEEQWISPANLSDESDTELDVKLFSQRRRLQPSELAGSWKVFEMSATPIYSEEDVTQETDDVPYVYLCTENLKKRSLPENPAYFGEEEMLDMQDVTVLWLPGGVTSYVDVNKDGILCIGVGWYSDEGVNLVMERDYEMDGKLKEVRWKSEMKRRWTNPPPMSLHL*

### NbL17g17560.1 peptidyl−prolyl_cis−trans_isomerase_FKBP20−2_chloroplastic [MW = 29.24 kDa]

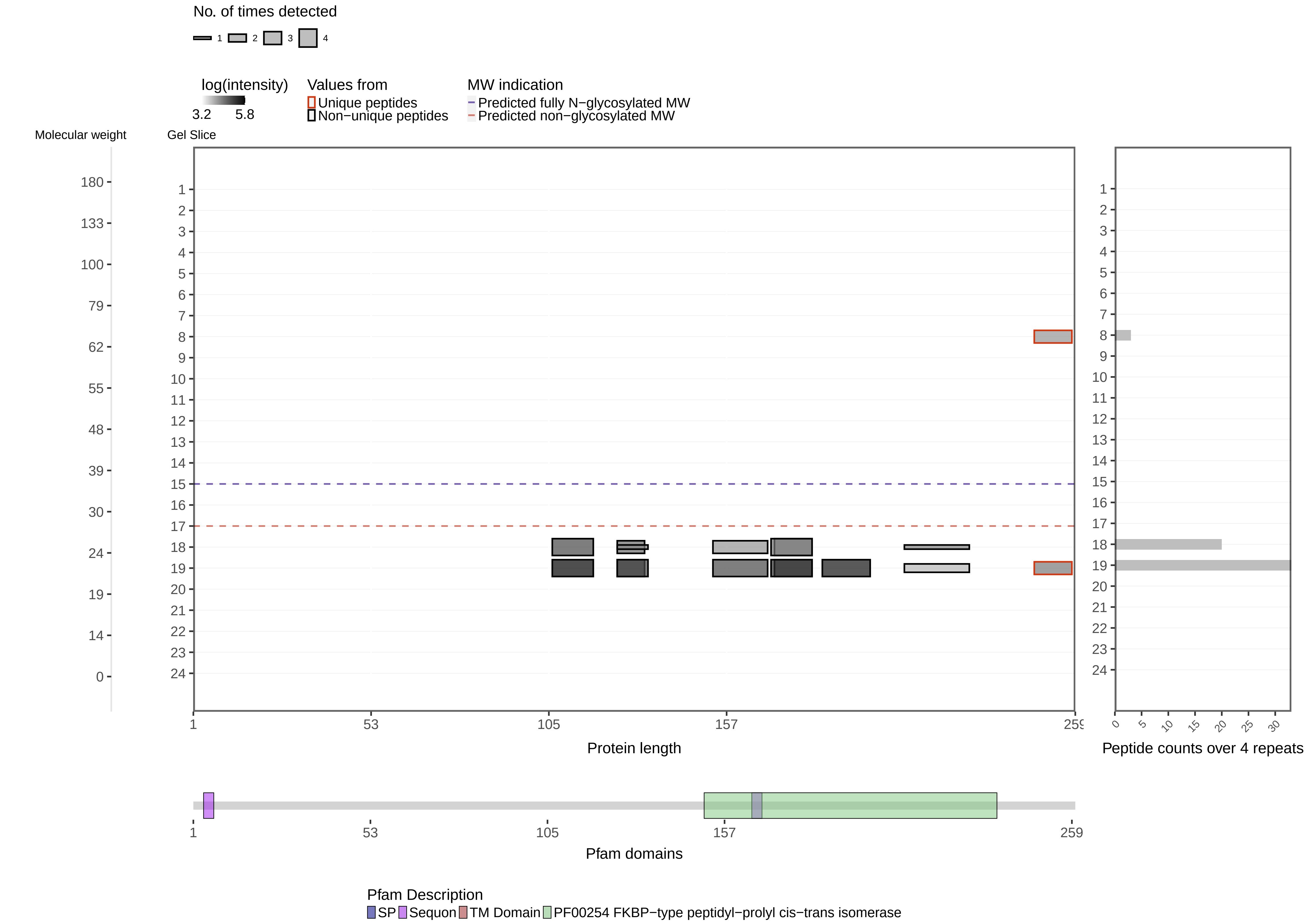


Protein sequence:

IREGFEVKVVTSDNYVKRDSGLILWDIAVGEGDCPKDGOQVTFHYIGYNESGRRIDSTYLQNAPAKIRMGTNALVPGFEEGIRDMRPGGRRRIIIPPELGPPVGPSTFFSSKQFEVFDVELLSIQDCKRRTIAGFYSDVVCN*

### NbL17g17380.1 ATP−dependent_Clp_protease_proteolytic_subunit_4_chloroplastic−like [MW = 34.06 kDa]

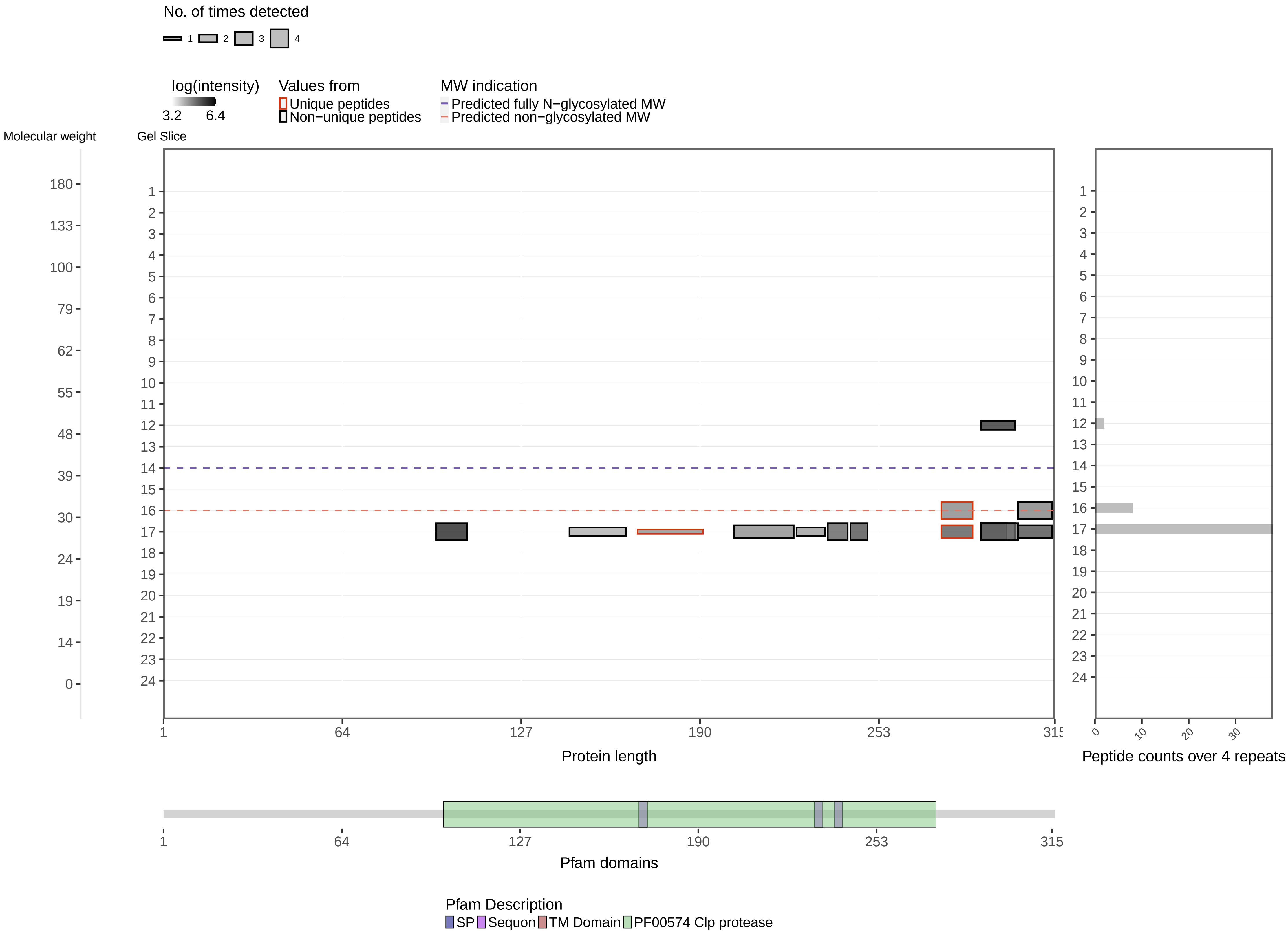


Protein sequence:

RLFINSPGGSLSATMAIFDVVKLVRANVSTVALGIAASTASIILGGGTKGKRFAMPNARIMIHOPLGGASGOAIDVEIOAREIMHNKDNVTKIISNCTGRSYEOVOKDIDRDRYMSPIEAVEYGLIDGVIDRDSIIPLEPVPERVKSTLNYOEISKDPKKFLNPEIPDDEIY*

### NbL17g17340.1 60S_ribosomal_protein_L13−2 [MW = 23.58 kDa]

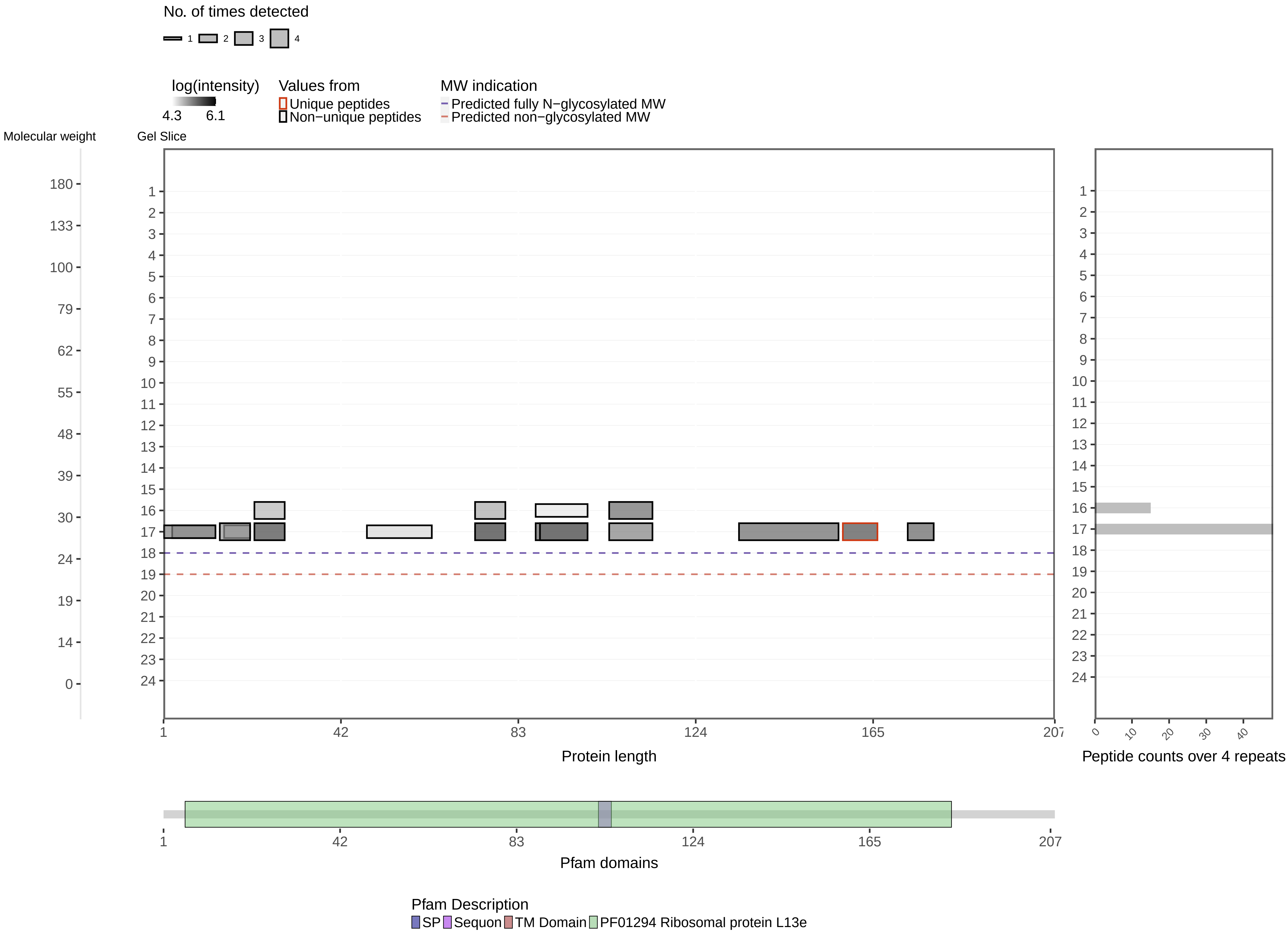


Protein sequence:

APTIGIAVDHRRRNRSLEGLQTNVQRLKTYKAKLVIFPRRAKKIKAGDSSAEELATATQVQGSYMPITREQPAVDLVRVTDEMKSFNAYGKLRIERTNARHIGARLKRAAEAEKEEKK*

### NbL17g17250.1 neutral_alpha−glucosidase_C [MW = 124.3 kDa]

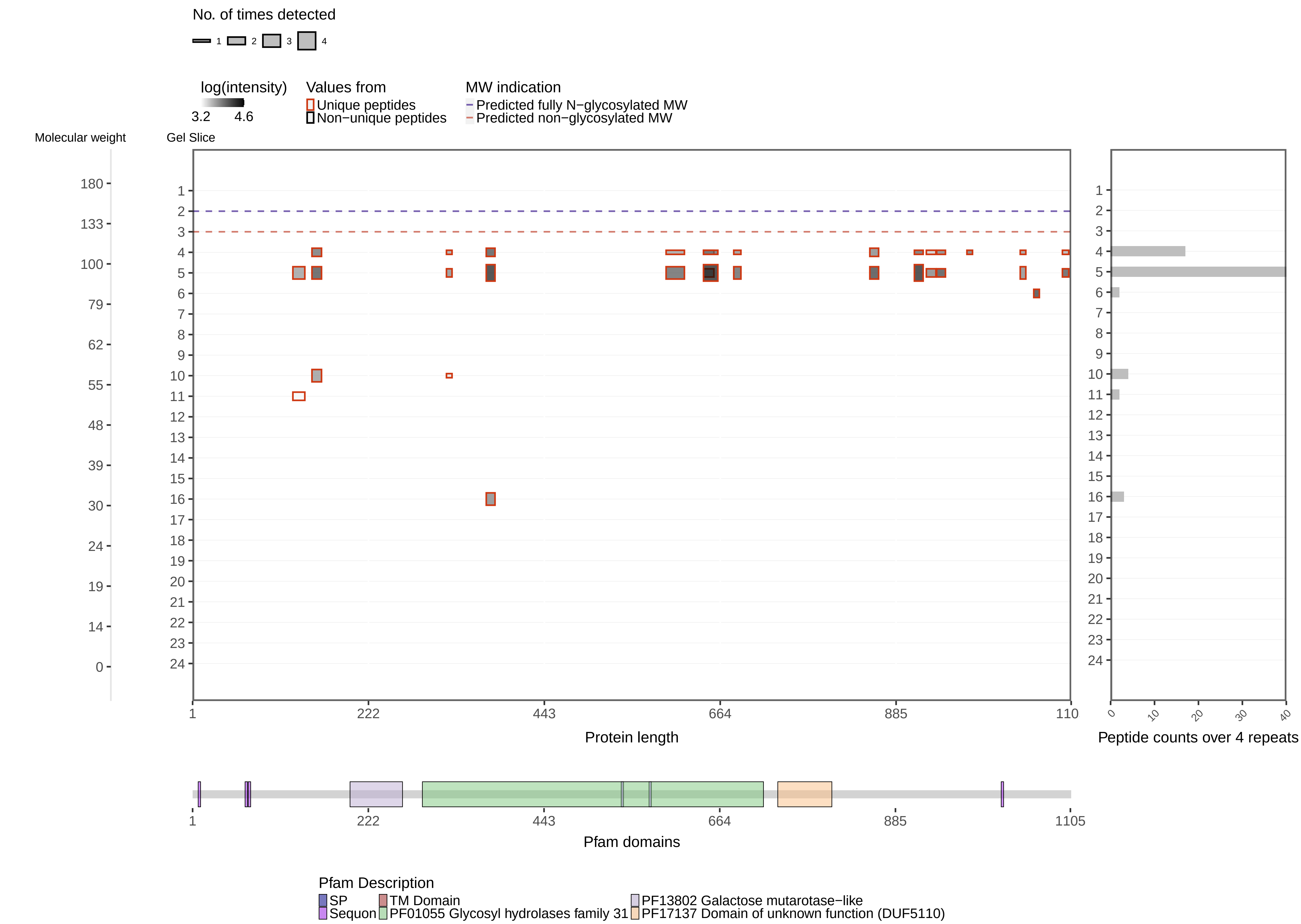


Protein sequence:

### NbL17g17190.1 probable_carotenoid_cleavage_dioxygenase_4_chloroplastic [MW = 65.96 kDa]

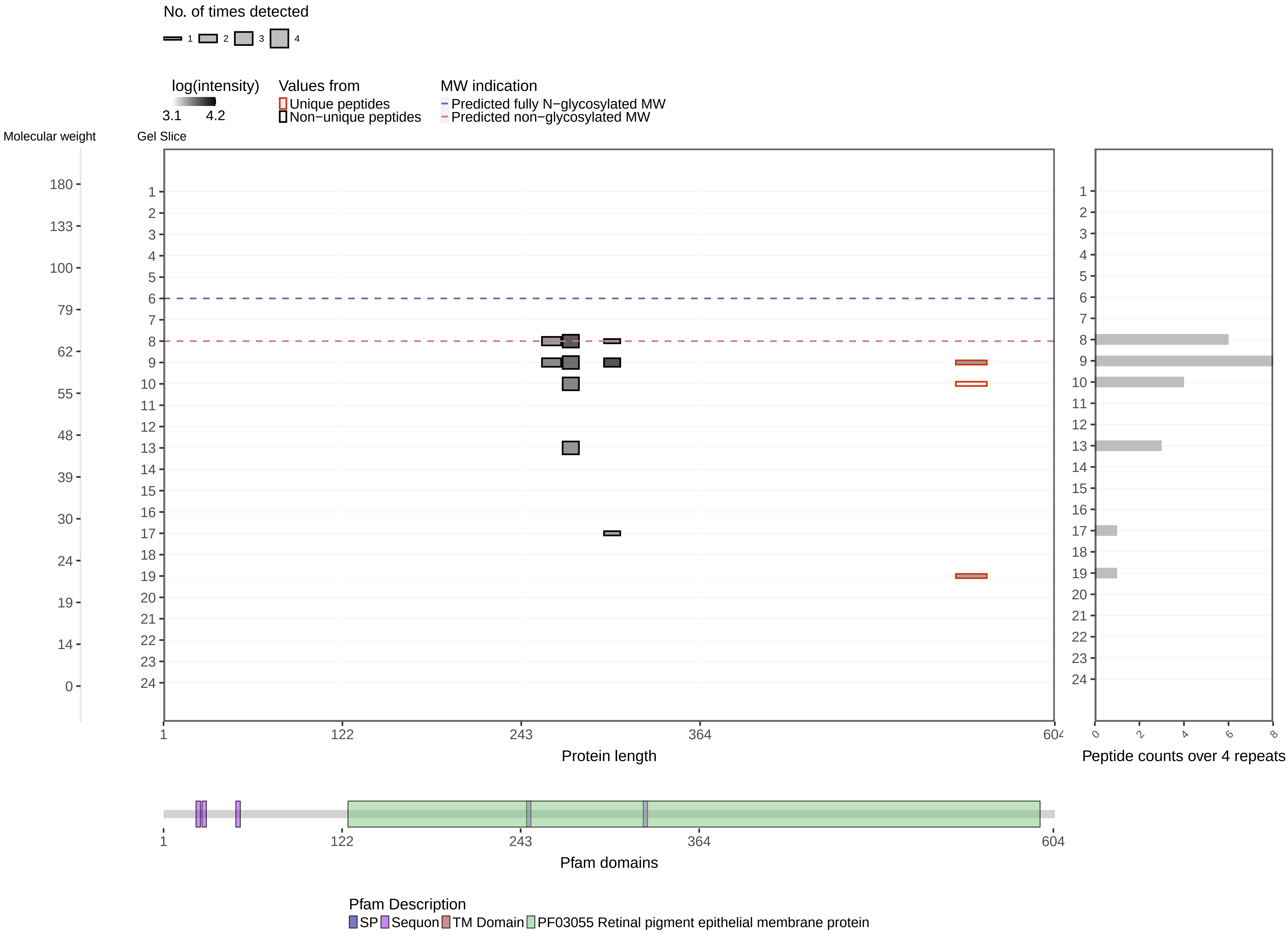


Protein sequence:

MSMTAHPKIDPDTNEAFAFRYGPMPPFLSYFRIEPNGTKTPDVPIFSMTRPSFLHDFAITKKYAIFSDIQIGMNPLEFITGGSPVSCDSGKIPRLGVIPRDAKDESKMKWFDVPGFNIVHAINAWDEDDGDTIVMVAPNILSVEHTLERVDMIHASVEKVKIDLKSGMVSRHPVSTRNLDFGVTNPAYVGKKNKYVYAAIGDPMPRIAGIAKLDVSVAETDRRDCIVGCRLFGEGCFGGEPYFVAKDPNNPTADEDDGYVVSYVHNEKTGESRFLVMDAKSPNLDIVAAVKLPRR

NbL17g17060.1 ABC_transporter_E_family_member_2 [MW = 68.21 kDa]

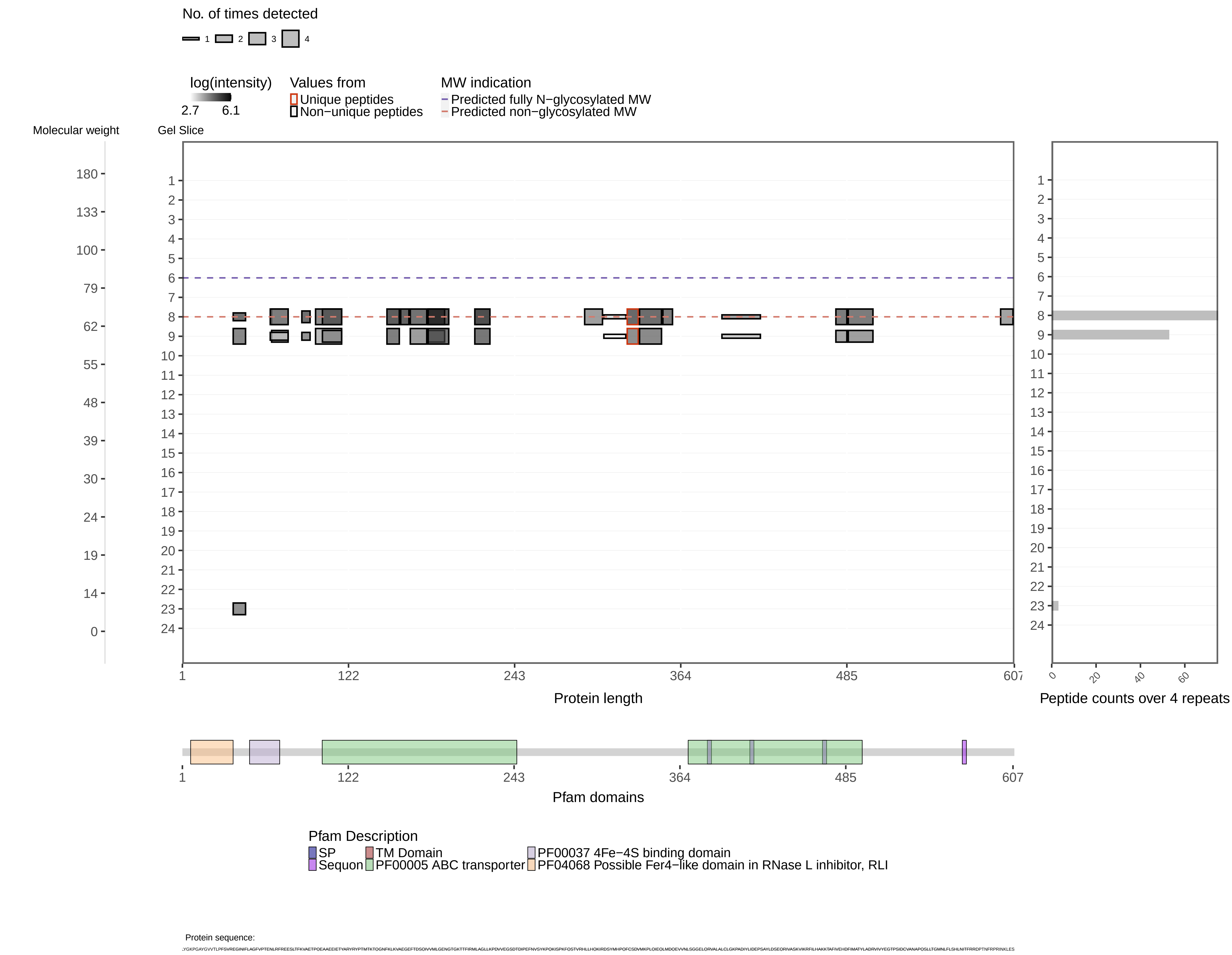

### NbL17g17000.1 AP−4_complex_subunit_epsilon [MW = 105.88 kDa]

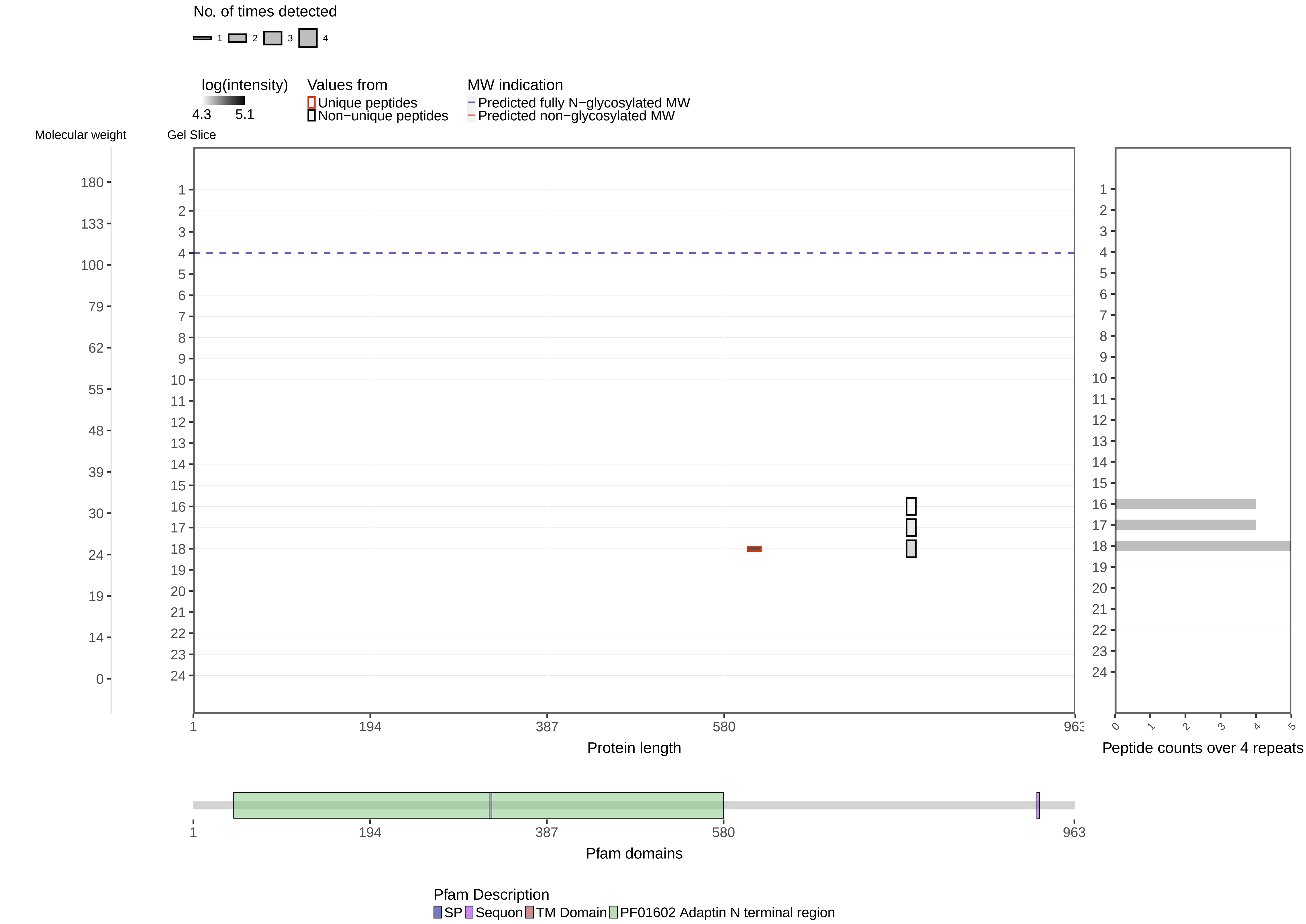


Protein sequence:

[illegible]

### NbL17g16960.1 long−chain−alcohol_oxidase_FAO4A−like [MW = 84.84 kDa]

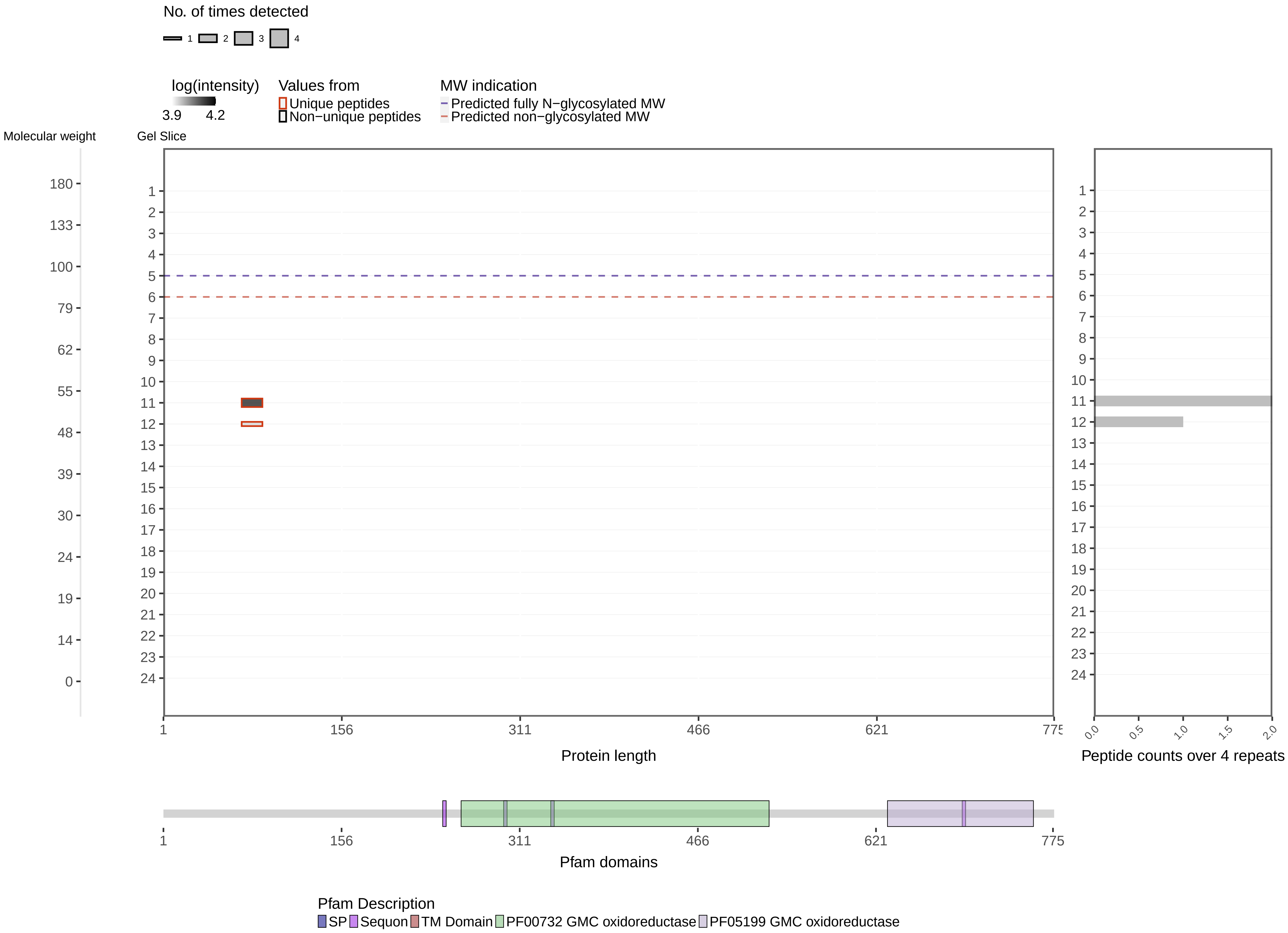


Protein sequence:

MGVQSEIEDEGFQNMILRKGCQELGYQVETIPRNAQSDHYCGWCSMGCKDGKKKGTAETWLVDLVKSGNGAILPECEALEVIHVKKNDKSGKSKAIGVAFAFQNGKGVREICMVKSKVTIVACGALSTPSLLKKSGLKNPNIGRNLHLHPVVIAWGYFPDTPSDSNVLWPEADKKSYEGGIMTAMSKVVANFEGSGYGAIIQTPGLHPGMFSALMPWVSGLDIKMRMCKYSRTAYLLALARDKGSGEAFSPYSVTYKMDGIDEENLKLGLEKMLRILAAAGAEEIGTQHGK

NbL17g16800.1 60S_ribosomal_protein_L7−2−like [MW = 28.74 kDa]

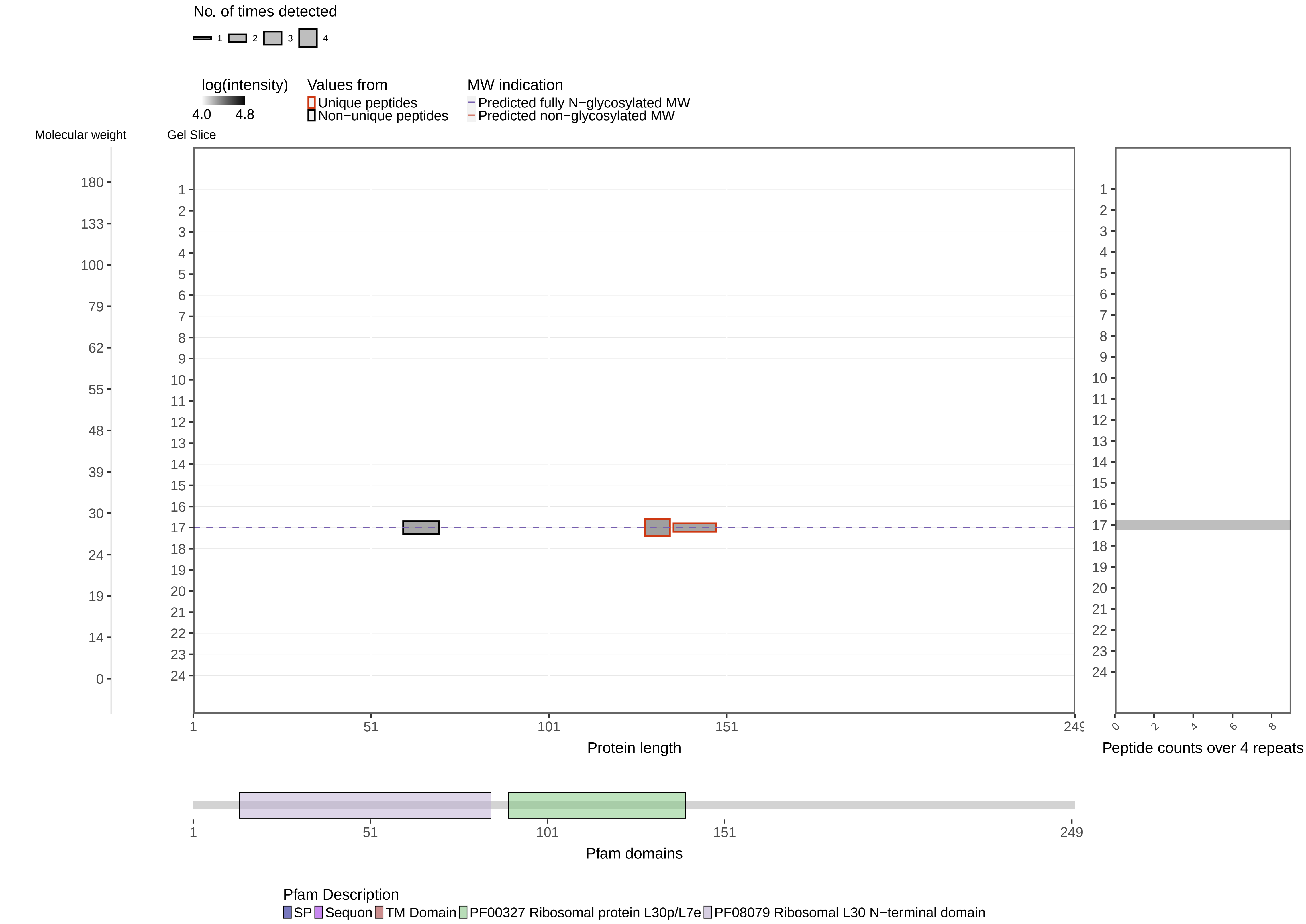


Protein sequence:

LLRLRQIFNGVFLKVNKATLNMLLRIEPYVTYGYPNLKSVRELIYKRGFGKVDKQRIALTDNSVIEQVLGKHGIICMEDLVHEILTVGPHFKEANNFLWPFQLKAPLGGLKKKRNHYVEGGDAGNRENYINELIRRMN*

NbL17g16680.1 carbon_catabolite_repressor_protein_4_homolog_4−like [MW = 43.19 kDa]

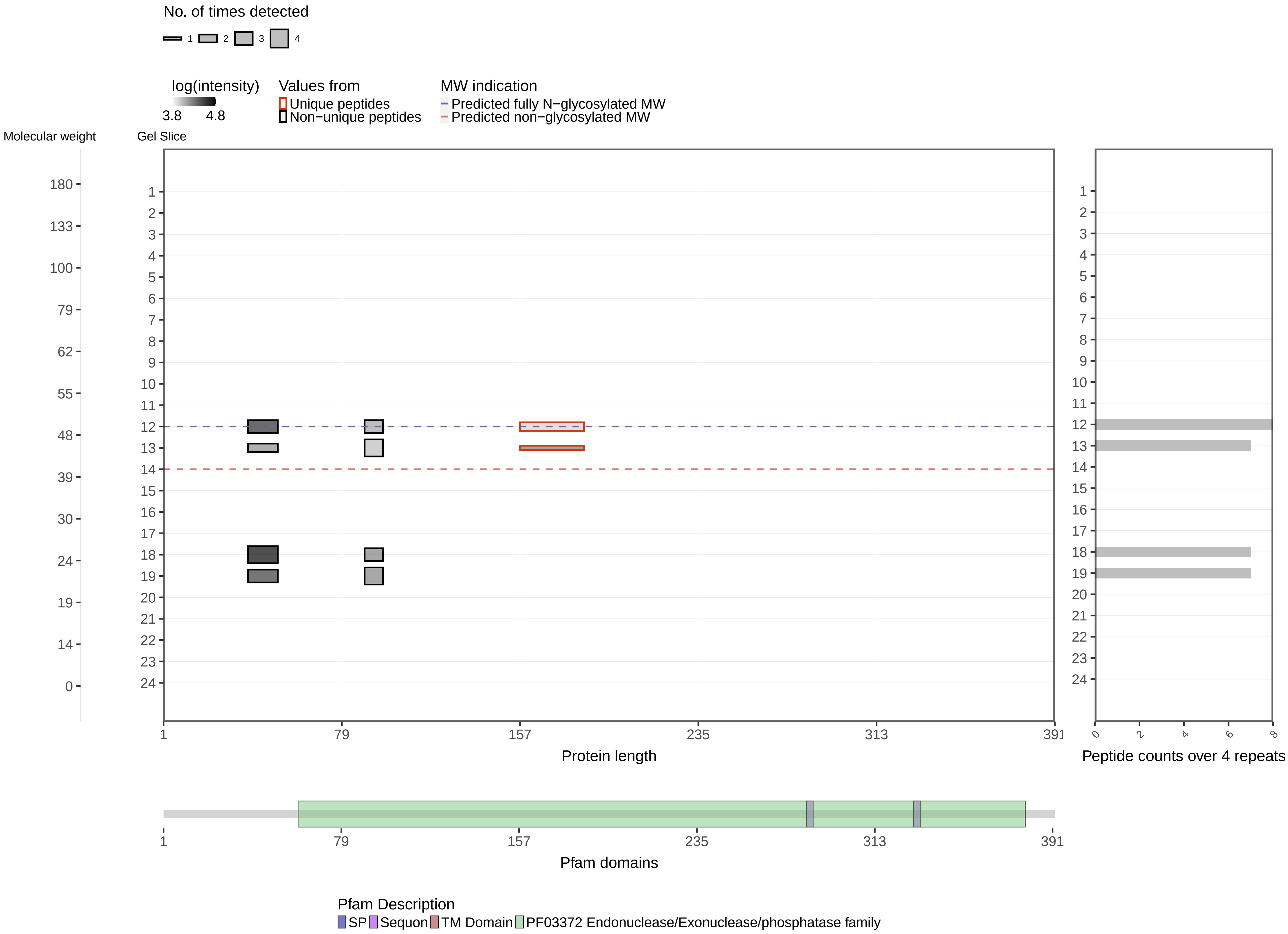


Protein sequence:

NKKLESKGVSSKNSQADHGDLNDPRIRLKRDCVGIMAAFRLKDPCQLIIVANTHIYWDPELADVKLAQARYLLSRLAQFKLLVSDKFDCSPSVVVAGDFNSTPGDQVYQYLISGSSLVETLPESSNDMPIPLCSVYASTKGEPHFTNCTPGFTGTLDYILFSPSGDIKPVSYLELPEPEASDVQGGLPNYYHPSDHLPIGAEFEIVH*

### NbL17g16660.1 pectin_acetylesterase_8−like [MW − SP = 42.04 kDa]

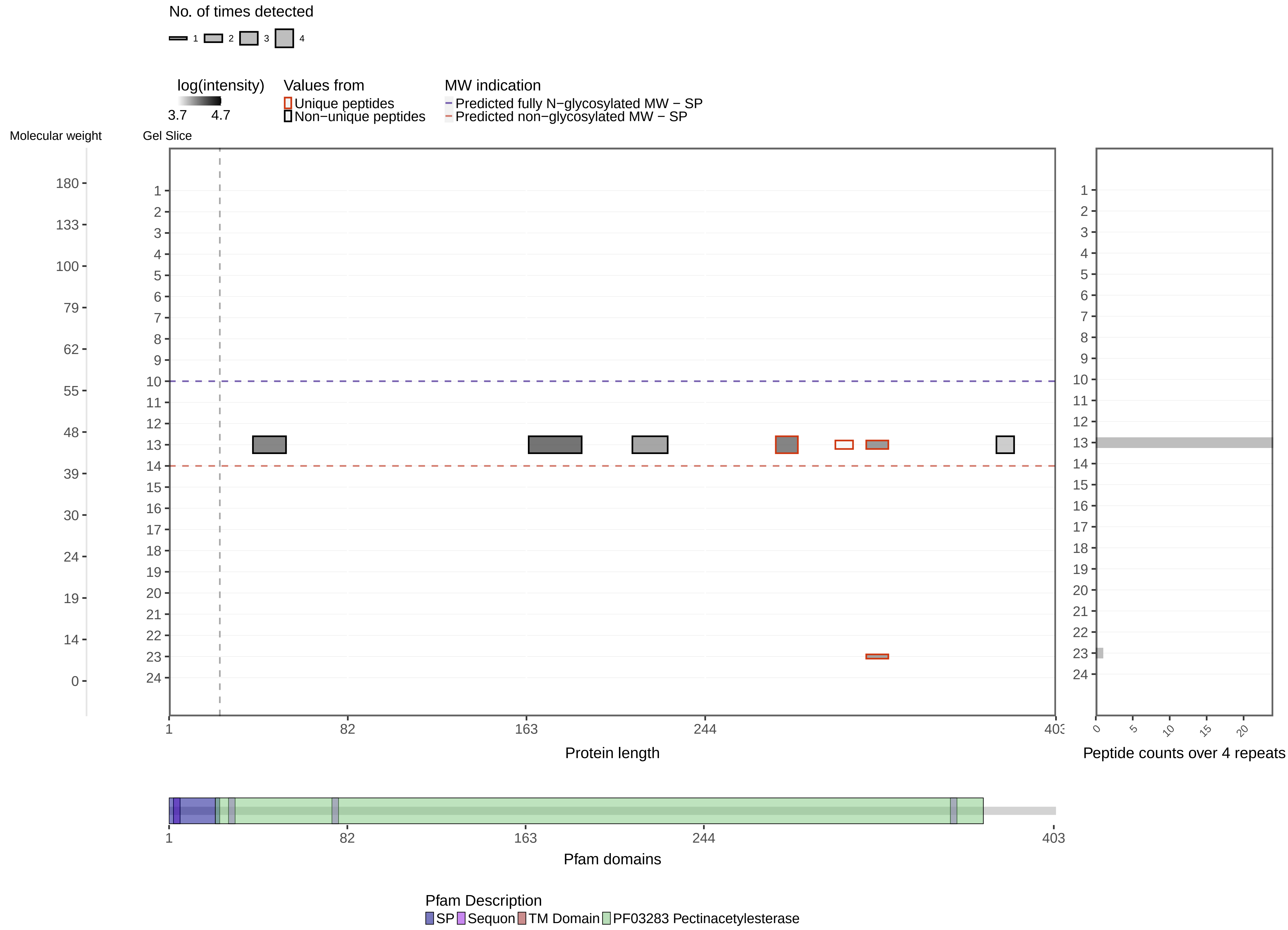


Protein sequence:

FKALLPMGAKVKCFSDAGYFINVKDISGAPHIEEFFGDVVRLHGSAKNLPLSCTSRLKPNLCFFPQNVVQHVRTPLFLVNAAYDSWQIKNILAPSVADPRGVWRNCKLDILRCSSKQLQTMQGFRLGFLRALYALGPSSSRGYFINSCYAHCQTEVQETWFRADSPKLANKTIAKALGDWFFDKNPFQKIDCPYPCDKTCHNRIFNPNVHPFDIDM*

NbL17g16590.1 eukaryotic_translation_initiation_factor_3_subunit_G−like [MW = 32.27 kDa]

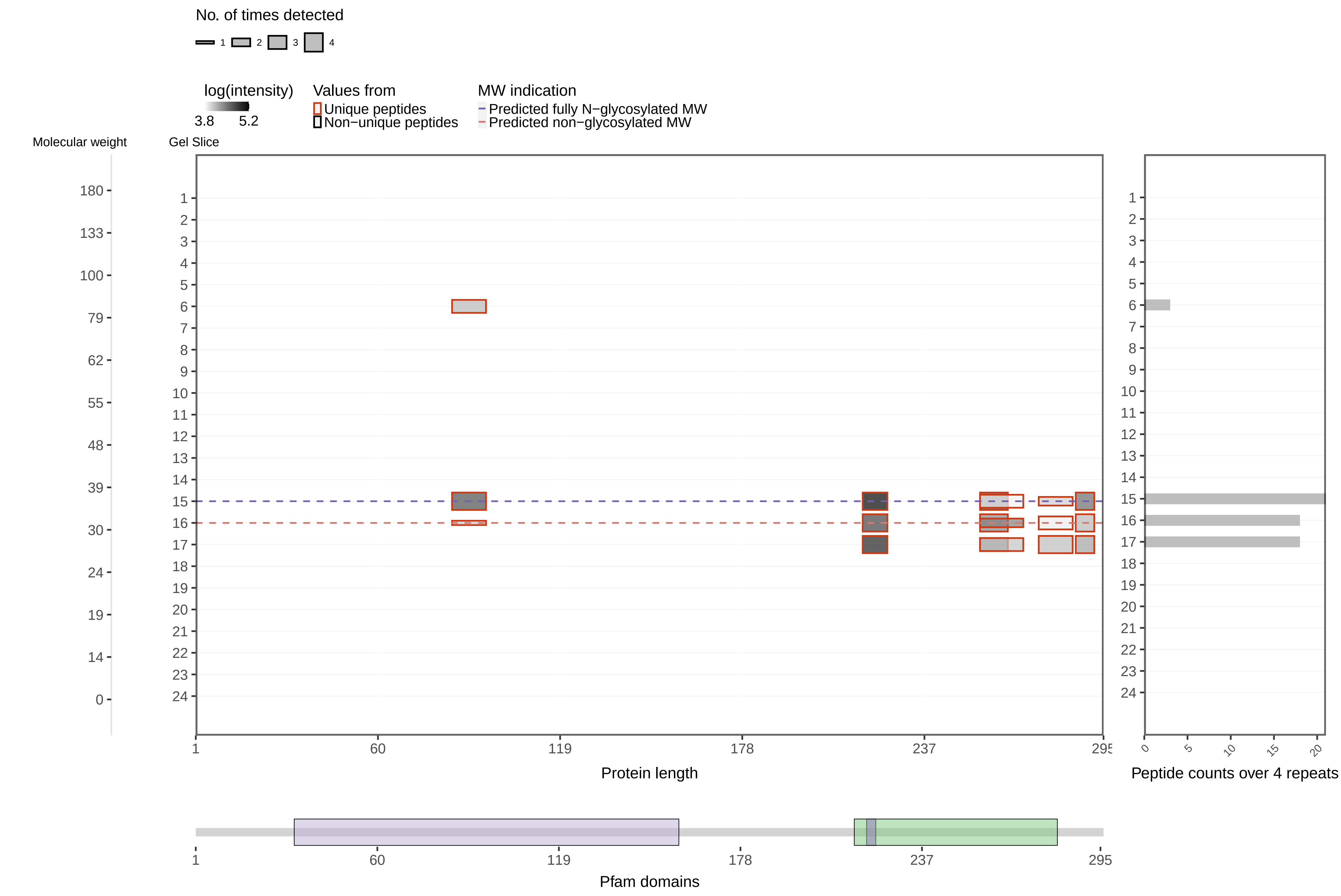


Pfam Description

SP Sequon TM Domain PF00076 RNA recognition motif. (a.k.a. RRM, RBD, or RNP domain) PF12353 Eukaryotic translation initiation factor 3 subunit G

Protein sequence:

IVCRTCGKKGDHWTSKCPFKDLAQPSDTFMDKPPSAEAPAAGAGPQKGAYVPPSMRGGGAERGSGGGTEMRRRNEENSVRVTNLSEDTREADLLELFHPFGHVSRVYVATDQKTGMSRGFGFVNFVNREDAERAINKLNGYGYDNLILRVEWAAPRTN*

### NbL17g16540.1 polyphenol_oxidase_E_chloroplastic−like [MW = 66.53 kDa]

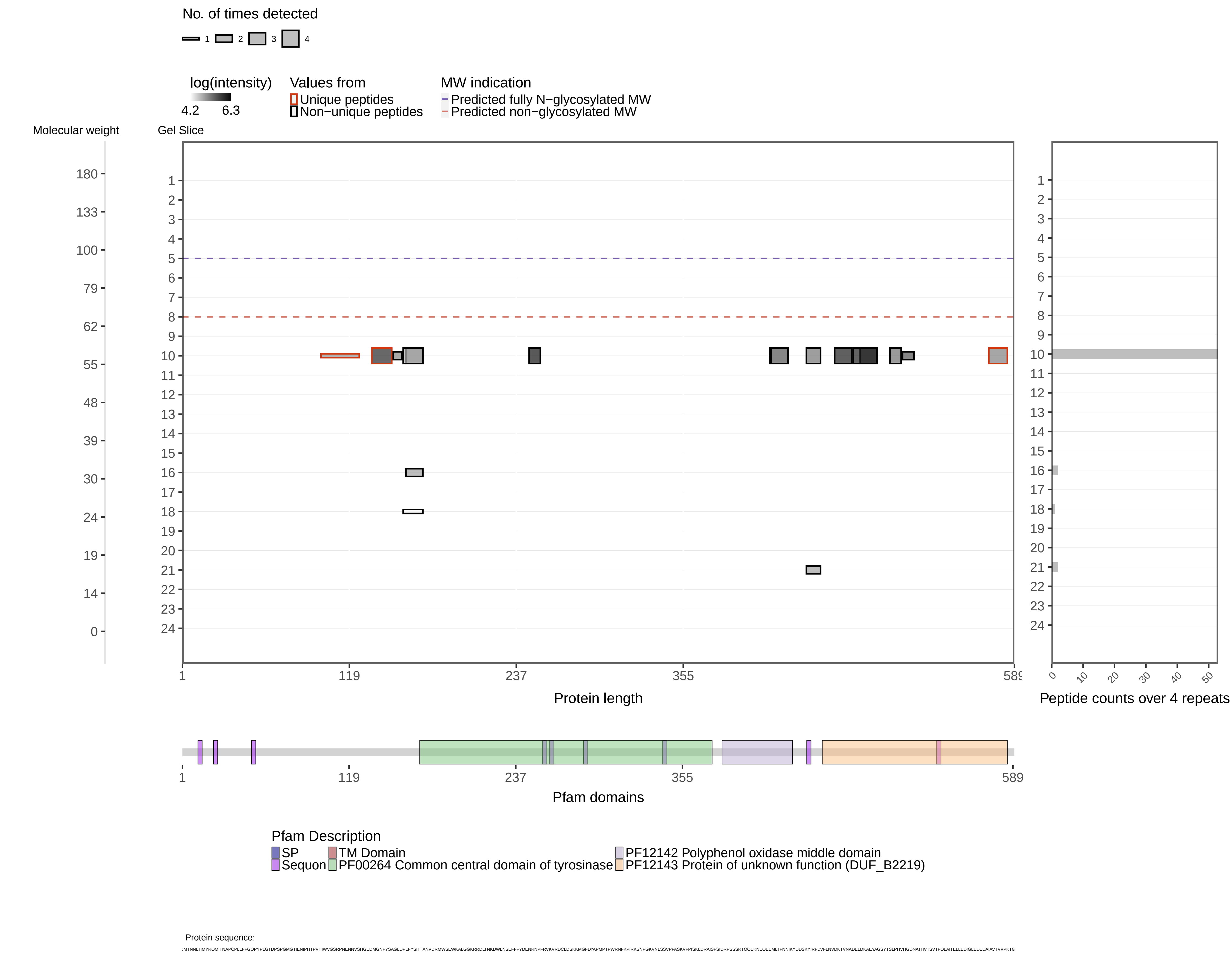

NbL17g16350.1 L−arabinokinase−like [MW = 108.33 kDa]

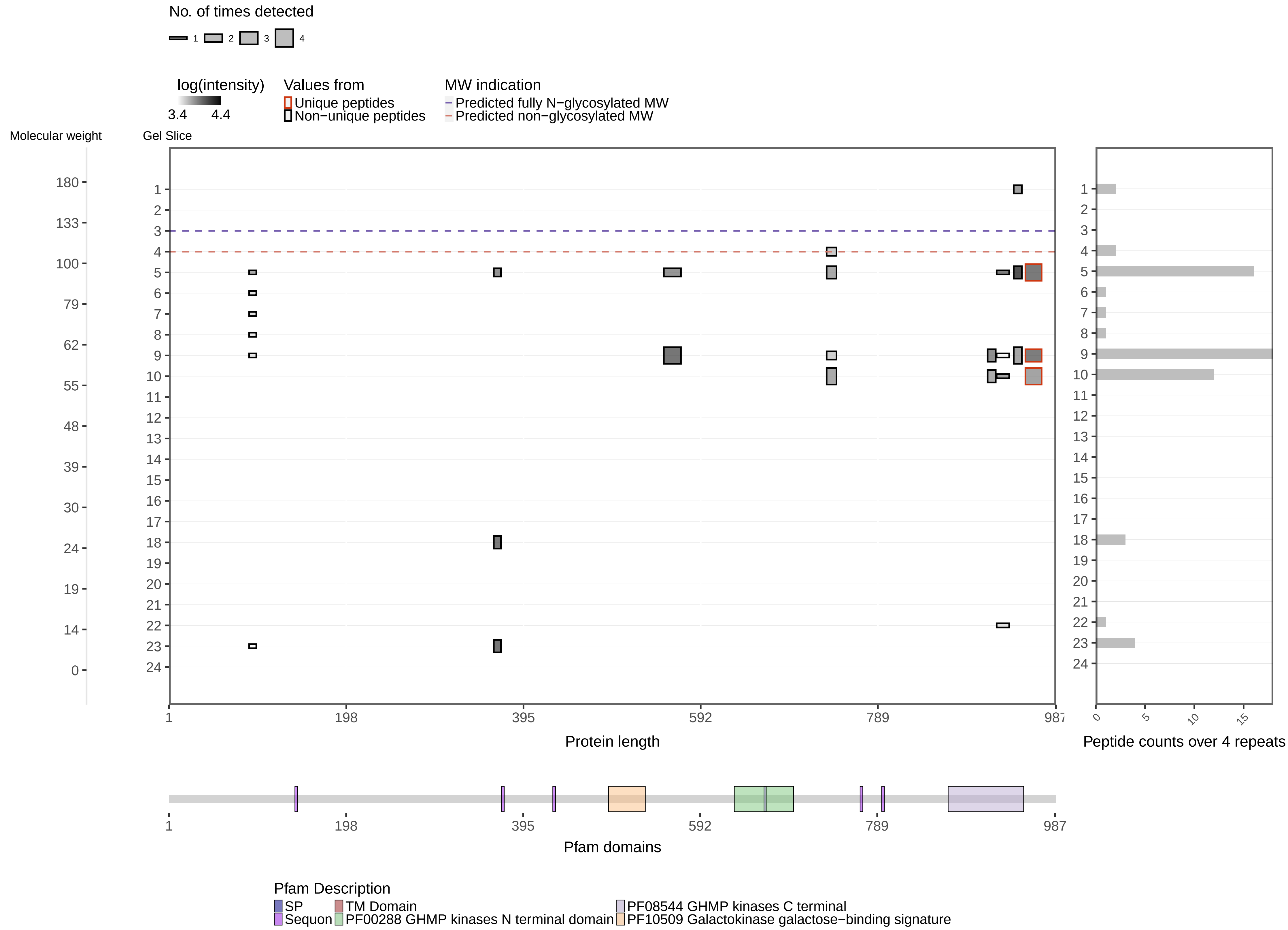


Protein sequence:

AAGLFNWEEDIFVARAPGRLDVMGGIADYSGSLVLQMPIREACHVAVQKIHPSKQRLWKHALARQQAKGQGPTPVLQIVSYGSELSNRGPTFDMDLSDFLEGDEPITYEKSRQYFARDPSQRWAAYVAGTVLVLMKELGIRFENSISLLVSSAVPEGKGVSSSASVEVASMSAIAAAHGLNISPRELALLCQKVENHVVGAPCGVMDQMTSACGEANKLLAMVCQPAEVLGLVDIPGHIRVWGIDSGIRHSVGGADYGSVRIGAFMGREIVKSIASKLLSQSLSTNGNHTDE

### NbL17g16320.1 40S_ribosomal_protein_S6−like [MW = 28.29 kDa]

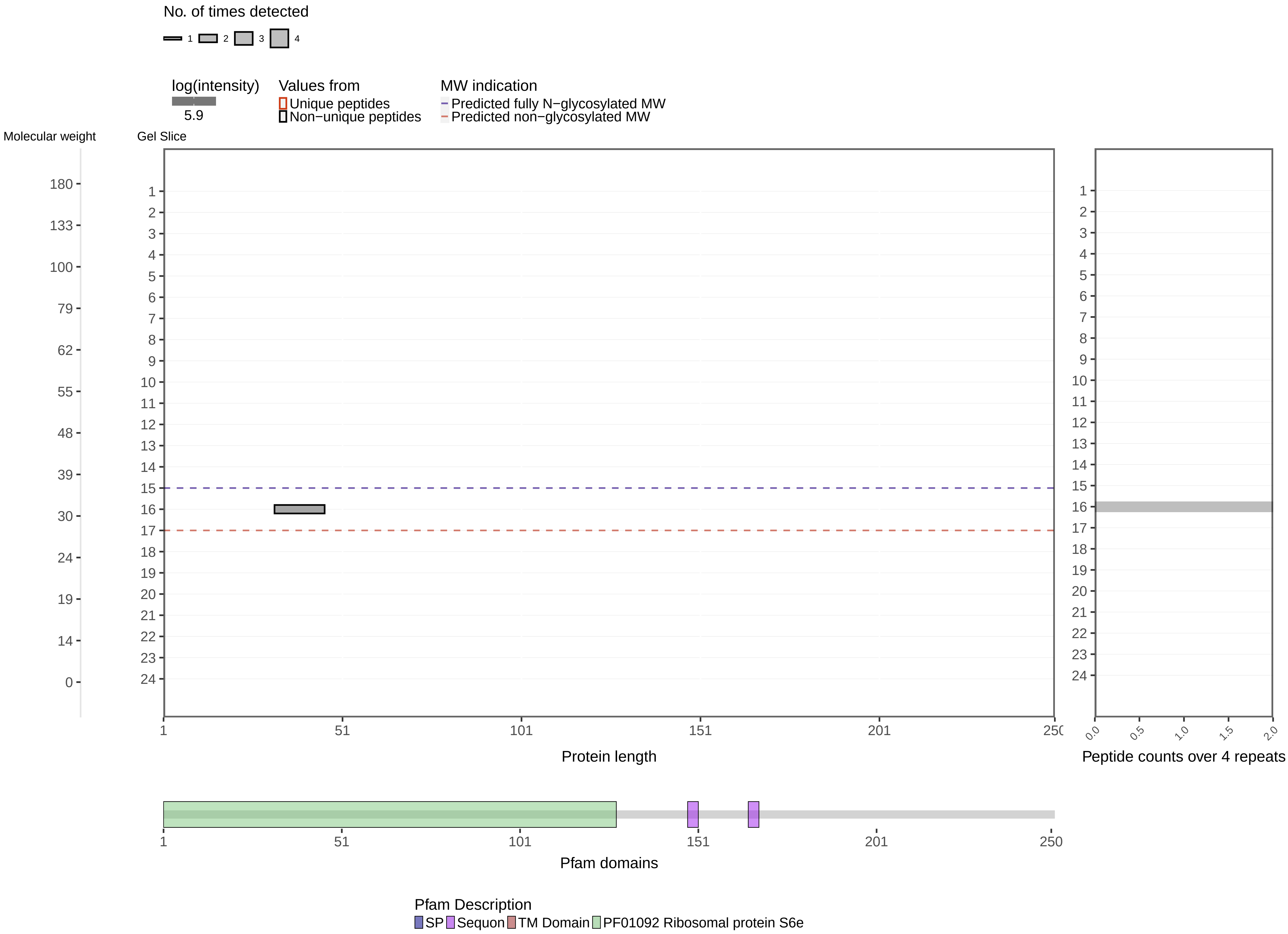


Protein sequence:

/LNLVIVKKGENDLPGLTDTEKPRMRGPKRASKIRKLFNLSKEDDVRKYVNTYRRNFTTKTGKKASKAPKIQRLVTPLTLQRKRARIADKKKRIAKAKSEAAEYQKLLASRLKEQREKRSESLAKKRSRLSAASKPSIAA*

### NbL17g16290.1 cryptochrome_DASH_chloroplastic/mitochondrial [MW = 66.82 kDa]

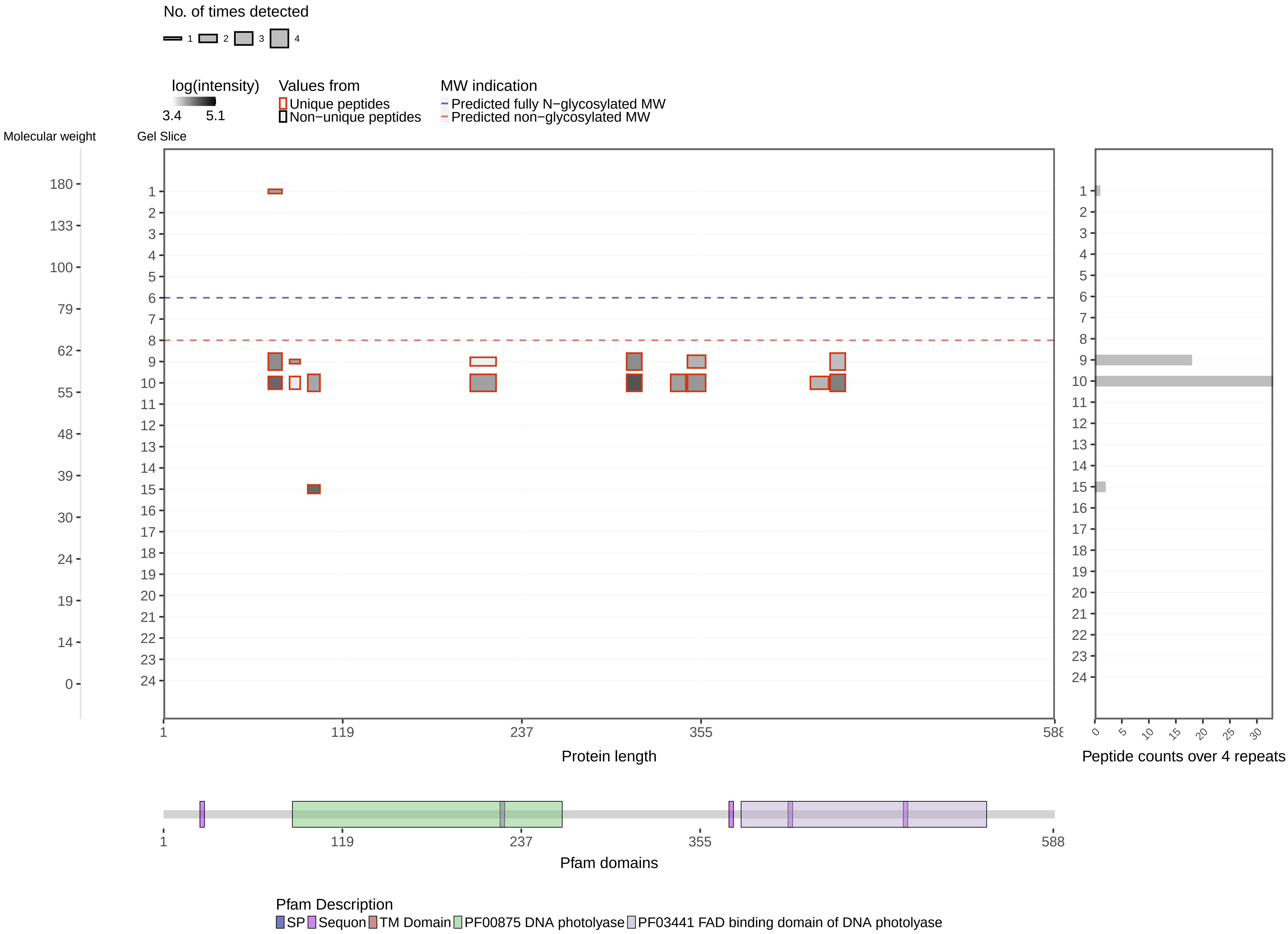


Protein sequence:

CVPQVTELGLQSEKVSKGMKFIGGESAALGRVHDYFWKKDLLKVYKVTRNGMLGPDYSTKFSPWLASGSLSPRFIYEEVKRYEKERQSNDSTYWVLFELIWRDYFRFLSIKQGNLLFHAGGPRKVDINWSQDQTMFDAWRRGQTGYPLIDANMKELDTTGFMSNRGRQIVCSFLVRDMGIDWRMGAEWFETCLLDYDPCSNYGNWTYGAGVGNDPREDRYFSIPKQAQNYDPEGEFVAYWLPELRALPREKRHSPGMMYLKPIVALKHGYTKKTGDSKMPYGSRR

### NbL17g16230.1 glucan_endo−13−beta−glucosidase_5−like [MW − SP = 50.27 kDa]

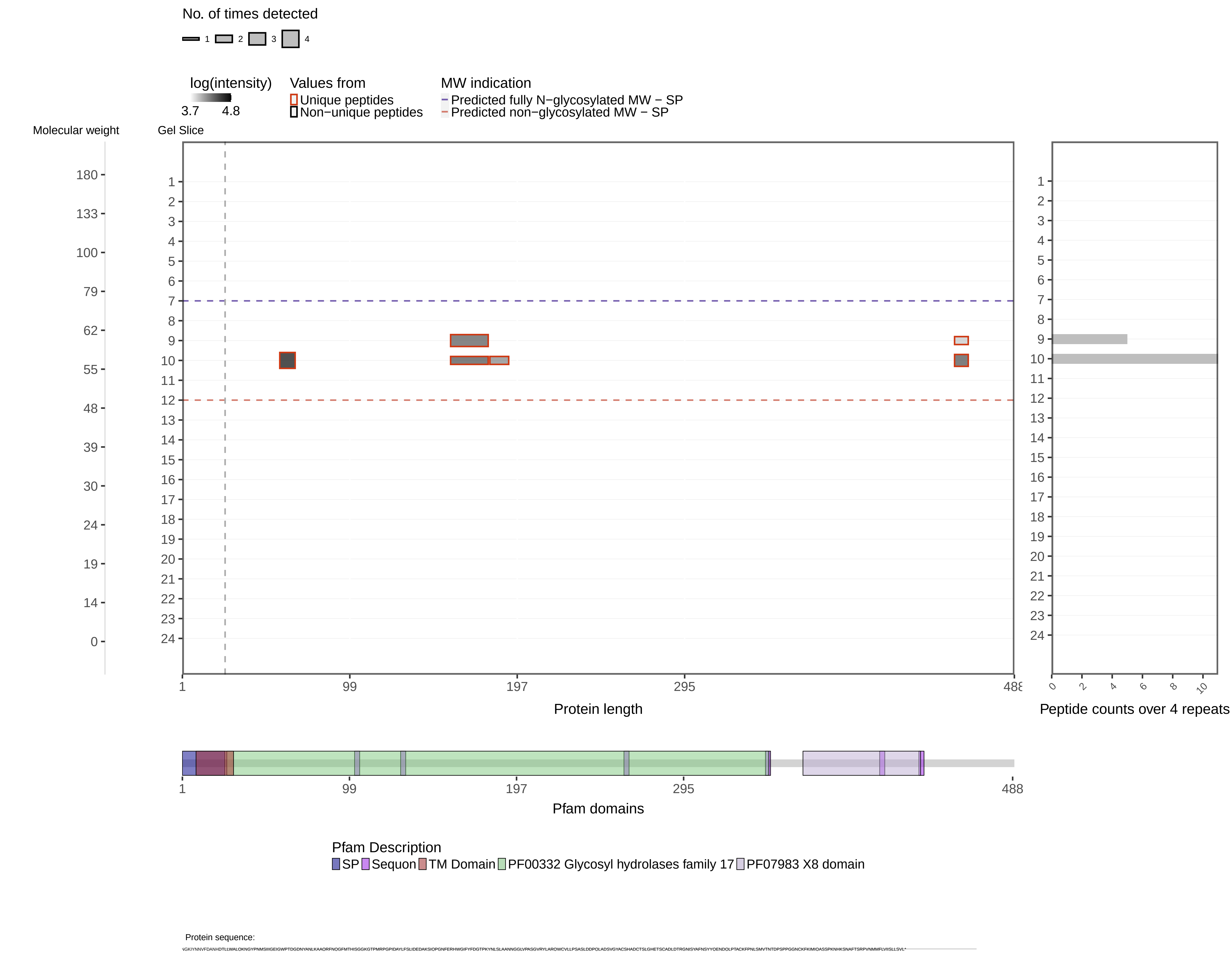

NbL17g16140.1 delta−aminolevulinic_acid_dehydratase_chloroplastic−like [MW = 46.71 kDa]

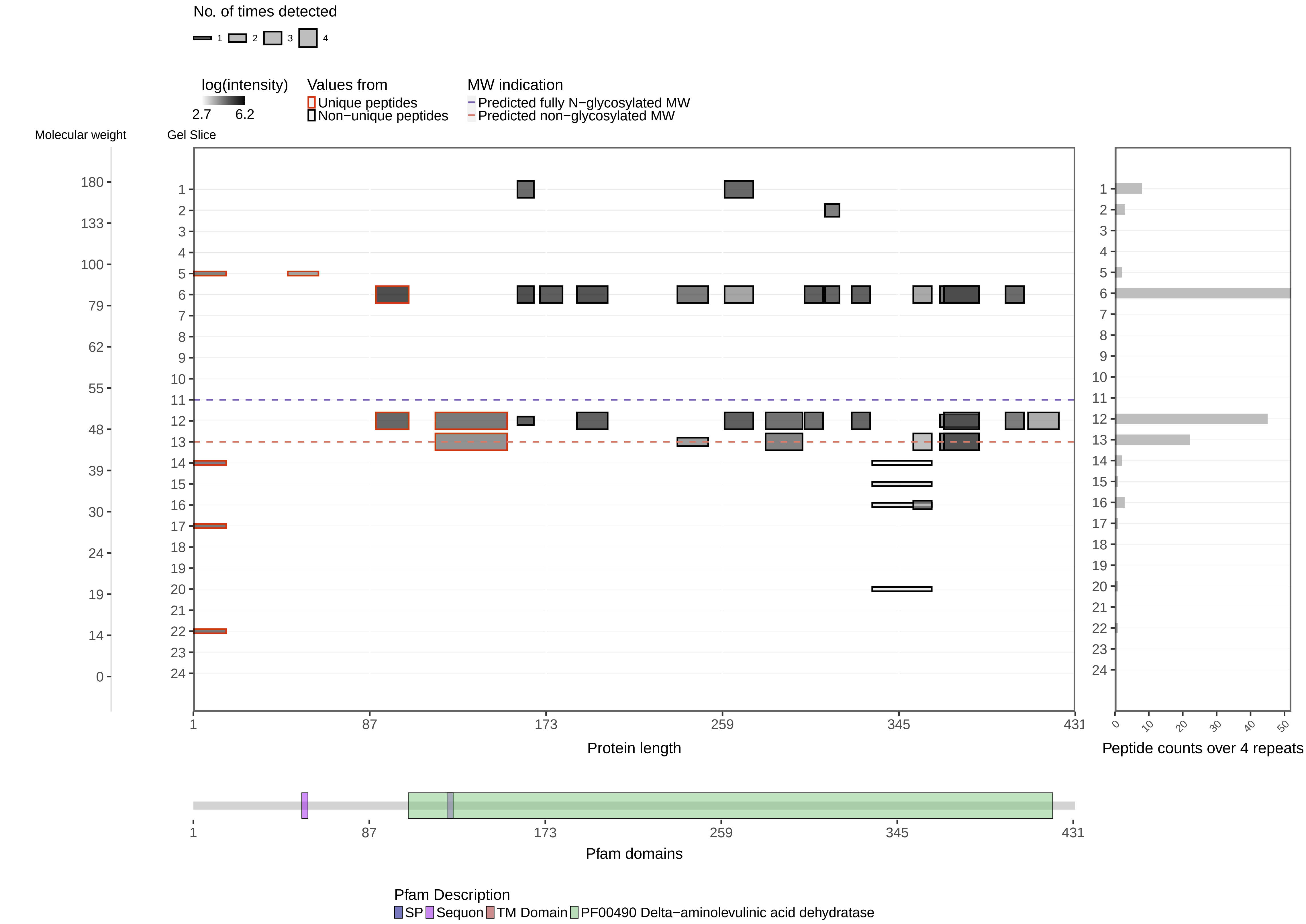


Protein sequence:

PRAIRFLKDKYPDLVIYTDVALDPYSSDGHDGIVREDGVIMNDETVHQLCKQAVAQARAGADVVSPSDMMDGRVGAIRAALDAEGFQHVSIMSYTAKYASSFYGPFREALDSNPRFGDKKTYQMNPANYREALVELQADESEGADILLVKPGLPYLDIIRLLRDKSALPIAAYQVSGEYSMIKAGGVLKMIDEERVMMESLMCLRRAGADIILTYFALQAGRCLCGEKR*

NbL17g16130.1 peroxidase_16−like [MW − SP = 33.46 kDa]

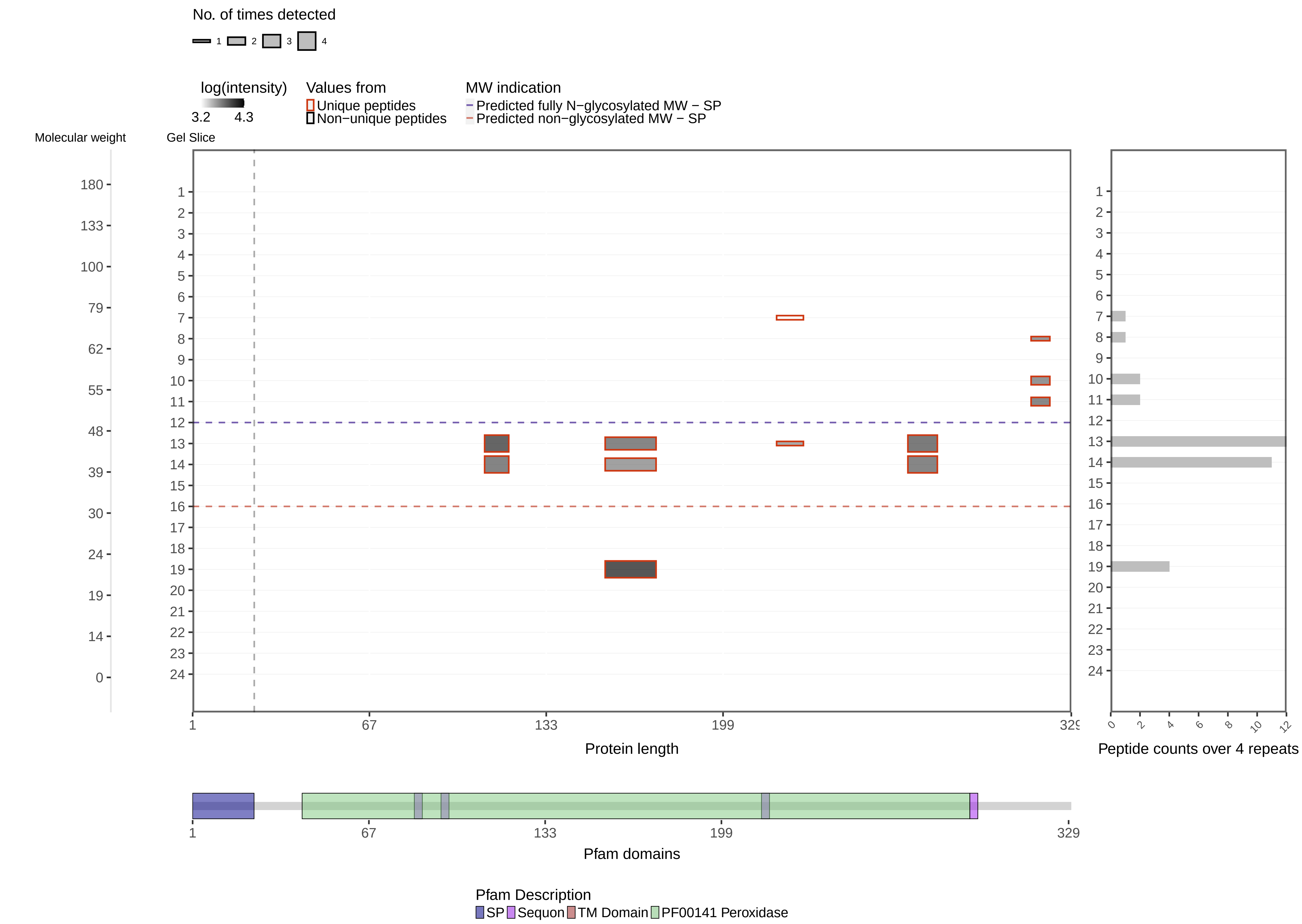


Protein sequence:

GRMSSQTSVQNNLPHSNFDLQKLLPMFASRGLSIRHLIALSGAHTLGFSHCNQFSSRLYNFNSTHKVDPTIDVTYAKKLQEMCPQDALPGVVIPMDPDTPQLFDNTYYKNLQRGKGLFTSDQTLVTKRGSRRIVNIFASNKTAFERVFIAAMSKLGRFGVKTGNLGEIRKDCAVVN*

### NbL17g15990.1 probable_plastid−lipid−associated_protein_10_chloroplastic [MW = 30.53 kDa]

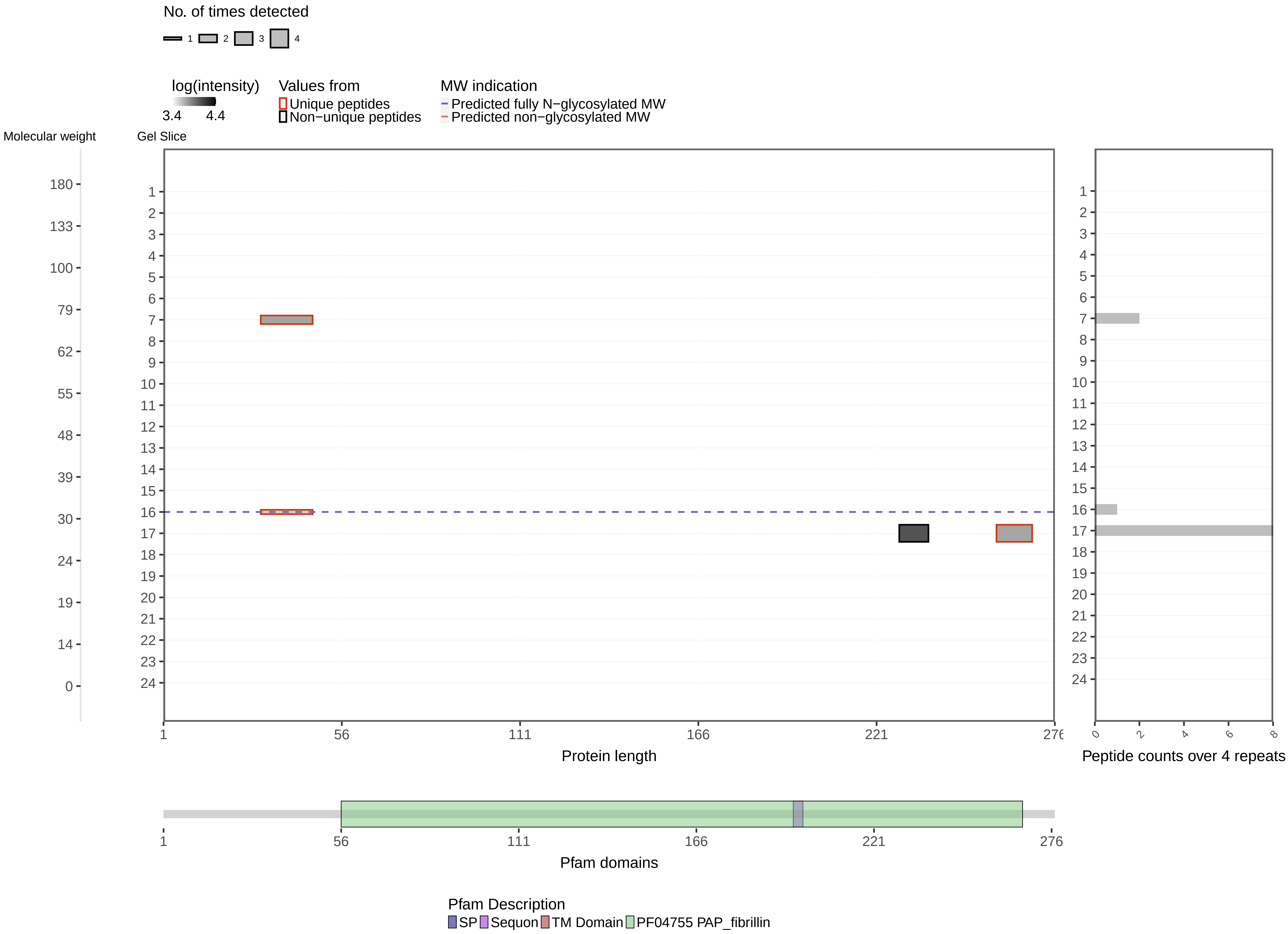


Protein sequence:

RLPFFQVGQIFQKFECKSESREGLVRNVIKWSVPRLLEENEGATLVVSARFSCVSARNIYLKFEEISLQNINISEDLQAVIAPAILPRSFLSLQVLQFIRSFKAQVPVTSPERRSVGGLYYLSYLDNNMLLGRAVGGGGVFVFTRAQSLTF*

### NbL17g15960.1 probable_tocopherol_cyclase_chloroplastic [MW = 57.53 kDa]

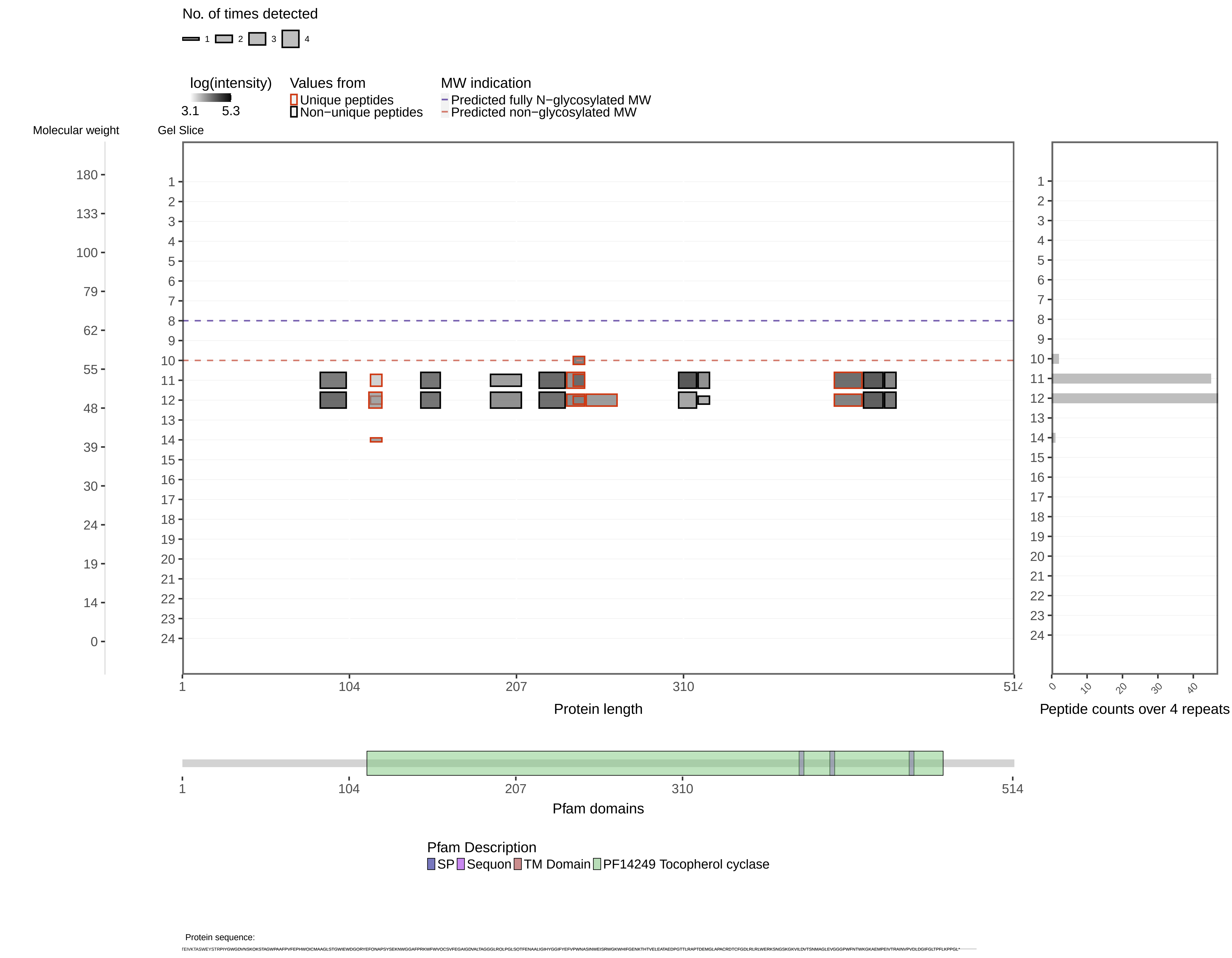

NbL17g15920.1 peroxisomal_fatty_acid_beta−oxidation_multifunctional_protein_AIM1−like [MW = 76.11 kDa]

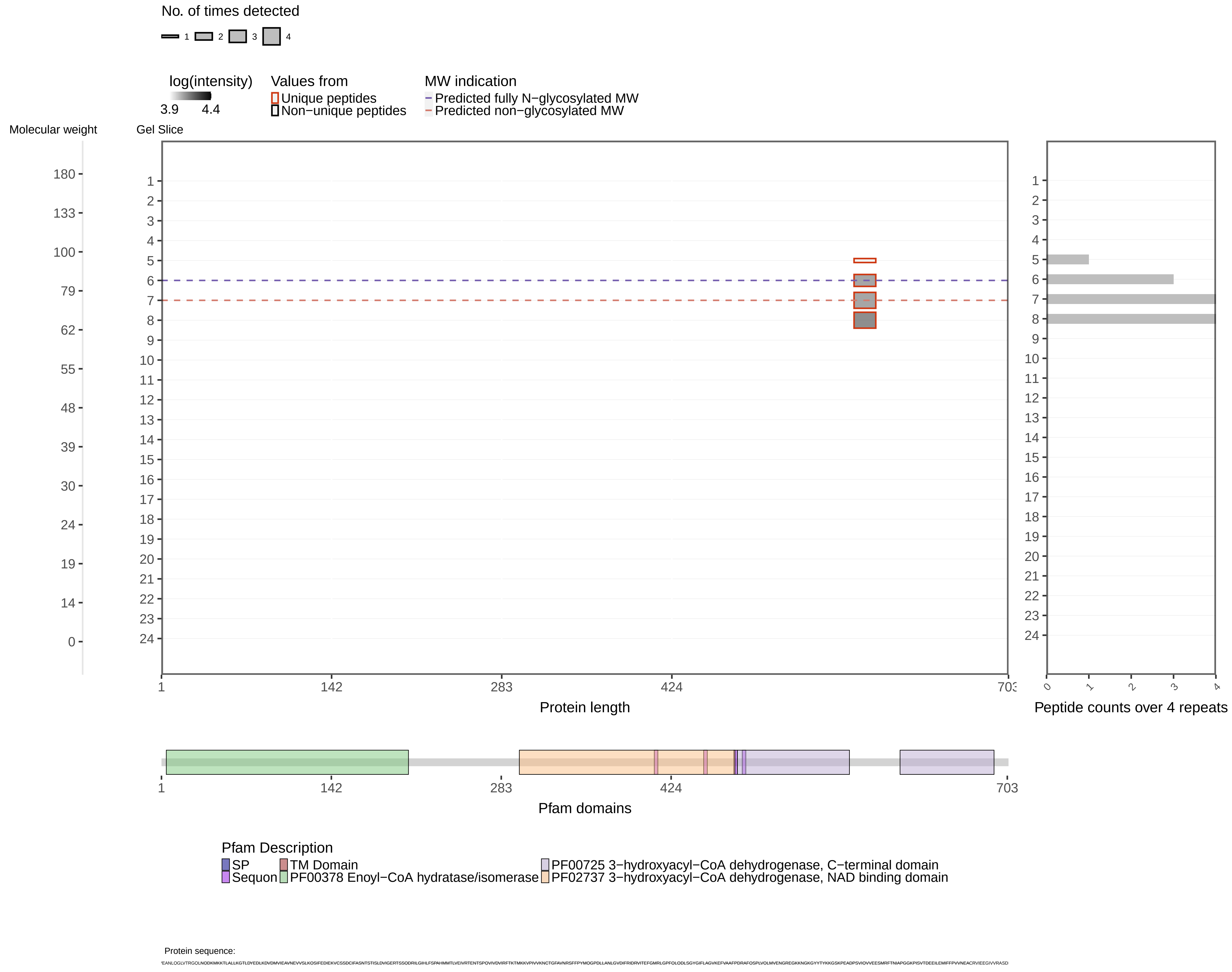

### NbL17g15860.1 eukaryotic_translation_initiation_factor_3_subunit_L−like [MW = 60.65 kDa]

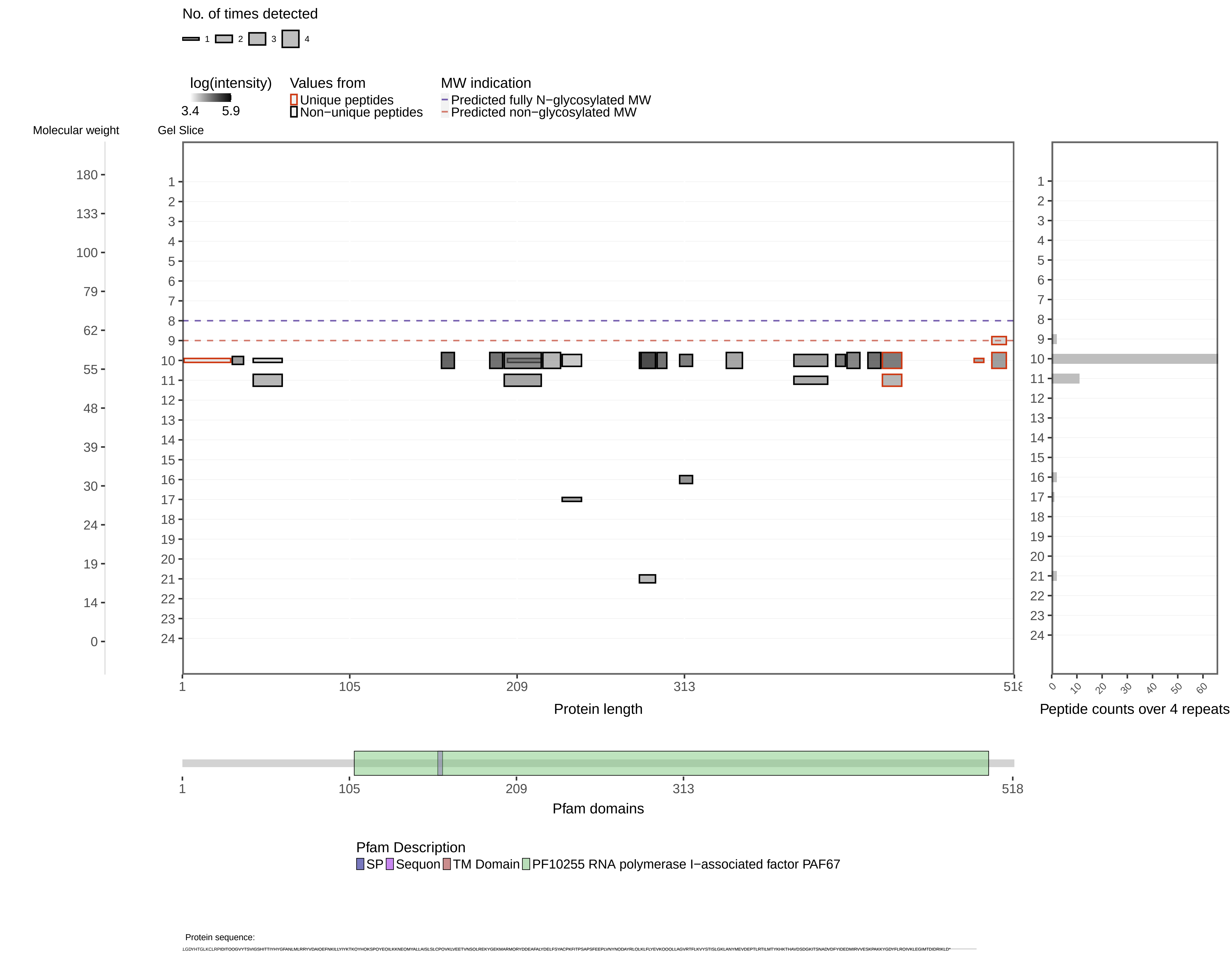

NbL17g15850.1 aspartate_aminotransferase_cytoplasmic [MW = 49.32 kDa]

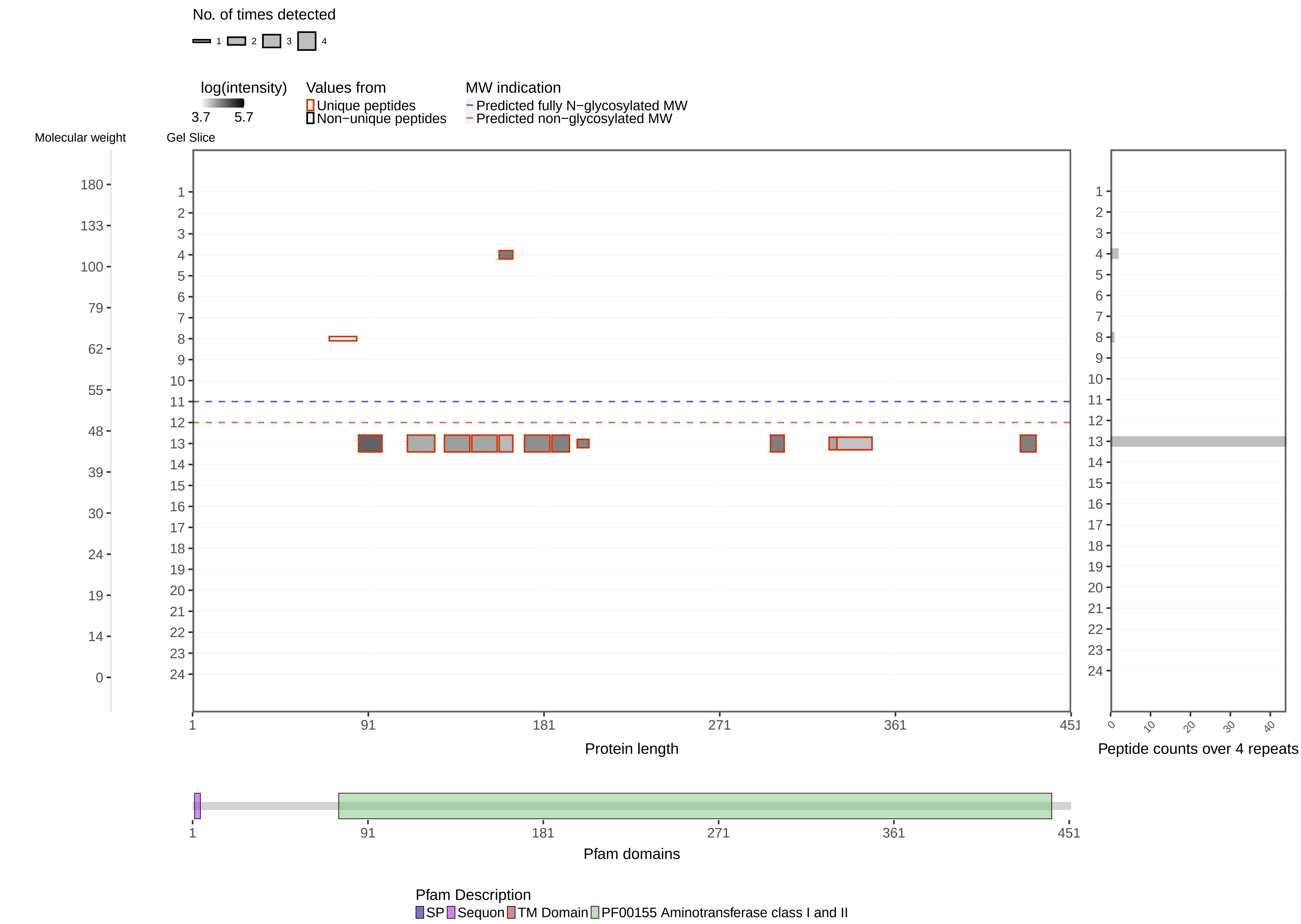


Protein sequence:

EDLGSAPSGAIVLLHACAHNPTGVDPTIDQWEQIRRLMRSRGLLPFFDSAYQGFASGSLDTDAQSVRMFVADGGEVLVAQSYAKNMGLYGERVGALSIVCRNADVASRVESQLKLVIRPMYSNPPIHGASIVATILKERNMYHEWTLELKAMADRIIRMRQQLFDALRARGTPGDWSHIIKQIGMFTFTGLNSEQVAFMTKEYHIYMTSDGRISMAGLSSRTVPHLADAIHAAVARAR*

### NbL17g15830.1 la_protein_1 [MW = 51.5 kDa]

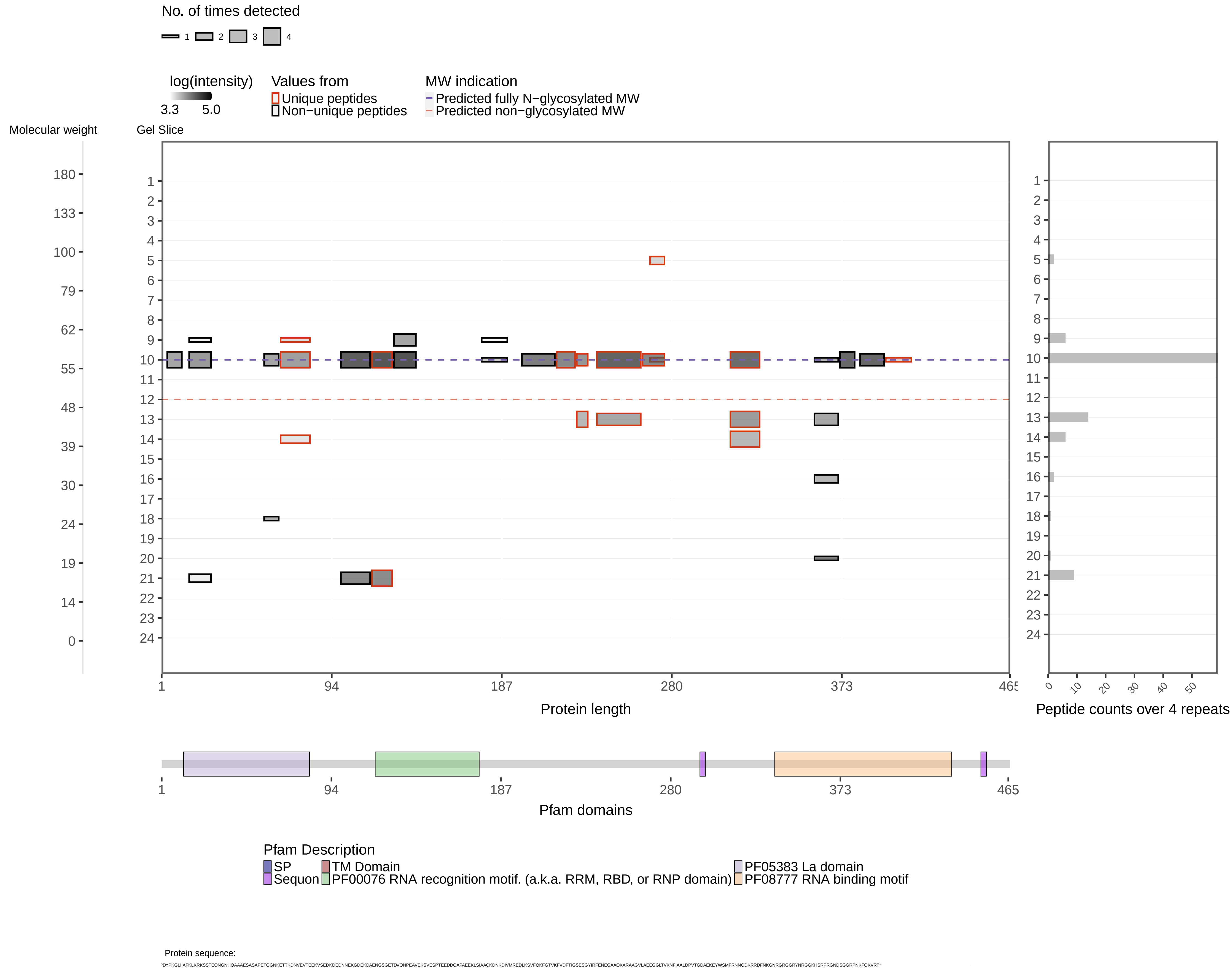

### NbL17g15800.1 coiled−coil_domain−containing_protein_25 [MW = 25.3 kDa]

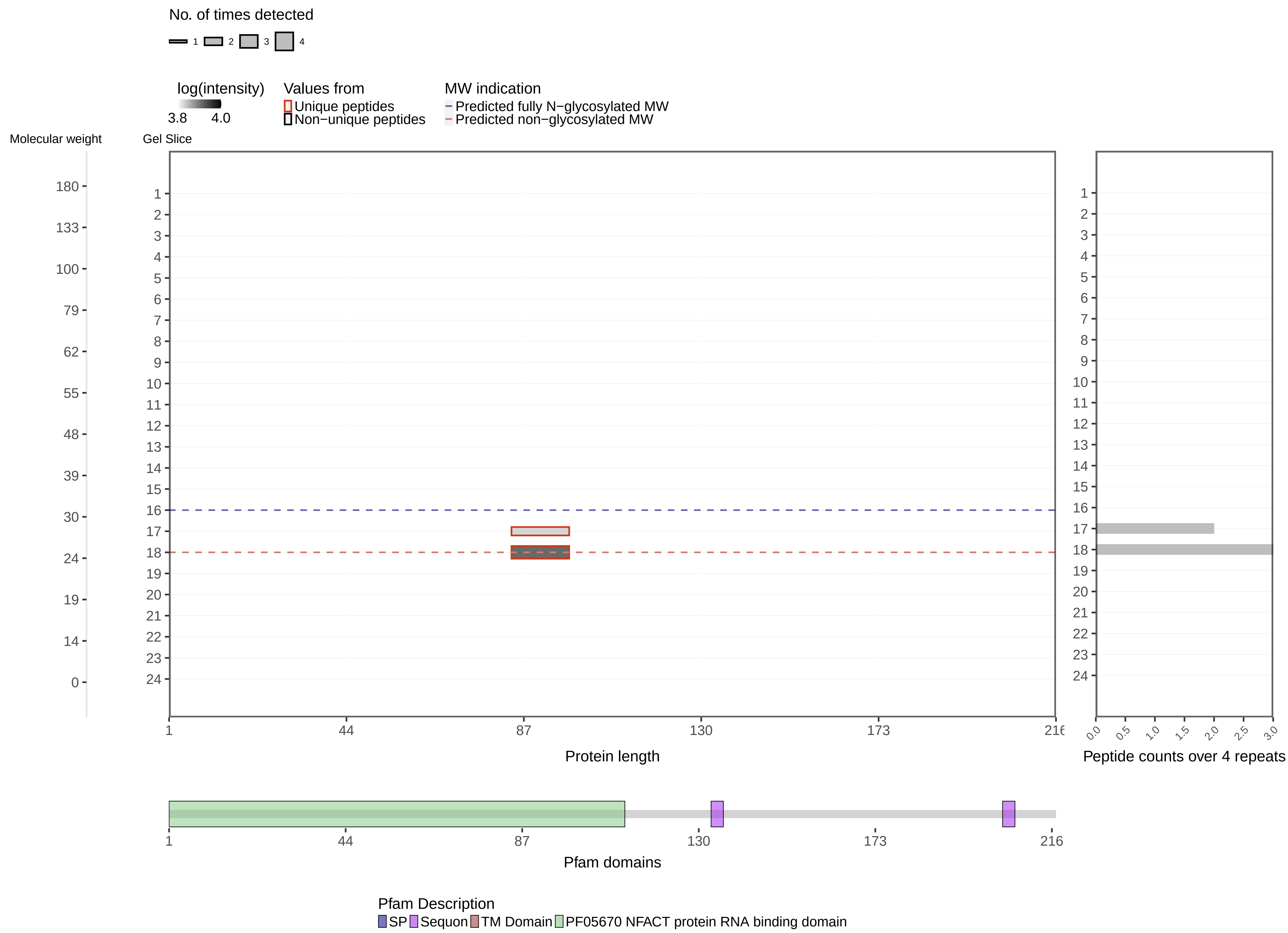


Protein sequence:

NLKKTASMDVGQVSFHNPKMVRTVKVEKRINEIVNRLNRTKVERKPDLKAEREAVNAAERAERKQQLREKKRREDMDRLEKERQAEMRSYKNLMVADKMTSNKDIASTNKSLQELEEDFM*

### NbL17g15610.1 54S_ribosomal_protein_L24_mitochondrial [MW = 24.37 kDa]

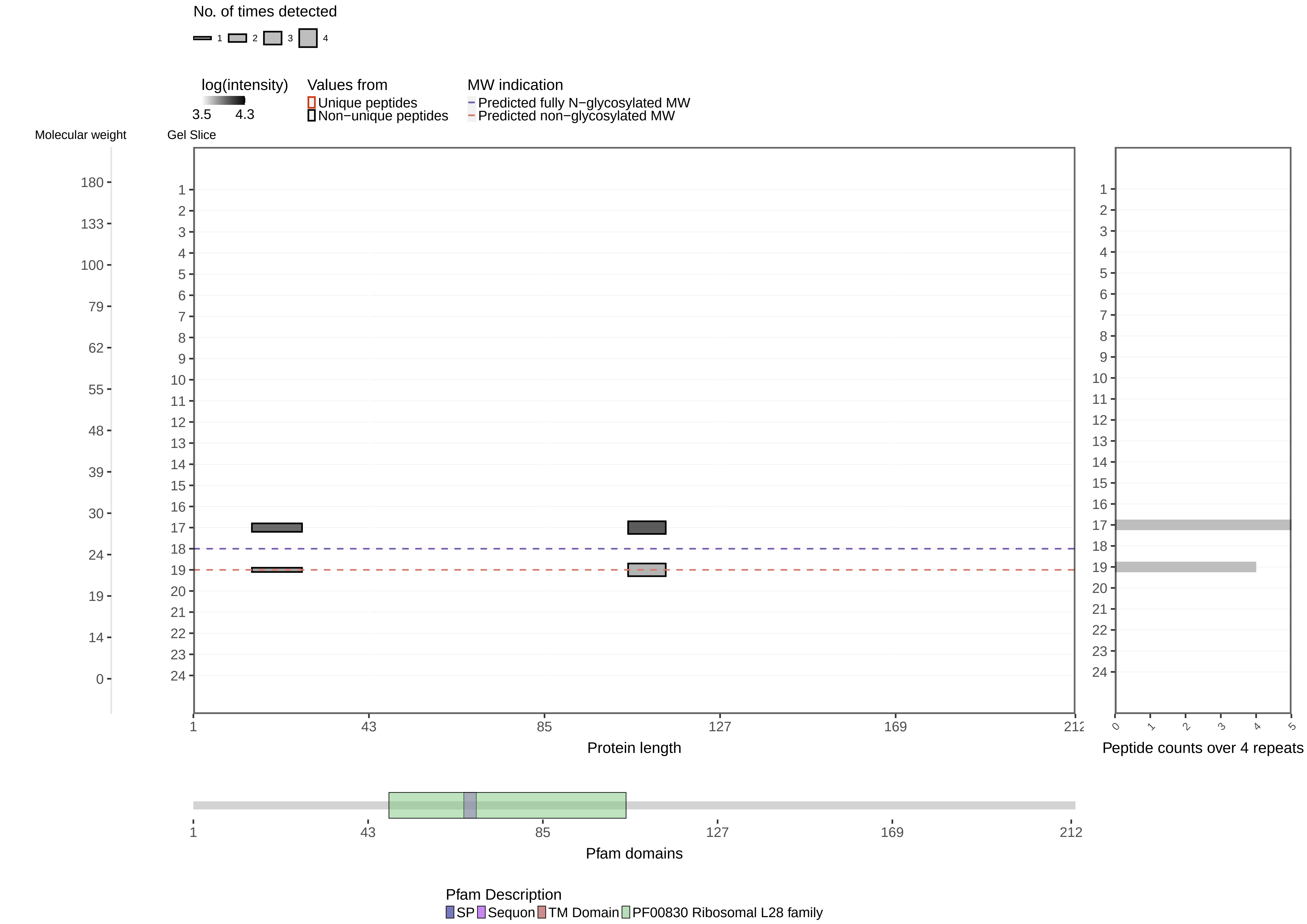


Protein sequence:

[illegible]VTTHALRCIDKAGGIDEYLLKTPYHEMDTELGLFWKAKIEKLYEELGKMEVVFFTPEDEAKLEEQFREVKLEERAARREARRKMYGWSPKSELKADSEGTDGEGYLQSDFHEGMVANA*

### NbL17g15510.1 50S_ribosomal_protein_L17−like [MW = 18.46 kDa]

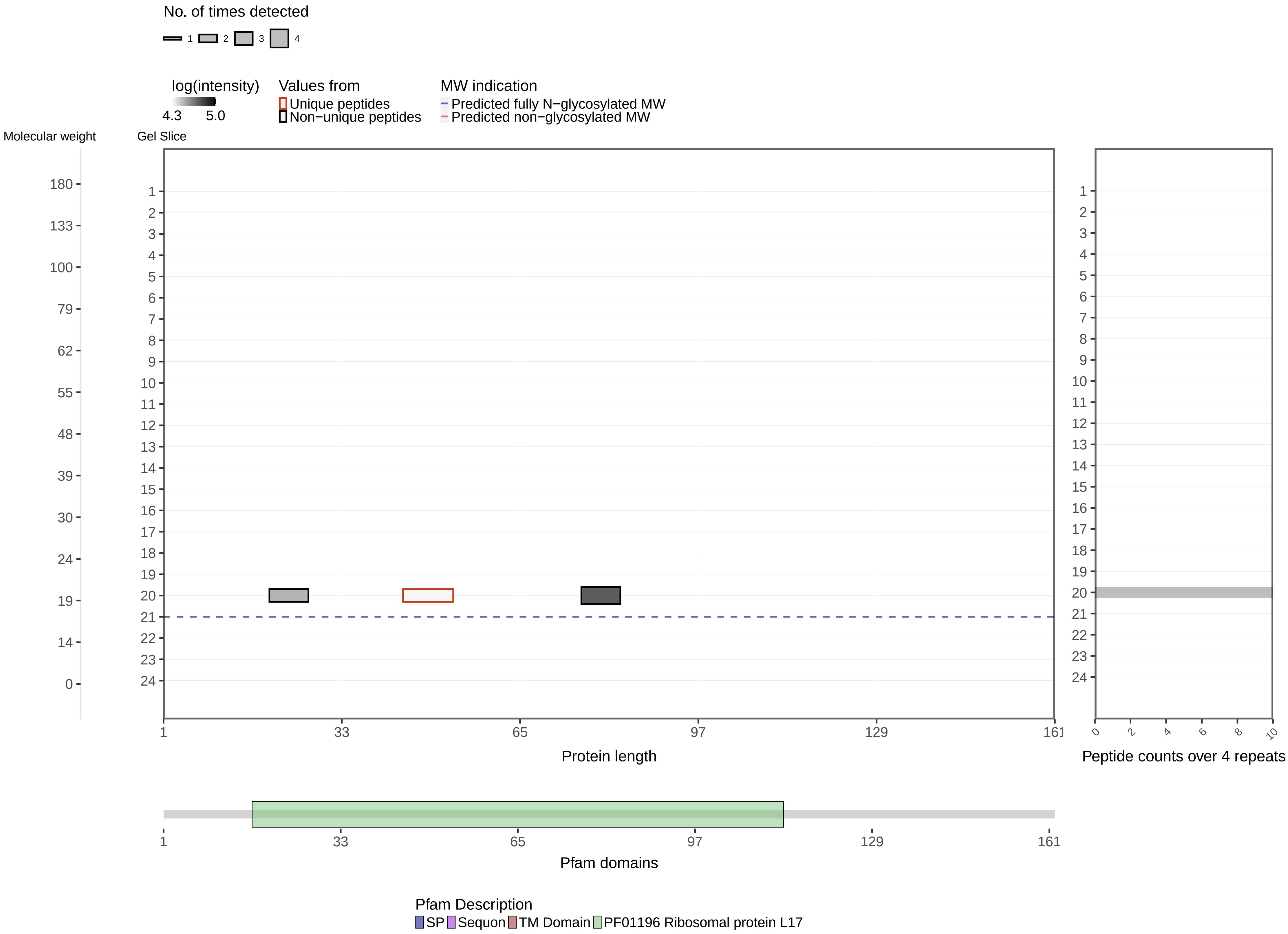


Protein sequence:

VRGDDVIHKLFTELAYRYKDRAGGYTRVLRTRIRVGDAAPMAYIEFIDRENELRQSKPPNPQPPQRPAMDPWTRSRLSRQFAPPKEEKSSDSDI*

### NbL17g15420.1 GDSL_esterase/lipase_CPRD49−like [MW = 29.06 kDa]

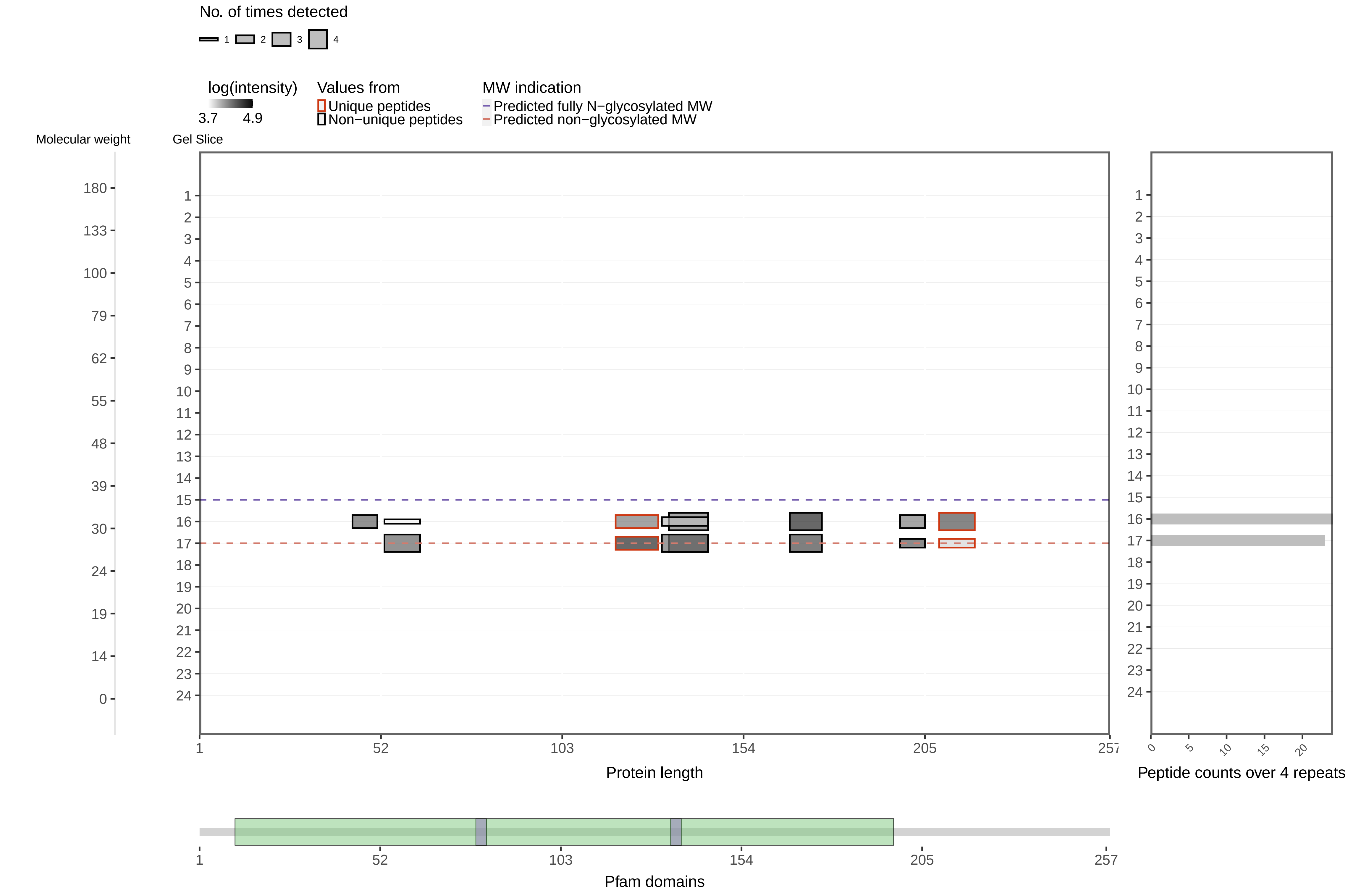


Protein sequence:

ENIRIIFLSCPPVDEARVRENTSSYFSELVRTNELCRQYSDACIELCNEMNLKVVDLWTALQKREDWLTSCFTDGIHLAEAGSKIVVEEIIKVLKEAEWTPSLYWKSMPTEFAEDSSYDLVLADGKTTINPSEWTYHREIQWD*

### NbL17g15310.1 30S_ribosomal_protein_S31_chloroplastic [MW = 12.09 kDa]

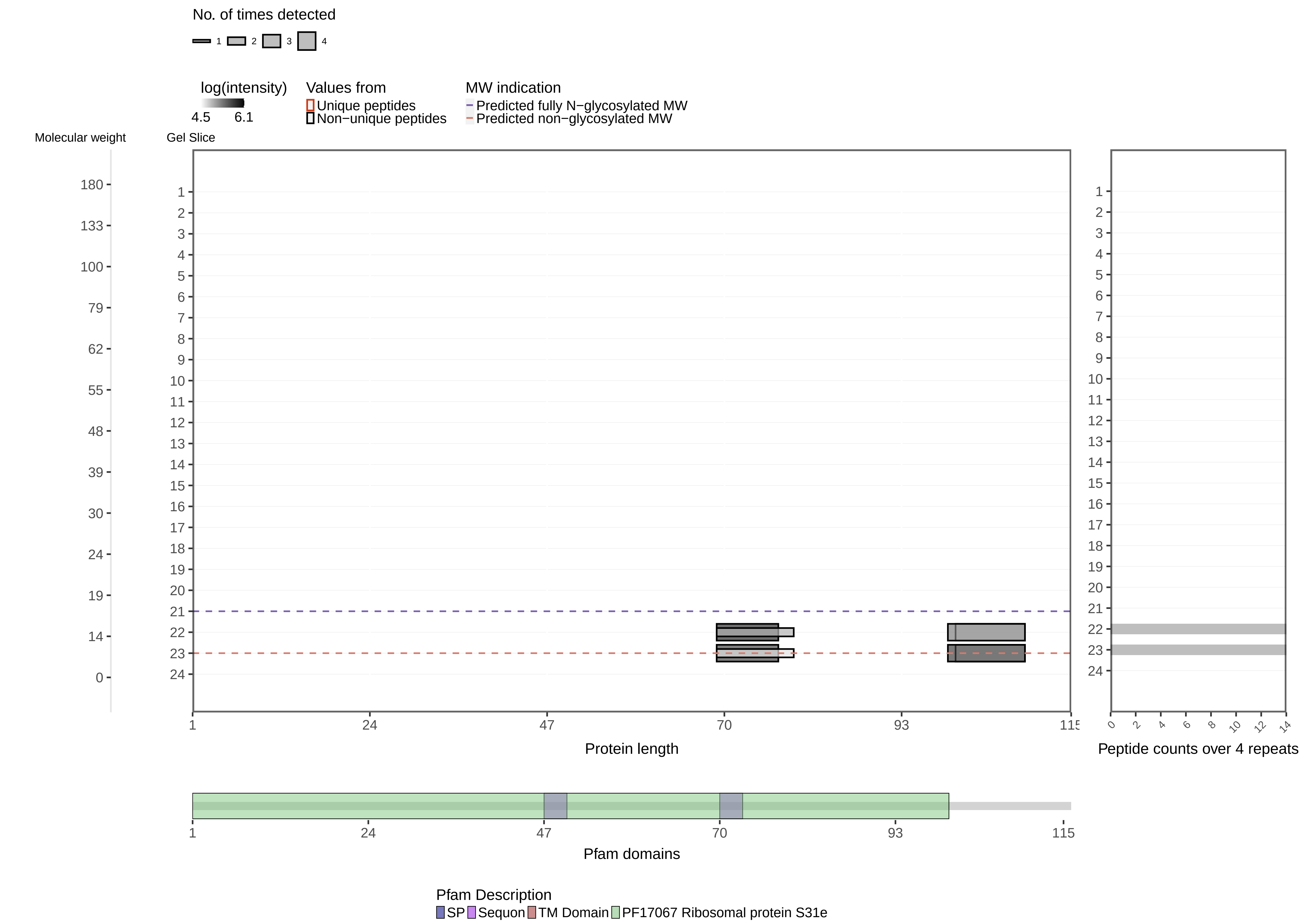


Protein sequence:

ANSSSLPLIYCGRGDKKTAKGKRFNHSFGNARPRNKKKGRGPPRVVAPPAAPKRDPFDDGQVIKIEINE*

NbL17g15230.1 60S_acidic_ribosomal_protein_P0 [MW = 34.16 kDa]

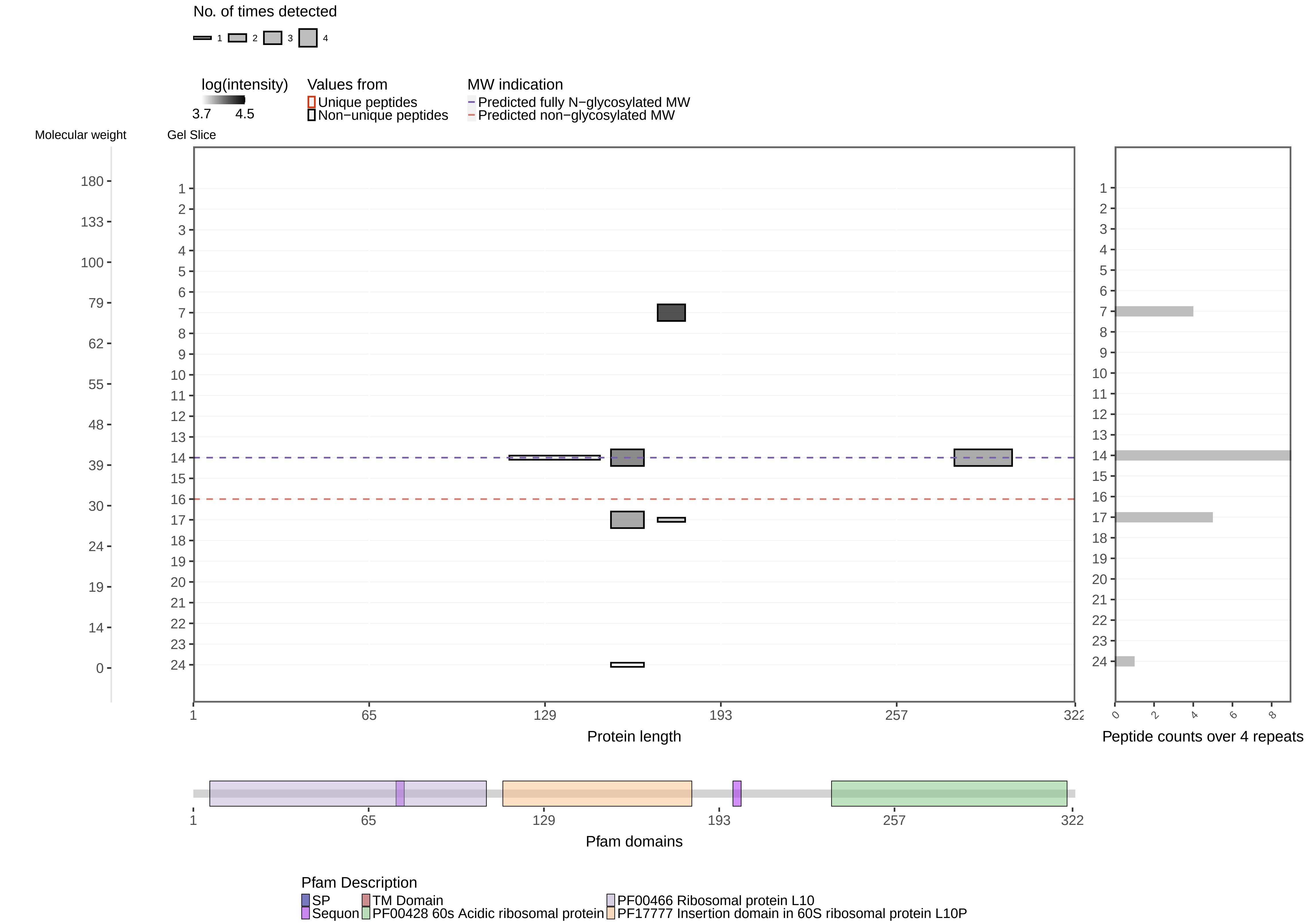


Protein sequence:

[illegible]TKINKGTVEIITPVELIKKGDKVGSSEAALLAKLGIRPFSYGLVVLSVYDNGSVFSPEVLDLTEDDLVEKFAMGVSMVTSLSLAISYPTLAAAPHMFTNAYKNVLAIAVETDYSFPLADKVKEYLEDPSKFAAVAAAPVAAAGSGAAPAAAKEEEKKDEPAEESDDDMGFSLFD*

### NbL17g15030.1 LIM_domain−containing_protein_WLIM2b−like [MW = 20.84 kDa]

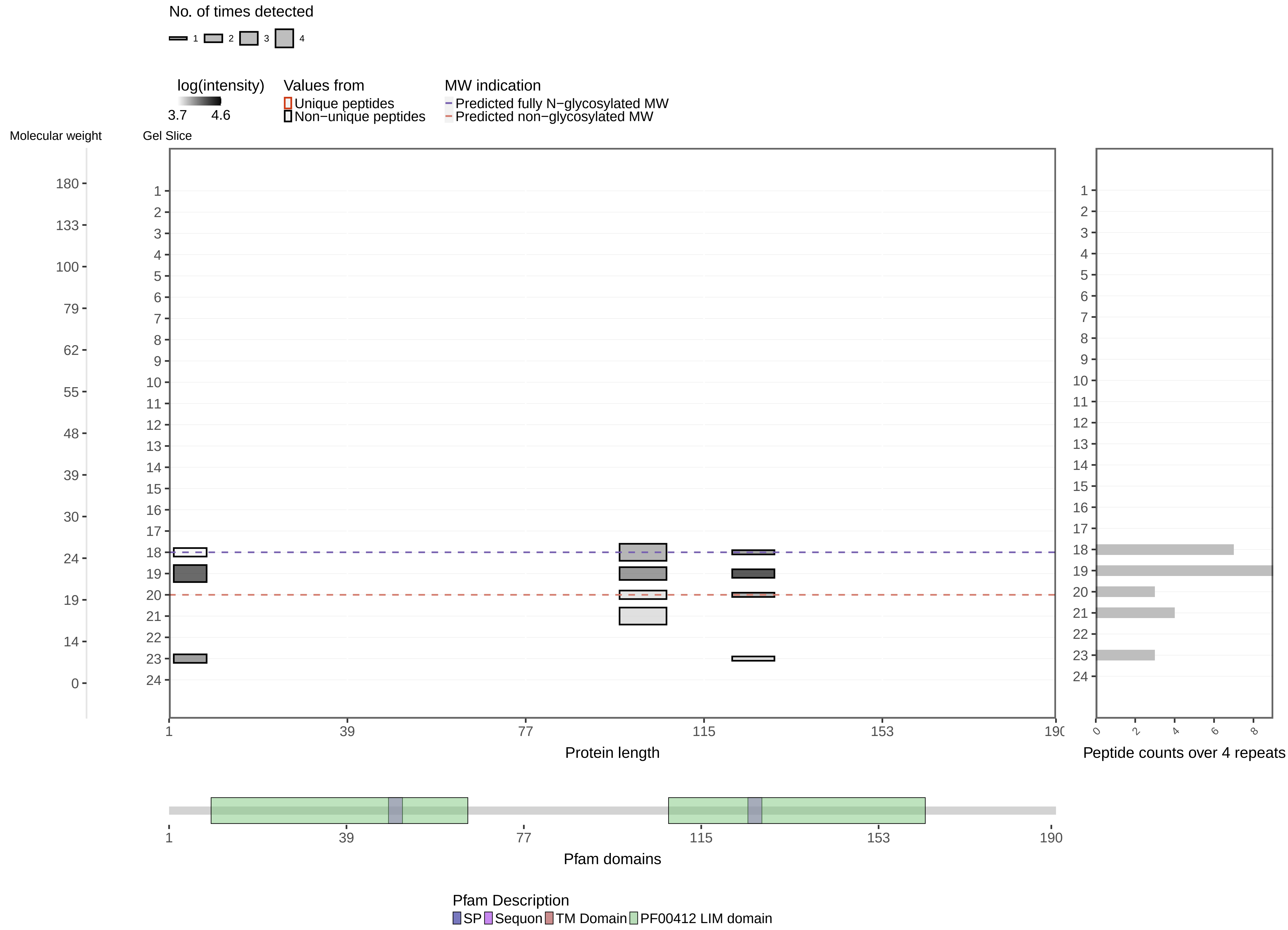


Protein sequence:

SAEKLTPELTRSPSKAAGMFSGTQEKCATCGKTAYPLEKVTVENQSYHKTCFKCSHGGCSLSPSNYAALNGILYCKPHFSQLFKEKGSYNHLIKSASMKRPAATVPDS*

NbL17g14540.1 uncharacterized_LOC104238289 [MW = 81.41 kDa]

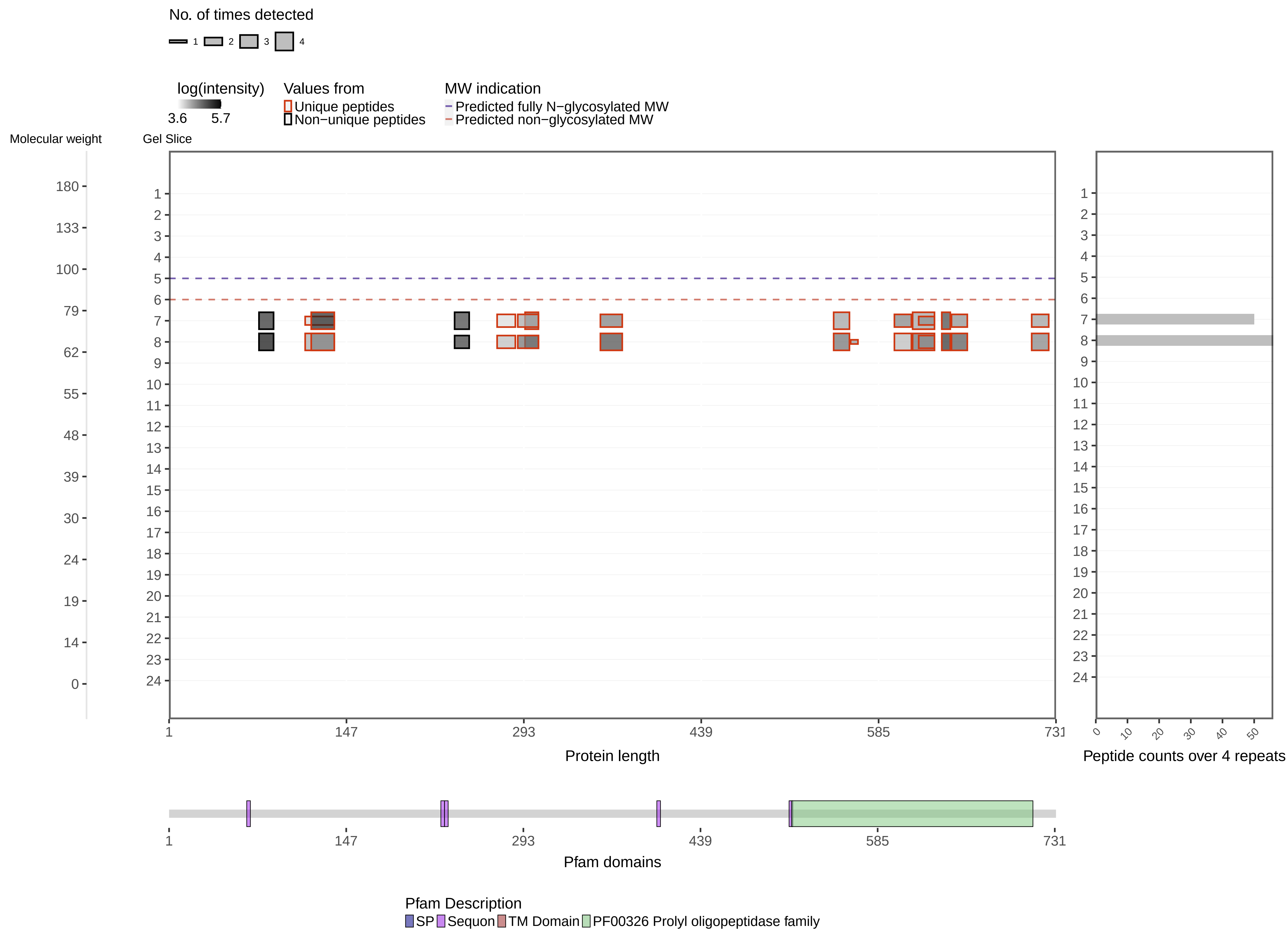


Protein sequence:

[illegible]

### NbL17g14340.1 jacalin−related_lectin_3 [MW = 71.29 kDa]

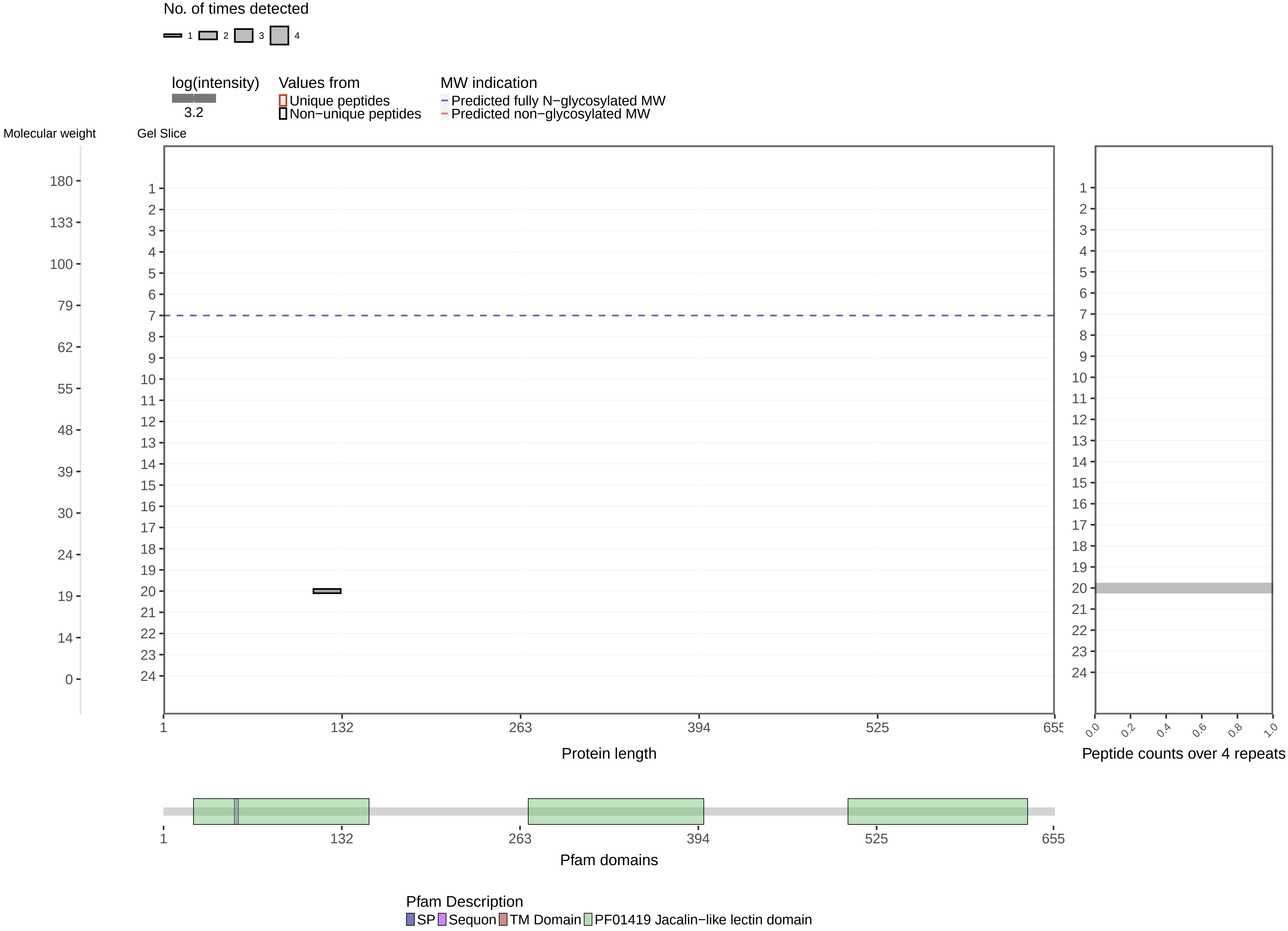


Protein sequence:

GGTGGFKTEKIIFDYPSEILTHITGYYGPAMIMGPNIIQSLTFHTTKGKHGPYGEEQGQQFSTKLKEGIIVGFHGRKGLFLDALGVHVLEGKVVSCLPPSPTQTSIKPSAASVPSLPAPAPPKSIKPDAAIVPSLPAPAPPKSIEPKPAFVTEVDDRPQWSFRLGKNGLTEEGVQRIVKDPAPYGPGPWGGDGGKPWDDGVFTGIKQIILTQSSEGICCIEIEYDRNGQSVWSVKHGANAGKCTNRVKLEYPHEVLTCITGFYGPISKDMGLKVIKSLTFHTTRRKFGPFGEELGTYI

NbL17g14280.1 coatomer_subunit_alpha−1−like [MW = 137.1 kDa]

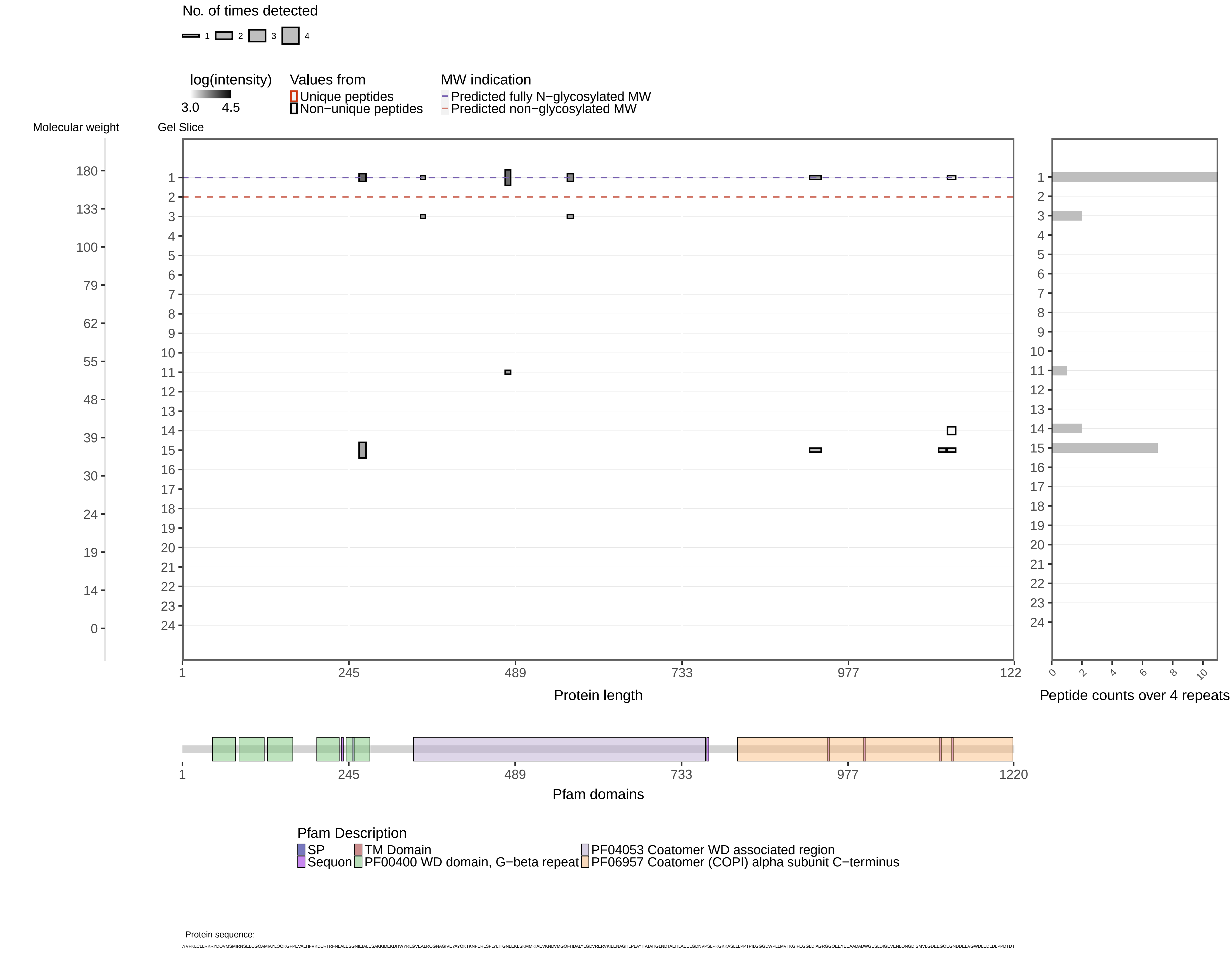

NbL17g14160.1 peroxisomal_24−dienoyl−CoA_reductase [MW = 31.88 kDa]

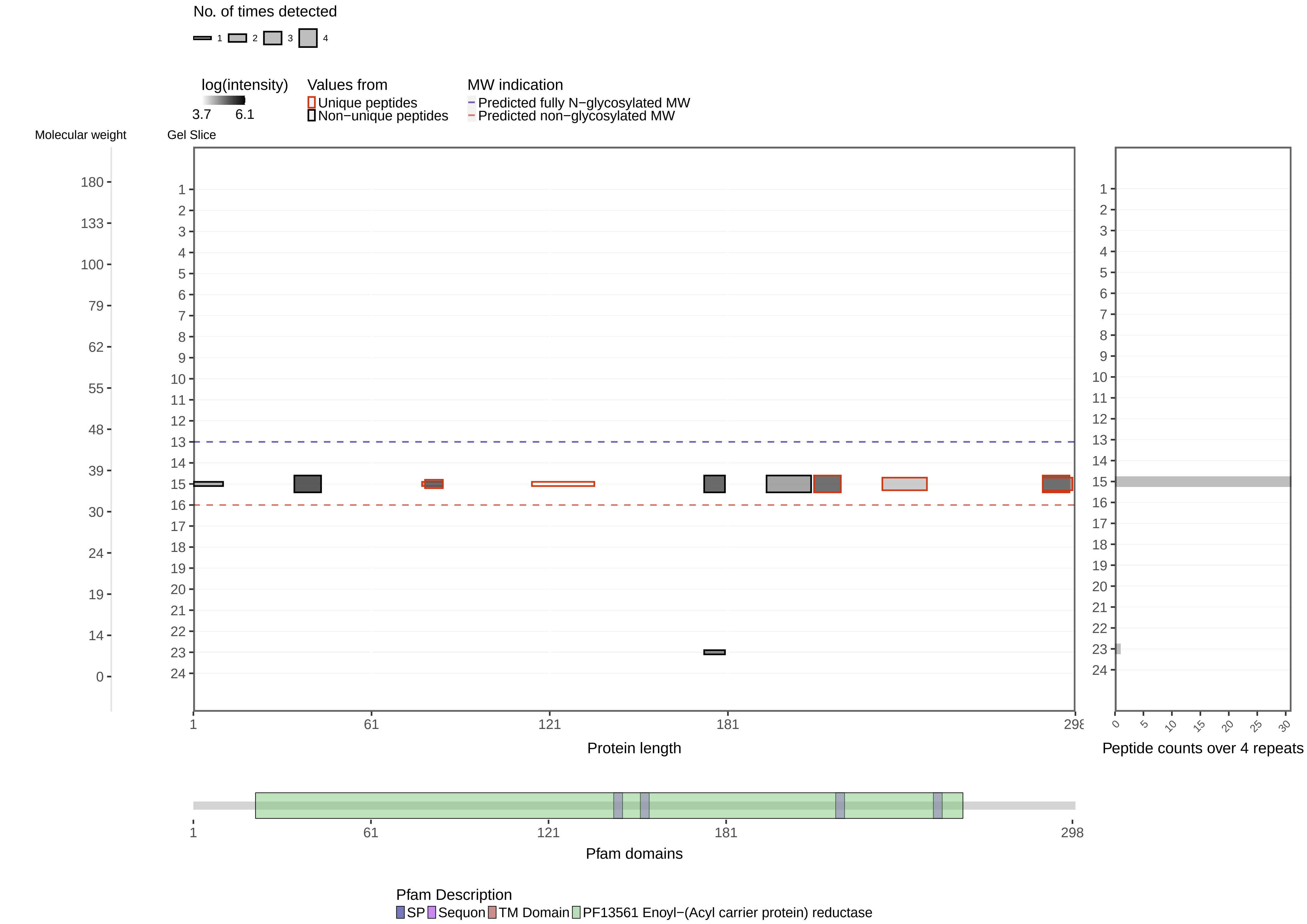


Protein sequence:

[illegible]KGGLGRNSTSGGIILNISATLHYTASWYOIHVAAAKAAVDAVTRNLALEWGTDYDIRVNGIAPGPIGDTAGMRKLVPDEINNKTREYMPLYKLGEKYDIAMAALYLASDAGKYINGTTIIVDGGLWLSRPRHLPKDAVKELSRTVEKRSRNAPVGVPSSKL*

### NbL17g14140.1 phosphoglycerate_kinase_chloroplastic [MW = 50.01 kDa]

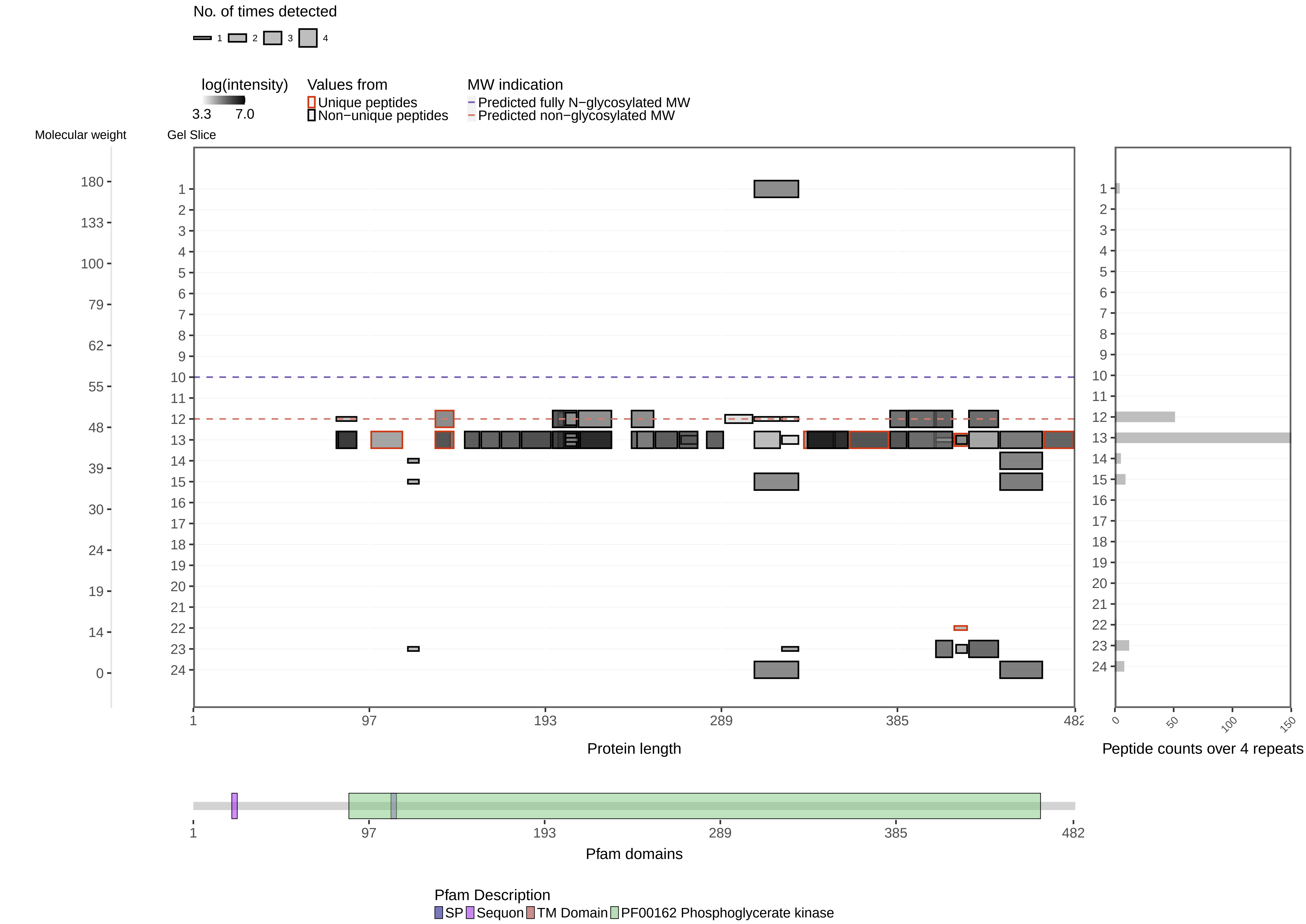


Protein sequence:

HRAHASTEGVTKFLKPSVAGFLLQKELDYLVGAVSNPKRPFAAIVGGSKVSSKIGVIESLLEKCDILLLGGGMIFTFYKAQGLSVGSSLVEEDKLELATSLLEKAKAKGVSLLLPSDVVIADKFAPDANSKIVPASAIPDGWMGLDIGPDSVKTFNDALDTTKTVIWNGPMGVFEFDKFAVGTEAIAKKLADLSGKGVTTIIGGGDSVAAVEKVGVASVMSHISTGGGASLELLEGKVLPGVVALDEADAPVAV*

NbL17g14120.1 thiosulfate/3−mercaptopyruvate_sulfurtransferase_1_mitochondrial−like [MW = 41.38 kDa]

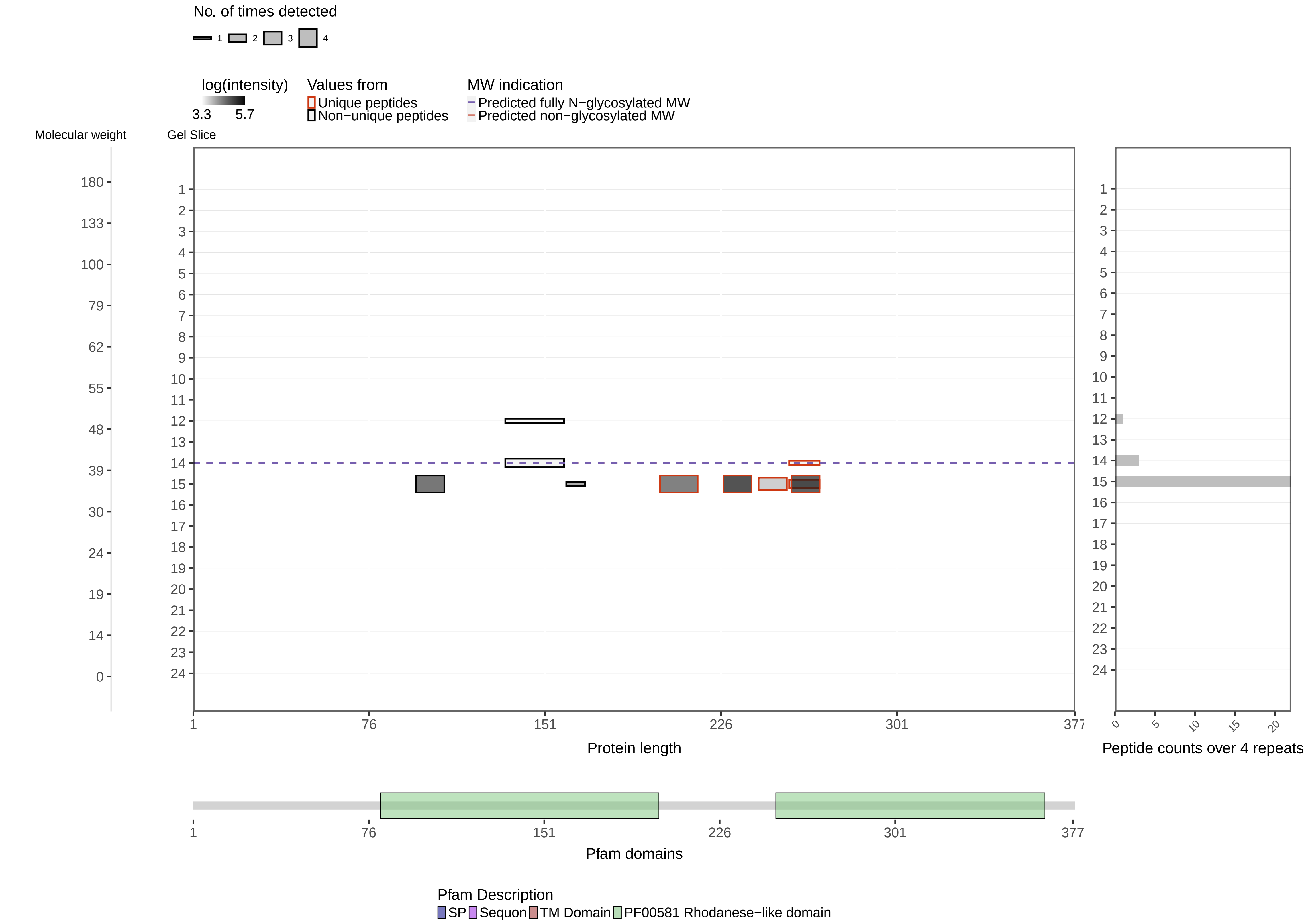


Protein sequence:

WWMFRVFGHDRVWVLDGGLPRWRASGYDVESSASGDAILKASAASEAIEKVYQGQTVAPITFRTKFQPNLVWTLDQVRKNIEETTYQHIDARSKARFDGVAPEPRKGIRSGHVPGSKCIPFAQMLDGSYTLLSTDELKKKFDQEGISLDKPVITSCGTGVTACILALGLHRLGKTDVPVYDGSWTEWGAHPDTPVSTSEA*

NbL17g14000.1 porphobilinogen_deaminase_chloroplastic−like [MW = 39.26 kDa]

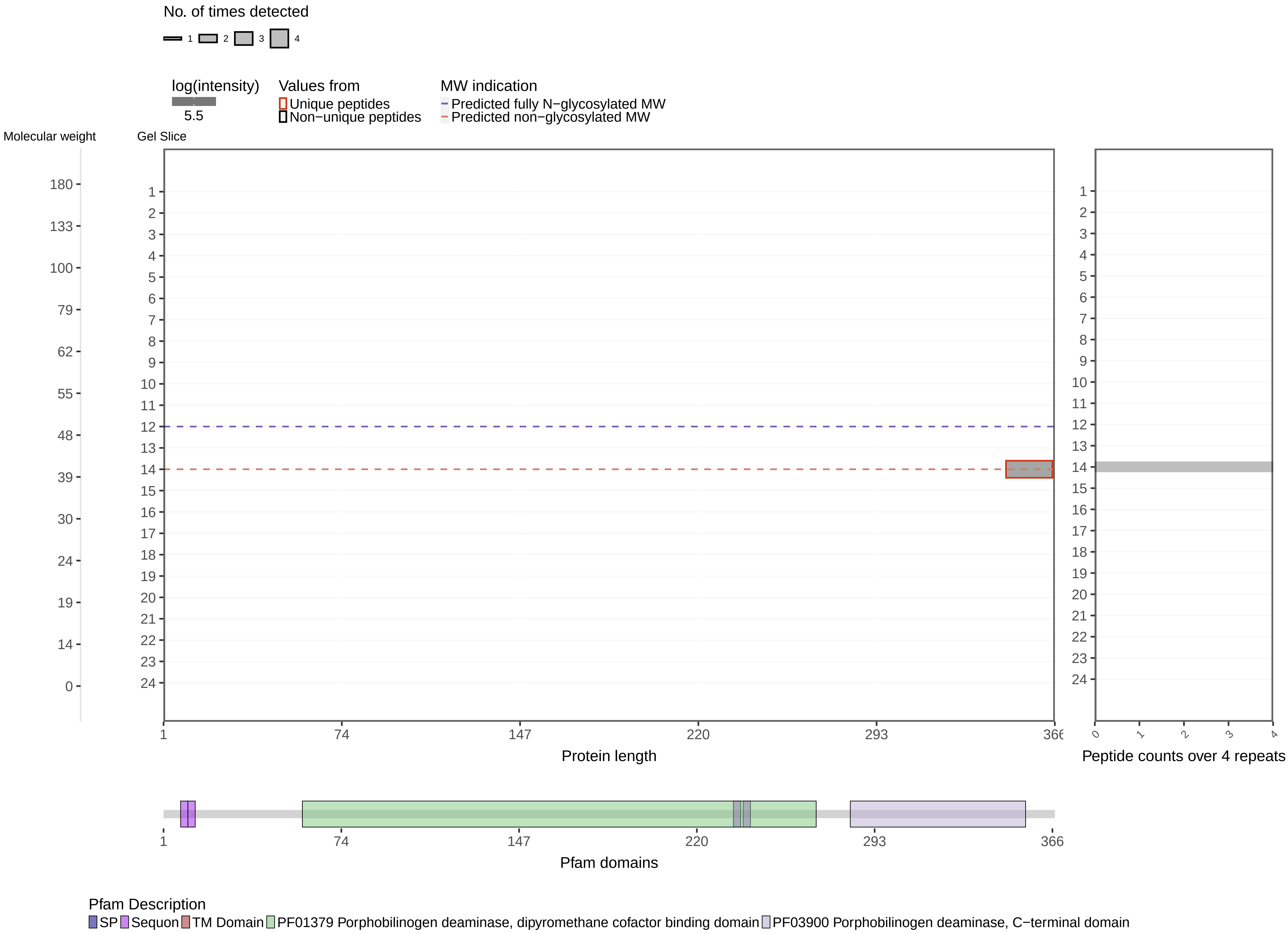


Protein sequence:

AQLPSGSTIGTASLRRKSQILHRYPSLNVLENFRGNVQTRLRKLNEGVVQATLLALAGLKRMNMTENVTSILPIEDMLPAVAQGAIGIACRTDDETMANYIASLNHEETRLAVACERAFLKTLDGSCRTPIAGYACRGEDGDCIFKGLVASPDGTRVIETSRKGPYTFEDMIRLGEDAGKELLAEAGPGFFGN*

NbL17g13990.1 protein_Iojap_chloroplastic [MW = 25.89 kDa]

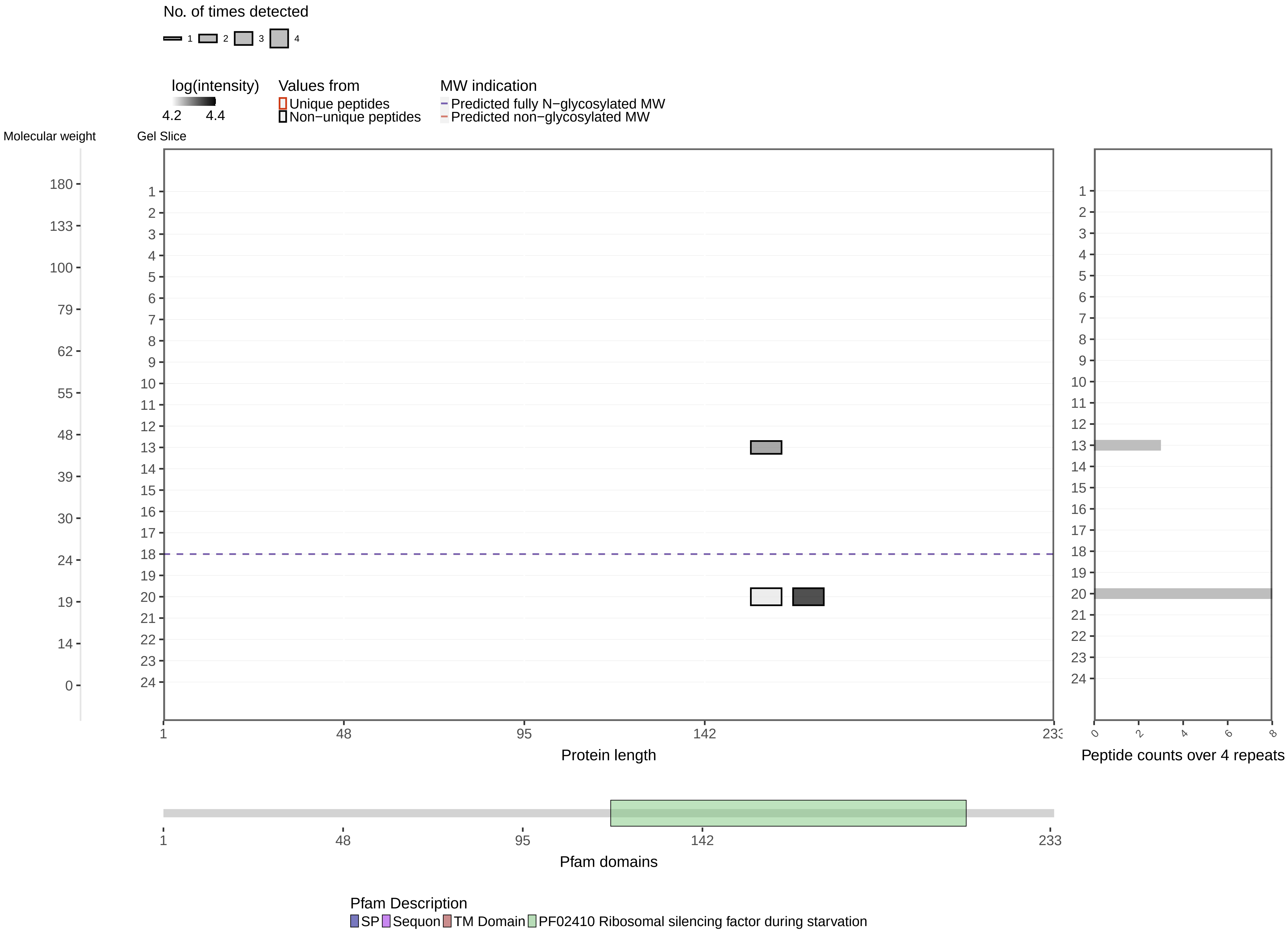


Protein sequence:

TEEIDDDAESLSFAVAVAKVASDVKAGDIKVLFVKPLVYWTRFFIIATAFSRPOIDAIGTRIRDMAEVQYGRVASGDFKPNSWTLLDFGDVVVHIFLPQQREYYNLEEFYGNAAPIELPFENQQLRGSKGY*

NbL17g13950.1 ribosome−binding_ATPase_YchF−like [MW = 45.68 kDa]

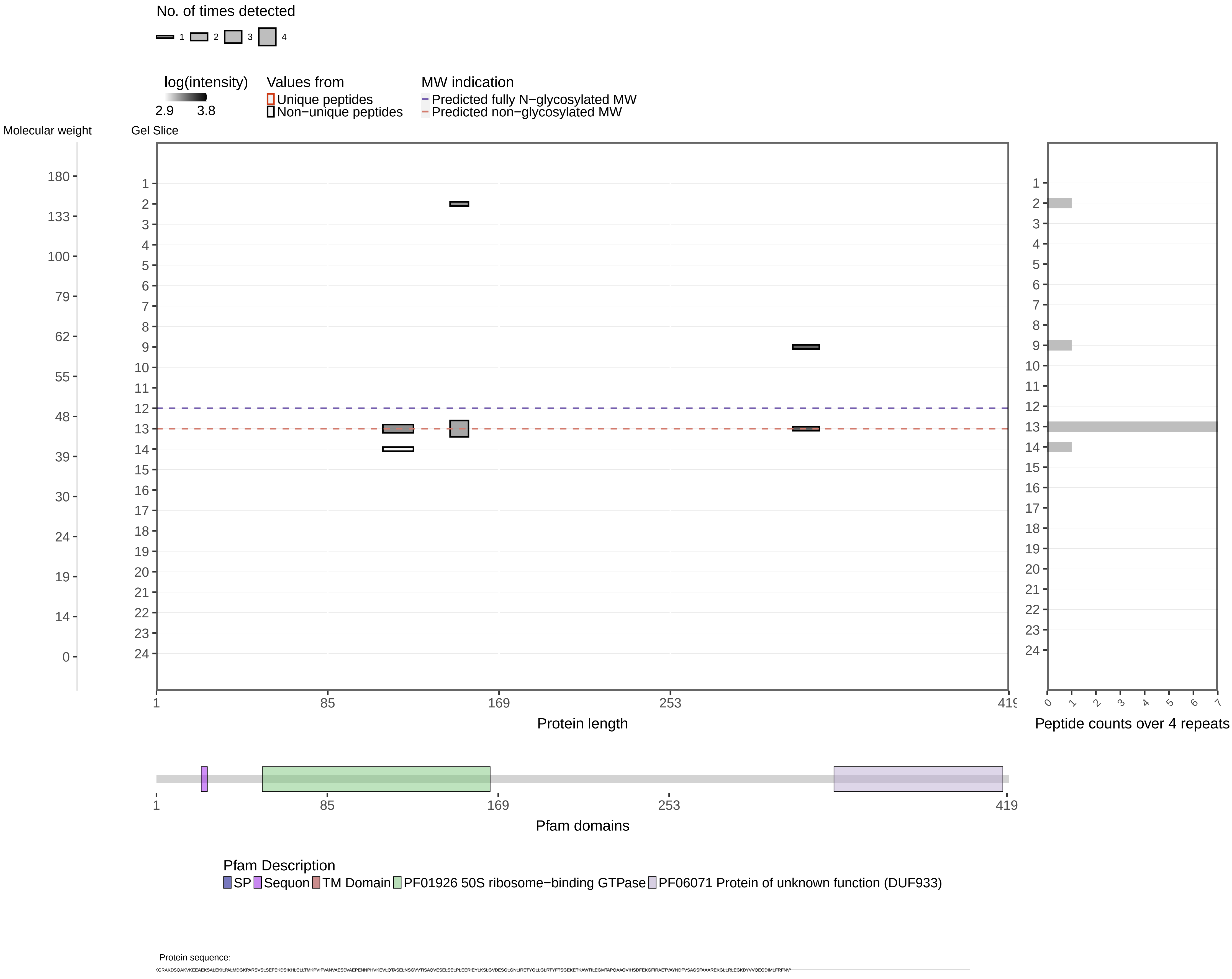

NbL17g13940.1 peptidyl−prolyl_cis−trans_isomerase_CYP59 [MW = 69.17 kDa]

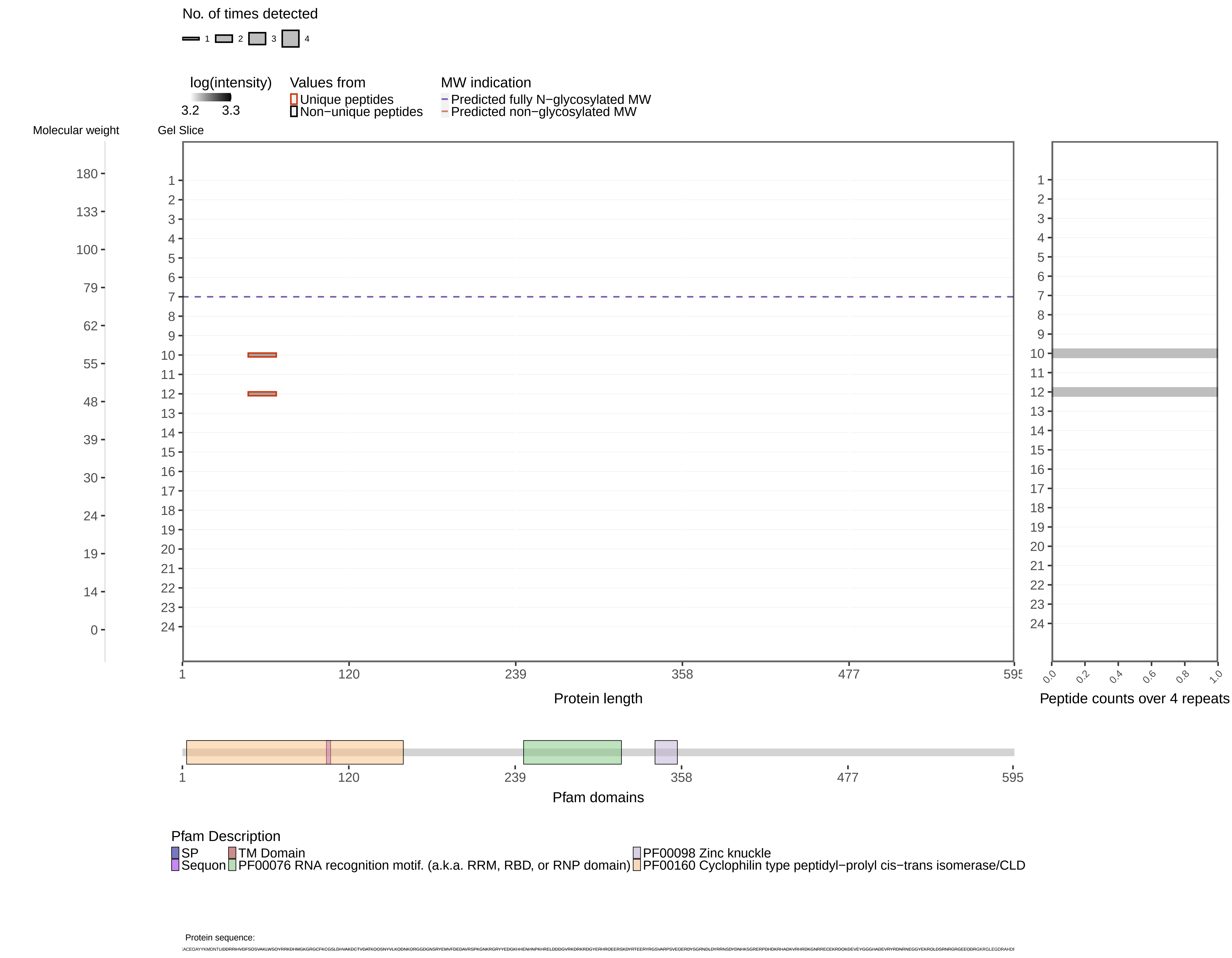

### NbL17g13920.1 uncharacterized_LOC107785455 [MW = 17.95 kDa]

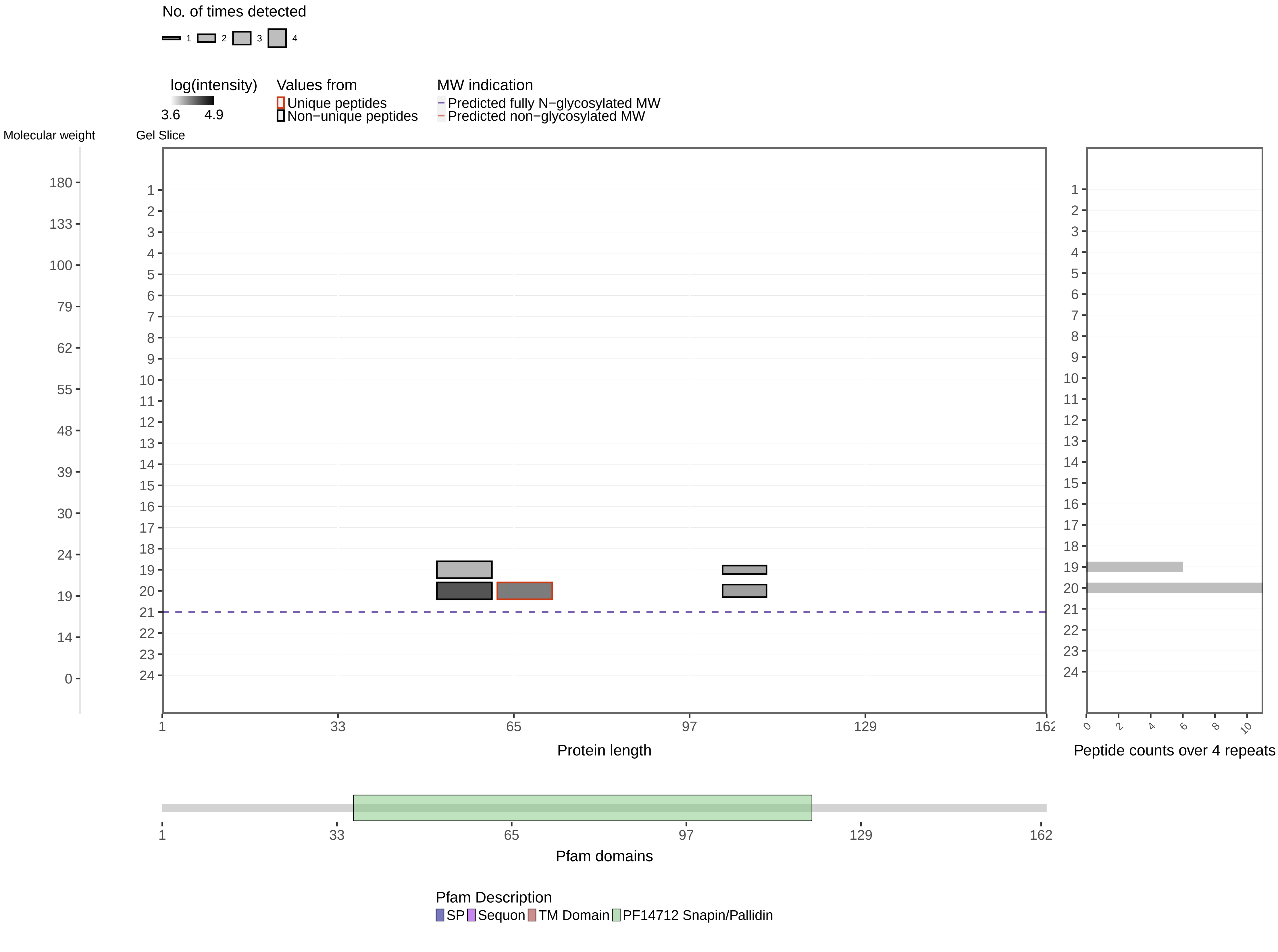


Protein sequence:

SLDRLTGELDKLLEDAPLPFIMQHAARISGVRKRITSLNSLLKSIQRRLDNIDHTLSAGLIHVNRGWWARPRESKQSSLTRSWRRFKSLSQCQ*

### NbL17g13870.1 photosystem_II_10_kDa_polypeptide_chloroplastic [MW = 14.11 kDa]

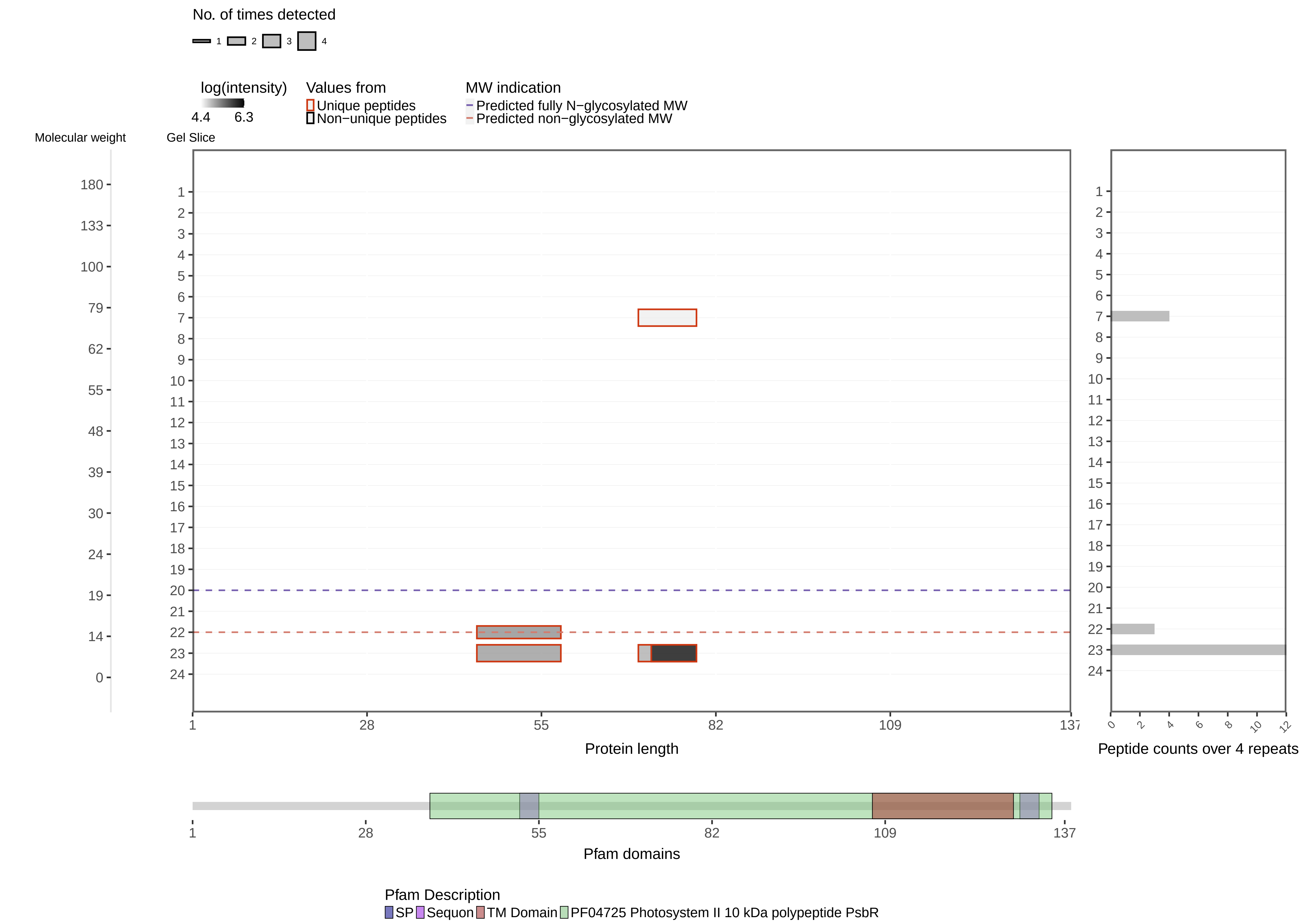


Protein sequence:

SLRDGVDASGRKPKGKGVYQFVDKYGANVDGYSPIYNTDDWSPSGDVYVGGTTGLAIWAVTLVGILAGGALLVFNTSALAQ*

### NbL17g13800.1 beta−ureidopropionase [MW = 46.58 kDa]

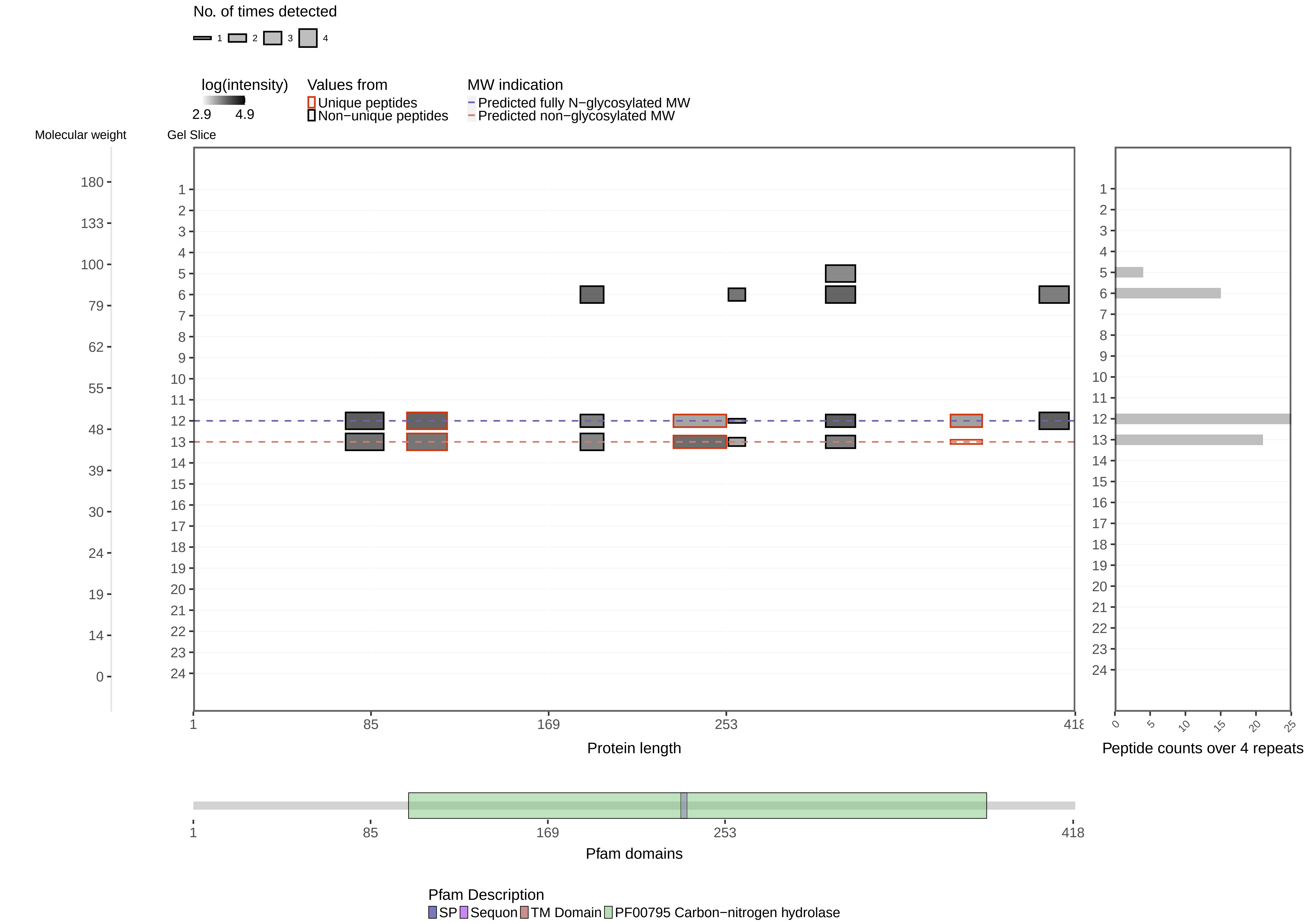


Protein sequence:

YHGETLWNTAVIIGNHGNIIGKHRKNHIPRVGDFNESTYYMEGNTGHPVFDTAYGKIAVNICYGRHHPLNWLAFGLNGAEIVFNPSATVGELSEPMWPIEARNAAIANSYFVGSINRVGTEVFPNPFTSGDGKPQHADFGHFYGSSHFSAPDSSCTPSLSRHKDGLLISEMDLNLCRQLKEKWGFRMTARYEVYADLLARYVKHDFEPQVISDPLLYKNA*

### NbL17g13780.1 uncharacterized_protein_C6C3.02c−like [MW = 14.27 kDa]

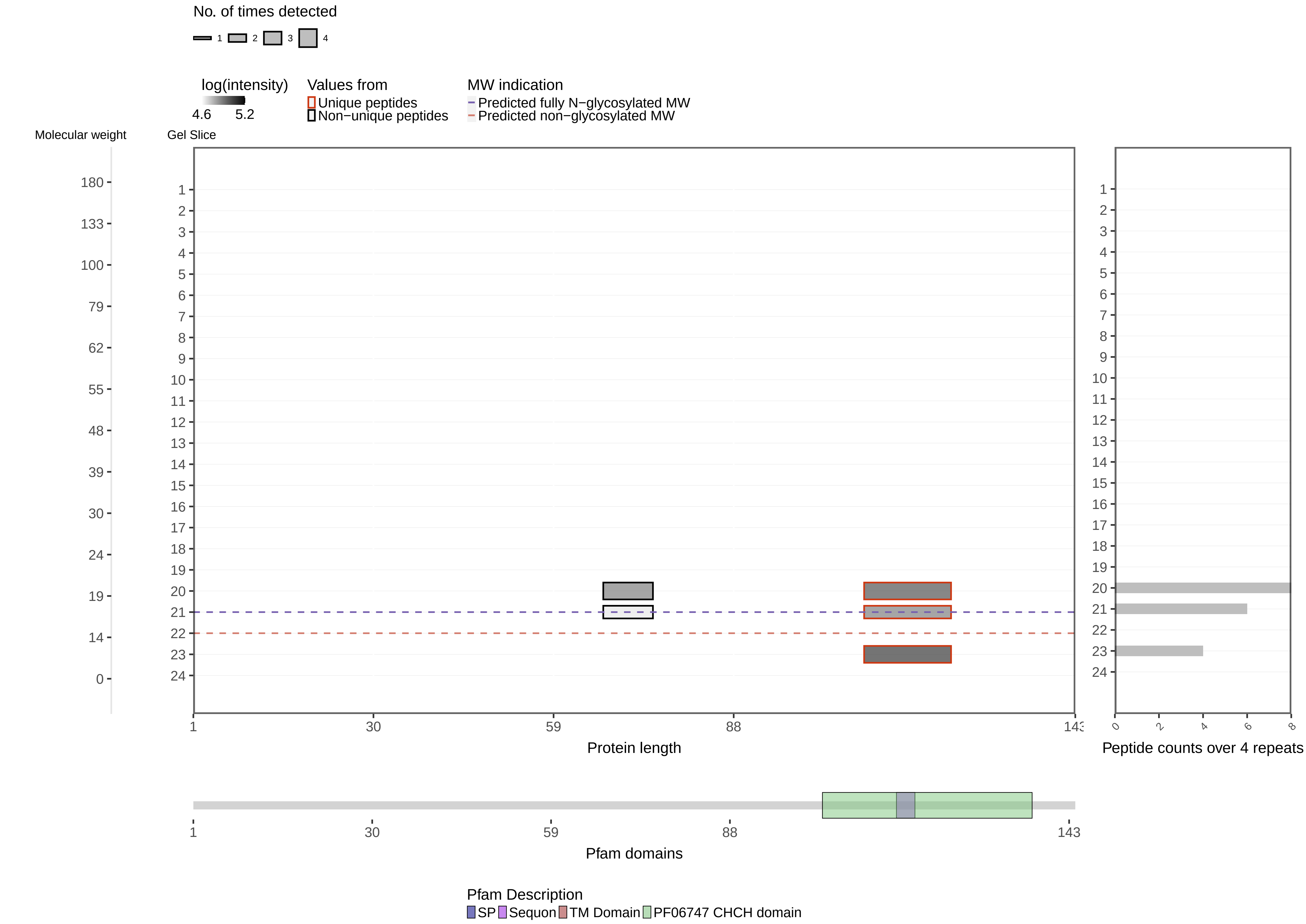


Protein sequence:

TGSAVAHRAVDAVMGPRTIQHETVASAVPAAAAAPTTVGAASDACGMHSKAFQDCINSSGSDIGKCQFYMDMLSECRRNSMLNA*

### NbL17g13730.1 3−oxo−Delta(45)−steroid_5−beta−reductase−like [MW = 44.37 kDa]

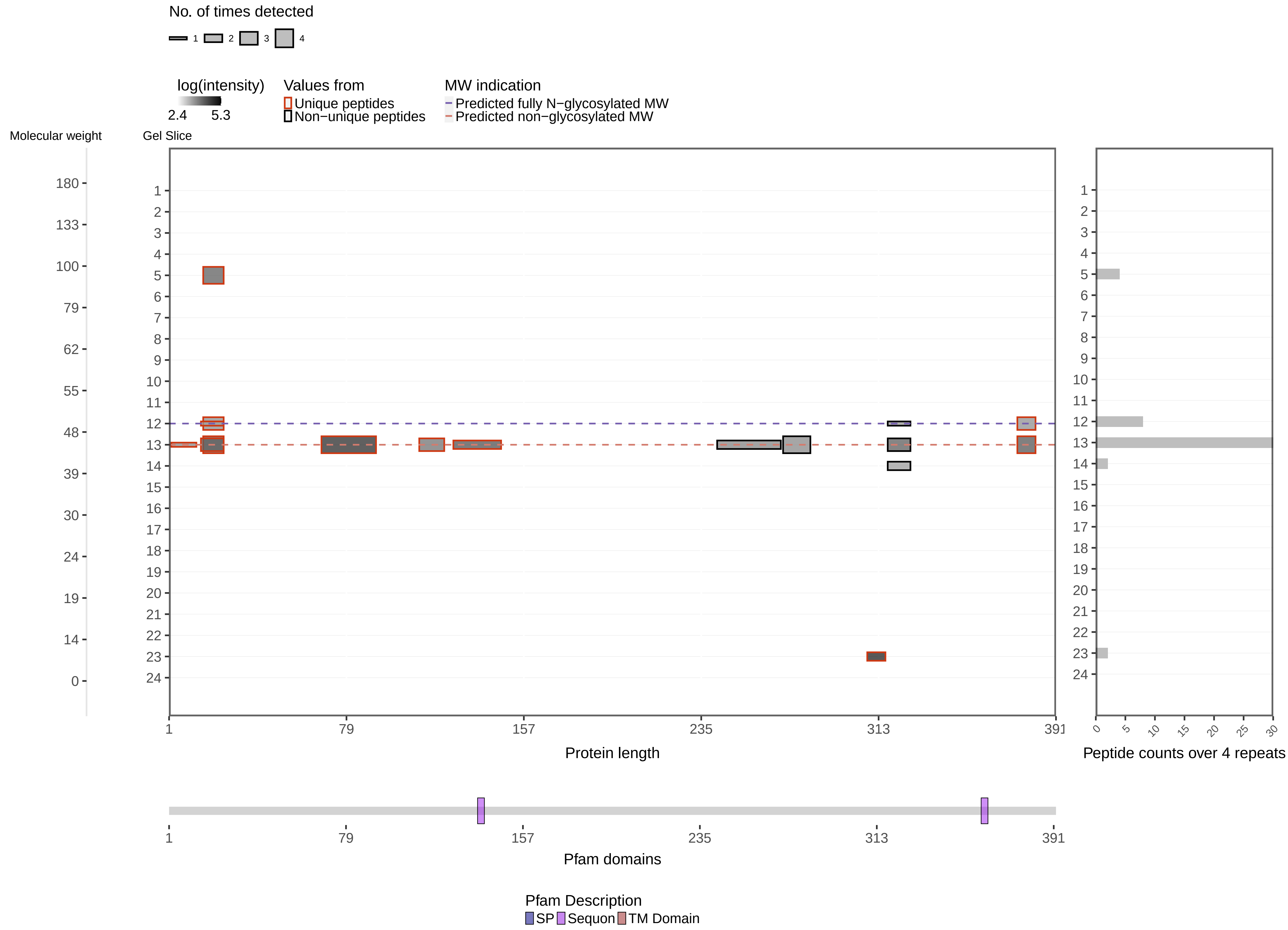


Protein sequence:
.EDILFEEVKKKEKLTWSVHRPGLIFGFSPYSLMNIIASLCVYAAICKHEGLKLKFPGVKSAWDGYSDCSDADLIAEHQIWASVDPNAKNEAFDVSNGDVFKWKHLWGILAEEFEVEAEEFDENKKGWRLEEMMKDKGEIWDEIVKENGLVPNKLEQVGGWWFADIVLNDECPLDTMNKSKEHGFLGFRNSKNAFISWIDKIKAYKIVP*

### NbL17g13720.1 3−oxo−Delta(45)−steroid_5−beta−reductase−like [MW = 43.71 kDa]

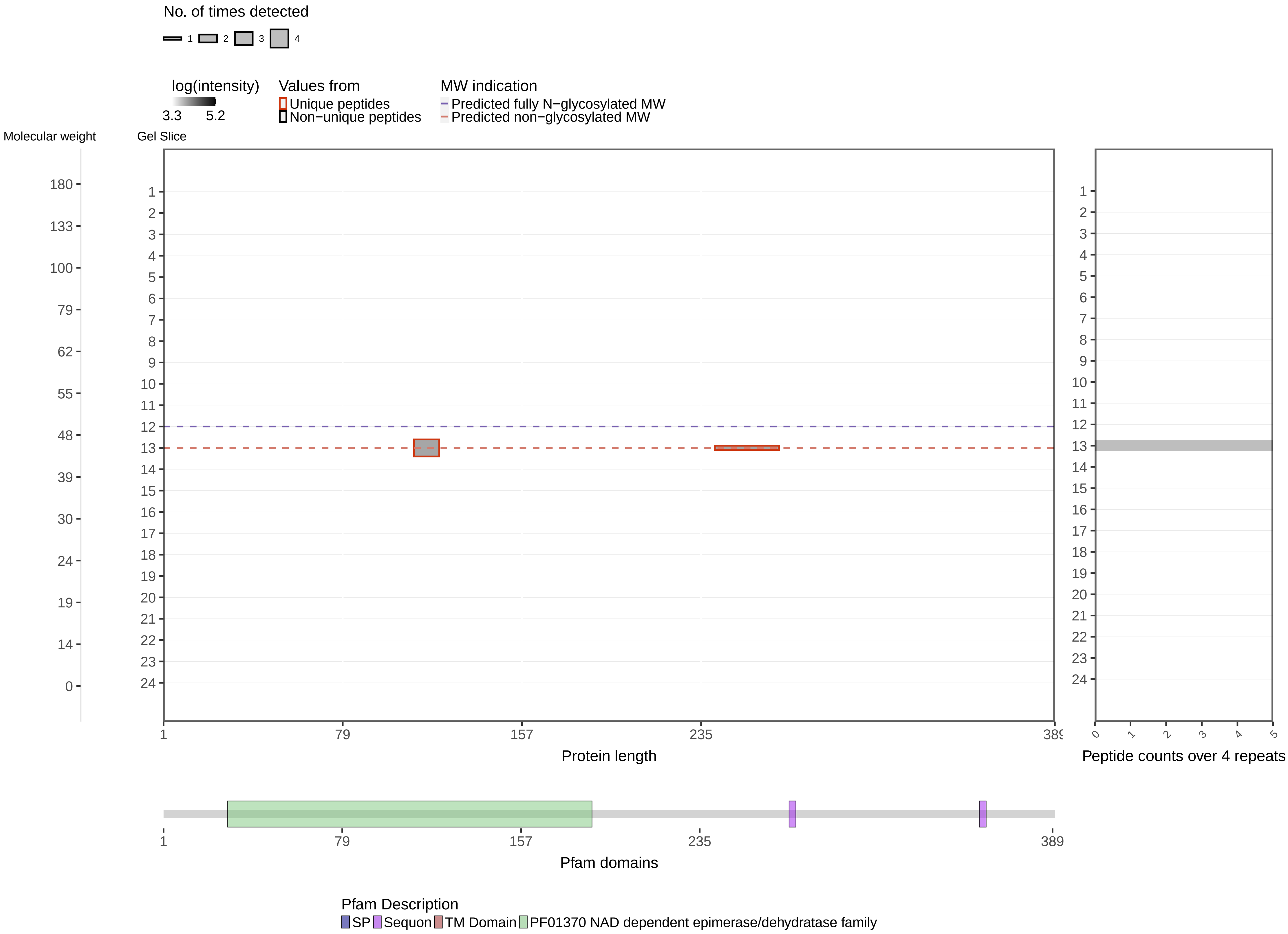


Protein sequence:

DILFKEVEKKEGLTWSVHRPGTIFGFSPYSLMNIVGTLCVYAAICKHEGLPLKFPGVKPAWGGYSDCSDADLIAEHQIWAAVDPYAKNEAFNVSNGDVFKWKHFWKVLAEQFGVEATEFDEVNRCTLAEMMKDKGPVWDEIVKENGLTATKLEDVGVWWFVDLILAGDCPLDTMNKSKEHGFLGFRNSQKAFISWIDKVKAYKVVP*

### NbL17g13670.1 ATP−dependent_Clp_protease_proteolytic_subunit−related_protein_1_chloroplastic [MW = 44.92 kDa]

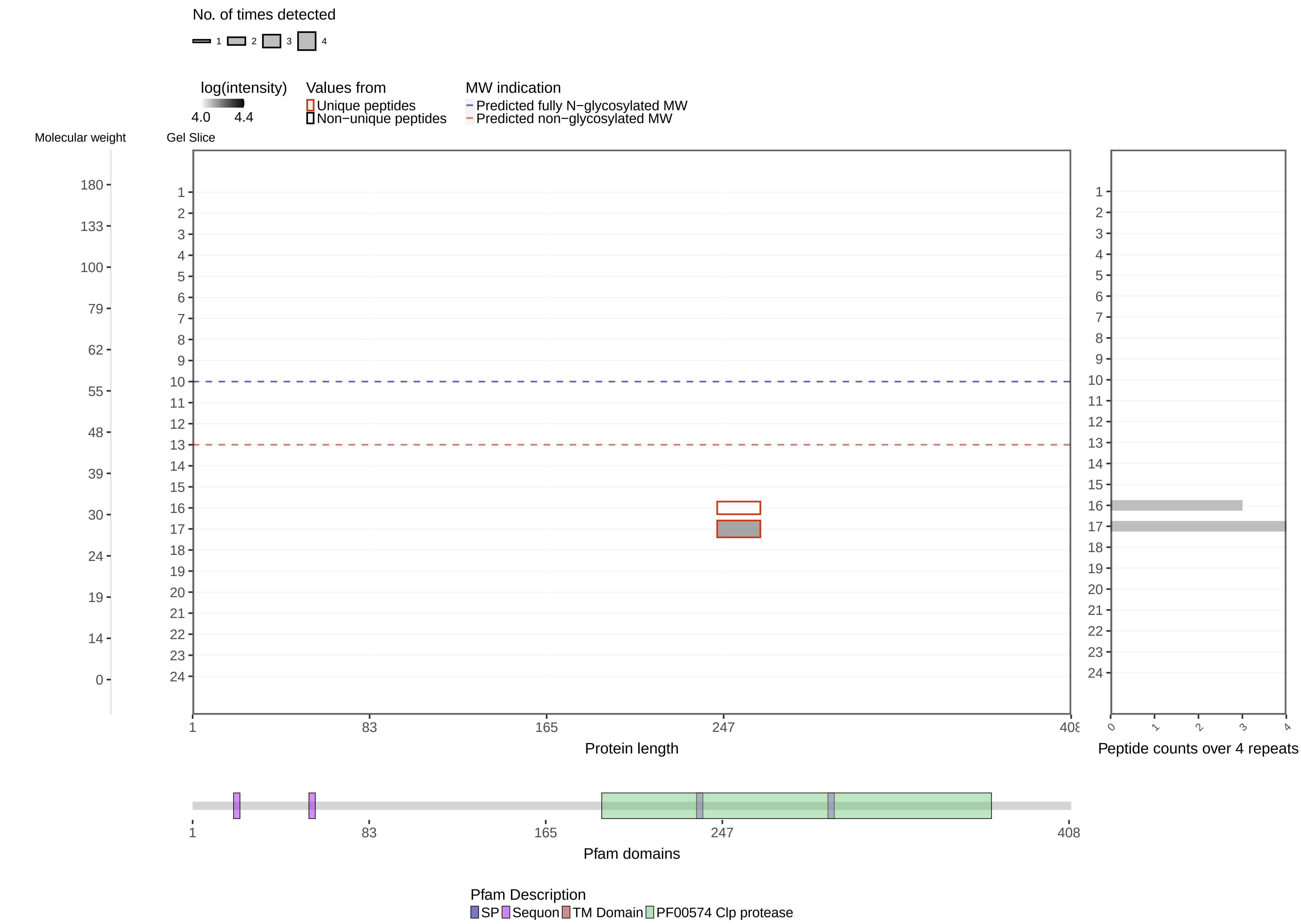


Protein sequence:

PPDLPSLLLDARIVYLGMPIVPAVTELLVAQFMWLDYDNPSKPIYLYINSSGTQNEKMETVGSETEAYAIADTMAYCKSDIYTVNCGMAYGQSAMLLSLGKRGFRAMQPNSSTKLYLPKVSKSSGAVIDMWIKAKELDANTEYYLELLAKGIGKPKEEIEKDIQRPKYLRAQEAIDYGIADKIIDSRDNAFEKRVYDEILAQSRAMRKAGPGAQAAPSGFR*

NbL17g13420.1 pectin_acetylesterase_10−like [MW − SP = 43.82 kDa]

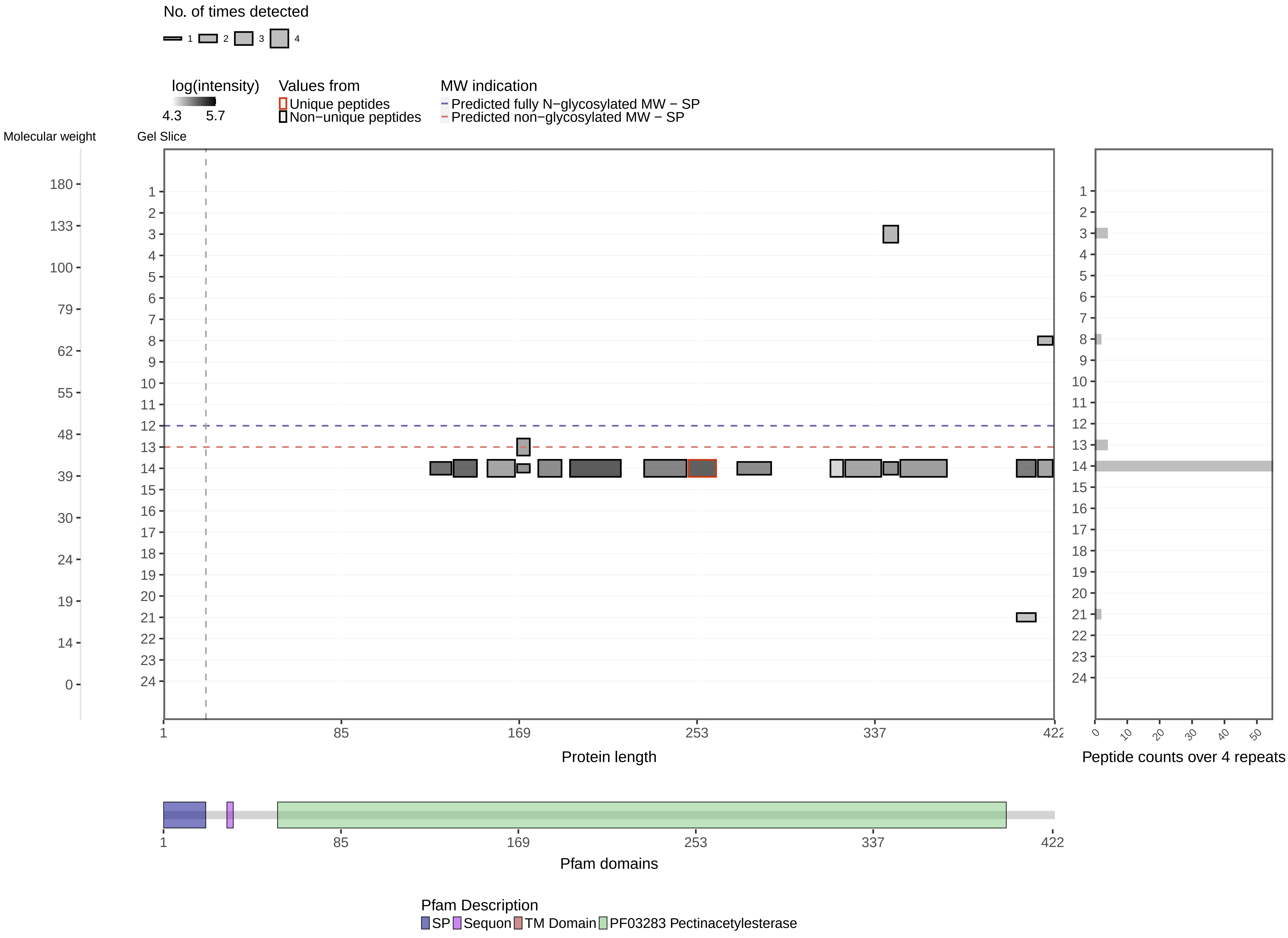


Protein sequence:

ALLSGCSAGGLASILHCDEFRTLFPSSTKVKCLSDAGLFMDATDVSGGHALRNLYQGVVNVQGLQKTLPSTCTNKLDPTSCFFPQNLIGNIKTPLFLLNAAYDSWQVQSSLAPPVADPKGLWHDCKLNNEQCSASQIQFLQGFRNDMLNAVKGFAASTQNGLFINSCFAHCQSERQDTWFADDSPLIDNKPVALAVGDWYFDRAGVKSIDCAYPCDKTCHNLVFK*

NbL17g13300.1 endoglucanase_25−like [MW = 76.89 kDa]

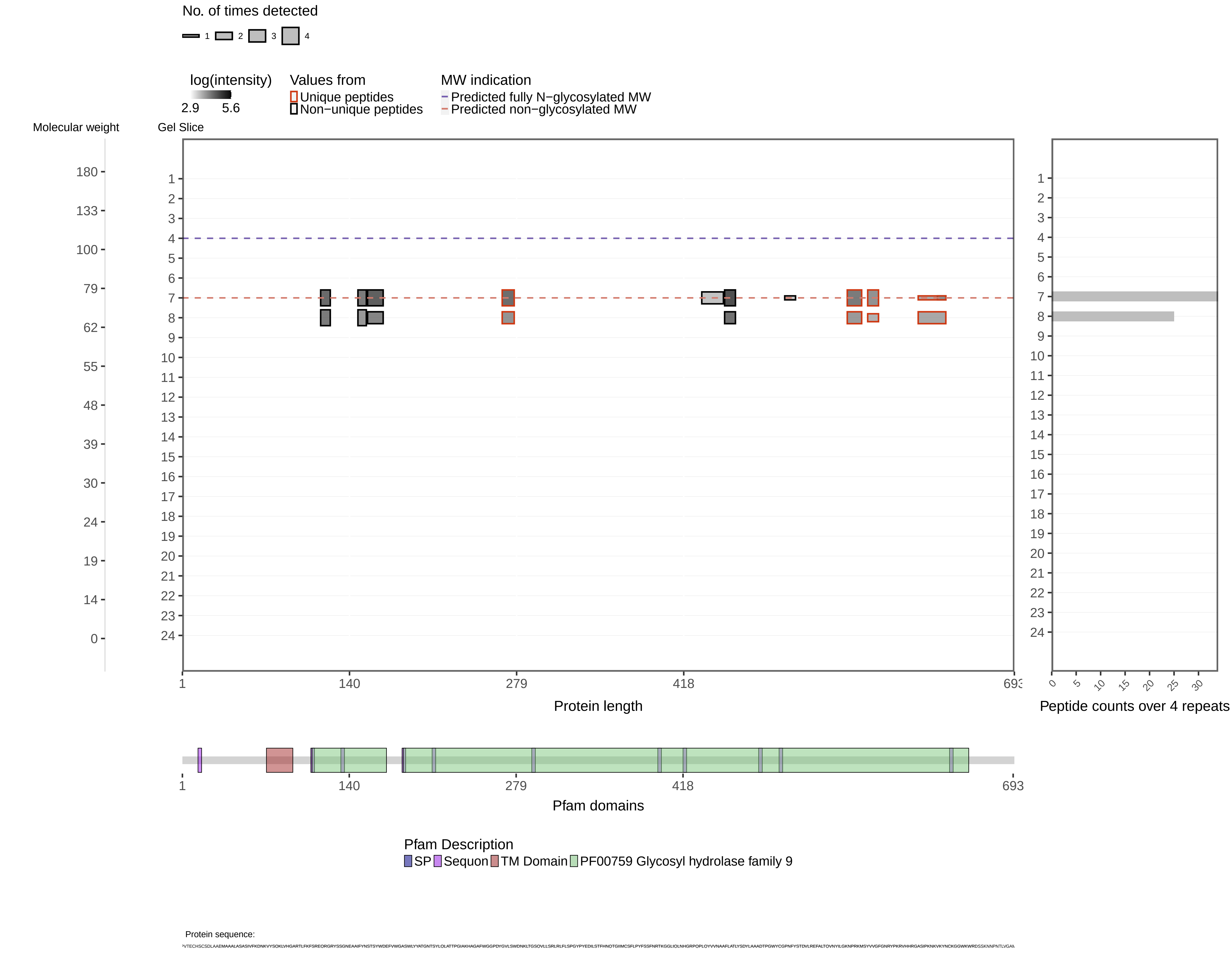

### NbL17g13110.1 microtubule−associated_protein_70−2−like [MW = 69.02 kDa]

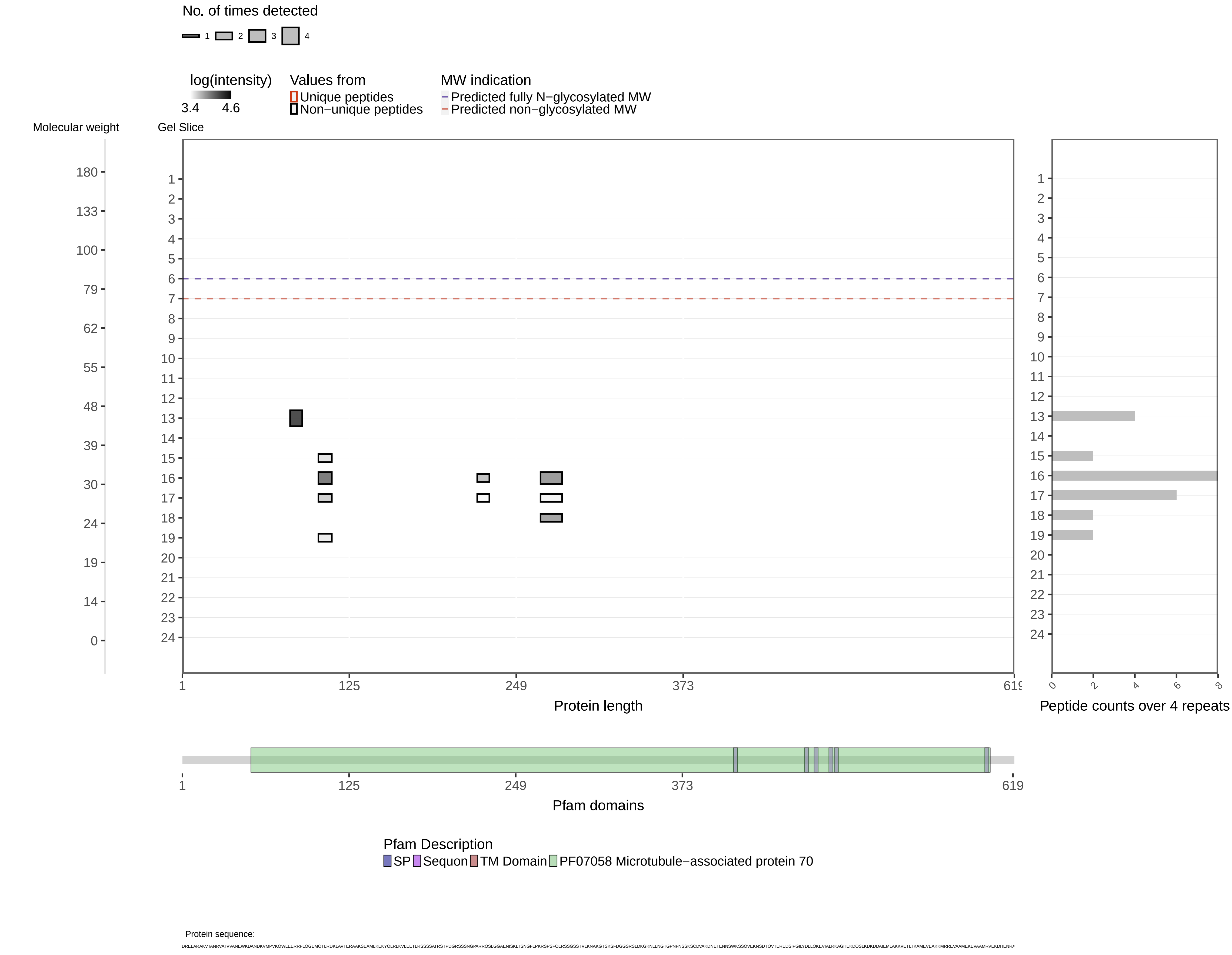

### NbL17g13050.1 aspartate−semialdehyde_dehydrogenase−like [MW = 44.42 kDa]

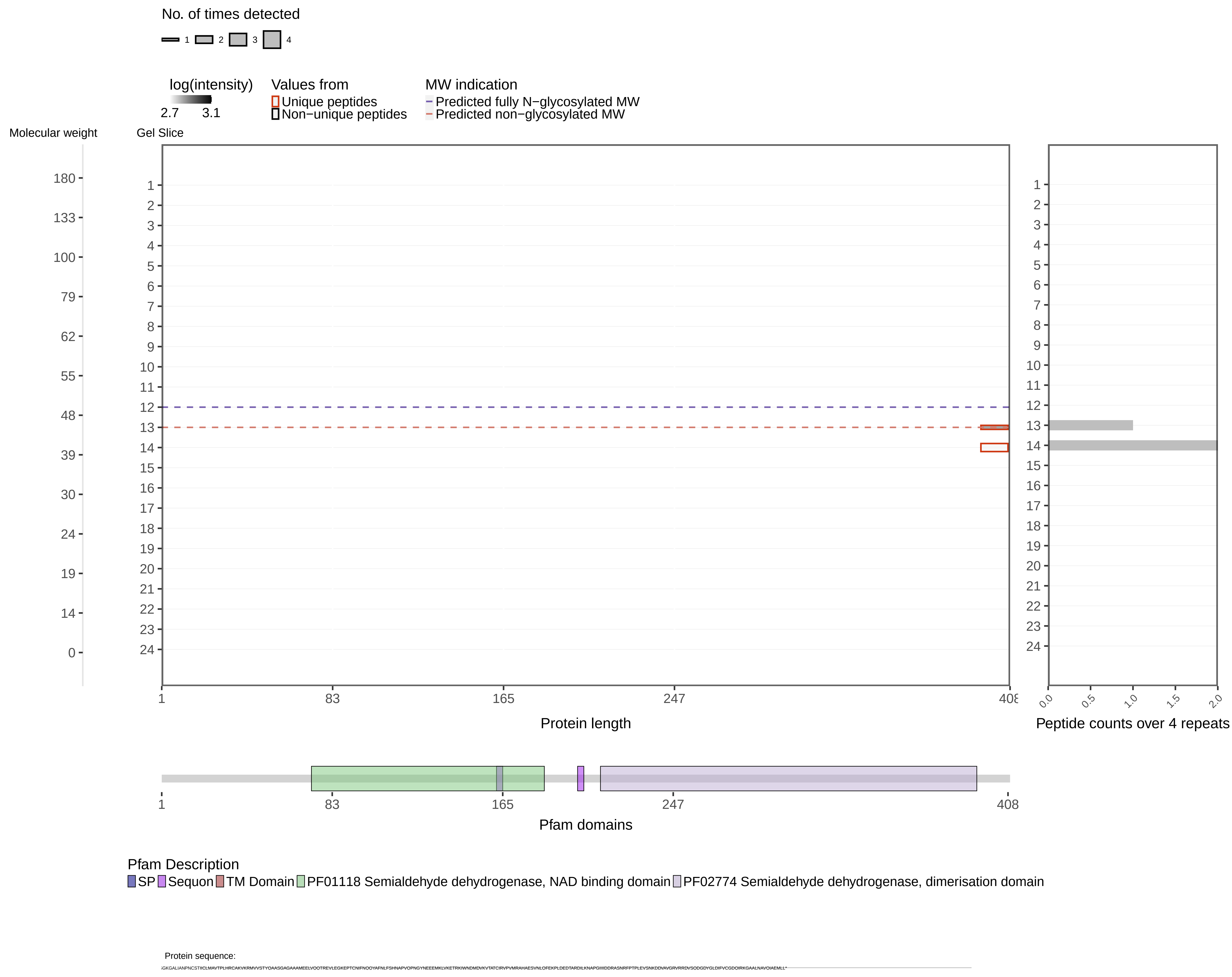

NbL17g12900.1 G−type_lectin_S−receptor−like_serine/threonine−protein_kinase_At1g11330 [MW = 94.54 kDa]

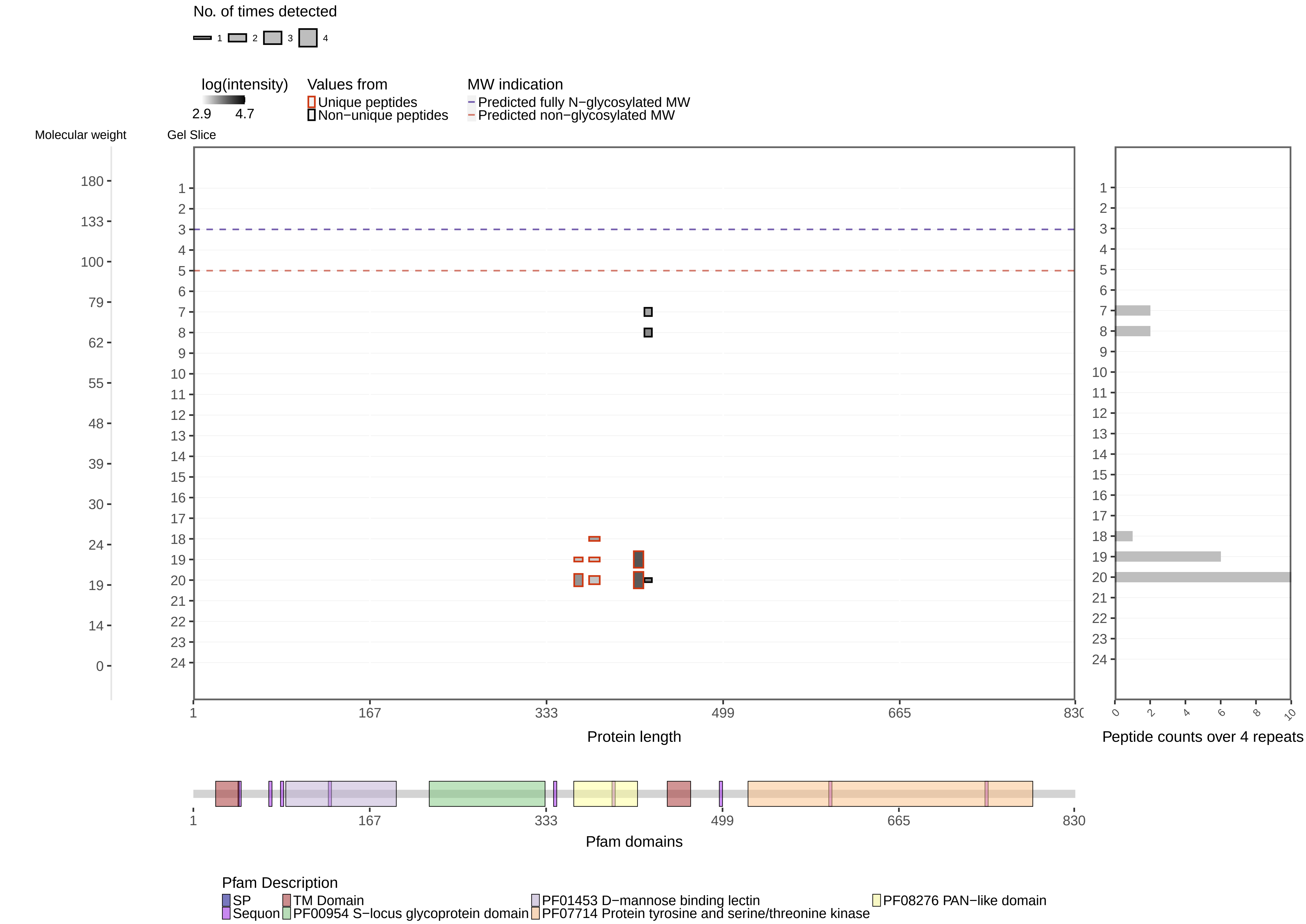


Protein sequence:

[illegible]

NbL17g12790.1 NADH_dehydrogenase_[ubiquinone]_1_beta_subcomplex_subunit_7−like [MW = 11.22 kDa]

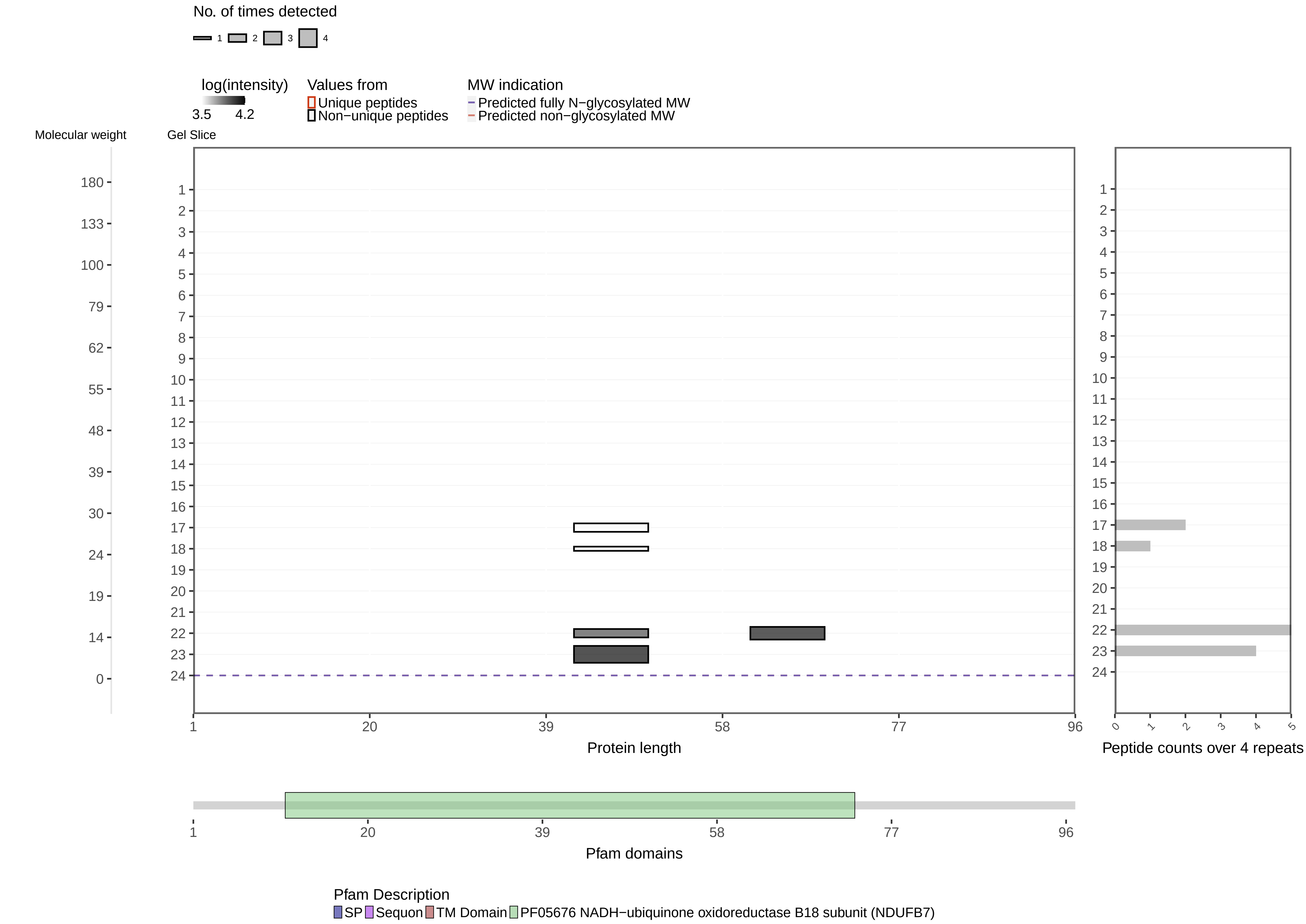


Protein sequence:

PLNKCRQSEFYLPWKCEDERHSYEKCEYELVMERMLQMQKIRQQQKGQQSIPLFPKTANA*

NbL17g12650.1 uncharacterized_LOC109243108 [MW = 31.09 kDa]

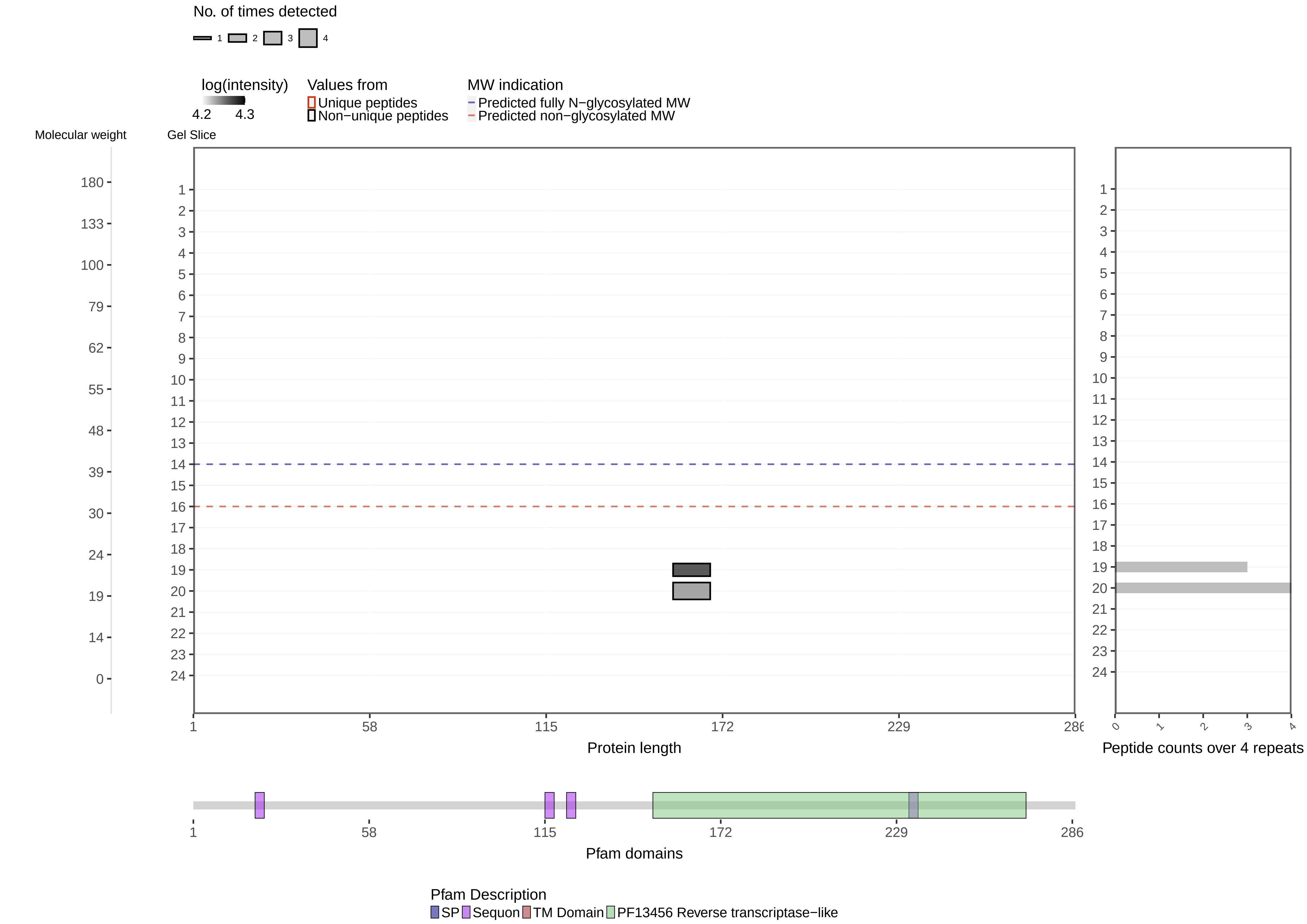


Protein sequence:

IIKLDLPKGDQALSSGRSCTLEFDGASKGNPGQAGAGAVVRADDGSLICRLREGLGIATTSVAEYRGFILGLKYAHSKGFTSIRAQGDSKLVCMQIQGLWKVKNKNISTLFQQAKQLKDRFLSFRIIHVLRESNCDADQQANLAVGLPDGHIQEEIEK*

NbL17g12600.1 ADP−ribosylation_factor_2 [MW = 19.5 kDa]

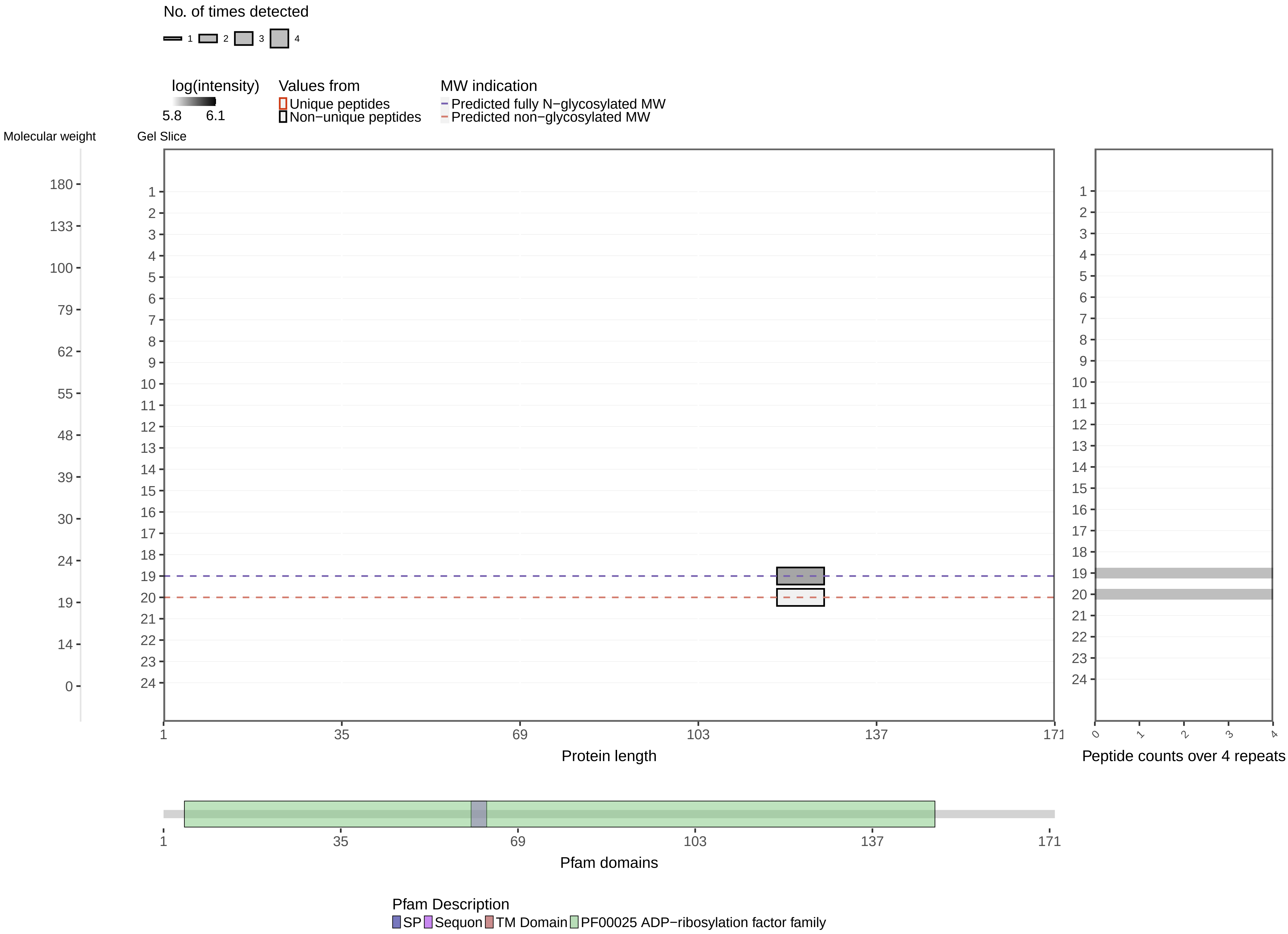


Protein sequence:

RPLWRHYFQNTQGLIFVVDSNDRDRVVEARDELHRMLNEDELRDAVLLVFANKQDLPNAMNAAEITDKLGLHSLRQLSLVHPEHLCNFWRGTLRGA*

NbL17g12510.1 succinate−−CoA_ligase_[ADP−forming]_subunit_alpha−2_mitochondrial−like [MW = 35.19 kDa]

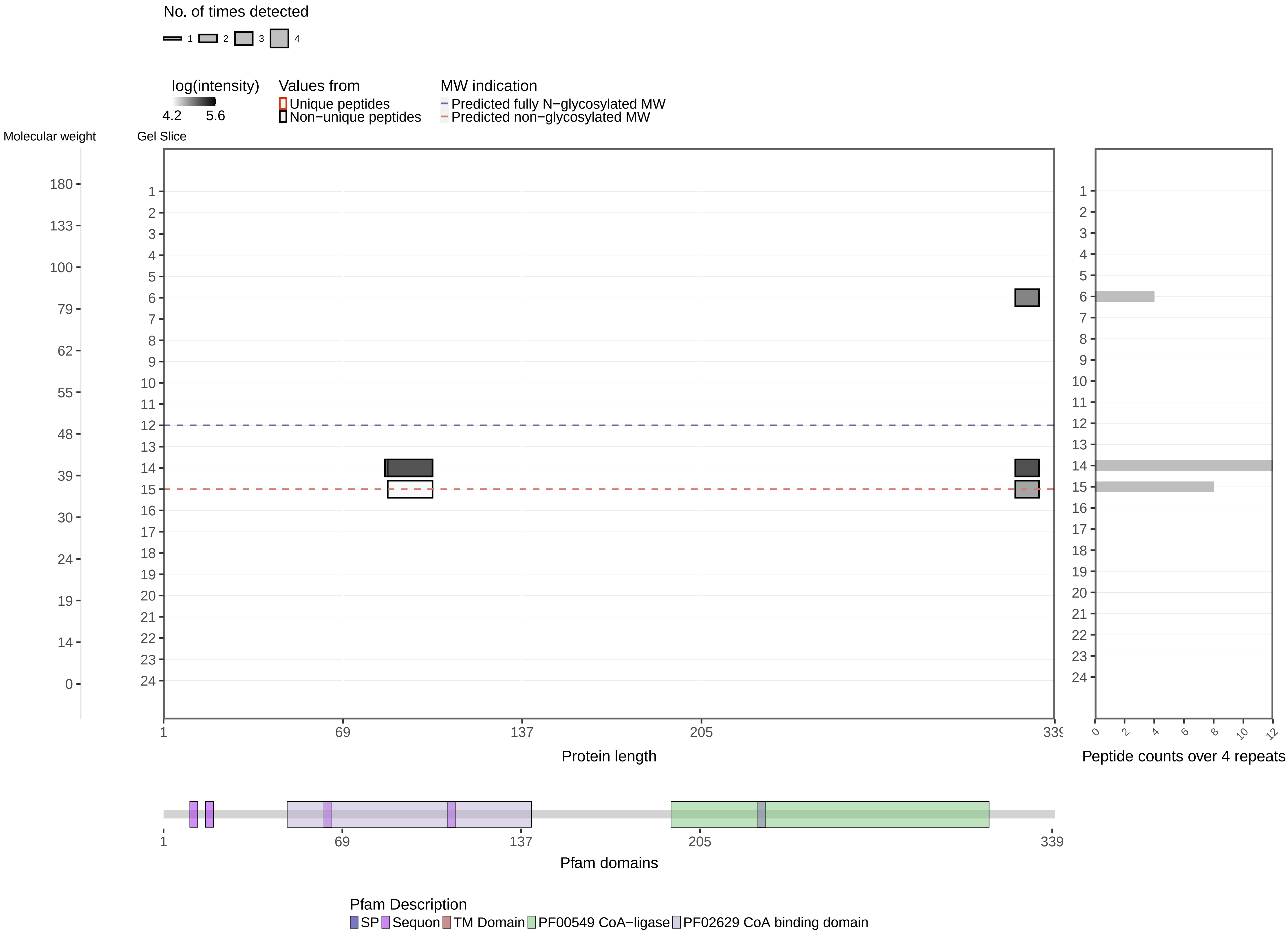


Protein sequence:

KKQSKTRLIGPNCPGIIKPGECKIGIMPGYIHKPGRIGIVSRSGTLTYEAVFQTTAVGLGQSTCVGIGGDPFNGTNFVDCLEKFVADPQTEGIVLIGEIGGTAEEDAAAFIKESGTQKPVVAFIAGLTAPPGRRMGHAGAIVSGGKGTAQDKIKALKEAGVTVCESPAKIGASMLDVFKQRGLV*

### NbL17g12450.1 thioredoxin_X_chloroplastic [MW = 20.73 kDa]

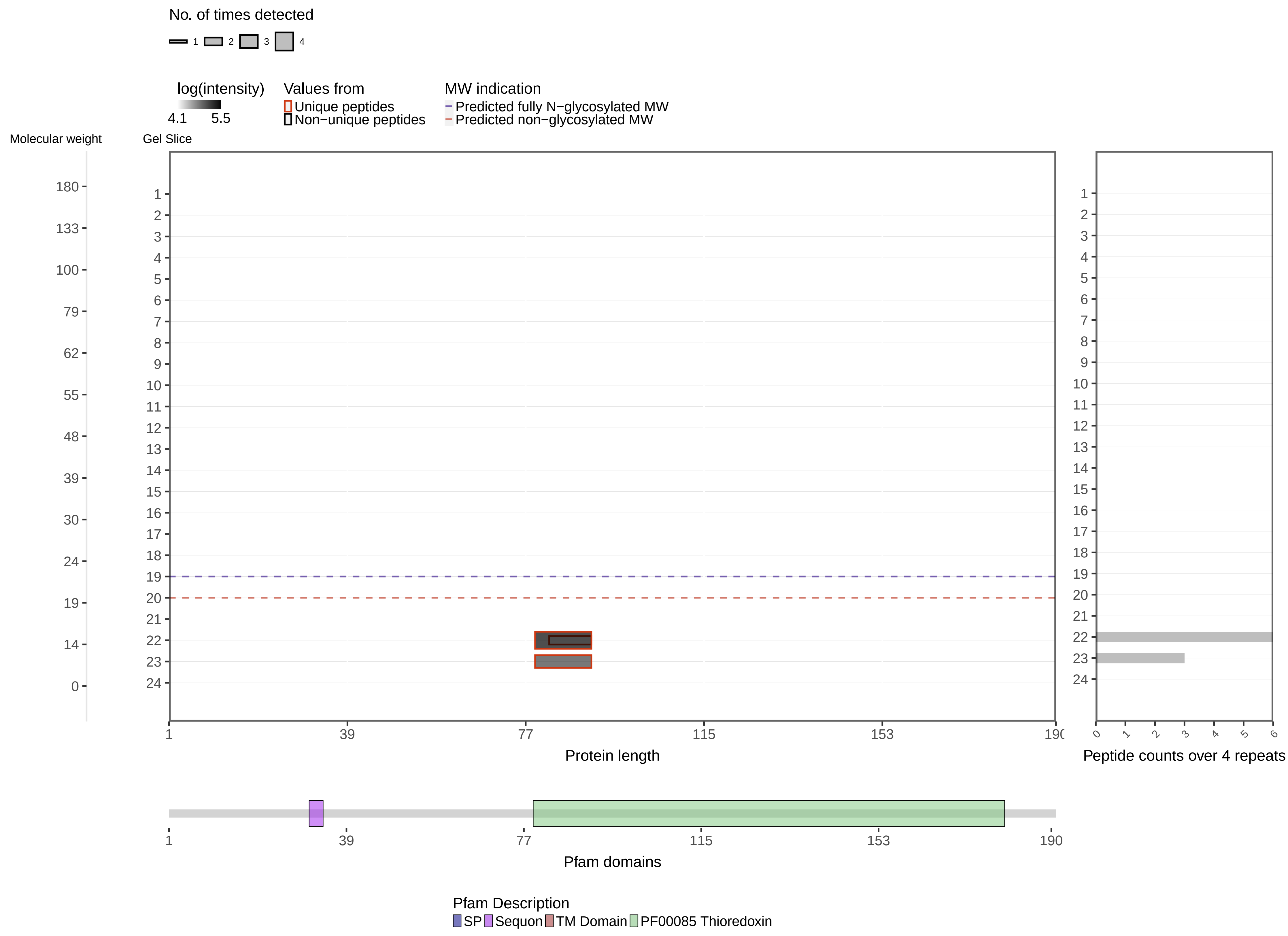


Protein sequence:

KEDEFSDVVLKSDRPVLVEFVATWCGPCRLIGPAMESVAEVEYKEKITVVKIDHDANPKLIEEFKVYGLPTLILFKDGKEVPYSKREGAITKVKLKEYLDGLLKTVSIA*

### NbL17g12420.1 probable_prefoldin_subunit_5 [MW = 16.62 kDa]

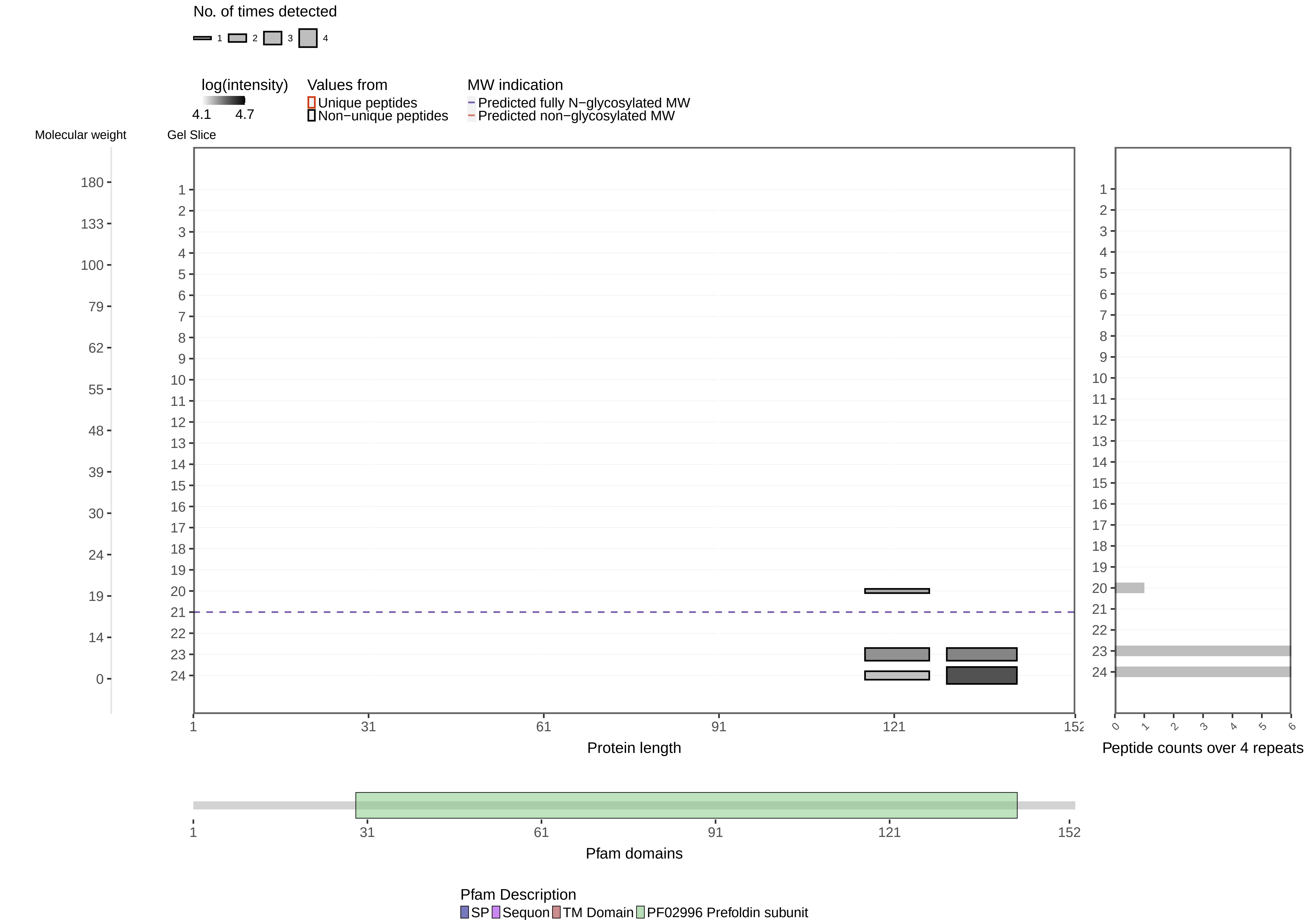


Protein sequence:

GKKMLVPLTASLYVPGTLDDADKVLVDVGTGYFIEKTMTEGKDYCERKINLLKSNYEQLLDVATKKKSIADETGVILQAKLRQLAPPAR*

### NbL17g12400.1 probable_leucine−rich_repeat_receptor−like_protein_kinase_At1g35710 [MW − SP = 34.6 kDa]

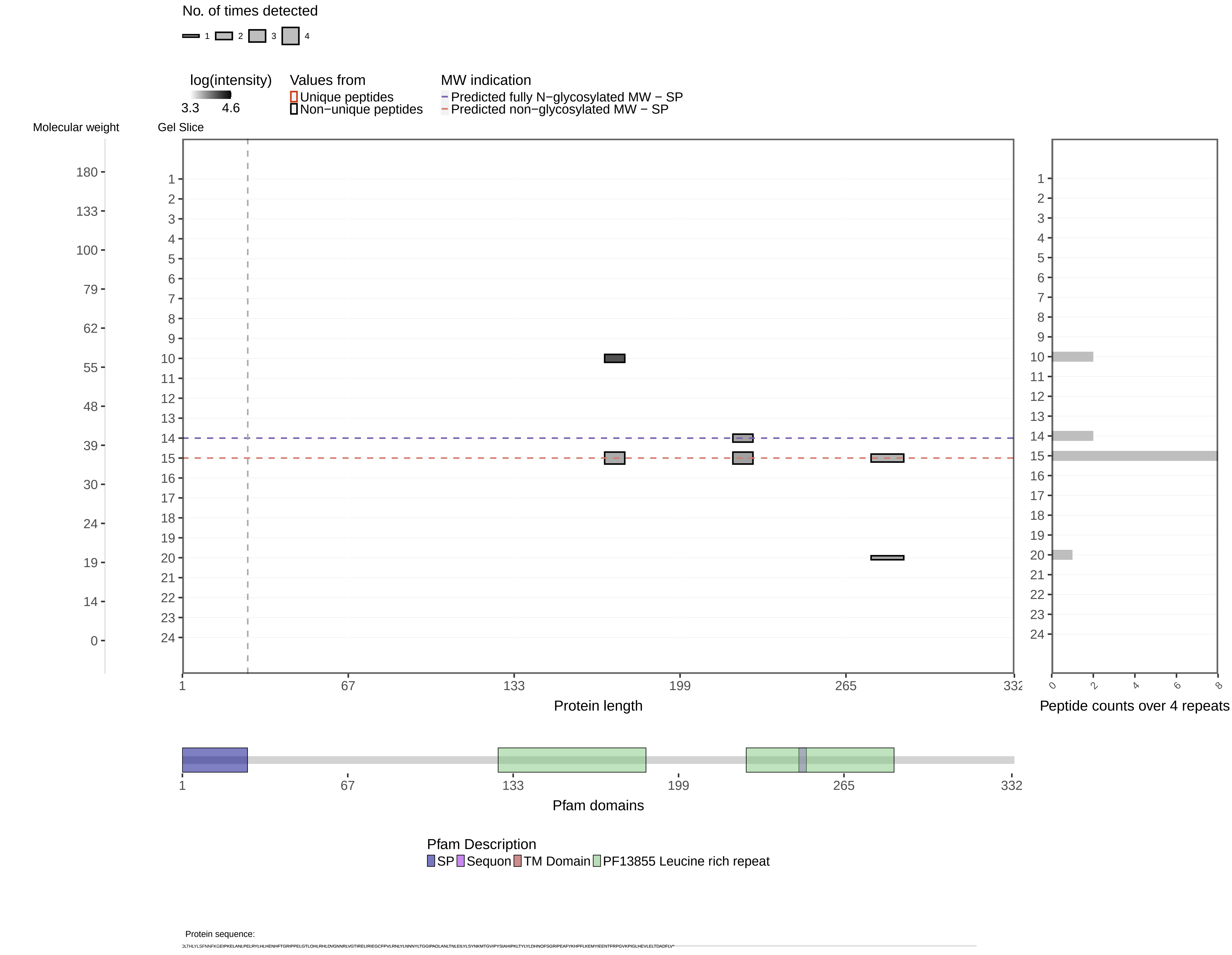

NbL17g12290.1 50S_ribosomal_protein_L19−like [MW = 29.01 kDa]

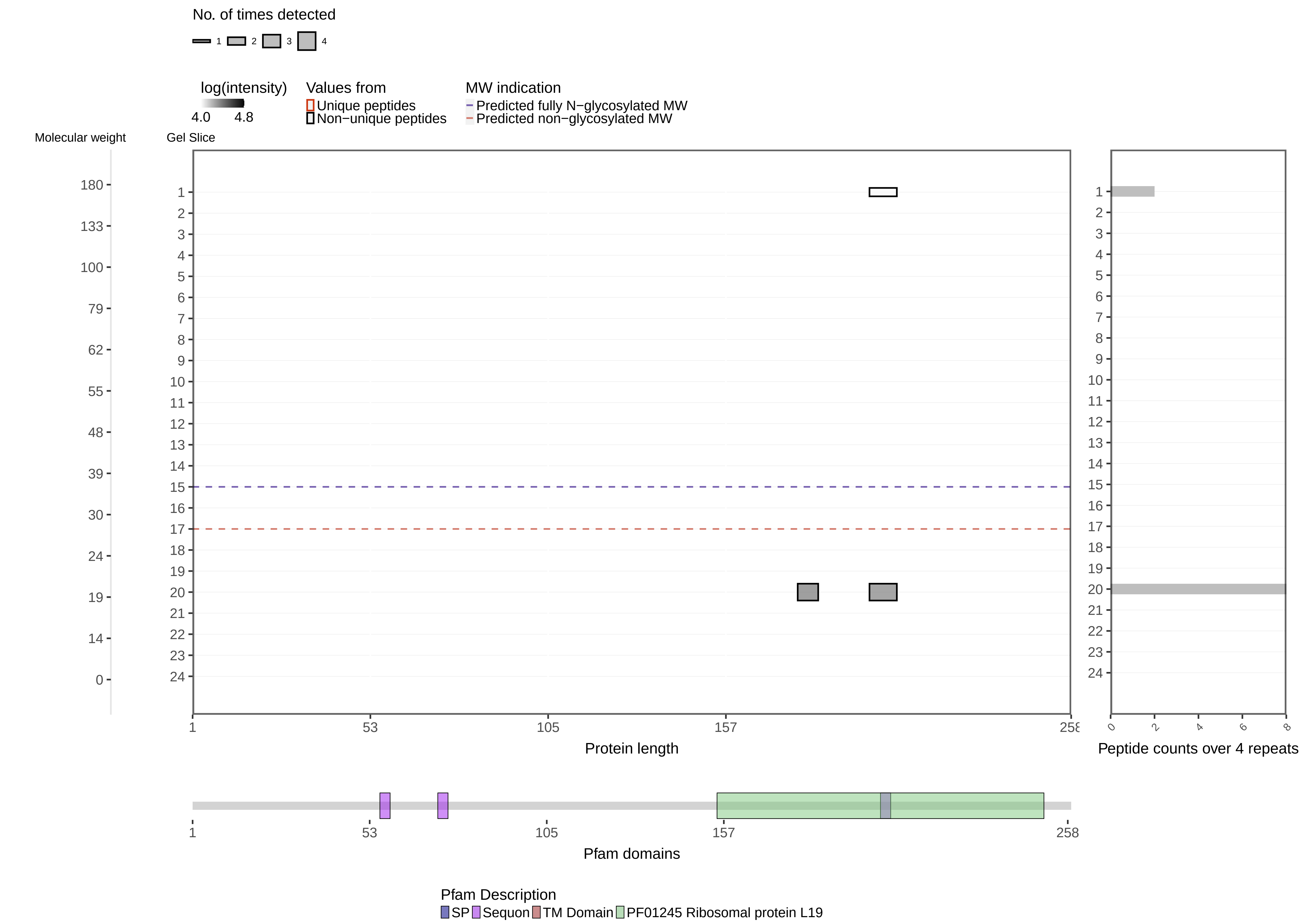


Protein sequence:

GATSVDKVPEDASSSLATPRIKFKRLDKQAAHIMQILDKEAVEEVKAQSPIPDIKPGYIVQLKVEVPENKRRVSIVKGIVIARRNAGLNTTFRLRRLVAGVGVENLFHLYSPNIKEMKVLDKKKVRRAKLYYLRDKMNALKKH*

### NbL17g12260.1 phenylalanine−−tRNA_ligase_beta_subunit_cytoplasmic [MW = 70.79 kDa]

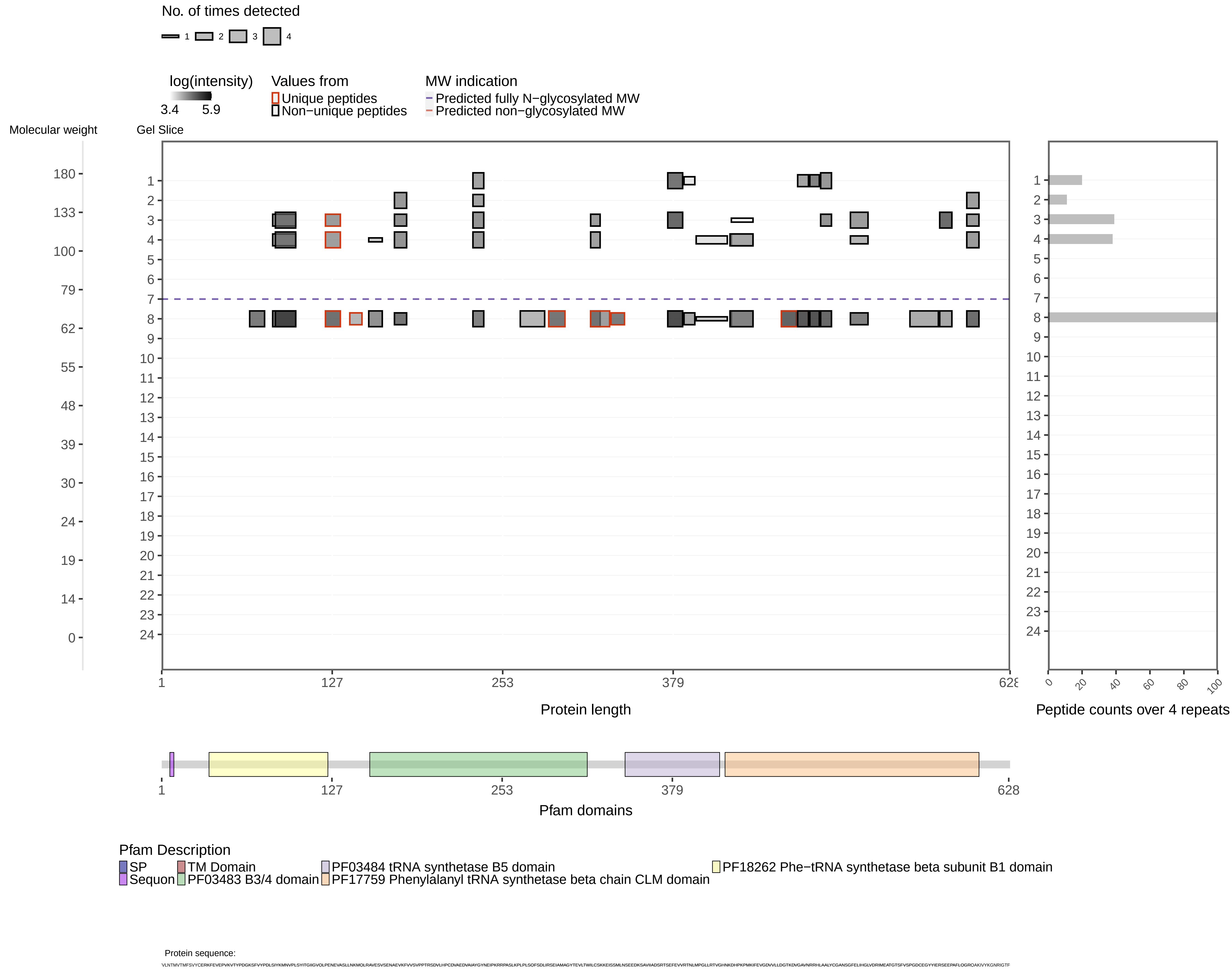

NbL17g12240.1 eukaryotic_translation_initiation_factor_5A−2 [MW = 17.36 kDa]

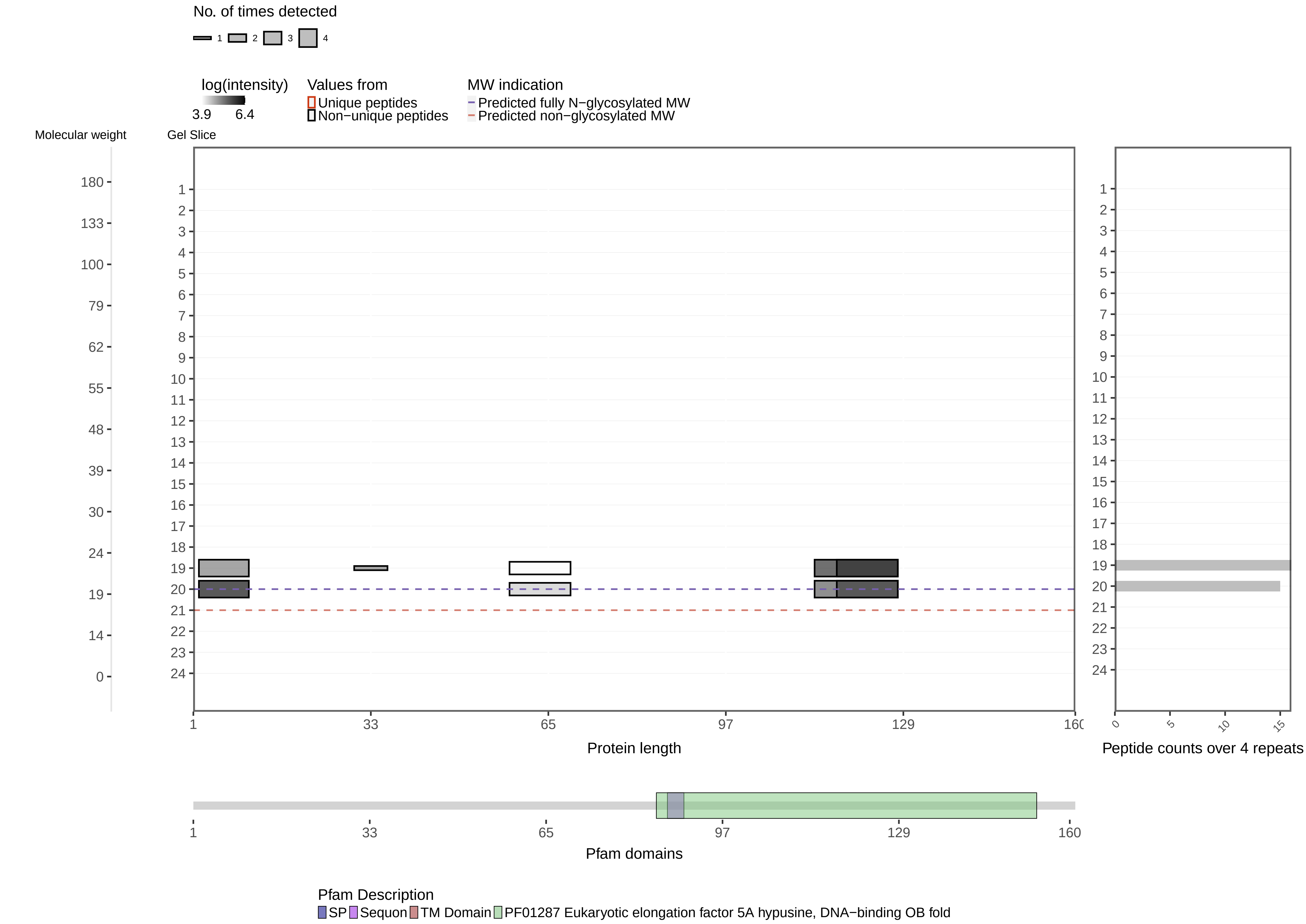


Protein sequence:

TGKKLEDIVPSSHNCDVPHVNRTDYQLIDISEDGFVSLLTENGNTKDDLRLPTDDNLLTQIKDGFAEGKDLVVSVMSAMGEEQICALKDIGPK*

### NbL17g12070.1 40S_ribosomal_protein_S9−2−like [MW = 23.06 kDa]

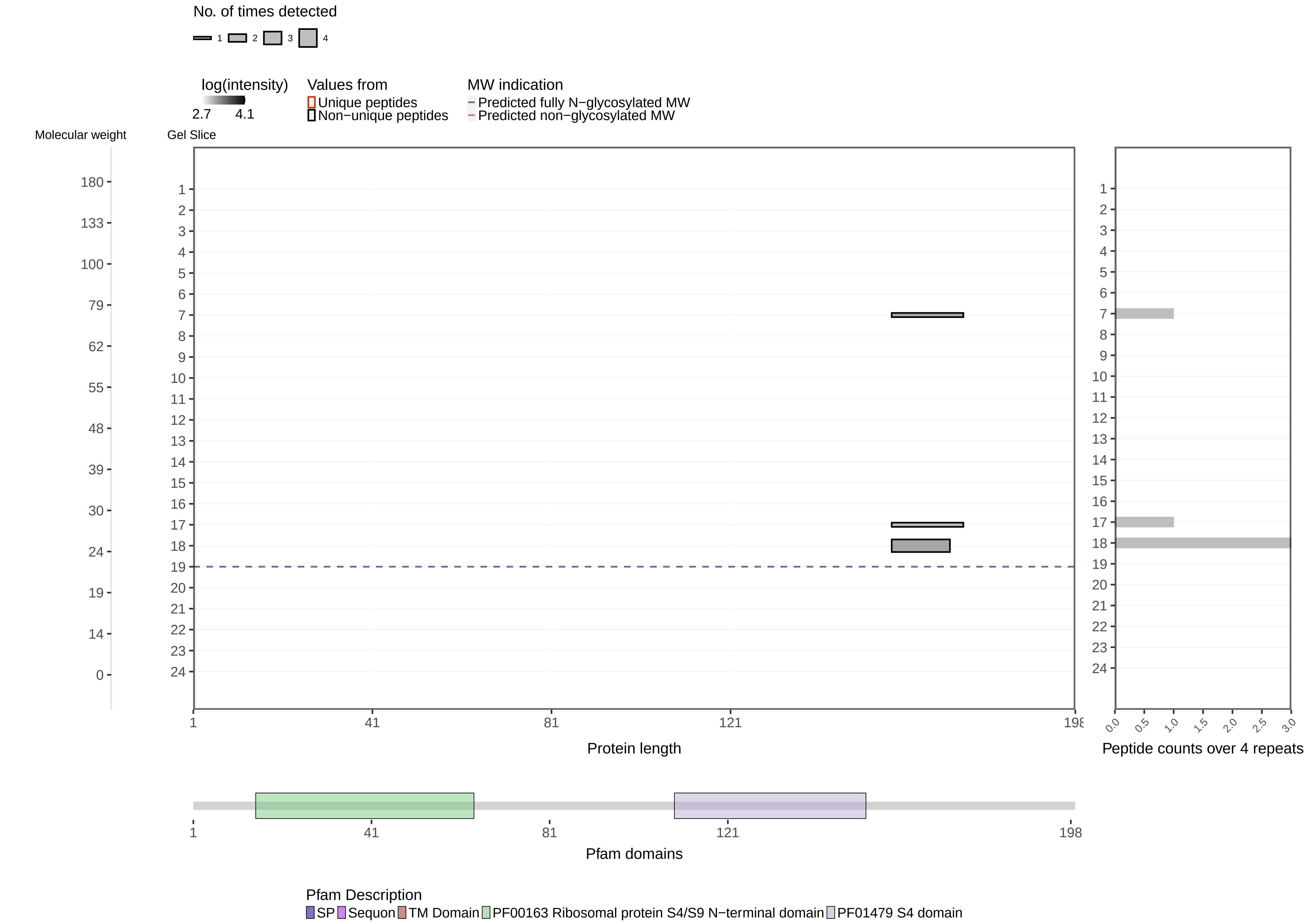


Protein sequence:

LLDESQNKLDYVLSLTVENFLERRLQTLVFKTGMAKSIHHARVLIRQRHIRVGRQVVNVPSFMVRLDSQKHIDFSLISPFGGGRPGRVKRKNQKAAAKKASGGDGDEEDEE*

### NbL17g12060.1 peptidyl−prolyl_cis−trans_isomerase_CYP23 [MW − SP = 22.9 kDa]

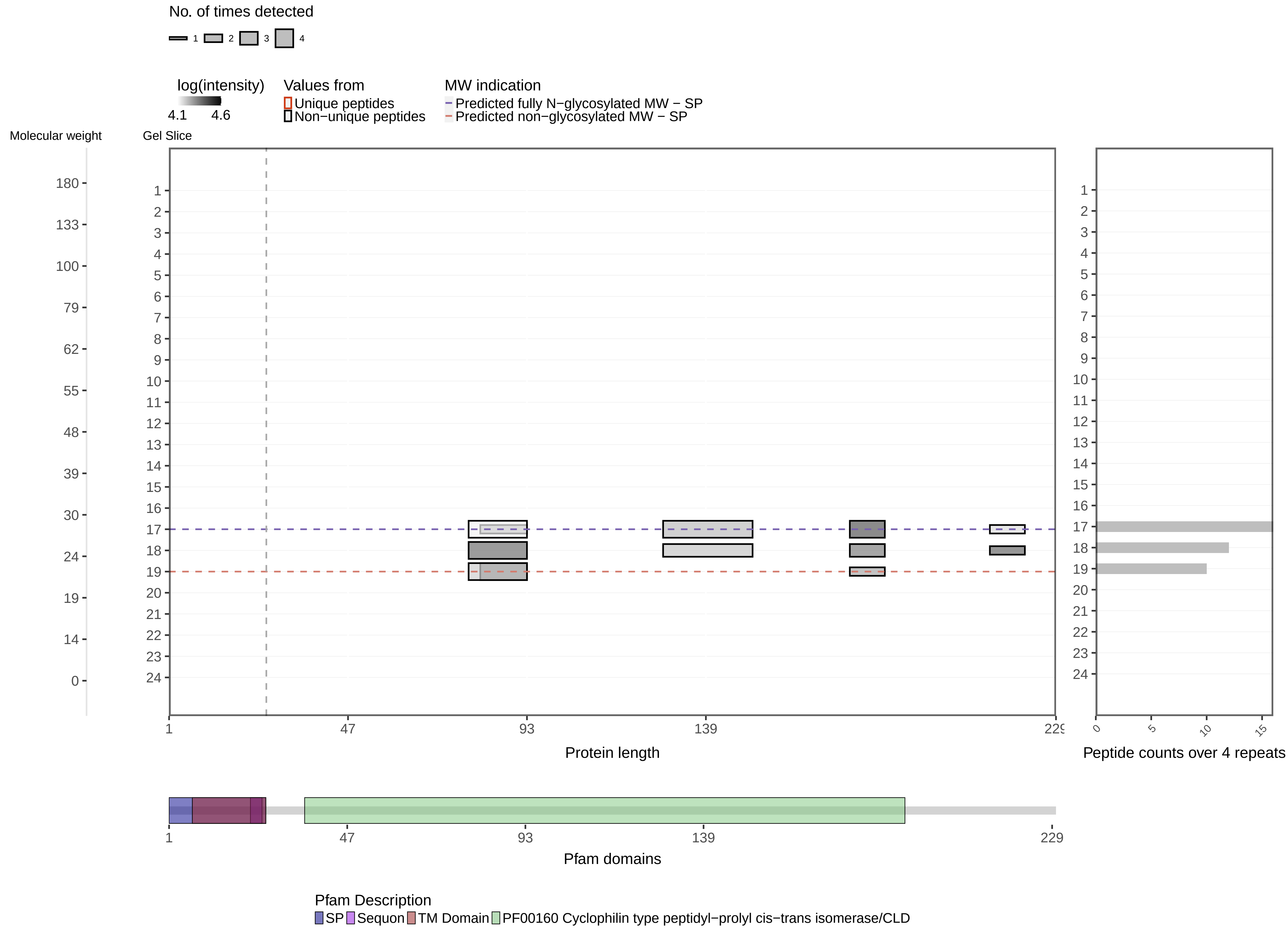


Protein sequence:

RAEAEKTVVGELSEVKHVRGILSMGRYDDPDSGTSSFSMLLGDAPHLDGKYAIFGKVTKGDETLRKLEQLPTRREGIFVMPTERITIQSTYYYDTNMEICEEERSLLKRKLSAAAVEIEKQRMKCFP*

NbL17g12000.1 mitochondrial_outer_membrane_protein_porin_of_36_kDa [MW = 29.18 kDa]

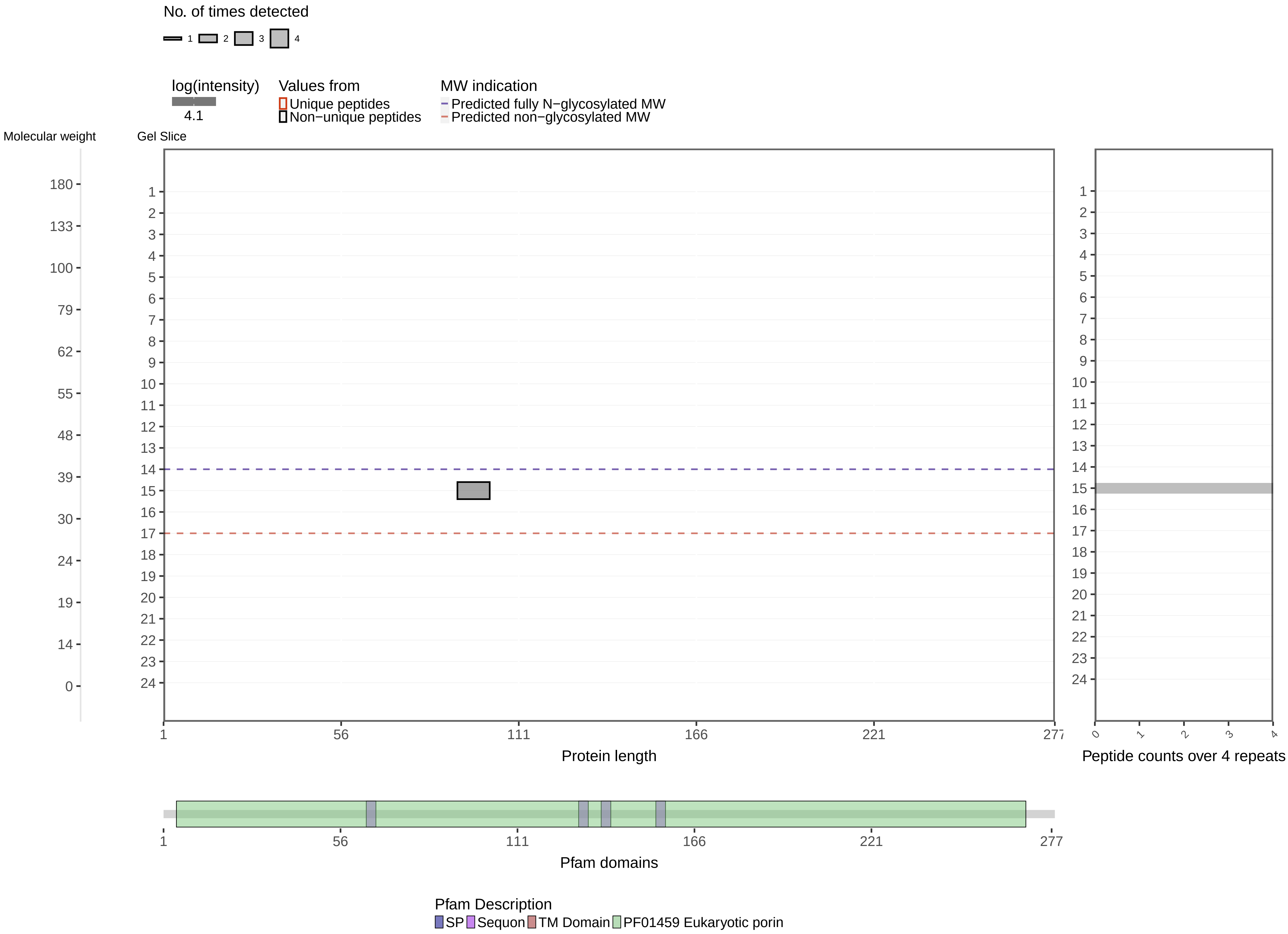


Protein sequence:

SPLVNFSGVAGNNTVALGTDLSFDTASGNFTKCNAGLSFSTSDLIASLALNDKGDTLSASYYHSVKPVTNTAVGAELTHSFSSNENTLTIGTQHLLDPLTTVKARVNSYGKASALIQHEWRPKSLFTVSGEVDTRAIEKSAKIGLAVALKP*

### NbL17g11850.1 protein_SPIRAL1−like_1 [MW = 10.57 kDa]

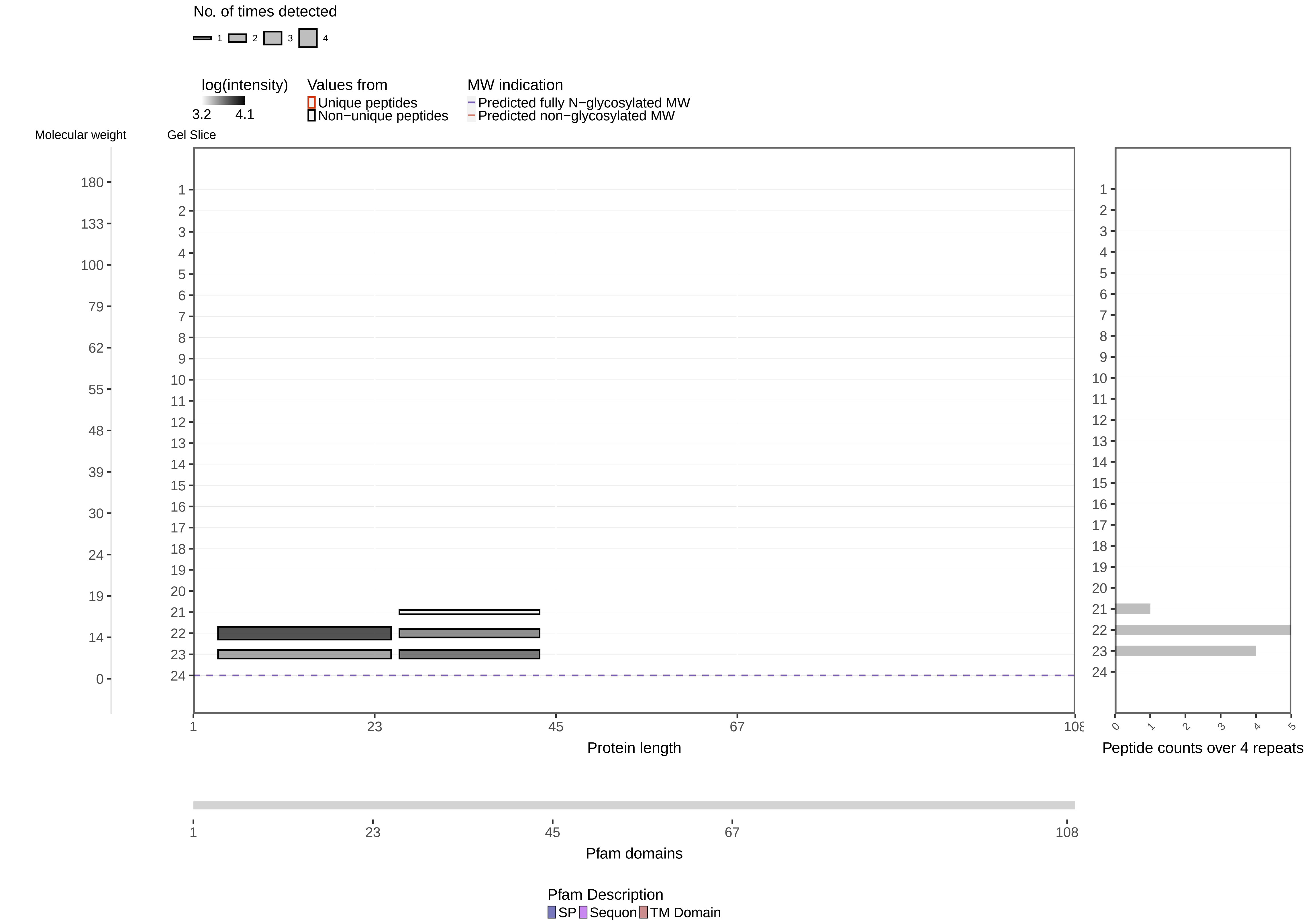


Protein sequence:

PINKEPSPKPVAAAPLADATKQIPAGIQSIRADGQNTGNFITDRPSTKIHAAPGGGSSLGYLFGGGSS*

### NbL17g11810.1 peptidyl−prolyl_cis−trans_isomerase_E [MW = 18.12 kDa]

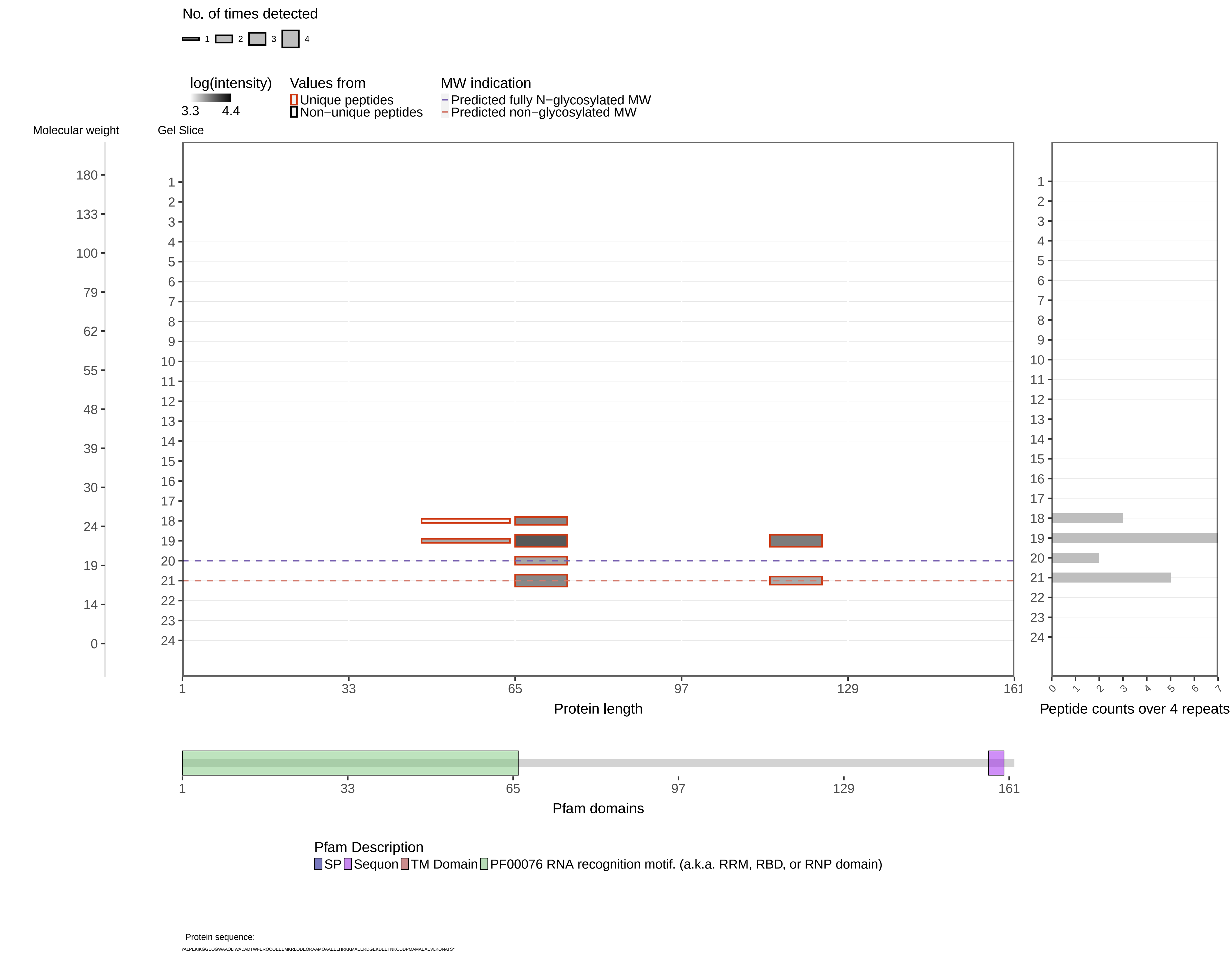

### NbL17g11800.1 alpha−14−glucan−protein_synthase_[UDP−forming]_2−like [MW = 40.92 kDa]

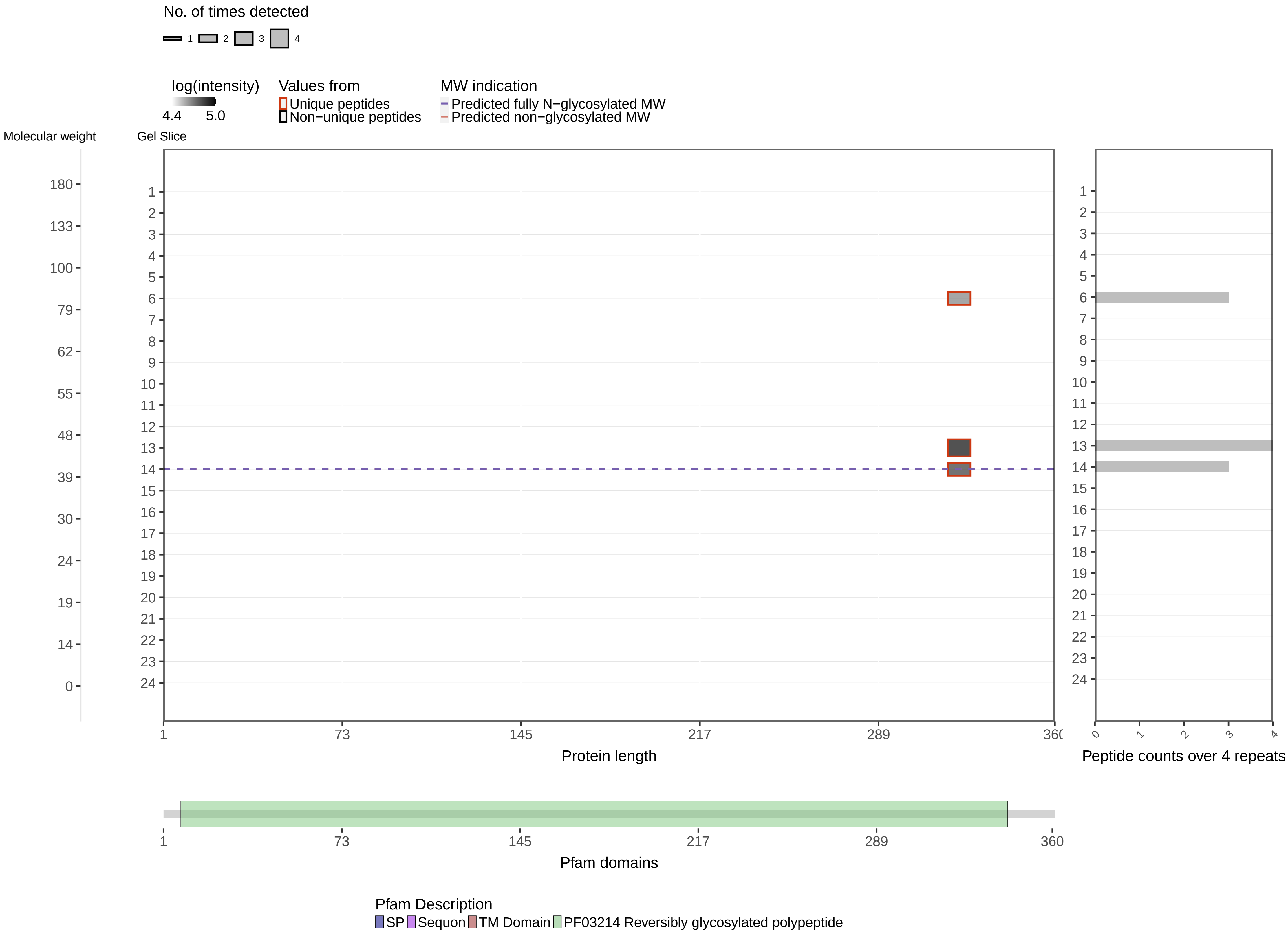


Protein sequence:

NIPDYDAPTQLVKPRERNTRYVDAVMTIPKGTLFPMCGMNLAFNRELIGPAMYFGLMGDGQPIGRYDDMWAGWCTKVICDHLGLGIKTGLPYIWHSKASNPFVNLKKEYKGIYWQEEIIPFFQAVTLPKECTTVQQCYVELAKQVKAKLSNVDSYFTKLAEAMATWIEAWDELNPTGENLAKLSISNDKTK*

### NbL17g11490.1 2−pyrone−46−dicarbaxylate_hydrolase−like [MW = 39.06 kDa]

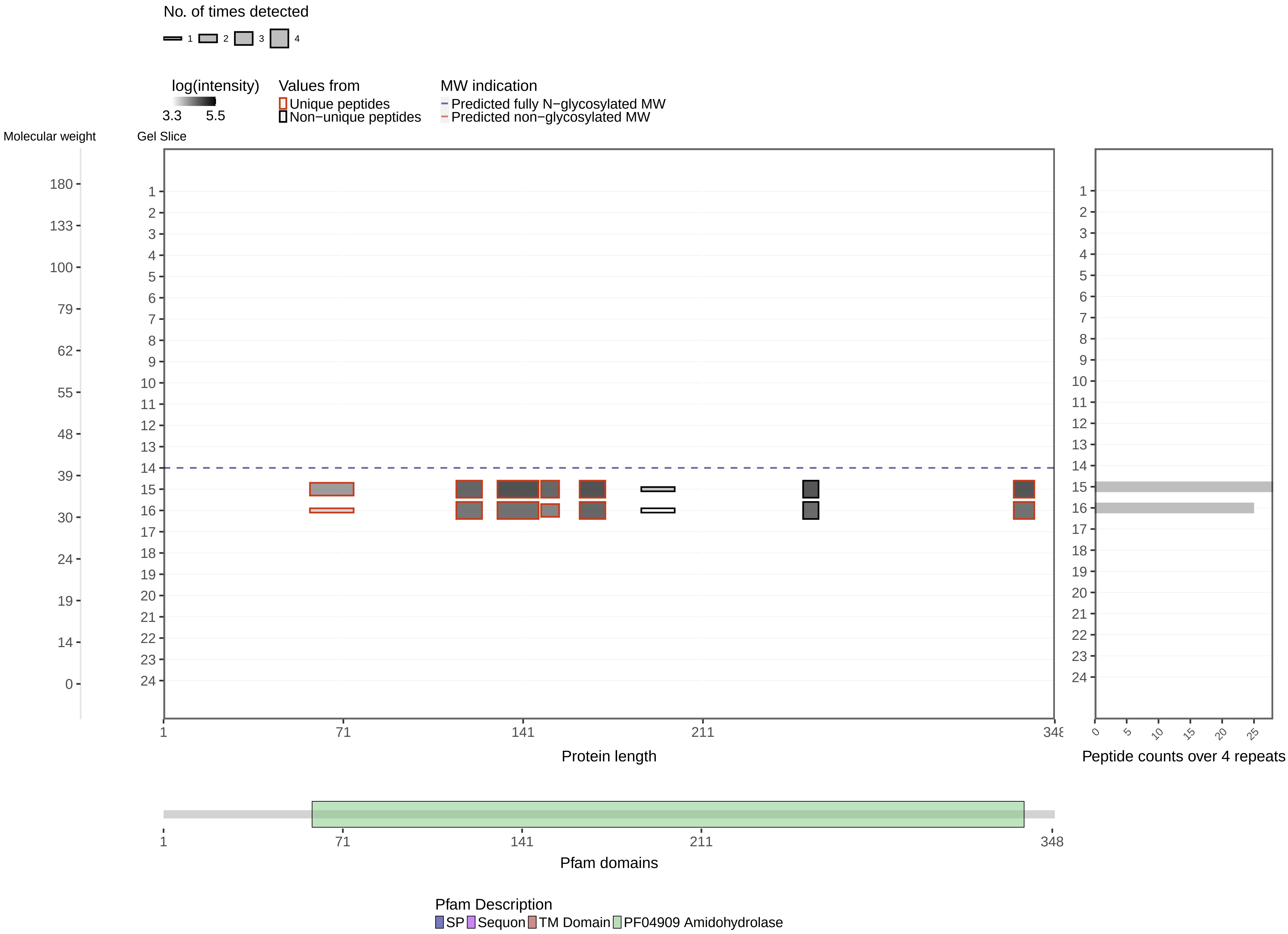


Protein sequence:

RFNPYLWPTGEKQMTNEVGKALFSKAGELGVPVGFMCMKGLDLHLQEIEELCTEFPSTVVLLDHLAFCKLPNNDEERPSFSELLKLSRFPQVYVKFSALFRVSRNRYPYEDLSQVLAQLVSSYGAHRVMWGSDFPYVVPECGYKEAREAVHLLANQGHLPPSATEWILGKTIMQLFDGKWSSVVTC*

### NbL17g11450.1 short−chain_dehydrogenase/reductase_2b−like [MW = 41.54 kDa]

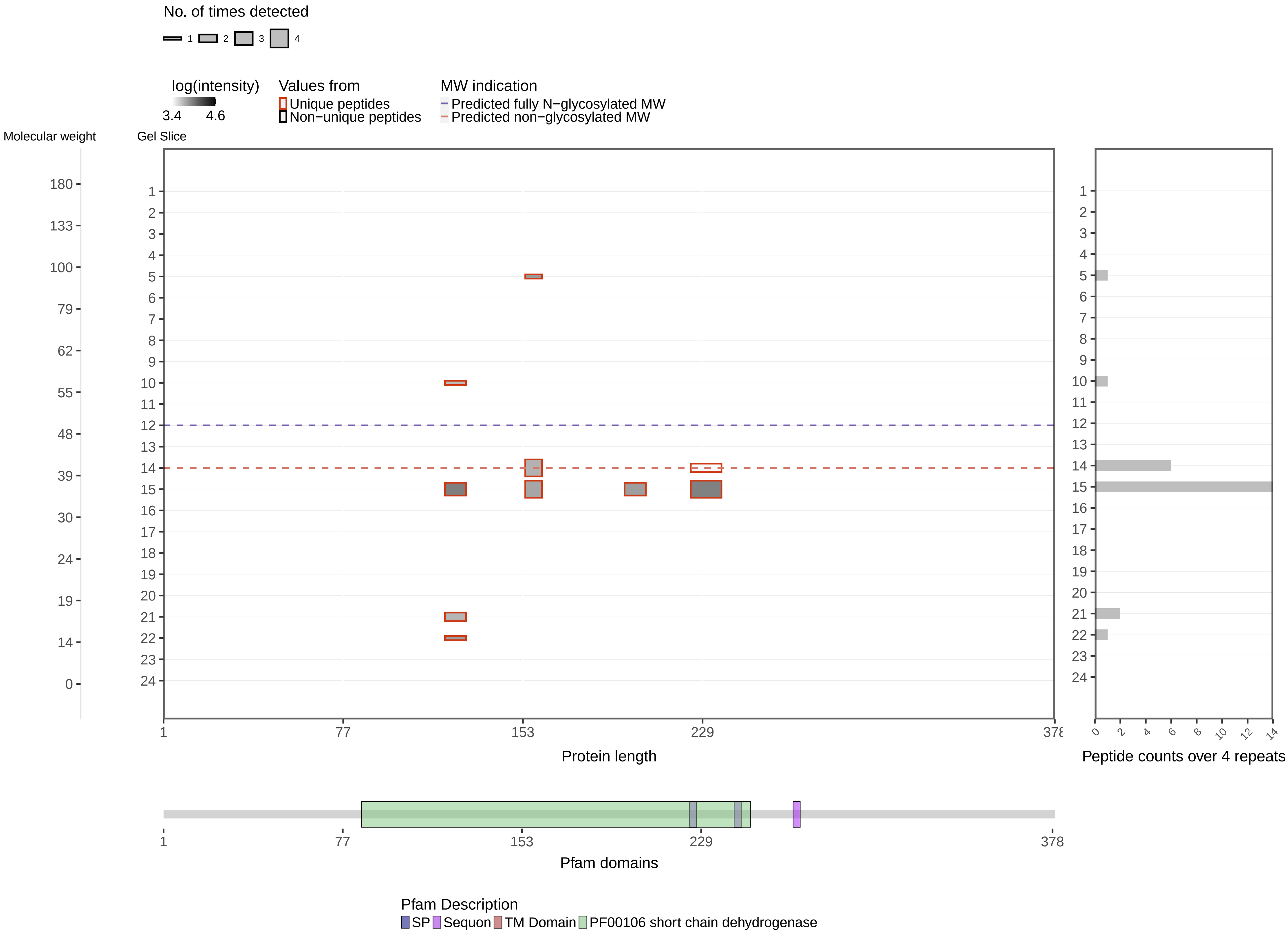


Protein sequence:

DEATLKALNIDPVDWLAGKAANVVQVAMKTTYESAKLCLDTNYYGVKNVTEALLPLLQNSRSAKIVNVSSLRSELKRVPNEDRRKELGDIENLTEDKIEKILQKFLHDLKQDALEVNGWQIMLPAYSISKVSLNAYTRVLARRYPKICINCVHPGYVNTDINWHTGTMPVEEGAEGPVMLALLPDGGPTGCYFDRTVVAQF*

NbL17g11440.1 non−specific_lipid_transfer_protein_GPI−anchored_2−like [MW − SP = 16.91 kDa]

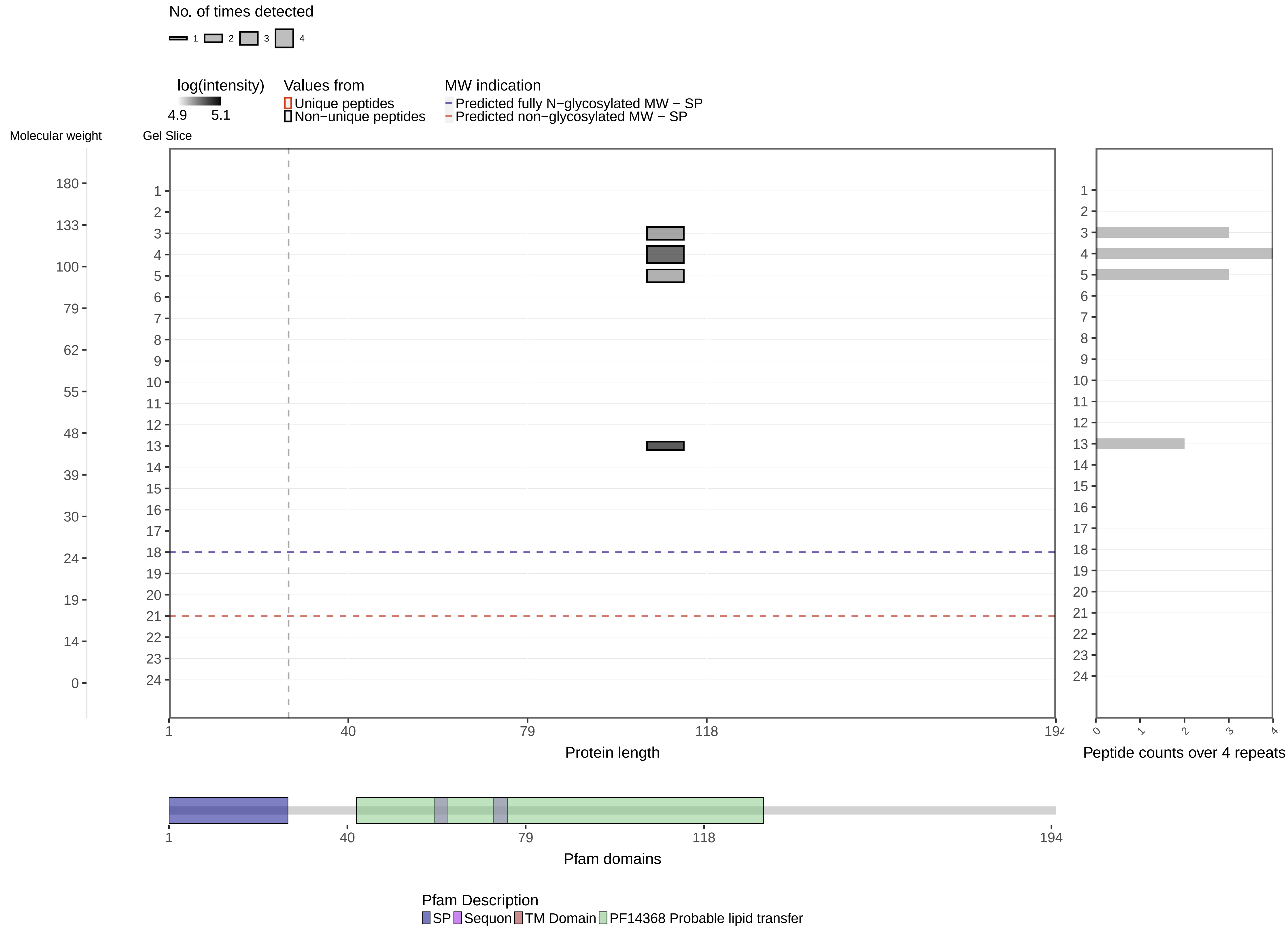


Protein sequence:

ELAGLVDSNPICLCQLLGDPDKLGVQIDISRALKLPNICSVETPPVSTCAALGIPVAAPTASDELAPALSPGGLASSPTTENNDKNAAQTMAFSKMNIVLGFGIMFFTTIF*

### NbL17g11330.1 elongation_factor_1−beta−like [MW = 24.49 kDa]

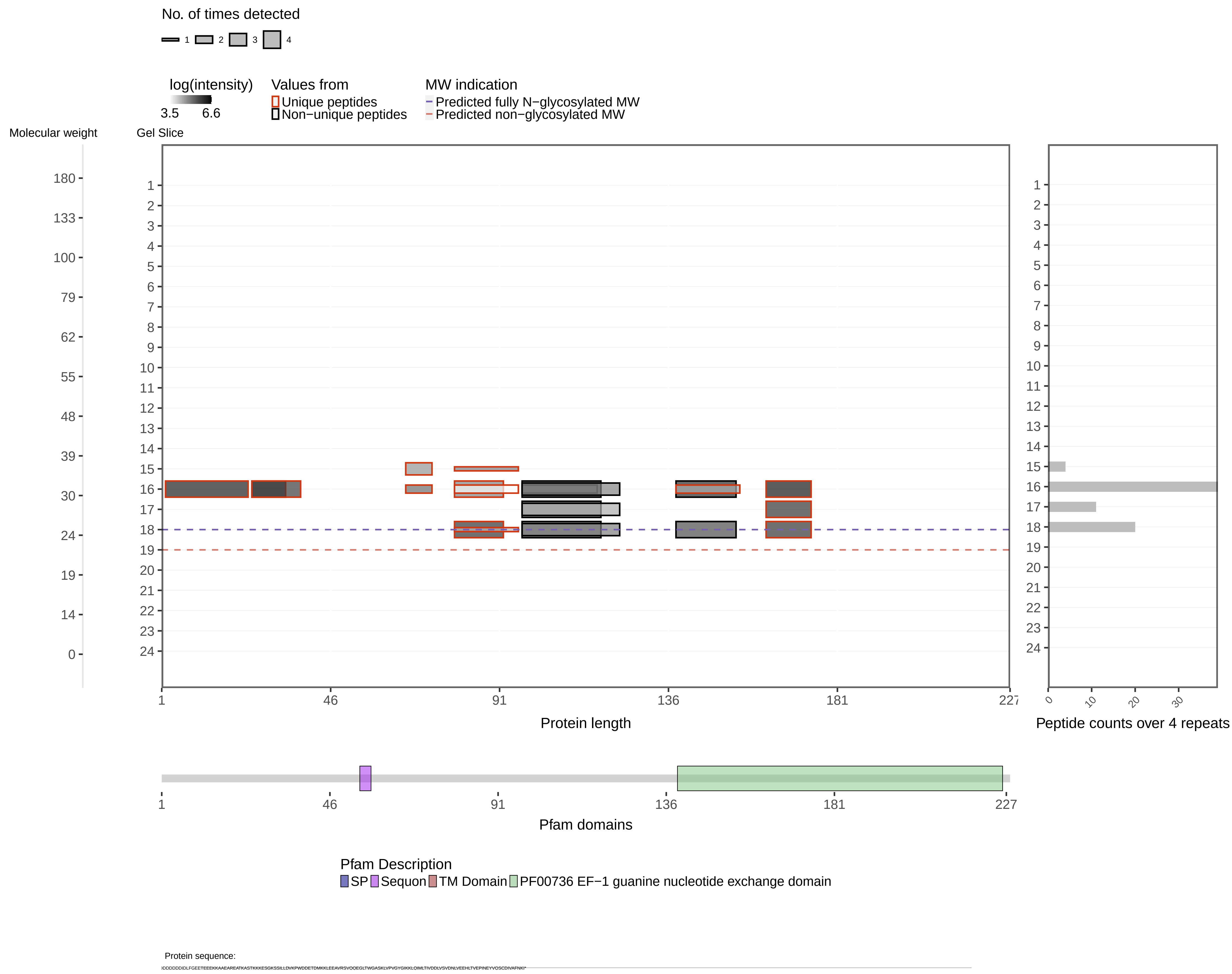

### NbL17g11260.1 calmodulin−7−like [MW = 20.76 kDa]

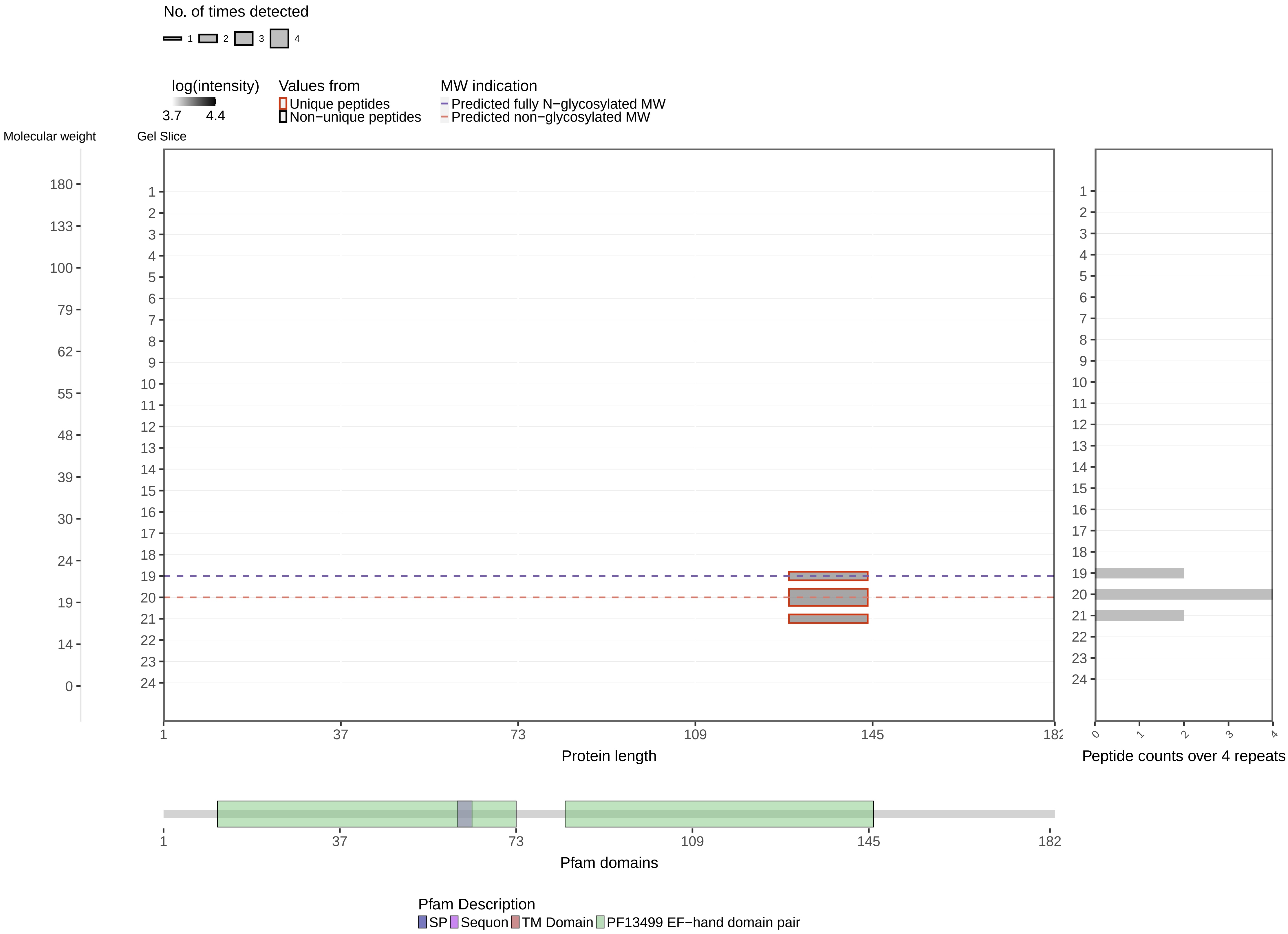


Protein sequence:

KDTDSEEELKEAFRVFDKDQNGFISAAELRHVMTNLGEKLTDEEVDEMIREADVDGNGQINYEEFVKVMMANRRRRRIEESKRGYSISQSNGRKSRKRDRCTIL*

NbL17g11170.1 uncharacterized_LOC104119948 [MW = 29.58 kDa]

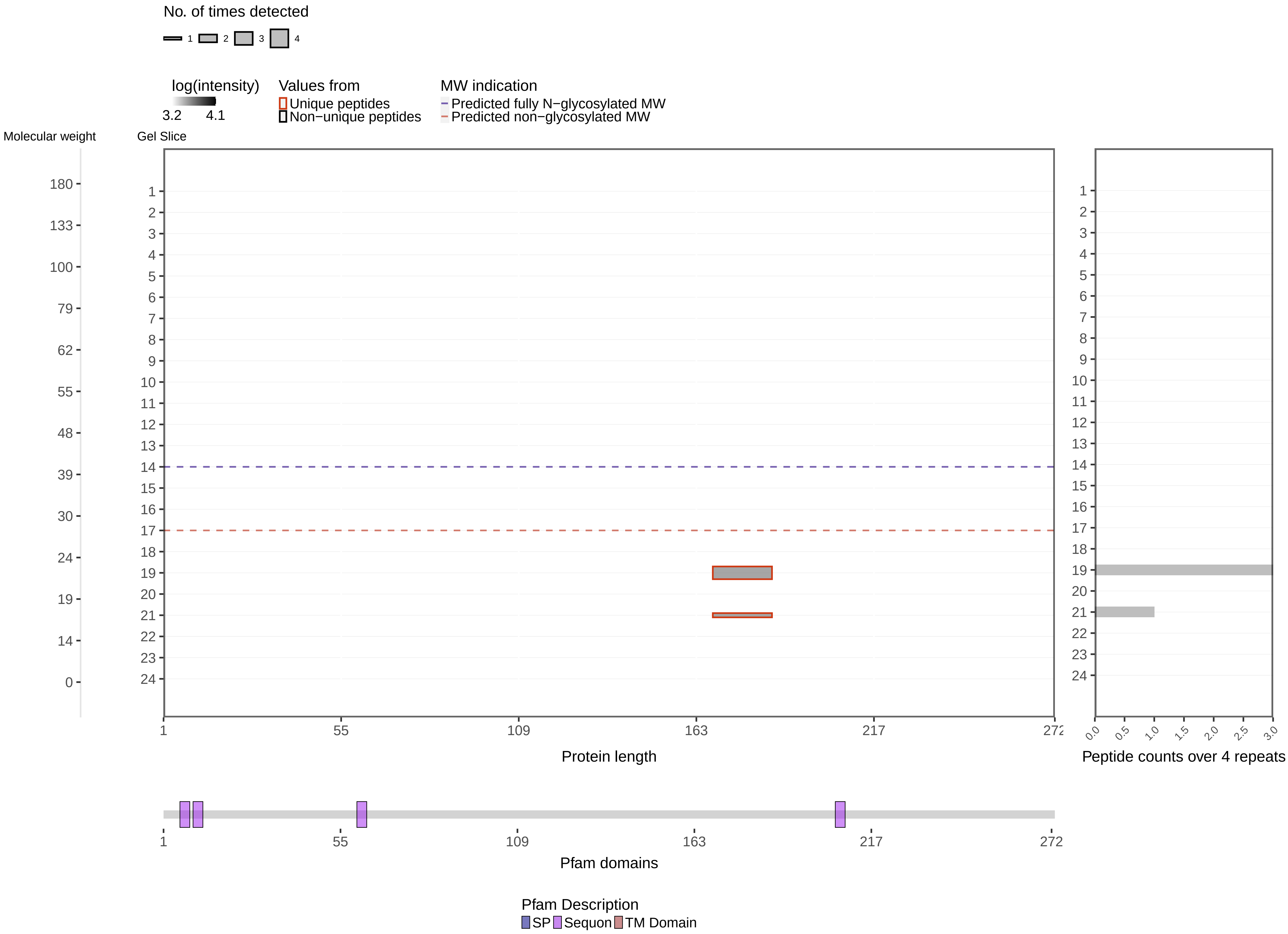


Protein sequence:

EEATKQKLCEDLNSLVLESSNSQLARLEELKRRLEAMNPSRASTAPAPDQNPVEPVQSSRTQGVSSVKDTQQPSSGSSGNTSKEGNDRSSGAENGQNQLPSLAEARVKKKINLQGRGKGIGIVPKGRGSTAPGWTGAGFDVDGRS*

NbL17g10900.1 short−chain_type_dehydrogenase/reductase−like [MW = 28.07 kDa]

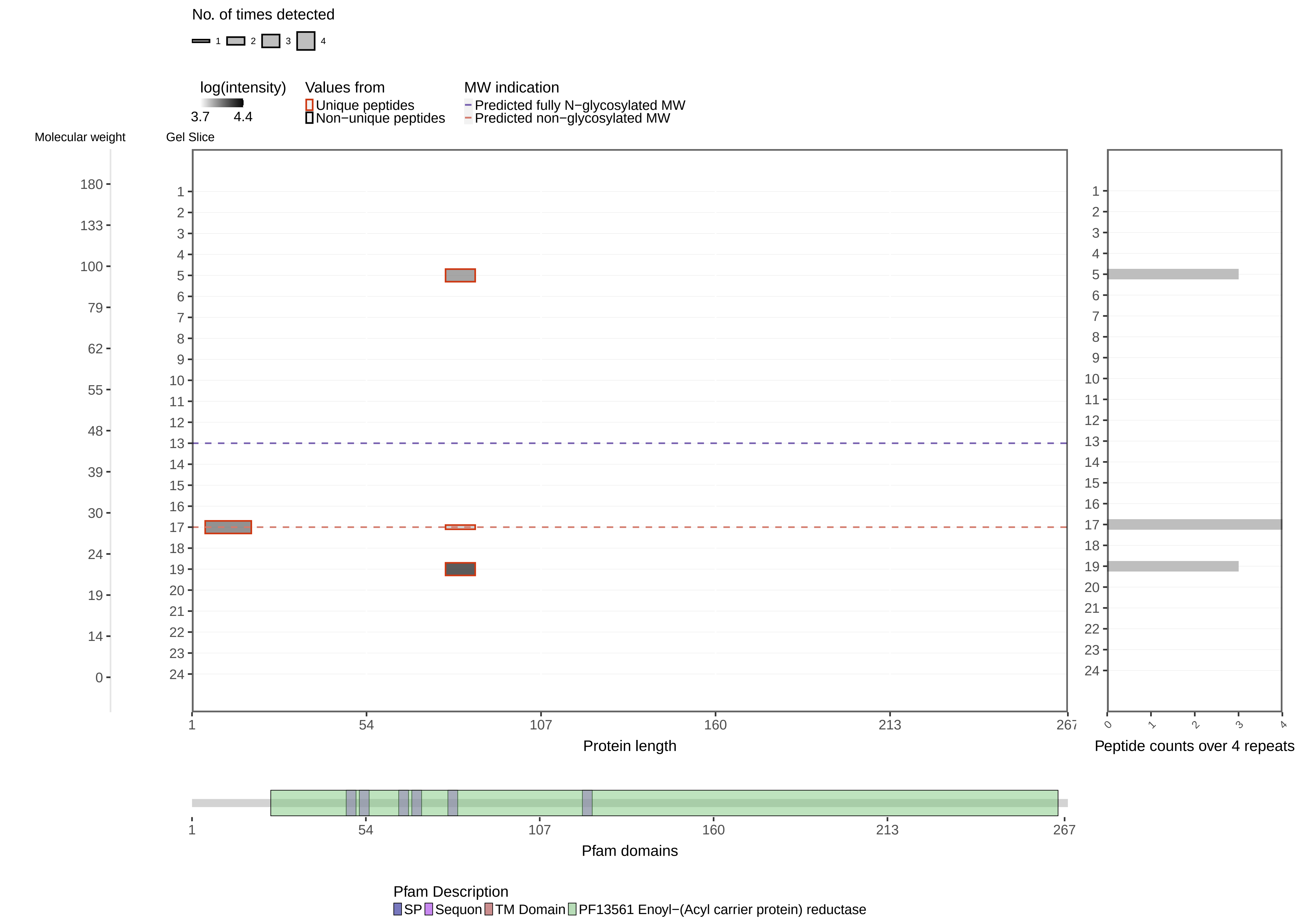


Protein sequence:

TTLEDFDRTFEVNARGAFICCKEGASRIKRGGGGRIICLTTSLAAAFRPGYGAYTASKAAVEAMVKILAKELKGTGITANCVAPGPIATEMFYDGKTEEMIKRVIDECPHGRLGQTEDVAPVVGFLAADASEWVNGQIIRVNGGYI*

NbL17g10840.1 lysosomal_beta_glucosidase−like [MW = 63.72 kDa]

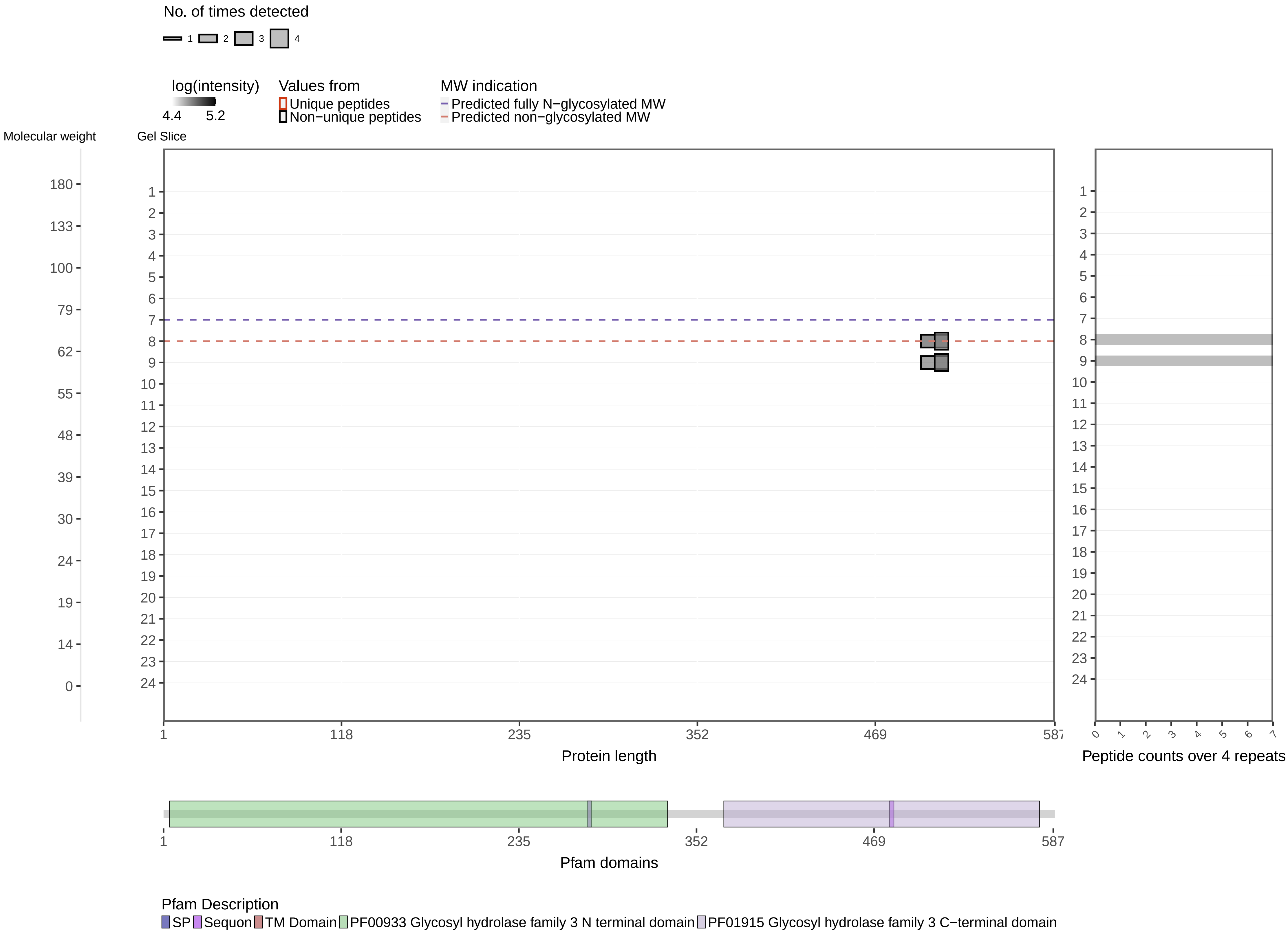


Protein sequence:

NYSYSVQAGIMAGIDMIMVPENYREFIDSLTSQVKANIIPMSRIDDAVKRILRVKFVMGLFENPMSDPSLANQLGSQEHRELAREAVRKSLVLLKNGKTPSQPLLPLPKKAPKILVAGTHADNLGYQCGGWTIEWQGVAGNDLTIGTTILTAIKKTVDPSTQVVYQQNPDANFVKSNKFSYAIVVVGEVPYAEMFGDSSNLTIAEPGPSTISNVCGAVKCVVVVVSGRPVVLEPYVSKMDALVAAWLPGTEGQGVADALFGDYGFTGKLARTWFKRVDQLPMNFDDAHVDPLFPI

NbL17g10520.1 GDP−mannose_46_dehydratase_1 [MW = 41.44 kDa]

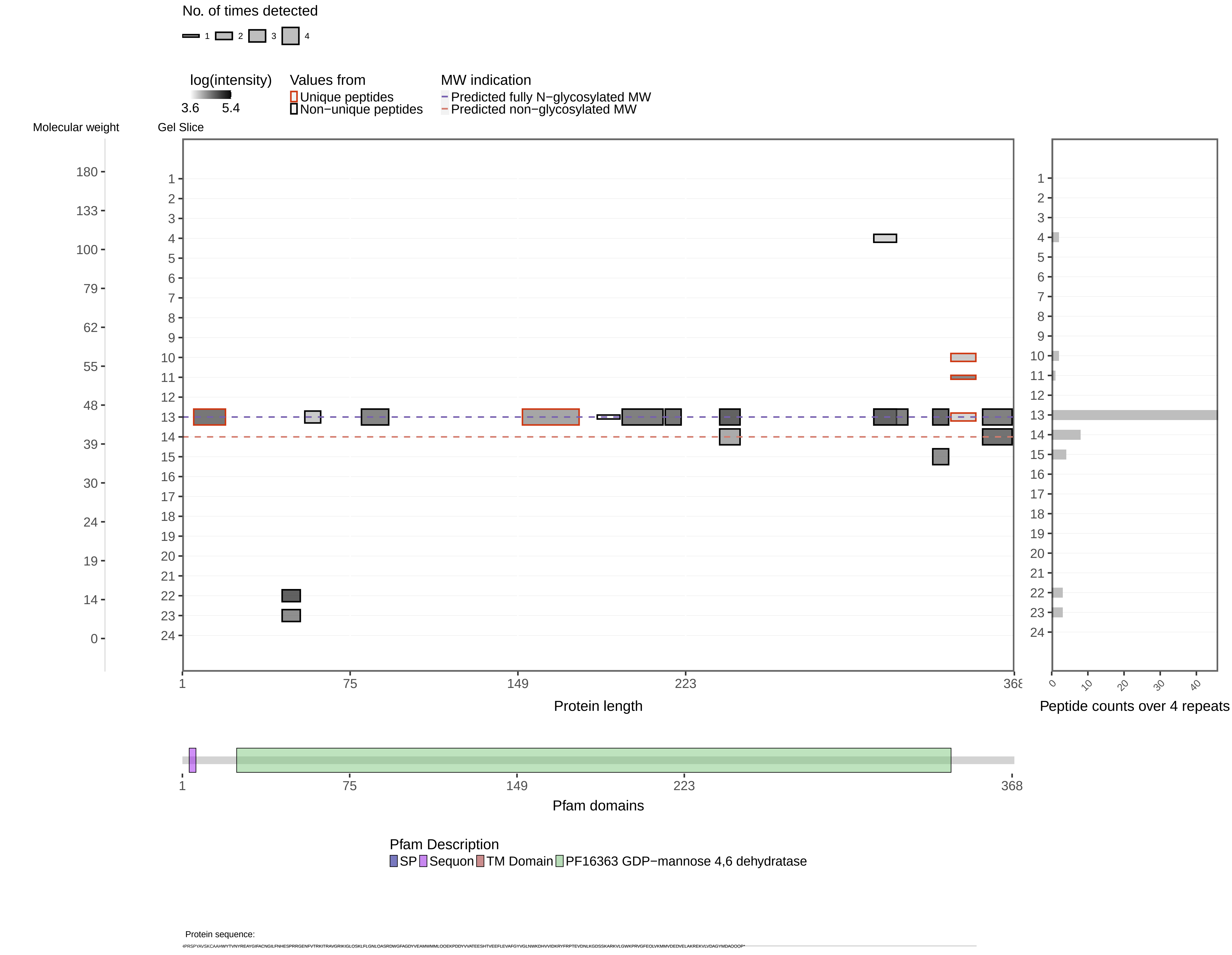

NbL17g10510.1 dihydropyrimidinase [MW − SP = 55.41 kDa]

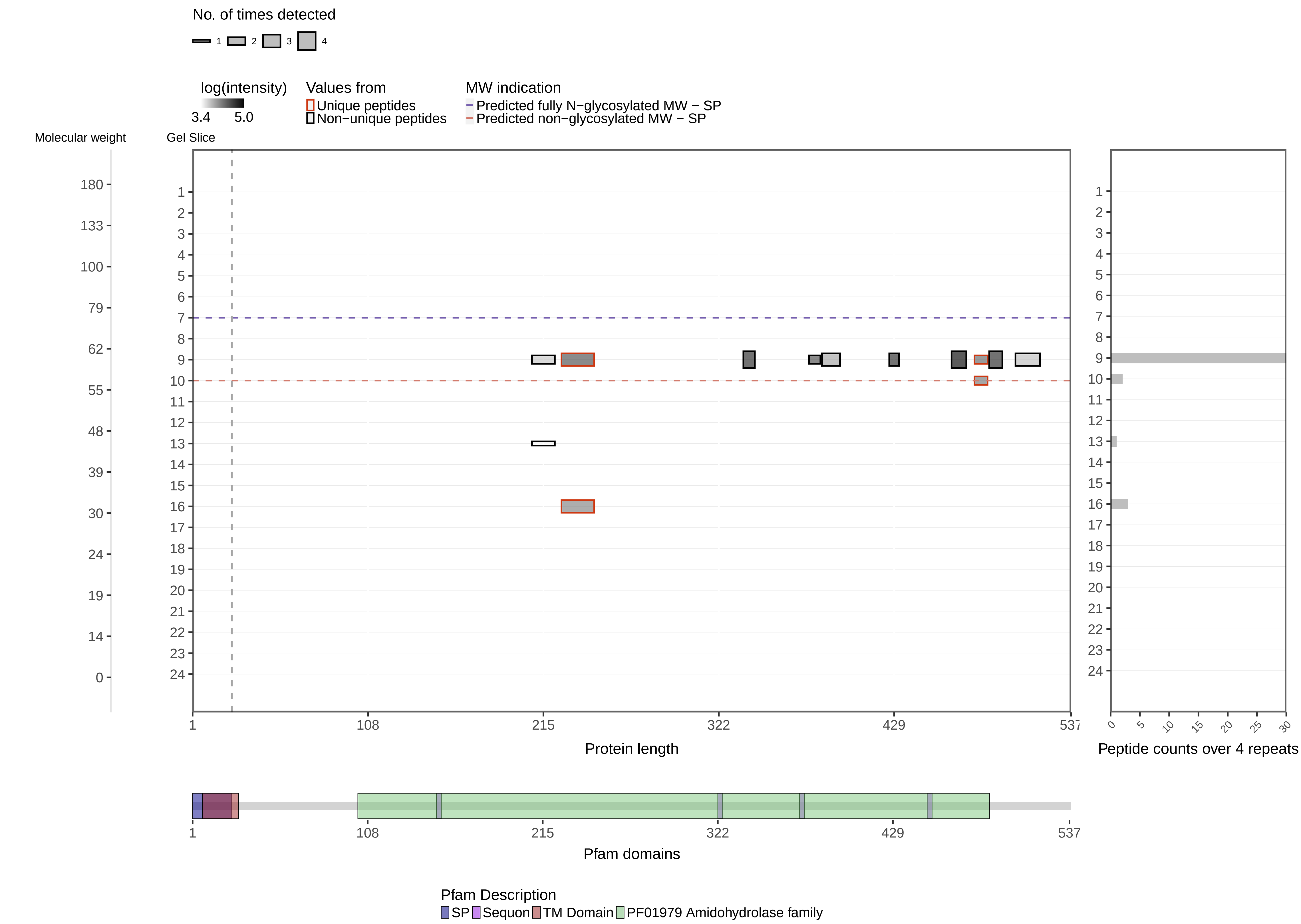


Protein sequence:

[illegible]EGHTLSRPPVLEGEATARAIRLAGFVNTPLYVVHVMSIDAMEEIARARKSGQRVVGEPVVSGLVLNDSVLWDPDFHFAARYVMSPPIRAAGHGTALQGALAMGLLQVVGTDHCTFNSTQKALGIGDFRKIPNGVNGMEERMHLVWDTMVASGQISVTDYVRITSTECARIFNIYPRKGAILVGSDADIIILNPNSSFEISSKSHHYRTDTNVYEGWRGKGKVEITIAGGRIVWENDELKVVPGAGKYIEMPPFSYLFDGIDKADAKYLASLRAPVKRSTT*

### NbL17g10120.1 glucosidase_2_subunit_beta−like [MW − SP = 64.28 kDa]

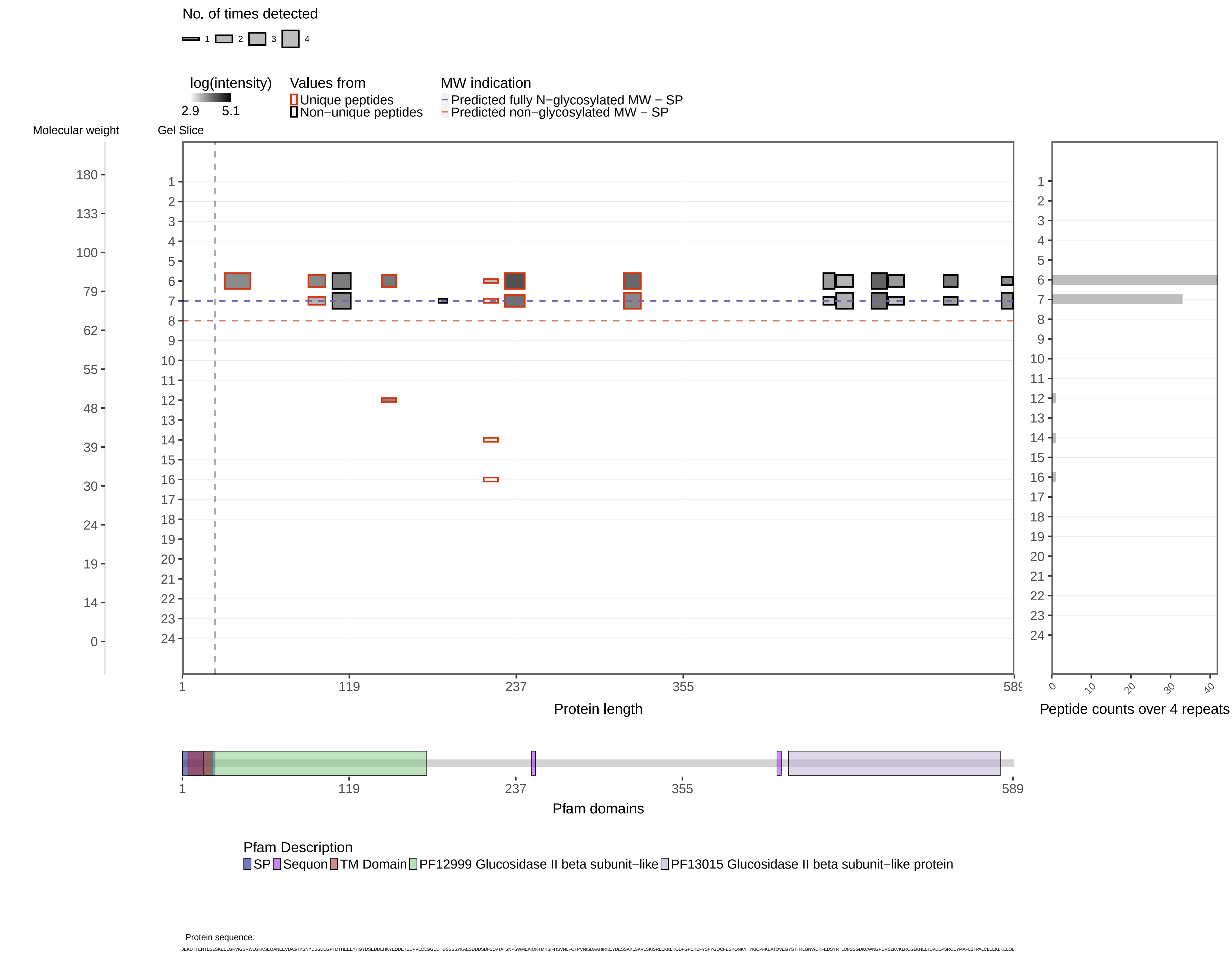

### NbL17g10110.1 sulfite_reductase_1_[ferredoxin]_chloroplastic [MW = 77.69 kDa]

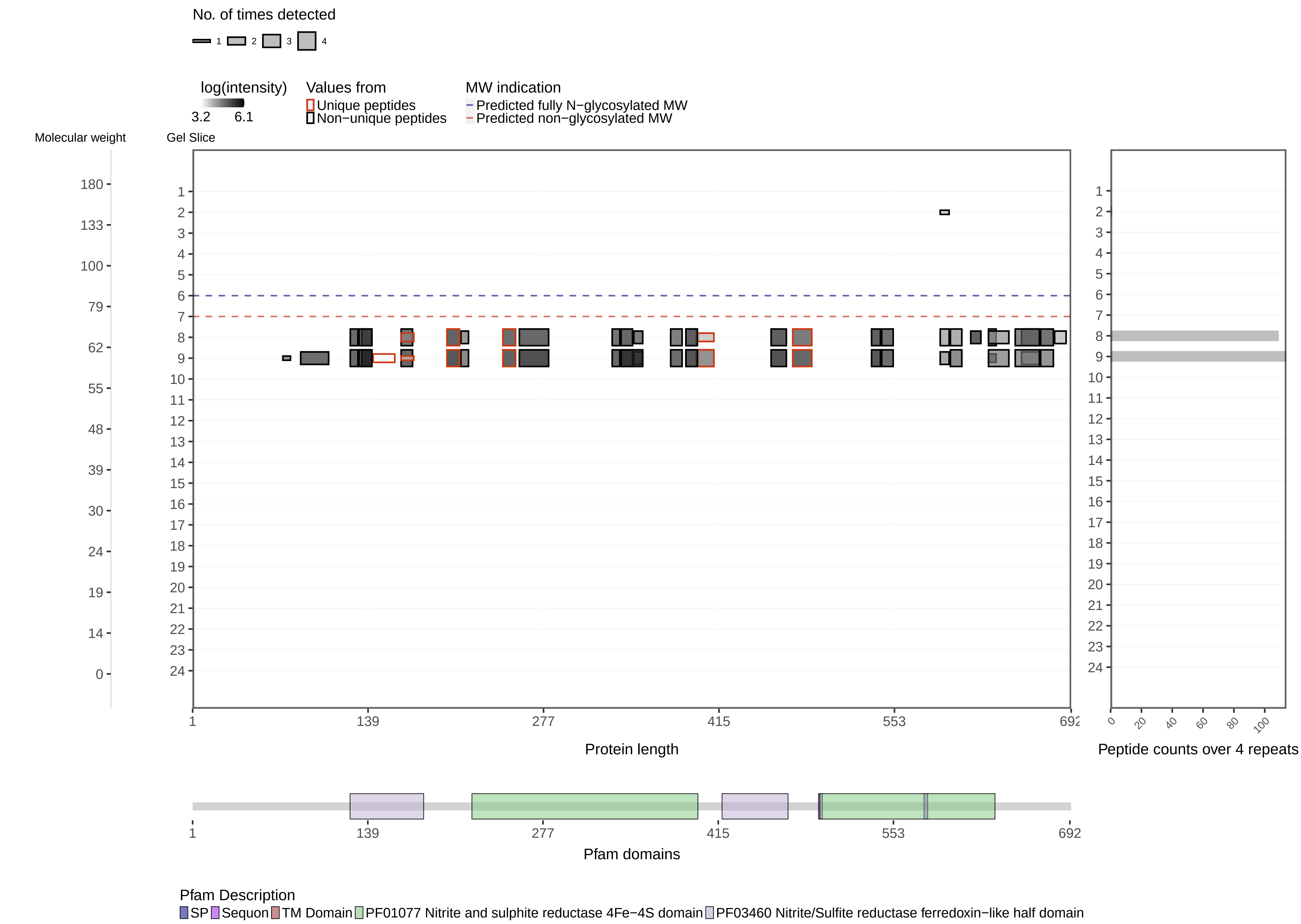


Protein sequence:

### NbL17g09950.1 THO_complex_subunit_4A−like [MW = 26.78 kDa]

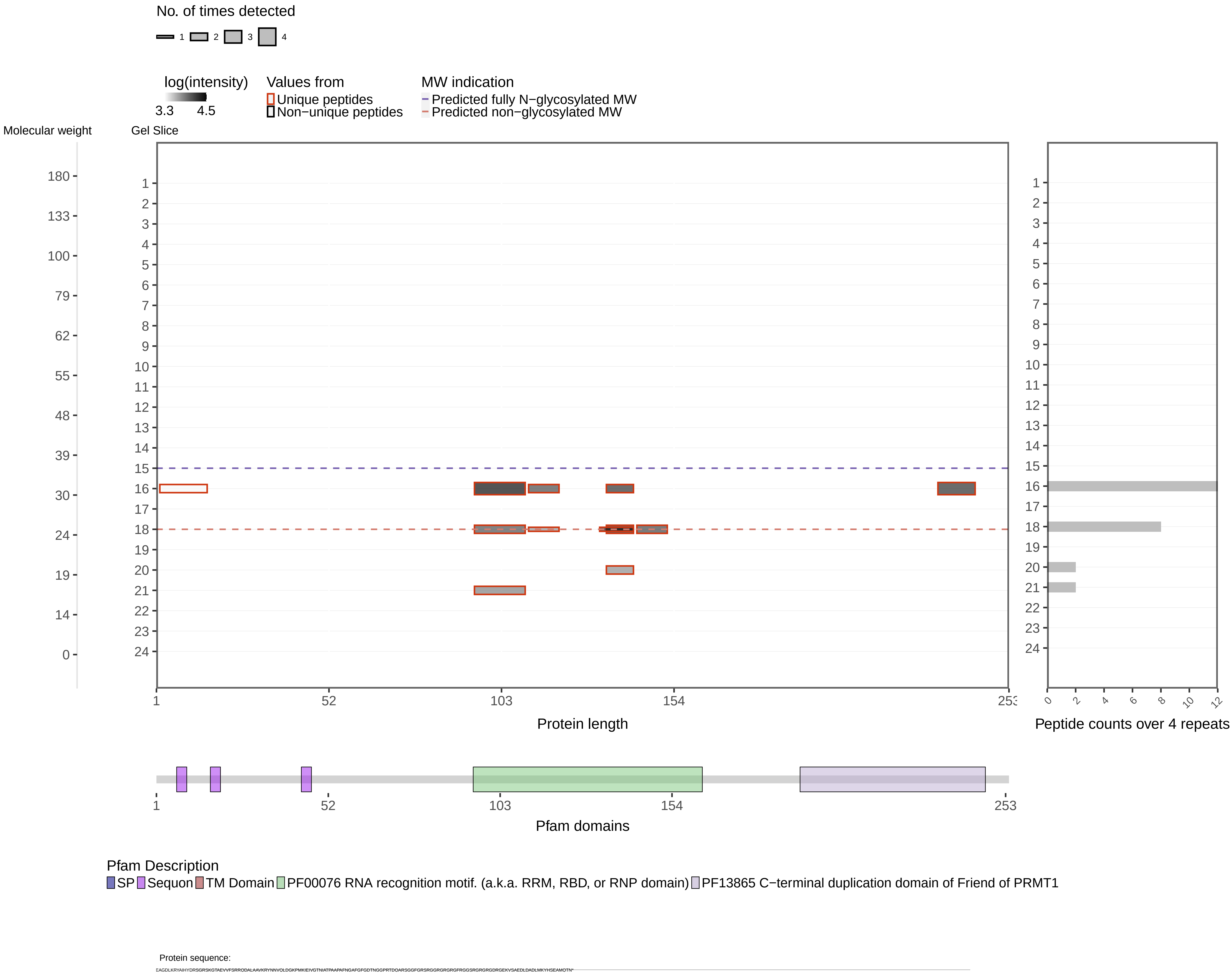

### NbL17g09920.1 histone_H4 [MW = 11.41 kDa]

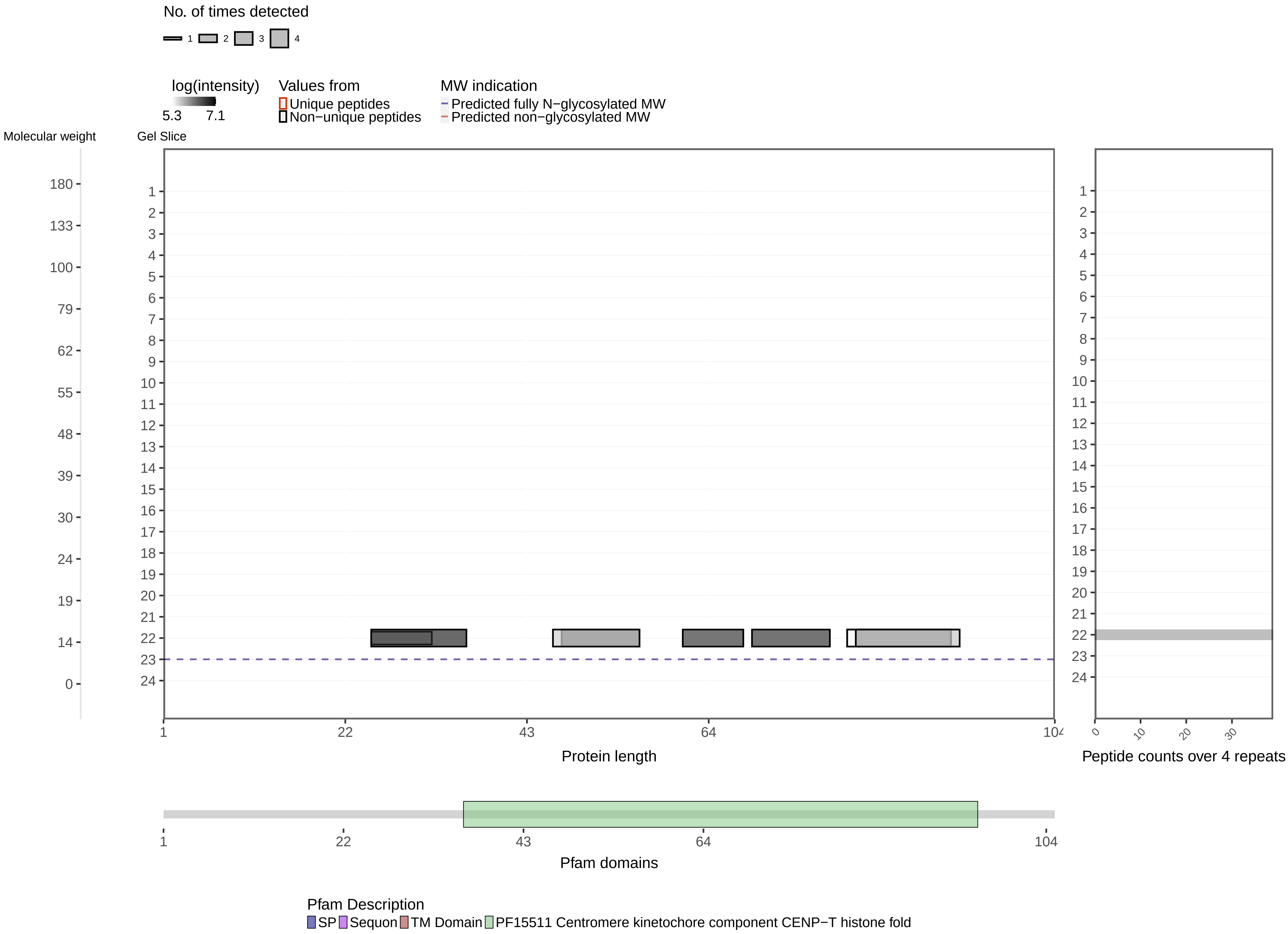


Protein sequence:

ARRGGVKRISGLIYEETRGVLKIFLENVIRDAVTYTEHARRKTVTAMDVVYALKRQGRTLYGFGG*

### NbL17g09880.1 osmotin−like_protein [MW − SP = 24.49 kDa]

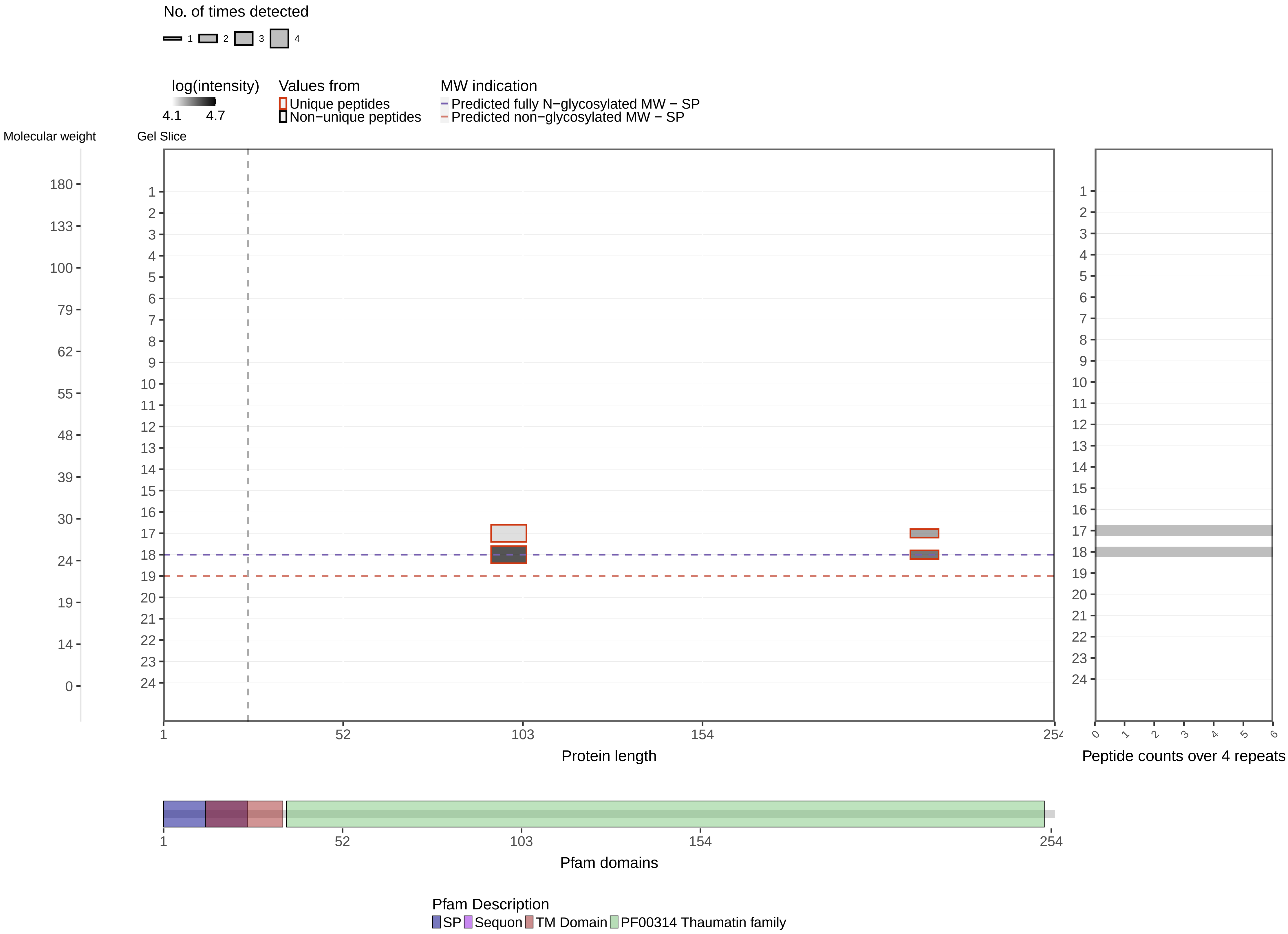


Protein sequence:

ATLAQFALHHGHADFSSYGVSLVDGFNIPLTVTPHEGKGQCPVVGCRAEILQTCPAVLQLRSHGGHAVVGCKSGCEAFKTDELCCRNHFNSPQTCKGSSYSQFFKHACPATFTYAHDRPSLMHECSSPRELKVIFCH*

### NbL17g09850.1 heat_shock_cognate_70_kDa_protein [MW = 52.43 kDa]

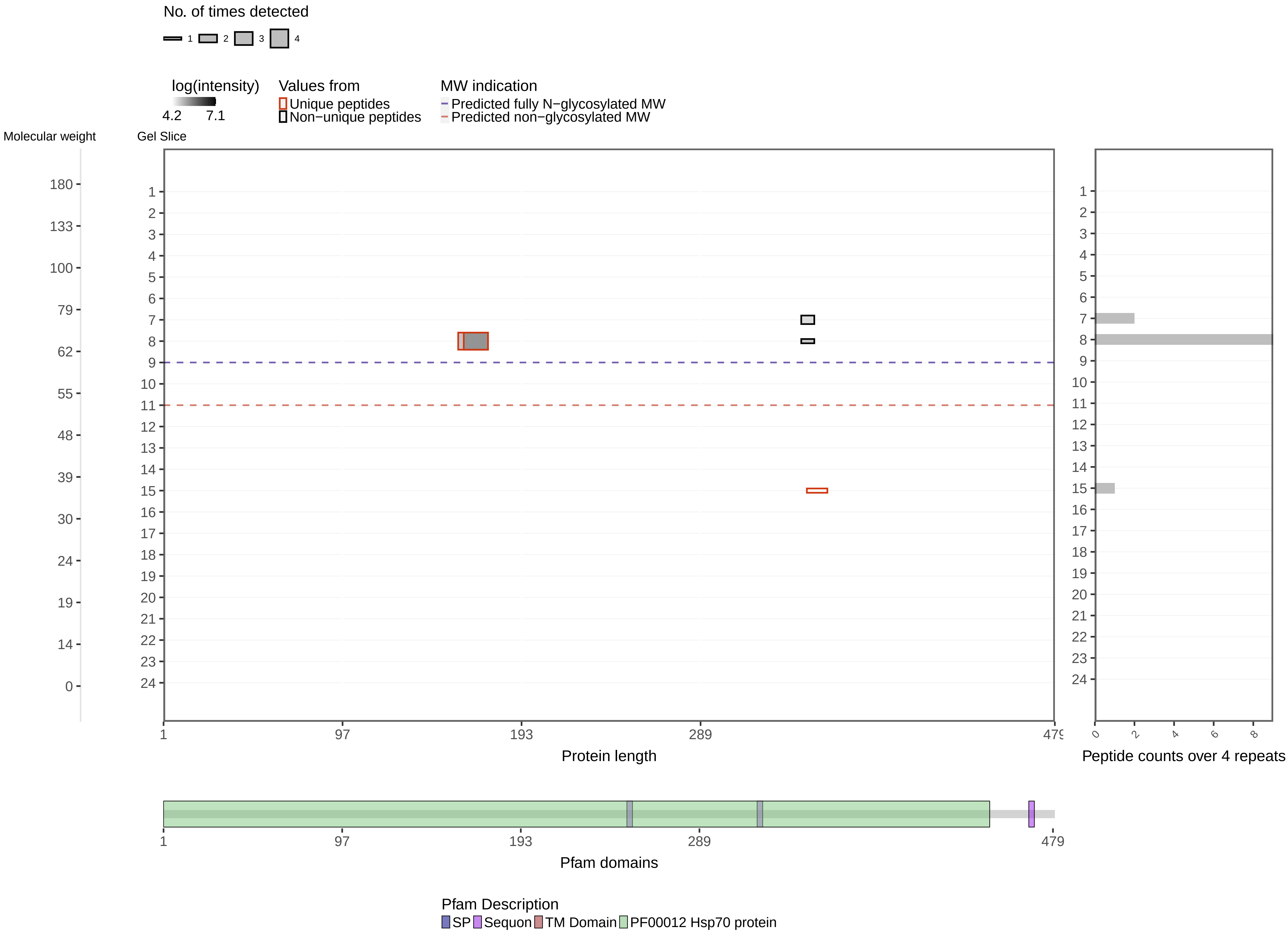


Protein sequence:

DVTPLSLGLETAGGVMTVLIPRNTTIPTKKEQVFSTYSDNQPGVLIQVYEGERARTKDNNLLGKFELSGIPPAPRGVPQINVCFDIDANGILNVSAEDKTTGQKNKITITNDKGRLSKEEIERMVQEAEKYKSEDEELKKKVEAKNALENYAYNMRNTIKDDKISSQLPAADRKKIEDAIDEAIKWLDSNQLAEVDEFEDKMKELEGICNPIIAKMYQGGAGGATMDEDGPSVAGGAGNGSGAGPKIEEVD*

NbL17g09630.1 60S_ribosomal_protein_L35−like [MW = 14.2 kDa]

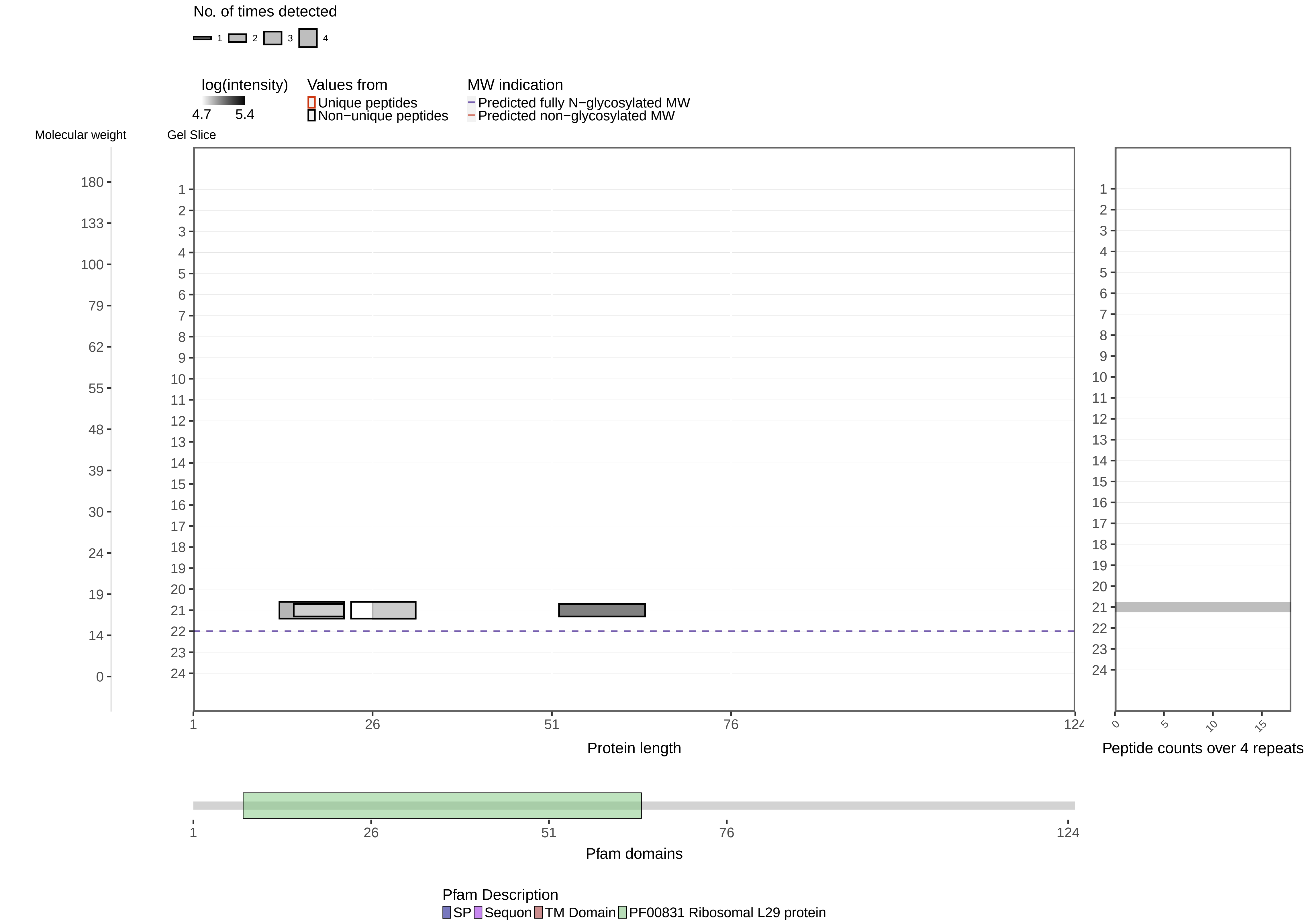


Protein sequence:
KVVRLSIAQVLTVISQKQKTALREAYKNKKYLPLDLRPKKTRAIRKRLTKHQASLKTEREKKKEMYFPIRKYAIKV*

### NbL17g09530.1 calcium−transporting_ATPase_4_endoplasmic_reticulum−type−like [MW = 116.18 kDa]

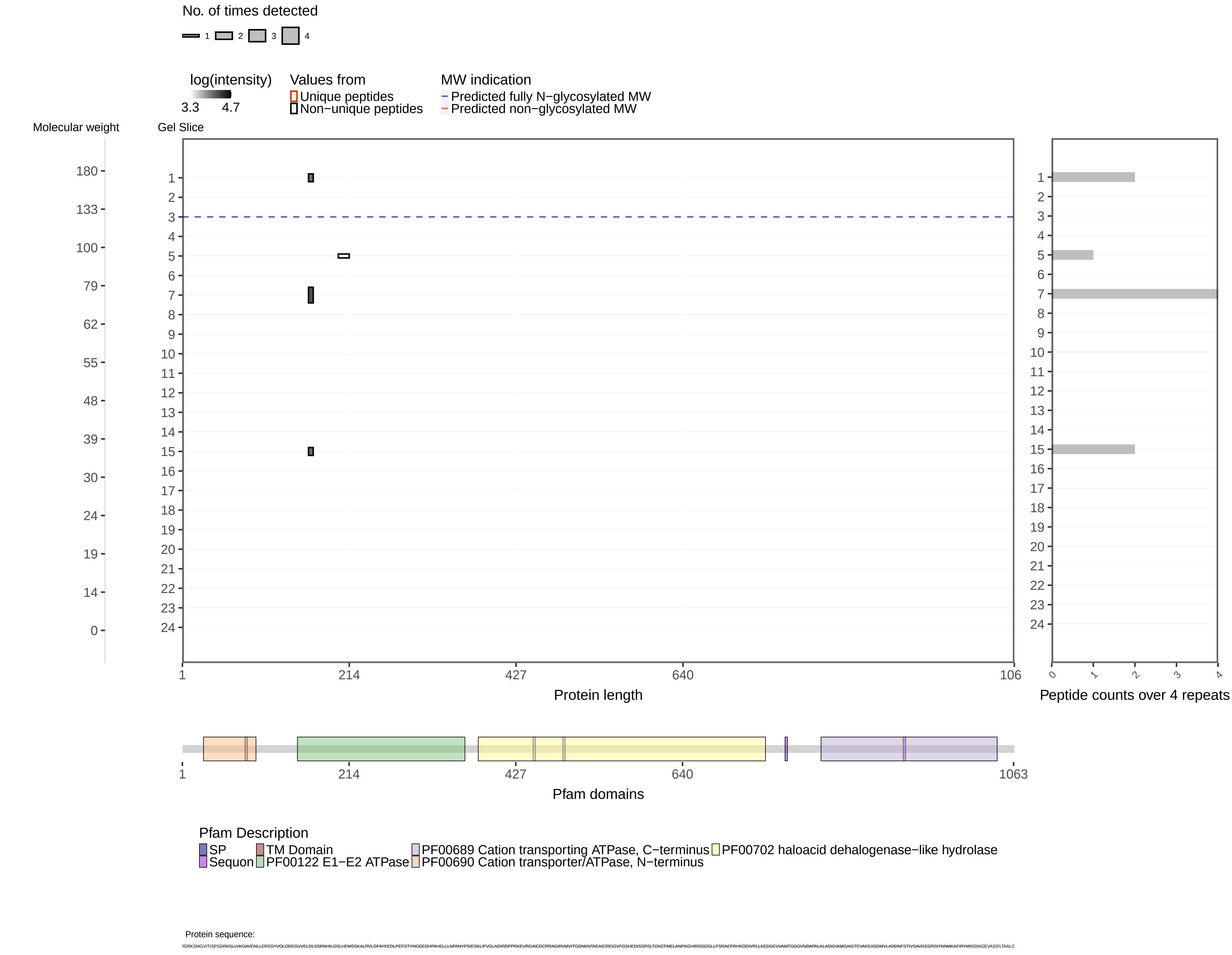

### NbL17g09400.1 phototropin−1−like [MW = 114.89 kDa]

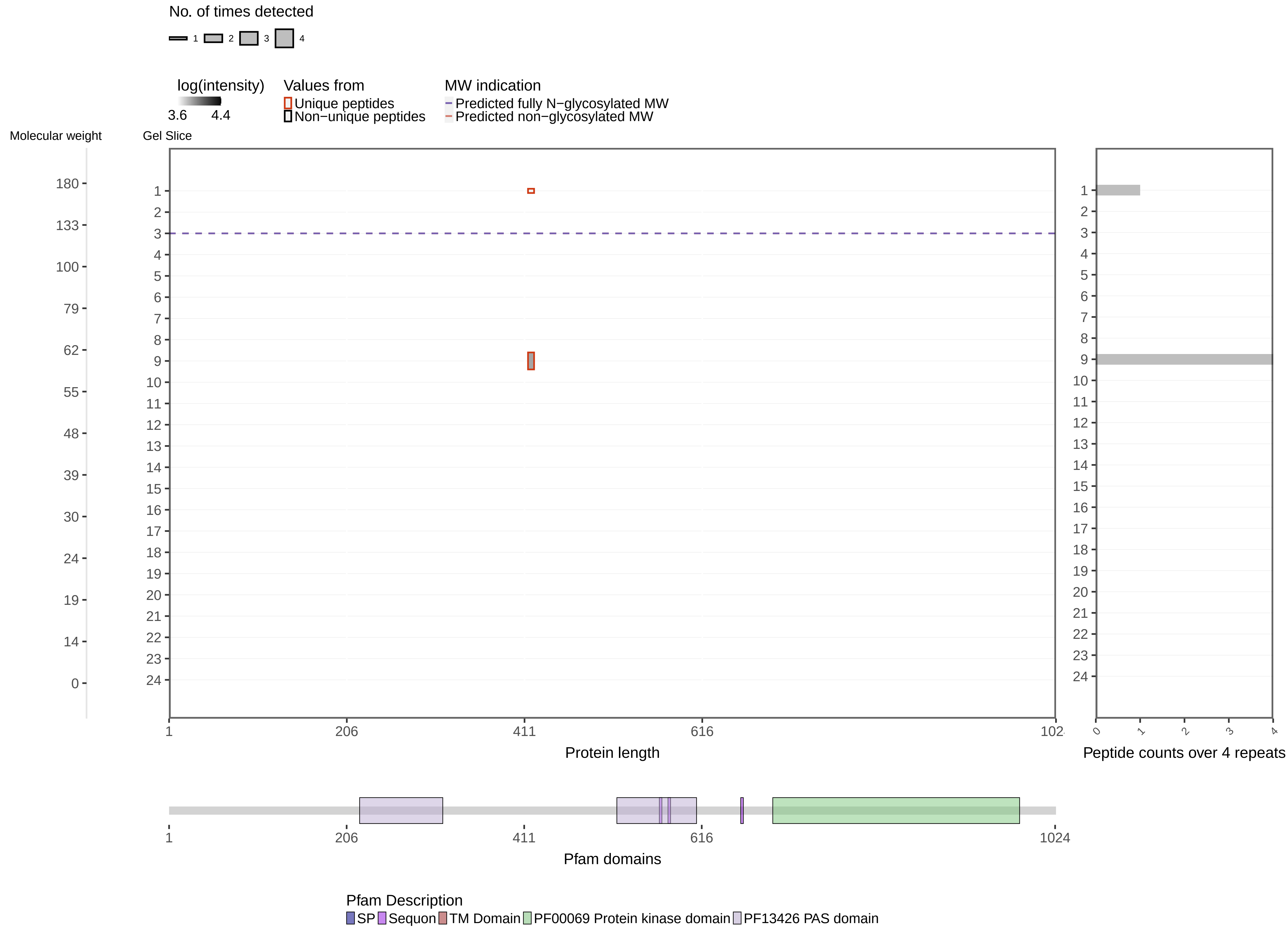


Protein sequence:

[illegible]GIDLATTLERIEKNFVITDPRLPDNPIIFASDSFLELTEYSREEILGRNCRFLQGPETDPATVKKIRQAIDNQTDVTVQLINYTKTGKKFWNLFHLQPMRDQKGEVQYFIGVQLDGSQHVEPLHNSIPEDKATESAKLVKETAENVDEAVRELPDANSKPEDLWRNHSKVVHAKPHRKDSPSWQAIQKVLDSGEPIGLKHFKPIKPLGSGDTGSVHLVELCGTGQCFAMKAMDKSIMLNRNKVHRACAEREILDMLDHPFLPALYASFQTKTHICLITDYYPGGELFLLLDRQPTKV[illegible]

NbL17g09360.1 2−dehydro−3−deoxyphosphooctonate_aldolase_1 [MW = 32.79 kDa]

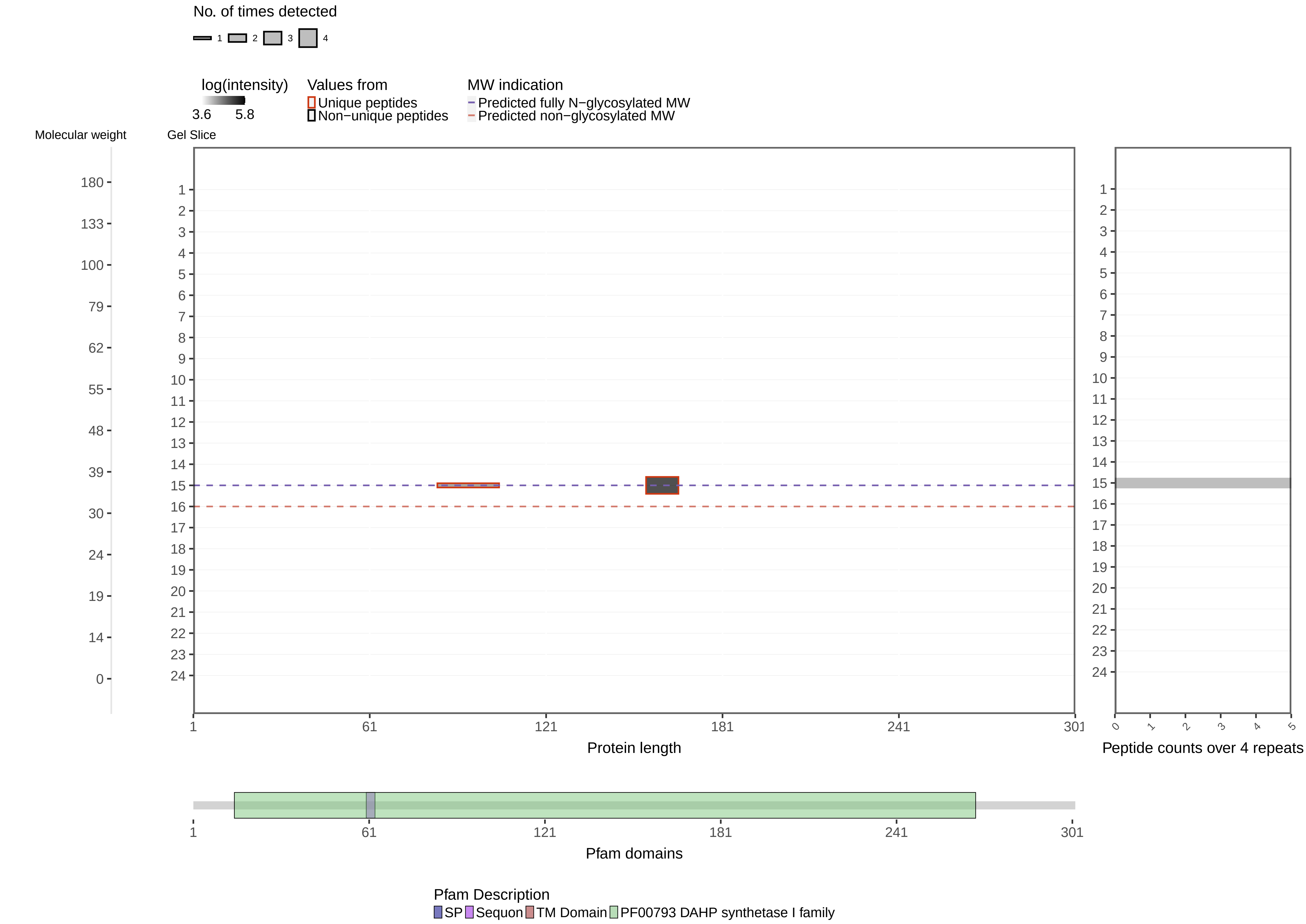


Protein sequence:

CAPSVMANSAEKVRLAGNQNVMVCERGTMFGYNDLIVDPRNFEWMREANCPVVADITHSLQQPAGKKLEGGGVASGGLRELIPCIARTAVAVGVDGIFMEVHNDPLSAPVDGPTQWPLRHLEELLEELVTLGVRELAKGSSSSKLISLHFVIRTQKFGYL*

### NbL17g09290.1 probable_trans−2−enoyl−CoA_reductase_mitochondrial [MW = 41.16 kDa]

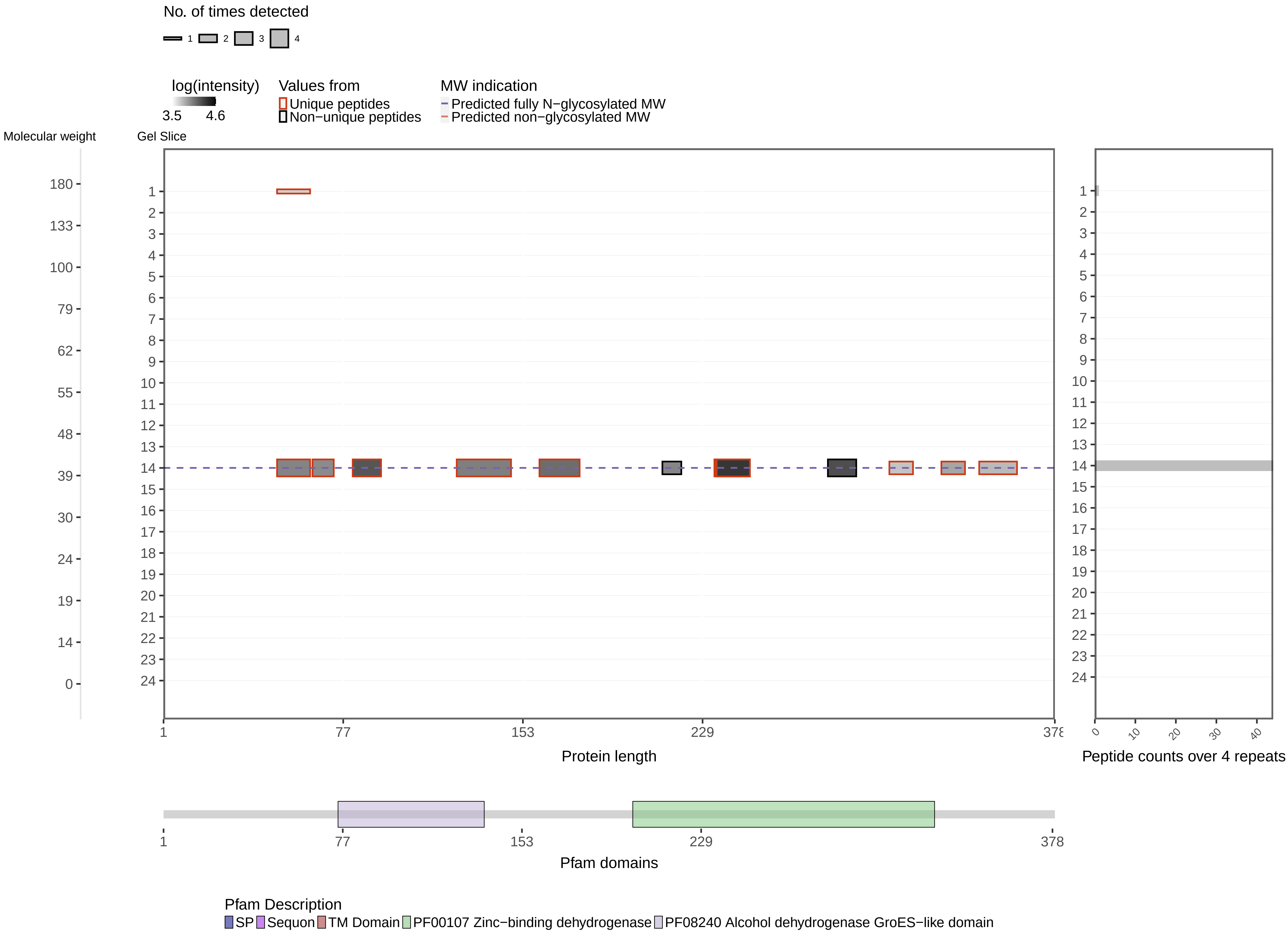


Protein sequence:

RMLEDFVTLKPGNSIVQNGATSMVGQCFIQLARLRGIHSINIIRDRAGSDEAKEYLKKLGADEVYTESQLEVKNVRGLLGSISEPALGFNCVGGNAALLVLKFLKQGGTMVTYGGMSKRPITVSTSSFIFKELTLTGFWLQRWMSSDKAEECRSMIDYLLGLCRDGKLKYELEVAPFDDFHSALEKAMGKRGSQPKQVLKF*

### NbL17g09070.1 ATP_synthase_subunit_d_mitochondrial−like [MW = 19.79 kDa]

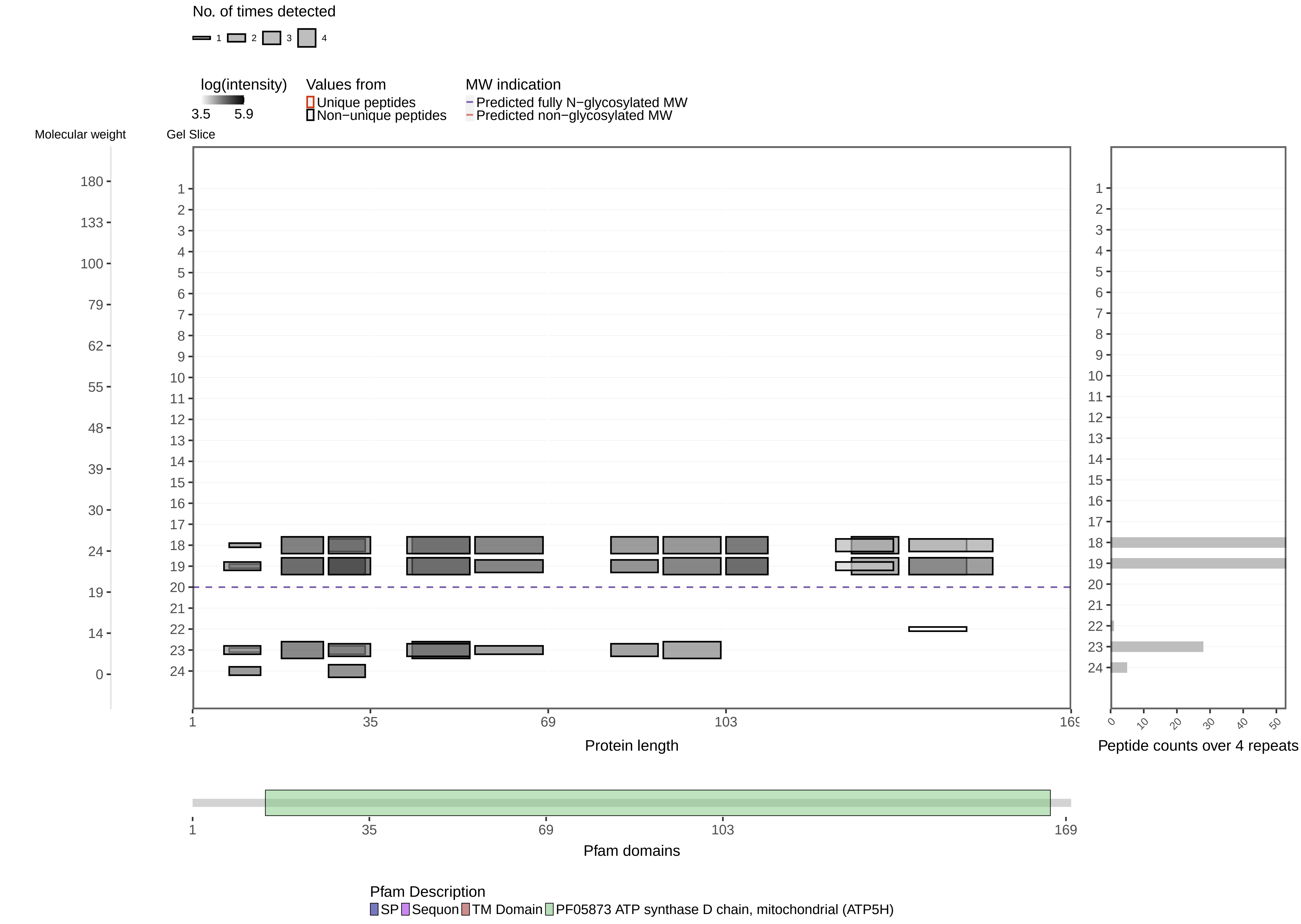


Protein sequence:

ːIGSRLVDMYKEAYESIEIPKFEDTVTPQYKPKFDALLVELKEAEQKSLKESERLEKEVAEVQELKKKLSTMTAEEYFEKHPELKKKFDDEIRNDYWGY*

NbL17g08990.1 elongation_factor_1−alpha [MW = 49.3 kDa]

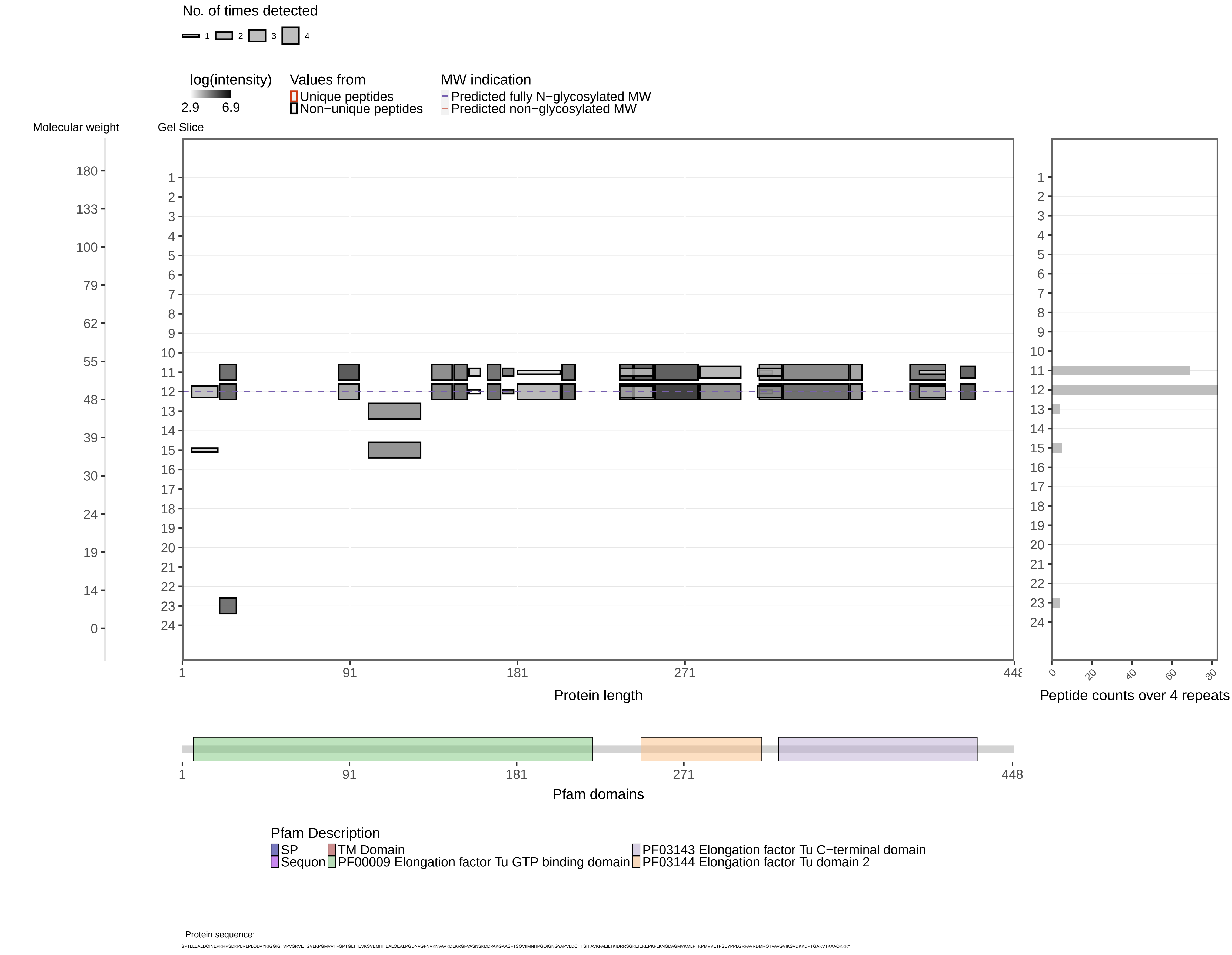

### NbL17g08980.1 protein_disulfide−isomerase_5−1 [MW − SP = 13.76 kDa]

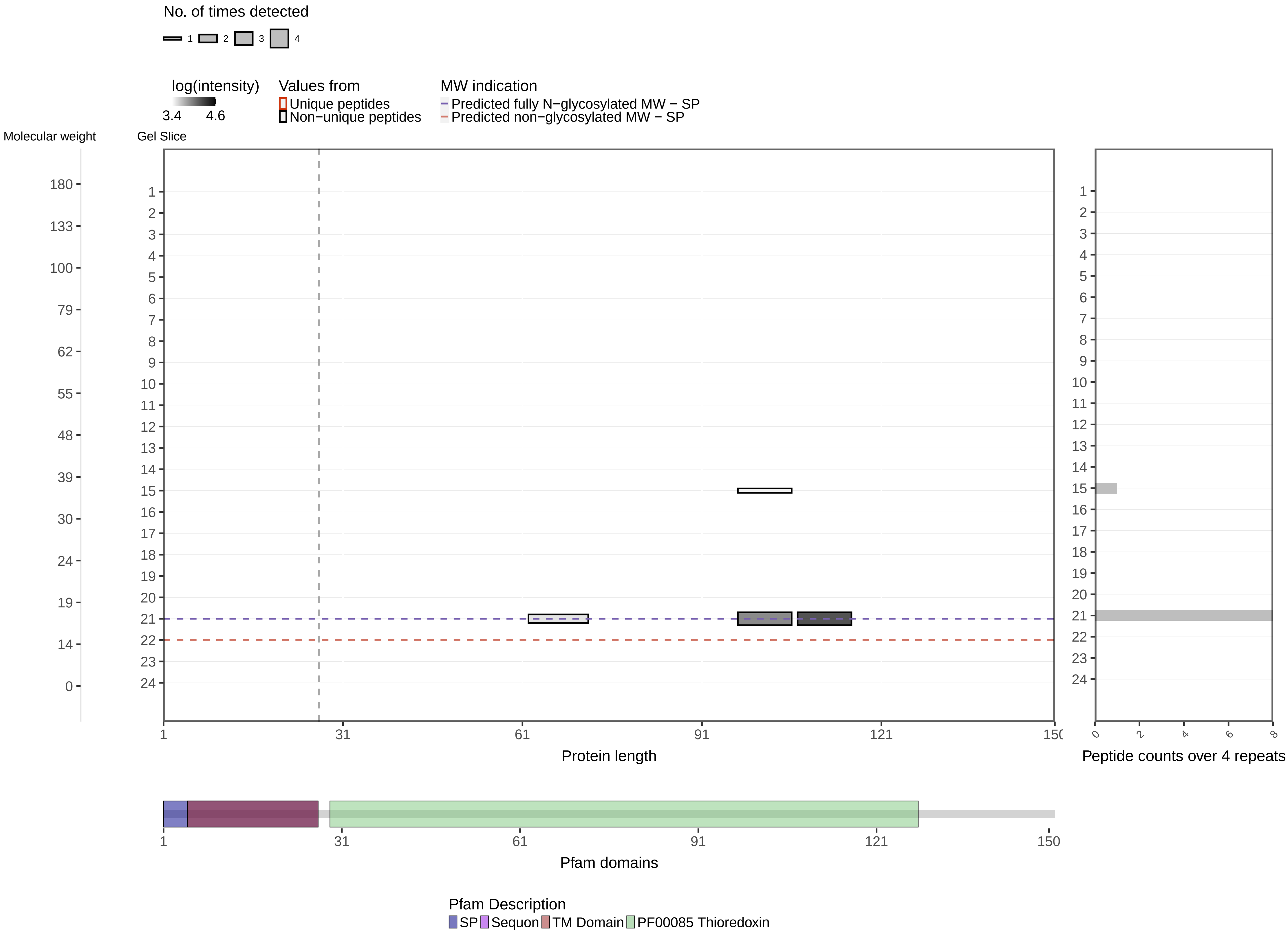


Protein sequence:

GTLWDDLGKAMEGEDEIEVGQVDCGTHKPVCNKVDIHSYPTFKLFYNGEEVAKYQGPRDIESLKIFALEEAEKAATKAEAGDDKEL*

NbL17g08910.1 ruBisCO_large_subunit−binding_protein_subunit_alpha [MW = 62.14 kDa]

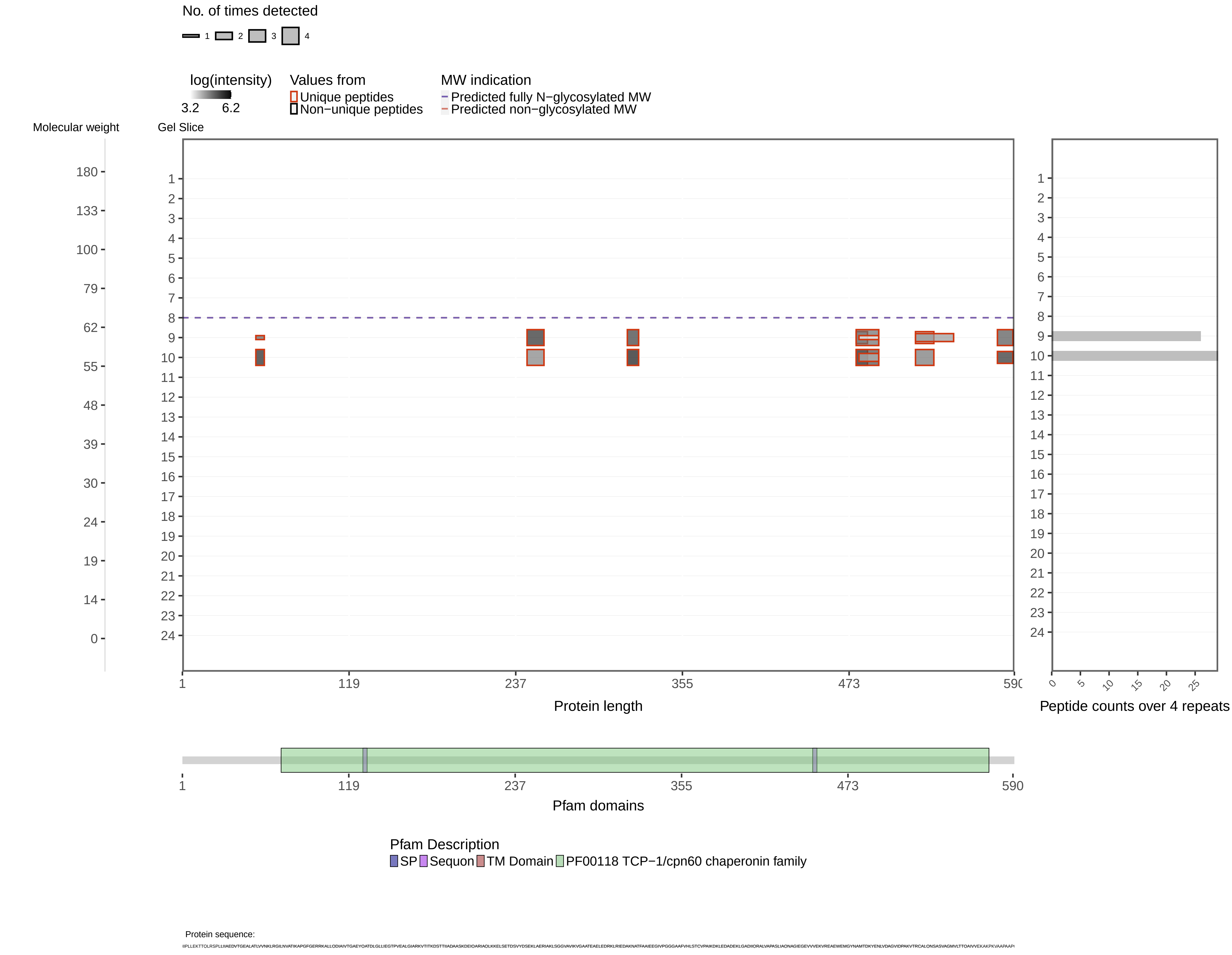

### NbL17g08830.1 glutaredoxin−C4 [MW − SP = 12.08 kDa]

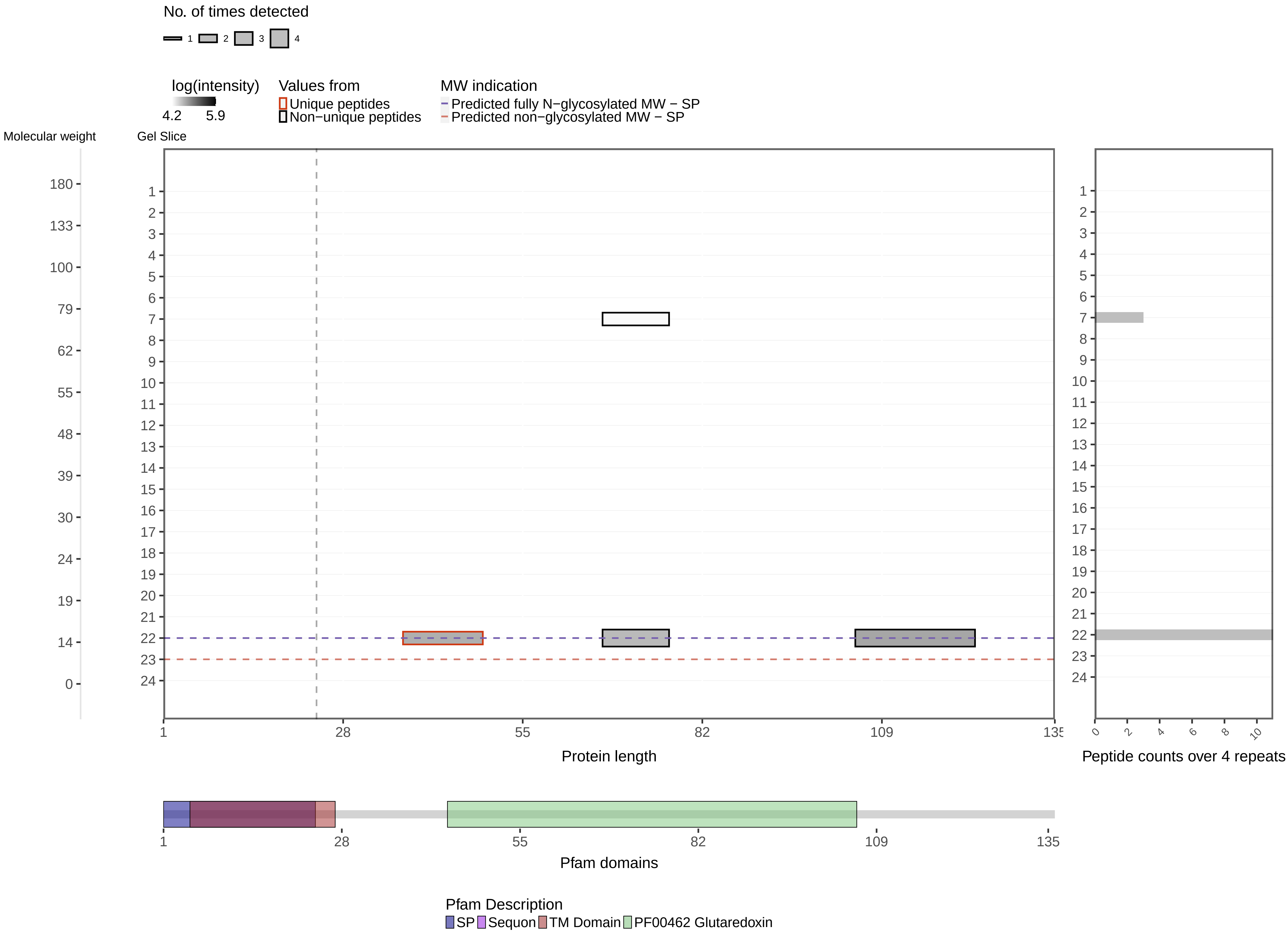


Protein sequence:
RKAKAVFKELKQKPYVVELDERDDGWSIQDALSEIVGRRTVPQVFINGKHIGGSDDTVEAYENGELAKLLGLSAKKDDL*

NbL17g08820.1 cysteine−rich_receptor−like_protein_kinase_10 [MW − SP = 78.14 kDa]

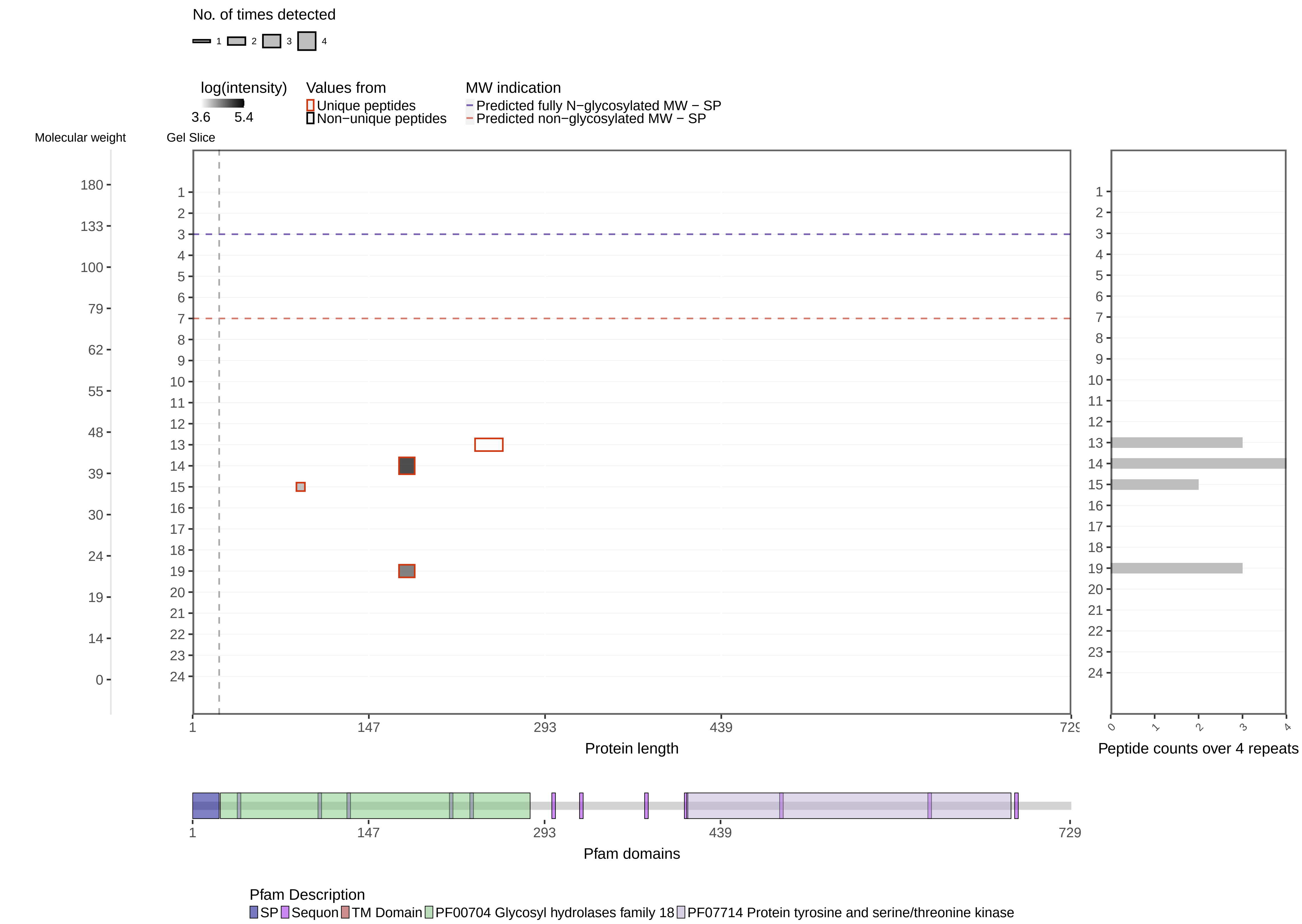


Protein sequence:

LSSLCYLLLRKGKAKVTALLLLNQTTNRAKRRKMDKKMSEEVQFYSFESLGIATNNFSLGNKLGEGGFGPVYQGKFSDGQEVAIKRLSTSSGQGVLEFKNEILLIAKLQHTNLVRLLGYCTQGEERVLVYEYMHNKSLDFFLFDTNKKERINWGTRFRIIEGVAQGLLYLHKYSRLTVVHRDLKASNILLDSNMNPKISDFGMARIFGQQESEANTKRIVGTRGYMSPEYALRGIVSTKTDVFSFGVLLLEIVSGKKNNSCYDSEHPLNLIGLAWELWREERALELLDGTLIESC

NbL17g08580.1 4−hydroxy−3−methylbut−2−en−1−yl_diphosphate_synthase_(ferredoxin)_chloroplastic [MW = 82.1 kDa]

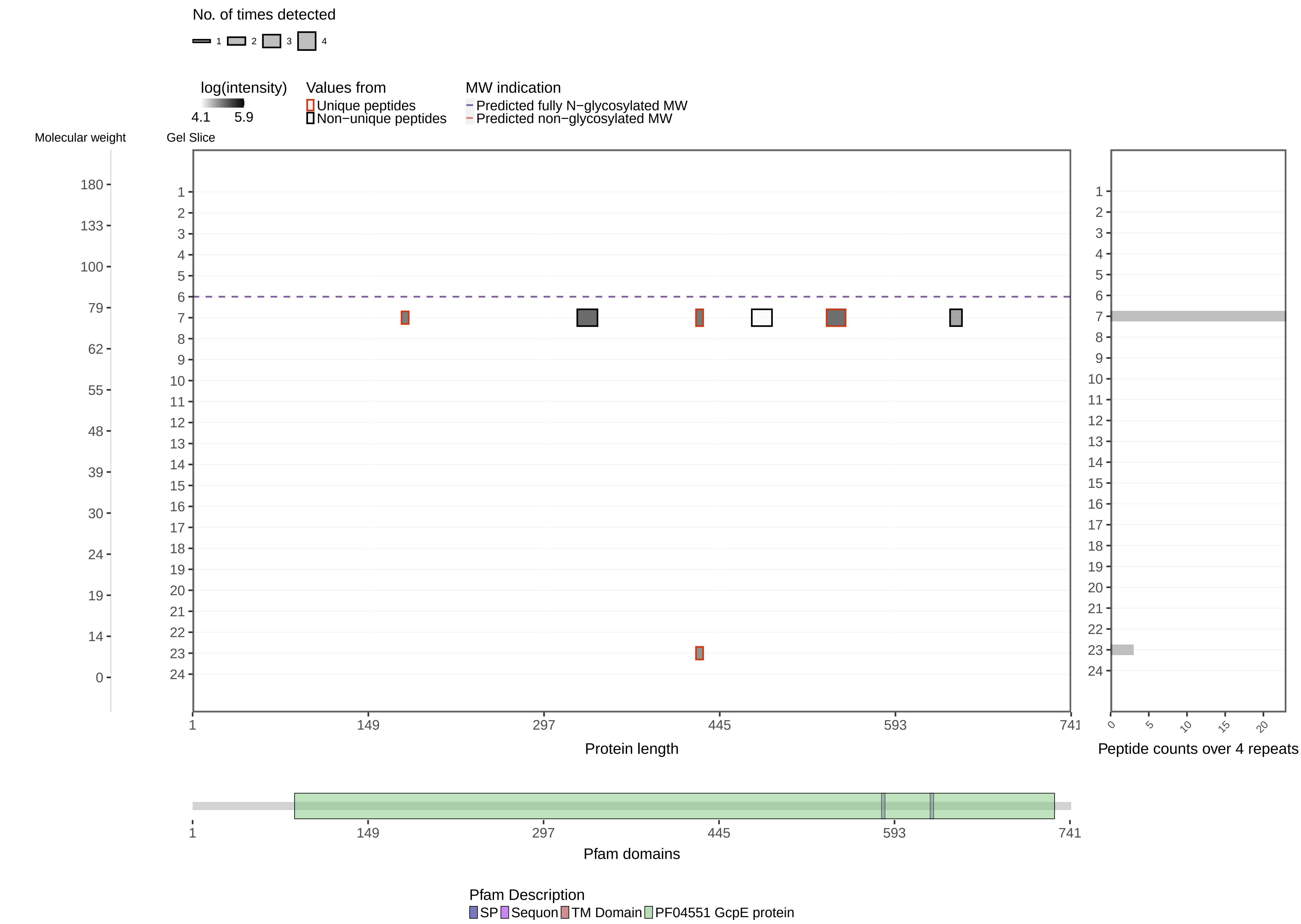


Protein sequence:

RLANLGKRAAEIQQGVAPFEEKNRRYFDFQRRSGELPAQKEGDEVDYRGVLHRDGSVLMSVSLDQLKAPELLYRSLAAKLVVGMPFKDLATVDSILLRELPPVDDKDSRLALKRLVDISMGVITPLSEQLTKPLTNVMVLVTLKELSGGAQKLLPEGTRLVVSVRGDESKDELEILKSTDVTMILHHLPYTEEKIGRVHAARRLFEYLSQNSLNFPVIHHIHFPNDTHRDDLVIGAGANAGALLVDGLGDGILLEAPDQDFDFLRNTSFNLLQGCRMRNTKTEYVSCPSCGRTLF

NbL17g08320.1 isovaleryl−CoA_dehydrogenase_mitochondrial [MW = 48.67 kDa]

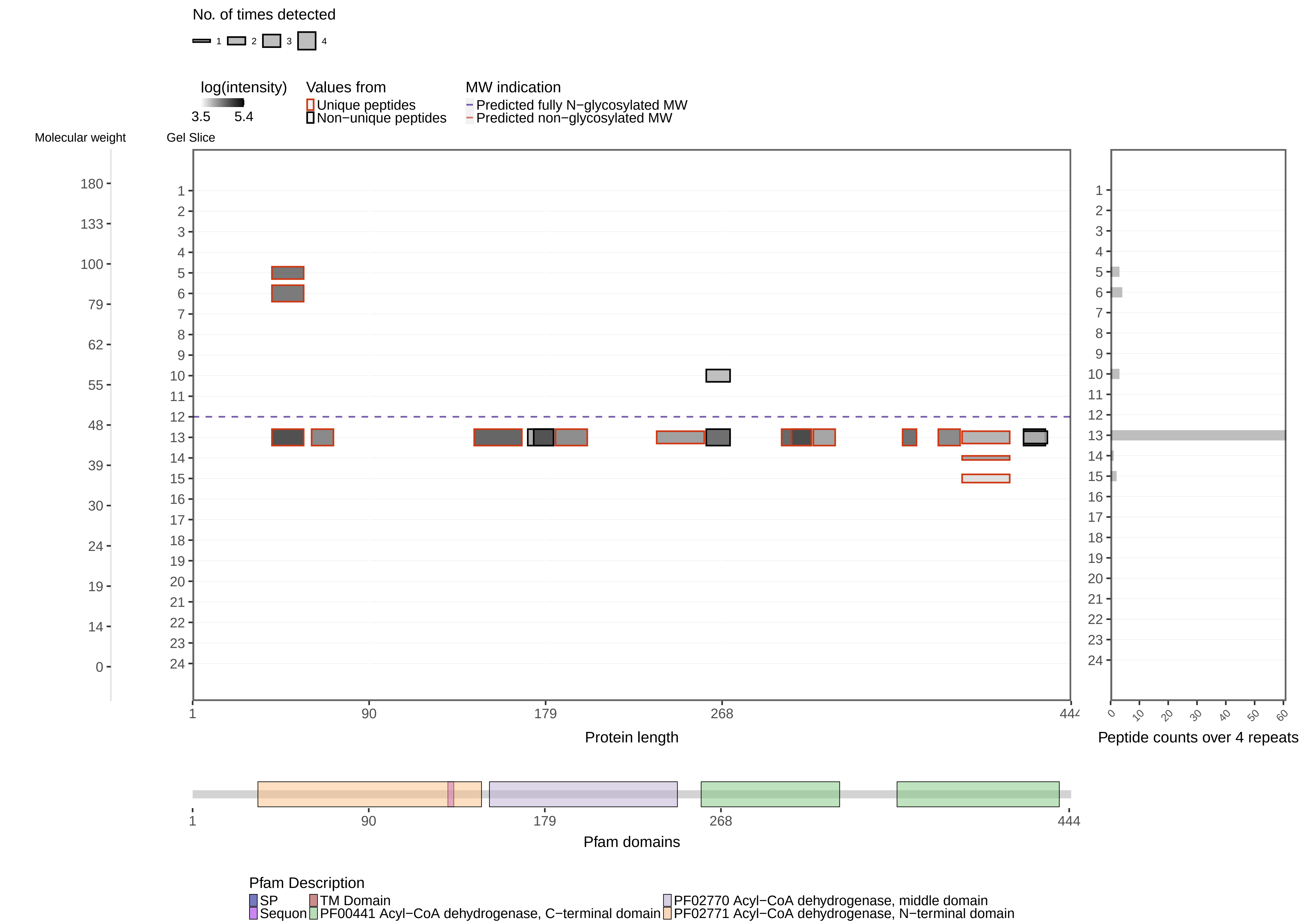


Protein sequence:

SKGITAFIIEKGMKGFSTAQKLDKLGMRGSDTCELVFENCFVPEENILGQVGKGVYVLMSGLDLERLVLSSGPVGLMQACLDVVLPYVKQREQFGRPIGEFQFIQGKVADMYTSLQSSRSGVLCCHTSSISFSLLTLINNFMYLFVFIVHNRAYVYSVARECDSGNINTKDCAGVILIAAERATQVALQAVQCLGGNGYVNEYPTGRLLRDAKLYEIGAGTSEIRRMIIGRELFKEQ*

### NbL17g08300.1 proteasome_subunit_beta_type−4−like [MW = 27.66 kDa]

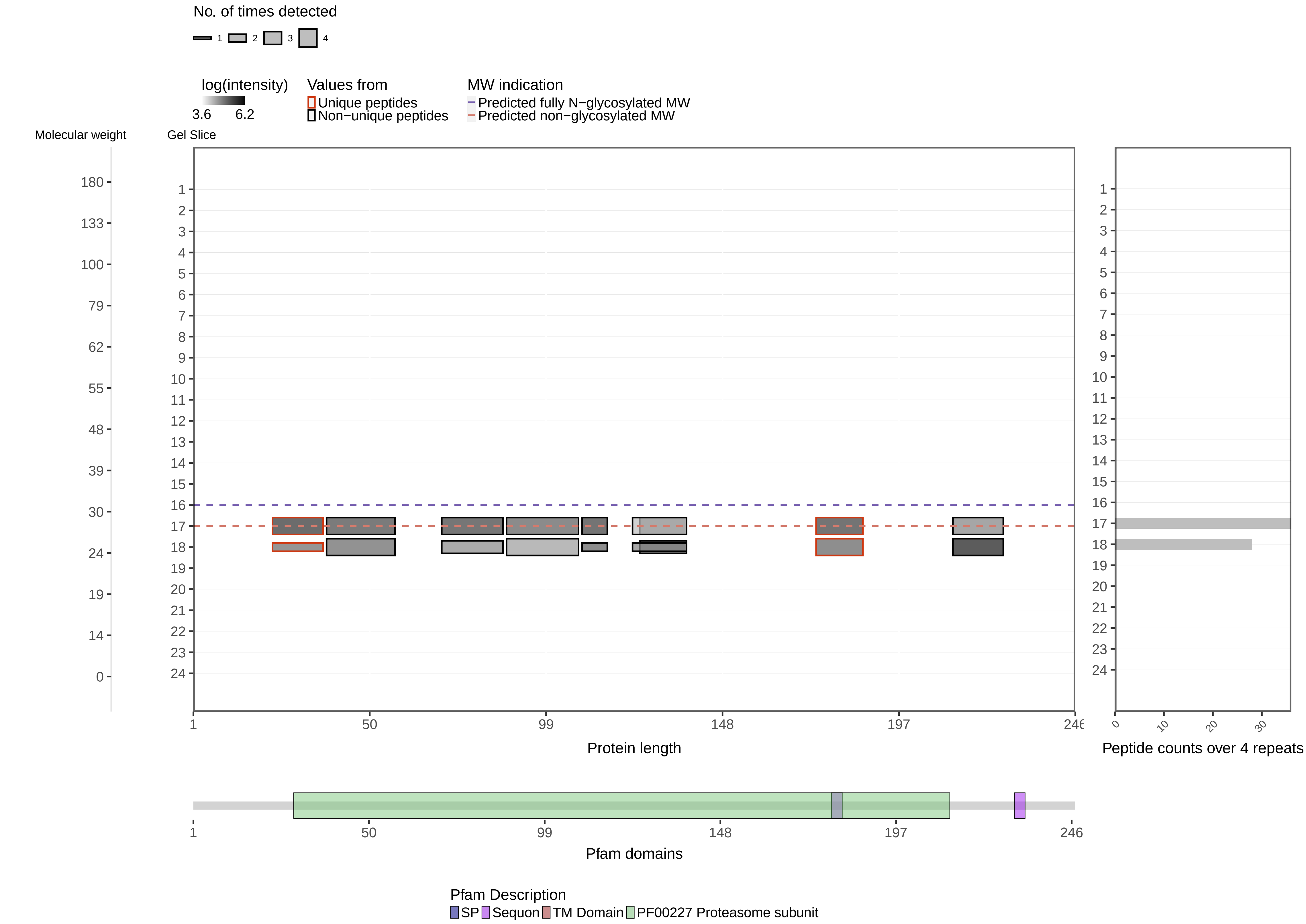


Protein sequence:

INYLTRMMYNRRNKFNPLWNSLVLGGVKNGQKYLGTVSMIGVHYEDDHVATGFGNHLARPILREEWHENLSYEEGVKLLEKCMRVLLYRDRSAVNKLQIAKITEEGVTVSQPYSLKTFWNFSAFQNPTVGAEGSW*

### NbL17g08290.1 ABC_transporter_F_family_member_1−like [MW = 67.48 kDa]

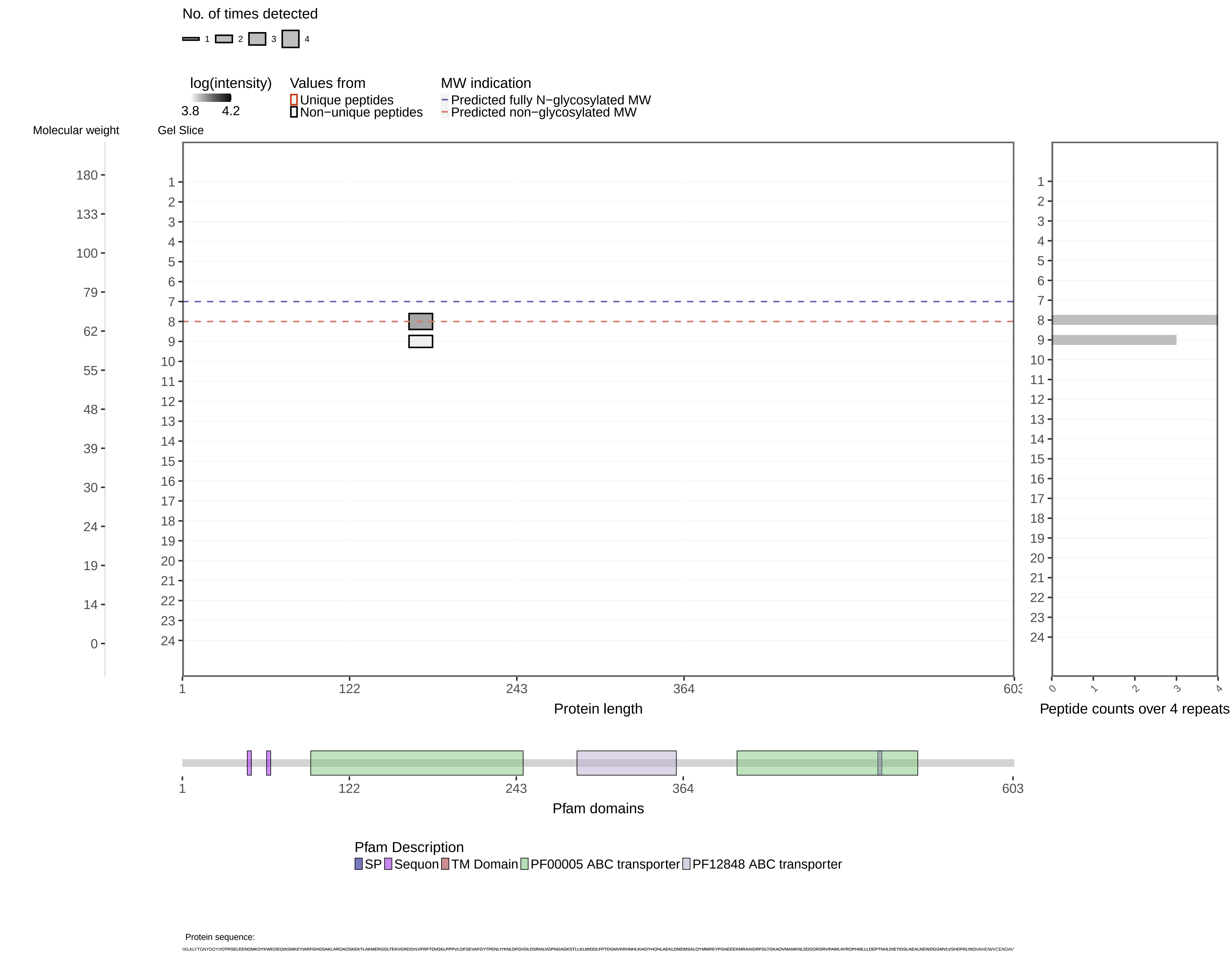

NbL17g08240.1 4−coumarate−−CoA_ligase−like_7 [MW = 58.9 kDa]

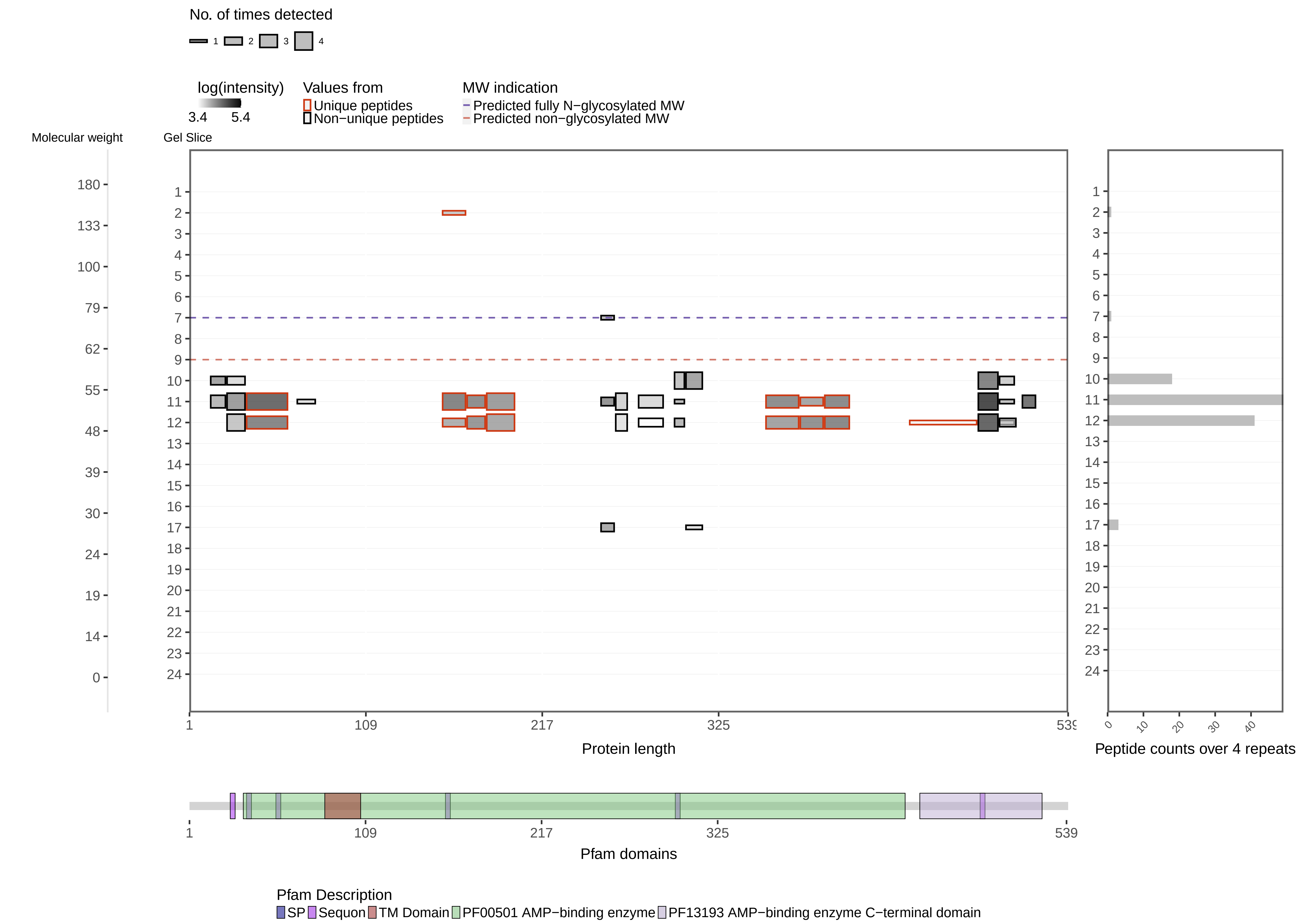


Protein sequence:

MAKFDLEMILKSVDKYRVTHLWIVPPVVLALAKNPVVKKYNLTSLRNIGSGAAPLGKELMEECAKNFPQVVVLQGYGMTETCGIVAVESPYVGPRHTGSAGTLVPGVEAQIVSVDKLKPLTPGQLGEIWVRGPNIMQGYFNNPQATKLTIDNQGWVHTGDLGYFDEEGMLYVVDRIKELIKYKGFQVAPAELEGLLVSHPEILDAVVIPFPDAEAGEVPIGYVVRSPNSSLTEEDVQKFIADQVAPFKRLRKVTFVNSVPKSASGKILRRELIEKVRSKL*

### NbL17g08190.1 T−complex_protein_1_subunit_beta−like [MW = 57.01 kDa]

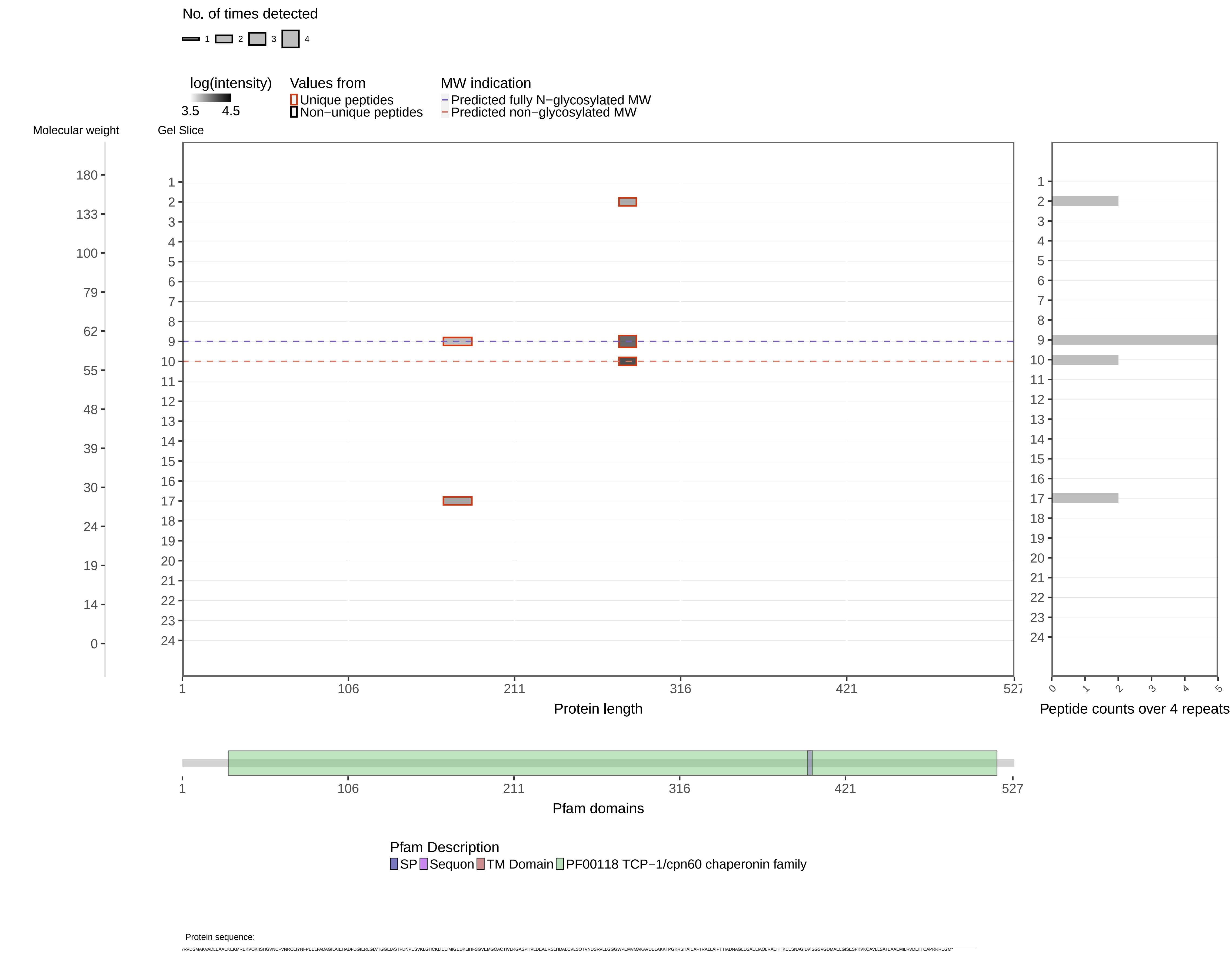

### NbL17g08130.1 xyloglucan_endotransglucosylase/hydrolase_protein_31−like [MW = 42.3 kDa]

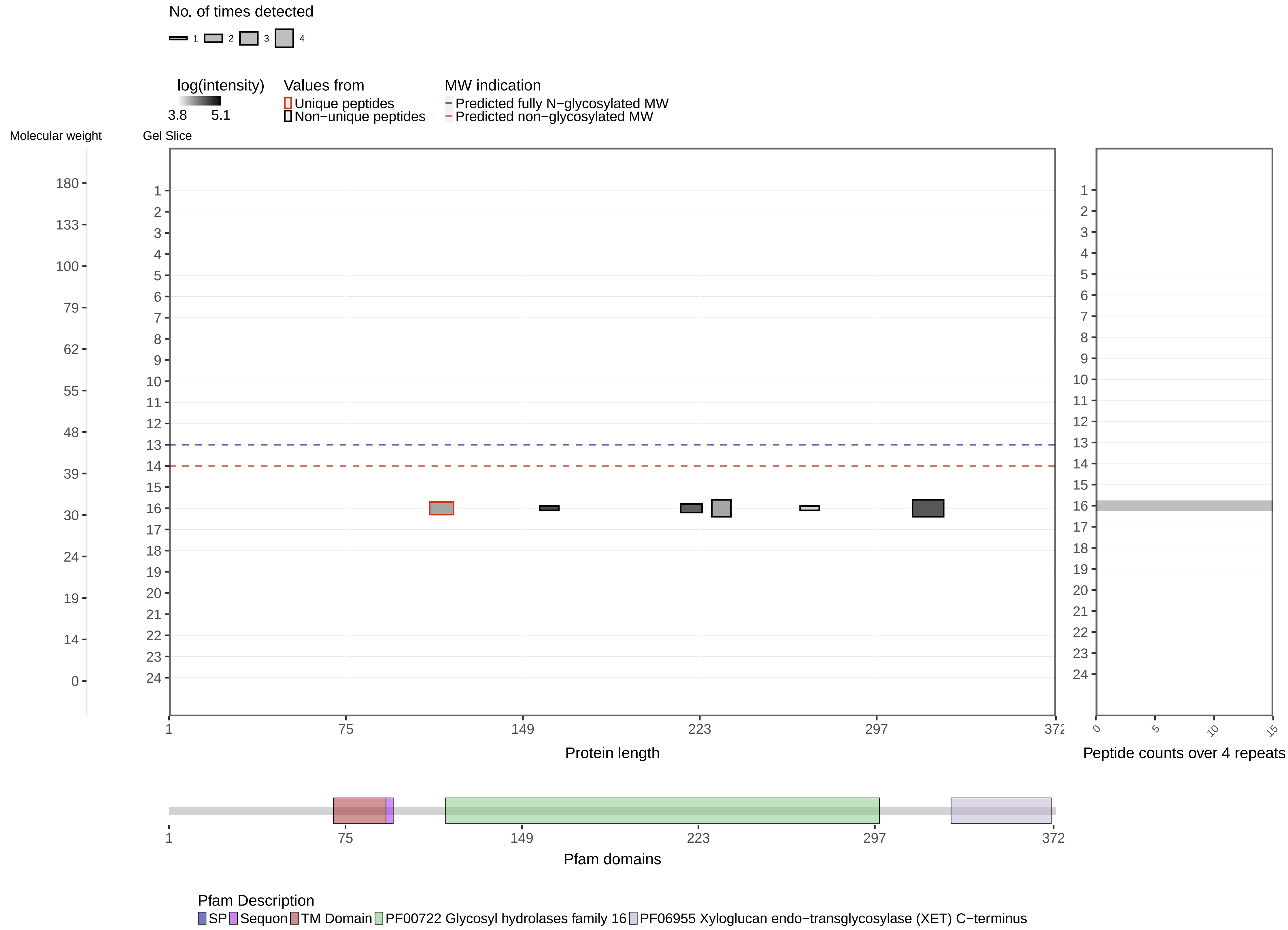


Protein sequence:

### NbL17g07970.1 superoxide_dismutase_[Cu−Zn]_chloroplastic [MW = 29.56 kDa]

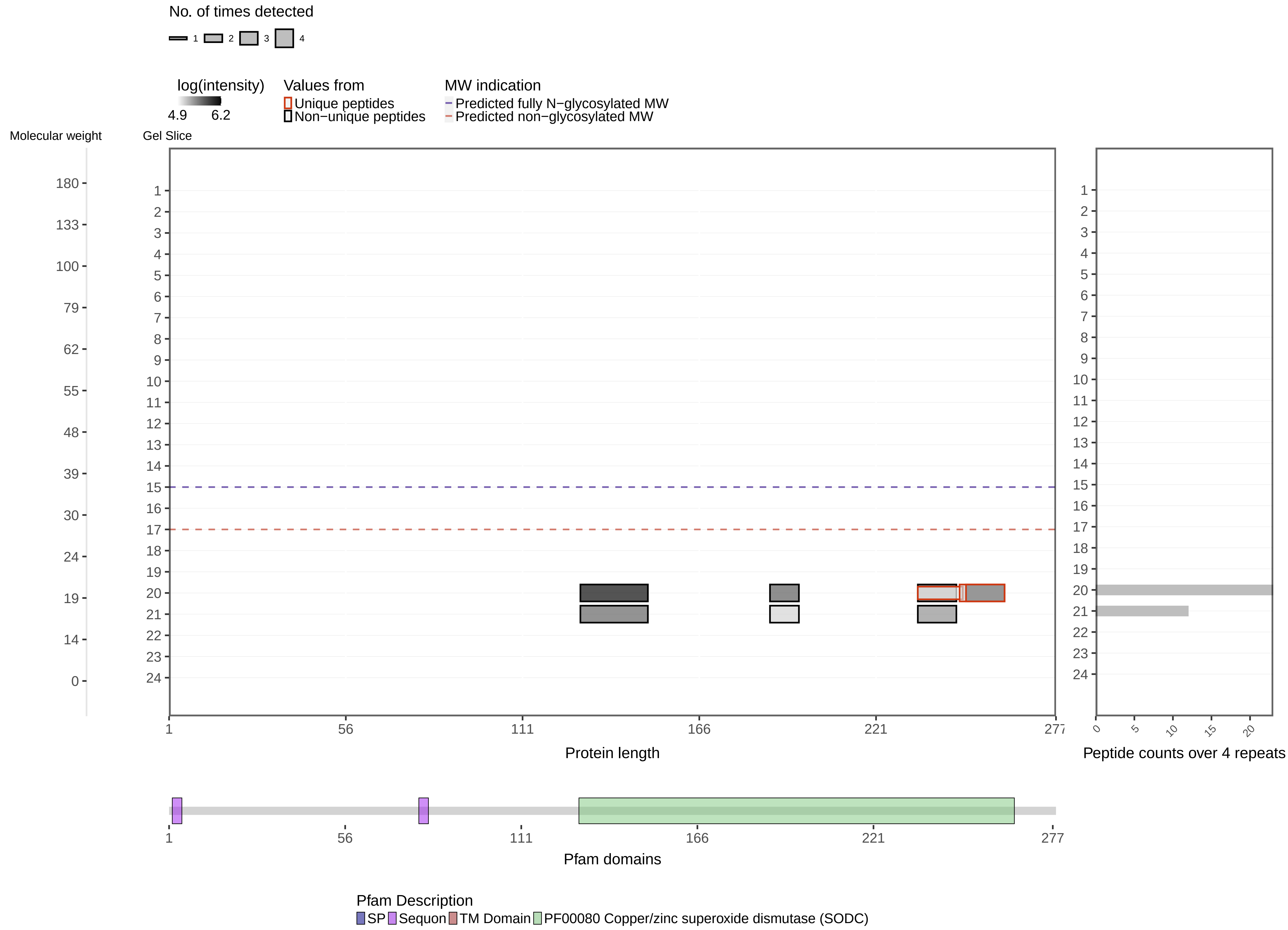


Protein sequence:

NSNVEGVVTLSQDDDGPTTVKVRITGLTPGLHGFHLHEYGDTTNGCMSTGPHFNPDGKTHGAPEDEIRHAGDLGNIVANADGVAEATIVDNQIPLYGPNSVVGRALVVHELEDDLGKGRHELSLTTGNAGGRLACGKKHTWFDSNMK*

### NbL17g07960.1 50S_ribosomal_protein_L9_chloroplastic [MW = 21.64 kDa]

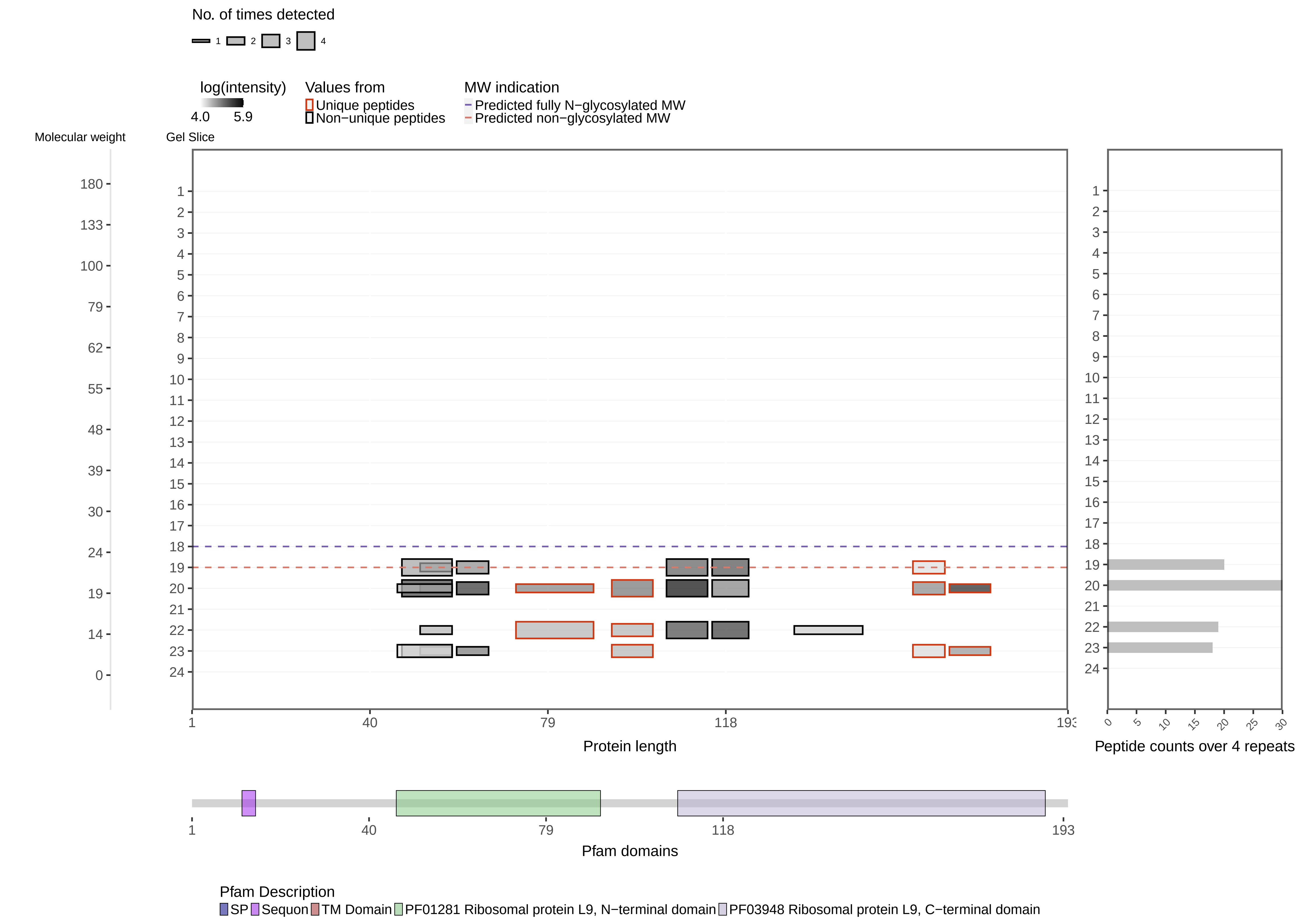


Protein sequence:

PTLLKEMKMEDERIEAEKQRVKEEALQLARIFETVGAFKVKRKGGKGKQIFGSVTAQDLVDIIKAQLQREVDKKIVSVPEIRETGEYVAELKLHQDVTARVRLTVFAN*

### NbL17g07940.1 histone_H2B−like [MW = 16.03 kDa]

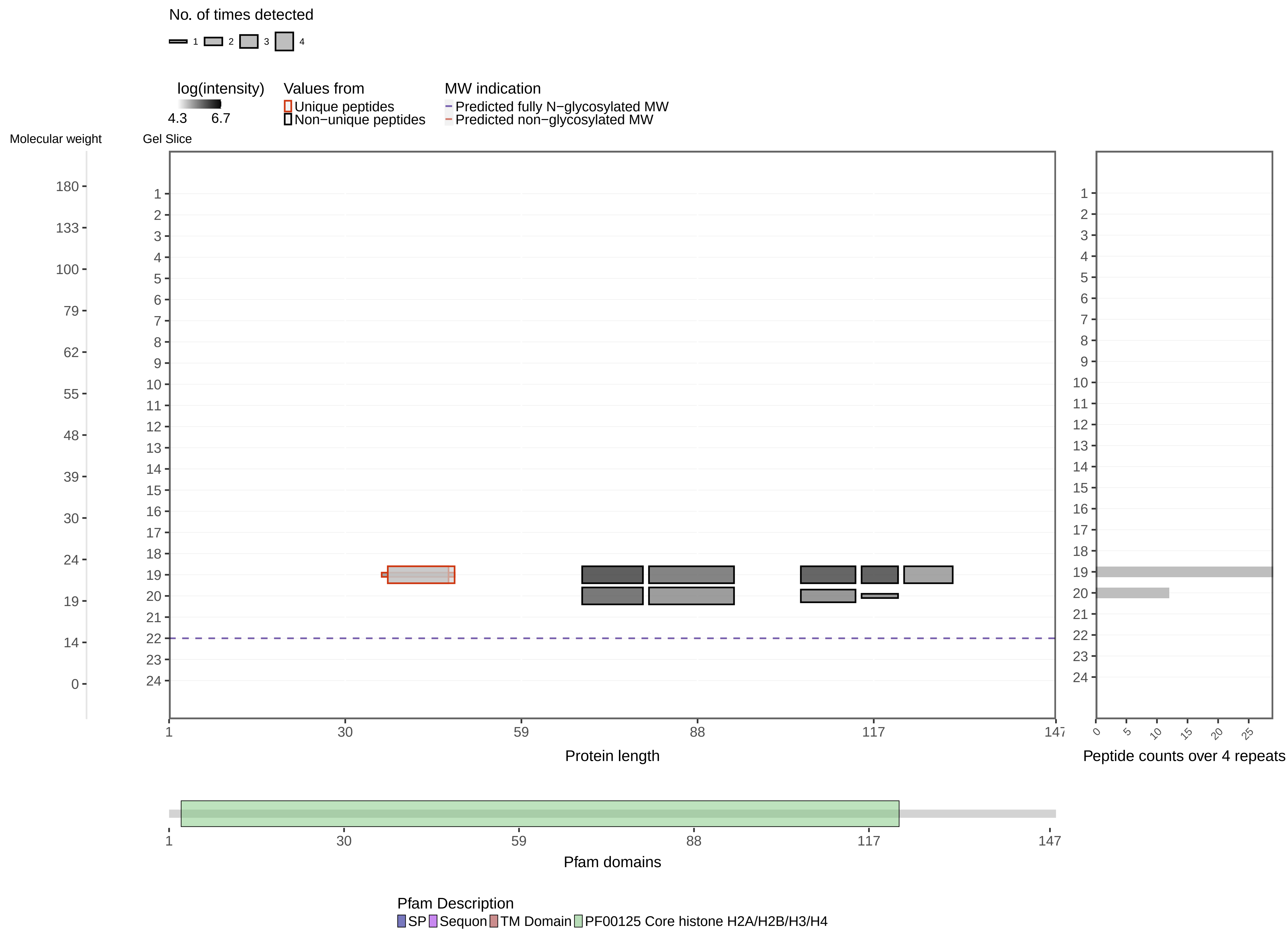


Protein sequence:

TYKIYIFKVLKQVHPDIGISSKAMGIMNSFINDIFEKLAQESSRLARYNKKPTITSREIQTAVRLVLPGELAKHAVSEGTKAVTKFTSS*

NbL17g07830.1 aspartic_proteinase−like_protein_2 [MW = 32.86 kDa]

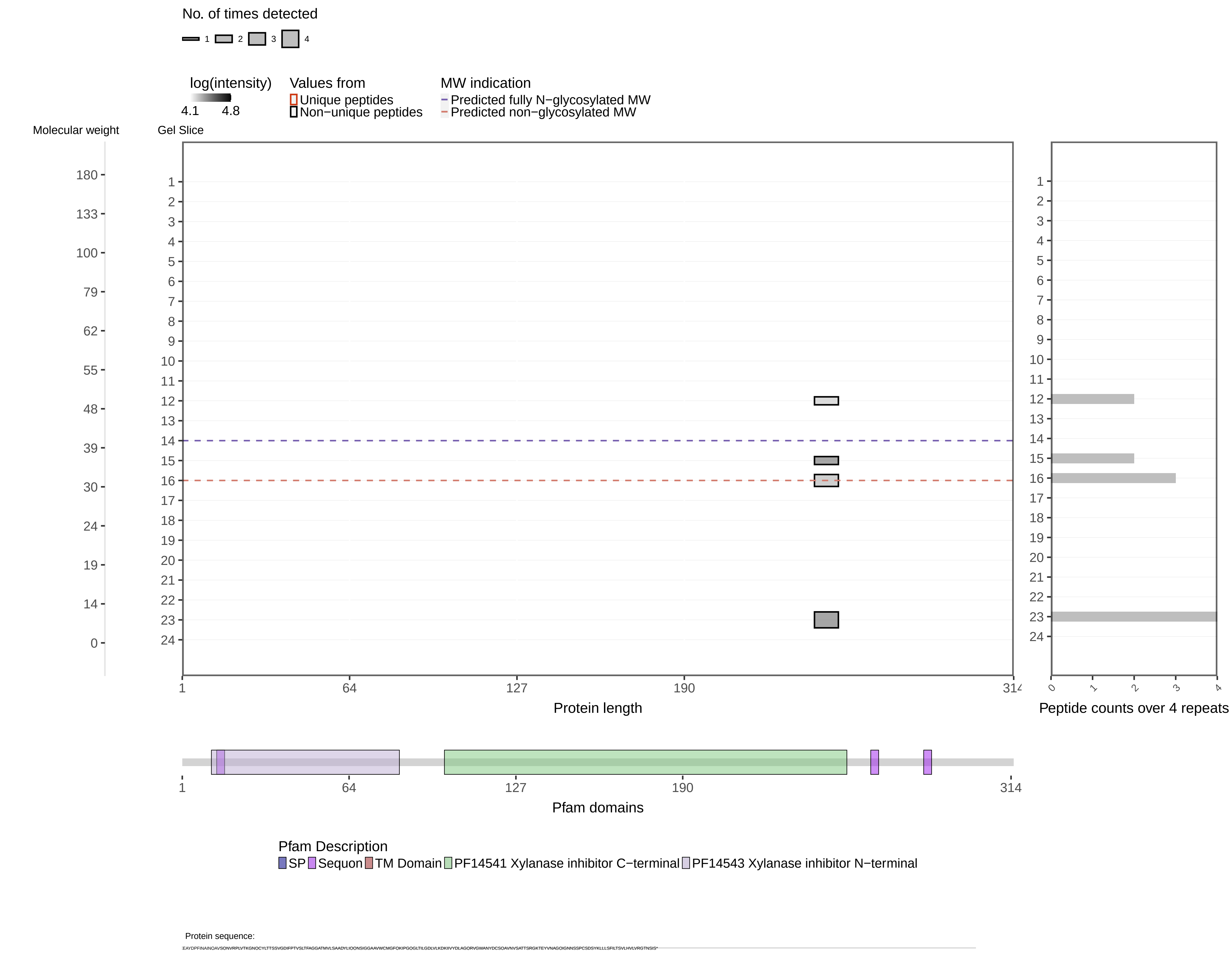

NbL17g07680.1 UDP−D−apiose/UDP−D−xylose_synthase_2 [MW = 43.48 kDa]

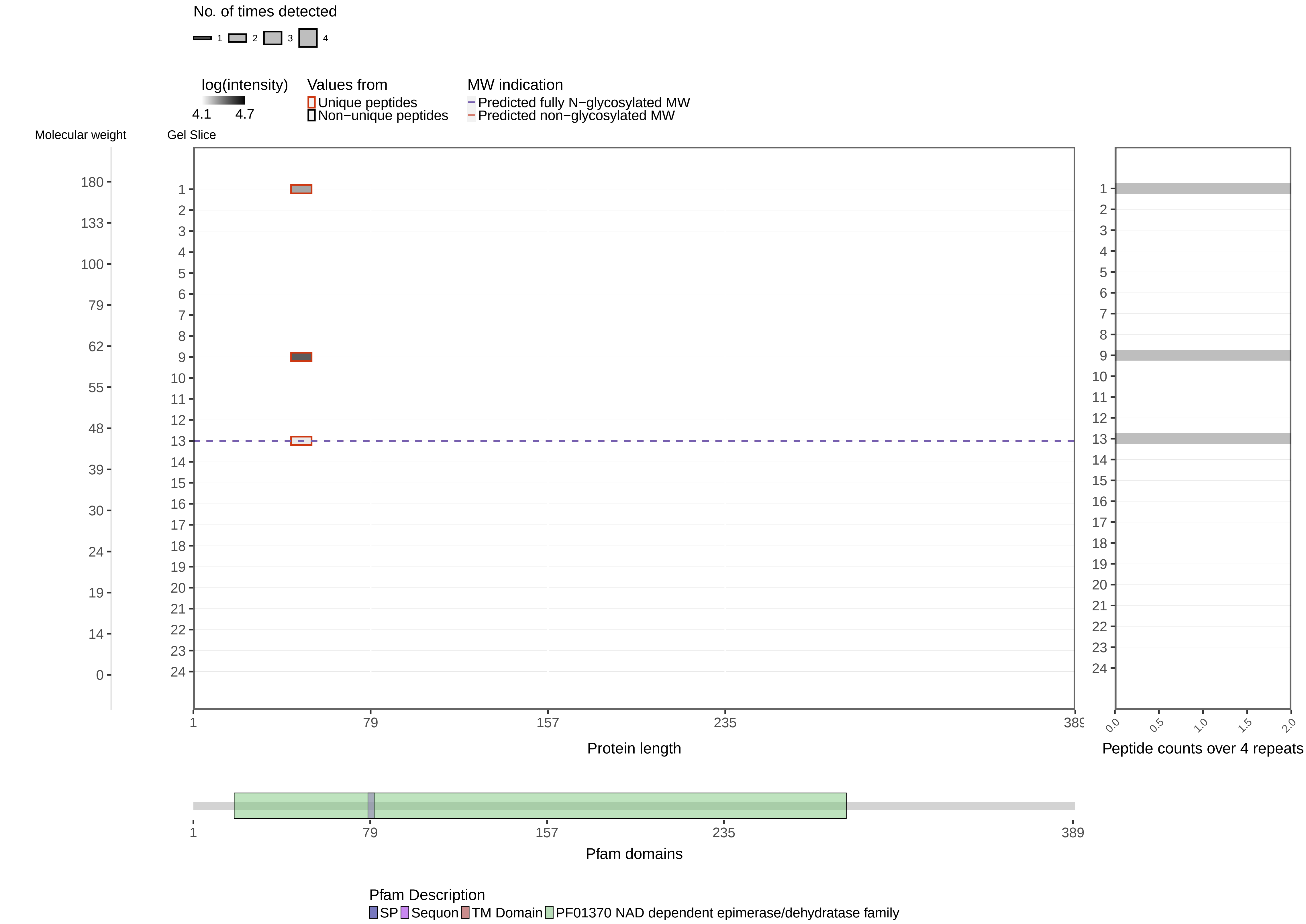


Protein sequence:
SYACAKQLIERLVFAEGAENGLEFTIVRPFNWIGPRMDFIPGIDGPSEGVPRVLACFSNNLFRREPLKLVDGGESQRTFIYIKDAIEAVLLMIENPARANGHIFNVGNPNNEVTVRQLAEMMTQVYSKVSGESSIETPTIDVSSKEFYGEGYDDSDKRIPDMTIINRQLGWNPKTSLWDLLESTLTYQHRTYAEAVKQAMSKTTAN*

### NbL17g07450.1 stress−response_A/B_barrel_domain−containing_protein_At5g22580 [MW = 12.23 kDa]

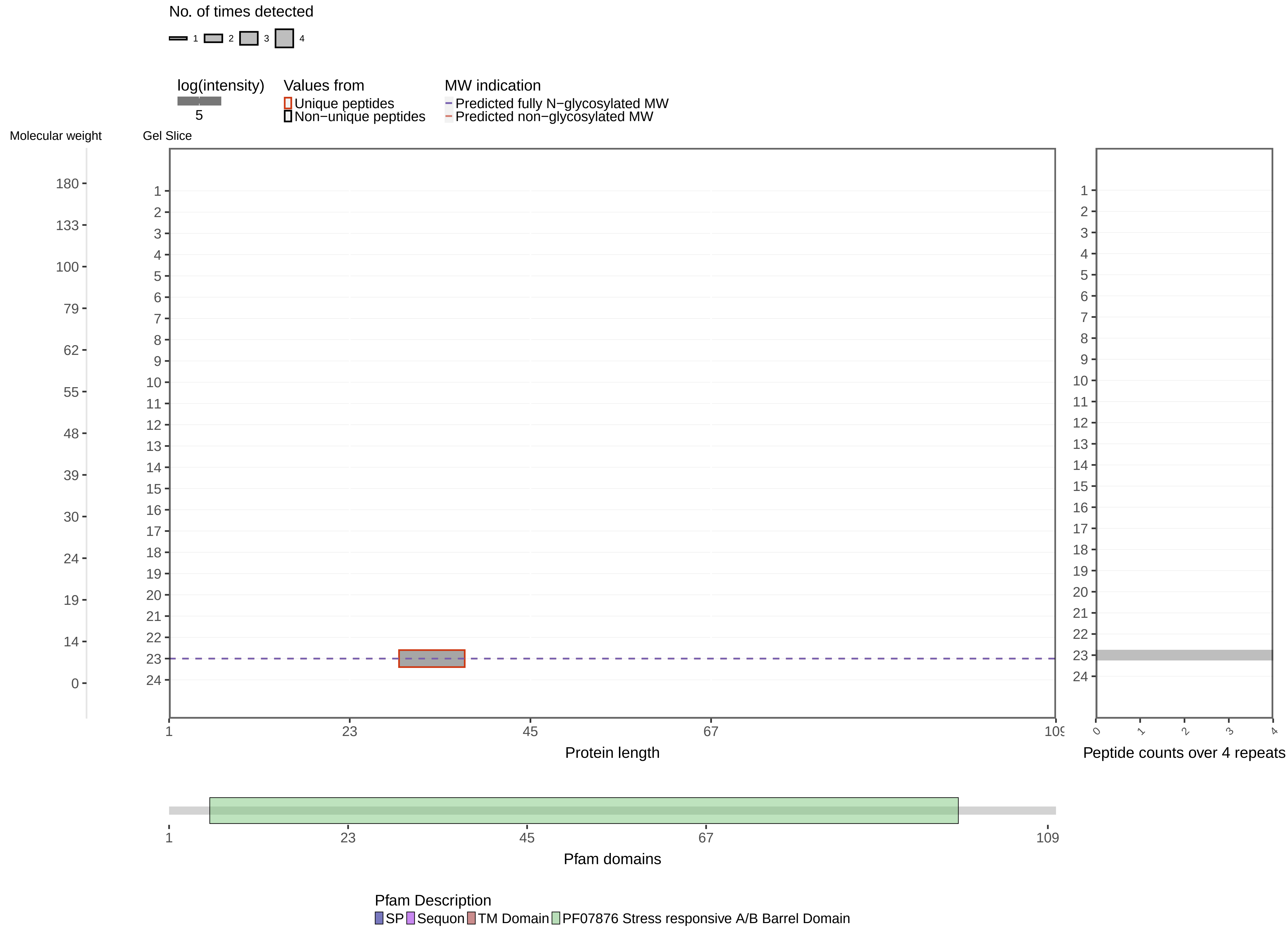


Protein sequence:

'GKDAESHEMLRQGFTHAFLMTFNNKDDYTTFVSHPNHVEFSATFSTVIDKAVLLDFPAISVKAPALA*

### NbL17g07340.1 peptidyl−prolyl_cis−trans_isomerase_CYP71 [MW = 70.48 kDa]

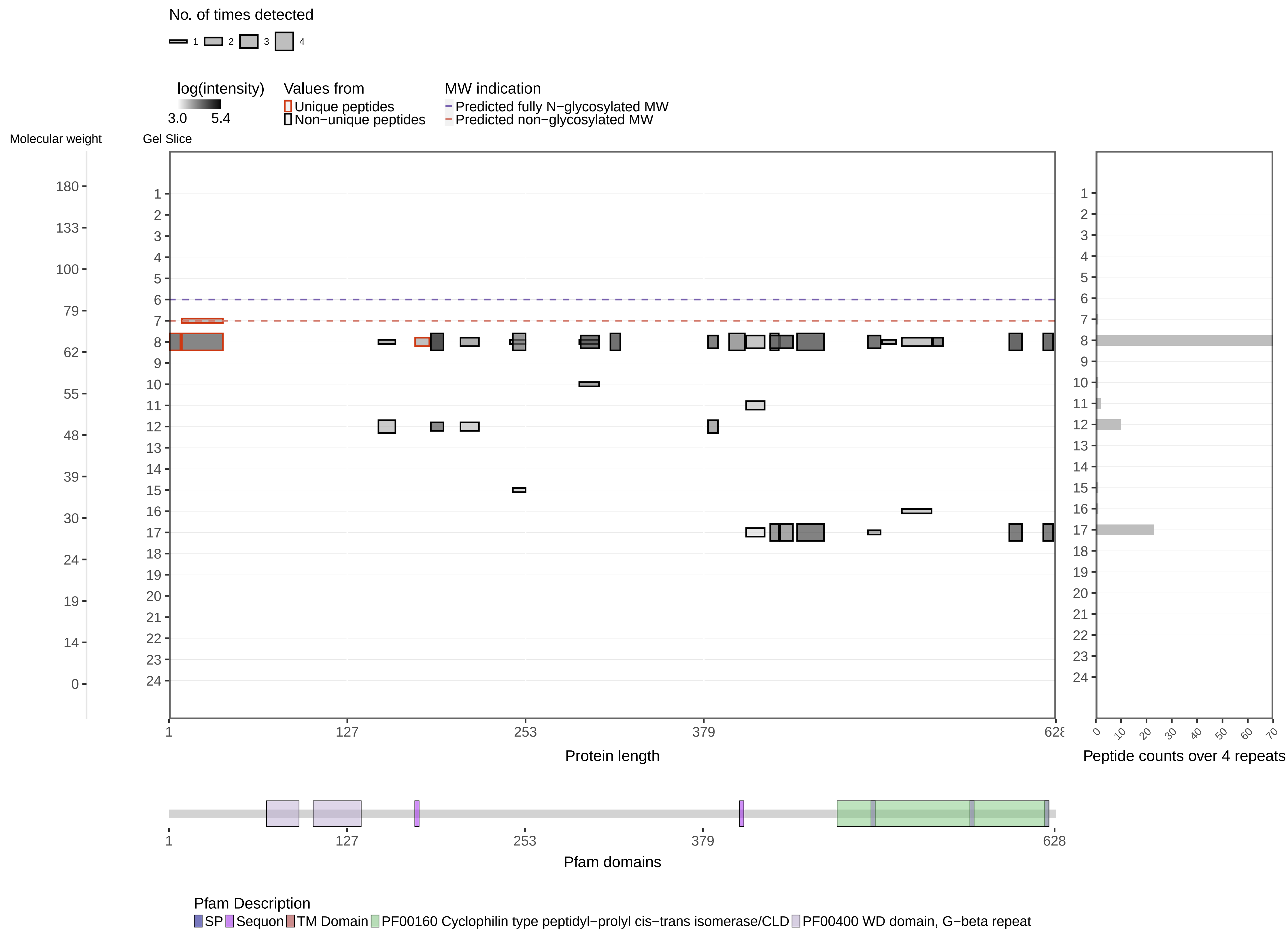


Protein sequence:
[illegible]

NbL17g07330.1 60S_acidic_ribosomal_protein_P2A−like [MW − SP = 9.72 kDa]

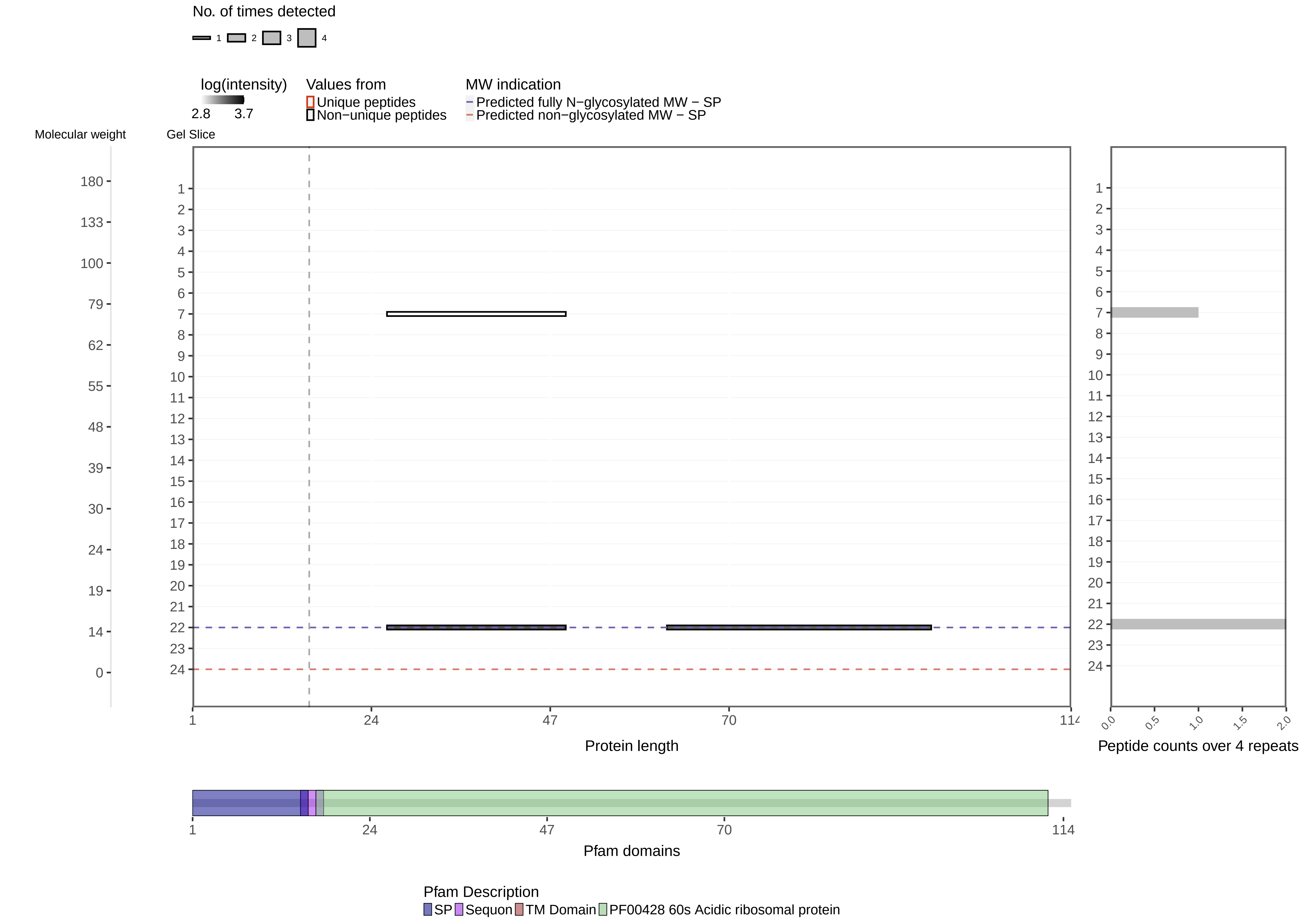


Protein sequence:

### NbL17g07320.1 flagellar_radial_spoke_protein_5 [MW = 51.22 kDa]

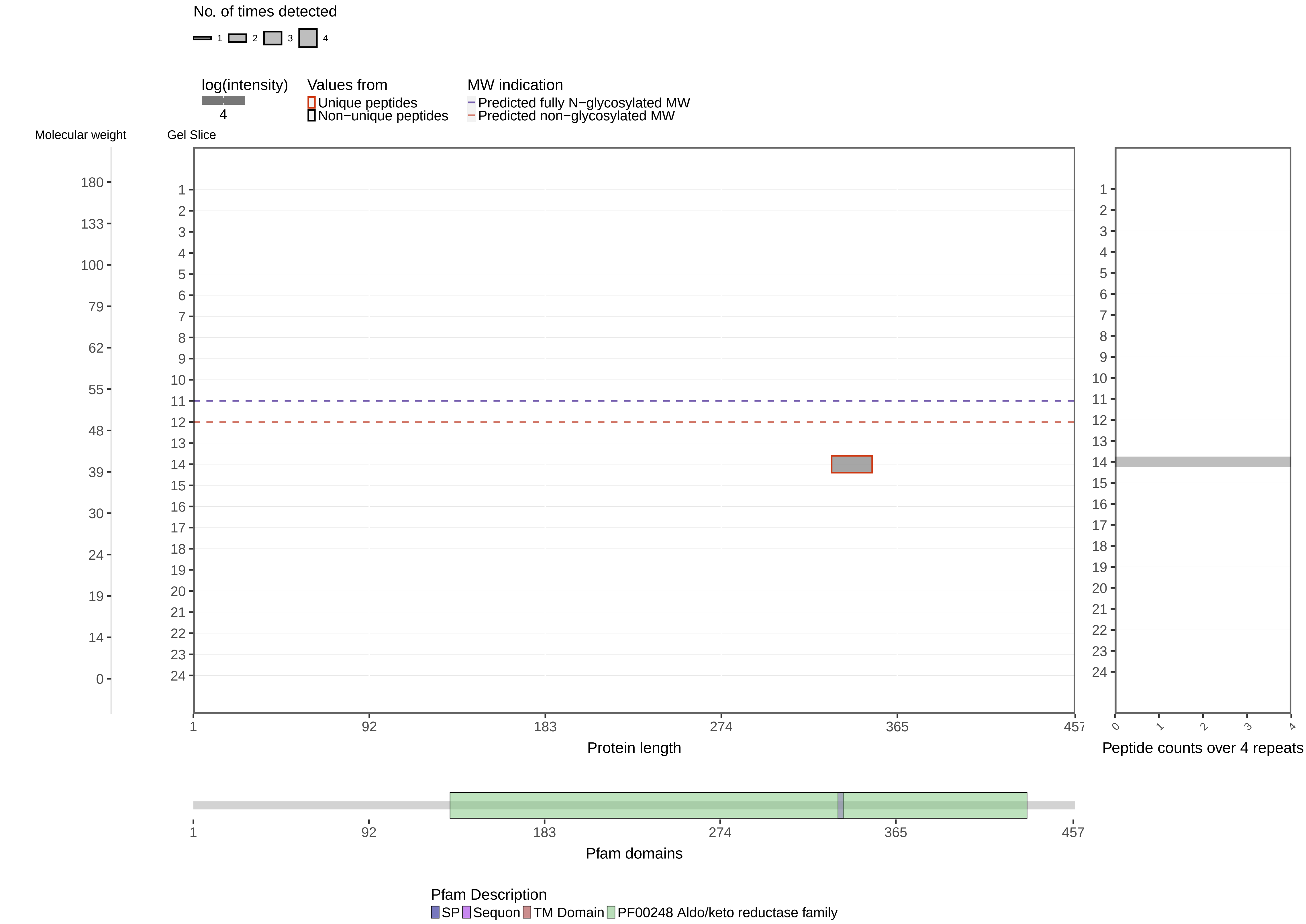


Protein sequence:

RSFVVQNIDVSRKRMDVASLDMLQFHWWDYTNSGYLDALKHLTDLKEEGKIKTIALTNFDTERLRIILENGIPVVSNQVQHSLVDMRPQQKMAELCQLTGVKLITYGTVMGGLLSEKFLGTNLSIPFAAPPLNTPSLQKYKRMVDAWGGWSLFQELLRTLKTVANKHGVSIPTVAVKYILDQPTVAGSMVGVRLGLSEHIKDVNAVFSLVLDEEDIRSILQVTNKGKDLQKIIGDCGDEYRRA*

NbL17g07310.1 uncharacterized_LOC109212815 [MW = 20.25 kDa]

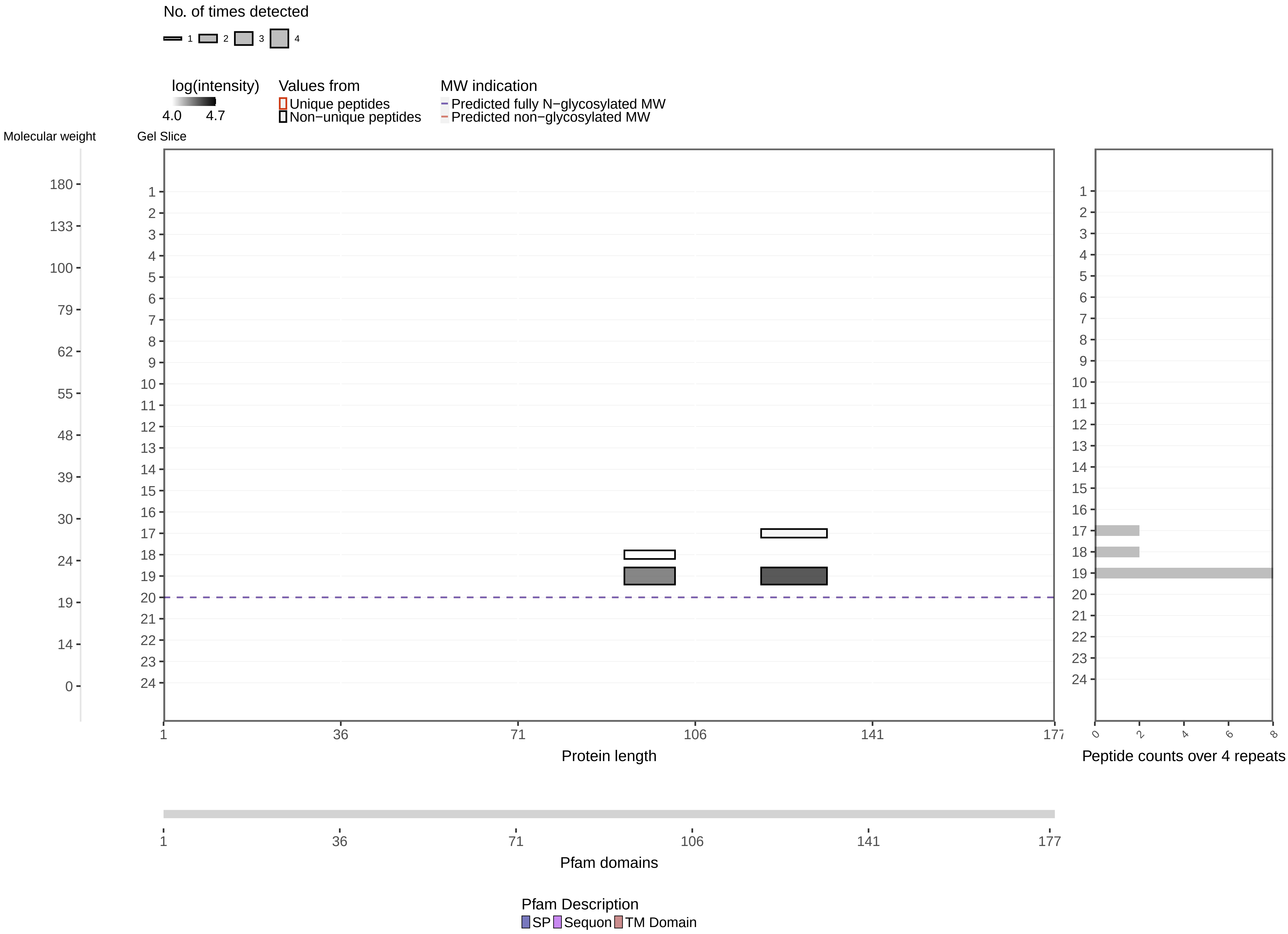


Protein sequence:

DKELRQRLQDSEQKLGLSMPLDQAKDRATQLESEVTLLERRLILASGAEGMDGFRQRWSLHGRLTDTKKRLEALKGGMENRKKDEPAEIEPAKKRWFFW*

### NbL17g07190.1 60S_ribosomal_protein_L10a−1−like [MW = 24.69 kDa]

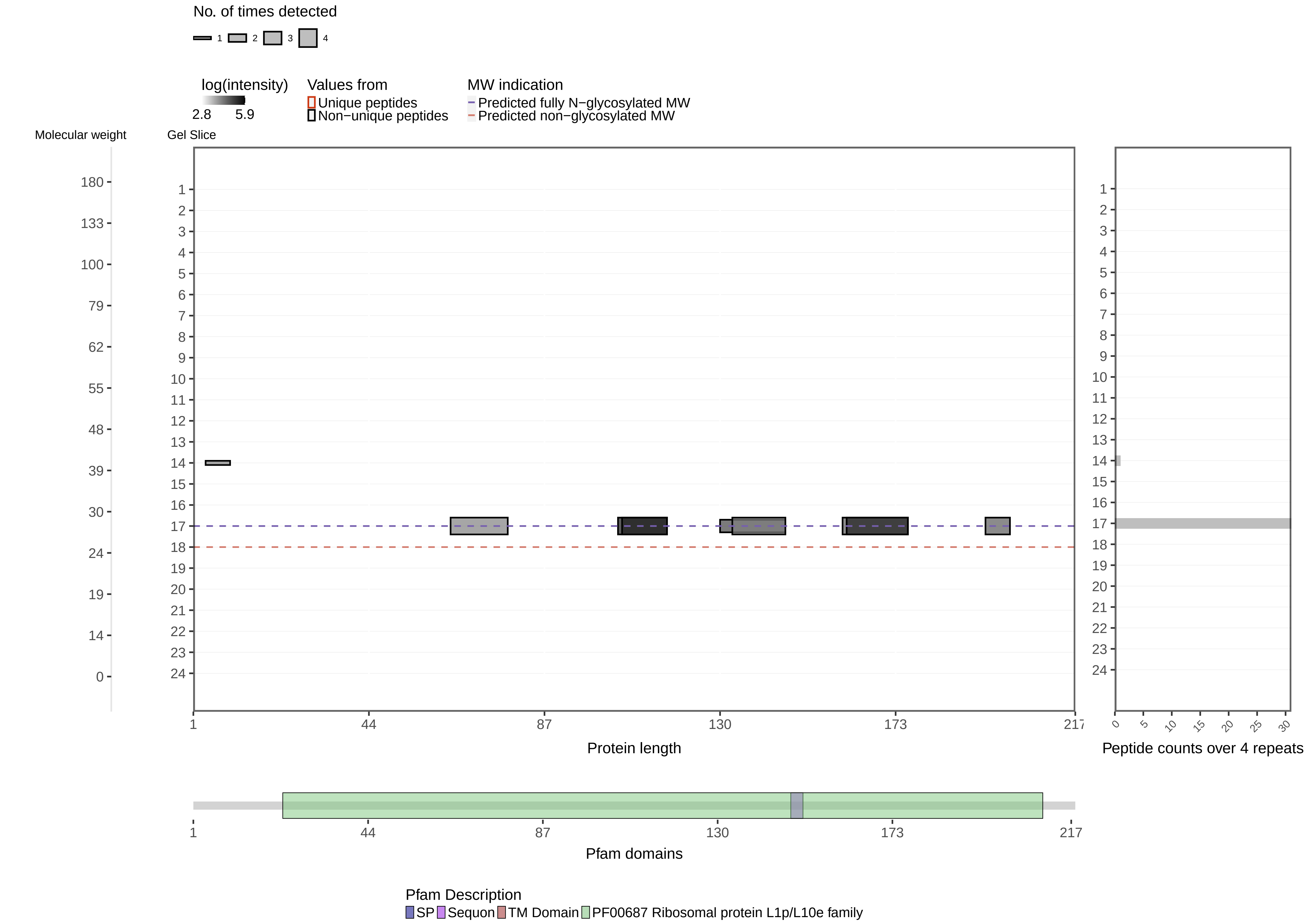


Protein sequence:

KKLVKKLAKKYHAFLASESVIKQIPRLLGPGLNKAGKFPSLVSHQESLESKVNETKATIKFQLKKVLCMGVAVGNLDMEEKQIFQNVQMSVNFLVSLLKKNWQNVRCLYLKSTMGKTQRVF*

### NbL17g07180.1 ferredoxin_root_R−B1−like [MW = 16.1 kDa]

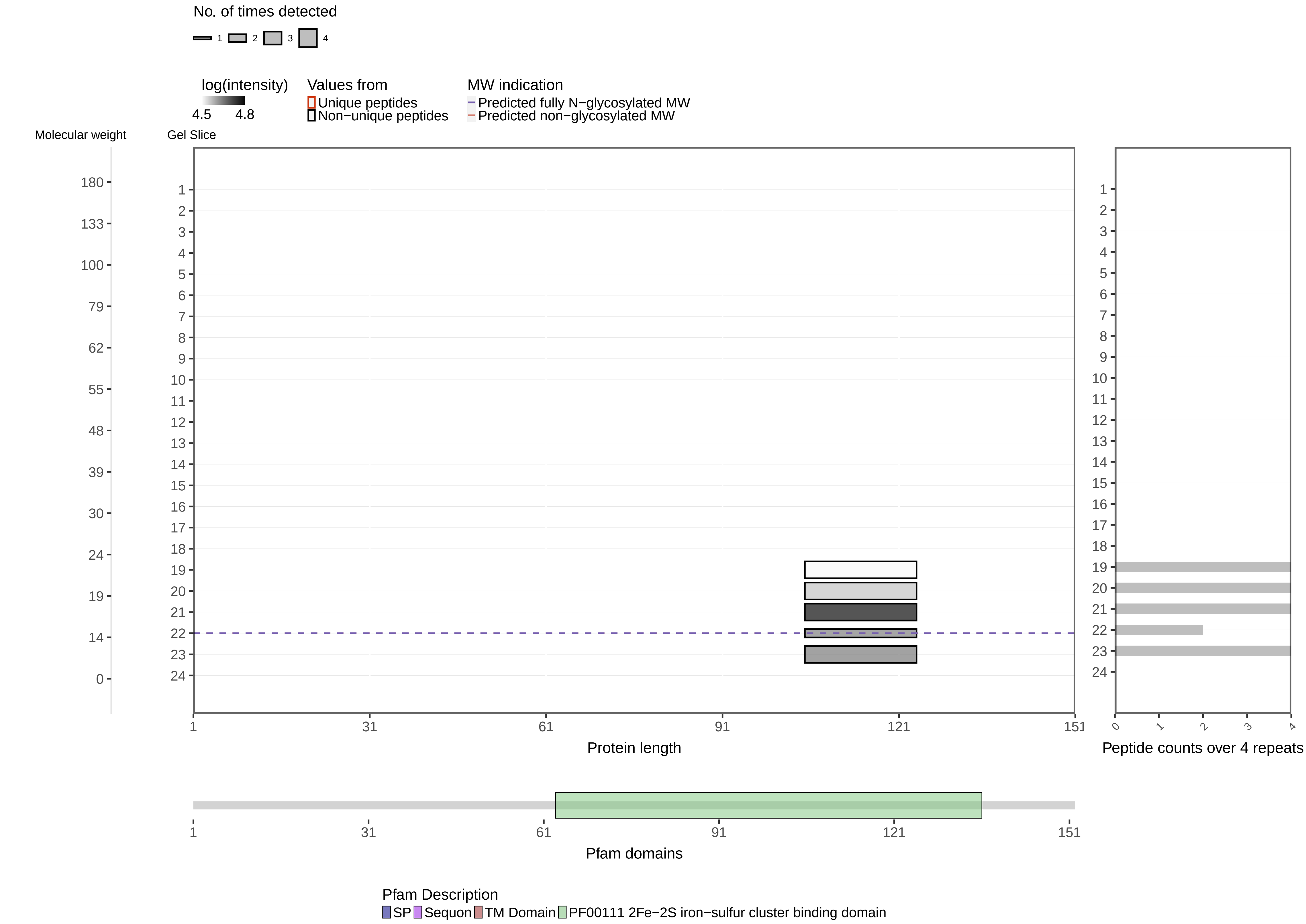


Protein sequence:

PDGTEHEFEAPSDTYILDAAEEAGVELPYSCRAGACSTCAGKIESGSVDQSDGSFLDDNQMKEGYLLTCVSYPTSDCVIYTHKEGDLY*

### NbL17g07110.1 protein_transport_protein_SEC24−like [MW = 81.99 kDa]

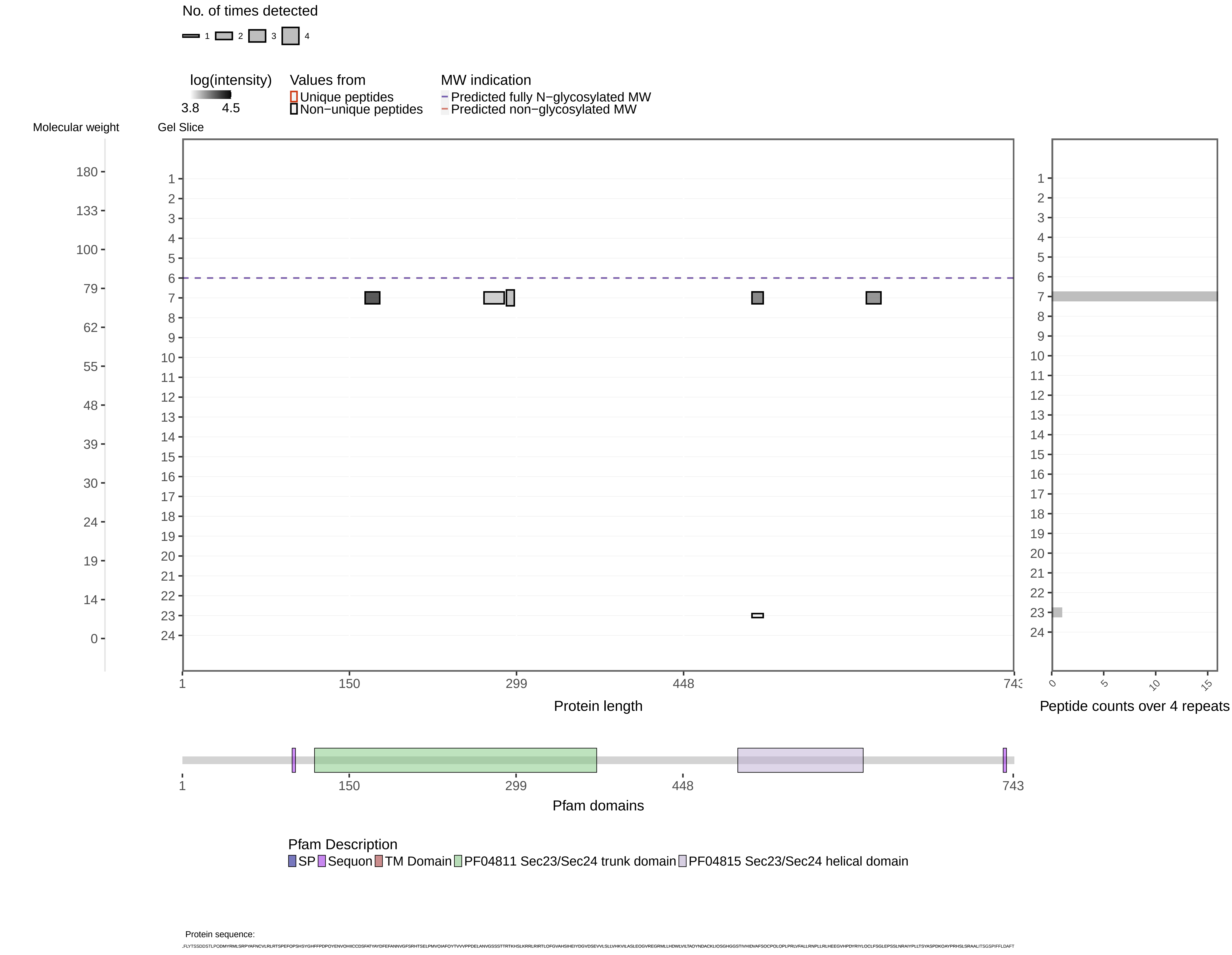

### NbL17g06950.1 DNA−damage−repair/toleration_protein_DRT100−like [MW − SP = 37.82 kDa]

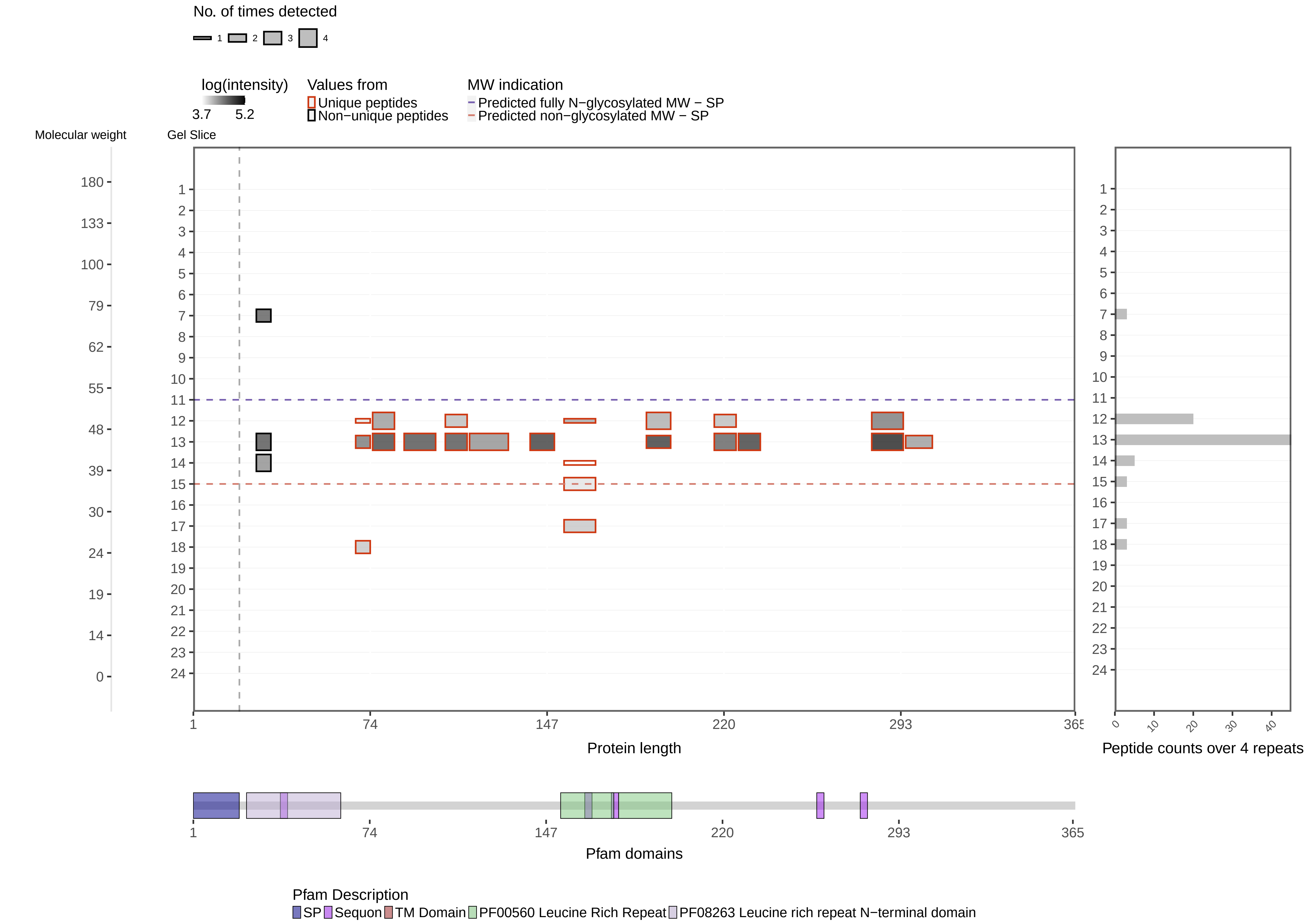


Protein sequence:

'RSLTNLSSLMHLDLRNNRIFGTLPTNFGKLRMLSRALLSGNRLIGKIPYSISYIYRLSDLDLSLNKLSGTIPPSLGKMRVLATLNLDGNNISGTIPPTLINSRINILNLSKNSIEGYIPDSFDERSYFMVMDLSYNKLRGKFPKSISSATFIGHFDVSHNHLCGOIPAGSPFDHLEASSFTYNDCLCGKPLKPCR*

### NbL17g06930.1 uncharacterized_LOC109227436 [MW − SP = 94.02 kDa]

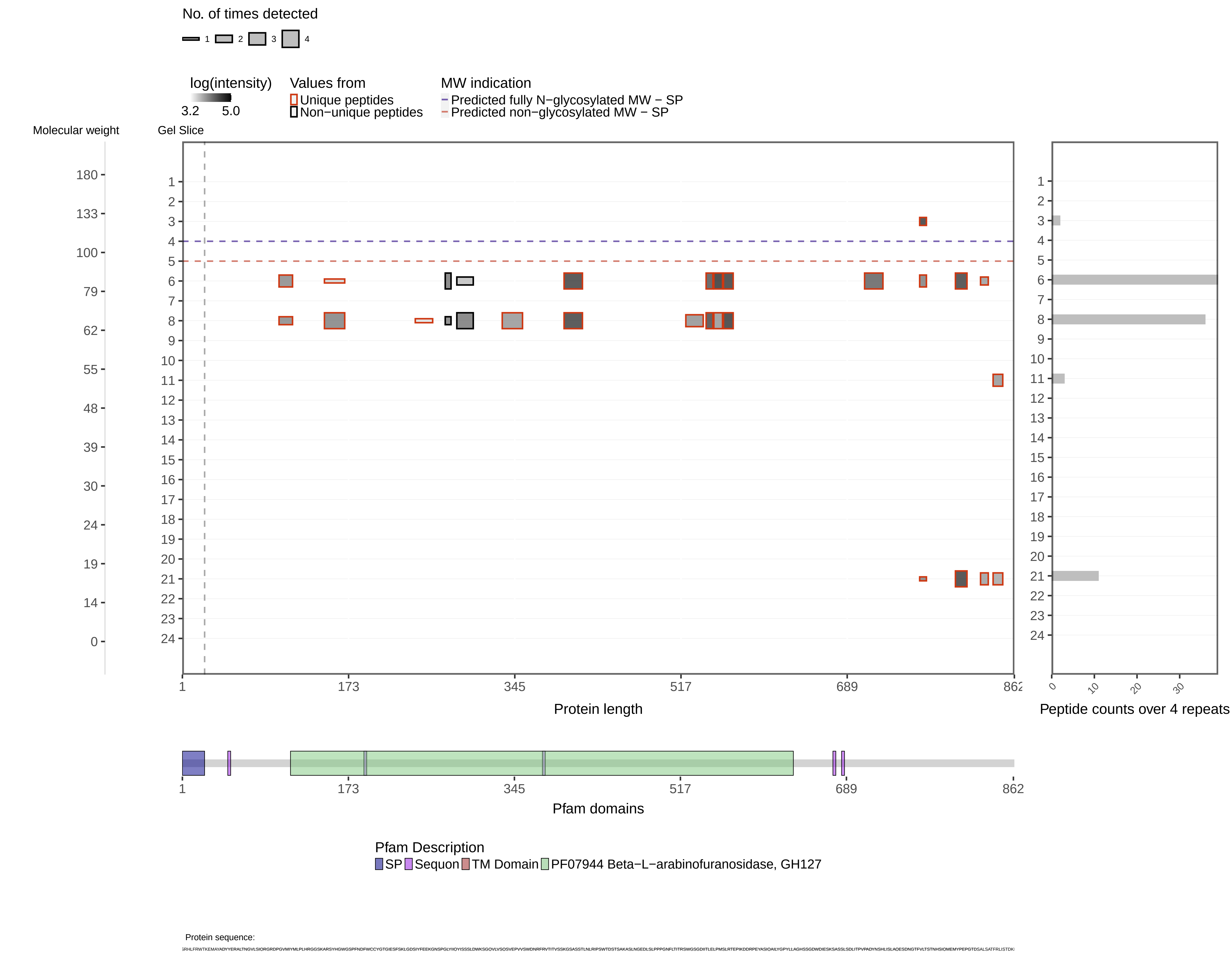

### NbL17g06820.1 actin−104 [MW = 41.72 kDa]

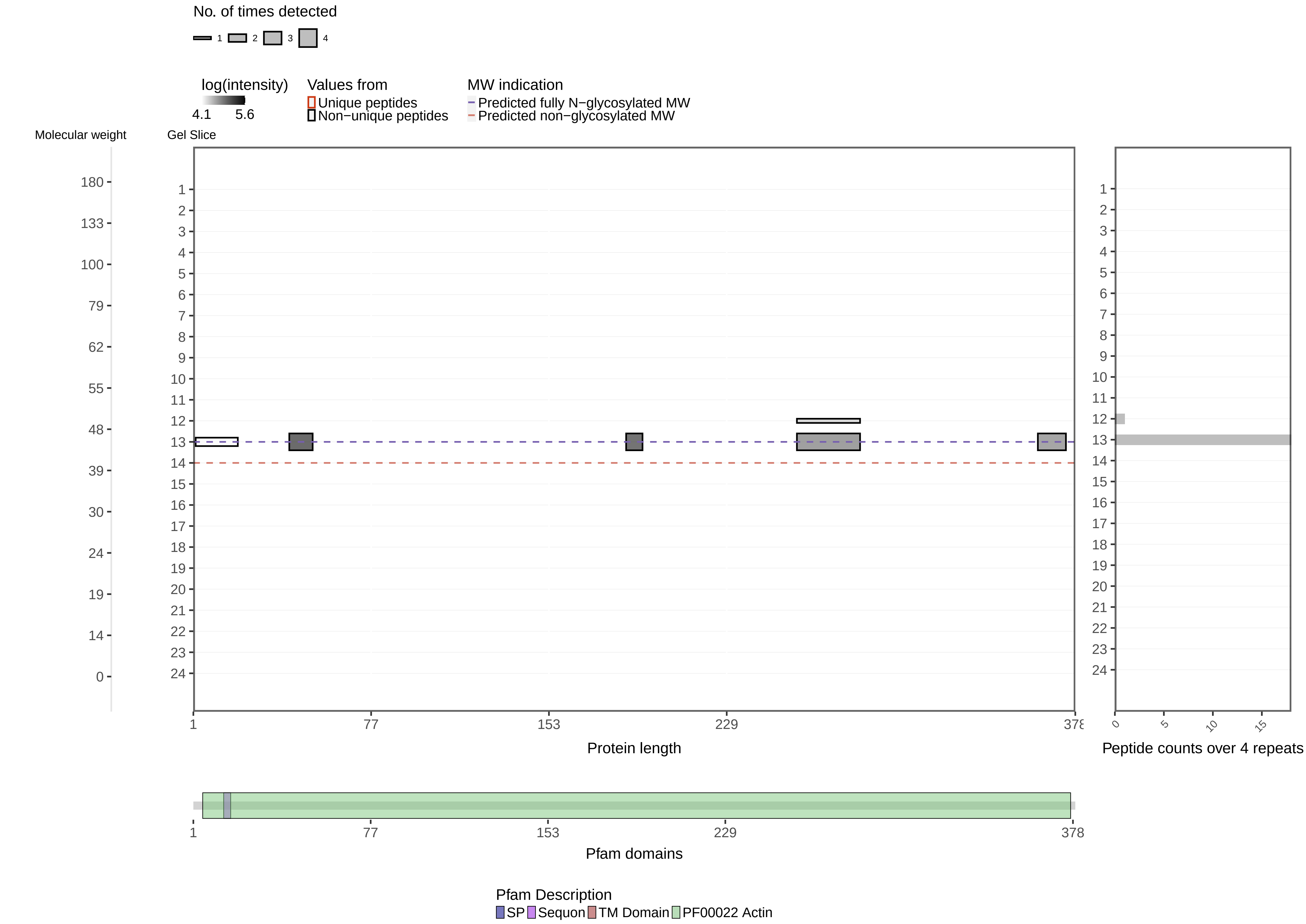


Protein sequence:

.PHAILRLDLAGRDLTDYLMKILTERGYMFTTSAEREIVRDVKEKLAYVALDFEQELETAKSSSSVEKSYELPDGQVITIGAERFRCPEVLFQPSLIGMEAAGIHETTYNSIMKCDVDIRKDLYGNIVLSGGSTMFPGIADRMSKEITALAPSSMKIKVVAPPERKYSVWIGGSILASLSTFQQMWISKAEYDESGPAIVHRKCF*

NbL17g06660.1 signal_recognition_particle_subunit_SRP72−like [MW = 77.98 kDa]

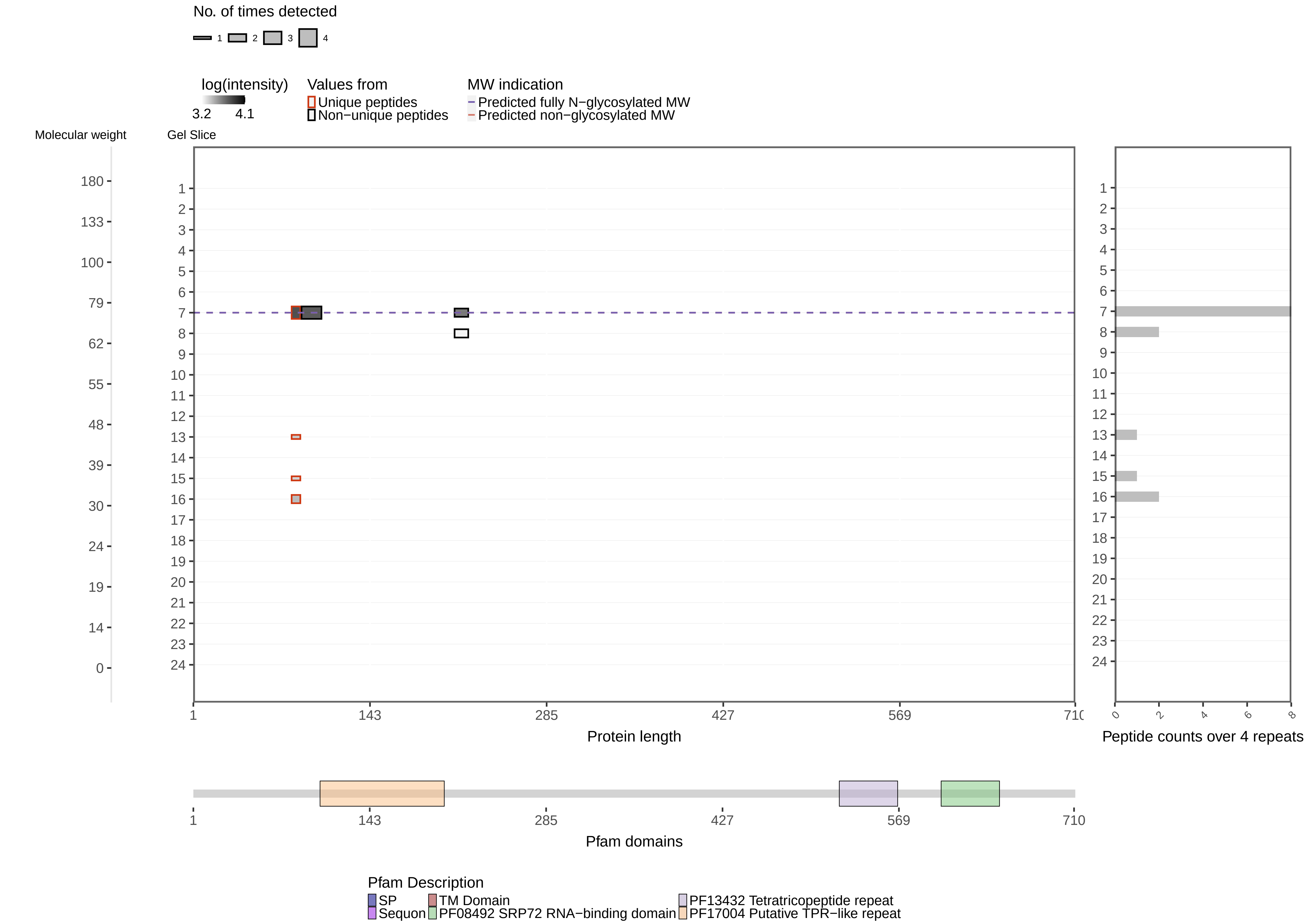


Protein sequence:

NbL17g06570.1 OTU_domain−containing_protein_6B−like [MW = 36.71 kDa]

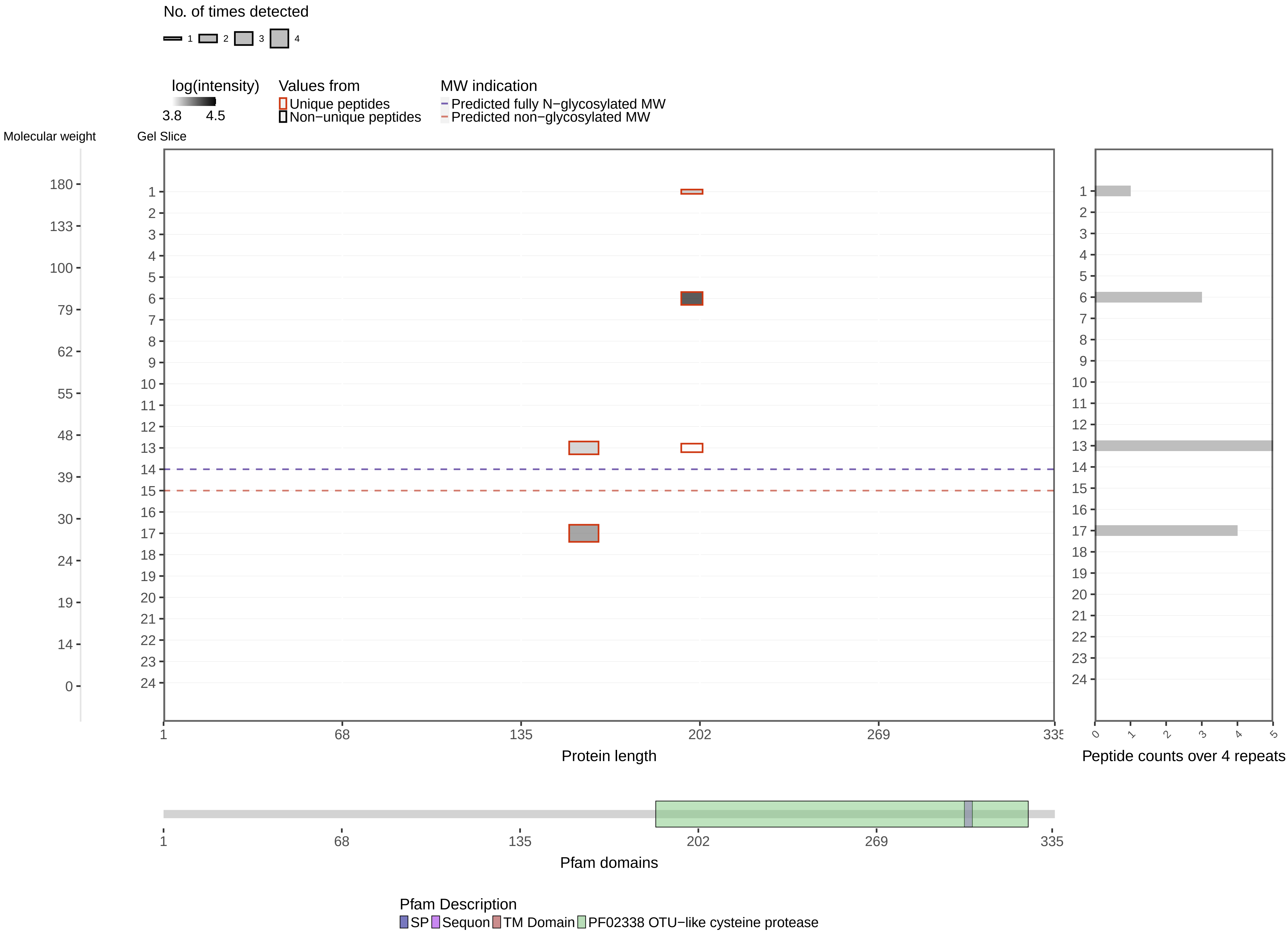


Protein sequence:

QEEQSSIVSDRVIENEKLDSKLEPLGLMVNEIKPDGHCLYRAVENQLAIRSGGSSPYTYLELRQMVAAYMRKHATNFLPFFLSENVEGGESDDSLVERFENYCGEVESTAAWGGQLELGALTHILKKHIMIFSGSFPDVEMGKEYKSDNGSGSSTSSIMLSYHMHAFGLGEHYNSVVPSSA*

NbL17g06500.1 probable_sucrose−phosphate_synthase_1 [MW = 117.24 kDa]

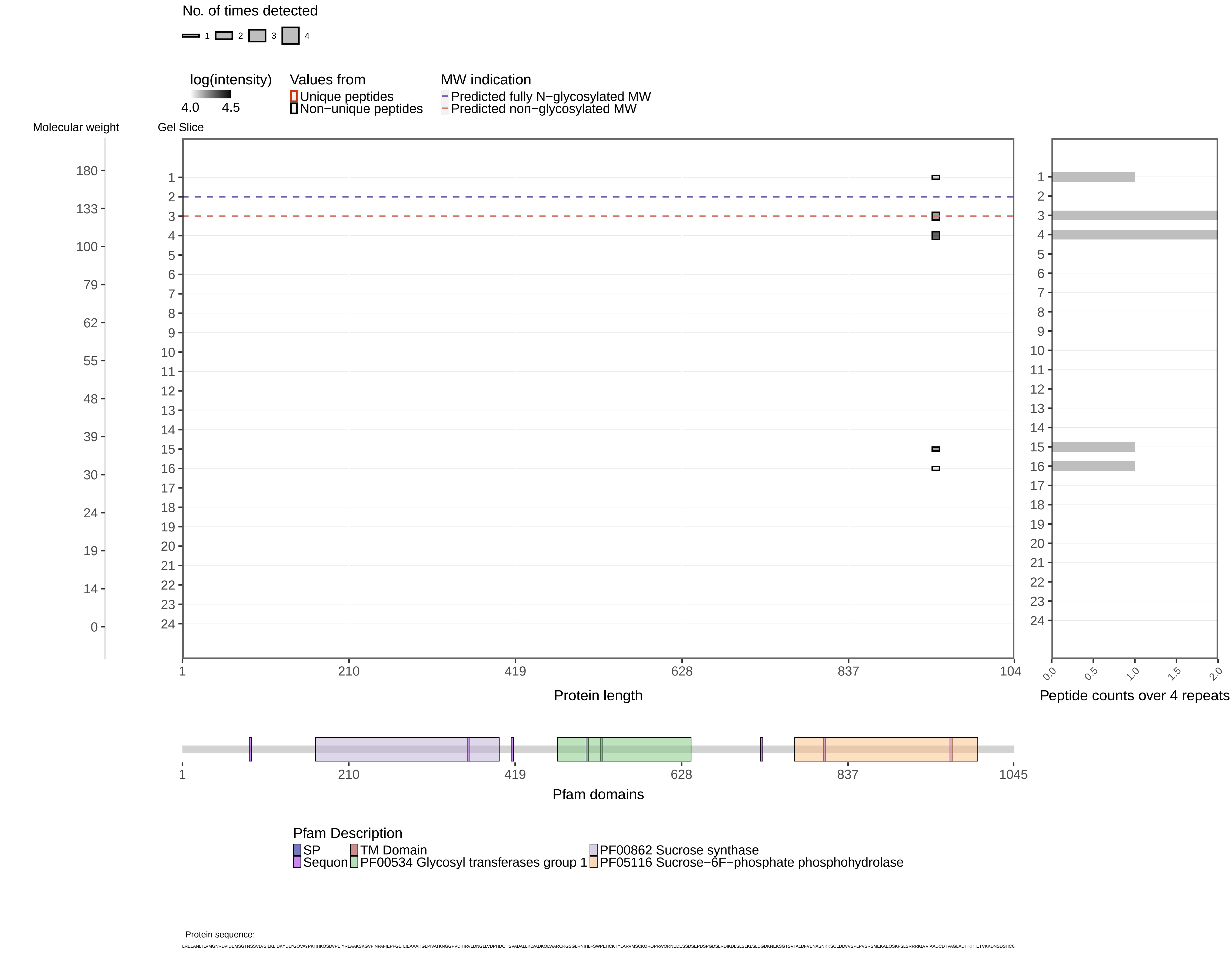

NbL17g06470.1 vesicle−associated_membrane_protein_711 [MW = 25.21 kDa]

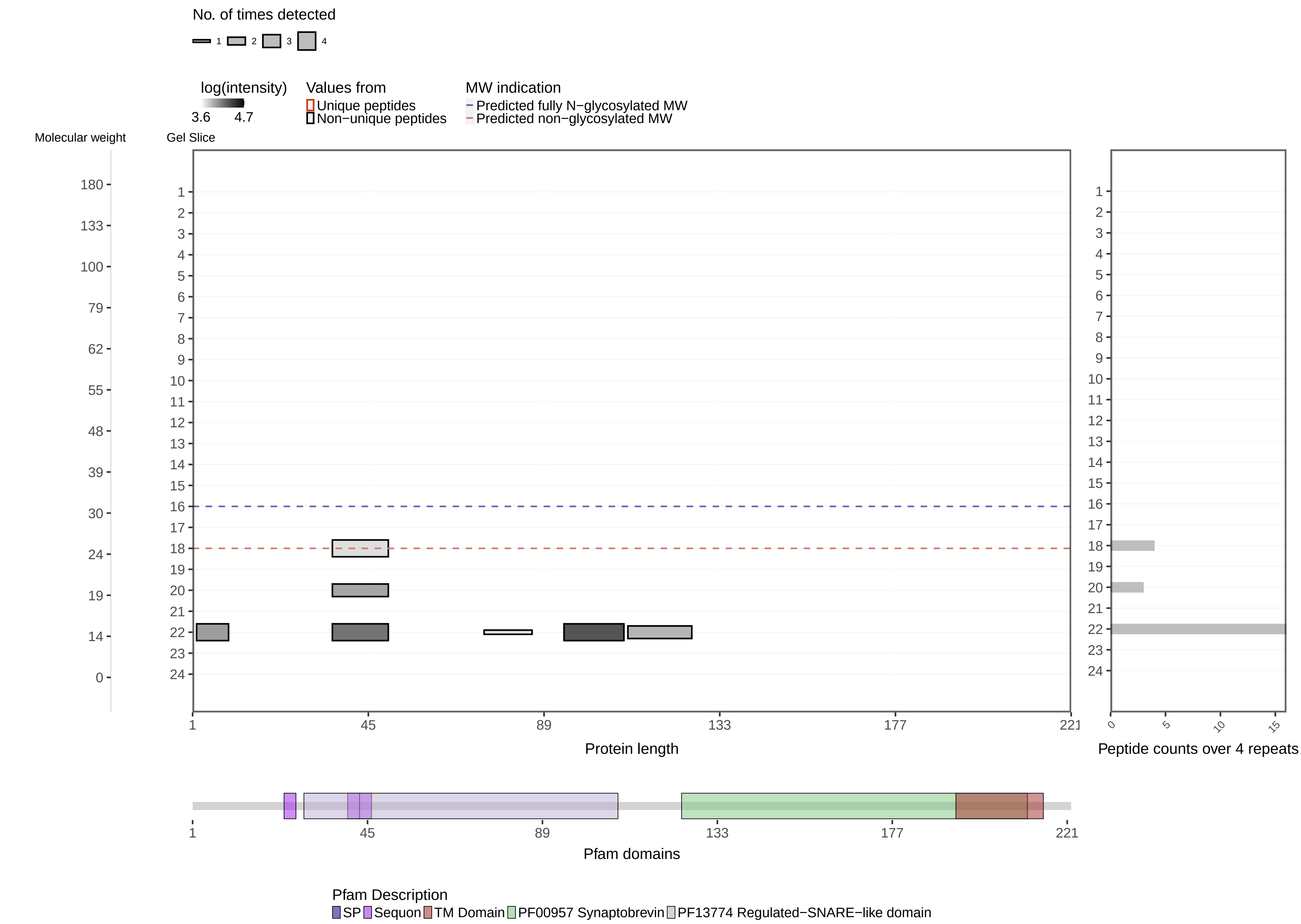


Protein sequence:

AQAYAMNDEFSRVLSQQIEYYSSDPNADRINRLKGEMSQVRNVMIENIDKVLERGDRLELLVDKTANMQGNTFRFRKQARRFRSTVWWRNVKLTVALIFLLLVIIYVILAFVCHGLTLPTCLK*

NbL17g06440.1 adenine_phosphoribosyltransferase_5−like [MW = 20.64 kDa]

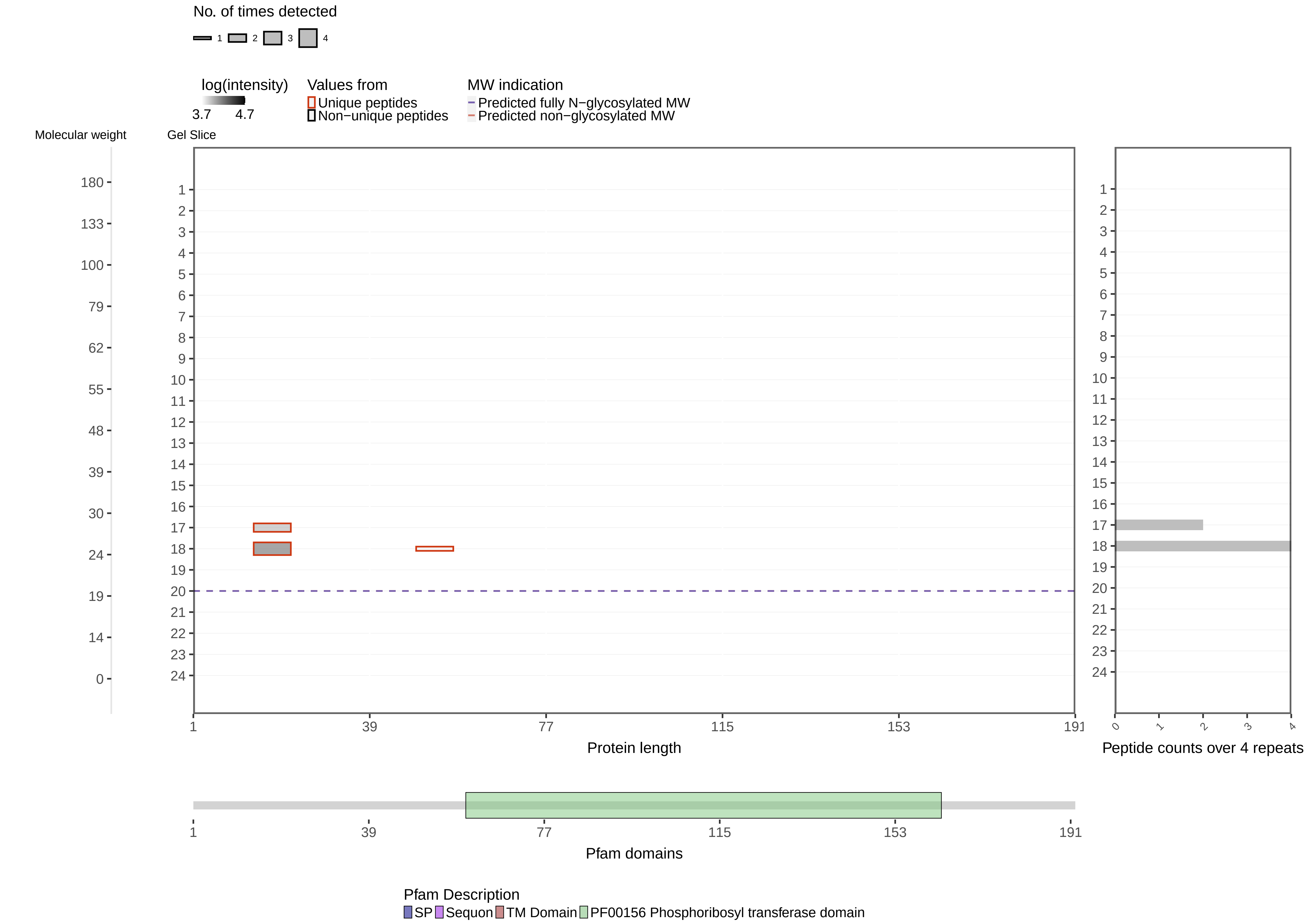


Protein sequence:

AIGAKFVPLRKPGKLPGKVIAESYELEYGKDCLEMHVGAVQQGDKAVILDDLVATGGTLSAAIKLLERMGAEVVECGCVIGLPEVKGQSRLKGKPLYILVEPRELEDF*

### NbL17g06410.1 ATP_synthase_subunit_b'_chloroplastic−like [MW = 23.98 kDa]

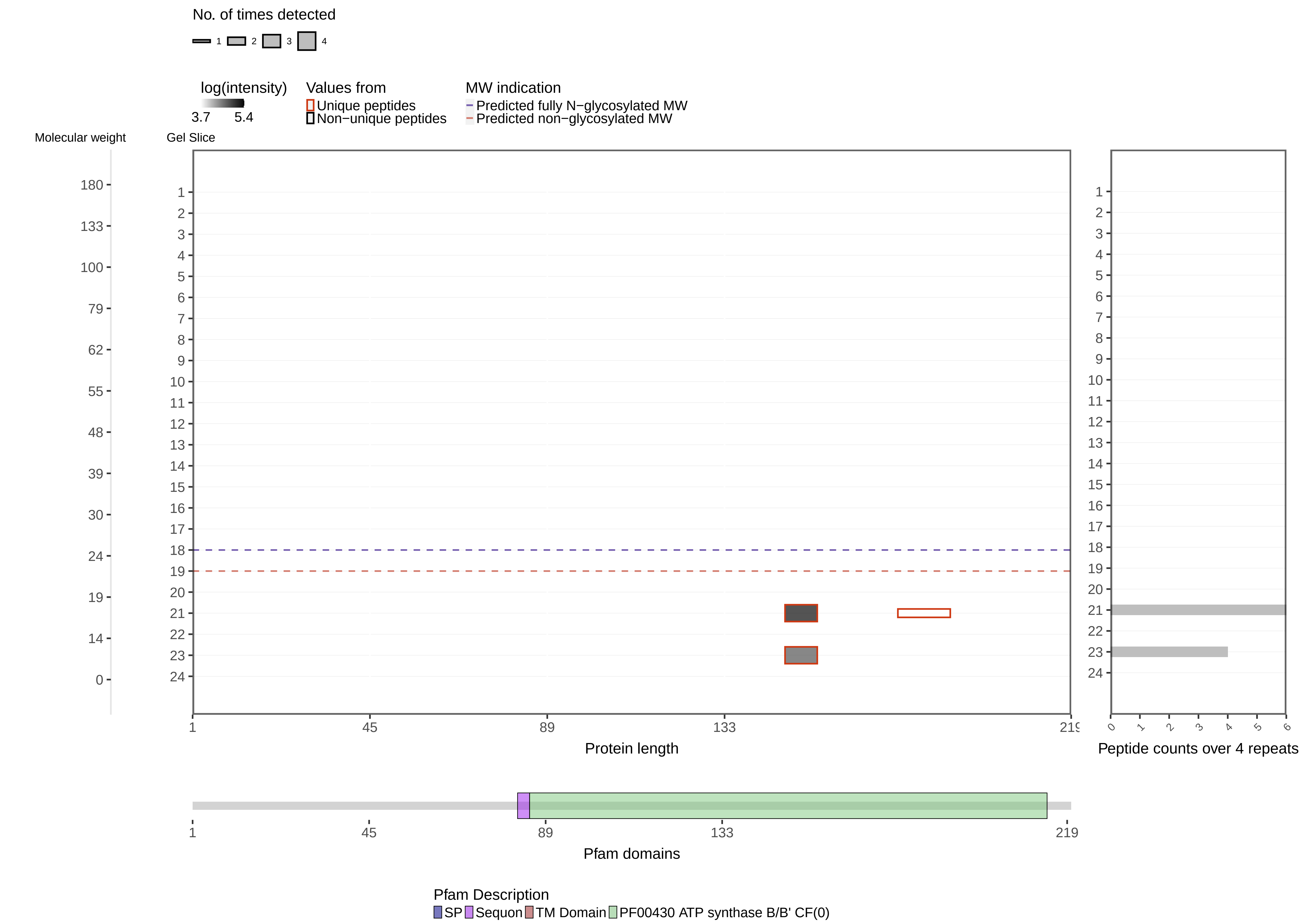


Protein sequence:

LDKLYFSPLGKFMDERDSAIKEKLNSVKDTSAEVKQLEDQAAAIMKAARAEISAALNKMKKETQLEVEQKIAEGRKKVEAELQEALASLDRQKEETIKSLDSQIAALSDEIVKKVLPVSN*

### NbL17g06350.1 putative_clathrin_assembly_protein_At2g25430 [MW = 70.4 kDa]

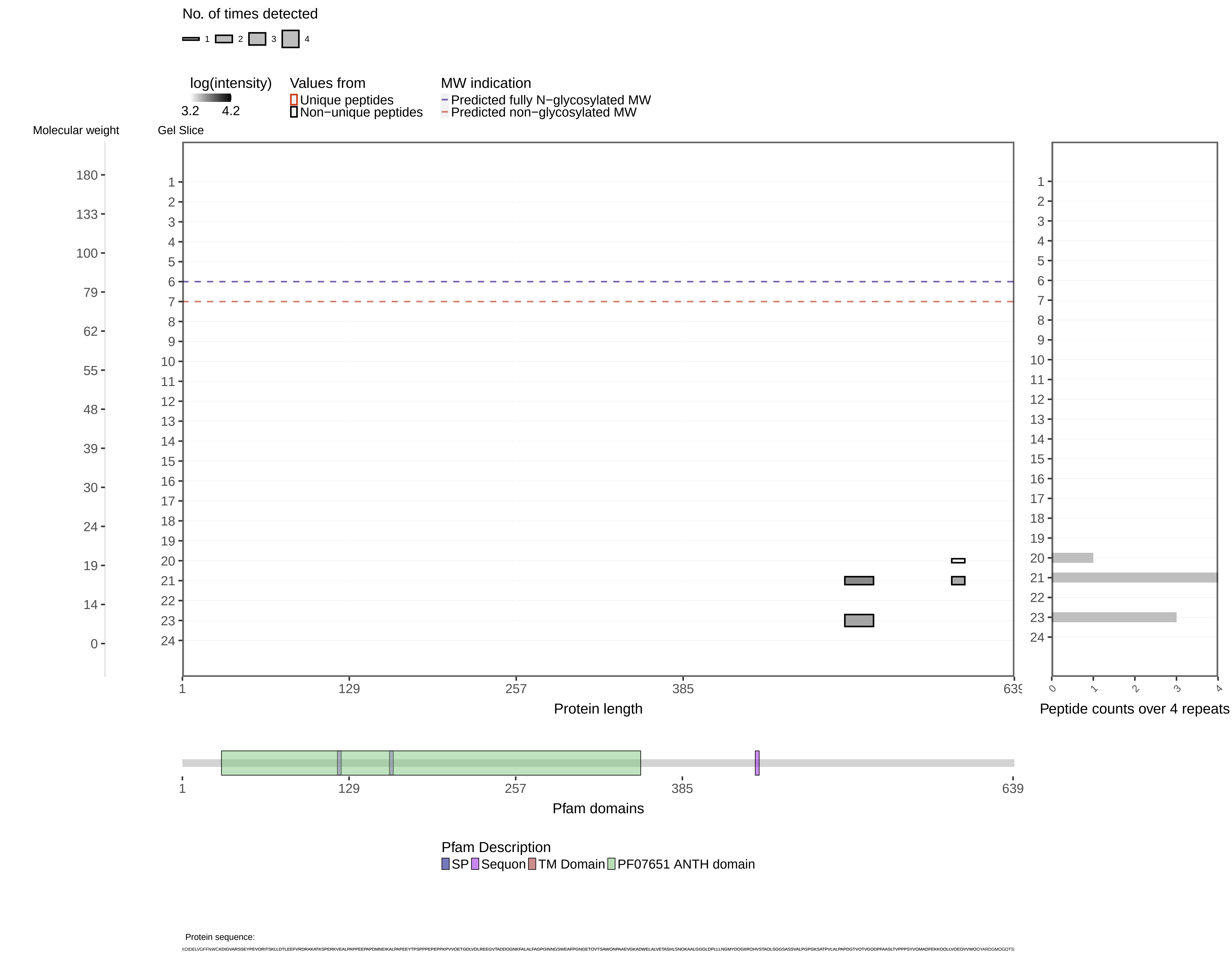

### NbL17g06280.1 putative_L−ascorbate_peroxidase_6 [MW = 36.73 kDa]

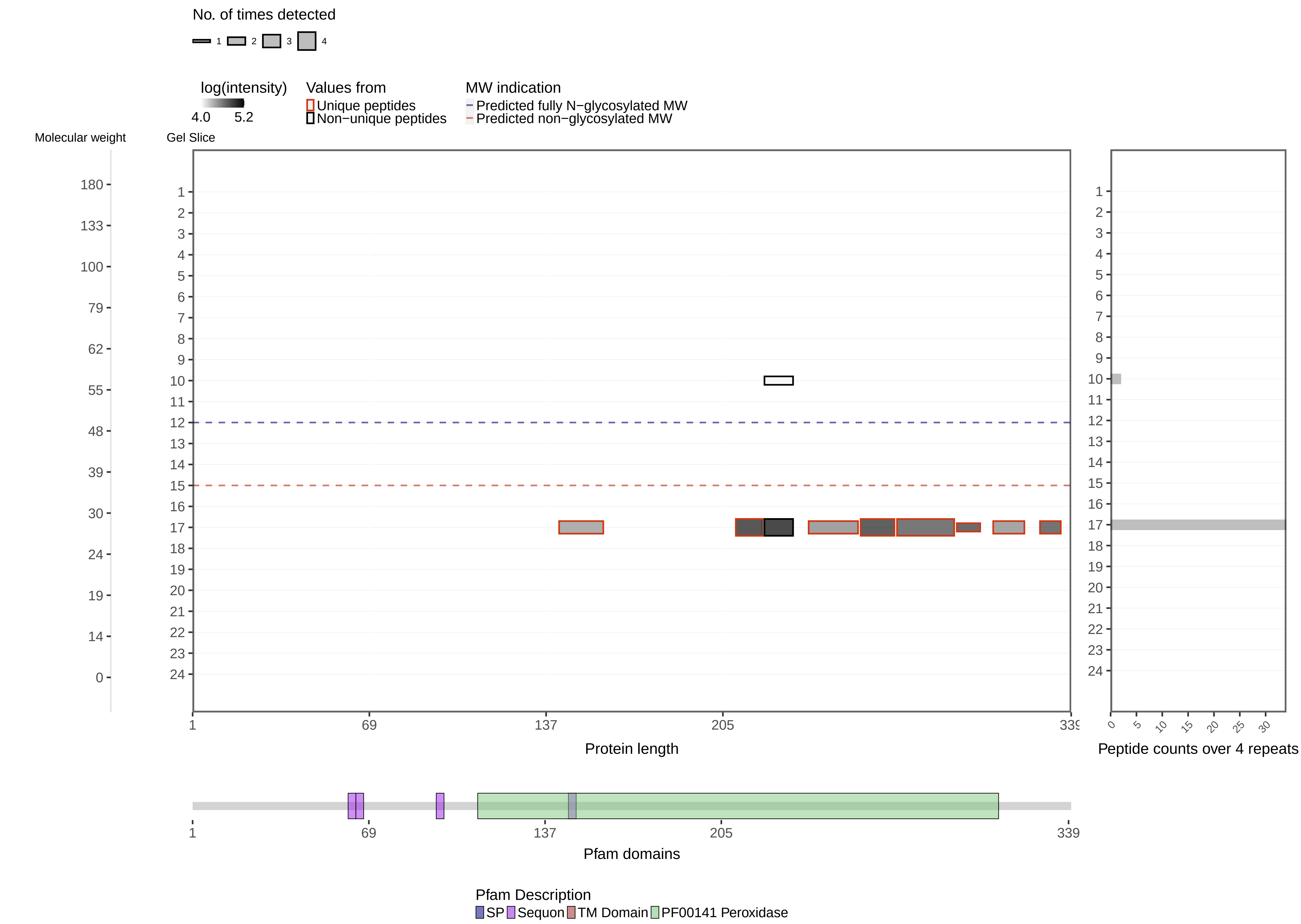


Protein sequence:

ENKGLKKSVKILEKAKSQIDVVQPVSWADIIALAGAEAVSLCGGPSIPIQLGRIDSMMPDPEGKLPEESLDATSLKQCFERKGFSTQELVALSGAHTLGSKGFGNPVVFDNAYFKILMEKPWLSSAGMTSMVGLPSDRALVEDDECVRWISKYAEDQSLFFEDFKNAYTKLVDTGATWKKSL*

### NbL17g06180.1 RNA−dependent_RNA_polymerase_6 [MW = 135.65 kDa]

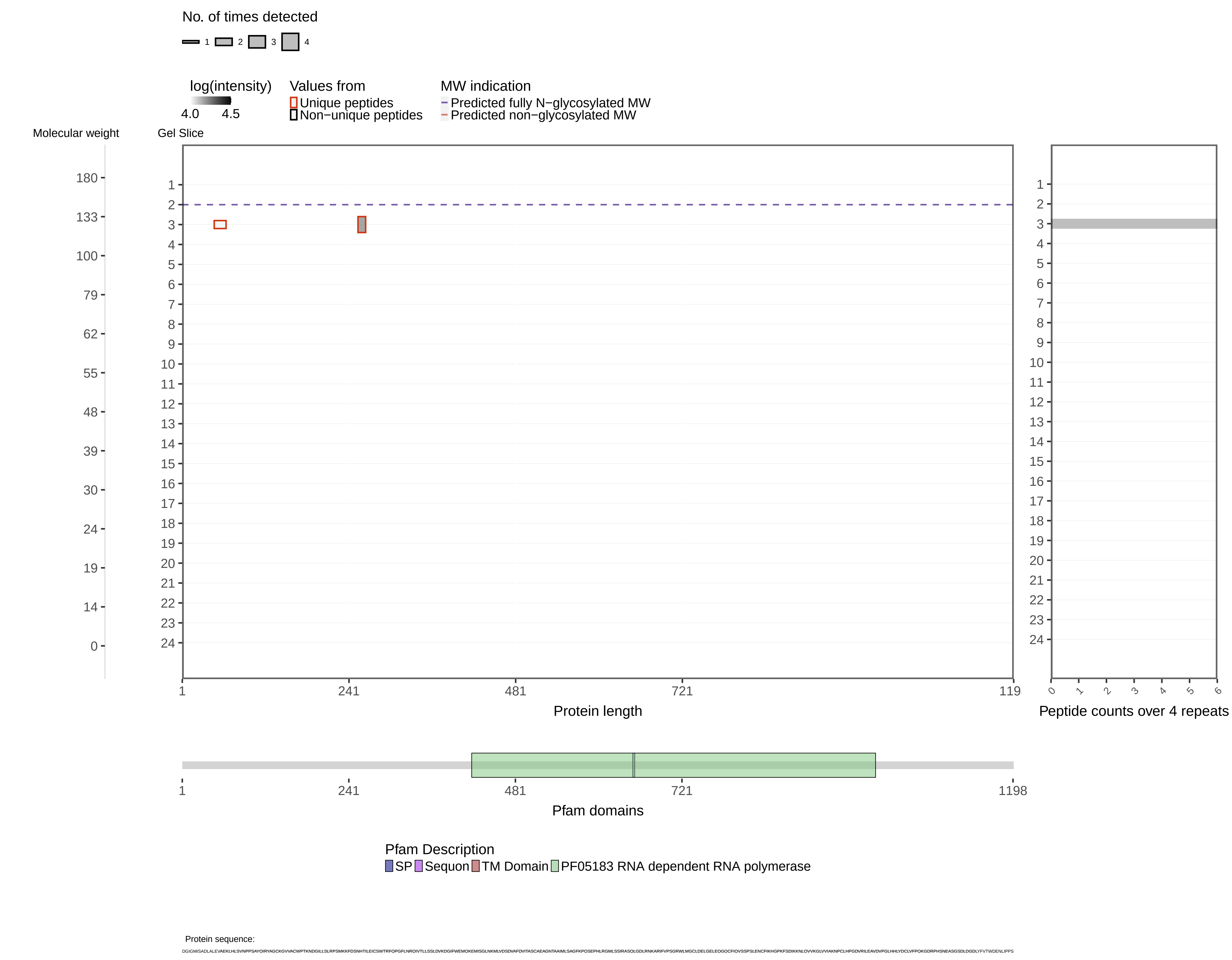

NbL17g06080.1 DEAD−box_ATP−dependent_RNA_helicase_56−like [MW = 48.43 kDa]

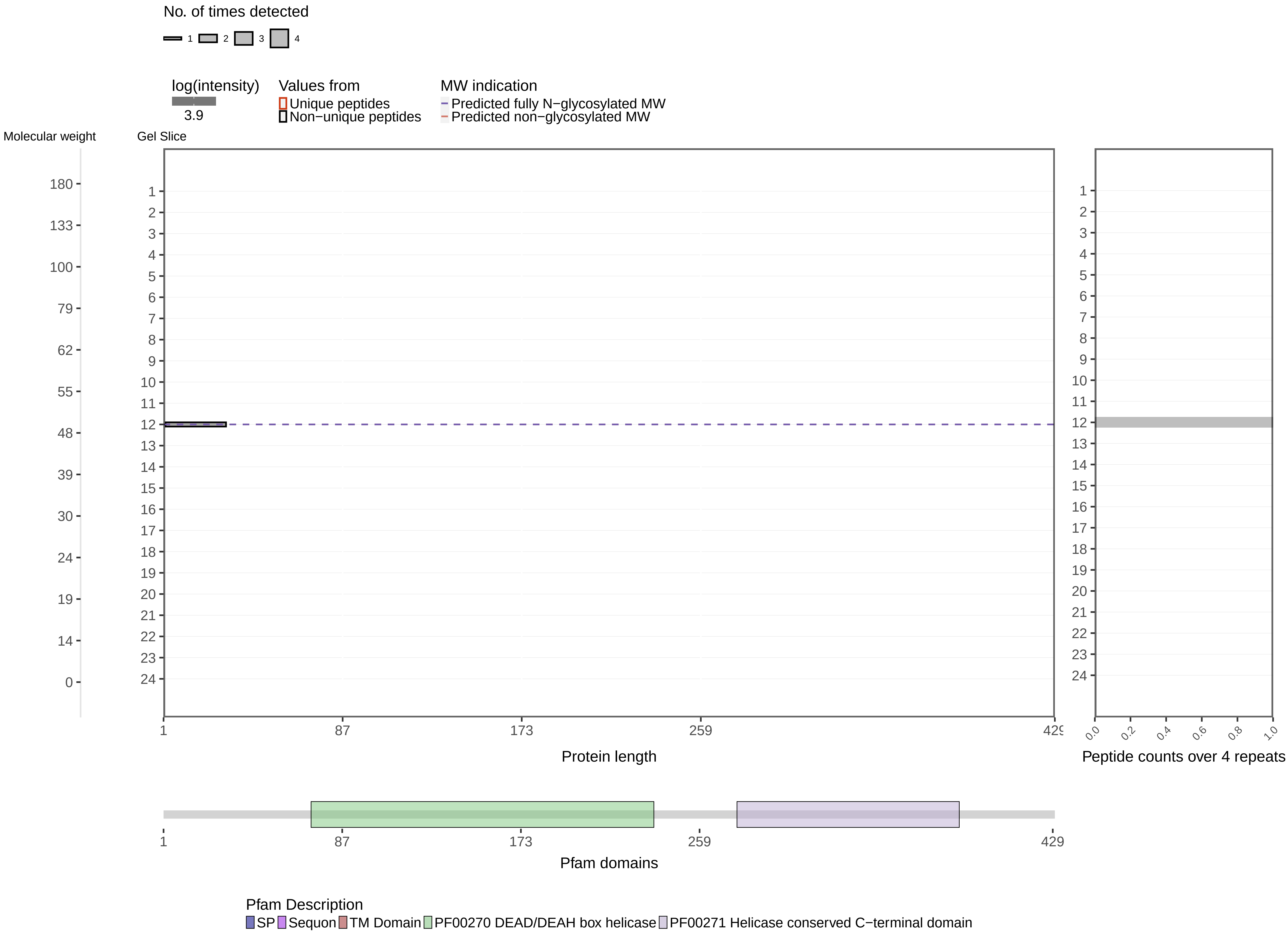


Protein sequence:

KMLESLDMRRDVQEIFKMTPHDKQVMMFSATLSKEIRPVCKKFMQDPMEIYVDDEAKLTLHGLVQHYIKLSETEKNRKLNDLLDALDFNQVVIFVKSVSRAAELNKLLVECNFPSICIHSGMTQEERLTRYKGFKEGHKRILVATDLVGRGIDIERVNIVINYDMPDSADTYLHRVGRAGRFGTKGLAITFVSSASDSDVLNQVQERFEVDIKELPEQIDTSTYMPS*

NbL17g05800.1 uncharacterized_LOC107805848 [MW − SP = 21.28 kDa]

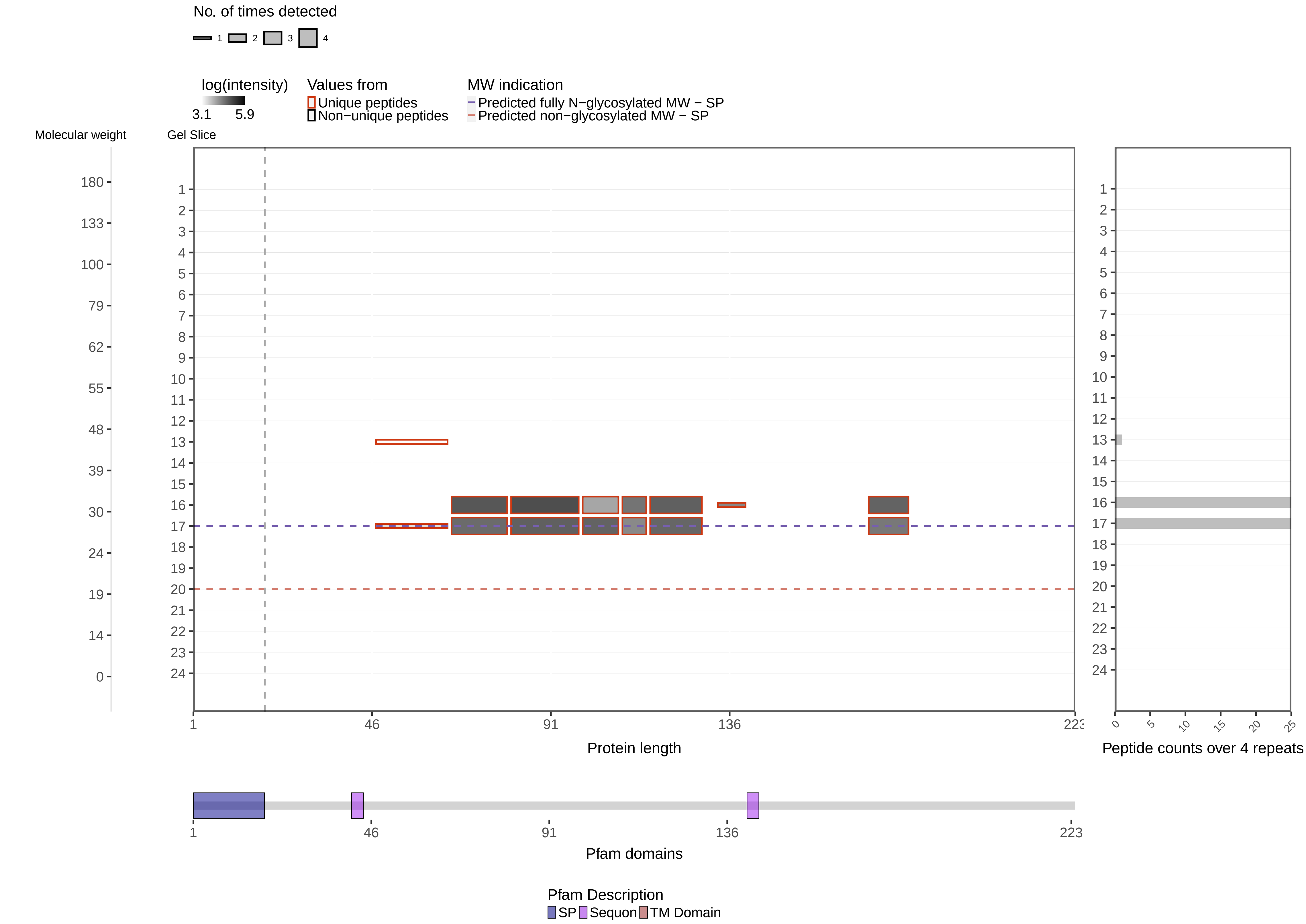


Protein sequence:

ASSLAQITGRFLACQSKYAEPSEAFNDILIRNSKSLDILNSKNHTEVGAAVSGSDGGGPYFWCVLFSNGKPKSSFVTEGGEPKVTRPGCFSGANDQCNGGNTLSRTIHLWPTALGSFAALLYAL*

### NbL17g05670.1 ADPATP_carrier_protein_mitochondrial [MW = 41.9 kDa]

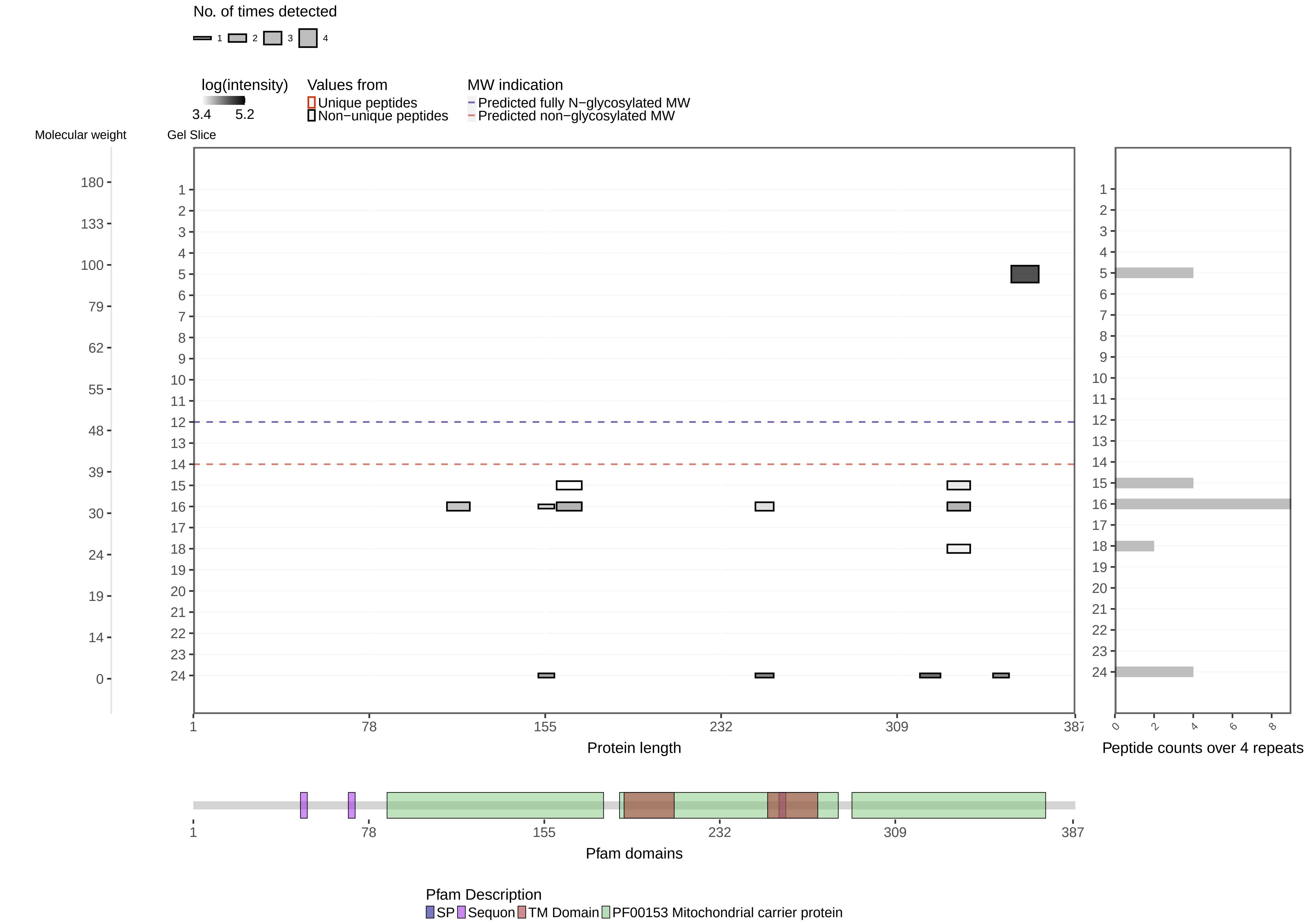


Protein sequence:

KKDRDGYWKWFAGNLASGGAAGASSLLFVYSLDYARTRLANDAKASKKGGERQFNGLVDVYRKTLKSDGIAGLYRGFNISCVGIIVYRGLYFGMYDSLKPVLLTGNLQDSFFASFGLGWLITNGAGLASYPIDTVRRRMMMTSGEAVKYKSSFDAFSQIVKNEGPKSLFKGAGANILRAVAGAGVLAGYDKLQVIVFGKKYGSGGA*

NbL17g05660.1 omega−amidasechloroplastic [MW = 40.96 kDa]

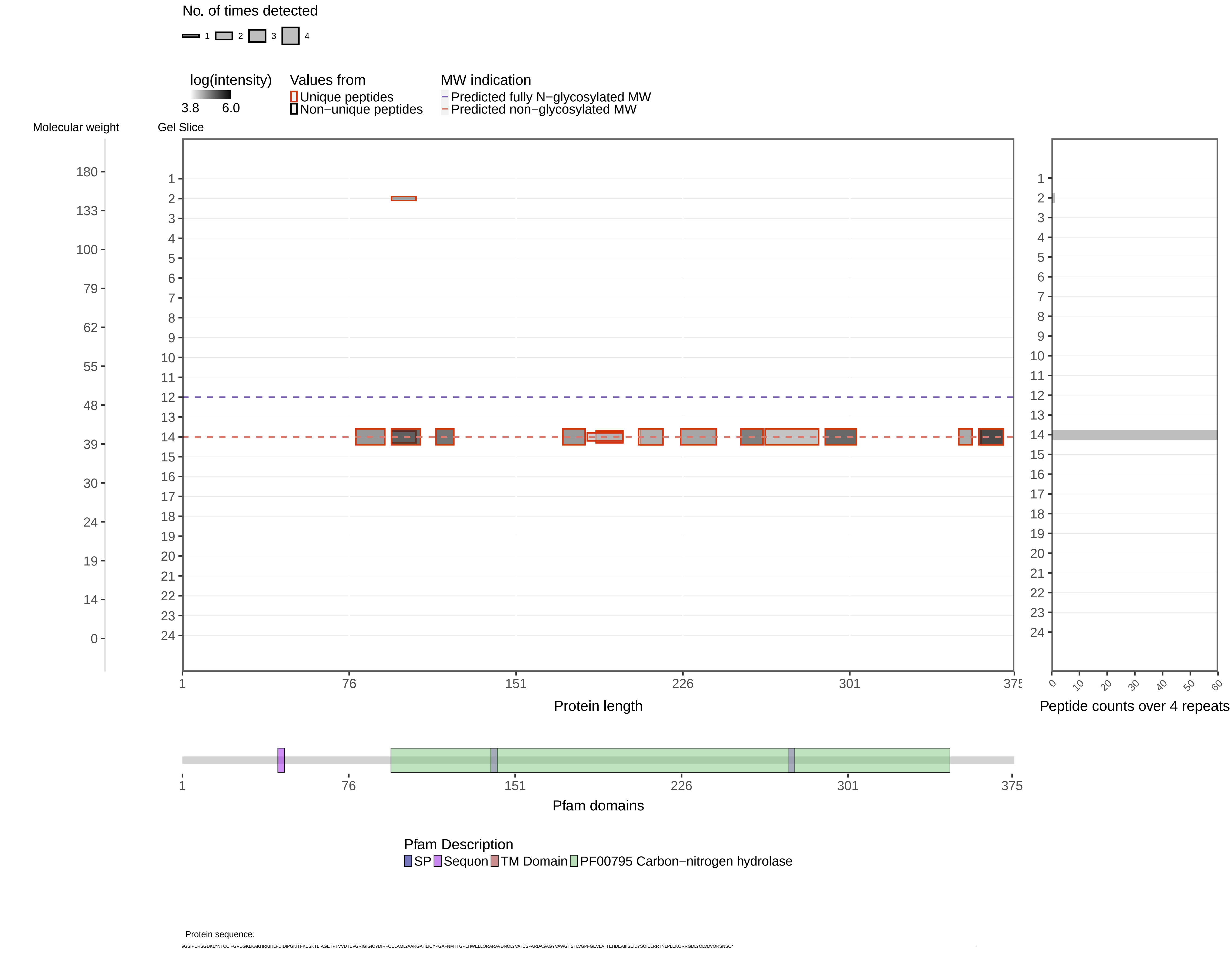

### NbL17g05460.1 aminopeptidase_M1 [MW = 99.18 kDa]

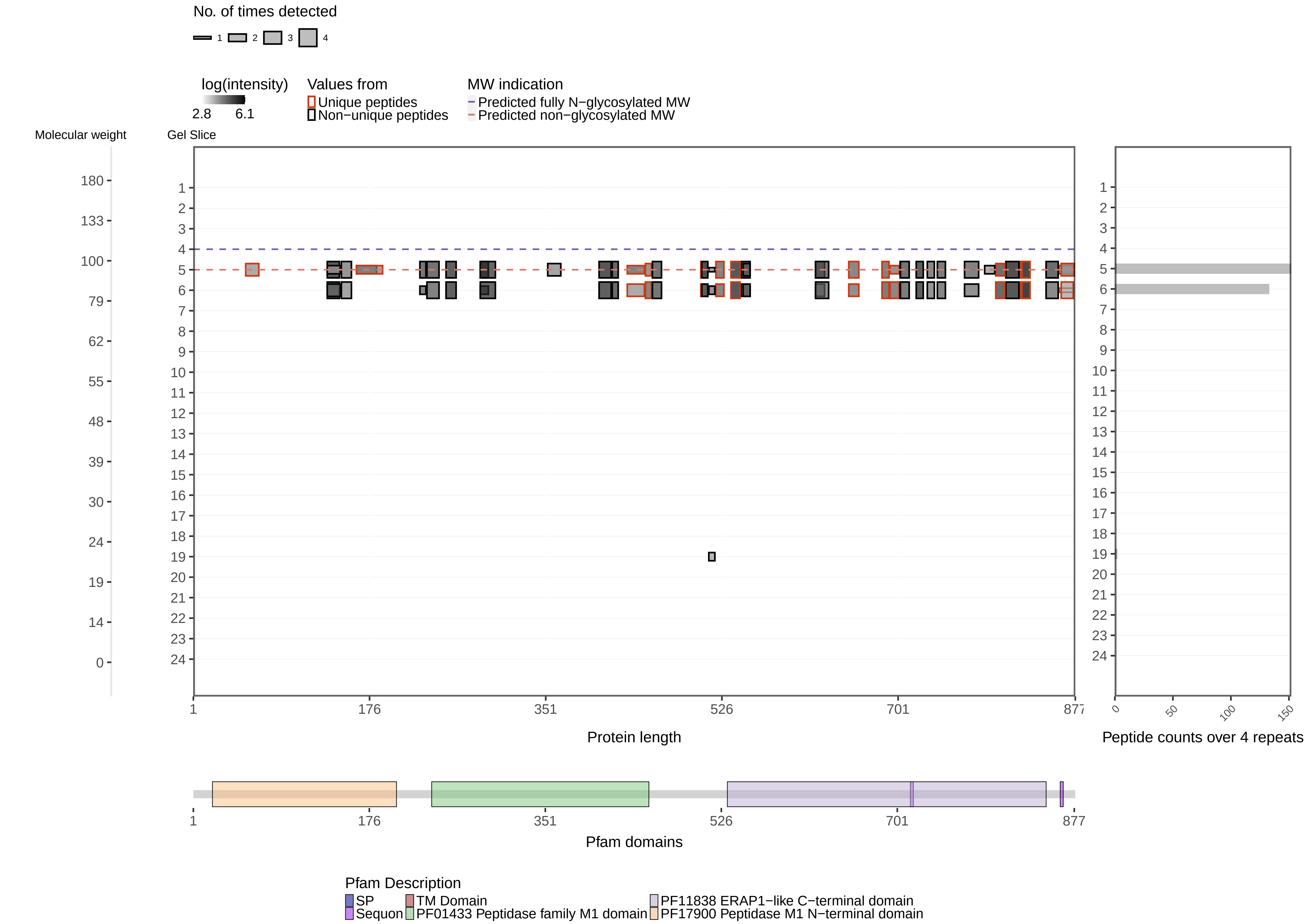


Protein sequence:

ACSNAKTEDLWSVLQEESGQPVNKLMNSWTKQQGYPVVSVKIKDQKLECDQTQFLLSGSHGDGQWIVPLTLCCGSYEARKSFLMQEKSEALDVKGLLGSSSSKGNPWIKVNVDQTGFYRVKYDDELSARLRYAIESKCLSTNDKYGILDDSYALSMACHQSLSSLLALMASFREELDYTVLSNLISISCKVSRIAADAVPDLKDHIKLFFINLFQFSAERLGWDPKQGESHLDAMLRGELLNALAAFGHDETINEAIRRFYIFLDDRNTPVLPPDLRRAVYVAVMQRVNKSDF

NbL17g05430.1 ran−binding_protein_1_homolog_c−like [MW = 24.49 kDa]

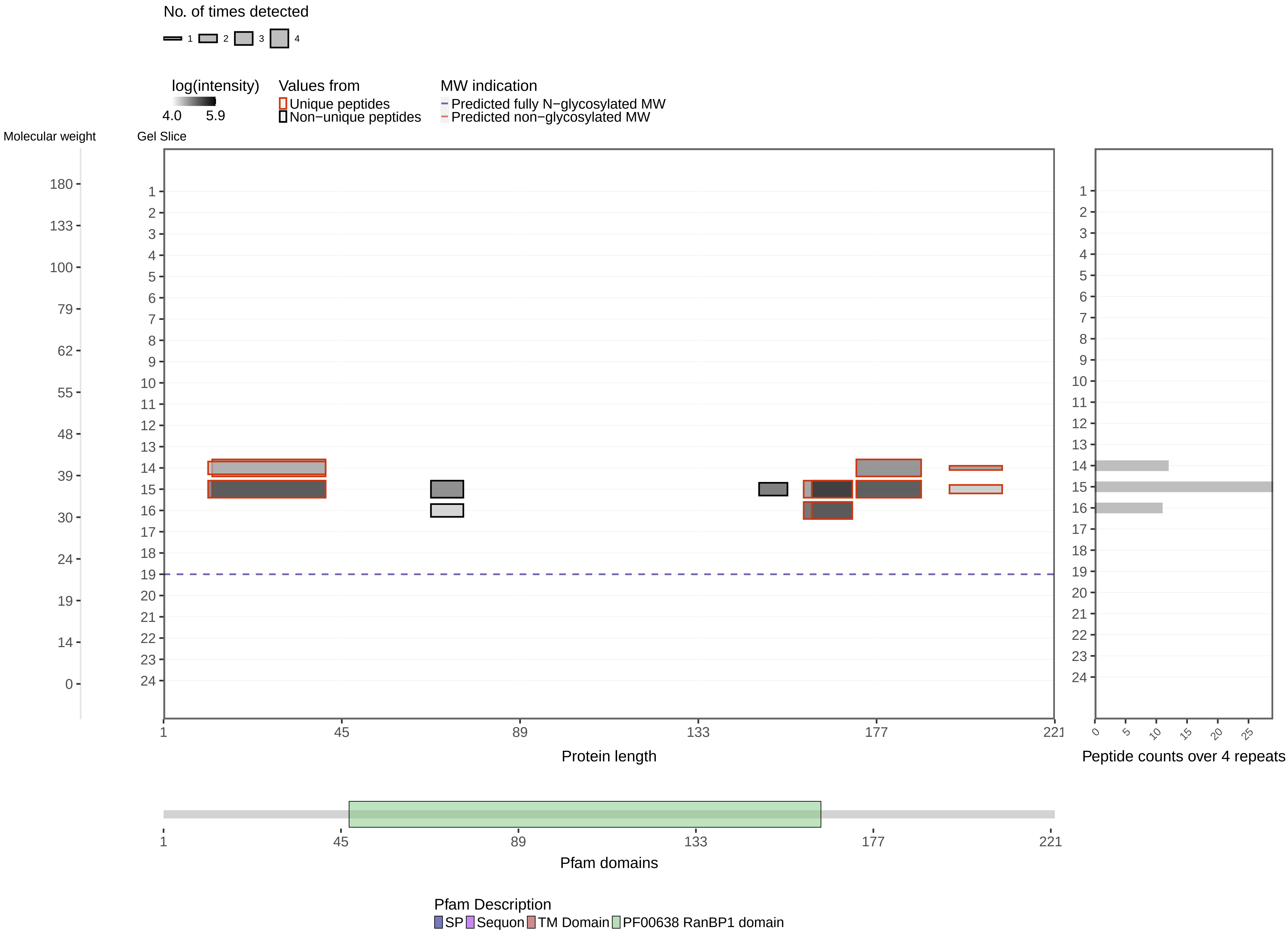


Protein sequence:

MRQSKTLKICANHLVLPTMTIQEHAGNEKSCVWYAADFADGELKEETFCIRFASIENCKSFKEKVEEIAESQQTKSEESEGAAAAATELIEKLSVESKDKEDKESPAPTEEKENKKEENADEKN*

### NbL17g05330.1 eukaryotic_initiation_factor_4A−14−like [MW = 46.93 kDa]

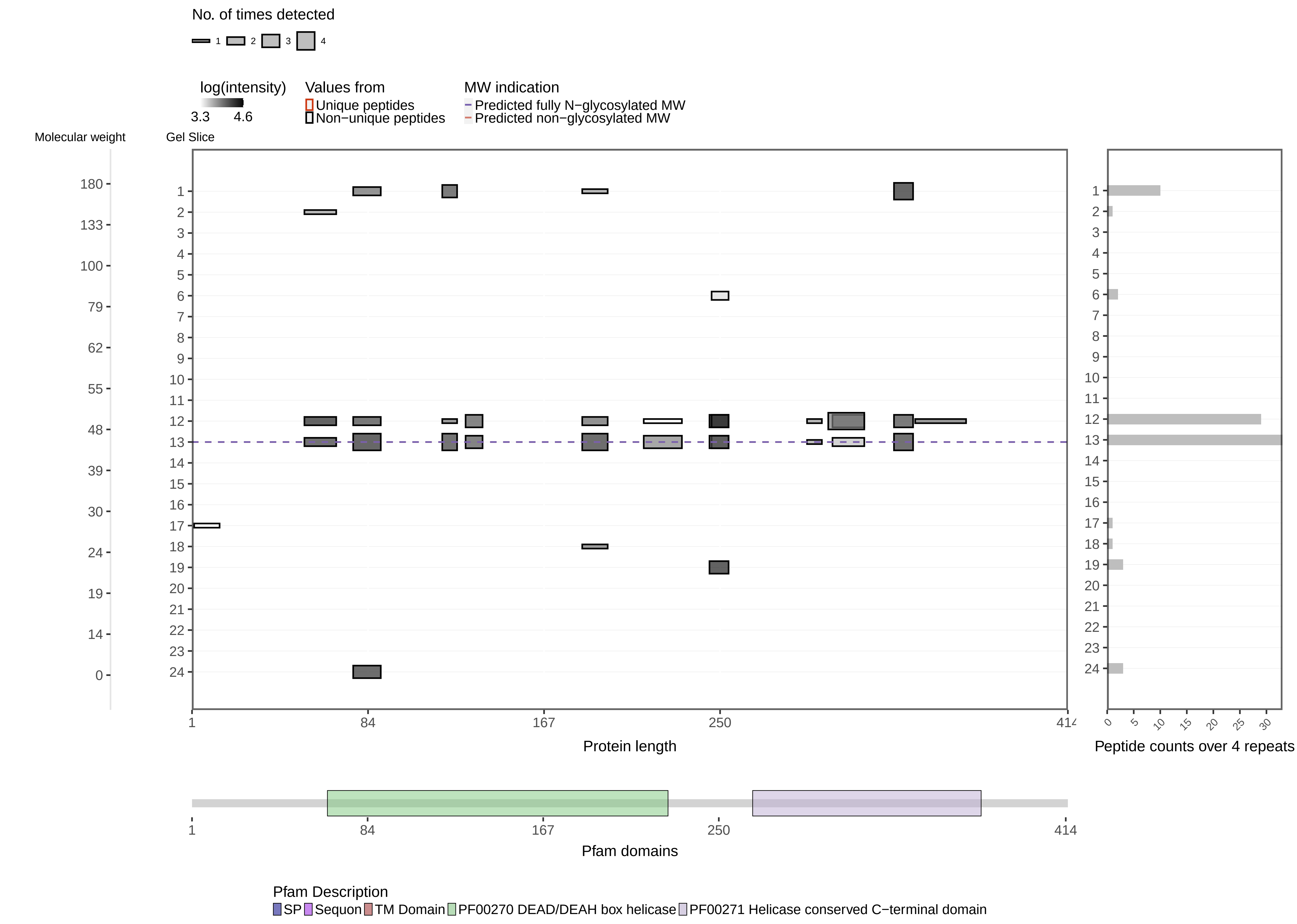


Protein sequence:

ADEMLSRGFKDOIYDIFOLLPPKIOVGVFSATMPPEALEITRKFMNKPVRILVKRDELTLEGIKOFYVNVDKEEWKLETLCDLYETLAITOSVIFVNTRRKVDWLTDKMRSRDHTVSATHGDMDONTRDIIMREFRSGSSRVLITTDLLARGIDVOOVSLVINYDLPTOPENYLHRIGRSGRFGRKGVAINFVTKDDERMLFDIOKFYNVVIEELPANVADLL*

NbL17g05300.1 cytokinin_riboside_5'−monophosphate_phosphoribohydrolase_LOG8−like [MW = 23.27 kDa]

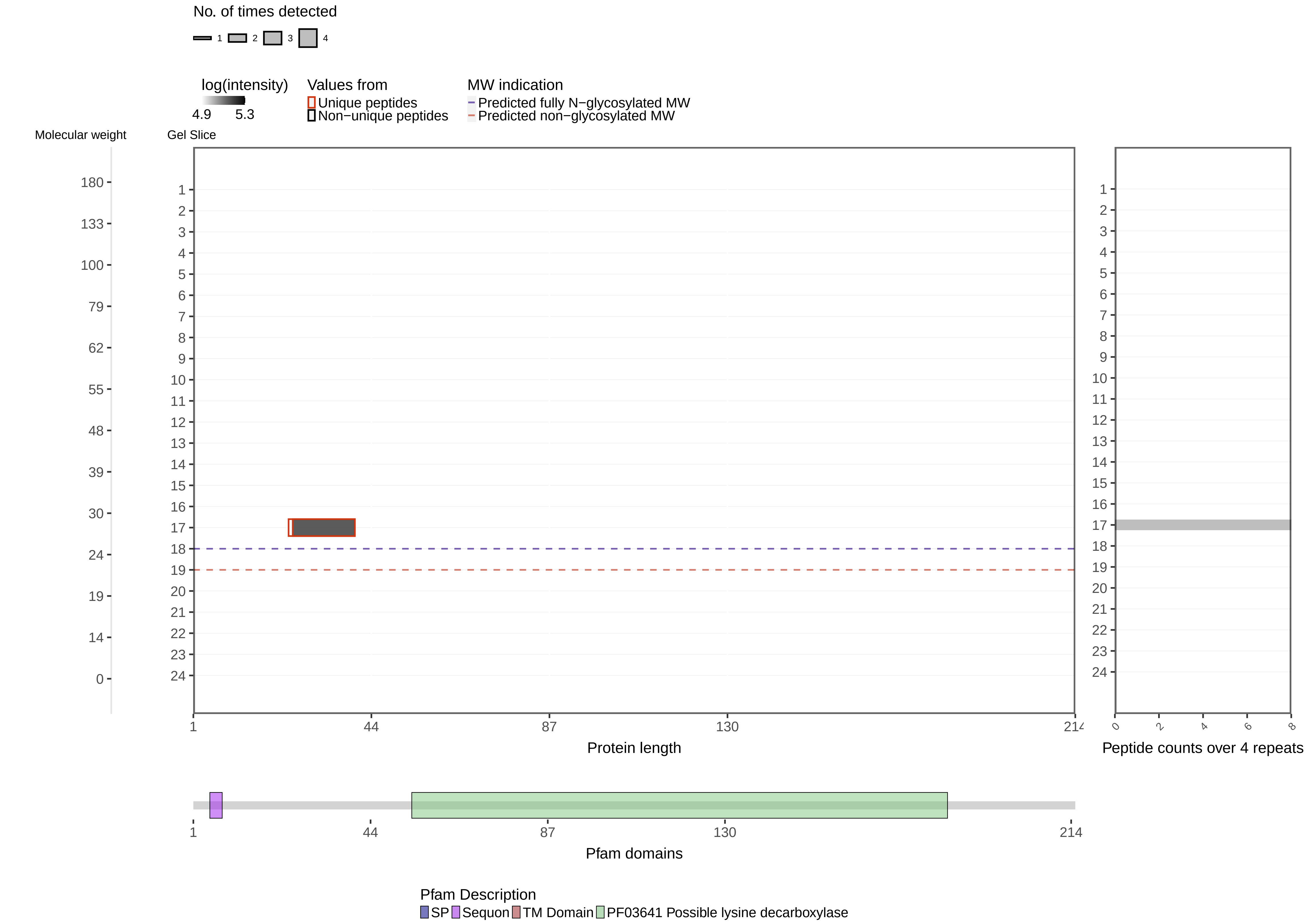


Protein sequence:

MHERKAEMASQADAFIALPGGYGTMEETLEMITWAQLGIHKKPVGLLNVEGYYNSLLTLFDNGVEEGFIKPGARDIVLAAPTARELLSKMEQYTPSHDHVAPHESWQMEELAYPKEQSP*

### NbL17g05280.1 elongation_factor_2 [MW = 93.97 kDa]

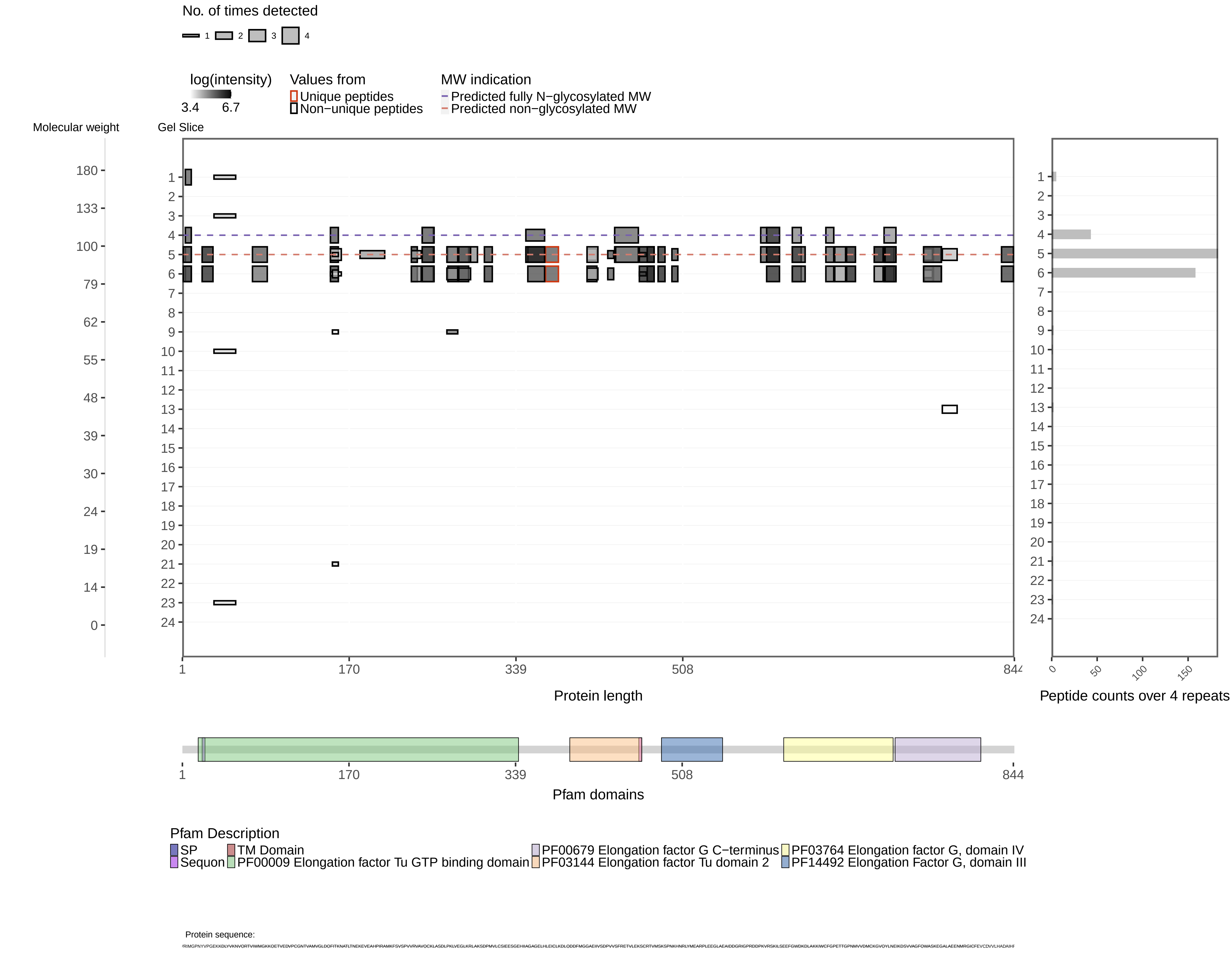

NbL17g05260.1 elongation_factor_2 [MW = 93.99 kDa]

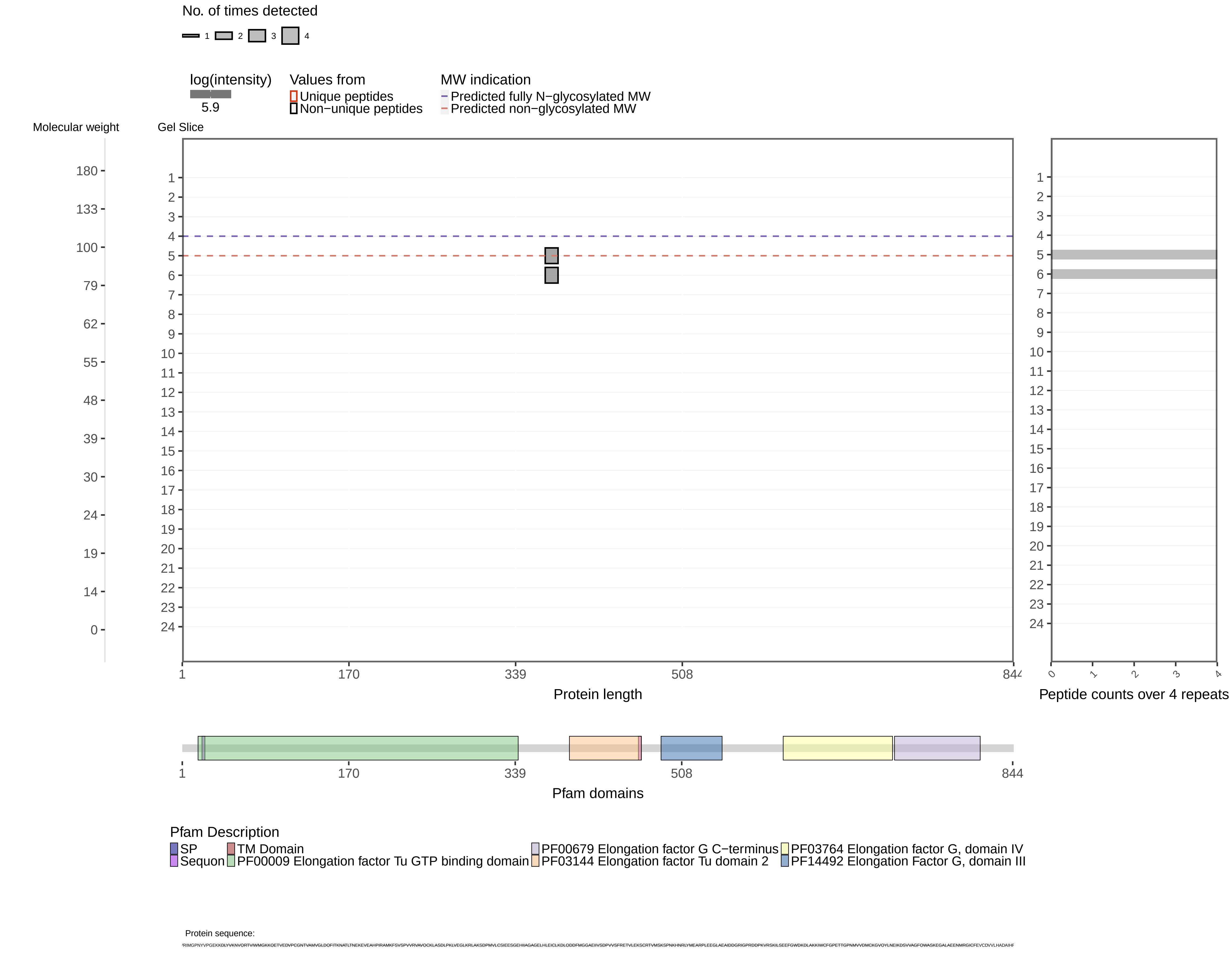

### NbL17g05150.1 UDP−sulfoquinovose_synthase_chloroplastic−like [MW = 53.39 kDa]

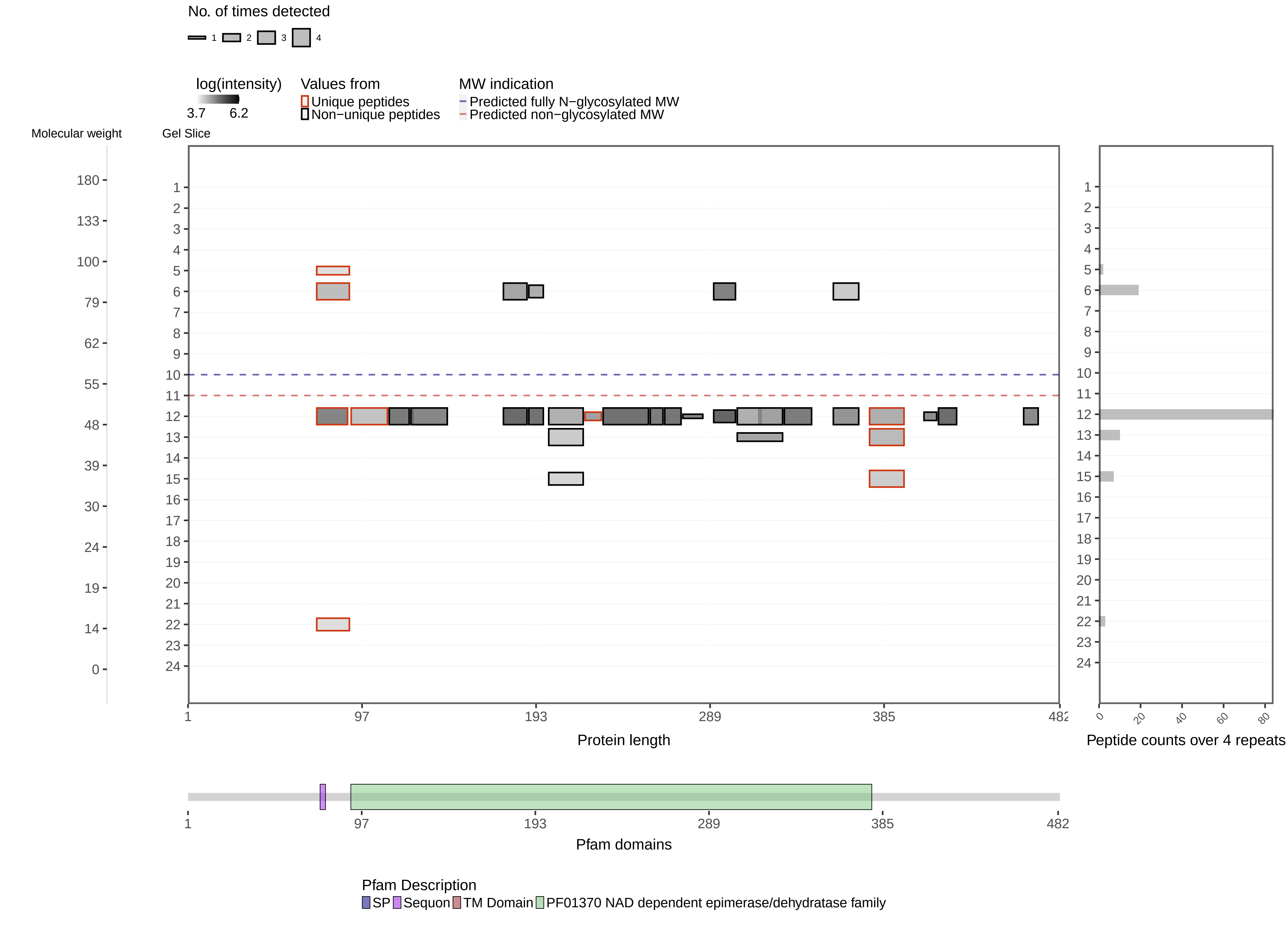


Protein sequence:

[illegible]

### NbL17g05050.1 glycine_dehydrogenase_(decarboxylating)_mitochondrial [MW = 113.92 kDa]

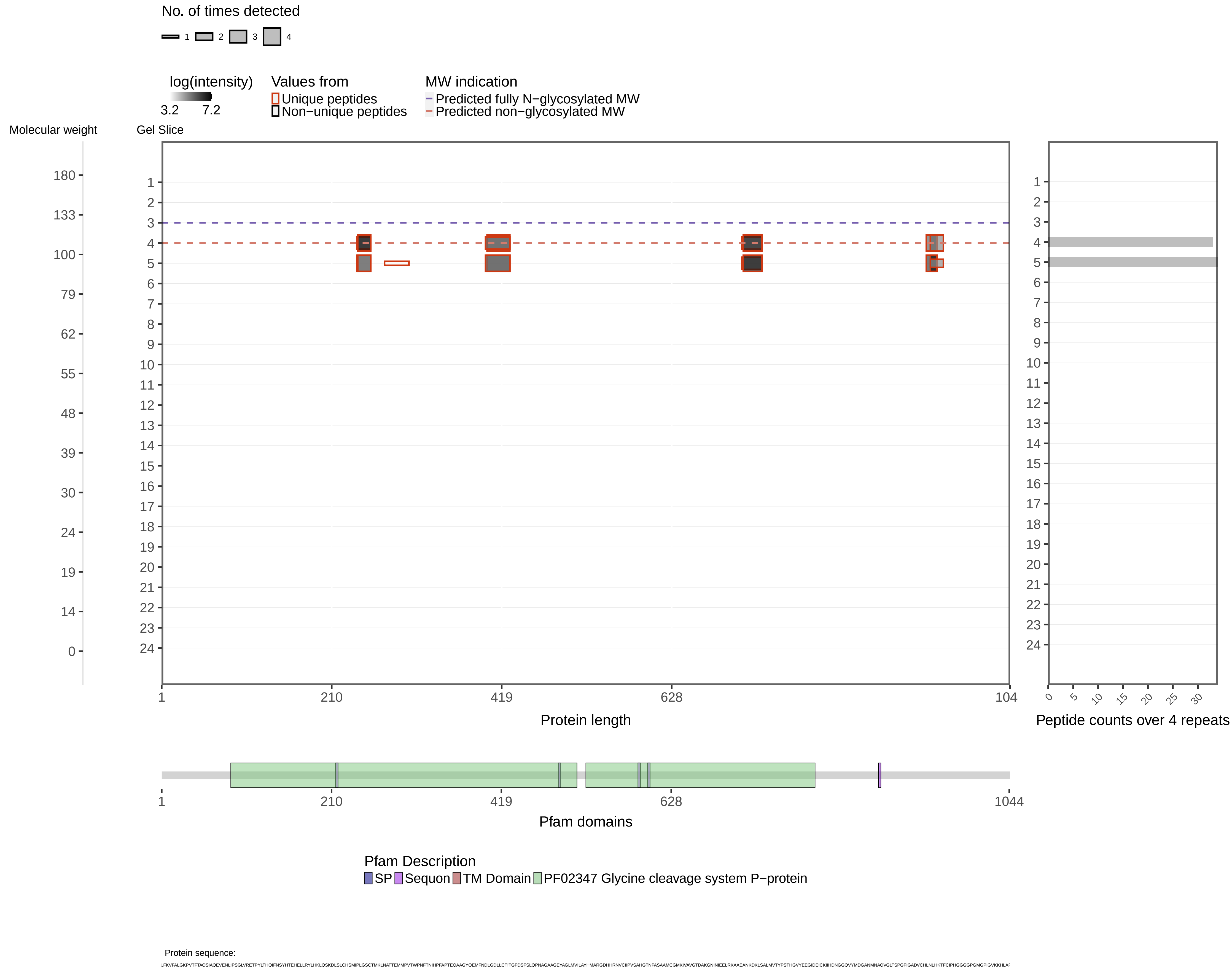

### NbL17g04950.1 abscisic_acid_receptor_PYL2−like [MW = 20.9 kDa]

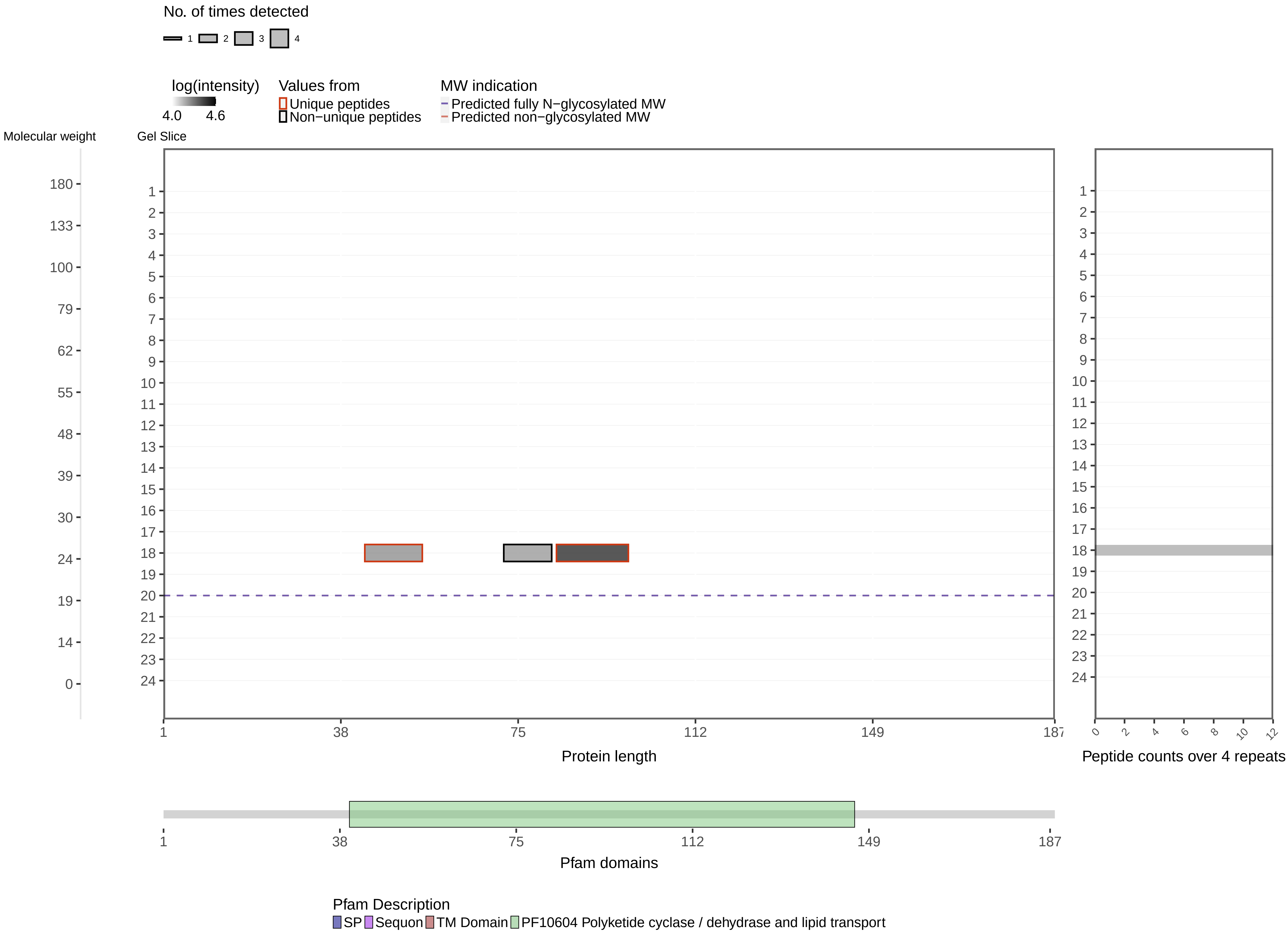


Protein sequence:

SMREVSVVSGIPASTSTERLEILDDEKHILSFKVVGGEHRLNNYRSVTSVNEFKKNGKAYTIVLESYIVDIPEGNTGEDTKMFTDTVVKLNLQKLGVVAMAALHGQE*

### NbL17g04900.1 NADH_dehydrogenase_[ubiquinone]_iron−sulfur_protein_7_mitochondrial−like [MW = 23.6 kDa]

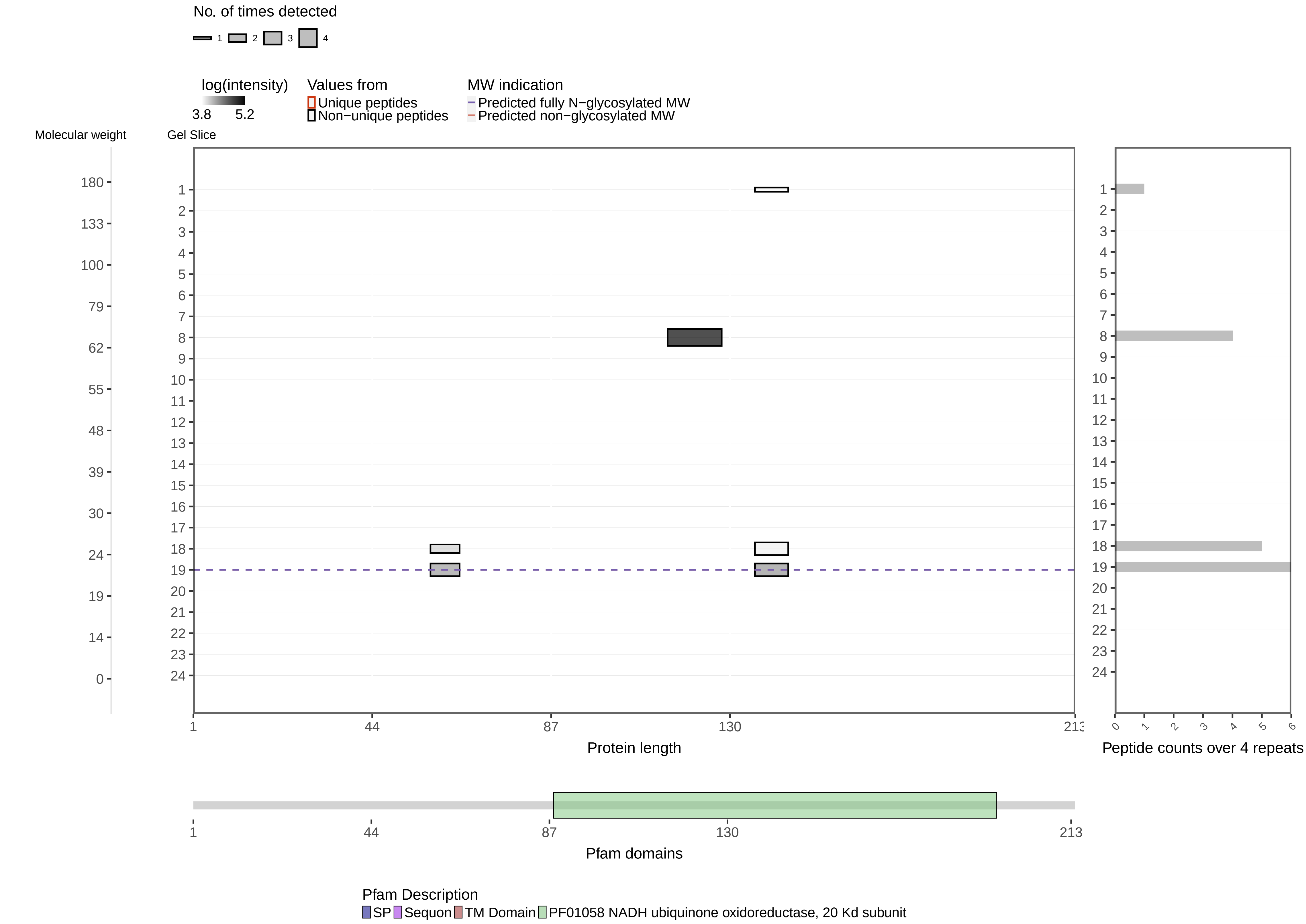


Protein sequence:

AGAARYDFDRFGVIFRPSPRQSDCMIVAGTLTNKMAPALRKVYDQMPEPRWVISMGSCANGGGYYHYSYSVVRGCDRIVPVDIYVPGCPPTAEALLYGVLQLQKKINRRKDFLMWWTK*

### NbL17g04850.1 serine_hydroxymethyltransferase_3_chloroplastic−like [MW = 56.36 kDa]

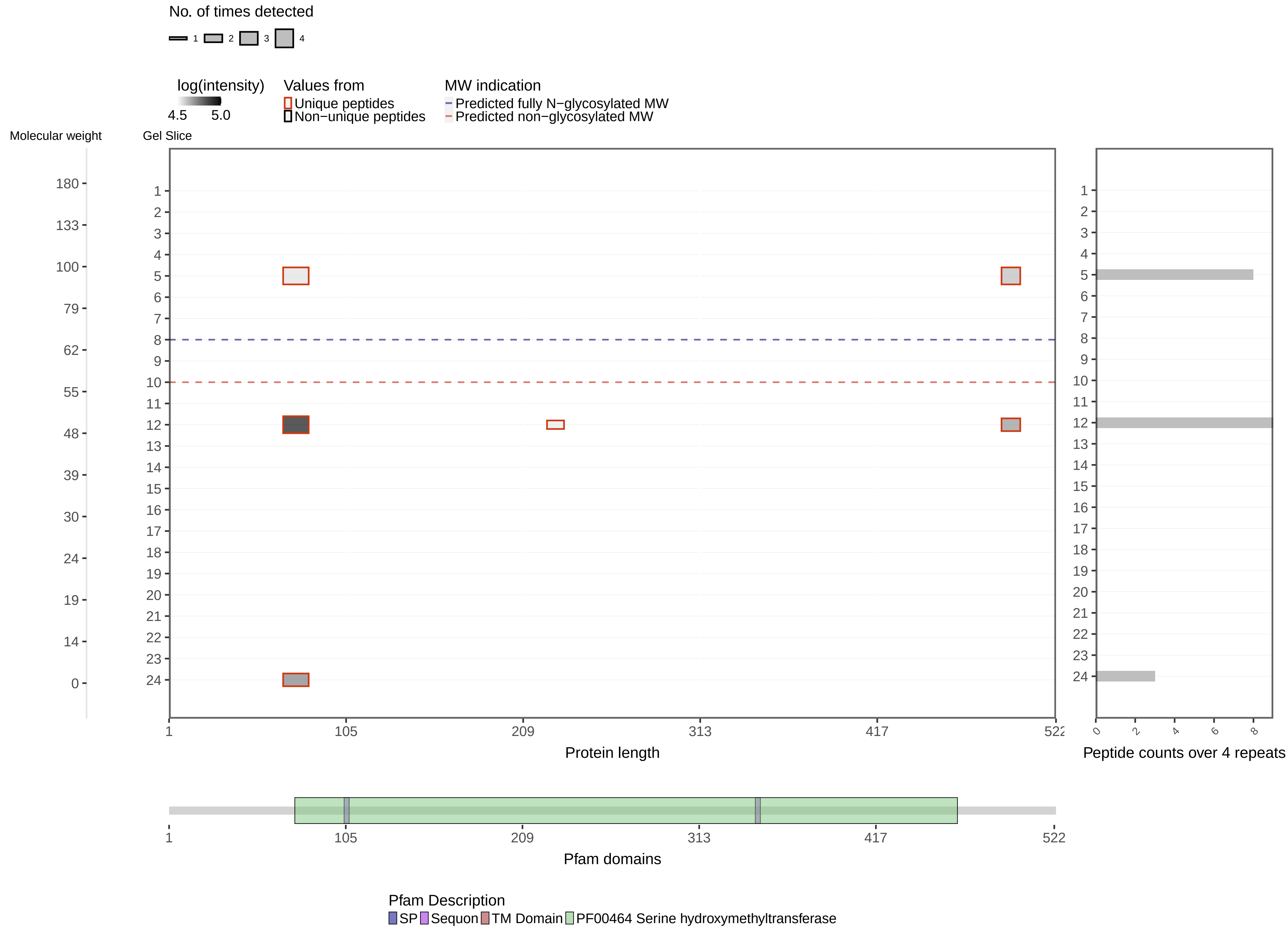


Protein sequence:

KLIIAGASAYPRDFDYPRLRKIADAVGAFLMMDMAHISGLVAASVLANPFEYCDIVTTTTHKSLRGPRGGMIFFKKDPVLGVDLESAINNAVFPGLQGGPHNHTIGGLAVCLKHAKSPEFKAYQNQVVSNCRALASRLTELGYKLVSGGSDNHLVLVDLRPLGTDGARVEKILDMASITLNKNSVPGDKSALVPGGIRIGSPAMTTRGFTEKEFVAVADFIHEGVQITLEAKKSVSSTKLQDFLKFVTAPDFNLIDRVADLQRRVEAFTSQYPLPGL*

### NbL17g04840.1 PITH_domain−containing_protein_1 [MW = 22.52 kDa]

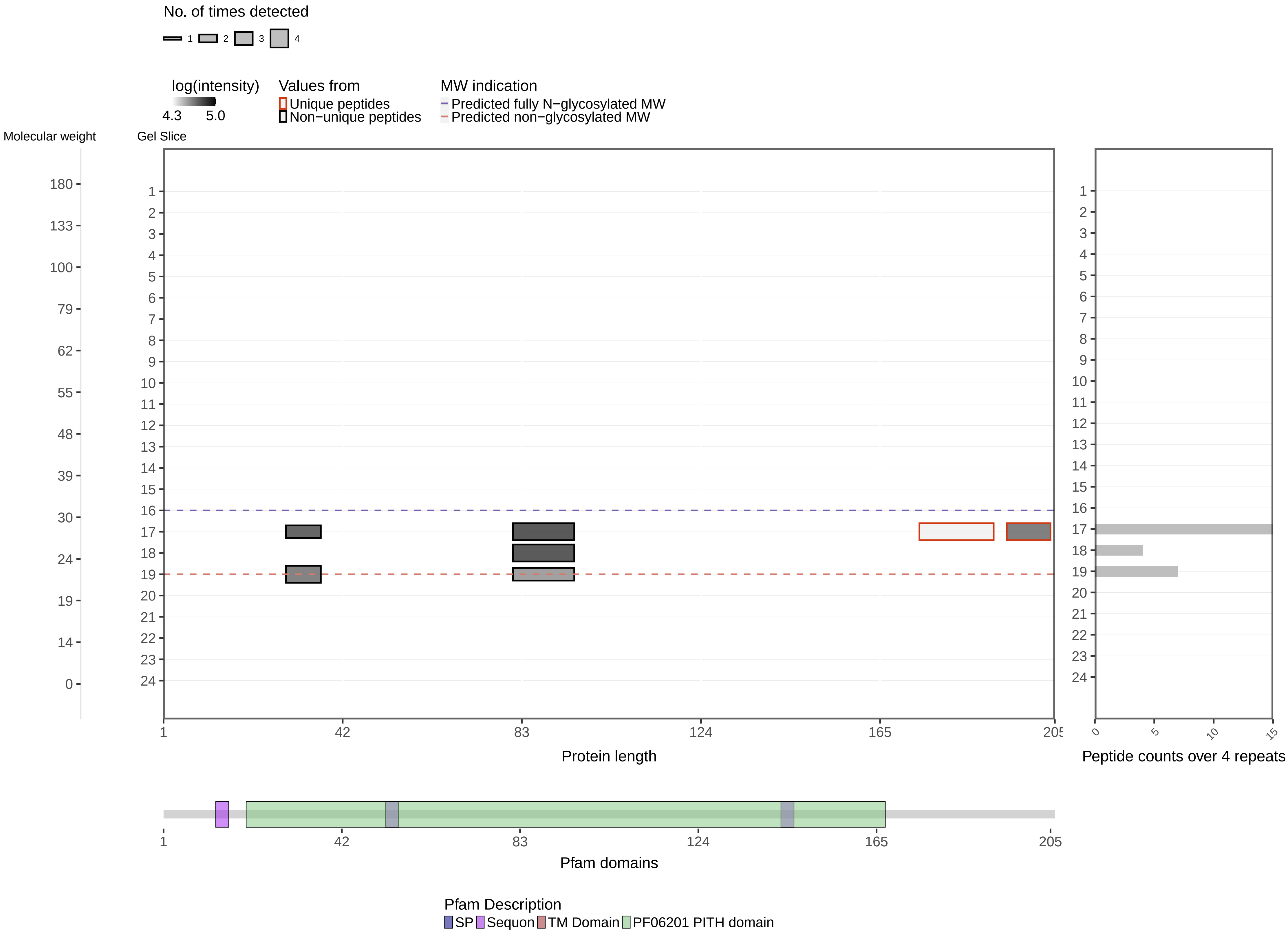


Protein sequence:

TSPAKMRAFINRDGIDFSDAQSMQPVQEWDLAENLQGVLEYQTRYSRFQSVGNITLHFPDNFGGDTTQIHYIGLKGEATQMKRDVVANIVYEVVPNPSDHKTRAEGGGGLSHVE*

### NbL17g04760.1 vacuolar−processing_enzyme−like [MW = 57.08 kDa]

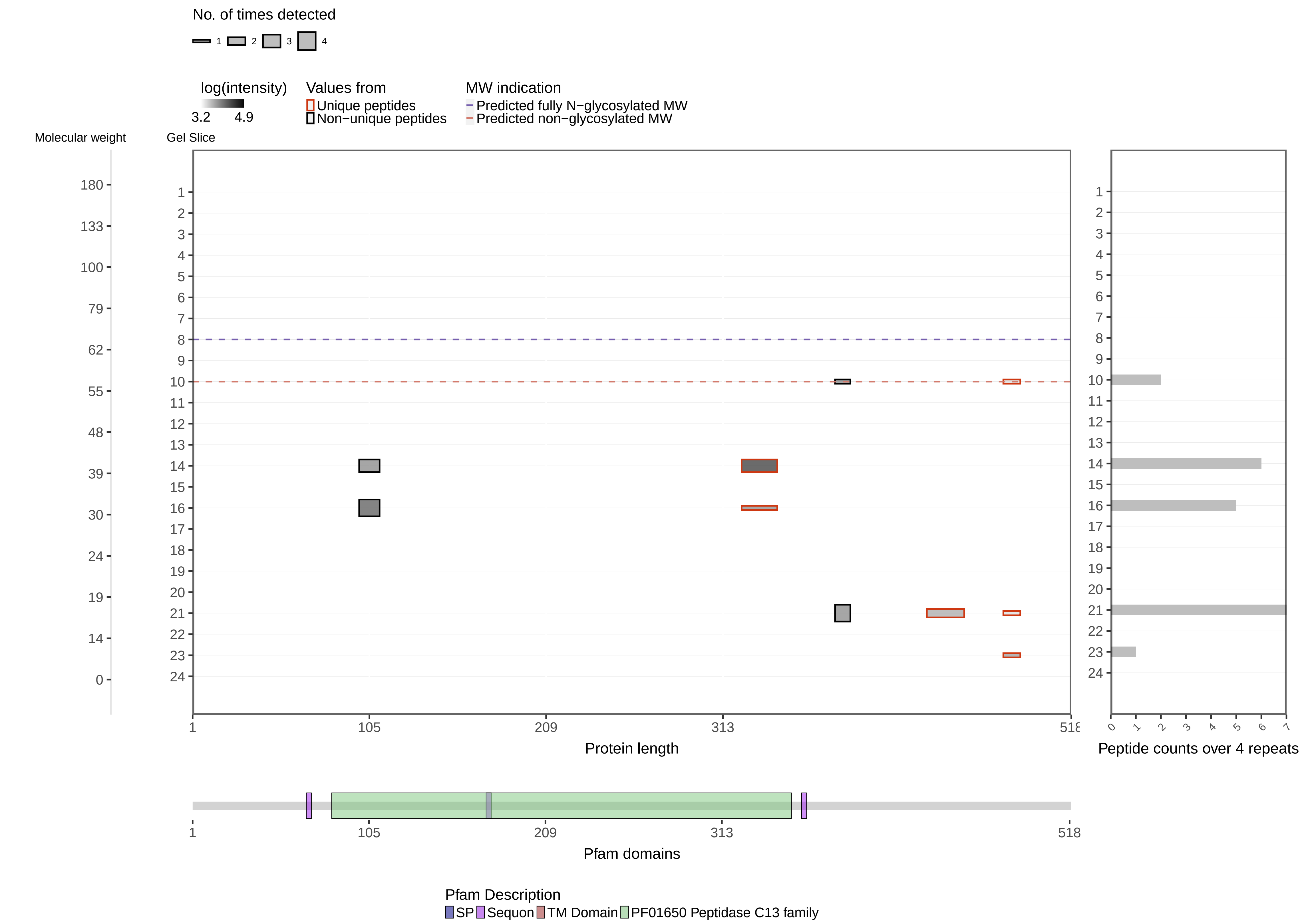


Protein sequence:

NbL17g04650.1 uncharacterized_LOC109241480 [MW − SP = 151.69 kDa]

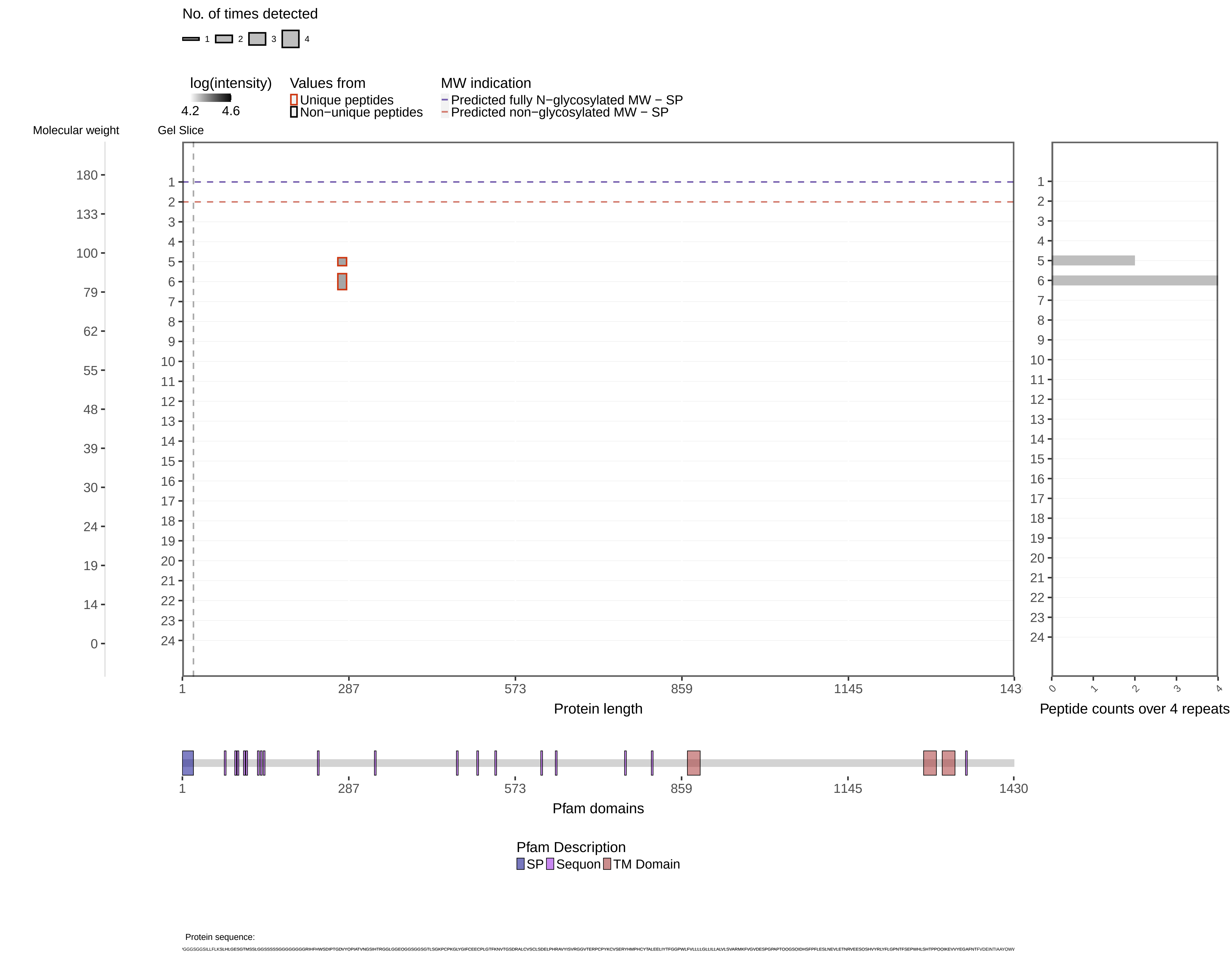

NbL17g04640.1 glutamyl−tRNA(Gln)_amidotransferase_subunit_C_chloroplastic/mitochondrial−like [MW = 15.82 kDa]

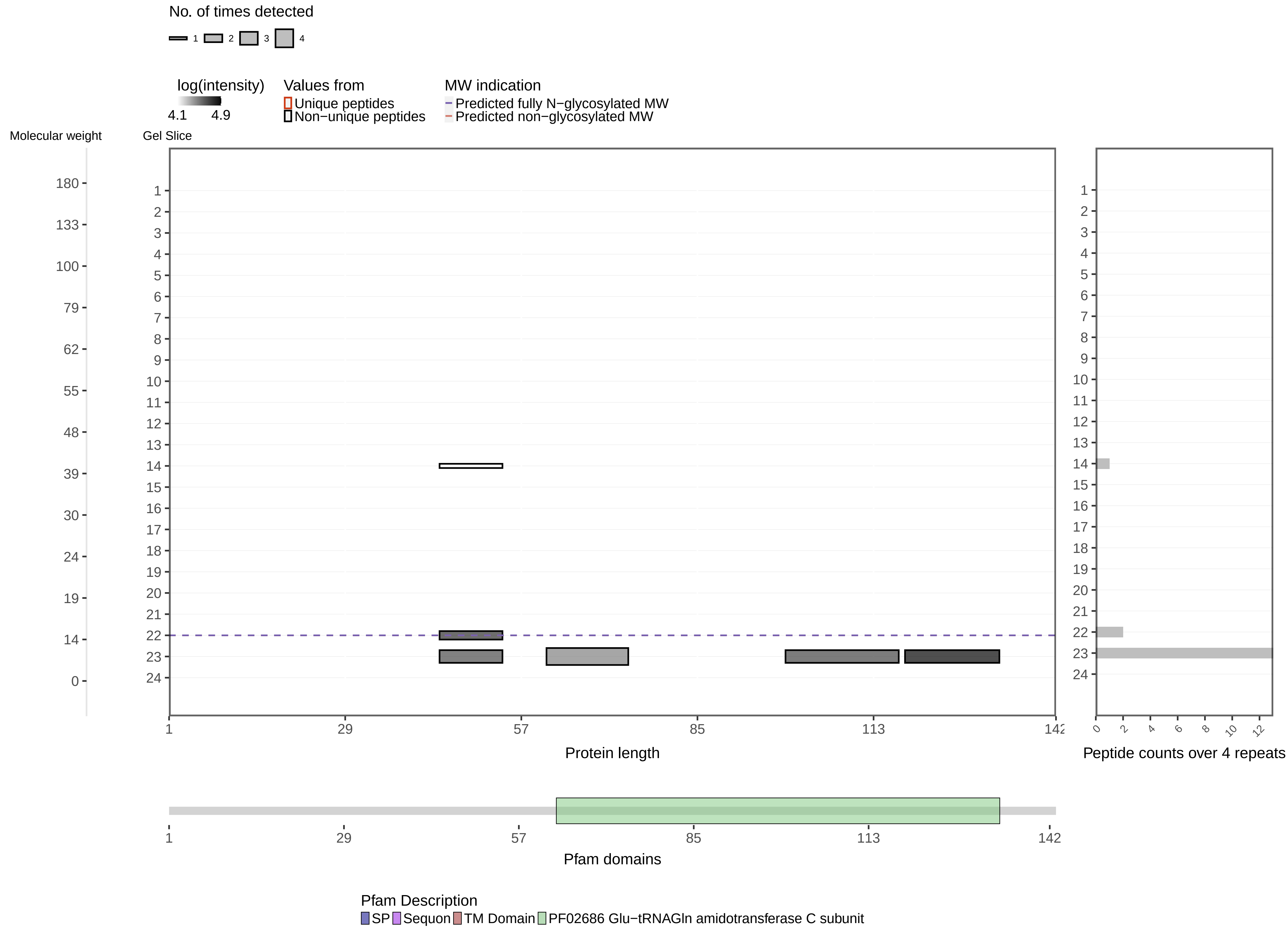


Protein sequence:
TARISLTPQQVEEFAPKIRQVIDWFGQLQAVDLQSIEPAIRADTEGDNLRDDVPEKFENREALISAVPSFEEPYIKVPKVLNKE*

### NbL17g04560.1 profilin−2 [MW = 14.14 kDa]

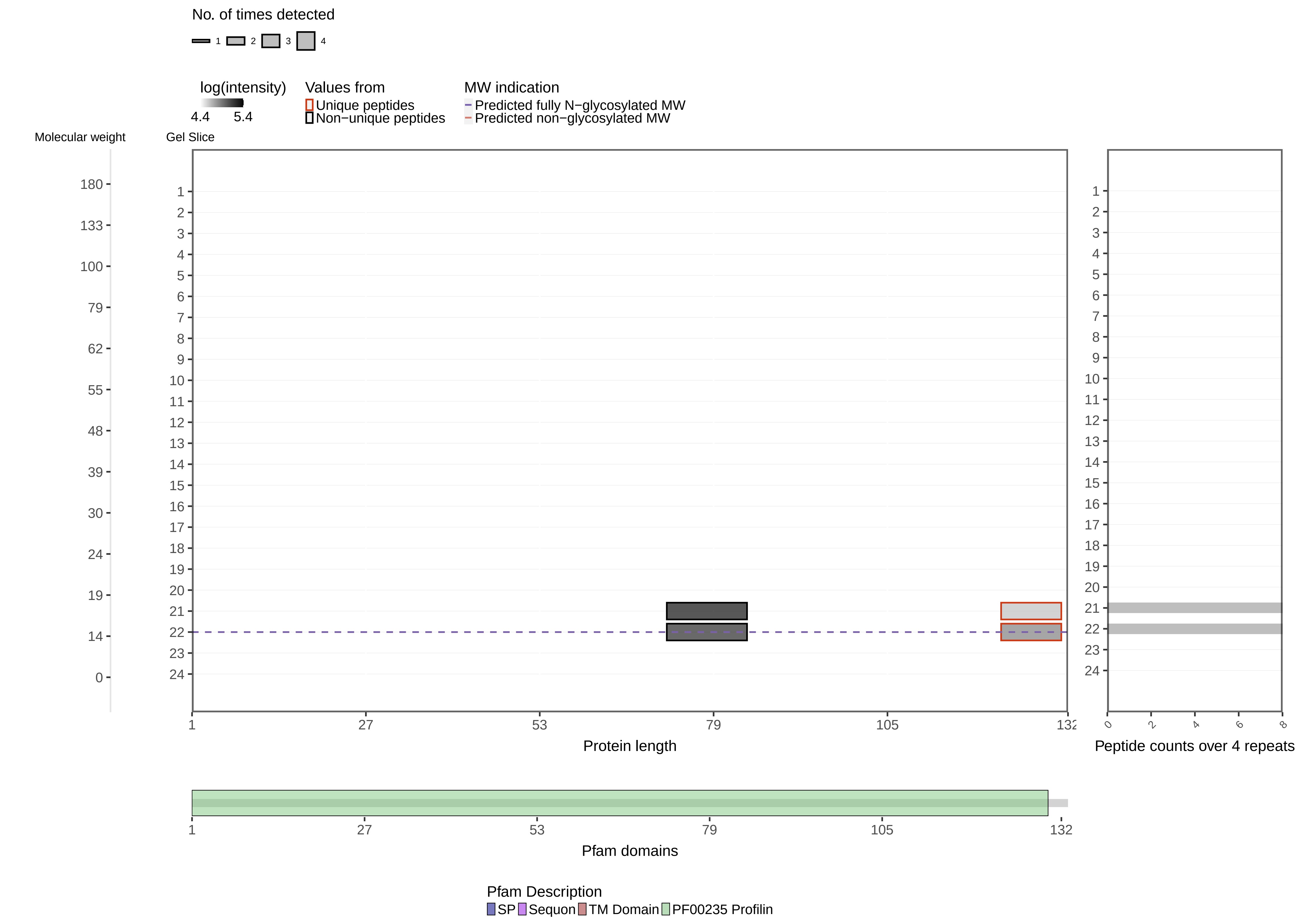


Protein sequence:

IDFAEPGTLAPTGLYLGGTKYMVIQGEPGAVIRGKKGPGGITIKKTNQALIIGIYDEPMTPGQCNMIVERLGDYLVDQGL*

### NbL17g04500.1 NADP−dependent_malic_enzyme−like [MW = 69.39 kDa]

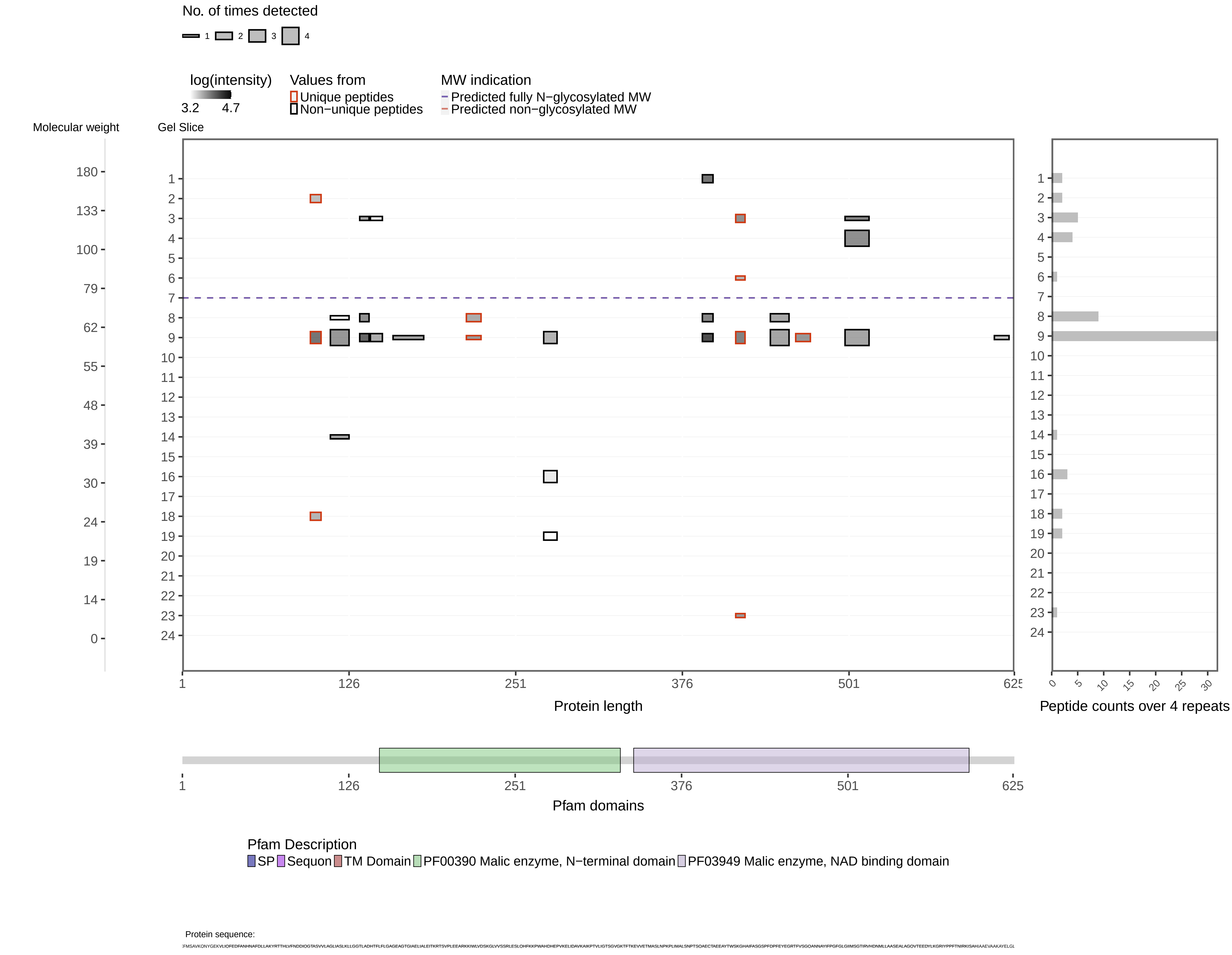

NbL17g04360.1 probable_transcriptional_regulatory_protein_At2g25830 [MW = 36.98 kDa]

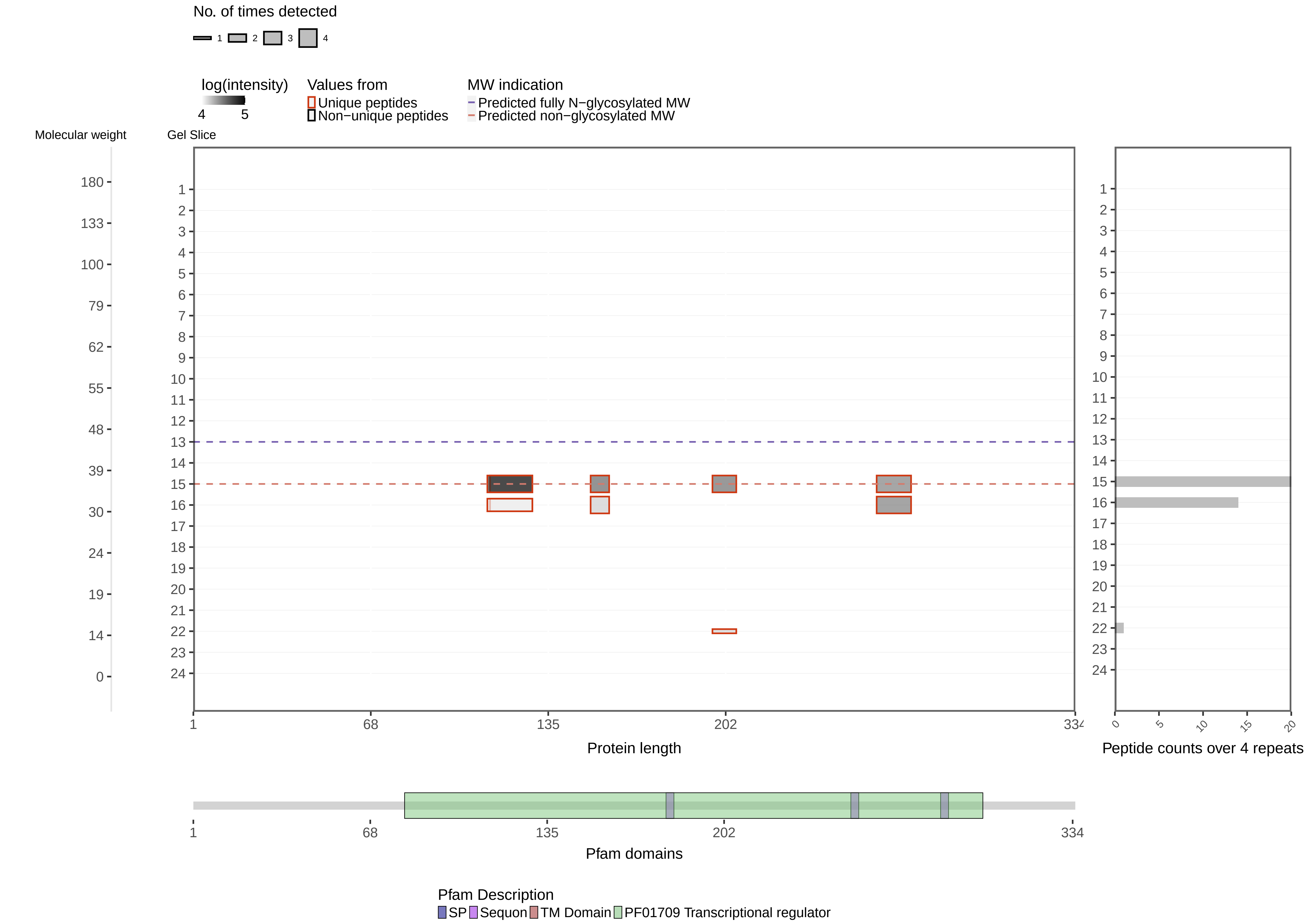


Protein sequence:

EAFMEKIYEVYGYGGVGIIVEVLTDKVNRSVAAVREVVKDCGGKMADSGSIMFKFRRARVANVKVTDIDRDQLLTIALDAGAEDIIEPSMDDYDTEANSSESRVYKVVSSADNYPVILSKLQEEGIKFEPDNGSELLPITPIEVRKHSISEFYSSNLVALVNLLRGEYYFNHEILQKEFNN*

### NbL17g04330.1 nuclear_pore_complex_protein_NUP85 [MW = 81.21 kDa]

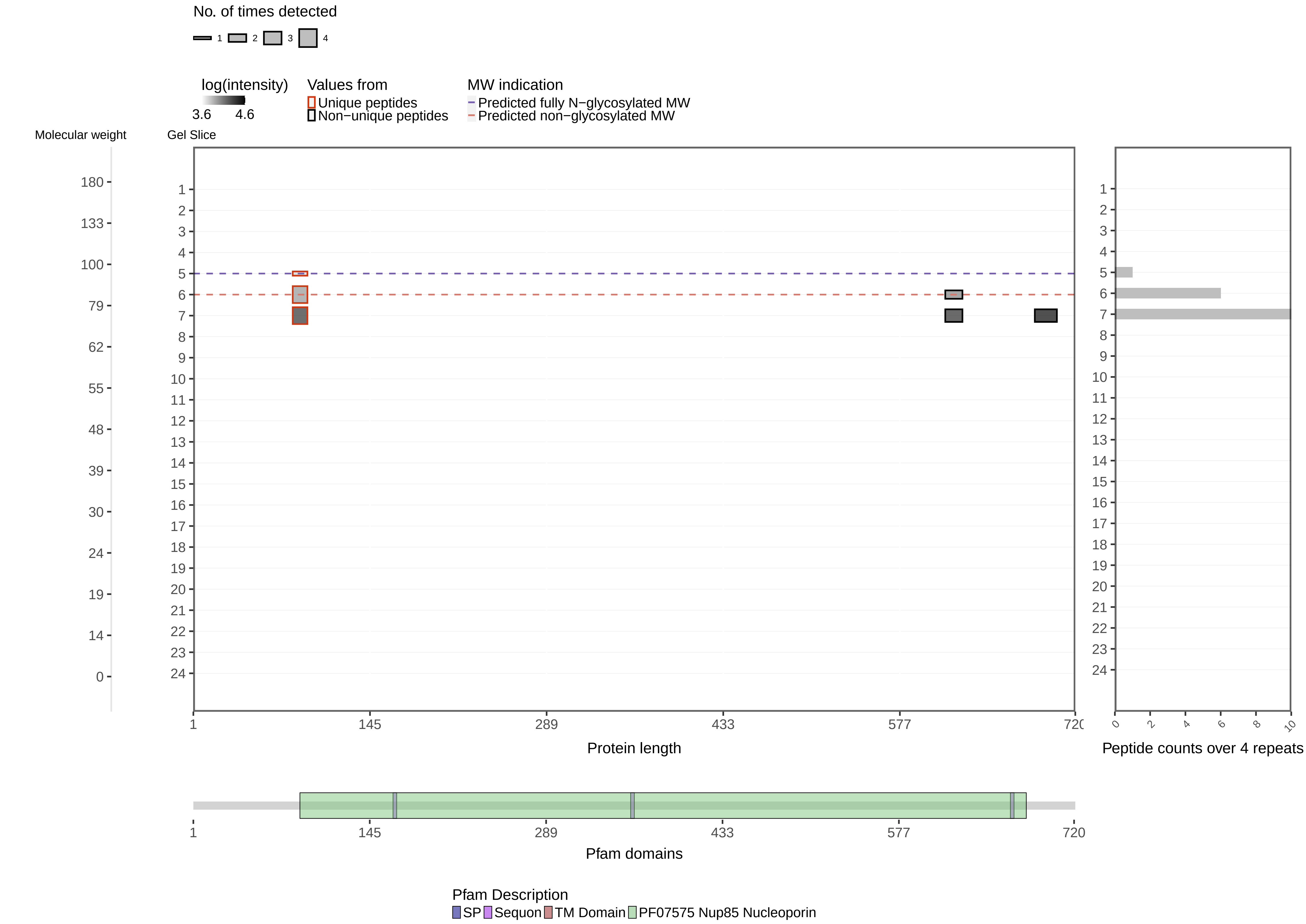


Protein sequence:

[illegible]

### NbL17g04140.1 probable_lactoylglutathione_lyase_chloroplast [MW = 40.12 kDa]

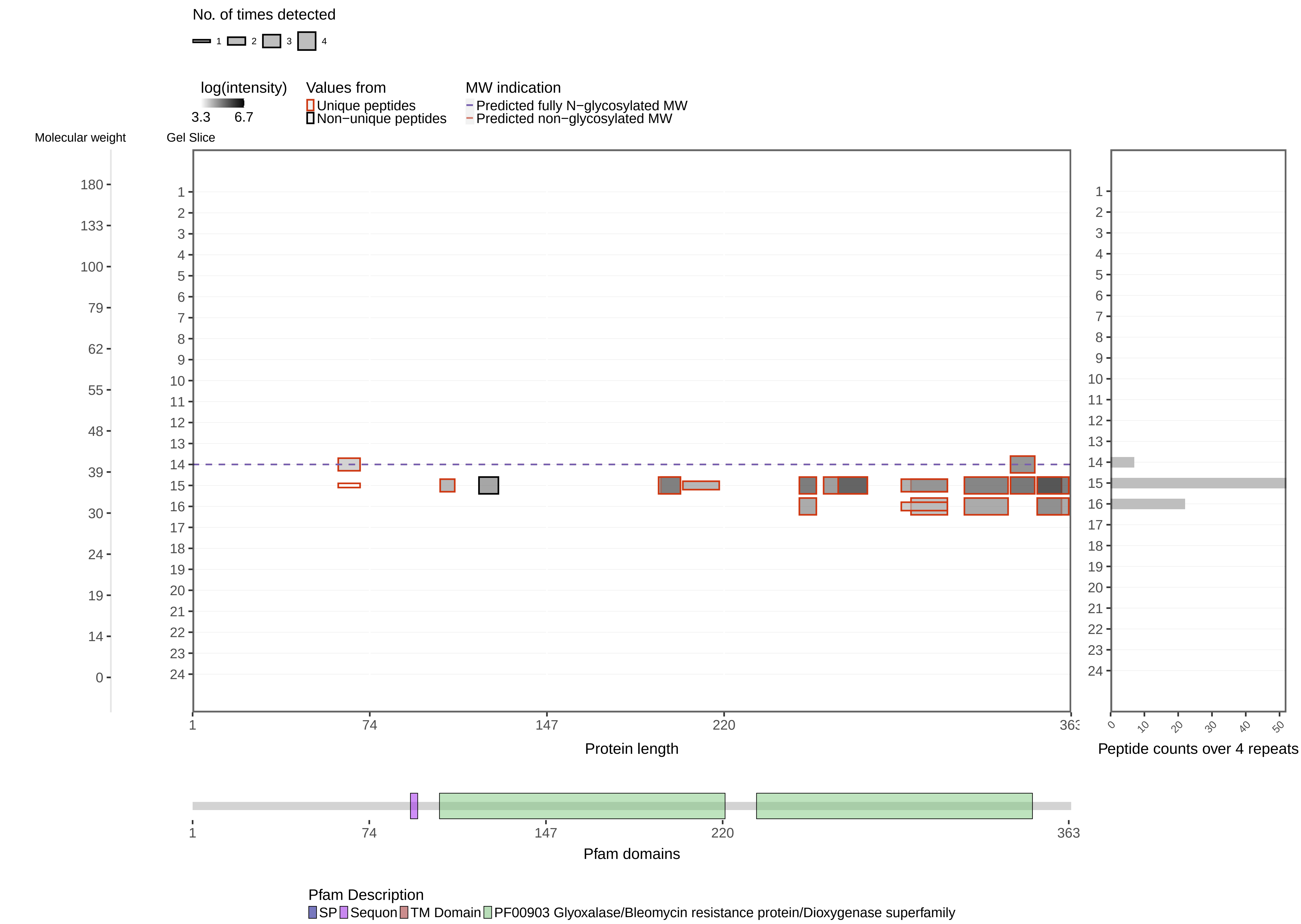


Protein sequence:

TGFGHFGIAVEDVAKTVDLVKAKRGTWNLIQSKVGGTIIALVEDPDGYKFELLQRGPTPEPFREVMLRVGDLDRSINFYKKAFGMELLRKRDNPDNKNTIAVLGYGPEDKNAVLVLTYTYGVKSYDKGNGYAQIAVGTDDVYKTAEAIKLYGGEIVLEPGPLPGINTKIMACVDPDGWKTVFVDNADFMKELE*

NbL17g04060.1 uncharacterized_LOC107794167 [MW − SP = 38.26 kDa]

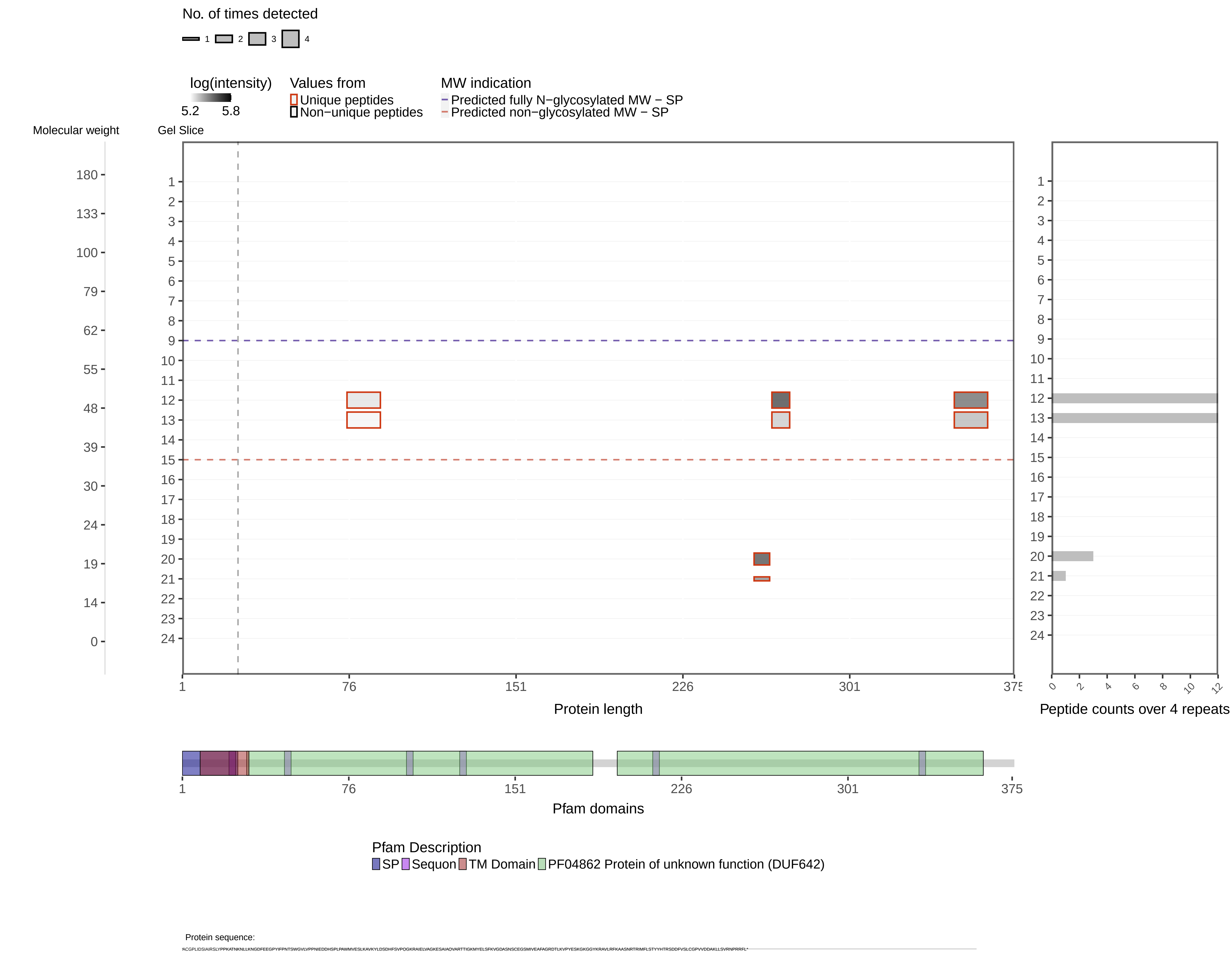

### NbL17g04020.1 GTP−binding_nuclear_protein_Ran−A1 [MW = 25.02 kDa]

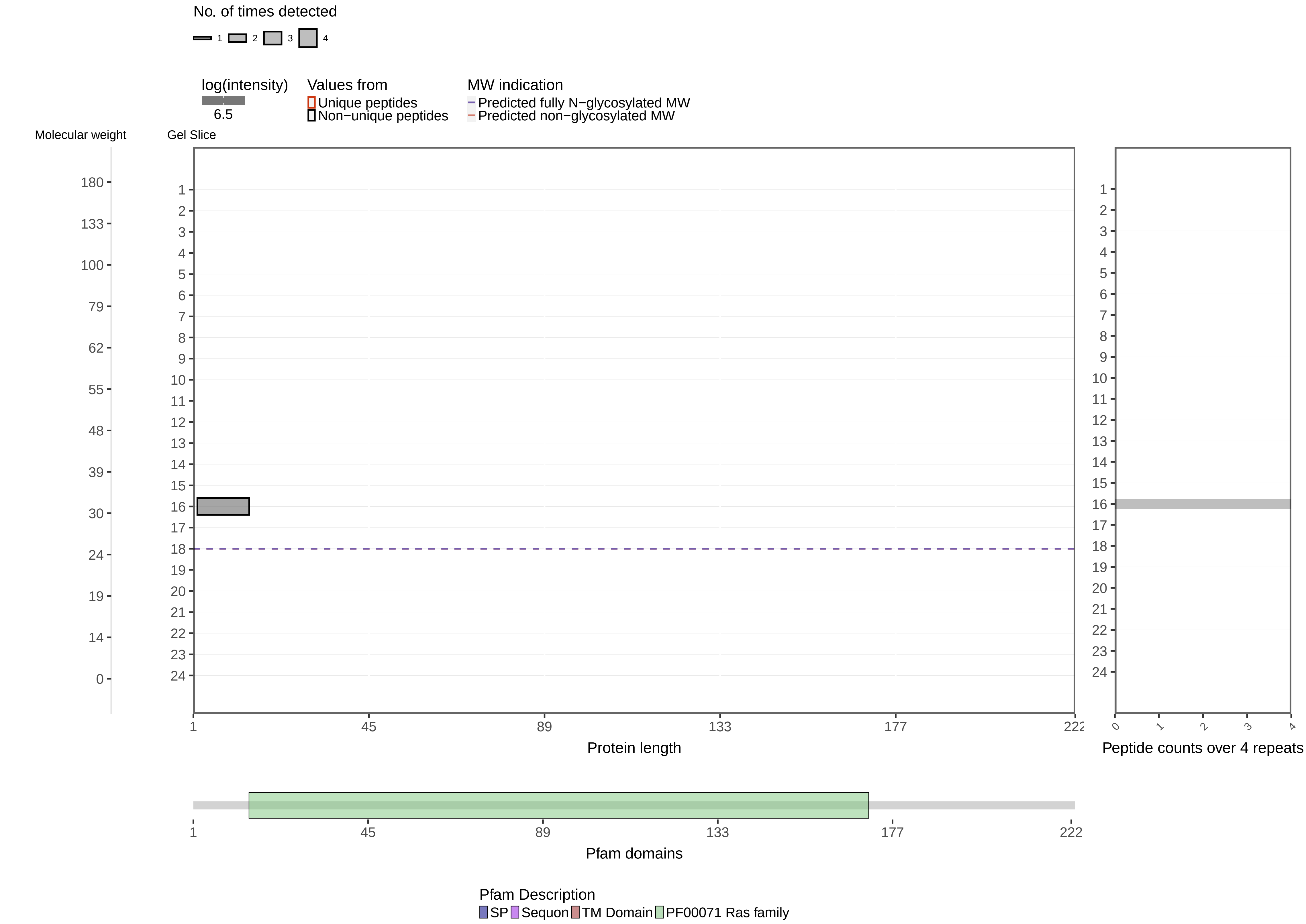


Protein sequence:

LTYKNVPTWHRDLCRVCENIPIVLCGNKVDVKNRQVKAKQVTFHRKKNLQYYEISAKSNYNFEKPFLYLARKLAGDPNLHFVESPALAPPEVQIDLAAQQQHEAELAAAASQPLPDDDDETFD*

NbL17g03950.1 aspartyl_protease_family_protein_At5g10770−like [MW = 55.63 kDa]

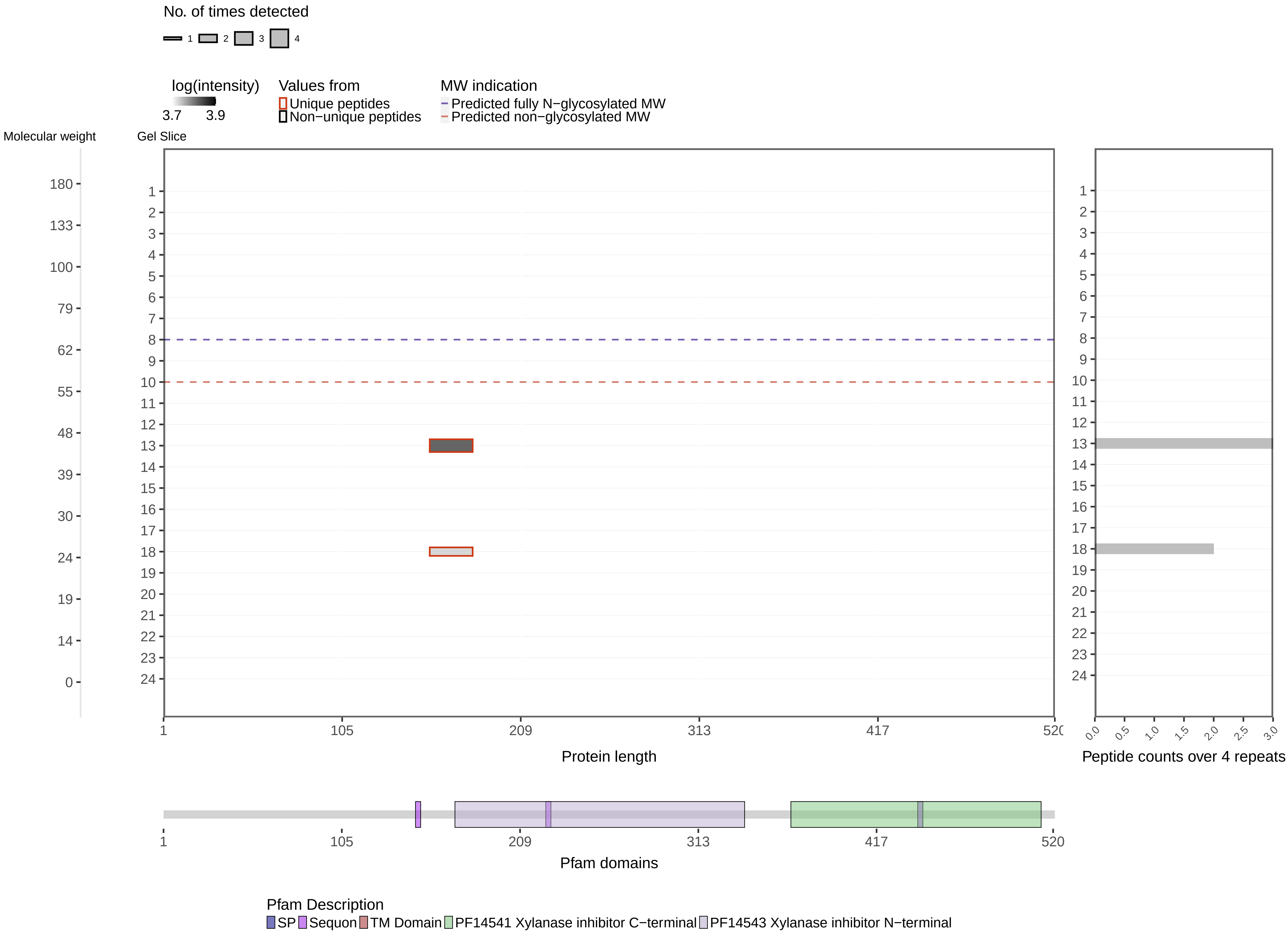


Protein sequence:

SSSNCVYGIQYGDTSFSVGFFAKDKLTLTPKDVFDGFMFGCGQNNRGLFGKTAGLIGLGRDPLSIVQQTAQKFGKYFSYCLPTSRGSNGHLTFGNGNGVKSSKAVKNGIIFTPFASSQGATFYFIDVLGISVGGKALSISPMLFQNAGTIIDSGTVITRLPSTAYSSLKSTFKQFMSKYPPAPALSLLDTCYDLSNYTSISIPKISFNFNSNANVDLDPNGILITNGASQVCLAFAGNGDDDSIGIFGNIQQQTLEVVYDVAGGQLGFGYRGCS*

### NbL17g03930.1 acyl−protein_thioesterase_2−like [MW = 28.43 kDa]

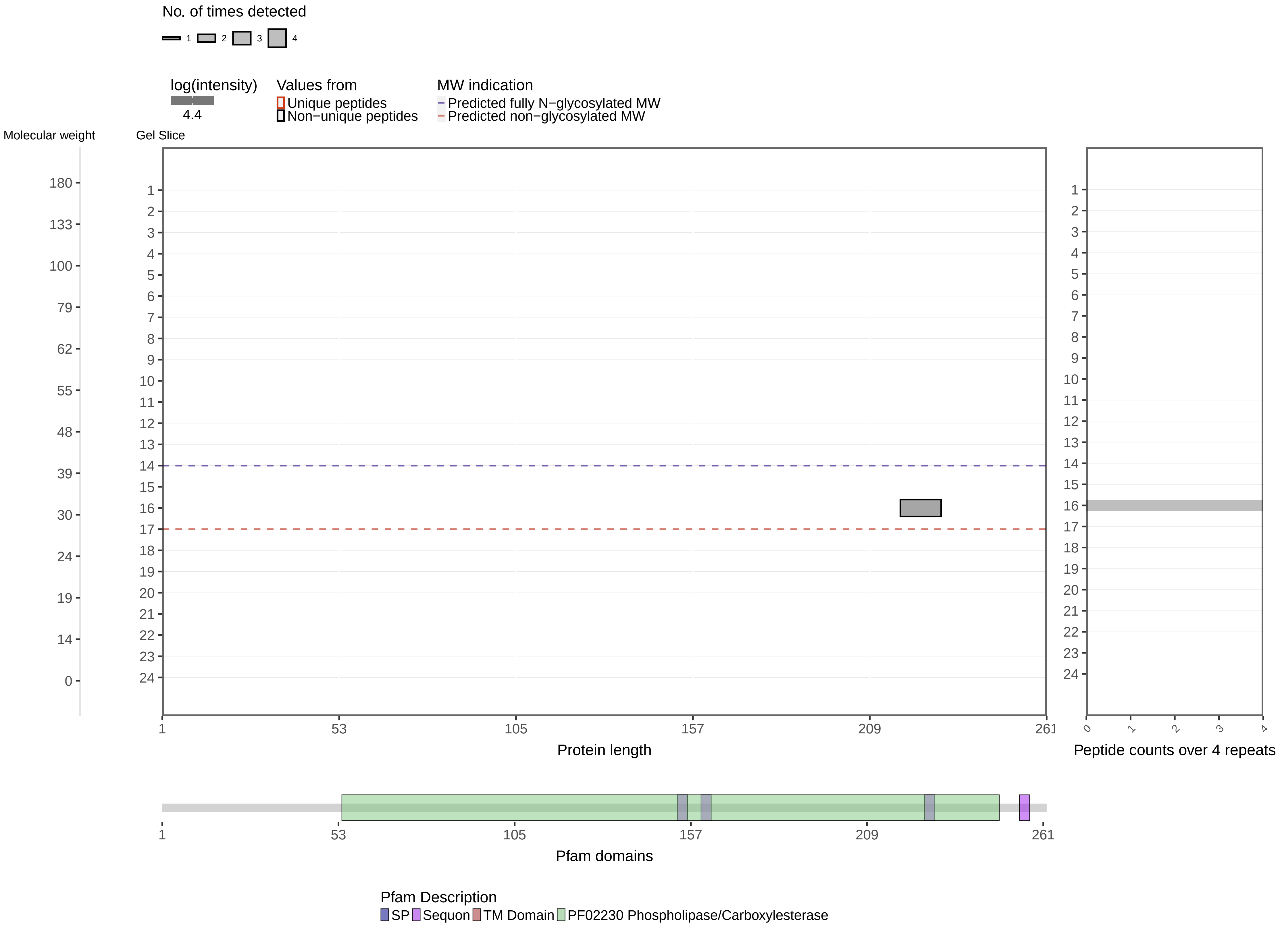


Protein sequence:

STEPADVKLCVGGFSMGAAAALYSATCHAFKQYGNGSPYPLNLSAVVGLSGWLPCSRTLRNRMQGMNDAGRRAASLPILLCHGTGDDVVAYQHGEKSARILSSSGFQNLTFRNYQGLGHYTIPEETDEVCCWLAANLSLGGT*

NbL17g03820.1 E3_ubiquitin−protein_ligase_AIP2−like [MW = 35.71 kDa]

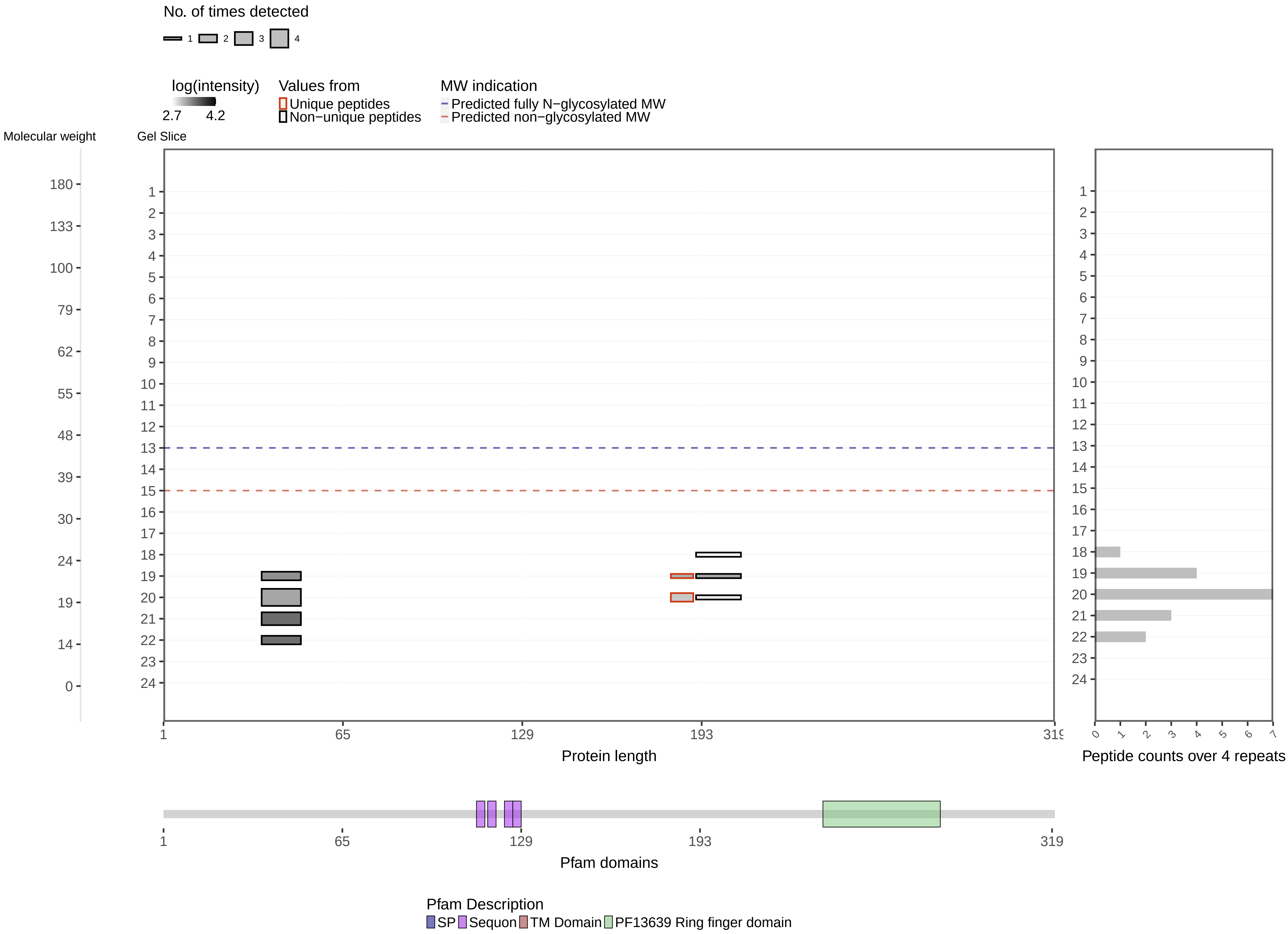


Protein sequence:

PGWLVQSNLLTAAATLFAGESSEGHLAGADASEGSRNILEELVARLDNIIPEILDDGPAAPRAPPASKKVVANLPVTTVTDEFLTKMGADAECAICREVLVVDDKMQELPCKHMFHPPCLKPWLDEHNSCPICRHELLTDDHHYESWKEREKEAEEERKGAANAVREGEYMYV*

### NbL17g03660.1 photosystem_II_protein_D1−like [MW = 38.95 kDa]

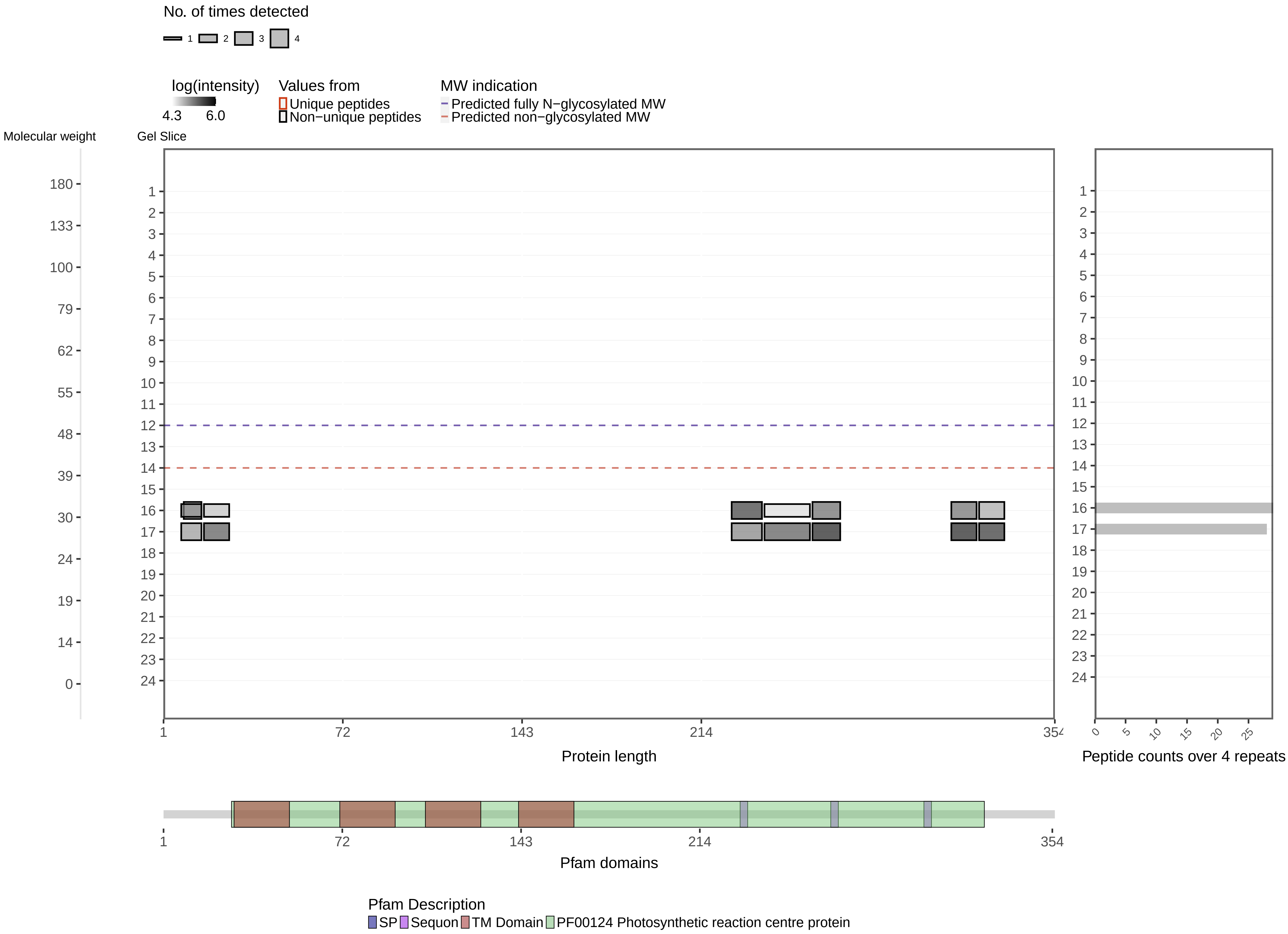


Protein sequence:

FSDGMPLGISGTFNFMIVFQAEHNILMHPFHMLGVAGVFGGSLFSAMHGSLVTSSLIRETTENESANEGYRFGQEEETYNIVAAHGYFGRLIFQYASFNNSRSLHFFLAAWPVVGIWFTALGISTMAFNLNGFNFNQSVVDSQGRVINTWADIINRANLGMEVMHERNAHNFPLDLAAIEAPSTNG*

### NbL17g03500.1 psbP_domain−containing_protein_5_chloroplastic−like [MW = 33.34 kDa]

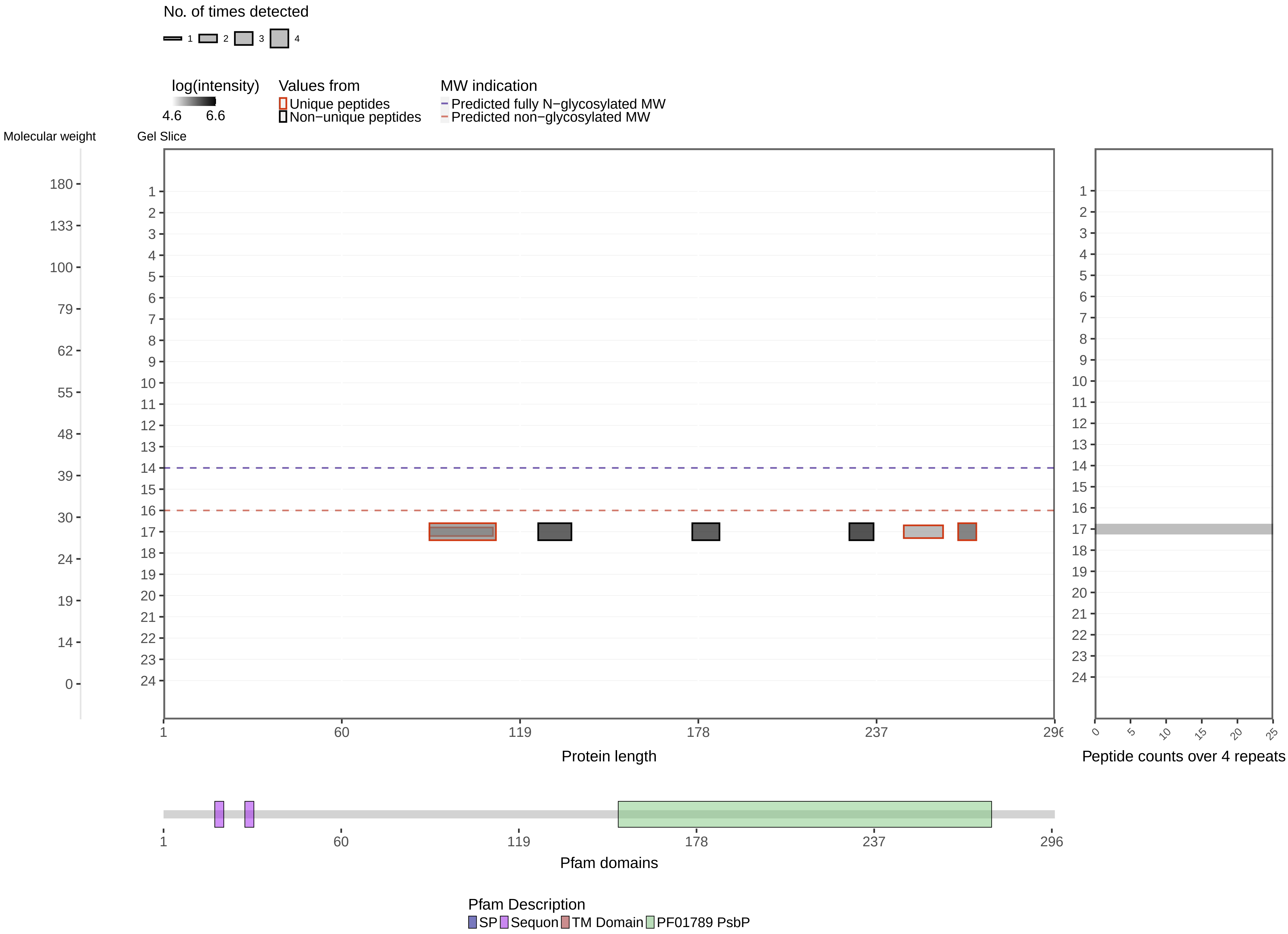


Protein sequence:

RLRIVSERVDIIDNLIISVSIGPPNSQFLKSKEKSTWTAKDVADSVLSDKSALRVTSTQRLAESSLLDAHTSEIDGEPYWYYEYLVRKSPTKTAQETNLFRHYAASTAERDGYLYTFSASALGKQWDKMGPFLEKSVASFRLLPPTESYVPPYKDPWRFW*

### NbL17g03090.1 peptide−N4−(N−acetyl−beta−glucosaminyl)asparagine_amidase_A−like [MW − SP = 62.44 kDa]

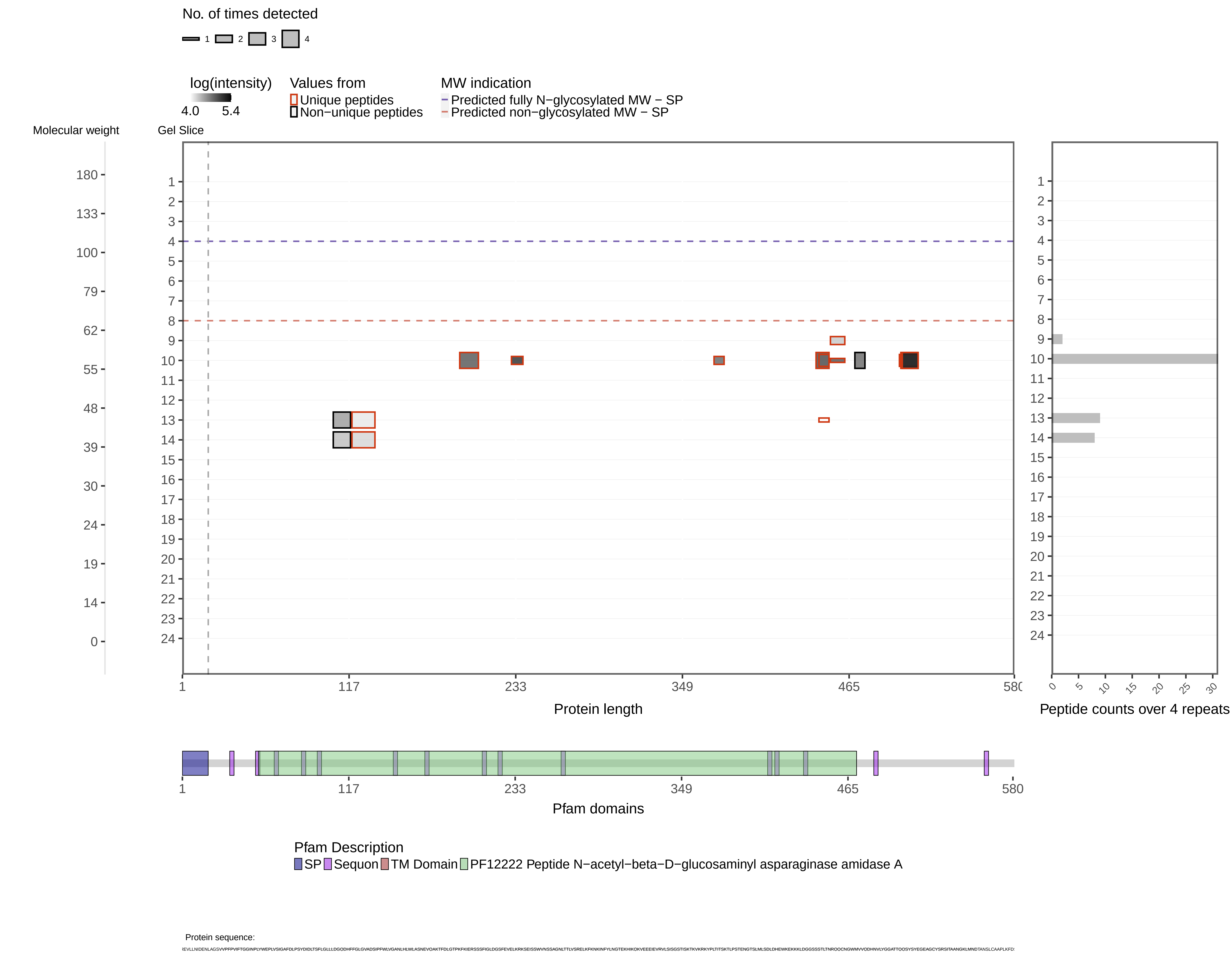

NbL17g03070.1 eukaryotic_translation_initiation_factor_2_subunit_alpha−like [MW = 38.95 kDa]

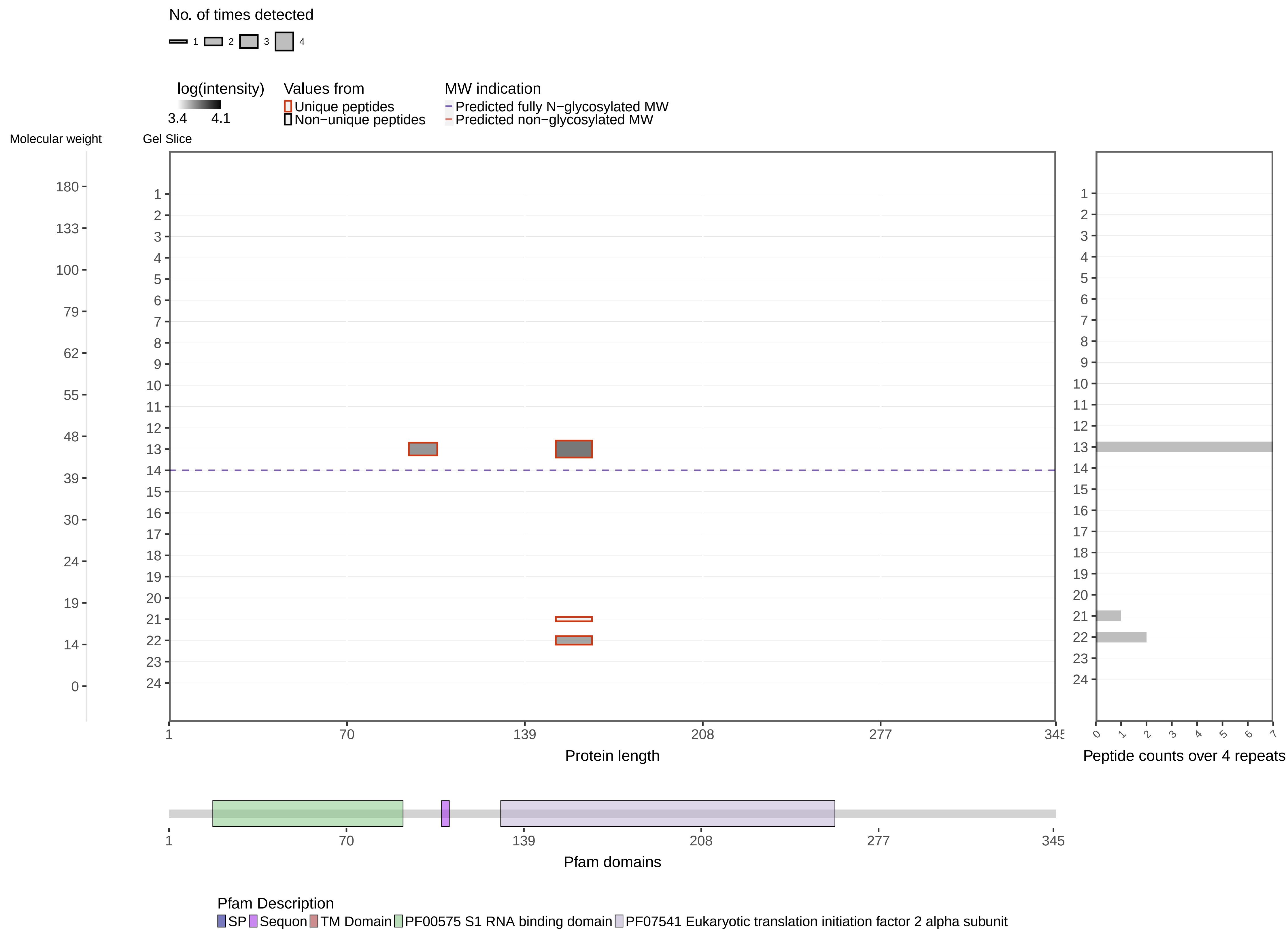


Protein sequence:

.NSLTREIKETGPDGQEVTKVVPALSEEVKDVLVKNIRRRMTPQPLKIRADIEMKCFQFDGVIHIKEAMRKAEAAGNQDCPVKIKLVAPPVYVVLNAQTLDKEQGIAVLSKAIAACTQEIERHKGKLTVKEAPRAVSEREDKLLAEQMAKLGRENEERSGDEDSEEEEDTGMGEIDLEKSGTGITD*

### NbL17g02910.1 PRA1_family_protein_B1−like [MW = 23.3 kDa]

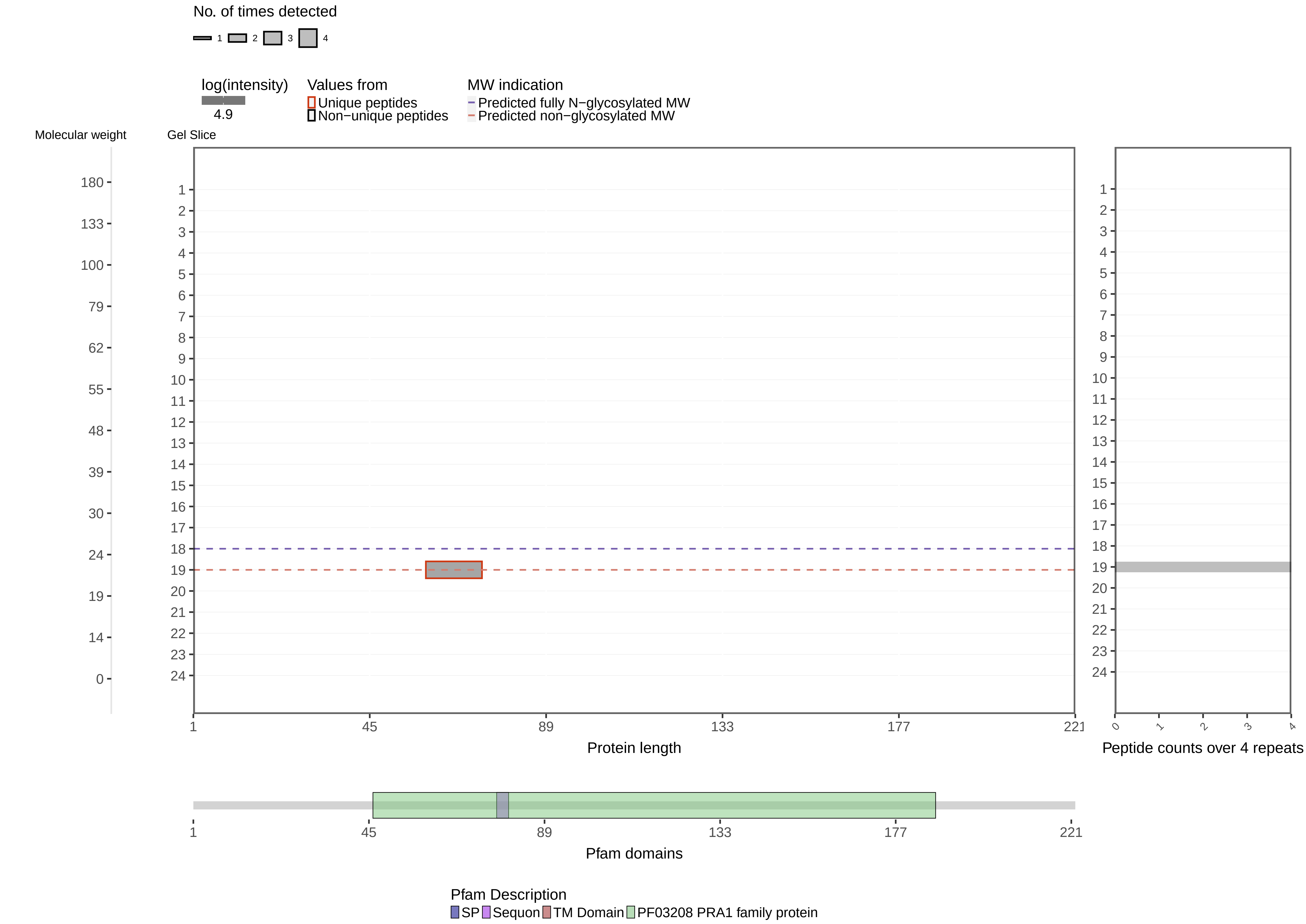


Protein sequence:

LLSHPFSLLVLLALLGGWFFLYLFRPSDQPVVILGRTFSDRETLGILVVSTIVVVFLTSVGSLLISALLVGLAIVCAHGAFRVPEDLFLDDQEPGNVGFLSFLGGAASSAAAAAAAAPAVAARV*

### NbL17g02890.1 uncharacterized_LOC104234041 [MW = 30.72 kDa]

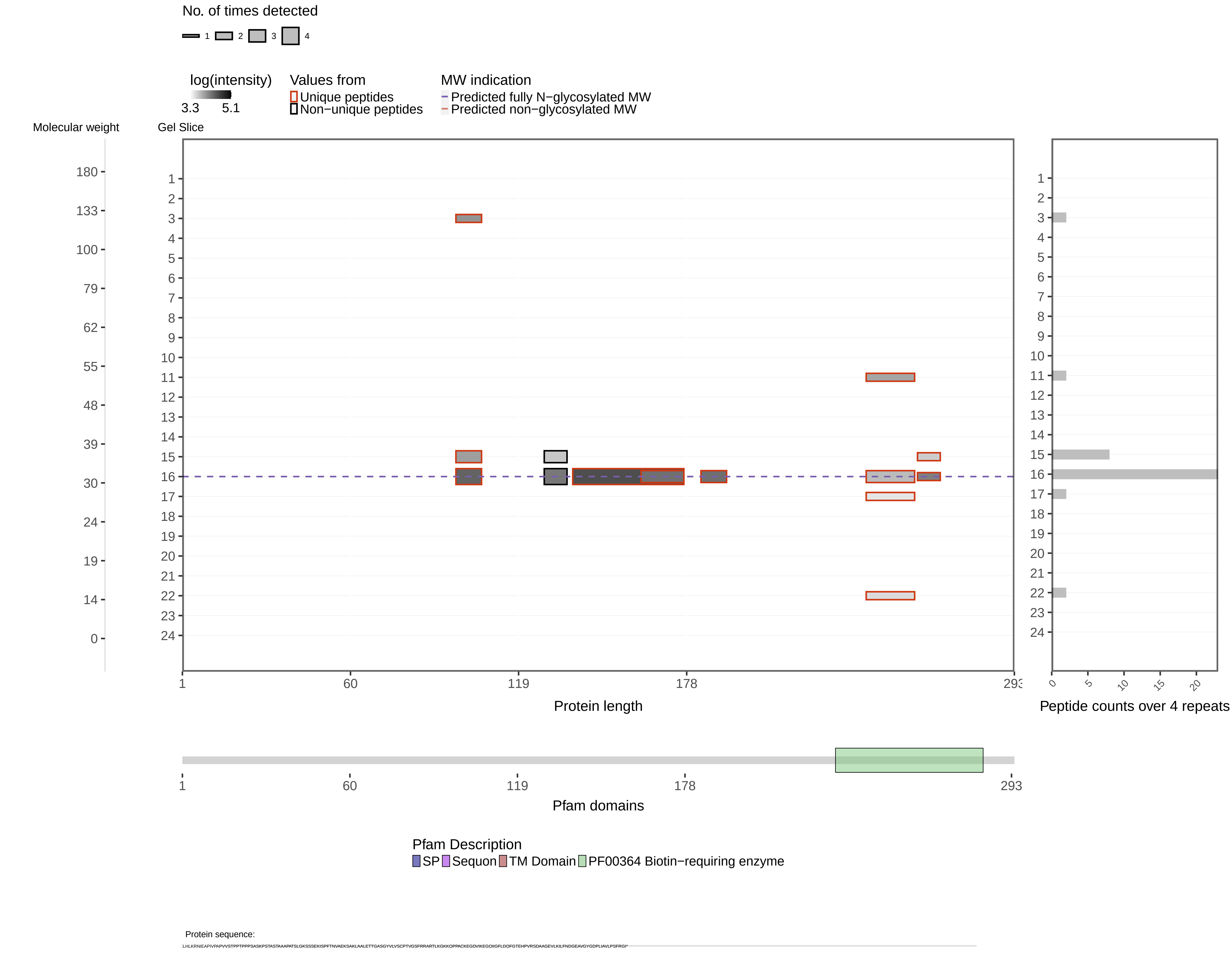

NbL17g02740.1 adenylosuccinate_synthetase_chloroplastic−like [MW = 55.02 kDa]

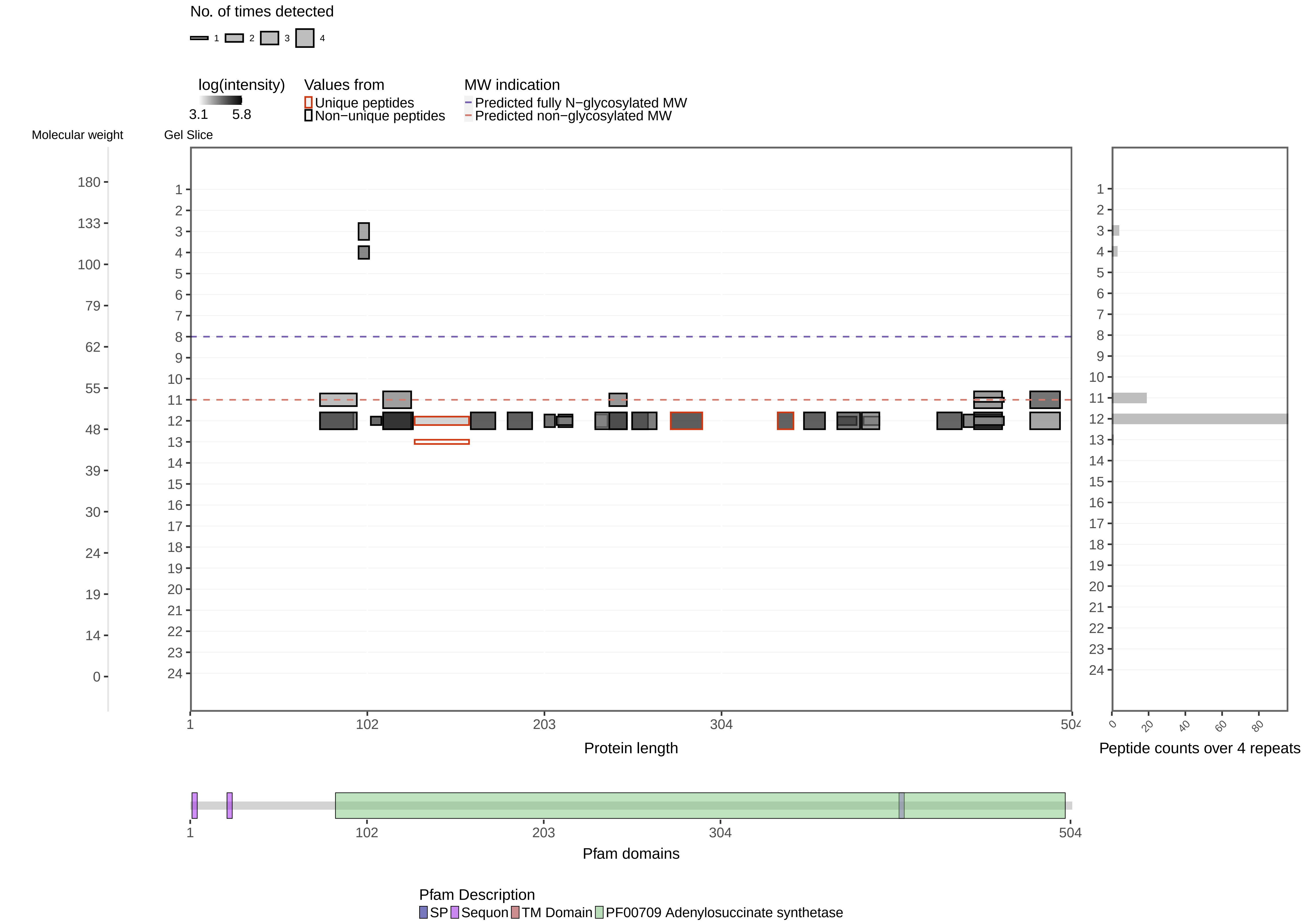


Protein sequence:
QKLDLLLSDAASRFKGFNYGPDMLREEVERYKKFAERLEPFITDTVHFMNDALSQKKKILVEGGQATMLDIDFGTYPFVTSSSPSAGGICTGLGIAPRVVGDLIGVVKAYTTRVGSGPFPTEILGKGGDLLRFAGQEFGTTTGRPRRCGWLDIVALRFCCQINGFASLNLTKLDVLSDLSEIELGVSYRQPDGTPVESFPSDLRLLDQIKVEYEVLPGWQTDISSIRKYSELPKAAREYVERIEELVGVPIHYIGVGPGRDALIYK*

### NbL17g02570.1 GTP−binding_protein_TypA/BipA_homolog [MW = 74.93 kDa]

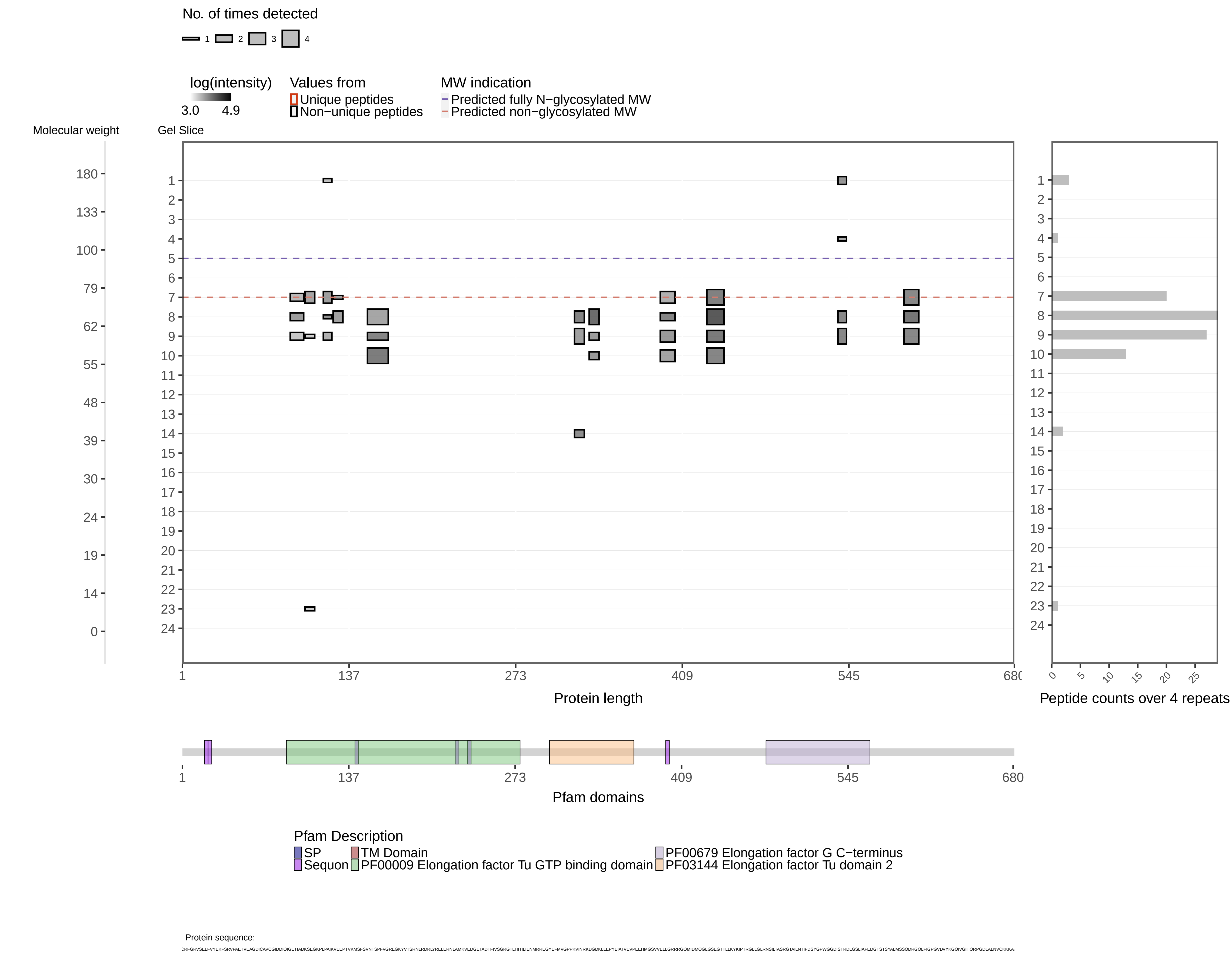

### NbL17g02560.1 aspartate_carbamoyltransferase_1_chloroplastic [MW = 45.42 kDa]

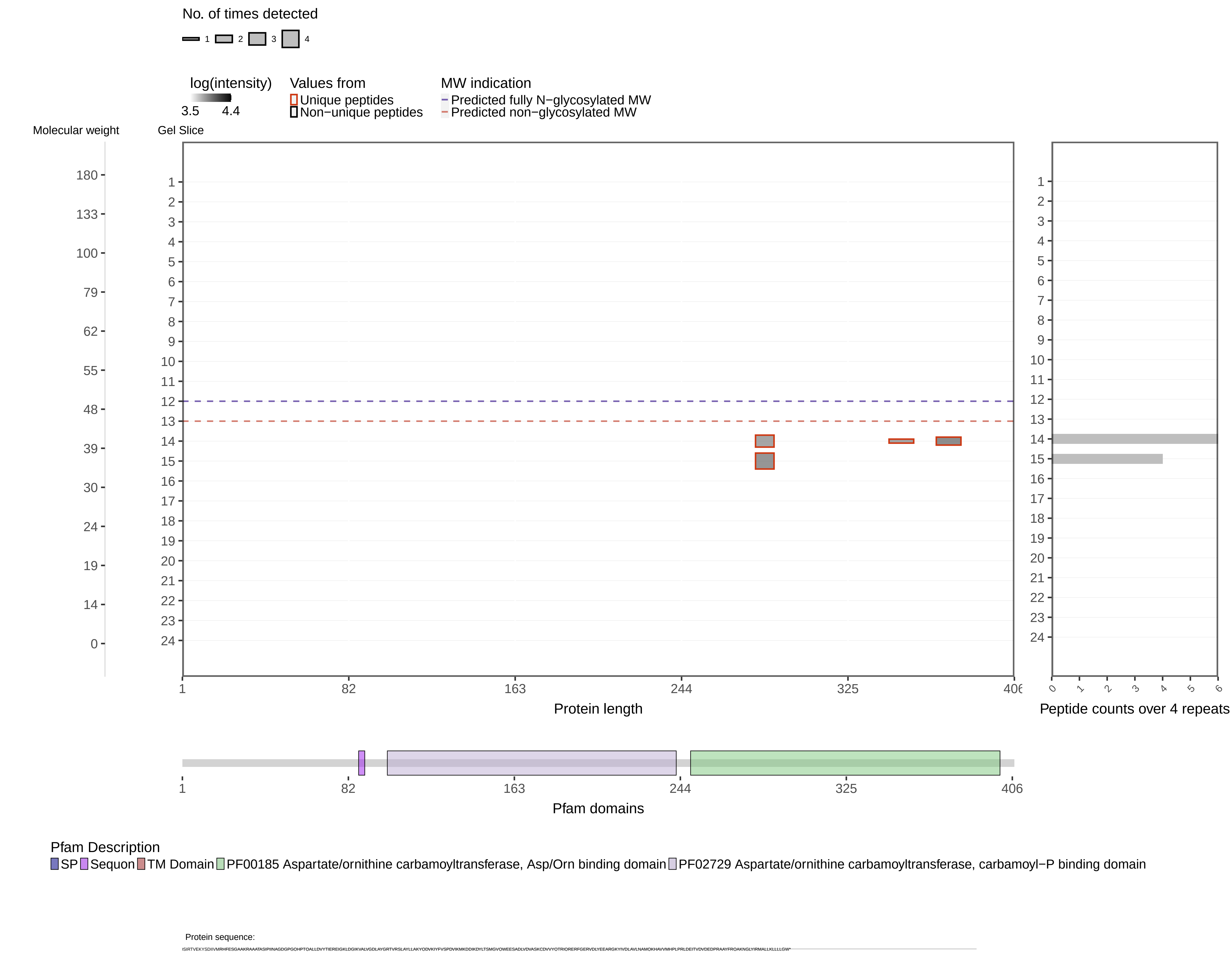

### NbL17g02410.1 stearoyl−[acyl−carrier−protein]_9−desaturase_chloroplastic [MW = 44.78 kDa]

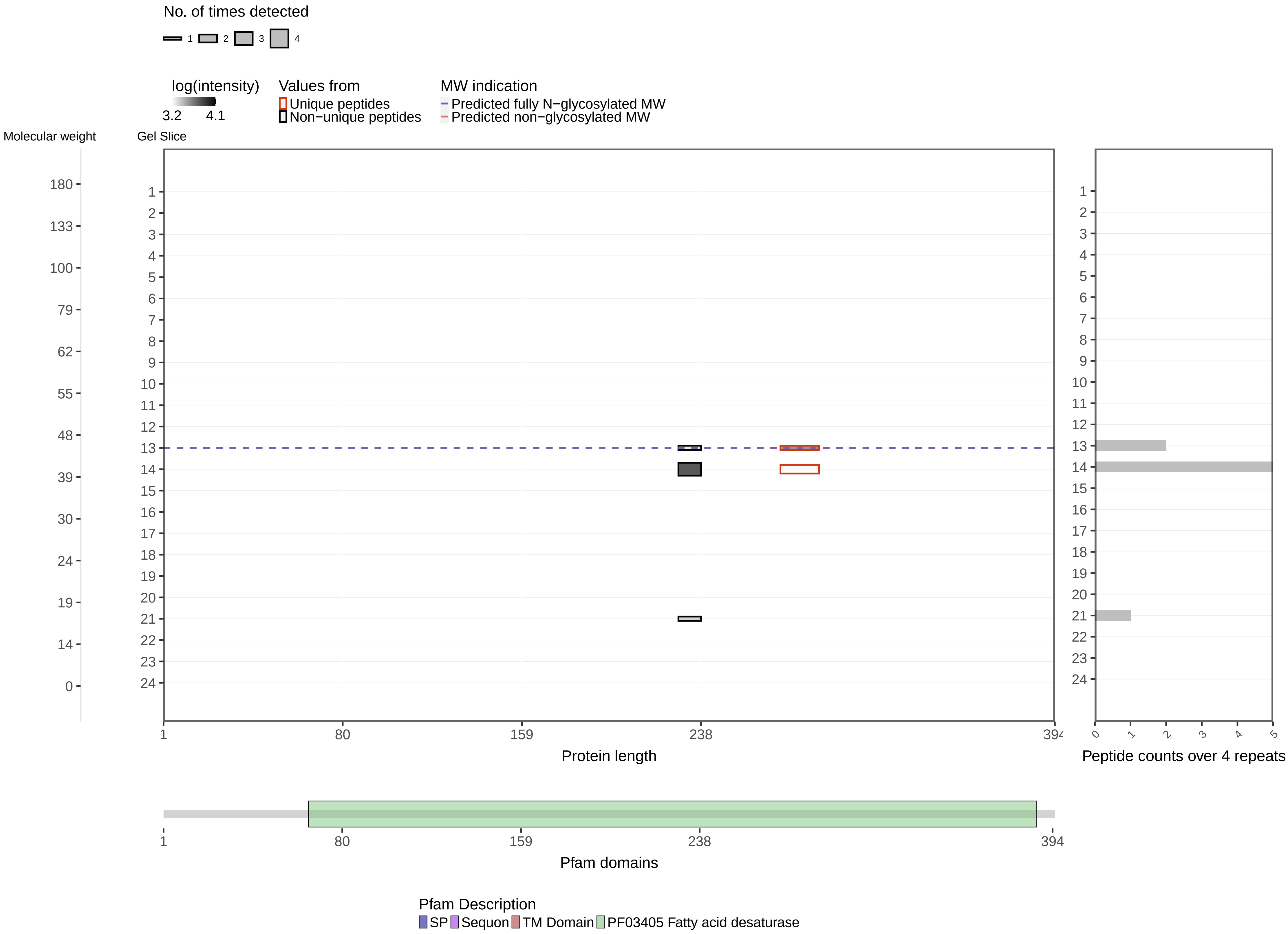


Protein sequence:

YLYLSGRVDMRQIEKTIQYLIGSGMDPRTENNPYLGFIYTSFQERATFVSHGNTARHAKEHGDLKLAQVCGIIAADEKRHETAYTKIVEKLFEVDPDGTVQAVADMMRKKISMPAHLMYDGRDDNLFEHFSAVAQRLGVYTAKDYADILEFLVGRWEMEKLTGLSGEGRIAQDYVCGLAPRIRKLEERAQARAKQKAPVPFSWVFGREIKL*

### NbL17g02110.1 reductase_1−like [MW = 42.03 kDa]

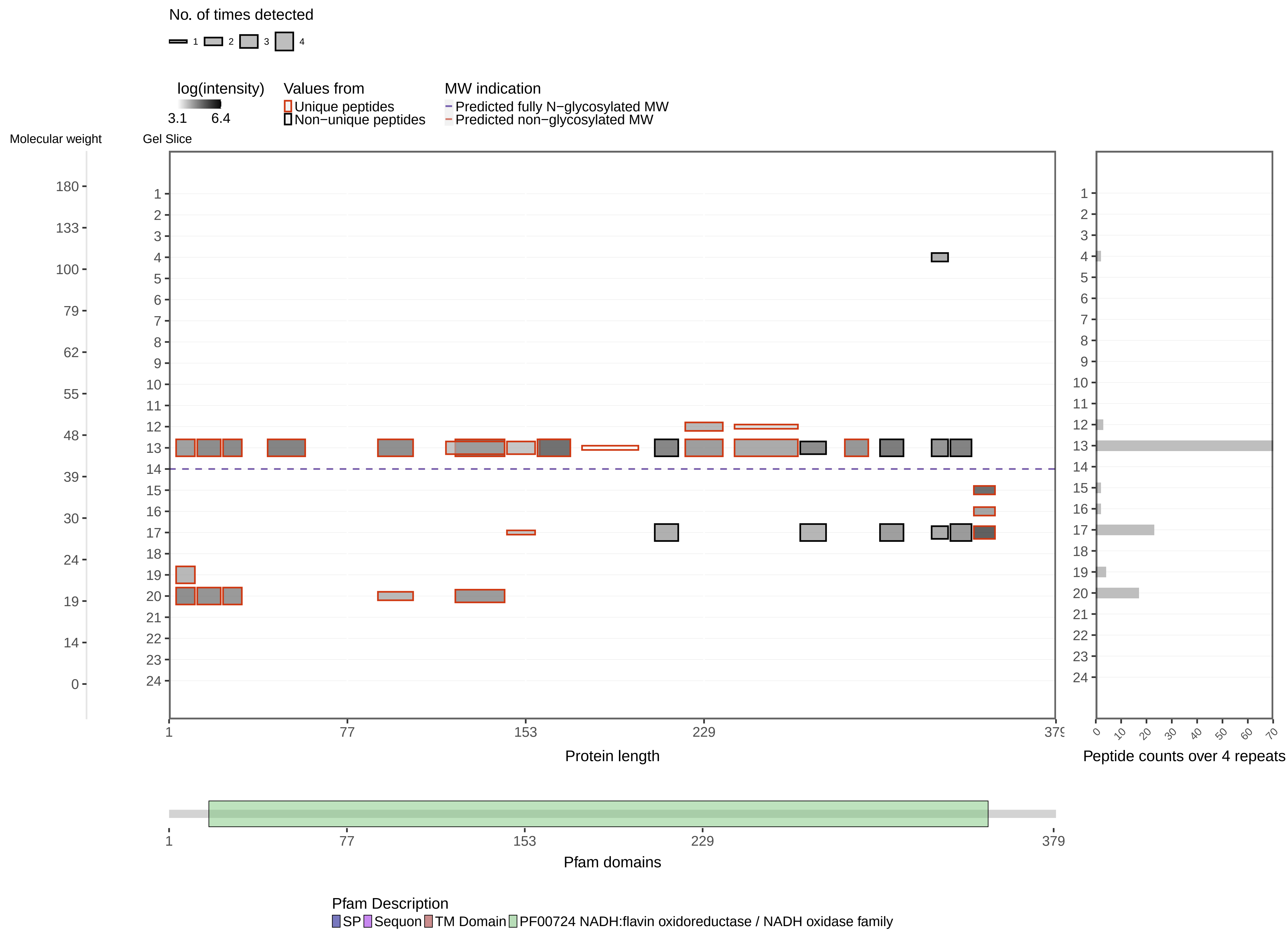


Protein sequence:

IAIEAGFDGVEIHGAHGYLIDQFMKDQINDRTDHYGGSLENRCRFAHEIVEAVVNEIGADRVGIRLSPFASYMESGDSNPSALGLYMAESLNKYGIAYCHMVEPRMKTVGEKVECSESLVPMRKAFKGTFMVAGGYDRGDGNKAVVEDRADLVAYGRLFLANPDLPRRFELDVPLNKYNRDTFYISDPVVGYSDYPFLETTA*

### NbL17g02080.1 phosphoglucan_phosphatase_LSF2_chloroplastic [MW = 34.59 kDa]

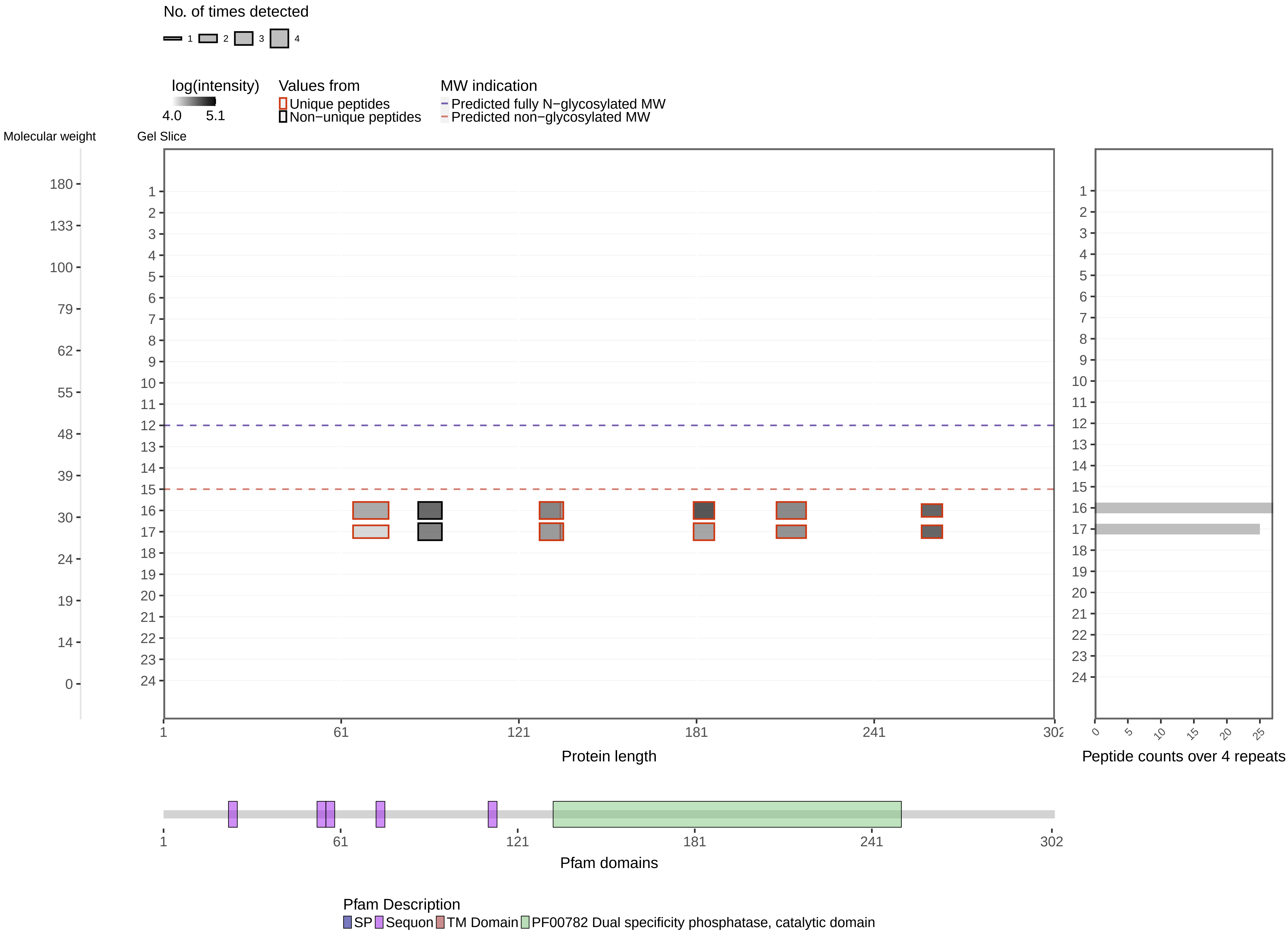


Protein sequence:

EENVAFILNLQQDKDIEFWGIDLQSIVRRCSELGIYHMRRPAKDFDPDSLRVVLPEAVSSLEWAISEGKGRVYVHCTAGLGRAPAVSIAYMFWFCGMDLNAAYDTLTSKRPCGPNKRSIRAATYDLAKNDHWKEPFENLPDYAFADVADWERKLIQDRVRGLRNT*

### NbL17g01880.1 protein_LST8_homolog [MW = 35.41 kDa]

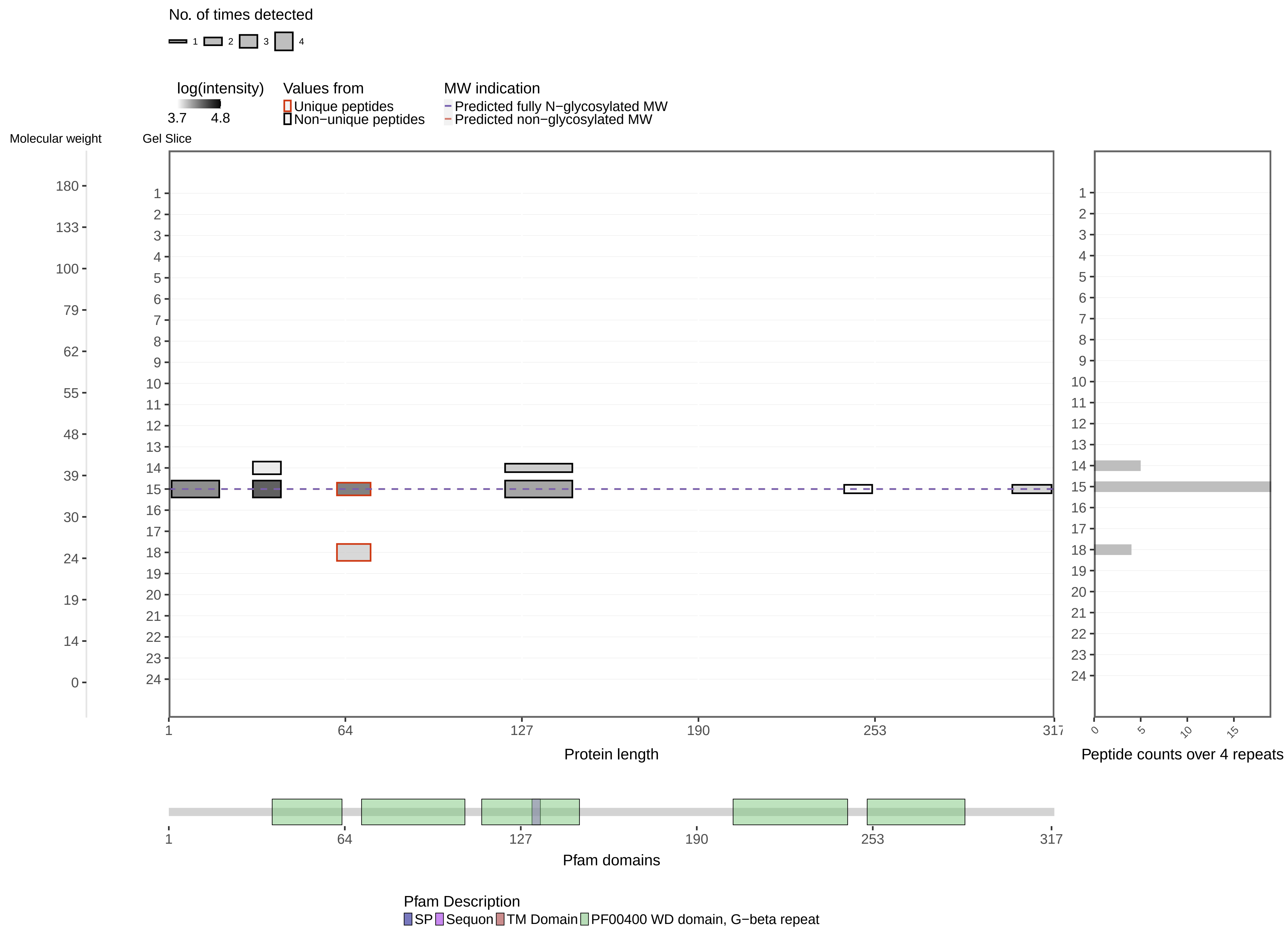


Protein sequence:

RVWDLRANSCSCELVPEVDTAVRSLTIMWDGSLIVAANNRGTCYVWRLLRGTETMTNFEPLHKLQAHDGYILKCLLSPELCEPHRYLATASSDHTVKIWNVDGFTLEKTLIGHERWVWDCVFSVDGAFLITASSDTTARLWSTSTGKDIKVYQGHHKATVCCALHDGPEPSS*

NbL17g01640.1 probable_polygalacturonase_At3g15720 [MW − SP = 50.29 kDa]

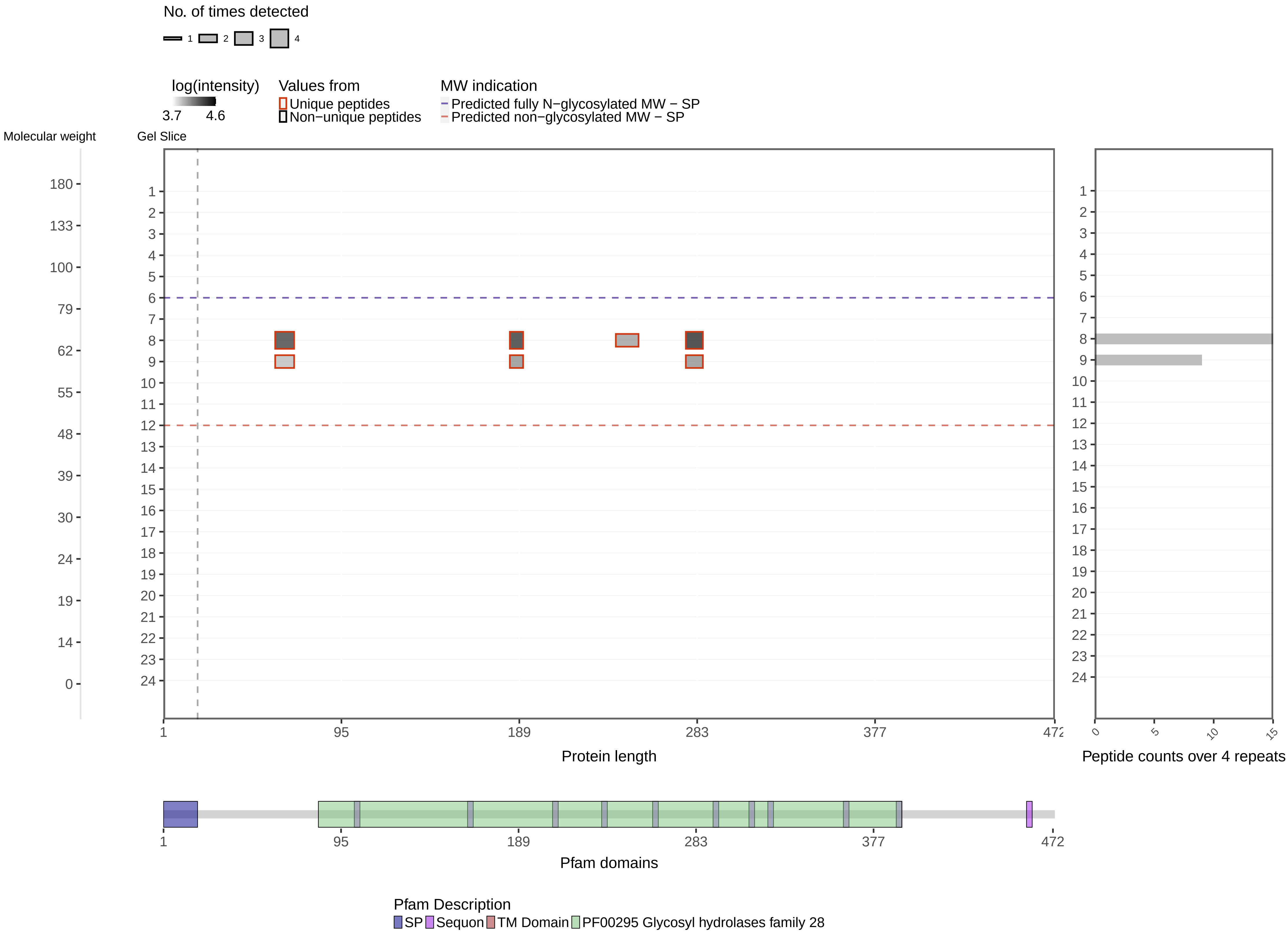


Protein sequence:

DGIDIEDSNNTVITRCNINTGDDAICPKSSTGPVHNLTATNCWIRTKSSAIKLGSASFYTFKNFLFENITIFESHRGLALQIRDGGNVSDITFSNINTSTRHYHSSWWGRAEPIYVTTCPRDATSKAGSISNLLFVNITSTSENGIFLSGSGGGVLSNLKFINMNLTYKRWTKYPDGWVDYRPGCQELVKHQTAGFMMEHIDGLVFENVVMKWSNDESMRWNNPLDFRPSTVNNISMLNFYSRSYQQ*

### NbL17g01460.1 elongation_factor_1−gamma_2−like [MW = 47.67 kDa]

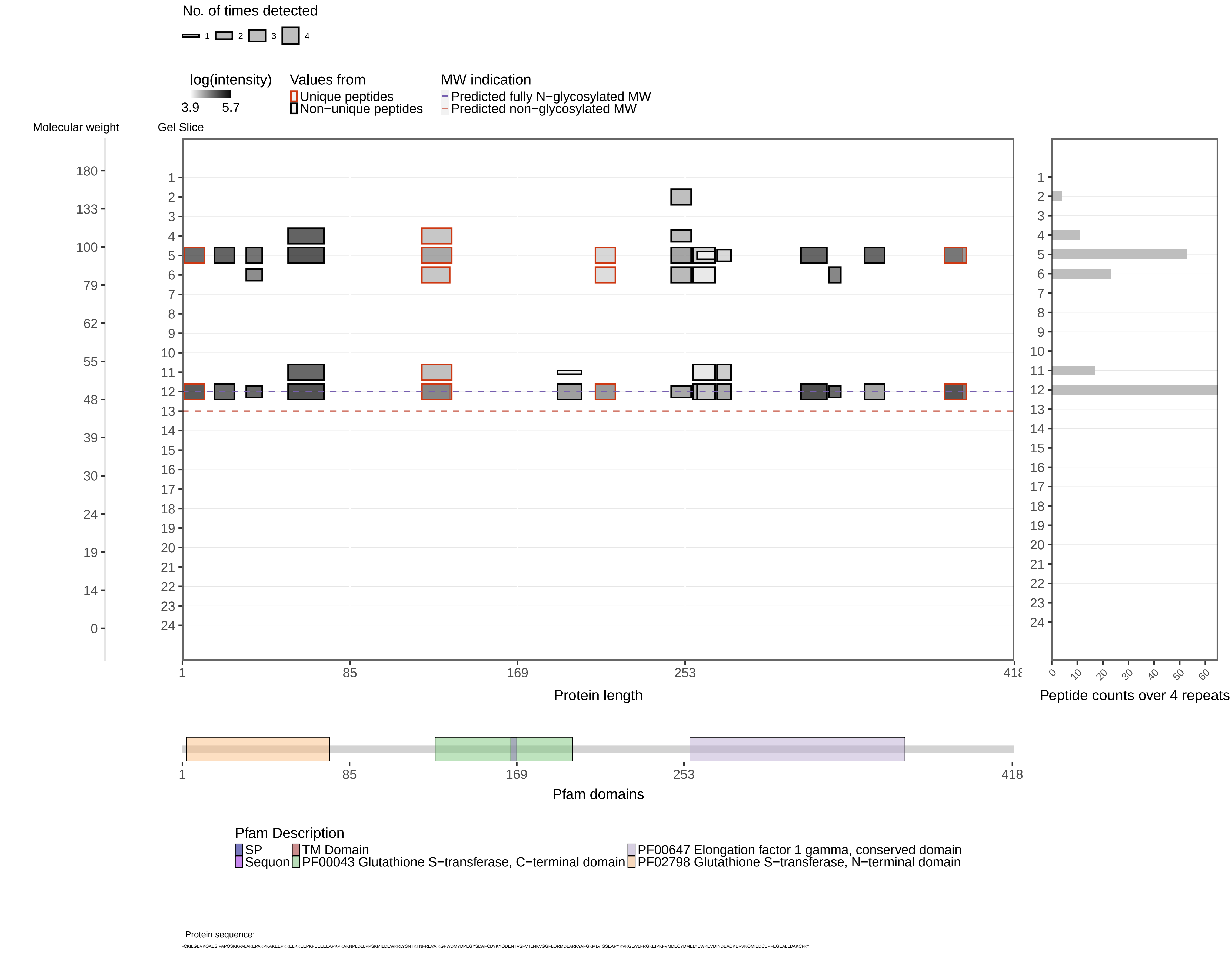

### NbL17g01390.1 uncharacterized_LOC107819606 [MW = 19.86 kDa]

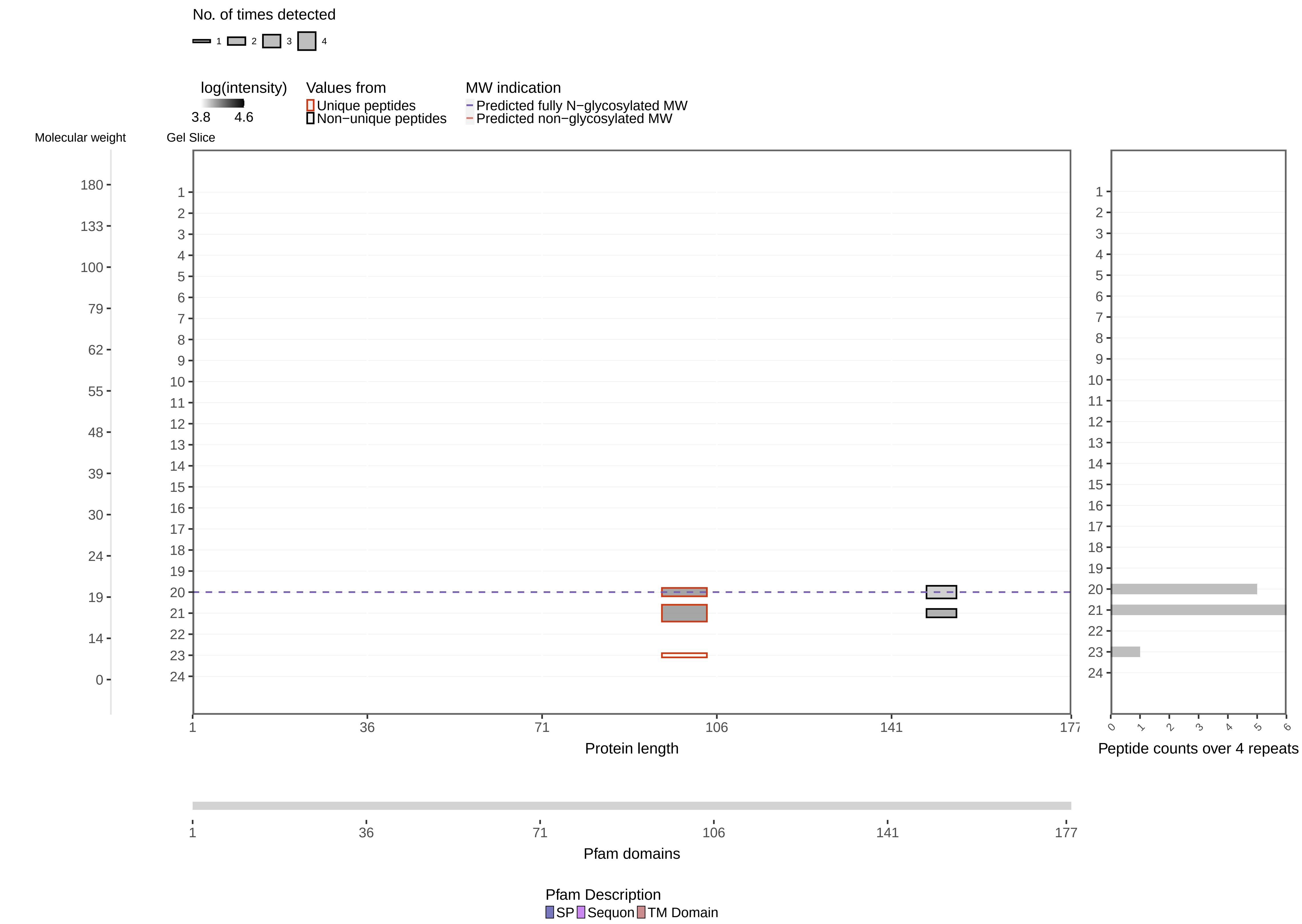


Protein sequence:

KDAELIDKAREATQSGERMSSEQYGALRRKIGGTYKDFFKSYVEVKGEYVEEGWIDKTCKVCKKDTRGEARQVDNFGRYAHVACLENSKSGNFFTNIFSR*

NbL17g00940.1 patellin−3−like [MW = 69.35 kDa]

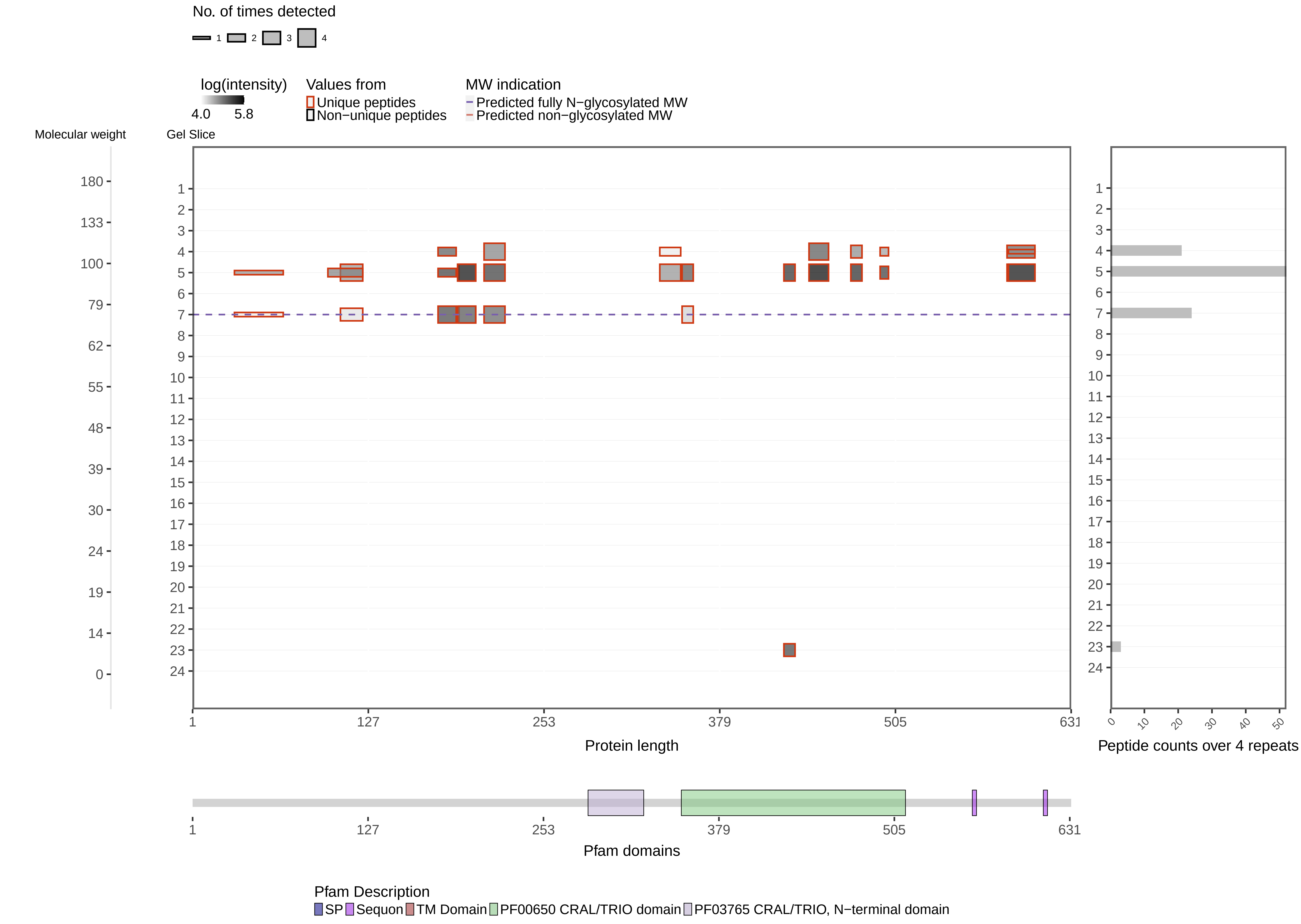


Protein sequence:

VILLKFLRARDFKVKEAFAMLKSVVAWRKEFKIDELLVEDLSGLGLEKVVYNHGVDKEGHPVCYNAFGAFQDNELYQNTFADKEKMNKFLRWRIQFMEKSIRNLDFSPDGICTFVQVIDLKNSPGLYLFKKELRQATNRAVQLLQDNYPEFVAKQVFINVPWWYPAYYRMTNALFTTRTKSKFVFAGPSRSAETLLKYIVPEQVPAQYGGLSREGEQEFTTAEAATEDTIKPTSKHTIEFPVTEKTSLVWEARVIGGNVSYGAEFVPSAEGGYTIIVEKSRKVAAADETVITNTYT

### NbL17g00930.1 patellin−3 [MW = 59.8 kDa]

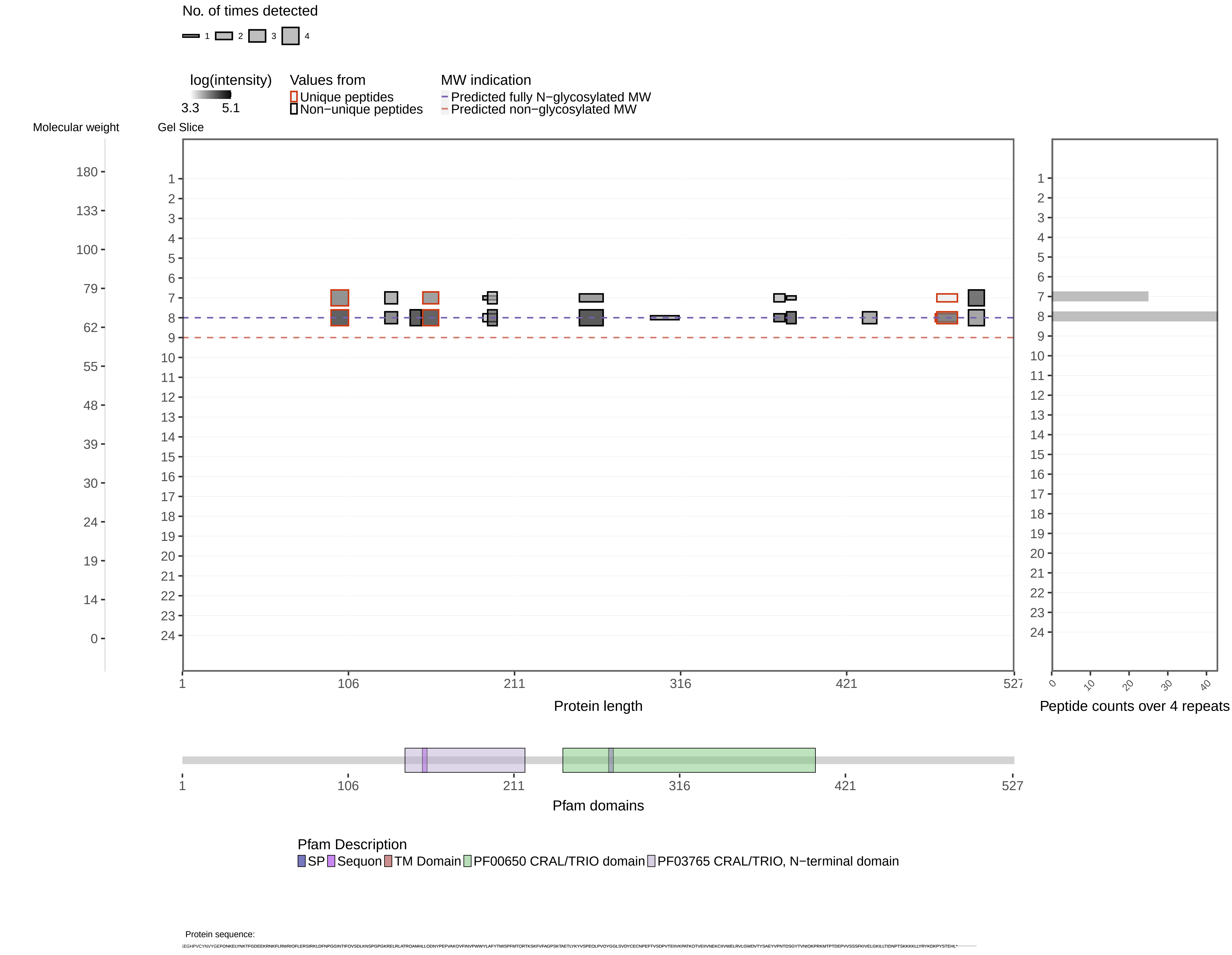

NbL17g00900.1 carboxymethylenebutenolidase_homolog [MW = 33.5 kDa]

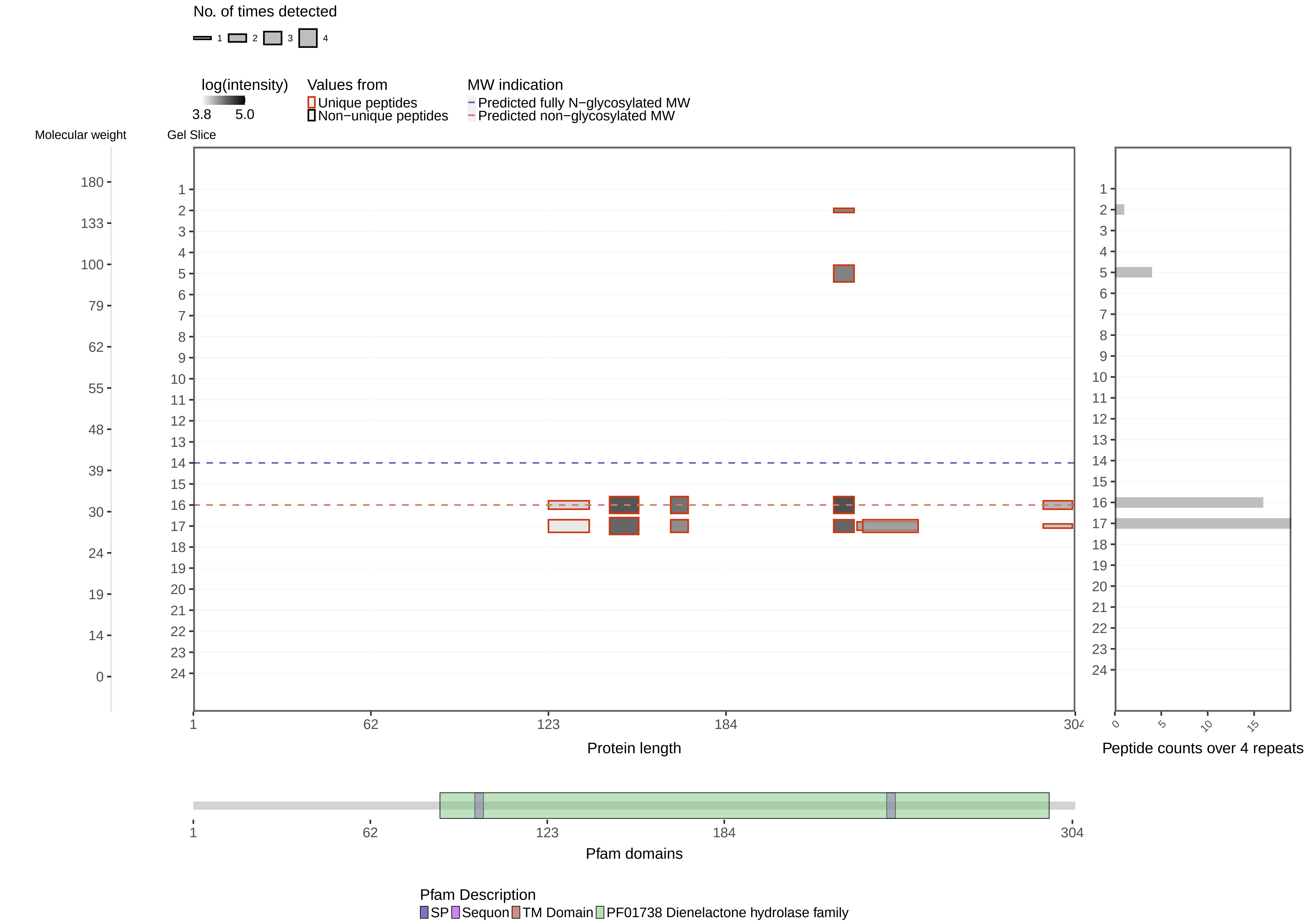


Protein sequence:

WGKDKPQASFEQWRGSLDKQRVVRDIFTSTKWMVDEFVAAGISKKLGVIGFCFGGGLLIDILAQDQGSQFGVGISFYGTRIDVSVASKIKVPLLLIAGDNDSLCPVNVLKEVDKNANGCKVVVFEGRGHGFAHRPQSLDDDKDAEEAFLMMRNWLHDGLVSEY*

### NbL17g00510.1 uncharacterized_LOC104227417 [MW − SP = 27.62 kDa]

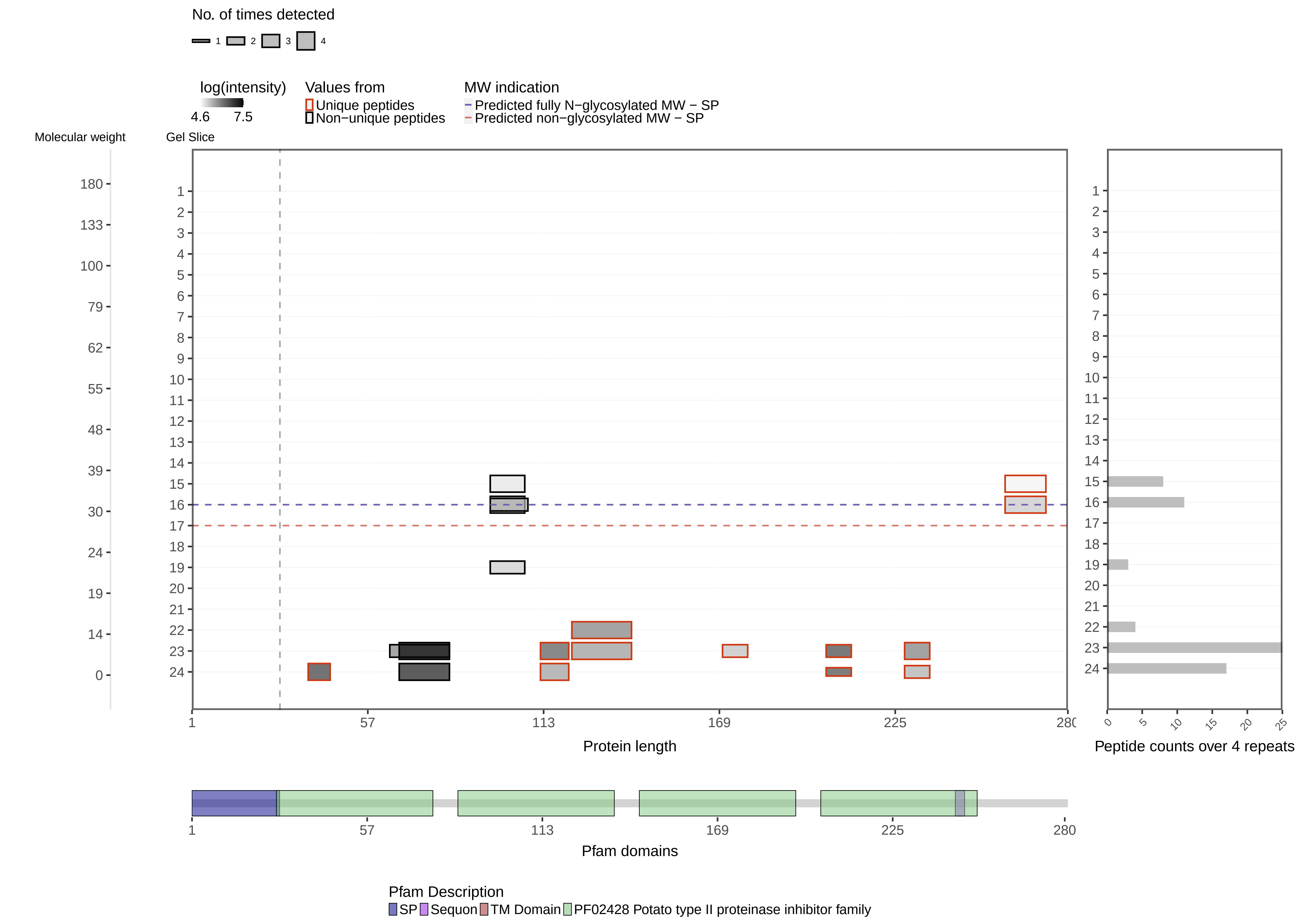


Protein sequence:

ADGTFVCEGESDPRNPKACPRNCDPRIDYRVCPRSEEKKNDRICTNCCSGKKGCKYFSDDGTFVCEGESDPRNPKGCTFECDPRIAYGICPLSEEKKNNRLCTNCCAGKKGCNYFSSNGTFICEGESEYGSKVVEYVGEVEDDLOKSKVAVS*

NbL17g00400.1 adenylate_kinase_chloroplastic [MW = 32.82 kDa]

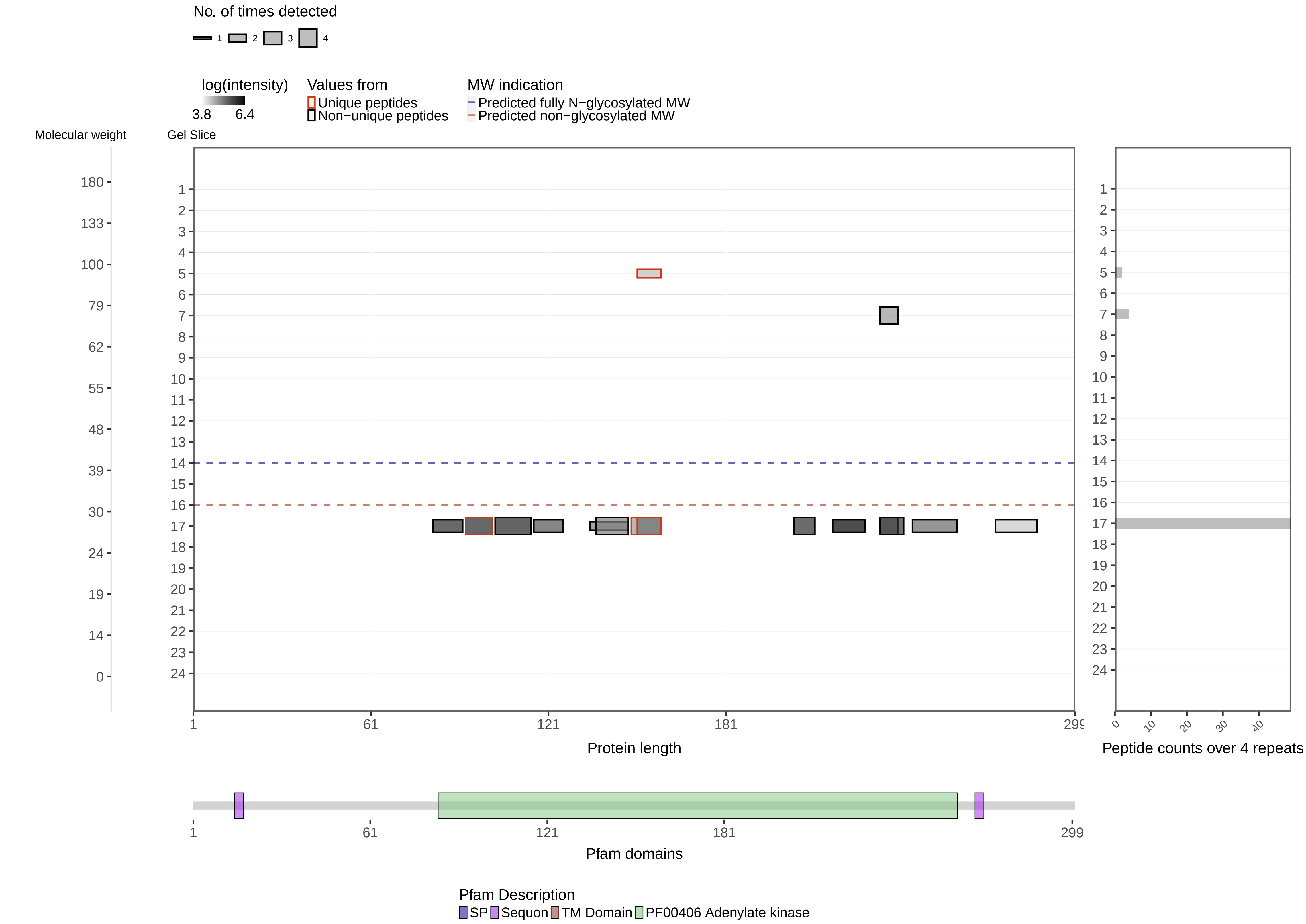


Protein sequence:

KLVPDEIVVTMVKERLNGPDSQEKGWLLDGYPRSSSQAKALKEFGFQPDLFILLEVPEEILVERVVGRRLDPVTGKIYHLKYSPPETEEIAARLTQRFDDTEEKVKLRLHTHHQNVEAVLSLYKDITVKVNGSASKQDVFAQIDDALTQLREQKQGKLGTVAA*

### NbL17g00380.1 protein_TIC_40_chloroplastic [MW = 49.49 kDa]

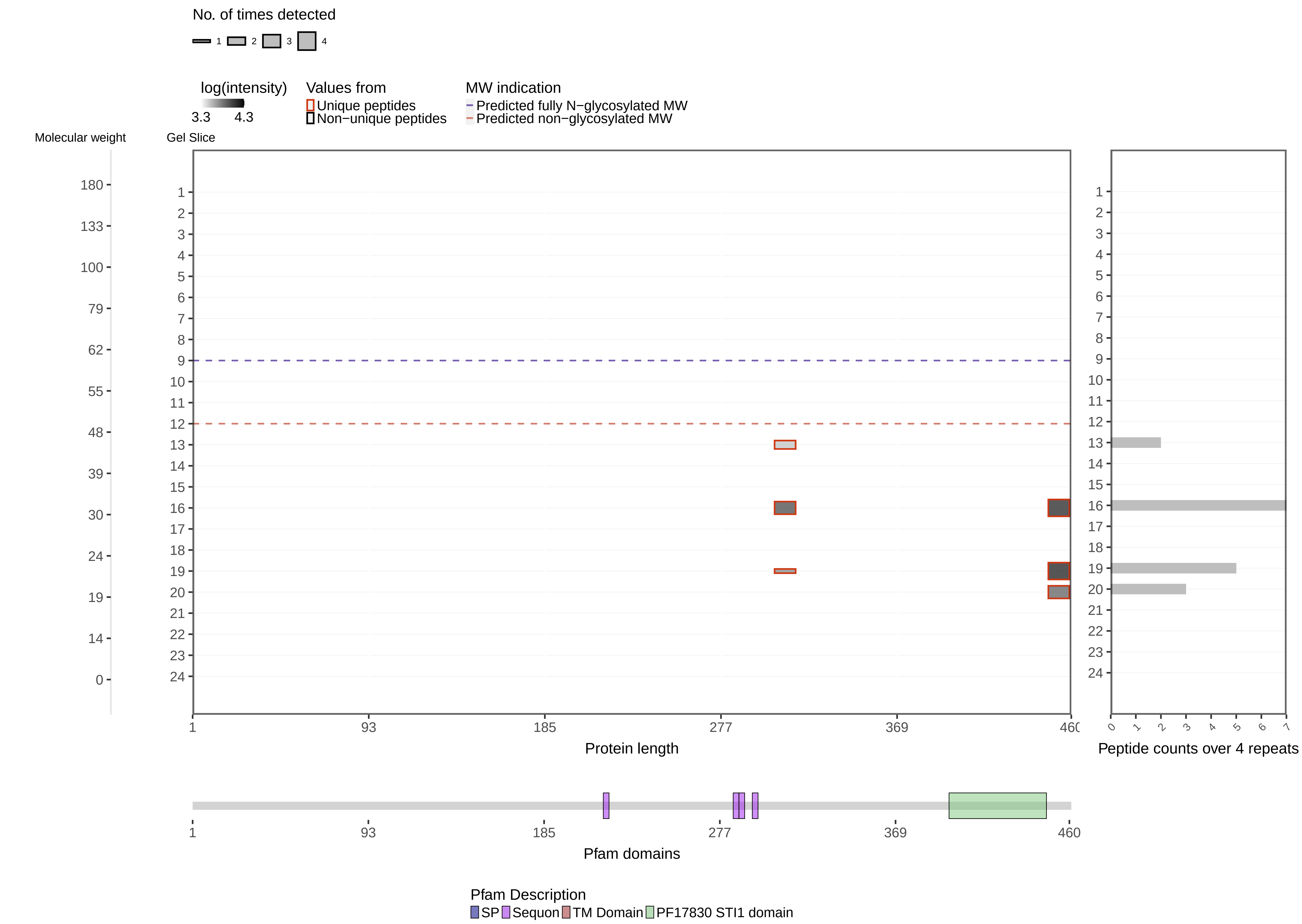


Protein sequence:

EAERVPKKNAFVDISPDETFQKGAFEDYKESTETASVTVDQVTQNGAASKSGFGASTSDSTGFGANTSNTTGFGANTSGSTSSTEKSNPLMSVEALEKMMEDPTVQKMVYPYLPEEMRNPTTFKWMLQNPQYRQQLQDMLNNMGGNPEWDNRMVDSLKNFDLSSPEIKQQFDQIGLTPEEVISKIMANPDVAMAFQNPRVQAAIMDCSQNPLSIAKYQNDKEVMDVFNKISELFPGVSGSP*

### NbL17g00350.1 alkaline/neutral_invertase_CINV2 [MW = 62.69 kDa]

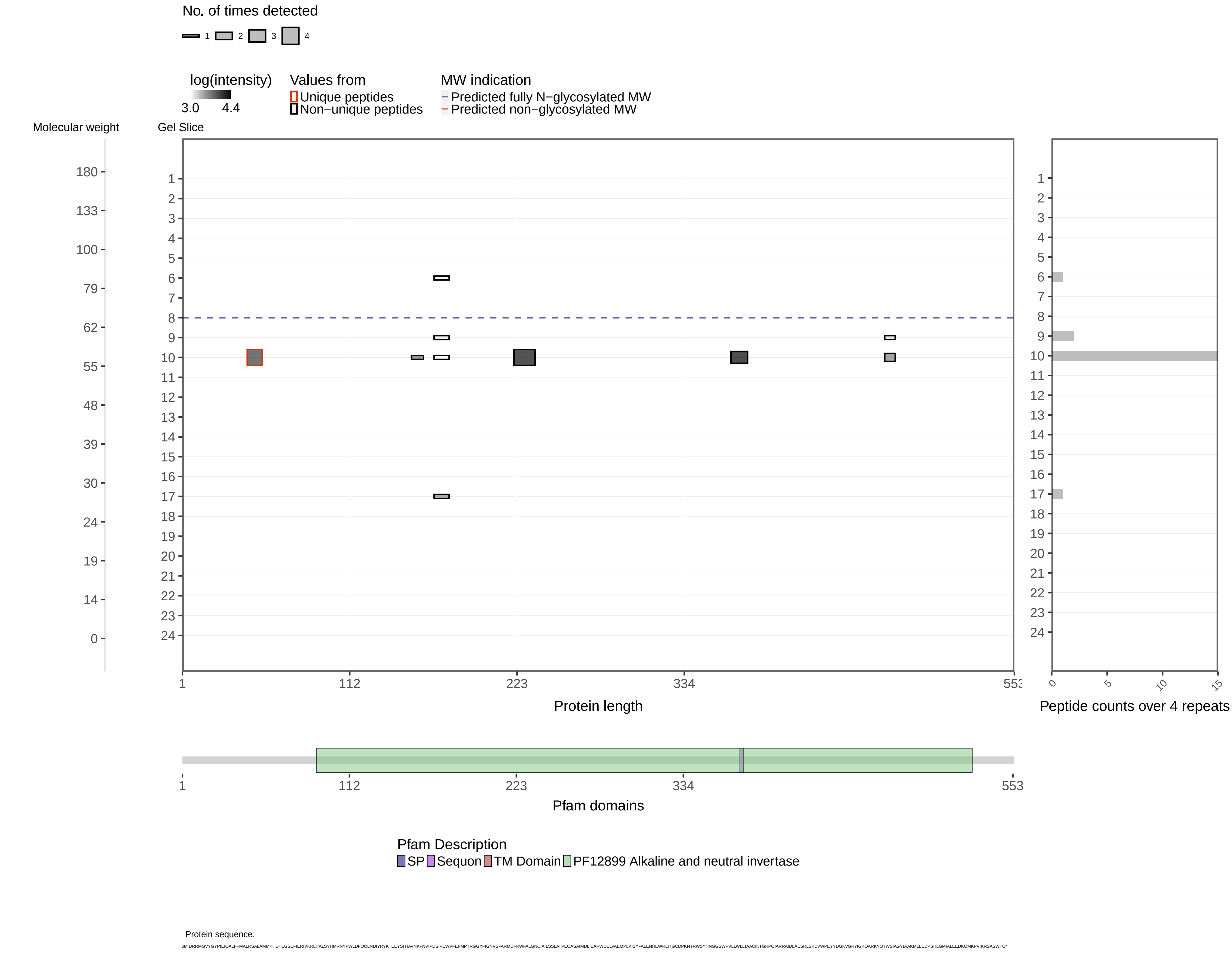

### NbL17g00240.1 UPF0160_protein [MW = 42.3 kDa]

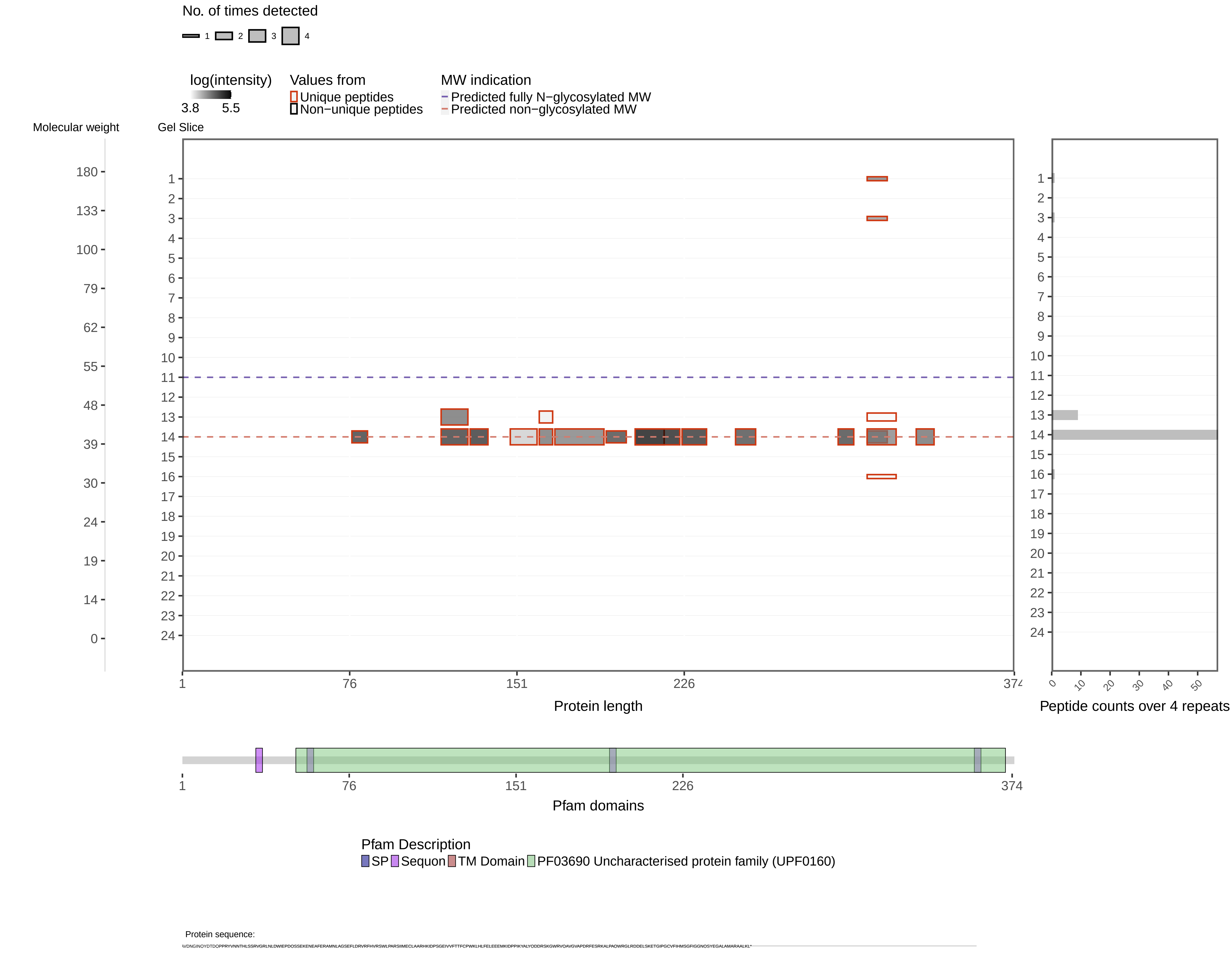

### NbL17g00170.1 hydroxypyruvate_reductase−like [MW = 43.93 kDa]

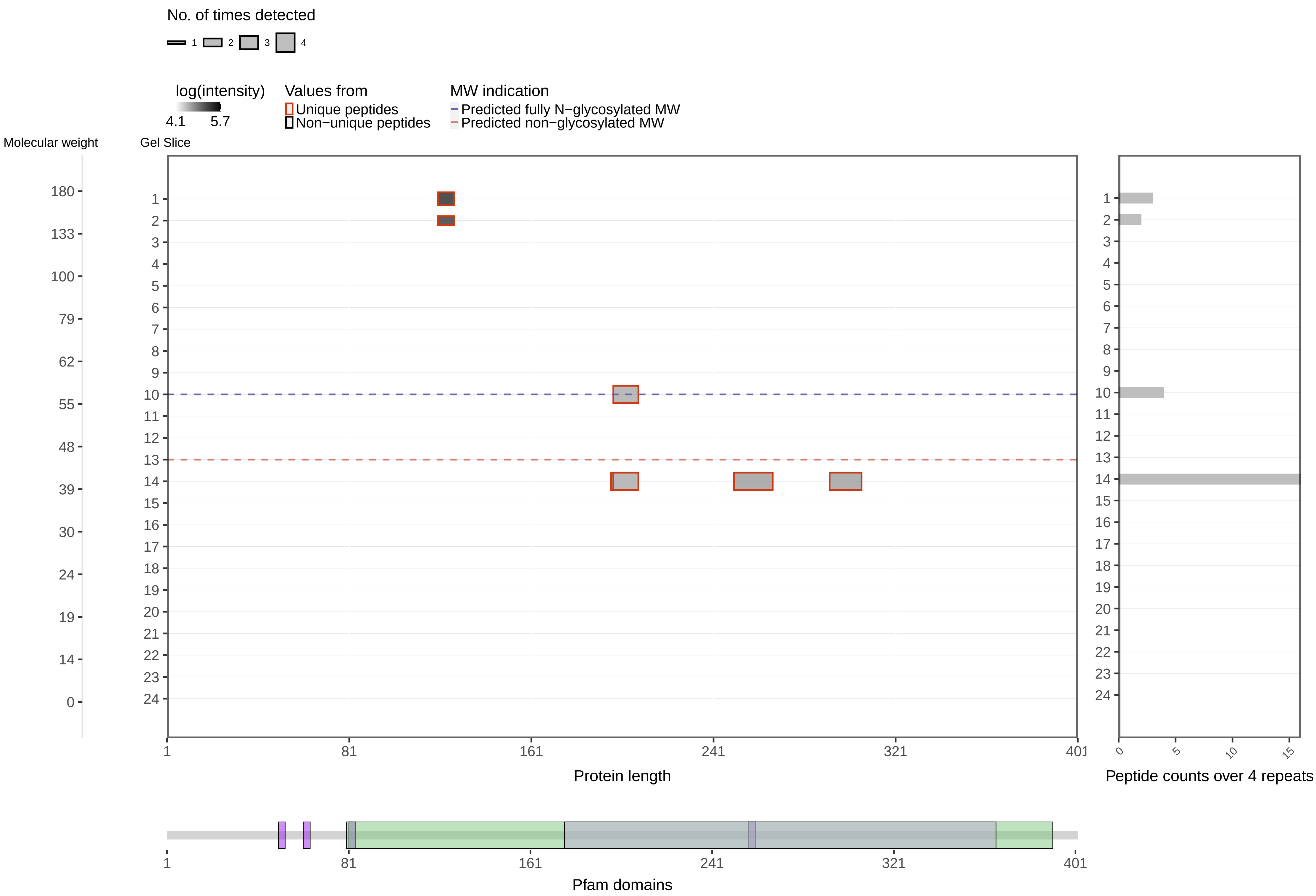


Pfam Description

SP Sequon TM Domain PF00389 D−isomer specific 2−hydroxyacid dehydrogenase, catalytic domain PF02826 D−isomer specific 2−hydroxyacid dehydrogenase, NAD binding domain

Protein sequence:

ƆHQLKISVEQRKLGEPIGDNLEGKTVFILGFGNIGIHLAKRLRPFDVKILATKRSWSRQAQDSSKSEAPYVENGSYDDLVDERGNHDDILKFASKADIVICCLAMNKETAGIVNNDFISAMKKGAILINIARGGLLDYDAVFHHLKSGHLGGLGIDVAWAEPFDPDDAILKFPEVIITPHVAGVTEKSYRYMAKVVGDVALQLHAGEPFTGIEIVN*

NbL17g00160.1 stromal_70_kDa_heat_shock−related_protein_chloroplastic−like [MW = 74.09 kDa]

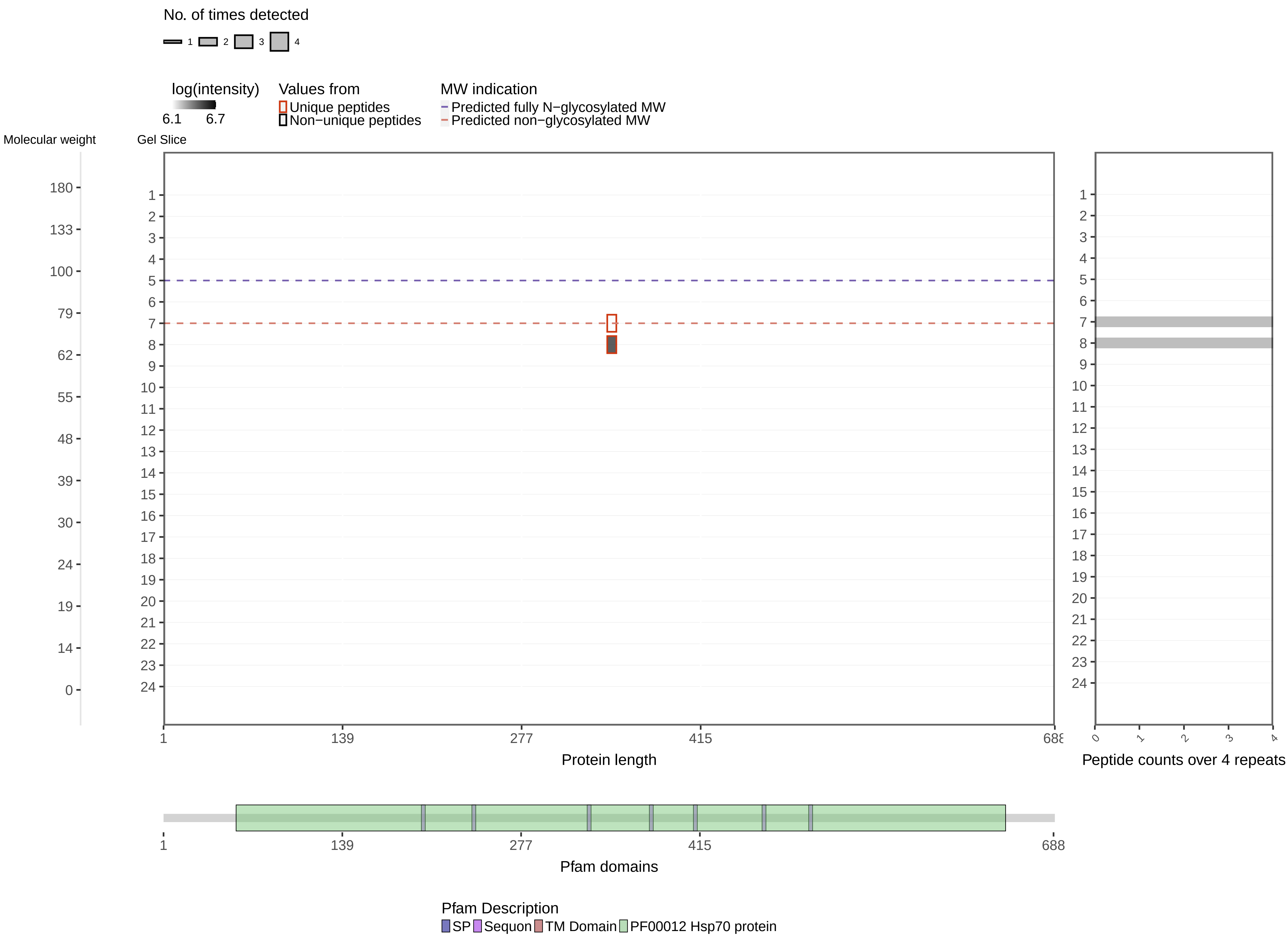


Protein sequence:

LPFITATADGPKHIETTLTRAKFEELCTDLLDRLKTPVQTALKDANLSFKDIDEVVLVGGSTRIPAVQNLVRKMTGKEPNVSVNPDEVVALGASVQAGVLAGDVSDIVLLDVTPLSLGLETLGGVMTKIIPRNTTLPTSKSEVFSTAADGQTSVEINVLQGEREFVKDNKSLGRFRLDGIPPAPRGVPQIEVKFDIDANGILSVTATDKGTGKKQDITITGASTLPKDEVDRMVQEAEKFAKEDKEKRDSIDTKNQAESVVYQTEKQLKELGDKVPADVKDKVESKLRELKDAISGGST

NbL16g27900.1 uncharacterized_LOC109206410 [MW − SP = 7.53 kDa]

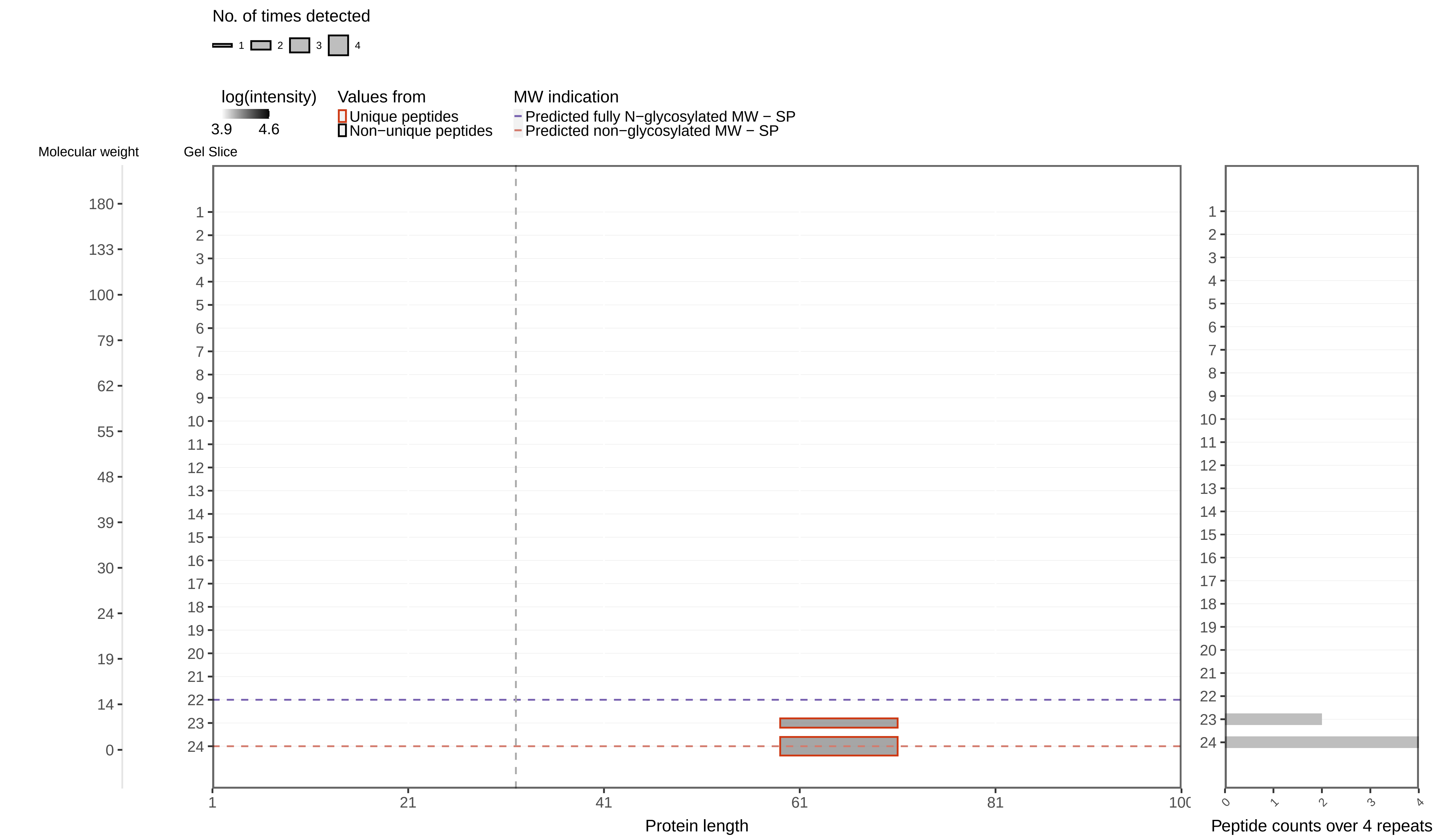


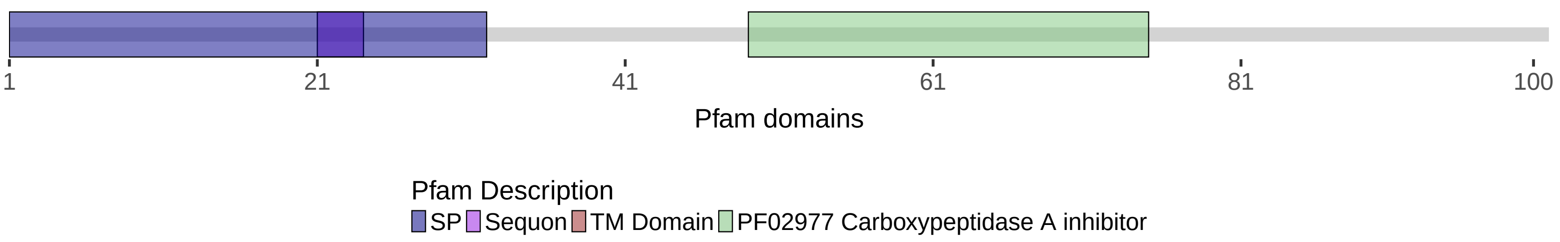


Protein sequence:

QEIAMVKTKLPPLTDFITCGKACSSDSDCSDGWRCTYCIYQNEVVIVSPVPIYRAAGYYCES*

### NbL16g27730.1 NADH_dehydrogenase_[ubiquinone]_flavoprotein_2_mitochondrial−like [MW = 28.59 kDa]

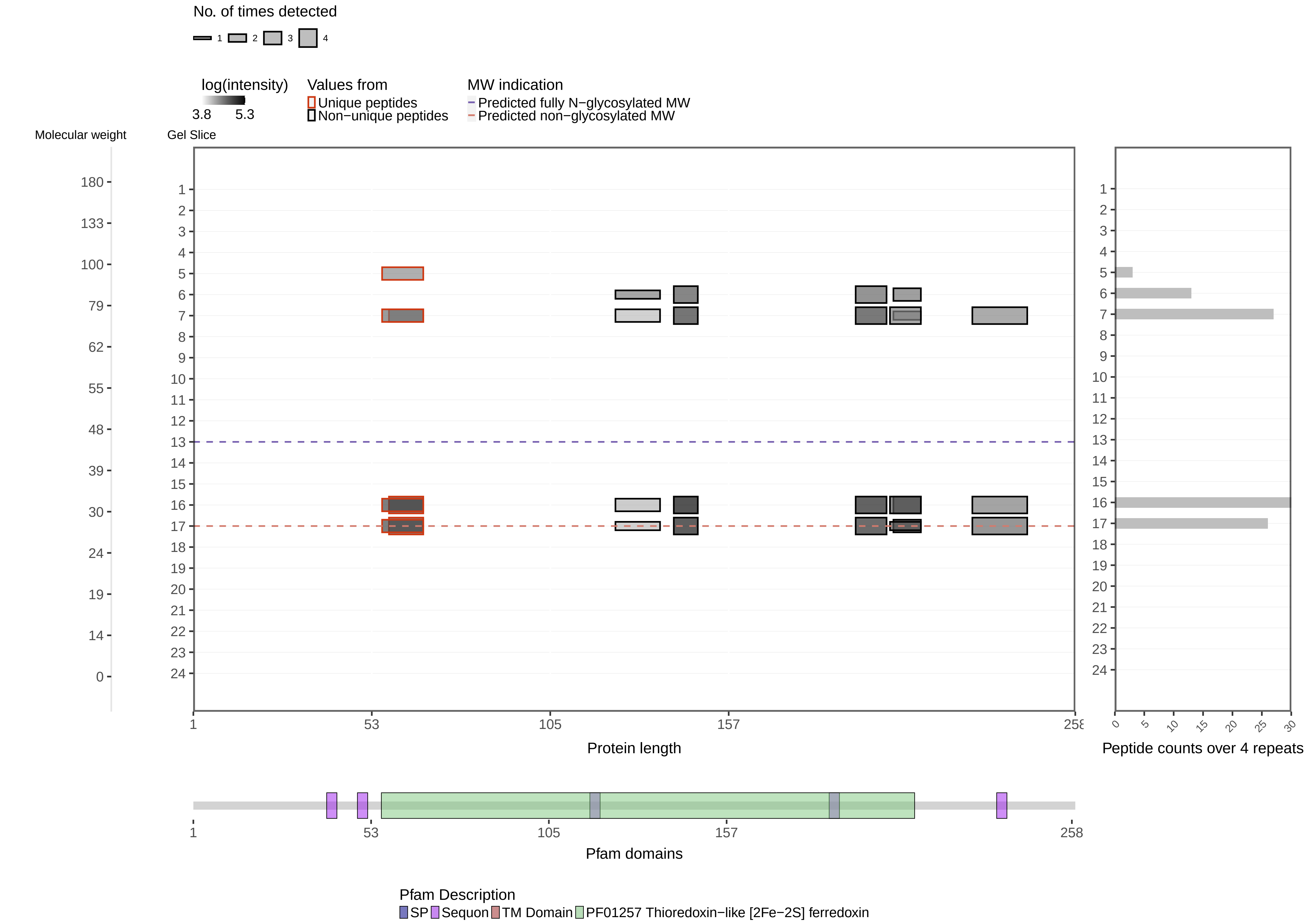


Protein sequence:

RTKVGKYHLLVCGTTPCMIRGSRGIEEAILKHLGVKRNEVTKDGLFSVGEMECMGCCVNAPMMTVADYSNGSEGYKYNYYEDLTPKRAVEIVEAFRKGEKPQRGTQNPGRINCGPEGGNTTLLGEPKAPPCRDLDACSAE*

### NbL16g27710.1 protein_OSB2_chloroplastic−like [MW = 35.24 kDa]

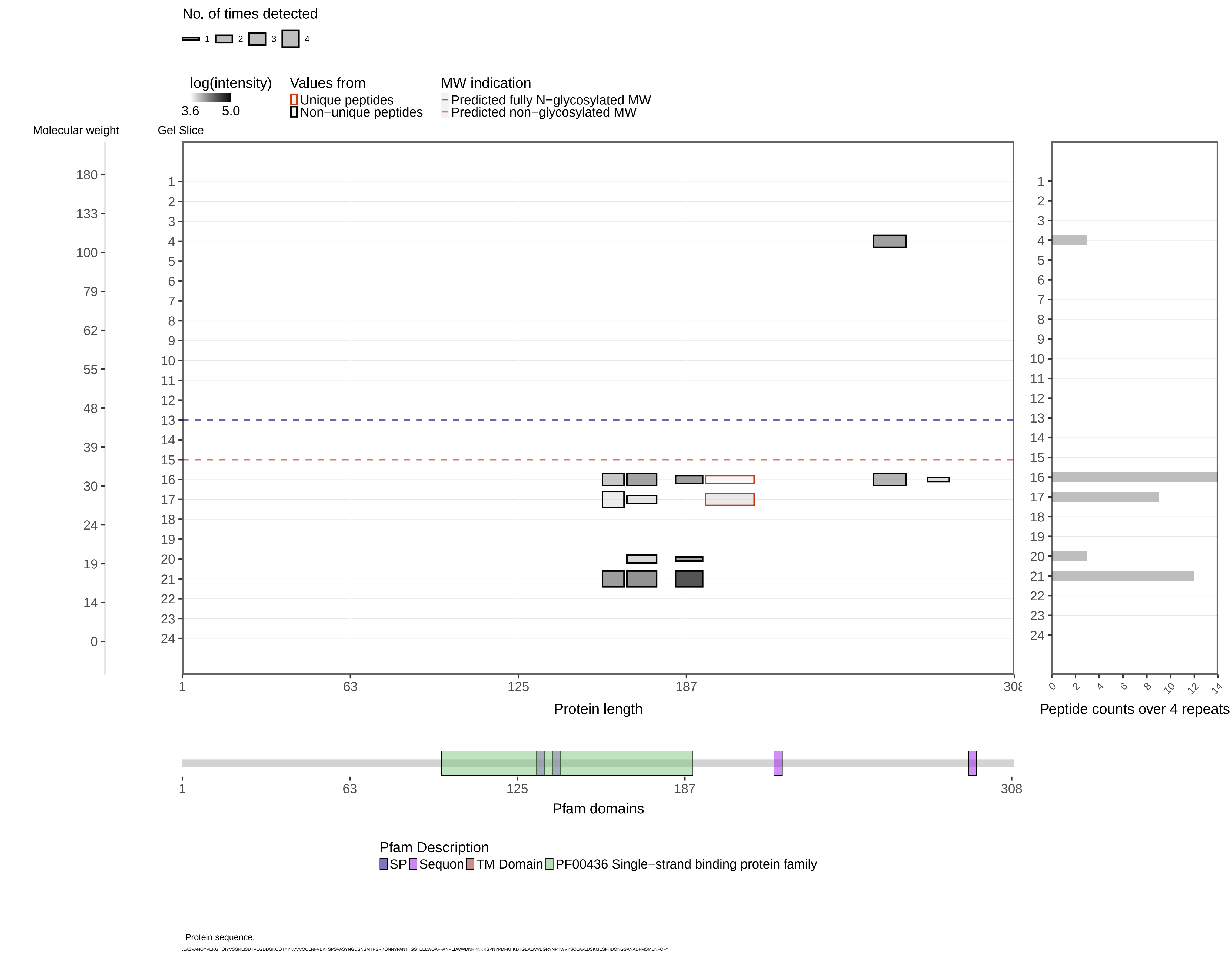

NbL16g27660.1 glutathione_reductase_chloroplastic [MW = 60.4 kDa]

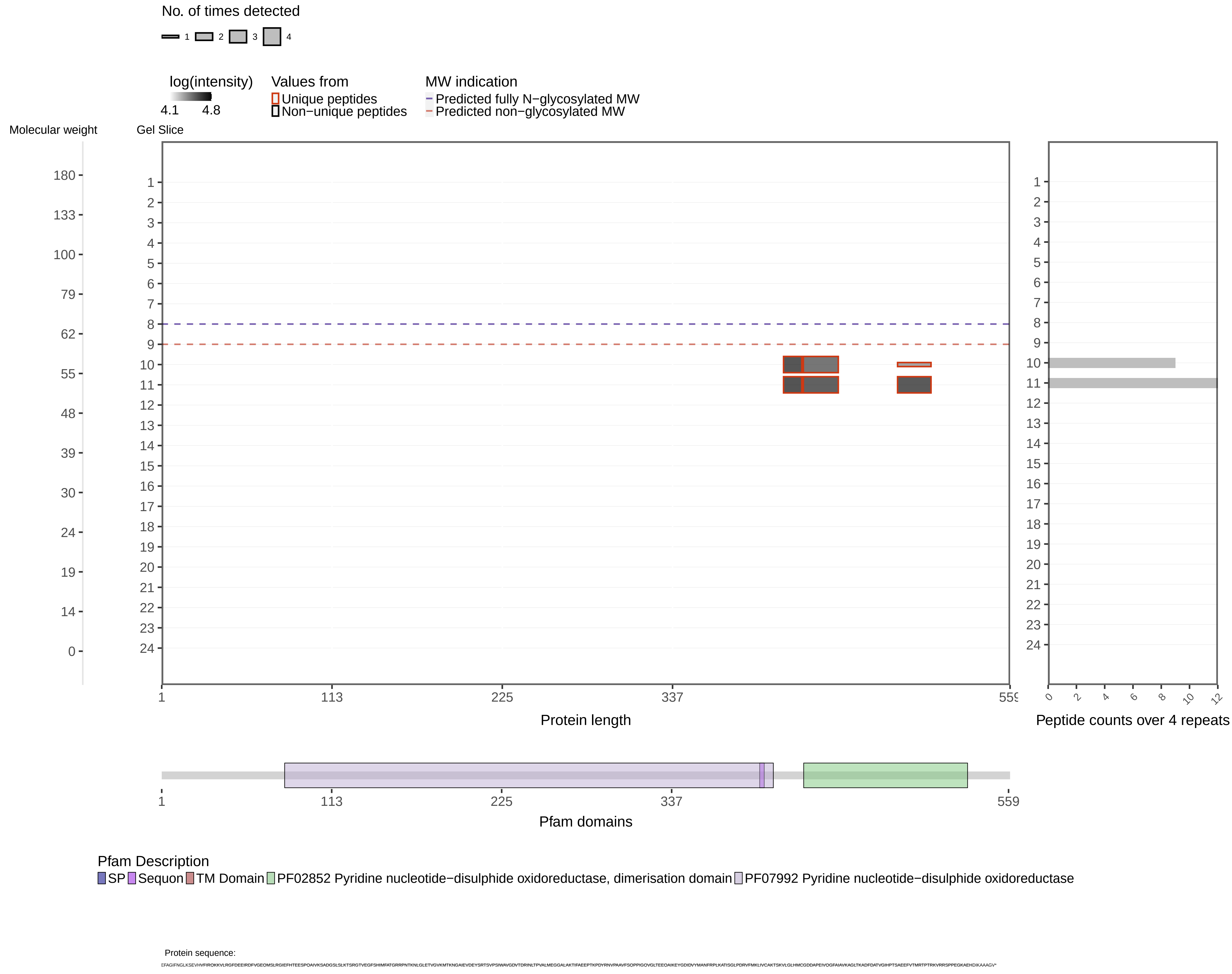

### NbL16g27590.1 histone_H1−like [MW = 30.63 kDa]

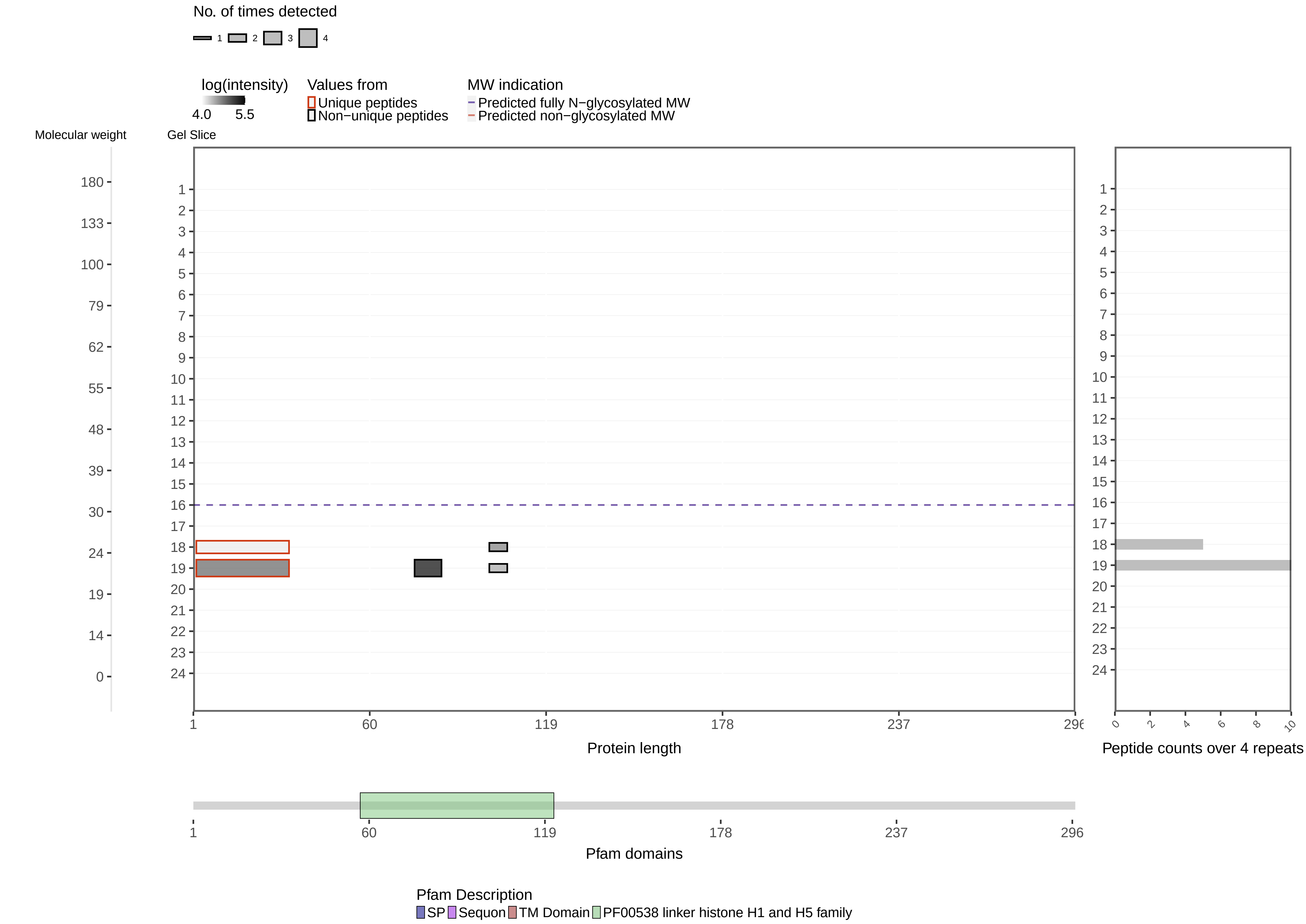


Protein sequence:

PADASAPAKKKTAVAKAAKAKPAAKSKAAAKPKAKPAAKAKPAAKAKPAAKAKPAAKAKPAAKAKPAANAAKSAAKSKPKPKPKATVAAKPKATPVKAKAAPKPKAKETKPAKVAKTATRTTPSRKAAPKAAPAKKEPAKKTPAKSVKSPAKKASTRRGRK*

### NbL16g27480.1 poly_[ADP−ribose]_polymerase_1 [MW = 112.03 kDa]

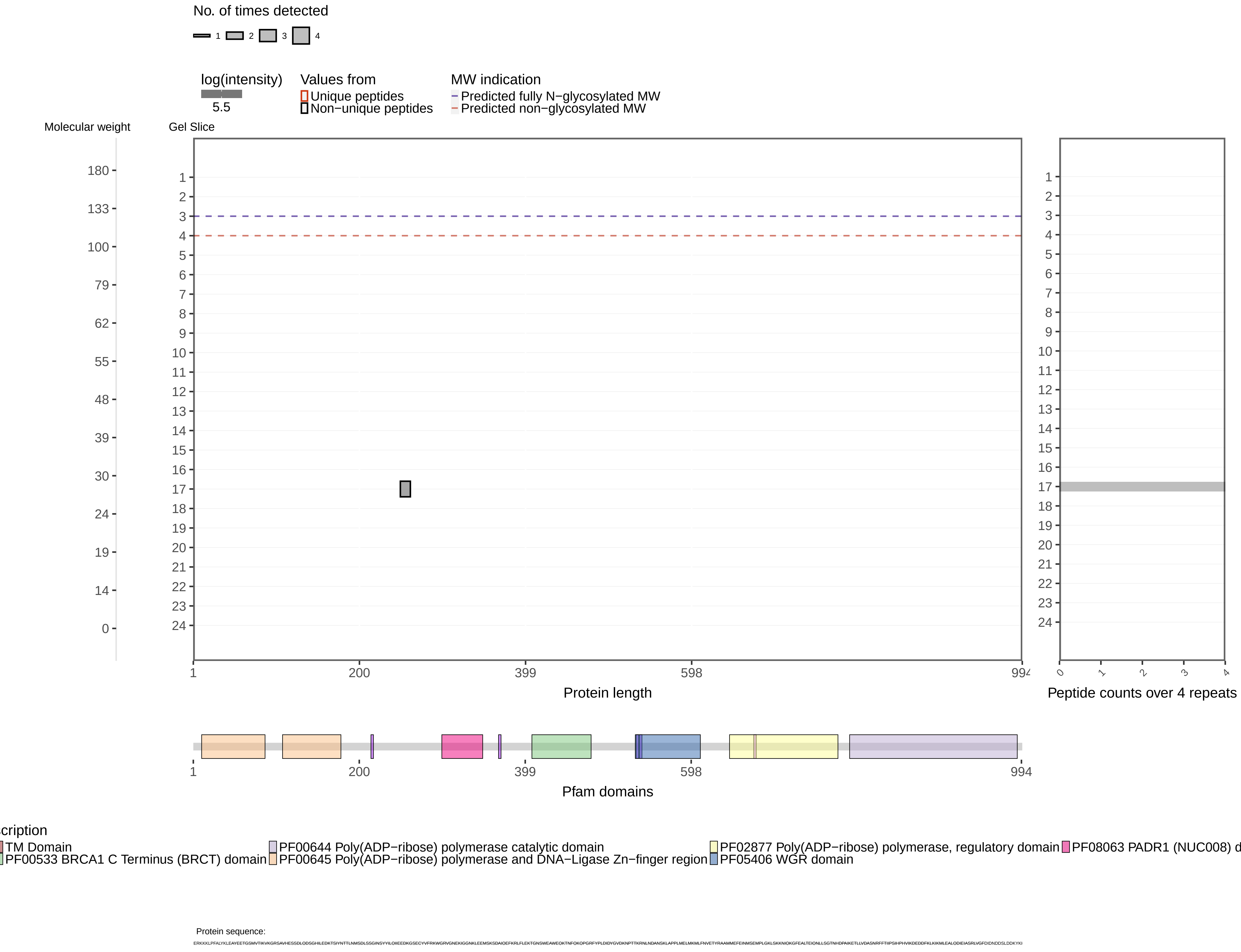

### NbL16g27250.1 TOM1−like_protein_2 [MW = 42.21 kDa]

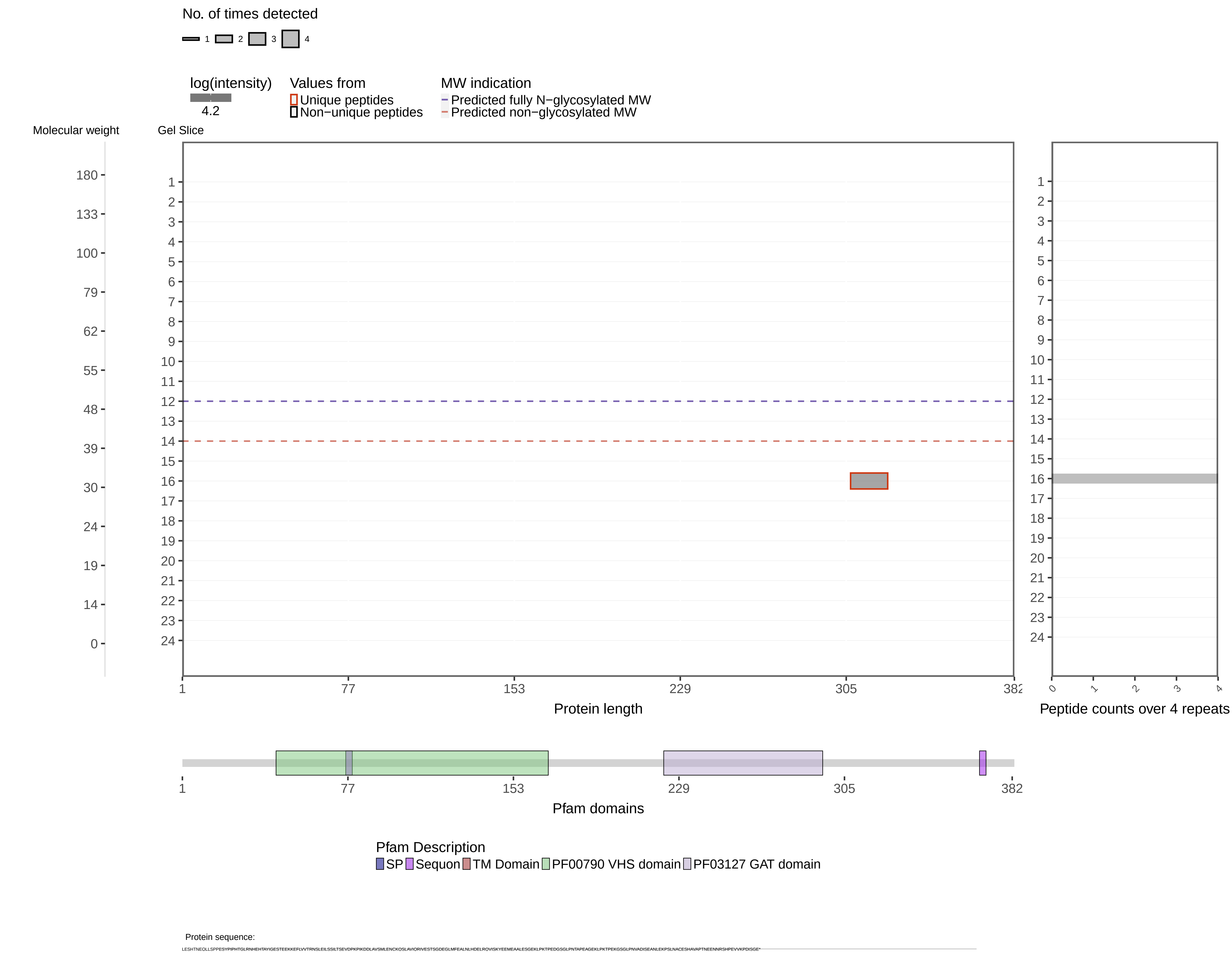

NbL16g27240.1 putative_exosome_complex_component_rrp40 [MW = 26.14 kDa]

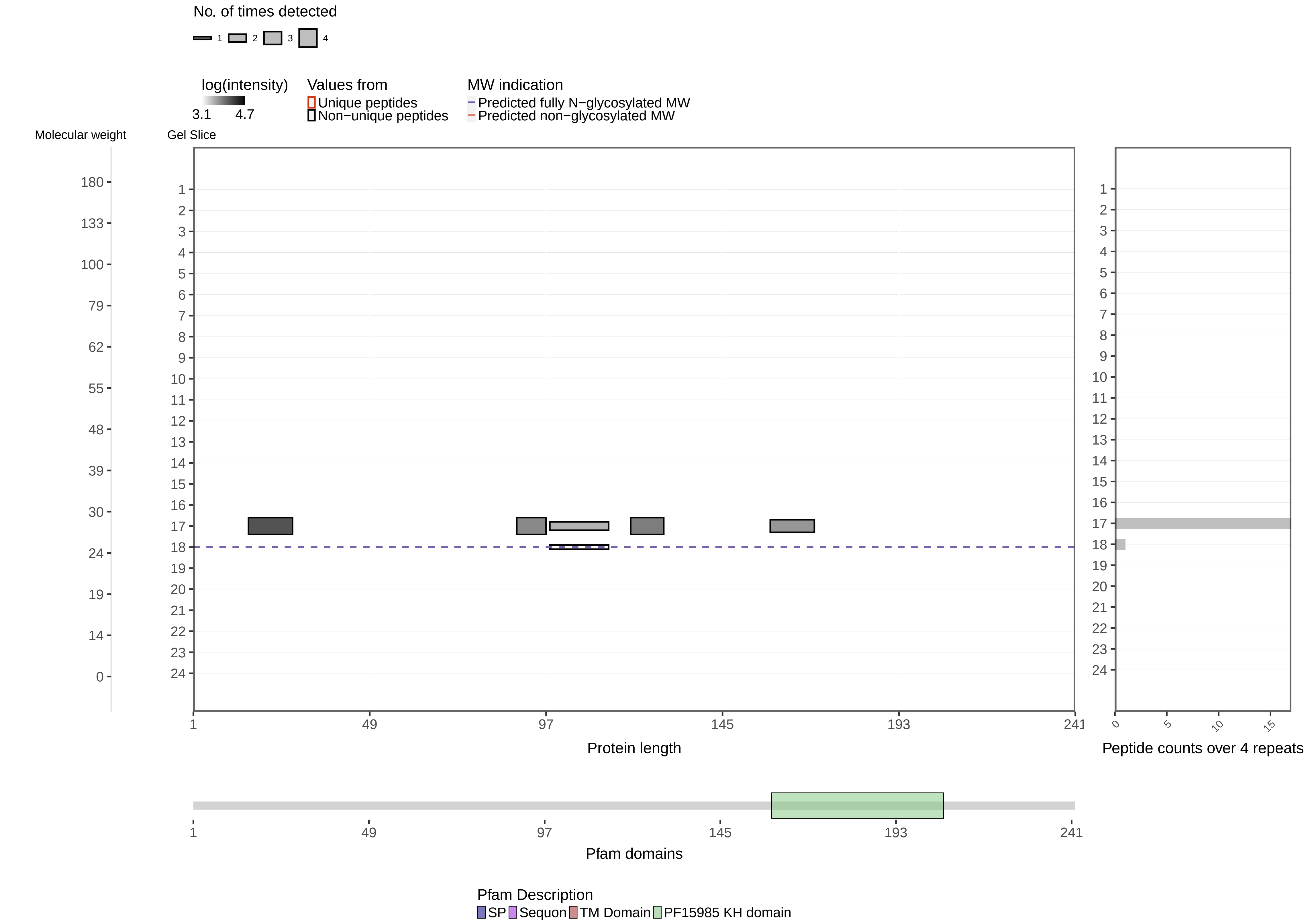


Protein sequence:

FEGGTRRNIPKFEVGTLIYARVVKANTGMNPELSCMDASGKAAEFGLIKDGYMFESSTGLSRMLLSSPTCPVLEGLGKKLAFEIAVGLNGRVWVNAEHOSTIILVANAIMNSESLTPVOOKIMVERLLDRVN*

### NbL16g27200.1 rho−N_domain−containing_protein_1_chloroplastic [MW = 42.71 kDa]

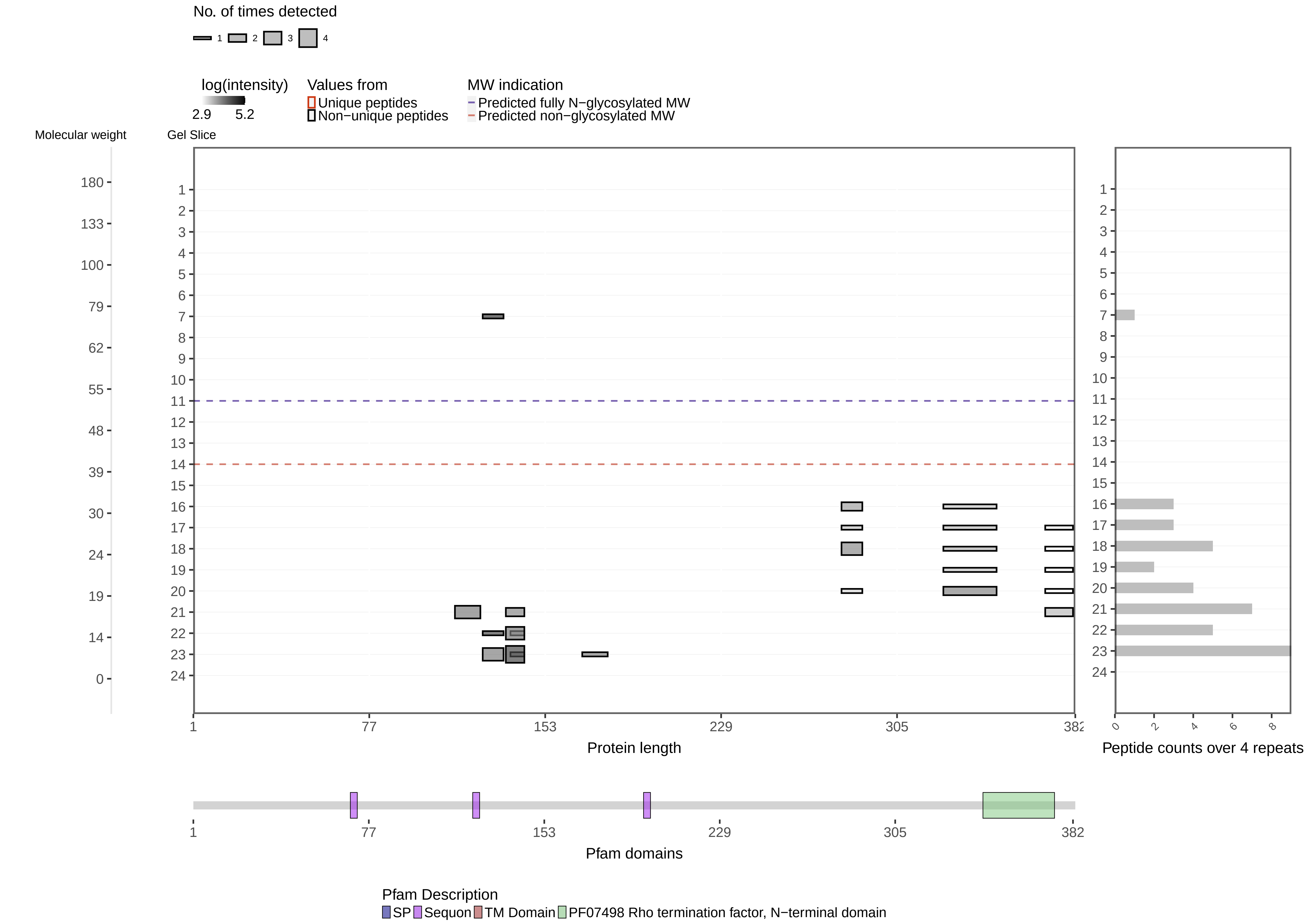


Protein sequence:

LKLLRKHSVQKGKKTNSNRSSNFVLDQPDQSNVFFEERAGSSTELNSDVNHVAQDSGTPVVNRPKSNFQRRSPVSRTKFQPVYHEGDTDNLVASTATDKTEKDIHLDPELGRTPKNKVDSTAHPDIEPIFSDEDVFDEMADDDTAKMFENVDANDDEEQHQVGEELAEMKLTELRAIAKSRGLKGYSKLKKVELIELLSADQV*

### NbL16g27140.1 uncharacterized_protein_At1g03900−like [MW = 30.14 kDa]

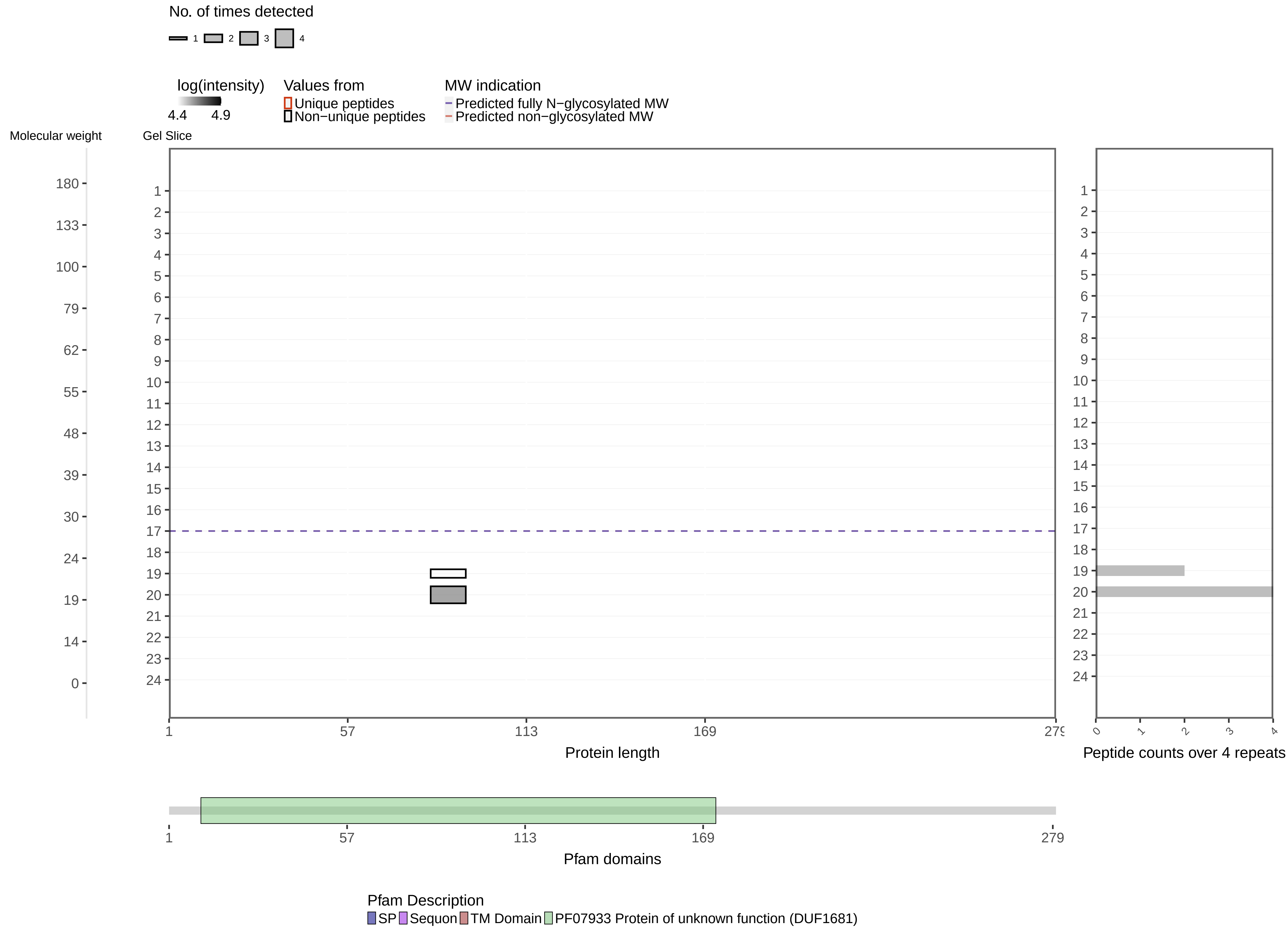


Protein sequence:

'ALSDHDKYVKRENEKEVGDGEAGDDNHIDIHPAVNHRLKEGETIRISVKNKPSSGTGMLSAVGLMAGTTGAVKKTPALAPPPSGGNKIKSPLPPPPNDPATARKTSTSPSIALKENTKHSTDPLSDLAEIERSLPSAAGSRSSKPTAAGWAAF*

NbL16g27120.1 ABC_transporter_I_family_member_21−like [MW = 32.06 kDa]

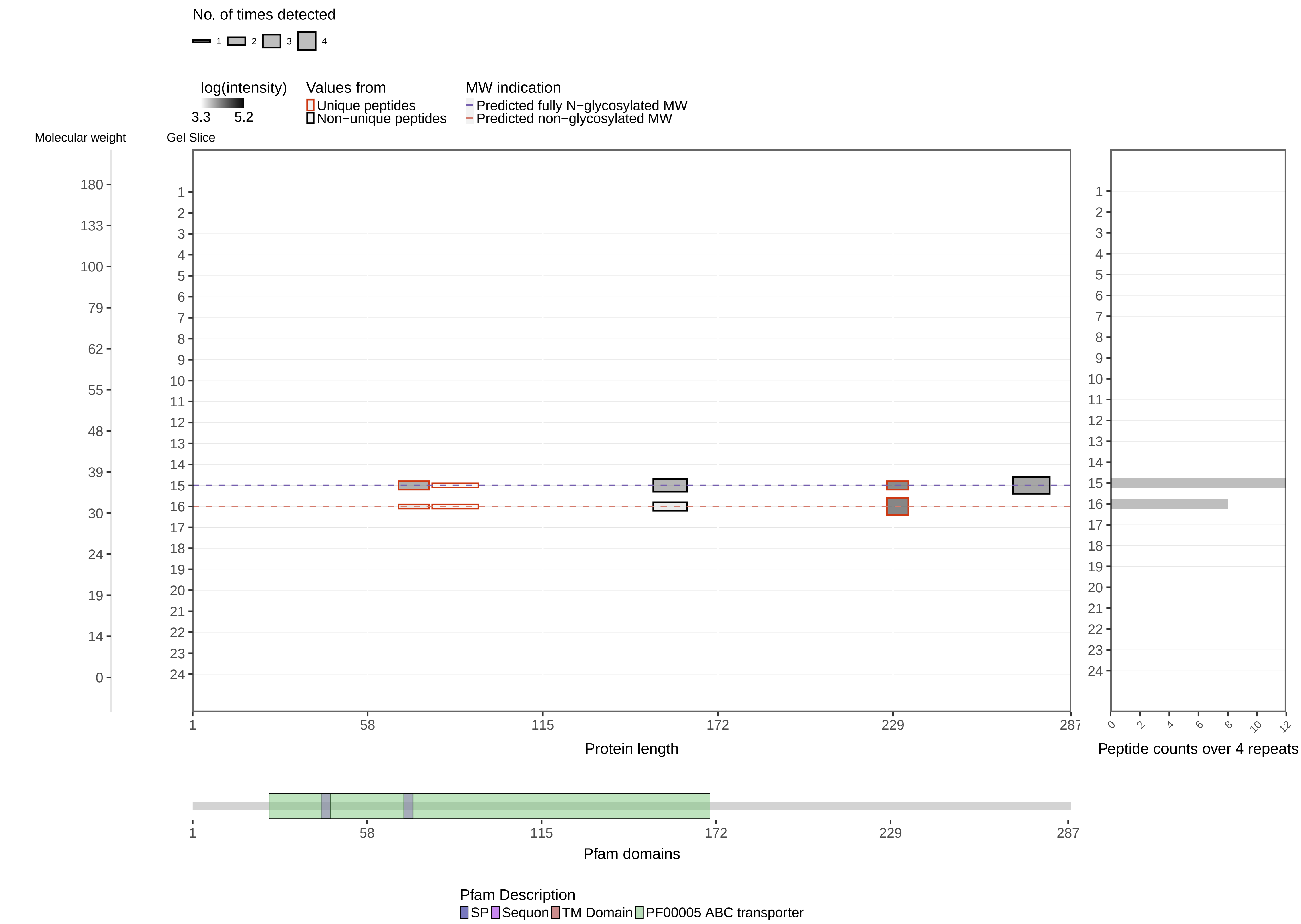


Protein sequence:

ELLDINLQWRMHKVSDGQRRRVQICMGLLHPYKVLLLDEVTVDLDVVARMDLLDFFKEECEQRGATIVYATHIFDGLESWATDLVYIQDGVLKRSEKLSELPDLKSSPNLLSVVENWLRSETTIEKKKPVIAPPKVQKSSPFGSSPFQSSRHMAYFR*

### NbL16g27090.1 probable_60S_ribosomal_protein_L14 [MW = 15.89 kDa]

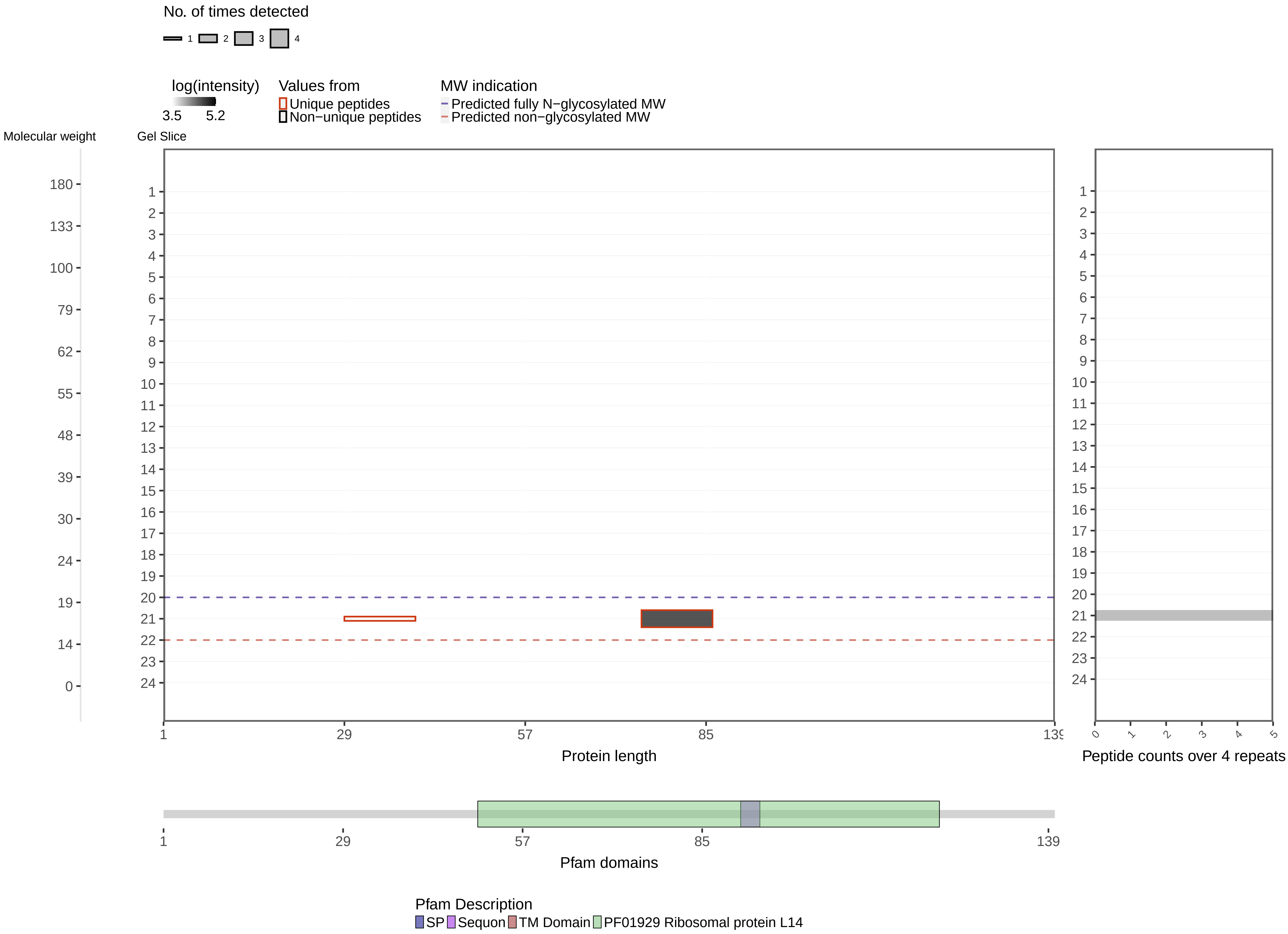


Protein sequence:

KRLSLTDIKIDIKRIPKKKTLVEAMEAADVKTKWENSSWGRKLIVQKRRAALNDFDRFKLMLAKIKRAGVVRQELAKLKKTAA*

NbL16g27000.1 rho_GDP−dissociation_inhibitor_1−like [MW = 27.82 kDa]

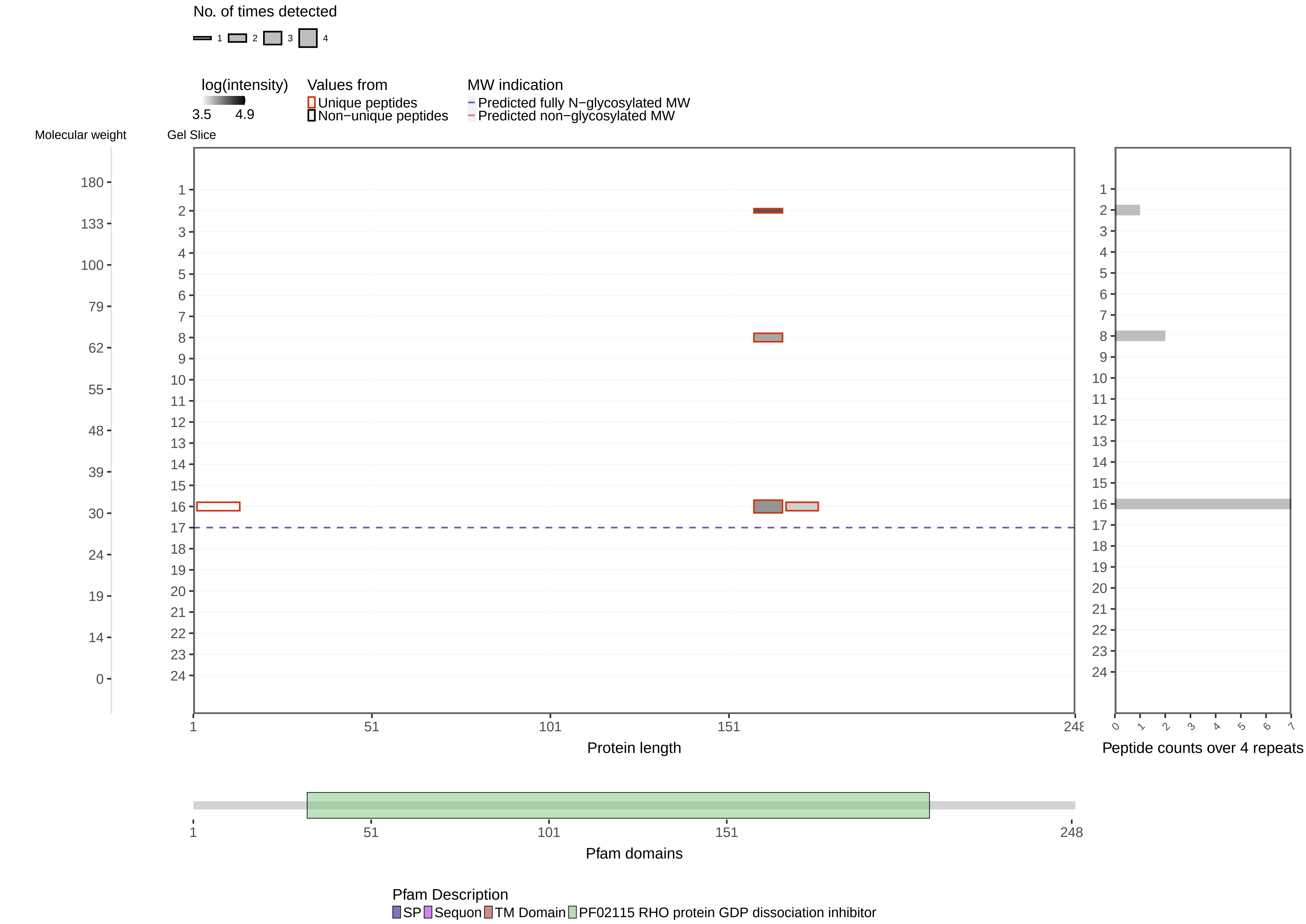


Protein sequence:

NTPLPIKEGQRKSVLFTLKEGSEYRLKLTFSVLHNIVSGLAYTNTVWKAGLQVDQSKGMVGTFAPQREPYIHMLEEETTPSGALARGTYTAKLMALDDIGETRIHILNKQNSHRLLIERVLVMIVHGVFTLSNYGDLQ*

### NbL16g26980.1 phospholipid−−sterol_O−acyltransferase−like [MW − SP = 69.67 kDa]

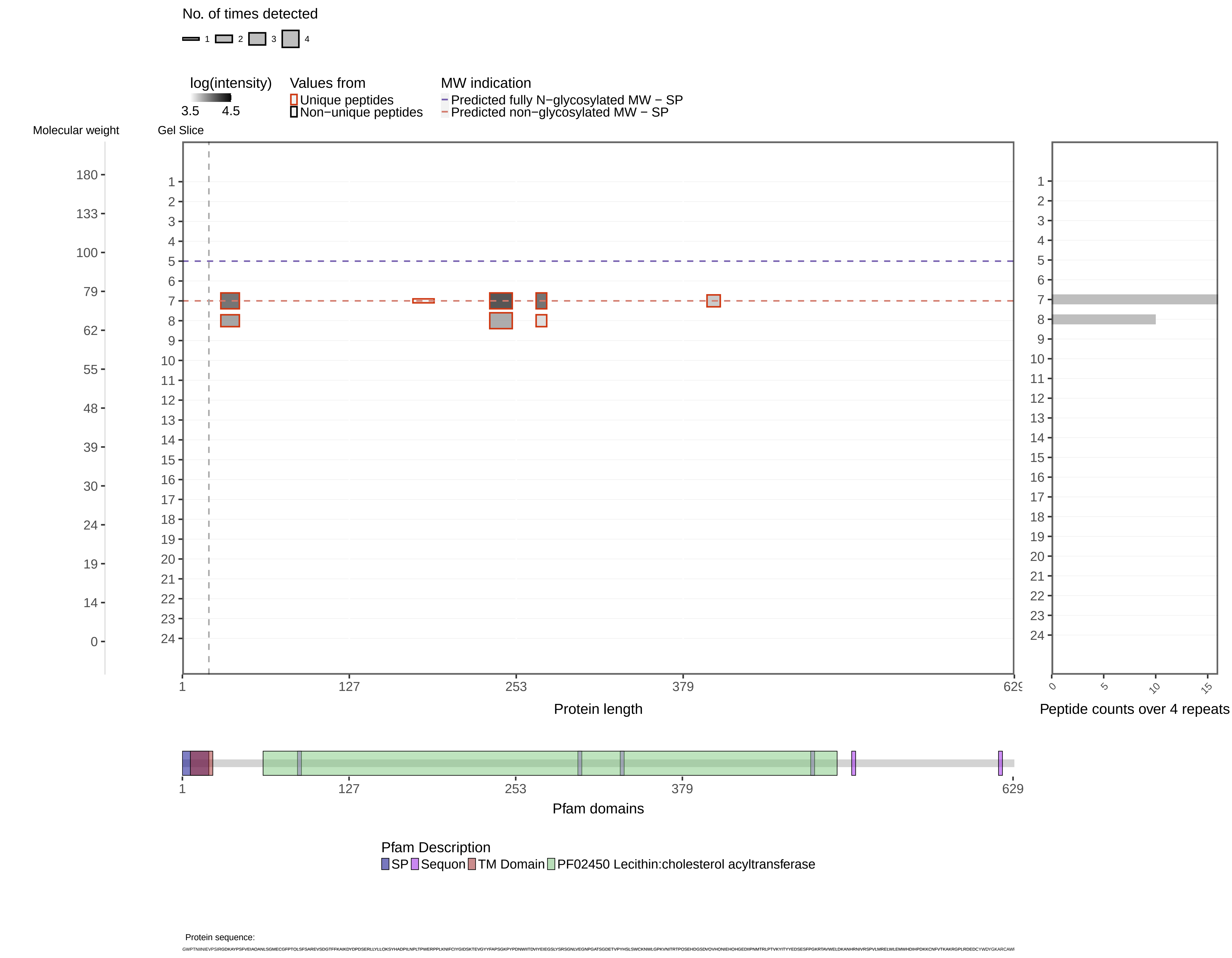

### NbL16g26880.1 60S_ribosomal_protein_L23−like [MW = 15.01 kDa]

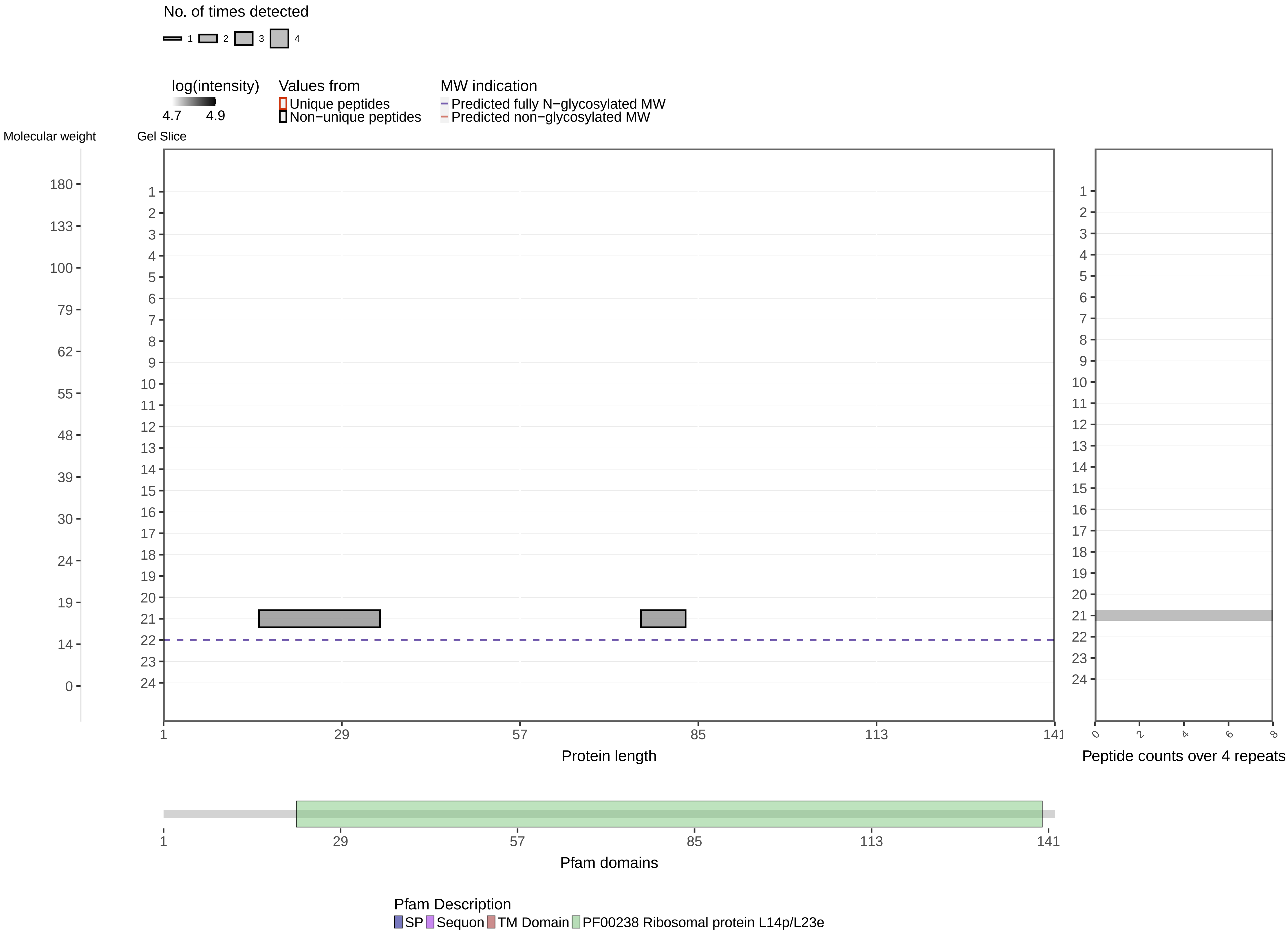


Protein sequence:

GDMVMATVKKGKPDLRKKVMPAVIVRQRKPWRRKDGVFMYFEDNAGVIVNPKGEMKGSAITGPIGKECADLWPRIASAANAIV*

NbL16g26760.1 NADH_dehydrogenase_[ubiquinone]_1_alpha_subcomplex_subunit_9_mitochondrial [MW = 43.45 kDa]

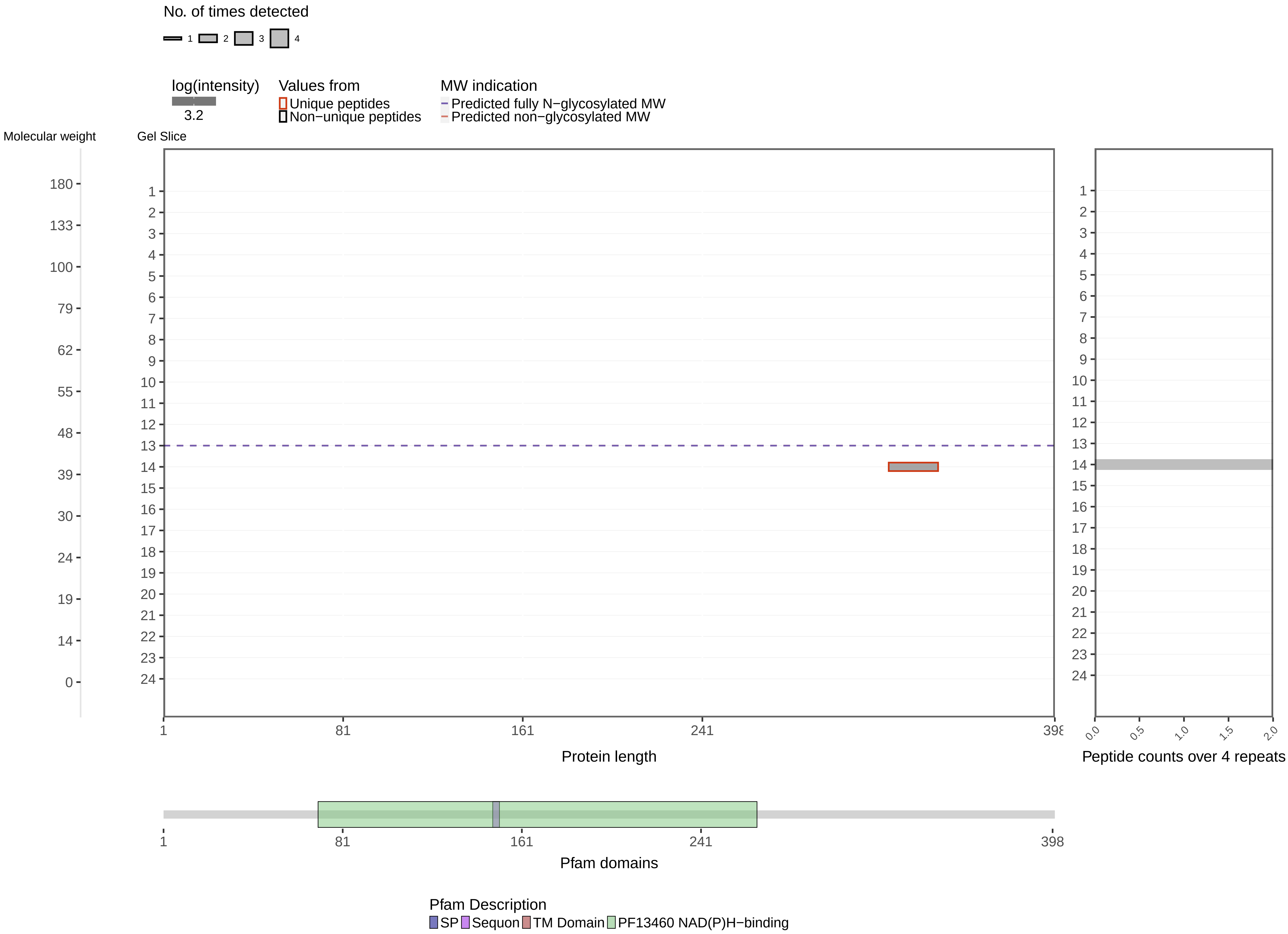


Protein sequence:

SPTSPSRMLRAKAAAEEAVAHELPEATIMRPAVMIGTEDRILNPWAHFAKKYGFIPLFGDGSTKIQPVFVADVASAIVASLKDNGISMGKVYELGGPDVYTMHELAELMFDMIREWPRYVNVPFPIAKALASPREFLLNKVPAPMPTPTIFNLDAIEALATDKIVSKDALTFEDLGLVPHKVKGYPVEFLIQYRKGGPNYGSTVSERVSPESYP*

### NbL16g26700.1 alcohol_dehydrogenase_class−3−like [MW = 25.78 kDa]

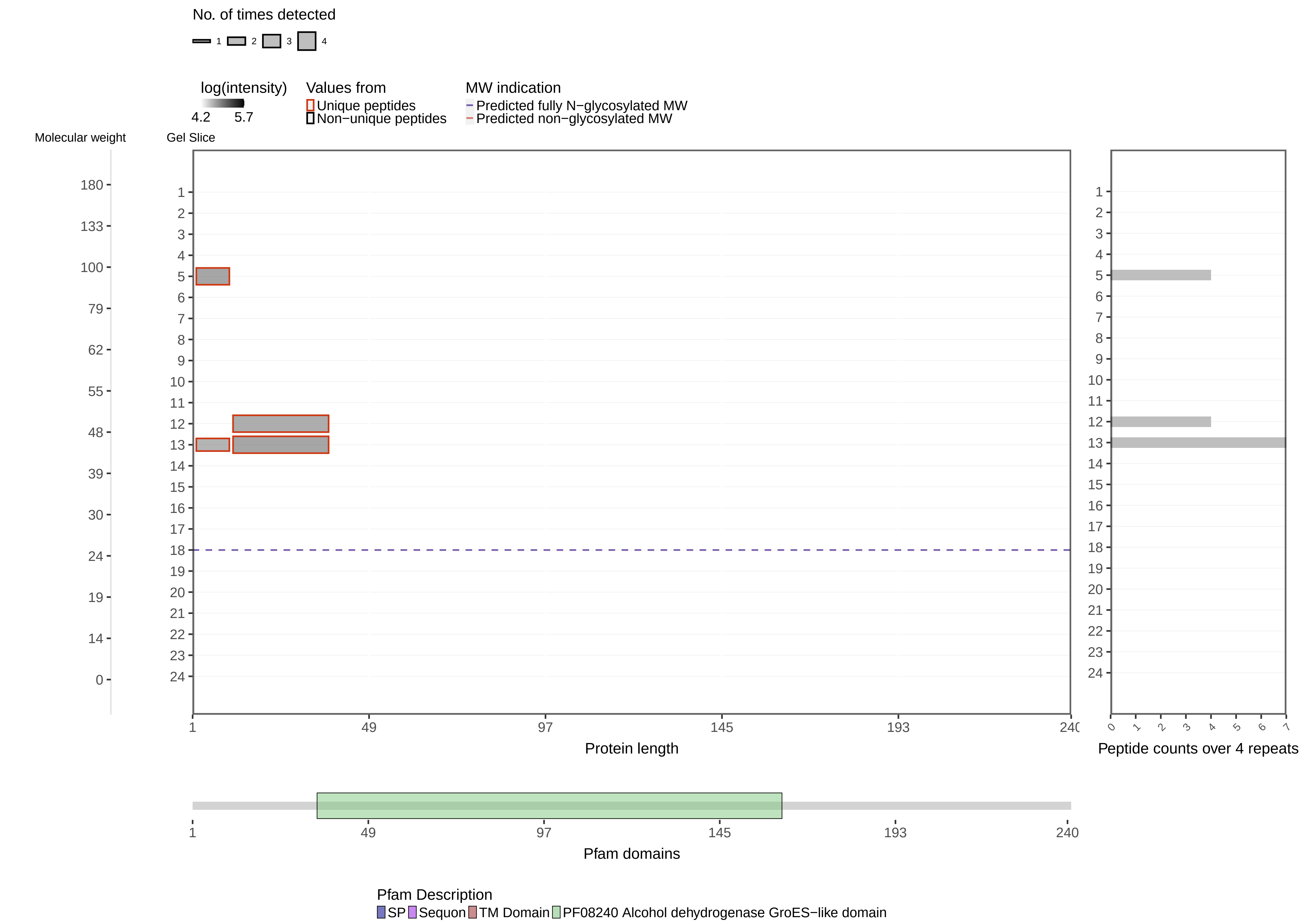


Protein sequence:

SGKTNLCGKVRAATGVGVMMNDRQSRFSINGKPIYHFMGTSTFSQYTVVHDVSVAKIDPVAPLEKVCLLGCGVPTGLGAVWNTAKVEPGPLLLSLAWAQLVLLWRRVQRLLVLRELLGLILTAKSLIEPKSLV*

NbL16g26680.1 ribokinase−like [MW = 43.47 kDa]

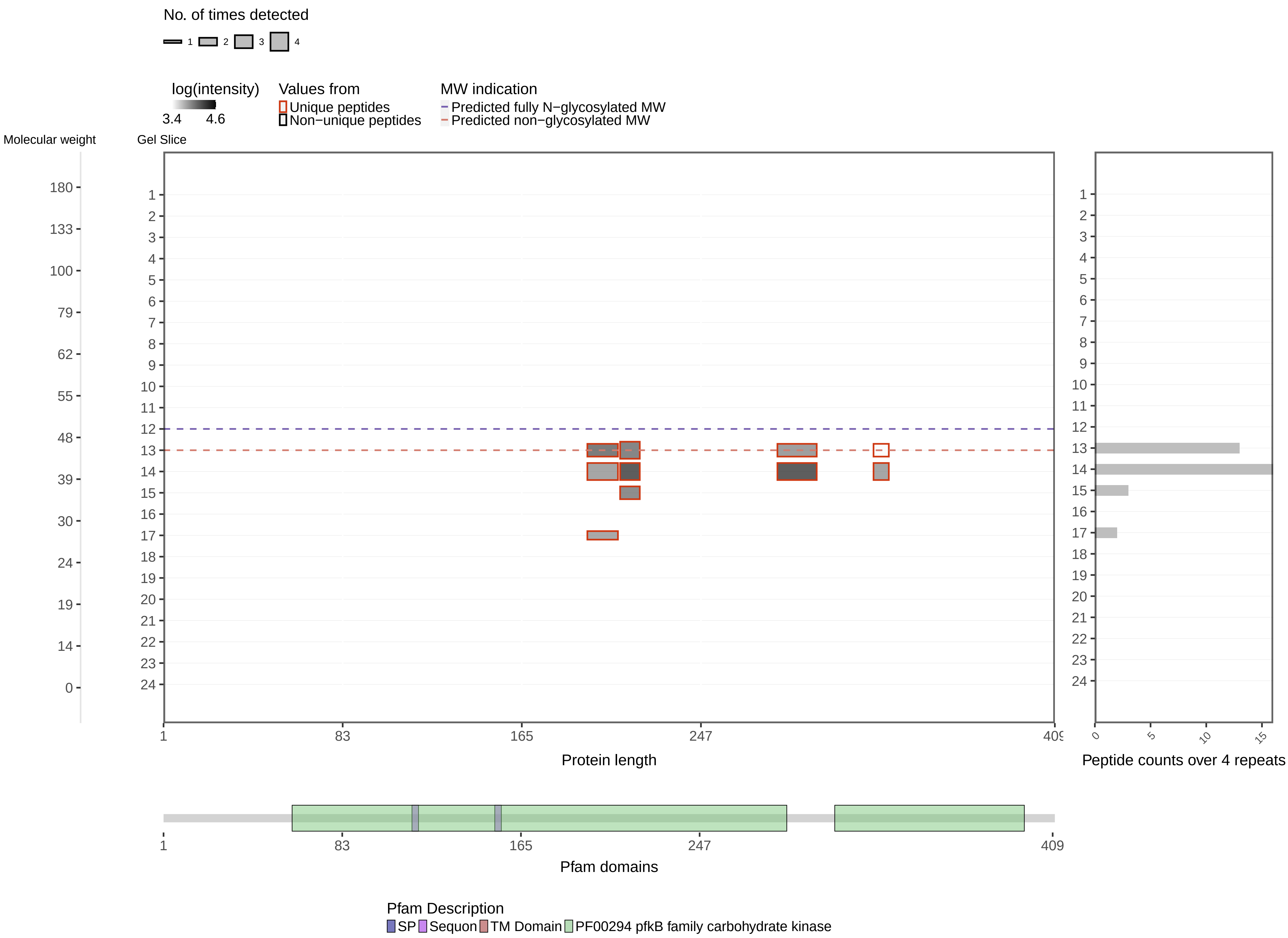


Protein sequence:

[illegible]FDGRLHETAAVVAEEANNRGIPILIDTERKREGLDHLLSFASYVVCSTKFPQEWTEASSIPSALVSMLLRLPKAKFVIVTLGKDGCIMLQRTEMENVHSEEMDVDNLFEKLNQEKDTSATVPACISSNVVKLHAKGIGTISGRLLVGTAESIPQTELIDTTGAGDAFIGAVLYSICANIPPEKMLPFAAQVAAIKCRALGARAGLPHYTDHRLTPFLV*

### NbL16g26420.1 ribonuclease_P_protein_subunit_p25−like_protein [MW = 26.92 kDa]

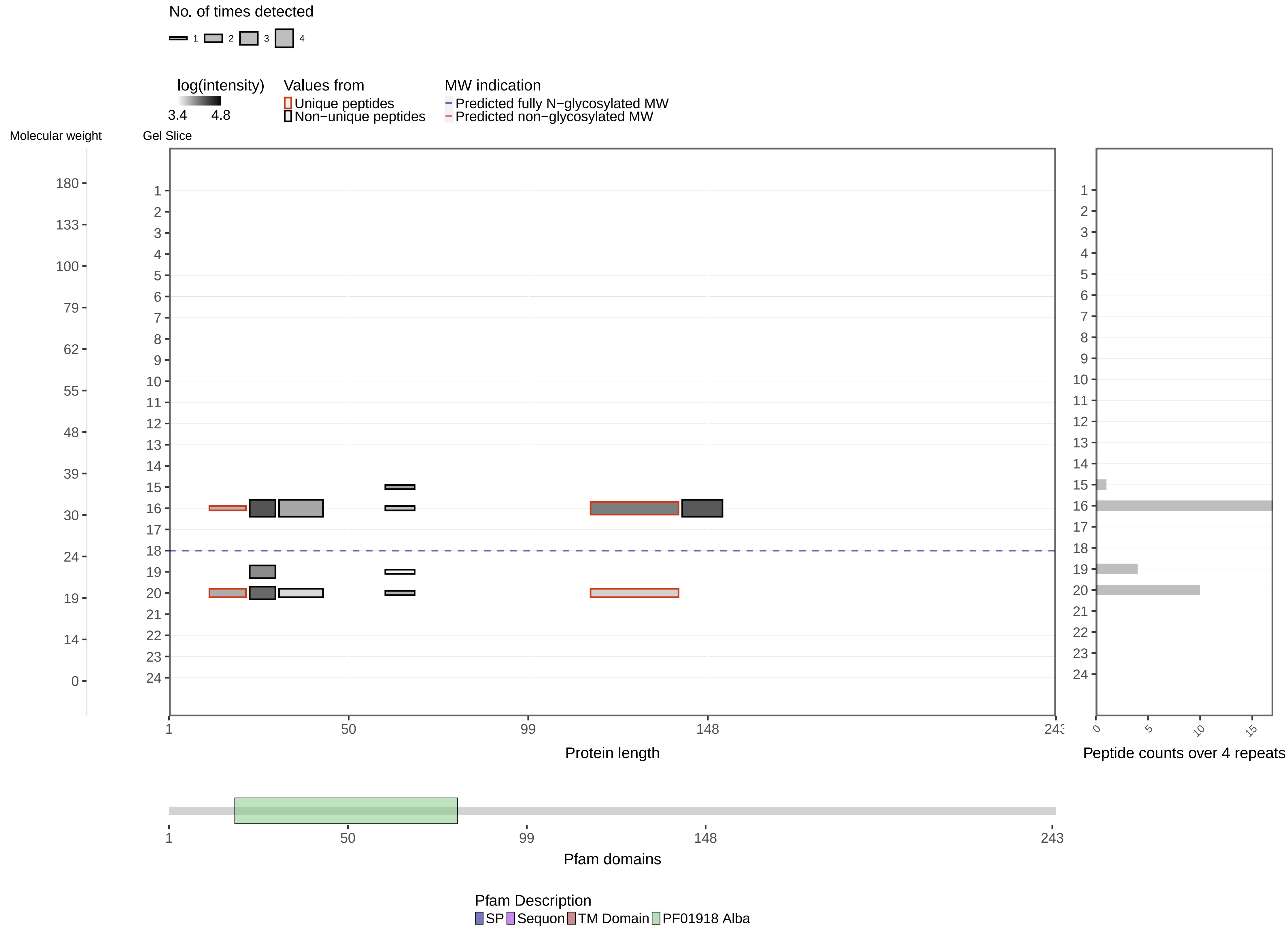


Protein sequence:

ANSPGYQAPSGVEVEQMKPQYRNYQPQLQQQPPRQTRAVYYAGNEDSYGRGRGRGRGRGRGWSRGGYGNYQENGDYSNWGRGGGRGGWGYPGSEYGRGRGGGGRGYGRGRGRMGPRPRGGGNQA*

NbL16g26410.1 pantoate−−beta−alanine_ligase−like [MW = 35.73 kDa]

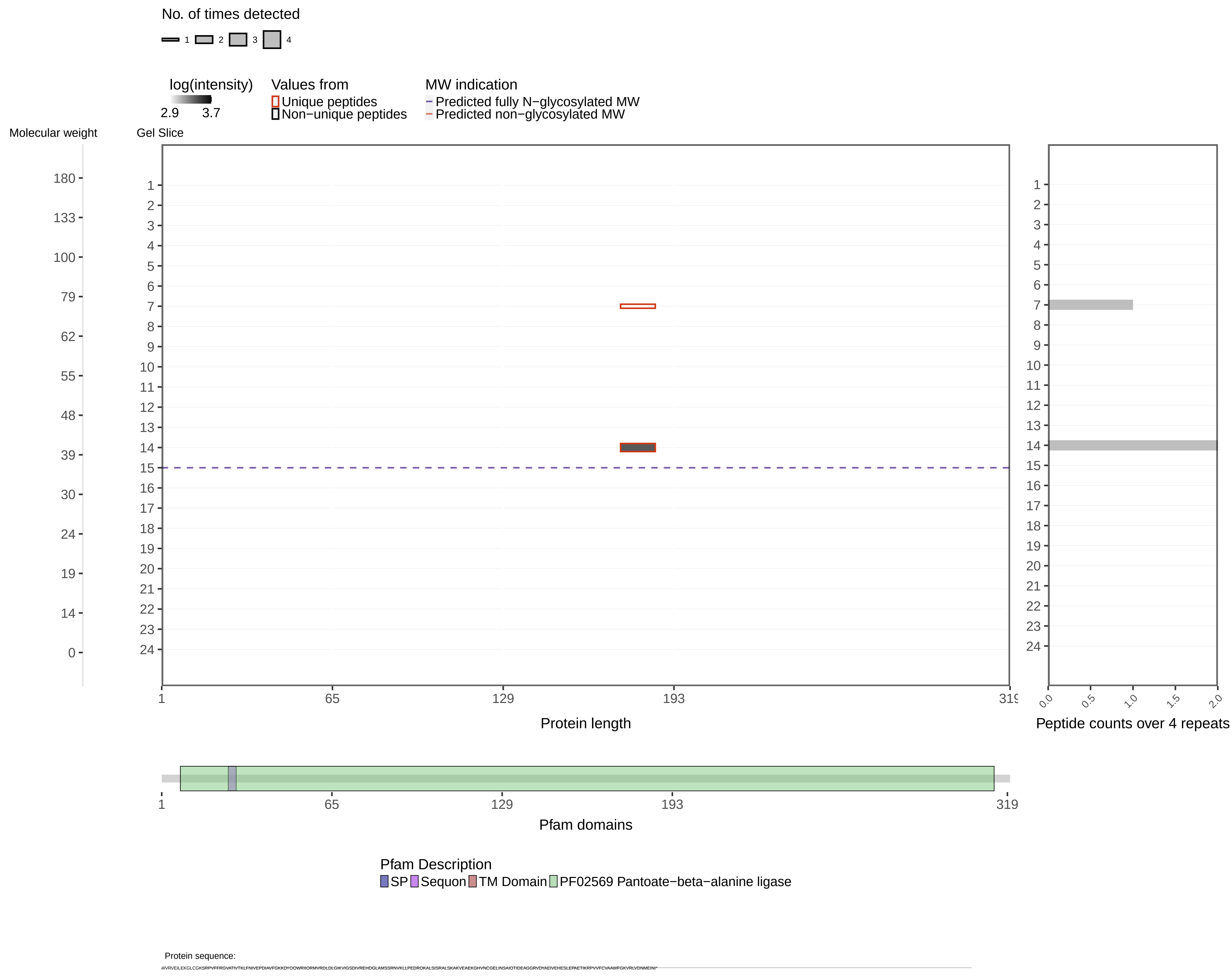

### NbL16g26400.1 vesicle−associated_membrane_protein_722 [MW = 24.77 kDa]

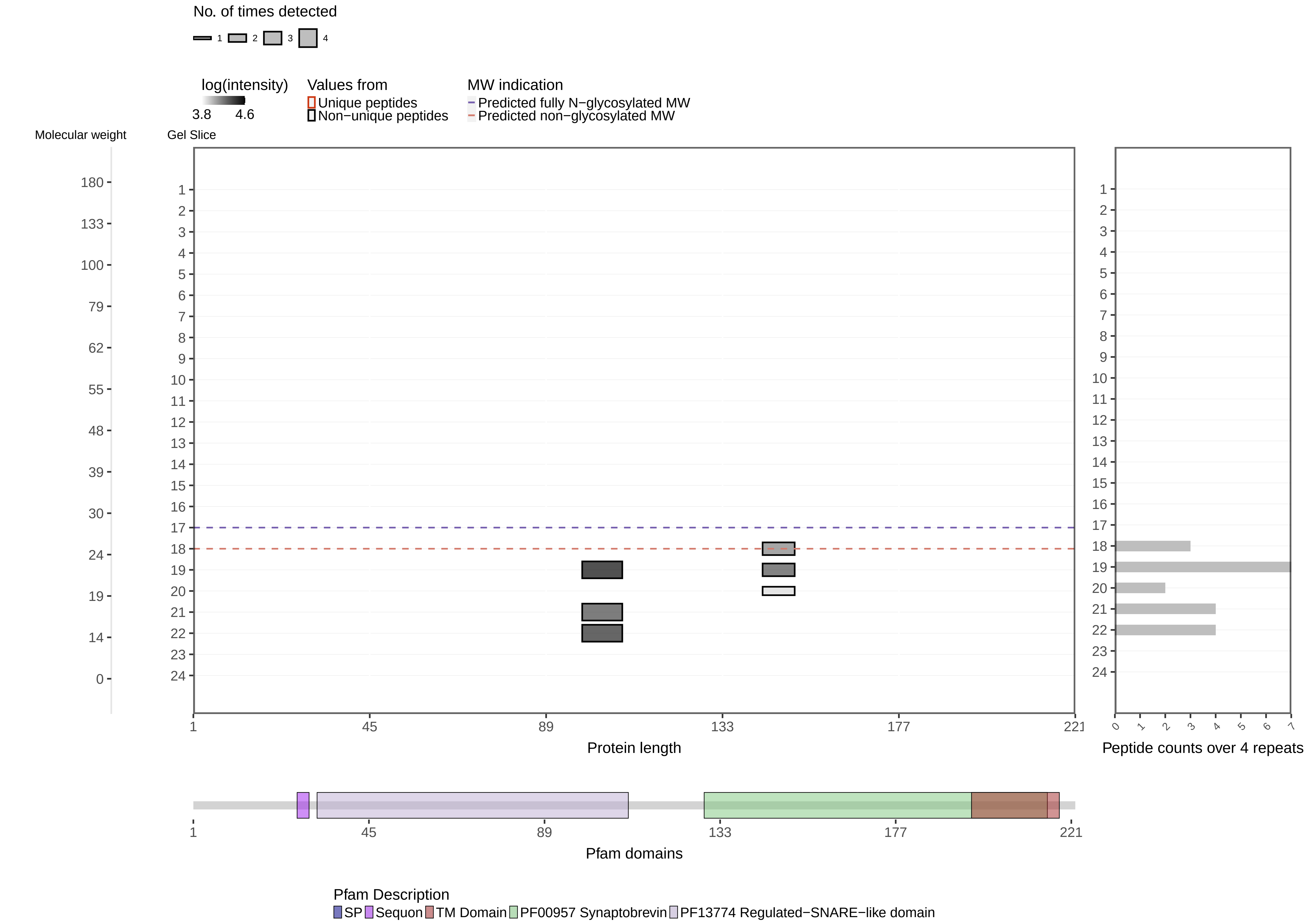


Protein sequence:

KAATAVANSLNREFGPKLKEQMQYCVDHPEEISKLAKVKAQVSEVKGVMMENIEKVLDRGEKIELLVDKTENLRSQAQDFRTQGTTMRRKMWLQNMKIKLIVLGIIIALILIIVLSVCGGFNCH*

### NbL16g26330.1 desumoylating_isopeptidase_1−like [MW = 28.82 kDa]

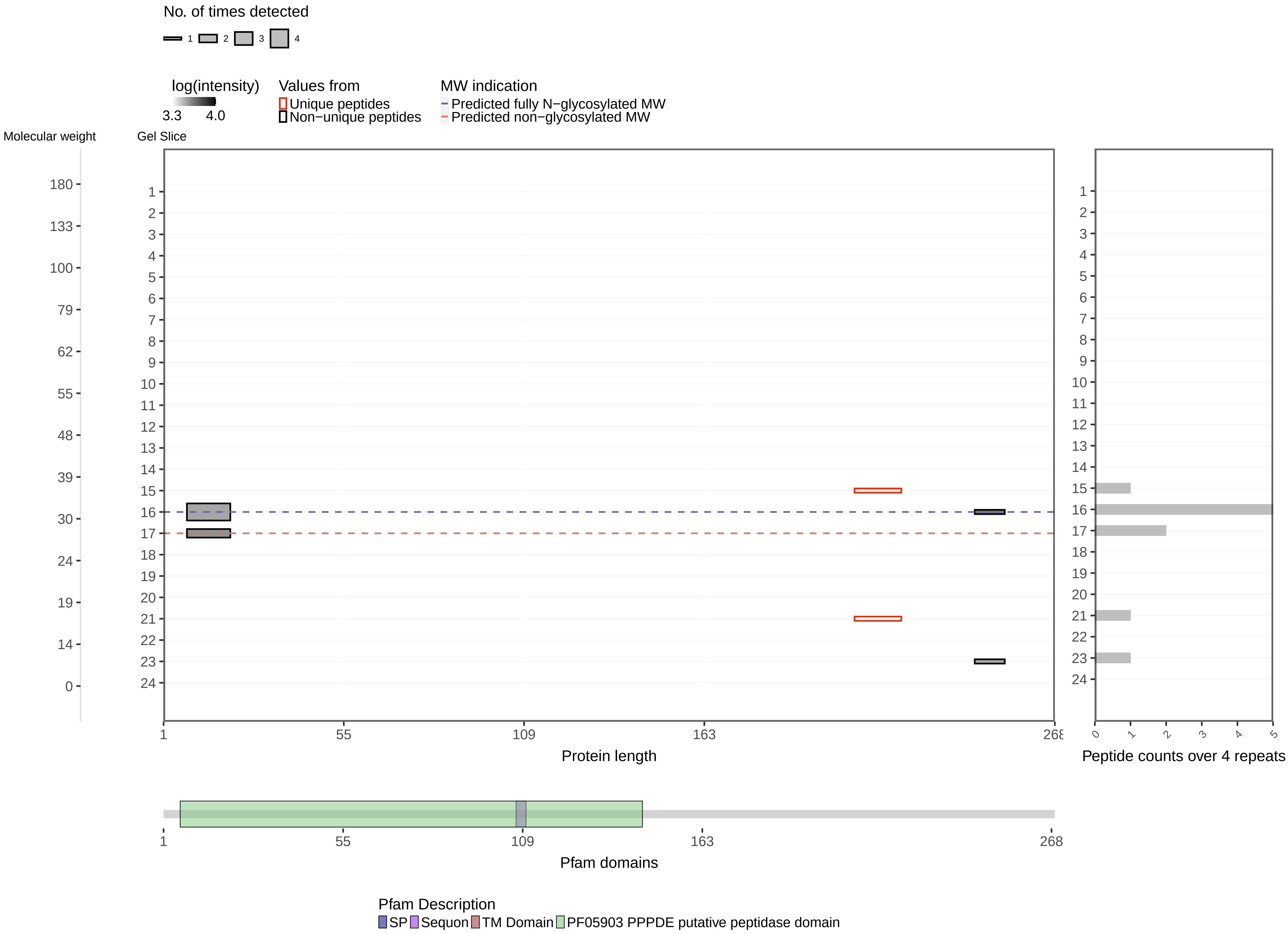


Protein sequence:

PDDILNLPNEVMNSPMGALIMPMIQQLESTLRANAVPQAPQFRPPTVASASQQMVSGGKSQVGEDKSGSVVKPAKREKKPVGEQEKTPANTAVSDPLGDSRSKVQEEITREFAAIMATGTLRASEAAALATKRVMQRYGHTTAALN*

### NbL16g26250.1 cytochrome_b5 [MW = 14.98 kDa]

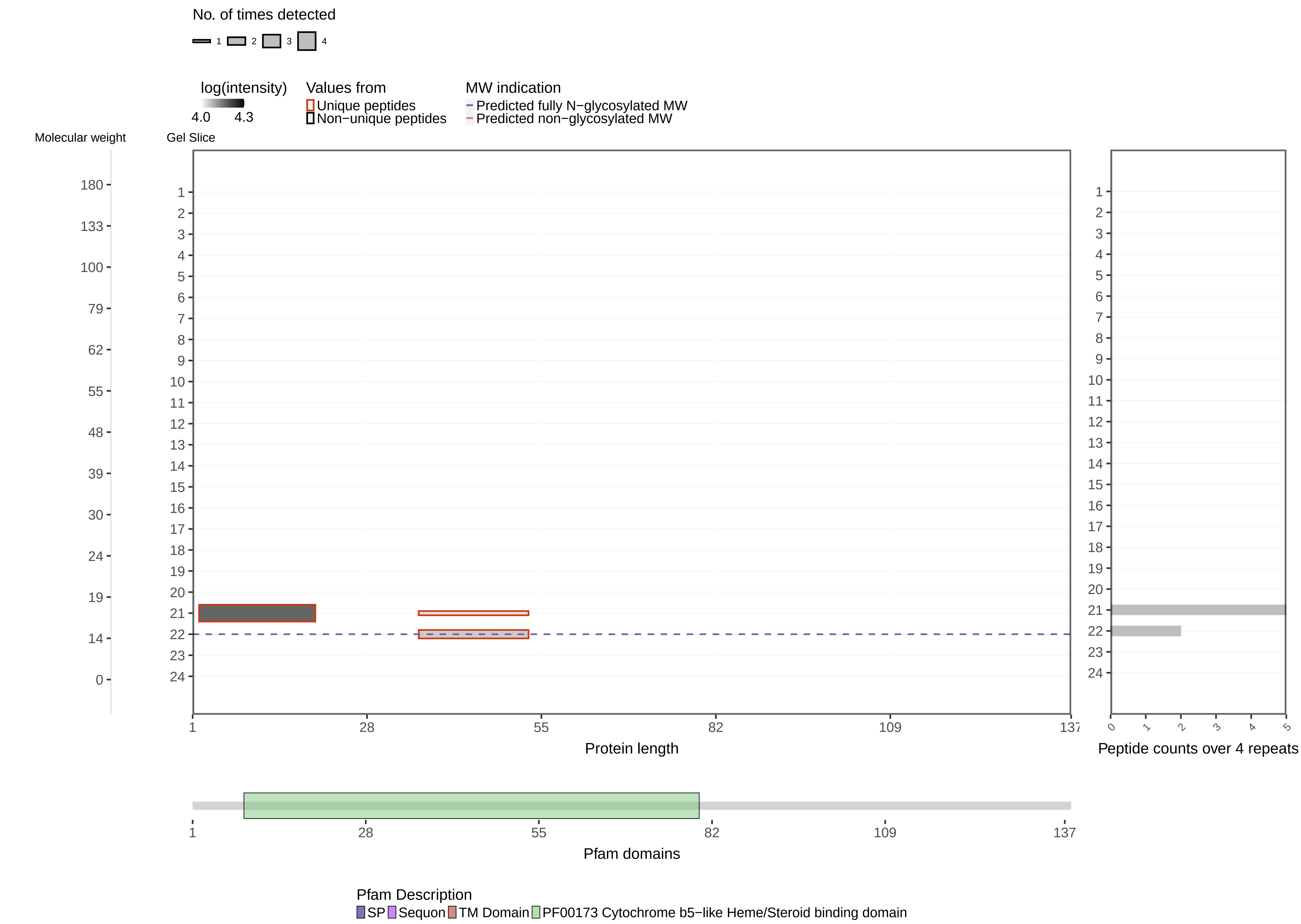


Protein sequence:

DATDDFEDVGHSSSARAMLDEYYVGDIDSATIPTKTKYTPPNQPHYNQDKTSEFVVKLLQFLVPLIILGVAFGIRFYIKQSSA*

### NbL16g25980.1 putative_G3BP−like_protein [MW = 50.95 kDa]

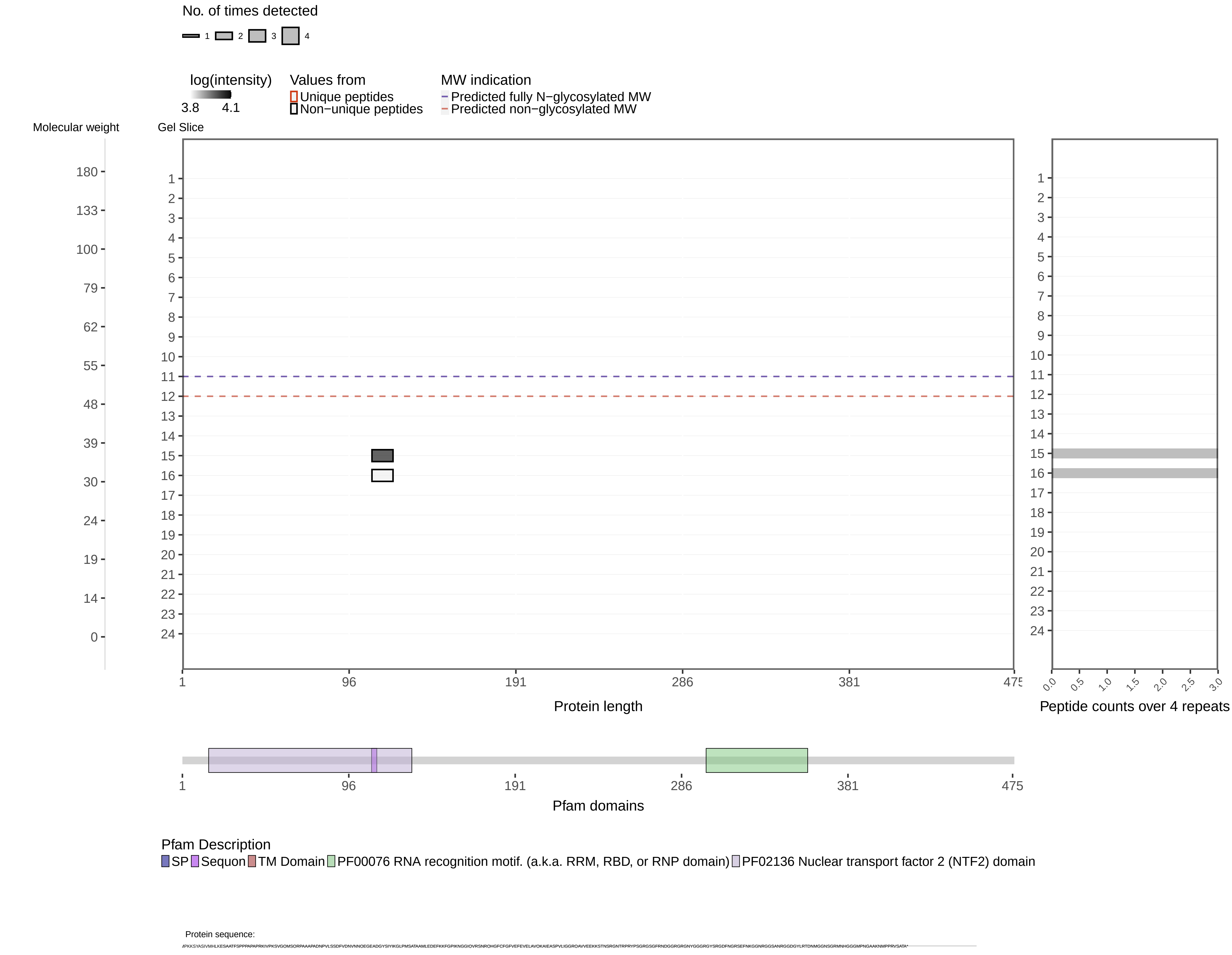

NbL16g25960.1 anthocyanidin_3−O−glucosyltransferase [MW = 52.54 kDa]

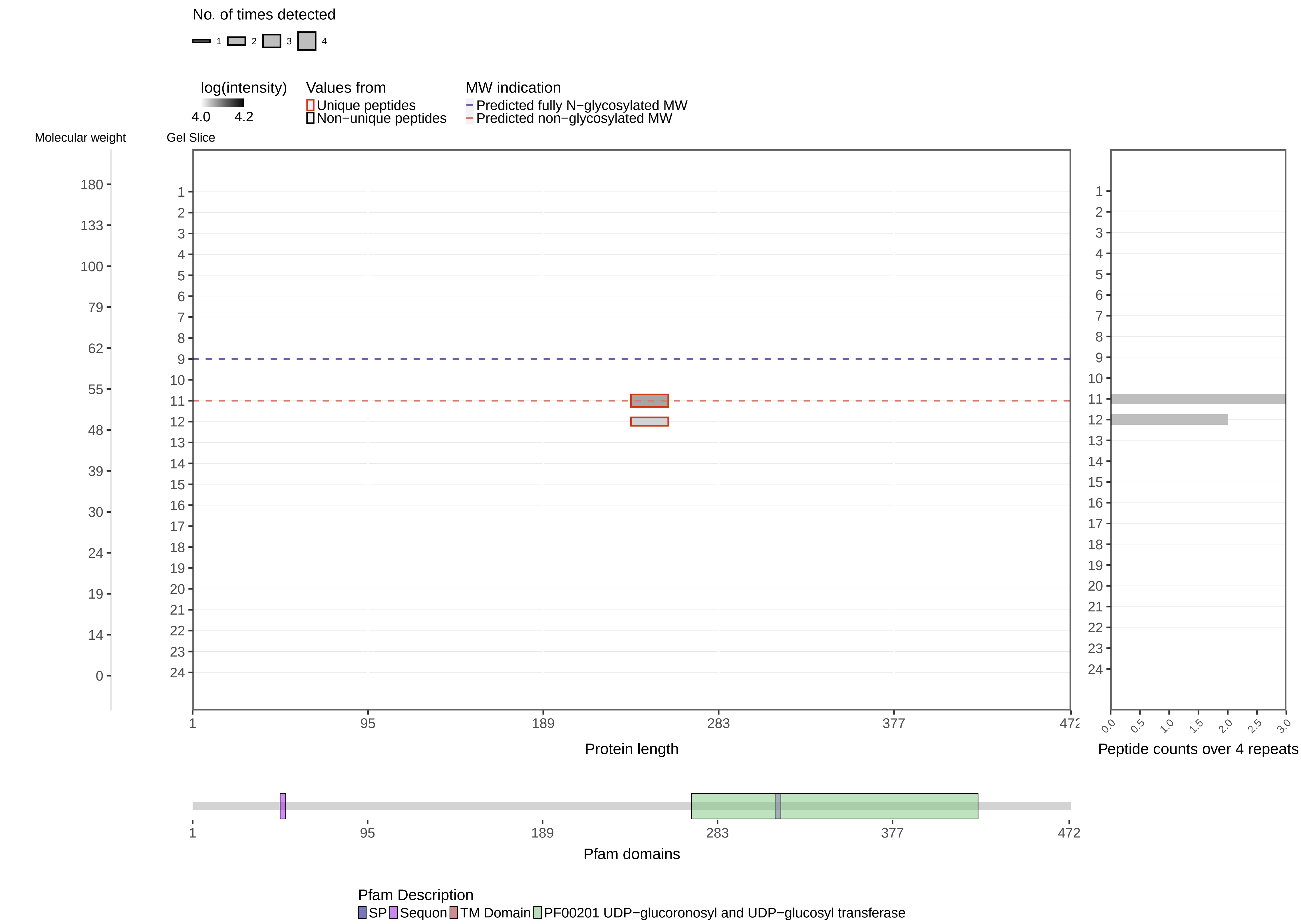


Protein sequence:

MEGPYIEYVKAQFNKPVFLVGPVVPDPPSGKLEERWSSWLNKFEAGTVIYCSFGSETFLKDEQIKELALGLEQTALPFFLVLNFPANVNFSVELNRALPEGFLERVKDKGIIHSGWVQQQHILAHSSVGCYVCHAGFSSVIEALVNDCQVVMLPQKGDQFLNAKLVSGDMKAGVEVNRRDEDGYYGKEDIKEAVEMVMVEIDKQPGELIRENQKKWKEFLLNKDIQCKFIEDLVQEMMAMTKVSST*

### NbL16g25940.1 putative_quinone−oxidoreductase_homolog_chloroplastic [MW = 34.65 kDa]

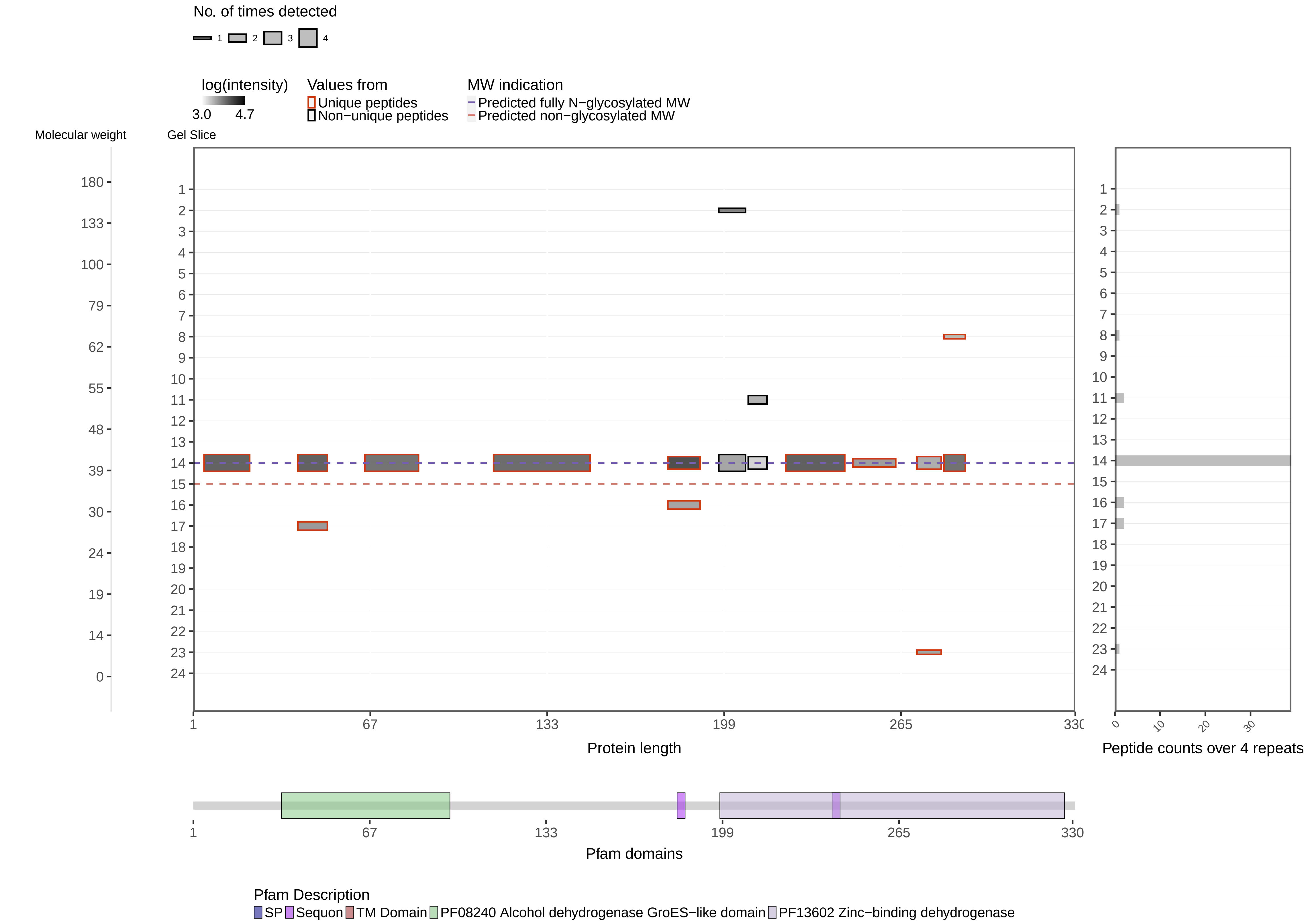


Protein sequence:

.DGSGPRKNILVTAASGGVGHYAVQLAKLGNTNVTATCGARNFDFVKSLGADEVLDYKTPEGAALKSPSGQKYDAVIHCTTGIPWSTFEPNLSAKGKVIDLTPGPSAMWTYAVKKLTFSKKQLVPLLLIPKKEDLDLVVRLVKEGKLKTVIDSKHPLSKAEDAWSRSIDGHATGKIIVEP*

NbL16g25880.1 peptidyl−prolyl_cis−trans_isomerase_FKBP15−1−like [MW = 21.89 kDa]

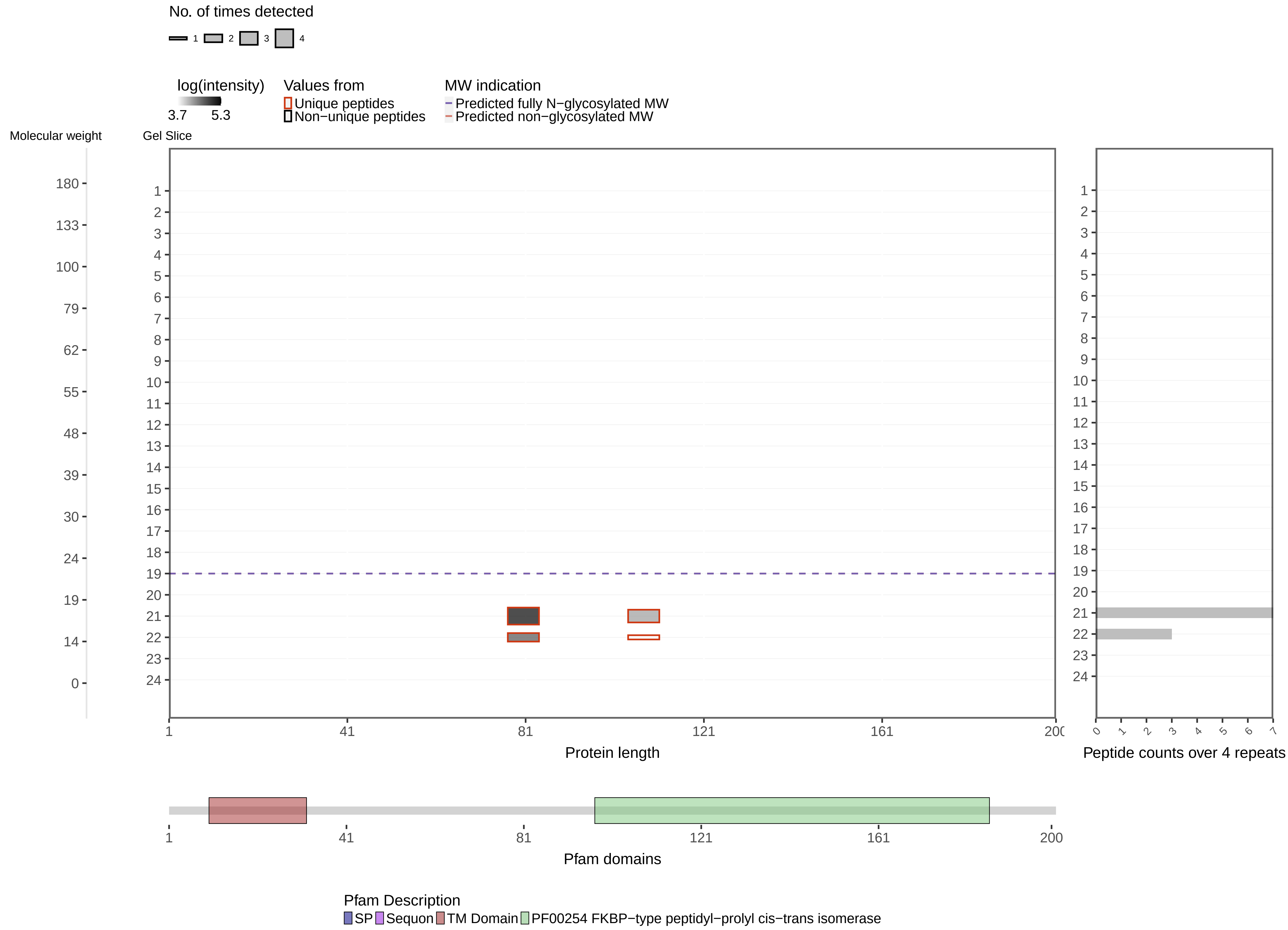


Protein sequence:

FKPKSCQLKAHKGDRVSVHYSGKLTDGTVFDSSYERNDPIEFELGSGQVIKGWDQGLLGMCVGEKRKLKIPAKLGYGESGSPPKIPGGATLVFDTELVAVNGKKSAADSEL*

### NbL16g25870.1 peptidyl−prolyl_cis−trans_isomerase_FKBP62−like [MW = 63.81 kDa]

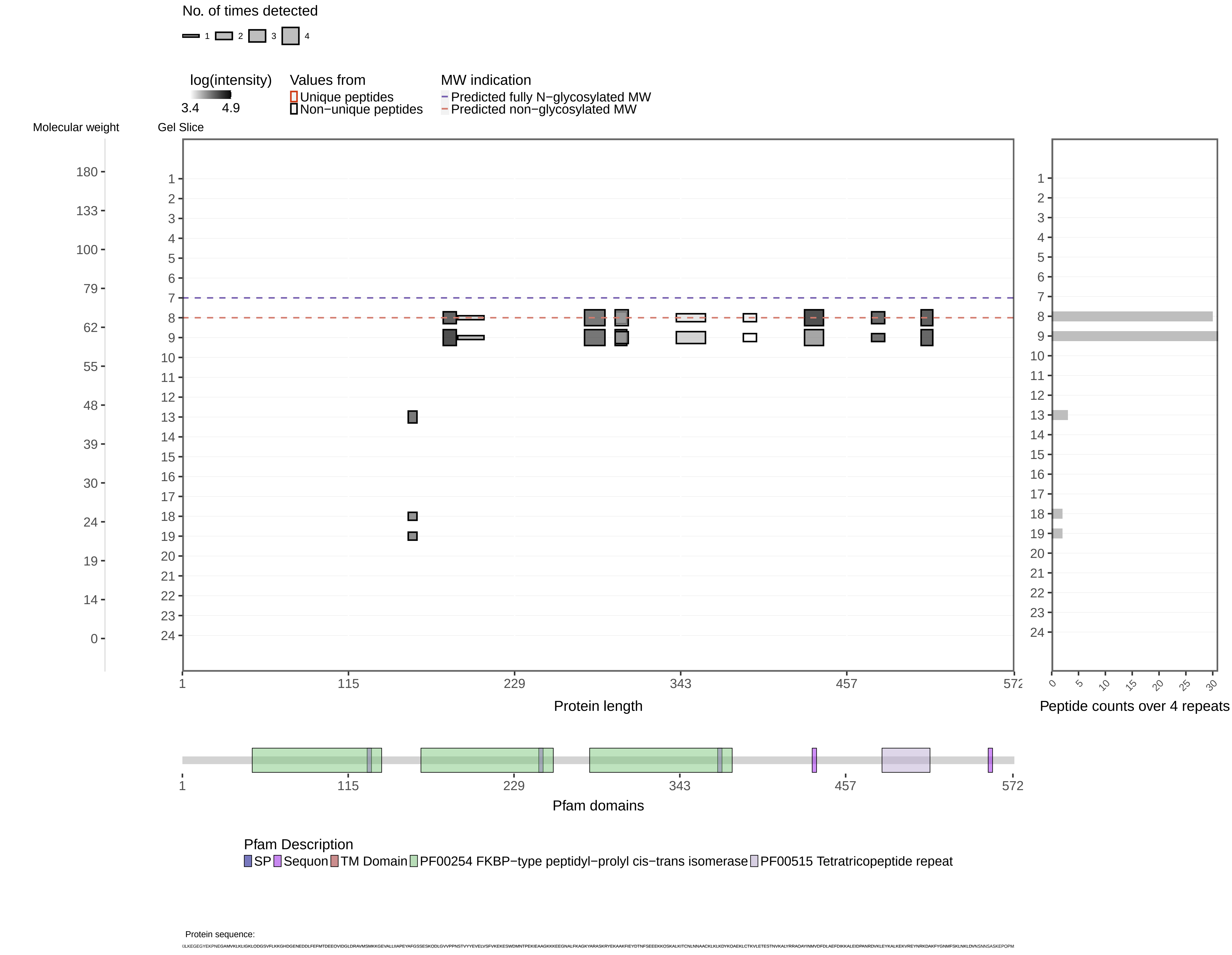

### NbL16g25810.1 adenylylsulfatase_HINT3 [MW = 22.35 kDa]

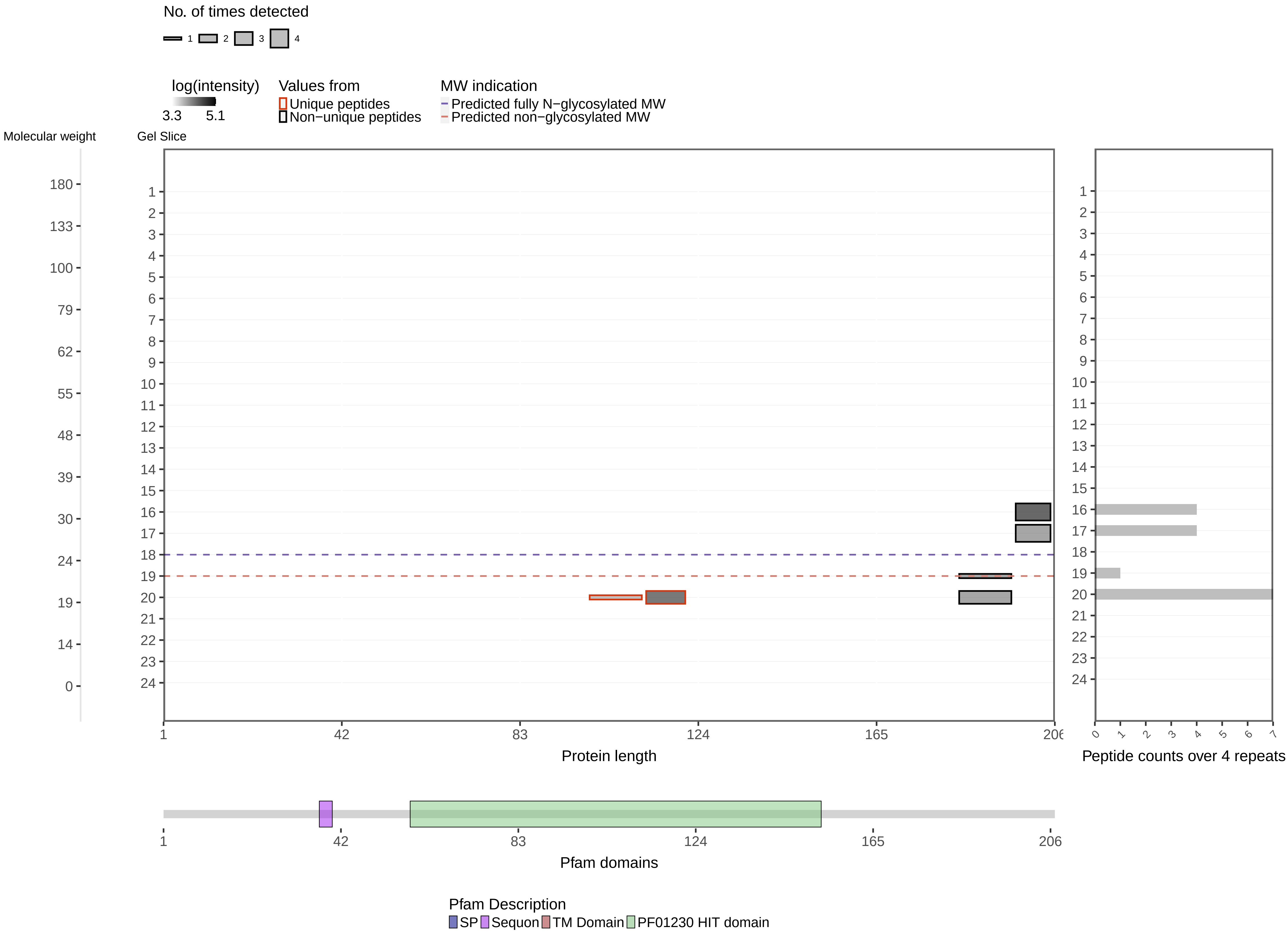


Protein sequence:

KCHFSSLKETPPSVVASMFSKVPAISSAVIKATGYDSFNLLVNNGEAAGQVVFHTHIHIIPRKASDCLWASESLRRRPLKLDQEAKKLGENIRENLPSLGNIEDSKGQVSDLIGN*

### NbL16g25790.1 cytochrome_b561_and_DOMON_domain−containing_protein_At3g25290−like [MW − SP = 23.39 kDa]

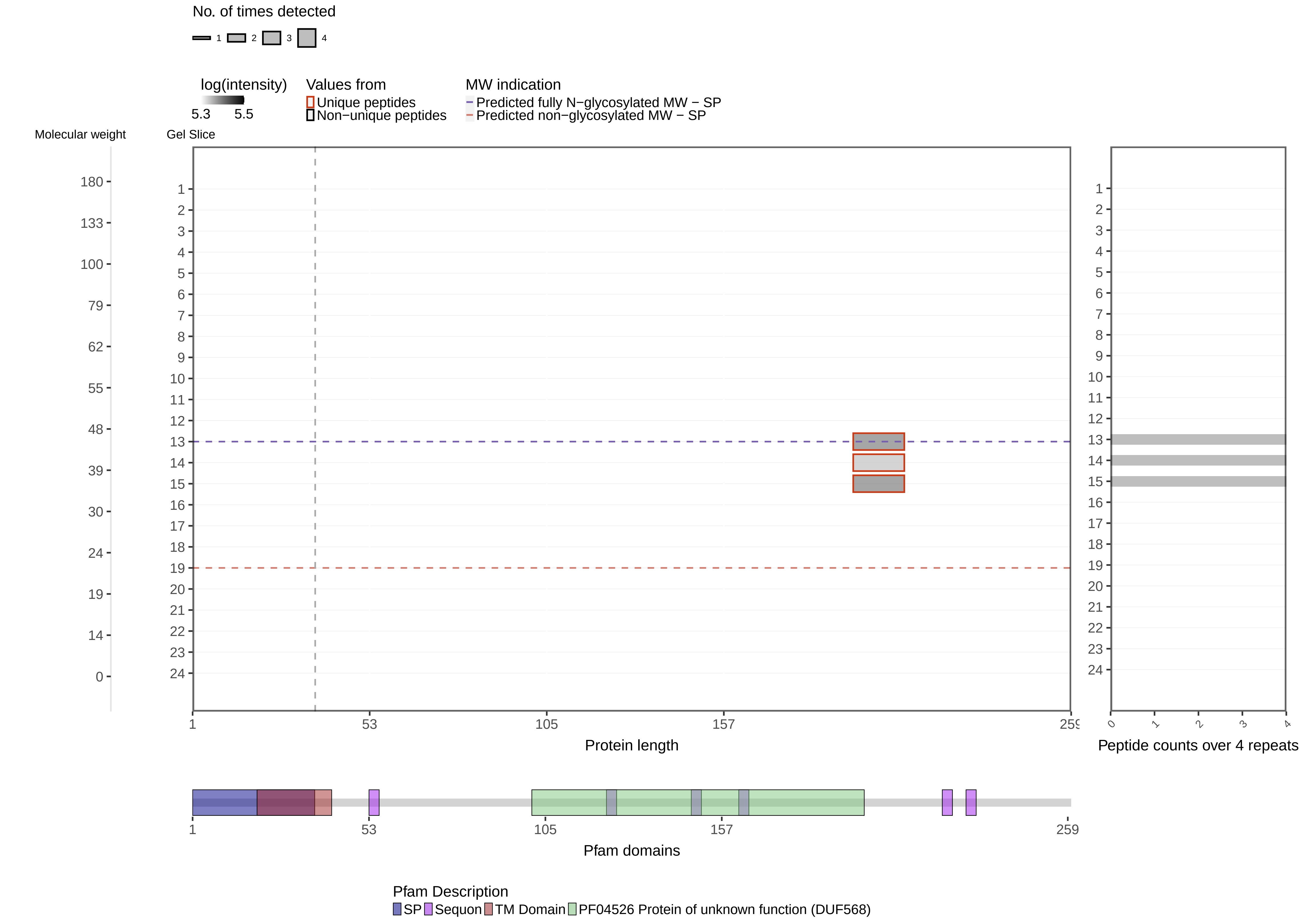


Protein sequence:

MVVKTYNLTSYKSITESKLLYNVLDSKAESSNGTMKIFATLELPENTTTVNQVWQVGPAVNDGKPVVHKFDTENLASKATLDLAATSSAGEGNKNAGSSASNSNSNSSSQSGNETGGSSRILKSENGIFAFLFFLGVLLLQL*

NbL16g25770.1 cytochrome_b561_and_DOMON_domain−containing_protein_At3g25290−like [MW − SP = 39.66 kDa]

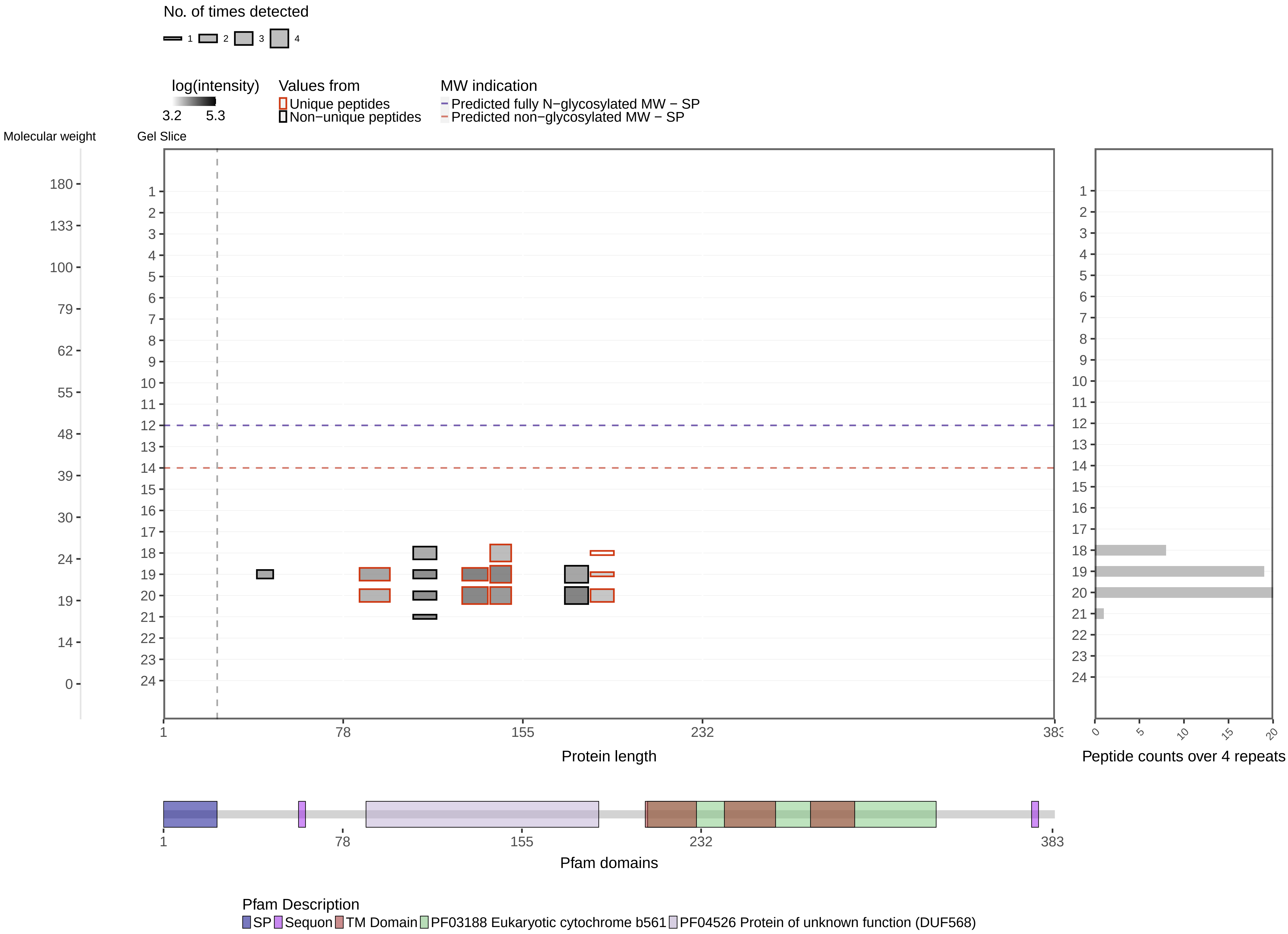


Protein sequence:

FQPANLNSKGTLDLVSGESKTGASGNSKVKRKNIHGILNAISWGILFPIGIIIARYMRTFKSADPAWFYLHVSCQCSAYIIGVAGWATGLKLGSESKGITYDTHRNIGIALFSFATLQVFALLLRPKKDNKYRFYWNIYHHGVGYTMLILSLINVFKGLDILDPATKWRSAYIGILVVLGAIAVLLEVITWIVVLKRKSNKSTKPYDG*

NbL16g25750.1 ras−related_protein_RABA5b−like [MW = 23.87 kDa]

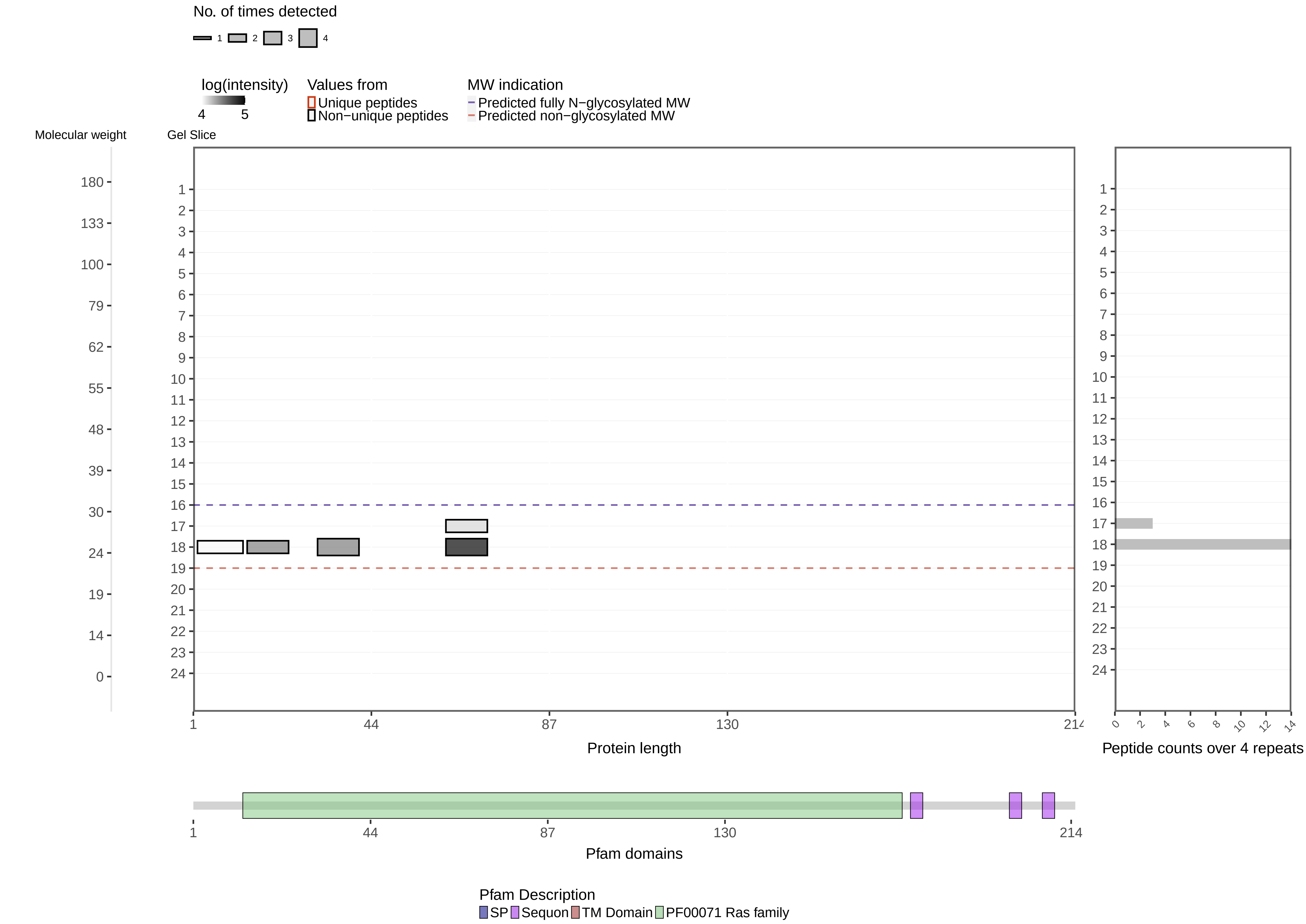


Protein sequence:

RKTTFENIKRWLDELNTHCDTTVARMLVGNKCDLENIRDVSVEEGKNLAEEEGLFFIETSALDSTNVNTAFEVVIREIYKNLSRKVLNSDSYKAELSVNRVSLANGTDMSKQNASCCSR*

### NbL16g25740.1 cysteine−rich_repeat_secretory_protein_55−like [MW − SP = 27.18 kDa]

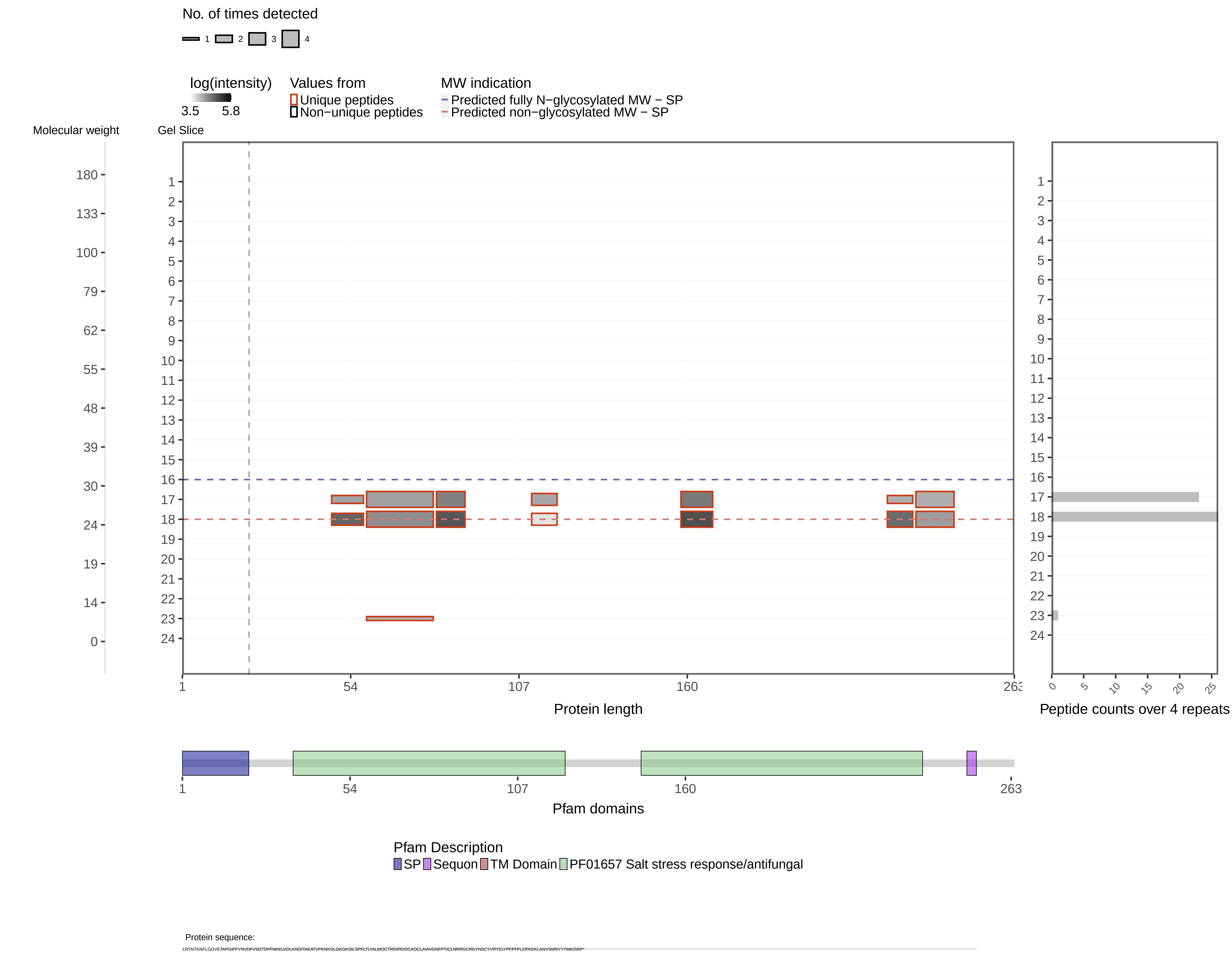

### NbL16g25690.1 putative_phosphatidylglycerol/phosphatidylinositol_transfer_protein_DDB_G0282179 [MW − SP = 15.84 kDa]

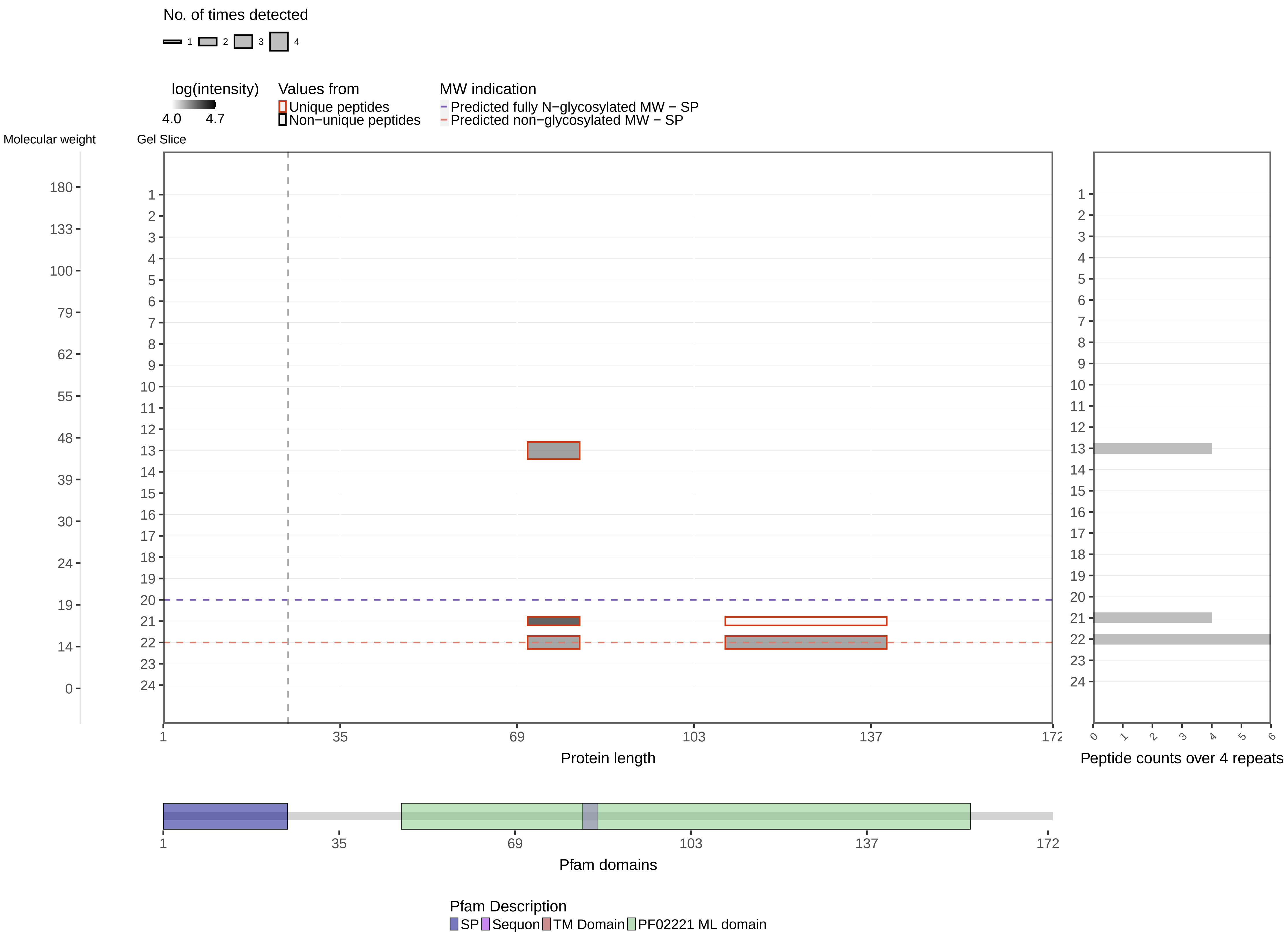


Protein sequence:

FSISASTGKNLTGGKLVIDVNYLFFHVHHEAIDLCKETSCPASGDFLISHSQELPGFTPPGSYTLTMKMVDDKNQQLSCISFGFSIGFIAESQALAADS*

NbL16g25650.1 33_kDa_ribonucleoprotein_chloroplastic [MW = 35.4 kDa]

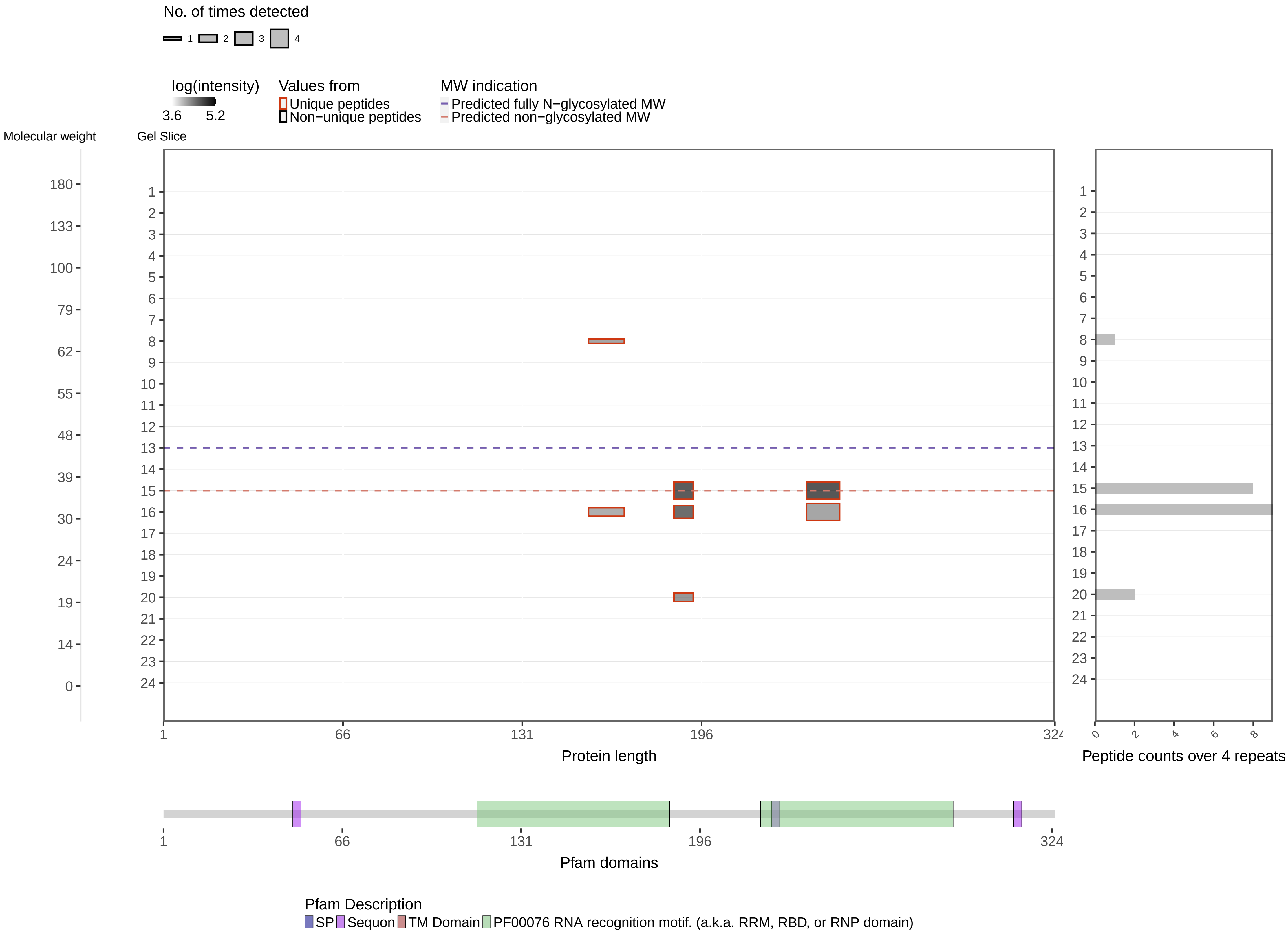


Protein sequence:

SRGFAFVTMGSVEEAREAIRLFDGSQVGGRTVKVNFPEVPRGGEREVMSAKIRSTYQGFVDSPHKLYVANLSWALTSQGLRDAFADQPGFMSAKVIYDRSSGRSRGFGFITFSSAEAMESALDTMNEVELEGRPLRLNVAGQKAPLSSPPVVETSPENDSENSELLSSLSS*

### NbL16g25540.1 epoxide_hydrolase_3 [MW = 44.25 kDa]

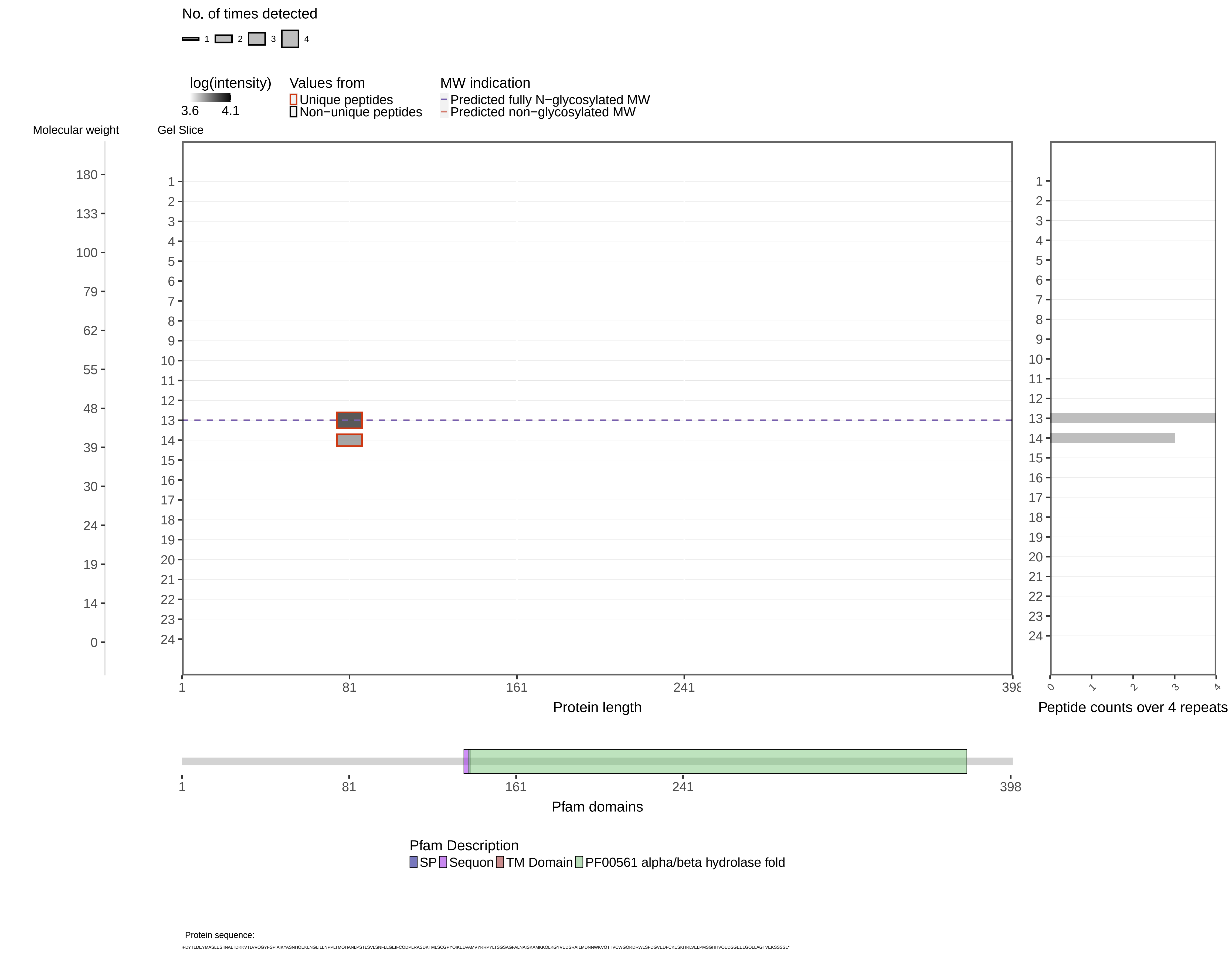

### NbL16g25440.1 monocopper_oxidase−like_protein_SKU5 [MW − SP = 64.22 kDa]

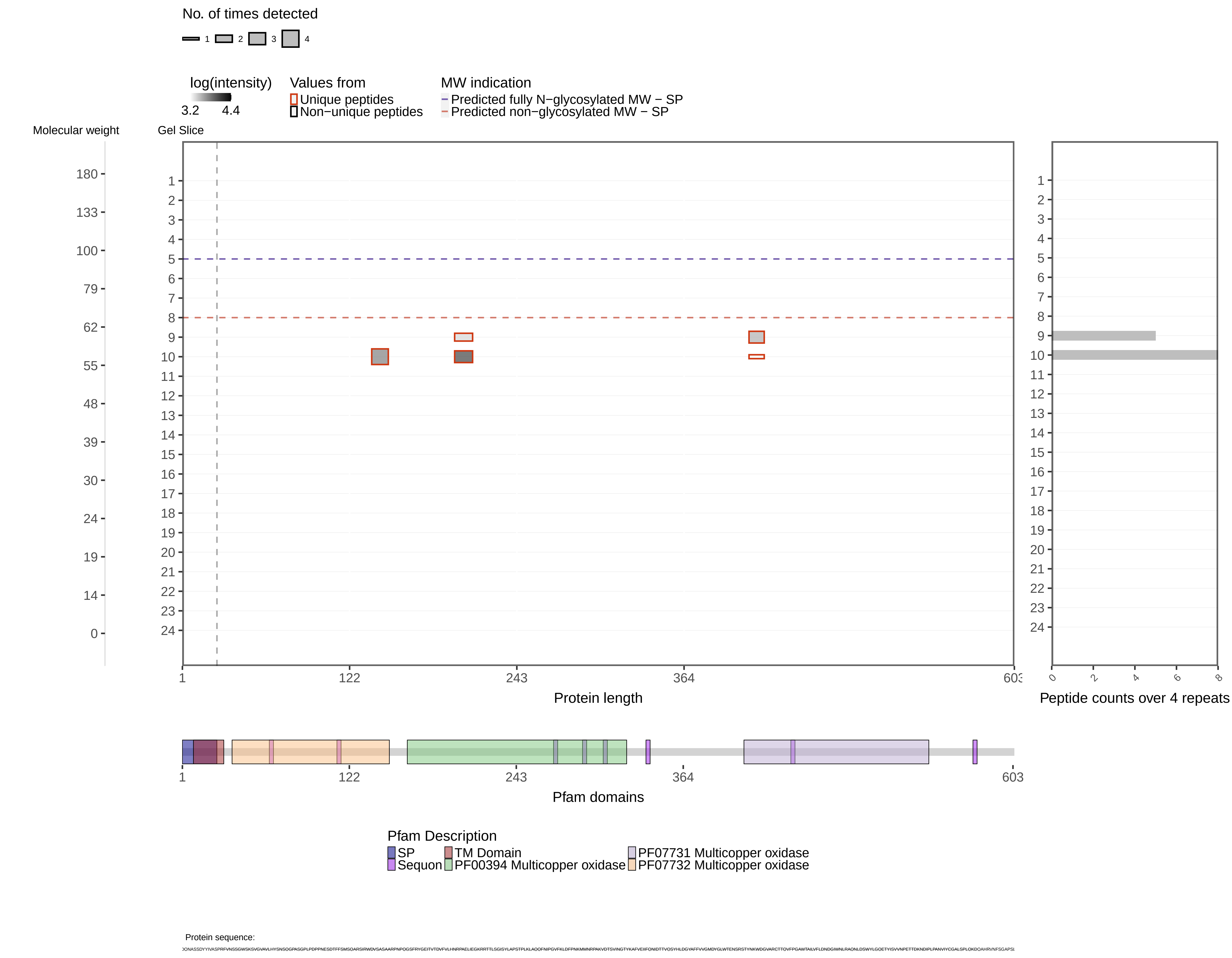

### NbL16g25360.1 uncharacterized_protein_At5g48480−like [MW = 16.93 kDa]

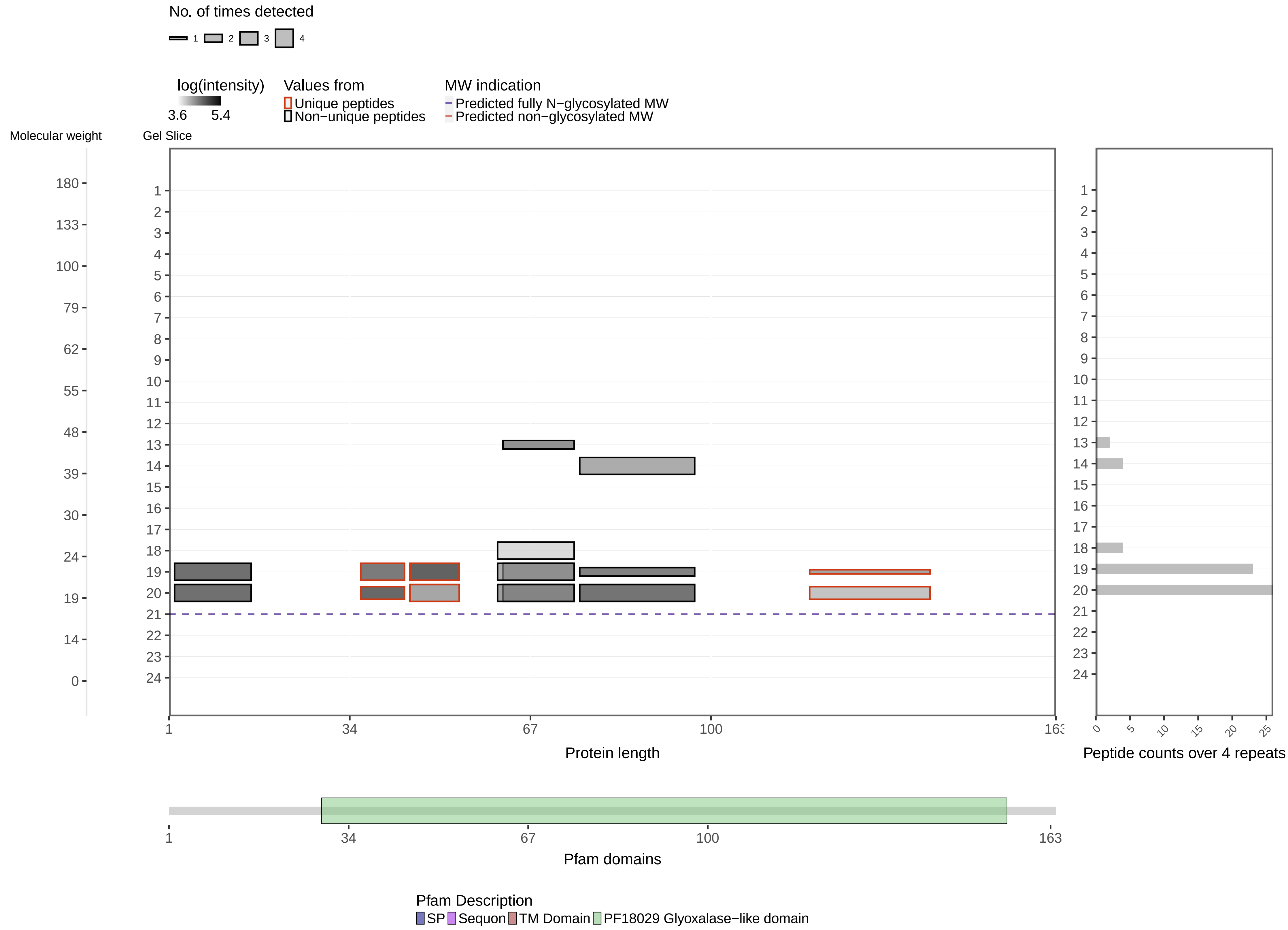


Protein sequence:

LILSAELKLGSTTFLVSDSDLTDDSSAPVKTASGCIFCLETEDVEGAVAKAVTAGAVSEGEIAEGDGACYGGRVGKLKDPYGNVWLVCSPSKKYE*

NbL16g25210.1 ubiquitin_carboxyl−terminal_hydrolase_12−like [MW = 83.99 kDa]

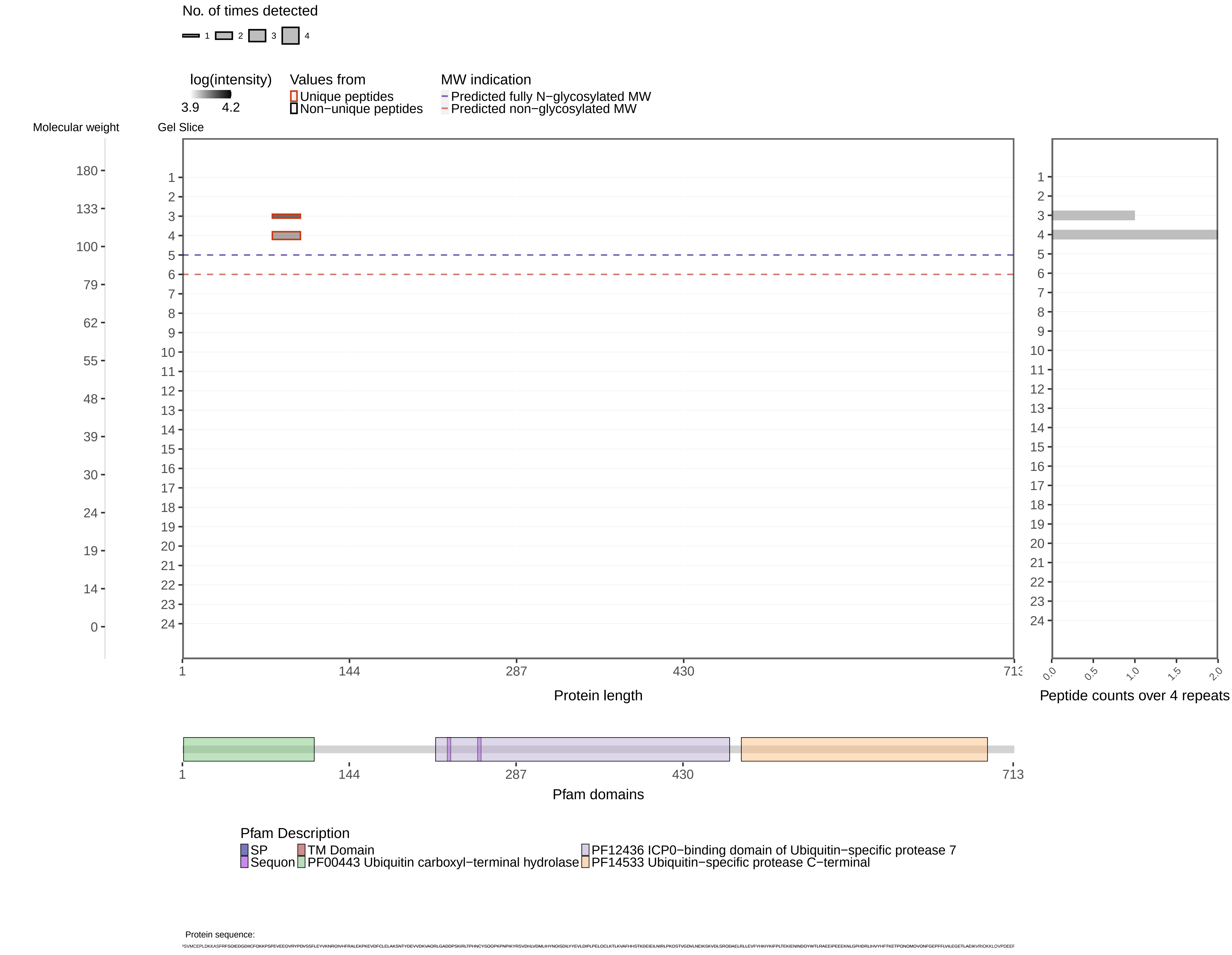

### NbL16g25200.1 ubiquitin_carboxyl−terminal_hydrolase_12−like [MW = 43.72 kDa]

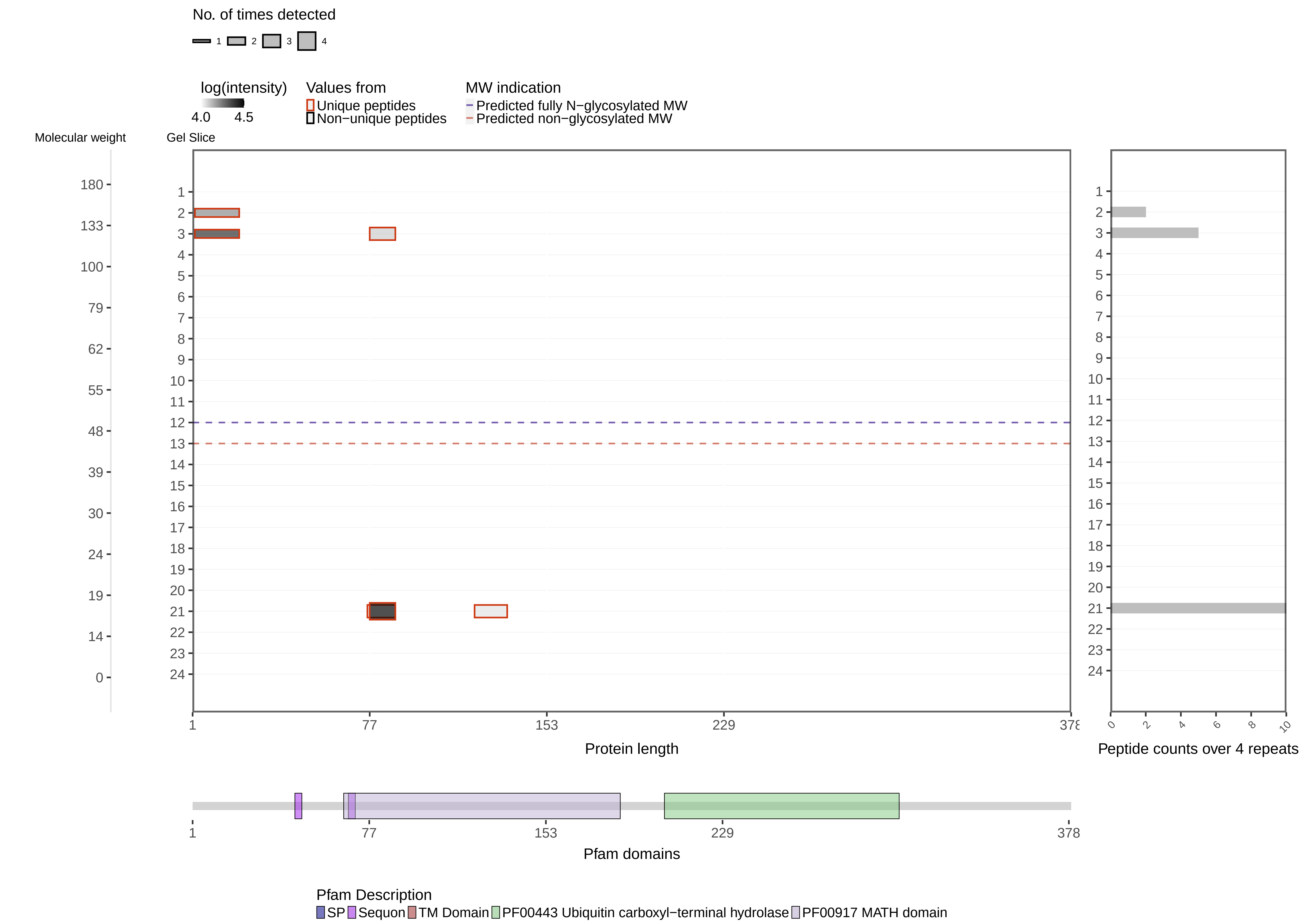


Protein sequence:

DRVVIEADVAVRKVIDYWTYDSKKETGYVGLKNQGATCYMNSLLQTQYHIPYFRKAVYHMPTTENDMPSGSIPLALQSLFYKLQYSDKSVATKELTKSFGWDTYDSFMQHDVQELNRVLCEKLEDKMKGTVVEGTIQKLFEGHHMNYIECINVDFKSTRKESFYDLQLDVKGCRDVYASFDKYVEVERLEGDNKYHAEEHGLQ

NbL16g25160.1 D−lactate_dehydrogenase_[cytochrome]_mitochondrial [MW = 62.77 kDa]

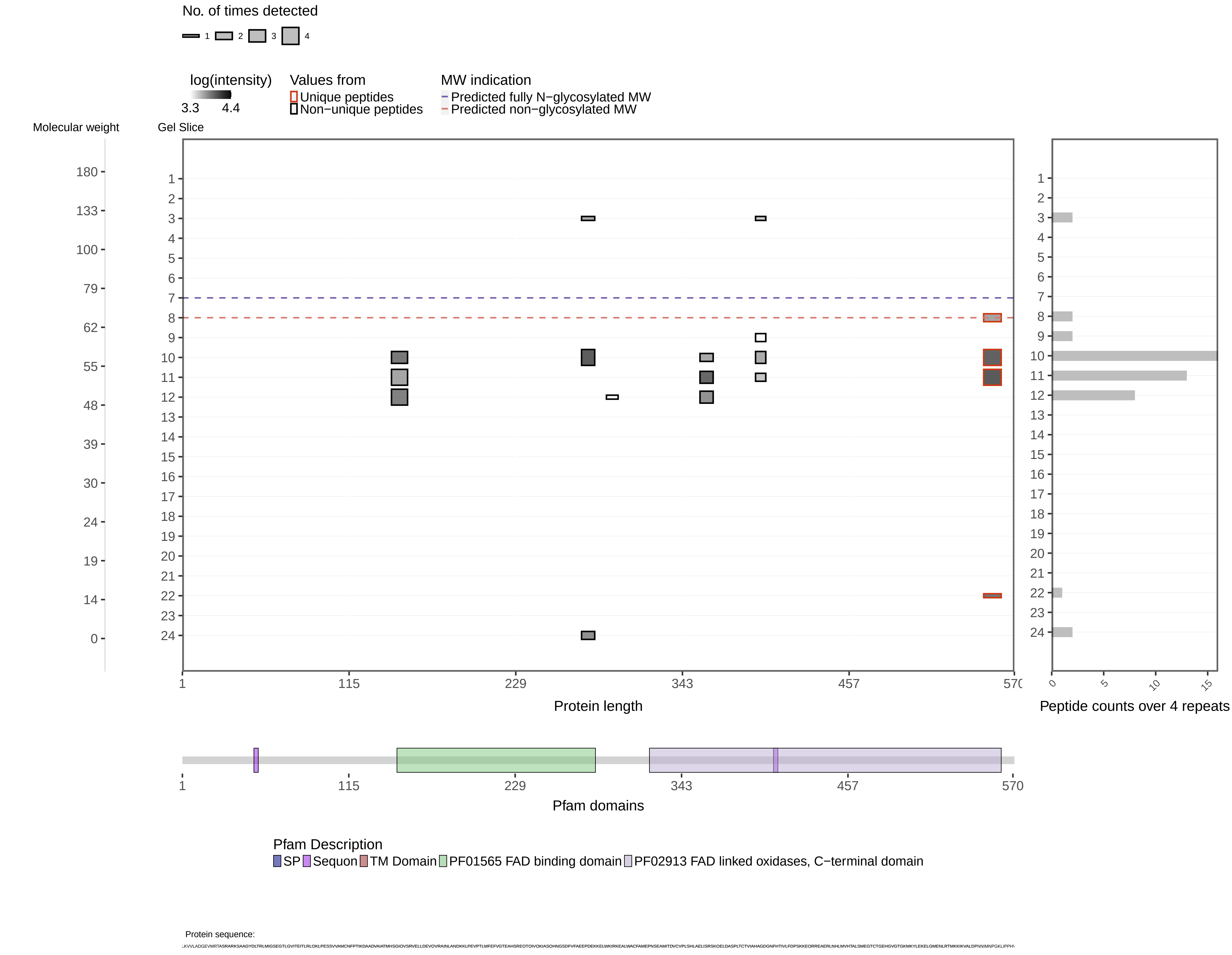

NbL16g25090.1 sucrose−phosphatase_1 [MW = 52.67 kDa]

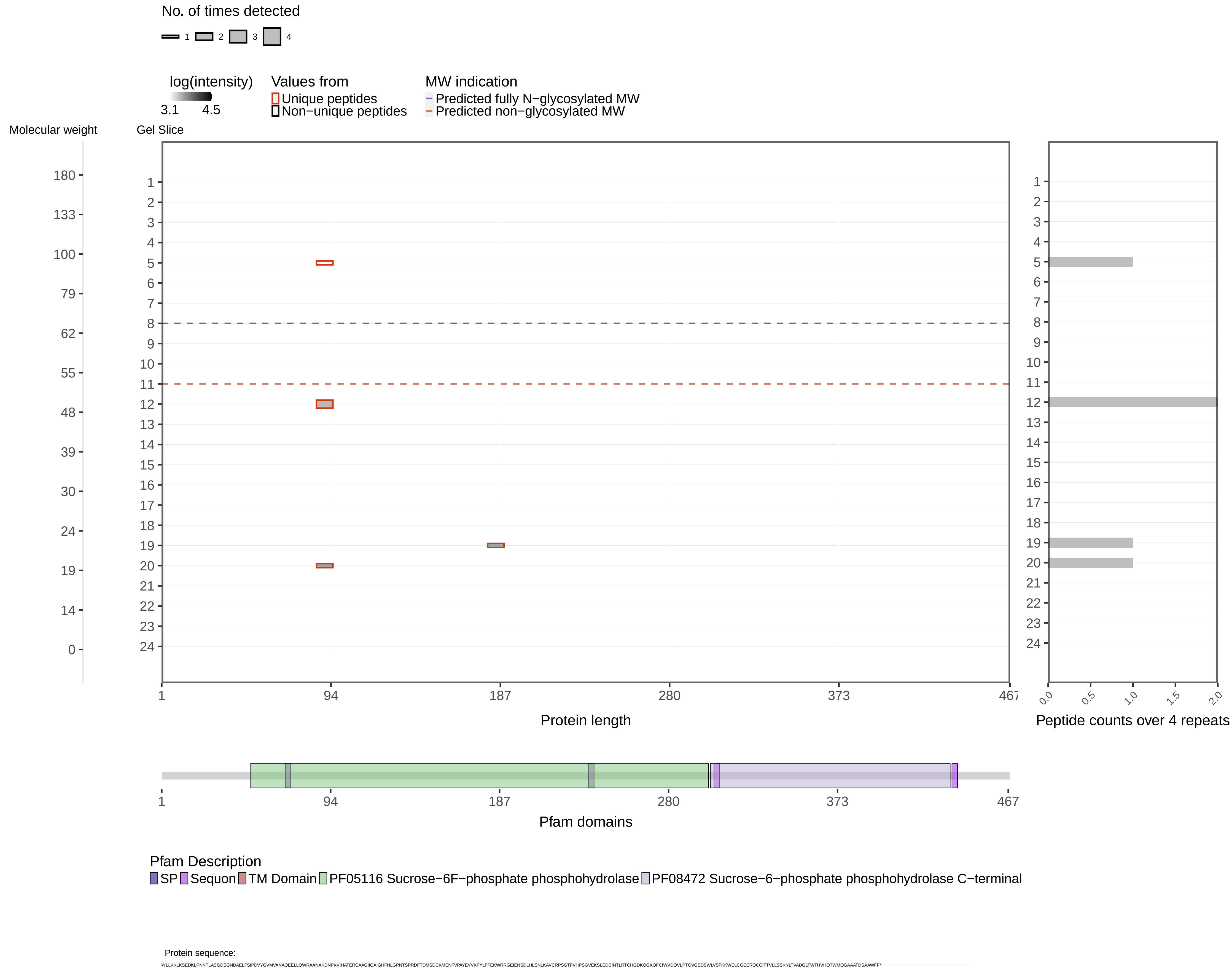

NbL16g25060.1 methionine_aminopeptidase_2B−like [MW = 48.7 kDa]

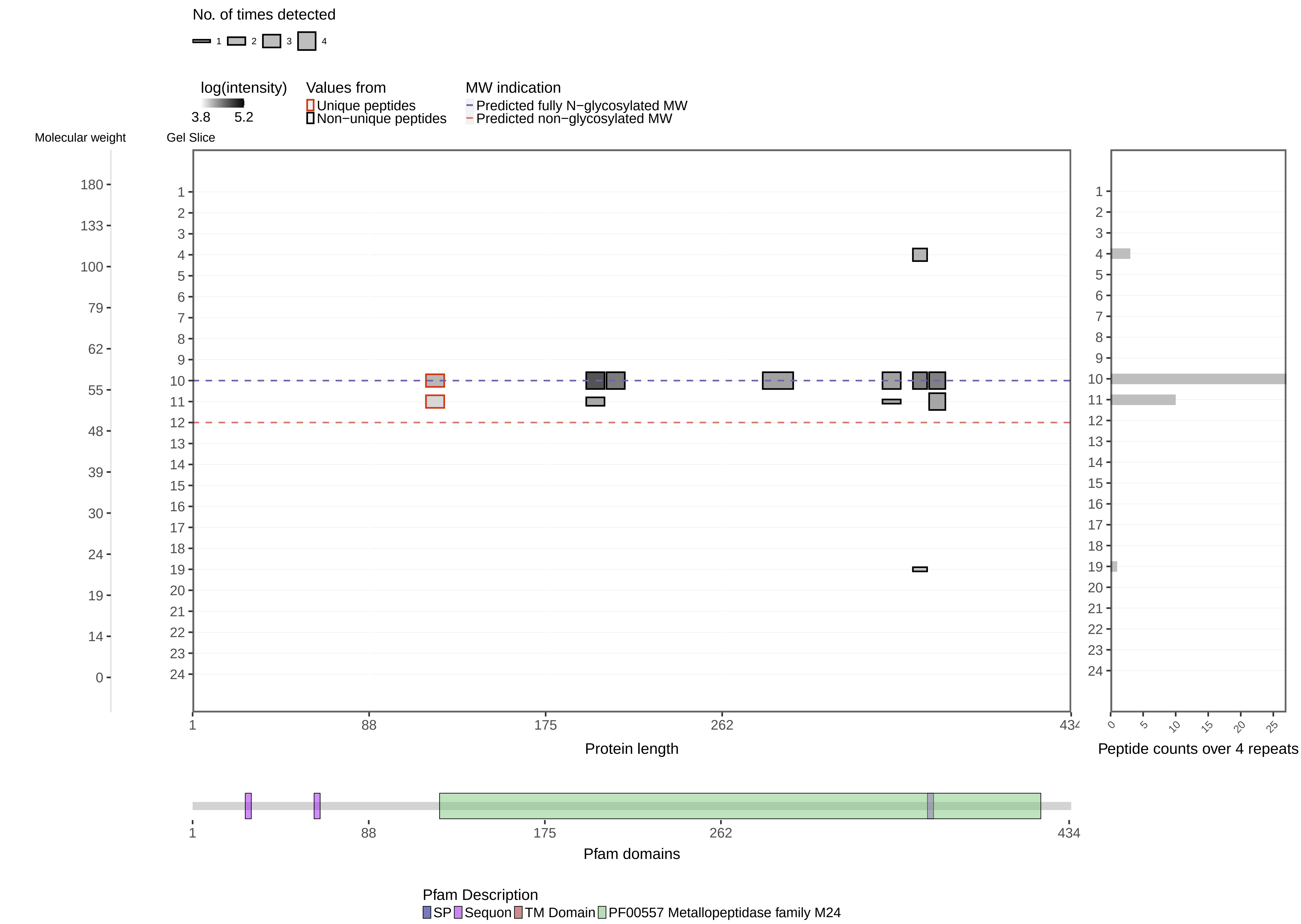


Protein sequence:

[illegible]VMKLDFGTHIDGRIVDCAFTVAFNPMFDPLLEASREATNTGIKEAGIDVRLCDVGAAIOEVMESYEVEINGKVFOVKSIRNLNGHSIGOYOIHAGKSVPIVKGGEOTKMEEGEFYAIETFGSTGKGYVREDLECSHYMKNFDIGHIPLRLPRAKOLLATINKNFSTLAFCRRYLDRLGETKYLMALKNLCDAGIVOPYPPLCDIKGSYVSOFEHTILLRPTCKEVVSRGDDY*

NbL16g25050.1 fasciclin−like_arabinogalactan_protein_17 [MW − SP = 47.11 kDa]

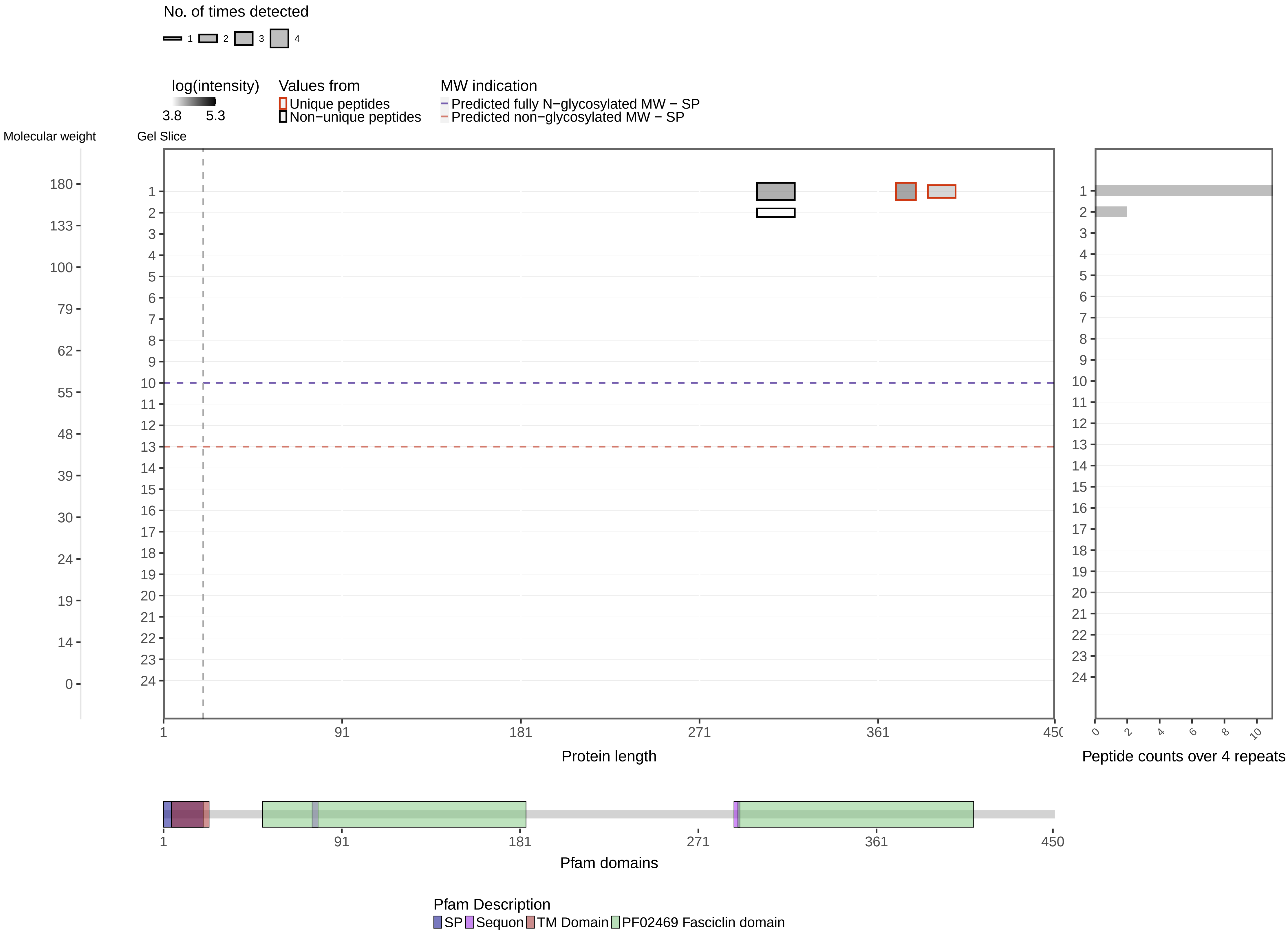


Protein sequence:

TNRLKKKPTILVPAGAPPVLPIYSAMAPGPSLAPAPAPGPGGPHHHFDGESQVKDFIQTLLHYGGYNEFADILVNLTSLATEMGRLVSEGYVLTVLAPNDEAMAKLTTDQLSEPGAPEQIMYYHLIPEYQTEESMYNAVRRFGKVKYDTLRLPHKVVAQEADGSVKFGVAKGSAYLFDPDIYTDGRISVQGIDGVLFPVEEIKAAPVAAPVAKVVAKPRRGKLMEVACSVLGSFC*

### NbL16g25020.1 iron−sulfur_cluster_co−chaperone_protein_HscB_mitochondrial [MW = 30.39 kDa]

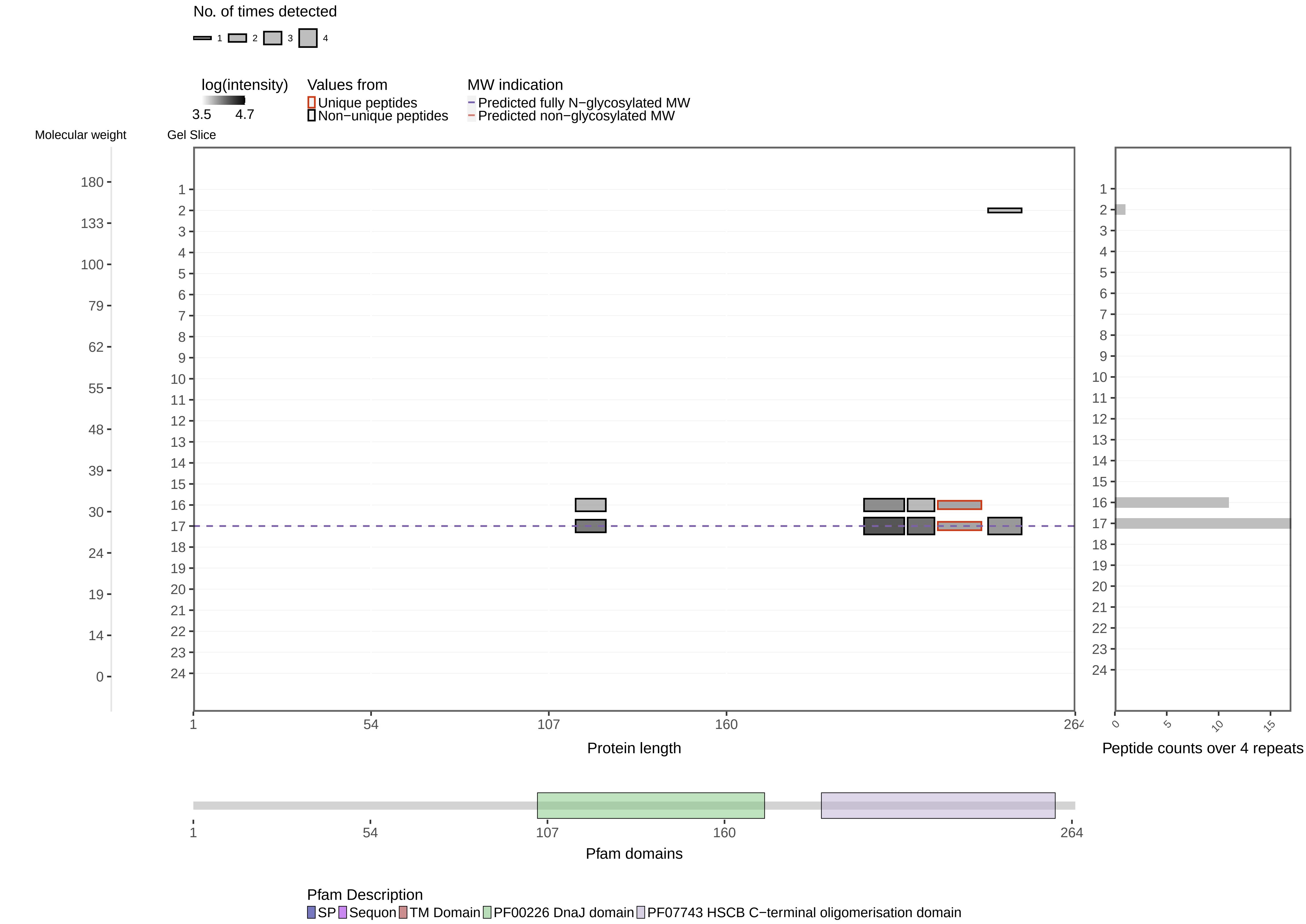


Protein sequence:

EGENLERKYKDWQRKLHPDLVHTKSEKEREYAAEQSARVIDAYRTLSDPLSRAIYILKLEGVHVDEEERIDDLELLAEIMELRETVEEAANAQALKQIQGEIQGKFEQSSISFANALQKRKYEEAVAATQRMTYYKRANEEIVKKL*

### NbL16g24920.1 5−methyltetrahydropteroyltriglutamate−−homocysteine_methyltransferase [MW = 84.58 kDa]

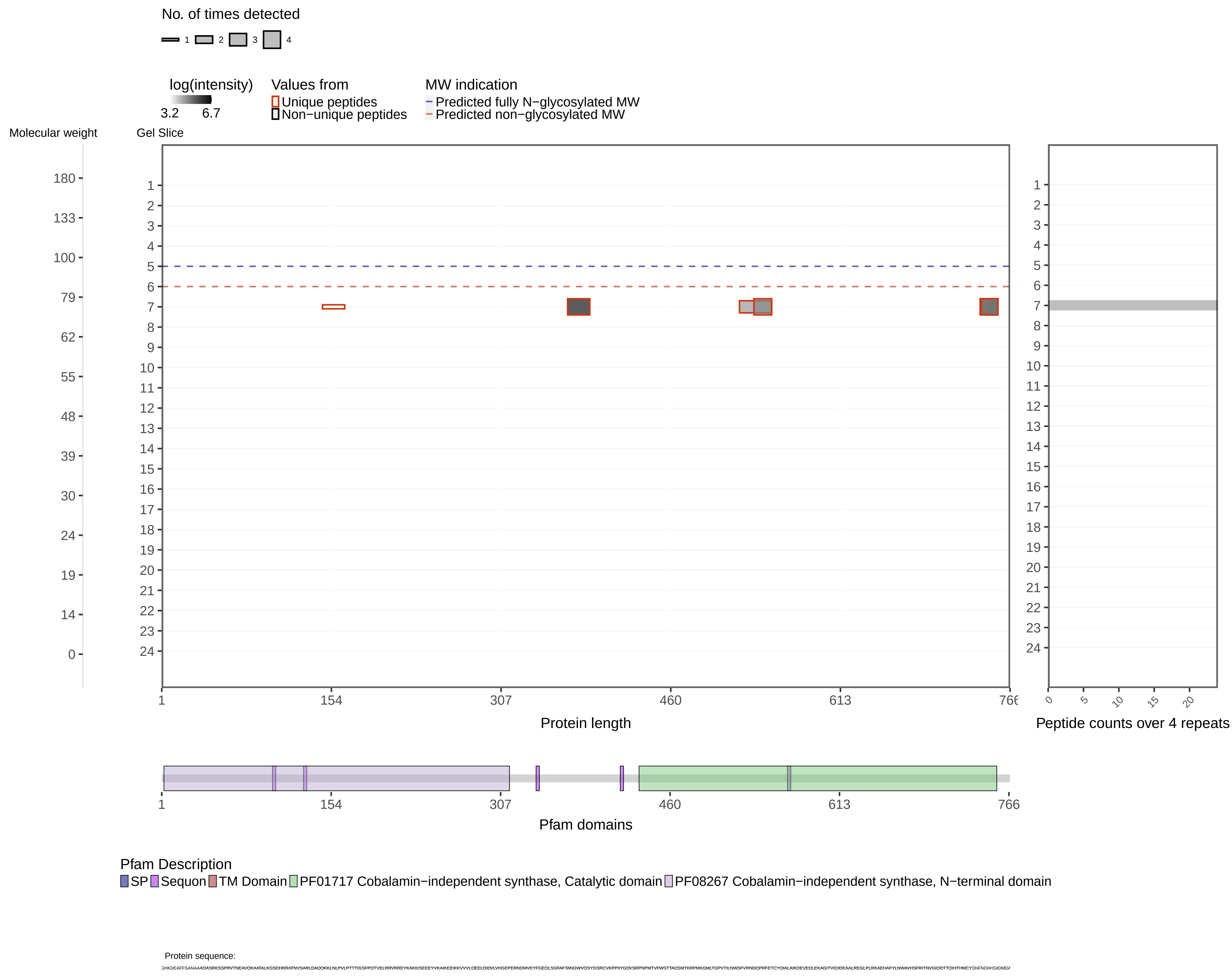

NbL16g24840.1 NADH−−cytochrome_b5_reductase_1−like [MW = 31.08 kDa]

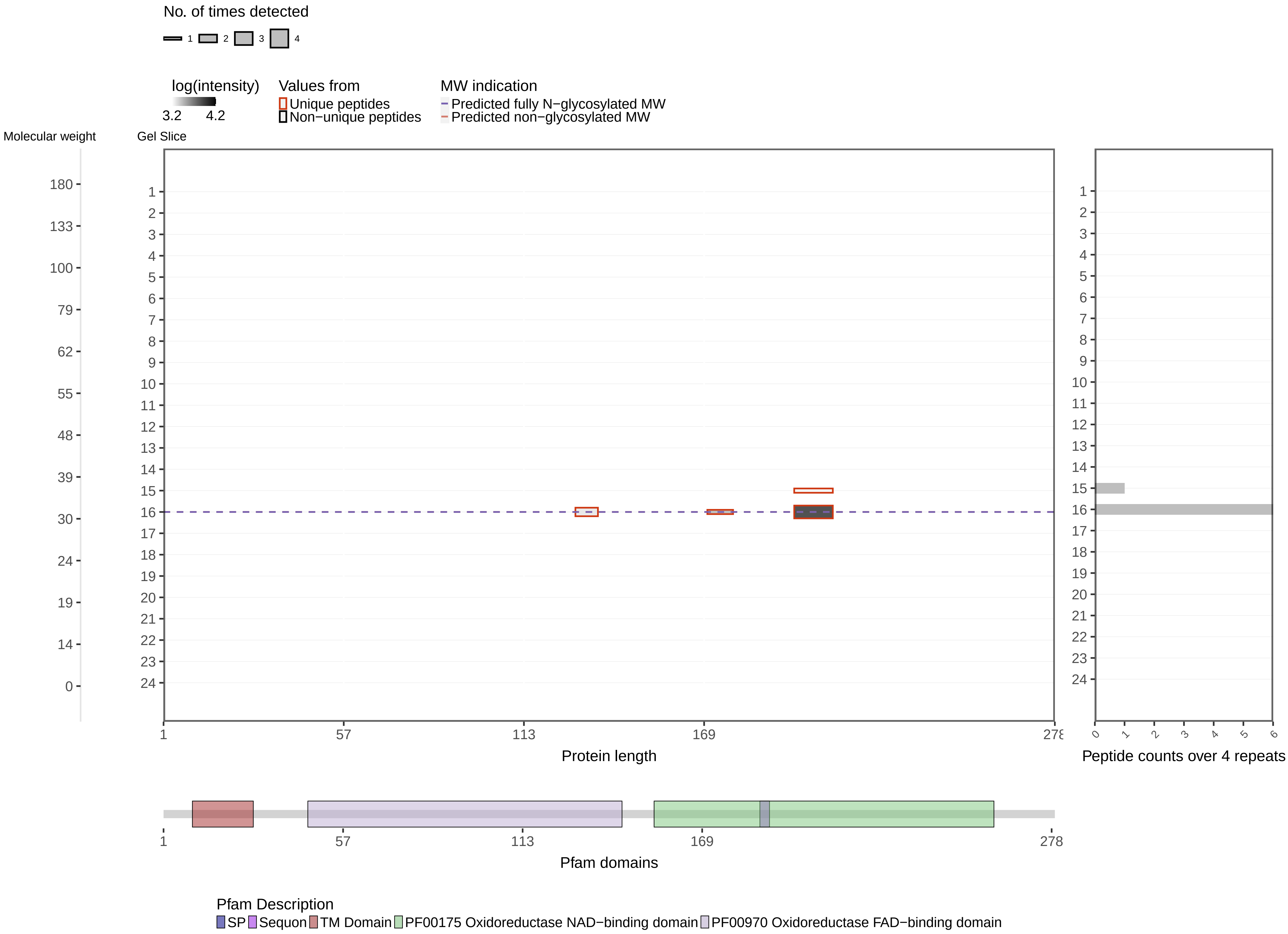


Protein sequence:

REGDYMAVKGPKGRFKYQPGQVRAFGMLAGGSGITPMFQVARAILENPNDKTKVHLIYANVTYEDILLKEQLDGLAANYPDRFKIYYVLNQPPEVWSGGVGFVSKEMIQNHCPAPASDIQILRCGPPPMNKAMAAHLEALGYTPEMQFQF*

NbL16g24700.1 4−alpha−glucanotransferase_DPE2 [MW = 81.21 kDa]

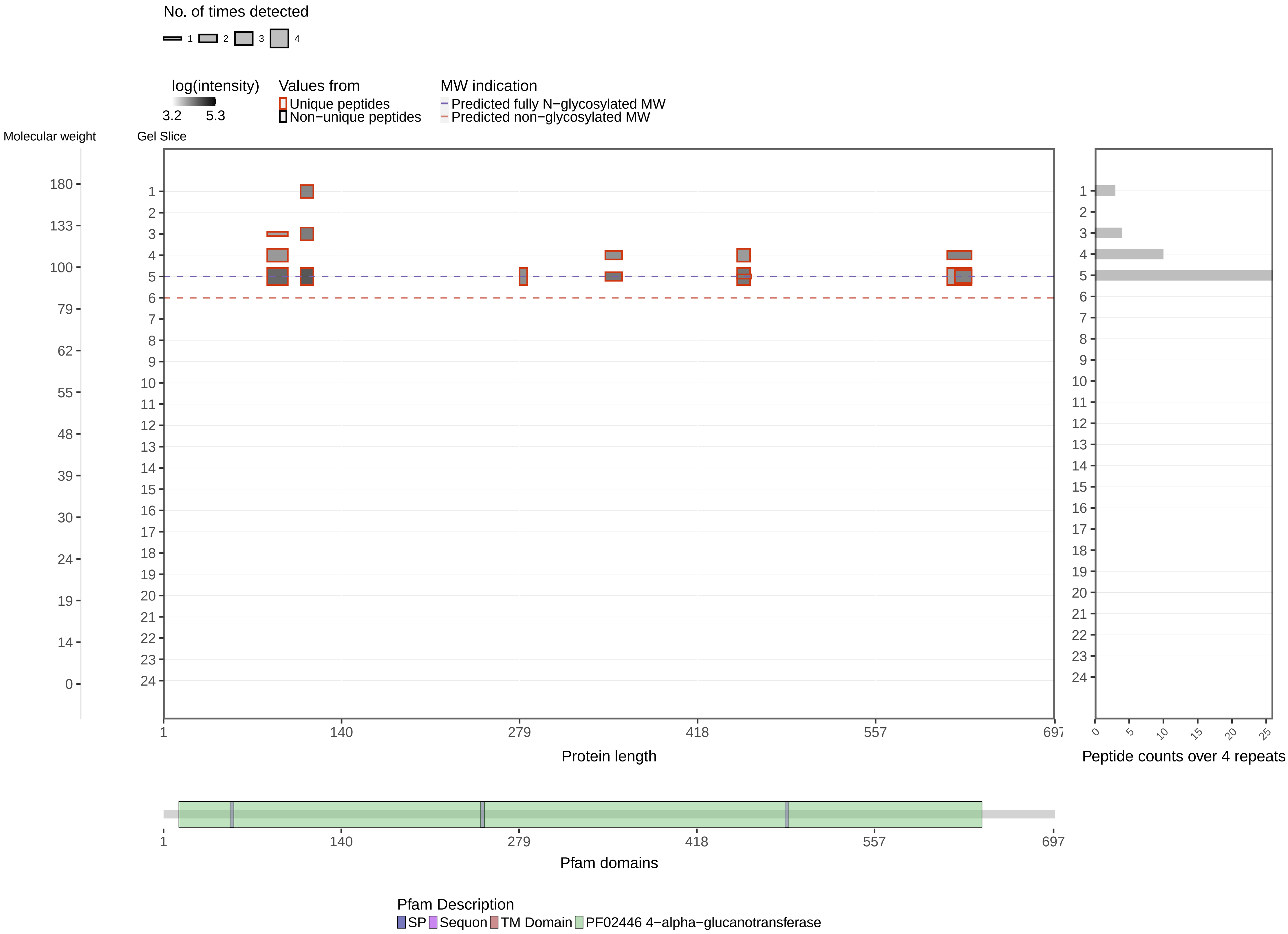


Protein sequence:

ESDGLWDFNRLTQPYIGHELLQEKFGASWTIIASTFLNEYRKGLYEFKEECNTEKKIASVLKSFLETSMLVESEEKLRRKLFDLLQNVALIKDPEDLRKFYPRFNIEDTTSFKDLDHHSQNVLKRLYYDYYFQRQEGLWRQNALKTLPVLLNSSNMLACGEDLGLIPSCVHPVMQELGLVGLRIQRMPSEPGLEFGIPSQYSYMTVCAPSCHDCSTLRAWWEEDEERRHRLFQAVVGSDELPPDQCTPEIVHFVLRQHVEAPSMWSIFPLQDLLALKEDYTTRPAVEETINDP

NbL16g24500.1 oxygen−dependent_coproporphyrinogen−III_oxidase_chloroplastic [MW = 34.51 kDa]

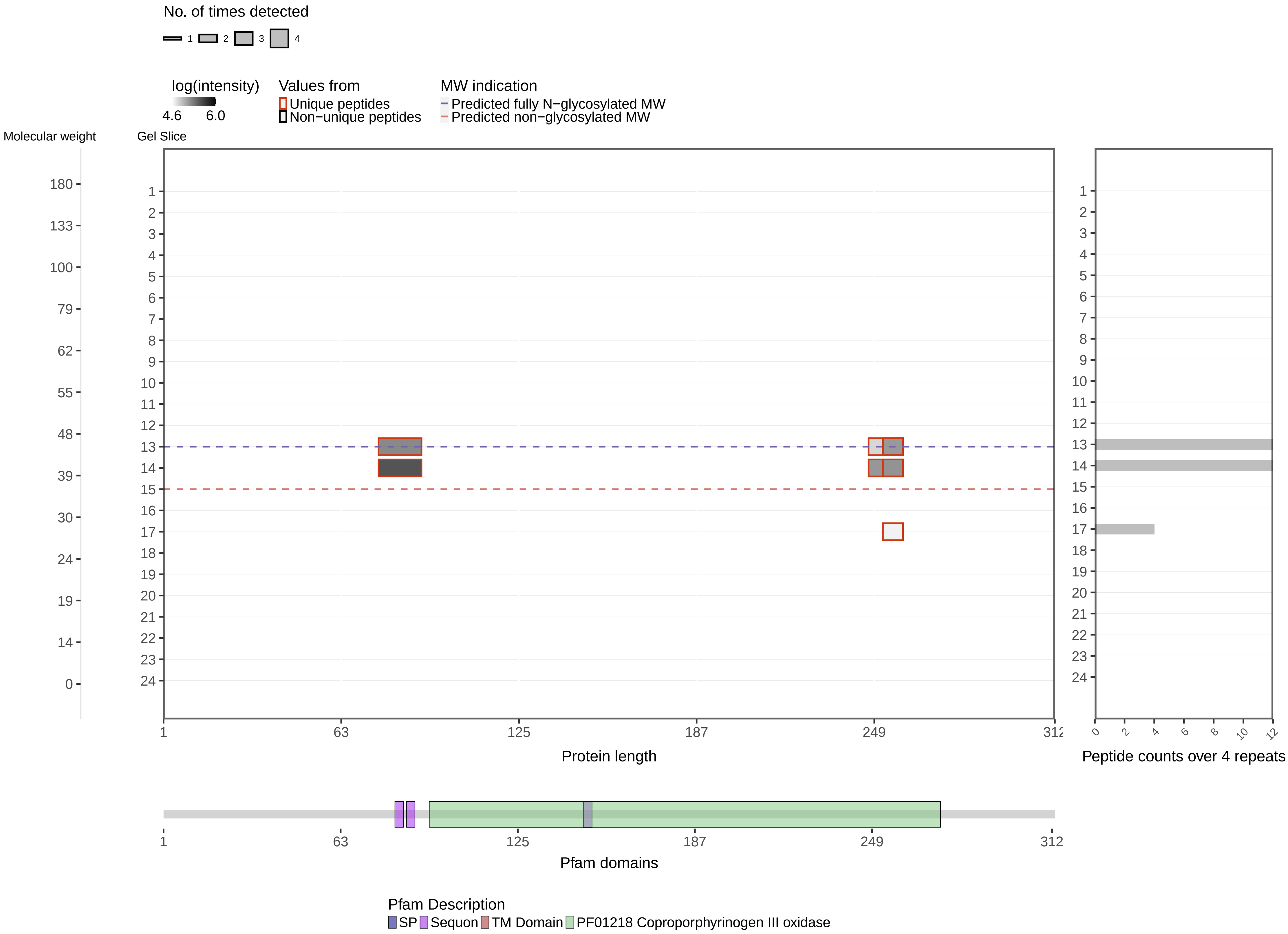


Protein sequence:

KAGVNVSVVYGVMPPEAYRAARPTDNGNVKPGPIPFFAAGVSSVLHPKNPFAPTLHFNYRYFETDAPKDAPGAPRQWWFGGGTDFTPAYIFEEDVKHFHSVQKAACDKFDASFYPRFKKWCDDYFYIKVKASVPLETCFCWLLELAIESSKILAFLLKYLLGLFTSHP*

### NbL16g24490.1 alpha−ketoglutarate−dependent_dioxygenase_alkB_homolog_6 [MW = 28.34 kDa]

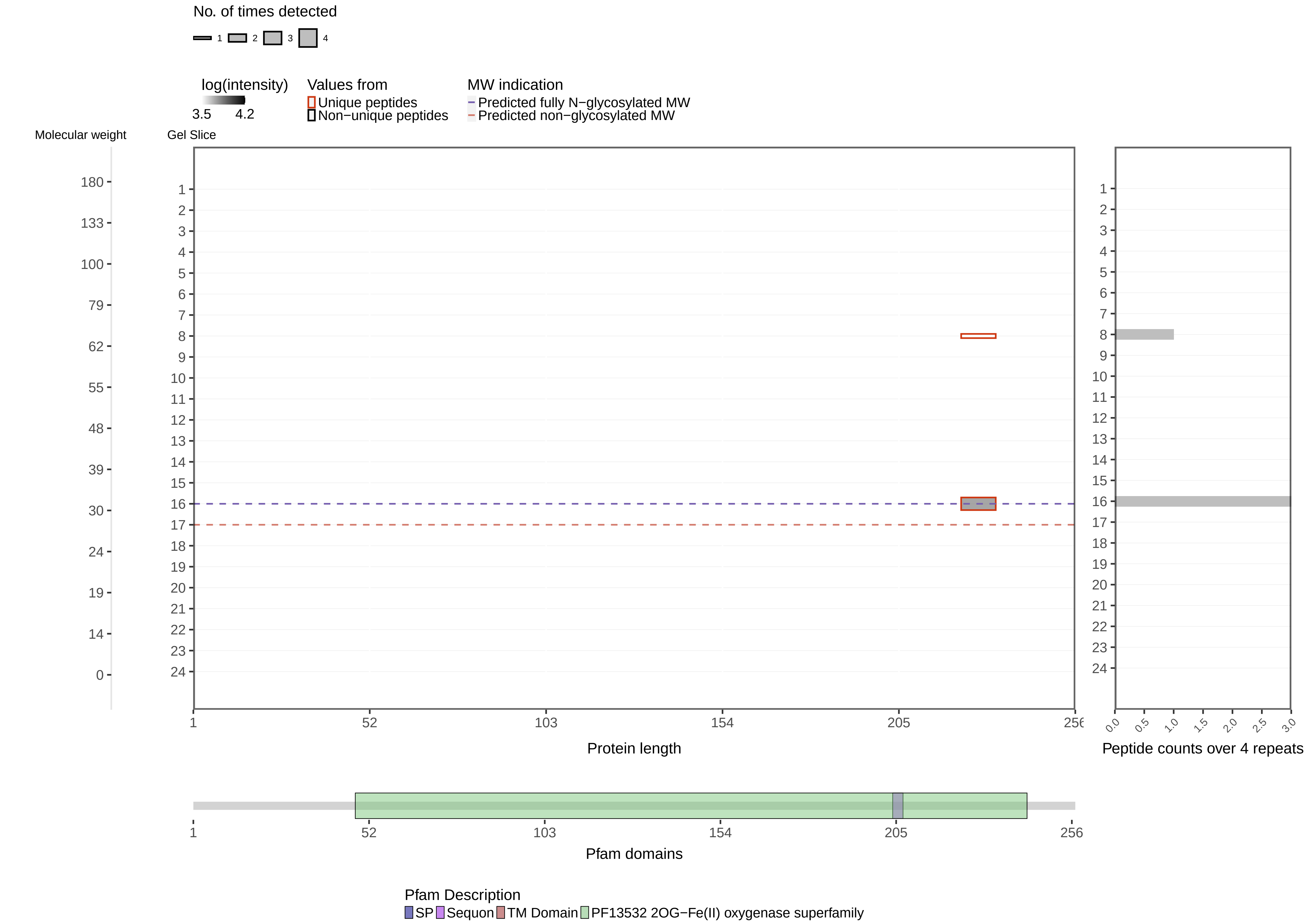


Protein sequence:

LSLGSPVVMDFTPRPNLRSCVGTHGNGVDEKISDPEAAVMNSGEQLDNCLPFSIVLMPRSLLIFKDLVYSDYLHGIKDSEVQRCNEAVNLTNAQNDGVVQHLSGSVKALDSDETVIYRANTRVSLTCRVVTKVCKNIFKL*

### NbL16g24470.1 transcription_factor_Pur−alpha_1−like [MW = 31.78 kDa]

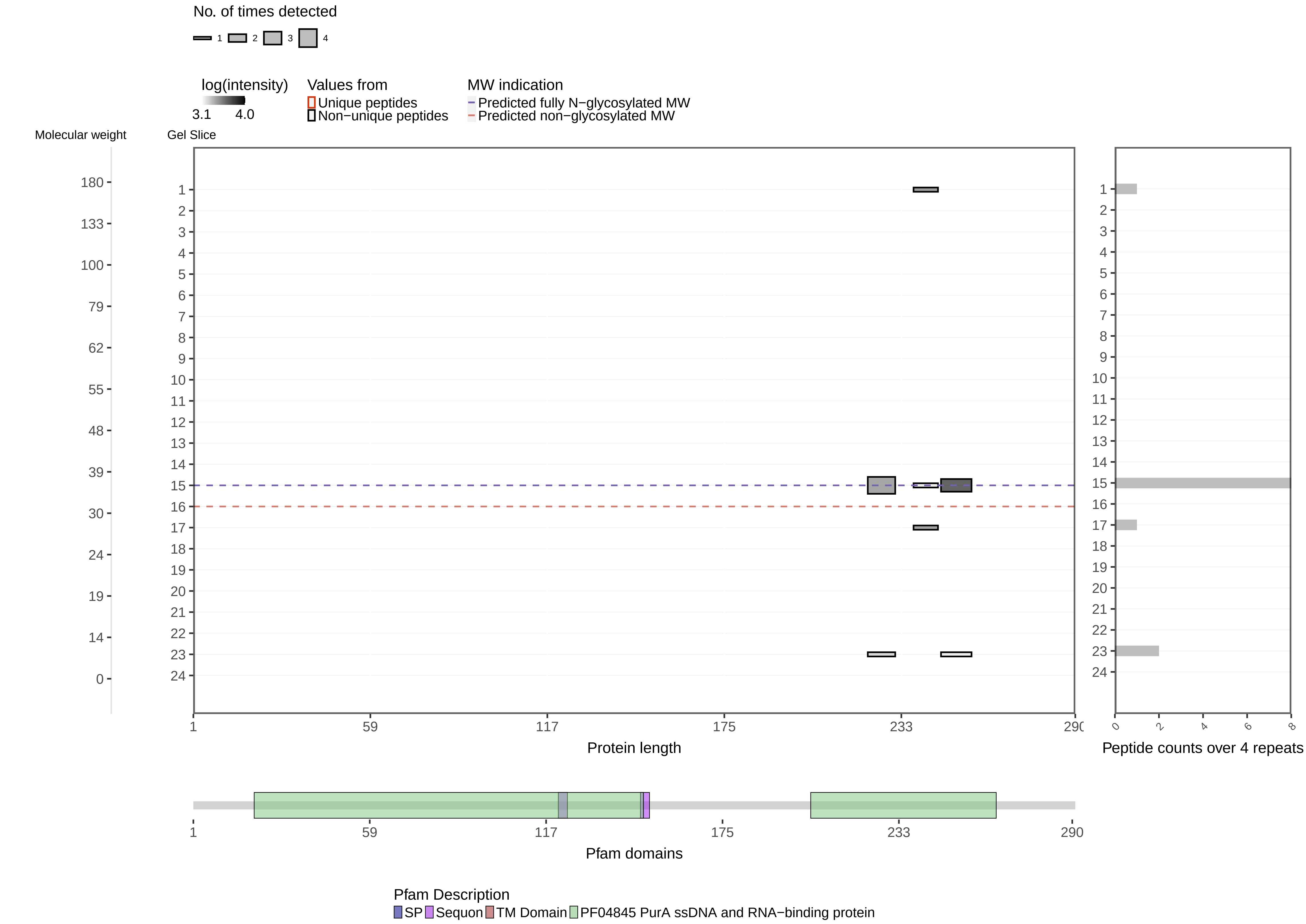


Protein sequence:

SAQDEGWAAFRNILAEINETSRLFISPNQQTSEPSDRLGLSDDVGAGFISSHSSQSGPTADLSVERTIDLPAADEVSNLGVSKVIRADQKRFFFDLGSNNRGHFLRISEVAGSDRSSIILPLSGLKQFHEMVGHFVEISKDRLEGITGANVRTIDSPQR*

NbL16g24360.1 pyruvate_kinase_cytosolic_isozyme [MW = 55.03 kDa]

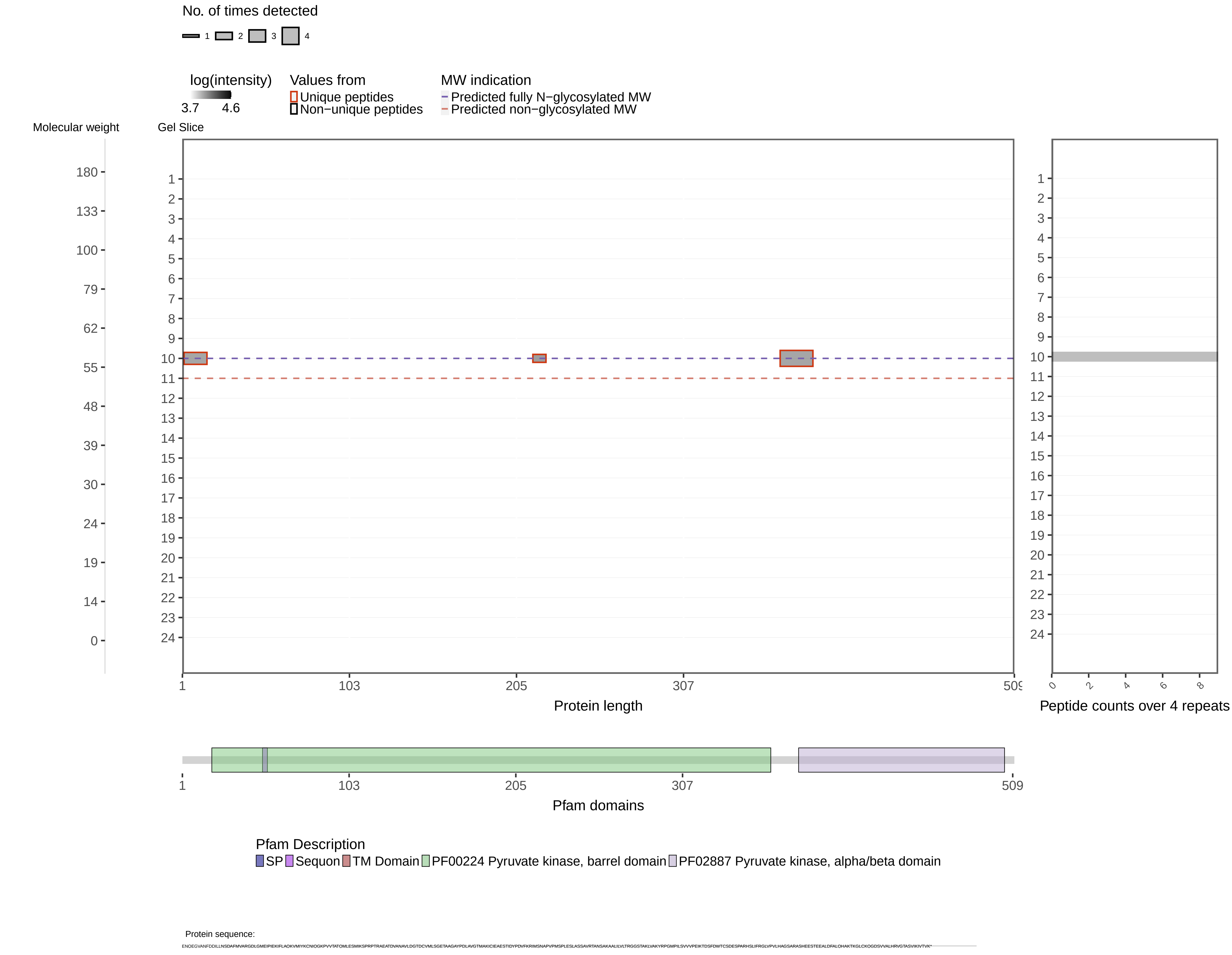

NbL16g24300.1 2−oxoisovalerate_dehydrogenase_subunit_beta_1_mitochondrial−like [MW = 39.6 kDa]

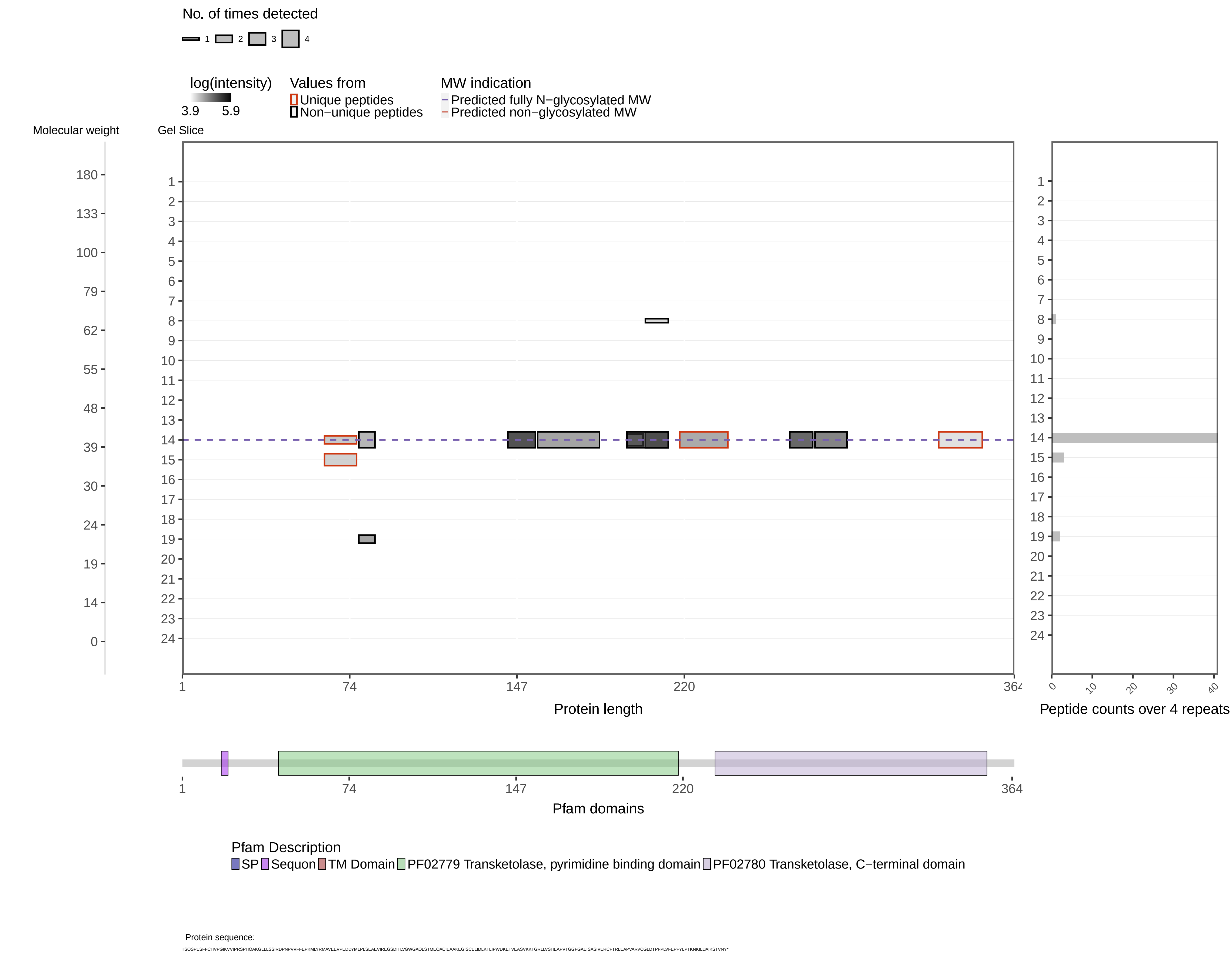

NbL16g24290.1 protein_kinase_and_PP2C−like_domain−containing_protein [MW = 72.07 kDa]

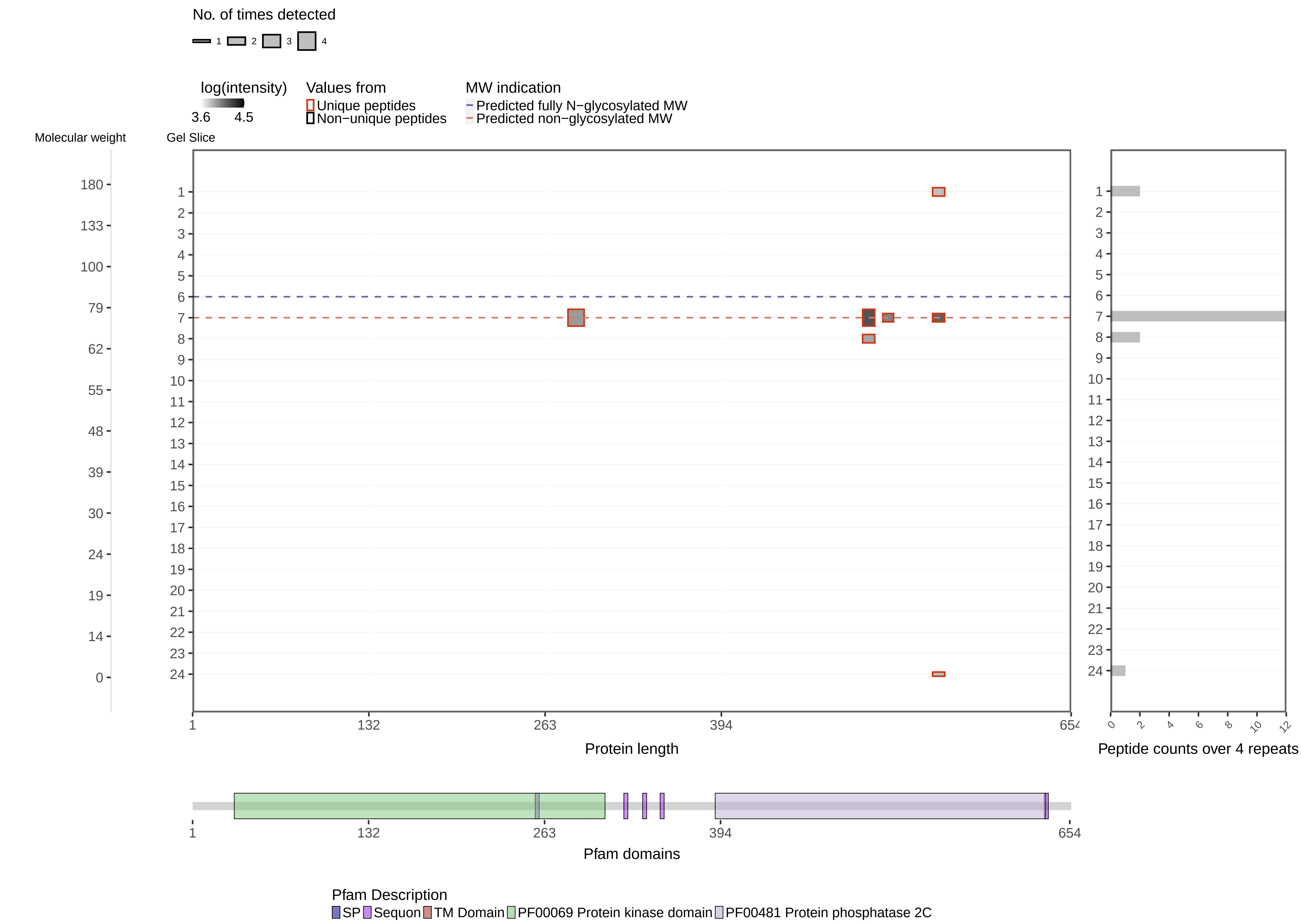


Protein sequence:

EDEIGTNHTEMASADSFISSNGSDGANIQSYQENISWFGQGKEFSKRVPTALAASAWLDYSDEQVYSPVLSWGSFASCGRRETMEDRHFLMPQLCDEKDIHVFAIFDGHRGSAAAEFSAGALPGFLQNLGSISSPSDALFEAFIKTDVAFRTQLDSCRKRKGAIQKDWHPGCTAIAALIVKNKLIVANAGDCRTILCRAGNPYALSRDHVASCHEERERIIRAGGLVKWQVDTWRVGDAALQVTRSIGDDDLKPAVTAEPEITLTTLSAEDEYIVMASDGLWDVVSETDVVNIIF

### NbL16g24250.1 chaperonin_60_subunit_beta_2_chloroplastic−like [MW = 63.31 kDa]

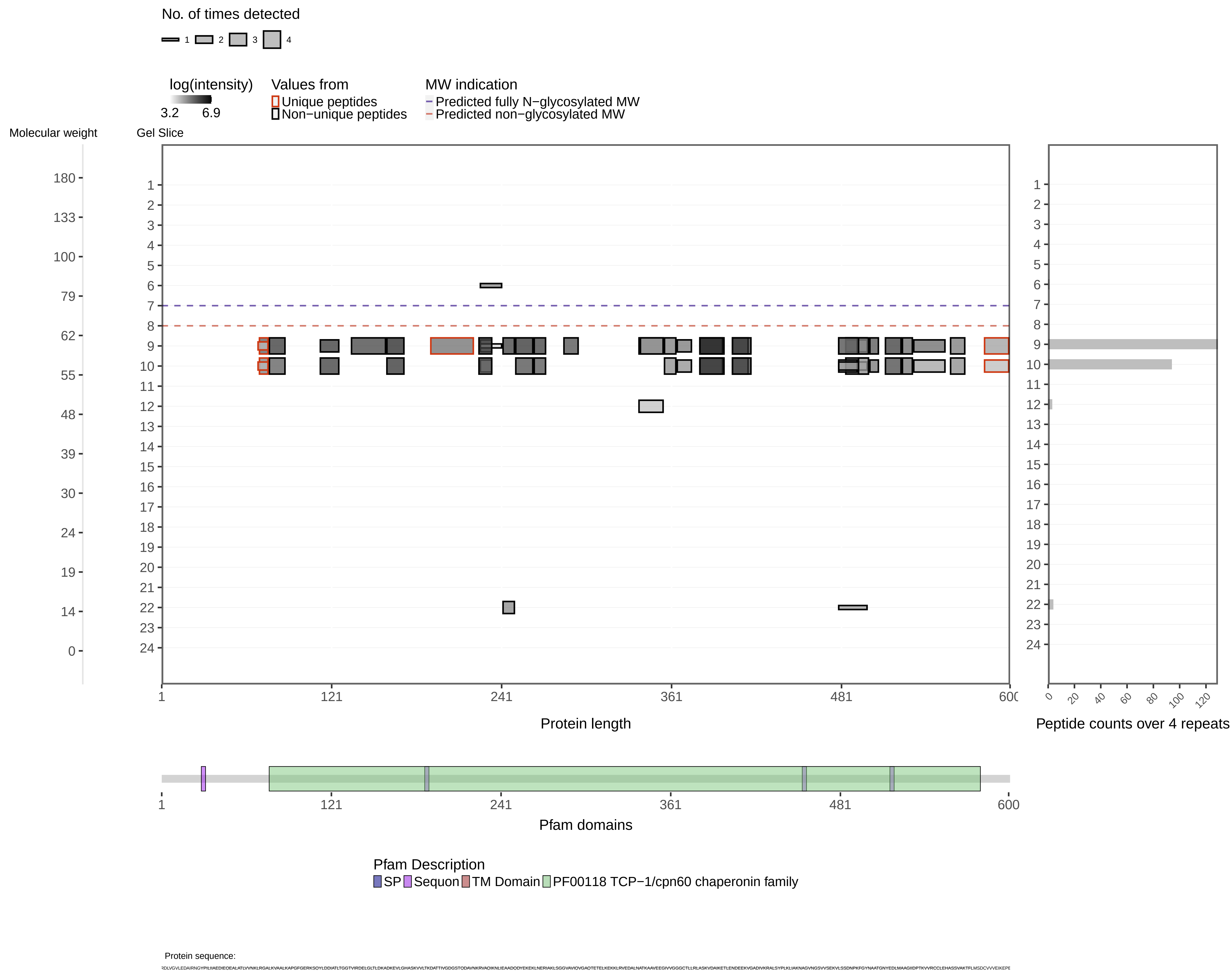

### NbL16g24010.1 uncharacterized_LOC107786789 [MW = 27.48 kDa]

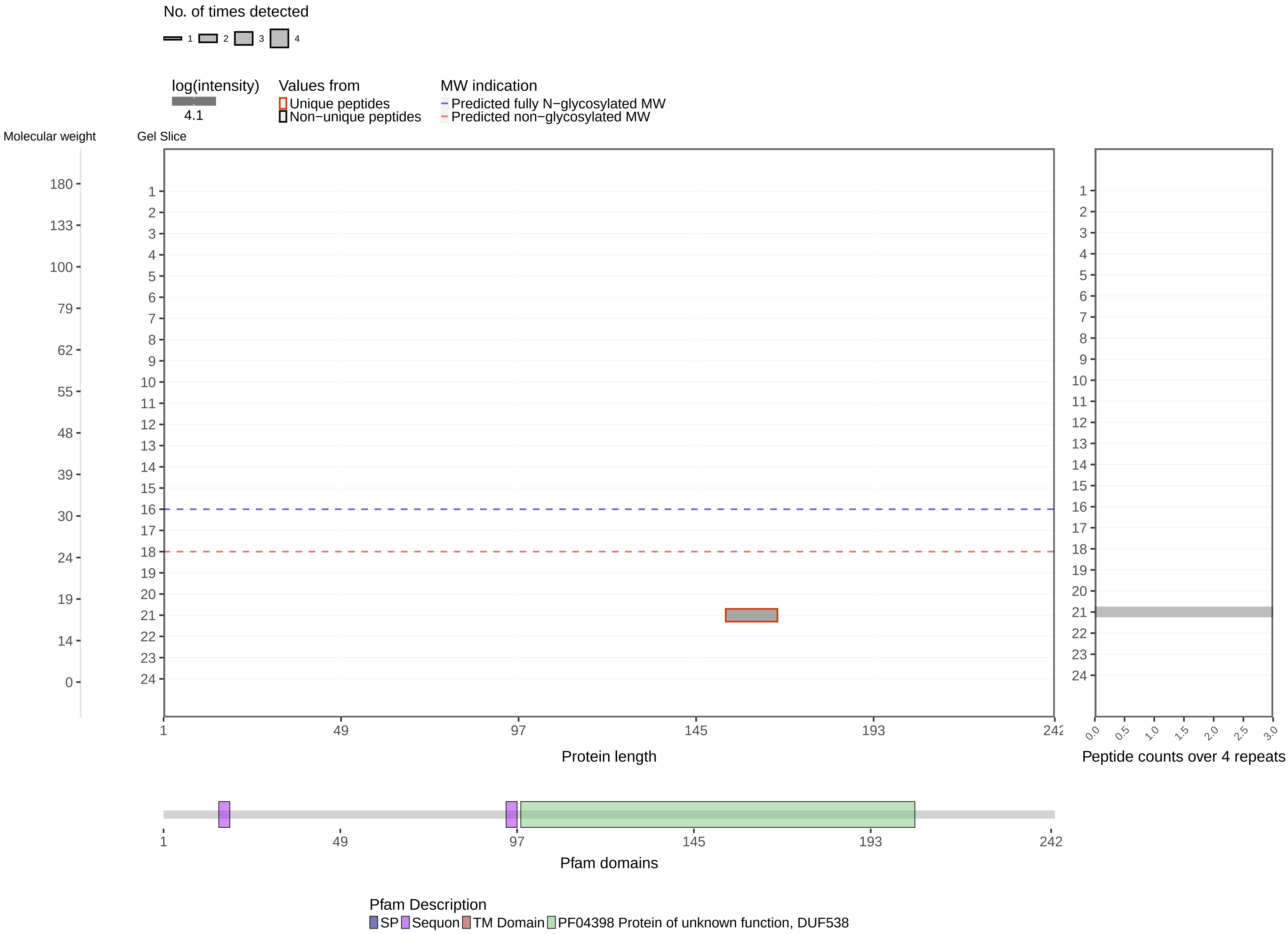


Protein sequence:

KGLLPDSVKSYSLSKDGNFEVELEKPCYVEFEYLVYYAEKITGKLSIGSITDLEGIQVKRFFLWLNVNEIRVDLPSSGSIYFQVGIINKKLDMNQFETVHSCRDNALALCGVSLRQVLQLPNPVNDMPMLITE*

### NbL16g23980.1 dirigent_protein_22−like [MW − SP = 18.57 kDa]

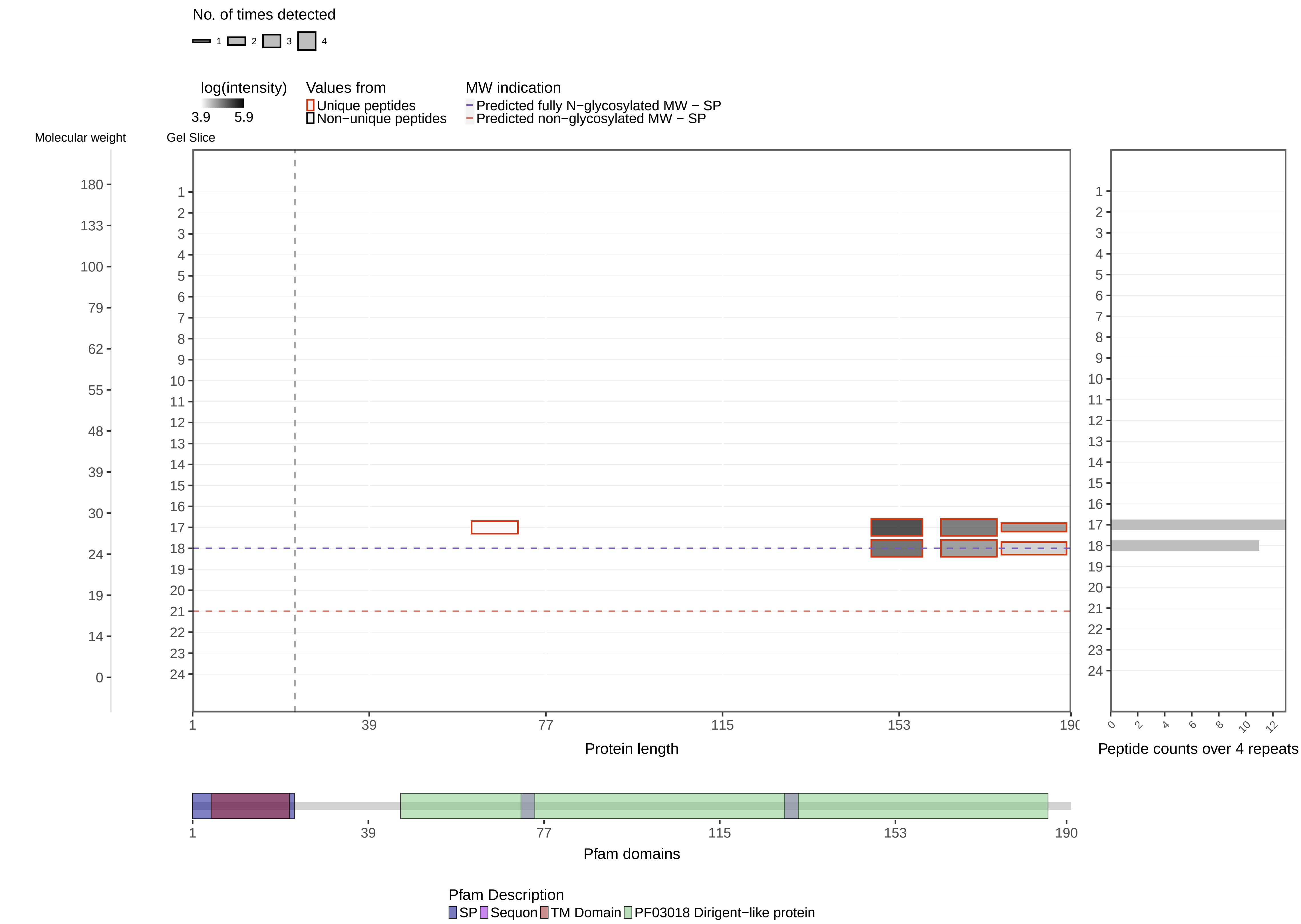


Protein sequence:

NALTLGPELSSKIVGRSQGFYASASQNDVGLMMVMNFAFVEGKYNGSTFTVLGRNSVFEKVREMAVIGGSGLFRFARGYVQASTHSWDMKTGDATVQYDAYVLHY*

NbL16g23540.1 glucan_endo−13−beta−glucosidase_acidic_isoform_GI9−like [MW = 38.99 kDa]

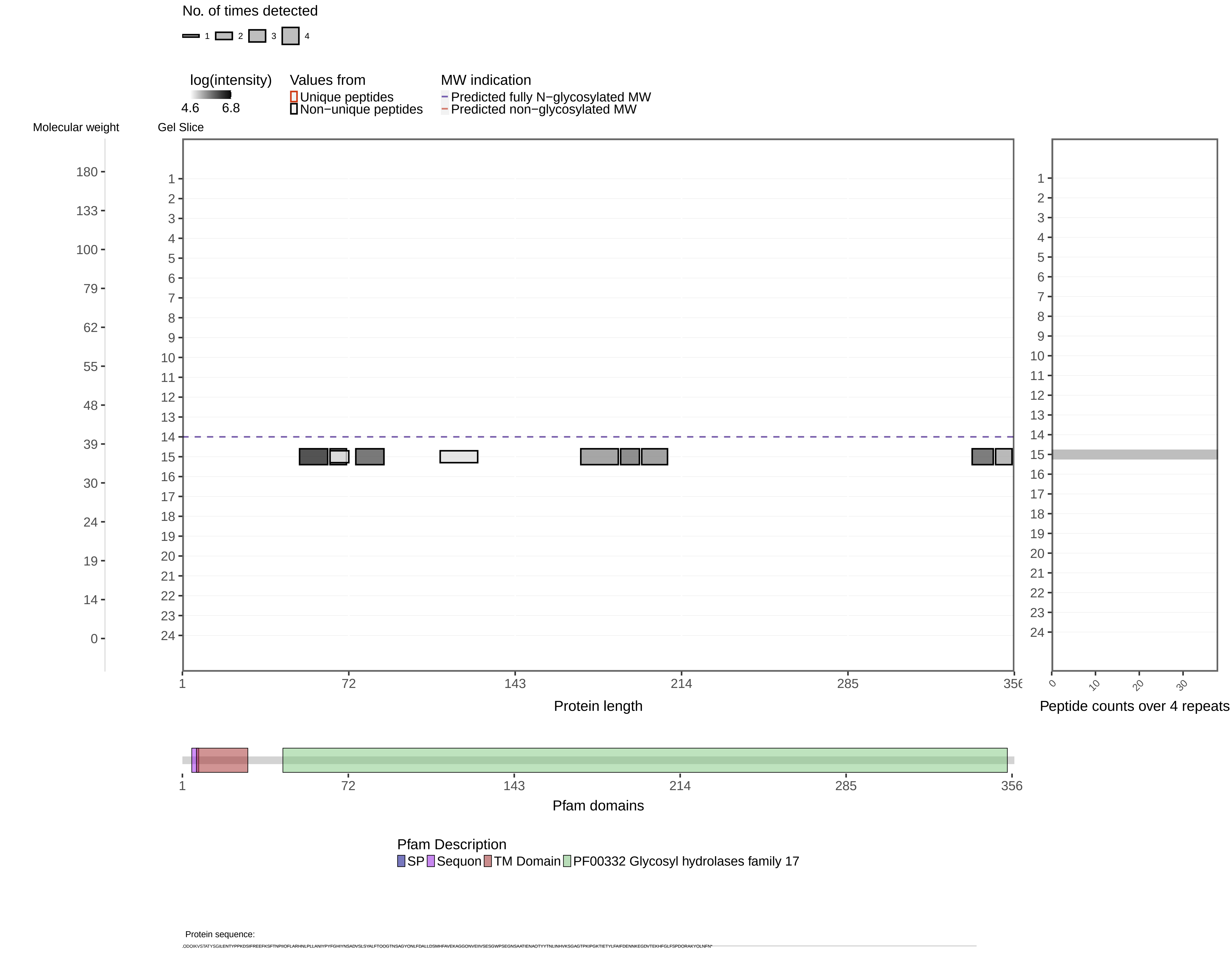

NbL16g23410.1 CBL−interacting_serine/threonine−protein_kinase_3−like [MW = 50.02 kDa]

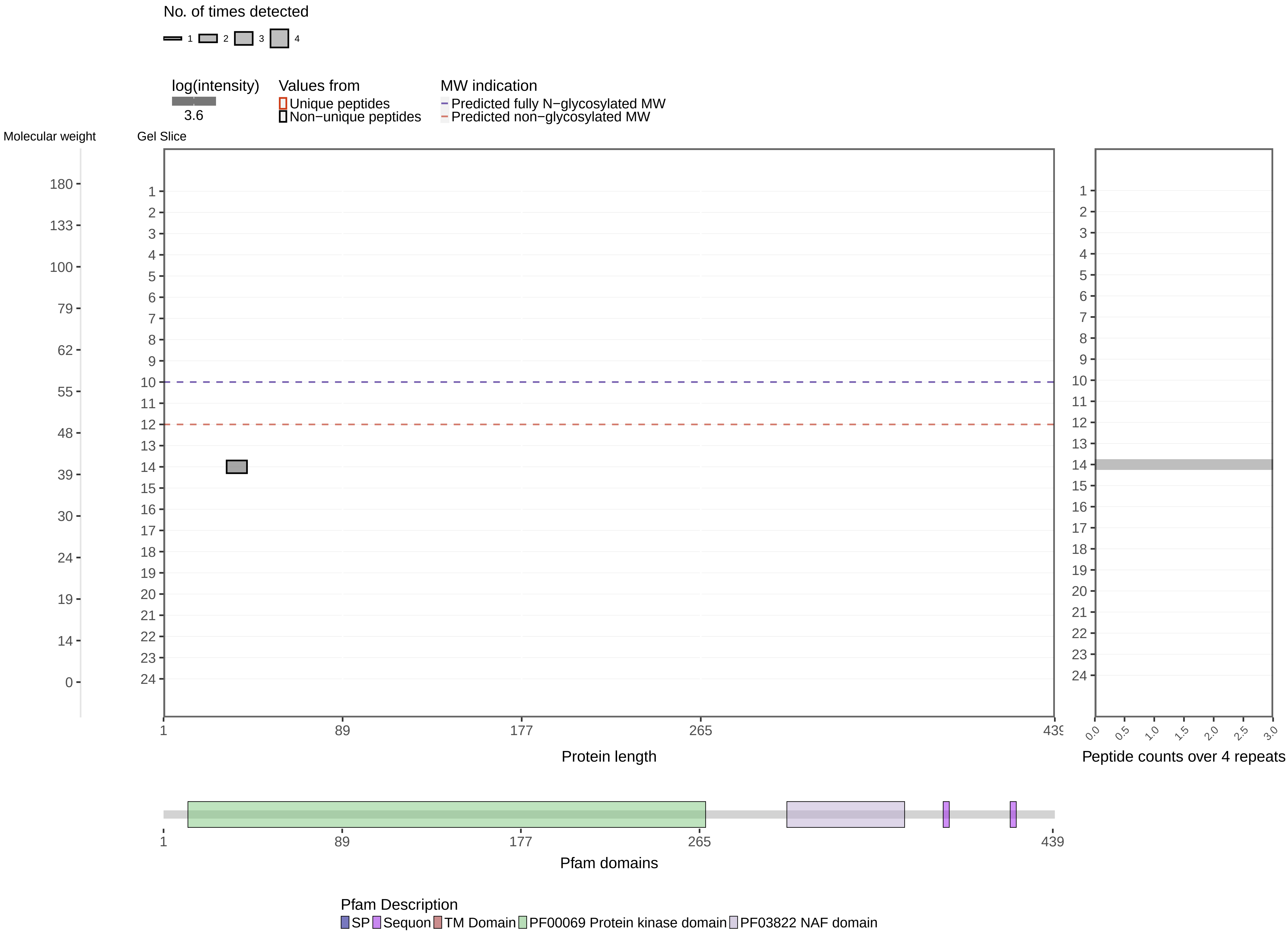


Protein sequence:

LLAGYLPFDDSNLMNLYNKISAAEFTCPPWISFAAMKLITRILDPNPTTRITAPEILEDEWFKKDYRPPVFDEIEDANLDDVEAVFKDSEEYHVTEKKEEKPTSMNAFELISMSQGLNLGNLFDEQGFKRETRFTSKCPASEIISKIEEAAKPLGFDVSKKNYKMRLQNLKAGRKGNLNVSTEVFQVAPSLHMVEVRKAKGDTLEFHKFYKNLSTSLDEVVWKTEENMEKKK*

NbL16g23370.1 uncharacterized_protein_At2g37660_chloroplastic−like [MW = 36.74 kDa]

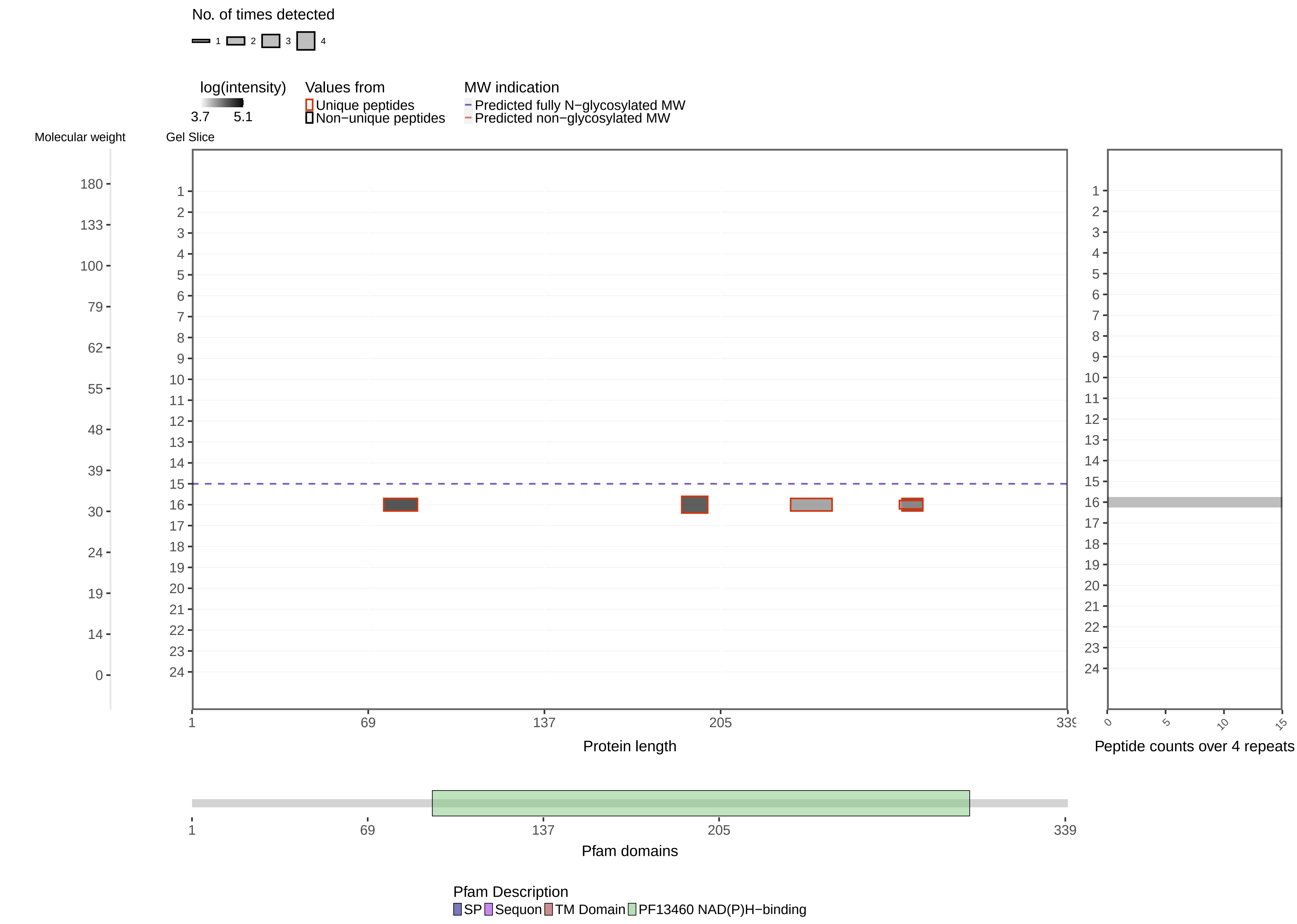


Protein sequence:

…GVTHVICCTGTTAFPSRRWDGDNTPERVDWEGVRNLVSALPQSLKRIVLVSSIGVTKFNELPWSIMNLFGVLKYKKMGEDFVCNSGLPFTIIRAGRLTDGPYTSYDLNTLLQATAGERRAVLIGQGDKLVGEVSRLVVAEACIQALDVDFTEGQTYEINSIPGDGPGTDTLKWQELFRASHNN*

### NbL16g23330.1 UBP1−associated_protein_2B−like [MW = 52.03 kDa]

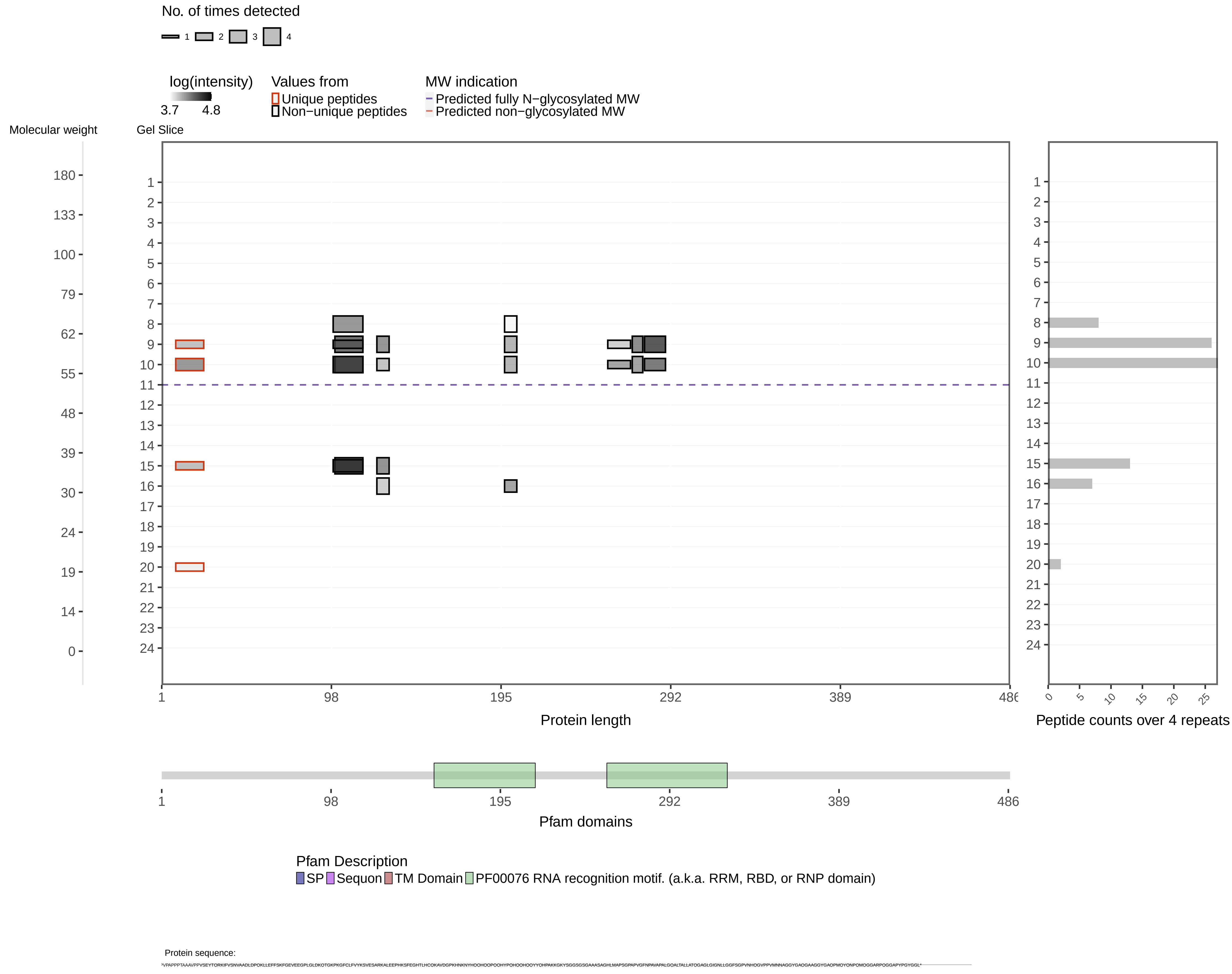

NbL16g23230.1 molybdopterin_biosynthesis_protein_CNX1−like [MW = 70.28 kDa]

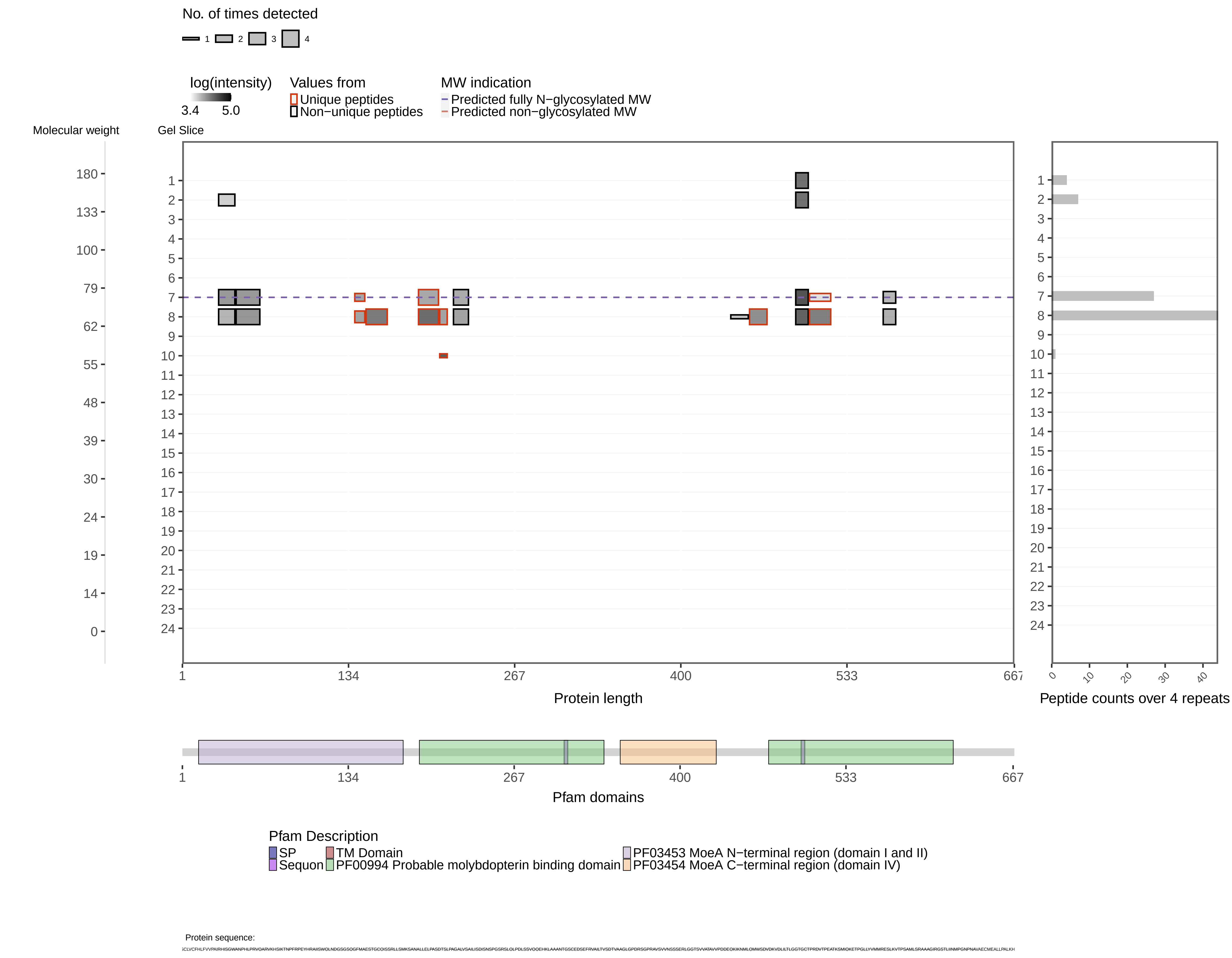

NbL16g23040.1 kirola−like [MW = 16.8 kDa]

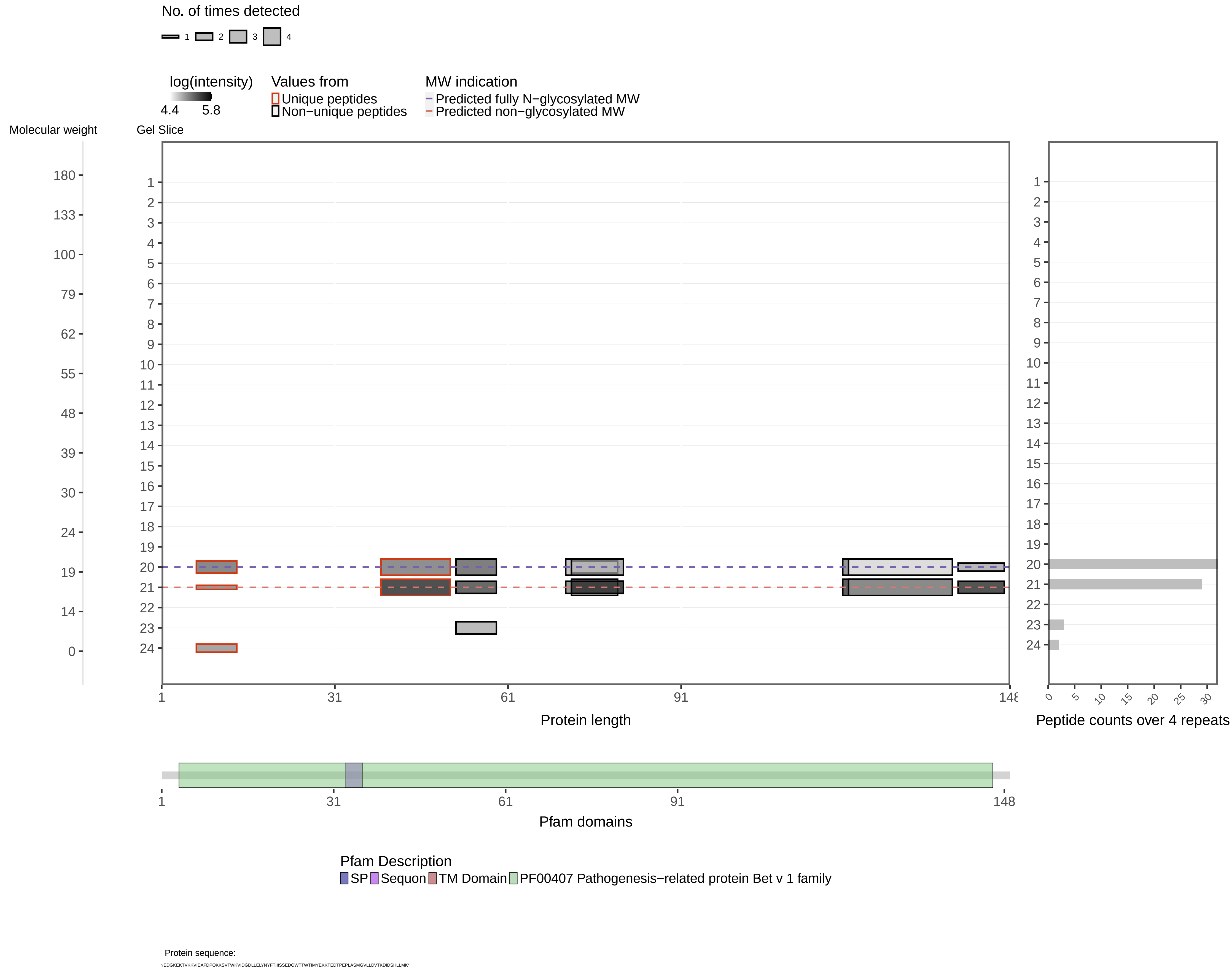

NbL16g23010.1 indole−3−acetaldehyde_oxidase−like [MW = 149.36 kDa]

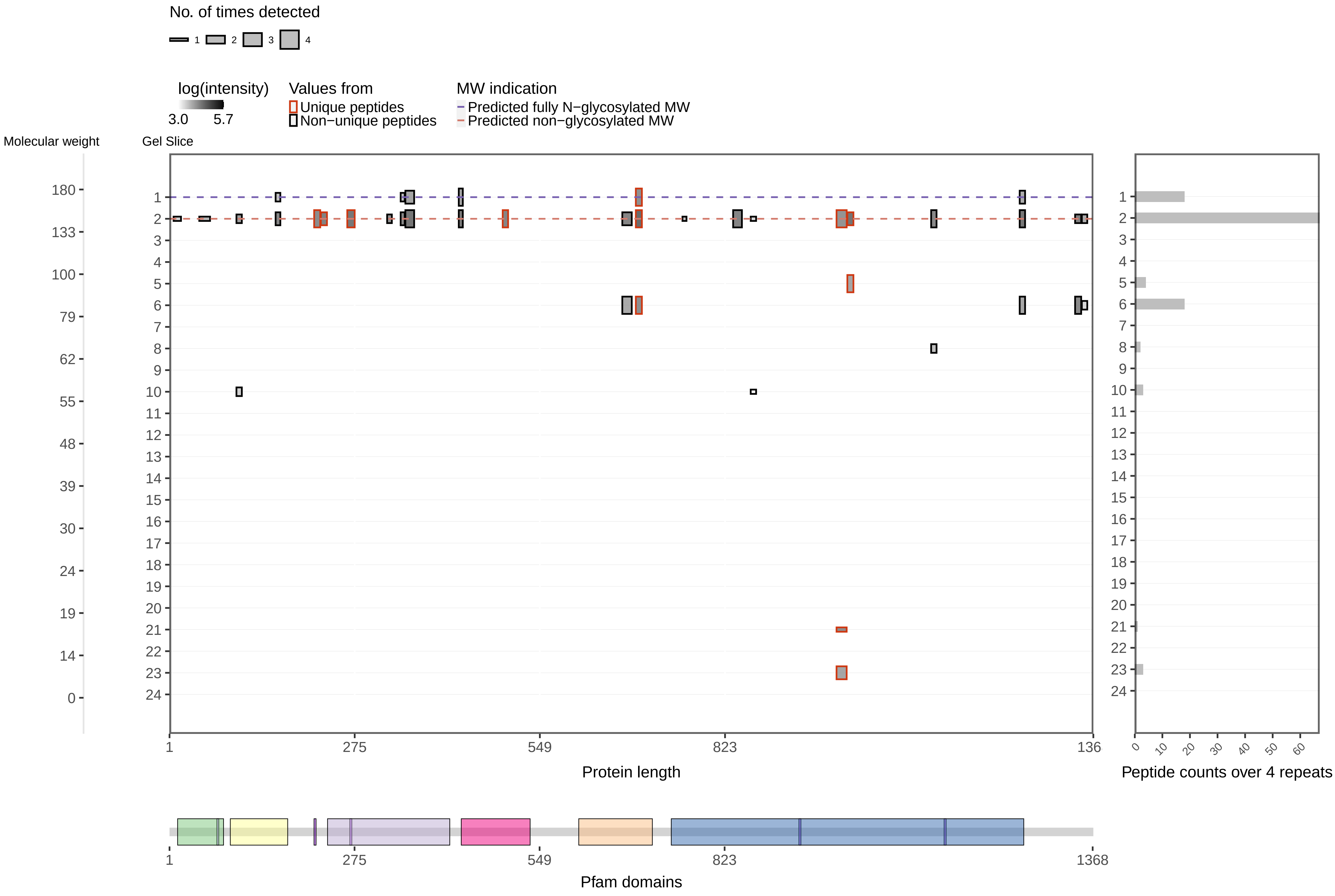


PF00941 FAD binding domain in molybdopterin dehydrogenase
PF01799 [2Fe−2S] binding domain
PF03450 CO dehydrogenase
sulfur cluster binding domain
PF01315 Aldehyde oxidase and xanthine dehydrogenase, a/b hammerhead domain
PF02738 Molybdopterin−binding domain of aldehyde dehydrogenase

Protein sequence:

### NbL16g22990.1 uncharacterized_LOC107782810 [MW = 13.57 kDa]

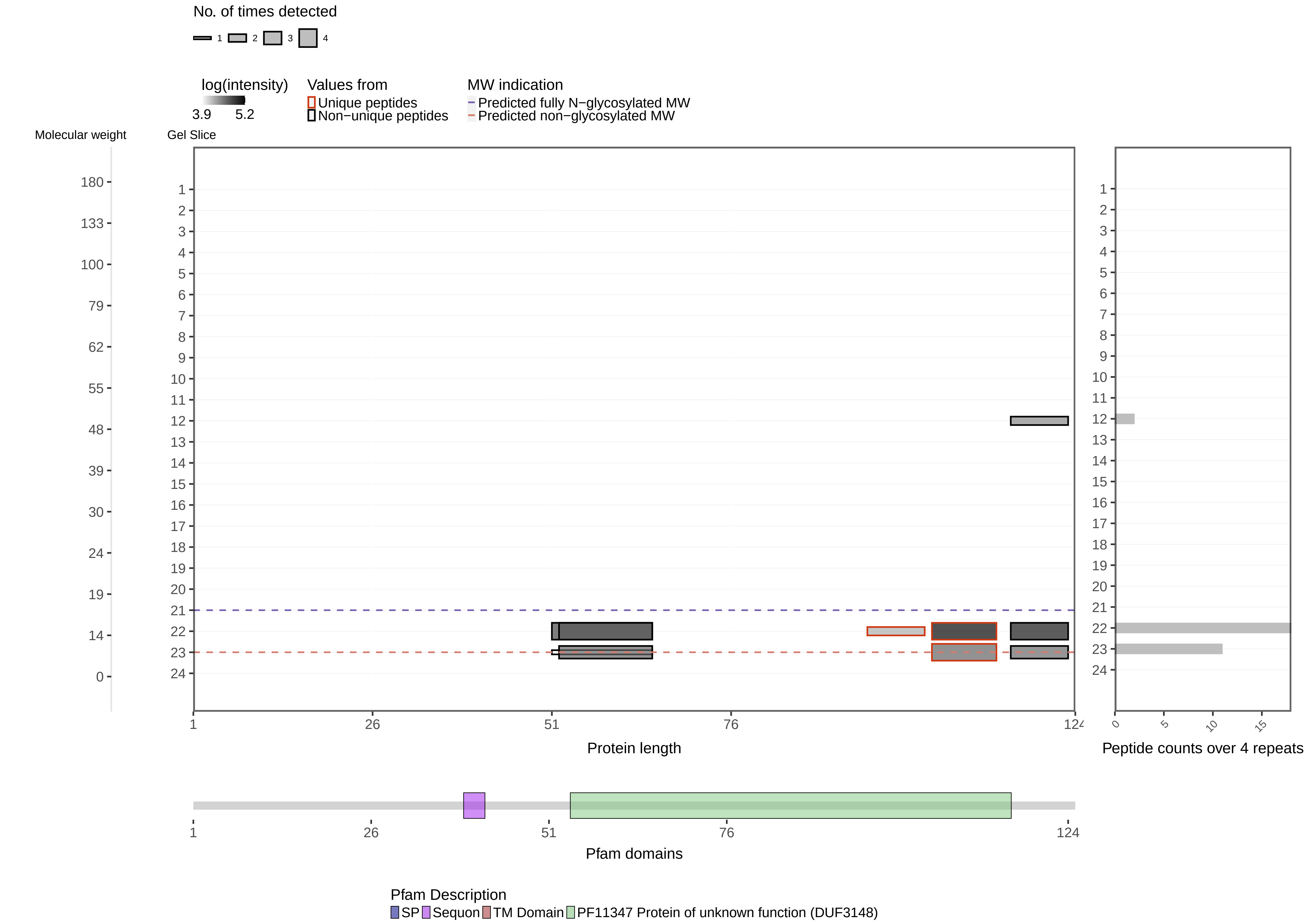


Protein sequence:

DSKKLGIGSPIIVIEAPKLLKTAASVPCLRPNDGLVKPGDVGRIVSRKPMDVWAIRLSIGTYLIDRKYFKPLELEN*

NbL16g22950.1 serine/threonine−protein_phosphatase_BSL3−like [MW = 104.34 kDa]

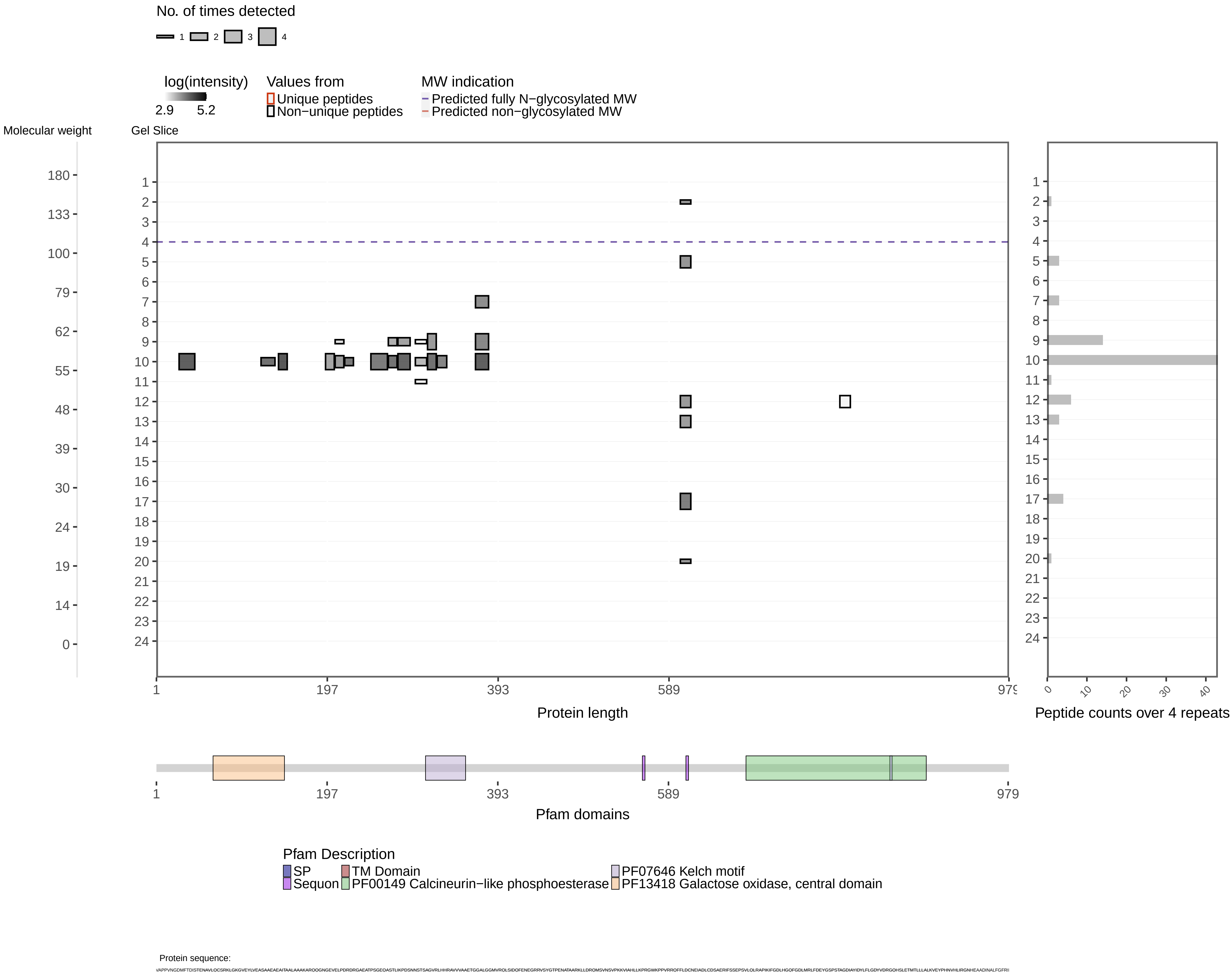

### NbL16g22940.1 protein_CLP1_homolog [MW = 48.01 kDa]

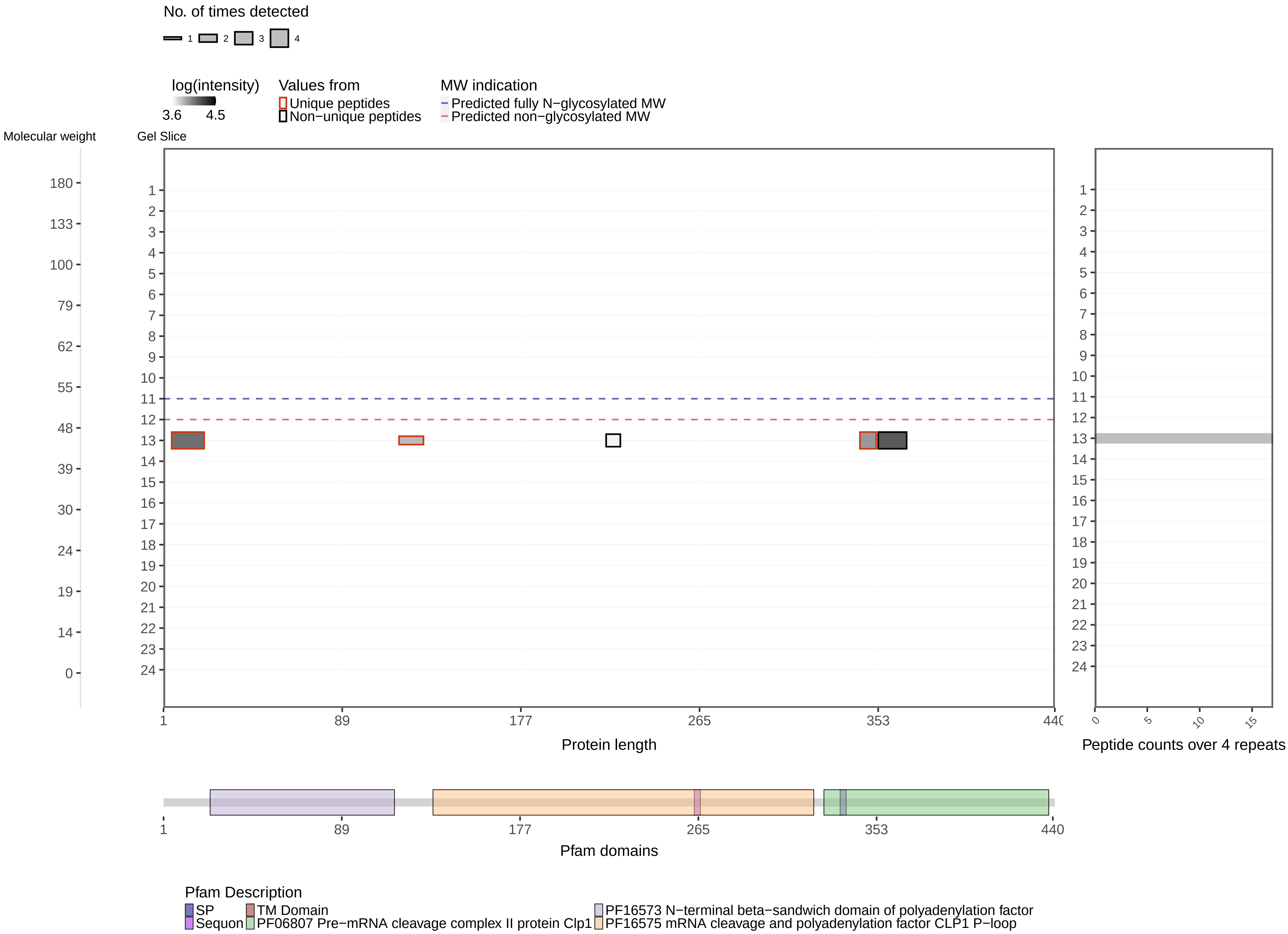


Protein sequence:

SANIDHYKVLVKELAQTLERQFSGNAESRASGMVINTMGWIEGVGYELLLHAIDTFNATVILVLGQEKLCSMLKDVLKNRPNVDVVKLQKSGGVVSRNAKVRQKARGYRIREYFYGPSNDLSPHSNVVNFSDLFIYRIGGGPQAPRSALPIGAEPAADPTRLVLVSINRDLLHLVLAVSYANEPDQIISSNVAGFIYITDVDMHRKKITYLAPCAGELPSKYLIVGTLTWIEN*

NbL16g22840.1 UDP−D−apiose/UDP−D−xylose_synthase_2−like [MW = 43.32 kDa]

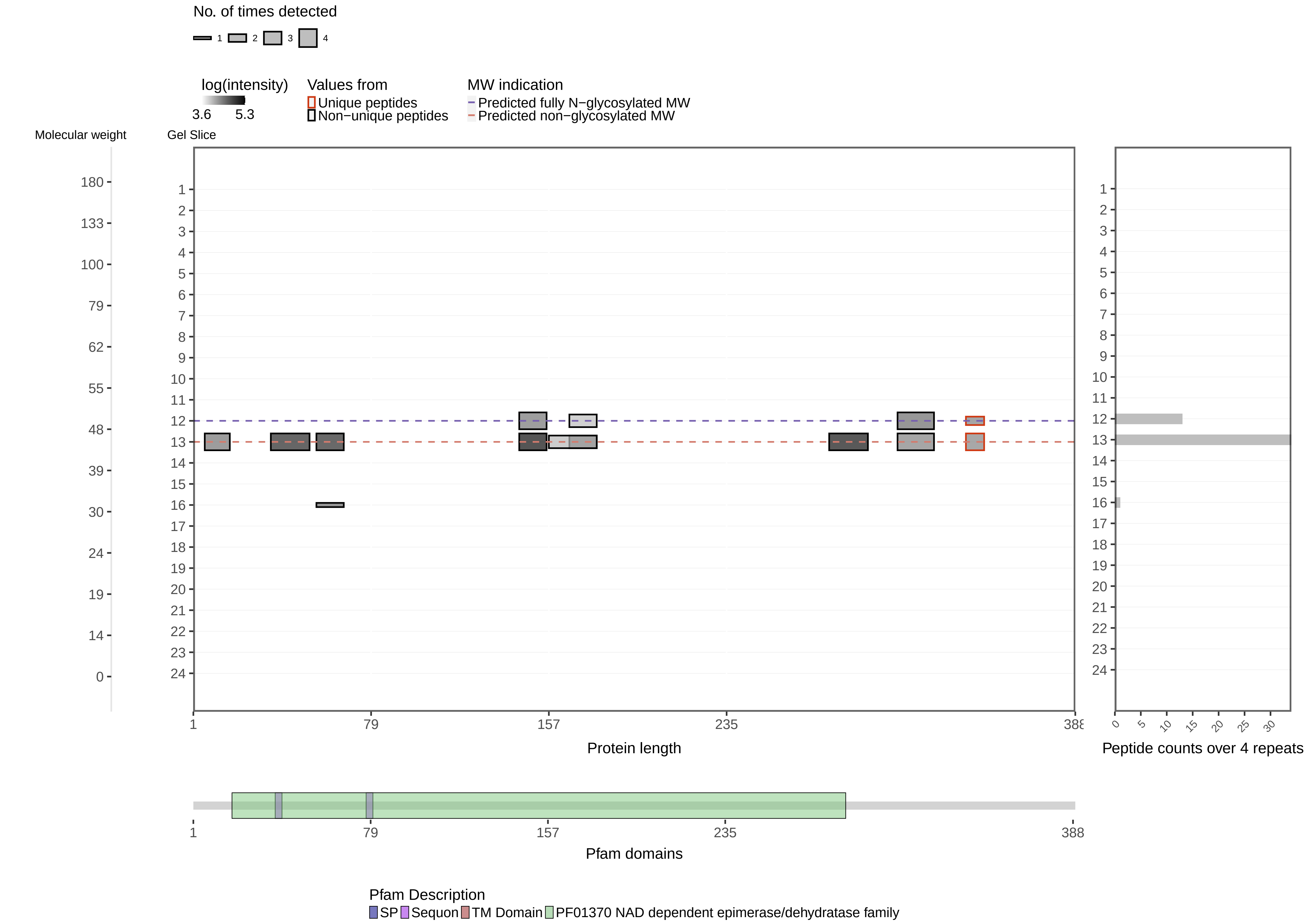


Protein sequence:

SYACAKQLIERLIYAEGAENGLEFTIVRPFNWIGPRMDFIPGIDGPSEGVPRVLACFSNNLLRHEPLKLVDGGHSQRTFIYIKDAIEAVLLMIENPARANGQIFNVGNPNNEVTVRQLAEMMTQVYSKVSGESPPETPTIDVSSKEFYGEGYDDSDKRIPDMTLINRQLGWNPKTSLWDLLESTLTYQHRTYAEAVKQAMSKTTAN*

NbL16g22720.1 60S_ribosomal_protein_L10a−1−like [MW = 24.61 kDa]

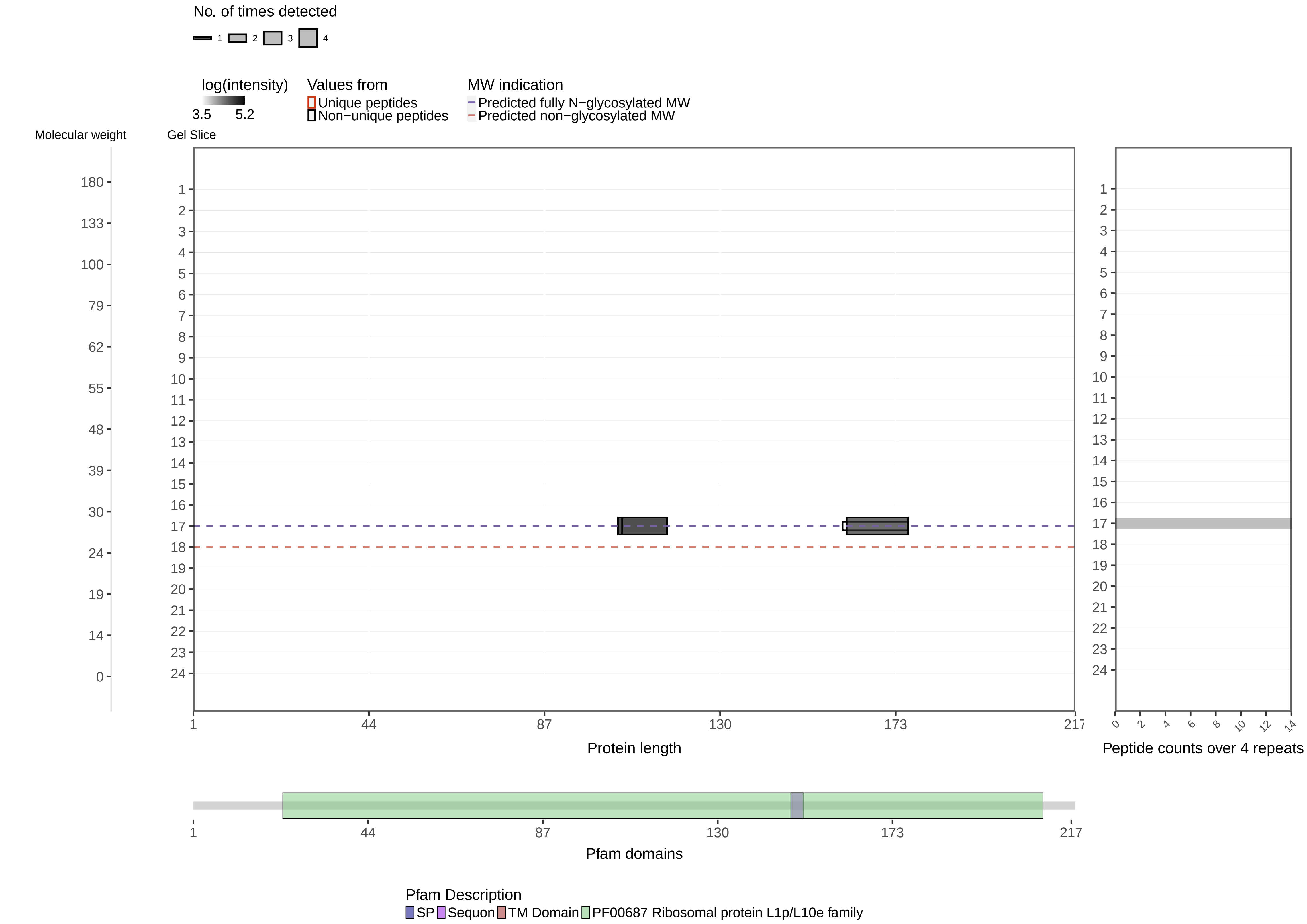


Protein sequence:

KKLVKKLAKKYHAFLASEAVIKQIPRLLGPGLNKAGKFPSLVSHQESLESKVNETKATIKFQLKKVLCMGVAVGNMDMEEKQIFQNVQMSVNFLVSLLKKNWQNVRCLYLKSTMGKTQRVF*

### NbL16g22580.1 proteasome_subunit_alpha_type−3−like [MW = 27.22 kDa]

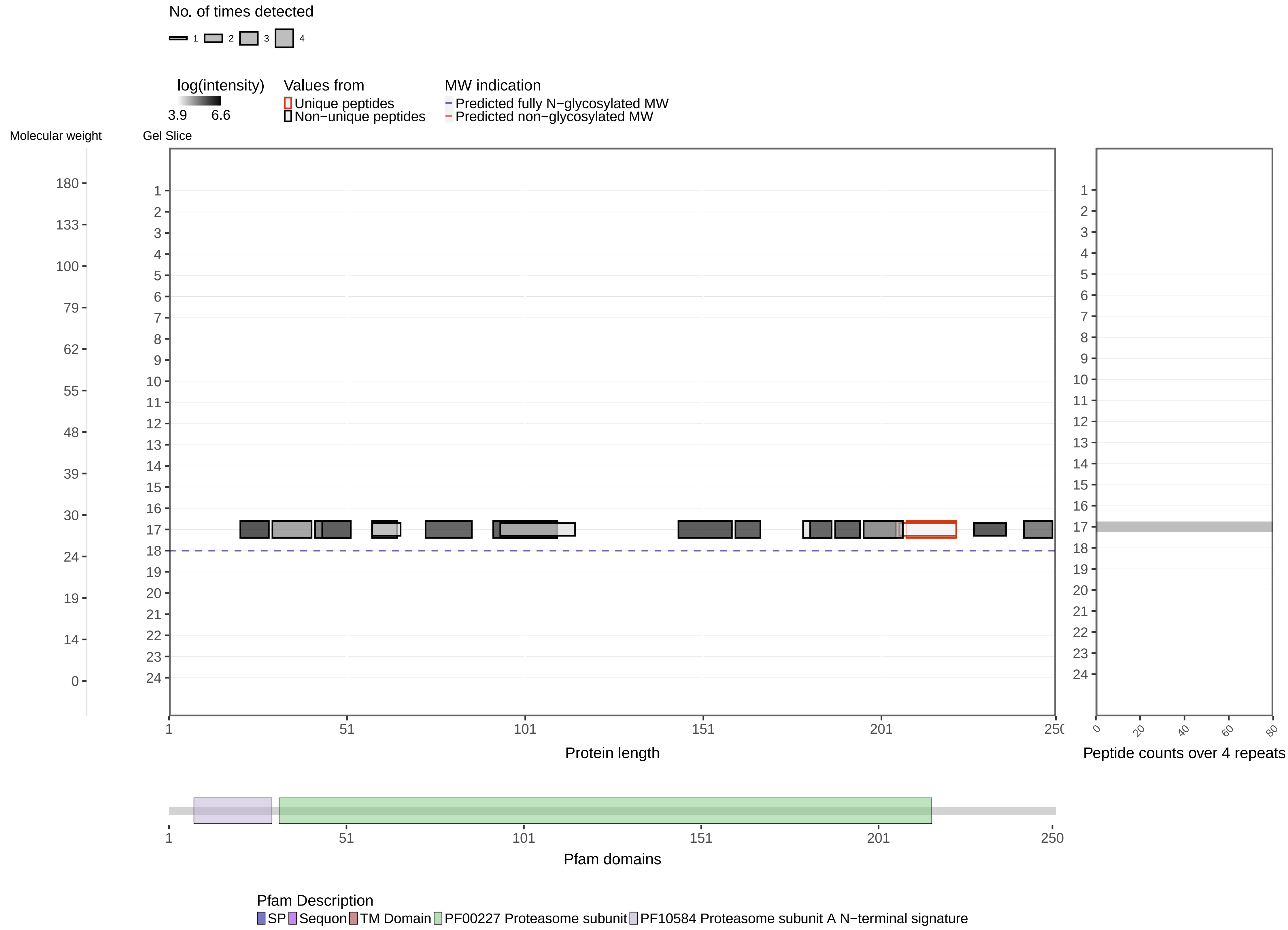


Protein sequence:

AERVASYVHLCTLYWWLRPFGCGVILGGYDRDGPQLYLVEPSGVSYRYFGAAIGKGRQAAKTEIEKLKLSEMTCRQGVIEVAKIIYGVHDEAKDKAFELEMSWICDESNRQHQKVPDDILEEAKTAAKAALEEMDAD*

### NbL16g22570.1 magnesium−protoporphyrin_IX_monomethyl_ester_[oxidative]_cyclase_chloroplastic−like [MW = 47.3 kDa]

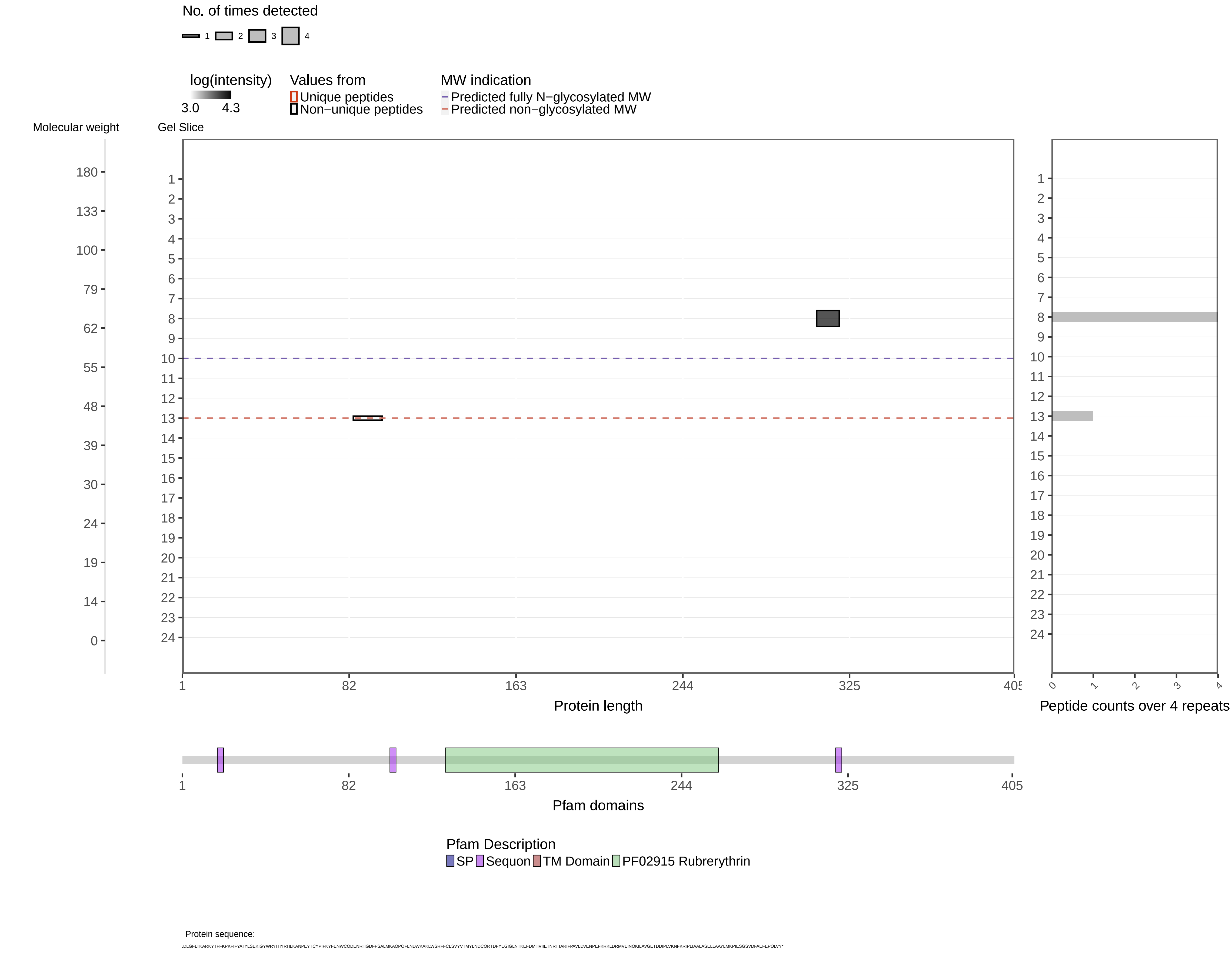

NbL16g22490.1 nascent_polypeptide−associated_complex_subunit_alpha−like_protein_1 [MW = 21.45 kDa]

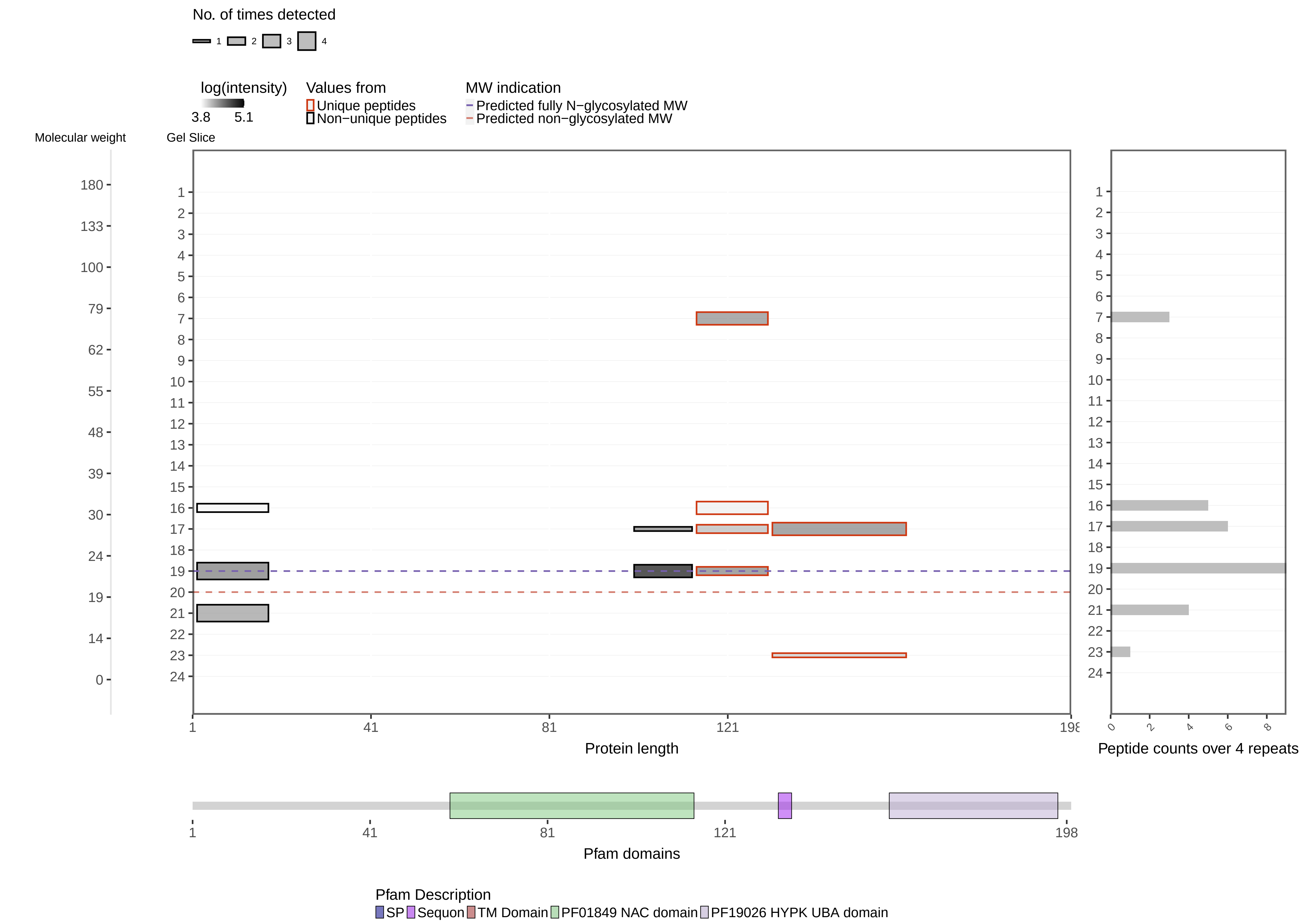


Protein sequence:

SKNILFVISKPDVFKSPASDTYVIFGEAKIEDLSSQLQSQAAEQFKAPNLSNVISKPEPSTVAQDDEDVDETGIEPKDIELVMTQAGVSRAKAVKALKGADGDIVSAIMELTN*

### NbL16g22420.1 vinorine_synthase−like [MW = 50.77 kDa]

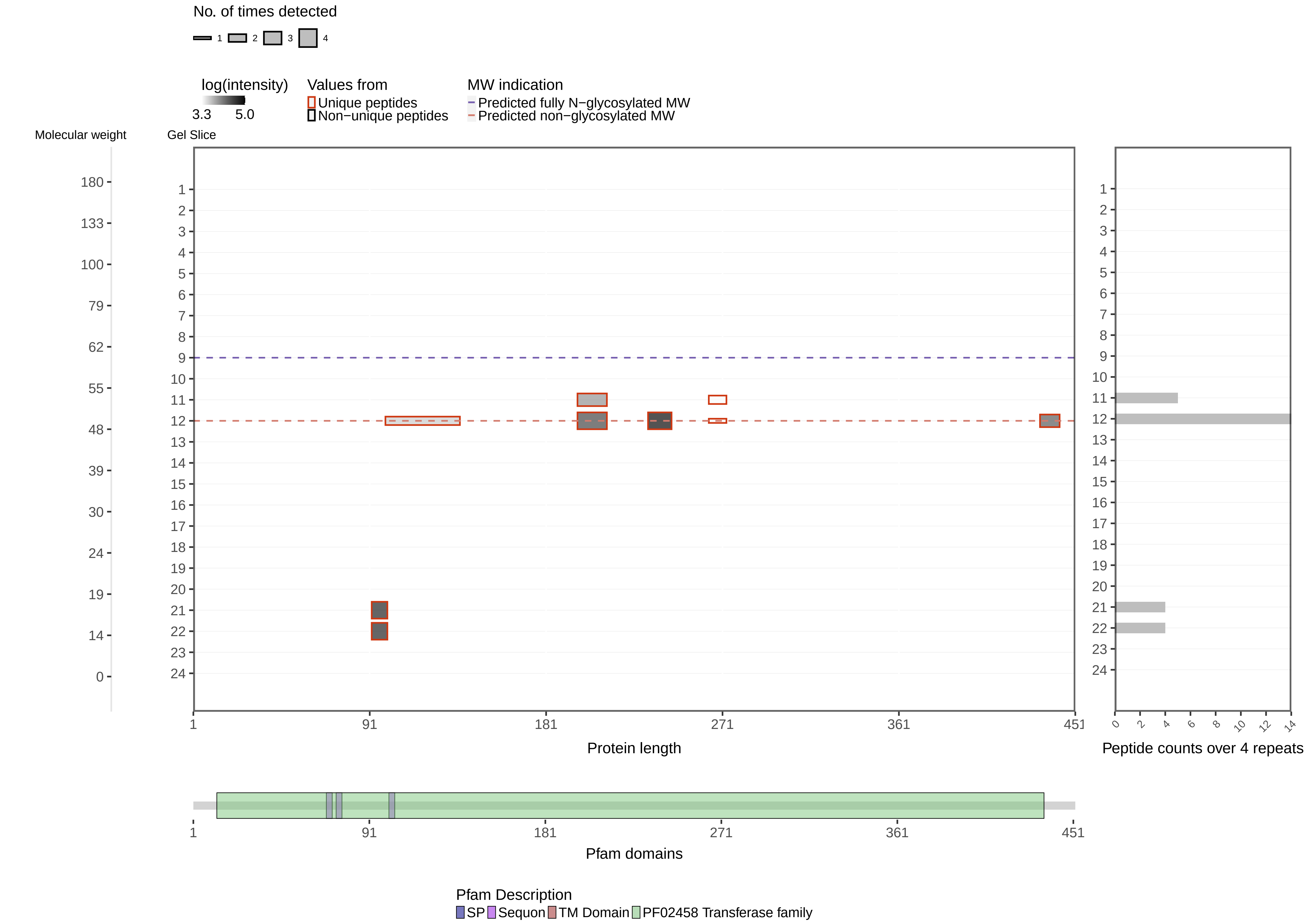


Protein sequence:

PRNECPPSHSTEGAFSRGALTNVYDFNPSRIAFLKTKAASENIPKPSRVNVVTSVIWKCLTAHSKRPSTLVHLVNMRKRVDPPMHDHYIGNFVGFSEASKDENEKEIANMVALMRKGLAKFDNDCVKRLKGDEAIGTILEHVKDAVDLNSRKGVDLYYFTSWCGYSFYDVNFGWGNPIKFSTTEGKFKNMIMLMDTKDGGVEAWVTLSEDDMEIFEKNTELLTYATLKASFPDRE*

NbL16g22270.1 tubulin_beta−2_chain [MW = 50.21 kDa]

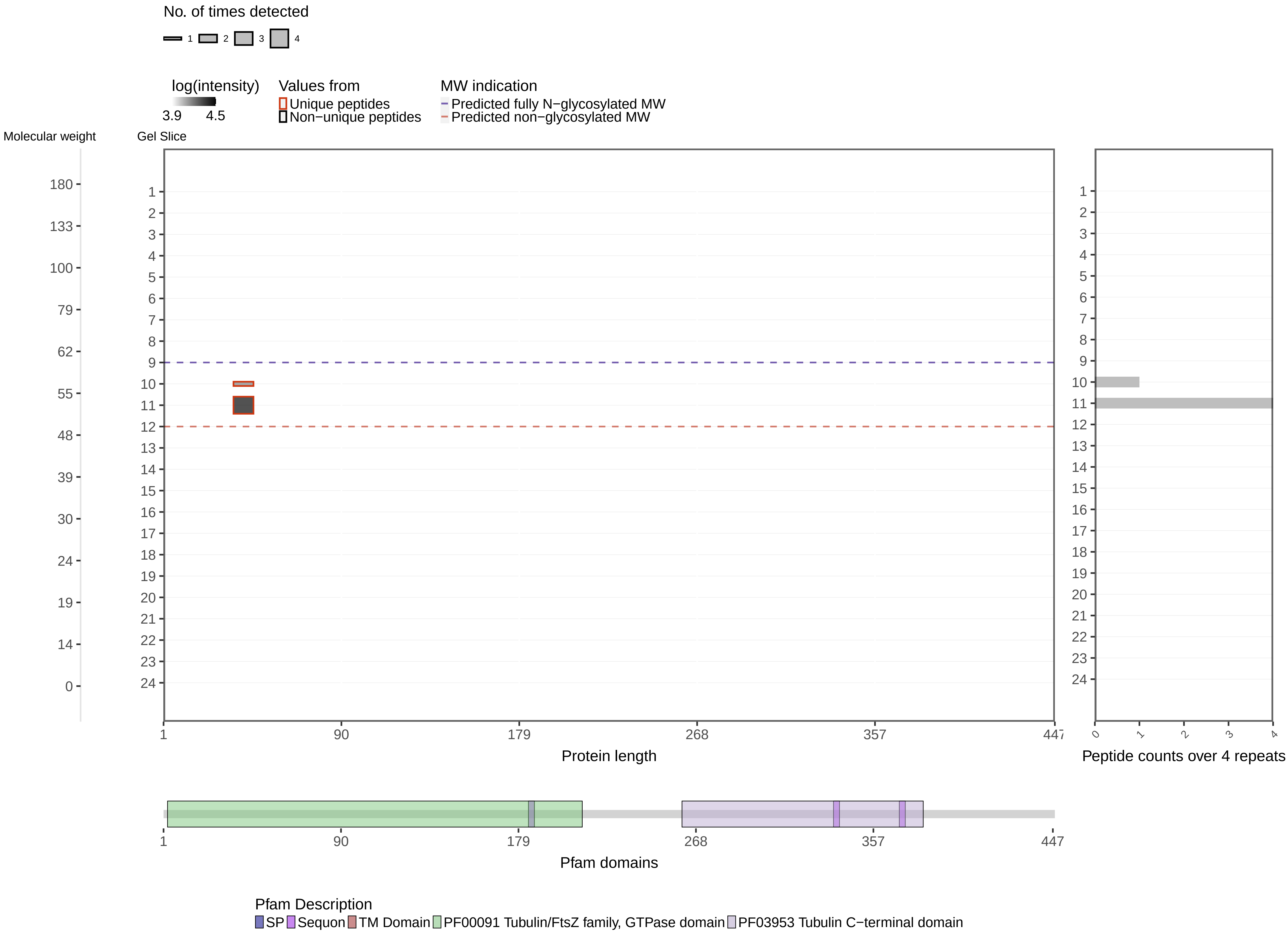


Protein sequence:

FRTLKLTTPSFGDLNHLISATMSGVTCCLRFPGOLNSDLRKLAVNLIPFPRLHFFMVGFAPLTSRGSOOYRALTVPELTOOMWDSKNMMCAADPRHGRYLTASAMFRGKMSTKEVDEOMLNVONKNSSYFVEWIPNNVKSTVCDIPPTGLKMASTFIGNSTSIOEMFRRVSEOFTAMFRRKAFLHWYTGEGMDEMEFTEAESNMNDLVSEYOOYODATADEEGEYYEDEEEAOED*

### NbL16g22210.1 laccase−4−like [MW − SP = 58.62 kDa]

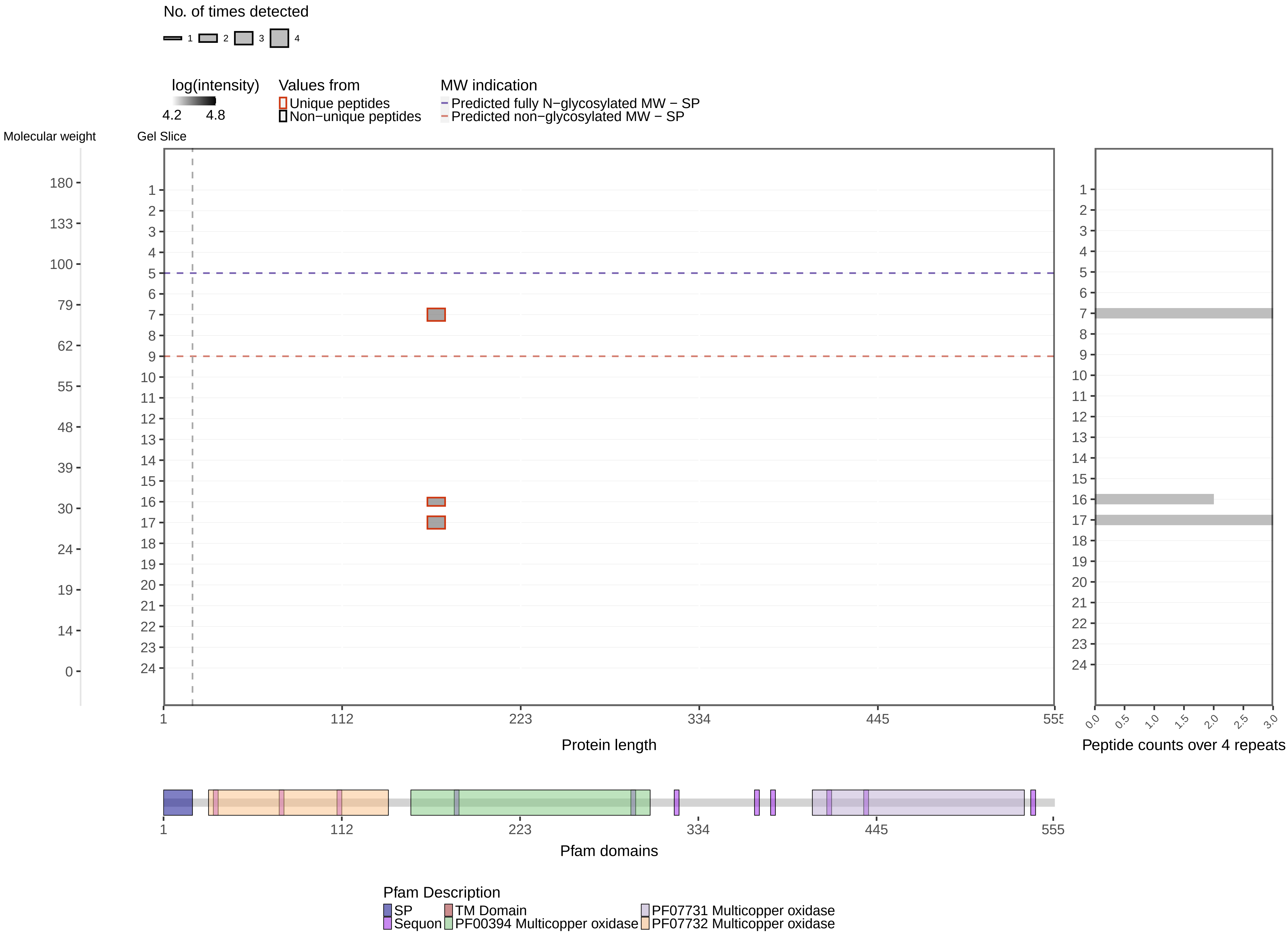


Protein sequence:

NVIVTADQSFGKYMVAASPFMDAPIAVDNITATATLHYSGALGSSPTTLTSTPPQNATSVANNFLDALKSLNSKKYPAKVPQTVDRSLFFTVGLGINPCPTCKQANGSRVVASVNNITFVMPTVALLQAHFFGINGVFTTDFPANPPFVFNYTGTSPTNLATTKGTKVYRLPYNATVQLVLQDTGIIAPENHPIHLHGFNFFLVGKGLGNFNSKTDPKNFNLIDPVERNTIGVPSGGWVAIRWRADNPGVWFMHCHLEVHTTWGLKMAFLVDNGKGPNESLLPPPKDLPKC*

NbL16g22110.1 oxysterol−binding_protein−related_protein_3A [MW = 52.17 kDa]

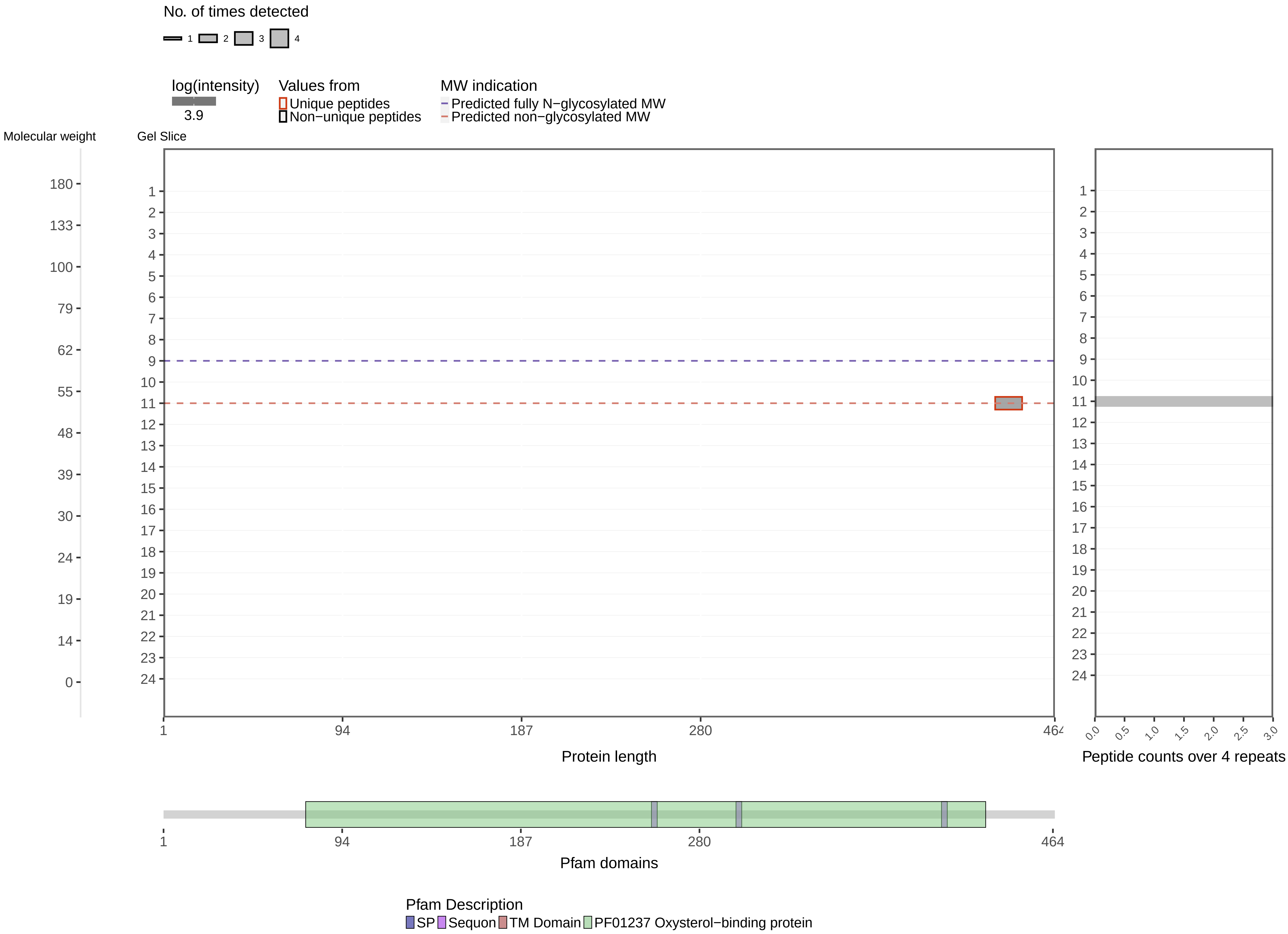


Protein sequence:

KKSGVVLDLVPPPTKVHNLIFGRTWIDSPGEMILTNLTTGDKVLLYFQPCGWFGAGRYEVDGYVVYNSEEEPKILMTGKWNESLSYQPCDLEGEPLPGSTMKEVWRTADAPKNDKFQYTYFAHKINSFDTAPKKLLASDSRLRPDRYALEKGDLSKASSEKSRLEERQRADRRTREAKGDEFKPKWFNLSNEVCPTPWGDLEVYEYNGKYHEHRAAIDSSGHAEEADVKTTEFNPWQYEDSVVSA*

### NbL16g22070.1 tyrosine_aminotransferase−like [MW = 46.52 kDa]

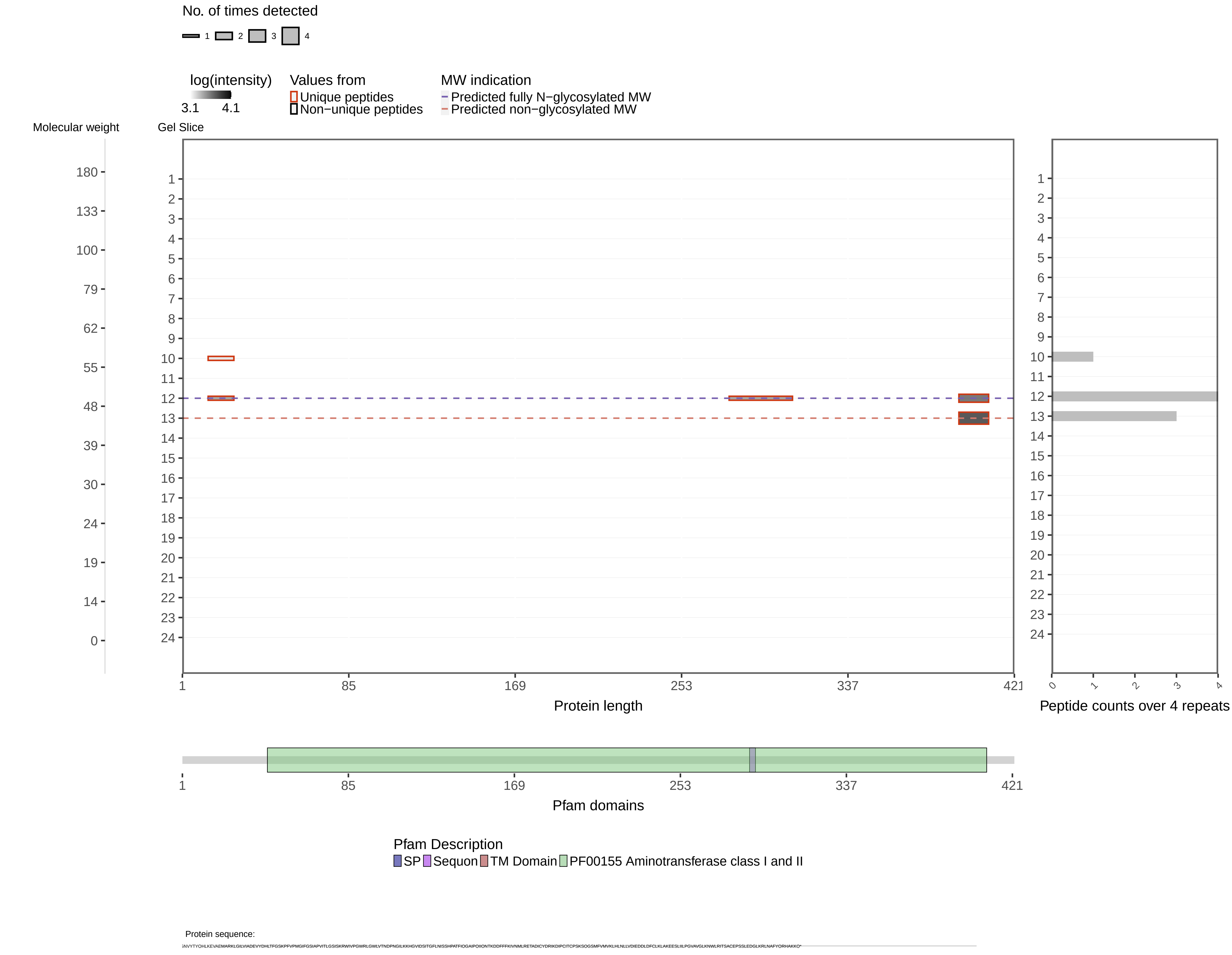

### NbL16g22030.1 probable_ADP−ribosylation_factor_GTPase−activating_protein_AGD6 [MW = 49.18 kDa]

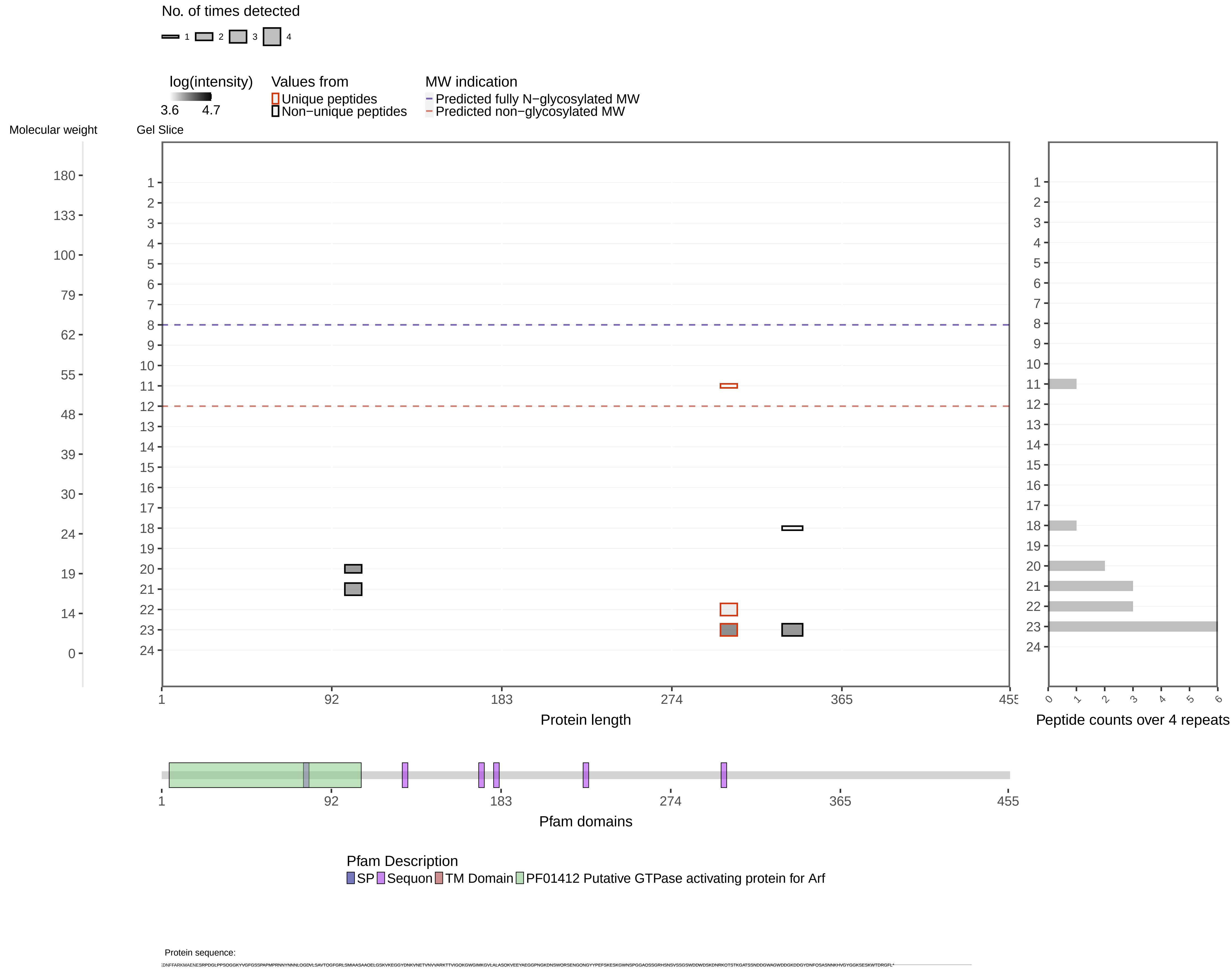

NbL16g21970.1 60S_ribosomal_protein_L23a−like [MW = 17.24 kDa]

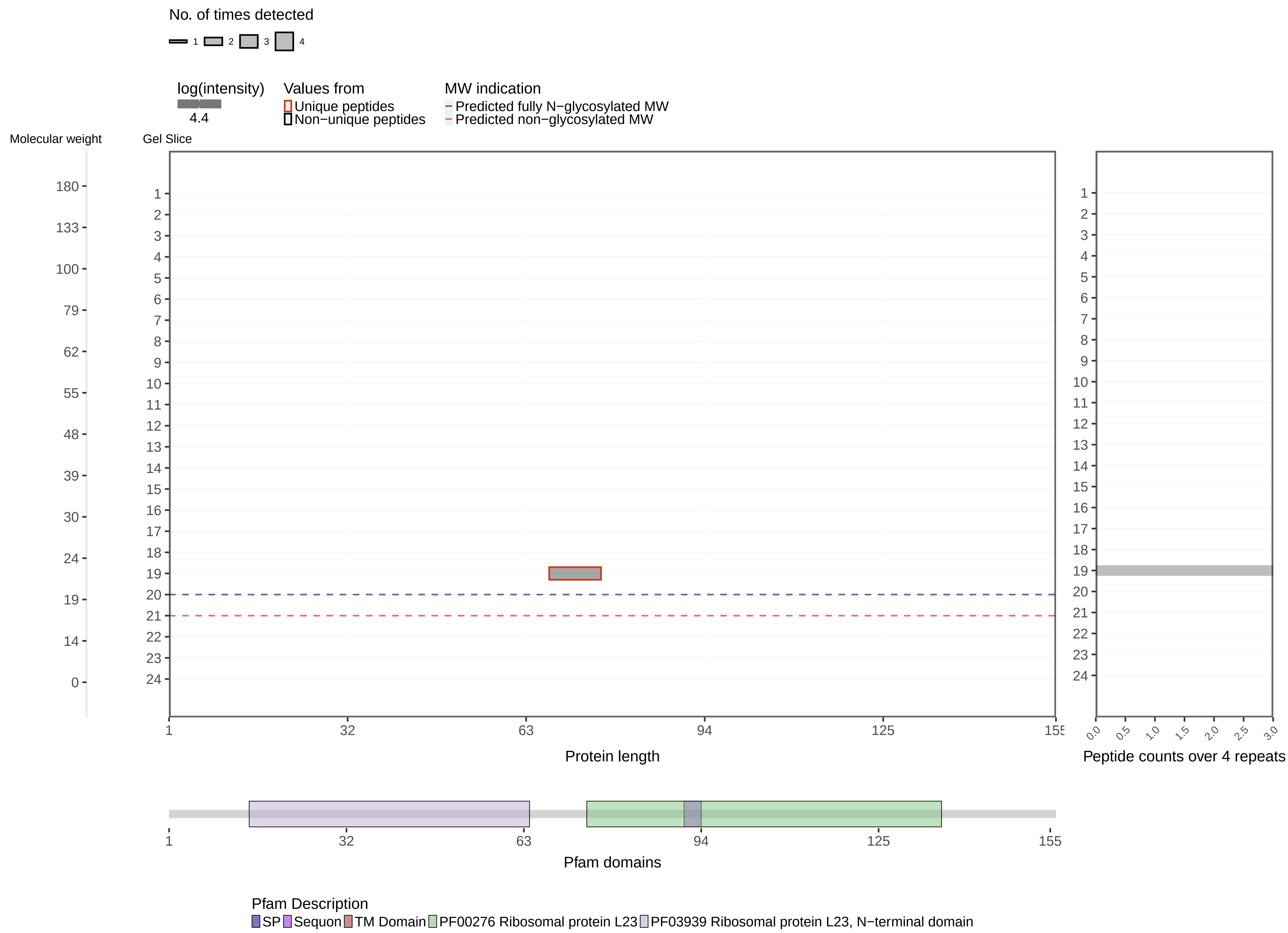


Protein sequence:

APGRNKLDHYGILKYPLTTESAMKKIEDNNTLVFIVDIKADKKKIKDAVKKMYDIQTKKVNTLIRPDGTKKAYVRLTPDYDALDVANKIGII*

### NbL16g21880.1 TIP2−1−like [MW = 10.04 kDa]

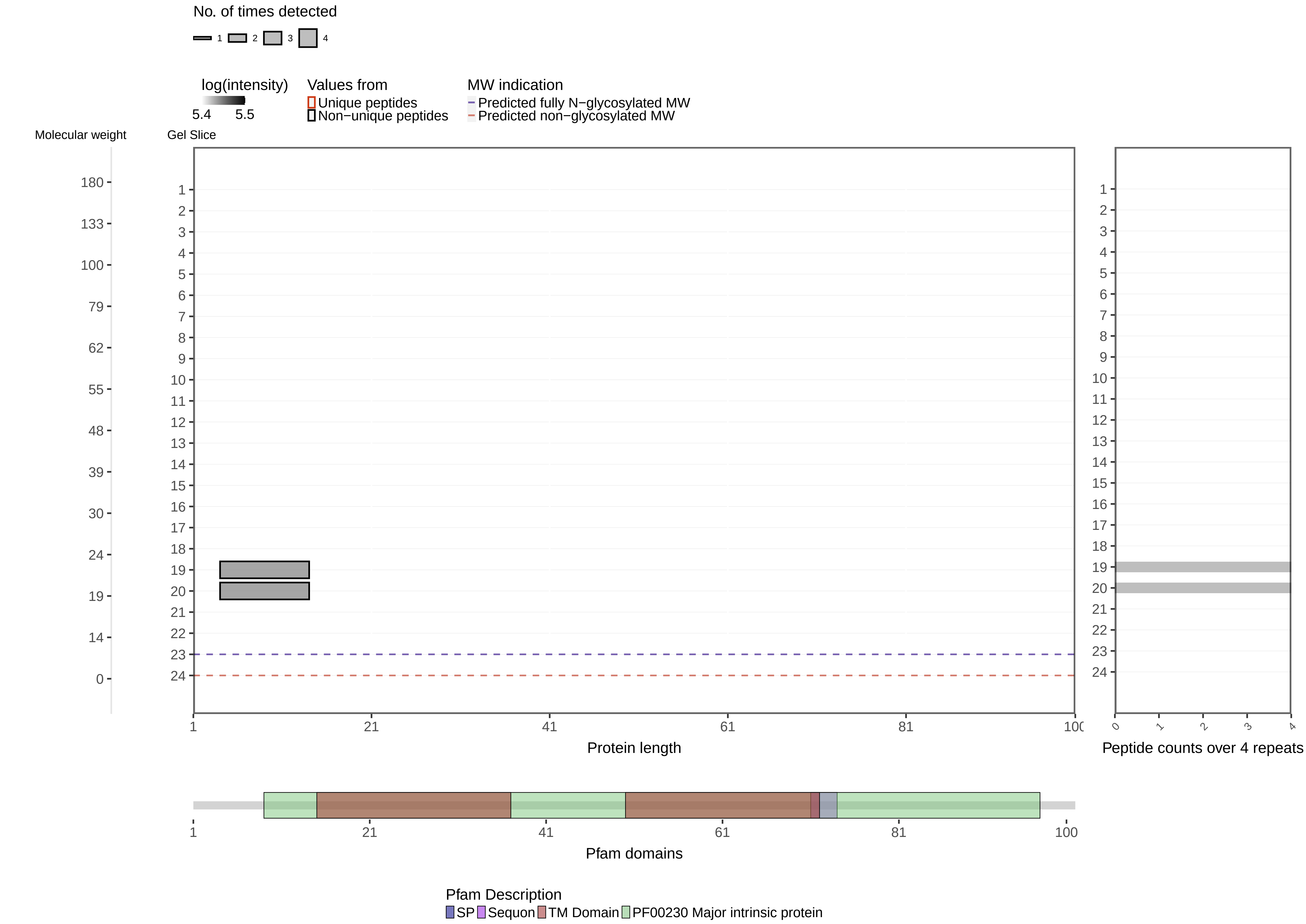


Protein sequence:

YNKLTADAALDPAGLVAVAVCHGFGLFVAVAVGANISGGHVNPAVTFGLAVGGQITILTGLFLH

NbL16g21800.1 60S_ribosomal_protein_L4−1−like [MW = 44.61 kDa]

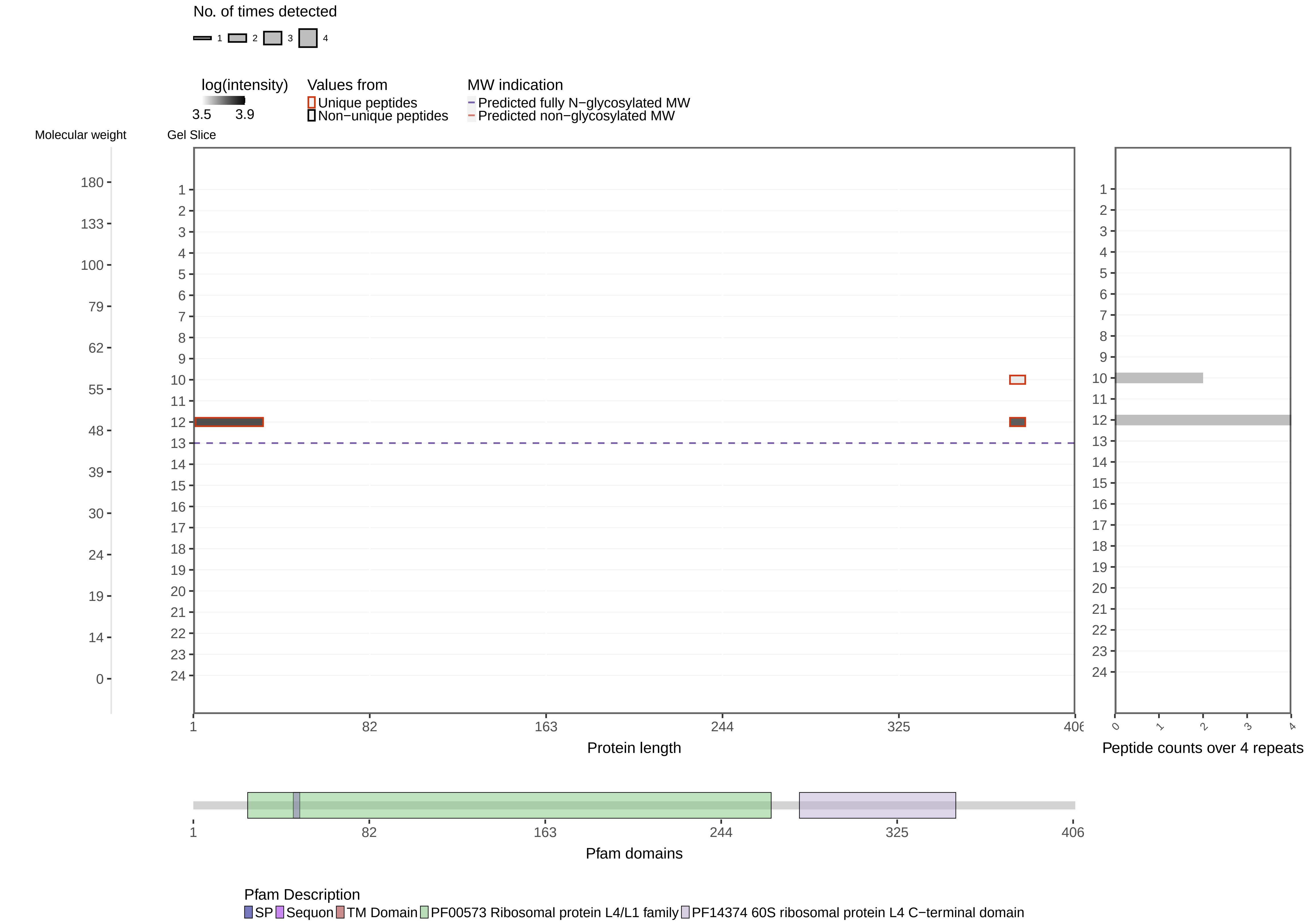


Protein sequence:

SHAIRPGKGKMRNRRYISRKGPLIVYGTEGAKLVKAFRNIPGVEICHVERLNLLKLAPGGHLGRFVVWTKSAYEKLDEIYGSFDKPSEKKKGYVLPRAKMVNADIARIINSDEVQSVVRPIKKDVKRATLKKNPLKNLNVLLKLNPYAKTARRMSLLAEAQRVKAKKEKLDKKRHQVTKEEASAIRSASSAWYKTMISDSDYAEFDVFTKWLGVSQ*

### NbL16g21520.1 tyramine_N−feruloyltransferase_10/30 [MW = 25.97 kDa]

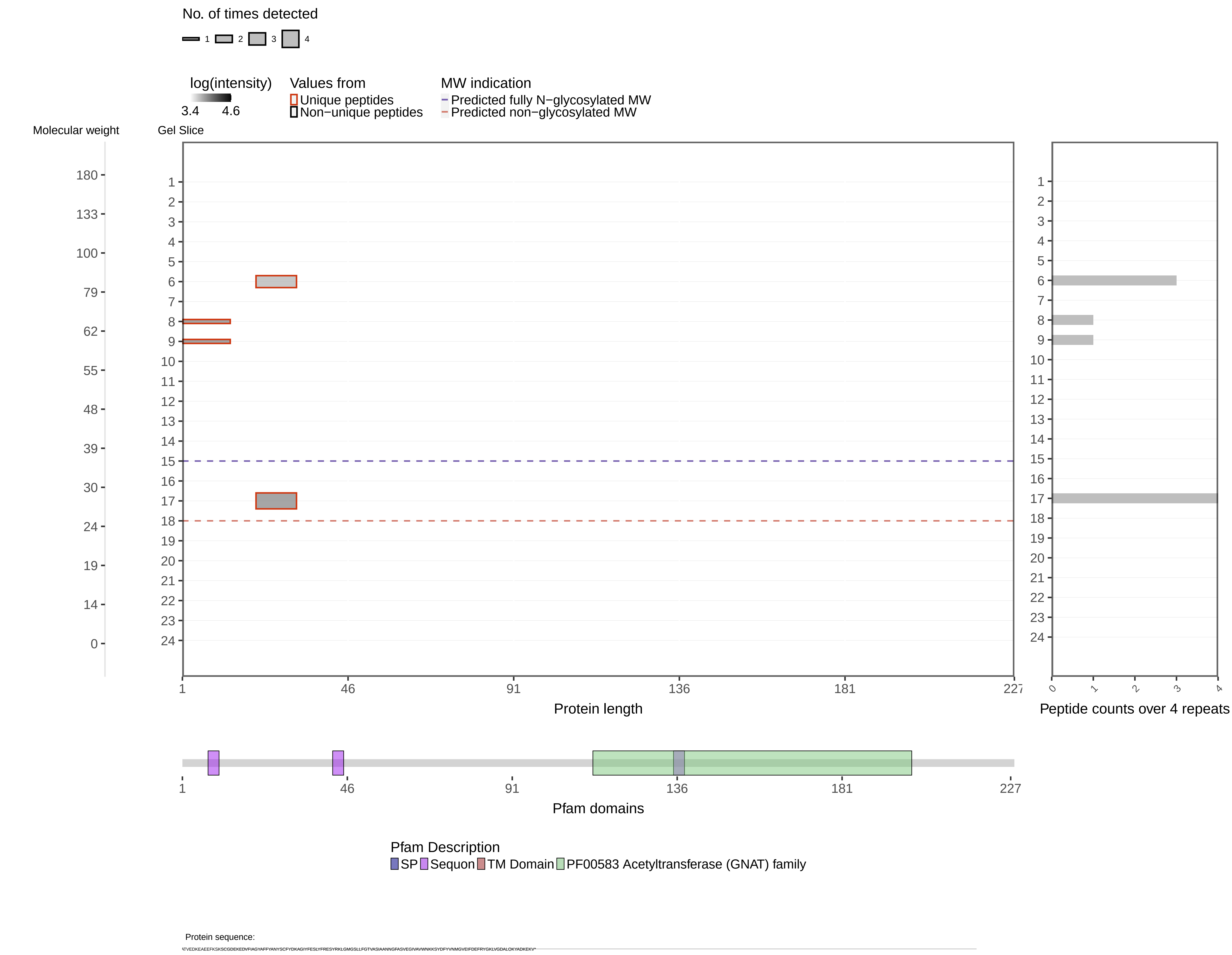

NbL16g21460.1 nucleoid−associated_protein_At4g30620_chloroplastic−like [MW = 21.48 kDa]

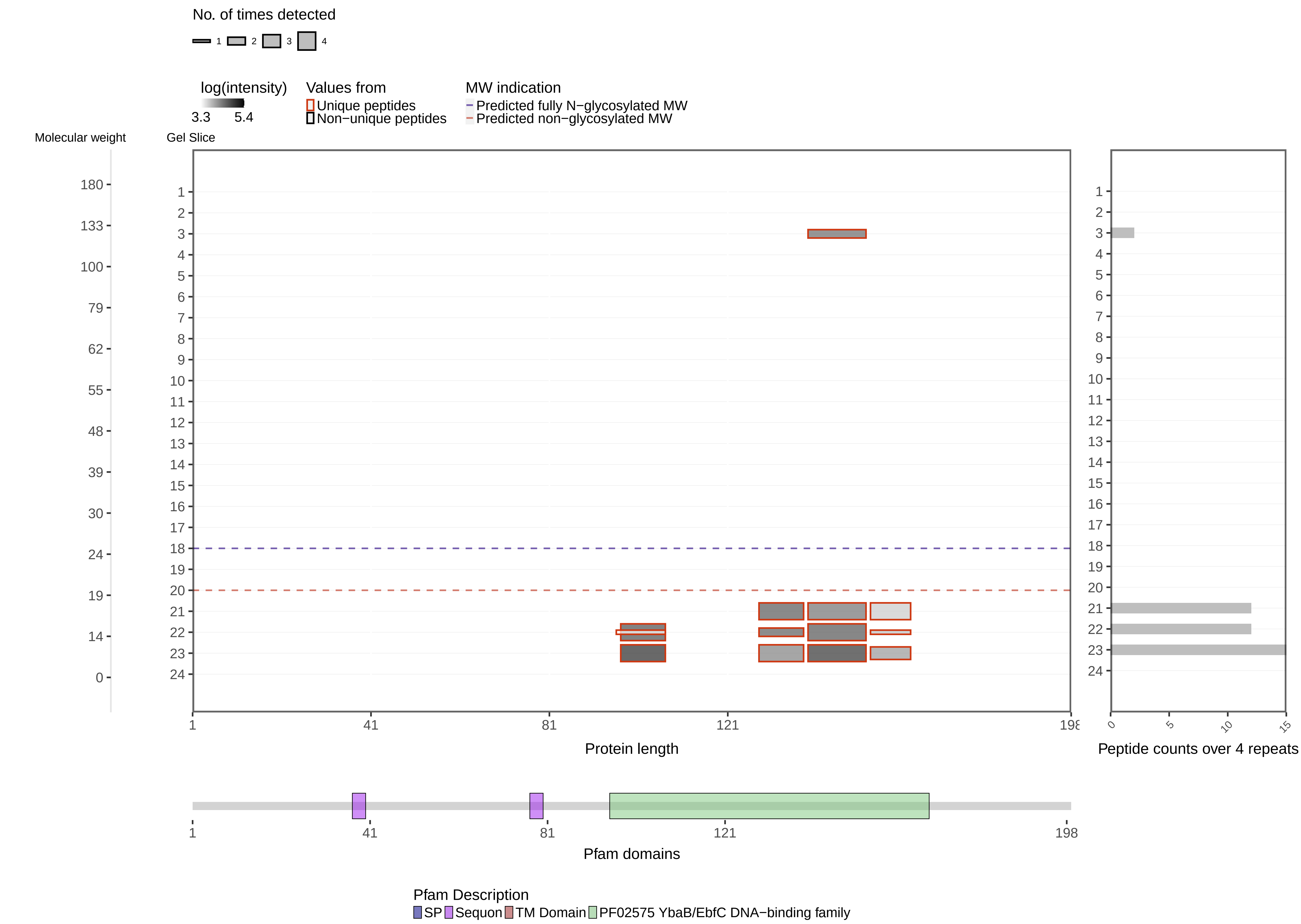


Protein sequence:

NMQNLYETVKKAQMVVQVEAVRVQKELAAAEFDGYCEGELIKATLSGNQQPVRIEITDAAMDLGPEKLSLLVTEAYKDAHQNSVLVRTIKLACRMVAFDHLVSTFLILRFYT*

### NbL16g21420.1 early_nodulin−like_protein_1 [MW − SP = 17.53 kDa]

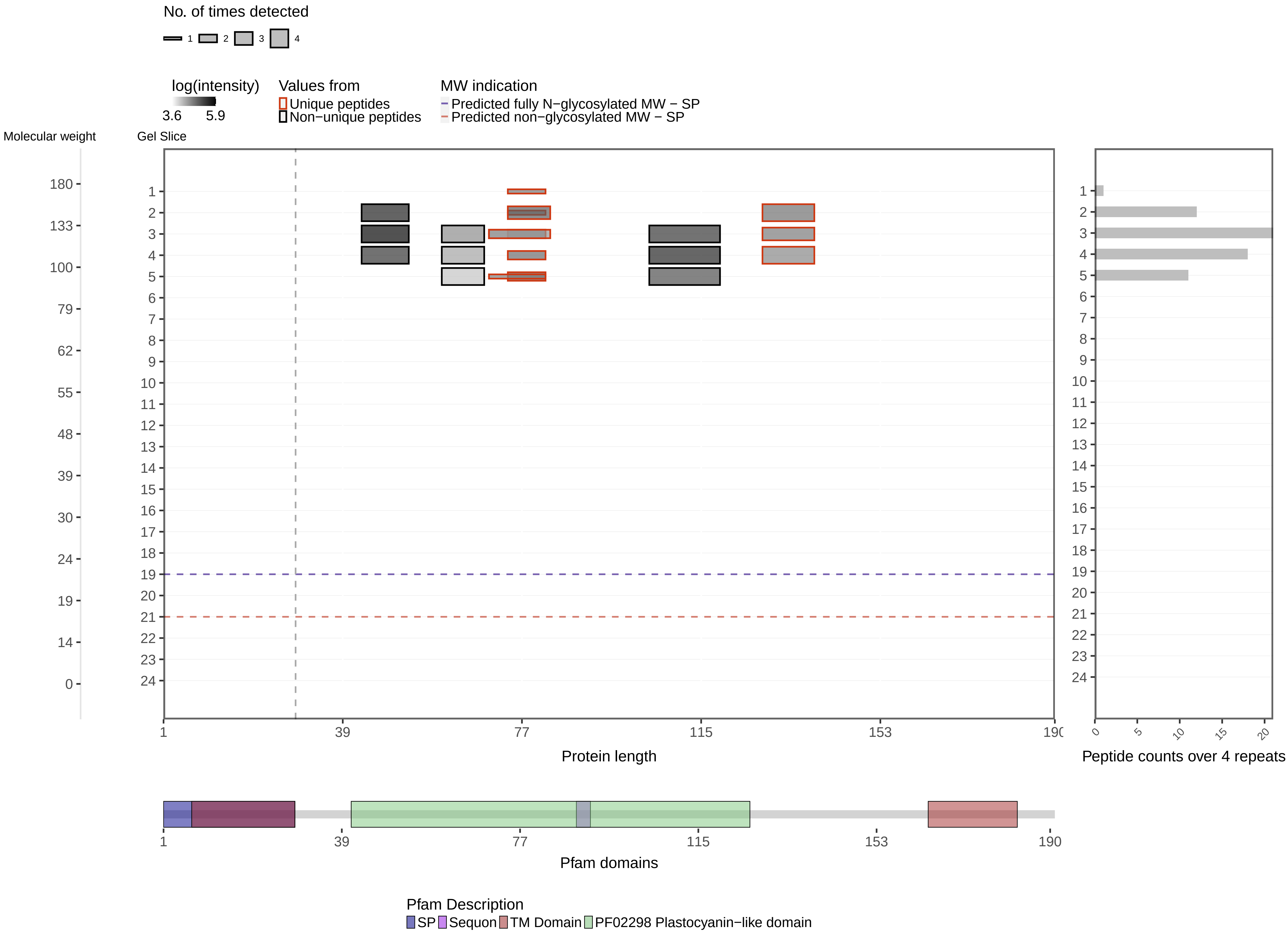


Protein sequence:

RDYVTCNTSSPIAVHNDGNTKIELTHSGAYYFISGAKGHCEQGQKLIVVTLSESNMRRFMGAPAPSQTEFDAPAMAPTSNSATLKAGLVMAFGVLMGFFFLNEGYSH*

### NbL16g21370.1 DNA−directed_RNA_polymerases_II_and_IV_subunit_5A−like [MW = 20.14 kDa]

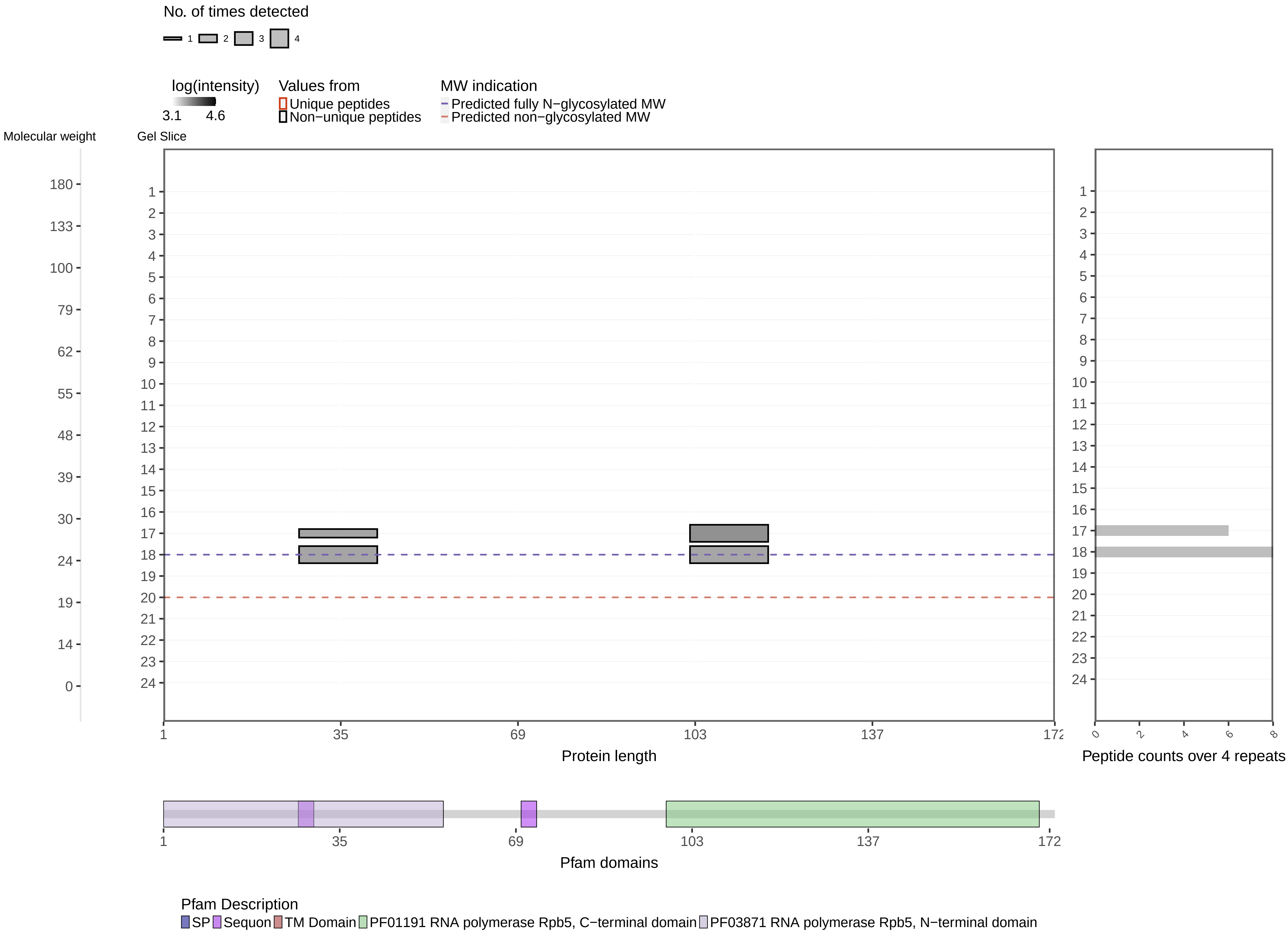


Protein sequence:

NLTPFARTCISEISTKFHLEVFQEAELLVNIKNHVLVPEHQPLTPEEKKTLLERYTVKETQLPRIQITDPIARYYGLKRGQVVKIIRPSETAGRYITYRYVV*

NbL16g21360.1 uncharacterized_LOC104225434 [MW = 48.02 kDa]

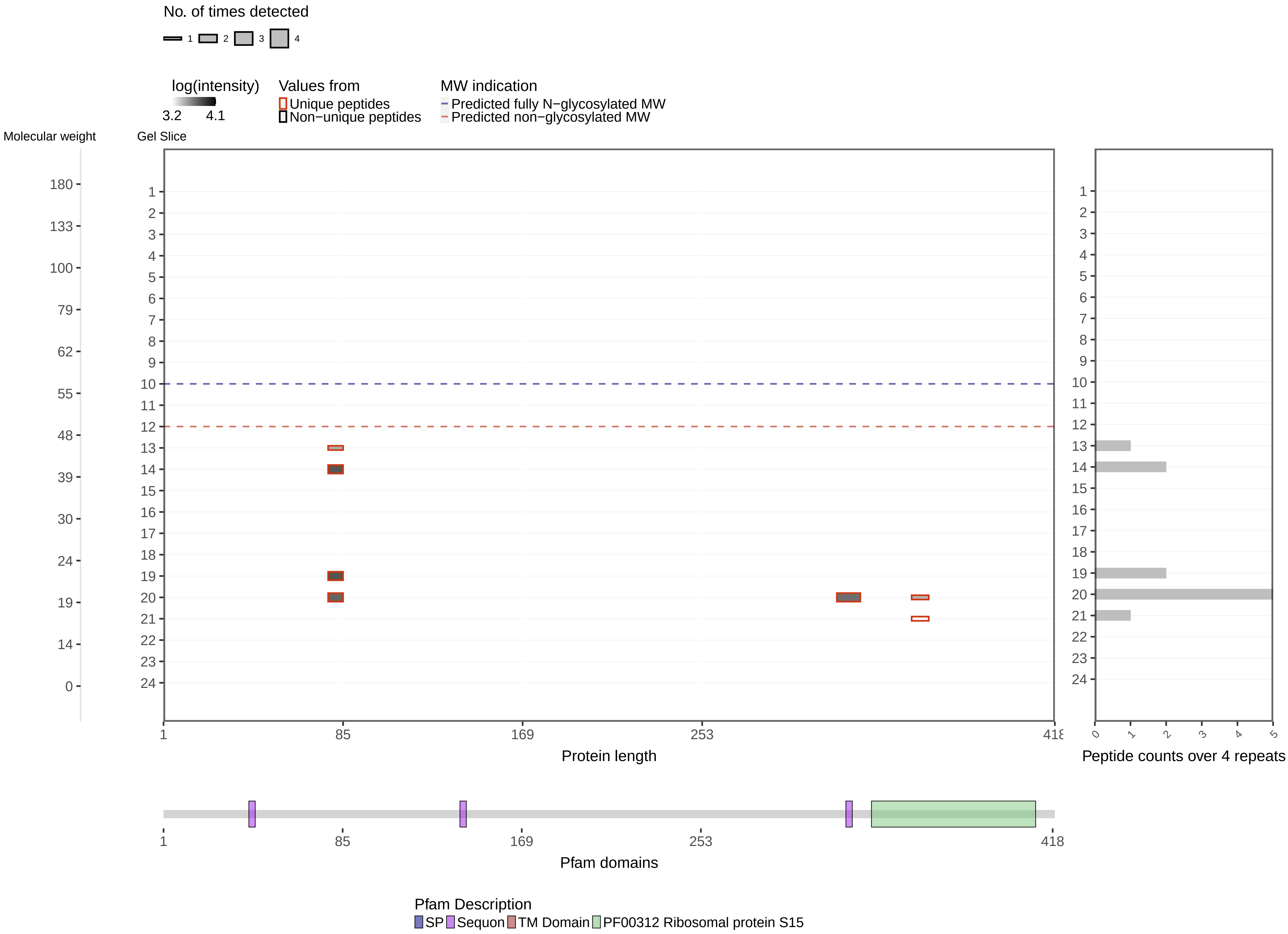


Protein sequence:

GDKEGMRPEFVRMYSHGELGEKLKMLRPAKKKEGEKGWFSLKELNDRLAKLRETEEKESESRMADSVFSEIKKSLVKMSTDDEEKAKRTMIQRLDVLGQLGGTPSFMLHPPKEELVEKYFHPDNMSSEEKMKLELQKVRDEFKMSESDCGSSRVQVAQLTTKIKHLSSVLHKKDKHSRKGLQAMVQKRKKLLKYLRRTDWDSYCLVLSKLGLRDNPDYKF*

### NbL16g21340.1 ABC_transporter_C_family_member_2−like [MW = 180.89 kDa]

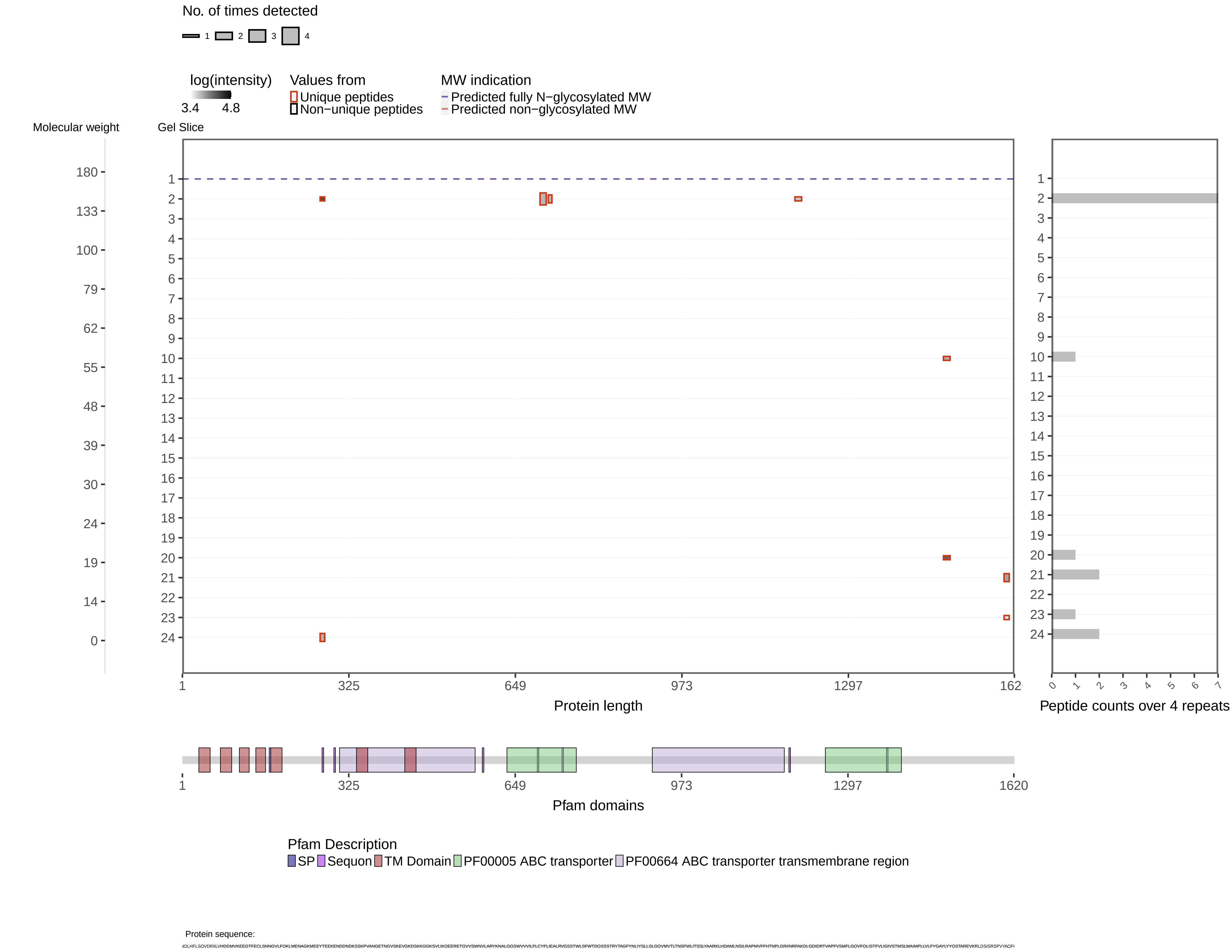

### NbL16g21330.1 tubulin_alpha−2_chain−like [MW = 49.77 kDa]

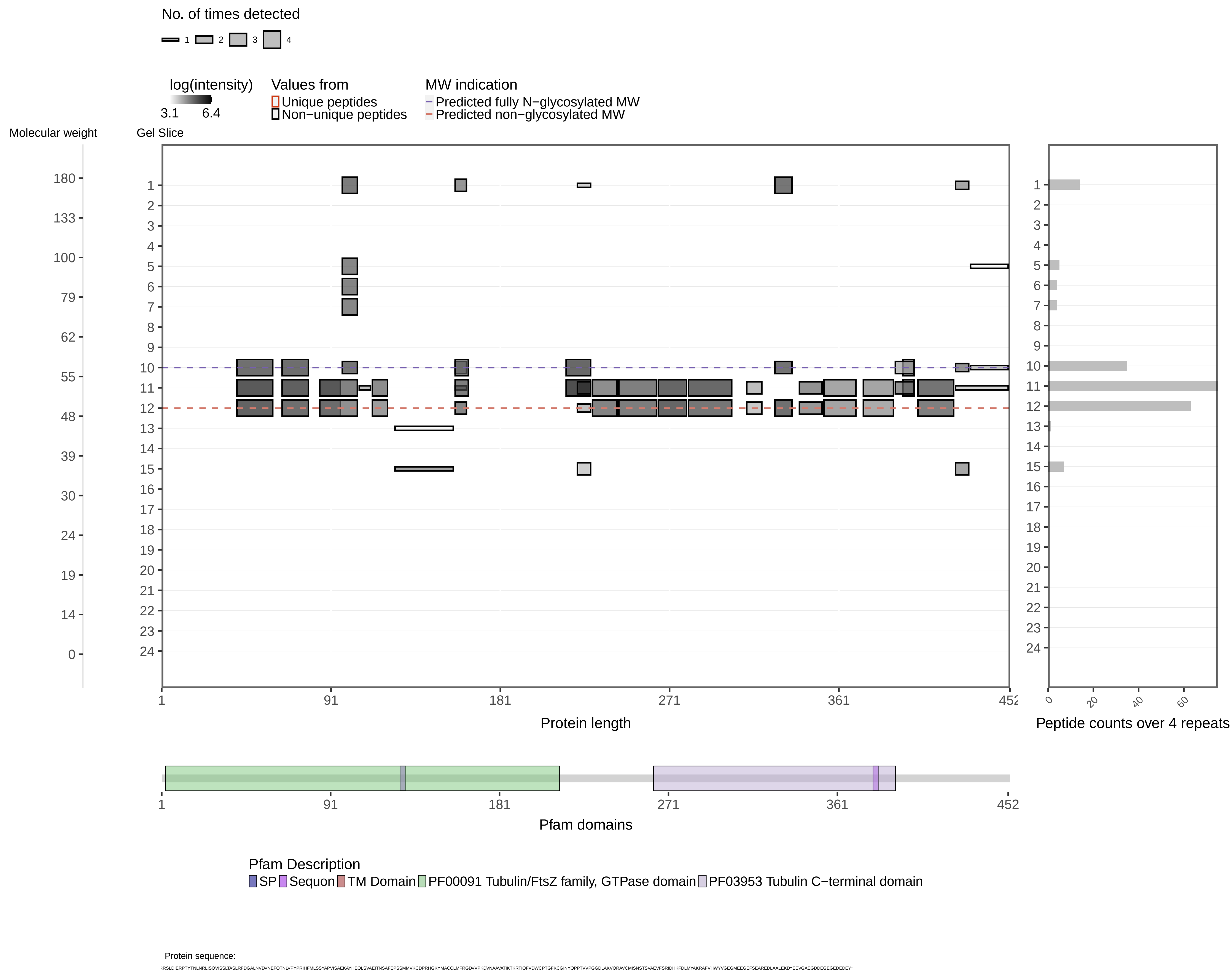

### NbL16g21320.1 60S_ribosomal_protein_L32−1−like [MW = 15.56 kDa]

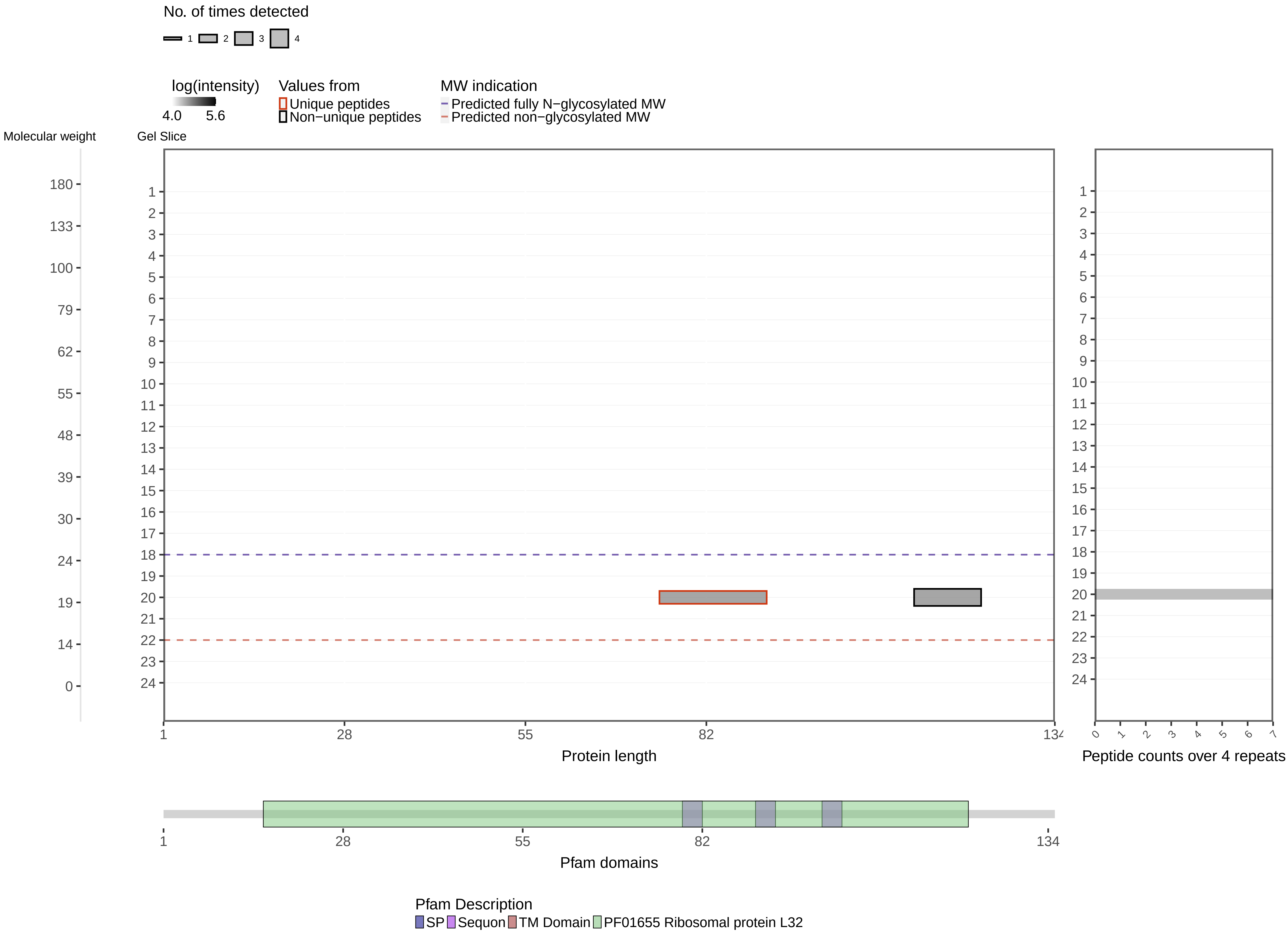


Protein sequence:

PNIGYGSDKKTRHYLPNGFKKFVVHNASELDVLMMHNRTYCAEIAHNVSTRKRKEIVERAAQLDVVVTNKLARLRSQEDE*

### NbL16g21290.1 photosystem_I_reaction_center_subunit_psaK_chloroplastic [MW = 13.4 kDa]

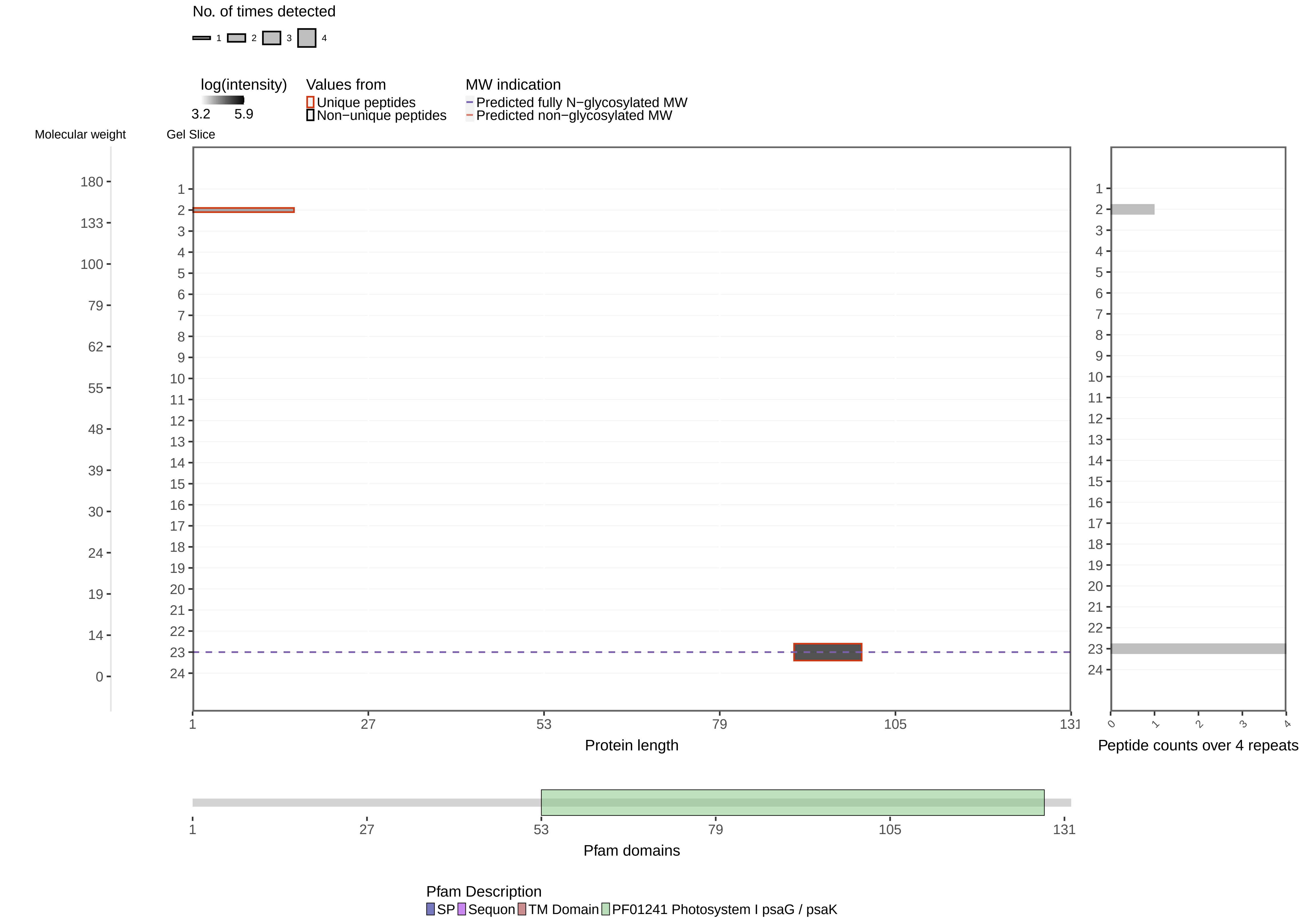


Protein sequence:

SPTNLIMVTSTSLMLFAGRFGLAPSANRKATAGLKLEVRDSGLQTGDPARFTLADTLACGVVGHIIGVGVVLGLKNIGAL*

### NbL16g21230.1 3−isopropylmalate_dehydrogenase_2_chloroplastic−like [MW = 43.43 kDa]

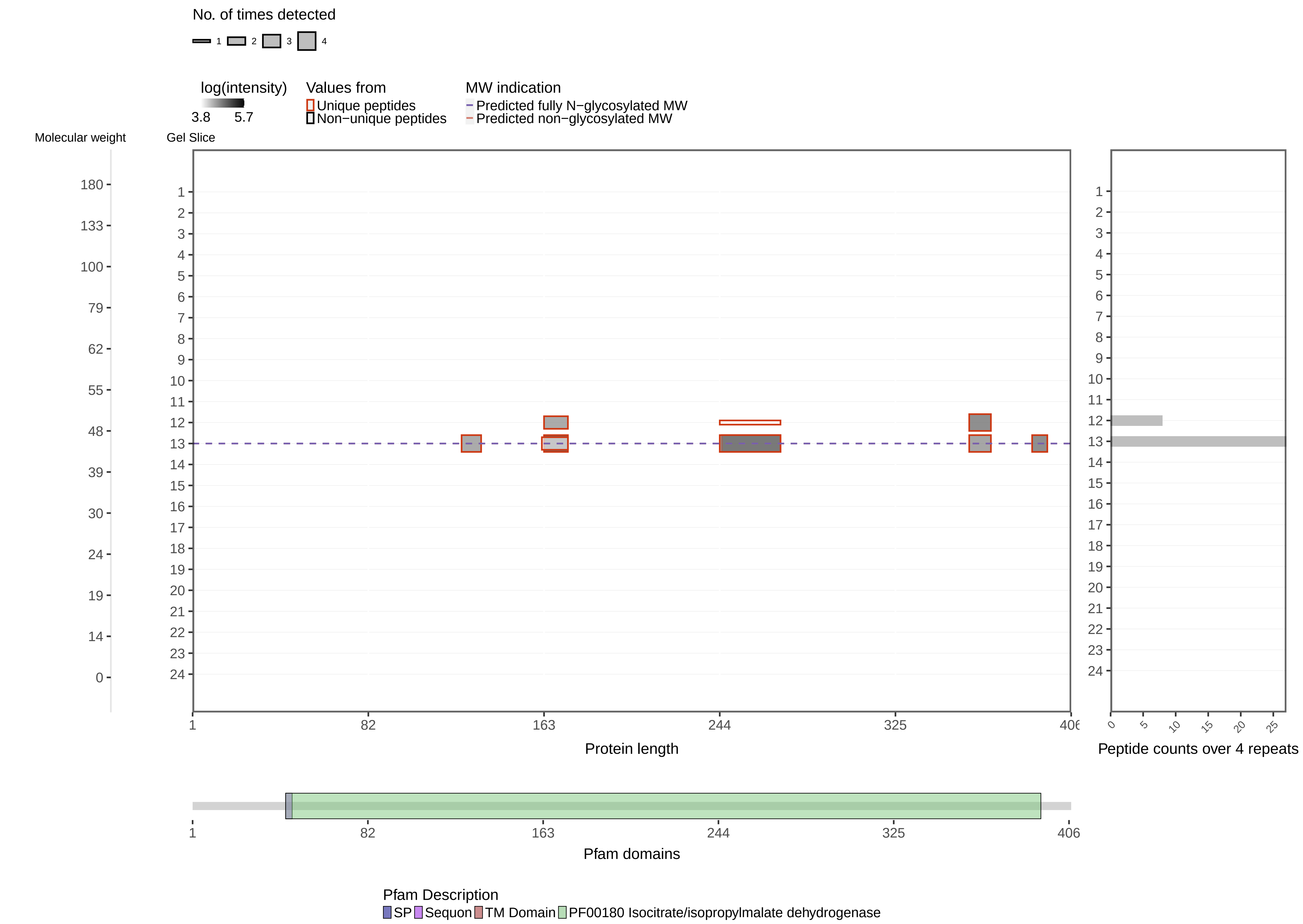


Protein sequence:

TNENGQEIGFNTEVYAVYEIERIARIAFETARKRRGKLCSVDKANVLEASMLWRKIVTALASEYPDVELSHMYVDNAAMQLVRNPKQFDTVVTNNIFGDILSDEASMITGSIGMLPSASLGETGPGLFEPIHGSAPDIAGQDKANPLATVLSAAMLLKYGLGEEKAAQRIEAAVLDALNRGFRTGDIHSAGHKLVGCKEMGEEVLKSIDSKTPAAV*

NbL16g21110.1 putative_hydrolase_C777.06c [MW = 35.9 kDa]

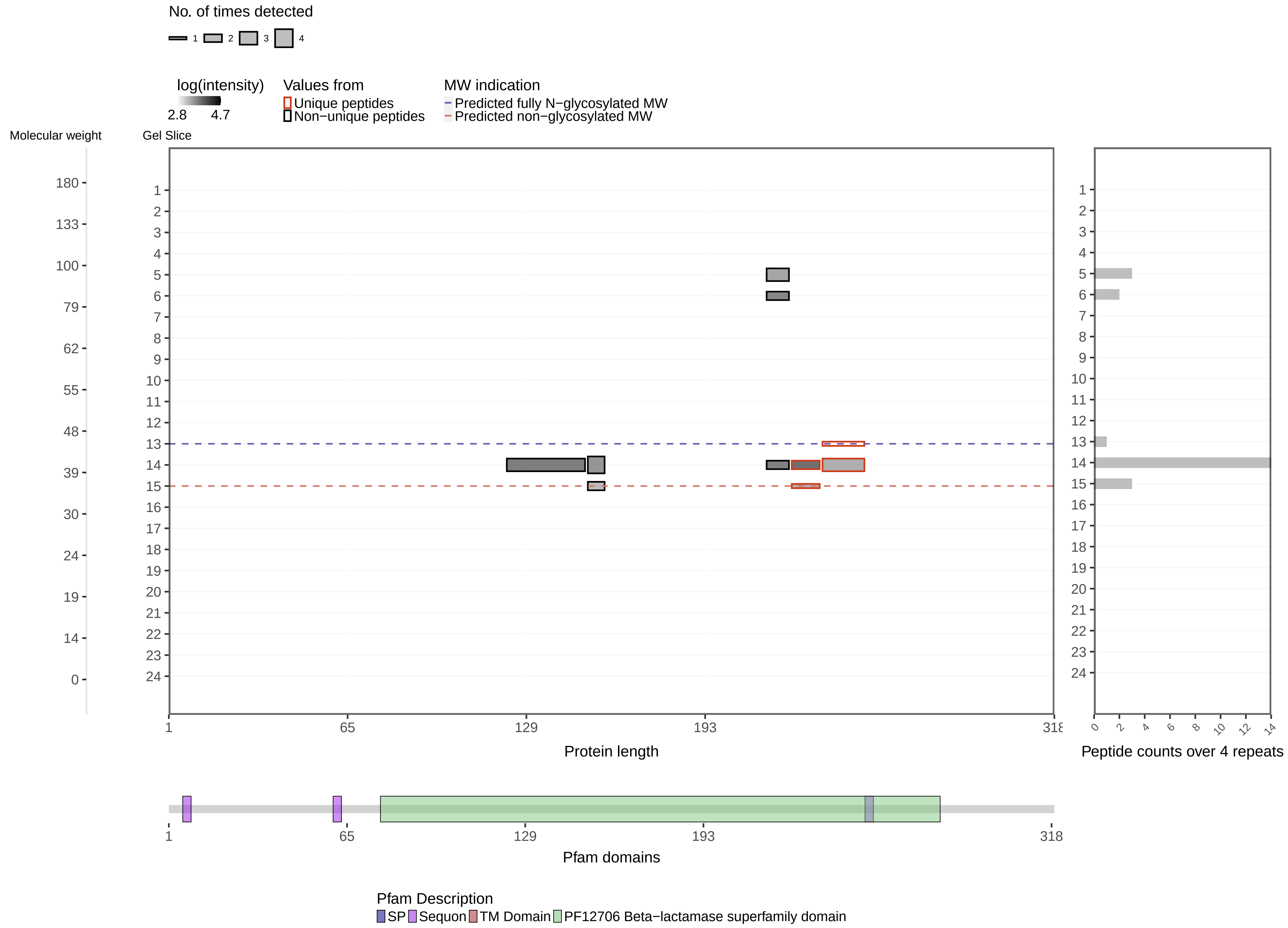


Protein sequence:

IDQKFPYLVQKKLKEGQEIRRVAQLDWQIIENDCETPFVASGLEFVPLPVMHGEDYVCLGFLFGKNFRVAYISDVSRFLPTTVSYISKHSGQQLDLLILDTLYKNGSHNVHLCLTQTLEALKMLCPKRALLIGMTHEFDHHKDNEFLVEWSRREGIEVQLAHDGLRVPVDL*

NbL16g21000.1 ornithine_aminotransferase_mitochondrial−like [MW = 53.43 kDa]

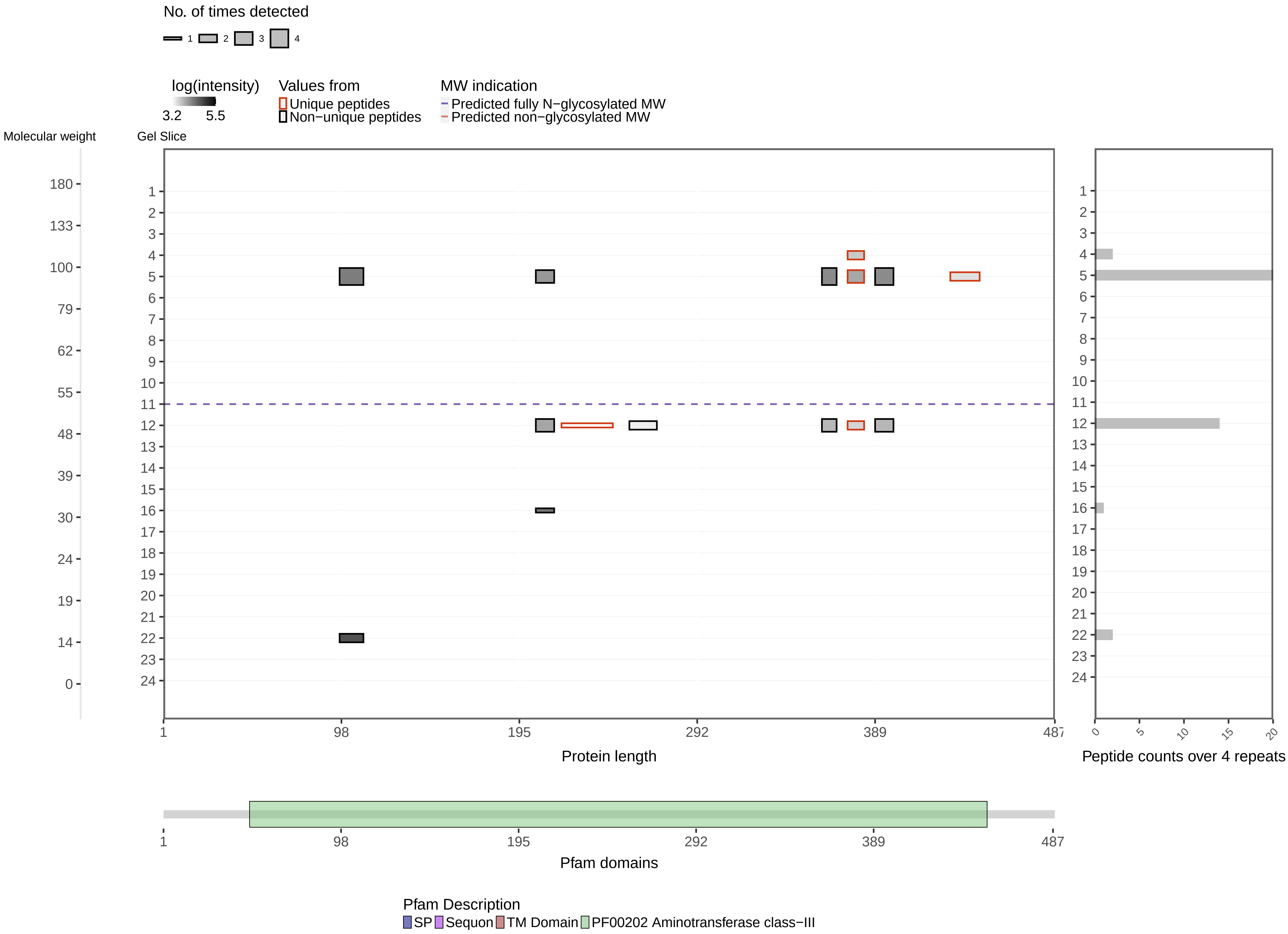


Protein sequence:

EPIQGEAGVIIPPEGYLKAVRDHCSKYNILMIADEIQSGLARSGRLLACDGEEVRPDIVILGKALGGGVLPVSAVLADKDVMLCIQAGEHGSTFGGNPLASAVAIASLDVIRDEGLAERYGLLWSKLDIERSAQMGEQLRHQLMKVQGQFPNFIKEVRGKGLFNAVELNSKALLPVTAYDICMKLKERGILAKPTHDSIIRLTPPLSMSLEELQEGSKALHDVFELDLPKMQKEKPATVSHATSSFCDRCGRNLYGSS*

NbL16g20970.1 protease_Do−like_5_chloroplastic [MW = 32.64 kDa]

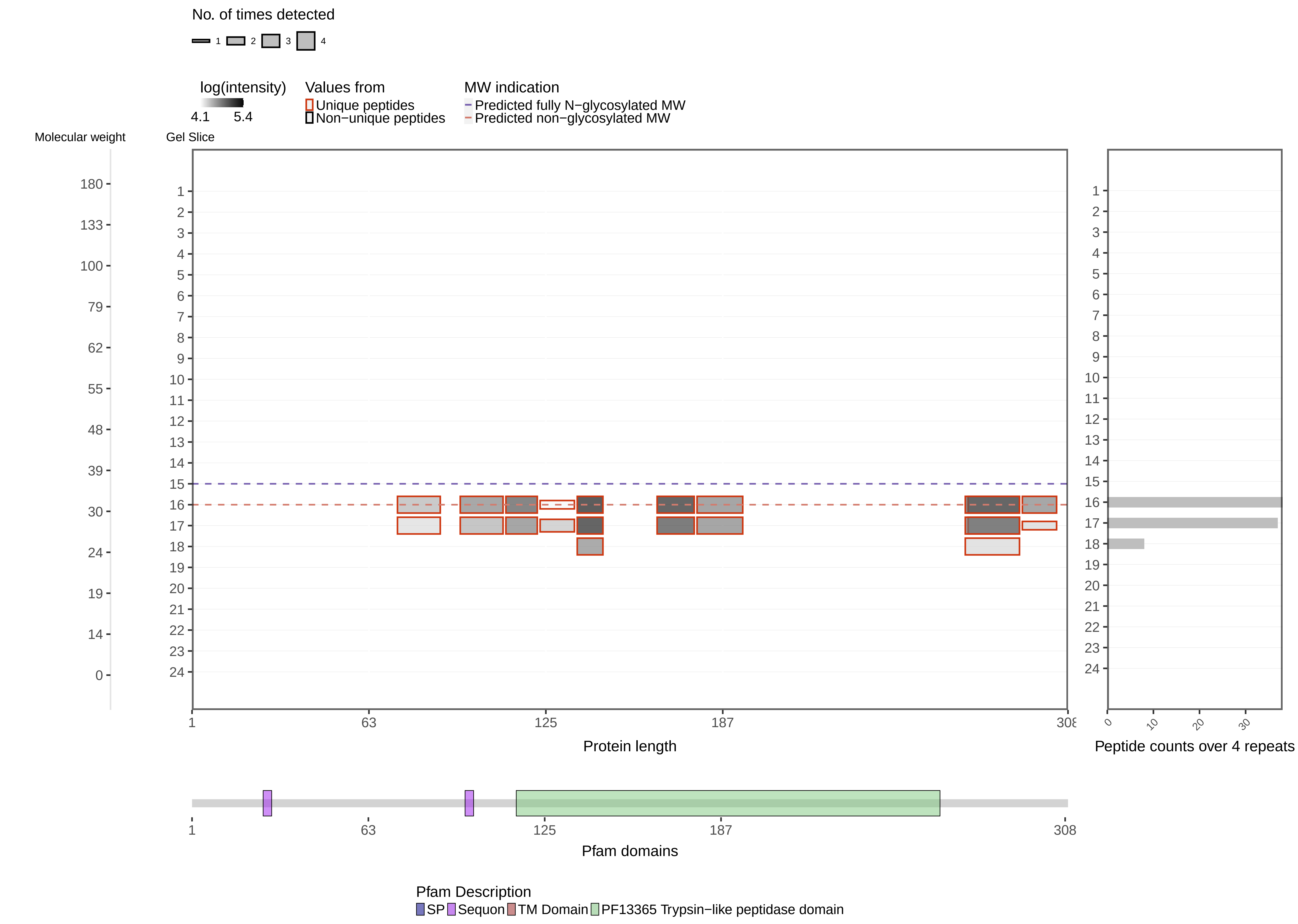


Protein sequence:

SGLQRCKVSLVNAKGESIAKEAKIVGVDPAYDLAVLKVDVEGDEVKPISVGTSRGLRVGQSCFAIGNPYGFENTLTTGVVSGLGREIPAPNGAAIKGAIQTDAAINAGNSGGPLIDSSGHVIGINTATFTRRGSGMSSGVNFAIPIDTVVRTIPYLIVYGTTYKDRY*

NbL16g20800.1 prefoldin_subunit_6−like [MW = 14.9 kDa]

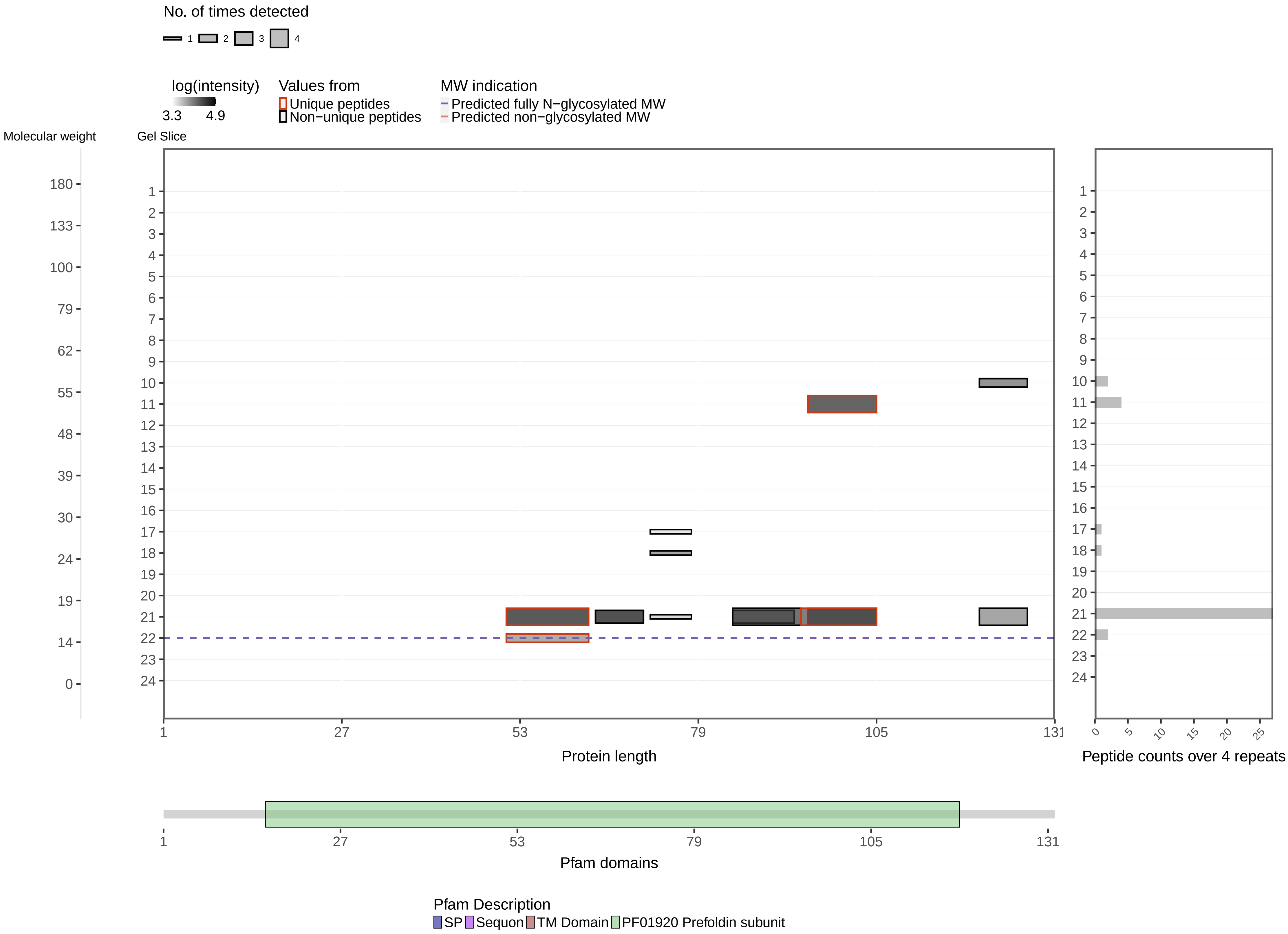


Protein sequence:

.DLLNEDANVYKLIGPVLVKQDLAEARANVKKRIEYISAELKRLDGTLQDLEEKQNSKRKRILKFQQKIQSLQAGKGKA*

### NbL16g20750.1 60S_ribosomal_protein_L18a−like [MW = 21.45 kDa]

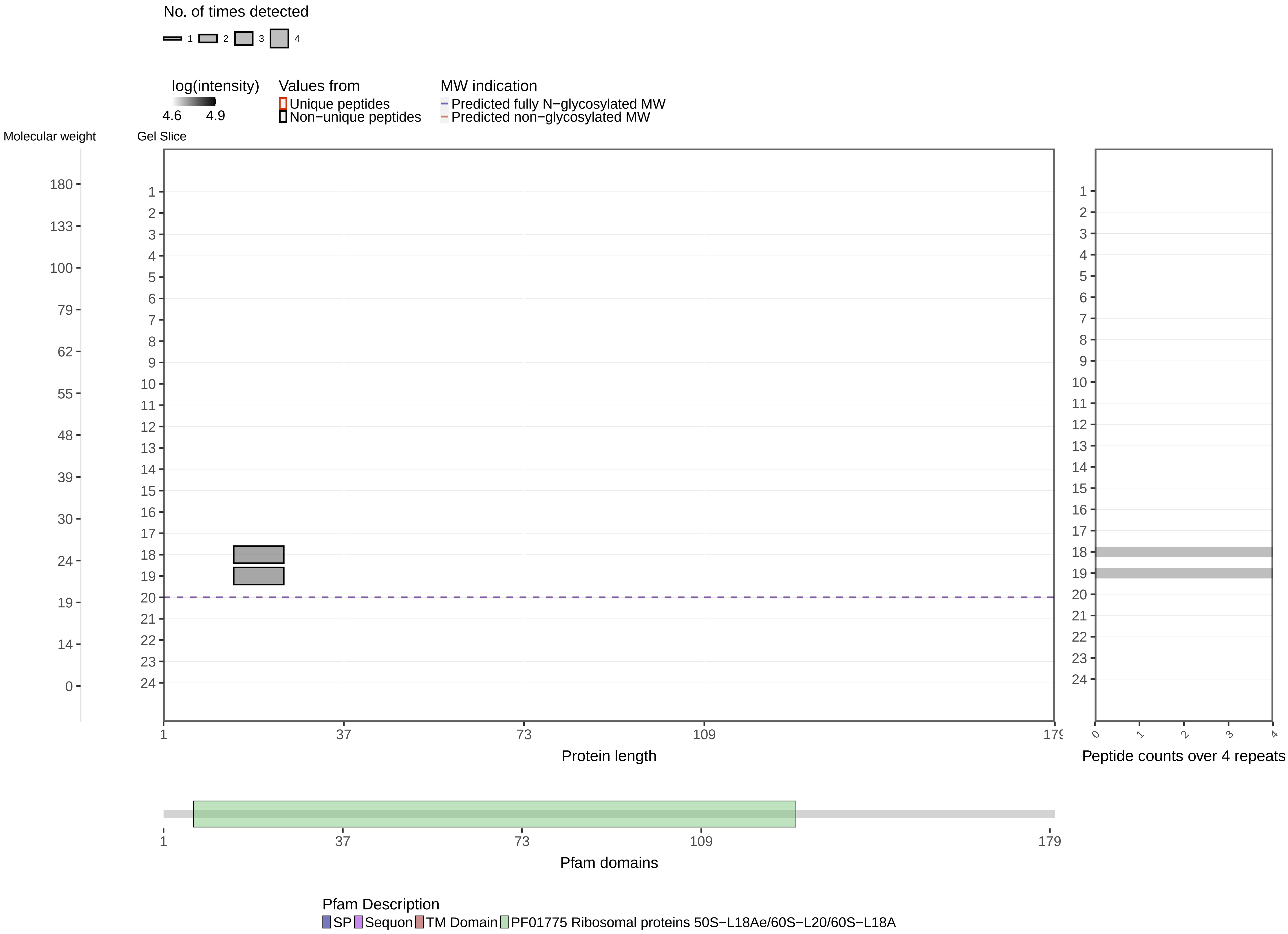


Protein sequence:

NYGIWLRYQSRTGYHNMYKEYRDTTLNGGVEQMYTEMASRHRVRHHCIQIIKTATIPAKLCKRESTKQFHDSKIKFPLVFKKVRPPSRKLKTTYKATRPNLFM*

### NbL16g20710.1 uncharacterized_LOC109214655 [MW = 66.18 kDa]

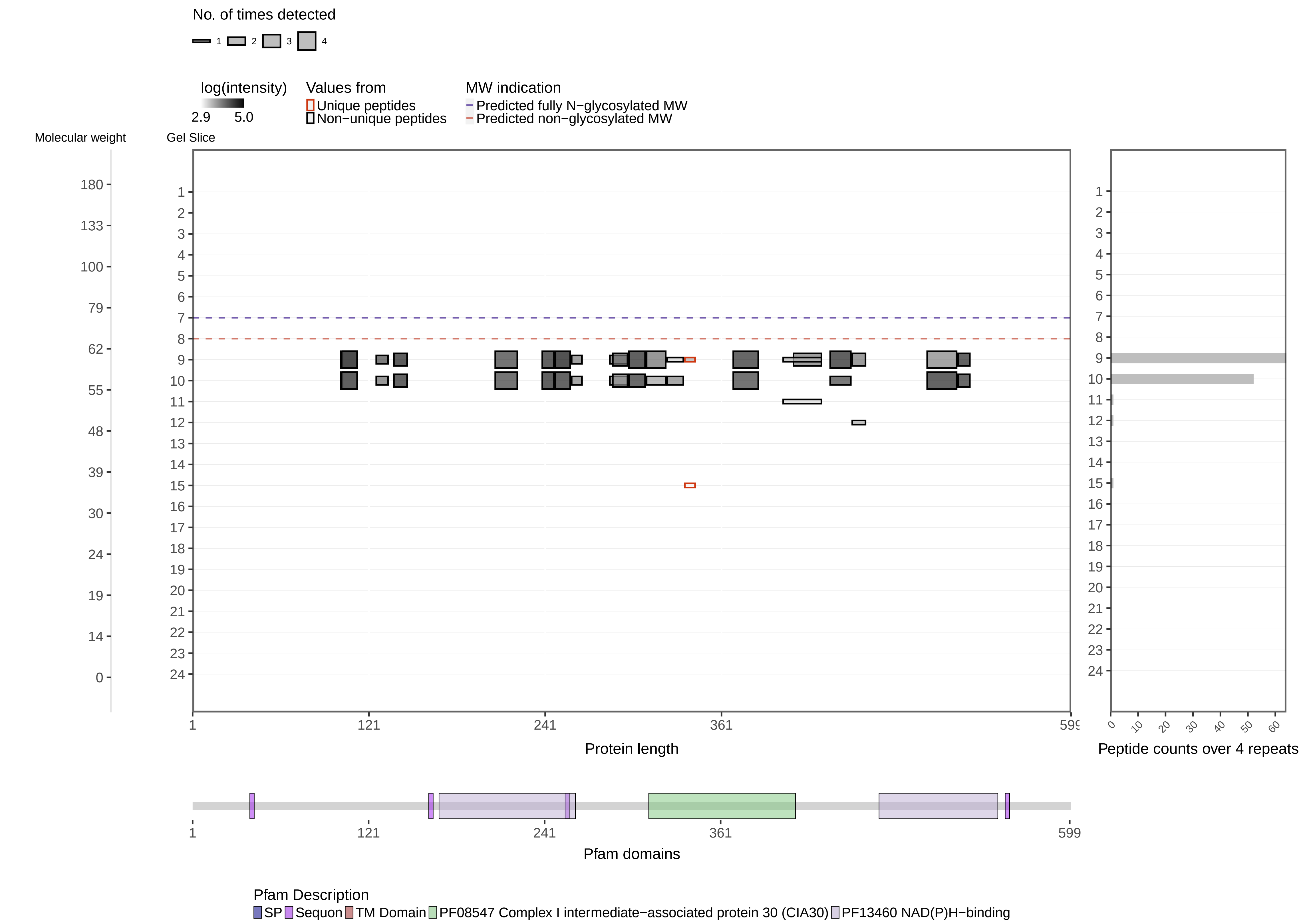


Protein sequence:

NbL16g20690.1 Golgi_to_ER_traffic_protein_4_homolog [MW = 37.07 kDa]

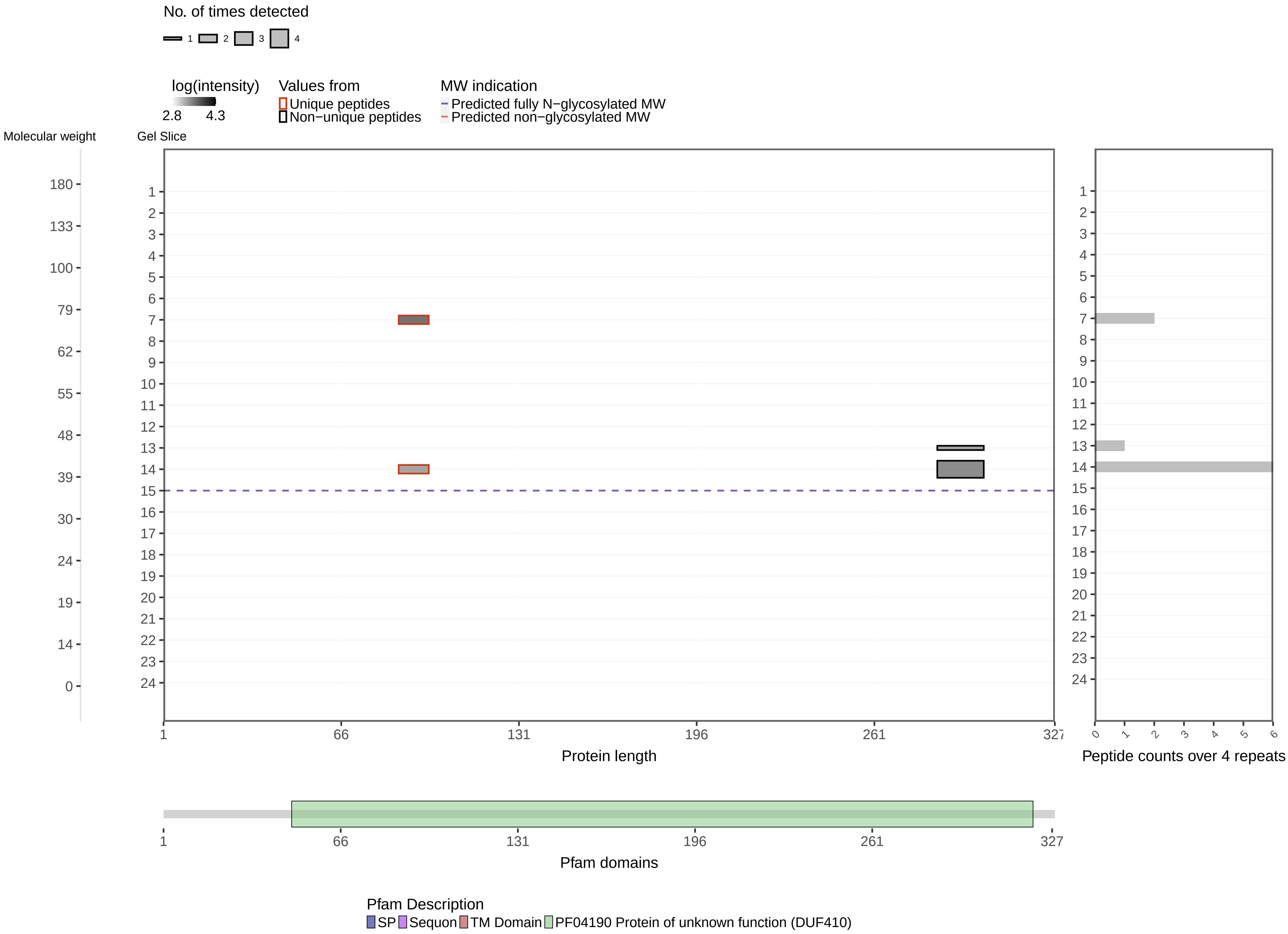


Protein sequence:

SAEFGAHKHGSPELHEMLADYIYSOSTEVDMVKVSFHFVRGRNLKKFASTIINFMGKCYPGEDDLAIARAILMYLSLGNLRDANKLMEEVEKEMEAKHLGFPOSELMOFVNYLLLTLORDALPLFNMLROSYKSSIDRDPLLNELLDEIAKKFYGVORKNPLOGIFGDIFKMMGGE*

NbL16g20680.1 aminoacyl_tRNA_synthase_complex−interacting_multifunctional_protein_1−like [MW = 30.46 kDa]

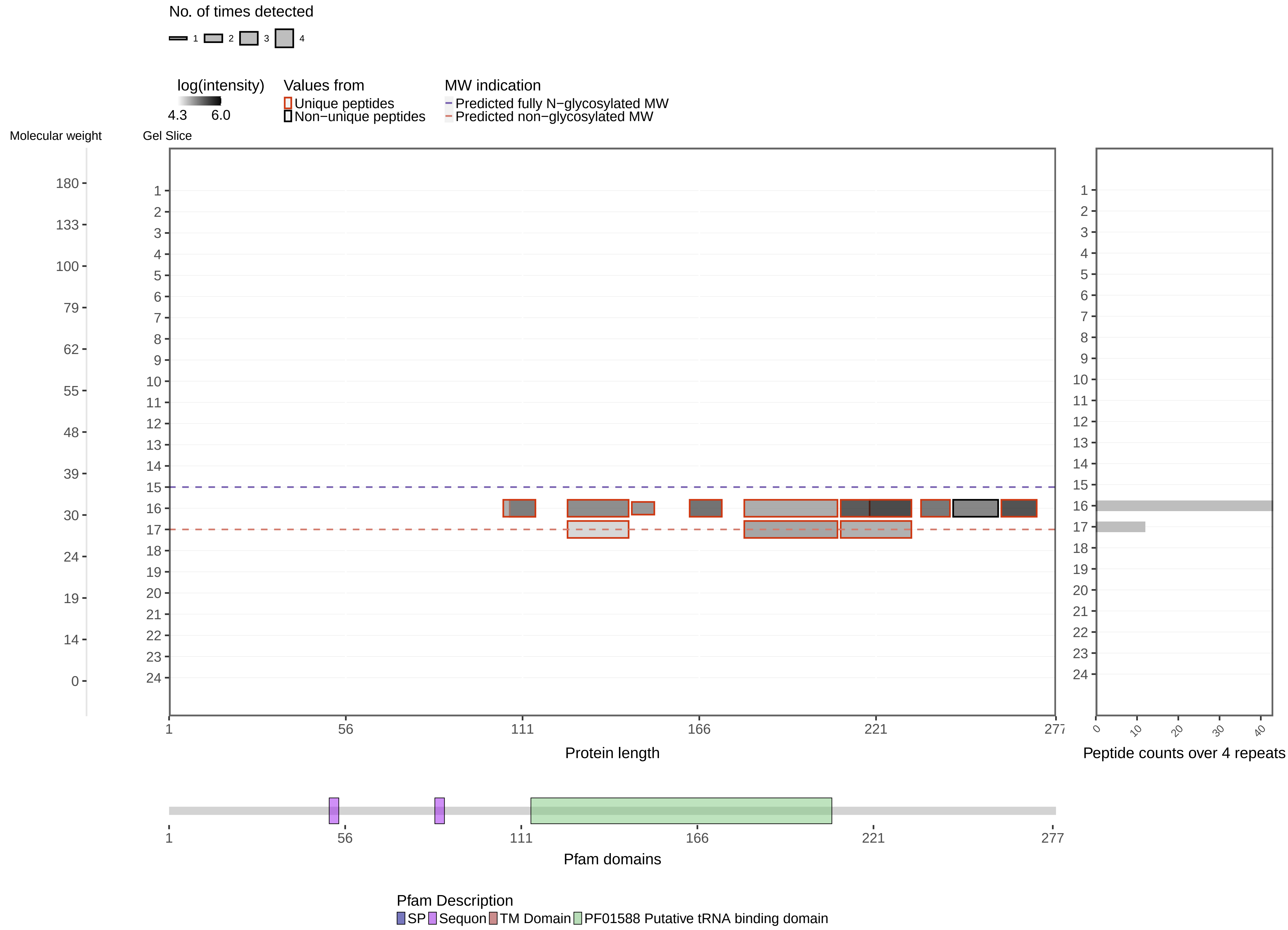


Protein sequence:

### NbL16g20660.1 guanosine_nucleotide_diphosphate_dissociation_inhibitor_2−like [MW = 49.68 kDa]

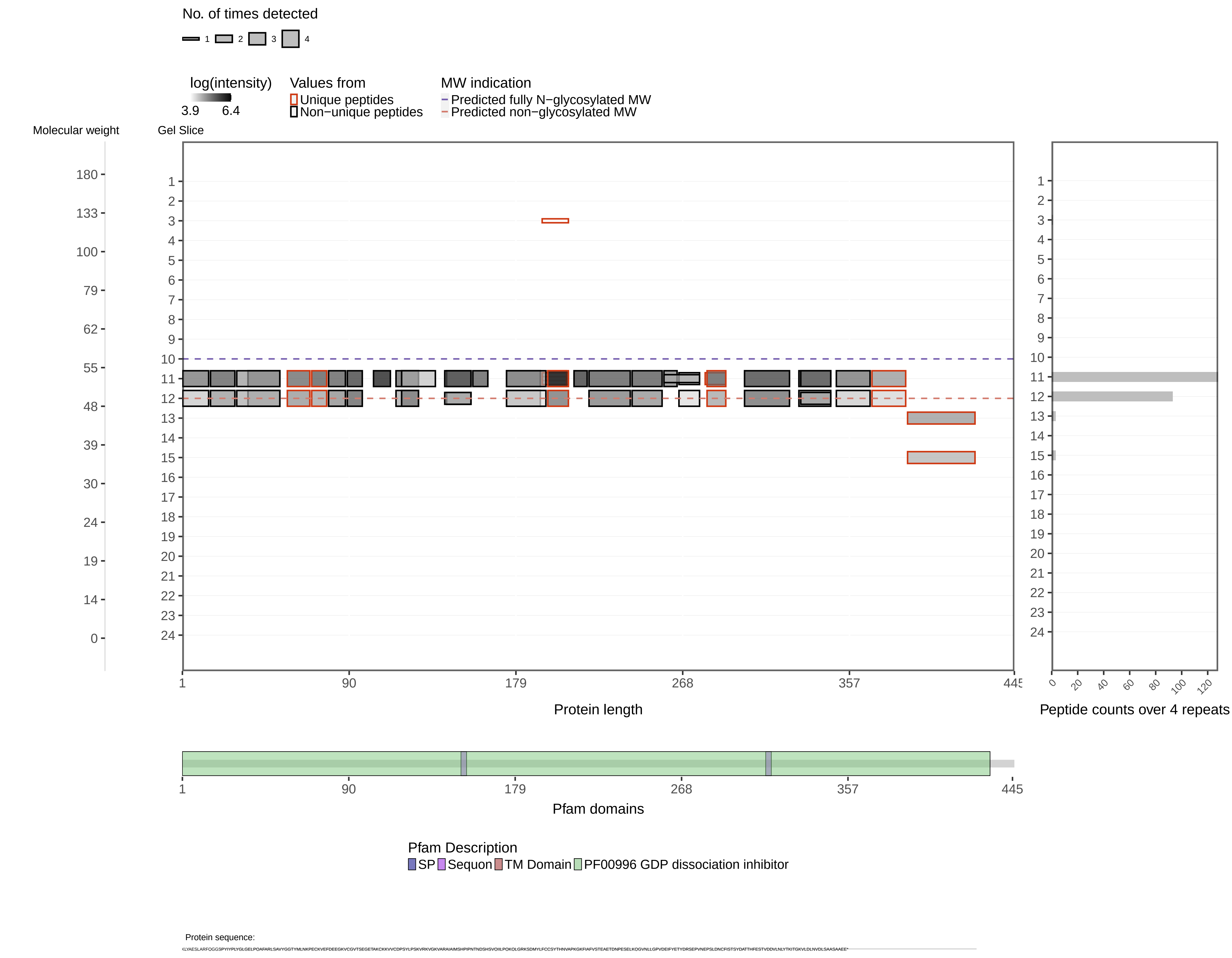

NbL16g20610.1 pyruvate_dehydrogenase_E1_component_subunit_beta−3_chloroplastic−like [MW = 44.46 kDa]

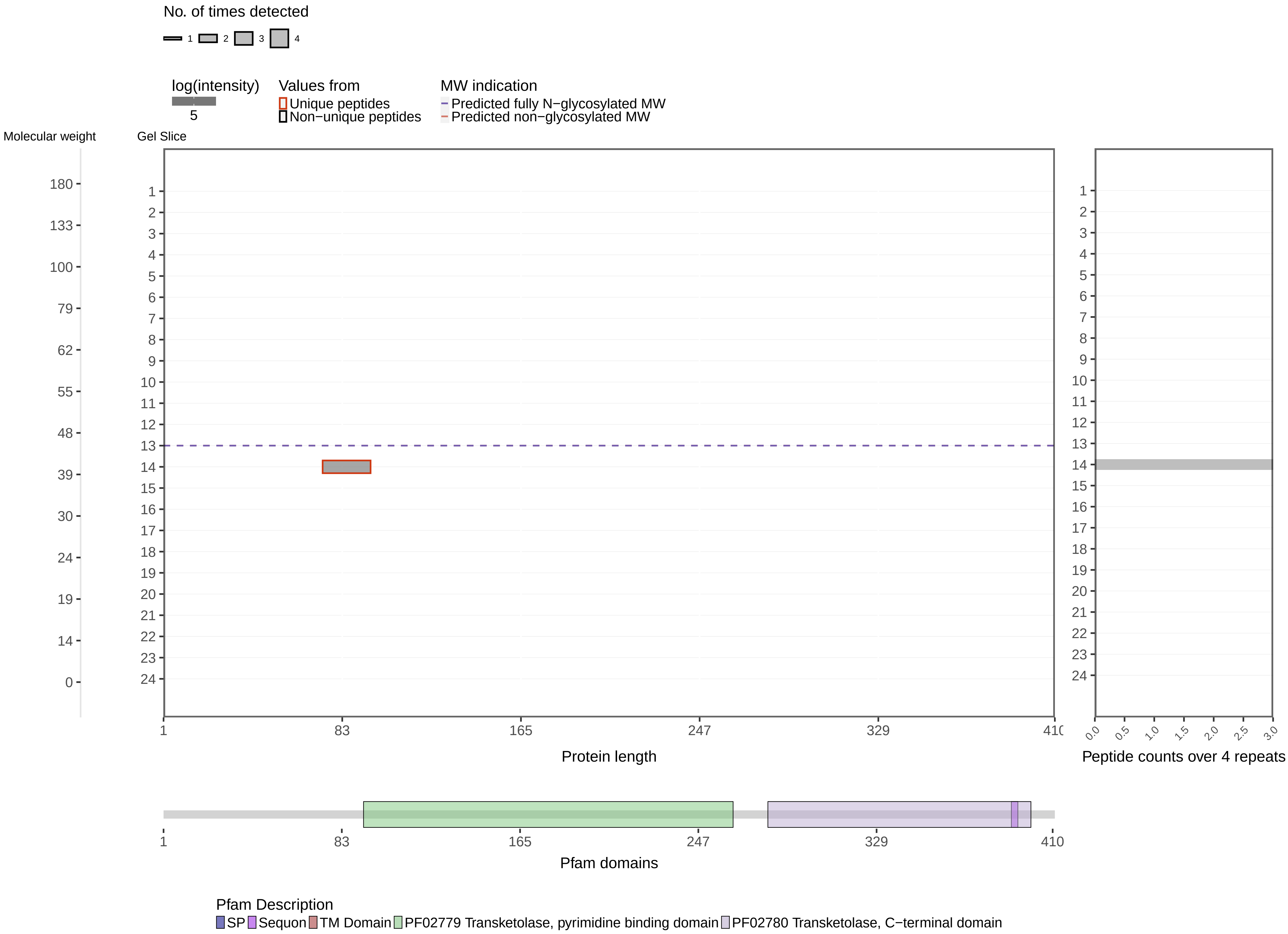


Protein sequence:

SGGQFKIPVVIRGPGGVGRQLGAEHSQRLESYFQSIPGIQMVACSTPYNAKGLMKAAIRSDNPVILFEHVLLYNLKERIPDEEYVLNLEEAEMVRPGEDVTILTYSRMRYHVMQAAKTLVDKGYDPEVIDIRSLKPFDLHTIGNSVKKTHRVVIVEECMRTGGIGASLTAAITENFHDYLDAPIVCLSSQDVPTPYAGTLENWTVVQPPQIVTAVEQLCQ*

NbL16g20560.1 vacuolar_protein_sorting−associated_protein_25−like [MW = 20.85 kDa]

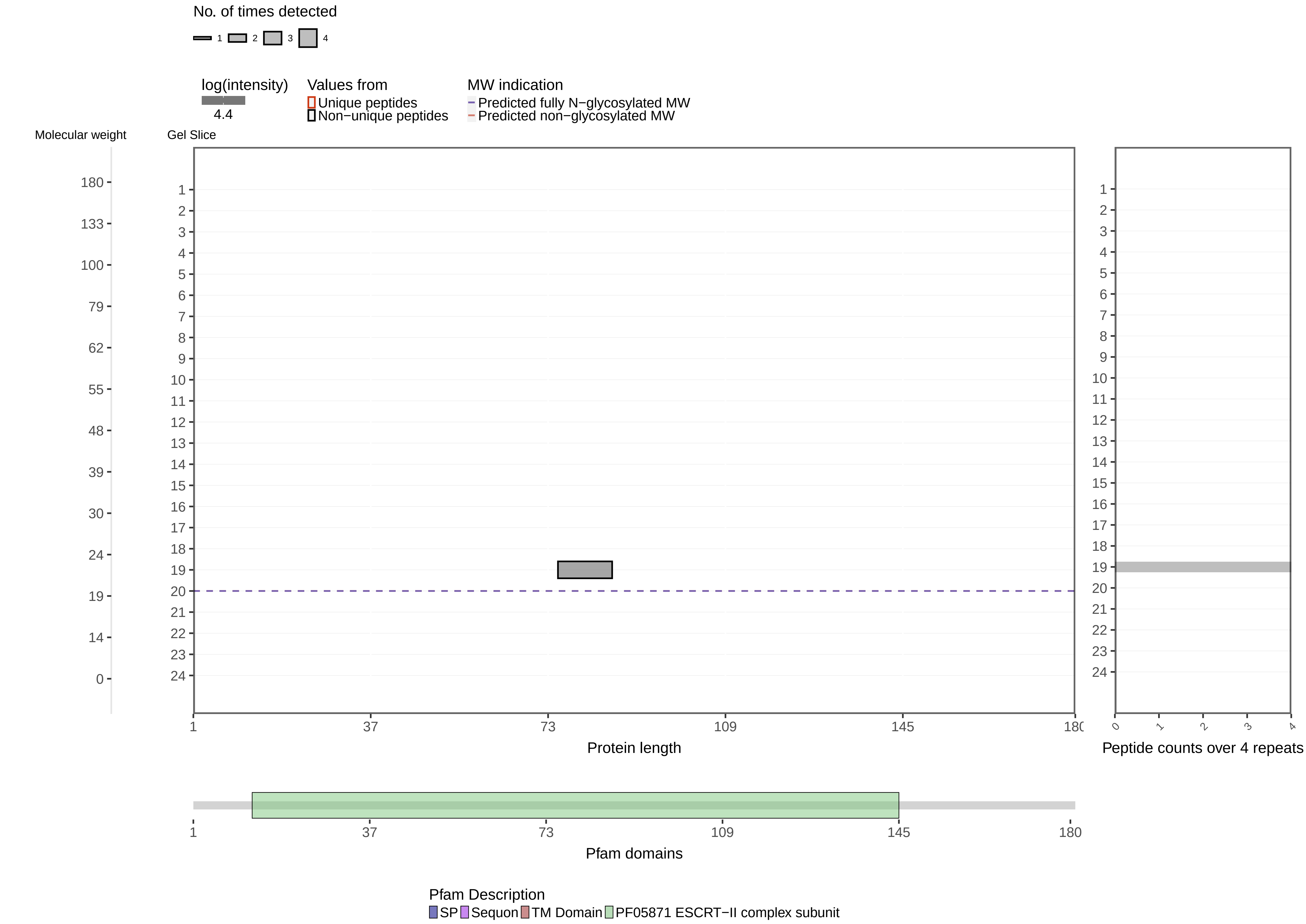


Protein sequence:

.SALVSDGRAEWMDKGHRKCLILWHRIQDWADLIVNFVKEYGLEDSVMTVEEICSGTESRGTELYGMDRTVLMRALKLLEHKGKLAIFKGTSADDEGVKFSL*

### NbL16g20540.1 inositol−tetrakisphosphate_1−kinase_4 [MW = 57.81 kDa]

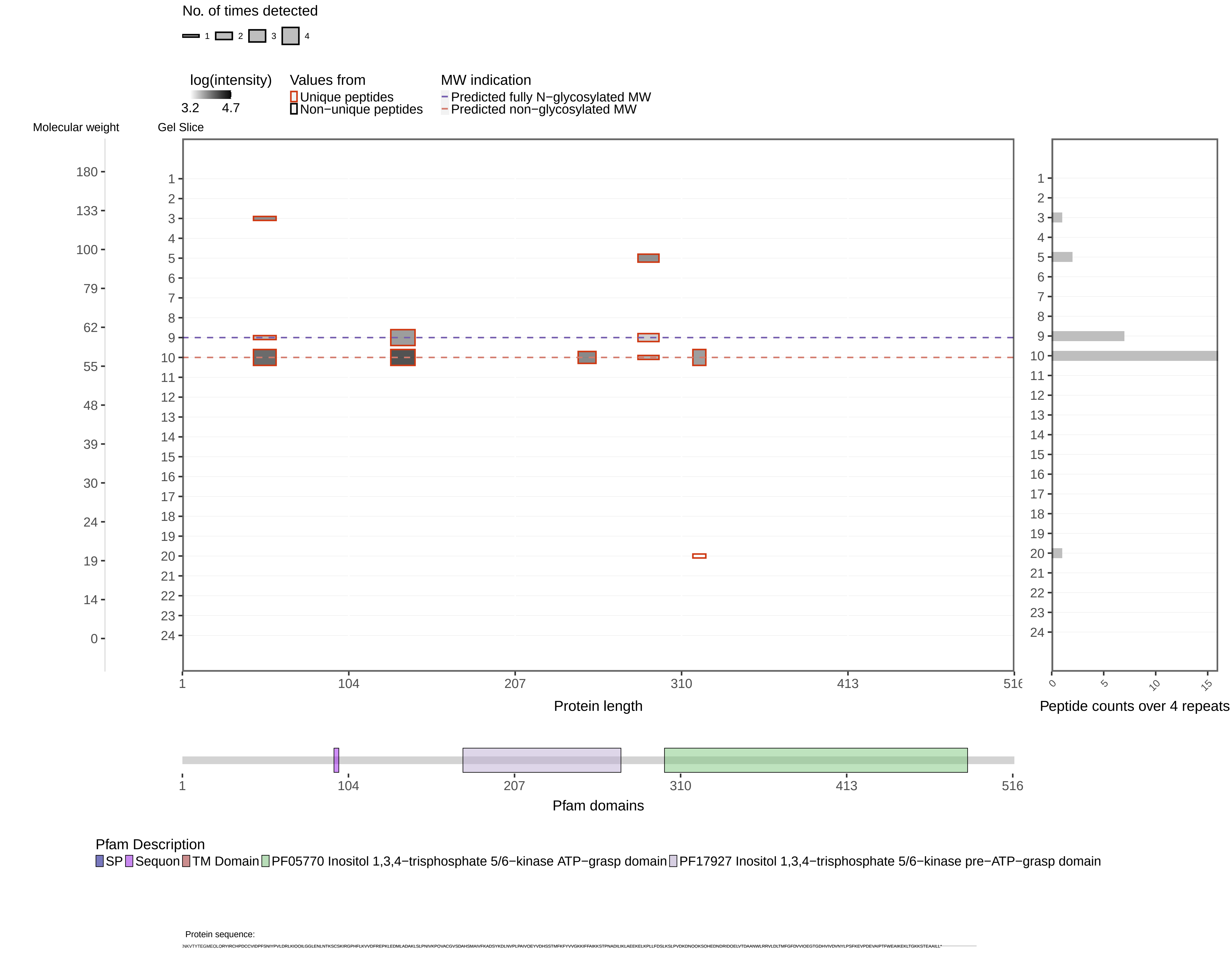

### NbL16g20470.1 60S_ribosomal_protein_L9−1−like [MW = 21.99 kDa]

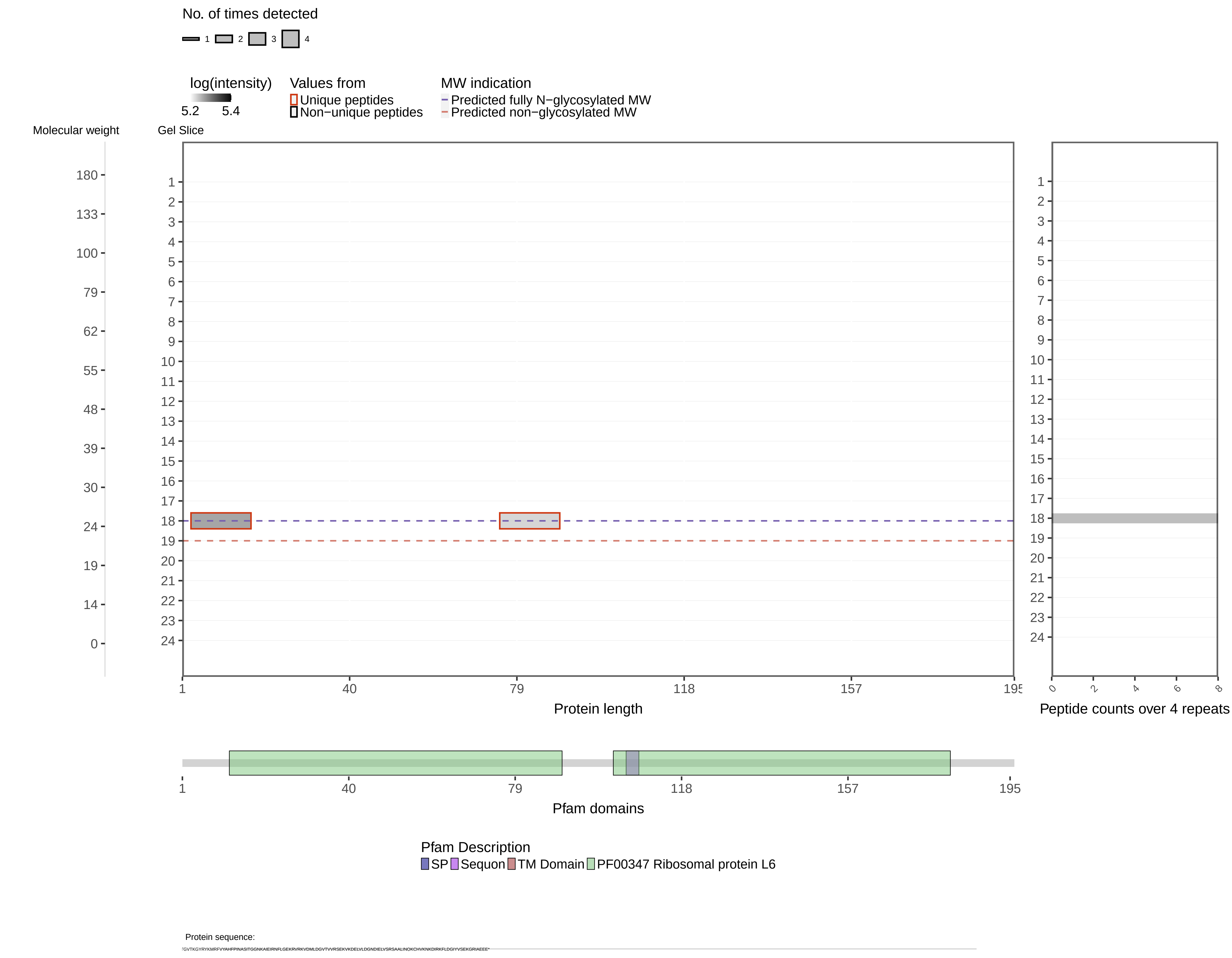

### NbL16g20450.1 beta−galactosidase−like [MW = 125.88 kDa]

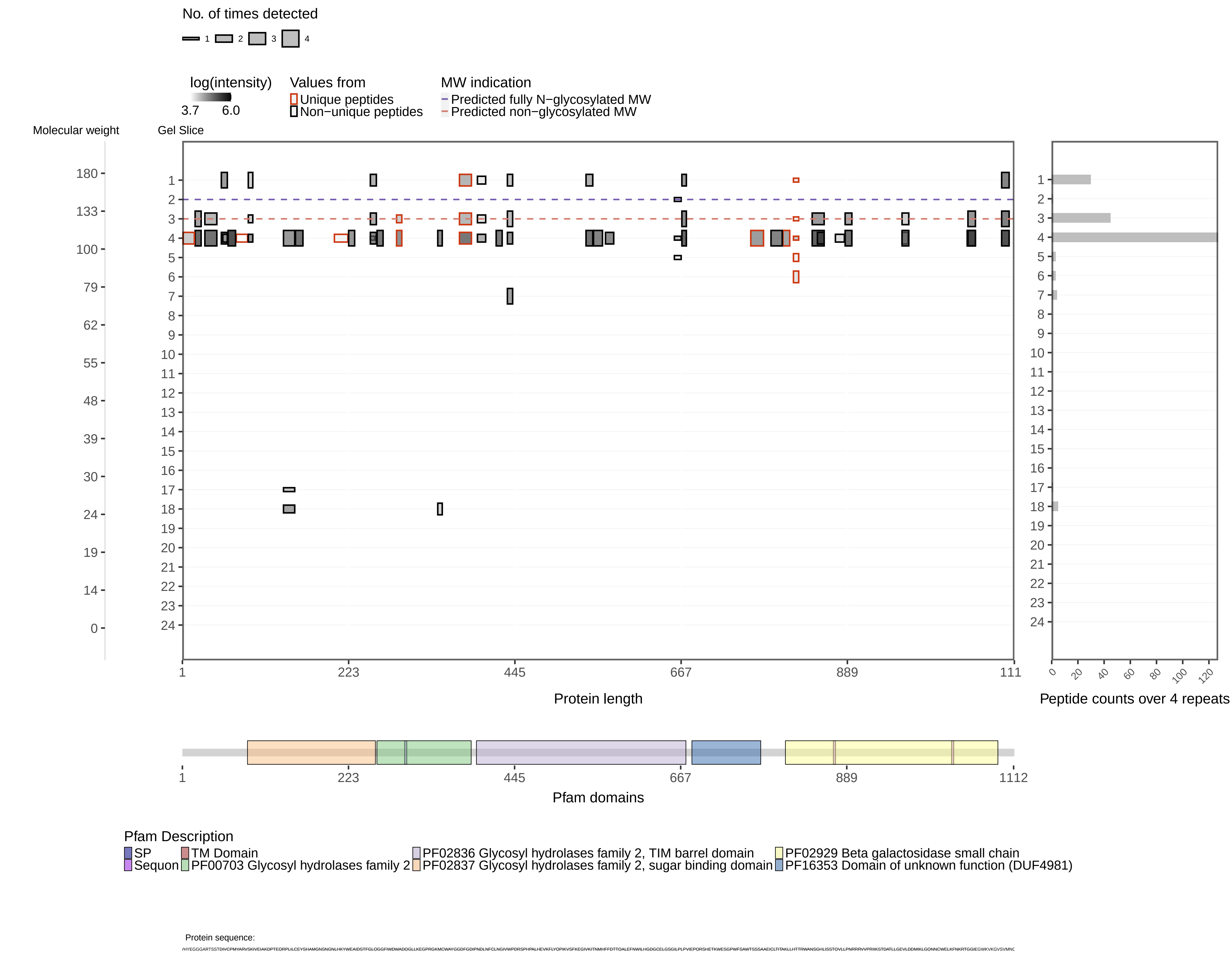

### NbL16g20410.1 mitogen−activated_protein_kinase_homolog_NTF4 [MW = 45.23 kDa]

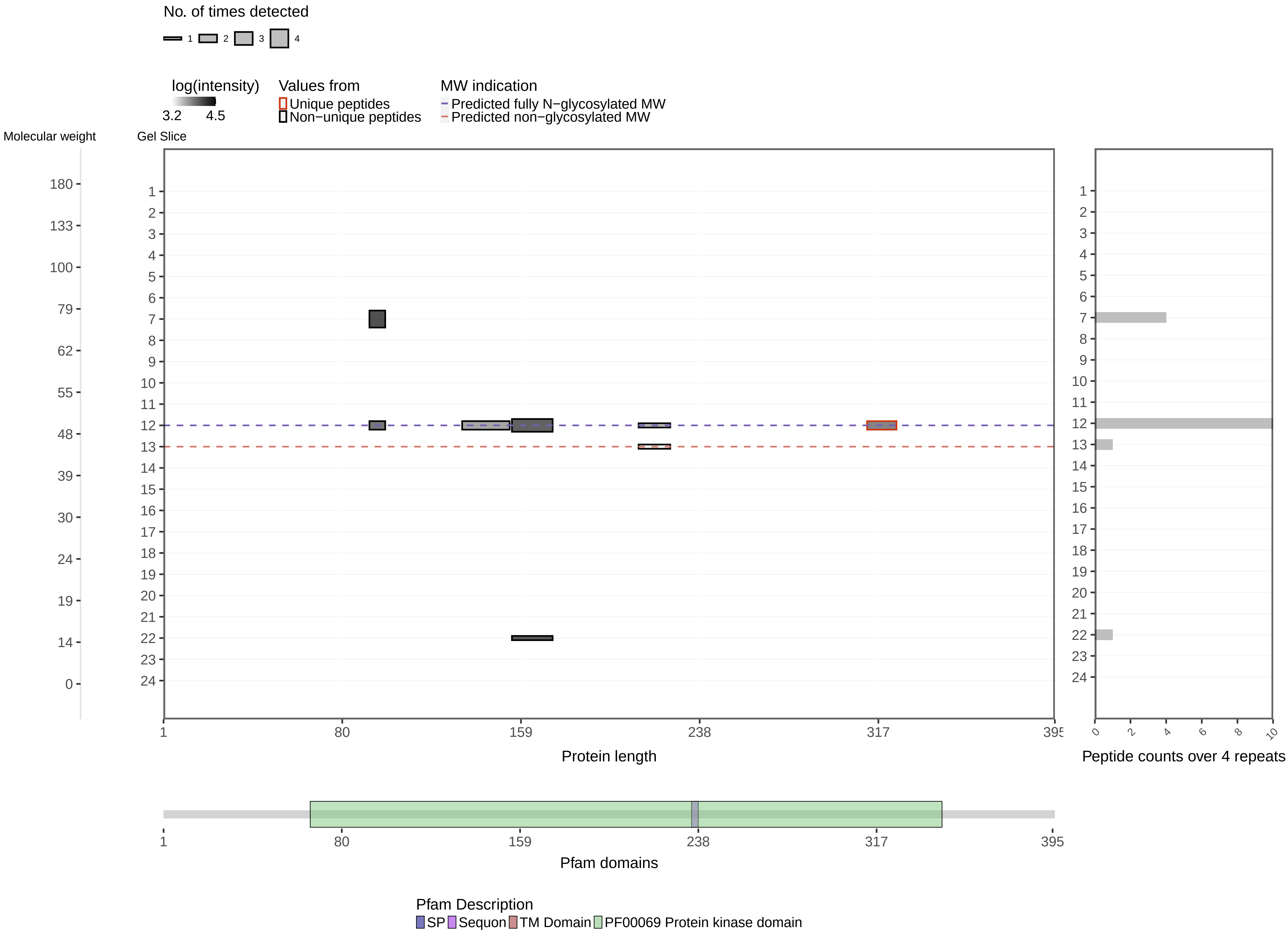


Protein sequence:

DLKPSNLLLNANCDLKICDFGLARVTSETDFMTEYVVTRWYRPPELLLNSSDYTAAIDVWSVGCIFMELMDRKPLFPGRDHVHQLRLLMELIGTPSEAEMEFLNENAKRYIRQLPLYRRQSFVEKFPHVNPAAIDLVEKMLTFDPRRRITVEDALAHPYLTSLHDISDEPVCMTPFNFDFEQHALTEEQMKELIYREGLAFNPEYQHM*

NbL16g20390.1 cysteine_synthase_chloroplastic/chromoplastic [MW = 41.26 kDa]

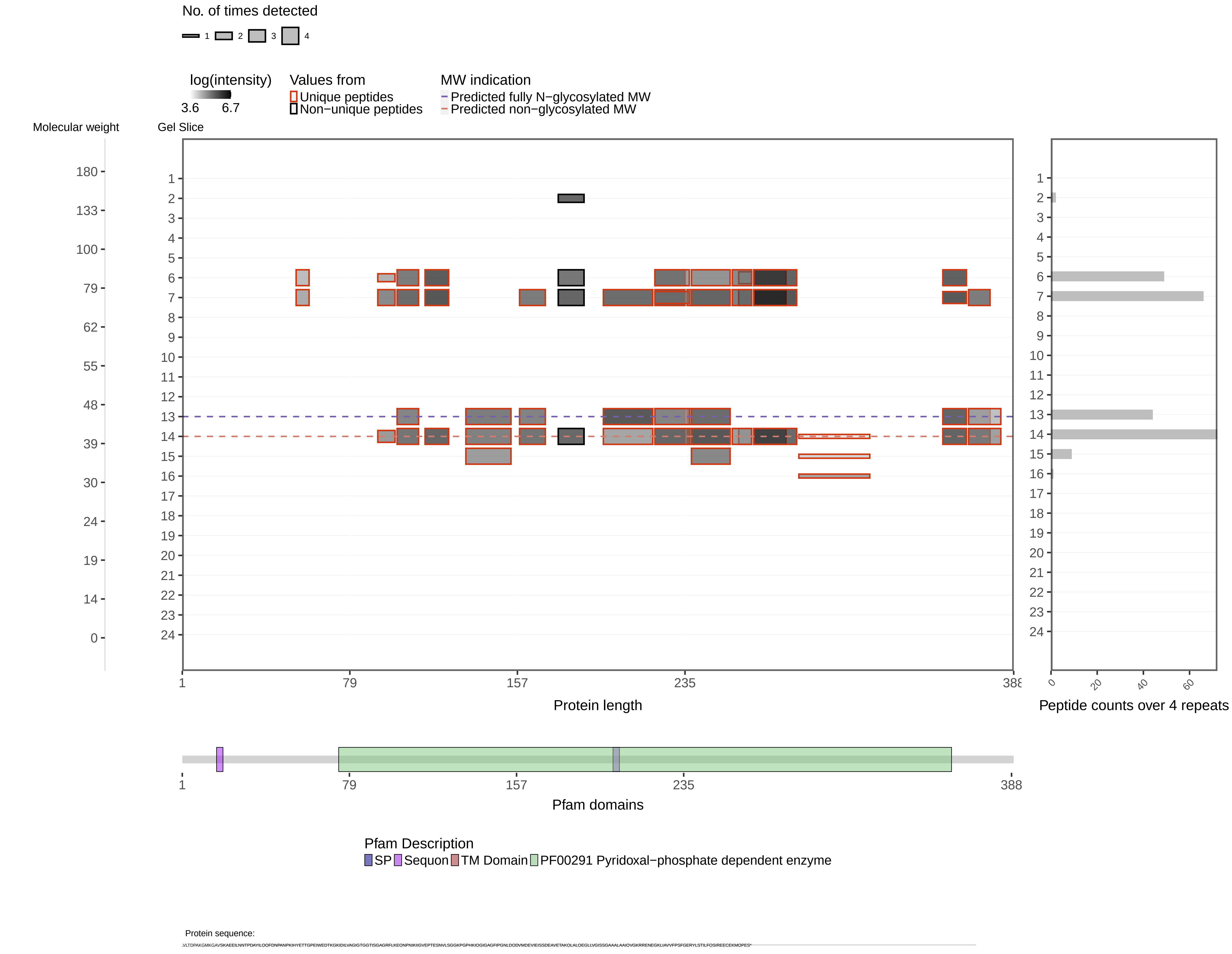

NbL16g20370.1 uncharacterized_LOC107775190 [MW = 16.8 kDa]

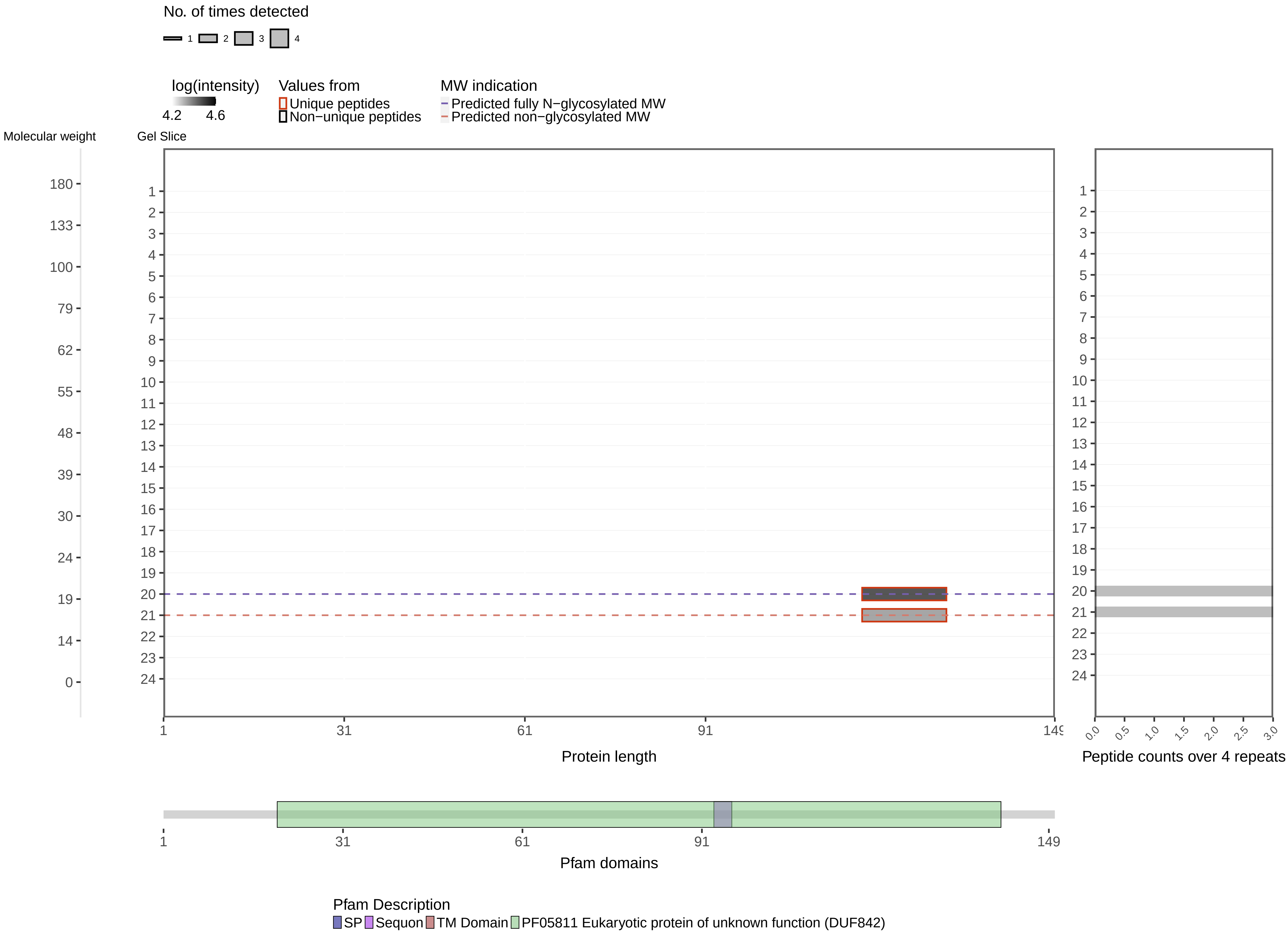


Protein sequence:

EIDNCVEYCSVPVLQAQNLVENEMANFQEKMNRSLRVCQDKYETAKLQKSSDATKVVESCVDQSVQDSIKTLPHLVGKLKVSLGVTD*

### NbL16g20300.1 2−isopropylmalate_synthase_A−like [MW = 59.88 kDa]

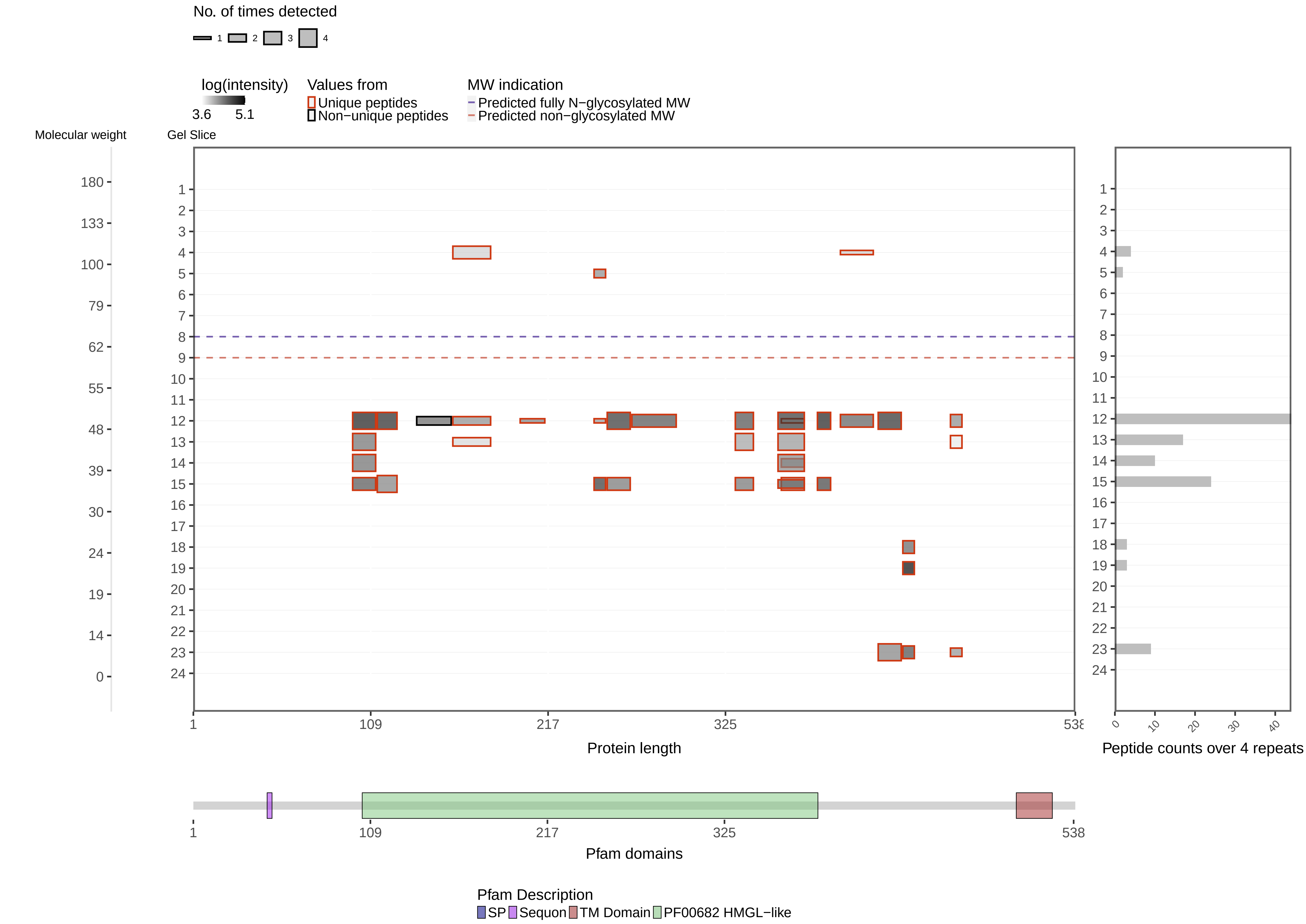


Protein sequence:

[illegible]

### NbL16g20000.1 probable_U3_small_nucleolar_RNA−associated_protein_7 [MW = 19.07 kDa]

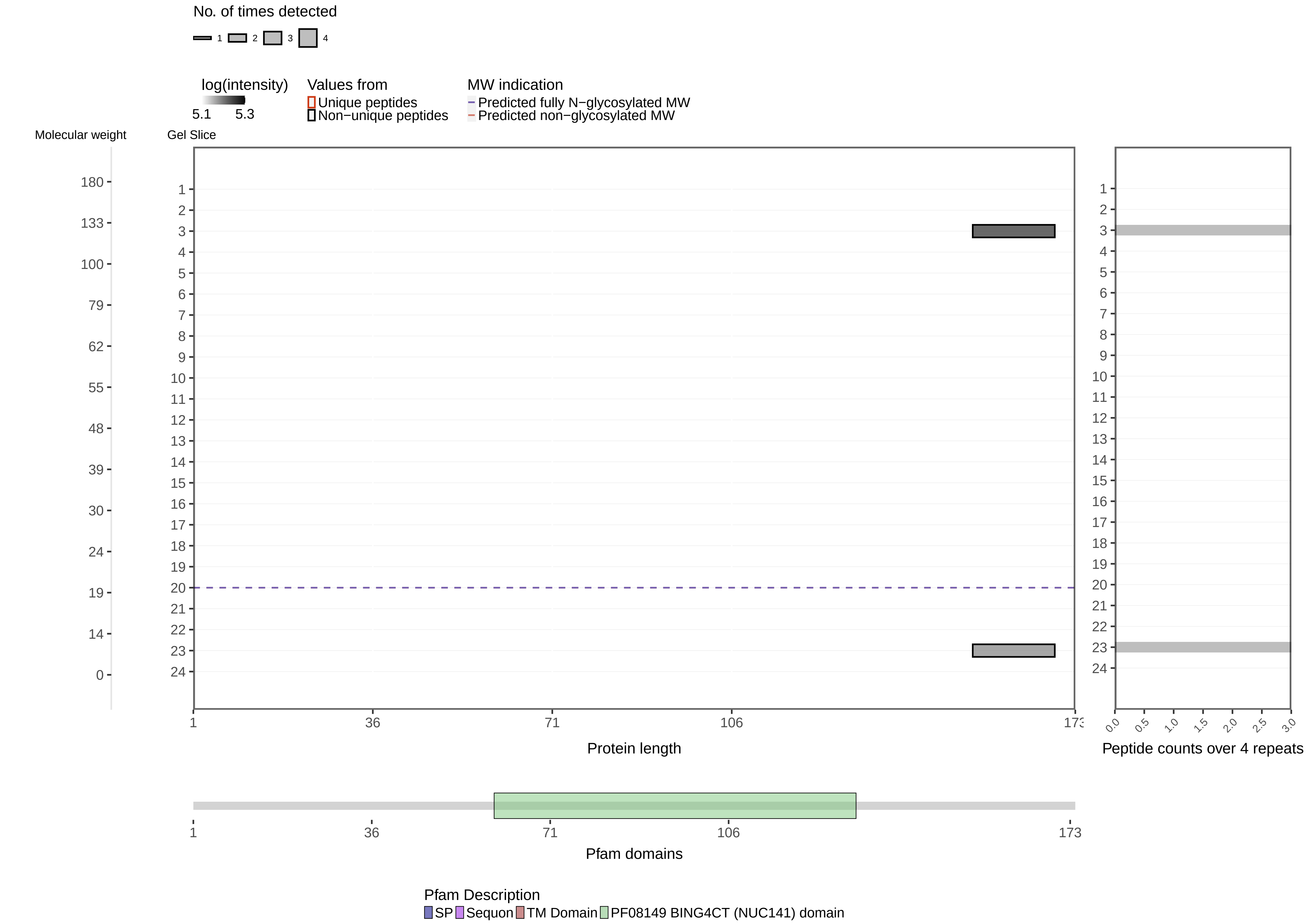


Protein sequence:

DVLGIGHSMGWSFIIIPGSGEPNFDSWVASPFETSKQRREKEVRSLLDKLPPETIMLDPTKIGTVRSTRKKEQPTKDDKEAEMEAAIEEAKSMPMKAVV*

### NbL16g19980.1 50S_ribosomal_protein_L3_chloroplastic [MW = 29.99 kDa]

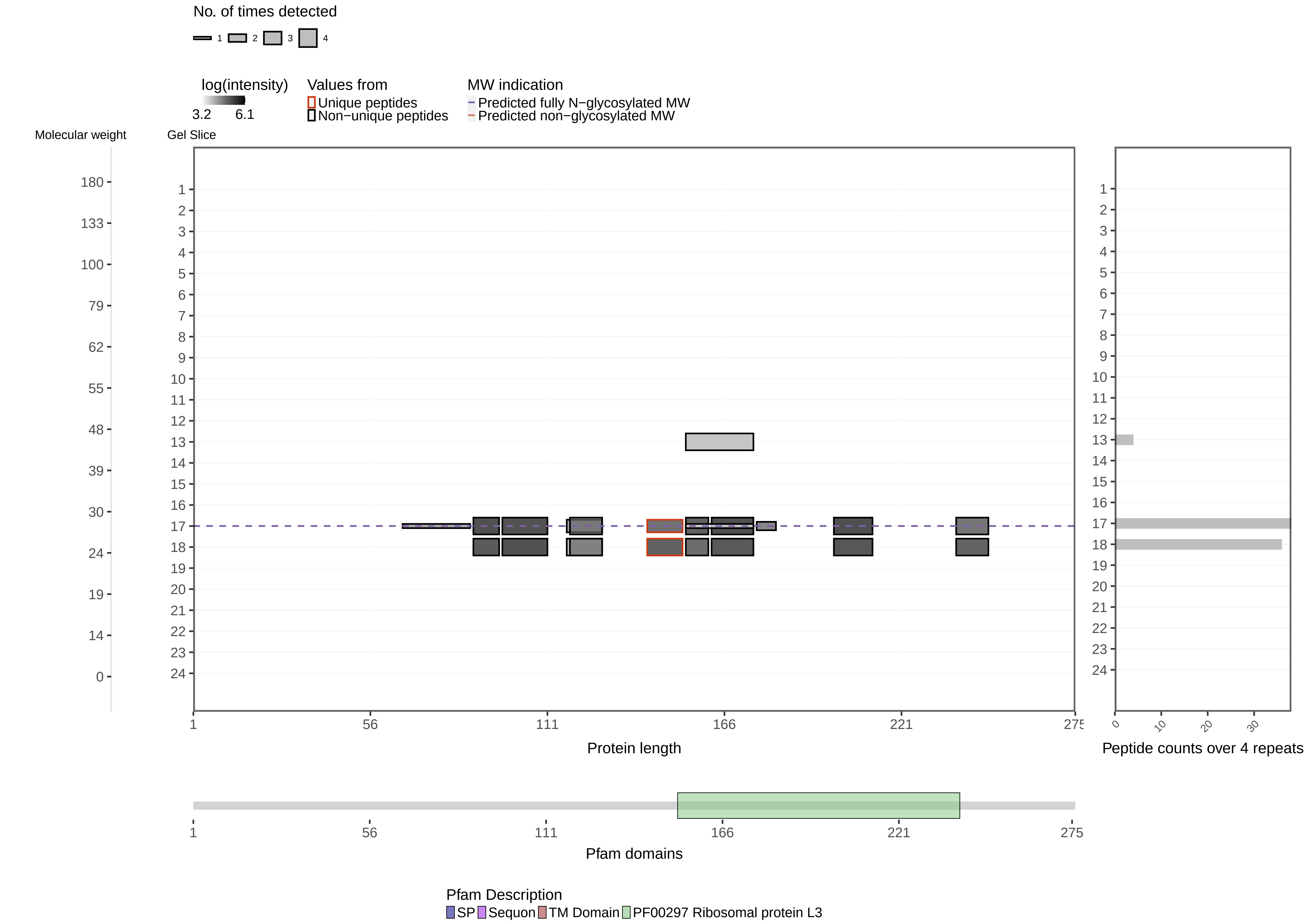


Protein sequence:

GHLEKSGIIPLRHLQEFRLQSIDGFEVTQKLDFGELFKEGDLVDVSGTTIGKGFQGGIKRHNFKRGQMTHGSKSHRQLGSIGAGTTPGRVYKGKKMPGRMGGTKRKIRKLKIVKIDDQLNIIMIKGALPGKPGNLLRIAPAKIVGKNIPKS*

NbL16g19800.1 DNA−damage−repair/toleration_protein_DRT100−like [MW − SP = 36.8 kDa]

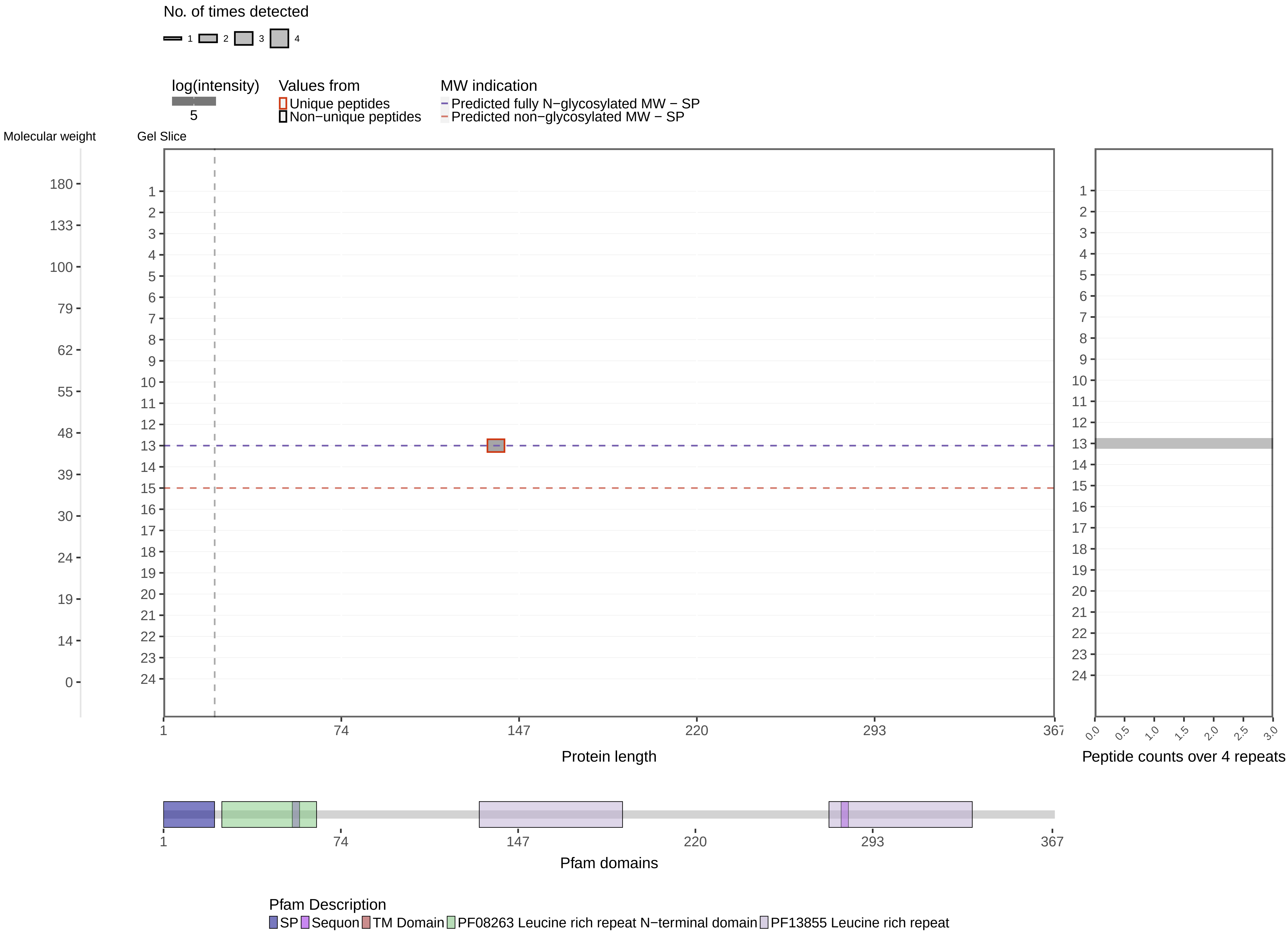


Protein sequence:

IPSSIVTLGRLIHLDLSNNMLTGEIPSDIGKMSMLSRAIMNNNQISGSIPTSIAKLYRLVDLDLSMNQISGSIPAELGSMPVLTVLNLDSNQISGSIPTNLLTSSGLNILNLSRNSLEGILPDIFVPTTYFTHLDLSYNNLGGSIPKTLSSAKYVGHLDLSNNHLCGSIPEGSFFNSLEASSFSNNDCLCGSPLLSC*

### NbL16g19660.1 40S_ribosomal_protein_S16−like [MW = 16.93 kDa]

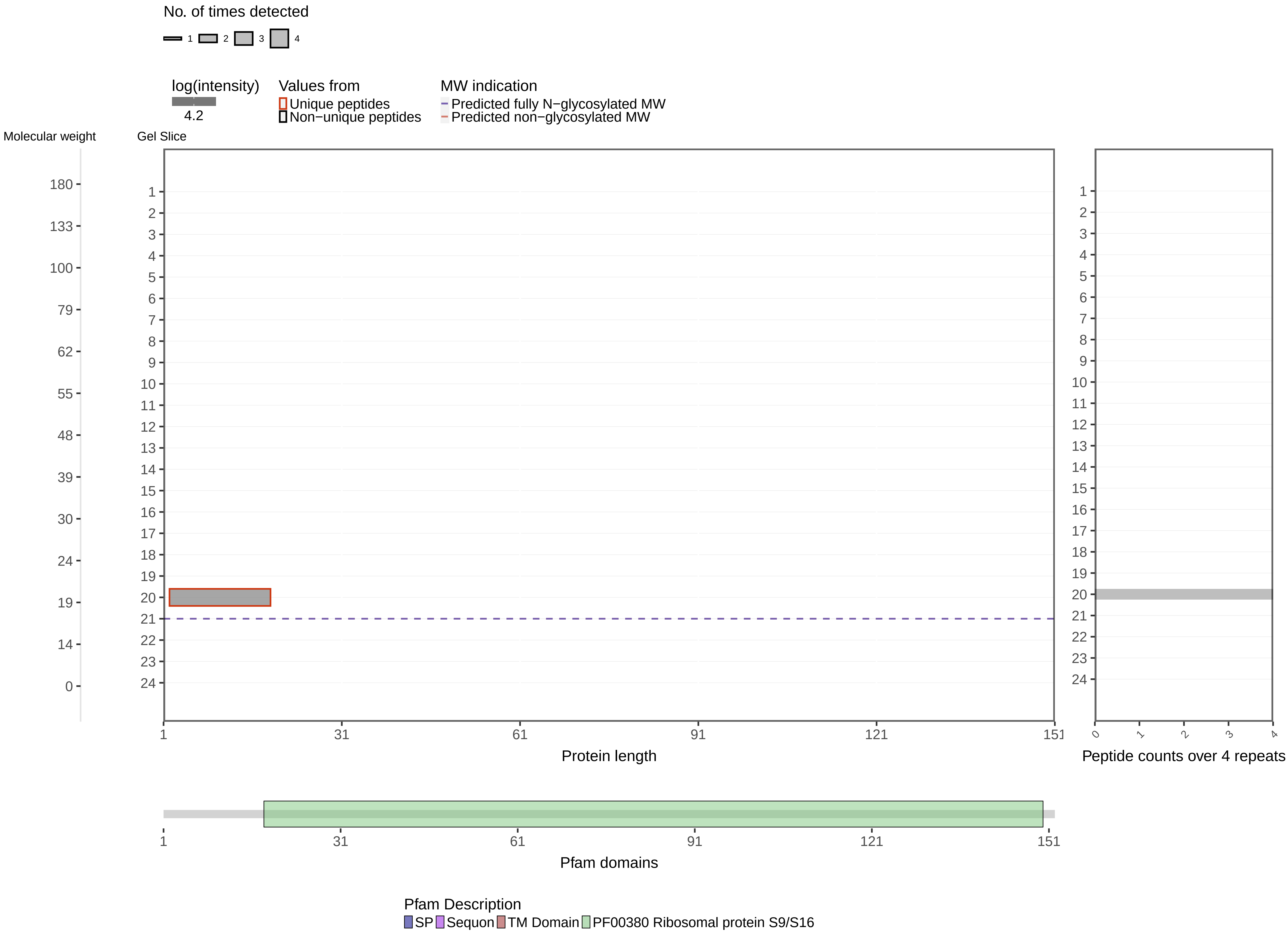


Protein sequence:

RHRFAGVDMRIRVKGGGHTSQIYAIRQSIAKALVAFYQKYVDEQSKKEIKDILVRYDRTLLVADPRRCEPKKFGGRGARARFQKSYR*

NbL16g19620.1 uncharacterized_LOC107811196 [MW = 25.56 kDa]

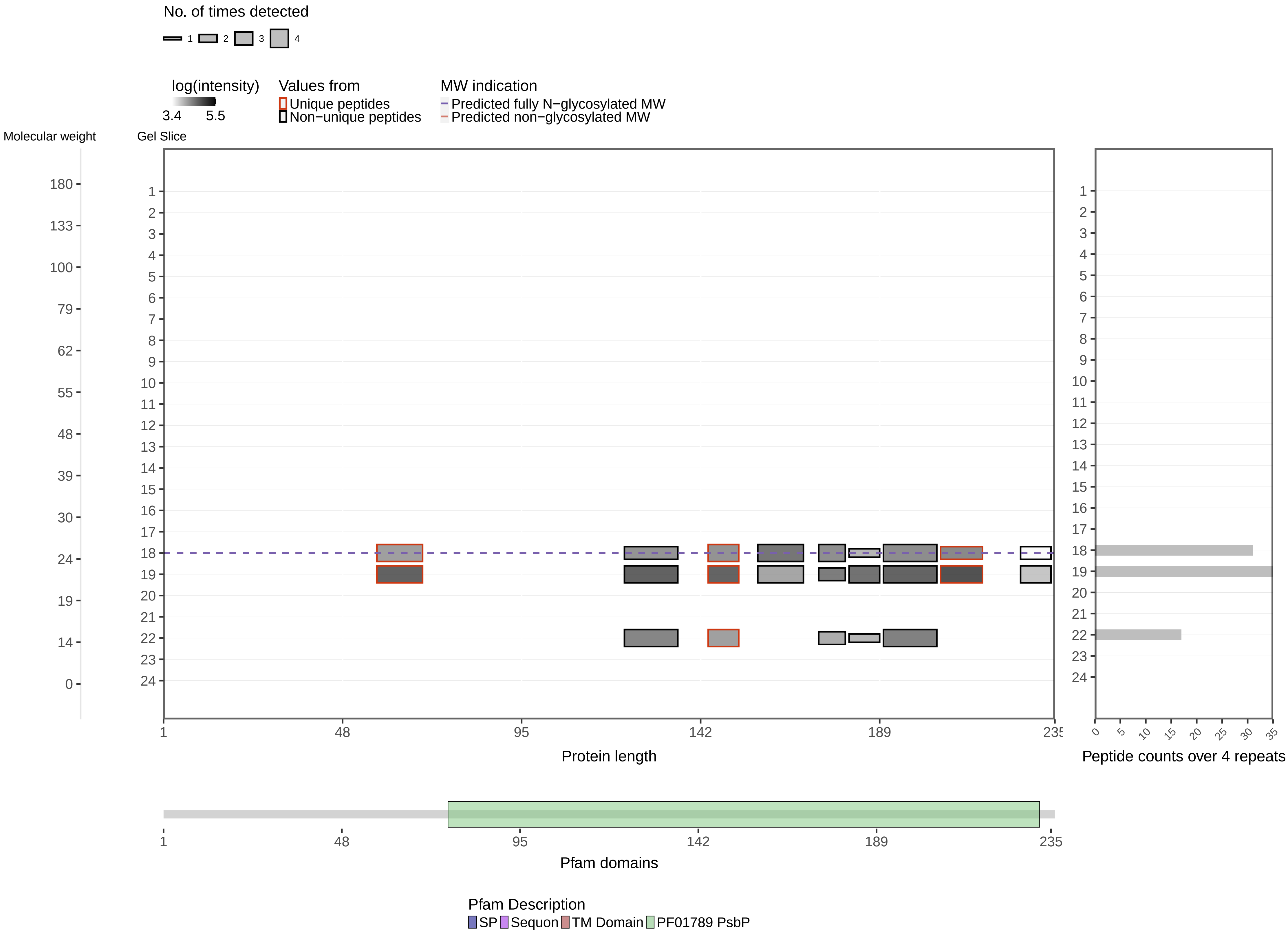


Protein sequence:

GDKAKPKKFAARFASSDRSEVLSVVIRPSNQLKITFLEAKDITDLGSLKEAAKIFVPAGATLYSGRSIKIKEDEGFRTYYFYEFARDEQHVALVAAVNSGKAYIAAATAPESKWNDDGLKLRSAAVSLMVL*

NbL16g19580.1 linoleate_9S−lipoxygenase_6 [MW = 97.25 kDa]

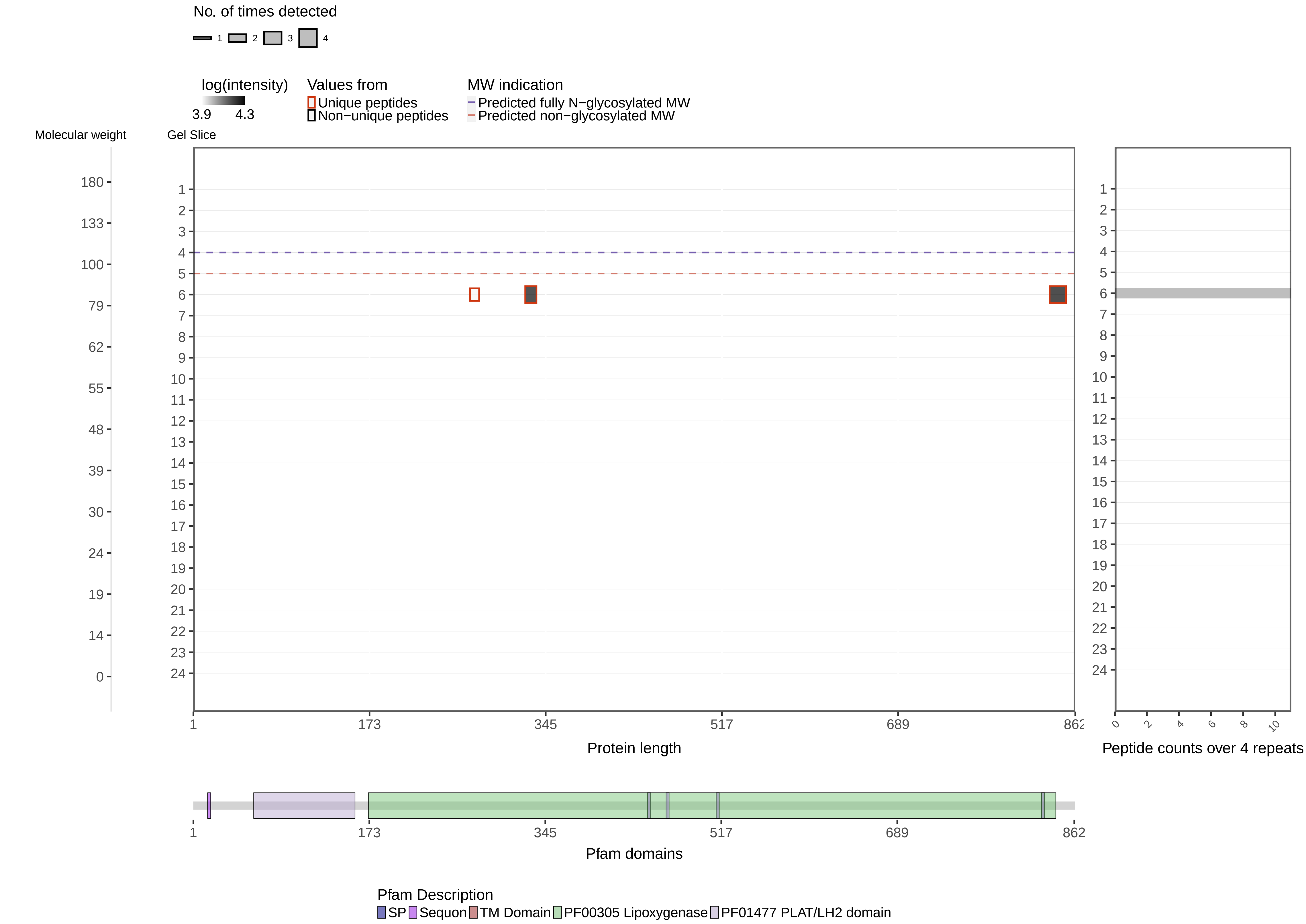


Protein sequence:

TIDEAIKSNRLFILNHHDTIMPYLRRINTTTTKTYASRTLLFLQDNGSLKPLAIELSLPHPDGDQFGAISKVYTPADEGVEGSIWELAKAYVAVNDSGVHQLISHWLNTHAVIEPFVIATNRQLSVLHPIHKLLHPHFRDTMNINAMARQILINAGGVLESTVFPSKYAMEMSAVVYKNWIFPDQALPTDLVKRGMAVEDSSSPHGIRLLIQDYPYAVDGLEIWSAIKSWVTEYCSFYYKSDDSILKDNELQAWWKELREEGHGDLKDEPWWPKMENSQELIDSCTIIIWTASALHAAV

NbL16g19540.1 delta−1−pyrroline−5−carboxylate_synthase [MW = 77.79 kDa]

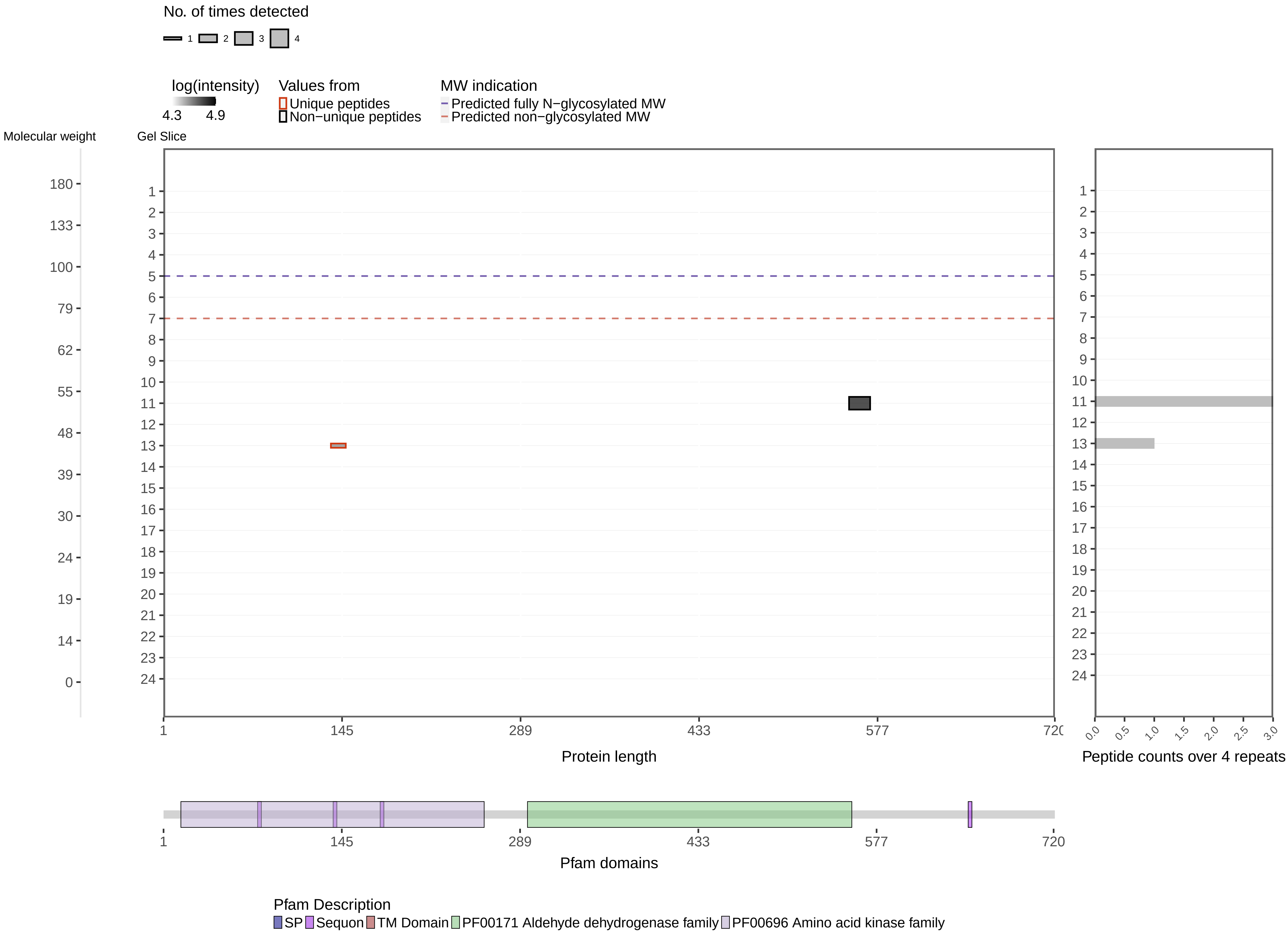


Protein sequence:

AAQQAGYEKSLISRLALKPGKISNLAKSVRVLANMDEPIGRILRRTEVADGLILEKSSSPLGVLLIIFESRPDALVQIASLAVRSGNGLLLKGGKEAKRSNAILHKVITSAIPGTVGERLIGLVTSREEIPELLKHDDVIDLVIPRGSNKLVSQIKASTKIPVLGHADGICHVYVDKSADMDMAKRVVVDAKTDYPAACNAMETLLVHKDLAQNGGLNDLIVELQTKGVSLYGGPKASSLLKIPEARTFHHEYGSLACTVEVVEDVYAAIDHIHQHGSAHTDSIITKDQEAAEIFLRQVDSA

### NbL16g19530.1 uncharacterized_protein_At2g34160−like [MW = 14.5 kDa]

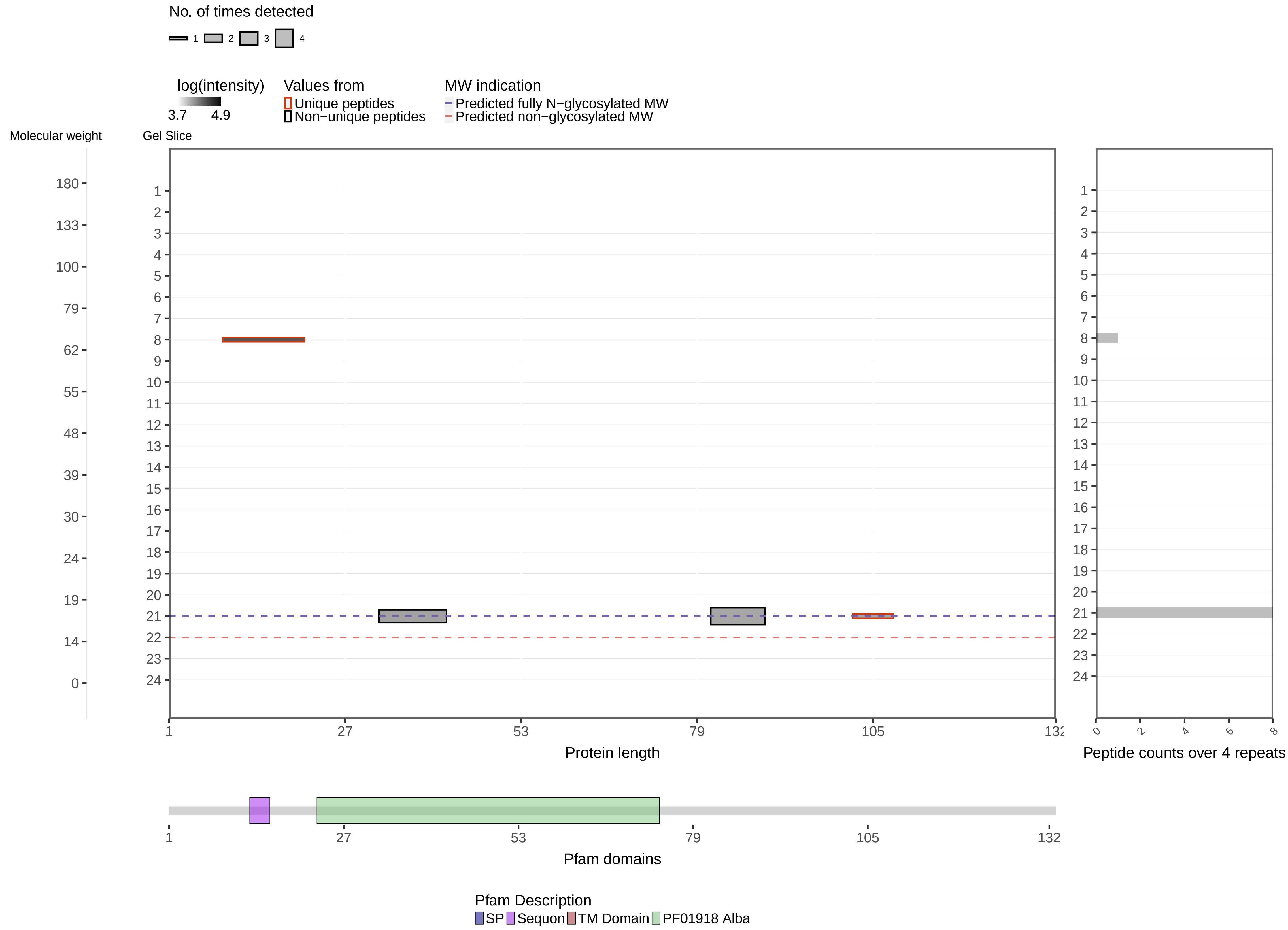


Protein sequence:

ELSALGMAIATVVSIAEILKSNGFAVEKKIMTSTVDVKDGSRGRPVSKAKIEVLLGKSEKFDNLMAAAAERELGDGEEOS*

NbL16g19450.1 importin_subunit_alpha−2−like [MW = 58.48 kDa]

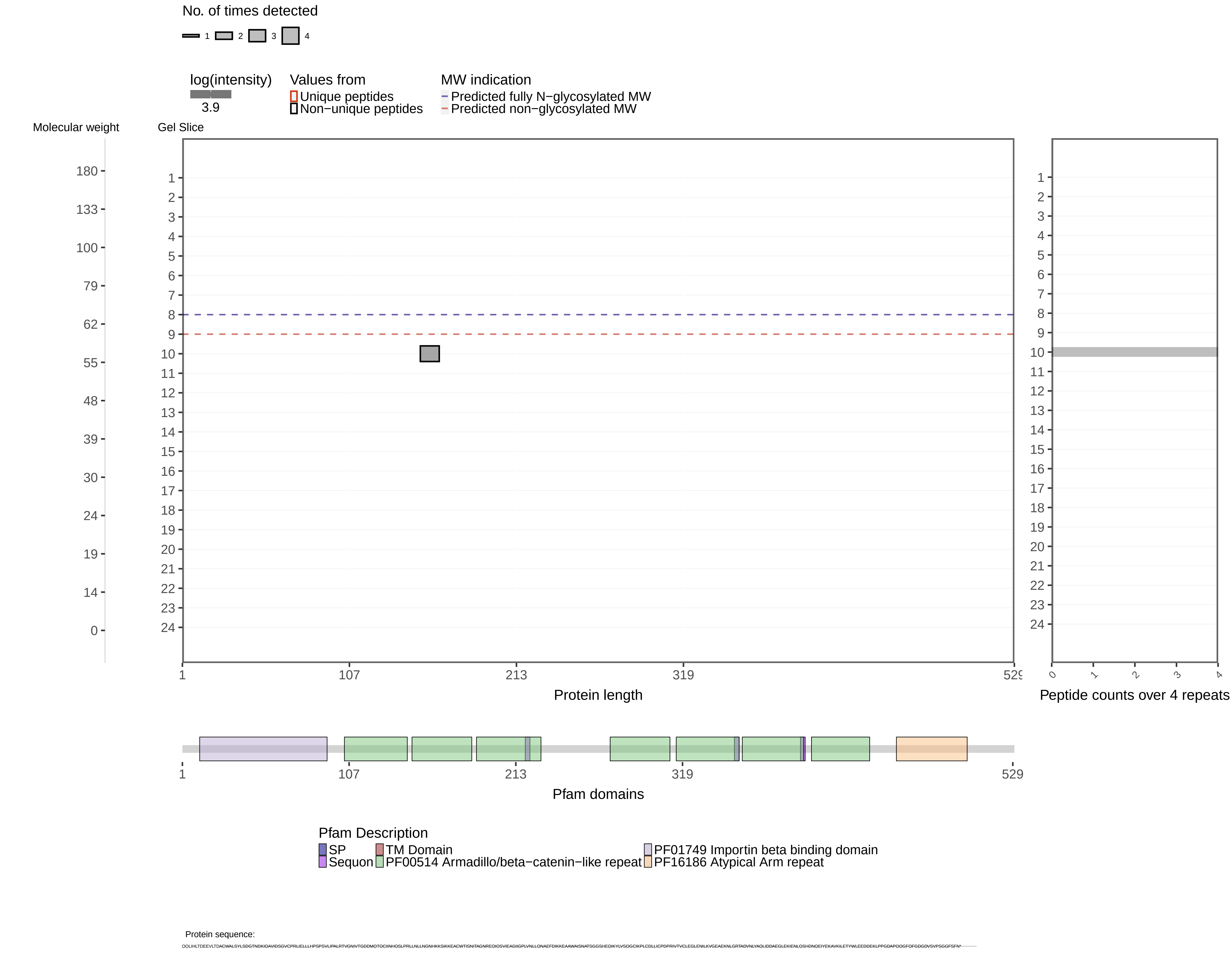

### NbL16g19390.1 40S_ribosomal_protein_S20−2 [MW = 13.63 kDa]

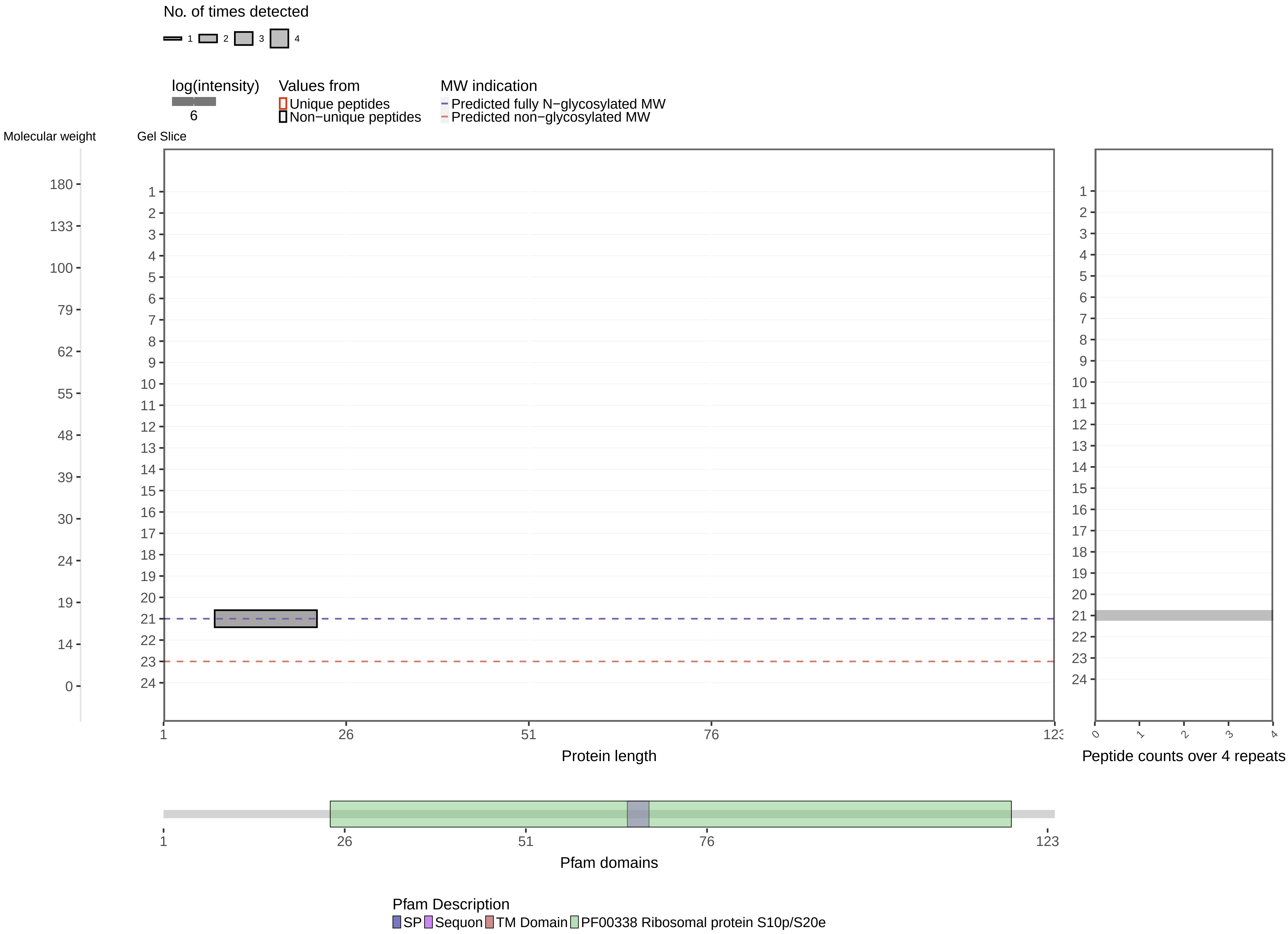


Protein sequence:

KRLRVKGPVRMPTKVLNITTRKSPCGEGTNTWDRFELRVHKRVIDLFSSADVVKQITSITIEPGVEVEVTIADS*

### NbL16g19360.1 nudix_hydrolase_16_mitochondrial−like [MW = 20.43 kDa]

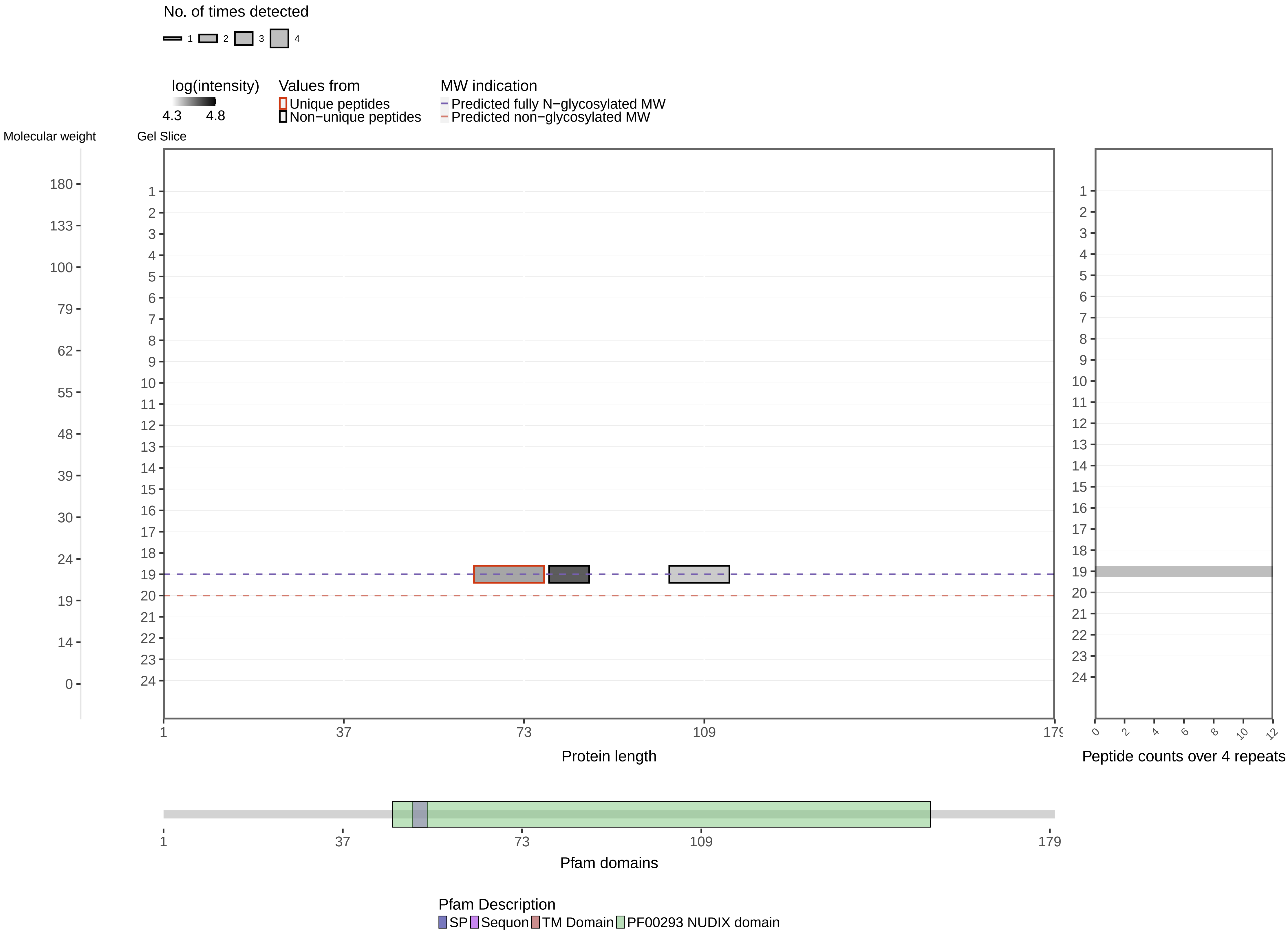


Protein sequence:

EAIEEAGVRGDLVHFLGYYFFKSKTLQDEYSPEGLCRAAMFALLVKEELECWPEQRLRRRSWLTIPEAIQCCRHPWMRQALEEGFSKWHDSGMASTMNNED*

NbL16g19330.1 ferredoxin−dependent_glutamate_synthase_chloroplastic [MW = 177.69 kDa]

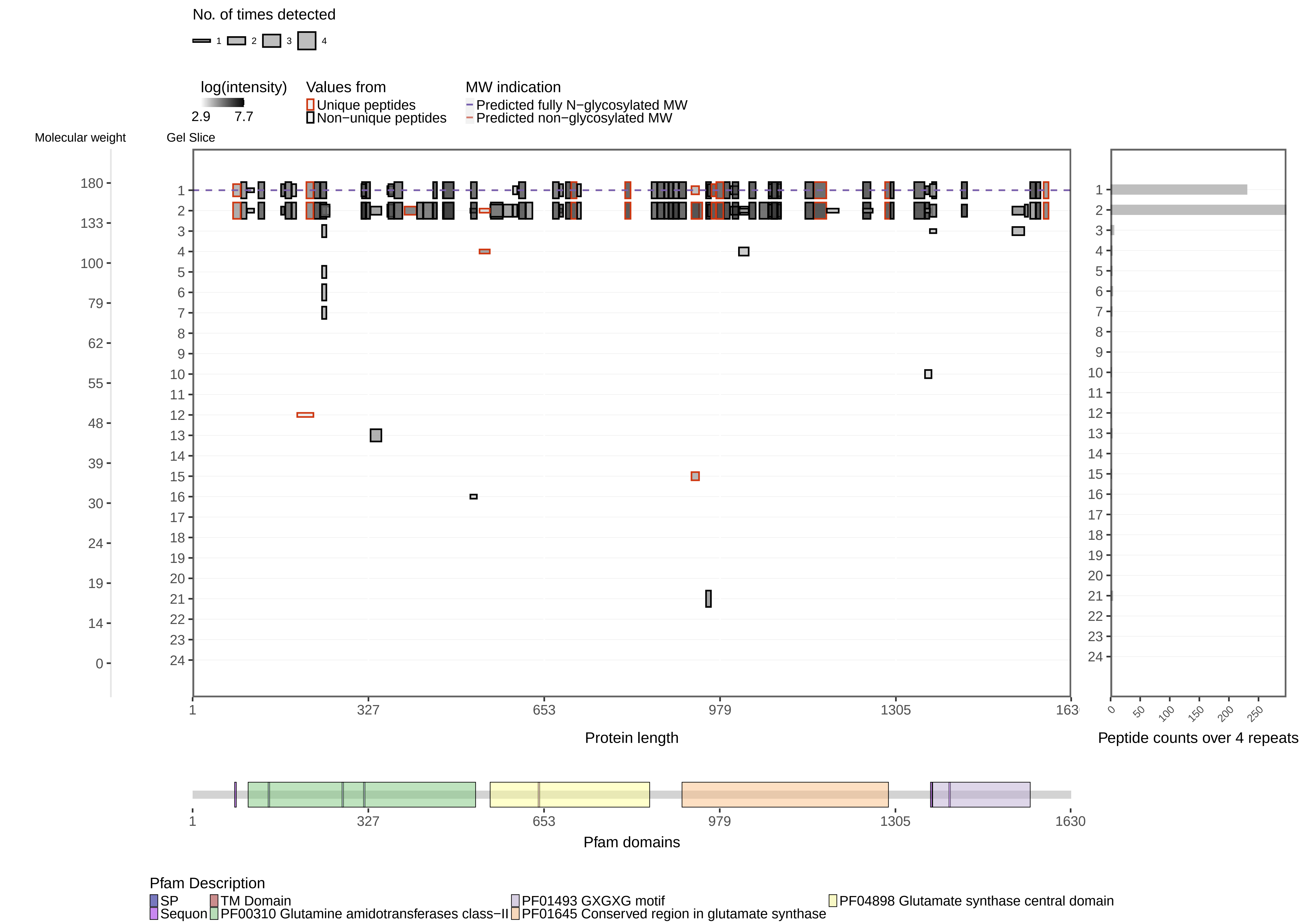


Protein sequence:

### NbL16g19310.1 ultraviolet−B_receptor_UVR8−like [MW = 47.44 kDa]

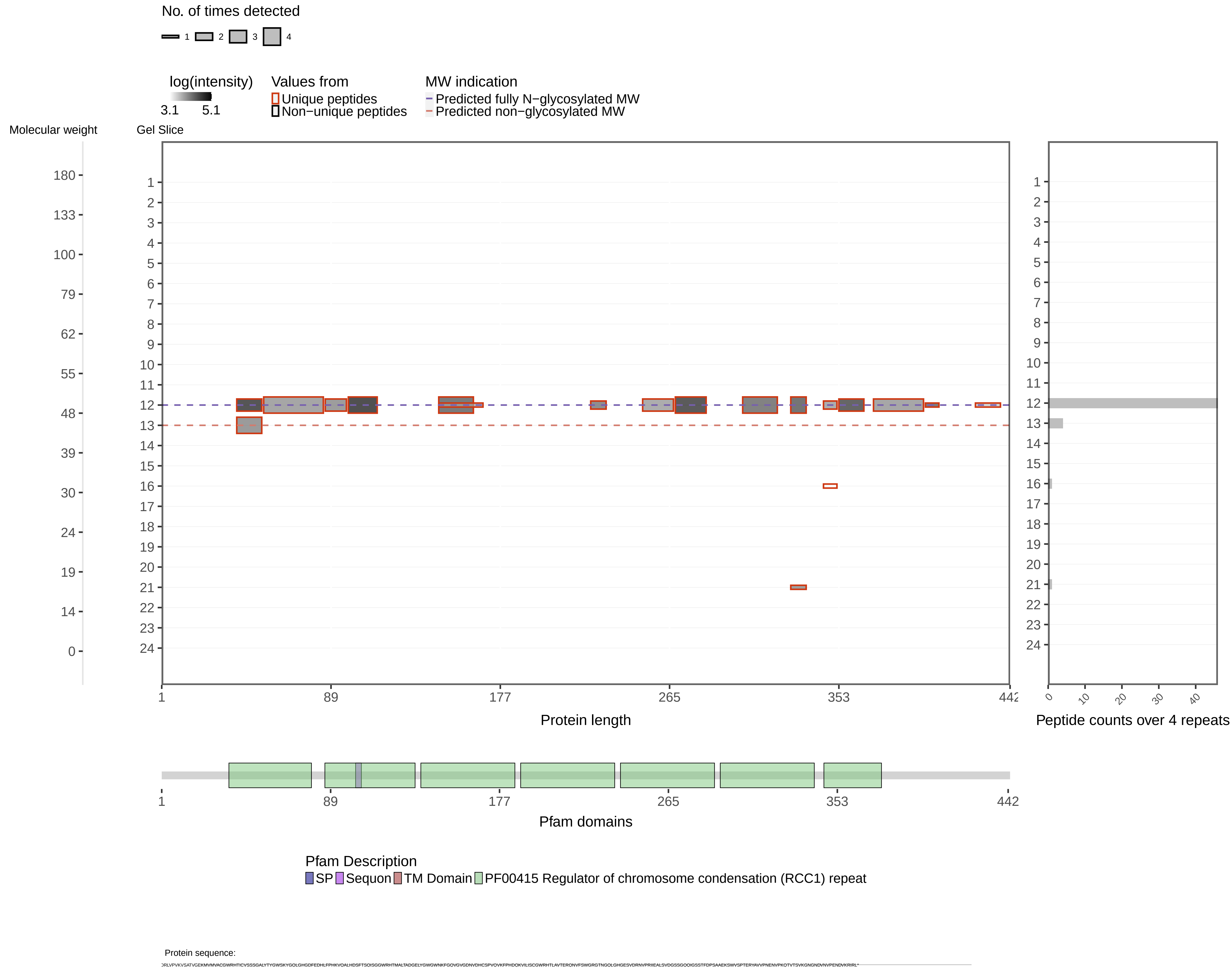

NbL16g19100.1 glucose−6−phosphate_1−dehydrogenase_chloroplastic [MW = 67.31 kDa]

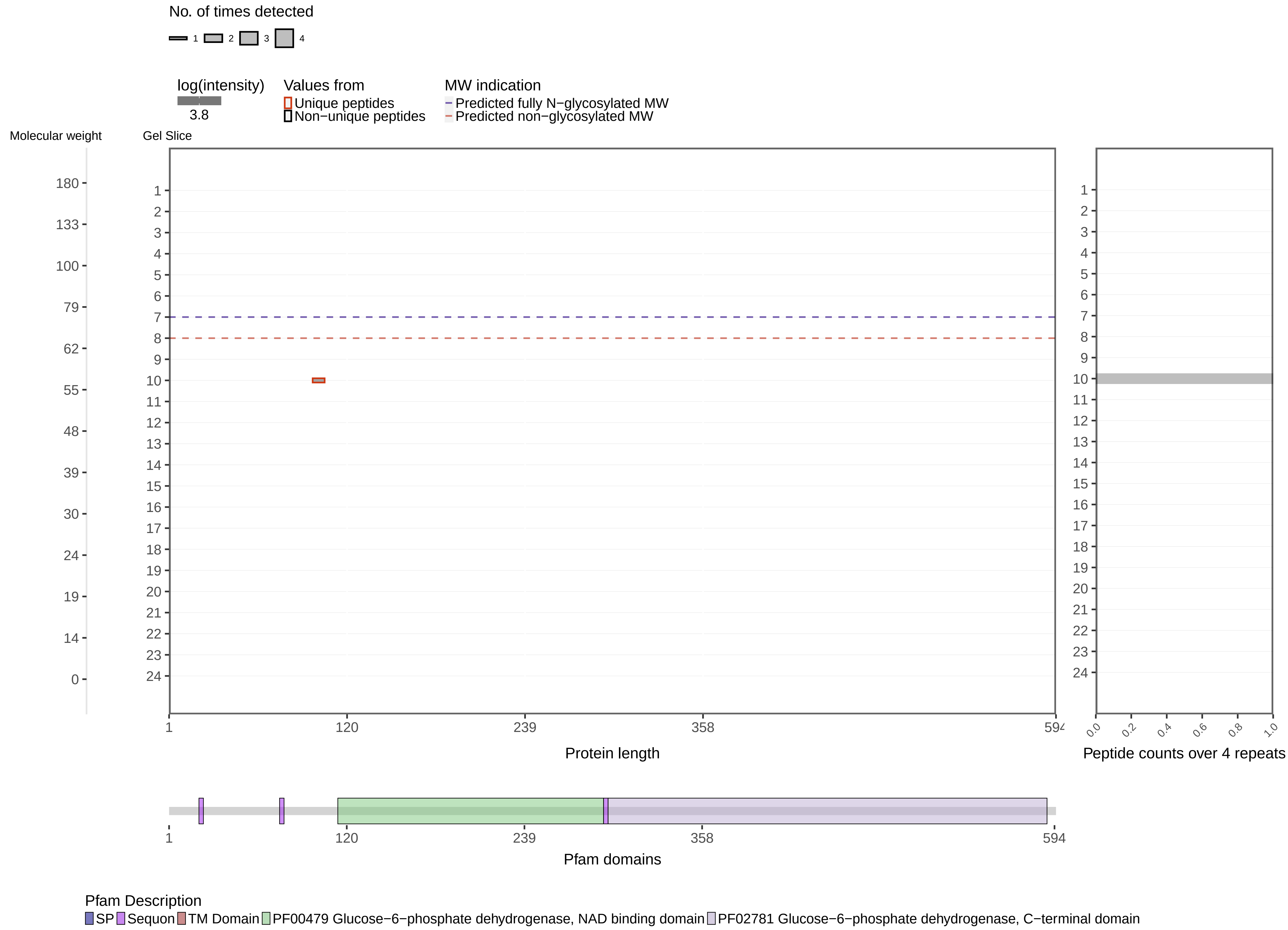


Protein sequence:

YLGKELVENLSVLRFSNLIFEPLWSRQYIRNVQFIFSEDFGTEGRGGYFDHYGIIRDIMQNHLLQILALFAMETPVSLDAEDIRNEKVKVLRSMRPLQLDDVIIGQYKSHTKGDVTYPGYTDDKTVLKDSLTPTFAAAALFIDNARWDGVPFLMKAGKALHTRSAEIRVQFRHVPGNLYNKNFGSDLDQATNELVIRVQPNEAIYLKINNKVPGLGMRLDRSNLNLLYSARYSKEIPDAYERLLLDAIEGERRLFIRSDELDAAWSLFTPVLKELEDKKIVPEYYPYGSRGPIGAHYL[illegible]

### NbL16g18660.1 NADH_dehydrogenase_[ubiquinone]_flavoprotein_1_mitochondrial [MW = 56.6 kDa]

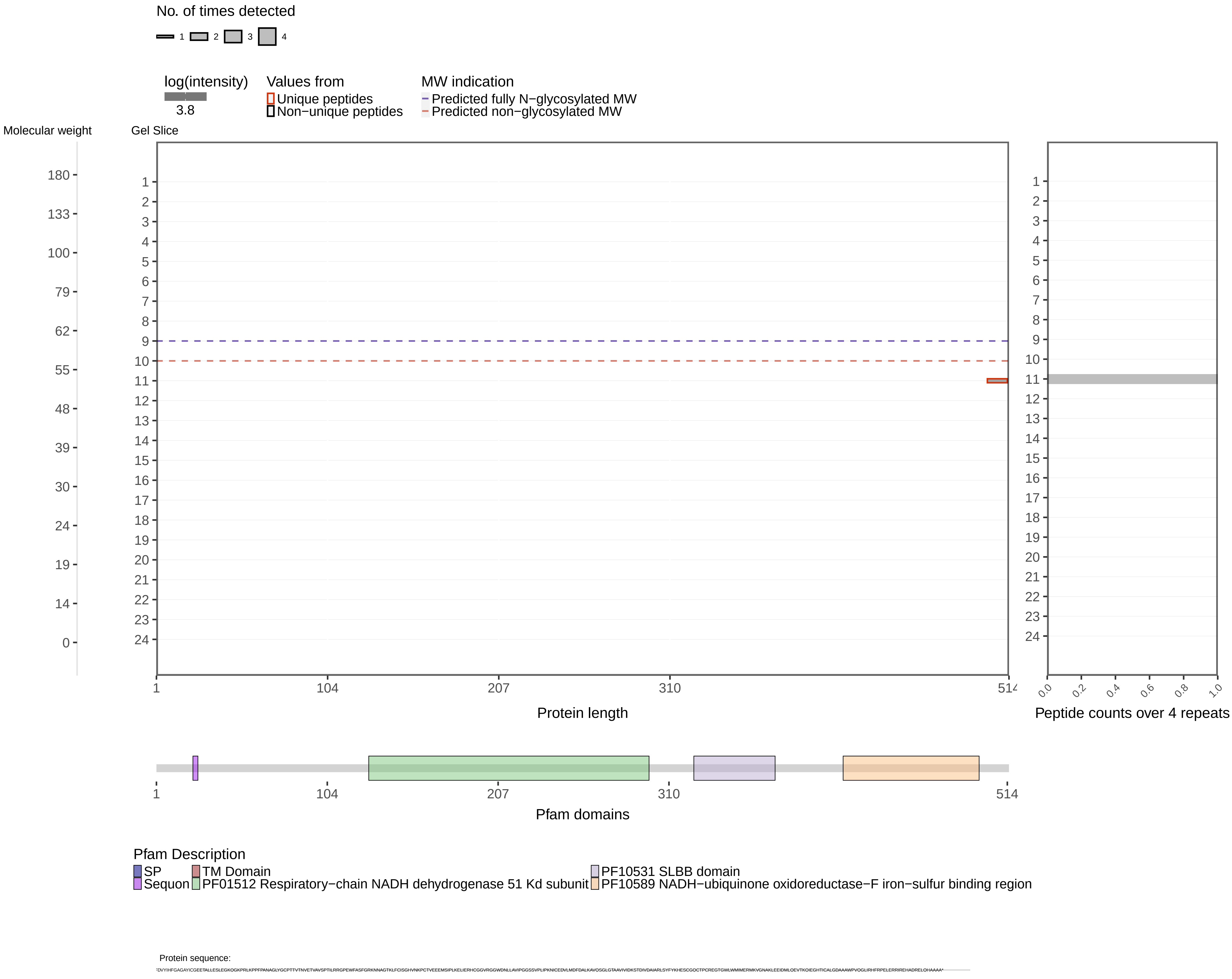

### NbL16g18640.1 adenylate_kinase_4 [MW = 26.49 kDa]

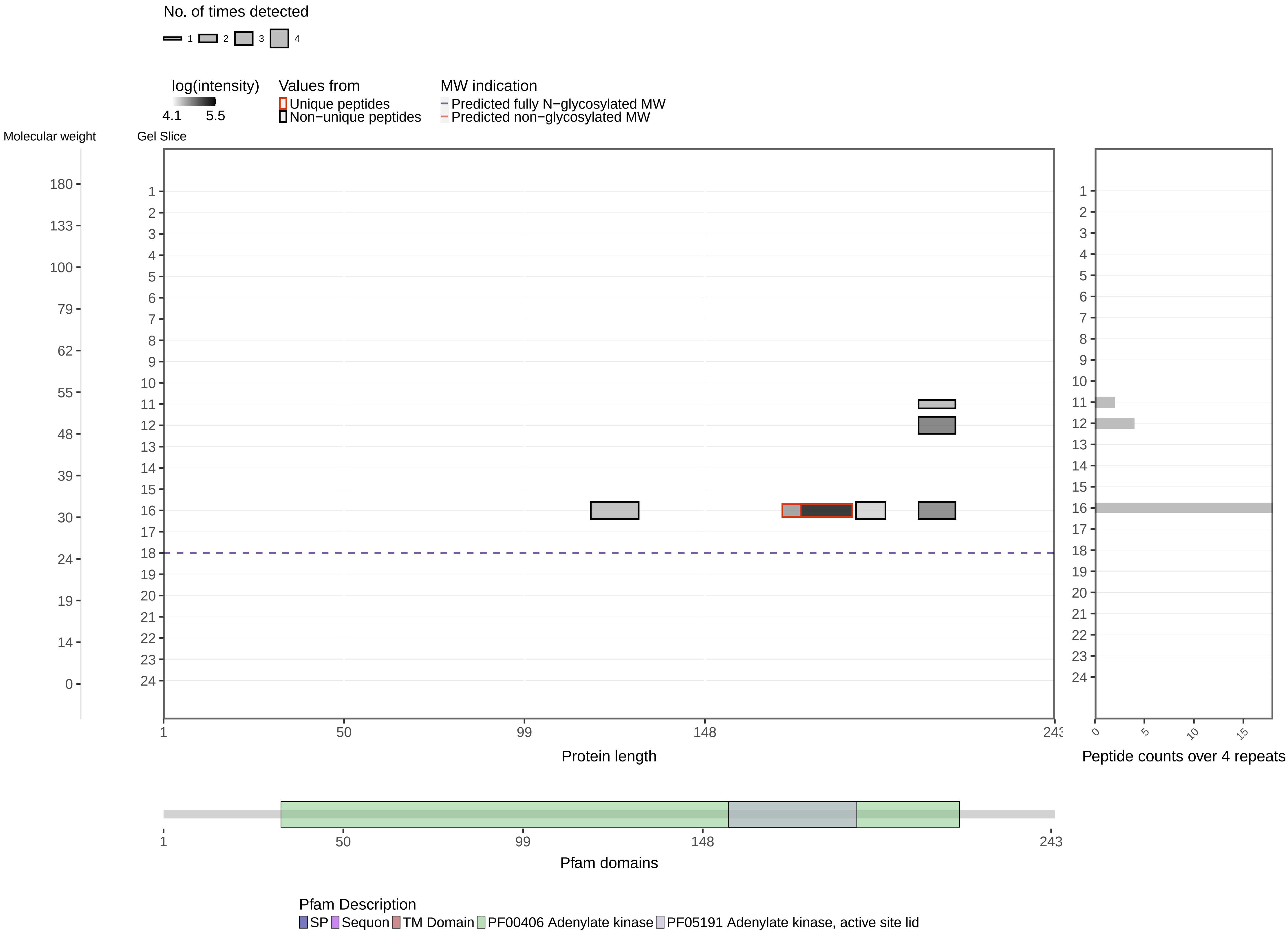


Protein sequence:

GFILDGFPRTVAQAEKLDEMLQKQGTKIDKVLNFAIDDAILEERITGRWIHPSSGRSYHTKFQPPKLPGVDDITGEPLIQRKDDTAEVLKSRLDAFHRQTEPVINYYSKKGVVASLHAEKPPKEVTSEVKNVLSS*

### NbL16g18420.1 glyceraldehyde−3−phosphate_dehydrogenase_cytosolic [MW = 36.99 kDa]

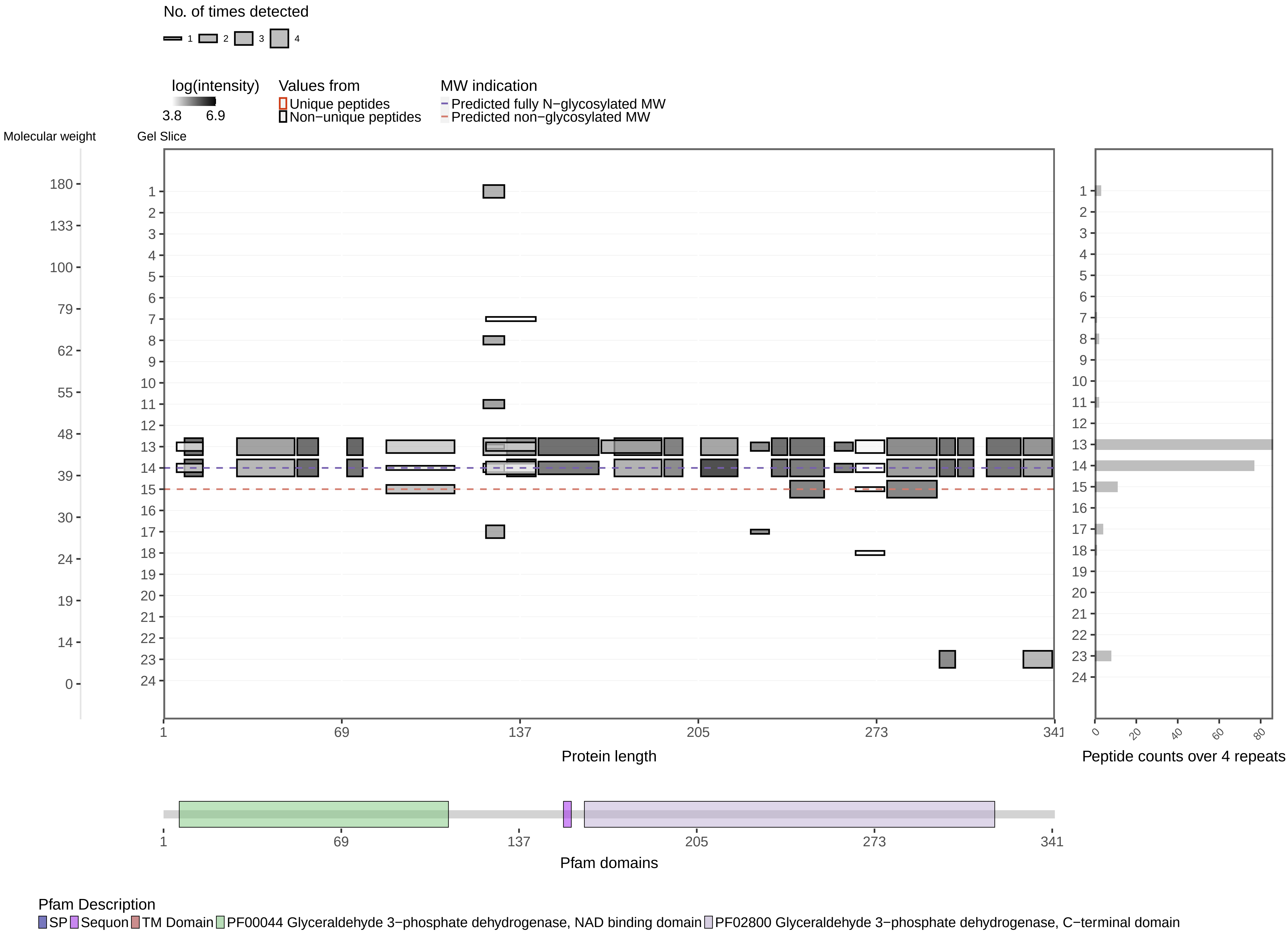


Protein sequence:

CTTNCLAPLAKVINDRFGIVEGLMTTVHSLTATQKTVDGPSMKDWRGGRAASFNIIPSSTGAAKAVGKVLPALNGKLTGMAFRVPTVDVSVVDLTVRLEKEASYDEIKAAIKEESEGKLKGILGFTEDDVVSTDFVGDSRSSIFDAKAGIALSKSFVKLVSWYDNEWGYSSRVIDLICHMASVA*

### NbL16g18340.1 protein_HHL1_chloroplastic [MW = 25.92 kDa]

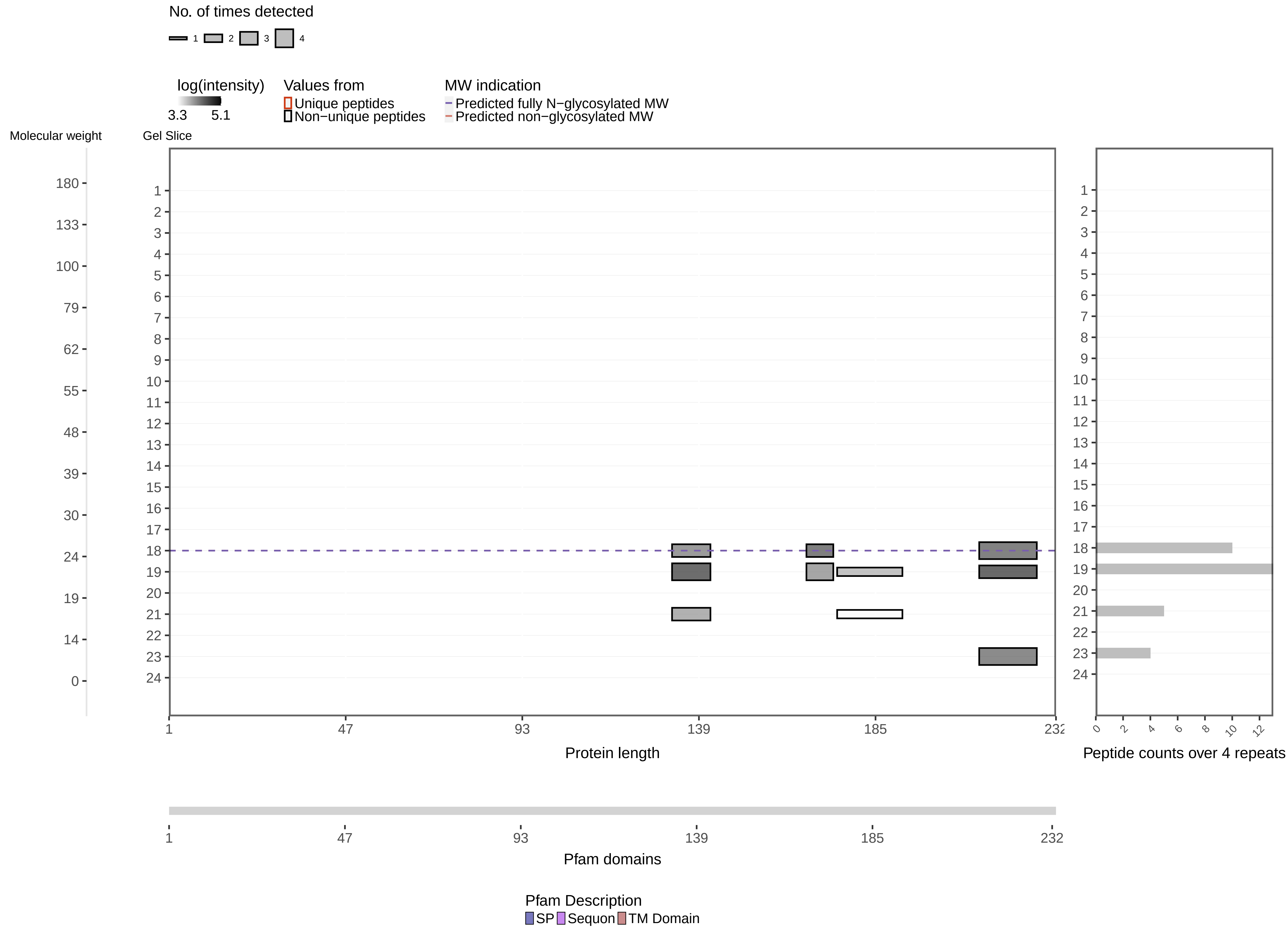


Protein sequence:

IIVTGGTTAKIMVAAKDNFLGKYVYKDTLARNLAAVIYNDEKNIKKLAMKQHRVLKSATEFRYGYKLVENNNLRAALSTTDVIELPTPDQLKTVVDKVKDFFGDAKESFGKLTALPTSESSEEDSPKEGPK*

### NbL16g18190.1 probable_pectate_lyase_5 [MW − SP = 42.53 kDa]

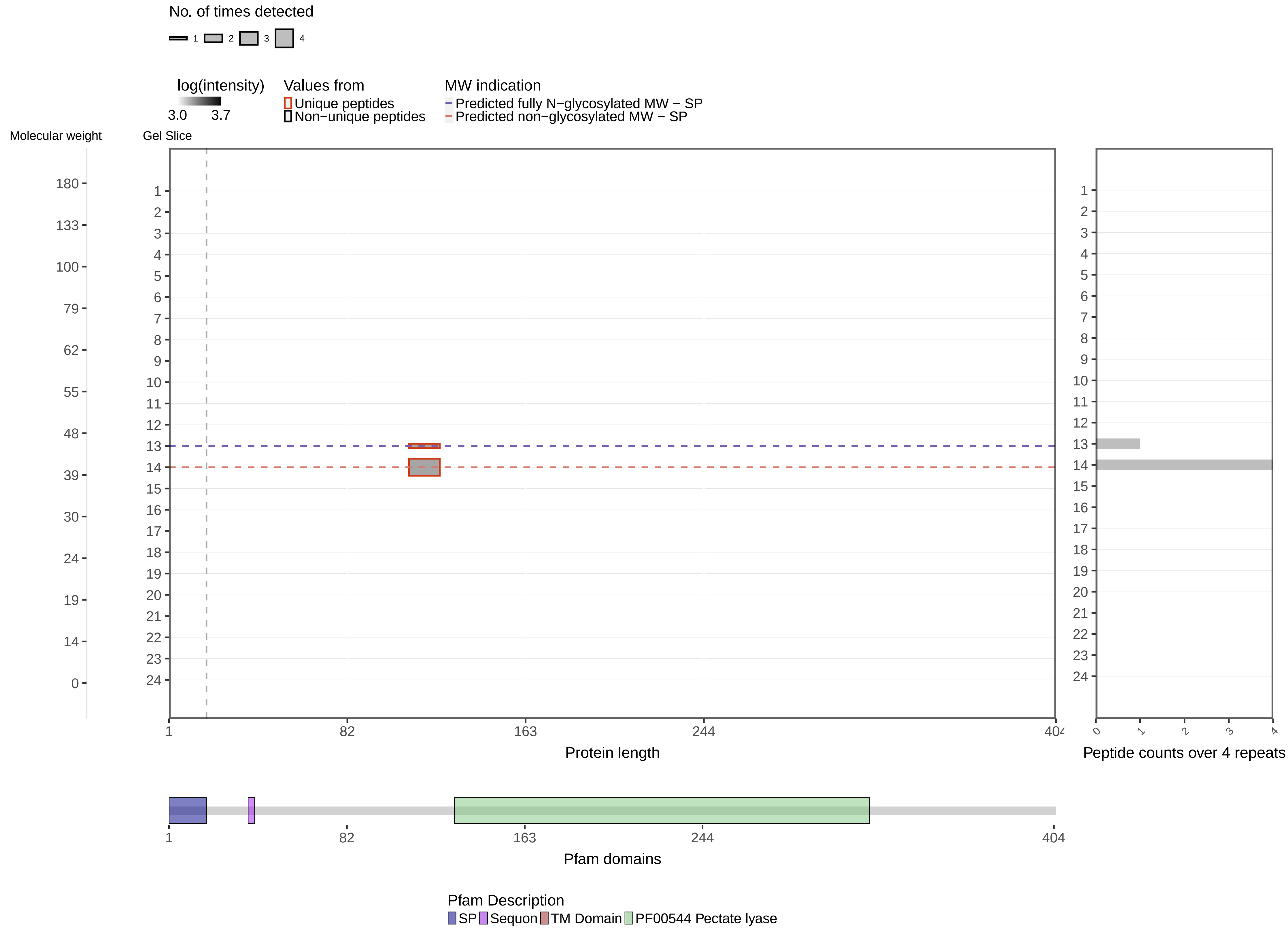


Protein sequence:

RTLSDGDGVSIFGGSHVWVDHCSLSNCRDGLIDAIRGSTAITISNNYMTHHNKVMLLGHSDSFTRDKNMQVTIAFNHFGEGLVQRMPSENCRCRHGYFHVVNNDYTHWEMYAVGGSASPTINSQGNRFLAPNDIFNKEVTKHEDAAESEWKNWNWRSEGDLMLNGAFFTRSGAGASKSYAKASSLSARPSTLVSSITLNAGALGCKKAKRC*

NbL16g18180.1 T−complex_protein_1_subunit_epsilon−like [MW = 59.04 kDa]

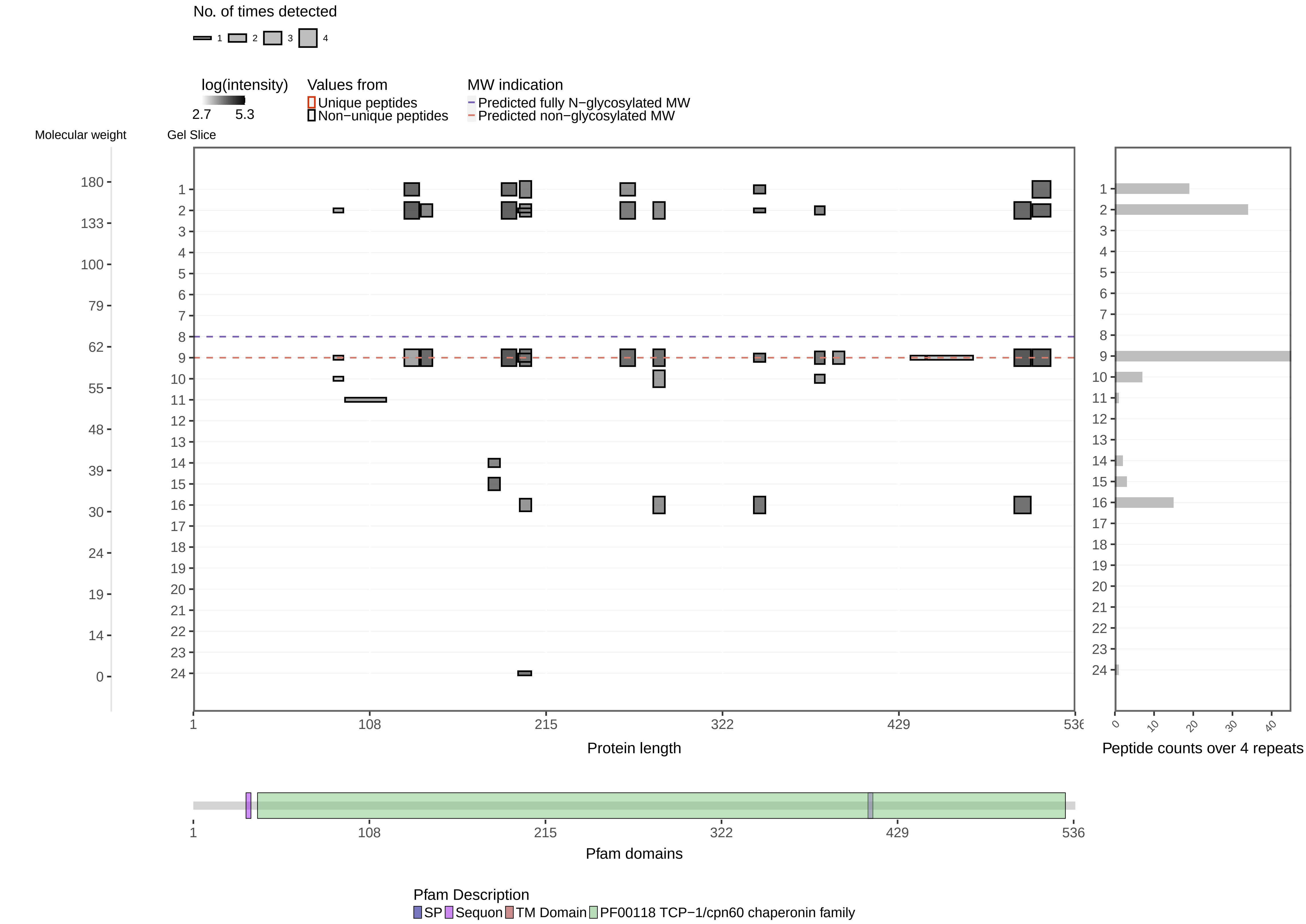


Protein sequence:

[illegible]

### NbL16g18060.1 60S_ribosomal_protein_L18−2−like [MW = 20.86 kDa]

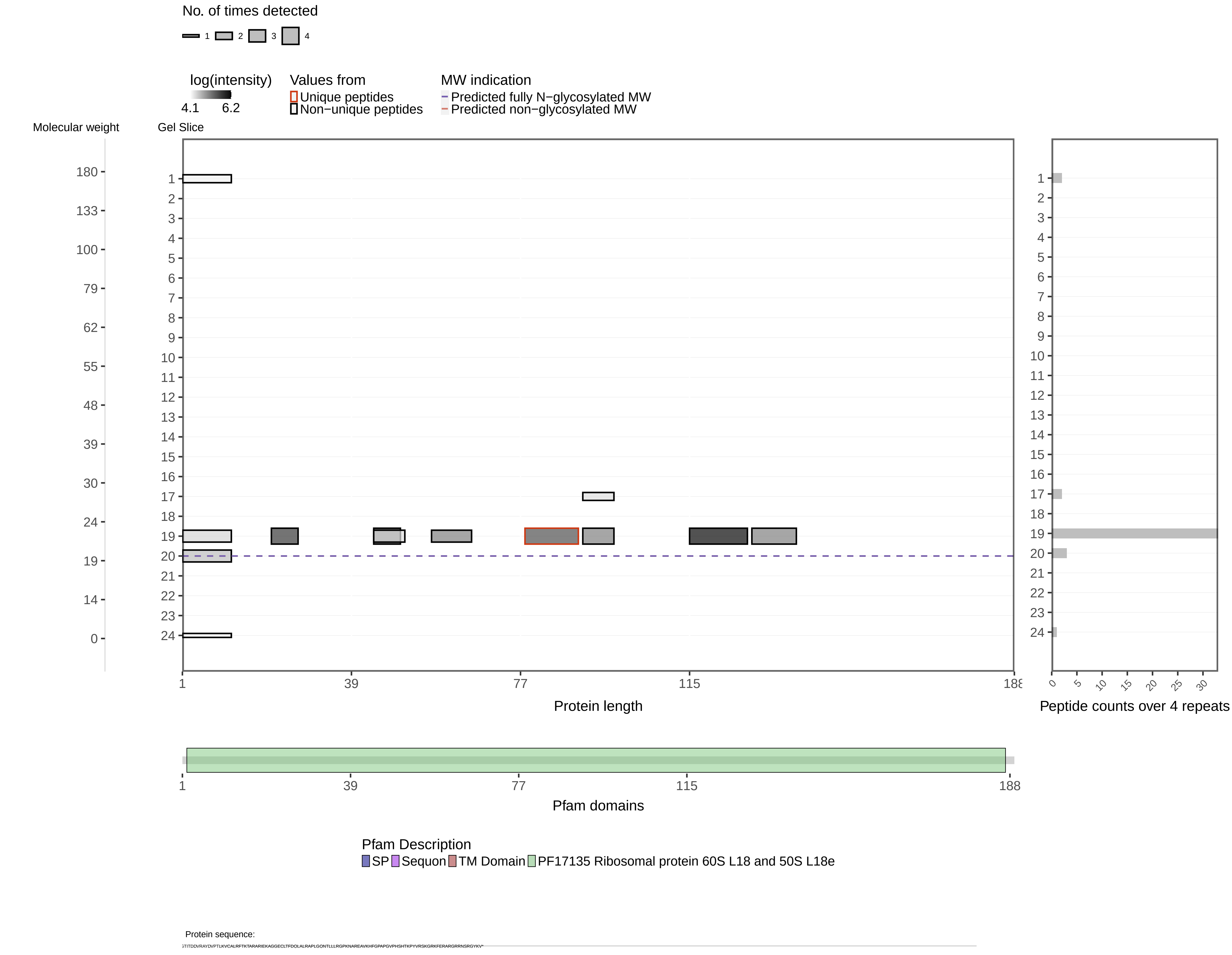

### NbL16g18040.1 probable_pectinesterase/pectinesterase_inhibitor_41 [MW − SP = 54.26 kDa]

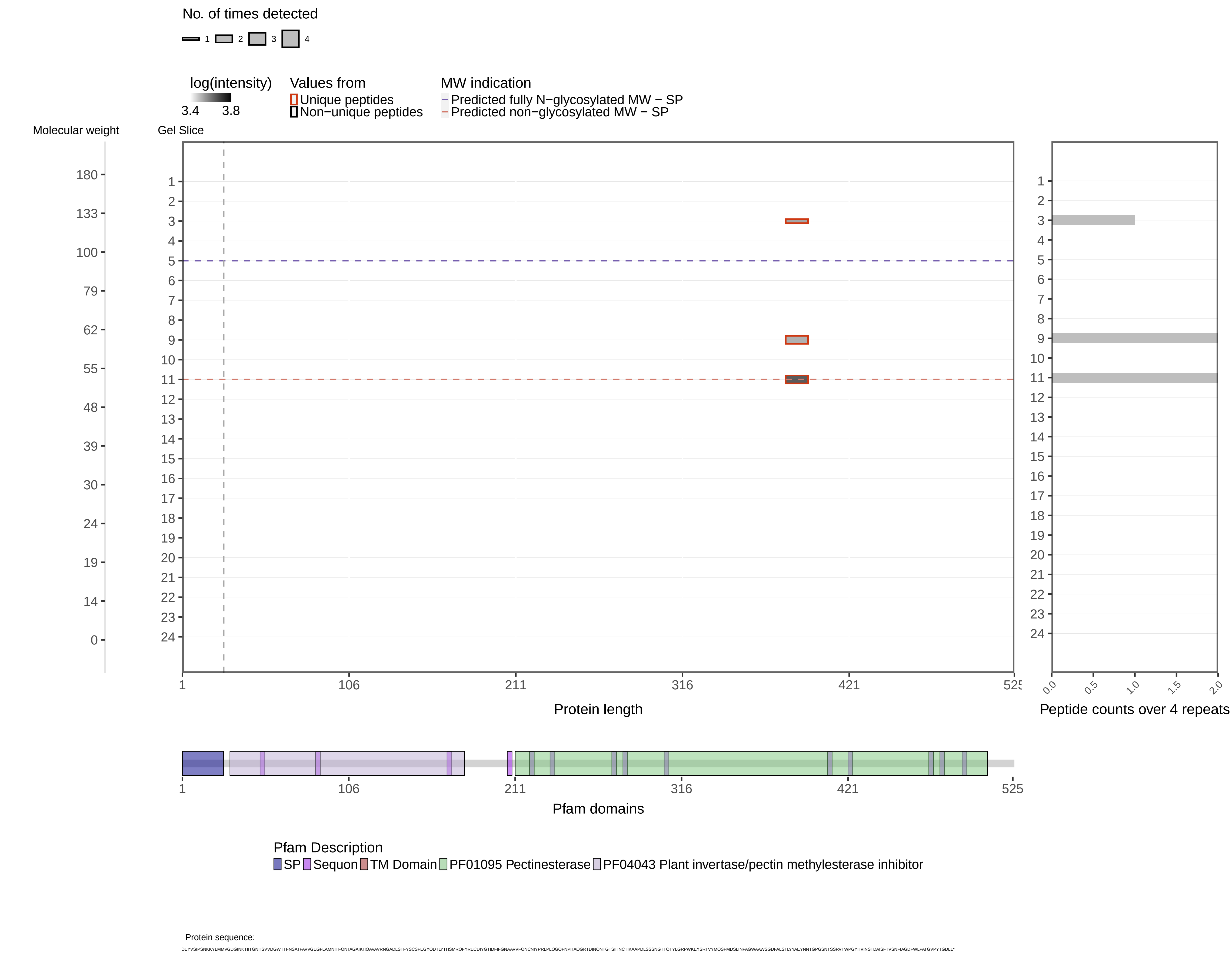

NbL16g18030.1 uncharacterized_LOC107803223 [MW = 33.57 kDa]

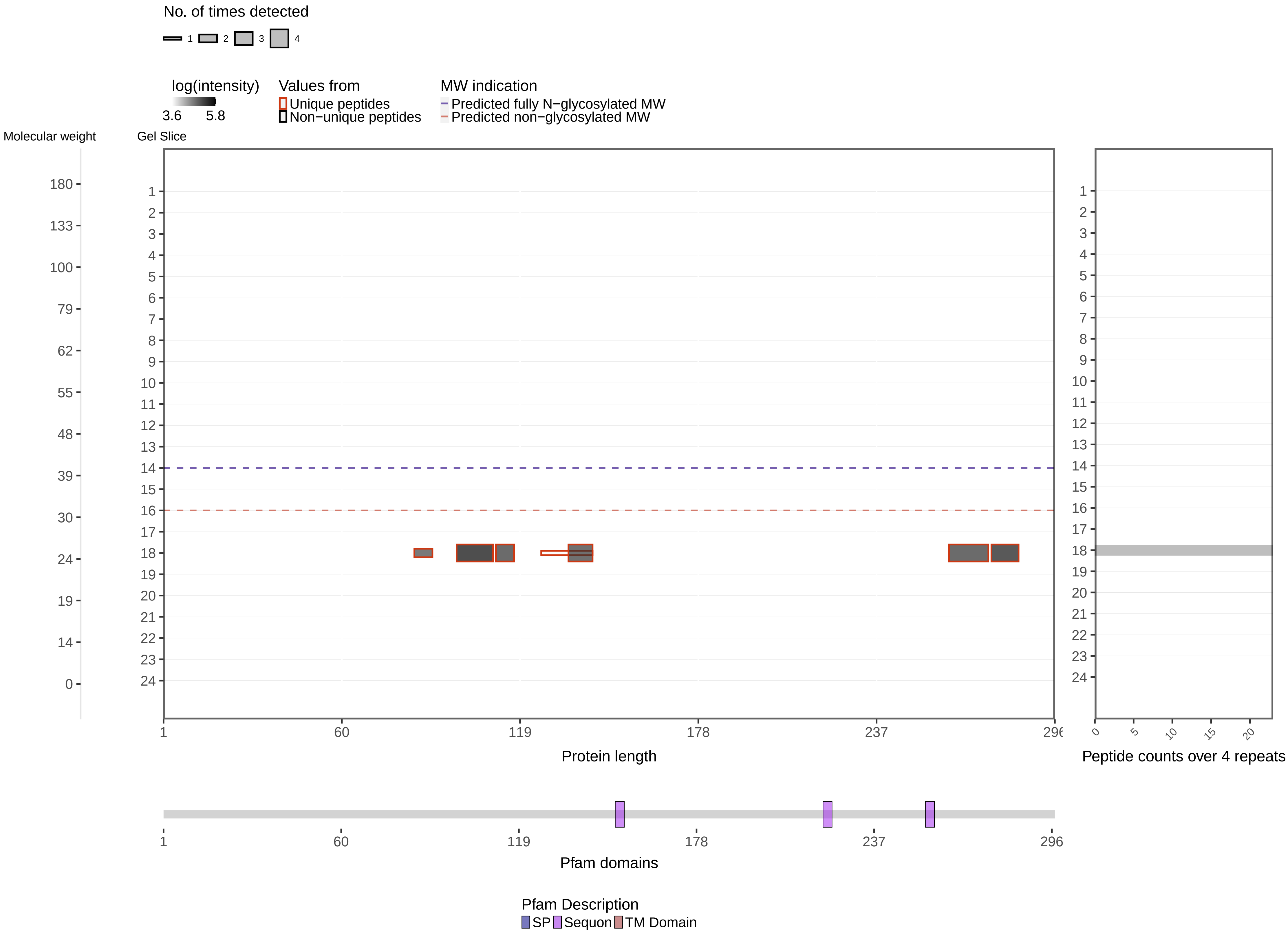


Protein sequence:
EGAEQFRLDVAQNPNDTEESIWCFLCEAQLYGVGEARKRYLELMLCVQVGRDSRPVMREAYNMFKDGGNPEKLWLVRIFSLTTNYSLILKLVAAFSSGQPNEYFYASLYAGLYYESQNESDAAKVHLIAACQSAYGSRSDDYMAALAKVHCKCRNWNLN*

NbL16g17960.1 dihydrolipoyl_dehydrogenase_2_chloroplastic−like [MW = 60.26 kDa]

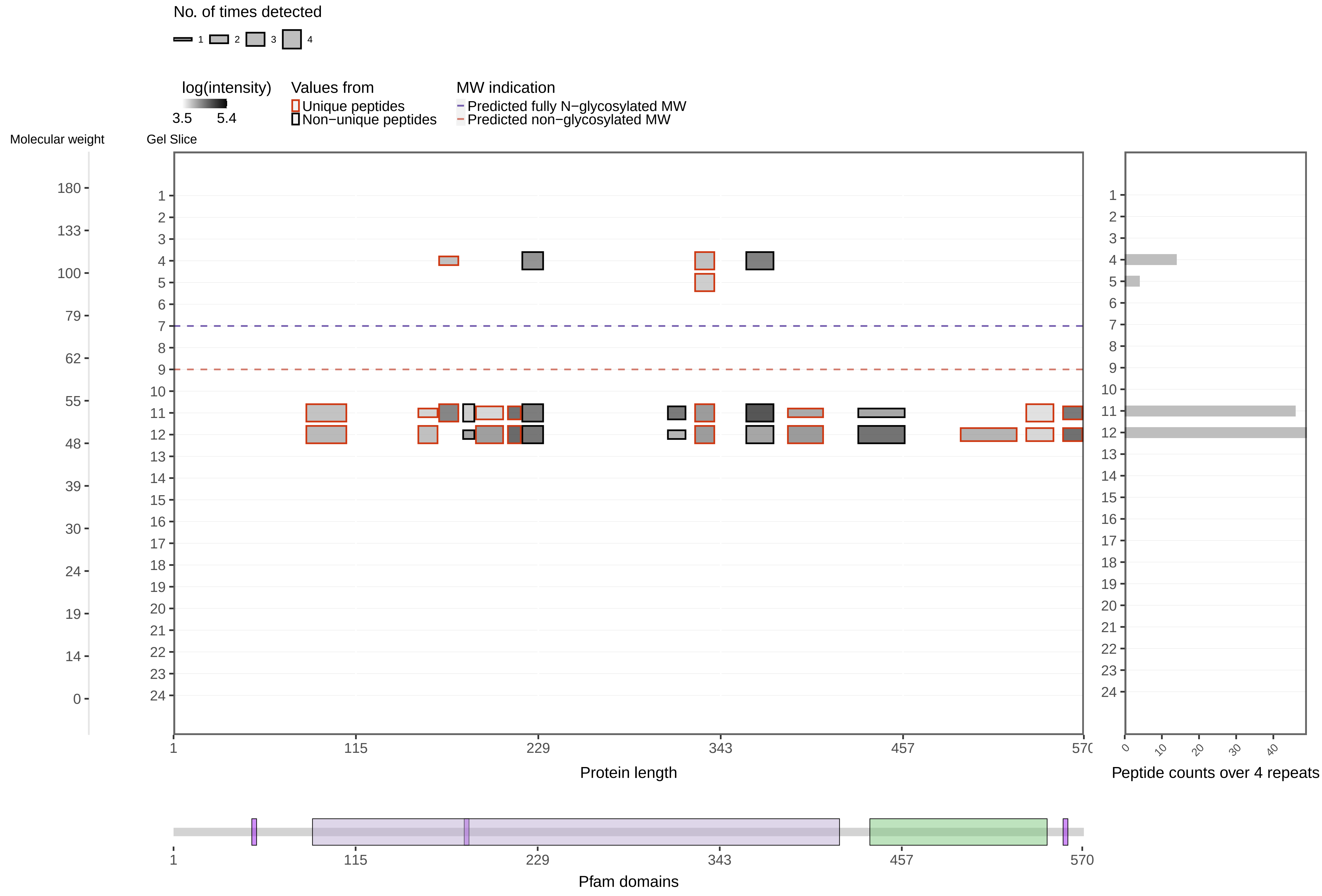


Pfam Description

SP Sequon TM Domain PF02852 Pyridine nucleotide−disulphide oxidoreductase, dimerisation domain PF07992 Pyridine nucleotide−disulphide oxidoreductase

Protein sequence:

DVYTALGSEVTFVEALDQLMPGFDPEIGKLAQRVLINPRKIDYHTGVFASKITPARDGKPVLIELIDAKTKEPKDTLEVDAALIATGRAPFTQGLGLENINVQTQRGFVPVDERMQVIDANGELVPHLYCIGDANGKMMLAHAASAQGISVVEHVTGKDHILNHLSIPAACFTHPEISMVGLTEPQAREKAEKEGFEISIAKTSFKANTKALAENEGEGLAKLIYRPDNGEILGVHIFGMHAADLIHEASNAIALGSRIQDIKYAVHAHPTLSEVIDELFKSAKVKTNASRPIAEPVAA*

### NbL16g17940.1 lon_protease_homolog_1_mitochondrial−like [MW = 108.13 kDa]

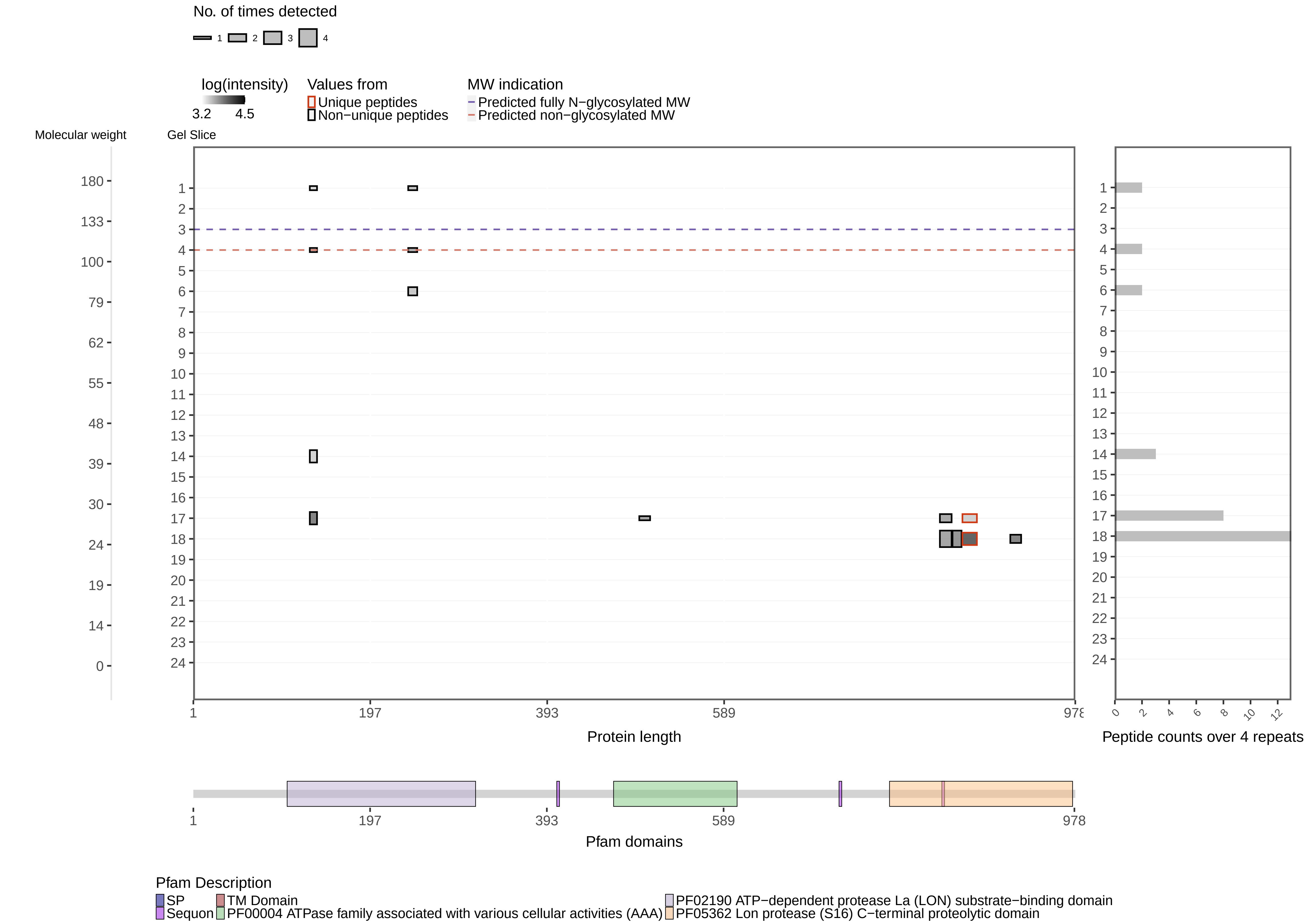


Protein sequence:

[illegible]

NbL16g17920.1 ATP−dependent_Clp_protease_proteolytic_subunit_5_chloroplastic−like [MW = 31.63 kDa]

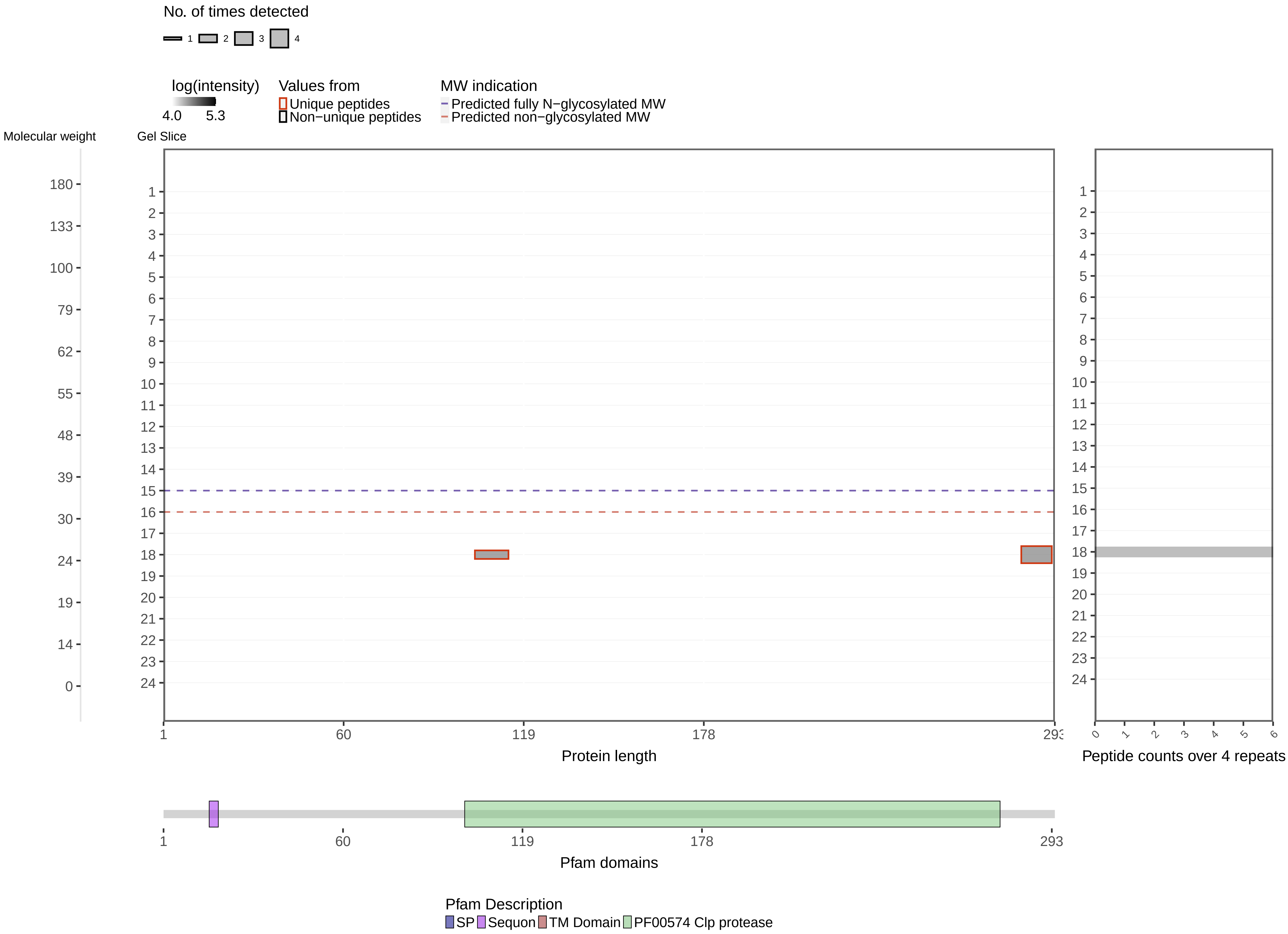


Protein sequence:
LYLDAVDPTKDIVMYVNSPGGSVTAGMAIFDTMRHIRPDVSTVCVGLAASMGAFLLSAGTKGKRYSLPNSRIMIHQPLGGAQGGQTDIDIQANEMLHHKANLNGYLAYHTGQSLEKINQDTDRDFFMSAKEAKEYGLIDGVILNPMKALQPIAAAAEQ*

NbL16g17830.1 uncharacterized_LOC109210602 [MW = 27.17 kDa]

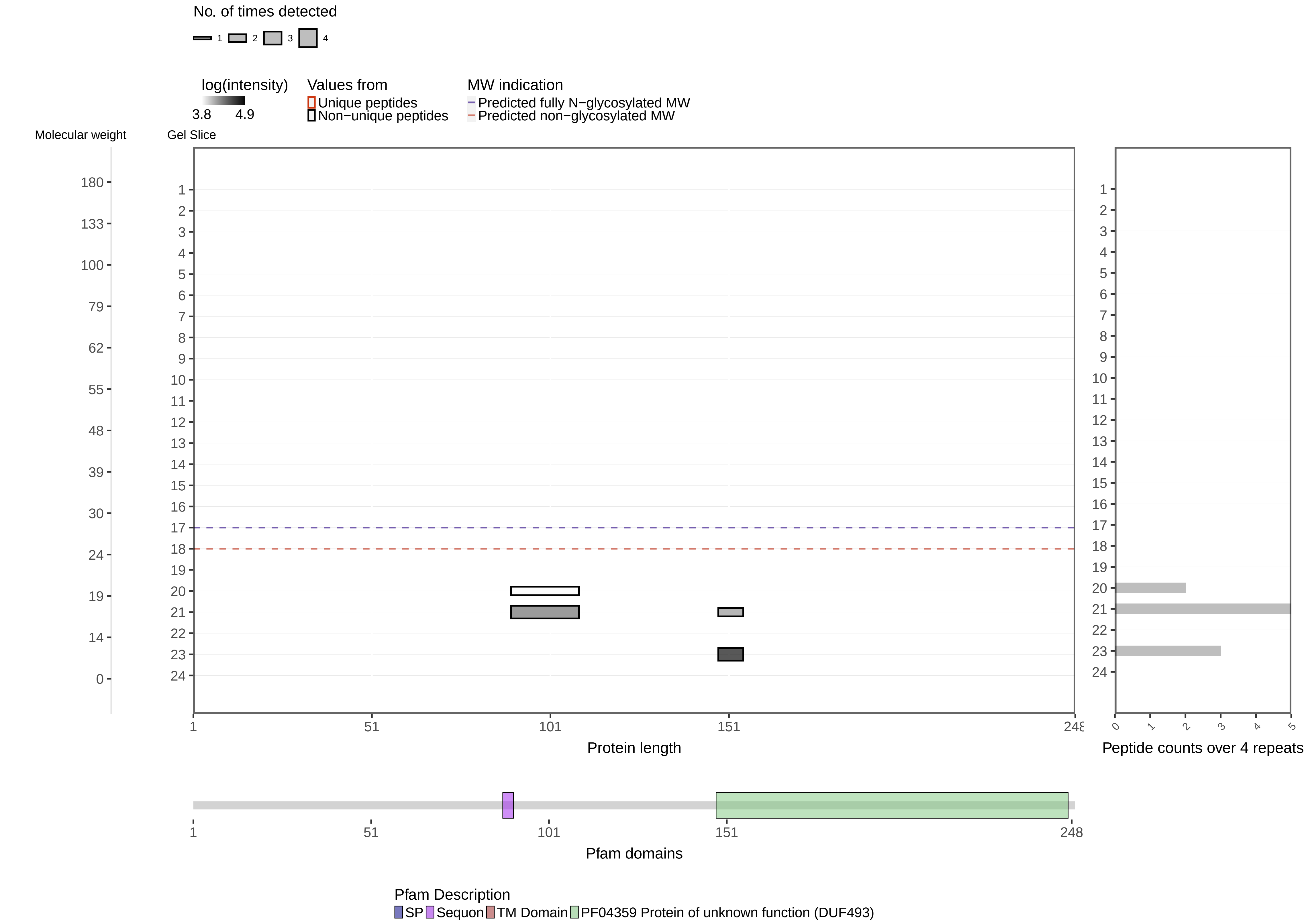


Protein sequence:

SEVSKSEGRVGQTTNVIIGGTVQDDATNEWLALDKKVNSYPTERGFTAIGTGGDDFVQAMVVAVESVIQQPISEASVDPPDVHDDVDGFVGKVRQKLSSGGKYVSVNIGPIEVVSSEQVQAVYNAMRKDDRMKYFL*

NbL16g17670.1 sm−like_protein_LSM36B [MW = 11.89 kDa]

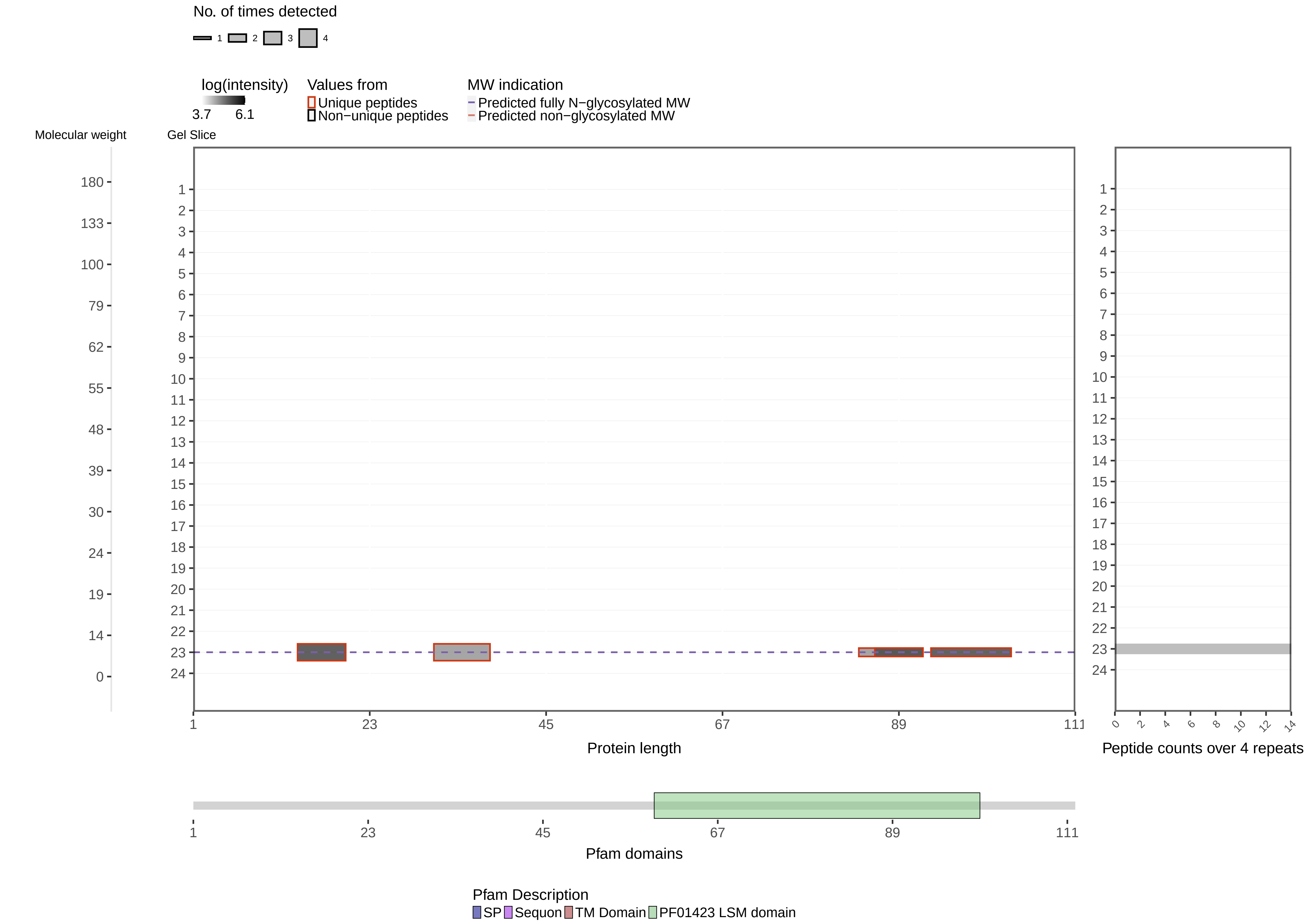


Protein sequence:

SATILLSHHIHGFSSSSILACLDGYMNIAMEQTEEYVNGQLKNKYGDAFIRGNNVLYISTSKRTQGEGA*

### NbL16g17620.1 glucan_endo−13−beta−glucosidase_8−like [MW − SP = 51.13 kDa]

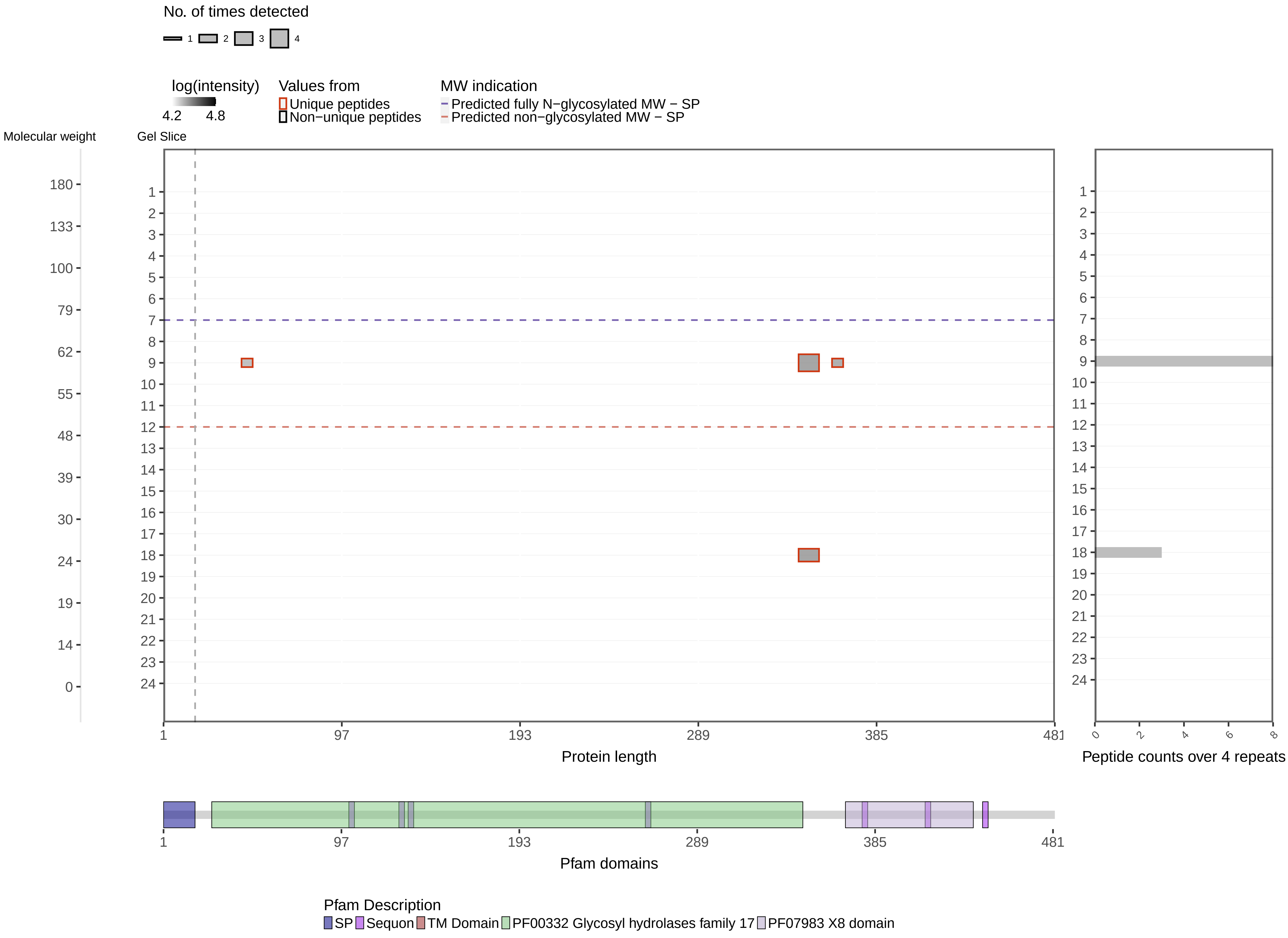


Protein sequence:

SNPVVDNGVEYTNVFDANFDTLYSALKAVGYGNMTILVGEVGWPTDGDKNANSNNAYRFYKGLFSRLAANKGTPLRPGYIEVYLFGLIDEDAKSIAPGNFERHWGIFRYDGQPKFAMDISGQGQDKHLVAAKNVQYLPKRWCMLNPNANNLSILANNIDYACTFSDCTALGYGSSCNNLDANGNASYAFNMYYQVQNQLDLSCDFQGLAMVTDKNISQGTCNFIIQTGIYSVSSKVLPGIVVLLSGFTFLLL*

### NbL16g17580.1 glutamate_dehydrogenase_B [MW = 44.52 kDa]

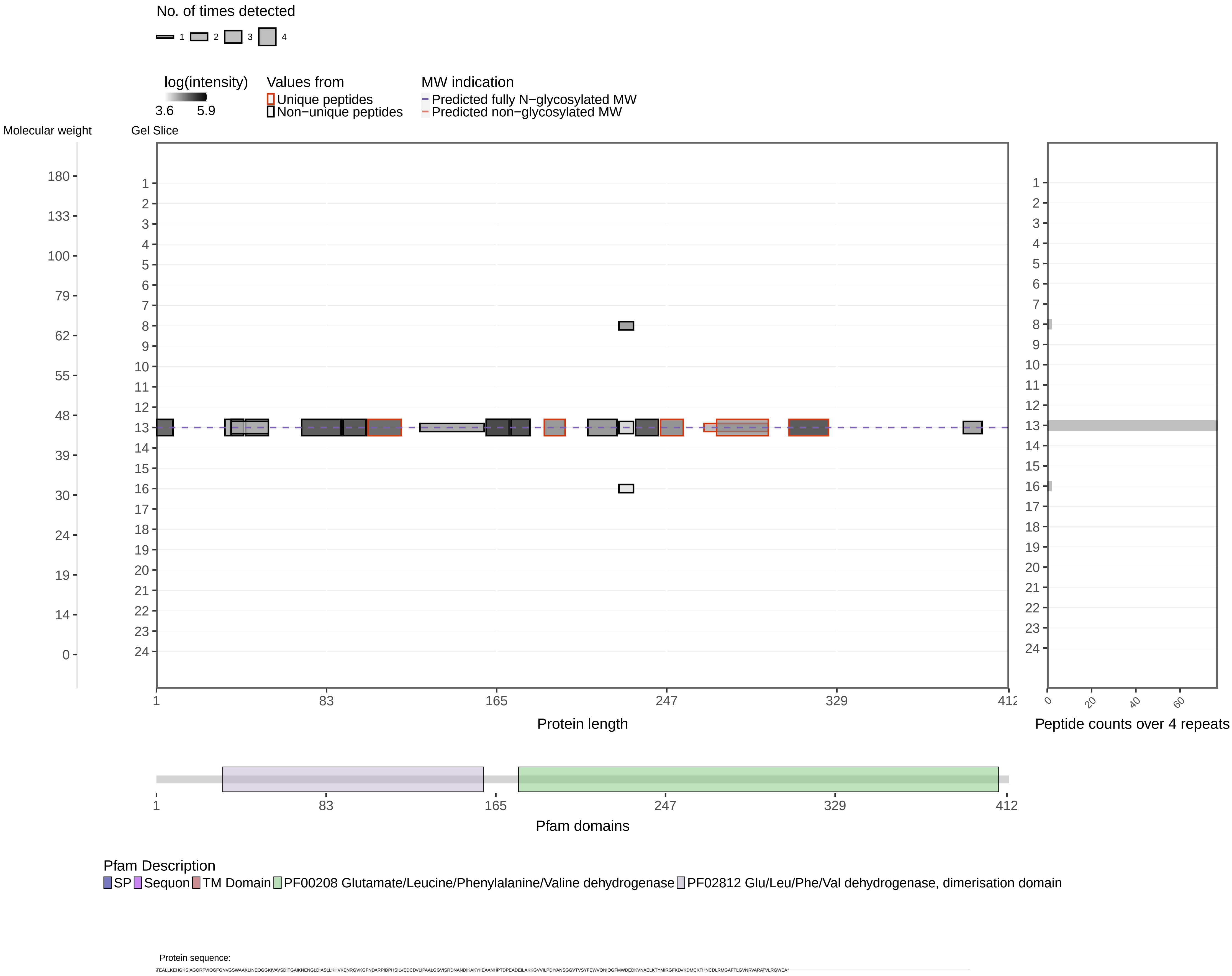

### NbL16g17540.1 40S_ribosomal_protein_S28 [MW = 11.97 kDa]

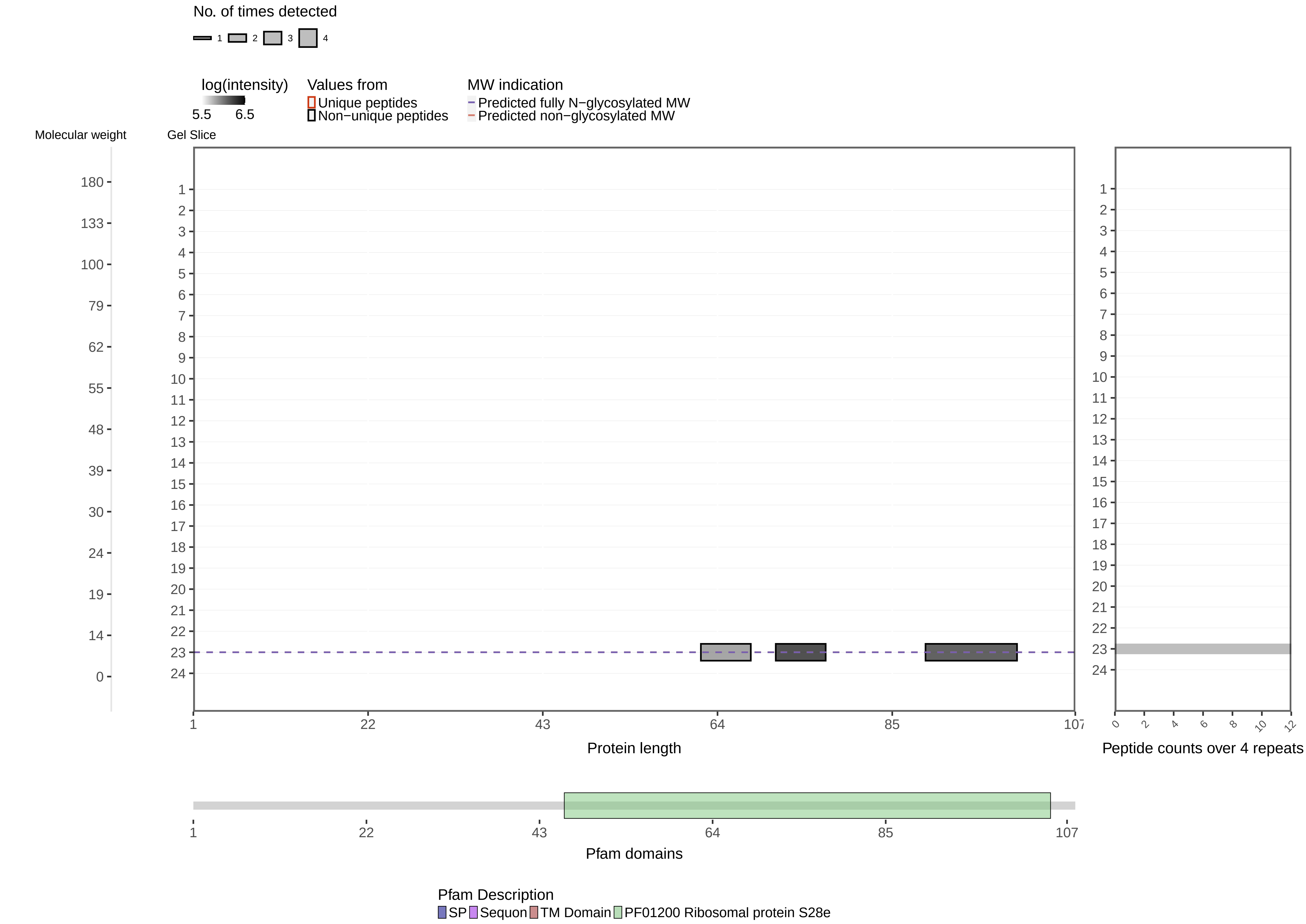


Protein sequence:

IESATKHAIVVKVMGRTGSRGQVTQVRVKFLDDQNRFIMRNVKGPVREGDVLTLLESEREARRLR*

### NbL16g17380.1 protein_trichome_birefringence−like [MW = 76.6 kDa]

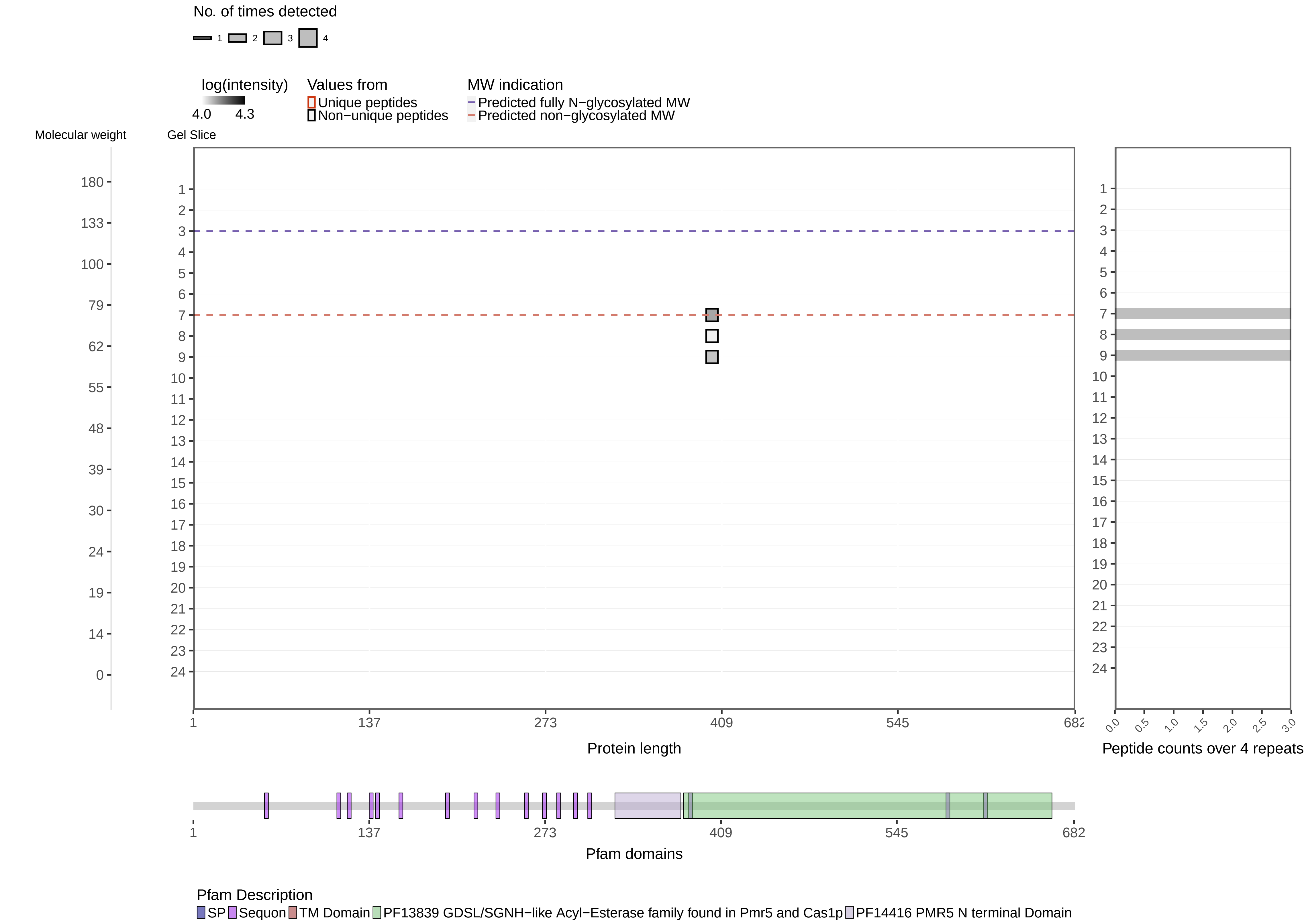


Protein sequence:

FFDGKWVKDESYPLYKPGSCSLIDEQFNCFLNGRPDSNYIKMKWKPNACTLPRLNGTHMLELLRGKRLVFVGDSLNRNMWESLVCILRNSVEDQKKVYEEFGRHHFRTEASYSFLFEEYNCRVEFFVSPFLVQEWEFADKKGAKKETLRLDLIGQSADQYKSADILIFNTGHWWTHEKTSLGKDYYQEGSHVYSELDVLEAFRKALTTWGRWIDSHVNPHKTFVLFRGYSASHFSGGQWNSGGACDHETEPIKNNTYLTPYPSKMNVLERVLRGMETQVSYLNVTRM

### NbL16g17260.1 DNA−directed_RNA_polymerase_III_subunit_2 [MW = 129.89 kDa]

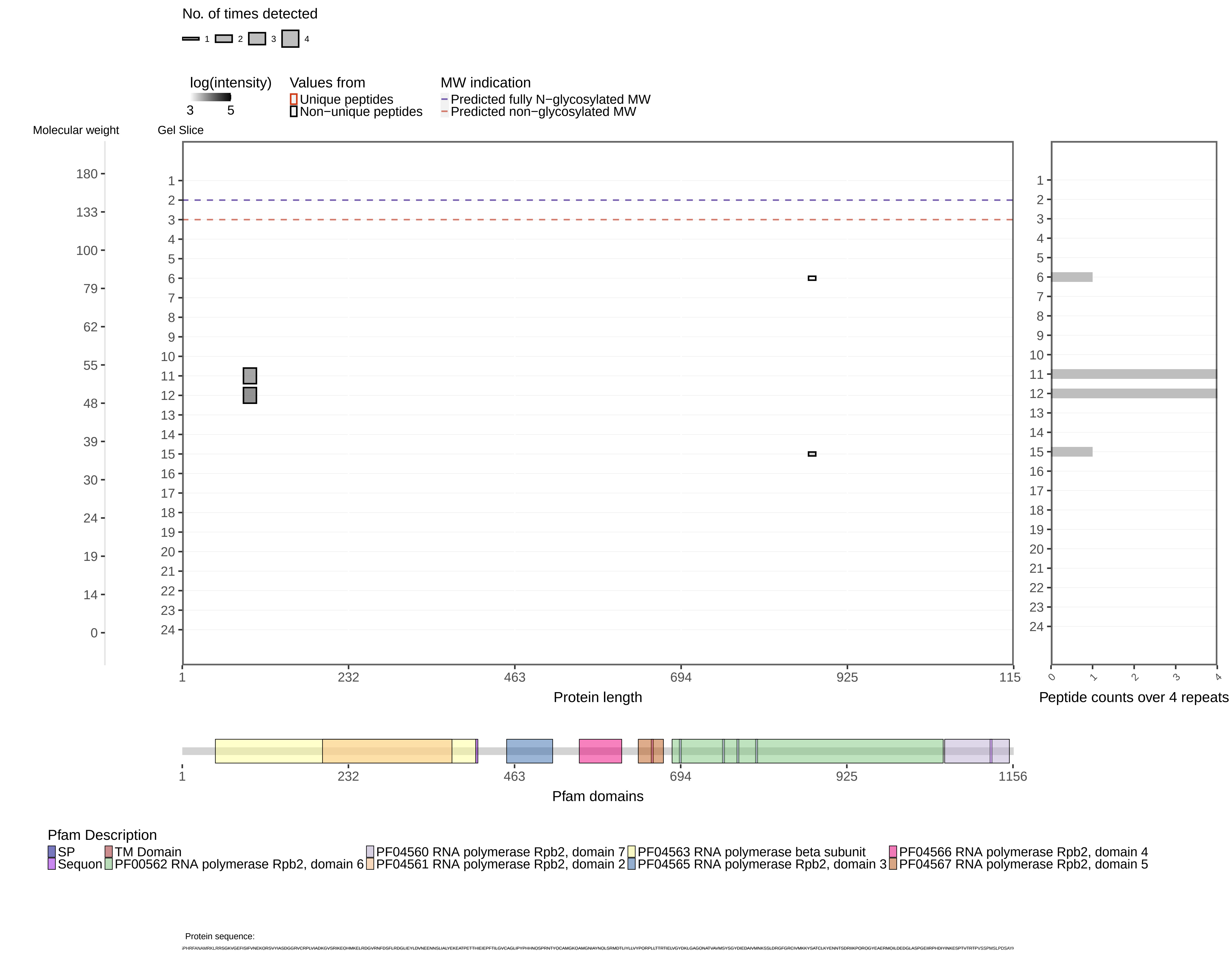

### NbL16g16760.1 glutamine_synthetase_chloroplastic [MW = 47.51 kDa]

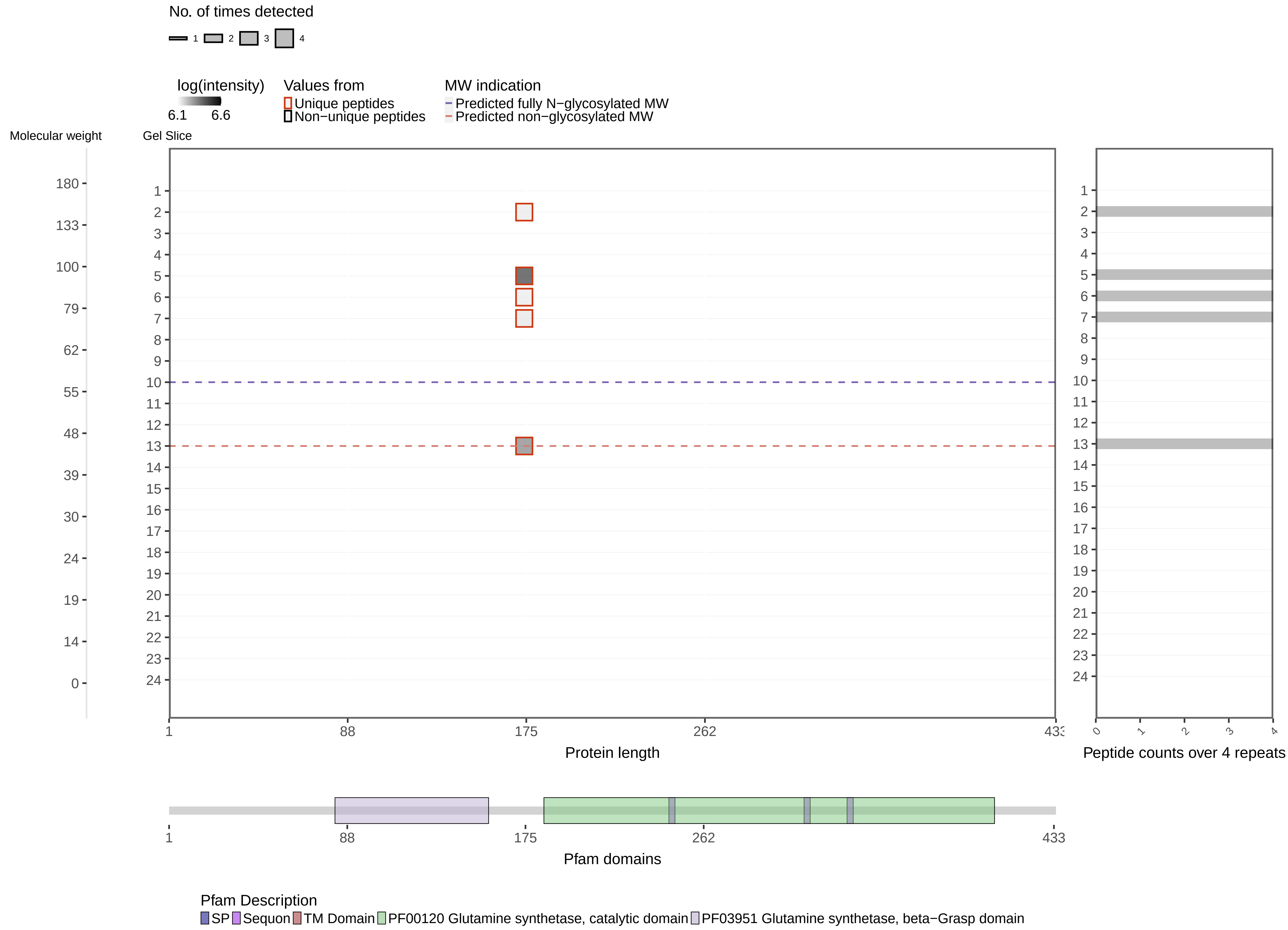


Protein sequence:

SWPVGGYPGPQGPYYCGAGADKSFGRDISDAHYKACLYAGINISGTNGEVMPGQWEFQVGPSVGIEAGDHIWCARYILERITEQAGVVLSLDPKPIEGDWNGAGCHTNYSTLSMREEGGFEVIKKAILNLSLRHKEHISAYGEGNERRLTGKHETASIDKFSWGVANRGASIRVGRDTEKQGKGYLEDRRPASNMDPYVVTGLLAETTILWEPTLEAEALAAQKLALNV*

### NbL16g16660.1 UPF0613_protein_PB24D3.06c [MW = 37.07 kDa]

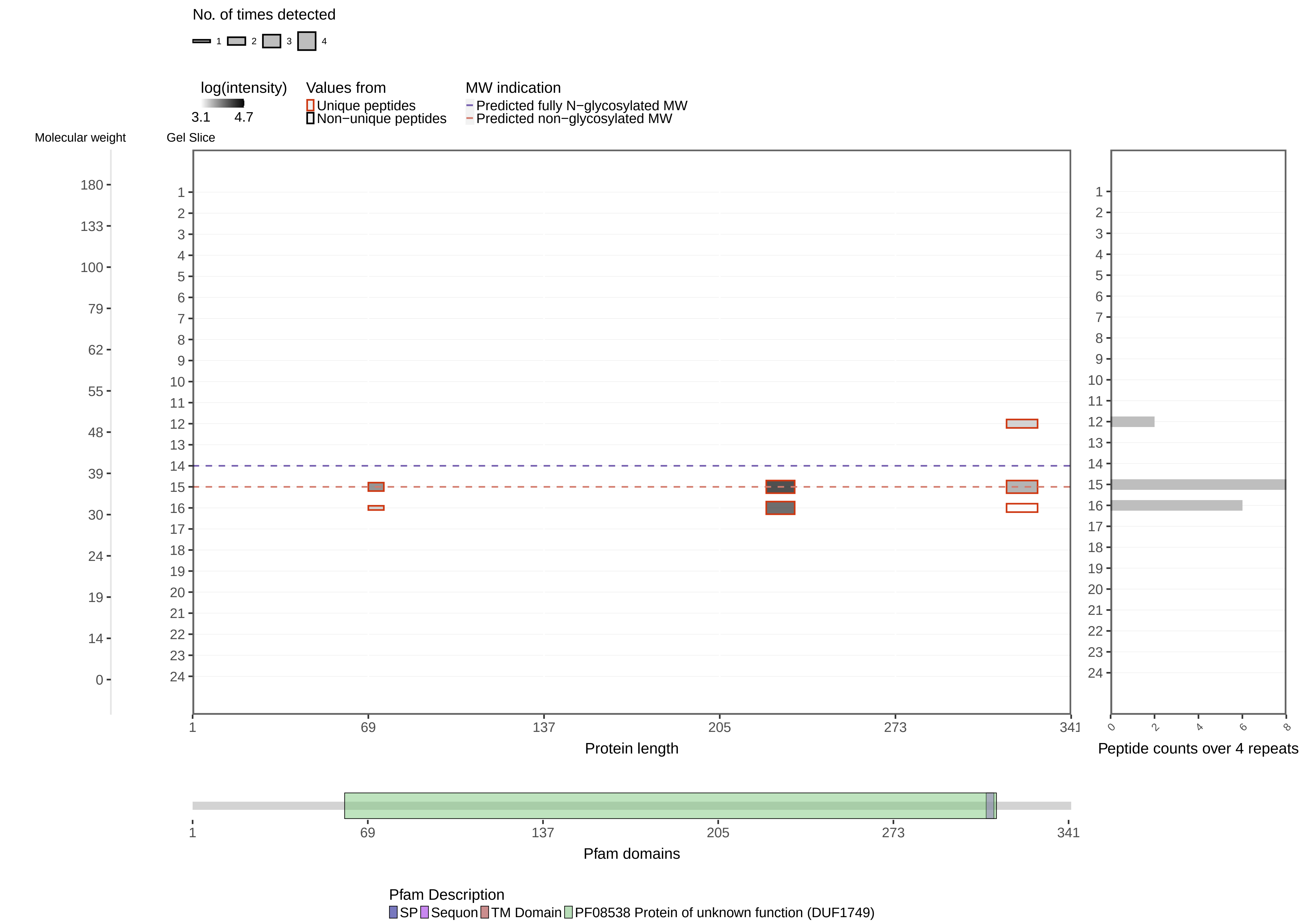


Protein sequence:

STGCQDIVYYMRTNAACSRAVRAAILQAPVSDREYKATLPDTASMIDLASNMISEGRESELMPRDANPDAPITAFRYHSLCAYNGEDDLFSSDFSDDQLKQRLGHMSNTPCQVIFSMGDEYVPDYVDKKALVDRLCRAMGGAEKVEIEHGNHSLSNRVAEAIEAIMSFVKREGPSGWDDPWN*

### NbL16g16530.1 basic_7S_globulin−like [MW − SP = 44.81 kDa]

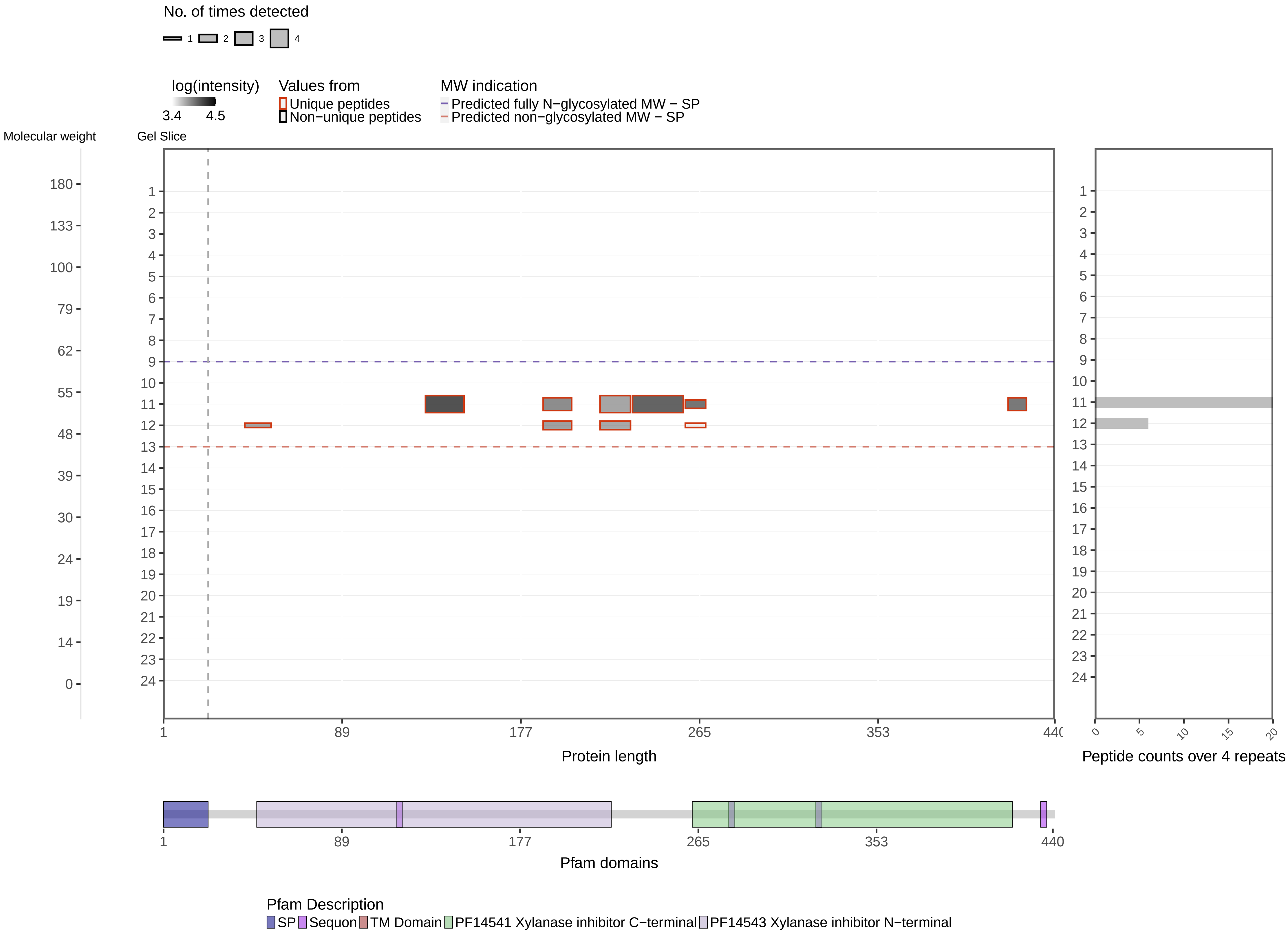


Protein sequence:

FAVCLSSSTNSKGVVLFGDGPYTFLPNREFANNDFSYTPLFINPVSTASAFSSREPSSEYFIGVKSIKINEKVVPINTTLLSIDNQGVGGTKISTVNPYTILETSIYNAVTNFFVKELVNITRVASVAPFRACFDSRNIASTRVGPAVPSIDLVLQNENVFWRIFGANSMVQVSENVLCLGFVDGGVSPRTSIVVGGYTIEDNLLQFDLARSRLGFTSSILFRQTTCANFNFTSIA*

### NbL16g16480.1 ammonium_transporter_1_member_3 [MW = 50.05 kDa]

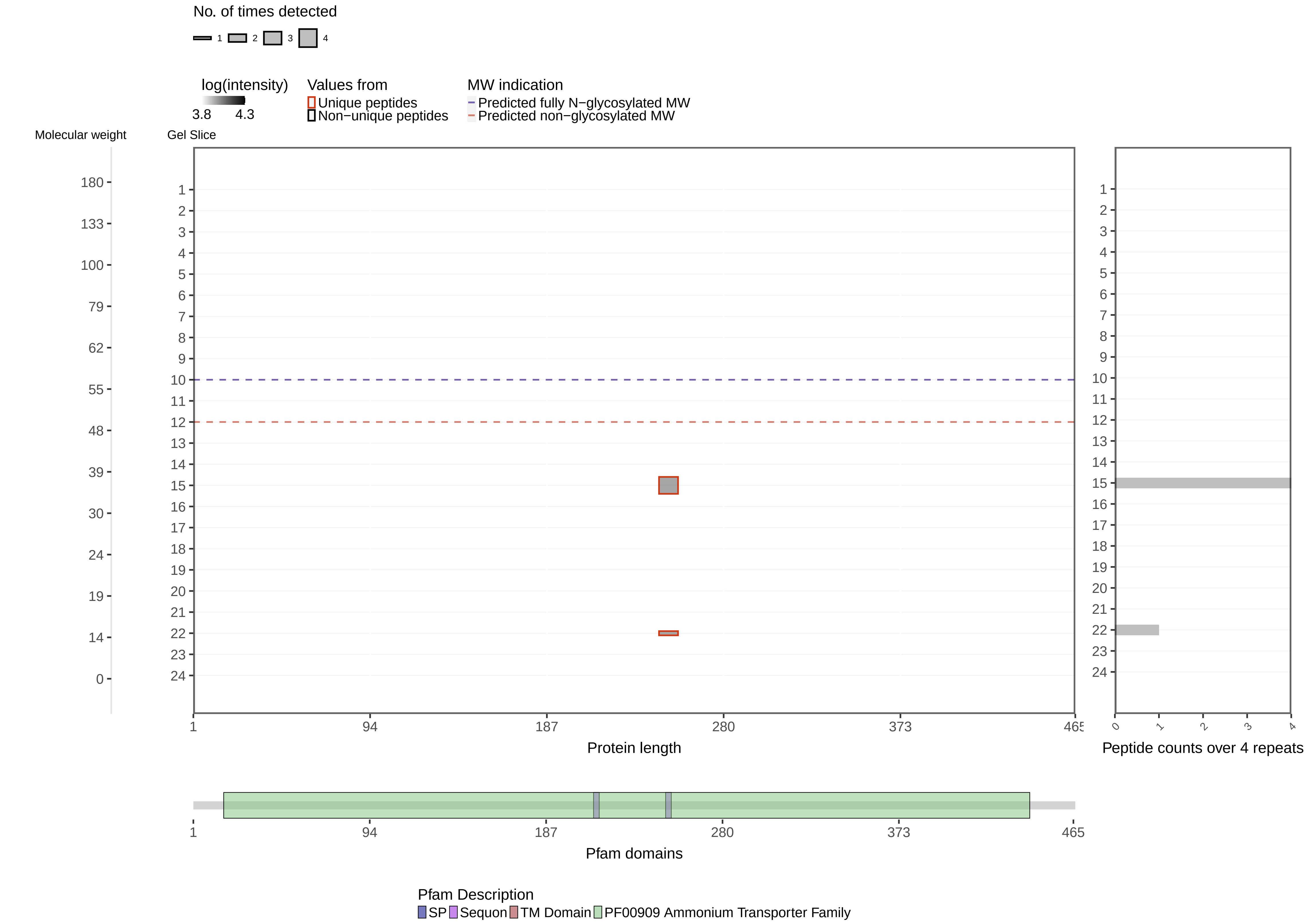


Protein sequence:

FLLWFGWFGFNPGSFDKILVPYPNKIDQGNWTSVGRTAVTTTLAGSTAGIVTLFGRRLLVGHWDATDVCNGVLGGFVAITSGCSVVEPWAAIVCGFCAAWVLIGLNILALKLKFDDPLEAAQLHGGCGAWGLIFTGLFAKEEFVLQAYNSGKTQVVRPYGLILGGGWGLFGAQVVELLSIVVWVSLTMGPLFYTLQKLGILRITADEEVAGLDISSHGGYAYEANQEESGPRFYGEYLRMQHQS*

### NbL16g16470.1 stem−specific_protein_TSJT1−like [MW = 28.11 kDa]

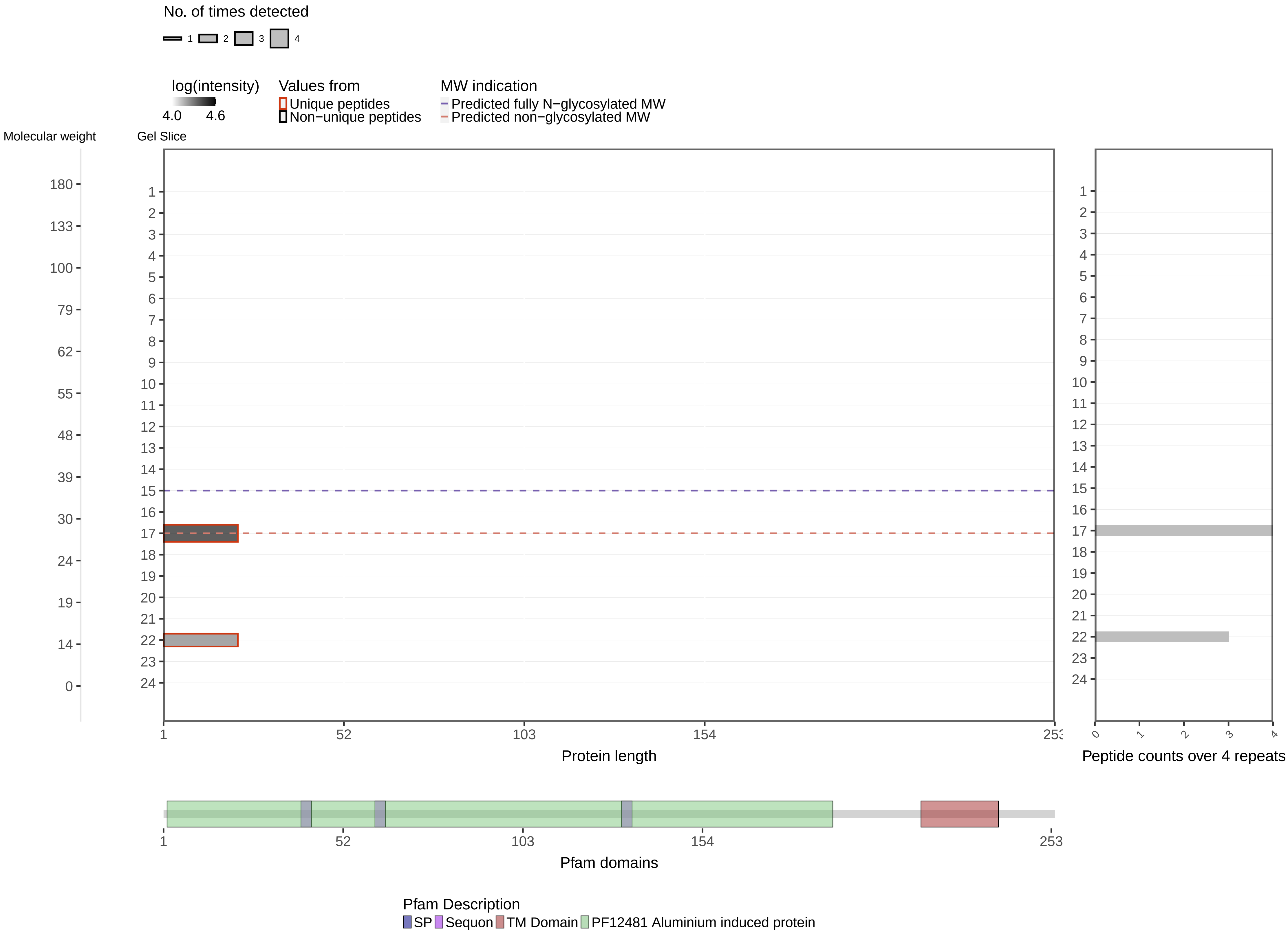


Protein sequence:

ALRDRAPYPPNHVVGHLEGNFTFIVFDKSTSTLFVATDQVGKVPLYWGITADGYVAFANDADLLKGACGKSLASFPQGWFDRLFFSFRQSDLSSGITIEGIMLIYFVLYILFLNSISFVCIFRVFLLYISWRIEKLREPQK*

### NbL16g16450.1 ATP−dependent_(S)−NAD(P)H−hydrate_dehydratase [MW = 34.29 kDa]

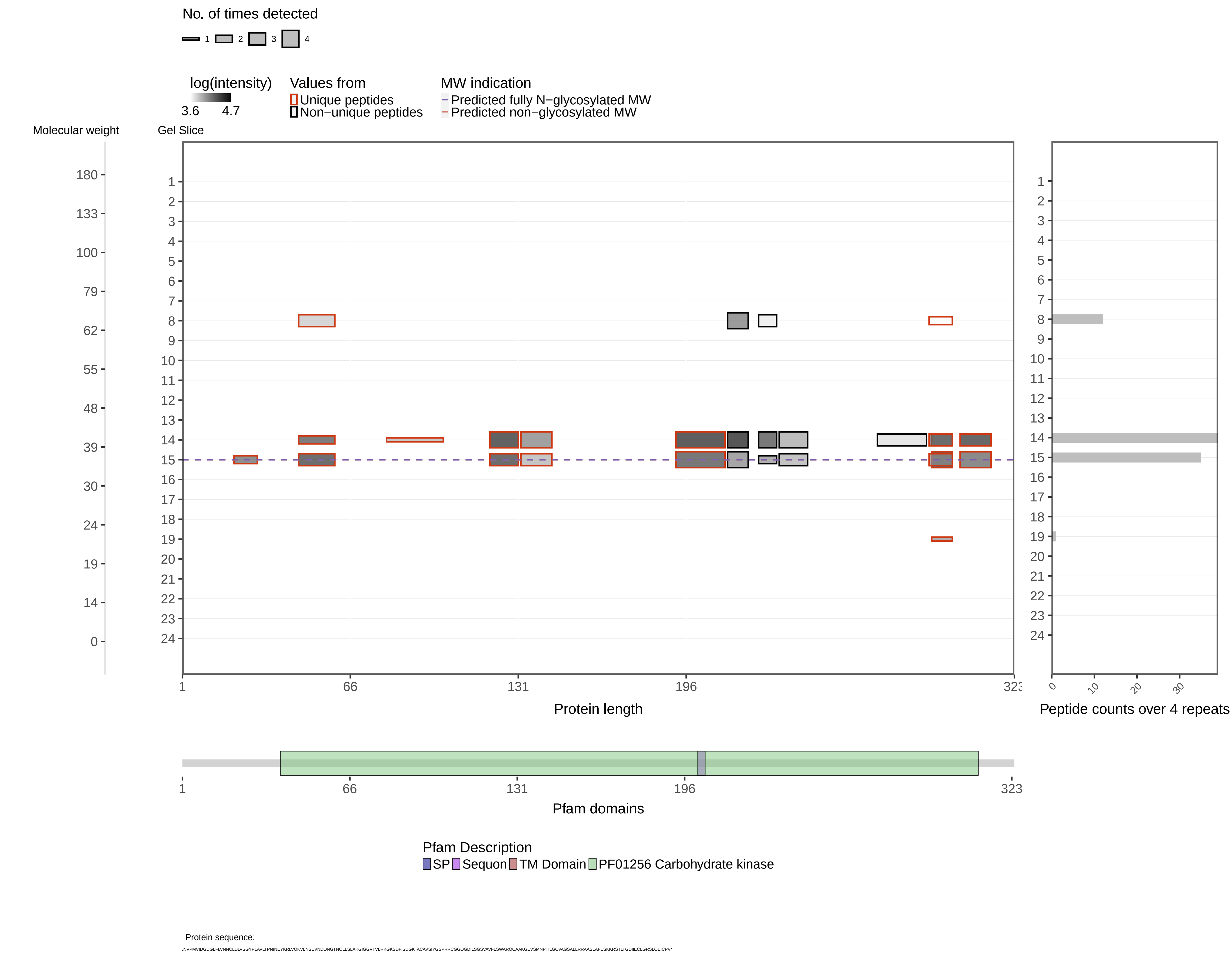

### NbL16g16350.1 COBRA−like_protein_7 [MW − SP = 68.92 kDa]

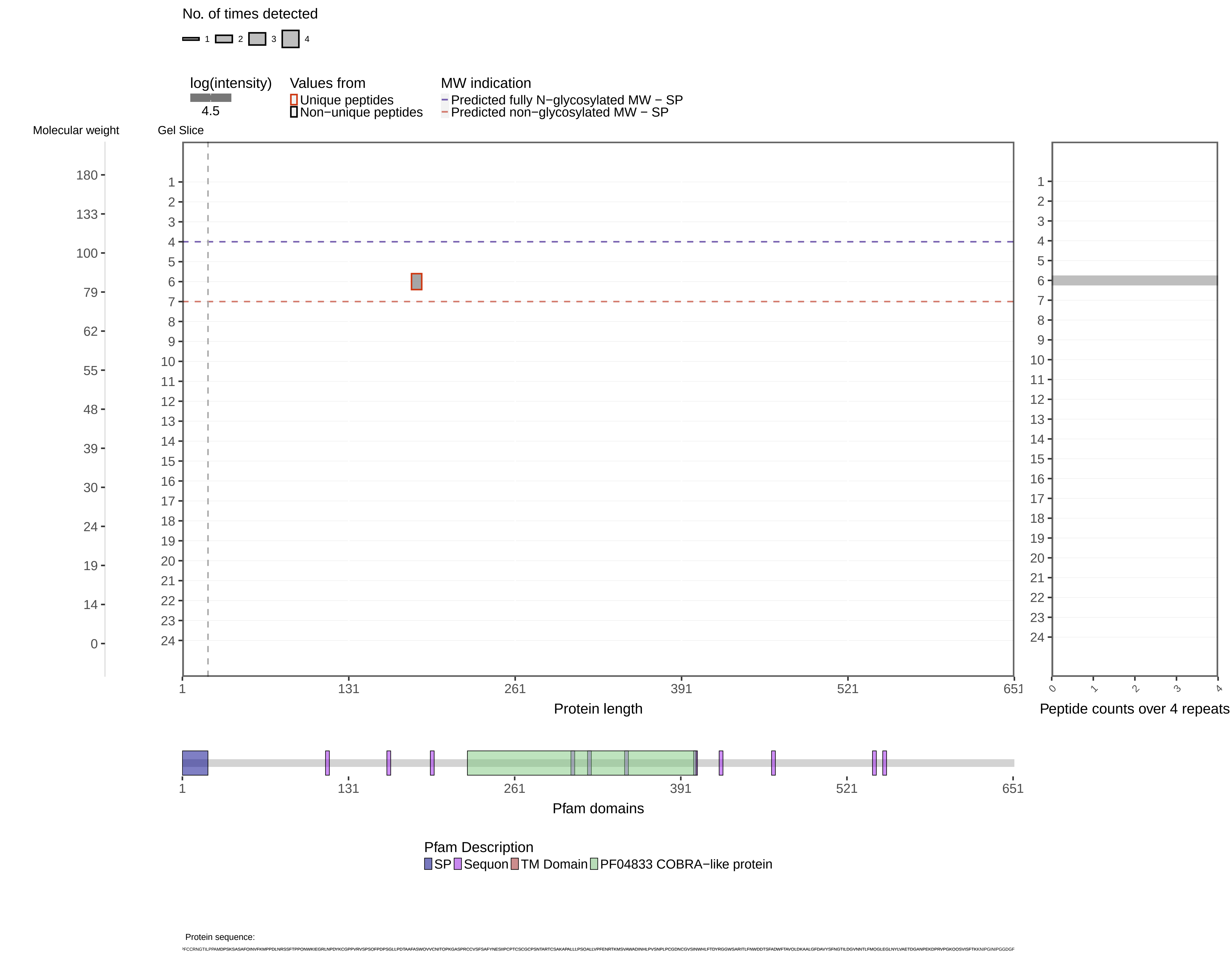

NbL16g16320.1 importin_subunit_alpha−2−like [MW = 58.5 kDa]

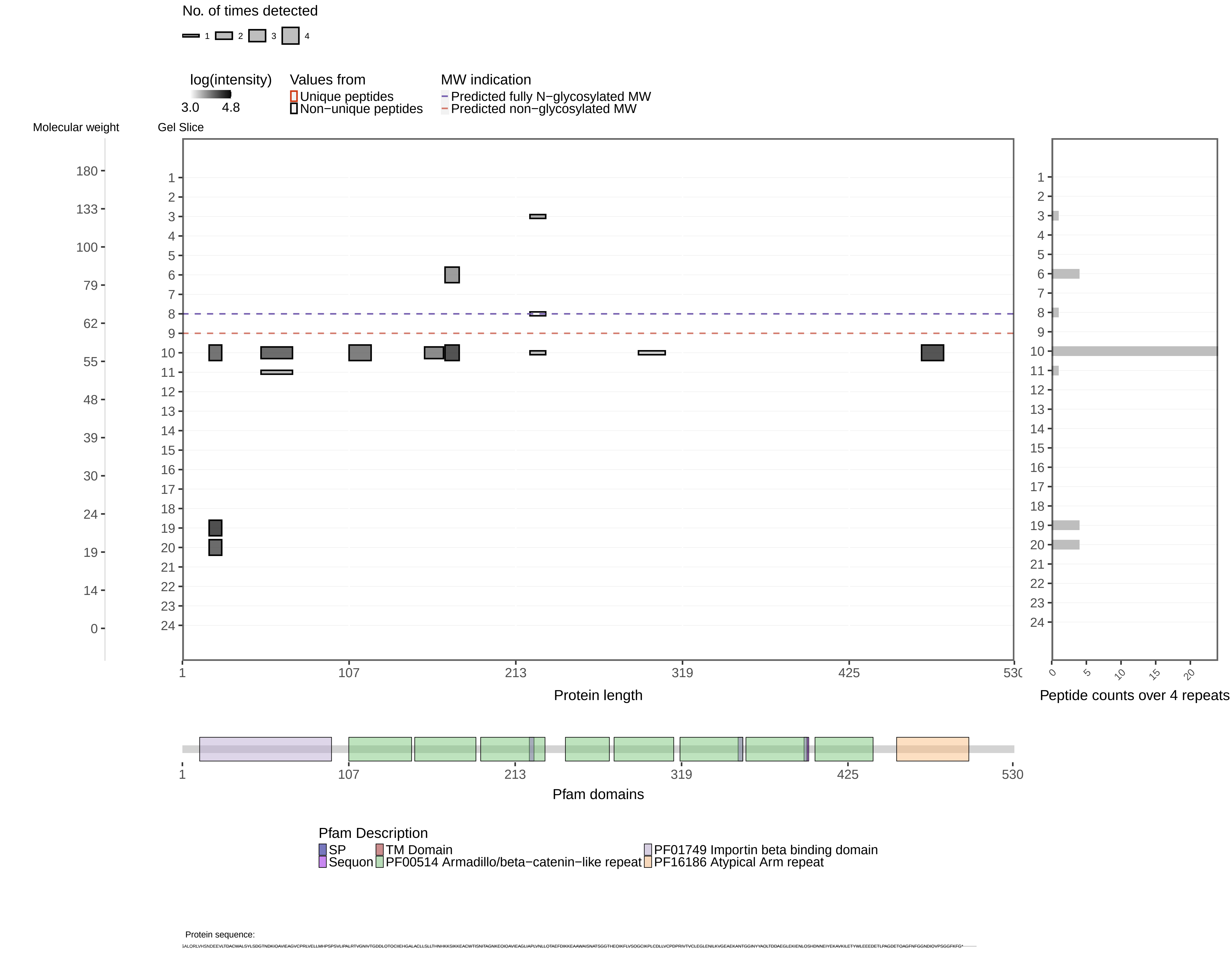

### NbL16g16250.1 acetate/butyrate−−CoA_ligase_AAE7_peroxisomal−like [MW = 62.53 kDa]

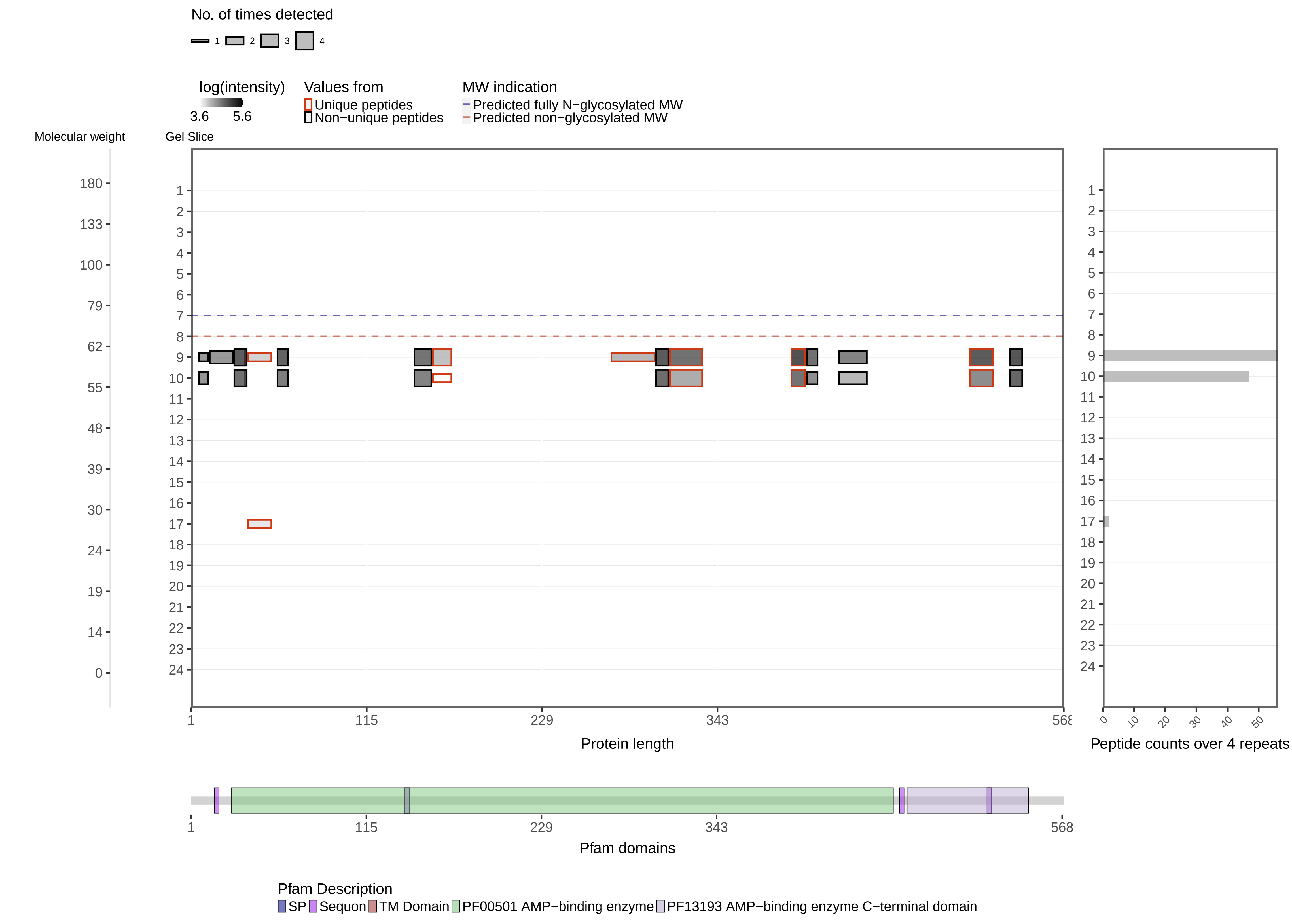


Protein sequence:

### NbL16g16230.1 GTP−binding_protein_SAR2 [MW = 21.94 kDa]

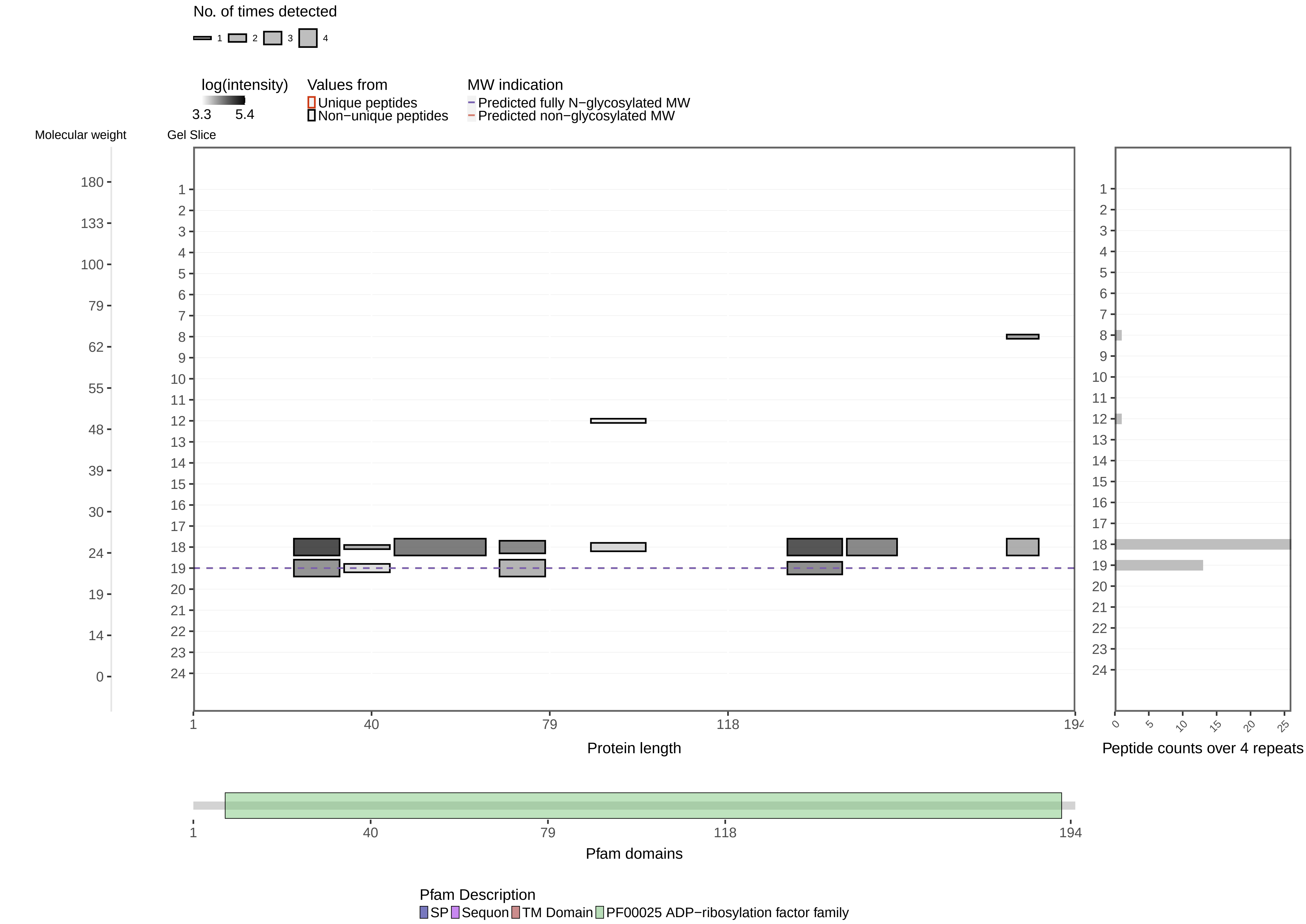


Protein sequence:

)YYAKVDAVVYLVDAYDRERFPESKKELDGLLSDESLANVPFLILGNKIDIPYAASEDELRYHLGLTGVTTGKGNVNLAGTNVRPIEVFMCSIVRKMGYGEGFKWMSQYIK*

NbL16g16190.1 uncharacterized_protein_C167.05−like [MW = 17.84 kDa]

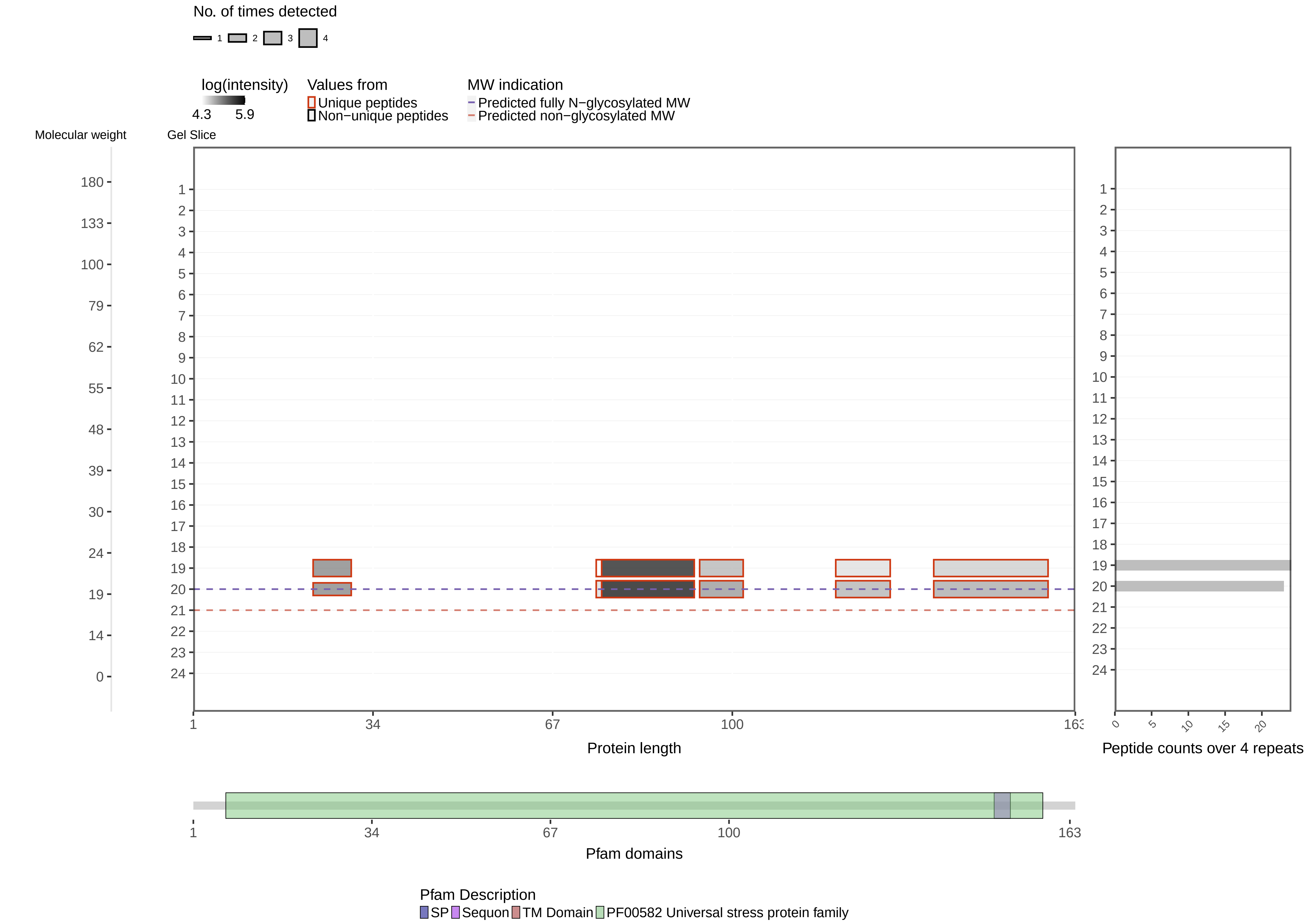


Protein sequence:

### NbL16g16150.1 phytochrome_B−like [MW = 125.82 kDa]

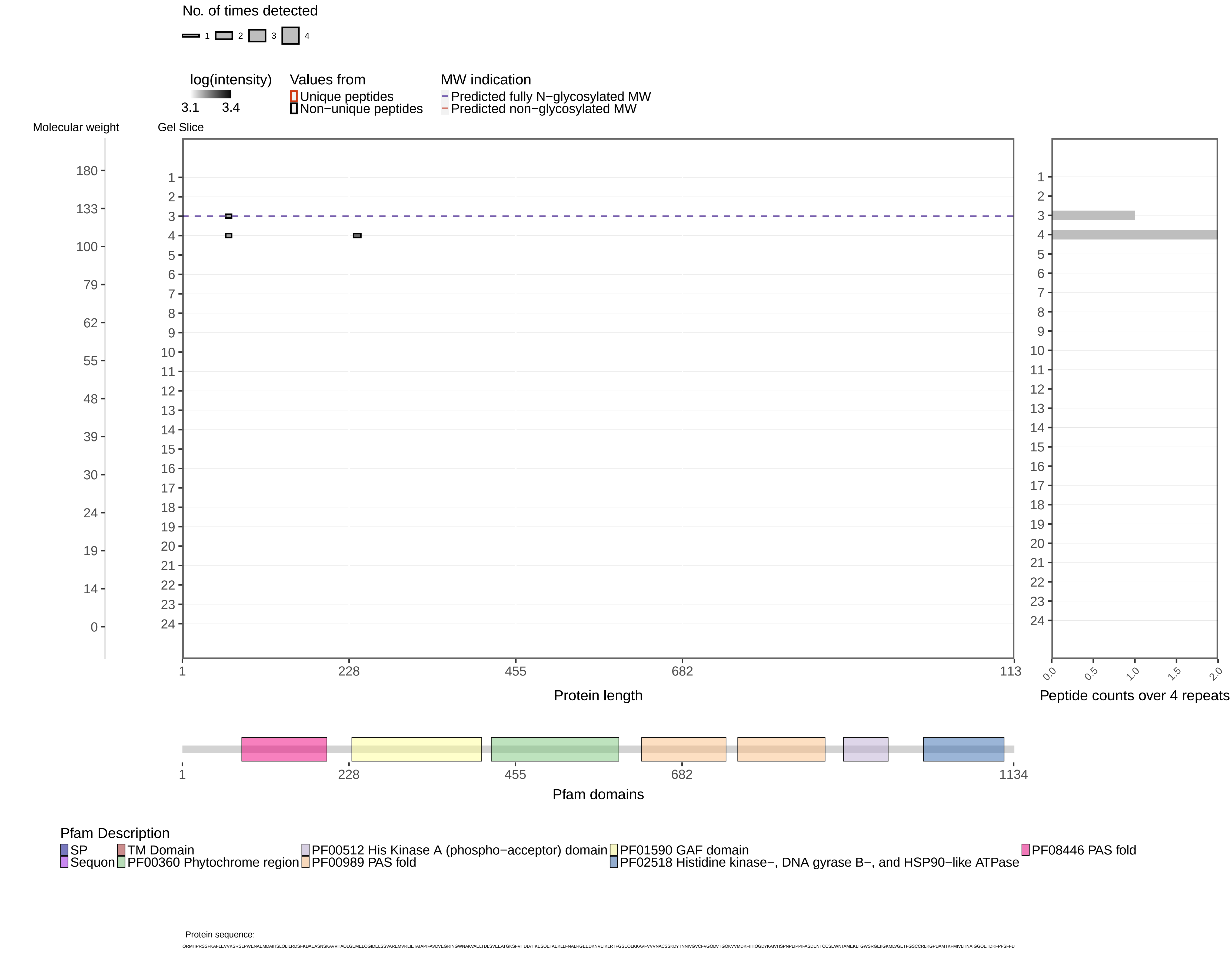

NbL16g16030.1 galactokinase−like [MW = 54.53 kDa]

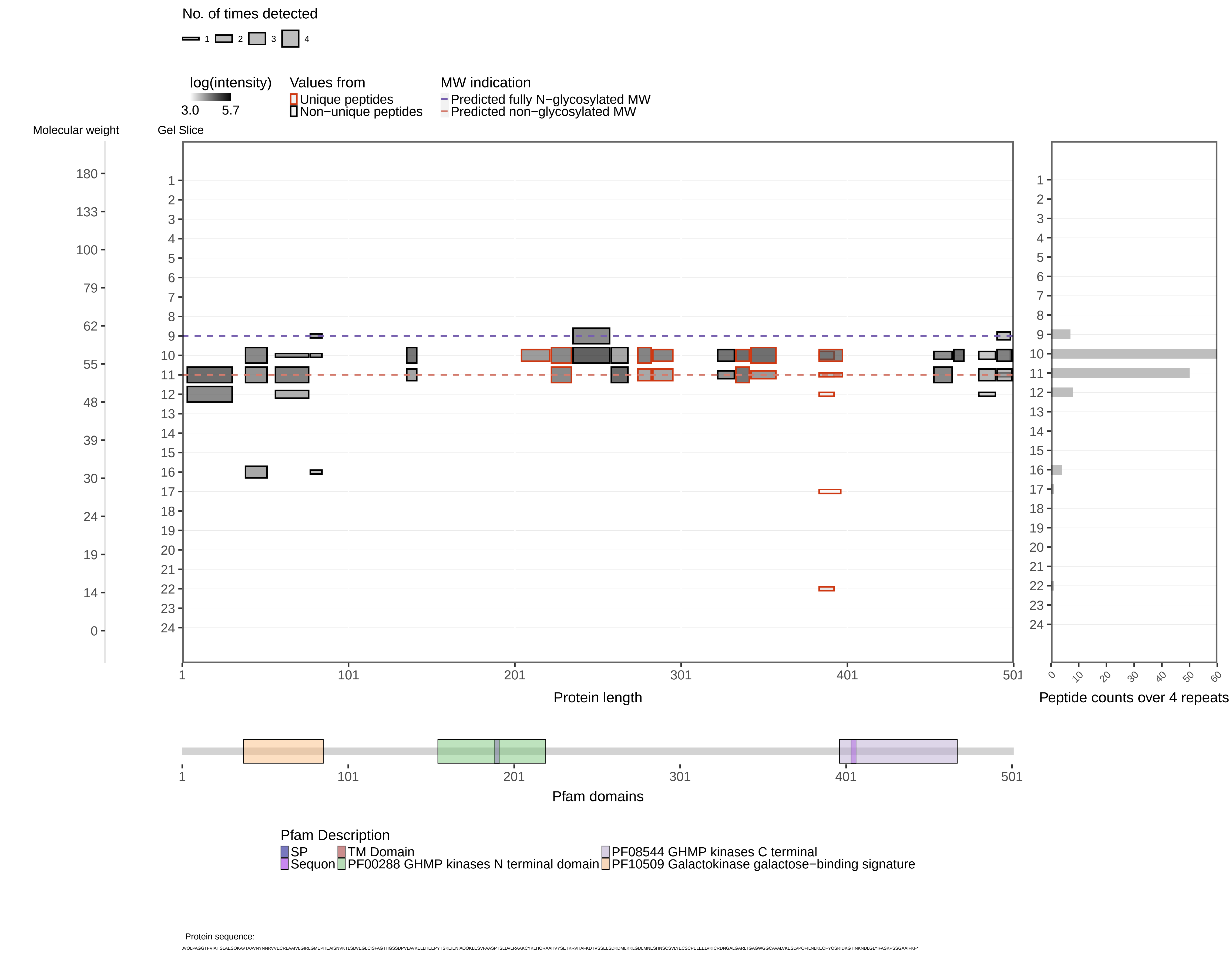

NbL16g15970.1 30S_ribosomal_protein_S1_chloroplastic−like [MW = 45.59 kDa]

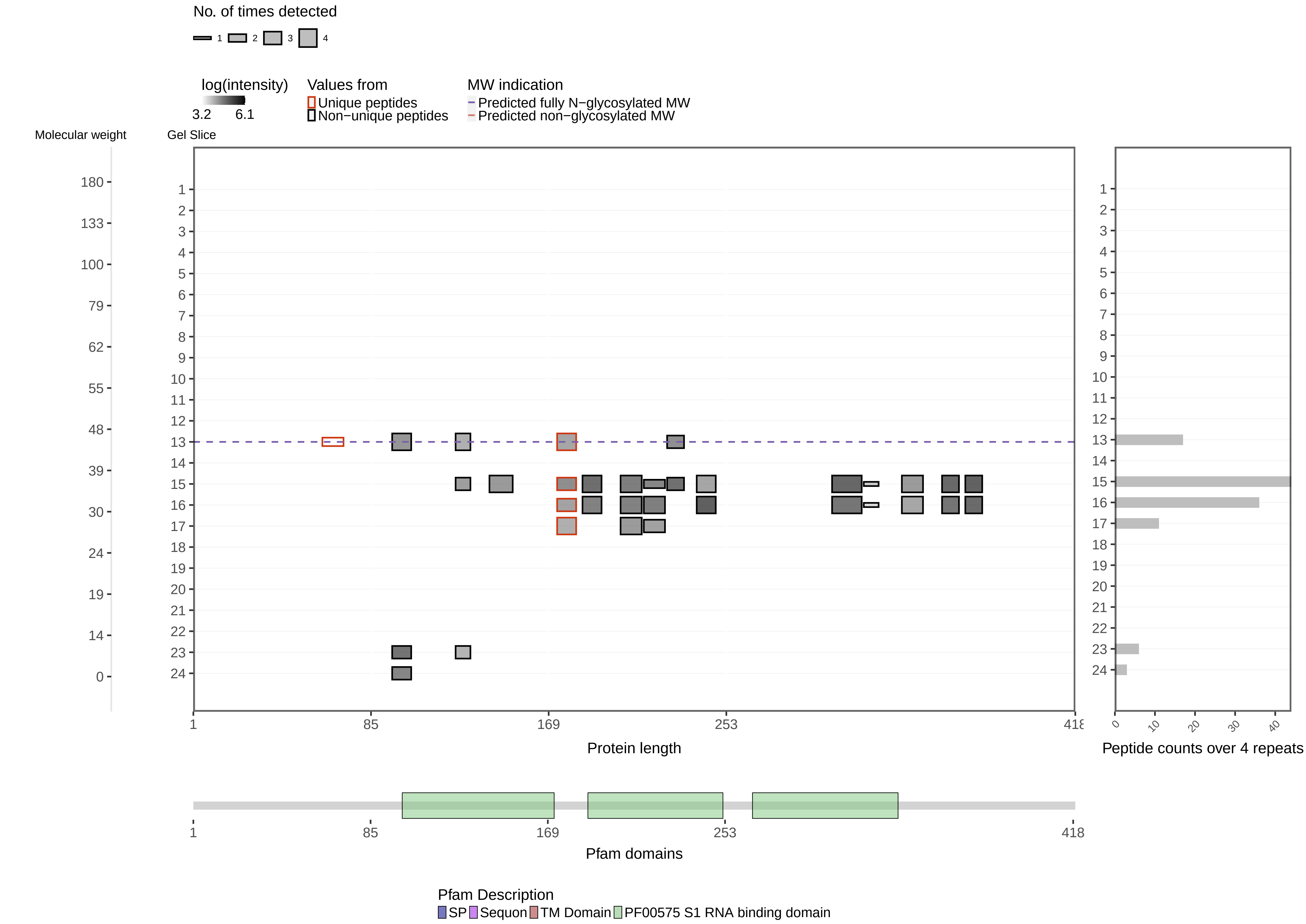


Protein sequence:

KVVGANKGGVVALVEGLRGFVPFSQISTKSTAEELLEKELPLKFVEVDEEQSRLVLSNRKAMADSQAQLGIGSVVLGTVQSLKPYGAFIDIGGINGLLHVSQISHDRVSDIATVLQPGDTLKVMILSHDRERGRVSLSTKKLEPTPGDMIRNPKLVFEKAEEMAQTFRQRIAQAEAMARADMLRFQPESGLTLNSDGILGPLTSELPEDGLDLSEIPPAEDL*

### NbL16g15950.1 DEAD−box_ATP−dependent_RNA_helicase_40−like [MW = 125.02 kDa]

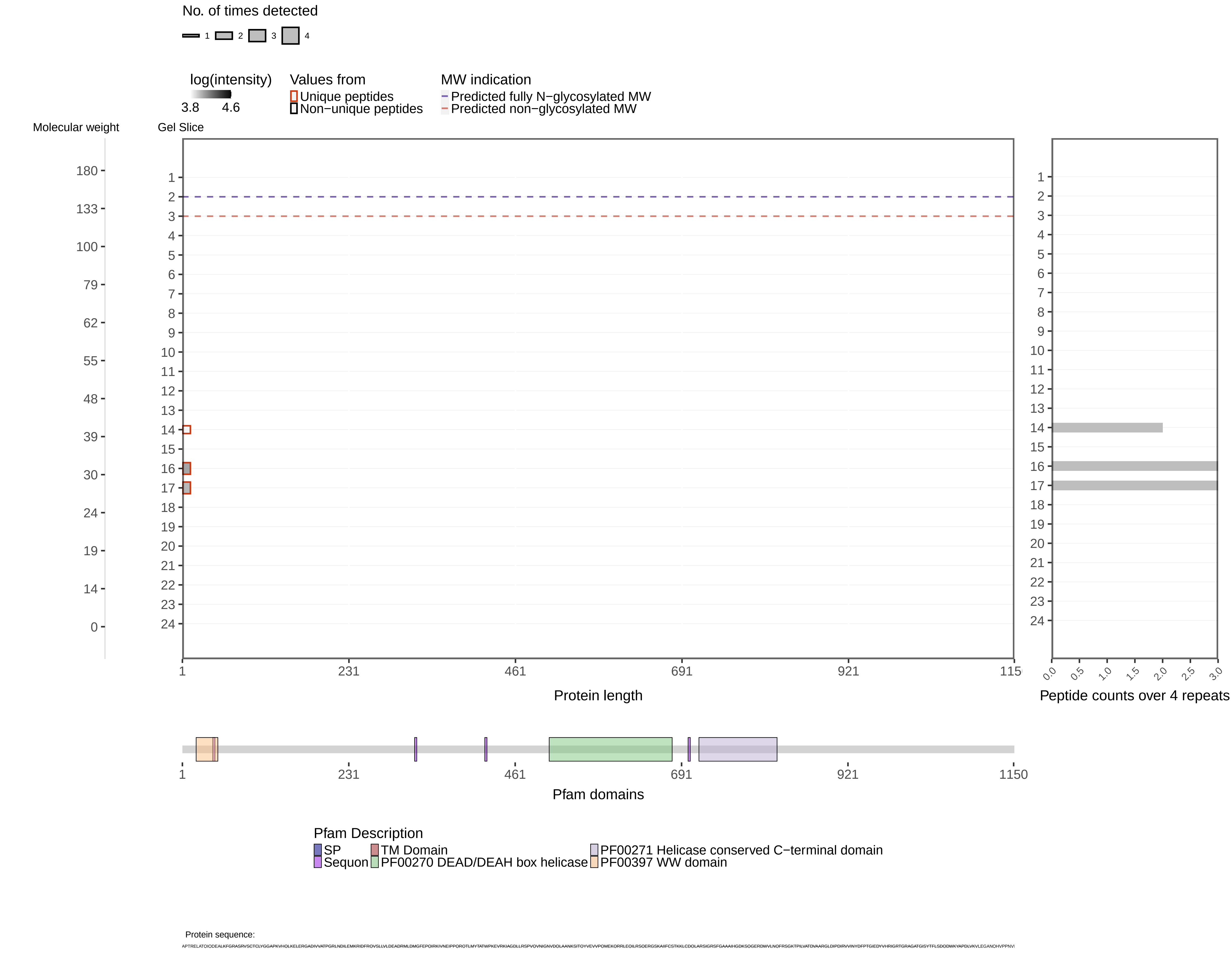

### NbL16g15820.1 pyruvate_dehydrogenase_(acetyl−transferring)_kinase_mitochondrial−like [MW = 42.26 kDa]

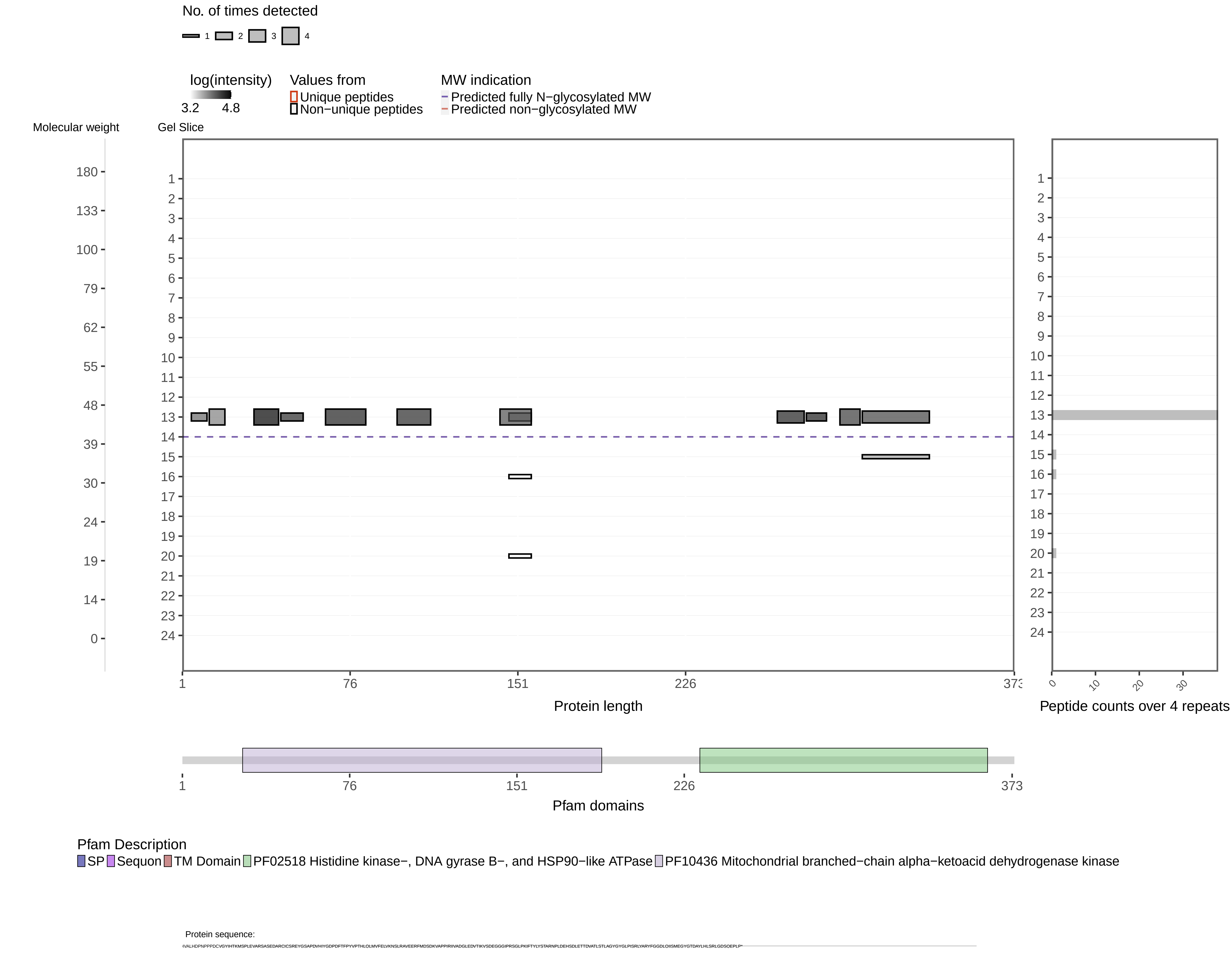

NbL16g15720.1 photosynthetic_NDH_subunit_of_lumenal_location_1_chloroplastic [MW = 25.81 kDa]

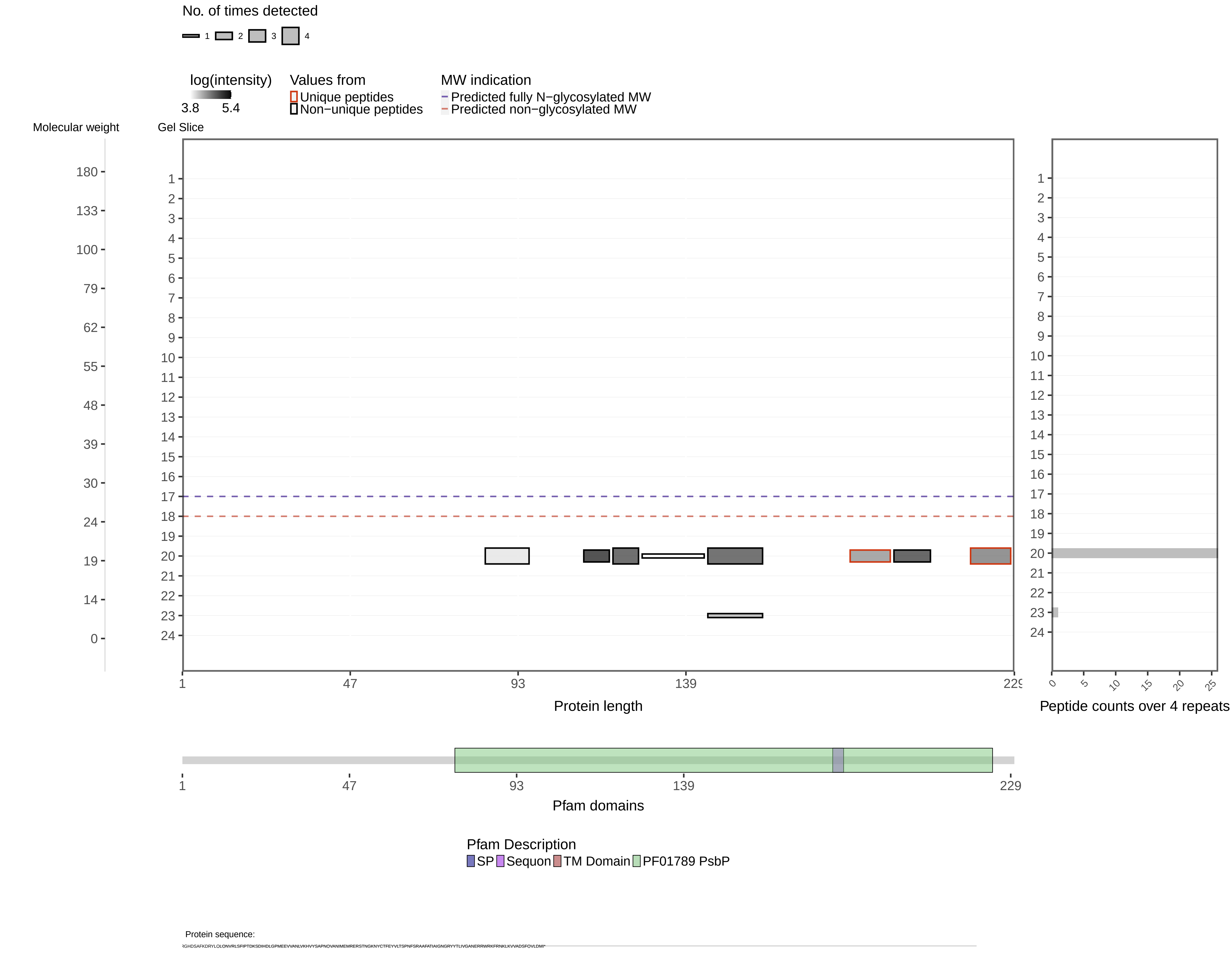

NbL16g15670.1 dihydrolipoyllysine−residue_acetyltransferase_component_4_of_pyruvate_dehydrogenase_complex_chloroplastic−like [MW = 48.99 kDa]

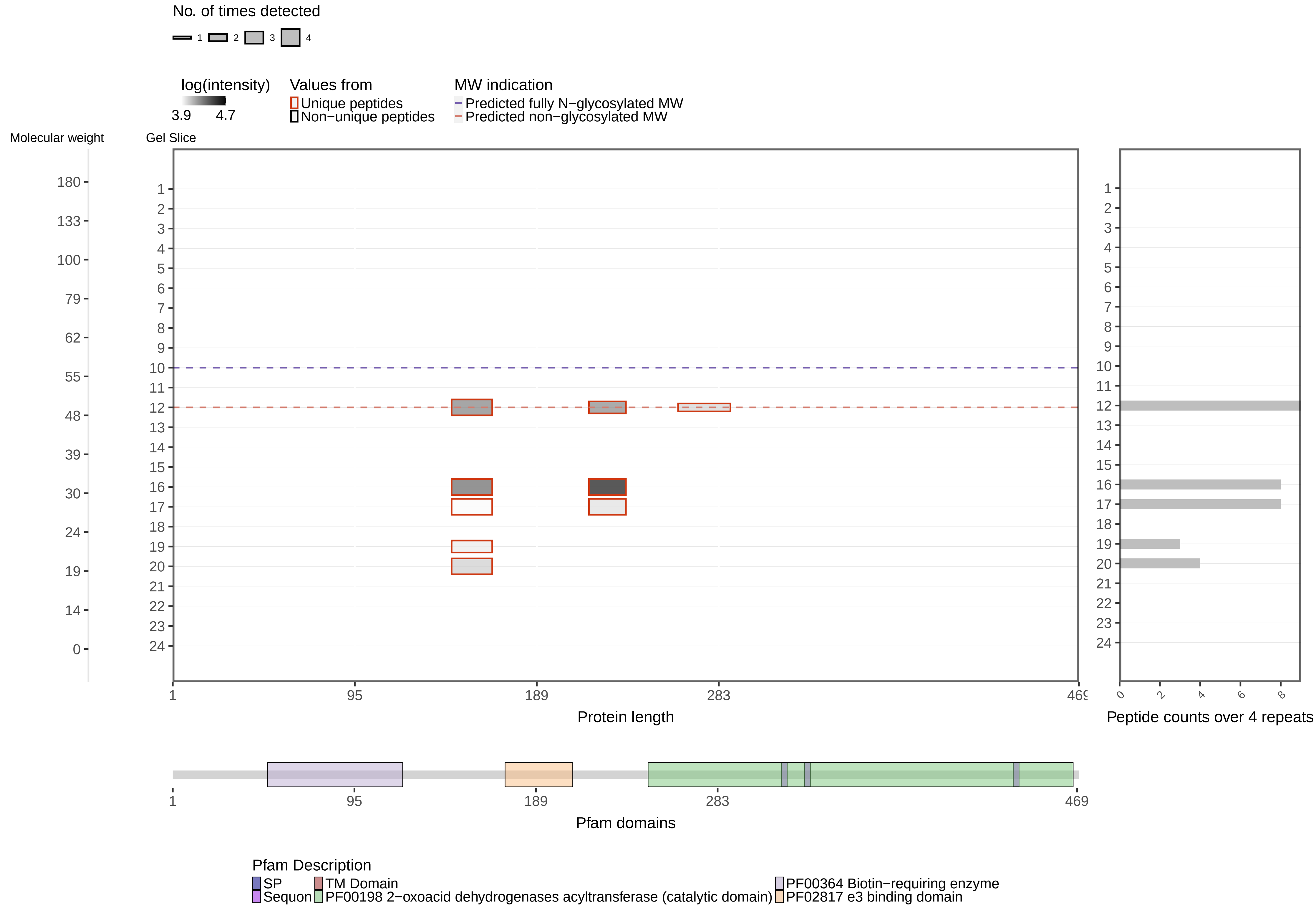


Protein sequence:

/AAPIPSAAAPAKASPTSFPDIPGSTVVPFTTMOIAVSKNMMESLSVPTFHVGYPVSTNSLDALYEKVKPKGVTMTALLAKAAAMALAOHPVVNATCKDGKSFTYNSSINIAVAVAINGGLITPVLODADKLDLYLLSOKWKELVDKARAKOLOPHEYNSGTFTLSNLGMFGVDRFDAILPPGOGAIMAVGASKPTVVSDSEGFFSVKNKMLVNVTADHRIIYGADLAAFLOTFSKIVENPESLTM*

NbL16g15340.1 anthranilate_phosphoribosyltransferase_chloroplastic−like [MW = 38.74 kDa]

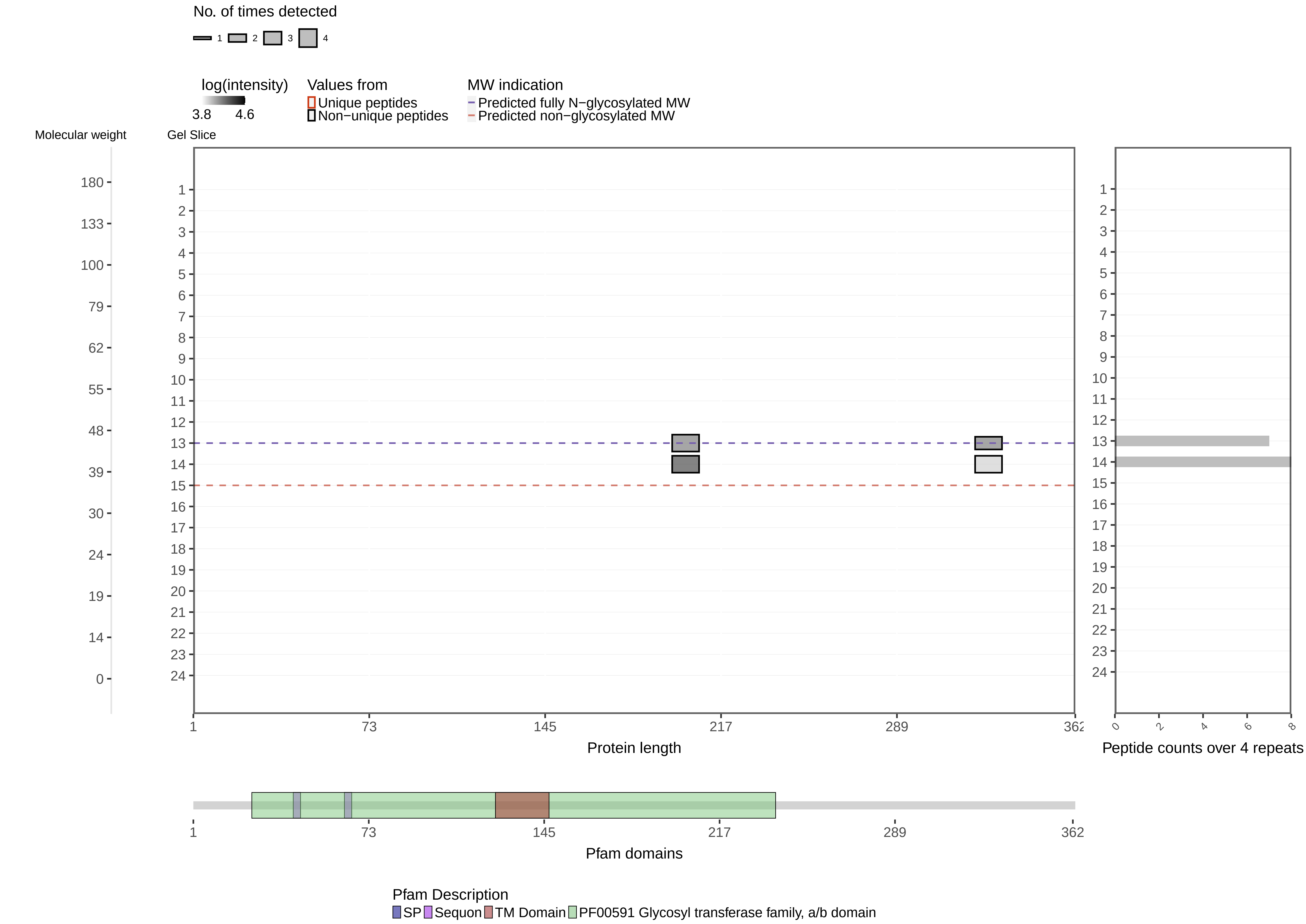


Protein sequence:

.FVHSEGLDEMSPLGPGVVLDVTAEKIEKFSFDPLDFGIPRCTLGDLRGGGPEHNAGVLRRVLSGETGPMADALDIATLVKTYDICHHSLSAGRWHKNMLLGLCLLHSFGCILASKDLVONILYSHSDVALHHYKHMPLMILNAAAALLVSRRVGNLAEGVAVARTTHOSGKALDTLNHWMMFOVHLMLLEYRG*

NbL16g15300.1 universal_stress_protein_A−like_protein [MW = 21.16 kDa]

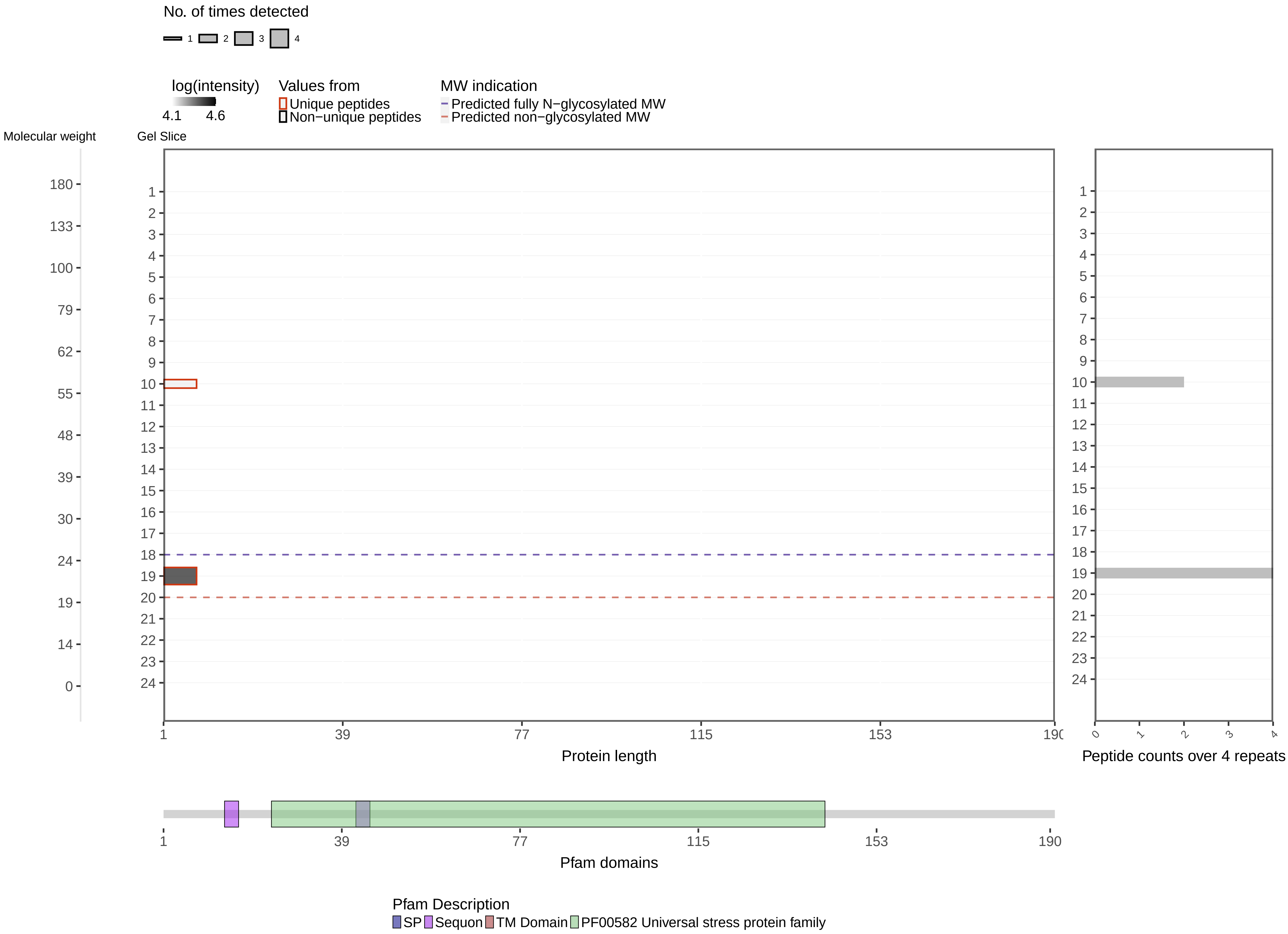


Protein sequence:

RDKIRGLHLLECFVNRCHEIGVPCEAWIKQGDPKEVVCLEVKRVQPDLLVVGCRGLGPFQRIAKRLPFPGRSLEFGSIGVSTGLQLVISVSSRISKAYFRCEITIPELD*

NbL16g15290.1 50S_ribosomal_protein_L12_chloroplastic−like [MW = 19.68 kDa]

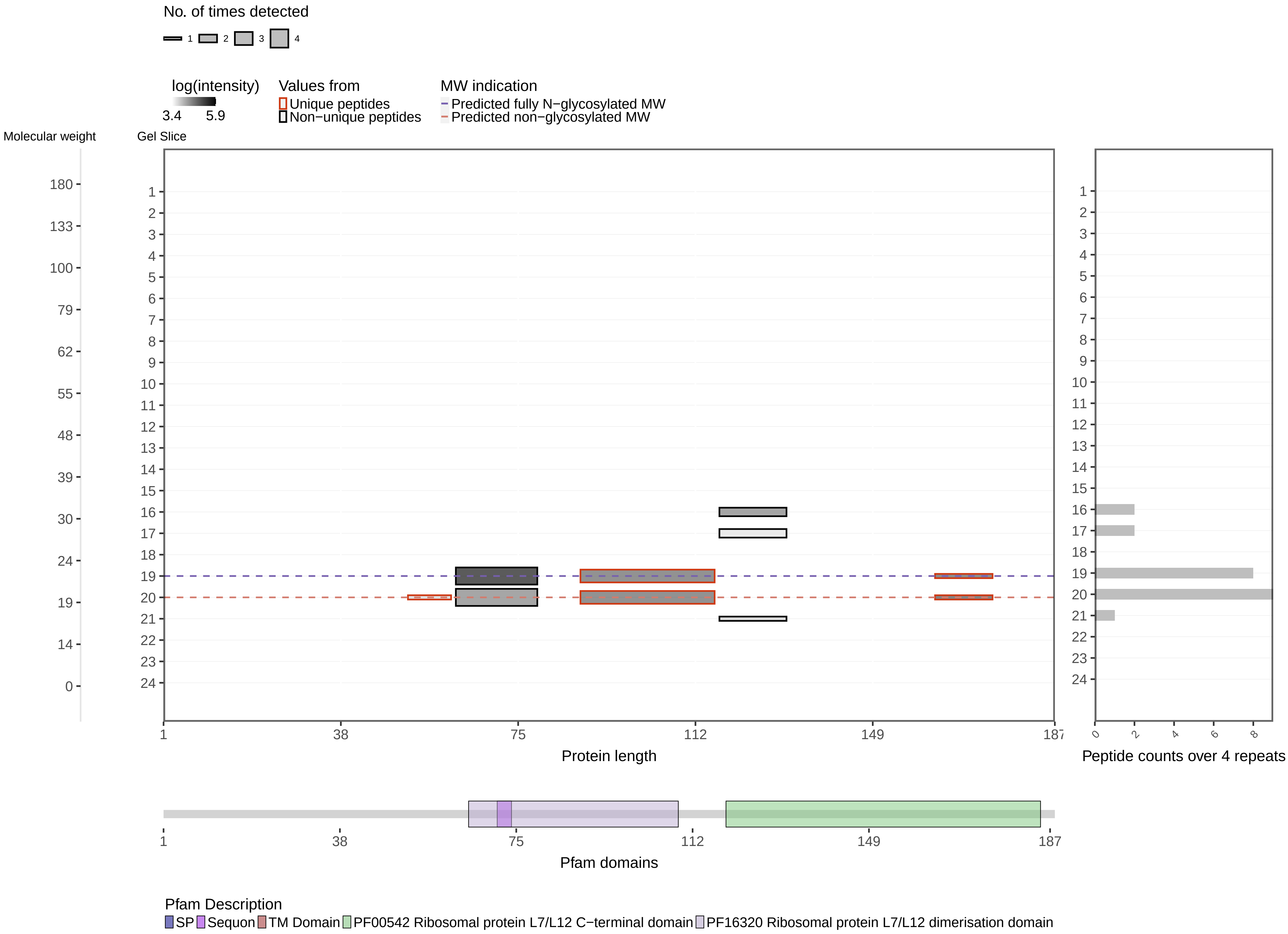


Protein sequence:

.VEYLQDKLGVTAASFAPAAVSAVPAAAEAPAVVEEKTEFDVIIEEVPSNARIAAIKAVRALTNLALKEAKELIEGLPKKFKEGISKDEAEDAKKQLEEAGAKVSIA*

NbL16g15050.1 3−dehydroquinate_synthase_homolog [MW = 43.91 kDa]

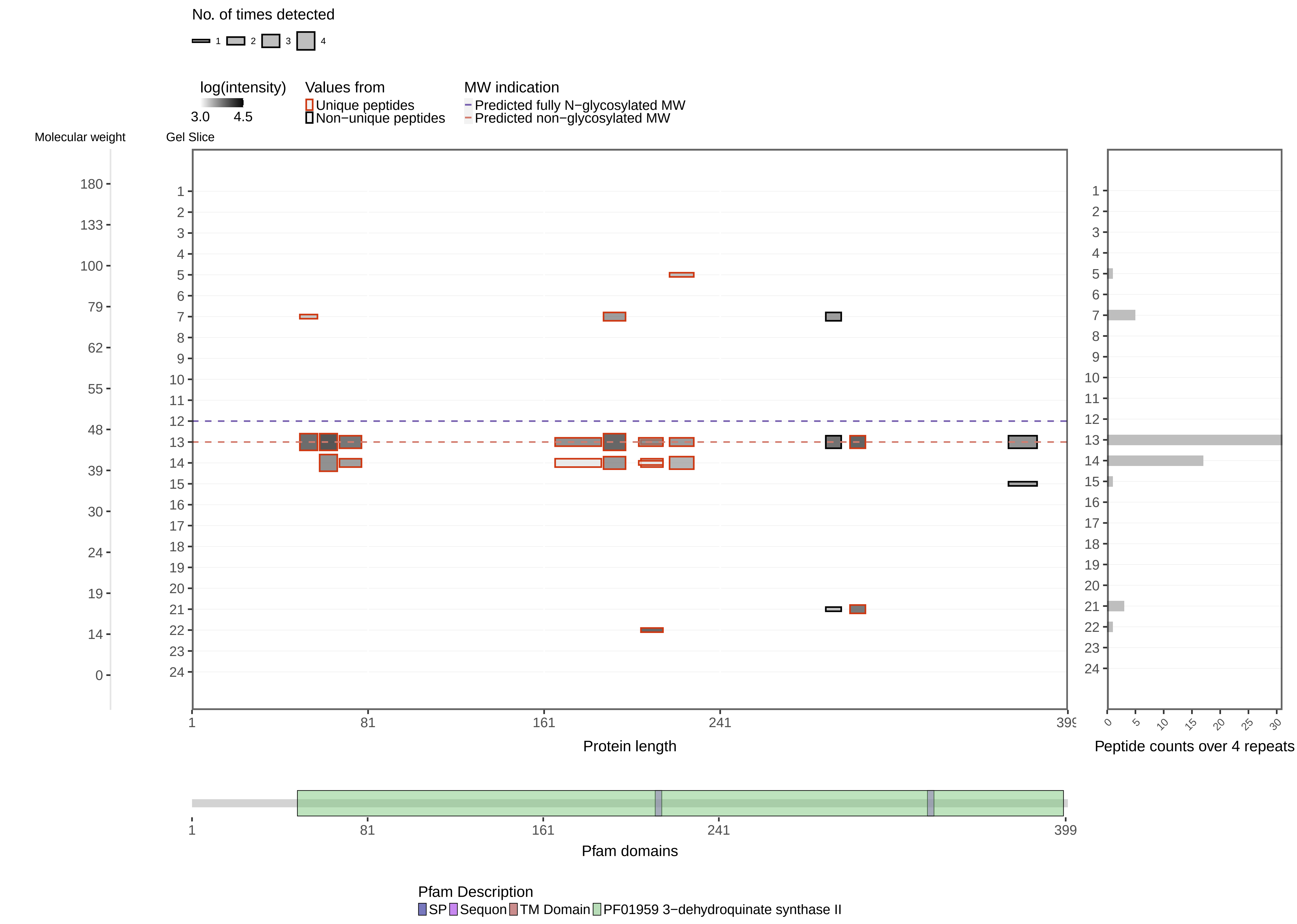


Protein sequence:

/MKVEDVGAILDLKGYFDRRNEEDSLLNLTKAKITHVQSTGMGDRVCVDICSLMRPGEGLLVGSFARGLFLVHSECLESNYISSRPFRVNAGPVHAYVAVPGGKTSYLSELKSGKEIIVVDQRGMQRTAIVGRVKIETRPLILVEAEVESKNESYSILLQNAETVGLVSPLQEEGYQRTTIPVTSLKVGDEVCLLVQGGARHTGIEIKEFIVEK*

### NbL16g14910.1 alpha−14_glucan_phosphorylase_L−1_isozyme_chloroplastic/amyloplastic [MW = 110.39 kDa]

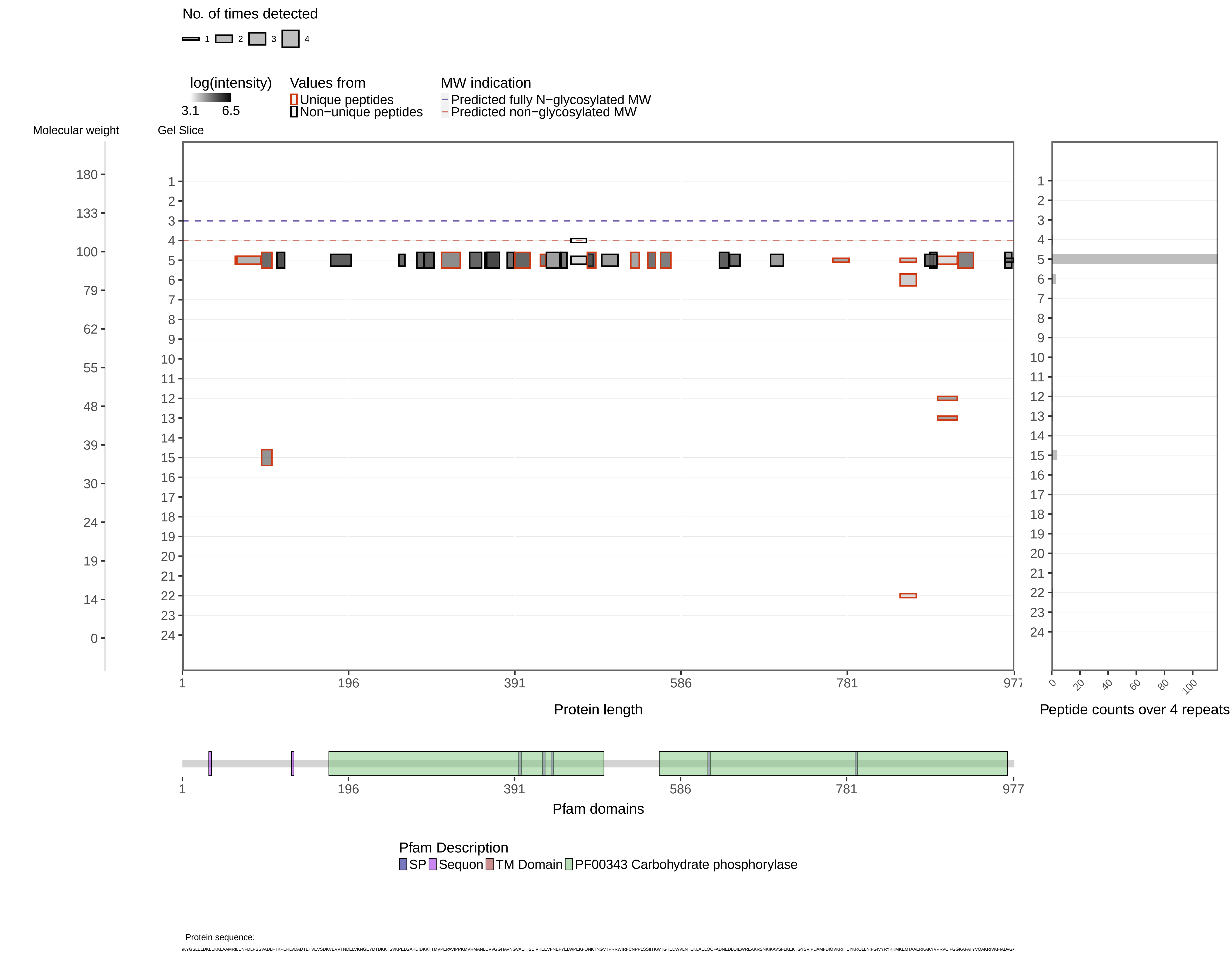

NbL16g14470.1 nicotinate_phosphoribosyltransferase−like [MW = 62.82 kDa]

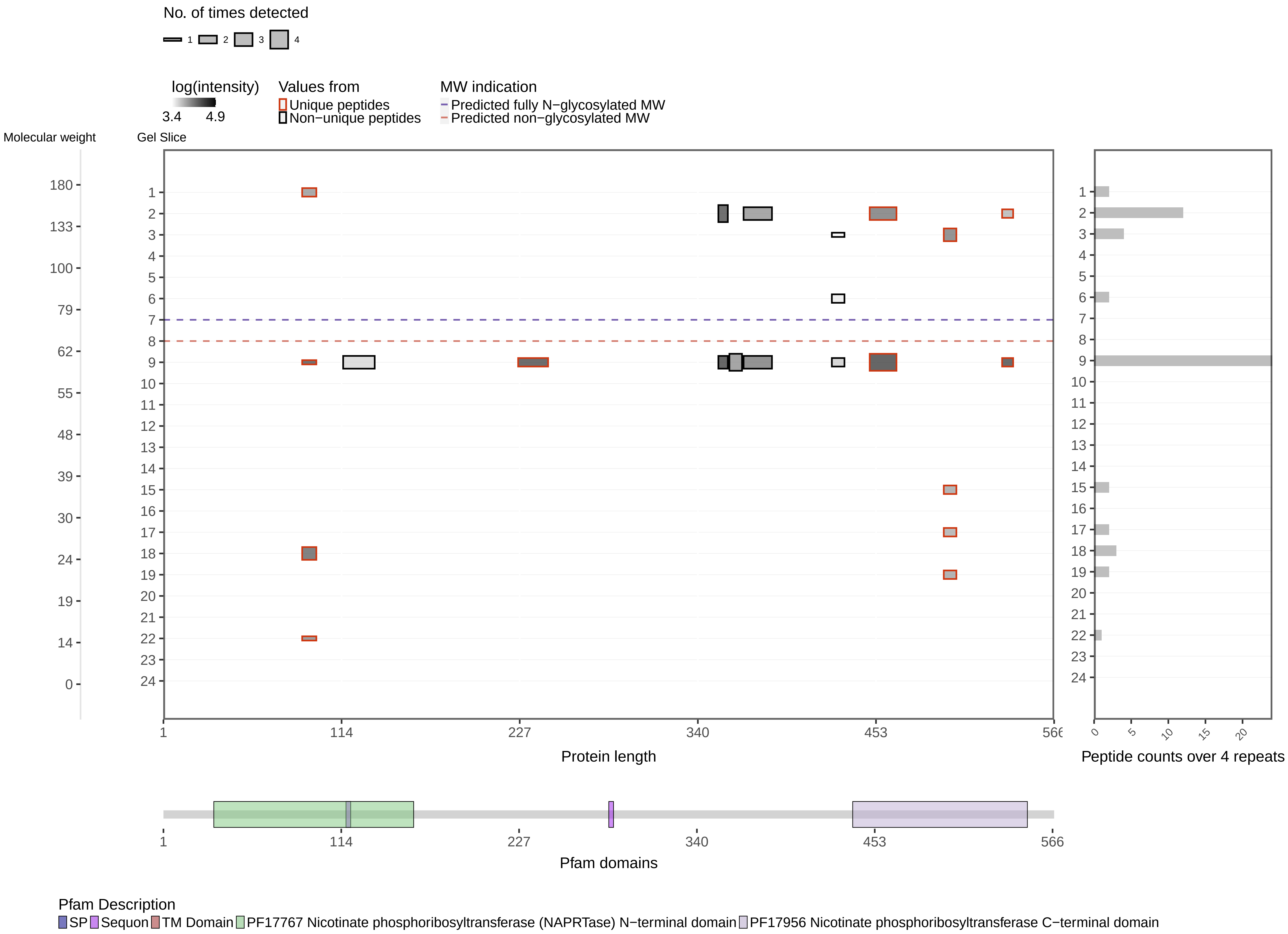


Protein sequence:

GSHILGGIFSETNQSELAAFTSYGLAFAGSFLALVDTYDVMRSGVPNFCAVALALNDLGYRAVGIRLDSGDLAYQSCEARKFFQTIEKEFGVSGFGKMSITASNDLNEETLDALNKQGHEVDSFGIGTHLVTCYAQPALGVVFKLVEINNQPRIKLSEDVTKVSIPCKKRCFRLYGKEGFPLVDIMAGENEPPPKVGERILCRHPFSESKRAYVVPQRVEELLKCFWPGSSDKKREDLPPLKEIRERCIKQLEQMRPDHMRKLNPTPYKVSVSAKLYDFIHFLWLNEAPVGELO*

NbL16g14460.1 probable_phosphoribosylformylglycinamidine_synthase_chloroplastic/mitochondrial [MW = 154.18 kDa]

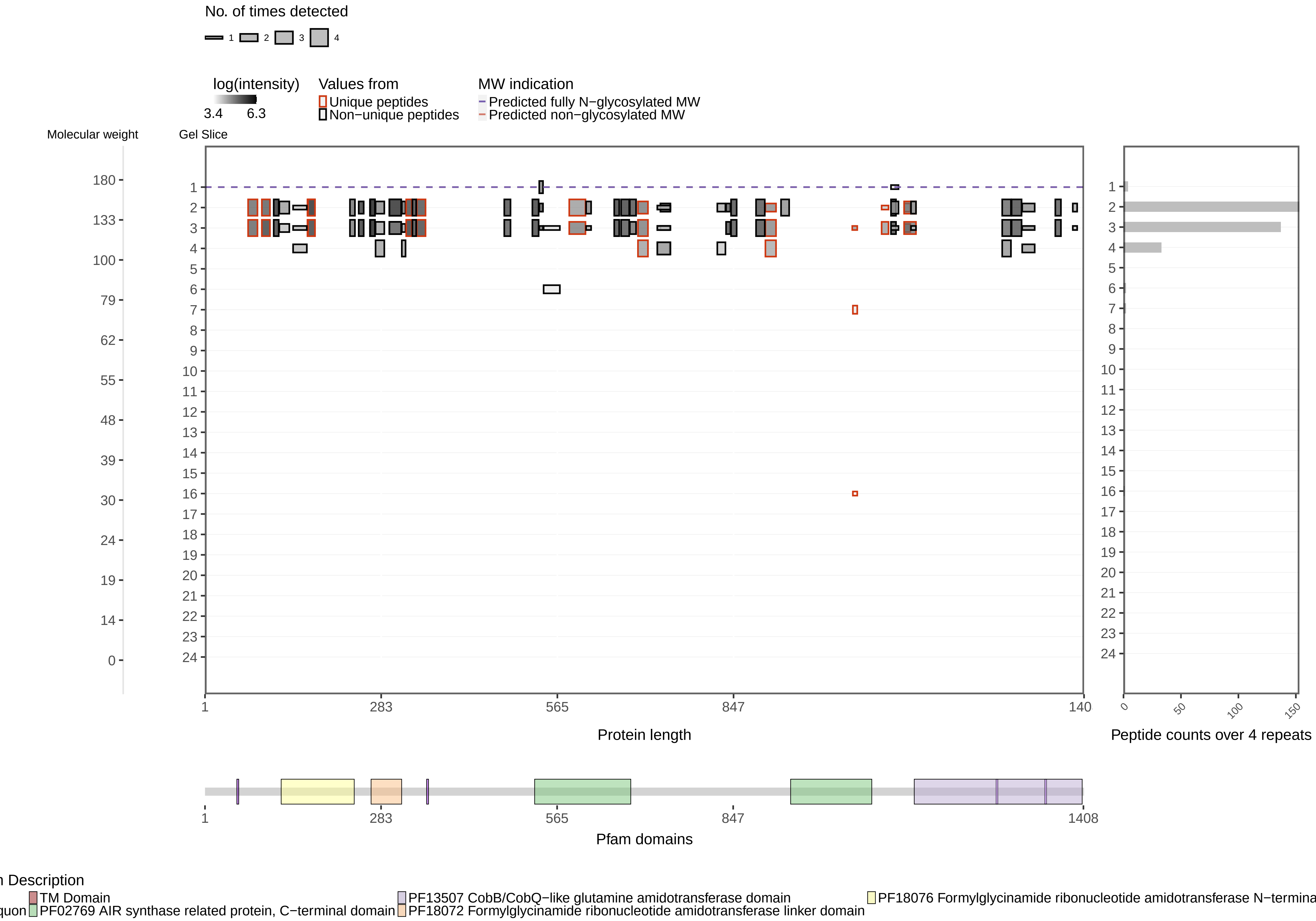


Protein sequence:

NbL16g14390.1 putative_oxidoreductase_TDA3 [MW = 44.35 kDa]

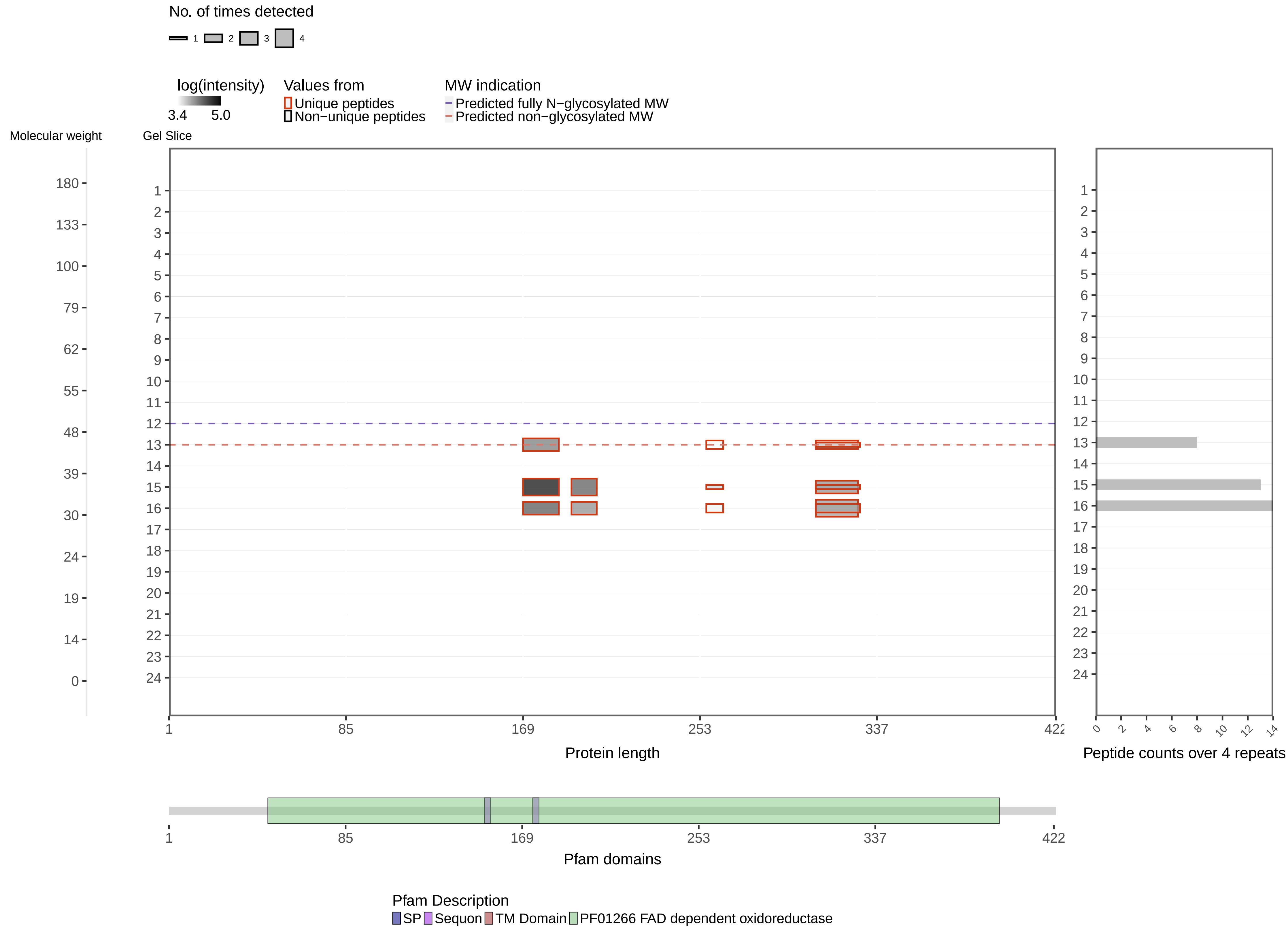


Protein sequence:

IGVEVVIGKAERVEMAEGKTKGIVLEDGREIVADAVVLALGPWTKRFSILGSLFGVYGIKAHSIVLEPKEPNRITPHALFLTYYPAHGGKPMDPEVYPRPTGEVYICGMSAREEVPDDPGQVIPVPESINALKRVAASVSSHLVEGEAAVTAEQACILPCSEDDVPIIGEVPGIKGCYVATGHSCWGILNGPATGAAMAELILDGQANIVDLSQFTPARFASSLKK*

NbL16g14350.1 subtilisin−like_protease_SBT1.7 [MW − SP = 78.77 kDa]

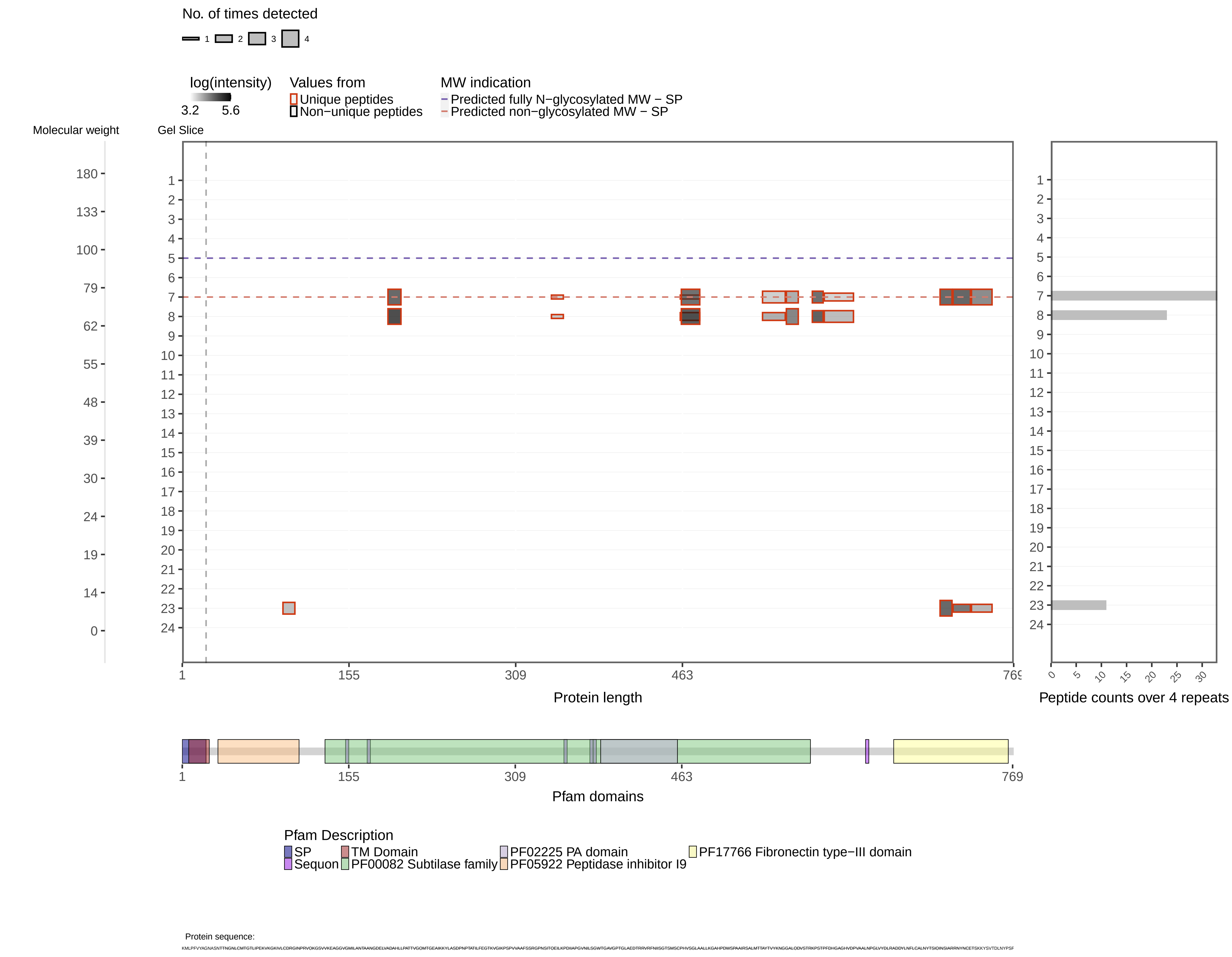

### NbL16g14340.1 casein_kinase_II_subunit_alpha−2 [MW = 39.35 kDa]

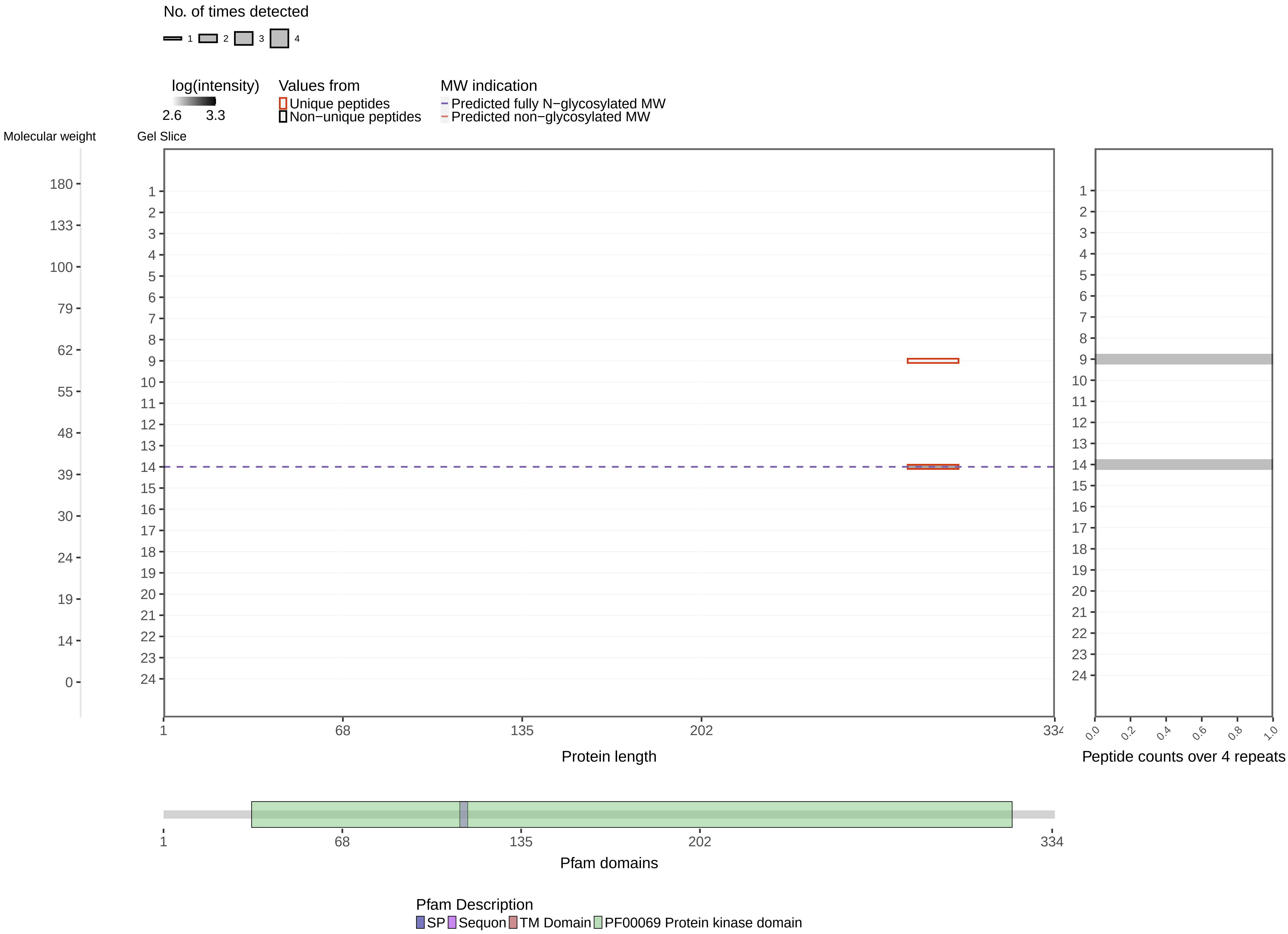


Protein sequence:

VMIDHELRKLRLIDWGLAEFYHPGKEYNVRVASRYFKGPELLVDLQDYDYSLDMWSLGCMFAGMIFRKEPFFYGHDNQDQLVKIAKVLGTDELNAYLHKYQLELDPQLEAMVGRHSRKPWSKFINADNQHLVLSEAIDFLDKLLRYDHQDRLTAREAMAHPYFLQVRAAENSRMRTQ*

NbL16g13950.1 protease_Do−like_7 [MW = 57.84 kDa]

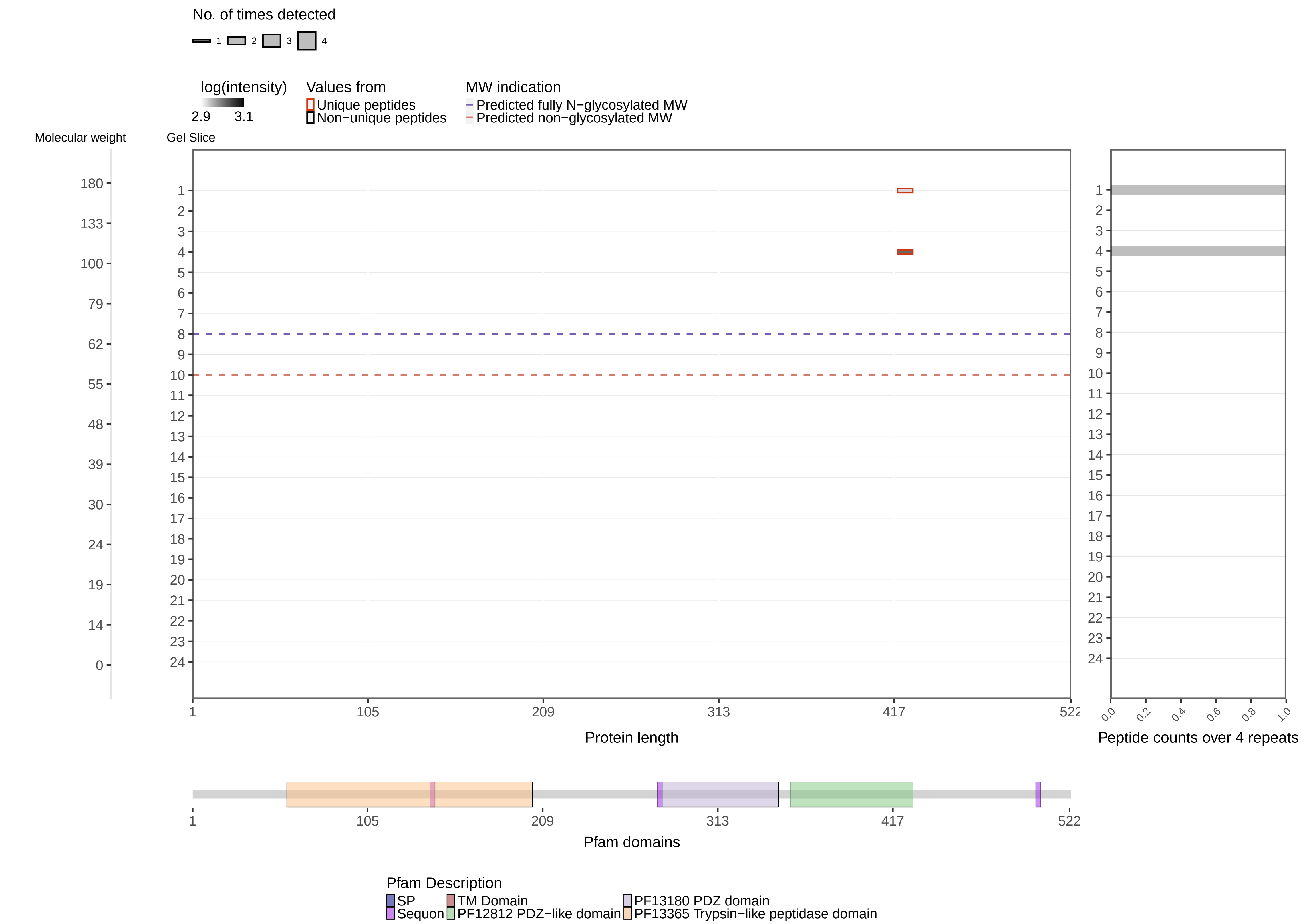


Protein sequence:

NbL16g13890.1 ribulose_bisphosphate_carboxylase_small_chain_chloroplastic [MW = 20.25 kDa]

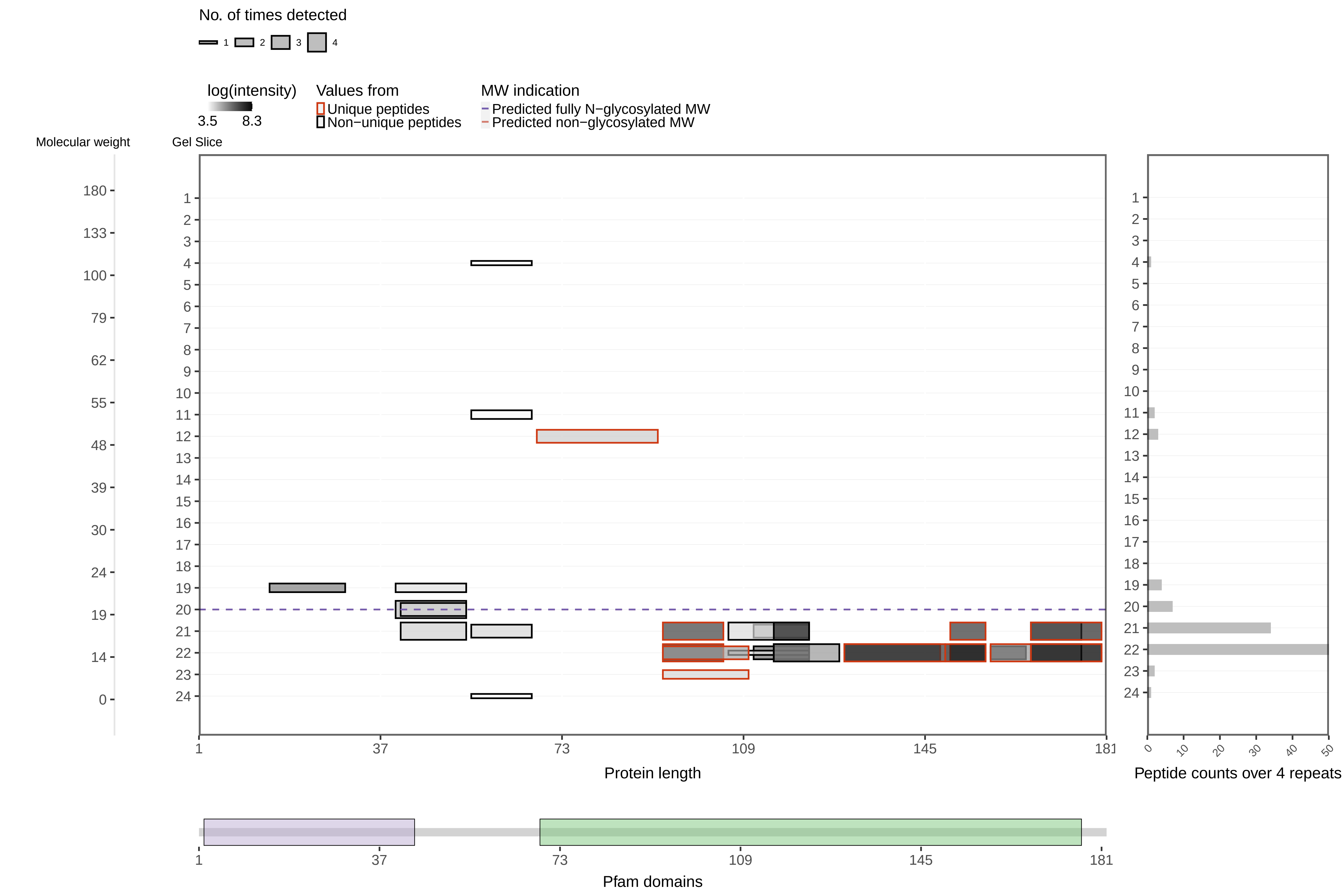


Pfam Description
SP Sequon TM Domain PF00101 Ribulose bisphosphate carboxylase, small chain PF12338 Ribulose−1,5−bisphosphate carboxylase small subunit

Protein sequence:
.SQEQLLSEIEYLLKNGWVPCLEFETEKGFVYREHHKSPGYYDGRYWTMWKLPMFGCTDATQVLAEVGEAKKEYPQAWVRIIGFDNVRQVQCISFIASKPDGY*

### NbL16g13840.1 14−3−3−like_protein_E [MW = 31.39 kDa]

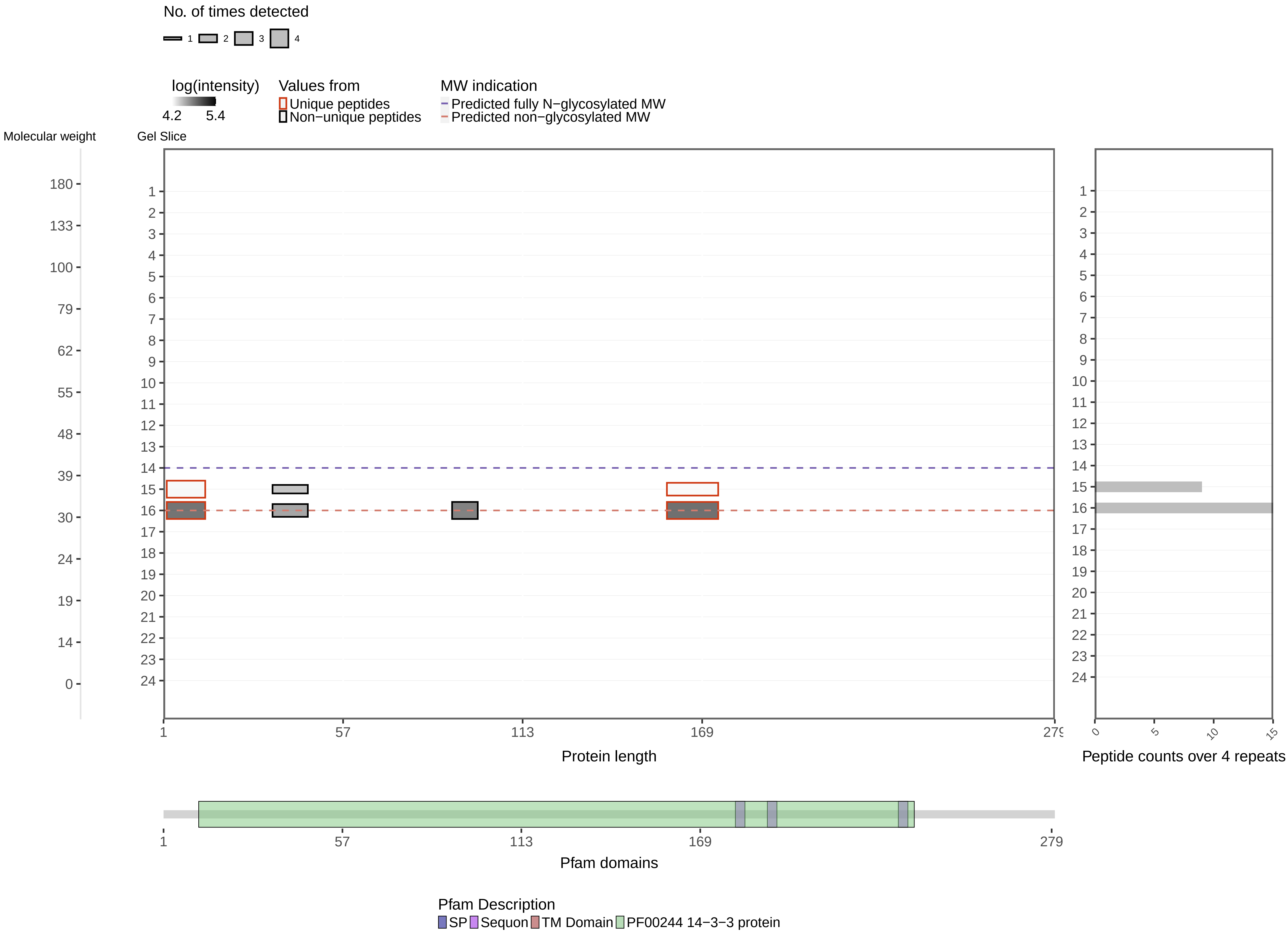


Protein sequence:

KMKGDYHRYLAEFKTGAERKEAAENTLLAYKSAQDIALAELAPTHPIRLGLALNFSVFYYEILNSSDRACNLAKQAFDDAIAELDTLGEESYKDSTLIMQLLRDNLTLWTSDTTVSLSFLPPSLPPSLALSHTEKYKPTNADAPTHKHCYAH*

### NbL16g13610.1 proteasome_subunit_beta_type−1−like [MW = 24.61 kDa]

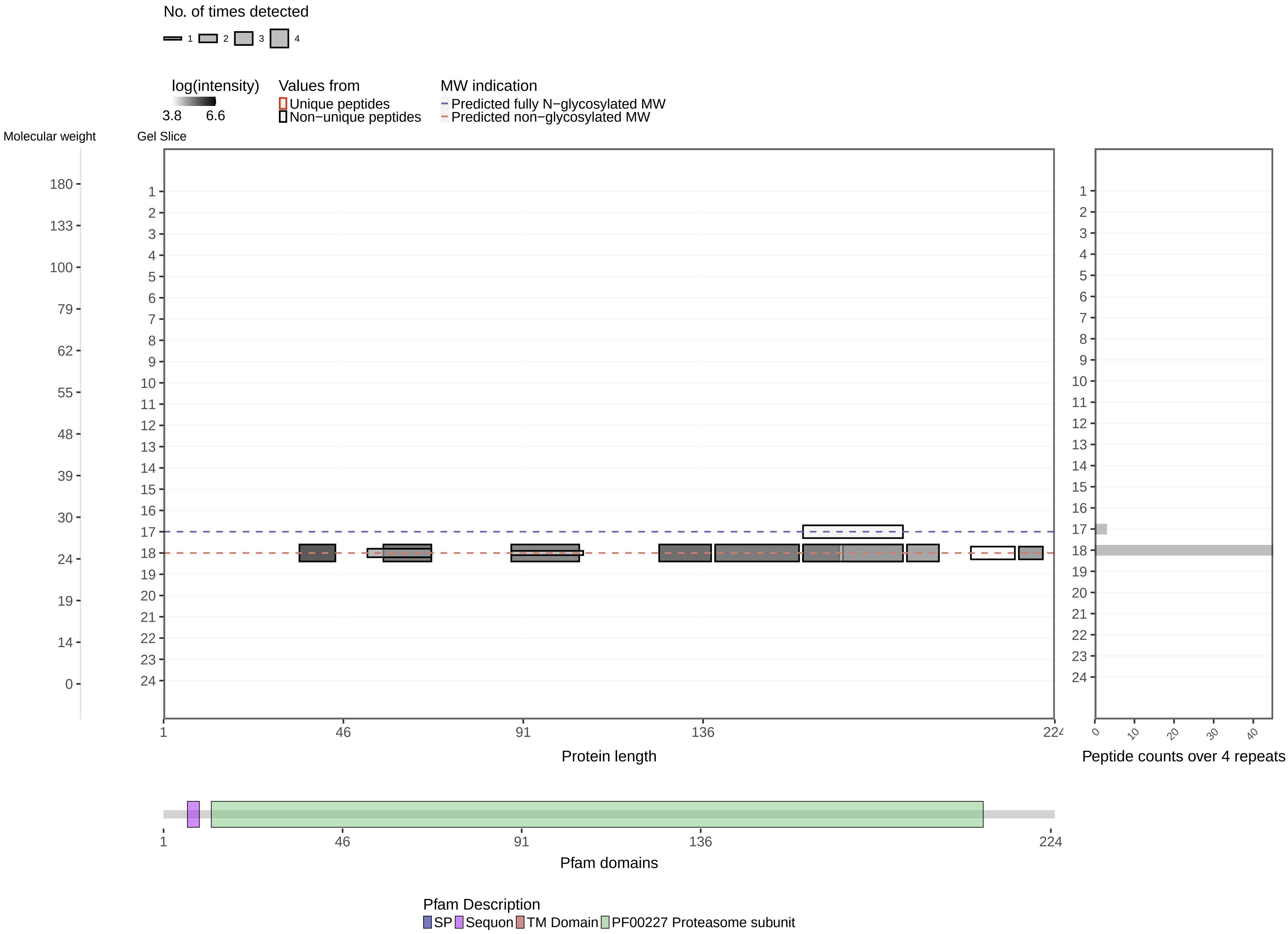


Protein sequence:

LSNTLYYKRFFPYYAFNVLGGLDNEGKGCVFTYDAVGSYERVGYSSQGSGSTLIMPFLDNQLKSPSPLLLPAKDAVTPLSESEAIDLVKTCFASATERDIYTGDKLEIVILNADGVRREEMDLRRD*

### NbL16g13590.1 inosine_triphosphate_pyrophosphatase [MW = 22.67 kDa]

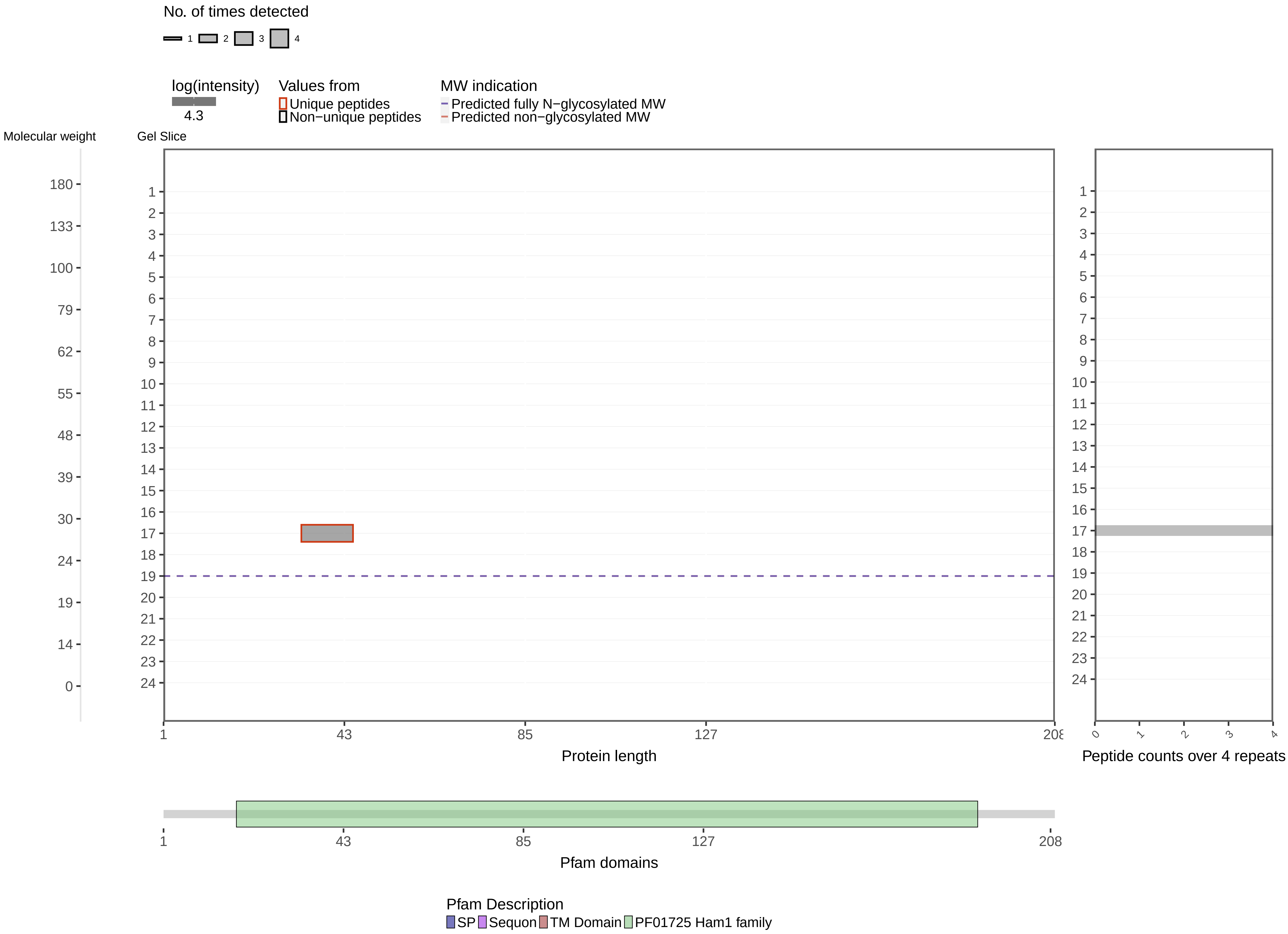


Protein sequence:

GPYIKWFLQKIGHEGLNNLLLAYEDKTAYAMCVFSLALGPNSEPLTFVGKTLGRIVPARGPNDFGWDPIFQPHGYDQTYAEMPKEGKNKISHRGKALELVKSHFAEARYTFQTDTTA*

NbL16g13290.1 chaperonin_CPN60−2_mitochondrial [MW = 61.38 kDa]

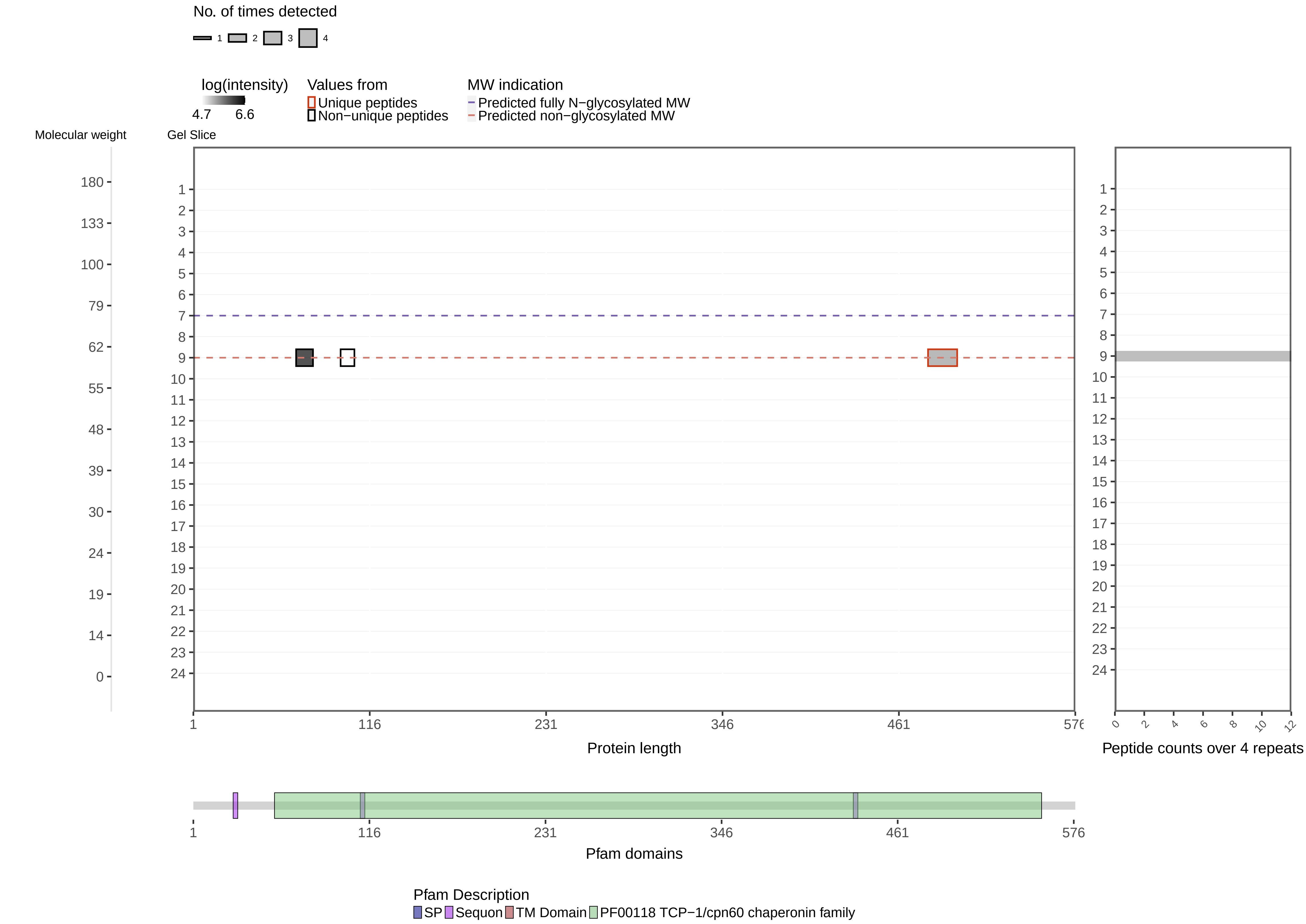


Protein sequence:

KRQRPLLIVAEDVDSEALATLILNKLRAGIKVCAIKAPGFGENRKANLQDLAILTGGQVVTEELGLNIENVELEMLGSCKKVAISKDDTVILDGAGEKKSIEERCEQIRSTIELSTSDYDKEKLQERLAKISGGVAVLKIGGASEAEVGEKKDRVTDALNATKAAVEEGIVPGGGVALLYAAKELDKLETANFDQKIGVQIIQNALKTPVHIIASNAGVEGAVVVGKLLEQDDTDLGYDAAKGEYVDMVKAGIIDPLKVIRTALVDAASVSSLLTTTEAVVVELPKDEKETPAMGGGMGGI

### NbL16g13240.1 7−hydroxymethyl_chlorophyll_a_reductase_chloroplastic [MW = 50.46 kDa]

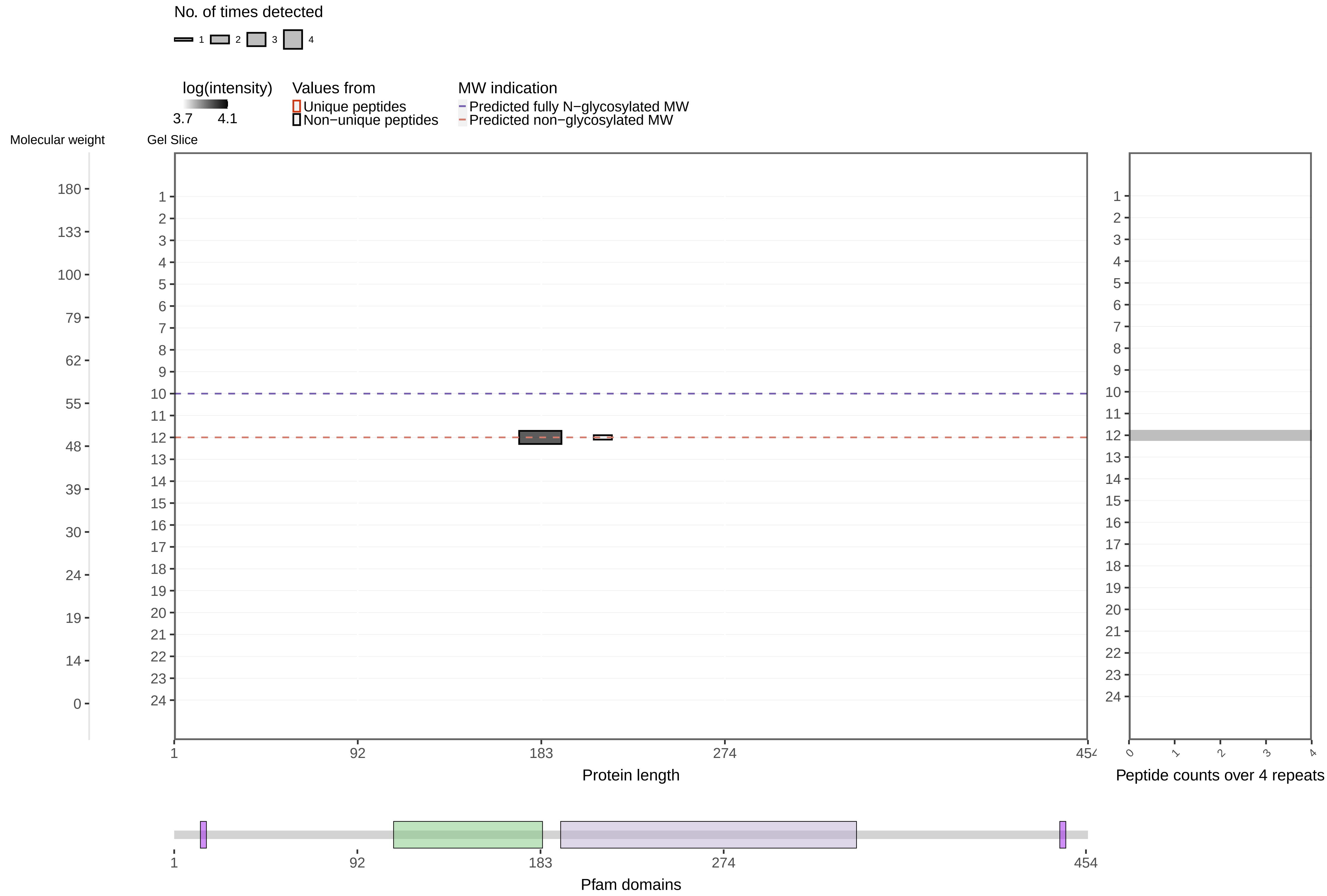


Pfam Description

SP Sequon TM Domain PF04422 Coenzyme F420 hydrogenase/dehydrogenase, beta subunit N−term PF04432 Coenzyme F420 hydrogenase/dehydrogenase, beta subunit C terminus

Protein sequence:

LNLEKLYVLGTNCVDNGKREGLDKFLKAASSEPETVLHYEFMQDYKVHLKHLDGHIEEVPYFCLPANELVDVIAPSCYSCFDYTNGLADLVVGYMGVPKYSGISMMQHPQYVTVRNERGREMLSLVENLLEITPTISSGNRRPFVVETVKADDEAKFGRGPAQPAPKFVGNLIAFFLNIVGPKGLEFARYSLDYHTIRNYLHVVRKWGKERADRHMPAYAKKIVSMYNQSGEIDQMLSSK*

NbL16g13200.1 malate_dehydrogenase_chloroplastic−like [MW = 43.93 kDa]

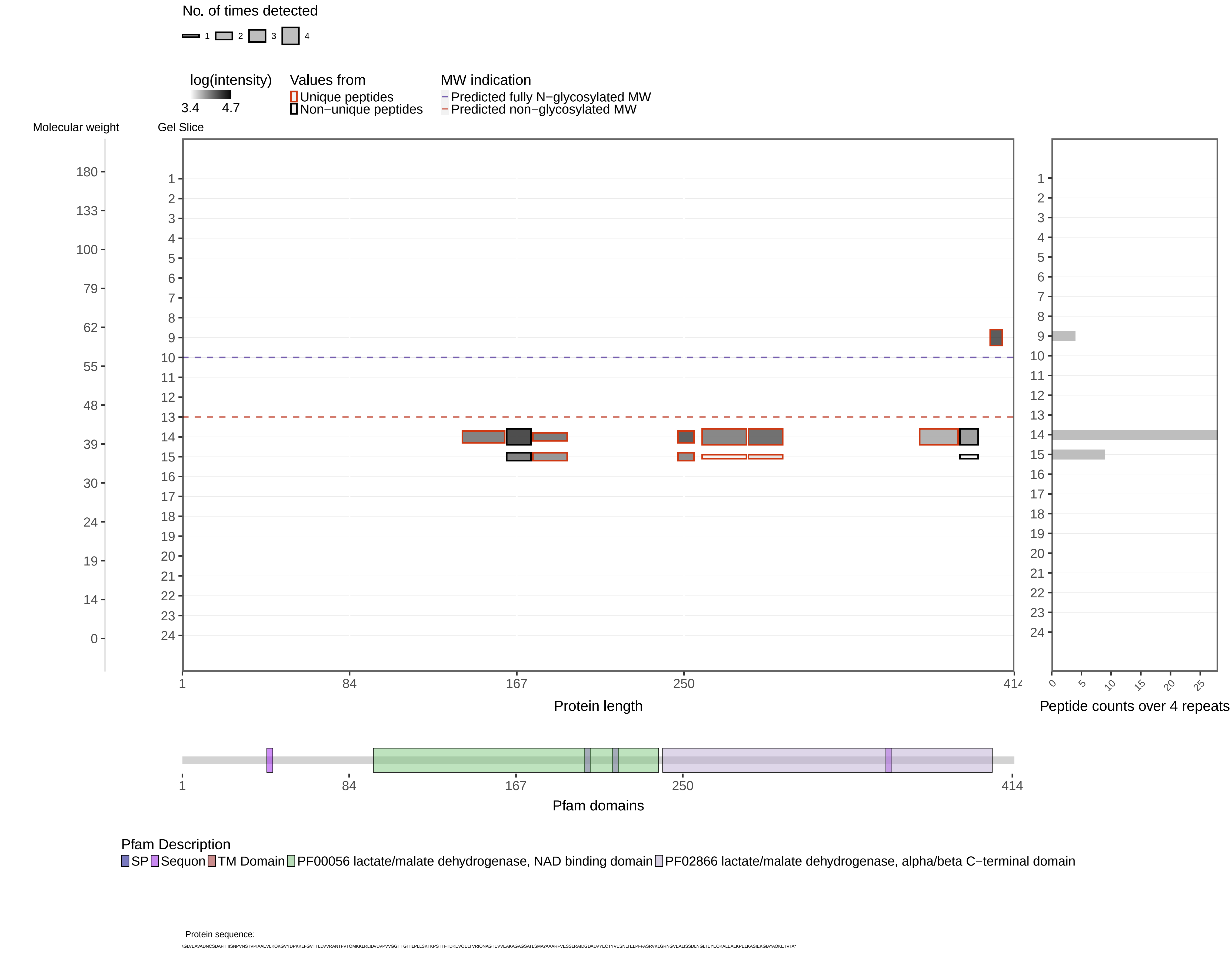

### NbL16g13130.1 haloacid_dehalogenase−like_hydrolase_domain−containing_protein_At2g33255 [MW = 28.46 kDa]

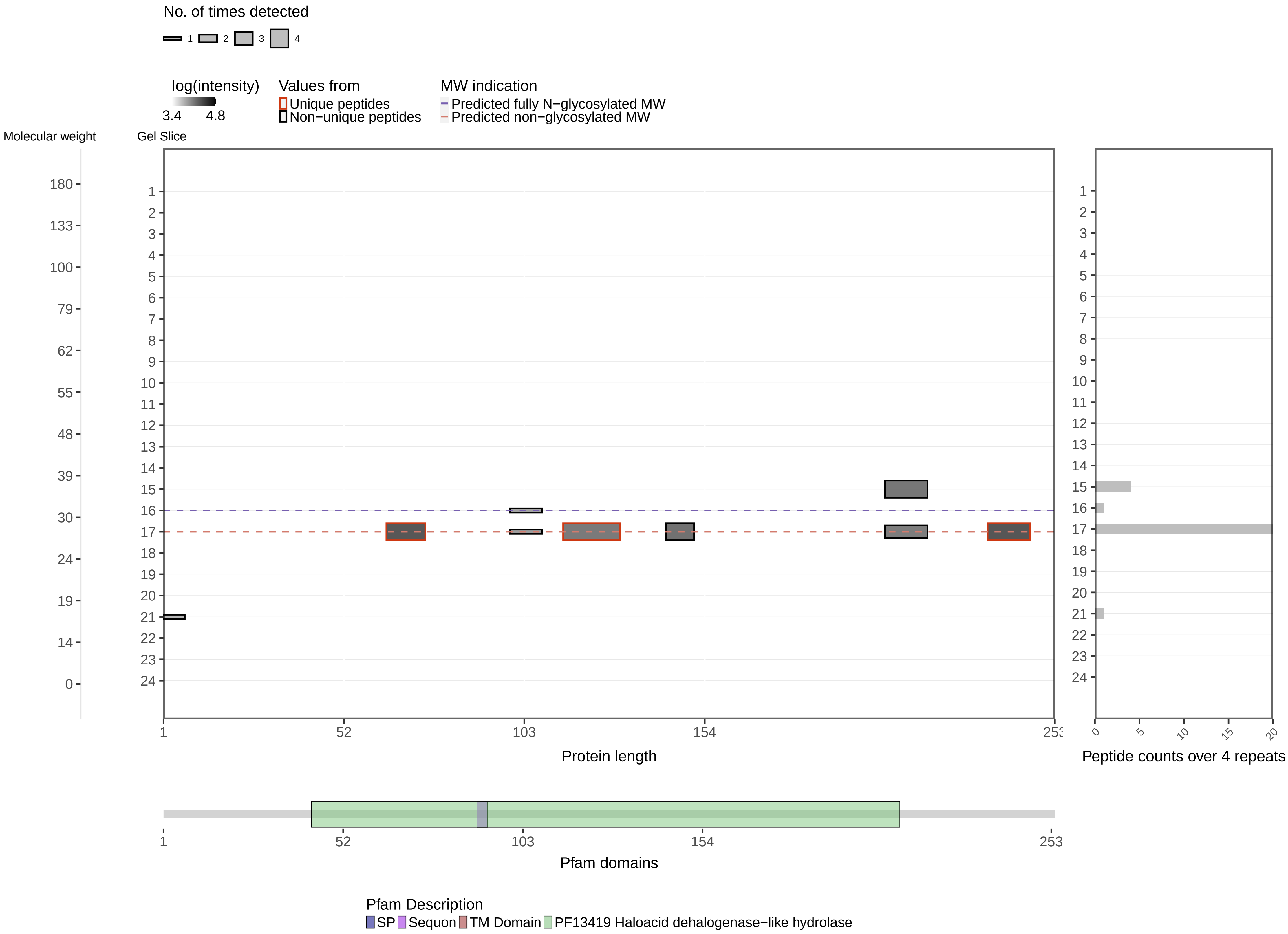


Protein sequence:

QIMPGTAELCSFLDSKNIRRGLITRNVKDAVDLFHVRFGVKFSPALSREFRPYKPDPAPLLHICSTWEVQPNEVMMIGDSLKDDVACGKRAGAFTCLLDETGRYDAPEYANVEHQPDYKVSSLVEIQSLLERHFELAS*

### NbL16g13010.1 glutamyl−tRNA(Gln)_amidotransferase_subunit_B_chloroplastic/mitochondrial [MW = 60.6 kDa]

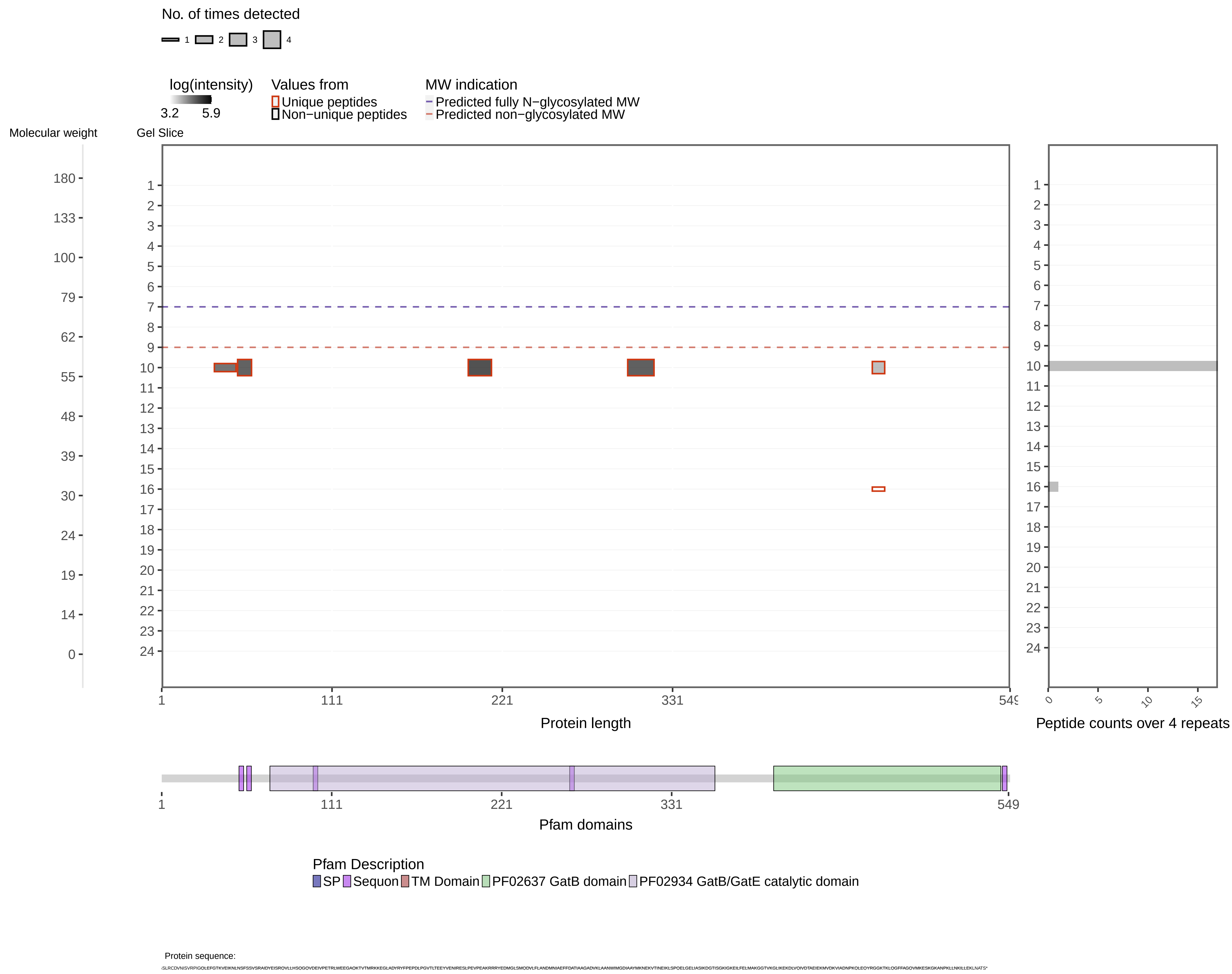

NbL16g12970.1 3−isopropylmalate_dehydratase_large_subunit_chloroplastic−like [MW = 55.43 kDa]

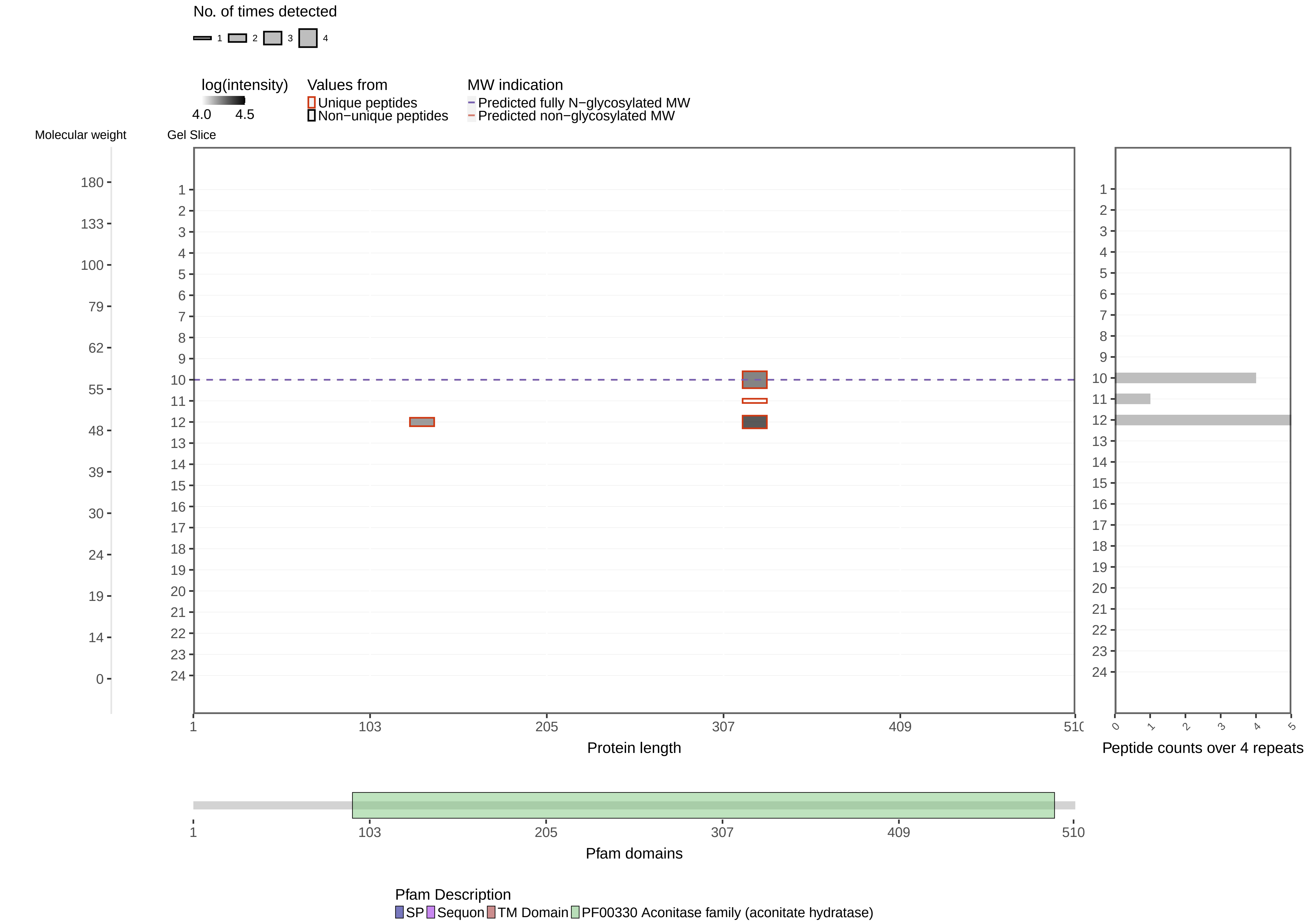


Protein sequence:

GEMPNYLLAKDLMLQIIGEISVAGATYKTMEFAGTTAESLNMEERMTLCNMVVEAGGKNGVIPADKTTYDYLKDKTTVPYEPVYSDGGARFLKEYRIDISKLEPLVAKPHSPDNRALARECKDVKIDRVYIGSCTGGKTEDFMSAAKLFQAHGKKVKVPTFLVPATREVWMDIYSLPVPGSGGKTCSQIFQEAGCYEPASPSCGACPGGLPDTYGRLNEPKVCVSTTNRNFPGRMGHIEGQIYLASPYTAAASALTGYVTDPREFLM*

### NbL16g12960.1 3−isopropylmalate_dehydratase_large_subunit_chloroplastic−like [MW = 55.75 kDa]

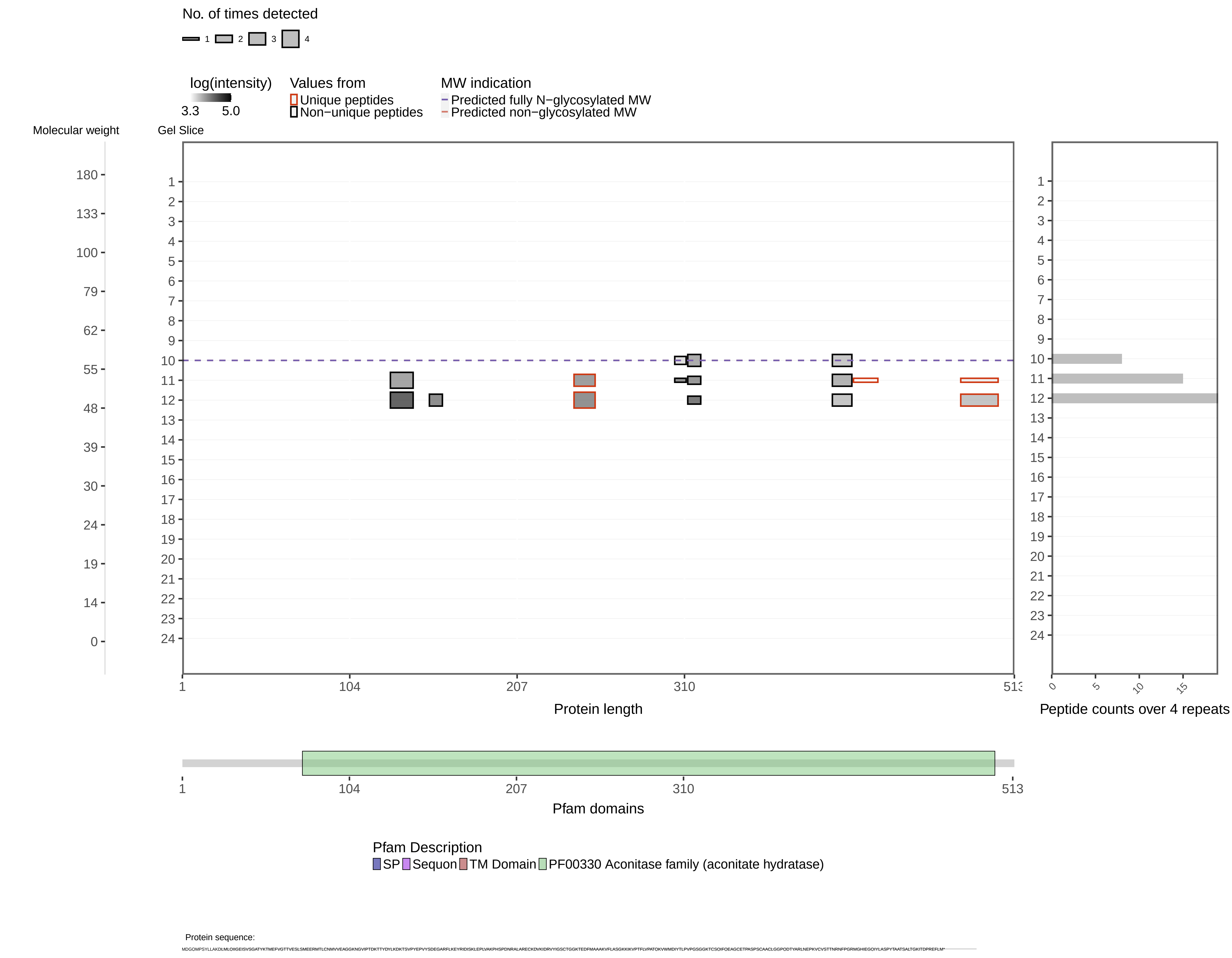

### NbL16g12910.1 putative_G3BP−like_protein [MW = 52.34 kDa]

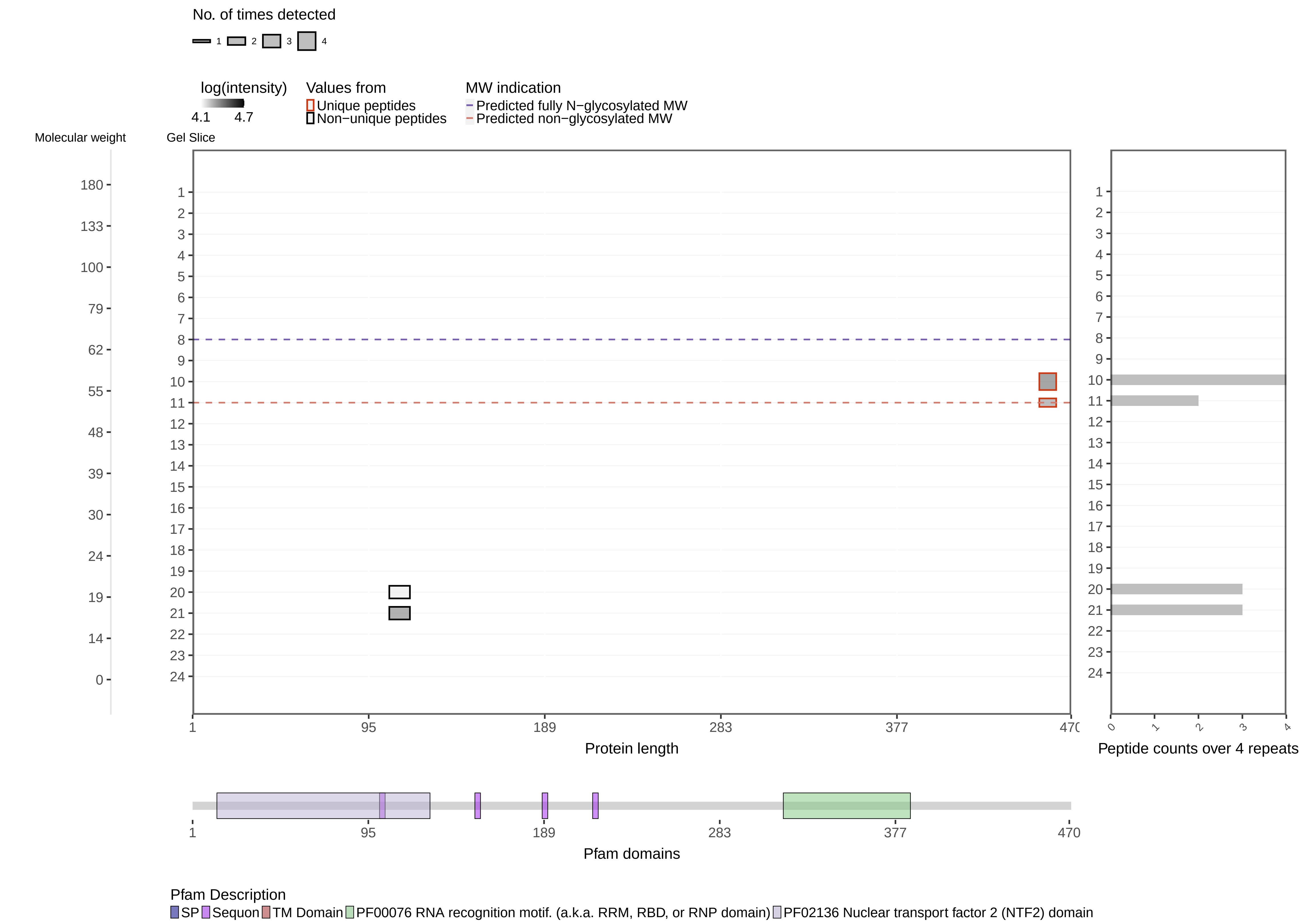


Protein sequence:

VQEHLSSPIEEPVAEPQKHTYASILQVTKGQPAQGGPAQSSFNKPTLPPSEWQHVSEPPSLSSVPSTNAIERSITEATEEASAVEDEVEVKSVYVKNVPTTMAAYEIEEEFKKFGKLKPDAVAIRTRKDIDVCYAFVEFEDVTGVQNAIEASTVQIGGHQLYIEGRRPNRNNLIRGRGRGRGRVSYSMDGRGRYGSRGFGRGGQDGGDRDYGRSRGNGYYRQAPRQERAYSGNQQSLRNEQYSWE*

NbL16g12900.1 RNA−binding_protein_CP33_chloroplastic [MW = 32.42 kDa]

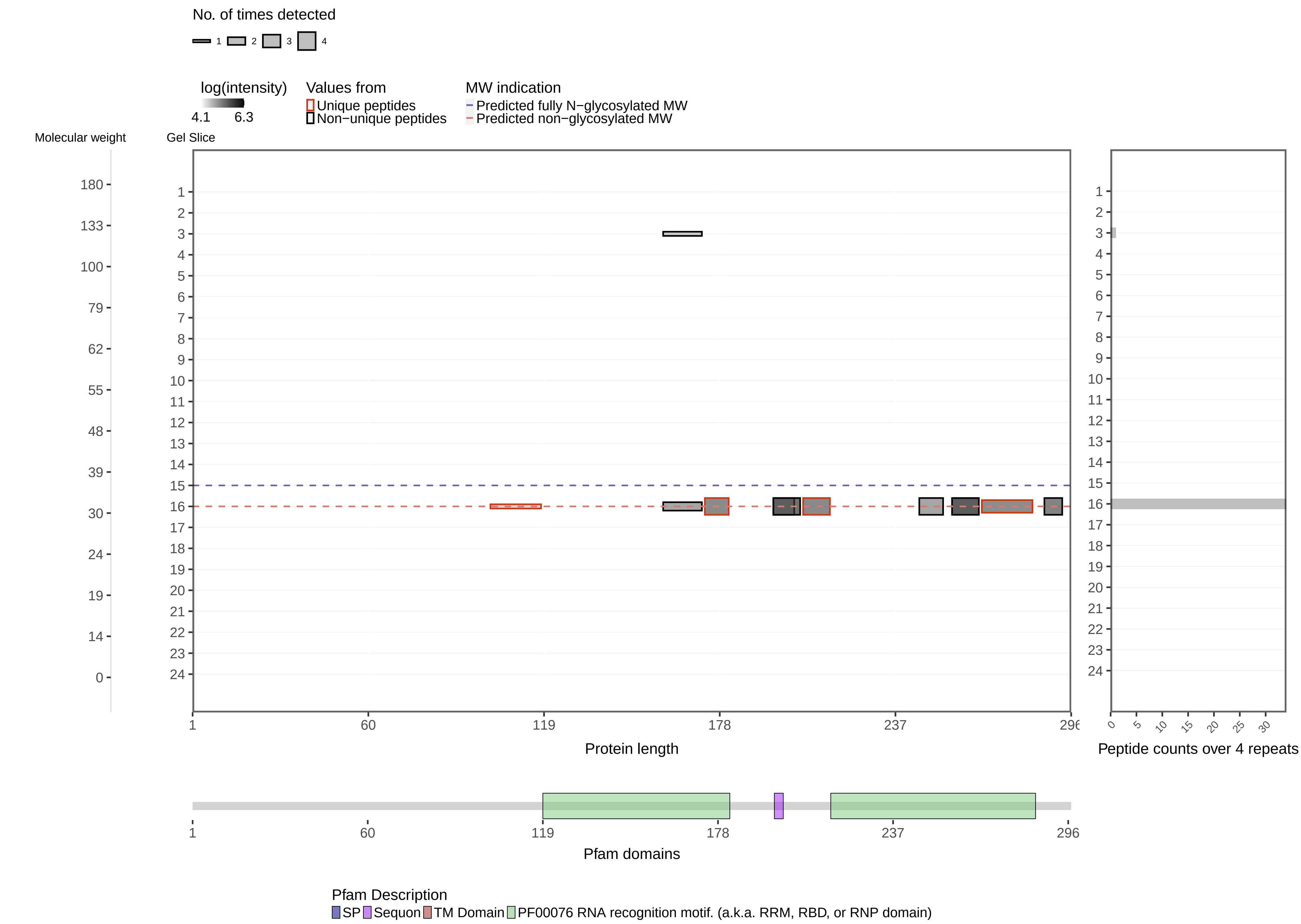


Protein sequence:

VQDYATPELVEVLYDRETGKSRGFAFVTMSSLEDCKTVIENLDGRQDLLTSGVWGRTLRVNFSDKPKPKEPLYPETEYKLFVGNLAWSVTSESLAQAFQEYGTVVGARVLYDGETGRSRGYGFVSYENREQQENALQNLNGVELDGRAMRISLAQGKKQ*

### NbL16g12820.1 glucose−6−phosphate_isomerase_cytosolic [MW = 70.28 kDa]

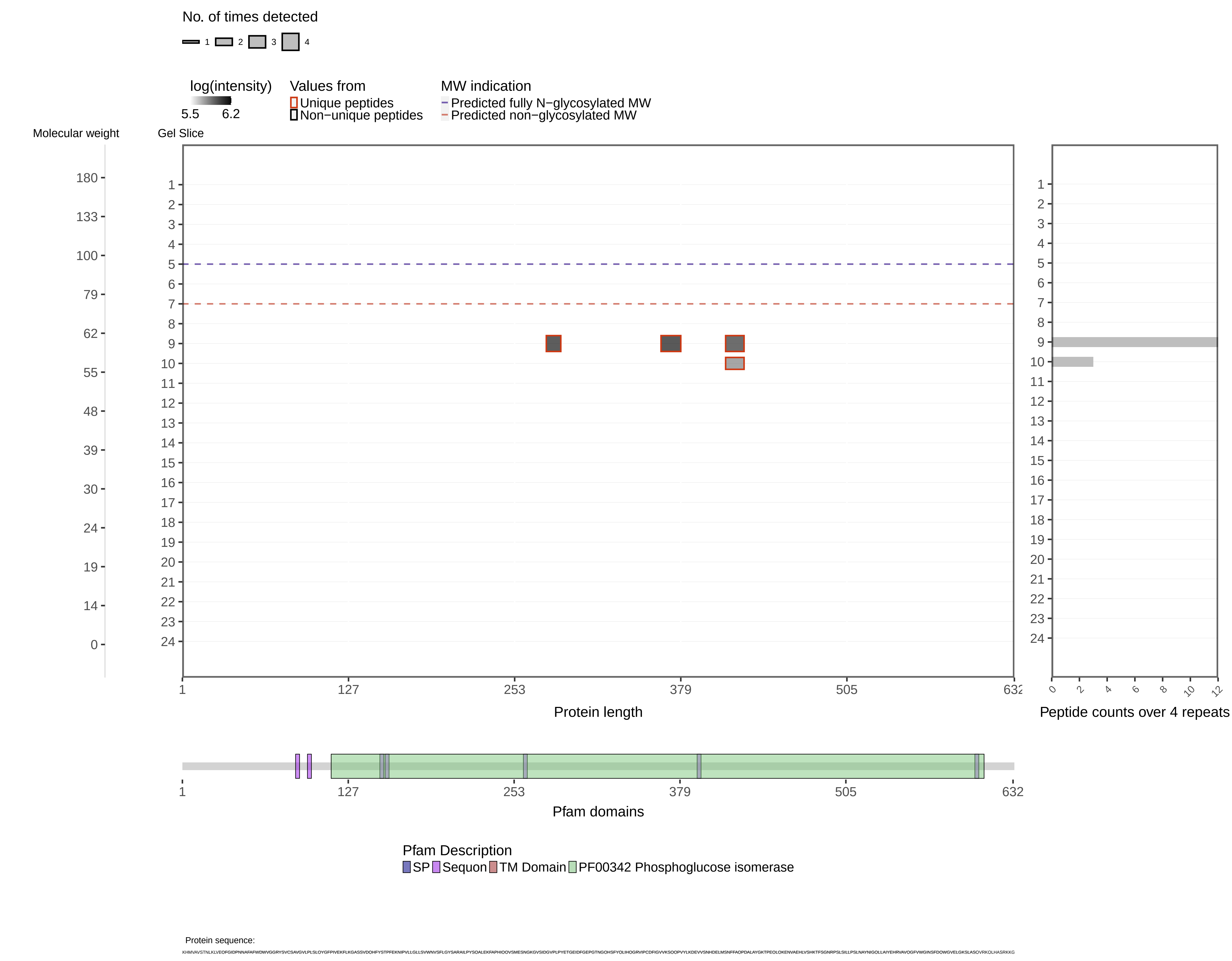

### NbL16g12780.1 pathogenesis−related_protein_STH−2−like [MW − SP = 16.93 kDa]

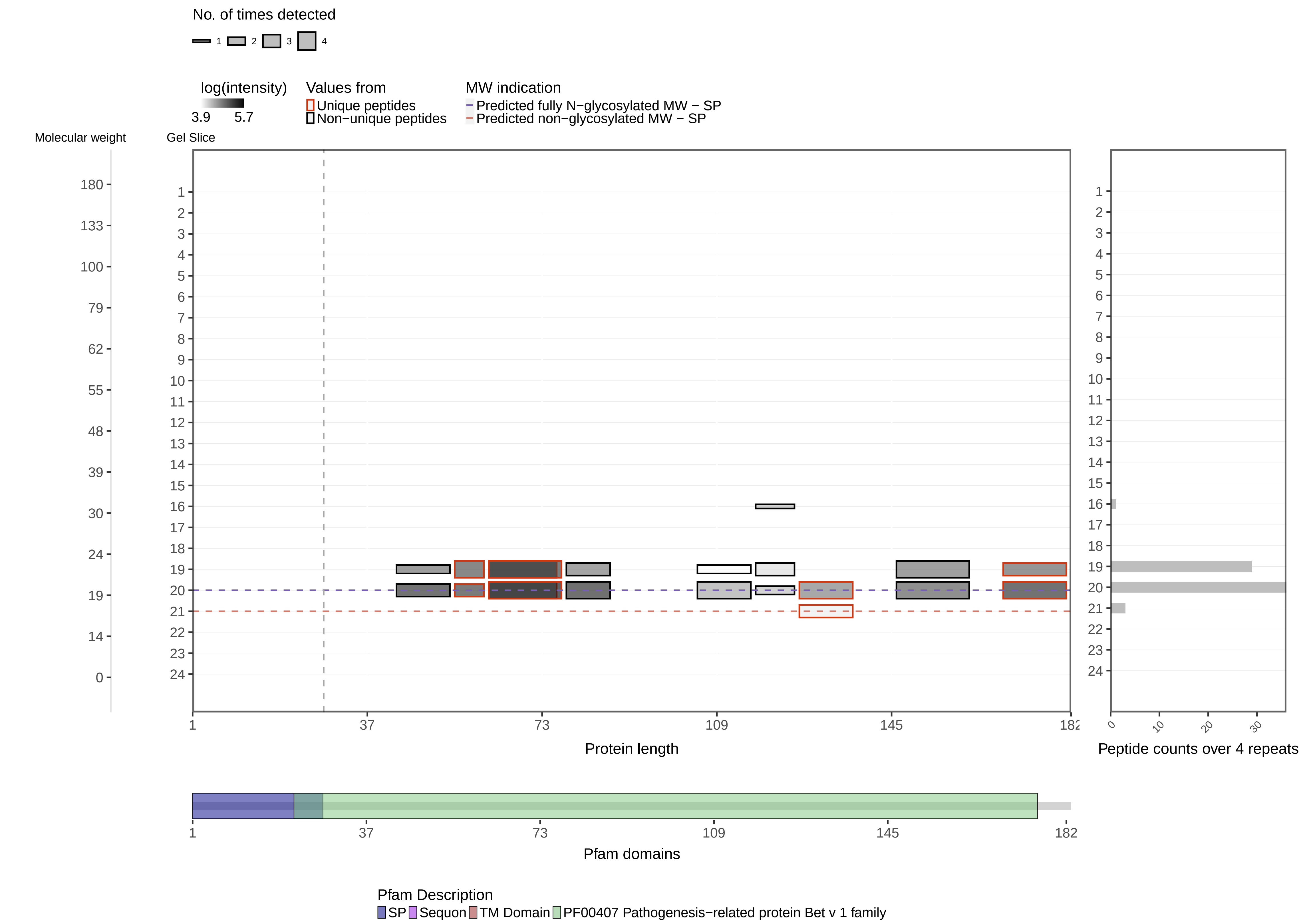


Protein sequence:

INFVEGGPIKYLKHKLHVIDNKNLVTKYSLIEGDVLGDKLESITYDVKFETSANGGCICKTSTEYHTKGDYVFKEEEHNEGKDKAMELFKAVEDYLLANPTVYA*

### NbL16g12770.1 pathogenesis−related_protein_STH−2−like [MW = 17.81 kDa]

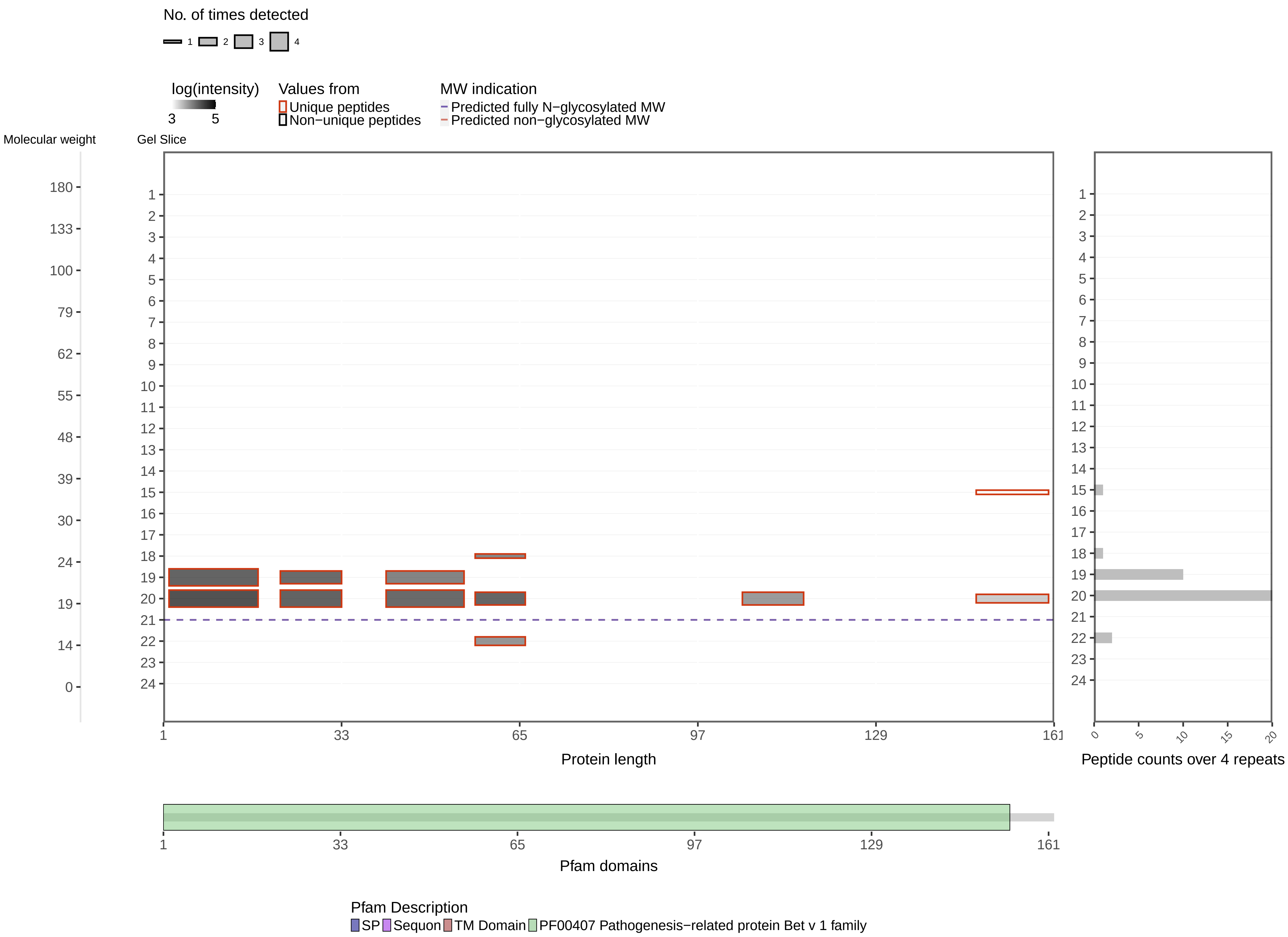


Protein sequence:

YLKHKIHVIDDKNLVTKYSLIEGDVLGDKLEFVTYDIKFEASGNGGCICKTSTEYHTKGDYVFKEEEHSEGKDQAMELFKTVEDYLIANPSLYV*

### NbL16g12620.1 cyanate_hydratase [MW = 18.58 kDa]

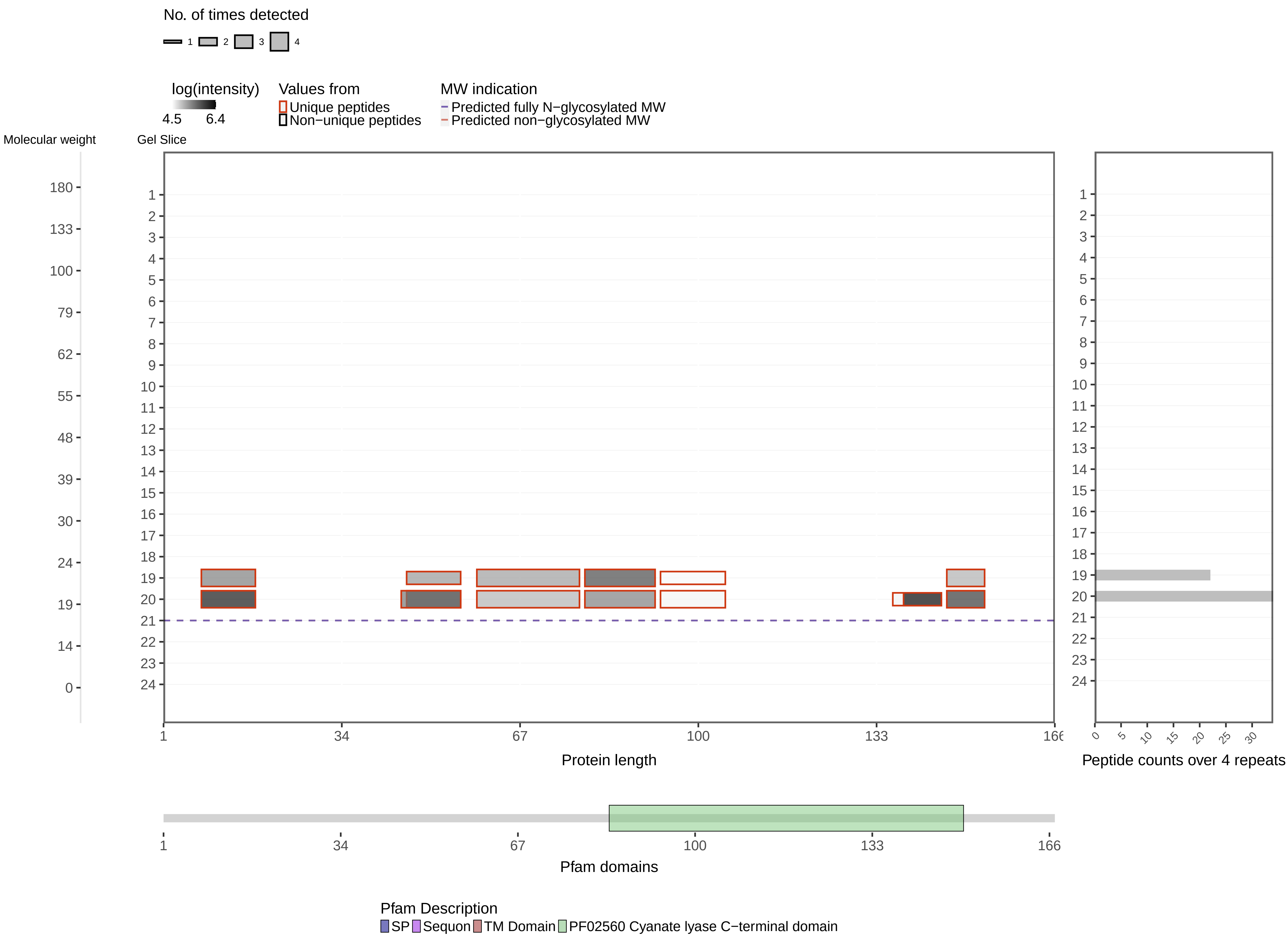


Protein sequence:

IDMMEAPLRSYDPNLIQDPTVYRLNEAVMHFGGSIKEIINEEFGDGIMSAIDFFCSVDKVKGVDGKDRVVVTFDGKYLPHTEQKSEHMVSRLMRKE*

### NbL16g12590.1 50S_ribosomal_protein_L28_chloroplastic [MW = 16.34 kDa]

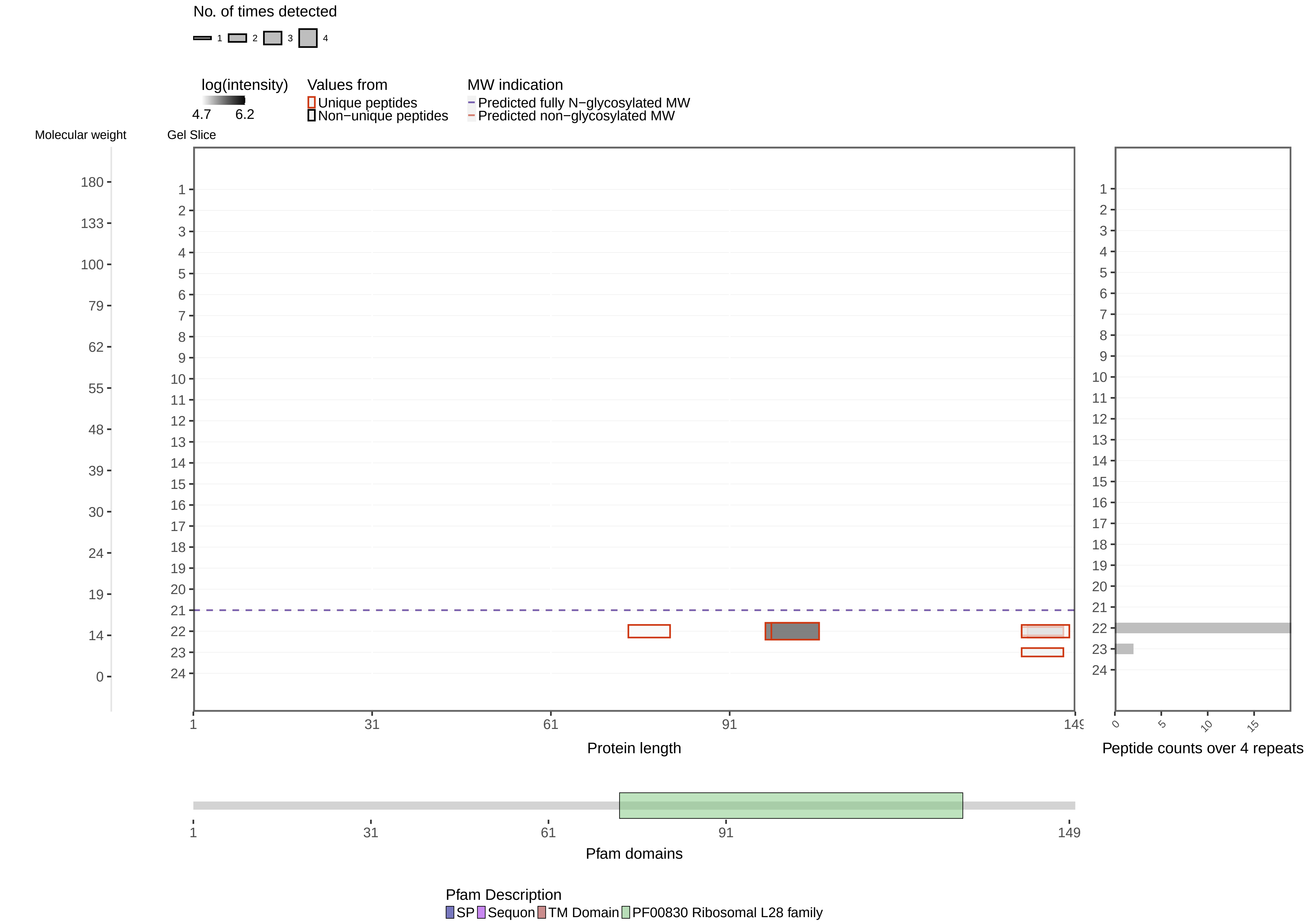


Protein sequence:

FKPSLQPVARRICPFTGKKSNRANKVSHSNHKTKKLQFVNLQYKRIWWEAGKRYVKLRLSTKALKTIEKNGLDAVAKKAGIDLSKK*

### NbL16g12530.1 probable_plastid−lipid−associated_protein_6_chloroplastic [MW = 30.17 kDa]

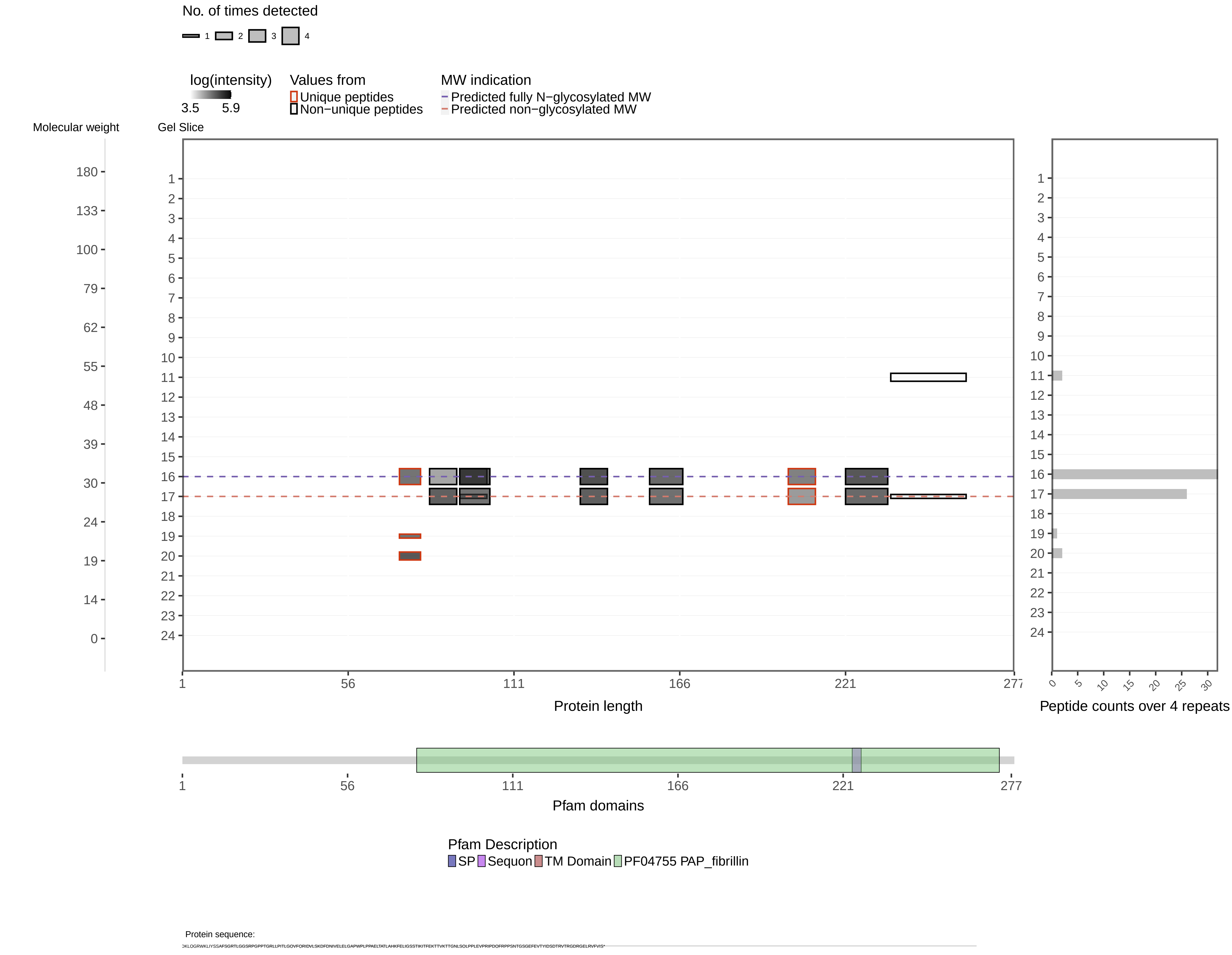

NbL16g12430.1 probable_methyltransferase_PMT3 [MW = 70.25 kDa]

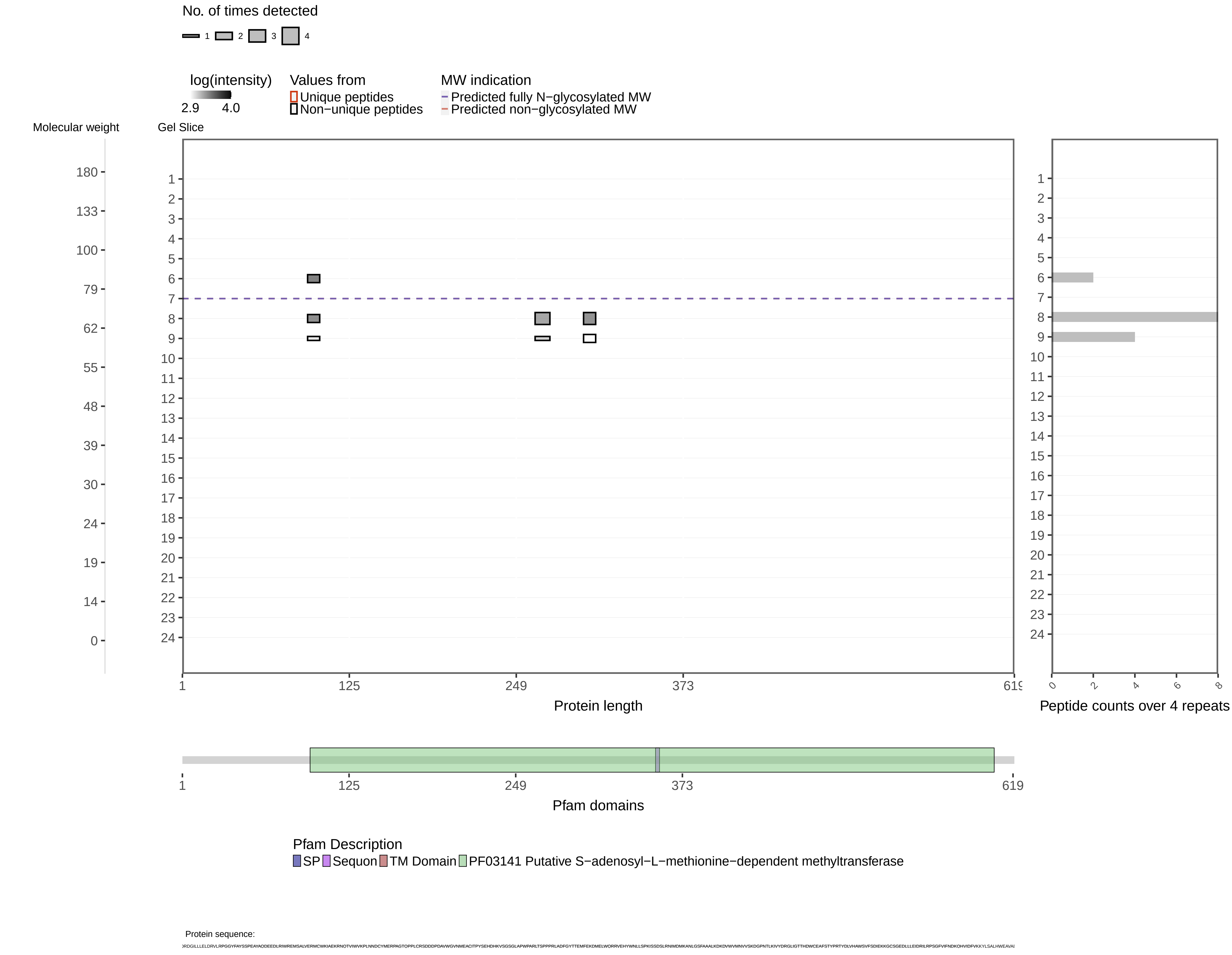

### NbL16g12270.1 methylcrotonoyl−CoA_carboxylase_subunit_alpha_mitochondrial [MW − SP = 82.29 kDa]

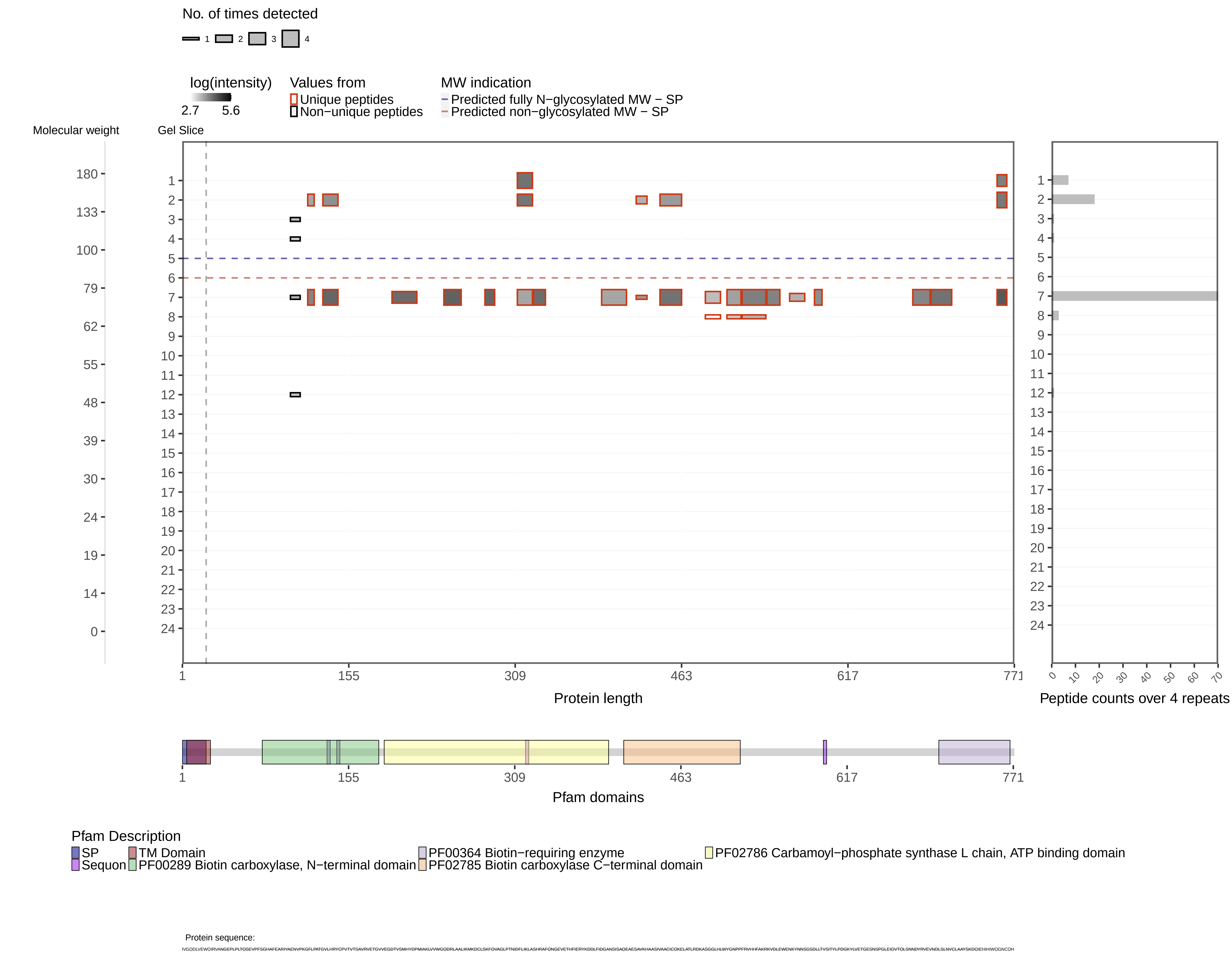

NbL16g12180.1 ETO1−like_protein_1 [MW = 100.22 kDa]

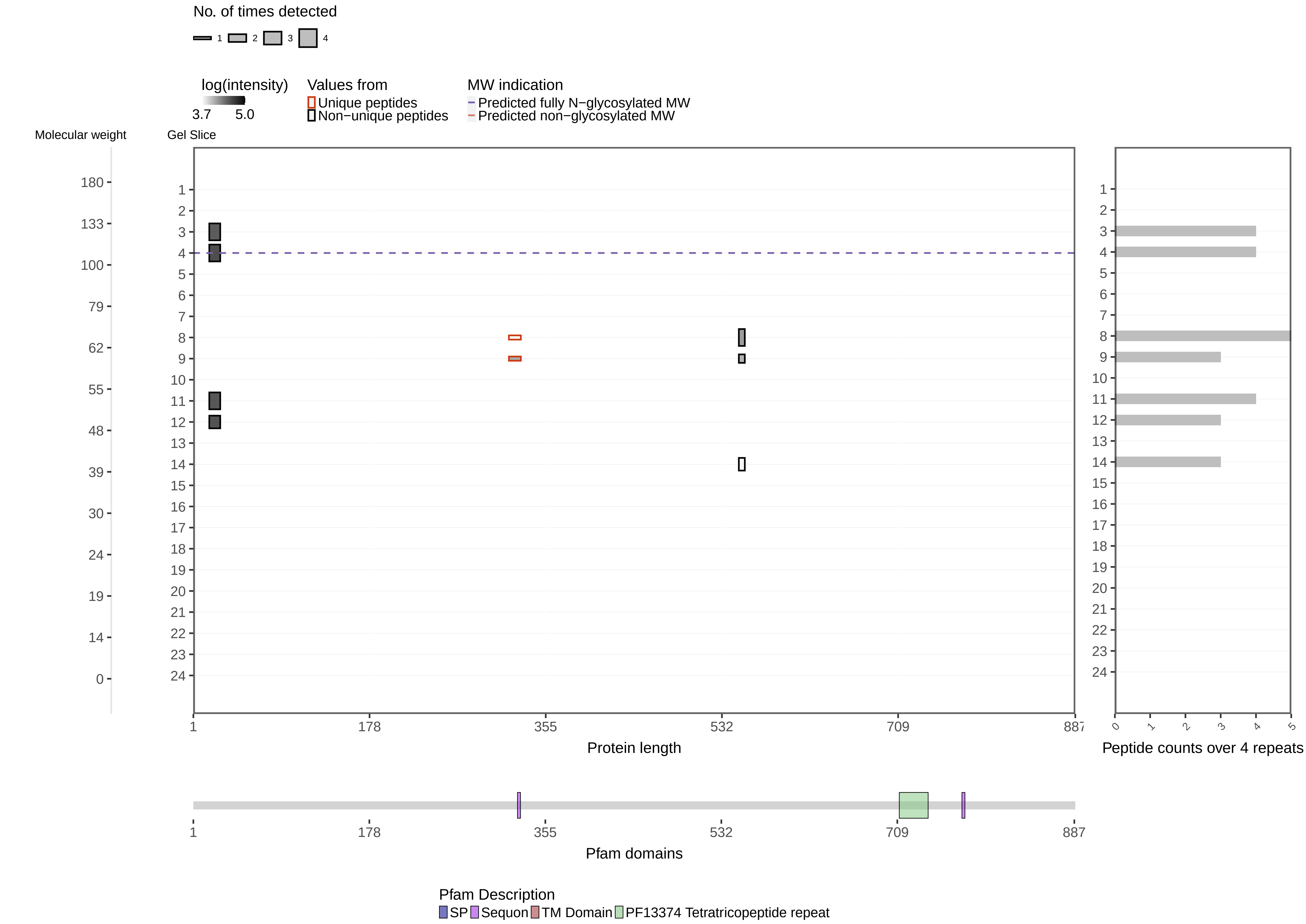


Protein sequence:

EKLSSVISSSTPLGWMYQESSLYCDGEKRLDDLGKATELDPTLTYPYMYRAASLMRKQNVQAALSEINRILGFKLALECLELRFCFYLALEDYRLAICDIQAILTLCPDYRVFEGRVAALQLRTLLREHVESWTEADCWLQLYDRWSSVDDIGSLSVIYQMLESDAAKGVLYFRQSLLLLRLNCPDAAMRSLHLARQHASSEHERLVYEGWILYDTGHCEEGLQKAEESISIKRSFEAYFLKAYALADSSPDASCSSTVISLLEEALRCPSDRLRKGQALNNLGSVYVDCGKLDA

### NbL16g12050.1 probable_histone_H2A_variant_1 [MW = 14.67 kDa]

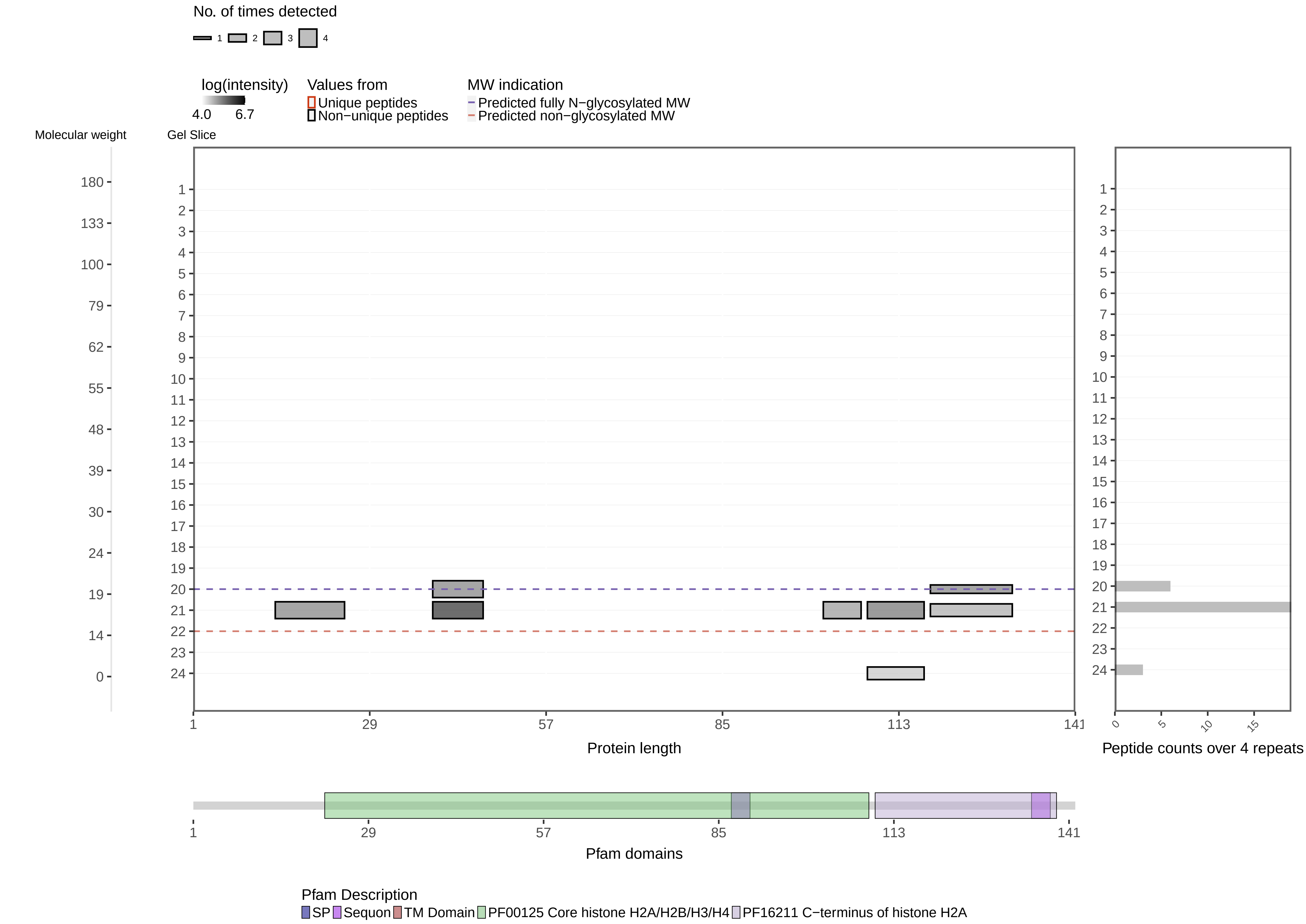


Protein sequence:

RTSAHGRVGATAAVYSAAILEYLTAEVLELAGNASKDLKVKRITPRHLQLAIRGDEELDTLIKGTIAGGGVIPHIHKSLINKTAKE*

NbL16g11980.1 T−complex_protein_1_subunit_theta−like [MW = 58.15 kDa]

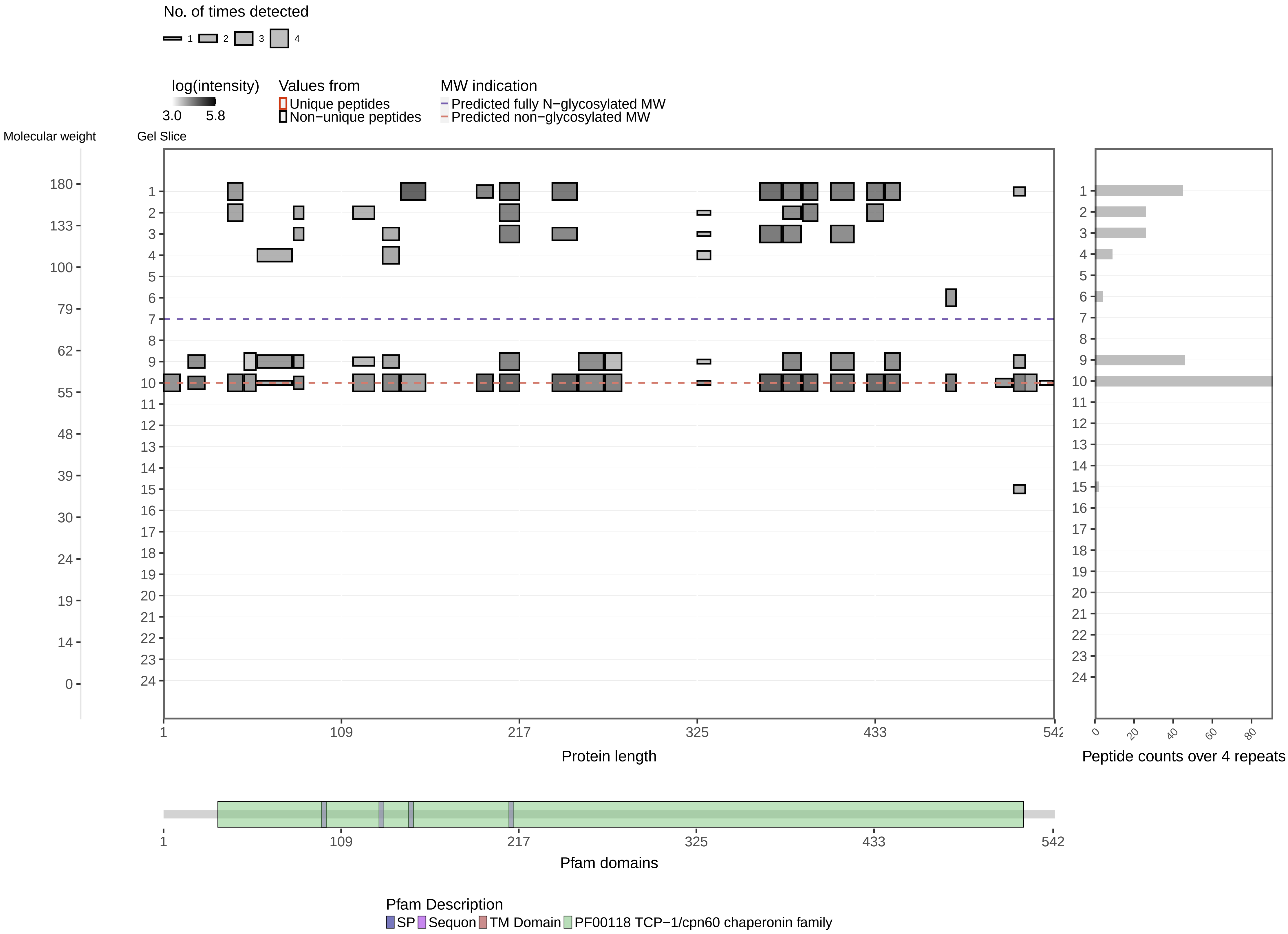


Protein sequence:

SAEQLENYAKTEEAKVEELIKAVADSGAKVIVSGGAVGEMALHFCERYKLMVLKISSKFELRRFCRTTGAVALLKLSQPNADDLGYVDSVSVEEIGGVRVTVVRNEEGGNSVSTVVLRGSTDSILDDLERAVDDGVNTYKAMCRDSRIVPGAAATEIELASKLKEFSFKETGLDQYAIAKFAESFEMVPKTLAENAGLNAMEIISSLYAEHASGNTKVGLNLEKGACEDISALSIWDLYVTKFFALKYAADAVCTVLRVDQIIMSKPAGGPSRRDQPAGMDED*

### NbL16g11750.1 zinc_finger_protein_598−like [MW = 105.8 kDa]

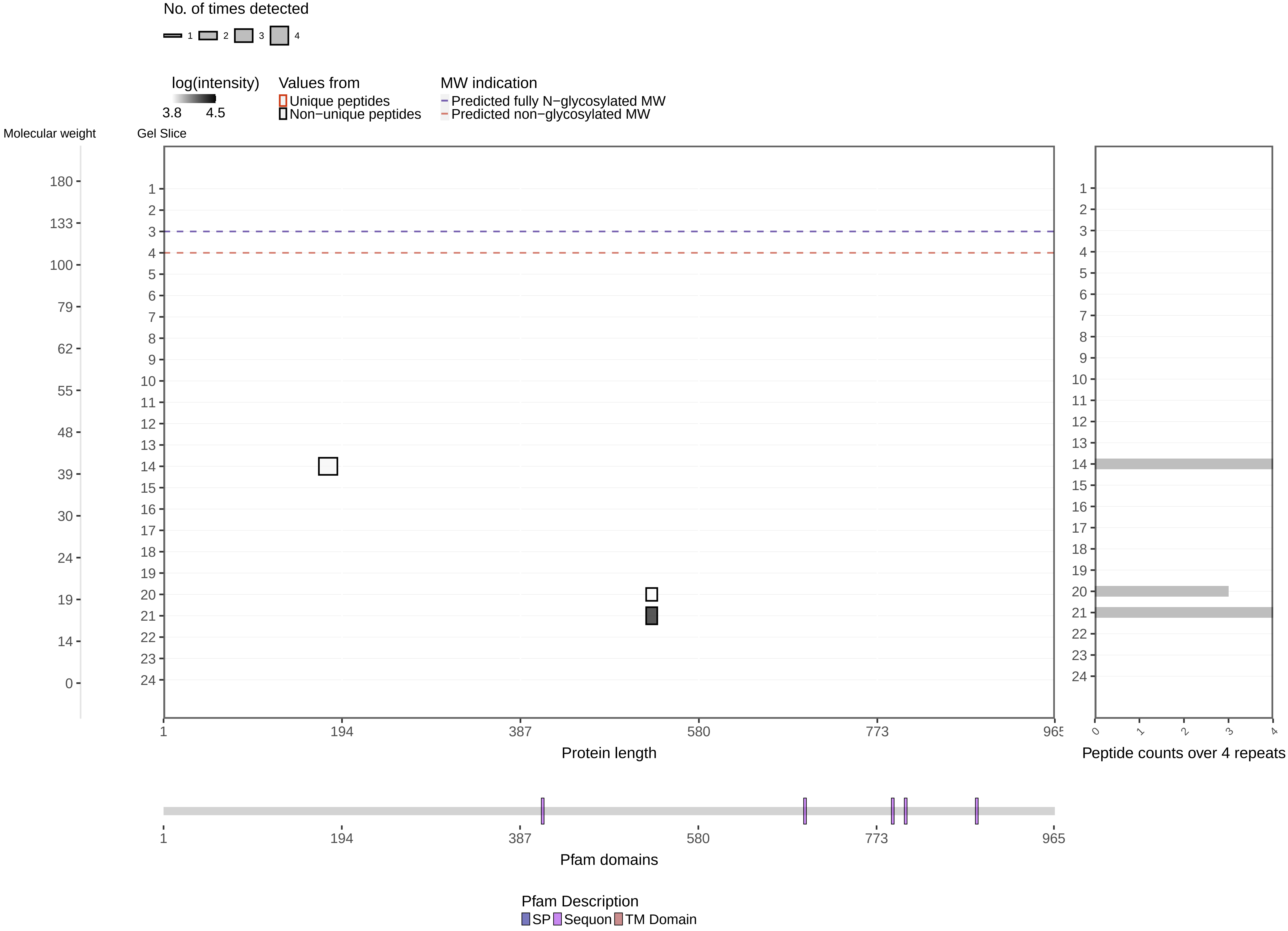


Protein sequence:

ASGSSSSSRHSKAATDDPSPPVITTQHAWPAVNGAIGSASGSSQIKPPSSSSRYSKAATDNPSAPVITREHAWPAVSGATGSTSGSSQIKLPSSSSRYSKAATDNPSAPVITREHAWPAVGGATGSTSGSSQIKPSTVDGPPPSSYLSSVAALSSLVHEASSSSVGSSKNWYQSNRISHSTSAPNLVQSGSFDSSVSDFPPVSAAQTRKLPARGQQAVTNVEDVQTANKSLVERMRLALQFDQDKFAAFKDISAEYRQGIIDAETYLAYVEQFGLSHLILELARLCPDAERQKE

### NbL16g11520.1 protoporphyrinogen_oxidase_chloroplastic [MW = 59.8 kDa]

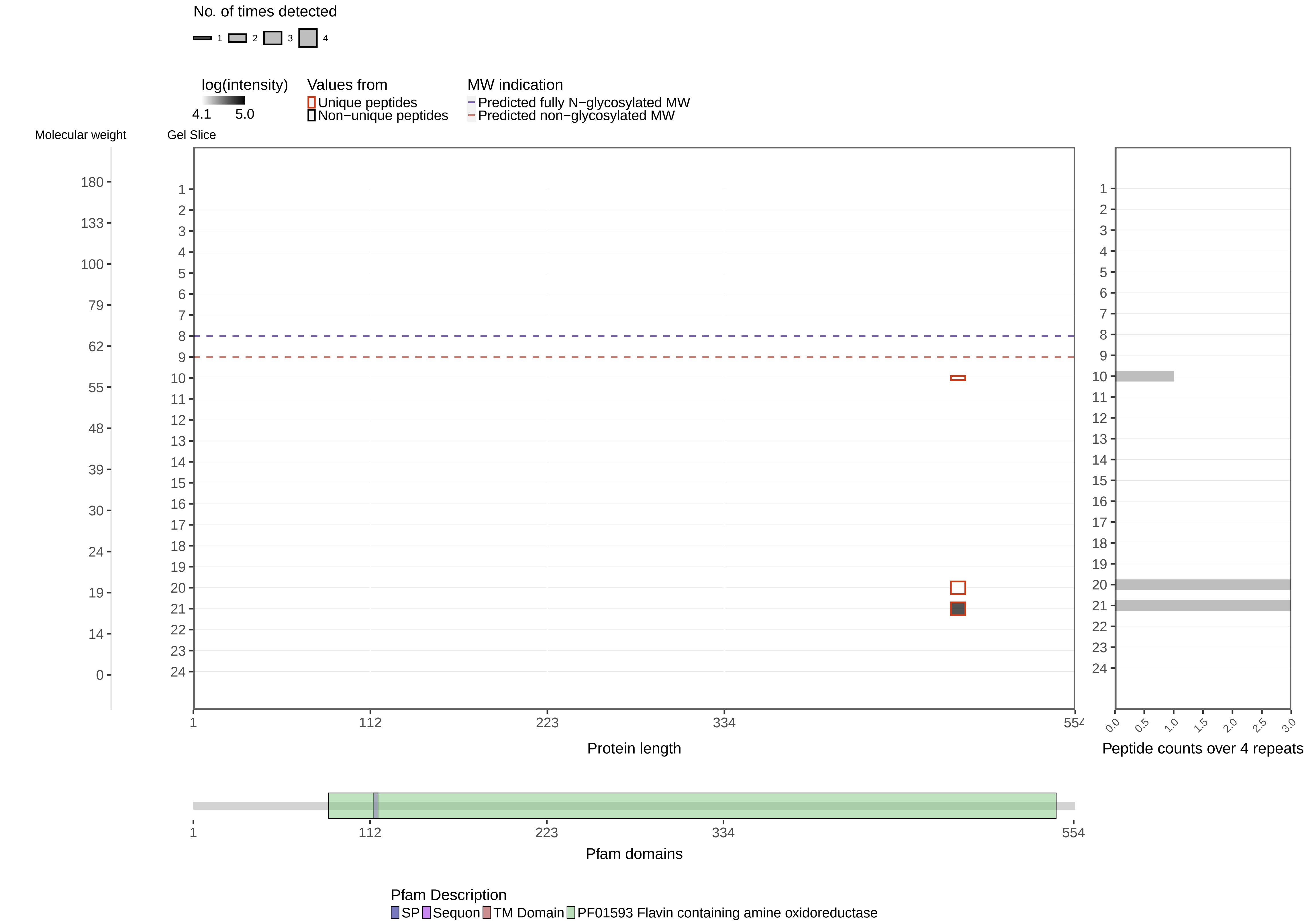


Protein sequence:

IGGTFKAIKERSSTPKAPRDPRLPKPKGQTVGSFRKGLRMLPDAISARLGSKLKLSWKLSSITKSEKGRYRLTYETPEGVVSLQSRSIVMTVPSYVASNILRPLSVAAADALSNFYYPPVGAVTISYPQEAIRDERLVDGELKGFGQLHPRTQGVETLGTIYSSSLFPNRAPKGRVLLLNYIGGAKNPEILSKTESQLVEVVDRDLRKMLIKPKAQDPVVVGVRVWPQAIPQFLVGHLDTLSTAKAAMSDNGLEGLFLGGNYVSGVALGRCVECAYEVASEVTGFLSRYAYK*

NbL16g11450.1 protein_trichome_birefringence−like_42 [MW = 42.53 kDa]

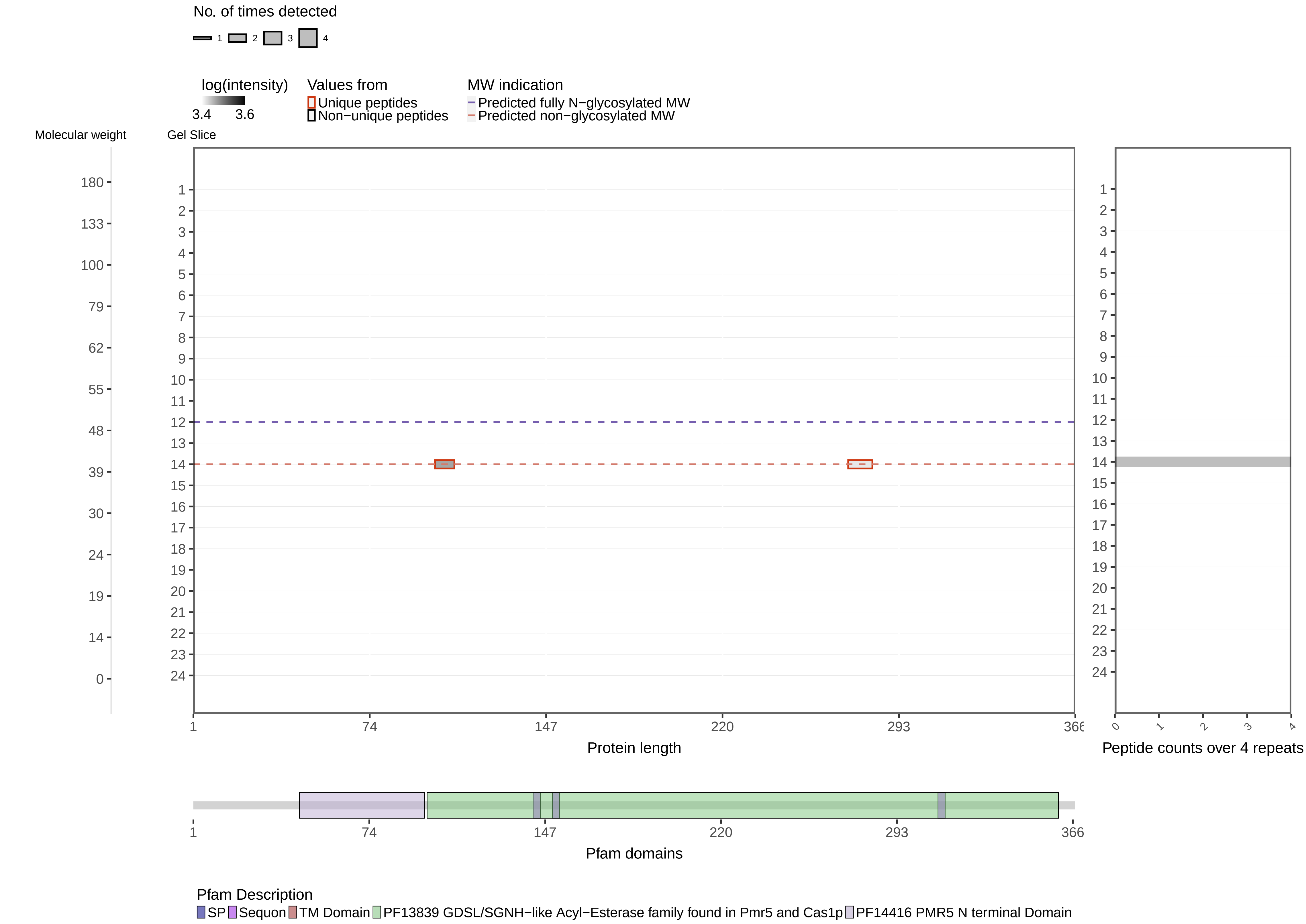


Protein sequence:

LIDLVVEETGRVLKLDSLSRSDQWKGVDFLIFNSYHWWIHTGNLQTWDYFQVGDKIYKEMDHMEAYKIALTTWANWVDSNIDPAMTKVFFQGISAVHYHGKDWNEPMVQDCRGQTKPIEGSNYPGDRYPGEAVVKNVLSNMTKSVYLLDITLLTQLRKDGHPSRYANGETDCSHWCVAGVPDAWNELLYTILLQK*

NbL16g11430.1
trifunctional_UDP−glucose_46−dehydratase/UDP−4−keto−6−deoxy−D−glucose_35−epimerase/UDP−4−keto−L−rhamnose−reductase_RHM1
[MW = 78.93 kDa]

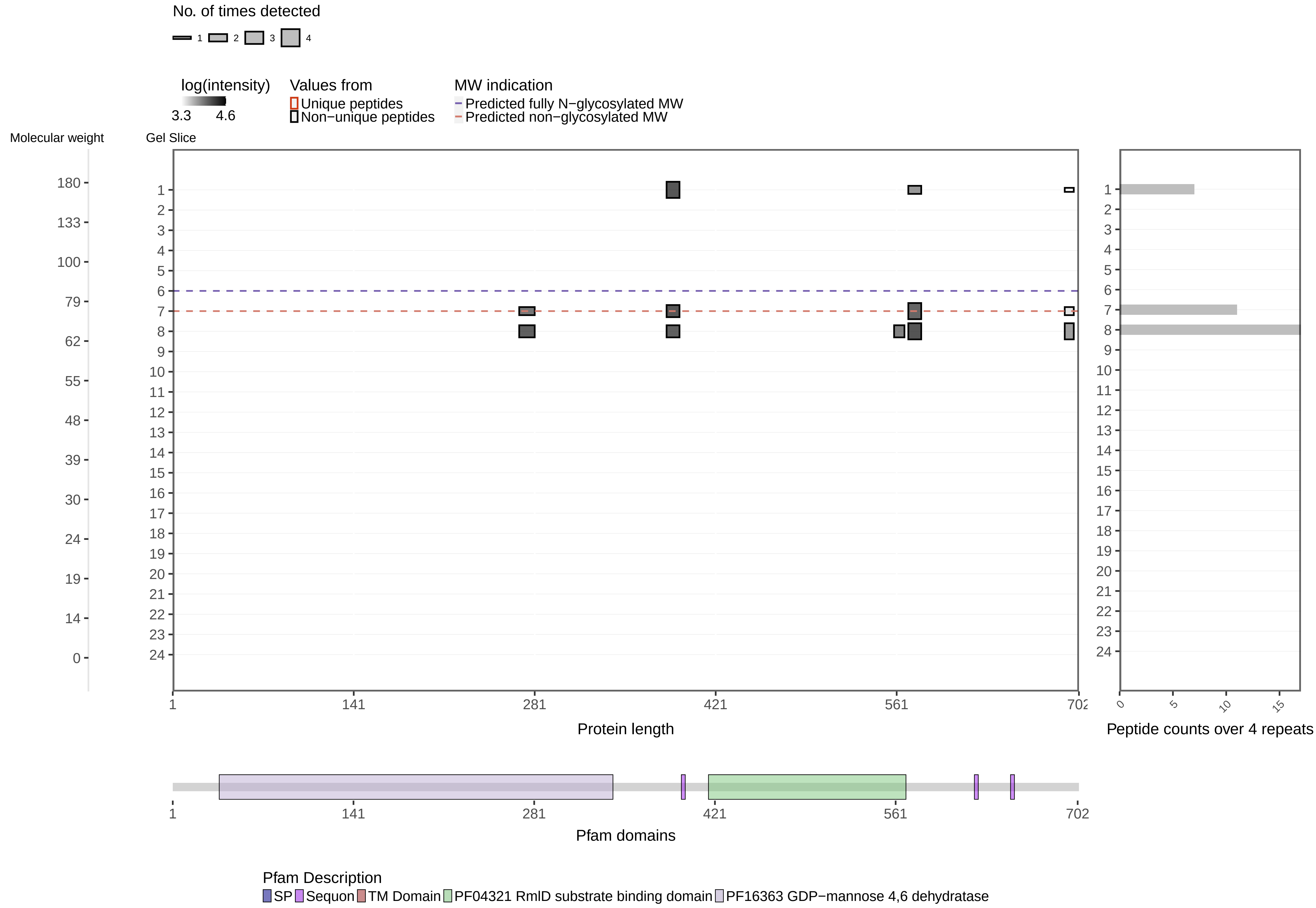


Protein sequence:
[illegible]

### NbL16g11330.1 peroxisomal_(S)−2−hydroxy−acid_oxidase [MW = 40.52 kDa]

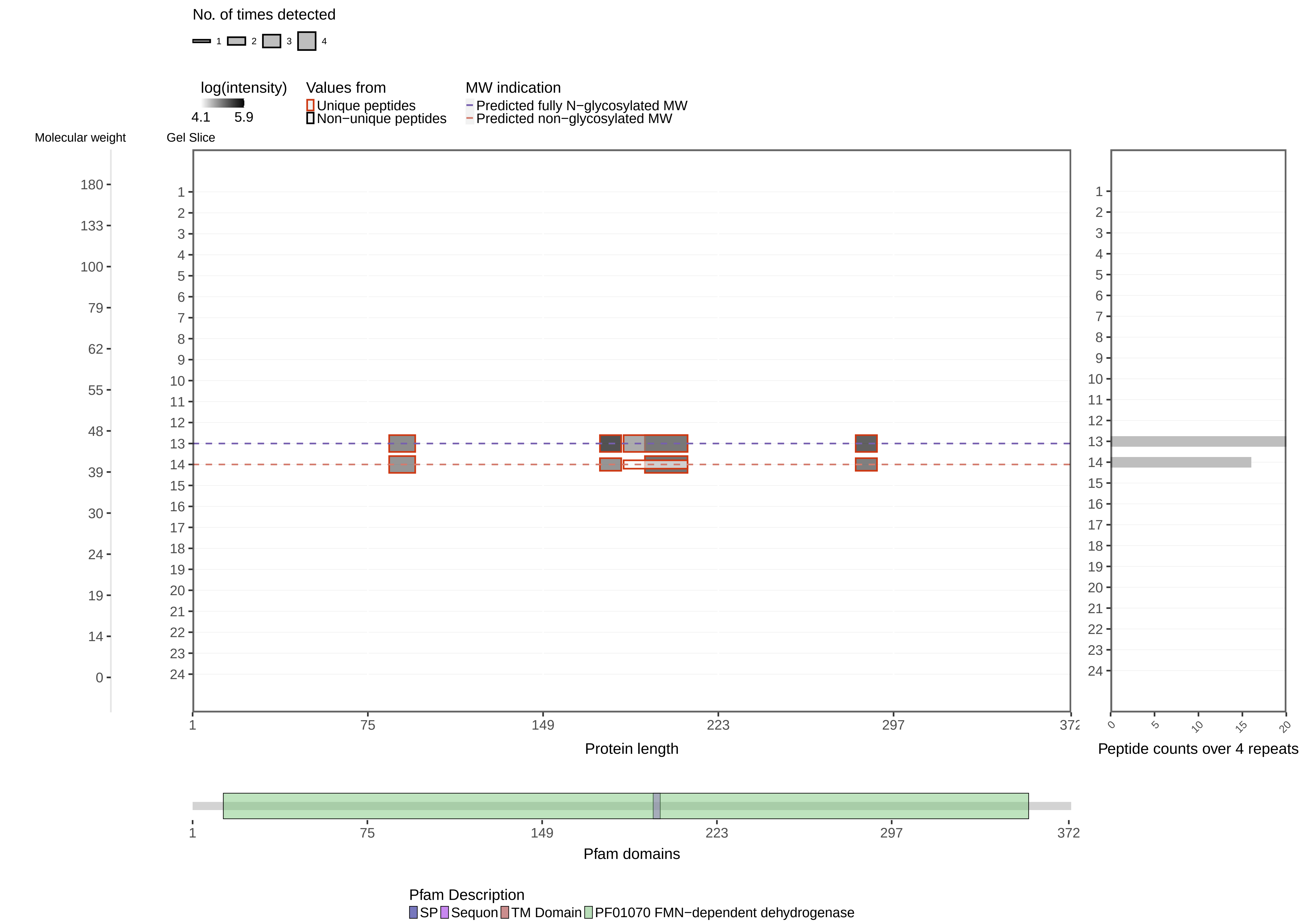


Protein sequence:

TLPPFLTLKNFEGLDLGKMDEANDSGLASYVAGOIDRSLSWKDVOWLOTITSLPILVKGVLTAEDARIAVOAGAAGIIVSNHGAROLDYVPATIMALEEVVKAAOGRVPVFLDGGVRRGTDVFKALALGASGVFIGRPVVFSLAAEGEAGIRKVLOMMRDEFELTMALSGCRSLKEITRNHIVADWDAPRAALPAPSL*

NbL16g11260.1 chlorophyll_a−b_binding_protein_8_chloroplastic [MW = 29.37 kDa]

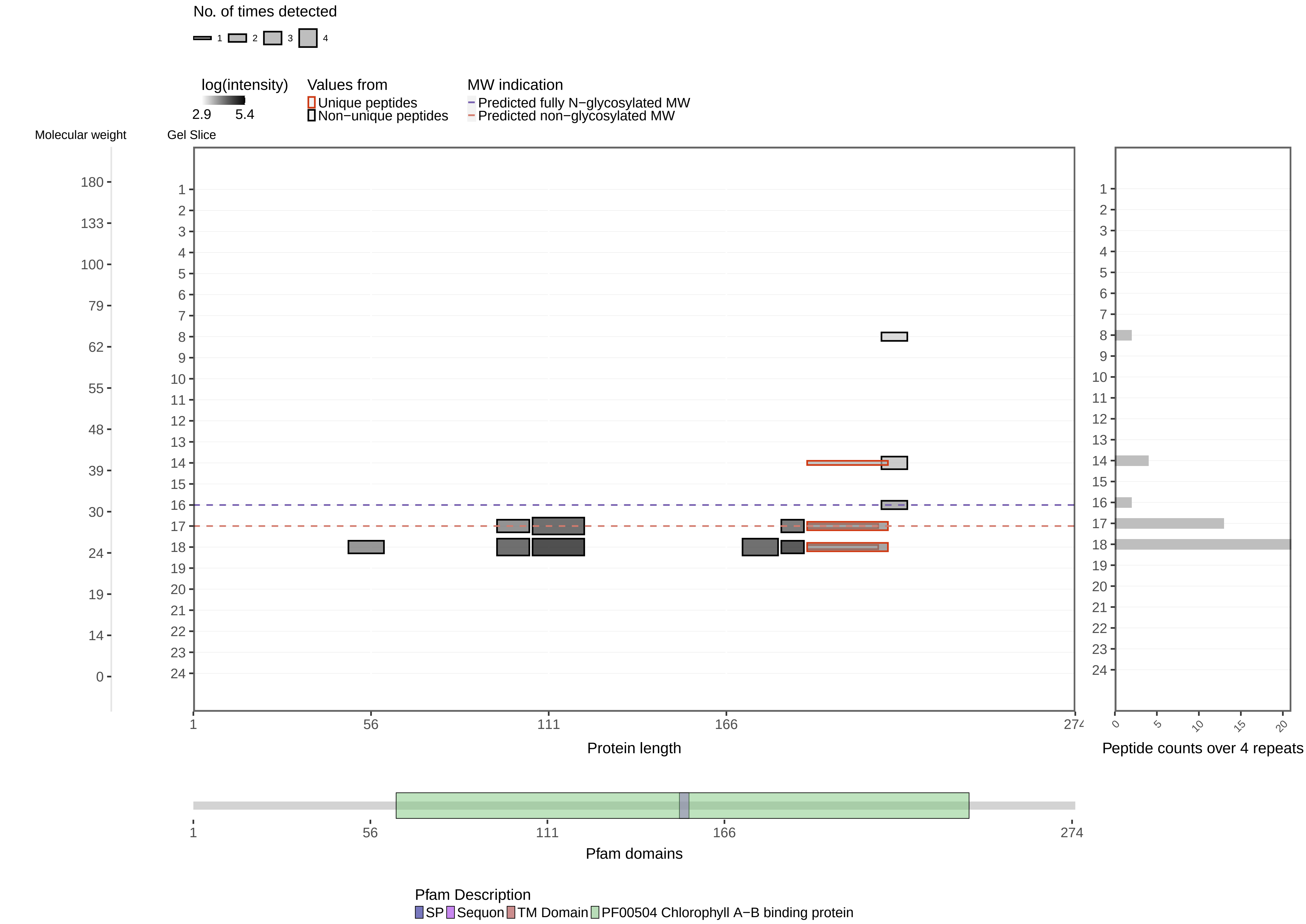


Protein sequence:

LIPPETALPWFOTGVIPPAGTYNYWADNYTLFVLEMALMGFAEHRRFODWAKPGSMGKOYFLGLEKGLGGSGEPAYPGGSFFNPLGFGKDEKSMKDLKLKEVKNGRLAMLAILGYFIOGLVTGVGPYONLLDHLADPVNNNILTNFKFH*

NbL16g11160.1 60S_ribosomal_protein_L11−1 [MW = 20.74 kDa]

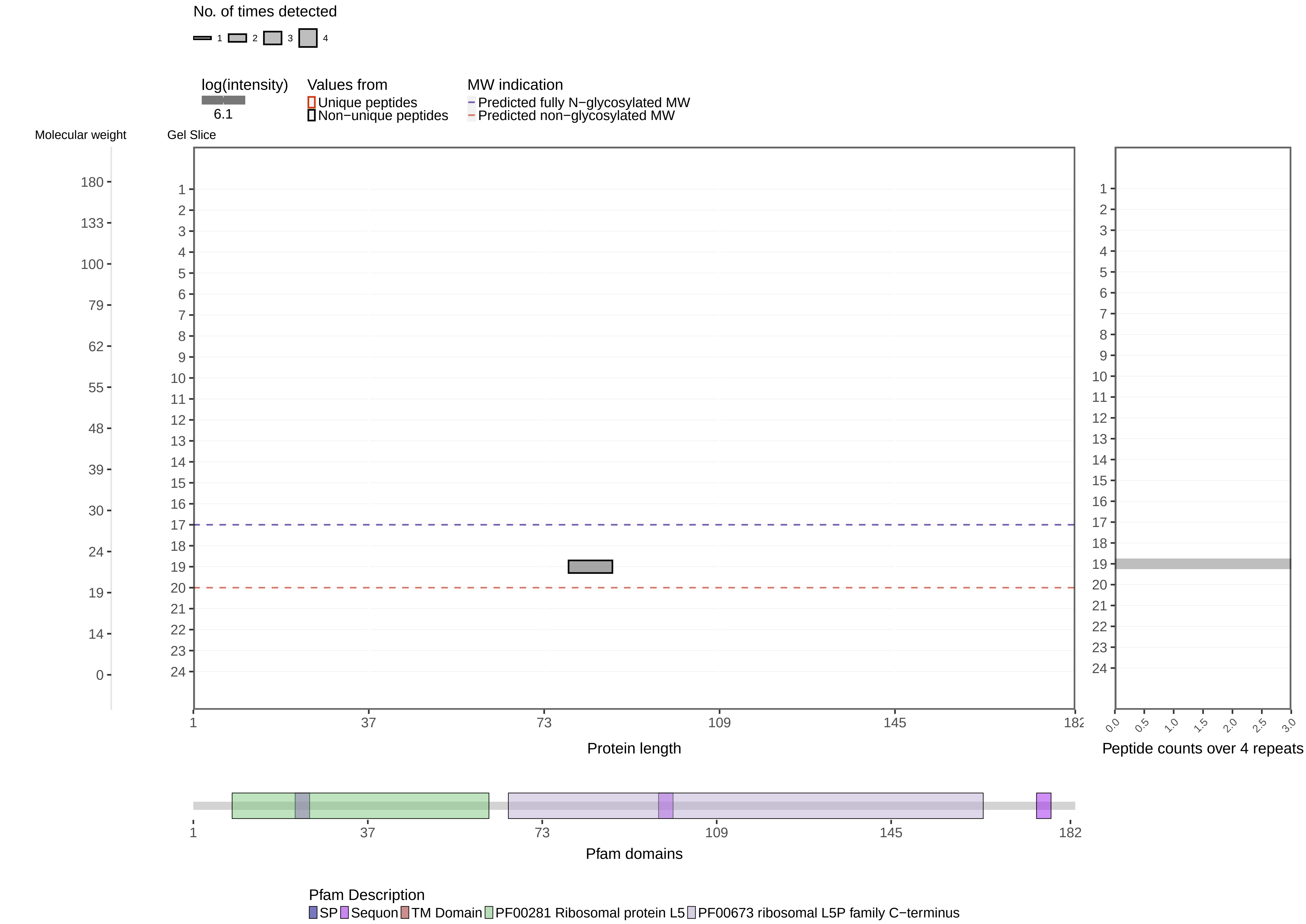


Protein sequence:

MQLLESGLKVKEYELLRRNFSETGCFGFGIQEHIDLGIKYDPSTGIYGMDFYVVLERPGYRVARRRRCKARVGIQHRVTKDDAMKWFQVKYEGVILNKSSNIQ*

NbL16g11070.1 uncharacterized_LOC107795773 [MW − SP = 9.99 kDa]

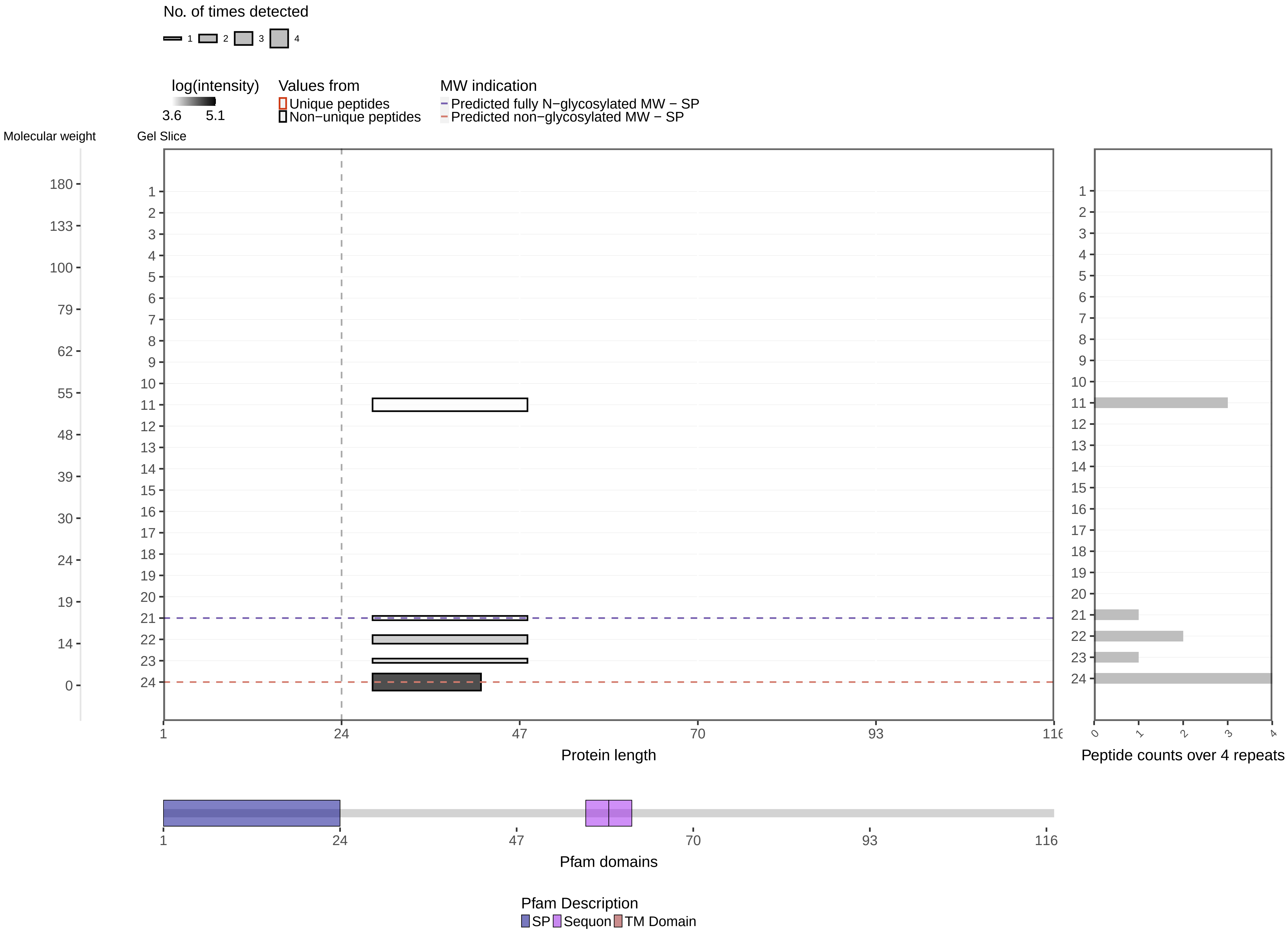


Protein sequence:

CLARCMARCFLNASNDTLDESHSTSLEARNRVCNVGCSLGHCSEFLFQYITRSLELACQTAVRTIAWATML*

### NbL16g11060.1 uncharacterized_LOC107795773 [MW − SP = 10.45 kDa]

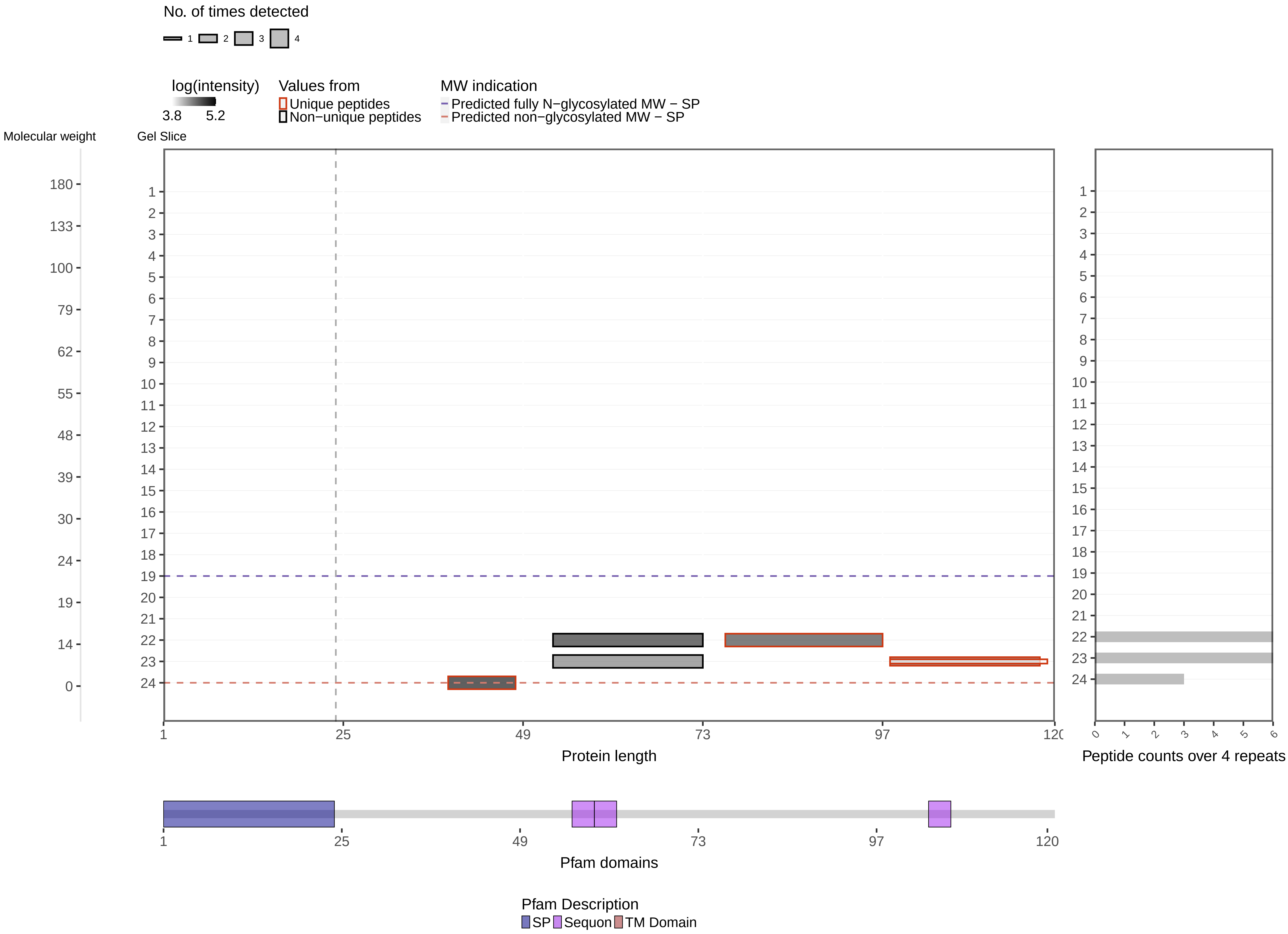


Protein sequence:

RCMARCFLNASNDTLDESHSTSLEARNRVCNVGCSLGHCSEFLFQYDNEKFGTCMSNCSENYCMGNNALEKA*

### NbL16g10770.1 probable_plastid−lipid−associated_protein_11_chloroplastic [MW = 19.66 kDa]

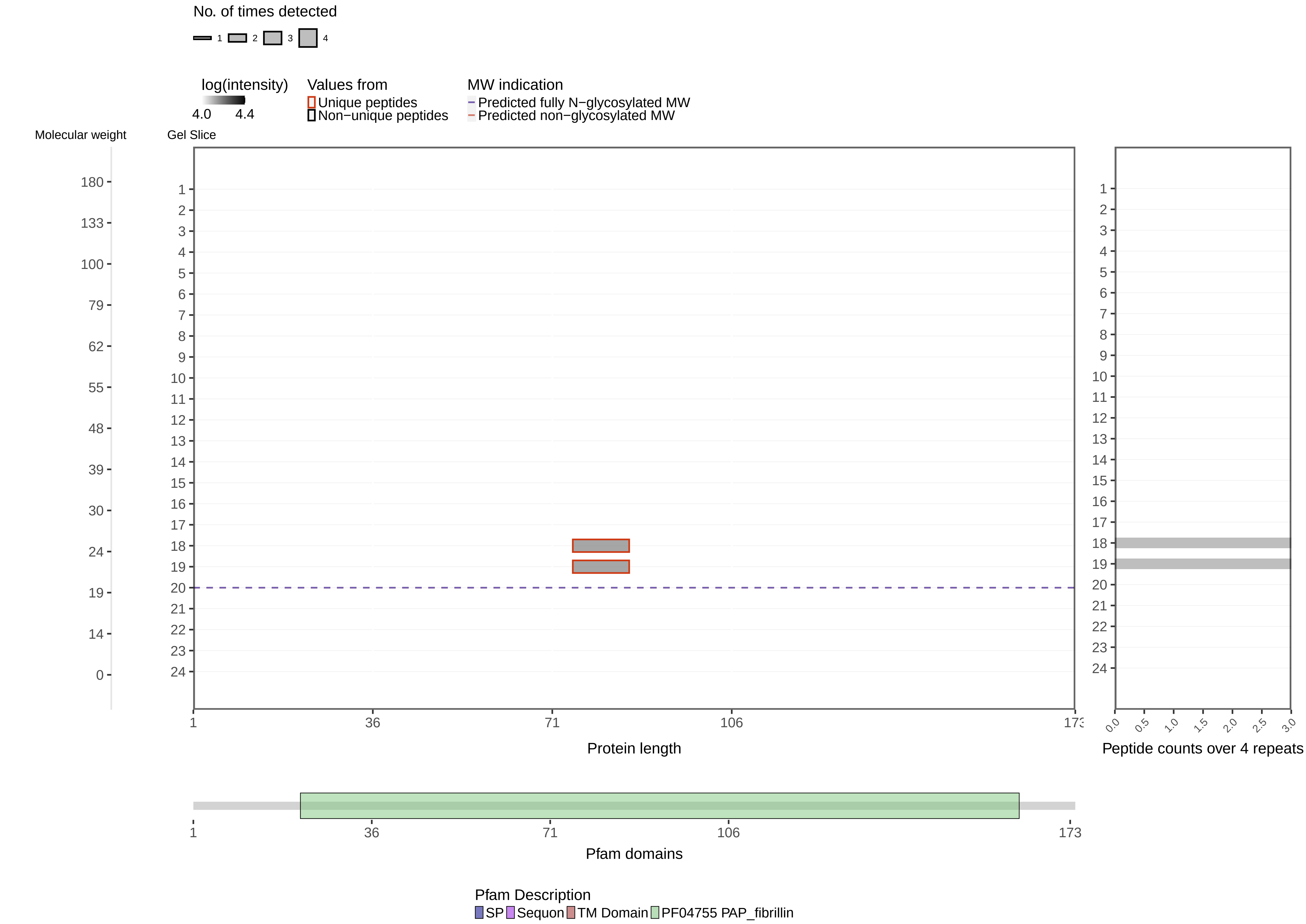


Protein sequence:

GDVLQVIDVEKGTLNNVITFPPDGVFFVRYTIDVASSQRVNFRFTSAVLRGKNWEFPLPPFGQGWFETVYLDDEIRVVKDIRQDYLVVERAPYTWKE*

### NbL16g10490.1 peroxisomal_membrane_protein_11C [MW = 25.73 kDa]

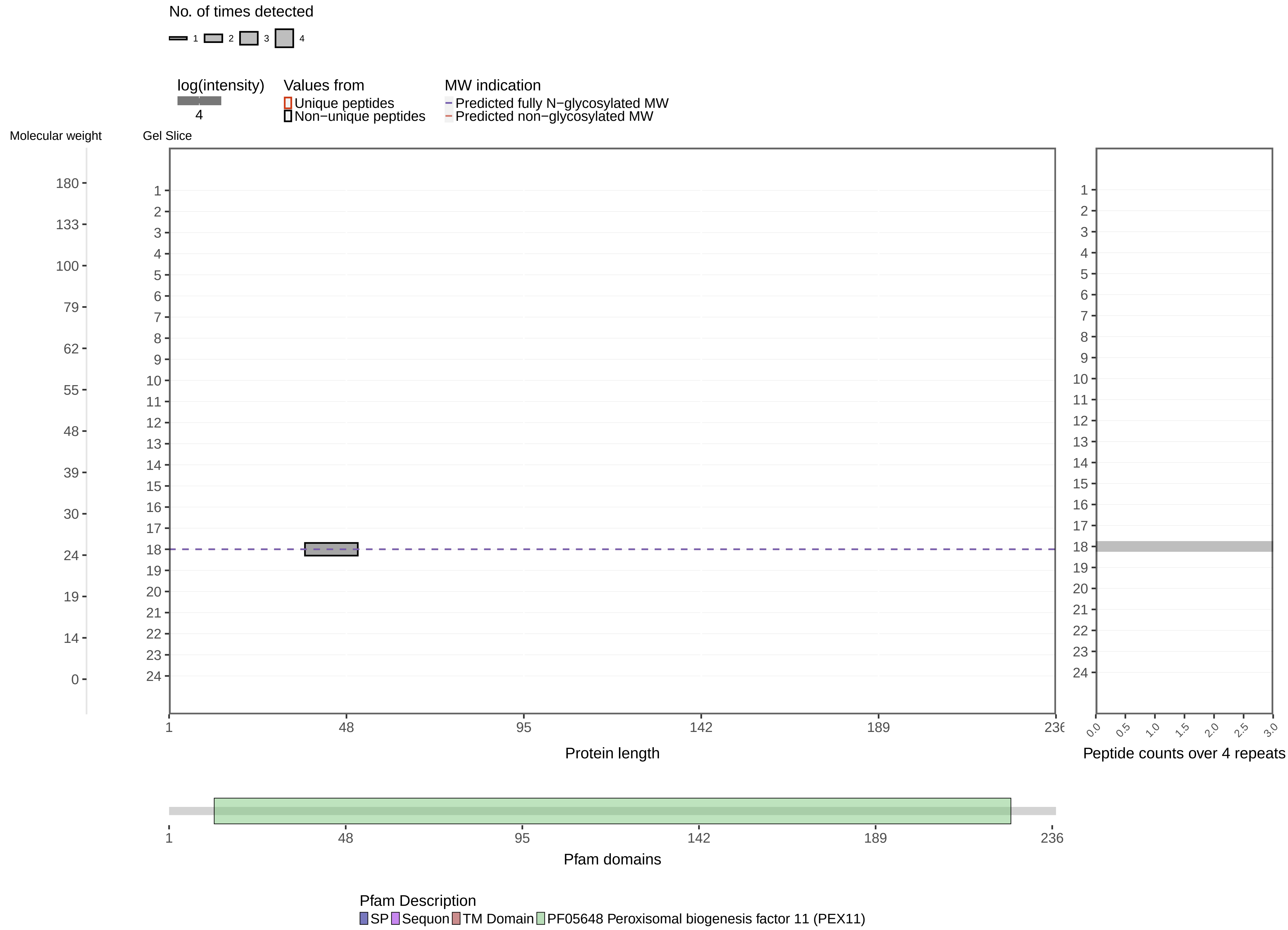


Protein sequence:

WLGRTGIYKDKERTELIGRISLFCWMGSSICTALVEIGELGRLSGSLKKLEKGLKNTDKHKNEQYQSKIQKSNERSLALIKAGMDIVVAVGLLQLAPKKVTPRVTGAFGFVSSLISCYQLLPSPPKIKAS*

NbL16g10300.1 uncharacterized_LOC109231095 [MW = 30.26 kDa]

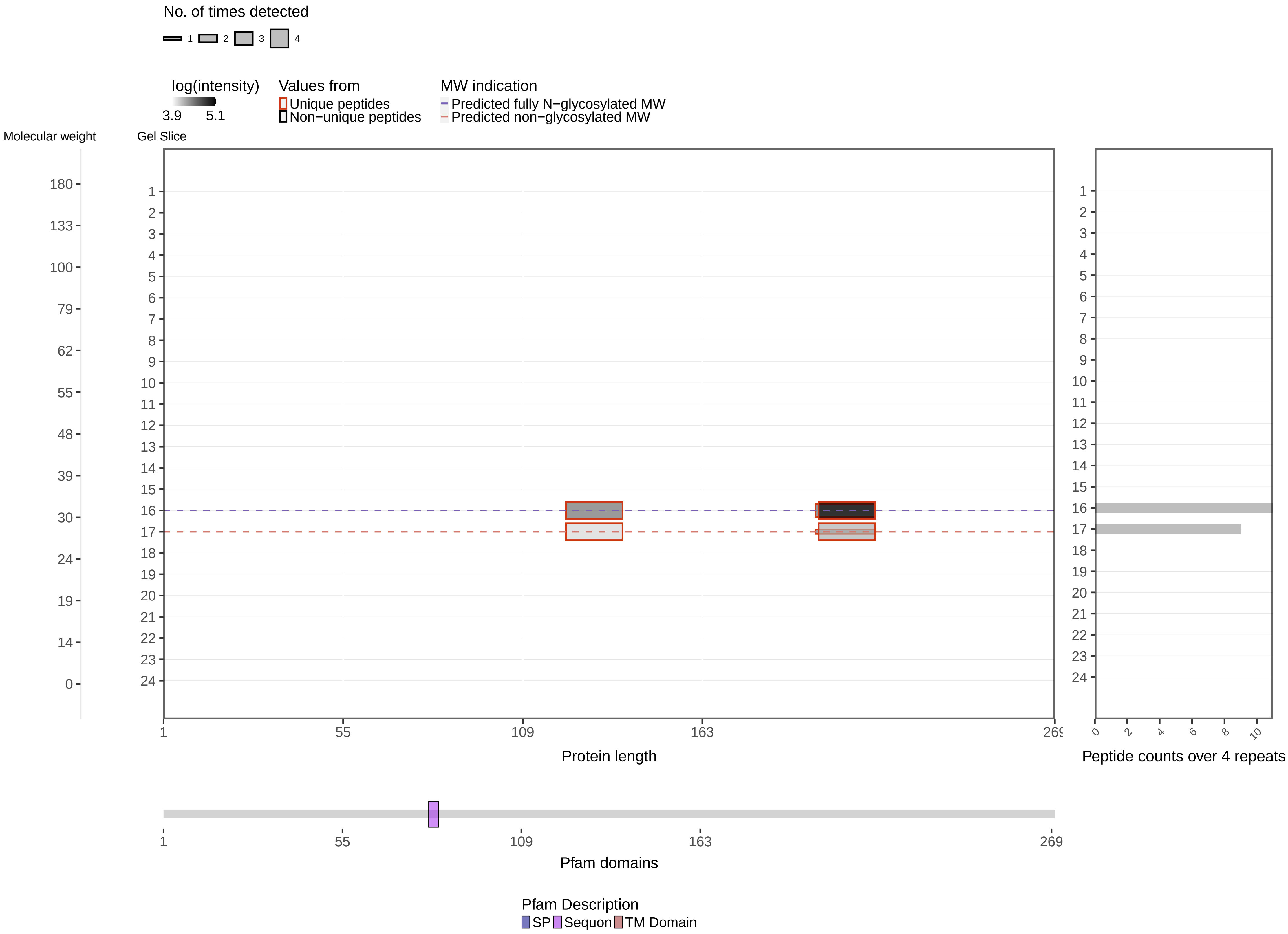


Protein sequence:

:VRDEWMDNDLATWIGANRFYPGVPDALKFASSKLYIVTTKQSRFADALLRELAGITIPAERIYGLGTGPKVKVLKKLQEMPEHQGLTLHFVEDRLATLKNVIKESELDGWNLYLGDWGYNTQKEREEAASIPRIRLLGLSDFSNKLK*

### NbL16g10200.1 putative_UDP−rhamnose:rhamnosyltransferase_1 [MW = 52.96 kDa]

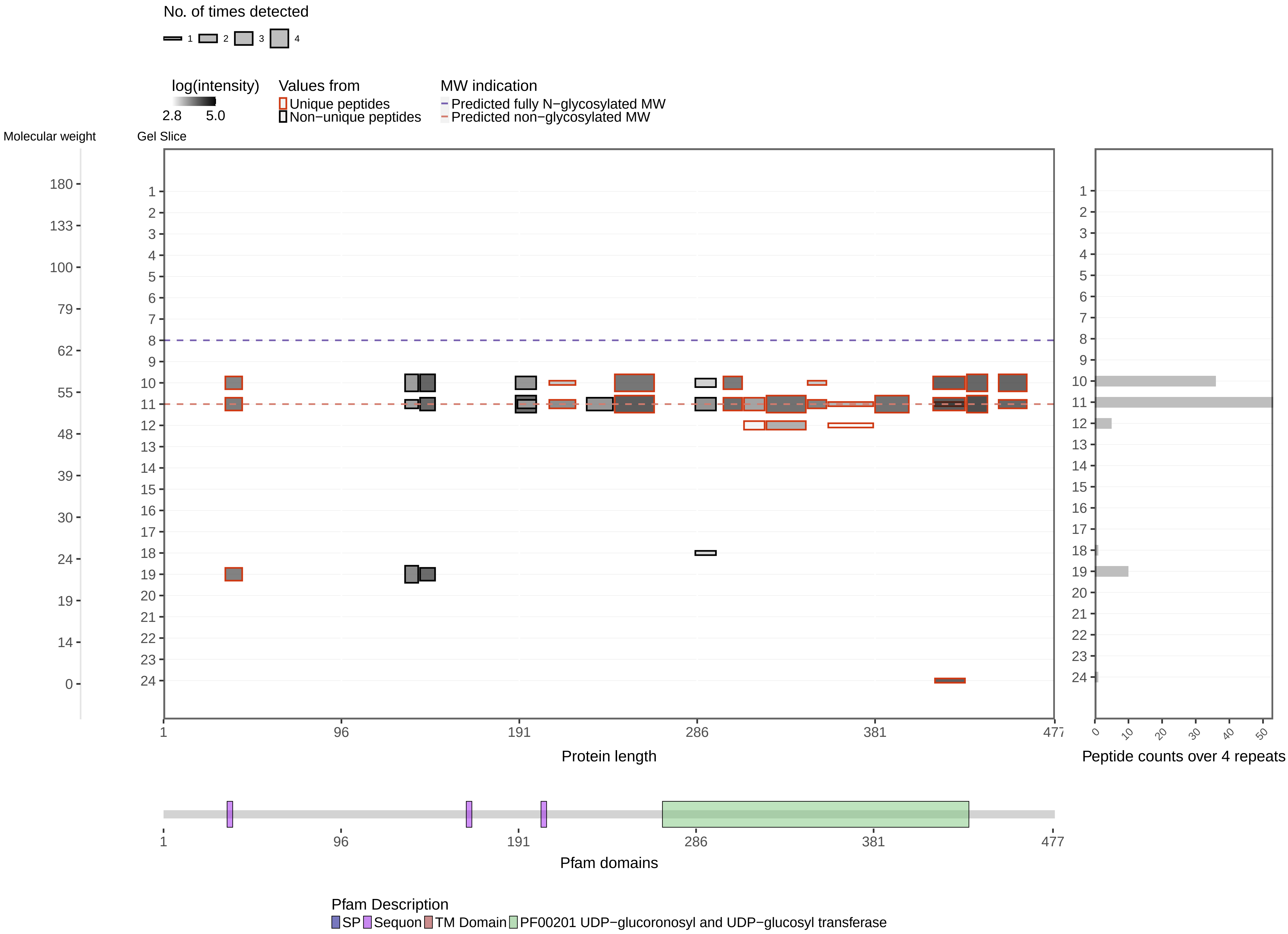


Protein sequence:

MEFEGEYLDTYNKVAGKPVIPIGLLPPEELPADERTLDCQPNWQKICKWLDEQKPRSVLFVGFGSECKLSKNQVYEIANGVKLSGLPFFWILQRPSWALNDIDALPSGFGASTEGRGLVHIGWAPQKEILAHPSIGGSLFHGGWGSTIETLRYGHVLVVLPFVFDQGLNARLLVEKGVAIEVKRNEEDGSFSGNDIATSLREAMVSEDGEELRARAGKADGIFGDQNLHDSYVKNFVEYMRINGEMKV*

### NbL16g10190.1 UDP−glycosyltransferase_91C1−like [MW = 52.56 kDa]

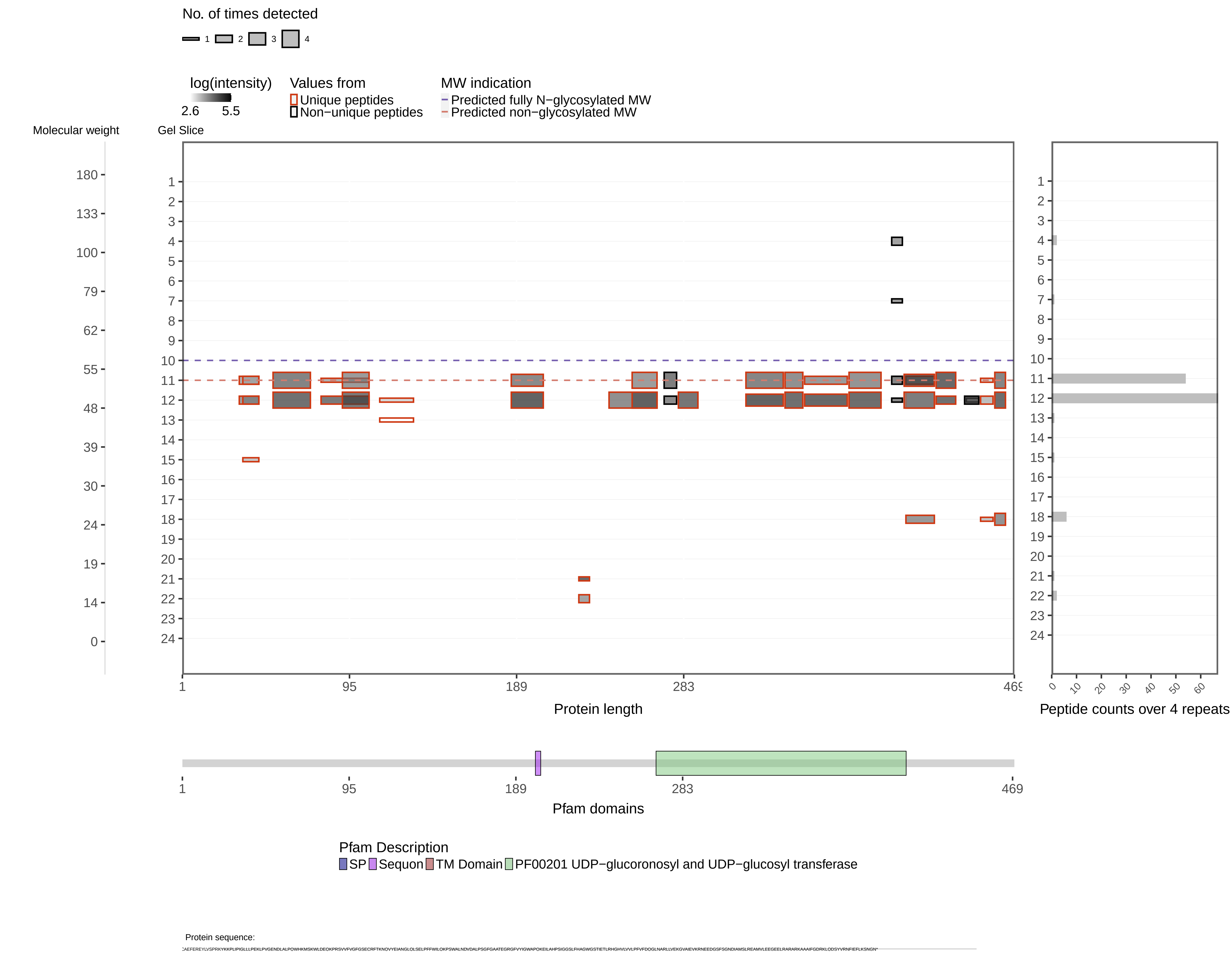

NbL16g10160.1 uncharacterized_LOC109218791 [MW = 12.48 kDa]

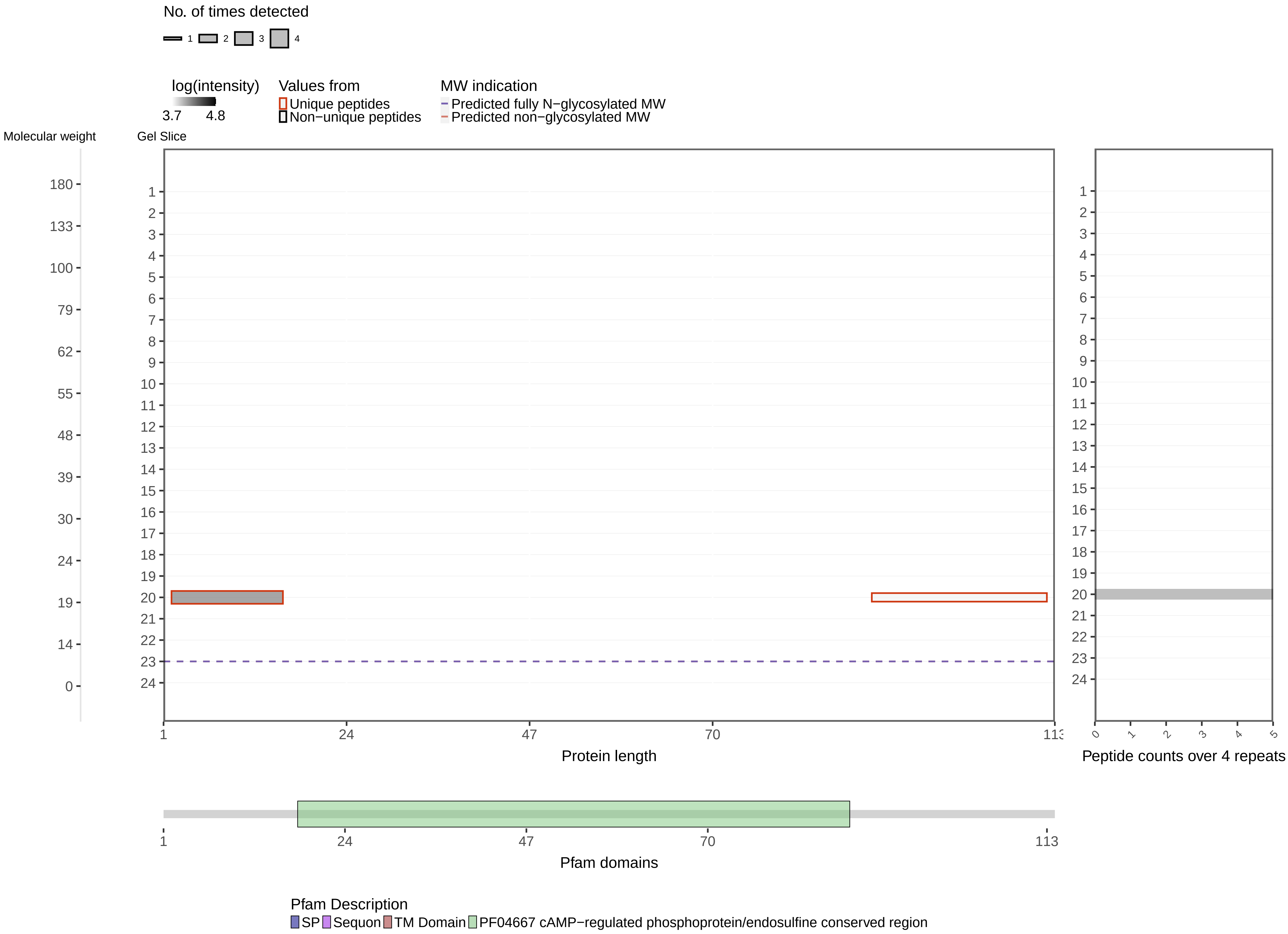


Protein sequence:

KDHERAYFDSADWALGKQGVAKPKGPLEALRPKLQPTNQQTRYRKSPYAPSEGEDGSNAPPEDPTANE*

NbL16g09900.1 rhodanese−like_domain−containing_protein_4_chloroplastic [MW = 50.87 kDa]

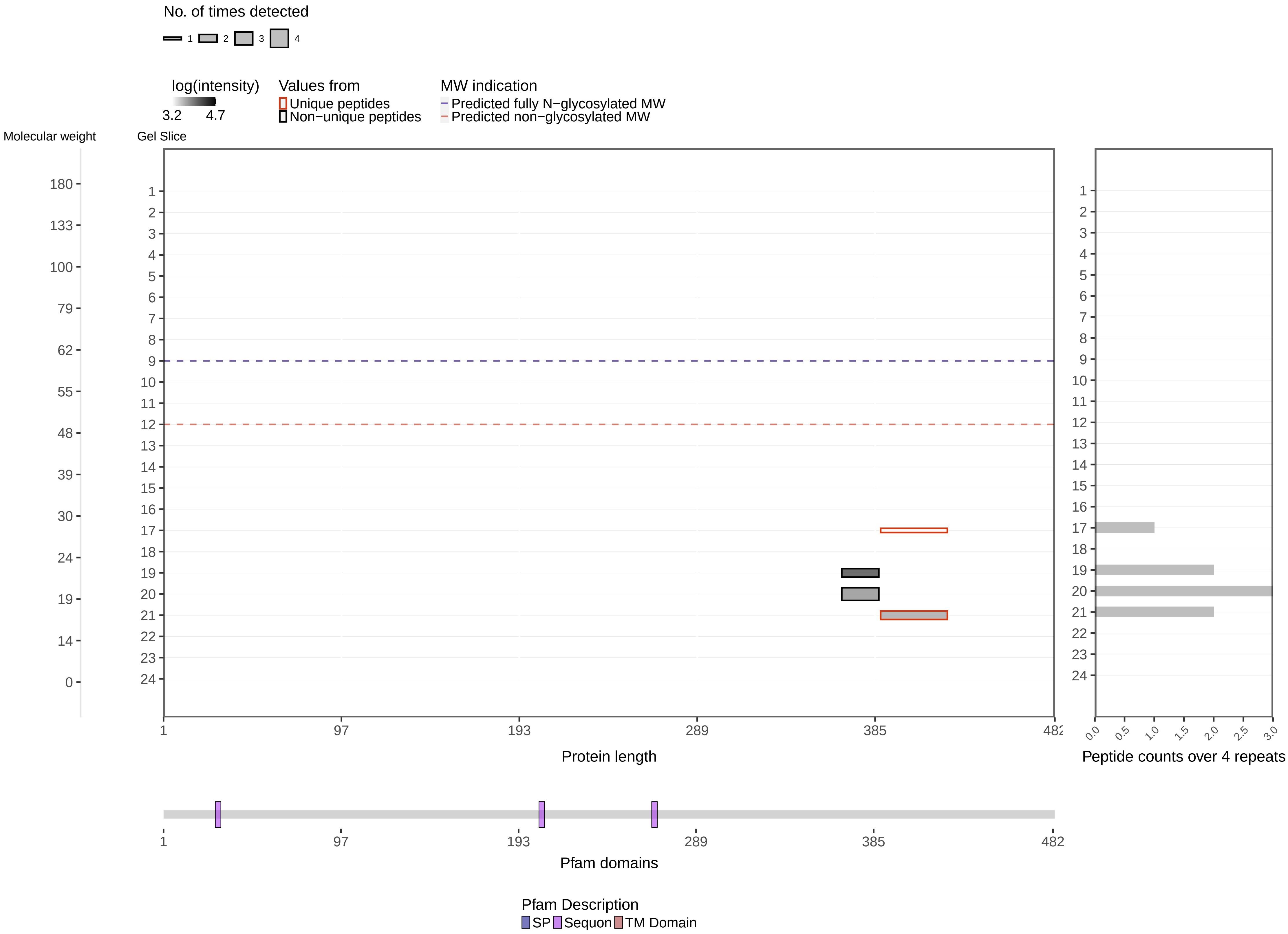


Protein sequence:

GFKAAYAIKDGAEGPRGWKNSGLPWILPKKTLSLDLGNLSDALDGVPVDGSDAIAVGLGVAAATVLGFLVFSEVETILQVLGSAALIQLVSTKLLFAEVRKKTLQQVDAFVTTNIAPKELVGDIKQIGMALLPTPVTSKSLPAPAETTPEINSVPKPEVKVEALTGISKPLSPYPSYSDLKPPTSPMPSQPSGSAEKTEAVSKVELSAESTPEINSVPKLEVKAEALAGSSKSLSPYPNYPDLKPPTSPTPSQA*

NbL16g09580.1 serine/threonine−protein_phosphatase_2A_65_kDa_regulatory_subunit_A_beta_isoform [MW = 65.57 kDa]

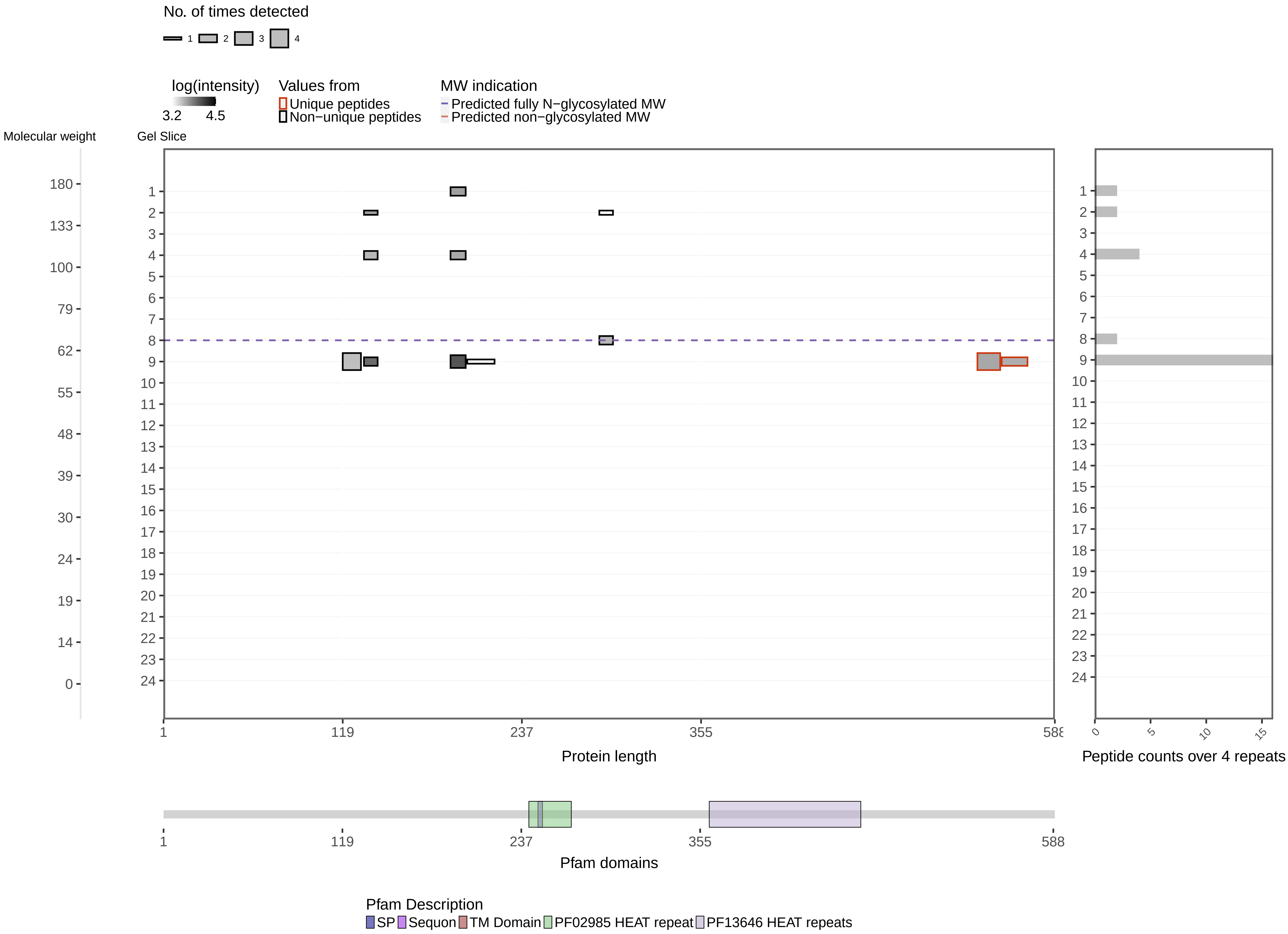


Protein sequence:

TRTDLVPAYVRLLRDNEAEVRIAAAGKVTKFCRILSPELAIQHILPCVKELSSDSSQHVRSALASVIMGMAPVLGKDATIEHLLPIFLSLLKDEFPDVRLNIISKLDQVNQVIGIDLLSQSLLPAIVELAEDRHWRVRLAIIEYIPLLASQLGIGFFDDKLGALCMQWLQDKVYSIRDAAANNLKRLAEEFGPEWAMQHIIPQVLDMTTSPHYLYRMTILRAISLLAPVMGSEITCSKLLPVVITATKDRVPNIKFNVAKVLQSLIPIVDHSMVEKTIRPSLVELAEDPDVDVRFYANQALQSID

### NbL16g09440.1 pyruvate_dehydrogenase_E1_component_subunit_alpha_mitochondrial [MW = 42.95 kDa]

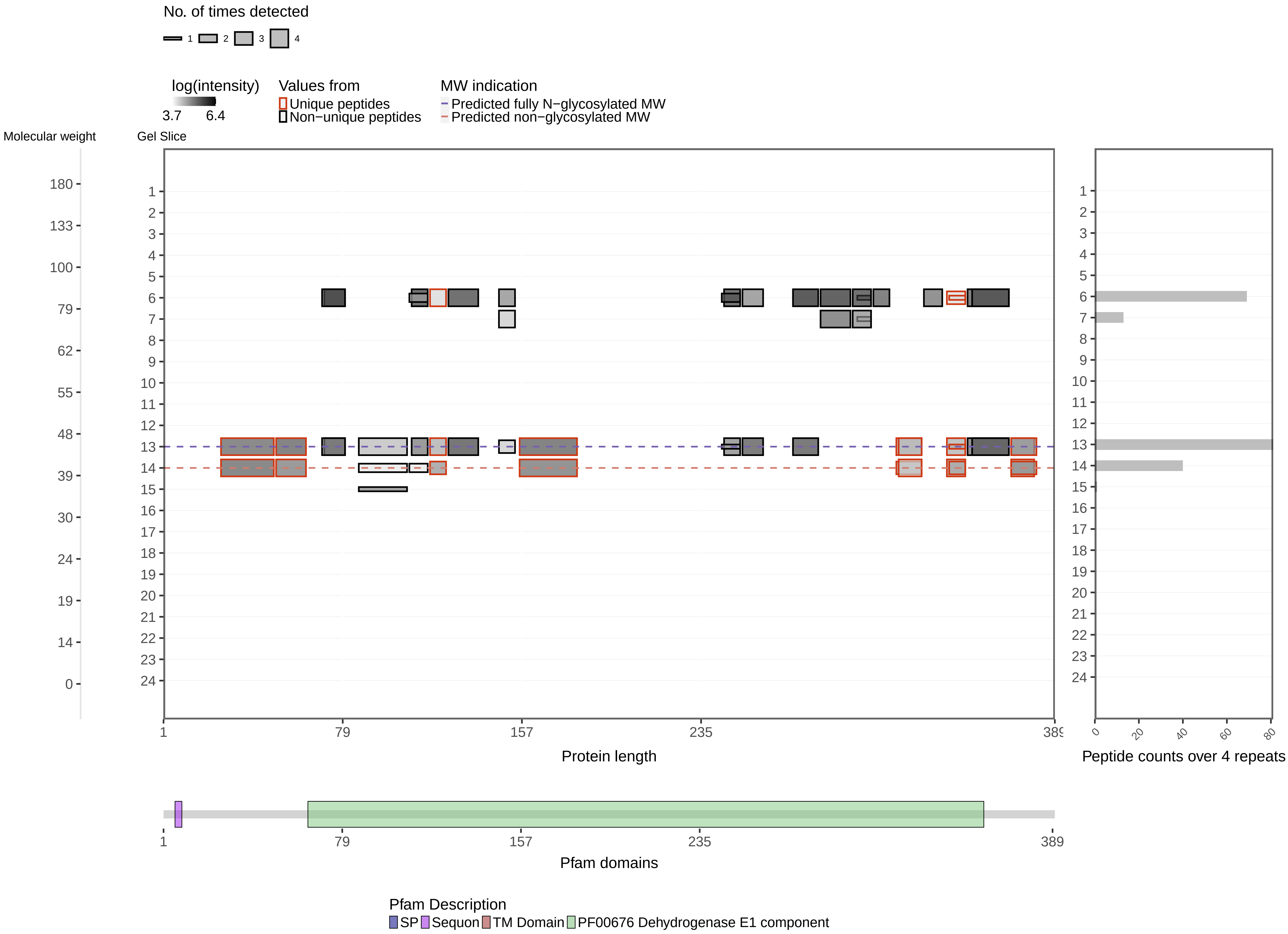


Protein sequence:

SKEEHVTFAMYGDGAANQGQLFEALNMAALWDLPAILVCENNHYGMGTAEWRAAKSPAYYKRGDYVPGLKVDGMDALAVKQACKFAKEHALKNGPIILEMDTYRYHGHSMSDPGSTYRTRDEISGVRQERDPIERIRKLVLAHDIATEKELKDIEKEKRKVVDEAIAKAKESPMPDPSELFTNVYVKGFGVESFGADRKEVRTALP*

### NbL16g09420.1 serine/threonine−protein_phosphatase_PP2A−2_catalytic_subunit−like [MW = 35 kDa]

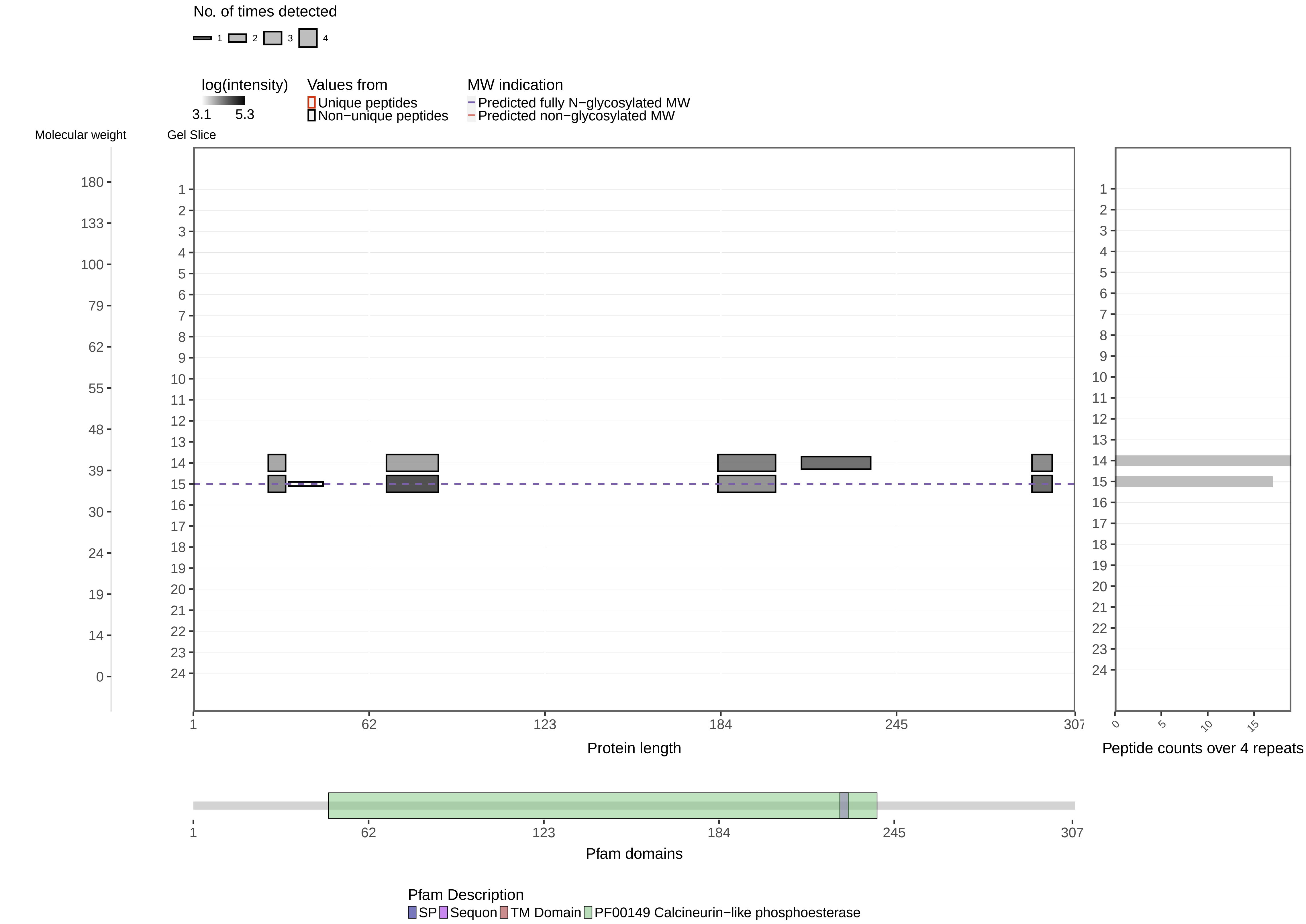


Protein sequence:

'KYFTDLFDYLPLTALIESQIFCLHGGLSPSLDTLDNIRSLDRIQEVPHEGPMCDLLWSDPDDRCGWGISPRGAGYTFGQDIASQFNHTNGLTLISRAHQLVMEGFNWCQDKNVVTVFSAPNYCYRCGNMAAILEIGENMEQNFLQFDPAPRQIEPDTTRKTPDYFL*

NbL16g09180.1 26S_proteasome_non−ATPase_regulatory_subunit_11_homolog [MW = 47.13 kDa]

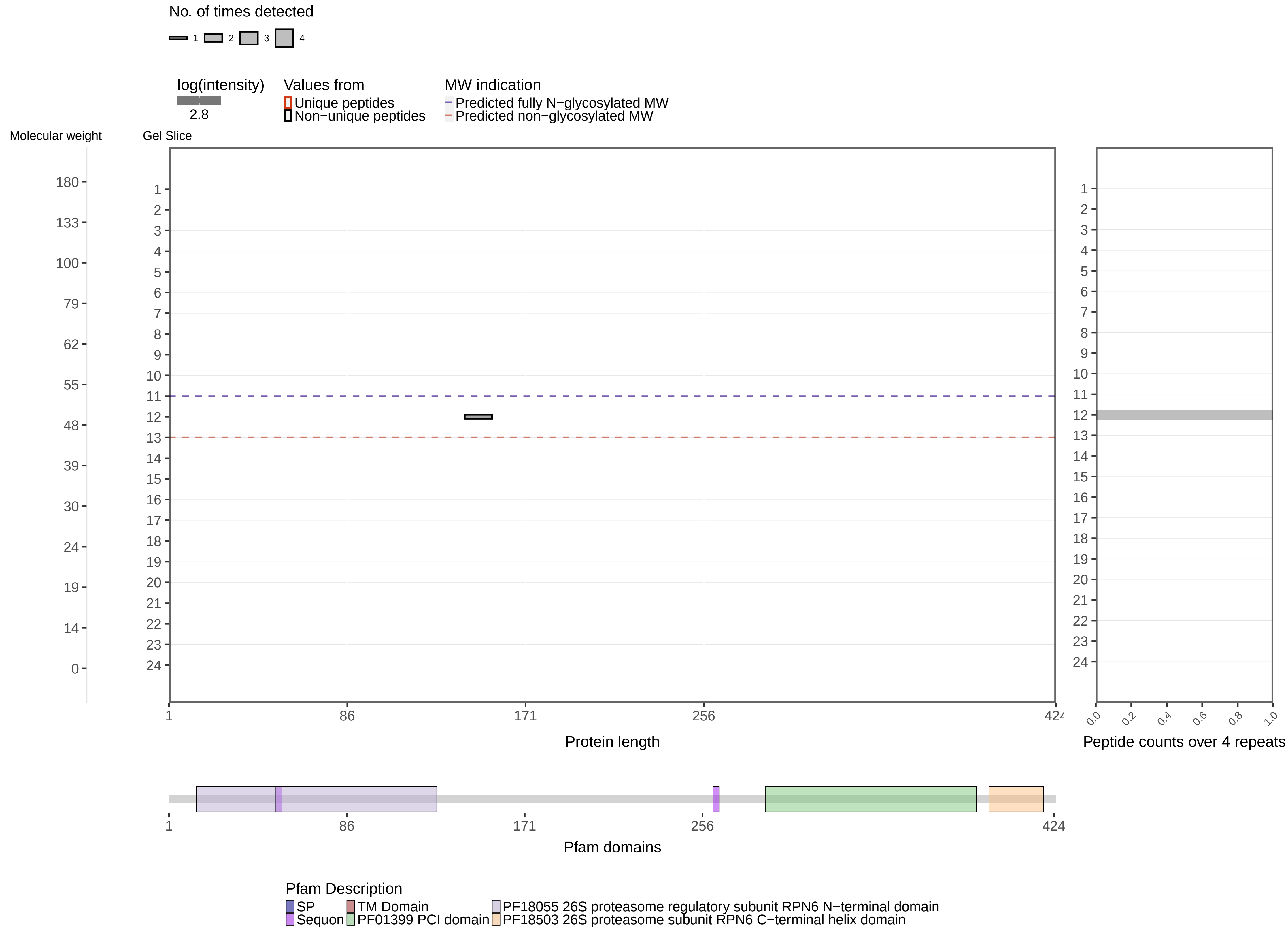


Protein sequence:

VPPAQQGSIDLQSGILHAEEKDYKTGYSYFYEAFEAFNALEDPQAIYSLKYMLLCKIMVNQSDDVAGITSSPKVGLQYQGPEVDAMKAVADAHSKRSLKLFETALRNFKAQLDEDPIVHRHLSSLYDTLLEQNLCRLIEPFSRVEIAHMAELIELPGDHVEKKLSQMILDKKFAGTLDQGAGCLIIFEDPKPDAIYPATLETVSNMGKVVDSLFVRSSRIIA*

### NbL16g09160.1 eukaryotic_translation_initiation_factor_5A−2−like [MW = 17.35 kDa]

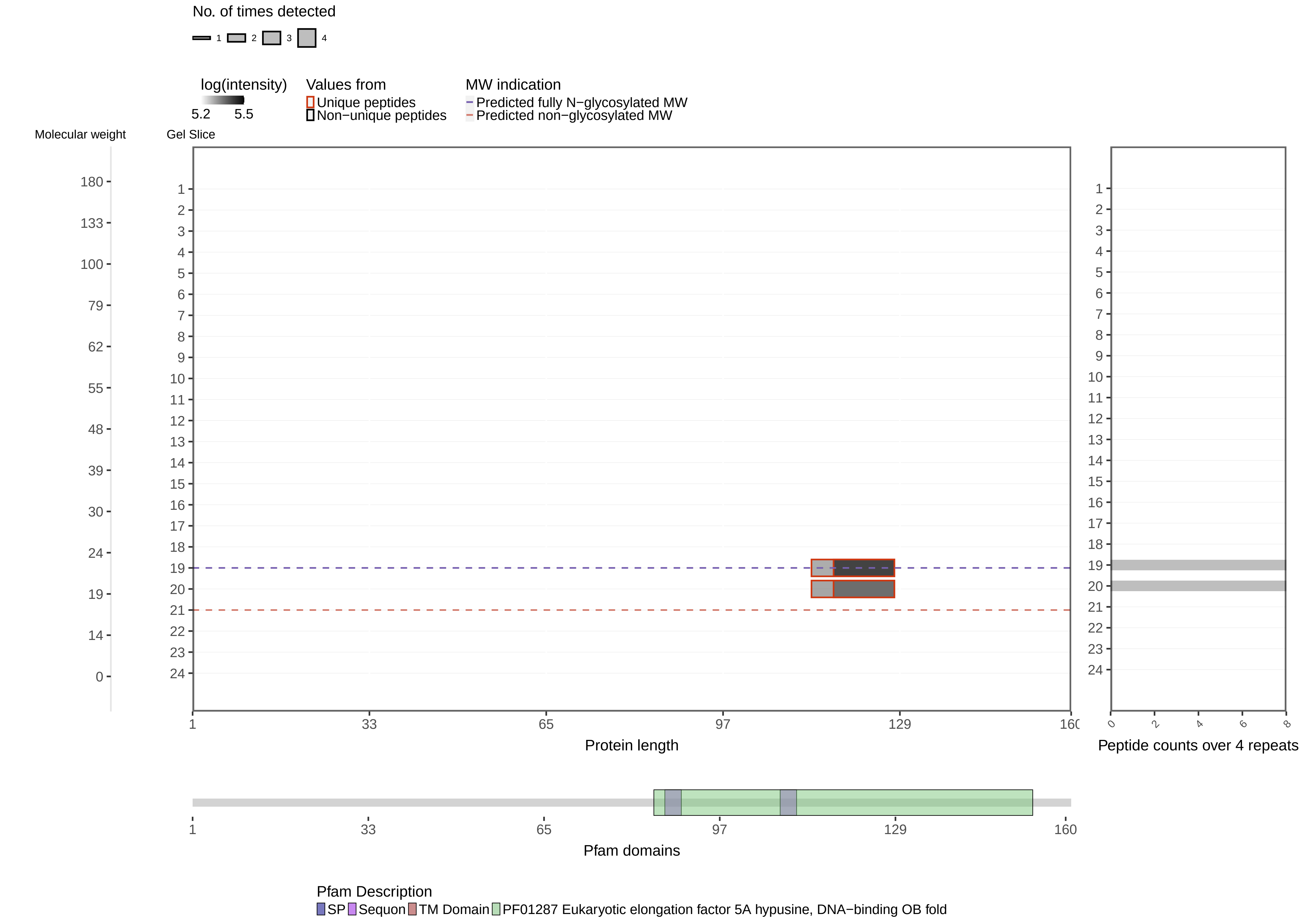


Protein sequence:

TGKKLEDIVPSSHNCDGPHVNRTDYQLIDISEDGFVSLLTENGSTKDDLRLPTDENLLTQIKDGFAEGKDIVVSVMSAMGEEQICALKDIGPK*

### NbL16g09080.1 PLASMODESMATA_CALLOSE−BINDING_PROTEIN_3−like [MW = 29.27 kDa]

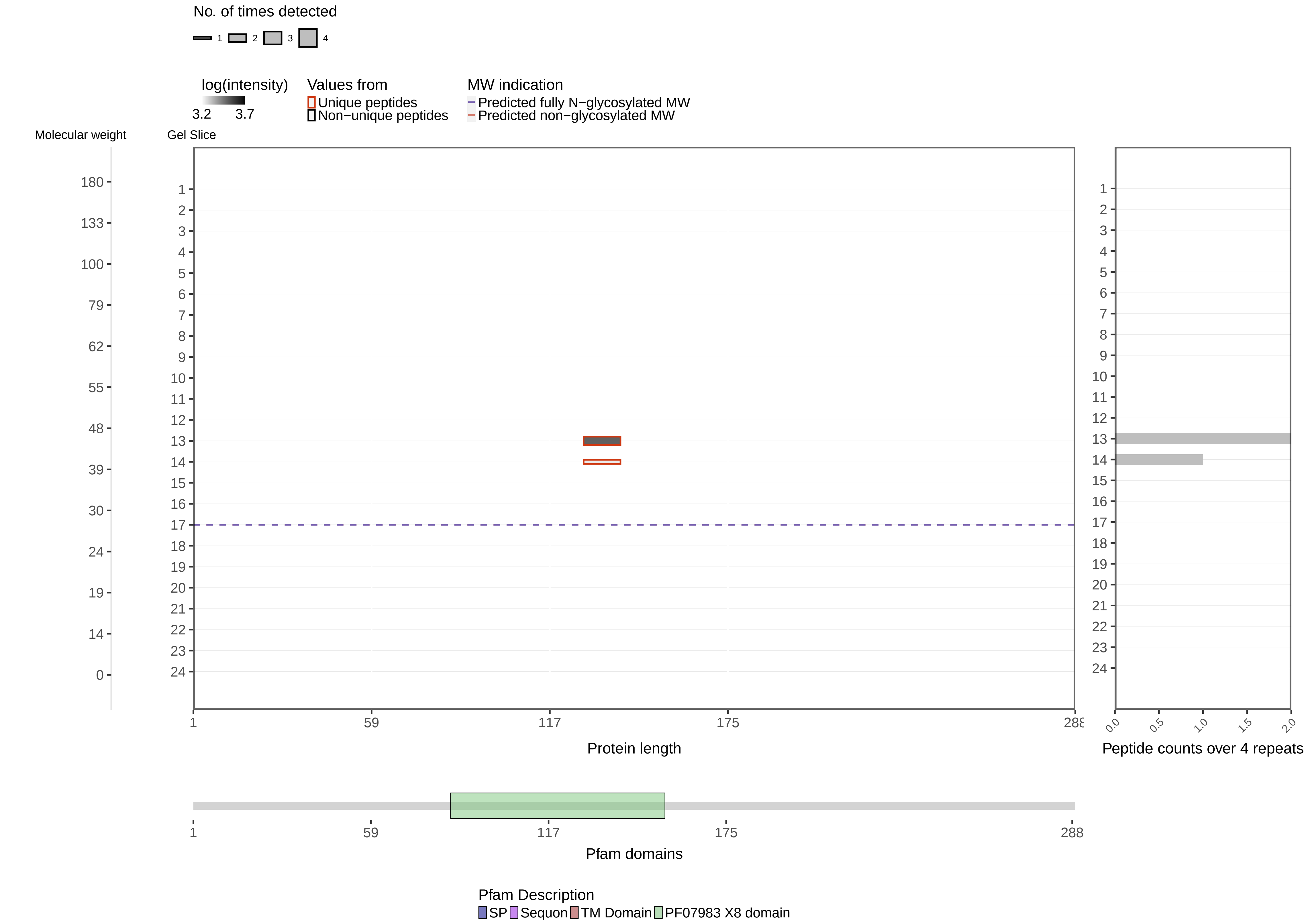


Protein sequence:

YAVNSYYQRKGAAGASCDFSGTATTTPNPPTTTGSGCVYQSTPGNTGGGTTTPTIAPPGSTTPTIAPPGTTTPTIAPPGTTIPTTAPPGTTTPGIGTGGTGTGIATTPGTNNQVFGFGPSGSGISSTDGNAAGHFKNSVFLTTALLAISLLICLRA*

### NbL16g09020.1 alpha−14−glucan−protein_synthase_[UDP−forming]_2 [MW = 41.36 kDa]

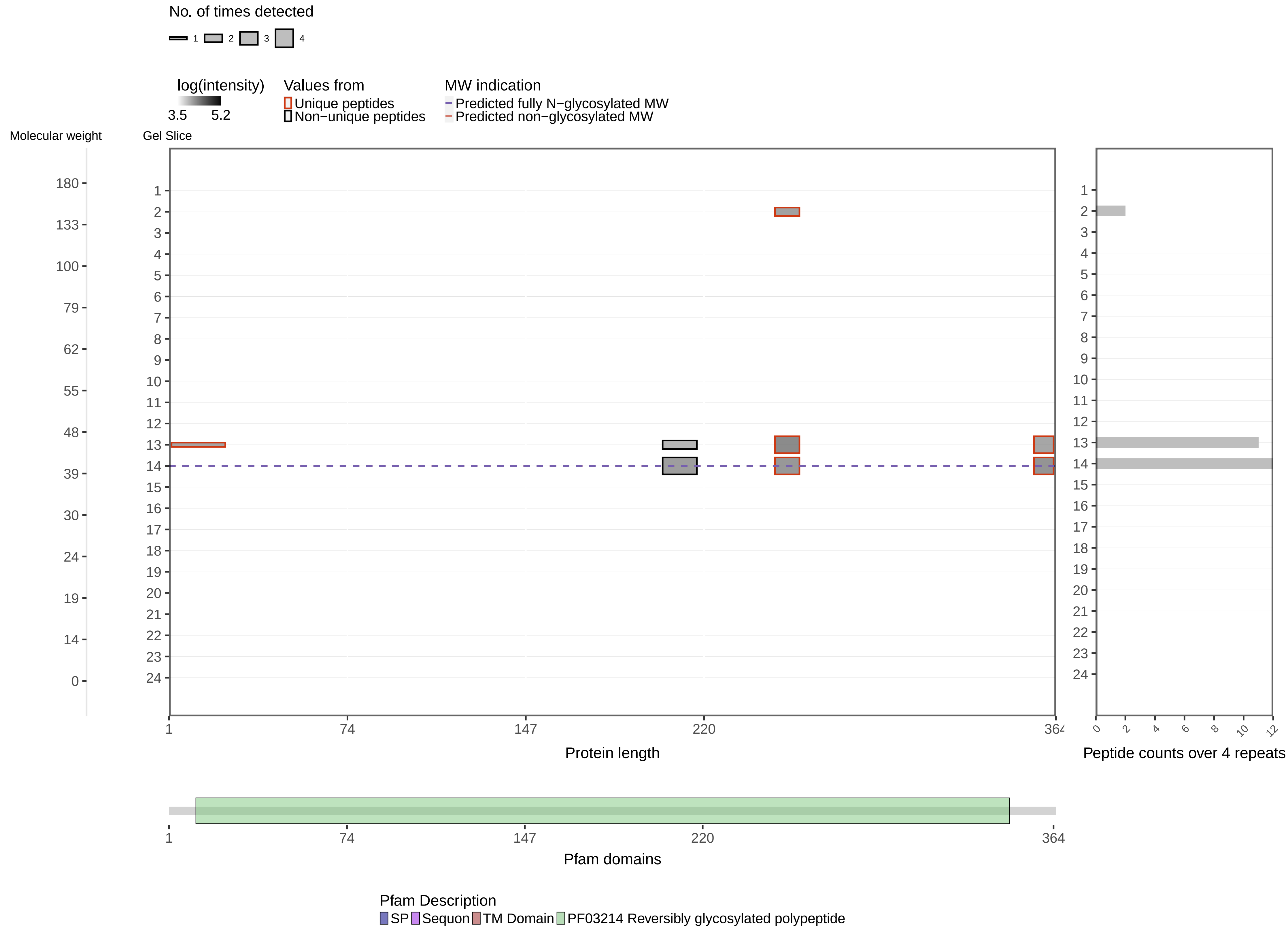


Protein sequence:

WLNIPDYDAPTQLVKPRERNTRYVDAVMTIPKGTLFPMCGMNLAFNRELIGPAMYFGLMGDGQPIGRYDDMWAGWCTKVICDHLGMGIKTGLPYIWHSKASNPFVNLKKEYKGIYWQEEIIPFFQSATLPKDCTSVQQCYIELSKQVKDKLSKVDPYFTKLADAMVTWIEAWDELNPTGEGLAKLSISNVPVK*

NbL16g08920.1 2−hydroxyisoflavanone_dehydratase−like [MW = 35.75 kDa]

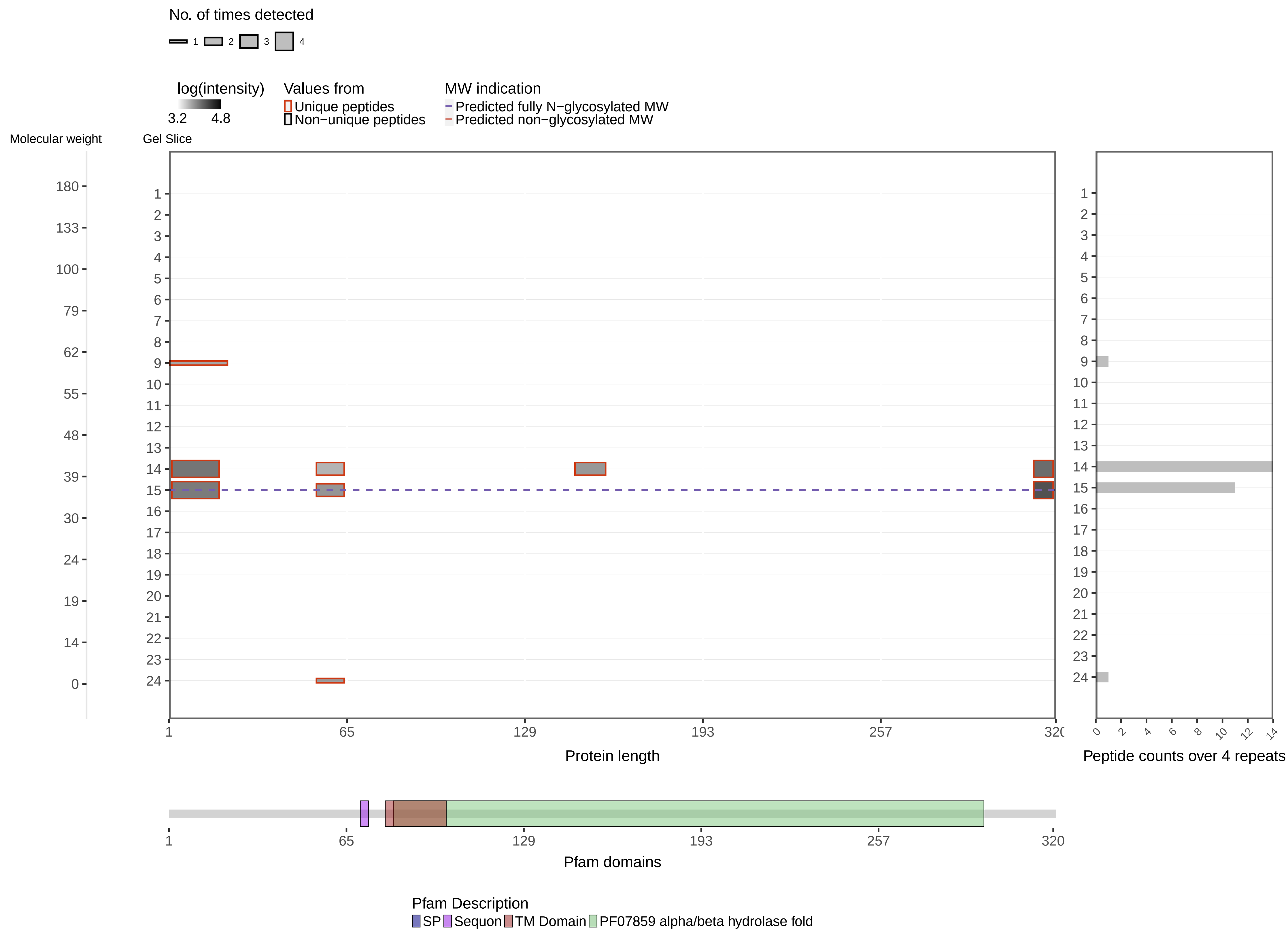


Protein sequence:

LWLTNYGDFNKIFLVGESGGANIVFNVVMRAGREKLNGDVKICGSILACPCFLIPHENVDVKDLMAYKLLTNVICPKIEPLDCPMINPLCKMAPSLSGLGCSKLFVCLAEKDEWGPGEMELQFVEGVKKSGWNGEFVFFVVEGEGHSFFILDPETEKARDLIKRFANFIQVK*

### NbL16g08900.1 protein_PHYLLO_chloroplastic [MW = 150.95 kDa]

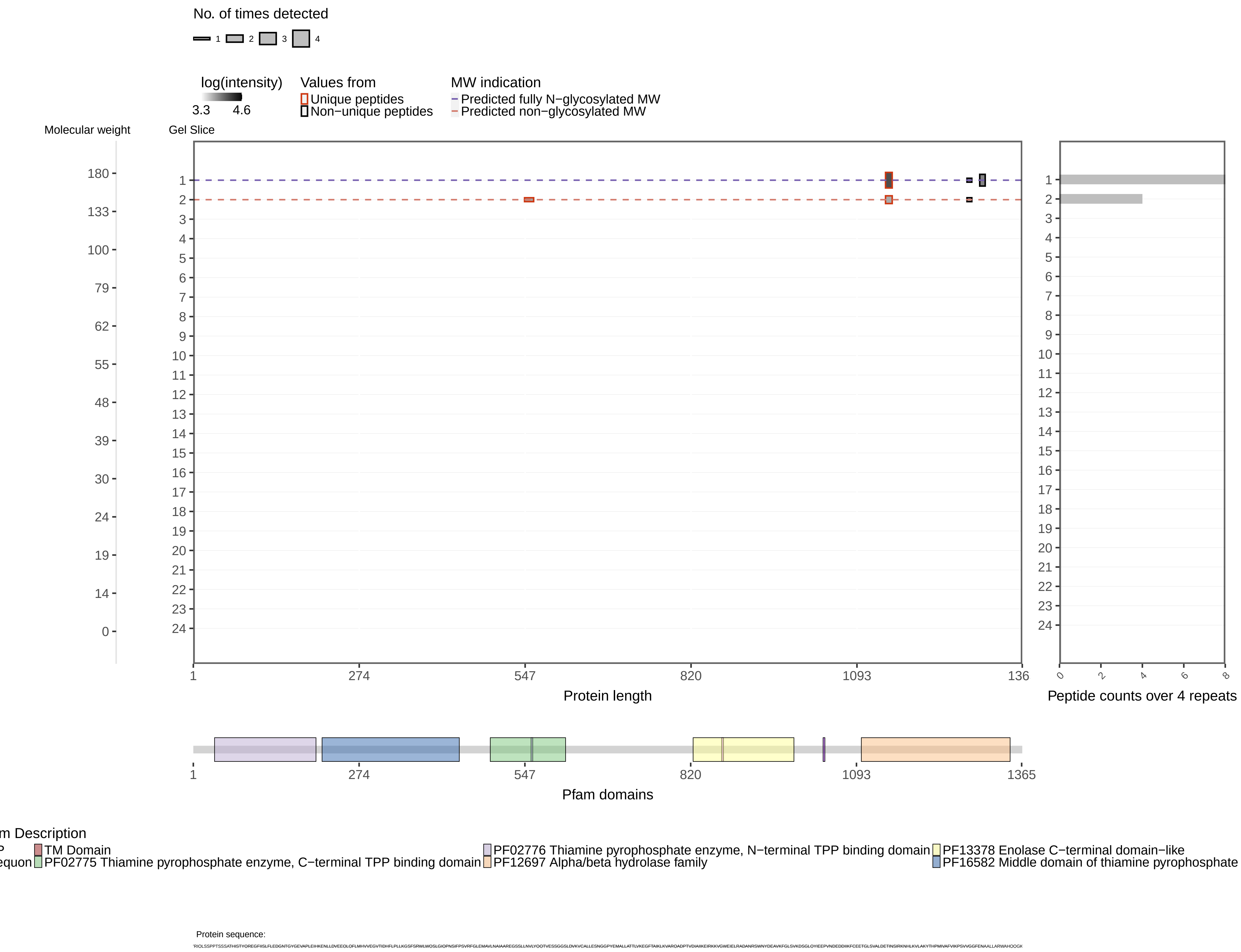

### NbL16g08880.1 6−phosphogluconate_dehydrogenase_decarboxylating_1 [MW = 53.31 kDa]

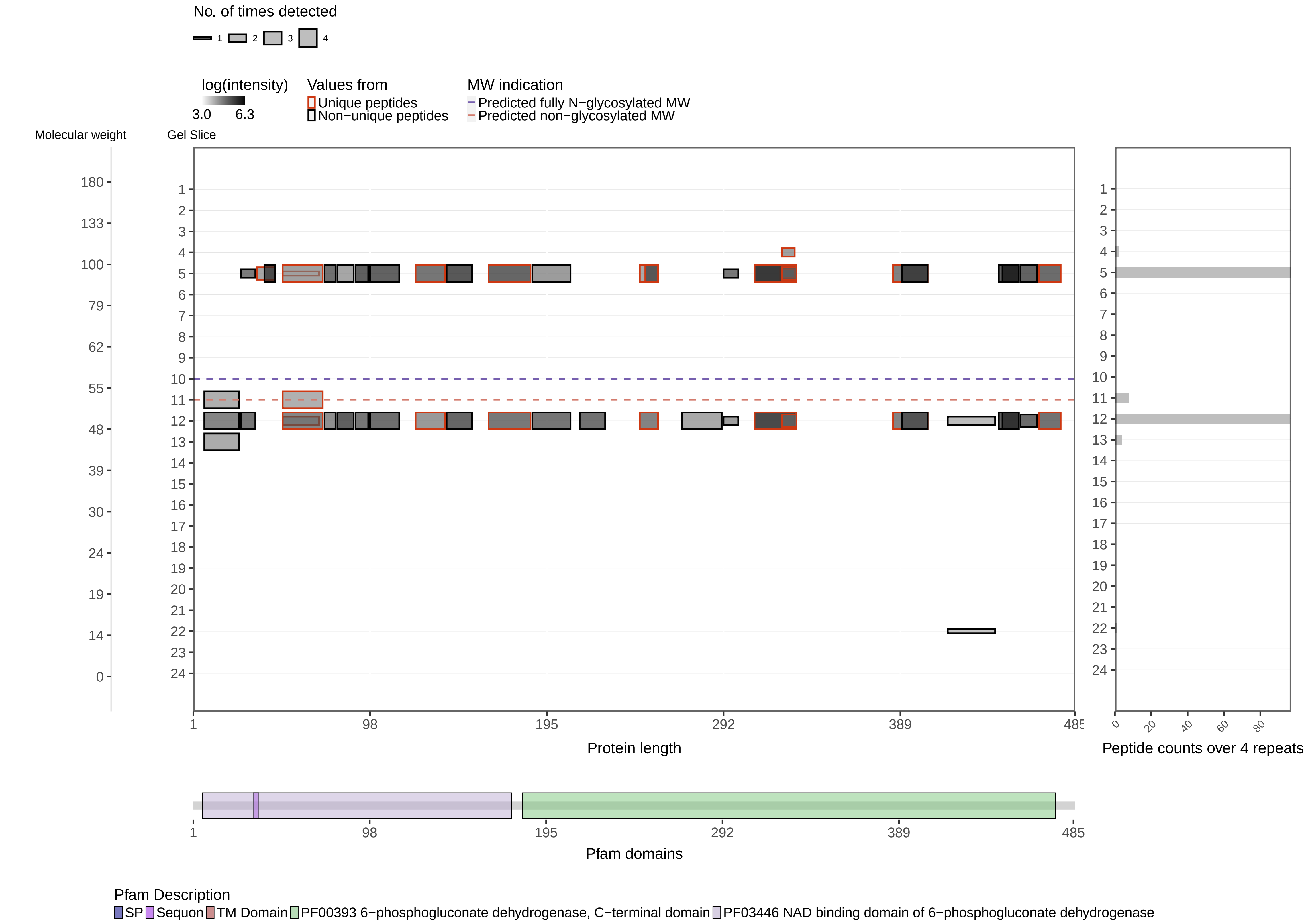


Protein sequence:

ELLSFLIEITADIFGIKDDKGDGYLVDKVLDKTGMKGTGKWTVQQAAELSVAAPTIASSLDSRFLSGLKDERVQAAKVFKASGVSDILVDQSVDKNQLIDDVRKALYASKICSYAQGMNLIRAKSVEKGWDLKLGELARIWKGGCIIRAIFLDRIKGAYDRNPDLANLLVDEEFAKEMVERQSAWRRVVCLAINSGISTPGMSSSLAYFDSYRRERLPANLVQAQRDYFGAHTYERTDVQGSYHTEWFKIARQSKN*

NbL16g08690.1 phosphoenolpyruvate_carboxylase_4 [MW = 117.38 kDa]

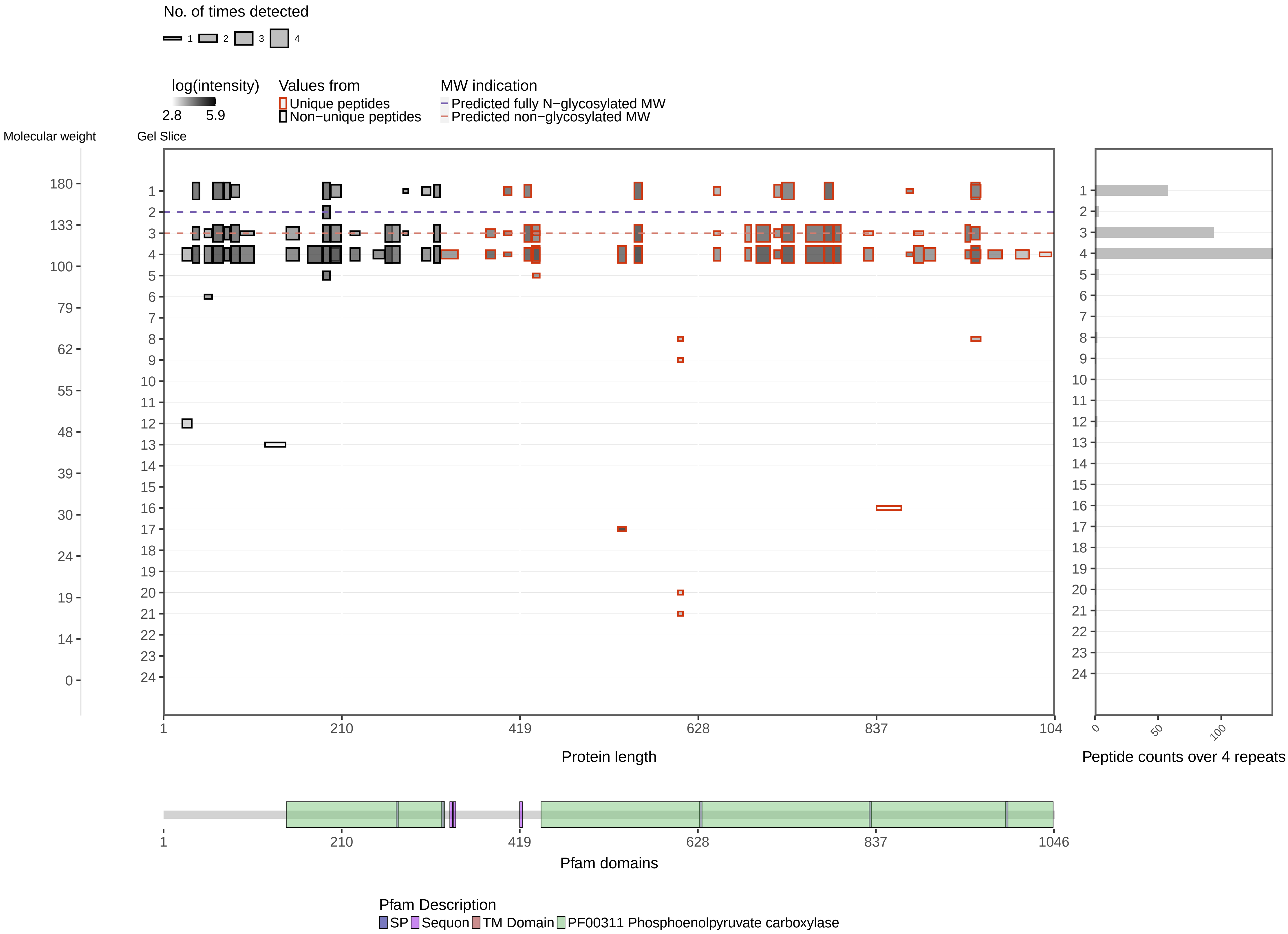


Protein sequence:

CYDSLQSCGSGVLADGRLADLIRRVATFGMVLMKLDLRQESGRHSEALDAITKYLDMGTYSEWDEEQKLDFLIKELKGKRPLVPPTIEVPPDVKEVLDTFKVAAELGSDSLGAYVISMASNASDVLAVELLQKDARLAAAGELGRPCPGGTLRVVPLFETVKDLREAGSVIRRLLSIDWYREHIIKNHDGHQEVMVGYSDSGKDAGRFTAAWELYKAQEDVVAACNEYGIKVTLFHGRGGSIGRGGGPTYLAIQSQPPGSVMGTLRSTEQGEMVQAKFGLPQMAVRQLEIYT

### NbL16g08610.1 ubiquitin_receptor_RAD23c−like [MW = 41.22 kDa]

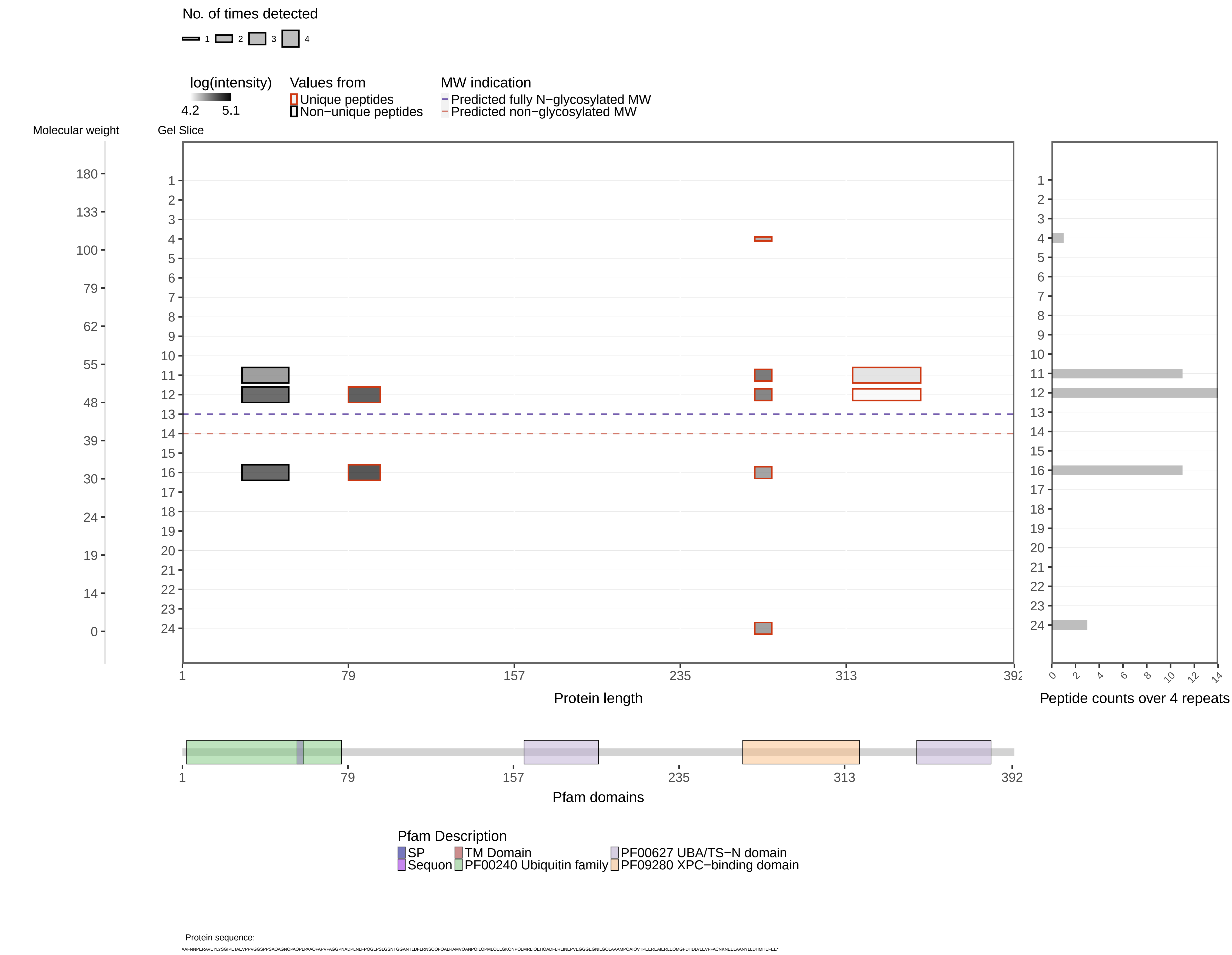

### NbL16g08600.1 alpha−xylosidase_1−like [MW − SP = 101.62 kDa]

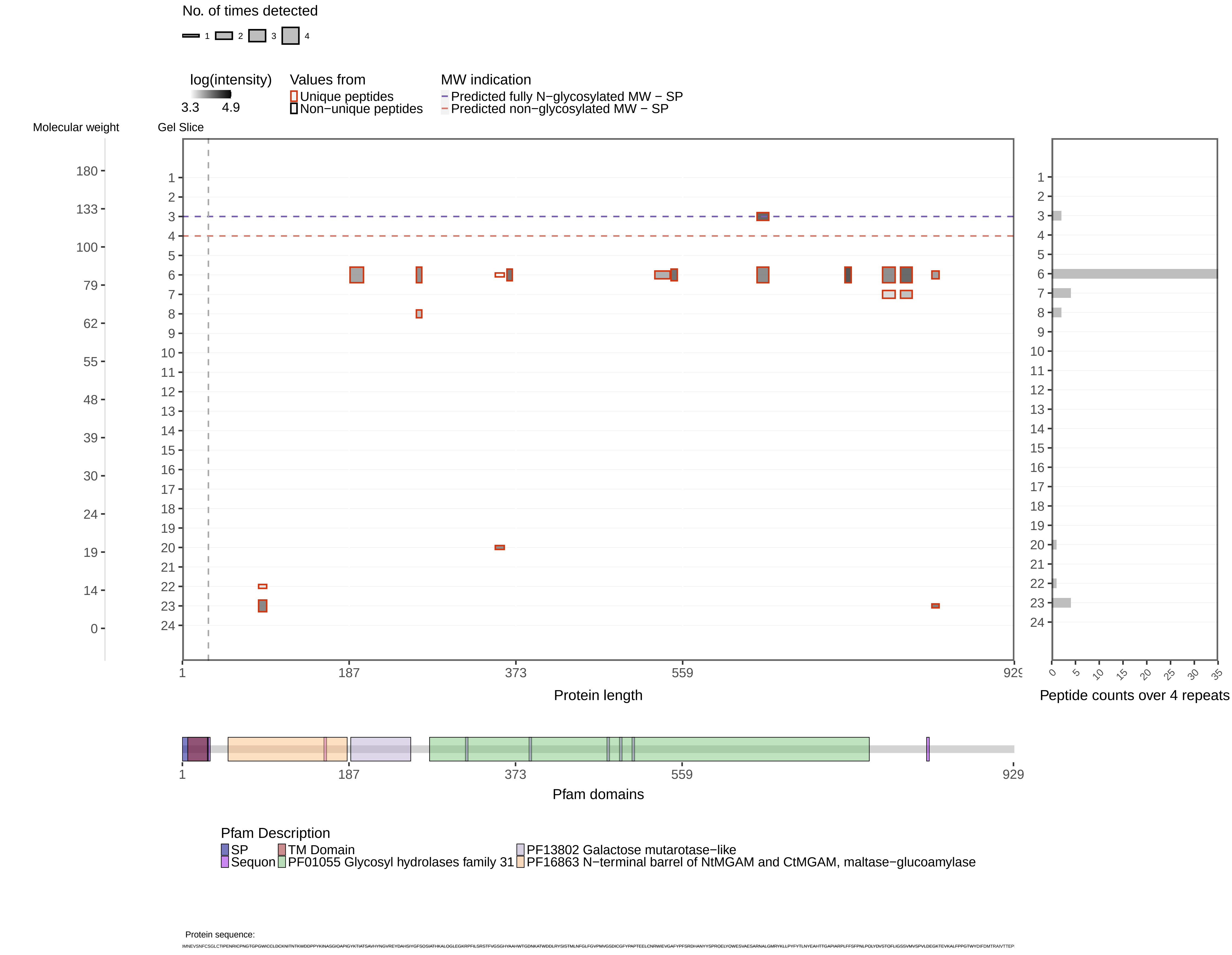

NbL16g08500.1 acyl−acyl_carrier_protein_thioesterase_ATL3_chloroplastic−like [MW = 24.14 kDa]

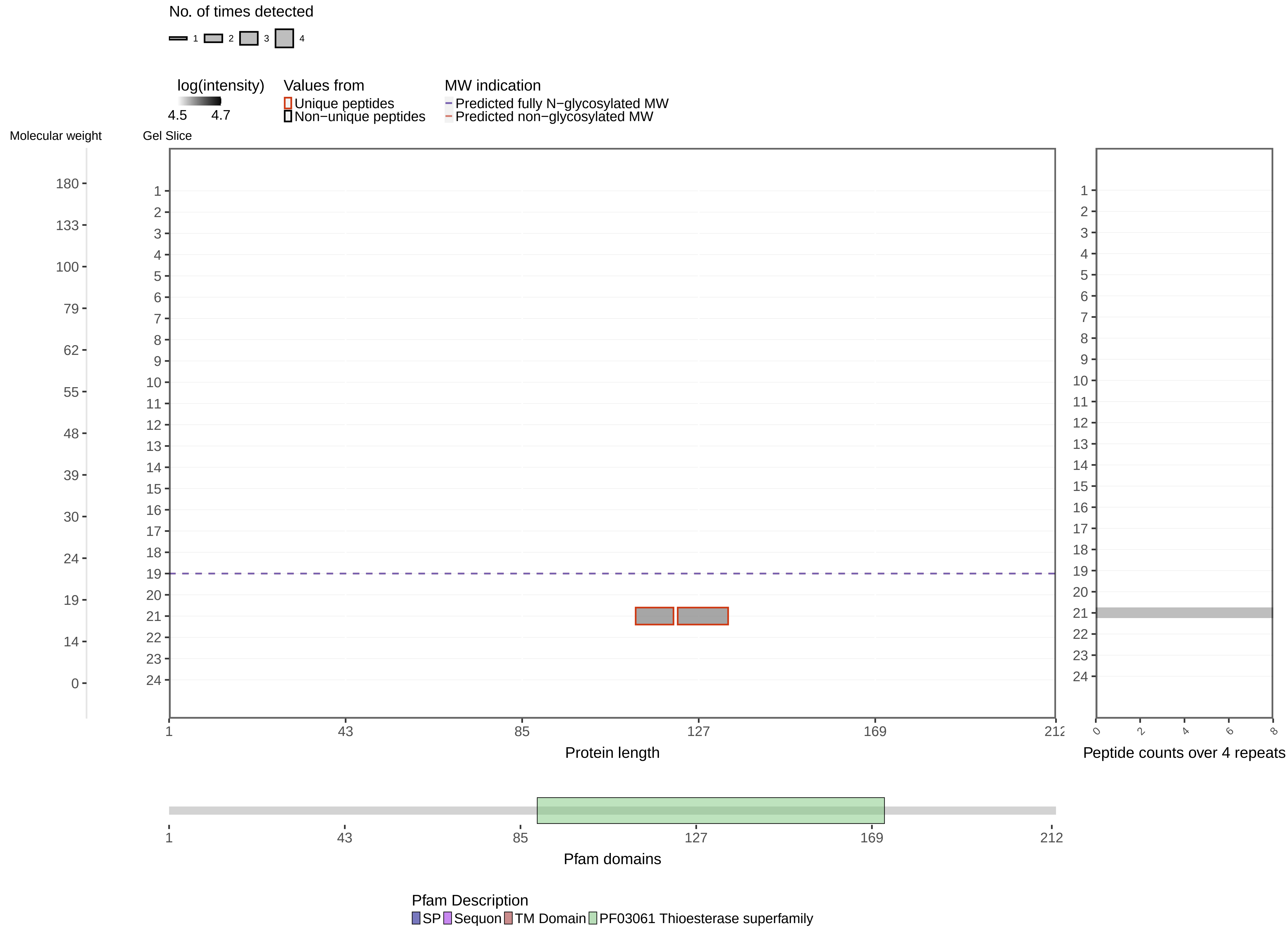


Protein sequence:

VNNAIYASYCQHCRHELLERLGISADTVARDGDALALTELSLKYLAPLRSGDRFVVKVRISDSSSVRLFFEHFIFKLPGQEPILEATGTAVWLNKNYRPVRIPSEFRSKYVQFLRQEASK*

### NbL16g08230.1 DNA−directed_RNA_polymerases_II_IV_and_V_subunit_10−like [MW = 11.84 kDa]

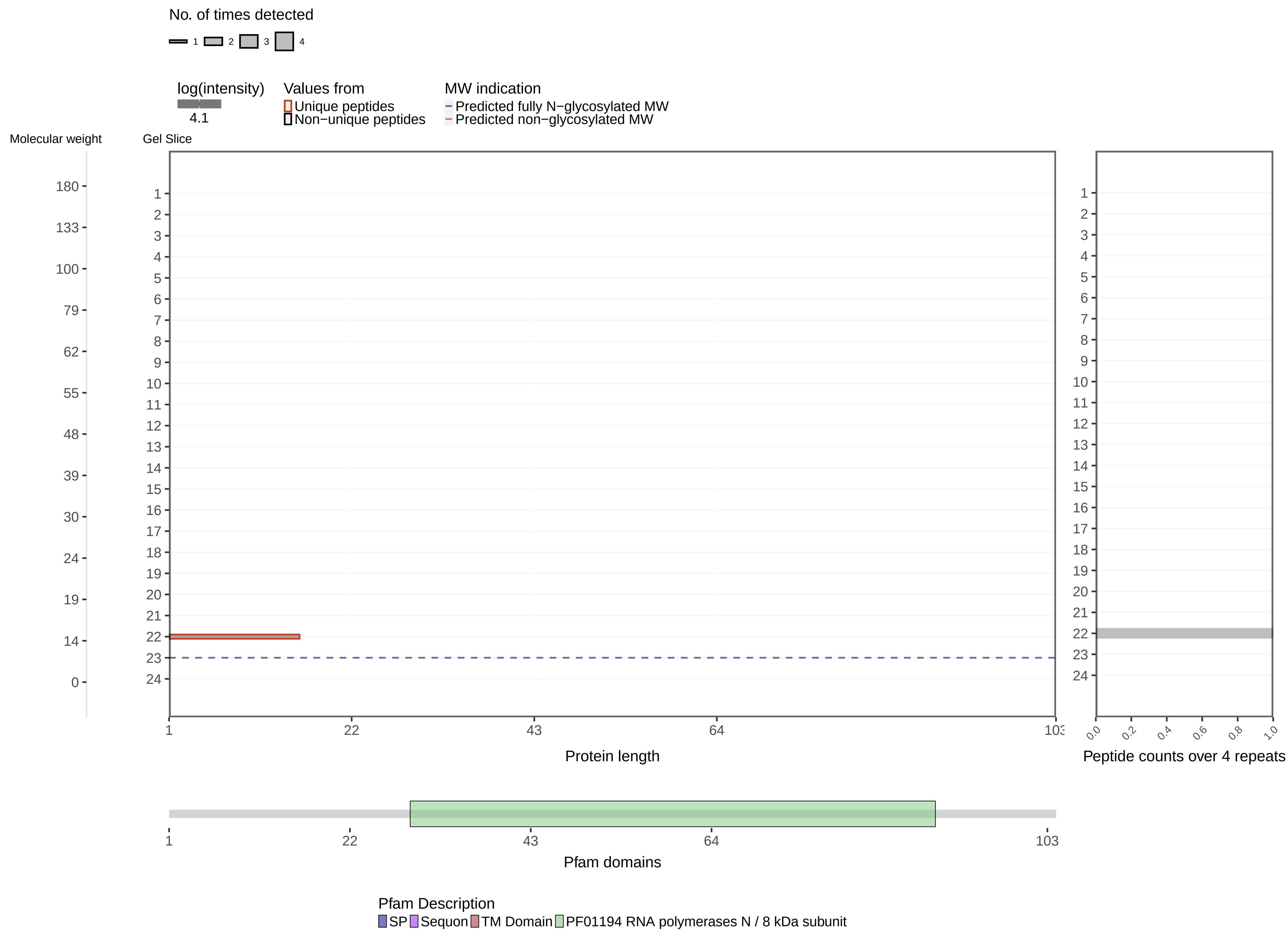


Protein sequence:

GNKWDQYLDLVLASYDKKDGQTAEAEALDALGLVRYCCRRMLMTHVDLIEKLLNYNTLGKSEGS*

### NbL16g08210.1 ALG−2_interacting_protein_X−like [MW = 97.97 kDa]

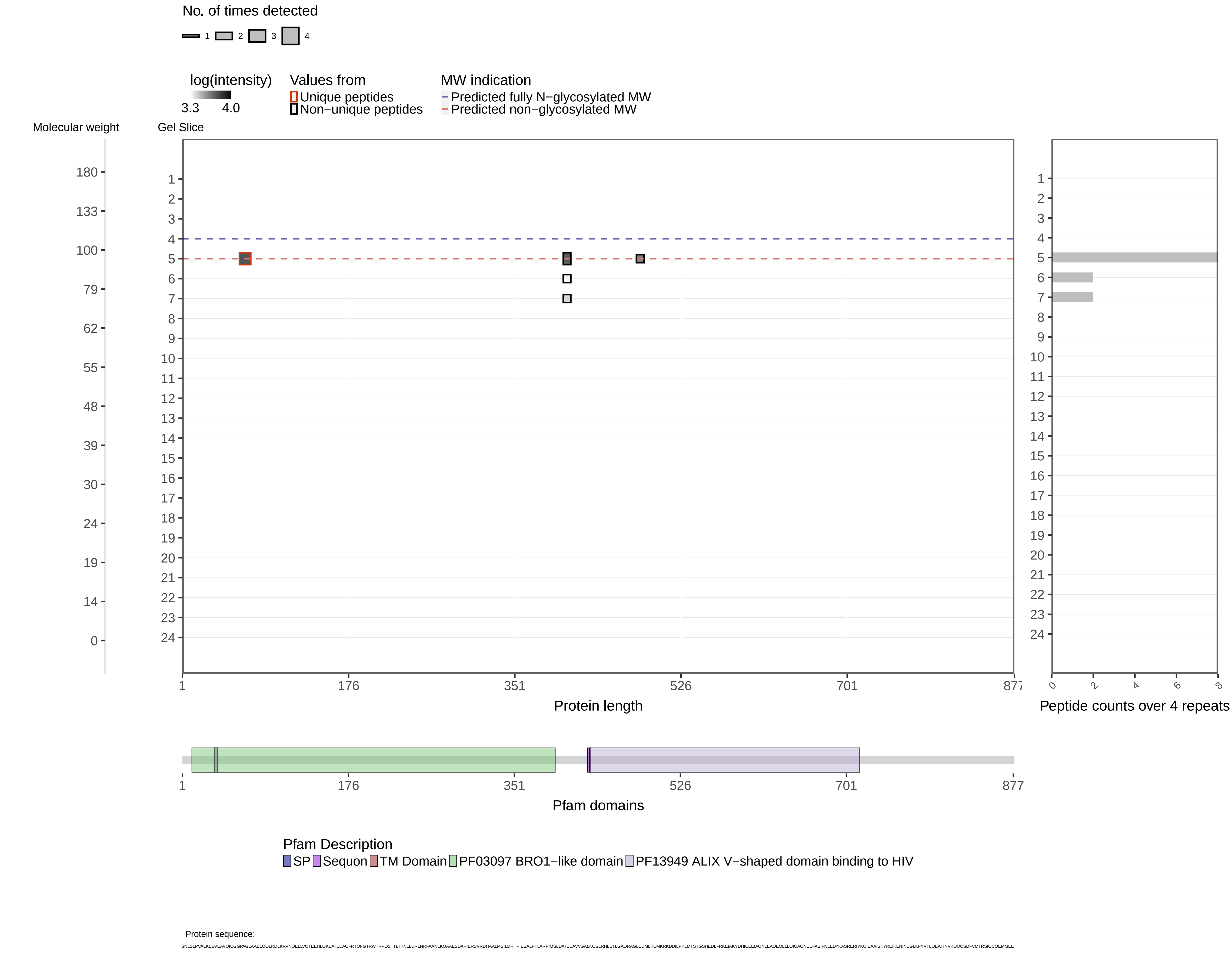

### NbL16g08100.1 neurofilament_heavy_polypeptide [MW = 101.32 kDa]

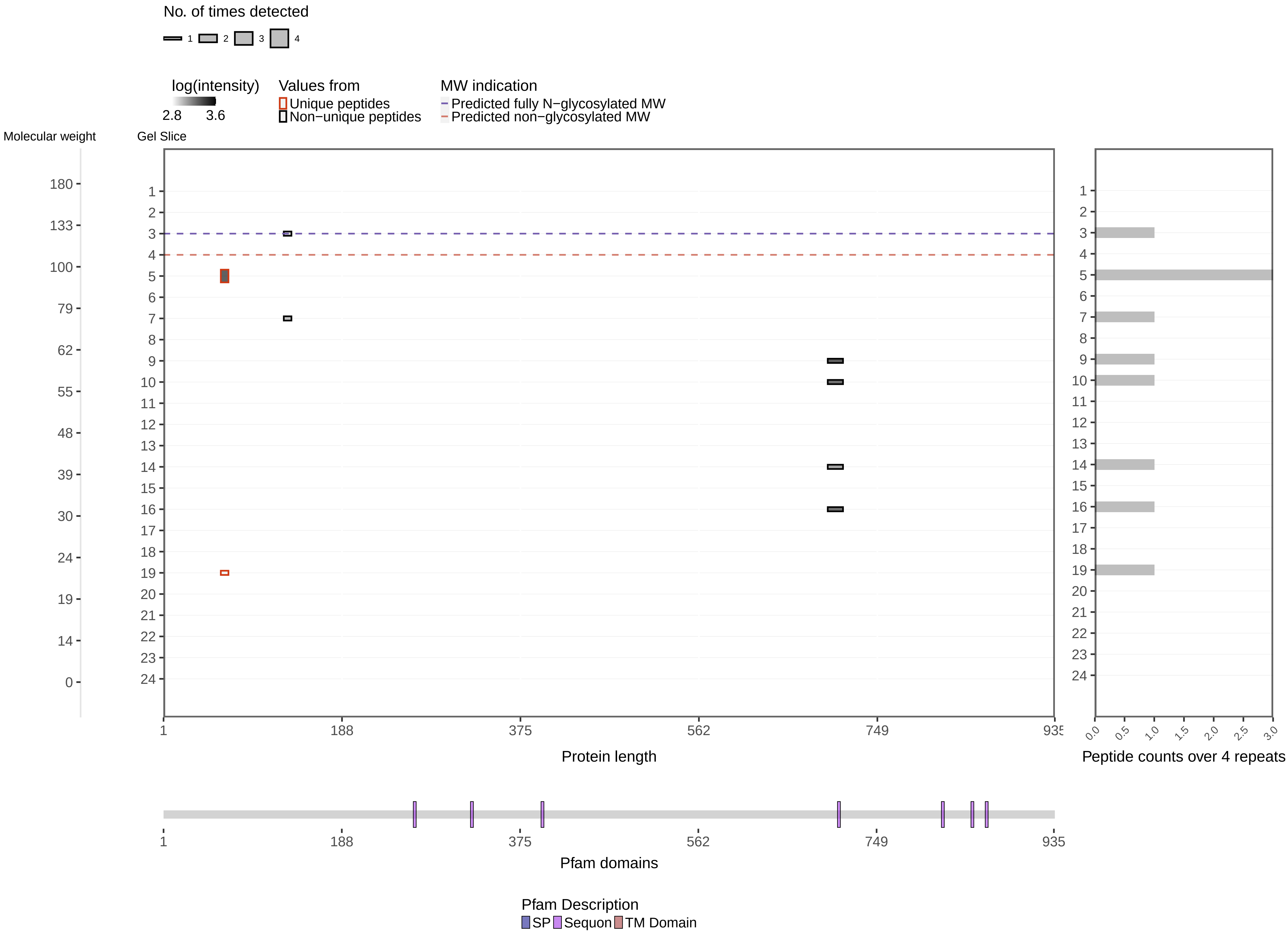


Protein sequence:

EAVNVAPSSTVLSLPGESAPKKGGRQKEDSLNQEECVSEKESEATSDLEVKQVRRPSKKAPAEPSHKENEGGSTSDVEAKKQKKSDKKIDTKNKNQVGPSVRNKEDSKKRGRGKASPETAPPQESPDDSAKHECNEEDIPKTSAKKKPSSGKGRERVPSPESPDNSAKHENNEEETPRTSAKRKQRSSGKDKVSGTVQHGESLVGKKIKVWWPLDKEFYEGVVEKFDSAKKKYRVAYTDGDVEIINLTEERWKLVEDDPMSEGEQIASADAASERQKKKKPRNAES[illegible]

### NbL16g07920.1 probable_N−acetyltransferase_san [MW = 18.41 kDa]

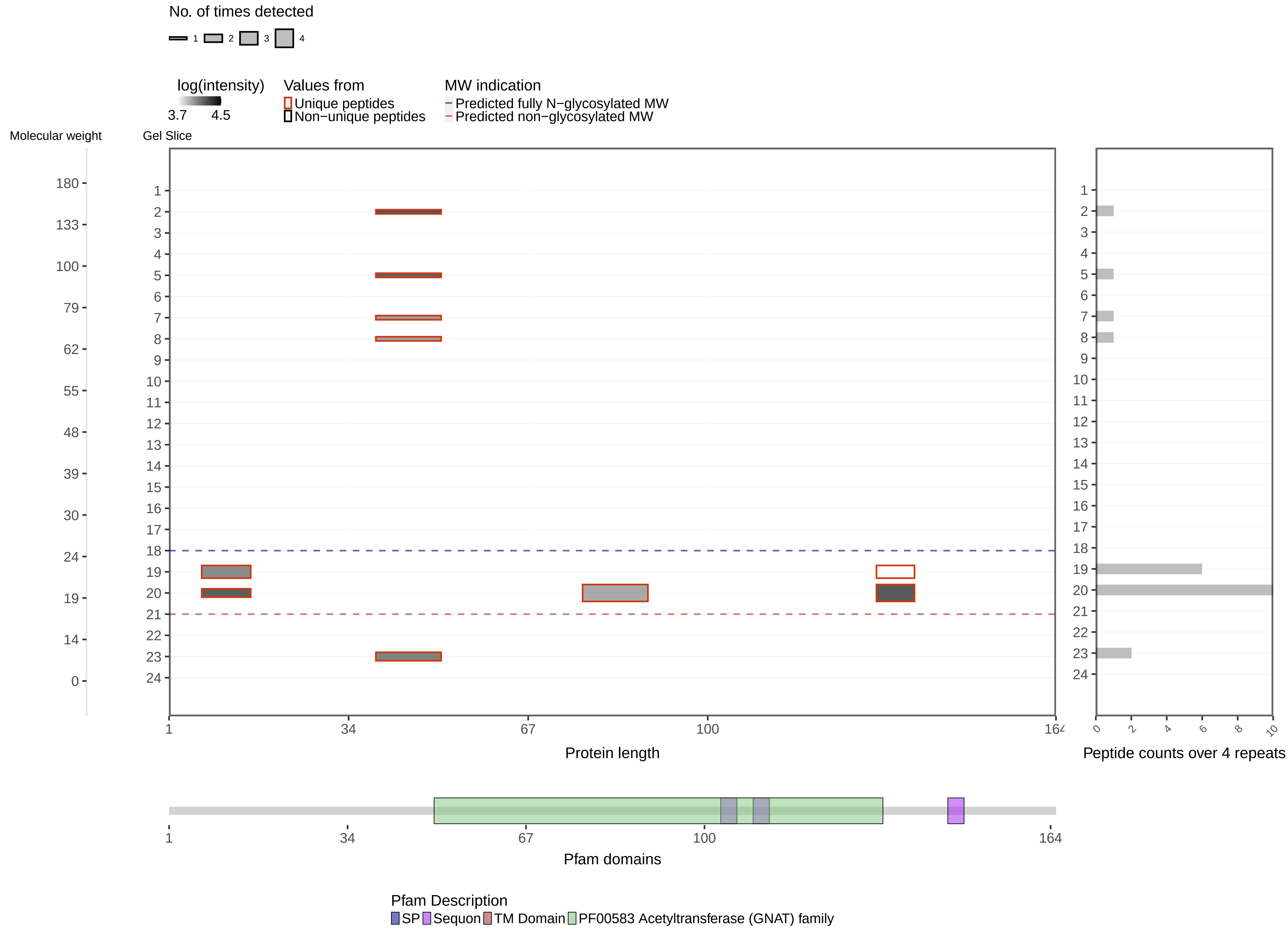


Protein sequence:

EKKEGGAVRVYIMTLGVLAPYRGLGIGKMLLNHILNLSAKQNVSEIYLHVHTINEDALNFYKKFGFEVTDKIQNYYTNITPPDCFVLTKFITQTKK*

NbL16g07840.1 psbP_domain−containing_protein_6_chloroplastic−like [MW = 28.21 kDa]

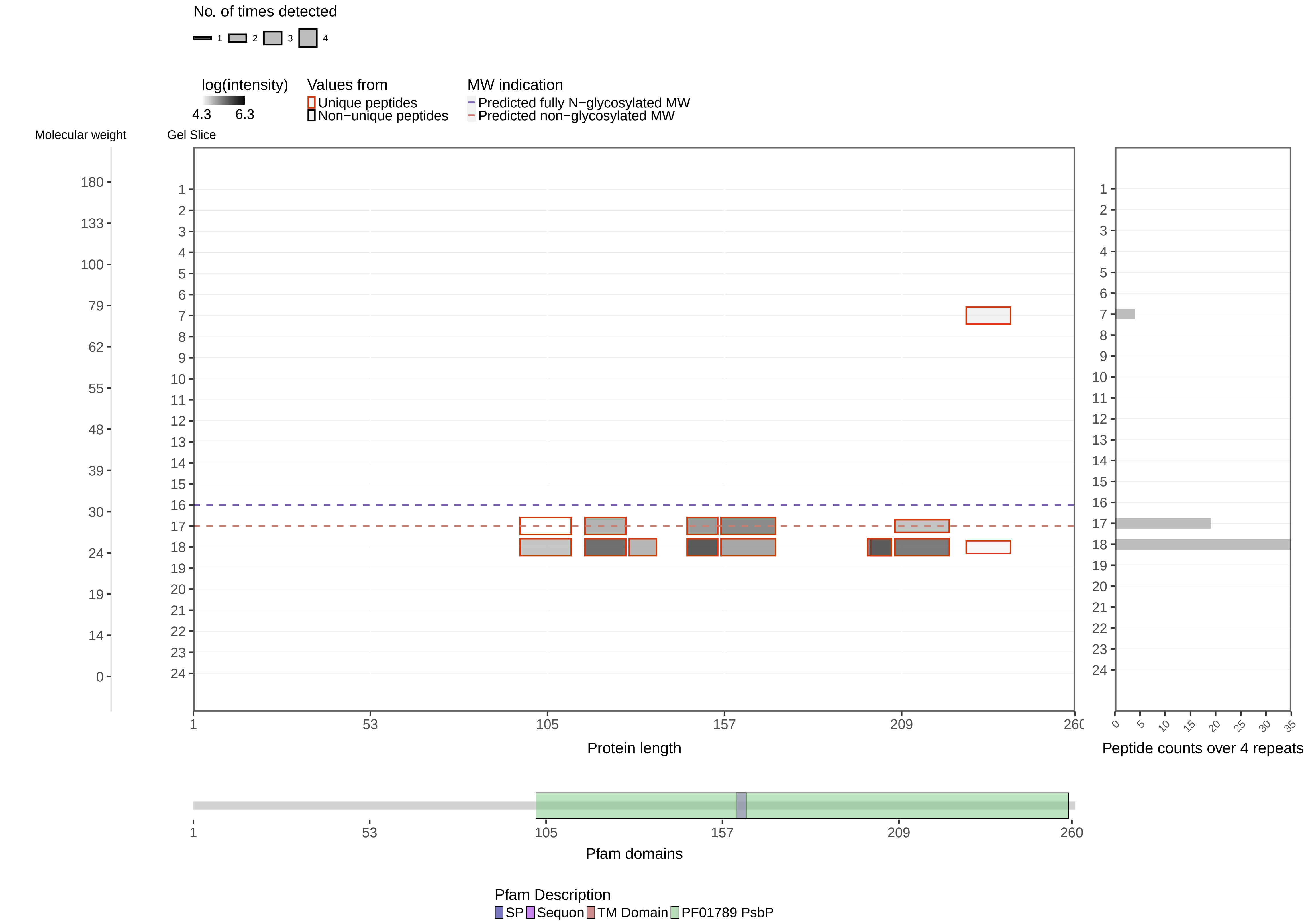


Protein sequence:

ANILSGNYCQPKCAEPWVEVKFEDEKQGKVQVVASPLIRLTNKPNATIEDIGTPEKVIASLGPFVTGNTLDPDELLQTAIEKRGDLTYYKYVLETPYALTGSHNLAKATAKGNTVVLFVVSANDKQWSSSQKTLEAILDSFEV*

NbL16g07410.1 poly(rC)−binding_protein_3−like [MW = 57.37 kDa]

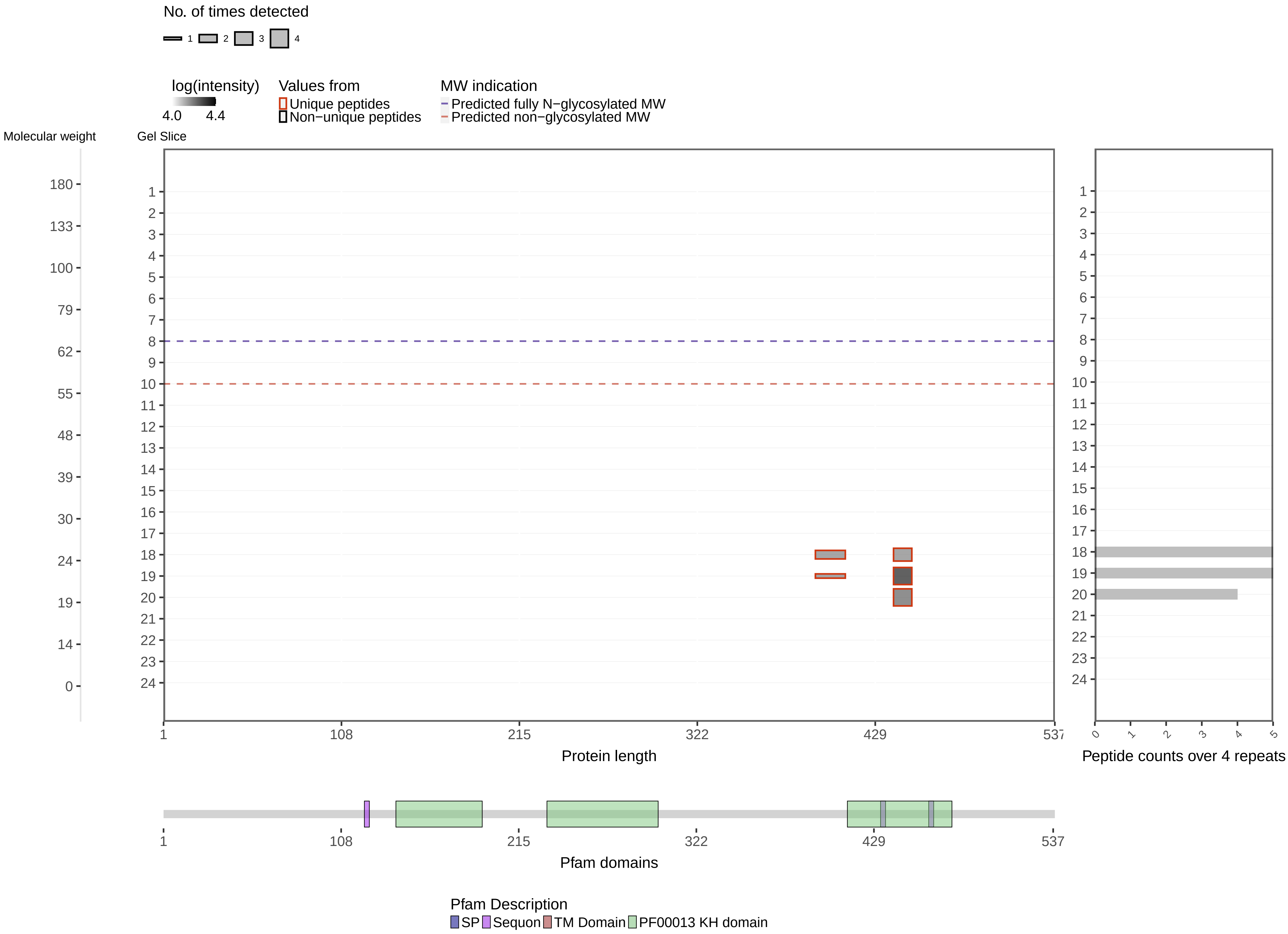


Protein sequence:

[illegible]

NbL16g07280.1 EKC/KEOPS_complex_subunit_bud32−like [MW = 33.54 kDa]

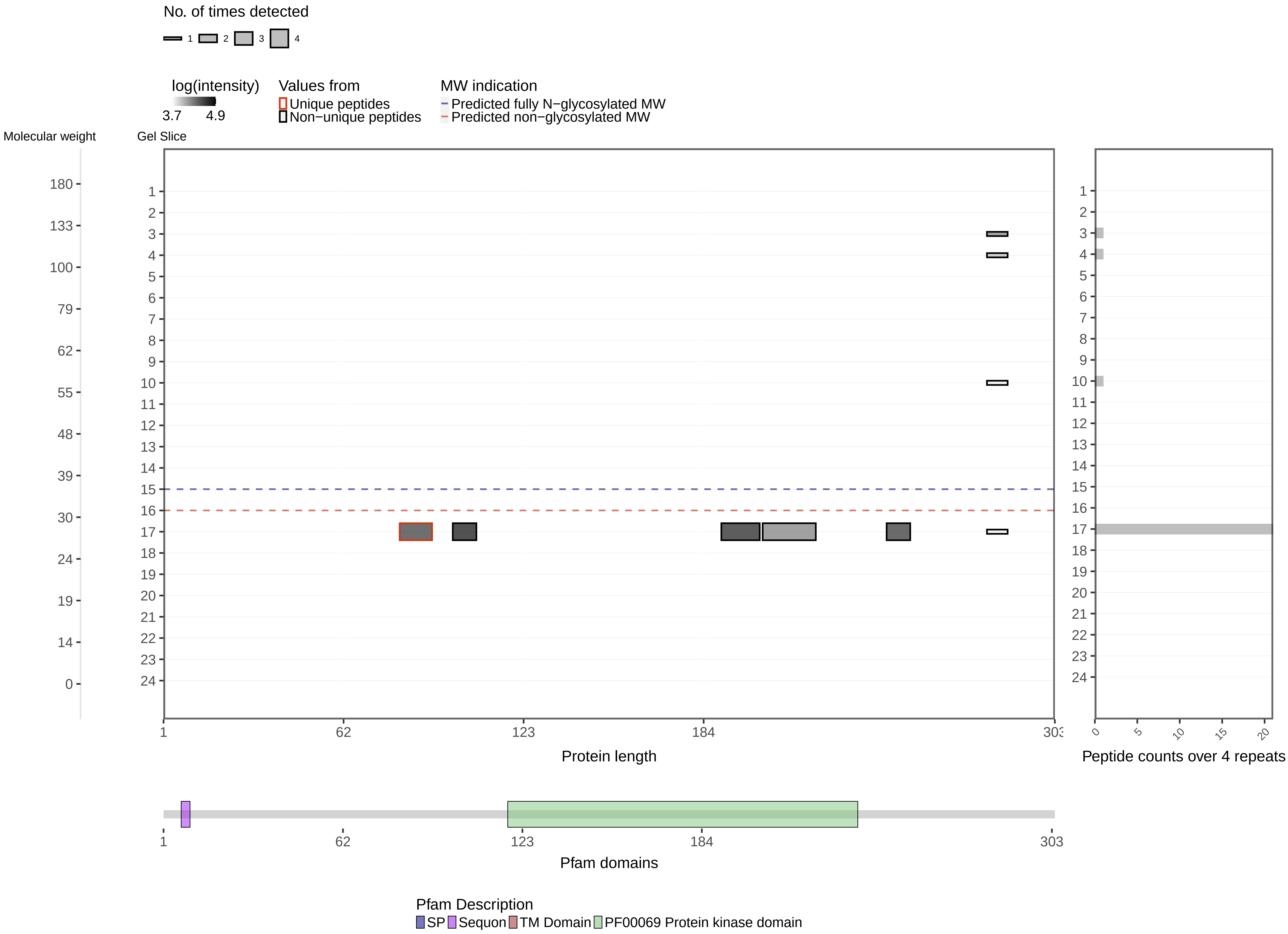


Protein sequence:

ARRLGVTTPVLYAVDPVTHTLTFEYVEGPSVKDIFLGFGLVGVDDERMTDIATQIGNAIGKLHDGGLVHGDLTTSNMLMRSSANQLVLIDFGLSFTSTLPEDKAVDLYVLERALLSMHSSCGNVMDQILAGYKKSSKQWSSTFNKLAQVRQRGRKRTMVG*

### NbL16g07220.1 40S_ribosomal_protein_S24−1 [MW = 15.39 kDa]

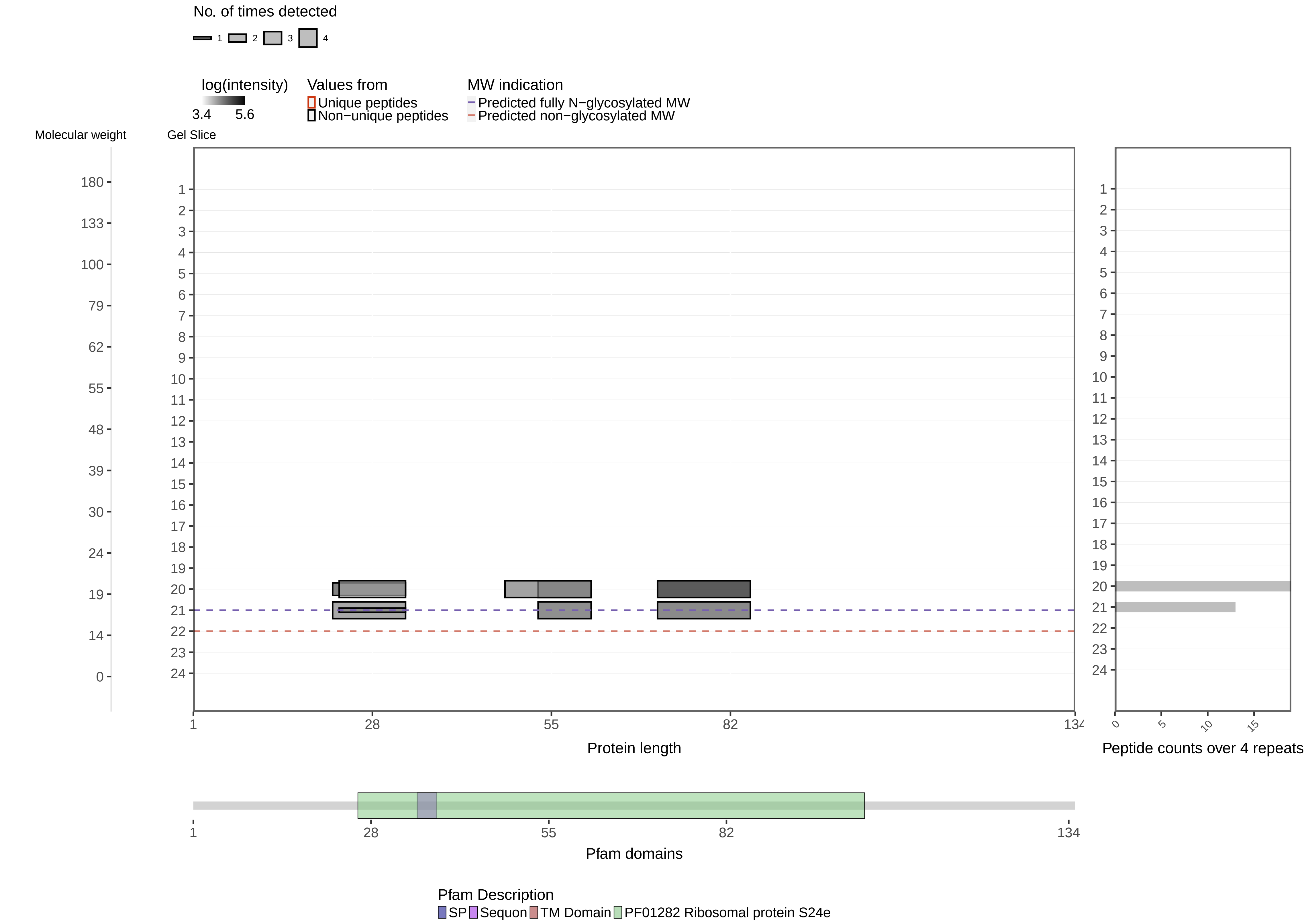


Protein sequence:

NAIFVFKFRTHFGGGKSTGFGLIYDSVENAKKYEPKYRLIRNGLDTKVEKSRKQMKERKNRAKKVRGVKKTKAGDAKKK*

### NbL16g07120.1 probable_glutathione_peroxidase_5 [MW = 18.91 kDa]

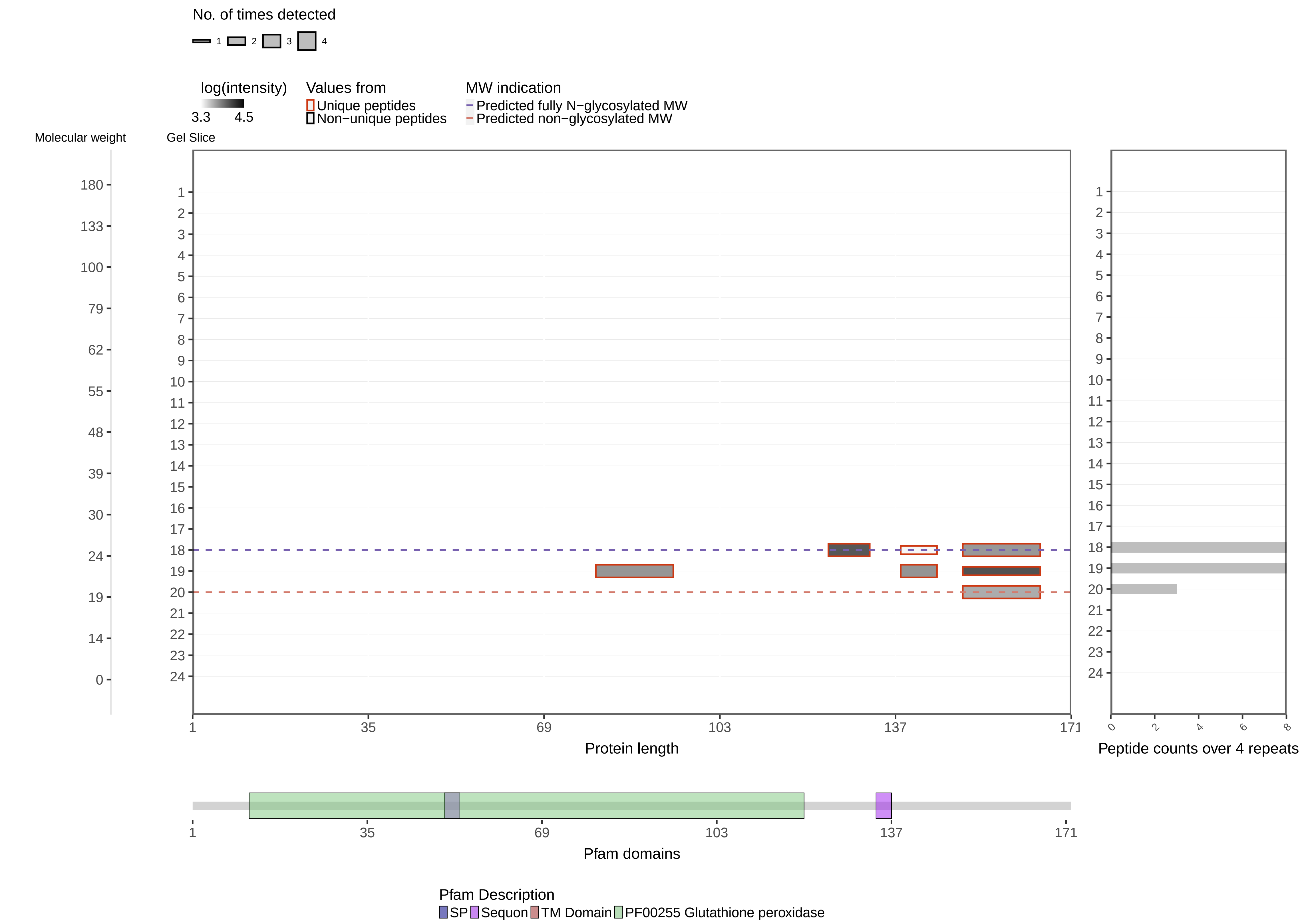


Protein sequence:

PCNQFLKQEPGTSEEAQQFACTRFKAEYPIFQKVRVNGPDAAPVYQFLKSSKGGFLGSSIKWNFTKFLIDKEGKVIKRYGSTTTPLAIEADIQKALGEV*

NbL16g07060.1 auxin−binding_protein_T92 [MW = 26.21 kDa]

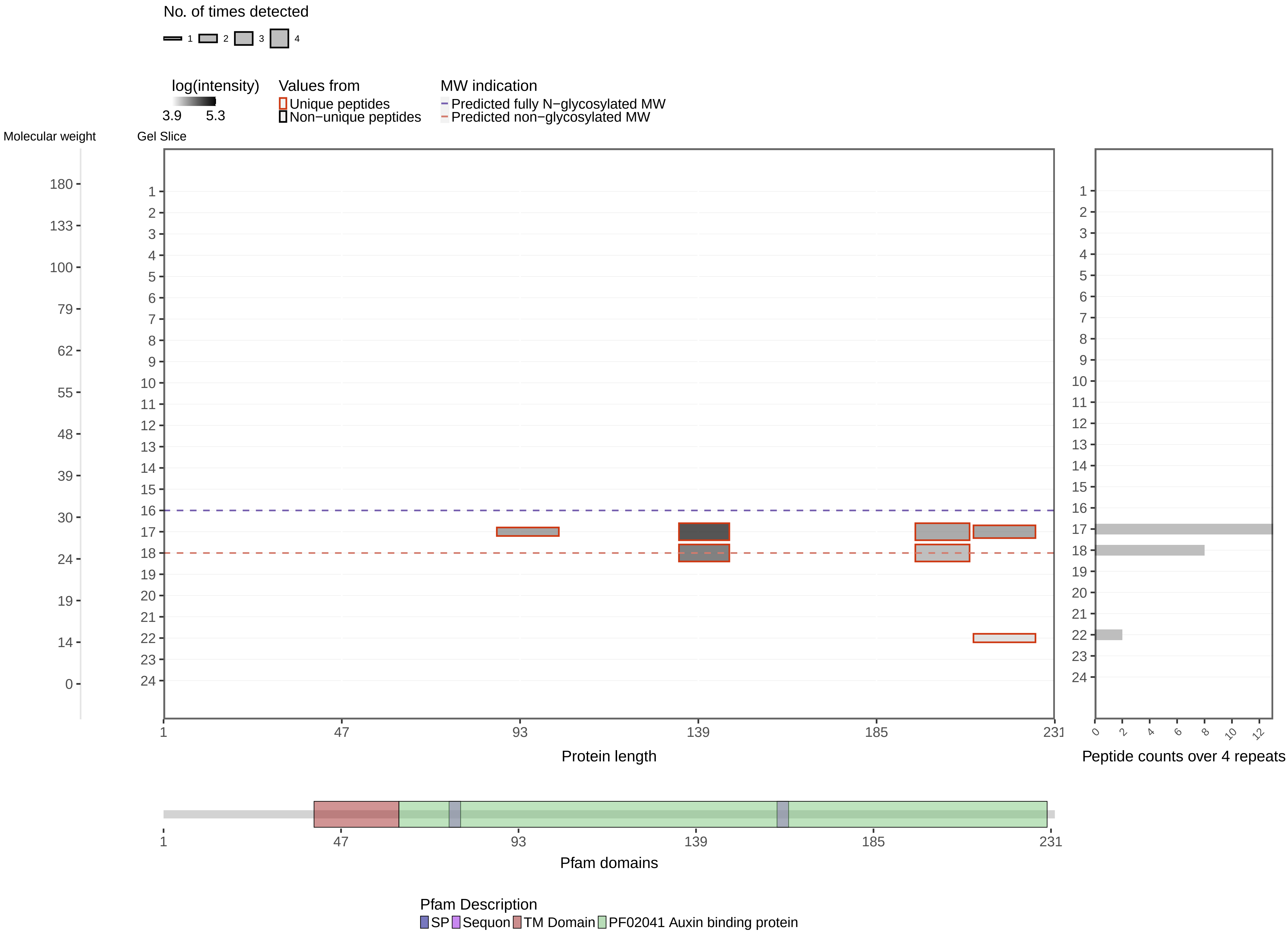


Protein sequence:

:EIEVWLQTFAPGSRTPIHRHSCEEILIVLKGQGTLYLTPSSHSKYPGNPQEFHIFPNSTVHIPVNDVHQVWNTGEHEDLQVLVVISRPPVKVFMYDDWSMPHTAAKLKFPYYWDEECYQTTSRKDEL*

### NbL16g06970.1 histone_H2AX−like [MW = 15.22 kDa]

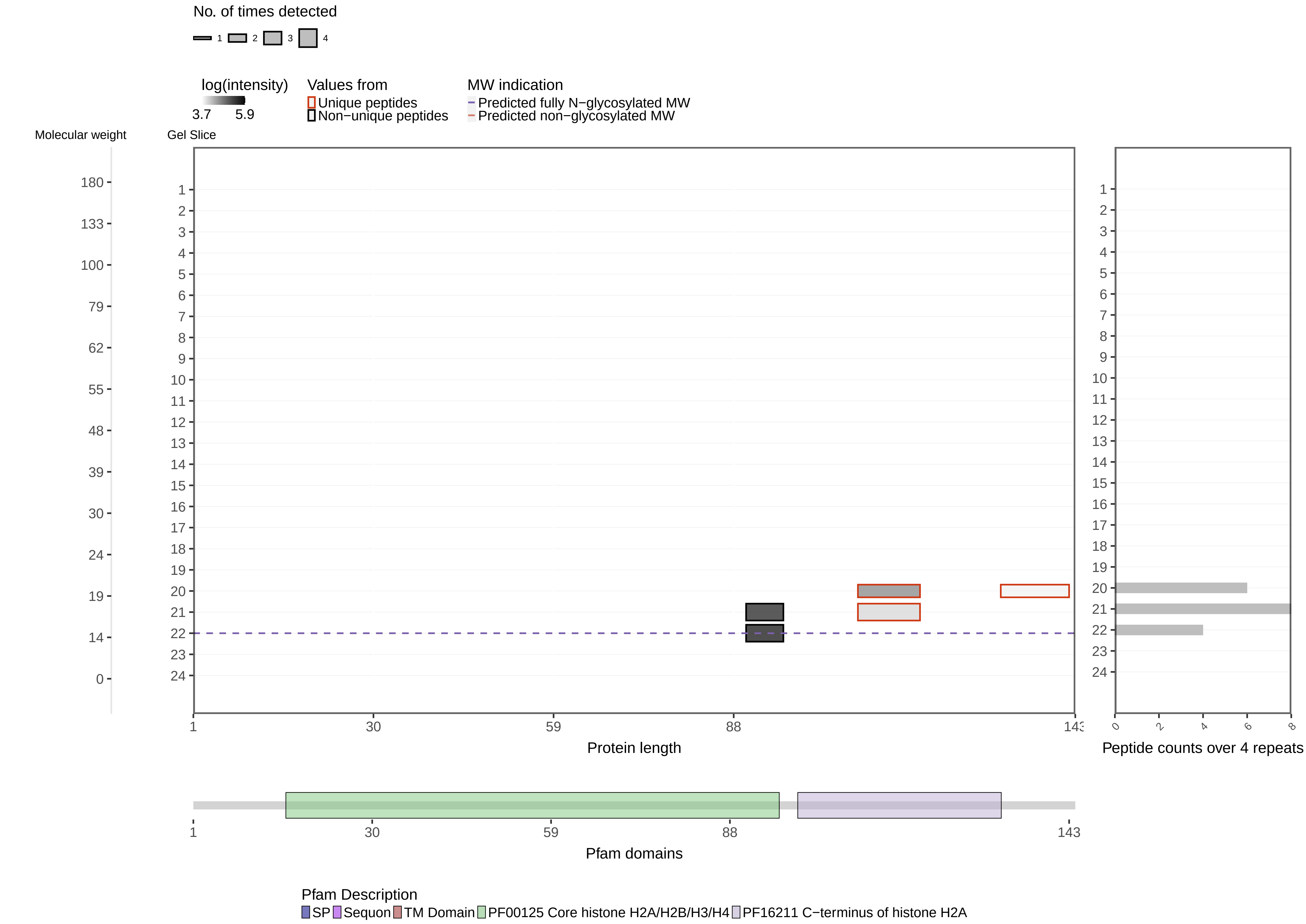


Protein sequence:

YLSAVLEYLAAEVLELAGNAARDNKKTRIVPRHIQLAVRNDEELSKLLGKVTIANGGVLPKIHPNLLPNKNGKGKLEAGTQSQEF*

### NbL16g06920.1 bifunctional_3−dehydroquinate_dehydratase/shikimate_dehydrogenase_chloroplastic−like [MW = 56.48 kDa]

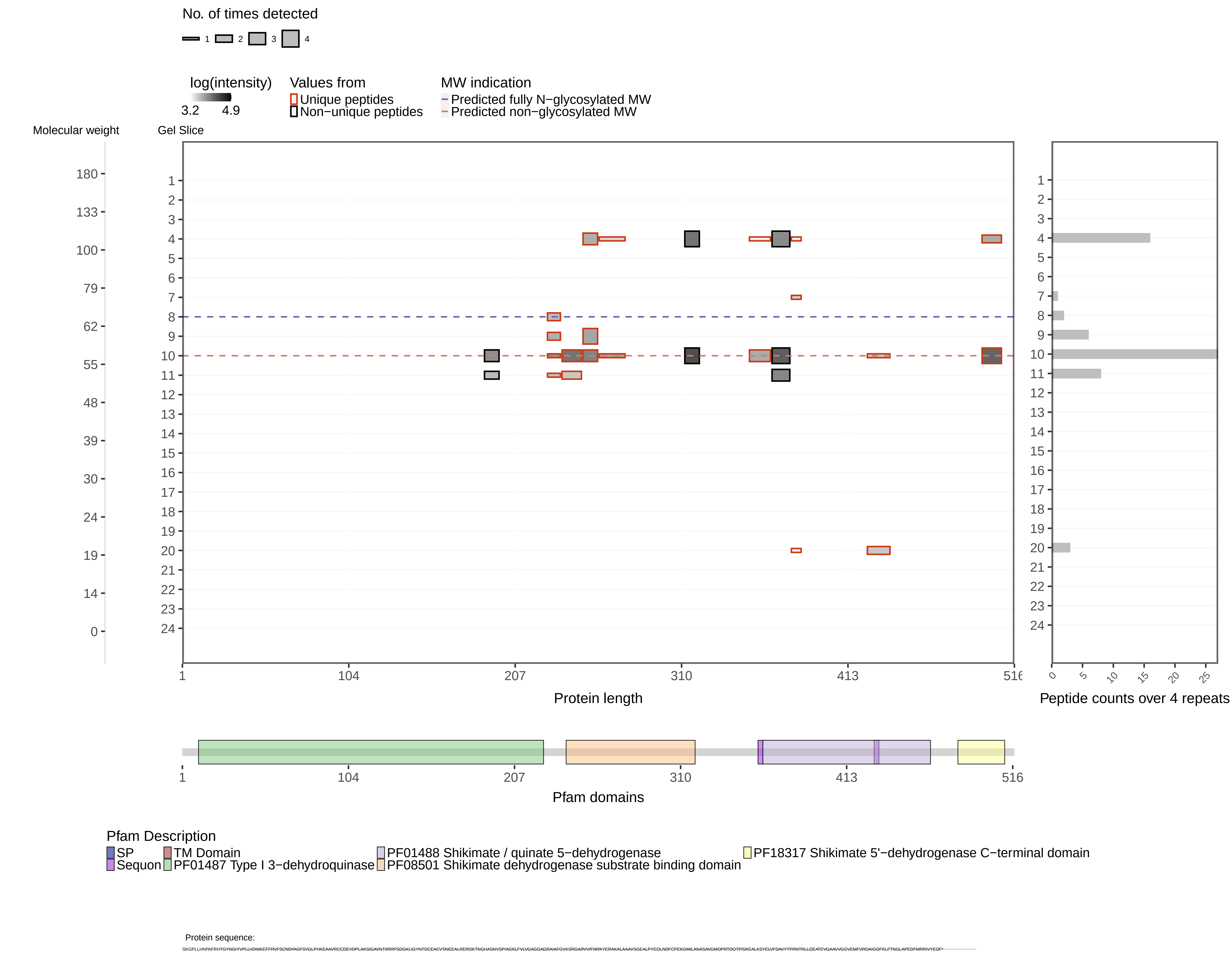

NbL16g06900.1 protease_Do−like_2_chloroplastic [MW = 66.9 kDa]

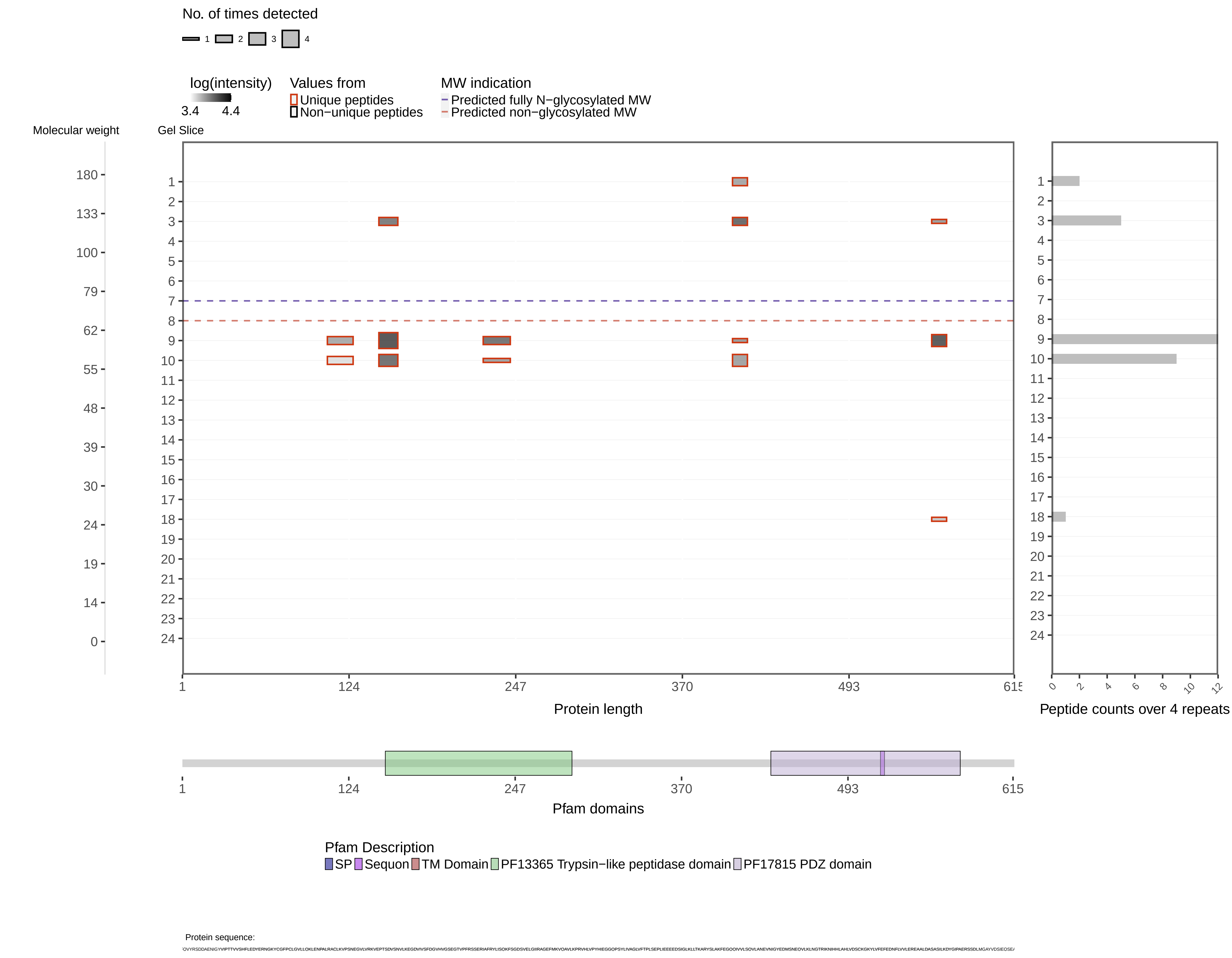

### NbL16g06810.1 universal_stress_protein_PHOS34−like [MW = 17.5 kDa]

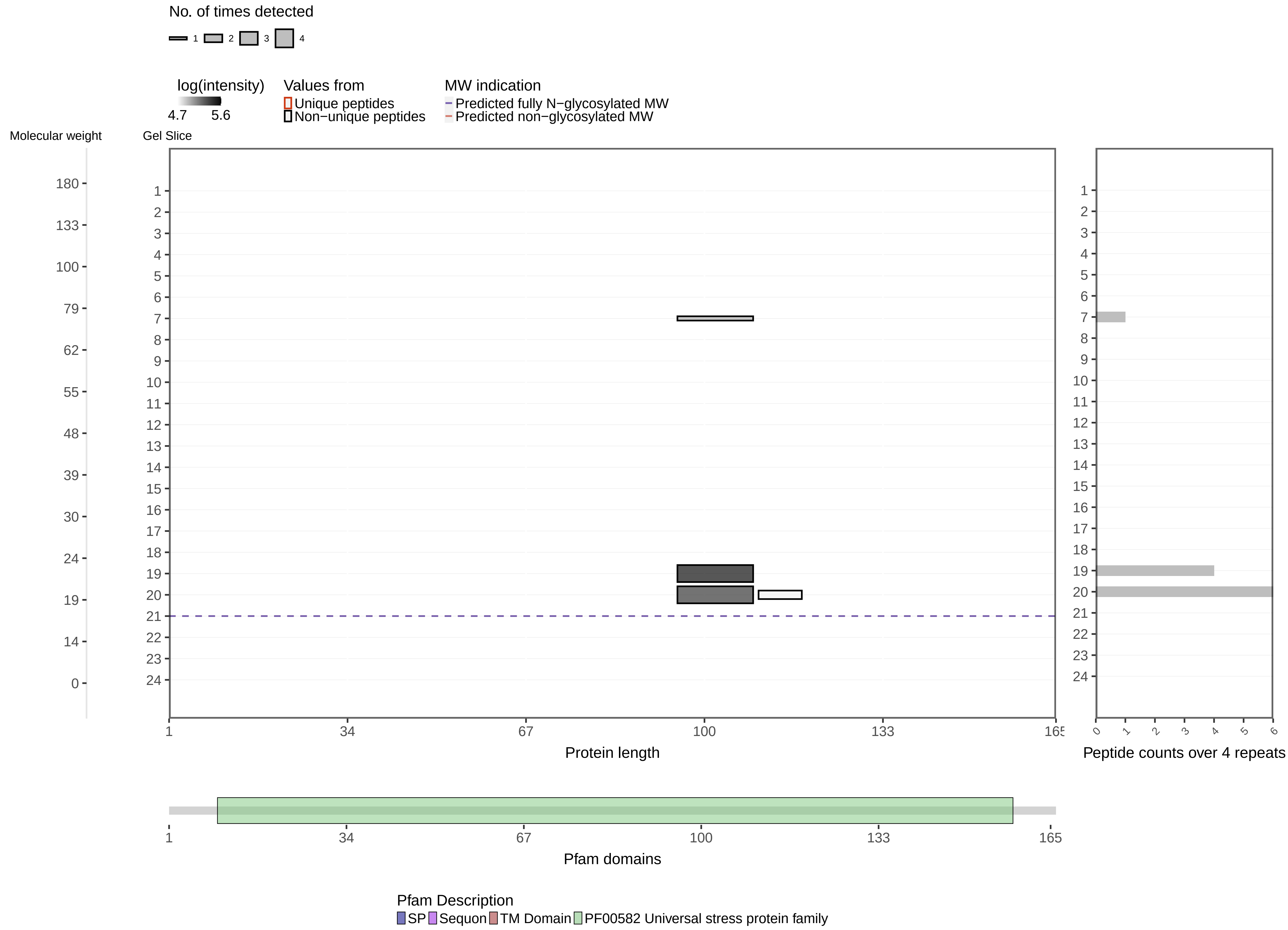


Protein sequence:

ᴾYVEADLRKISSRVAEKAKQICSAKSVNDVVVEVMEGDARNVLCDAVEKHHASILVVGSHGYGVLKRTVLGSVSDYCAHHAHCSVMIVKRPKVKA*

### NbL16g06800.1 beta−glucosidase_11−like [MW − SP = 55.64 kDa]

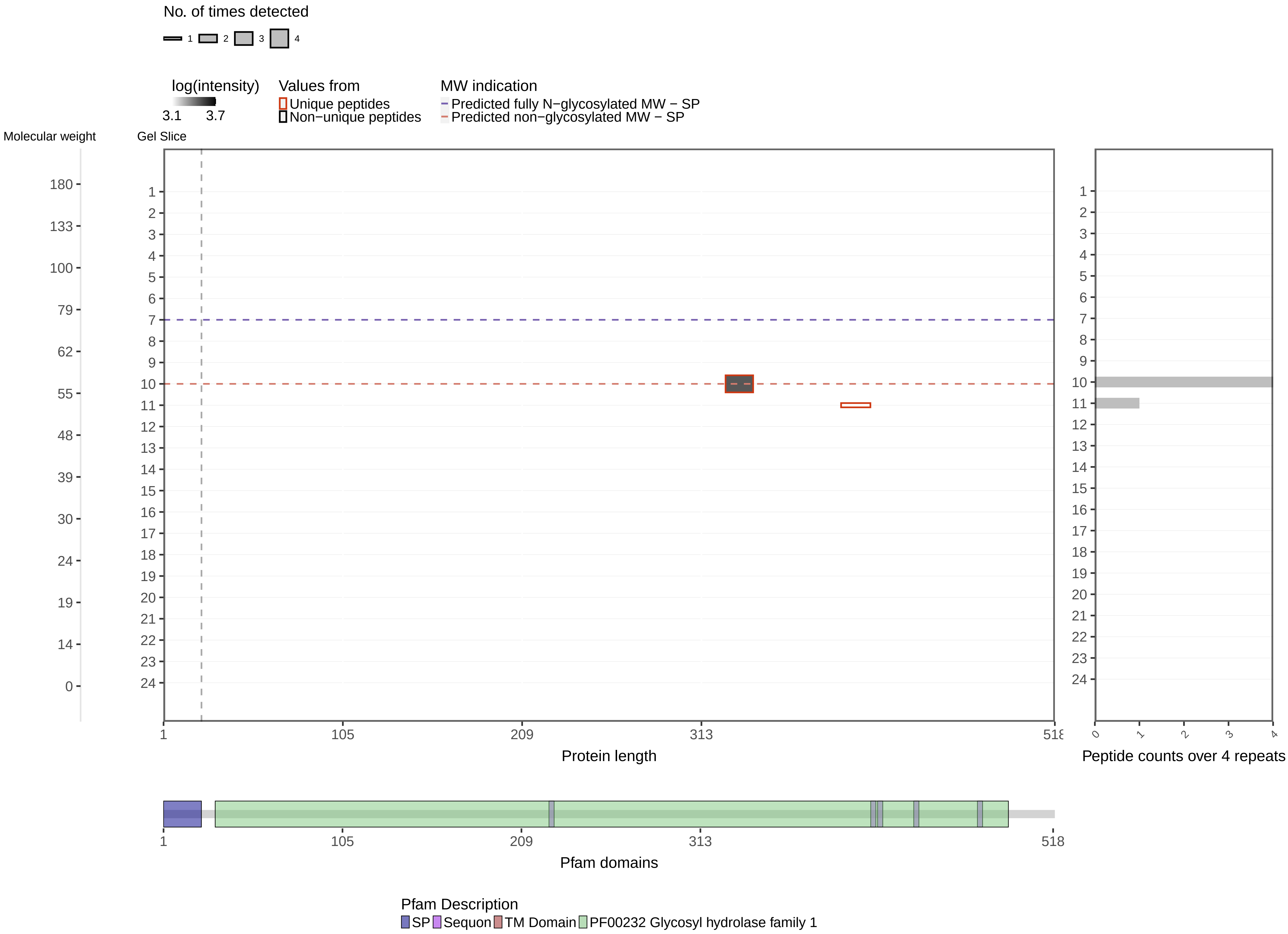


Protein sequence:

RLYKRKYKSIQHGFVGLNIFAYRFIPHTNAKADEGATLRAYAFYLGWFMSPLIFGDYPVTMKDRVGSRMPTFTRQESEQVKGAIDFIGLNHYATLPVKDRPSAFERDYRDFSTDAAIQLVFEQGGVTPKGEYPWTQPGLRAVLEYFKKTYGNLPIYIHENGQTTPRNATLNDTGRVEYLHAYIRNLLDAMRNGSNVRGYFSWSFLDGLELLDGYKSAYGLYYVDLDDENLTRYPKLSQHWYSNFLKGKSRKSGVDLVIGKEVSHSSESKFSS*

### NbL16g06720.1 60S_ribosomal_protein_L19−3−like [MW = 27.35 kDa]

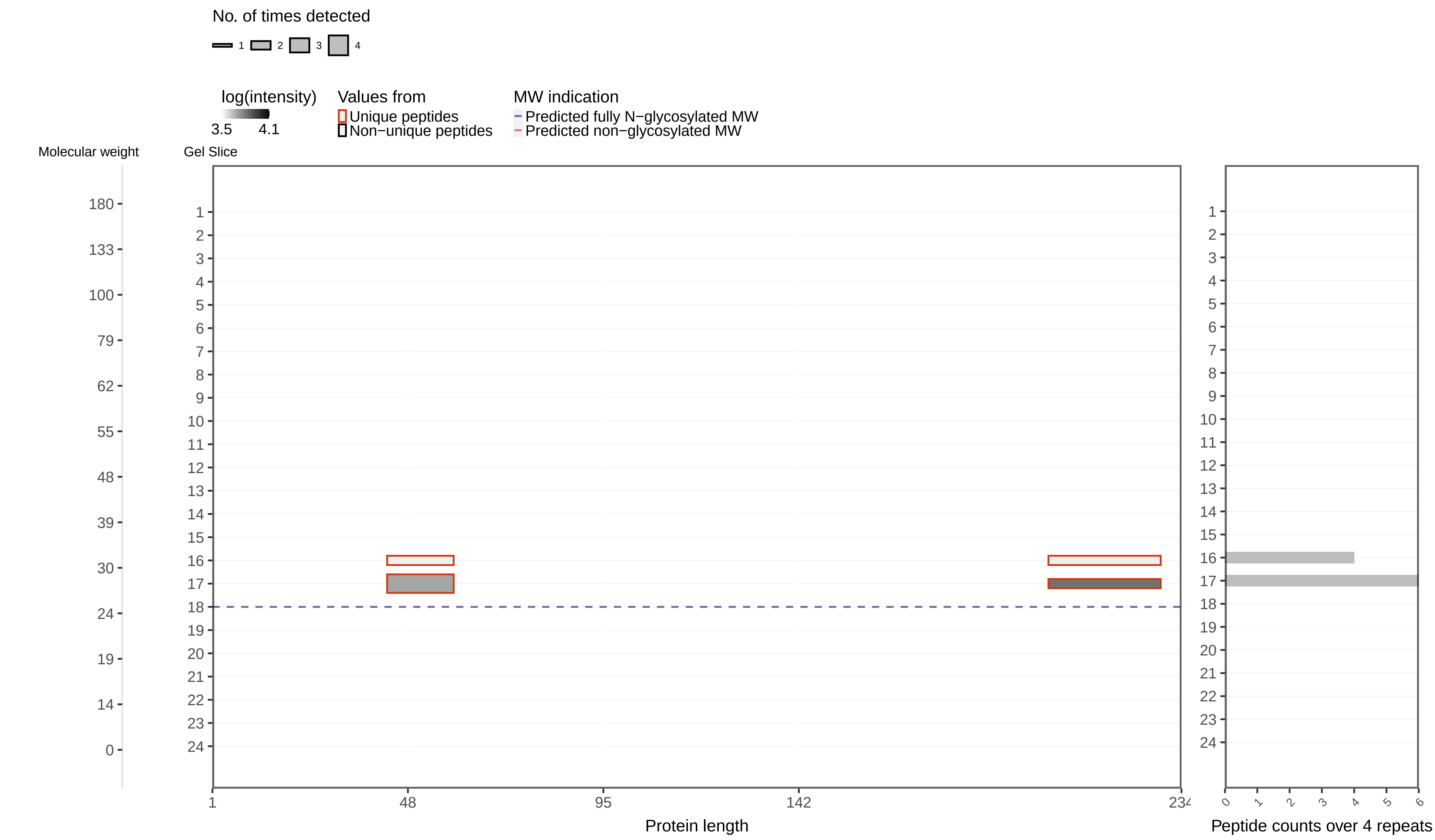


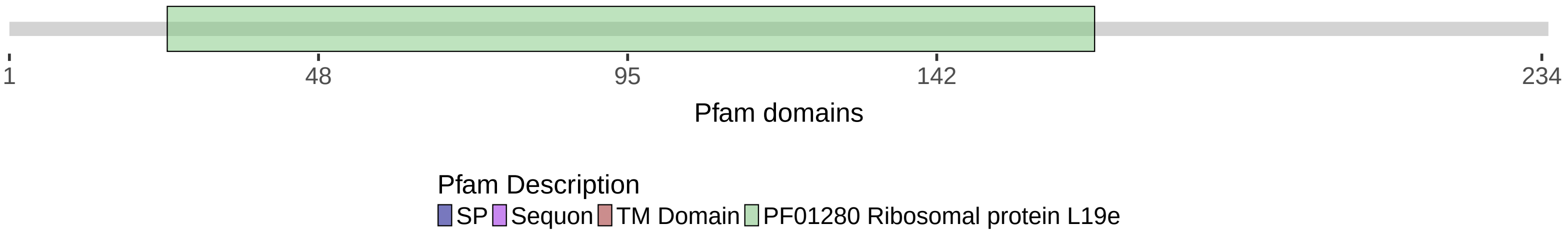


Protein sequence:

TREARLPTKVLWMRRMRVLRRLLRKYRESKKIDRHMYHDMYMKVKGNVFKNKRVLMENIHKTKAEKAREKTLSDOFEARRAKNKASRERKFARREERLAOGPGERPIOPAAATATPOAOPAOGSKKSKK*

### NbL16g06700.1 methionine_aminotransferase−like [MW = 44.94 kDa]

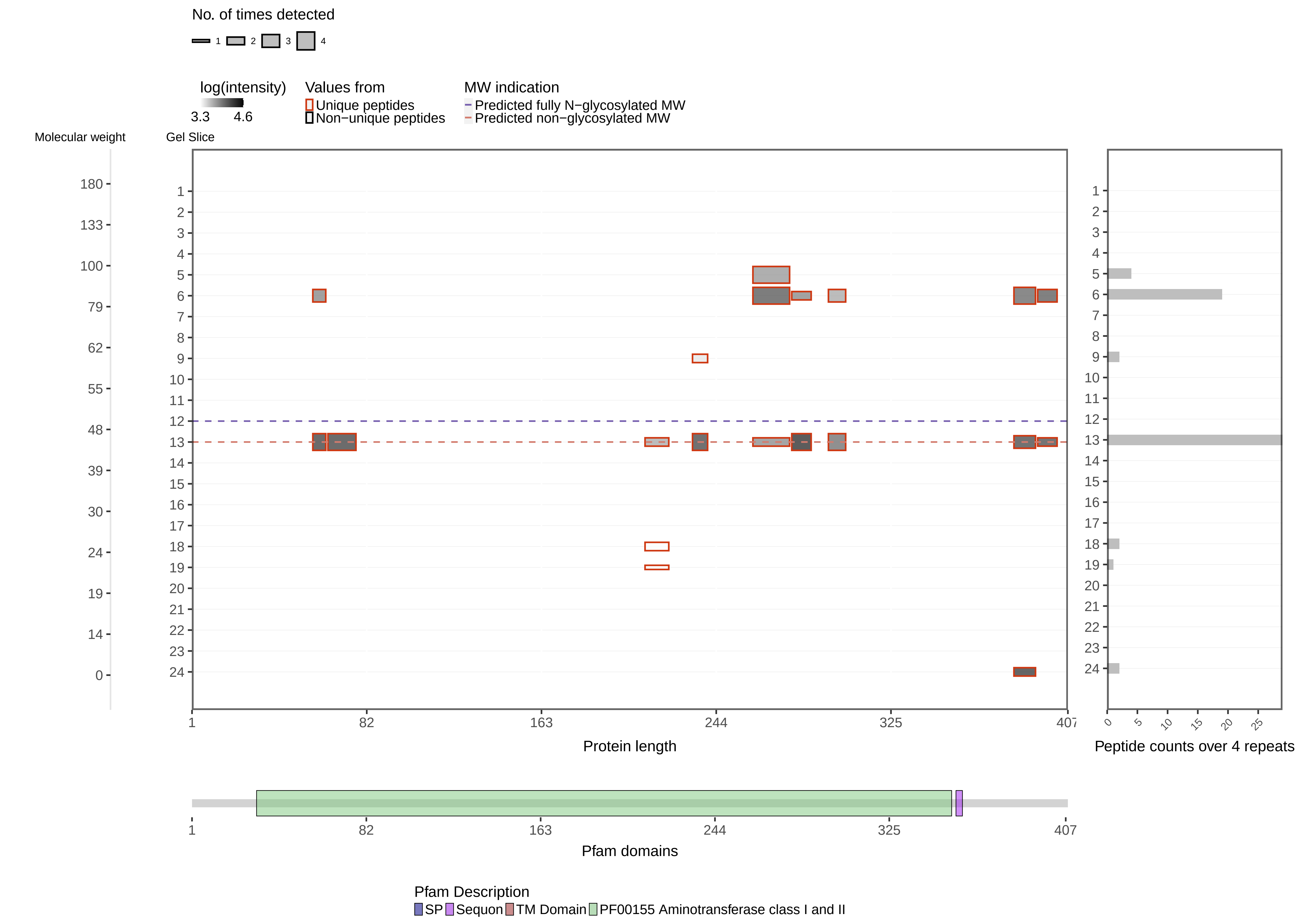


Protein sequence:

RTLDILAVTDEVYEHITFDGERHTSIATFPGMQKRTIITSSLSKSFSVTGWRIGWAIGPACFASAIRNIHVKITDSAPAPFQEAALTALRSSPEYFDALRQDYESKRDYLTQVLTKVGFQVPFMPKGSFFLFMELPGTCTLSDVEFVEELIKQAGVVAVPGHVFFHTNSSLLEDFSSTAISYHRRYIRFAFCKSEETLAAASQKLTELVDGSGRLKLF*

### NbL16g06670.1 CTP_synthase−like [MW = 67.33 kDa]

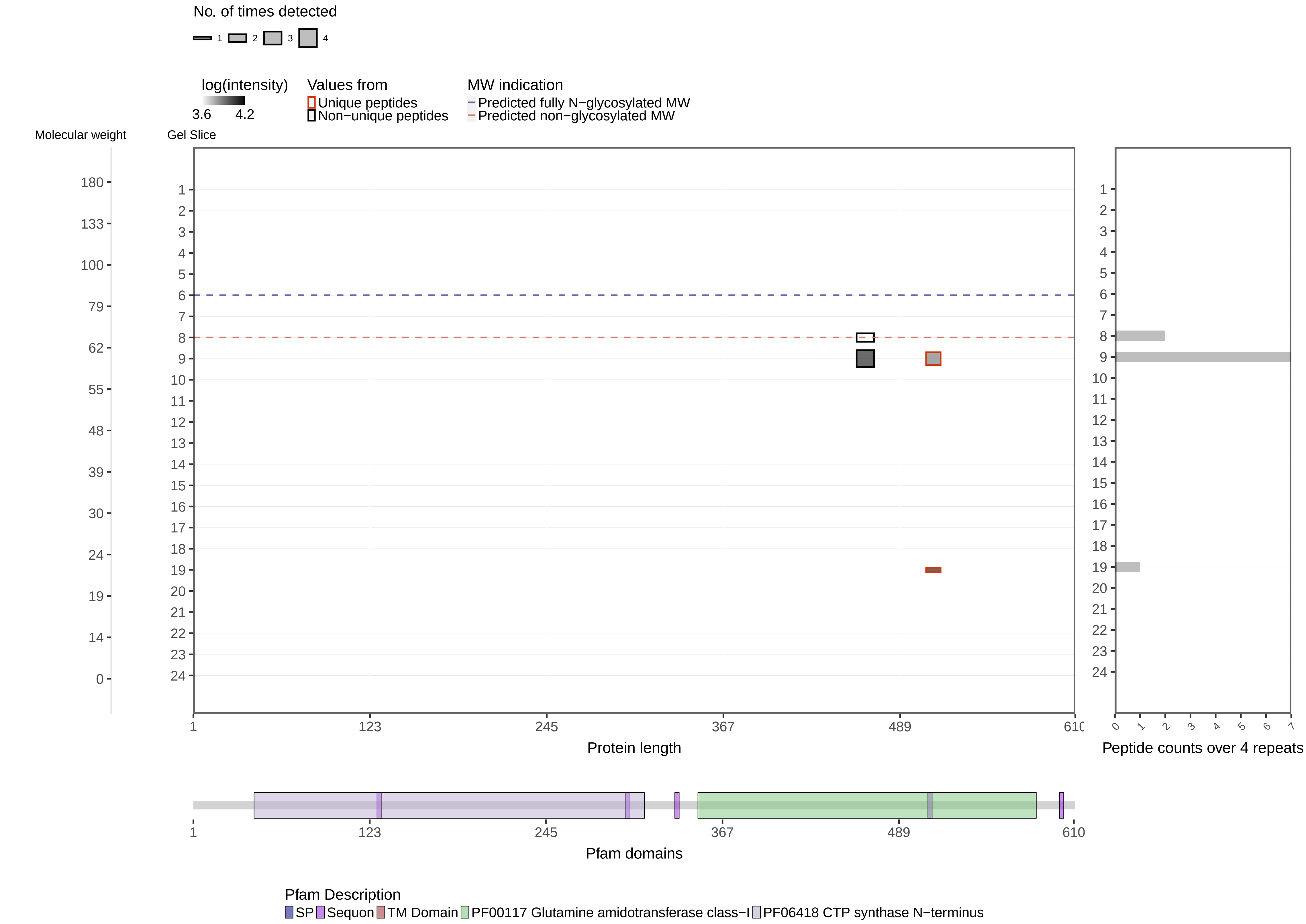


Protein sequence:

IIPLLLQNQSAHDAILKQLDLLSVARHVDLQEWTRRAETSDNITKSVSIAMVGKYVGLTDSYLSVVKALMHACIACSLKPSIDWIAASDLEDDTANLNPEAHAAAWKTLRGAACVLVPGGFGDRGVKGMMLAARYARENSVPYLGICLGMQISVIEFARSVLHLEKANSEEFDPQTPERVVIFMPEGSKTHMGSTMRLGSRRTLFRTPDCITAKLYNNSEYVDERHRHRYEVNPEIVGSLEEAGLKFVGRDESGNRMEILELPDHPFYIGVQFHPEFKSRPGRPSAPFLGLVLA

NbL16g06550.1 glycolipid_transfer_protein_1 [MW = 22.34 kDa]

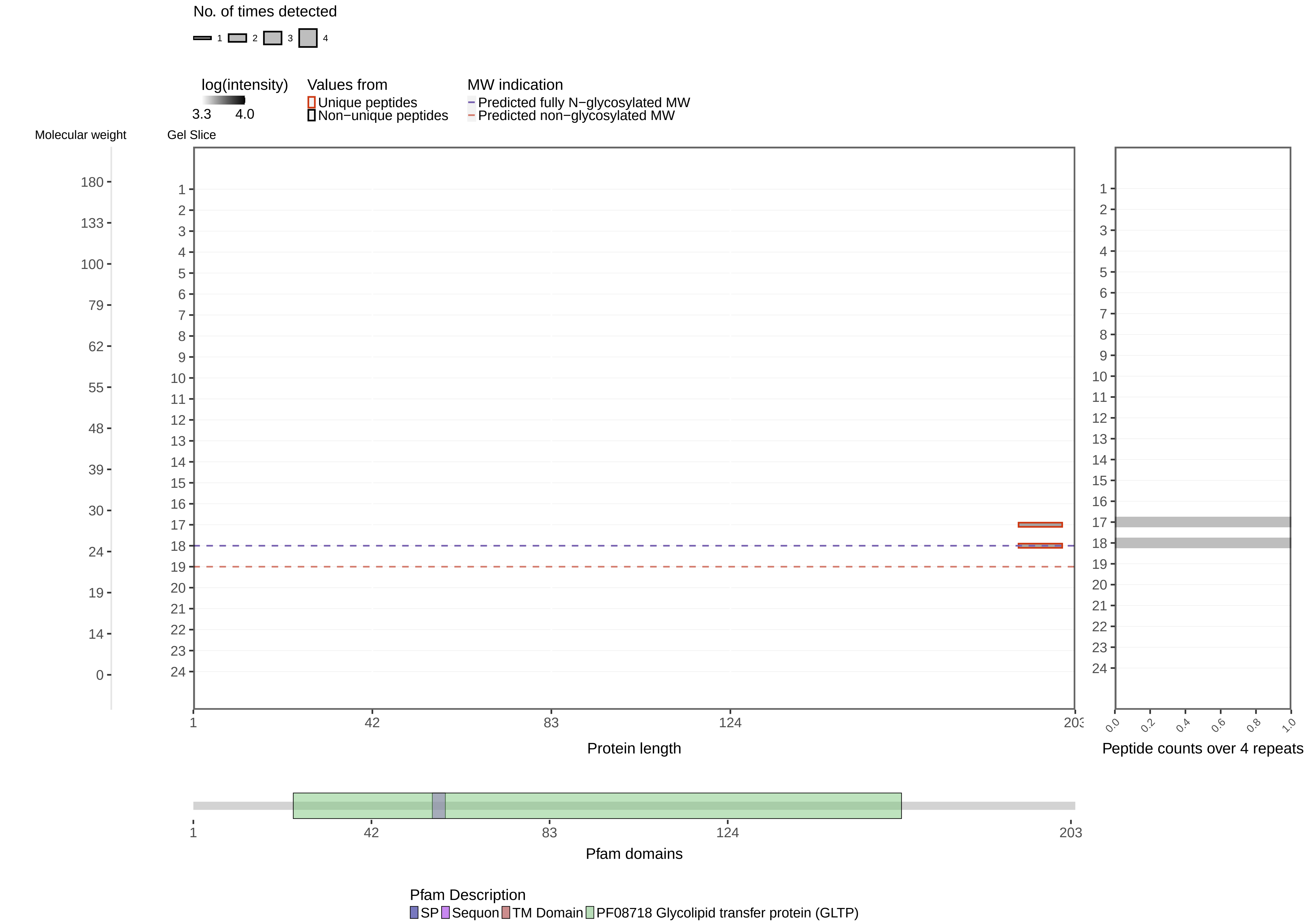


Protein sequence:

SSCTNGLLWLARAMDFLVELFRNLLEHGDWAMSQACSDSYSKTLKKFHGWLASSSFSVAMKLAPDRKKFMDVICGTGDIDSDMGQFCTTFSPLLEENHKFLASVGMDDLKAS*

NbL16g06530.1 succinyl−CoA_ligase_[ADP−forming]_subunit_beta_mitochondrial [MW = 45.15 kDa]

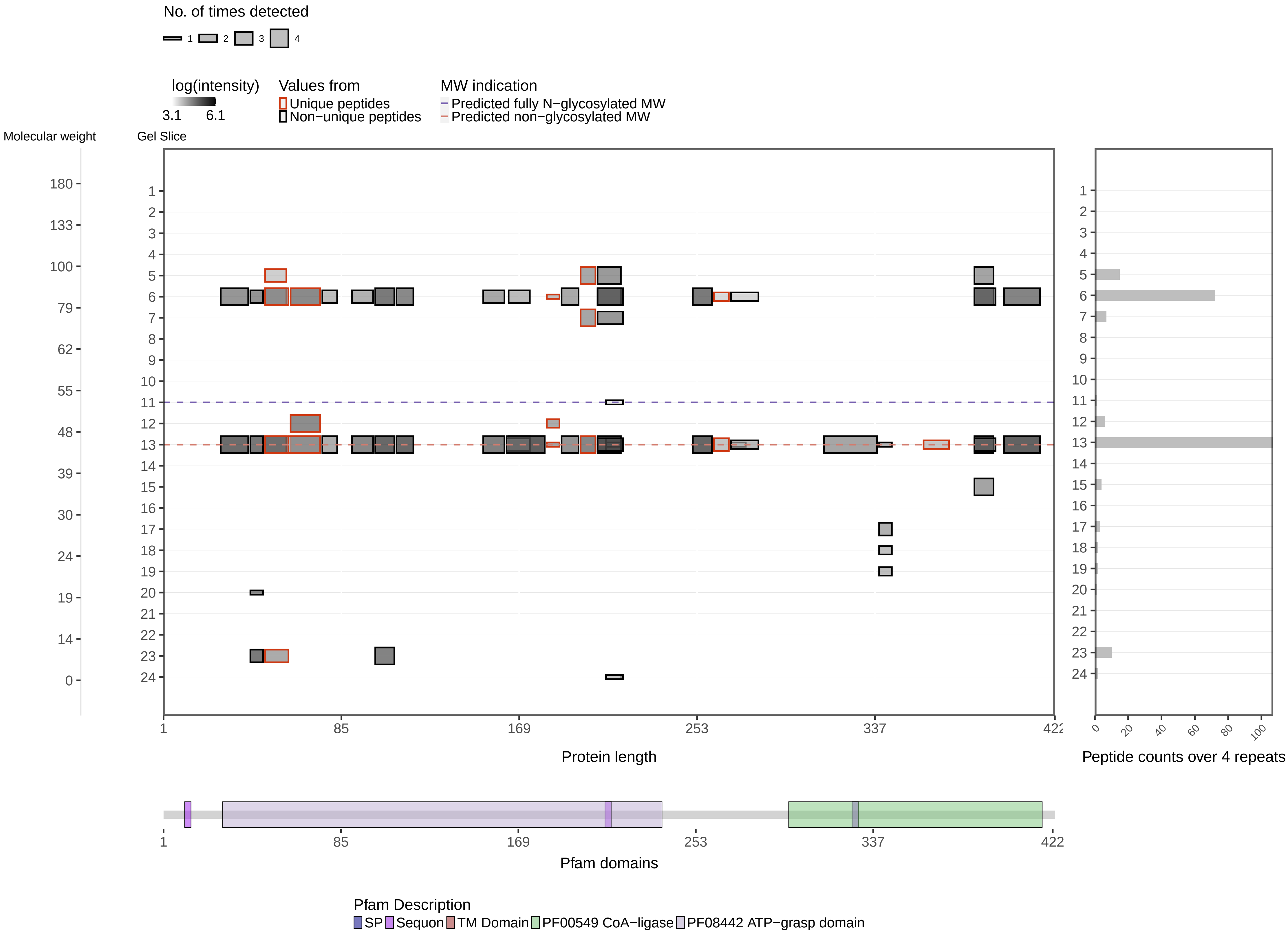


Protein sequence:

KVVDGLSPKVADRNDSIEQVKKLYKLFCESDCTMLEINPLAETSDNKLVAADAKLNFDDNAAYRQKDIFSLRDSSQEDPREVAAAKADLNYIGLDGEIGCMVNGAGLAMATMDIIKLHGGTPANFLDVGGNATEGQVVEAFKILTADEKVKAILVNIFGGIMKCDVIASGIANAAKQAQLKVPVVVRLEGTNVEQGKRILKESGMTLITAEDLDDAAEKAVKALA*

### NbL16g06510.1 60S_ribosomal_protein_L23 [MW = 15.01 kDa]

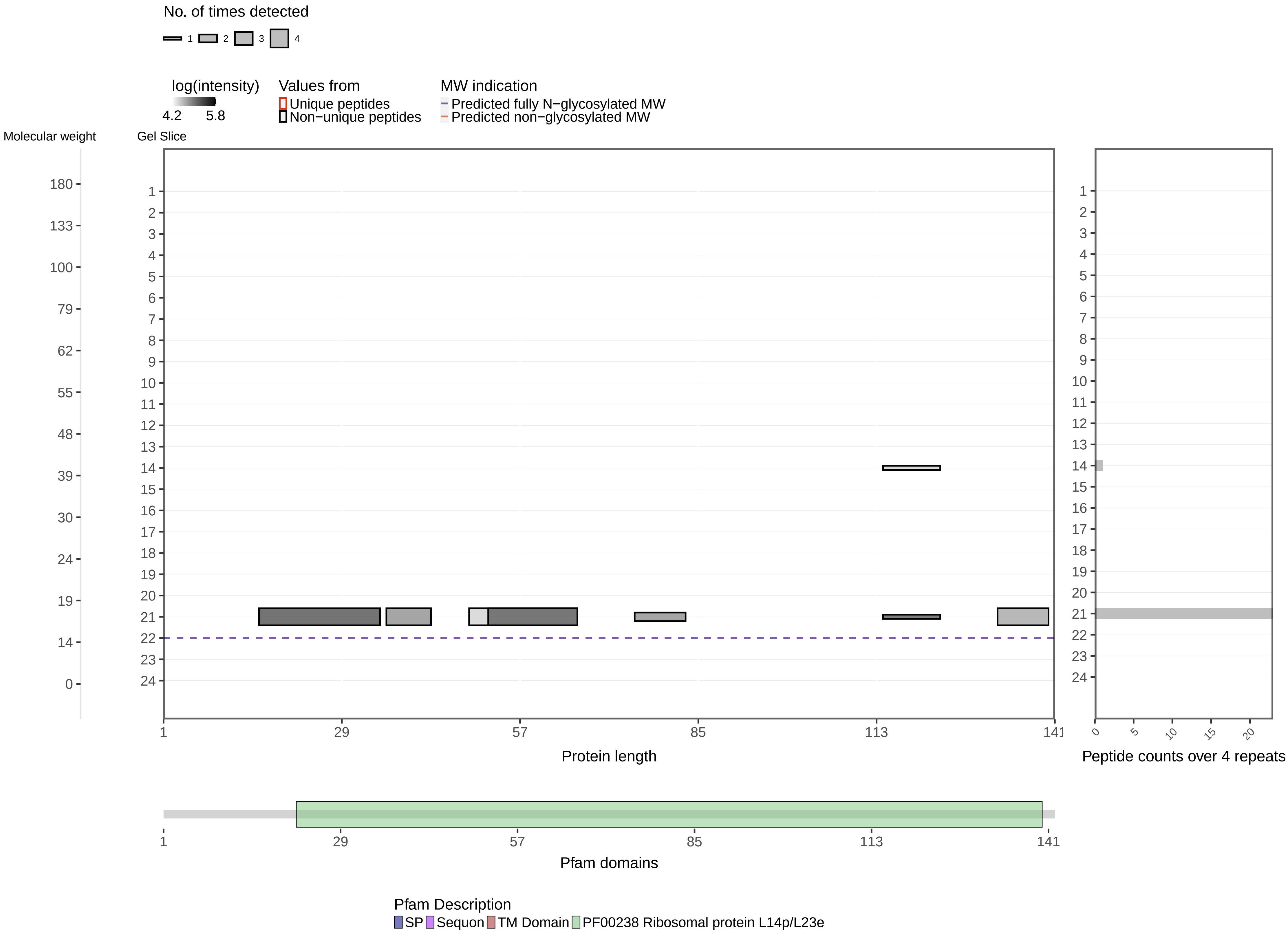


Protein sequence:

DMVMATVKKGKPDLRKKVMPAVVVRQRKPWRRKDGVFMYFEDNAGVIVNPKGEMKGSAITGPIGKECADLWPRIASAANAIV*

### NbL16g06460.1 putative_G3BP−like_protein [MW = 49.47 kDa]

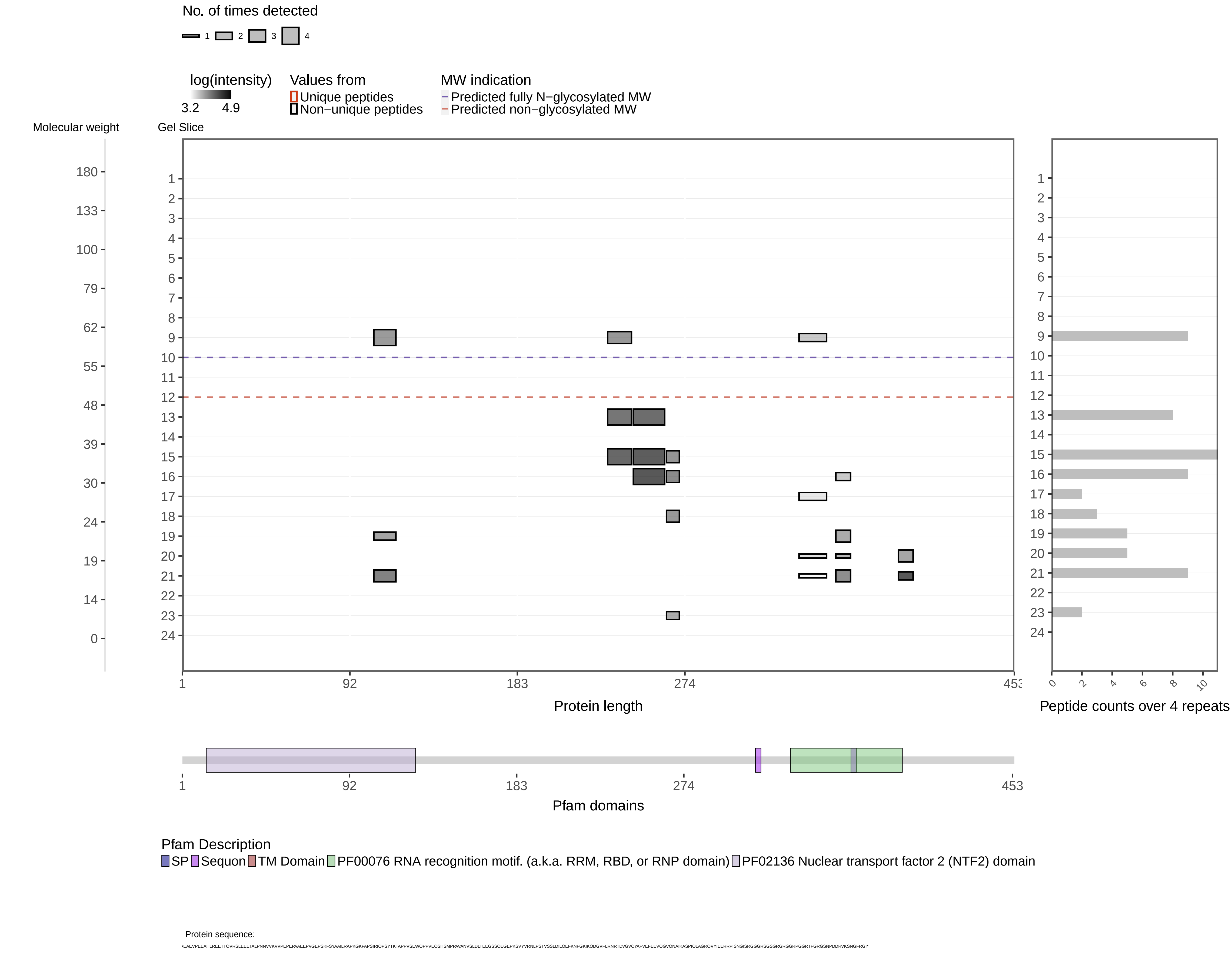

### NbL16g06390.1 photosystem_I_reaction_center_subunit_IV_A_chloroplastic [MW = 15.05 kDa]

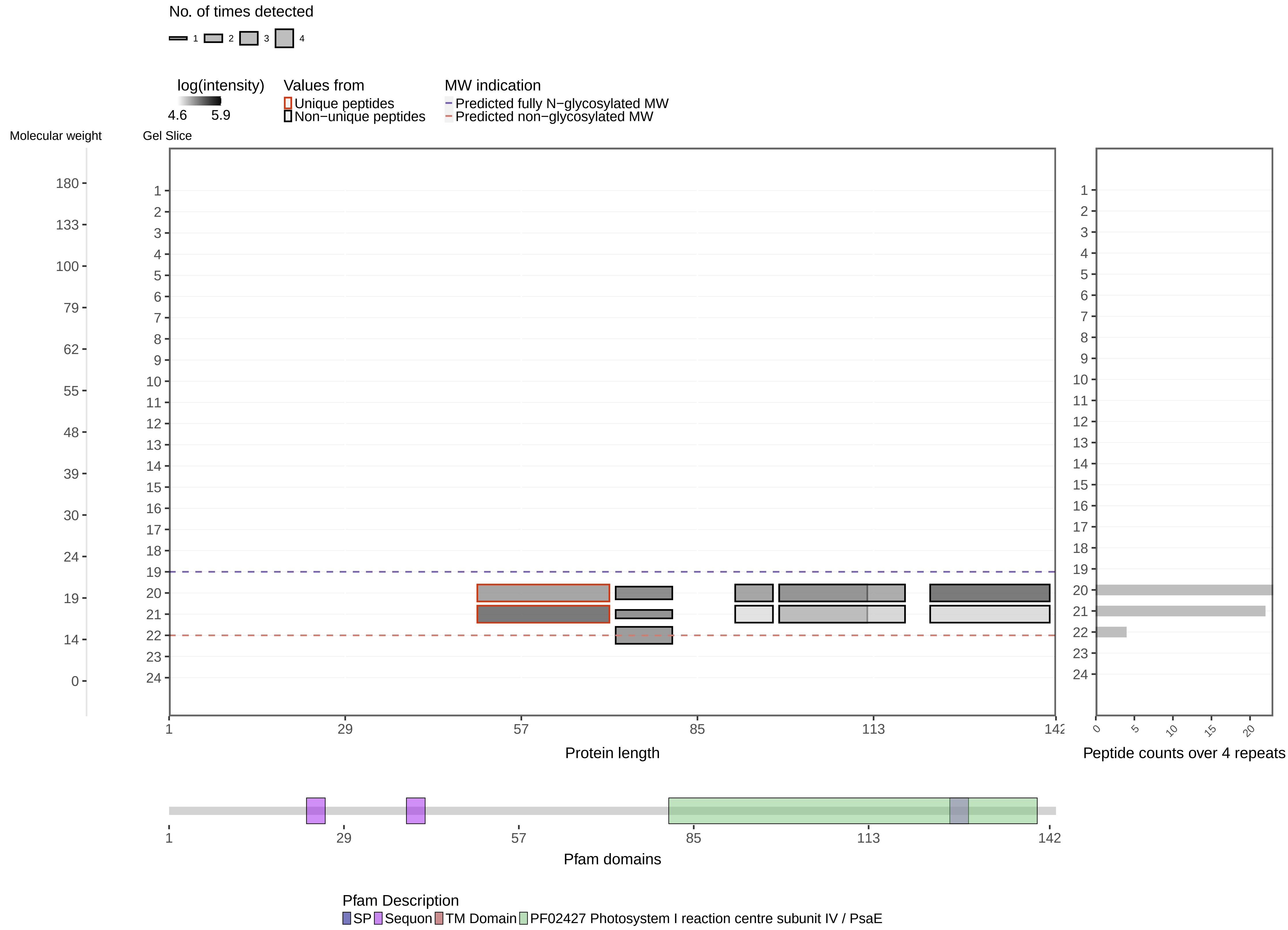


Protein sequence:

AATTAEPAEAPVKAAKPPPIGPKRGTKVRILRKESYWYKGTGSVVACDQDPNTRYPVVVRFNKVNYANVSTNNYALDEIEEVK*

### NbL16g06360.1 GDSL_esterase/lipase_LTL1−like [MW − SP = 36.96 kDa]

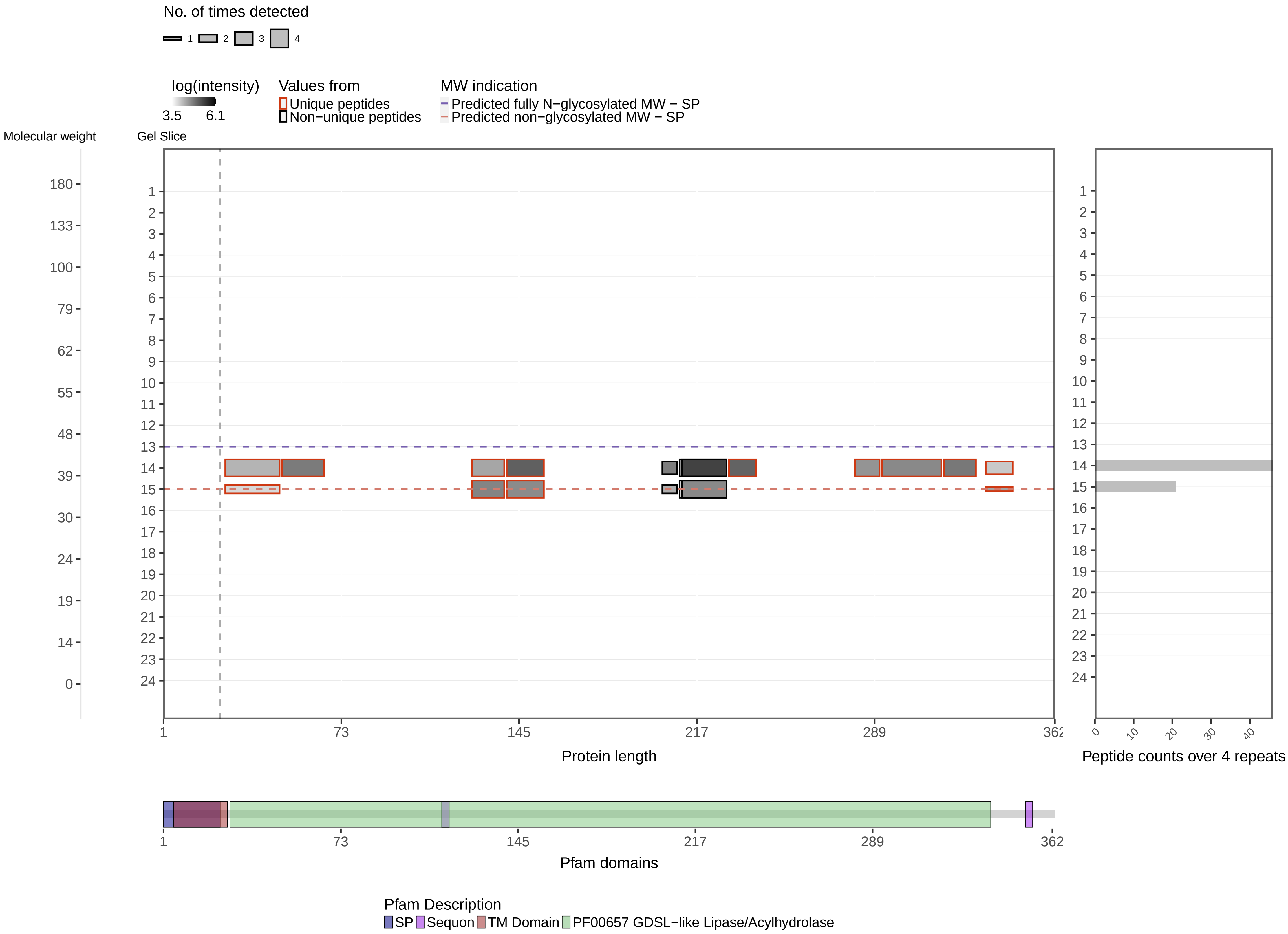


Protein sequence:

NYFFTPVSARRLQYNLEDFSVYIISEYRKILMRLYDLGARRVLVTGSGPLGCVPSSLASRSSNGECAEEPQRASAIFNPLLVEMIQSLNQELGSDVFVAANAVDMQNDFIRDPQAFGFITSKVACCGQGPYNGVGICSAASNLCPNRDVYAFWDPFHPTERANKIIVQTIFTGSDKYMTPMNLSTIMAIDSL*

### NbL16g06330.1 26S_proteasome_regulatory_subunit_4_homolog_A [MW = 49.5 kDa]

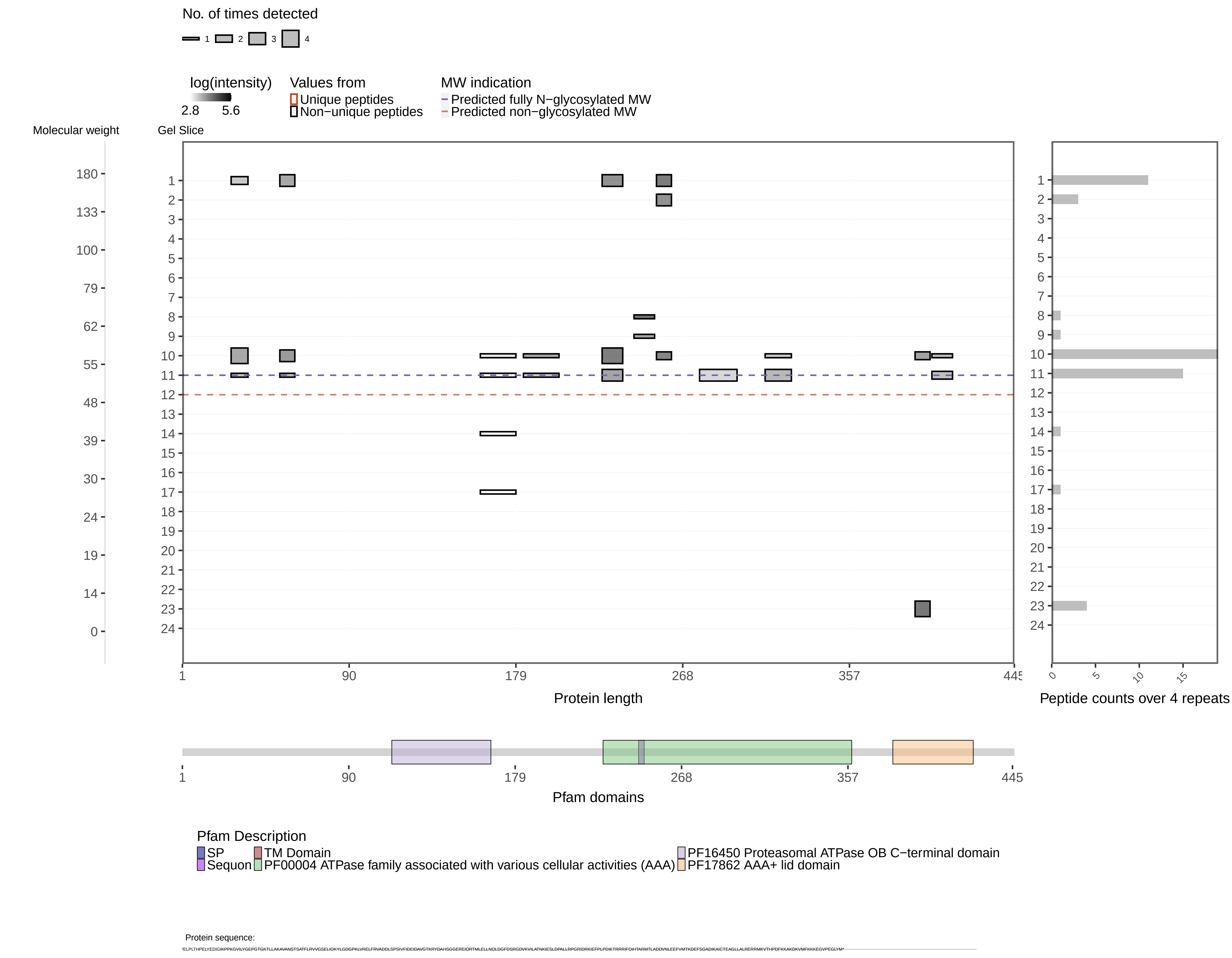

### NbL16g06310.1 probable_pectate_lyase_8 [MW − SP = 46.38 kDa]

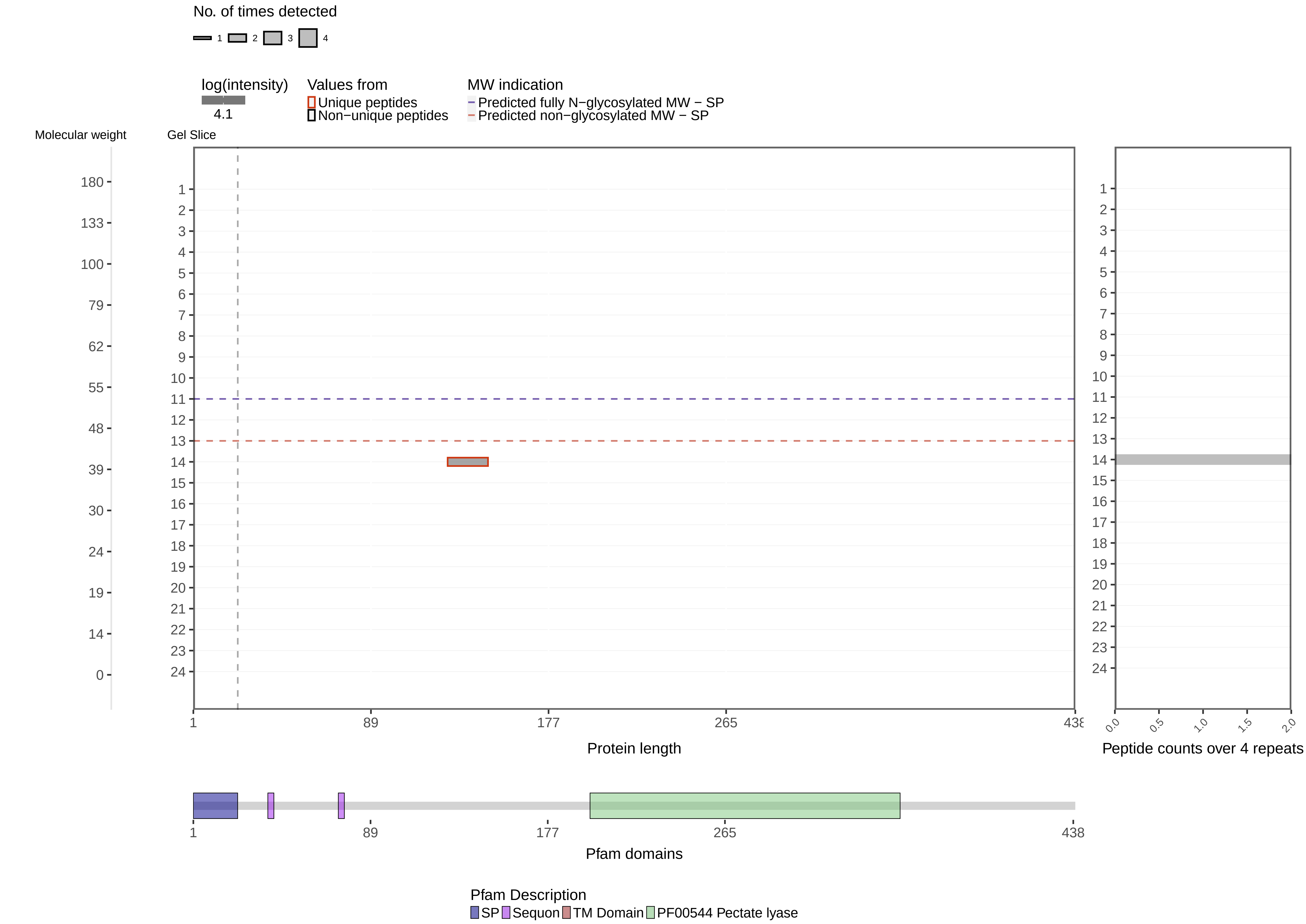


Protein sequence:

HIHDCKPTGNAMVRNSPSHYGWRTMADGDGISIFGSSHIWIDHNSLSNCADGLIDAIMGSTAITISNNYFTHHNEVILLGHSDSYIRDKVMQVTIAFNHFGEGLIQRMPRCRHGYFHVVNNDYTHWEMYAIGGSADPTINSQGNRYLAPVNPFAKEVTKRVEPWEGSWKHWNWRSEGDLMLNGAYFTASGRAAPRSYARASSLAAKSSSMVGTITSNAGALSCHIGFQC*

### NbL16g06060.1 oleoyl−acyl_carrier_protein_thioesterase_1_chloroplastic−like [MW = 42.06 kDa]

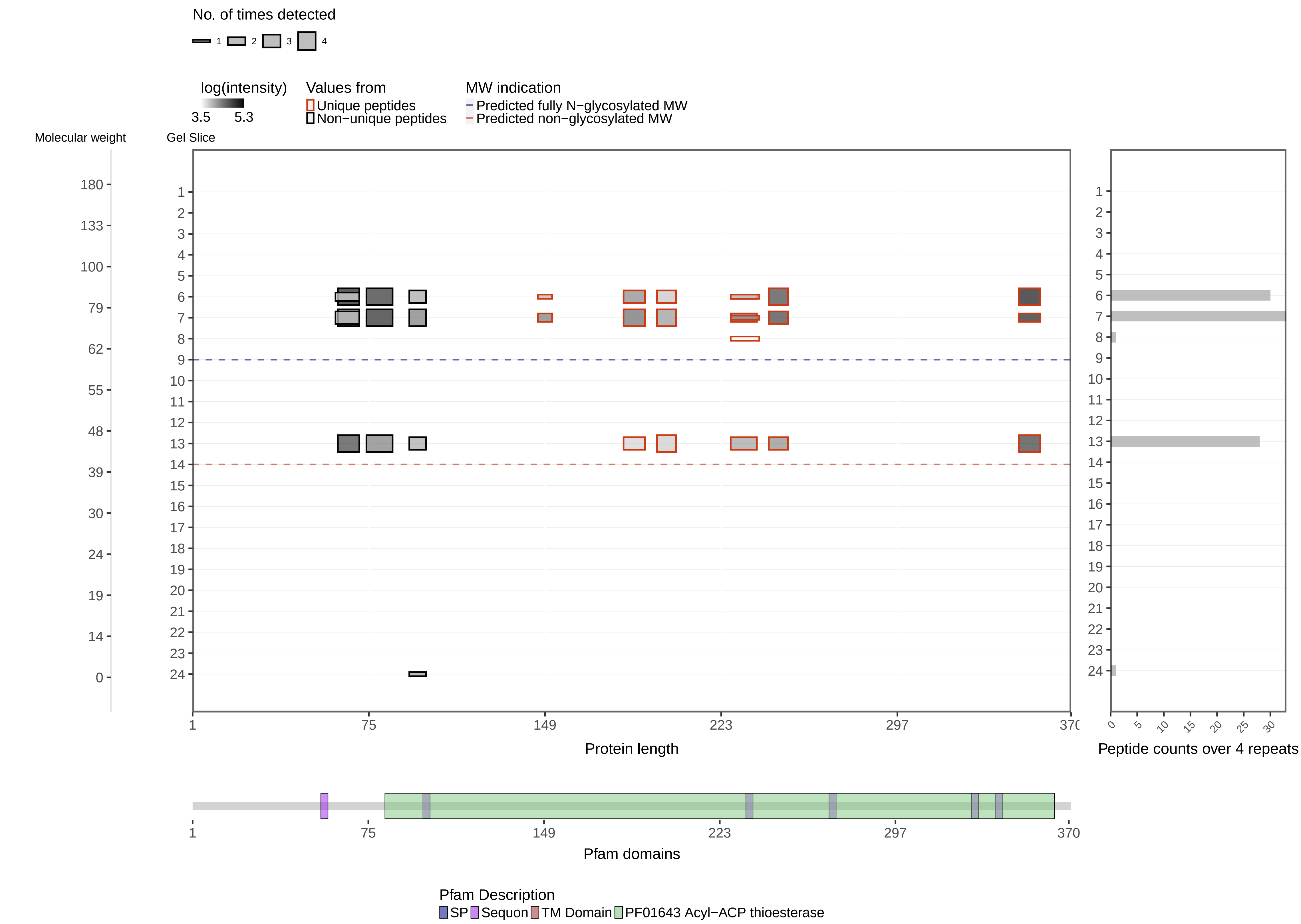


Protein sequence:

/GTRRDWILRDCATGDVIGRATSKWIMMNODTRRLOKVADEVRDELSIYFPKELRLAFPEENNGSLKKIAKLEDPAEYSKLGLVPRRADLDMNOHVNNVTYIGWVLESMPOEIIDTHELETITIDYRRECOHDDVVDSLTSYEPIEDAASLEIEGAANGSGTAPKDLNKSFLHLLRLSSDGLEINRCRTEWRKKPARI*

NbL16g06030.1 galacturonosyltransferase_8 [MW = 65.39 kDa]

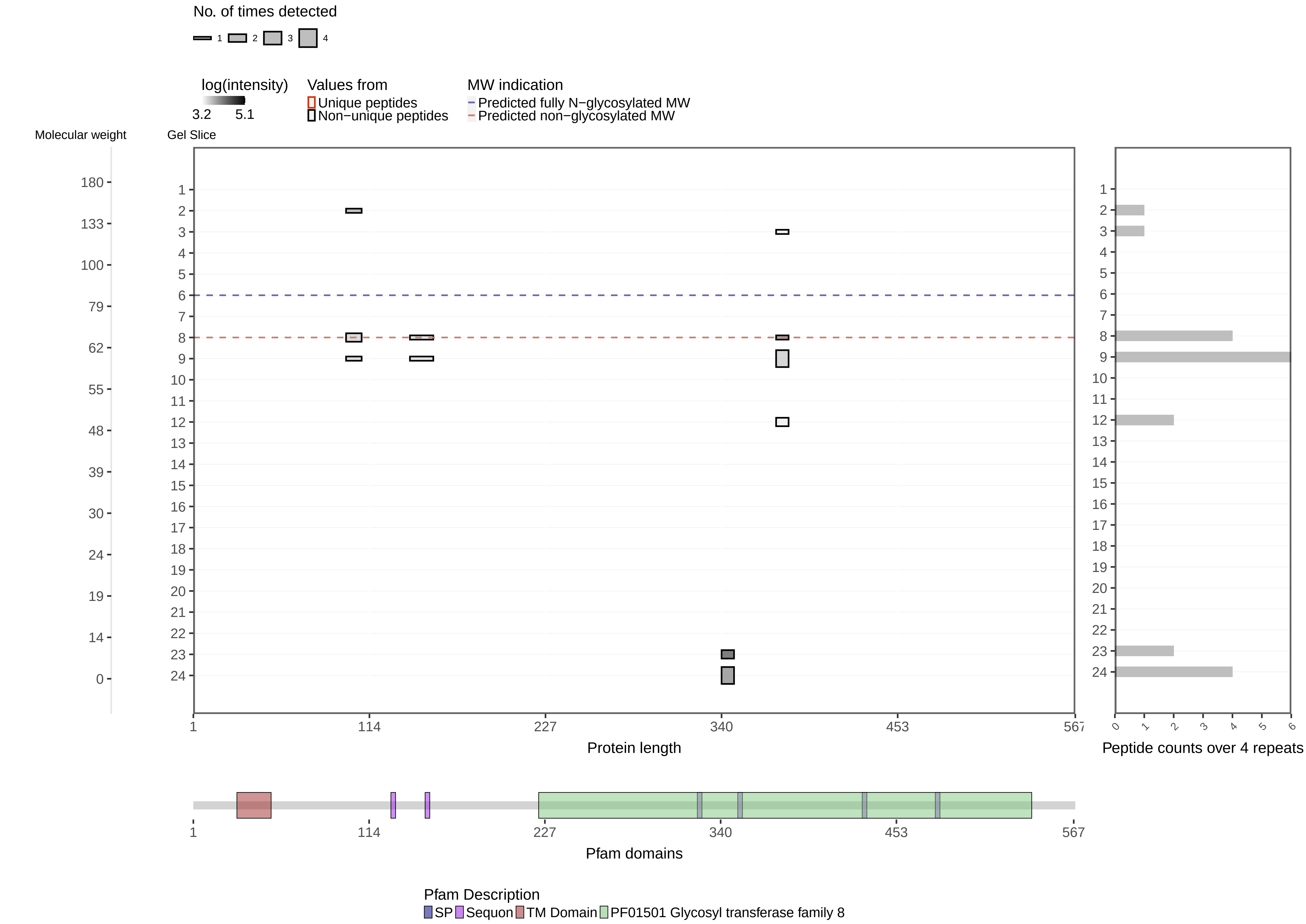


Protein sequence:

AVKHSKDPSKHVFHVVTDKMNLGAMQVMFKMKDYNGAFIEVKSVEDYKFLNSSYVPVLKQLESAKLQQFYFENKLENATKDTTNMKFRNPKYLSILNHIRFYLPELYPKLHRILFLDDDIVVQKDLTGLWKIDMDGKVNGAVETCFGSFHRYAQYMNFSHPLIKEKFNPKACAWAYGMNFFDLDAWRREKLTEEYHYWQSLNENRTLWKLGTLPPGLITFYSTTKPLDKSWHVLGLGYNPSISMDEINNAAVVHFNGNMKPWLDIAMNQFRPLWTKYVDYENEYVQACNF

### NbL16g06020.1 putative_G3BP−like_protein [MW = 50.43 kDa]

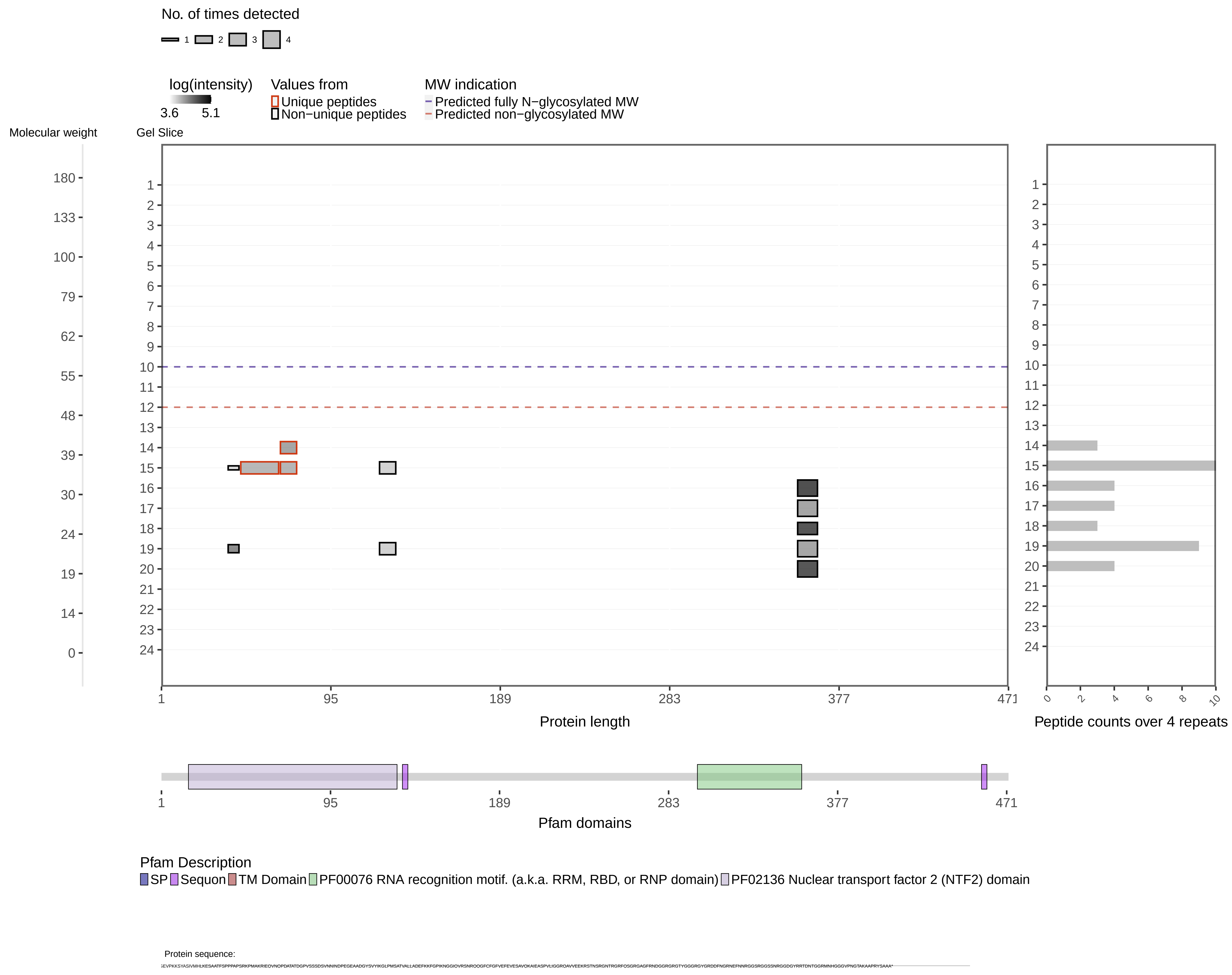

NbL16g05940.1 peptidyl−prolyl_cis−trans_isomerase_FKBP62−like [MW = 64.08 kDa]

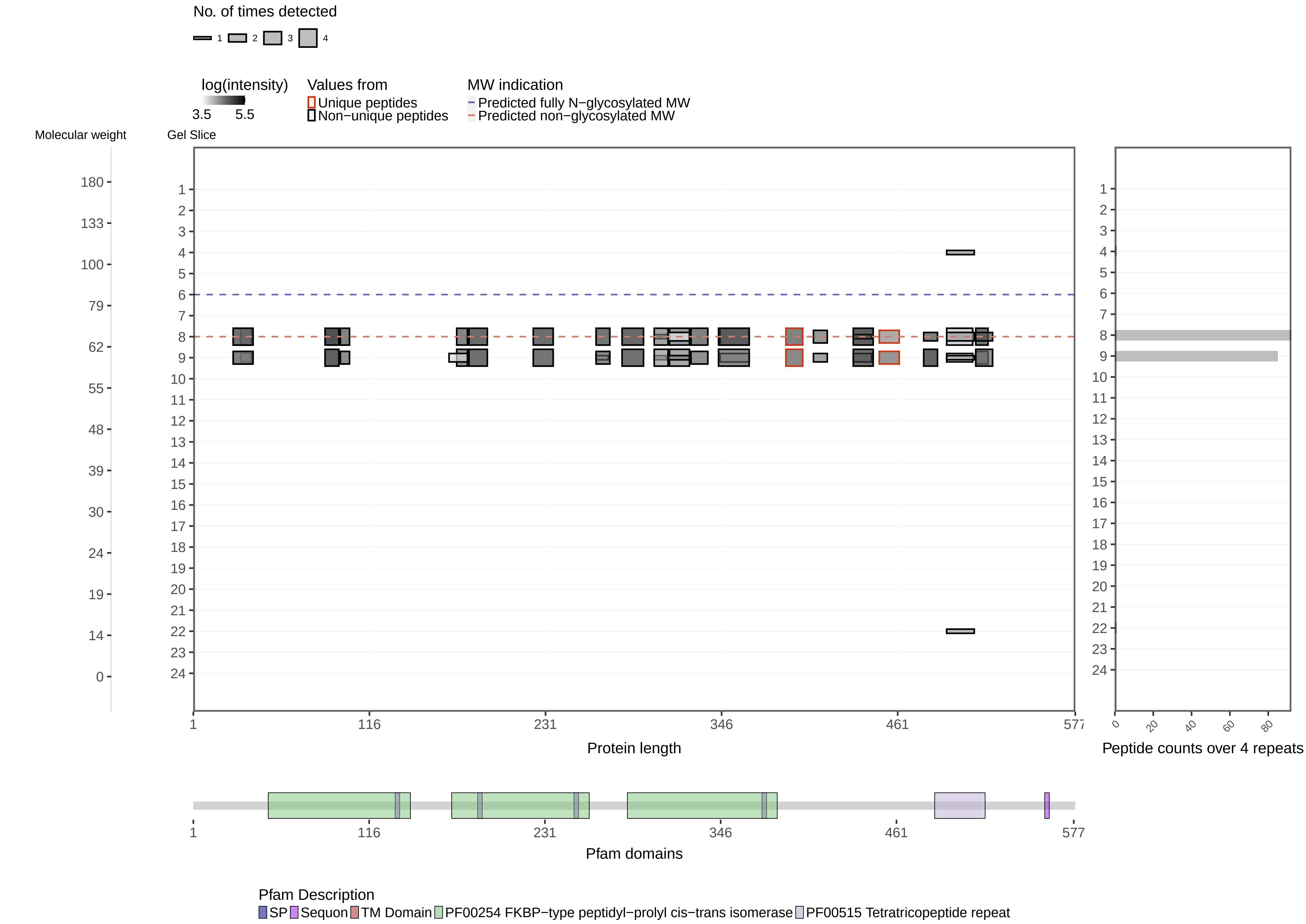


Protein sequence:

VVKKILKEGEGYEKPNEGAVVKLKLIGKLQDGTVFIKKGHDGENEDELFEFKTDEEQVIDGLDRAVMTMKKGEVALLTIAPEYAFGSSESKQDLAVIPPNSTVHYEVELVSFVKDKESWDMNTQEKIEAAGKKKEEGNALFKAGKYTRASKRYEKAAKFIEYDTSFSEEEKKQSKALTISCNLNNAACKLKLKDYKQAEKLCTKVLELESTNVKALYRRAQAYMNMADLDLAEFDIKKALEIDPDNRDVKLEYKALKDKVKEFNKKDAKFYGNMFAKLNKLETANSSKSAPKEPE

### NbL16g05930.1 40S_ribosomal_protein_S8 [MW = 25.02 kDa]

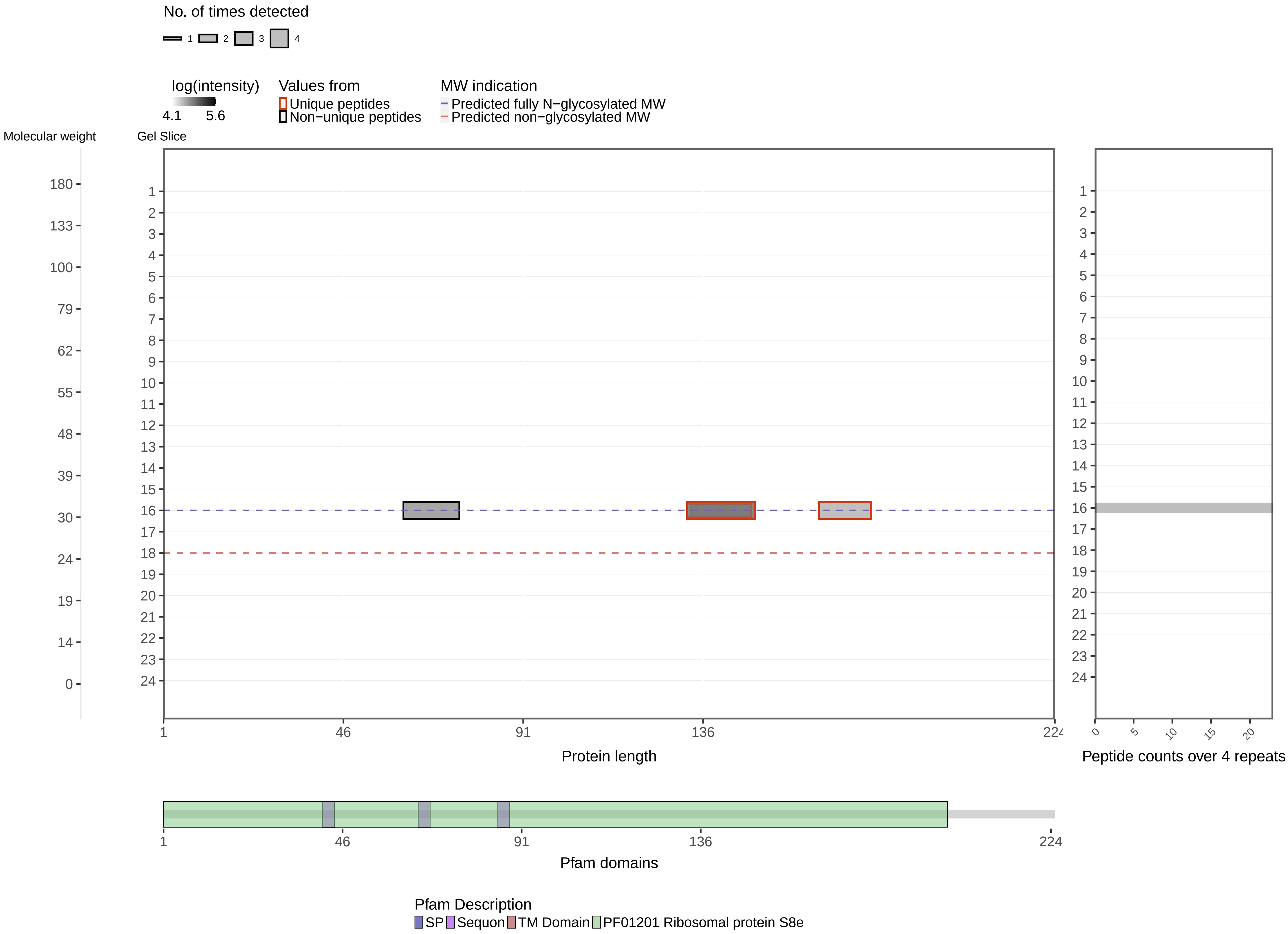


Protein sequence:

SAIVQVDAAPFKQWYLQHYGVDIGRKKKGAAKKETAEEGEGAAAAEETKKSNHVARKLEKRQKDRKLDAHLEEQFGAGRLLACISSRPGQRGRADGYILEGKELEFYMKKLQKKKGKAGAGGAA*

### NbL16g05790.1 protein_AE7−like [MW = 17.72 kDa]

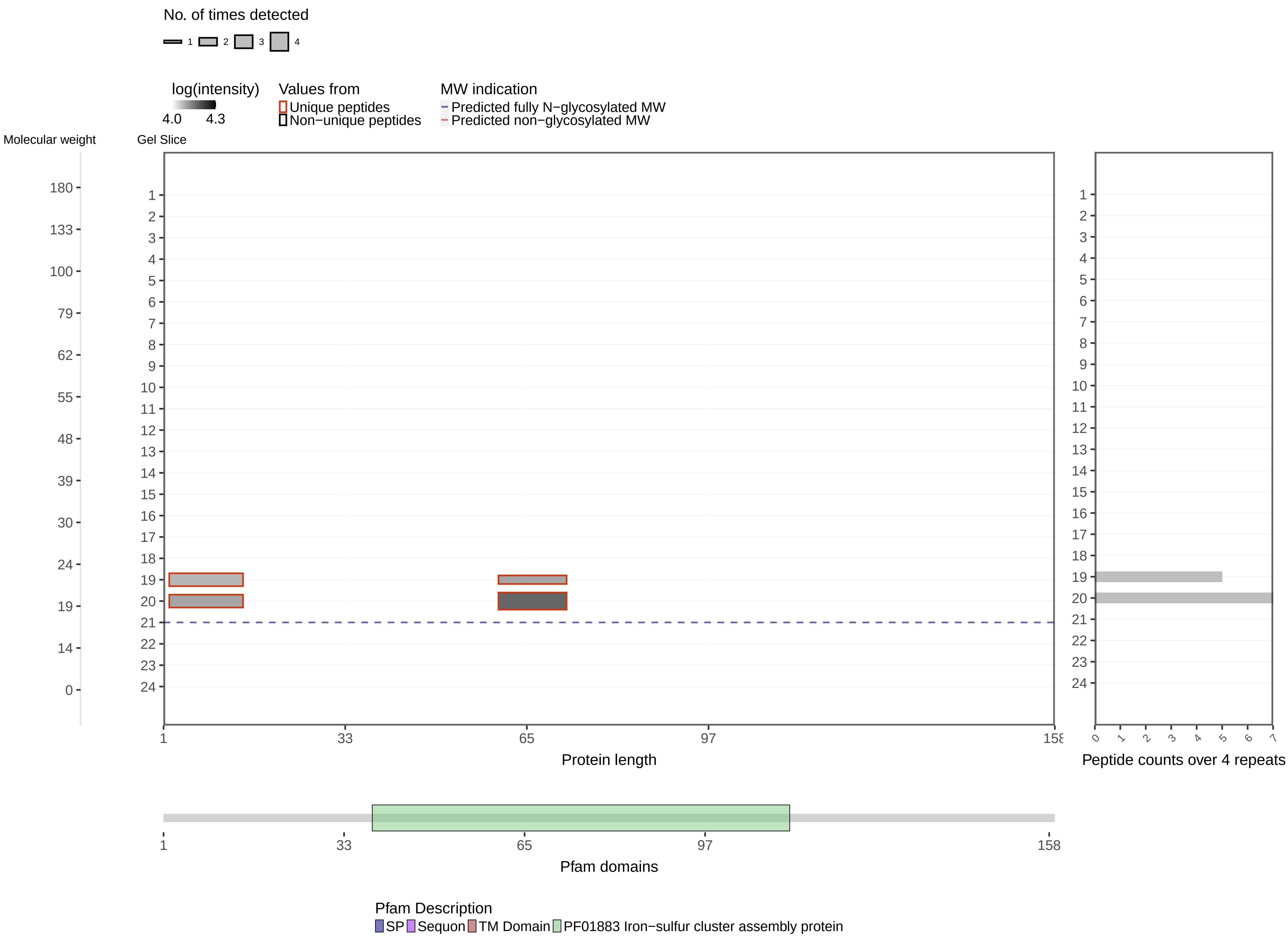


Protein sequence:

VDDERSYVRVTFTPTVEHCSMATVIGLCLRVKLMRSLPSRYKVDIRVAPGTHATEAAVNKQLNDKERVAAALENPNLVDMVDGCLAPSFE*

### NbL16g05780.1 serine_carboxypeptidase−like_20 [MW − SP = 53.01 kDa]

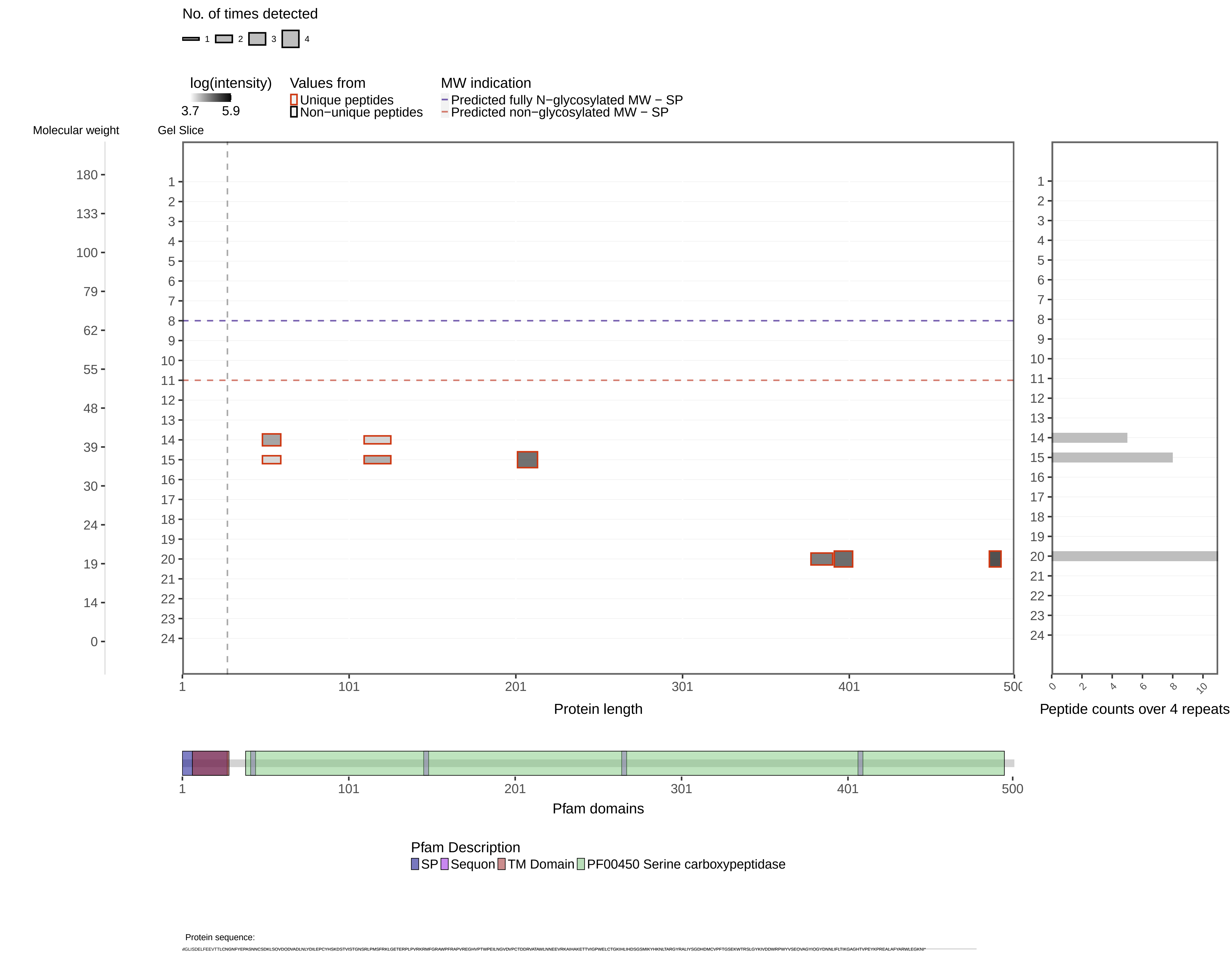

### NbL16g05660.1 60S_ribosomal_protein_L5 [MW = 34.67 kDa]

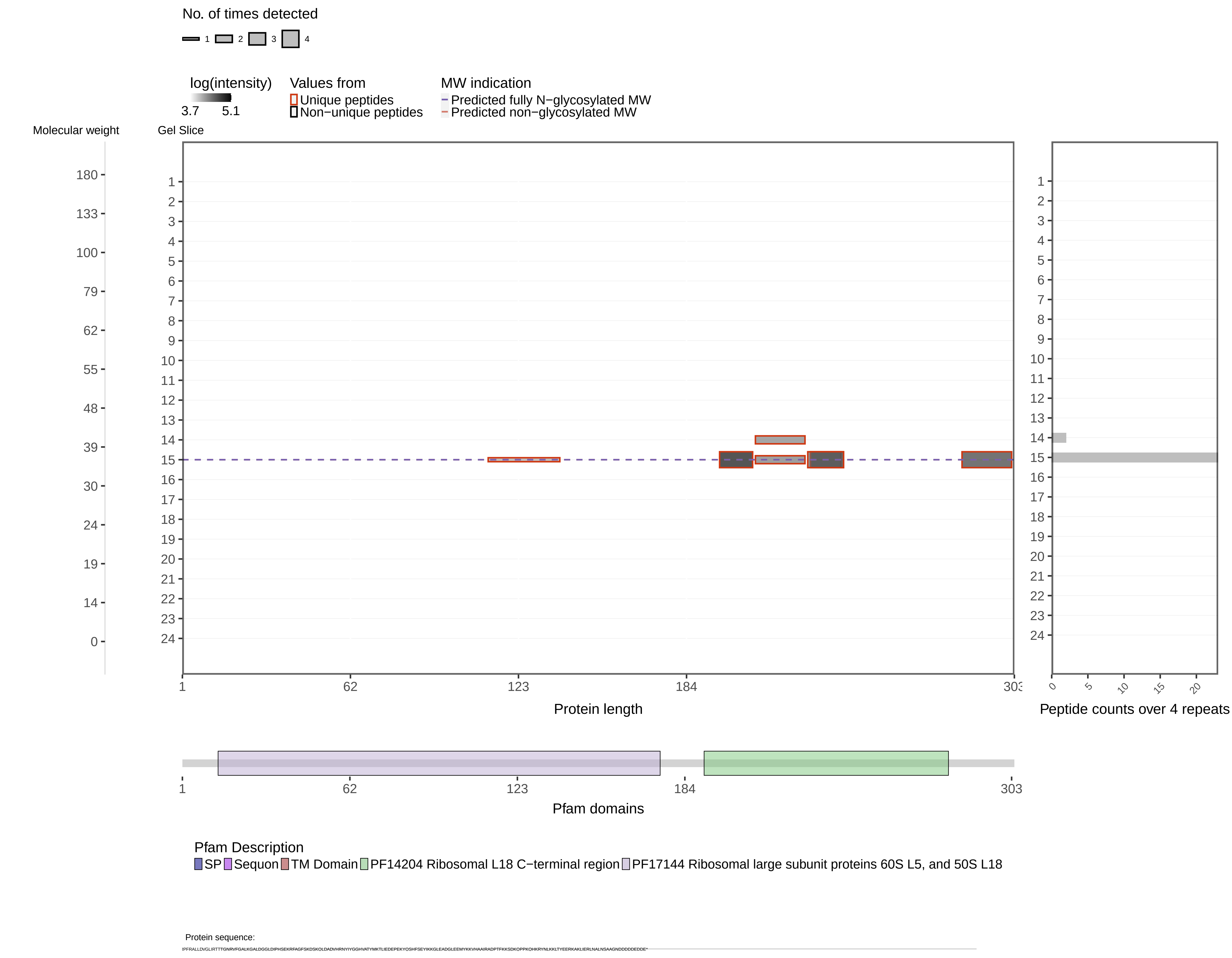

### NbL16g05560.1 50S_ribosomal_protein_L17_chloroplastic [MW = 23.16 kDa]

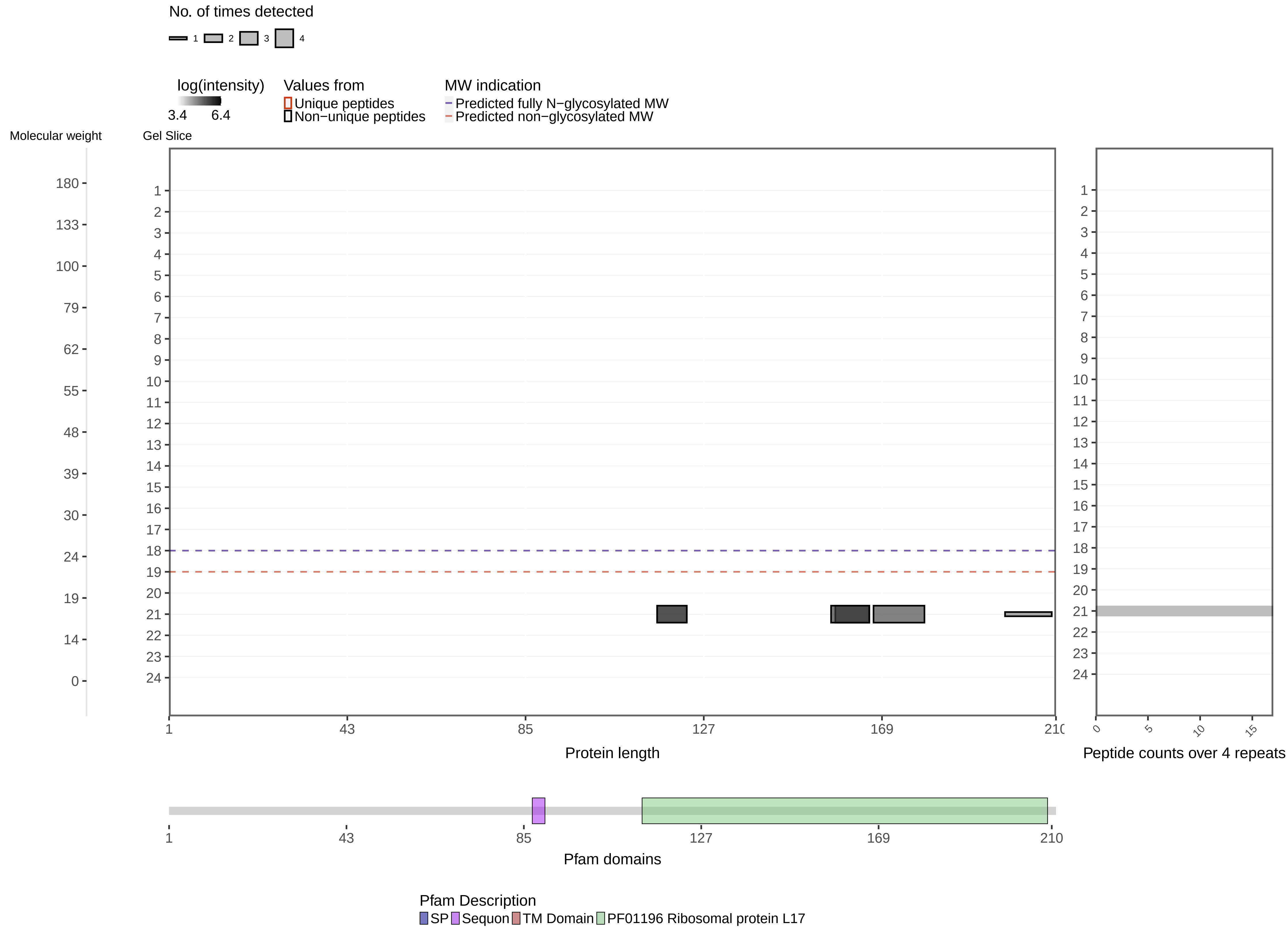


Protein sequence:

MRHGRKVPKLNRPPDQRRALLRGLTTQLLKHGRIKTTKARARAVRKYVDKMITMAKDGSLHKRRQALGFIYEKQIVHALFAEVPDRYGERNGGYTRIIRTLPRRGDNAPMAYIELV*

### NbL16g05490.1 pathogenesis−related_genes_transcriptional_activator_PTI6−like [MW = 29.68 kDa]

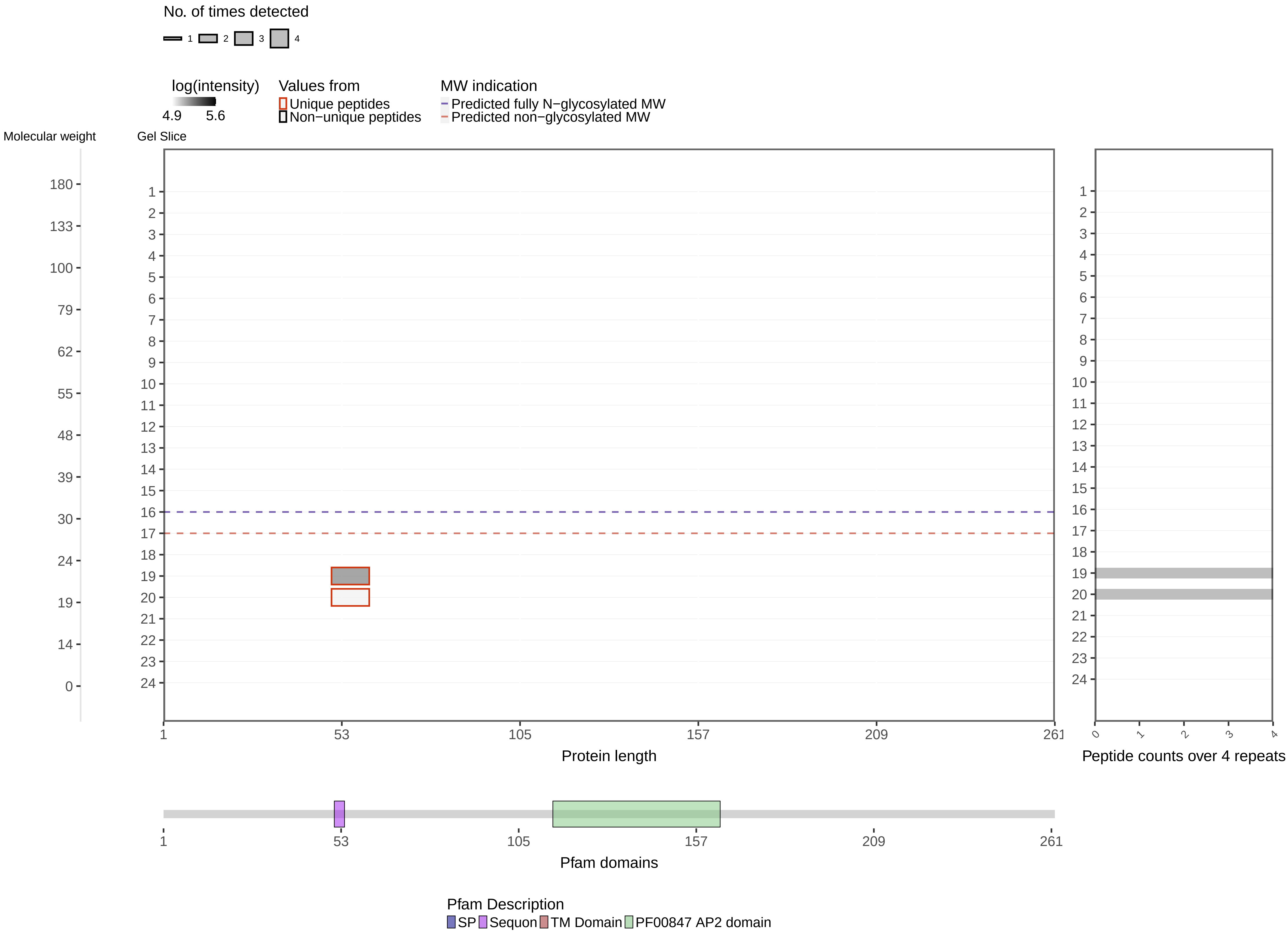


Protein sequence:

/RORPWGRWAAEIRDPTRGKRVWLGTYDTPEEAATVYDEAAVKLKGPDAVTNFPKVITADITETESVTDGGEKSENDFTLSPTSVLRNDYFTPFSDLGFCEVDAFGFDVDSLFROPEFGMSEKYYGEEFGEFDFDDFALEAR*

### NbL16g05480.1 eukaryotic_translation_initiation_factor_2_subunit_beta−like [MW = 35.55 kDa]

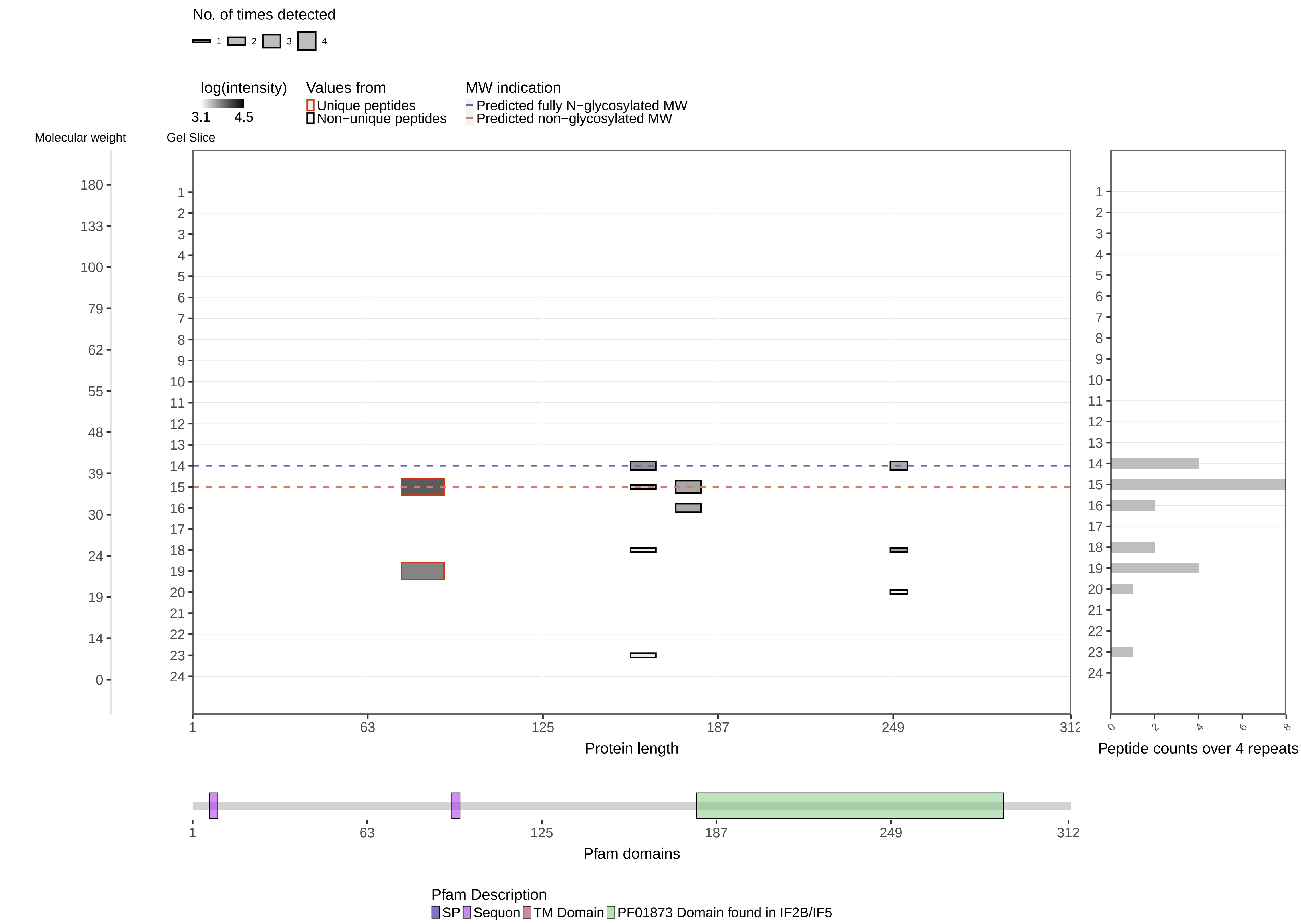


Protein sequence:

VLQQYPWEGTDRDYEYEELLGRVFNILRENNPELAGDRRRTVMRPPQVLREGTKKTVFVNFMDLCKTMHRQPEHVMTFLLAEMGTSGSLDGQQRLVIKGRFAPKNFEGILRRYVNEYVICNGCKSPDTILSKENRLFFLRCEKCGSGRSVAPIKAGFVARVGRRKAGT*

### NbL16g05410.1 40S_ribosomal_protein_S26−3−like [MW = 14.59 kDa]

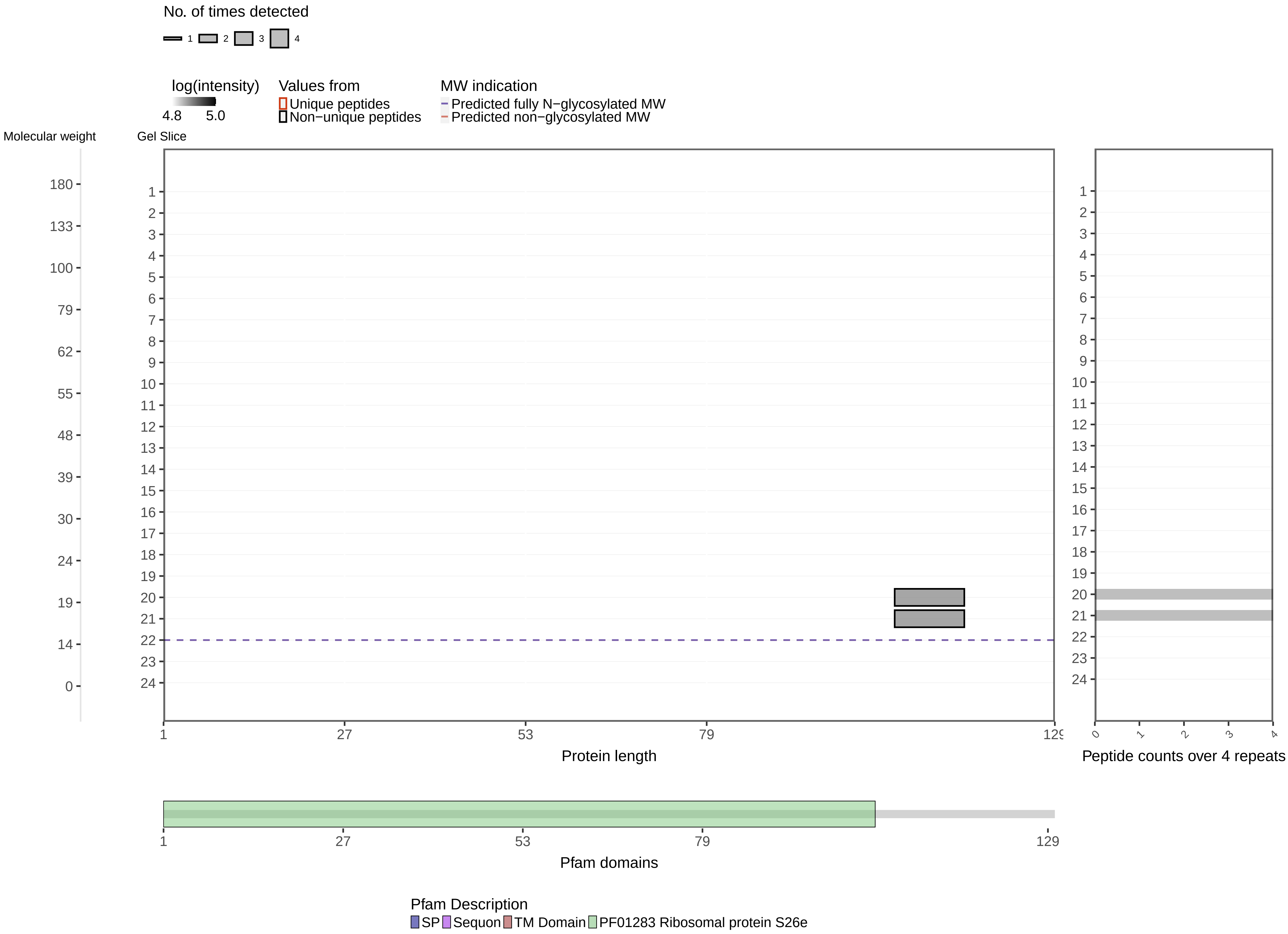


Protein sequence:

DVQEACAFETYTLPKLYLKMQYCVSCAIHSKVVRVRSRTDRRVREPPQRFRRPRDDAPKPGQAPRVPGAAPTAAART*

NbL16g05260.1 alpha−galactosidase_3−like [MW − SP = 48.93 kDa]

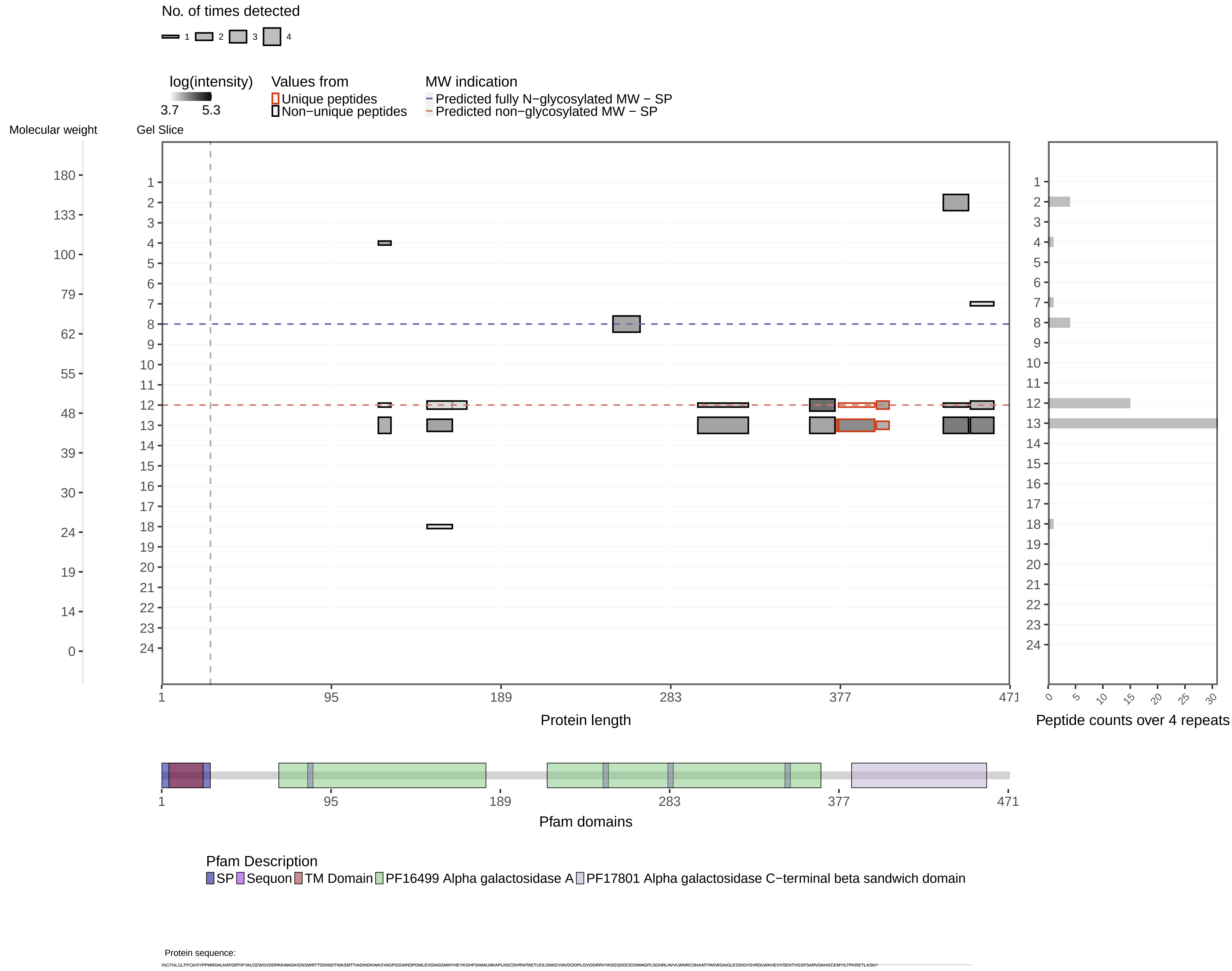

NbL16g05170.1 transcription_initiation_factor_TFIID_subunit_15b [MW = 44.63 kDa]

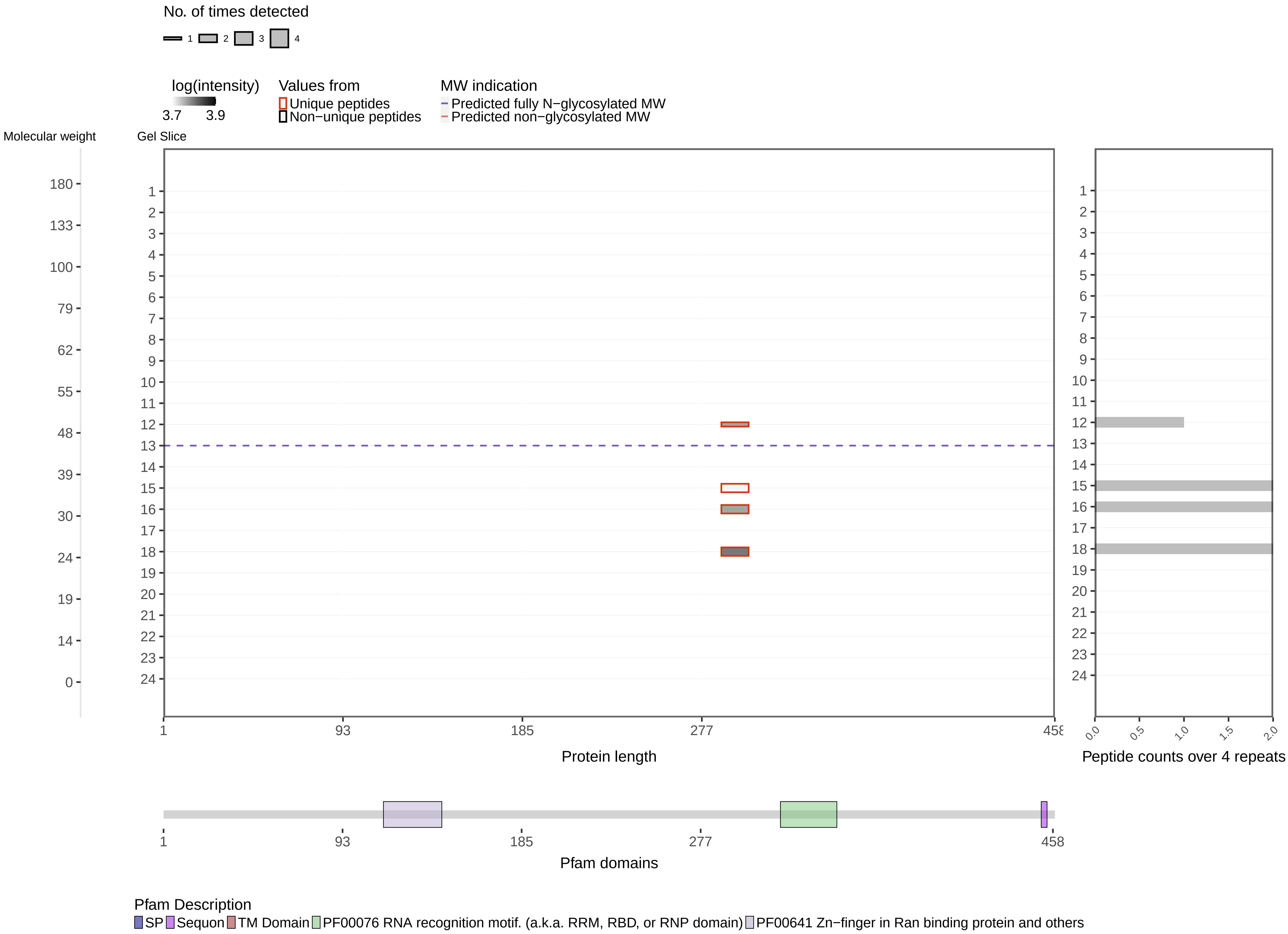


Protein sequence:

PPSYGAADSNYPPPSNTYGRNPDAVPPPTSYVGGPASYPPSYGAPGGYGGDKTANARGGGRGGPPGGYDGGYGGGQRNAGGYDISPAEAPEKVKQCDETCGDFCDNARIYISNLPPDVTIEELRELFGSIGQVARIKQKRGYKDQWPWSIKLYTDEQGKNKGDAVLSYEDPSAAHSAGGFFNNHDFRGYKISVAMAEKSAPKPAPAFGSGRGGRGGYGGGGGRRDNYRDSGGSGPDRQSHGGNRSRPY*

NbL16g05080.1 protein_DCL_chloroplastic−like [MW = 26.02 kDa]

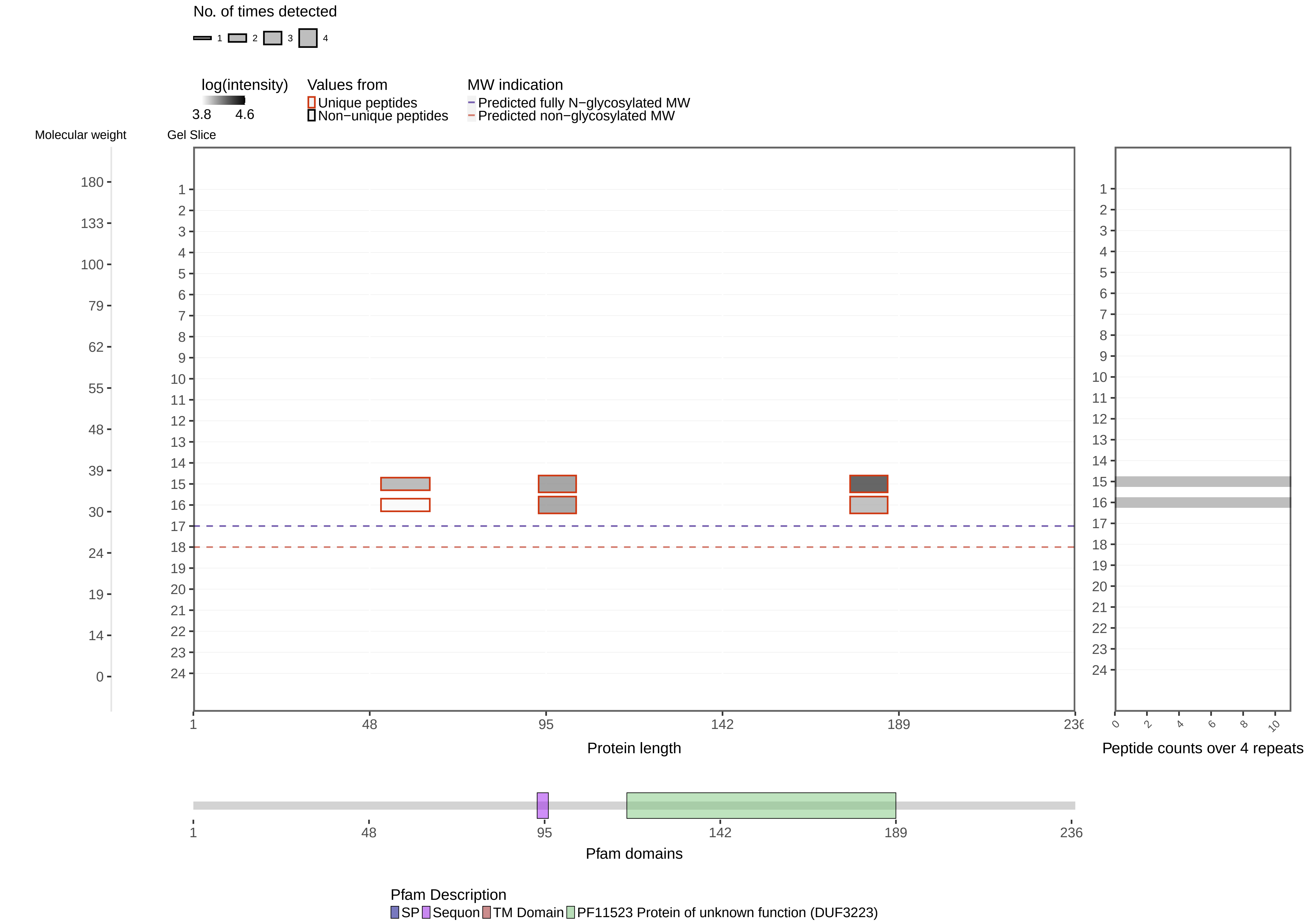


Protein sequence:

FGSSVEMFDYFYKLLHSWSPNLNLNKYEHLVLLDLLKKGHLEPERKIGTGIRAFQIRFHPQFKSRCFFVIREDGSVDDFSFRKCVDHILPLPENMQVKHEANKGLAHSGKGGGRKGGGHGRDRGNGGKSRY*

NbL16g05060.1 chlorophyll_a−b_binding_protein_4_chloroplastic−like [MW = 27.6 kDa]

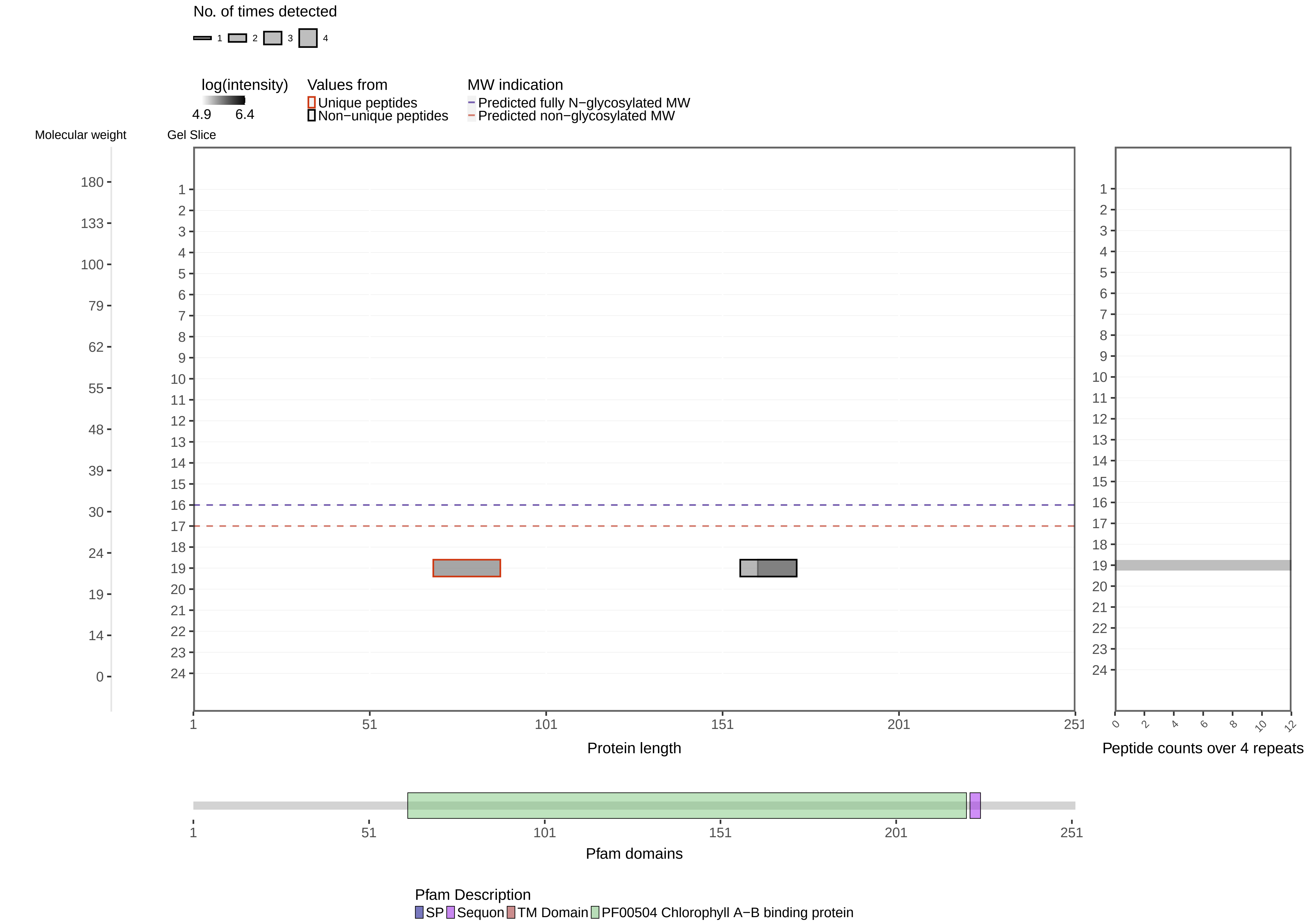


Protein sequence:

.PEVLTSAGLLNVPKWYDAGKSEYFASSSTLFVIEFILFHYVEIRRWQDIKNPGSVNQDPIFKSYSLPPNEVGYPGGIFNPLNFAPTLEAKEKEIANGRLAMLAFLGFLVQHNVTGKGPFDNLLQHLSDPWHNTIIQTFAN*

NbL16g04760.1 glycerophosphodiester_phosphodiesterase_GDPD6 [MW − SP = 43.5 kDa]

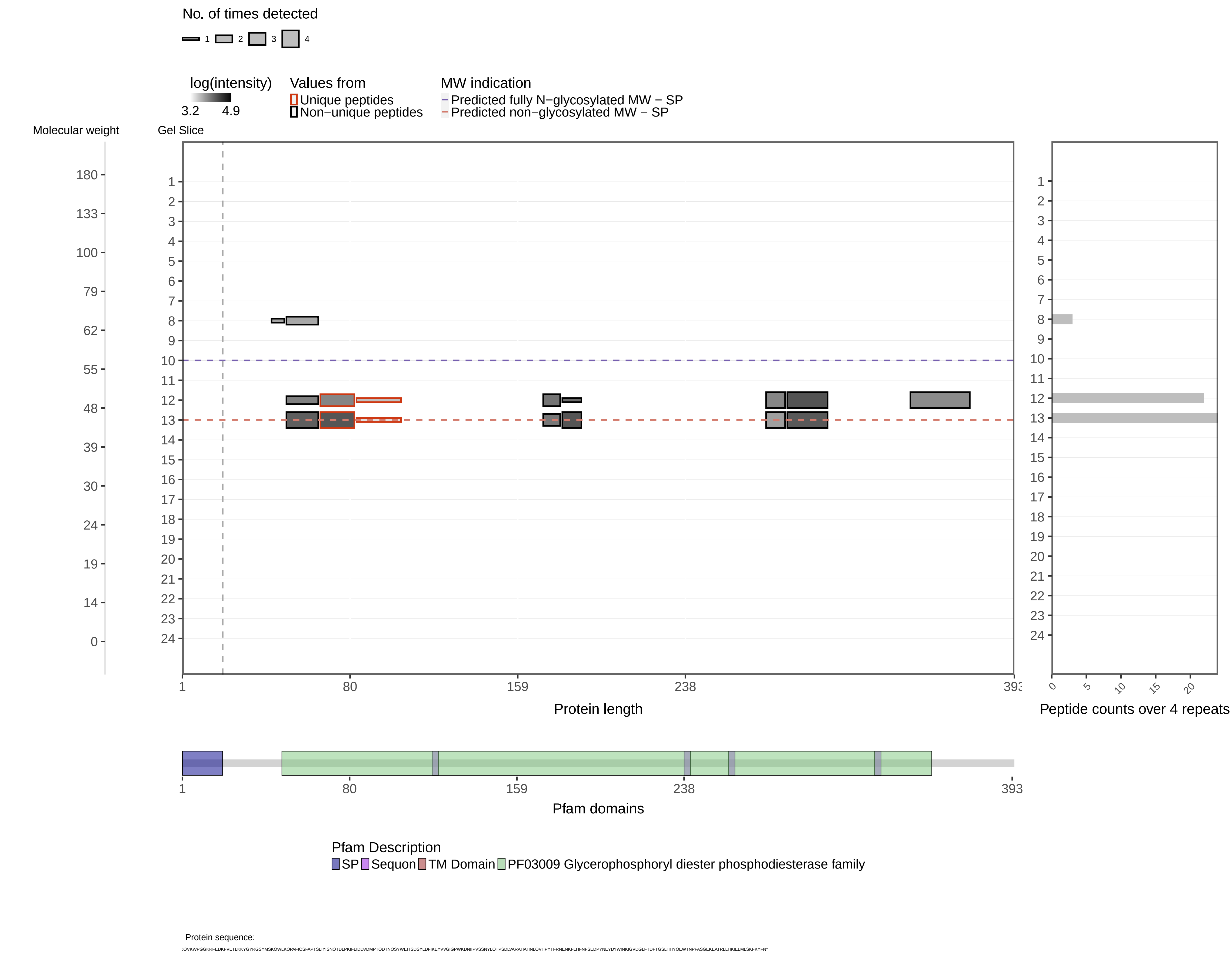

### NbL16g04660.1 D−aminoacyl−tRNA_deacylase−like [MW = 40.79 kDa]

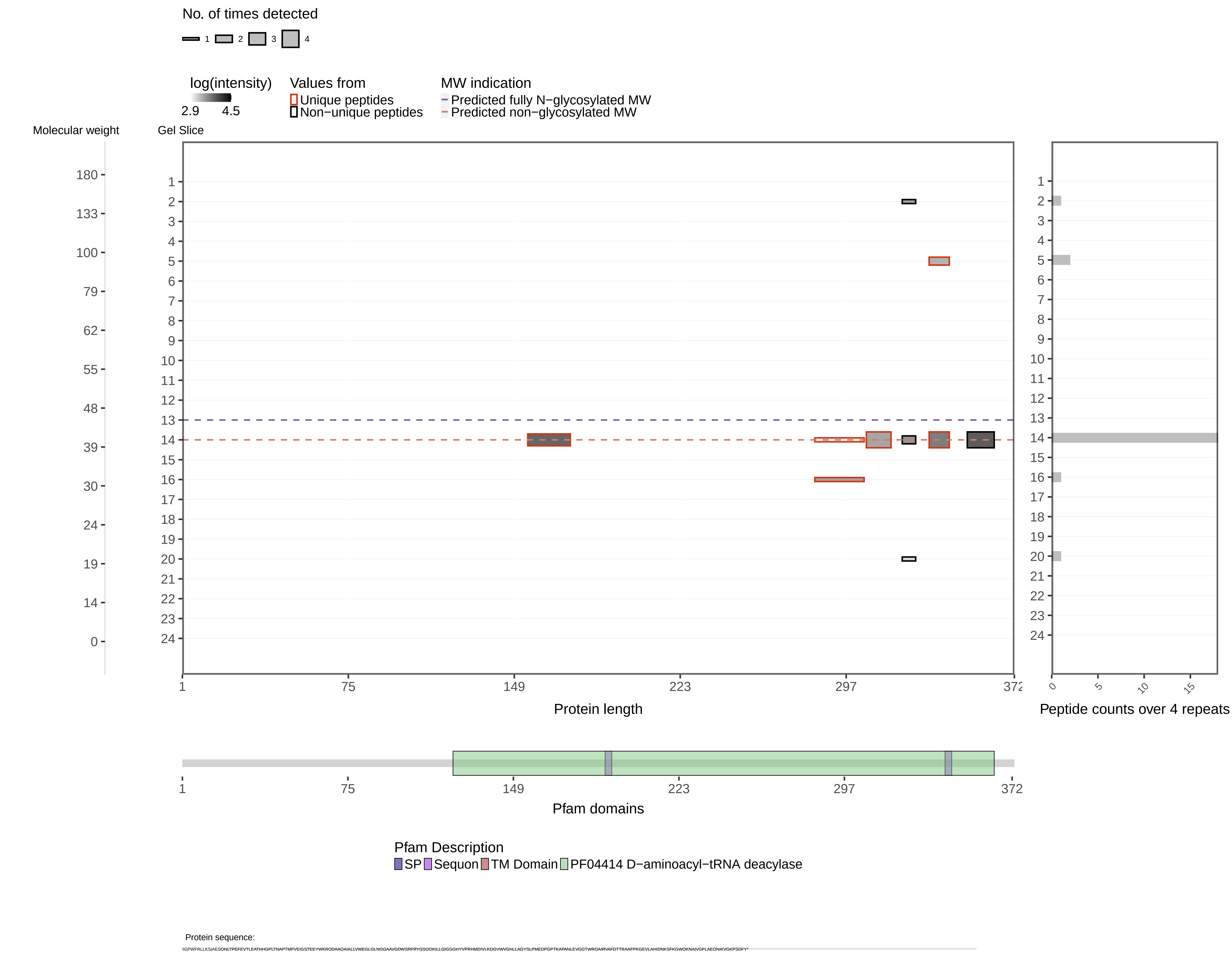

### NbL16g04550.1 ribulose_bisphosphate_carboxylase_large_chain [MW = 53.68 kDa]

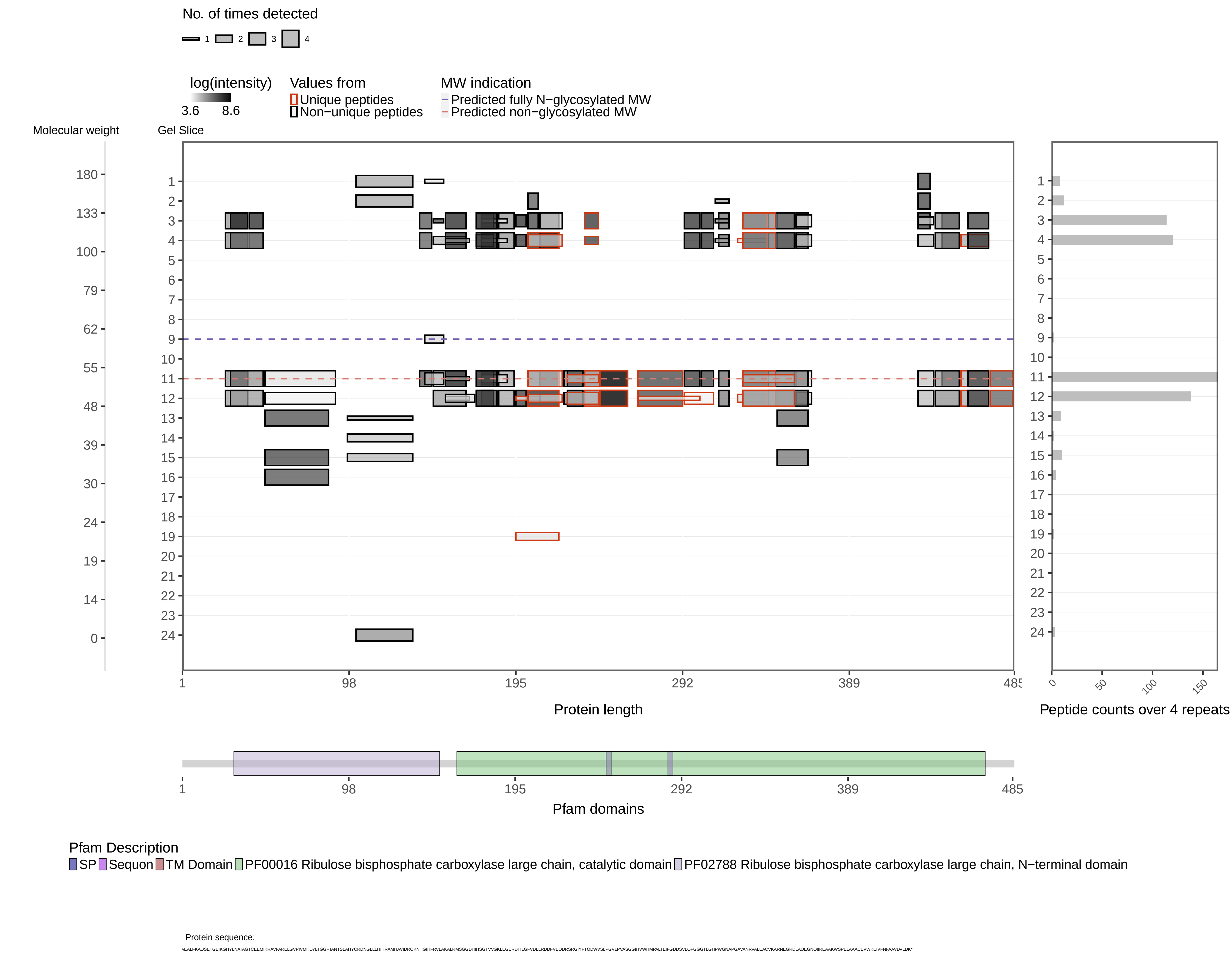

### NbL16g04540.1 ATP_synthase_subunit_beta_chloroplastic [MW = 25.73 kDa]

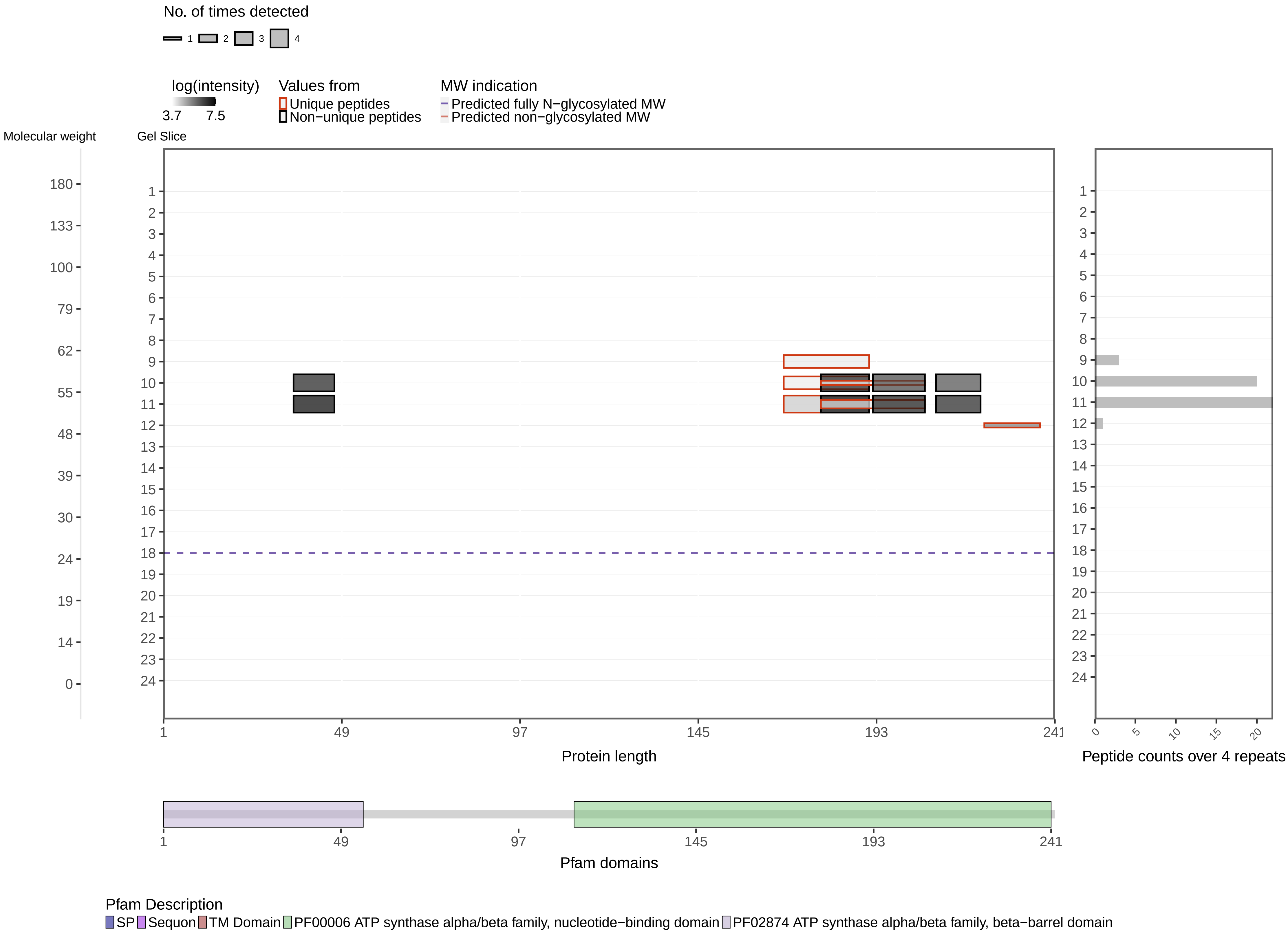


Protein sequence:

IFETGIKVVDLLAPYRRGGKIGLFGGAGVGKTVLIMELINNIAKAHGGVSVFGGVGERTREGNDLYMEMKESGVINEENIAESKVALVYGQMNEPPGARMRVGLTALTMAEYFRDVNEQDVLLFIDNIFRFVQA

NbL16g04530.1 ATP_synthase_epsilon_chain_chloroplastic−like [MW = 14.61 kDa]

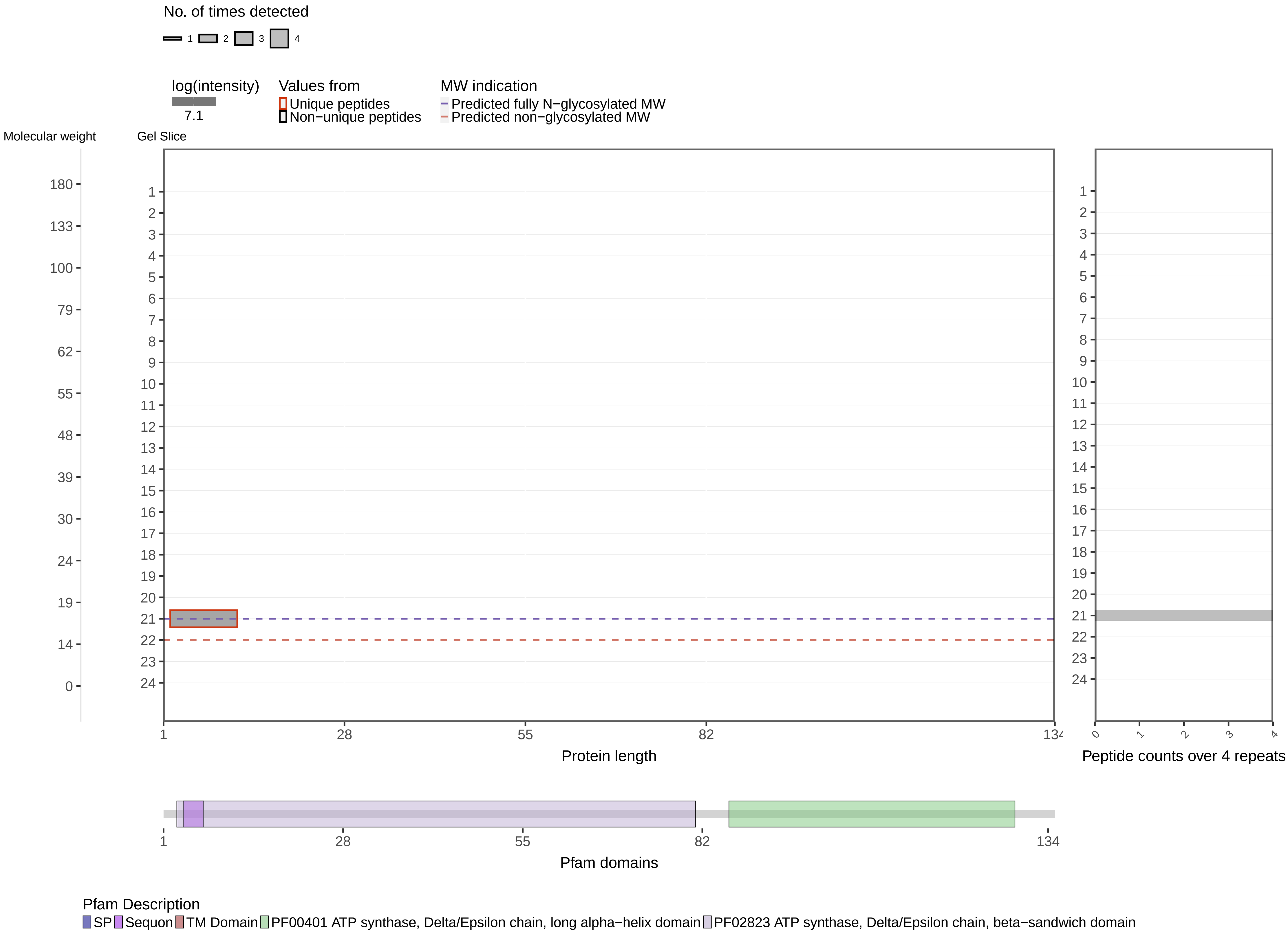


Protein sequence:

QWLTMALMGGFARIGNNEITVLVNDAEKGSDIDPQEAQQTLELAEANVKKAEGRRQKIEANLALRRARTRVEAINPIS*

NbL16g04380.1 ribosomal_RNA_small_subunit_methyltransferase_NEP1−like [MW = 31.39 kDa]

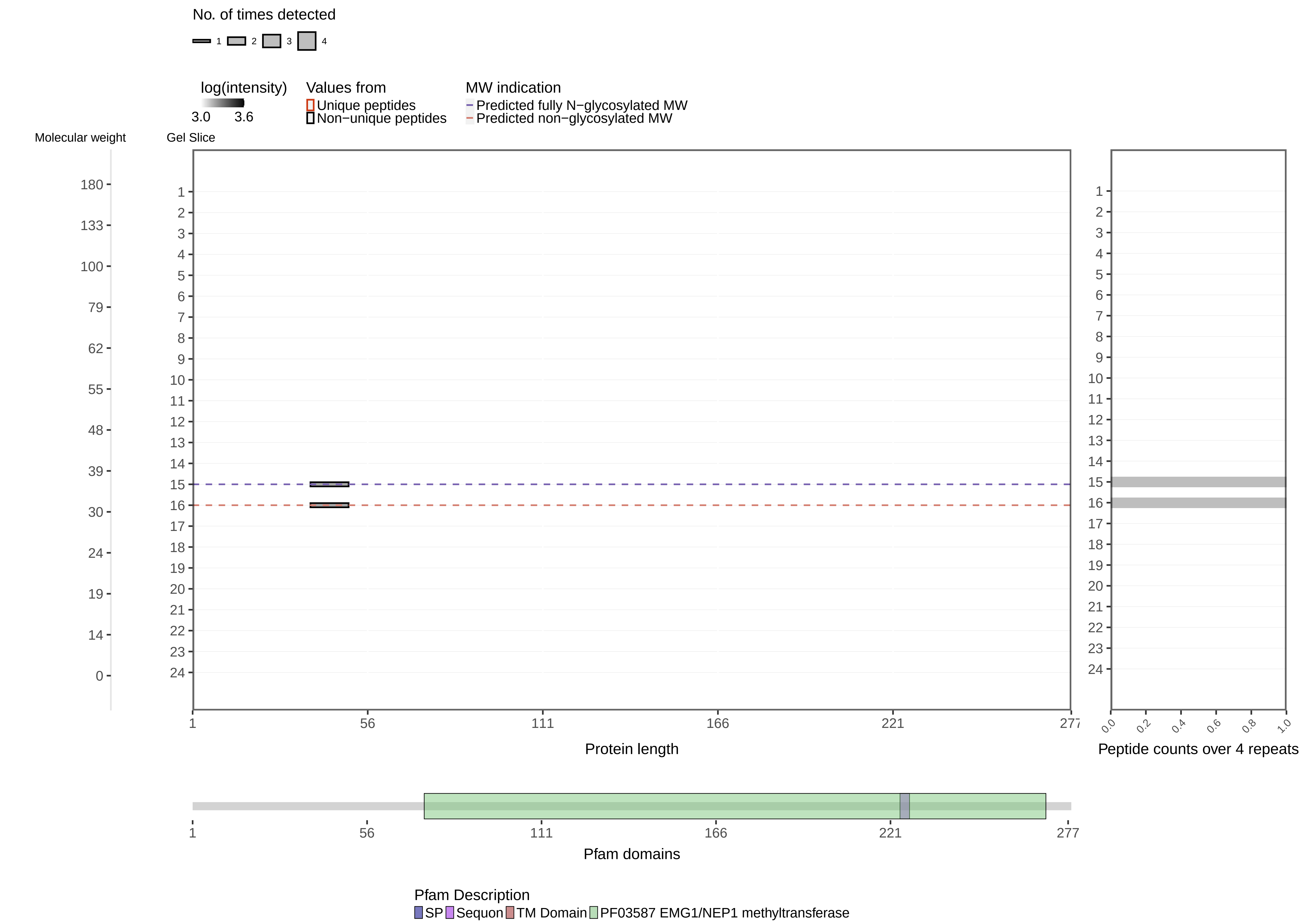


Protein sequence:

DSRVNKARRLKALYVKTEKGVLFEVKPHVRIPRTFKRFAGIMLQLLQKLSITAVGKREKLLRVIKNPVSQYLPIDCRKIGFSHSSEKLVDIQDYVNGINNETNLVFVVGAMAHGKIEKDYVEDYISVSDYPLSAAYCISMVTNAVERKWKIL*

NbL16g04330.1 40S_ribosomal_protein_S7−like [MW = 22.06 kDa]

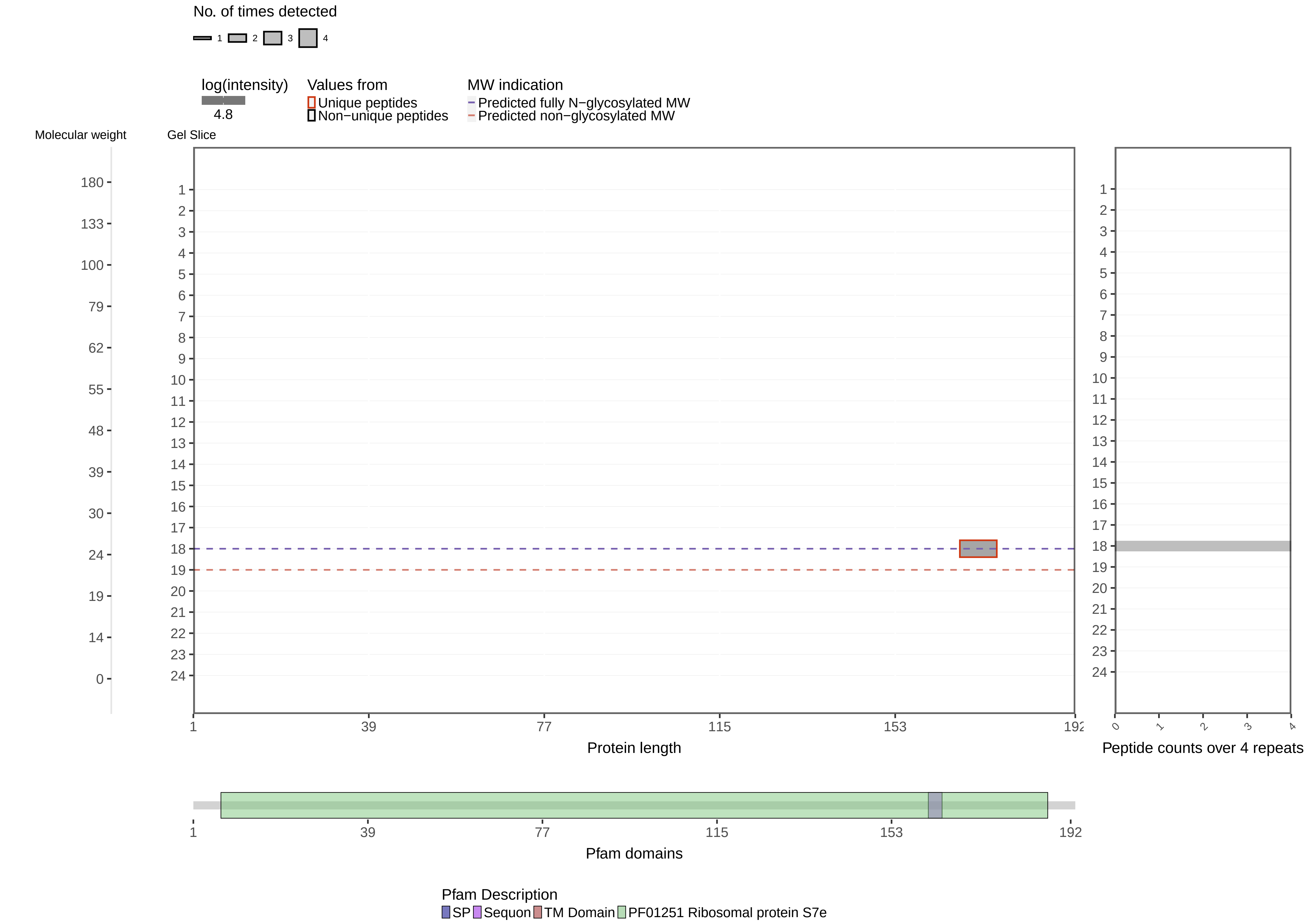


Protein sequence:

RELEKKFSGKDVIFIATRRIARPPKRGSAAQRPRSRTLTSVHDAILEDLVVPAEIVGKRTRYRIDGSKIMKVFLDPKERNNTEYKLEIFSAVYRKLSGKDVVFEYPITEA*

NbL16g04290.1 eukaryotic_peptide_chain_release_factor_GTP−binding_subunit_ERF3A−like [MW = 59.34 kDa]

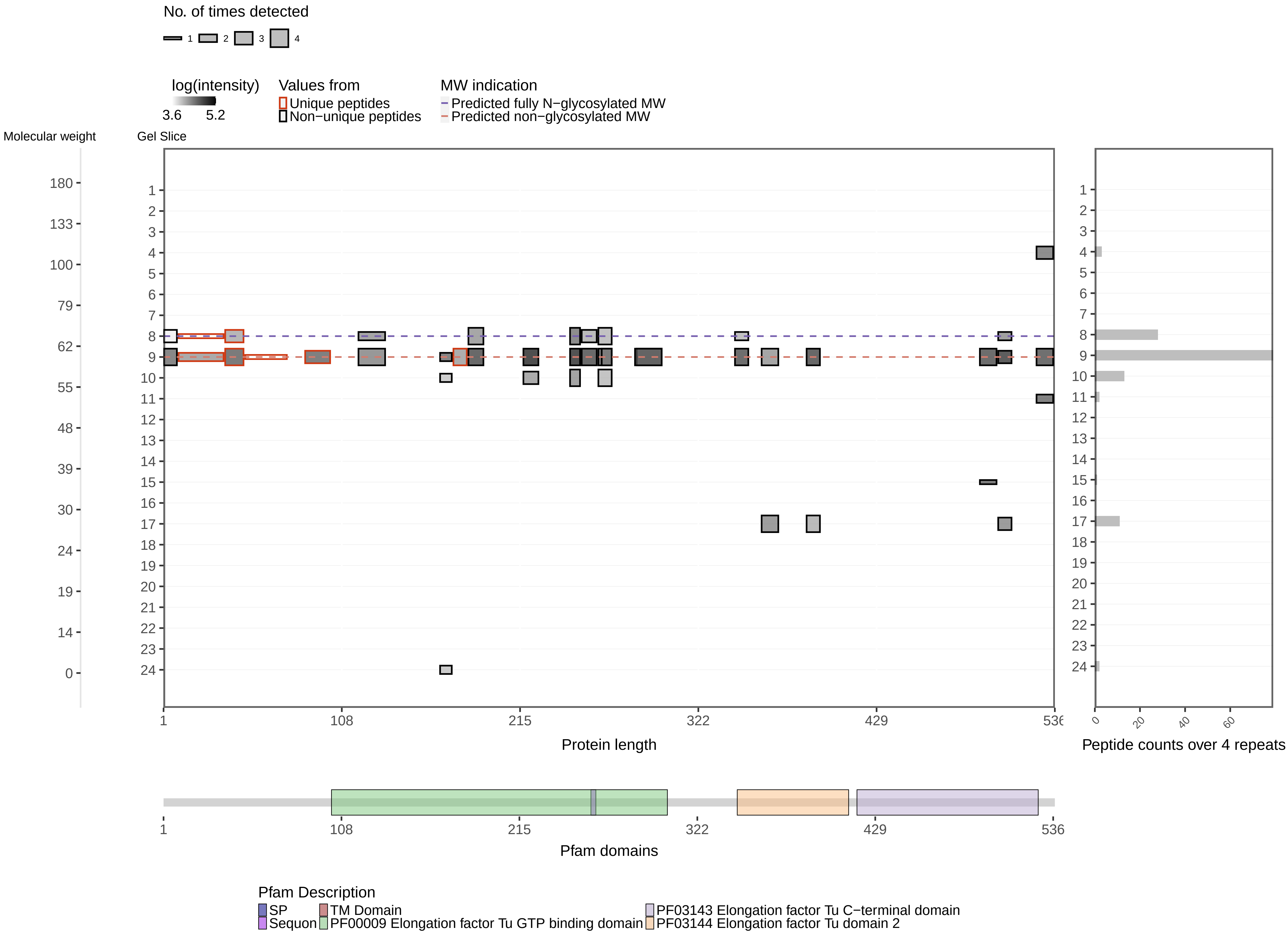


Protein sequence:

DDPTVNWSKERYDEIESKMVPFLRSSGYNVKKDVQFLPISGLLGSNLKTRMEKSVCPWWNGPCLFETLDAVEVPPRDPNGPLRMPIIDKFKDMGTVVMGKIESGSIREGDNLLVMPNKAAVKVLAIFCDEDKVRHVGPGENVRVRLSGVEEDDILSGFVLCSVAKPIHAVTEFVAQLQILELLDNAIFTAGYKAVLHIHSVVEECEIVELMQQIDLKTKKPMKKKPLFVKNGAIVLCRVQVNNLICVEKFSDFAQLGRFTLRTEGKTVAVGKITALPTVADNA*

NbL16g04260.1 guanine_nucleotide−binding_protein_subunit_beta−like_protein [MW = 35.72 kDa]

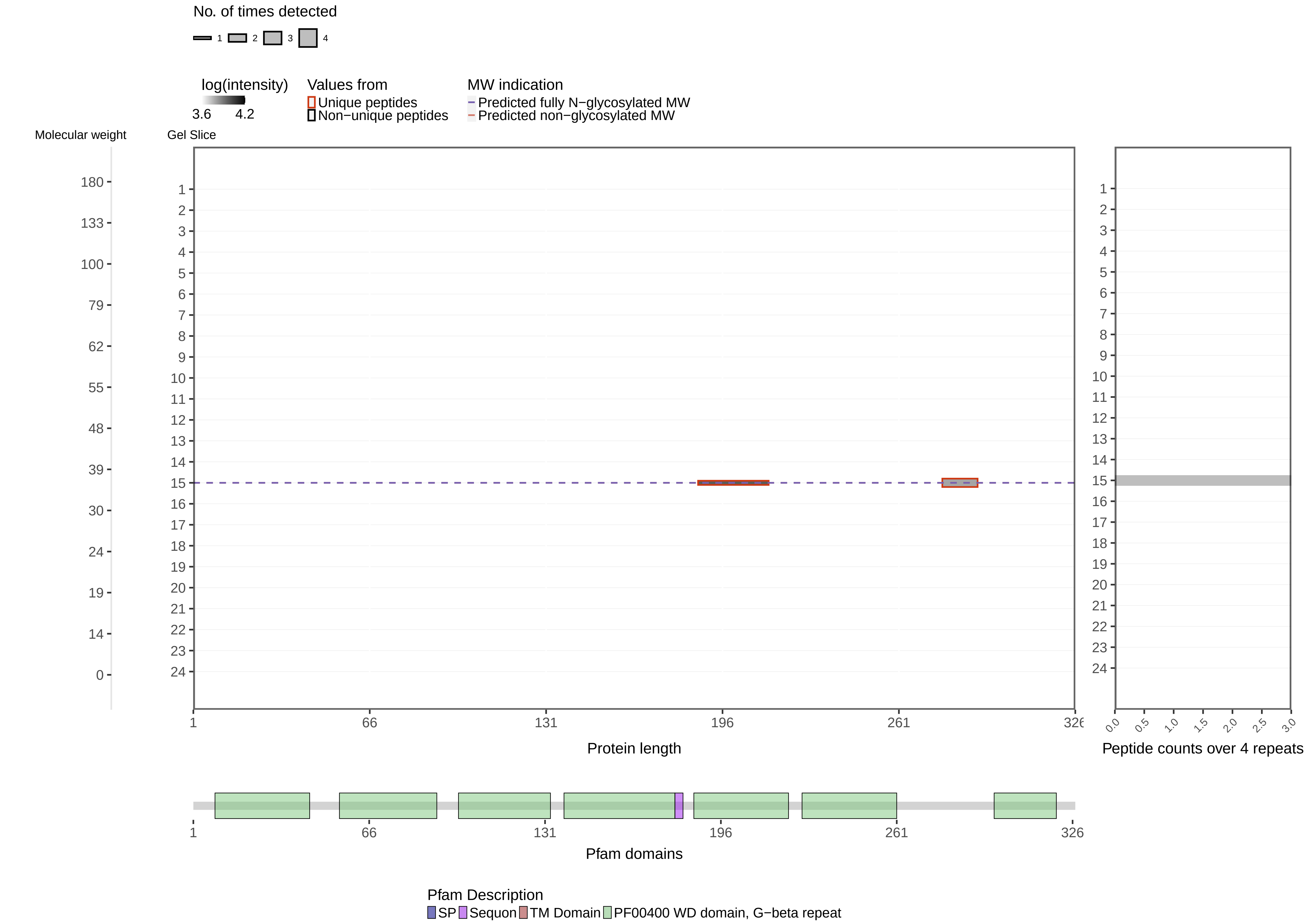


Protein sequence:

DWVSCVRFSPNNLQPTIVSGSWDRTVKIWNLTNCKLRSTLAGHSGYVNTVAVSPDGSLCASGGKDGVILLWDLAEGKKLYSLEAGSIIHSLCFSPNRYWLCAATESCIKIWDLESKSIVVDLKVDLKQESEMCAEGTAAAKNKIIYCTCLSWSADGSTLFGGYTDGLIRVWGIGRY*

### NbL16g04140.1 phosphoethanolamine_N−methyltransferase_1−like [MW = 56.19 kDa]

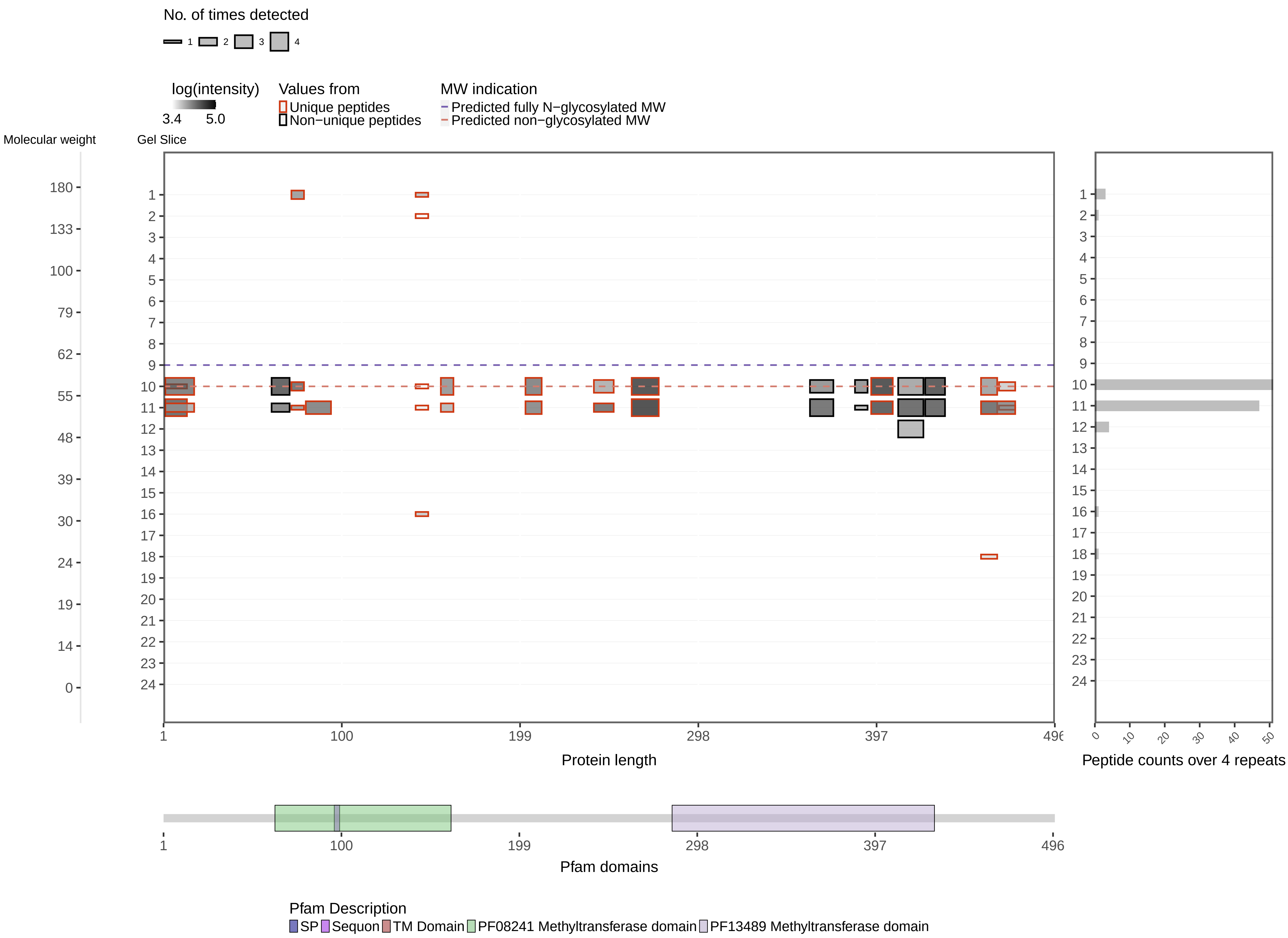


Protein sequence:

SADDRGFQHFLDTVQYKSNGILRYERVFGHSFVSTGGIDTTREFVAMLDLQPGQKVLDVGCGIGGGDFYMAEKYEVYVVGIDLSVNMISLAFERAVGRKCAVEFEVADCTTKTFPEGTFDVIYSRDTILHIQDKPALFRSFYKWLKPGGKVLISDYCKKAGPASEEFAEYIKQRGYDLHDVEAYGQMLRDAGFNDVVAEDRTEQFIKVLQKELDTVEKERESFIQEFSEKDYNDIVGGWKAKLIRSTSSGEQRWGLFIAKK*

### NbL16g04130.1 calcium−binding_allergen_Ole_e_8−like [MW = 17.28 kDa]

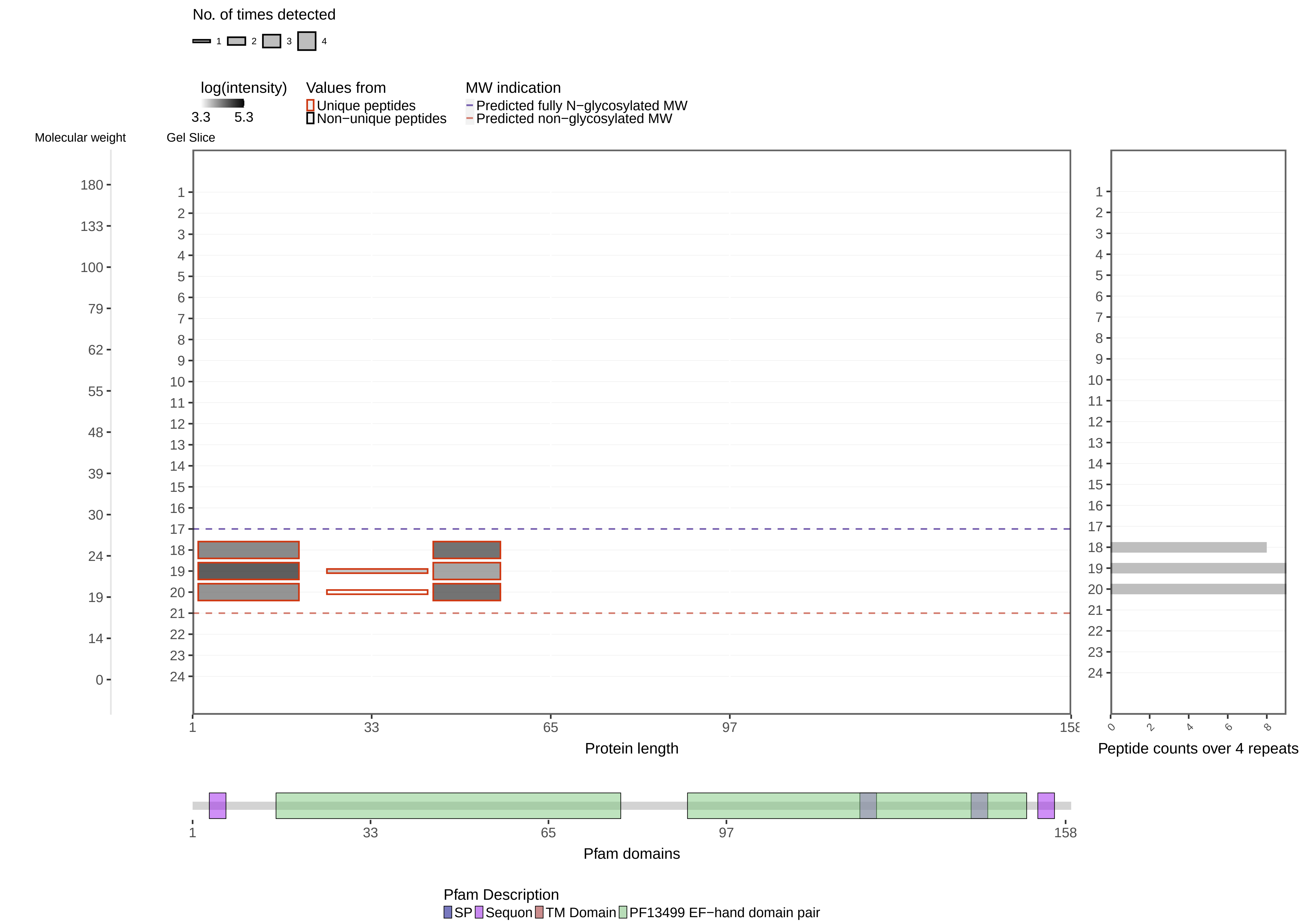


Protein sequence:

GCINLEEFADFCKSDDSVDGGAKELKEAFDLYDQNHNGLISAAELHQILNRLGQNCTVQDCTRMITSVDADGDGNVSFEEFKQMMCNNSKI*

NbL16g03910.1 60S_acidic_ribosomal_protein_P2A−like [MW − SP = 9.84 kDa]

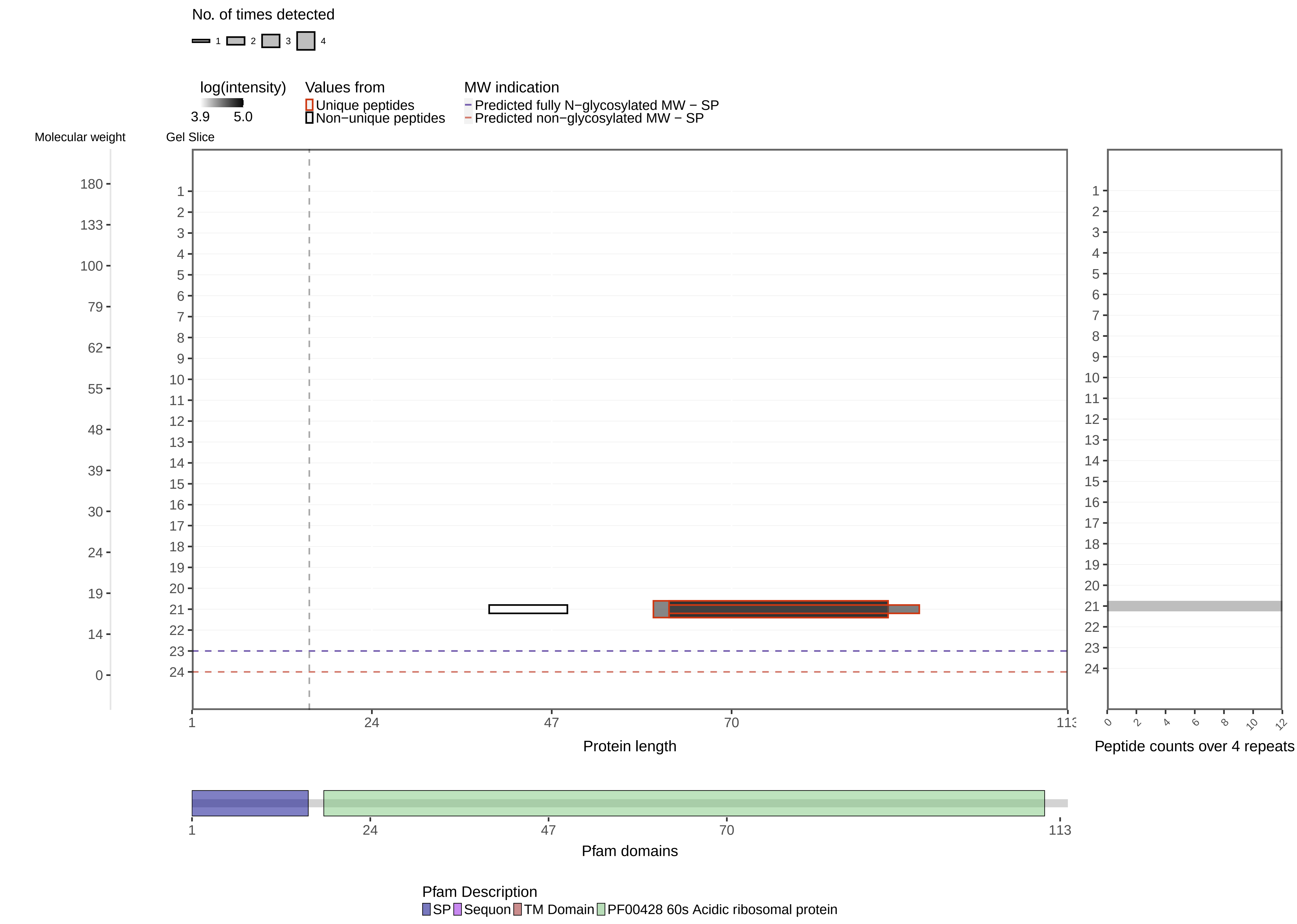


Protein sequence:

SQVEGKDITELIAAGREKLASVPAGGAGVAVAAPTGGAAAAPAAEEKKEEKKVEEKEESDDDMGFSLFD*

### NbL16g03870.1 60S_ribosomal_protein_L27−like [MW = 15.69 kDa]

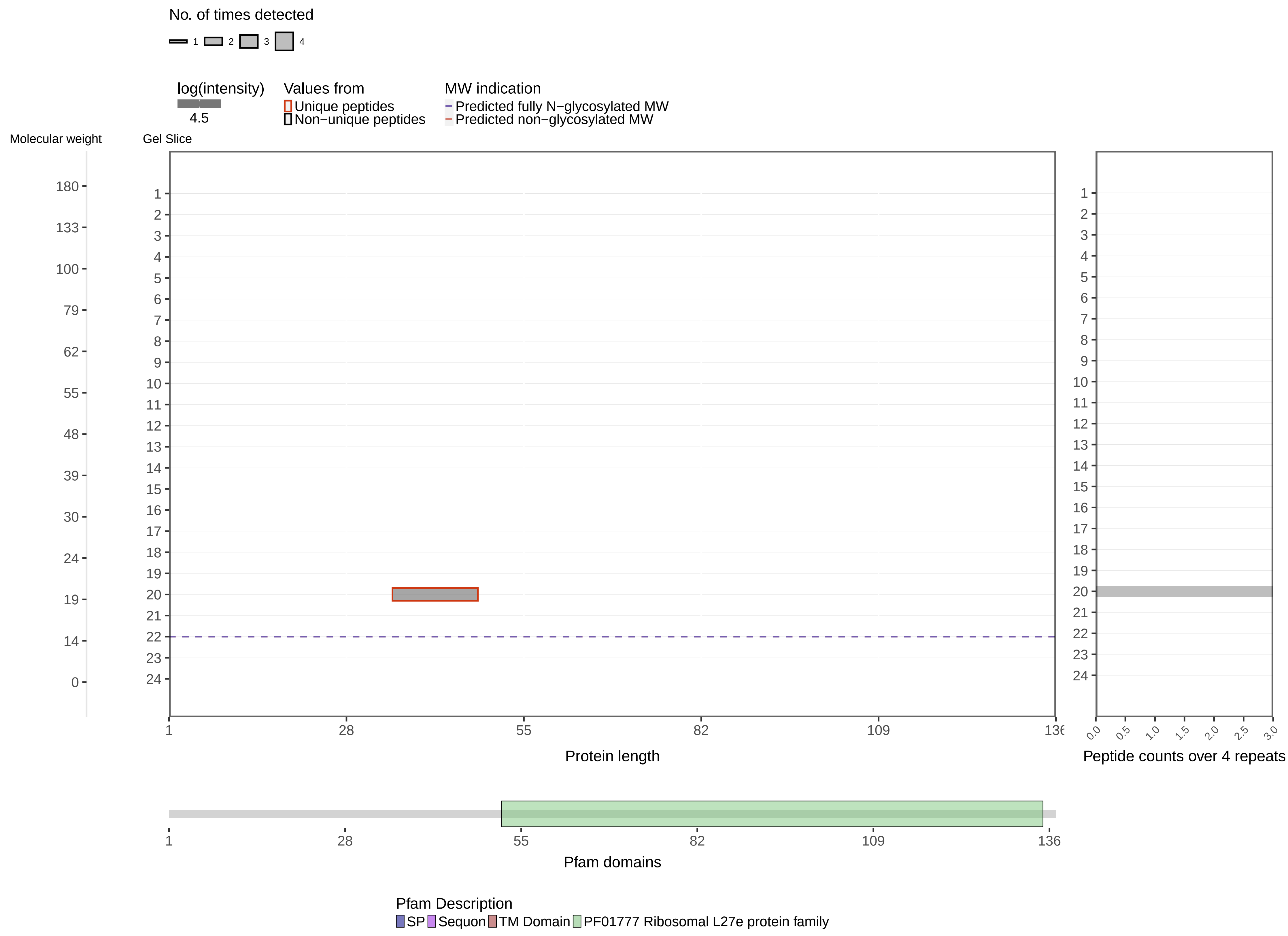


Protein sequence:

KDSAKKQAKKSRVKAFIKLVNYNHIMPTRYTLDVDLKDVVNVDVLQSRDKKVTAAKEAKARLEERFKTGKNRWFFTKLRF*

### NbL16g03830.1 chloroplast_stem−loop_binding_protein_of_41_kDa_b_chloroplastic [MW = 42.14 kDa]

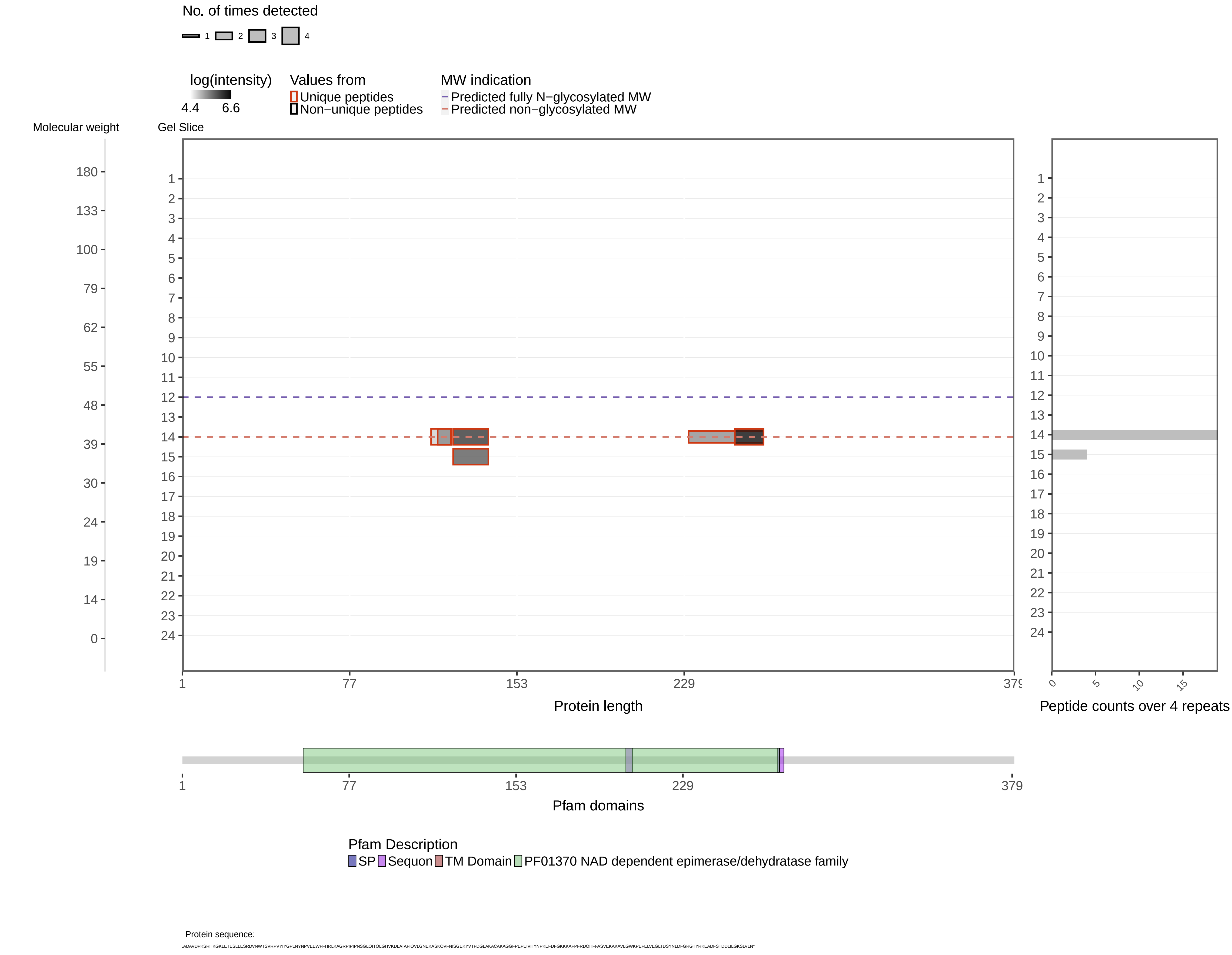

### NbL16g03810.1 probable_carbohydrate_esterase_At4g34215 [MW = 27.62 kDa]

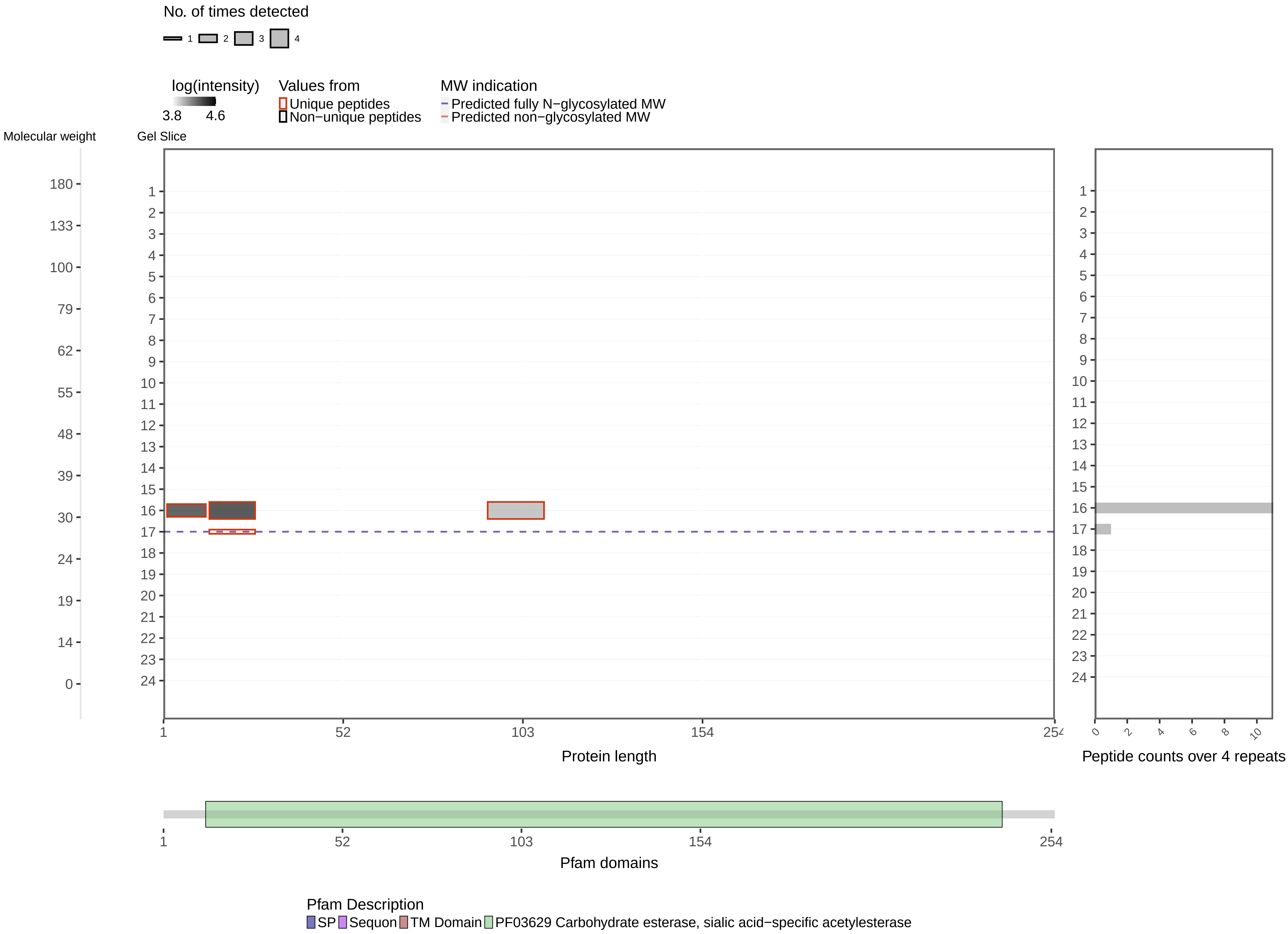


Protein sequence:

IGEHLYENMVKRARASVHQDGEIKALLWYQGESDTLSQHCAETYKANIEKLIHNVRADLDLPSLPIIQVVIASGDDKYIEKIREAQMGVDLPNVVCVDAMGLQLKEDNLHLCTEAQVKLGQMLADAYLTHFAPQQPSVASS*

### NbL16g03800.1 tRNA−dihydrouridine(20)_synthase_[NAD(P)+]−like [MW = 36.06 kDa]

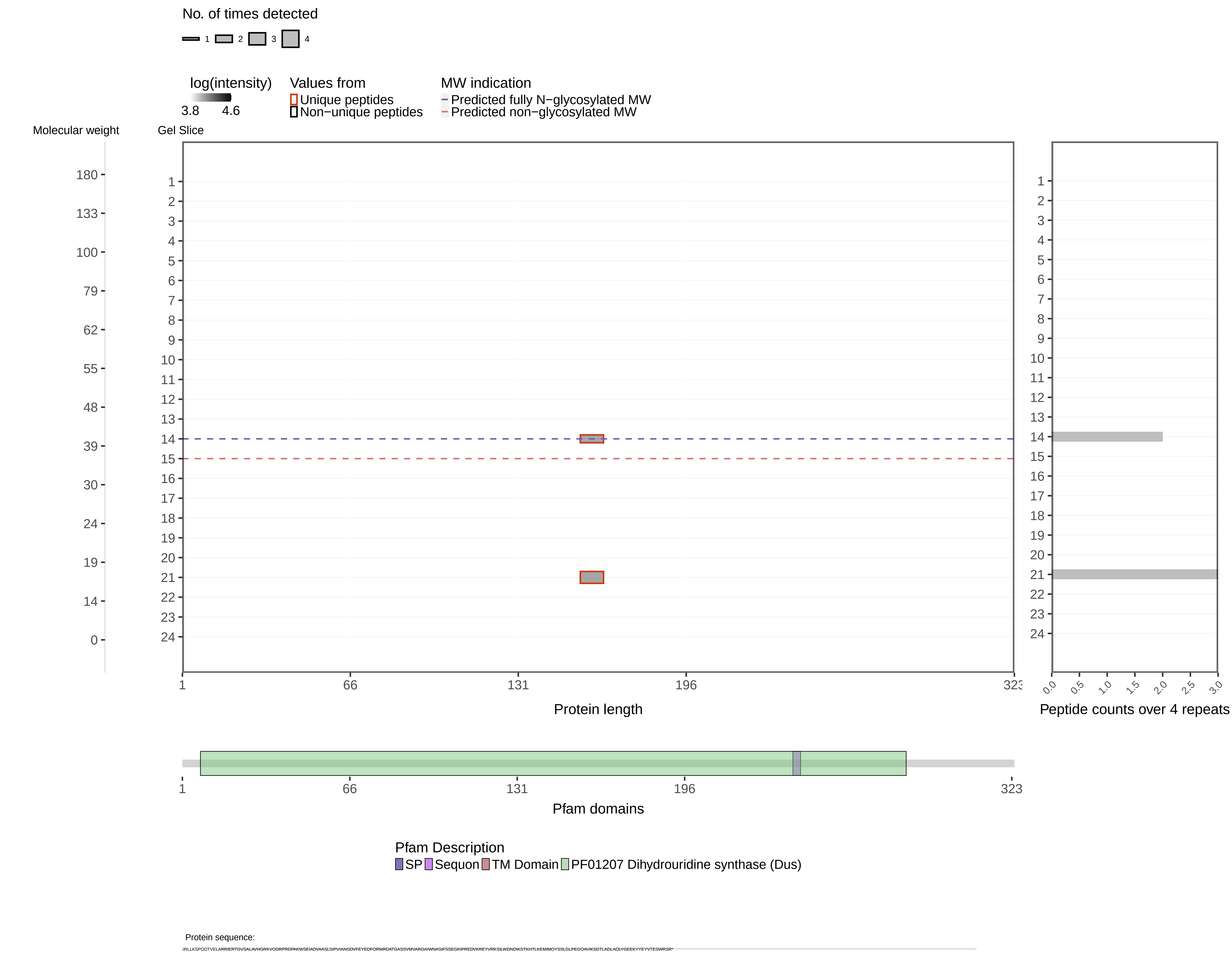

NbL16g03740.1 fructokinase−2 [MW = 34.78 kDa]

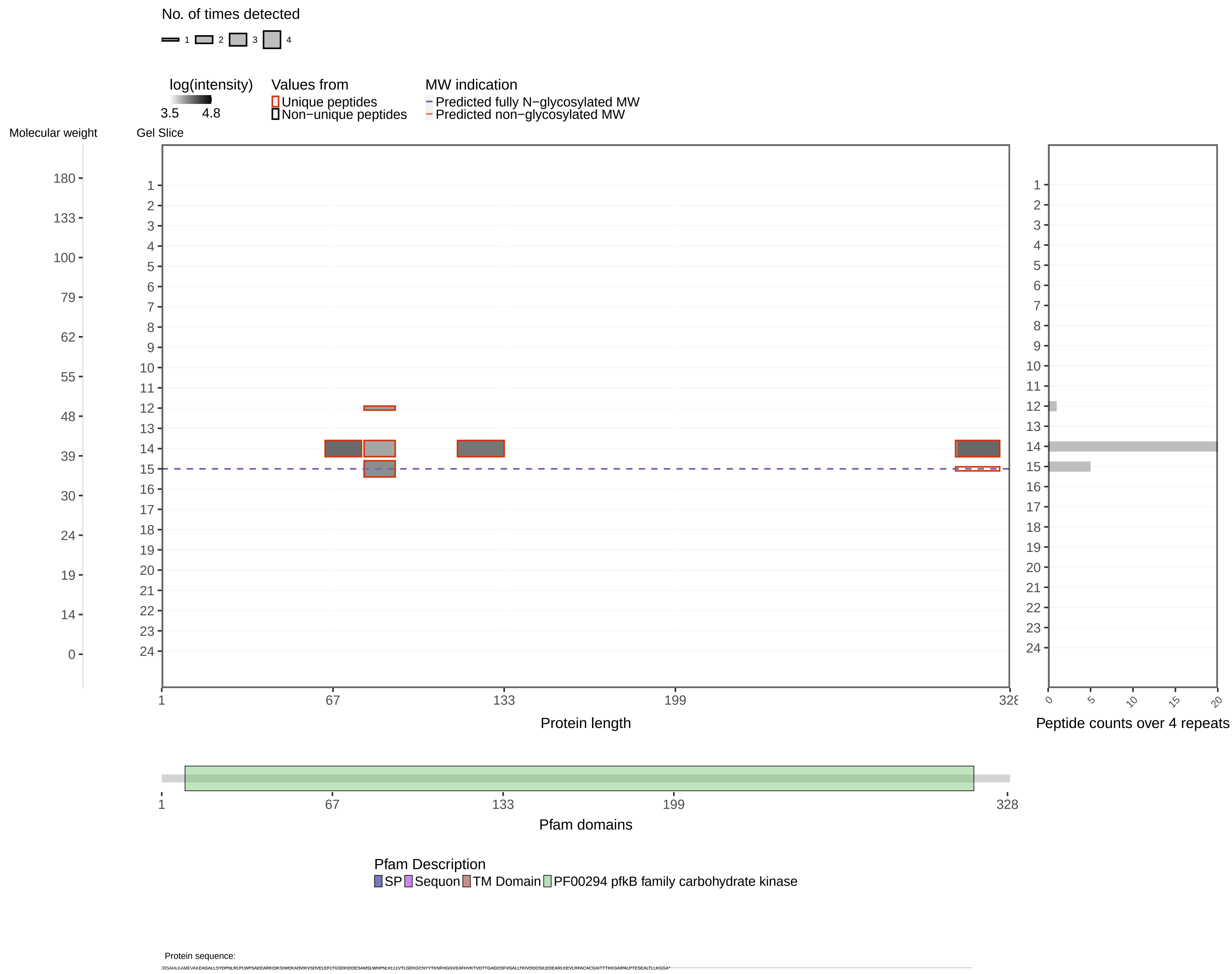

### NbL16g03630.1 haloalkane_dehalogenase−like [MW = 47.36 kDa]

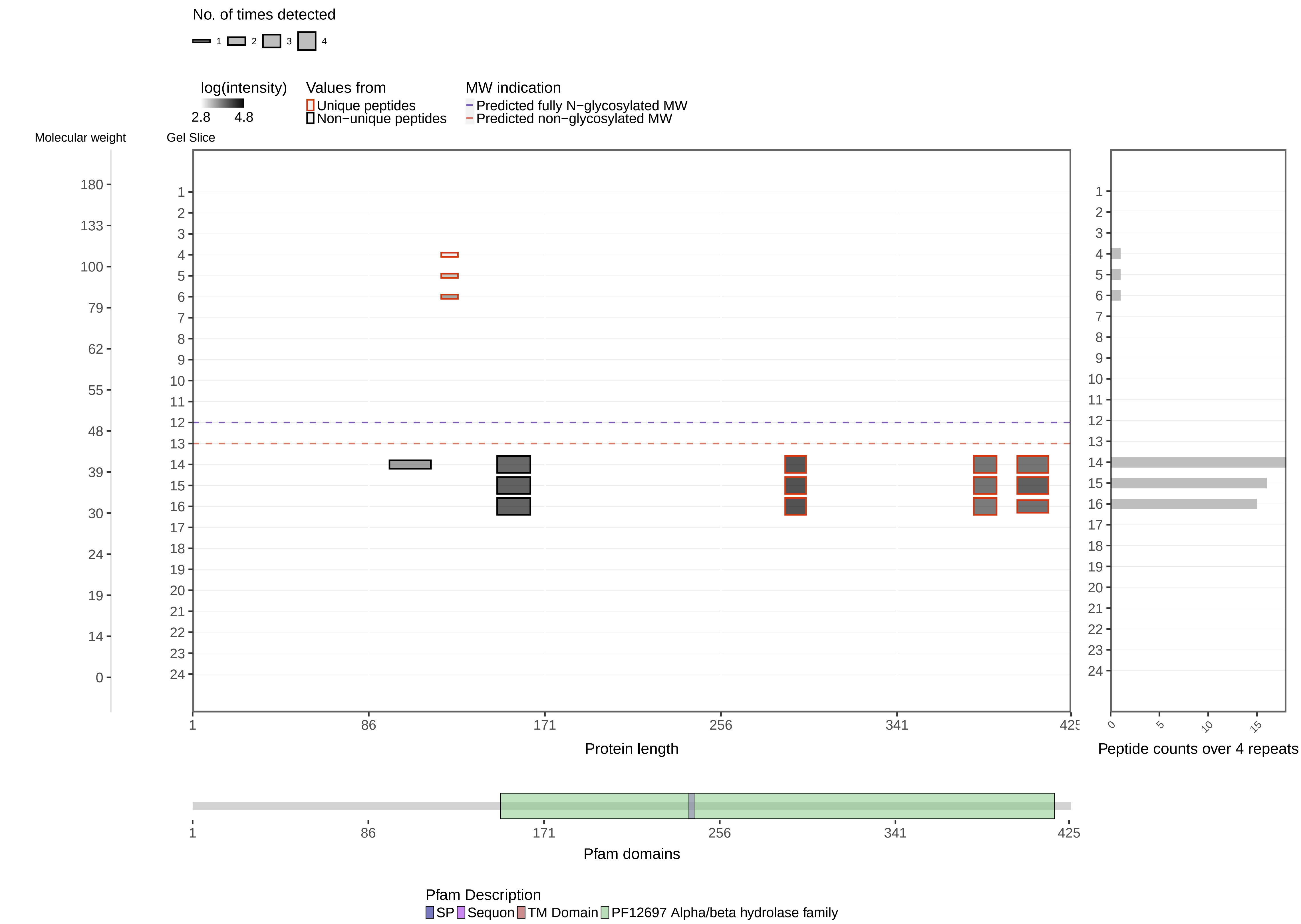


Protein sequence:

FDYTEKEFHEEFEKLLEVLGVTSPFFLVVQGFLVGSYGLTWTLKNTSKISKLAILNTPLTVSSPIPGLFQQLRIPFFGEFTCQNAVMAERFIEAGSTYVLKLEKADVYRLPYLASSGPGFGSVYRILERPCSPPHLSDLPRALLETARKAKFNEIASQISSGFASGSWDKPILVAWGISDKYLPQSIAEEFQKANPDSVKLKLIEGAGHMPQEDWPEKVVDALRMFFR*

### NbL16g03620.1 clustered_mitochondria_protein_homolog [MW = 188.94 kDa]

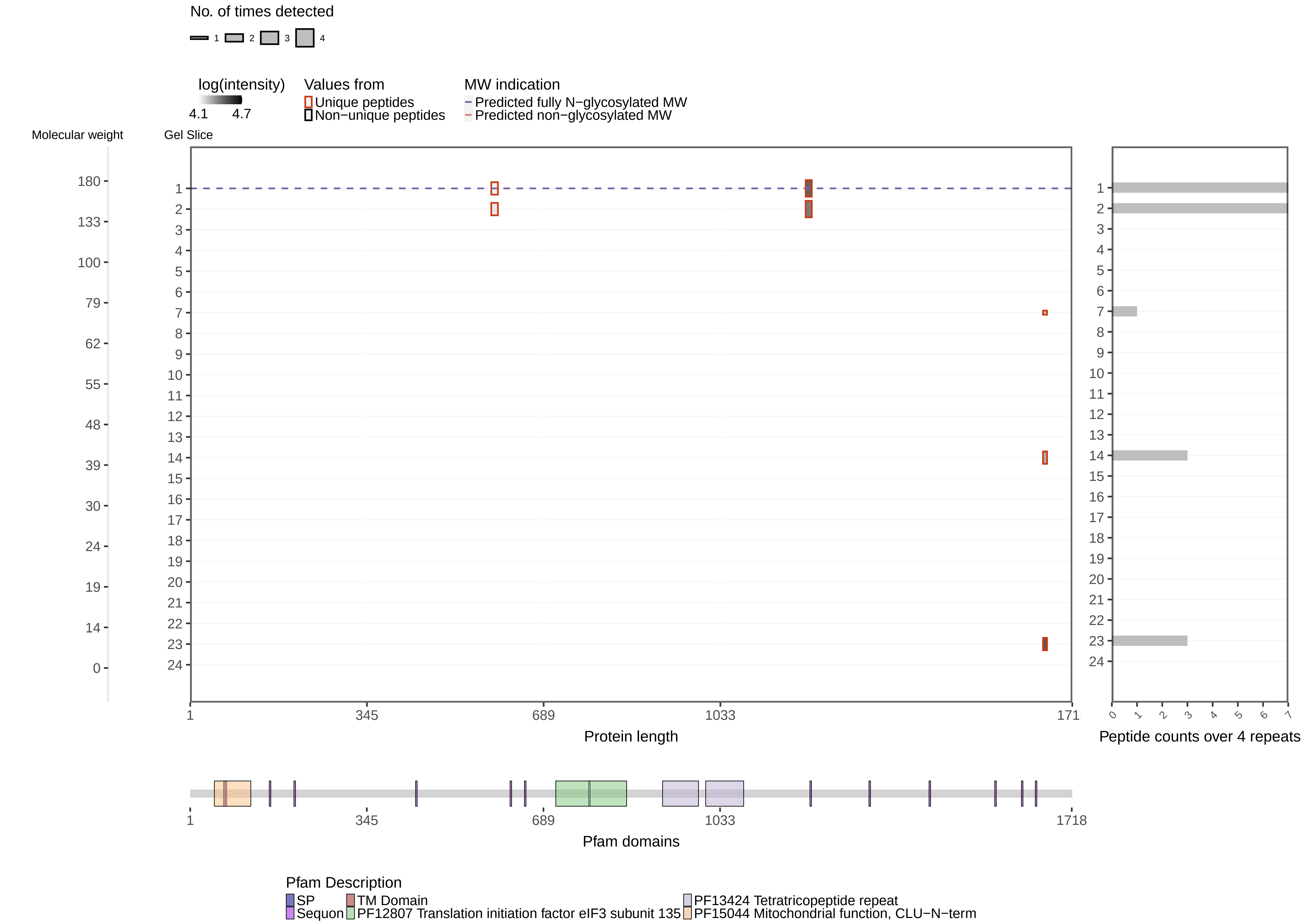


Protein sequence:

PKDYDMESPYPFSKTDVTSIVPLCKHVGCSSADGRTLLESSKVALDKGKLEDAVMYGTKALAKLIAVCGPYHRATASAYSLLAVVLYHTGDFNQATIYQQKALDINERELGLDHPDTMKSYGDLSVFYYRLQHIELALKYVNRALFLLHFTCGLSHPNTAATYINVAMMEEGMGNVHIALRYLHEALKCNQRLLGVDHIQTAASYHAIAIALSLMEAYSLSVQHEQTTLQILQAKLGAEDLRTQDAAAWLEYFESKALEQQEAARTGAPRLDATIASKGHLSVSDLLDYISPSQGSI

### NbL16g03600.1 triphosphate_tunel_metalloenzyme_3−like [MW = 28.08 kDa]

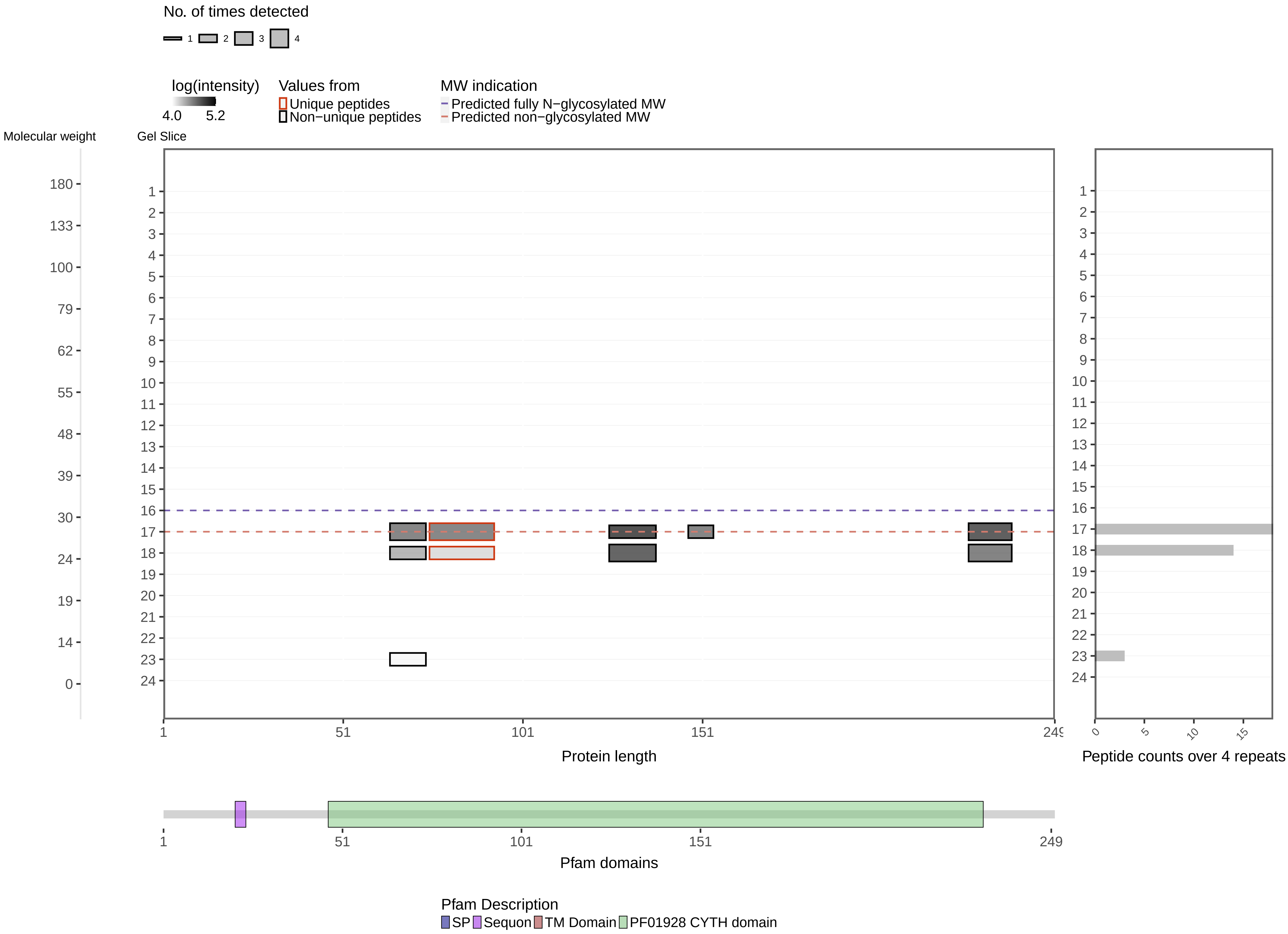


Protein sequence:
KAKPVISNGISRIEEDEEPIDPSIGRACVSDPWRLLLIDSSRVIQRVRDEYGIGDKGLVCLGGFRNVRAVYEWKGLKLELDETHYEFGMNYEIECESSDPERAKDMLEELLKSHGIEYSYSTVSKFAIFRSGKLPQ*

NbL16g03580.1 proline−−tRNA_ligase_cytoplasmic−like [MW = 57.03 kDa]

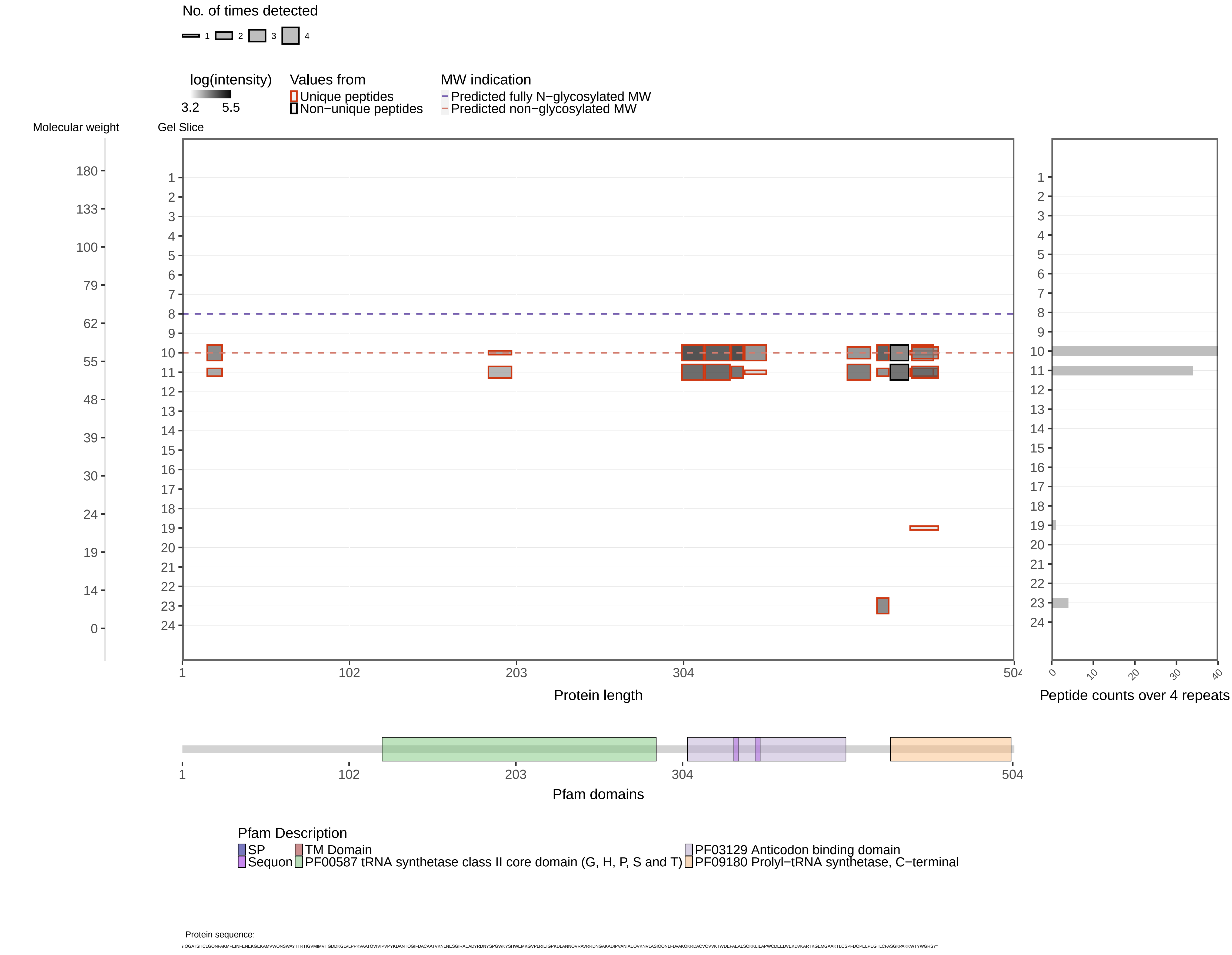

### NbL16g03320.1 aconitate_hydratase_cytoplasmic−like [MW = 105.48 kDa]

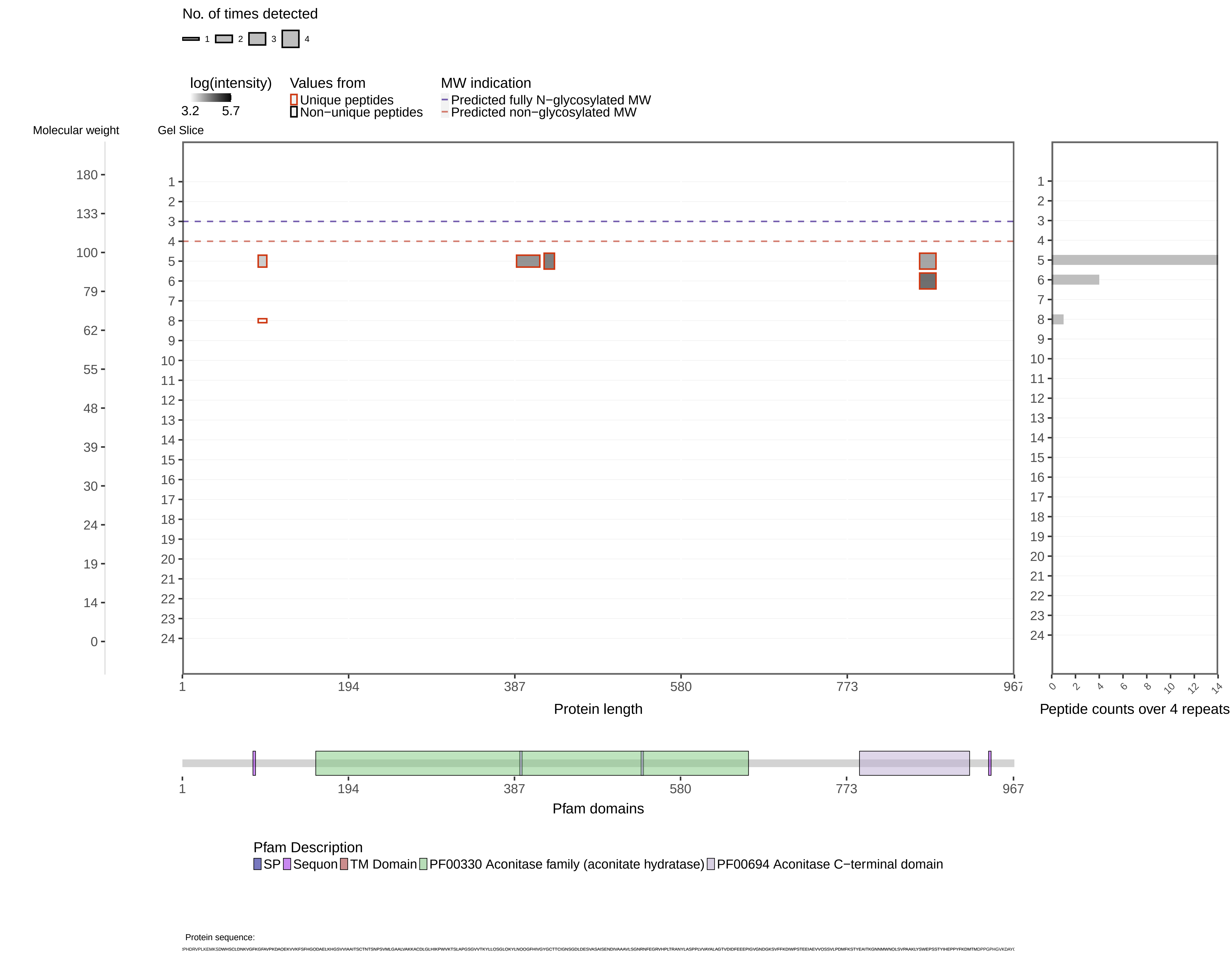

### NbL16g03310.1 NEDD8−activating_enzyme_E1_regulatory_subunit_AXR1−like [MW − SP = 60.43 kDa]

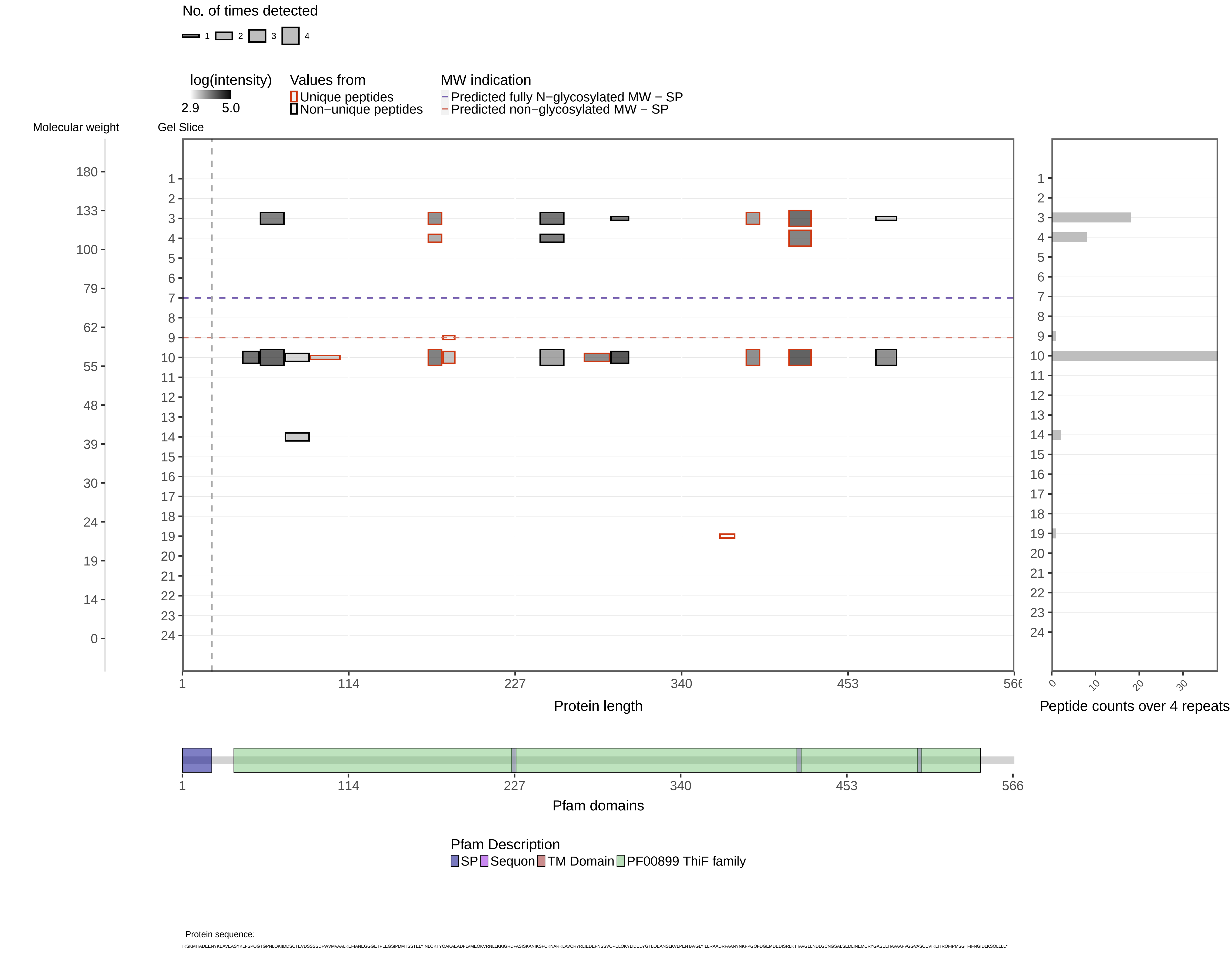

NbL16g03230.1 1−aminocyclopropane−1−carboxylate_oxidase_1 [MW = 36.05 kDa]

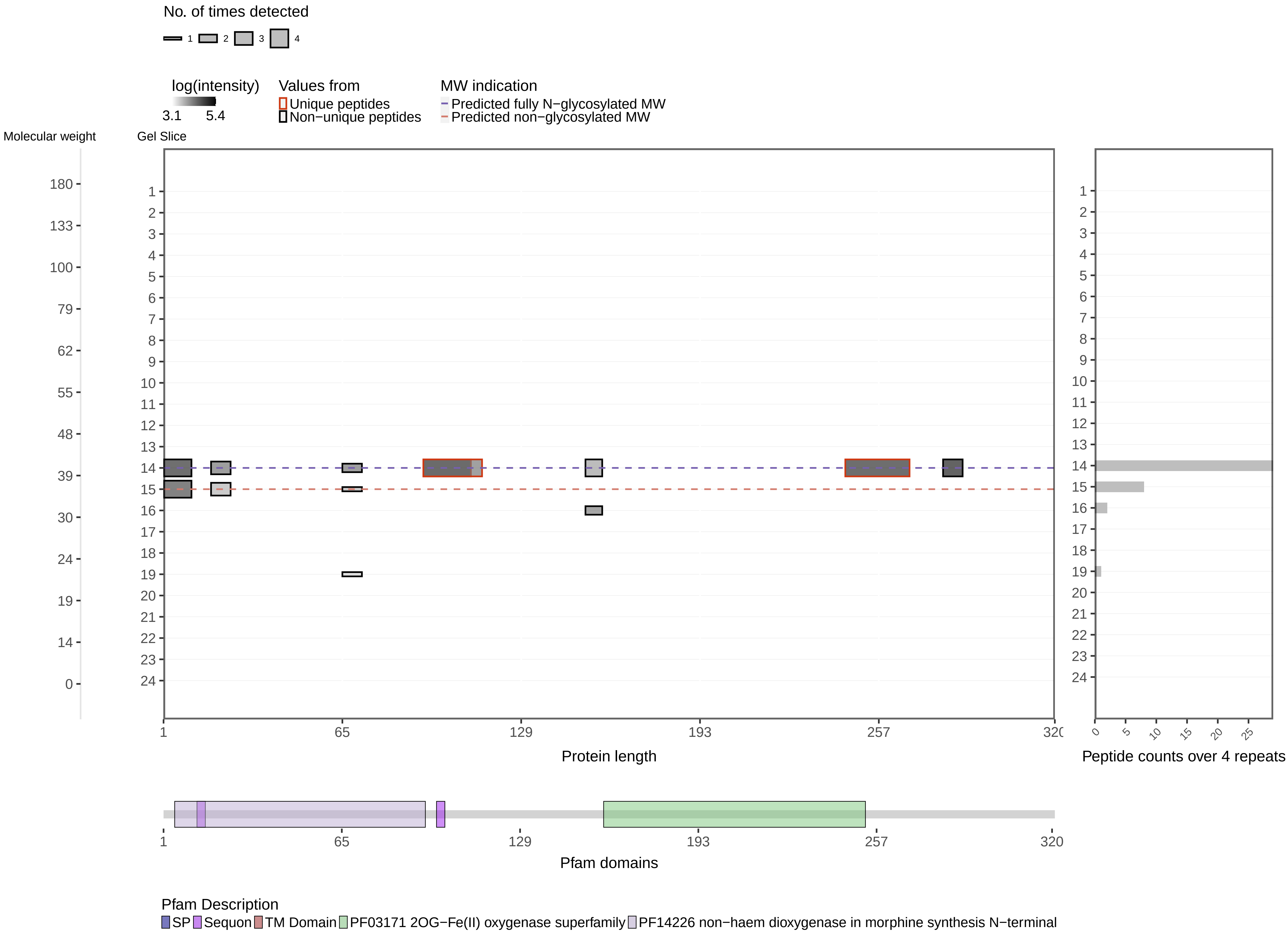


Protein sequence:

VFYGSKGPNFGTKVSNYPPCPKPDLIKGLRAHTDAGGIILLFQDDKVSGLQLLKDGQWIDVPPTRHSIVVNLGDQLEVITNGKYKSVMHRVIAQKDGTRMSLASFYNPGSDAVIYPAPALVEKEAVESKQVYPKFVFDDYMKLYAGLKFQAKEPRFEAMKSIESDVKMDAIATA*

NbL16g02880.1 acylsugar_acyltransferase_3−like [MW = 46.42 kDa]

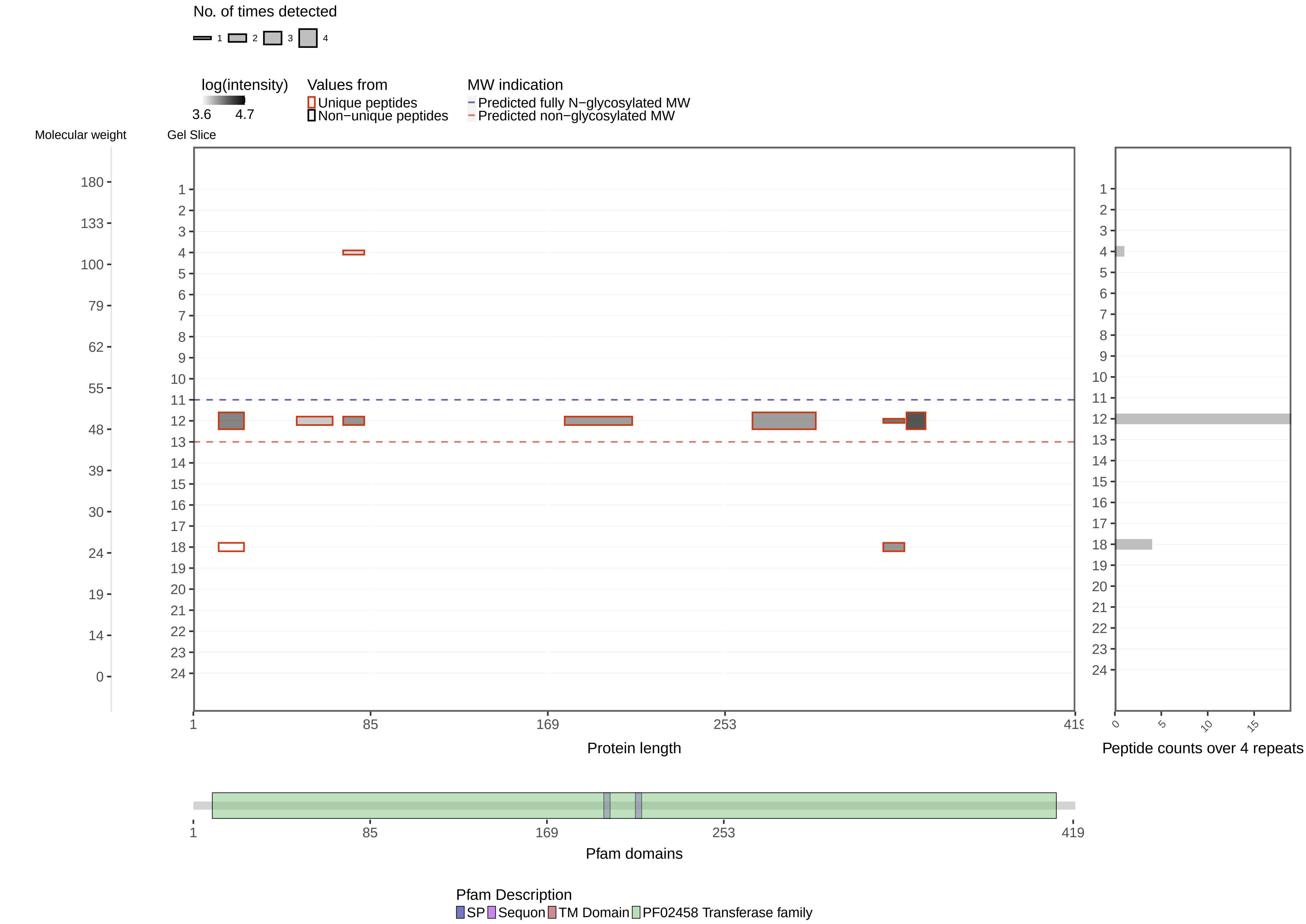


Protein sequence:

[illegible]

### NbL16g02770.1 pyruvate_dehydrogenase_E1_component_subunit_alpha−3_chloroplastic−like [MW − SP = 51.45 kDa]

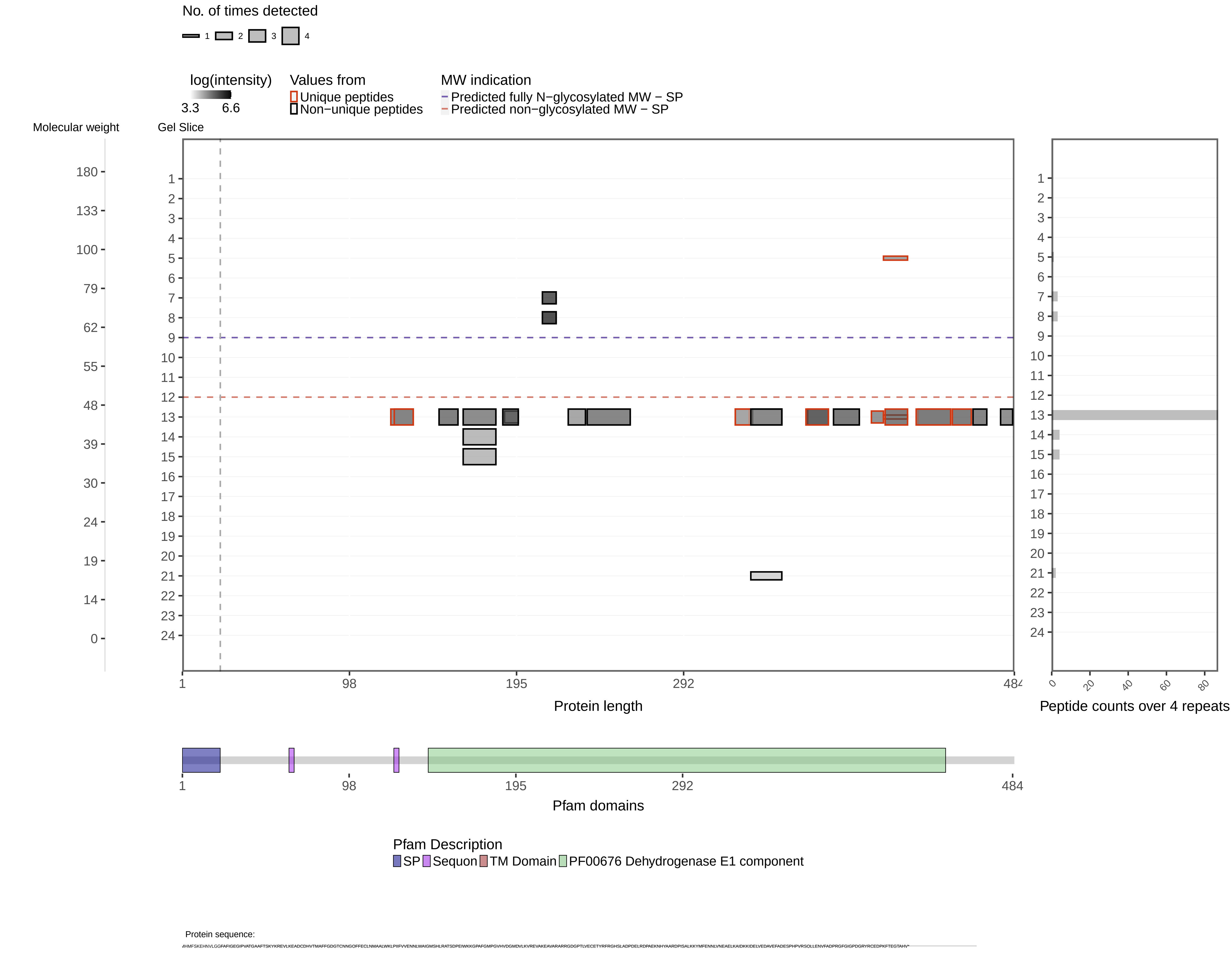

### NbL16g02690.1 20_kDa_chaperonin_chloroplastic−like [MW = 31.85 kDa]

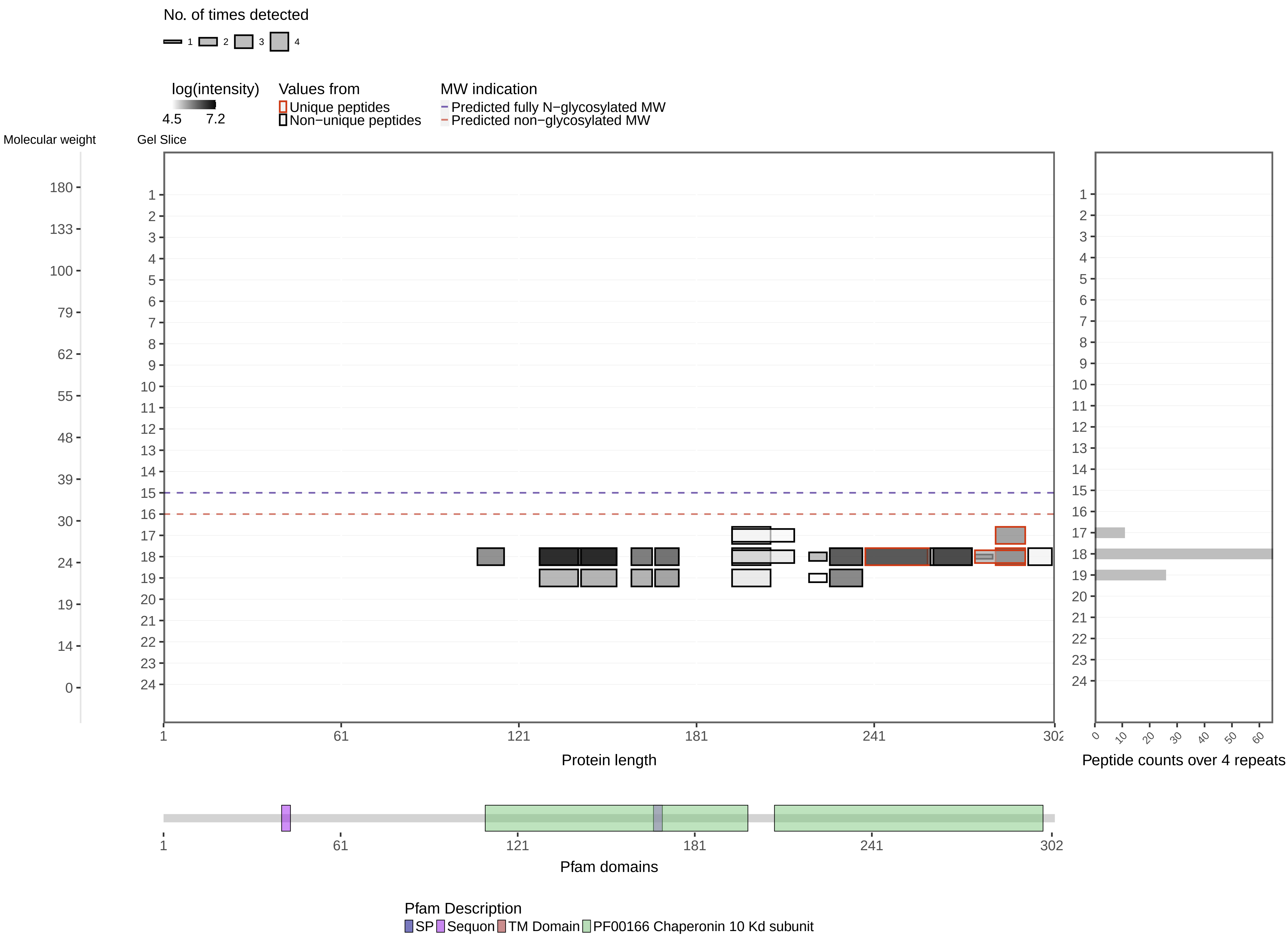


Protein sequence:

AOSKPOGGEIVAVGEGRSIGKTKVEISVKNGTOVLYSKYAGTEVLFDGSTHLILKEDDIVGILETDDIKDLOPLNDRILIKVAEVEEKTAGGLFLSEAAKEKPSIGSVIAAGPGPLDEEGNRKSLSVSPGNTVLYSKYAGNDFKGADGSDYVTLRASDVIAVLS*

NbL16g02640.1 probable_polygalacturonase [MW = 55.1 kDa]

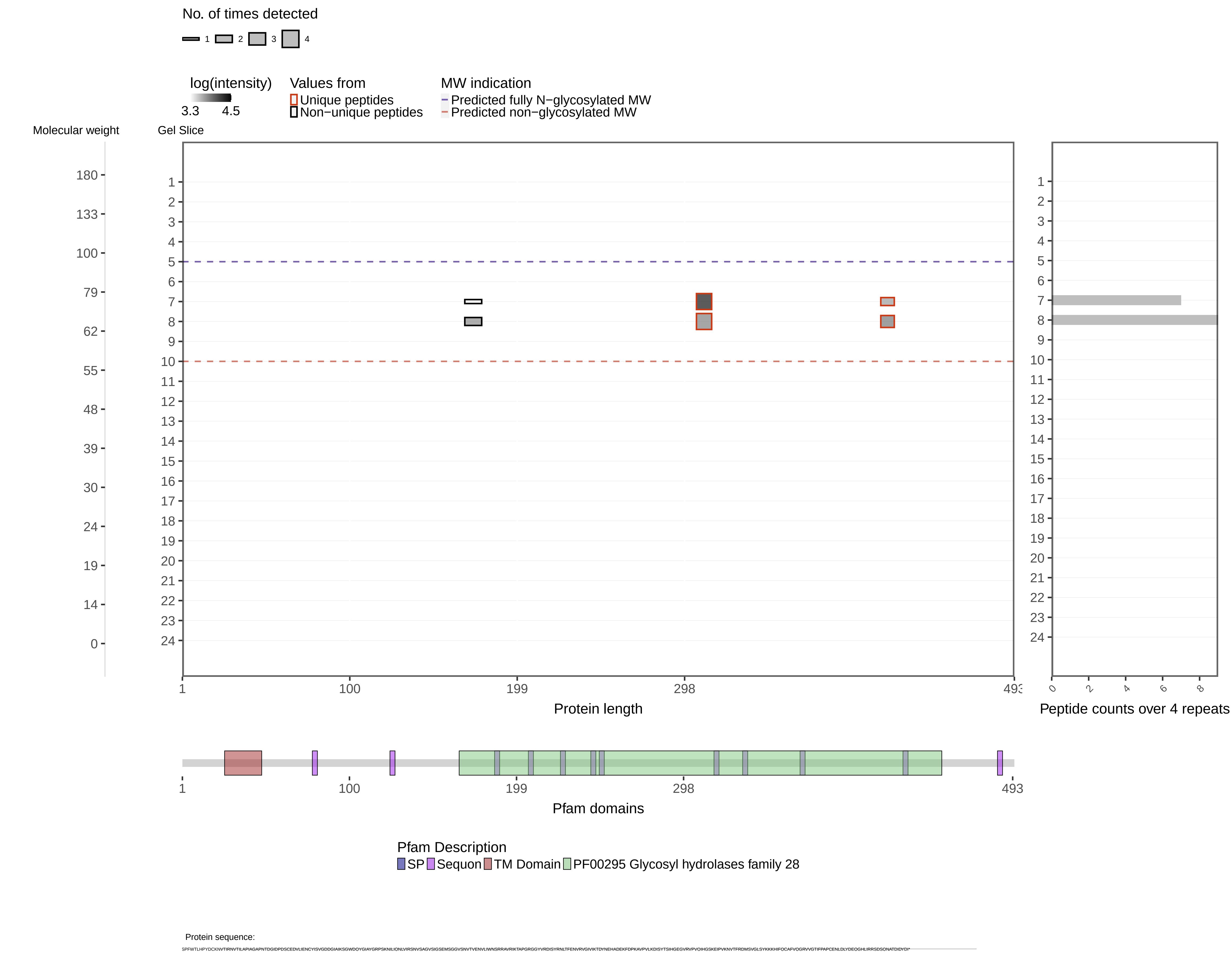

NbL16g02600.1 uncharacterized_LOC107785378 [MW = 21.73 kDa]

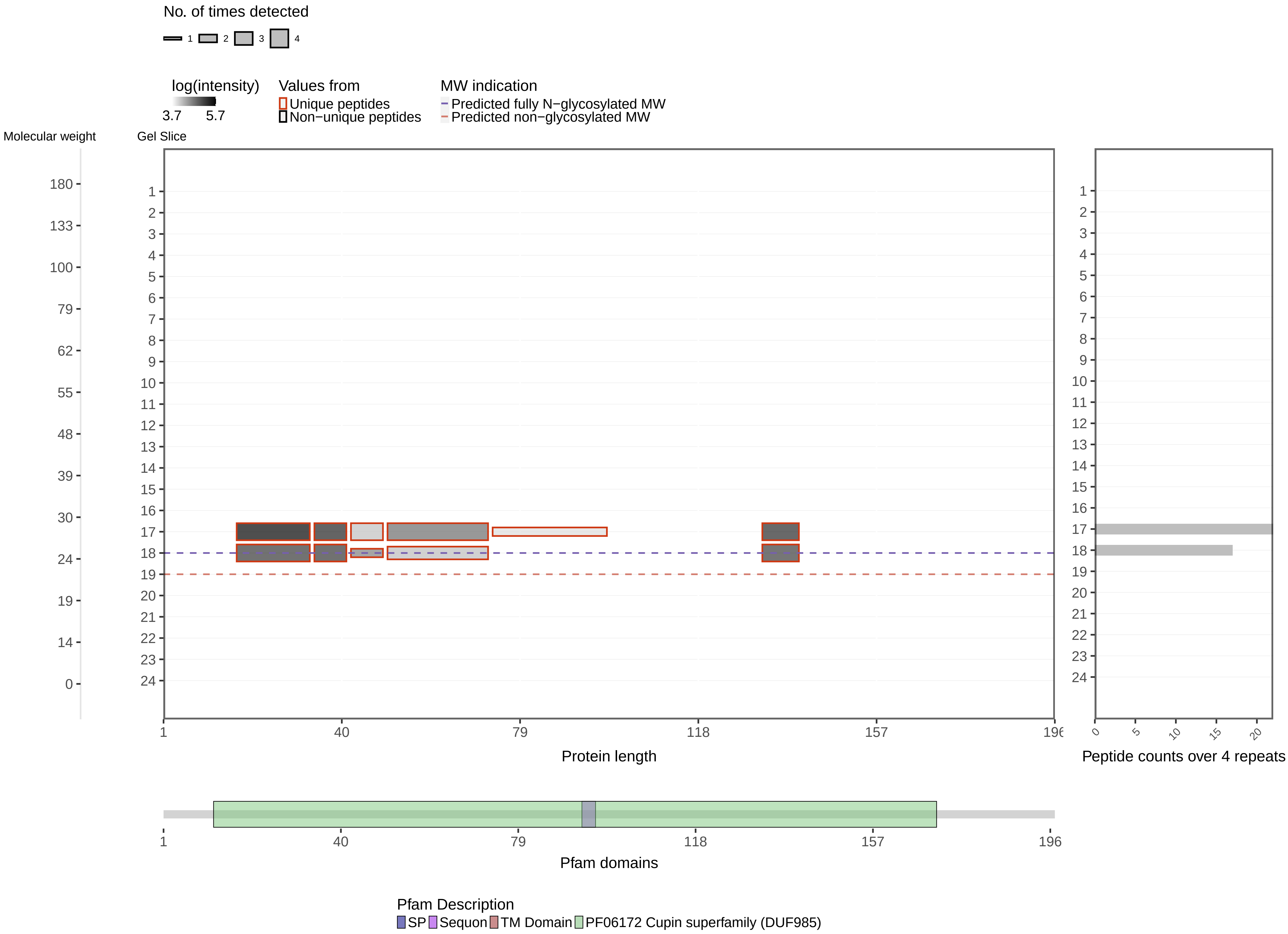


Protein sequence:

EPLTVMELNETDGKVKLTCIGPDPLADDQLVQYTVPPNVWFGAFPTRDFSISADMKVTKAPARDGEKHFSLVGCTCAPAFQFEDFELAKRSELVSRFPAHESLISLLTFLE*

### NbL16g02550.1 subunit_alpha_type−6 [MW = 27.29 kDa]

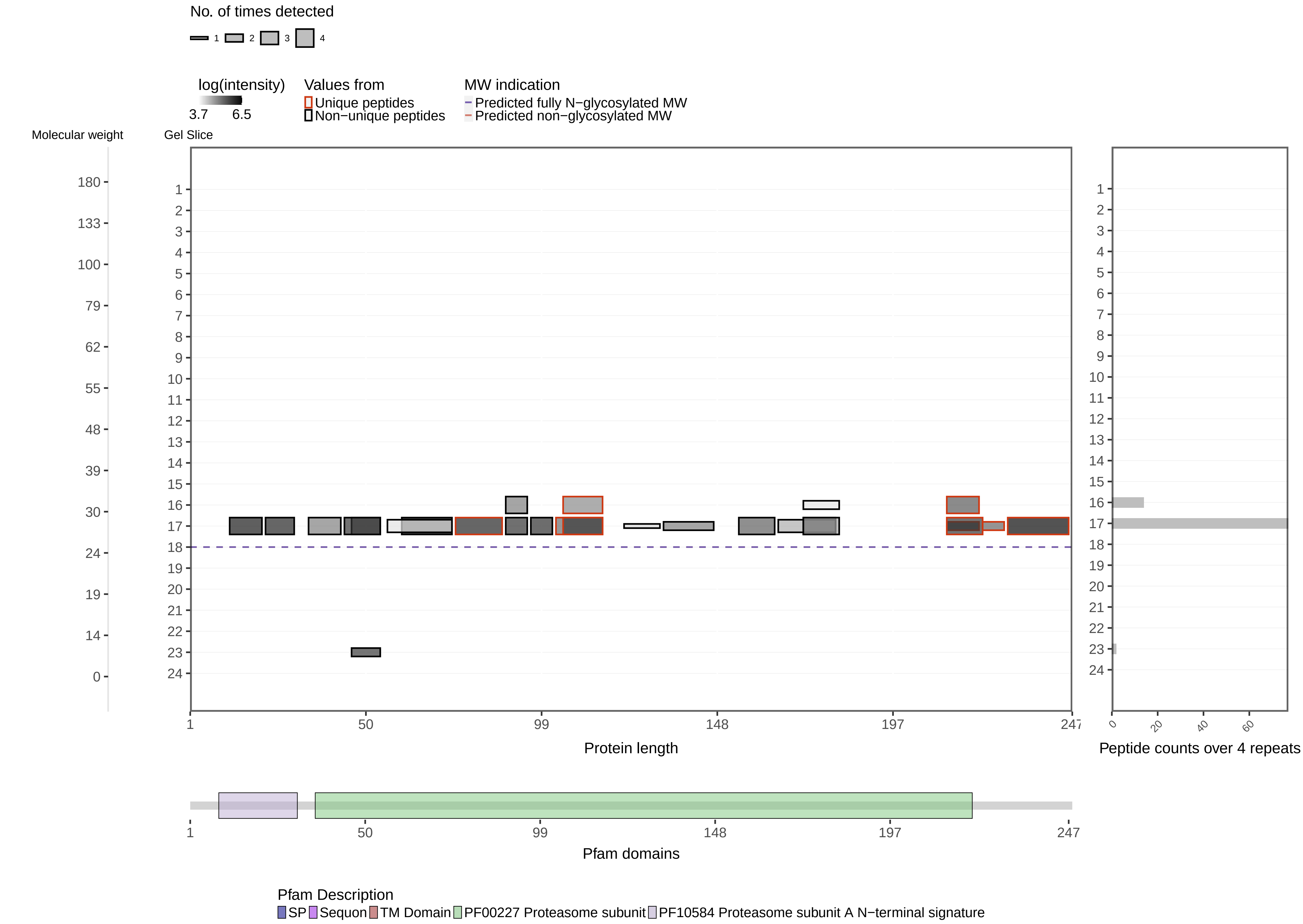


Protein sequence:

MPVDVLSKWIADKSQVYTQHAYMRPLGVVAMILGIDEEKGPQLFKCDPAGHFFGHKATSAGSKEQEAINFLEKKMKNDPAFSYEETVQTAISALQSVLQEDFKASEIEVGVVKKEDPIFRVLTTEEIDEHLTAISERD*

### NbL16g02480.1 40S_ribosomal_protein_S3−2−like [MW = 31.13 kDa]

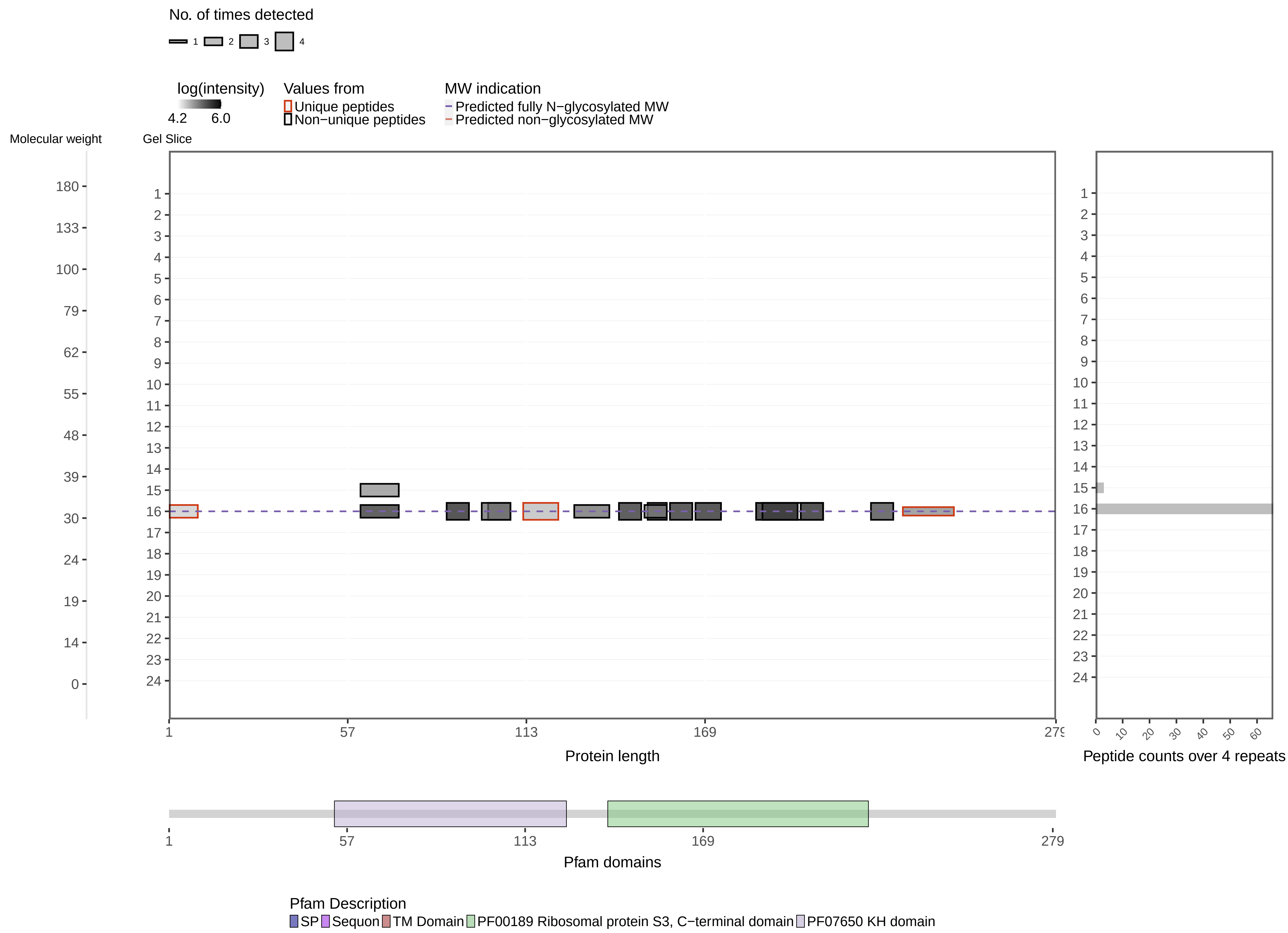


Protein sequence:

GLCAVAQAESLRYKLLGGLAVRRACYGVLRFIMESGAKGCEVIVSGKLRAQRAKSMKFKDGYMISSGQPVKEYIDSAVRHILMRQGVLGIKVKIMLDWDPKGKQGPTTPLPDLVTIHSPKEEKDYARTSVVVPDVVPTMLPTDIPIPVPVV*

NbL16g02460.1 vacuolar_protein_sorting−associated_protein_2_homolog_1−like [MW = 25.12 kDa]

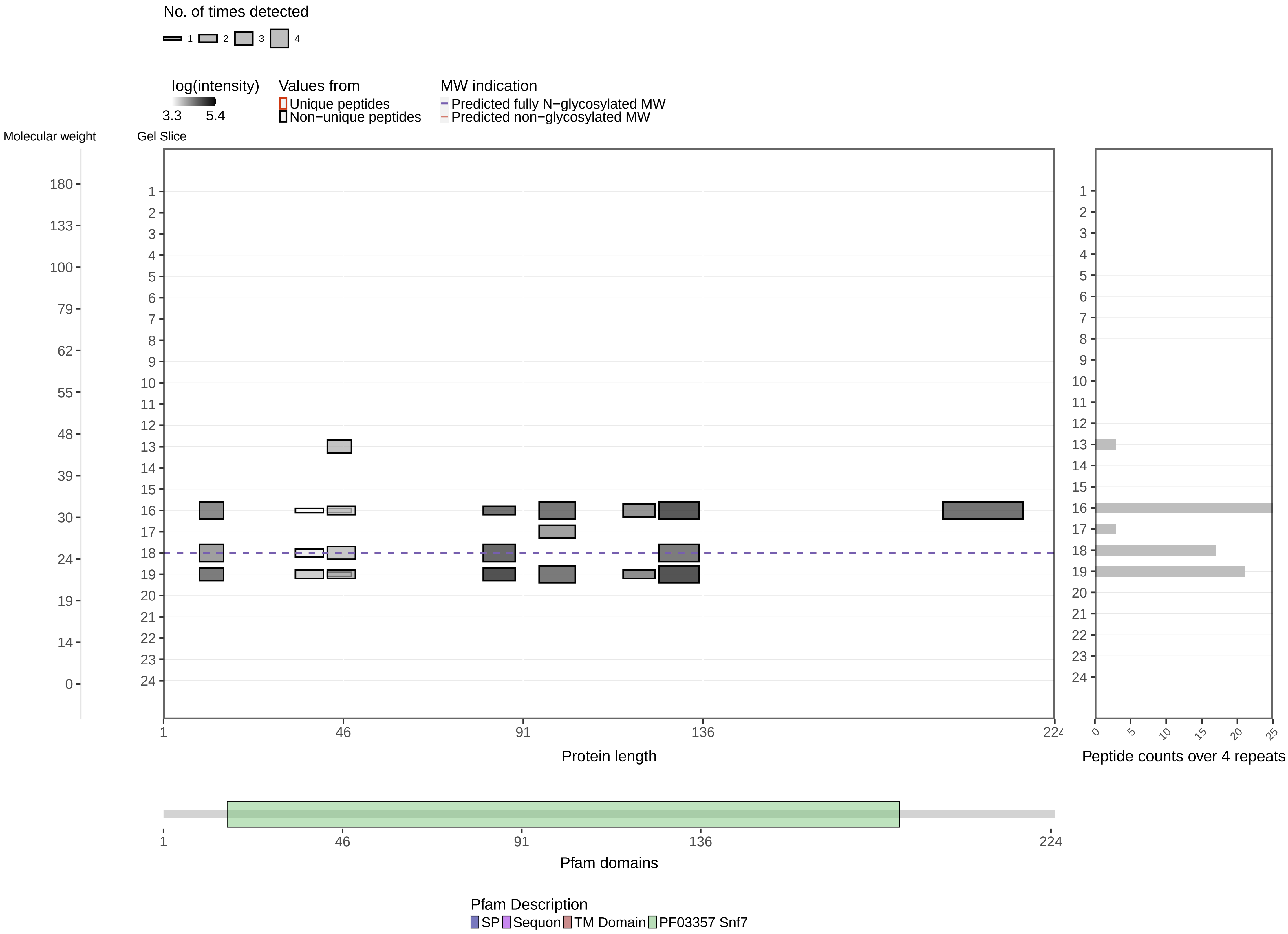


Protein sequence:

EAMKGVTKAMGQMNRQMNLPALQRIMQEFEMQNEKMEMVSEVMGDAIDDALEGDEEEEETEELVNQVLDEIGISMNNELVNAPSSAVAAPAAKNKVAQAEASEHDDGGIDSDLQARLDNLRKM*

### NbL16g02390.1 probable_inactive_receptor_kinase_At2g26730 [MW = 77.45 kDa]

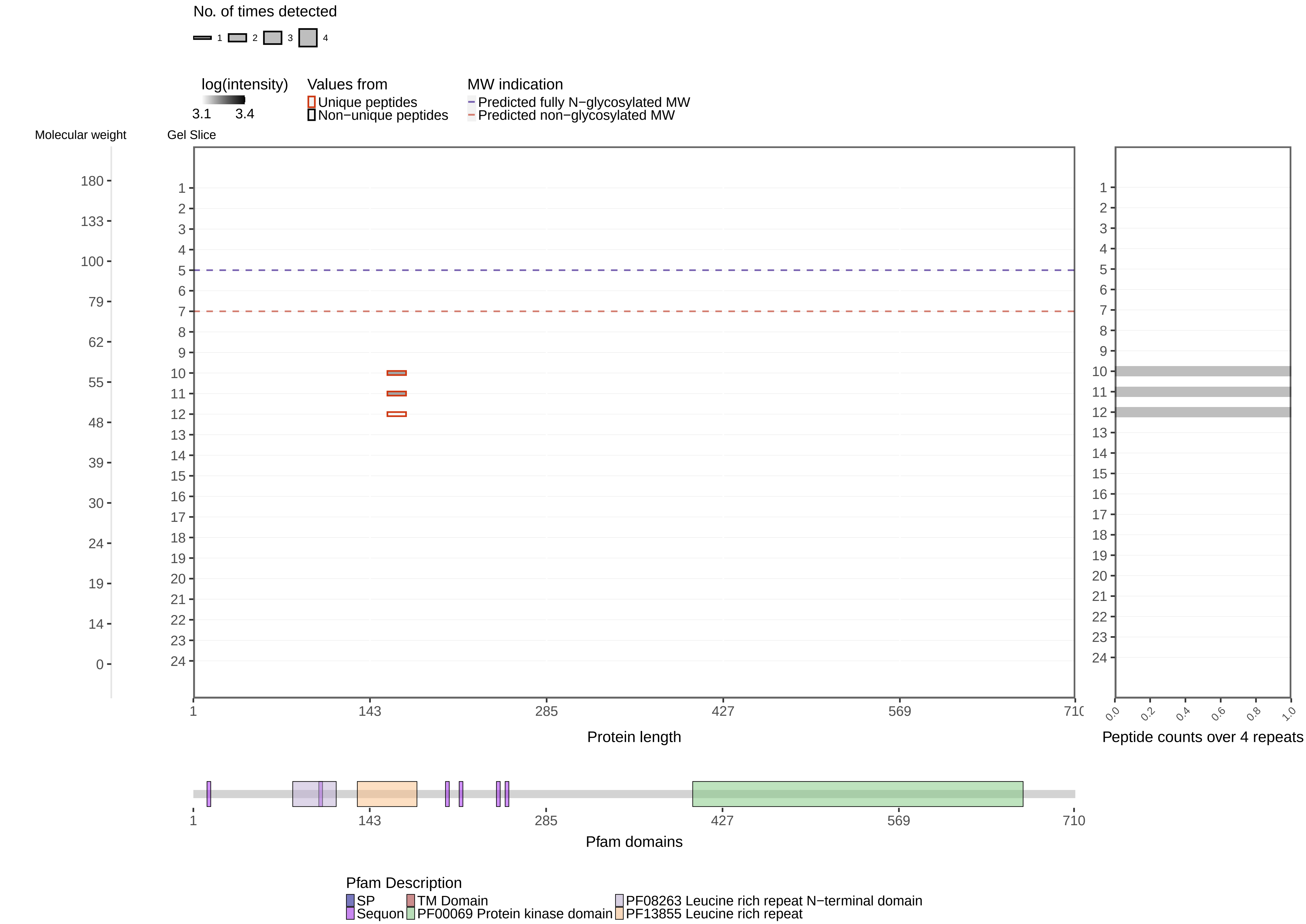


Protein sequence:

KPPTTTAAAFTGAGEAGTSSSKDDLTAEGERNKLVFFNGGGGYSFDLEDLLRASAEVLGKGSVGTSYKAVLEEGTTVVVKRLKDVVVTRKEFEQQMDVLGKMKHENVLPLRAFYYSKDEKLLVSDYVPAGSLSSLLHGSRGSGRTPLDWDSRMRIVLSAARGISYLHISGKIVHGNIKASNVLLKQDNNQDYACVSDYGLNPLFSTTATPVNNHRVAGYRAPEVLETRKVTFKSDVYSFGVLILELLTGKAPNQASLGEEGIDLPRWVQSVVREEWTAEVFDVELMRYHNVE

### NbL16g02310.1 translation_machinery−associated_protein_22−like [MW = 20.71 kDa]

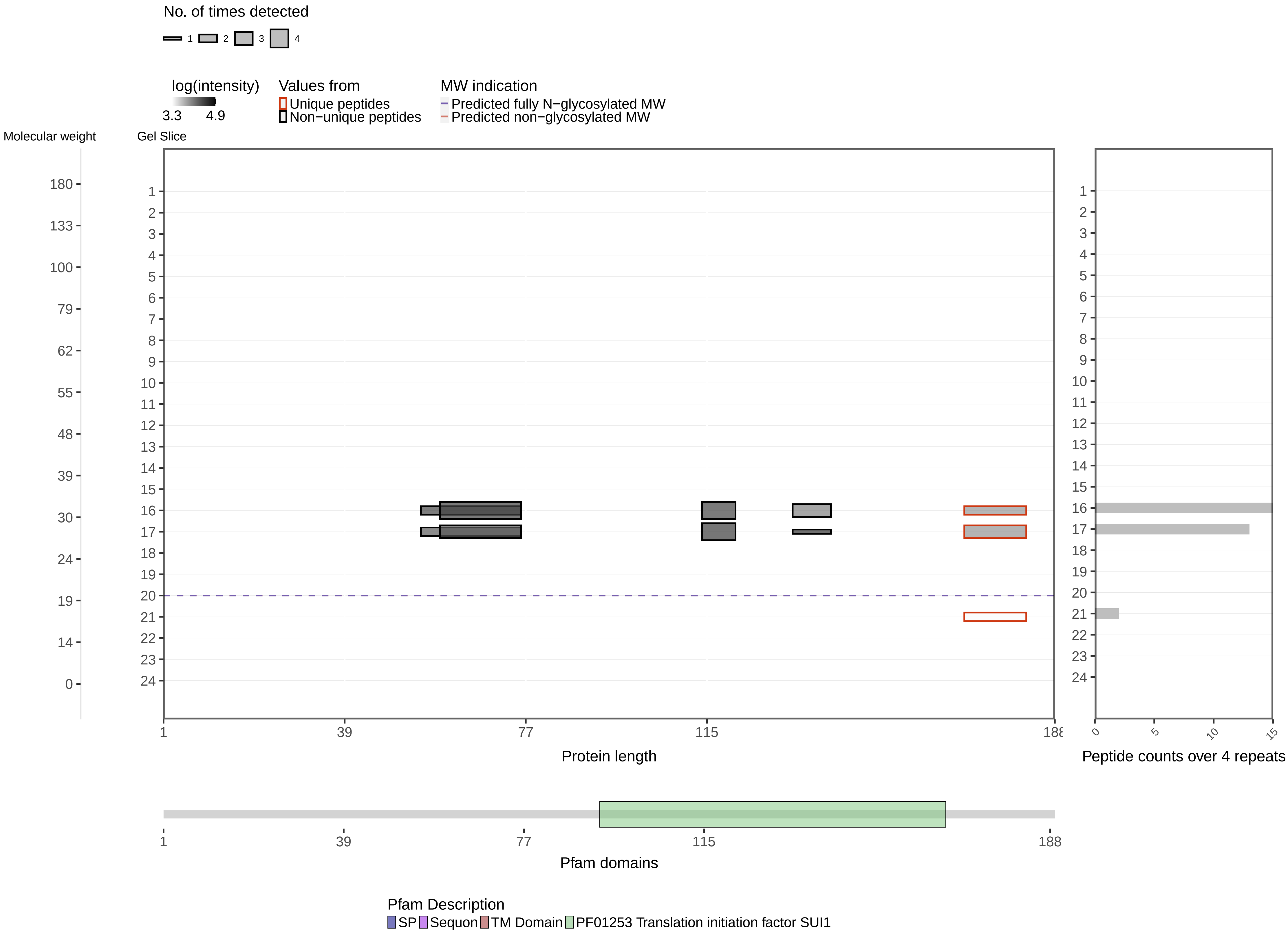


Protein sequence:

EVKRLPGGKIKKKERQEIVIEKVTRNRRKSITTIKGLDLFGVKLSDASKKLGKKFATGASVVKGPTEKEQIDVQGDISYDIVDFITETWRDVPASAVFFIEDGKKVPAM*

NbL16g02290.1 carbamoyl−phosphate_synthase_small_chain_chloroplastic [MW = 47.18 kDa]

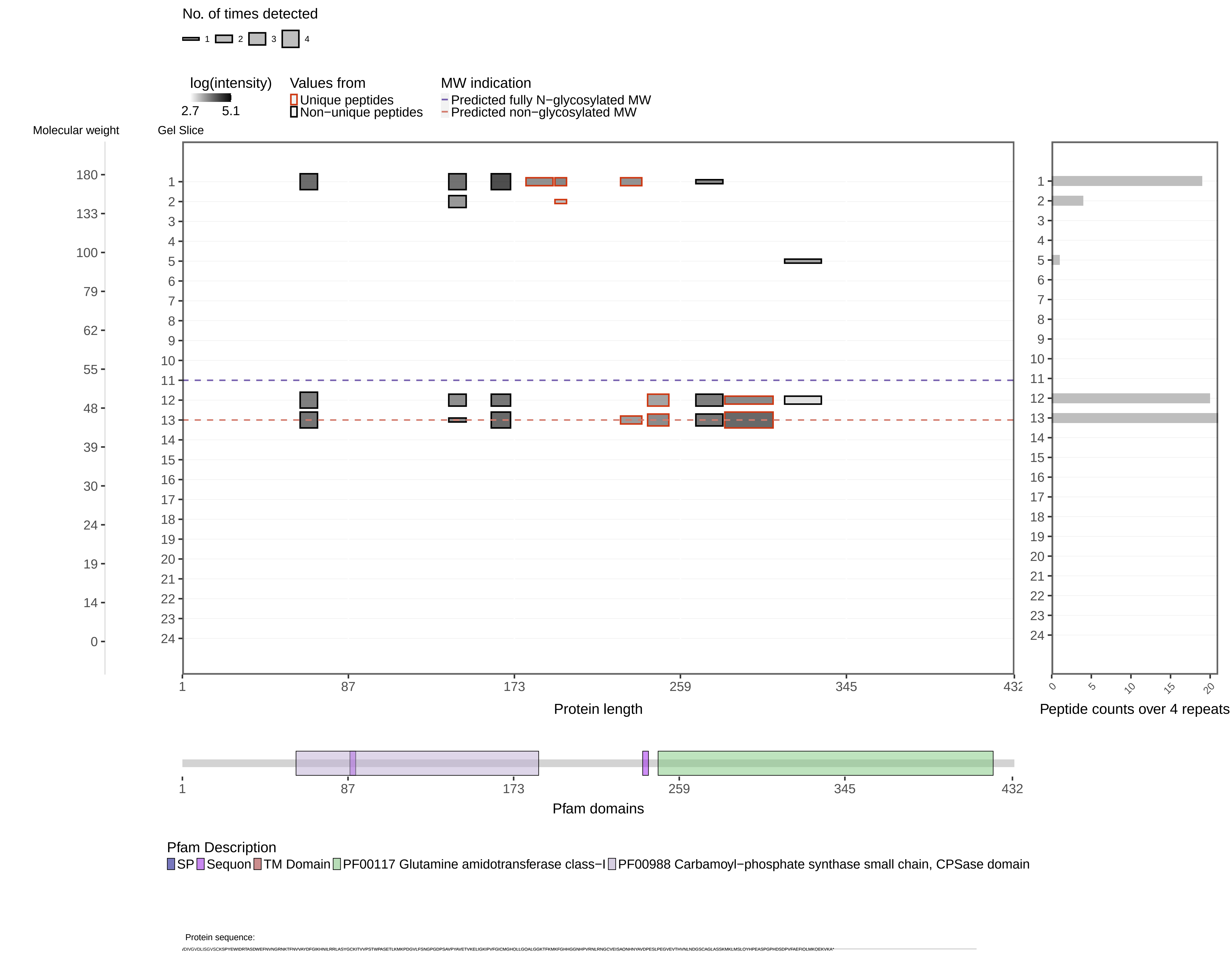

### NbL16g02270.1 beta−galactosidase−like [MW − SP = 78.3 kDa]

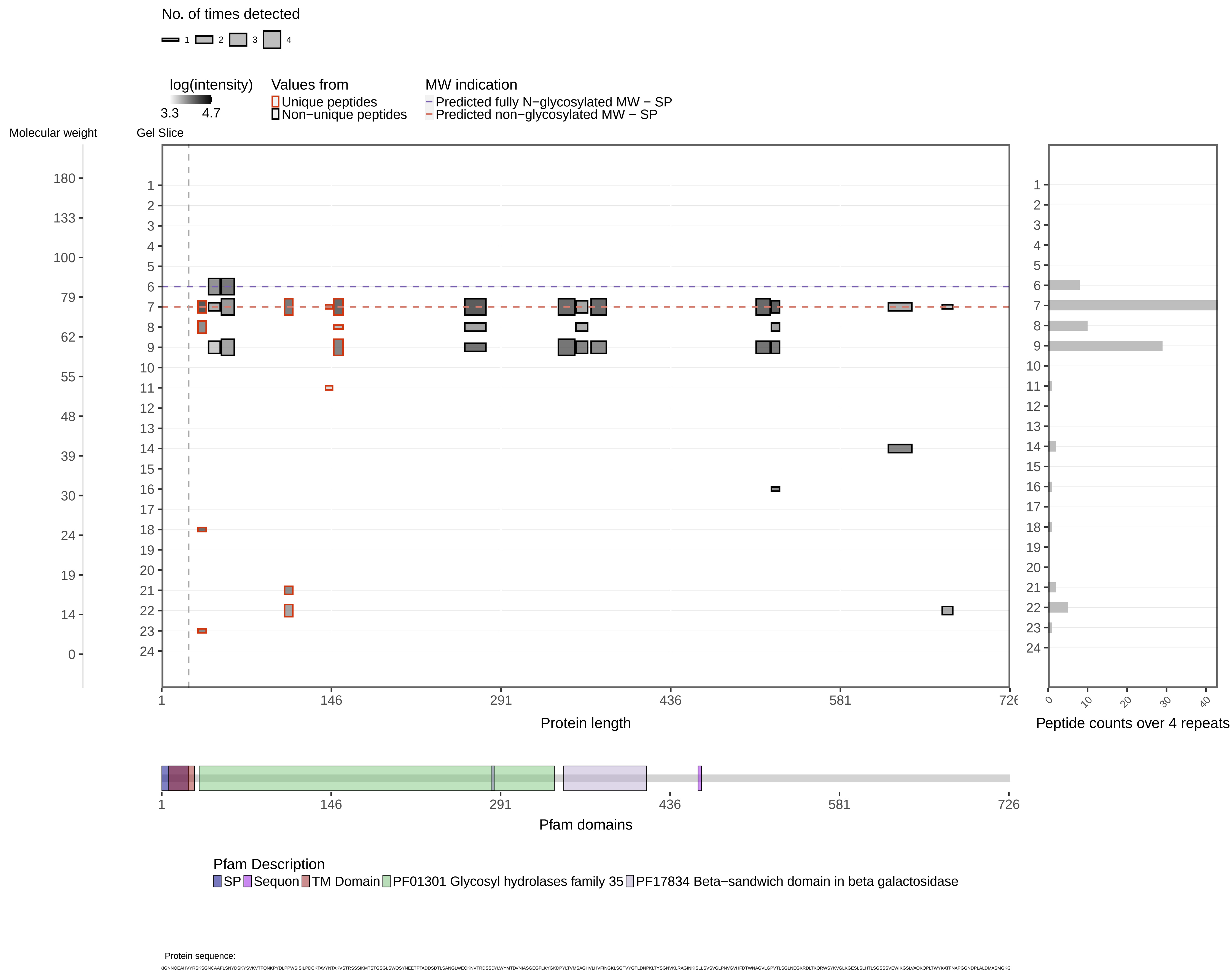

NbL16g02040.1 IAA−amino_acid_hydrolase_ILR1−like_4 [MW − SP = 46.43 kDa]

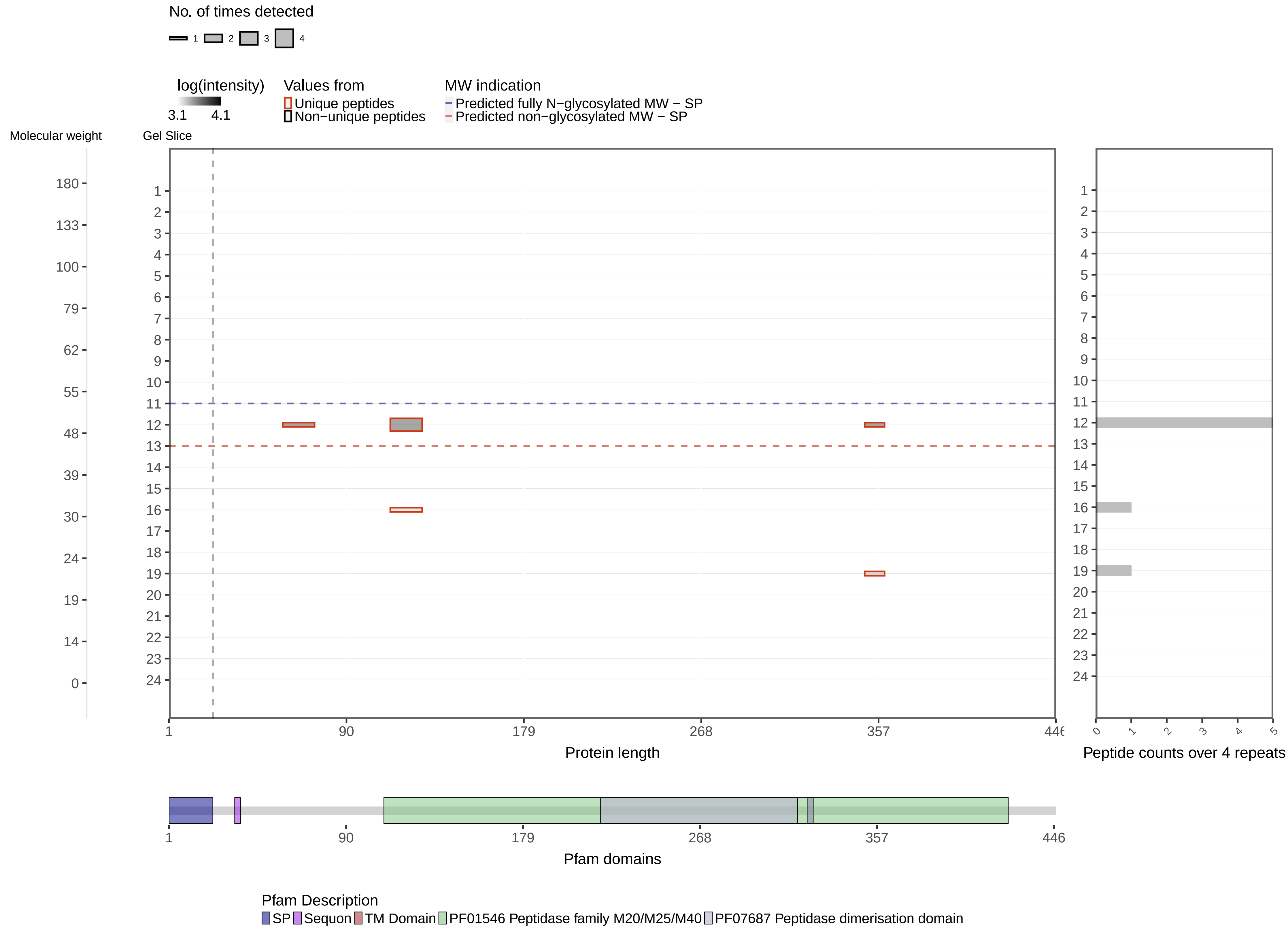


Protein sequence:

AKSGPILAGSGFFEAVINGKGGHAAIPQHTIDPILAASNIIVSLQHLVSREADPLDSQVVTVGKFKGGGAFNVIPDSVTIGGTFRAFSKESFTQLKQRIVEVITRQAAVQRCNATVNFDTNNKPFYPVTVNDKTLHEHFKNVAGQMLGTEKIRELKPLMGTEDFSFFAEAIPGYFYFVGMTDETKGRFESGHNPLYRVNEDALPYGVALHASLATTYLLDHQLKVTTQDRNVHDEL*

NbL16g01990.1 mitochondrial−processing_peptidase_subunit_alpha [MW = 54.81 kDa]

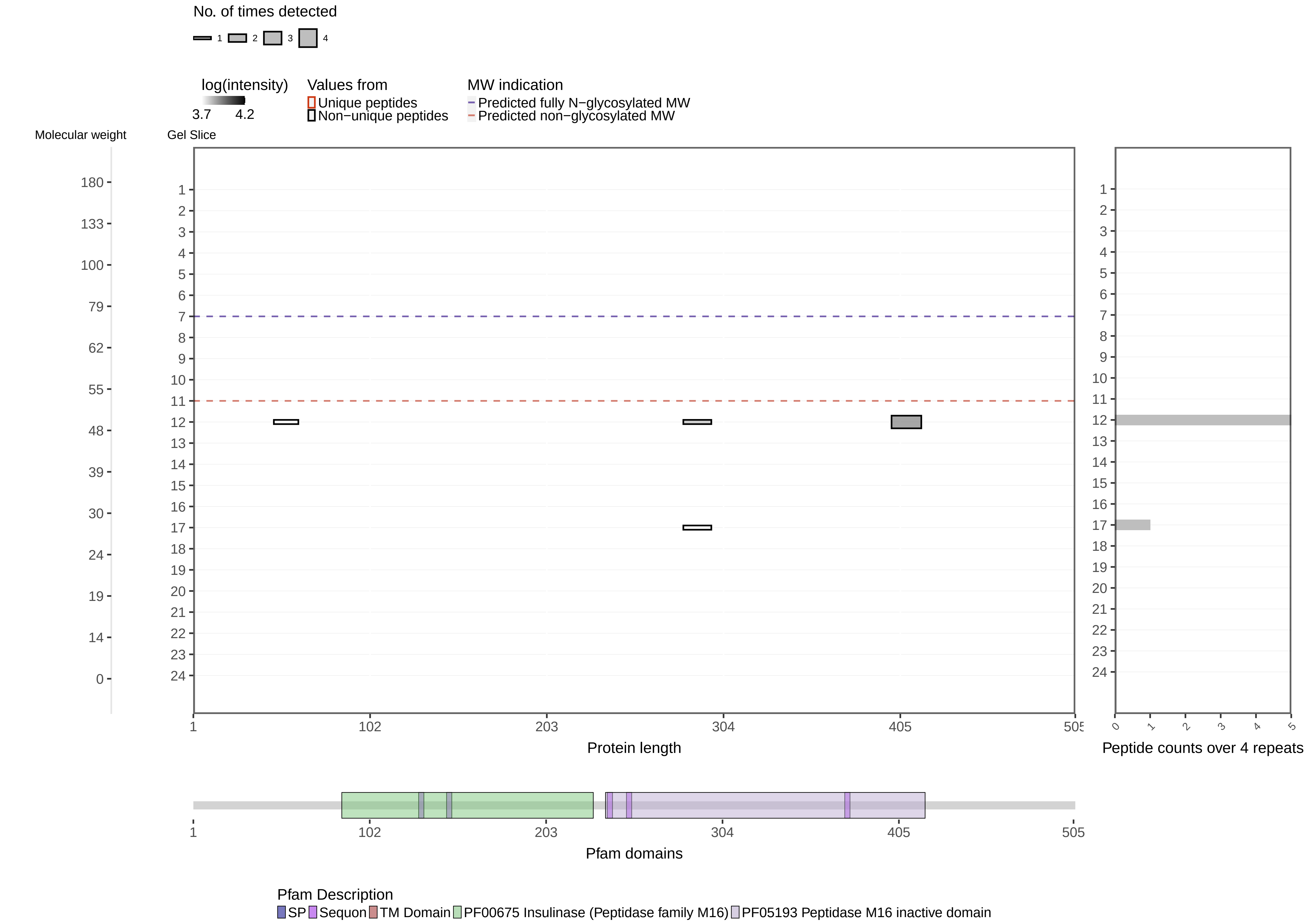


Protein sequence:

VLEEFVAENYTAPRMVLAASGVEHGELLKIAEPLLSDLPKVATSEVPKPVYVGGDYRRQADAEMTHFALAFEVPGGWMSEKESMTLTVLQMLMGGGGSFSAGGPGKGMYSRLYLRVLNQHPQIQAFSAFSSIYNDTGLFGIQATTTSDFGPQAVDVAVKELIAIANPGEVDQVQLDRAKQSTKSAILMNLESRMVASEDIGRQLLTYGERKPVEHFLKAIDAVSAKDIASVAQKLISSPLTMASYGDVLSLPSYDAVSNRFHSK*

NbL16g01960.1 profilin−1−like [MW = 17.27 kDa]

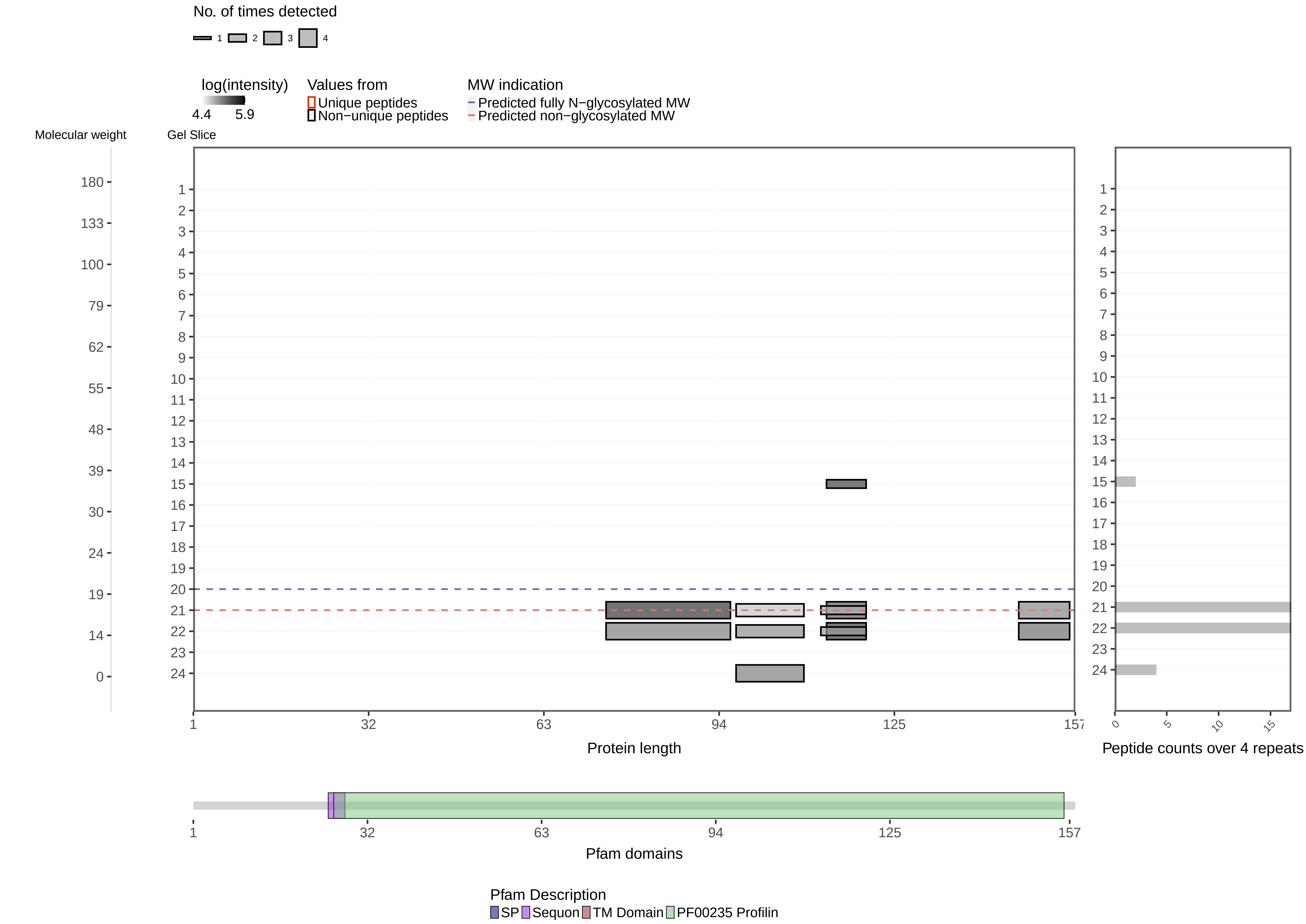


Protein sequence:

QFKPEEIKAIMNDFEEPGTLAPTGLHLGGTKYMVLQGEAGVVIRGKKGPGGITIKKTNMAILIGIYDEPMTPGQCNLVVERLGDYLLEQGF*

NbL16g01950.1 glucan_endo−13−beta−glucosidase_13−like [MW − SP = 51.37 kDa]

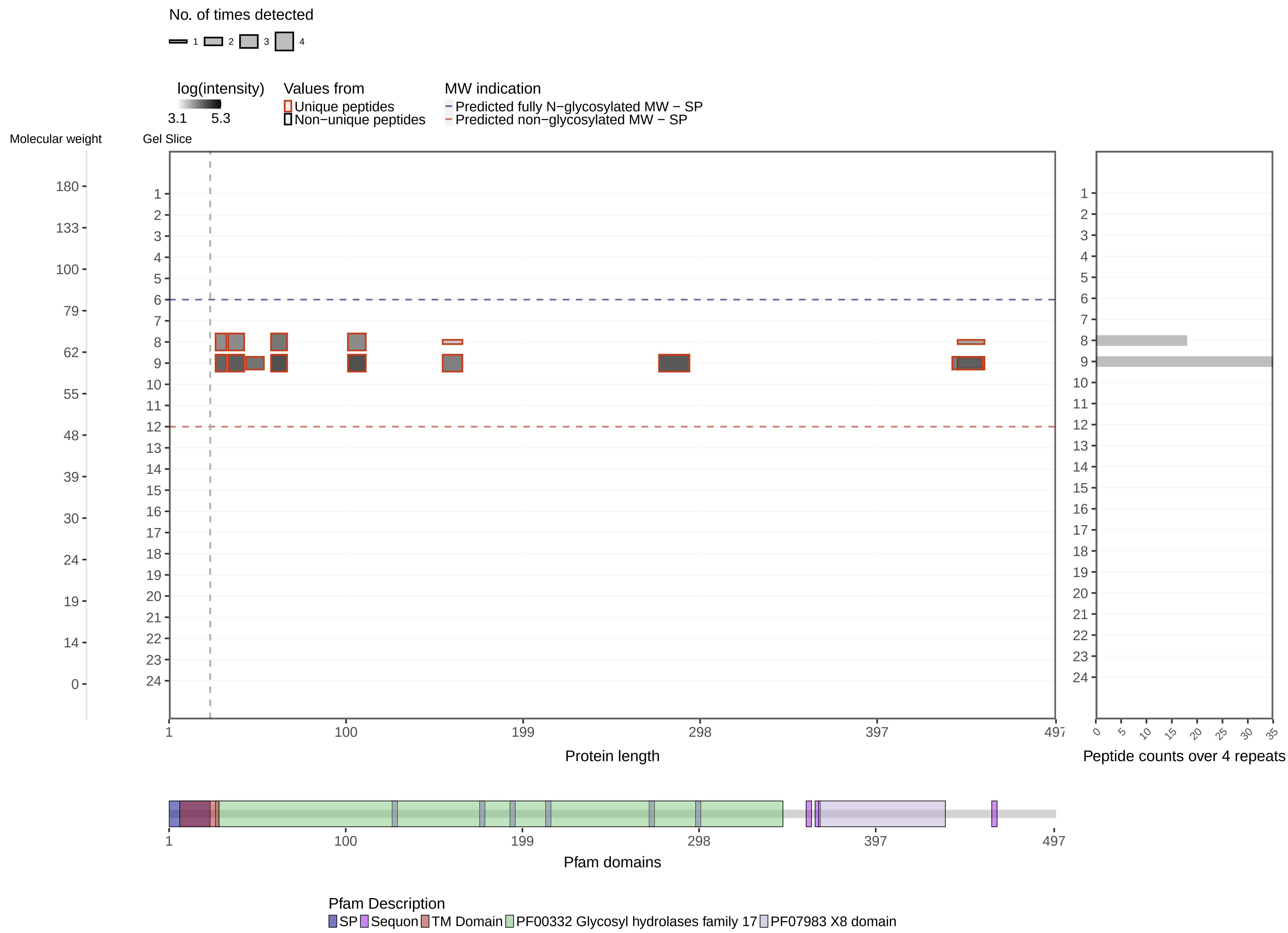


Protein sequence:

### NbL16g01930.1 arginine−−tRNA_ligase_chloroplastic/mitochondrial−like [MW = 73.29 kDa]

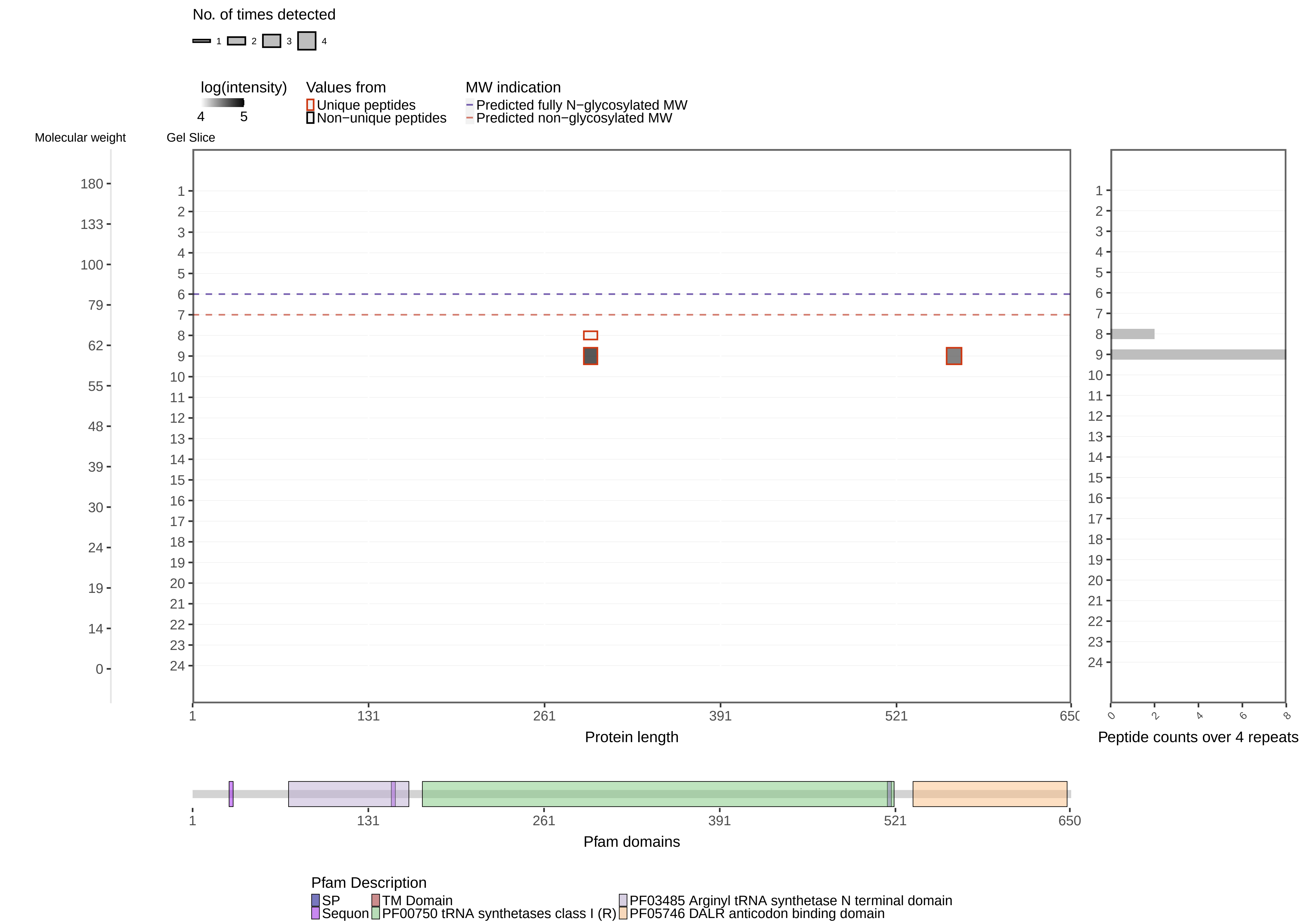


Protein sequence:

RKEFQKVYERLGVHLEEKGESFYNPYIPKALELLNEKGLIEESEGARVIFIERKKIPLIVVKRDGGFNYASTDLTALWYRLSEEKADWMIYVTDVGQREHFEMFFSAAKRAGWLPSNENDYPKASHVGFGLVLGEDGKRFRTRSTEVVKLVDLLDEAKSRCKLALEERGKAKEWTAEELDQTAEAVGYGAVKYADLKNNRLTNYTFNFDQMLSDKGNTAVYLLYAHARICSIIRKSSKDIEELKKSGAMGLAHQDERMLGLHLLQFPEIVEEACTNLLPNLLCEYLYKLSEDFTK[illegible]

NbL16g01910.1 probable_glutamate_carboxypeptidase_2 [MW = 79.91 kDa]

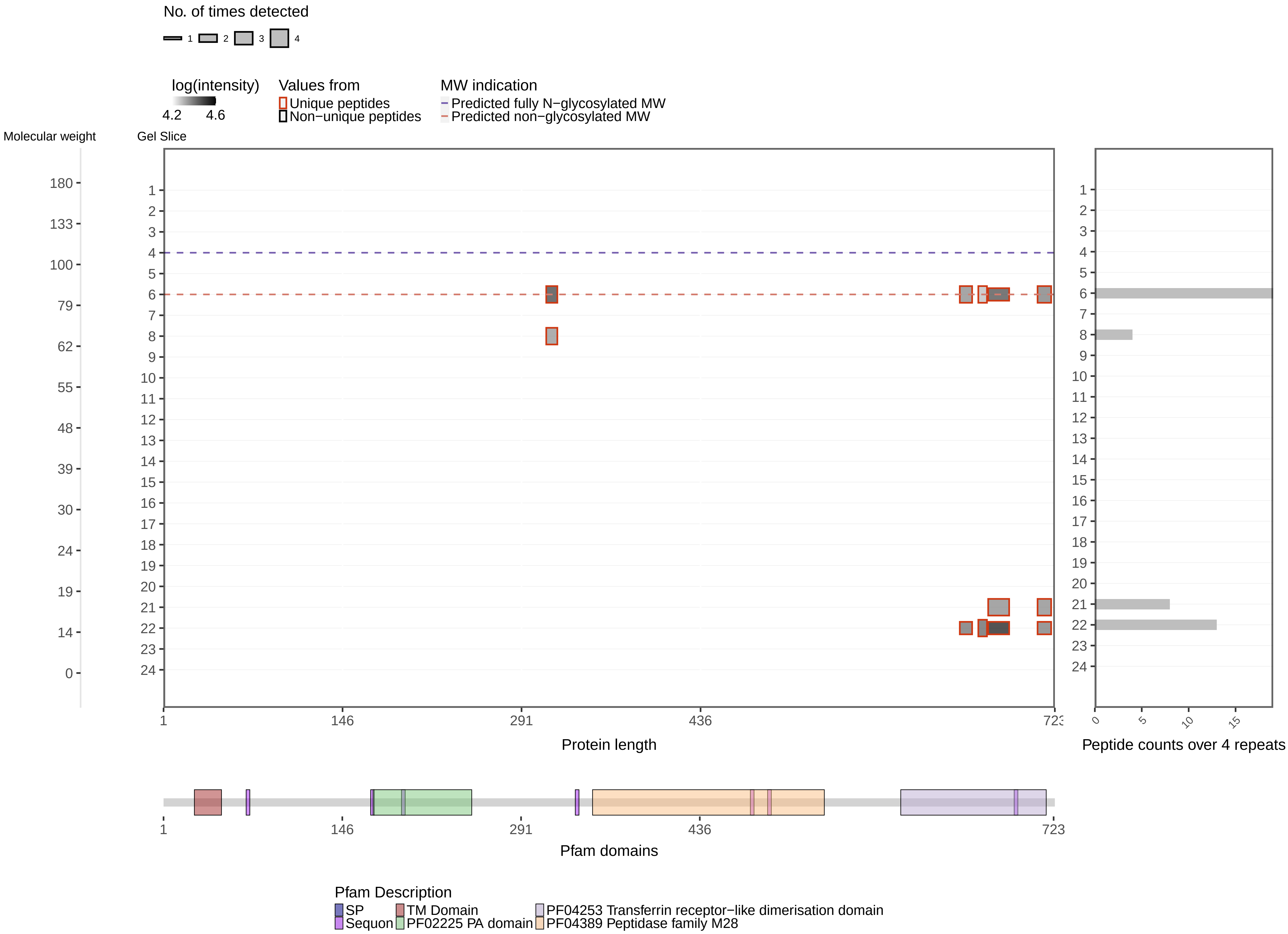


Protein sequence:

VIEGEEEPDRFVILGNHRDAWTFGAVDPNSGTAALLEIAORLEKLORKGWRPRRTIVLCNWDAEEYGLIGSTEWVEENREMLASRVVAYLNVDCAVOGGHFRASANPOLDELIIOVSOOIKDPDNSSOTIYOSWLDSGNDTTVKLGRLGGAGSDYAAFVOHIGIPALDLSFGDGYPVYHSMYDDFVWMKKFGDPMFHRHVAVASFWGLLALRLADDEILPFNYLTYAYELOKSAVOLEAEISENGIKLVPLYVSIEKFRKAAIRIEDDVKAVKAKRSWESLRELNDRLIMTERA

### NbL16g01720.1 probable_lactoylglutathione_lyase_chloroplastic [MW = 40.2 kDa]

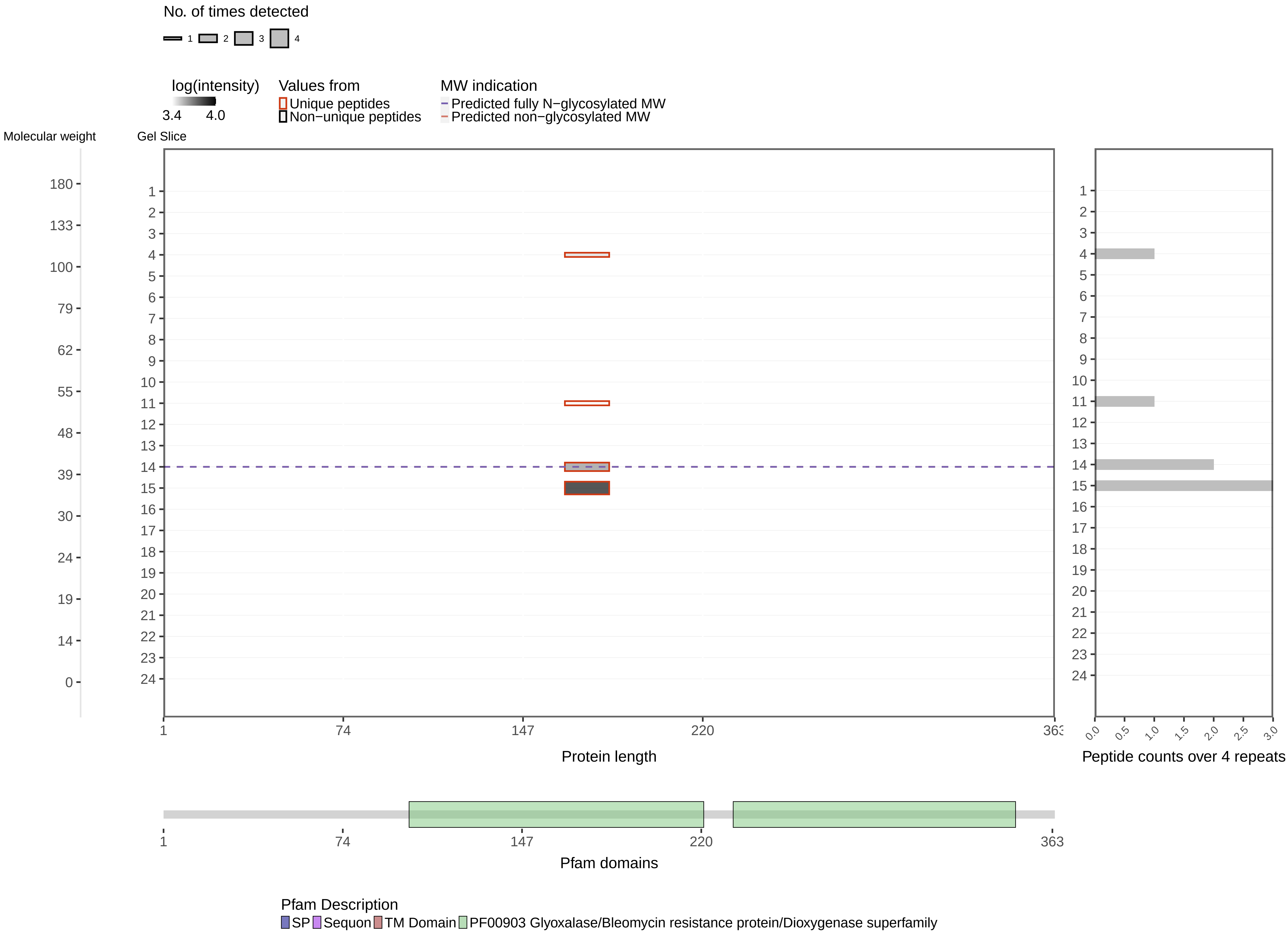


Protein sequence:

SGFGHFGIAVEDVSKTVELVKAKGGKVSREPGPVKGGKTVIAFIEDPDGYKFELLERGPTPEPLCQVMLRVGDLDRAINFYEKAYGMRLLRTRDNPEYKYTIAMLGYGPEDSSAVMELTYNYGVTEYDKGNAYAQIAIGTDDVYKTAEAIRLCGGKITREPGPLPGISTKIVACLDPDGWKSVFVDNVDFLKELE*

NbL16g01600.1 peroxisomal_fatty_acid_beta−oxidation_multifunctional_protein_AIM1 [MW = 82.66 kDa]

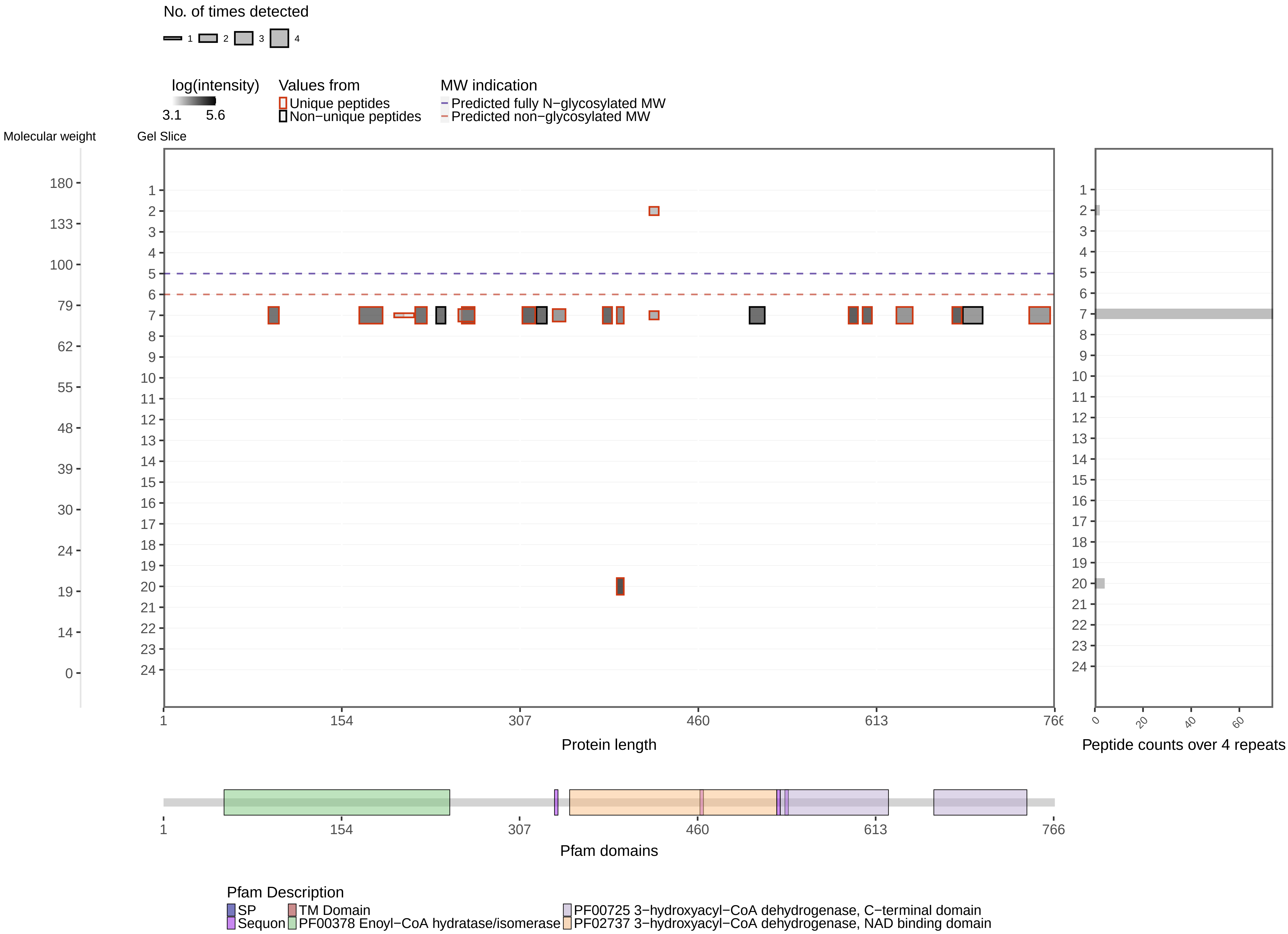


Protein sequence:

LVLSNIFVILKEINPEYLQKGIKAVEANIRGLVARKKLQQDKASKALSIVKGVLDYSEFKDVDMVIEAVIENVPLKQQIFSDIEKVCPPHCILATNTSTIDLNLIGEKTRSQDRIIGAHFFSPAHVMPLLEIVRTEKTSAQAILNLMAVGKSIKKVPVVVGNCTGFAVNRTFFPYAQSAHILANLGVDVYRIDSQITSFGLPIGPFQLQDLAGYGVAVATGKEFASAFPDRTFKSPLLDLLIKSGRNGKNNGKGYYIYKKGEKPKPDPSVLPIIEESRRLTNIMPGGKPISVTDQEIVEMILFPV

NbL16g01450.1 aldehyde_dehydrogenase_family_2_member_C4 [MW = 54.59 kDa]

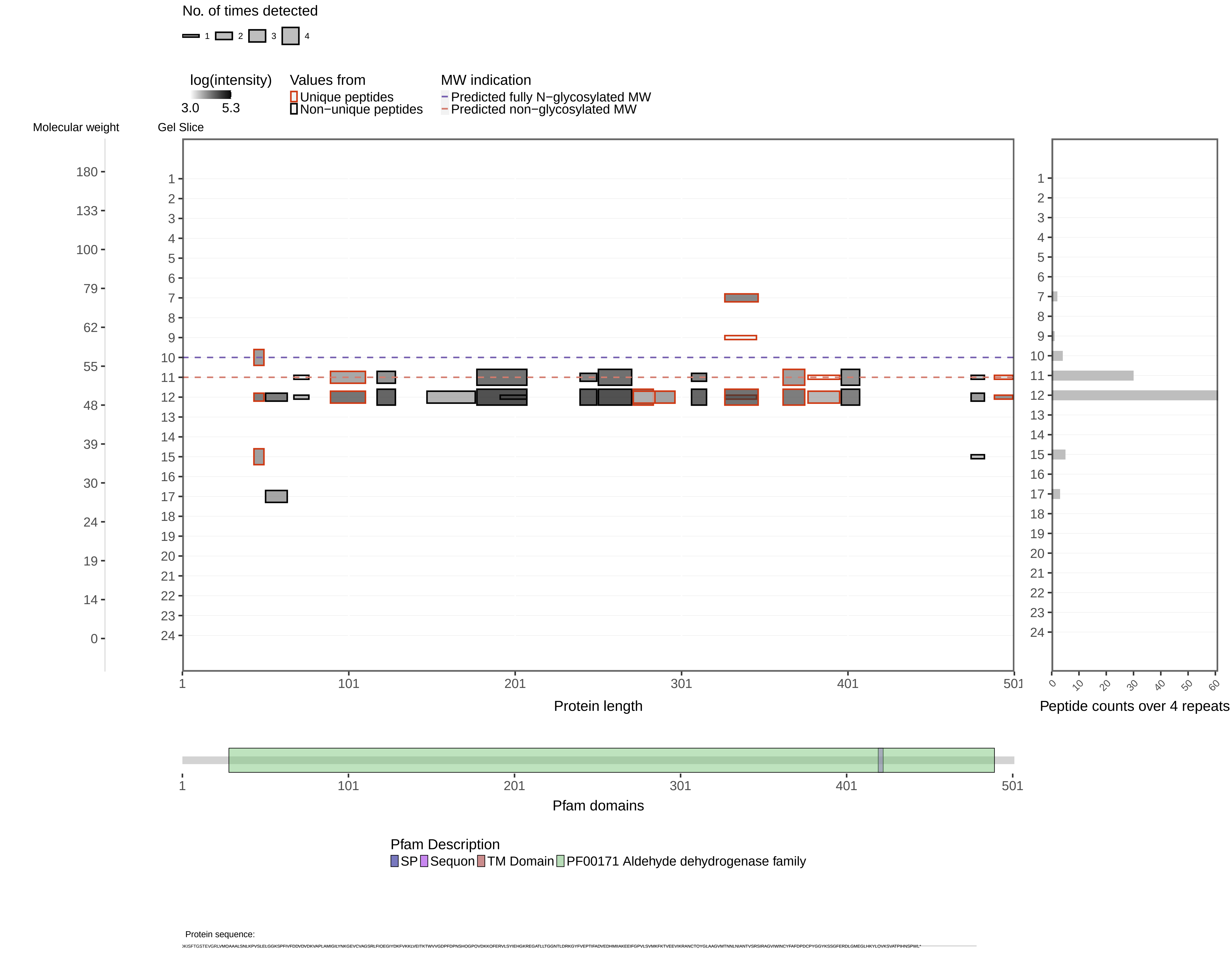

NbL16g01430.1 fe−S_cluster_assembly_factor_HCF101_chloroplastic−like [MW = 57.03 kDa]

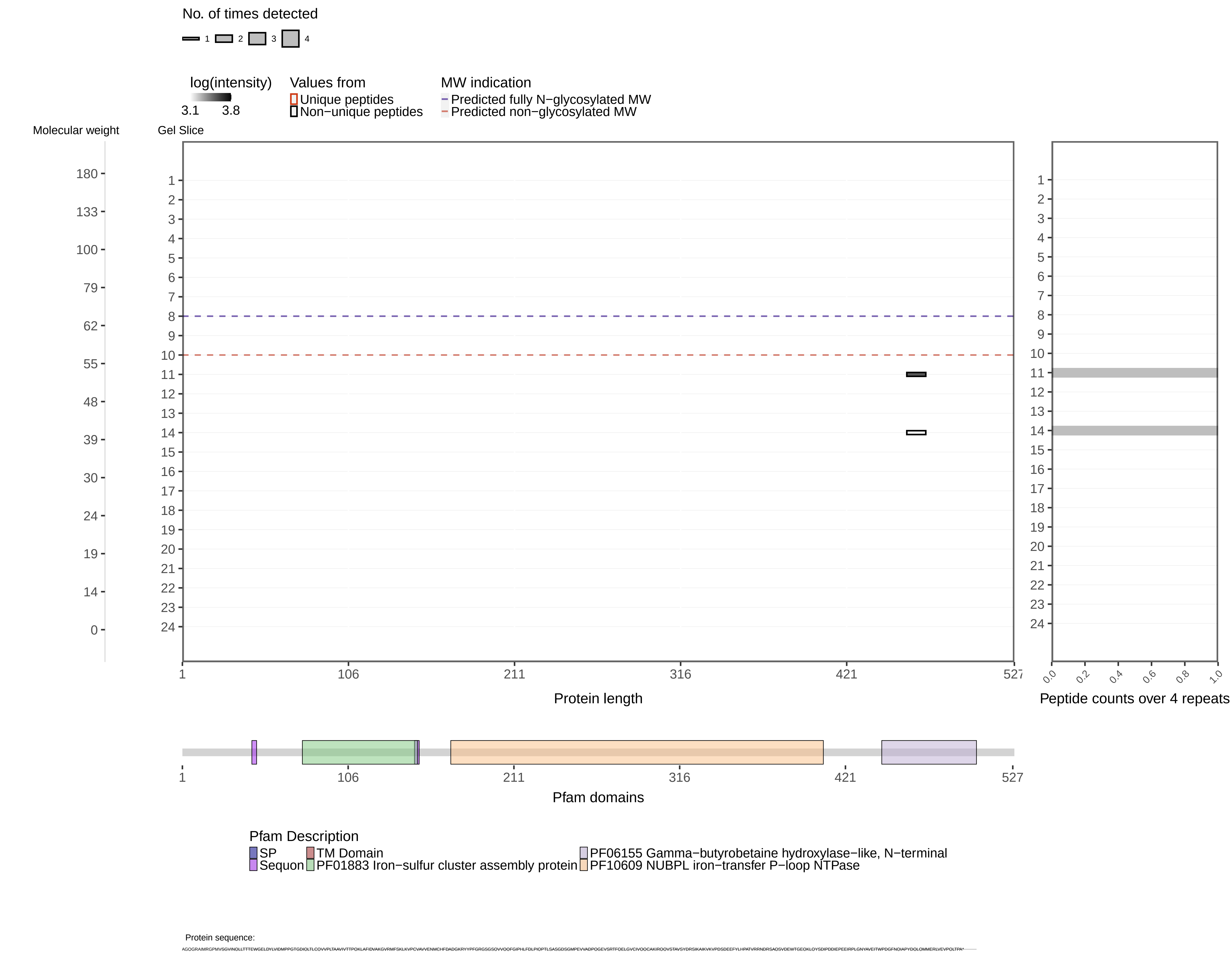

### NbL16g01420.1 uncharacterized_LOC107801219 [MW − SP = 14.23 kDa]

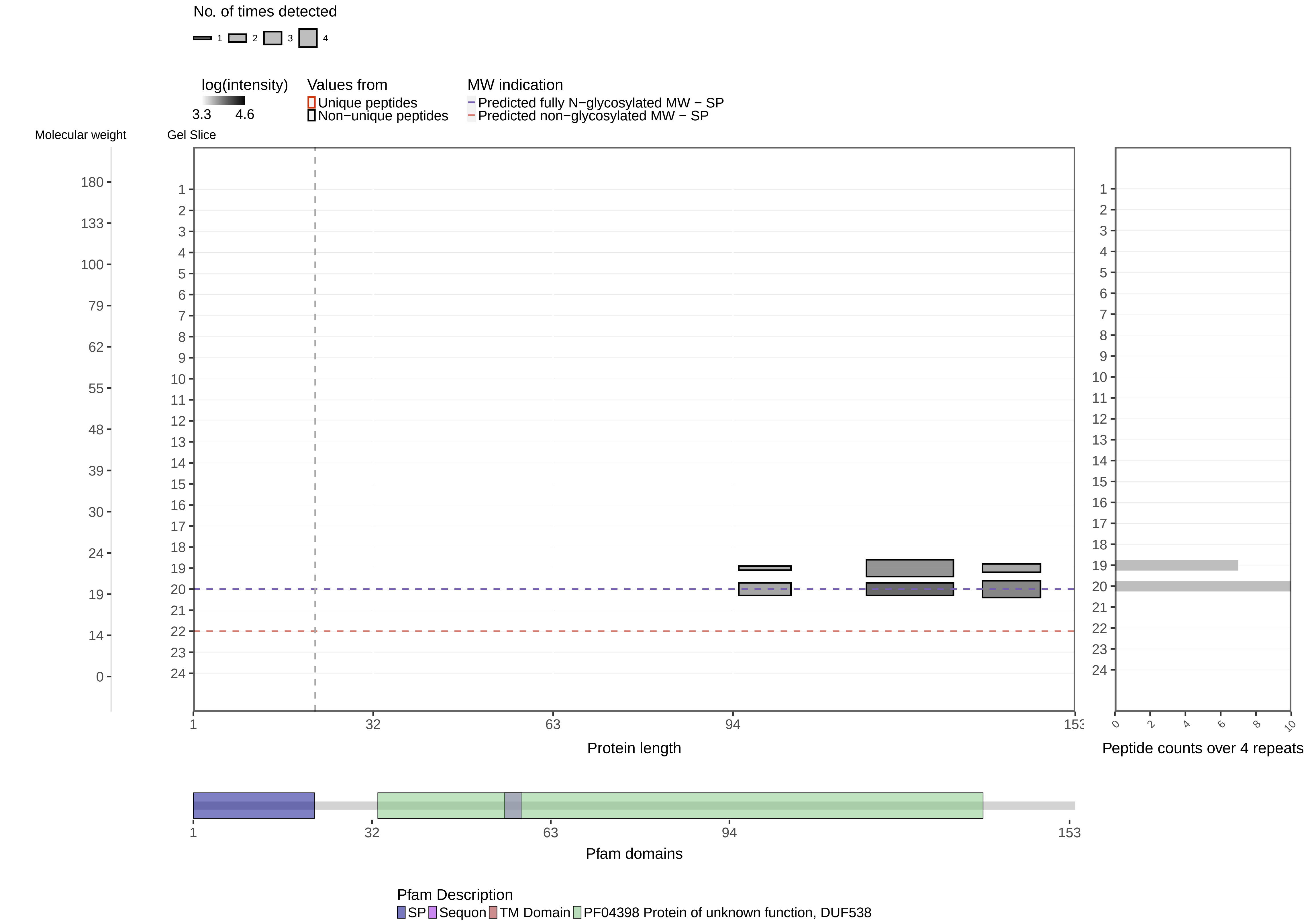


Protein sequence:

SLGATCKITLPPDNYLATYSKKITGKIVENRIAELDGISVRAFFKWWGITGIRSSGENLVFEVGMVTAKYPSKNFDESPECEGKKHSSS*

### NbL16g01390.1 leucine−rich_repeat_extensin−like_protein_4 [MW − SP = 47.34 kDa]

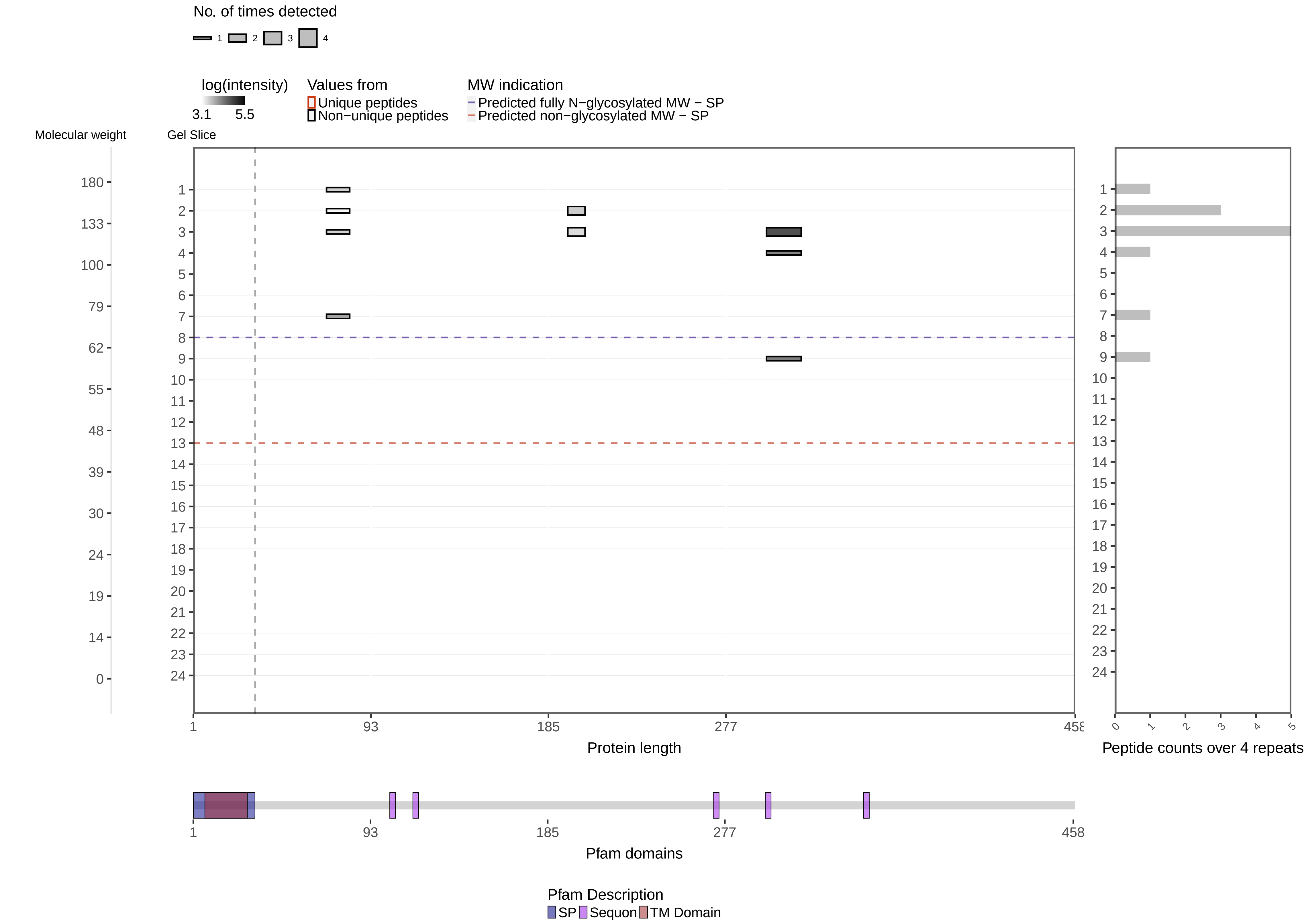


Protein sequence:

/PSQLFDKPLDAIFINHNRFMFELPDNFGNSPVSVIVLANNKFHGCLPASIGNMSNLNEVIMMNNGLRSCLPTEIGMLKNLTVFDVSFNQLMGPLPEKFGELVNLEQLNVAHNMLSGSIPKSICQLPKLENFTYSYNFFTGEPPVCLGLPEFDDQRNCLPNRPVQRSPAQCKAFLSKRIHCGAFKCHKFIPALPPPPPLSPPLPAPPPPVVVPISPSPPVLSPPPPPPSPPPPVYSPPPP

### NbL16g01280.1 GTP−binding_nuclear_protein_Ran−B1 [MW = 25.19 kDa]

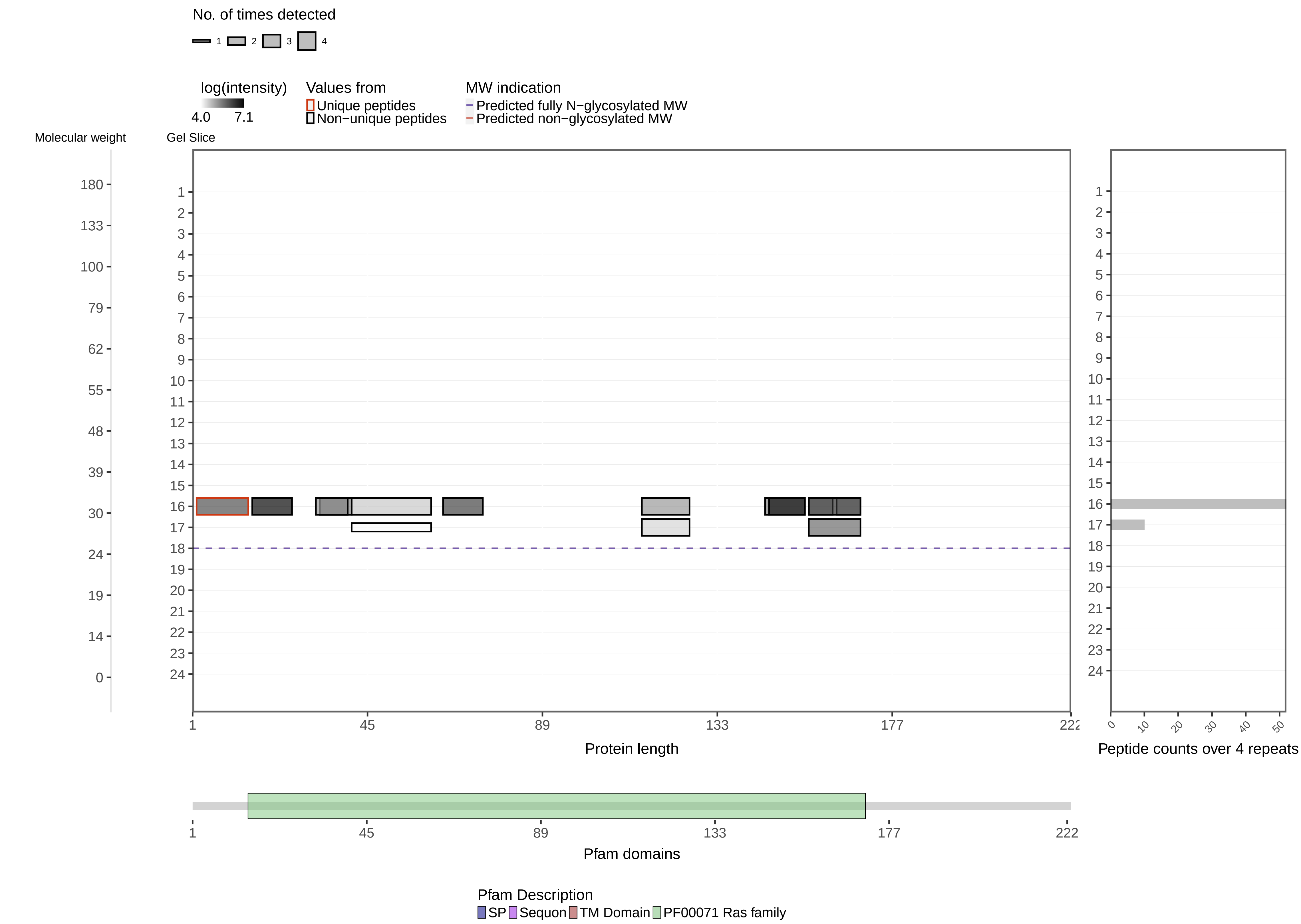


Protein sequence:

LTYKNVPTWHRDLCRVCENIPIVLCGNKVDVKNRQVKAKQVTFHRKKNLQYYEISAKSNYNFEKPFLYLARKLAGDANLHFVESPALAPPEVQIDLAAQQLHEQELLQAAAQPLPDDDDEAFE*

NbL16g00840.1 oxygenase_1_chloroplastic−like_(LOC107792958)_mRNA_HQ690822.1_Nicotiana_tabacum_heme_oxygenase_1 [MW = 32.13 kDa]

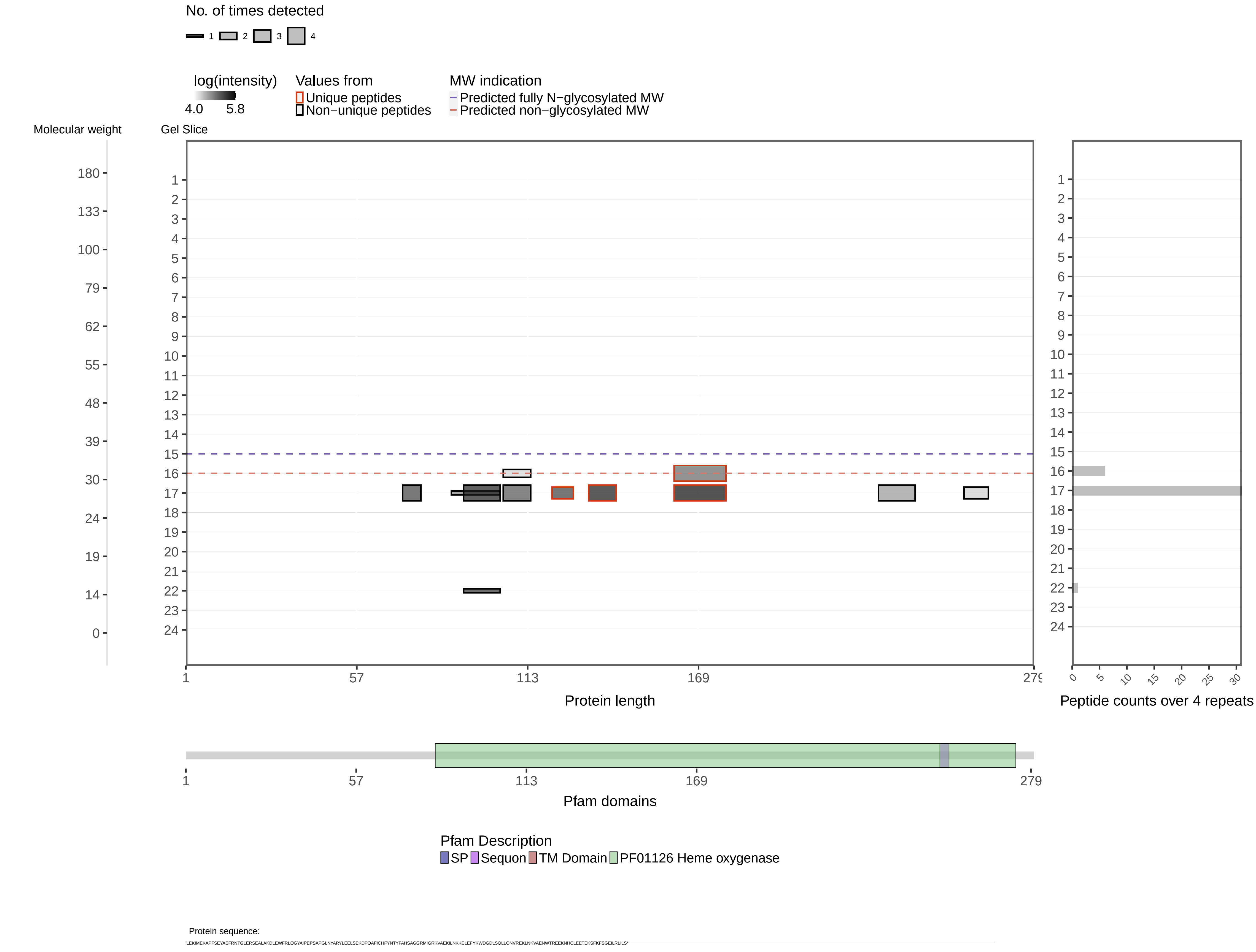

NbL16g00780.1 methyl−CpG−binding_domain−containing_protein_11−like [MW − SP = 28.99 kDa]

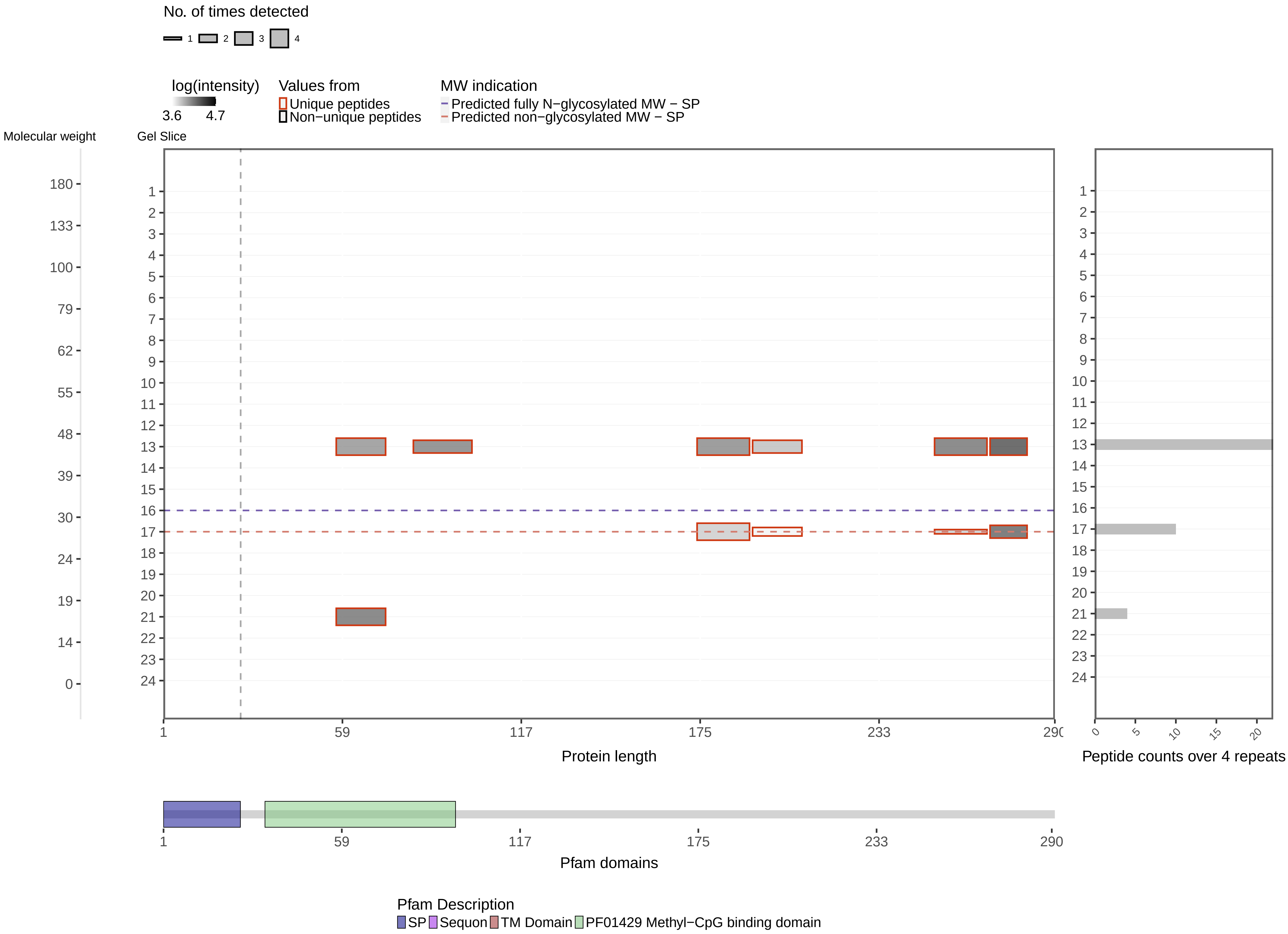


Protein sequence:

KDSNEVPEETEAAKDDDMEEAEEHEKDTVAVEAEKDVEQKQDENQDETQDGESELEKKEEAEATEKDVSEKDEIKSSENDGKKDESQNADGKVEDAPSEETQVEKDVEMADNVGHSKDVEEAPGDKVADDLEATKTNEEEKDVPVEEKVANSLLKQ*

NbL16g00770.1 thylakoid_lumenal_16.5_kDa_protein_chloroplastic−like [MW = 24.27 kDa]

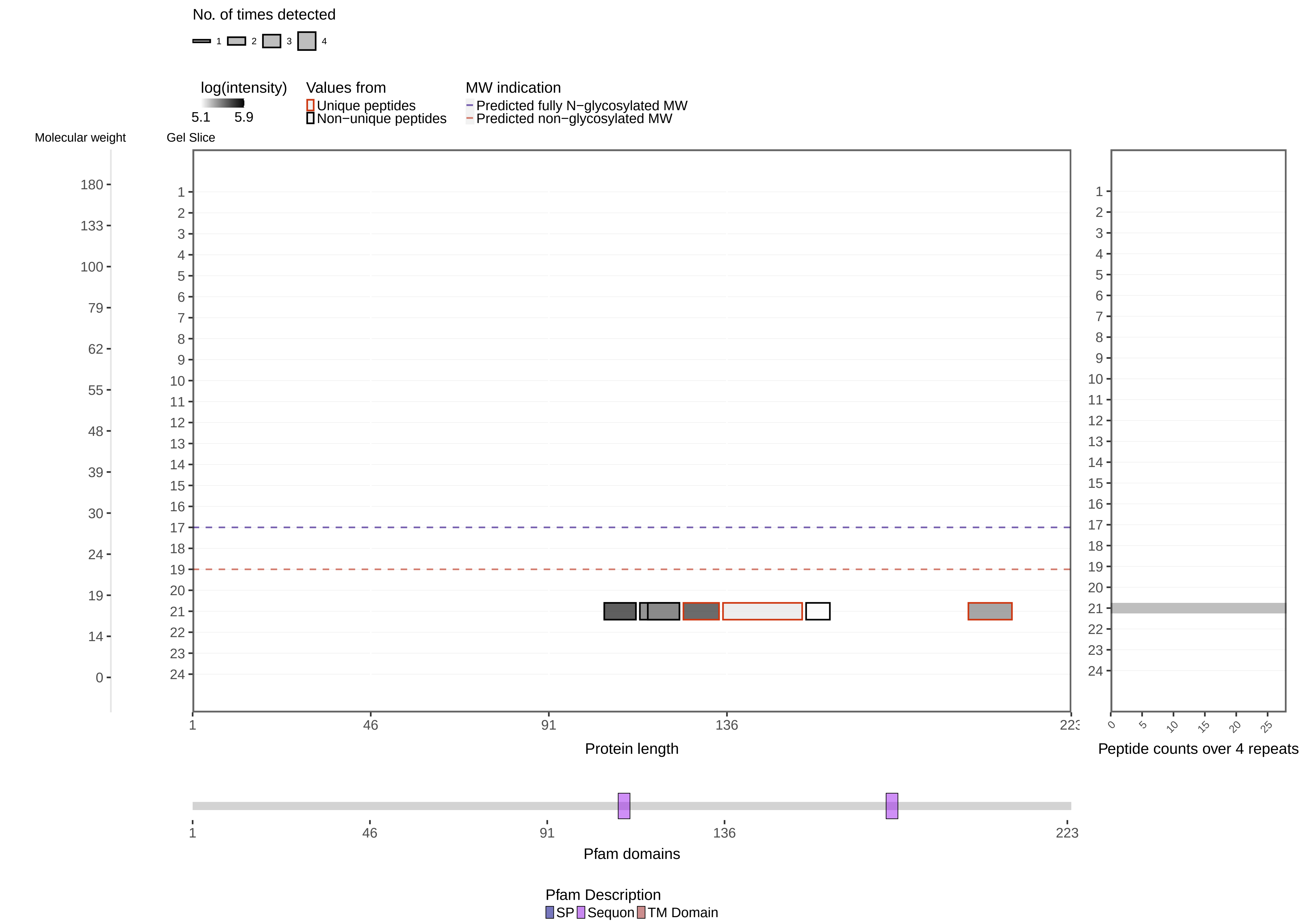


Protein sequence:

KKRLERQGIINSSTKEKGYLQDLVYKLSIVGQAIEKNDLSAASSVLGQSTDADWVQKVNSAFNKFSGSDEEKTEIDSFNSSLSFLVSSVTKNDIEGTKTAFLASASAFEKWTTLTGLIEELKGL*

### NbL16g00730.1 repetitive_proline−rich_cell_wall_protein_1−like [MW = 32.45 kDa]

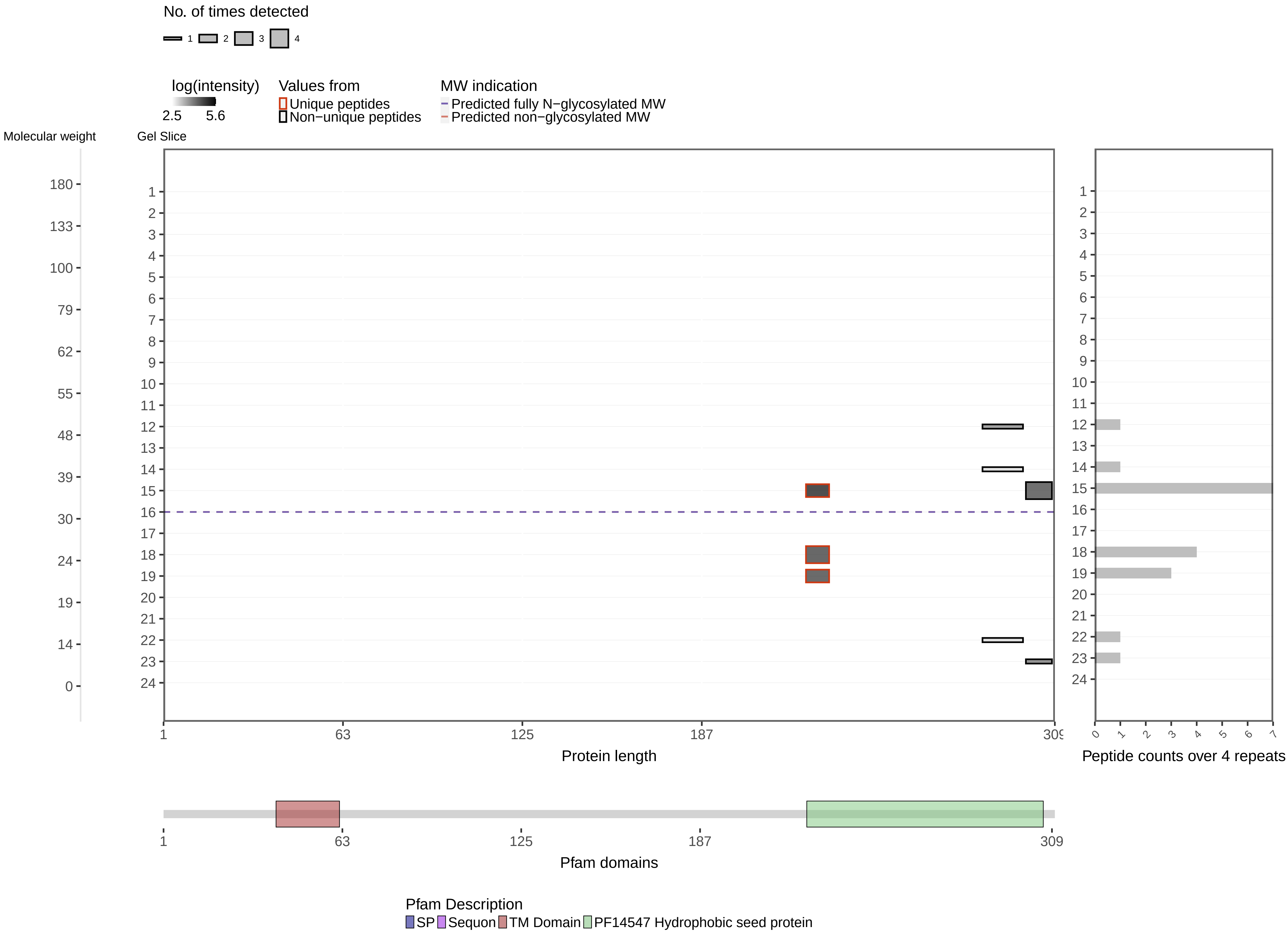


Protein sequence:

IGIPPVVKPPINFPPVVPPVIKPPIGIPPVTVPPVLKPPIGLPPIVKPPINLPPIGIPPVTVPPVITPSPKGKPCPSPPTTKATCPINTLKLGACVDLLGGLVHIGIGDPVVNECCPILHGLVELEAAACLCTTLKIKLLNLKIFVPLALOLLVTCGKTPPPGYTCSL*

NbL16g00670.1 bifunctional_purple_acid_phosphatase_26−like [MW − SP = 52.61 kDa]

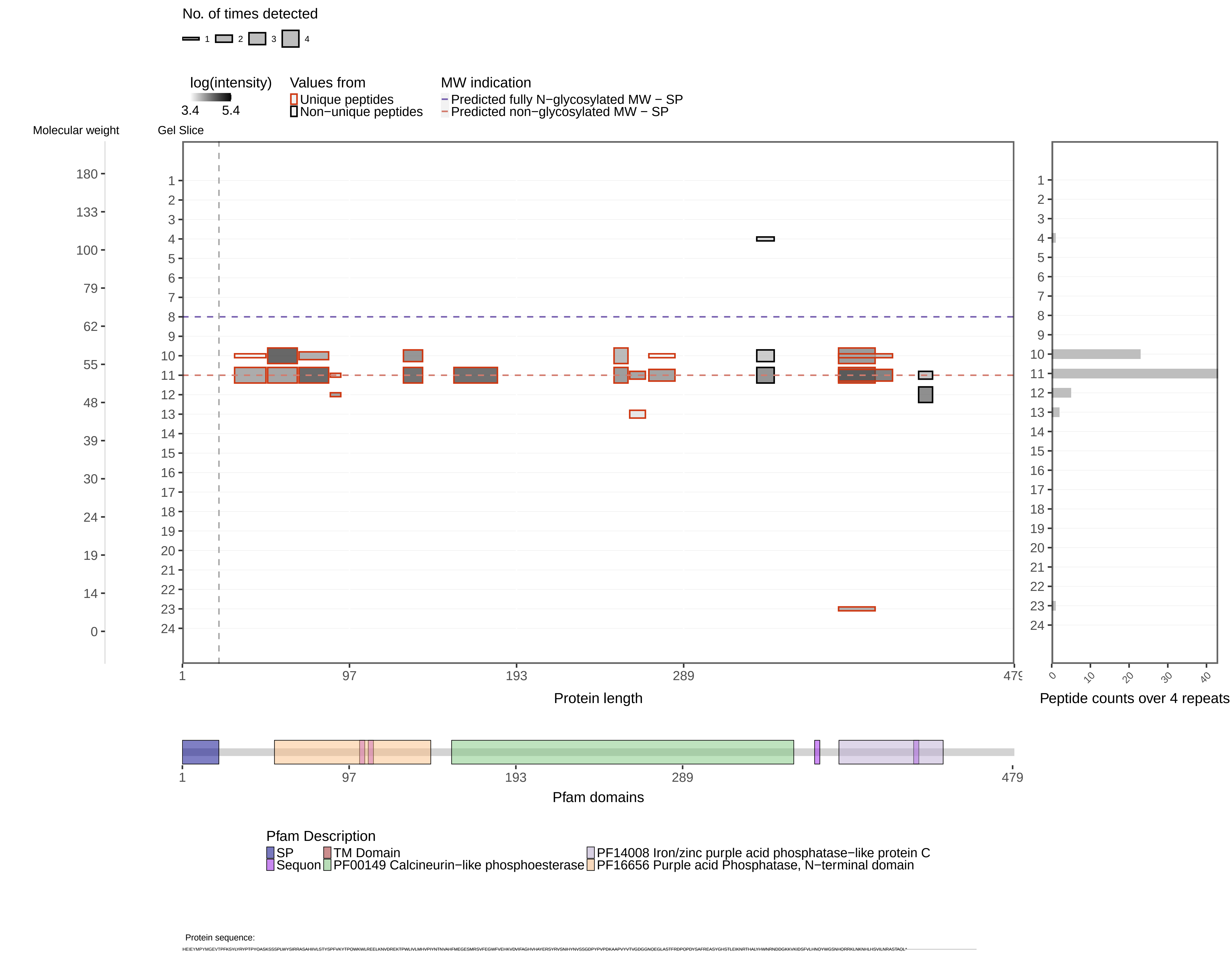

### NbL16g00610.1 pyrophosphate−energized_vacuolar_membrane_proton_pump [MW = 80.02 kDa]

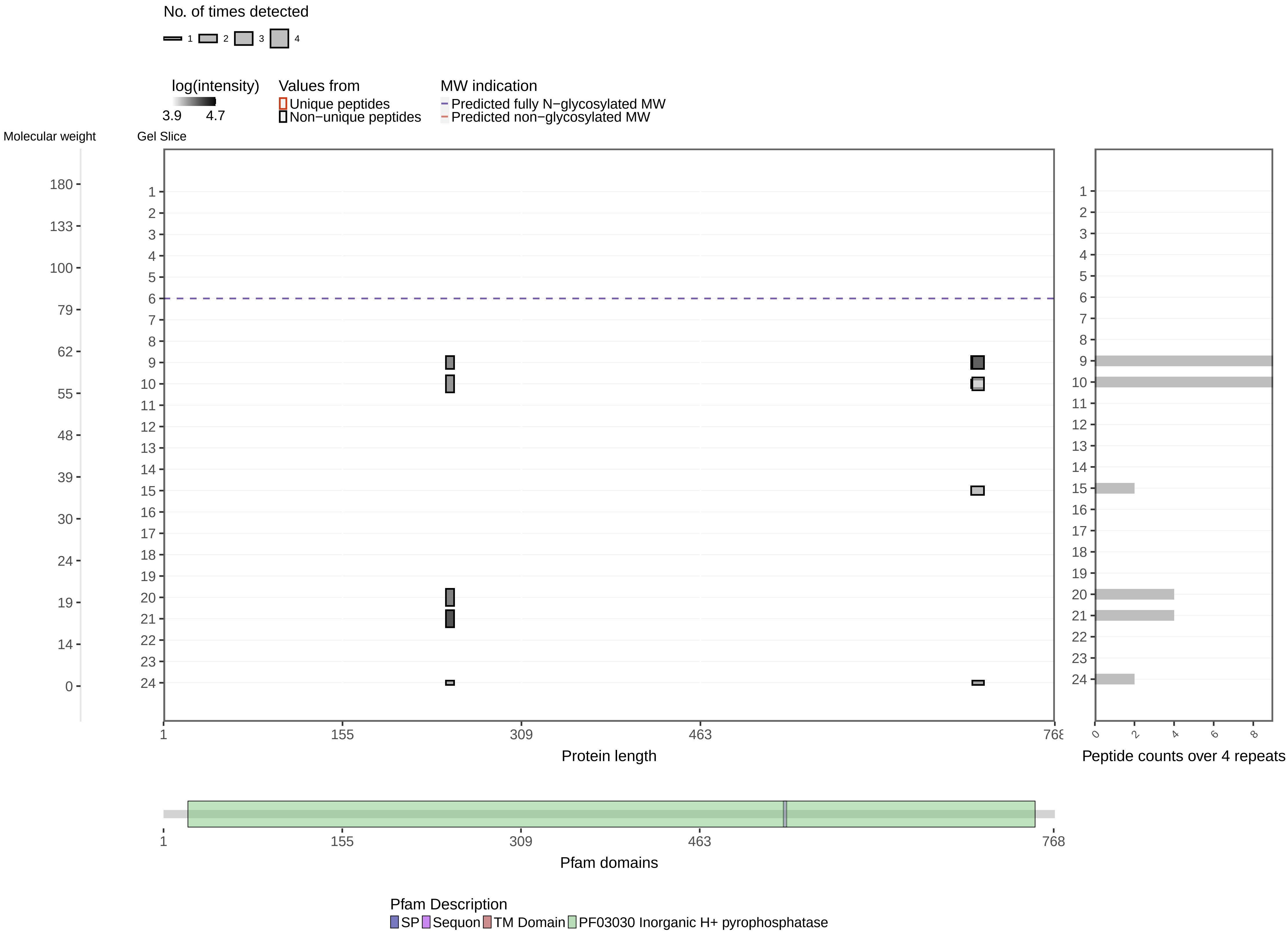


Protein sequence:

[illegible]

### NbL16g00550.1 eukaryotic_translation_initiation_factor−like [MW = 86.03 kDa]

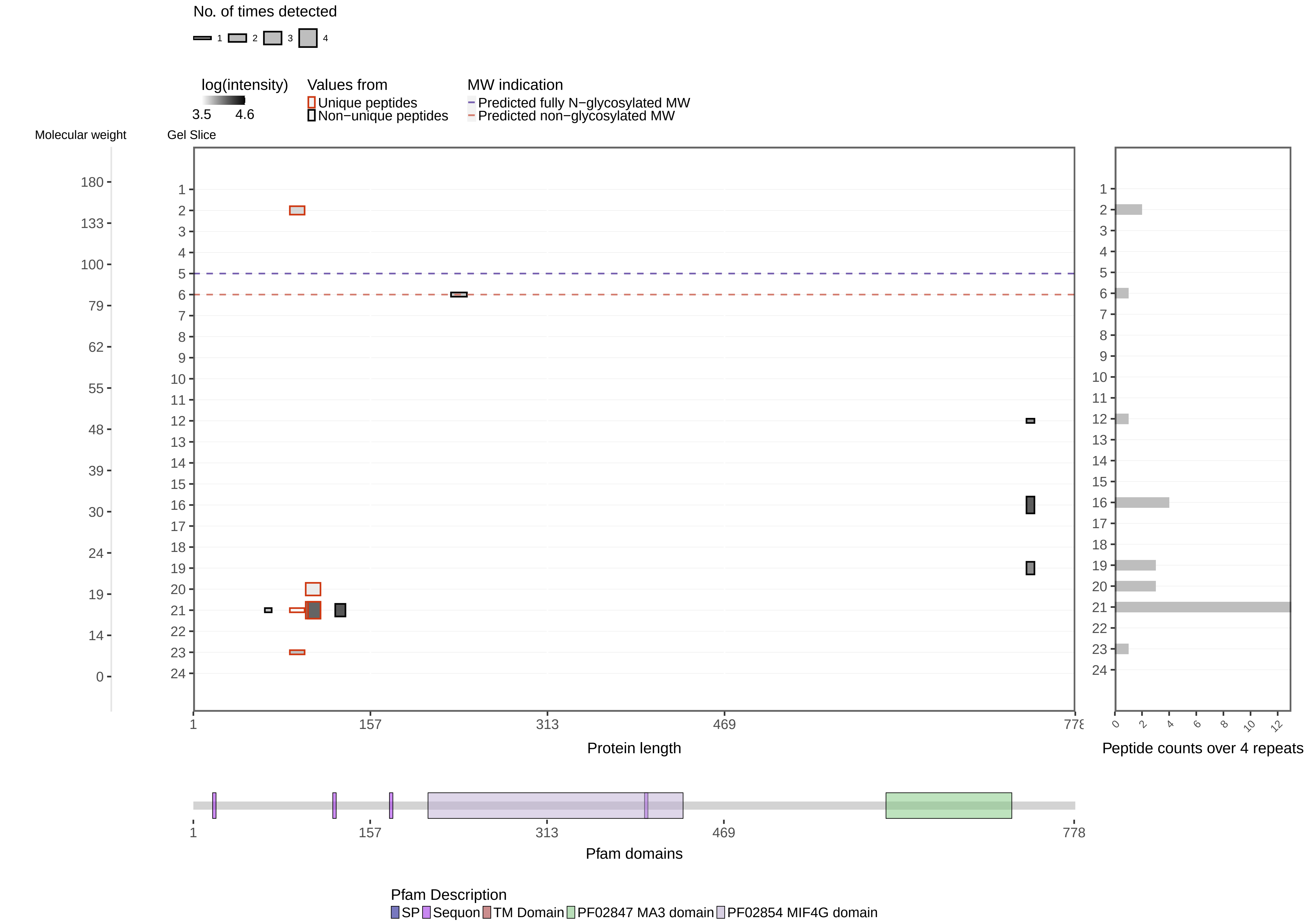


Protein sequence:

VEAICQFFNTIGKQLDEKQISRHINDTYFNRLKELSTNPQLASRLRFMVRDVLDLRANNWVPRREEVKAKTINEIHWEAEKSMGLRPGATASIRNARAIPSGAQGNLAAVGFRPGTGGMMPGMPGVRKMPGMPGMDNDNWEVQRSRSMPRGVQPPLIGKSPSLNTRQLPQGSGGIISGRSSALLQGSGGPPIRPSGYASAMEPVPQVPGPVRPSPATTVPSVAEKPRAPAARSNPEELKRKTVSLLEEYFNVRLLDEALQCIEELKSPDFYPEVVKETISLGLDKSPPRV

### NbL16g00540.1 shikimate_O−hydroxycinnamoyltransferase−like [MW = 49.28 kDa]

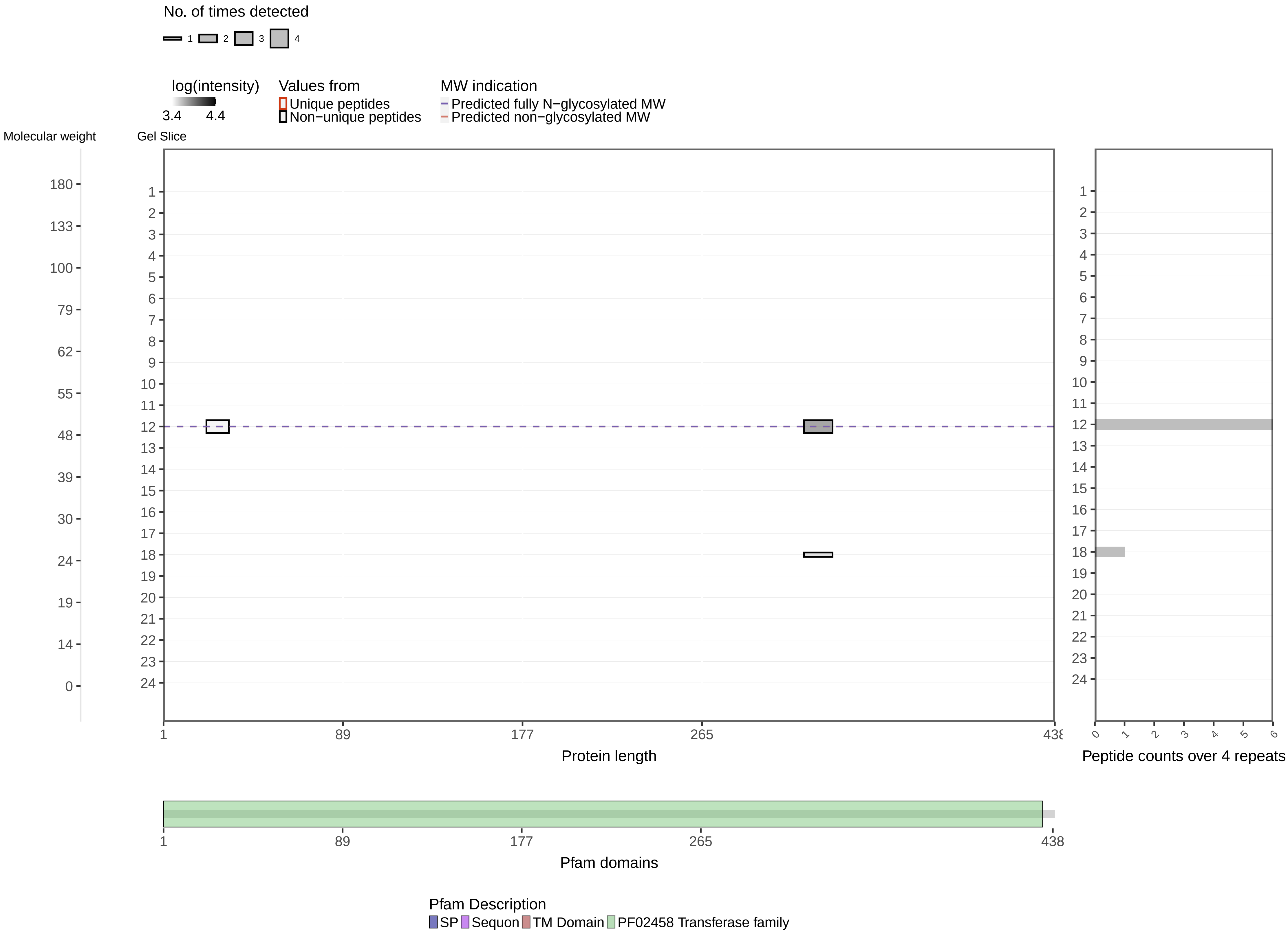


Protein sequence:

YHPPPTLKSSSEHVHGQVNGPKFSTTAMFKITSDQLTLLKTKSKHEGSRYEVLAAHIWRCTCKARALDDDQLTKLHVATDGRSRLFPPLPPGYLGNVVFTATPIAKSGEILSEPLMNTAQRIHTALARMDDEYLRSAIDYLELQTDLSKLIRGPTYFASPNLNINSWTRLPVHDSDFGWGKPVHMGPACILYEGTVYILPSPSDDKNLRLAVCLDADHMPLFEKYLYDF*

NbL16g00490.1 leucine_aminopeptidase_2_chloroplastic [MW = 60.87 kDa]

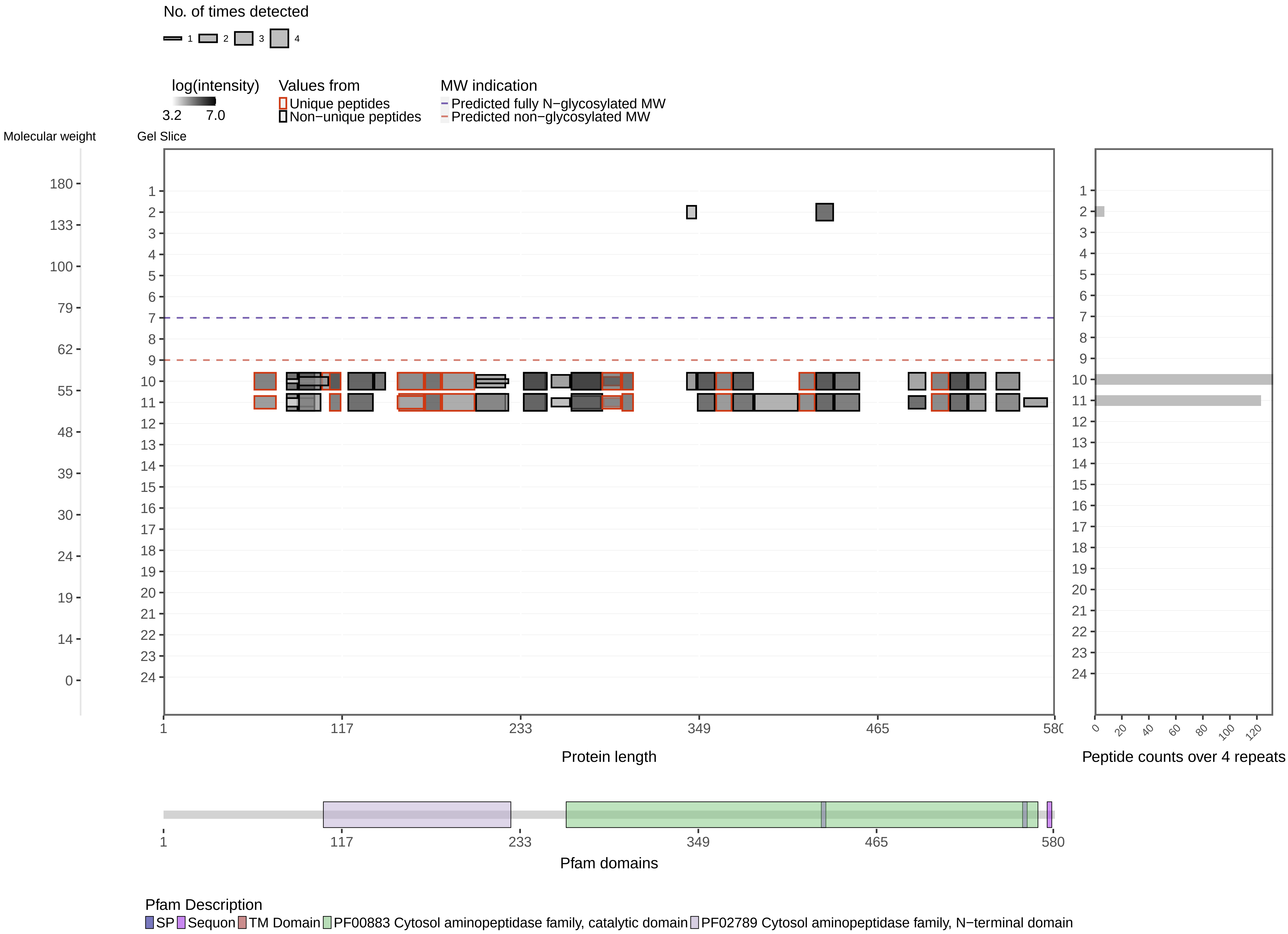


Protein sequence:

LAEEAKKIASTYSDVLSAKILNVEQCKELKMGSYLAVGEASANPPHFIHLCYKPTGGEIKKKLALVGKGLTFDSGGYNIKTGPGCLIELMKFDMGGSAAVLGAAKALGQIKPAGVEVHFIVAACENMISGTGMRPGDIVTASNGKTIEVNNTDAEGRLTLADALVYACNQGVEKIVDLATLTGACIVALGPSIAGVFTPSDDLAKEVAAASEVSGEKLWRMPLEESYWETMKSGVADMVNTGGRQGGAIVAALFLKQFVDEKVQWMHIDLAGPVWNDKKKNATGFGVSTLVEWV

### NbL16g00460.1 eukaryotic_translation_initiation_factor_5A−1/2 [MW = 17.48 kDa]

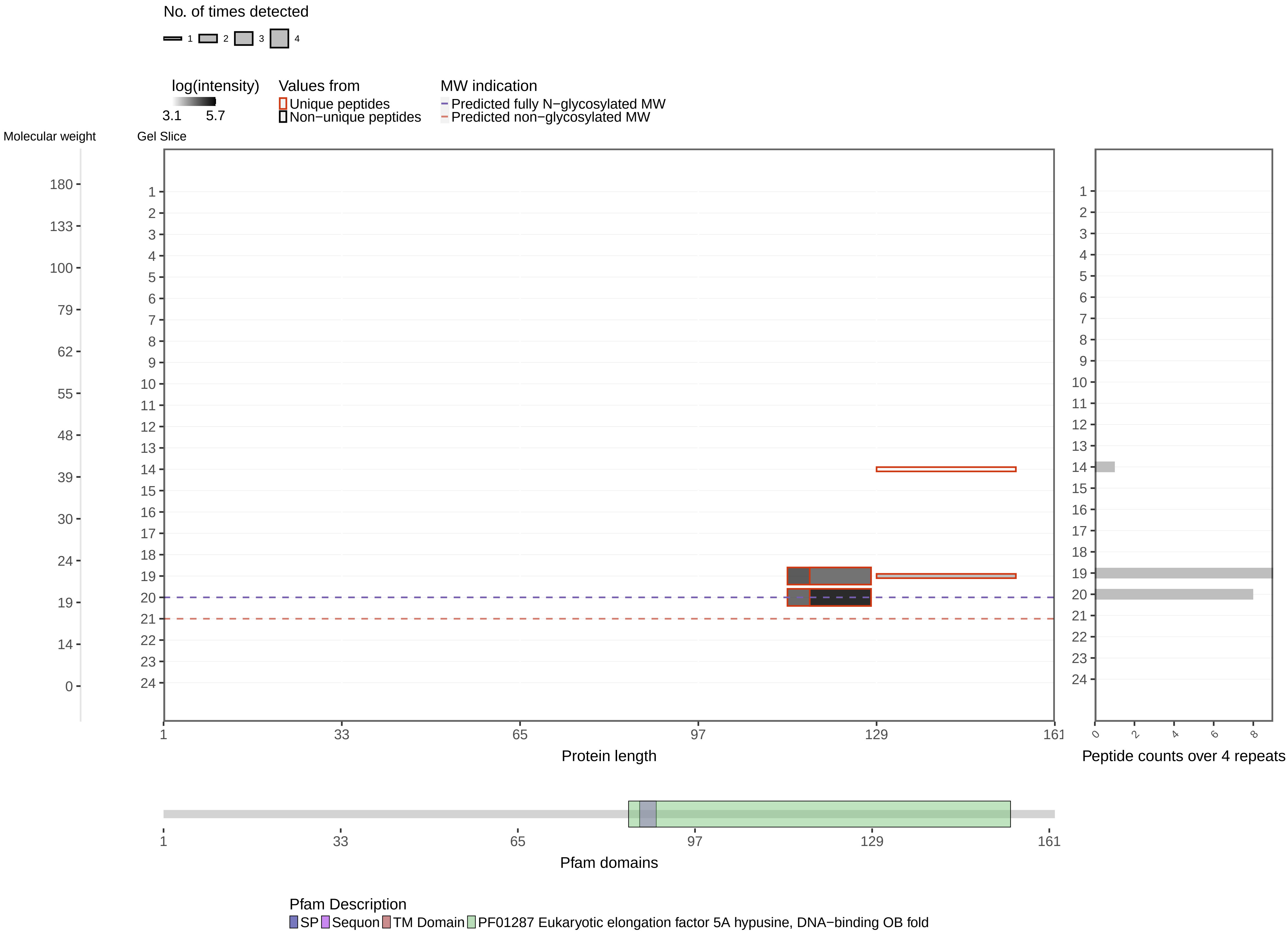


Protein sequence:

IGKKLEDIVPSSHNCDVPHVNRTDYQLIDISEDGFVSLLTENGNTKDDLRLPTDEALLSQVKGGFEEGKDLVLSVMSAMGEEQINAVKDIGTKN*

### NbL16g00370.1 anthranilate_synthase_beta_subunit_1_chloroplastic−like [MW = 31.26 kDa]

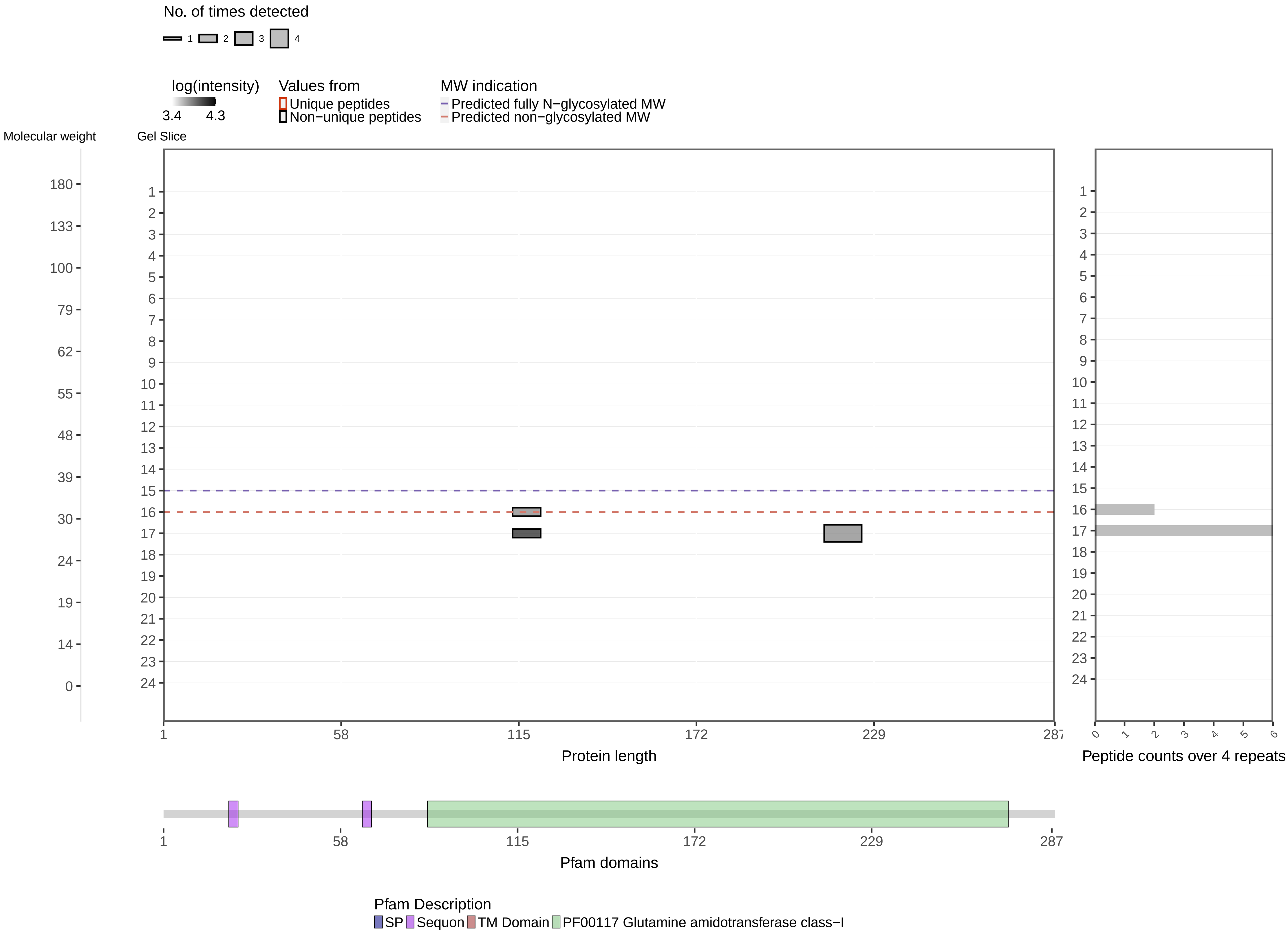


Protein sequence:

PGPGAPQDSGISLQTVLELGPTVPLFGVCMGLQCIGEAFGGKIVRAPYGVMHGKSSPVYYNEGGEDGLFAGLSNPFTAGRYHSLVIDKDTFPKGALEITAWTEDGLIMAARHKVYRHLQGVQFHPESIITTEGMTIVRNFVKMIERKEEAEAQD*

NbL16g00260.1 temperature−induced_lipocalin−1−like [MW = 21.48 kDa]

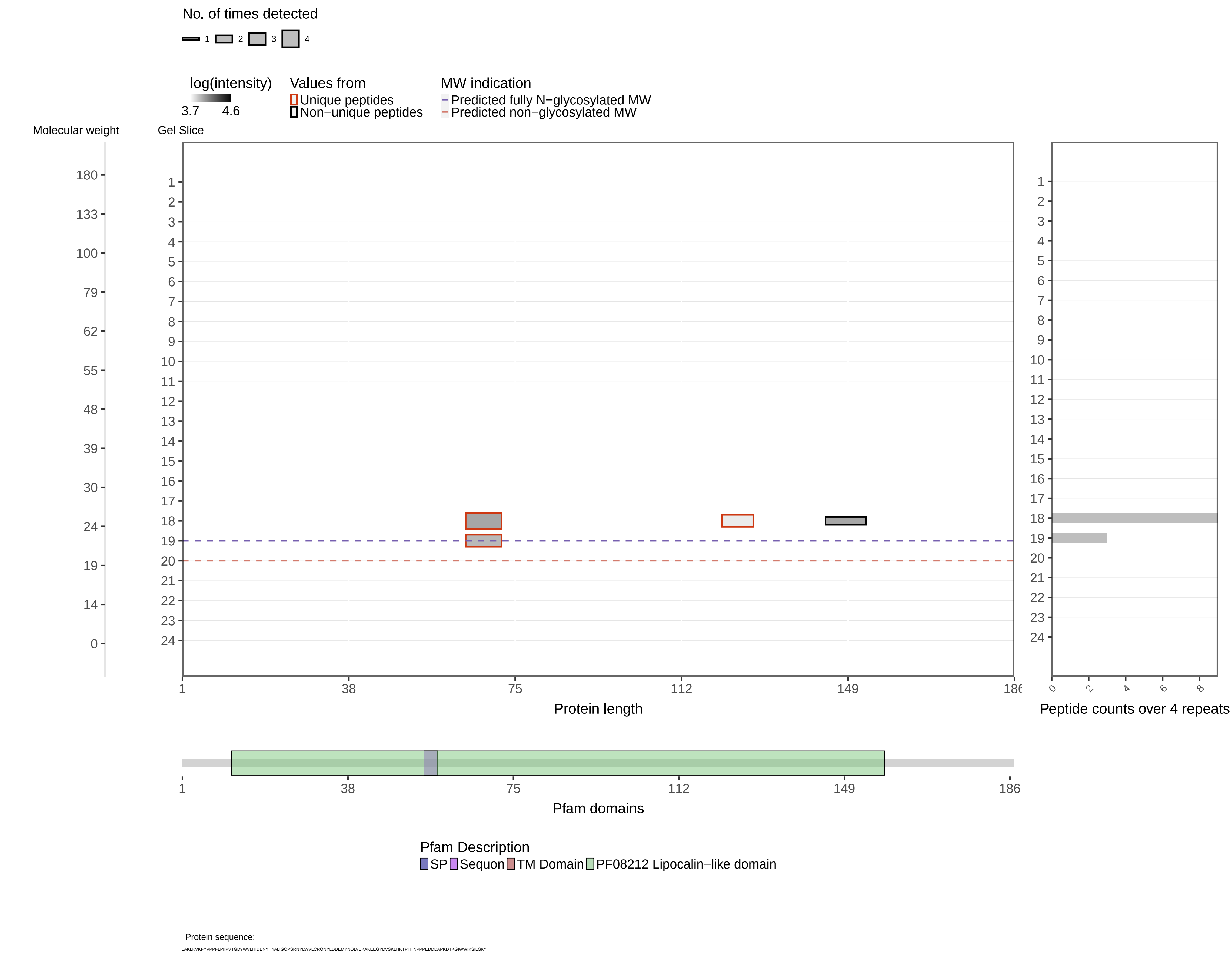

### NbL16g00210.1 adenylate_kinase_5_chloroplastic−like [MW = 70.68 kDa]

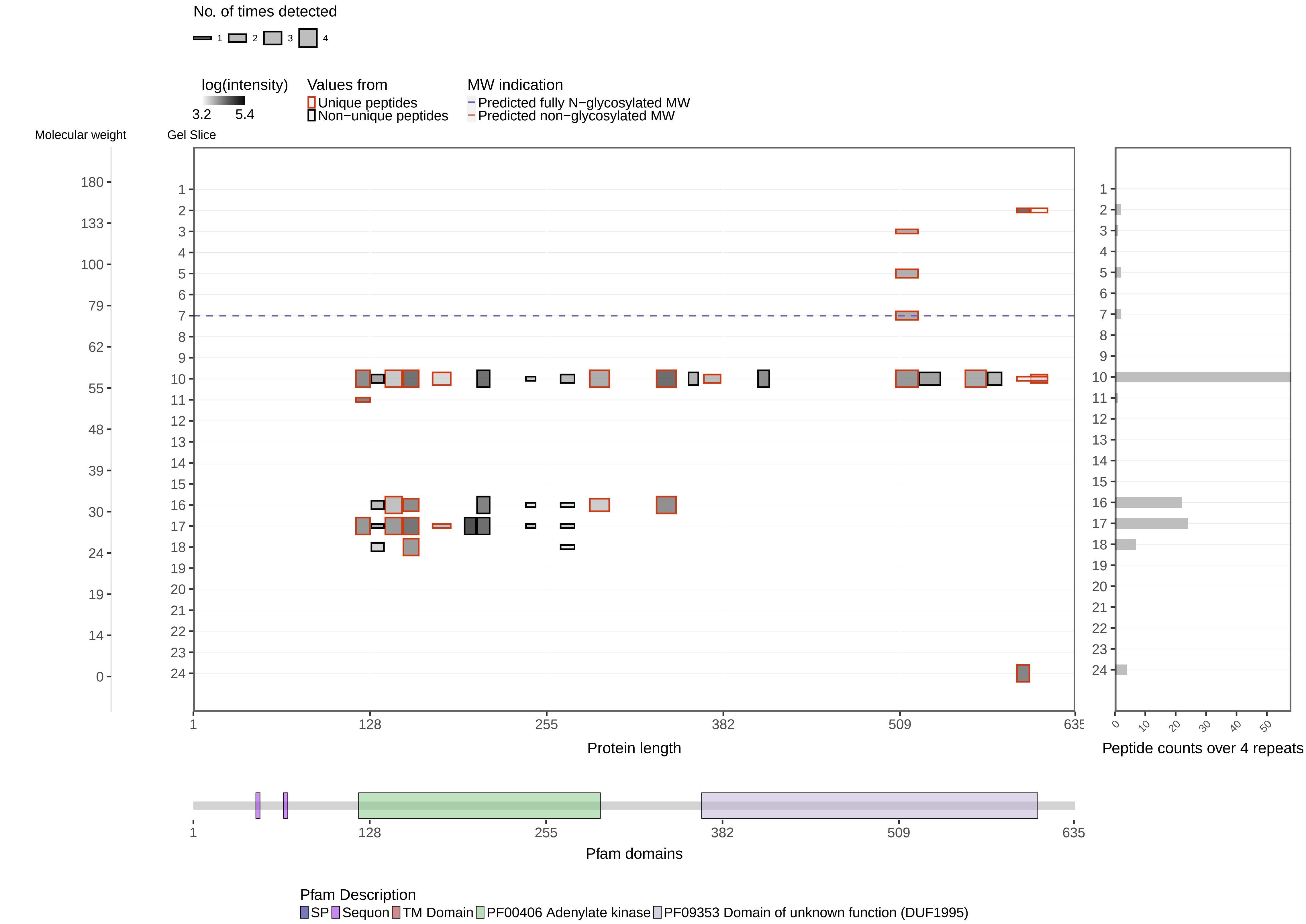


Protein sequence:

NbL15g25260.1 zinc_finger_CCCH_domain−containing_protein_11−like [MW = 42.98 kDa]

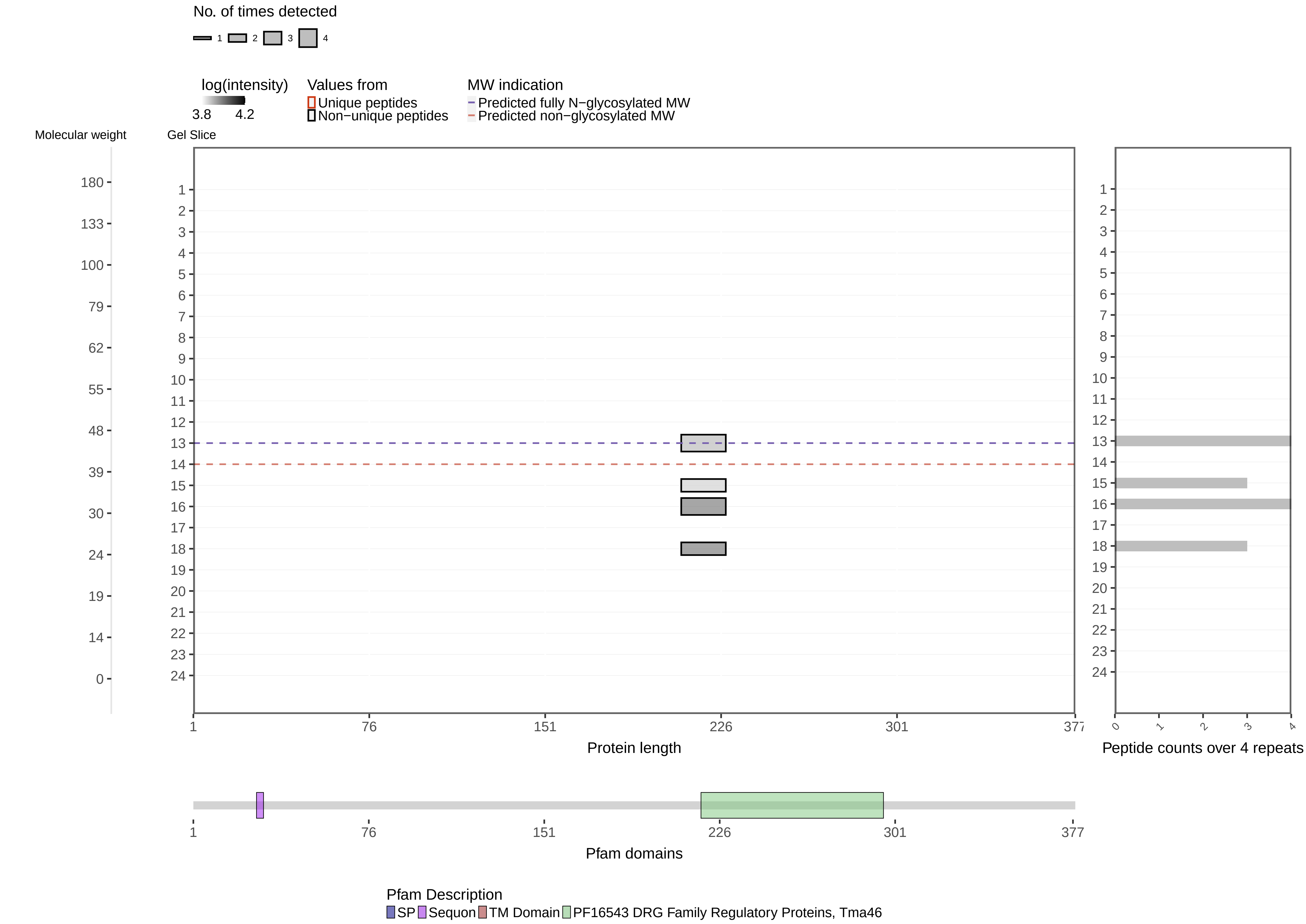


Protein sequence:

YGWFWVCPNGGKECHYRHALPPGYILKSQMKALLQEEADKMPIEEEIDEQRAKLTASTPLTTELFMEWKKKKMEEREASLAKQRADRAKNDRMSGRELFMSDASWFVDDVGAYDKYERDEESGEPPKDTARDEASSSMPARNIDDGGEASHNNEVNDNYDDDDDDDDDDDDDLDVDELNELEATLSKTSLQINEPGGHA*

NbL15g25240.1 60S_ribosomal_protein_L4−like [MW = 44.83 kDa]

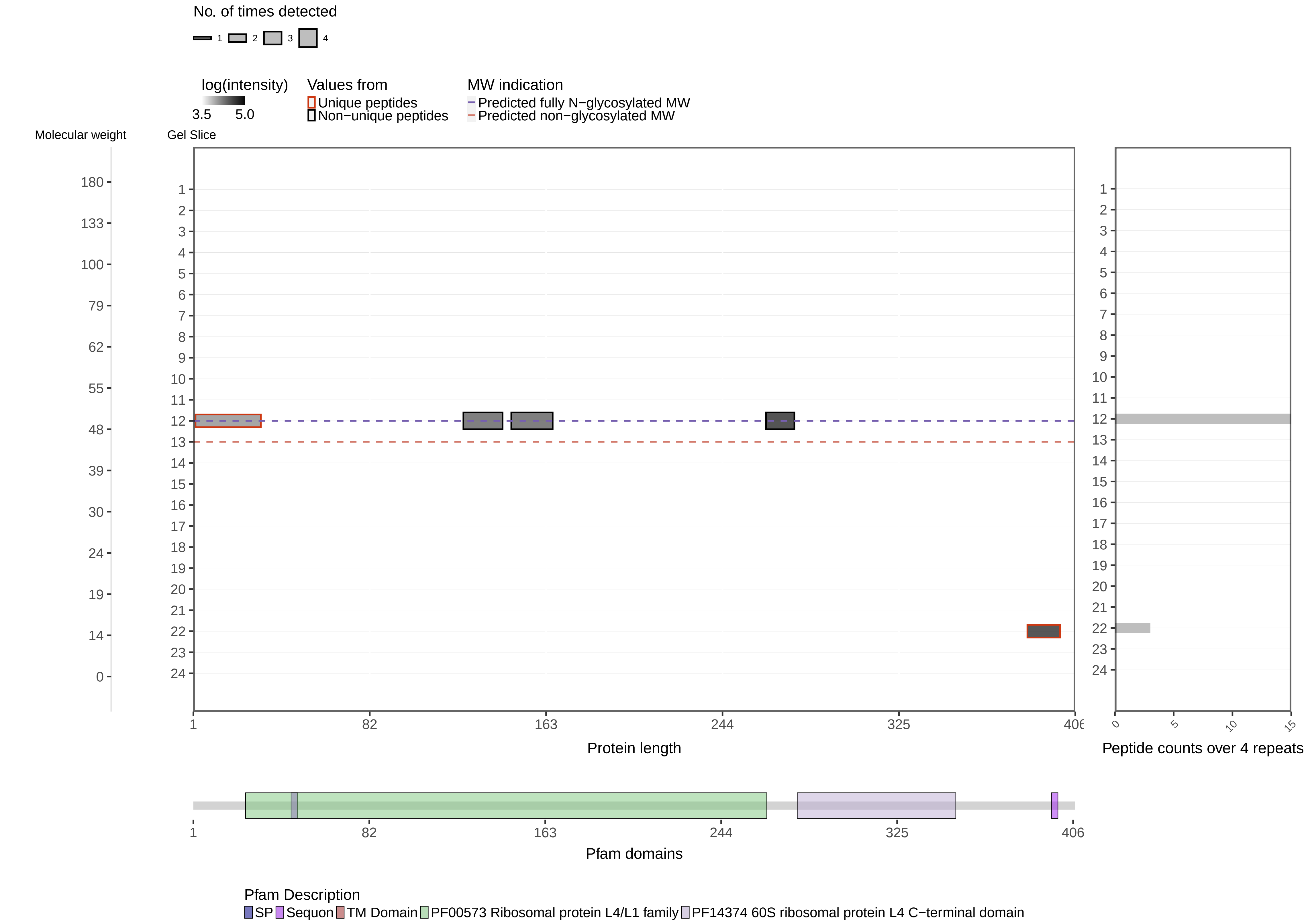


Protein sequence:

HAIRPGKGKMRNRRYFCRKGPLIVYGTEGAKLVKAFRNIPGVEICHVDRLNLLKLAPGGHLGRFVIWTKSAYEKLDSIYGSFDKPSEKKKGYLLPRPKMVNADLARIINSDEVQSVVRPIKKDVNKRATLKKNPLKNLNVLLKLNPYAKTARRMSLLAEAQRVKAKKEKLDKKRHQITKEEASAIRSASHSWYKTMISDSDYAEFDNFTKWLGVSQ*

NbL15g25210.1 probable_LRR_receptor−like_serine/threonine−protein_kinase_At1g67720 [MW − SP = 100.36 kDa]

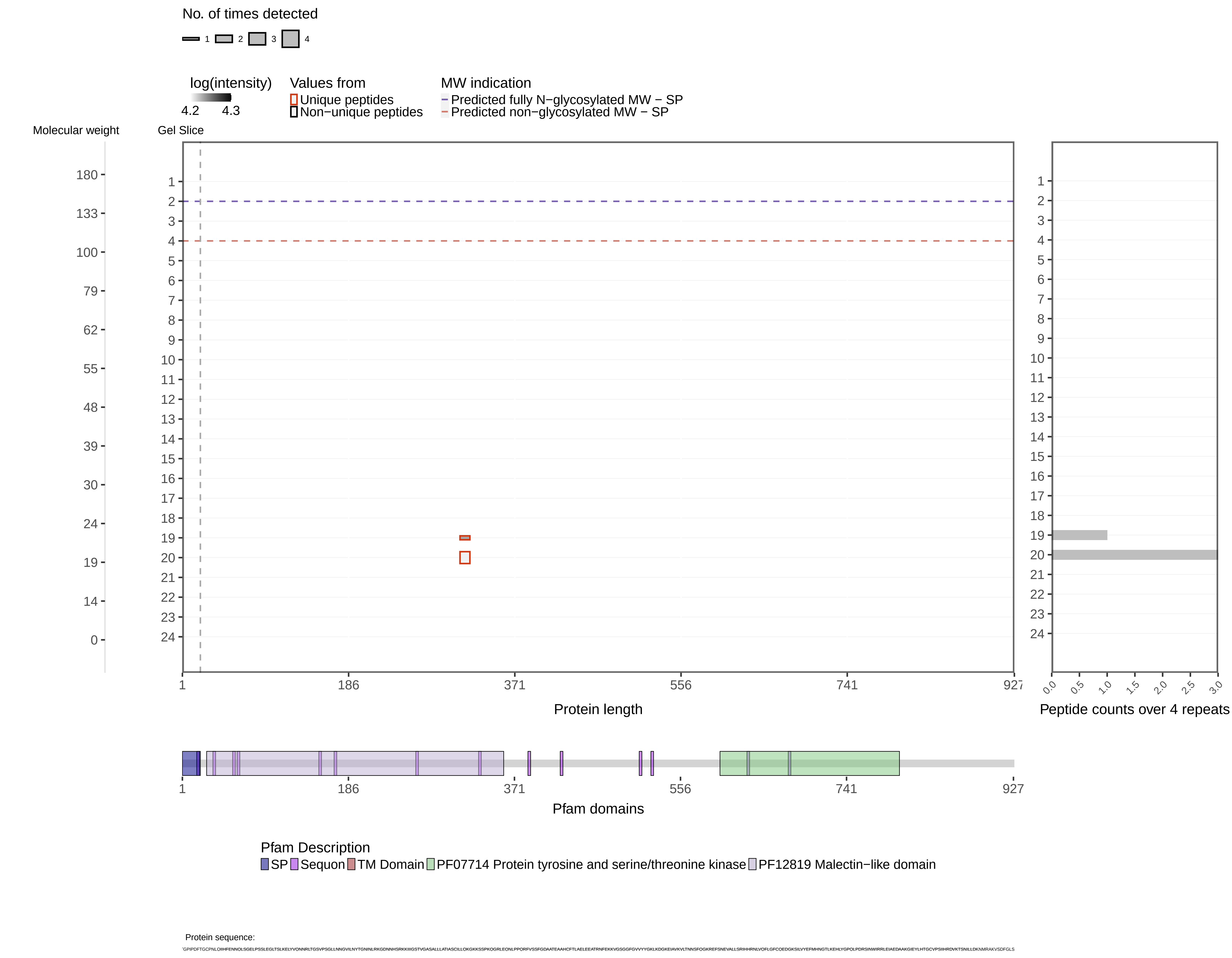

### NbL15g25120.1 40S_ribosomal_protein_S23 [MW = 15.73 kDa]

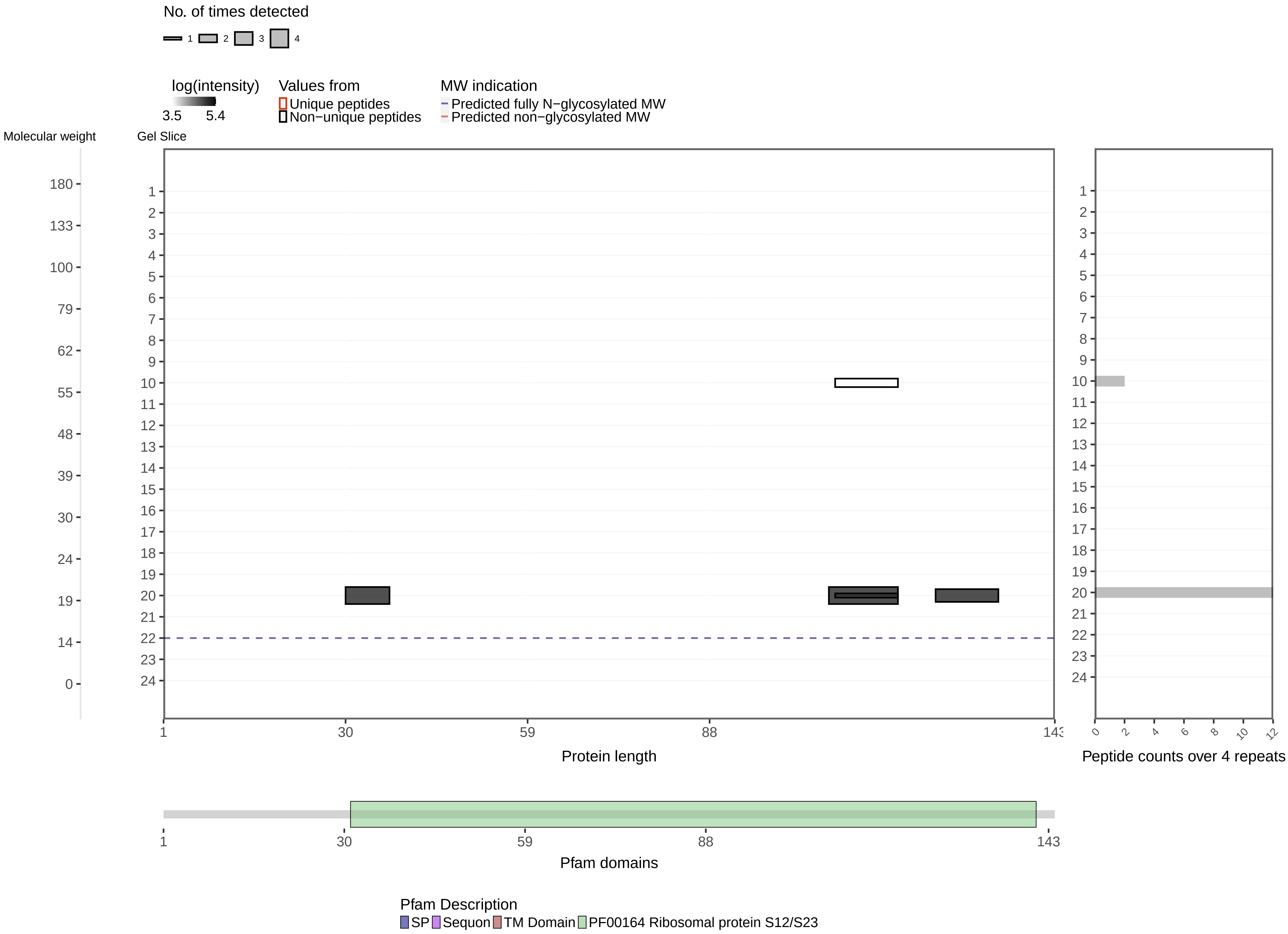


Protein sequence:

EAKQPNSAIRKCARVQLIKNGKKIAAFVPNDGCLNYIEENDEVLIAGFGRKGHAVGDIPGVRFKVVKVSGVSLLALFKEKKEKPRS*

### NbL15g24940.1 peroxidase_21 [MW − SP = 33.75 kDa]

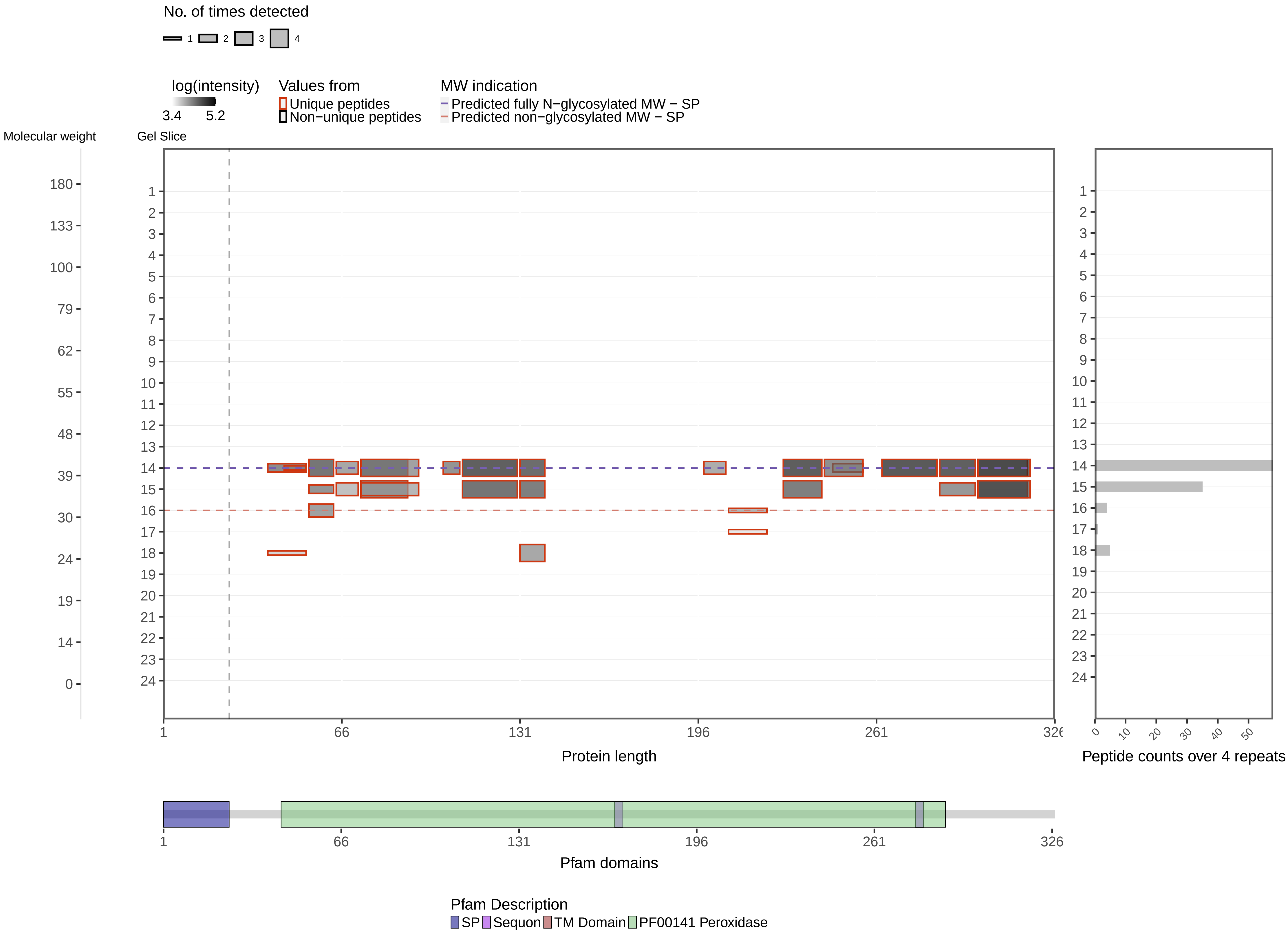


Protein sequence:

SKESYLAEVENSIPNHNDSMSSVLSRFOSIGVDTEGTVALFGAHSVGRVHCVNLVHRLYPTVDPTLDPDYAKYLKGRCPTPNPDPEAVLYSRNDRETTMILDNMYYKNILNHKGLLIVDOELVSDPNTSPFVEKMAADNGYFHDOFARALVILSENNPLTGDOGEIRKVCRYVNSA*

### NbL15g24920.1 valine−−tRNA_ligase_mitochondrial_1−like [MW = 123.32 kDa]

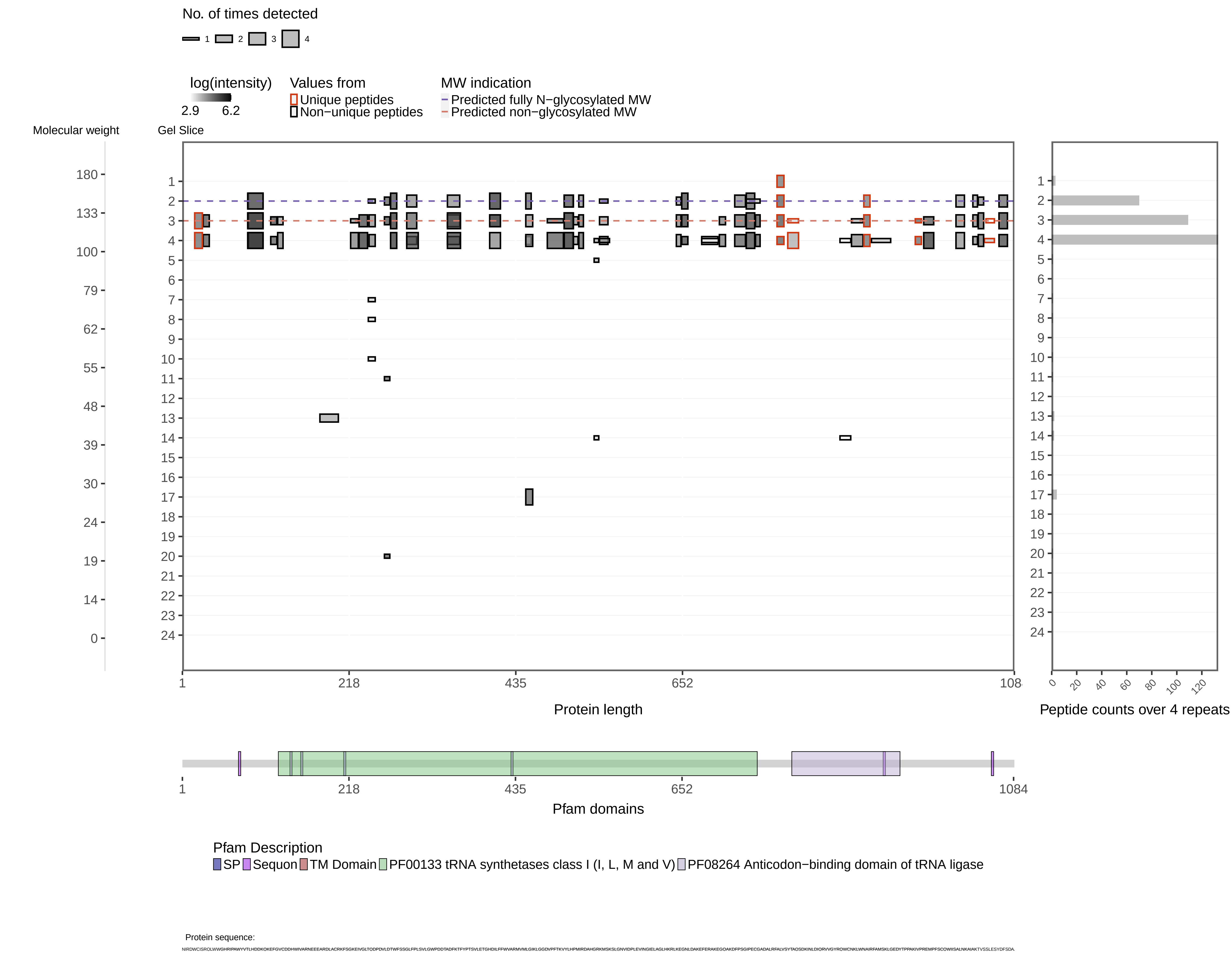

### NbL15g24890.1 dnaJ_protein_P58IPK_homolog [MW = 53.82 kDa]

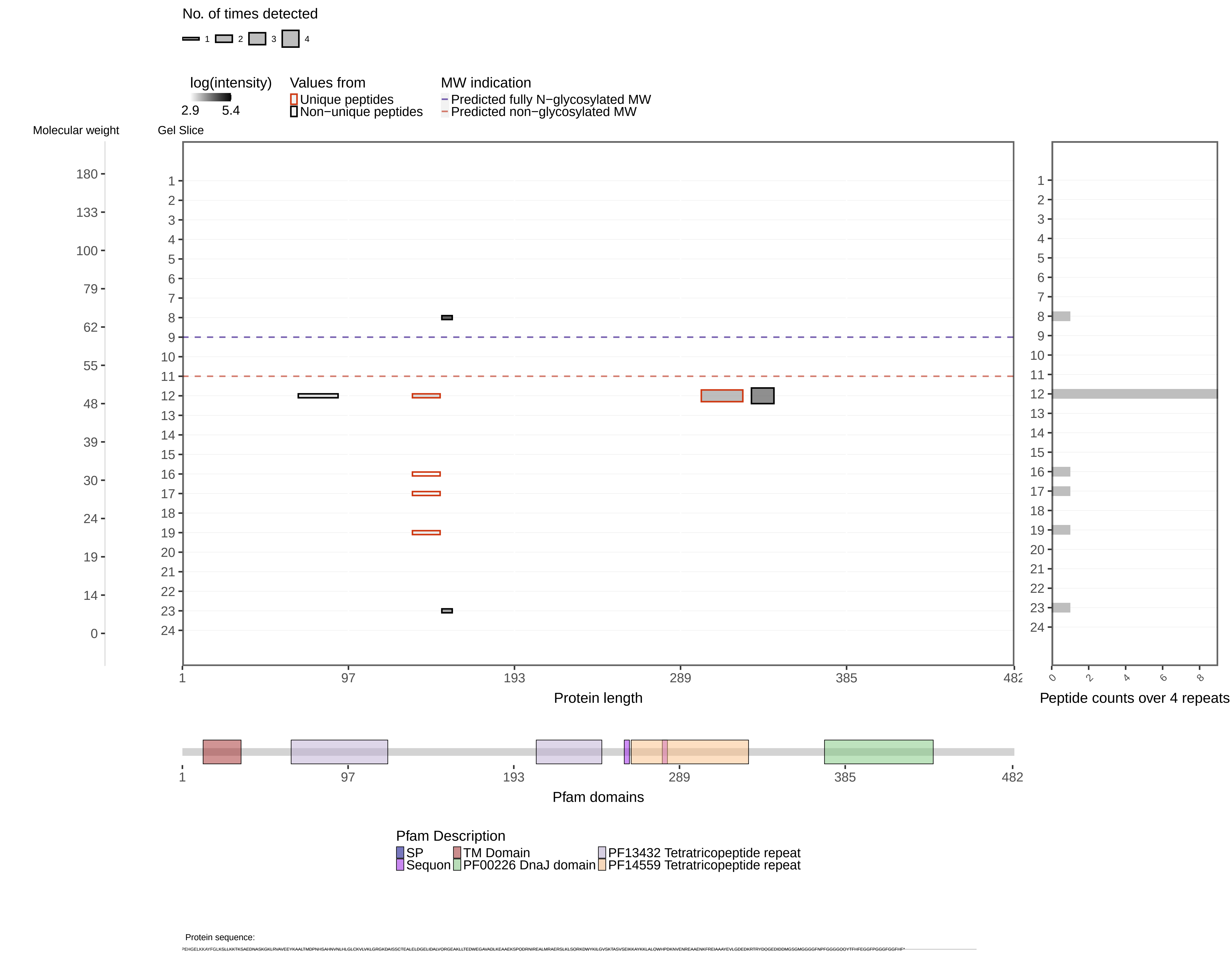

NbL15g24880.1 serine_carboxypeptidase−like_50 [MW − SP = 47.61 kDa]

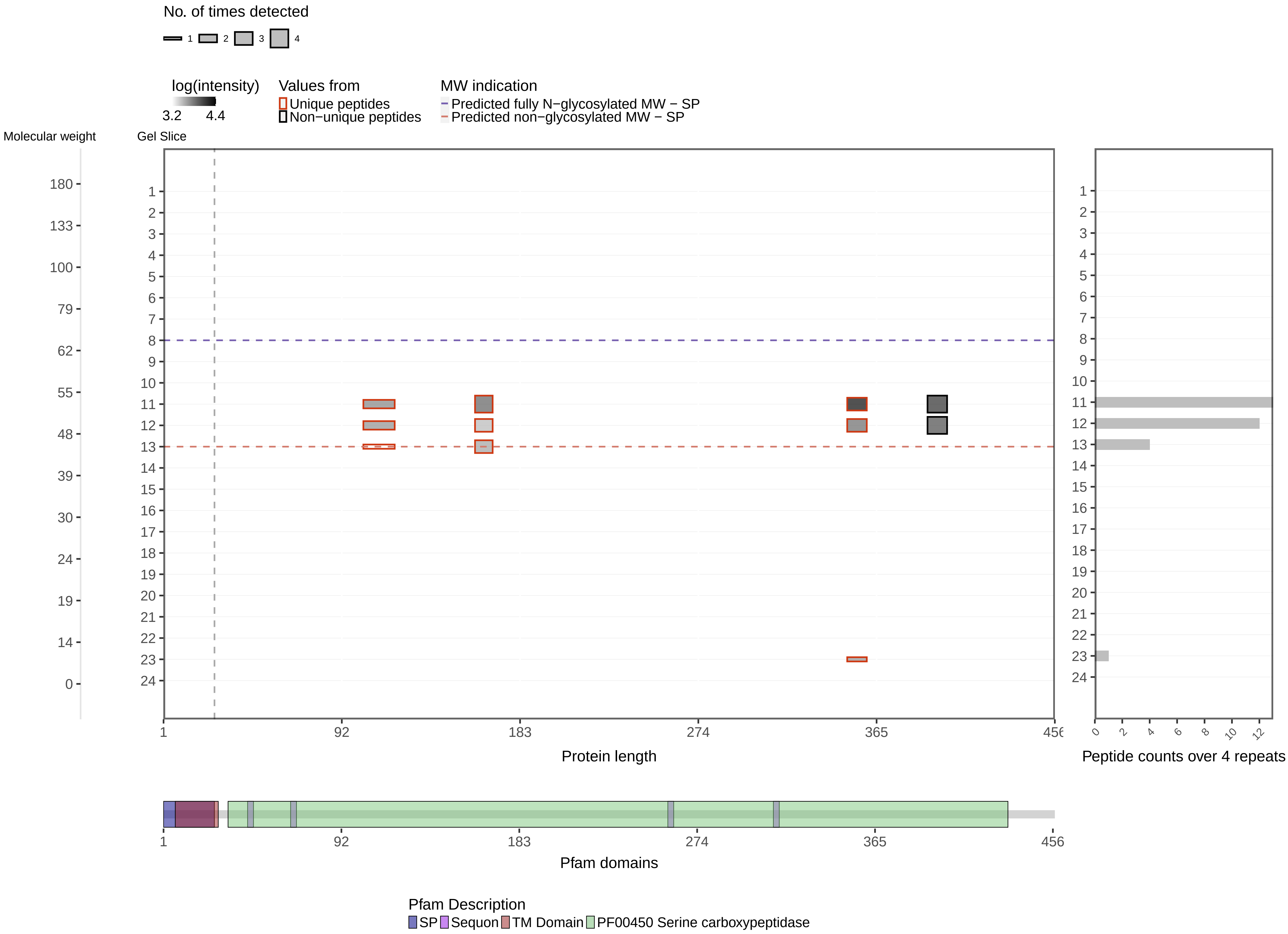


Protein sequence:

PETQVGTHALSAYYSGLINEKQKTQMESLQKEAIQLAKNGNWSEATSARSKVLGTLTNMAGLATLYDFRRLKPYEDELVAKFLSNVEIKRALKANESIVFEVCSDLVGKALHEDVMKSSRYMVEFLVKNTKVLLYQGQCDLRDGVVSTEAWMKKMKWDGIEKFLEAERKVWRVNDGLAGYVQKWESLSHVVVMNAGHLVPTDQAVNSQVMIEDWVLEKGLFATGQIKQKPSNIGKSL*

### NbL15g24820.1 PIP2−1−like [MW = 31.22 kDa]

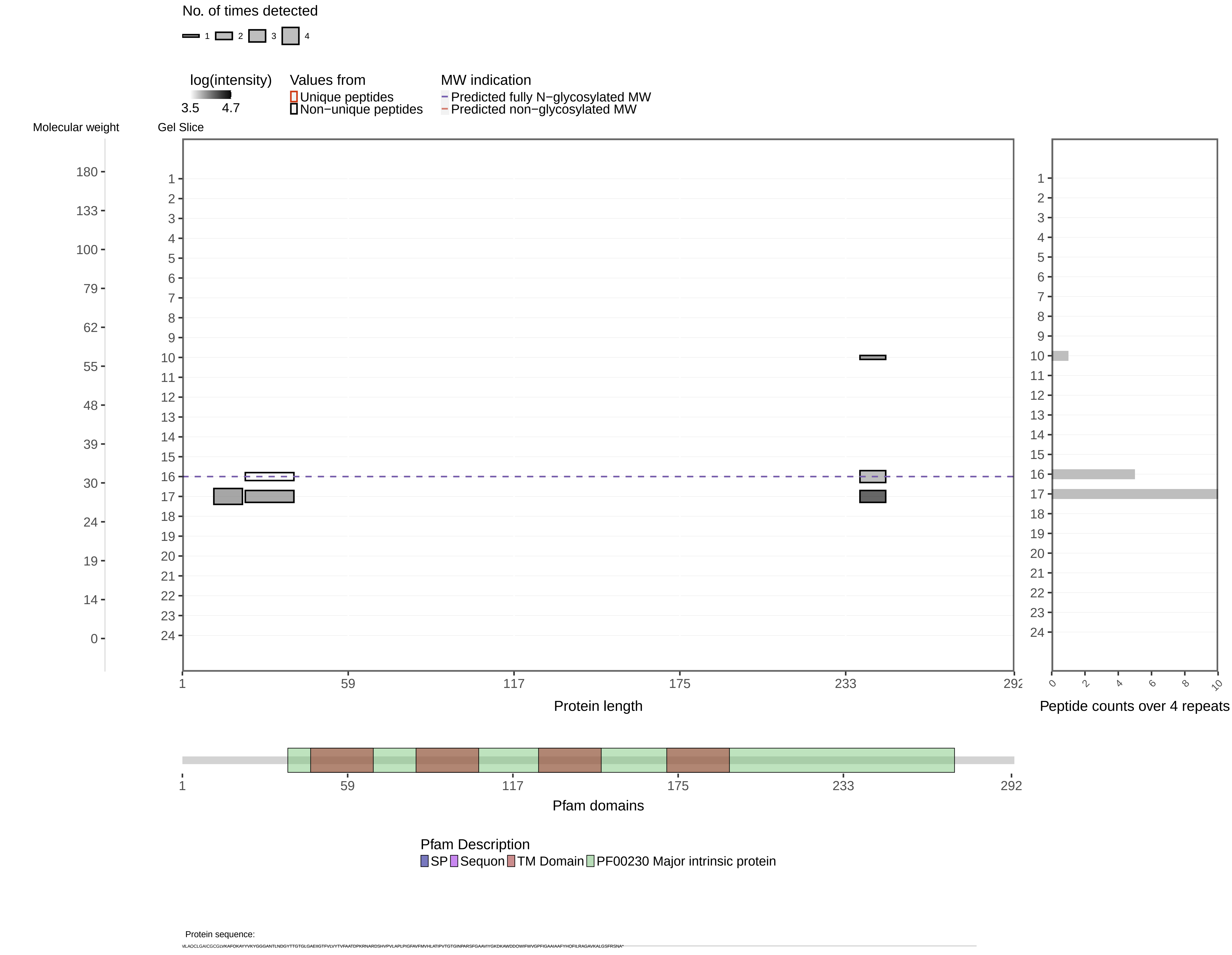

NbL15g24670.1 29_kDa_ribonucleoprotein_A_chloroplastic [MW = 35.71 kDa]

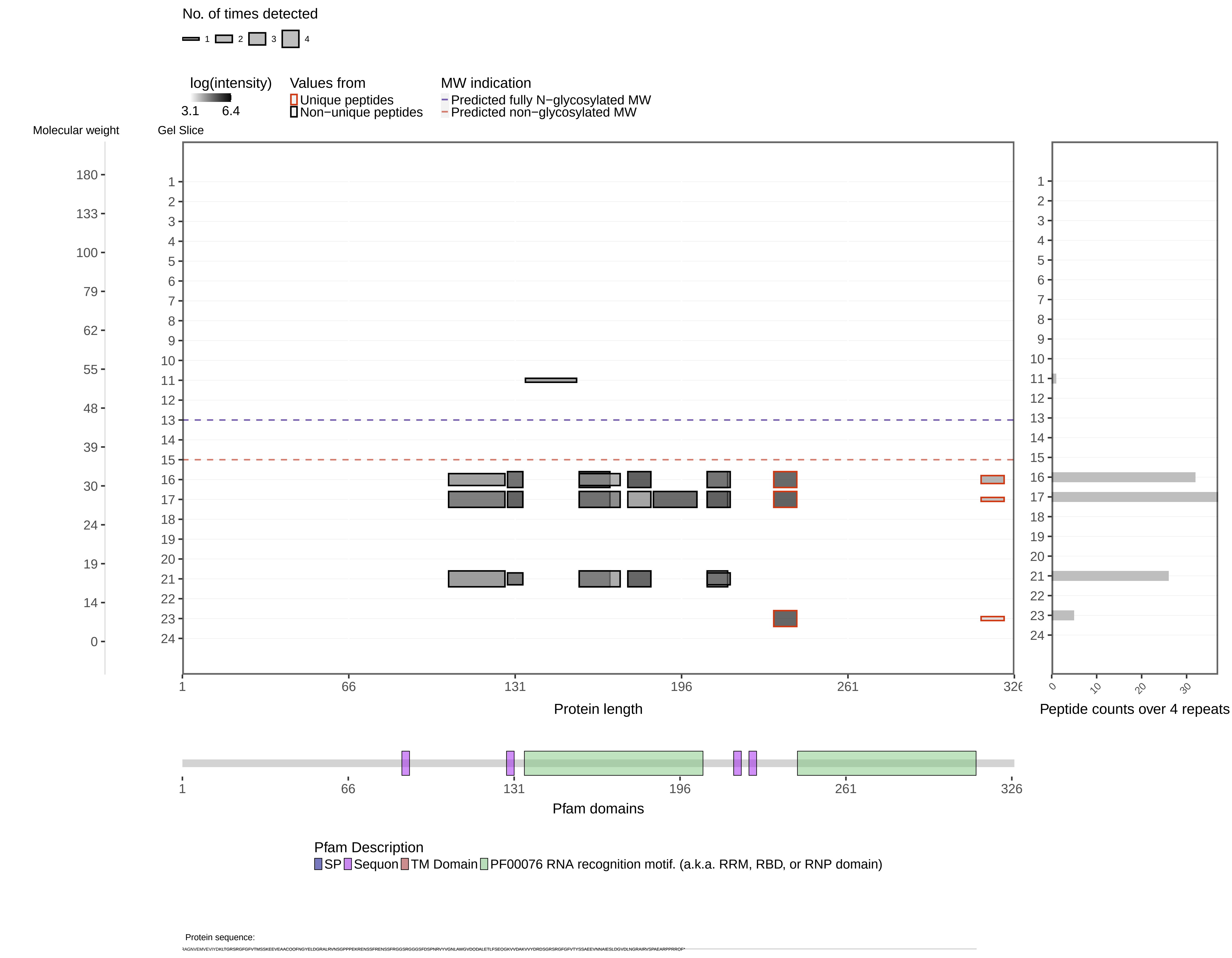

### NbL15g24630.1 ammonia−lyase−like [MW = 78.15 kDa]

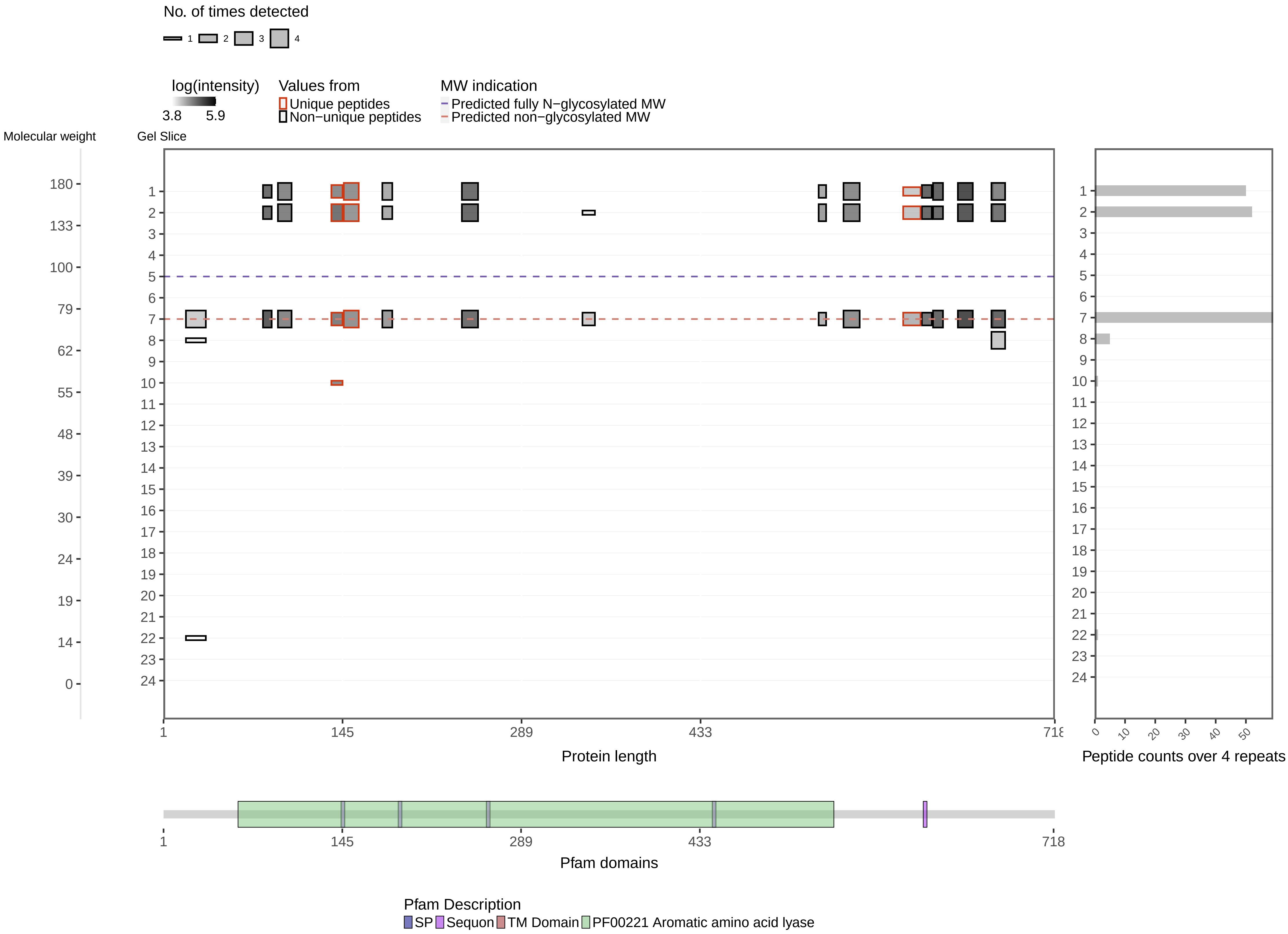


Protein sequence:

KPKQDRYALRTSPQWLGPQIEVIRSATKMIEREINSVNDNPLIDVSRNKALHGGNFQGTPIGVSMDNARLALASIGKLMFAQFSELVNDYYNNGLPSNLTASRNPSLDYGFKGAEIAMASYCSELQFLANPVTNHVQSAEQHNQDVNSLGLISARKTAEAVDILKLMSSTYLVALCQAIDLRHLEENLKNAVKNTVSQVAKRTLTMGANGELHPARFCEKELLRVVDREYLFAYADDPCSCNYPLMQKLRQVLVDHAMNNGESEKNVNSSIFQKIGAFEDELKAVLPKEVESAR

NbL15g24610.1 adenosine_kinase_2−like [MW = 37.48 kDa]

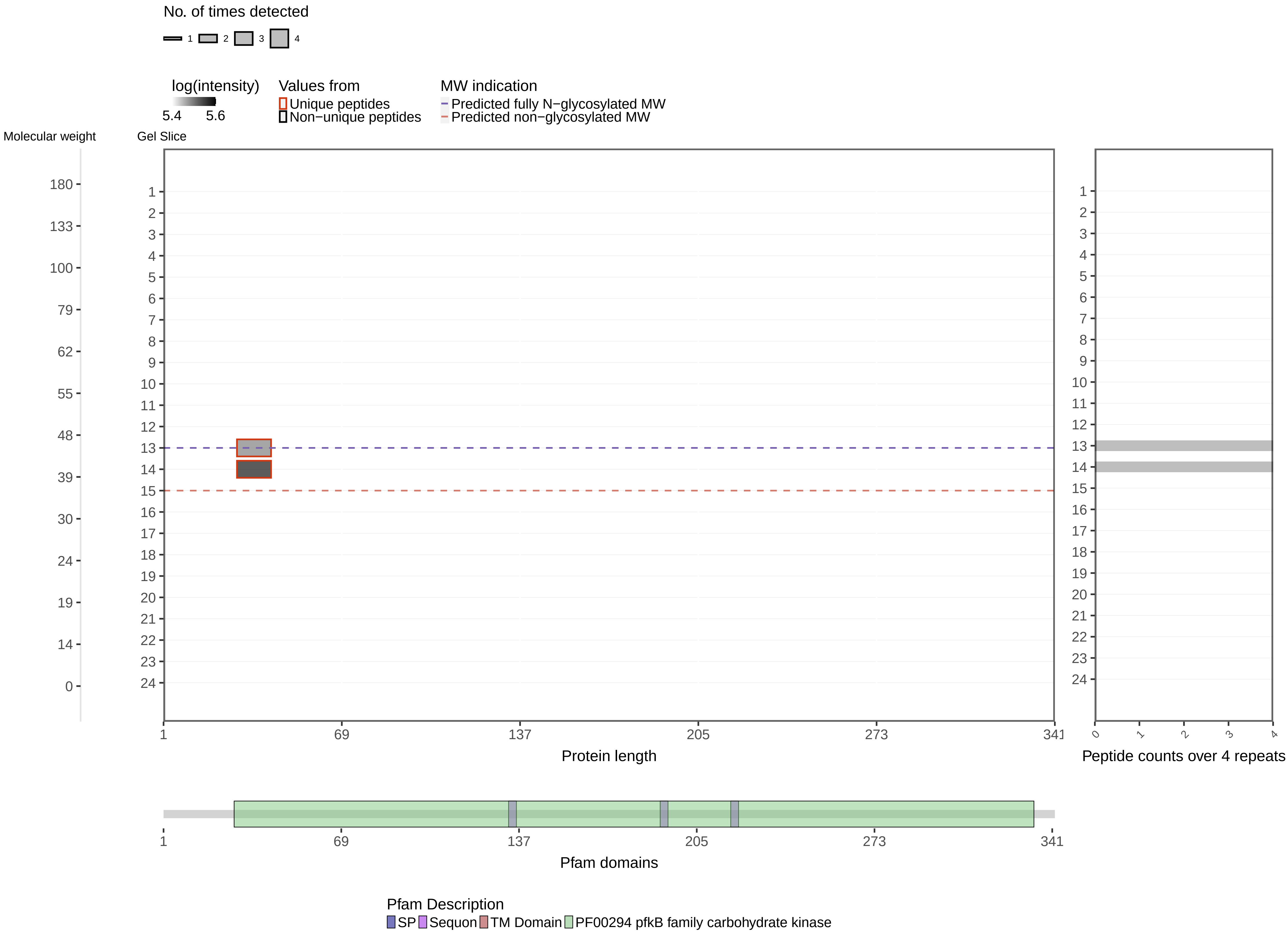


Protein sequence:

KAKFYYIAGFFLTVSPESIQLVAEHAAANNKIFSMNLSAPFICEFFRDPQEKALPYMDFVFGNETEARTFSKVHGWETDNVEEIALKISEWPKASETHKRITVITQGADPVVVAENGKVKLFPVIPLPKEKLVDTNGAGDAFVGGFLSQLVQGKPVEDCVRAGCYASNVIIQRSGCTYPEKPDFA*

### NbL15g24490.1 cell_division_cycle_protein_48_homolog [MW = 91.74 kDa]

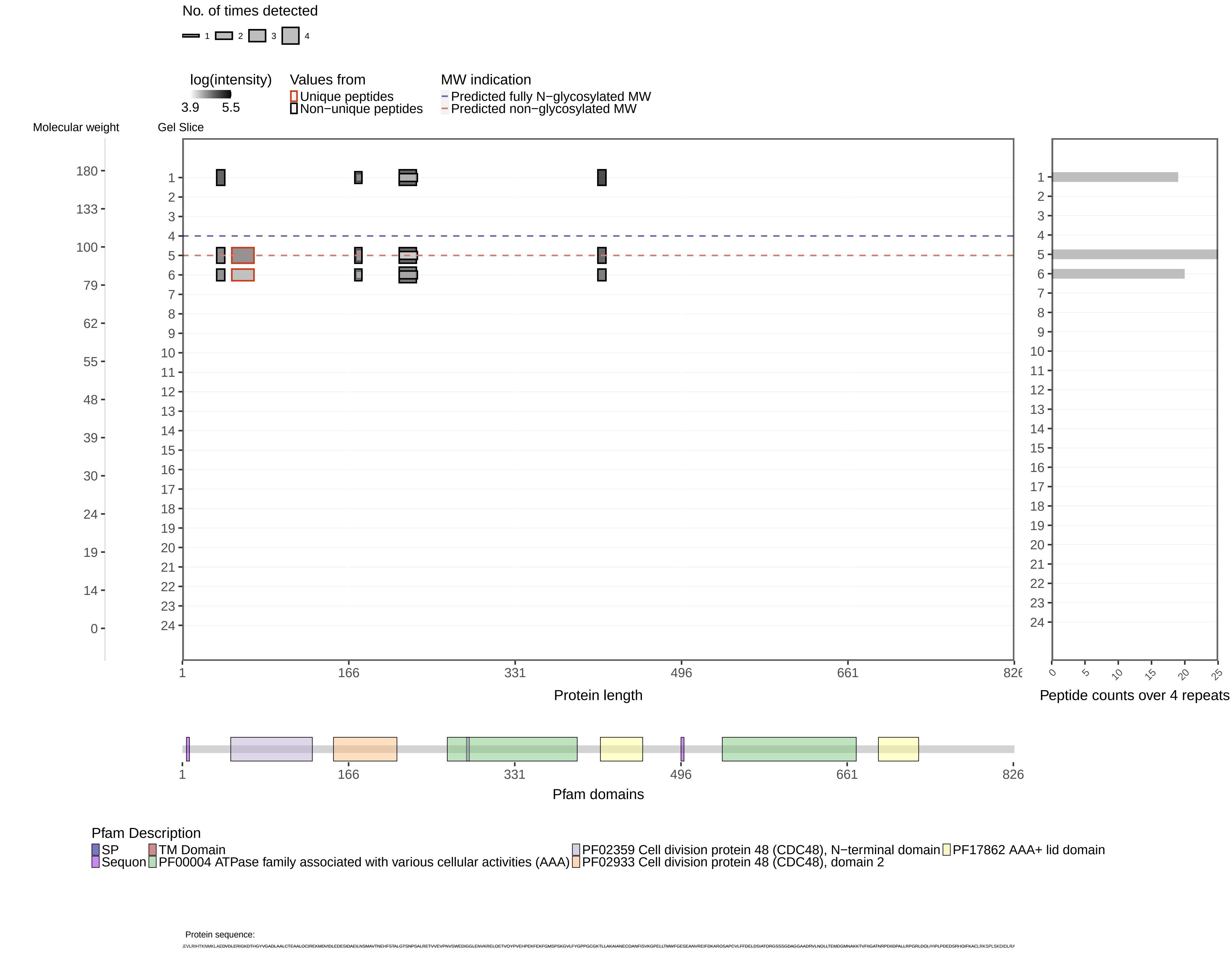

### NbL15g24390.1 uncharacterized_LOC109229140 [MW = 23.47 kDa]

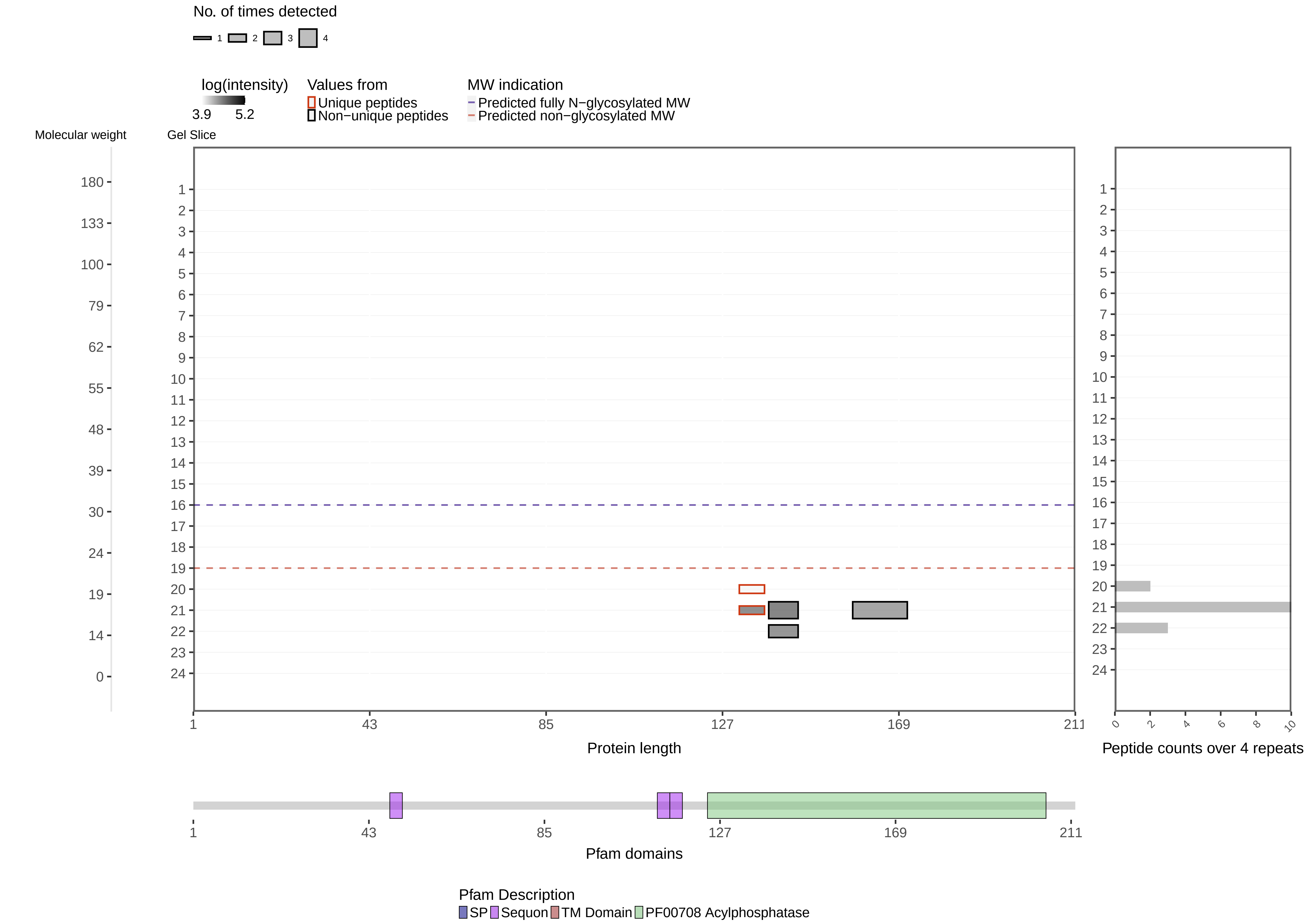


Protein sequence:

FRSPYFSSPMSSLTSDNNSNATPDVKTVRAVIKGRVQGVFYRDWTVENAKELGLKGWVRNRRDGSVEALFSGSPEKVQEMEQRCRRGPPAAIVTGLDVFPCDDDPGTGFECKQSG*

### NbL15g24320.1 glutamate_formimidoyltransferase [MW = 35.83 kDa]

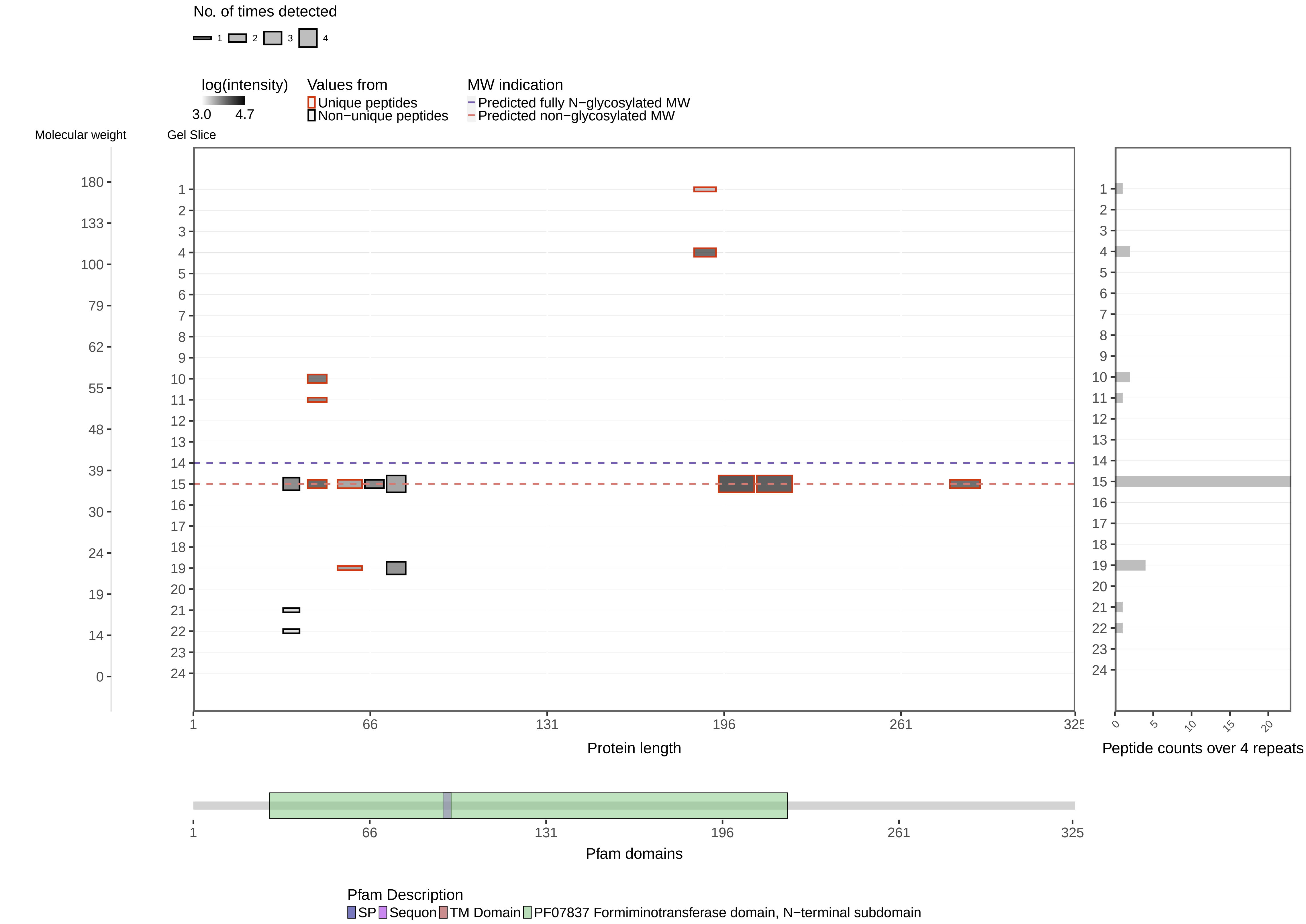


Protein sequence:

NLQVPTYLYGAAHQEGRSLDSIRRELGYFRPDSKENQWAGGPKWQTLQLKPDEGPSRATQATGVITIGATRWVDNYNIPVFTNDISIVRRIAKRVSGRGGGFPSVQSMALTHGGGTIEVACNLLEPARNGGDQVQLEVERLAREEGIAVGKGYYTDFSEDKIIESYLKLVQHSD*

NbL15g24260.1 photosystem_II_repair_protein_PSB27−H1_chloroplastic−like [MW = 19 kDa]

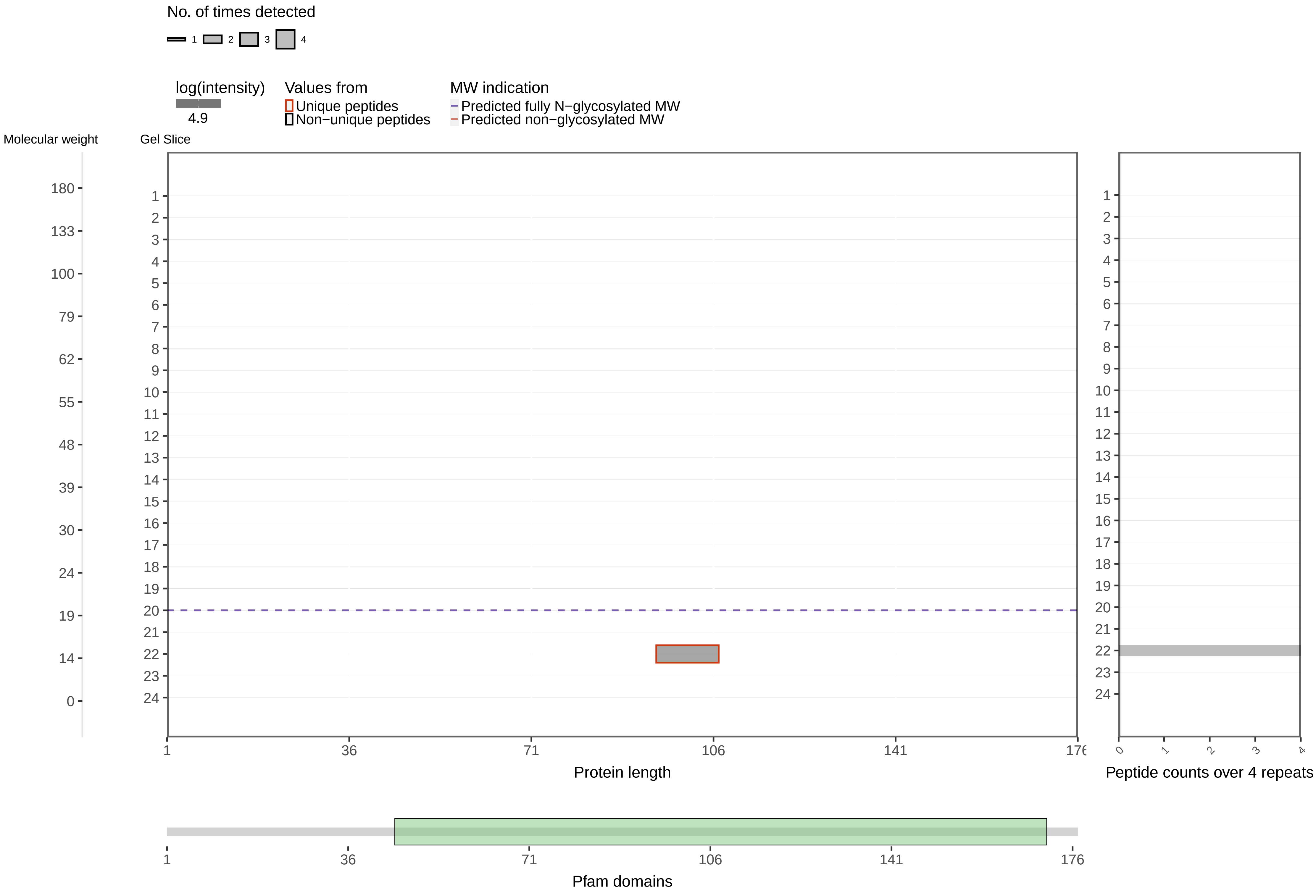


Pfam Description

SP Sequon TM Domain PF13326 Photosystem II Pbs27

Protein sequence:

REASEVIKKVRSTLSMEKGDPNIADSVTDLREASNYWVAKYRREKALLGRASFRDIYSALNAVSGHYVSFGPTTPIPAKRKQRILEEMDTAEKALQRGR*

NbL15g24190.1 protochlorophyllide_reductase_chloroplastic−like [MW = 43.25 kDa]

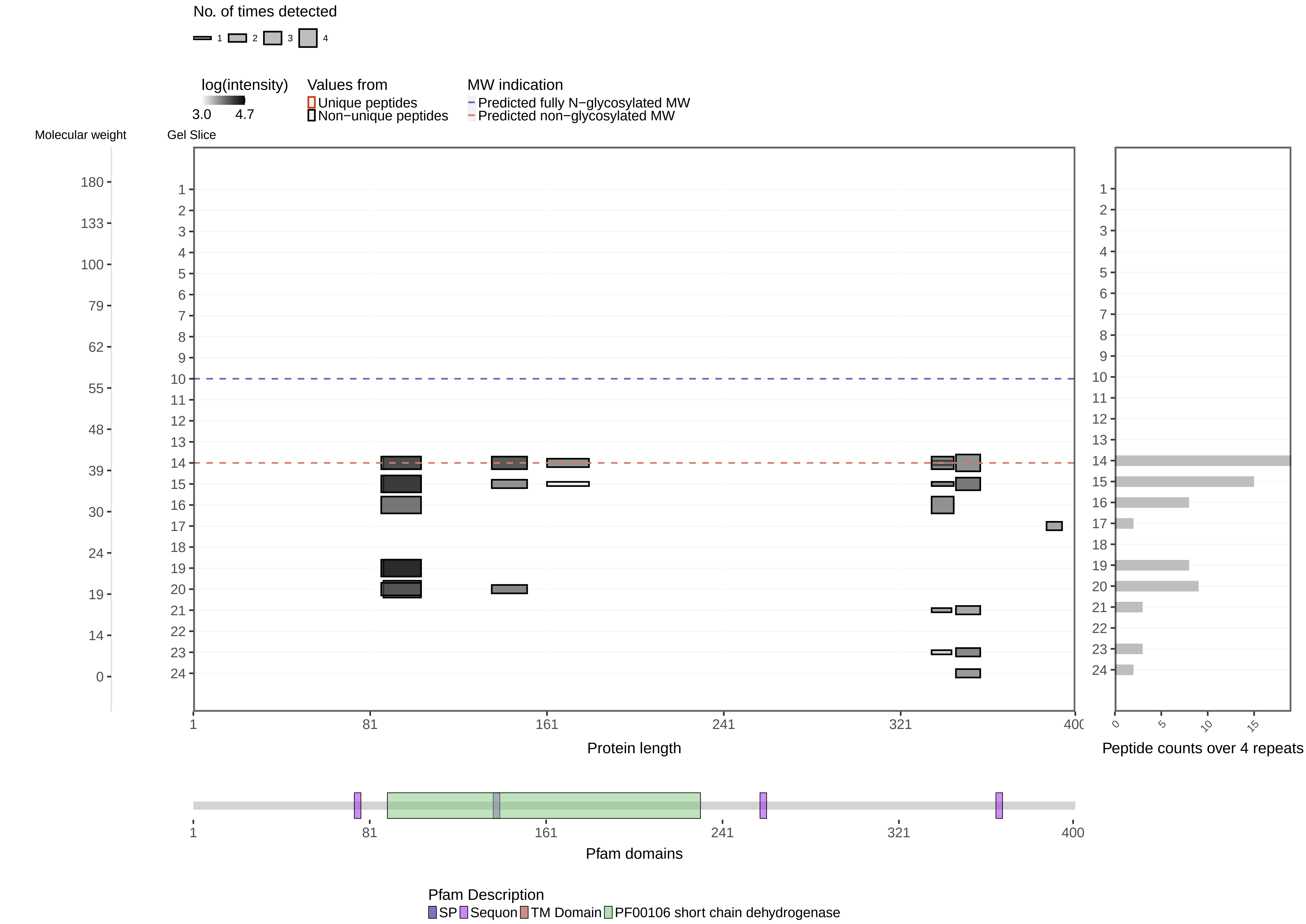


Protein sequence:

FELSVGTNHLGHFLLSRLLLDDLKQSDYPQKRLIIVGSITGNTNTLAGNVPPKANLGDLRGLSGGLNSLNCSPMIDGGEFDGAKAYKDSKVCNMLTMQEFHRRFHEENGIAFASLYPGCIAETGLFRNHIPLFRALFPPFQKYITKGYVSEAEAGKRLAQVVRDPSLSKSGVYWSWNSTSSSFENQLSKEASDAEKARKLWEVSEKLVGLA*

NbL15g24160.1 malate_dehydrogenase_glyoxysomal [MW = 19.31 kDa]

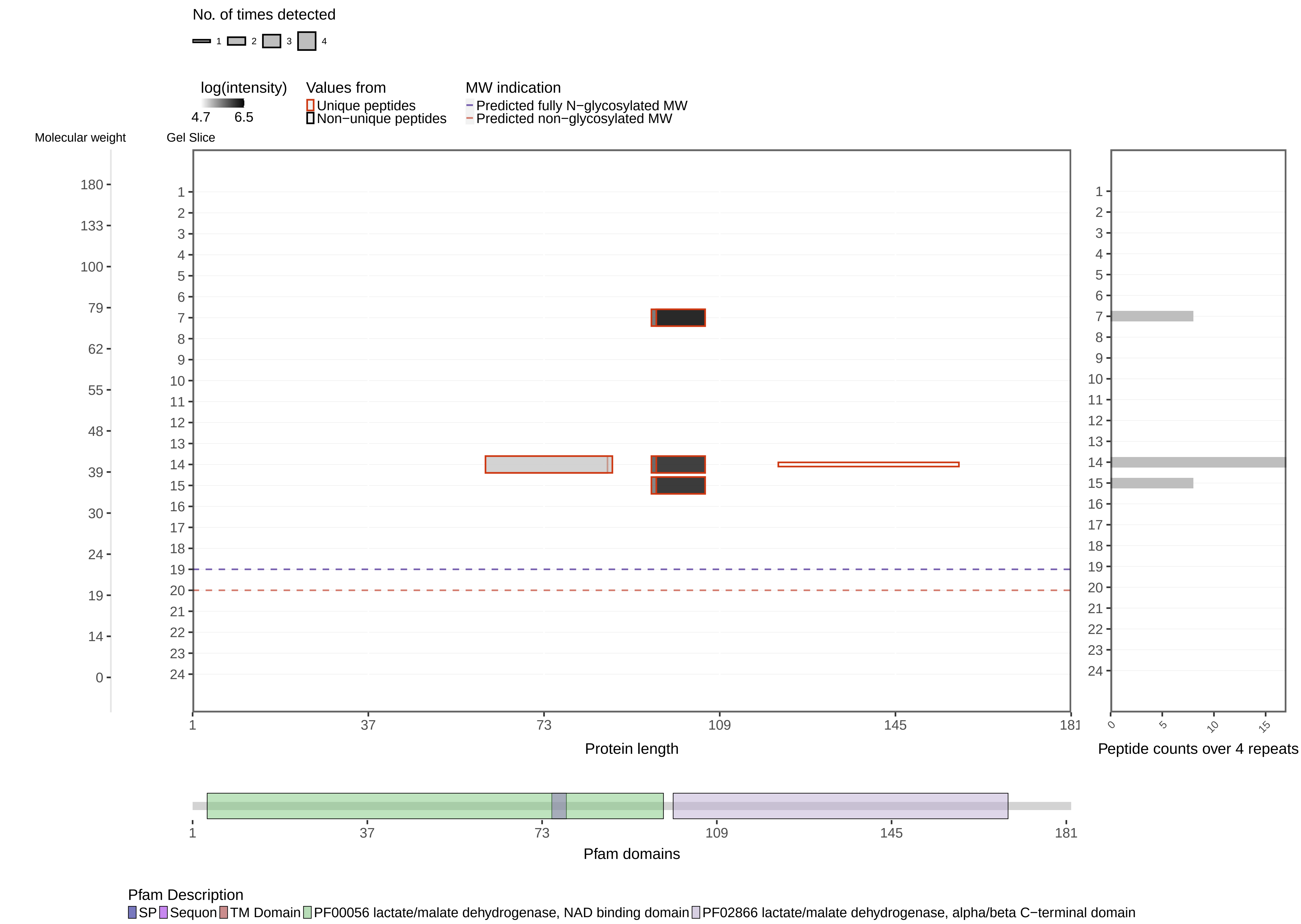


Protein sequence:

VPIAAEVFKKAGTYDPKKLLGVTSLDVVRTNTFVAEVLGLDPREVDVPVVGGHAGVTILPLLSQVKPPCSFTQEETEYLTKRIQDGGTEVVEVCVHYLQFSIY*

NbL15g24070.1 riboflavin_synthase−like [MW = 30.83 kDa]

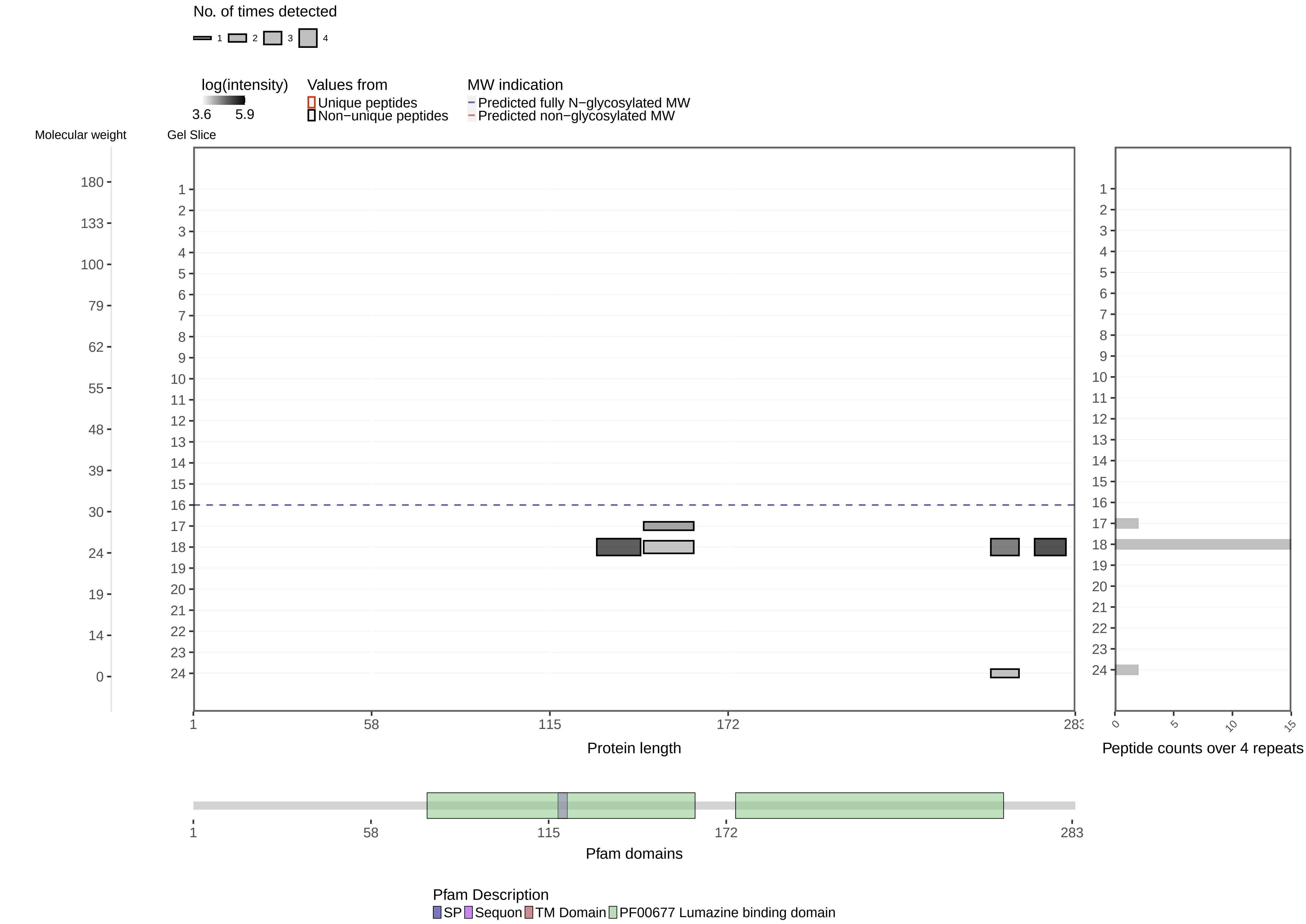


Protein sequence:

LSEFSVGLAPETLRKTSLIELEQGSIVNLERALKPSTRMGGHFVQGHVDGTGQIVELKPEGDSLWVKVKSSKEILRYIVPKGFIAVDGTSLTVVDVSEEEDCFNFMLVAYTQQNVVIPMKKVGQKVNLEVDILGKYVERLLSSGFVNAIKSS*

### NbL15g24000.1 tyrosine_aminotransferase−like [MW = 46.44 kDa]

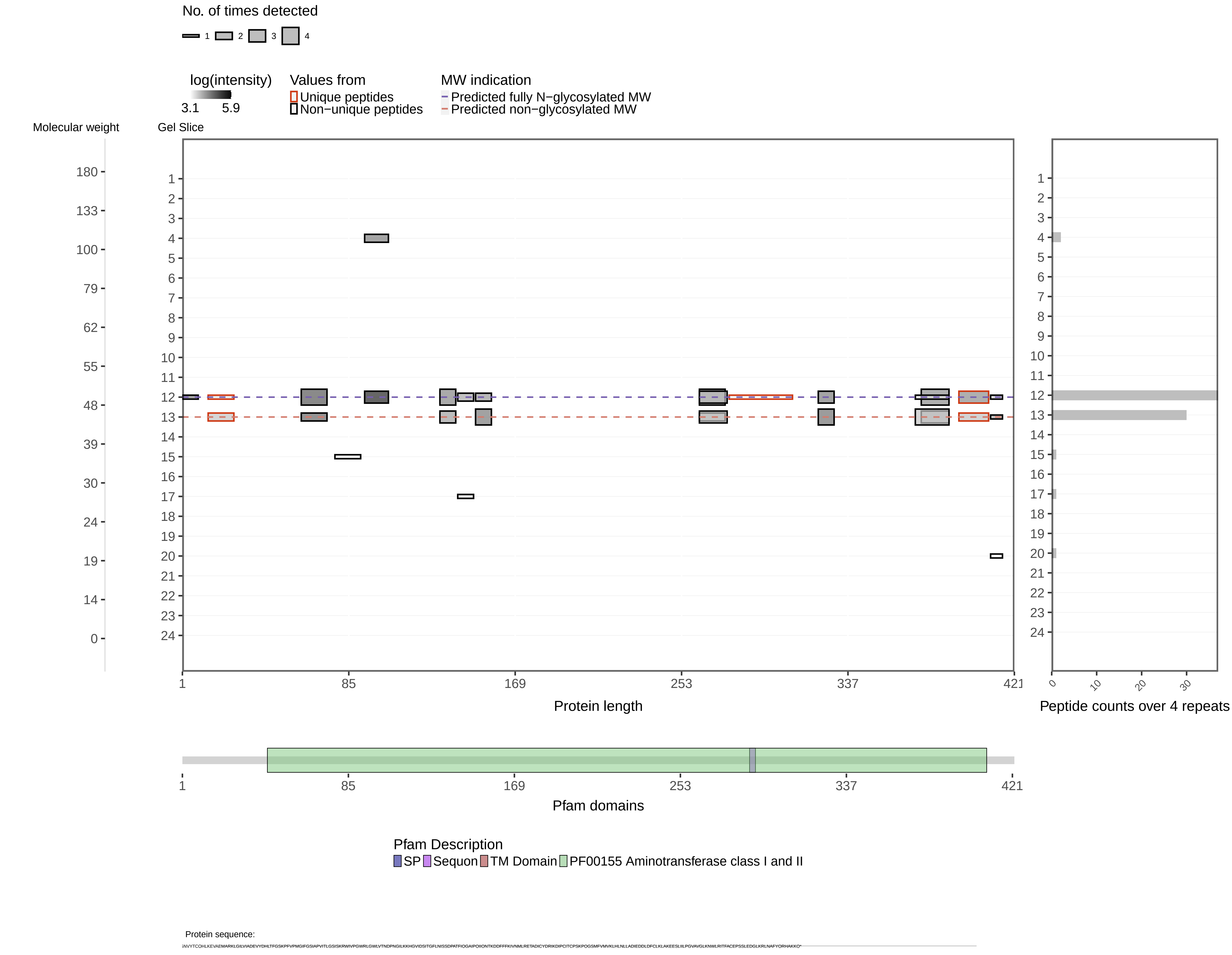

NbL15g23810.1 uncharacterized_protein_At4g28440−like [MW = 16.17 kDa]

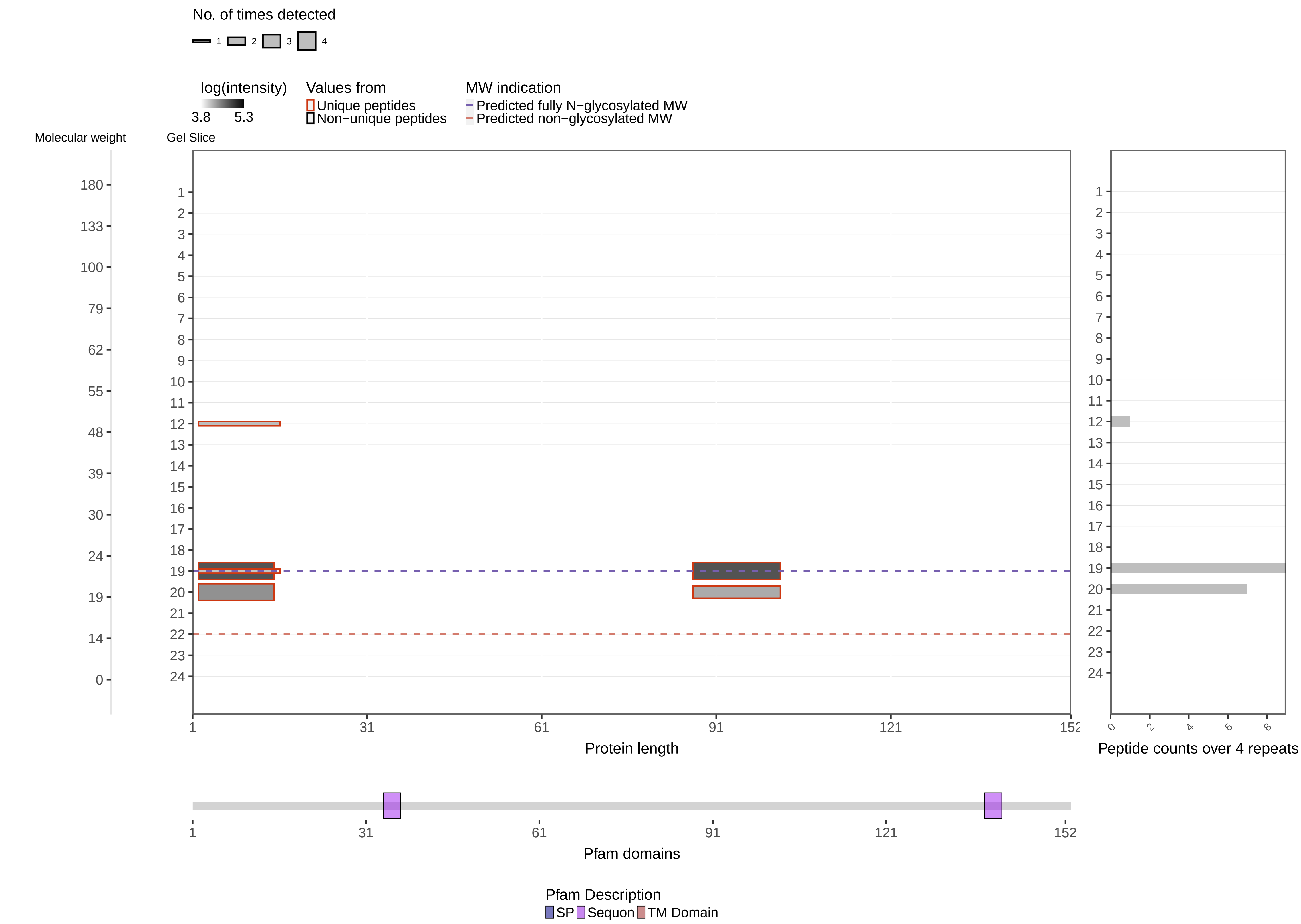


Protein sequence:

PRGPARLAECLVGDDTGSILFTARNEQVDMMKPDATVILRNAKIDMFKGSMRLAVDKWGRVEVTEPADFGVNEENNLSLVEYELVNVEE*

### NbL15g23780.1 26S_proteasome_non−ATPase_regulatory_subunit_2_homolog_A−like [MW = 78.02 kDa]

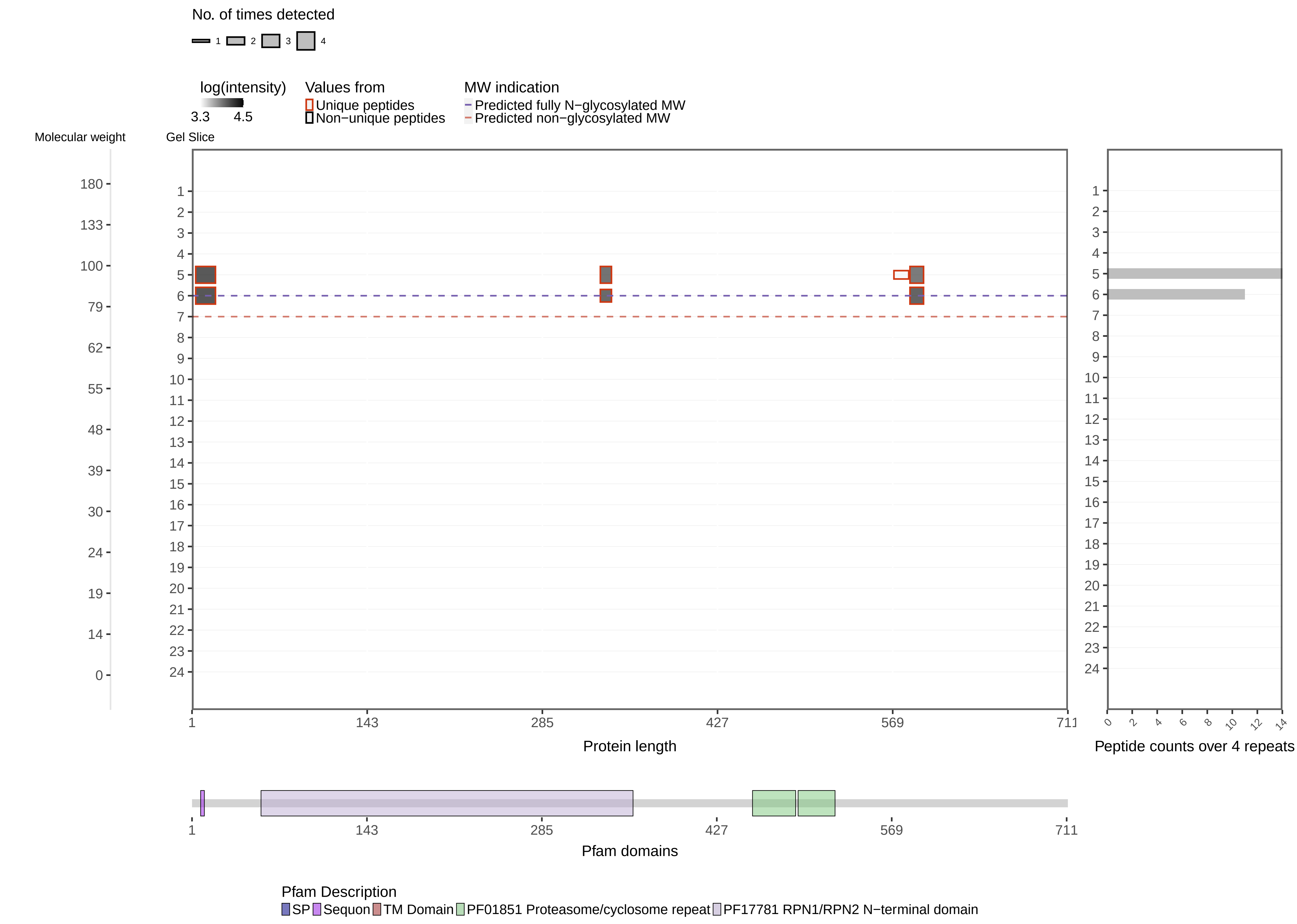


Protein sequence:

[illegible]

NbL15g23630.1 fasciclin−like_arabinogalactan_protein_9 [MW − SP = 23.98 kDa]

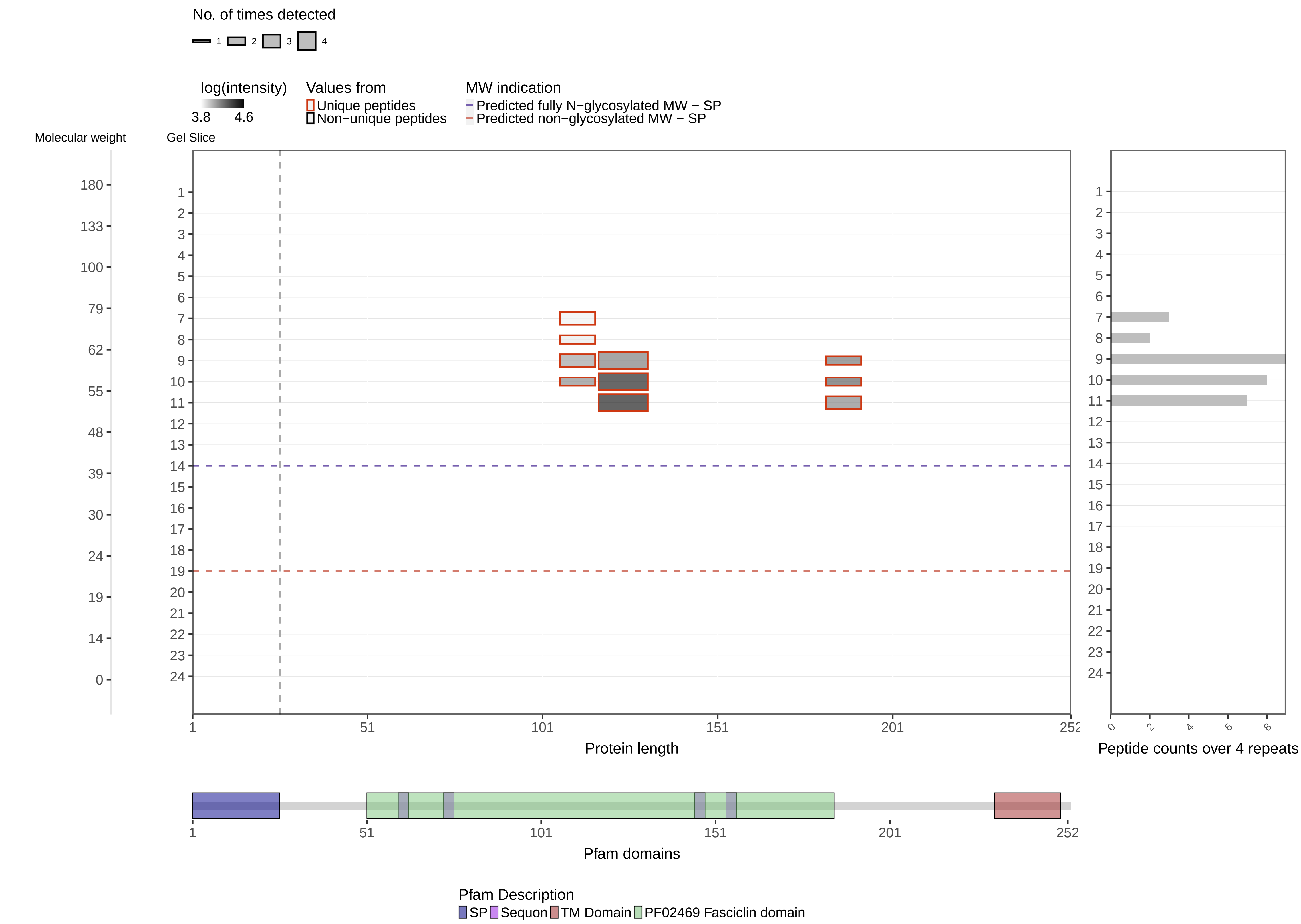


Protein sequence:

PKFYSFDDLQTVSNPVRTQATGQKGEPFGLNFTGQNNQVNVSSGTVETNIYNAIRKDPPLAVYQLDKVLIPLAFSDAKTPSSDDAPAPTDDSAKSGSTADKTKAQEPSPAPNGAKTINVGVFGLISGVFLFFMGALS*

NbL15g23580.1 GTP−binding_protein_TypA/BipA_homolog [MW = 72.05 kDa]

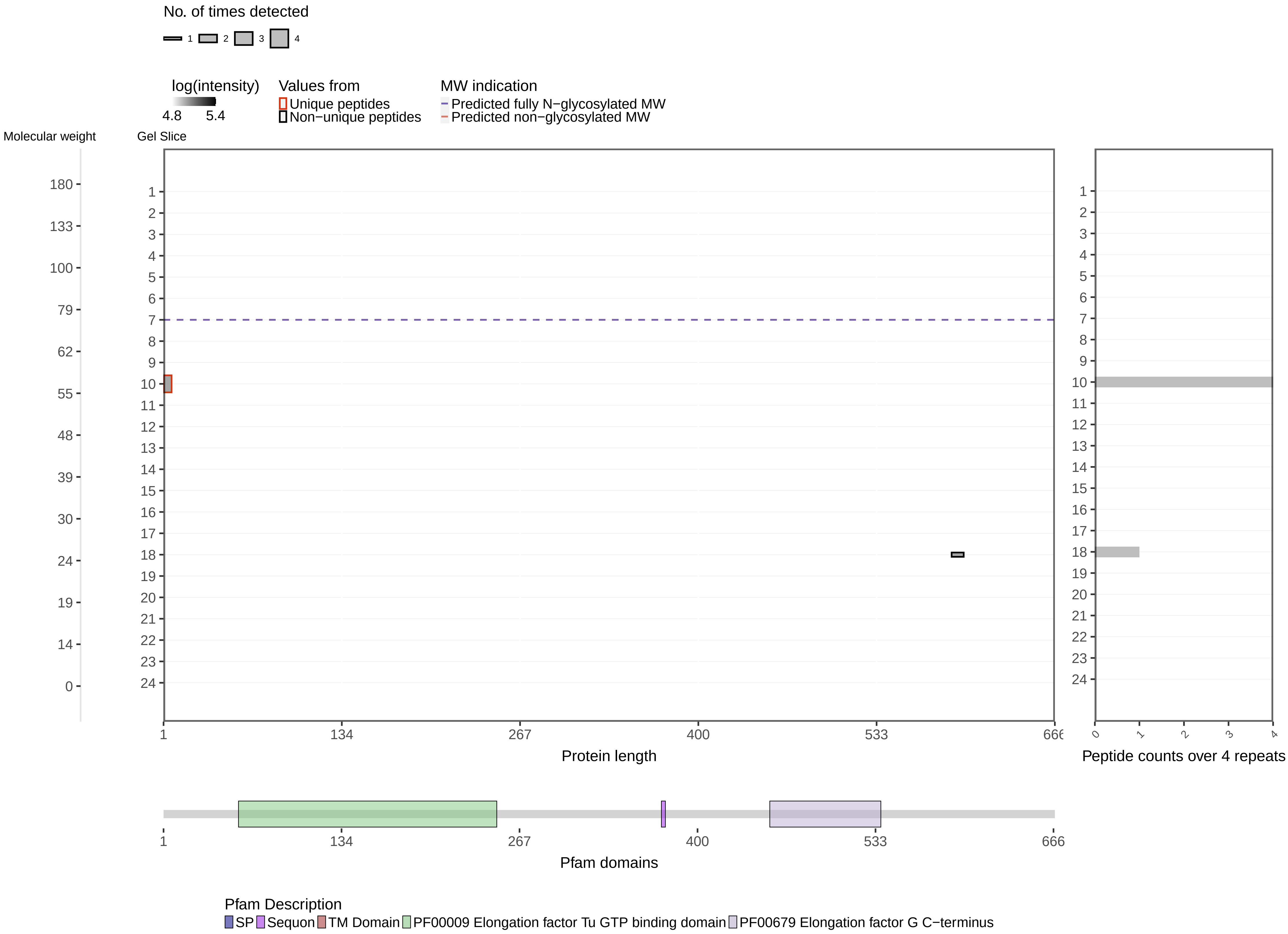


Protein sequence:

ADPVDSAGAGDIVSMAGLSSPTIGHTVANVEVMTALPTVVLDPPTISMTFGVNDSPLAGSDGIHLTGGKIGDRLLAEAETNLAINVLPSTSESYEVQGRGELQLGILIENMRREGFELSVSPPKVMYKVEKGVKLEPIEEVTIEVDDEHVGLVMEALSHRRGEVTDMGPVPGNTGRTRMCLTCPSRGLVGYRSVFSSDTRGAGFMHRAFLTYEKYRGPLGNVRKGVLVSMGRGLITAHALMNLEARGILFVAPGMETYDGMIIGEHSRDTDLDVNPVRTKELTNVRAACKDE

NbL15g23440.1 uncharacterized_protein_At1g03900−like [MW = 32.47 kDa]

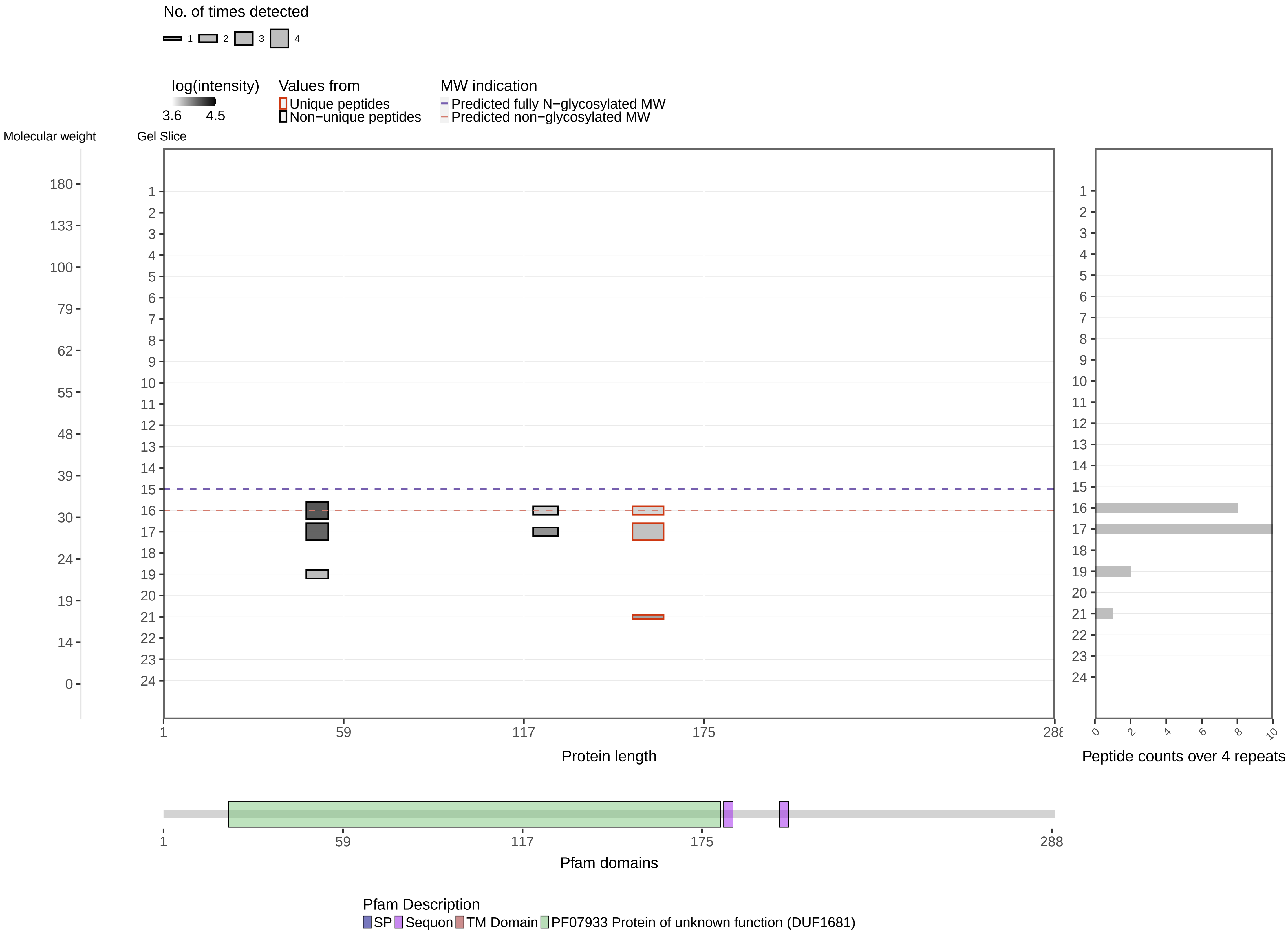


Protein sequence:

AYDFQAALHDHMKYLNKKKTAEEMEQQYQKSSSVDYSLKEGETLKLQIKNKTGGSTIRSKFFEQGLNNLSLEEKANRKDSTILIKPPPPPPAPLSPVDTAAMSPPVSPSKFSLEQSAETKDSASAKEQTNTPESLKDQGPQDVEDDDFGDFQAAG*

### NbL15g23370.1 ketol−acid_reductoisomerase_chloroplastic [MW = 63.33 kDa]

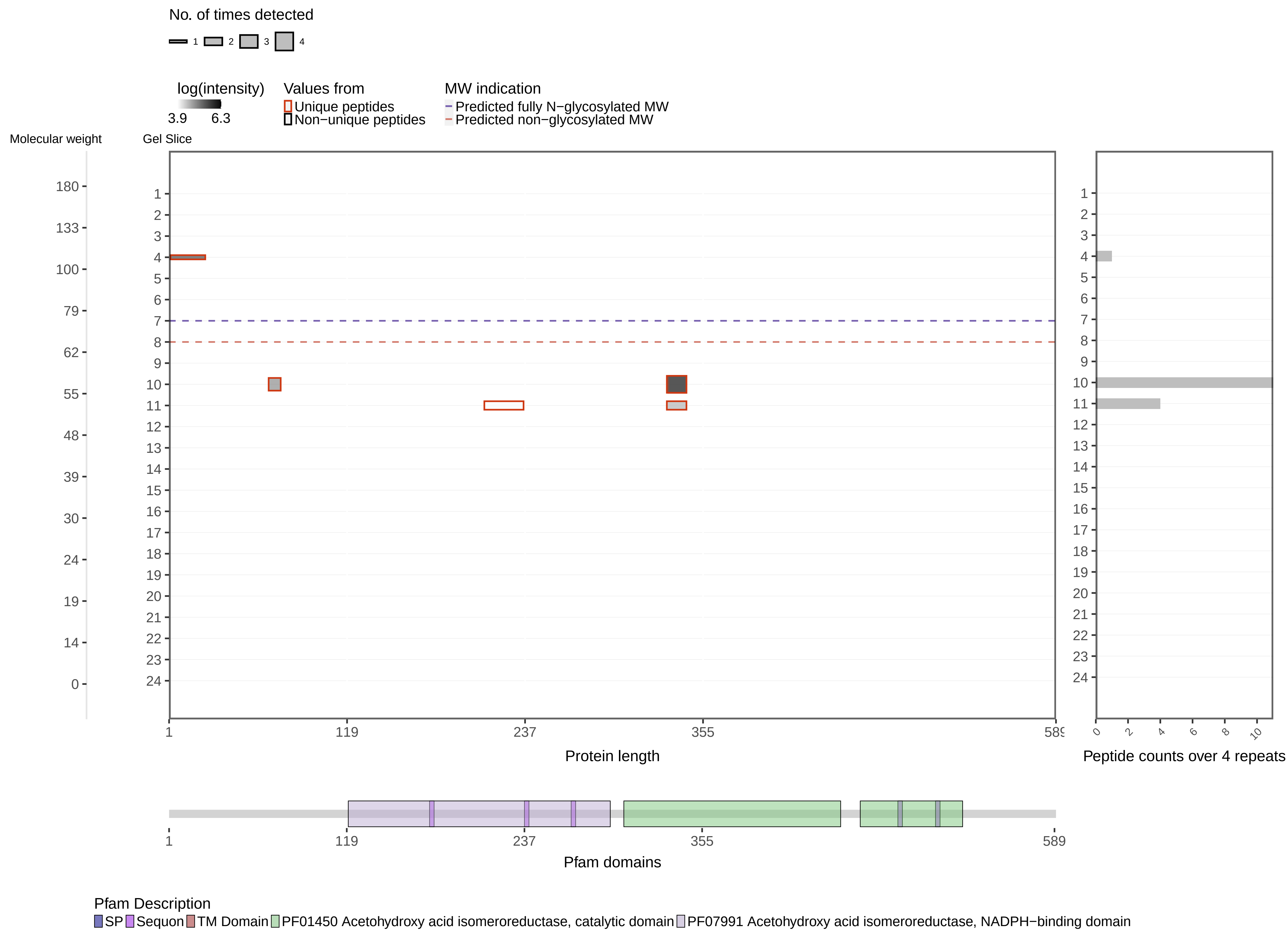


Protein sequence:

ATDVALGWSVALGSPFTFATTLEOEYKSDIFGERGILLGAVHGVVESLFRRFTVNGMNDELAYKNTVECITGIISKTISTKGMLAVYNSLTEEGKKEFEAAYSASYYPCMEILYECYEDVATGSEIRSVVLAGRRFYEKEGLPAFPMGKIDOTRMWKVGERVRAKRPAGDLGPMYPFTAGVFVALMMAOIEILRKKGHSYSEIINESVIESVDSLNPFMHARGVSFMVDNCSTTARLGSRKWAPRFDYILTOOALVAVDNNAAINODLISNFLSDPIHGAIEVCAOLRPTVDISVTAD

### NbL15g23360.1 bifunctional_riboflavin_kinase/FMN_phosphatase−like [MW = 41.42 kDa]

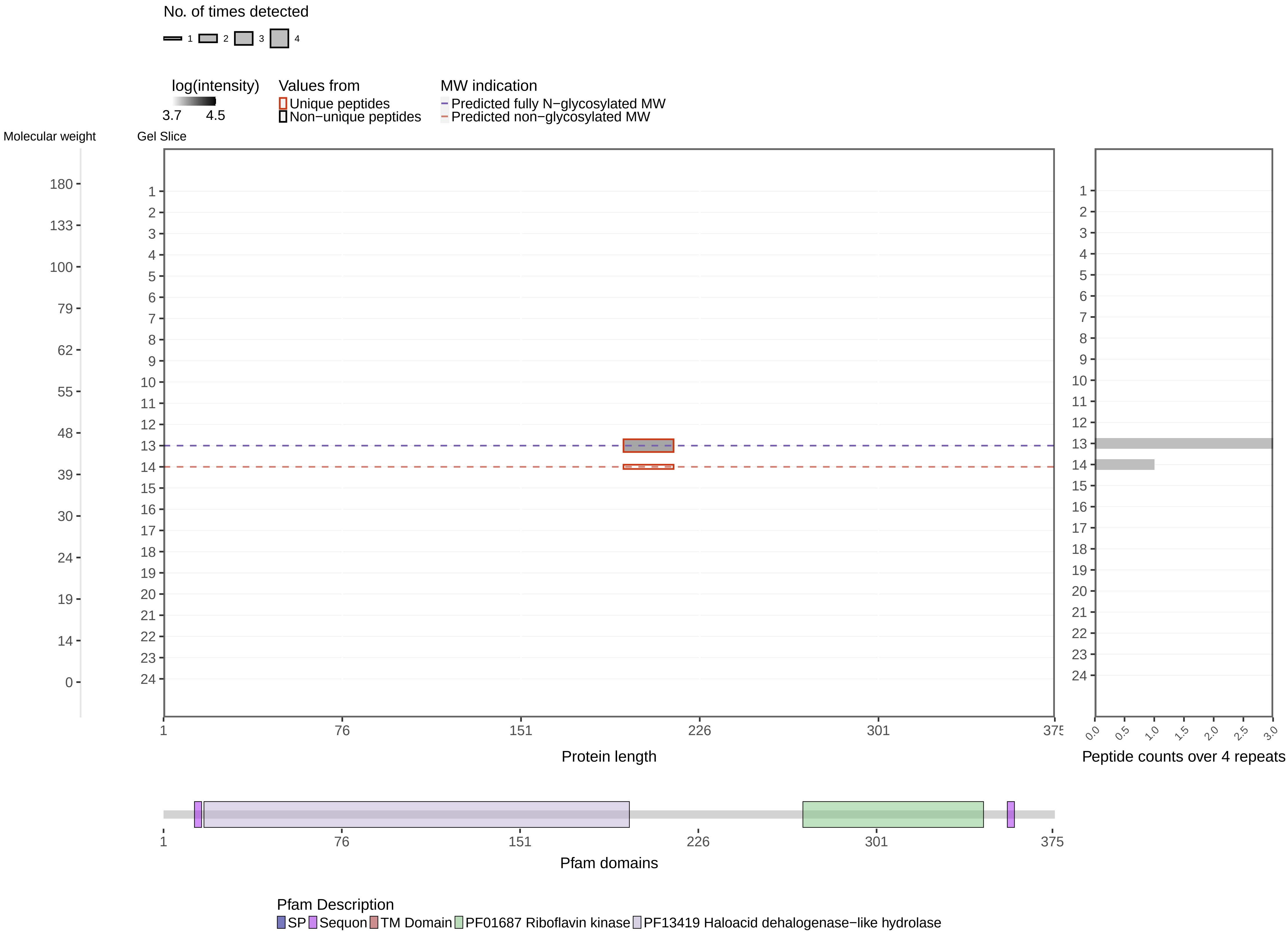


Protein sequence:

VIEDSVIGVKAGKAARMKVVAVPSFHSEFDOYSIADSVLRSLLELKPEVWGLPPFEDWVDNALLVEPVSFRGLYRNGLLHDFTDDGPSALPDOVFGLYFGWGRLEAHKFIKIVLGSGWGHGCCSSKRKMOACIVDSSDEOIOECKMEVVIVGYIRGSRDEGKWDKIEILEEDKSIASAALNRSEFSLGAFOSLFSEVAL*

NbL15g23350.1 14−3−3_protein_9 [MW = 29.54 kDa]

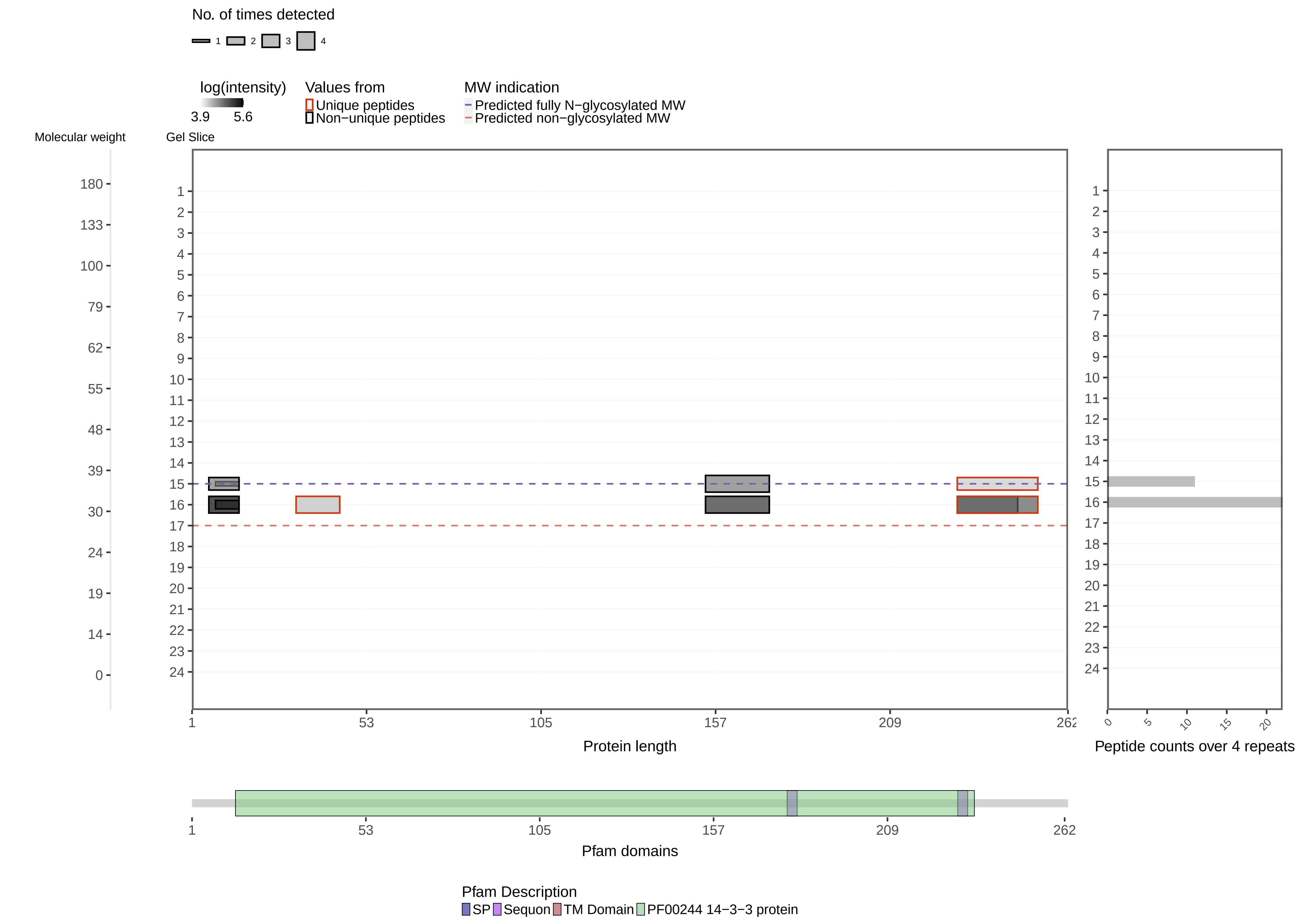


Protein sequence:

GESTVFYHKMKGDYYRYLAEFKSGNDKKEVAELSLKAYQSATTAAEAELPPTHPIRLGLALNFSVFYYEIMNSPERACHLAKQAFDEAISELDSLNEDSYKDSTLIMQLLRDNLTLWTSDLPEDAEDAQKGDATNKASGGEDAE*

### NbL15g23300.1 phosphoenolpyruvate_carboxylase−like [MW = 110.04 kDa]

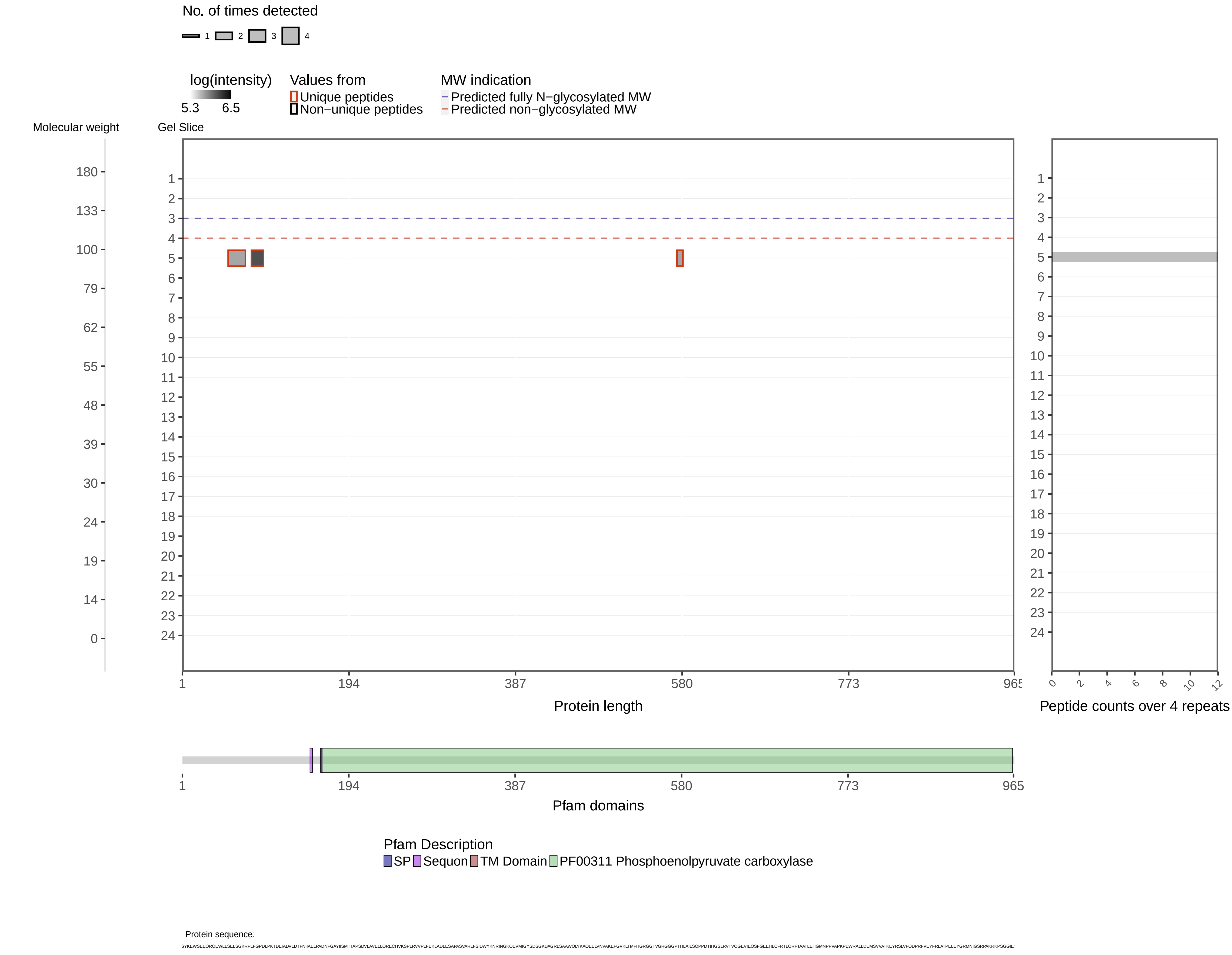

### NbL15g23290.1 protein_LIGHT−DEPENDENT_SHORT_HYPOCOTYLS_10−like [MW = 19.8 kDa]

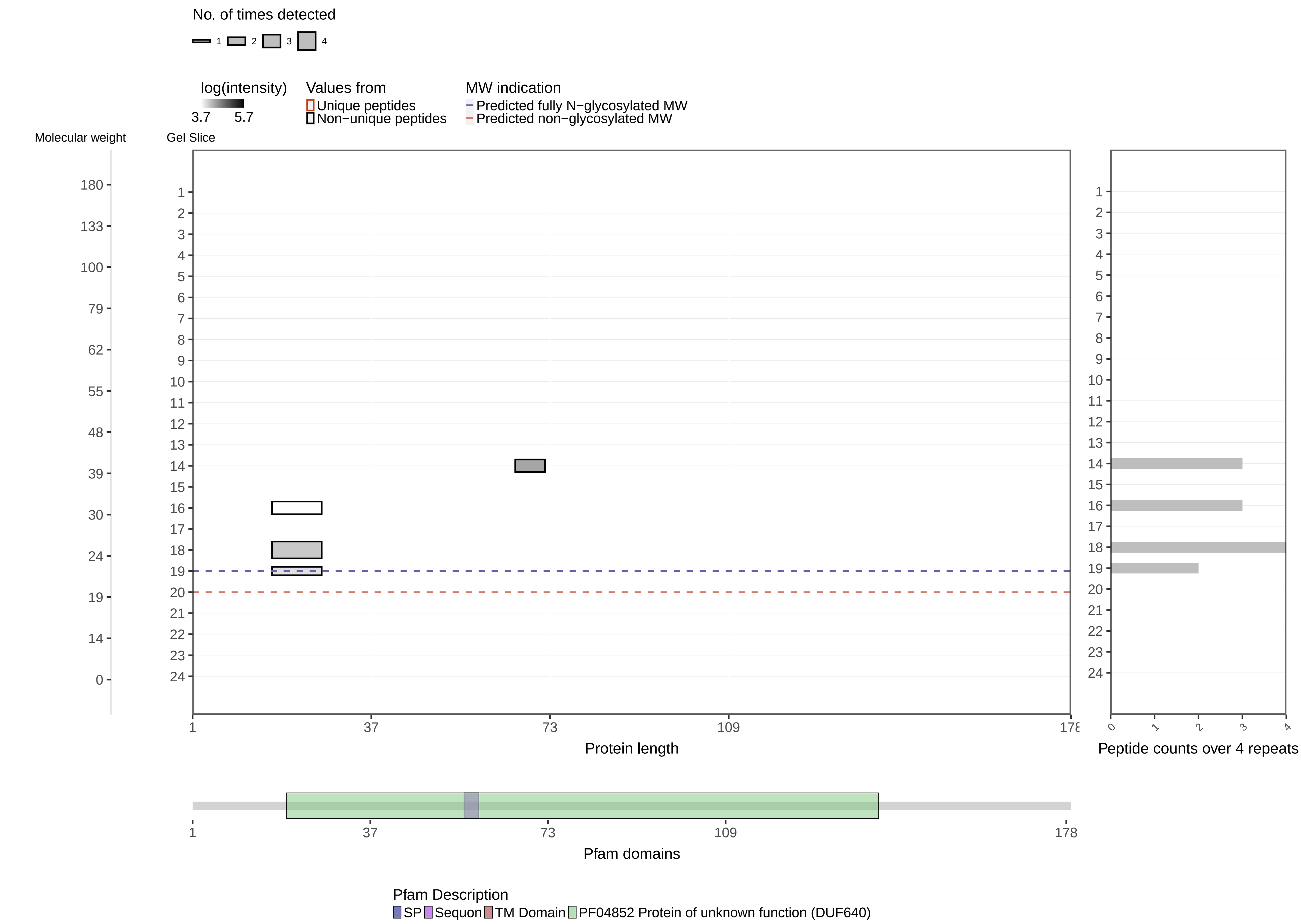


Protein sequence:

'HLQGCIFYGQPEPPAPCTCPLRQAWGSLDALIGRLRAAYEENGGSPETNPFASGAIRVYLREVKECQSKARGIPYKKKKKKIGGTSDSKGDDDSTSSHHPFS*

NbL15g23210.1 peroxidase_P7−like [MW − SP = 31.54 kDa]

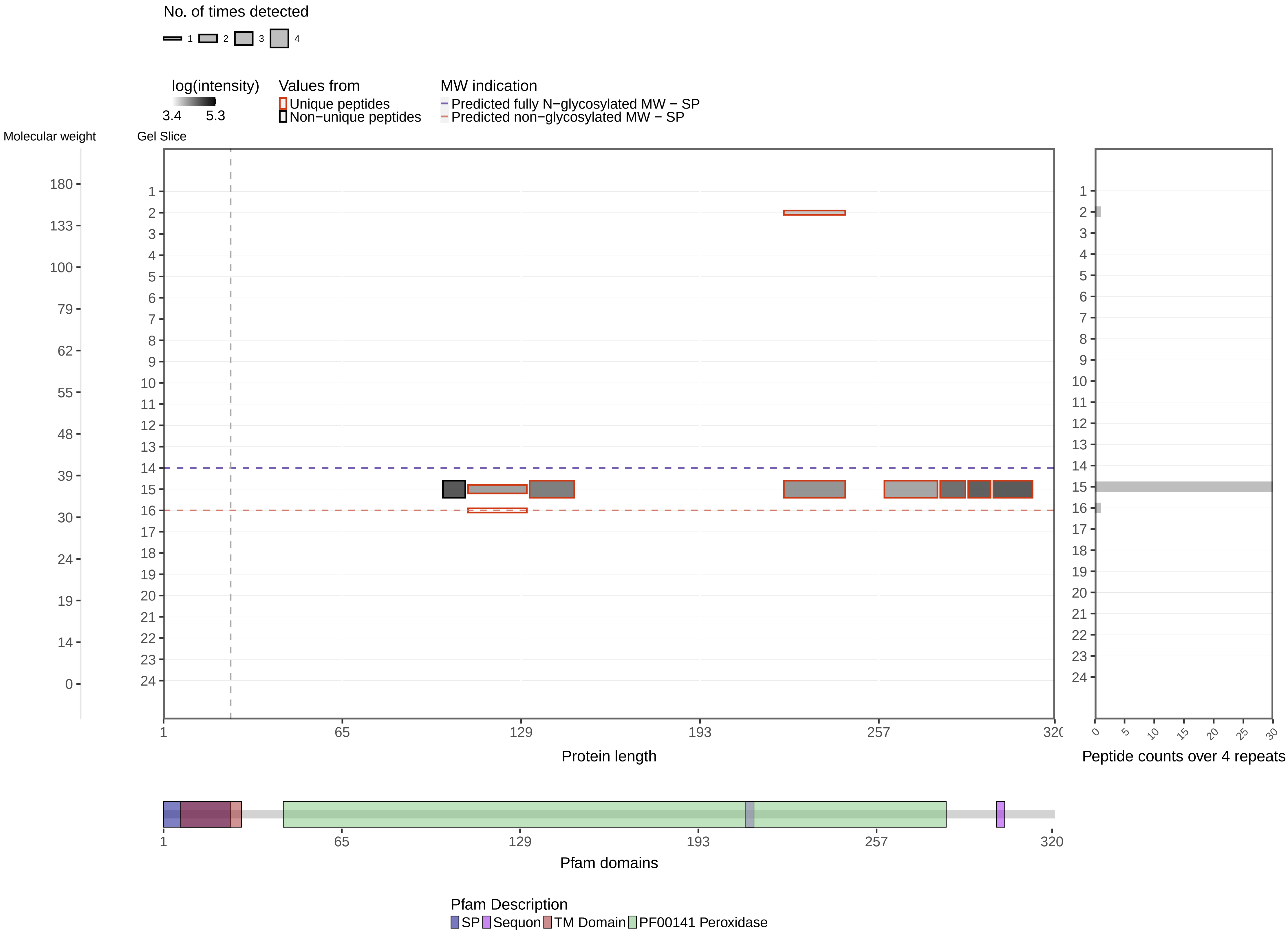


Protein sequence:

RRDARTASQSAANTQIPAPTSSLSTLLSMFSAKGLNARDMTALSGGHTIGQARCTTFRTRIHNDTNIDAQFATKRKATCPSSGGGANLAPLDIQTPNRFDNDYYQNLVARRGLLHSDQELFNDGSQDALVRSYSTNGAAFRSDFAAAMVRMGNISPLTGNNGEIRRNCRAIN*

### NbL15g23130.1 60S_ribosomal_protein_L11−1−like [MW = 20.76 kDa]

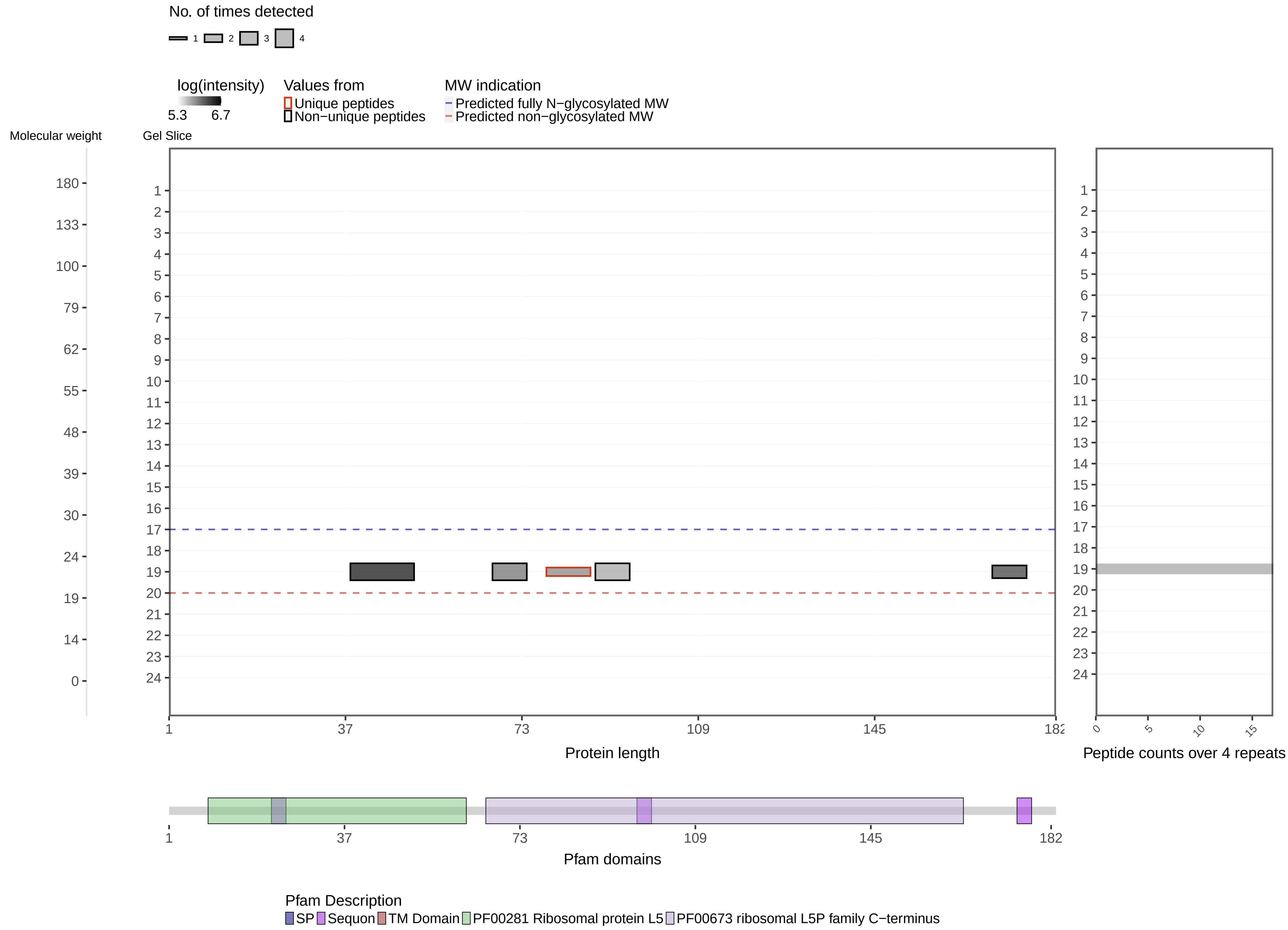


Protein sequence:

ATQLLESGLKVKEYELLRRNFSETGCFGFGIQEHIDLGIKYDPSTGIYGMDFYVVLERPGYRVARRRRCKSRVGIQHRVTKEDAMKWFQVKYEGVILNKSSNIQ*

### NbL15g23060.1 ATP−dependent_zinc_metalloprotease_FTSH_2_chloroplastic [MW = 52.72 kDa]

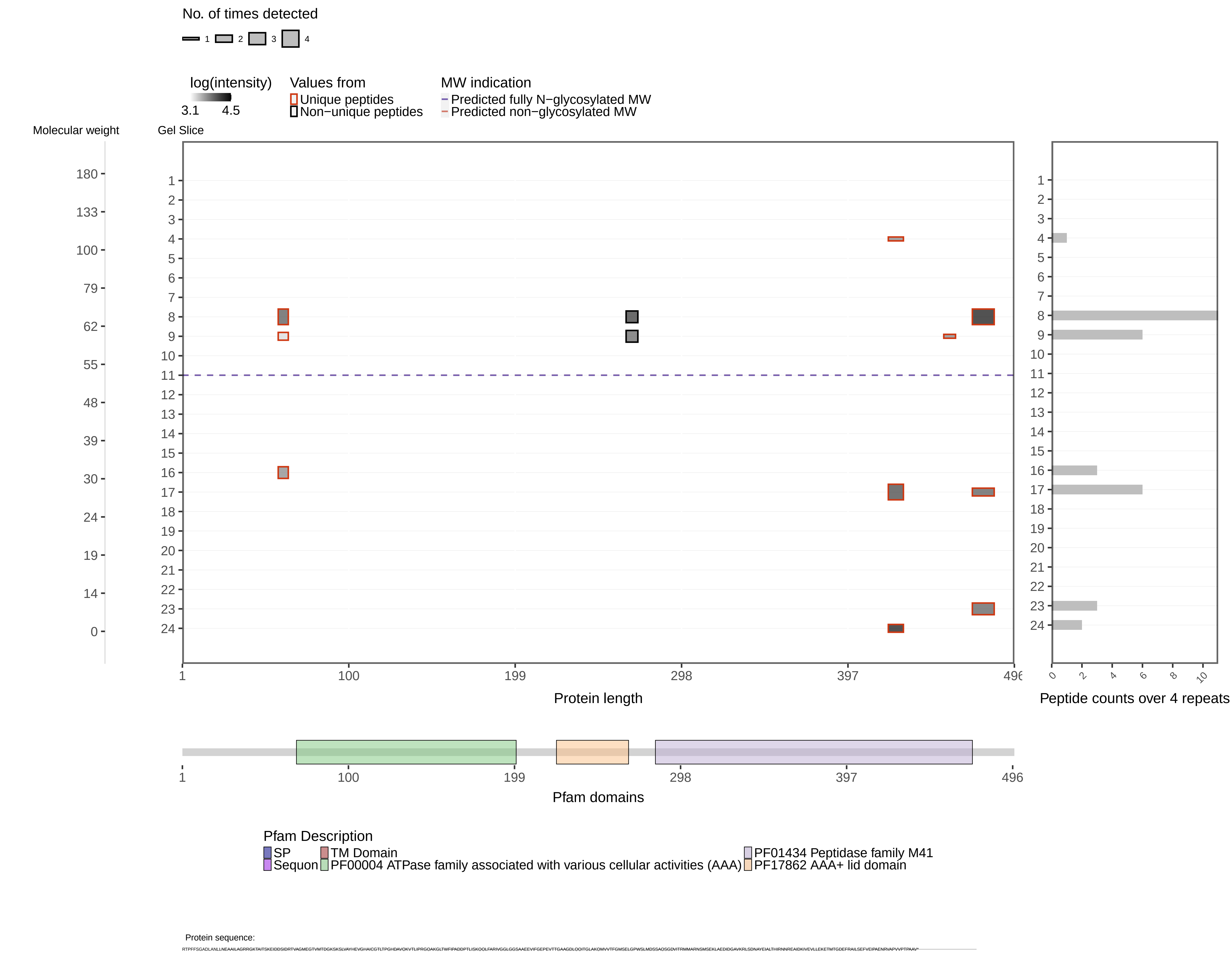

NbL15g22940.1 protein_STRICTOSIDINE_SYNTHASE−LIKE_10−like [MW − SP = 31.52 kDa]

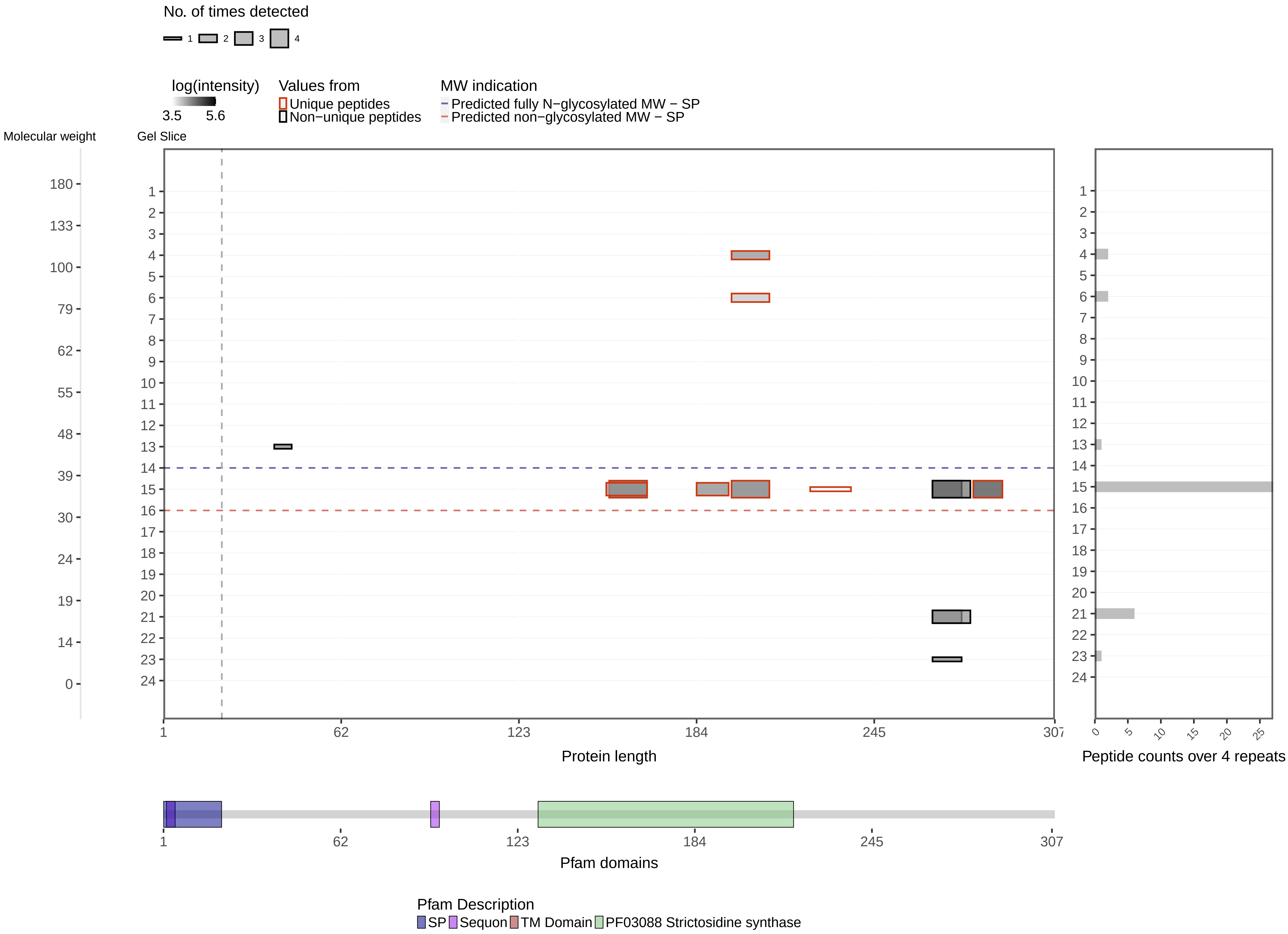


Protein sequence:

IYFTDSSTKYQRRQFSAAFVSGDTTGRLMKYDKSTKKVAVLLRGLALANGVALNKDGSFLLVSETGNCRIIRYWIKGPYAGRHDIFADVPGLPDNIRINSRGEFWVSLNAKTSATAIKLSAEGRMLEVLEDEEGKTLRYISEVEEQYGKLWFGSVMMPFLGVYGLS*

### NbL15g22840.1 translation_initiation_factor_IF−2_chloroplastic−like [MW = 108.26 kDa]

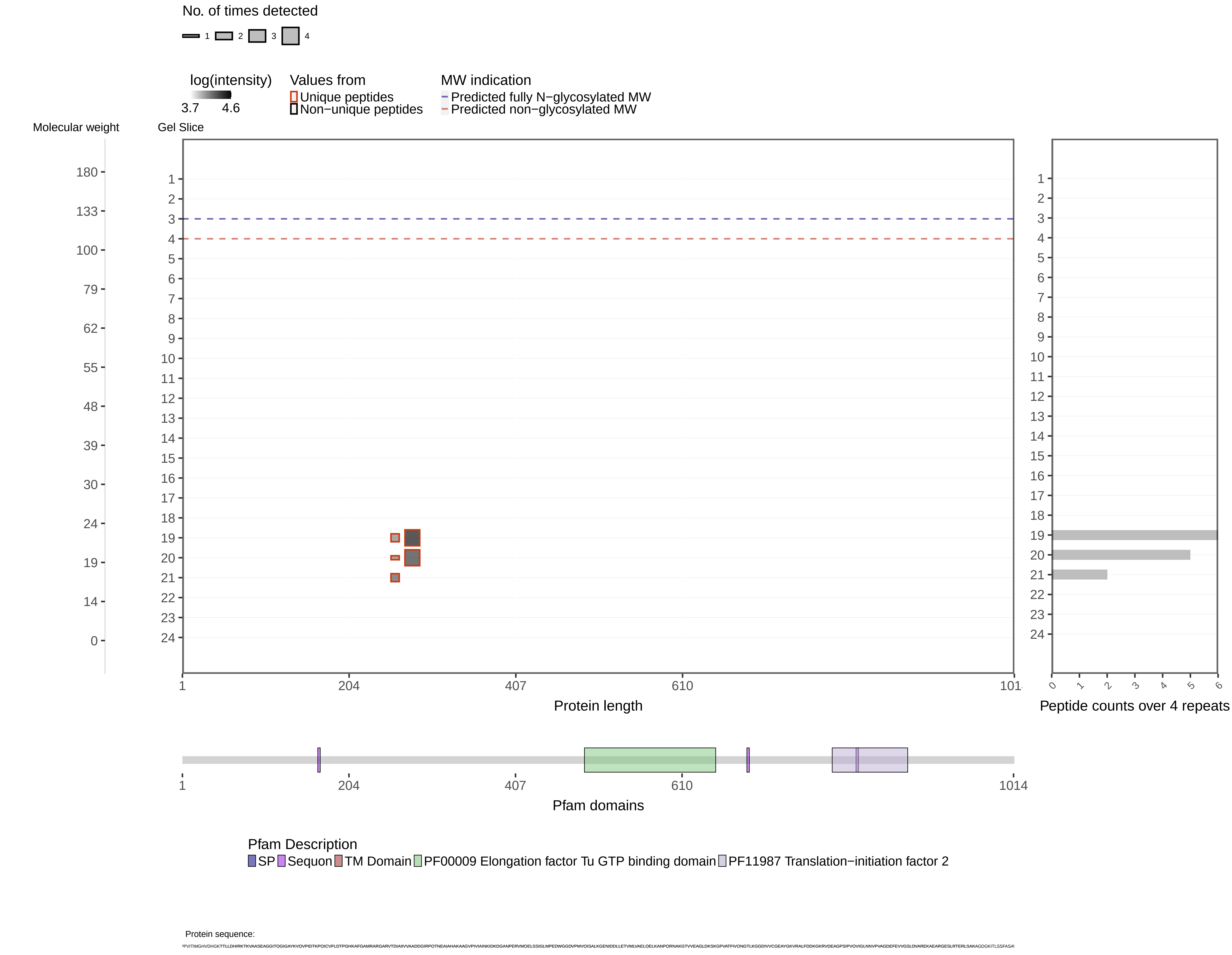

NbL15g22830.1 uncharacterized_LOC104238148 [MW = 102.38 kDa]

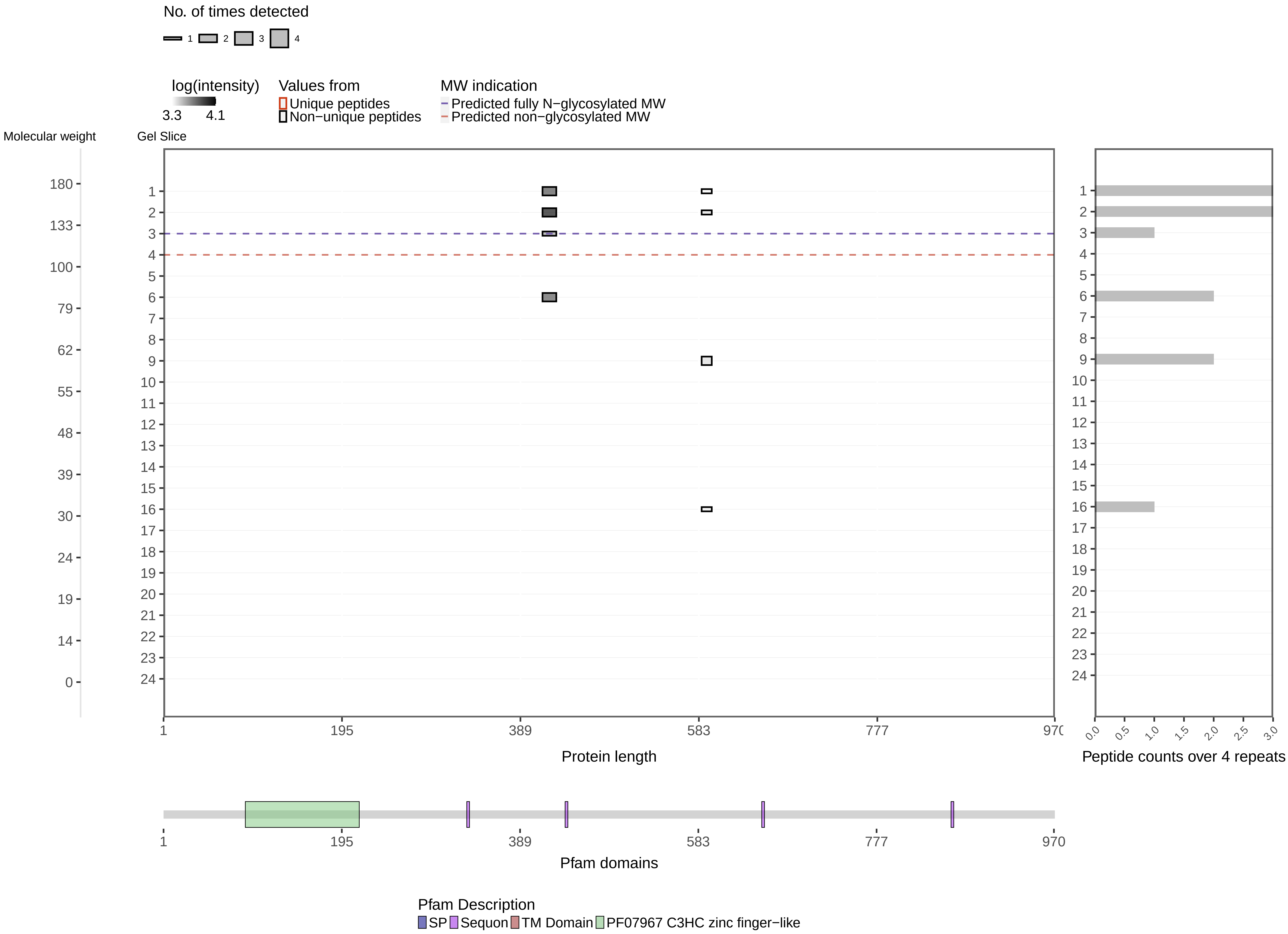


Protein sequence:

EVGGHAASYESRGPSSRKRNLEEGGSTVDRPQLPVQPADSVEGTVIDRDGDEVNDGSQYSAGPSKRACQSDAFGPQLTSYGKDSSGAGPSLSLGFEIGRDASRDDAFGRGQEQLTGMPSTRDSTHVSSVIAMDTVHSADDDSMESVENLPGDFDDVNFPSTSVLRSADPVETSELNYSNQAQQSTCPAVVRSAGEMGVSSTNDEEVVNTDTATGNVRDGPSVGISGGSIGMGASHEAEIHGTDASVHRADSVAGDVEAVAEITENQGQTGEFAPDPGLMGDYVPE

### NbL15g22750.1 glucose−1−phosphate_adenylyltransferase_small_subunit_chloroplastic/amyloplastic [MW = 58.89 kDa]

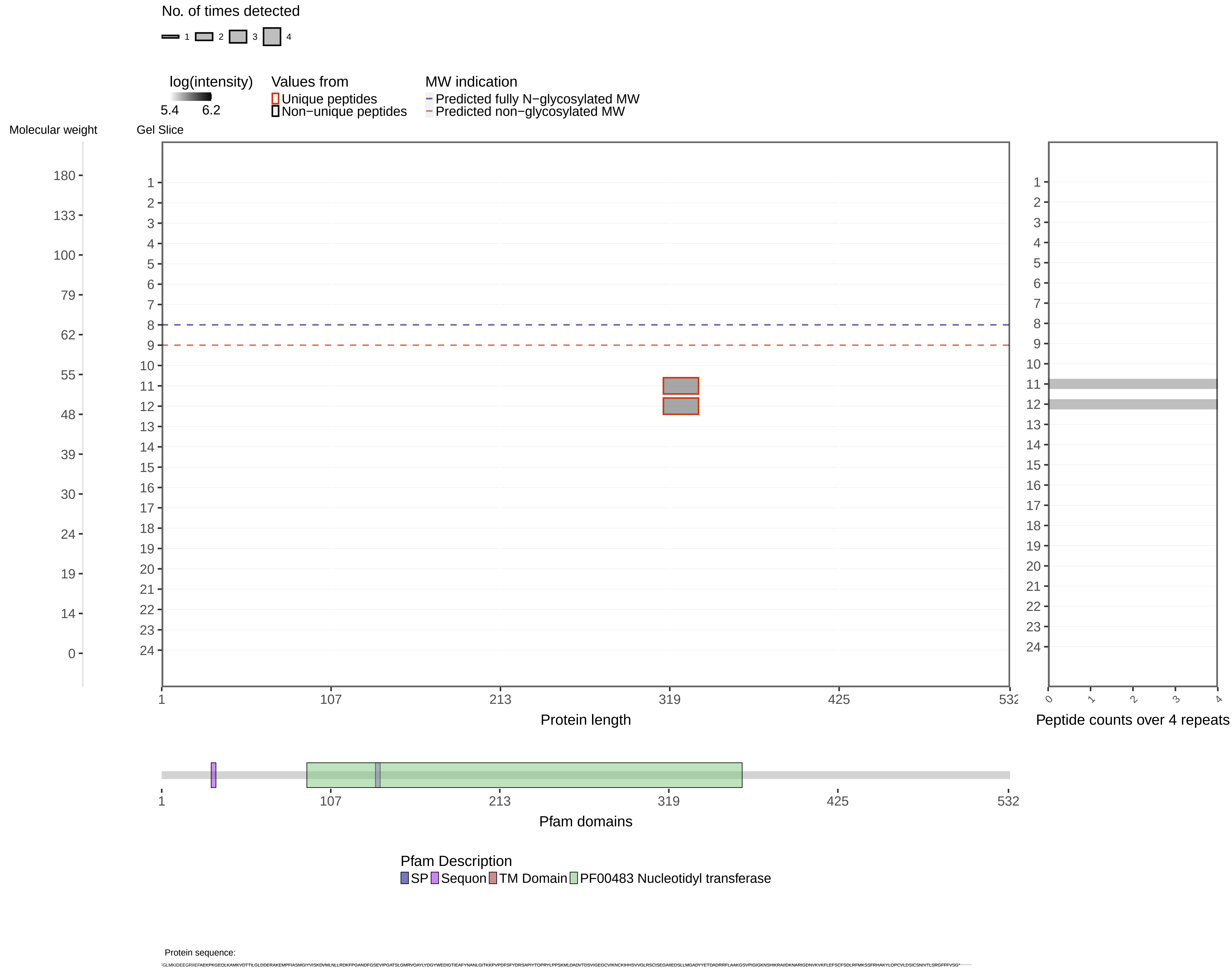

### NbL15g22710.1 subtilisin−like_protease_SBT5.3 [MW − SP = 80.06 kDa]

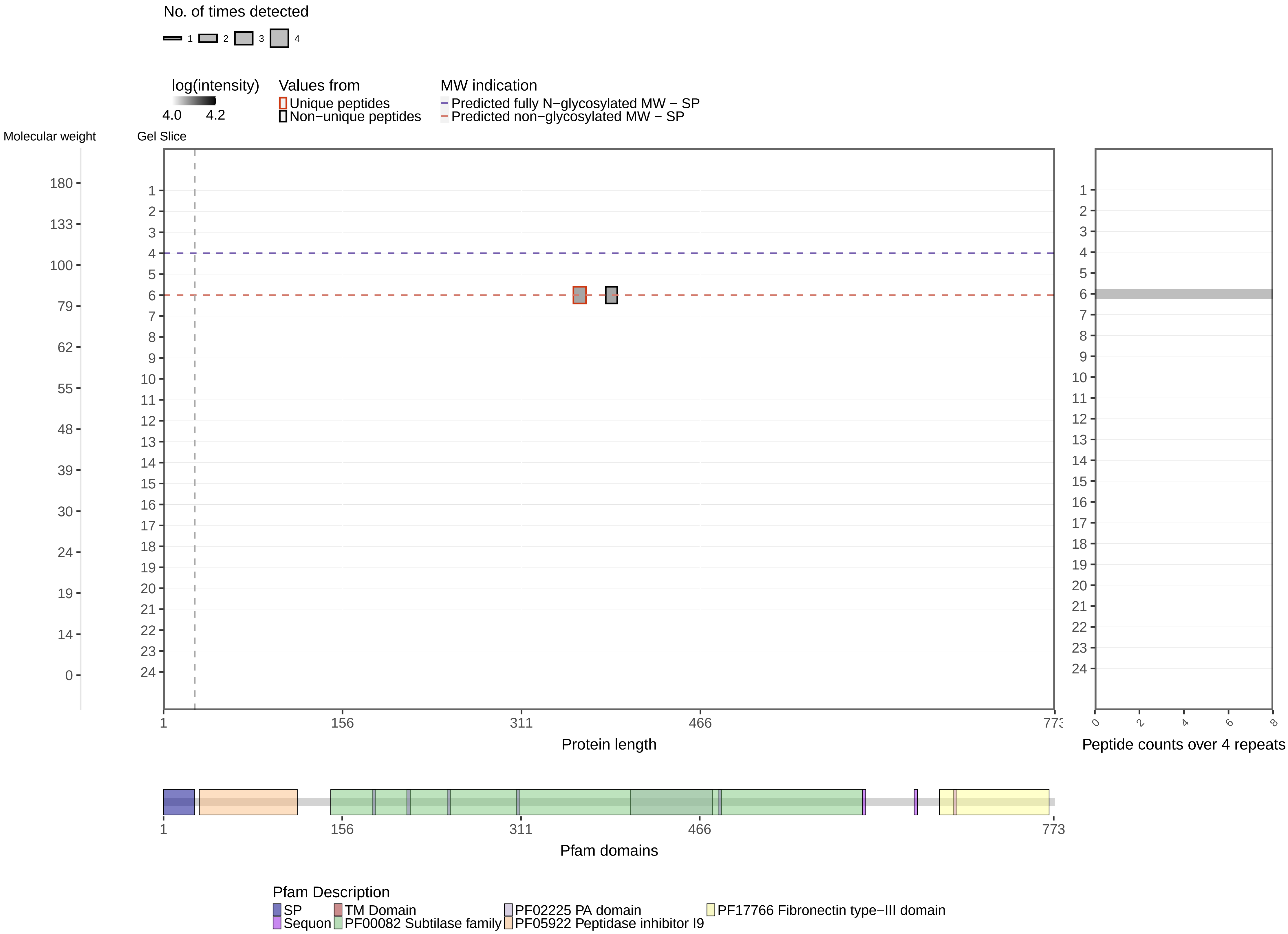


Protein sequence:

YKGESLSVETLPKGKFFPIINAASAKAPHASTDDARLCKAGALDPKKVKGTILVCLRGDNARVDKGQQAALAGAVGMVLANDYASGNEIIADPHVLPATQISYTDGLELFAYVNSTSTPTASITHPKTQLGTKPAPVMAAFSSIGPNTVTPEFLKPDITAPGVSIIAAYTGAEGPTNQDFDKRRVKFNSVSGTSMSCPHVSGIVGLLKTLYPTWSPSAIKSAIMTTARTRDNAIEPVLNASYIKTSPFAYGAGHVRPNRAMDPGLVYDLTIDDYLSFLYTQGYNETQIKTFTEGPFKCA

### NbL15g22480.1 peroxisomal_and_mitochondrial_division_factor_2−like [MW = 37.63 kDa]

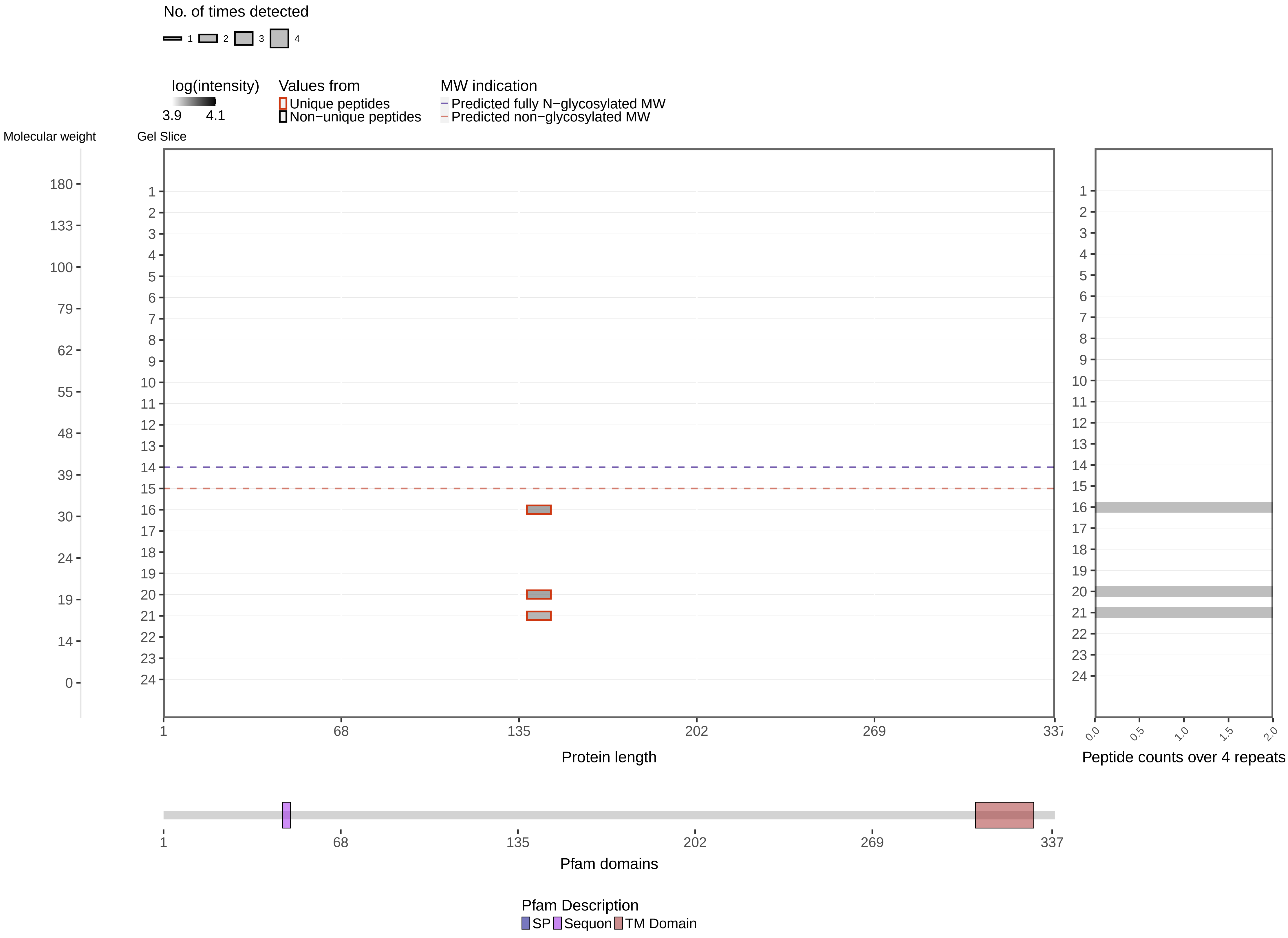


Protein sequence:

NDLHGSNSELSGVKSTLEGLKSSENEKTMKLDAIESERNLLLSKLEKLEASGNNQRGEVEVKEEEIRGLKKHIEELEGTVVNNEEWEKEKKELHMVKEELEKRVKEMISRAAELEKKLEEKEKVITERLVASNINGIPVGDFDKVGYFGAEVNLPVVAASSVVAVAVIGVVCYLKYGRKA*

NbL15g22260.1 probable_LRR_receptor−like_serine/threonine−protein_kinase_At1g07650 [MW − SP = 109.37 kDa]

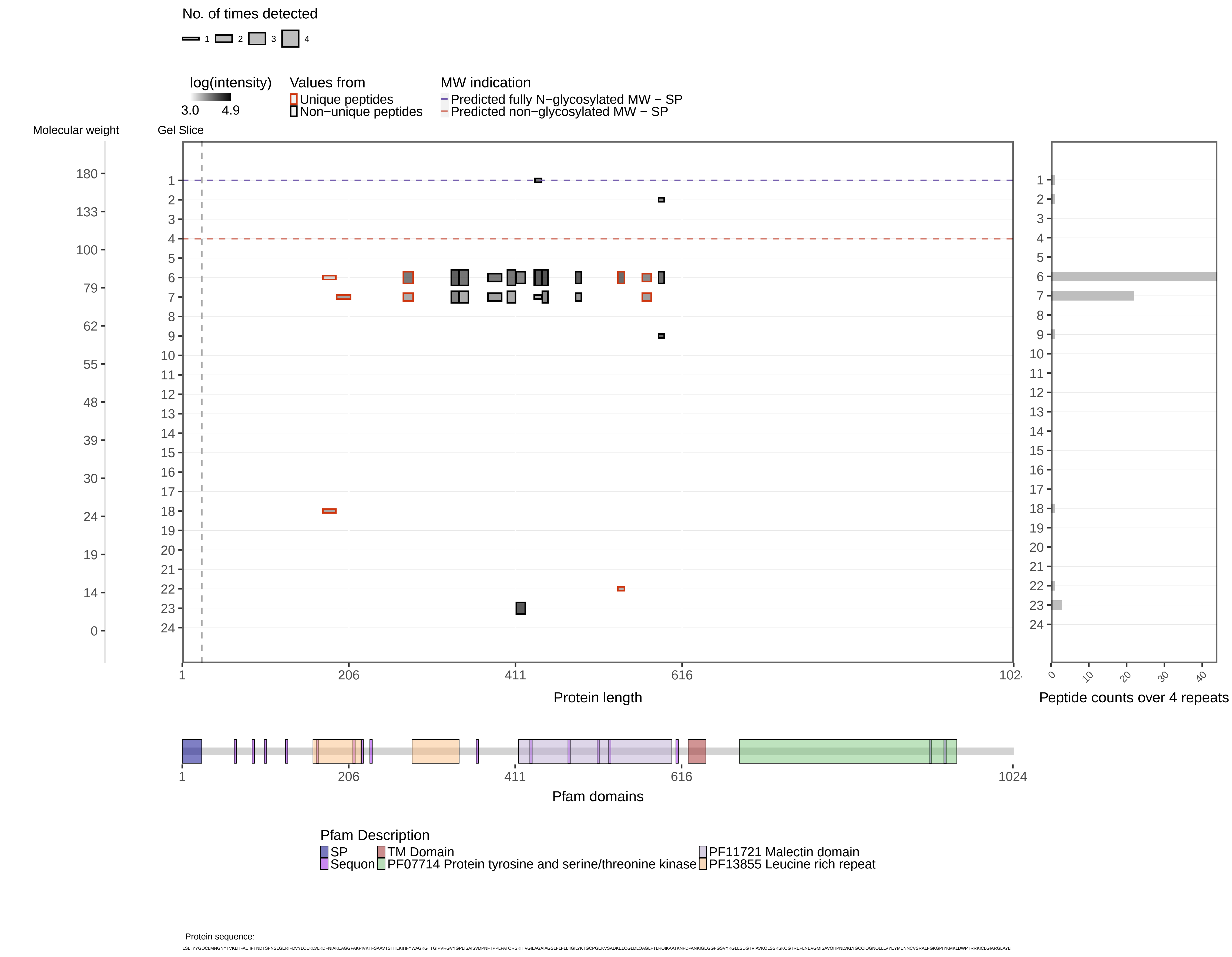

NbL15g22250.1 protein_STRICTOSIDINE_SYNTHASE−LIKE_10−like [MW − SP = 35.46 kDa]

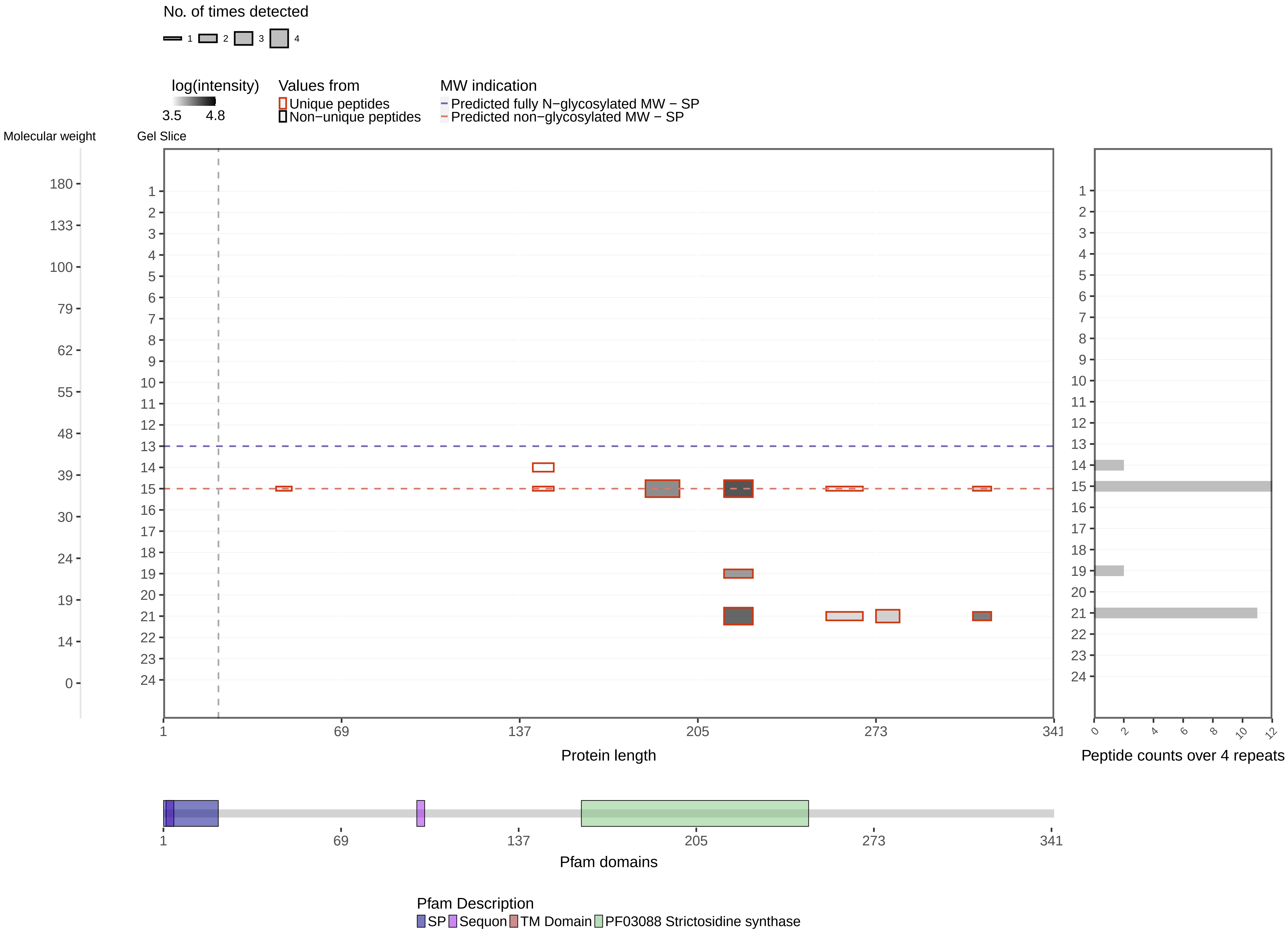


Protein sequence:

MFTNDIDIDNQNNVIYFTDSSTKYQRREFFAAFENGDATGRLMKYDKSTKKVTVLLRGLAFANGVALNKDRSFLLVSETANFRIIRYWIKGQYAGRHDIFADVPGFPDNIRINSRGEFWVALNAKTSATAIKLSTEGRMLEVLEDEEGKTLRDISQVEEKFGKLWFGSVTMPFLGVYGLFSSL*

### NbL15g22220.1 probable_glutathione_S−transferase [MW = 25.6 kDa]

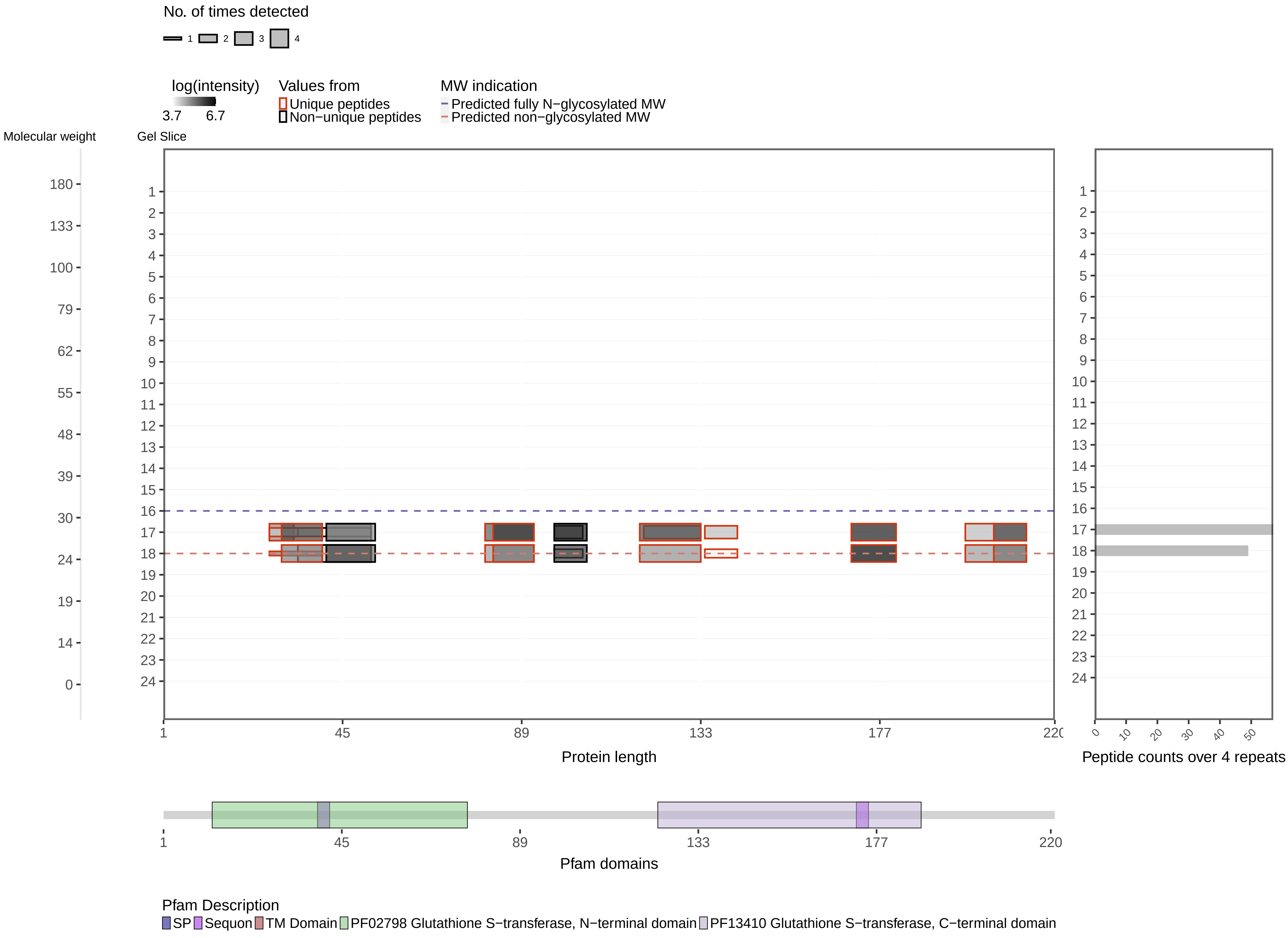


Protein sequence:

VADYIDKKIYDAGRKIWTTKKEDQEAANKEFIECWKLLEGELGDKPYFGGESFGFVDMALIPYYCWFPTYEKFGNFSIEAECPKIVAWAKRCMQKESVSKSLADPDKVYDFVVMLRQRFGIV*

### NbL15g22010.1 heme−binding_protein_2−like [MW = 28.61 kDa]

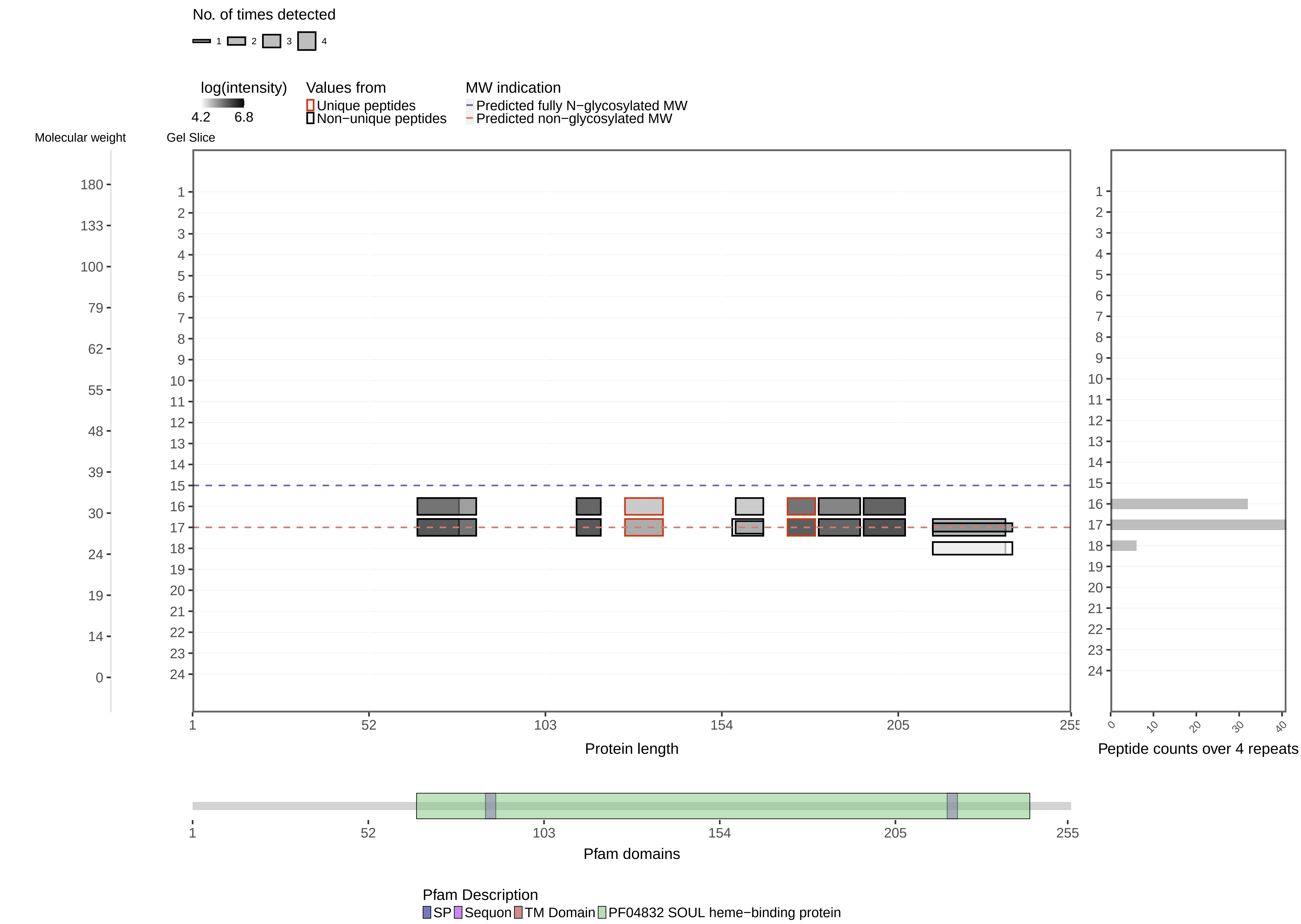


Protein sequence:

IQGKNGYQEKIEMTAPVITEVKPSDGPFCASSFVVSFYVPKKNQPNPPPAKGLHVQKWSNTYVAVRQFGGFVADVDVAKEAAALSASIADTKWAAAVEKSHAADNTTMYTVAGYNSPFEFKDRVNEIWFTFDLEKASAI*

### NbL15g21940.1 persulfide_dioxygenase_ETHE1_homolog_mitochondrial−like [MW = 32.36 kDa]

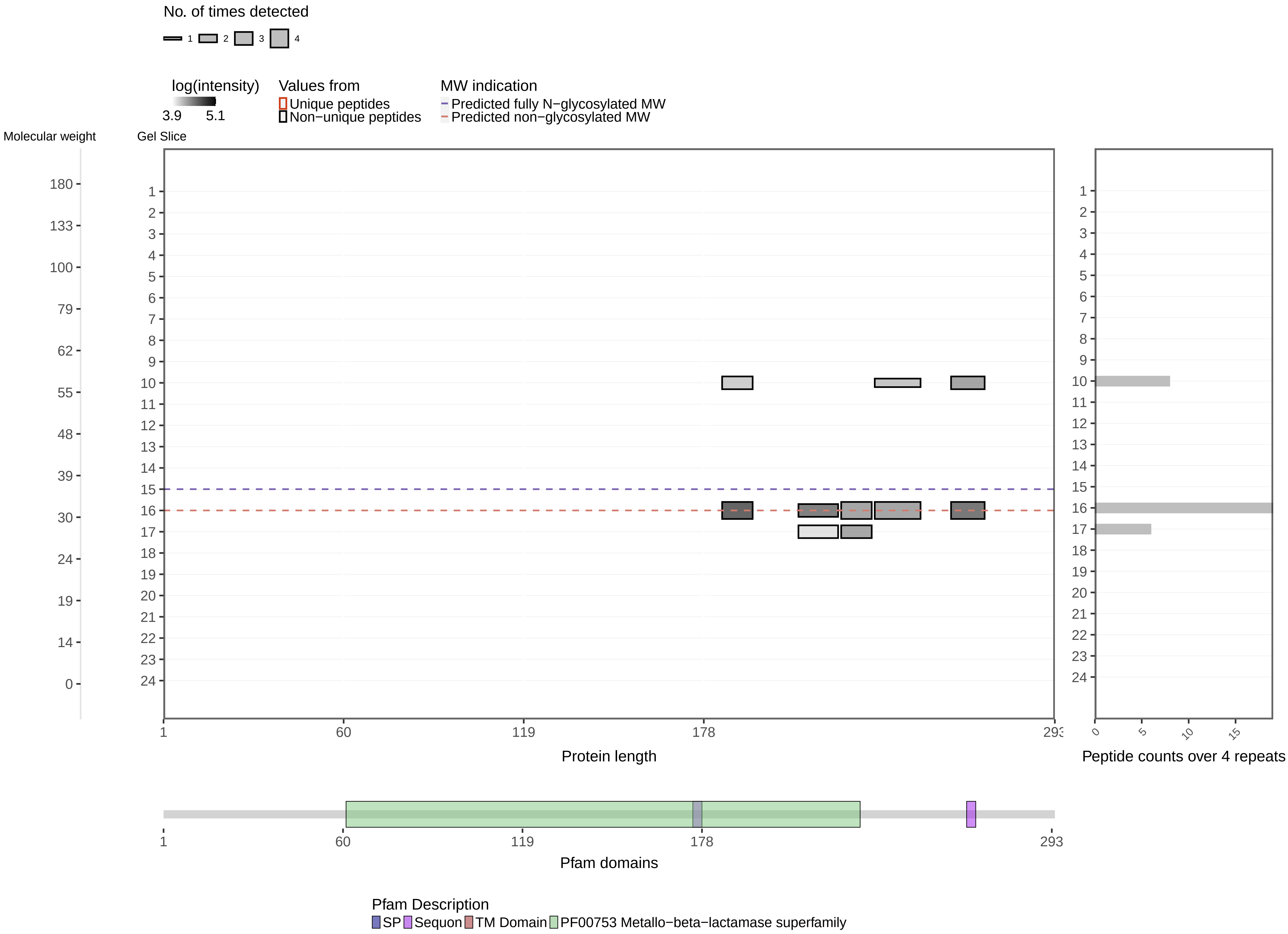


Protein sequence:

AKADLLVEPGDKIHFGDIFLEVRATPGHTLGCLTYITGNGSNQPHPRMAFTGDALLIRGCGRTDFQGGNSDKLYDSVHSQIFTLPKDTLVYPAHDYKGFTVSTVGEEMQYNPRLSKDKETFKNIMQNLNLSYPKMMDVAVPANMVCGLQETKSEAC*

### NbL15g21900.1 solanesyl−diphosphate_synthase_3_chloroplastic [MW = 44.01 kDa]

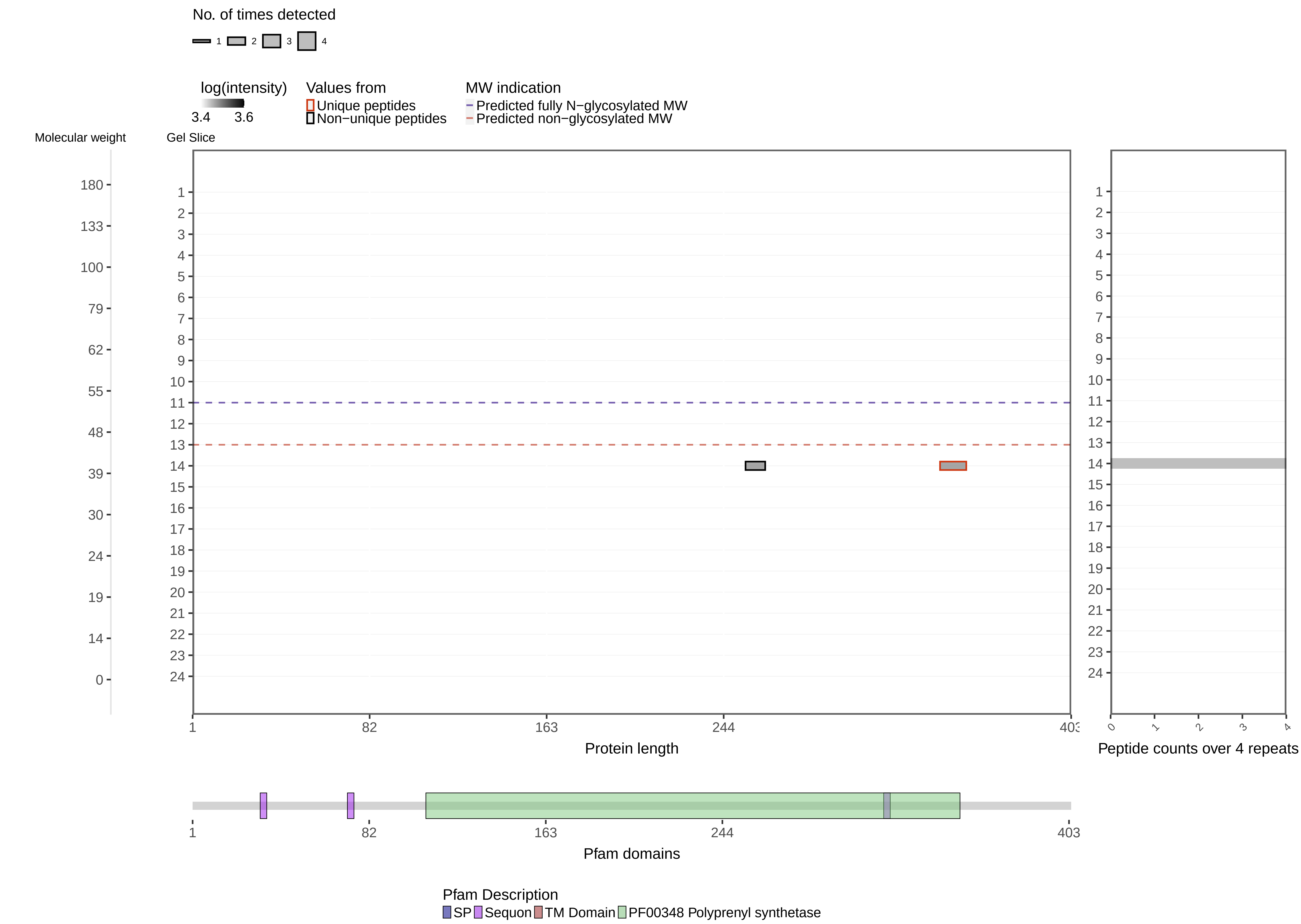


Protein sequence:

TRVAVLAGDFMFAQSSWYLANLENLEVIKLISQVIKDFASGEIKQASNLFDCDVELEEYLLKSYYKTASLIAASTKGAAIFSGVDRDISEQMFQYGKNLGLSFQVVDDILDFTQSAEQLGKPAGSDLAKGNLTAPVLFALEKEPKLRNIIESEFRDADSLEEAINLVKTTGGIQRAQDLAKEKADLALQSLKCLPSSPFRAALEEIVKYNLERIE*

NbL15g21850.1 peptide_methionine_sulfoxide_reductase_B1_chloroplastic−like [MW = 25.75 kDa]

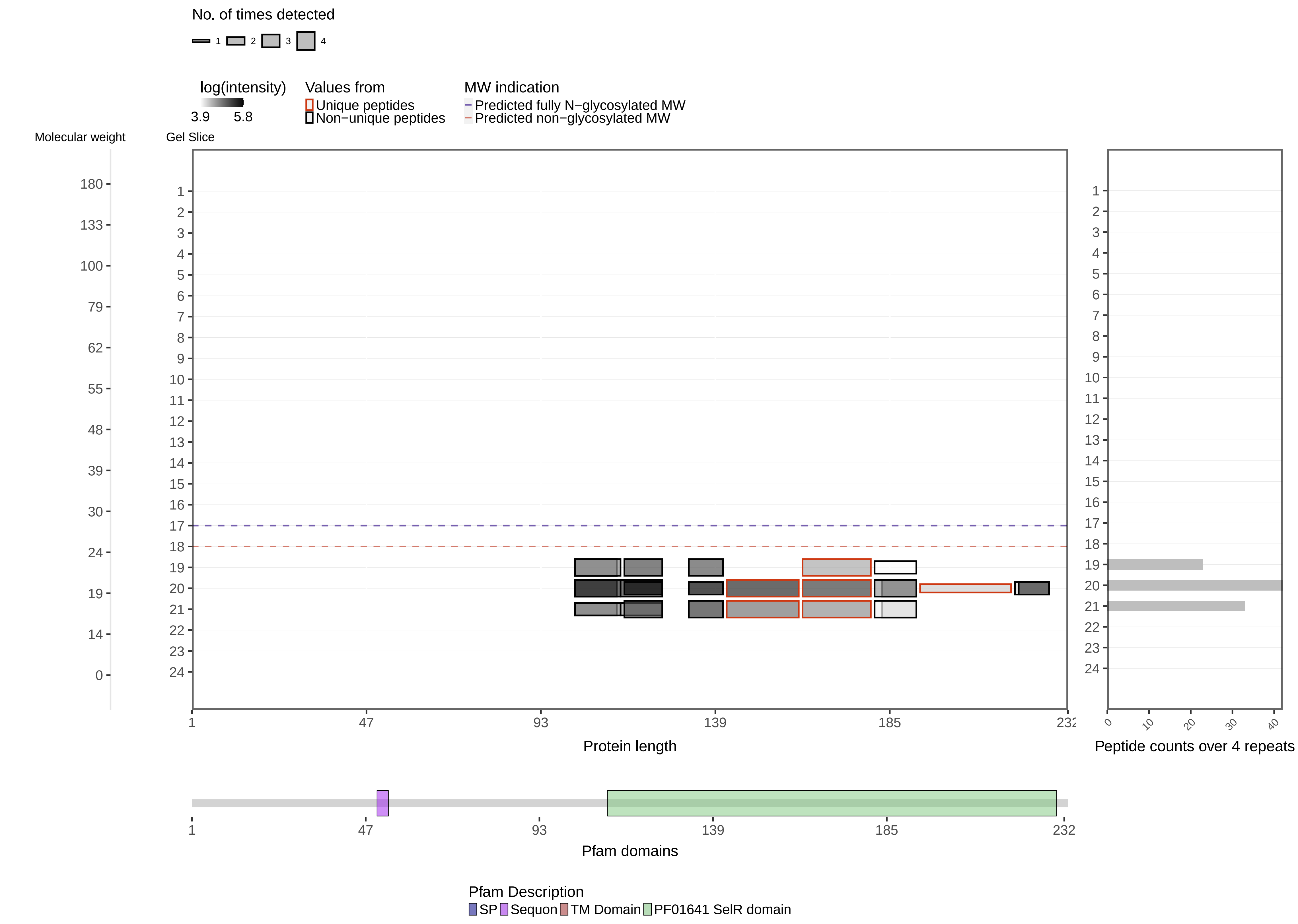


Protein sequence:

SSVSDEEWRKQLTNEQFYITRQKGTERAFTGEYWNTKTPGTYHCVCCDTPLFESSTKFDSGTGWPSYYQPVDNNVKSKMDLSIIFMPRQEVVCAVCDAHLGHVFDDGPPPTGKRYCINSAALKLKPK*

### NbL15g21780.1 probable_rhamnose_biosynthetic_enzyme_1 [MW = 75.11 kDa]

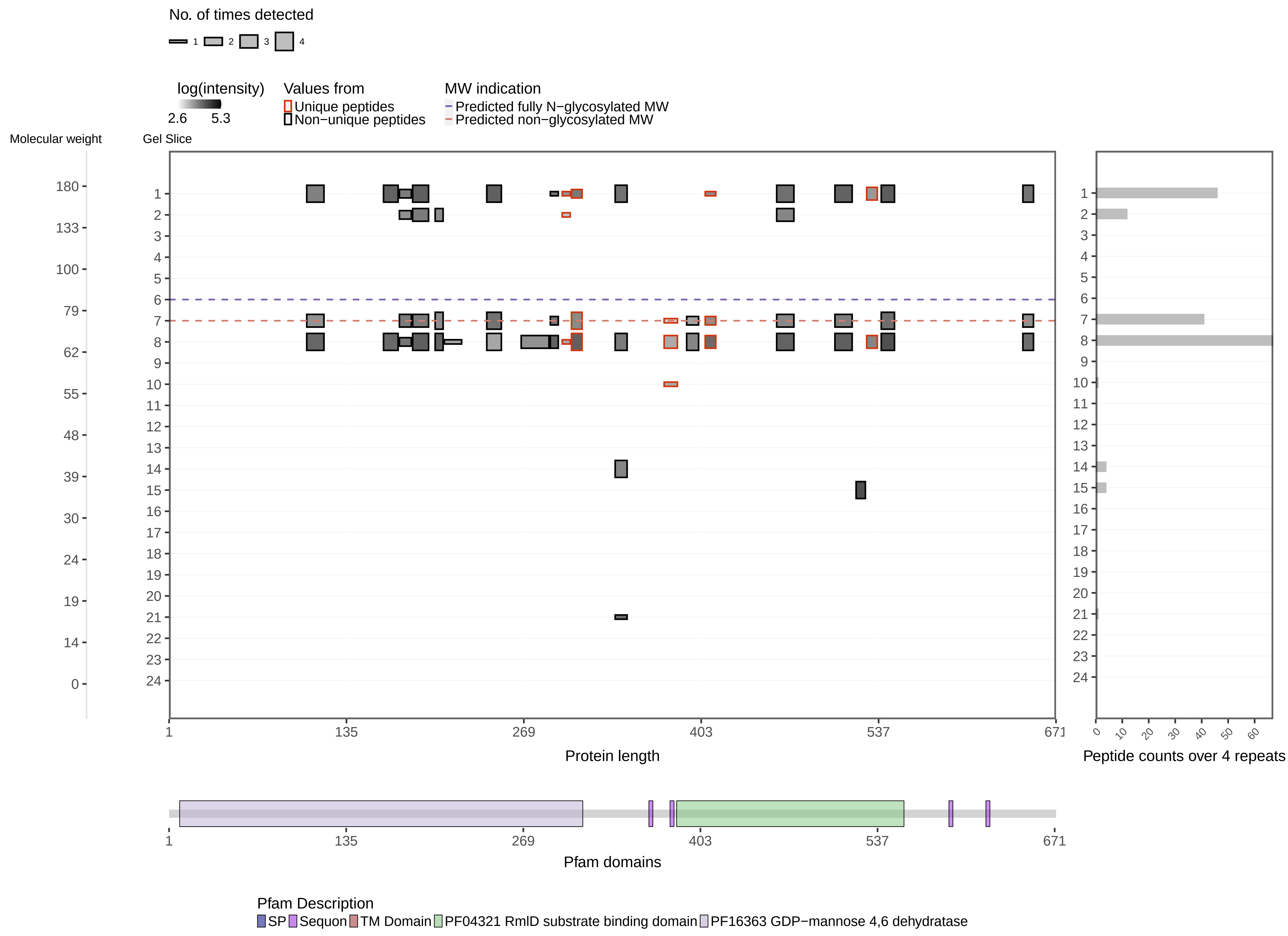


Protein sequence:

WWGDVSGALLPHPRMLMMPGGVERGAEKDDSGSSECSGNPNQTRMAVPASKIGNPPNKSPYKFLIYGRTGWIGGLLGKLCEKQGIPYDYGRGRLQDRSQLLADIHAVKPTHVFNAAGVTGRPNVDWCESHKTETIRTNVAGTLNLADVCKEHGLLMMNFATGCIFEYDAAHPEGSGIGFKEEDTPNFAGSFYSKTKAMVEELLKECENVCTLRVRMPISSDLSNPRNFITKISRYNKVVNIPNSMTILDELLPISIEMAKRNLKGIWNFTNPGVVSHNEILEMYKKYINPDF

NbL15g21490.1 secreted_glycoprotein_EP1−like [MW − SP = 31.36 kDa]

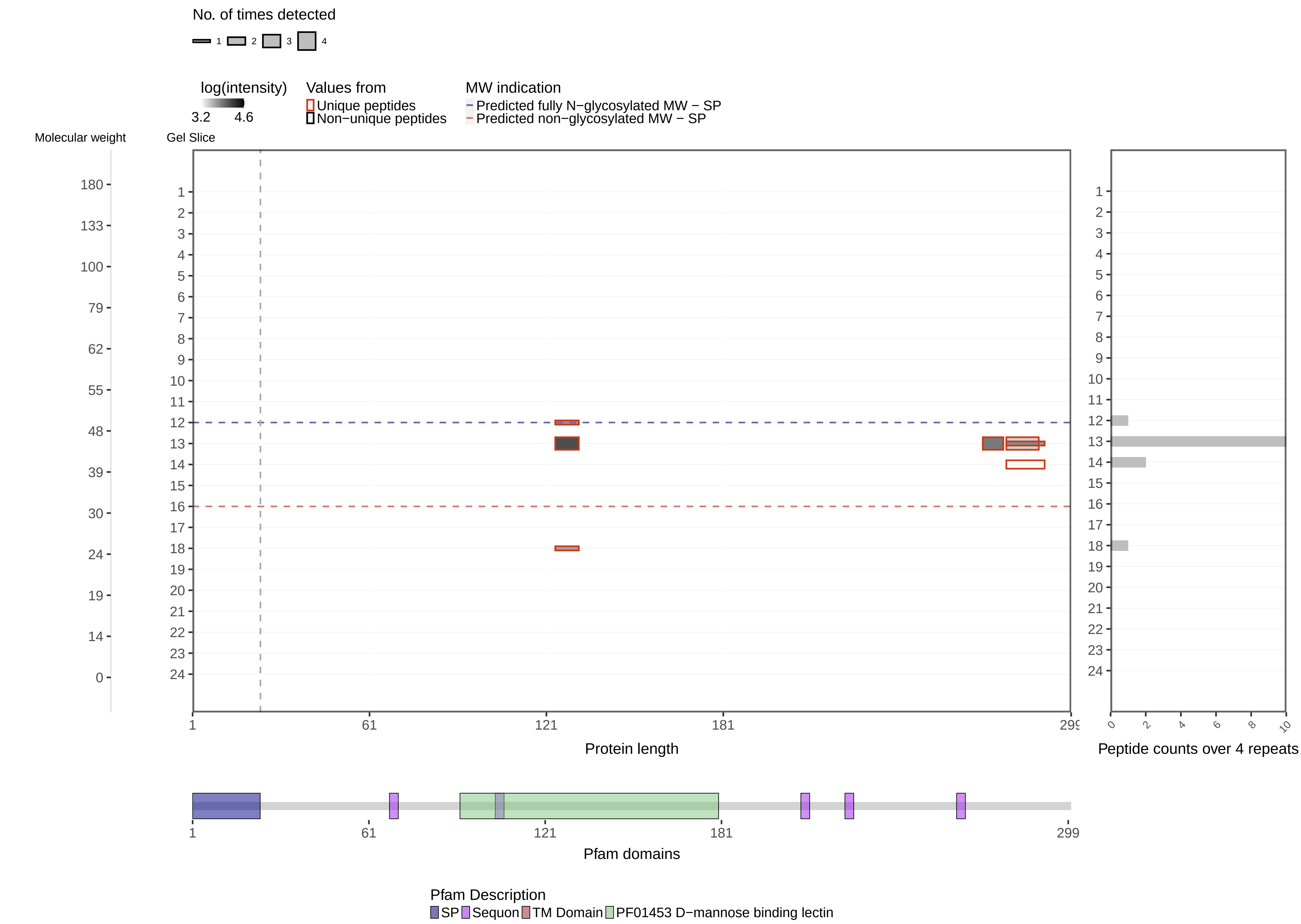


Protein sequence:

SGFKLLPNGNFVLHDSKGKFIWQSFDHPTDTLLVGQSLRLSGPNKLVSRASVKKNANGPYSLEVKPKIFGIYNKTKLHVELAWFDFKNSTLESVKLNSGNQRVKLDYRLAKSTTRNSHVMAFTEINTTLTYLRLEIDGNLRAYTFFEDKEEDRFRWRVTYSML*

### NbL15g21470.1 phosphoenolpyruvate_carboxylase−like [MW = 110.32 kDa]

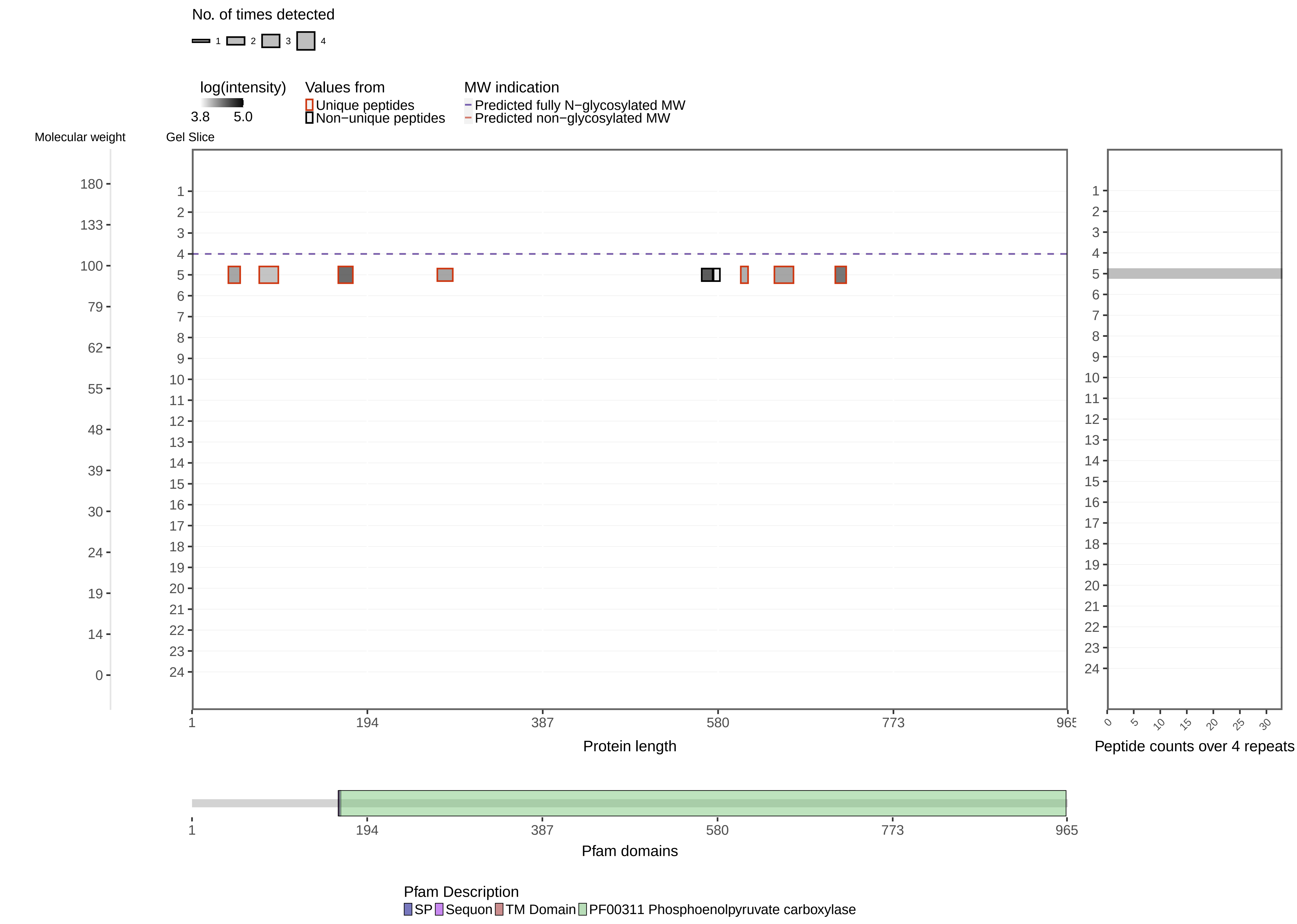


Protein sequence:

[illegible]

### NbL15g21320.1 uncharacterized_exonuclease_domain−containing_protein_At3g15140−like [MW = 36.71 kDa]

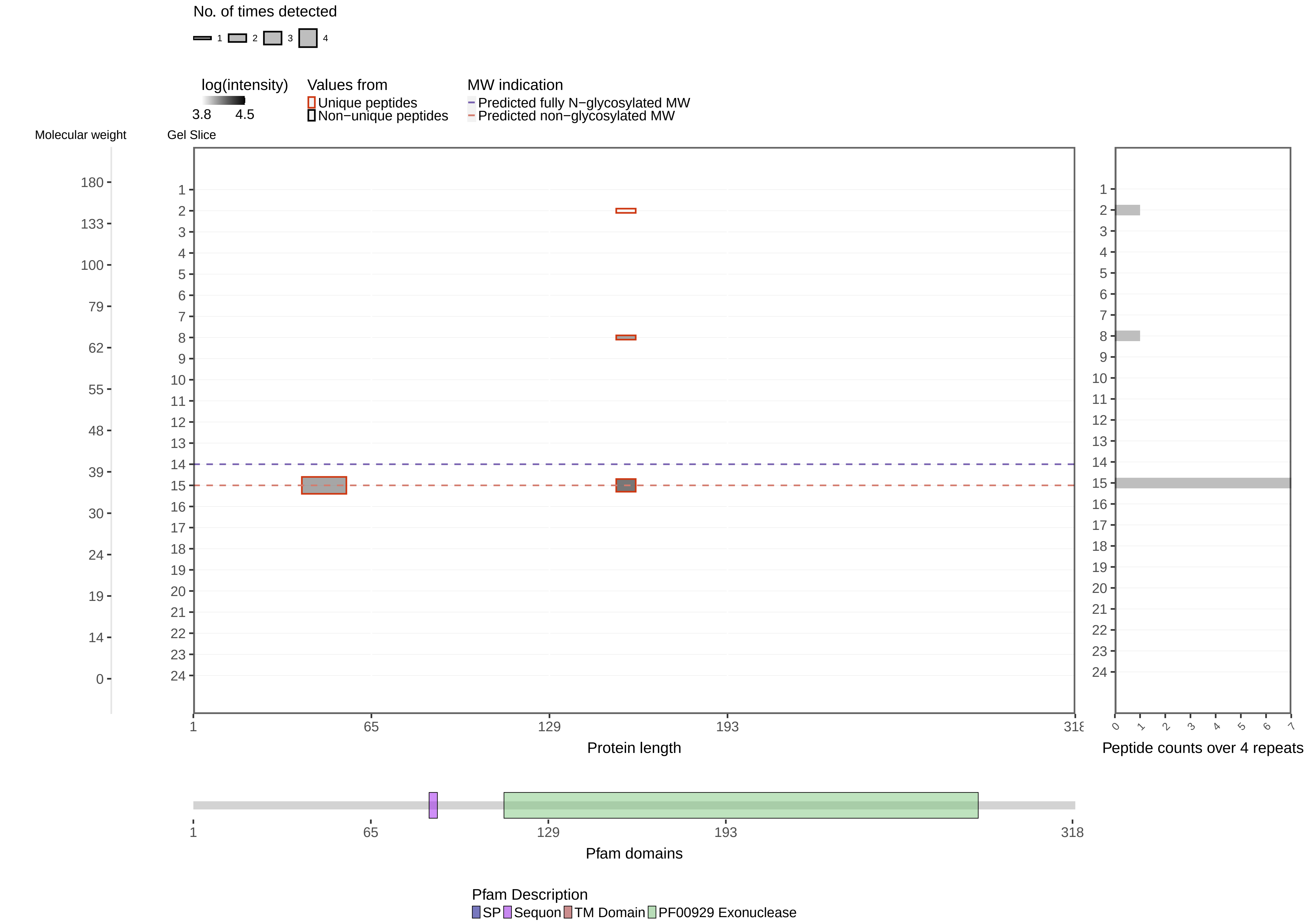


Protein sequence:

MHEDRINEYIEGKYGKLGVDRVWHDTAIPFGEVIEQFEVWLGERQLWRNEPGGCLNKAAFVTCGNWDLKTKVPQQCKVAGTKLPPYFMEWINLKDVFLNFYKRRAKGMLSMMRELQMPLLGSHHLGIDDAKNIARVLQHMLSDGALVQITARRNPHSPEKVEYLFEDRIV*

NbL15g21250.1 30S_ribosomal_protein_S20_chloroplastic−like [MW = 20.44 kDa]

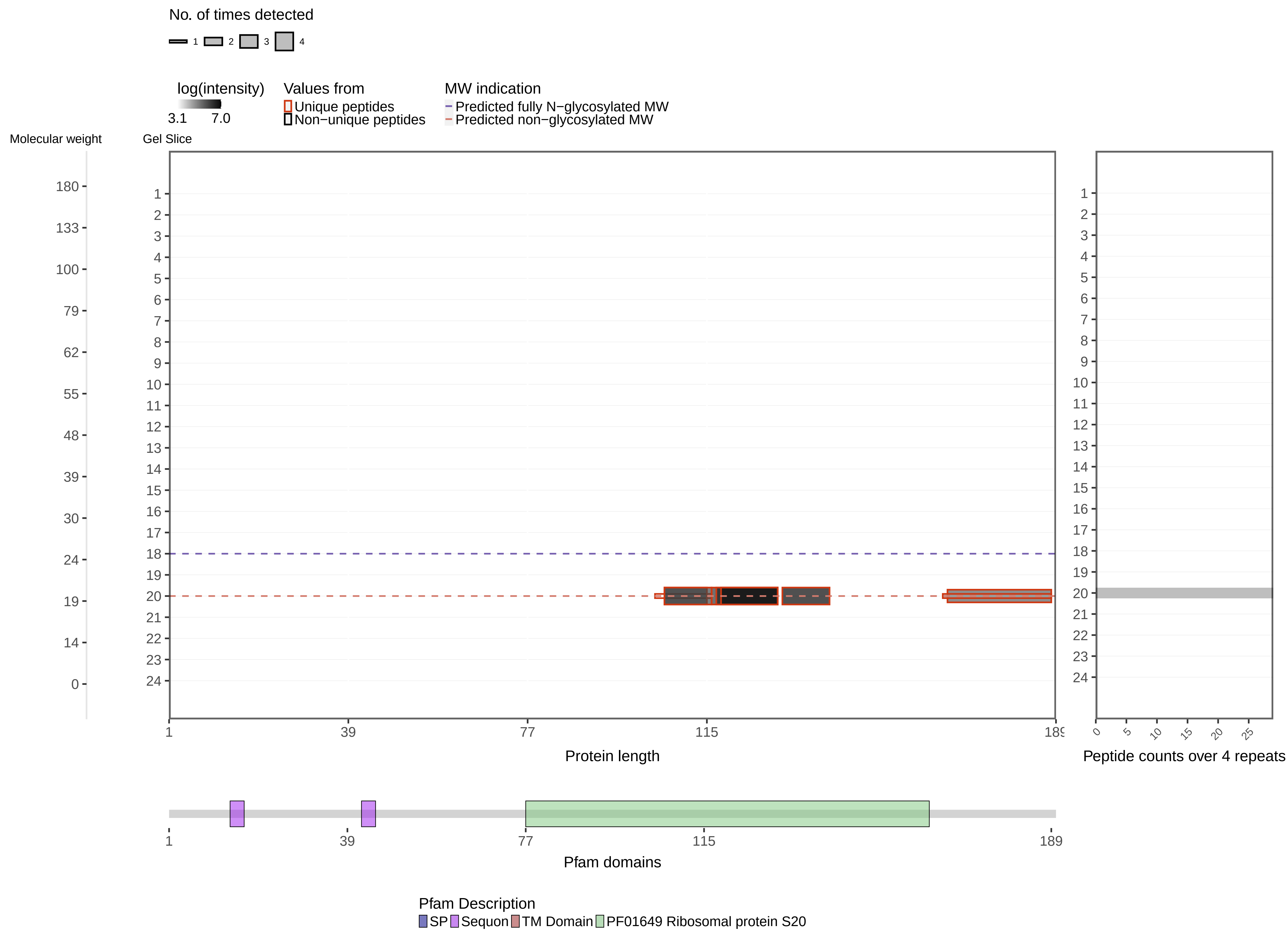


Protein sequence:

KRARQAEKRRLYNKAKKSEVKTRMKTVLEALDTLKKKTDAQSEEVISVEKLIAEAYSAIDKAVKAGSLHRNTGARRKSRLARRKKAVEIHHGWYVPAPVEEPALVATA*

NbL15g21170.1 probable_protein_phosphatase_2C_39 [MW = 31.01 kDa]

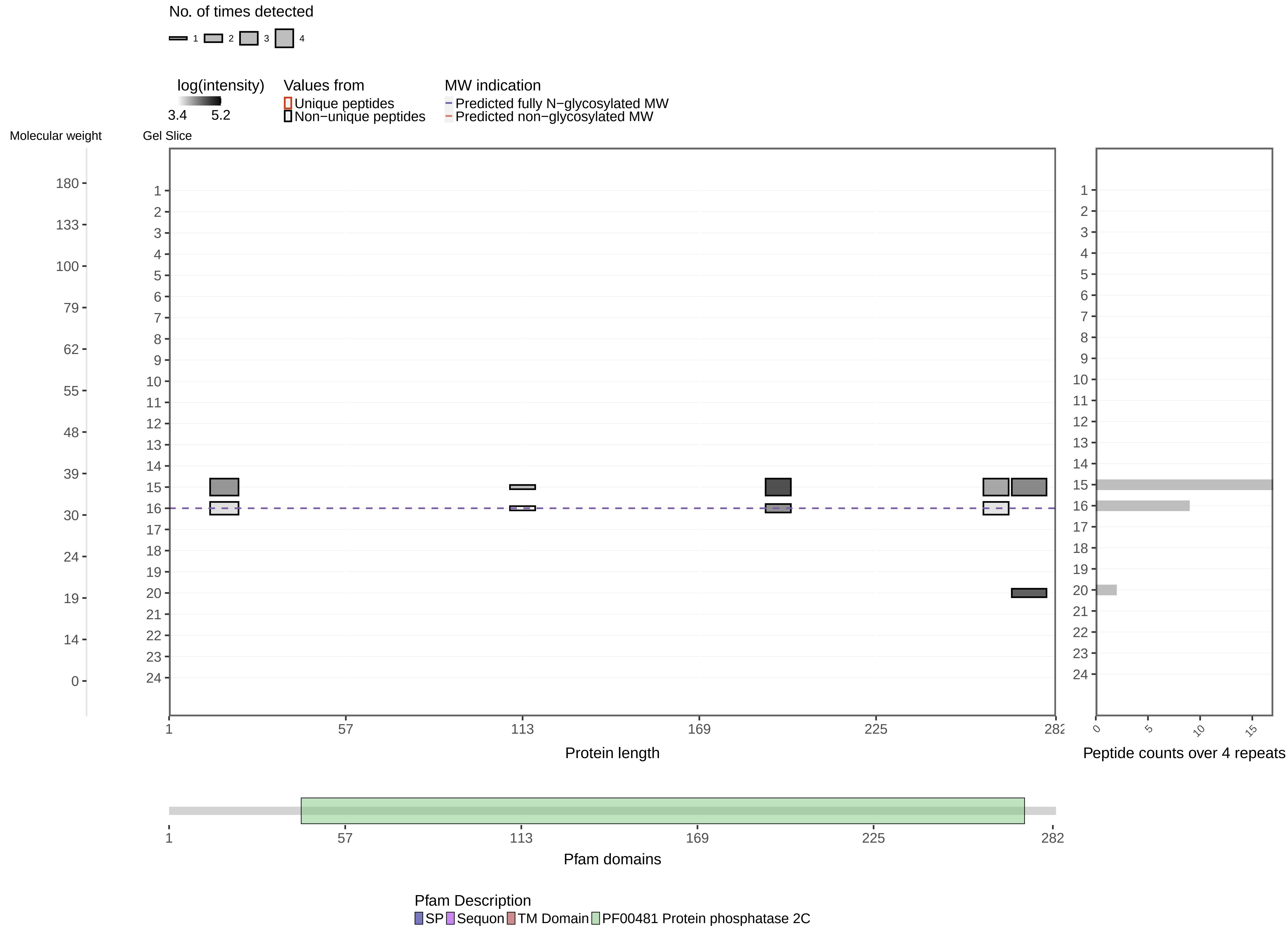


Protein sequence:

AVTAILINGQKLVVANVGDSRAVICENGVAKQLSIDHEPGRERKIIEDRGGFVSNFPGDVPRVDGQLAVARAFGDKSLKMHLSSEPDVMVELIDENTEFLILASDGVWKVLSNQEAADCIKSIKDARSAAKHLNEQALAKGSTDDISCVVVRFQ*

### NbL15g21150.1 3−hydroxybutyryl−CoA_dehydrogenase [MW = 32.05 kDa]

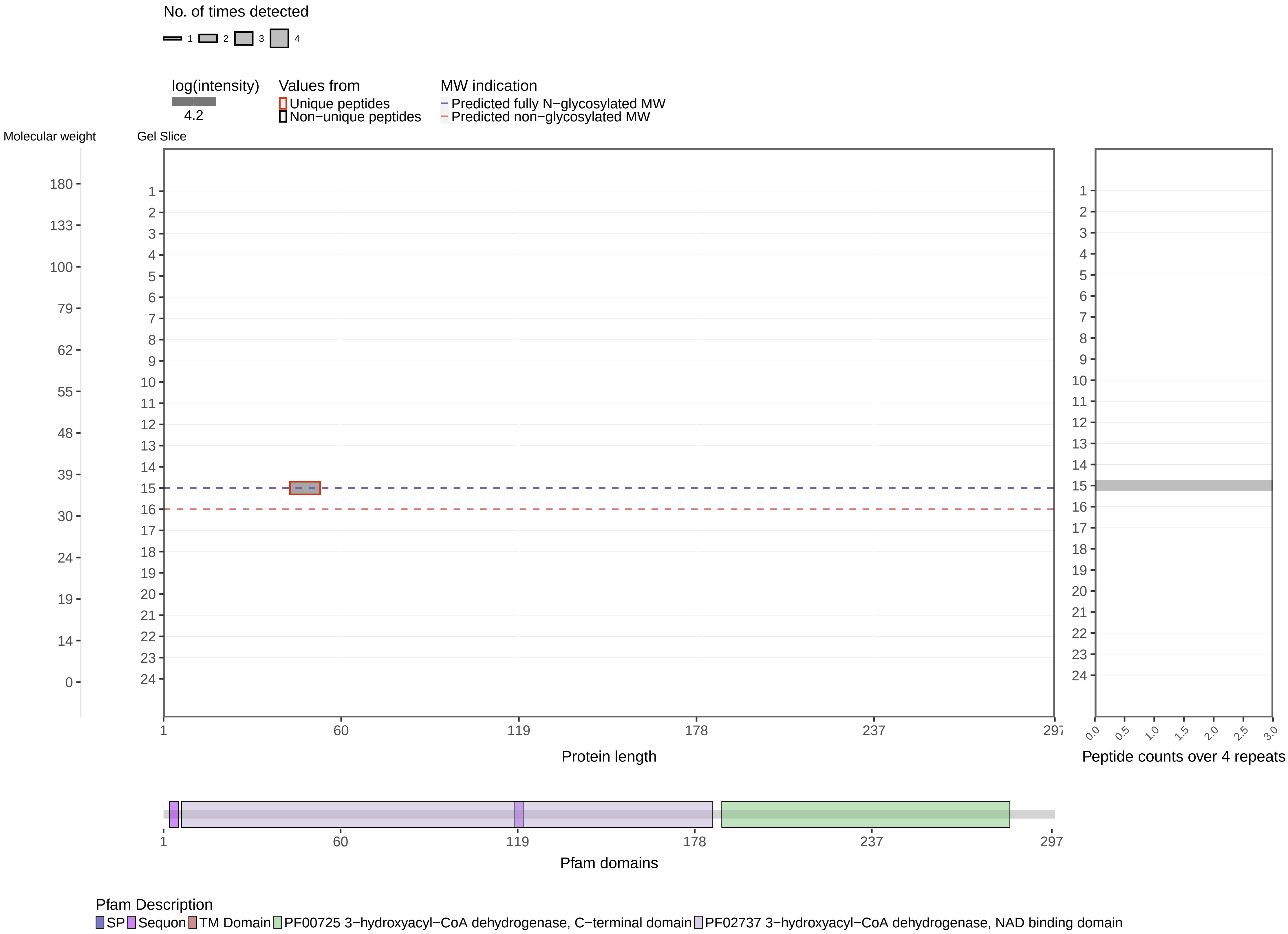


Protein sequence:

VIGMHFMNPPPIMKLVEIIRGADTSEDTYCATKCLAERFGKTVICSODYSGFIVNRILMPMINEAFHTLYTGVATKEDIDTGMKLGTNHPMGPLELADFIGLDVCLSIMKVLHVGLGDDKYAPCPLLVOYVDAGRLGKKRGIGVYDYRSAAADTGKSPRL*

### NbL15g21060.1 proton_pump−interactor_1−like [MW = 71.95 kDa]

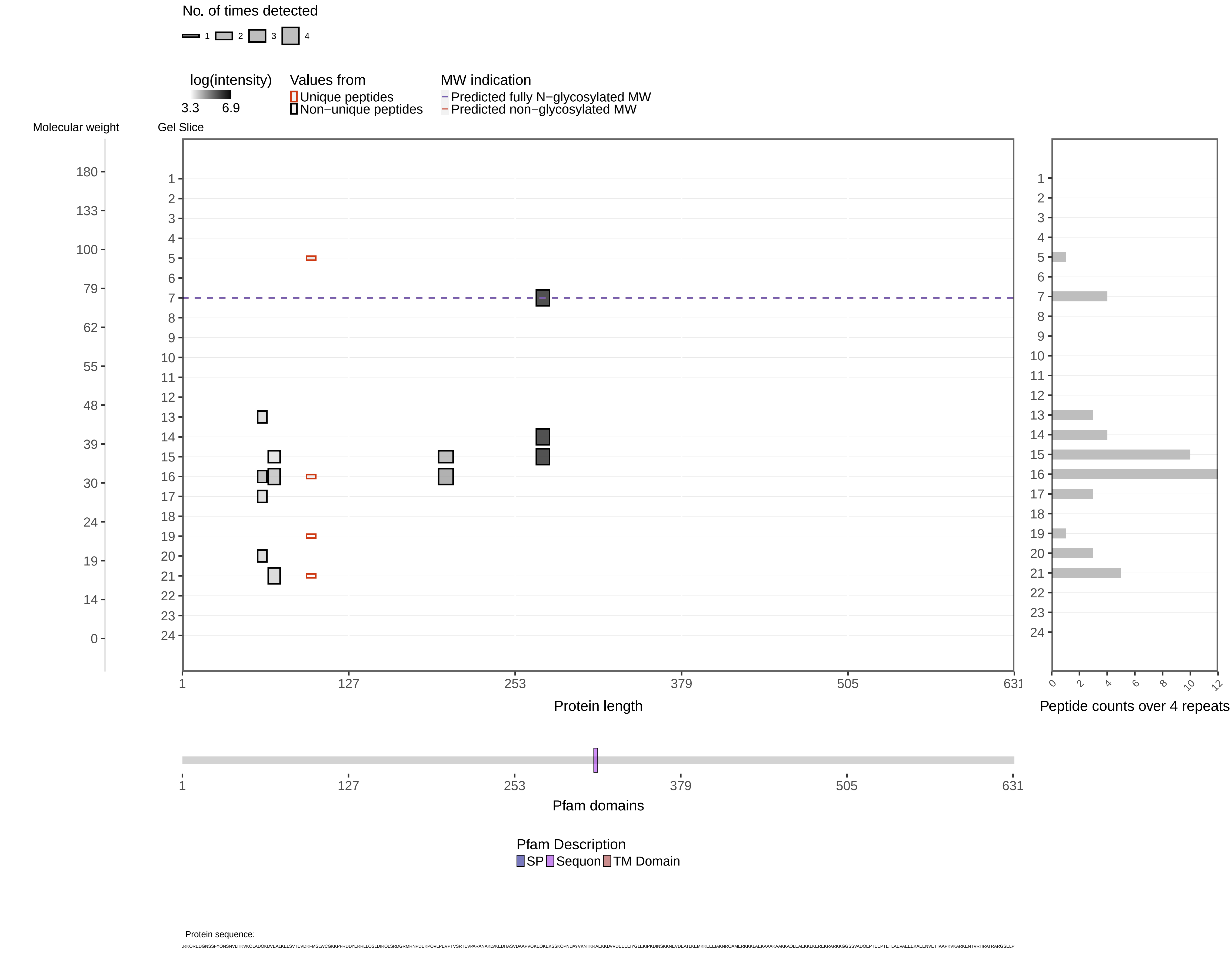

### NbL15g20960.1 thioredoxin−1−like [MW = 19.74 kDa]

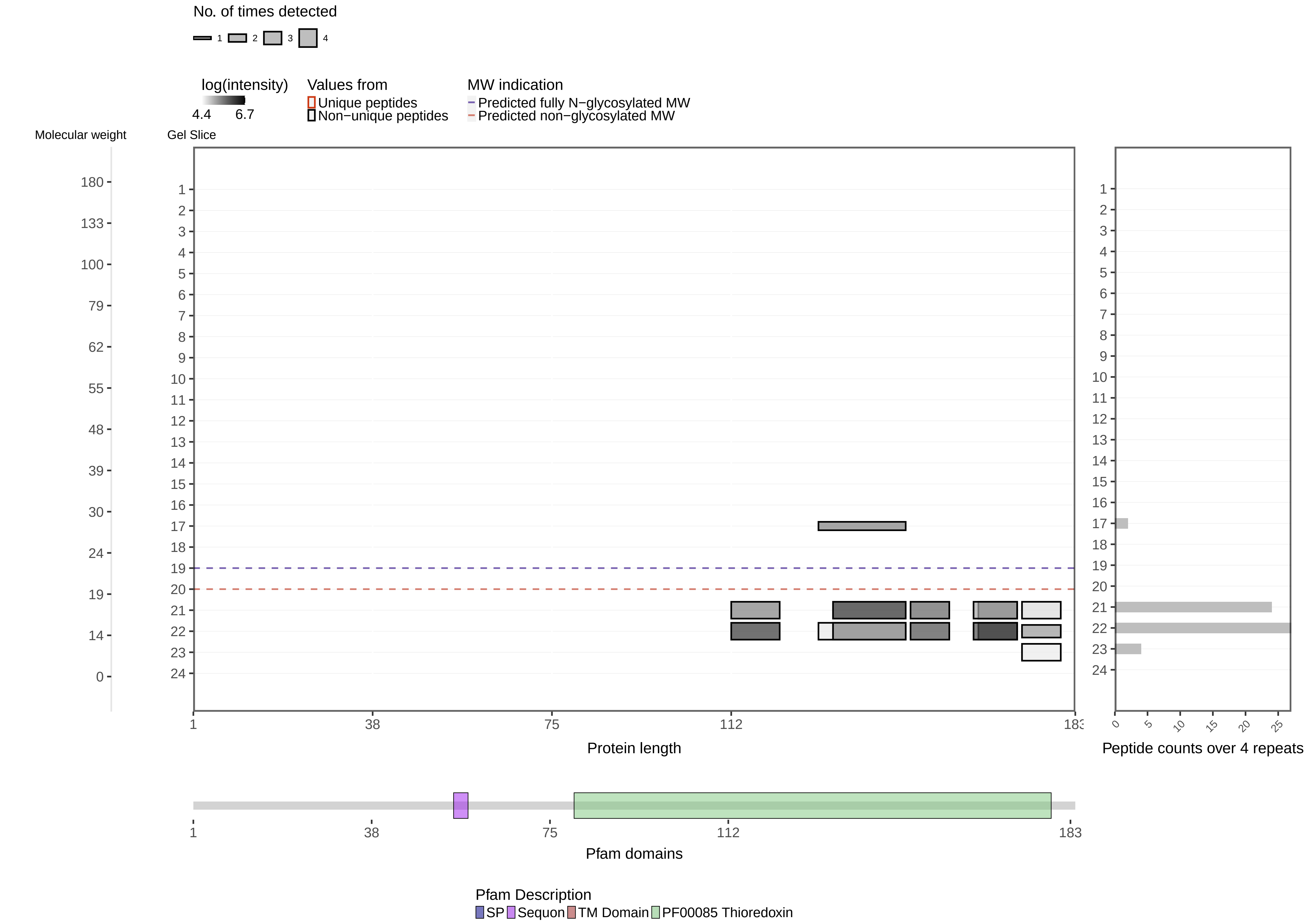


Protein sequence:

.EVGAVNDKTWKSLVVESDIPVLVEFWAPWCGPCRMIHPVIDELAKEYAGKLKFFKLNTDESPSTATELGIRSIPTVMIFKNGEKKDAVIGAVPKSTLTTCIEKFL*

### NbL15g20620.1 uncharacterized_protein_YnbB−like [MW = 48.13 kDa]

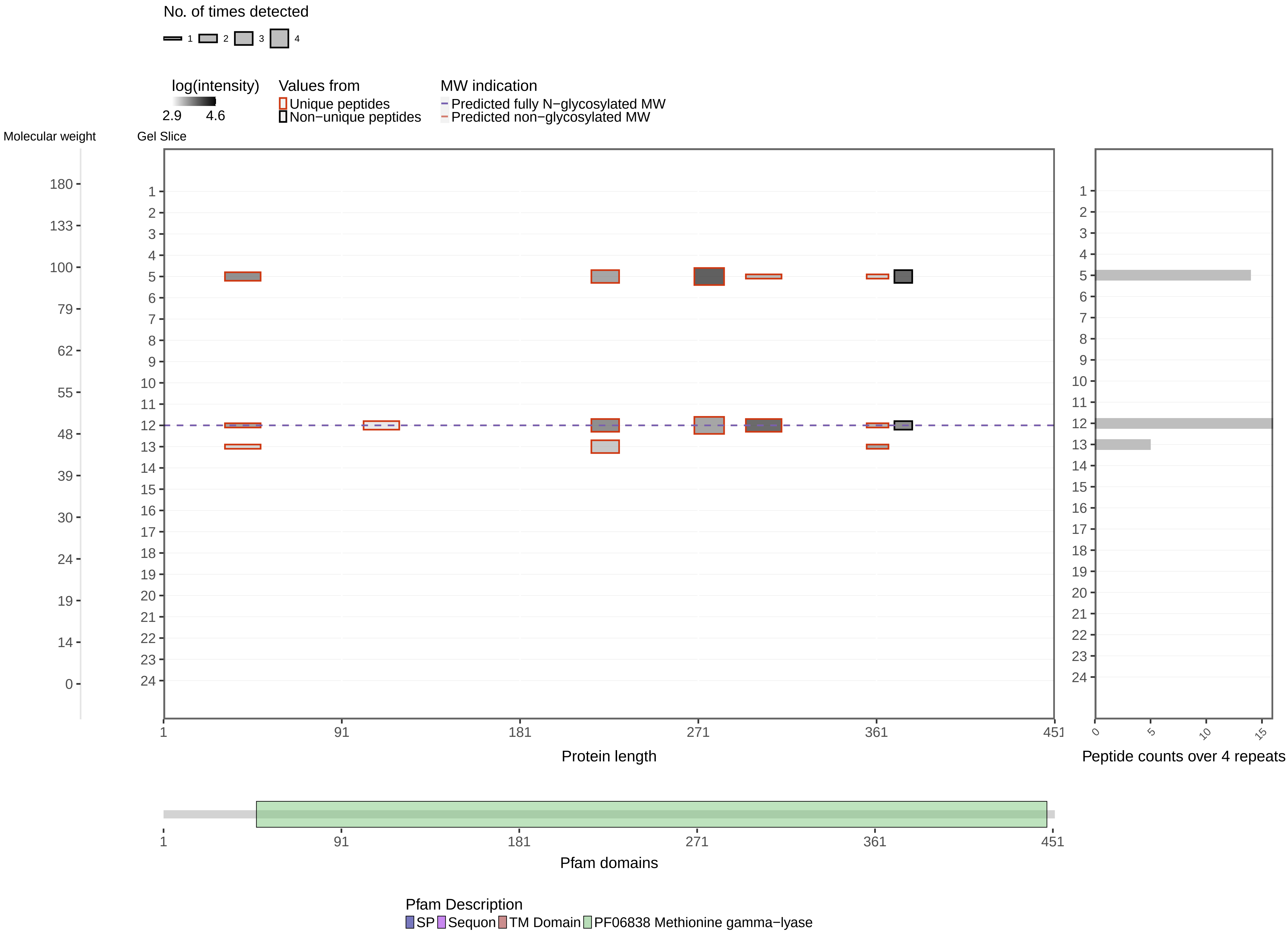


Protein sequence:

SWRRSLSVTEIGQAISIIKMQNPSCLVMVDNCYGEFVDDIEPPMVGADLIAGSLIKNPGGTIAPCGGYVAGRKRWVEAAAARLSAPGLGVDCGSTPGDIMRTFFQGLYLSPQMVGEAIKGSFLIAEVMAARGYKVQPLCRVKRHDTVQAVQLGSRERLLSFCEAVQRSSPVSSFIRPVAGATAGYASEVIFADGTFIDGSTSELSCDGPLREPFSVFCQGGTHWTQWGLVLGEVLKSL*

### NbL15g20500.1 vesicle−associated_protein_4−1−like [MW = 29.39 kDa]

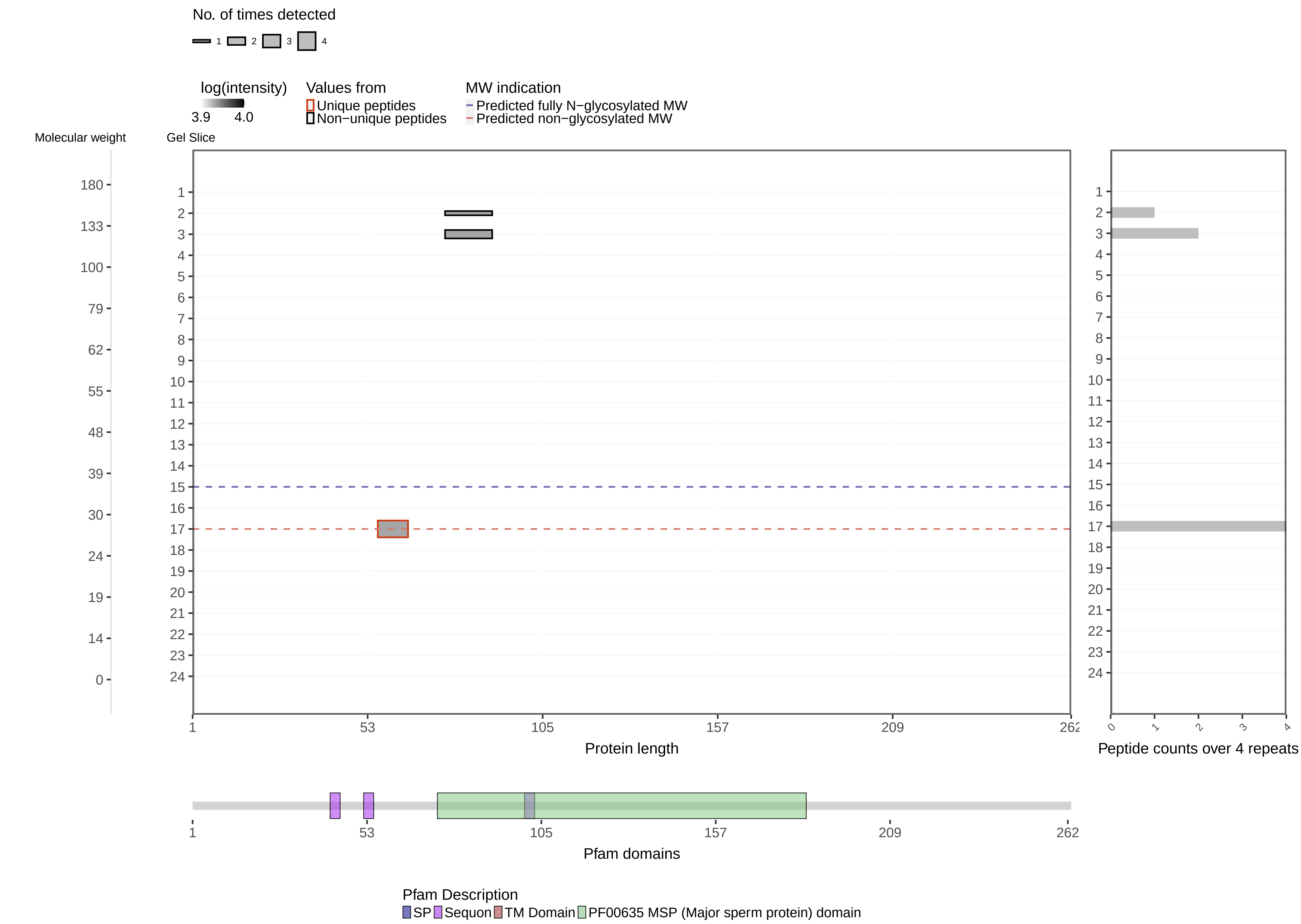


Protein sequence:

SCYMRPPGGILEPGESVIATVFKFVEHPENNEKSVDOKSKVKFKIMSLKVKEGTDYVPELFDEOKDOVTIERVLRVVFLDPERPSPALEKLKROLAEAEAALESRKKPPVETGPKVVGEGLVIDEWKERREKYLAROOVEAVDSV*

### NbL15g20480.1 ubiquitin_carboxyl−terminal_hydrolase_14−like [MW = 88.94 kDa]

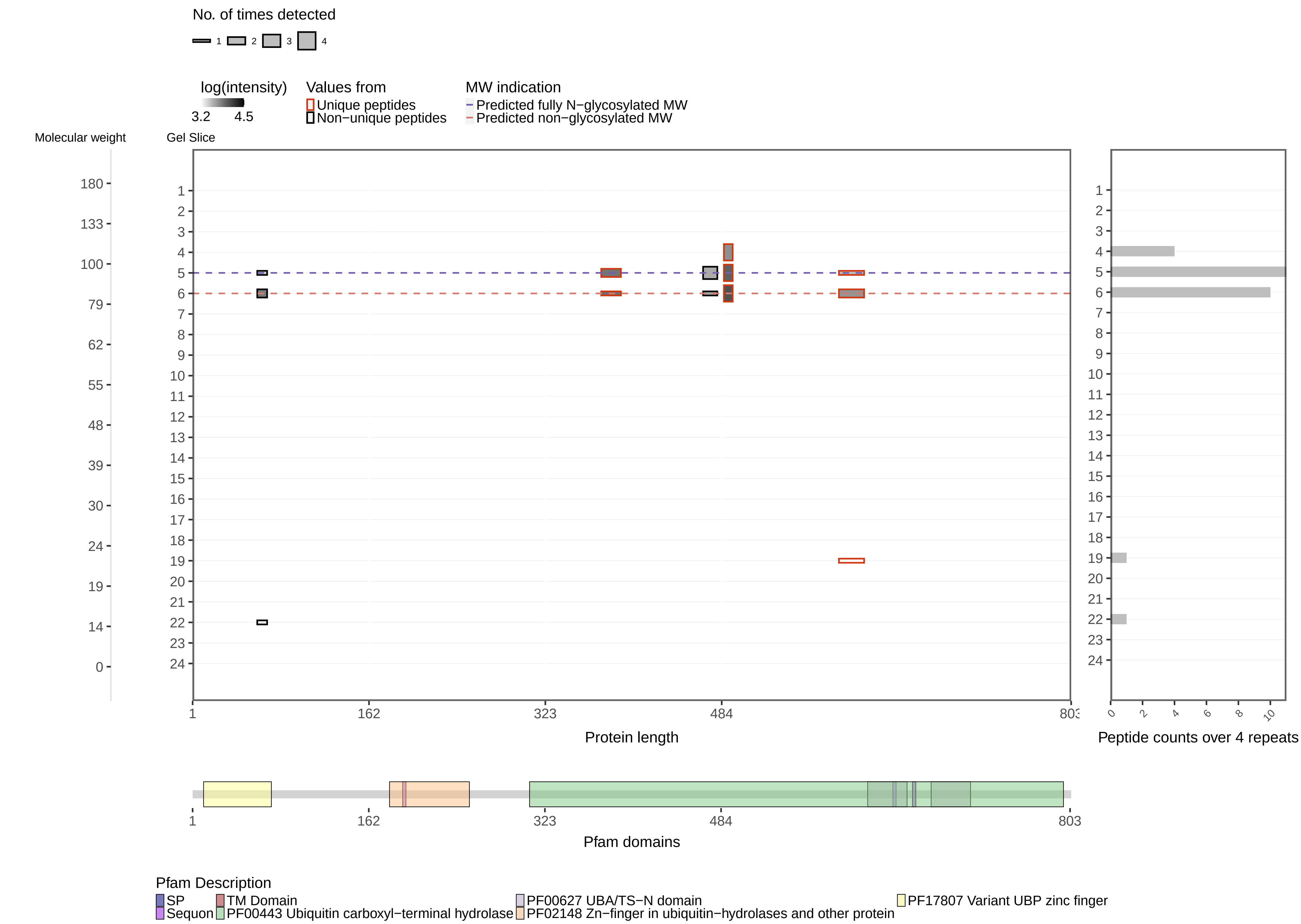


Protein sequence:

NbL15g20400.1 G−type_lectin_S−receptor−like_serine/threonine−protein_kinase_At4g27290 [MW − SP = 58.93 kDa]

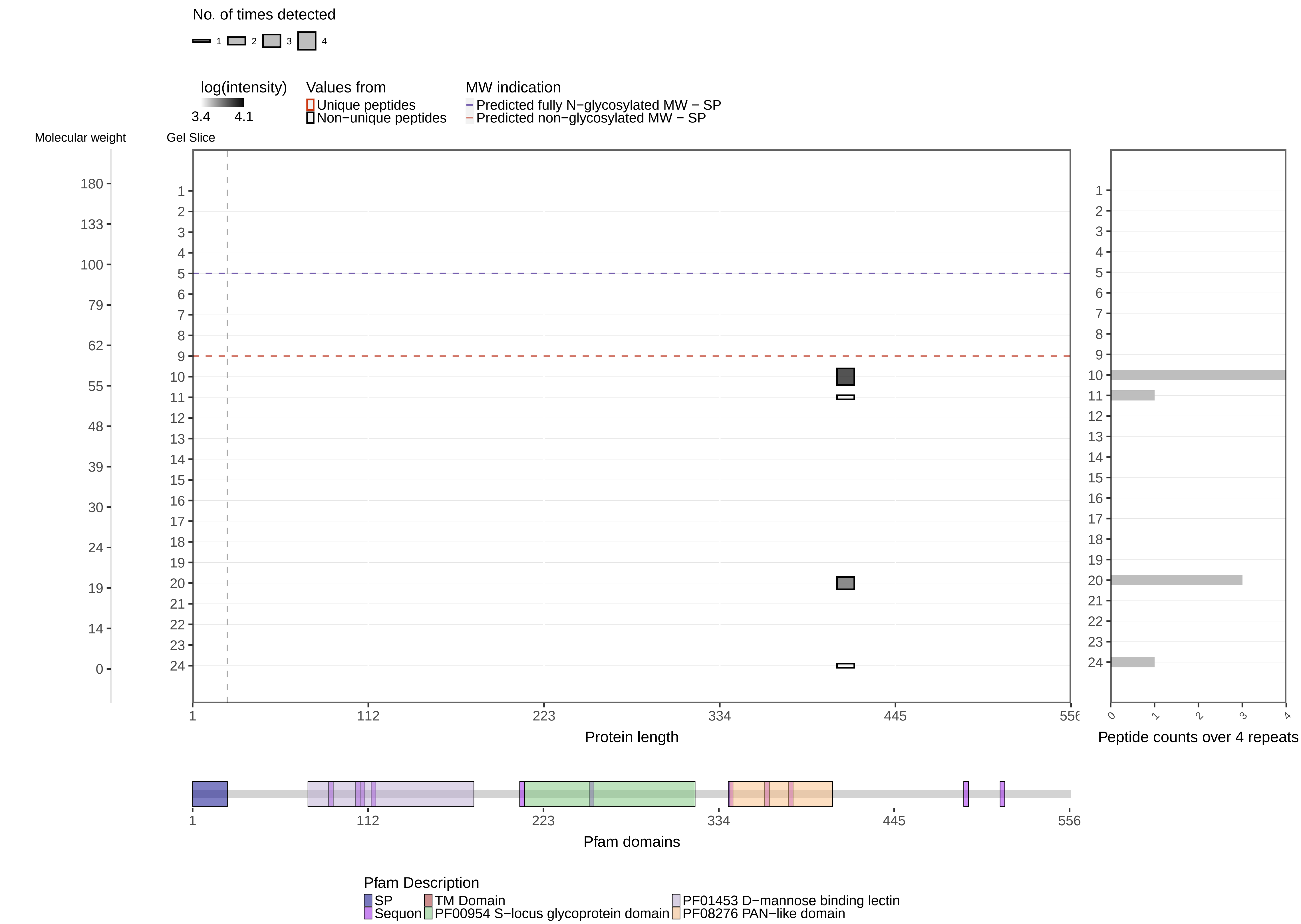


Protein sequence:

GKTMCIIRIEKTQSWVNYHSADADDCDTYKLCGAYSTCNILSDPFCQCLDKFEPKHPDDWIRSDWSSGCVRKRSLNCTADGFLKYSGVKLPDTRFSWFNETMTLDECRAVCLRNCSCMGYTNLDIRNGGSGCLLWIGELVDIRQLSQSGQDIYIRMSASDIDVAGSAGSNGKKAFILAVALPLLVALILLGLGVGLILYKRRREDPVITTKGRLDGHSNKNDNSNQSHSEDFELPLFDLFTLTKATDNFSFENKIGEGGFGQVYKGVLEDGQEVAVKRLSKIRARTSRVQE*

### NbL15g20240.1 probable_histone_H2A.1 [MW = 13.95 kDa]

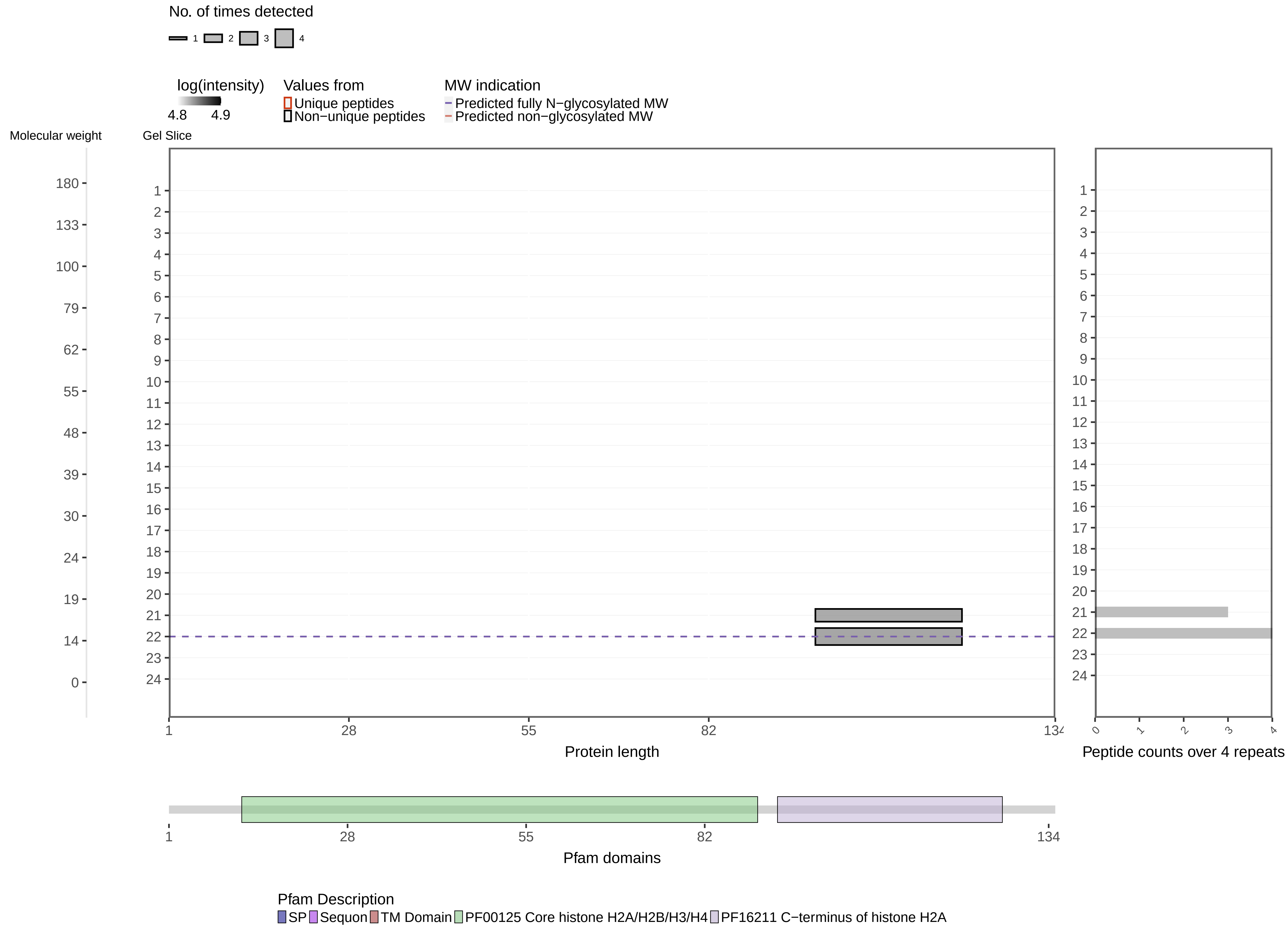


Protein sequence:

.AAVLEYLAAEVLELAGNAARDNKKTRIVPRHIQLAVRNDEELSKLLGDVTIANGGVMPNIHNLLLPKKTGGSSKPSADED*

NbL15g20110.1 protein_translation_factor_SUI1_homolog [MW = 25.14 kDa]

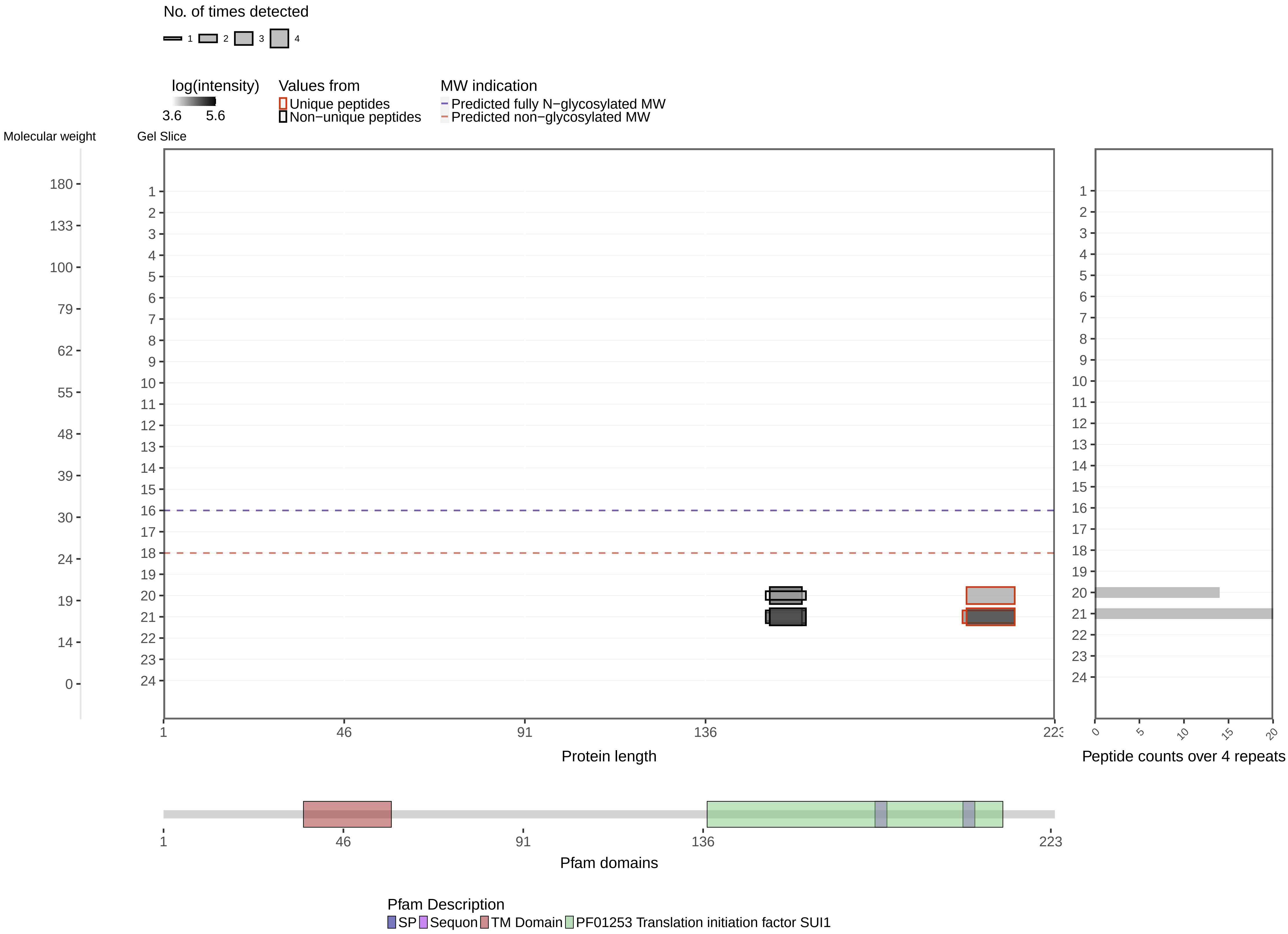


Protein sequence:

RLLSSTTQPSFMSDLDAQIPTAFDPFAEANADNSGAGSKDYVHIRIQQRNGRKSLTTVQGLKKEFSYNKILKDLKKDFCCNGTVVQDPELGQVIQLQGDQRKNVSAFLVQAGIVKKEHIKIHGF*

### NbL15g20100.1 thiamine_thiazole_synthase_chloroplastic [MW = 41.76 kDa]

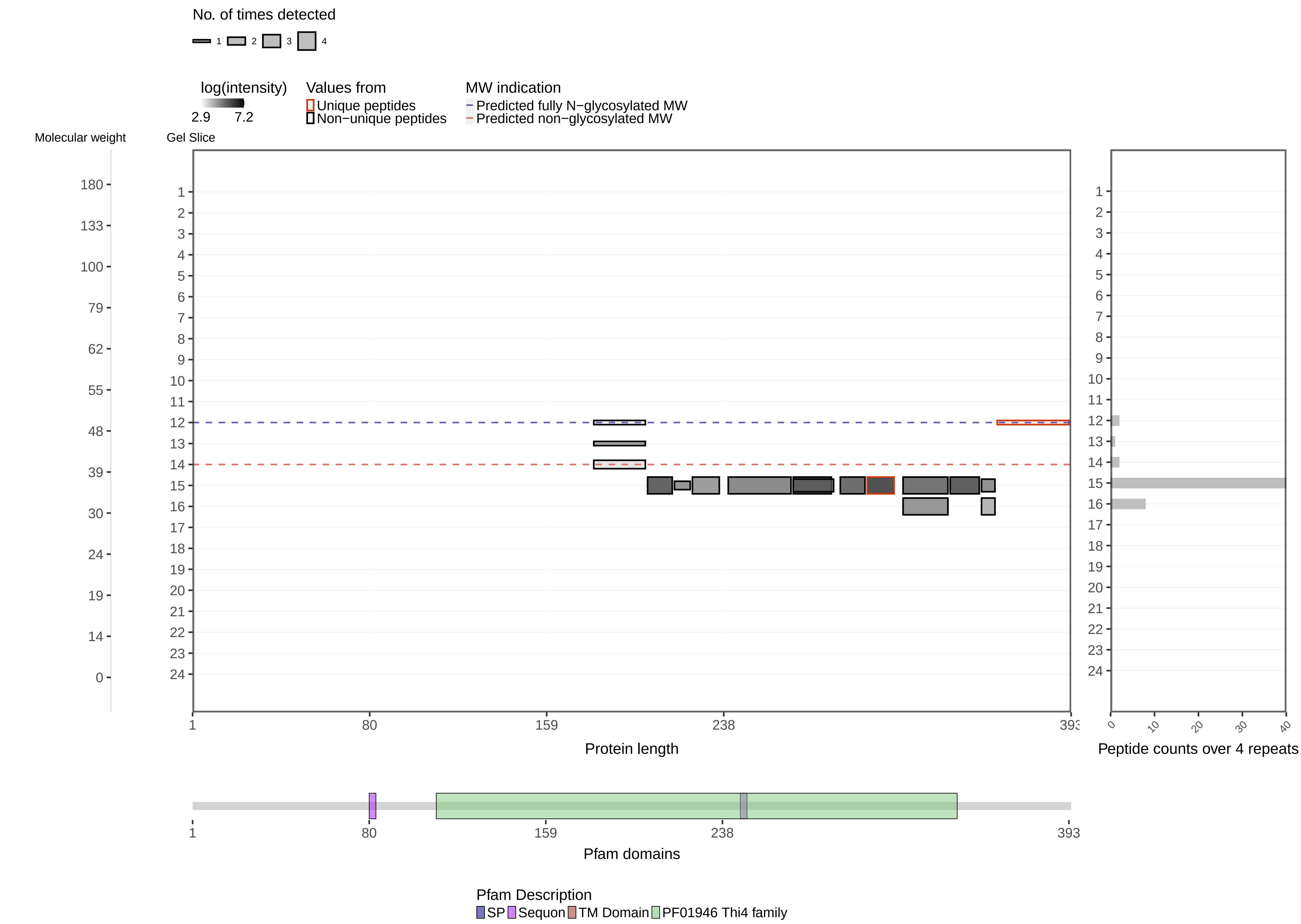


Protein sequence:

LFLNELGIDYDEQDNYVVIKHAALFTSTIMSKLLARPNVKLFNAVATEDLIVKNGRVGGVVTNWSLVSQNHDTQSCMDPNVMEAKIVVSSCGHDGPMGATGVKRLKSIGMINHVPGMKALDMNTAEDAIVRLTREVVPGMIVTGMEVAEIDGAPRMGPTFGAMMISGQKAAHLALRALGLPNALDGTAESSIHPELILAAADDAETADA*

NbL15g20030.1 protein_translation_factor_SUI1_homolog [MW = 12.63 kDa]

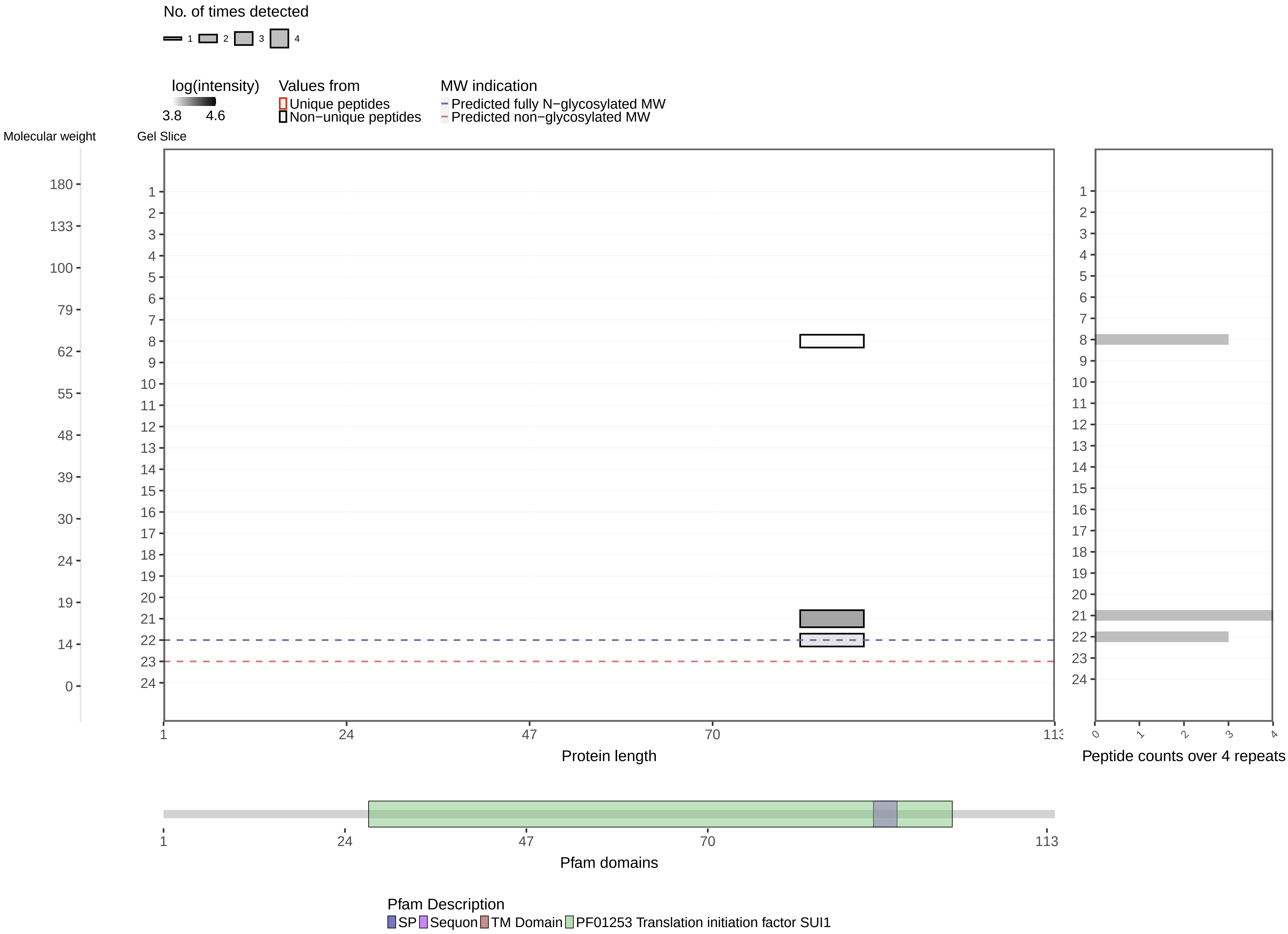


Protein sequence:

TTVQGLRKQFSYEKILKDLKKEFCCNGNVVQDKELGKVIQLQGDQRKNVSHFLVTAGVVKKDQIKIHGF*

NbL15g19800.1 DNA_polymerase_I_B_chloroplastic/mitochondrial−like [MW = 128.57 kDa]

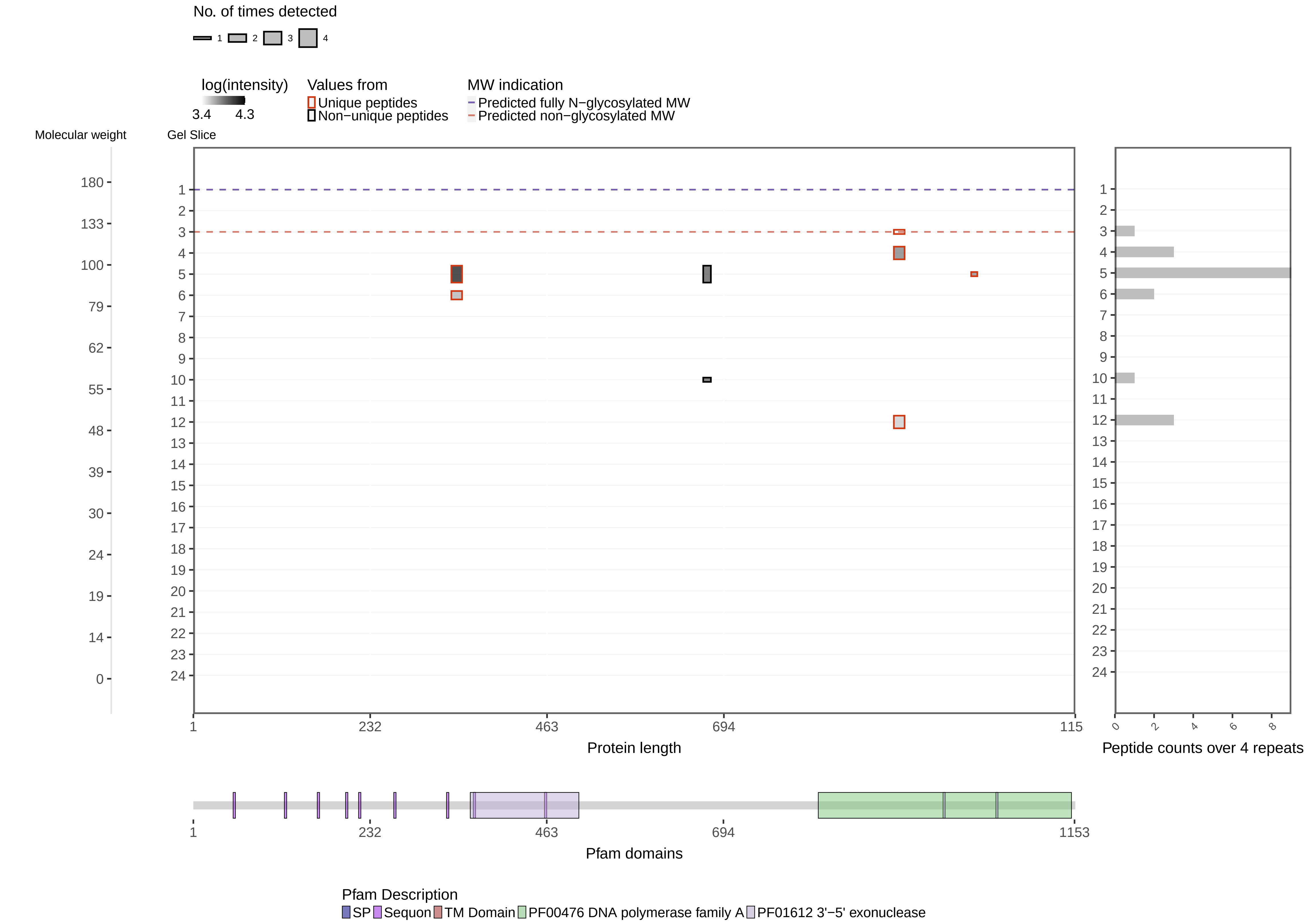


Protein sequence:

KTERELWICYSALDSISTLMLYESLKNKLSKRIWTFDGVSKGSMYEFYEKYWRPFGELLVQMETEGVLVDRSYLAEIEKVAKAEQQVAANRFRNWAAKYCPDAKYMNVGSDTQLRQLFFGGIQNRKNSDESLPYEKEFKVPNVDKVIEEGKKAPTKFRKIRLHRICDLIDTEMYTASGWPSVSGDALKALSGKVSADFDILDEADDNAEEDPETSIDEALATNNEVPSQEPEVSIYGSAYNAFGGGQKGIEACHAIAALCEMCSIDSLISNFILPLQGQDVSGENGRIHCSLN

### NbL15g19580.1 GDSL_esterase/lipase_At5g55050−like [MW − SP = 36.61 kDa]

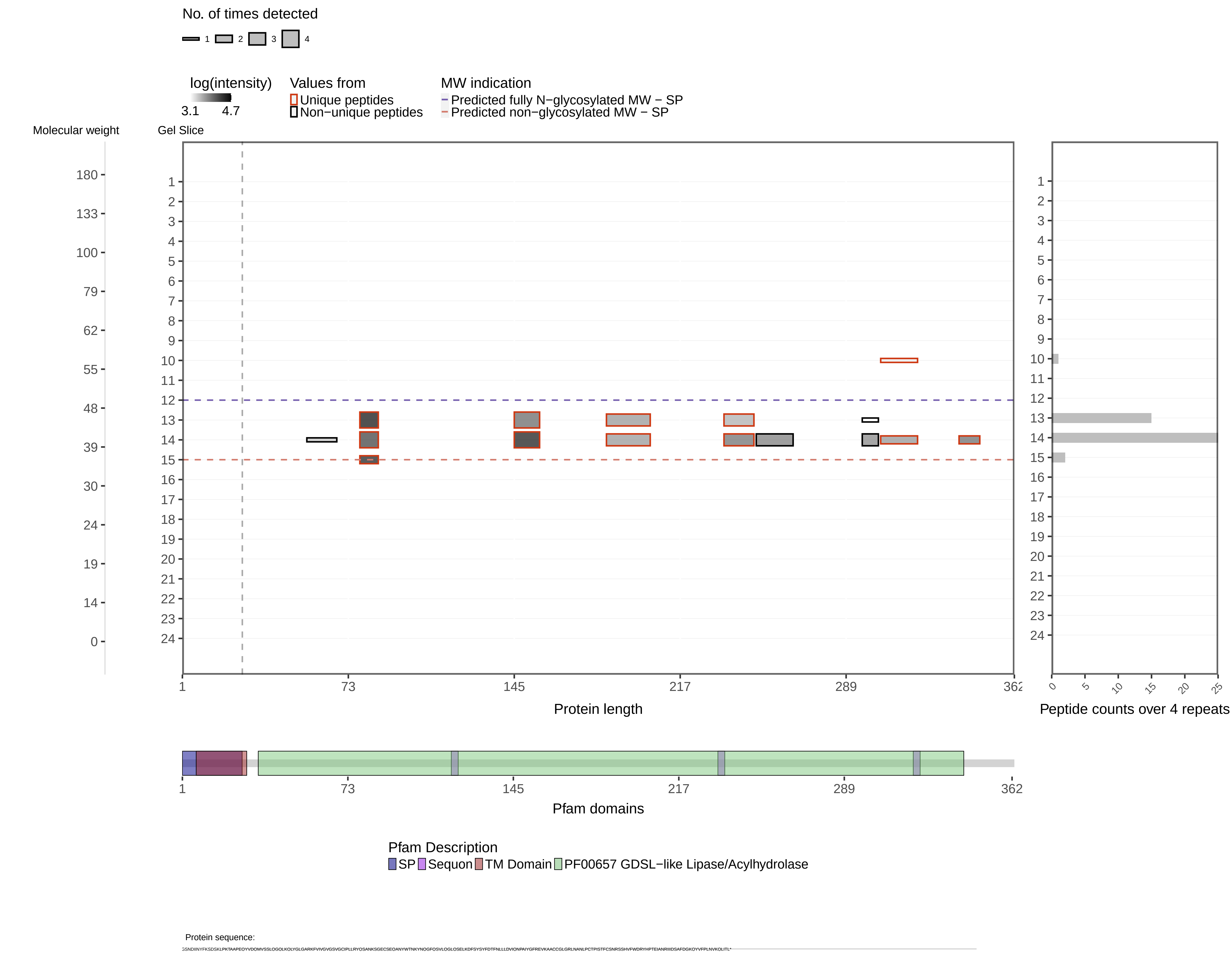

NbL15g19490.1
dihydrolipoyllysine−residue_succinyltransferase_component_of_2−oxoglutarate_dehydrogenase_complex_2_mitochondrial [MW = 54.9 kDa]

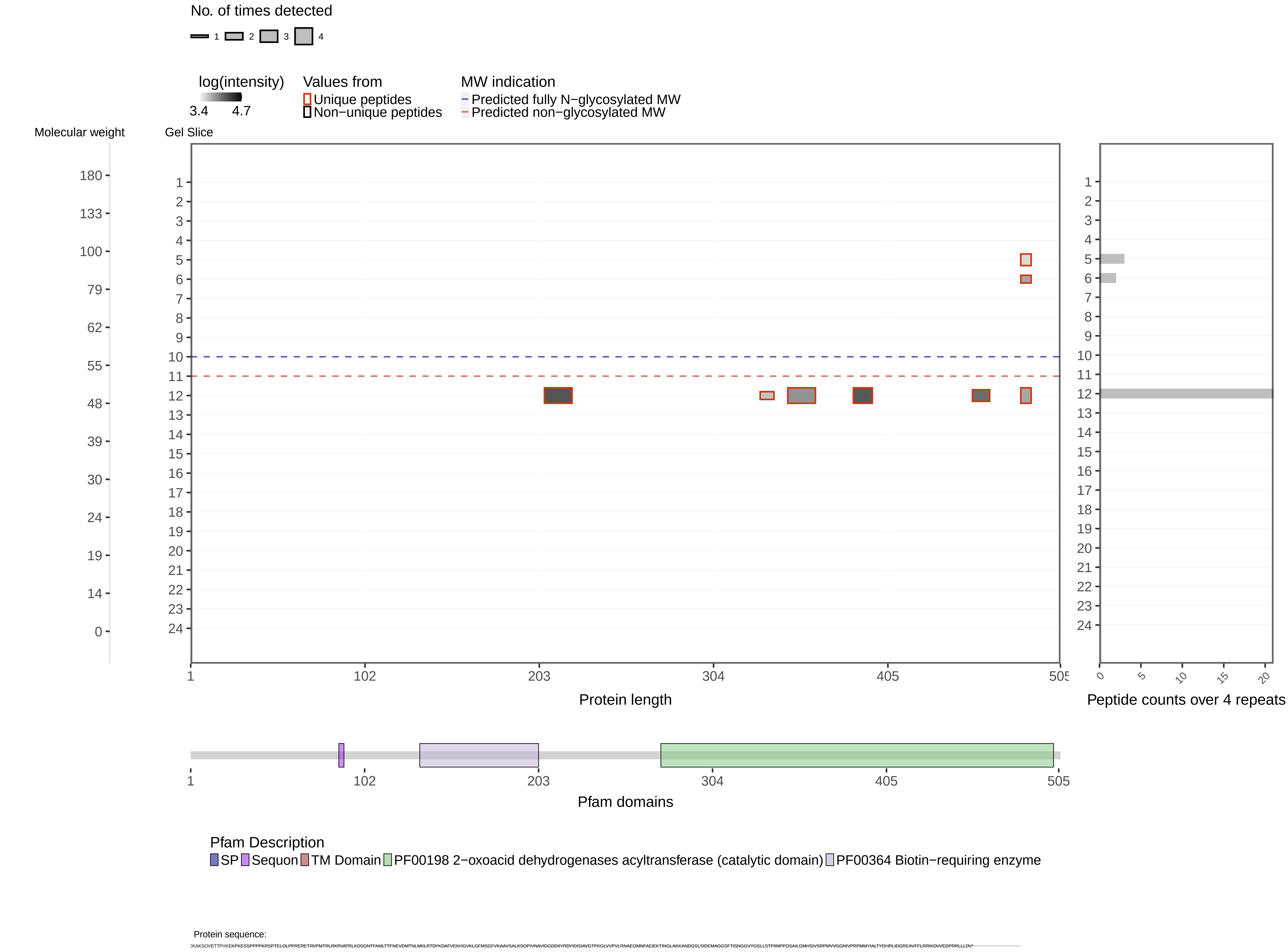

### NbL15g19480.1 imidazole_glycerol_phosphate_synthase_hisHF_chloroplastic [MW = 65.01 kDa]

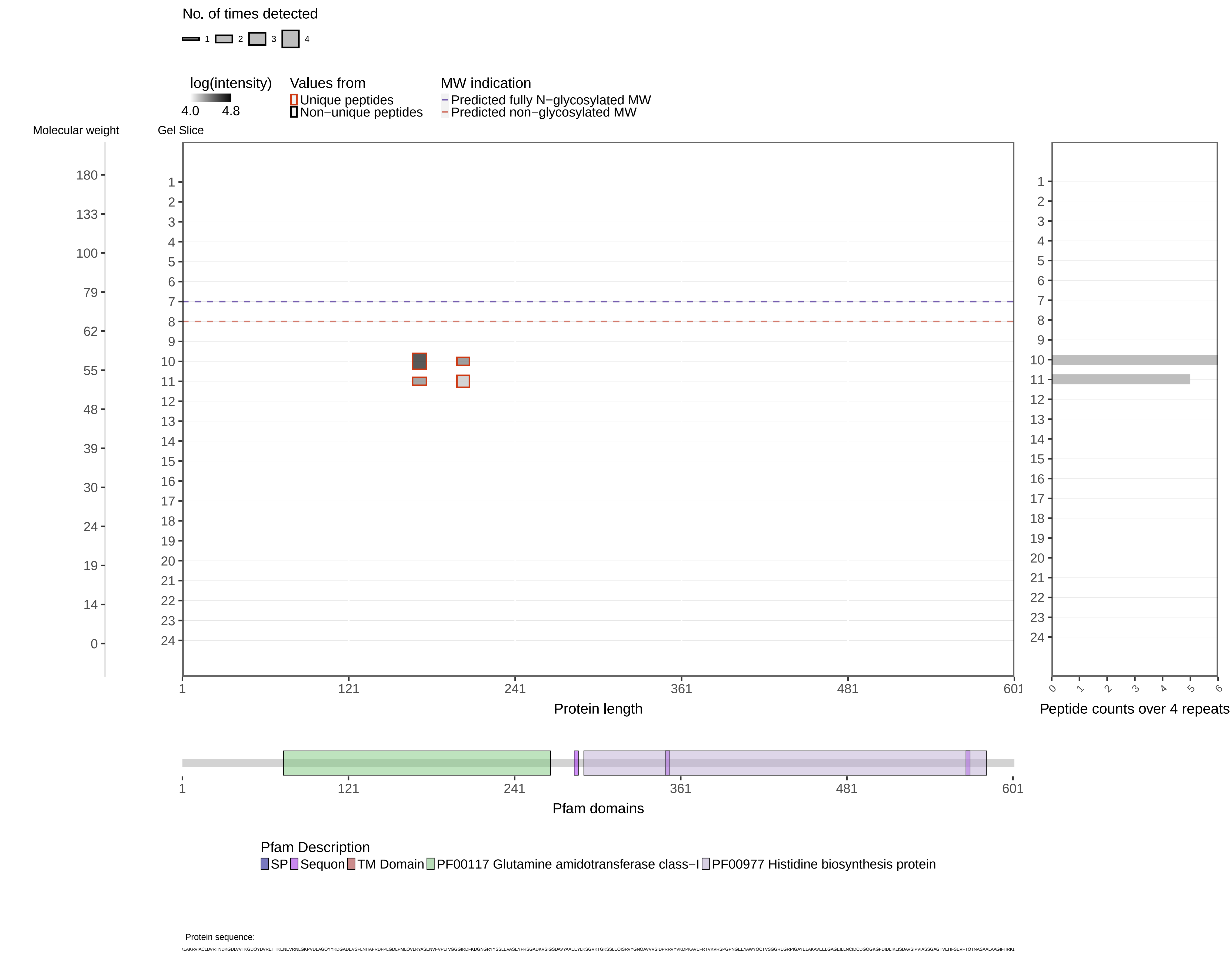

### NbL15g19410.1 small_ubiquitin−related_modifier_1−like [MW = 19.42 kDa]

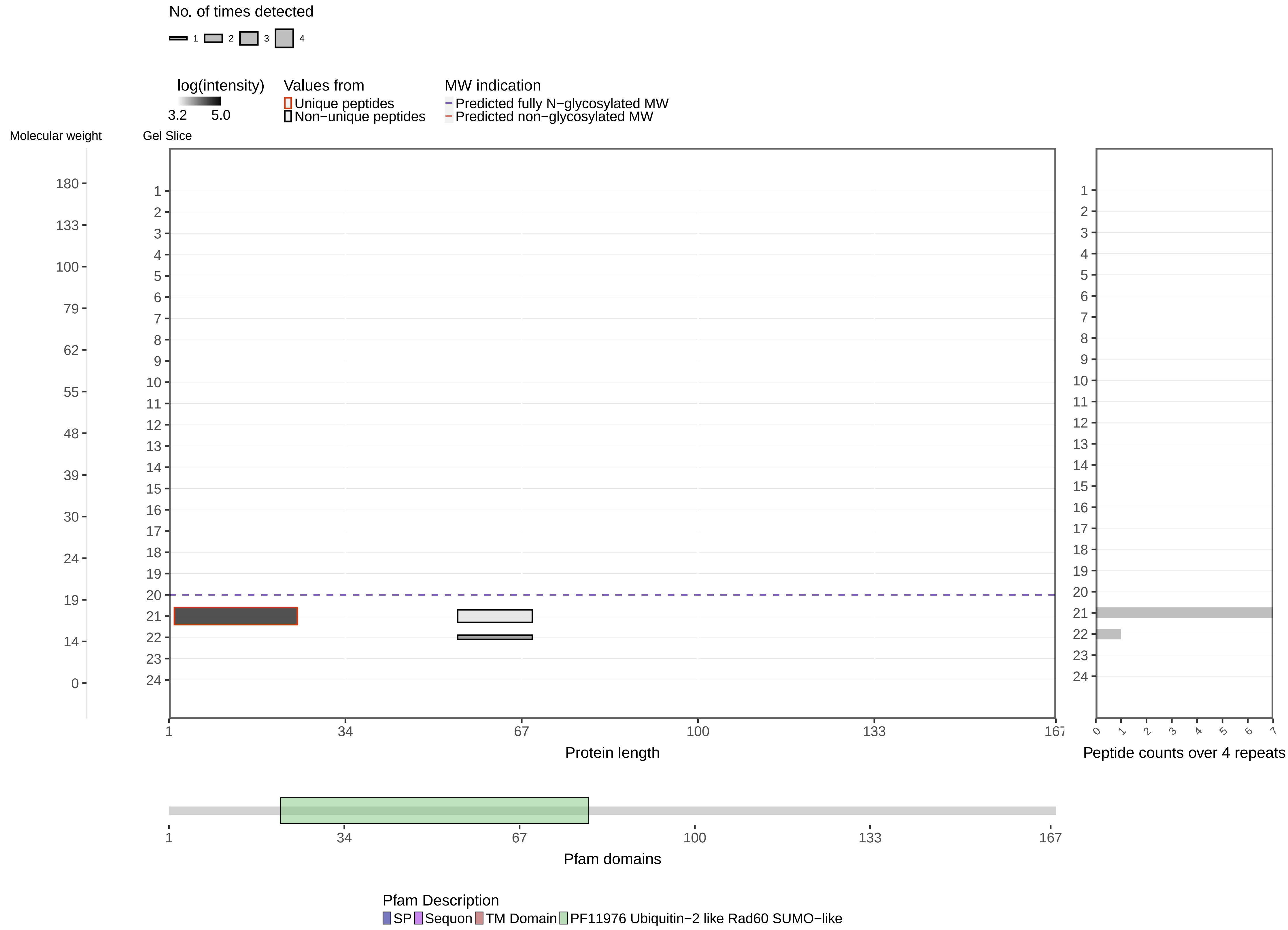


Protein sequence:

RRLRSEOTPDEVSVSPFFFTNPHSFHIONIFPCVFKCORICSVLSRYFFHILFGSLRWRMGMKLMLCCIRLVDTWVSFISEDFGLNYLRYILAELLRV*

### NbL15g19340.1 GrpE−like [MW = 33.76 kDa]

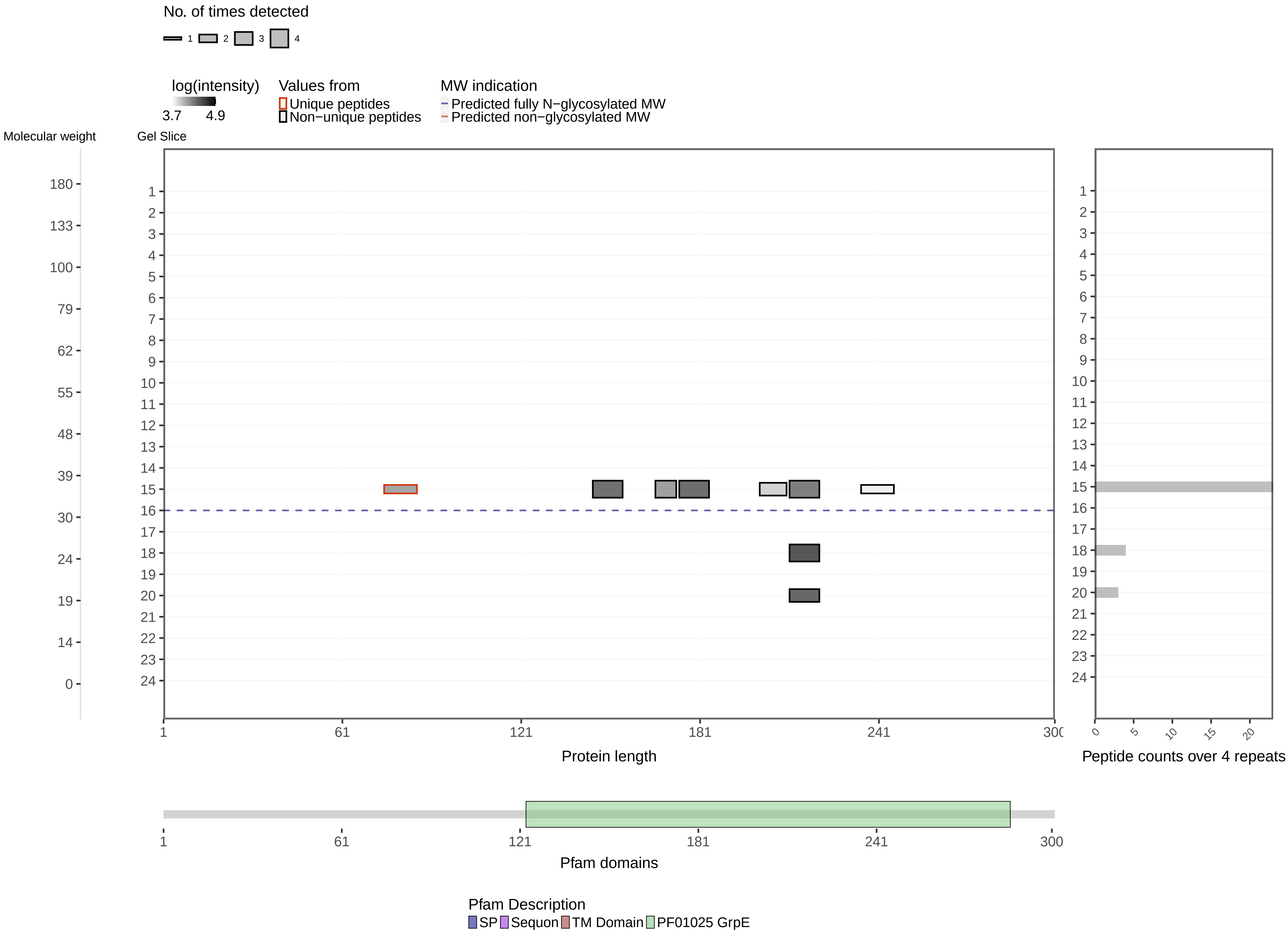


Protein sequence:

KLQDKFLRSYAEMENVMERTKREAENSKKFAIQNFVKALLDVPDNLGRASSVVKESFSKIDASKDTVGAMPLLKTLLEGVEMTDKQLAEVFKKFGVEKYDPTNEQFDPNKHNAVFQVPDPEKAPGVIAVCLKPGYTLHDRIIRPAEVGVTVAVETEADQSTNS*

### NbL15g19330.1 trigger_factor−like_protein_TIG_Chloroplastic [MW = 60.48 kDa]

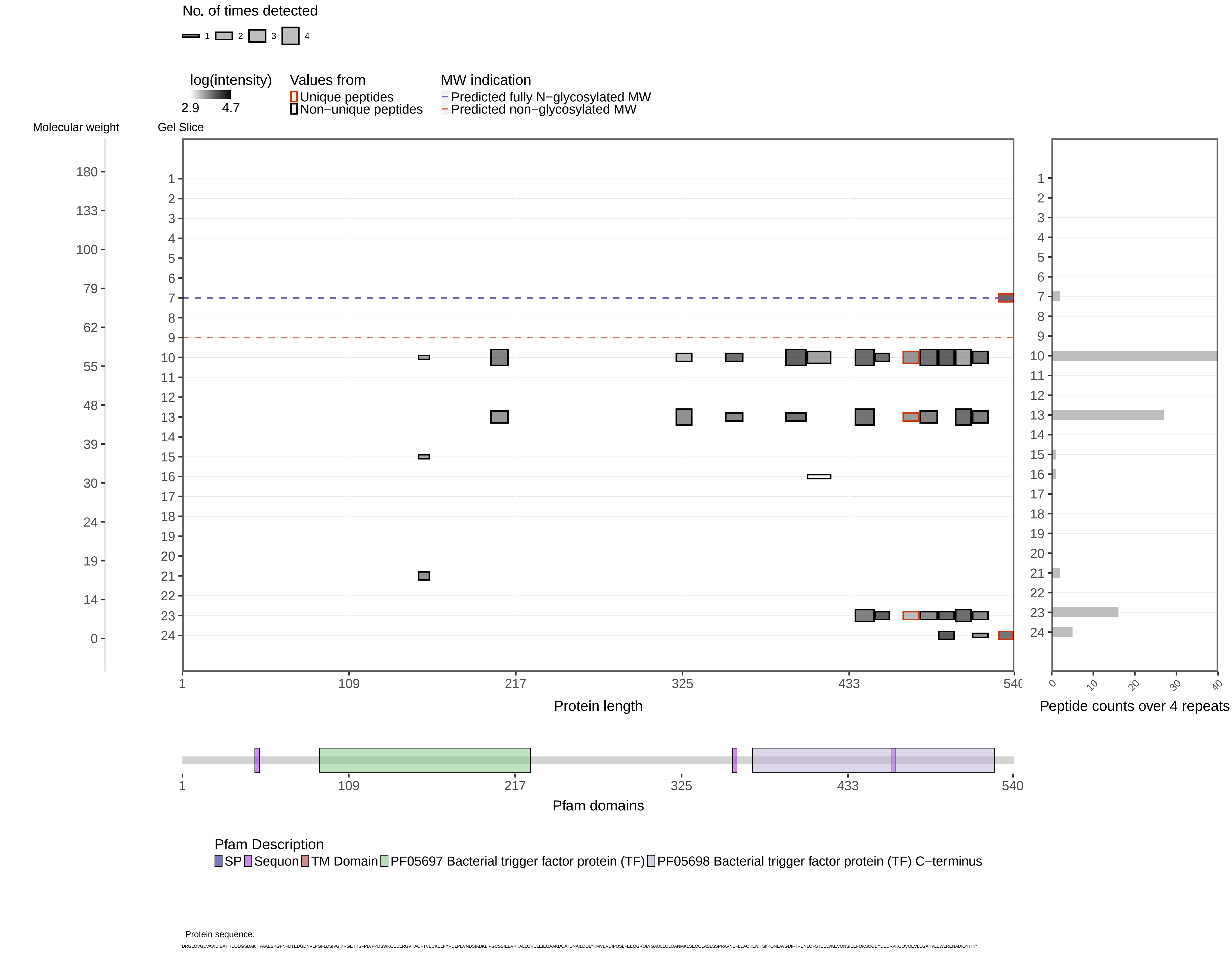

### NbL15g19320.1 65−kDa_microtubule−associated_protein_1−like [MW = 65.32 kDa]

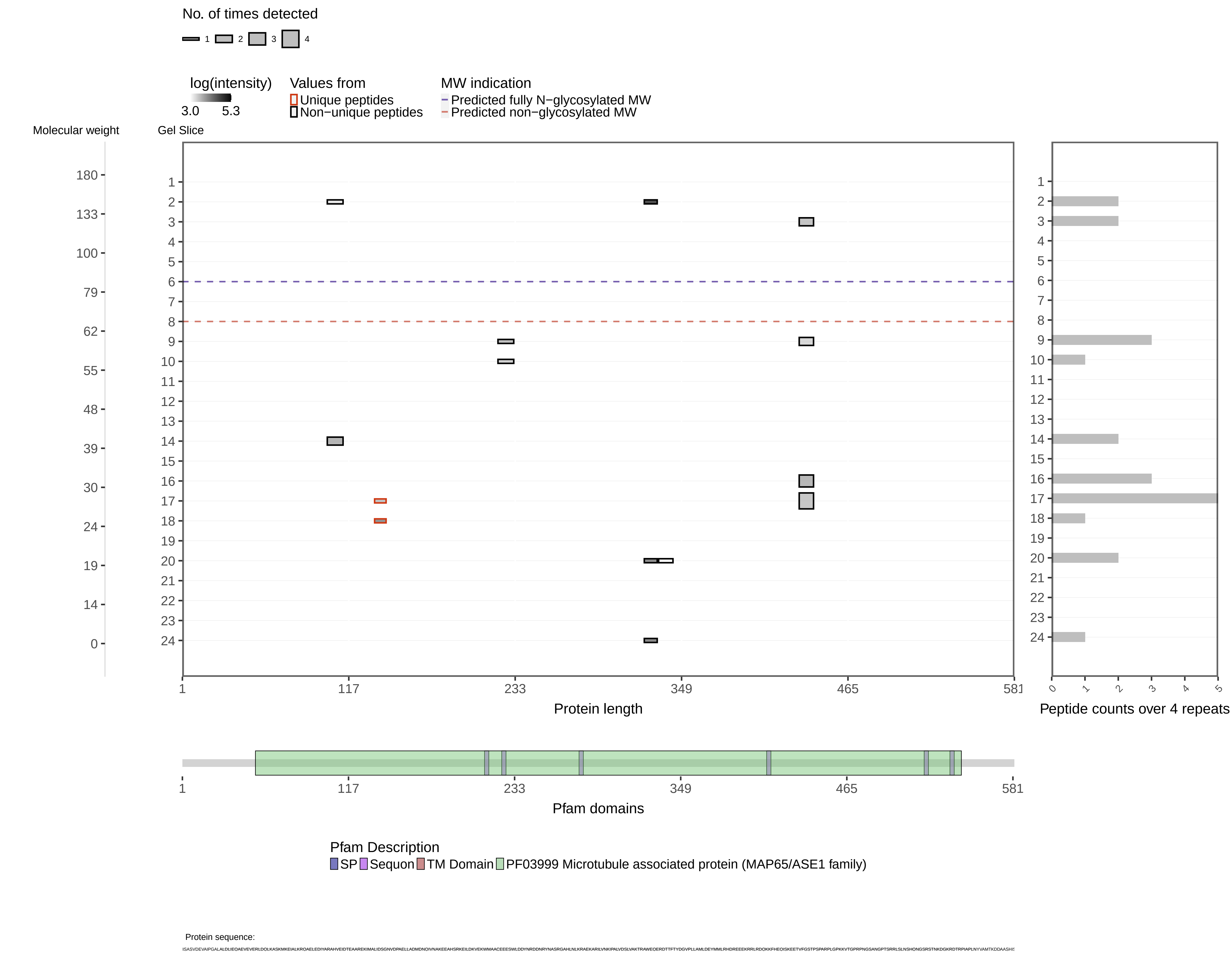

NbL15g19270.1 serine/threonine−protein_phosphatase_PP−X_isozyme_2 [MW = 34.87 kDa]

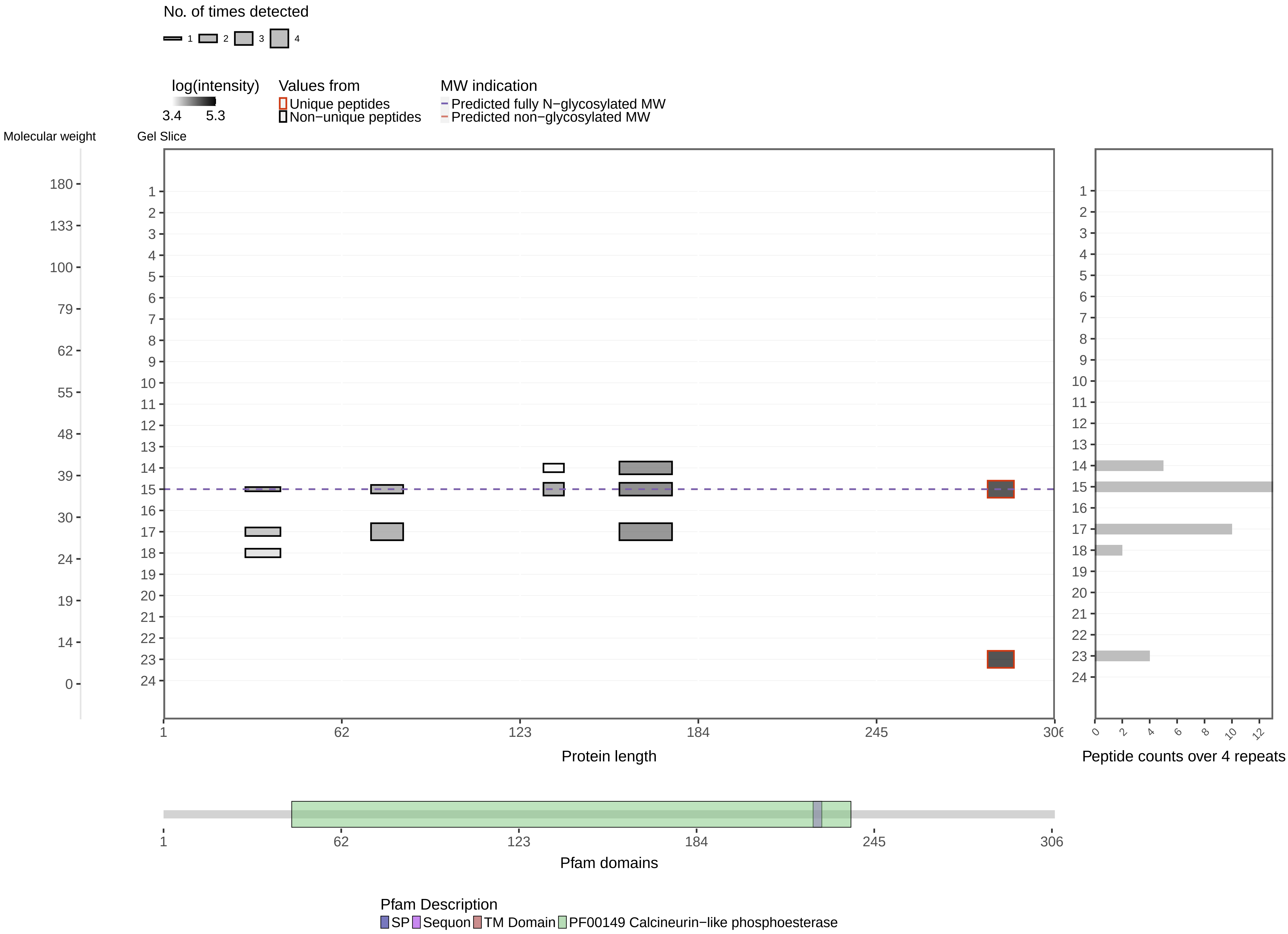


Protein sequence:

CTDIFDYLSLSALIENKIFSVHGGLSPTISTLDQIRTIDRKQEVPHEGAMCDLLWSDPEDIVDGWGLSPRGAGFLFGGSVVTSFNHTNNIDYICRAHQLVMEGFKWMFNNQIVTVWSAPNYCYRCGNVAAILELDENLNKQFRVFDAAPQESRGAPAKRPPPDYFL*

### NbL15g19240.1 fimbrin−1−like [MW = 58.46 kDa]

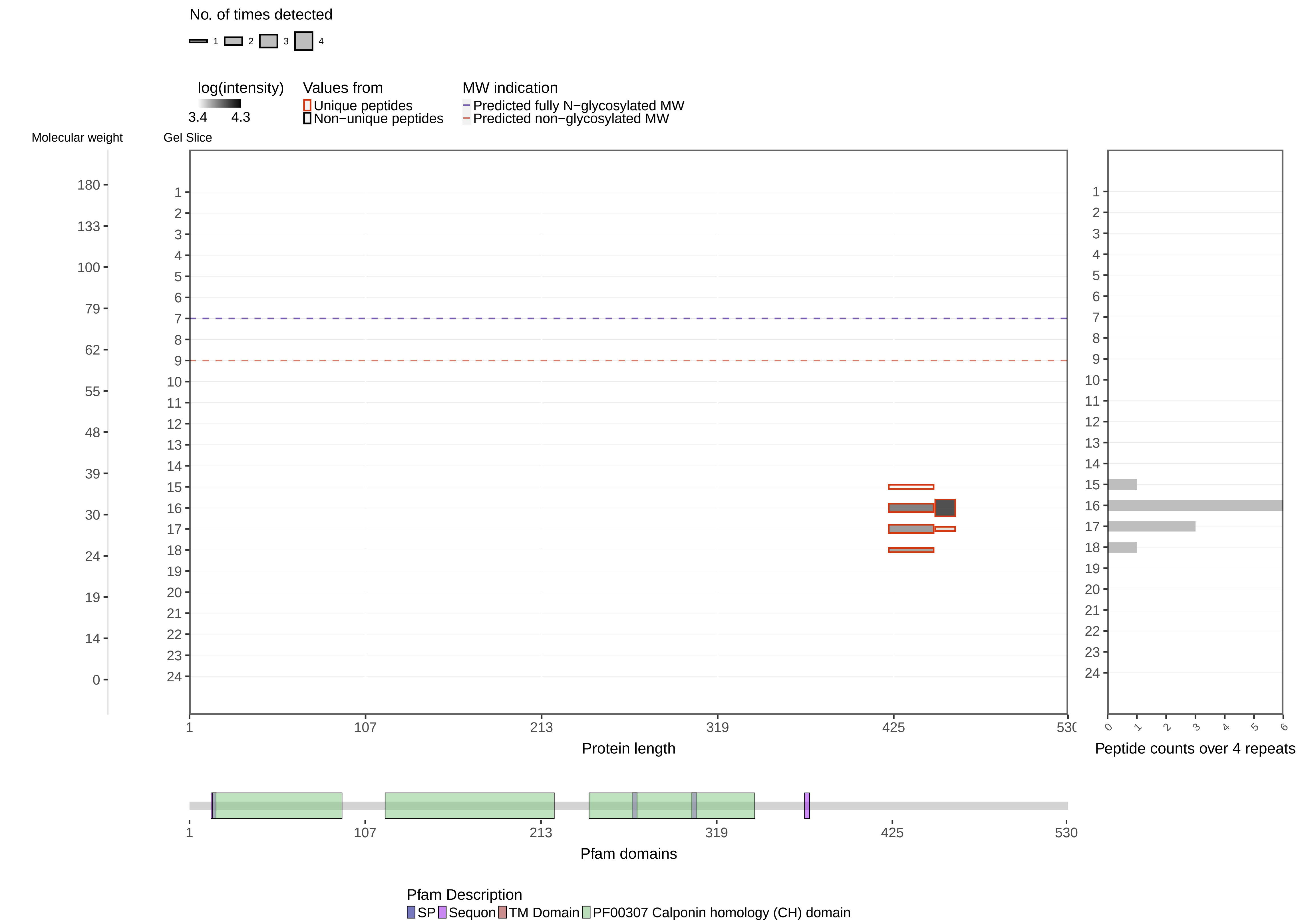


Protein sequence:

KKVKSTGRTSKMESFKDKNLSSGLFFLELLSAVEPRVVNWNLVTKGESDEAKKLNATYIISIARKLGCSIFLLPEDIMEVNQKMILTLTASIMYWSLQQTAEEAESTSPASTITDASPVRSTNGSMSPLIAASPDASPAPSISGASSPVVDASPAPSVNGDEESLLIAEVSKLADDDASSDVLASPEQAENVEIPSDVHRDAENAEIPSDVHRDAENAEIPTDVHRDDENAEIPTDVCTDADIAEIPSDTPSSPQLEDTEQQSGLSHEVENEGTKPDNEQ*

NbL15g19220.1 glycerophosphodiester_phosphodiesterase_GDPDL3−like [MW − SP = 79.45 kDa]

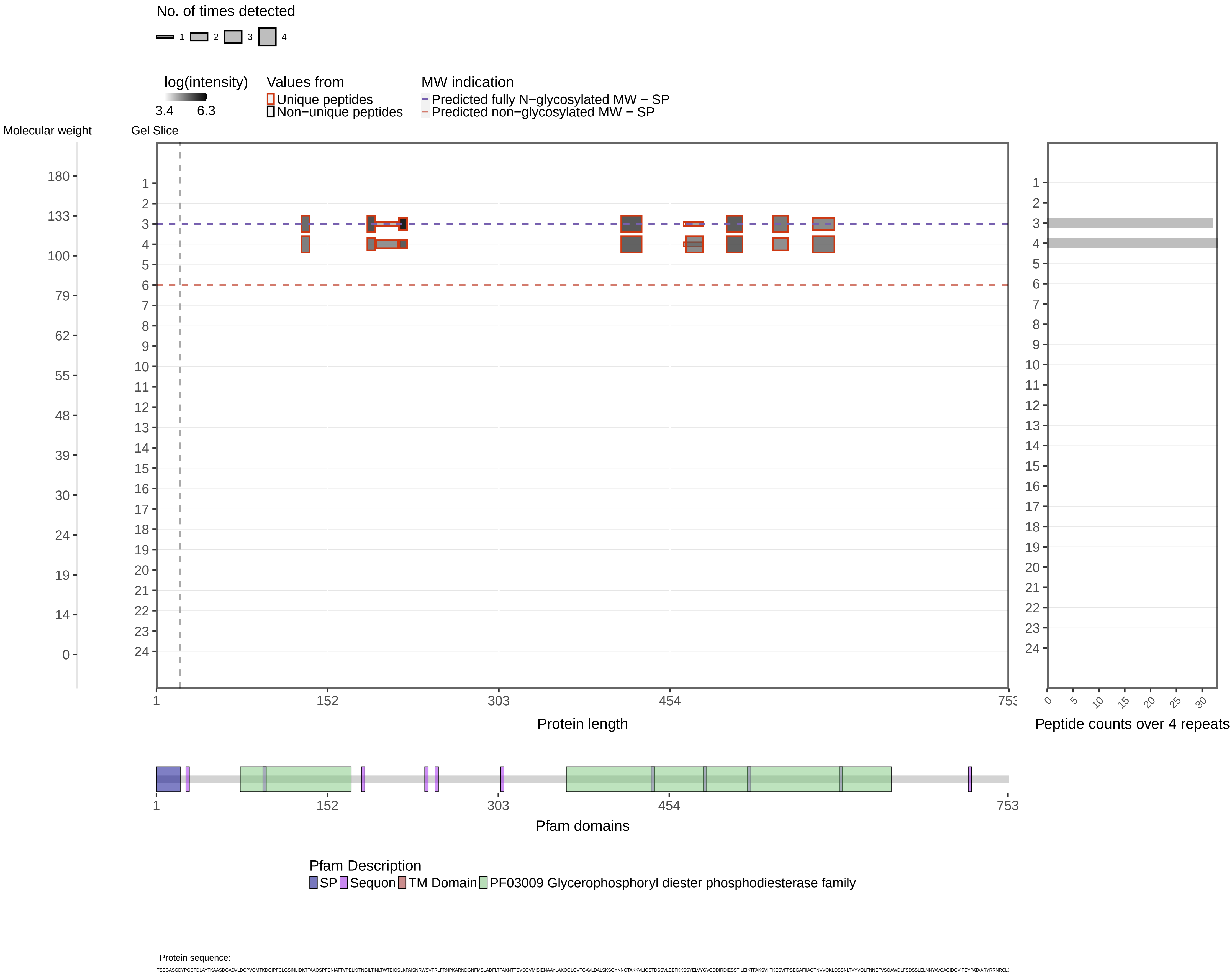

### NbL15g19070.1 uncharacterized_LOC109210041 [MW − SP = 46.5 kDa]

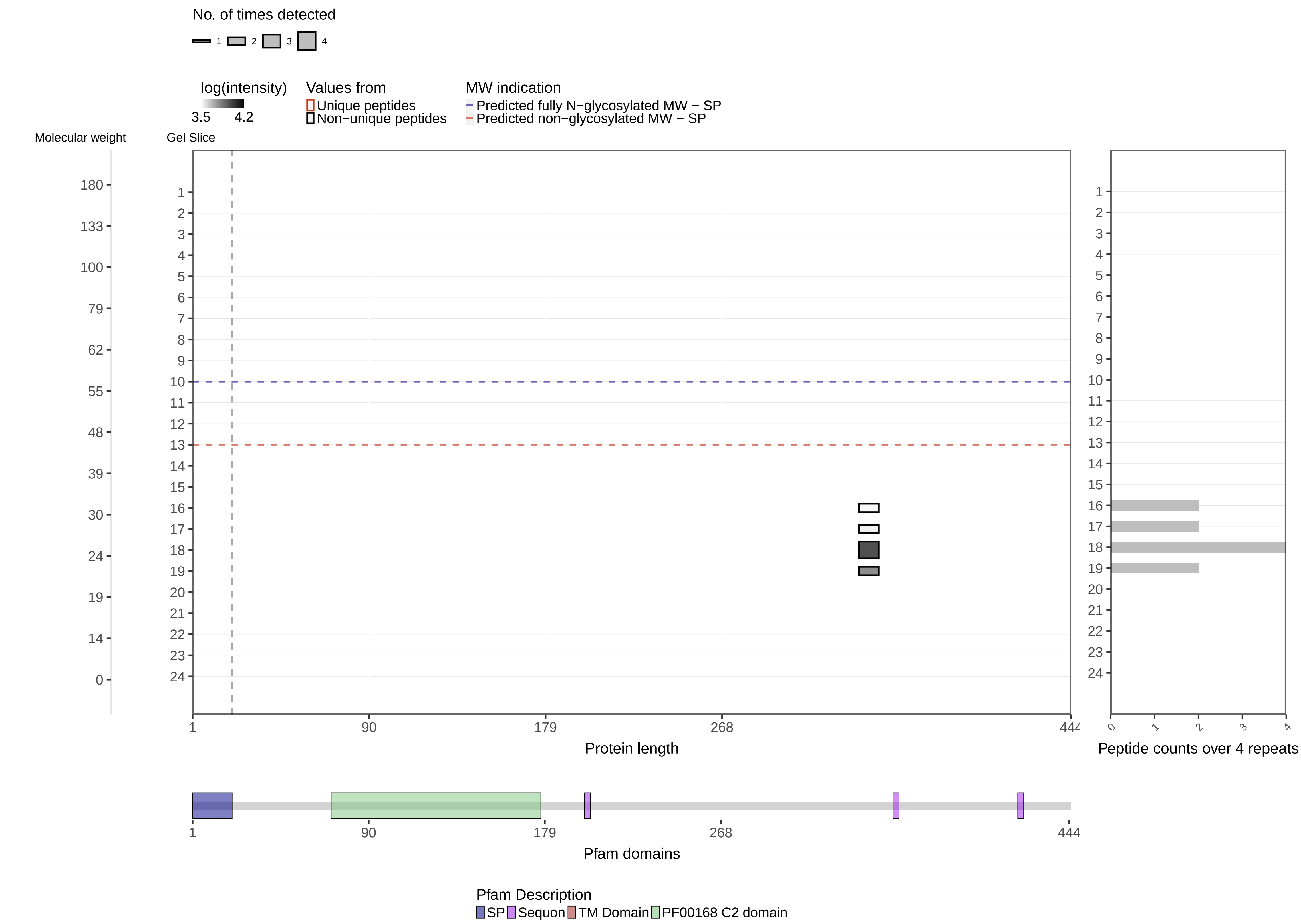


Protein sequence:

.PVSVDTDASRPPDSETPESLLKEFDKIEFPDPKILNENNLMVSEYFNIPGTNLDSQSSDSFVSSDTENQLSSDVDVHMENFPSGVANFHRYPKREFPPTSVSTIDSPSTLHPATSQSSDTQGASKSPGEEYASAPKASRDAERNISETKPSSAFPKPVVAVNIEPEQKVVQQDIVDMYMKSMQQFTESLAKMKLPLEMETRPTTSGNSSSDQTSQTPKSTGSRVFYGSRAFF*

### NbL15g18980.1 ABC_transporter_C_family_member_3 [MW = 174.35 kDa]

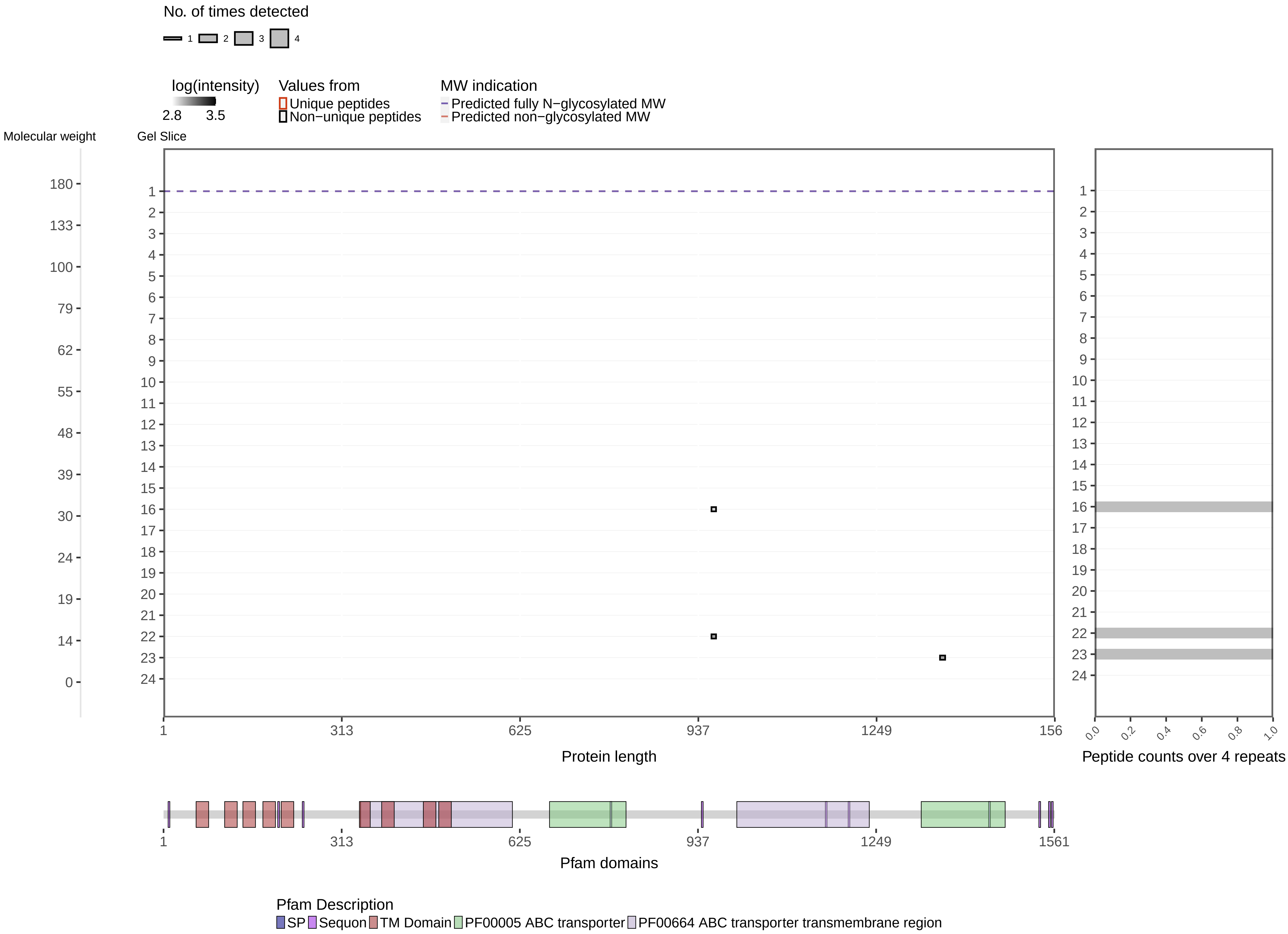


Protein sequence:

NbL15g18830.1 DNA_ligase_1−like [MW = 73.68 kDa]

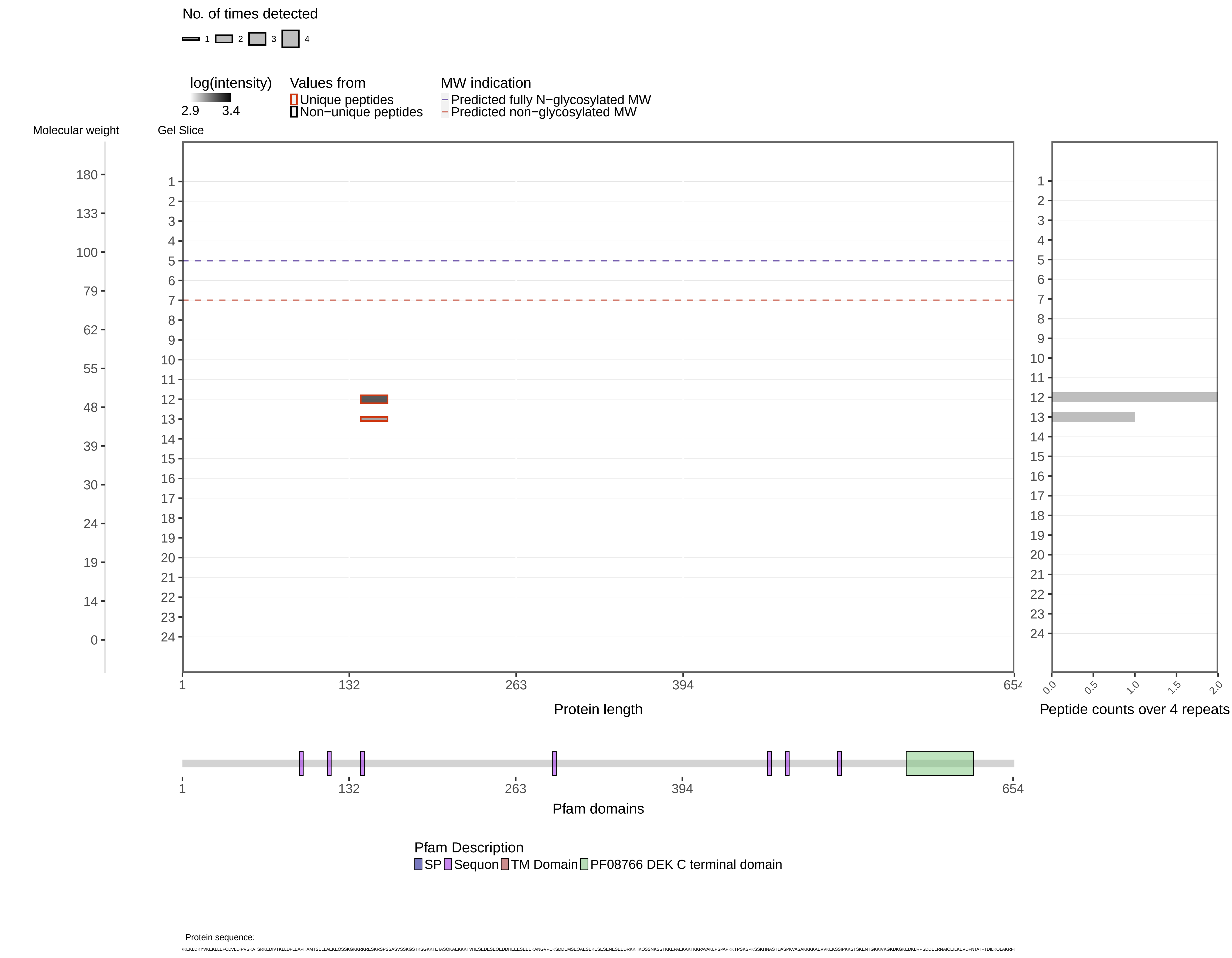

NbL15g18820.1 nudix_hydrolase_10−like [MW = 32.04 kDa]

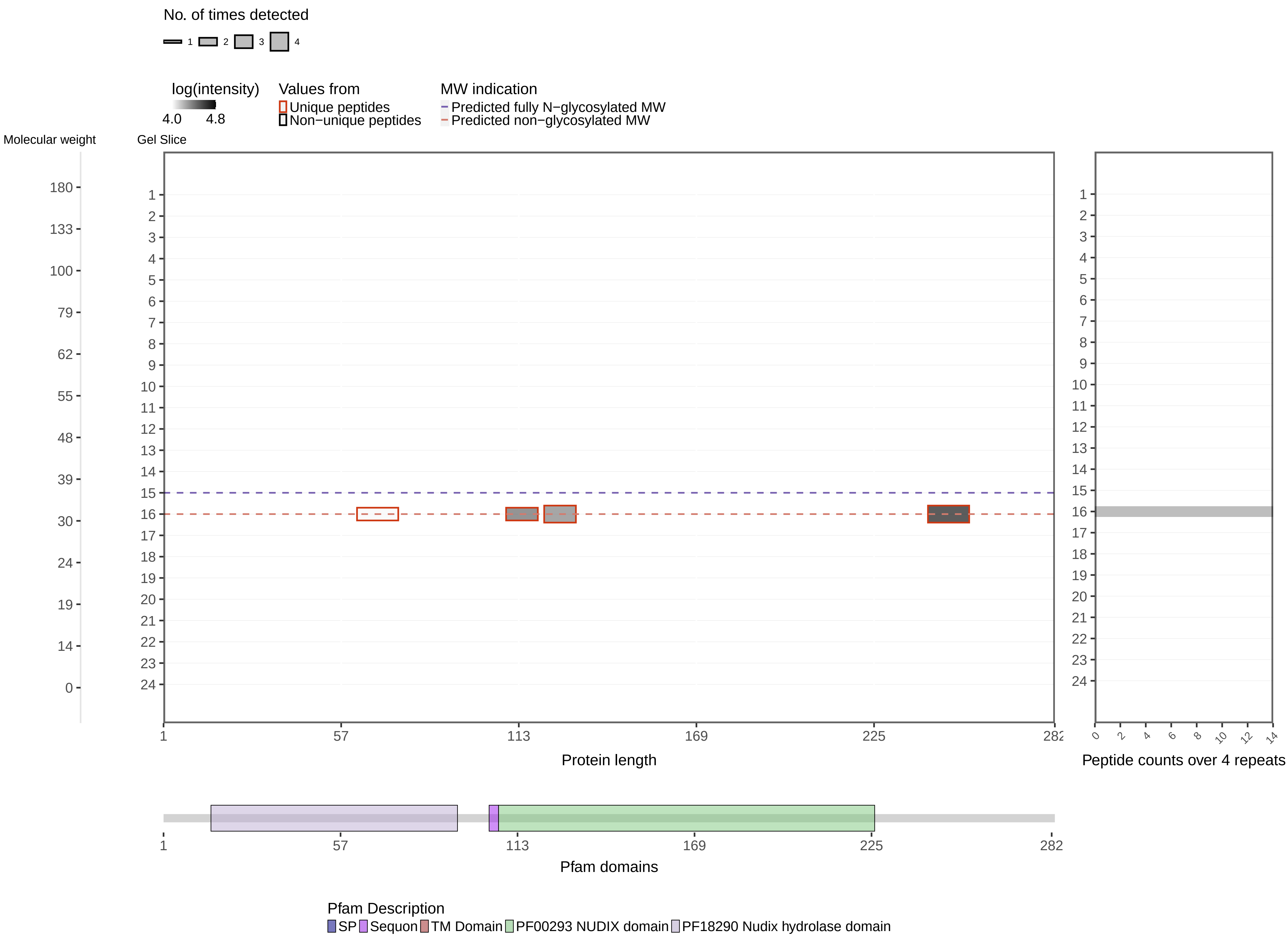


Protein sequence:

ESSGRMKGTGIWKIPTGVVEEGEDVFEAAIREVKEETGIDTEFLEVLAFRQVHKSFFGKSDLFFICMMHPLTFNLQKQDLEIEAVQWMSLEEYAAQPFVLRHGIFKYIKDLCLAKAEGGYLGFTPVPIKSYFDDPTSYLYFNEDGLVQENSAIHL*

### NbL15g18490.1 putative_late_blight_resistance_protein_homolog_R1A−3 [MW = 106.61 kDa]

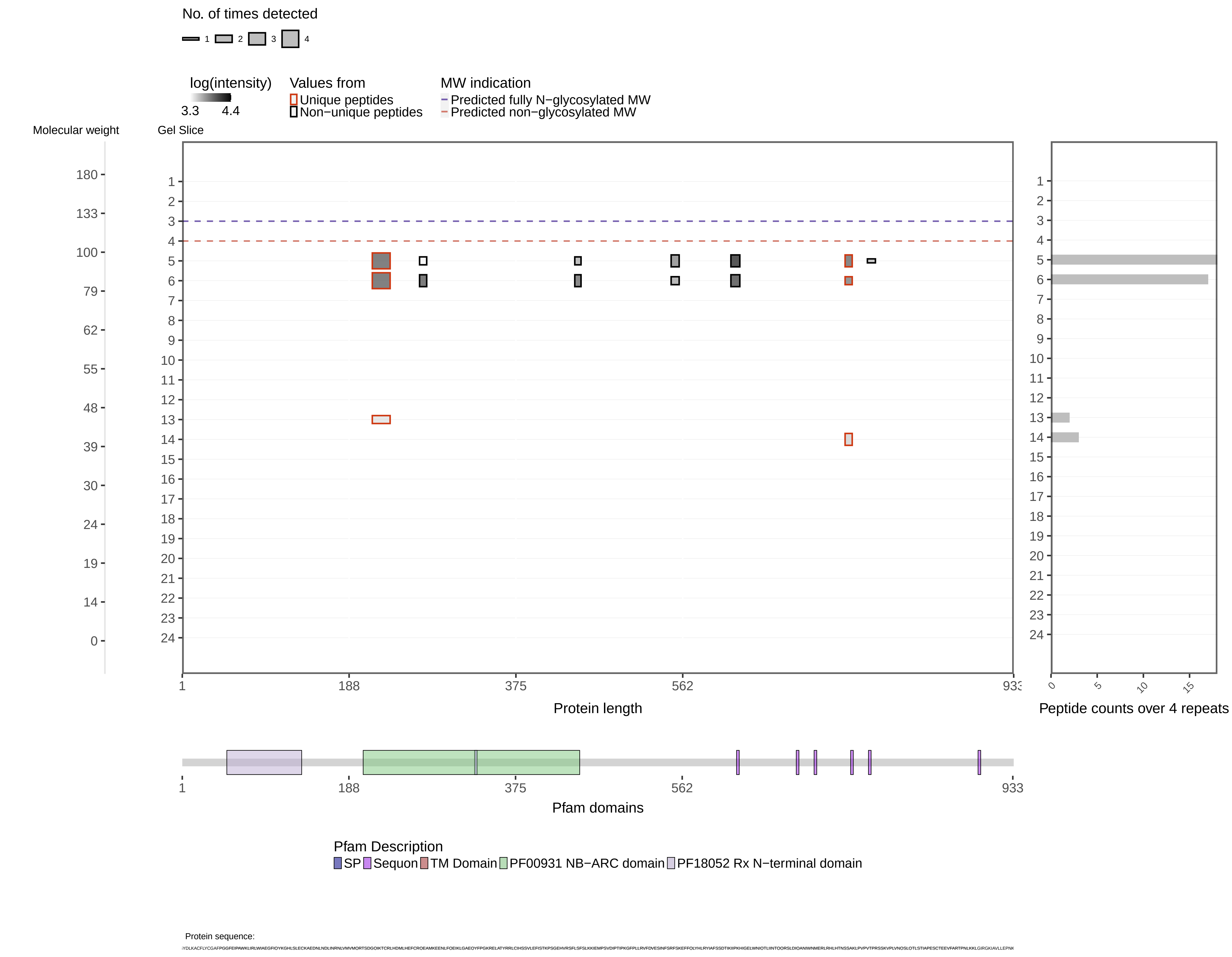

### NbL15g18400.1 heat_shock_cognate_protein_80 [MW = 80.19 kDa]

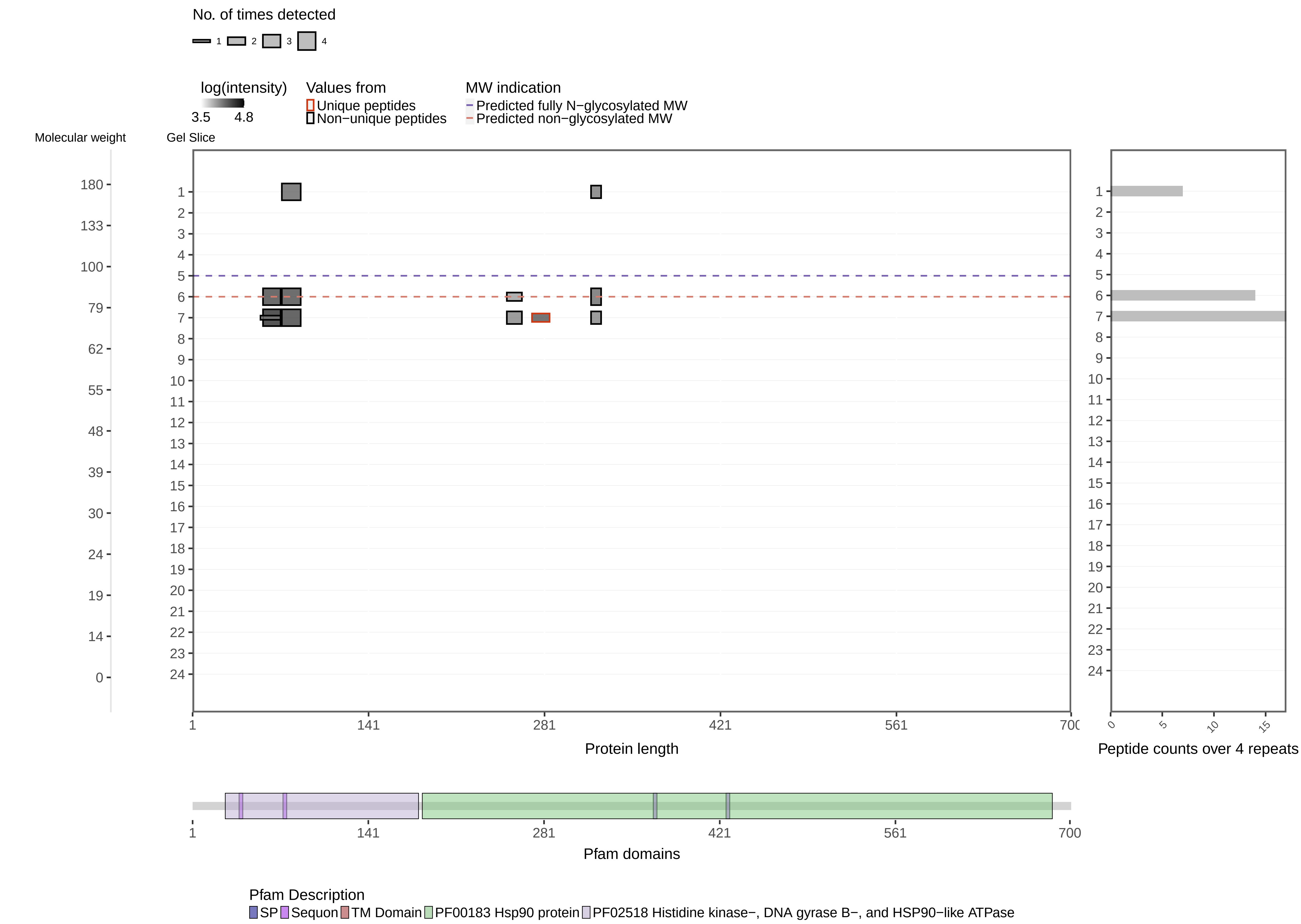


Protein sequence:

FIMDNCEELIPEYLSFVKGIVDSEDLPLNISREMLQQNKILKVIRKNLVKKCVELFFEIAENKEDYNKFYEAFSKNLKLGIHEDSQNRSKFAELLRYHSTKSGDEMTSLKDYVTRMKEGQNDIYYITGESKKAVENSPFLEKLKKKGYEVLYMVDAIDEYSIGQLKEFEGKKLVSATKEGLKLDESEDEKKKQEELKEKFEGLCKVIKDVLGDKVEKVVVSDRVVDSPCCLVTGEYGWTANMERIMKAQALRDSSMAGYMSSKKTMEINPENAIMEELRKRADADKNDKSVKDLV

NbL15g17860.1 protein_RAE1 [MW = 38.28 kDa]

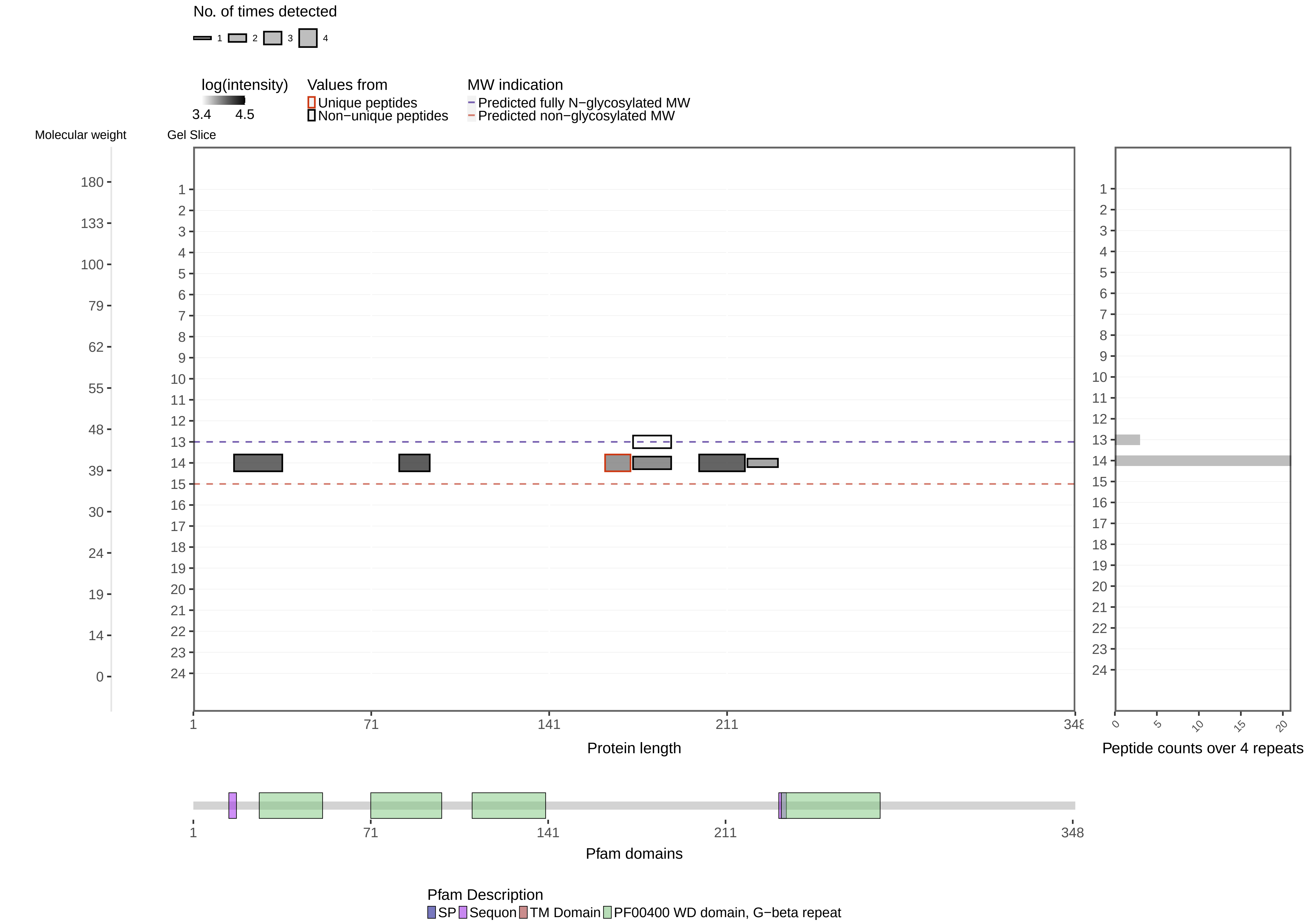


Protein sequence:

.TVKHPLMVAGTADRNLVVFNLQNPQTEFKRITSPLKYQTRCLAAFPDQQGFLVGSIEGRVGVHHLDDSQQSKNFTFKCHREGNEIYSVNSLNFHPVHGTFATAGSDGAFNFWDKDSKQRLKAMSRCSQPIPCSAFNHDGSIYAYAVCYDWSKGAENHNPSTAKTYIYLHFPQESEVKGKPRIGSSGRK*

### NbL15g17830.1 beta−xylosidase/alpha−L−arabinofuranosidase_2 [MW = 85.59 kDa]

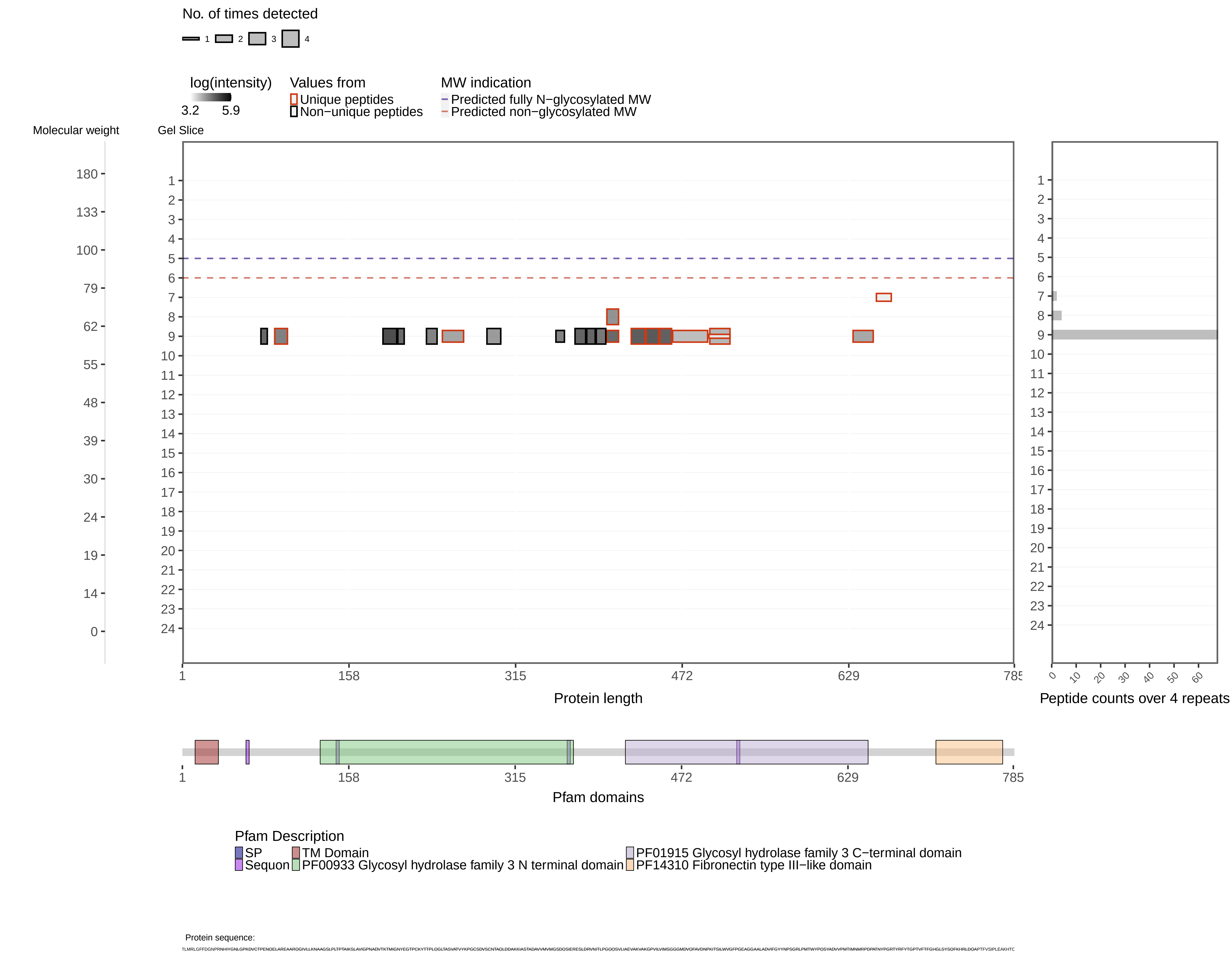

NbL15g17790.1 dynamin−like_protein_ARC5 [MW = 86.95 kDa]

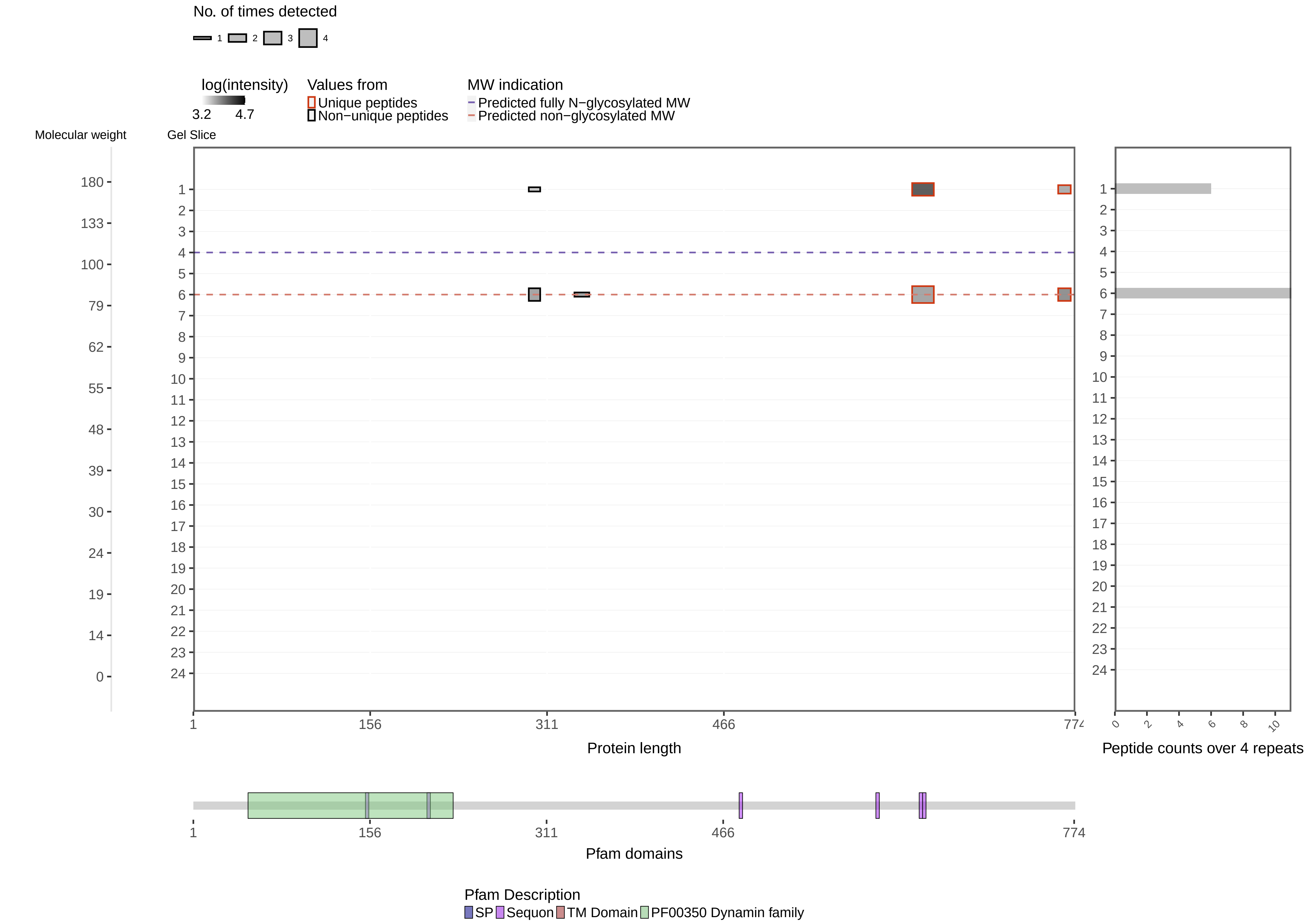


Protein sequence:

KGRNFHDLFLTKLSLLLKGTVIAPPDKFGETLQDERVNGGAFIGSDGLQFPHKLIPNAGMRLYGGAQYHRAMAEFRFVVGGIKCPPITREEIVNACGVEDIHDGTNYSRTACVIAVAKARDTFEPFLHQLGCRLLHILKRLLPIAVYLLQKEGEYLSGHDVFLRRVADAFNNFAESTEKSCREKCMEDLVSTTRYVTWSLHNKNRAGLRQFLDSFGGSEQSAIIGNSTSTGLSQDLSTASVASDNKQDVKPRTDVKLSHLASGNDSNTSTQTTETRLADLLDSTLWNRRLAPSS

NbL15g17720.1 GDSL_esterase/lipase_At1g28590−like [MW − SP = 38.02 kDa]

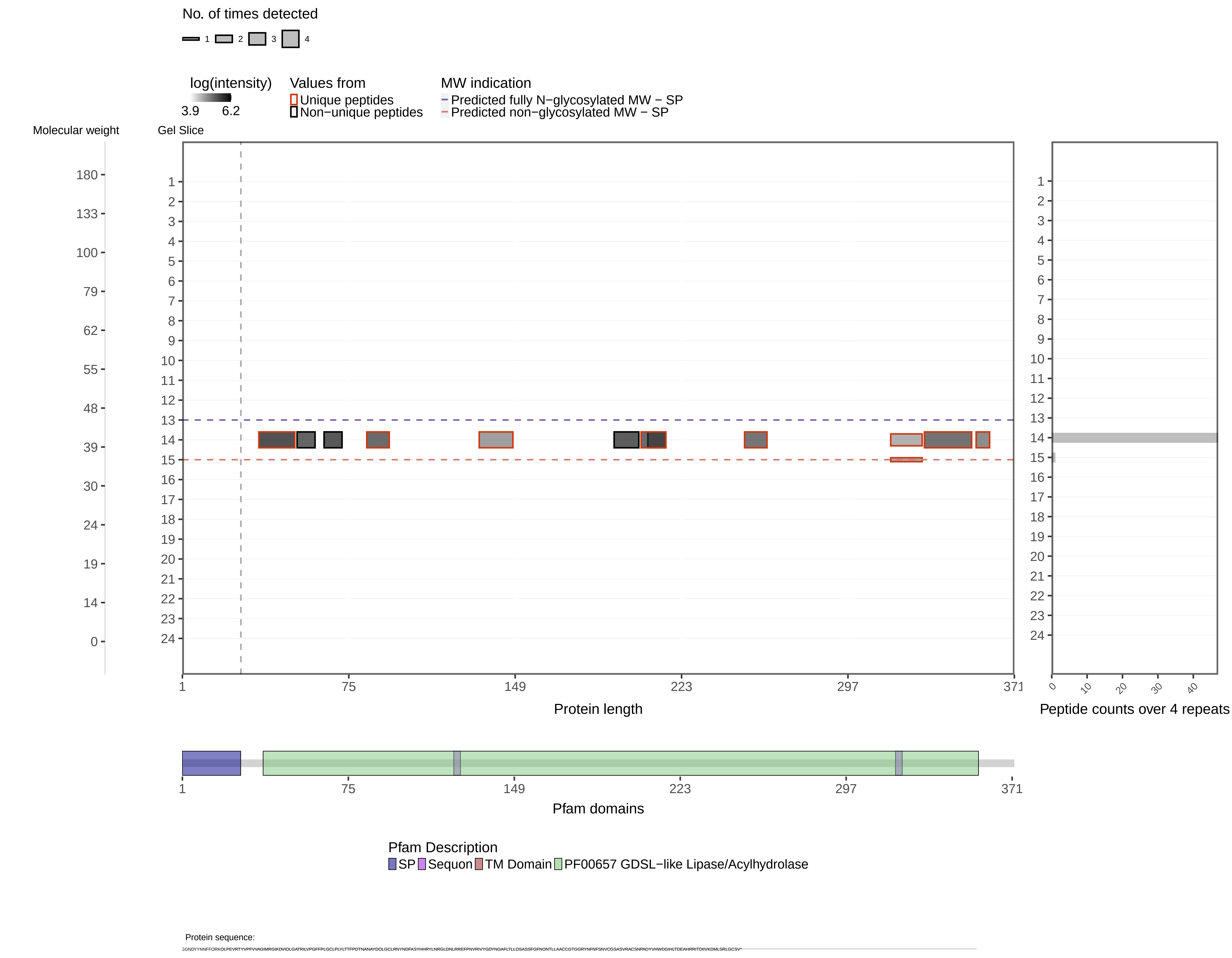

NbL15g17660.1 60S_ribosomal_protein_L3−like [MW = 48.51 kDa]

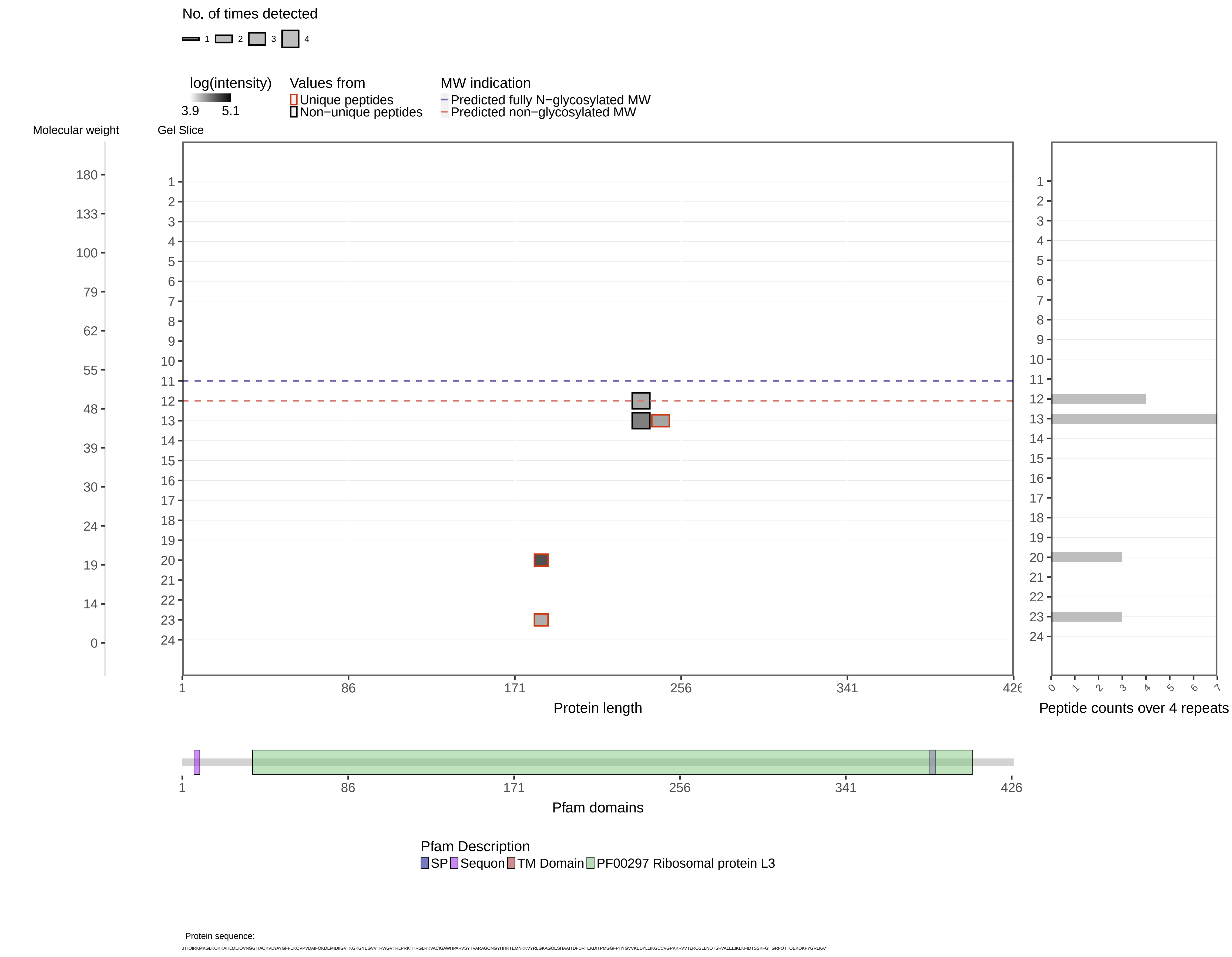

NbL15g17540.1 ankyrin_repeat_domain−containing_protein_2B−like [MW = 48.74 kDa]

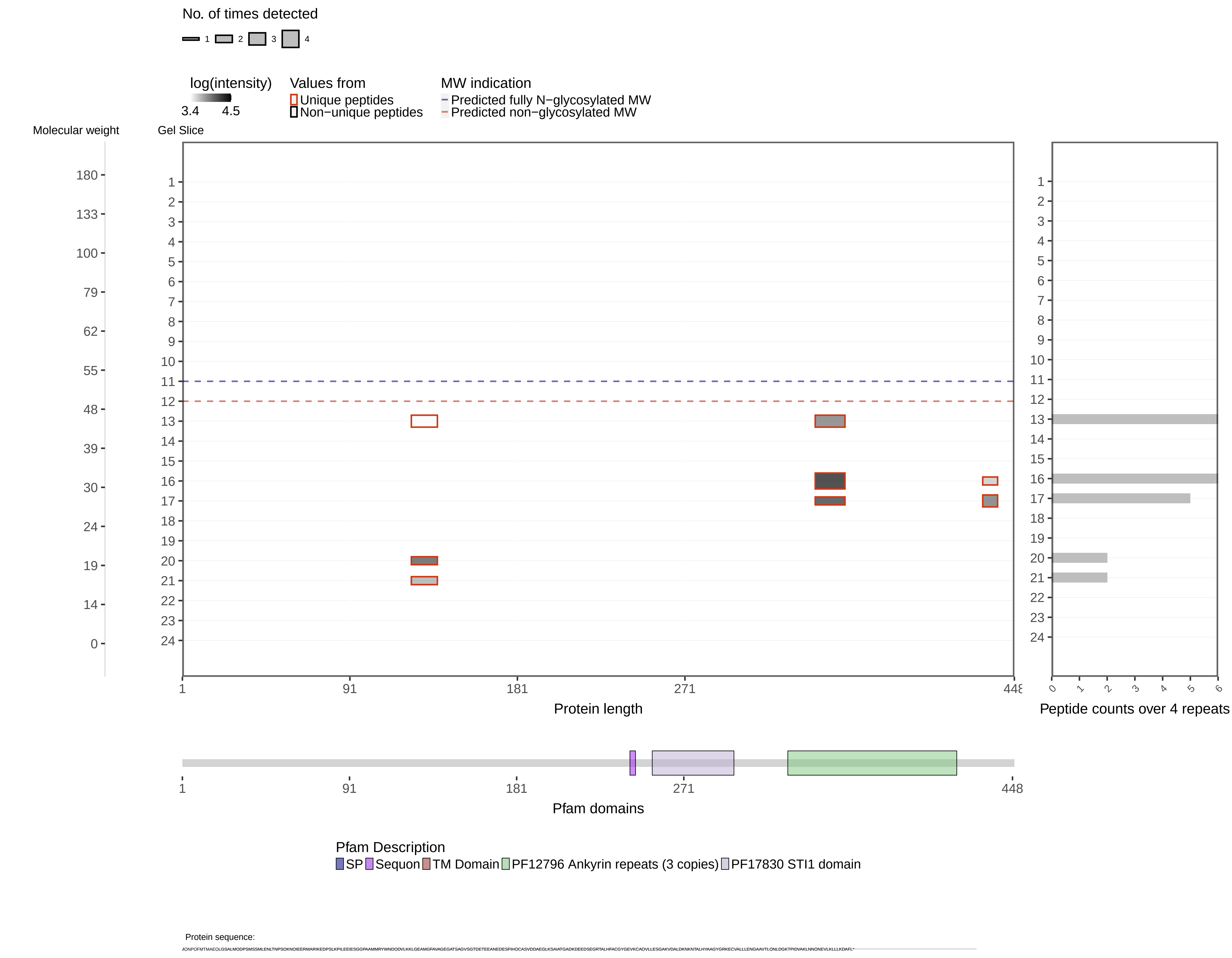

### NbL15g17440.1 60S_ribosomal_protein_L18−2−like [MW = 20.91 kDa]

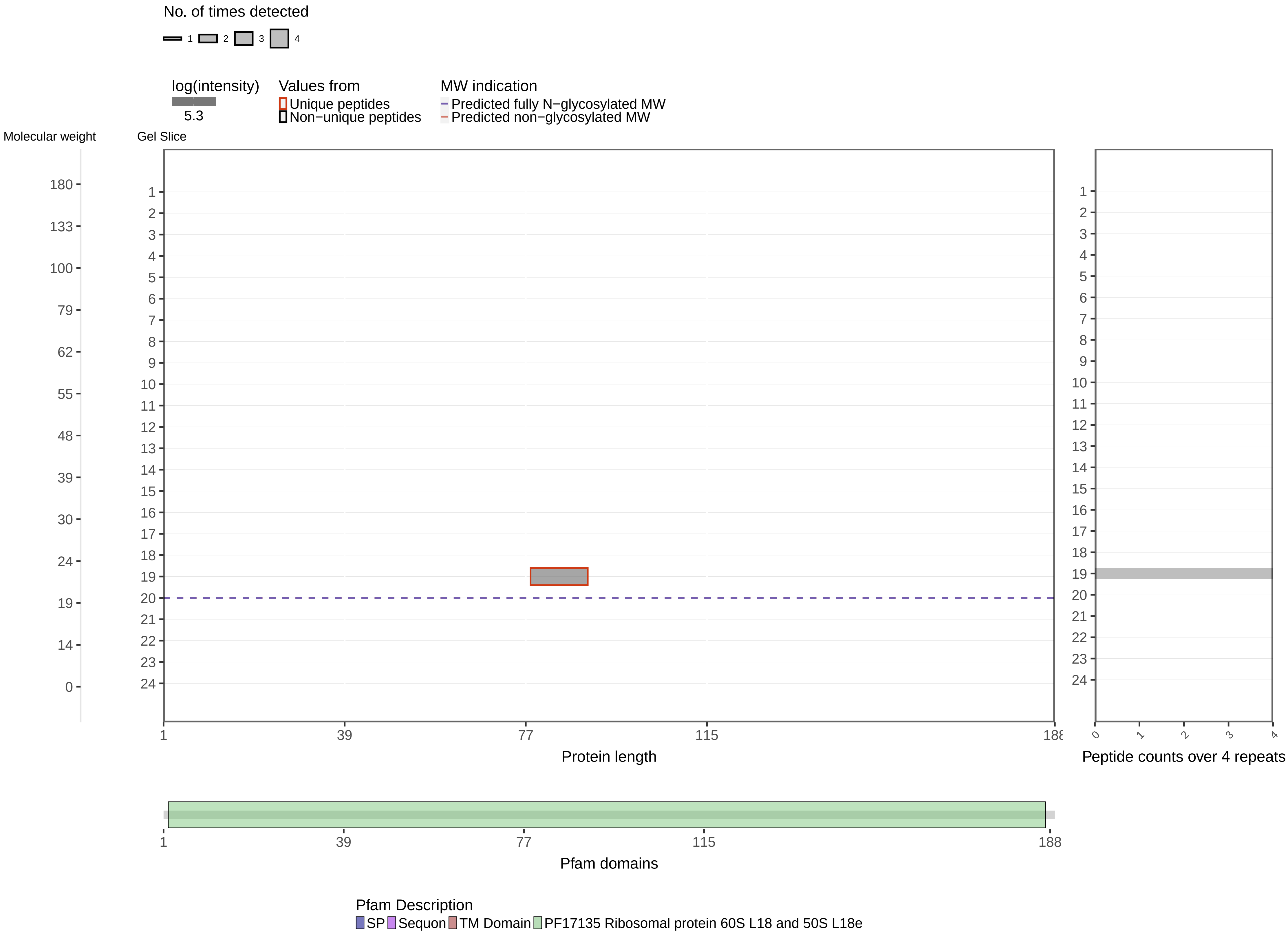


Protein sequence:

GTITDDVRAYDVPTLKVCALRFTKTARARIEKAGGECLTFDQLALRAPLGQNTLLLRGPKNAREAVKHFGPAPGVPHSHTKPYVRSKGRKFERARGRRKSRGYKV*

### NbL15g17410.1 60S_ribosomal_protein_L22−3−like [MW = 14 kDa]

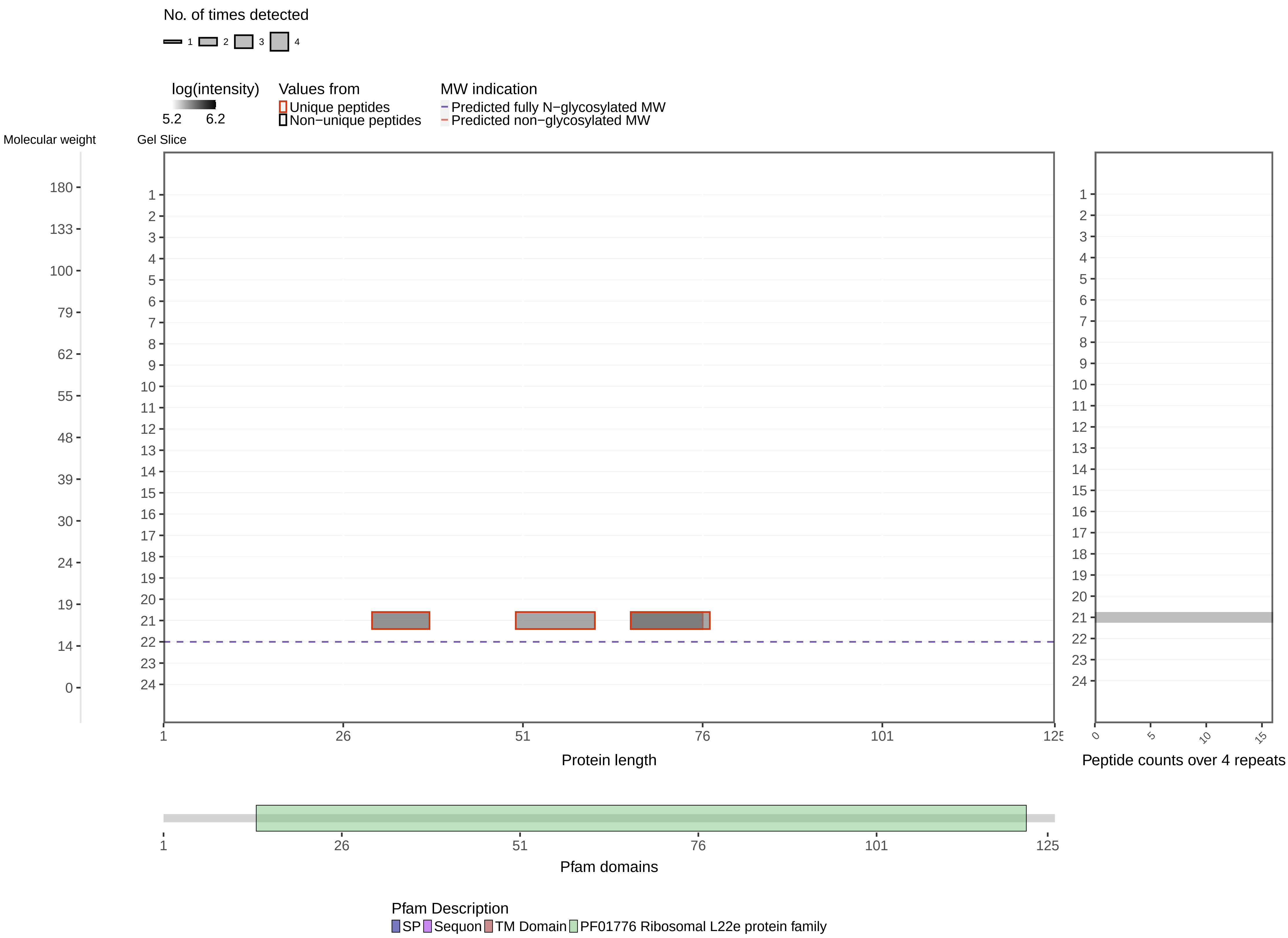


Protein sequence:

:AGALGDTVTVTRDKTKITVTSDSTFSKRYLKYLTKKYLKKNNVRDWLRVISSNKDRNVYELRYFNIAENEAEEED*

### NbL15g17080.1 40S_ribosomal_protein_S5−like [MW = 23.37 kDa]

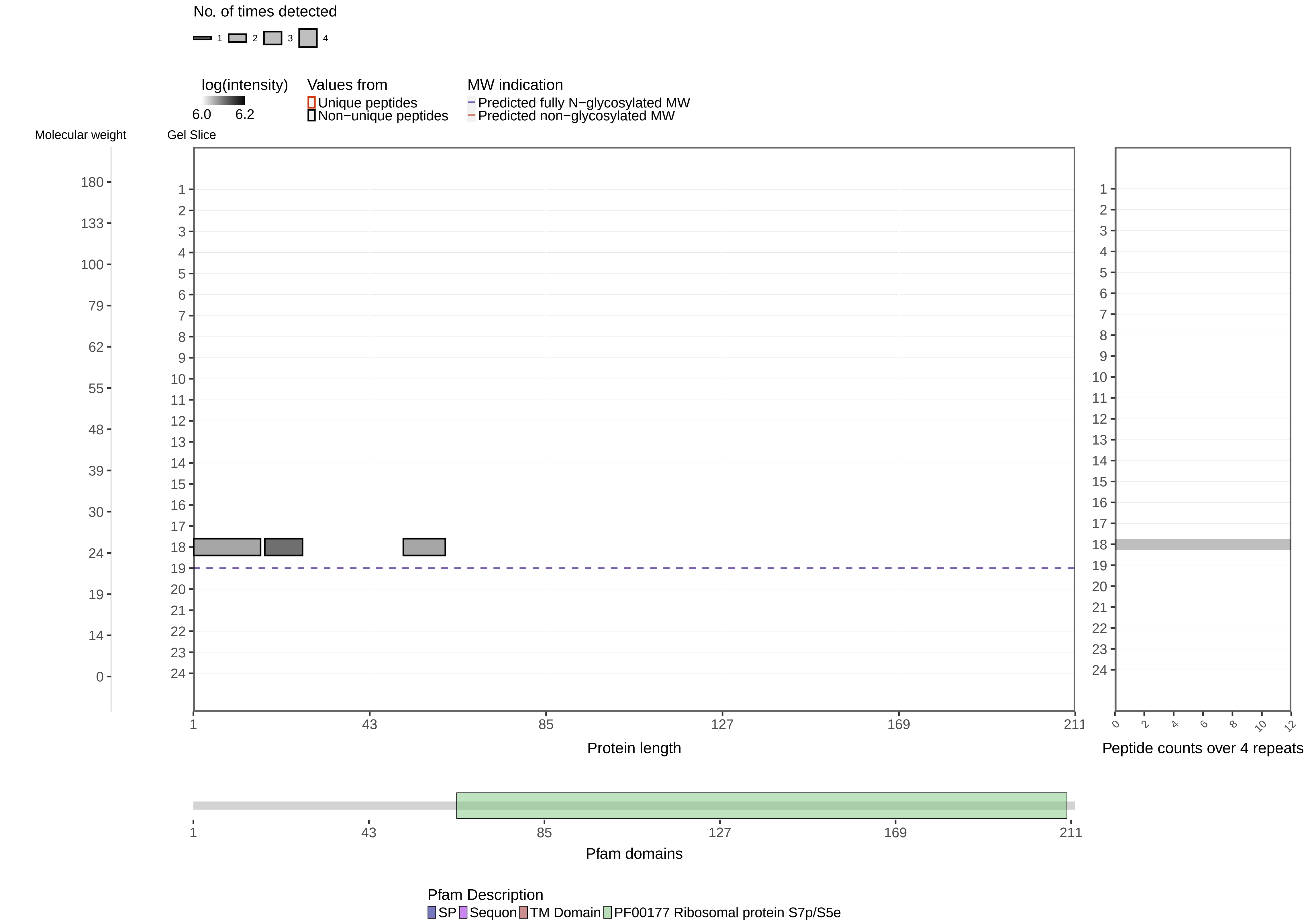


Protein sequence:

GKKLMAVRIIKHAMEIIHLLTDQNPIQVIVDAVINSGPREDATRIGSAGVVRRQAVDISPLRRVNQAIYLLTTGARESAFRNIKTIAECLADELINAAKGSSNSYAIKKKDEIERVAKANR*

NbL15g17000.1 carbon_catabolite_repressor_protein_4_homolog_1−like [MW = 66.98 kDa]

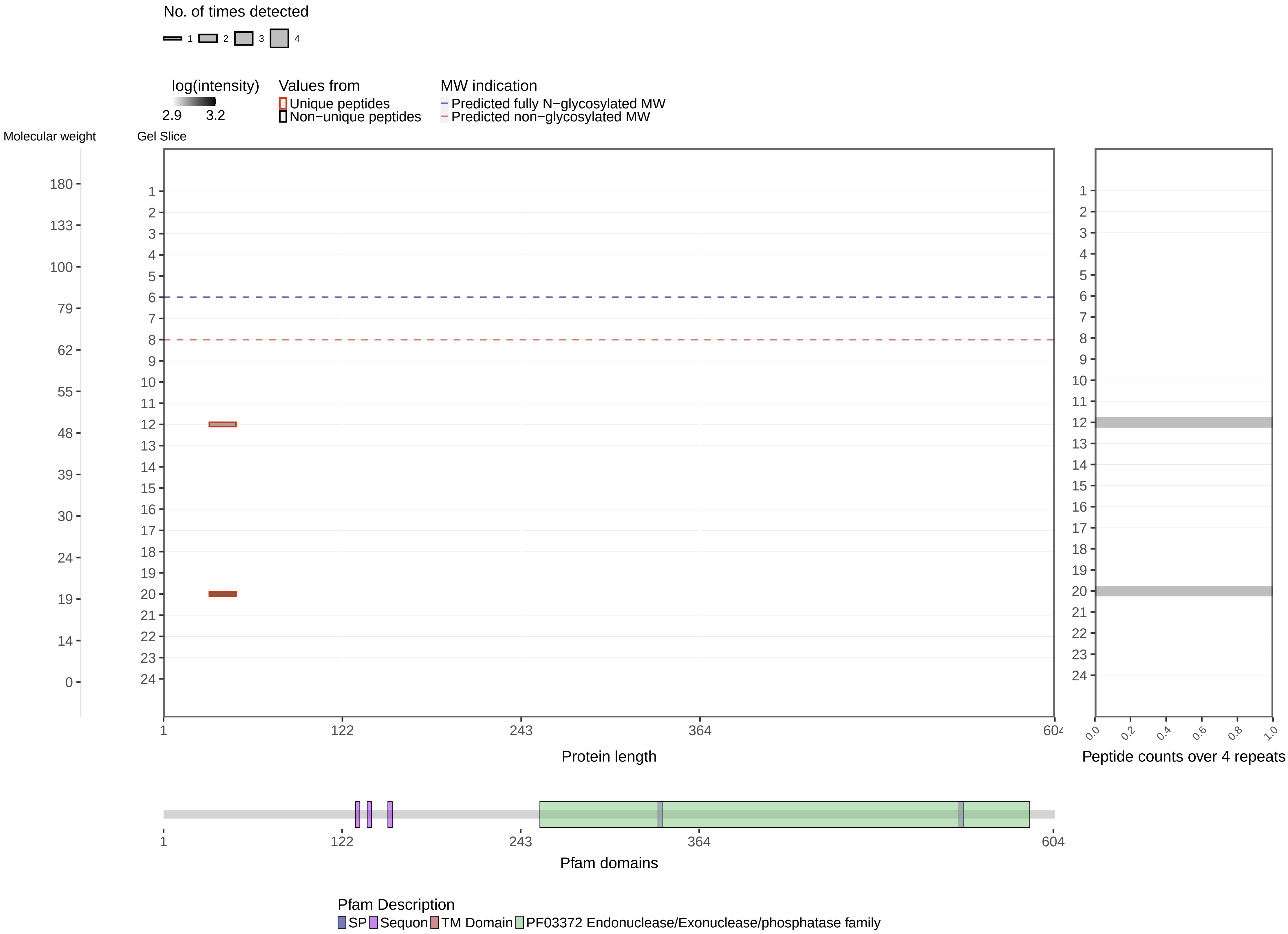


Protein sequence:

.REIVGYRADIVCLQEVQSNHFEEFFAPELDKHGYQALFKRKTAEVFNGSINNIDGCATFFRRDRFSHVKKYEVEFNKAAQSLTDAVVPIAQKKTALNRLVKDNVALIVVLEAKFSNQGVDNPGKRQLVCVANTHVNVQQELRDVKLWQVHTLLKGLEKIAASADIPMLVCGDFNSVPGSAPHTLLALGKVDPMHPDLAVDPLGILRPPTKLTHQLPLVSAYSSFARMGVGLGLEQQRRRMDPNTNEPLFTNCTRDFIGTHDYIFYSADSLTVESLLELLDEESLRKDTALPSPEW

NbL15g16830.1 enoyl−[acyl−carrier−protein]_reductase_[NADH]_2_chloroplastic−like [MW = 41.67 kDa]

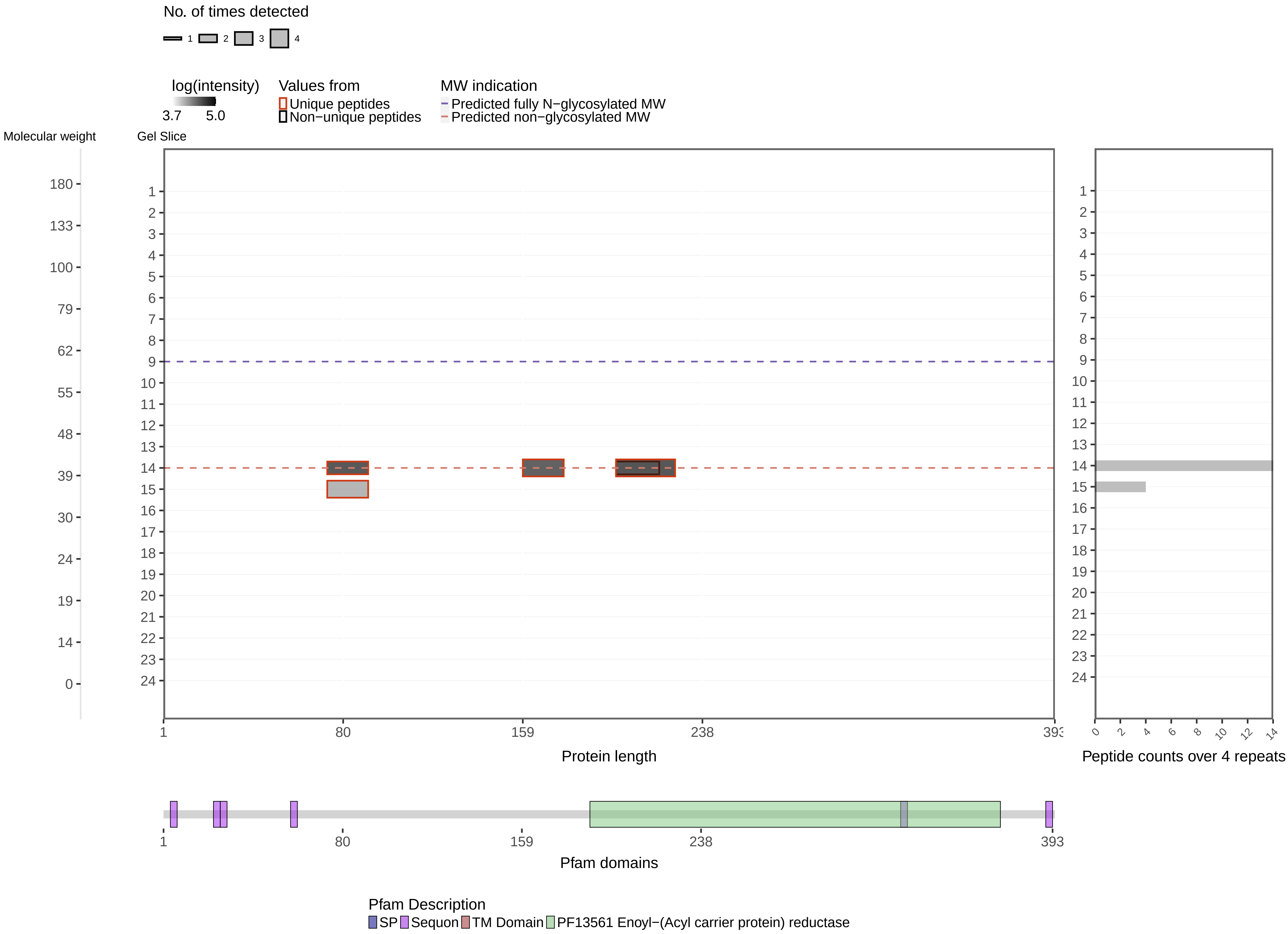


Protein sequence:

GSSKWTVKEVAECVKRDFGTIDILVHSLANGPEVSKPLSETSRKGYLAAVSASSYSFISLLRHFLPIINPGGATISLTYIASERIIPGYGGGMSSAKAALESDTRVLAFEAGRKHKVRVNTISAGPLGSRAAKAIGFIDMMINYSLENAPLQKELYAEEVGNTAAFLASPLASAITGATVYYVDNGLNAMGVGVDSPAFSGLDIPKDNKS*

### NbL15g16410.1 L−galactono−14−lactone_dehydrogenase_mitochondrial [MW = 71.75 kDa]

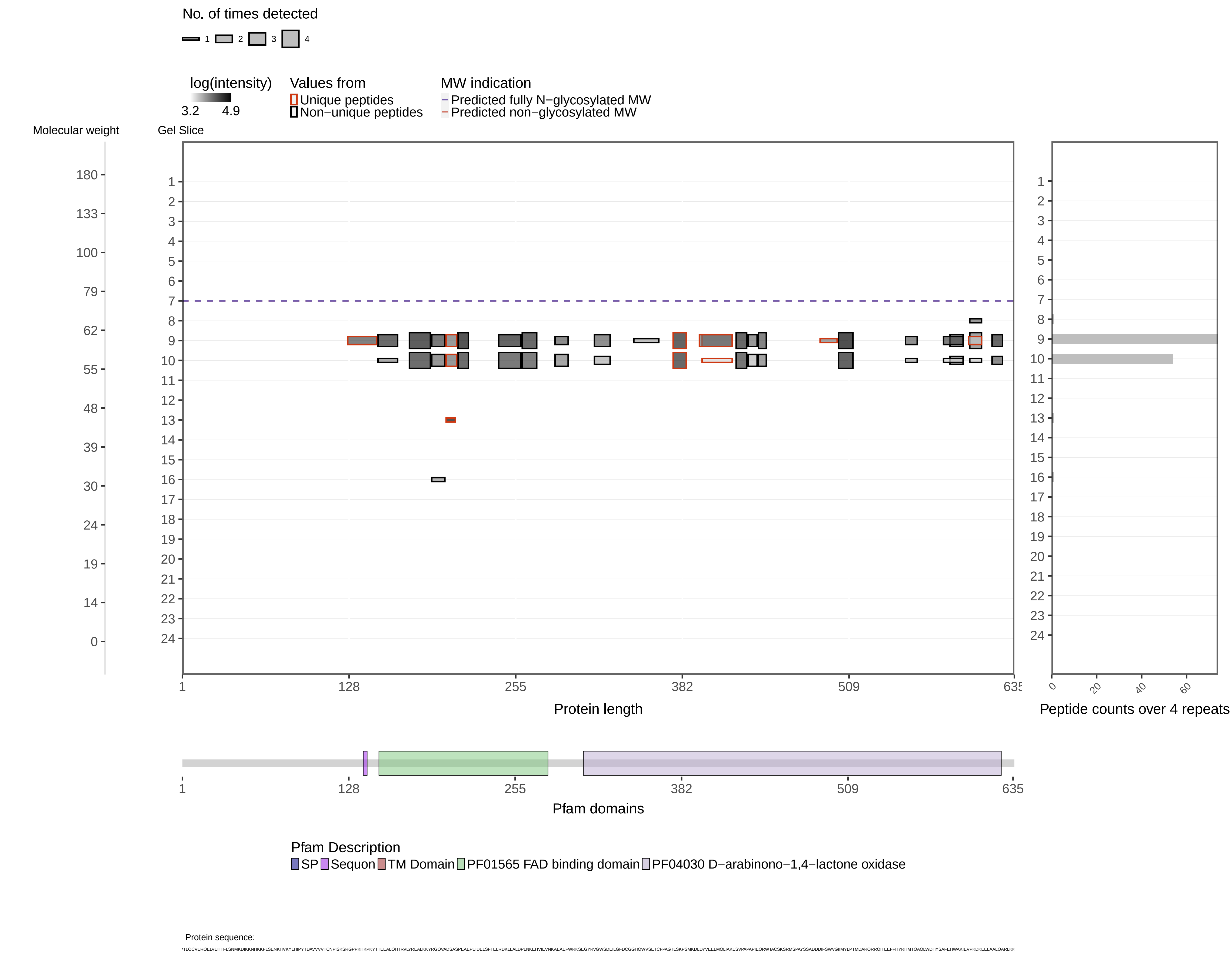

NbL15g16300.1 putative_glucose−6−phosphate_1−epimerase [MW = 34.76 kDa]

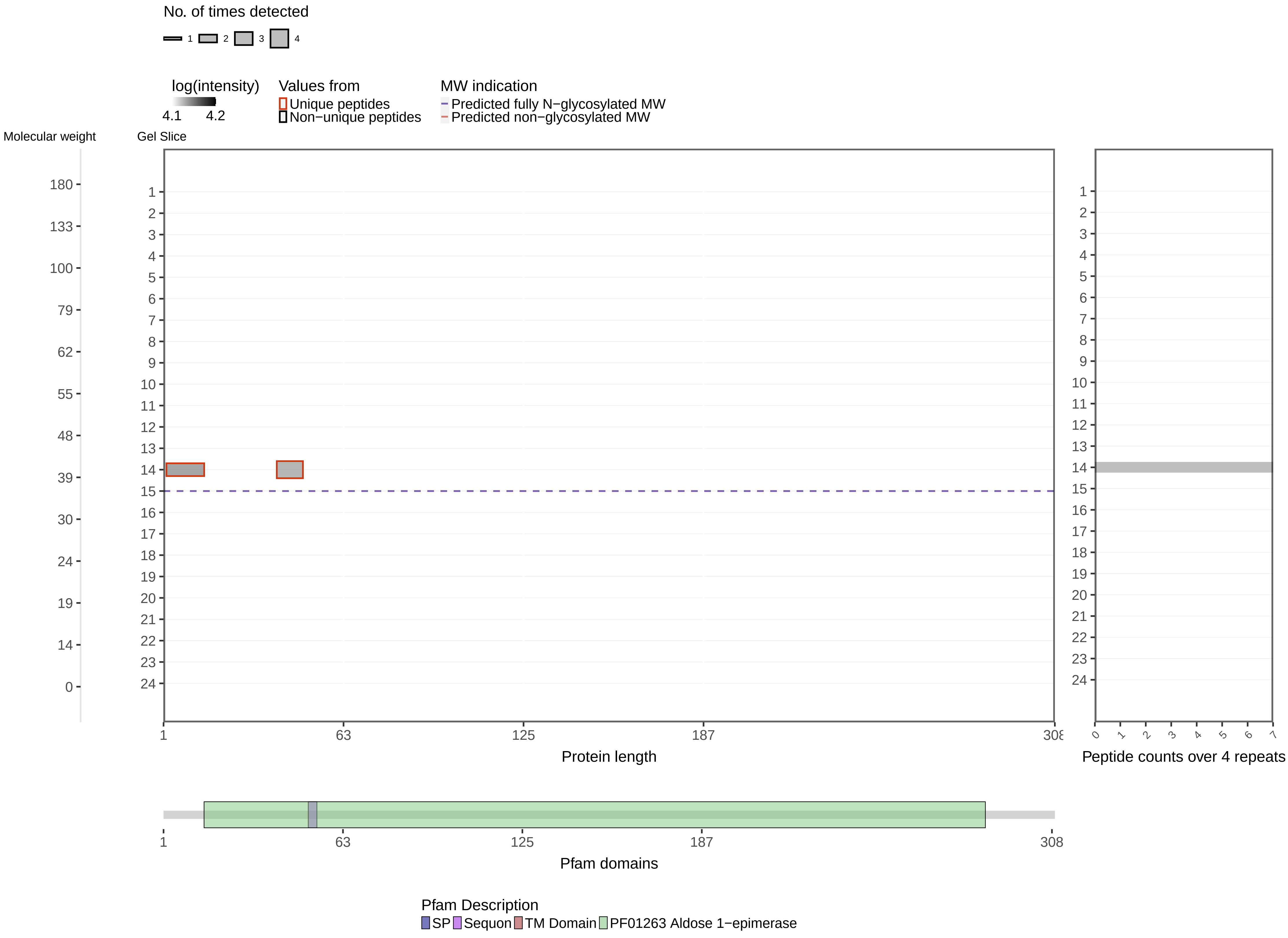


Protein sequence:

PRVRNTDNKPFSFTISLRNYLSVSDISEVRVEGLETLDYFDNLQQRERYTEQADAITFDSEIDRVYLSTPKKIAIIDHEKKRTYVIQKEGMLDAVVWNPWDKKAKSLPDLGDEDYKKMLCVDSAAVETPIVLKPSEEWKGRQELSTVSSSYCSGQLDPRKVLYGLA*

NbL15g16260.1 BAG_family_molecular_chaperone_regulator_4−like [MW = 26.07 kDa]

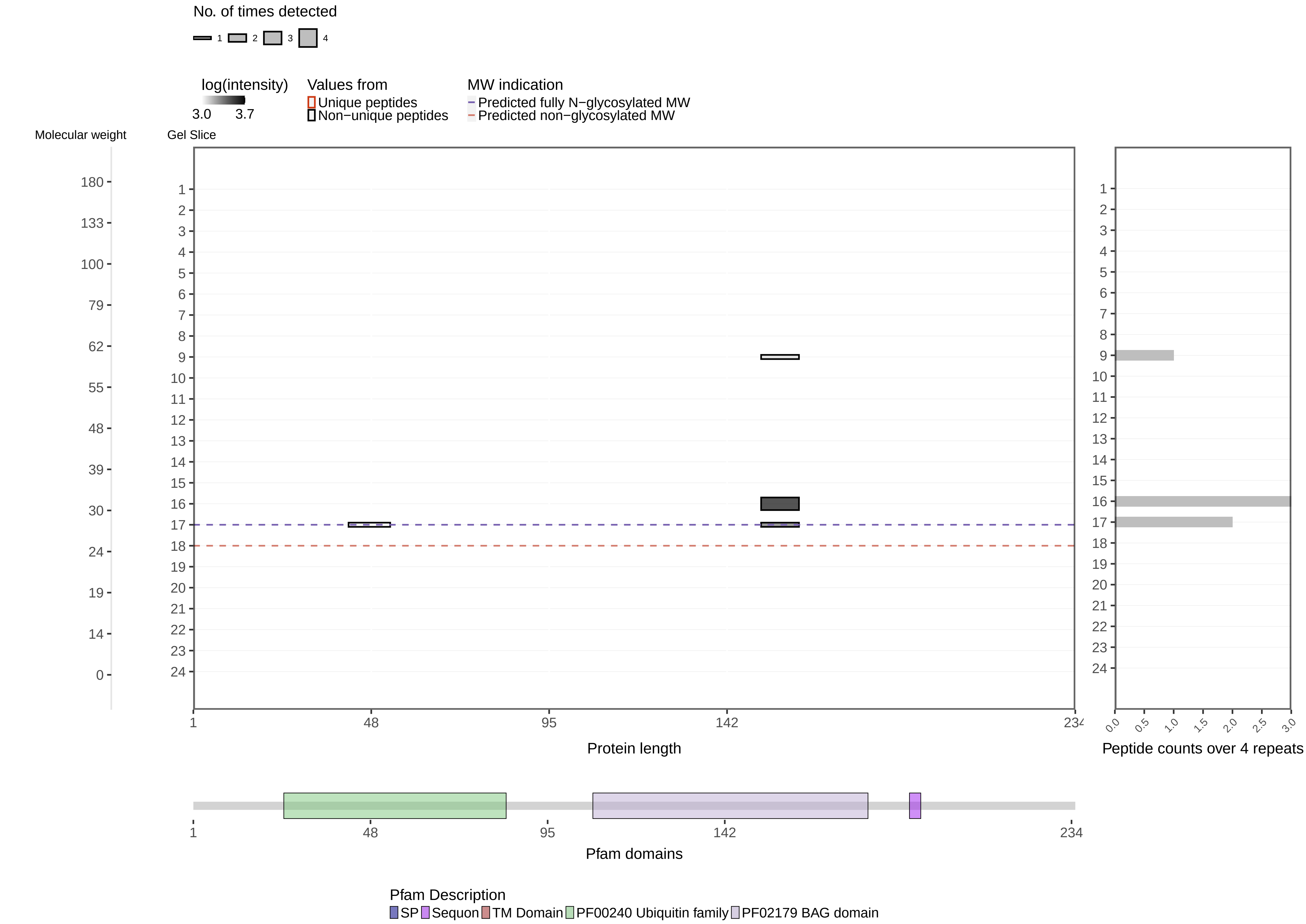


Protein sequence:

VAEVRTEVDKLSEQVSAVQAVVFGGSNVEEKDIICLTEMLMRQLLKLDEIDAEGEGKIQRKMEVRRVQSLVETMDAVKARNANPFSNNSNAVSVTTQWETFESGVGSLNAPPPPRPSSTVVTEDWEQFE*

NbL15g15910.1 mitochondrial_amidoxime_reducing_component_2−like [MW = 34.14 kDa]

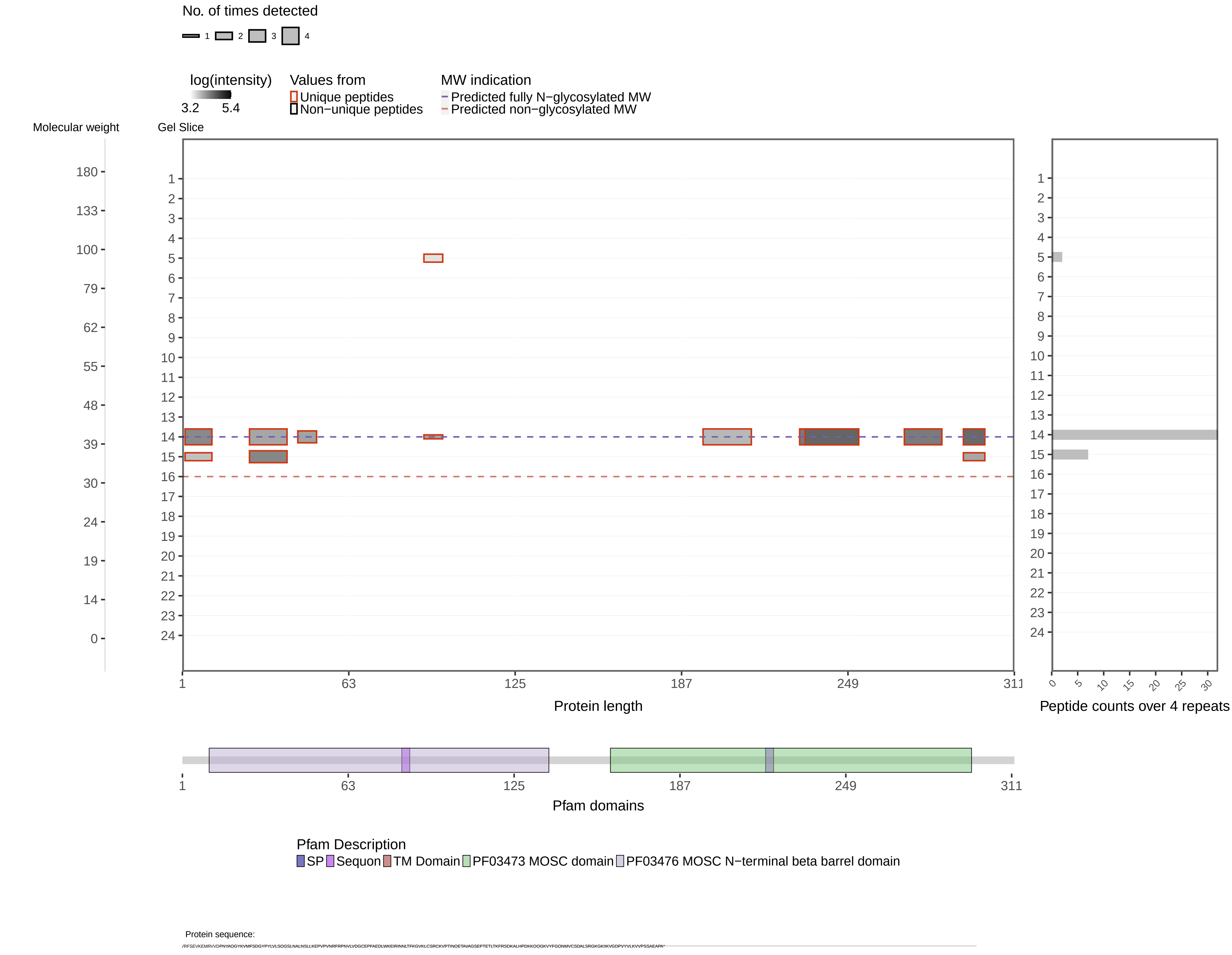

### NbL15g15850.1 uncharacterized_LOC107826751 [MW = 20.62 kDa]

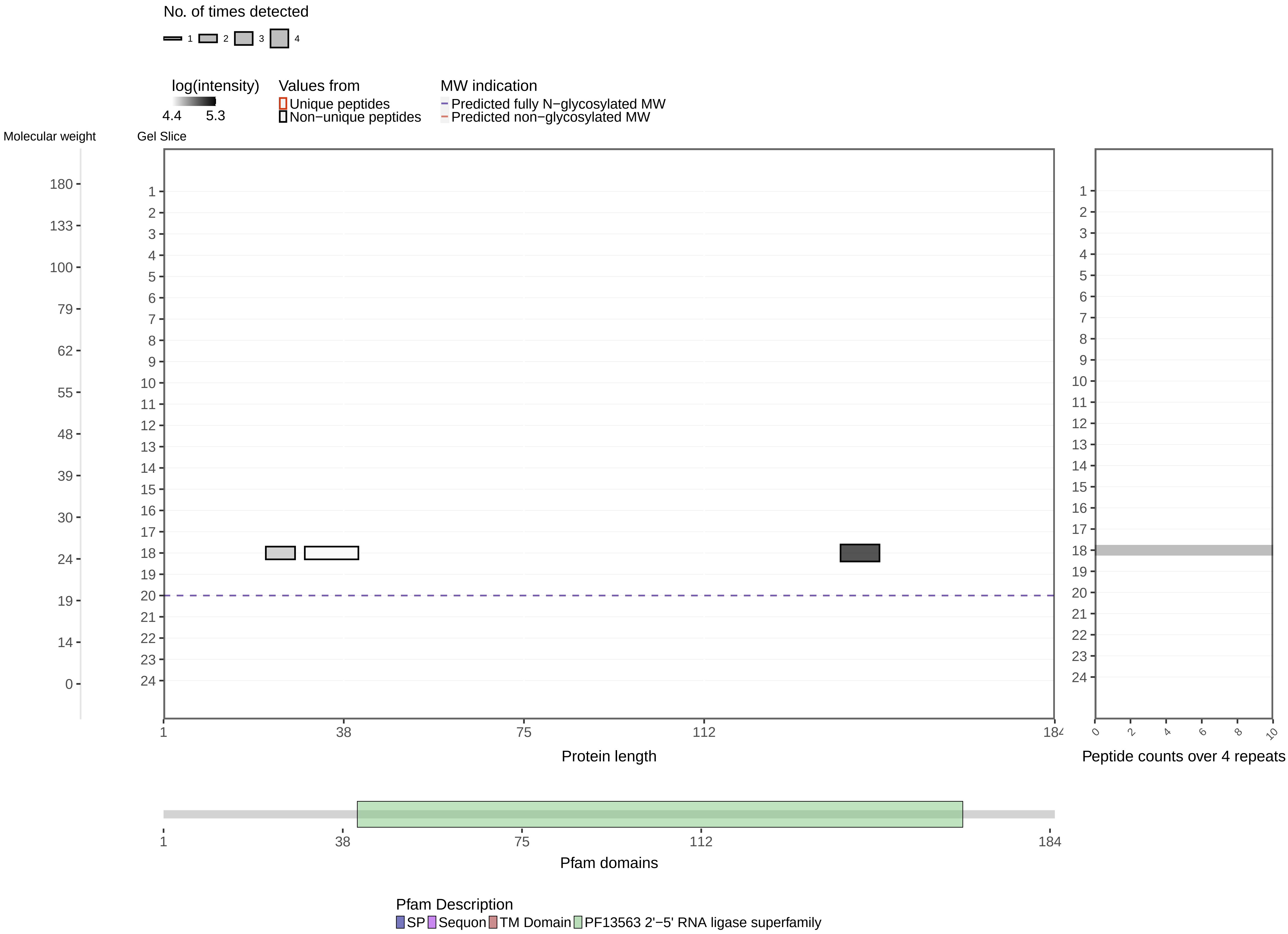


Protein sequence:

SLPNDDNILFLAPTPTLSLLQFHSQLFEAMKKEGIEIGEEHRPDTWIPYCPVAEEVPKTRMGEAFTVLRDLKLPVAGYAMEIALVEYPPVREVFSFVLGNTVEP*

### NbL15g15800.1 40S_ribosomal_protein_S15−like [MW = 17.24 kDa]

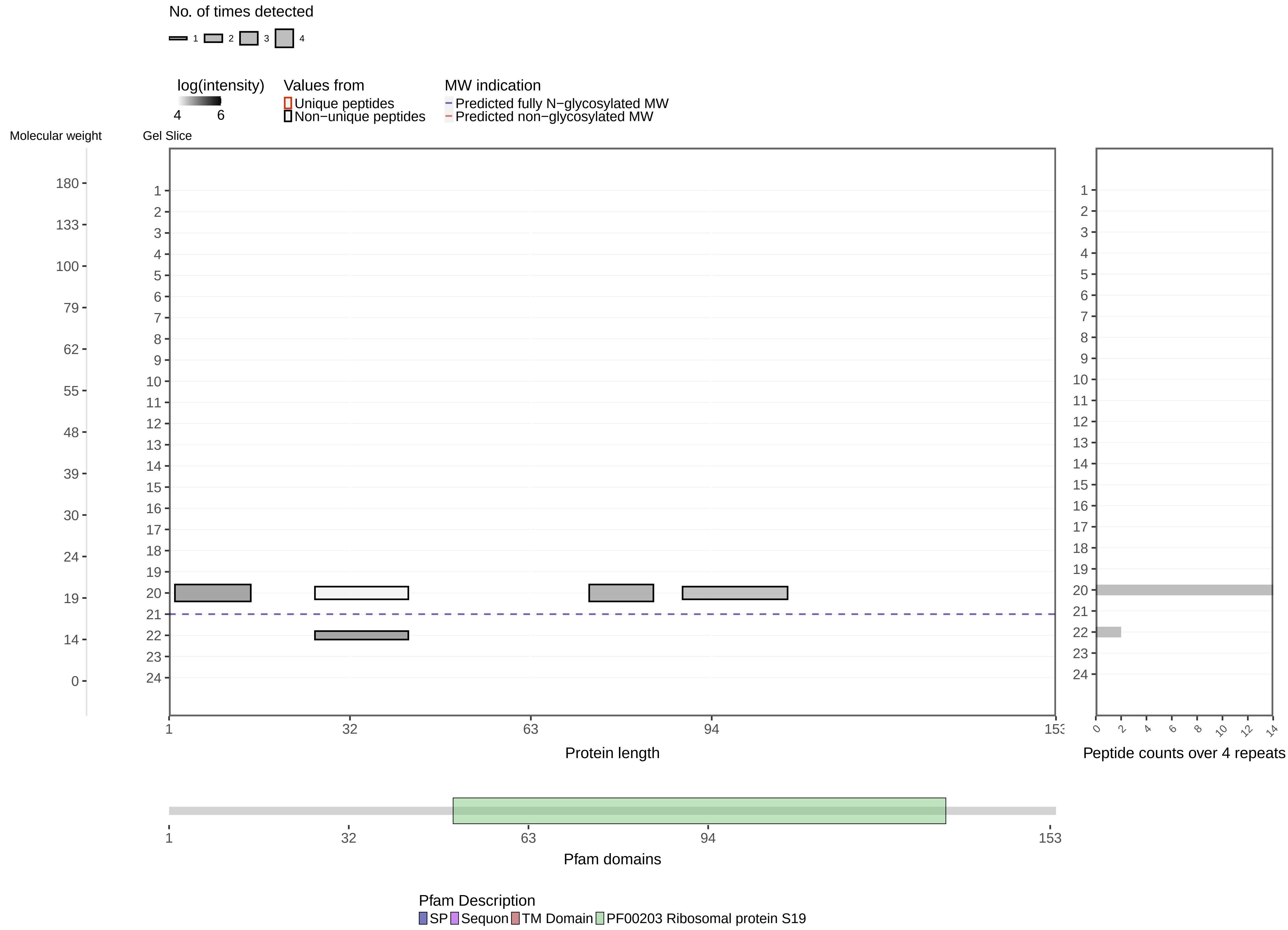


Protein sequence:

MALIKKLRKAKREAPPGEKPEVVRTHLRNMIIVPEMIGSVIGIYNGKTFNQIEVKPEMISHYLAEFSISYKPVKHGRPGIGATHSSRFIPLK*

NbL15g15540.1 haloacid_dehalogenase−like_hydrolase_domain−containing_protein_3 [MW = 29.08 kDa]

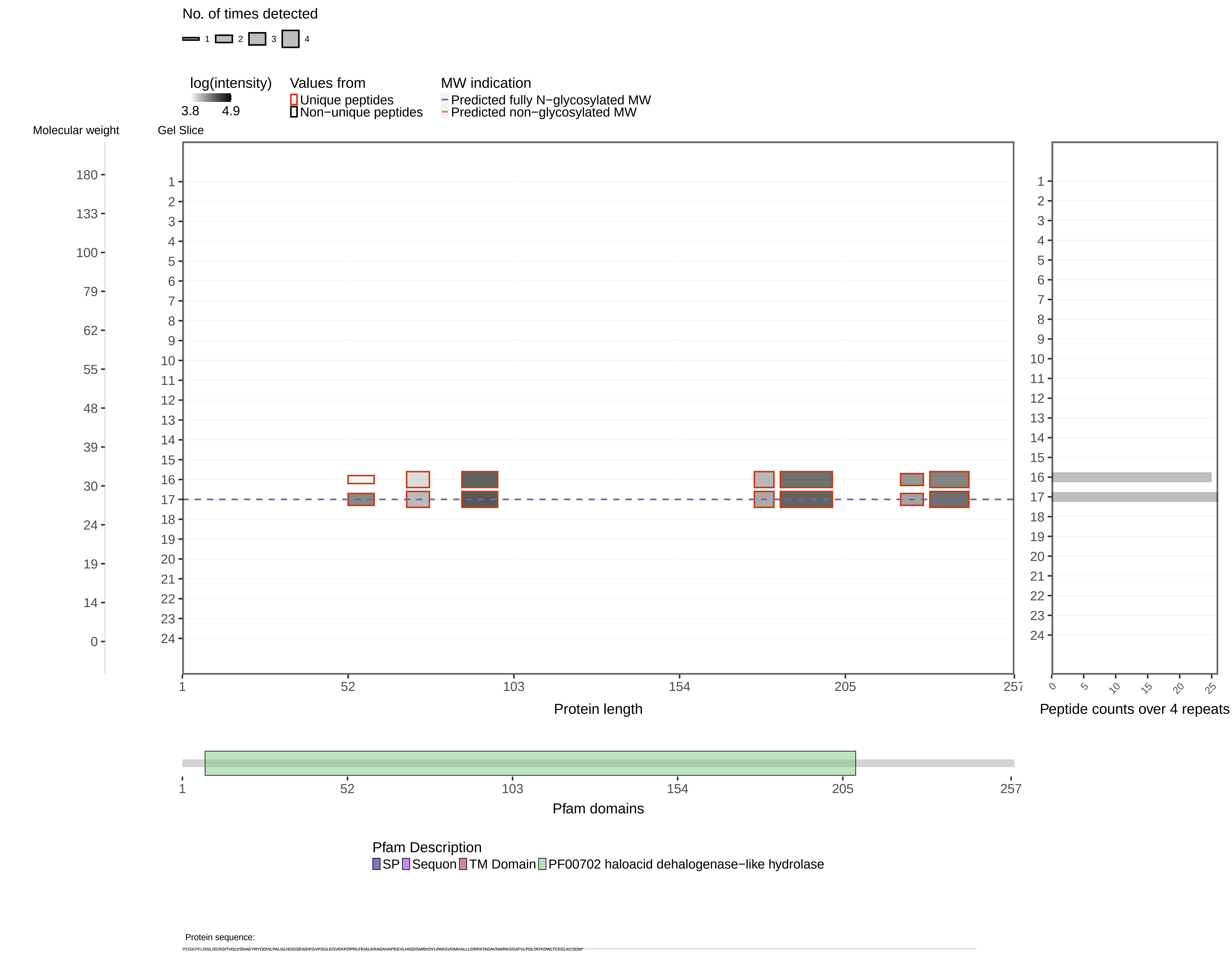

### NbL15g15510.1 proteasome_subunit_alpha_type−4 [MW = 27.15 kDa]

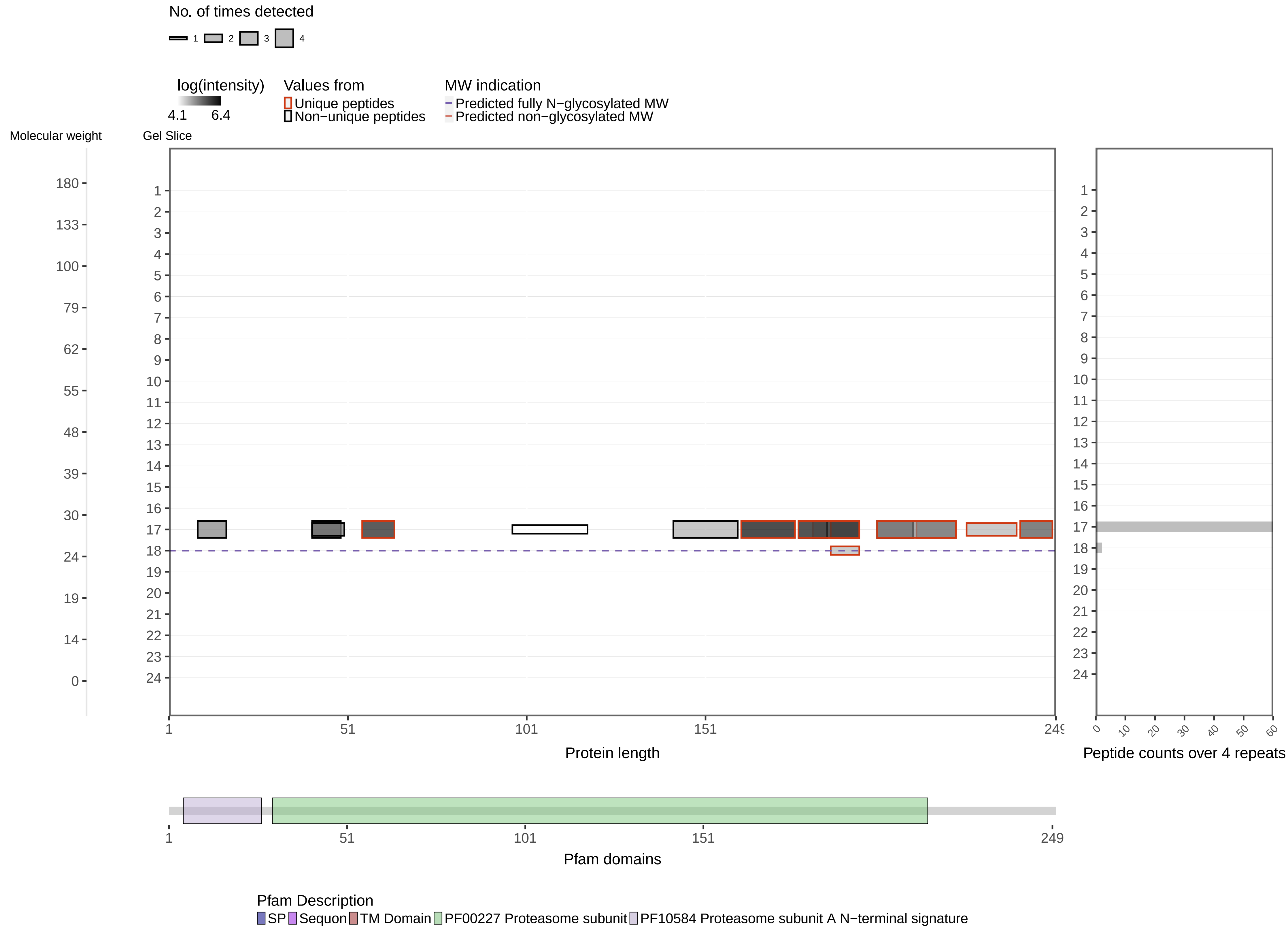


Protein sequence:

LCDTKQGYTQFGGLRPFGVSFLFAGWDKNYGFQLYMSDPSGNYGGWKAAAIGANNQAAQSILKQDYKDDITREEAVQLALKVLSKTMDSTSLTSEKLELAEVFLSNGKVKYQACSPESLNAMLVRSGLTQPAAEE*

### NbL15g15100.1 L−ascorbate_oxidase_homolog [MW − SP = 57.57 kDa]

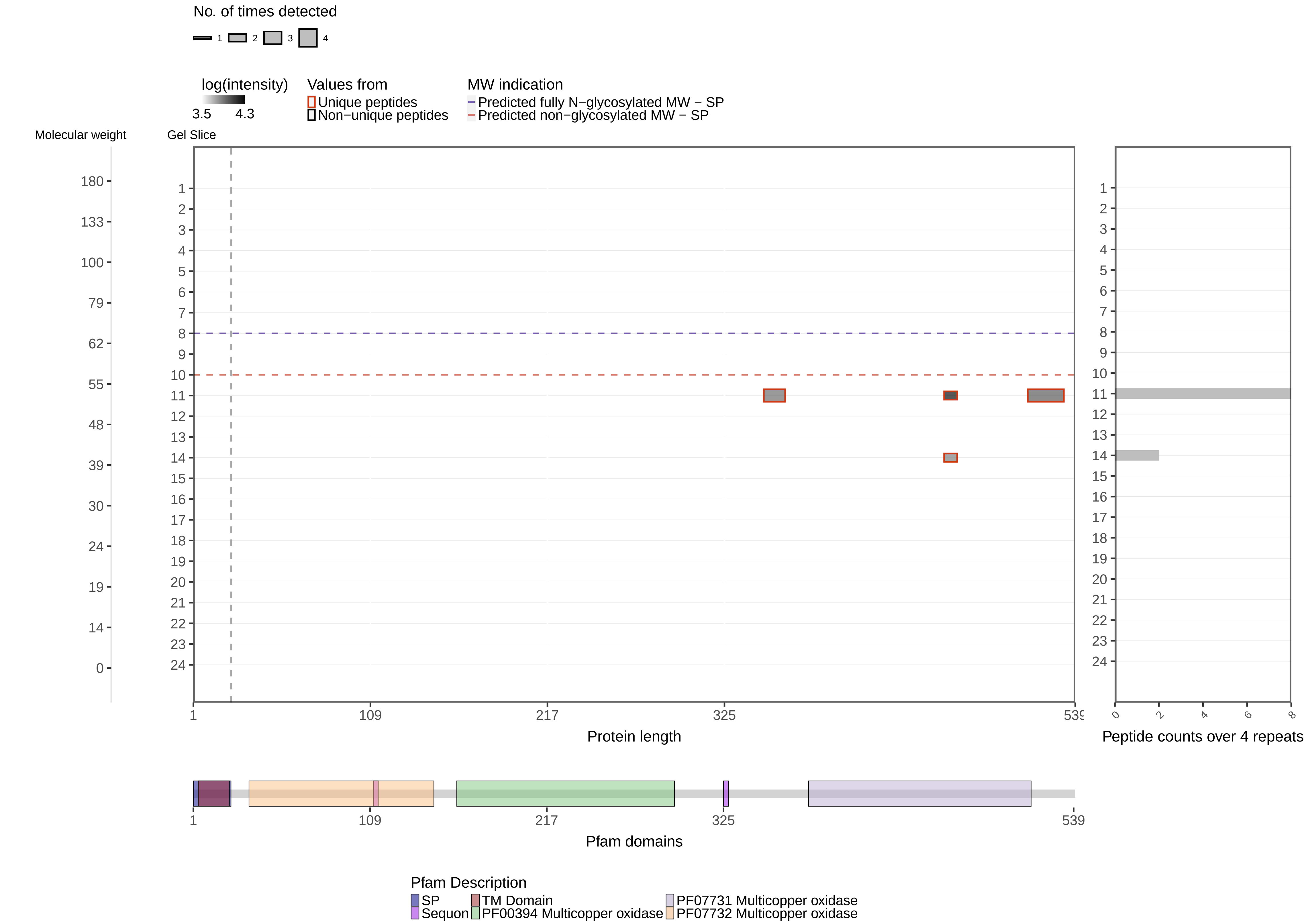


Protein sequence:

### NbL15g14860.1 ABC_transporter_F_family_member_5−like [MW = 78.54 kDa]

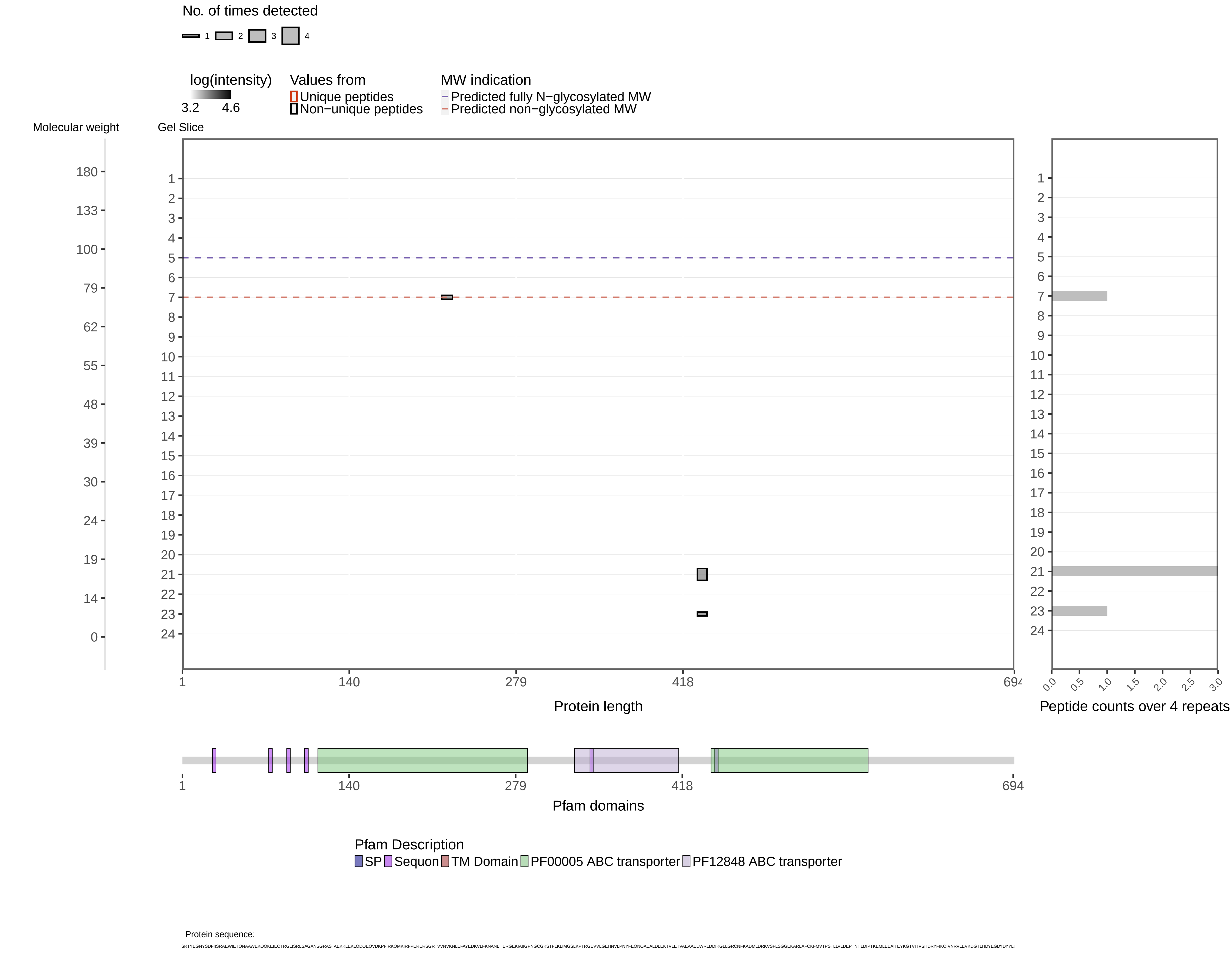

### NbL15g14840.1 developmentally−regulated_G−protein_3 [MW = 41.25 kDa]

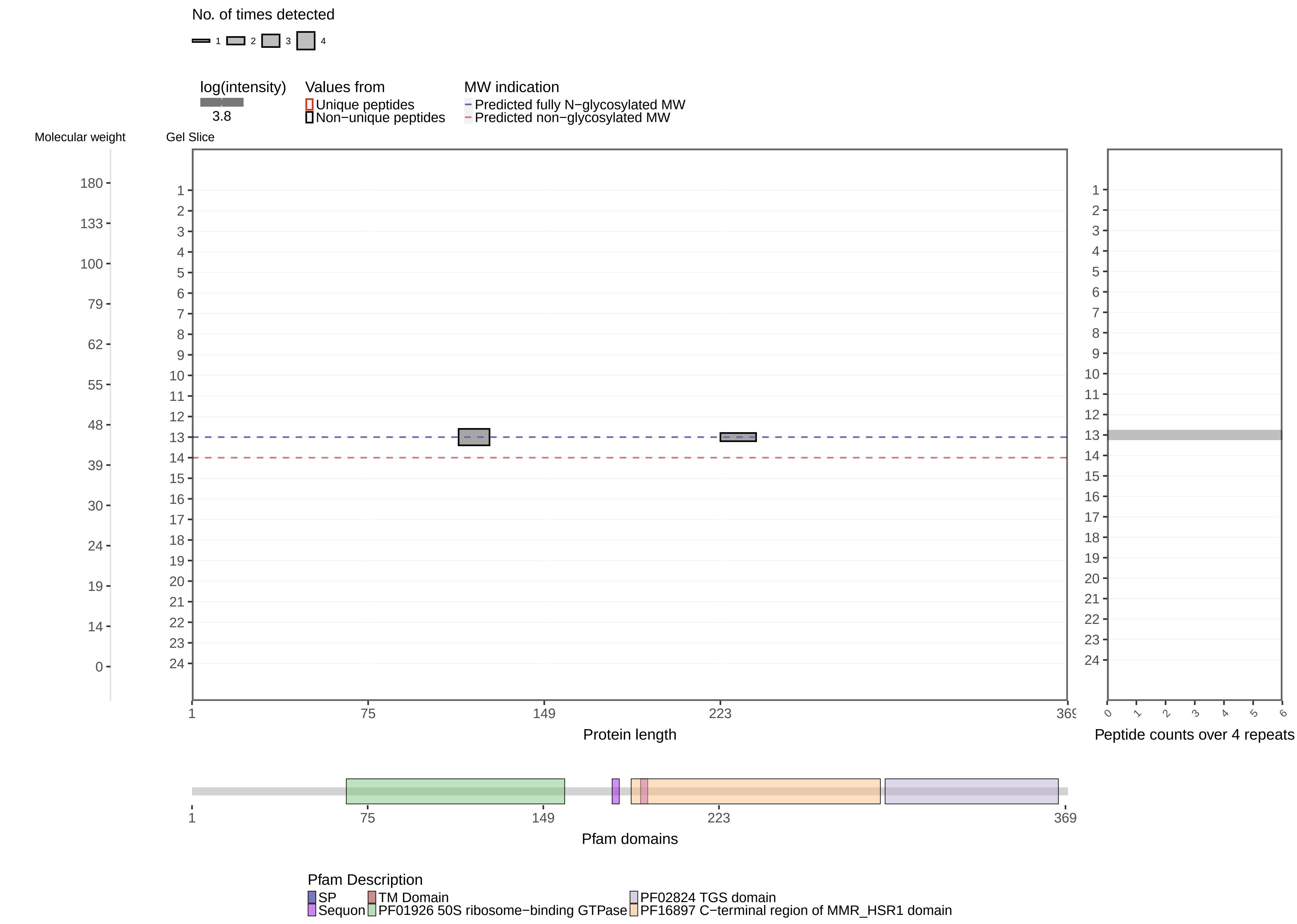


Protein sequence:

LNKEPPNLTFRKKEKGGINLTSTVTNTHLDLDTVKAICSEYRIHNADVHLRYDATADDLIDVIEGSRVYTPCIYVVNKIDQITMEELEILDKLPHYCPISAHLEWNLDGLLEKIWEYLSLTRIYTKPKGMNPDYEDPVILSSKRRTVEDFCDRIHKDMVKQFKYALVWGSSAKHKPQRVGREHELEDEDVVQIIKKV*

NbL15g14820.1 shikimate_kinase_chloroplastic [MW = 33.22 kDa]

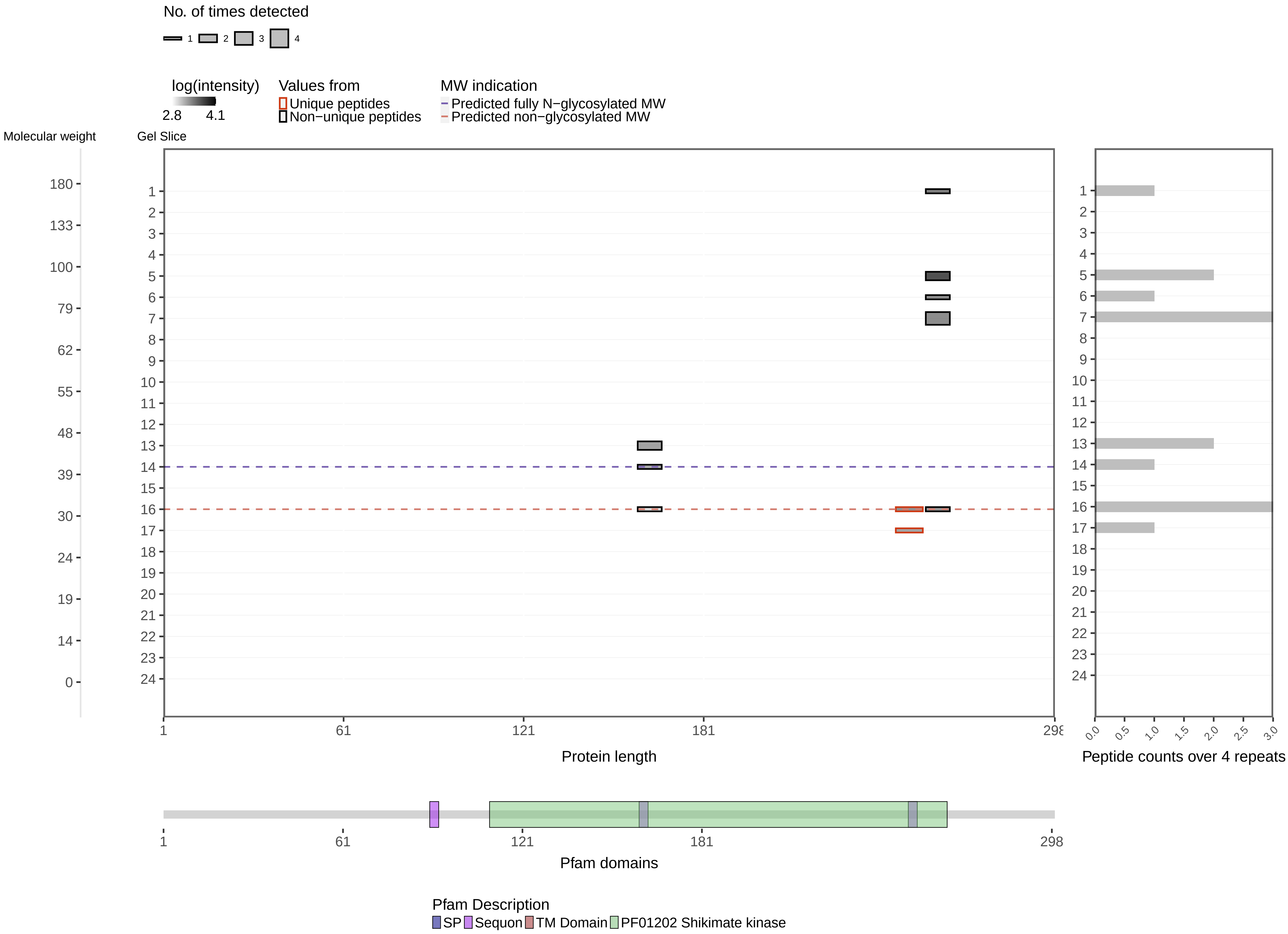


Protein sequence:

RLIEQAVGGTTVAEIFKLRGESFFRDNETEVLHKLSAMHRLVVSTGGGAVVRPINWRHMHKGISVWLDVPLDALAKRVTAEGTKSRPLLHEESGDIYDKTLKLRLTTLMETRGENYANASVRVSLENIAVKRGKDVCHITPTEITLEVLIQIQNFLKKQESVVVL*

NbL15g14720.1 4−alpha−glucanotransferase_chloroplastic/amyloplastic [MW = 65.37 kDa]

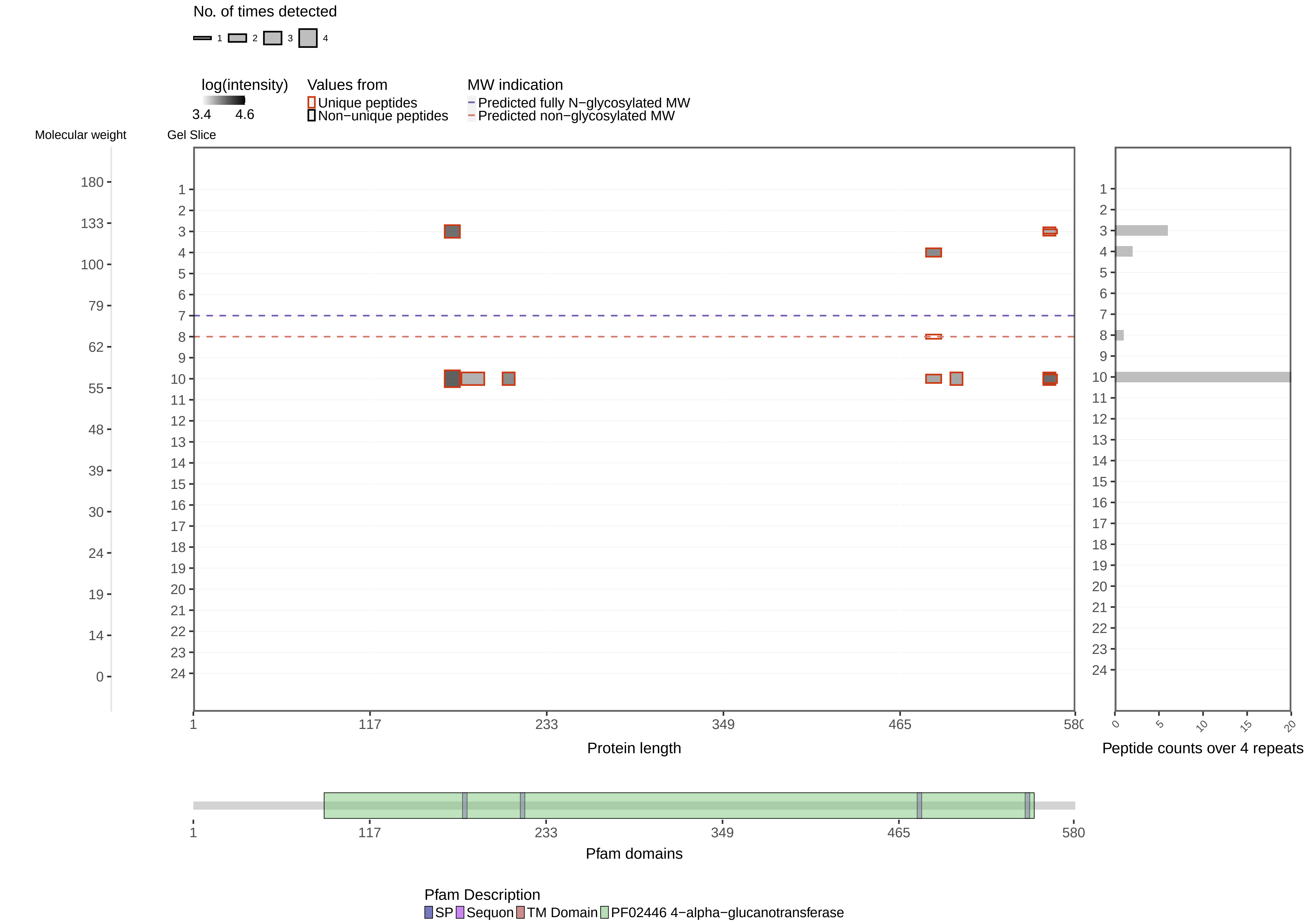


Protein sequence:

[illegible]WQKVRDYARSKGISIMGDMPIYVGYHSADVWANKKHFLLNRKGFPLIVSGVPPDAFSETGQLWGSPLYDWKAMEKDGFSWWIRRIRRATDLFDEFRIDHFRGFAGFWAVPSEAKVAMLGRWKVGPGKPLFDAIFQAVGKINIIAEDLGVITEDVVQLRKSIEAPGMAVLQFGFGSDAENPHLPHNHEQNQVVYTGTHDNDTTRGWWDILPQEEKSNVLKYLSNIEEEKISWGLIEAAISSVARIAIVPMQDVLGLGSESRMNIPATQFGNWSWRIPSSTRFDSLDAEAKKLR

NbL15g14640.1 2−hydroxyisoflavanone_dehydratase−like [MW = 35.73 kDa]

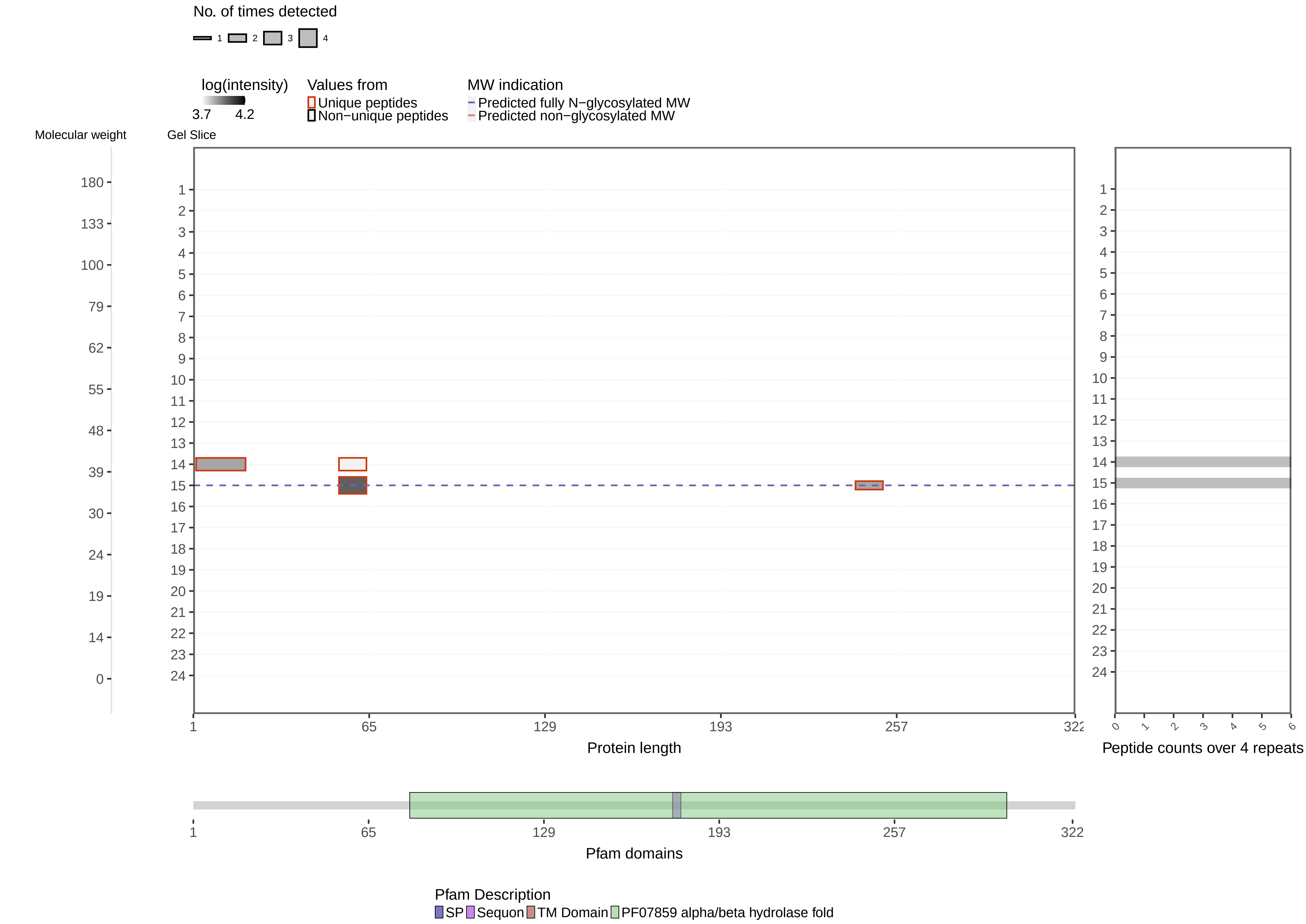


Protein sequence:

DAEPWITNHGDFNRLFIAGDSAGGNIVFNMTMRAGKESLIGGVKLVGAIFAFPFLLIPSVENIEGSLSYKLWNTICPPSEOGINSPMVNPVSEKGOSLSELGCSRIFLCAGEKDEHVPSEIVTRFAEAVKNSGWKGEVEFIEIEGEVHCFOAANPEAEKSKDLIKRMASFIORK*

### NbL15g14540.1 uncharacterized_protein_At2g39795_mitochondrial [MW = 31.65 kDa]

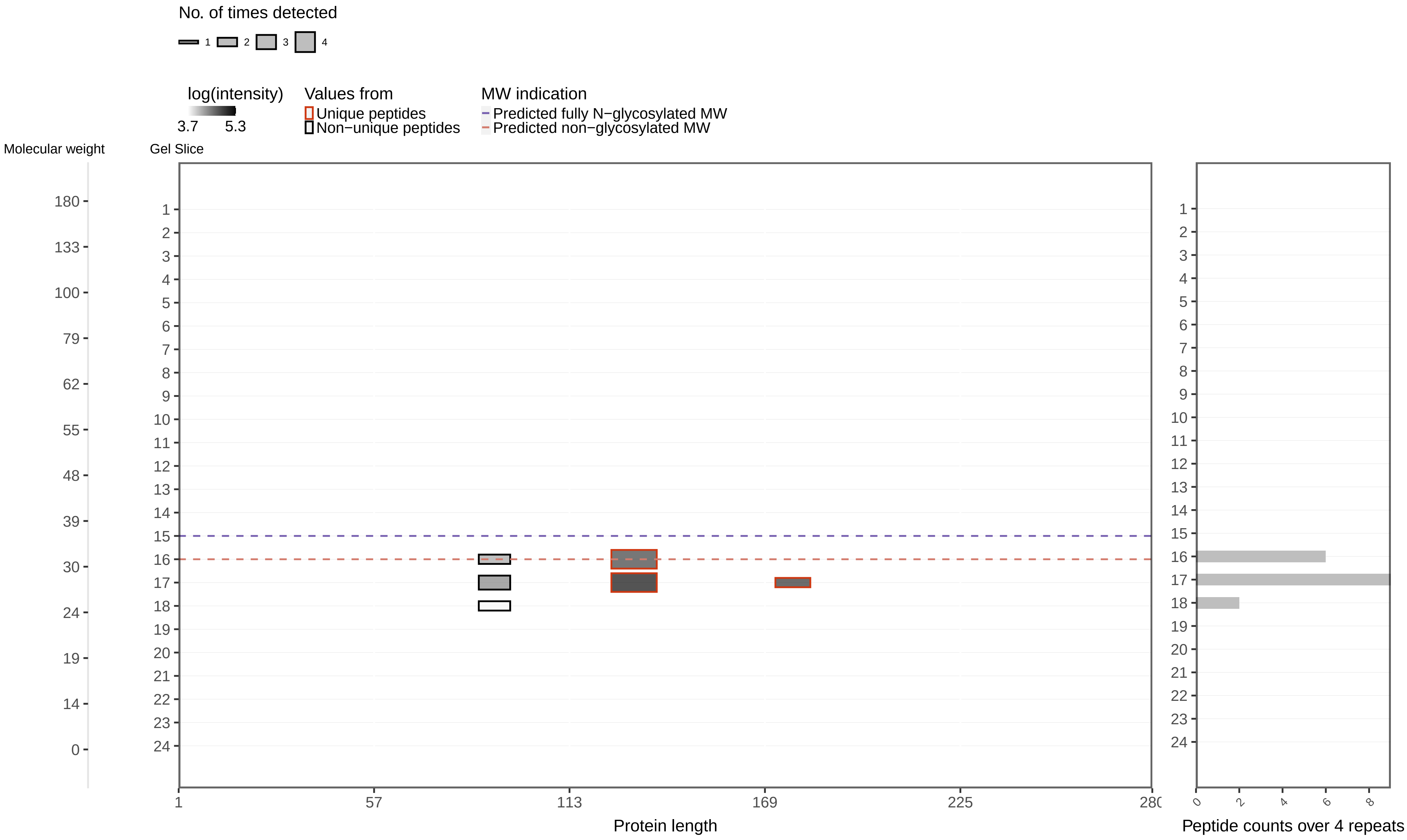


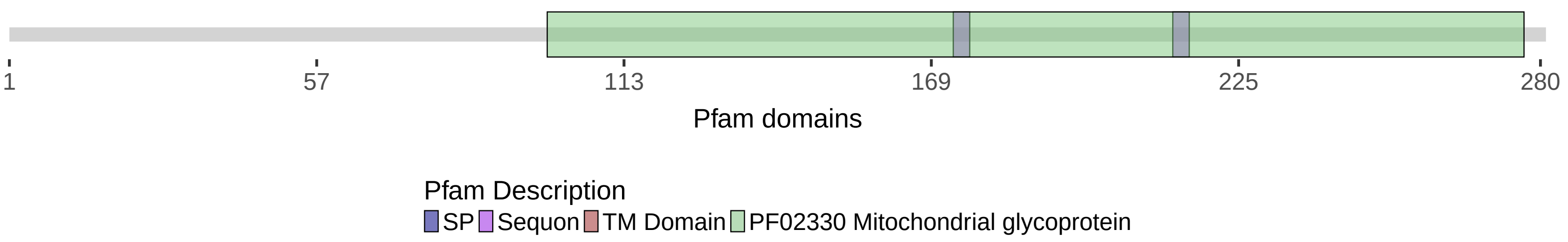


Protein sequence:

HPGQQTVTLIREYQGESIYVEVHMPDLVTGDEDENDNDDNDRERANQSQIPLVVRVSKKNGPSLEFGCTAFADEIAIETLSIKDPNASEDQIAYEGPDFSDLDENLQKAFHKYLEIRGIKPSATNFLHEYMINKDGREYVMWLKNLKKFIEA*

NbL15g14460.1 7−deoxyloganetic_acid_glucosyltransferase−like [MW = 53.48 kDa]

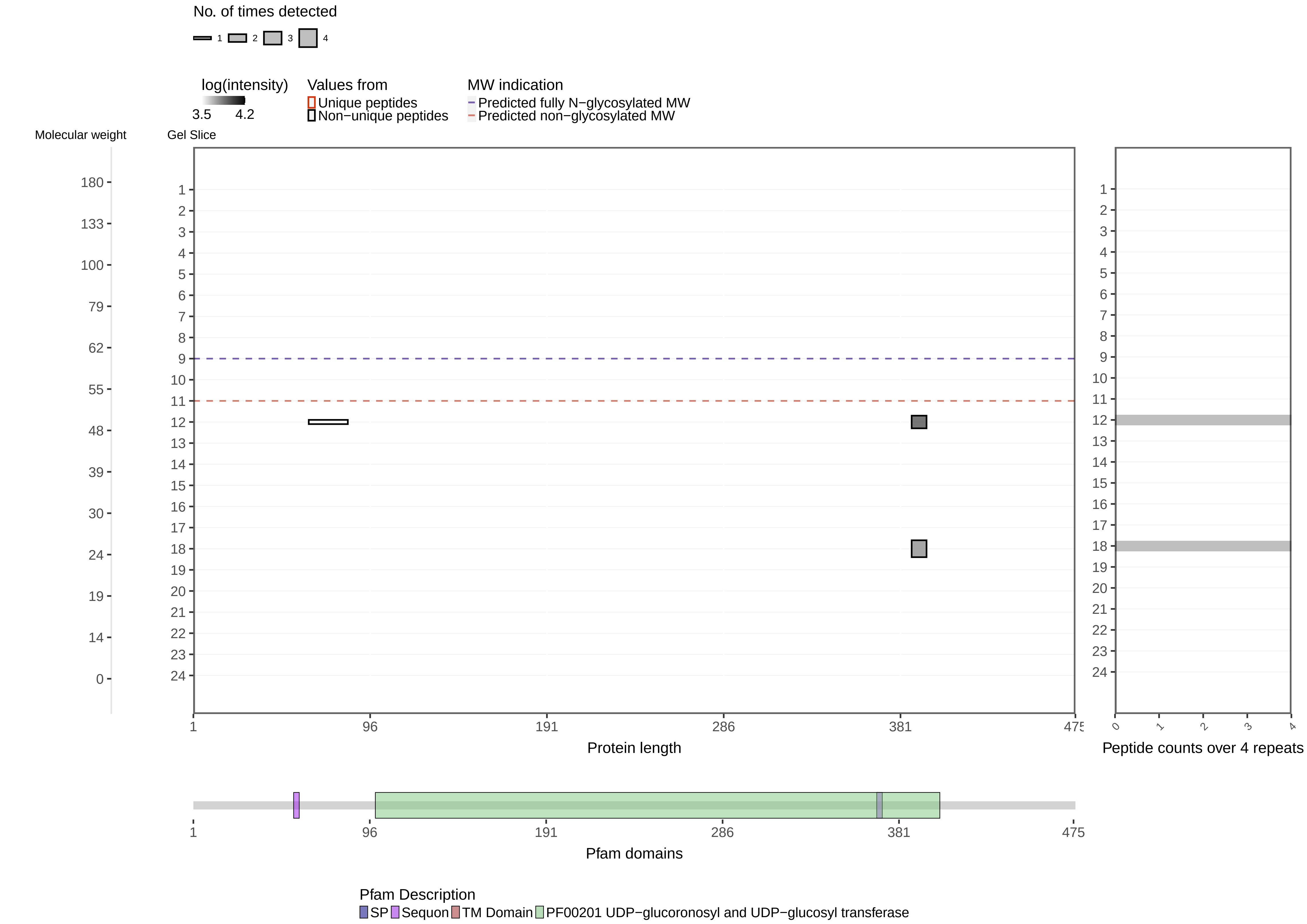


Protein sequence:

PILSQIRTVCPNVYTIGPVHAHLKTRLATKCTSSNSLWQEDENCINWLDTHPPKSVLYVSFGSIAGVTREELLEFWYGLVNSDQNFLWVMRADLIIGQEGKHEILEELEQGTKARGYMADWVPQEKVLAHTAIGGFLTHCGWNSTLESIVEGVPMICWPRFADQQVNSRFIGEVWKMGLDIKDTCDRDIIGKSIRDLMEKRRGEFLQRTEQMASMARRTVNEGGSSYINLDRLIQDIRLMCLPLKQFS*

### NbL15g14420.1 uncharacterized_LOC107802020 [MW = 32.76 kDa]

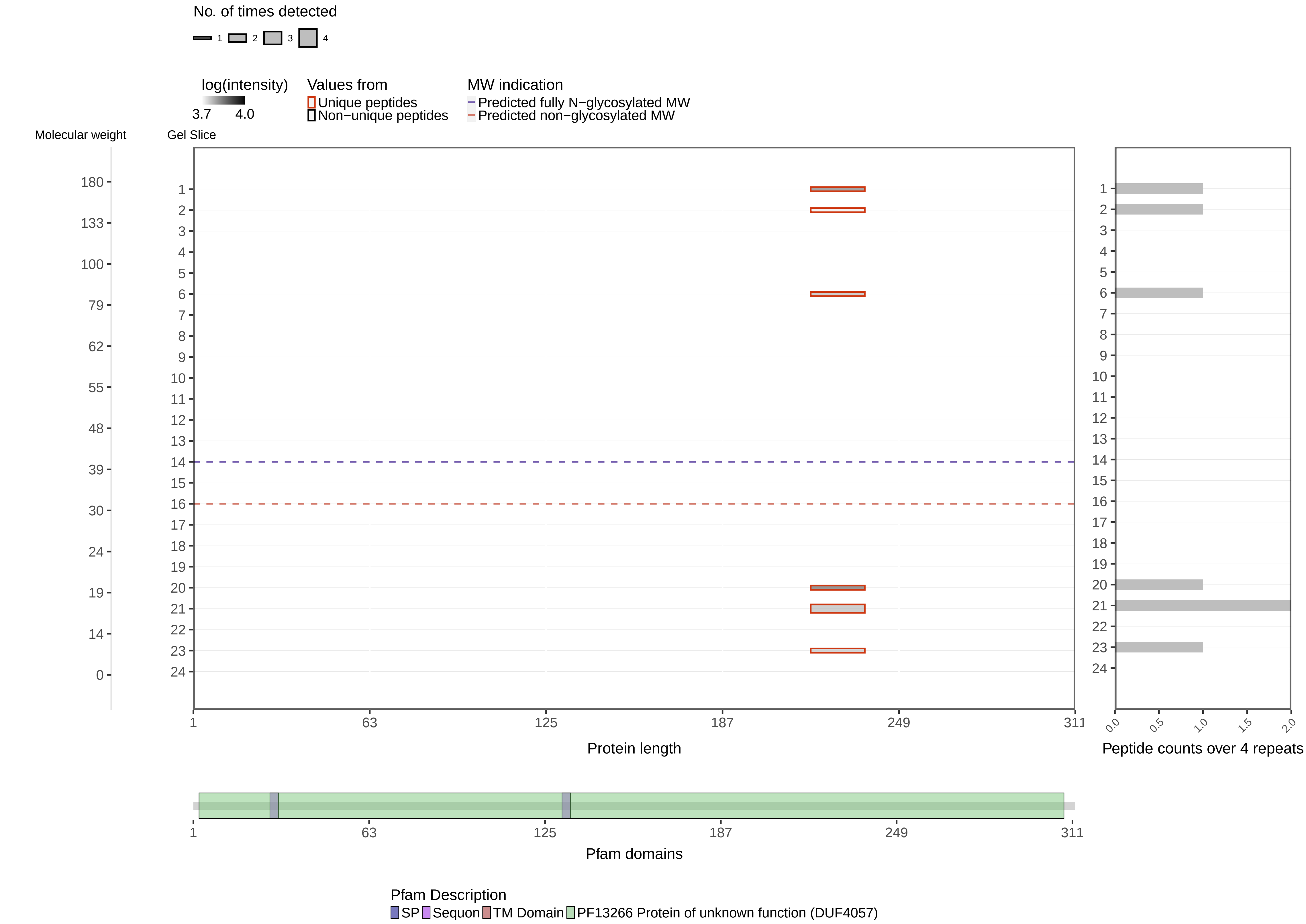


Protein sequence:

VAKQRELSGNLESESDSKIKKHLSDAKTKELSGNDIFGPPPEVPTRSLAAARSLESKESKDMGEPAPRTVRTSVRVSNPAGGQSSILFGDEPVGKTTKKIHNQKFAELTGNDIFKGDIPPGSADKPLSSAKLKEMSGSDIFSDGKIESRDHLGGVRKPPGGESTISLV*

NbL15g14380.1 tropinone_reductase_homolog_At5g06060−like [MW = 31.6 kDa]

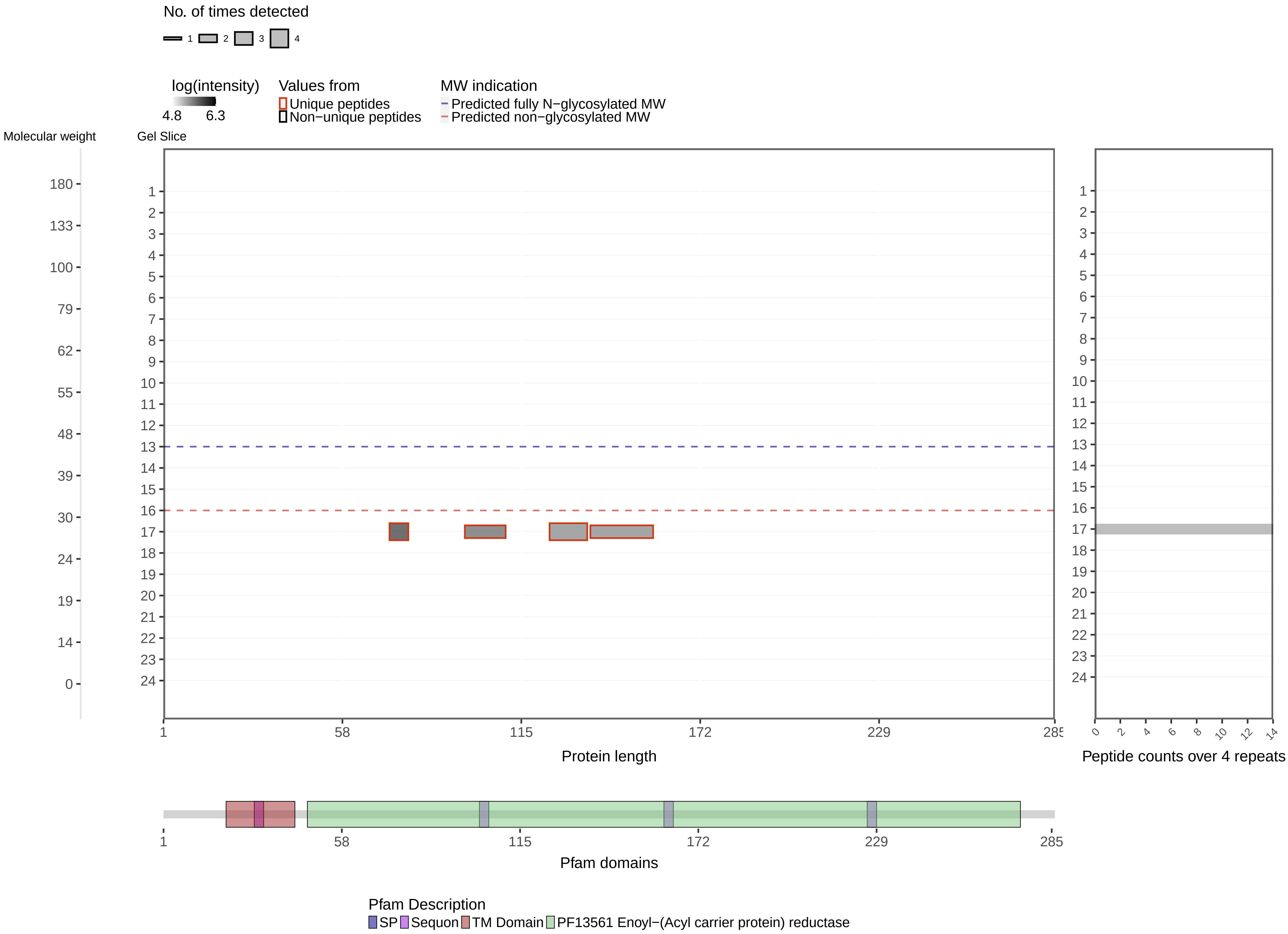


Protein sequence:

TDYAAEEYARLLGTNLESAYHLCQLAHPLLKASGNGSVVFISSVAGLVHISSGSIYGATKGAMNQLTRNLACEWAKDSIRVNGVAPWYIRTSLVEHLLENKTFLDGVISRTPLRRPGESQEVSSLVAYLCLPGASYITGQIIAVDGGFTVNGFEMTSY*

NbL15g14330.1 actin−7 [MW = 41.76 kDa]

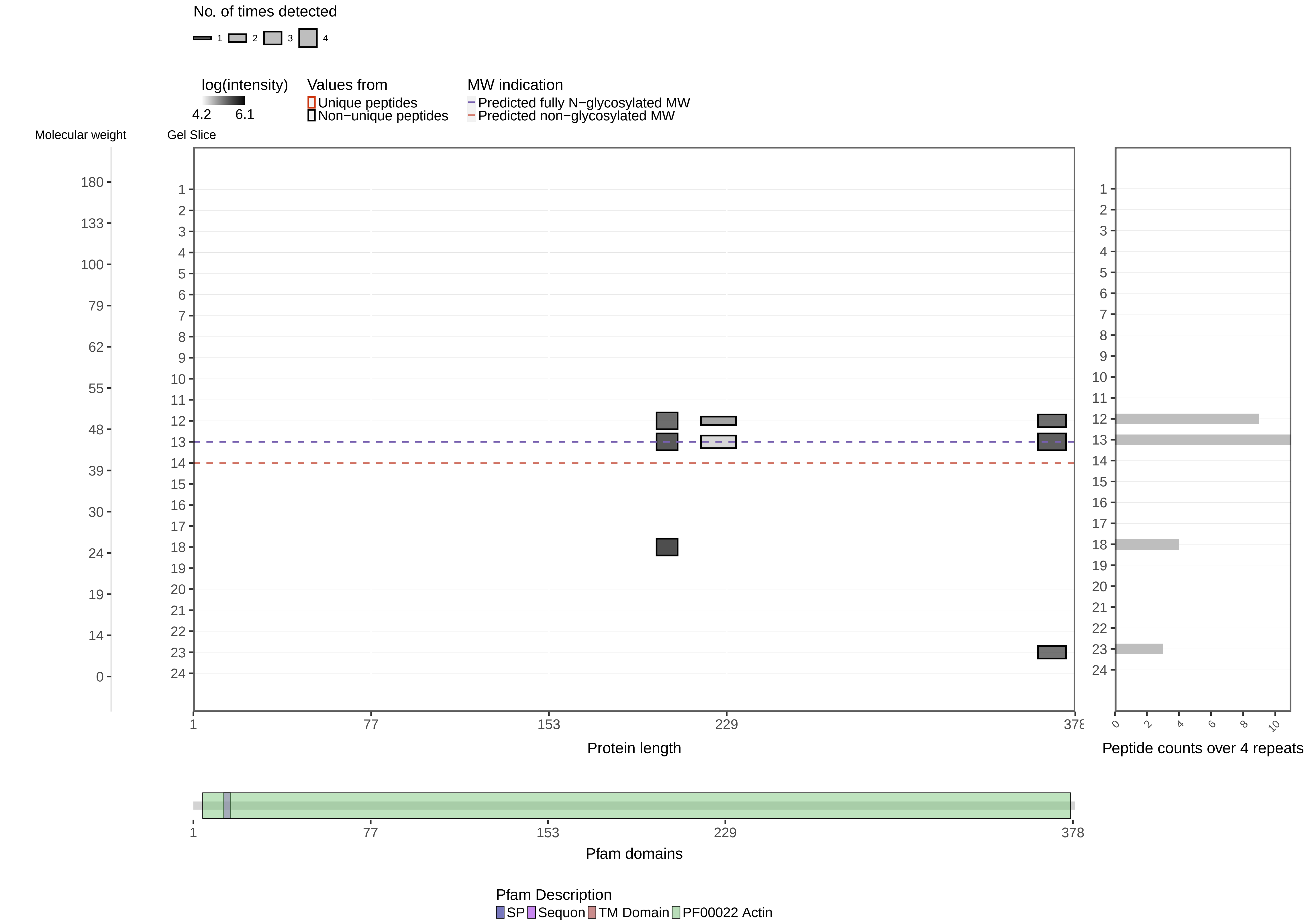


Protein sequence:

PHAILRLDLAGRDLTDSLMKILTERGYMFTTTAEREIVRDMKEKLAYVALDYEOELETARSSSSIEKNYELPDGOVITIGAERFRCPEVLFOPSMIGMEAAGIHETTYNSIMKCDVDIRKDLYGNIVLSGGSTMFPGIADRMSKEITALAPSSMKIKVVAPPERKYSVWIGGSILASLSTFOOMWISKGEYDESGPSIVHRKCF*

### NbL15g14290.1 40S_ribosomal_protein_S18−like [MW = 17.52 kDa]

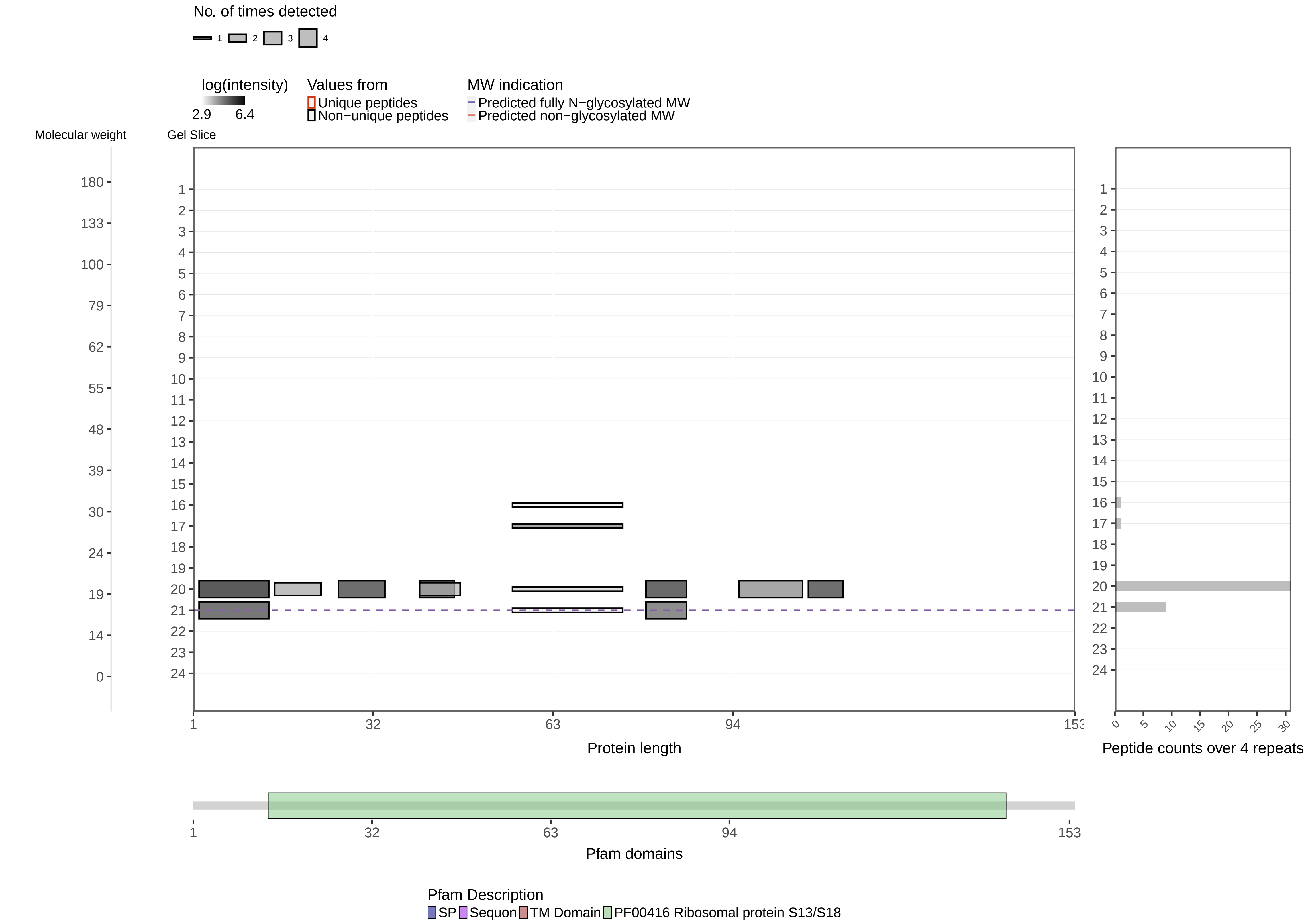


Protein sequence:

SVMVVVANPRQFKIPDWFLNRQKDYKDGKFSQVTSNALDMKLRDDLERLKKIRNHRGLRHYWGLRVRGQHTKTTGRRGKTVGVSKKR*

### NbL15g14280.1 uridylate_kinase−like [MW = 35.72 kDa]

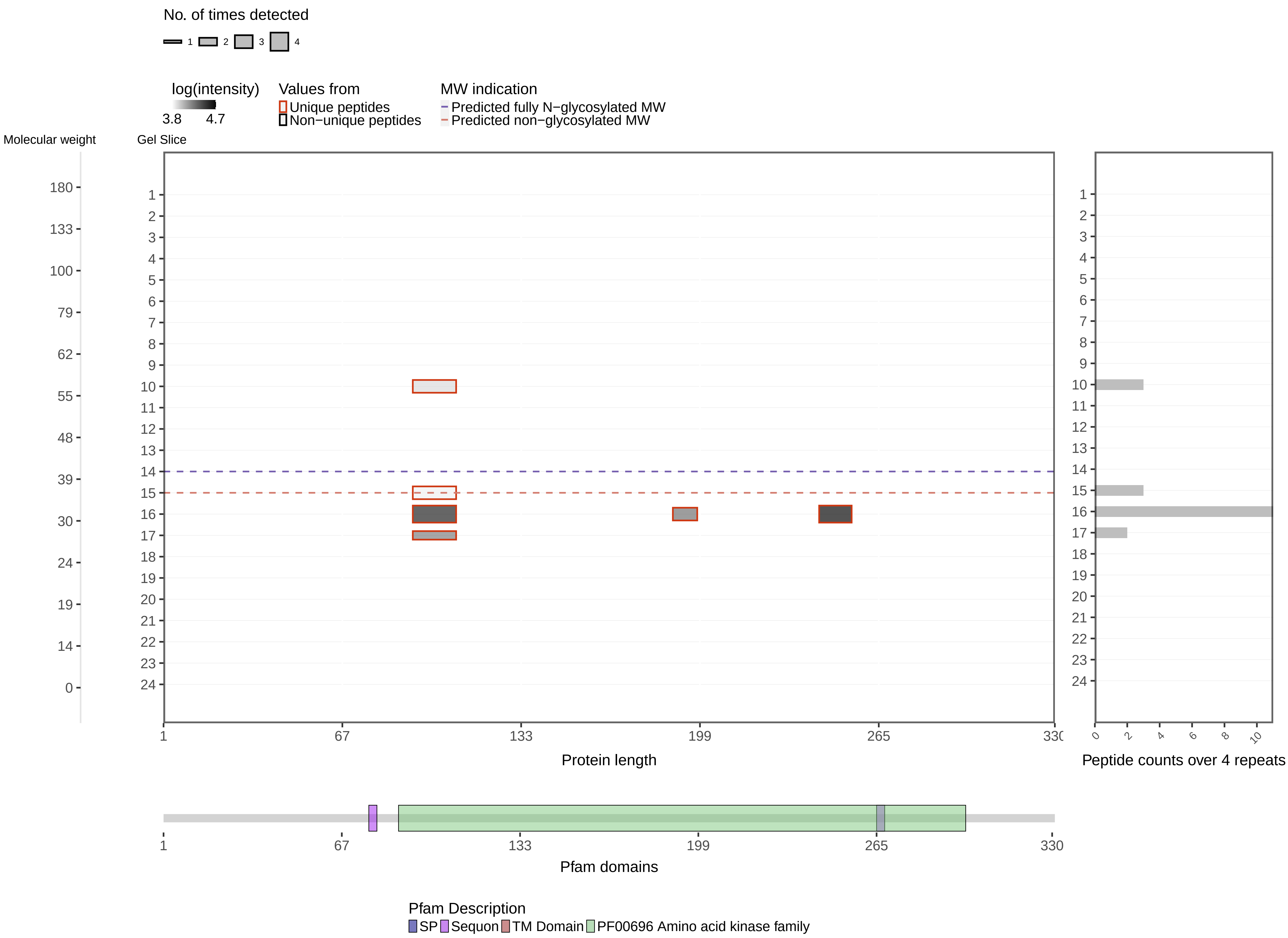


Protein sequence:

DRSSADYIGMLATVMNAIFLQATMESIGIPTRVQTAFRMSEVAEPYIRRRAVRHLEKGRVVIFAAGTGNPFFTTDTAAALRCAEINAEVVLKATNVDGVYDDNPRHNPDARLQDNLTYHDVISKELSVMDLTAITLCQENNIPVVVFNLNKPGNIAKAIKGERVGTLIGDTWNTEAAVS*

### NbL15g14200.1 thylakoid_lumenal_15_kDa_protein_1_chloroplastic−like [MW = 23.31 kDa]

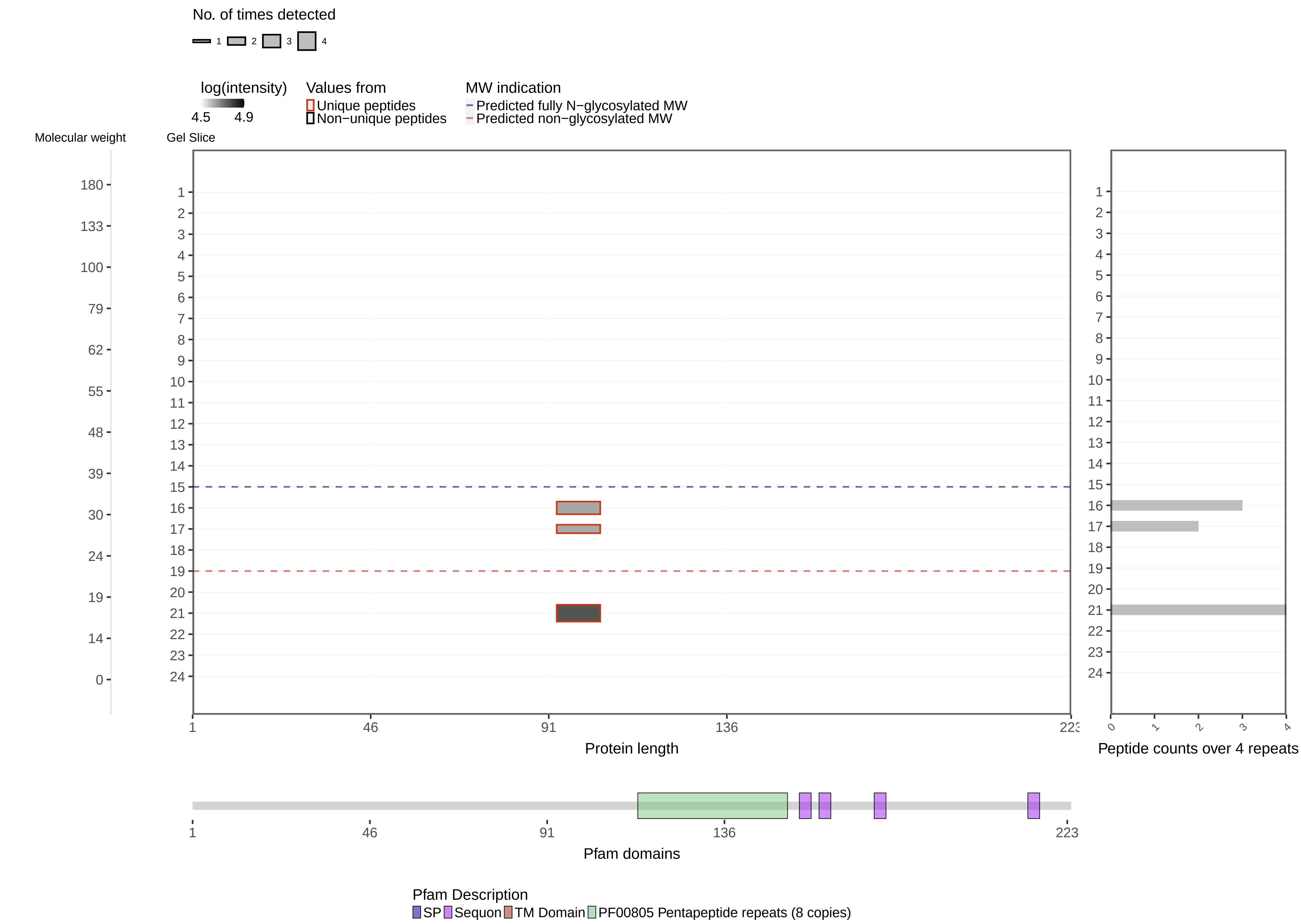


Protein sequence:

GKDFSGKTLIKQDFKTSILRQANFKGANLVGASFFDADLTGADLSDADLRGADFSLANVTKVNLSNANLEGALTTGNTSFKGSNIEGADFTDVPLRDDQREYLCKIADGVNPVTGNATRDTLFCK*

### NbL15g14020.1 3−hydroxyisobutyryl−CoA_hydrolase−like_protein_3_mitochondrial [MW = 46.8 kDa]

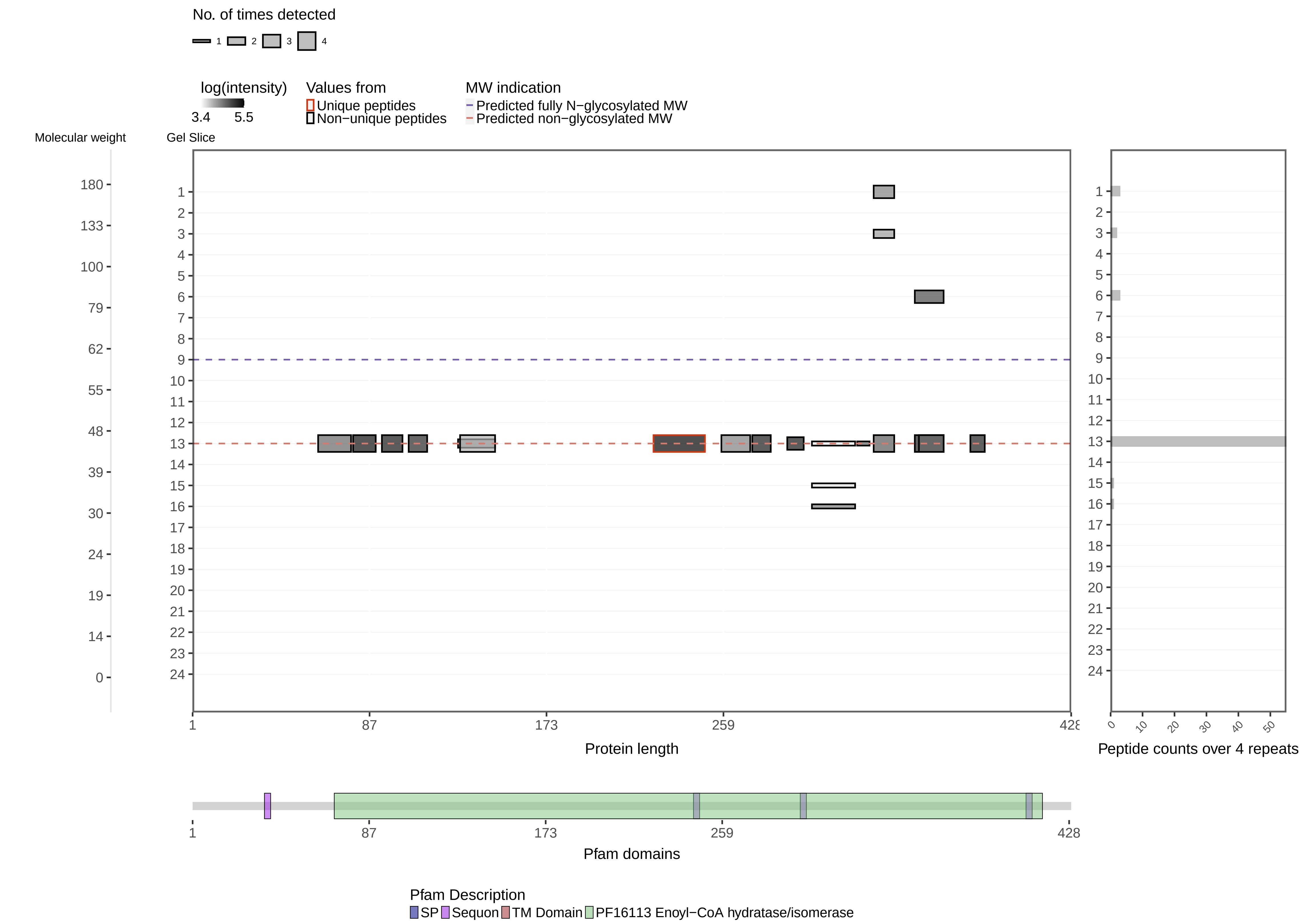


Protein sequence:

FSYIAAHGPGEGTVGAYLALTGNRISTPADALYVGLGTHYVPSANLSTLKEALLATKFSEDPDEEIKELLAKYSSNPDSEPRLKSLLPRIISTFAGNKSIREIIEELENHQQSADTLVVEWAKDALQGLRKGAPFSLCLTQKLFSRVASARGKNENDLSSLNGVMRTEYRIAIRSALRSDFAEGVRAVLVEKDQSPKWNPSCLEDVNLSEVEALFEPLSPDVGELNV*

NbL15g13940.1 serine−threonine_kinase_receptor−associated_protein−like [MW = 37.76 kDa]

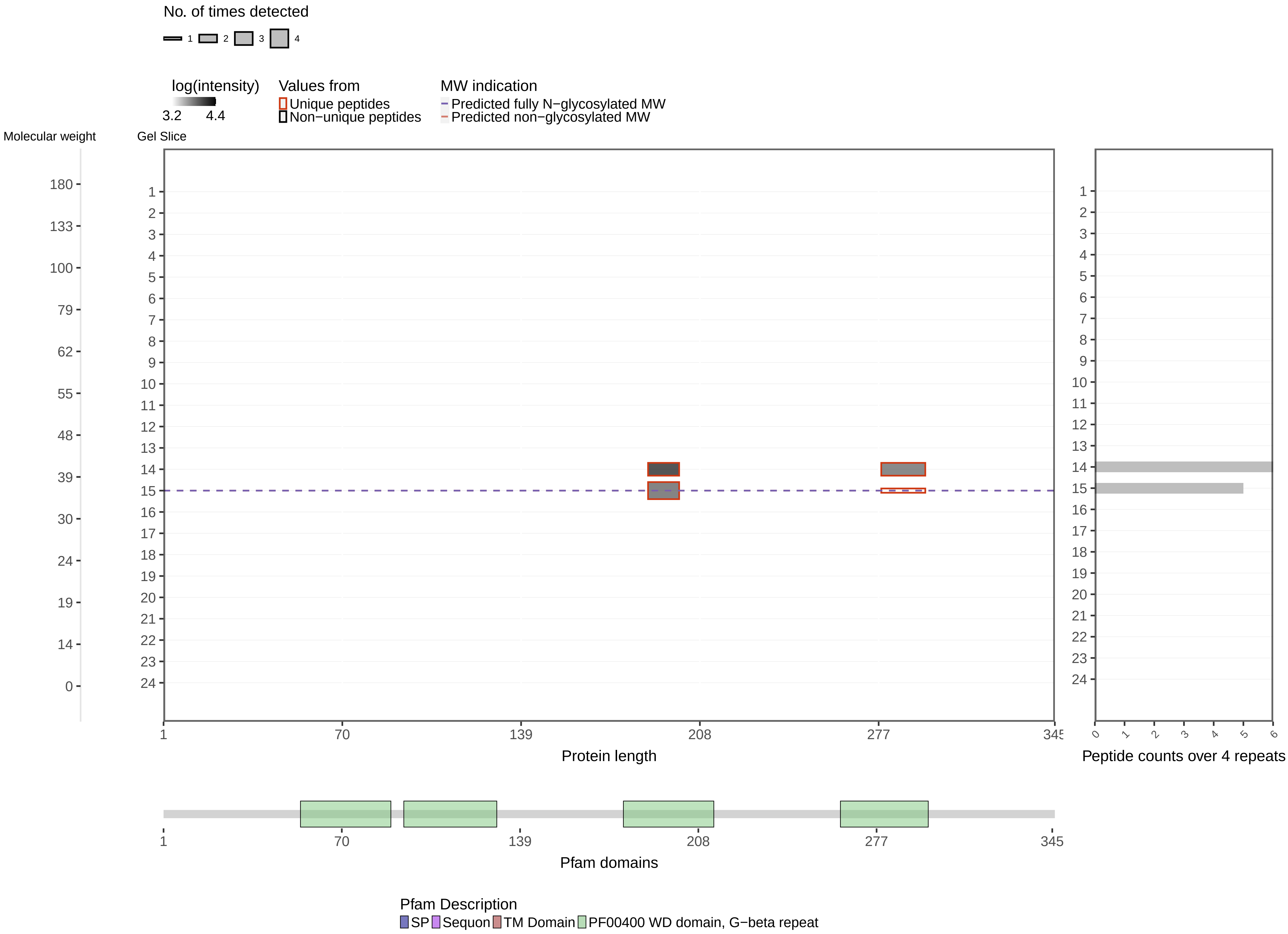


Protein sequence:

TILSSCTDSGGVRLWDVRTGDIVRTLETKSPVTSAEVSQDGRYITTCDGFSVKFWDANHFGLVKSYNMPCTVESASLEPKYGLKFVAGGEDMWIRVFNFHTGEELACNKGHHGPVHCVRFAPSGESYASGSEDGTIRIWQTGPLTNDDVDMTTNGKVKVSAEEVSRKIEGFQISDERKIKEEDLAD*

### NbL15g13870.1 protein_arginine_N−methyltransferase_PRMT10 [MW = 42 kDa]

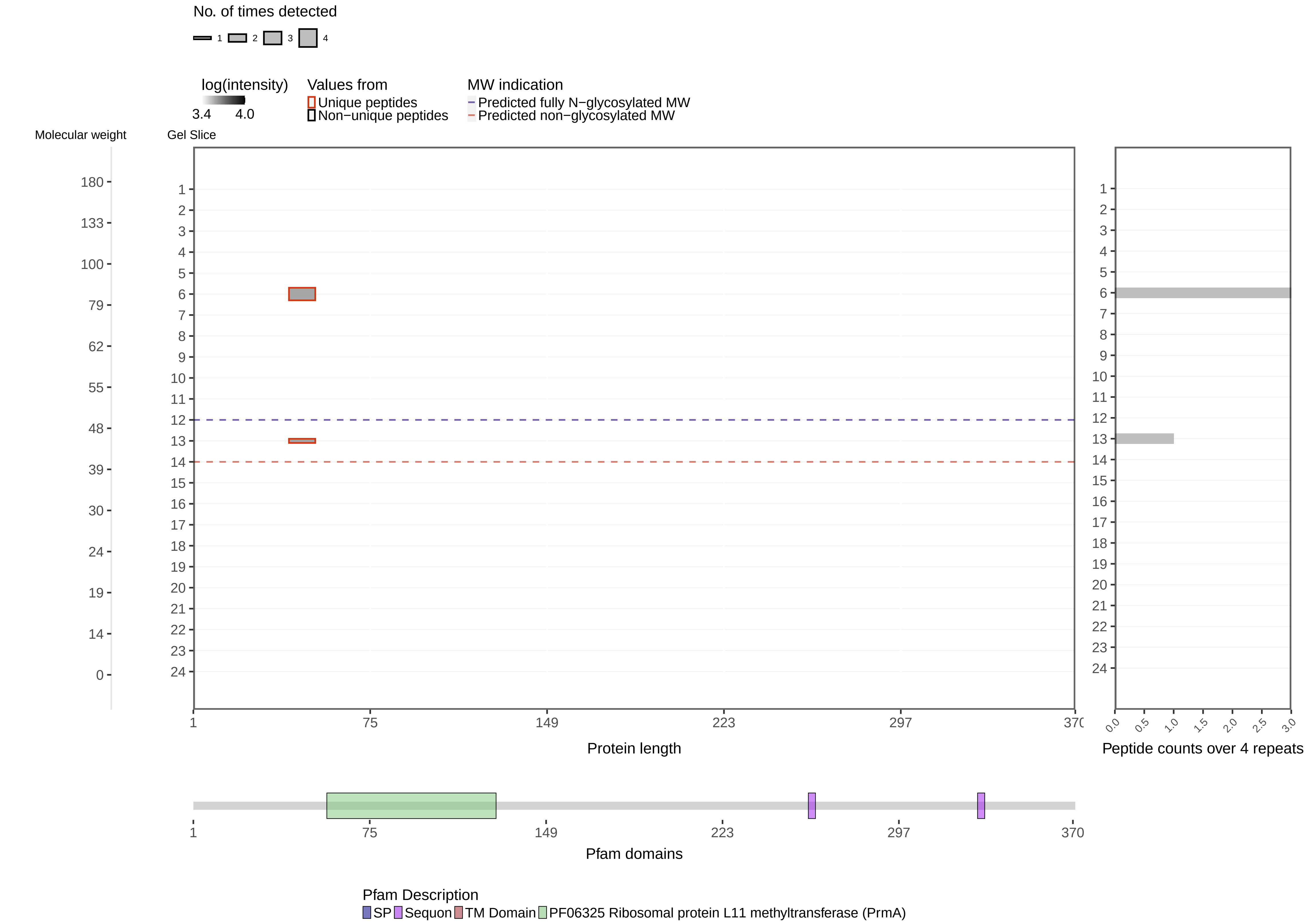


Protein sequence:

SGLVDOKKIDYDRAMDDWYHFVDETOTFYGVDMSSLSKPFTEEORKYYLETSLWNNLHPNOVIGKPAVIMEIDCLTSSVNDILSLOANISSSITAENSRLCGFGGWFDVHFRGRKESPAKHEIELTTAPSEDLGTHWGOOVFLLHPSIRVSOGDEMTINFSMYRSKENHRLLEVEFGCELROSSGKSLPSFSKKFHIE*

NbL15g13740.1 protein_phosphatase_1_regulatory_subunit_pprA [MW = 35.98 kDa]

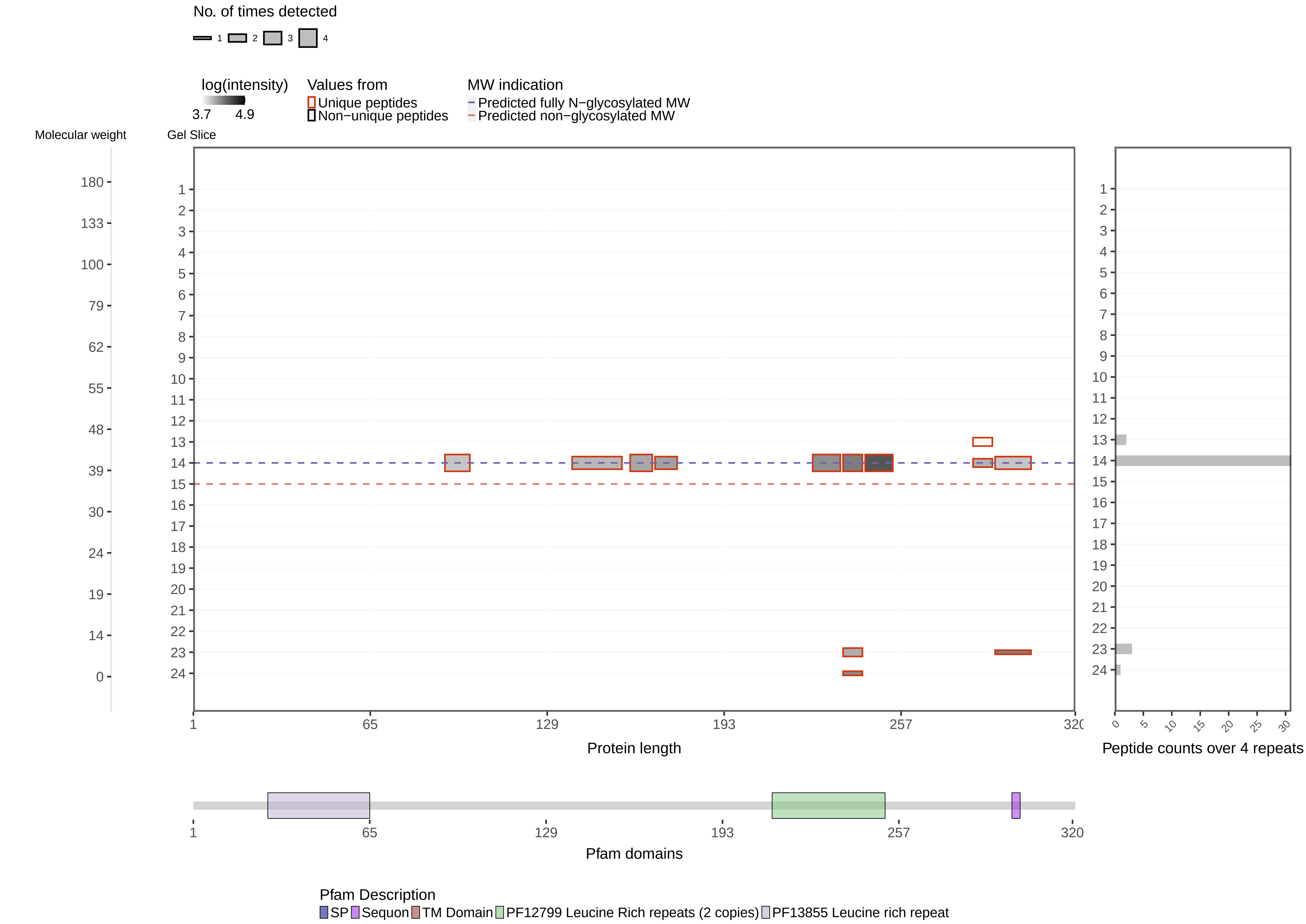


Protein sequence:

LQILELGSNRLRVMELLENLKNLQELWLGRNRIRTVNLCGLKCIKKISLQSNRLTSMIGFQECVALEELYLSHNGIAKMEGLSTLVNLRVLDVSANKLTEINDIENLRKLEDLWLNDNNIASLVGLAEAVAGAKEKLTTIYLERNPCAQSPNYTSSLRQIFPNIEQIDSEVYA*

### NbL15g13600.1 vacuolar_protein_sorting−associated_protein_32_homolog_2−like [MW = 24.32 kDa]

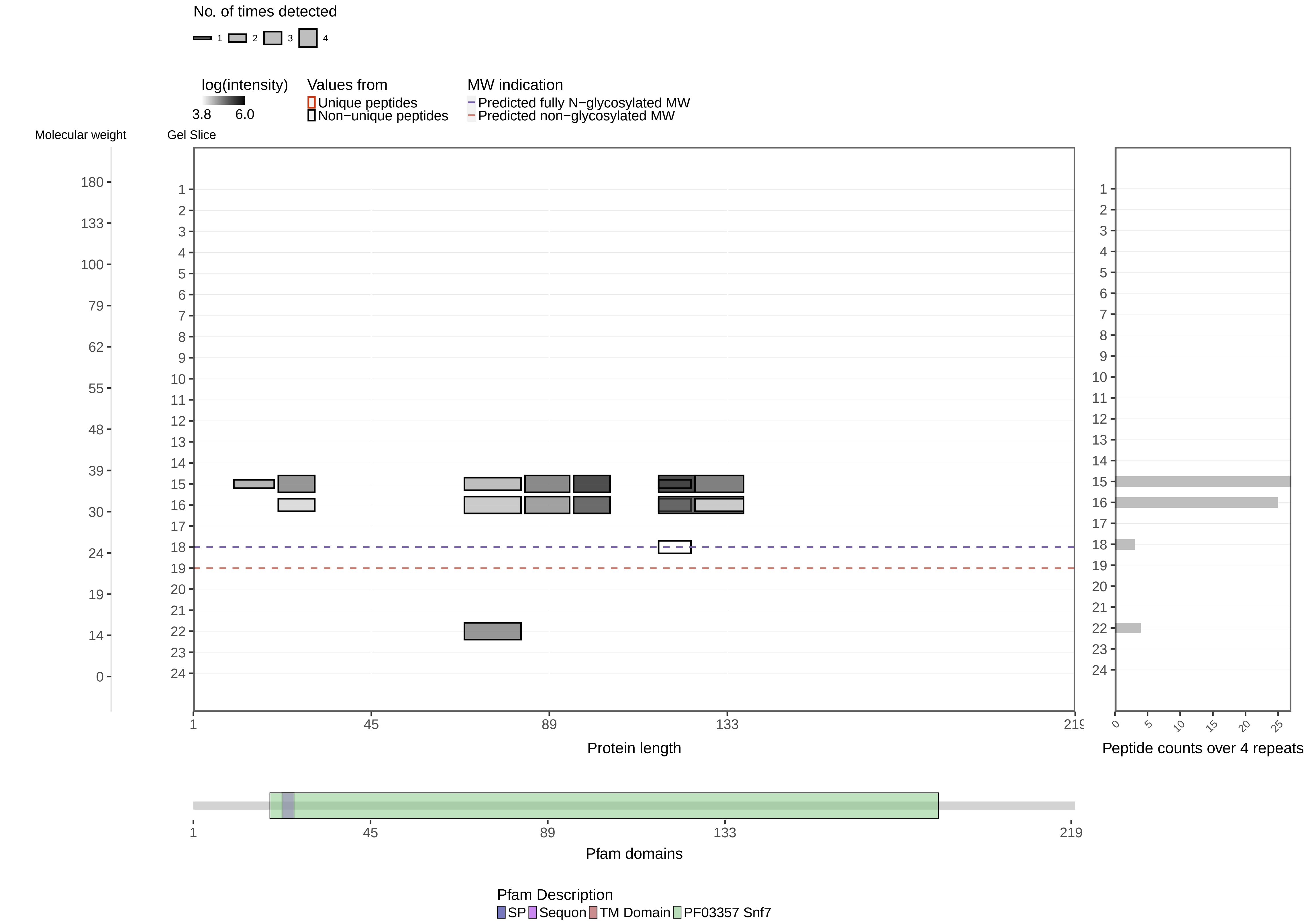


Protein sequence:

TETVDALRTGASAMKAMQKATNIDDVDKTMDEINEQTENMKQIQEALATPIGAAADFDEDELEAELEELEGAELEEQLLQPATTAPAAPVHVPAGRQPARPIPQRRTAEEDELAALQAEMAL*

NbL15g13580.1 mitochondrial−processing_peptidase_subunit_alpha−like [MW = 54.62 kDa]

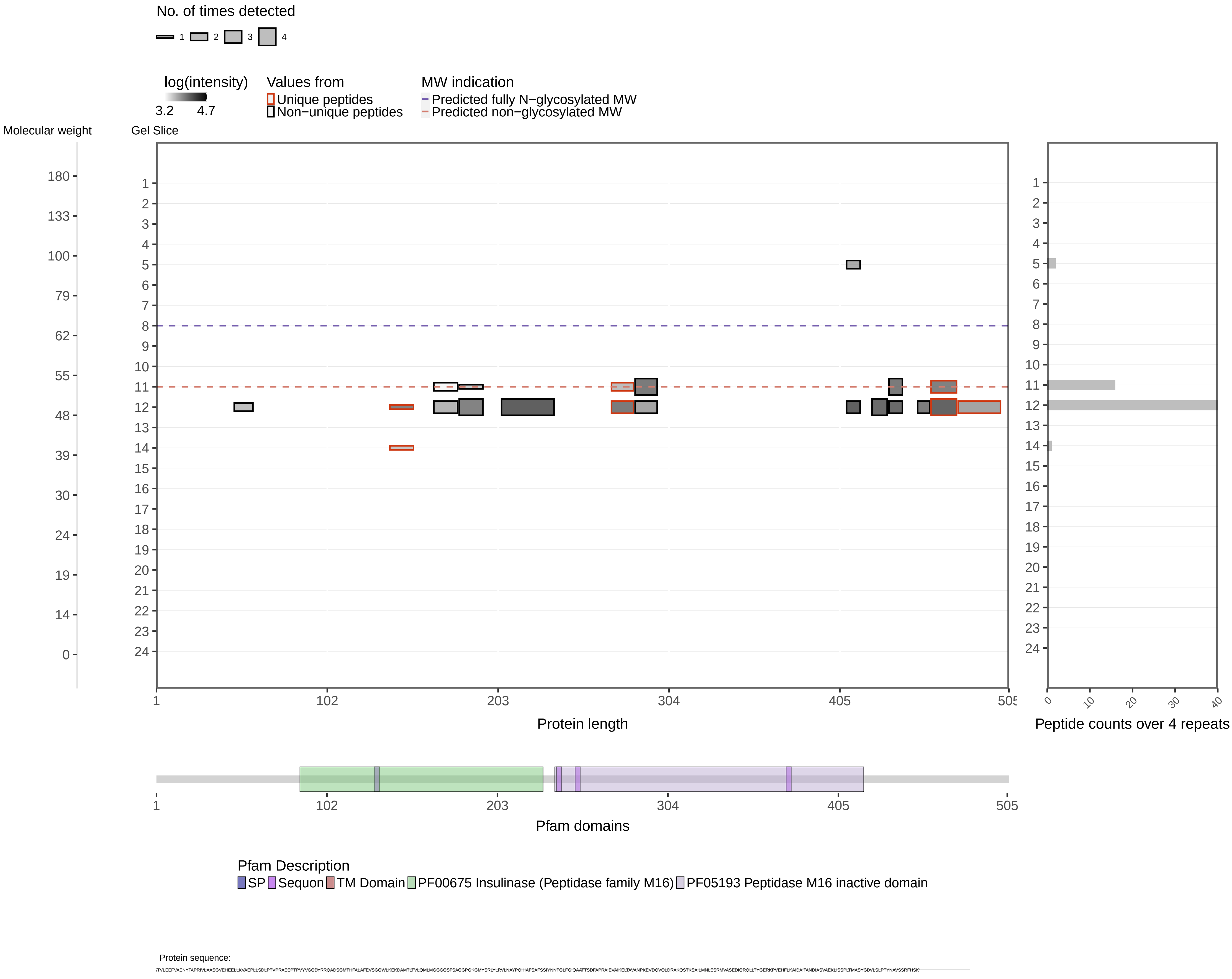

### NbL15g13530.1 probable_methionine−−tRNA_ligase [MW = 90.64 kDa]

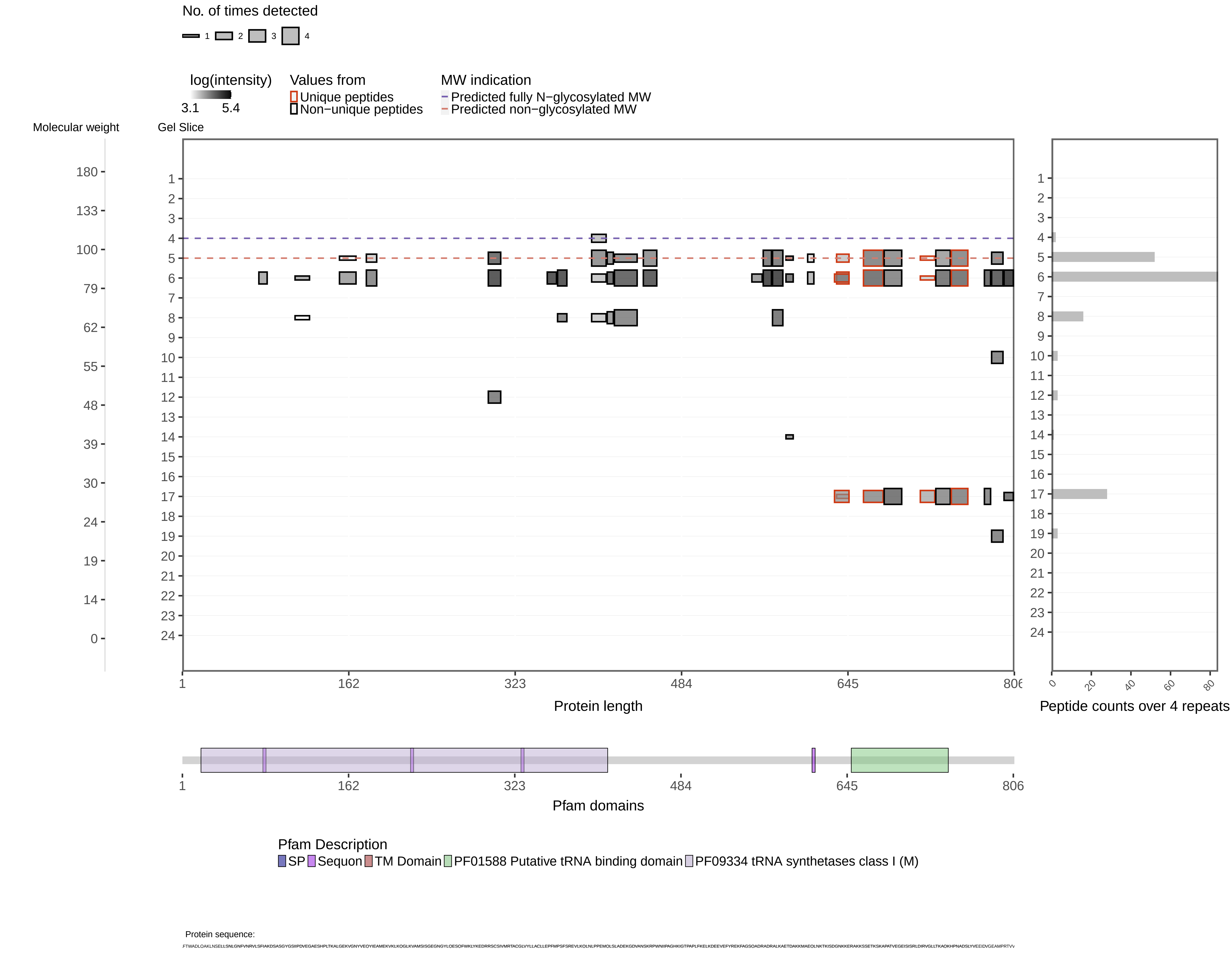

NbL15g13490.1 chalcone_synthase_2 [MW = 42.64 kDa]

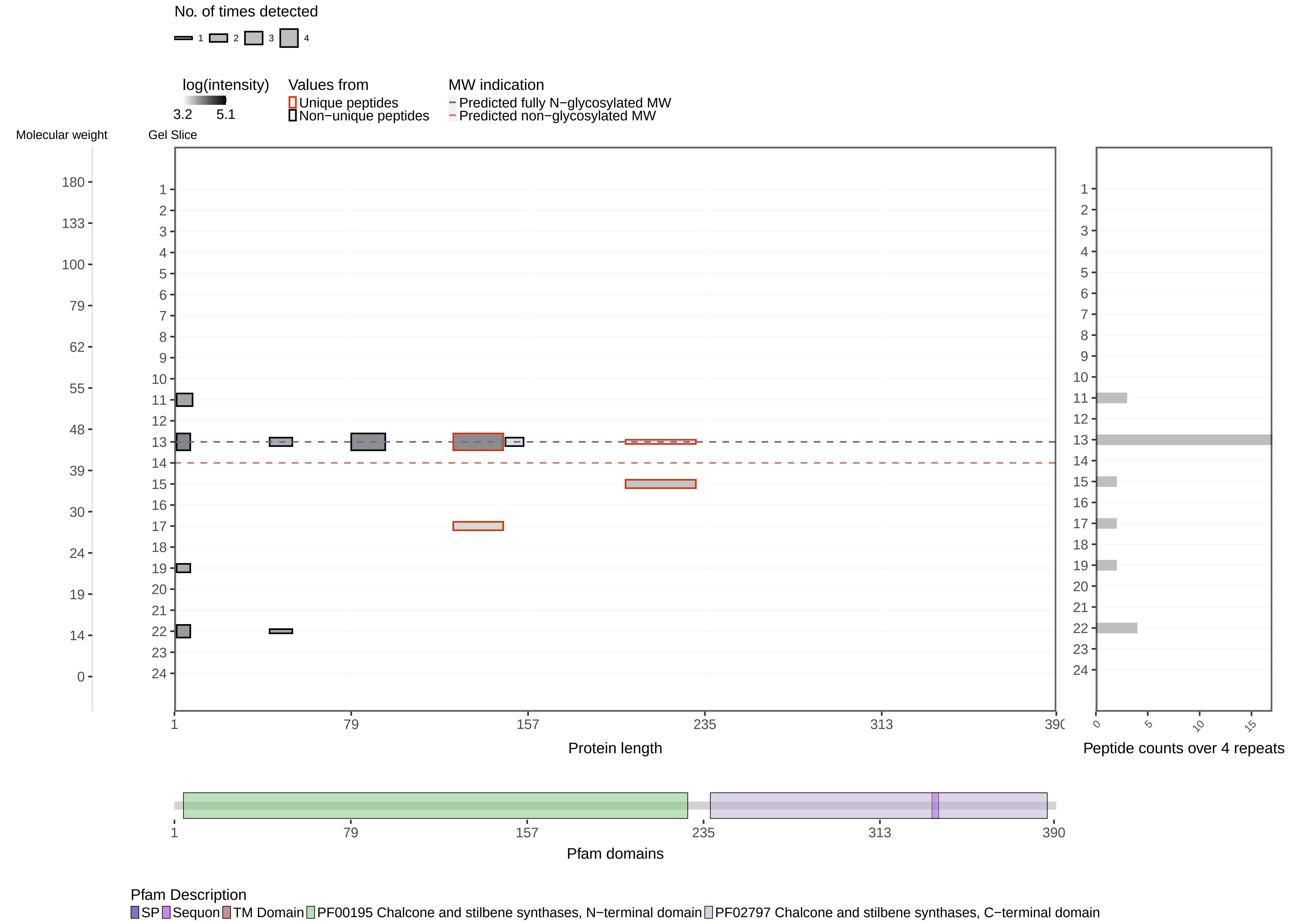


Protein sequence:

NKGARVLVVCSEITAVTFRGPSDTHLDSMVGQALFGDGAAAIIIGSDPVPRVERPLFELVSTAQTLLPDSEGAIDGHLREVGLTFHLLKDVPGLISKNIEKSLMEAFQPLGLSDWNSIFWIAHPGGPAILDQVELKLGLKPEKLRATRQVLSDYGNMSSACVLFILDEMRKASAKEGLGTTGEGLDWGVLFGFGPGLTVETVVLHSVST*

NbL15g13470.1 dihydroxy−acid_dehydratase_chloroplastic−like [MW = 72.72 kDa]

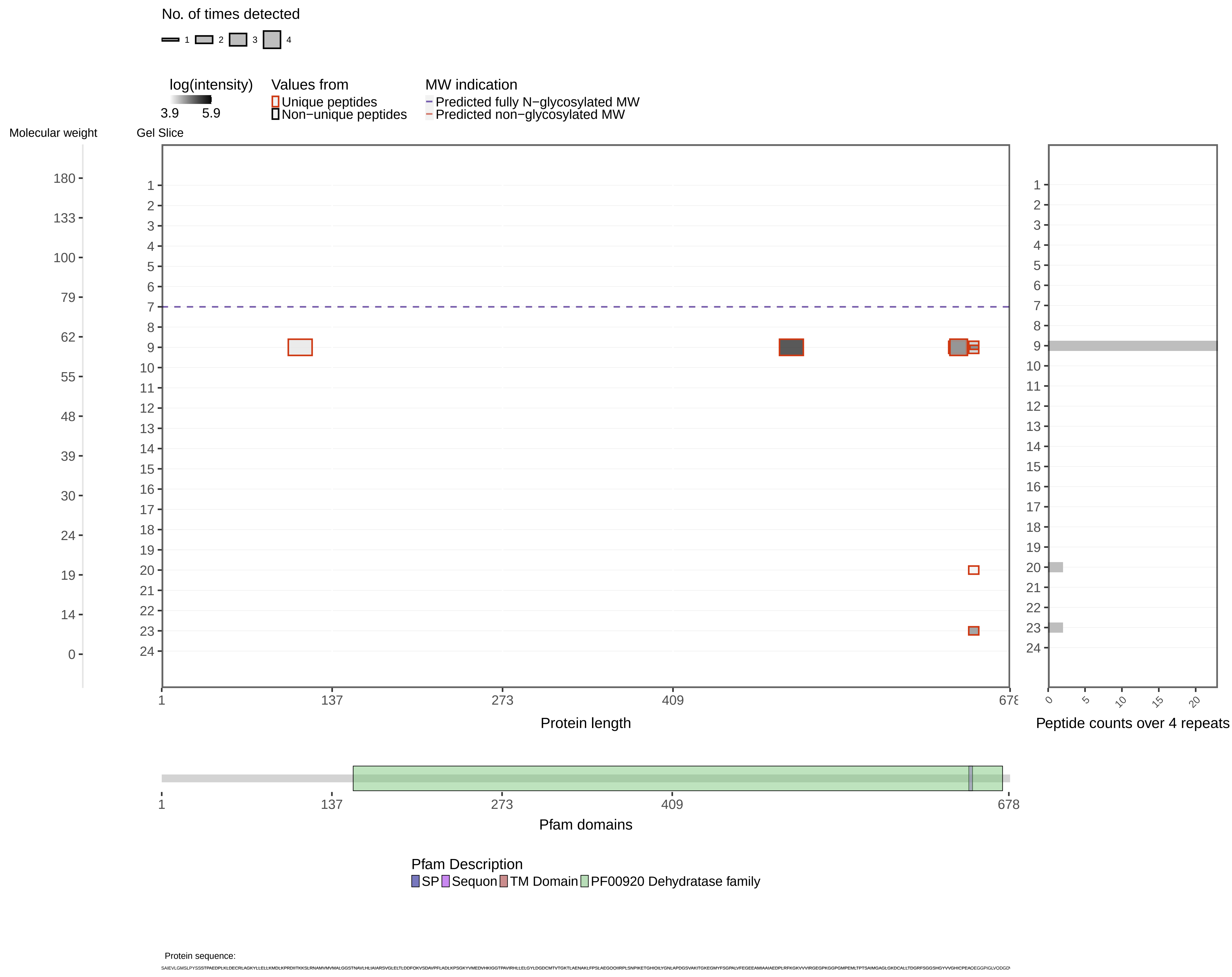

### NbL15g13400.1 chaperonin_CPN60−2_mitochondrial−like [MW = 61.09 kDa]

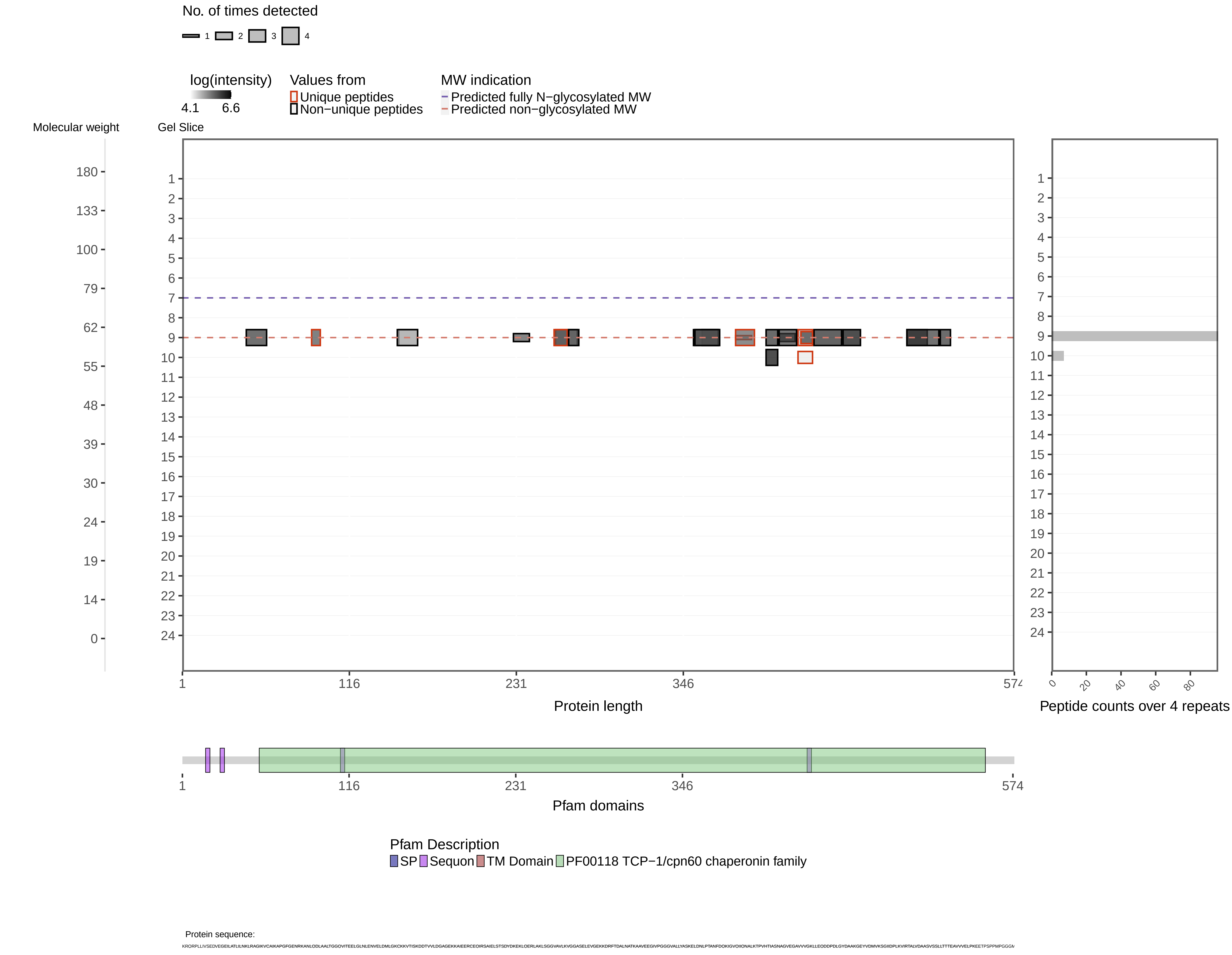

NbL15g13280.1 pto−interacting_protein_1−like [MW = 39.03 kDa]

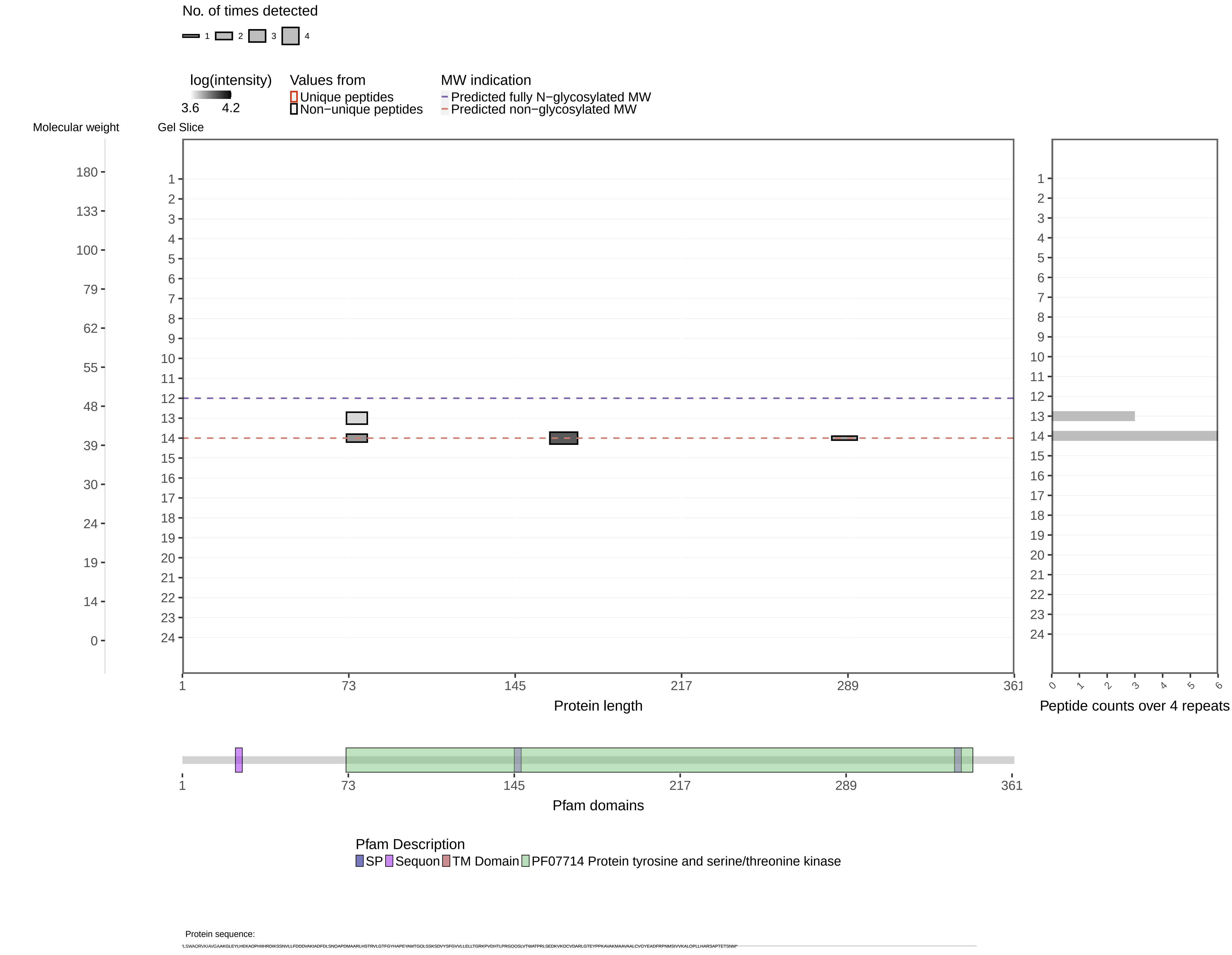

### NbL15g13180.1 dihydrolipoyl_dehydrogenase_mitochondrial [MW = 53.53 kDa]

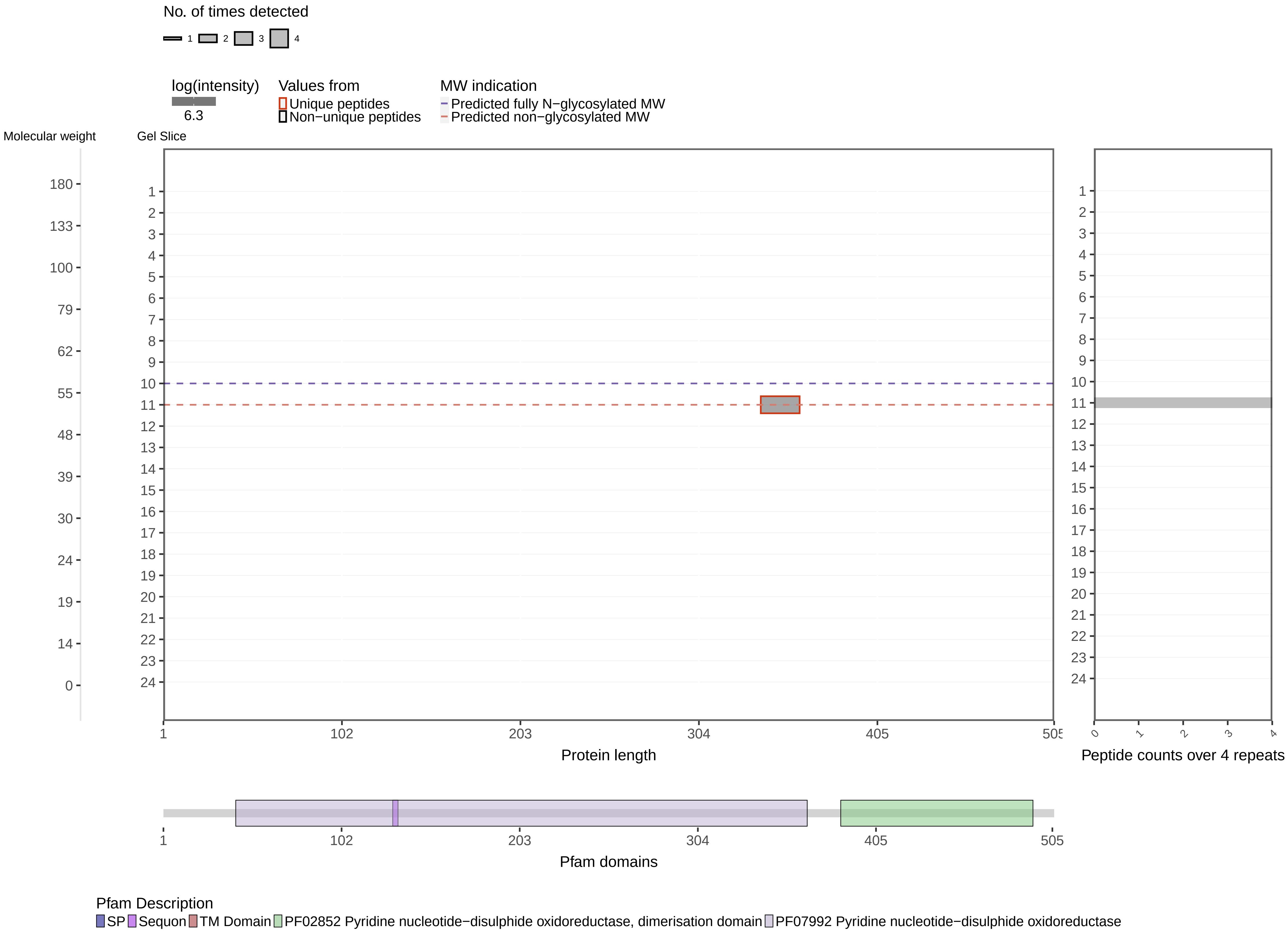


Protein sequence:

'VEFGPDIVPSMDGEVRKQFQRSLEKQKMKFILKTKVVSVETVGDGVKLTLEPAAGGDQTTLEADVVLVSAGRIPFTSGLGLDKIGVETDKAGRILVNERFVTNVPGVHAIGDVIPGPMLAHKAEEDGVACVEFIAGKEGHVDYDLVPGVVYTHPEVASVGKTEEQVKARGVDYRVGKFPFLANSRAKAIDDAEGIVKVLAEKETDKILGVHIMSPNAGELIHEAVLALHYGASSEDIARTCHAHPTMSEALKEAAMATYDKPIHI*

### NbL15g13110.1 cell_wall_/_vacuolar_inhibitor_of_fructosidase_1−like [MW − SP = 15.67 kDa]

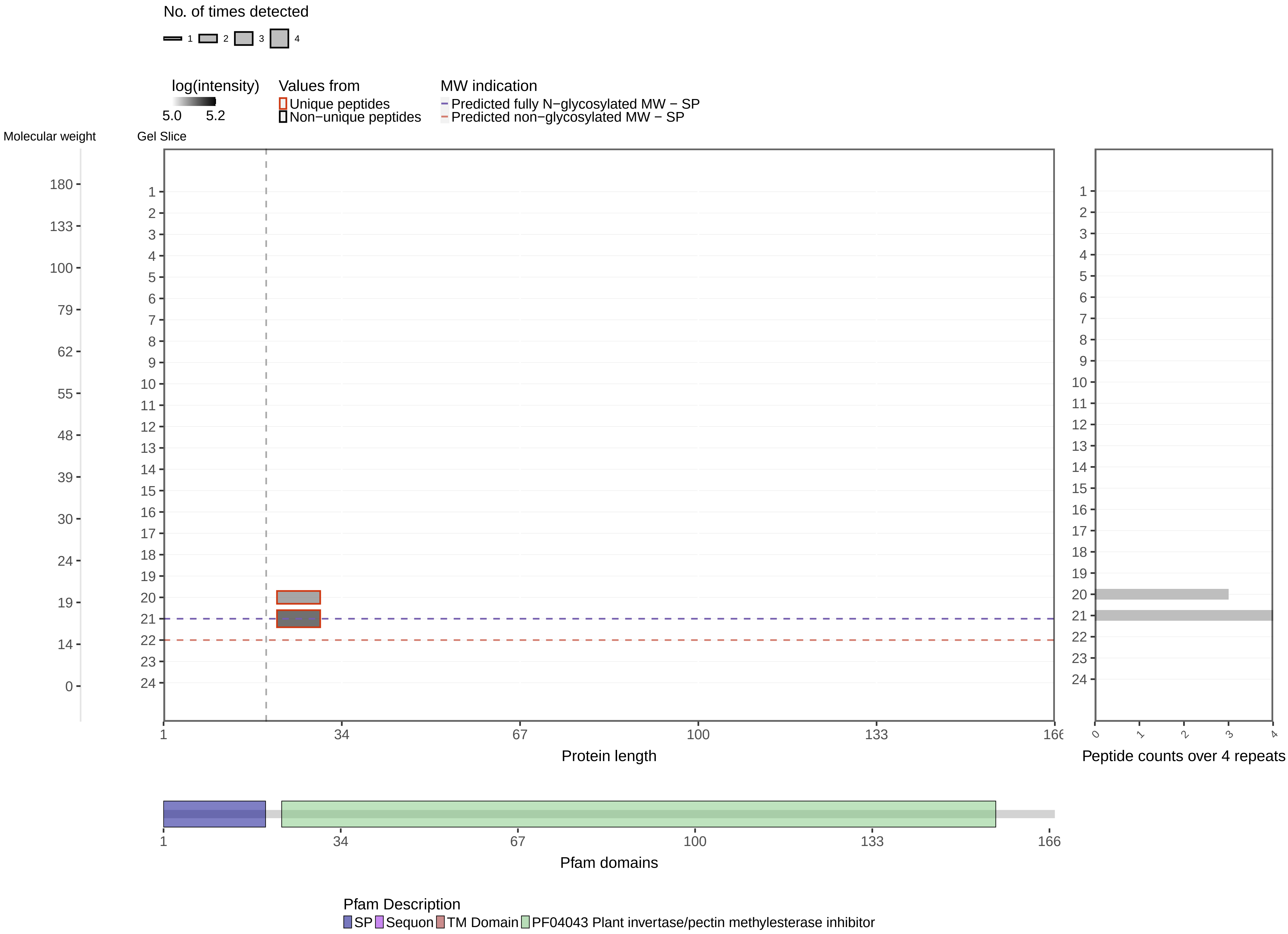


Protein sequence:

EIISKLRQSNPPAAWSVPLKKCAFSYKVILTVSLPEAIEALTKGNPKFAEDGMVGCFGDSQDCEENFKSCFSPLTGLNTAVLELSVVGRAIIRNLL*

### NbL15g13040.1 DNA−directed_RNA_polymerases_II_IV_and_V_subunit_9A [MW = 13.11 kDa]

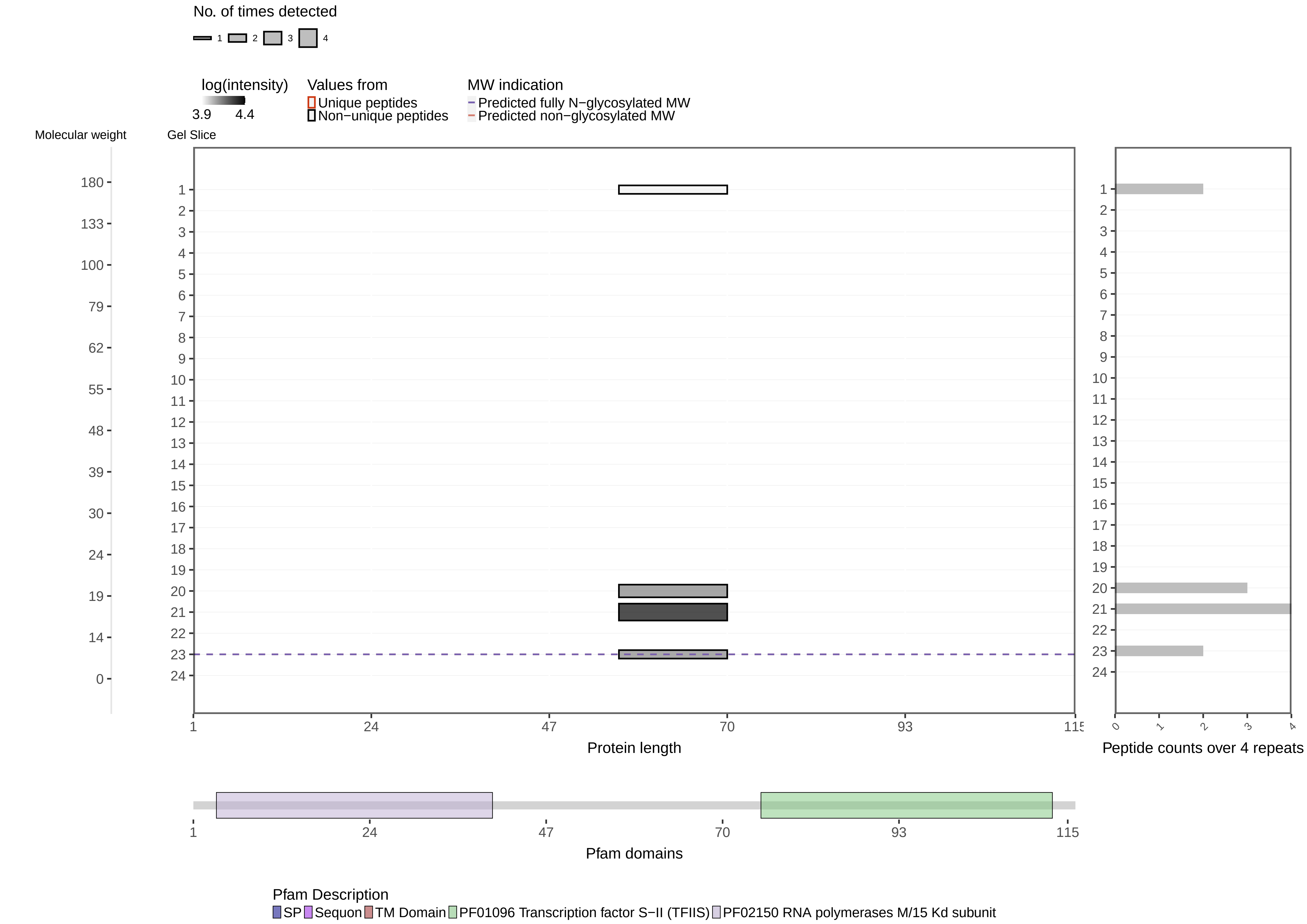


Protein sequence:

NEIHHSAAERTQVLQDVAADPTLPRTKSVRCSQCGHGEAVFFQATARGEEGMTLFFVCCNPNCGHRWRD*

NbL15g12970.1 L−type_lectin−domain_containing_receptor_kinase_S.4−like [MW − SP = 74.31 kDa]

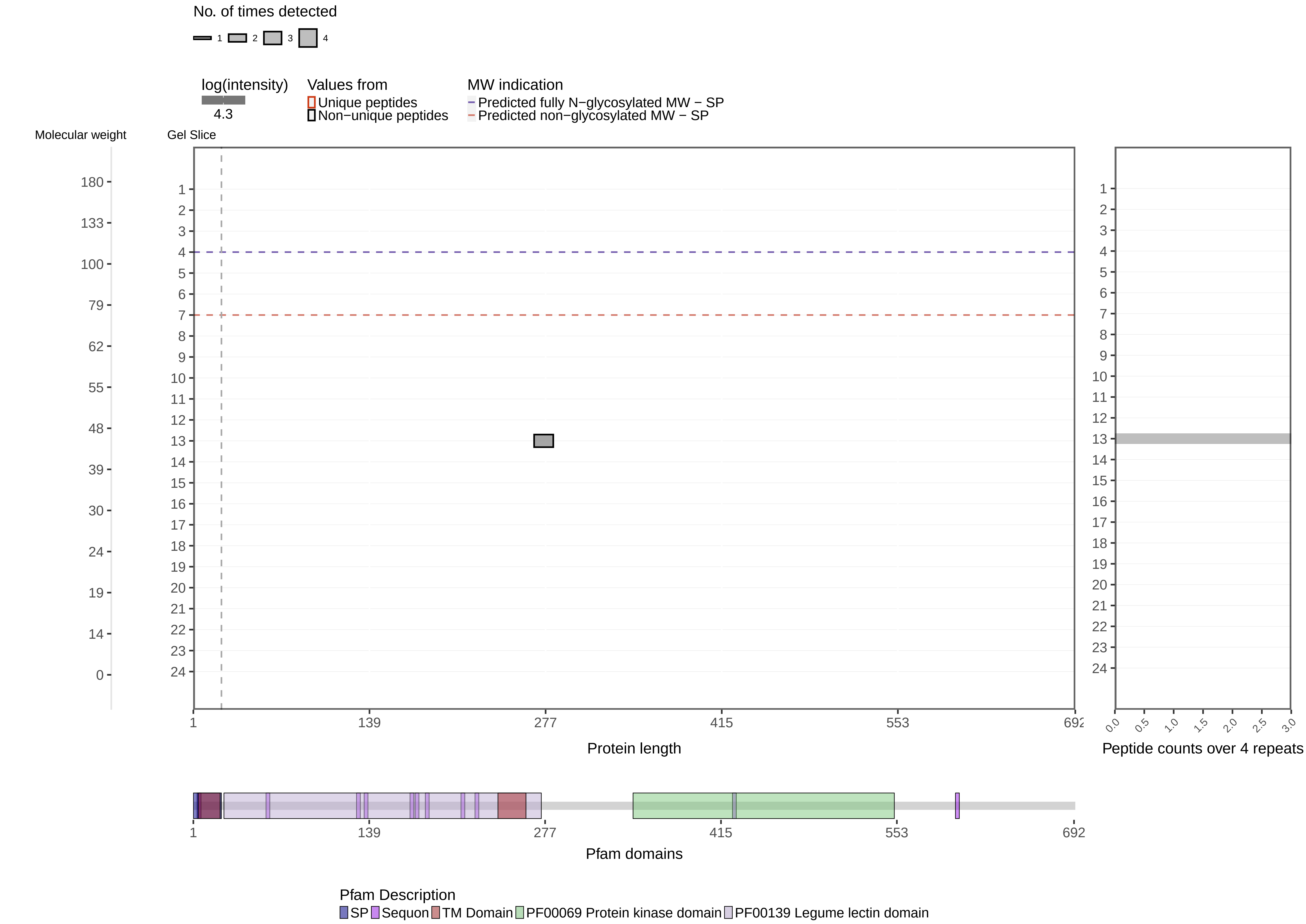


Protein sequence:

ELKKATRDFRDKELLGFGGFGRVYKGTLPKTSTEAAVKRINHDAKQGLQEFVSEIATIGRLRHRNLVQLIGWCRRKGDLLLVYDFMPNGSLDKYIYDEPRIILTWDQRFKIIKGVASGLLYLHEEWEQTVIHRDIKAGNVLLDAEMNGRLGDFGLAKLYEHGANPSTTRVVGTLGYLAPELTKTGKPTTSSDVFAFGALLLEVACGRRPIEAKALPEELILVDWVWDKWKEGAILEVVDPRLNSKYDEMEAVVVLKLGLMCSNNTPSKRPSMRLVVRYLEGEVALPEMLAAPDEY

### NbL15g12940.1 probable_methyltransferase_PMT11 [MW = 76.2 kDa]

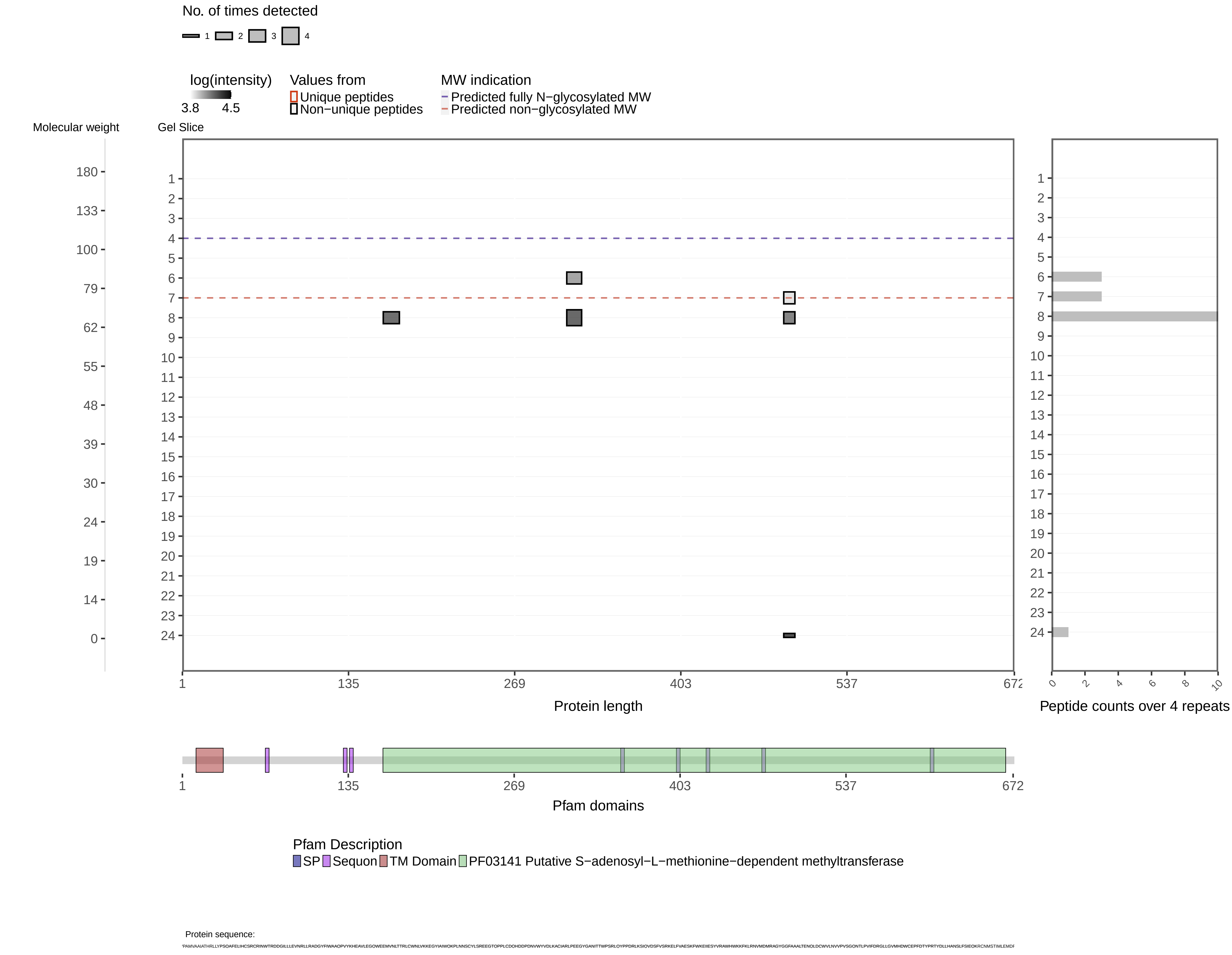

NbL15g12860.1 protease_regulatory_subunit_8_homolog_A−like [MW = 47.18 kDa]

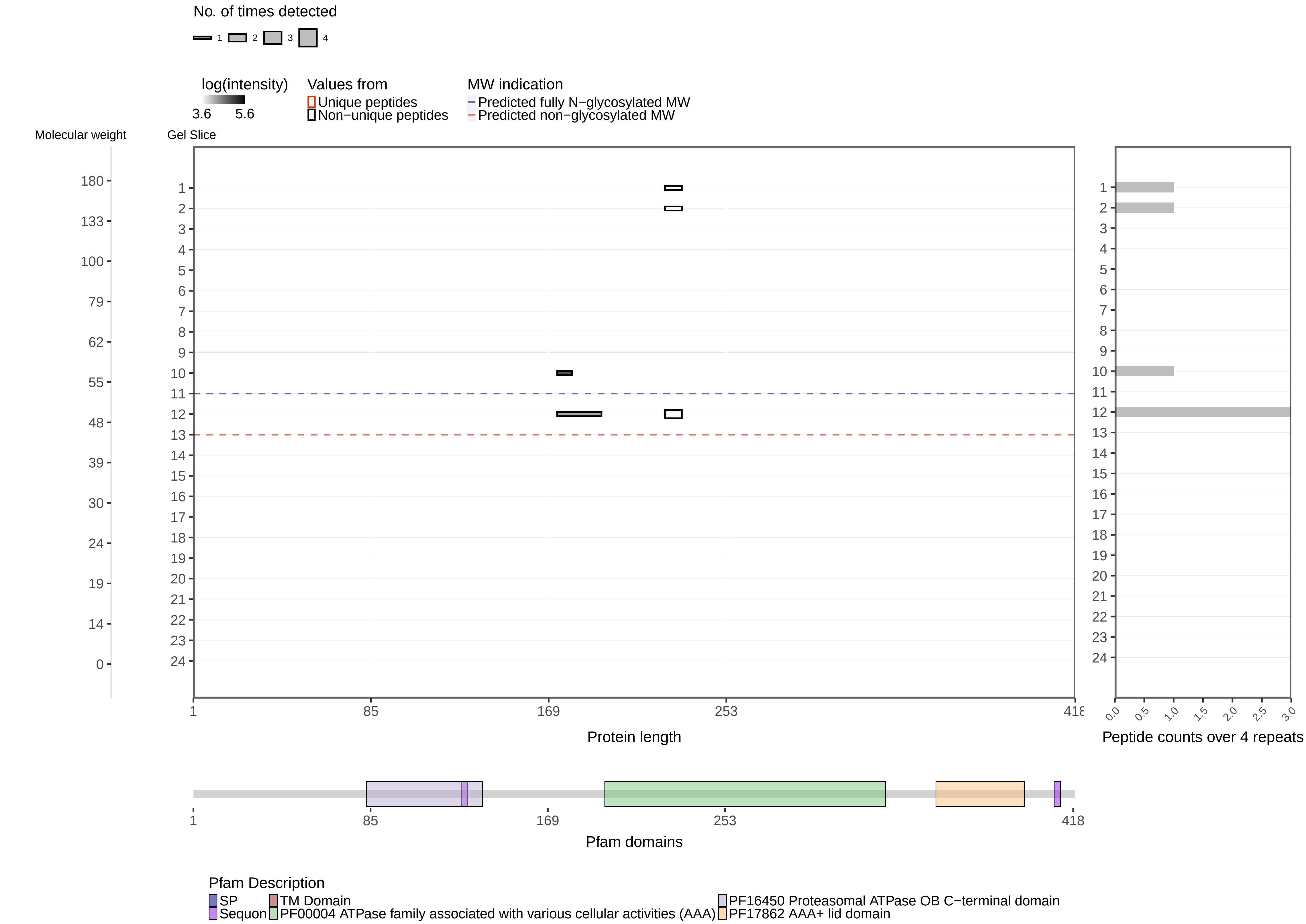


Protein sequence:

'LLYGPPGTGKTLLARAVAHHTDCTFIRVSGSELVQKYIGEGSRMVRELFVMAREHAPSIIFMDEIDSIGSARMESGSGNGDSEVQRTMLELLNQLDGFEASNKIKVLMATNRIDILDQALLRPGRIDRKIEFPNPNEESRLDILKIHSRKMNLMRGIDLKKIAEKMNGASGAELKAVCTEAGMFALRERRVHVTQEDFEMAVAKVMKKETEKNMSLRKLWK*

NbL15g12790.1 ras−related_protein_RABD2a [MW = 30.95 kDa]

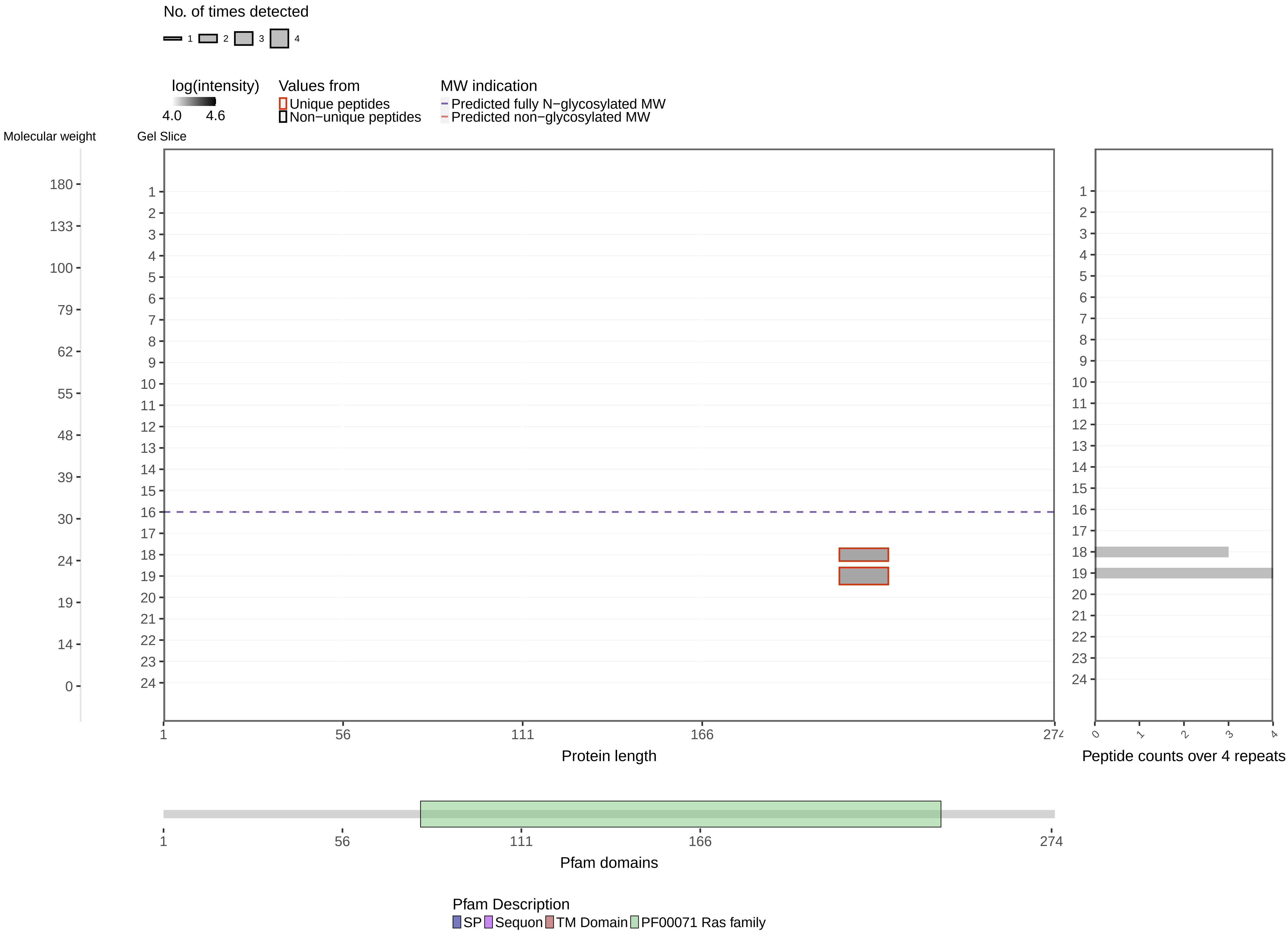


Protein sequence:

TIKLQIWDTAGQERFRTITSSYYRGAHGIIITYDITDQESFNNVKQWLSEIDRYASENVNKLLVGNKCDLTDNRAVSYDTAKAFADEIGIPFMETSAKSATNVEQAFMAMAAEIKNRMASQPTSNNARPPTVQIRGQPVNQKSGCCST*

NbL15g12760.1 glutamate_dehydrogenase_A−like [MW = 44.7 kDa]

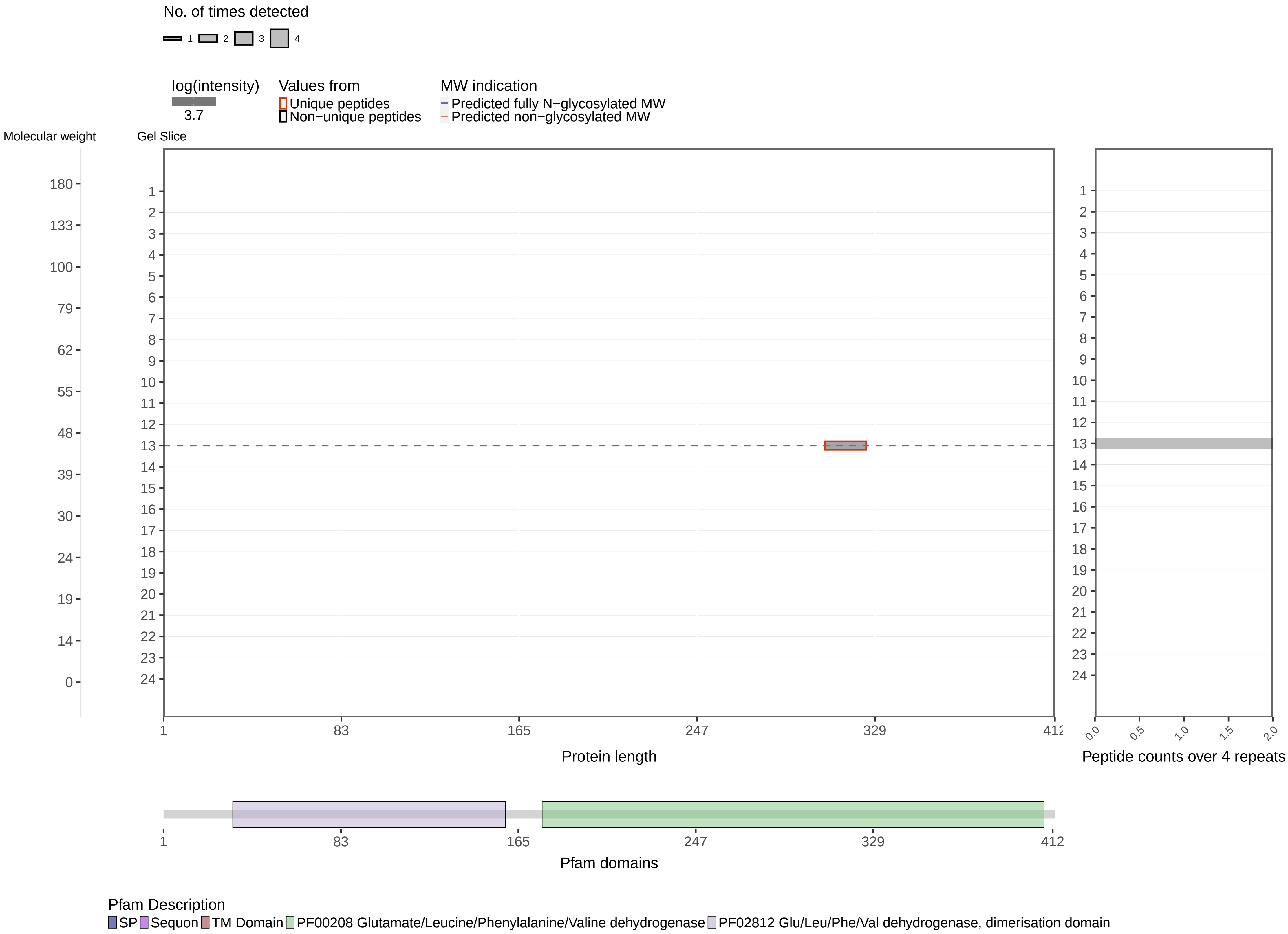


Protein sequence:

ALLAEHGKHIKDMTFAIQGFGNVGAWAARIIHERGGKVVAVSDITGAVKNPNGLDIPALLNHKEATGTLAGFSGSDAMSSDELLTHDCDVLIPCALGGVLNRENADSVKAKYIVEAANHPTDPDADEILSKKGVVILPDIYANAGGVTVSYFEWVQNIQGFMWDEEHVNRELKKYMTRAFHNLKNMCKSHNCNLRMGAFTLGVNRVARATQLRGWEA*

NbL15g12740.1 ATP_synthase_subunit_delta'_mitochondrial [MW = 27.92 kDa]

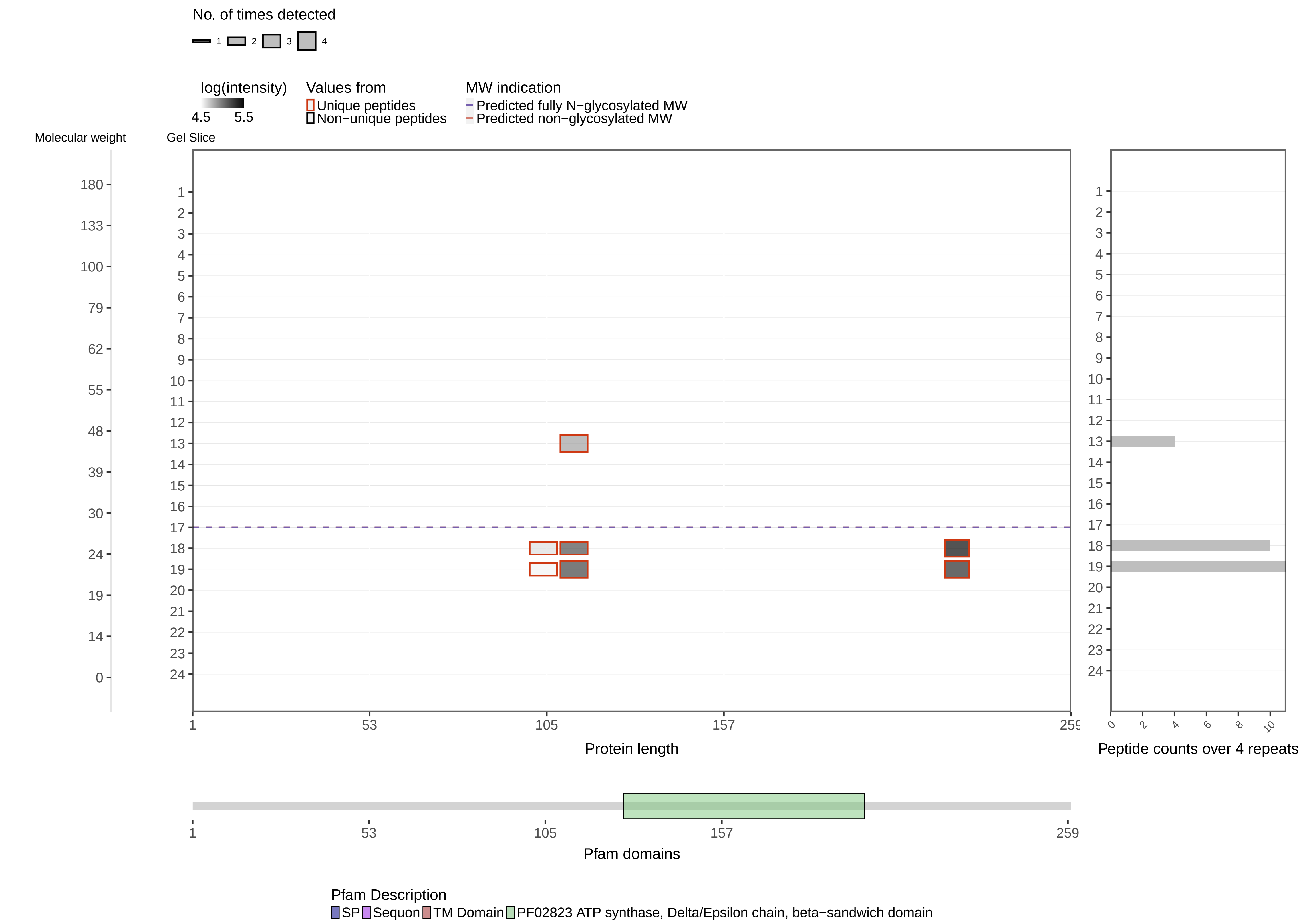


Protein sequence:

ARPATPSSIPSKLTVNFVLPYSSELSGKEVDMVIIPATTGQMGVLPGHVATIAELKPGVLSVHEGNDVTKYFVSGGFAFVHANSFADIIAVEAVPLDRIDPNLVQKGLTDFTQKLSTASTDVEKAEAQIGVDVHSALNAALTG*

NbL15g12660.1 macro_domain−containing_protein_VPA0103−like [MW = 30.74 kDa]

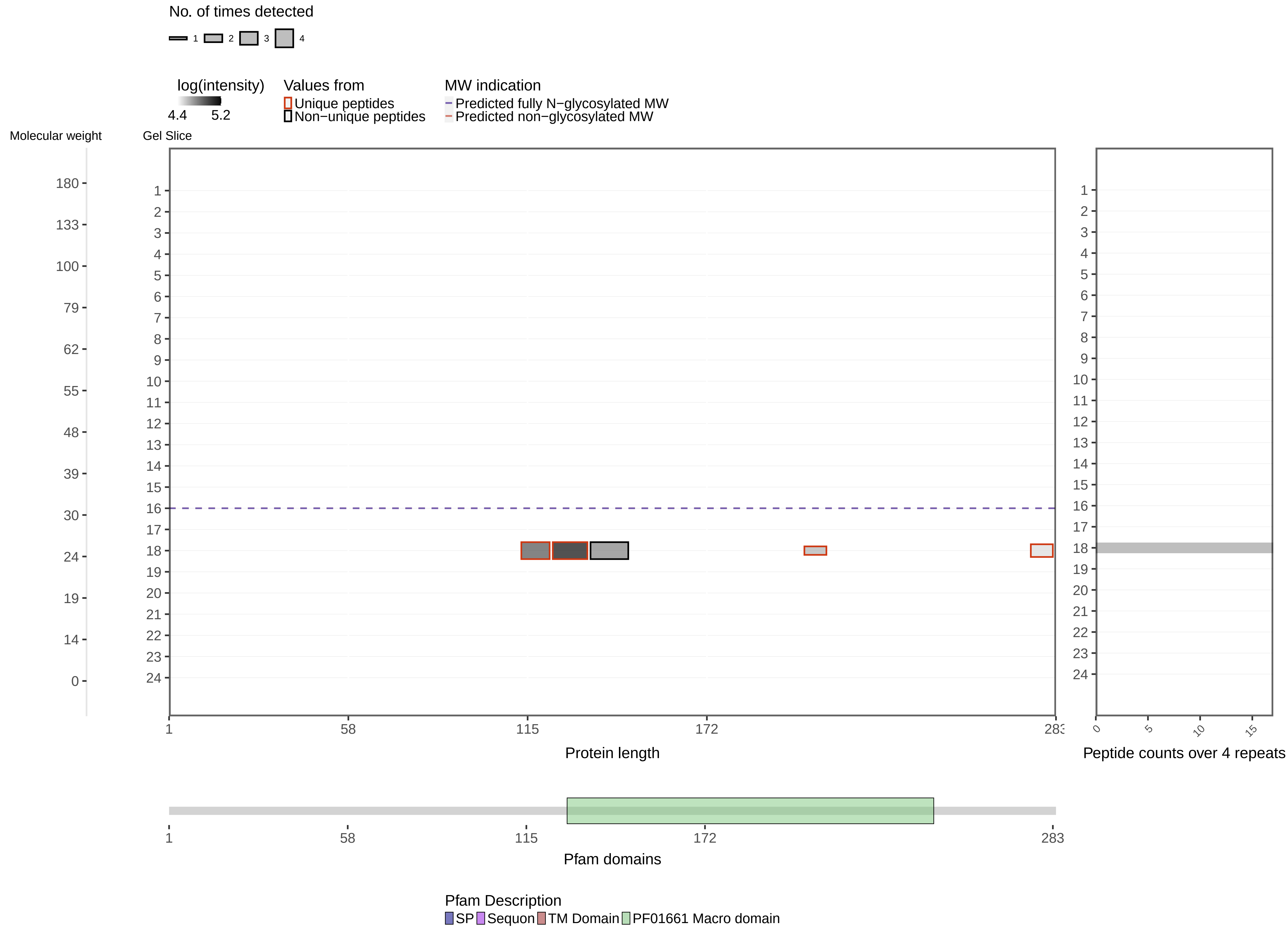


Protein sequence:

'NPANERMLGGGGADGAIHRAAGPELRDACYKVPEVRPGVRCPTGEARITLGFRLPTSHVIHTVGPIYDAVPNPKAYLTNSYRNSLRVAKENNIQYIAFPAISCGVFGYPFDEAATVAISTVKEFGSDLKEVHFVLFSDEIYDAWLKAANELLQN*

### NbL15g12410.1 clathrin_heavy_chain_1−like [MW = 192.69 kDa]

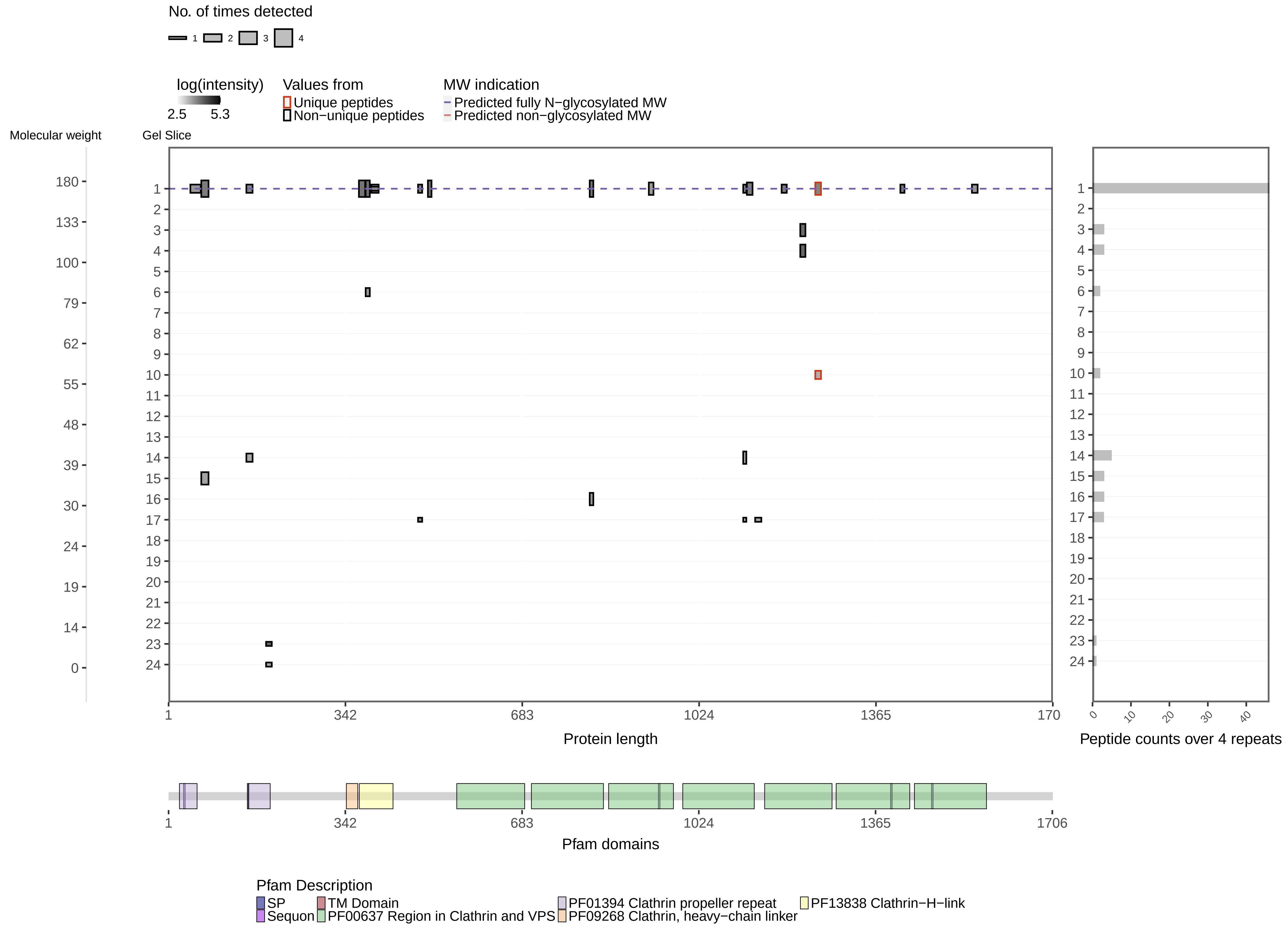


Protein sequence:

FIKGLILSVRSLLPVEPLVAECEKRNRLRLLTQFLEHLVSEGSQDVHVHNALGKIIIDSNNNPEHFLTTNPYYDSRVVGKYCEKRDPTLAVVAYRRGQCDEELINVTNKNSLFKLQARYVVERMDGDLWEKVLTPENEFRRQLIDQVVSTALPESKSPEQVSAAVKAFMTADLPHELIELLEKIVLQNSAFSGNFNLQNLLILTAIKADPSRVMDYINRLDNFDGPAVGEVAVEAQLYEEAFAIFKKFNLNVQAVNVLLDNIPDINRAVEFAFRVEEDAVWSQVAKAQLREGLVSDAIE

NbL15g12340.1 dual_specificity_protein_phosphatase_12−like [MW = 40.24 kDa]

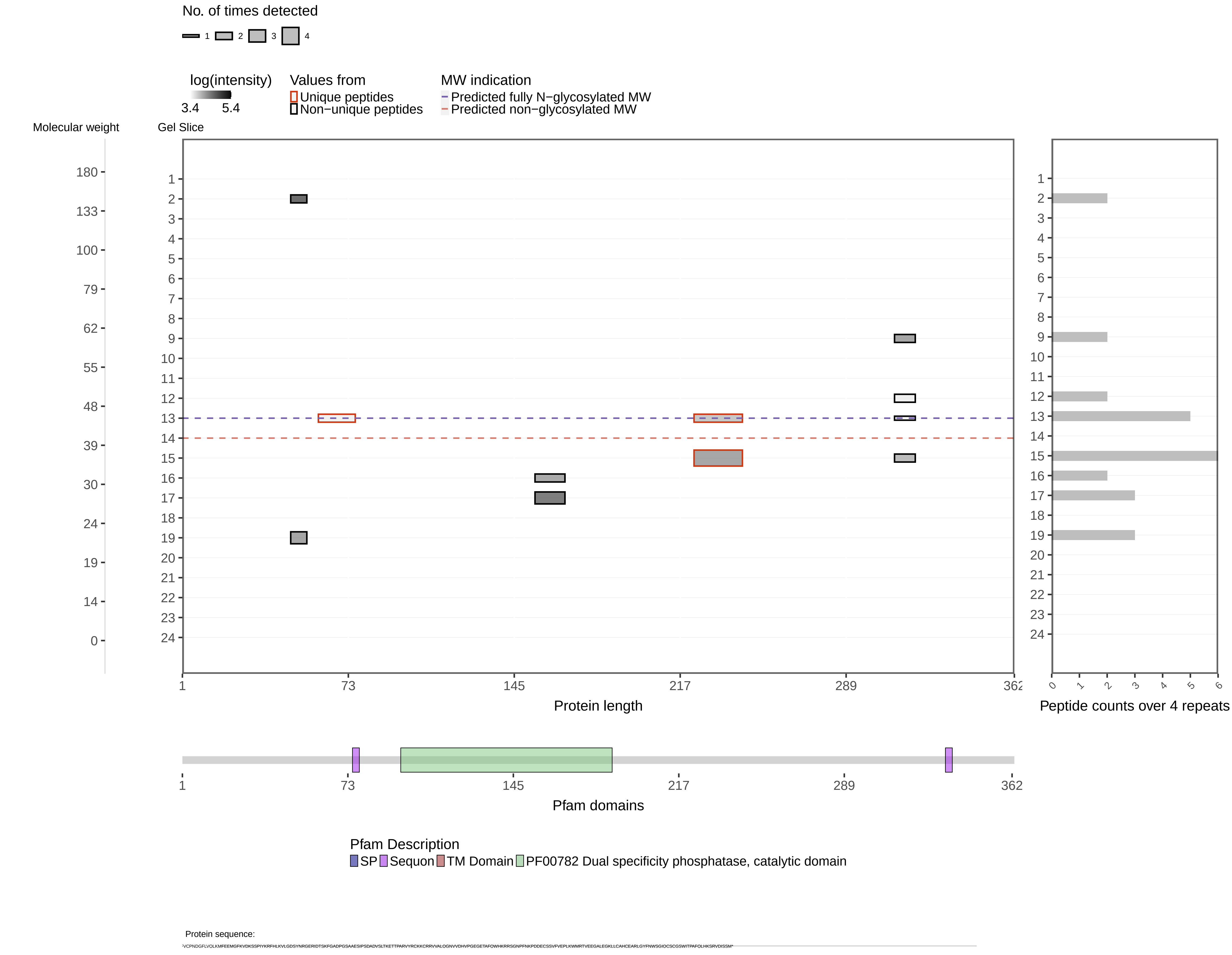

### NbL15g12310.1 sedoheptulose−17−bisphosphatase_chloroplastic−like [MW = 42.55 kDa]

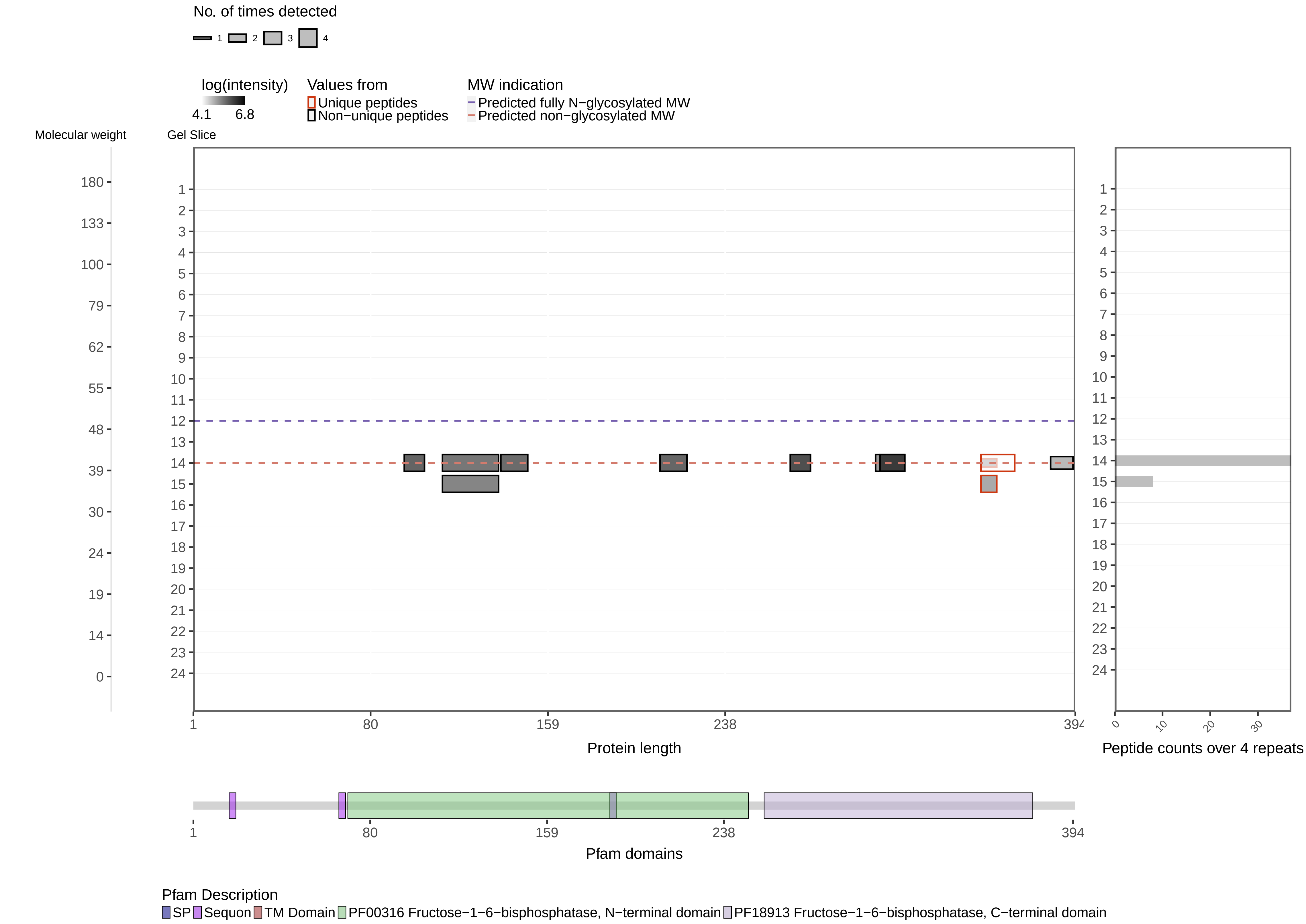


Protein sequence:

IVDTNFTVGTIFGVWPGDKLTGITGRDQVAAAMGIYGPRTTYVLALKDVPGTHEFLLLDEGKWQHVKDTTEIGEGKMFSPGNLRATFDNPDYAKLIDYYVKEKYTLRYTGGMVPDVNQIIVKEKGIFTNVISPTTKAKLRLLFEVAPLGFLIEKAGGYSSDGKQSVLDRVVVNLDDRTQVAYGSKNEIIRFEETLYGSSRLKSAEAVGAAA*

### NbL15g12220.1 30S_ribosomal_protein_S31_chloroplastic−like [MW = 11.06 kDa]

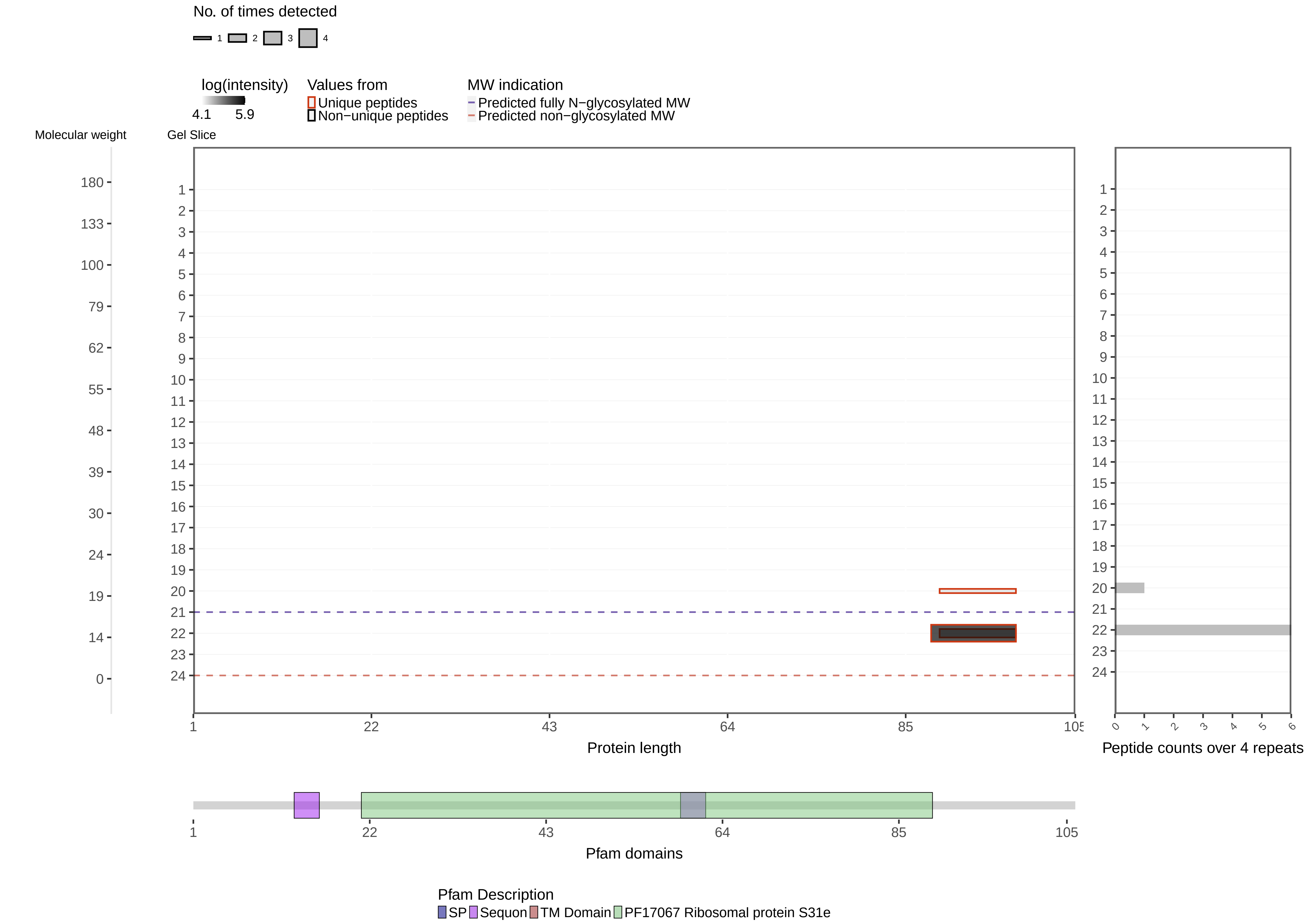


Protein sequence:

FPLIYCGRGDKKTAKGKRFNHSFGNARPRNKKKGRGPPRVVAPPAAPKRDPFDDGQVVKIEIVES*

### NbL15g12210.1 pleckstrin_homology_domain−containing_protein_1−like [MW = 16.88 kDa]

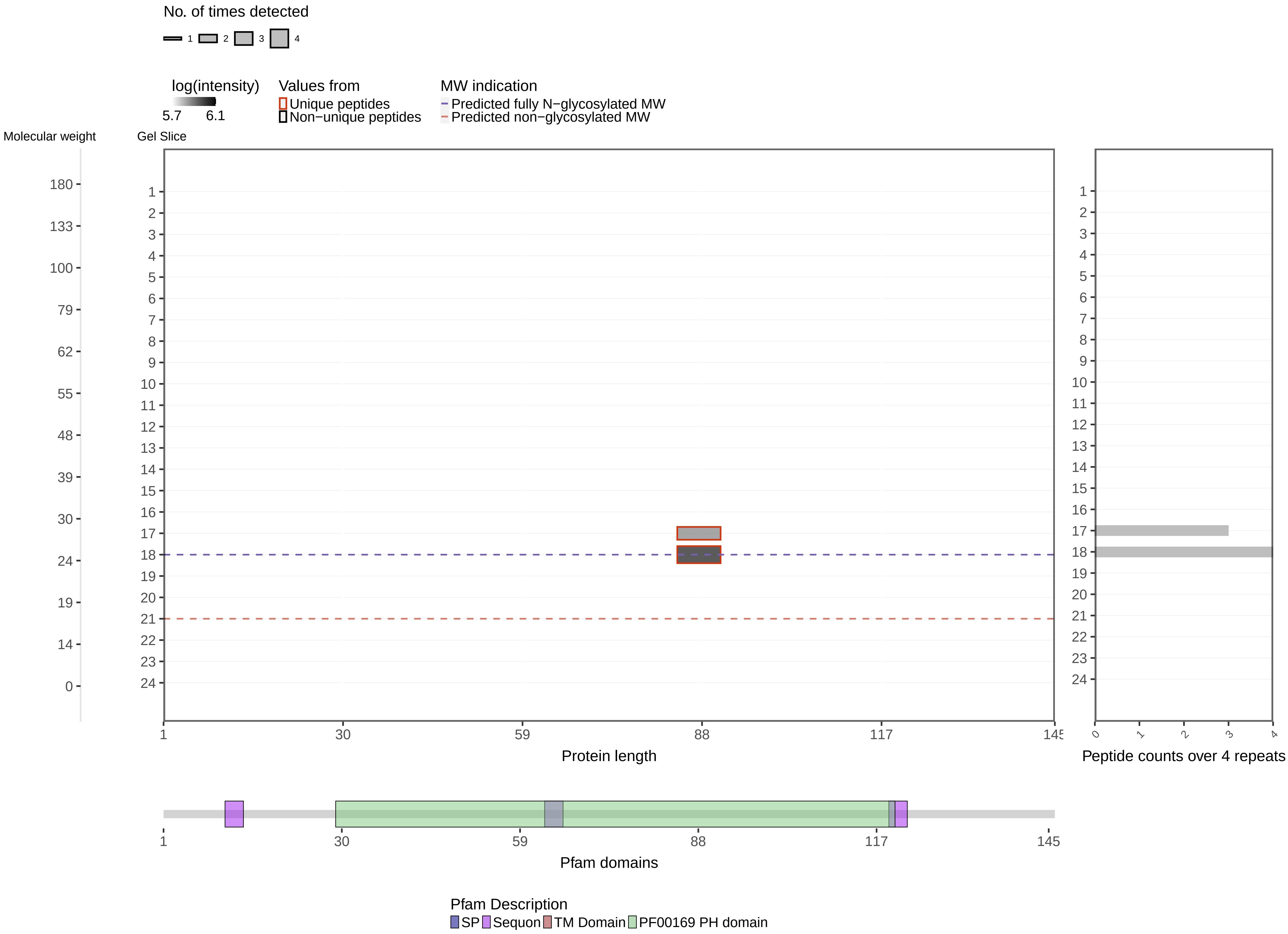


Protein sequence:

[illegible]WFKEANITRGSKPRGVIPVADCLTVKGAEDILNKQFAFELSTRADTMYFIADSEKEKEDWINSTGRCIVQHSRSVTHDEILDYDSRK*

NbL15g12090.1 7−deoxyloganetic_acid_glucosyltransferase−like [MW = 60.48 kDa]

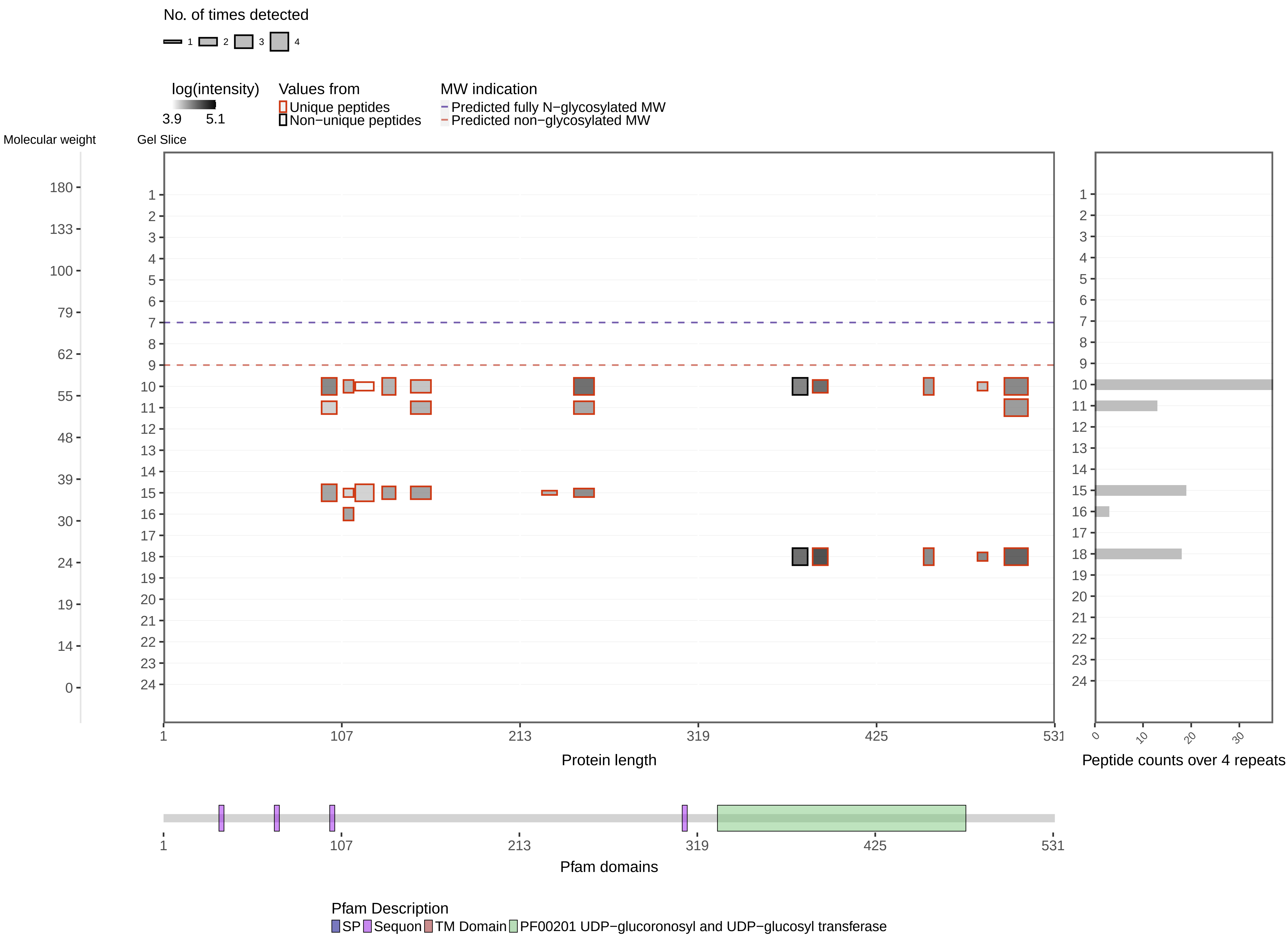


Protein sequence:

EGQENSRAYGLILNTFEELDGHLLSQMRNVCPNIYPIGPLHAHLKNKLSEENNSLPPSSNSIFEEDKSCINWLNNQPPNSVIYVSFGSIATMTKEQLMEFWYGLVNSKQRFLWVIRLDSVAGDDWKNKISVDLAEGTKERGYIVGWAPQEEVLAHPSVGGFWTHCGWNSILETVVYEGKPMICWPYFMDQQVNSRFVEAAWKLGLDMKDTCDREIVEKMVKDLMVVRKDEFLERANQMSKFAKECLVEGGSSYSNFERLVNDIKSLSDKVRHY*

### NbL15g12020.1 ribose−phosphate_pyrophosphokinase_1−like [MW = 45.67 kDa]

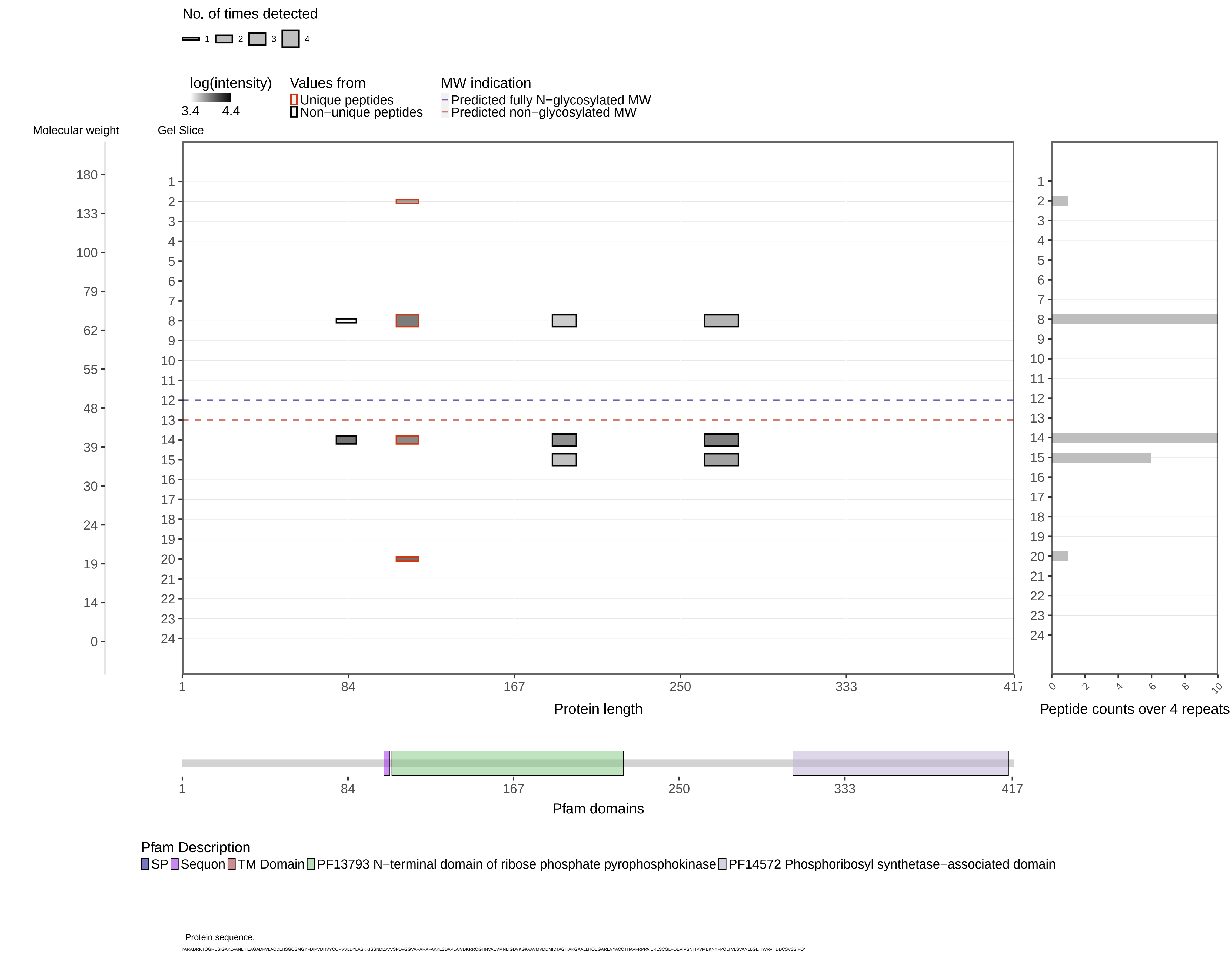

NbL15g12010.1 pyruvate_kinase_isozyme_G_chloroplastic [MW = 68.12 kDa]

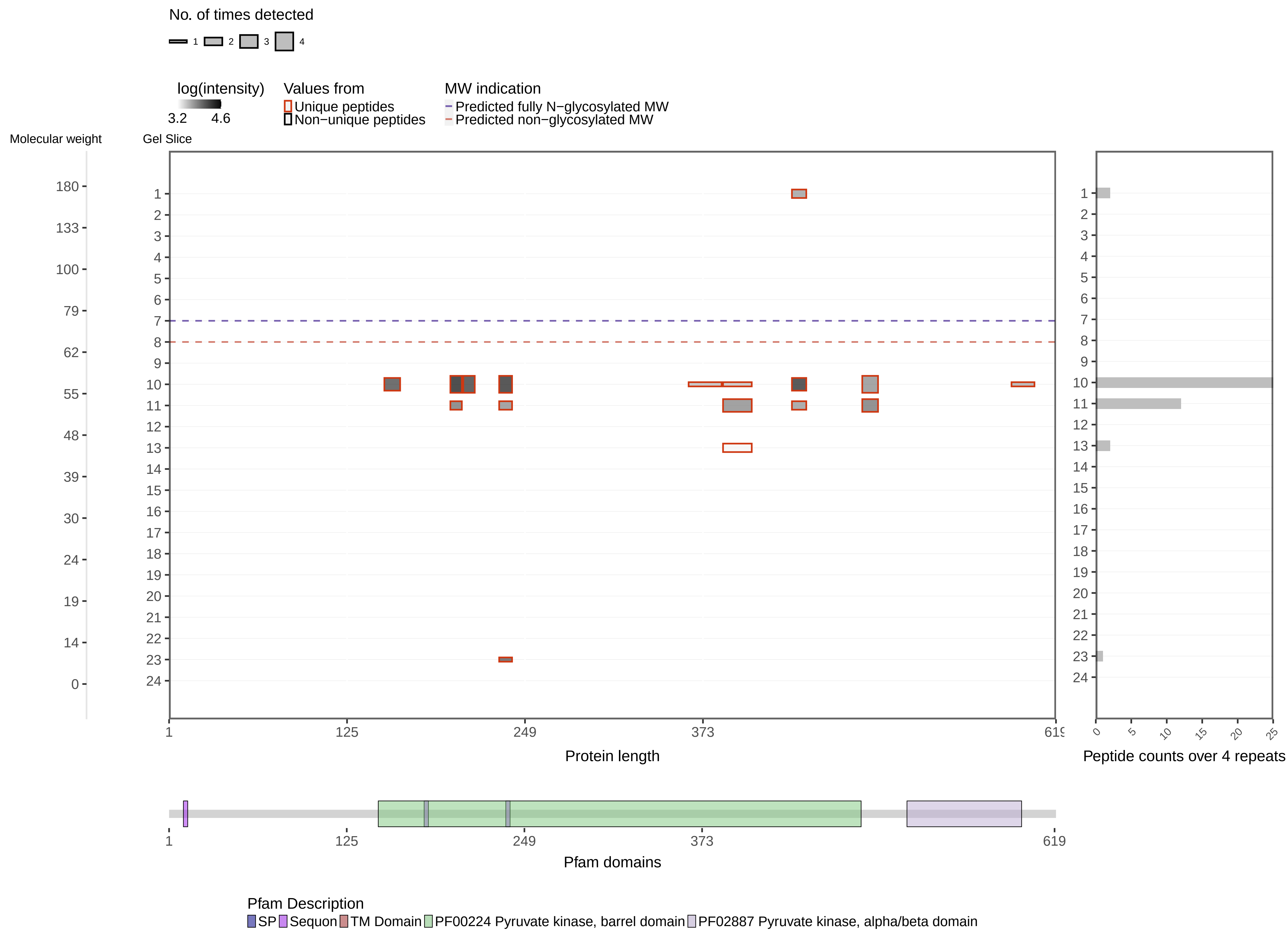


Protein sequence:

LKSRRHLNVRGKSATLPSITEKDWDDIKFGVDNQVDFYAVSFVKDAKVVHELKDYLKSCNADIHVIVKIESADSIPNLHSIISASDGAMVARGDLGAELPIEEVPLLQEDIIRRCQSMQKPVIVATNMLESMIDHPTPTRAEVSDISIAVREGADAVMLSGETAHGKYPLKAVKVMHIVALRTESSLQKSTSSPSQSAAYKSHMGEMFAFHSSSMANTLSTPIIVFTRTGSMAIILSHNRPSSTVFAFTNNERVKQRLALYHGVVPIYMEFSSDAEETFSRAIKLLLSKSLVKDGQYLTL

### NbL15g11960.1 syntaxin−22−like [MW = 30.04 kDa]

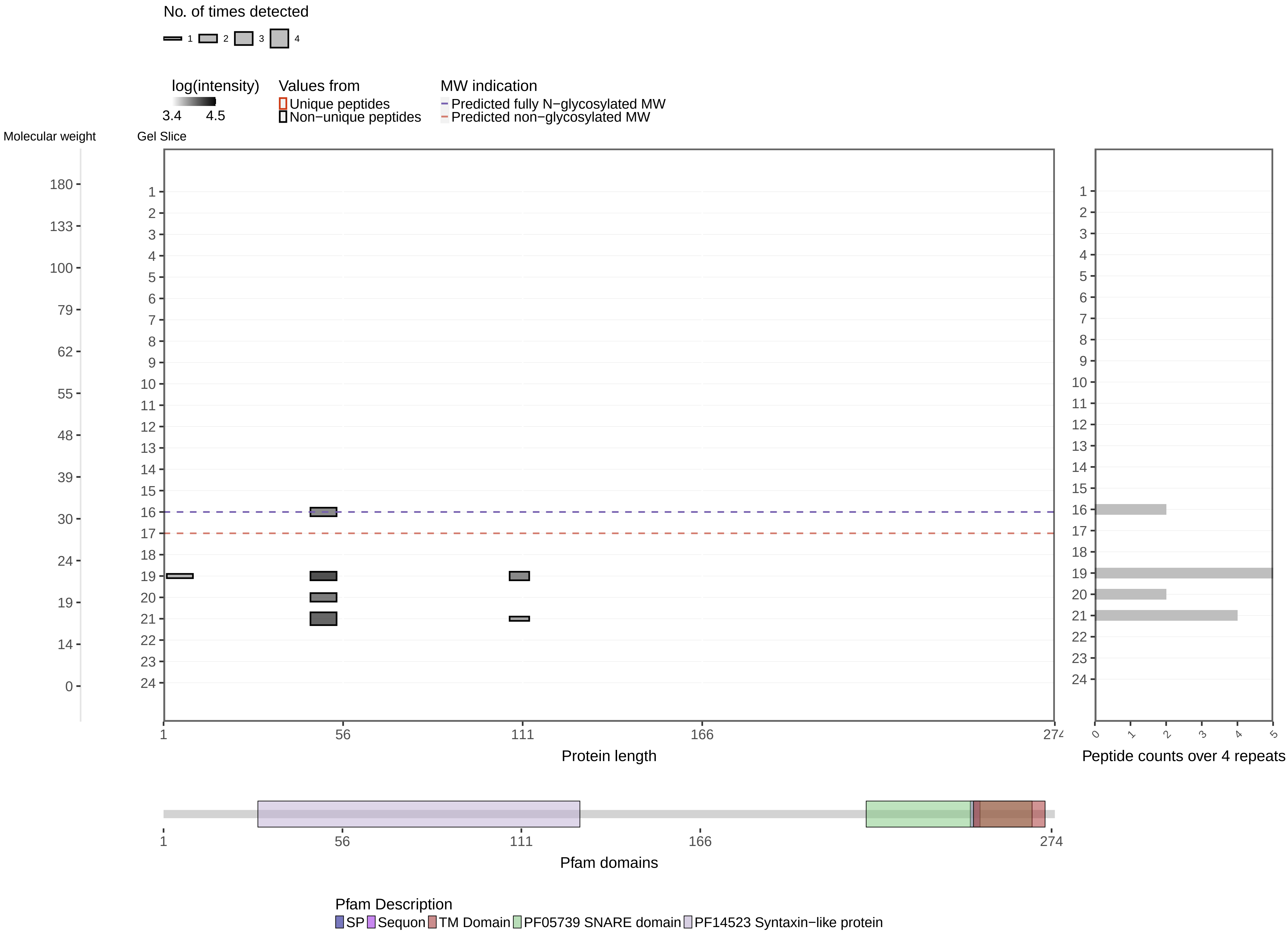


Protein sequence:

RLAAERETAYTPFVPQAVLPSSYTASEIDVSTEKSPEQRALLVESRRQEVLLLDSEIAFNEAIIEEREQGIQEIQQQIGEVNEIFKDLAVLVHEQGAMIDDIGSNVENAHAATAQGRSLLAKAAKTQRSNSSLTCLLLVIFGIVLLIVIIVLAA*

### NbL15g11810.1 uncharacterized_protein_At1g32220_chloroplastic [MW = 32.74 kDa]

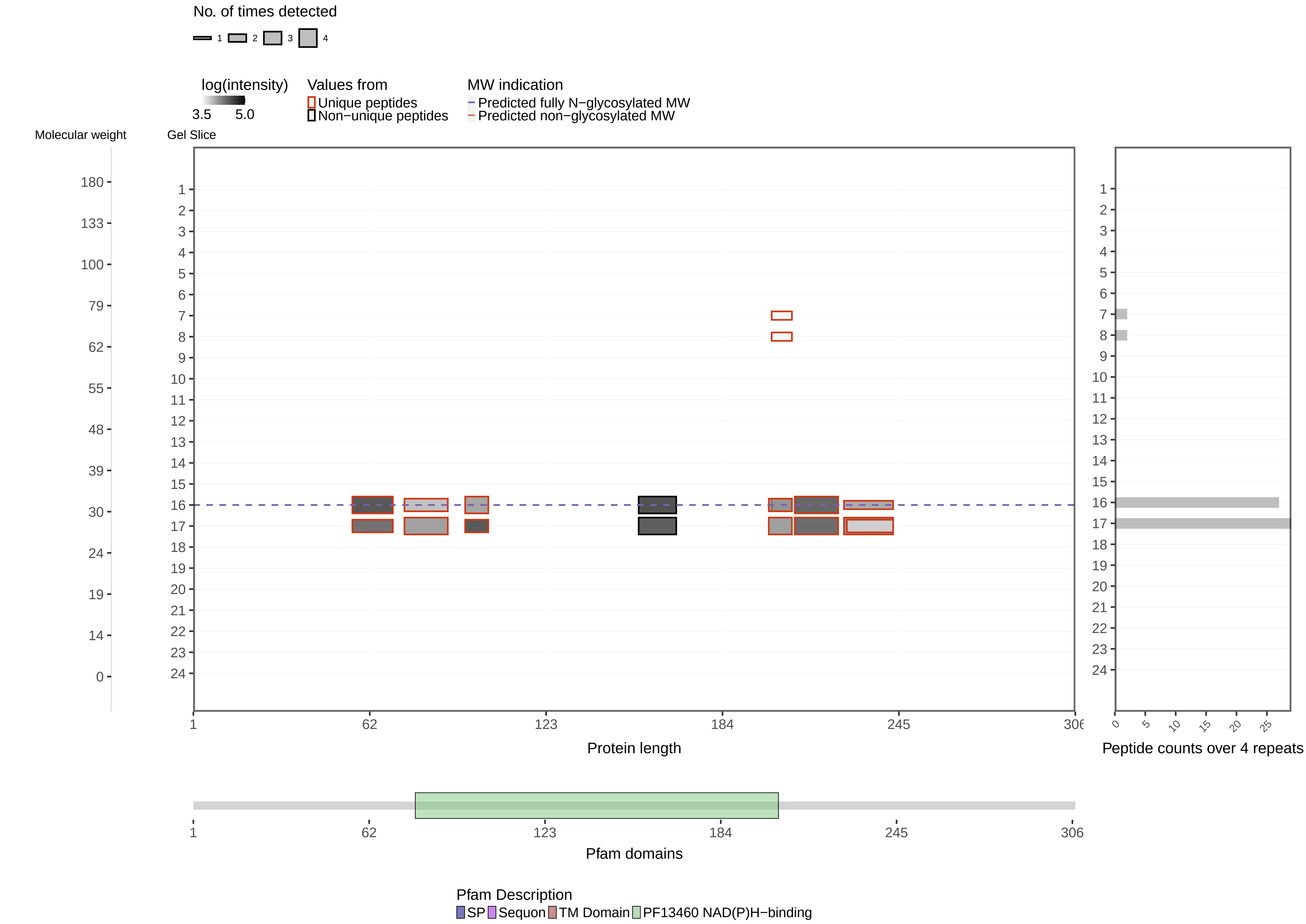


Protein sequence:

AVISTLGGFGSDEQMQRINGEANIVAVNAAKDYGIPKFILISVHDYNLPSFLLQSGYFTGKRKAESEVLSKYPNSGVVLRPAFIYGKRKVDGFEIPLDLIGEPLEKILRATESFTKPLSSLPASDLLLAPPVSVDDVAYAVINAVKDDDCFGIFTIGQIKEAAAGVKV*

### NbL15g11780.1 glycerol−3−phosphate_acyltransferase_chloroplastic [MW = 51.37 kDa]

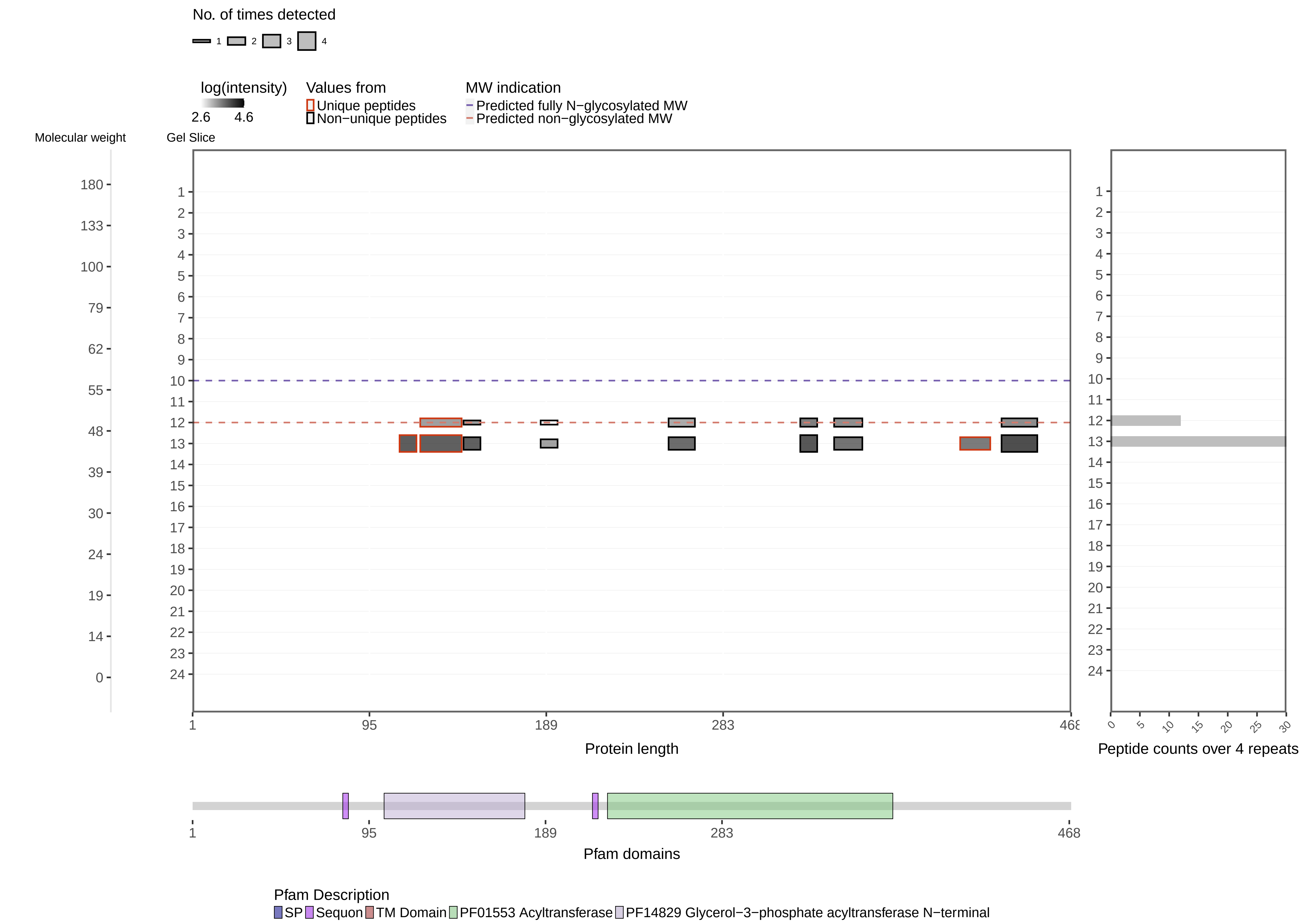


Protein sequence:

KLKQGVNVVLMSNHQSEADPAVIALLLESRHPYIAENIIYVAGDRVITDPLCKPFSMGRNLLCVYSKKHMGDDPKLVEKKRRANTRSLKEMAVLLRGGSKLIWIAPSGGRDRPNPVTKEWYPAPFDASSTDNMRRLVEHAGVPGHIYPLAILCYDIMPPPPQVEKNIGEKRIISFHGAGISVGPKIDFHEVAGALEDPEEAKMVYTKALYDSVNQQYNVLNSAIHGKQGQEASTRSVSLSQPWQ*

### NbL15g11460.1 3−isopropylmalate_dehydrogenase_2_chloroplastic−like [MW = 48.03 kDa]

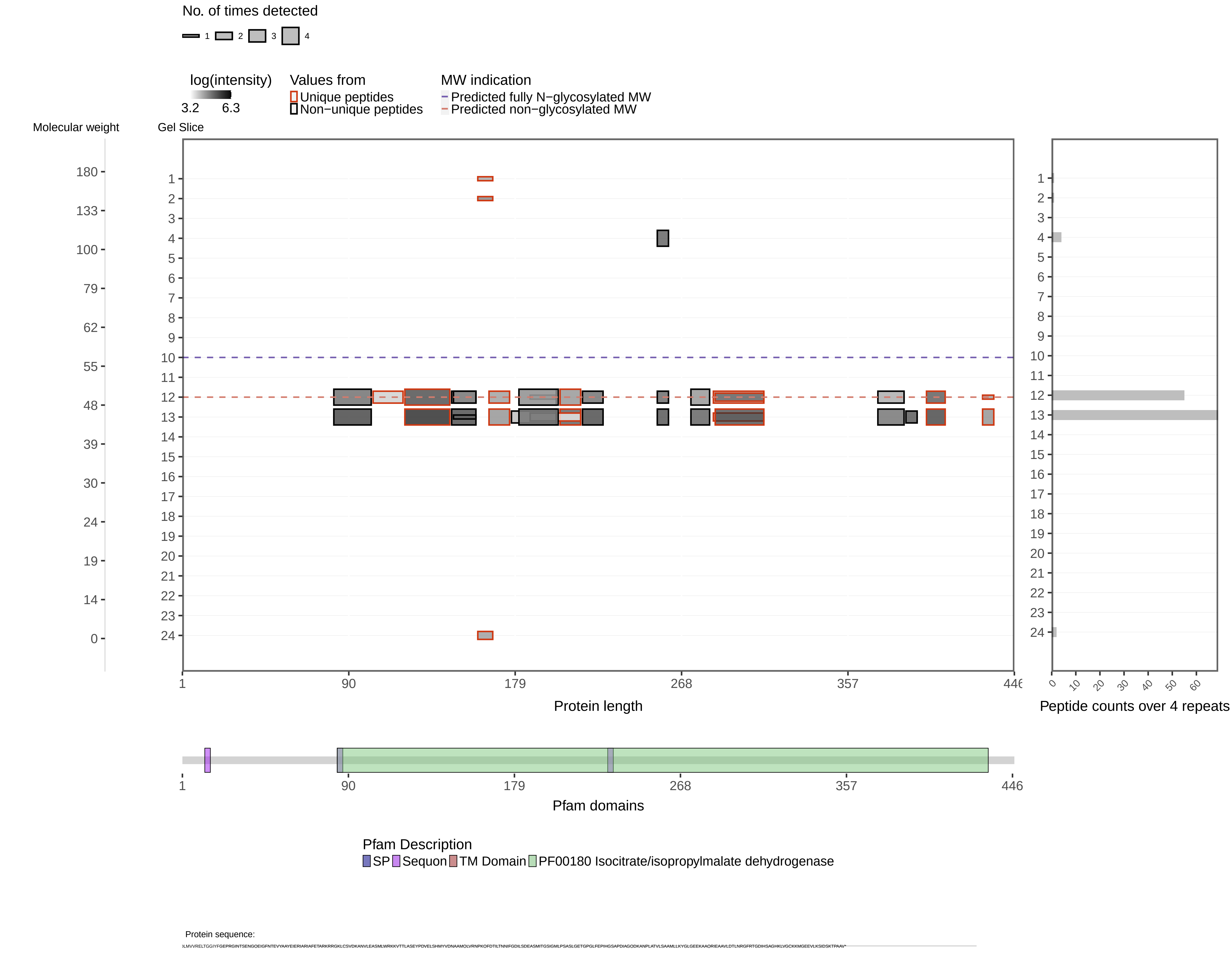

### NbL15g11380.1 O−methyltransferase_MdmC−like [MW = 32.12 kDa]

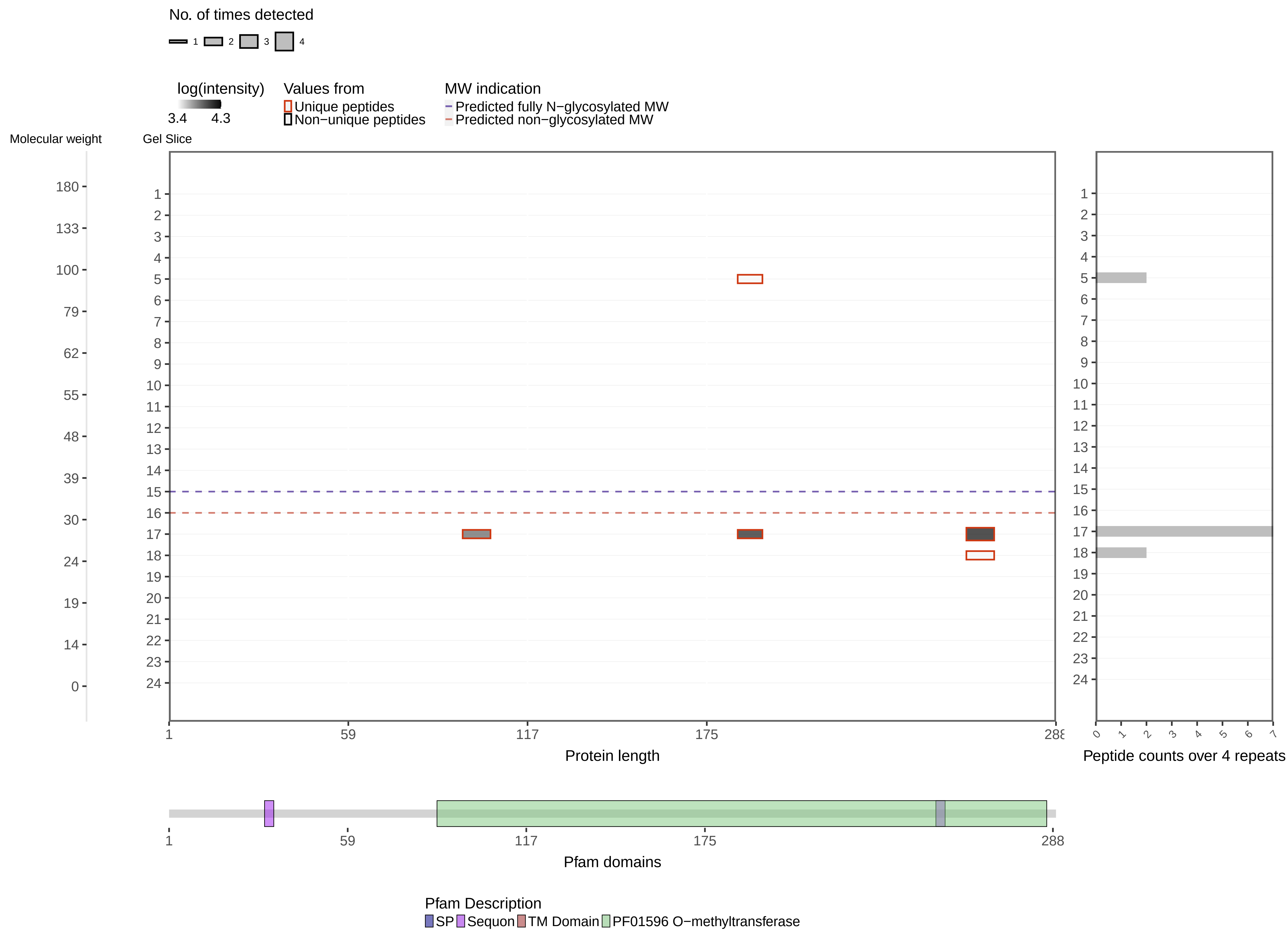


Protein sequence:

EVGTYTGYSSLAVALVLPEEGQLVACERDGKSIEVAKRYYDQAGVLHKVDVRHGLAADTLKSLIQNGESCRYDFAFVDAEKRMYQDYFELLLQLVRVGGLIVVDNVLWHGRVTDPLVNDSKTLSIRNFNQNLMEDKRVDISMVPIGDGMTICRKR*

### NbL15g11350.1 early_nodulin−like_protein_1 [MW − SP = 16.64 kDa]

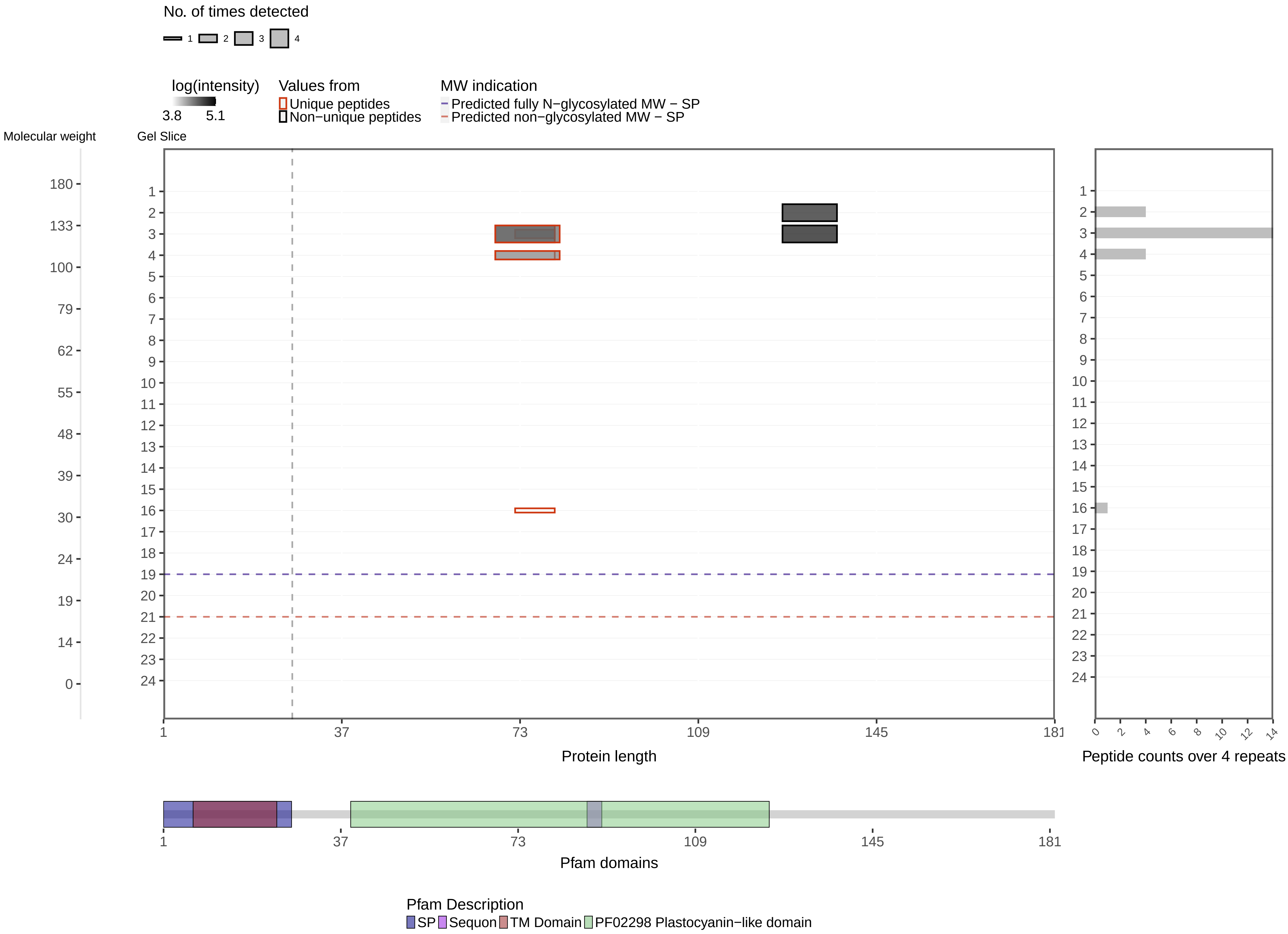


Protein sequence:

VSKRDYVTCNSSSPVAVHNDGNTKIELTHSGAYYFISGAKGHCEQGQKLIVVTLSENNMRRFMAAPAPSPAEFDGPAMAPTSSSASLKPSLFMAFGVLIGFLF*

NbL15g11290.1 probable_phospholipid_hydroperoxide_glutathione_peroxidase [MW = 25.74 kDa]

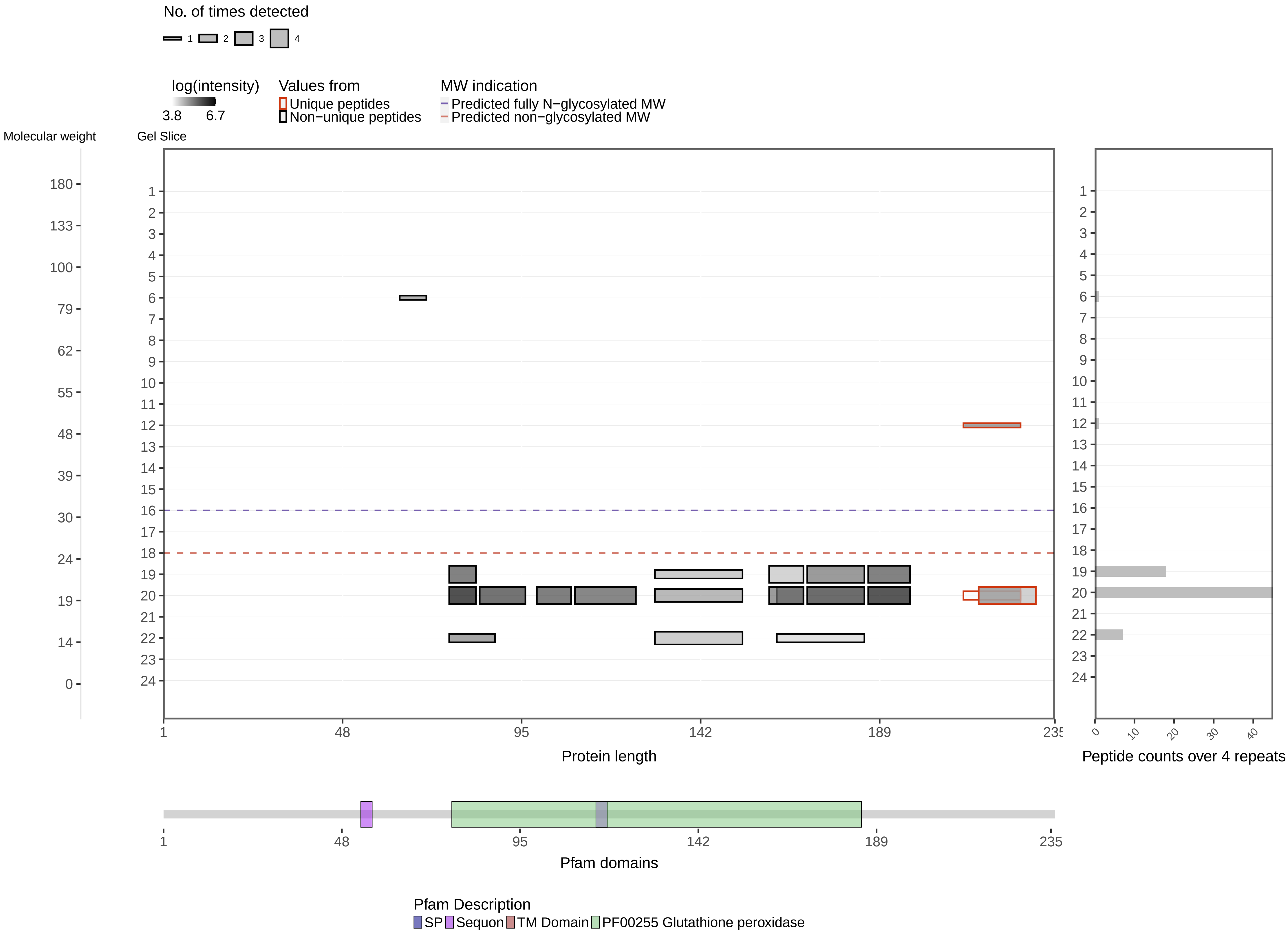


Protein sequence:

ASRCGLTTSNYTELSQVYEKYKTKGLEILAFPCNQFGAQEPGSNPEIKQFACTRFKAEFPIFDKVDVNGPNTAPVYQFLKSSAGGFLGDLVKWNFEKFLVDKNGKVVERYPPTTSPVQIEKDIQKLLAA*

### NbL15g11220.1 luminal−binding_protein_4−like [MW − SP = 70.34 kDa]

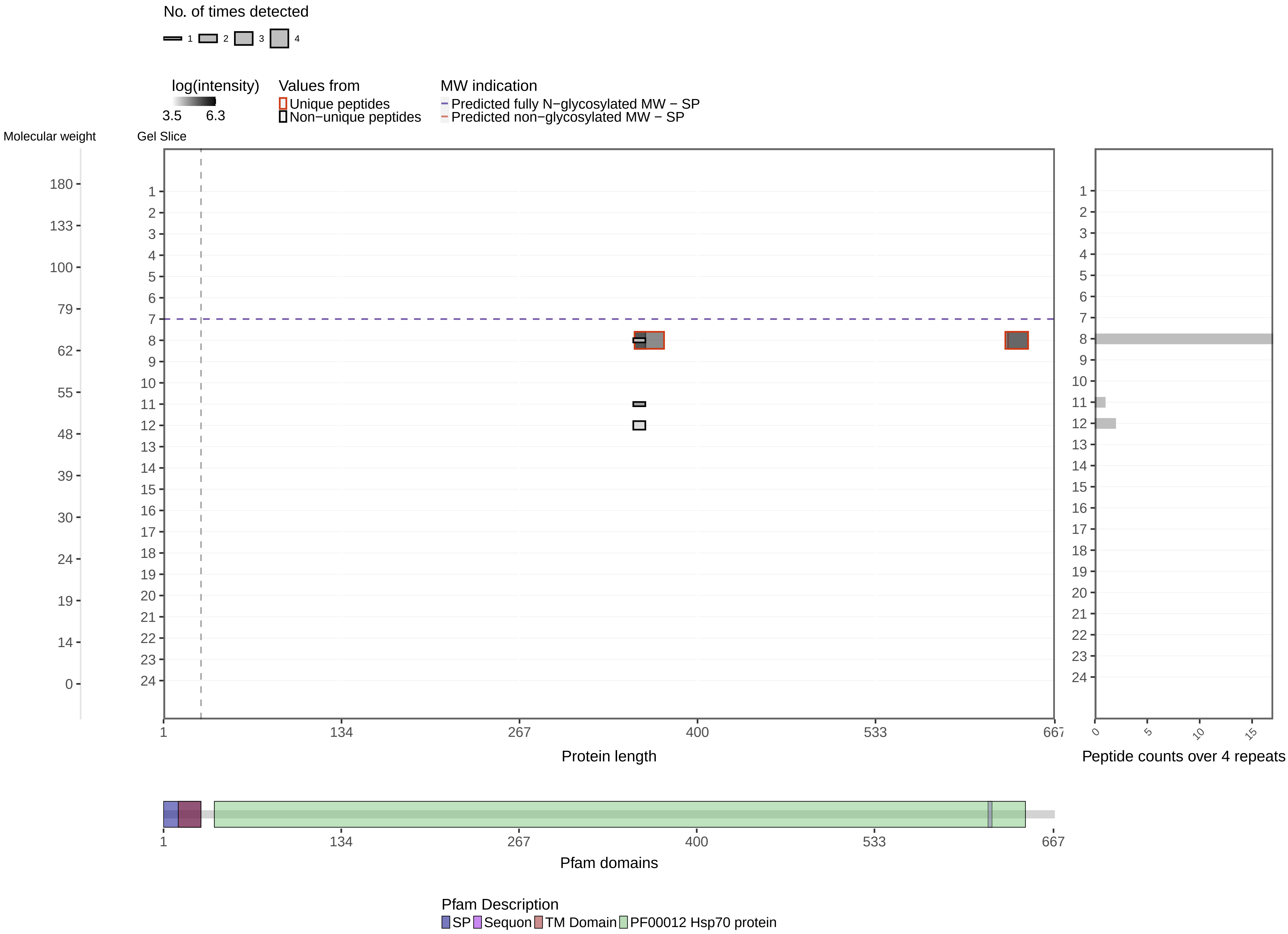


Protein sequence:

GTDFSEPLTRARFEELNNDLFRKTMGPVKKAMDDAGLEKNQIDEIVLVGGSTRIPKVQQLLKDYFEGKEPNKGVNPDEAVAYGAAVQGGILSGEGGDETKDILLLDVAPLTLGIETVGGVMTKLIPRNTVIPTKKSQVFTTYQDQQTTVTISVYEGERSLTKDCRLLGNFDLTGIPPAPRGTPQIEVTFEVDANGILNVKAEDKGTGKSEKITITNDKGRLSQEEIERMVREAEEFAEEDKKVKDRIDARNGLETYVYNMKNQINDKDKLADKLESDEKEKIDTAVKEALEWLDDN

NbL15g11190.1 nuclear_pore_complex_protein_NUP50A−like [MW = 47.78 kDa]

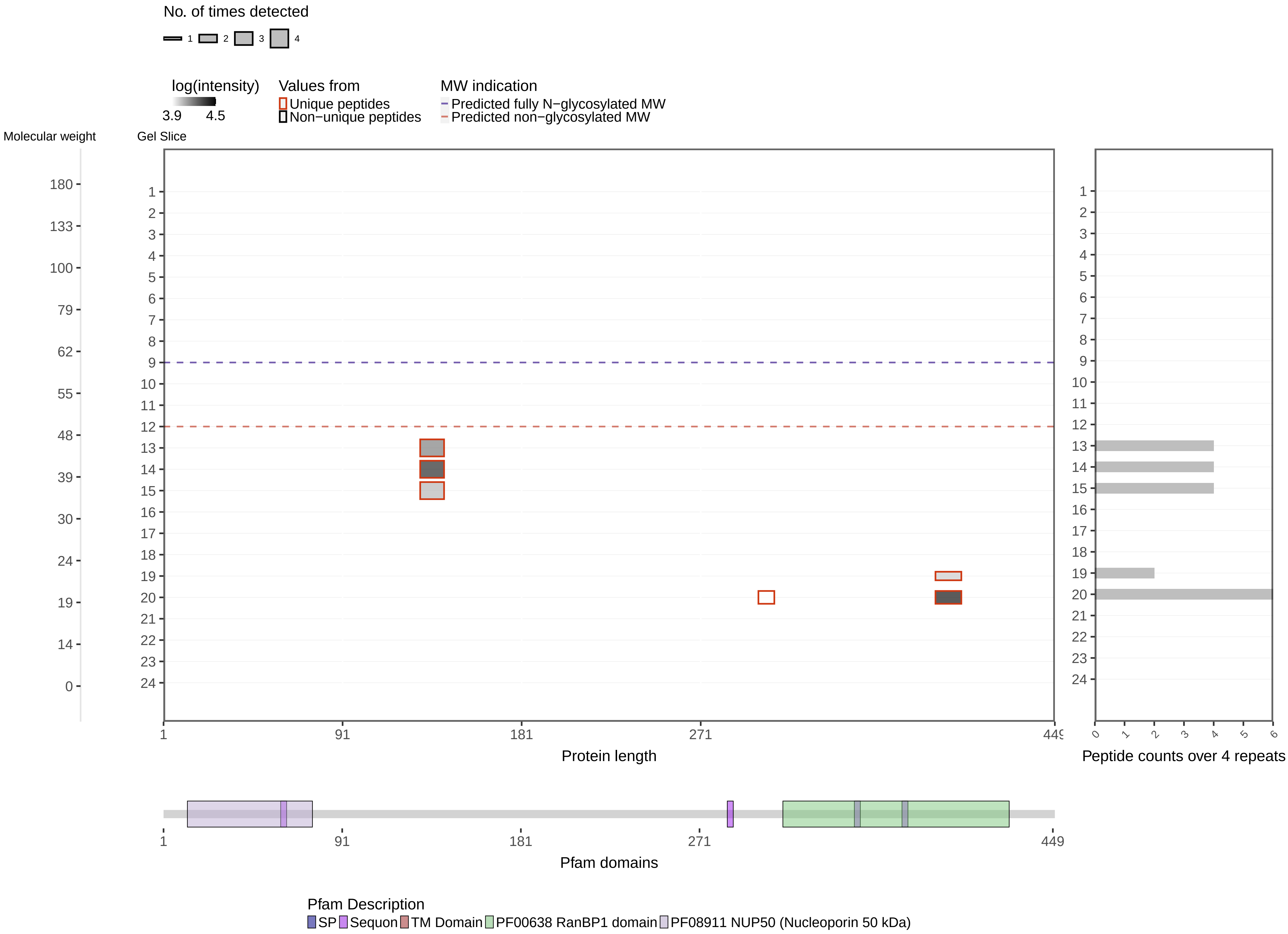


Protein sequence:

ETASFSSFQQLSSSQNAFTGFAGTGFTSTTFSFGAISKEGSPLGFGSESGAVSGSLFGAKSDQPSFGLTLPTNGNNSLFGNSGSSIVNKGEGTGFPSKEEVSVETGEENEKPVFTADSVLFEFLDGGWKERGKGELKVNVSTAREGKARLVMRTKGNYRLILNASLYPEMKLTNMDKRGLSFACLNSAGDGKEGLSTIALKFKDASIVEEFRAAVAKHKGTSTGSLKTPENSPKASDD*

### NbL15g11140.1 MAR−binding_filament−like_protein_1−1 [MW = 81.9 kDa]

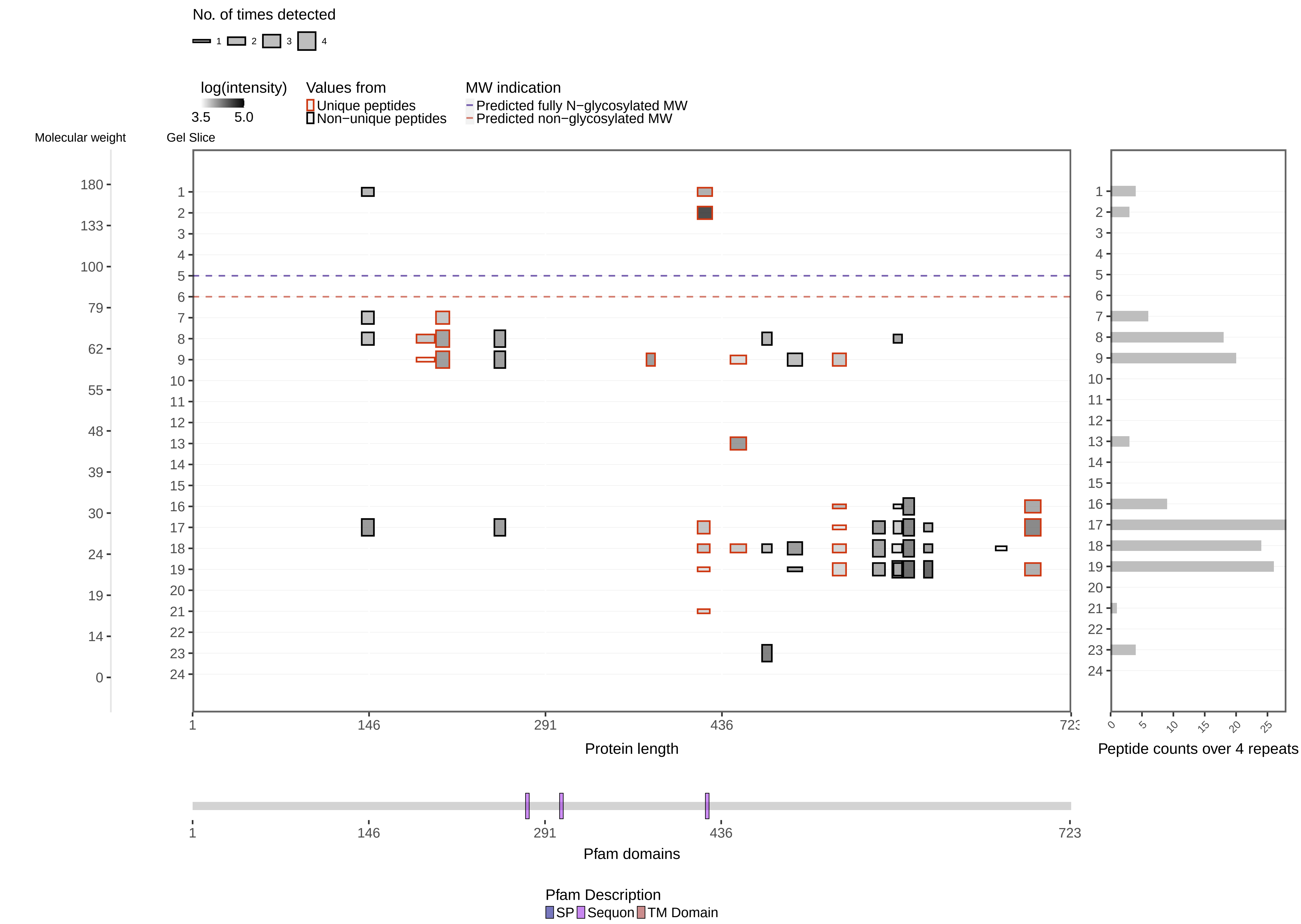


Protein sequence:
[illegible]

### NbL15g11020.1 uncharacterized_LOC107817663 [MW = 45.8 kDa]

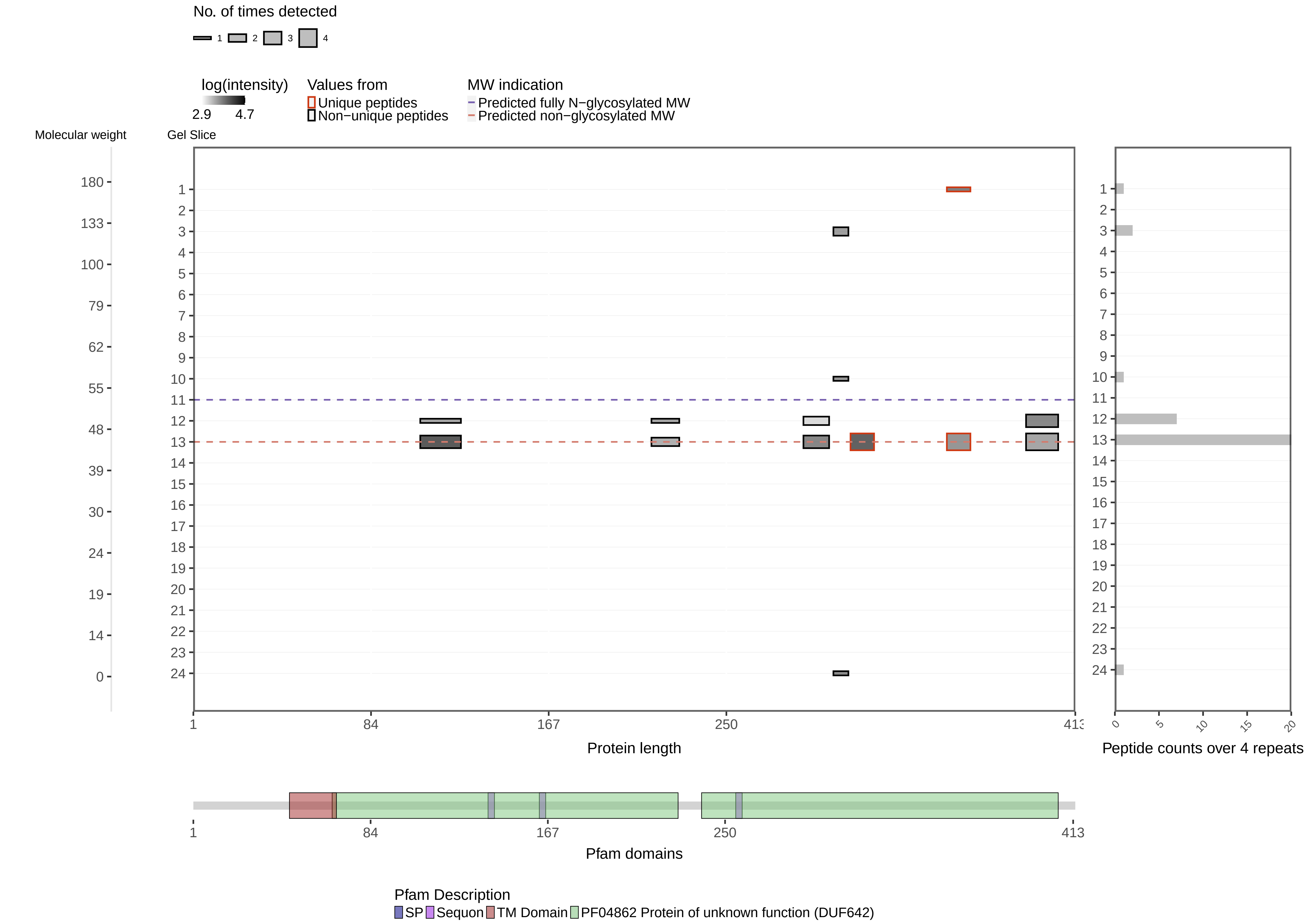


Protein sequence:

YSWGFLAQANEIDIMLHNPAIEKDPACGPLIDFIALKALKNPHRPKGNMLKNGNFEEGPYIFPNTSWGVLIPPNIEDDHSPLPGWMIESLKAVKYIDAAHFTVPEGKRAIELVAGRESAIAQQVITTKGKVYDLMFSVGDASNSCQGSMLIEAFAGQITLRVPYESTGNGGFKRAKLRFKAVSERTRVRFLSTYYHMKSDHSGSLCGPVVDDVRLVGVRYP*

### NbL15g11010.1 serine−threonine_kinase_receptor−associated_protein−like [MW = 38.45 kDa]

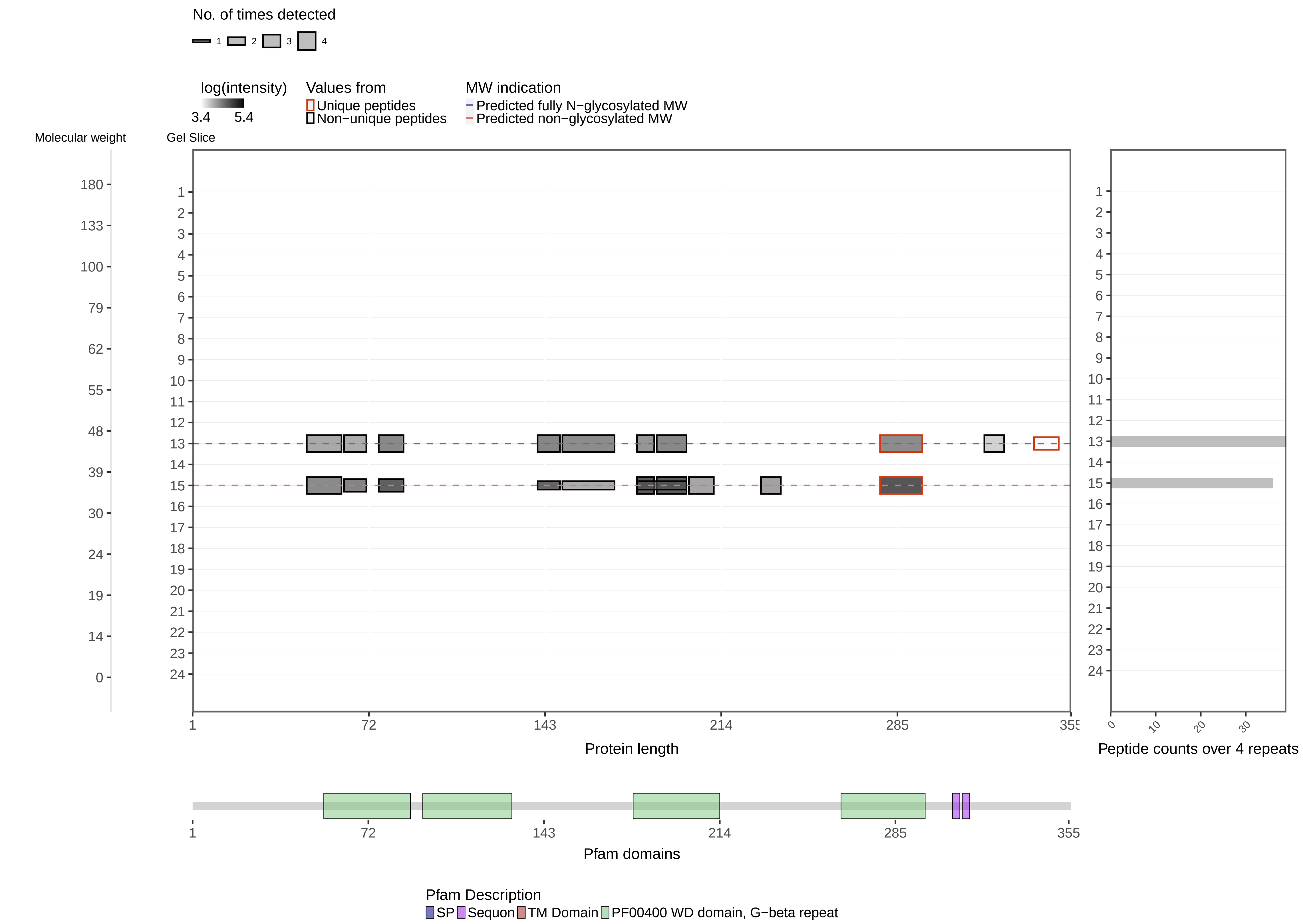


Protein sequence:

AGGLRLWDVRSGKVVQTLETKFPVTSAEVSQDGRYITTADGSSVKFWDANHFGLVKSHDLPCKVESASLEPKFGNRFIAGGEDMWVHVFDFHTGEEIGCNKGHHGPVHCLRFSPVGDTYASGSEDGTVRIWQLGPLGQNEDNATANGSIANAKDDMGEVTQKIEELDVSEAKKTEETQMDGAVKVADA*

NbL15g10750.1 ATP−dependent_Clp_protease_adapter_protein_CLPS1_chloroplastic−like [MW = 17.49 kDa]

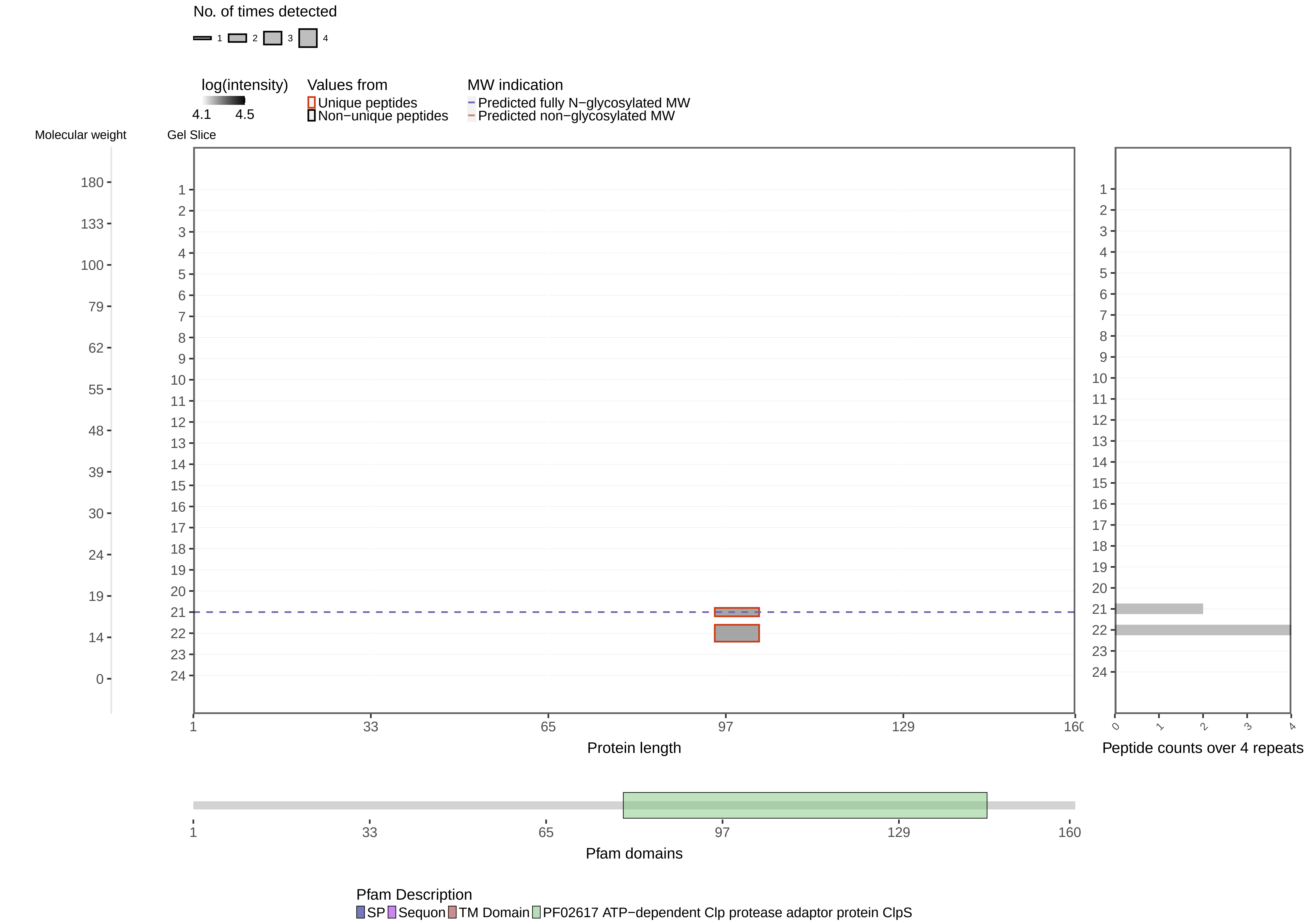


Protein sequence:

ESEFDLRKSRKMSPPYRVMLHNDNYNKREYVVQVLMKVIPGMTVDNAVNIMQEAHYNGLAVVIICAQADAEEHCTQLRGNGLLSSIEPASGGC*

NbL15g10740.1 protein_ASPARTIC_PROTEASE_IN_GUARD_CELL_1 [MW − SP = 49.22 kDa]

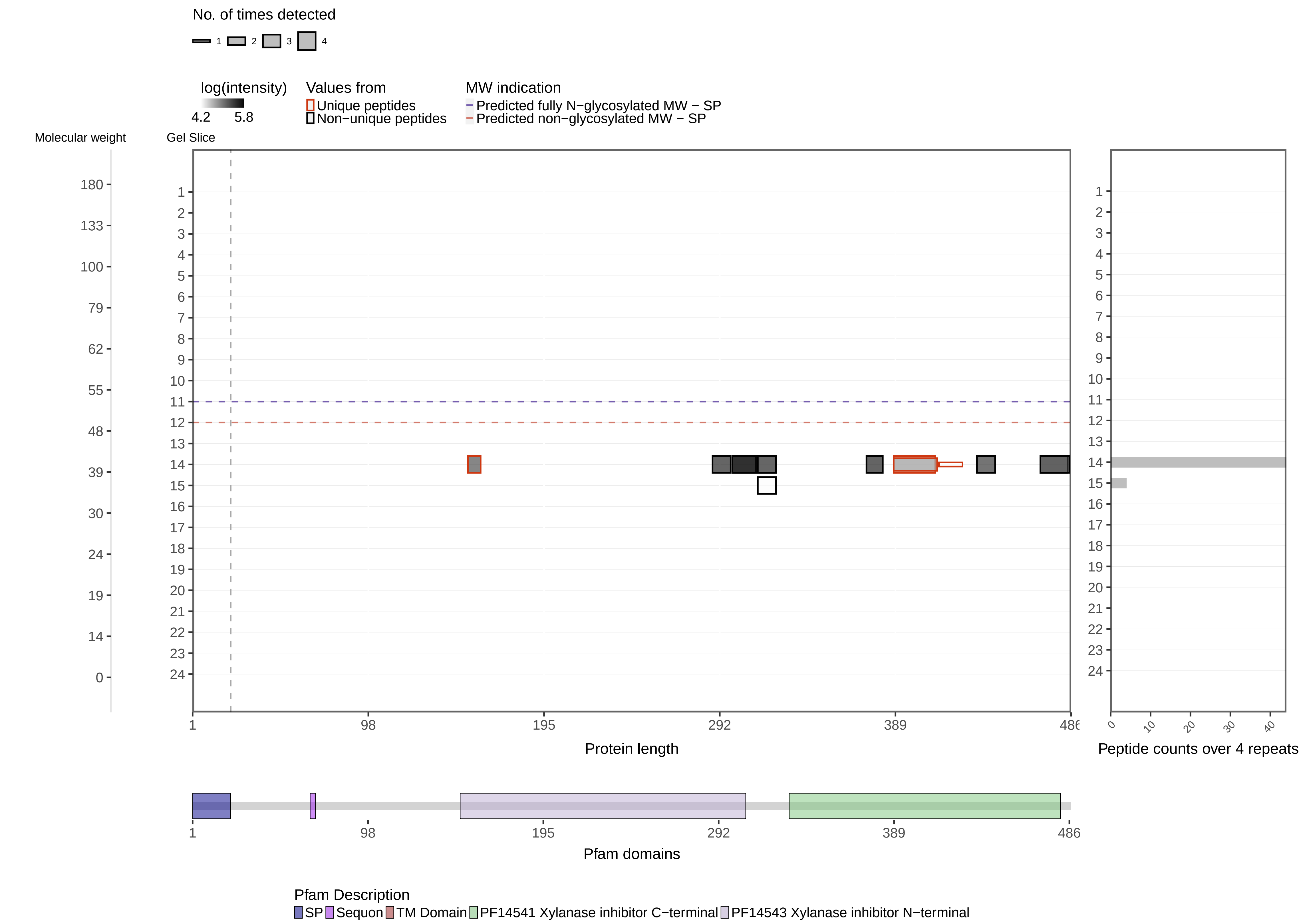


Protein sequence:

DGSFTVGEFATETVSFGNSGAFPKVAIGCGHDNEGLFVGAAGLIALGGGTLSLPSQIKASSFSYCFVDRDSASSSTLDFNSARPGDSVIAPLLRNSRRSTFFYVGLTGISVGGQMLSIPASVFQVDGSGNGGIIVDSGTAVTRLQTTAYNALRDSFVKFTQNLPSSGQFALFDTCYDLSSMKKVSVPTVALHFNGDKTLSLHPKNYLIPVDSSGKFCLAFAPTEGSLSIIGNIQQQGTRVSYDLANNLVGFSPDKC*

### NbL15g10620.1 phenylalanine−−tRNA_ligase_alpha_subunit_cytoplasmic−like [MW = 56.3 kDa]

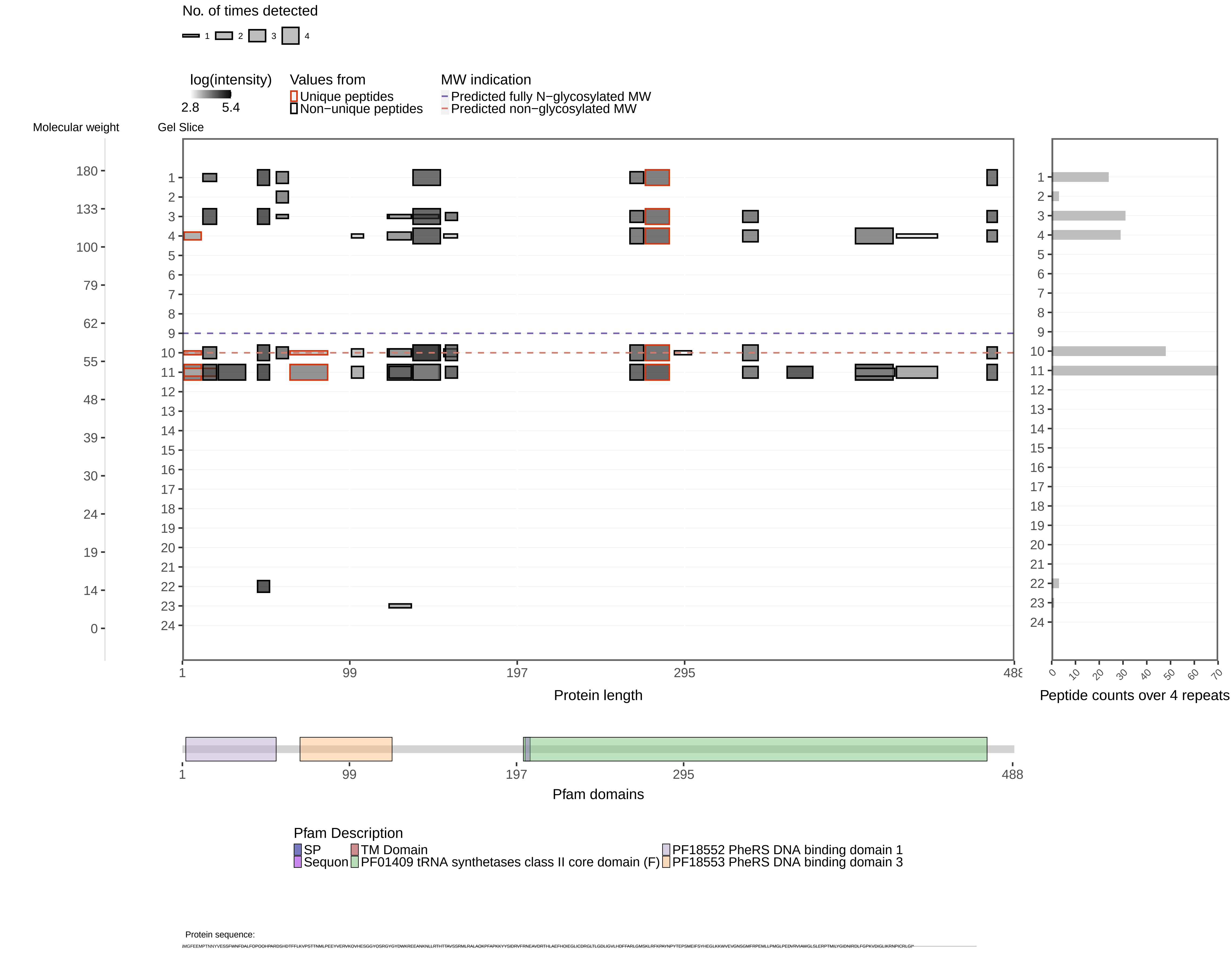

NbL15g10590.1 lysM_domain−containing_GPI−anchored_protein_1−like [MW − SP = 39.78 kDa]

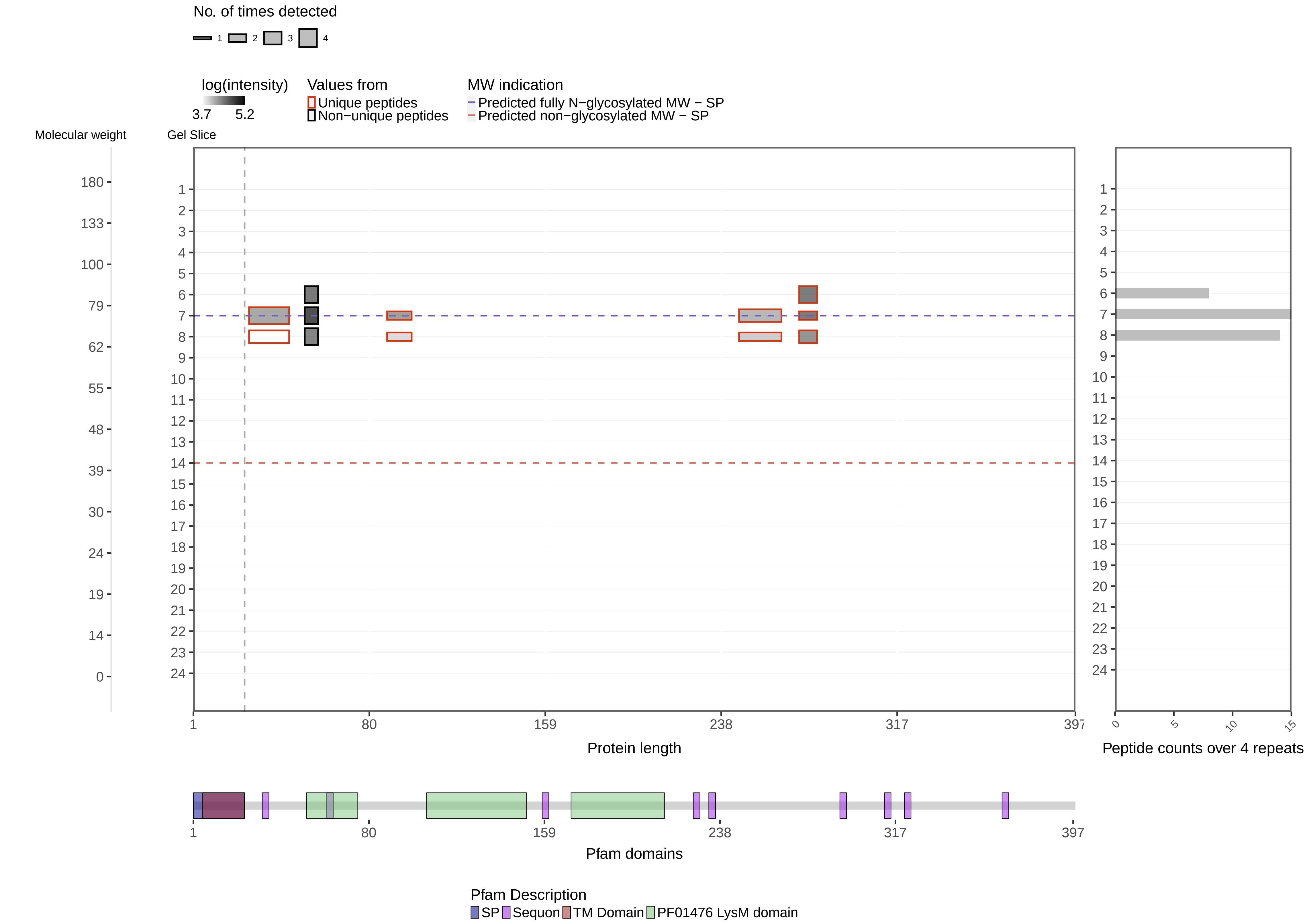


Protein sequence:

SVTIGNLGIINGLGQPQVDPGDILAIPLAACSSANLNWYNESLIVPNGSYALTANNCIKCGCRPTDLSLECSASGIVDKCSHLOCKDSNLFIGERREKHTAFGNCNVTACIYRGHLGGKTFRSLVNSSDVKCLGNOSOHTGLSMASPAADLALSPSPSPSPSPSYFSHSAKSHTSELMNYSPAHRHSKKSLSLALHFNLMVLELILEIFL*

### NbL15g10490.1 40S_ribosomal_protein_S7 [MW = 25.17 kDa]

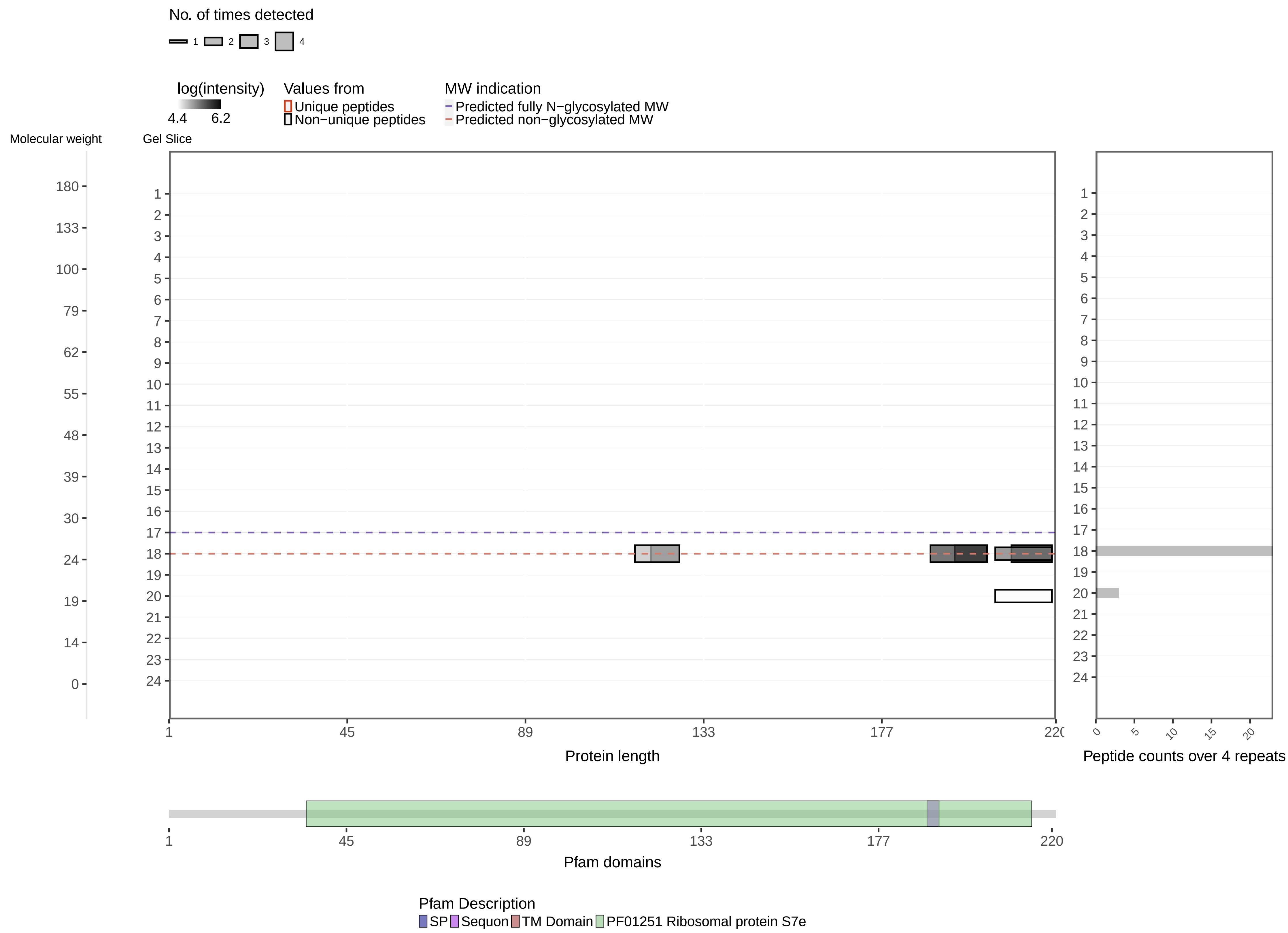


Protein sequence:

RLRKAFRKVHVRLVRELEKKFSGKDVIFIATRRIVRPPKRGSAAQRPRSRTLTSVHDAILEDLVVPAEIVGKRTRYRIDGSKIMKVFLDPKERNNTEYKLETFSAVYRKLSGKDVVFEYPITEA*

### NbL15g10410.1 histidine−−tRNA_ligase_chloroplastic/mitochondrial [MW = 56.7 kDa]

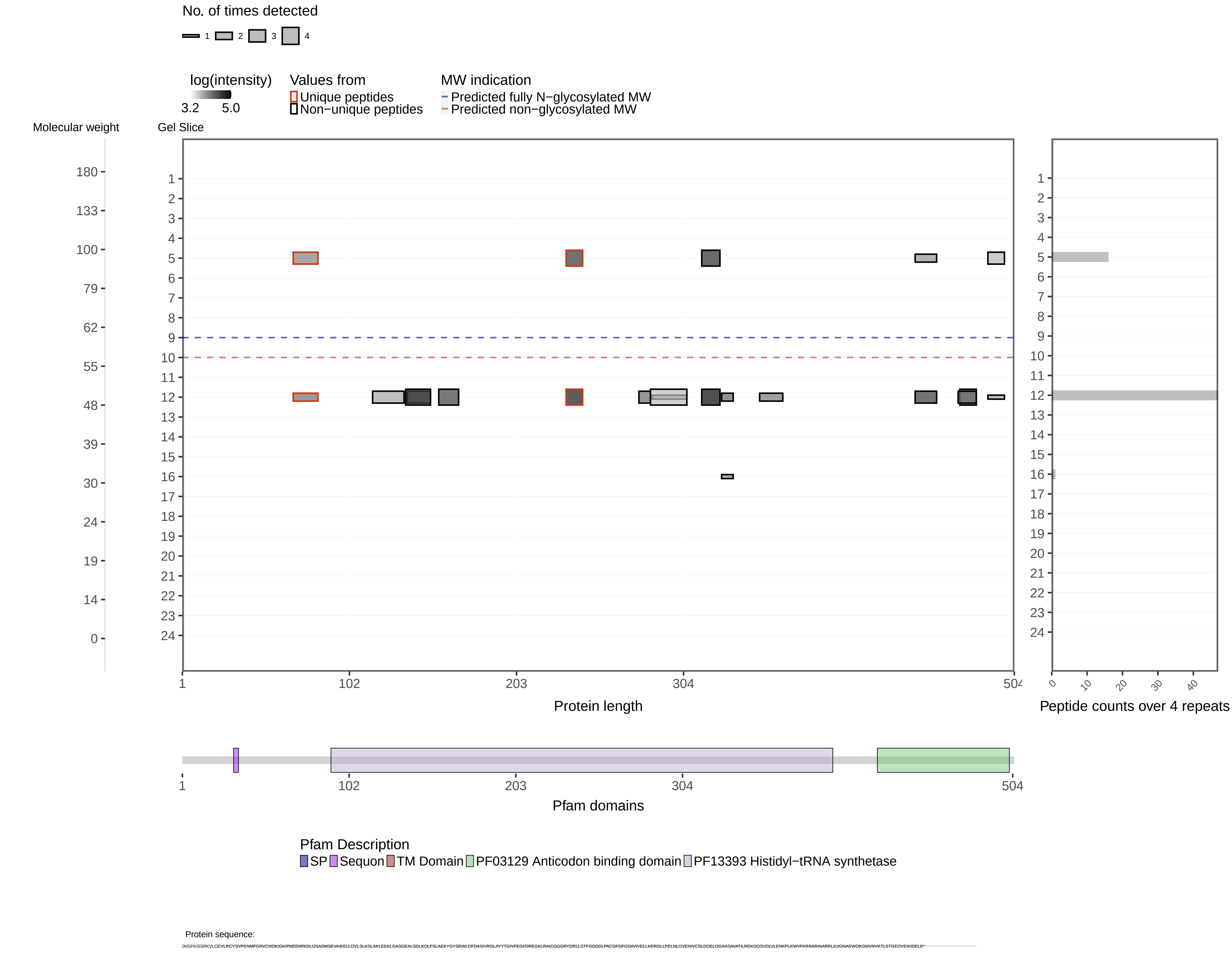

NbL15g10300.1 palmitoyl−protein_thioesterase_1−like [MW − SP = 36 kDa]

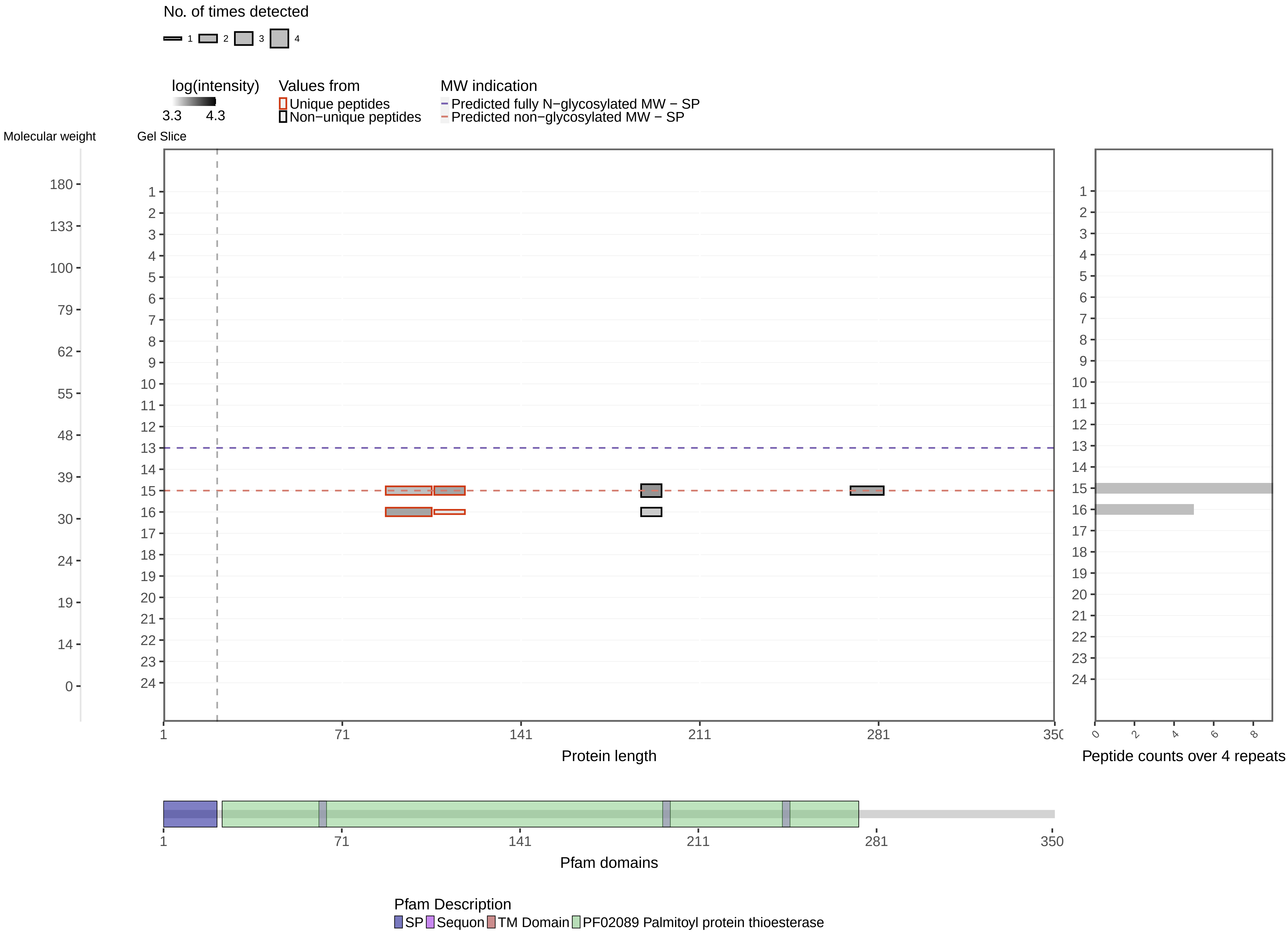


Protein sequence:
ILAPSGYLKLPNNIAGYMKACHFLPKLNNEIPSHRNSTYKKRFSSLENLVLIMFEHDTVLIPRETSWFGYYPDGAFDPILPPNKTELYTEDWIGLKTLDDSGKVOYVKVAGNHLOISSSDMRKHVVPYLAENAADGSSVDVESMAMHGLODESSTIFTFEGSSSYNWPAPVKSFFADLLGLTIDESVS*

NbL15g10190.1 peroxisomal_acyl−coenzyme_A_oxidase_1 [MW = 74.28 kDa]

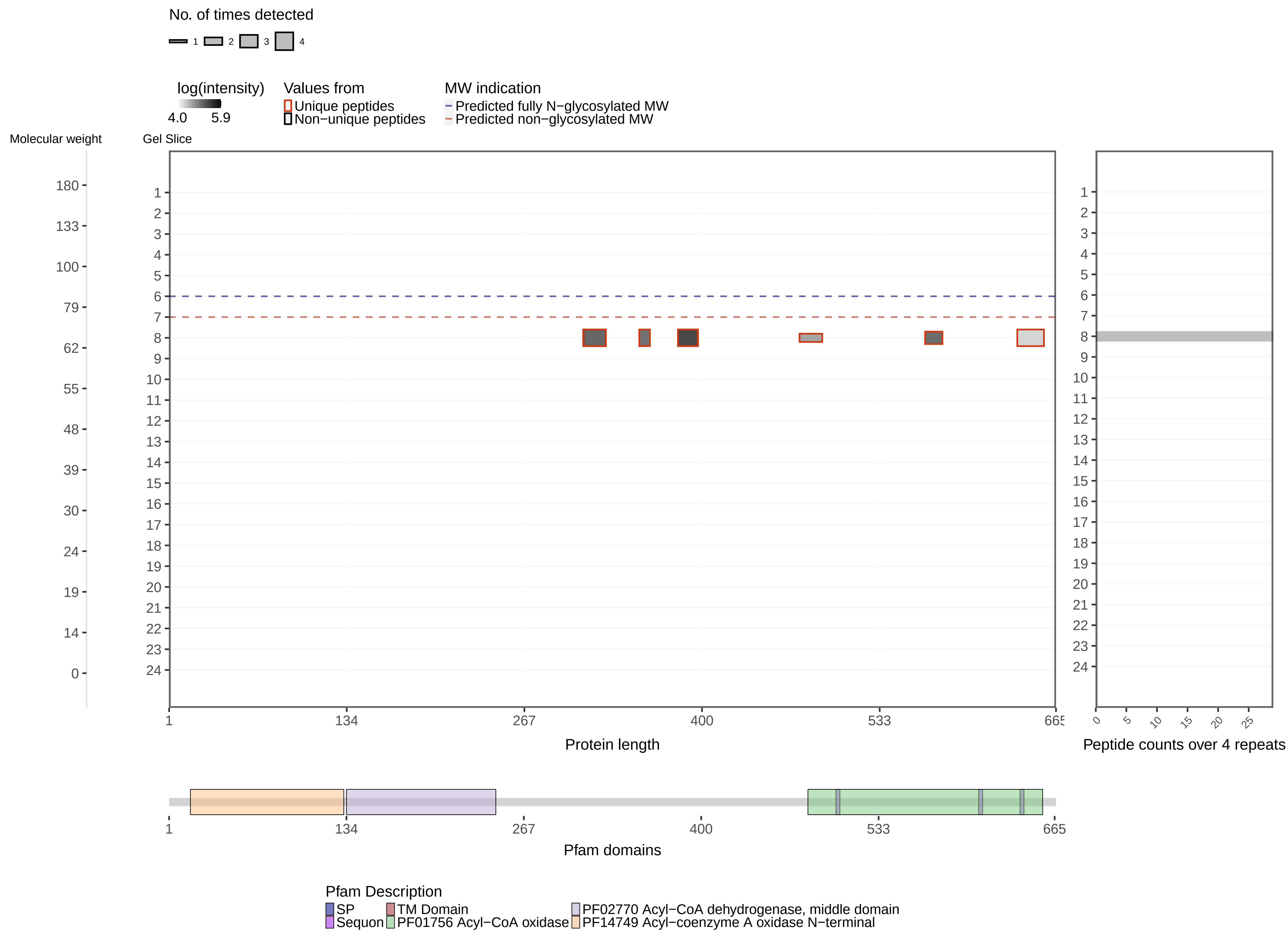


Protein sequence:

[illegible]

NbL15g10070.1 uncharacterized_LOC109231245 [MW = 231.86 kDa]

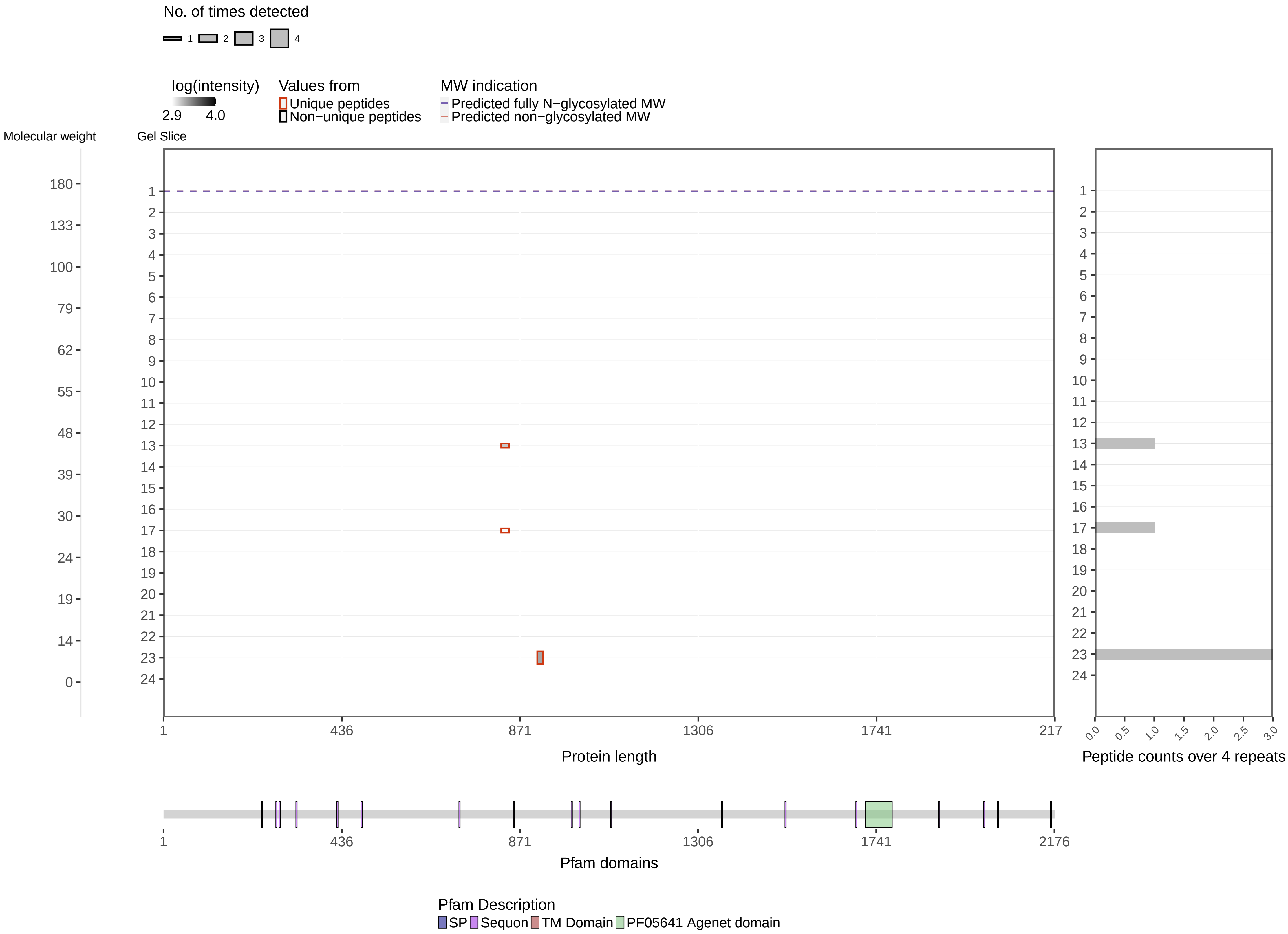


Protein sequence:

DRIRGQRSHTGNNETPSHPRSGPRTPDQANKQVMHQNKVTTPAAGRASGKAINSRVVSPIIPLSSPLWNIPTPSCDGLPSSGMARGAVIDYKALSPMHPYQTPPVRNFVGHTASWLPQAPFPGSWVASPQISAFPTLPVTEPVKLTPVKESSLSISAGAKHATPVSVAHAGESGILAGASPRDEKKASLLSADQKSRKRKKASGTEDRAHKSLRGTSSESVPAPAVCTQLSSKAPASDNFGQSSSVVVAPLVAQSQTGPAAAPILGHFSTSVVIAPLSSSAPINNSDIPIISTPS

### NbL15g09790.1 autophagy−related_protein_8f−like [MW = 14 kDa]

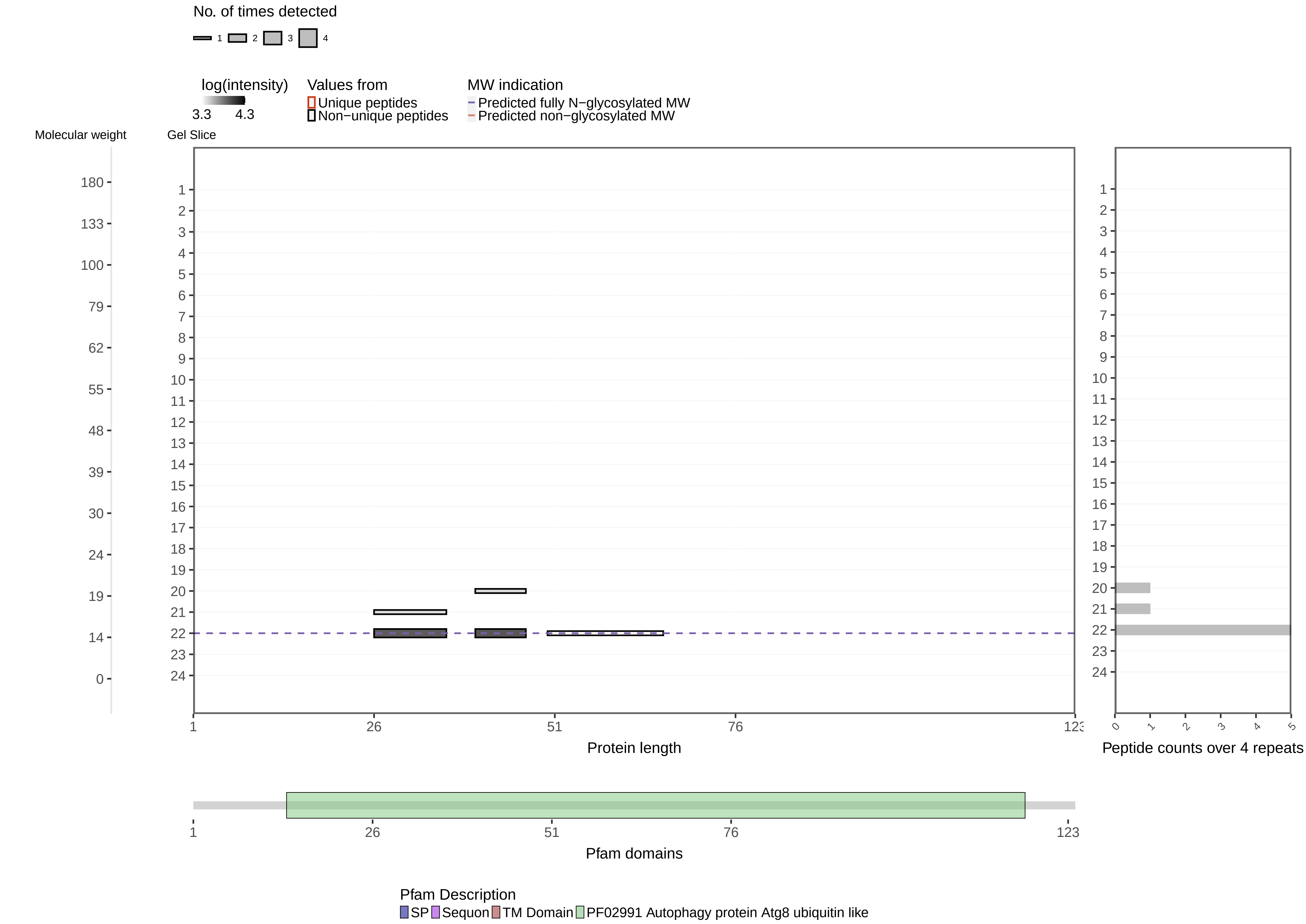


Protein sequence:

KKYLVPADLTVGQFVYVIRKRIKLSAEKAIFIFIDNVLPPTGAIISAIYDEKKDEDGFLYVTYSGENTFGDPNEL*

NbL15g09670.1 adenine_phosphoribosyltransferase_3−like [MW = 20.03 kDa]

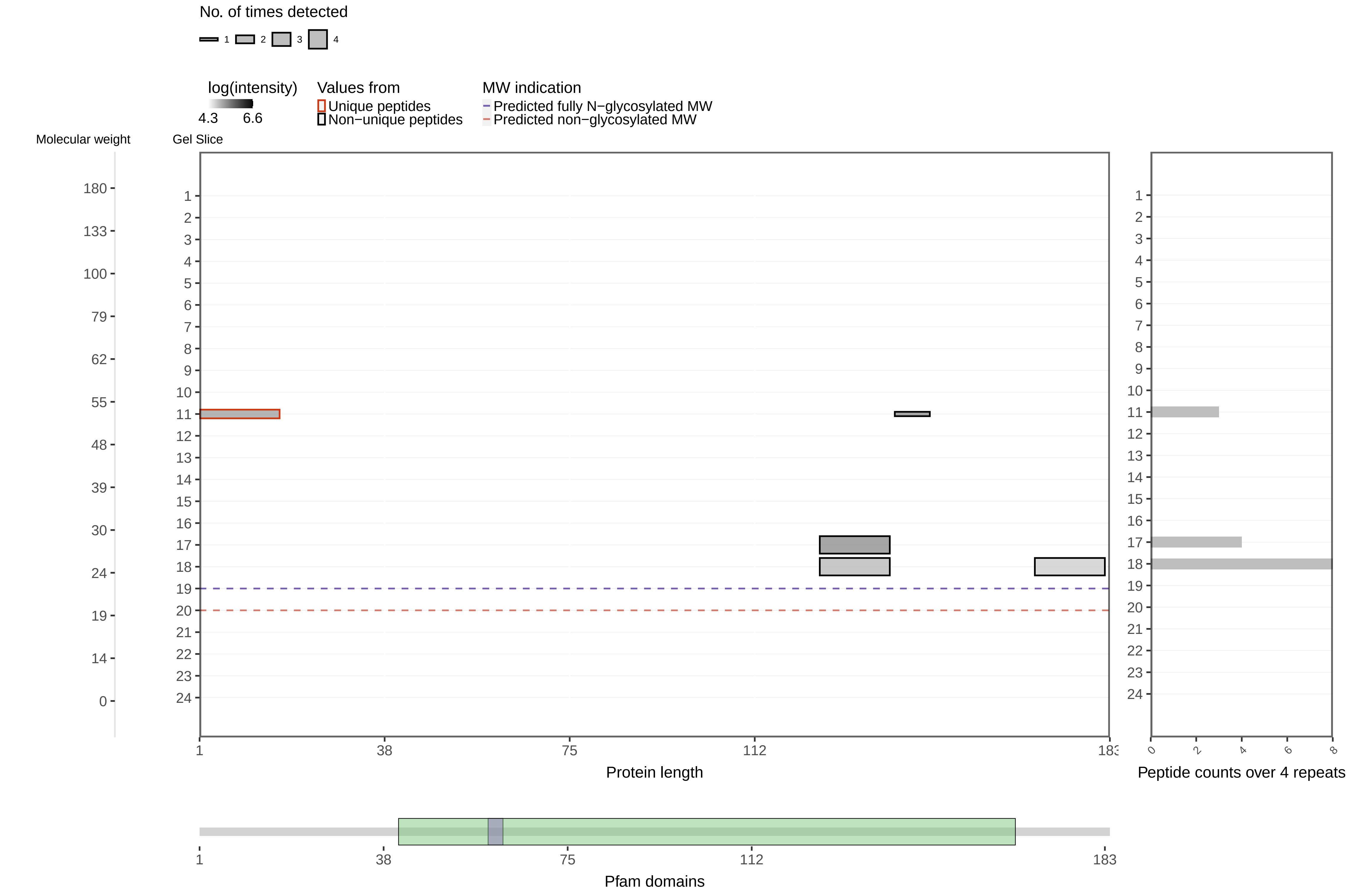


Protein sequence:

AIGAKYVPLRKPNKLPGKVFKQEYDLEYGSDCLEMHIGAVEAGERALVVDDLIATGGTLRAAMNLLERAGAEVVECACVIEIPDLKGKEQLNGKPLYVLVESQ*

### NbL15g09610.1 transaldolase [MW = 48.02 kDa]

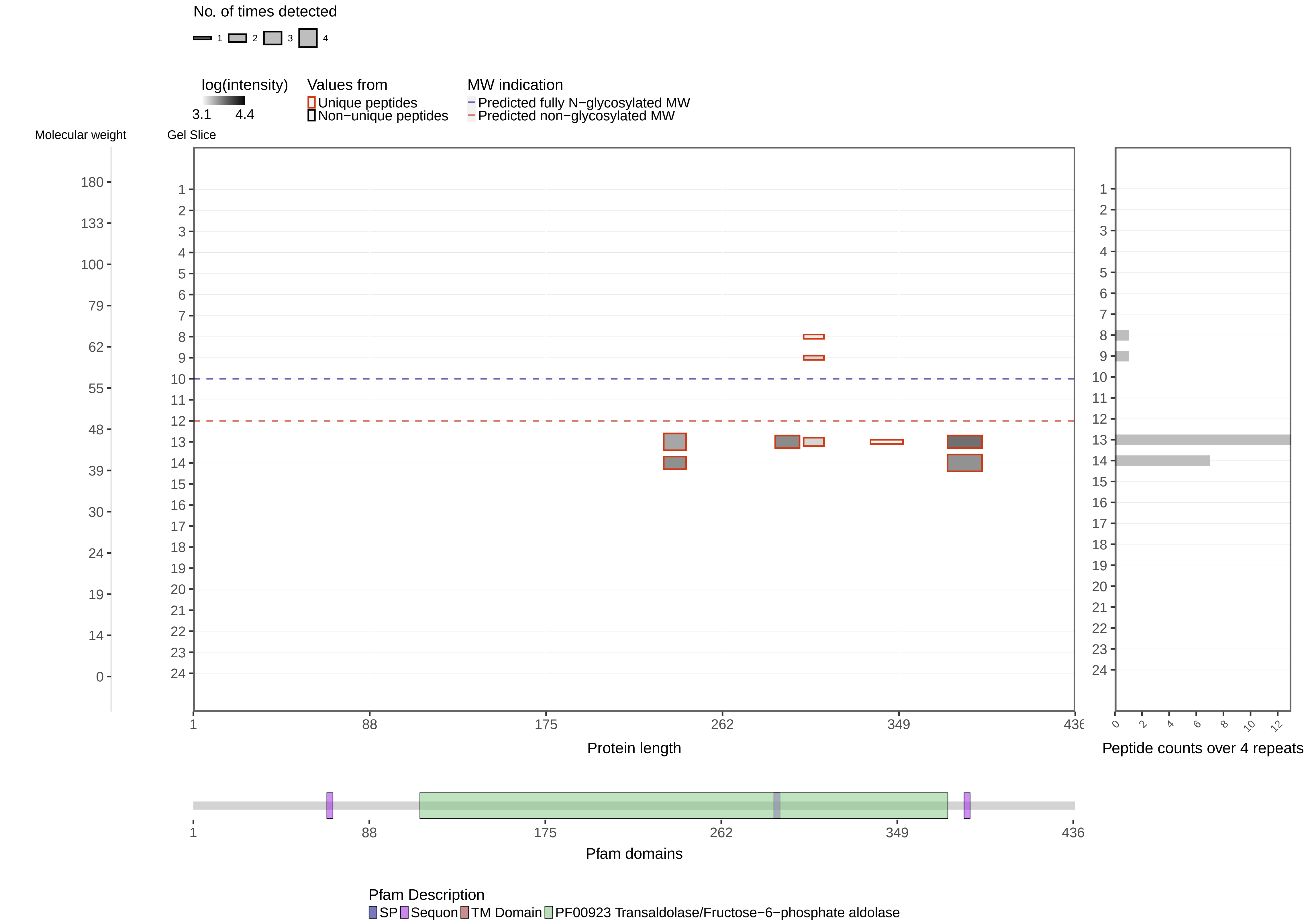


Protein sequence:

HGIVRKVHDLLKAYNEIEVPPERLLFKIPATWQGIEASRVLEAEGIQTHLTFVYSFCQAAAAAQAGASVIQIFVGRLRDWSRNHTGDSEIESALKRGEDPGLALVTKAYNYIHKYGHKSKLMAAAVRNKQDVFNLLGVDYIITPLKVLQSLKESVTPPDEKYSFIRRLSPQSAAAFNFTQDELQKWDQYSFSSAMGPAAVELLTIGMDGYTNQAKRVEEFFGKIWPPPNV*

NbL15g09590.1 monocopper_oxidase−like_protein_SKU5 [MW − SP = 63.74 kDa]

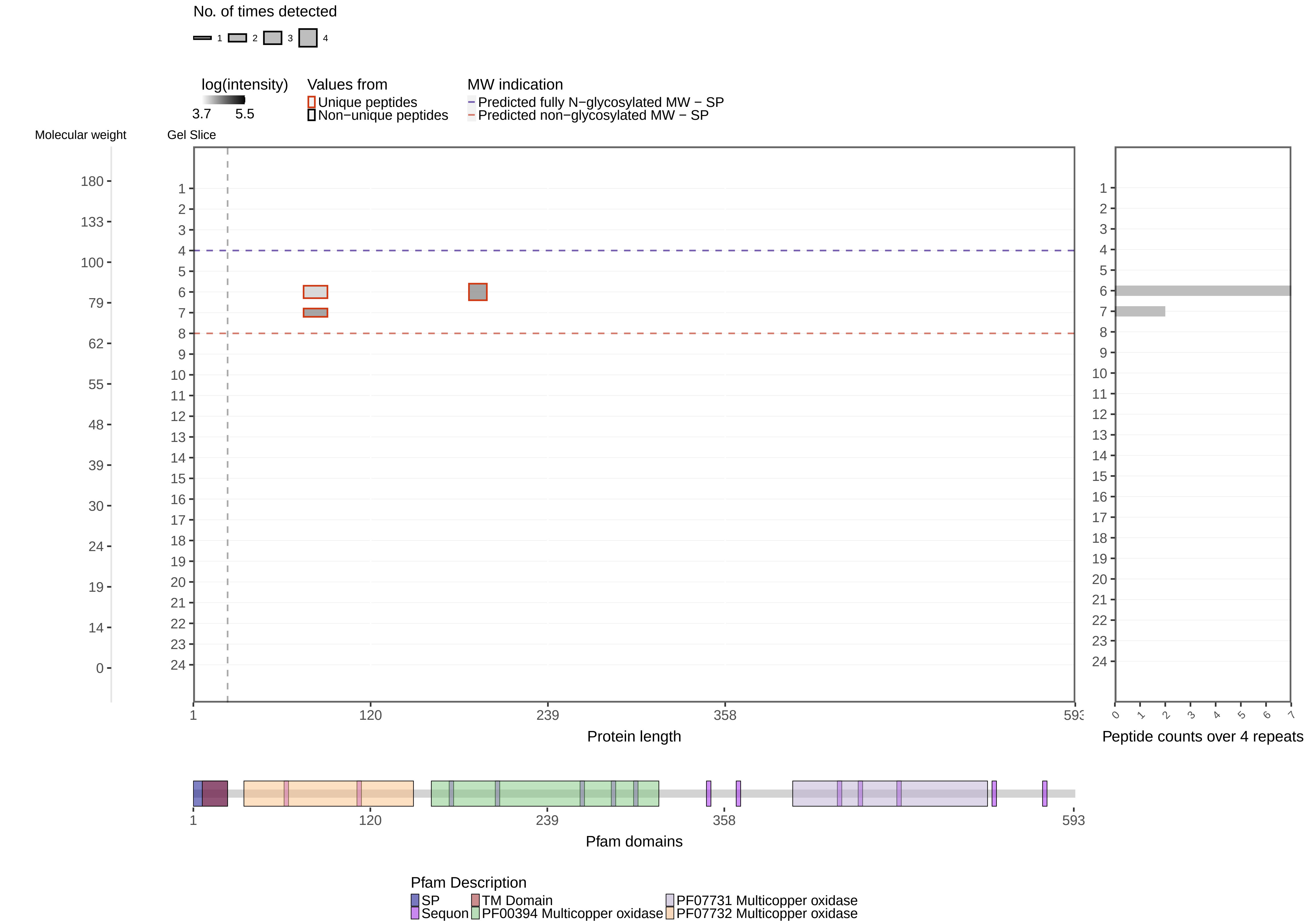


Protein sequence:

SDYYIVASARFVNQSQWQRVTGVGILKYTNSKGKAHGPLPEPPQDEFDKTYSMNQARSIRWNVTASGARPNPQGSFRYGSINVTEIYVIQNKPPTMVDGKRRTTLNGISFVNPKTPIRLADKYKLKGVYKLDFPTAPLTGPPKMETSVINGTYRGFMEIILQNNDTKVHTYHMSGYAFFVVGMDYGVWSNSSRGTYNKWDGIARTTTQVFPGAWTAILVSLDNVGIWNLRTENLDSWFLGQETYIRVVNPEPSNKTEMPMPDNALFCGQLSKLQKPQDISSATSIRGNGS[illegible]

### NbL15g09570.1 thylakoid_lumenal_protein_At1g12250_chloroplastic [MW = 28.5 kDa]

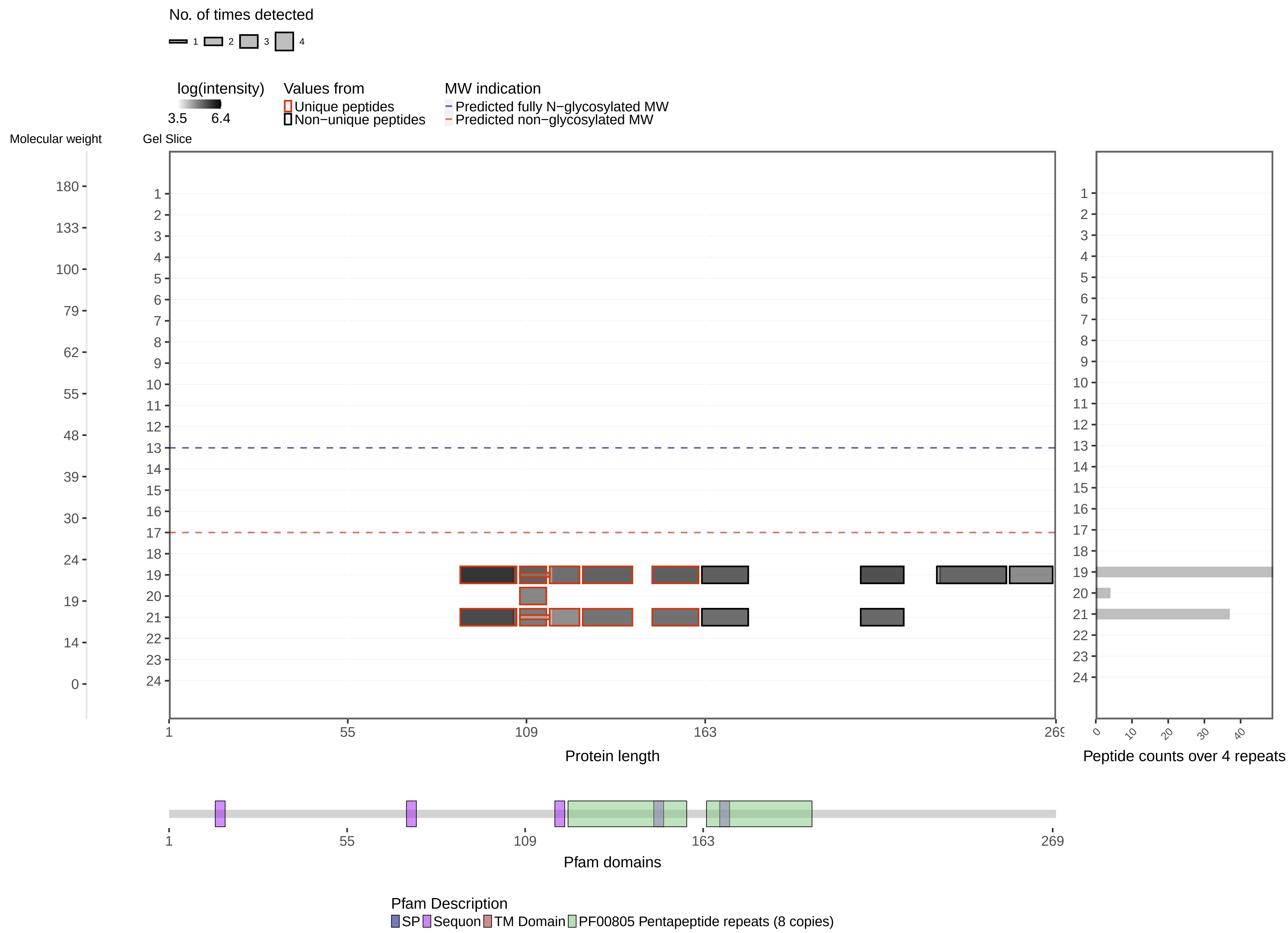


Protein sequence:

[illegible]MRESDFSGSTFNGAYLEKAVAYKANFSGADLSDTLMDRMVLNEANLTNAVLVRSVLTRSDLGGAIIEGADFSDAVIDLLQKQALCKYASGTNPVTGVSTRKSLGCGNSRRNAYGSPSSPLLSAPPQQLLDRDGFCDPATGLCDAS*

NbL15g09520.1 hsp70−Hsp90_organizing_protein_2 [MW = 67.64 kDa]

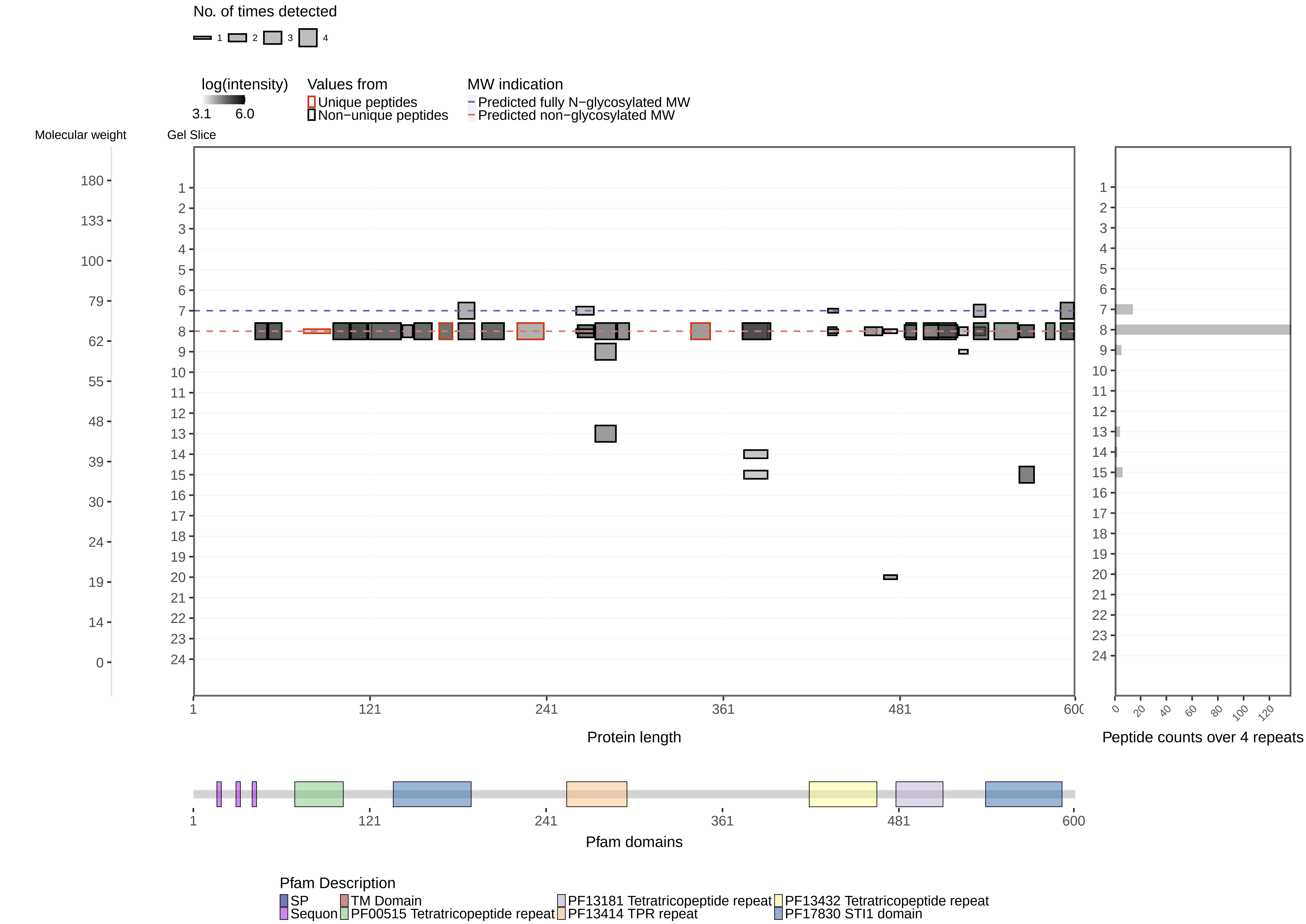


Protein sequence:

### NbL15g09510.1 elongation_factor_G−2_chloroplastic [MW = 86.04 kDa]

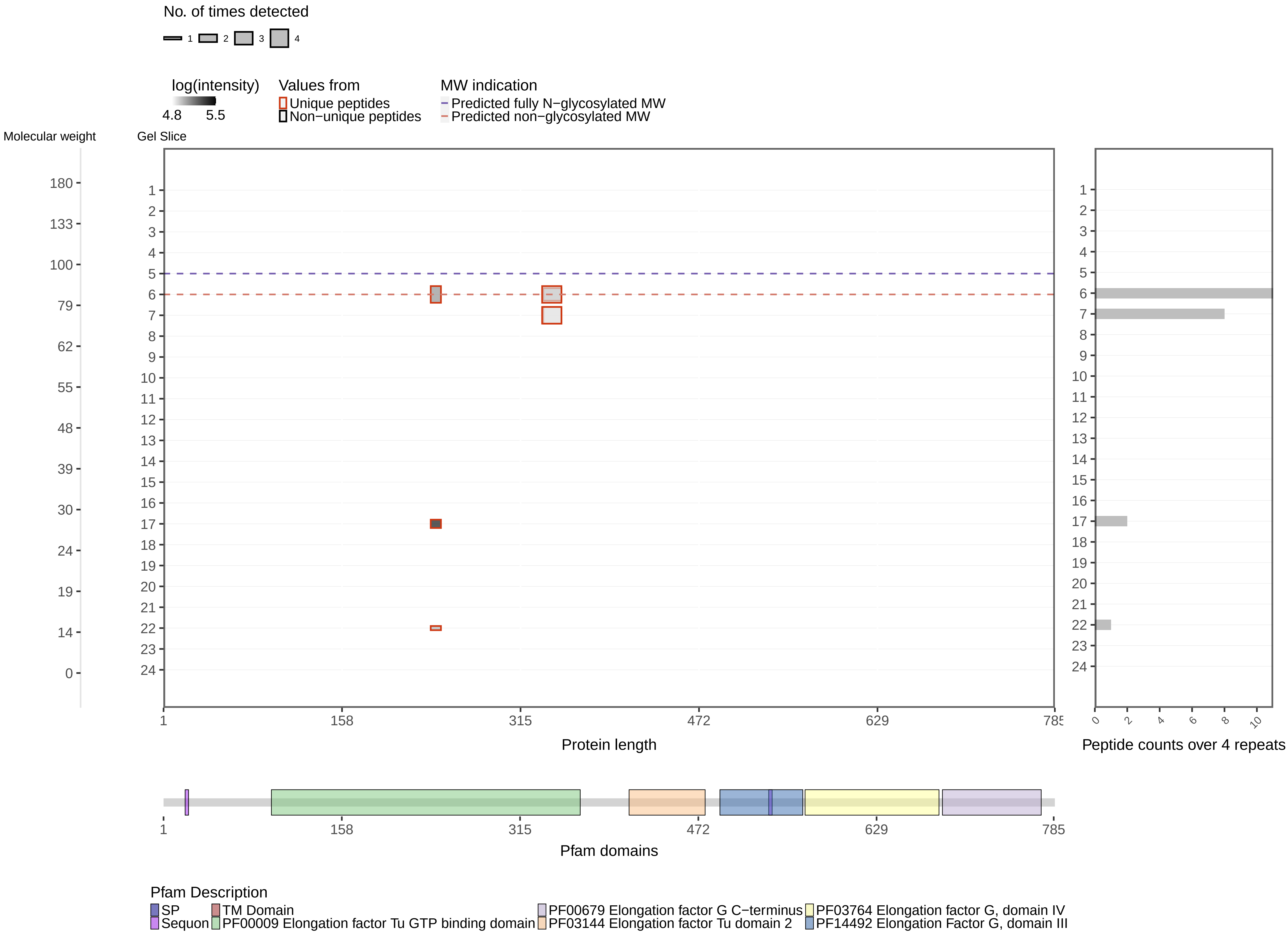


Protein sequence:

PDNPEVTIERAASDDEPFAGLAFKIMNDPFVGSLTFVRVYSGKLSAGSYVVNANKGRKERIGRLLEMHANSREDIKTALTGDIVALAGLKDTITGETLADPDKPVVLERMDFPDPVIKVAIEPKTKADIDKMATGLIKLAQEDPSFHFSRDEEINQTVIEGMGELHLEIIVDRLKREFKVEANVGAPQVNYRESISKISEVKYVHKKQSGGSGQFADITVRFEPMETGGGYEFKSEIKGGAVPKEYIPGVMKGLEECMSNGVLAGFPVVDVRAVLVDGSYHDVDSSVLAFQLAARG

### NbL15g09260.1 purple_acid_phosphatase_2−like [MW − SP = 51.36 kDa]

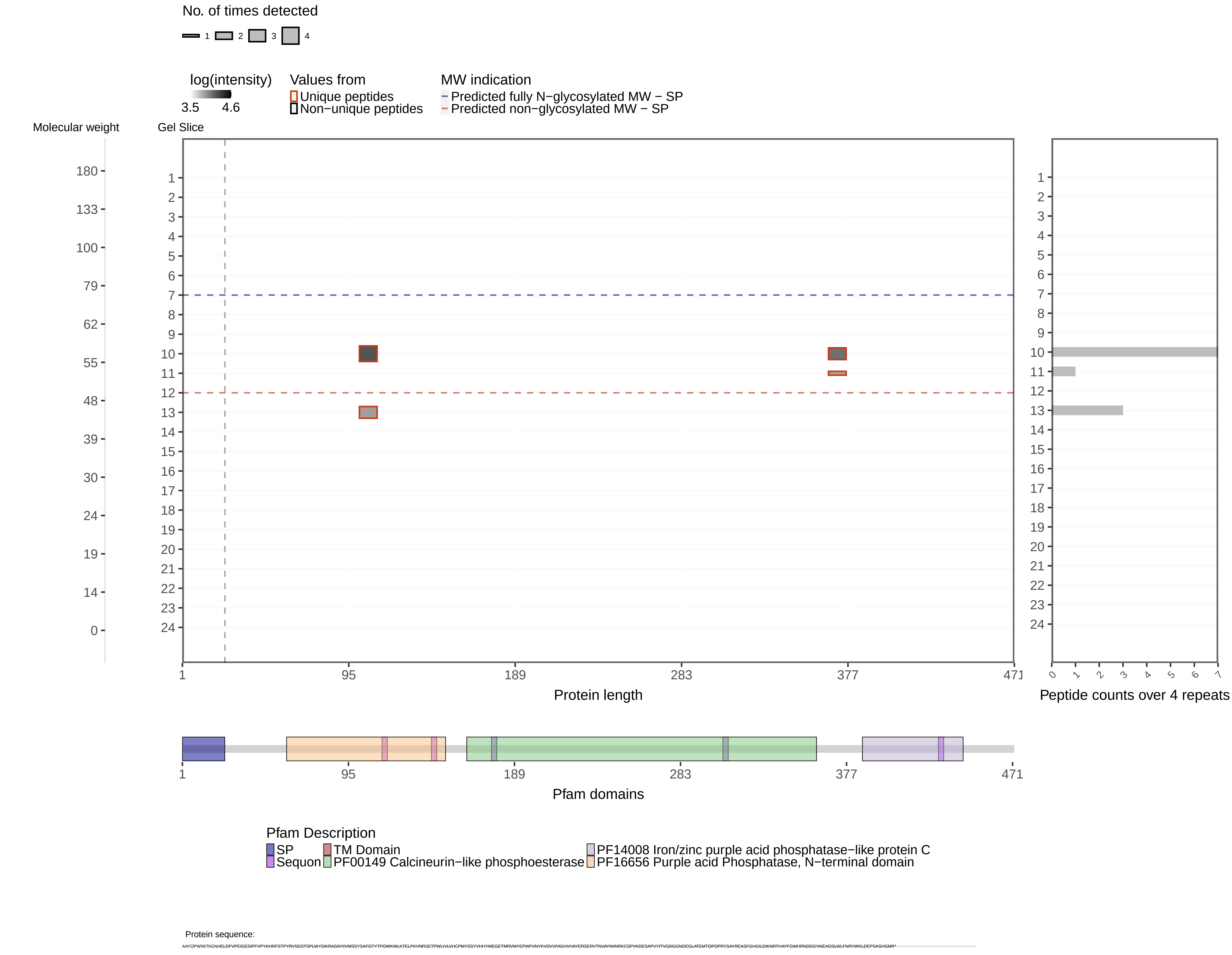

### NbL15g09210.1 zingipain−2−like [MW − SP = 45.33 kDa]

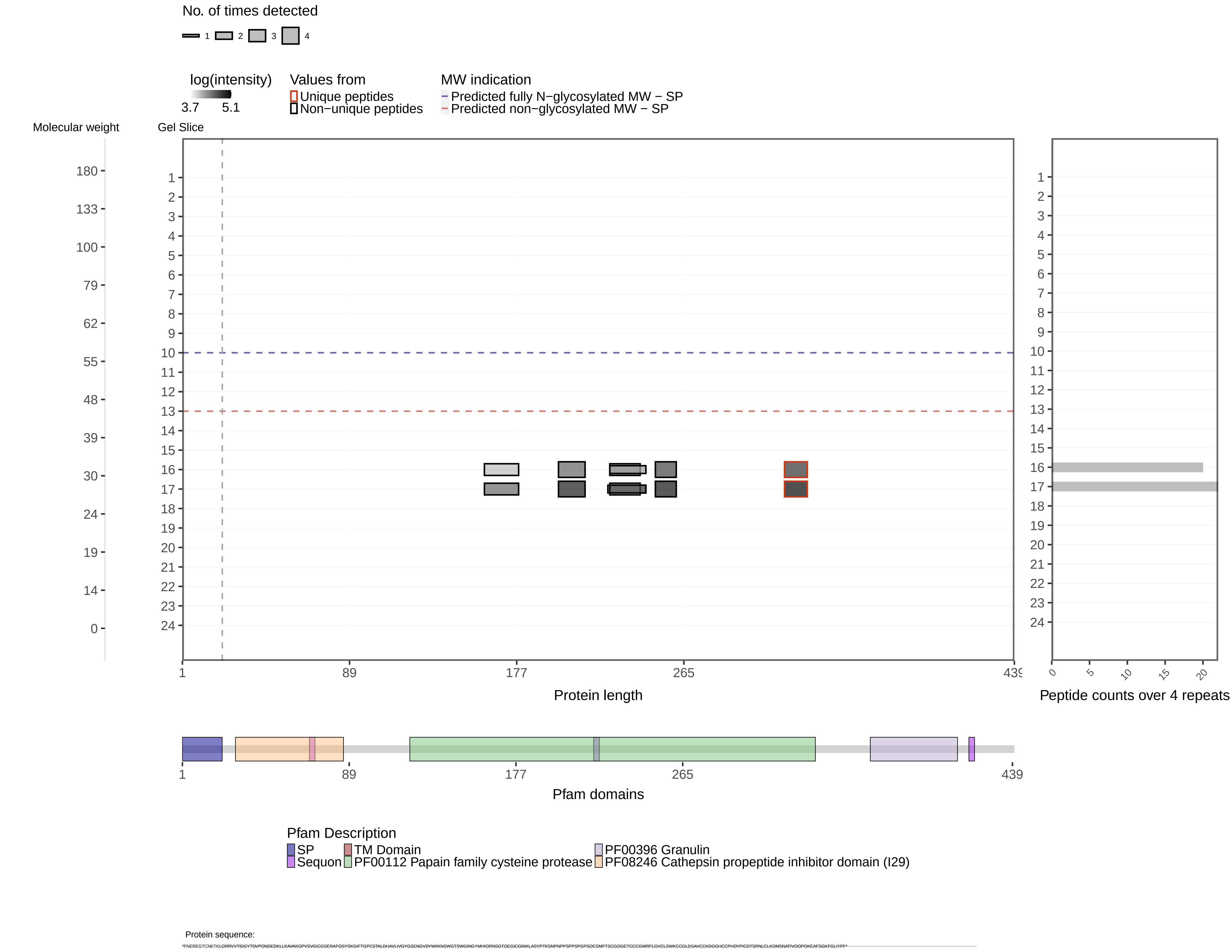

### NbL15g09200.1 lon_protease_2−like [MW = 32.55 kDa]

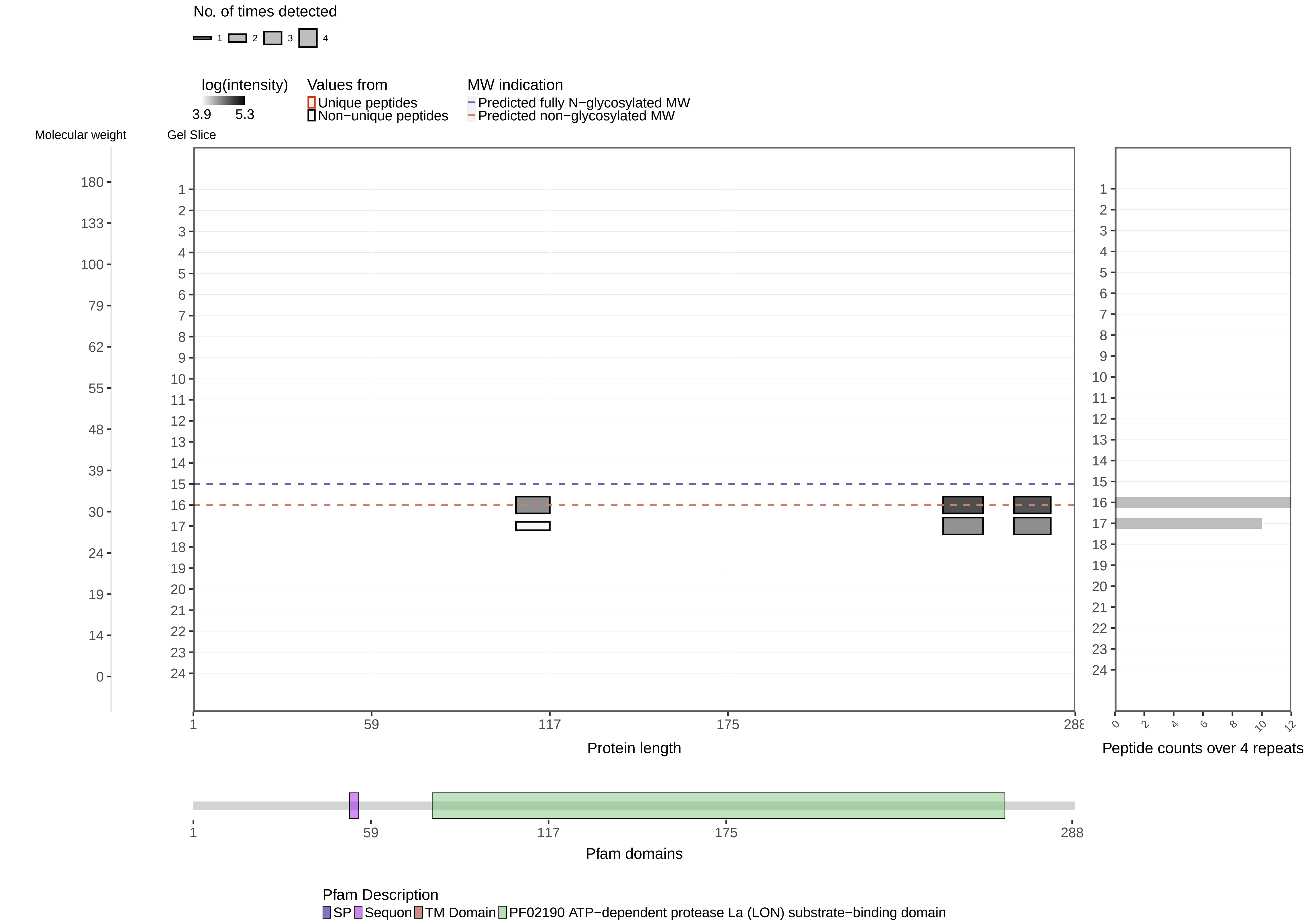


Protein sequence:

GCVGEVVKHERLVDDRFFLICKGQERFRVSKVVRSKPYLVAEVTWLEDRPSANGEDDVEGLANEVENYMKDVIRLSNRLNGKAEKEATDLRRNLFPTPFSFFVGSTFEGAPREQQALLELEDTAVRLKREKETLRNTLNYLTAASAVKDVFPSSG*

### NbL15g09000.1 50S_ribosomal_protein_L31_chloroplastic [MW = 14.55 kDa]

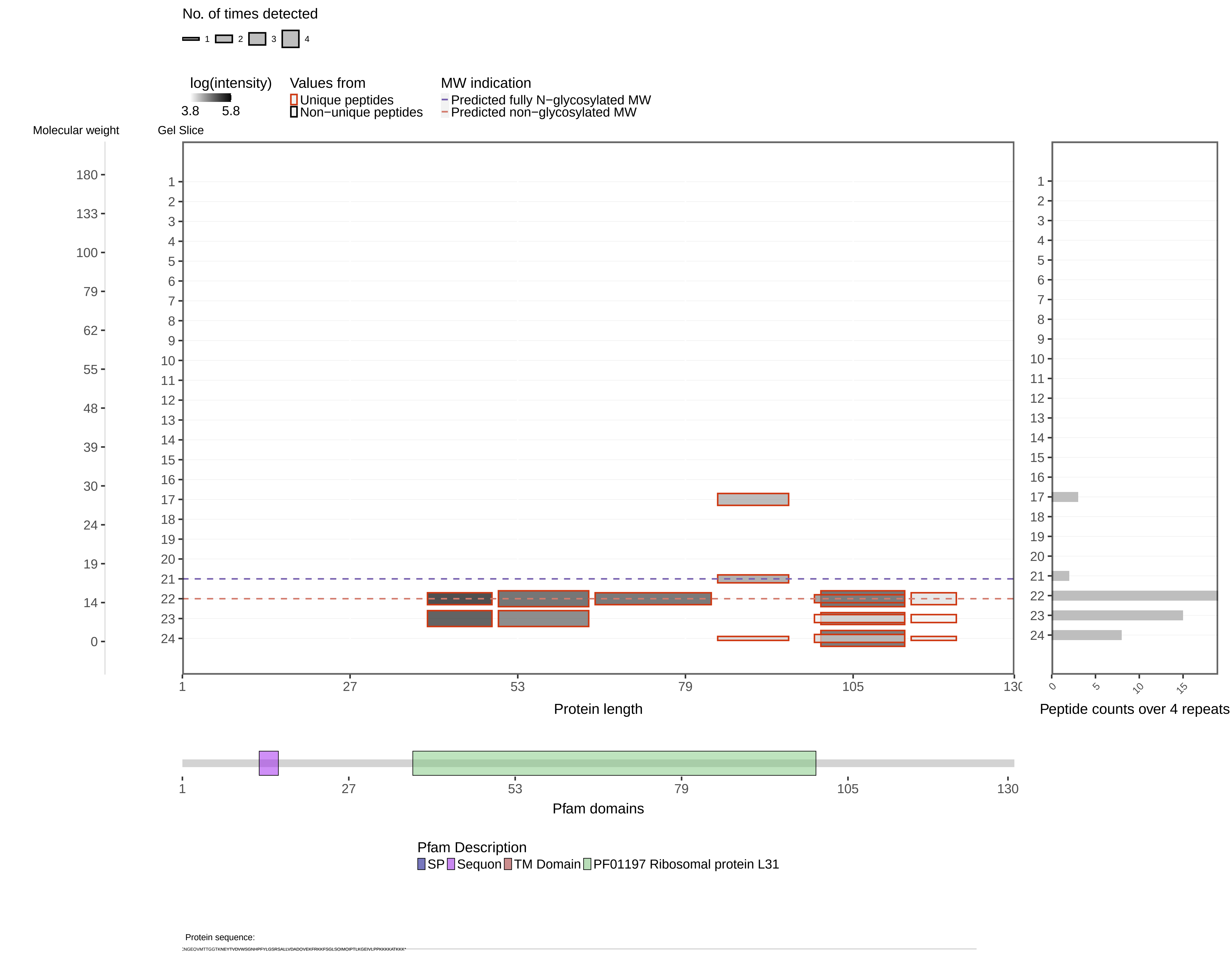

### NbL15g08950.1 proteasome_subunit_alpha_type−1−B−like [MW = 29.86 kDa]

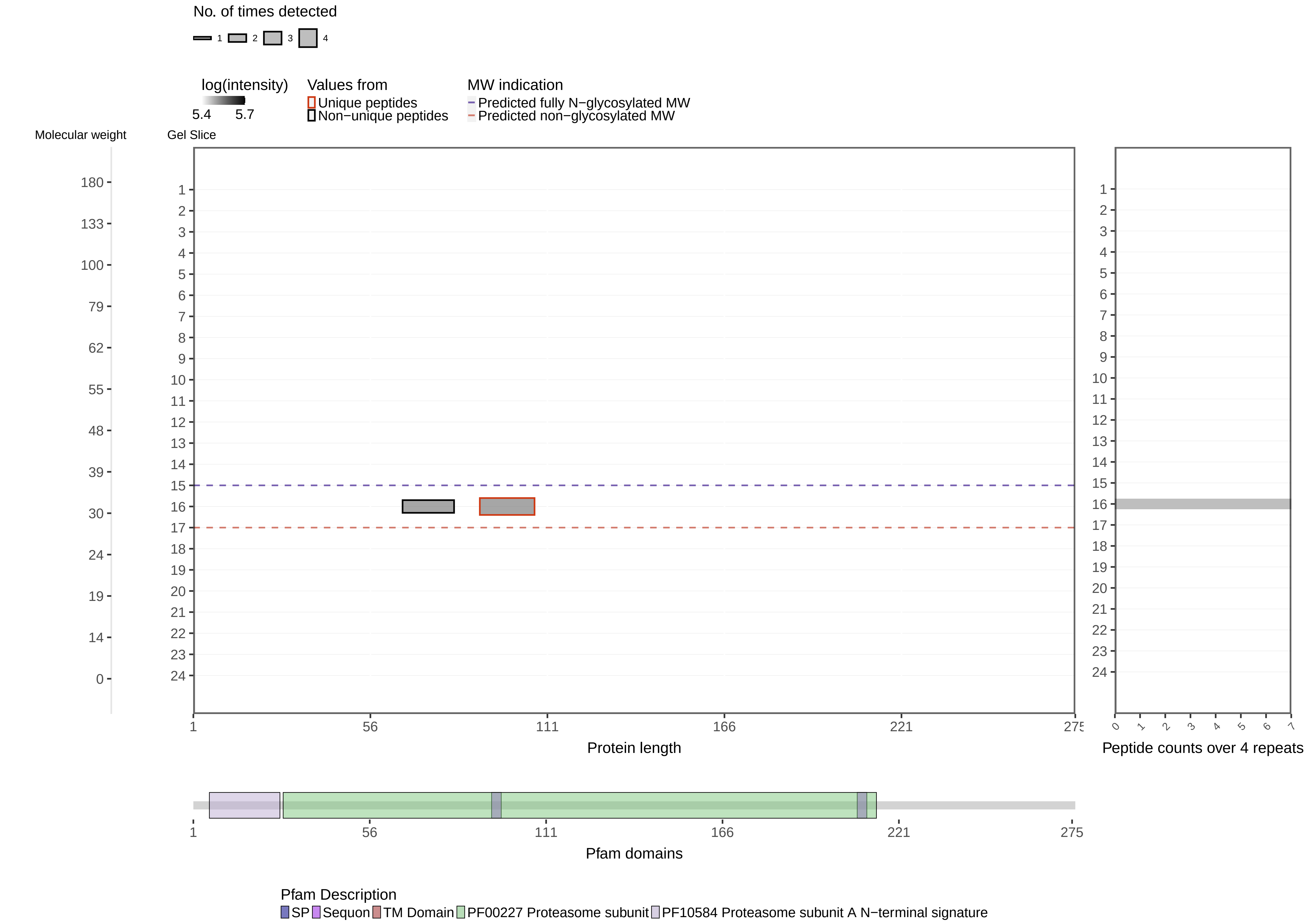


Protein sequence:

KRPYGVGLLVGGLDESGAHLYYNCPSGNYFEYQAFAIGSRSQAAKTYLERRFETFMDSSRESLLKDALFALRETLQGEKLTSTNCTVAVVGVGEAFHMLDKETIQALINEFEIAGEEAPAAAPDEAGSDEPAAAPEEGAPADQGAAPMDI*

### NbL15g08920.1 isoflavone_reductase−like_protein [MW = 37.79 kDa]

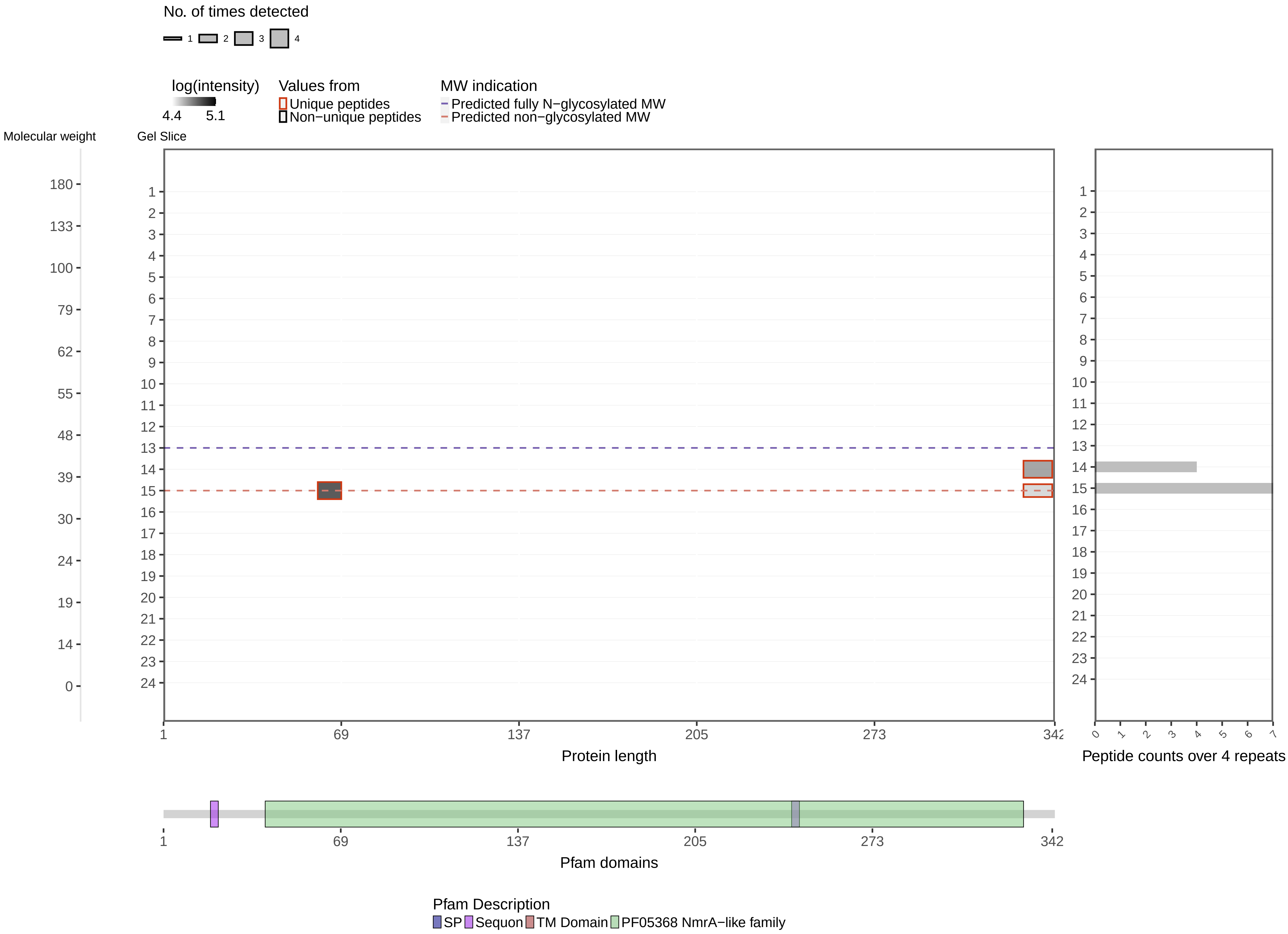


Protein sequence:

EPAKSAFAVKVQIRRAIEAEGIPYTYVSCNCFAGYFLPTMVQPGATAPPRDKVIIPGDGNVKAVFNEEHDIGTYTIKVVDDPRTLNKTLYIKPPKNTLSFNELVVVWEKMIGKTLEKIYIPEEQILKDIETSPMPLPVLLAINHATFVNGDQTNFKIEPSFGVEASELYPDVKYTTVEEYLGHFV*

### NbL15g08860.1 flocculation_protein_FLO11 [MW = 83.19 kDa]

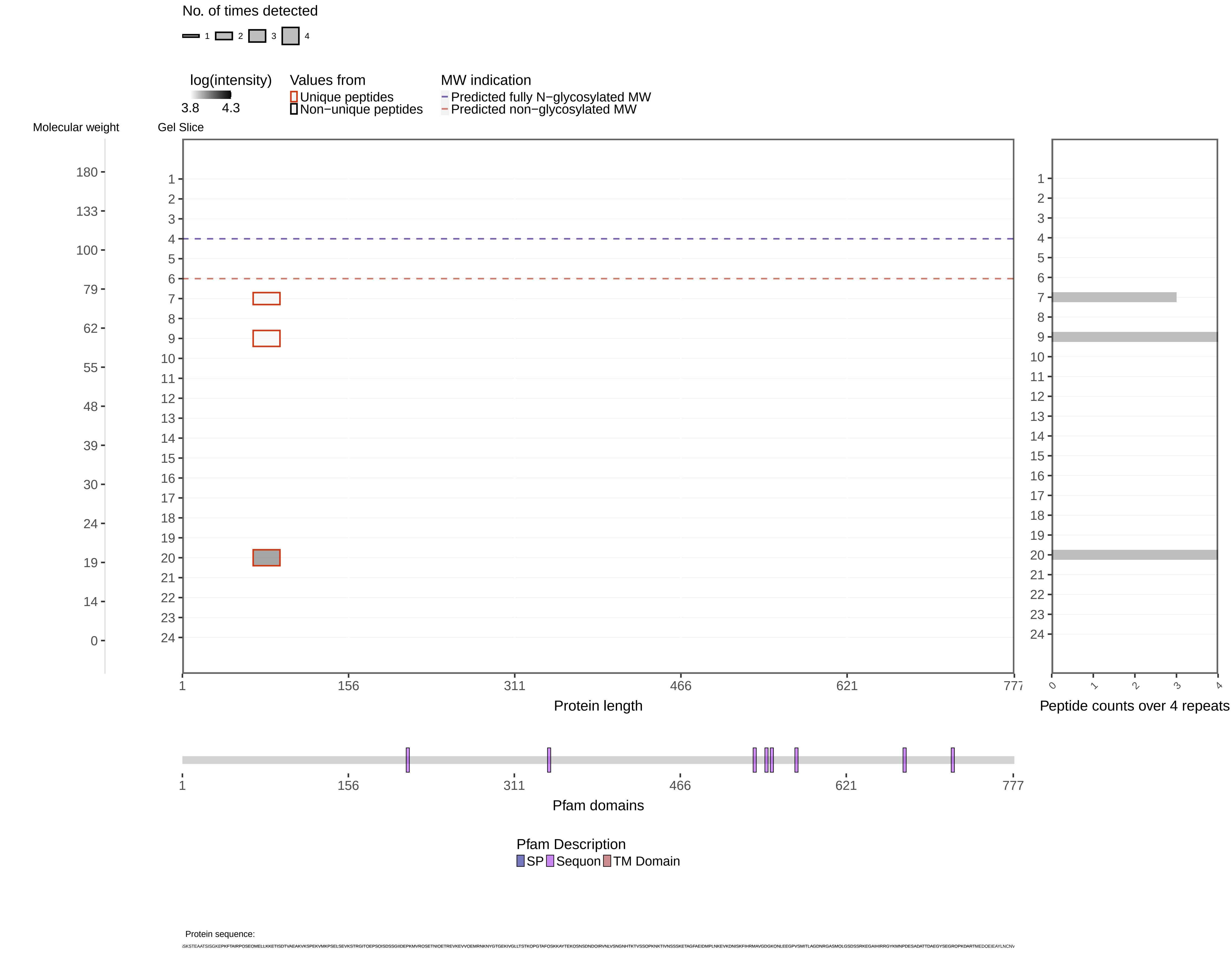

### NbL15g08780.1 maf−like_protein_DDB_G0281937 [MW = 26.92 kDa]

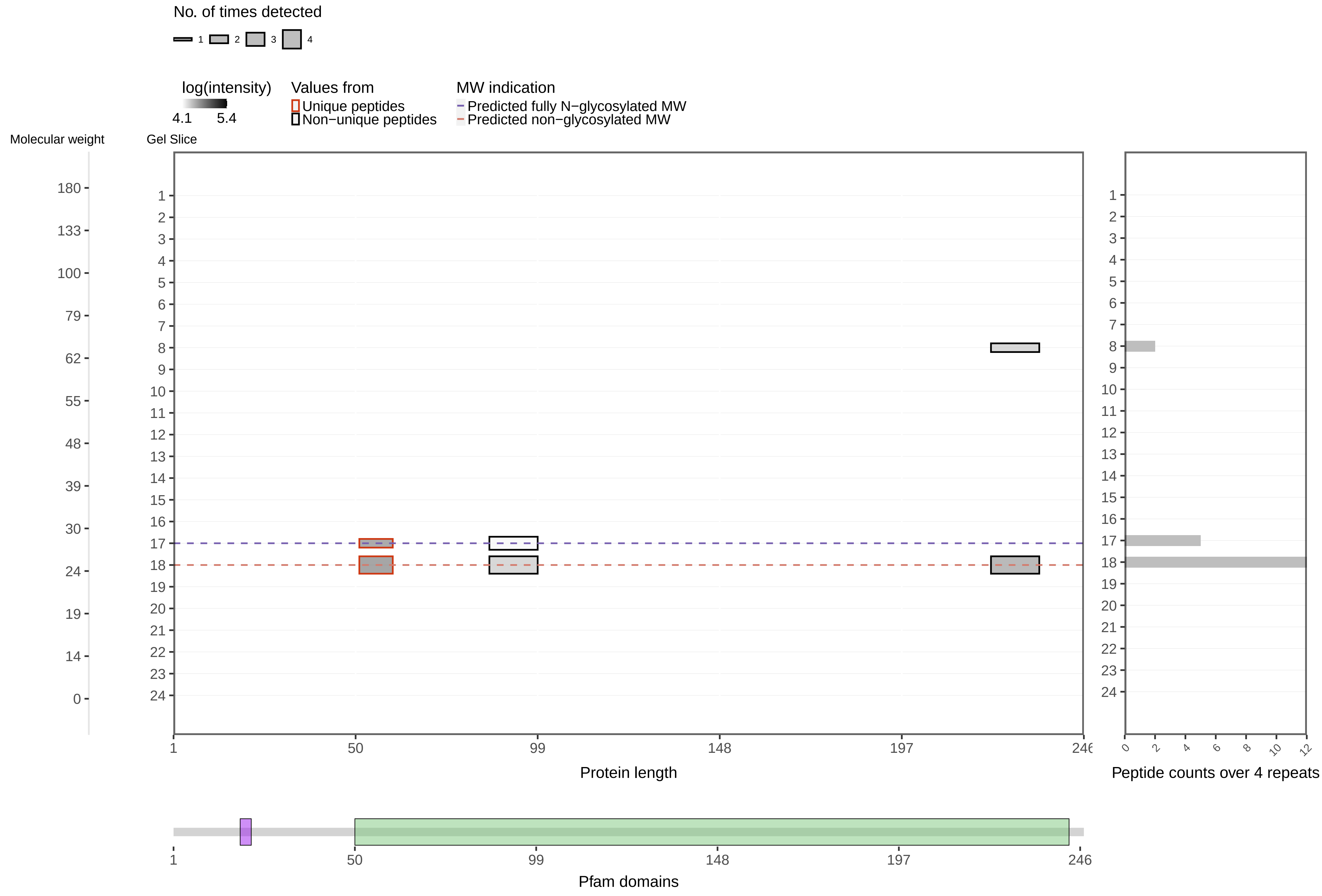


Pfam Description
SP Sequon TM Domain PF02545 Maf−like protein

Protein sequence:

SENLEKHSNPTLLVAADTVVVYEGIVREKPSSEAEARQFMKDYANGHAATISSVLVTNLVTGCRRGEWDKVEIYFHDIPDRVIDKLIEERIVLYAAGGLIIEHPLILPCIKEVVGSTDSVMGLPKALTERLIKEVL*

NbL15g08590.1 glucan_endo−13−beta−glucosidase_7 [MW − SP = 46.83 kDa]

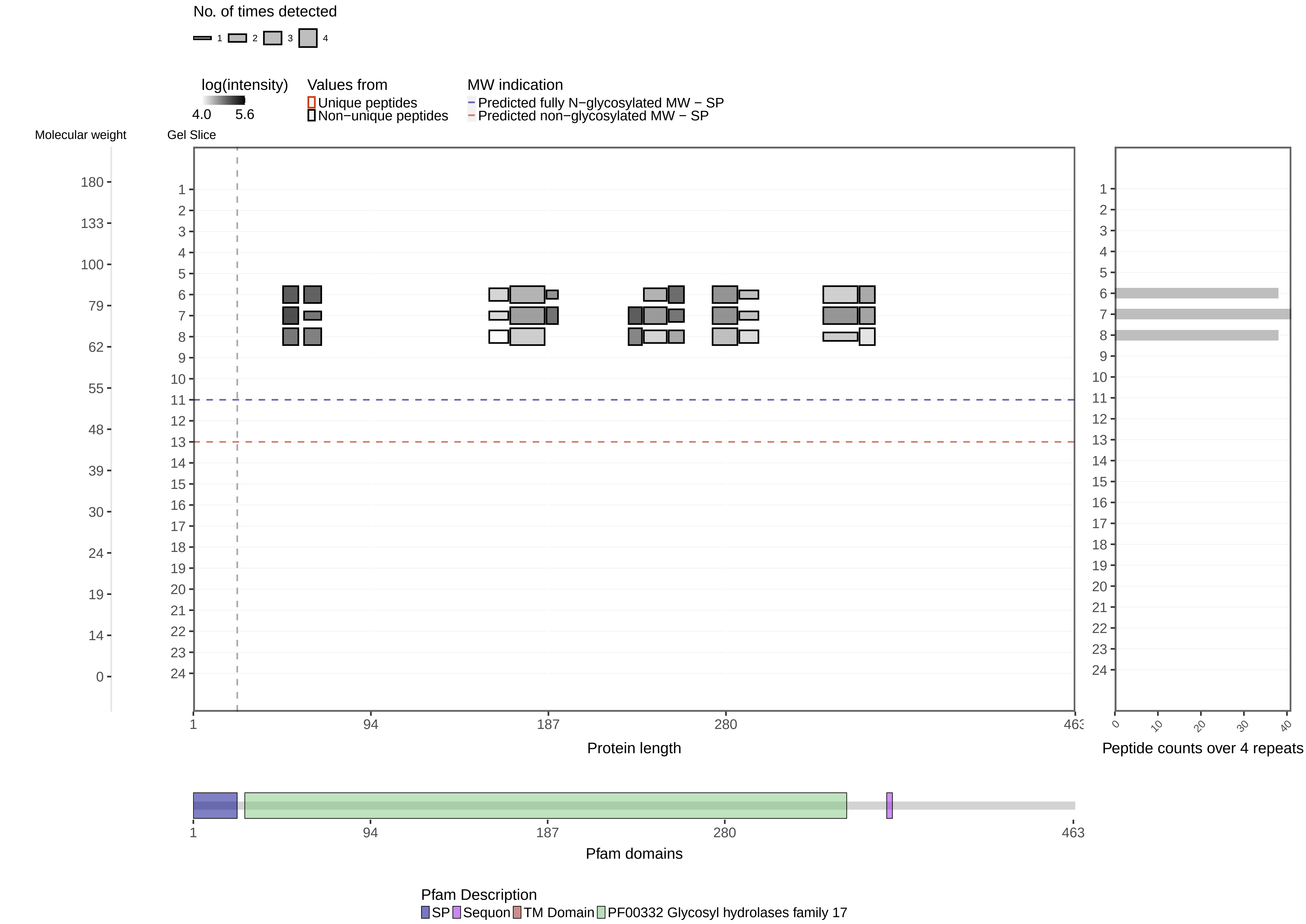


Protein sequence:

-FQPNSGRFDAGTNIKYTSMFDAQVDAVRSALNALGFKEVEIVIAETGWPYKGDSNEVGPSIENAKAYNGNLISHLRSMVGTPLMPGISVDTYLFALYDENLKPGPTSERSFGLFKPDLTMTYDAGLSKANSQDPMPKTLVTPSNTSMPPSSPKTPGTSSSPETPVTPSLPKTSMPPSSPETPATSSLPKPPVLSSSPKTKSVSKTDVSDARLQANIVYAVQFNLGVLVLSQILWHPMLHVL*

NbL15g08560.1 uncharacterized_LOC107783365 [MW = 200.02 kDa]

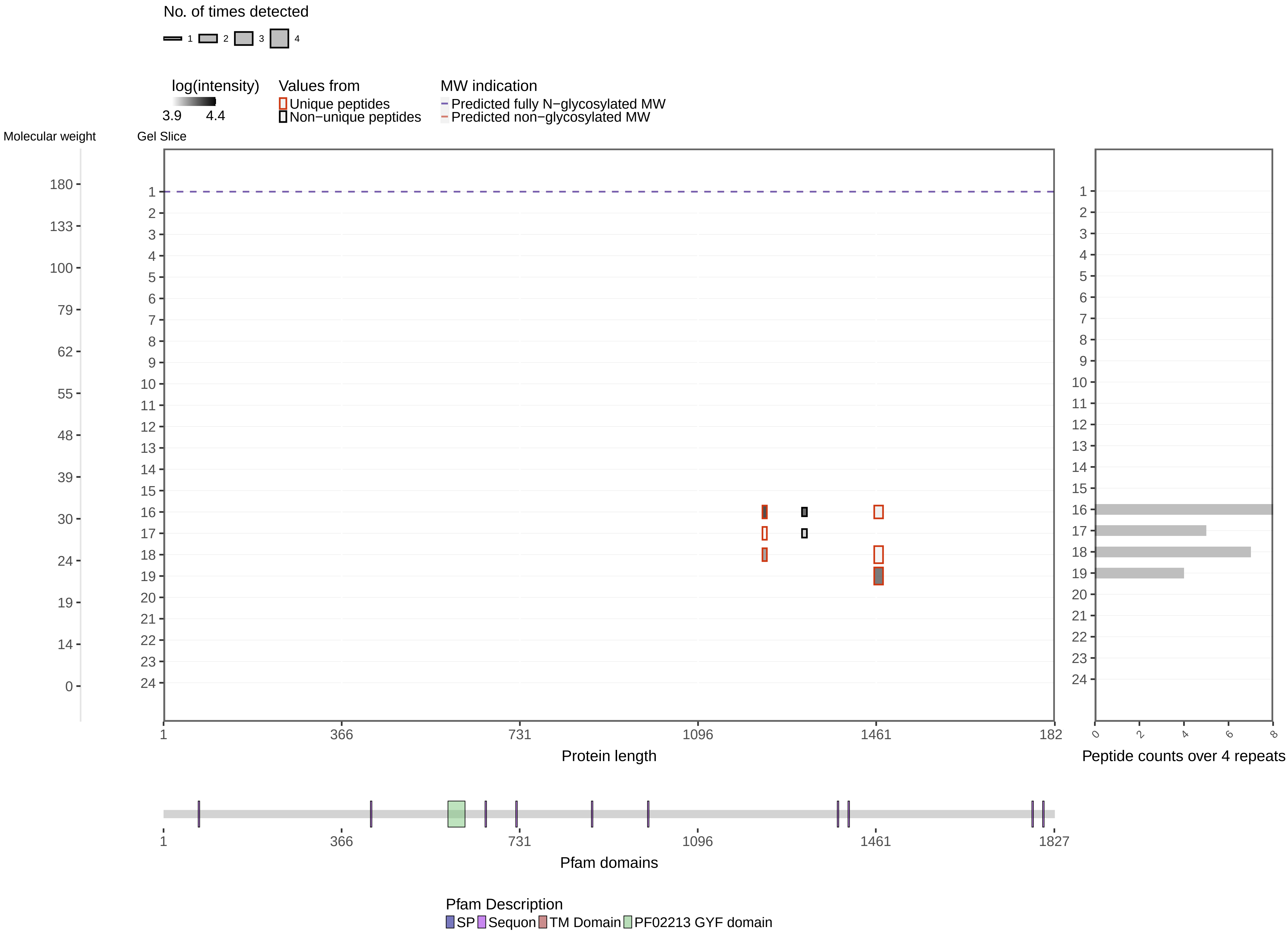


Protein sequence:

POLLNLLOOOYLMOLOSOAALGGPOOLSLLDKFLMLKOOOOOOOOOOOOOOOKOEEHOLMLLOOOOLLSOVLSDPHPHORFGEOPYGKLPTPGISAGNATVDPTRFOPSPNLFSMNTOVOHPVMEEAHASNFVLPPSISOGVTOIGSFETSSLHLPHOMFKDASSORSWGFAEOIDDIOOKVPRMATAMVDPSSHTEIASKHPLEGTENNEPLAVTTPEIASHFPLVEOVEKSVLLPSPLAVDNDLHOKNKLESPPAAISSEPOVERDPHNGLSVTKELKSVEPREV

### NbL15g08470.1 uncharacterized_LOC104247649 [MW = 72.82 kDa]

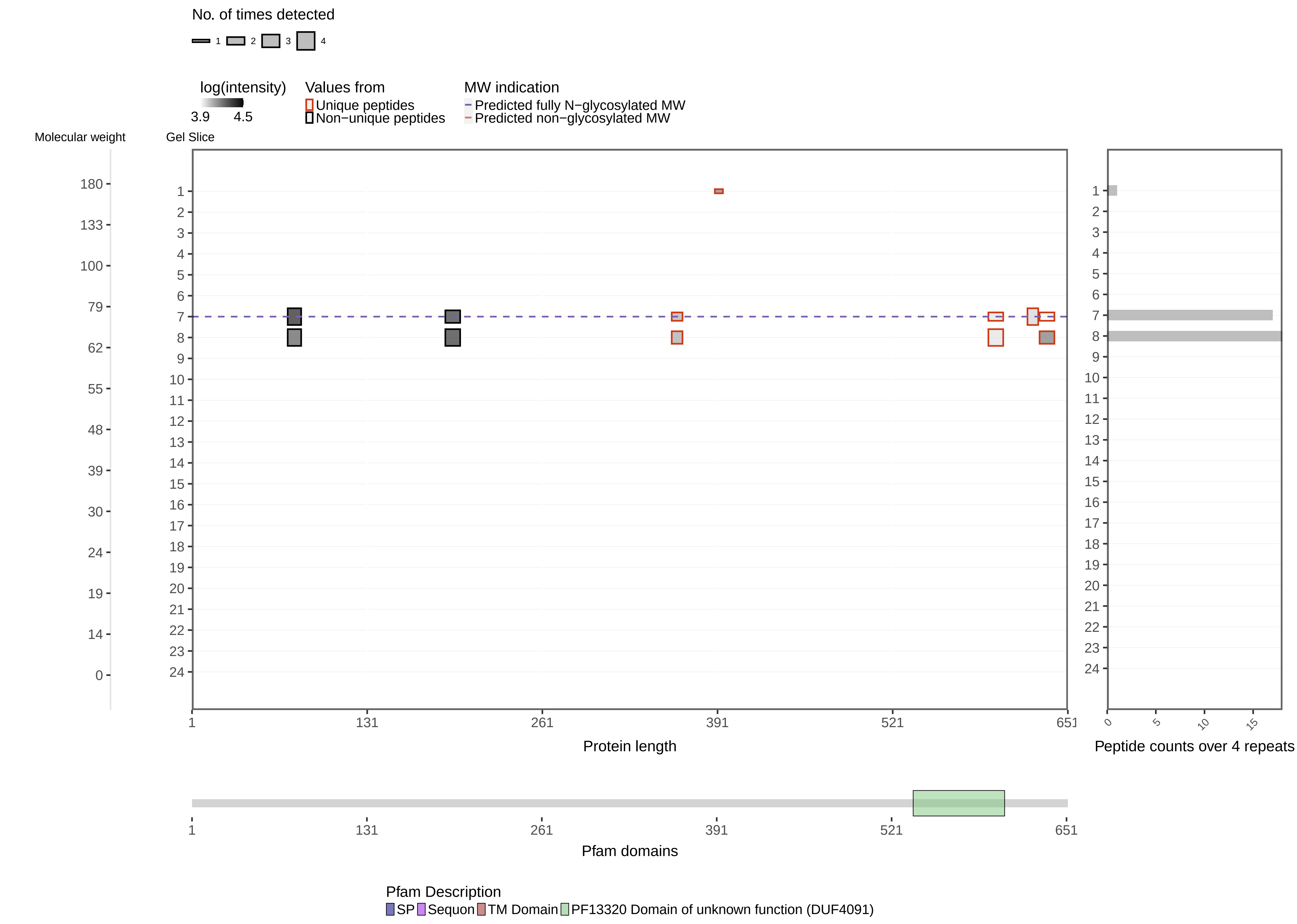


Protein sequence:

[illegible]

### NbL15g08380.1 histidine−−tRNA_ligase_cytoplasmic [MW = 94.21 kDa]

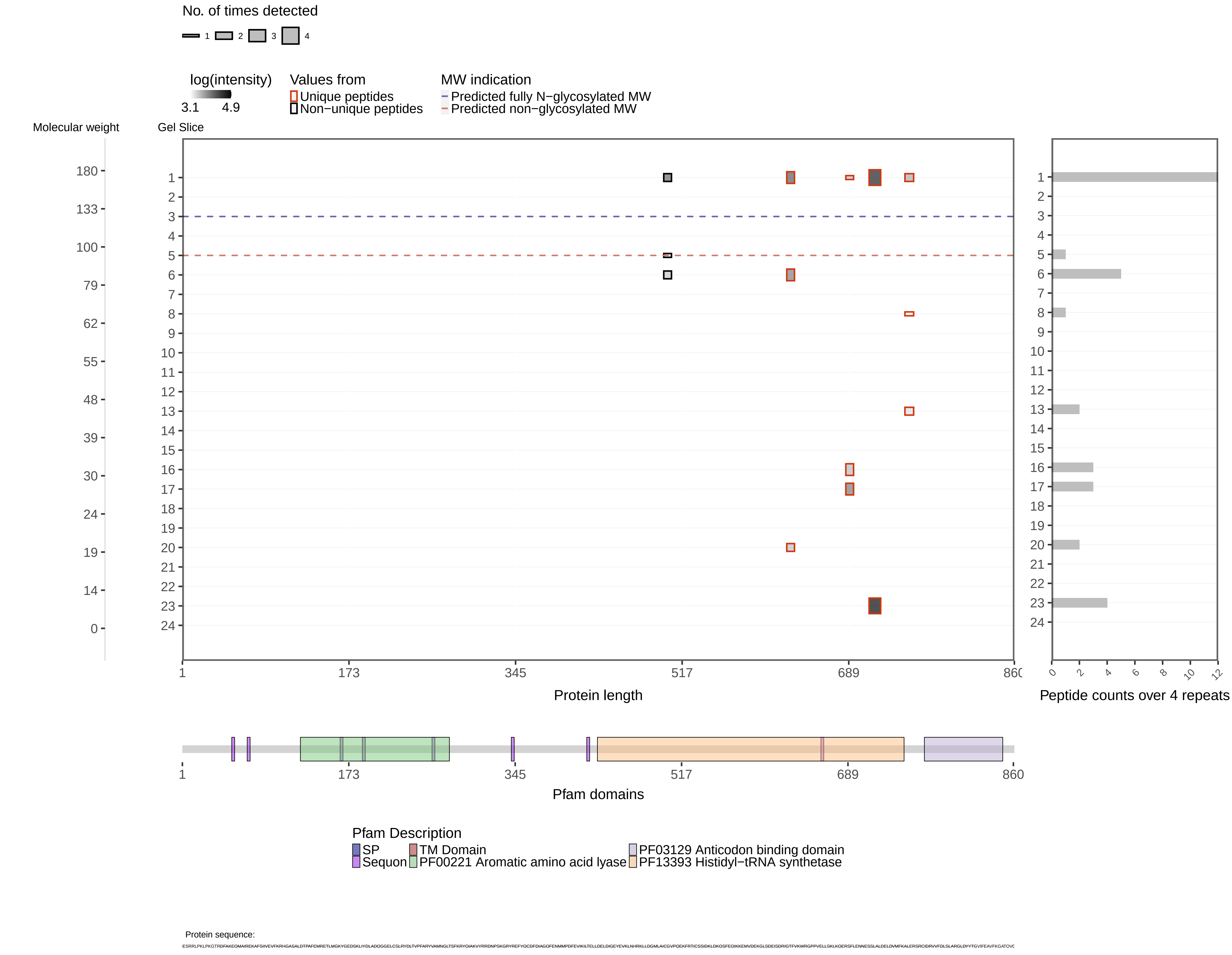

NbL15g08340.1 hydroquinone_glucosyltransferase−like [MW = 55.78 kDa]

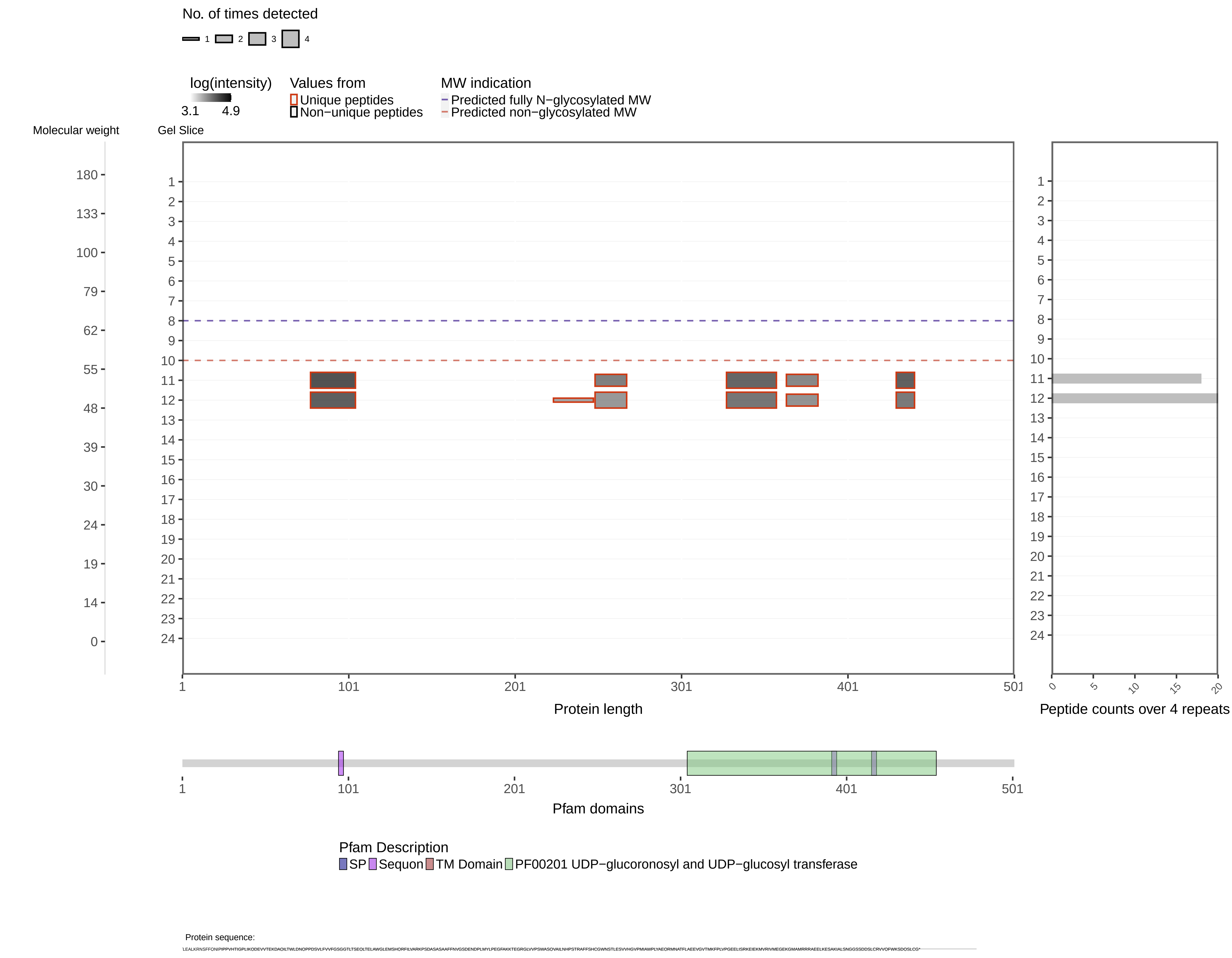

### NbL15g08300.1 NEDD8−conjugating_enzyme_Ubc12 [MW = 20.98 kDa]

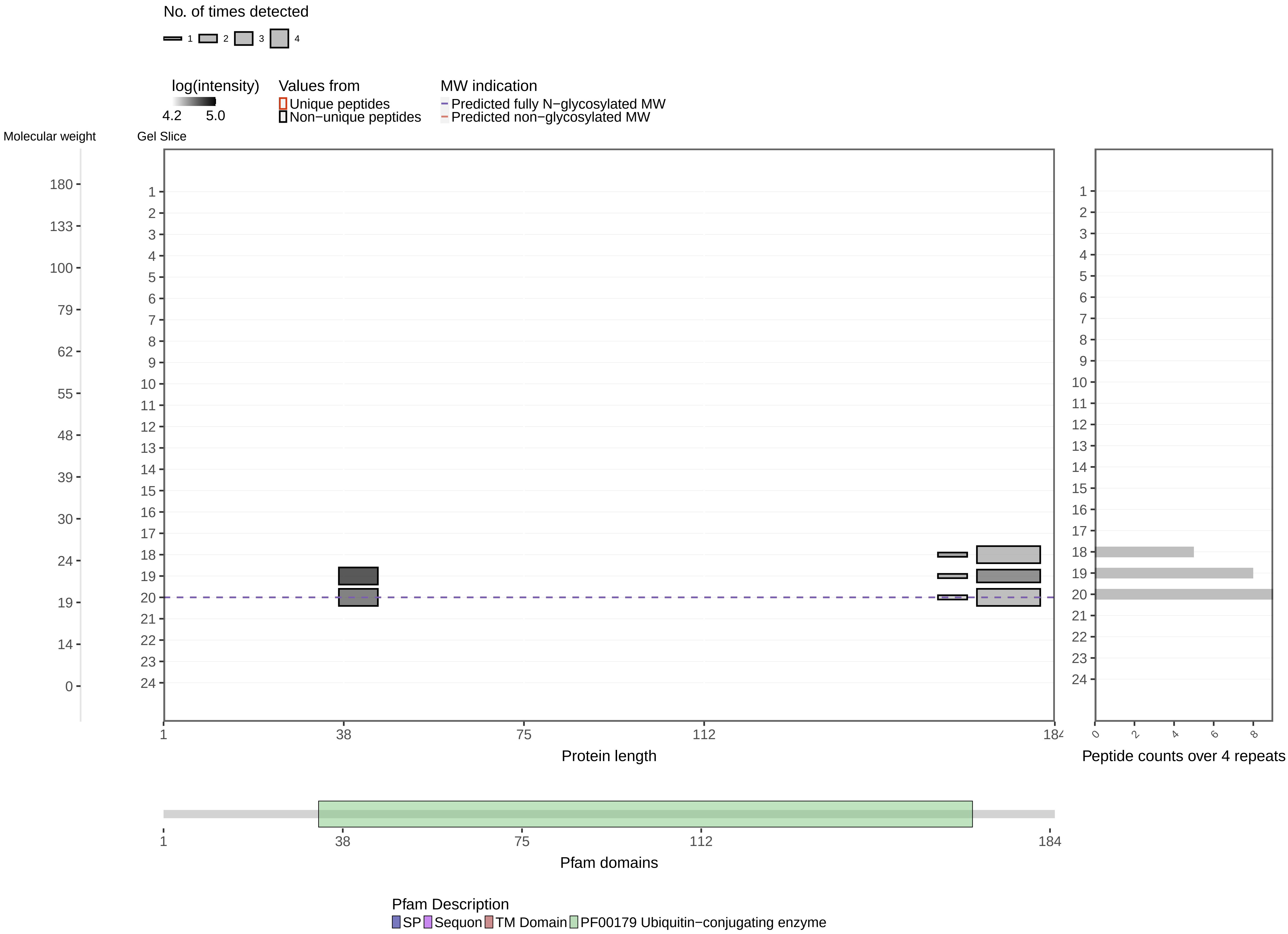


Protein sequence:

FSFSISPMYPHEAPKVKCKTKVYHPNIDLEGNVCLNILREDWKPVLNINTIIYGLYHLFTEPNHEDPLNHDAAAVLRDNPKLFESNVRRAMHGGYVGQTFFPRCM*

NbL15g08290.1 geranylgeranyl_pyrophosphate_synthase_chloroplastic−like [MW = 39.56 kDa]

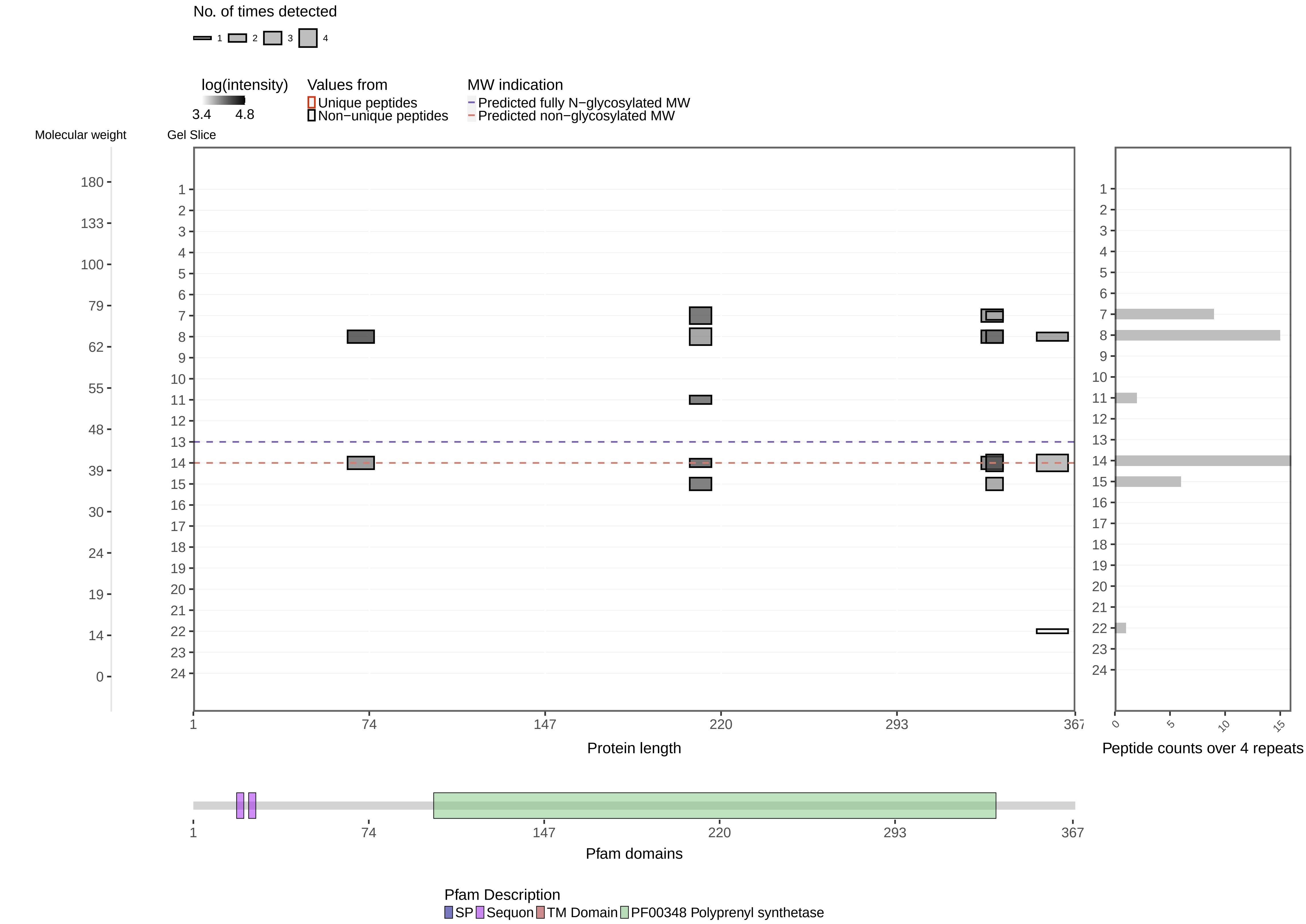


Protein sequence:

GKPTNHKVYGEDVAVLAGDSLLAFAFEYVATATAGVSPSRILAAIGELAKSIGTEGLVAGQVADIACTGNPNVGLDTLEFIHIHKTAALLEASVVLGAILGGGTDEEVEKLRRFARCIGLLFQVVDDILDVTKSSEELGKTAGKDLVVDKTTYPKLLGLEKAKEFAAELNRDAKQQLADFDPHKAAPLIALADYIAYREN*

### NbL15g08100.1 photosystem_II_CP47_reaction_center_protein−like [MW = 56.02 kDa]

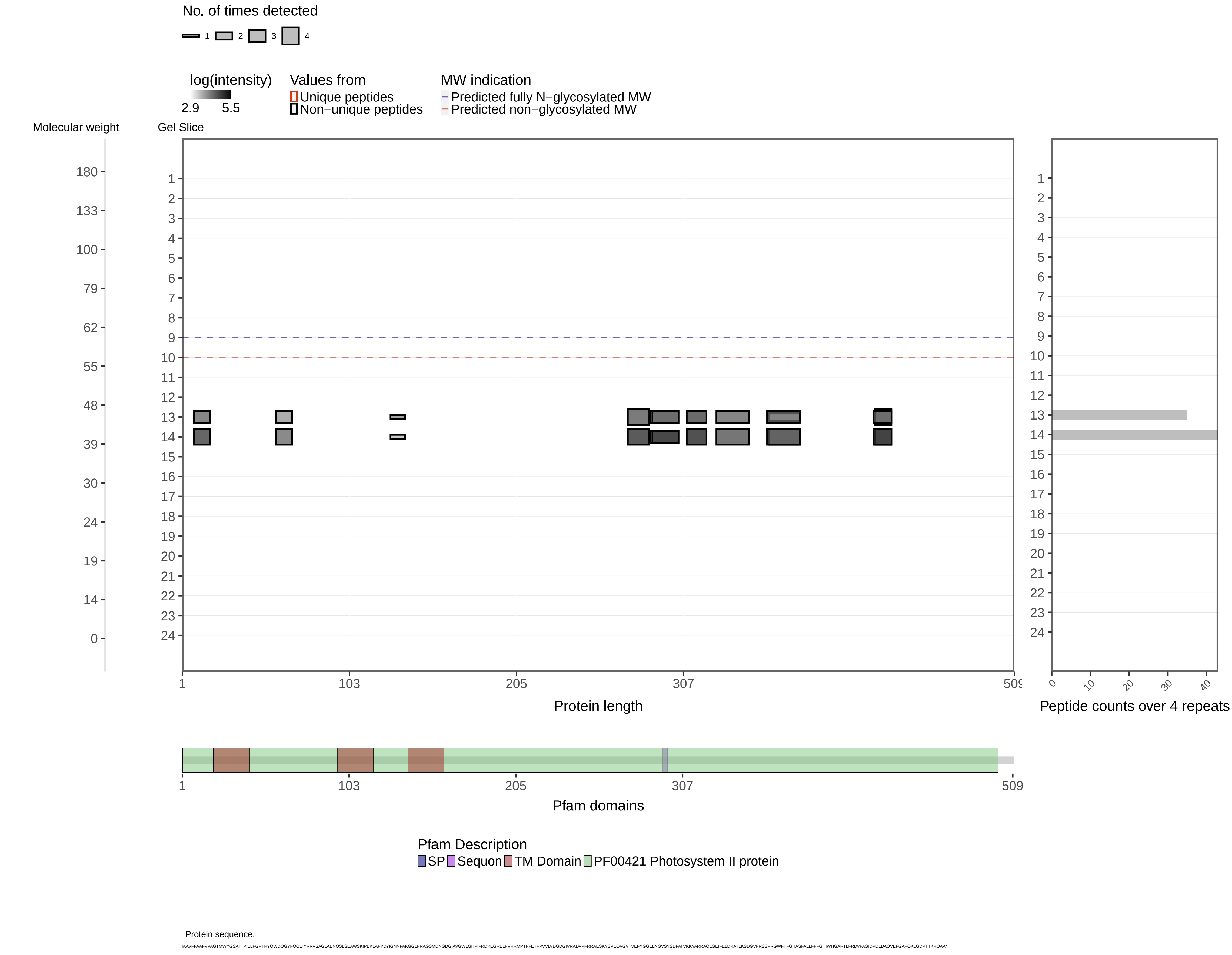

### NbL15g08090.1 ATP−dependent_Clp_protease_proteolytic_subunit−like [MW = 16.4 kDa]

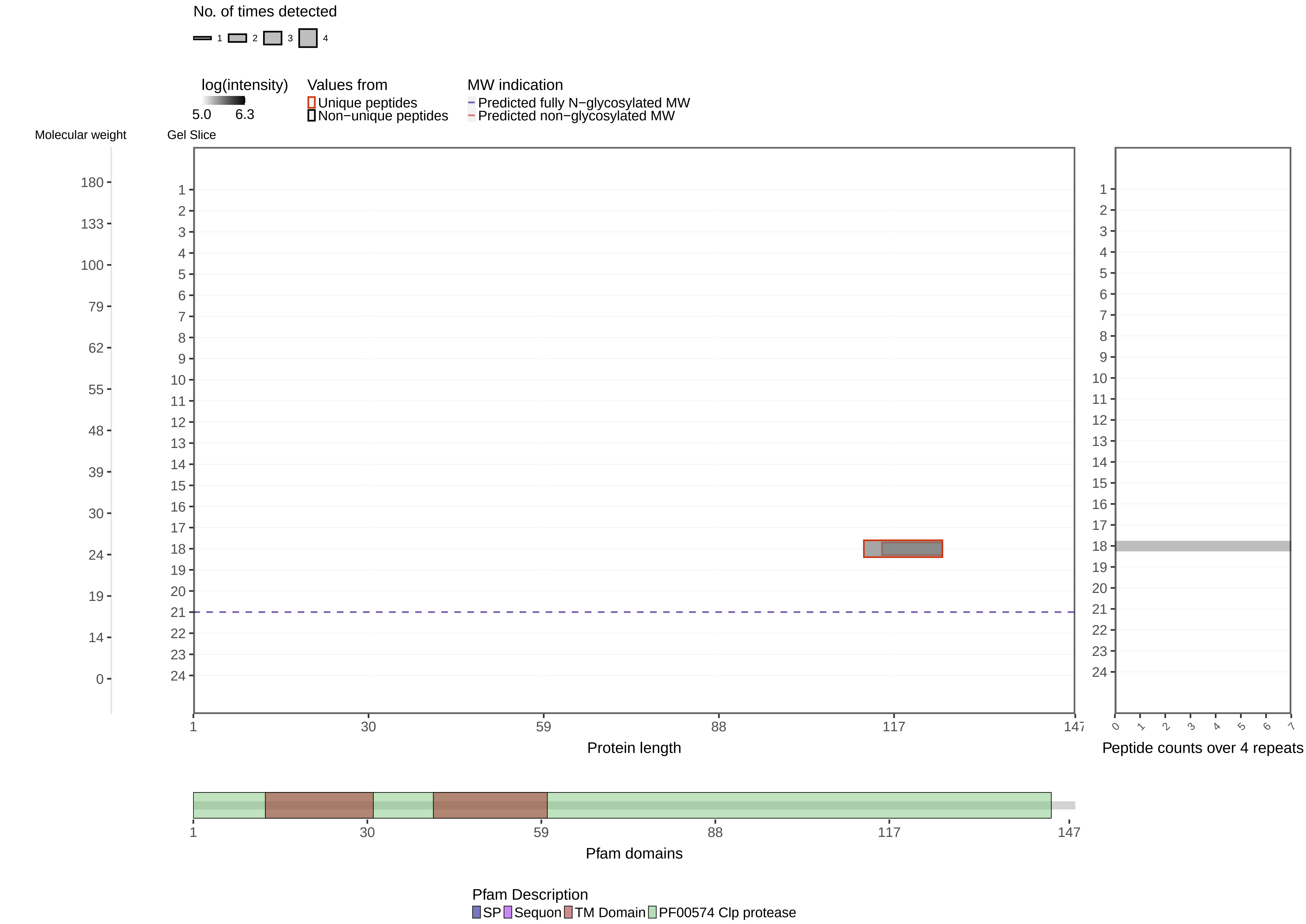


Protein sequence:

EITKRLAFPHAWVMIHQPASSFYEAQTGEFVLEAEELLKLRETLTRVYVQRTGKPLWVVSEDMERDVFMSATEAQAYGIVDLVAVE*

NbL15g07870.1 probable_phenylalanine−−tRNA_ligase_beta_subunit [MW = 66.8 kDa]

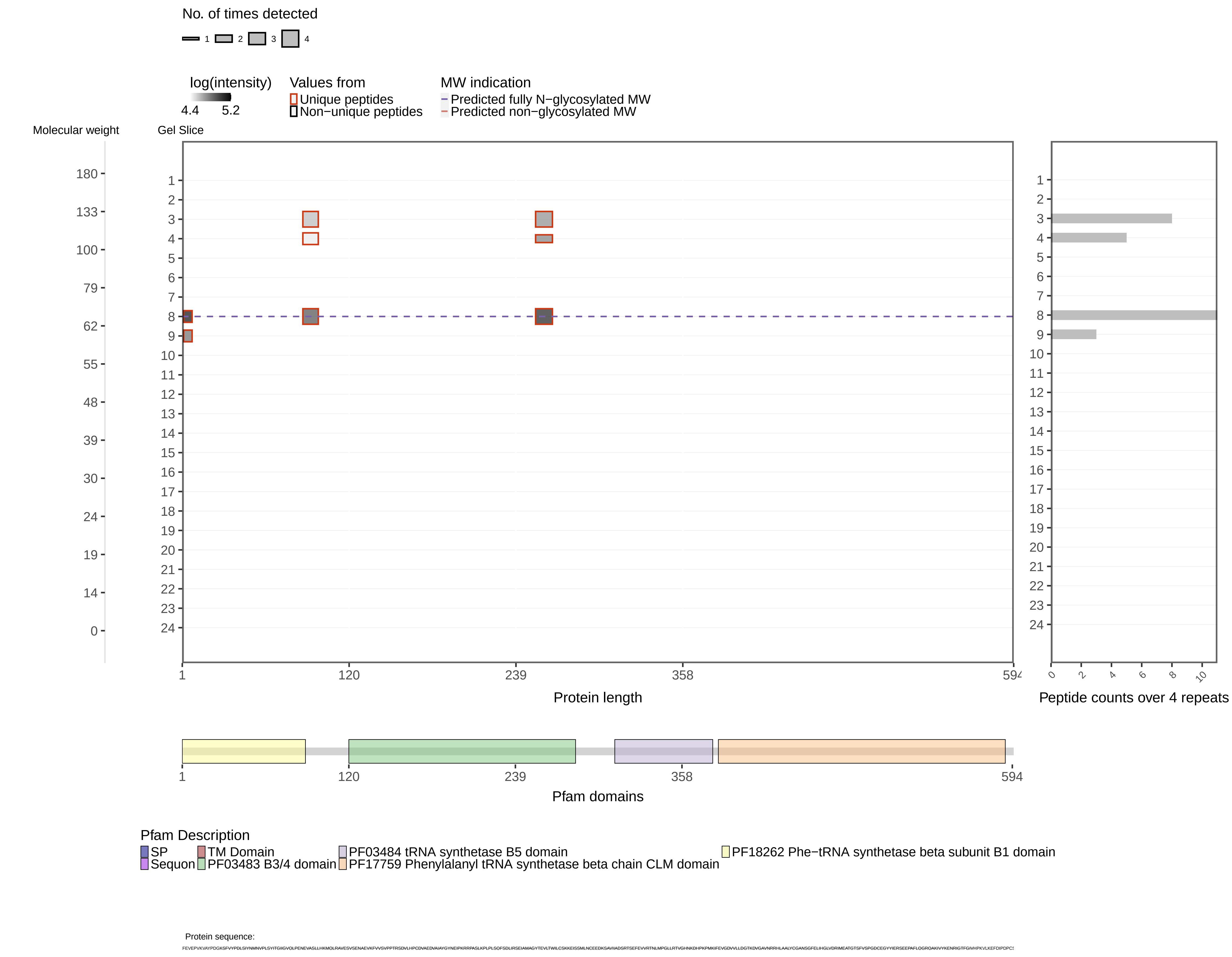

NbL15g07320.1 protein_TIC_22_chloroplastic [MW = 33.21 kDa]

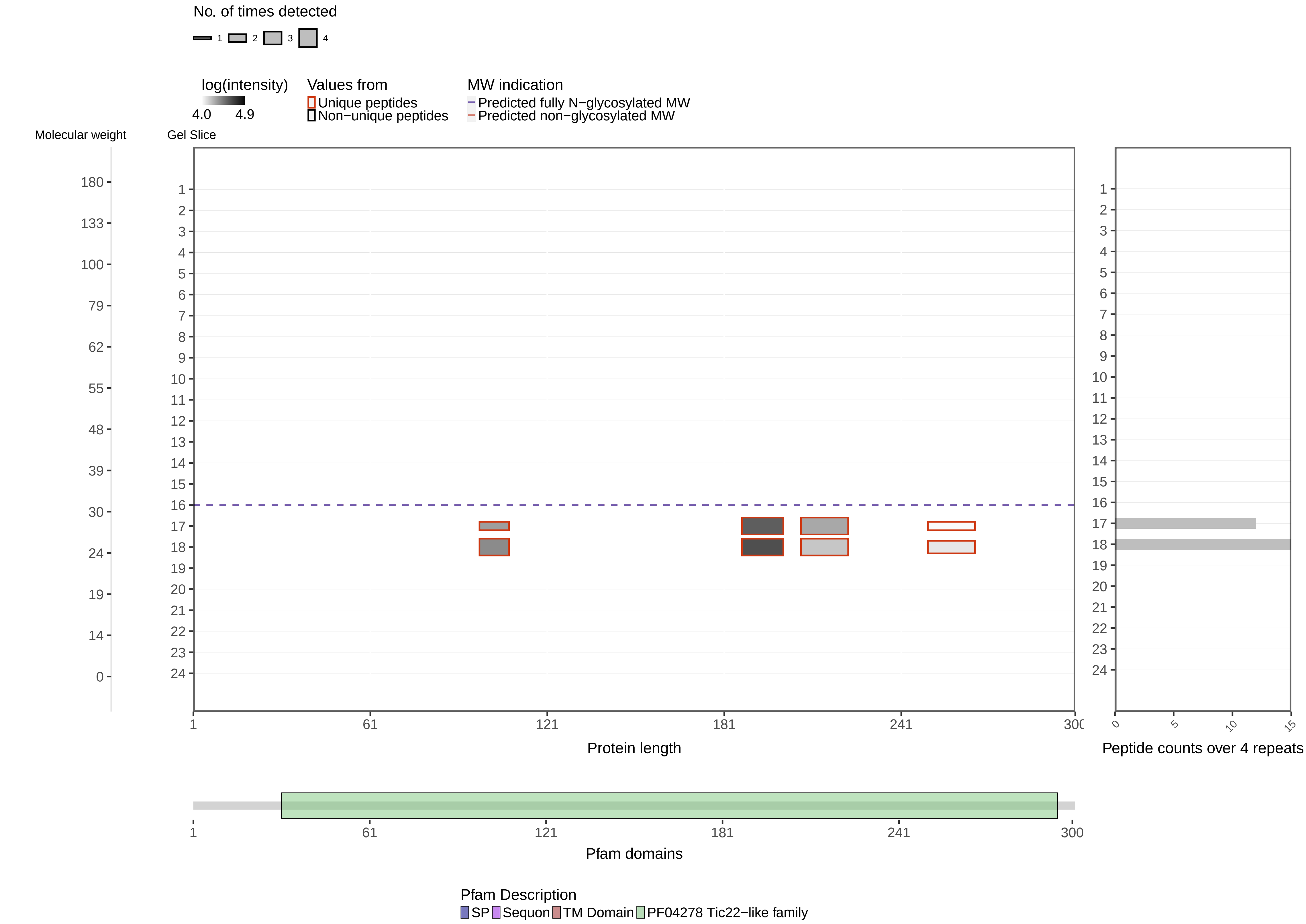


Protein sequence:
GLLCFRQEDAEAFLAQVRLRKGEVRGGAKVVPLTLDQVYMLKVEGIAFRFLPDPVQVQNAIESKLSDVKTEFNGVPVFQSDLLVVKKRNRRYCPIYFRKEDIEKELSSRASRGGVSQHIMVGSLEDVLKKMEISQRHSGWEDLIFIPPGKSHSQHIQDVTKA*

NbL15g07200.1 non−specific_lipid−transfer_protein−like [MW = 13.79 kDa]

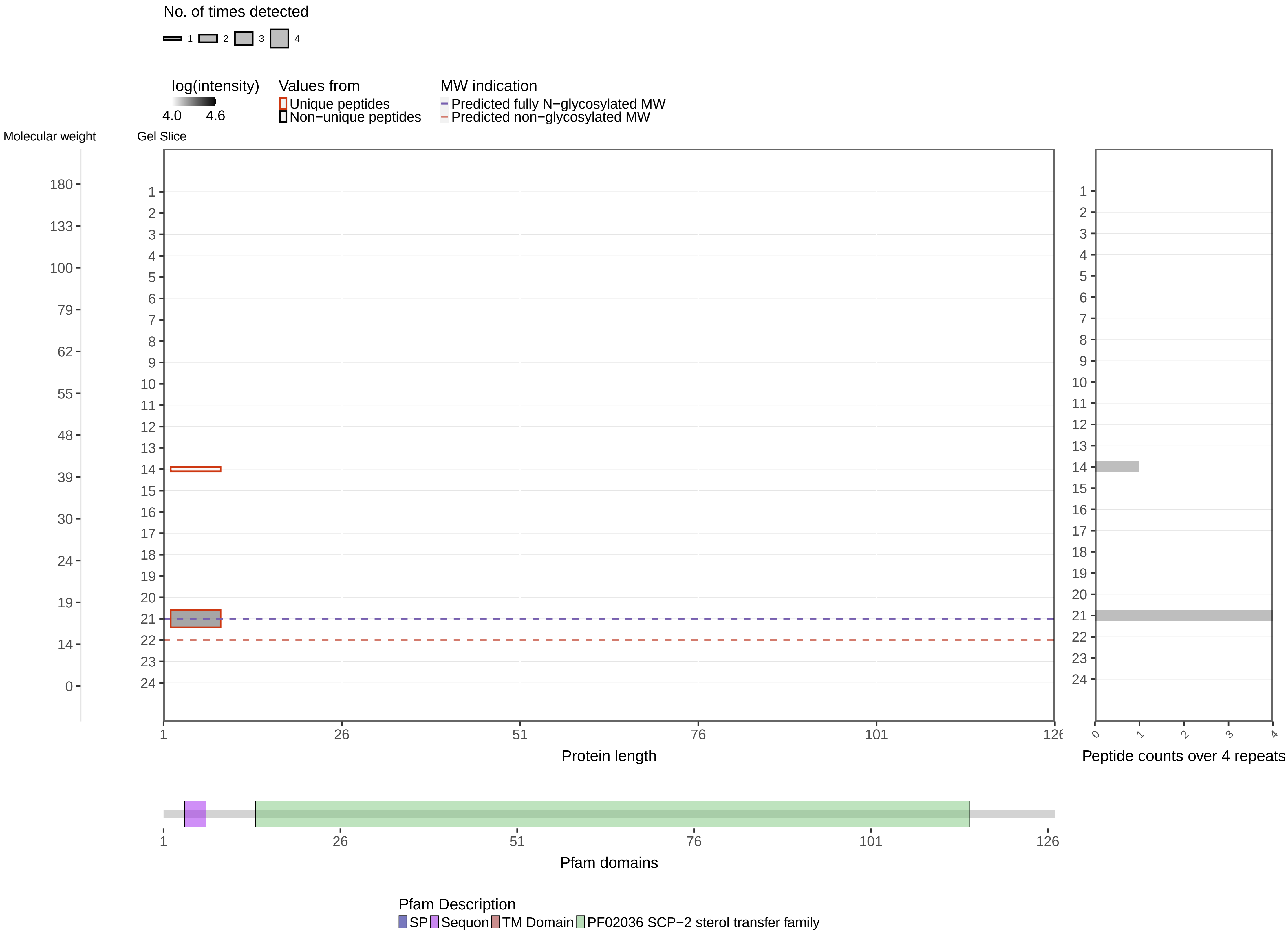


Protein sequence:

KVYVVDLKKGETKEGPYEGGKPDATFSFKDDDFVKIATGKMNPQIAFMRGAMKIKGSISAAQKFTPDIFPKPSKM*

### NbL15g07080.1 annexin_D5 [MW = 35.56 kDa]

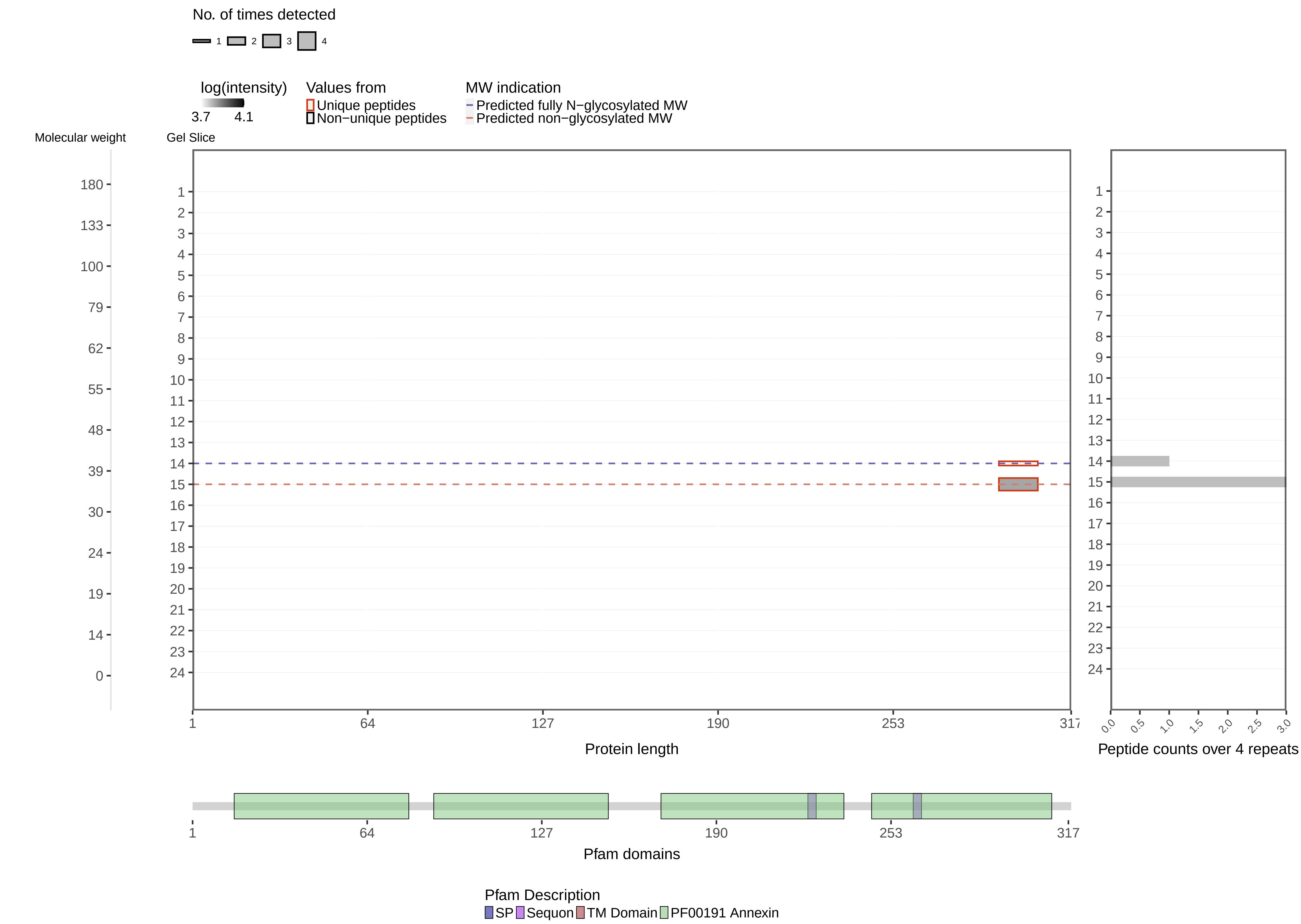


Protein sequence:

KKLLLAFVSTMRYEGPEVDRATVDYDAKALFKAGEKKLGTDEKTFIRIFSERSRAHLAAVSAAYHSMYGRELKKAVKNETSGHFEFGLLTILLCAENPAKYFAKELHKAMKGLGTNDTTLIRIVVTRTEIDMOYIKAEYOKKHKKSLNDAVHSETSGDYRTFLLSLLGPGH*

### NbL15g07050.1 MFP1_attachment_factor_1−like [MW = 23.63 kDa]

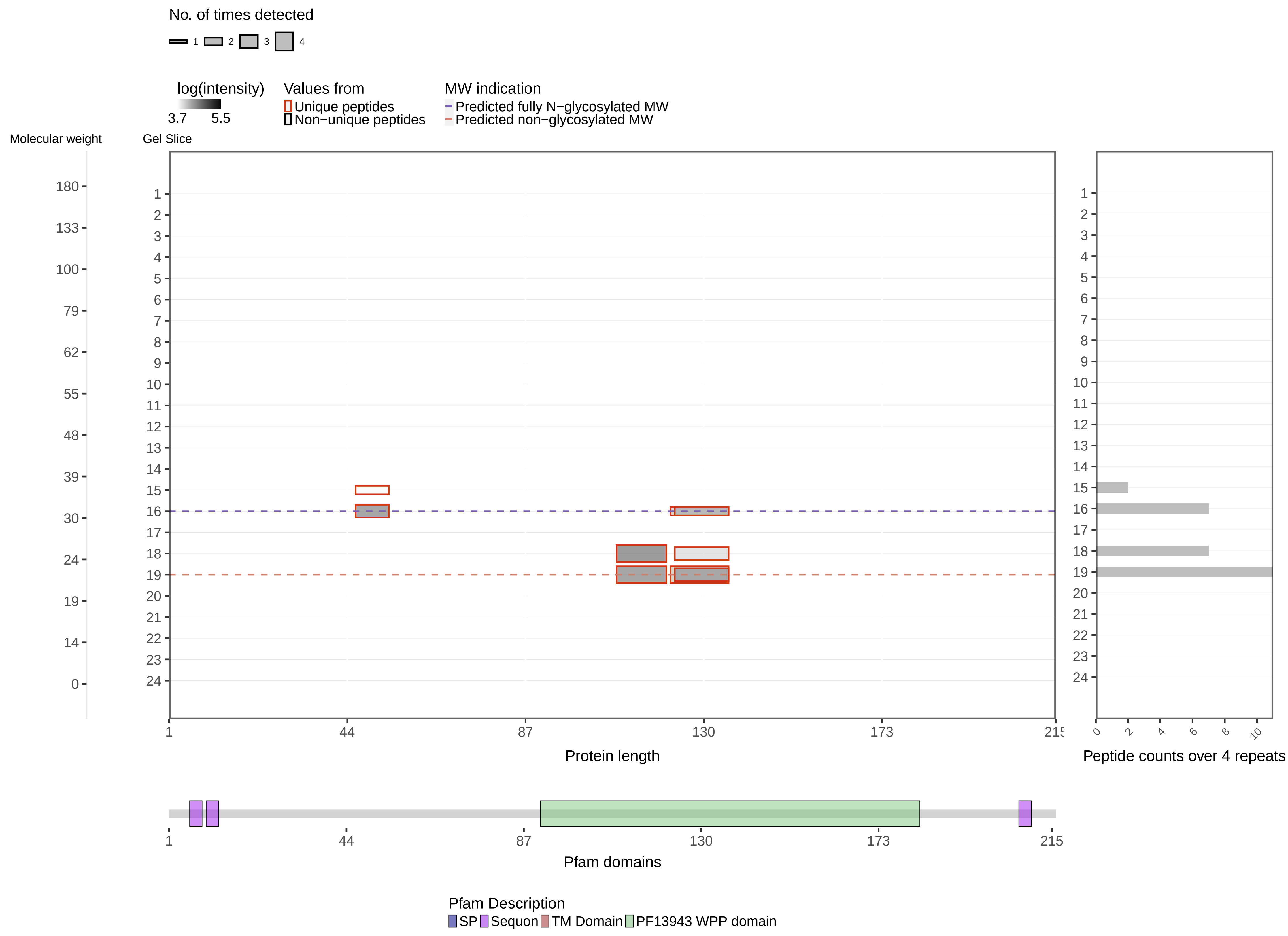


Protein sequence:

FSIWPPTQRTRDAVISRLIESLSTPSILSKRYGTIPHDEASDIAKLIEEEAFTTAGSTAIGDDDDIEILQVYSKEISKRMLDAVKSRSTPSADDSAALTDTETREPISDSAVTGENSSVVTKF*

### NbL15g06830.1 low−temperature−induced_cysteine_proteinase−like [MW − SP = 49.15 kDa]

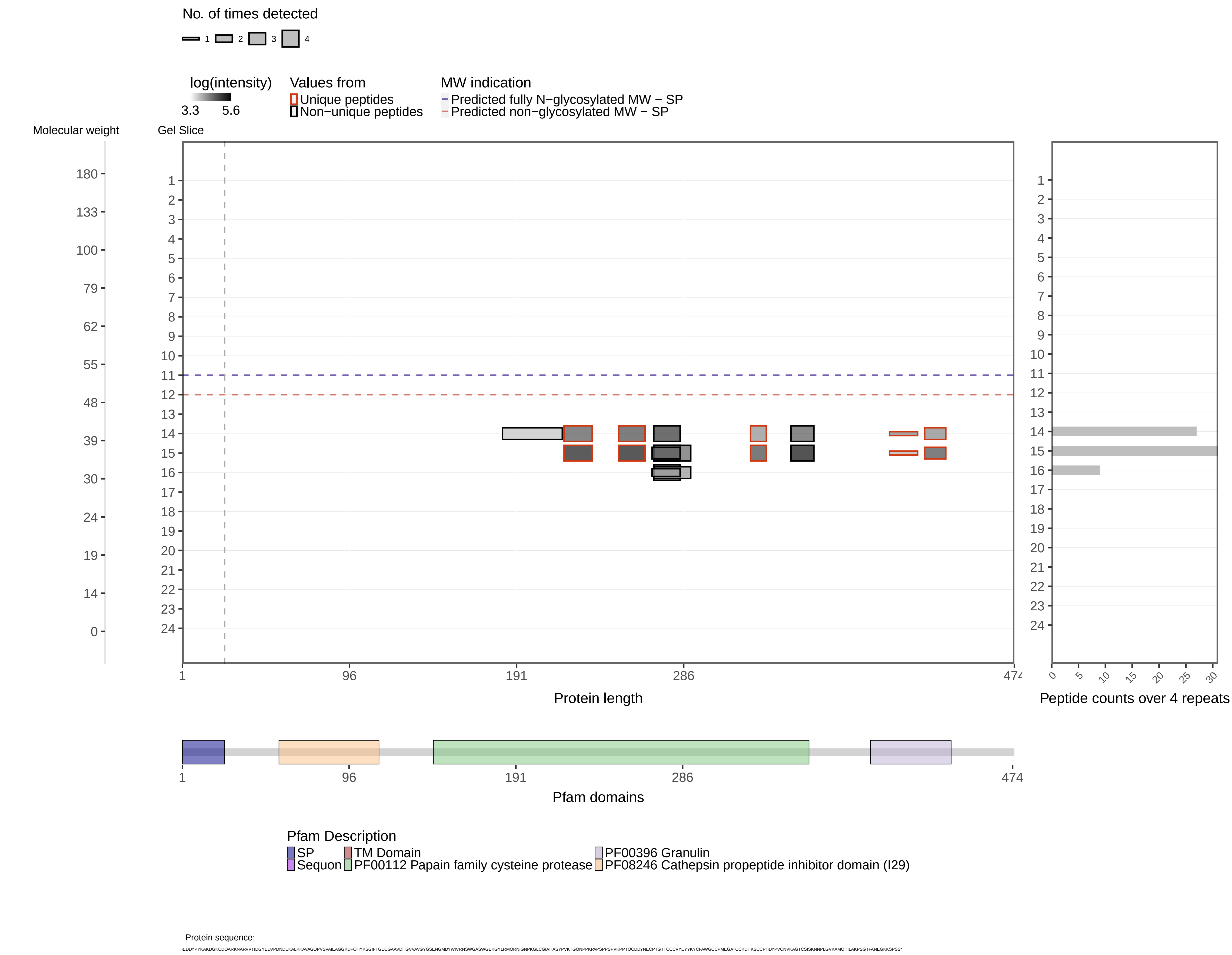

### NbL15g06780.1 uncharacterized_LOC104223854 [MW = 58.49 kDa]

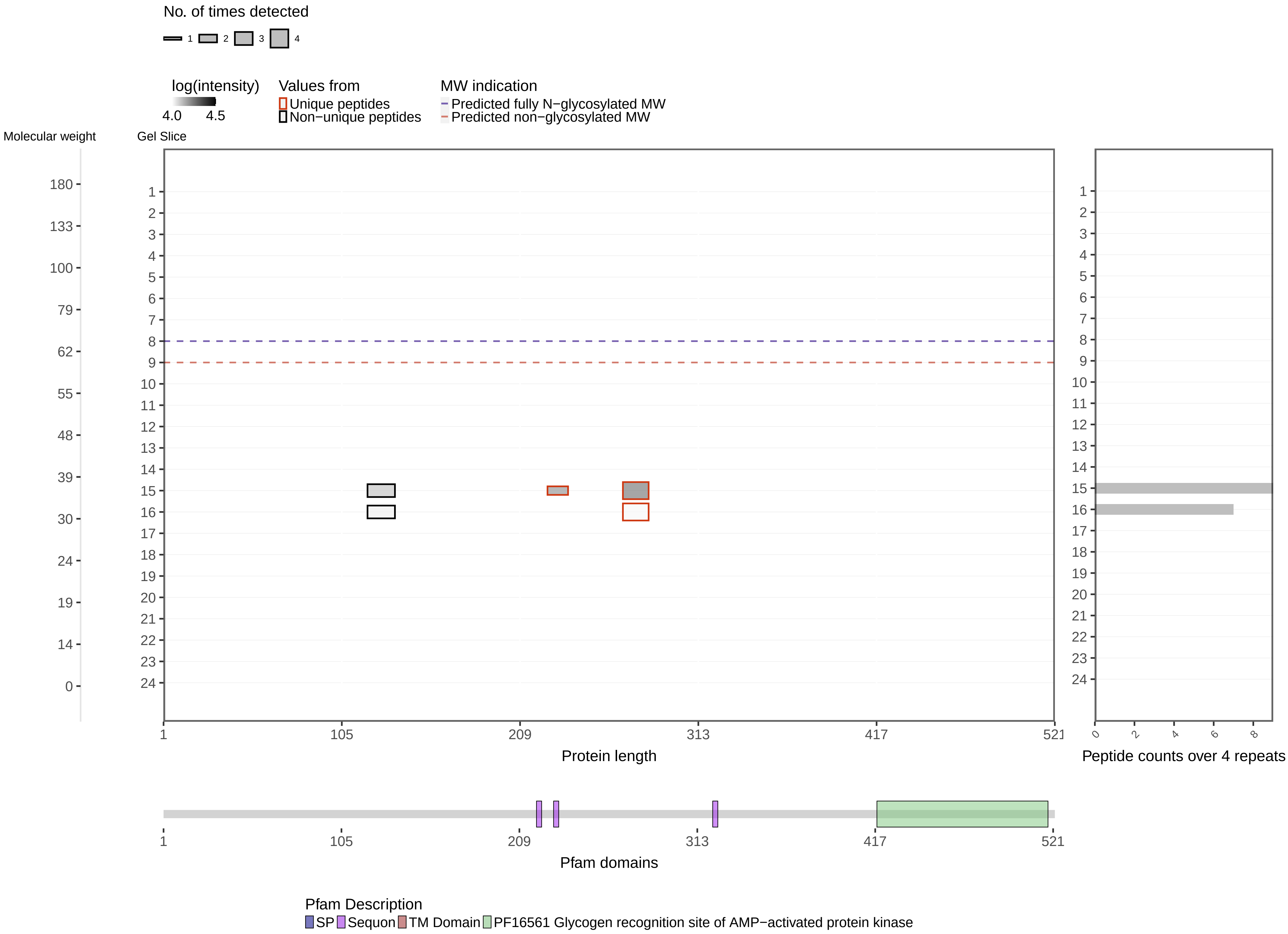


Protein sequence:

[illegible]IQNGELDSVEDSGFQIFNERGSEQVEGSVVGQNVVKTESTTPNGNSAEIFNGNSMPRSPQHVEHPCGLSFQIRNNSLSTEELTMAEGRNSADVEPQKVENQAEINRLKFMLHQKELELTQLKQQIEEEKRLLSILQIKAEREISEAQKLISEKDAELNAAEDSLSGLKEVELQYLADGESVEVAGSFNGWHHHKIKMDLESSSTIIDPIDQELSSDIVDPIGTRKPRLWKTMLWLYPGMYEIKFVVDGHWTTDPQMESVIRGAVHNNVLRVDR*

### NbL15g06760.1 S−adenosylmethionine_synthase_3−like [MW = 43.66 kDa]

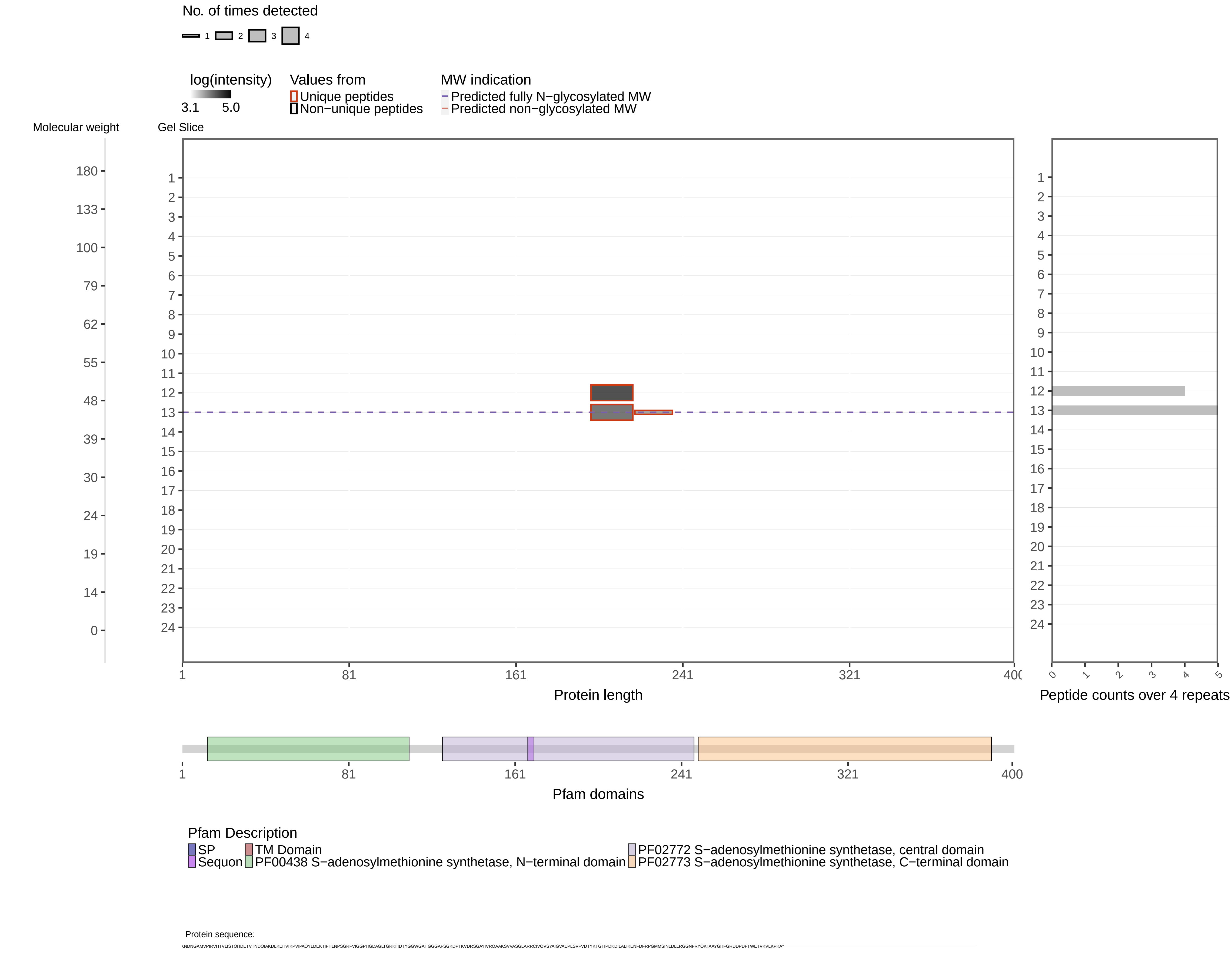

NbL15g06610.1 E3_ubiquitin−protein_ligase_MIEL1−like [MW = 34.18 kDa]

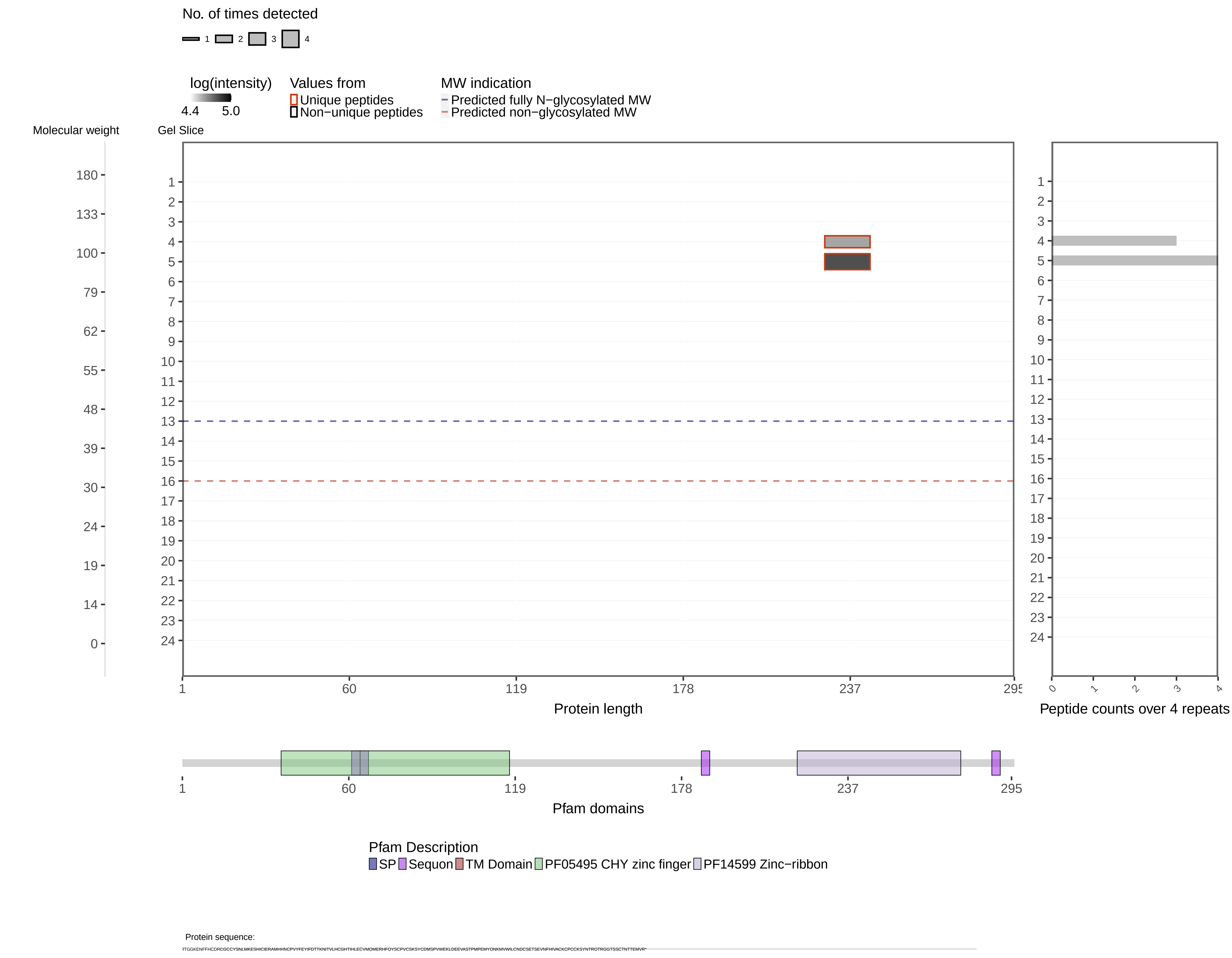

### NbL15g06460.1 UDP−glycosyltransferase_73C3−like [MW = 55.83 kDa]

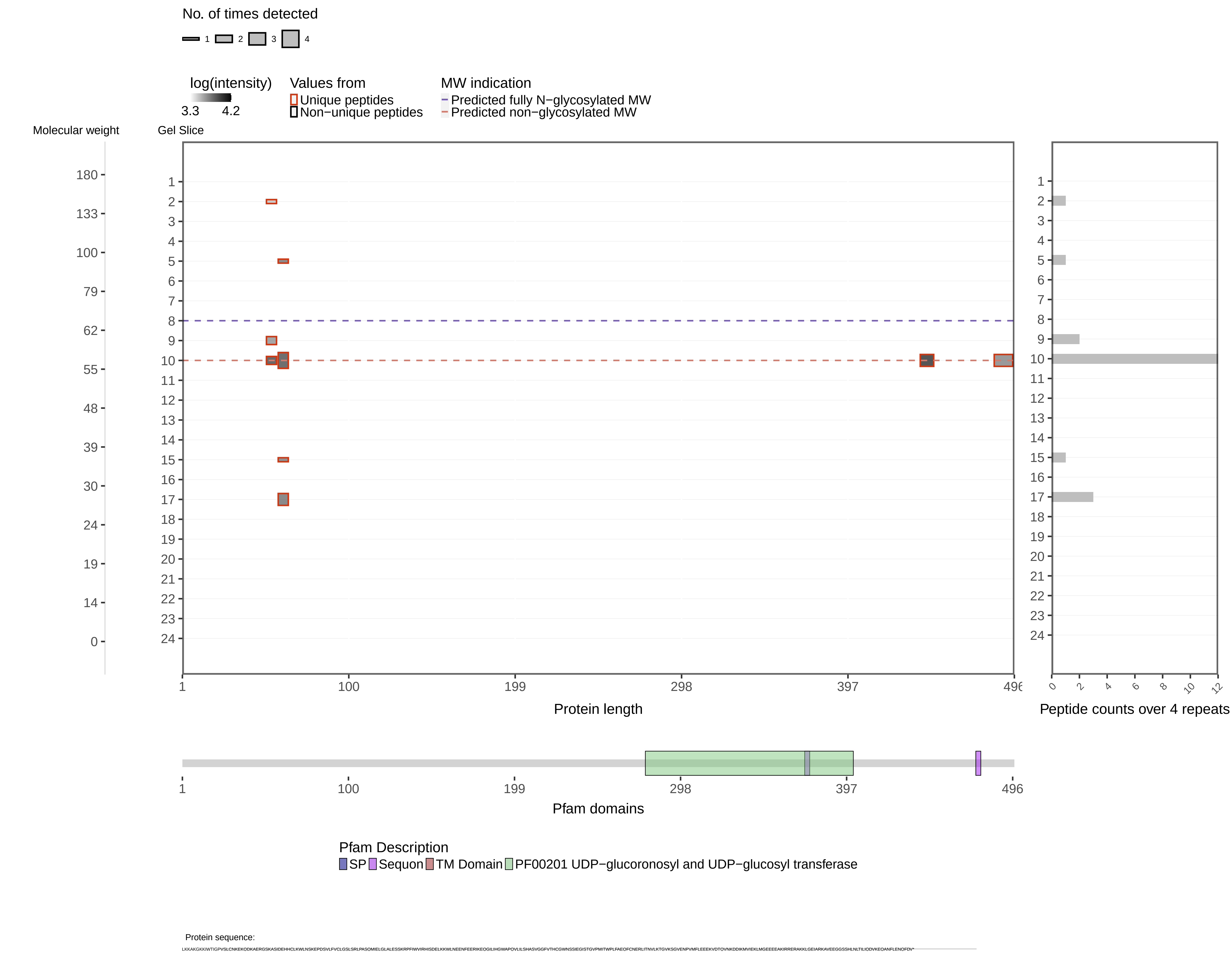

### NbL15g06380.1 threonine_dehydratase_biosynthetic−like [MW = 60.67 kDa]

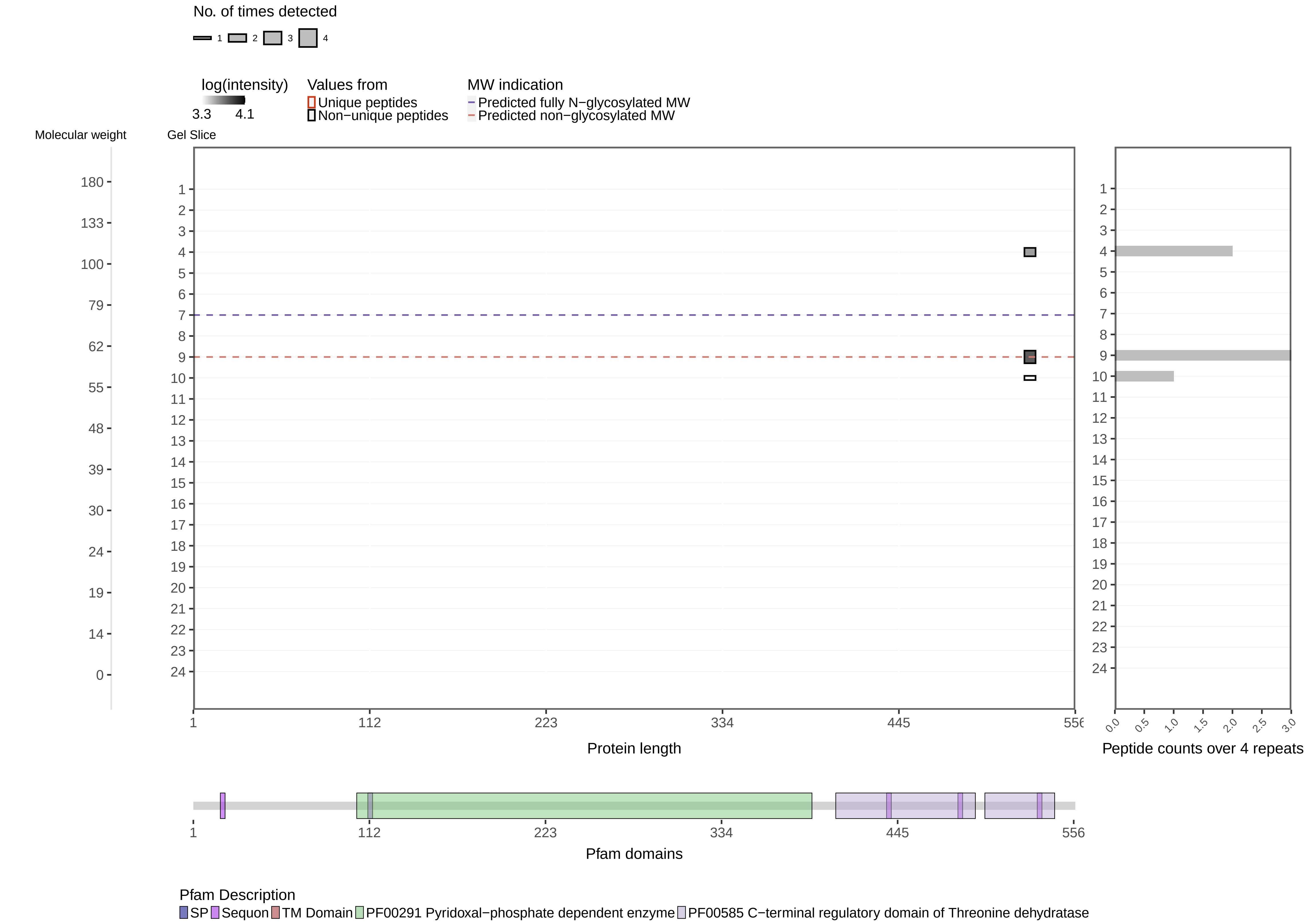


Protein sequence:

LIAGVATYFKRVAPHTKIIGVEPFGASSMTQSLYRGERVKLEQVDNFADGVAVALVGEETFRLCKDLIDGMVLVSNDAISAAVKDVYDEGRNILETSGALAIAGAEAYCKYYDIKGESVVVIASGANMDFSKLKLVVDLADIGGQREALLATFMPEEPGSFKKFCELAGLMNITEFKYRYNSGRKQALVLYSVGVTTKSDLESMLERMKSSQLNTVNLTNNNLVKEHIRHLMGGRSEPSNEIFCQFIFPEKPGALRKFLDAFSPRWNISLFHYREQGELCVSWIPSSKR*

NbL15g06370.1 peptidyl−prolyl_cis−trans_isomerase_FKBP16−4_chloroplastic [MW = 25.06 kDa]

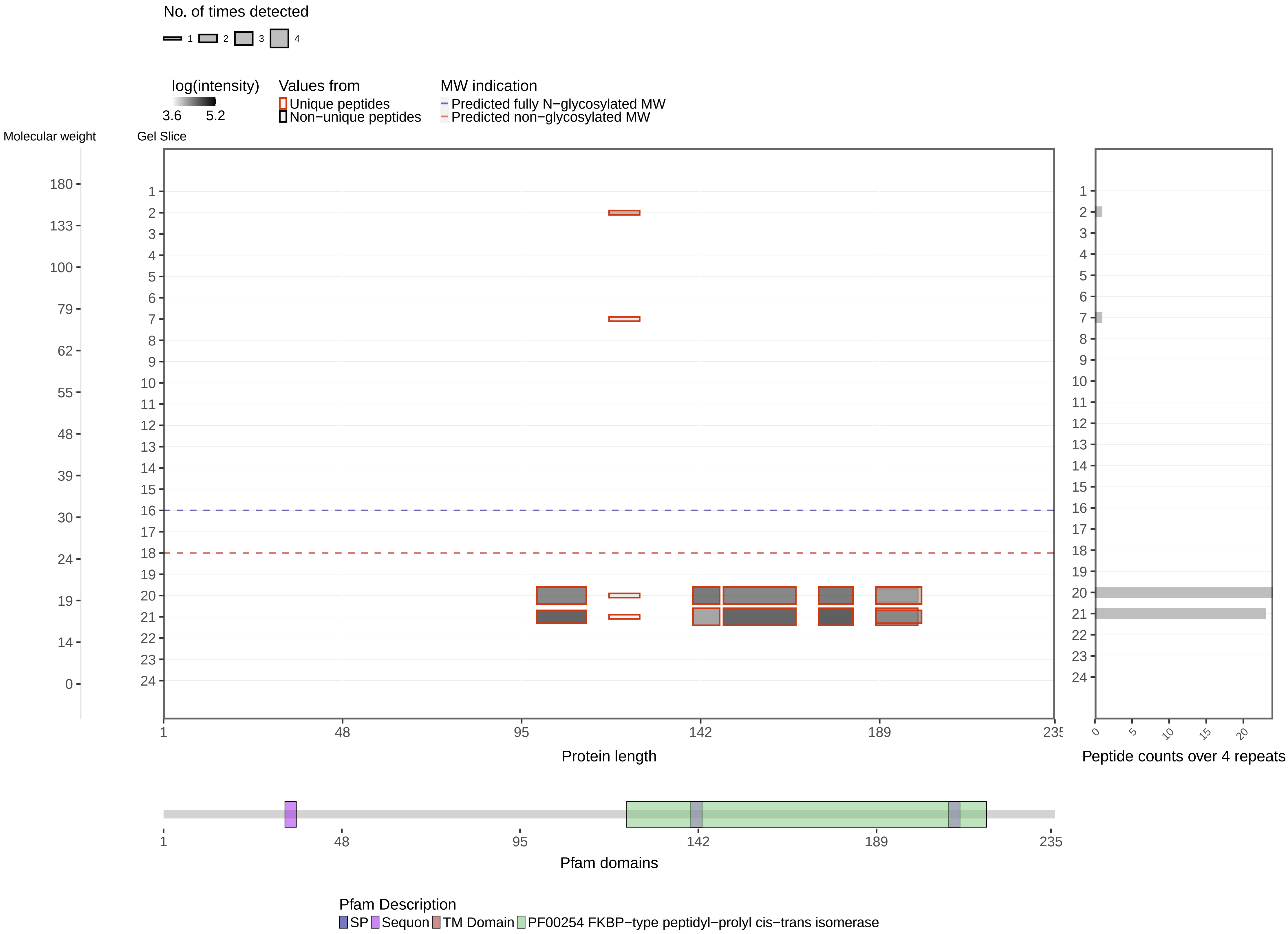


Protein sequence:

TLPNGLKYYDLKVGNGAEAVKGSRVAVHYVAKWRNITFMTSRQGMGVGGGTPYGFDVGQSERGTVLKGLDLGVQGMRVGGQRLLIVPPELAYGKKGVQEIPPNATIEIDIELLSIKQSPFGTAVKVVEG*

NbL15g06150.1 geranylgeranyl_pyrophosphate_synthase_small_subunit_chloroplastic−like [MW = 36.05 kDa]

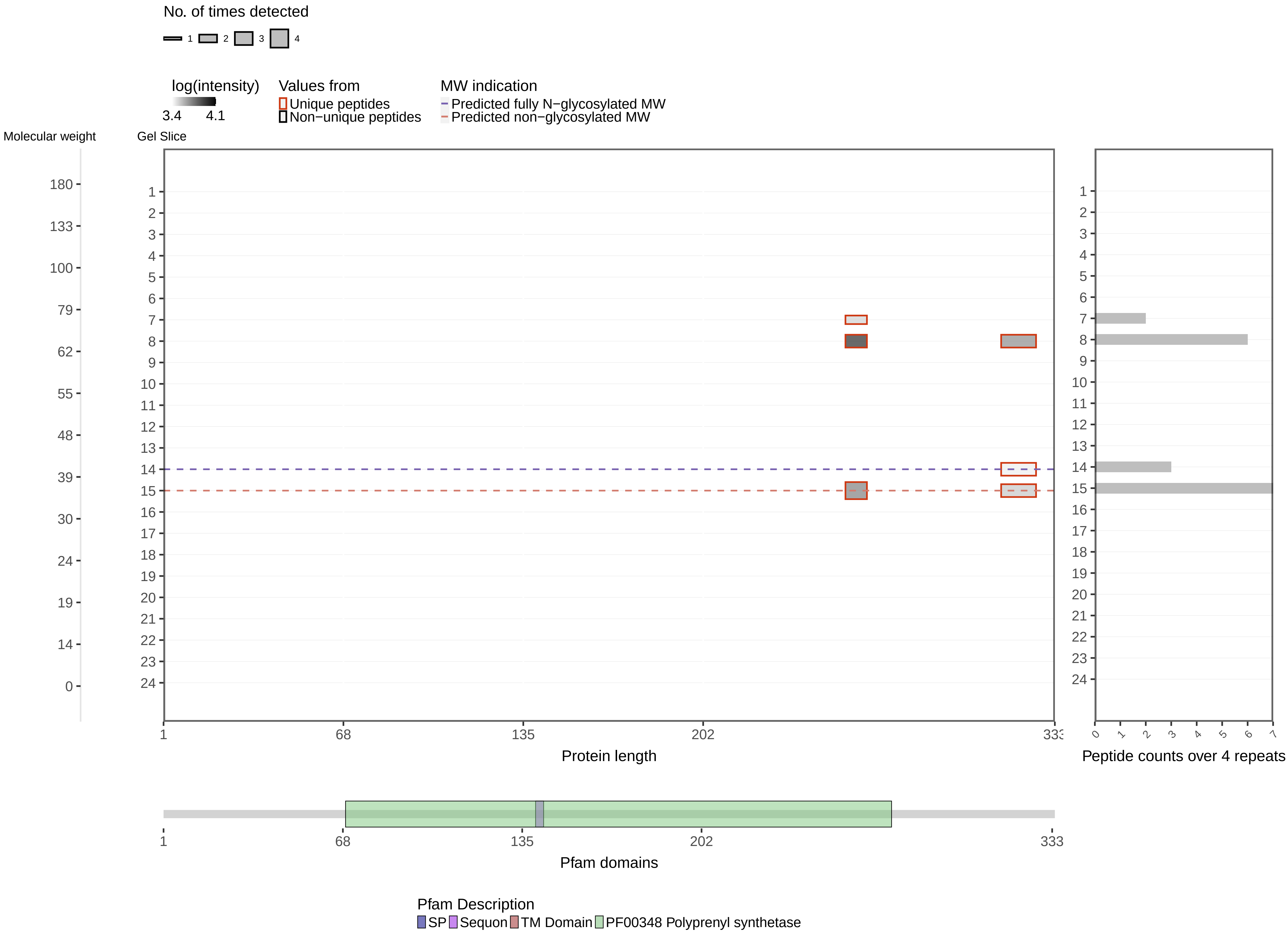


Protein sequence:

DALFPLGFQHIVSHTPSDLVPEDRVLKVITEIARAVGSTGMAAGQFLGLEGGPNAVDSVQEKKYGEMGECSAVCGGLLAGASDEEIQHLRRYGRAVGVLYRVVEDILEAKETGDKSEGKKKKGKSYVSLYGIEKAKKVAEDLRAQAKRELDGLGKYGDKVVPLYSFVDYAADRGFSLDN*

### NbL15g06100.1 peroxiredoxin−2E−2_chloroplastic [MW = 32.8 kDa]

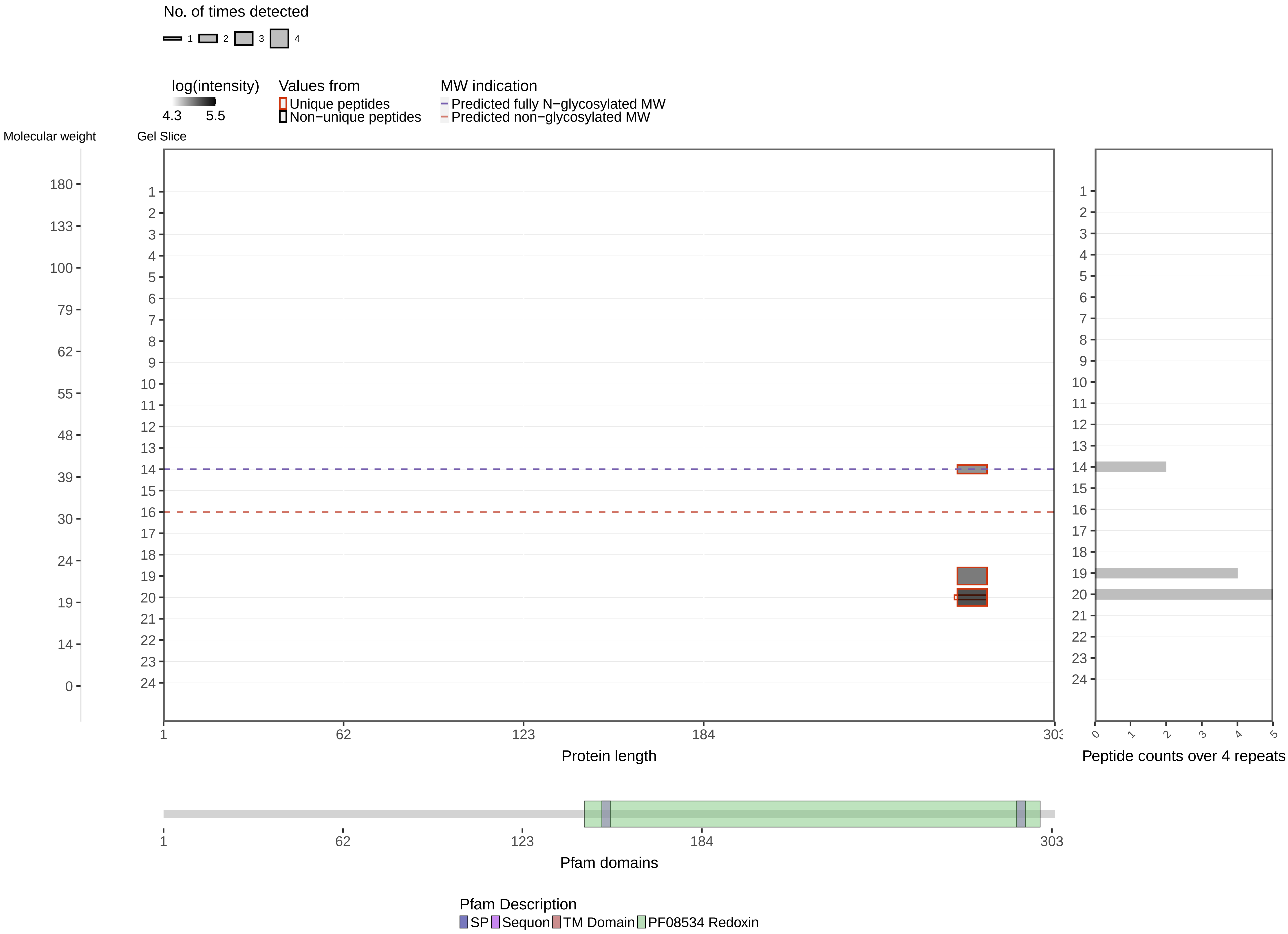


Protein sequence:

ATISVGDKLPNCTLSYFDSSDELKTLLISDLTNGKKVVLLAVPGAFTPTCSQKHLPGFVEKSKELKSKGVDTIACISVNDAFVMKYWKENLGINDEVMMLSDGNLEFTKAIGCELDLSDKPVGLGIRSRRYSMLVDDGVVKILNLEEGGAFNVSSAEDILKAL*

### NbL15g06080.1 NA [MW = 47.87 kDa]

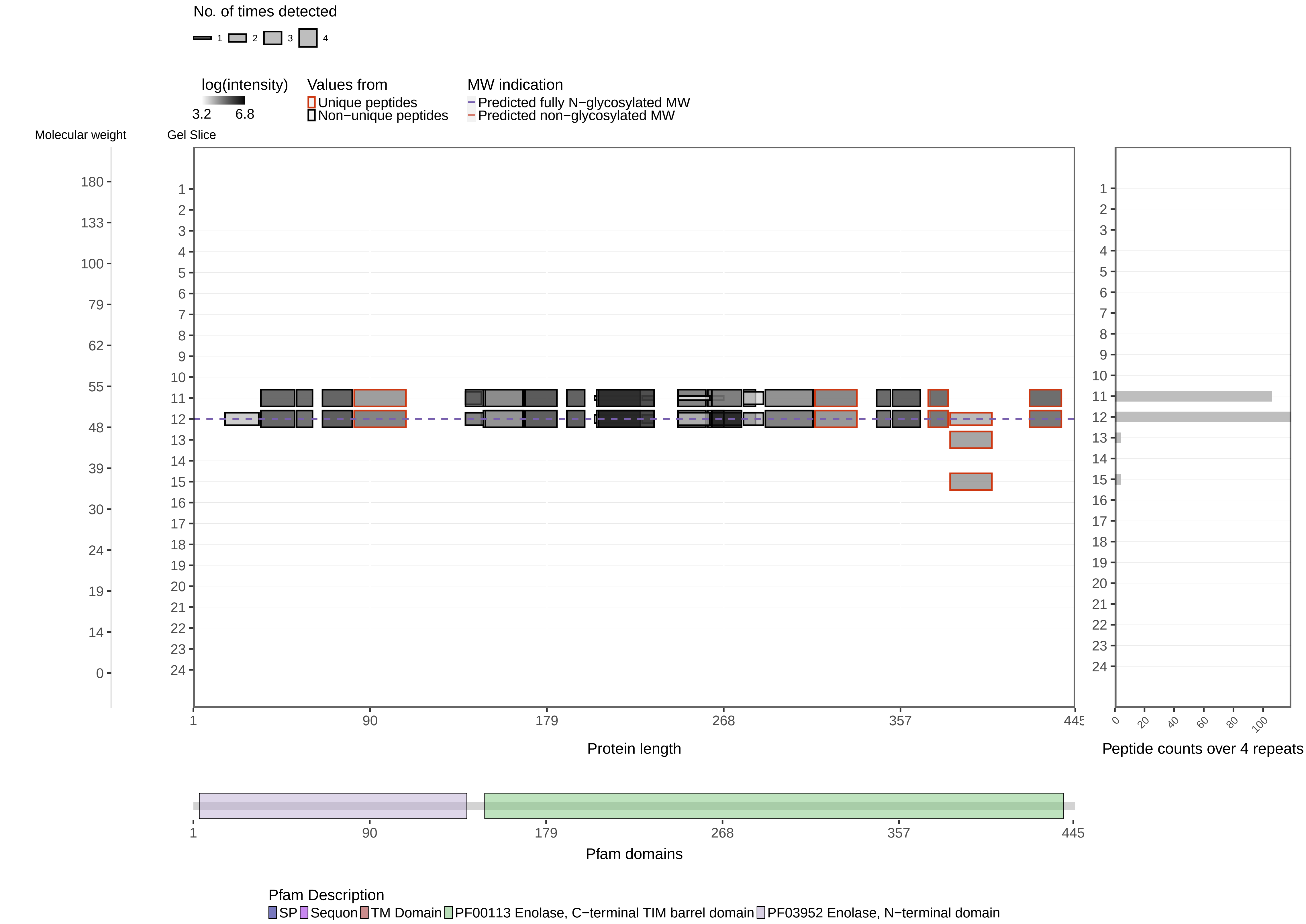


Protein sequence:

ATNVGDEGGFAPNIQENKEGLELLKTAIEKAGYTGKVVIGMDVAASEFFLKDKSYDLNFKEENNDGSQKISGDQLKDLYKTFVSEYPIVSIEDPFDQDDWETYAKLTTEMGEQVQIVGDDLLVTNPKRVAKAIAEKTCNALLLKVNQIGSVTESIEAVKMSKRAGWGVMTSHRSGETEDTFIADLAVGLSTGQIKTGAPCRSERLAKYNQLLRIEEELGSDAVYAGASFRKPVEPY*

NbL15g05910.1 14−alpha−glucan−branching_enzyme_1_chloroplastic/amyloplastic−like [MW = 98.76 kDa]

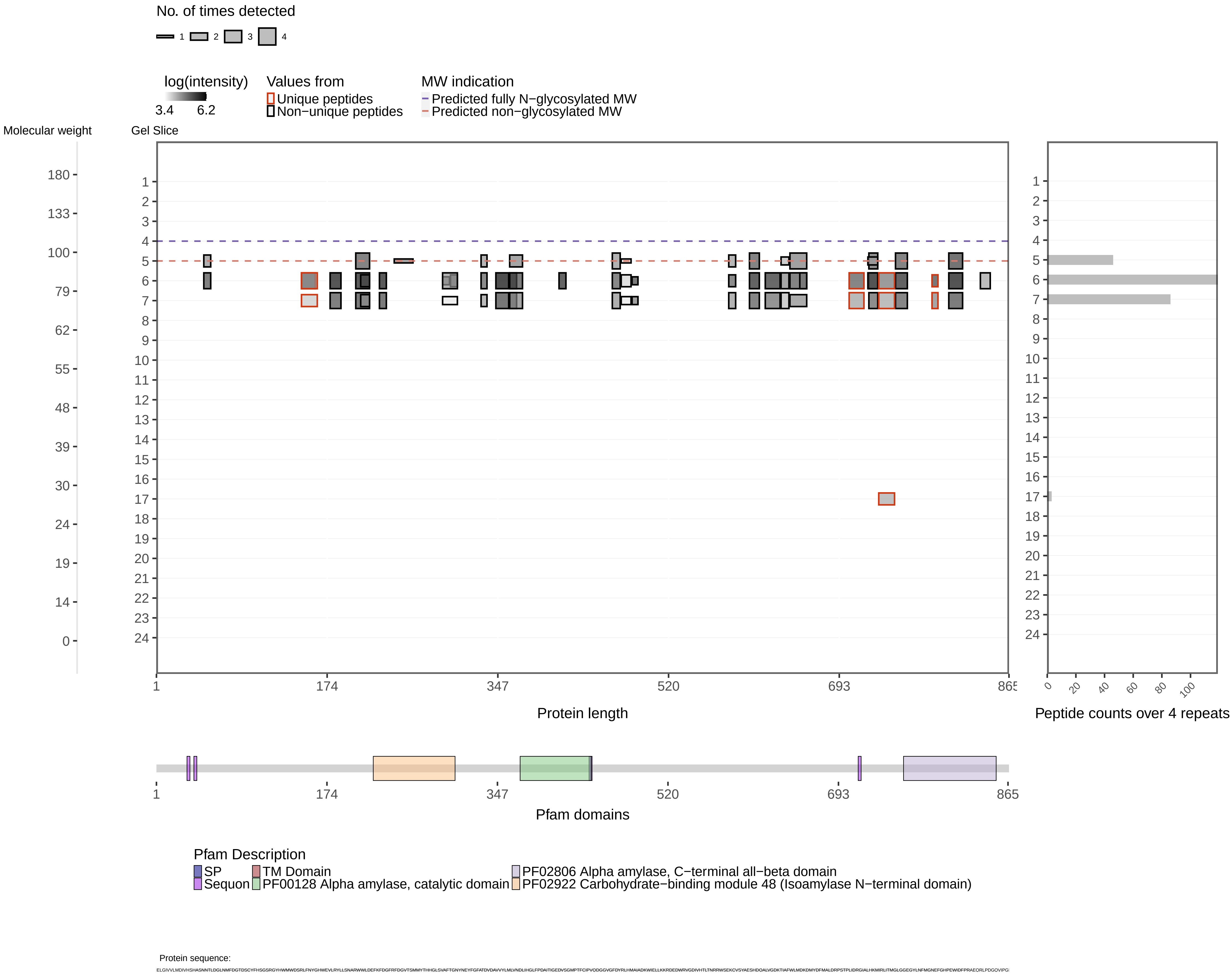

### NbL15g05890.1 probable_DNA_double−strand_break_repair_Rad50_ATPase [MW = 19.8 kDa]

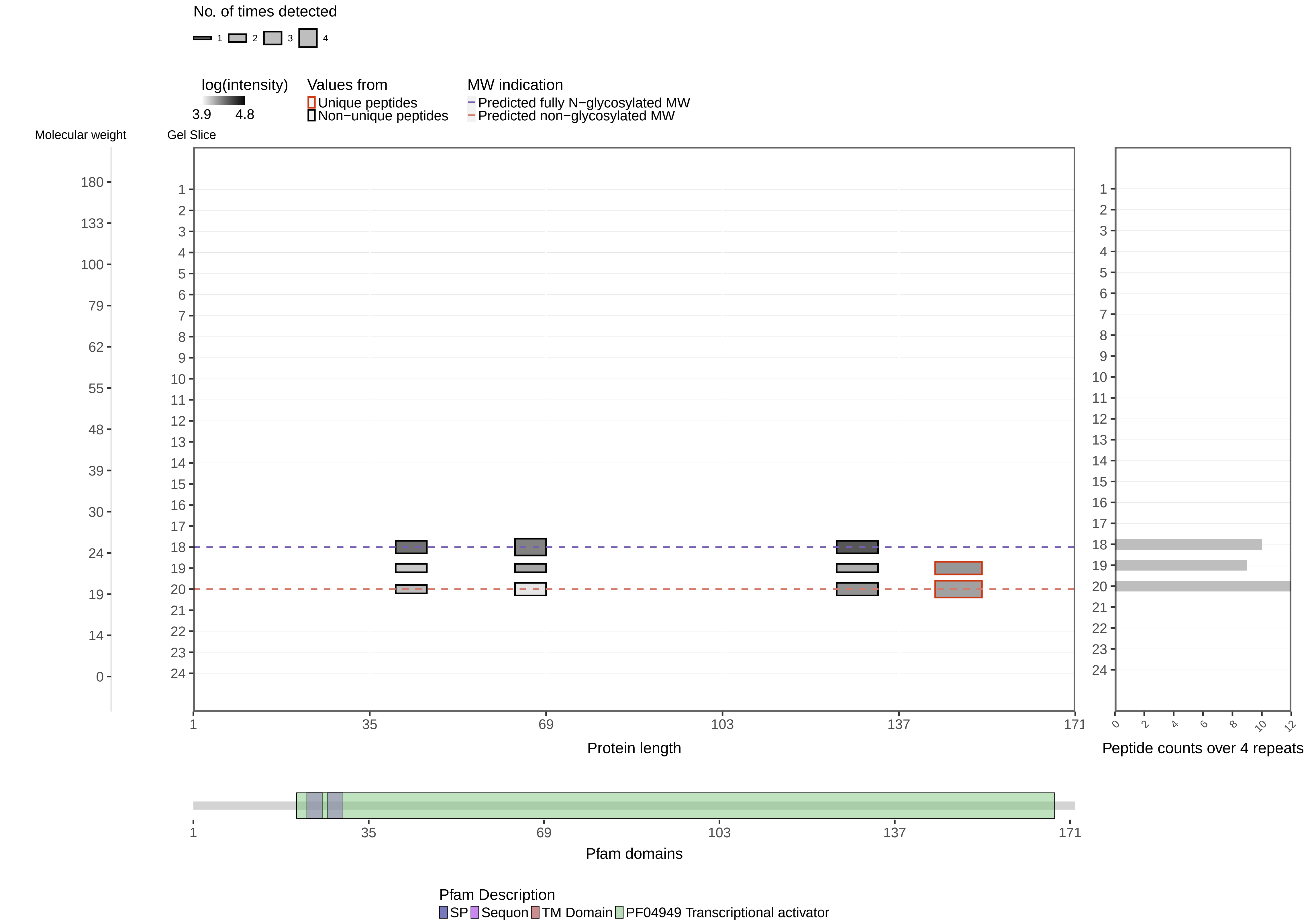


Protein sequence:
EETRRLAIIREELEALADPMKKEVSTVRKRIDAVNKELKPLGQTCQKKEREYKEALEAFNEKHKEKVLLISRLMELVGESEKMRMKKLEELSKSIETIQ*

NbL15g05850.1 fructose−bisphosphate_aldolase_cytoplasmic_isozyme [MW = 38.55 kDa]

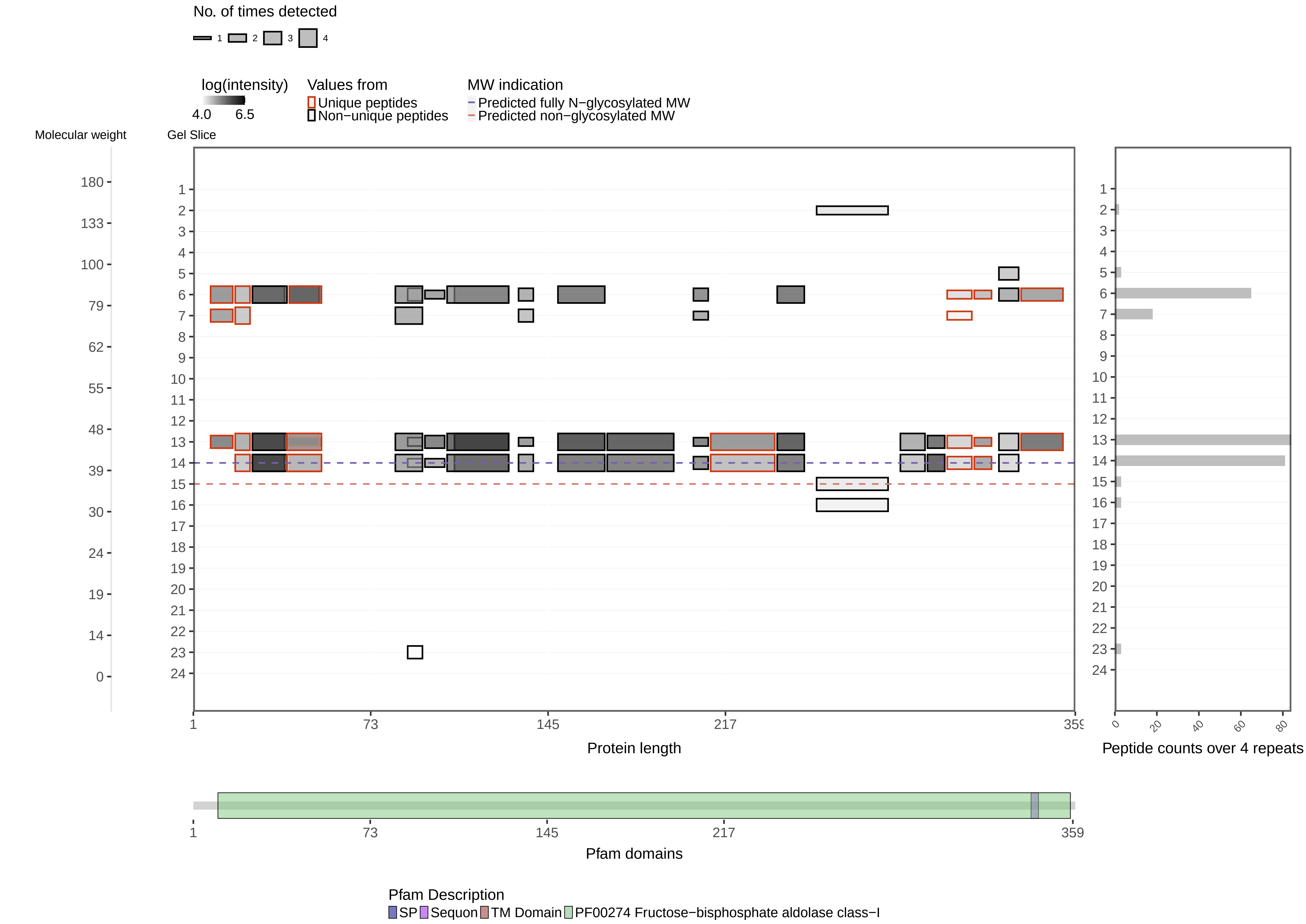


Protein sequence:

ARYAIICOONGLVPIVEPEILVDGSHDINKCADVTERVLAACYKALNDHHVLLEGTLLKPNMVTPGSEAAKVAPEVIAEYTIRALORTMPAAVPAVVFLSGGOSEEEATVNLNAMNKLOTKKPWSLSFSFGRALOOSTLKAWAGKEENVOKAOAAFLTRCKANSEATLGKYAGATNLSDGASESLHVKDYKY*

### NbL15g05680.1 ADP−ribosylation_factor [MW = 22.9 kDa]

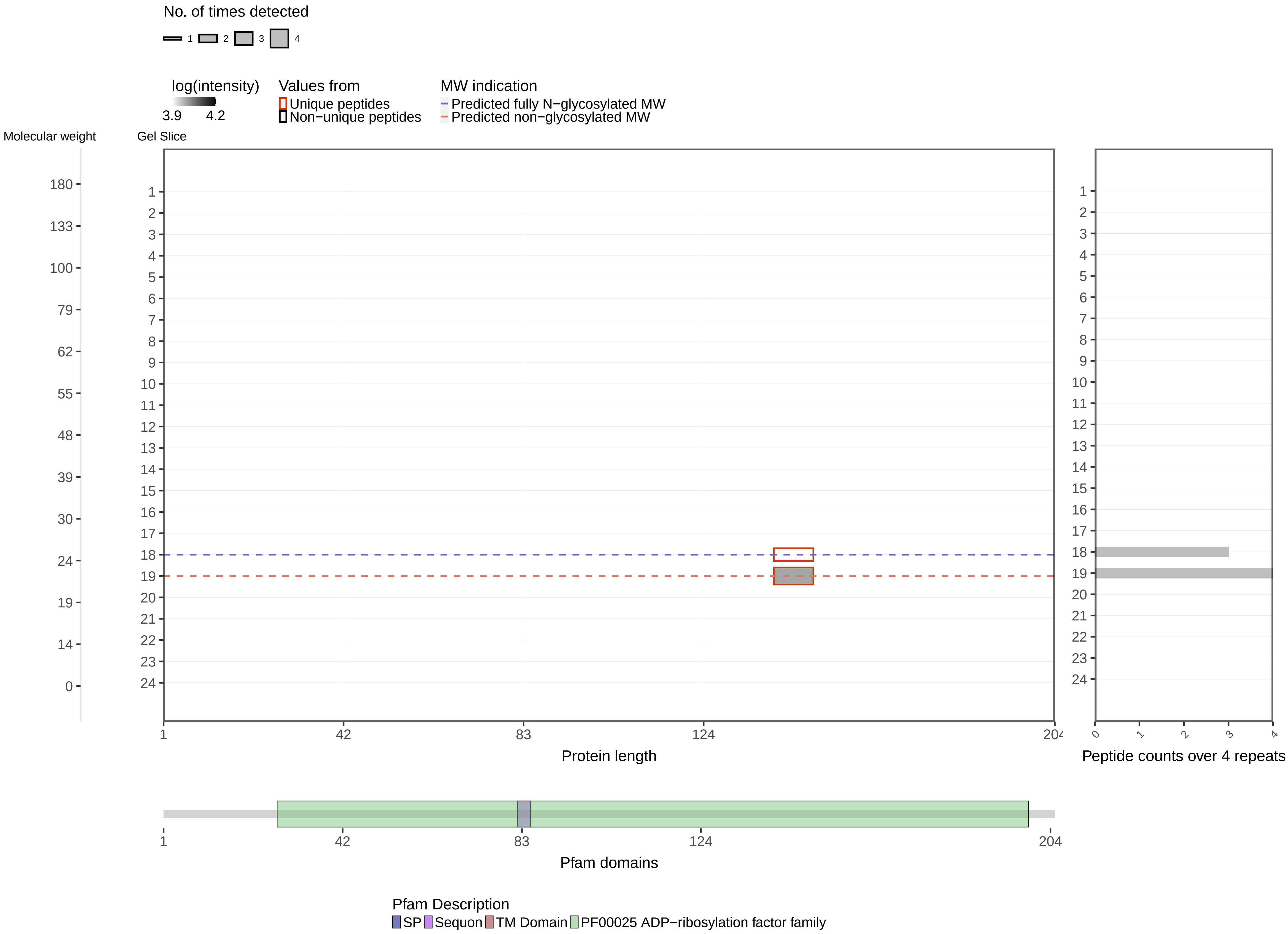


Protein sequence:
GQDKIRPLWRHYFQNTQGLIFVVDSNDRDRVVEARDELHRMLNEDELRDAVLLLFANKQDLPNAMNAAEITDKLGLHSLRQRHWYIQSTCATSGEGLYEGLDWLSNNIASKA*

NbL15g05480.1 probable_pectate_lyase_5 [MW − SP = 41.95 kDa]

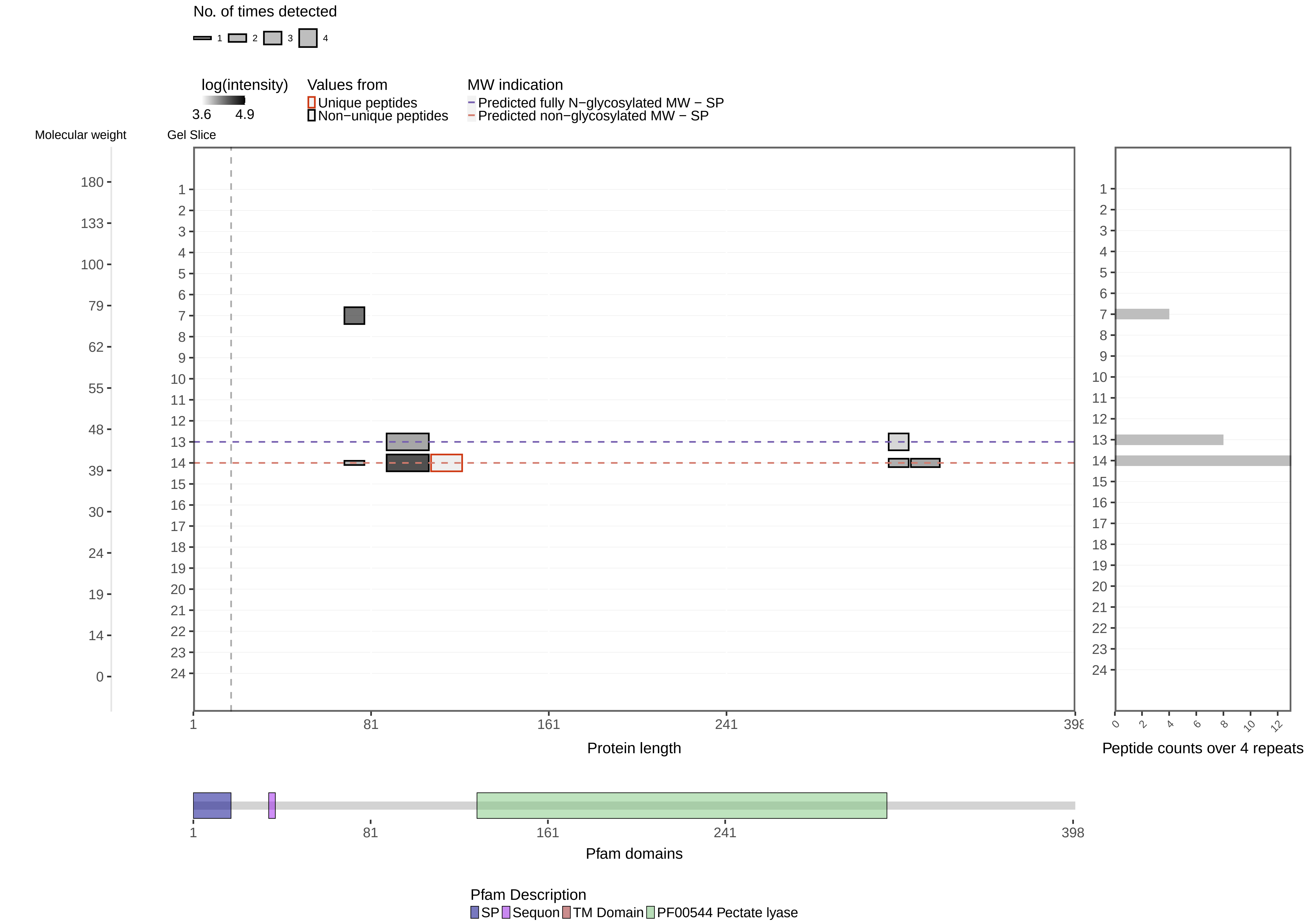


Protein sequence:
VRTLSDGDGVSIFGGSHVWVDHCSLSNCRDGLIDAIRGSTAITISNNYMTHHNKVMLLGHSDSFTRDKNMQVTIAFNHFGEGLVQRMPRCRHGYFHVVNNDYTHWEMYAVGGSASPTINSQGNRFLAPNDIFNKEVTKHEDAAESEWKKWNWRSEGDLMLNGAFFTRSGAGASKSYAKASSLSARPSTLVSSITLNAGALGCKKAKRC*

### NbL15g05280.1 glyceraldehyde−3−phosphate_dehydrogenase_cytosolic [MW = 37.02 kDa]

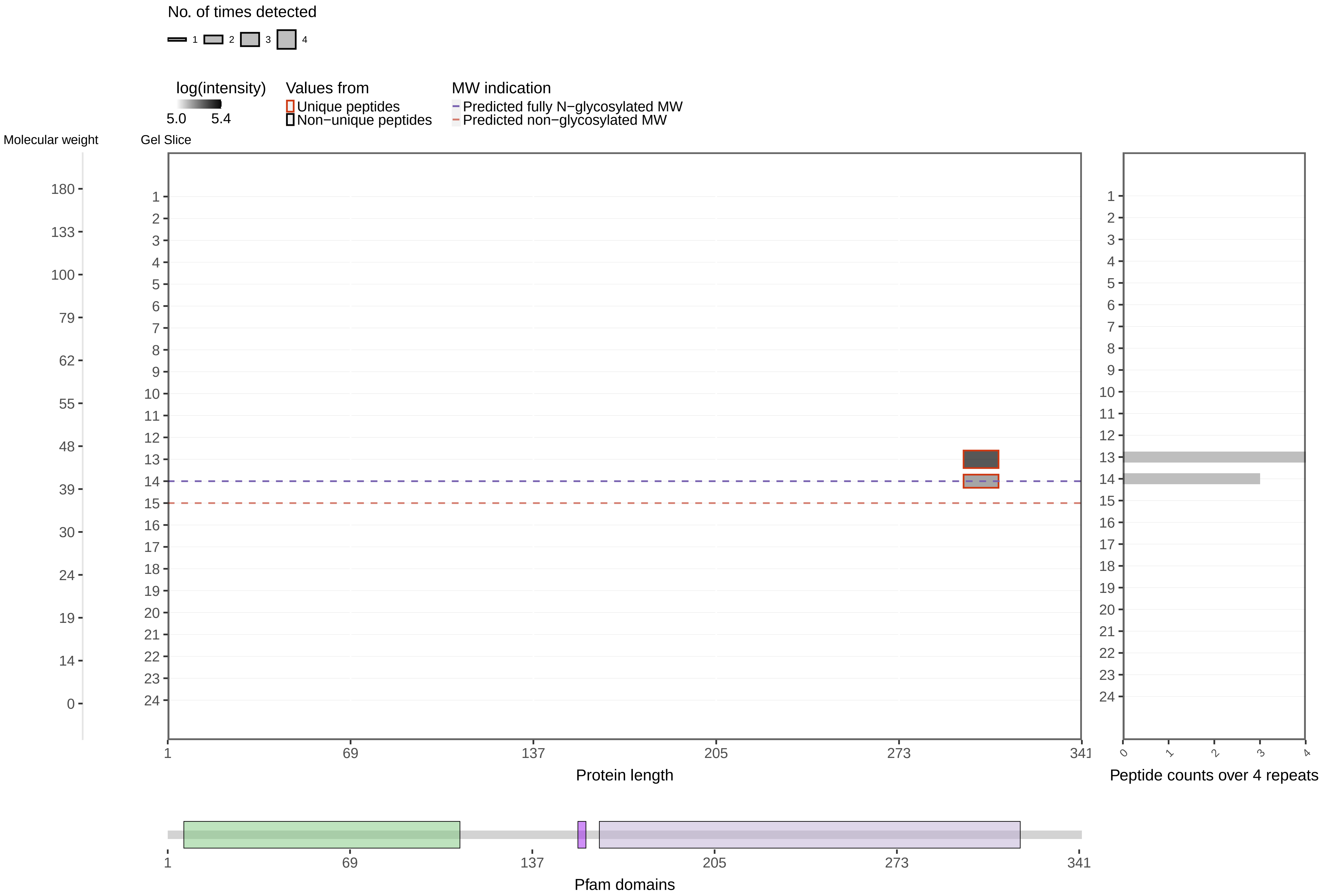


Pfam Description

SP Sequon TM Domain PF00044 Glyceraldehyde 3−phosphate dehydrogenase, NAD binding domain PF02800 Glyceraldehyde 3−phosphate dehydrogenase, C−terminal domain

Protein sequence:

CTTNCLAPLAKVINDRFGIVEGLMTTVHSLTATQKTVDGPSMKDWRGGRAASFNIIPSSTGAAKAVGKVLPALNGKLTGMAFRVPTVDVSVVDLTVRLEKEASYDEIKAAIKEESEGKLKGILGFTEDDVVSTDFVGDSRSSIFDAQAGIALSKNFVKLVSWYDNEWGYSSRVIDLICHMASVA*

### NbL15g05130.1 urease [MW = 89.74 kDa]

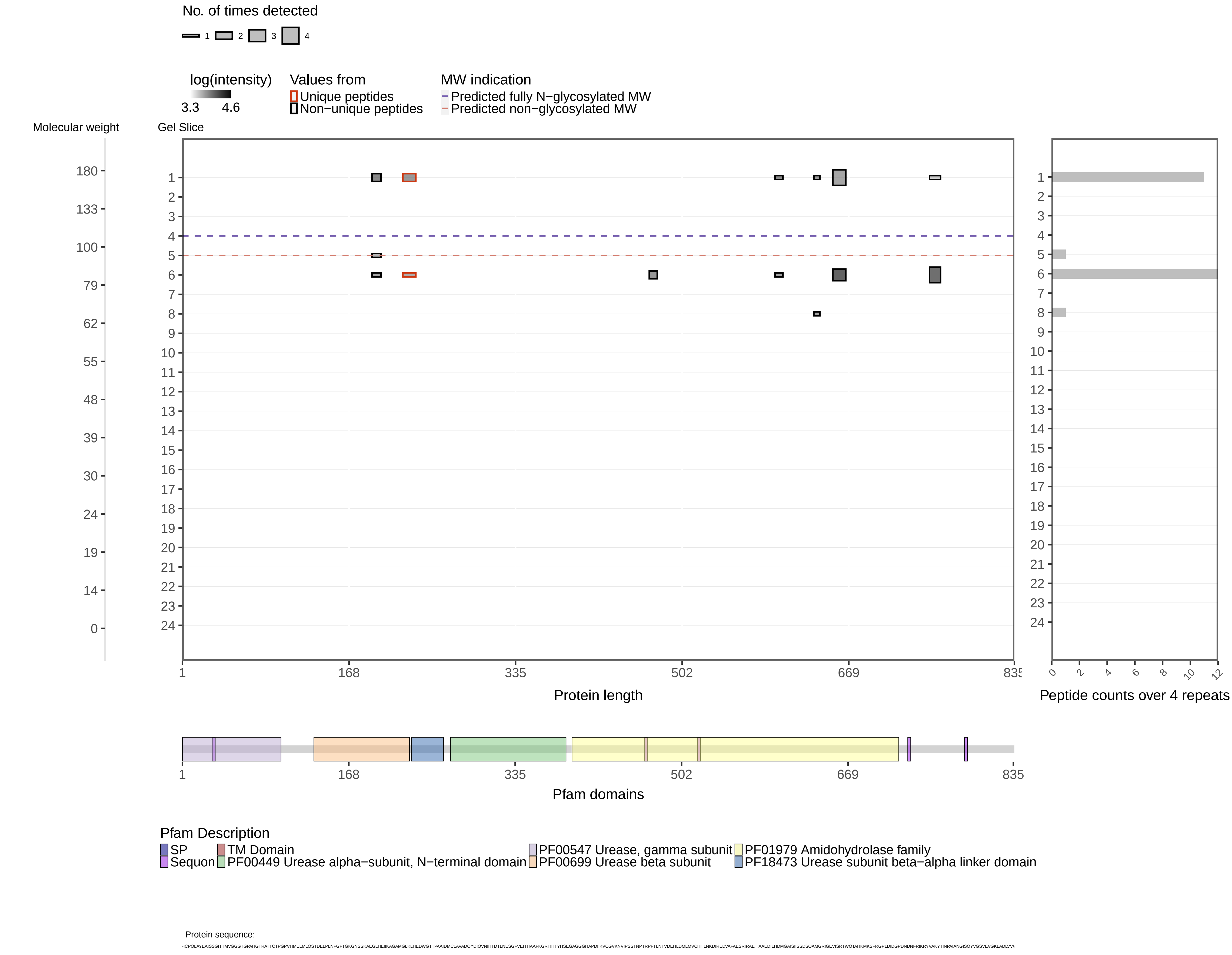

### NbL15g05110.1 adenylate_kinase_4 [MW = 26.51 kDa]

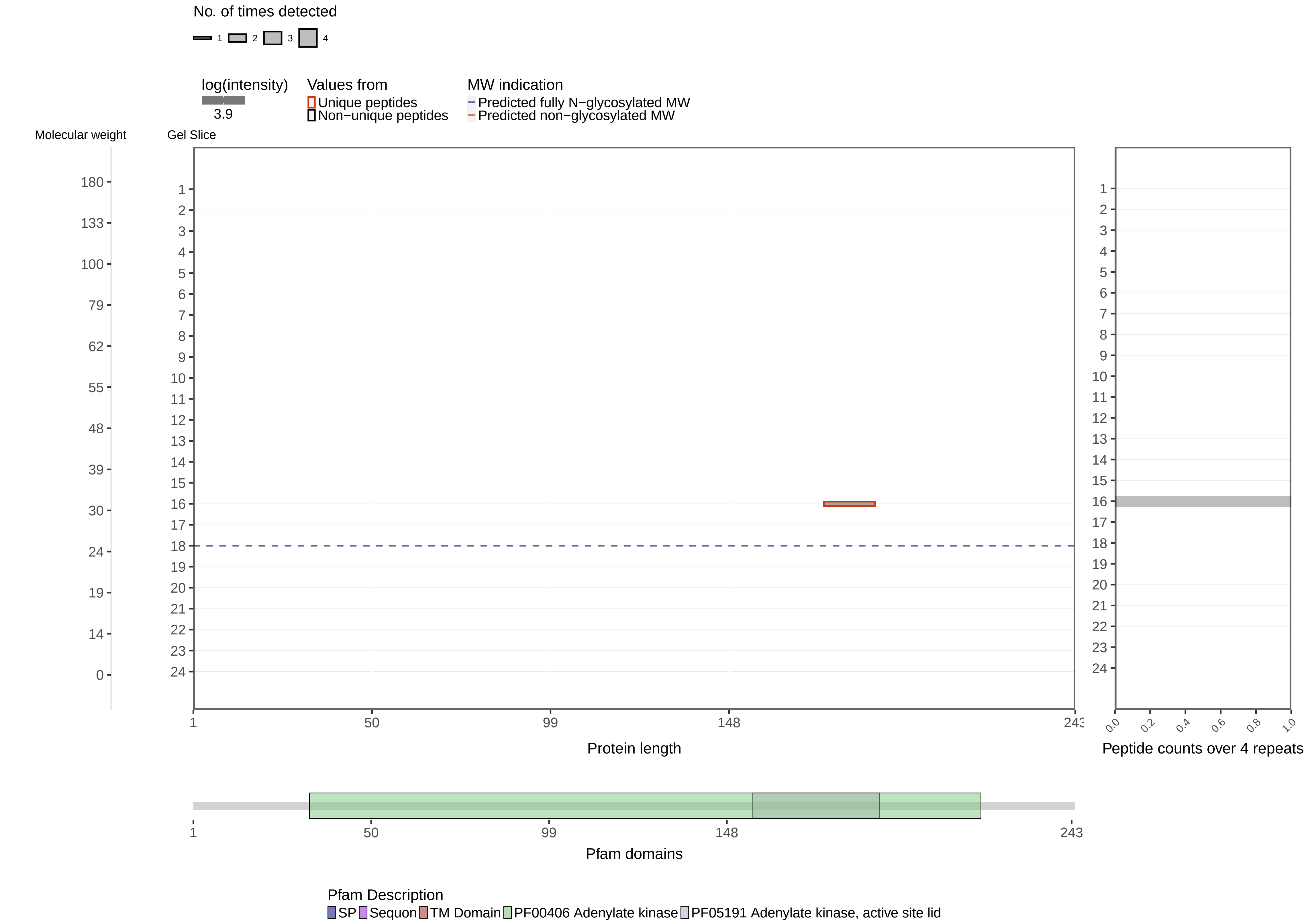


Protein sequence:

SFILDGFPRTVAQAEKLDEMLQKQGTKIDKVLNFAIDDAILEERITGRWIHPSSGRSYHTKFQPPKVPGVDDITGEPLIQRKDDTAEVLKSRLDAFHRQTEPVINYYSKKGVVASLHAEKPPKQVTSEVKNVLSS*

### NbL15g05100.1 probable_pre−mRNA−splicing_factor_ATP−dependent_RNA_helicase_DEAH4 [MW = 16.11 kDa]

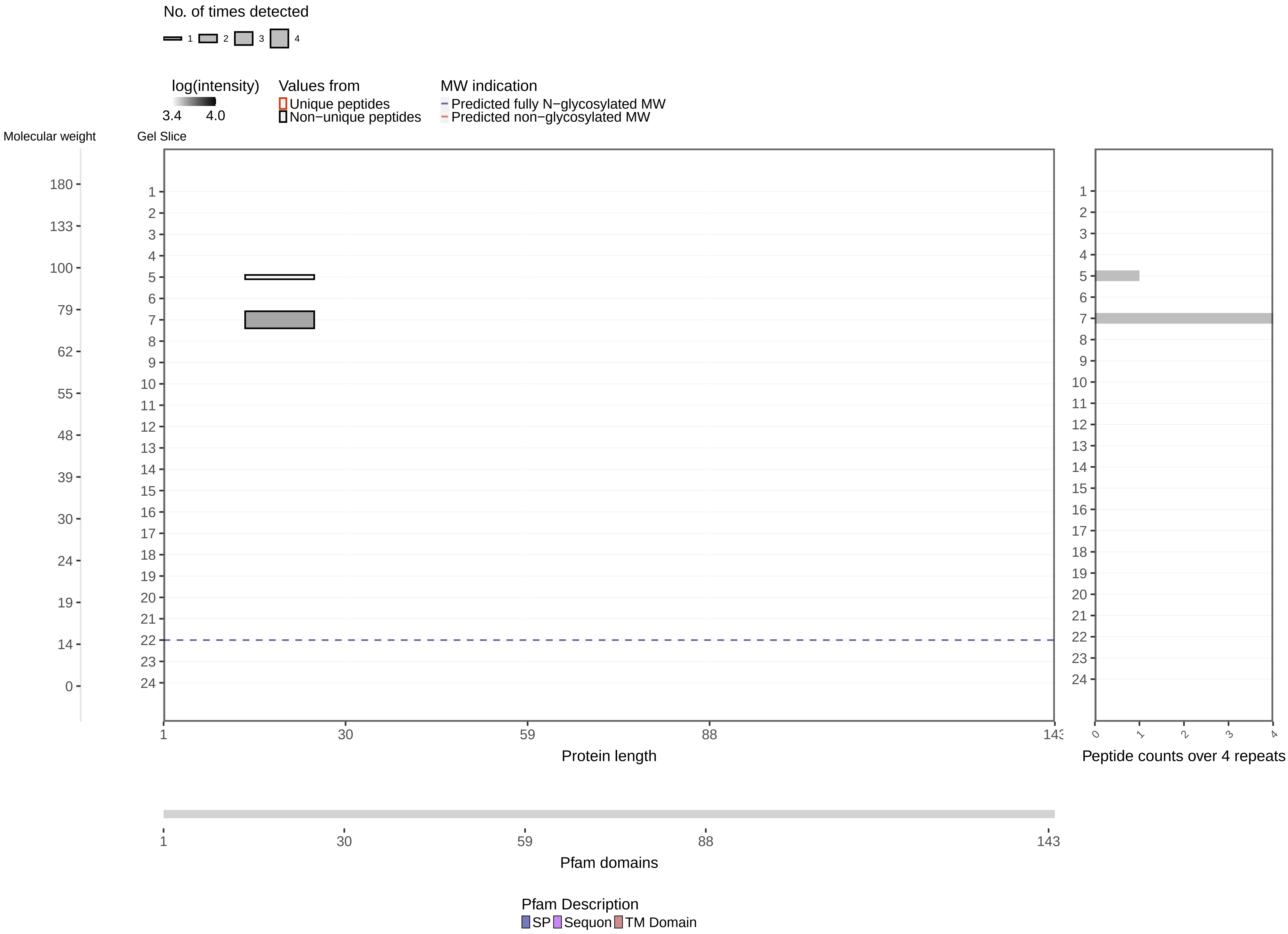


Protein sequence:

EASLKTAIDIHVREPEGDILIFLTGQVRVFSPPPPNCRRFTVSTNIAETSLTVDGVVWRLFNCKPSFQKERVAPPNVINPRGGKK

NbL15g05080.1 NADH_dehydrogenase_[ubiquinone]_flavoprotein_1_mitochondrial−like [MW = 55.46 kDa]

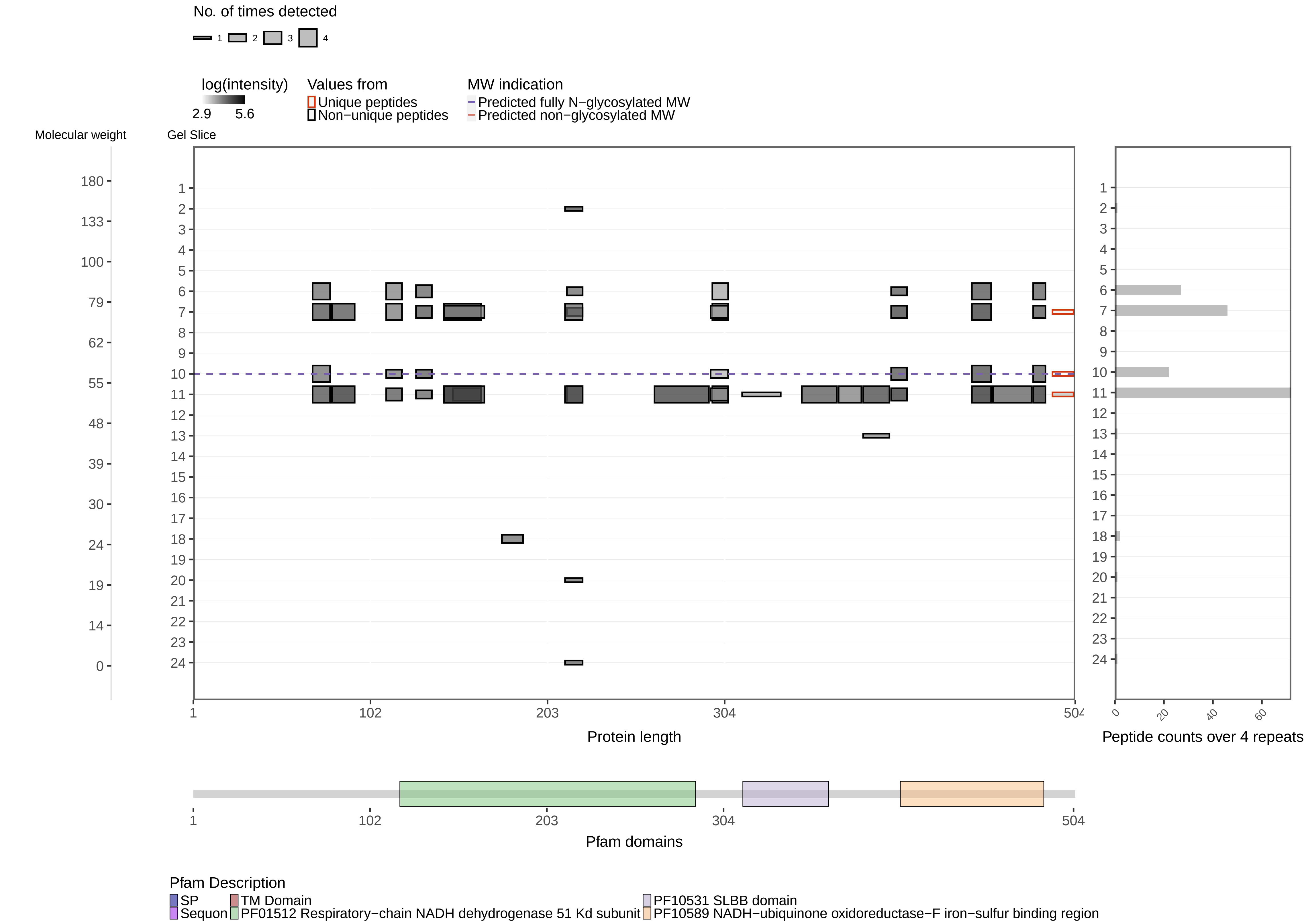


Protein sequence:

GAGAYICGEETALLESLEGKQGKPRLKPPFPANAGLYGCPTTVTNVETVAVSPTILRRGPEWFASFGRKNNAGTKLFCISGHVNKPCTVEEEMSIPLKELIERHCGGVRGGWDNLLAVIPGGSSVPLIPKNICEDVLMDFDALKAVQSGLGTAAVIVMDKSTDIVDAIARLSYFYKHESCGQCTPCREGTGWLWMIMERMKVGNAKLEEIDMLQEVTKQIEGHTICALGDAAAWPVQGLIRHFRPELERRIREHADRELQQAAAA*

### NbL15g04880.1 non−specific_lipid_transfer_protein_GPI−anchored_1 [MW − SP = 18.17 kDa]

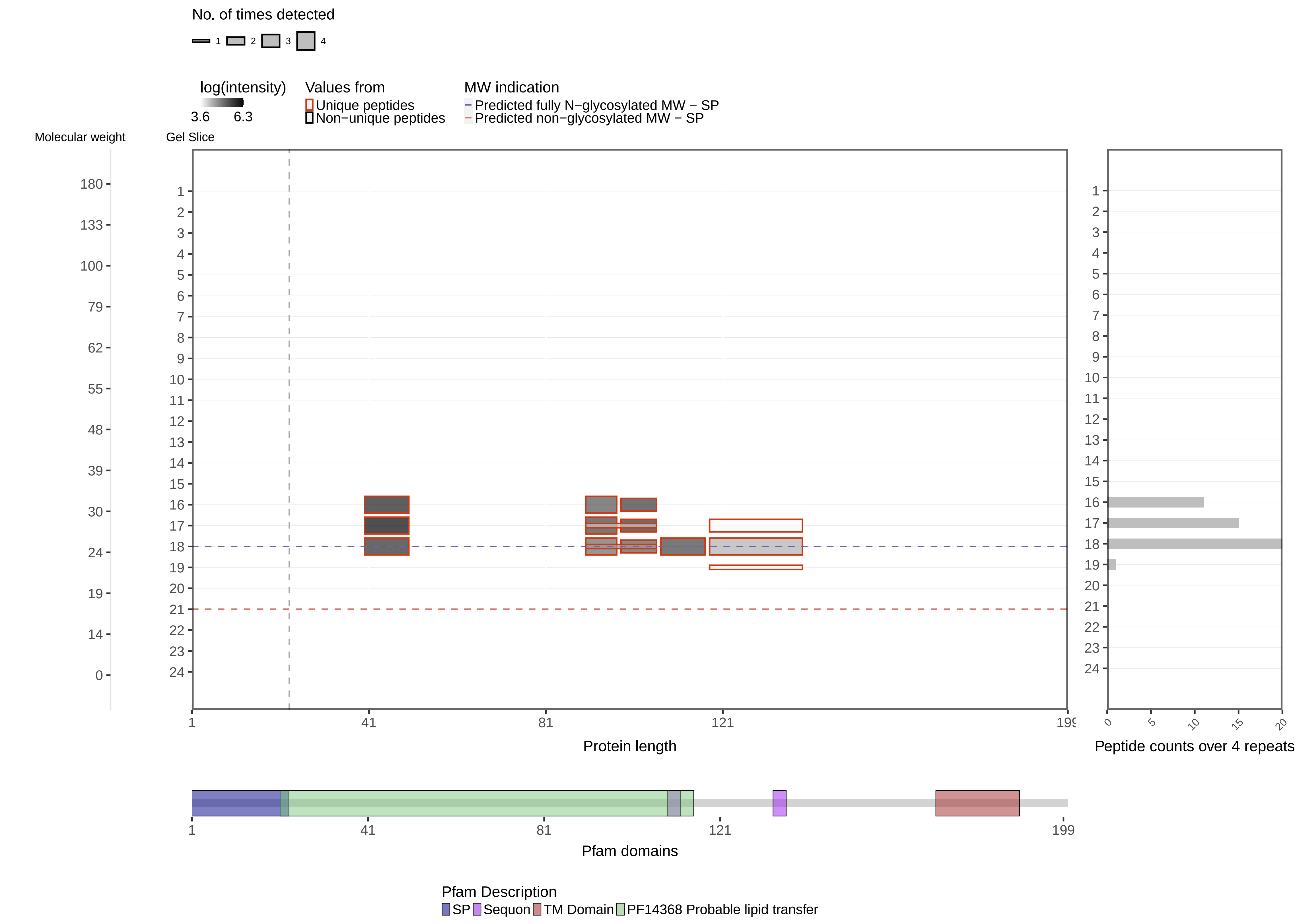


Protein sequence:

SPELKSMGIQEDKLLQLPSACKLTNASISNCPKLLNIPSTSPDYAIFTNASTASKTPAATPSGASSSSSPDTTKDSSNGFKNGPQLSIFGTIVAAAVVAVCSSVIPRRTTSVLM*

NbL15g04800.1 2−Cys_peroxiredoxin_BAS1_chloroplastic [MW = 29.9 kDa]

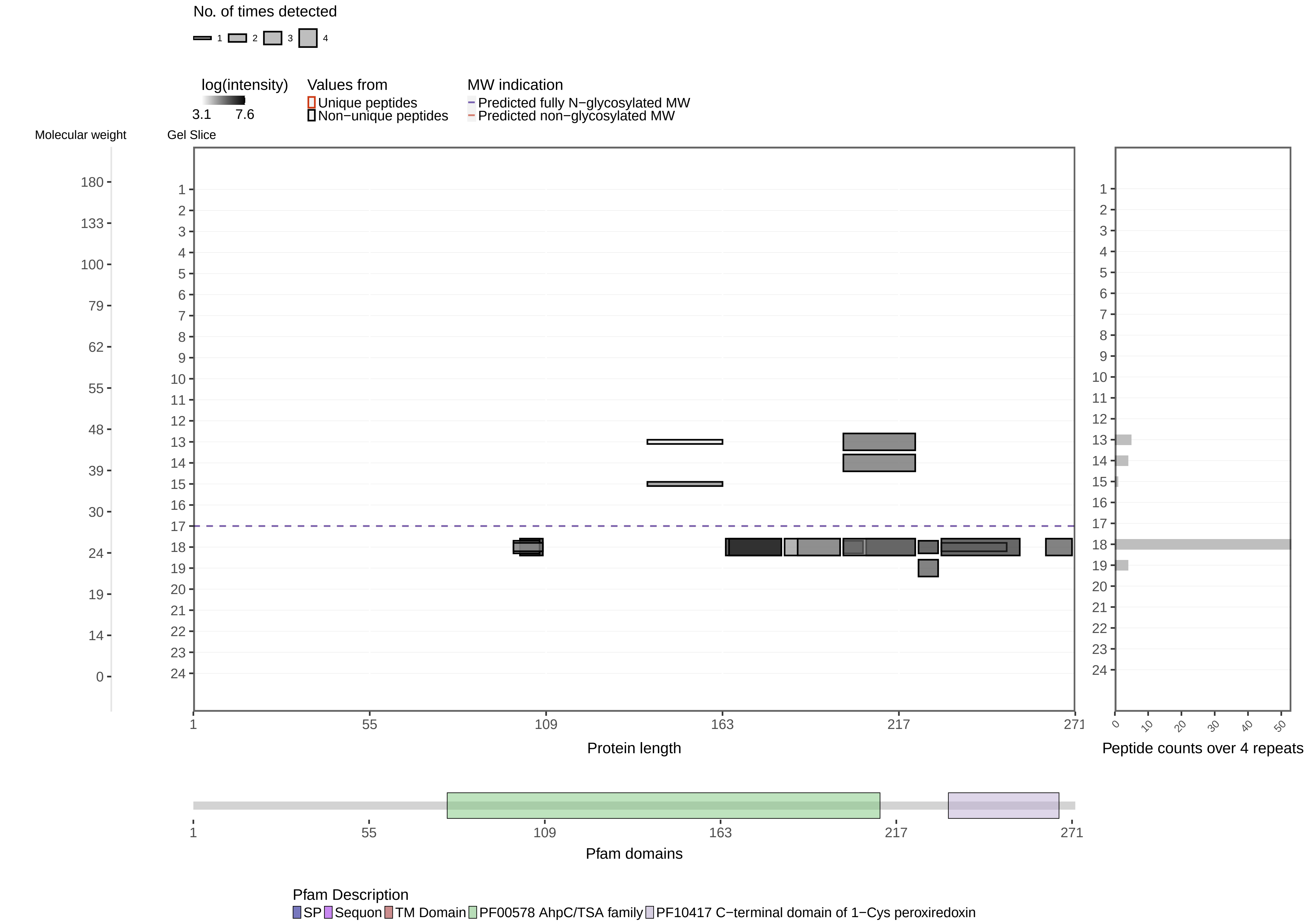


Protein sequence:

/CPTEITAFSDRYEEFEKLNTEILGVSVDSVFSHLAWVQTERKSGGLGDLNYPLISDVTKSISKSYNVLIPDQGIALRGLFIIDKEGVIQHSTINNLGIGRSVDETLRTLQALQYVQDNPDEVCPAGWKPGEKSMKPDPKGSKEYFASI*

NbL15g04580.1 putative_phosphatidylglycerol/phosphatidylinositol_transfer_protein_DDB_G0282179 [MW − SP = 14.7 kDa]

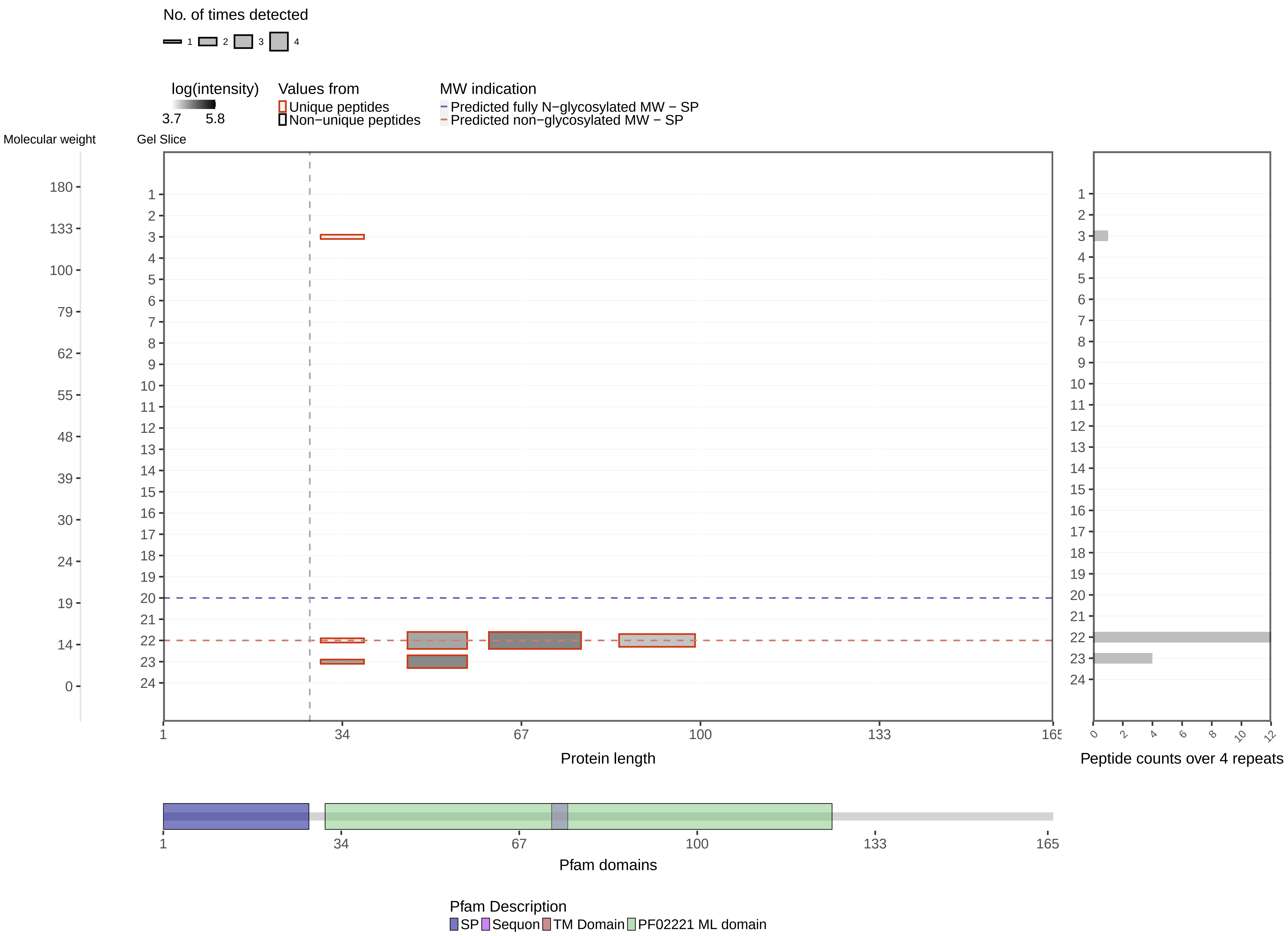


Protein sequence:
TDENISGGKLVIDVKYLFLHVHQESHDICKETSCPVSGDFVLSHSQALPGITPPVSHLSSLSSYFVMISFSGPLFCIASFIKRCSSASFLDFSDS*

NbL15g04420.1 sucrose−phosphatase_2−like [MW = 47.96 kDa]

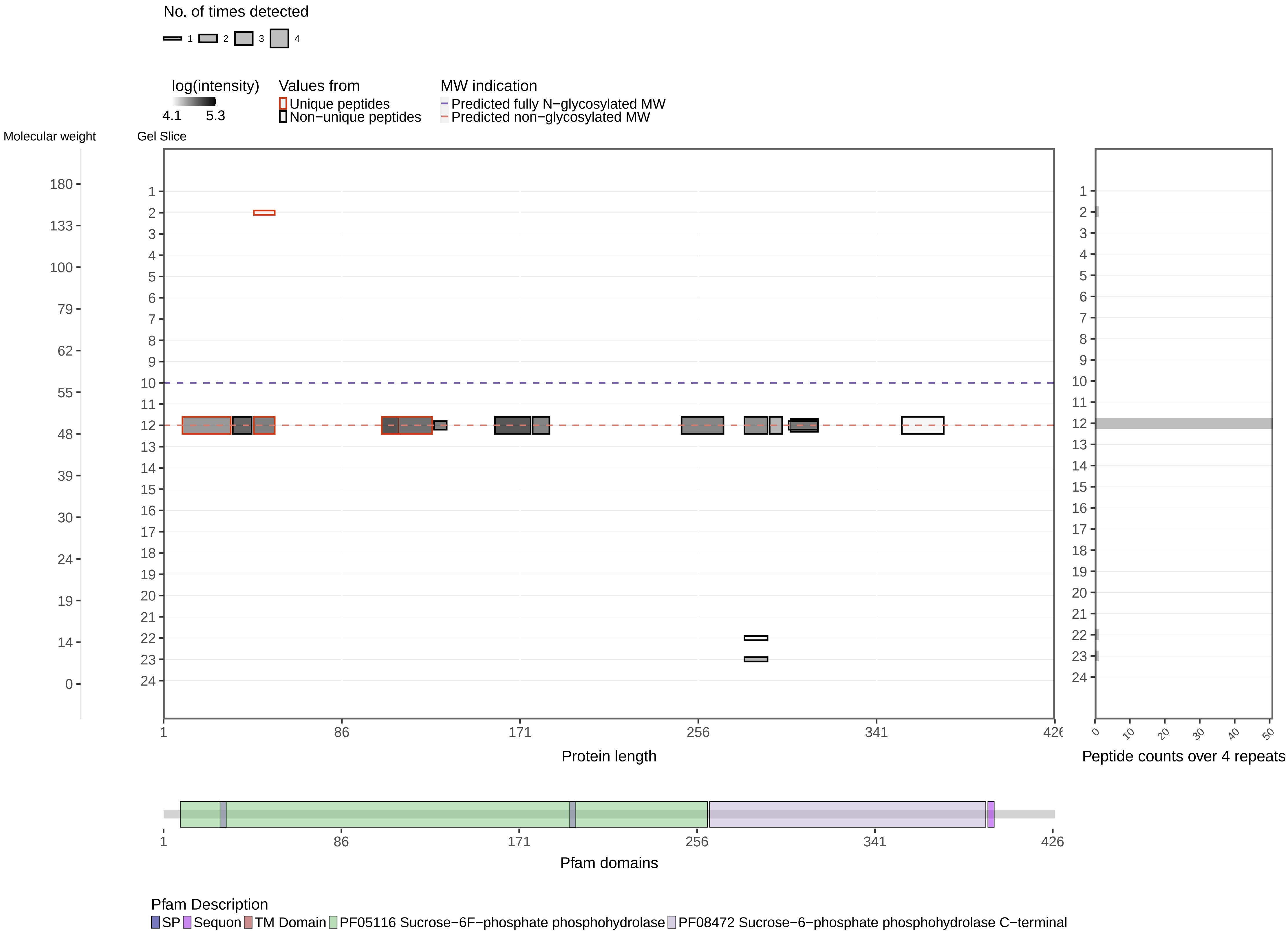


Protein sequence:

NbL15g04370.1 ubiquitin_carboxyl−terminal_hydrolase_12 [MW = 131.51 kDa]

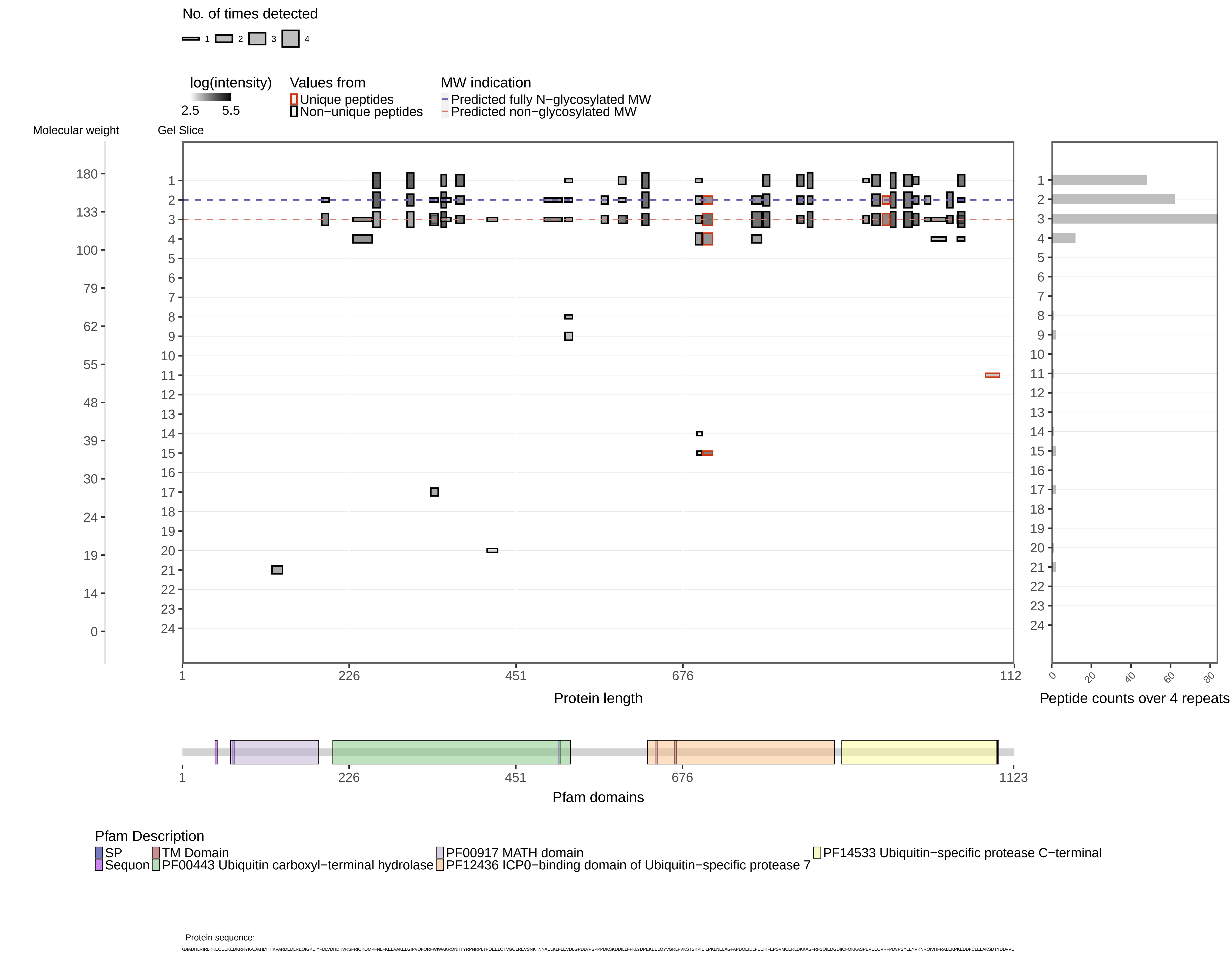

NbL15g04300.1 5−methyltetrahydropteroyltriglutamate−−homocysteine_methyltransferase−like [MW = 84.52 kDa]

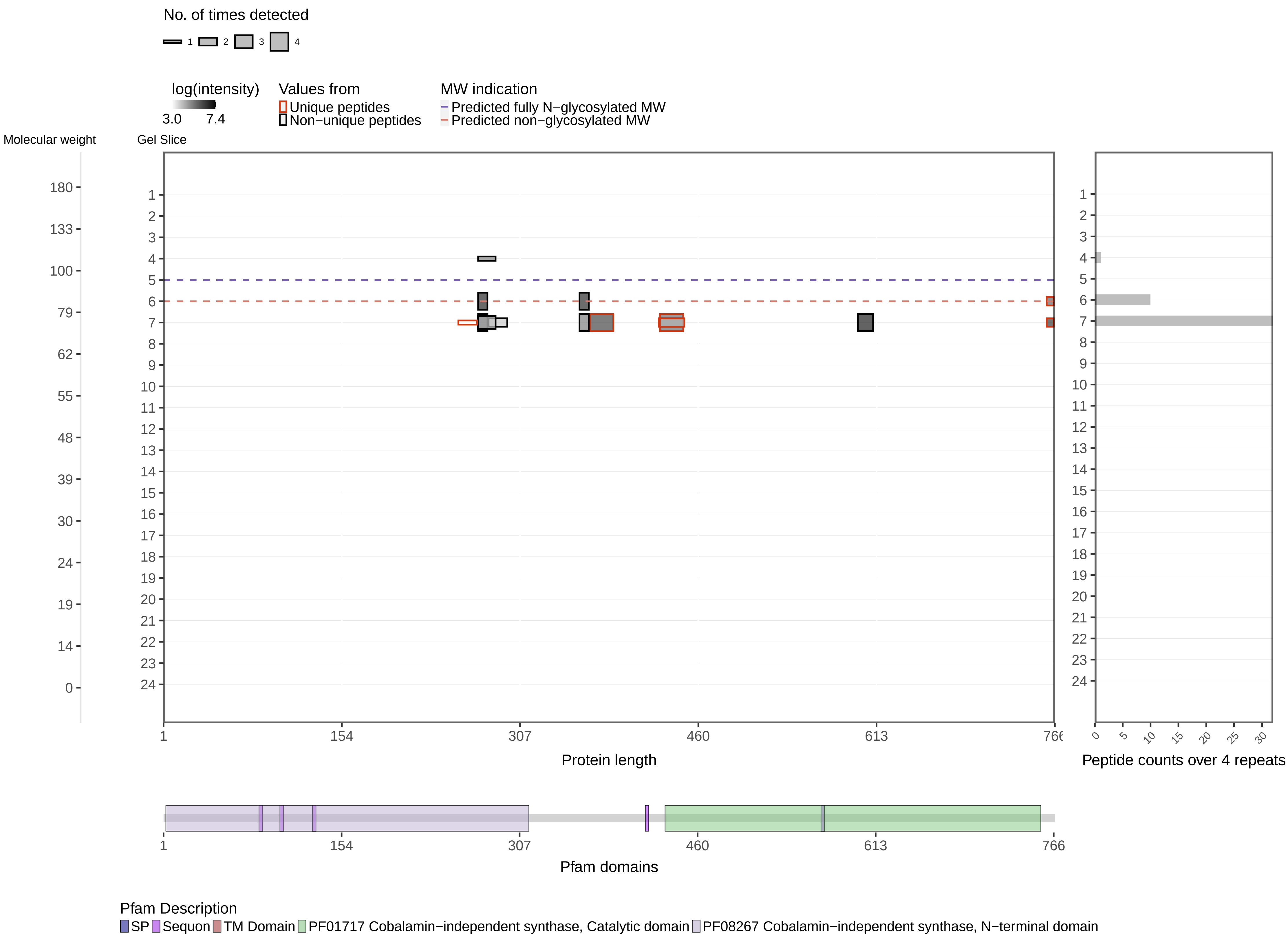


Protein sequence:

GAKDEAFFSANAAAQASRKSSPRVTNEAVQKASAALQGSDHRRATNVSARLDAQQKKLNLPILPTTTIGSFPQTVELRRVRREYKAKKISEEEYVKAIKEEIKKVVDLQEELDIDVLVHGEPERNDMVEYFGEQLSGFAFTANGWVQSYGSRCVKPPIIYGDVSRPKPMTVFWSSAAQSMTKRPMKGMLTGPVTILNWSFVRNDQPRFETCYQIALAIKDEVEDLEKAGITVIQIDEAALREGLPLRKSEHAFYLNWAVHSFRITNVGIQDTTQIHTHMCYSNFNDIIHSIIDMD

### NbL15g04170.1 probable_small_nuclear_ribonucleoprotein_F [MW = 12.01 kDa]

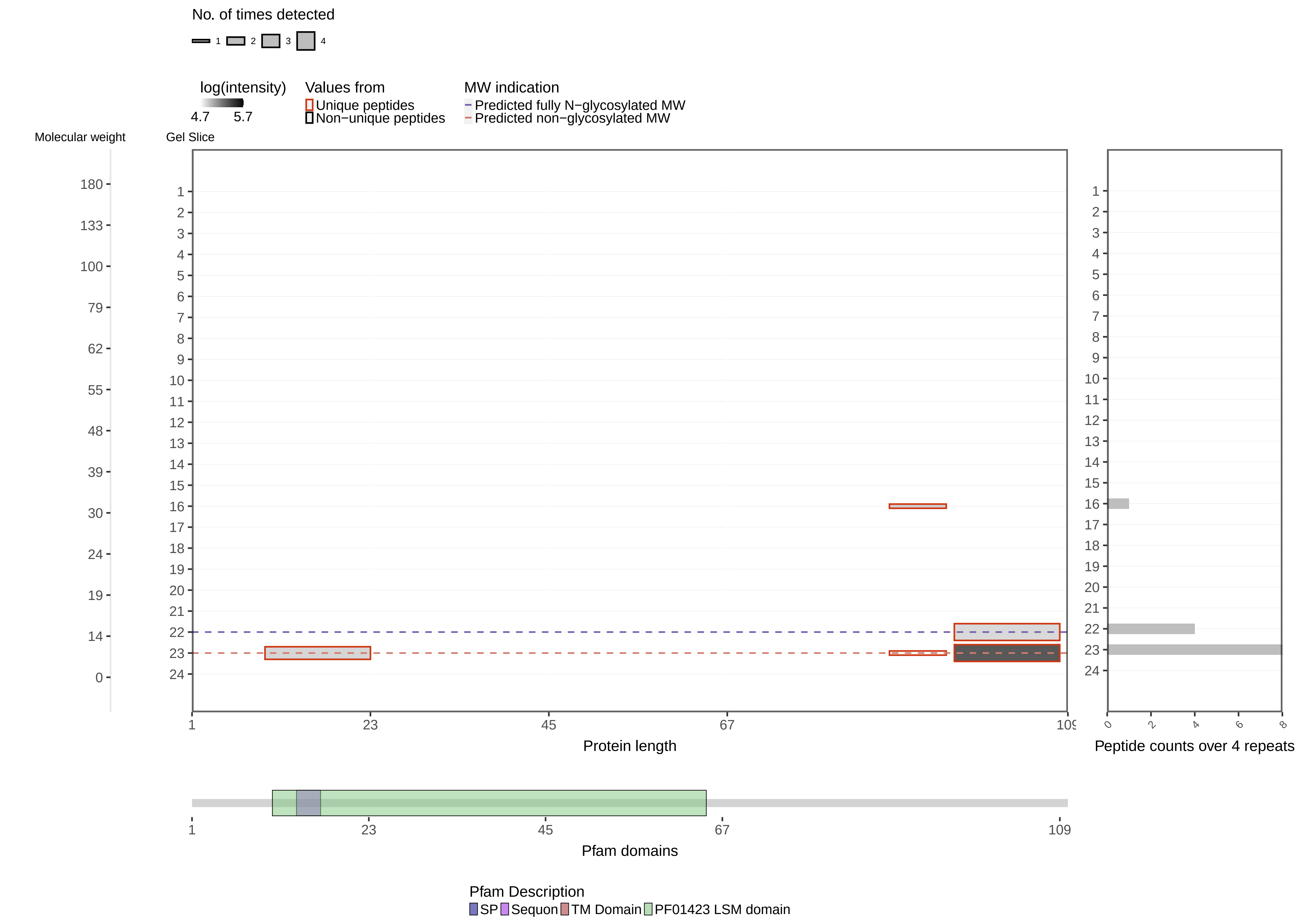


Protein sequence:

MNLQLANAEEYIDGQCTGSLGEILISSKATSIFFLSRLAPFSACIRCNNVLYLRGVPEDEELDDADRD*

### NbL15g03880.1 40S_ribosomal_protein_S16−like [MW = 16.75 kDa]

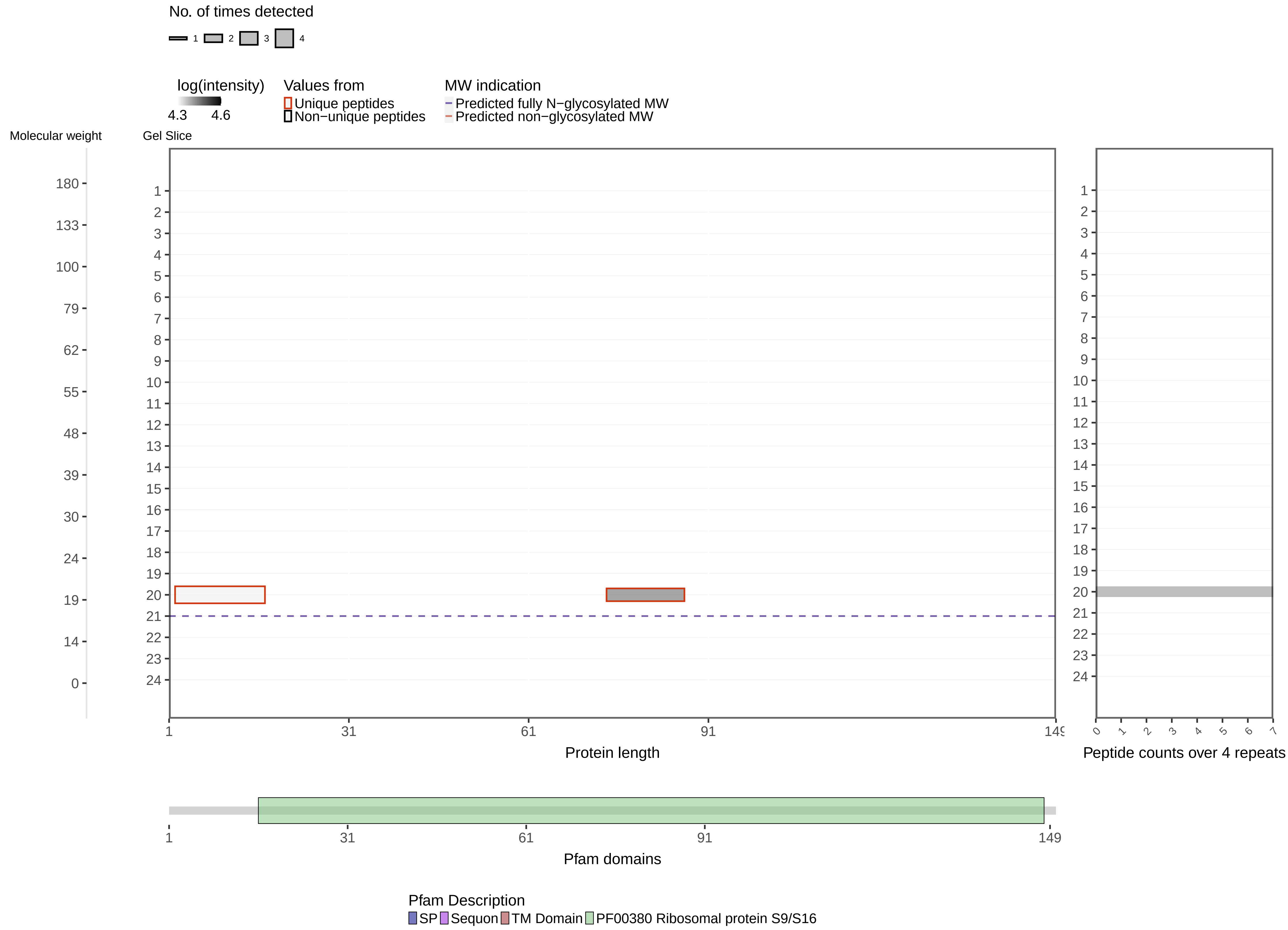


Protein sequence:

HRFAGVDMRIRVNGGGHTSQIYAIRQSIAKALVAFYQKYVDEQSKKEIKDILVRYDRTLLVADPRRCEPKKFGGRGARARFQKSYR*

### NbL15g03860.1 40S_ribosomal_protein_S10−1−like [MW = 20.01 kDa]

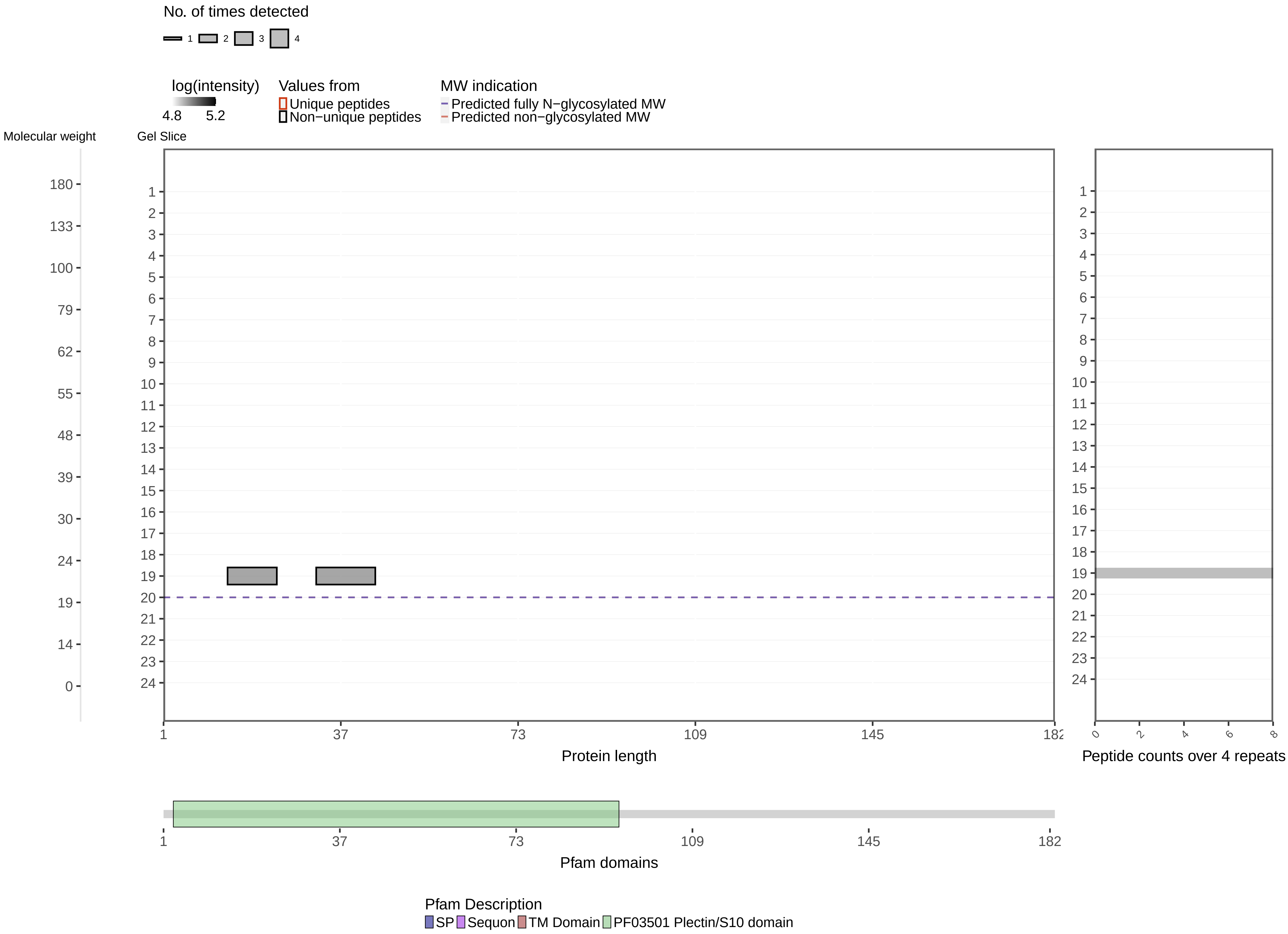


Protein sequence:

LPSEIVPATLKKSAKPLGRPMGGPPGDRPRGPPRFEGDRPRFGDREGYRAGPRGPPGEFGGEKGGAPADYQPAFRGGGGRPGFGRGAGGFGGAPPSSSFS*

### NbL15g03820.1 uncharacterized_LOC107811196 [MW = 28.71 kDa]

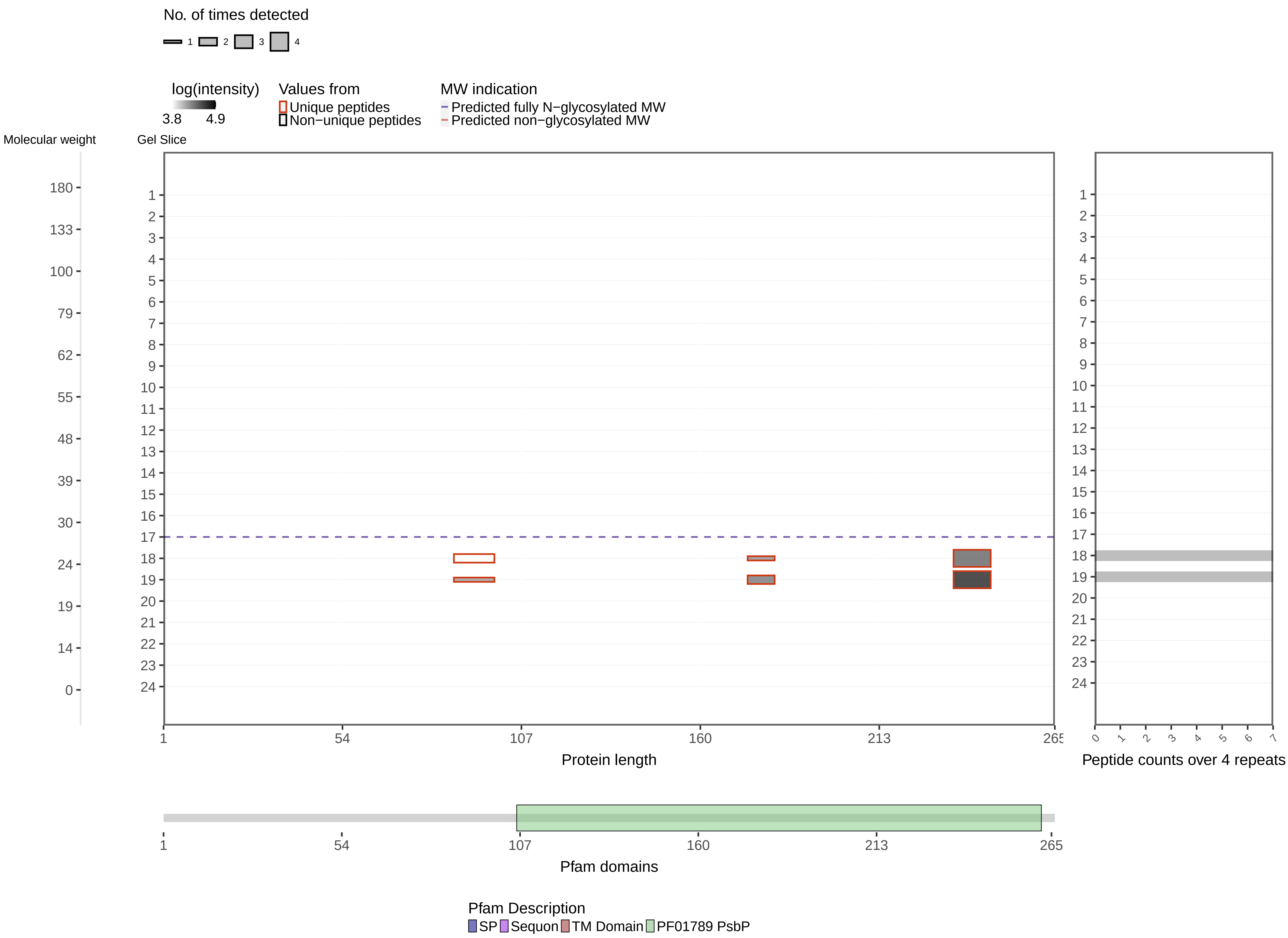


Protein sequence:

LTEPEDYNAGLSLYGDKAKPKKFAARFASSDRSEVLSVVIRPSNQLKITFLEAKDVTDLGSLKEAAKIFVPAGATLYSGRSIKIKEDEGFRTYYFYEFARDEQHVALVAAVNSGKAYIAGATAPESKWNDDGLKLRSAAVSLMVL*

### NbL15g03800.1 uncharacterized_protein_At2g34160−like [MW = 11.34 kDa]

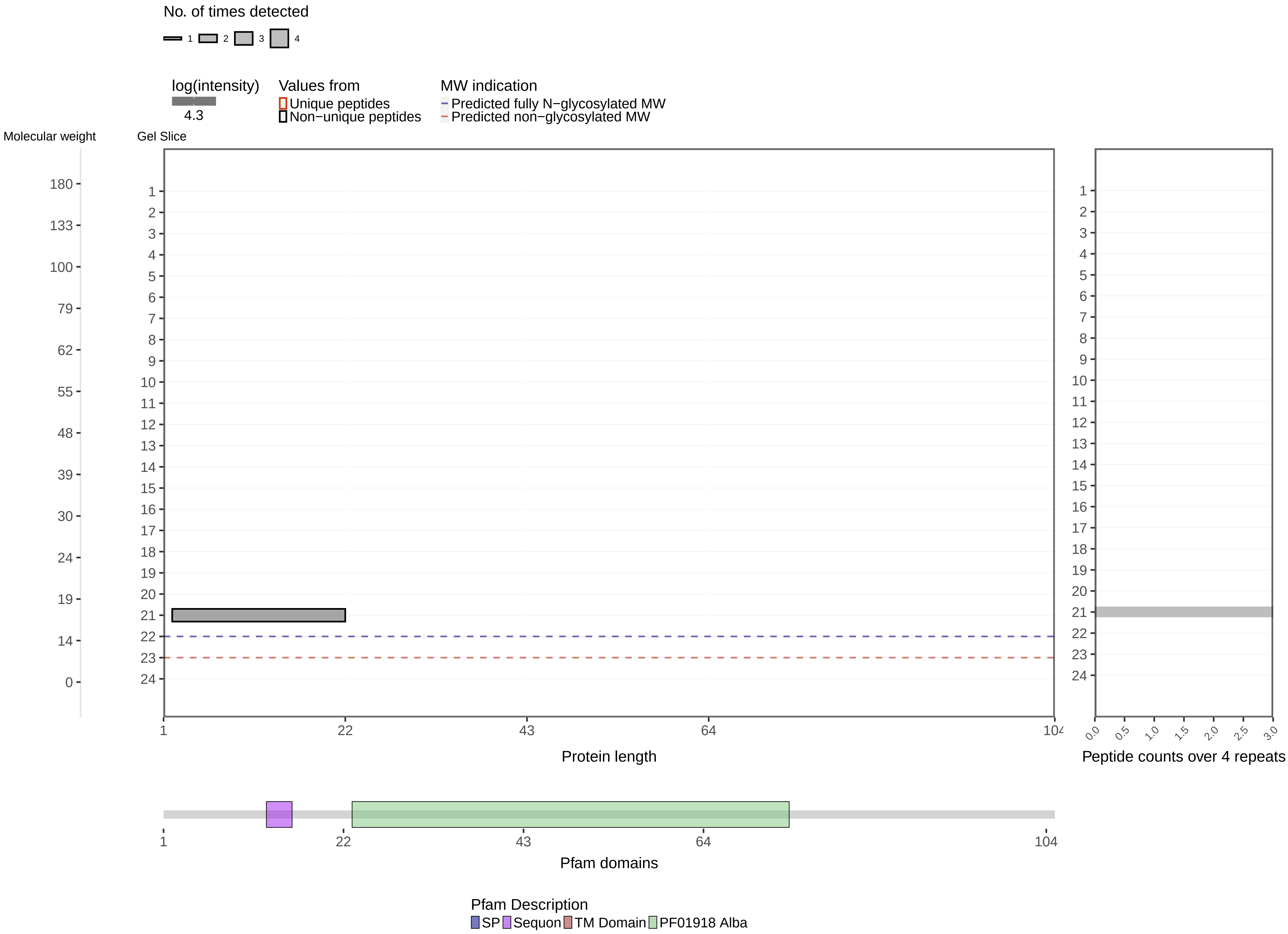


Protein sequence:

NLAKRYMQQYNDVELSALGMAIATVVSIAEILKSNGFAVEKKIMTSTVDVKDGSRGRPVSKAKLA*

### NbL15g03770.1 26S_protease_regulatory_subunit_6A_homolog [MW = 47.47 kDa]

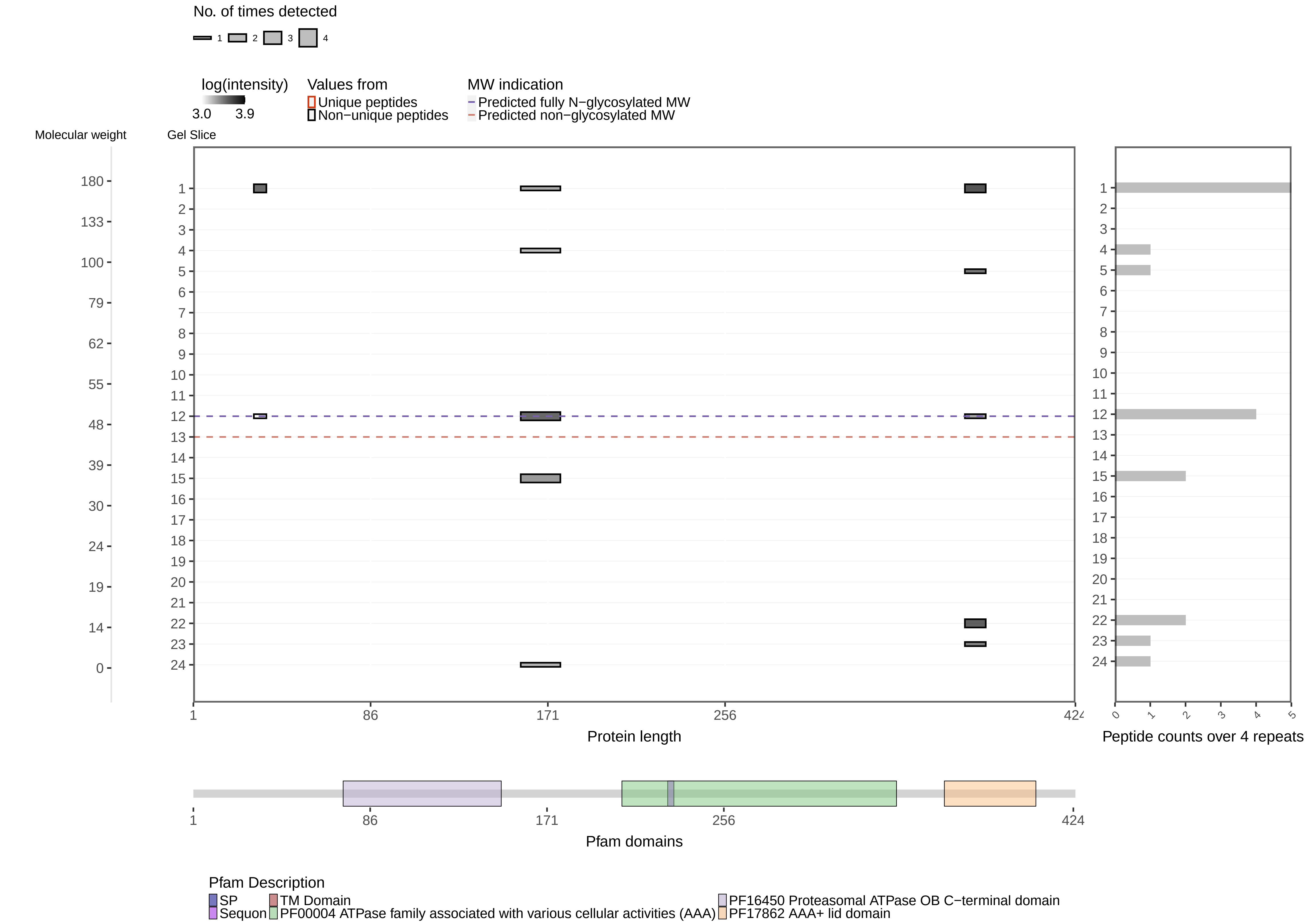


Protein sequence:

GVRPPKGVLLYGPPGTGKTLMARACAAQTNATFLKLAGPQLVQMFIGDGAKLVRDAFLLAKEKSPCIIFIDEIDAIGTKRFDSEVSGDREVQRTMLELLNQLDGFSSDDRIKVIAATNRADILDPALMRSGRLDRKIEFPHPTEEARARILQIHSRKMNVHPDVNFEELARSTDDFNGAQLKAVCVEAGMLALRRDATEVTHEDFNEGIIQVQAKKKASLNYYA*

NbL15g03560.1 ferredoxin−dependent_glutamate_synthase_chloroplastic [MW = 177.03 kDa]

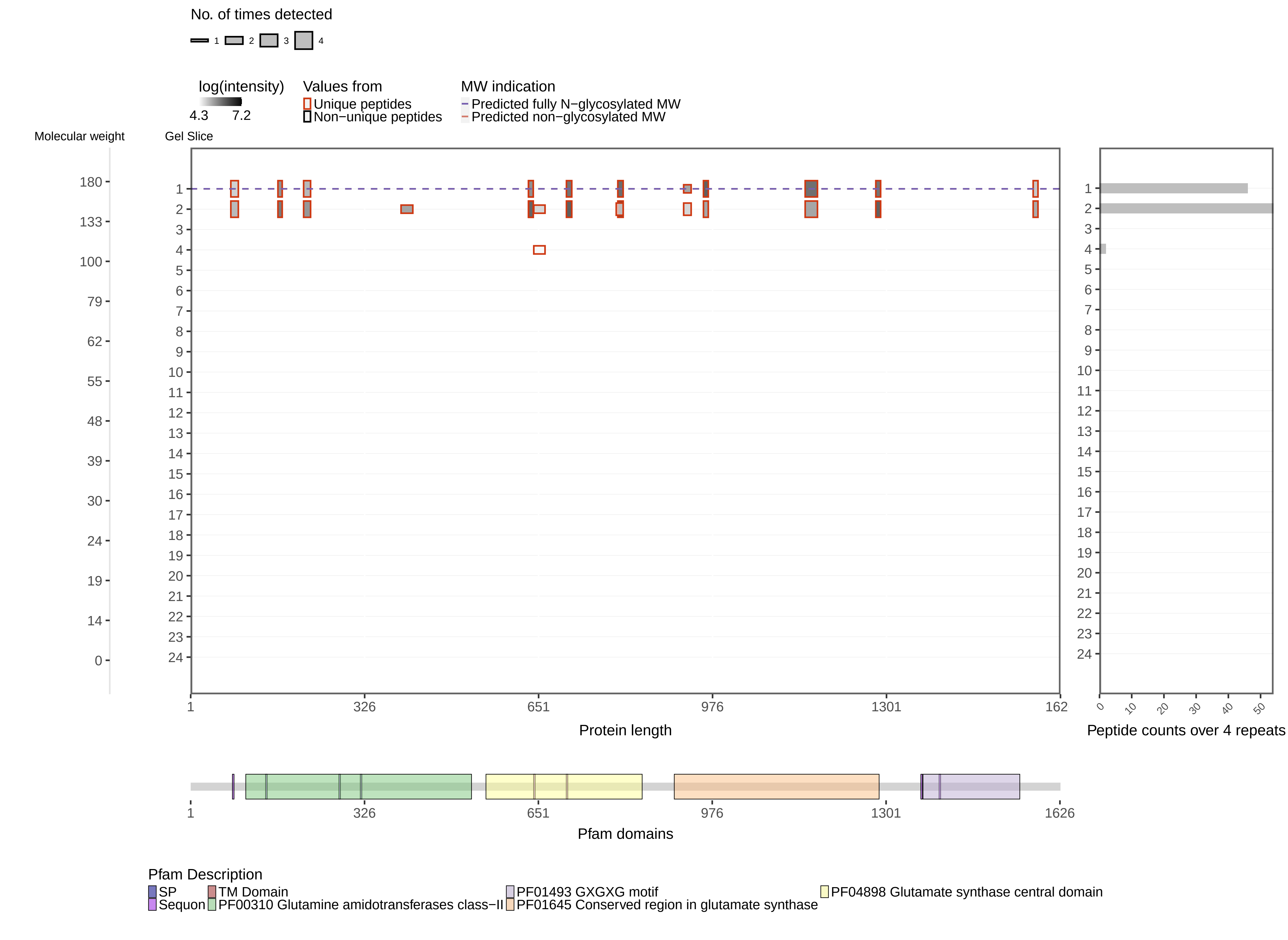


Protein sequence:

### NbL15g03310.1 glucose−6−phosphate_1−dehydrogenase_chloroplastic [MW = 70.75 kDa]

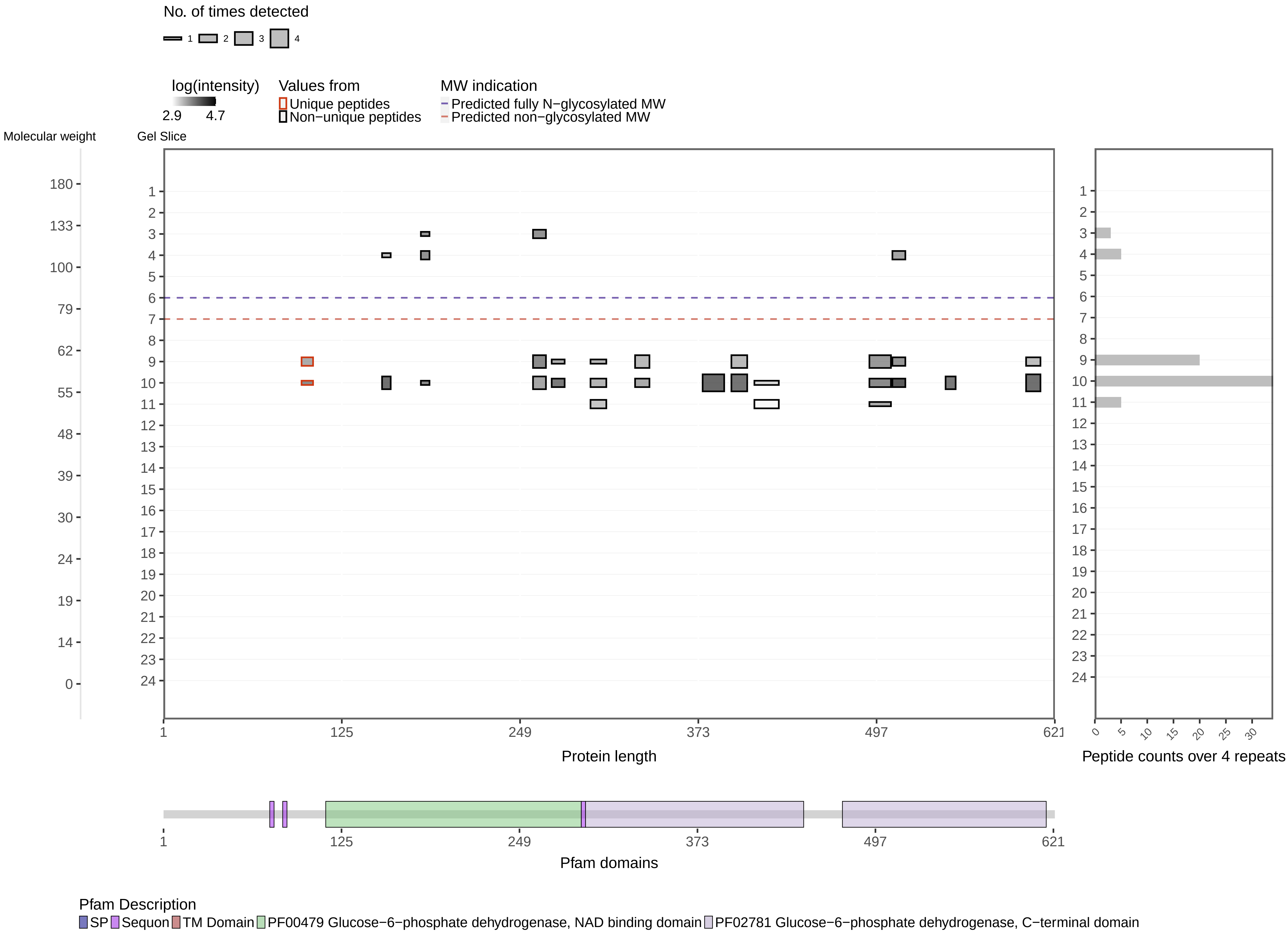


Protein sequence:

[illegible]

NbL15g03120.1 ubiquitin−conjugating_enzyme_E2_variant_1D [MW = 16.59 kDa]

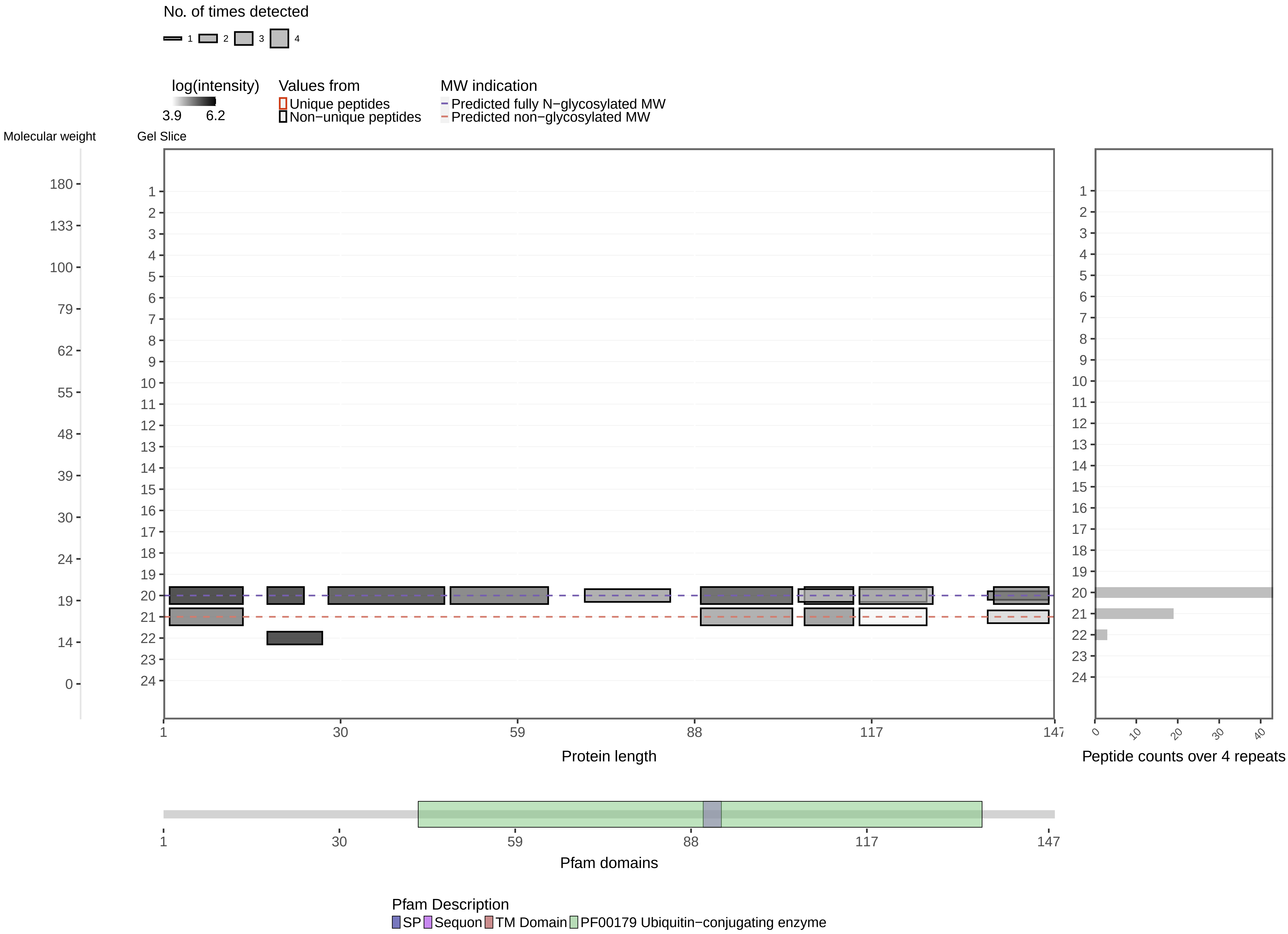


Protein sequence:

HEGRIYQLKLFCDKDYPEKPPSVRFHSRVNMTCVNHETGVVEPKKFALLANWQREYTMEDILVQLKKEMAAPHNRKLVQPPEGTYF*

NbL15g03090.1 TLD_domain−containing_protein_1 [MW = 46.98 kDa]

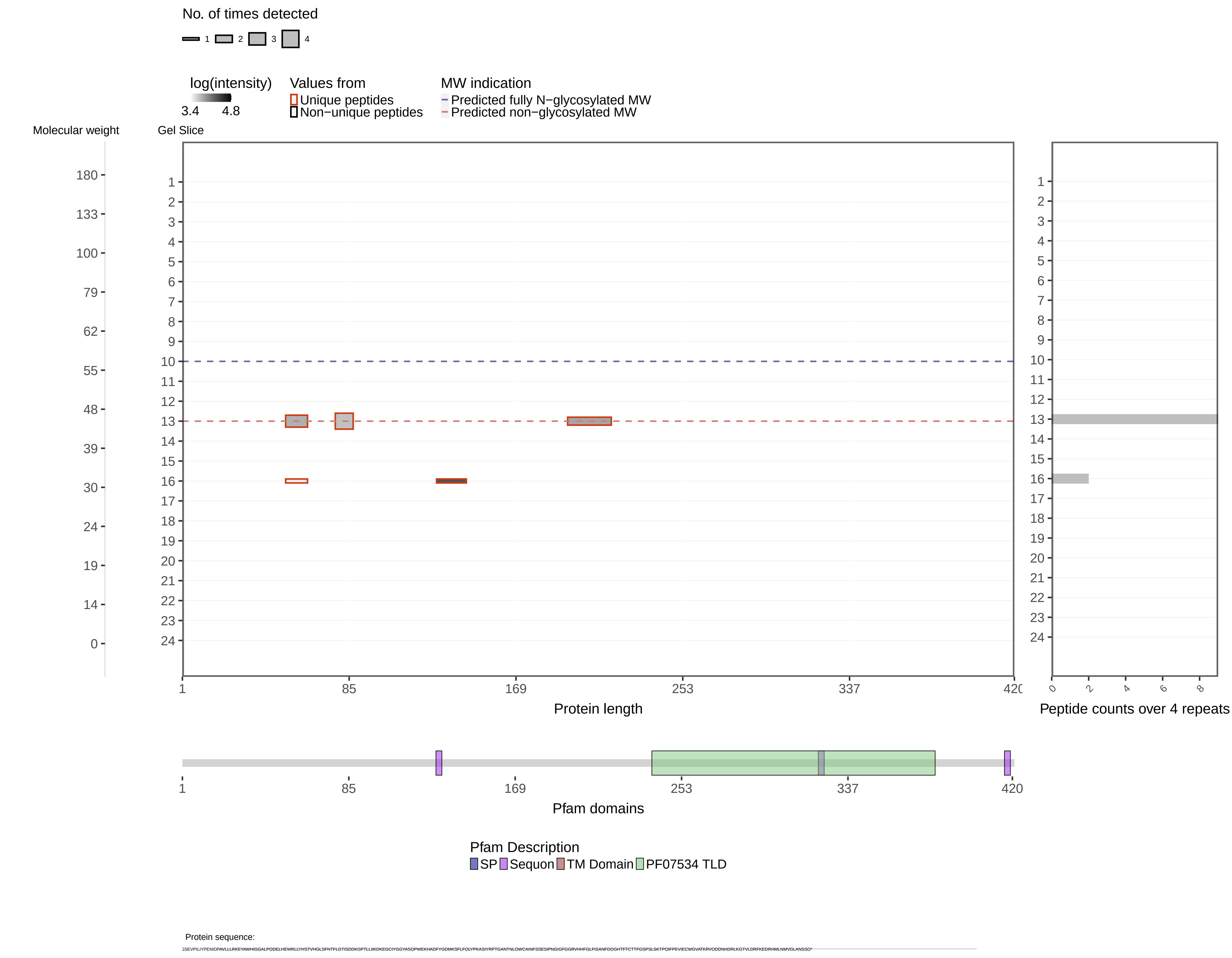

### NbL15g03070.1 secoisolariciresinol_dehydrogenase−like [MW = 29.48 kDa]

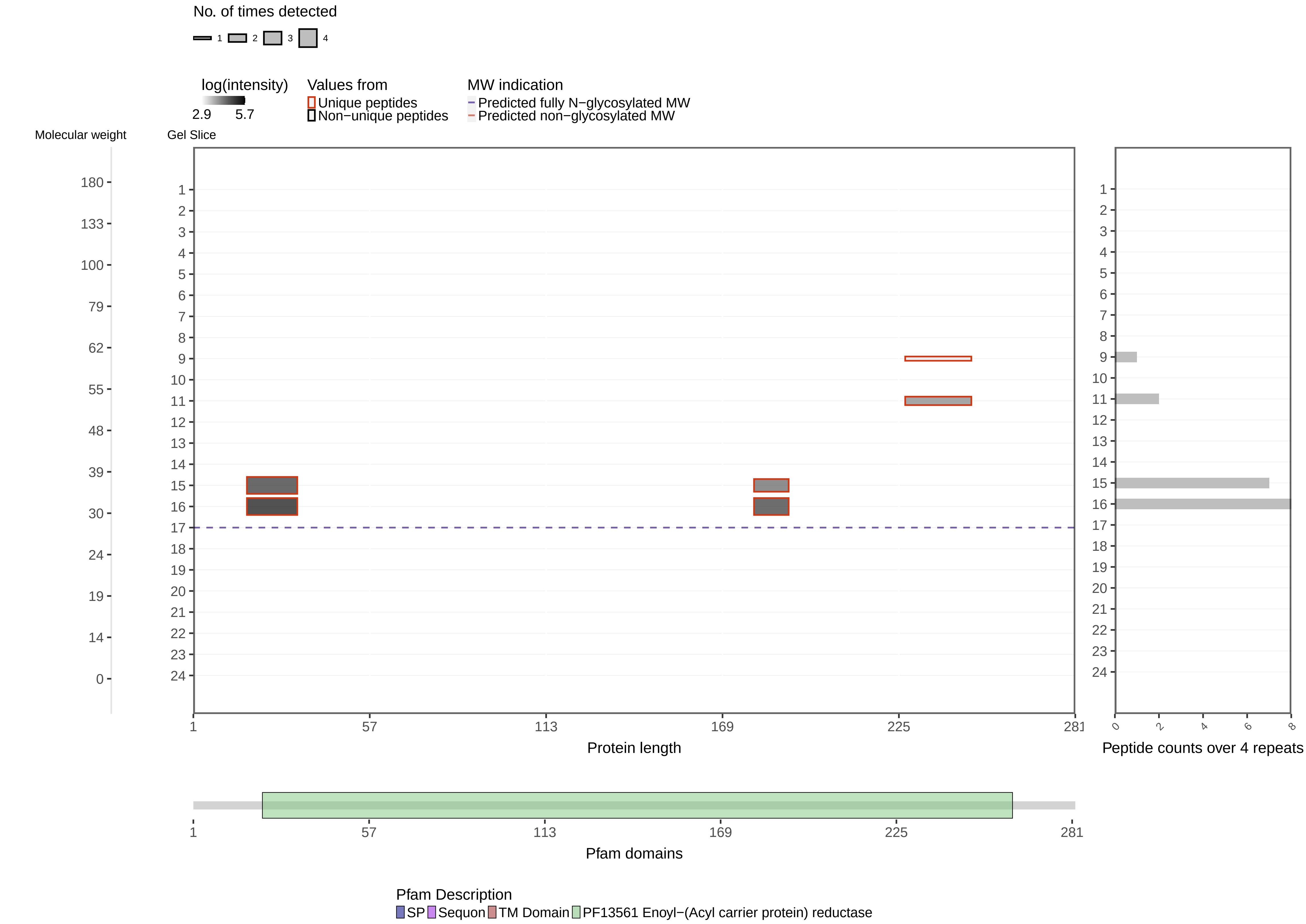


Protein sequence:

YGSFLGAKYAARVMIPAKKGVILFTSSVAAVNAGESPHSYTVSKHAVVGLTKNLCAELGQYGIRVNCVSPCAVATPLLVKAMGAEKSVVDGIICSSANLKGVVPTAEDVAEAALYLASDESKFVSGLNLVIDGGYSTTNQAYMKTIQSVLFSTV*

NbL15g03000.1 glycine−rich_RNA−binding_protein_4_mitochondrial−like [MW = 14 kDa]

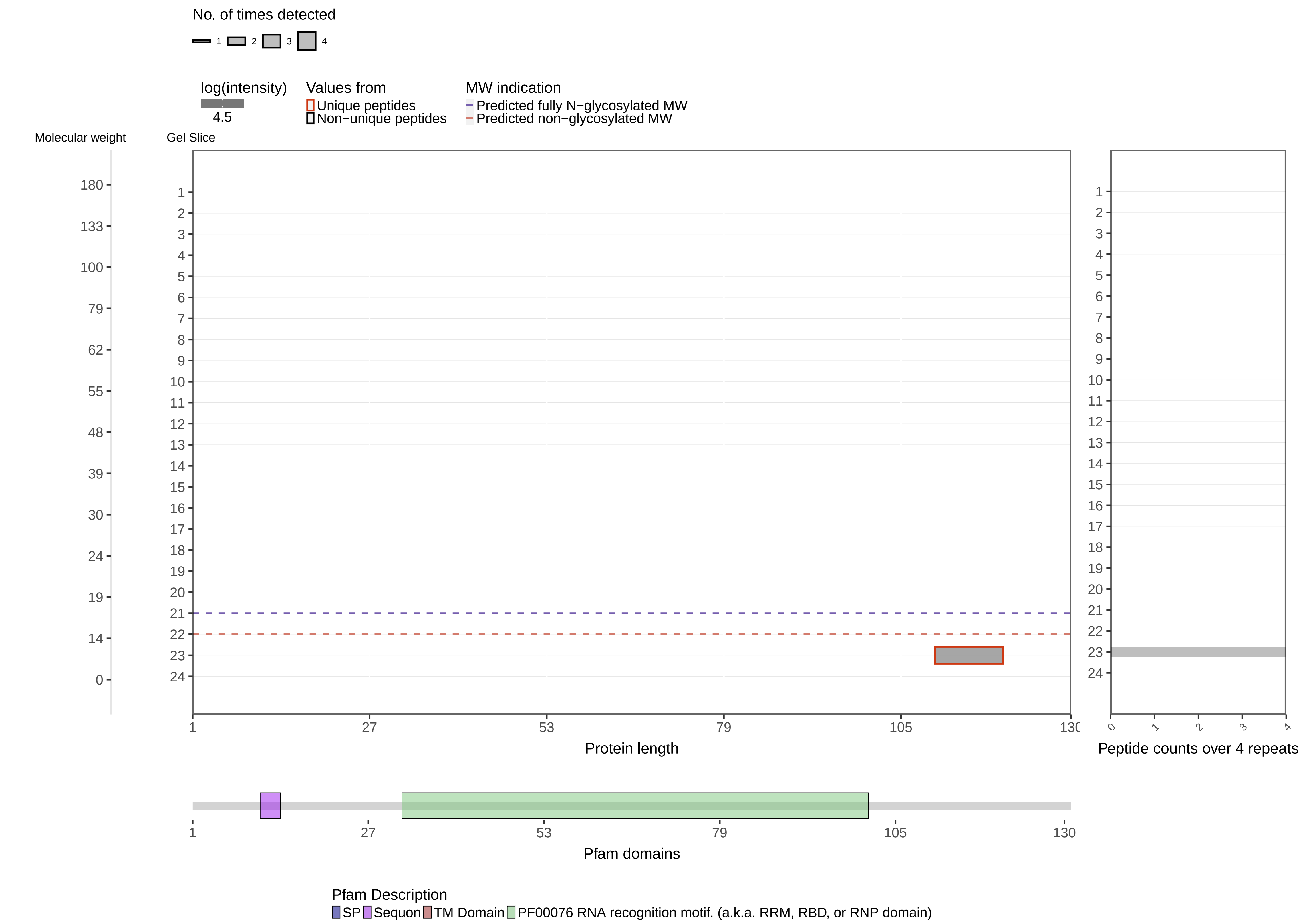


Protein sequence:

IGQVIEAKIVMDRVSDKSKGFGFVTYASEDDAKKALQEMNGKPLNGRVIFVDYAKPKADFGGAMPIARGPPEPPTEQ*

### NbL15g02960.1 protein_ABHD11−like [MW = 36.84 kDa]

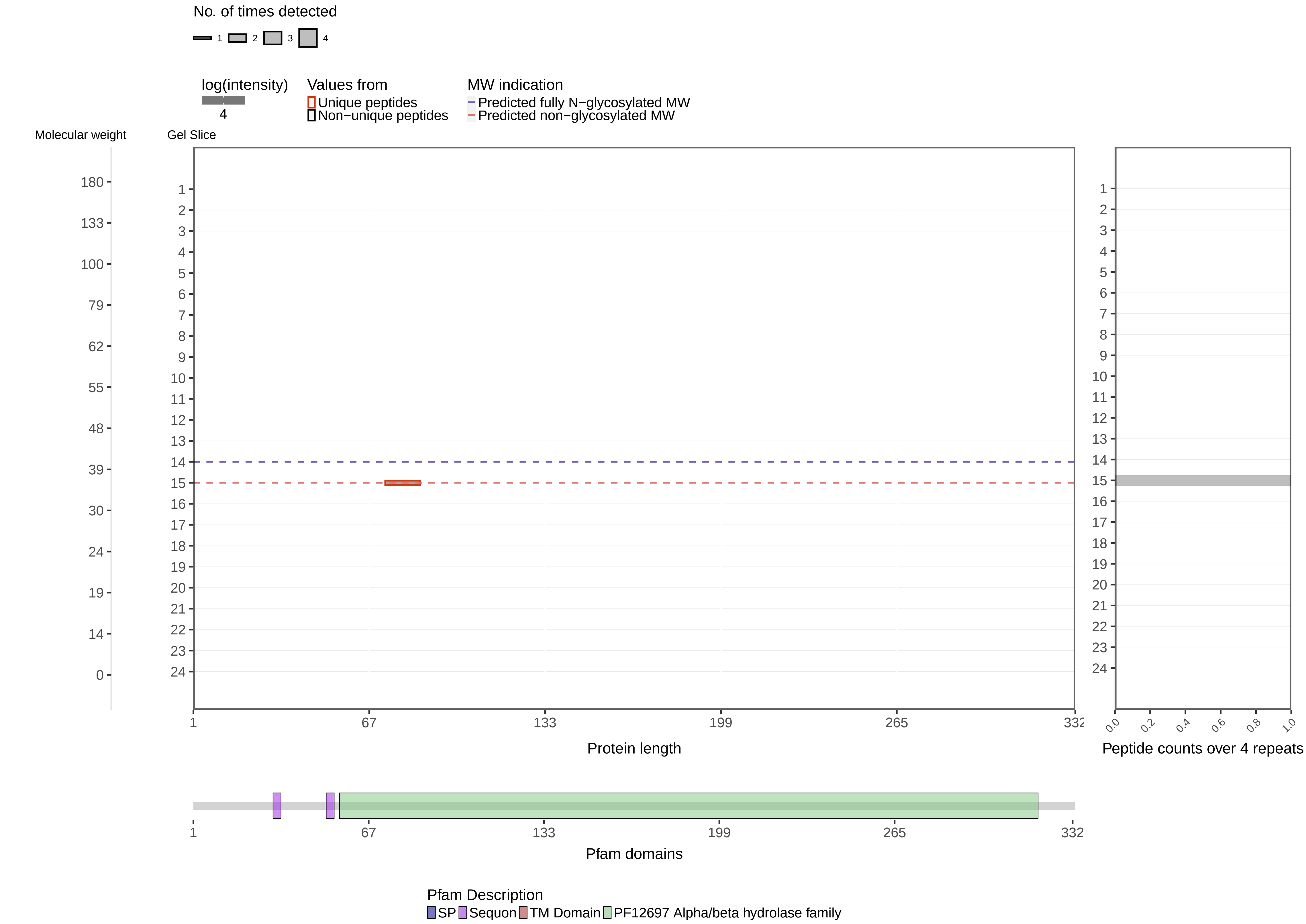


Protein sequence:

/YGQSARLPKQLWVLDSVPGKVDPEDSSREVEKVMQTLQSLPSSIPSRKWLVDHMLKLGFSKALSEWLGSNLQKSGDQMIWTFNIEAAIDMFNSYREKDYCPLLEHPPKGTEIAIVRAERSDRWDPETVQKLKSFASNGGGESEGKMSYHILPNSGHWVHVENPRGLLEIITPKLASLV*

NbL15g02940.1 beta−fructofuranosidase_insoluble_isoenzyme_1 [MW − SP = 62.37 kDa]

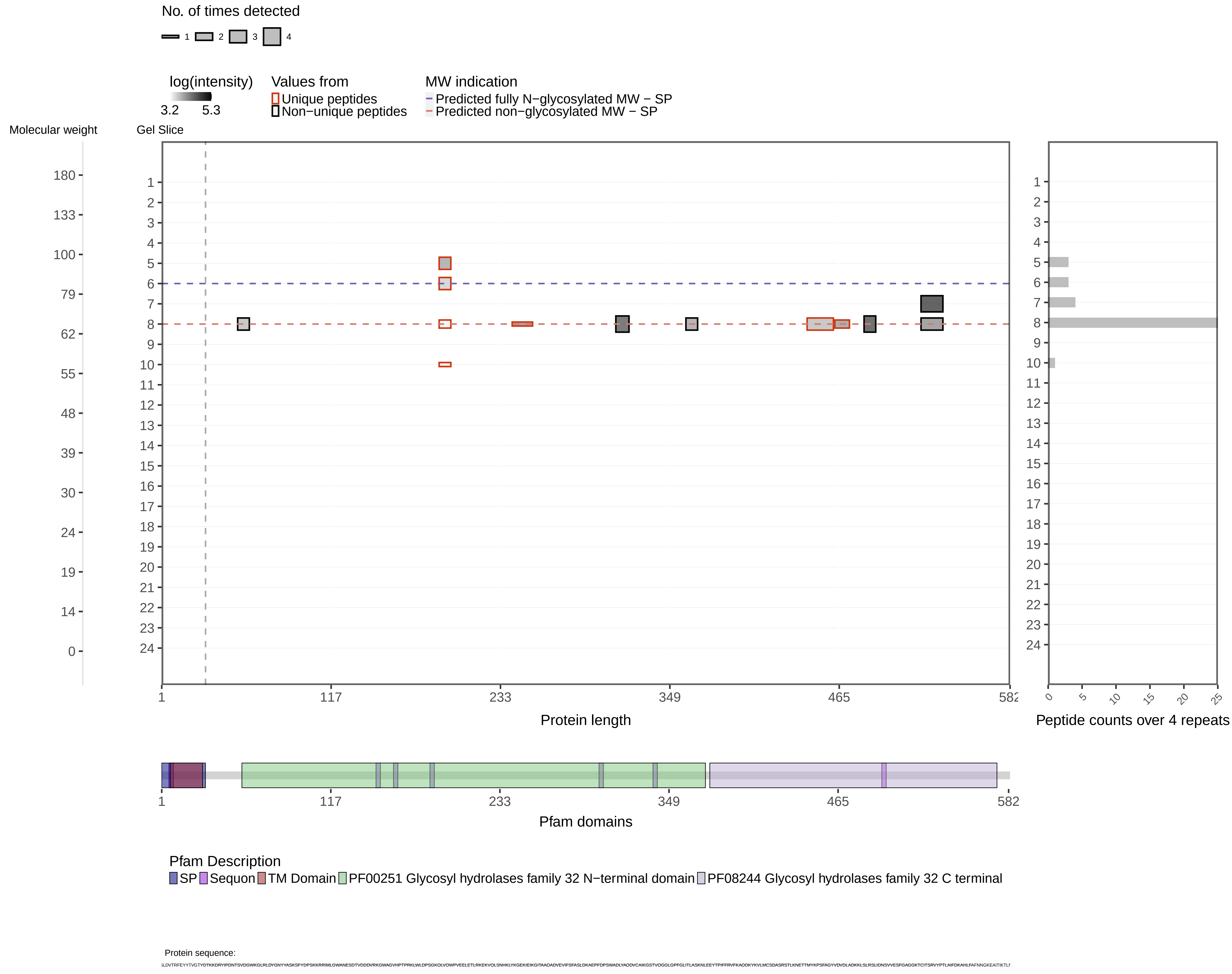

### NbL15g02790.1 pyruvate_decarboxylase_1−like [MW = 65.25 kDa]

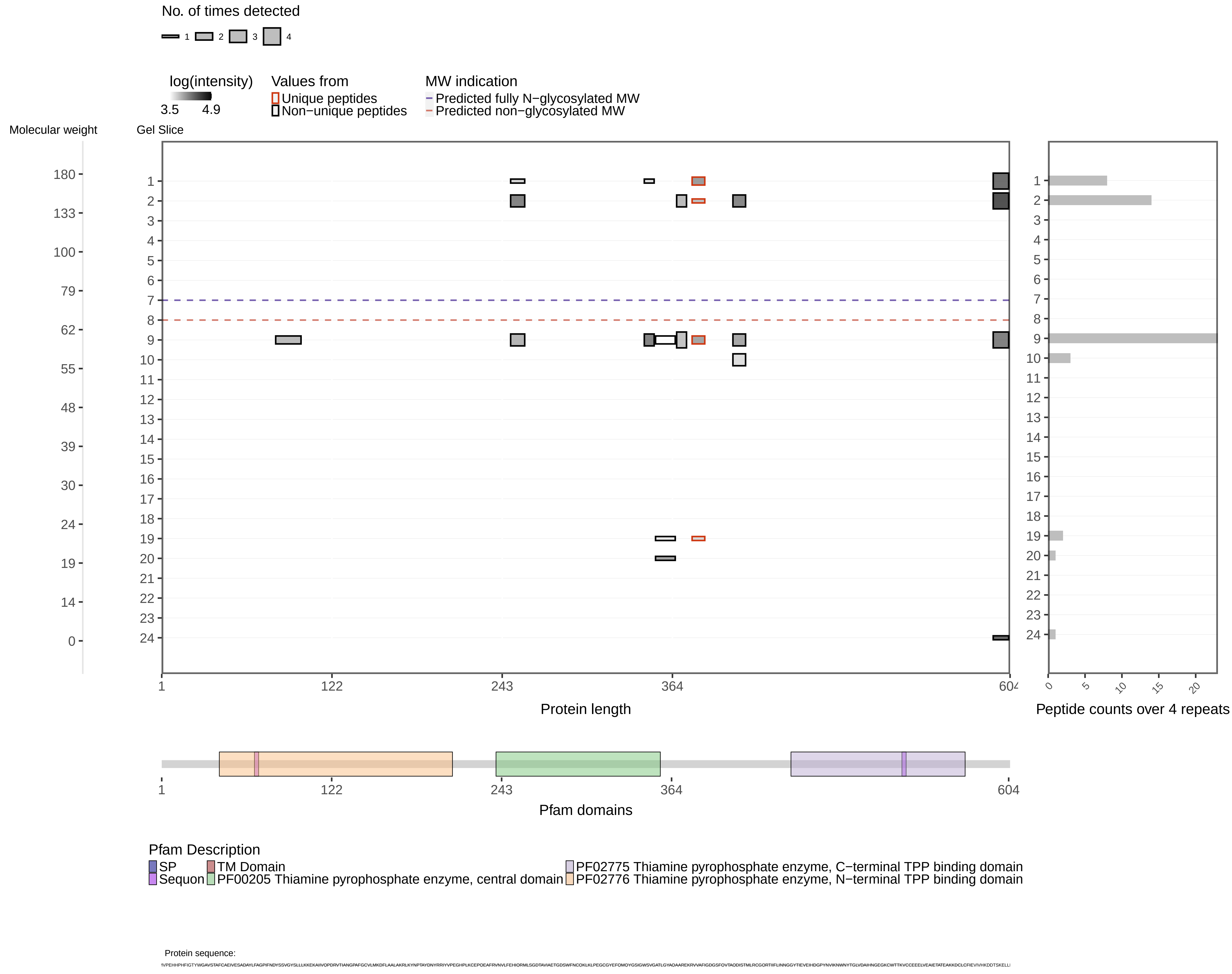

### NbL15g02760.1 acyl−coenzyme_A_oxidase_4_peroxisomal [MW = 48.4 kDa]

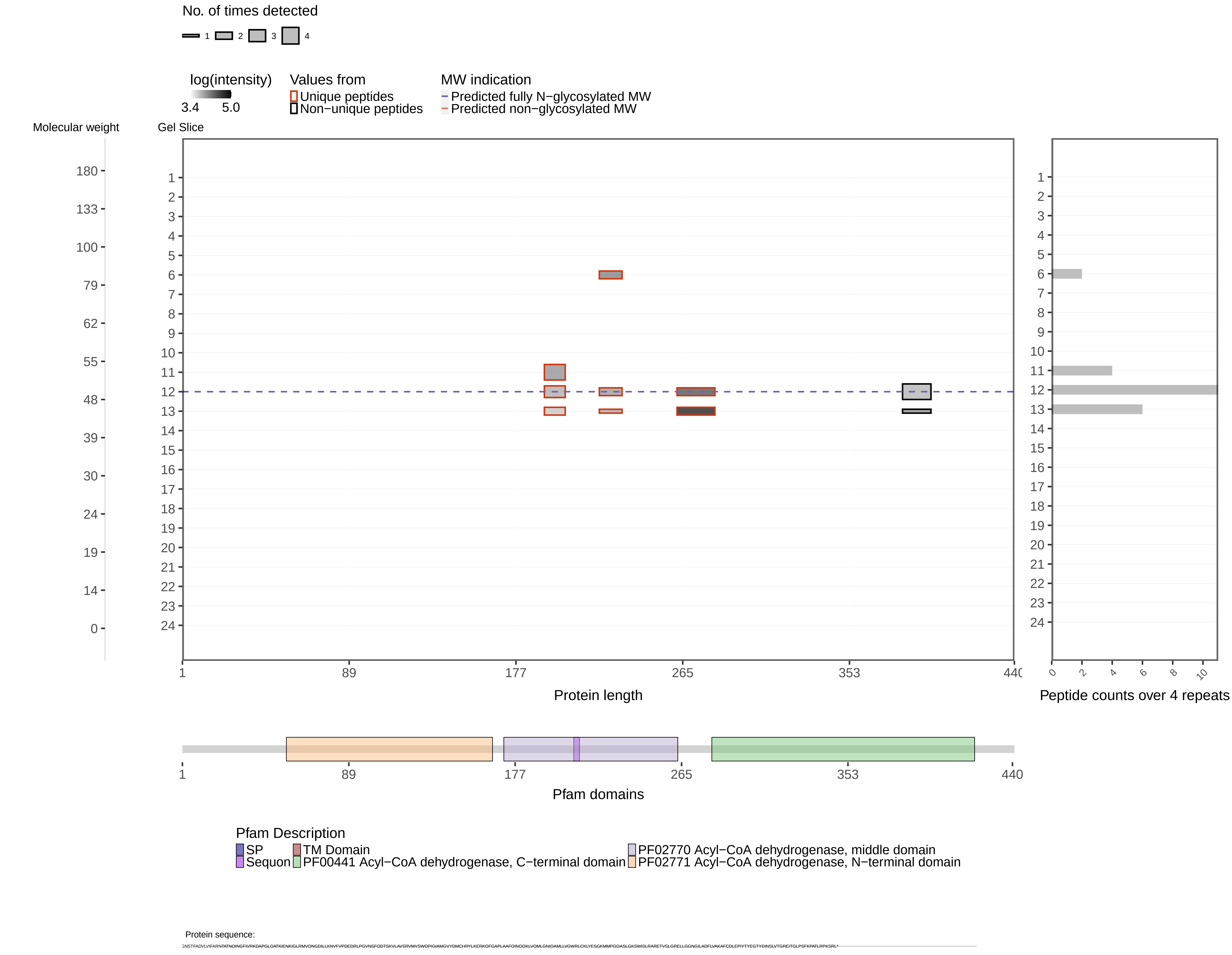

### NbL15g02730.1 rhodanese−like_domain−containing_protein_10 [MW = 24.28 kDa]

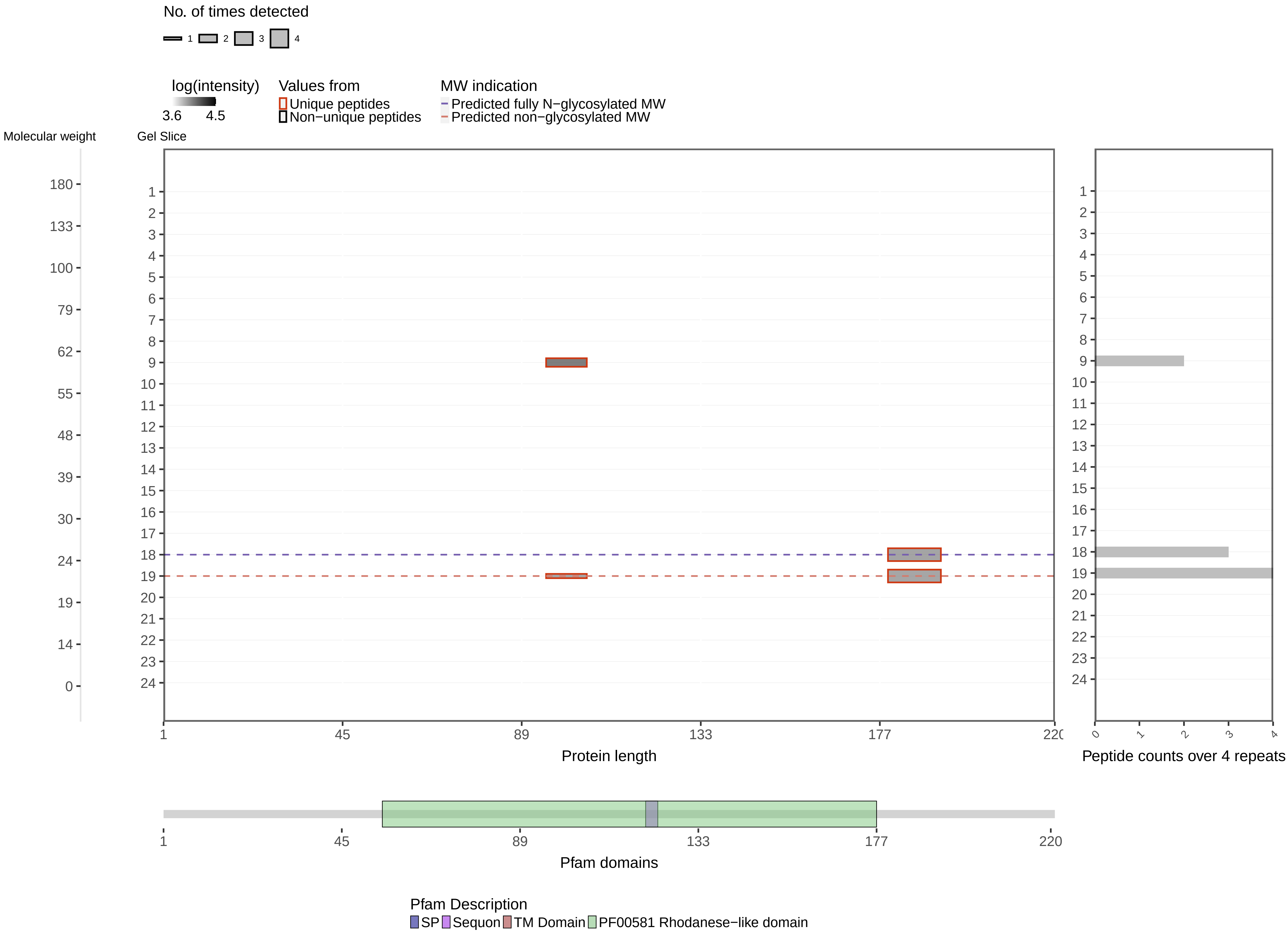


Protein sequence:

ISPITLLKKWVHFGYIGAWTGQNFTMINNDFVKQVEQQIPDKNNAKVLVACGEGLRSLMAISKLHKGEYKNLAWLVGGFTRASDSDFPAVEGPEKLQYATIGGVSYYLLQLLVLLKVVGKES*

NbL15g02610.1 fructose−16−bisphosphatase_chloroplastic−like [MW = 45.75 kDa]

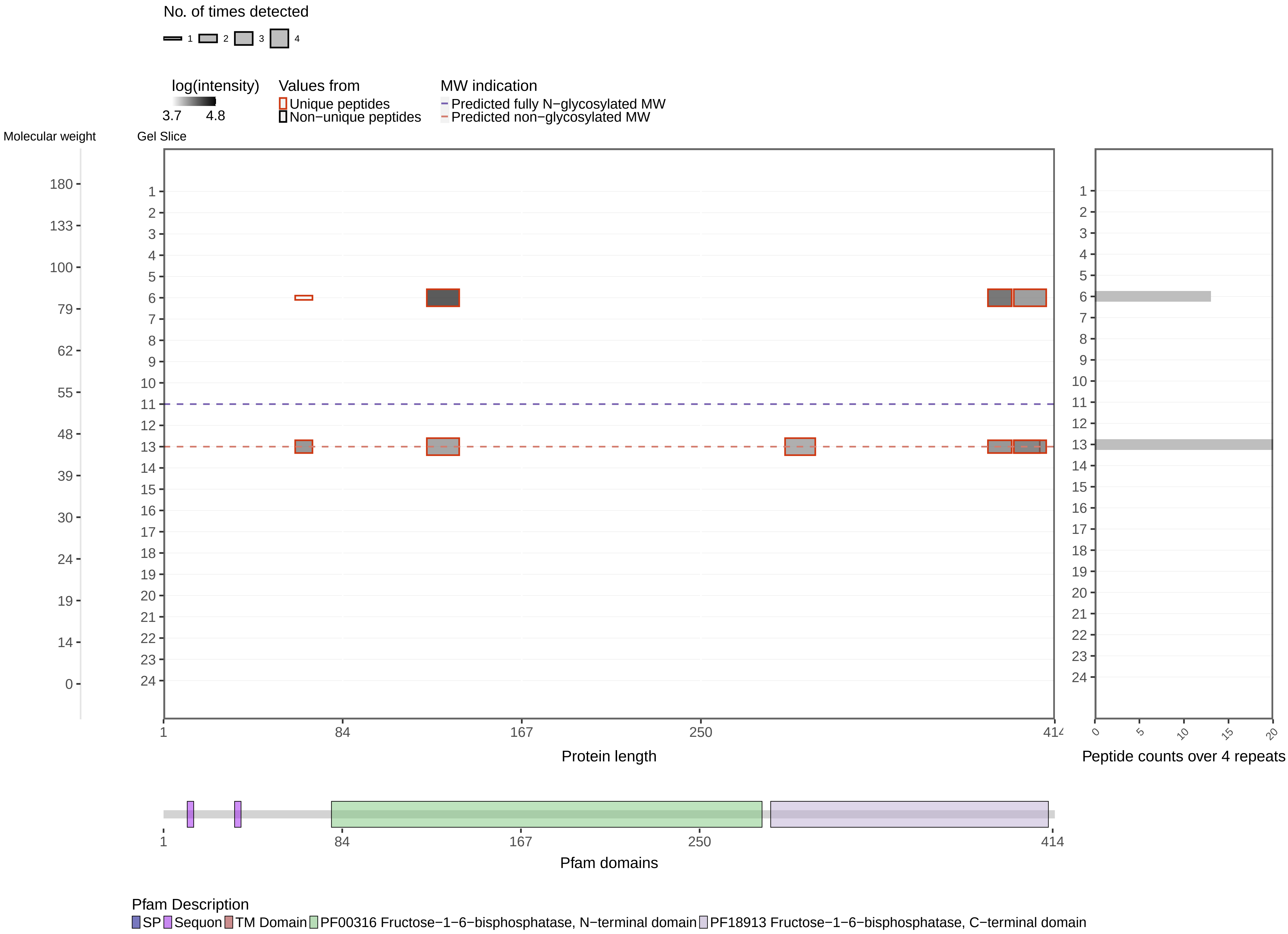


Protein sequence:

DAAVSTGSIFGIYSPNDDCLVDLEDTTMLDNVEQRCIVNVCQPGNNLLAAGYCMYSSSVIFVLTLGKGVFSFTLDPMFGEFILTYENIQIPKTGKIYAFNEGNYQLWDDRLKNYIDHLKEPGPNCKPYSARYIGSLVGDFHRTLLYGGIYGYPRDKKSRNGKLRLLYECAPMSFIVEQAGGKGSDGQQRILDIQPVEIHQRVPLFIGSTEEVEKLEKYLS*

NbL15g02590.1 protein_PLASTID_MOVEMENT_IMPAIRED_1 [MW = 105.09 kDa]

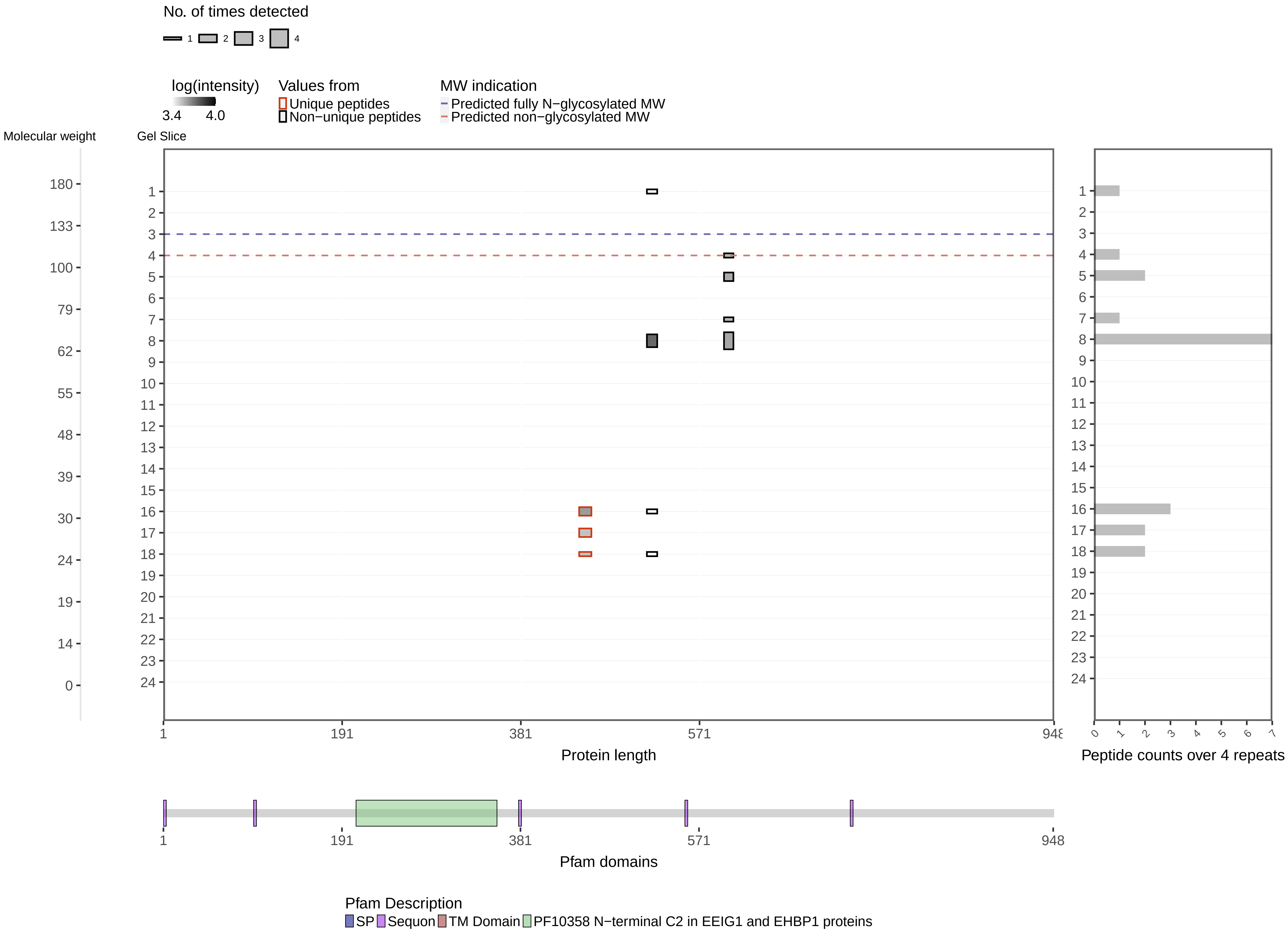


Protein sequence:

DEVKKEVEVKEEEEEAEEEEEEEQSEGNSDKRSVSSHEVVKEIVHDQHHLTRLSELDSIAQQIKALESMFKDENQVKTEDDDSESQRLDADEENVTREFLQMLEDSGGVNQFKTDQQETPTFKLQGGDEDSKKRESVIFIPDLGKGLGCVVQTRNGGFLAAMNPLNTVVSRKDTPKLAMQISKPFVFPSILSSMNGFEIFQRMAAIGMEEFTSKILSMMPMEELVGKTAEQIAFEGIASAIIQGRNKEVGASSTAARTVAAVKSMATAMNTSRNERISTGIWNICDEPLTVDEI

### NbL15g02550.1 NHP2−like_protein_1 [MW = 13.89 kDa]

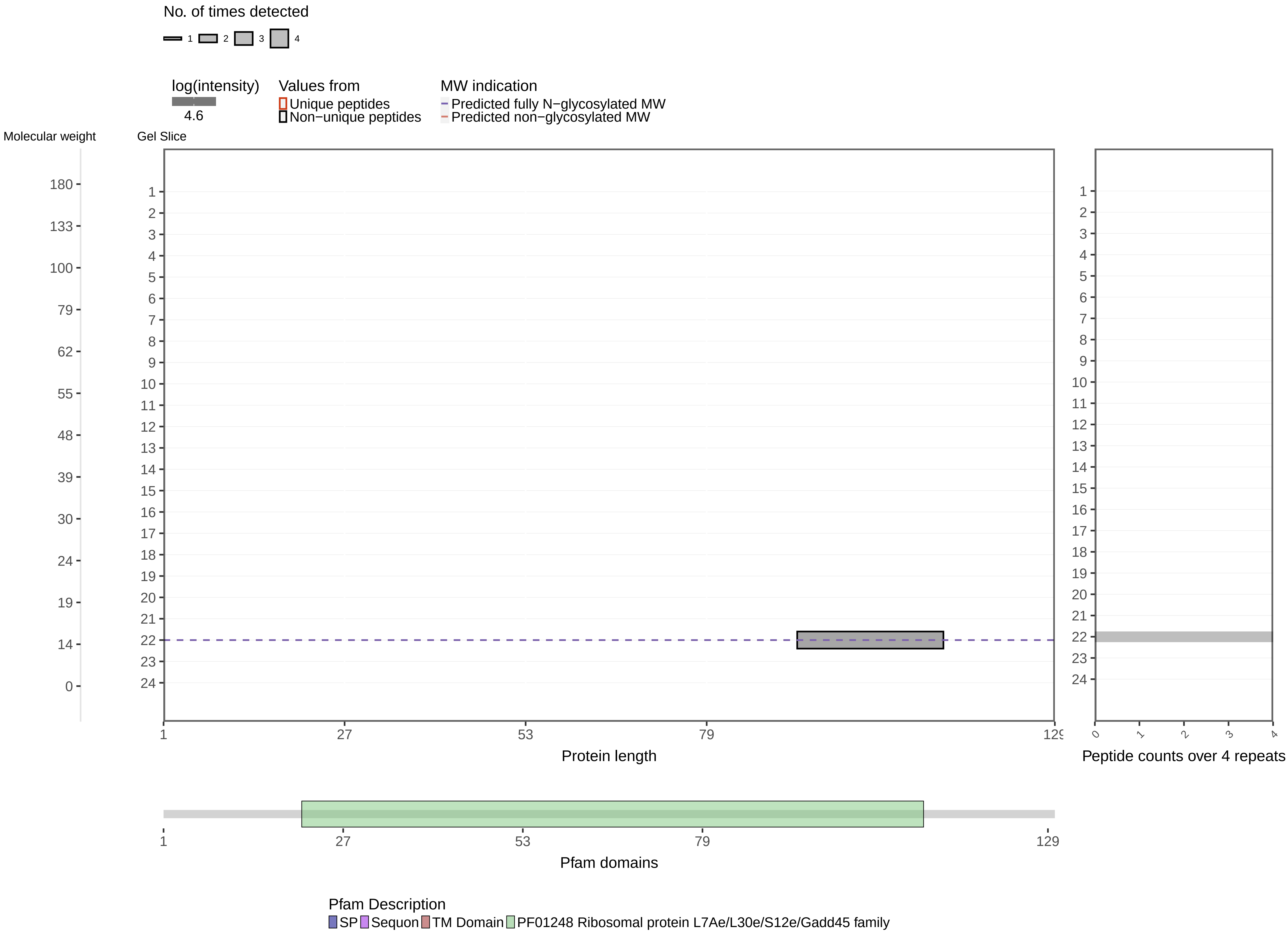


Protein sequence:

AEFIVMAADTEPLEILLHLPLLAEDKNVPYVFVPSKQALGRACGVTRPVIAASVTSNEGSQLKSQIQQLKDAIEKLLI*

NbL15g02500.1 ribulose_bisphosphate_carboxylase/oxygenase_activase_2_chloroplastic [MW = 48.33 kDa]

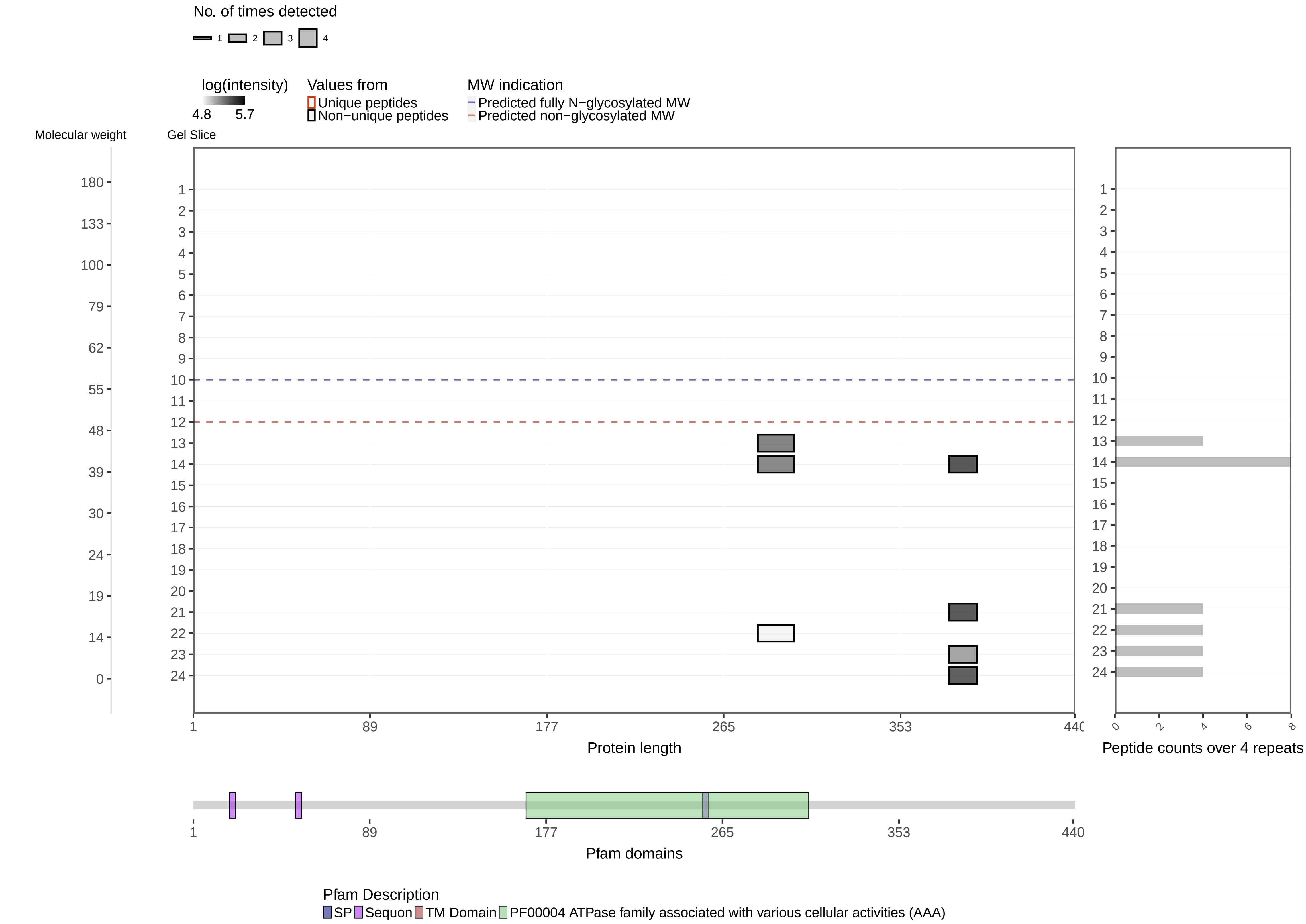


Protein sequence:

PAKLIRQRYREAAEIIRKGNMCCLFINDLDAGAGRMGGTTQYTVNNQMVNATLMNIADNPTNVQLPGMYNKQENARVPIIVTGNDFSTLYAPLIRDGRMEKFYWAPTREDRIGVCKGIFRTDNVPEEAVIKIVDTFPGQSIDFFGALRARVYDDEVRKWVSGTGIEAIGDKLLNSFDGPPTFEQPKMTVEKLLEYGNMLVQEQENVKRVQLAETYLKEAALGDANADAINTGNF*

NbL15g02490.1 uncharacterized_LOC104104784 [MW = 16.88 kDa]

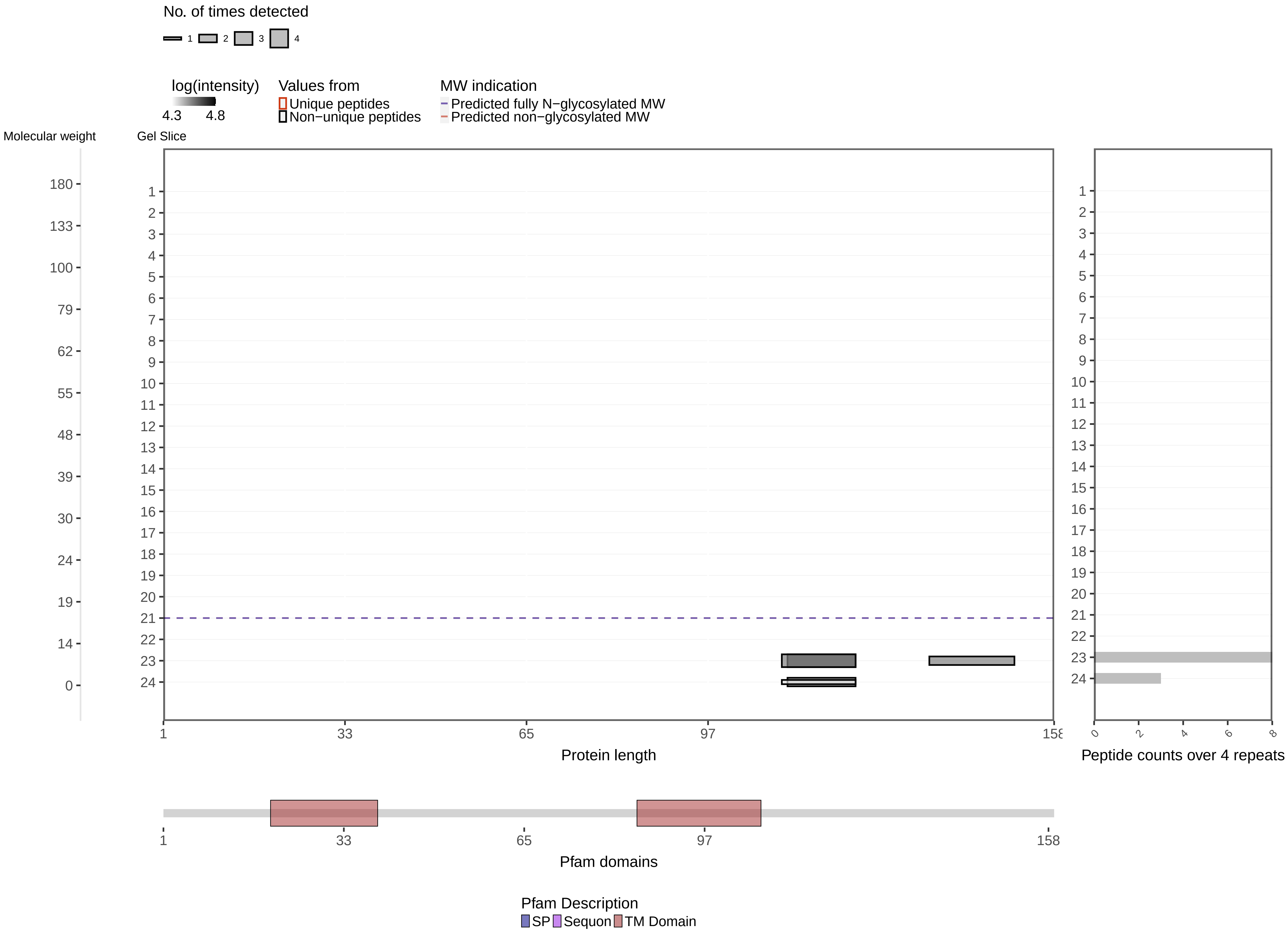


Protein sequence:

KHRSLKVVAAVGDVSSEGTTYLIAGAAAVALLGTAFPILFSRKDTCPECDGAGFVRKGGATLRANAARKDQVQIVCANCNGLGKLNQVDK*

NbL15g02430.1 mitochondrial_import_inner_membrane_translocase_subunit_Tim9 [MW = 13.93 kDa]

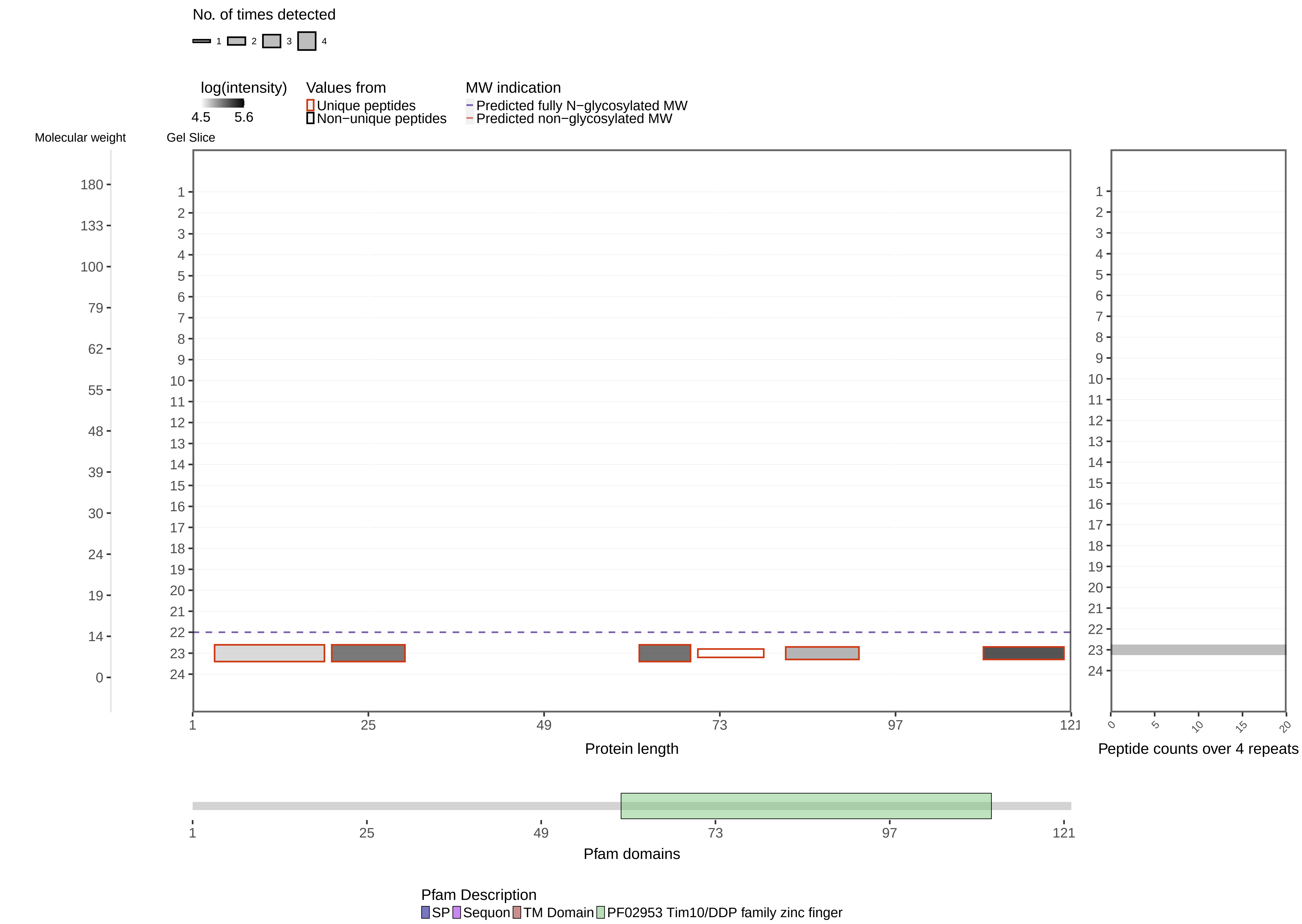


Protein sequence:

SDTDTQGKGHSLRMYNSLVERCFTDCVDTFKRKTLDKQEETCVRRCAEKFLKHSMRVGMRFAELNQGAPTPD*

### NbL15g02390.1 uncharacterized_LOC109223268 [MW = 20.95 kDa]

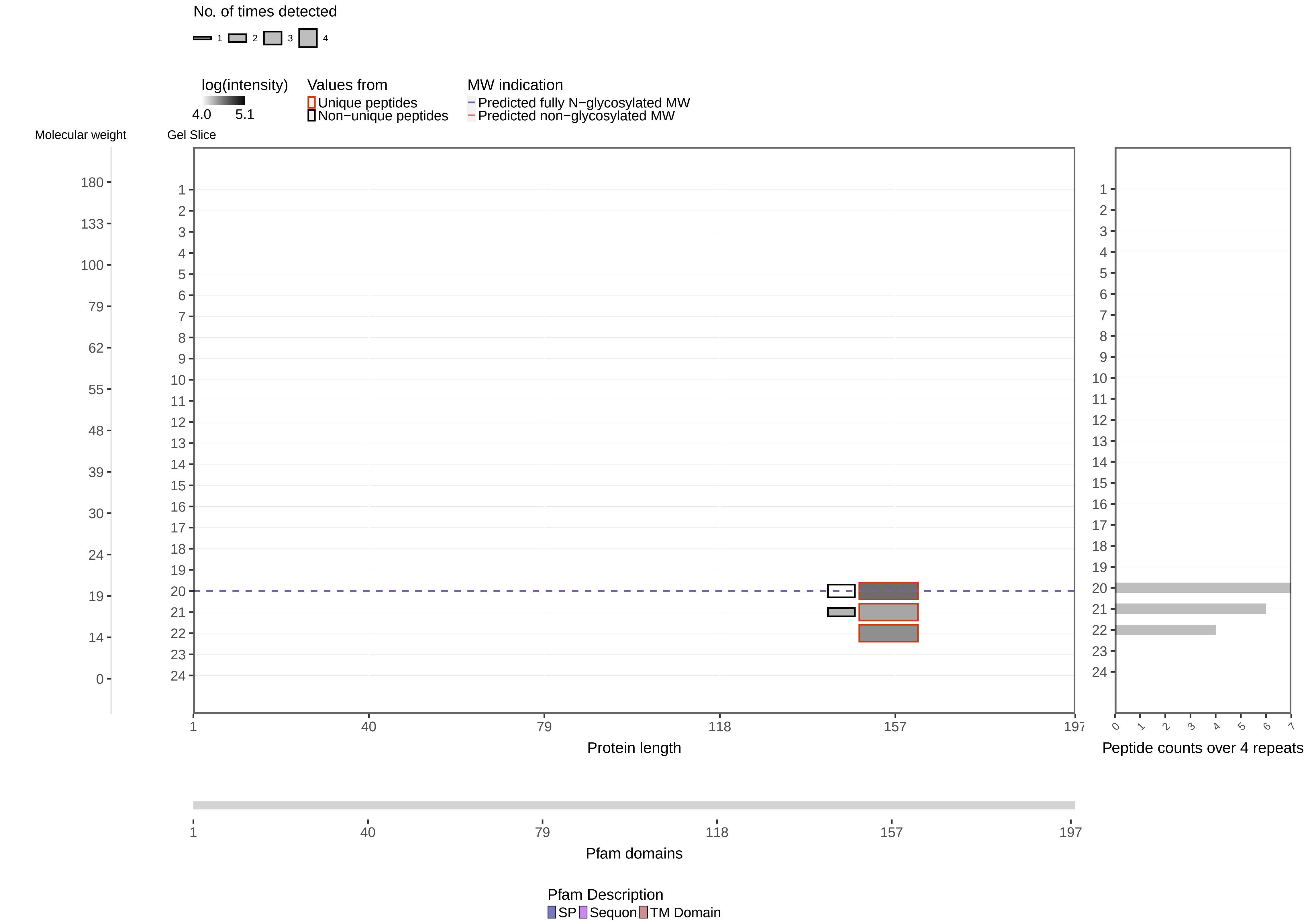


Protein sequence:

GAEAAEQLTQQEINGETFSNVPQTLSGECVEGQKDCKKARIQRPKSRQAETCTVKCVNTCIRGGLGSPGEGPLNVRRPLVVFKQGFRSRQYCLVECSDICNLIRDGEDGP*

NbL15g02340.1 heat_shock_cognate_70_kDa_protein_2−like [MW = 71 kDa]

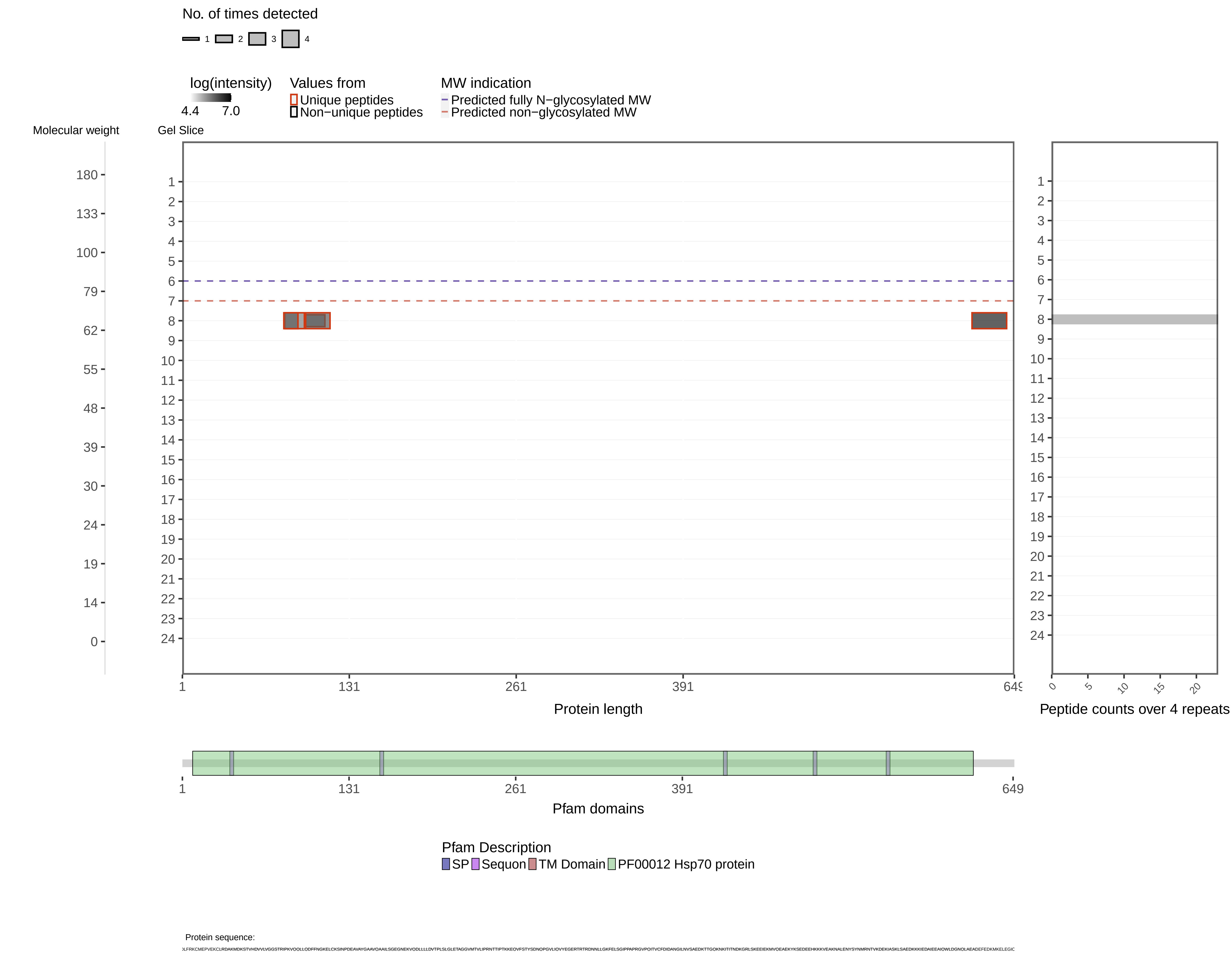

### NbL15g02330.1 THO_complex_subunit_4A−like [MW = 27.08 kDa]

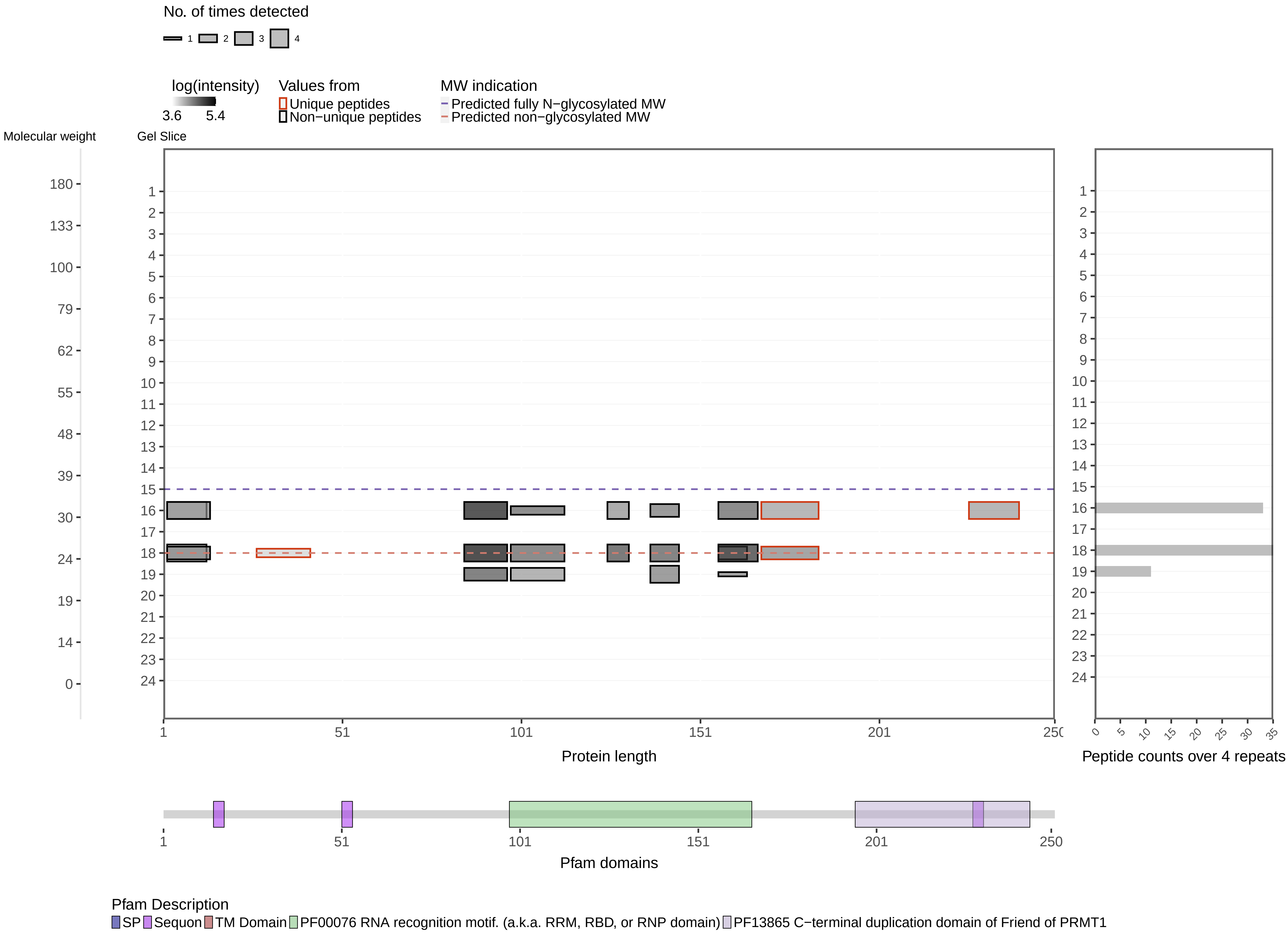


Protein sequence:

IKELFSEAGDIKRYSIHYDKSGRSKGTAEVIFSRRRDAEAAIKKYNNVQLDGKPMKIEFAGPNIGAPTLPPLRNRLYRNPNPAPRSQQRGGGFRRPPRGGRGSMRKEGGRGRGRVENITAEDLDADLEKYHSEAMETN*

### NbL15g02200.1 translin [MW = 34.53 kDa]

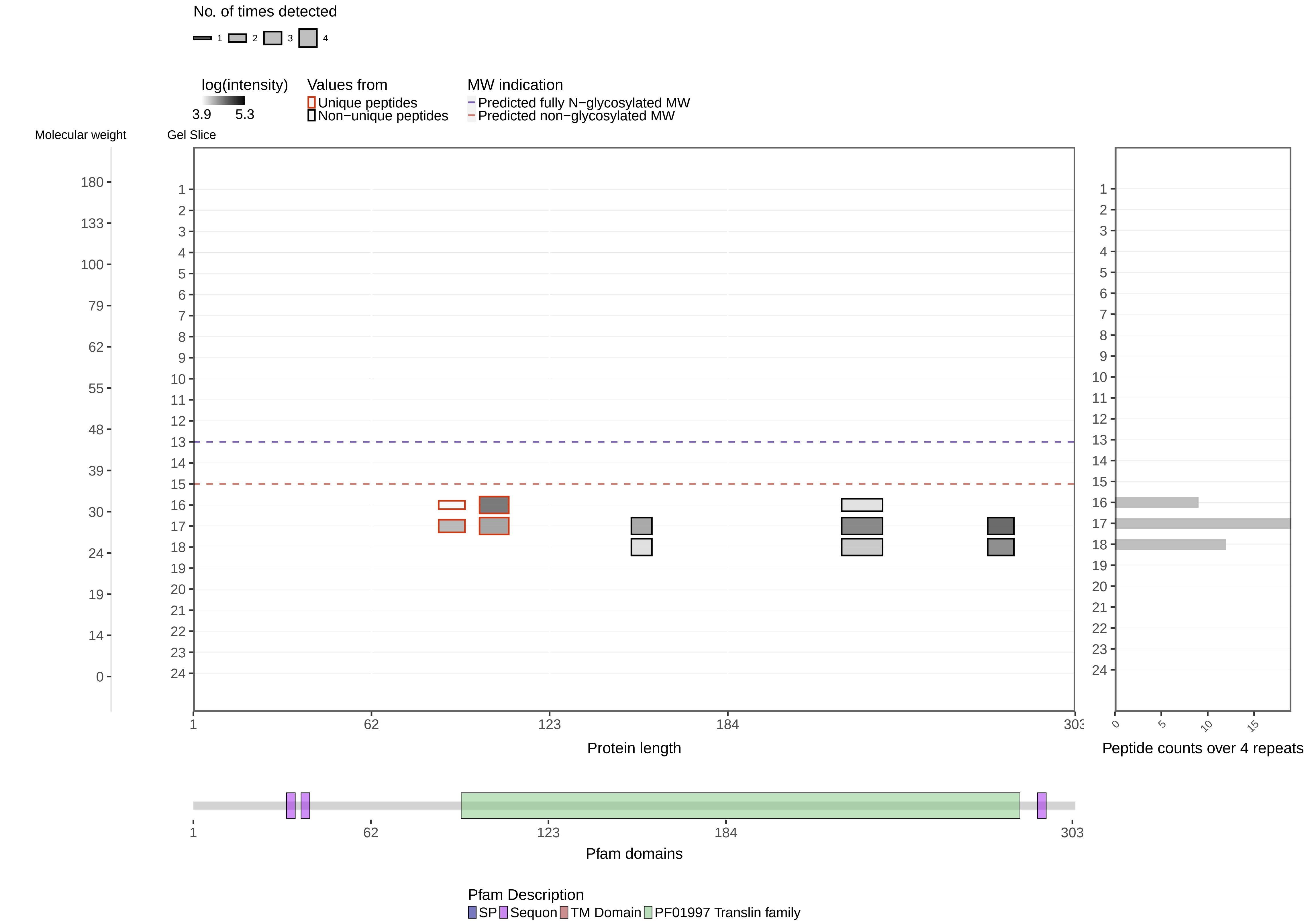


Protein sequence:

LYSRLSEIVRECPGOYFRYHGDWKSETOTVVSLLAFMHWLETGDLVLHNEVEEKLGLTPSEFGLDVEDYLIGICFMSNELPRYVVNOVTAGDYDCPRKVLKFLTDLHAAFRMLNLRNDFLRKKFDGMKYDLRKVEEVFYDVKIRGLAANGNSSGEKOVOGOS*

### NbL15g02180.1 adenosine_kinase_2 [MW = 37.51 kDa]

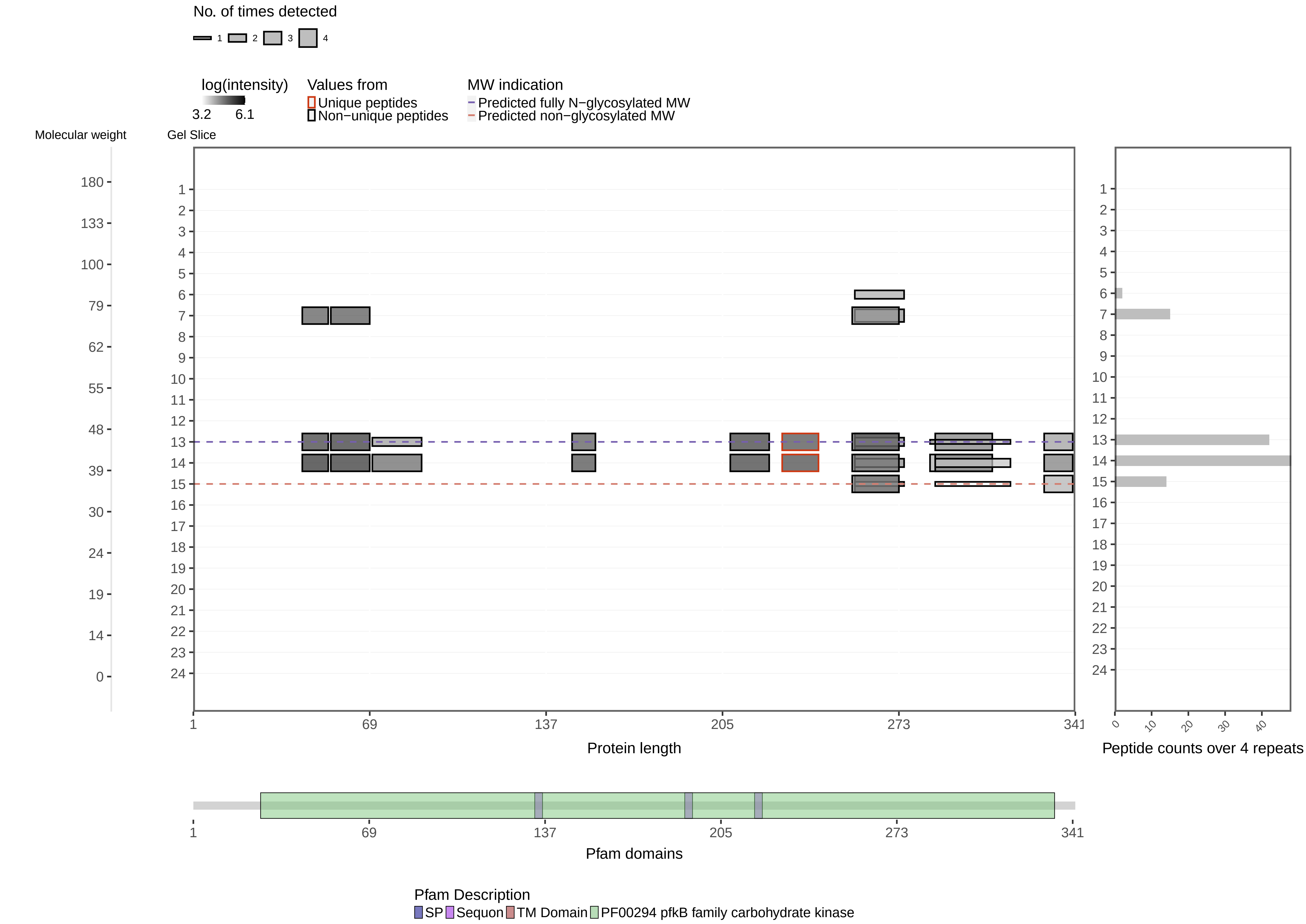


Protein sequence:

AKYYYIAGFFLTVSPESIQLVAEHAAAKNKVFSMNLSAPFICEFFKDQQEKVLPYMDFVFGNETEARTFSRVHGWETDNVEEIALRISQWPKASGTHKRITVITQGADPVVVAEDGKVKLFPVIPLPKEKLVDTNGAGDAFVGGFLAQLVQGKPIADCVKAGCYASNVIIQRSGCTYPEKPDFE*

### NbL15g02170.1 phenylalanine_ammonia−lyase [MW = 77.44 kDa]

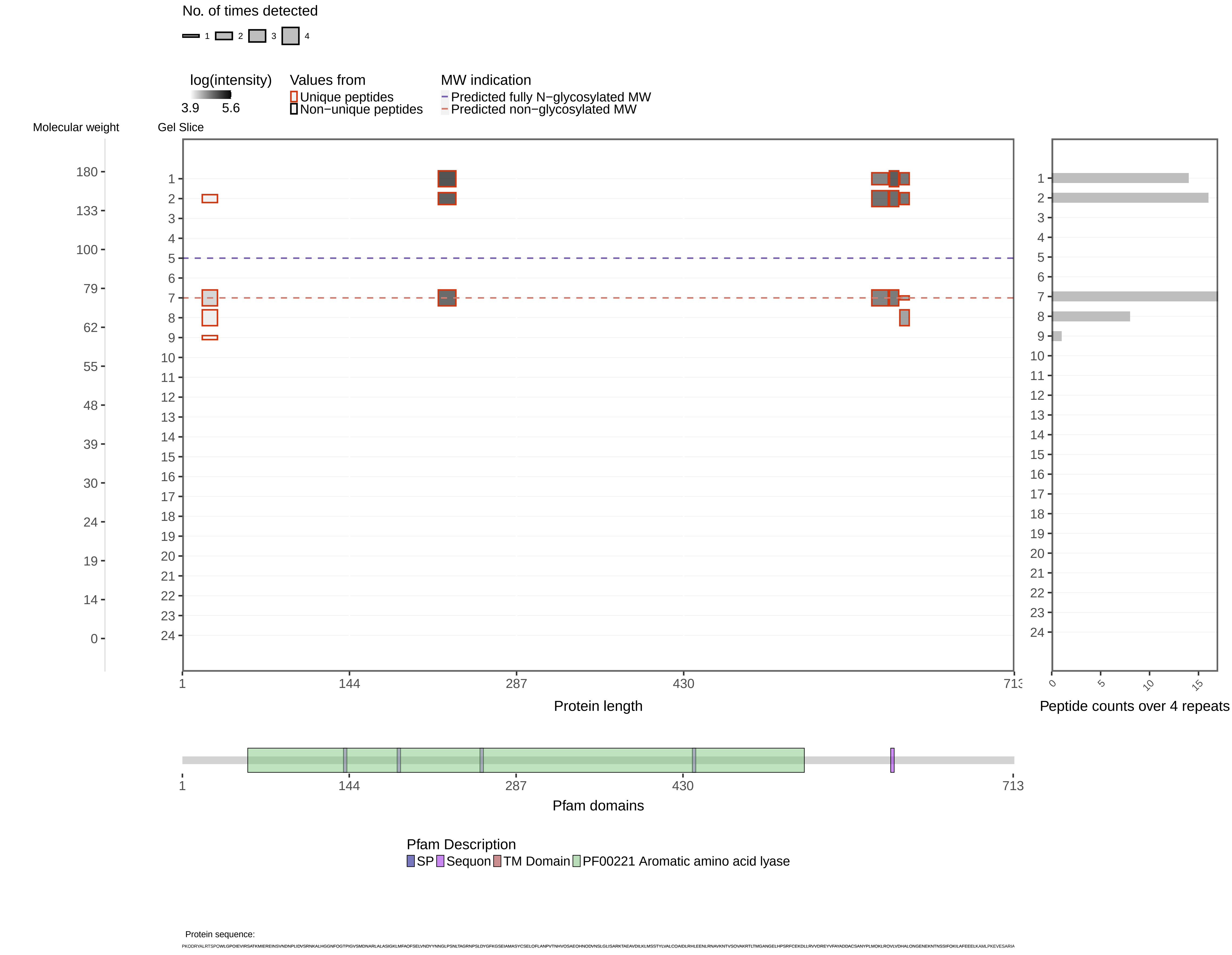

### NbL15g02160.1 29_kDa_ribonucleoprotein_B_chloroplastic [MW = 31.36 kDa]

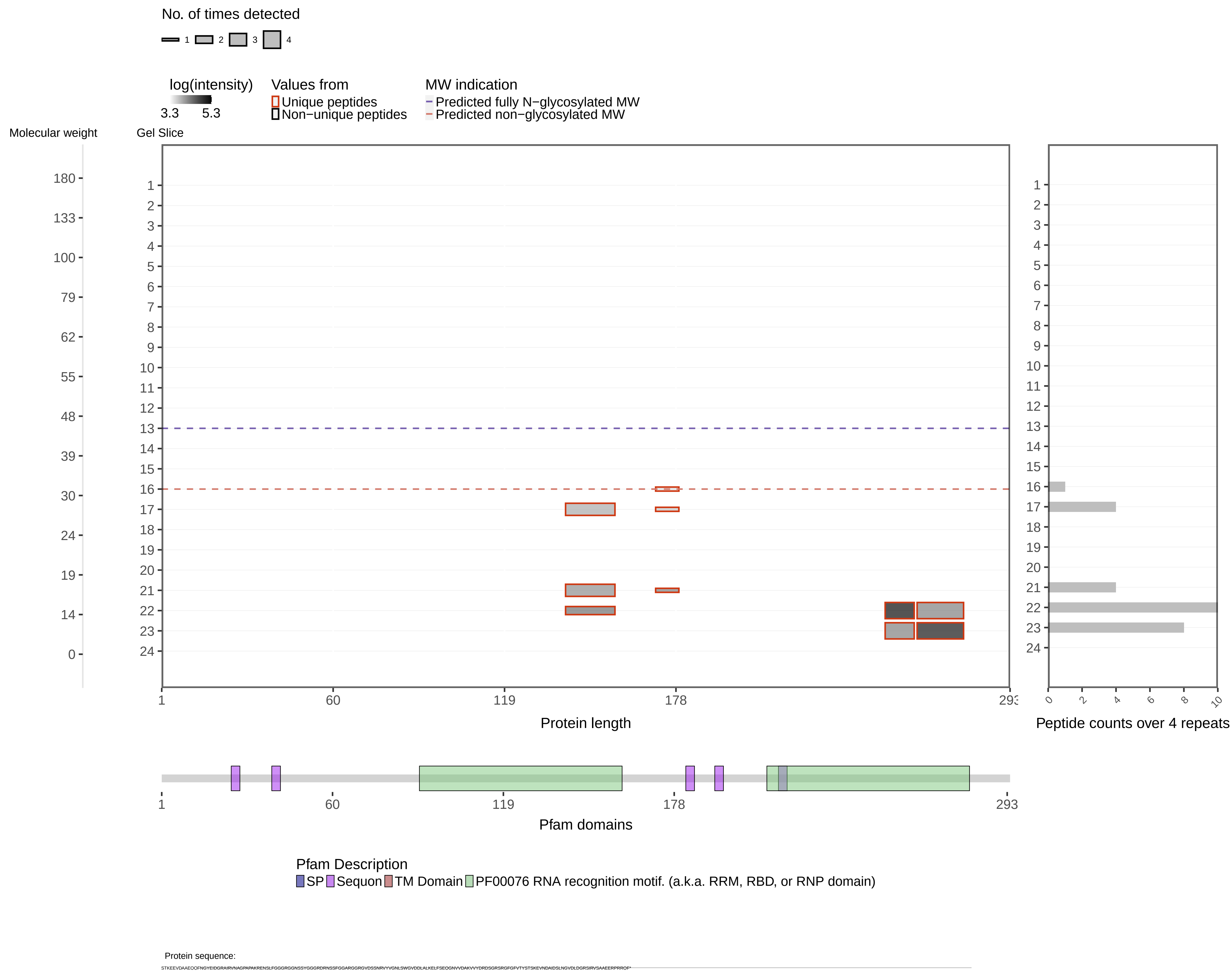

### NbL15g02100.1 WD_repeat−containing_protein_20−like [MW = 59.42 kDa]

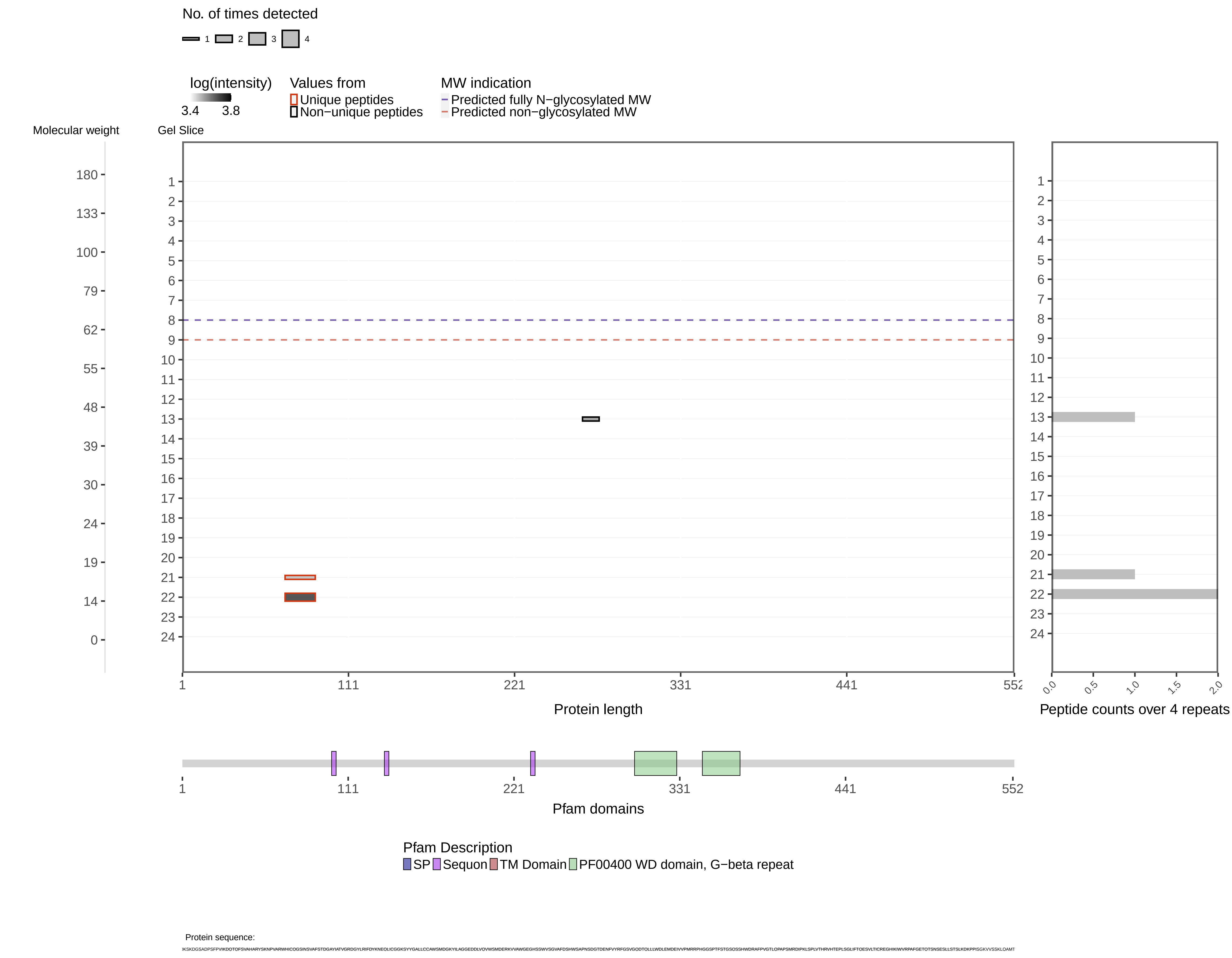

NbL15g02020.1 60S_ribosomal_protein_L4−like [MW = 44.7 kDa]

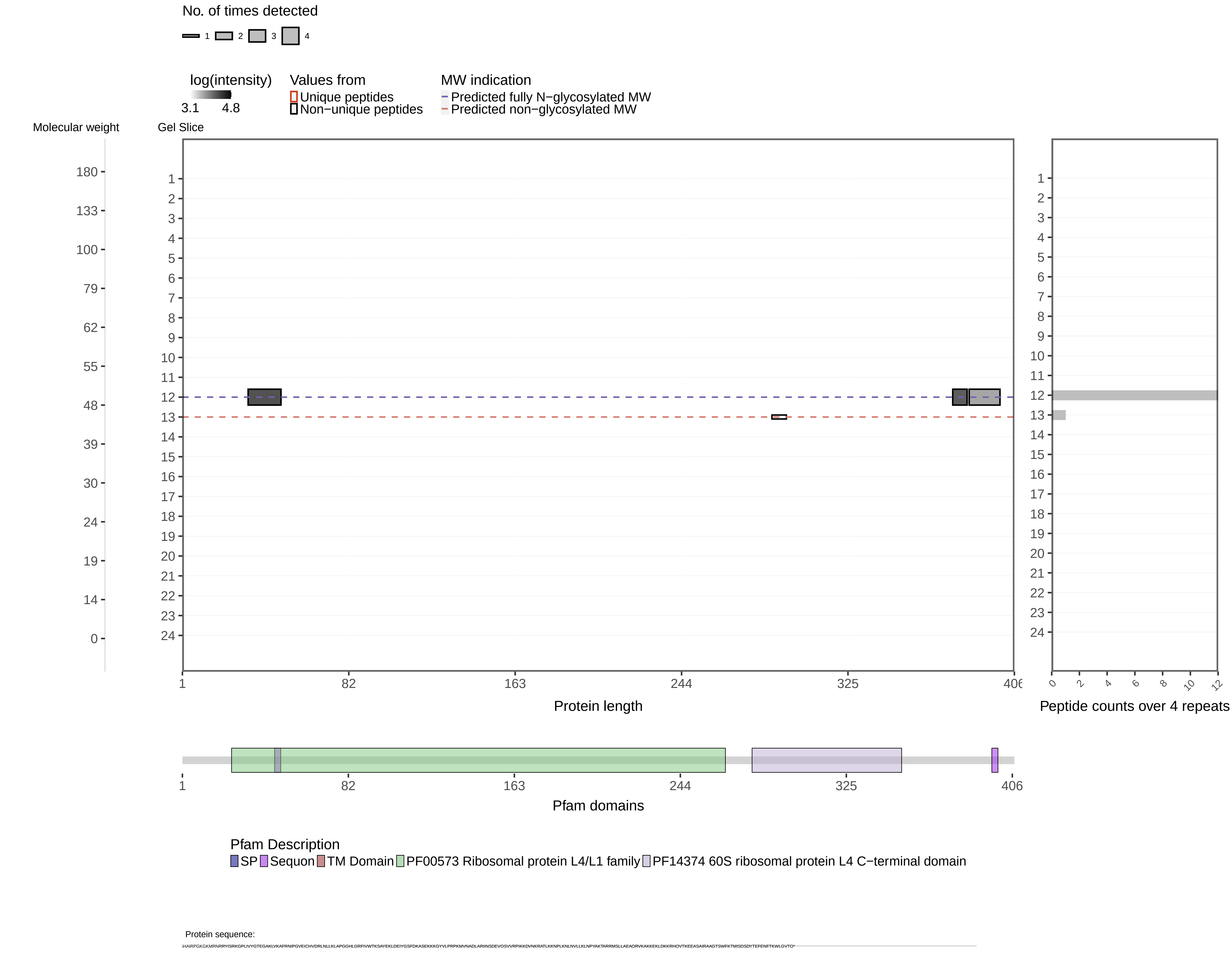

### NbL15g01970.1 protein_MODIFIER_OF_SNC1_11−like [MW = 30.03 kDa]

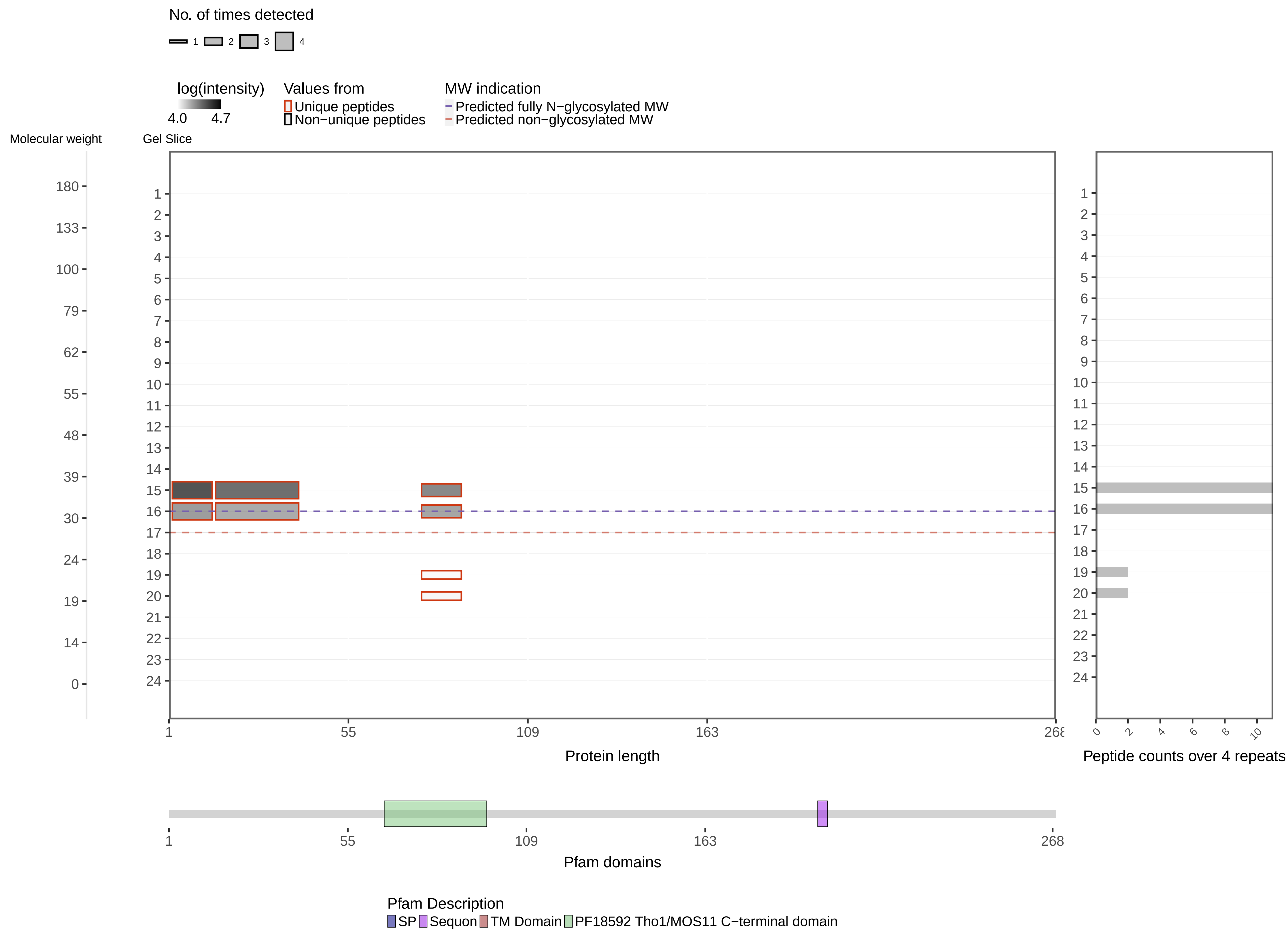


Protein sequence:

AERFGLVKSDTTEEEAKKKARLTRFAPPVKTDPVEEDKRKARALSFHSPSLLHNHKQMARETLSRMMLLWAKLERNLNDSYLTWLKSRWARILMFLSSPVDFRIQTFKFTAKTFSSLLMYHAVFSGNTRIIDNHLLGVKLPEMTCDQY*

### NbL15g01900.1 UDP−glycosyltransferase_73C5−like [MW = 55.45 kDa]

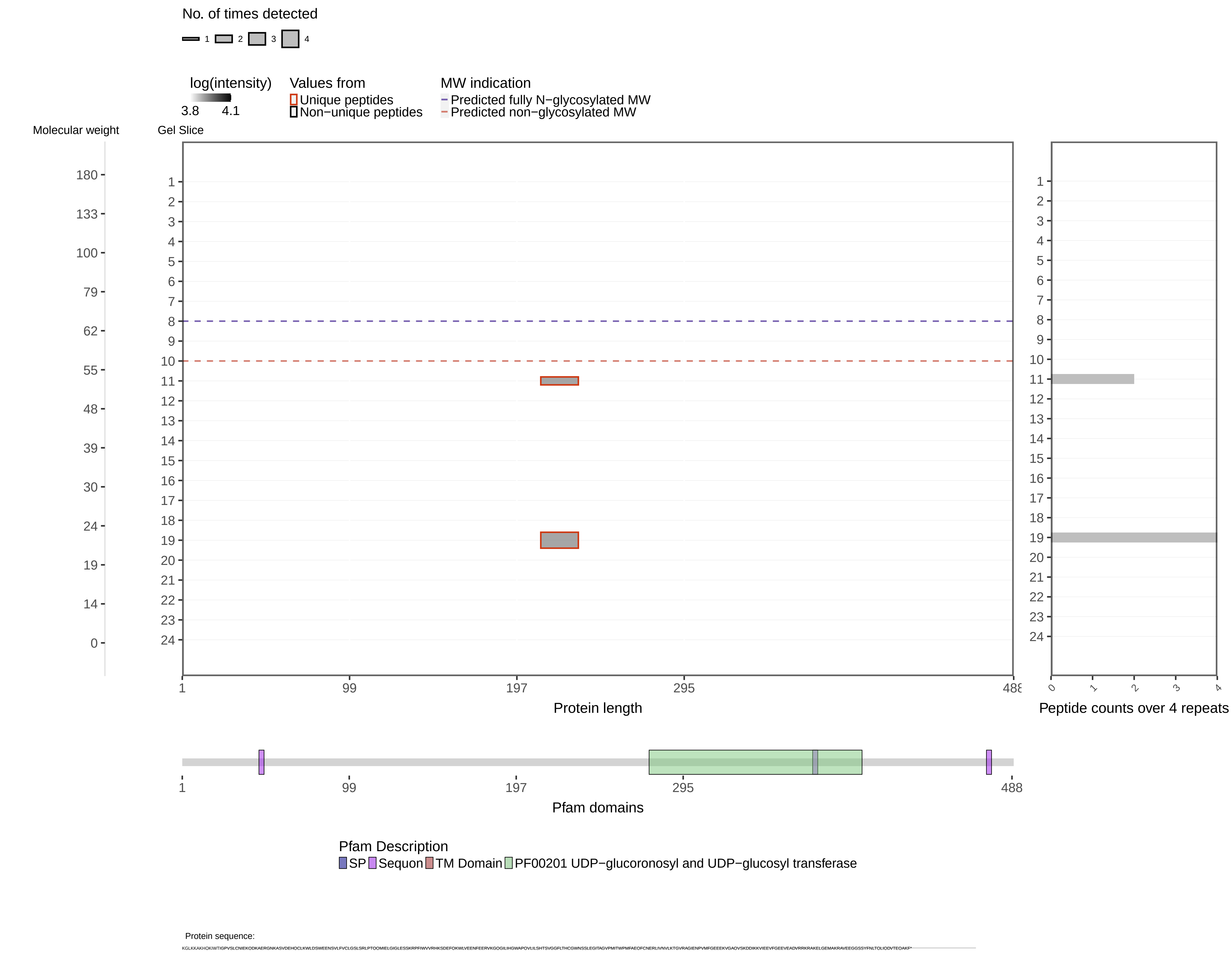

### NbL15g01610.1 protein_LURP−one−related_10−like [MW = 23.83 kDa]

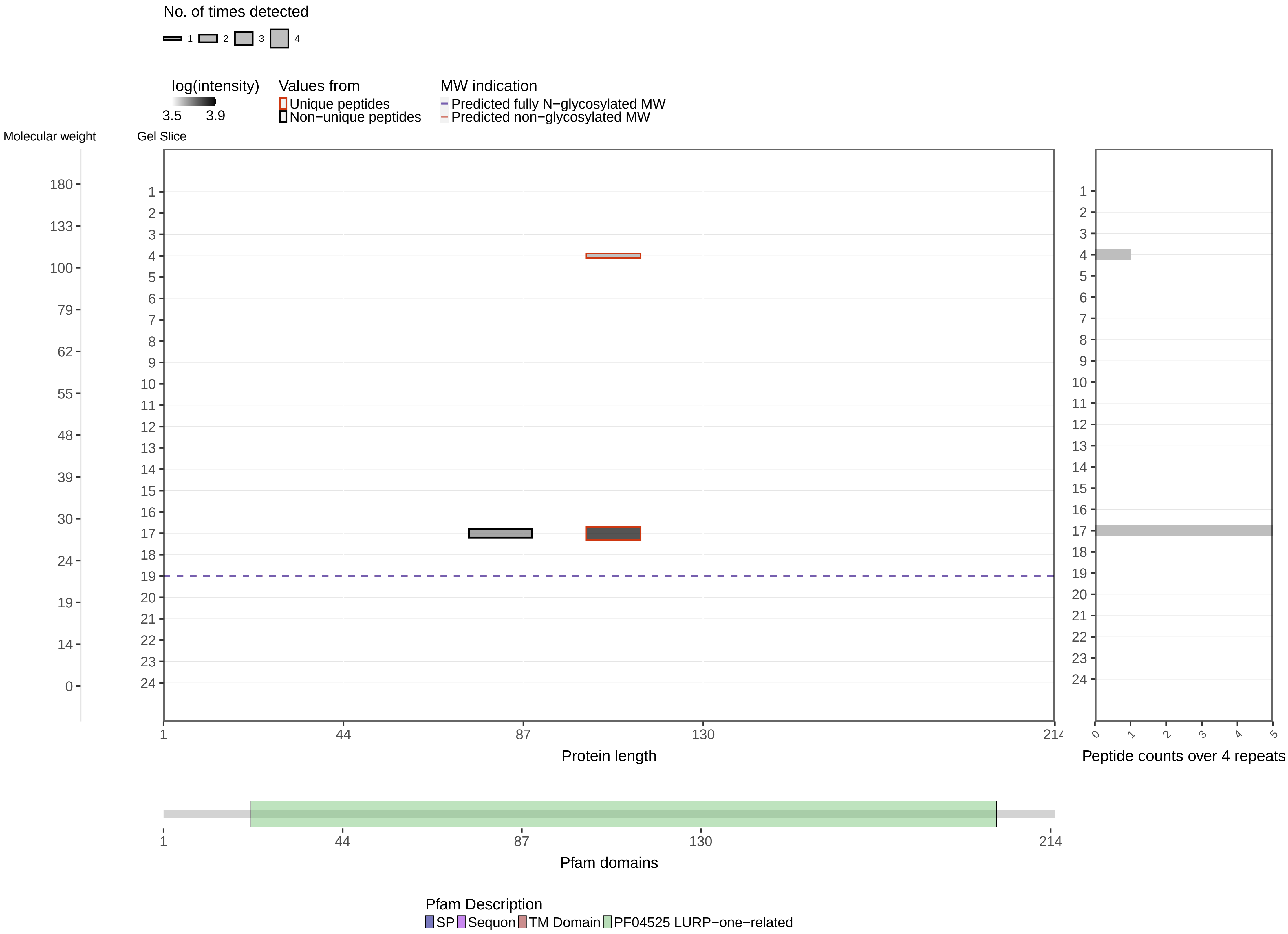


Protein sequence:

HRRWQVYRGESTDNENFLFSVKKSSLVQFKTKLDVFLAHNTKEDVCDFRIEGSWFERCCVIYAGNTNTIIAQMHRKHTAQSILLGKDNFGVTVYPNVDYAFVVALVVILEEINEDRSGAD*

### NbL15g01430.1 laccase−17−like [MW = 67.29 kDa]

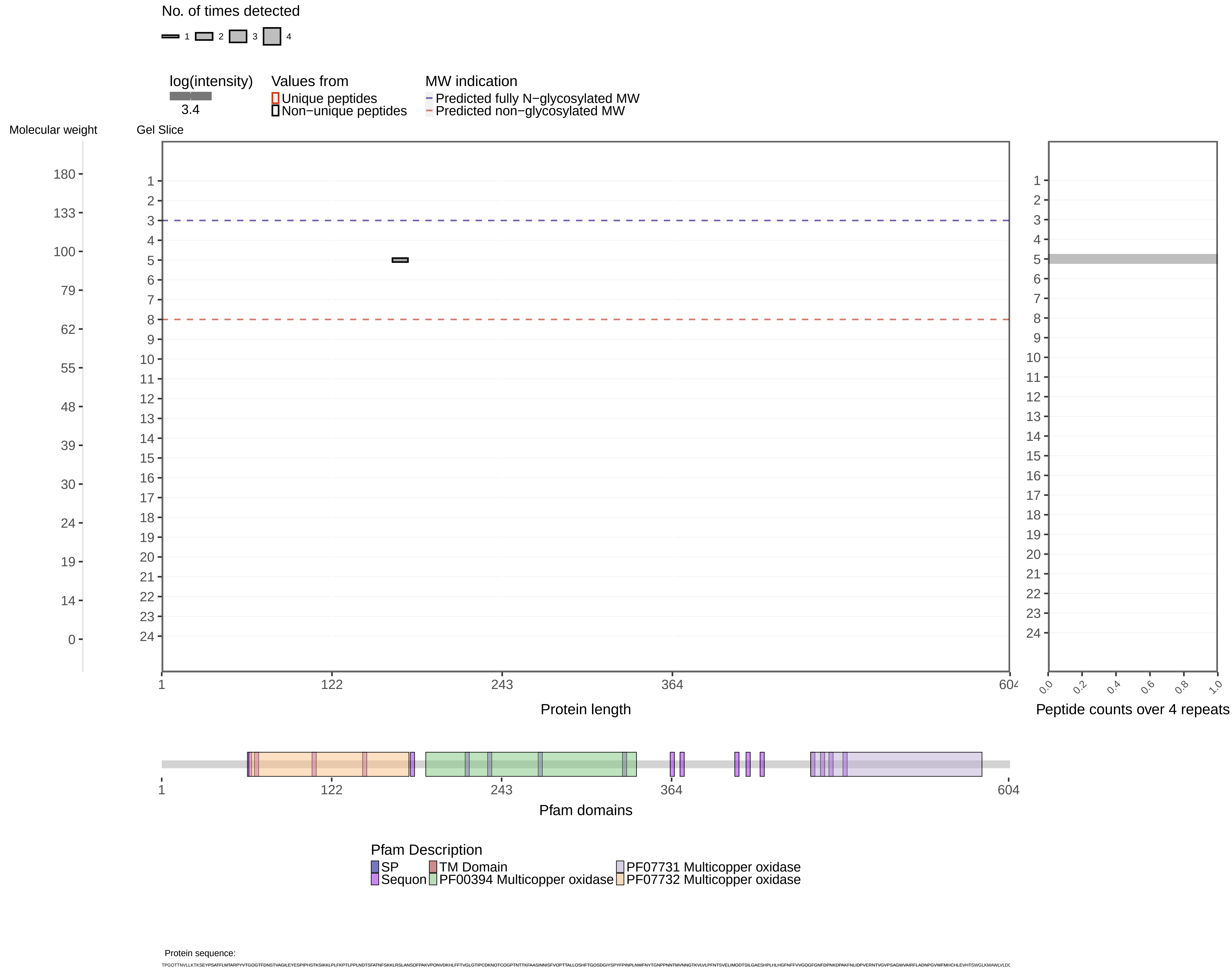

NbL15g01250.1 serine_hydroxymethyltransferase_mitochondrial−like [MW = 59.75 kDa]

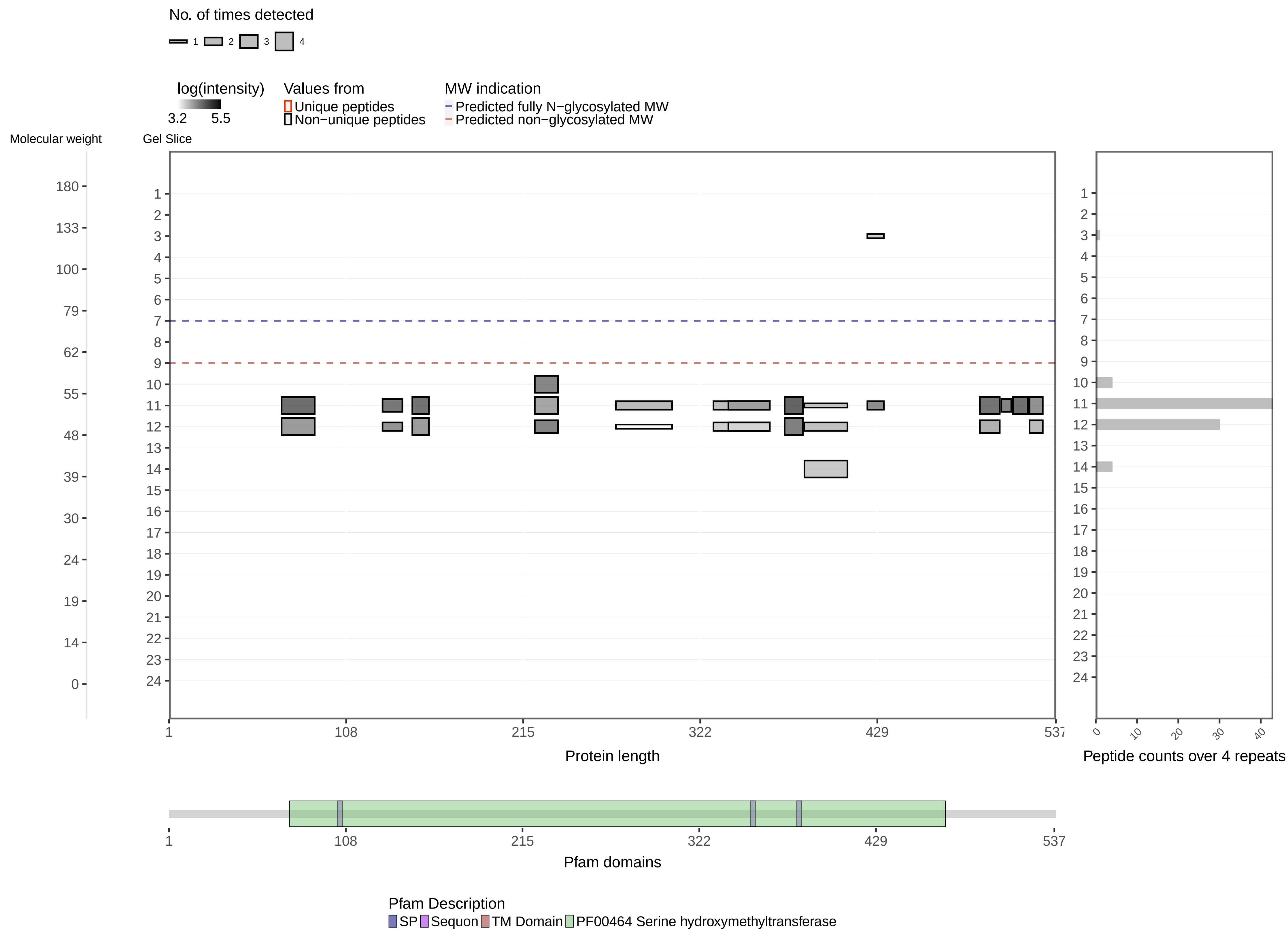


Protein sequence:

LYDYARIRKVCDKQKAVLLADMAHISGLVAAGVIPSPFEHADVVTTTTHKSLRGPRGAMIFFRKGLKEMNKQGKEVMYDYEDKINQAVFPGLQGGPHNHTISGLAVALKQVMTPEYKAYQEQVLSNCSKFAESLLANGYDLVSGGTENHLVLVNLRNKGIDGSRVEKVLESVHIAANKNTVPGDVSAMVPGGIRMGTPALTSRGFIEEDFVKVAEFFDAAVKLALKVKAETQGTKLKDFVETLSSDSKIQSEIARLRQDVEDYAKQFPTVGFEKETMKYKD*

### NbL15g01150.1 tubulin_beta−2_chain [MW = 50.18 kDa]

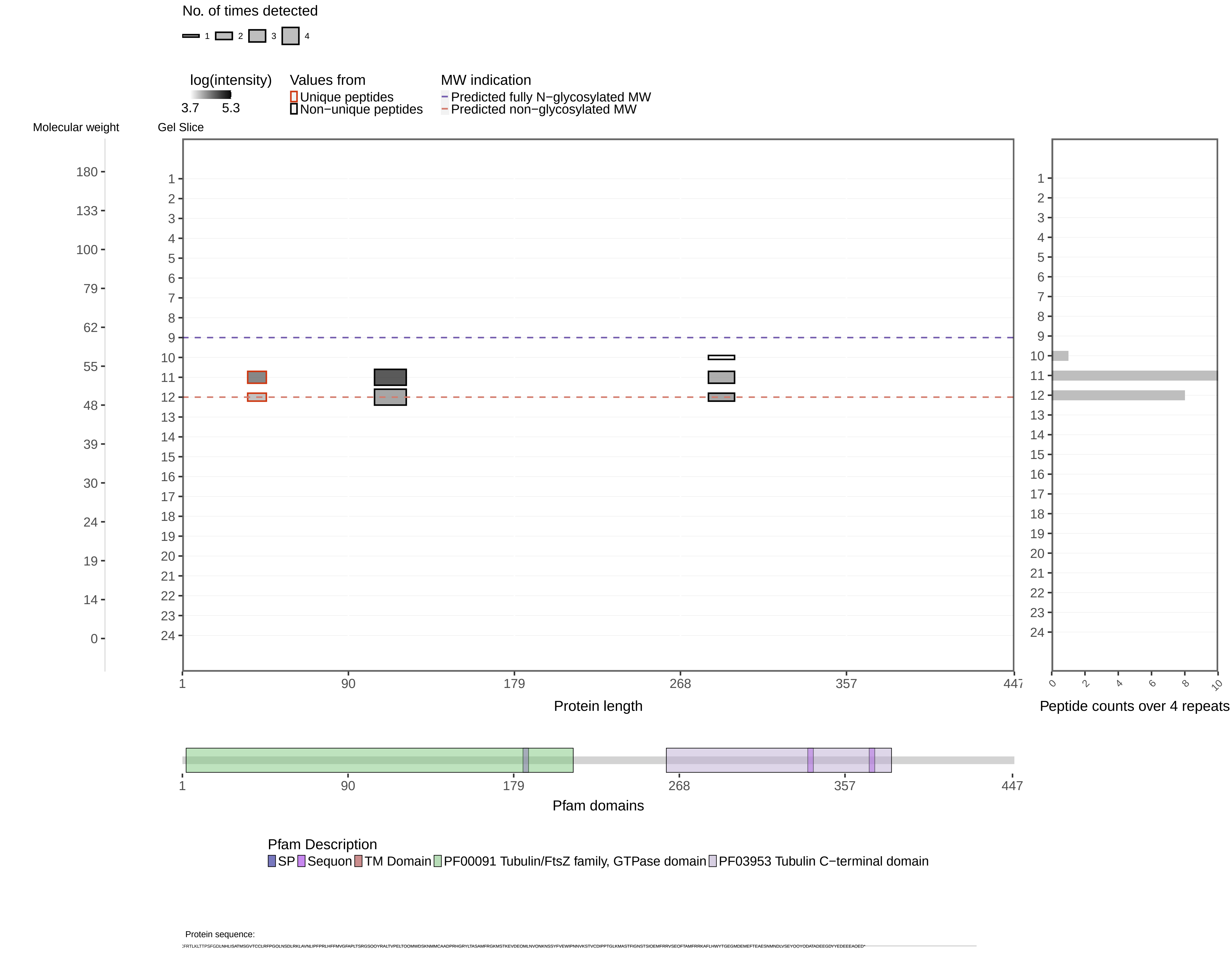

NbL15g01090.1 GDSL_esterase/lipase_At2g38180−like [MW = 27.78 kDa]

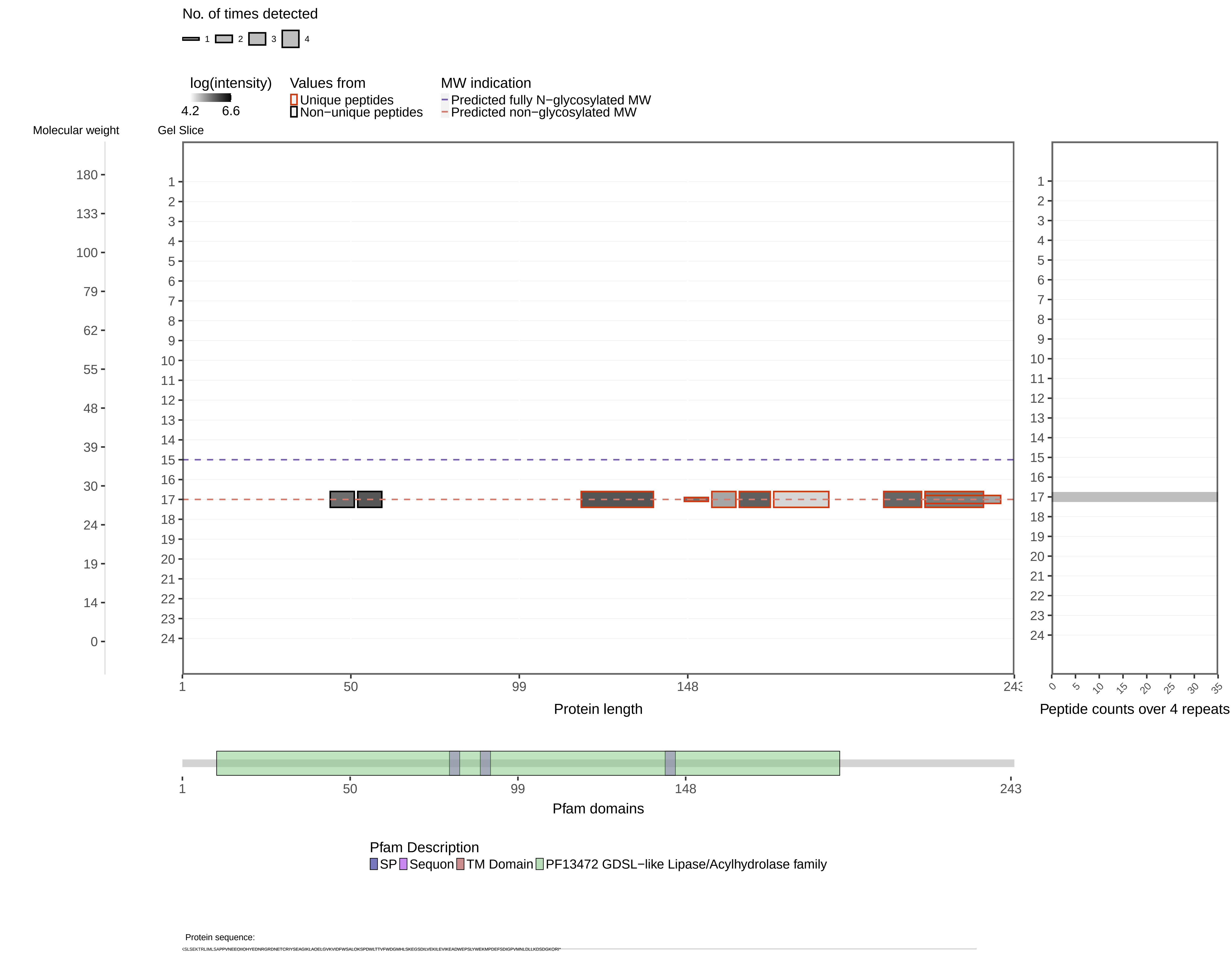

### NbL15g01020.1 fructose−bisphosphate_aldolase_1_chloroplastic [MW = 42.95 kDa]

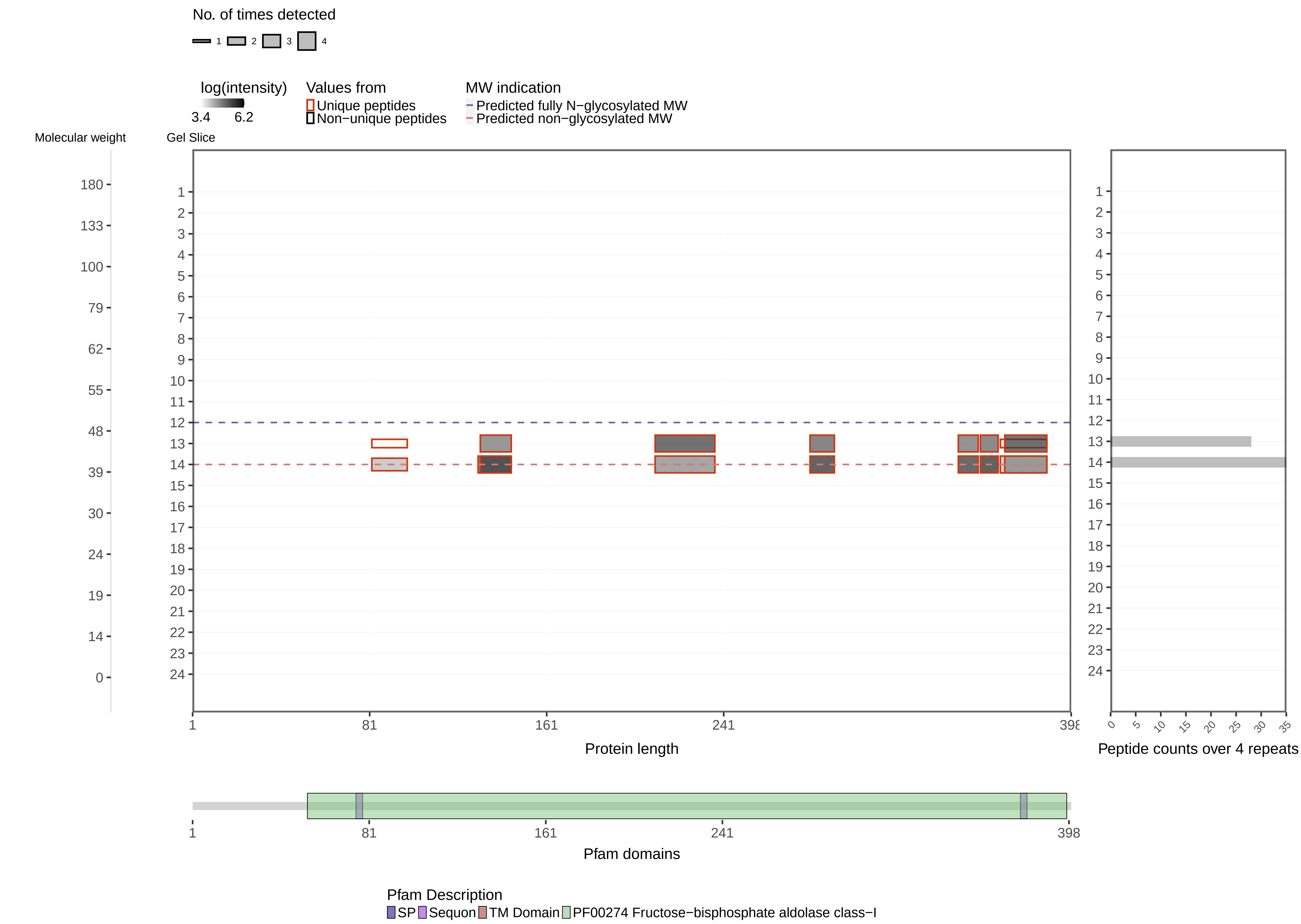


Protein sequence:

VVSIPNGPSALAVKEAAWGLARYAAIAQDSGLVPIVEPEILLDGEHGIDRTFEVAQQVWAEVFFYLAENNVMFEGILLKPSMVTPGAECKDRATPEQVADYTLKLLYRRIPHAVPGIMFLSGGQSEVEATYNLNAMNQAPNPWHVSFSYARALQNTCLKTWGGLPENVKAAQDALLVRAKANSLAQLGNYTGEGETDEAKQGMFVKGYVY*

### NbL15g00990.1 arginine_decarboxylase−like [MW = 79.12 kDa]

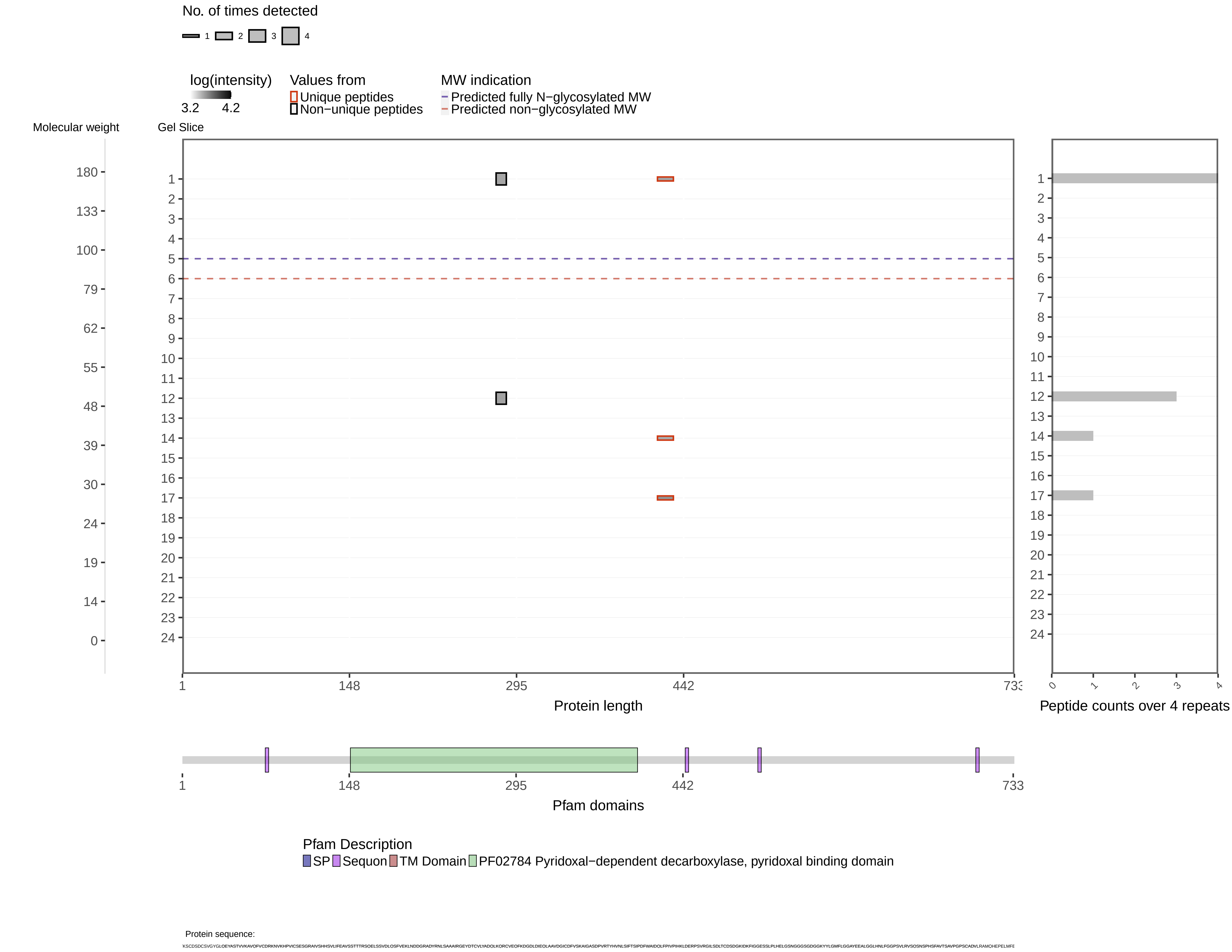

### NbL15g00930.1 eukaryotic_translation_initiation_factor_3_subunit_A−like [MW = 111.44 kDa]

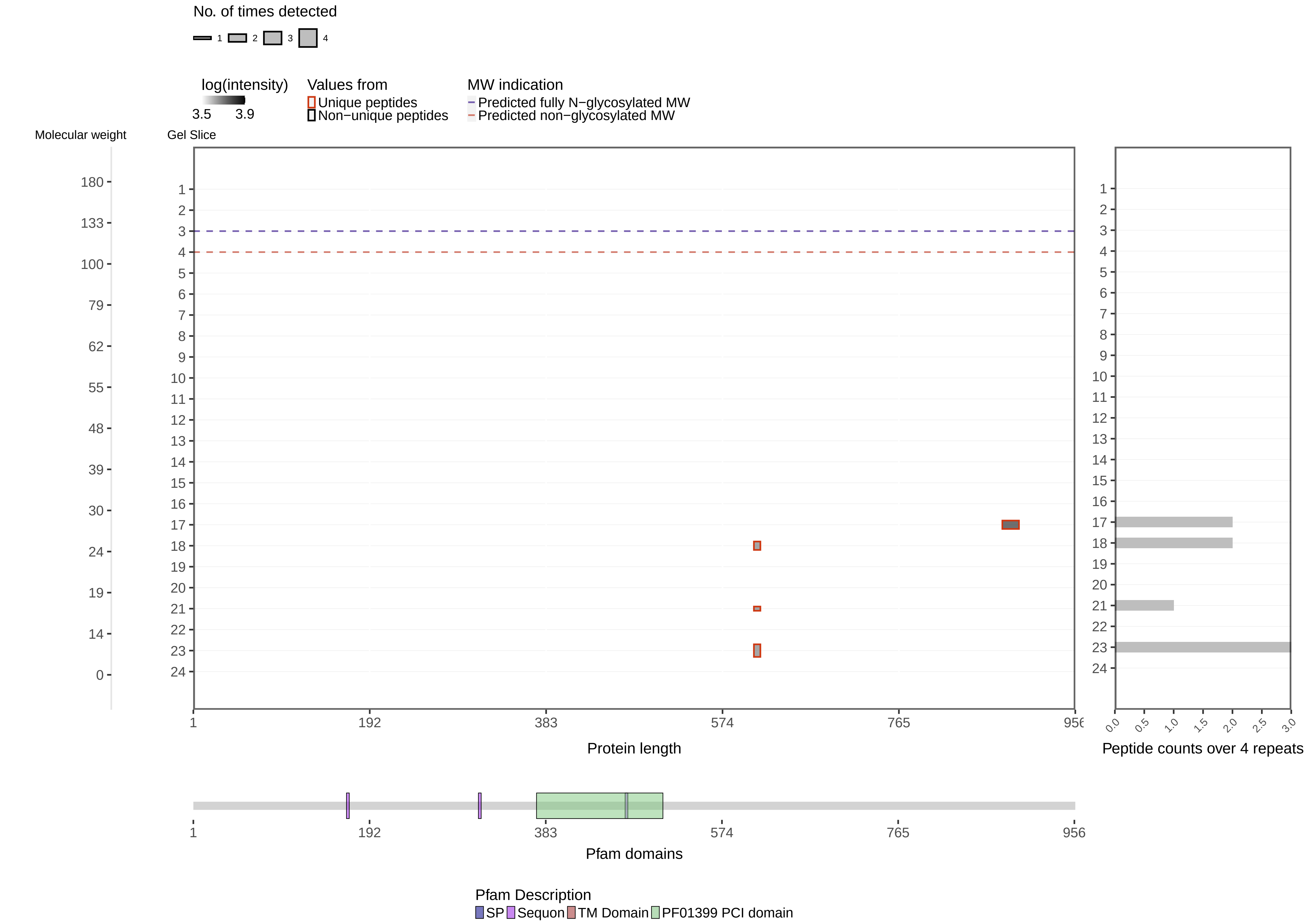


Protein sequence:

[illegible]

### NbL15g00860.1 oxysterol−binding_protein−related_protein_3A [MW = 52.19 kDa]

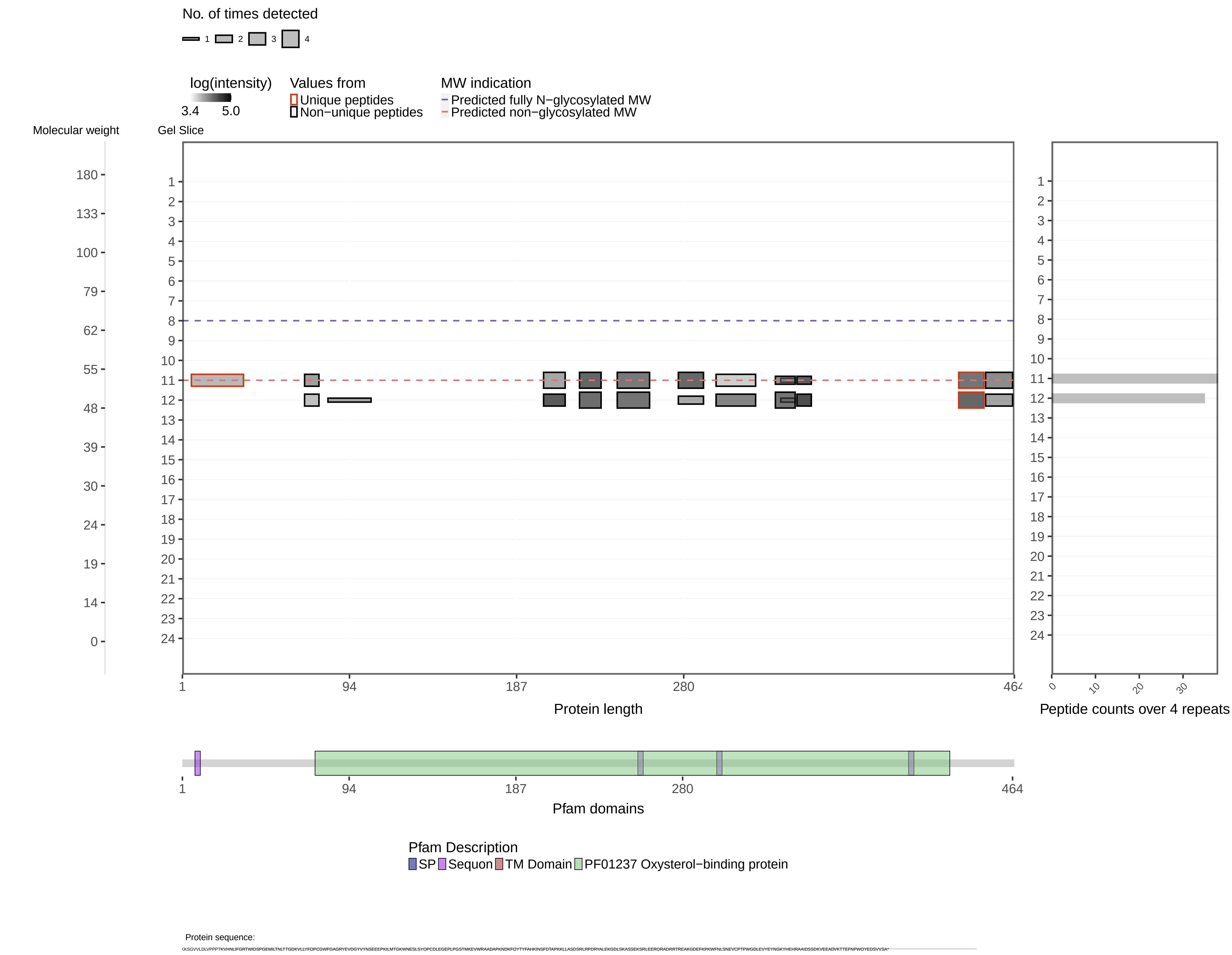

### NbL15g00710.1 glycine−rich_RNA−binding_protein−like [MW = 16.89 kDa]

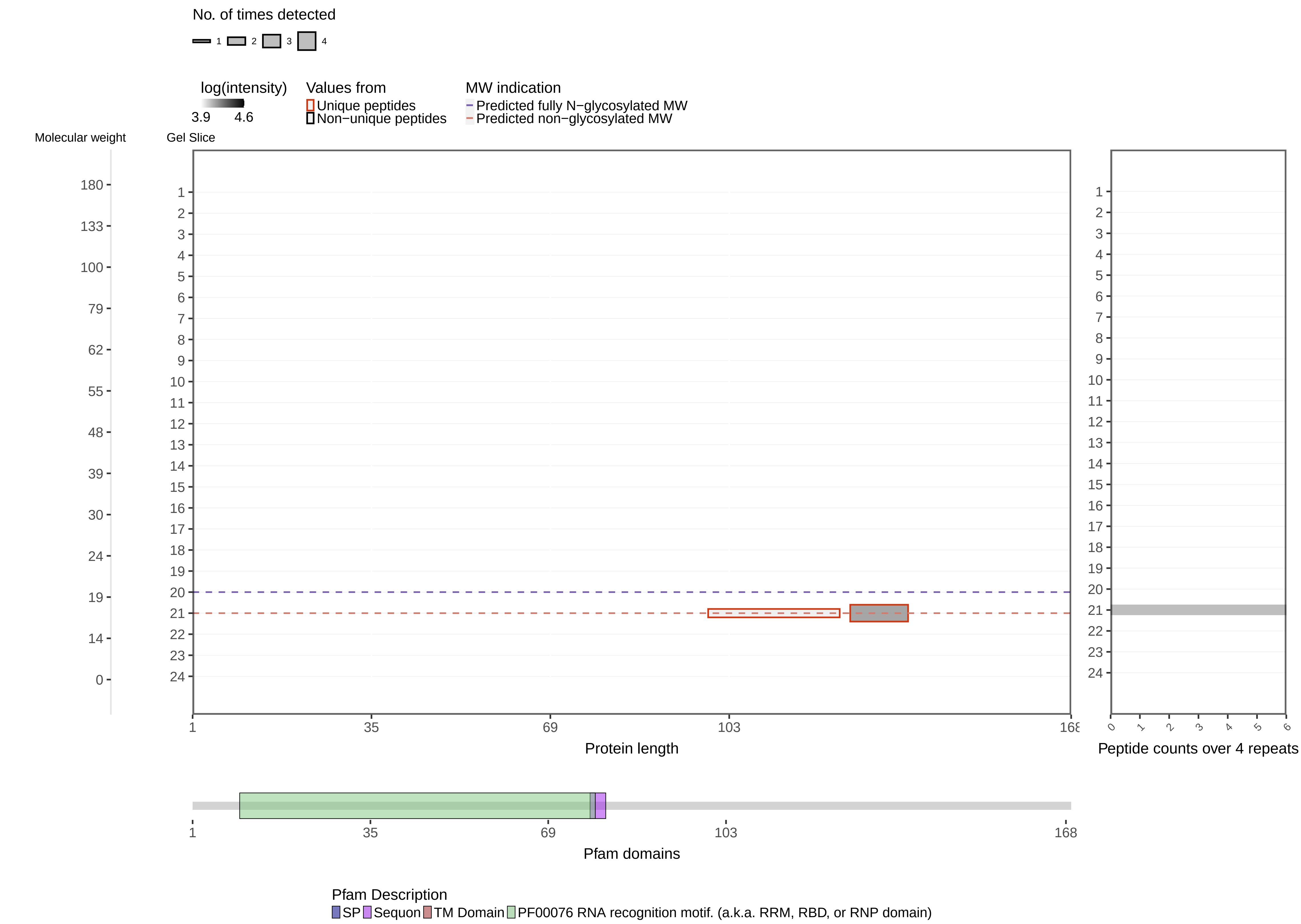


Protein sequence:

RNITVNEAQSRGGGGGFGGGRRREGGYSGGGGYSGGGGYSGGGGYGGGRREGGYGGGYGGGRDRGYGGGYGGGDGGSRYSRGGGASEGSWRN*

### NbL15g00670.1 ATP−dependent_Clp_protease_proteolytic_subunit_6_chloroplastic [MW = 30.88 kDa]

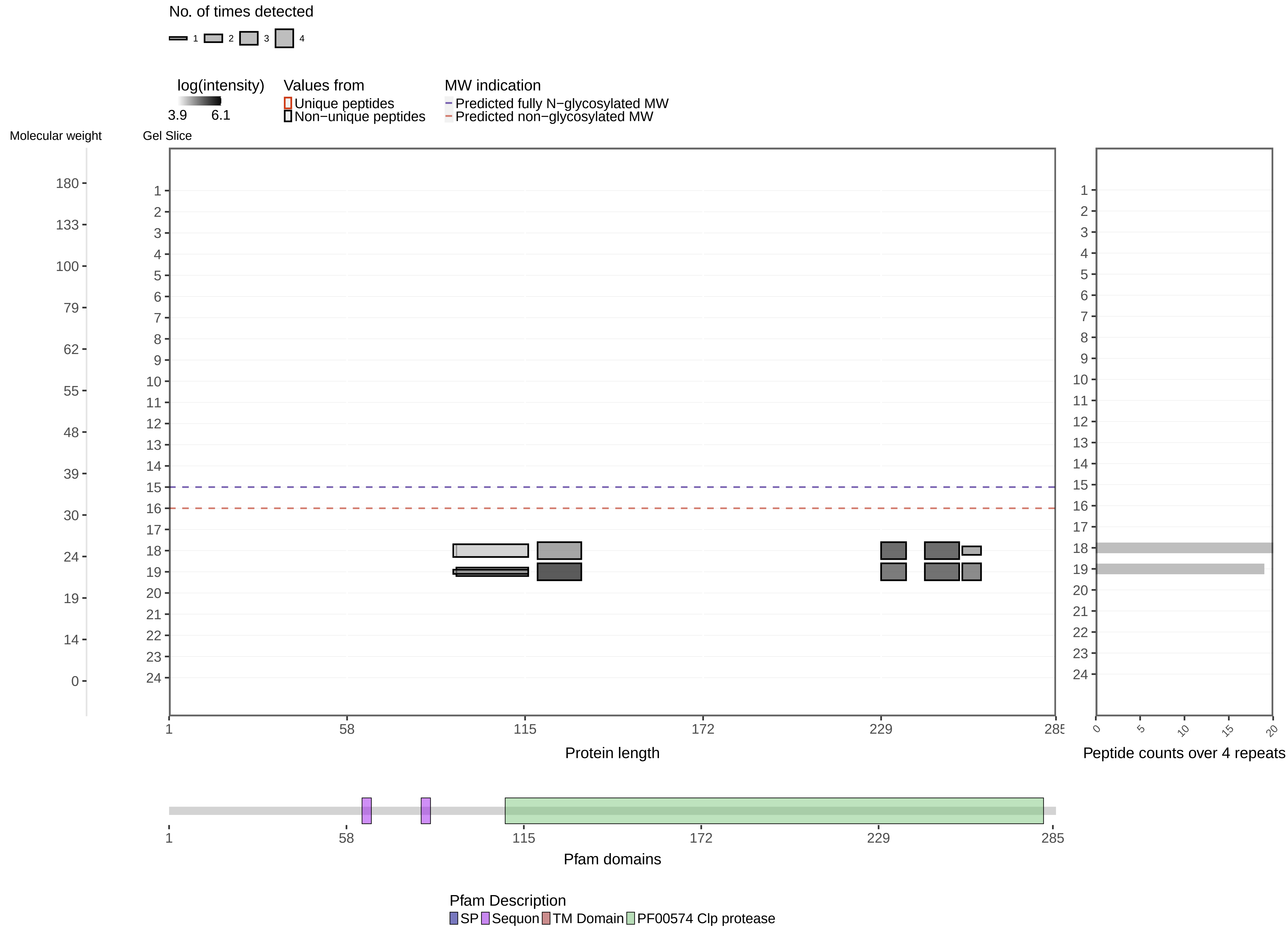


Protein sequence:

AQRVISQLVTLATIDENADILIYLNCPGGSTYSVLAIYDCMSWIKPKVGTVCFGVAASQGALLLAGGEKGMRYAMPNARIMIHQPQSGCGGHVEDVRRQVNEAVQSRHKIDKMYVAFTGQSIEKVQQYTERDRFLSVSEAMEFGLIDGVLETEY*

NbL15g00650.1 protein_TIC_62_chloroplastic [MW = 68.55 kDa]

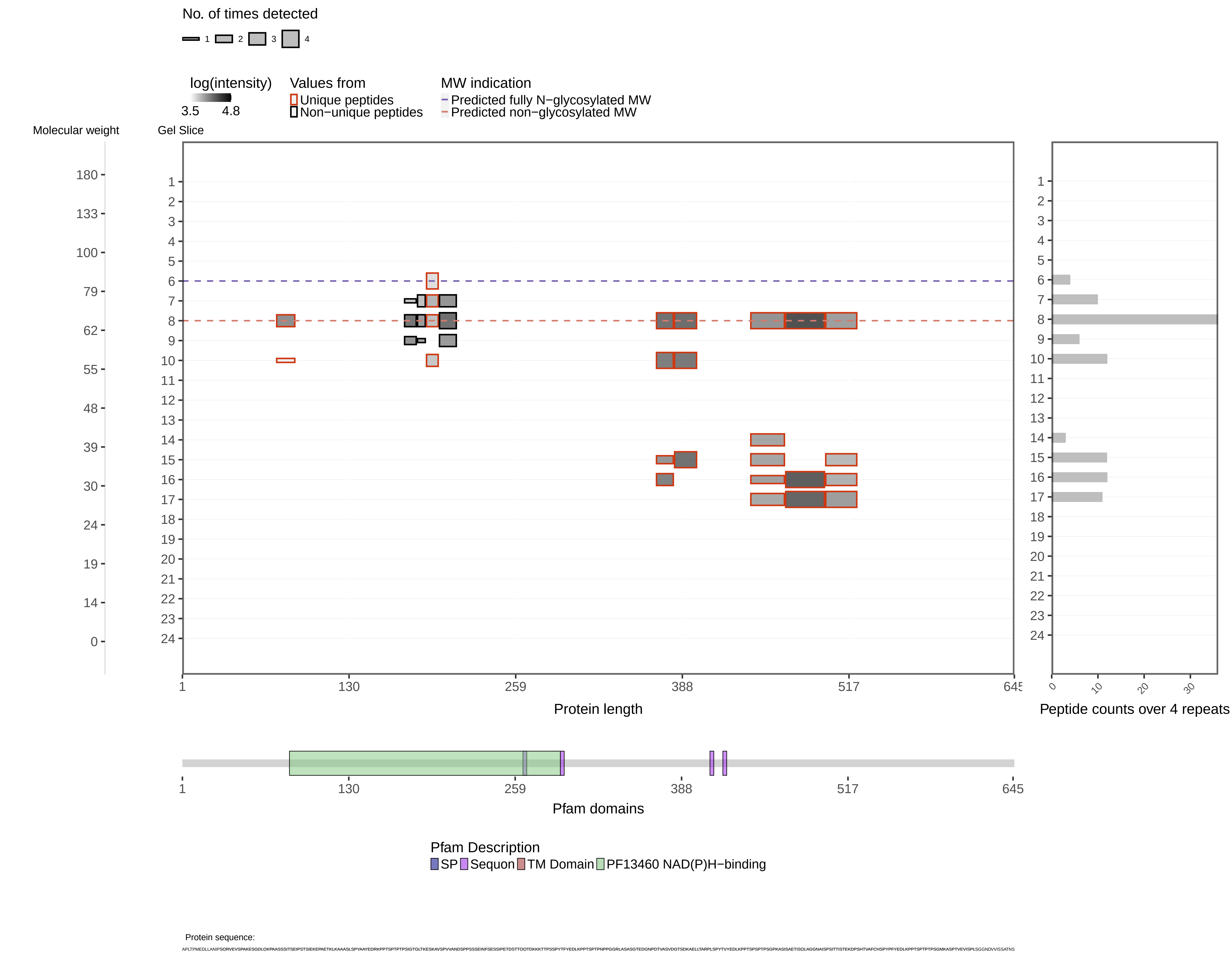

### NbL15g00590.1 ferredoxin−−nitrite_reductase_chloroplastic−like [MW = 65.6 kDa]

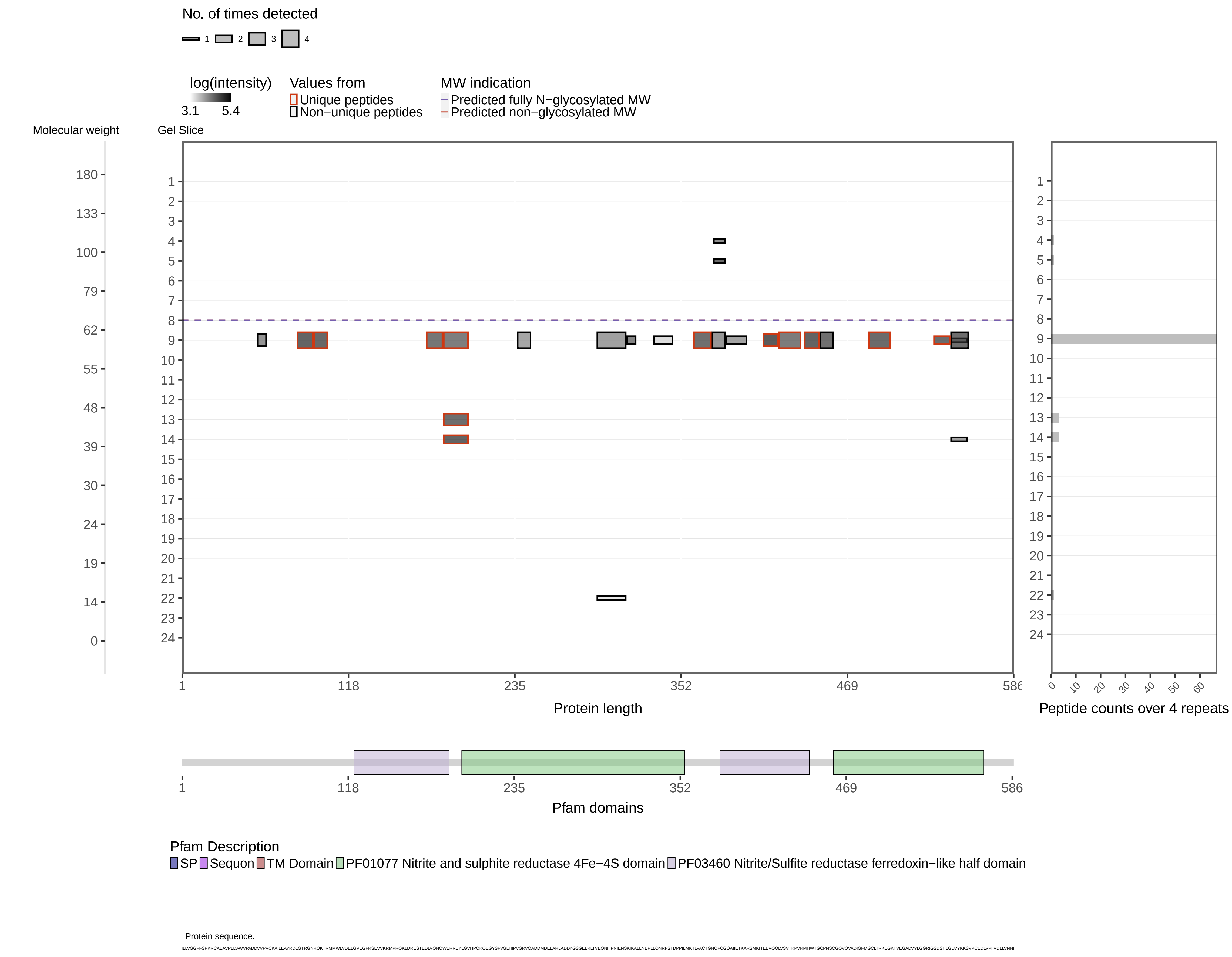

### NbL15g00470.1 uncharacterized_protein_At5g01610−like [MW = 19.27 kDa]

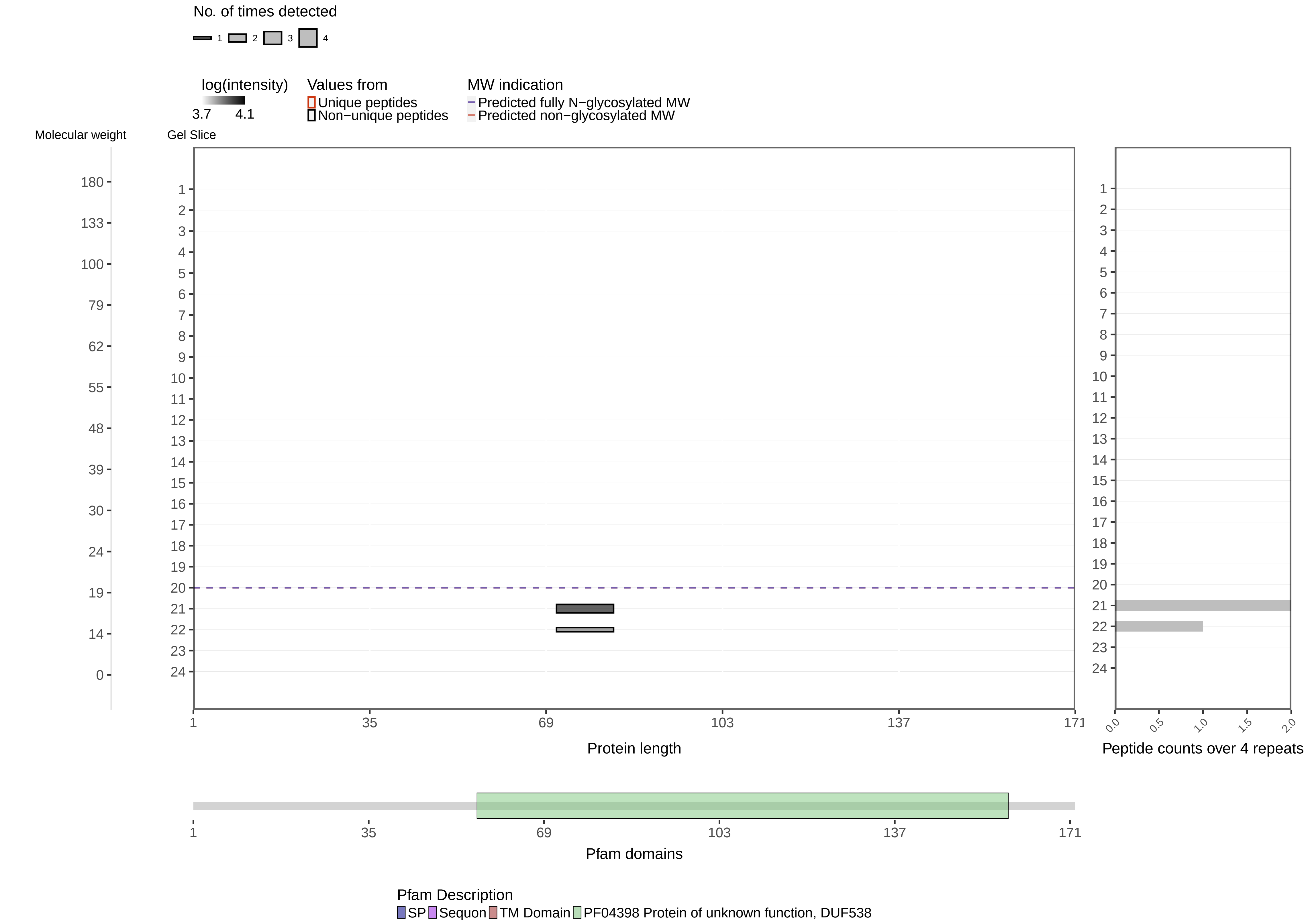


Protein sequence:

ATNYEFNEETRKLTVYIPSVCEVGYKDSSVLRFSTAVTGYLEKGKLVDIEGMKTKVMMWVKVTSISSEKSKIHFTAGLKKTRSREAYEVLRDGVAMEKF*

NbL15g00360.1 fructose−bisphosphate_aldolase_6_cytosolic−like [MW = 44.5 kDa]

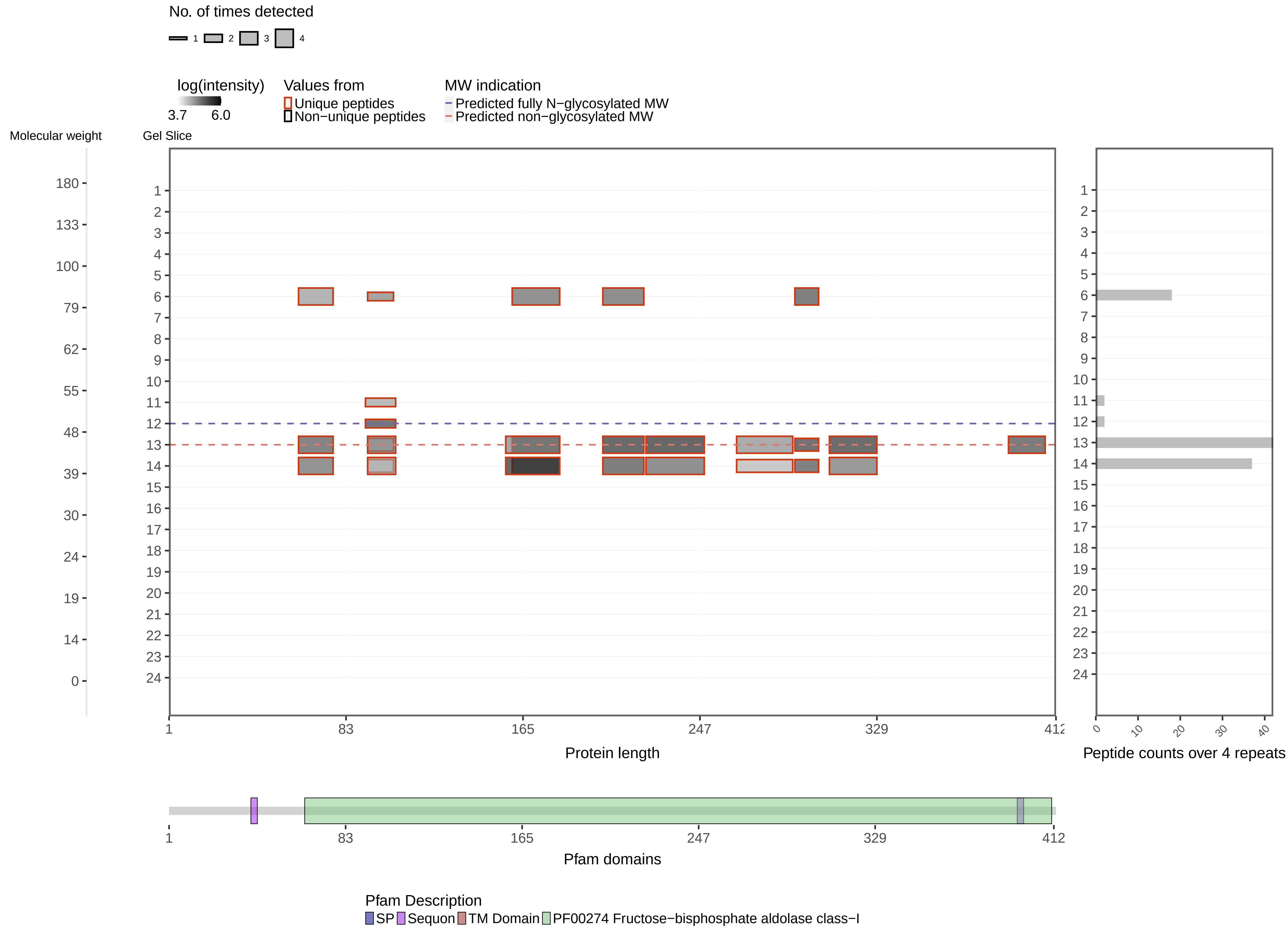


Protein sequence:

VRAVLKIGANEPSQLAINDNANGLARYAIICQQNGLVPIVEPEILVDGGHDINKCADVTERVLAACYKALNDHHVLLEGTLLKPNMVTPGSESPKVAPEVIAEYTVRALQRTMPAAVPAVVFLSGGQSEEEATRNLNAMNKLQSKRPWTLSFSFGRALQQSTLKAWGGKVENVEKARAAFLTRCKANSEATLGKYLGSSNLSEGASESLHVKDYKY*

### NbL15g00210.1 uncharacterized_LOC104247352 [MW = 26.34 kDa]

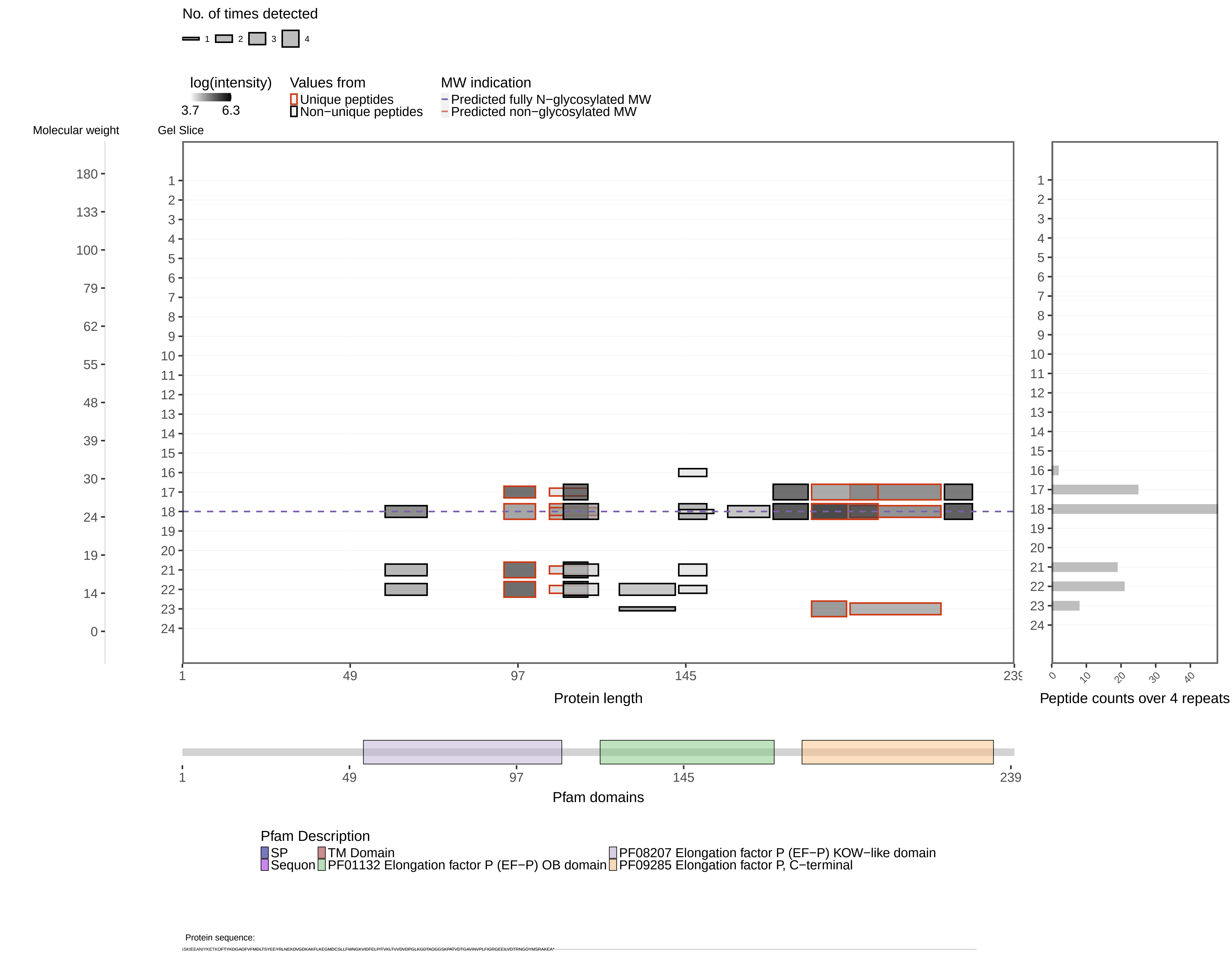

### NbL15g00200.1 apyrase_1−like [MW = 51.8 kDa]

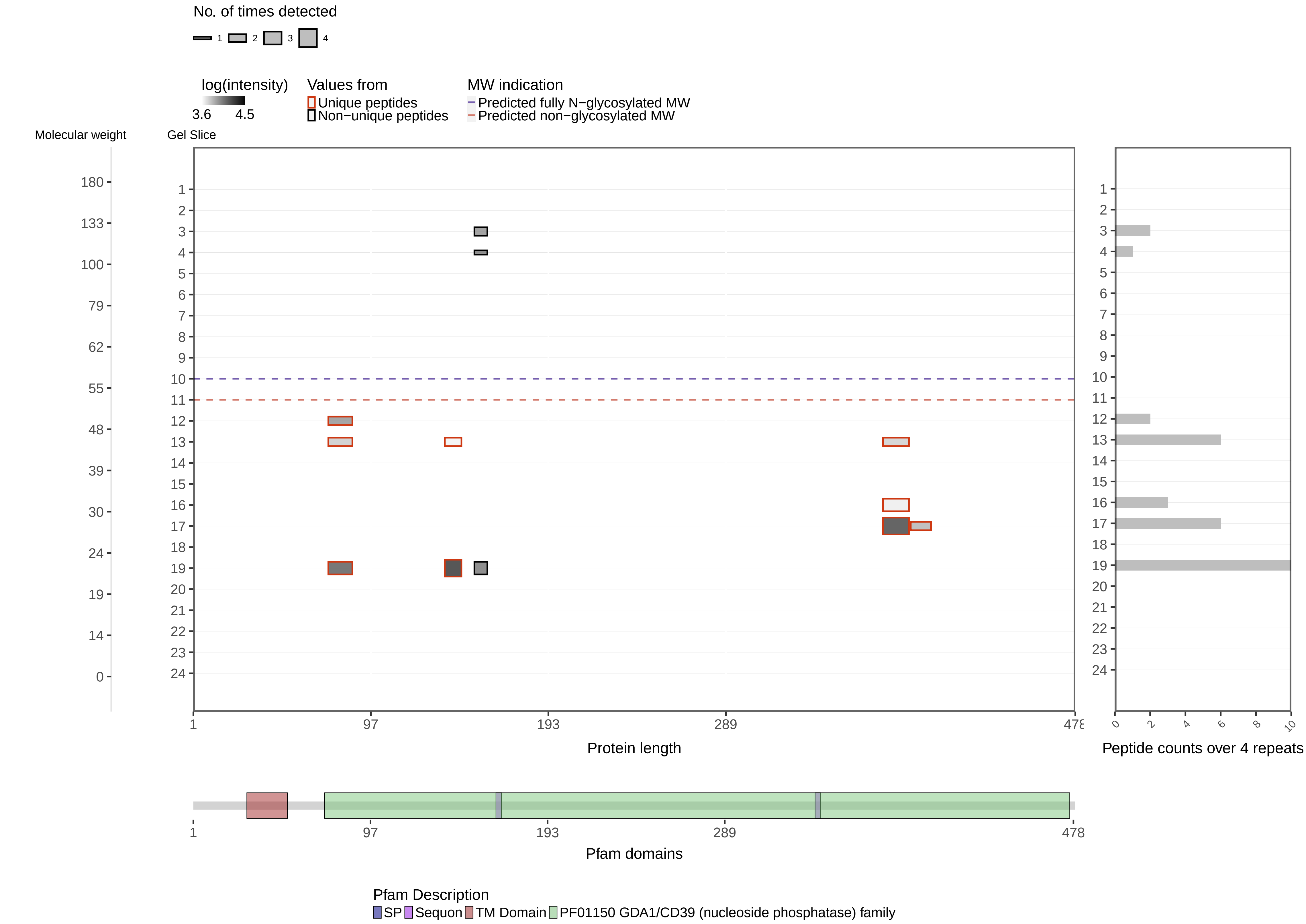


Protein sequence:

GGGSVQMAYAISEADAEKAPKLSGGEDTYVQEMYLKGAKYYLYVHSYLHYGLLAARAEILKVTKESGSPCILTGHHGSYKYGGKTYTASAMSQGSSLTNCRMVALKALKVNESACTHMKCTFGGVWNGGGGDGQKNMFVASFFFDRAVEVGFVDPNAAVAKVRPIDFESAAGRACDTRIDDAKATFPRVDPDNLPFLCMDLVYQYTLLVDGFGLDARQEMTLVKKVKYKNSLVEAAWPLGSAIEVASSMN*

NbL15g00180.1 uncharacterized_LOC104223818 [MW = 66.21 kDa]

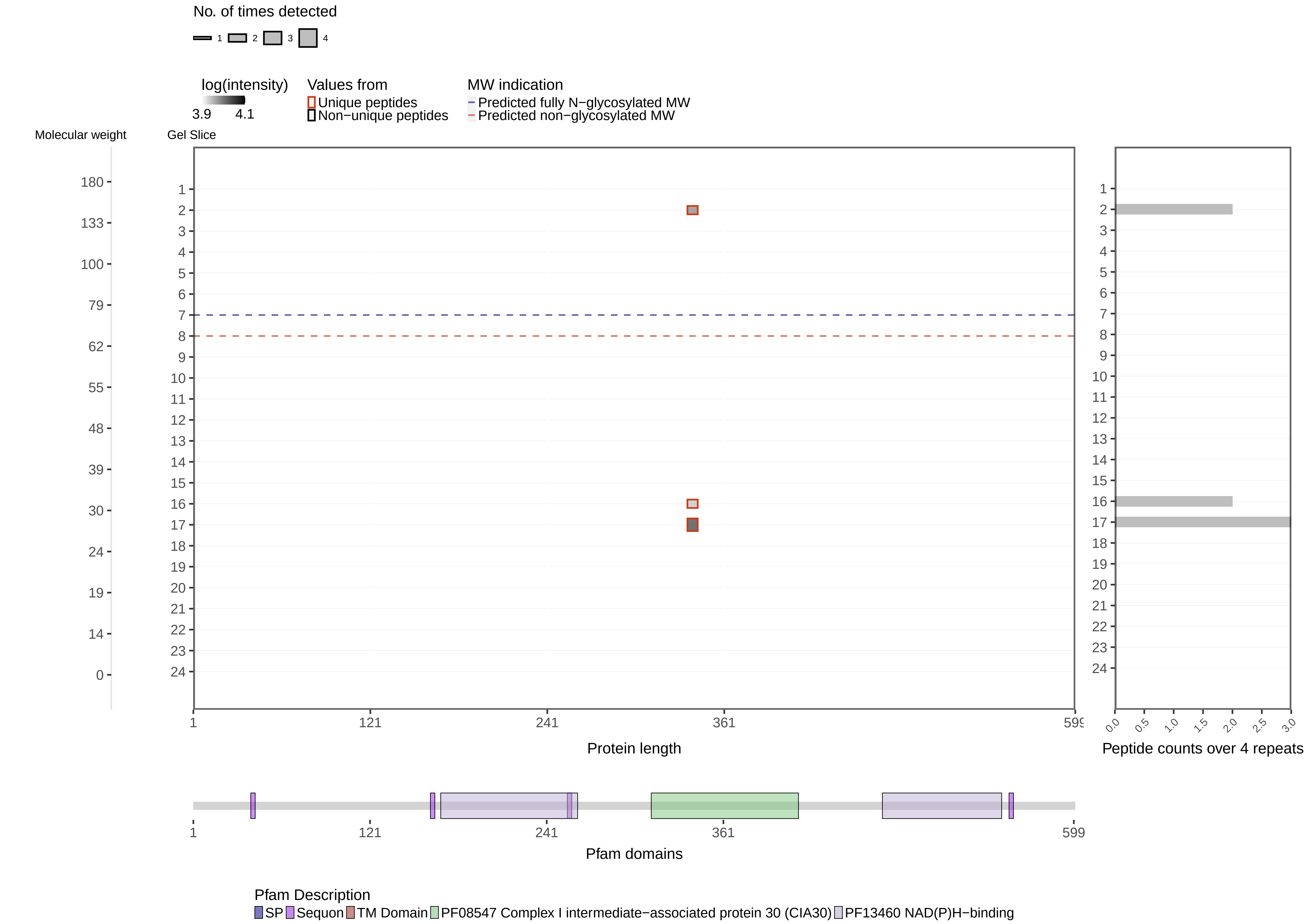


Protein sequence:

SEDSLNGWEVRQGTYFQDVVASKYDGGMDATFEFTERGEAIFSGYVFTRGGYVELTRKLSLPLGYTLDRYEGLVFSVGGNGRSYVVILEAGPSADTTQSKLYFARISTKAGFCRVRVPFSSFRPVSPDDPPLDPFLVHTLTIRFEPRRQRPIDGPSGVNQDLRSFQLIMEYIKALPTGQETDFVLVSCTGSGIEPTRREQVLRAKRAGEDSLRRSGLGYTIIRPGPLMEEPGGQRALIFDQGNRISQGISCADVADICVKALHDSTARNKSFDVCYEYVAEPGKELYELVAHLPI

NbL15g00090.1 26S_proteasome_non−ATPase_regulatory_subunit_12_homolog_A−like [MW = 39.02 kDa]

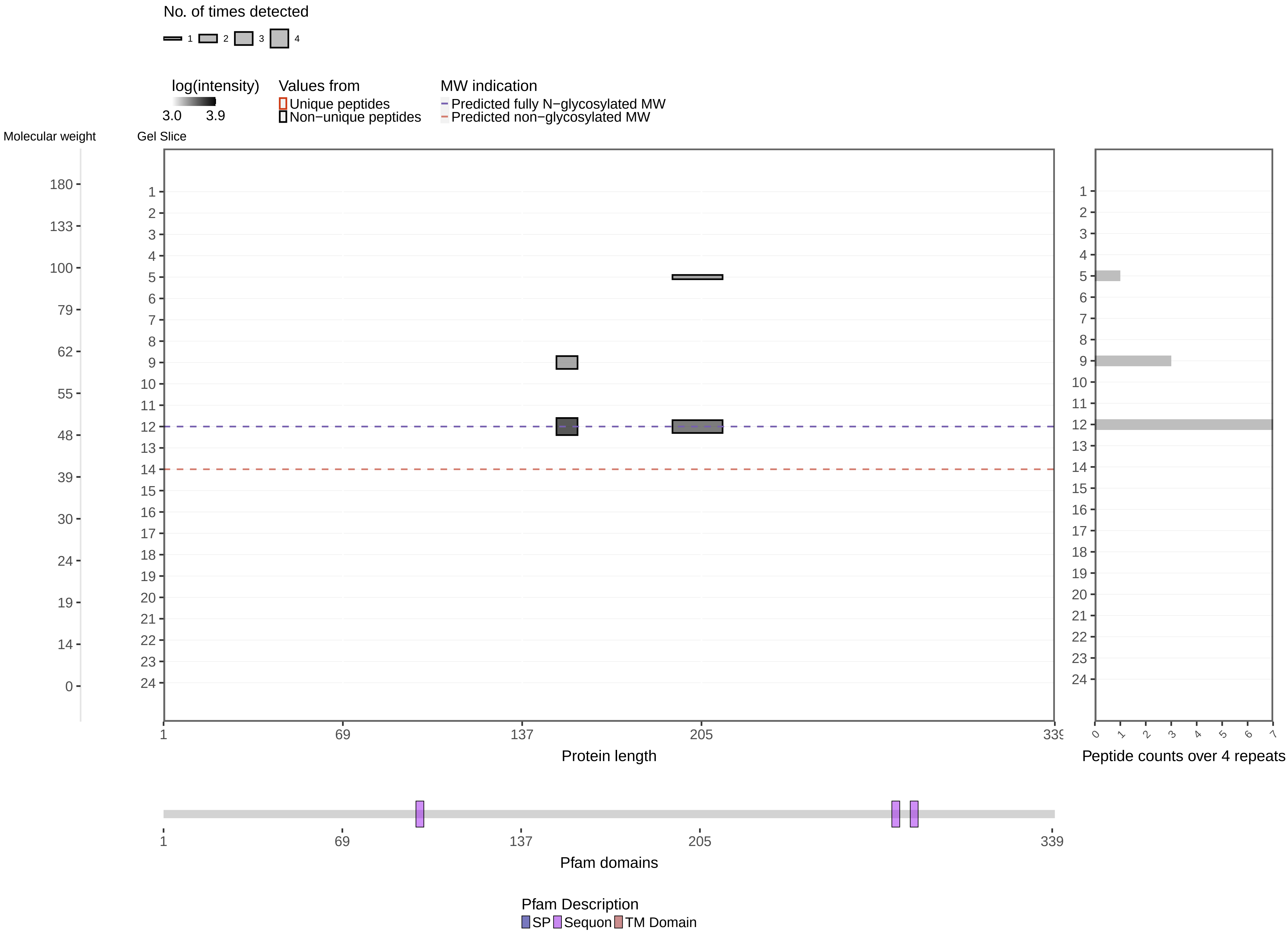


Protein sequence:

QVRLCLDRQDYVRAQILSRKINPRVFEAVPSKEKKKPKEGENVVEEPAADIPSLLELKQIYYQLMIRYYSHSNDYLEICRCYKAIYEIPSVKEDPAQWIPVLRKICWYLVLSPHDPMQSSLHNSTLEDKNLSEIPHFRLLLKQLITMEVIQWTALWNTFKMSLKMKRICLVALWVTRLPKISD*

### NbL15g00030.1 systemin_receptor_SR160 [MW = 132.88 kDa]

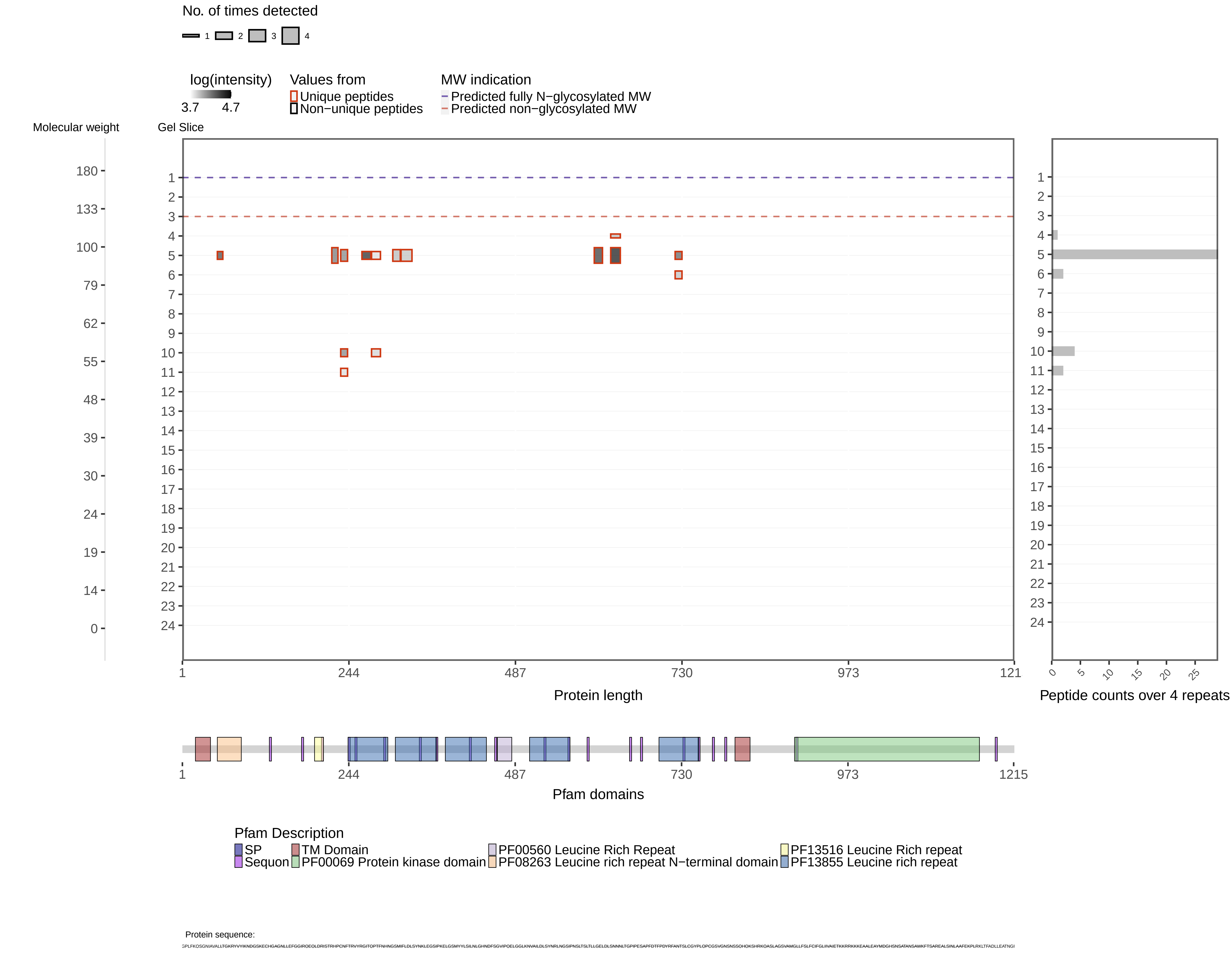

### NbL14g22030.1 fumarate_hydratase_1_mitochondrial−like [MW = 53.3 kDa]

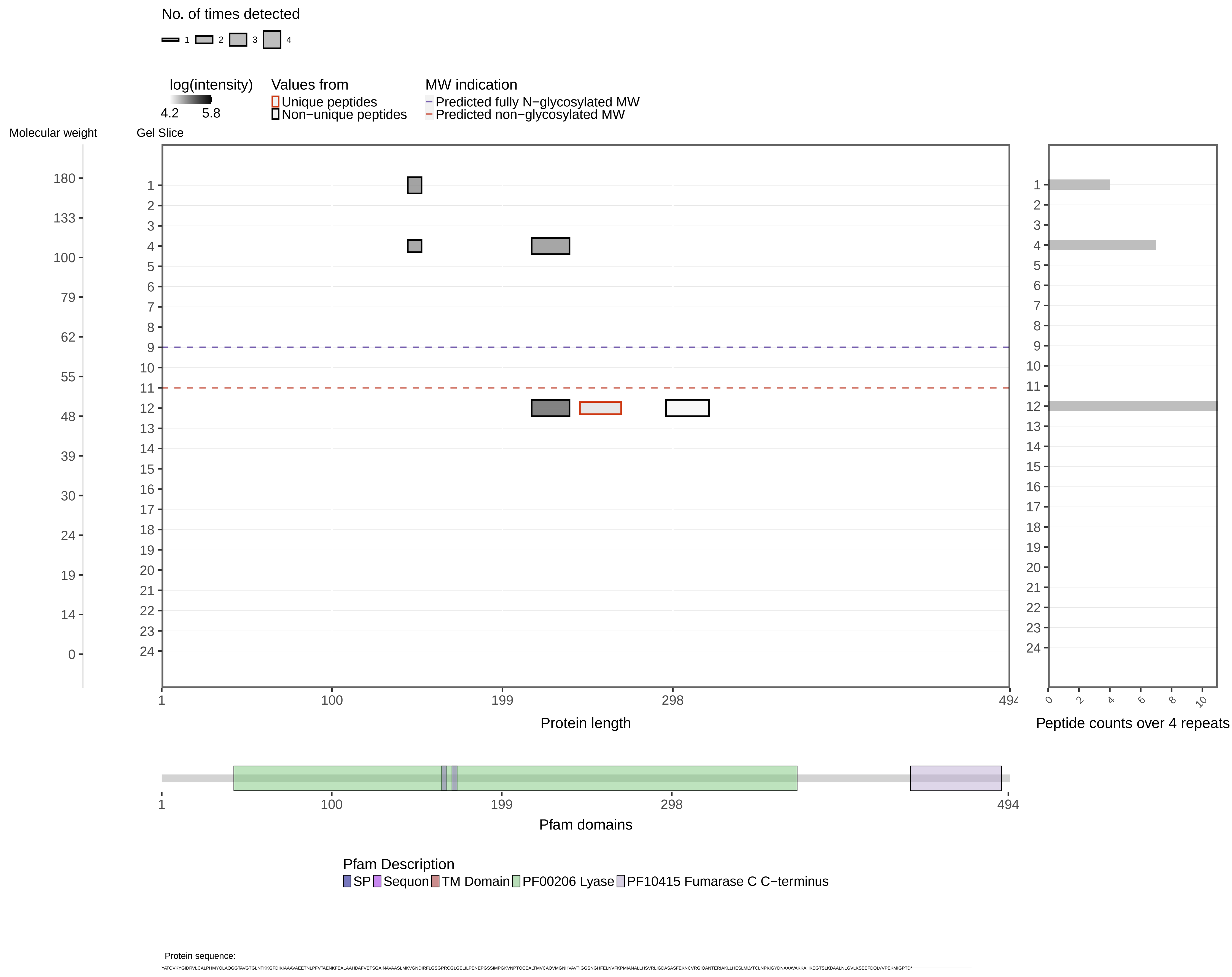

### NbL14g21940.1 methionine_aminopeptidase_1B_chloroplastic [MW = 39.53 kDa]

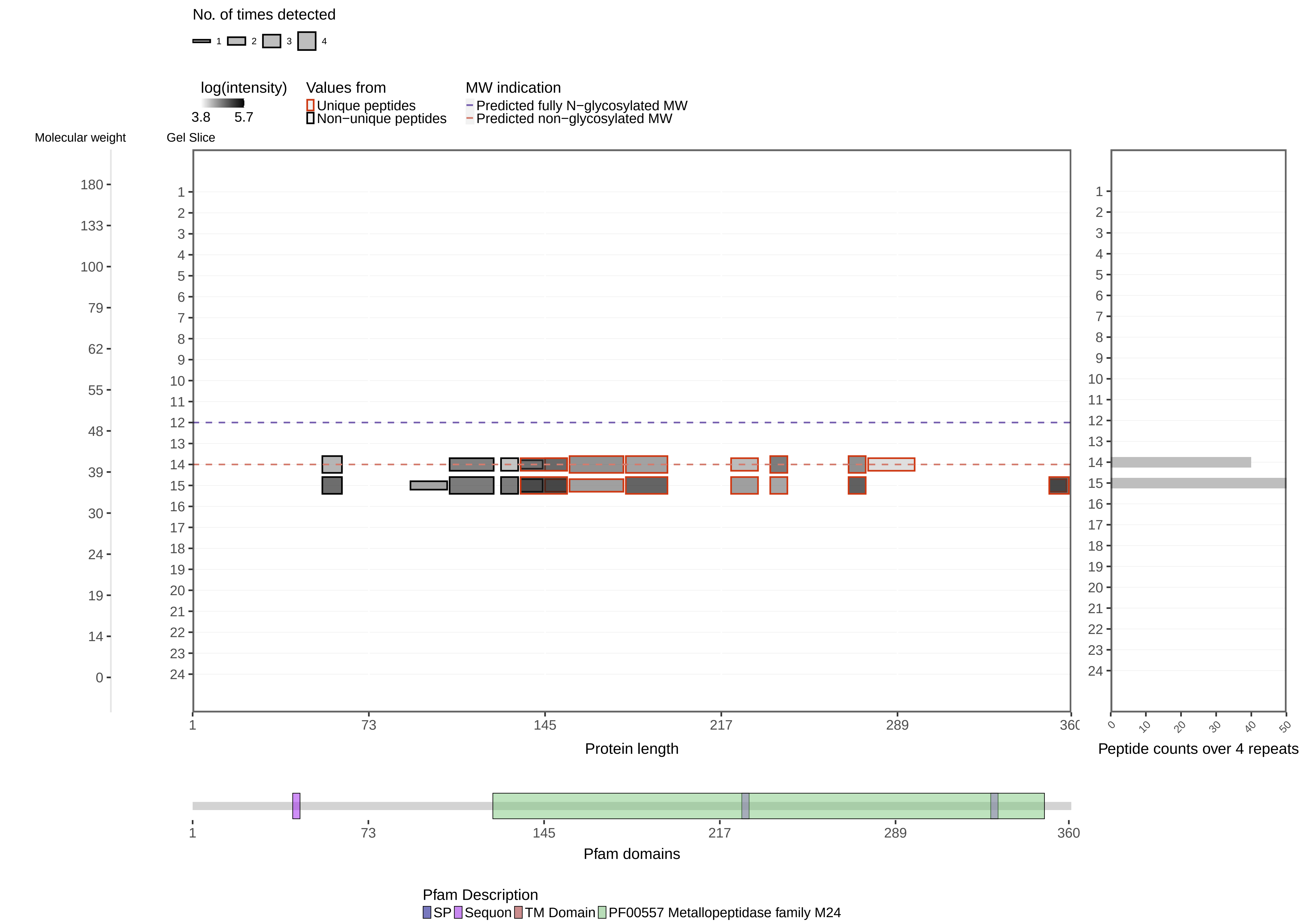


Protein sequence:

### NbL14g21860.1 HD_domain−containing_protein_2 [MW = 37.33 kDa]

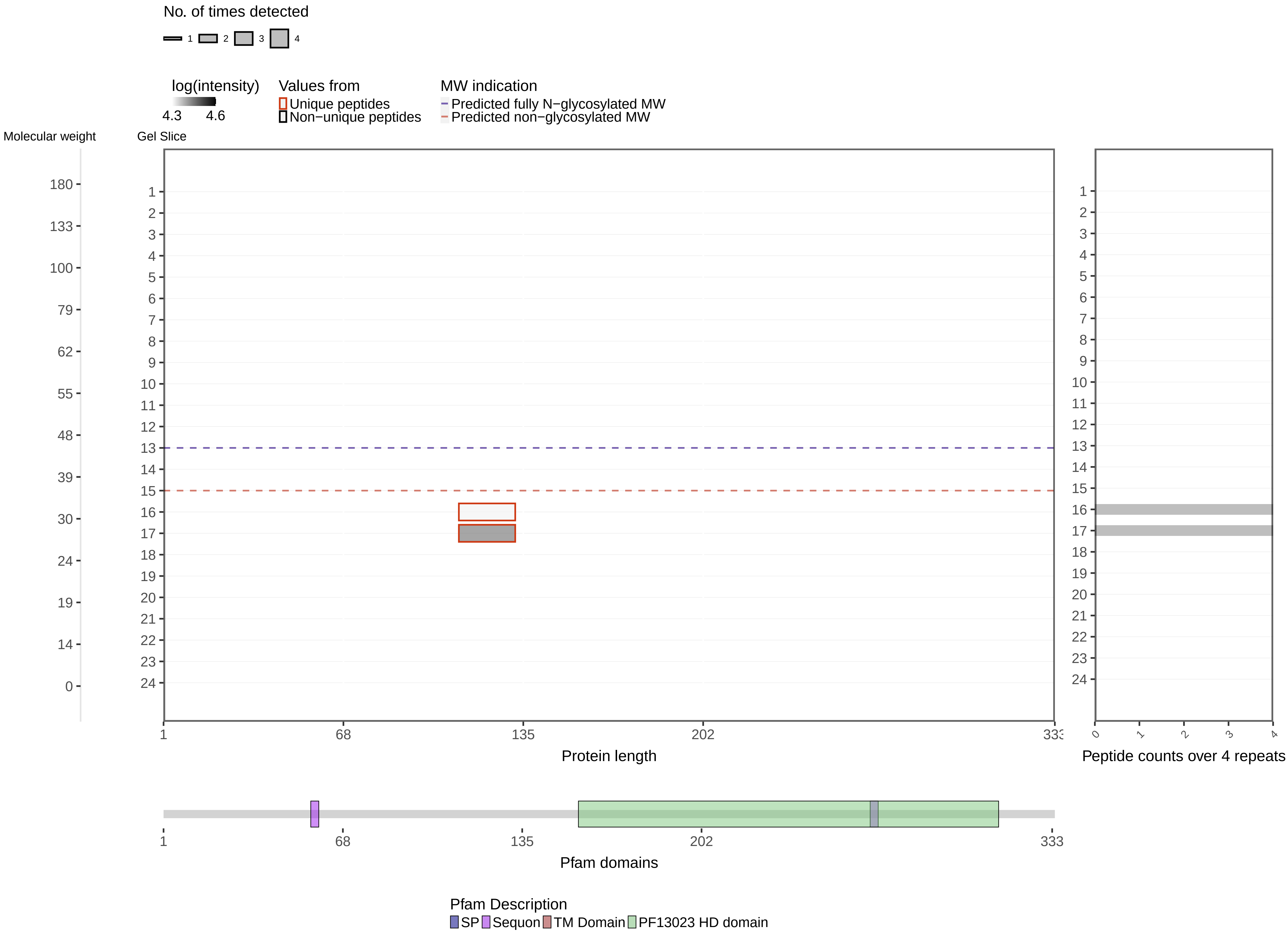


Protein sequence:

.CHSLKTQKRKGWVNHGIKGPESIADHMYLMSVMALIASGLPDVNRERCIKMAIVHDIAEAIVGDITPSDGVPKAEKSRREQAALNEMCEILGGGMRAEEIKELWQDYENNASLEANLVKDFDKVEMILQAFEYESEHGKVLDEFFLSTAGKFQTEIGKSWAAEIHSRRNSRLENRVN*

### NbL14g21720.1 protein_decapping_5 [MW = 63.98 kDa]

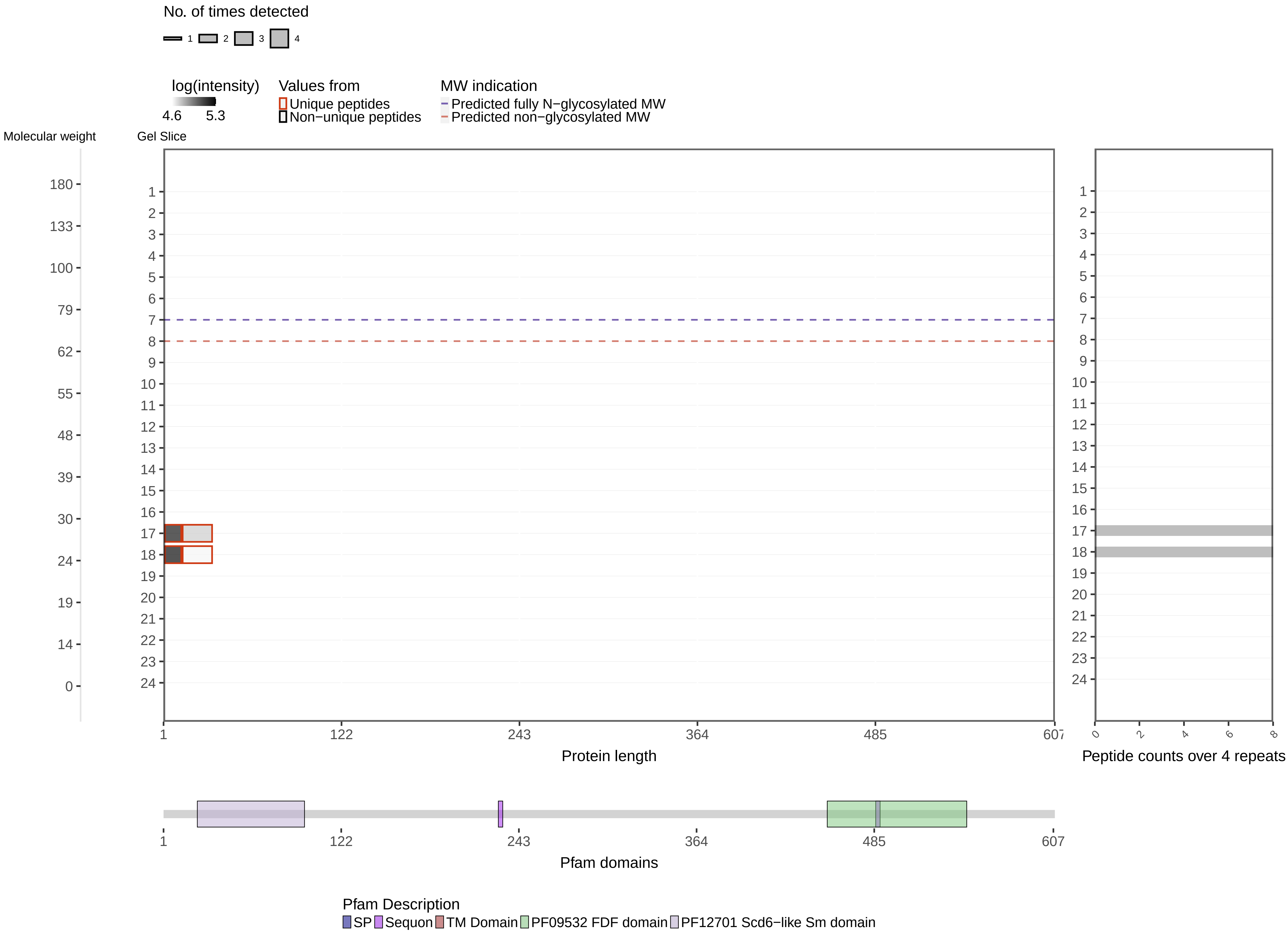


Protein sequence:

VSSIAPSSLSASLPTLPPLTTSPDVNPVVPPIPIKPNPVPSSALSQSVSTVMGPSSSNLVETPTPSLVTPGQLLQFGPIDVPSTQSTQTAQKDVEVVQVFPAPSSETPAPVKTEAQPPILPLPPQTRVQKTNGAPYQARYNNYRGRGGRGMGVSRPVTKFEEDFDFMAMNEKFKKDEVWGHLGKSNREGDGNGSDEDISFNKYDDDLPKIDVKPVYNKDDFFDSLSSNALDNDPSHGRTRFSEQRKIDVETFGDFSRYRGGRGGRGPGRGGRSRGSYYGRGGYGGYG[illegible]

### NbL14g21630.1 protein_CHUP1_chloroplastic−like [MW = 112.92 kDa]

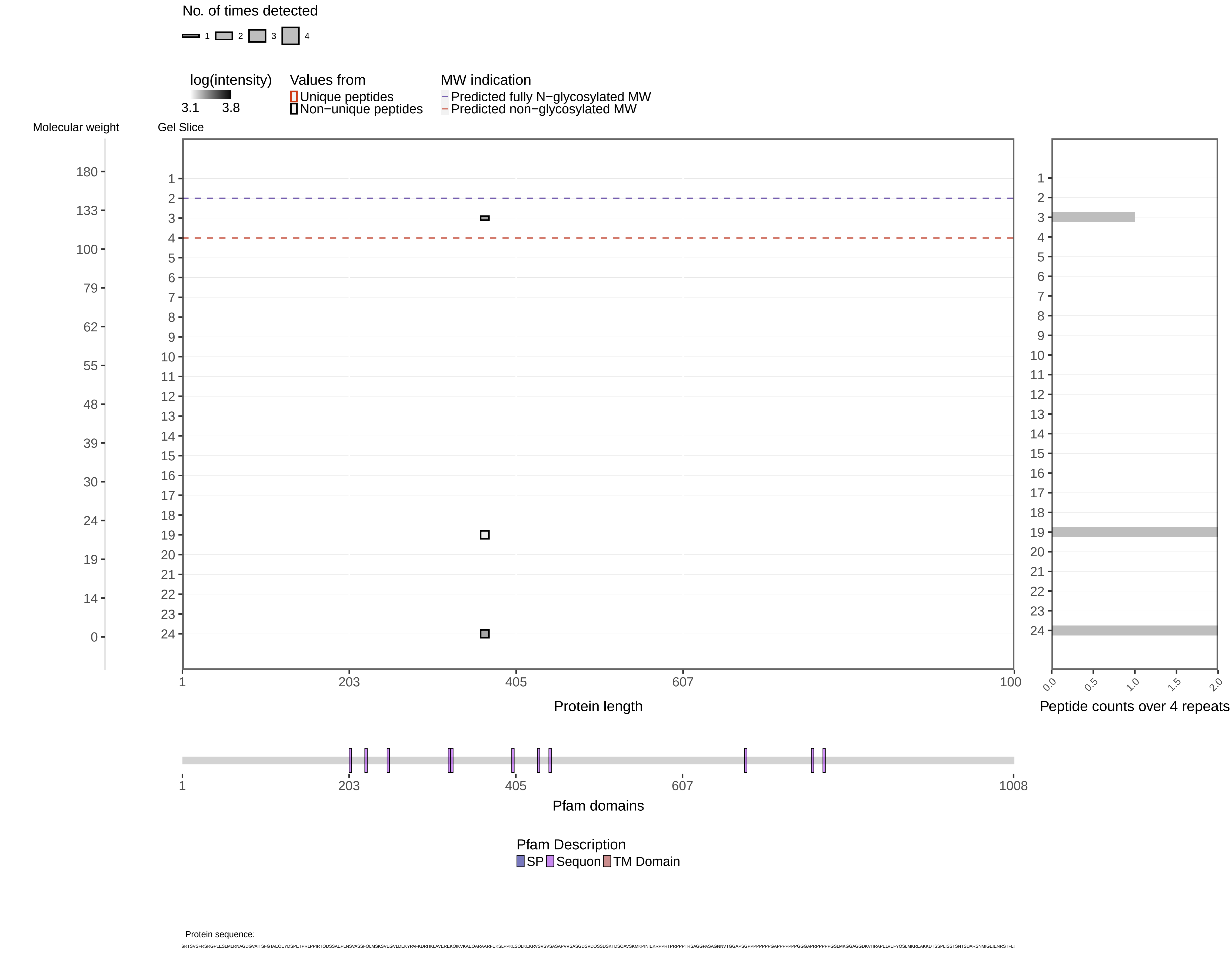

### NbL14g21620.1 uncharacterized_protein_At1g26090_chloroplastic−like [MW = 12.16 kDa]

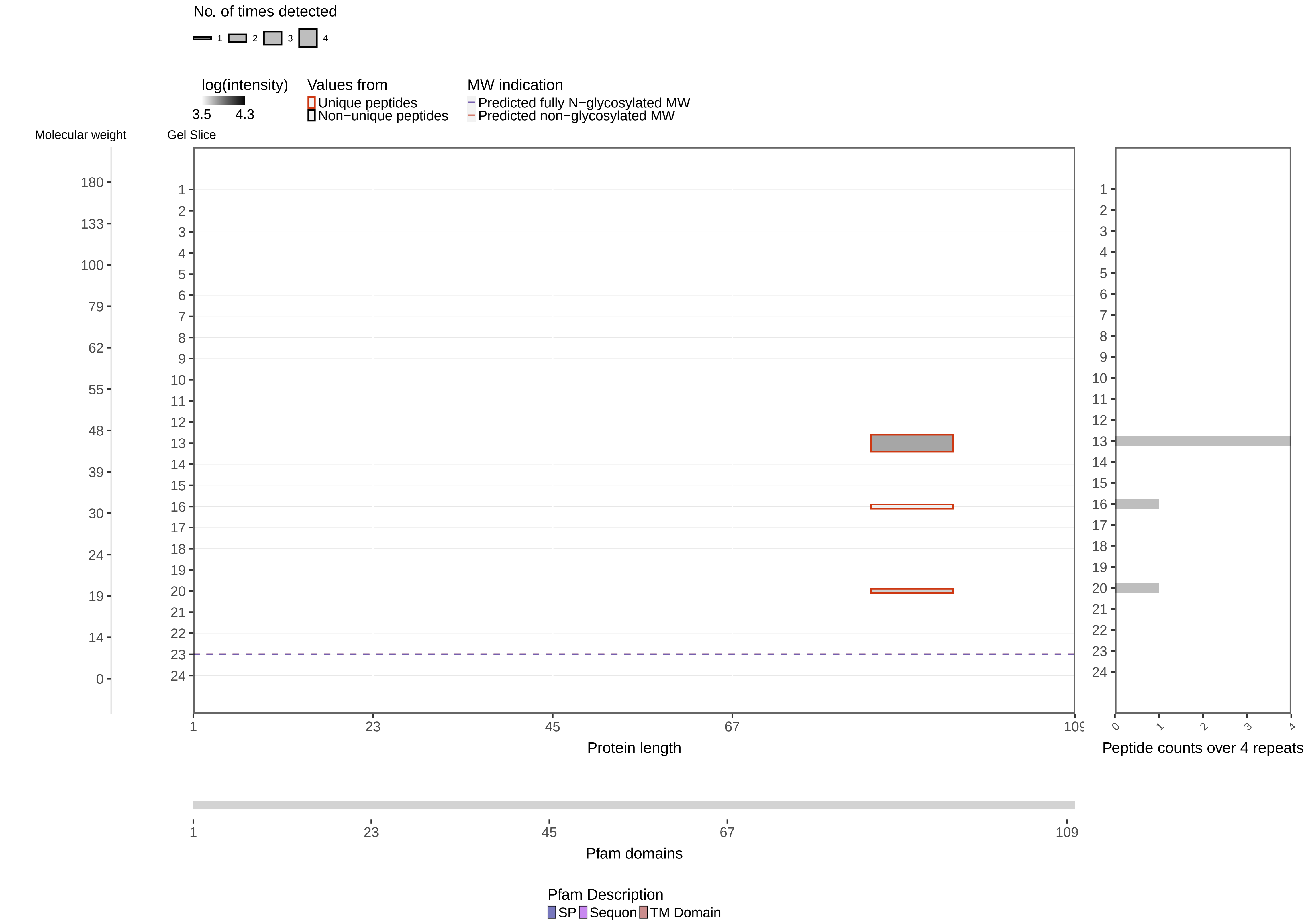


Protein sequence:

GISTEEMIRMIGAISKARLYLKYLRNVAEKTDLGRLASPSLLRLVDEAMTLSGRKLNLNGKTSSEIWD*

### NbL14g21380.1 MFP1_attachment_factor_1−like [MW = 16.36 kDa]

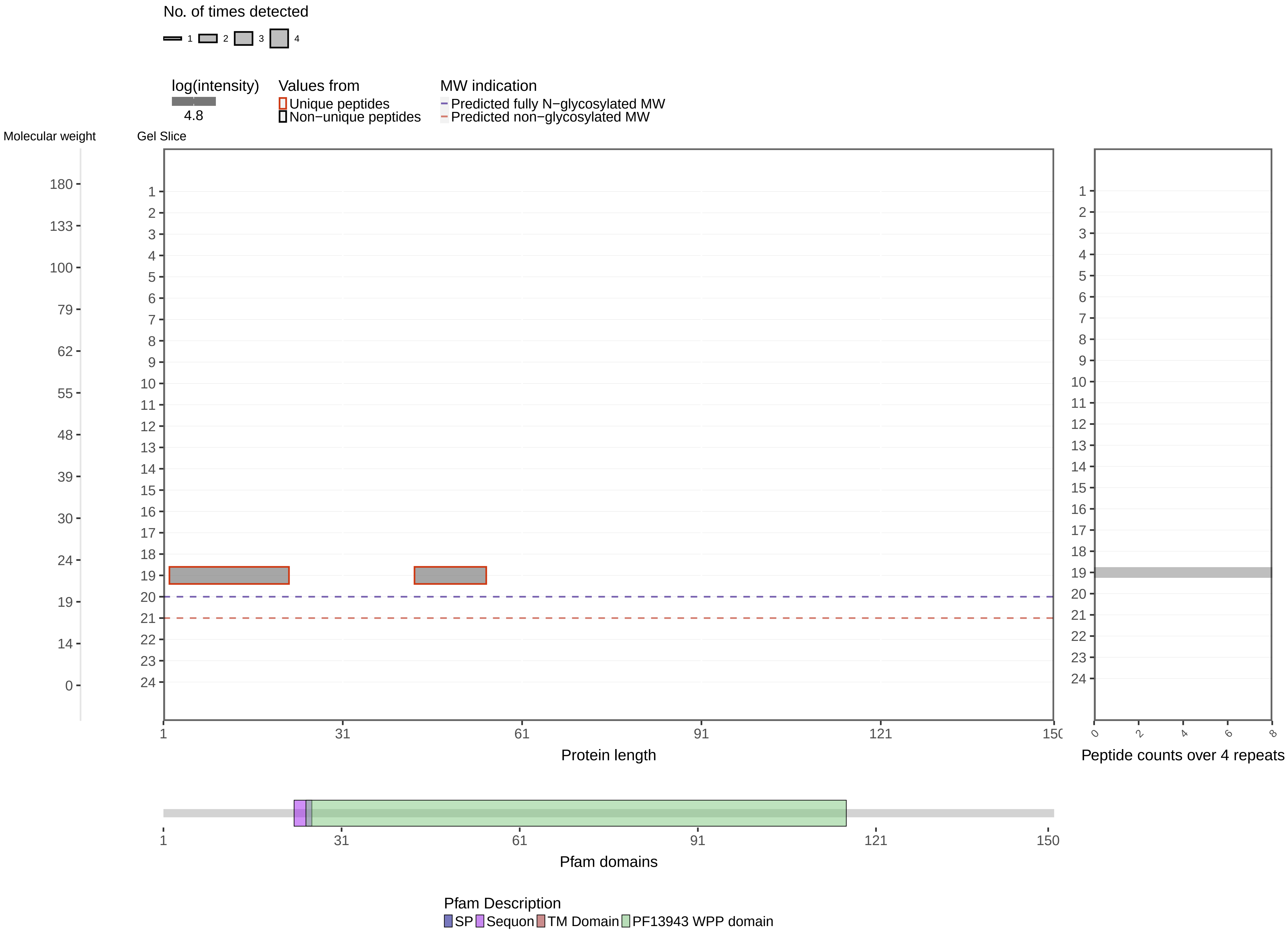


Protein sequence:

KNEEASDIARSIEEEAFTFAGSSVSPDDDGIEILQVYTKEISKRMLDAVKSRSAATTSPEAQTKPSESLVDASESSAHTGDISSIESES*

NbL14g21330.1 polyadenylate−binding_protein_2−like [MW = 72.04 kDa]

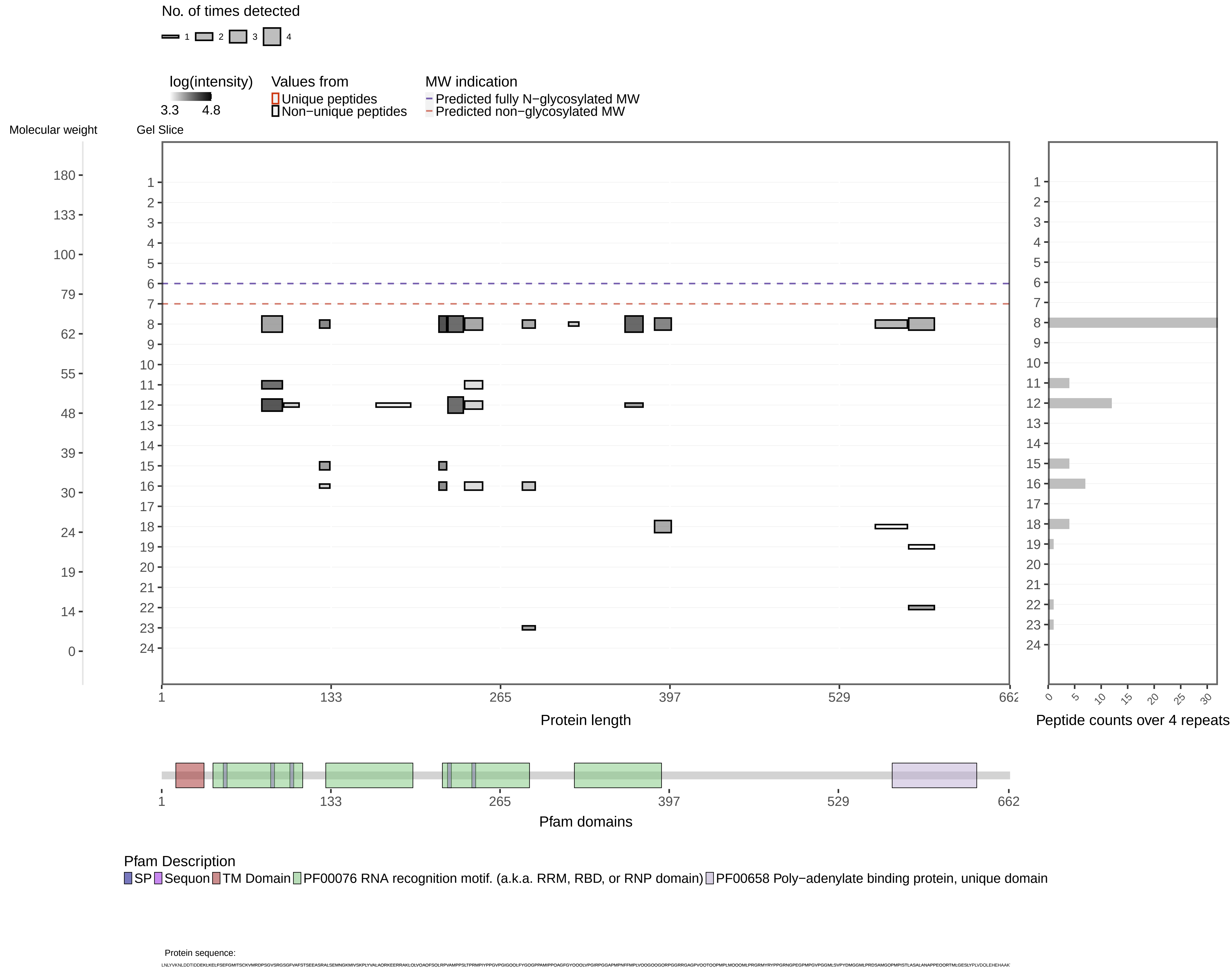

### NbL14g21320.1 50S_ribosomal_protein_L4_chloroplastic [MW = 34.4 kDa]

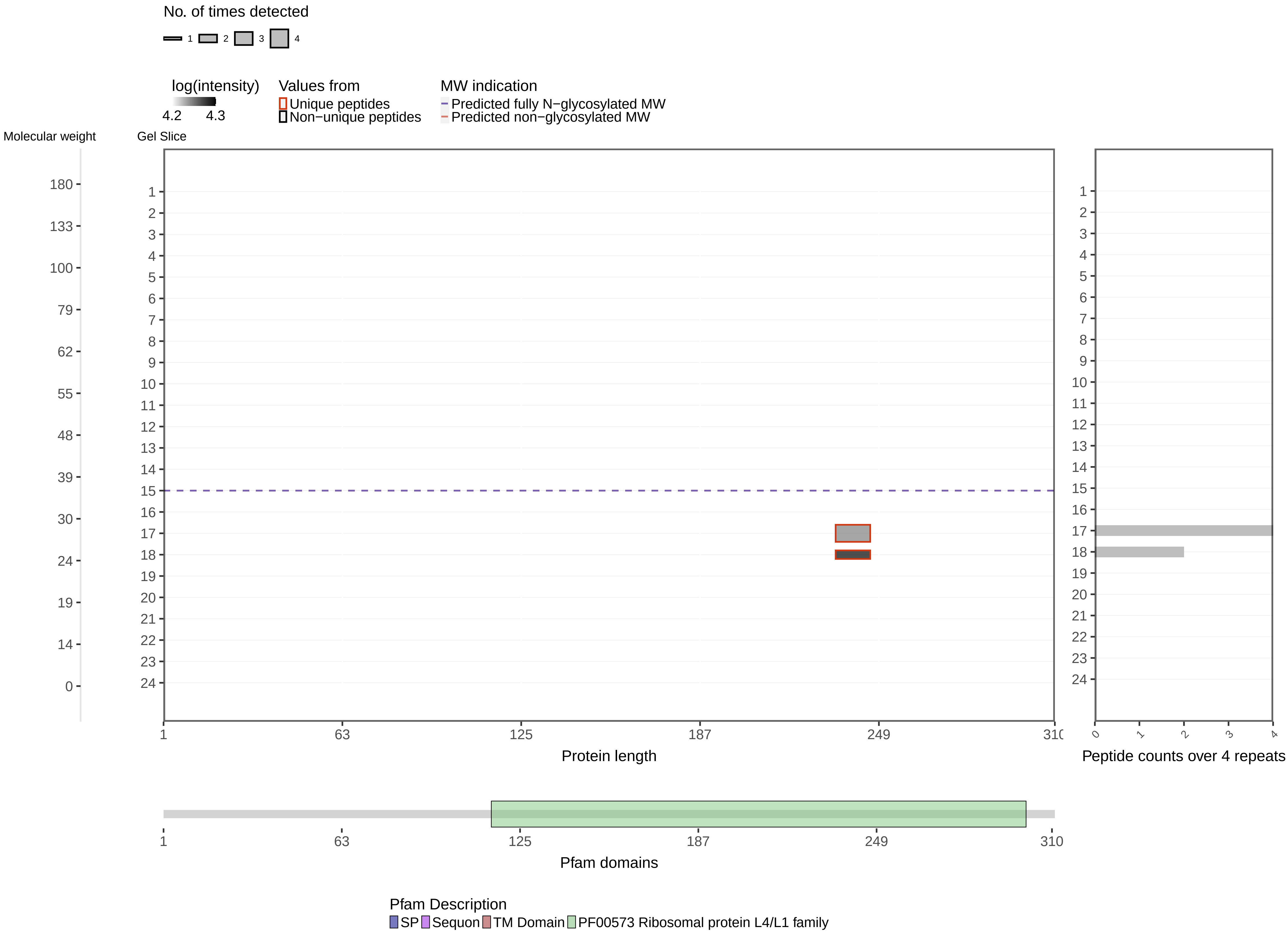


Protein sequence:

GTHSTKTISEVSGTGRKPWRQKGTGRARHGTLRGAQFRGGAAMHGPKPRSHAIKLNKKVRRLGLKIALSARTAEGKLMVFEDLELSSHKTKNIVNYFNQLENTKKLLLVDGGPISEKLKLATQNVHYVNVLPSIGLNVYSILLHDTLVMSRDAVNRIVERMHTPINR*

### NbL14g21060.1 protein_EXORDIUM−like [MW − SP = 31.13 kDa]

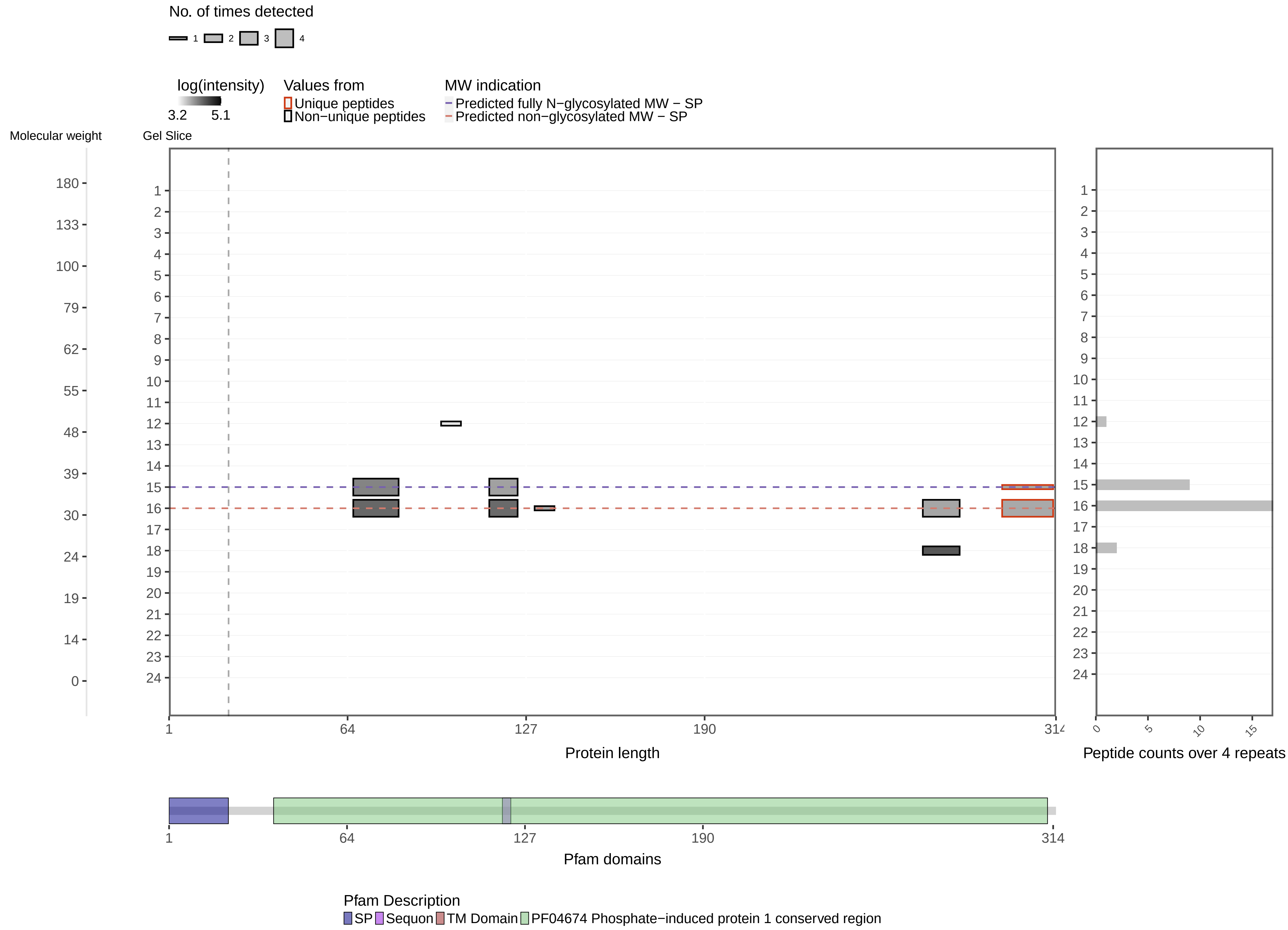


Protein sequence:

VLTASDVAVDGFCVNRCGTHGSSKGATIRGKTYKFAYIWVGNSETQCAGYCAWPFHQPIYGPQSPPLVAPNNDVGVDGMVINLASLLAGTATNPFGNGYYQGEADAPLEAASACPGVYAKGAYPGYAGDLLVDKTTGASYNAHGANGRKYVLPALYDPSTSTCSTLV*

NbL14g20920.1 cryptochrome−1−like [MW = 77.06 kDa]

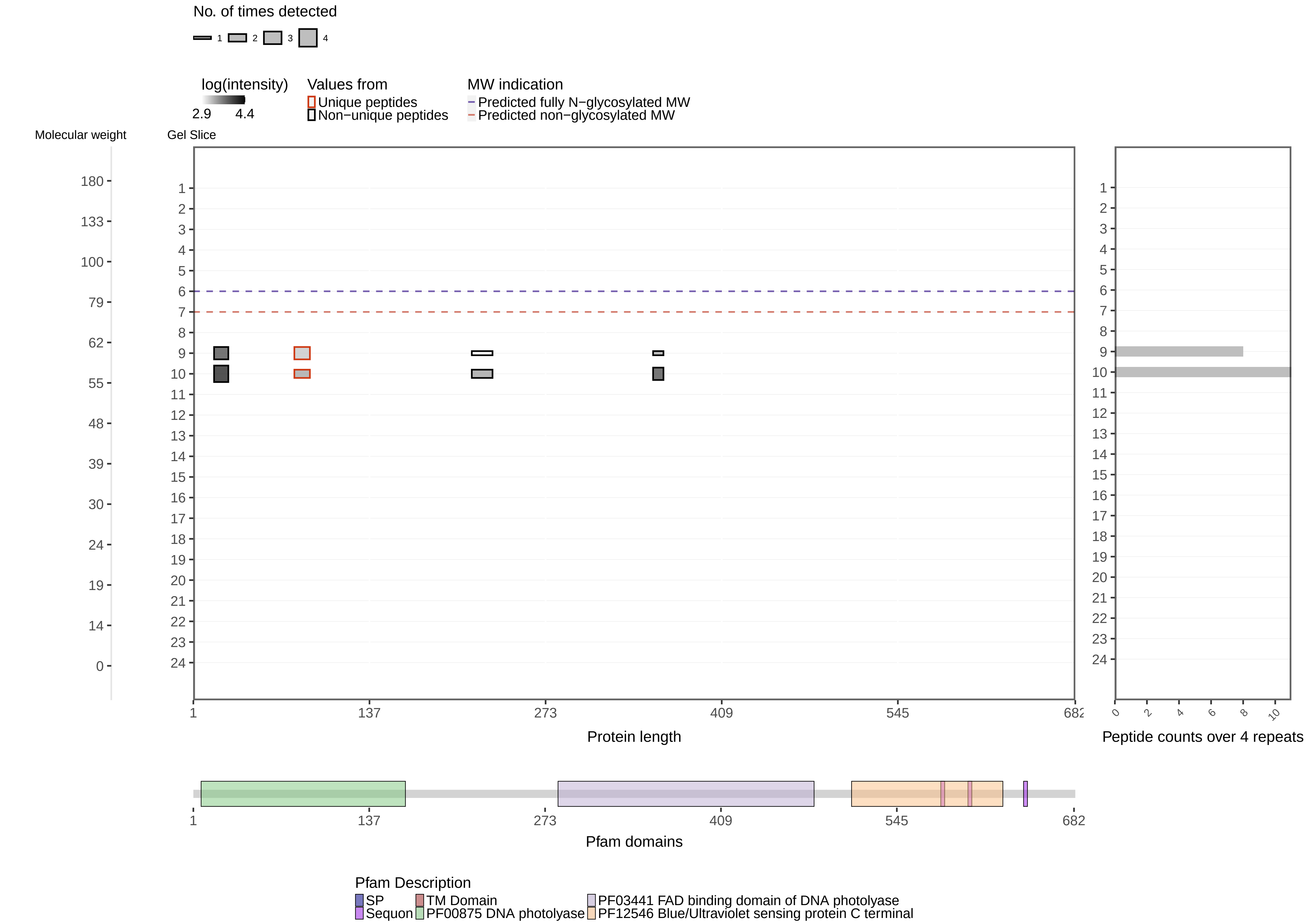


Protein sequence:

GYPLVDAGMRELWATGWLHDRIRVVVSSFFVKVLQLPWRWGMKYFWDTLLDADLESDALGWQYISGTLPDGREFDRIDNPQFVGYKCDPHGEYVRRWLPELSRLPTEWIHHPWNAPESVLEAAGIELGSNYPLPIVEIDAAKVRLEQALSQMWQHEAASRAAIENGMEEGHGDSADSPIAFPQAMHMEMDHEPVRNNNPVIVTVRRYEDQMVPSMTSSLFRVDDEETSVNIRNSVVDTRAEVPNDVNVTEGPRRDTLDQAVTQRARTNATPPQFNFVVGRRNSEDS

### NbL14g20810.1 pumilio_homolog_9−like [MW = 74.75 kDa]

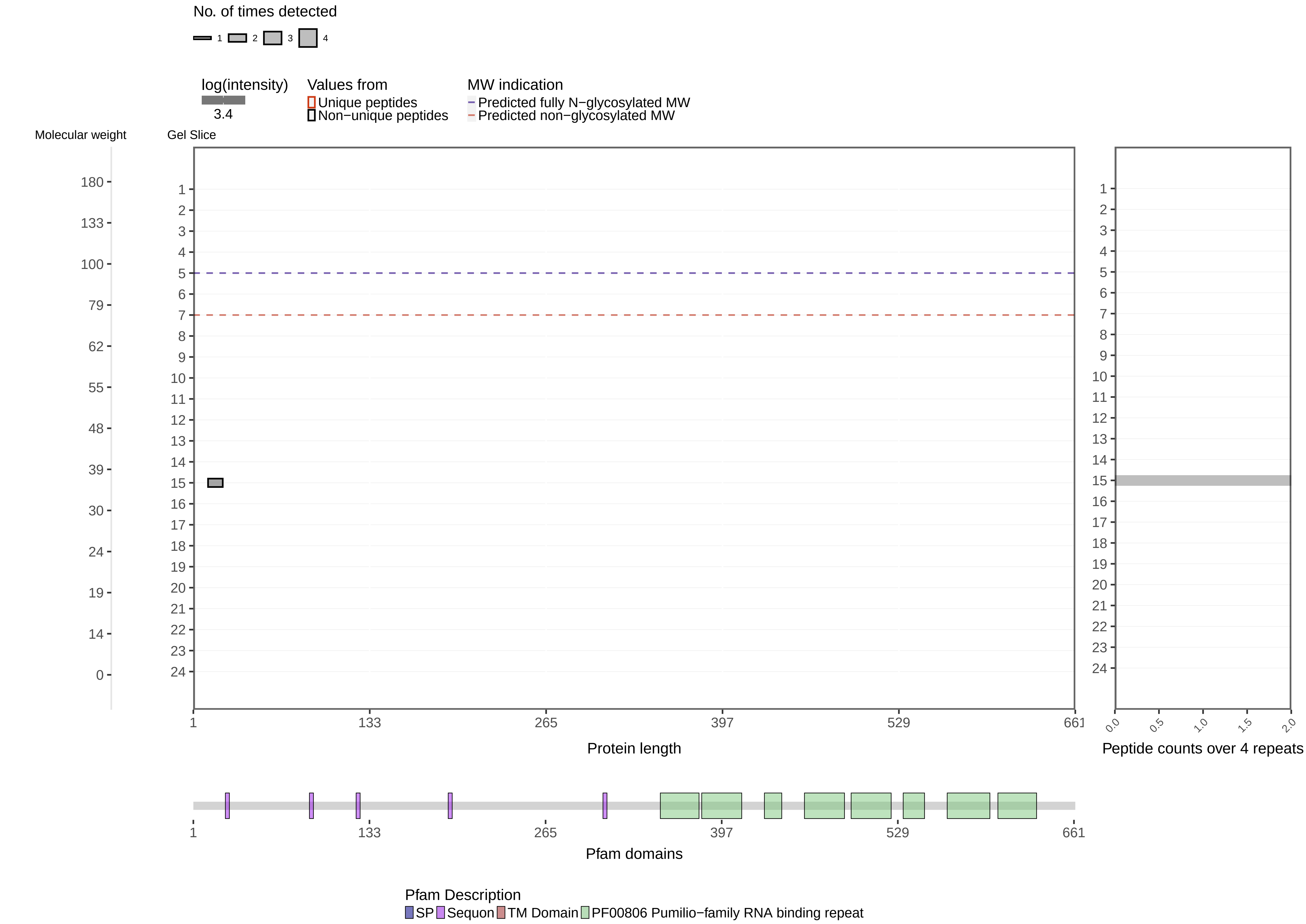


Protein sequence:

[illegible]

NbL14g20800.1 uncharacterized_LOC104112265 [MW = 30.44 kDa]

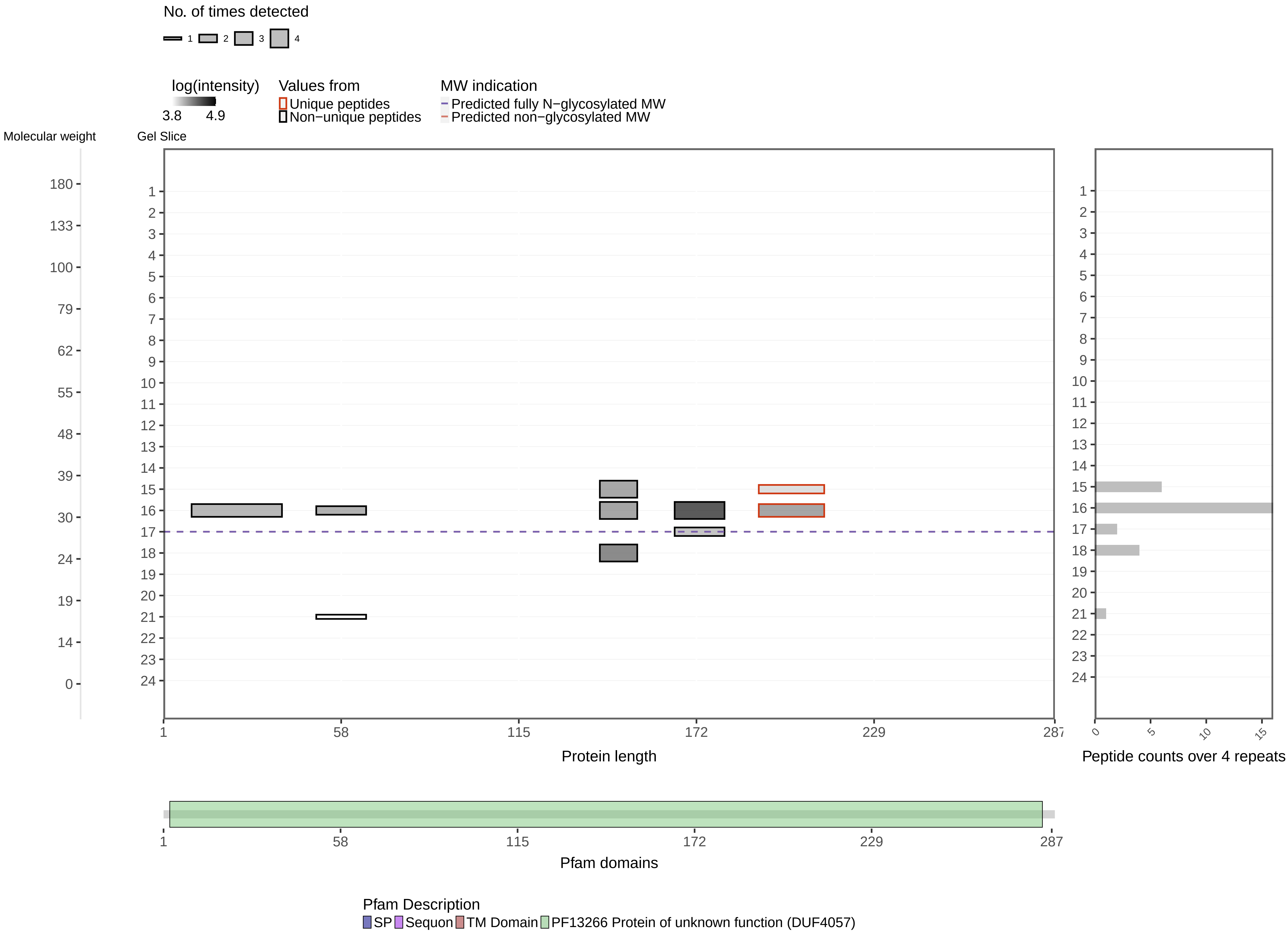


Protein sequence:

KPVTIPEVAKQRELSGTLESEAEMKLKKQLSDAKTKELVGHDIFAPPPEIQPRPLAARALALRESIAIGEPANNGDVSGDEQAVKTAKKIPEKKLAELTGNDIFKGDTPPASAEKHLSSAKLREMSGSNIFADGKVESRDFYGGVRKPPGGESSIALV*

### NbL14g20650.1 probable_beta−D−xylosidase_7 [MW − SP = 82.7 kDa]

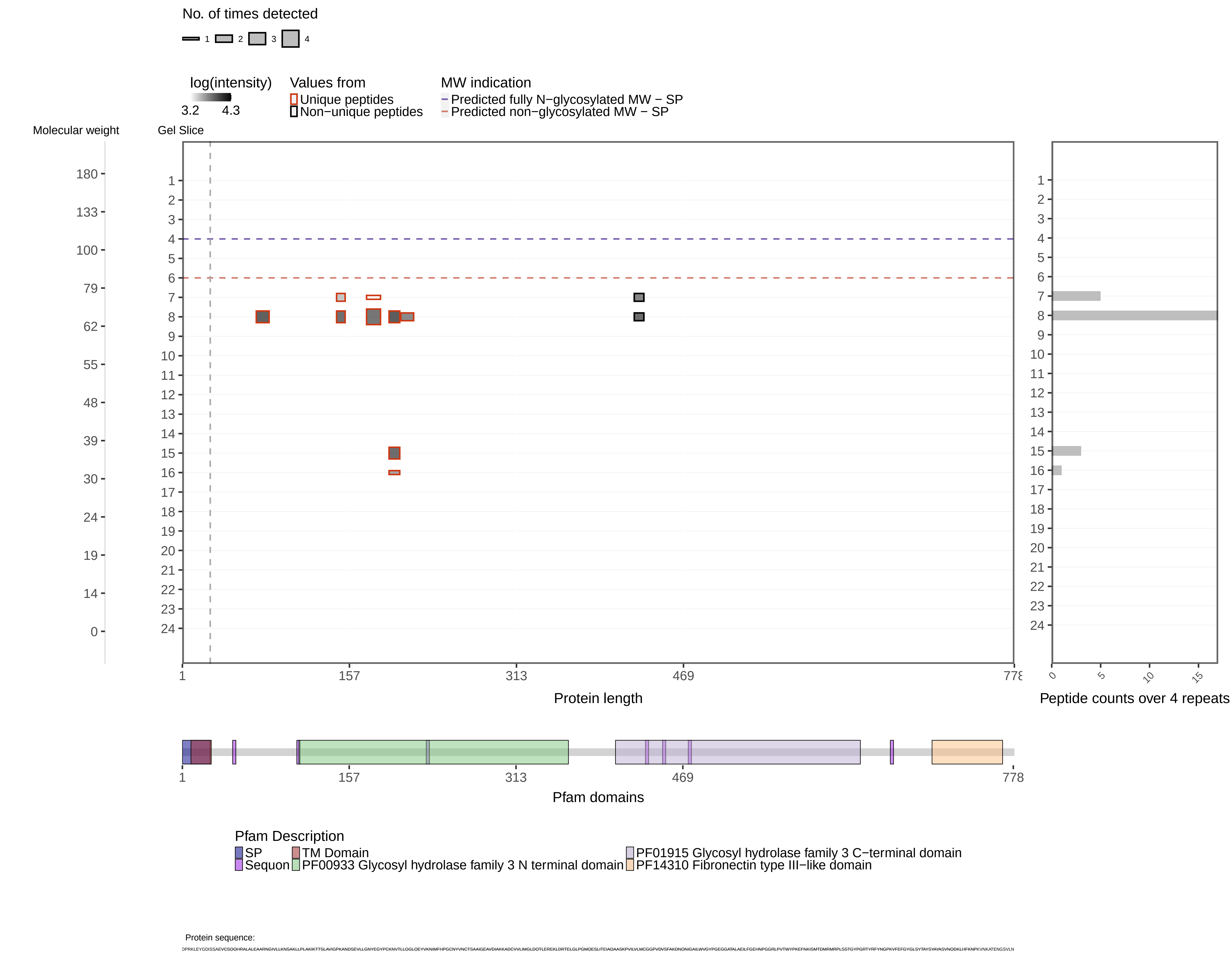

NbL14g20610.1 23−bisphosphoglycerate−dependent_phosphoglycerate_mutase [MW = 38.18 kDa]

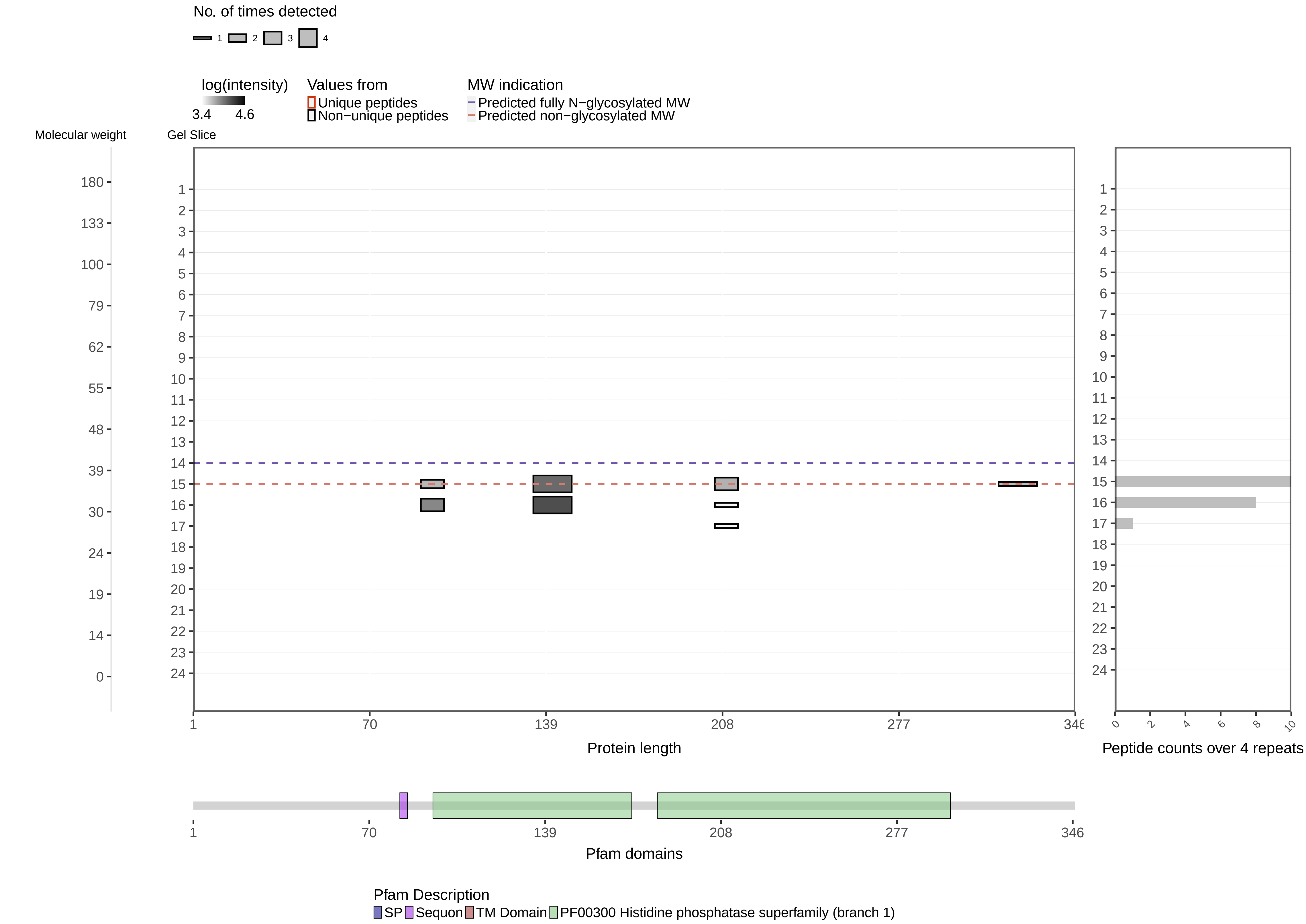


Protein sequence:

TOHRRKKVPTIIHEESEOAGAWSOIFSEETNKOCIPVVTAWOLNERMYGELOGLNKOETADRYGKEOVHEWRRSYDIPPPNGESLEMCAERAVAYFKEHIEPOLAGGKNIMIAAHGNSLRSIIMYLDKLTSOEVISLELSTGIPMLYIFKEGKFIRRGSPAAPTEAGVYAYTKTLAOYROKLDNMVO*

NbL14g20010.1 uncharacterized_oxidoreductase_At4g09670−like [MW = 39.81 kDa]

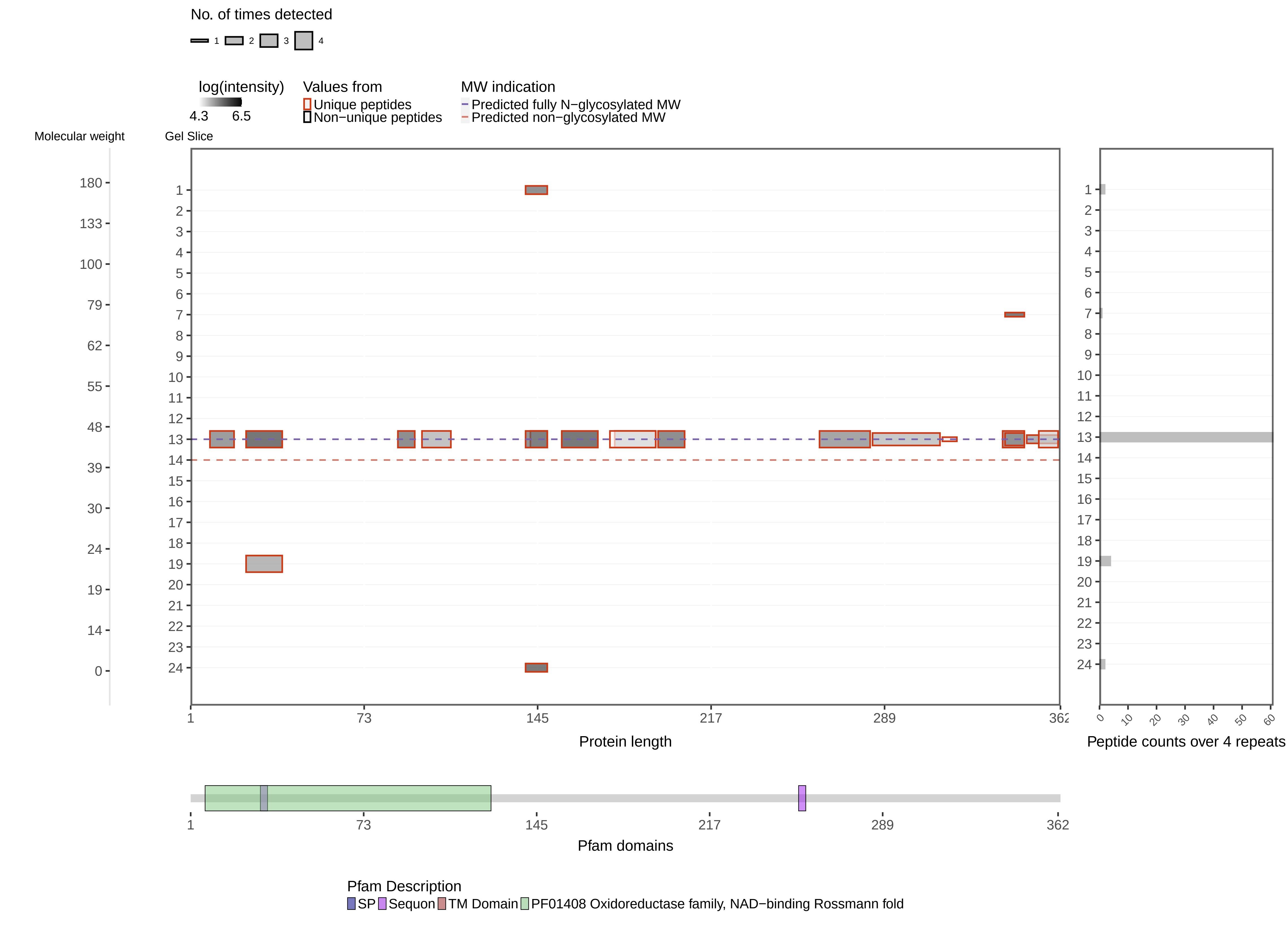


Protein sequence:

FLKNDIRVKADLDALGALGDAGWYSIRAILWTTDYELPKTVTALPDPELNEAGVILSCGASLSWKDGRVATFYCSFLANLVMDIAANGSKGNLRVHDFVIPFOENVAPFYTVESSRFGELSISIHPAPNEOIVSTDLPOEALMVKEFSNLVRSIKGEGCKPEKKWPTISRKTOLVVDAVKASIDKGFEPVEVVY*

NbL14g19890.1 ras−related_protein_Rab7 [MW = 22.91 kDa]

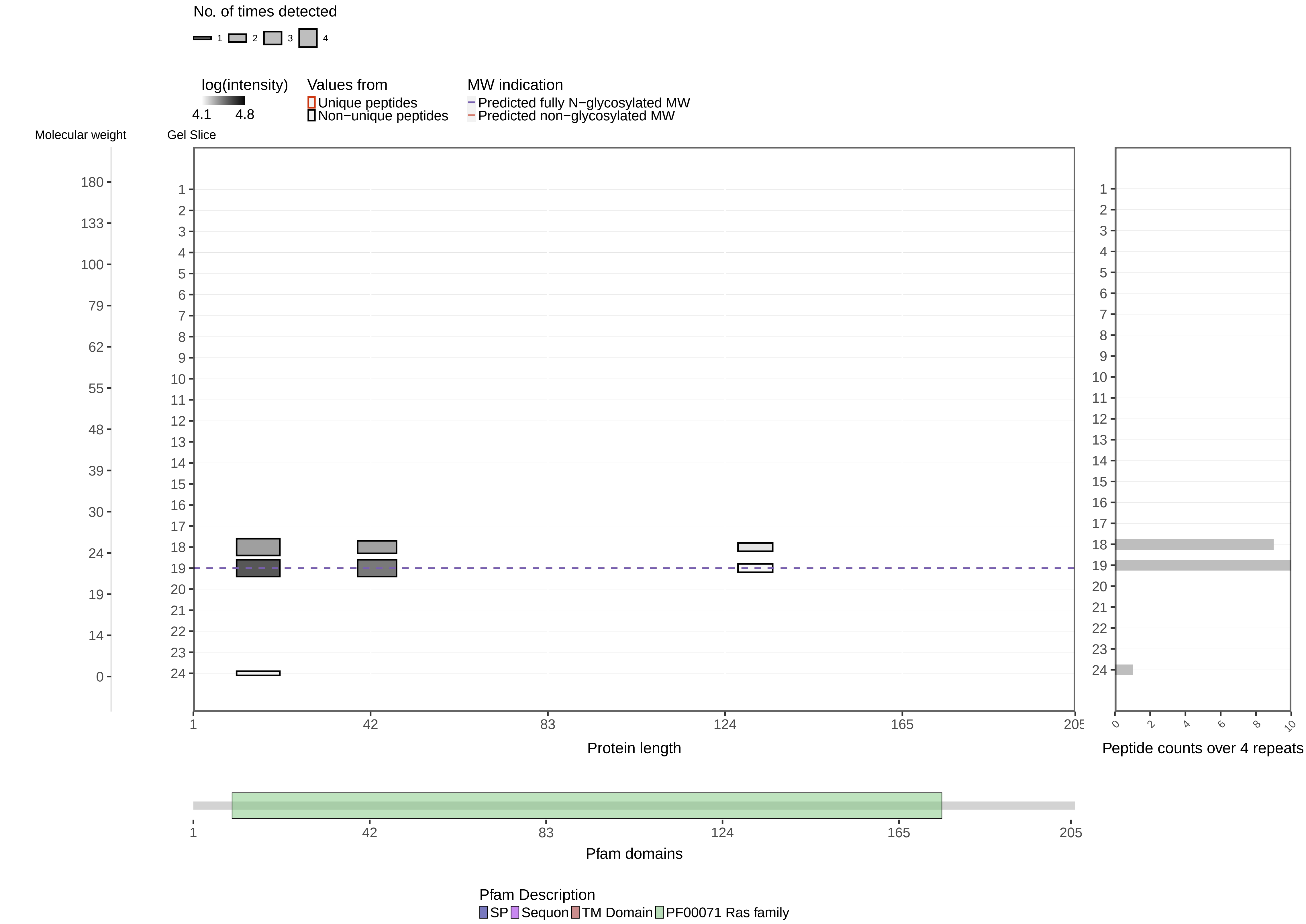


Protein sequence:

VNVMRSFDNLDNWHEEFLKQANPPDPKTFPFILLGNKIDIDGGNSRVVSEKKAKEWCASKGIPYFETSAKEDINVDAAFLSIAKTALANEHEQDIYFQGIPEAVSETEQRGGCAC*

### NbL14g19870.1 probable_ATP_synthase_24_kDa_subunit_mitochondrial [MW = 27.58 kDa]

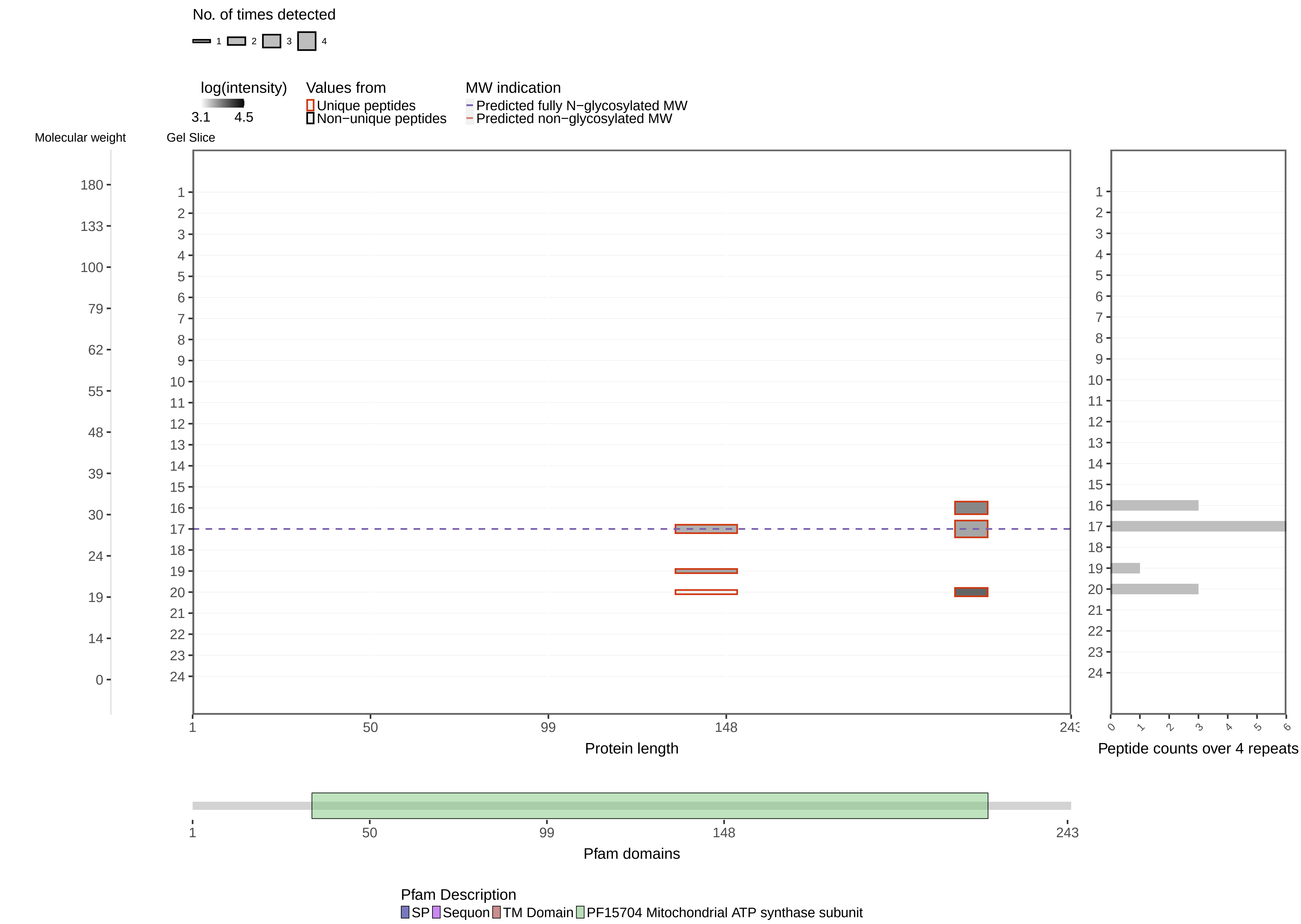


Protein sequence:

NIEVETHEIPDARTYLLKLKEMRVKRGLIDEQGIEDLQMAALDKVEKEIKKPLMRNDKKGIALLTAEFDKINQKLGIRKEDLPKYEEQLELKIAKAQLEELKKDALEAMETQKKREEFKDEEMPSVKSLDIRNFI*

### NbL14g19730.1 30S_ribosomal_protein_S1−like [MW = 59.43 kDa]

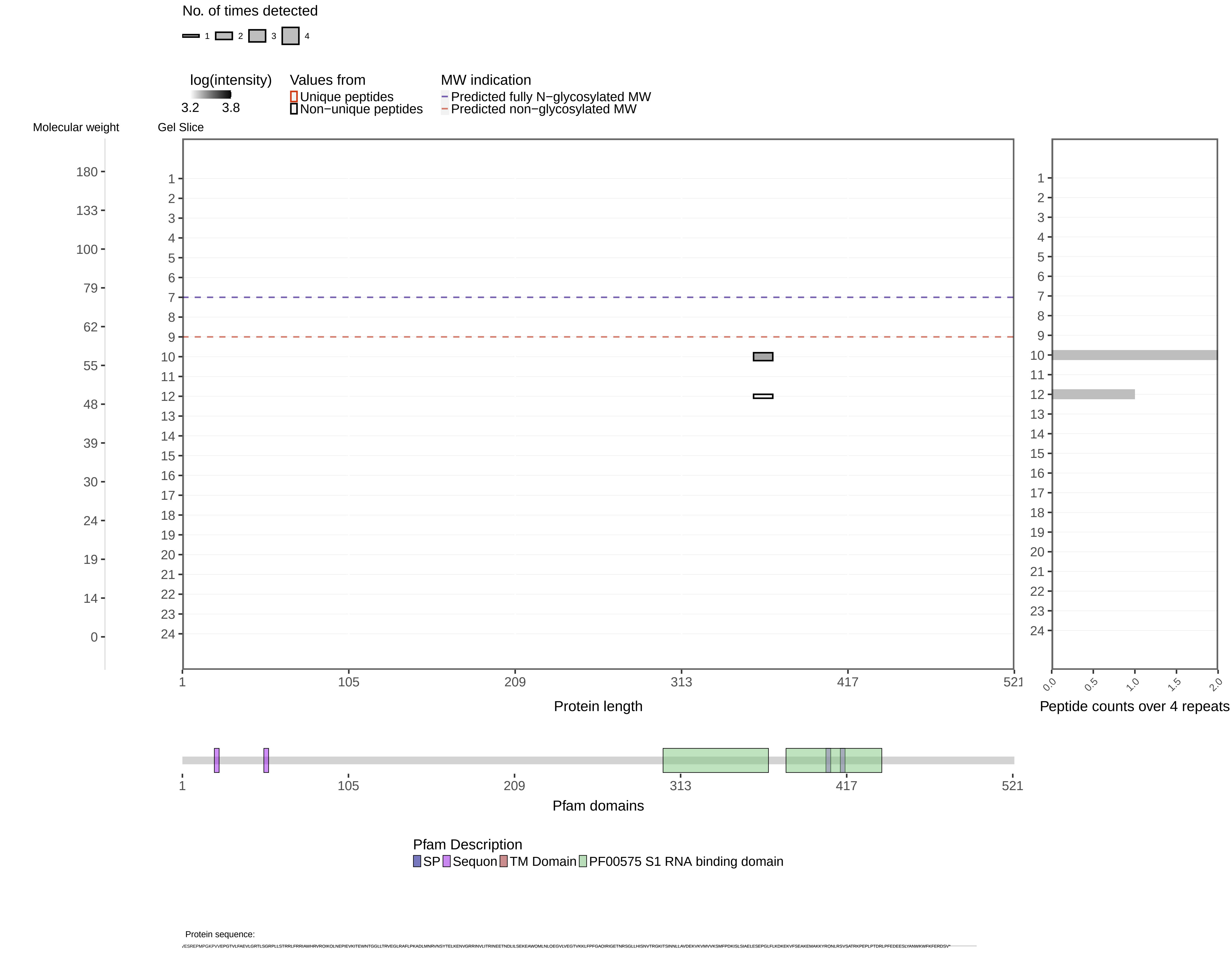

### NbL14g19580.1 xanthoxin_dehydrogenase−like [MW = 29.42 kDa]

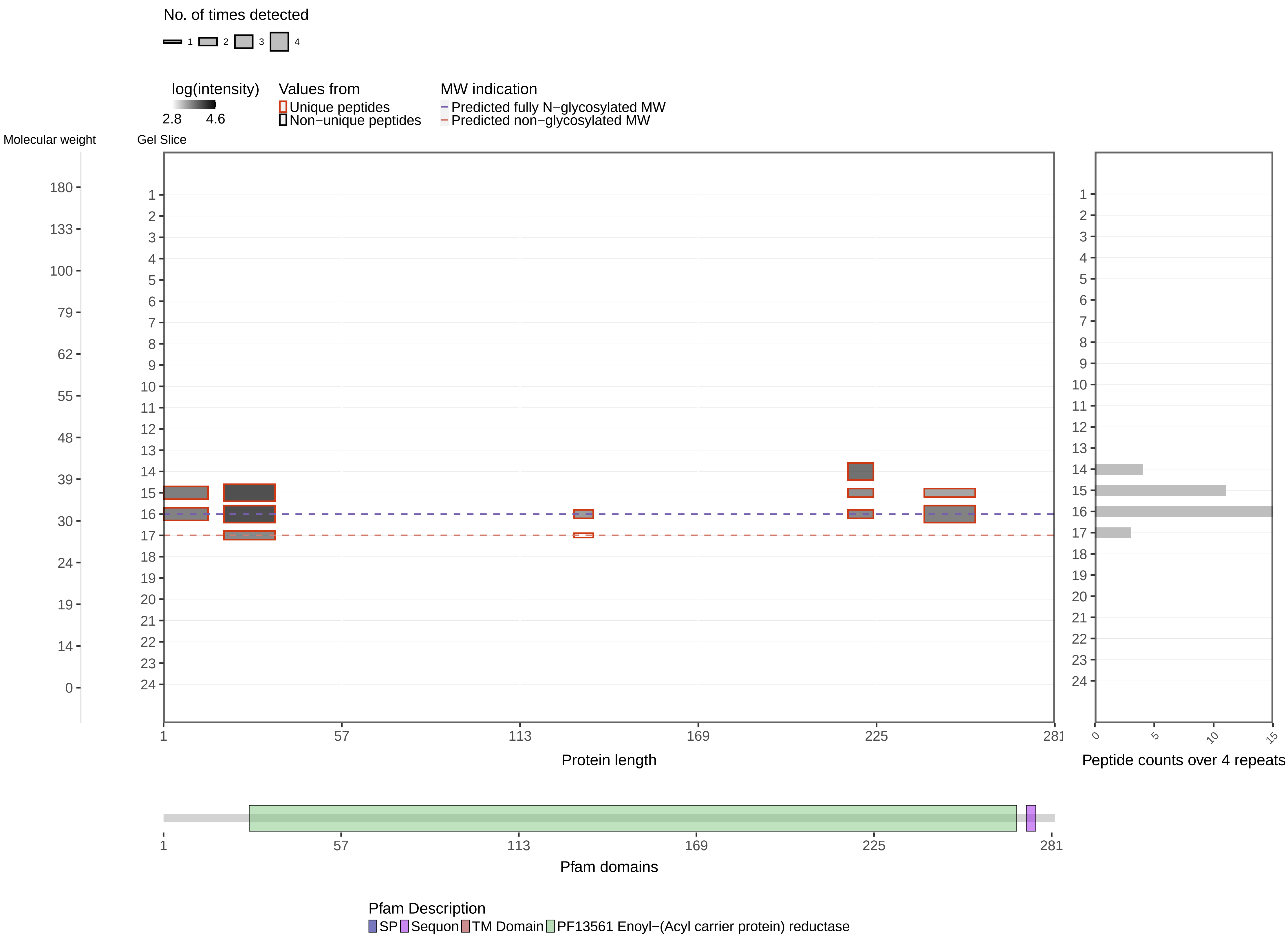


Protein sequence:

ILKGTFLGIKHAARIMIPQKKGSIVSLCSVGSAIAGIGPHAYIASKHAVLGLTQNVAAEMGKHGVRVNCVSPYAVATGLALAHLPEDERTNDALAGFRAFTGKNANLQGVELKTQDVANAVLFLASDEARYISGHNLMVDGGFSCVNHSLRVFR*

### NbL14g19570.1 uncharacterized_protein_DDB_G0284459−like [MW = 18.42 kDa]

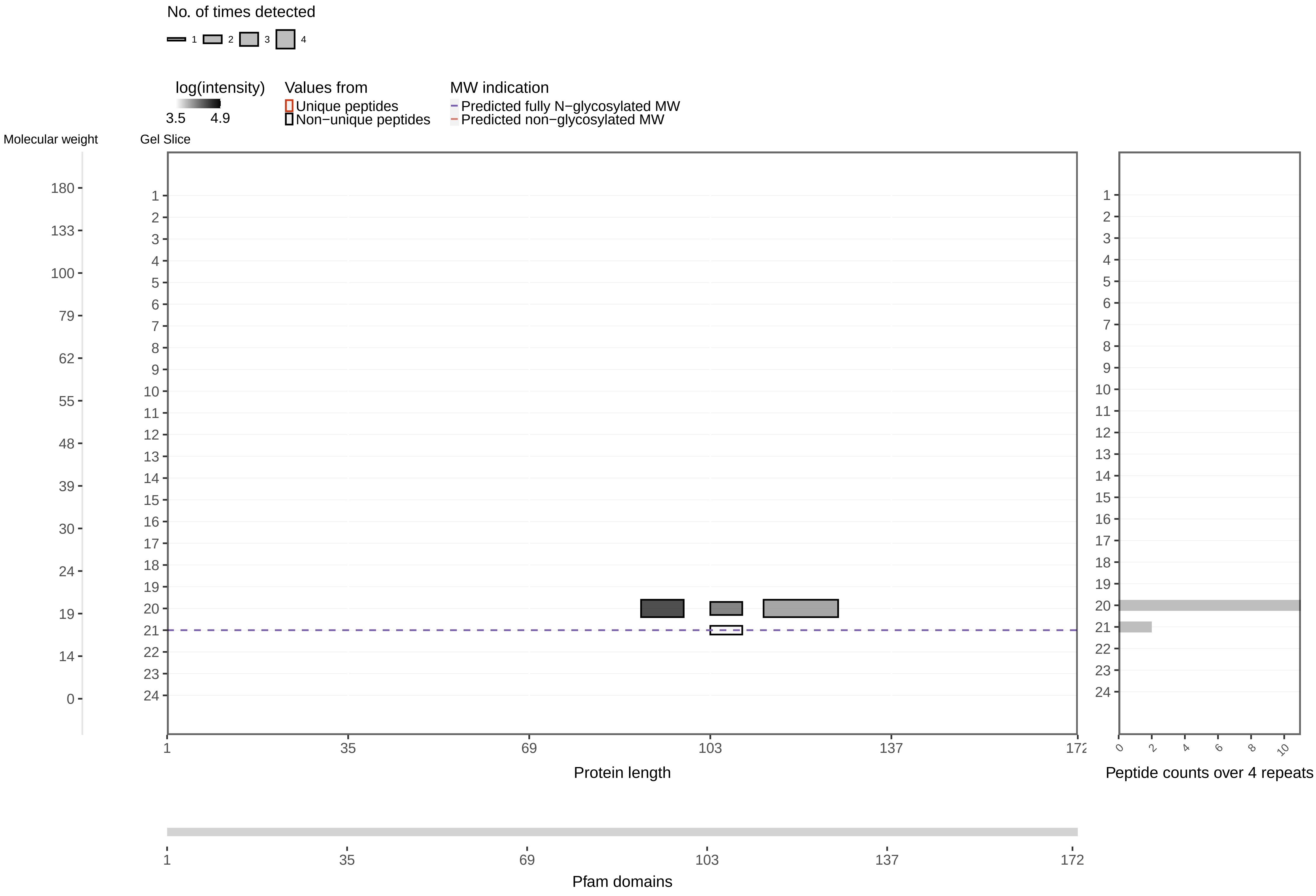


Pfam Description
SP Sequon TM Domain

Protein sequence:

PPSAPPKQVTVAALDGKNVITMEFQRQKAKELQEYFKQKKLEEANQGPFFGFIGKNEISNGRWAMFGFAVGMLTEYATGSDFVDQVKILLSNFGIVDLE*

NbL14g19390.1 aminoacylase−1 [MW − SP = 47.7 kDa]

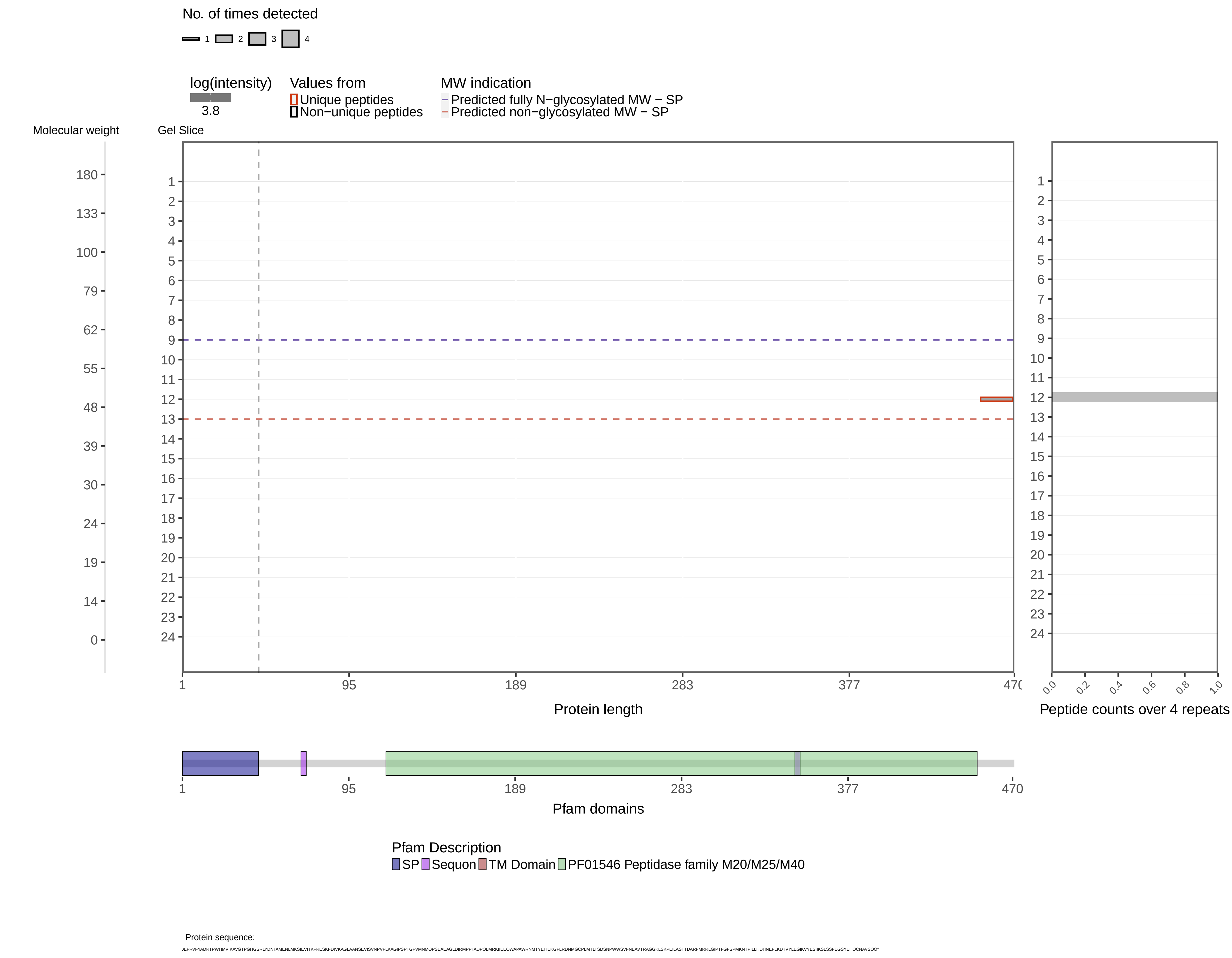

NbL14g19280.1 probable_N−succinyldiaminopimelate_aminotransferase_DapC [MW = 50.92 kDa]

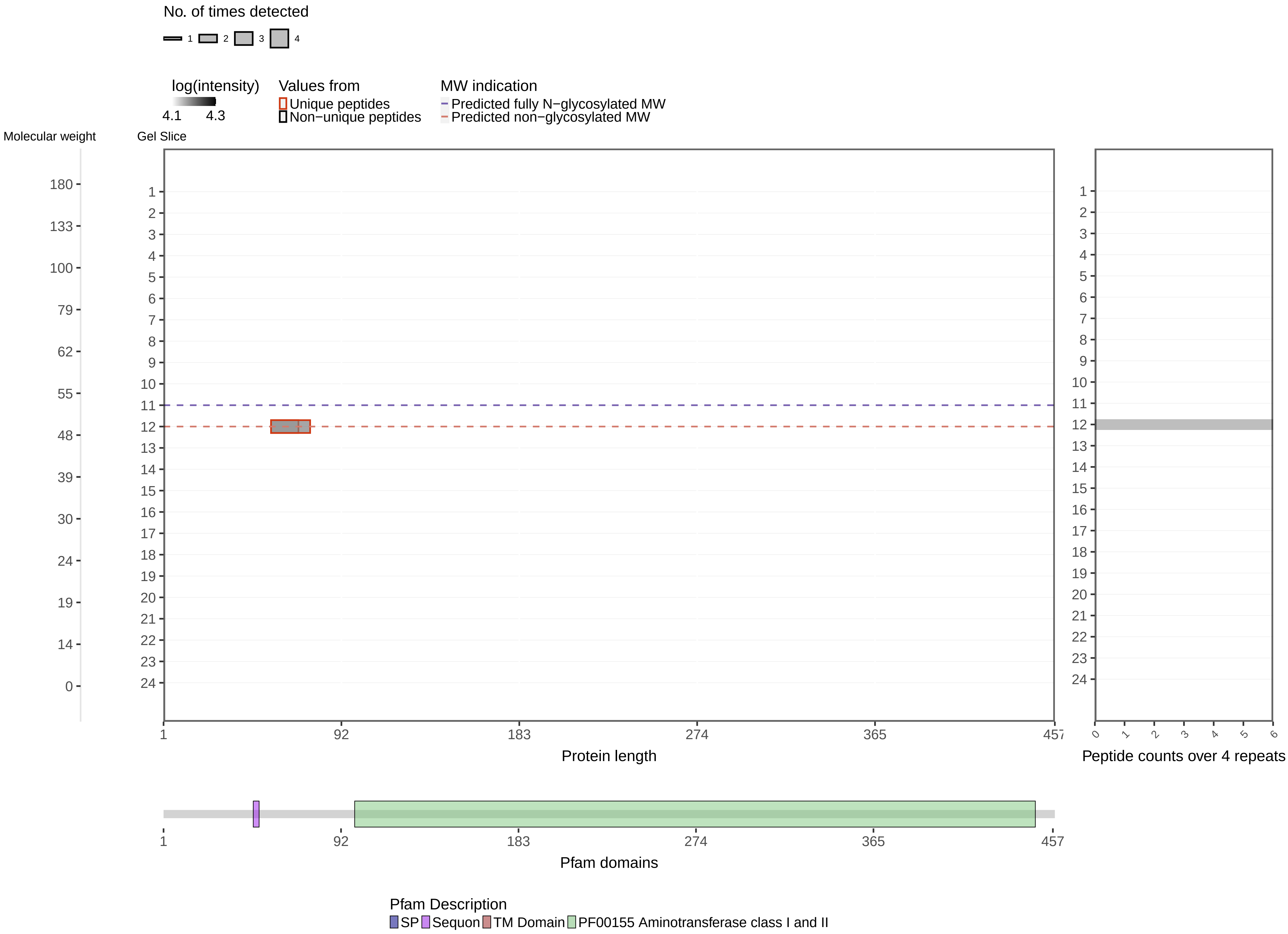


Protein sequence:

SLPIDELKSVVSKNTRAILLNTPHNPTGKMFNREELNVIASLCIENDILVFADEVYDKLAFEMEHISIASLPGMYERTVTMNSLGKTFSLTGWKIGWAIAPPHLTWGVRQAHSYLTFATSTPMQYAAATALRAPDSYYEELKRDYSSKKAILVEGLKAAGFTVYPSSGTYFVVVDHTPFGLENDIAFCEHLIKEVGVVAIPTSVFYLNPEEGKNLVRFTFCKDDDTLKNAVQRMKQKLSKK*

NbL14g19240.1 biotin−−protein_ligase_2 [MW = 41 kDa]

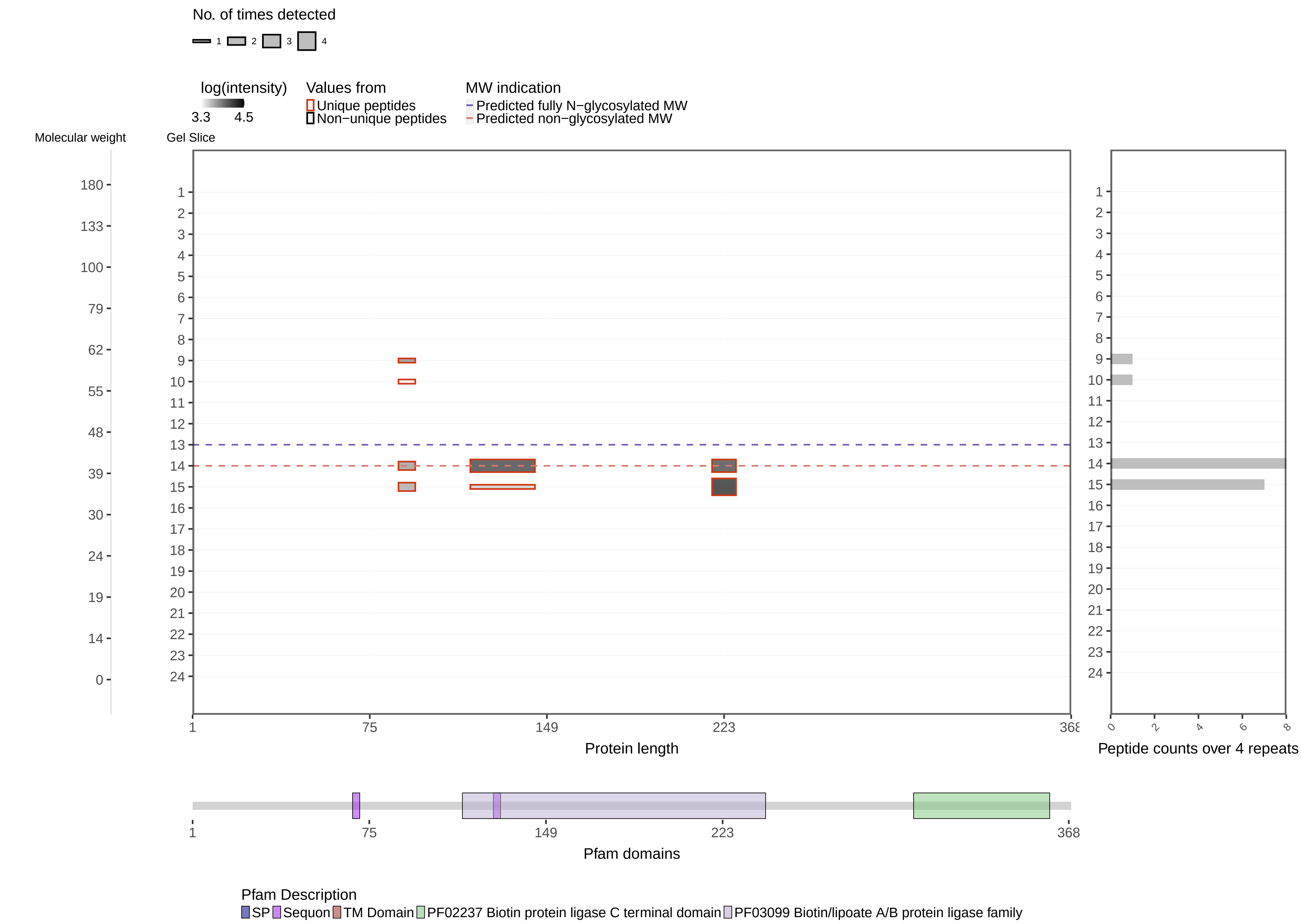


Protein sequence:

SRVVPHIQYVVCLAMTEAIKAVCLEKGTPELDVRIKWPNDLYLGGVKVGGILSTSVYRSKKFYVSAGVGLNVGNEKPTTCLNAVLKKVNPLSRELKREDVIAAFFNKFENLYDVFSEQGFQALEGLYYKTWLHSGQRVIVQERSGDQDYFFETVVTIQGLTSSGYLLGITDDGQMCELHPDGNSFDFFKGLIKRKMS*

### NbL14g19030.1 probable_eukaryotic_translation_initiation_factor_5−1 [MW = 50.84 kDa]

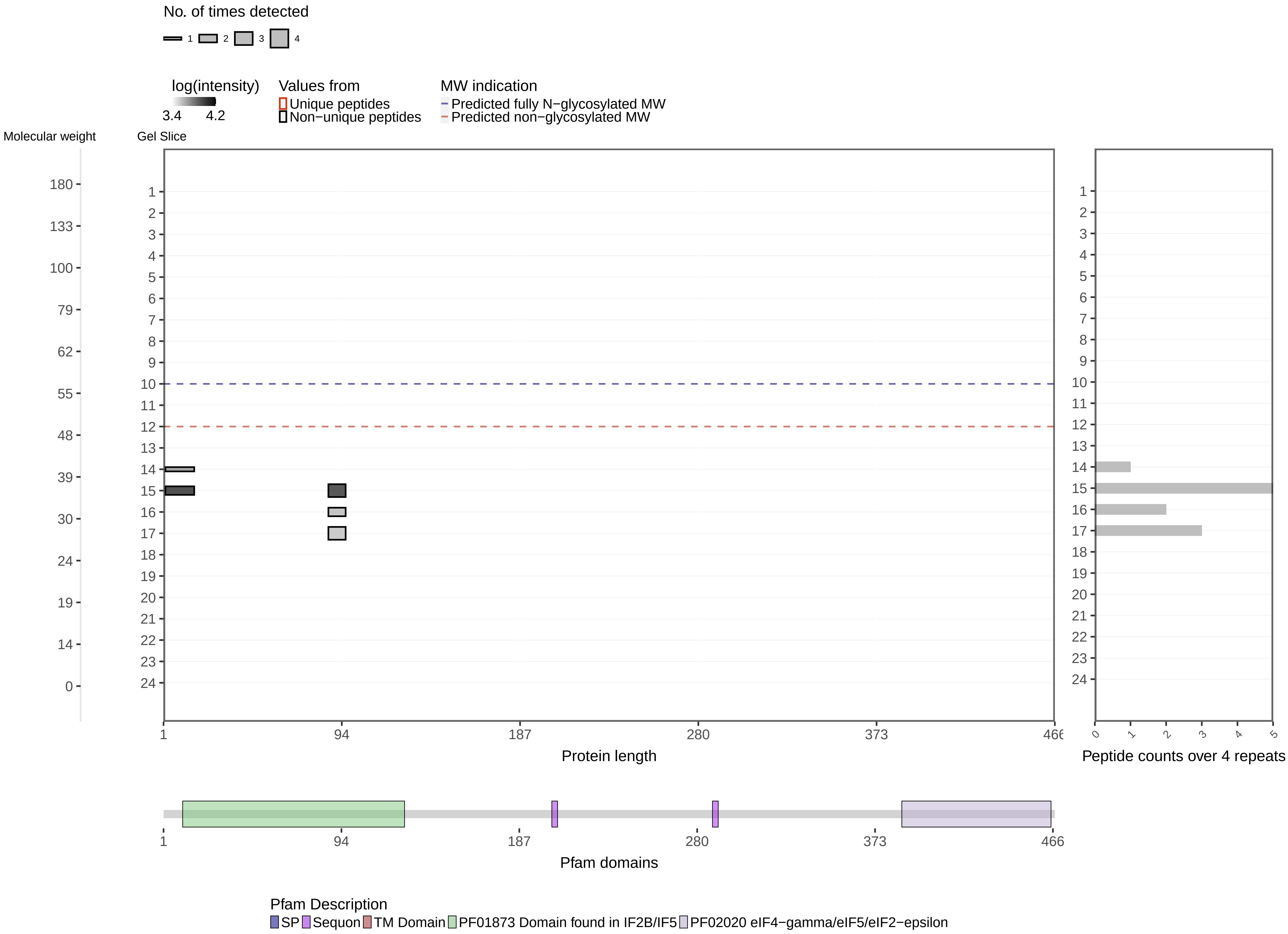


Protein sequence:

EAPPADSDDDVEWQTDTSMAAAQQRIQEQLNAVTASMVMLSATEEKSGKSSPAREEKPKANGQGTGVNCSNFKESLVAQTNGVERSNSKDKLVDAKERLADEIKEHLNKGSSVAQLKSFLGSISGTHEDIVDALFEALFGGVGKGFSKEVTKKKNYLAAVLQEDGSQLALLHAIESFCGKSSSEAVKEVALVLKALYDVDMVEEEFIVEWYEKGSKGSNKLSPIWKNVKPFVEWLQSAESETEDE*

### NbL14g18940.1 nitrate_reductase_[NADH]_2 [MW = 101.96 kDa]

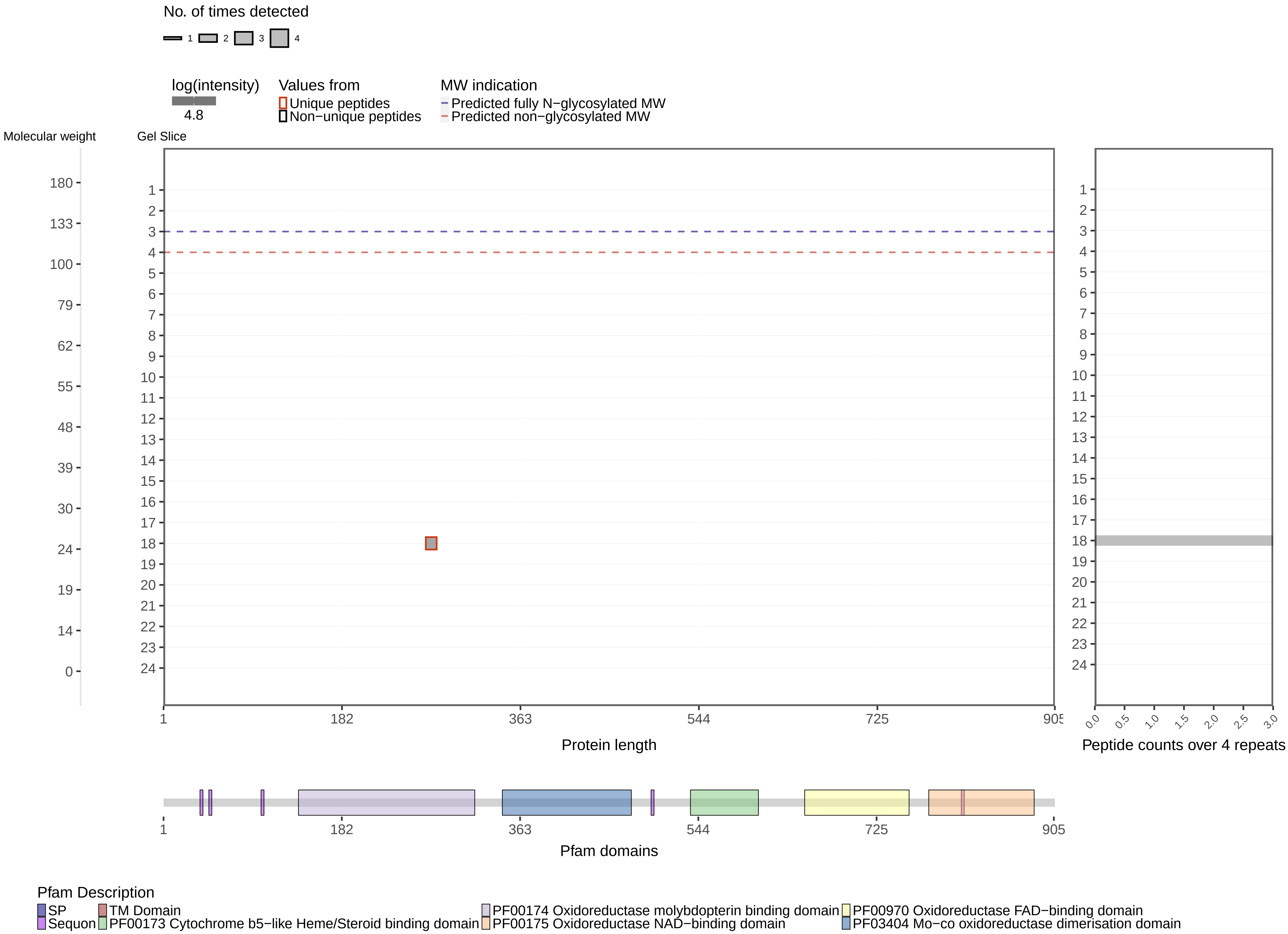


Protein sequence:

.DLLSAKEIAVRAWDETLNTQPEKLIWNVMGMMNNCWFRVKMNVCKPHKGEIGIVFEHPTLPGNQSGGWMAKERHLEISAEAPPTLKKSISTPFMNTASKMYSMSEVRKHSSADSAWIIVHGHIYDATRFLKDHPGGTDSILINAGTDCTEEFDAIHSDKAKKLLEDFRIGELITTGYTSDSPGNSVHGSSSFSSFLAPIKELVPAQRSVALIPREKIPCKLIDKQSISHDVRKFRFALPSEDQVLGLPVGKHIFLCAVIDDKLCMRAYTPTSTIDEVGYFELVVKIYFKGIHPKFPNG

NbL14g18880.1 G−type_lectin_S−receptor−like_serine/threonine−protein_kinase_At1g34300 [MW − SP = 89.96 kDa]

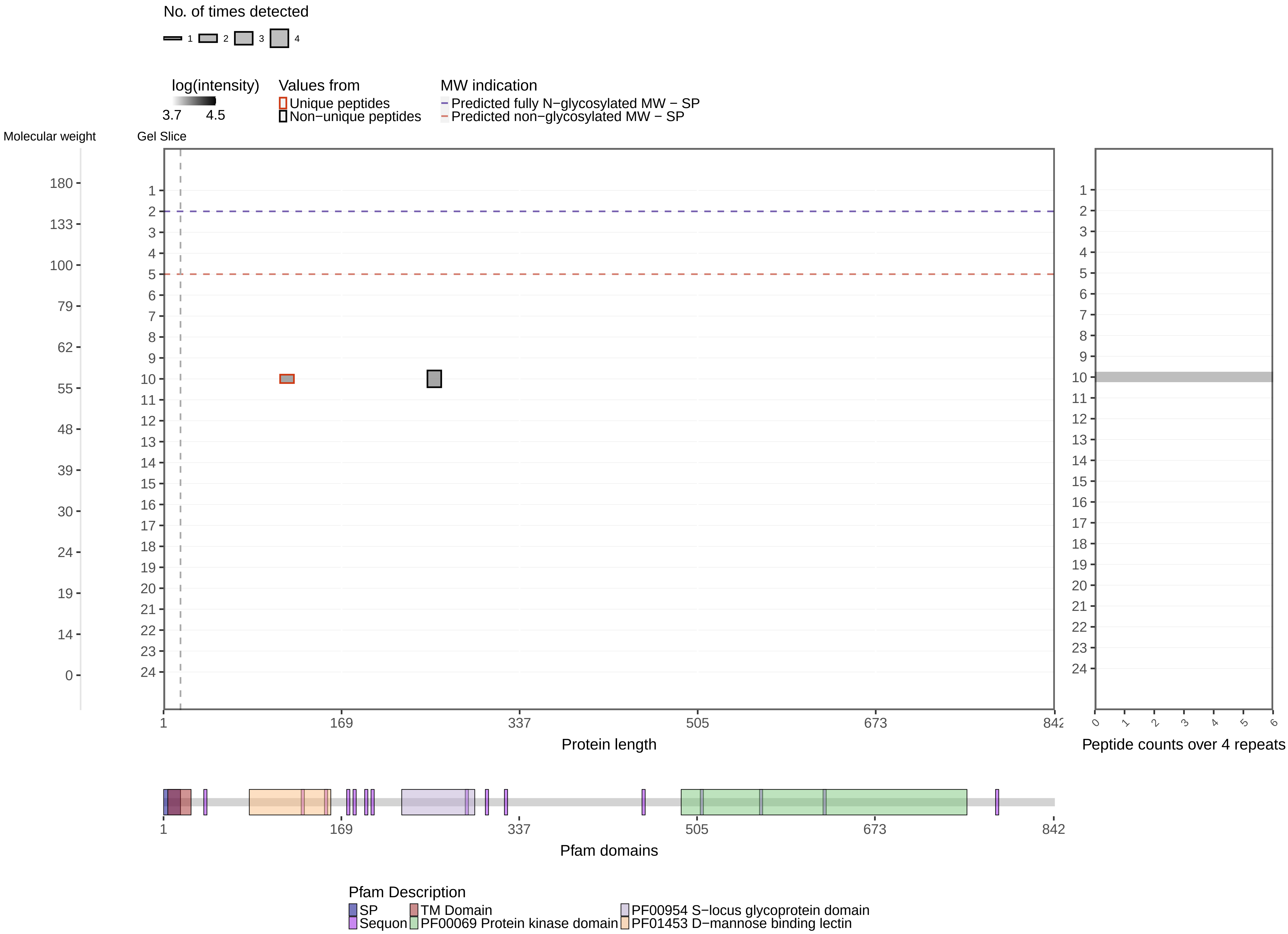


Protein sequence:

### NbL14g18810.1 RNA−binding_protein_8A−like [MW = 24.79 kDa]

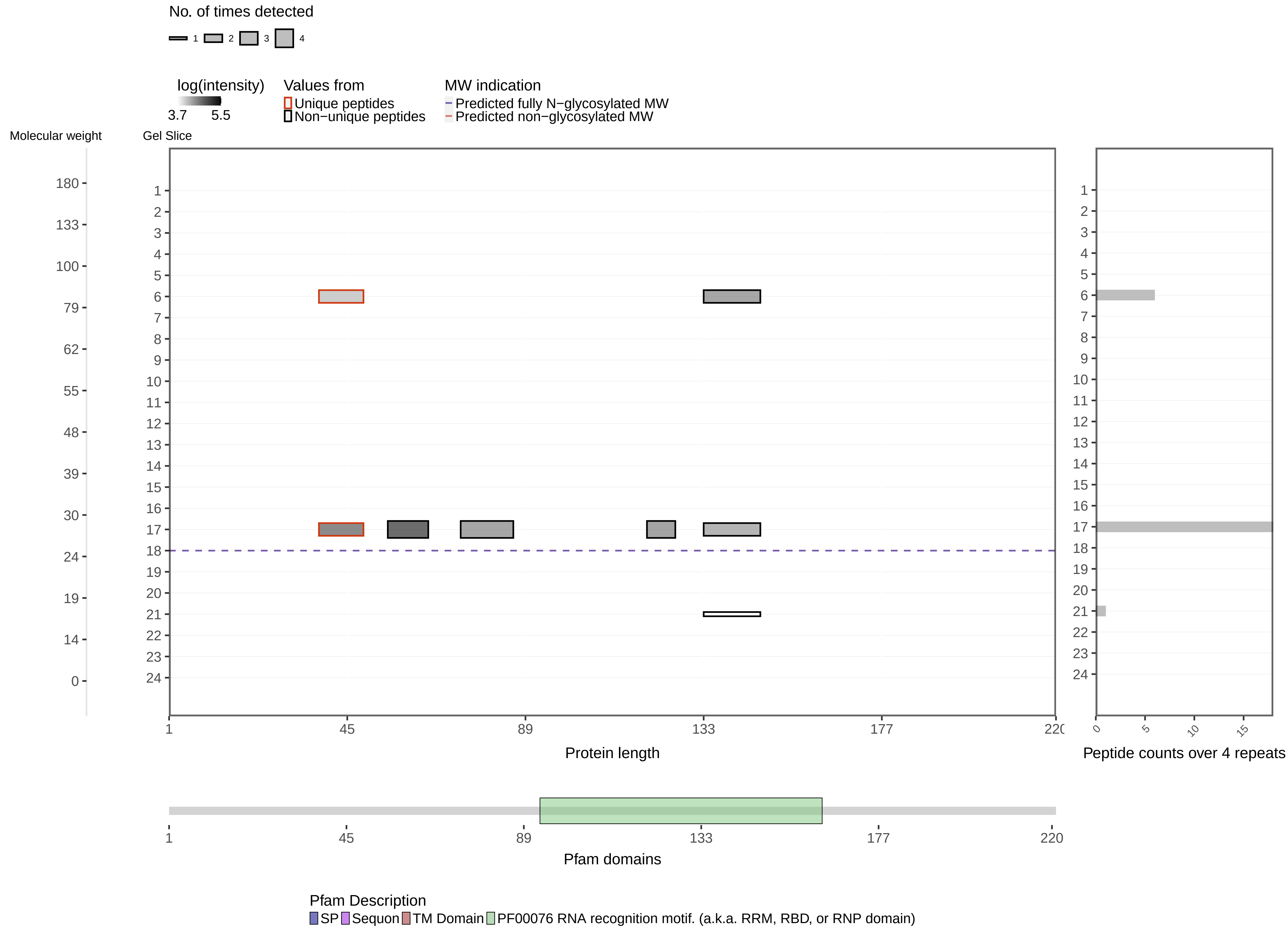


Protein sequence:
VNEETQEDDLHNEFGEFGEIKNVHLNLDRRTGFVKGYALIEYENFEEAQRAINEKDGAEFLTQTIHVDWAFSRGPFKRRNARRRYVHSLSMLLYLHMRFMENEAESSEEMMPAHWKDHSFSL*

### NbL14g18760.1 eukaryotic_translation_initiation_factor_3_subunit_I−like [MW = 35.91 kDa]

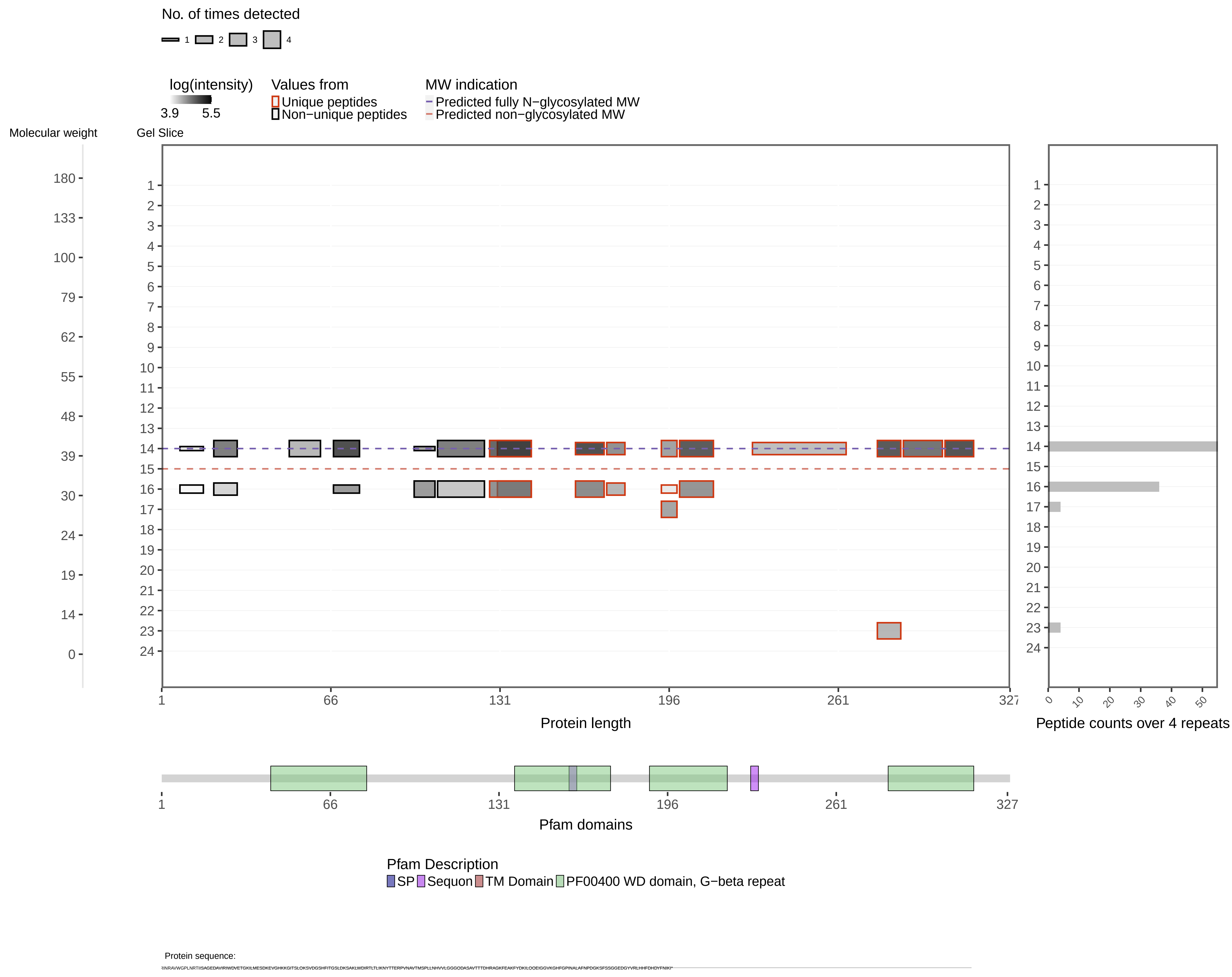

### NbL14g18660.1 uncharacterized_protein_ycf37 [MW = 34.46 kDa]

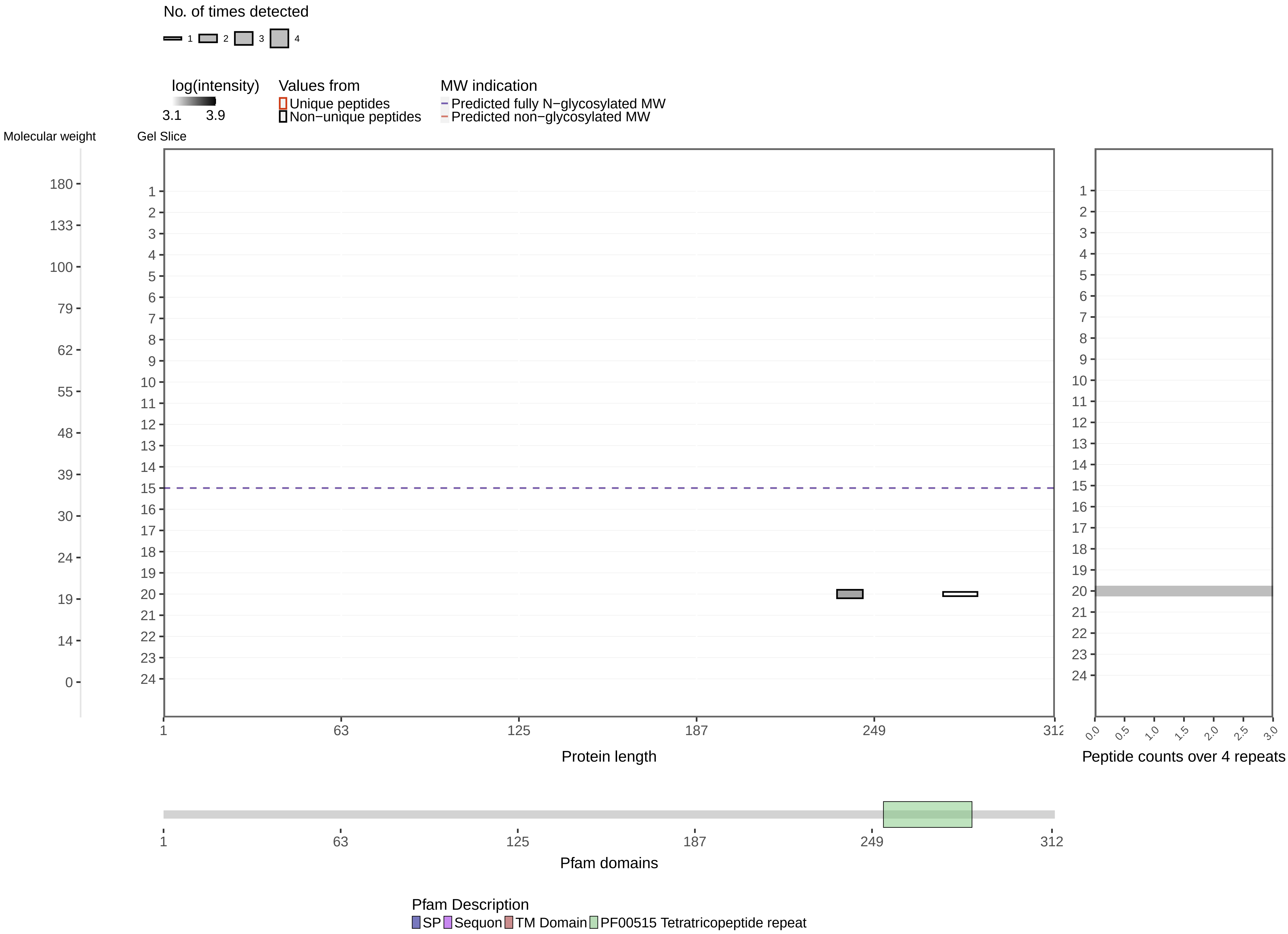


Protein sequence:

AGSFFVIRQVLVRRELDLSAKELQEQVRSGDASATELFELGAVMLRRKFYPAATKYLLQAVQKWDGDDQDLAQVYNALGVSYVLDEKIDKGIAQFENAVKLQPGYVTAWNNLGDAYEKTKDLTSALKAFEEVLLFDPNNKIARPRRDALKEKVQMYKGVPVKSKKS*

### NbL14g18650.1 chorismate_mutase_2 [MW = 29.38 kDa]

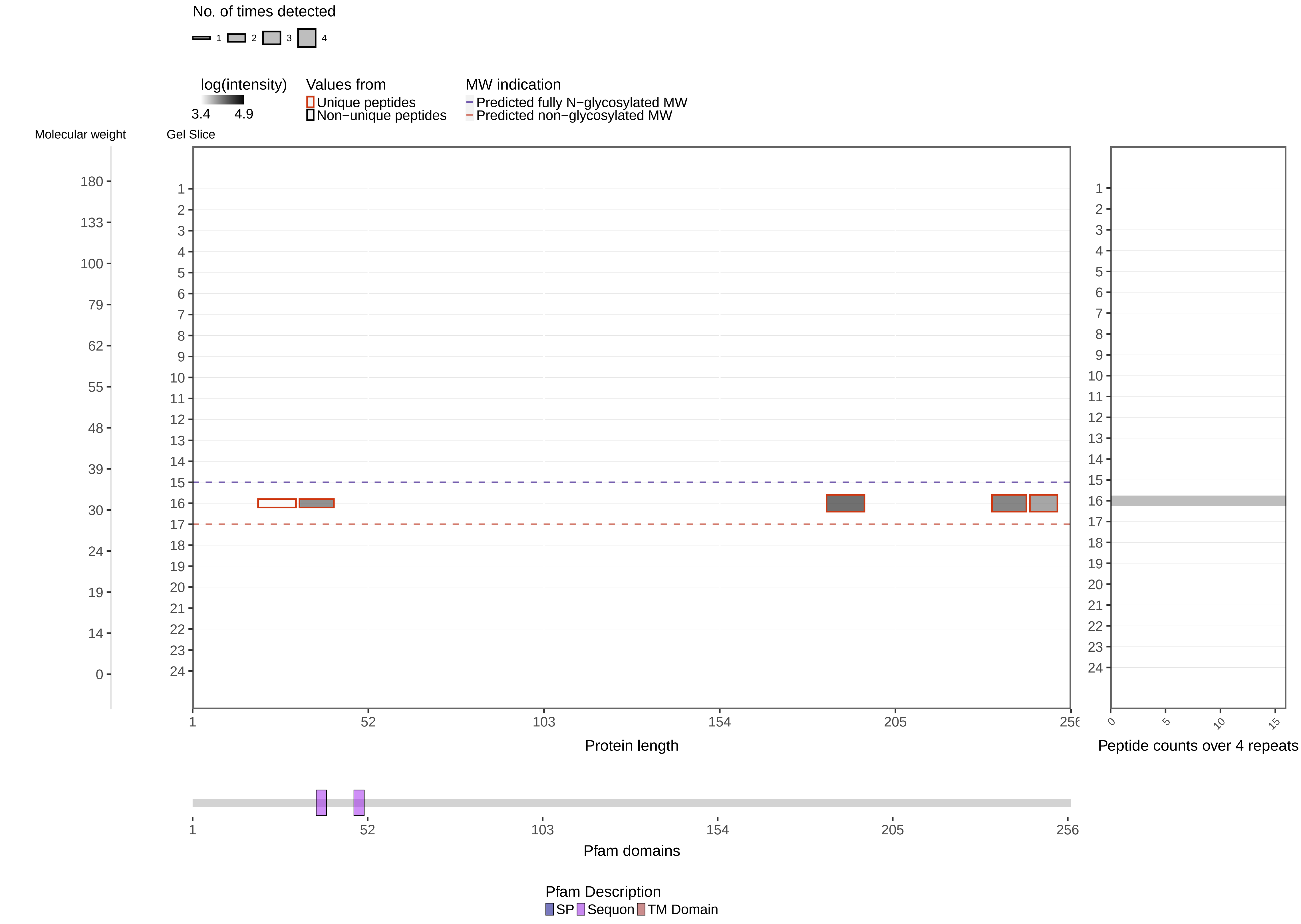


Protein sequence:

'IKQLLPLFCTDQADDEANFATTASCDIHLLQALSRRIHYGKFVAEVKFRDCTDQYKPFILAKDRDALMKLLTFEAVEEMVKKRVAKKALTFGQQVTLNIDDSTKEVKCKVDPSLVSRLYDEWIMPLTKLVEVEYLLRRLD*

NbL14g18640.1 dihydrolipoyllysine−residue_acetyltransferase_component_5_of_pyruvate_dehydrogenase_complex_chloroplastic [MW = 49.6 kDa]

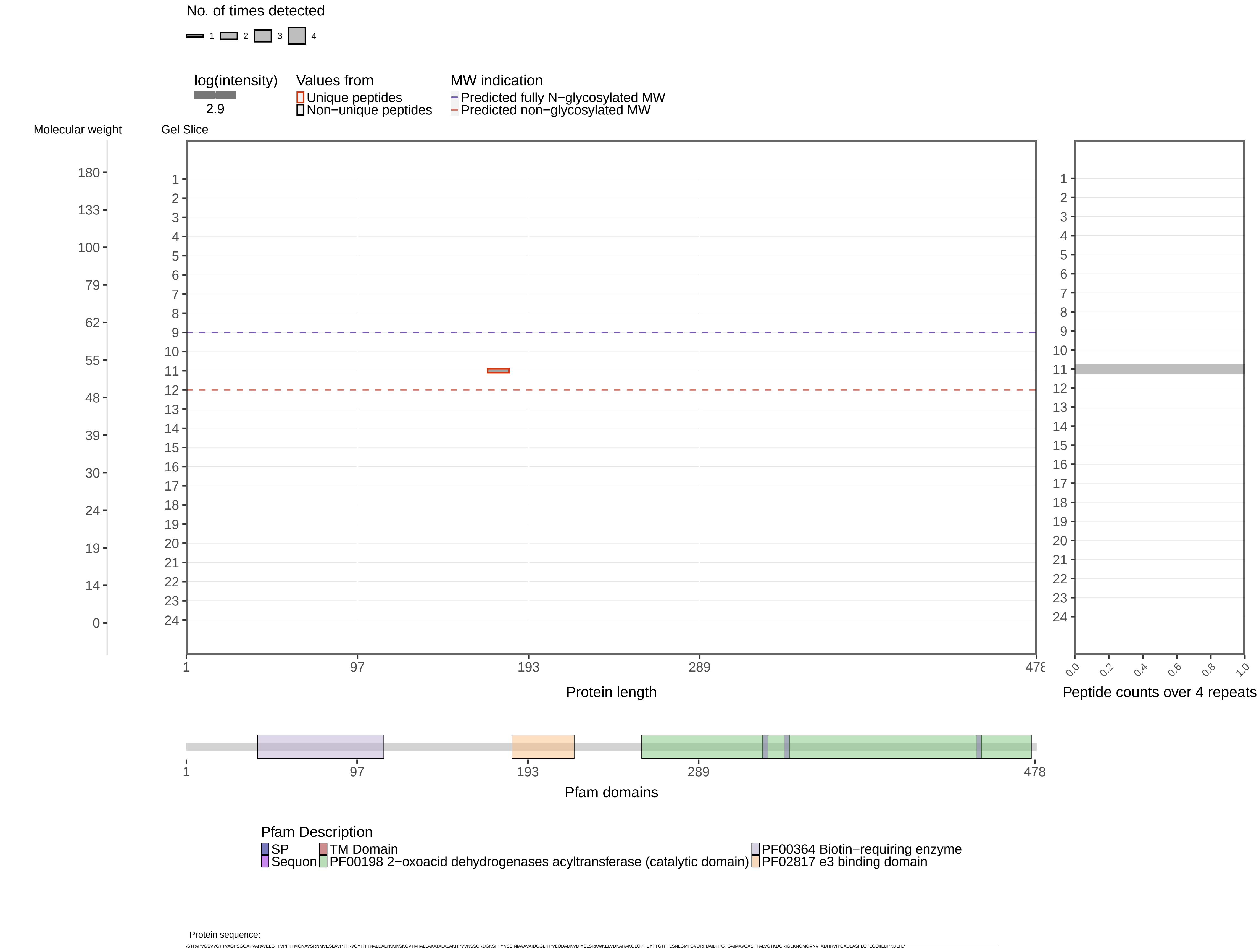

NbL14g18480.1 beta−galactosidase_16−like [MW − SP = 77.88 kDa]

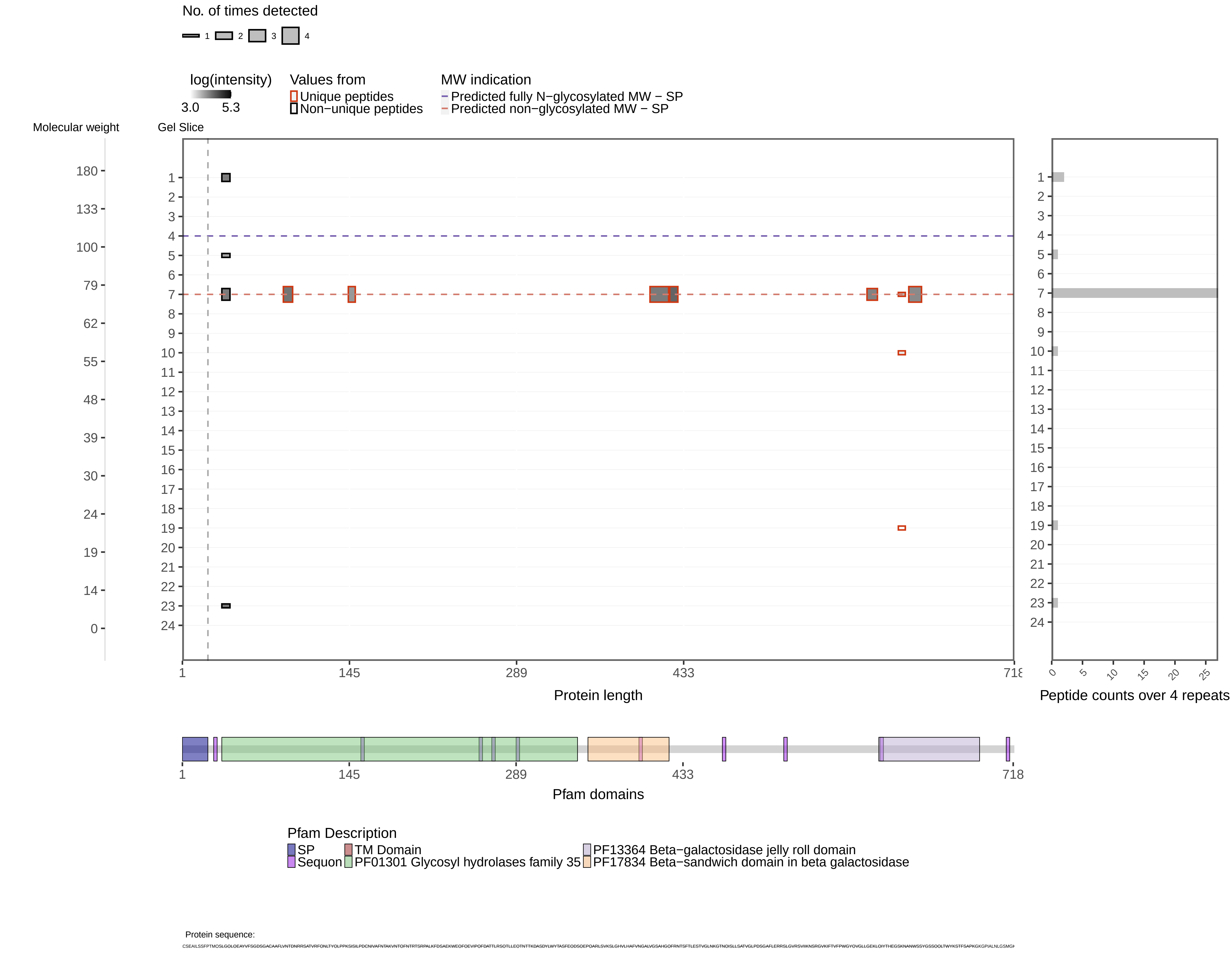

### NbL14g18470.1 beta−galactosidase_16−like [MW − SP = 76.47 kDa]

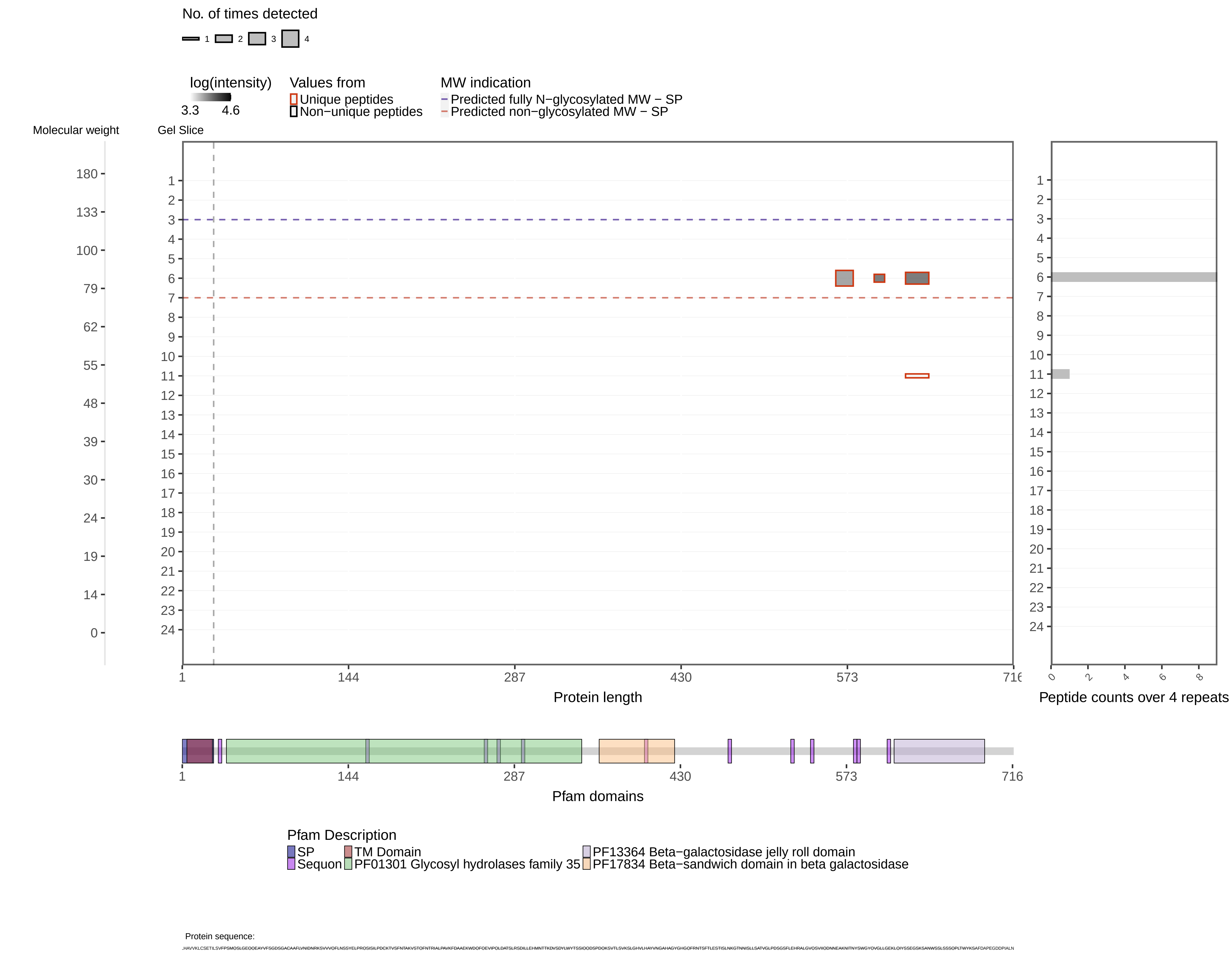

NbL14g18290.1 uncharacterized_LOC104240053 [MW = 38.14 kDa]

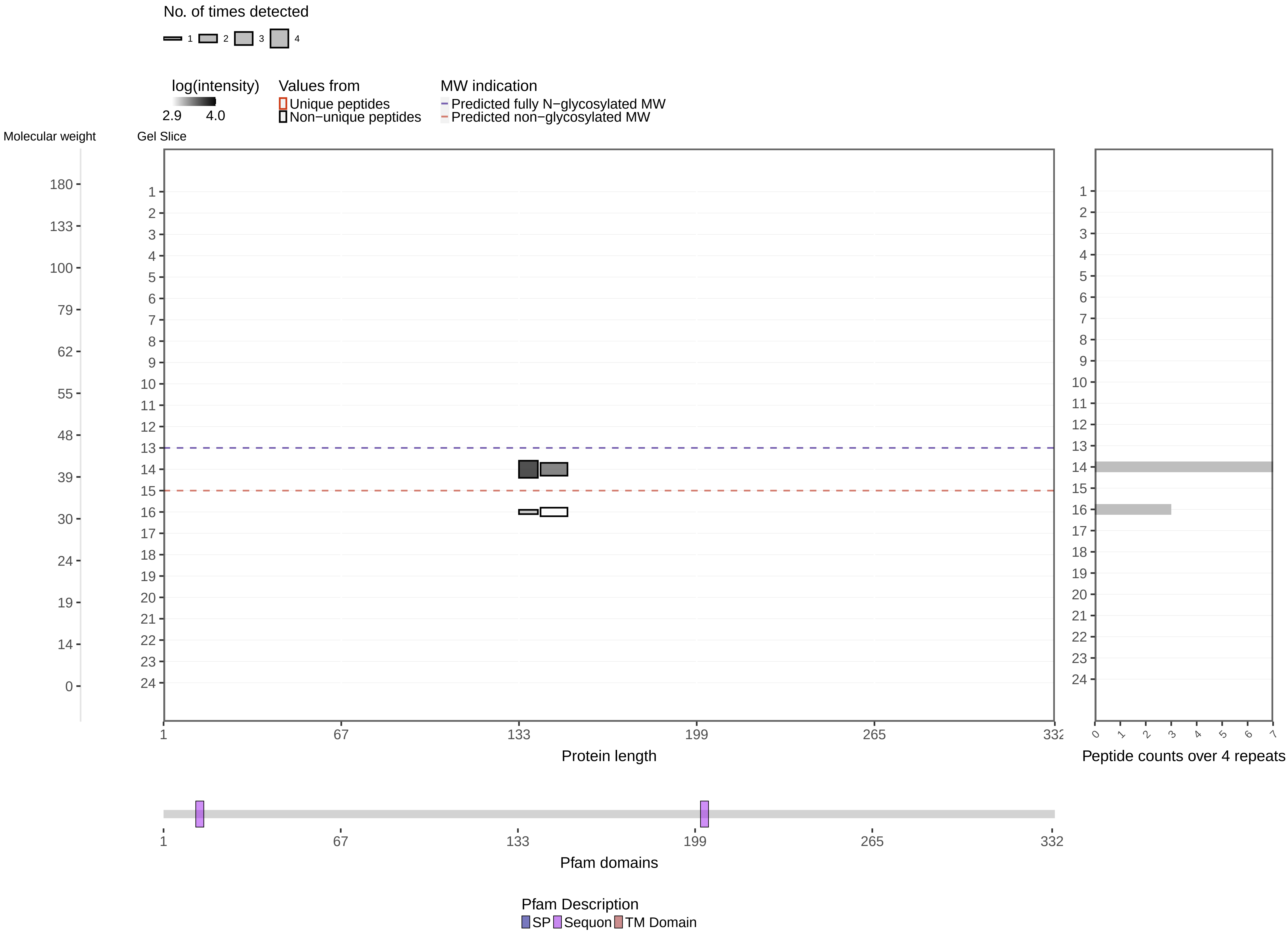


Protein sequence:

PDPDNDVYDFREMYVTPPDTDIYAIPRVLAPMPQKYIRCAMSDYGCYNVTEPPIDAPRDPMYKSEREVSKVFLTKHYRNRRVGDPEFALDFEEIYVIDSKTKSITRVKVVVTVPGGRNRDRKNDLLVIRDNGNSFKIIPSEERDDPTTVIEKEEWKKSRQDMERHLRKLRDFSVSNWF*

NbL14g18210.1 40S_ribosomal_protein_S15a−1 [MW = 14.78 kDa]

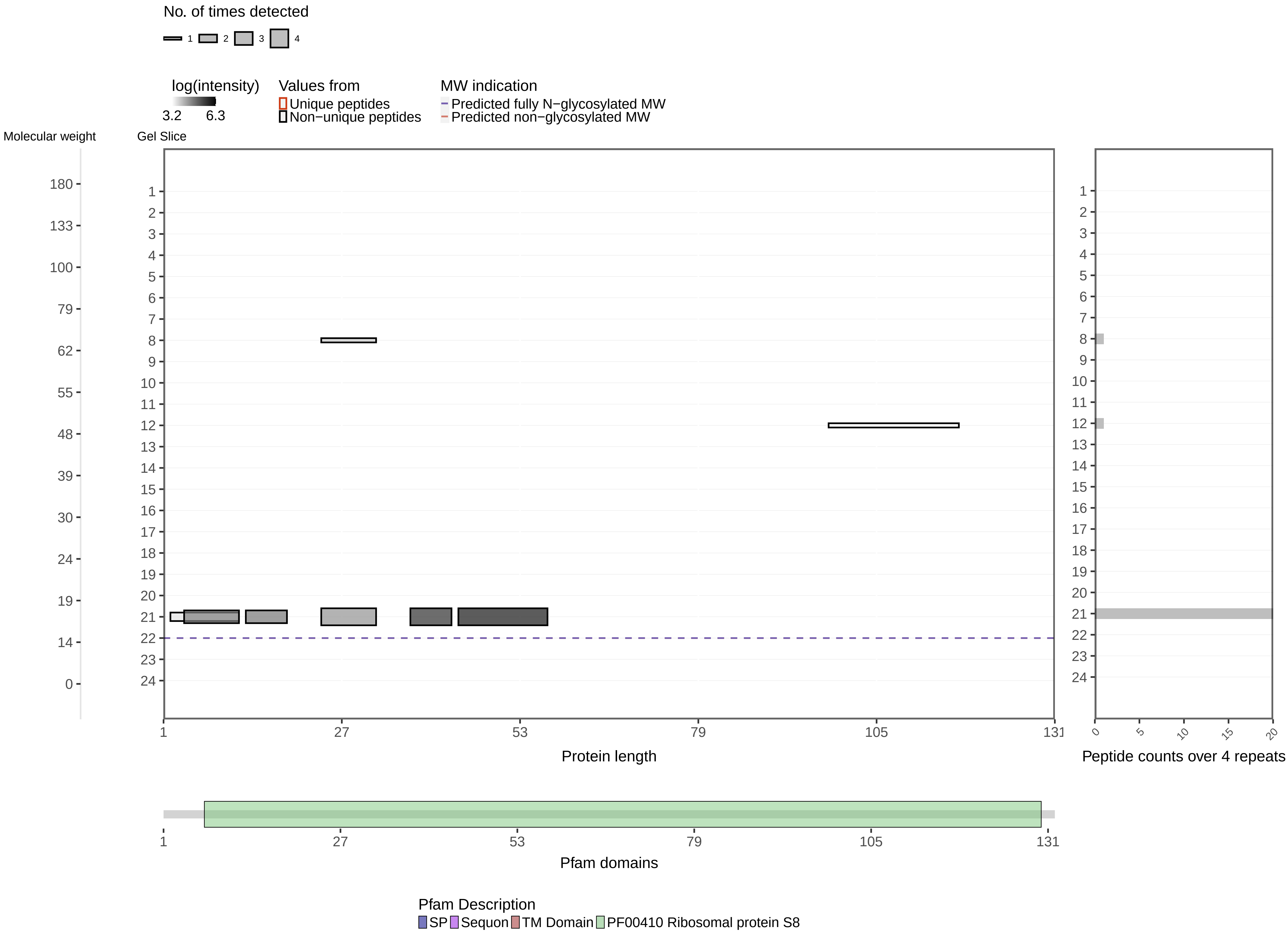


Protein sequence:

VDDHRSGKIVVELNGRLNKCGVISPRFDVGVKEIEGWTARLLPSRQFGYIVLTTSAGIMDHEEARRKNVGGKVLGFFY*

### NbL14g18160.1 UDP−sugar_pyrophosphorylase−like [MW = 68.76 kDa]

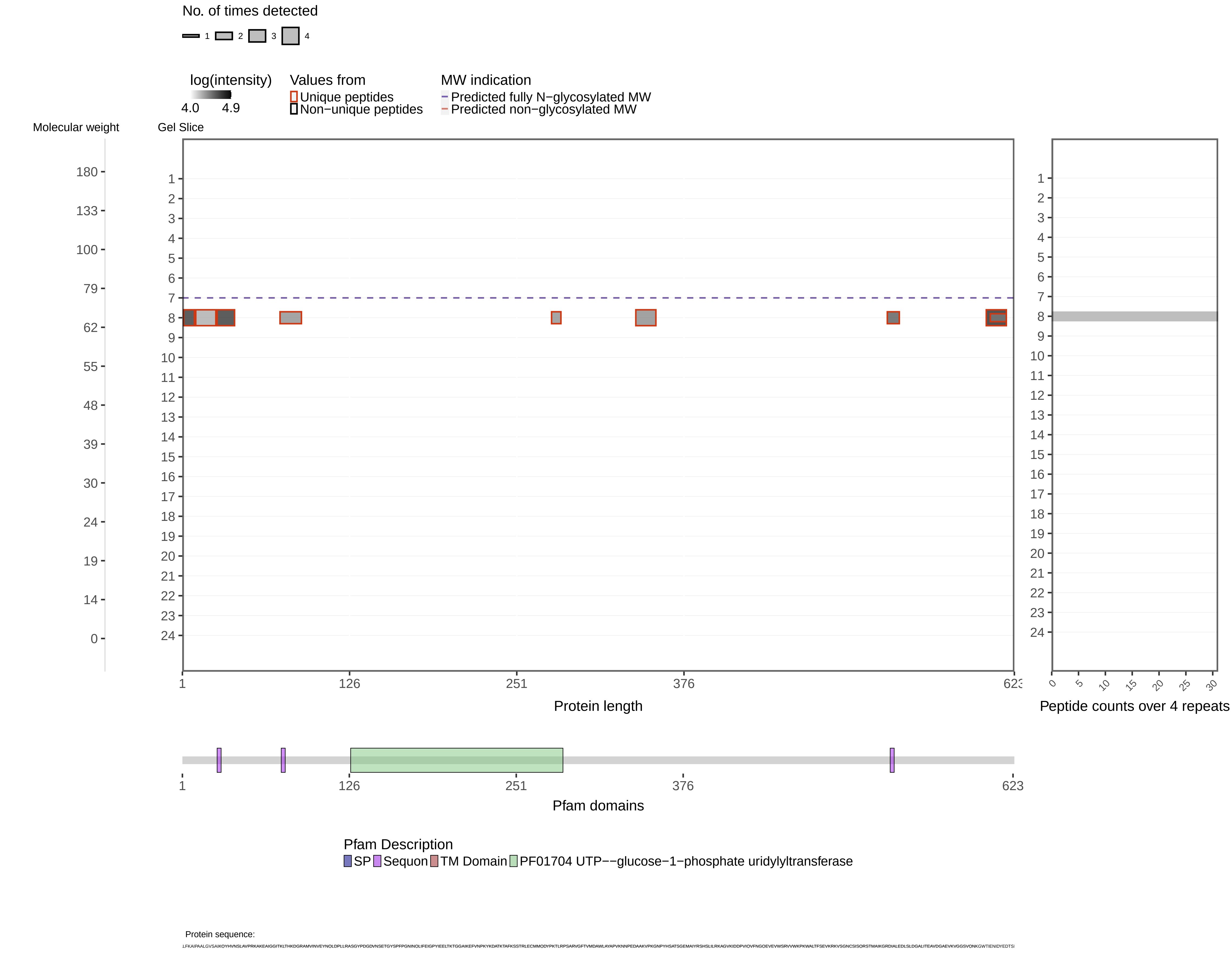

### NbL14g18030.1 TOM1−like_protein_2 [MW = 56.92 kDa]

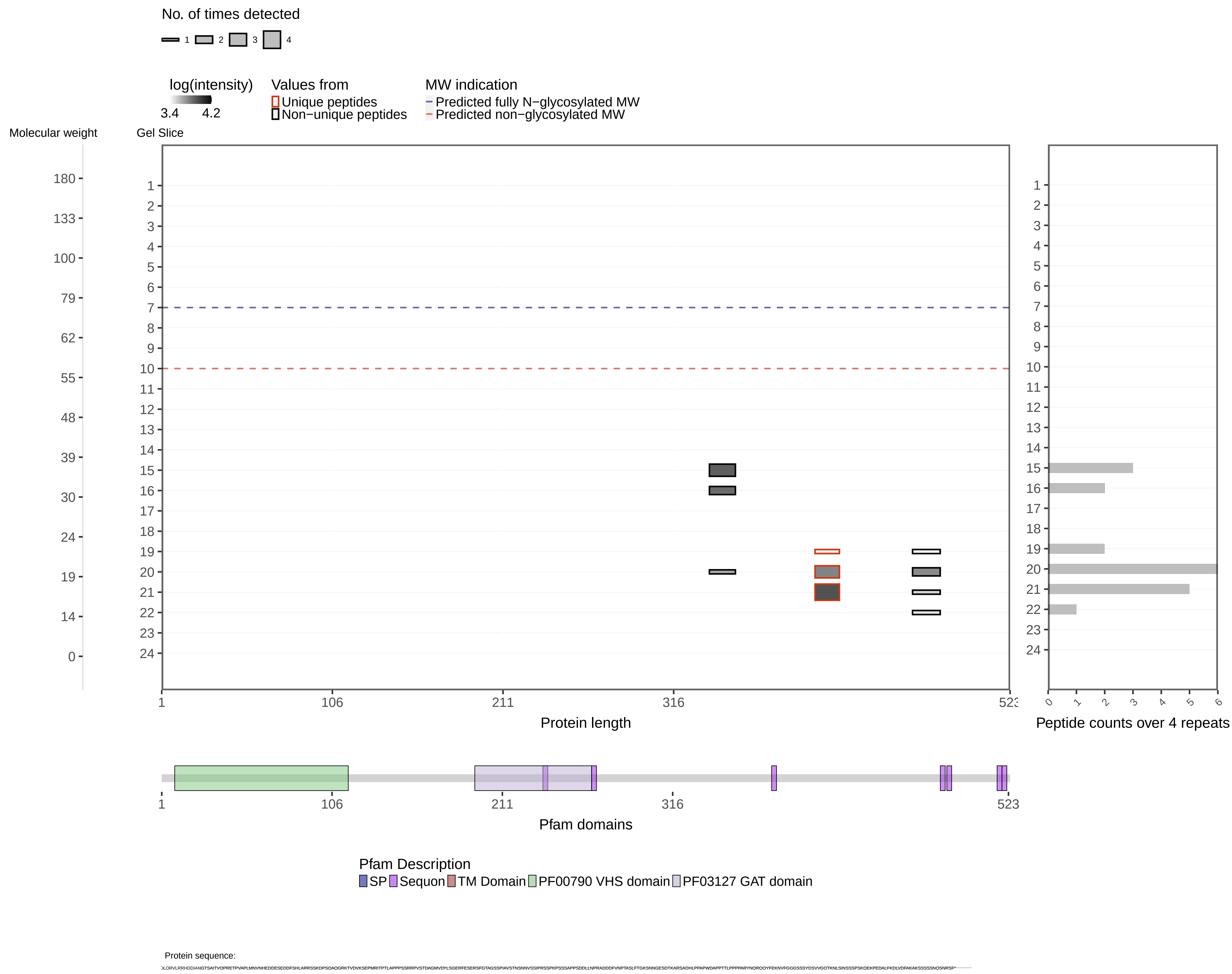

NbL14g17990.1 40S_ribosomal_protein_S2−3−like [MW = 30.24 kDa]

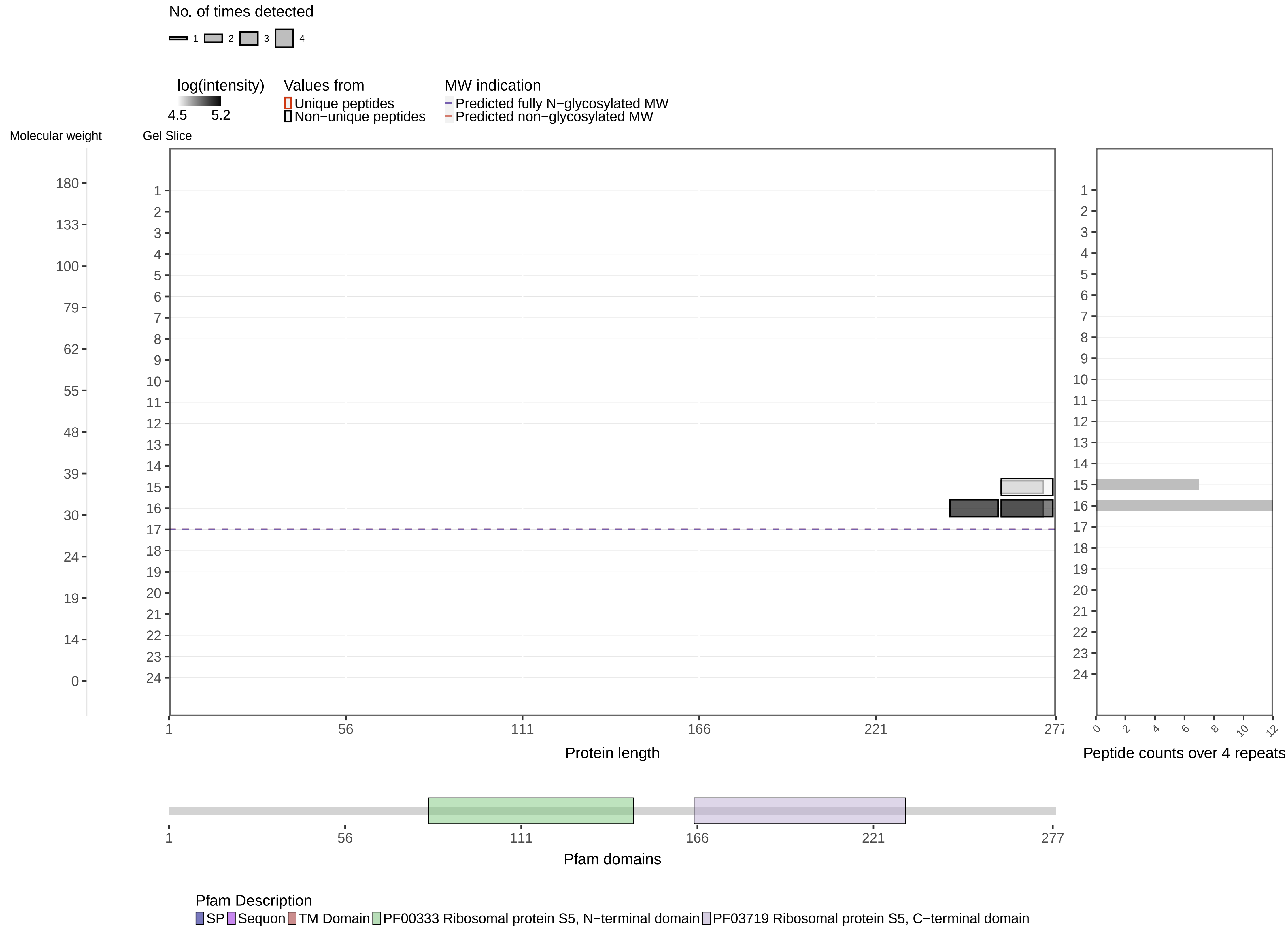


Protein sequence:

CSKEVATAIRGAIILAKLSVIPVRRGYWGNKIGKPHTVPCKVTGKCGSVTVRMVPAPRGAGIVAARVPKKVLQFAGIEDVFTSSRGSTKTLGNFVKATFDCLMKTYGFLTPDFWKETRFTKSPFQEYADILSKPANKVIVYSTEEAPAPERVEA*

NbL14g17950.1 2−oxoisovalerate_dehydrogenase_subunit_alpha_1_mitochondrial−like [MW = 13.77 kDa]

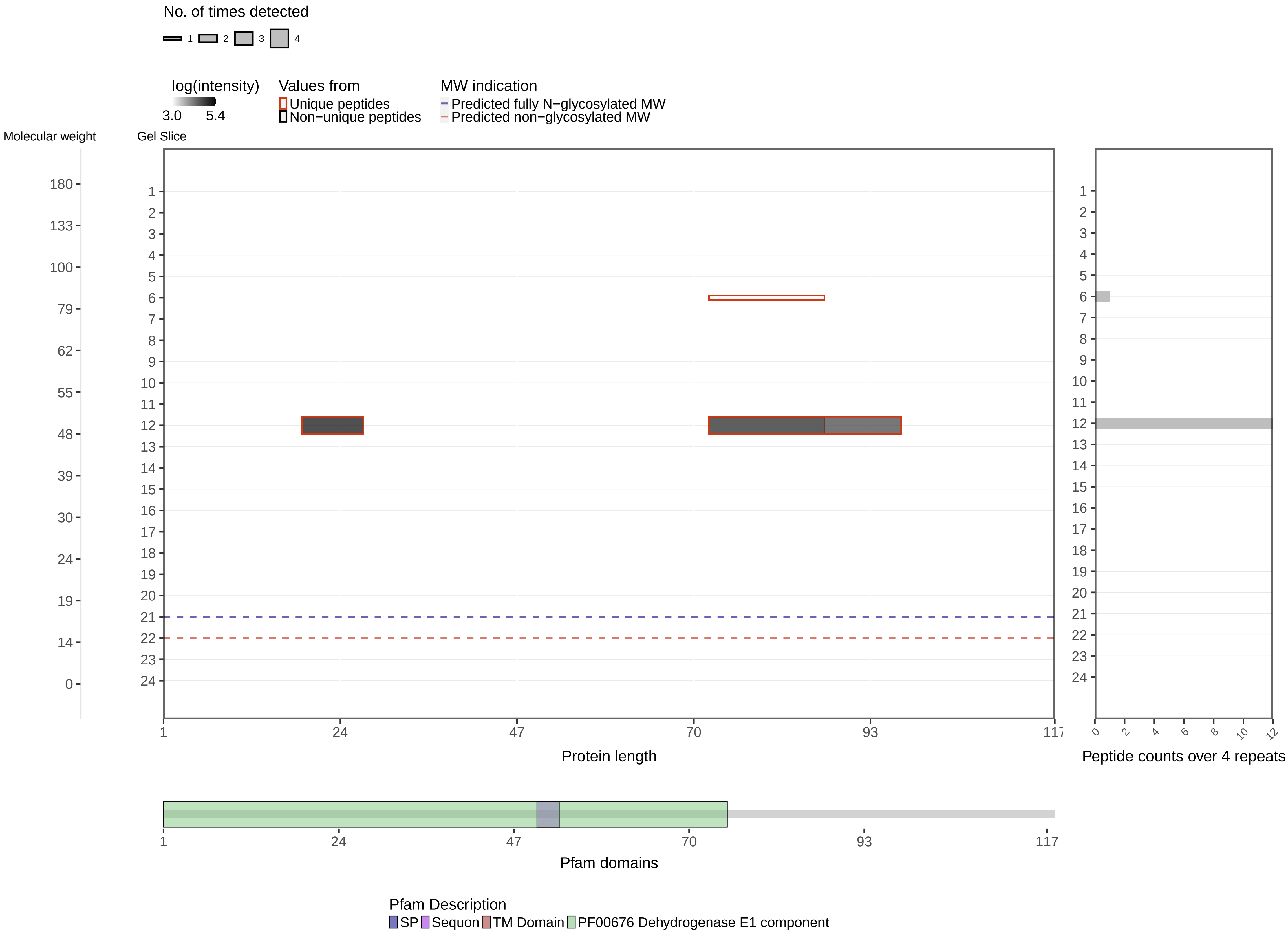


Protein sequence:

'WNDENESELRGDTRNKVLQAMQAAEKVEKPPLTDLFTDVYDKVPLNLQEQHKFIRDAVKRSPKEYPSDVPI*

### NbL14g17870.1 probable_protein_phosphatase_2C_11 [MW = 46.48 kDa]

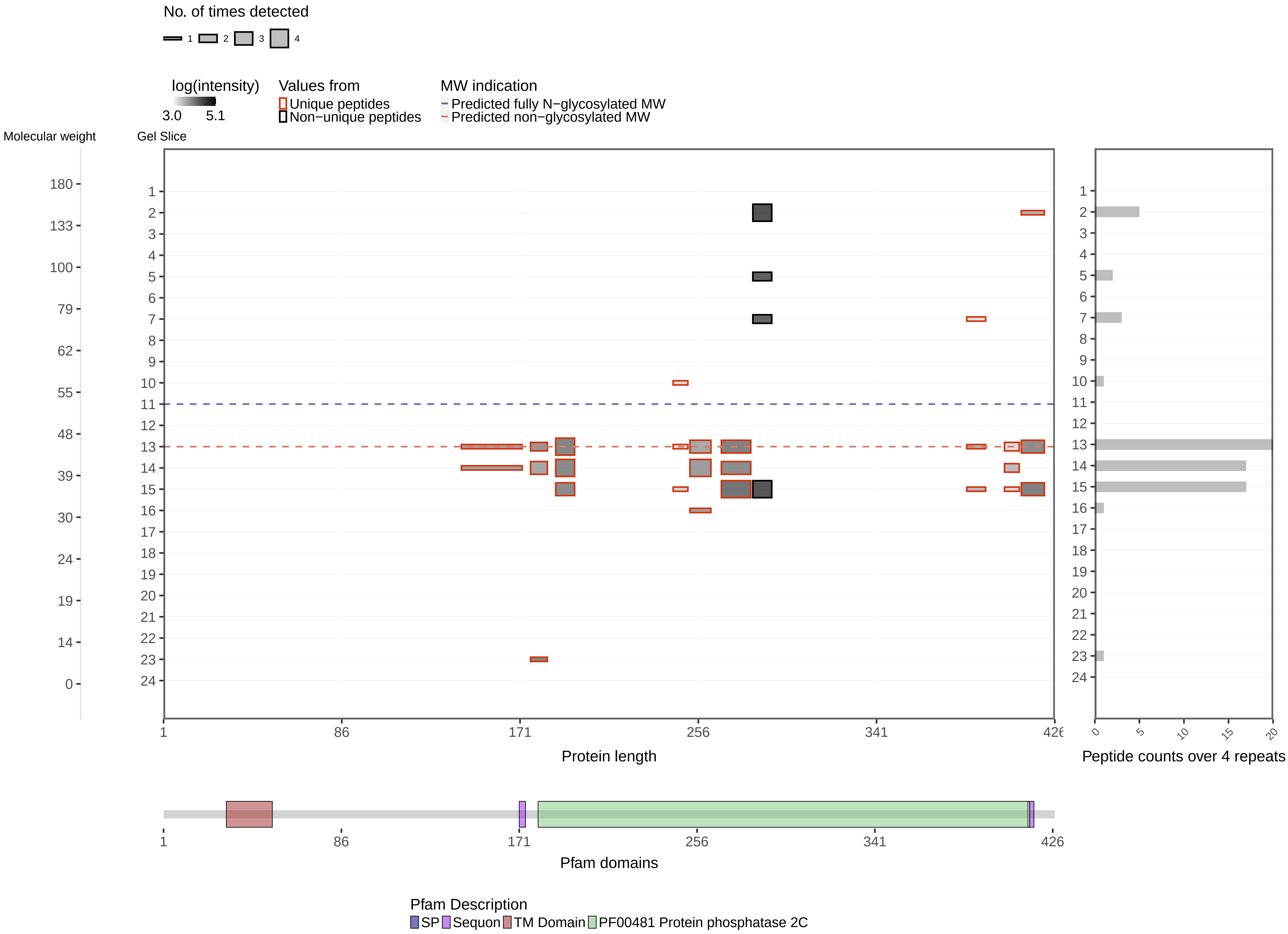


Protein sequence:

SEVDGQMVAFFGVFDGHGGSRTAEYLKNNLFKNLIGHPDFIKDTKSAIVEAFKQTDADYLNEEKGQQKDAGSTASTAVLLGDRLVVANVGDSRVVACRDGSAIPLSIDHKPDRSDERERIEQAGGFIIWAGTWRVGGVLAVSRAFGDKLLKPYVVADPEIQEEEIDGVDFIIIASDGLWNVLSNKDAVAIVQAIKDAEAASRKLIEEAYSRGSSDNITCVVVRFESA*

### NbL14g17630.1 serine_carboxypeptidase−like_42 [MW − SP = 50.43 kDa]

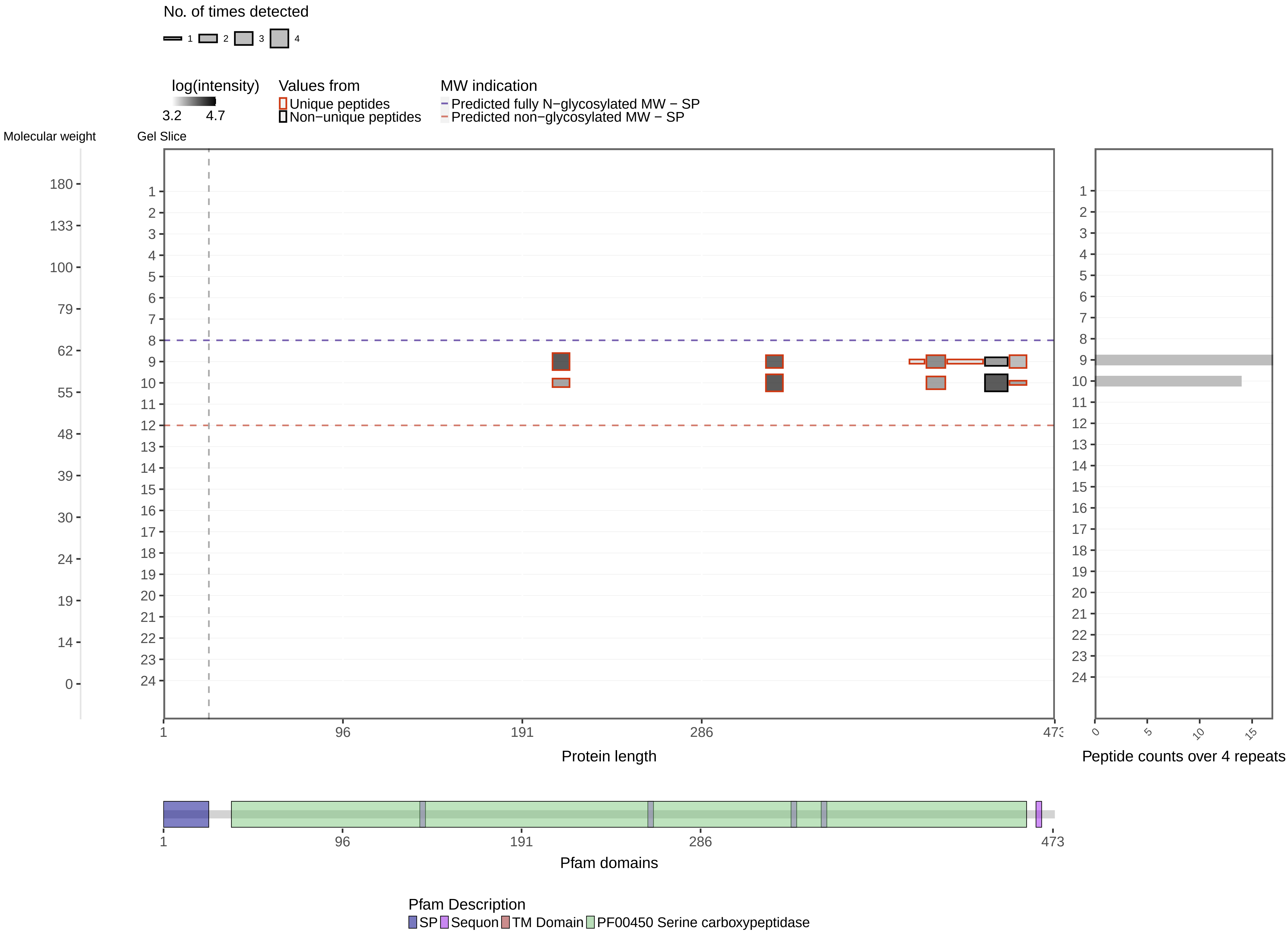


Protein sequence:

PATYEYFWSHGMISDEVGLAIMNQCDFEDYTFGSPHNVSHECNNAISEANDIIGEYINNYDVILDVCYPSIVEQELRLRKMATKMSVGVDVCMSYERRFYFNLPEVQKALHANRTNLAYSWSMCSSILNYSDVDGNINILPLFKRIIQNHIPVWIFSGDQDSVVPLLGSRTLVRELAHDMGFKITVPYGVWFHKGQVGGWQTEYGNILTFATVRGAAHMVPYAQPSRALHLFSSFIRGRRLPNNTRPSIED*

### NbL14g17590.1 uncharacterized_protein_At2g24330−like [MW = 44.61 kDa]

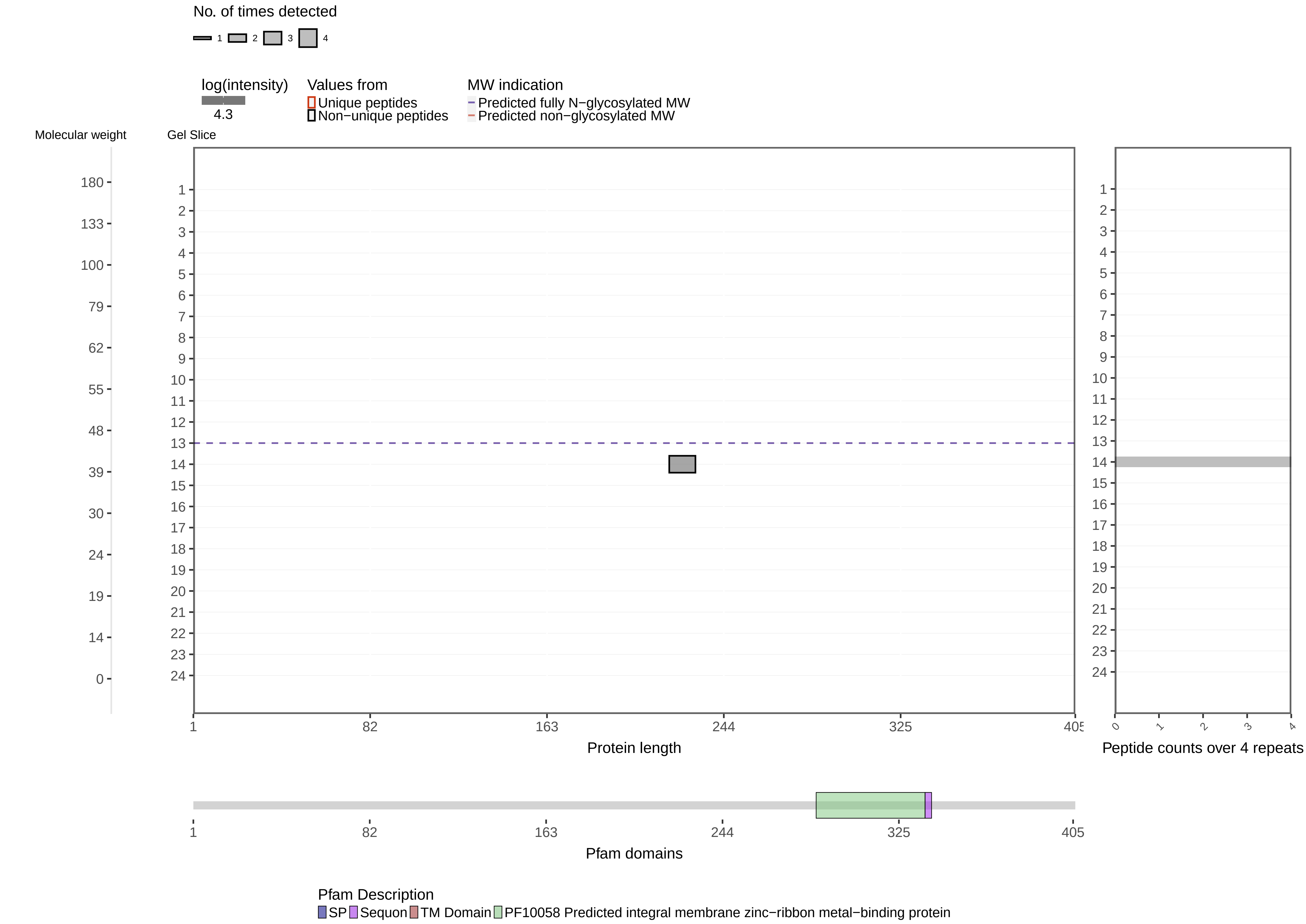


Protein sequence:

ASKLGADTGLKFHVGDDTNLNVPTGKSNDVEVVOSTGLRNRKOGRSSSPESAVLDHHEAEMLOOAOLEGSGTMOHOOTVVEHYNPTGSSTODGGWLSKIAALLVGEDPTOSYALICGNCHMHNGLARKEDFPYITYYCPHCHALNKSKOVDDRGSGTSTPNLGSTTVVADIDLVKOGGGSIPDKISASASGSPVAPPVATEGENIVSSTSNS*

### NbL14g17580.1 vacuolar_protein_sorting−associated_protein_45_homolog [MW = 65.21 kDa]

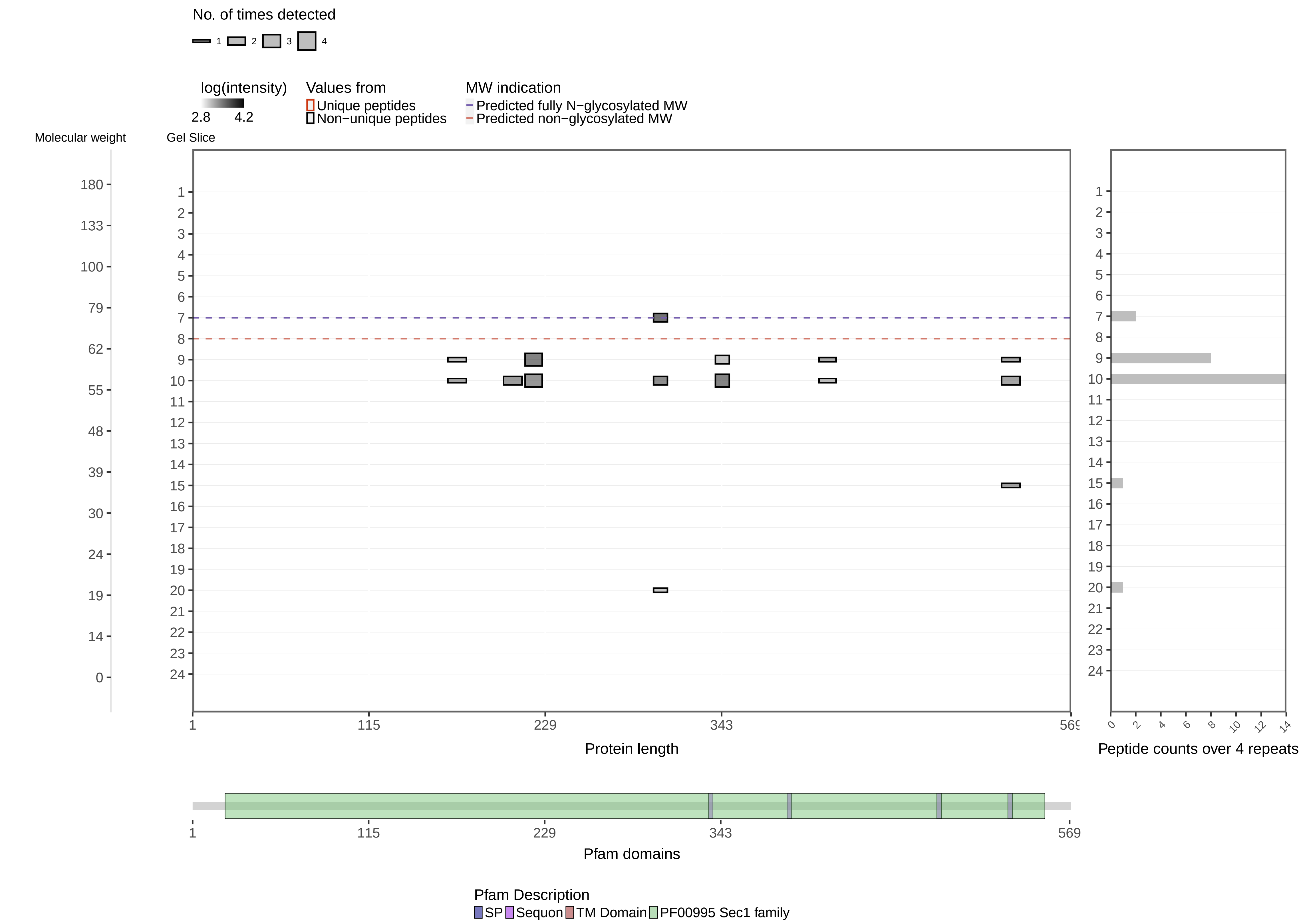


Protein sequence:

SSEQDAFFKTNMYDNFGDIGMNIKKMVDDFQQVAKSNQNIQTIEDMAKFVDNYPEYRKMQGNVSKHVTLVTEMSKIVEERKLMLVSQTEQELACNGGQGAAFEAVTTLLNNDNISDMDRLRLVMLYALRYEKESPVQLMQLFNKLASRSPKYKPGLVQFLLKQAGVDKRTGDLYGNRDLLNIARNMARGLKGVENVYTQHQPLLFQTMENITRGRLRDVDYPYVGNHFQQARPQEVVIFTVGGTTYEESRSVALQNSTNSGIRFILGGSALLNSKRFLRDLEEAQRIARTS

NbL14g17520.1 uncharacterized_LOC107768657 [MW = 29.09 kDa]

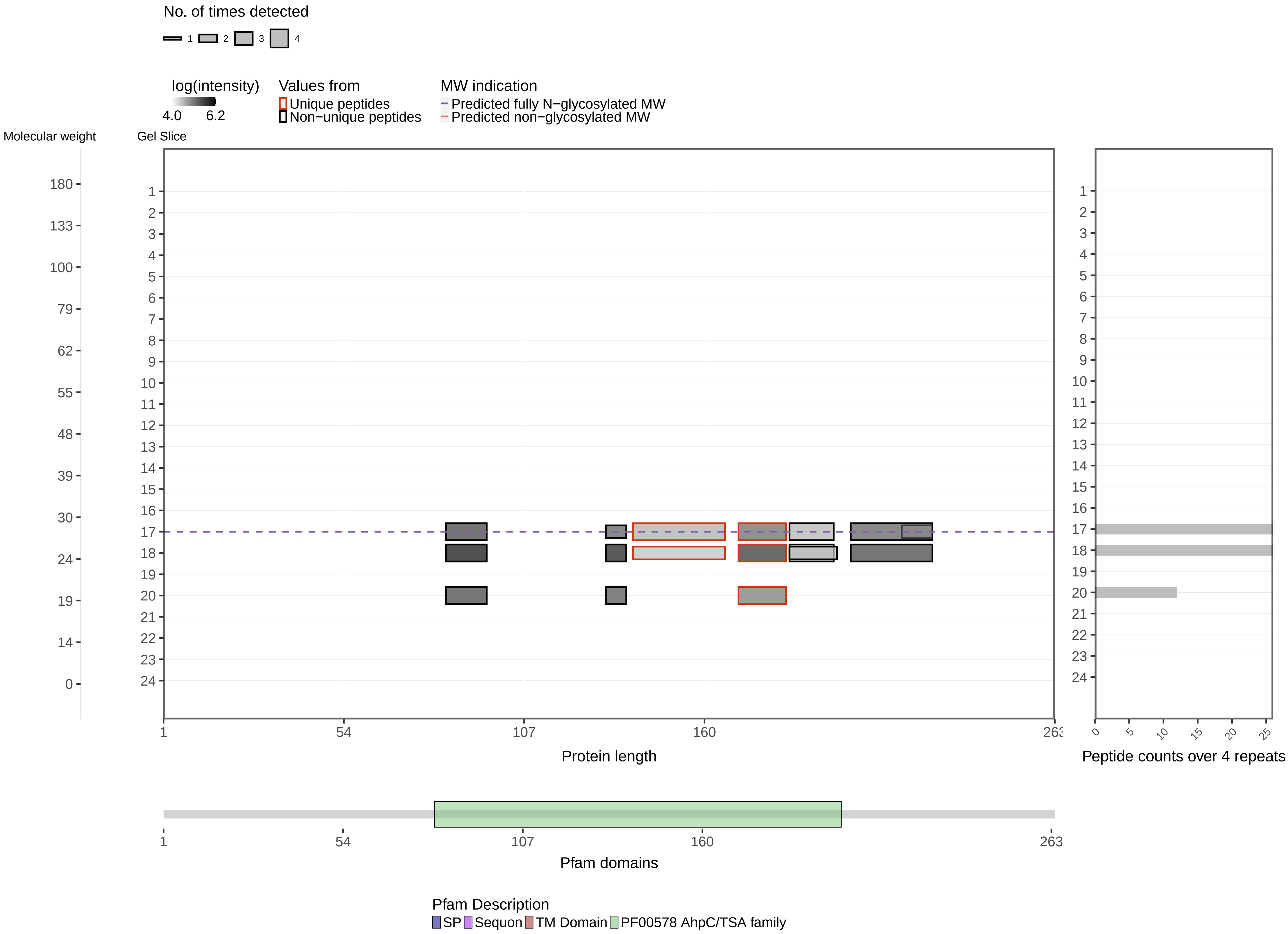


Protein sequence:

VIHLKKDIVKLSNFYMKKGLAVVAISSNSVVTHPQDGPEFMAEDAKLFKYPFPYLYDESQEVAKDFGAVCTPEFFLFKKDGRRPFELVYHGQFDDSRPSNNAPVTGRDLSLAIDCVLSGQPVSSNQKPSIGCSIKWHPGAKQ*

### NbL14g17390.1 RNA−binding_protein_2−like [MW = 33.27 kDa]

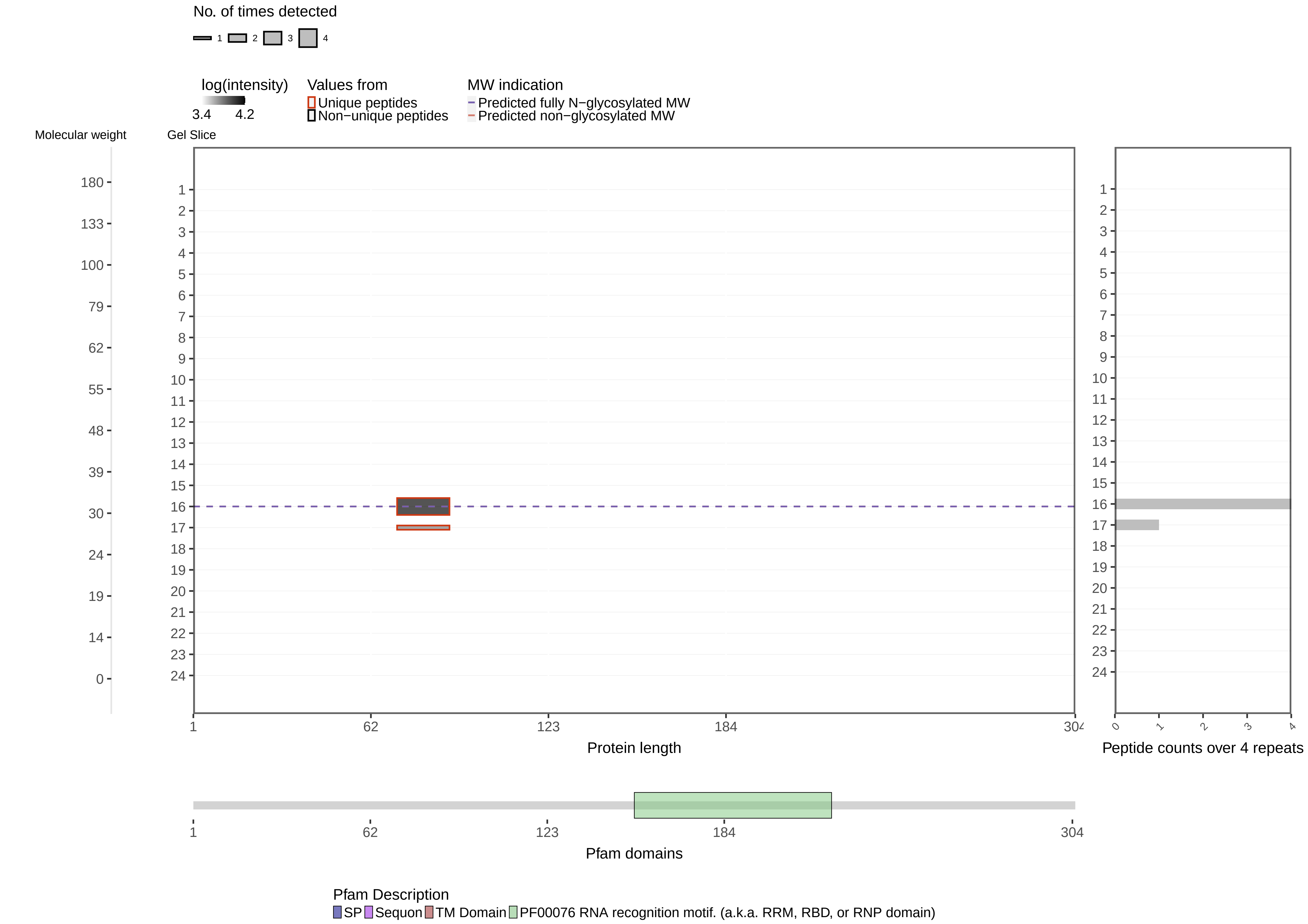


Protein sequence:

SLPRPRETLPLPPDASNTLYIEGLPADSSRREVAHIFRPFVGYKEVRLVRKESKHRGGDPLILCFVDFVDPACAATALSALOVLEVSRSEVWWLGESWEAINSIHOLFIYYFLLLRLNRLLLCTVTGICNSOLAOORAVSKVTVLCTLLVFEVGGFSLOOPLKPVKLSC*

NbL14g17090.1 protein_FAM188A−like [MW = 68.19 kDa]

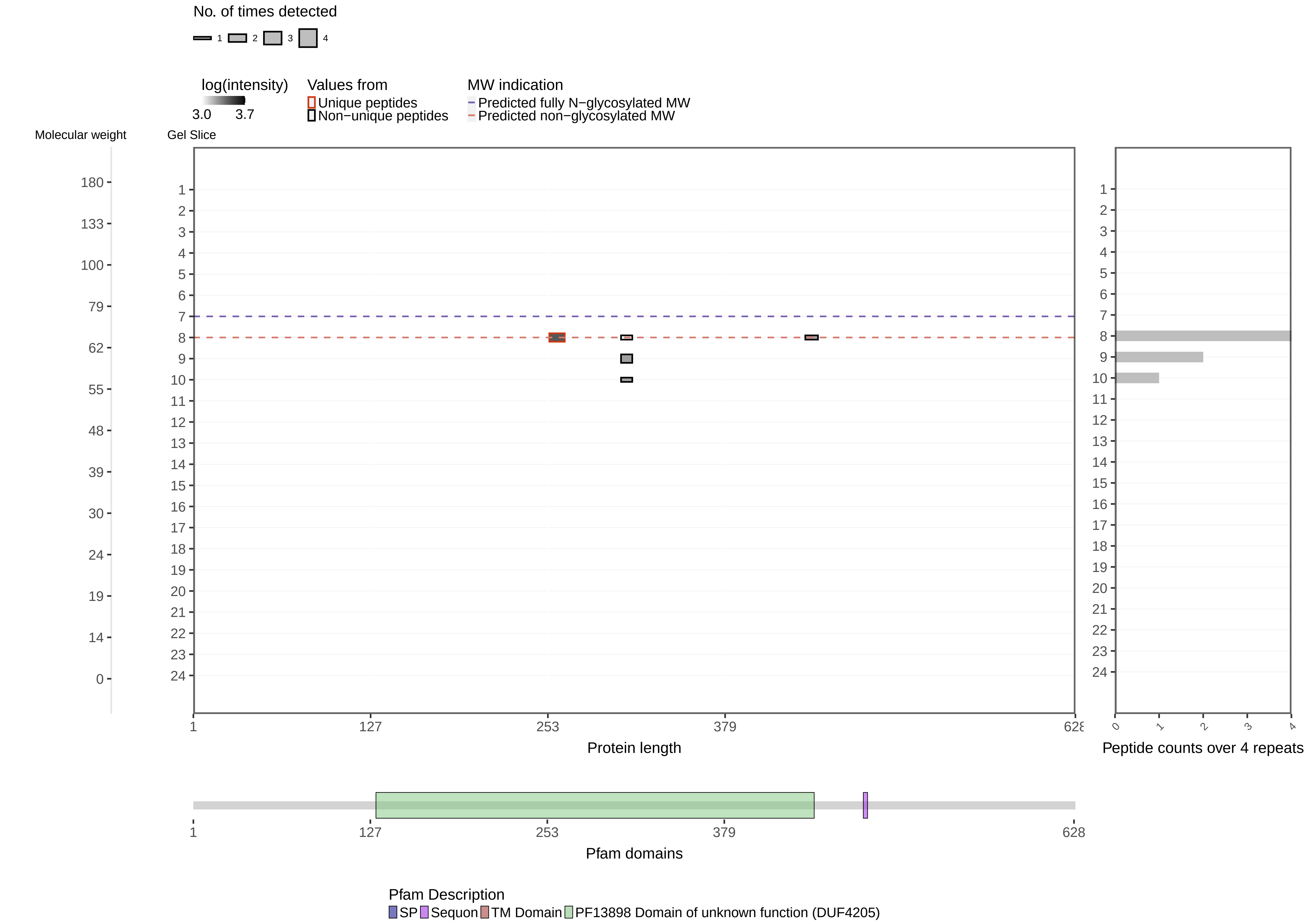


Protein sequence:

RLEALLPVFRSRMGAMLFLISALLSRGMDSVQADRDDPSQPLVTAPFGHASQEIVNLLLSGEAVANVFDGRMDLGGGMFVKGISTTVEIGFLTLLESLNFCKVGLHLKCPKWPIWVVGSESHYTVLFALDTKVQEENELEGRETKIRRAFDAQDQSGGGGFISVEGFHQVLRETNVSLPADKLQNLCSSGYIVWSEFWQVLLDLDKSMGGLKDPSGLMGKKVFDLYHFNGIAKSVMNGNQVSSVNDFPVQRPRLTKLRVSVPPRWTPEEFMSSGPGTSDPTAGKDSVT

### NbL14g17080.1 fructose−16−bisphosphatase_cytosolic [MW = 37.22 kDa]

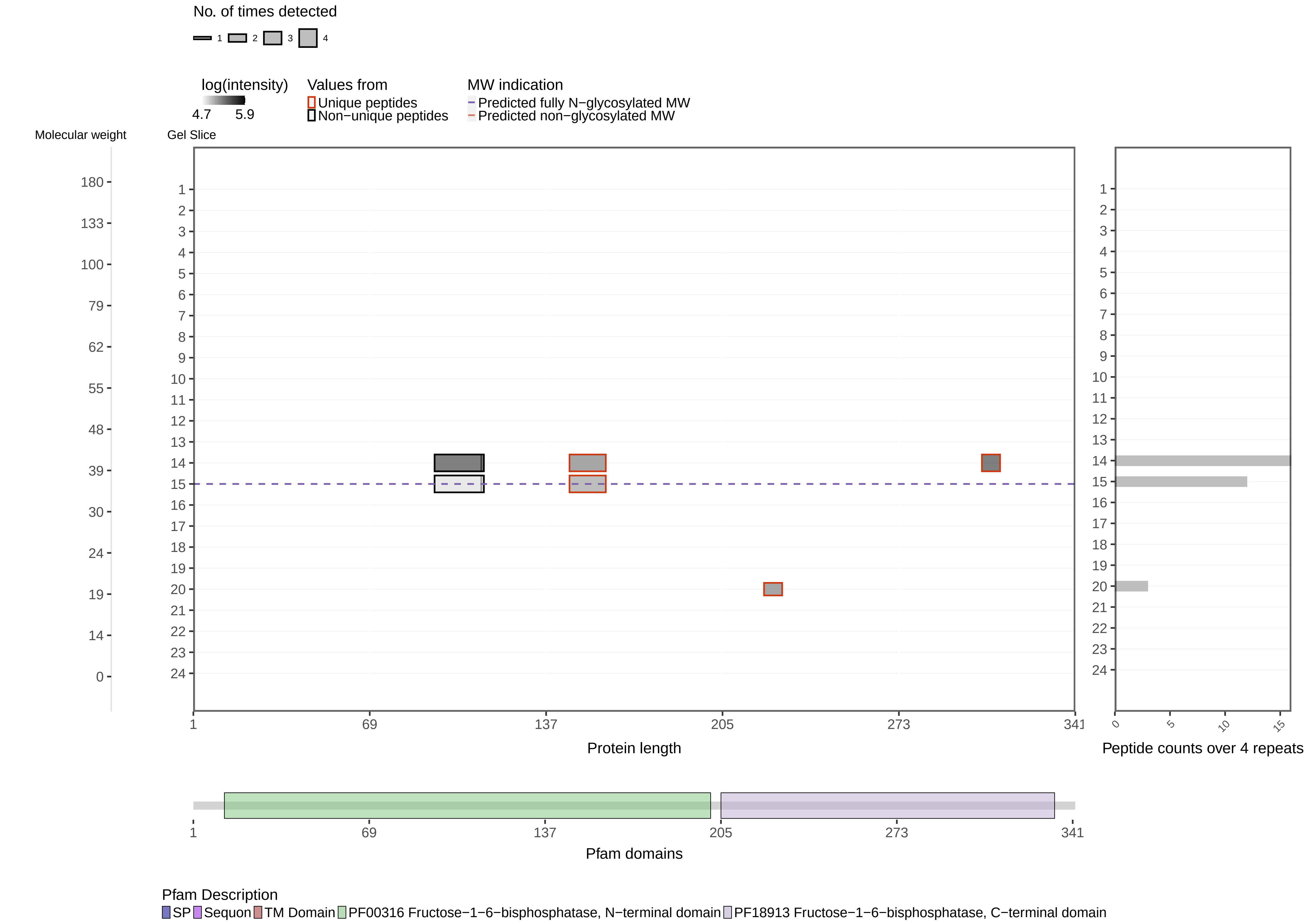


Protein sequence:

'GKNMLAAGYCMYGSSCTLVLSTGSGVNGFTLDPSLGEFILTHPDIKIPKKGKIYSVNEGNAKNWDAPTSKYVQNCKYPTDGSSPKSLRYIGSMVADVHRTLLYGGIFLYPADKKSPNGKLRVLYEVFPMSFLMEQAGGQSFTGKQRALDLVPEKIHERAPIFLGSYDDVEEIKKLYAAEEQN*

NbL14g16880.1 thioredoxin_Y1_chloroplastic−like [MW = 19.08 kDa]

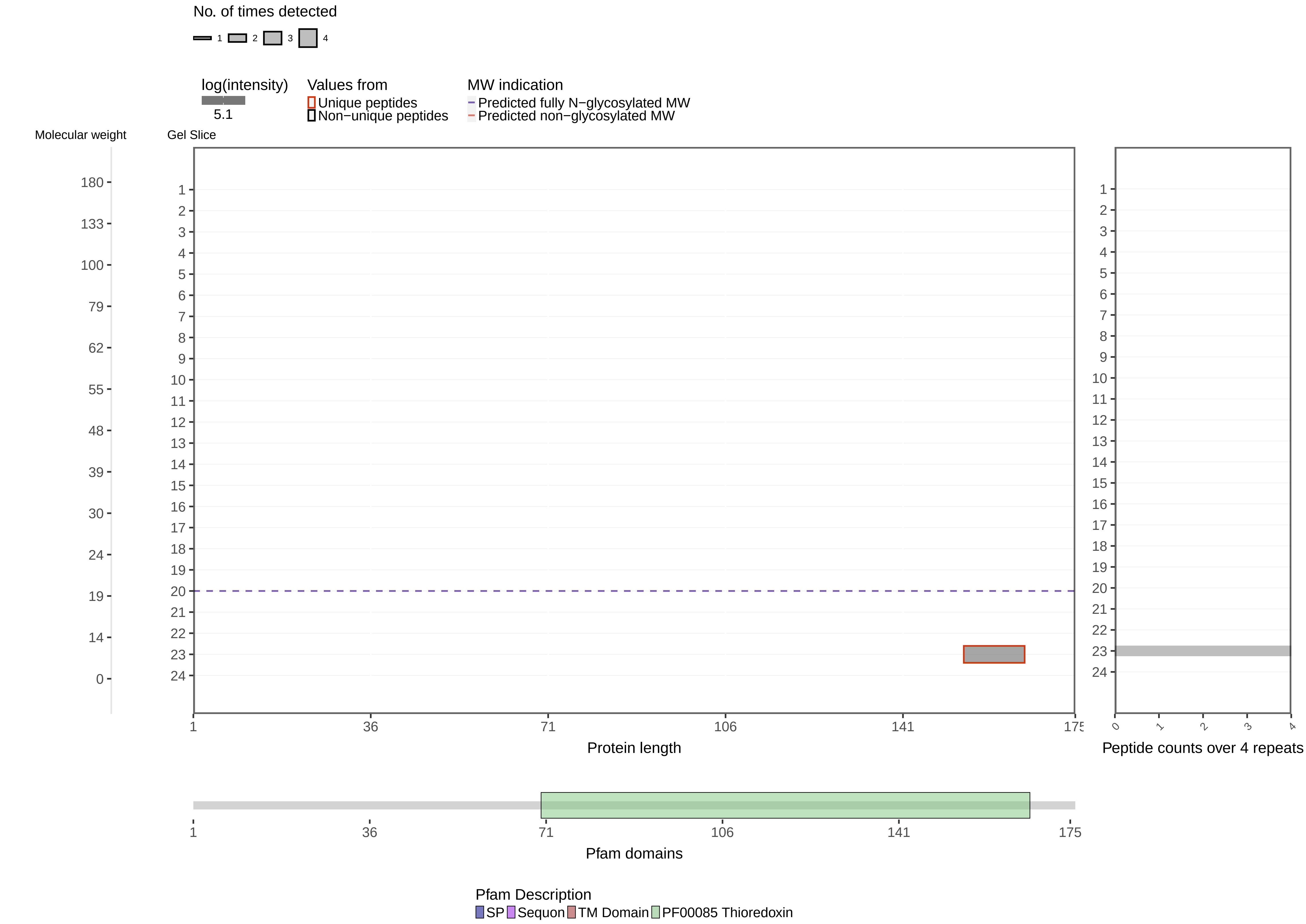


Protein sequence:

SFEELLEKSEKPVLVDFYATWCGPCQFMVPILNEVGESMKDKIQVVKIDSEKYPALADKYKIQALPTFILFKDGEVCDRFEGALNAPSLIQRIESALEVKQ*

NbL14g16810.1 stress_protein_DDR48−like [MW = 28.51 kDa]

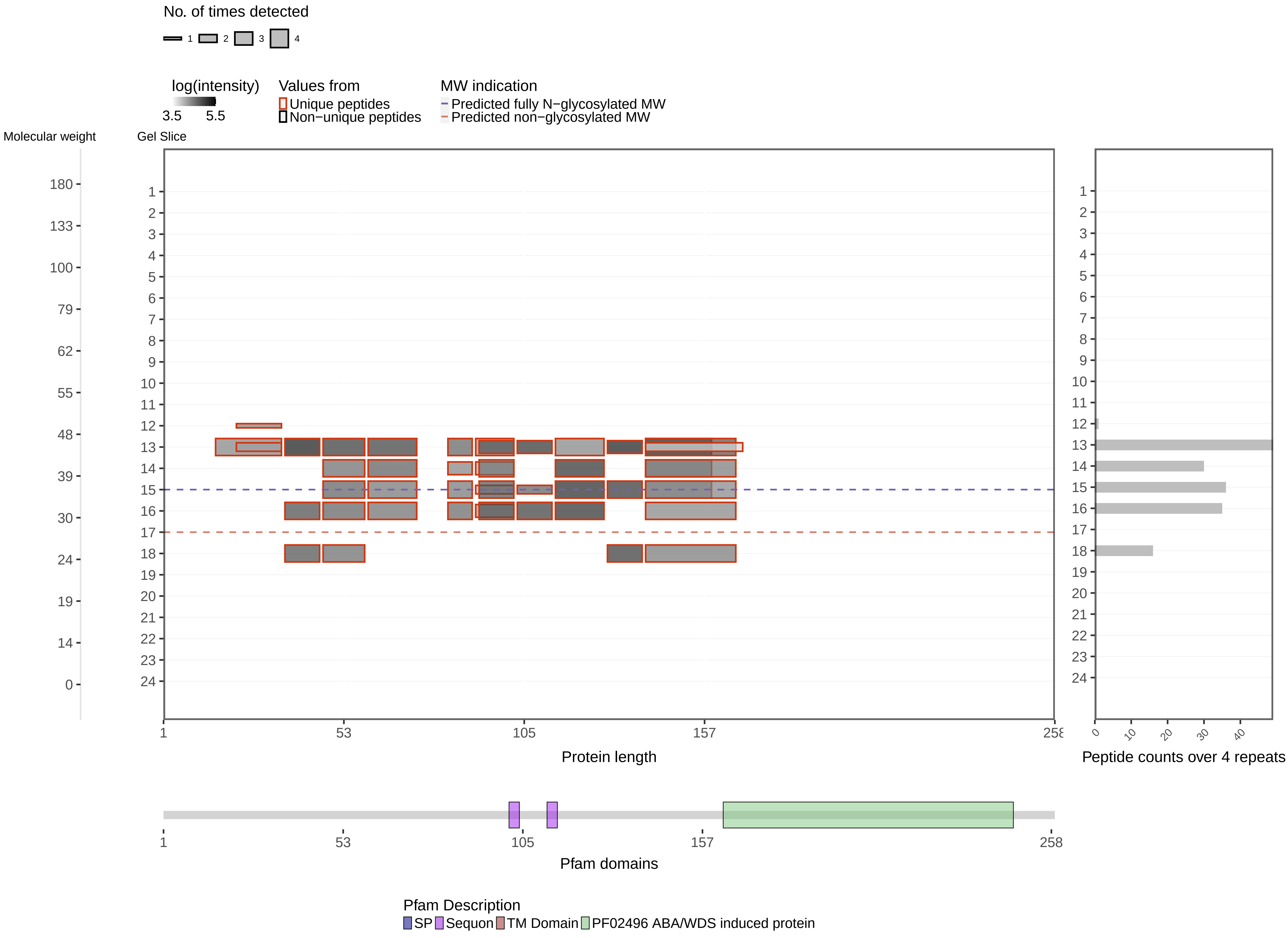


Protein sequence:
GYGDIPSETTYGEKTNYGEGDTYGKAAAHSSETTNFEENTDDEGKTGEDYEKEKRHHKHLEHVGGLGAAAAGAFALILAWKFQHEKHKAEKDPEHAHKHKIEEEIAAAAAVGAGGFAFHEHHEKKEAKEEEEEAEGKKKHHFF*

### NbL14g16790.1 uncharacterized_LOC107797753 [MW = 15.74 kDa]

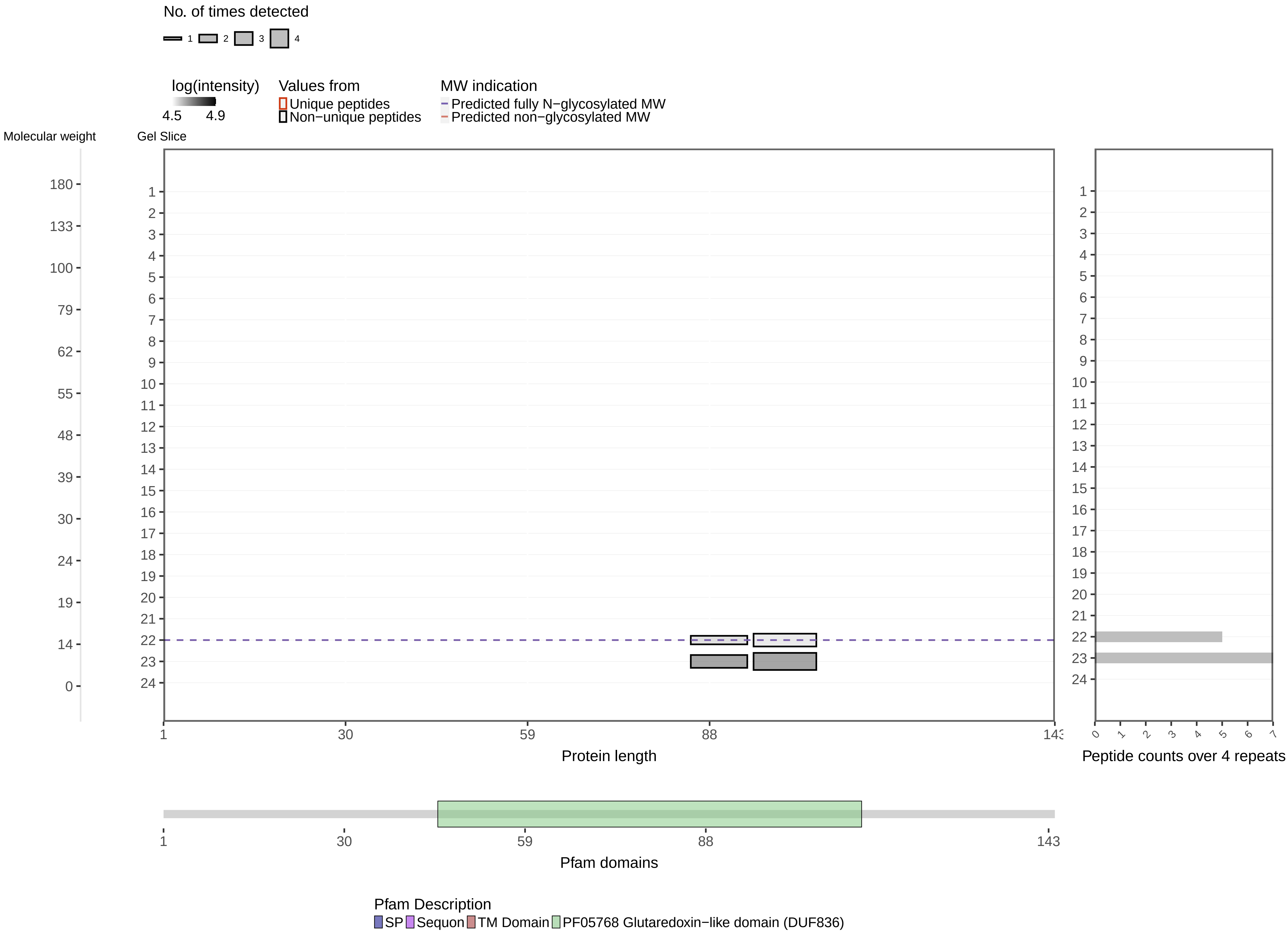


Protein sequence:

GLKEKLNAAFSLSTPHSLHDVQLQVRDITSNPEWEKAYQYEIPVLARVRPDGTEWSRLAVYSKARYGKGKRLENLEAAPRLYIV*

NbL14g16730.1 gamma−glutamylcyclotransferase_2−3 [MW = 21.1 kDa]

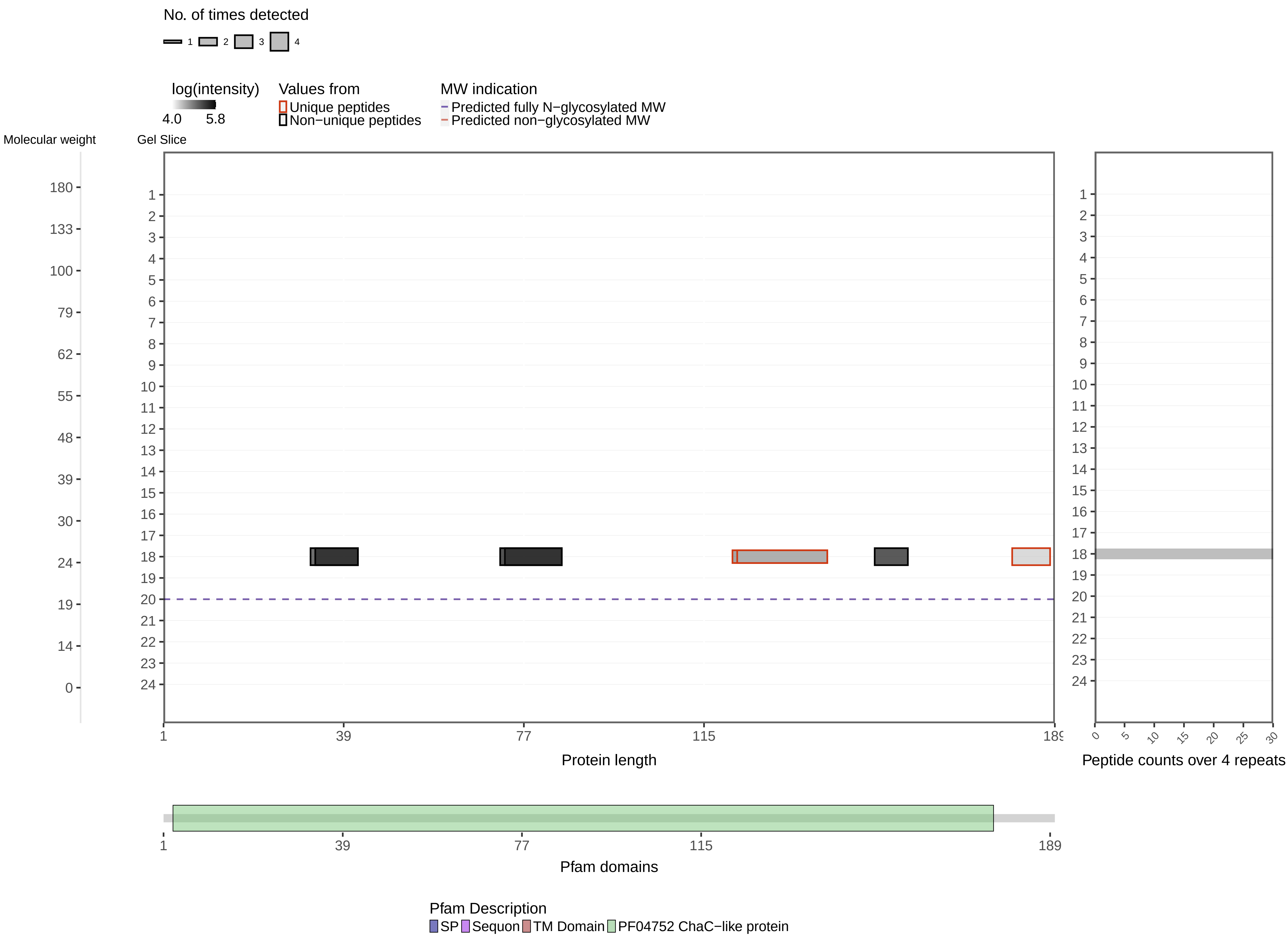


Protein sequence:

YLEVREKQYDKKAYVDLFTEPAASEPAVSGVLVYIGSSDKKVNQNYLGPASVEEIASQIVRAKGPSGPNRDYLFQLEKSLGLLGCEDKHVSDIAKEARRILSEEVTIS*

NbL14g16700.1 uncharacterized_LOC104218264 [MW = 47.65 kDa]

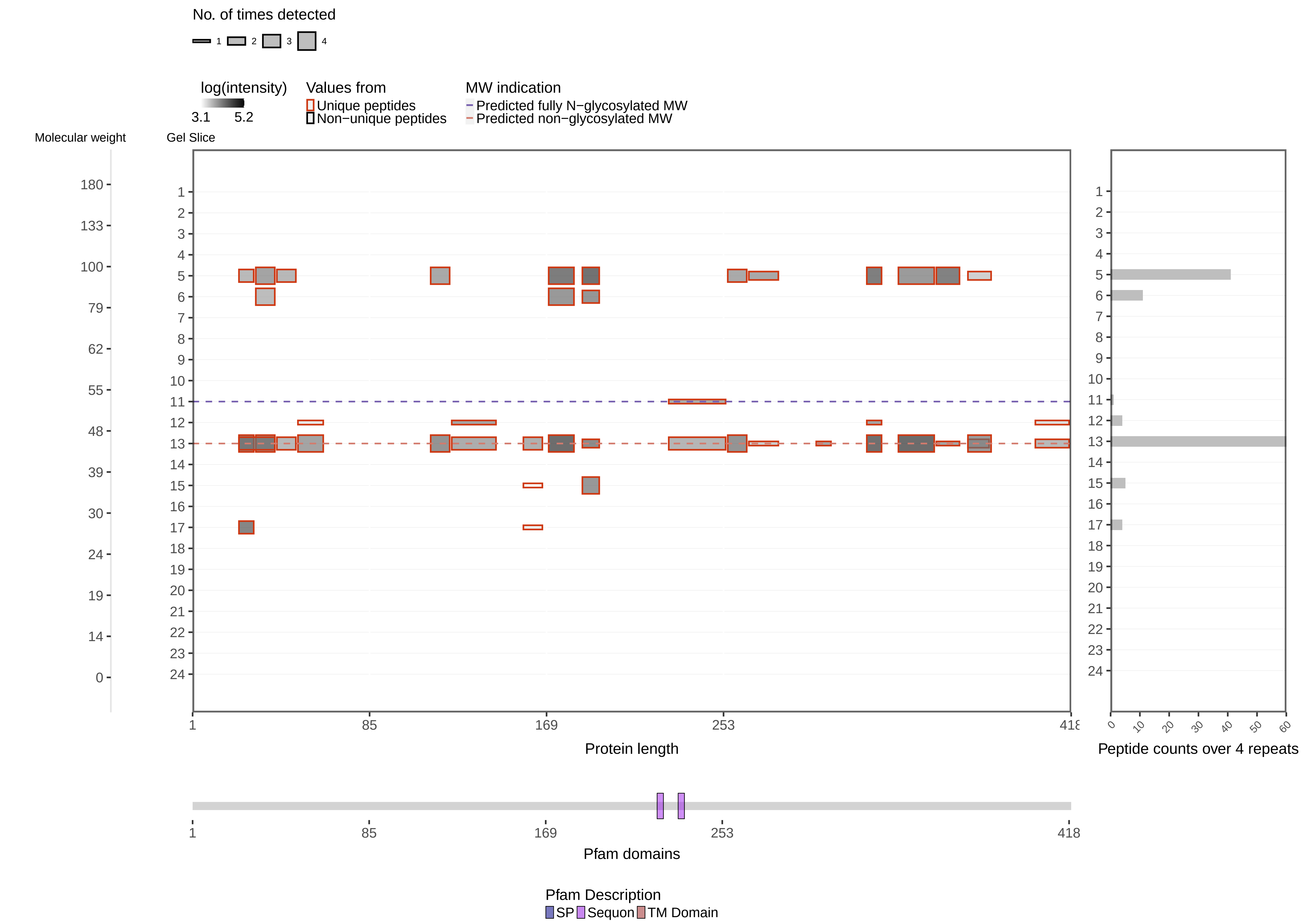


Protein sequence:

FLLMTMADFSDQLFNFQDILFDNSSGRLEFSGNNSQITLWPGDGKPGLWMNSISRMGALYSLIVREEDIFIEQNGHVQGRDEEIELVIPPVFDTCRRVLDAELQKEARELYWEGVCEASKRGLDWAEERLLTSVEKNPFVGEPHVVLGQIYLSKGKYEEAEKEAEKGLRLILEWGSPWDKRMSWEGWVAWTRVLLMKAKEKSWPQHSWGILNLGLVTK*

NbL14g16690.1 lysM_domain−containing_GPI−anchored_protein_1−like [MW − SP = 40.75 kDa]

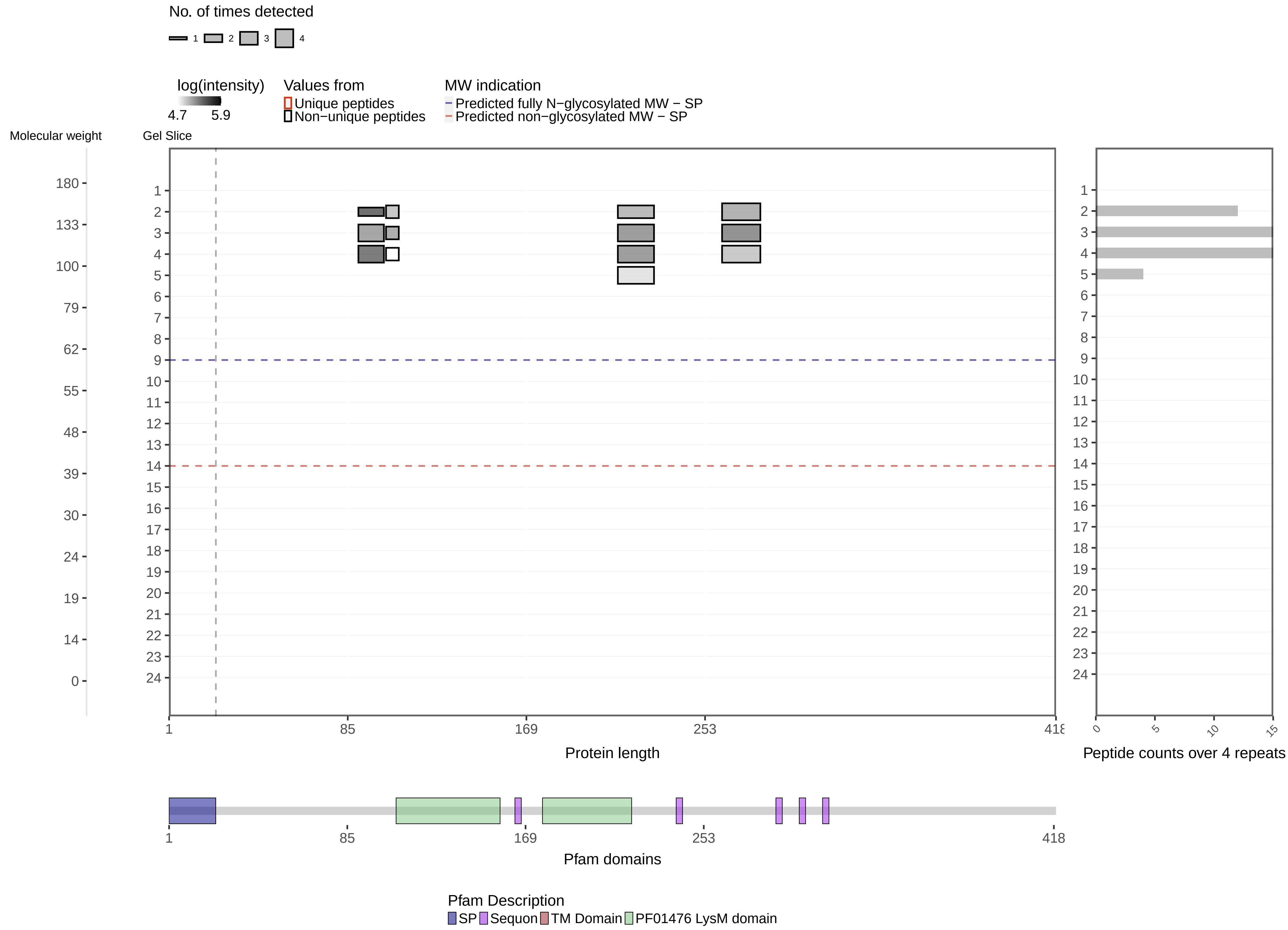


Protein sequence:

MNVNALGSPAIKEGDILAIPLSACASSFPRFASDYALSVANGSYAITASHCVOCSCGPGSRNLYCMPASLAVSCSSMOCKNSNLMLGNVTVOOTGGGCNVTSCNYGGFVNGTIITTLSSSLOPRCPGPOOFPPLVVPPTSVGPDSMFAPAPSPSESGGAPMDGPHSSVVPASGSVIAFPPSGGPSGSASSAWSLNPLVSFPIAILLYLCVKYVISLPL*

NbL14g16510.1 uncharacterized_LOC109217160 [MW = 47.31 kDa]

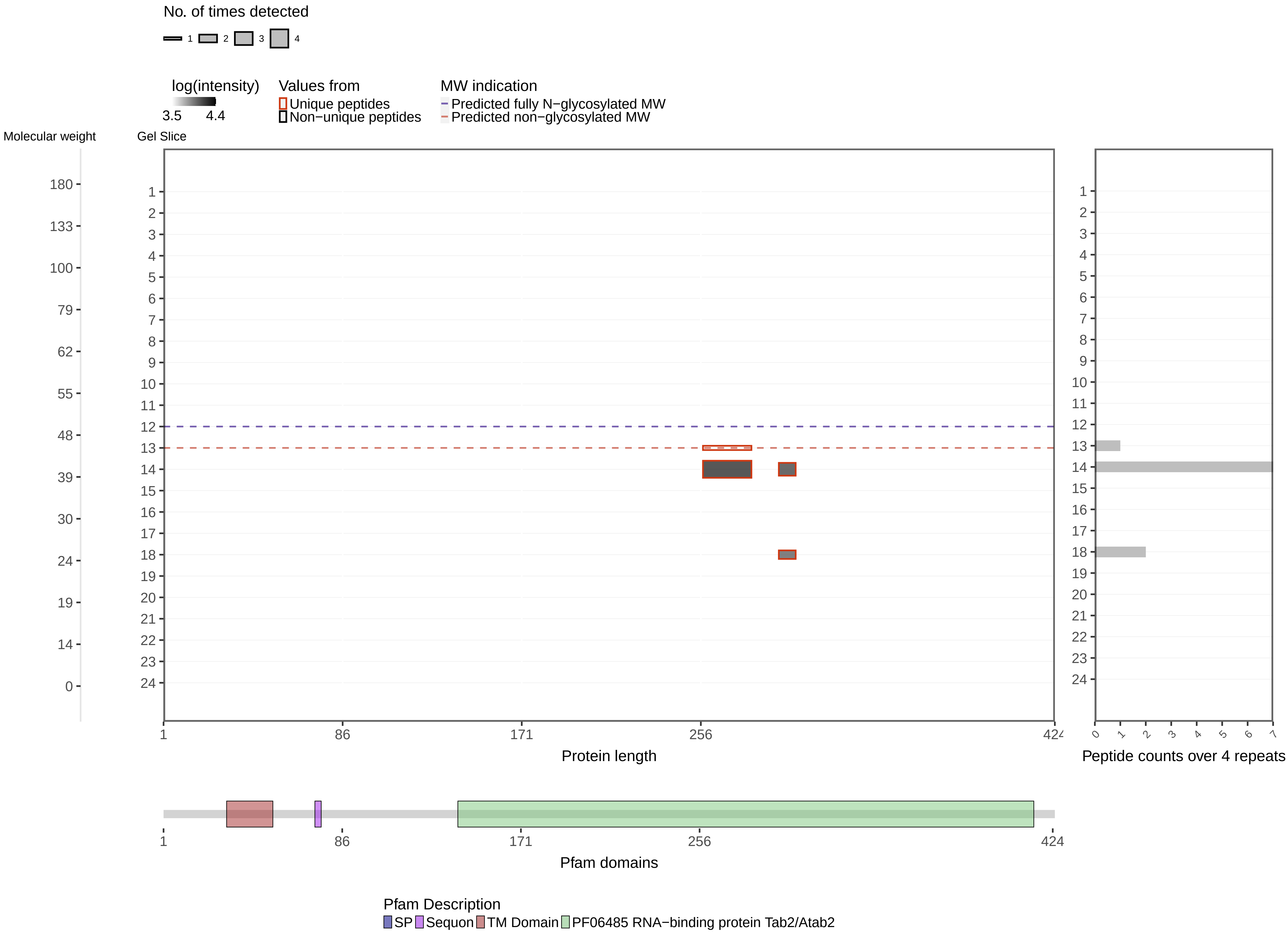


Protein sequence:

PDKIRFFRSQMQTIISRACNELGIKPVPSKRCLSLVLWLEDRYETVYTRHPGFQKGTKPLLALDNPFPMELPENLYGEKWAFVQLPFSAVREEVSNLETRLVFGASLDLDLLGIEIEDKTLIPGLAVATSRAKPLAAWMNGLEVCSIEADVARASLILSVGISTRYIYATYKKNPVSTSEAEAWEAAKKACGGLHFLAIQNDLNSDDCVGFWLLLDLPPPPV*

### NbL14g16250.1 photosystem_II_stability/assembly_factor_HCF136_chloroplastic [MW = 43.48 kDa]

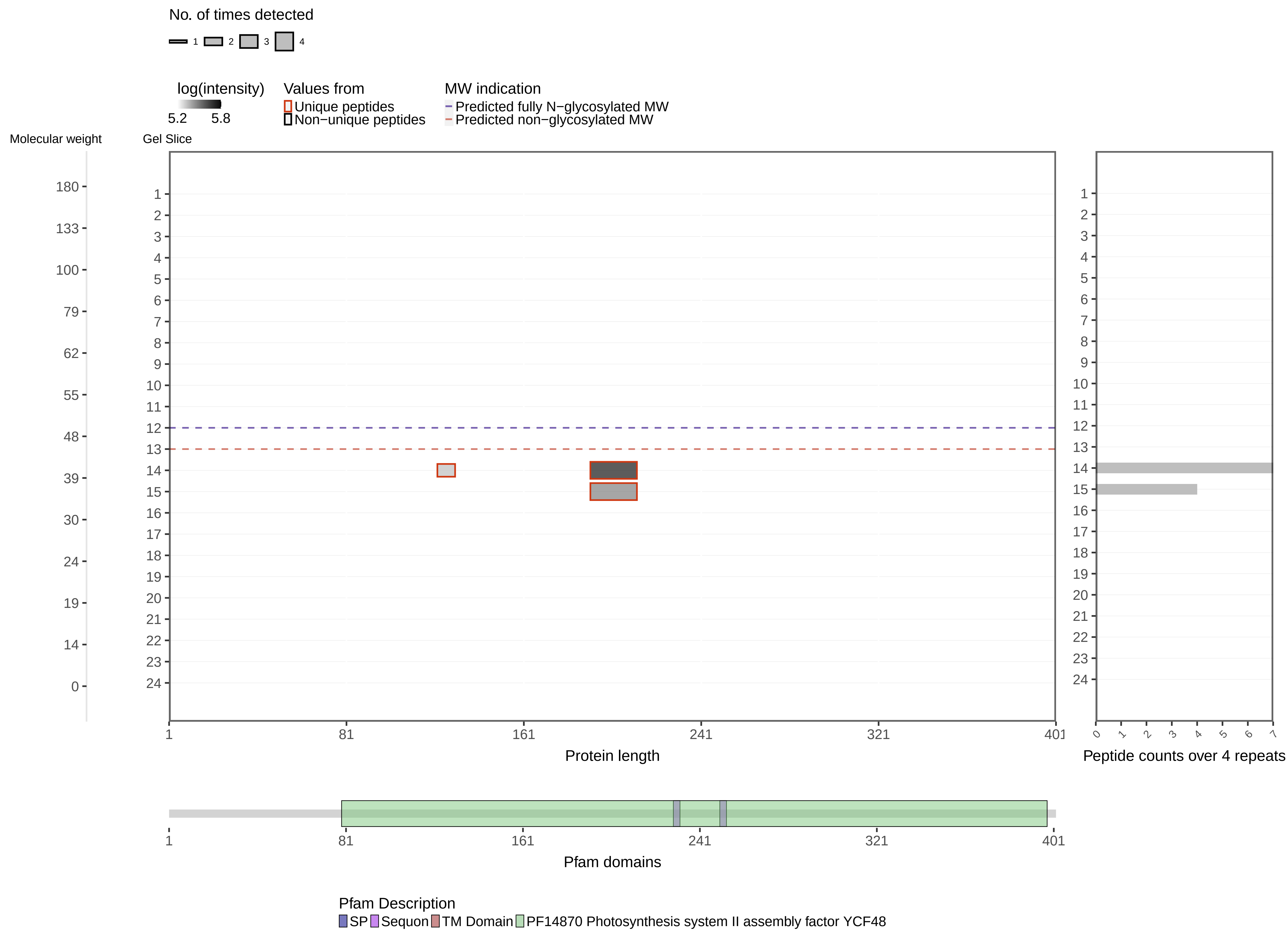


Protein sequence:

TGEESAEMVTDEGAIYVTSNKGYNWKAAIQETVSATLNRTVSSGISGASYYTGTFSTVNRSPDGRYVAVSSRGNFYLTWEPGQAYWQPHNRAVARRIQNMGWRADGGLWLLVRGGGLYLSKGTGLSEDFEEVSVQSRGFGILDVGYRSQDEAWAAGGSGILLKTNNGGKTWIRDKAADNIAANLYSVKFINDNQGFVLGNDGVLLKYLG*

NbL14g16220.1 ATP−dependent_Clp_protease_proteolytic_subunit_2_mitochondrial−like [MW = 26.41 kDa]

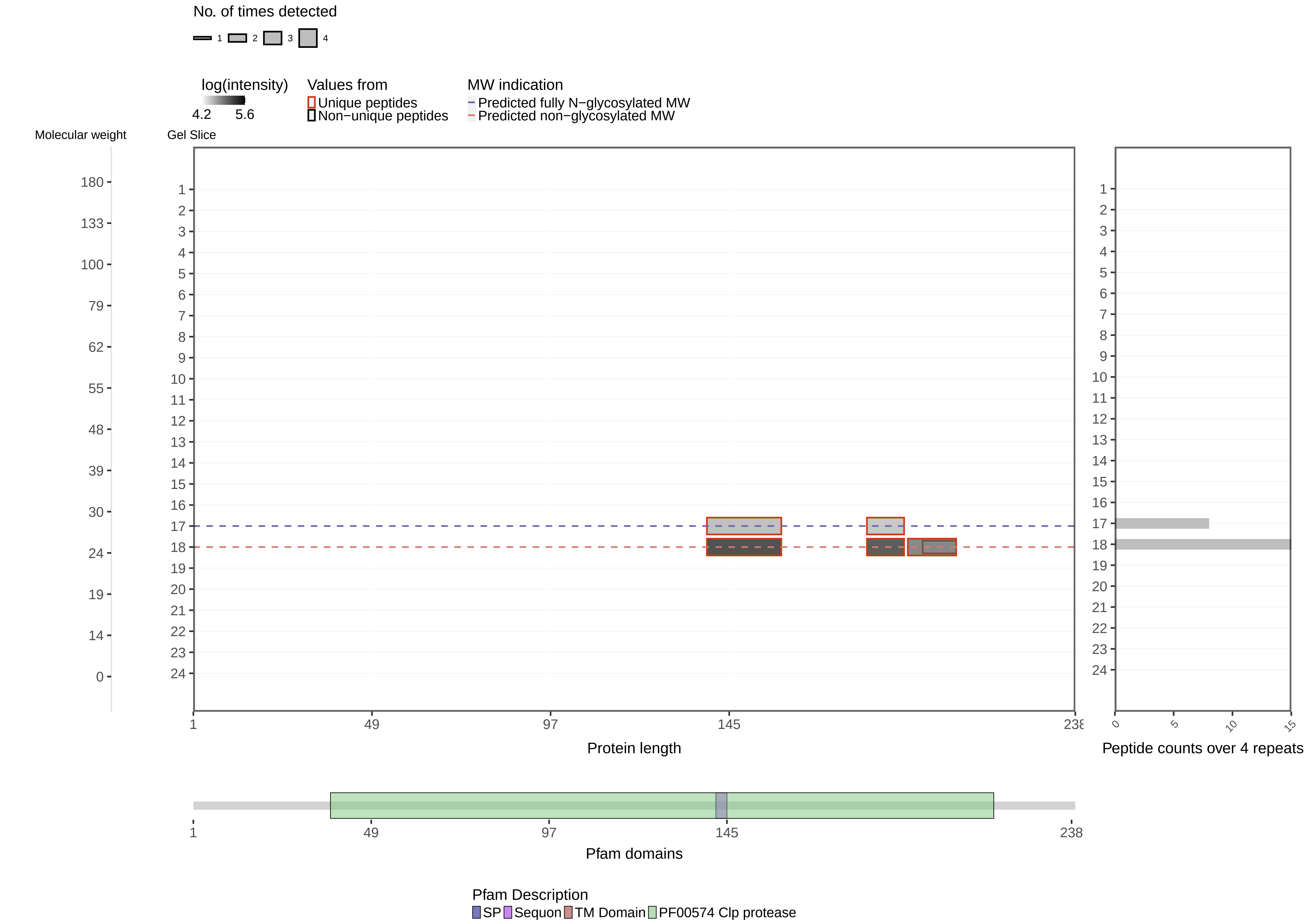


Protein sequence:

[illegible]QYIRSPINTICLGQAASMASLLLAAGAKGERRSLPNATIMIHQPSGGYSGQAKDMTIHTKQIVRVWDALNQLYSKHTGQPVEIIQKNMDRDYFMTAEEAKEFGIIDEVIDQRPMALVTDAIANEAKDKSSS*

### NbL14g16170.1 succinate−−CoA_ligase_[ADP−forming]_subunit_alpha−2_mitochondrial [MW = 35.58 kDa]

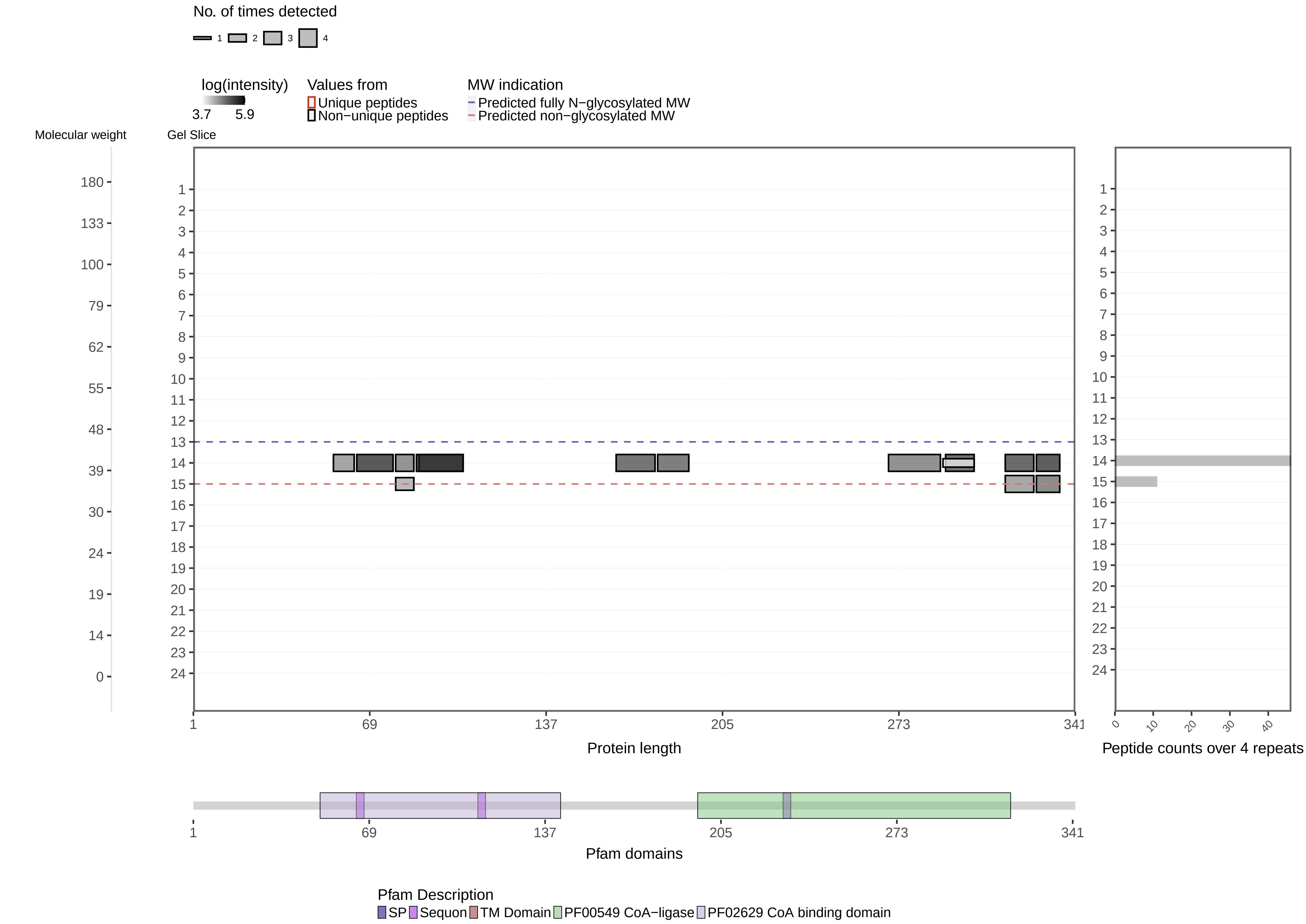


Protein sequence:

LKKQSKTRLIGPNCPGIIKPGECKIGIMPGYIHKPGRIGIVSRSGTLTYEAVFQTTAVGLGQSTCVGIGGDPFNGTNFVDCLEKFIADPQTEGIVLIGEIGGTAEEDAAALIKESGTQKPVVAFIAGLTAPPGRRMGHAGAIVSGGKGTAQDKIKALKEAGVTVCESPAKIGVTMLDVFKQRGLA*

### NbL14g16110.1 glutamate−1−semialdehyde_21−aminomutase_chloroplastic−like [MW = 59.69 kDa]

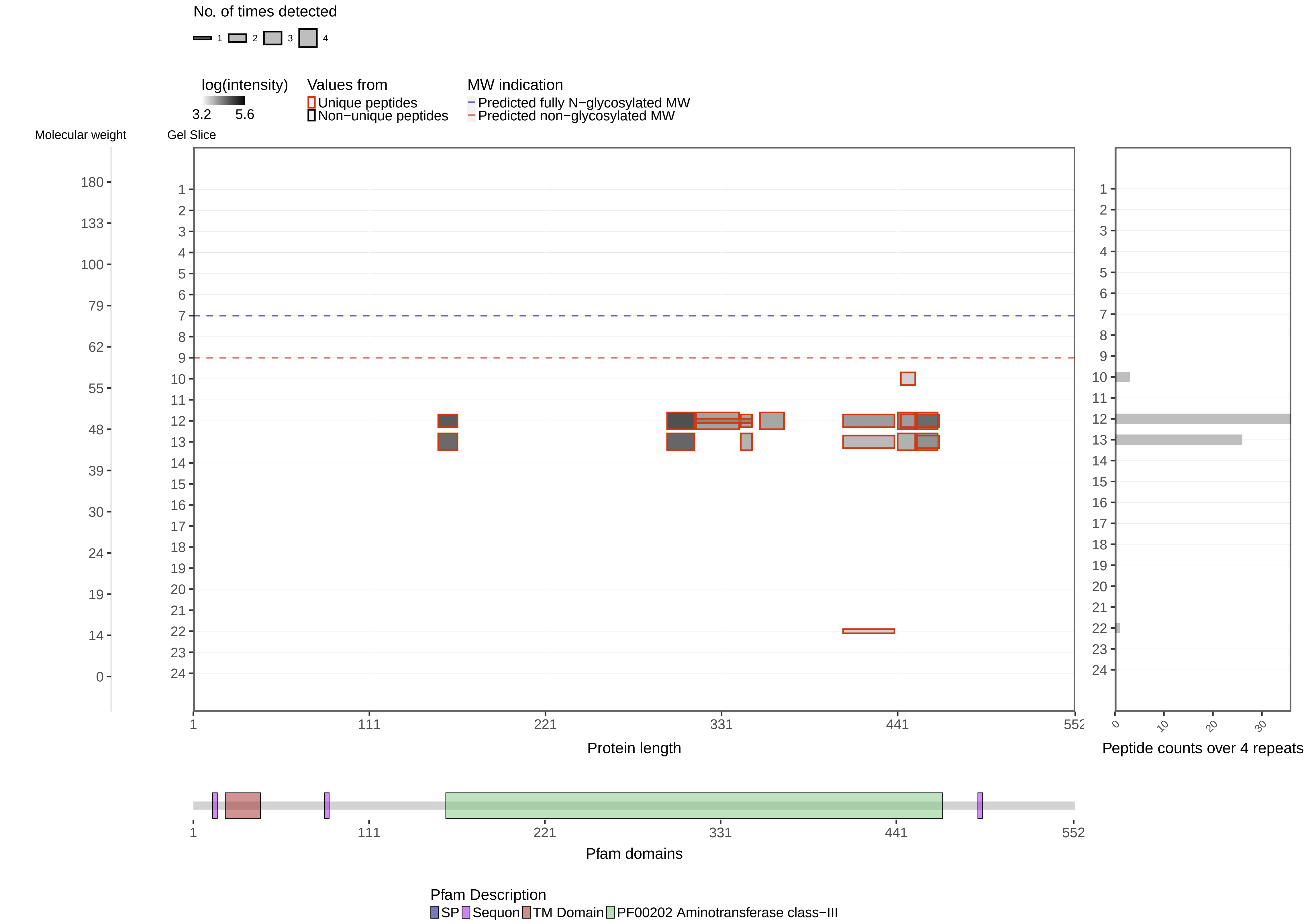


Protein sequence:

### NbL14g16020.1 protease_2 [MW = 86.34 kDa]

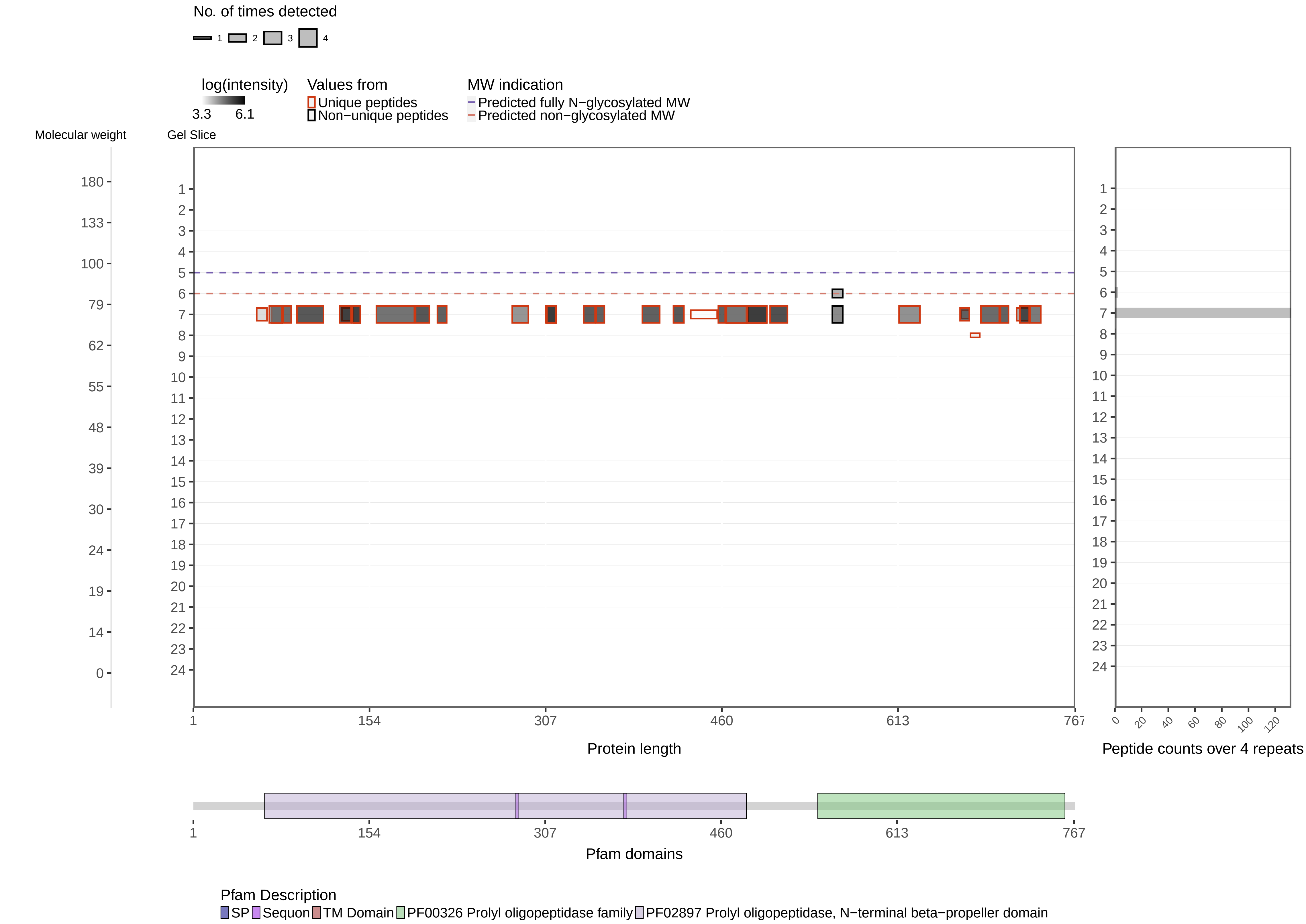


Protein sequence:

NbL14g15810.1 proteasome_subunit_beta_type−7−A−like [MW = 35.43 kDa]

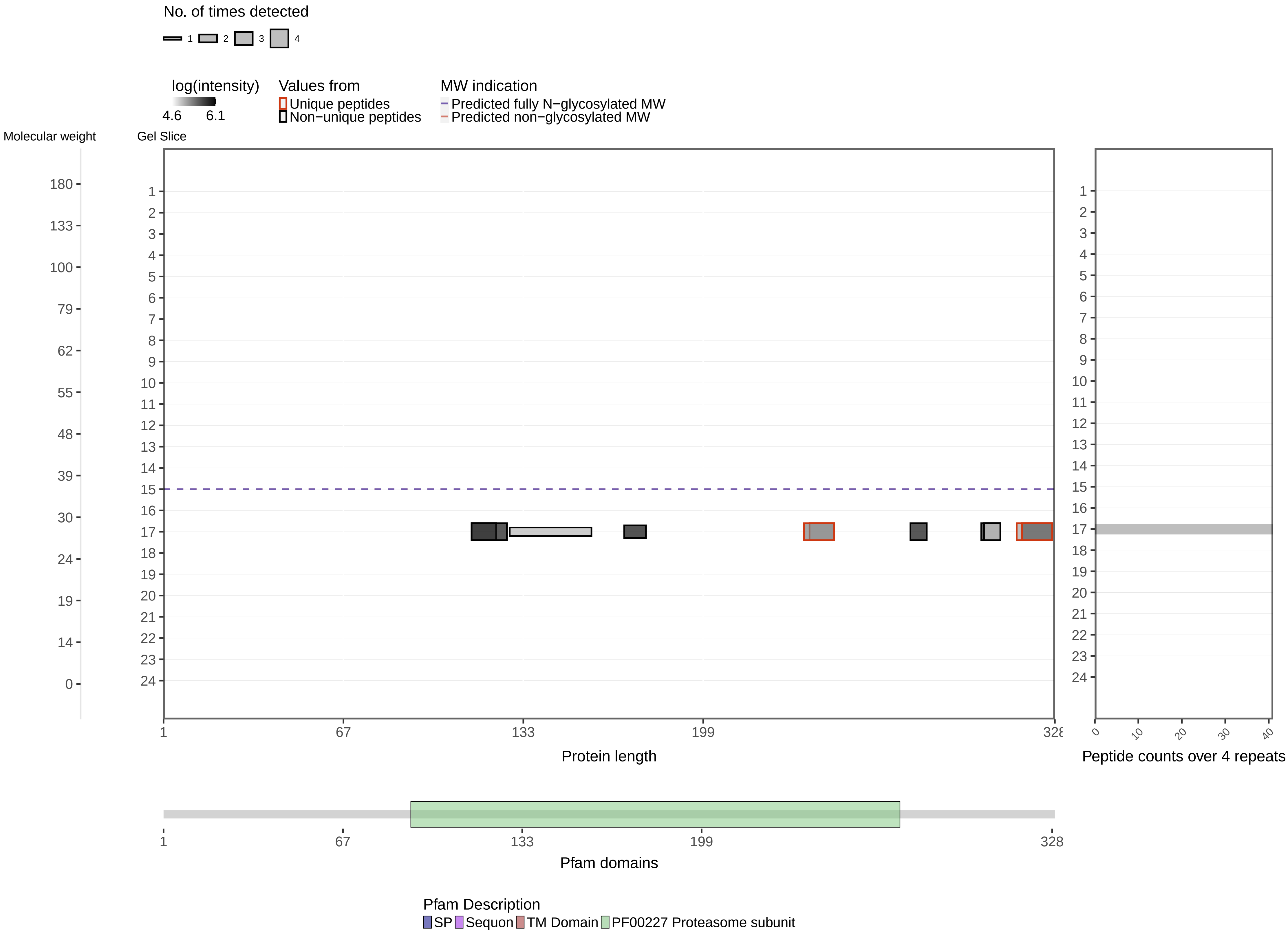


Protein sequence:

MVSSQLKLHRYHTGRESRVVTALTLLKTHLFSYQGYVSAALVLGGVDVTGPHLHTIYPHGSTDTLPYATMGSGSLAAMAIFESKYREGLSKDEGIKLVAEAILSGVFNDLGSGSNVDICIITKGNTEYLRNHMLPNPRTYPQKGYSFPKRTEVLLTKITPLRERVEVIEGGDAMEE*

### NbL14g15730.1 ribose−phosphate_pyrophosphokinase_4−like [MW = 44.73 kDa]

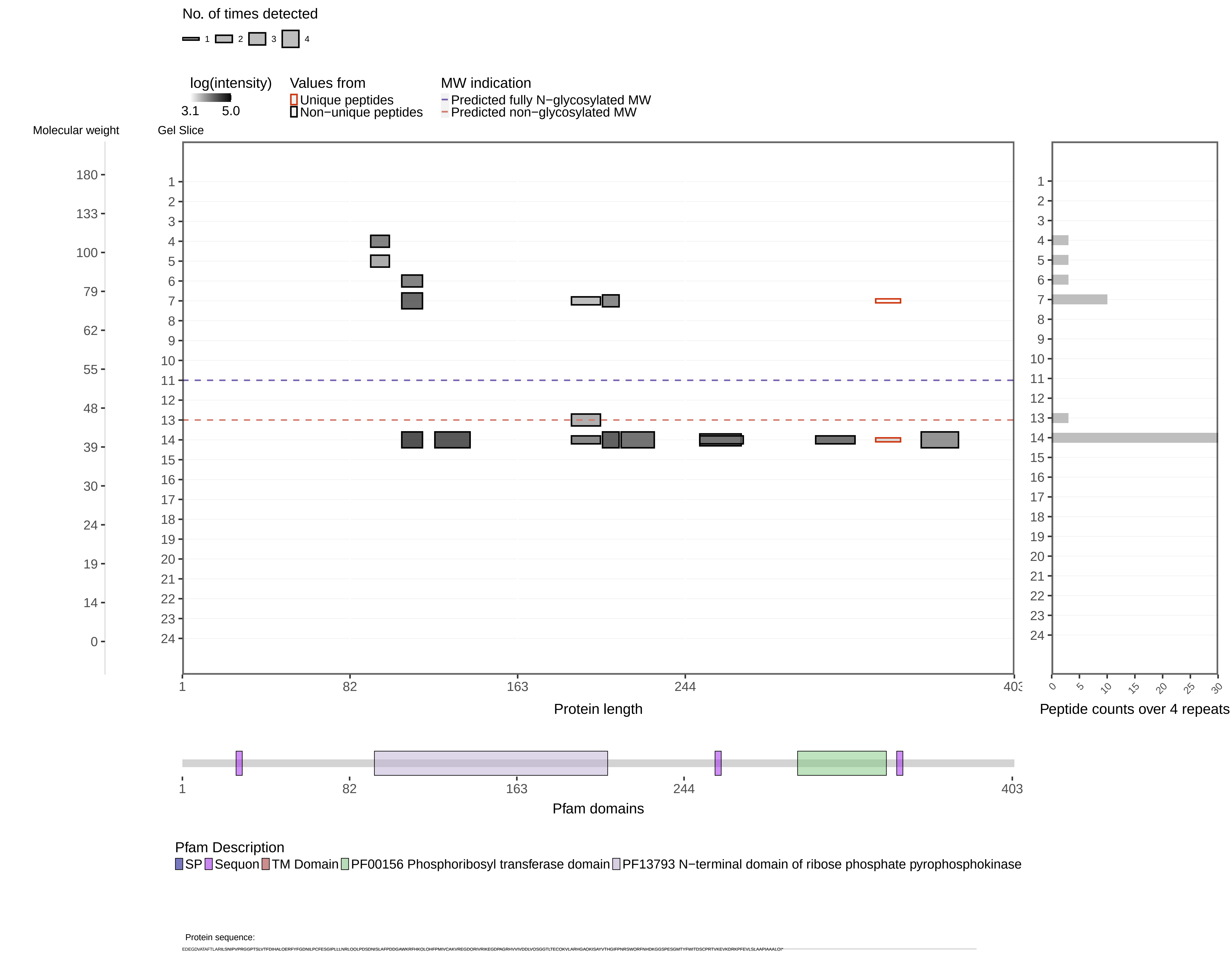

### NbL14g15720.1 uncharacterized_LOC104221143 [MW = 64.93 kDa]

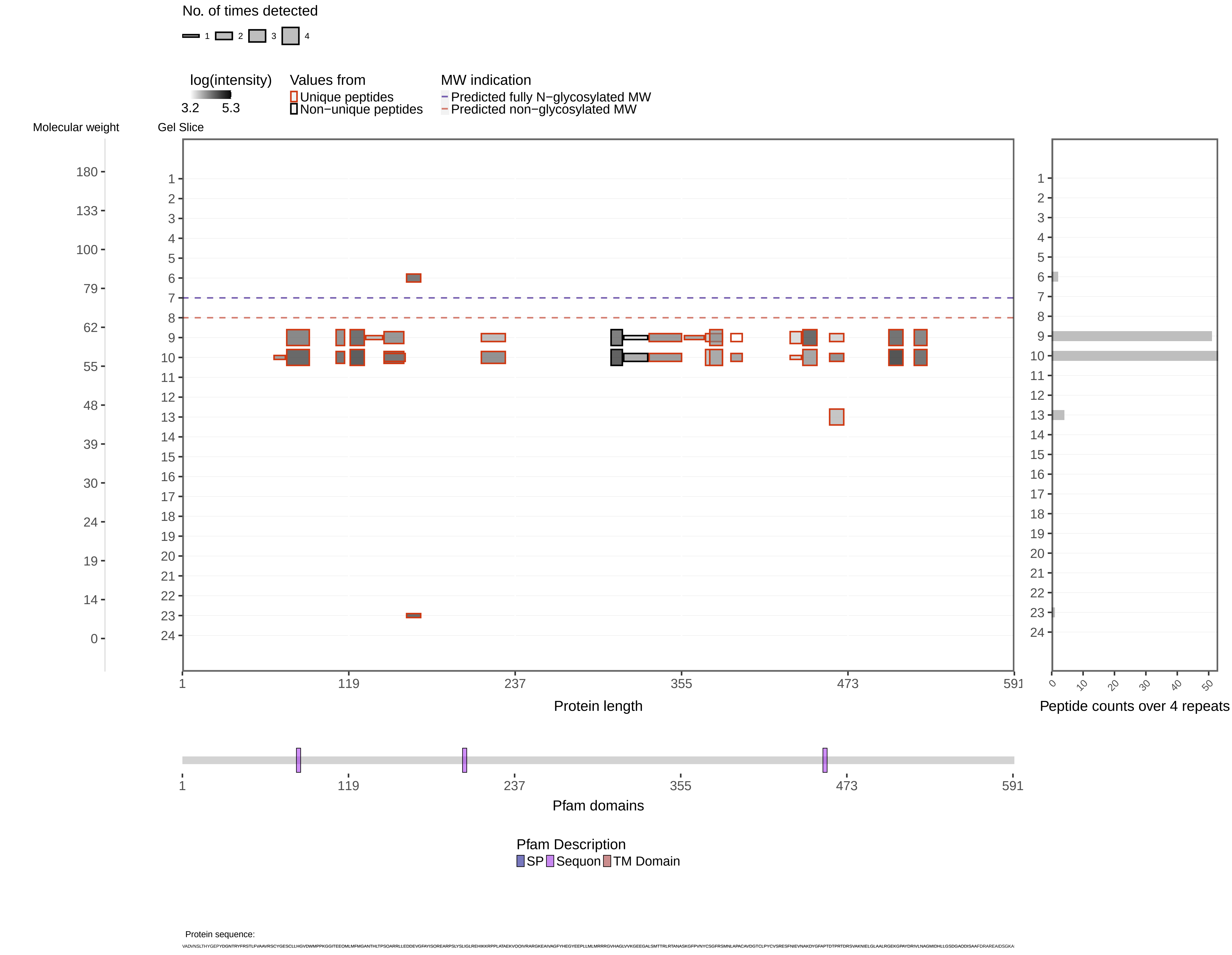

NbL14g15690.1 ATP−citrate_synthase_alpha_chain_protein_2 [MW = 46.61 kDa]

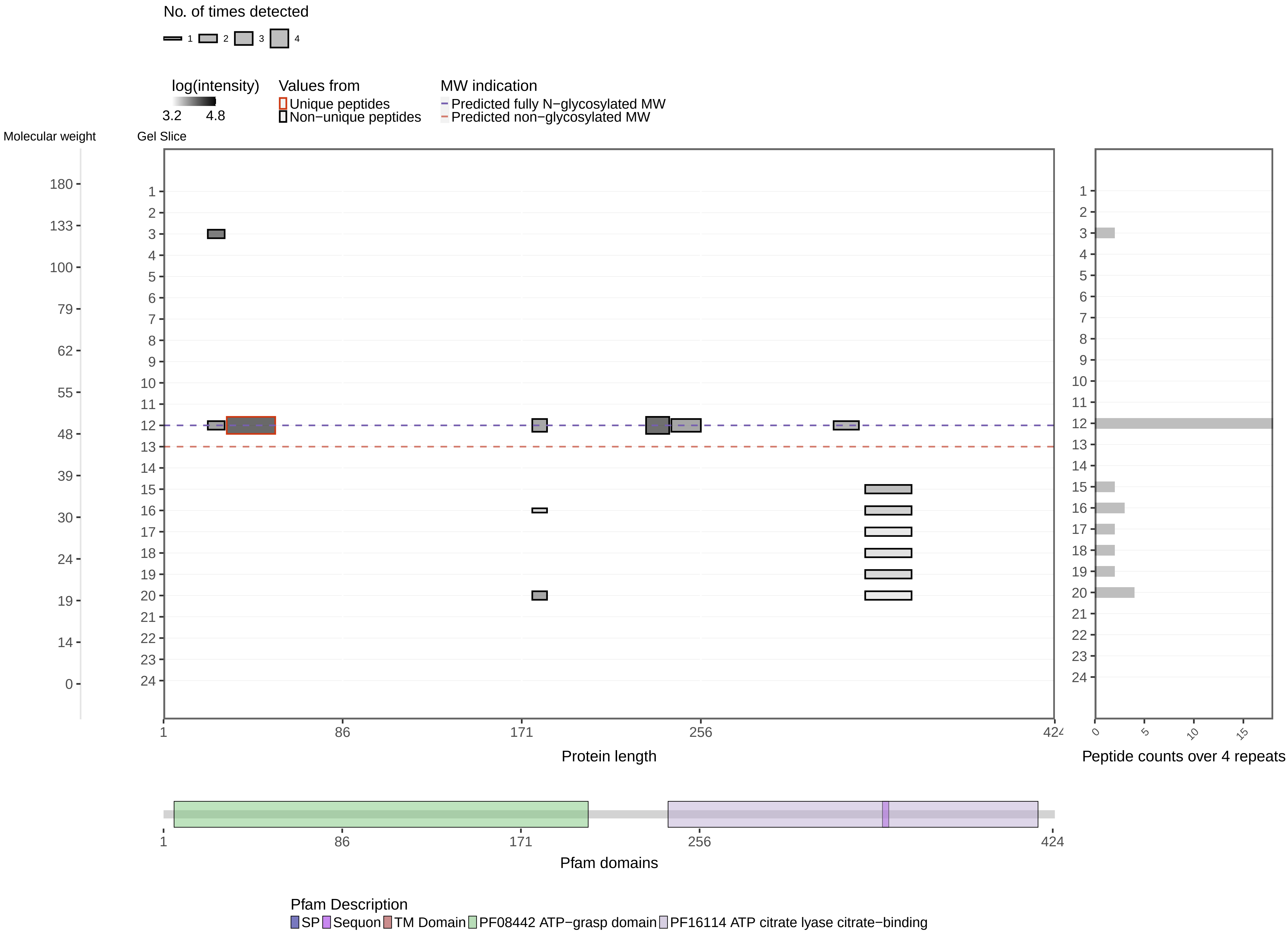


Protein sequence:

INPFTLVEGKPYPLDMRGELDDTAAFKNFKKWGNIEFPLPFGRVMSATESFIHGLDEKTSASLKFTVLNPKGRIWTMVAGGGASVIYADTVGDLGYASELGNYAEYSGAPNEEEVLQYARVVIDCATADPDGRKRALVIGGGIANFTDVAATFSGIIRALKEKESKLKAARMHIYVRRGGPNYQKGLTKMRSLGEEIGIPIEVYGPEATMTGICKQAIECITAAA*

### NbL14g15560.1 probable_splicing_factor_3A_subunit_1 [MW = 89.33 kDa]

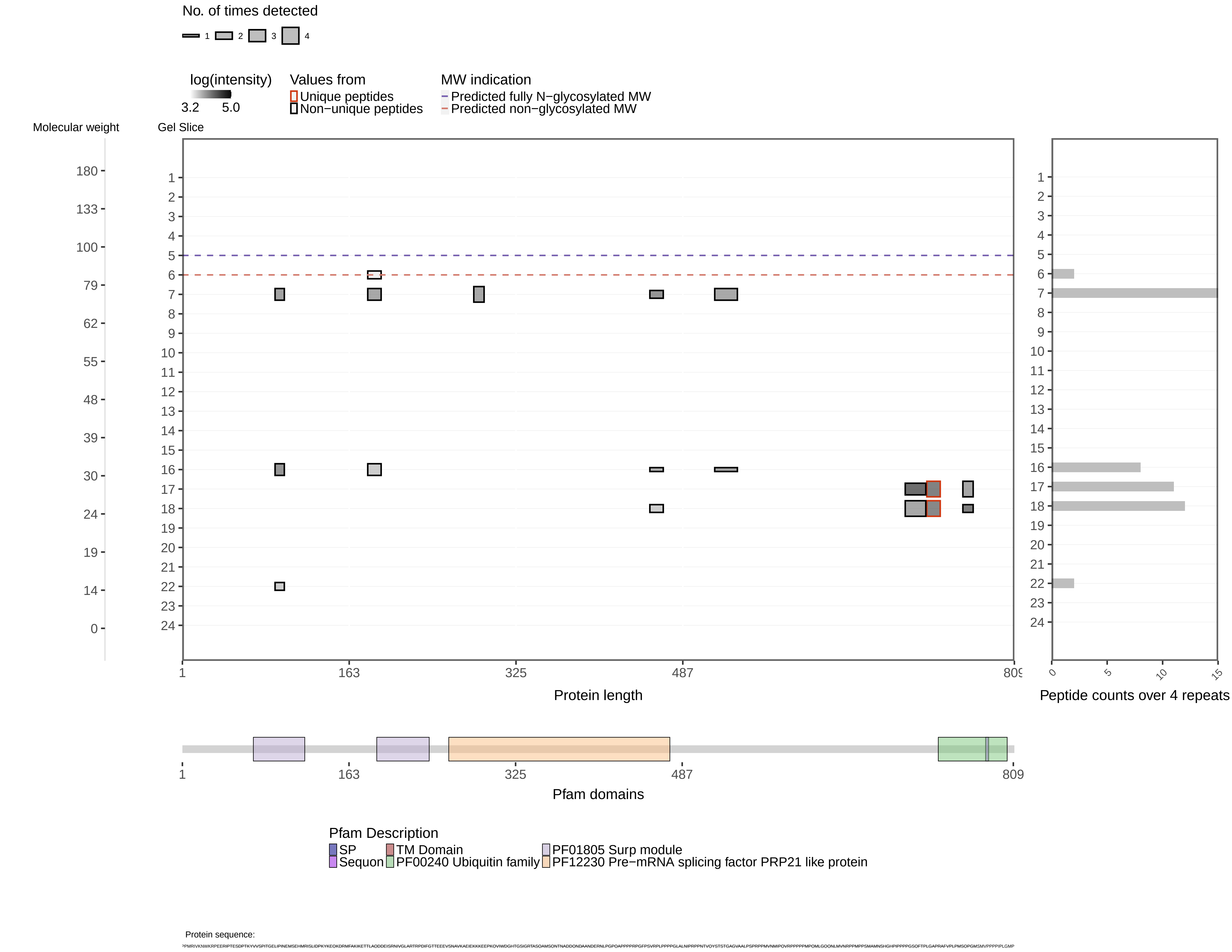

### NbL14g15350.1 protein_GAST1 [MW = 13.45 kDa]

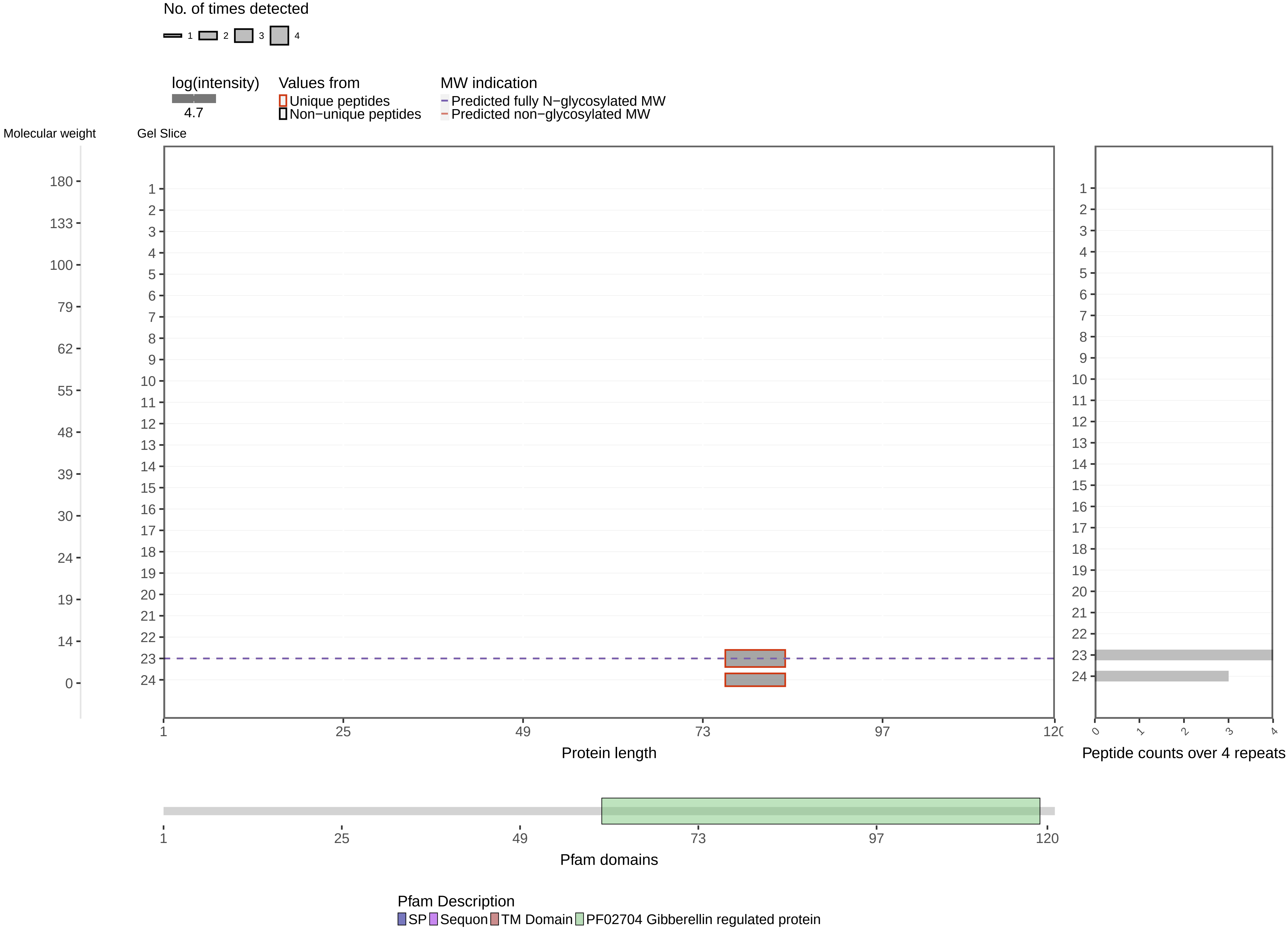


Protein sequence:
GTTPGSLHPQECLPKCTYRCSKTQYKKPCMFFCQKCCSKCLCVPPGTYGNKQFCPCYNNWKTKRGGPKCP*

### NbL14g15310.1 40S_ribosomal_protein_S19−3−like [MW = 17.99 kDa]

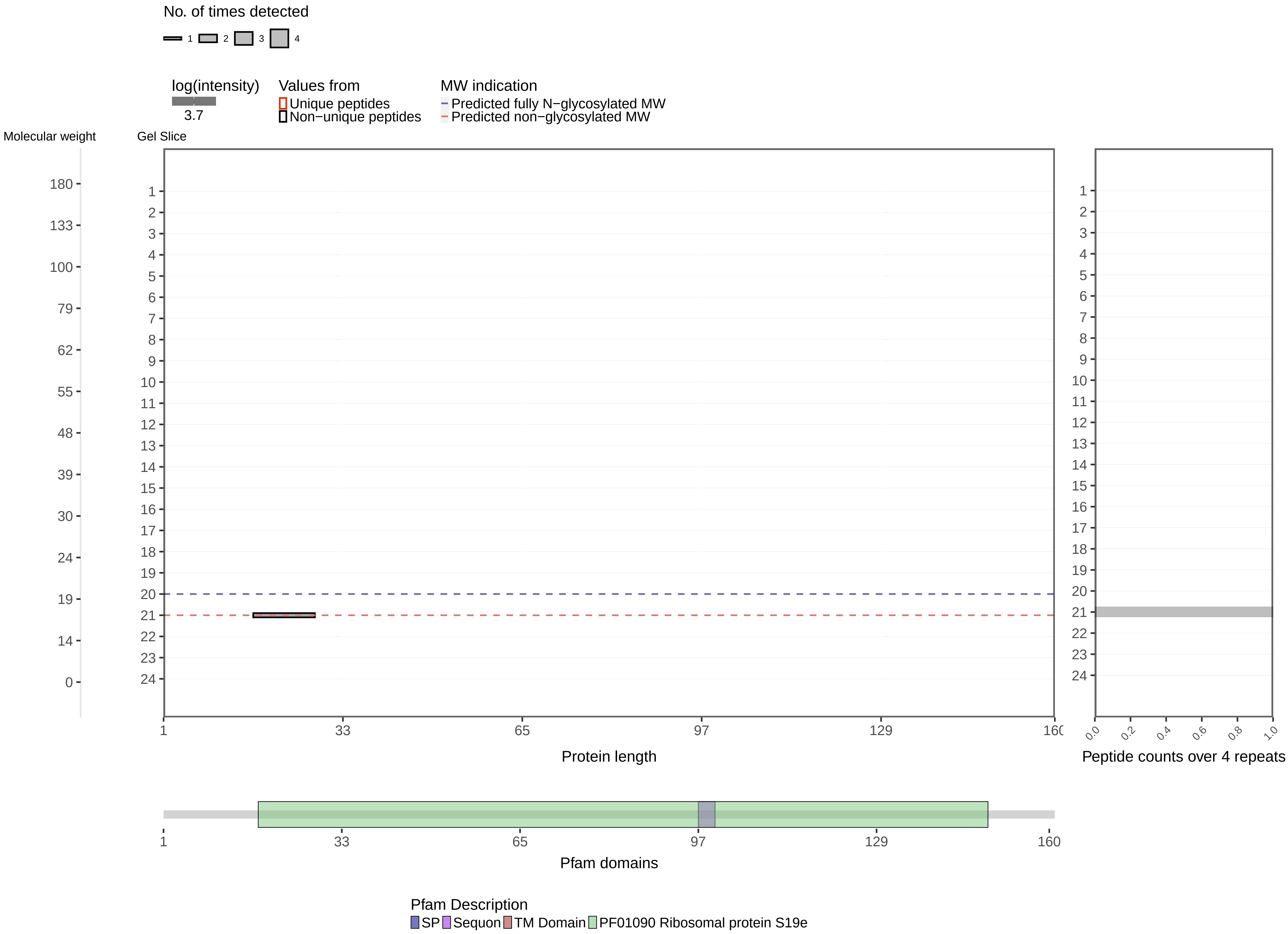


Protein sequence:

RAASMARKIYLRGGLGVGGFRRIYGGNQRNGSRPRHFCKSSGSVARNILQQLQNMNIIDFDPKGGRRITSNGQRDLDHCIRQDQDRRLNLKP*

### NbL14g15290.1 S−formylglutathione_hydrolase [MW = 31.9 kDa]

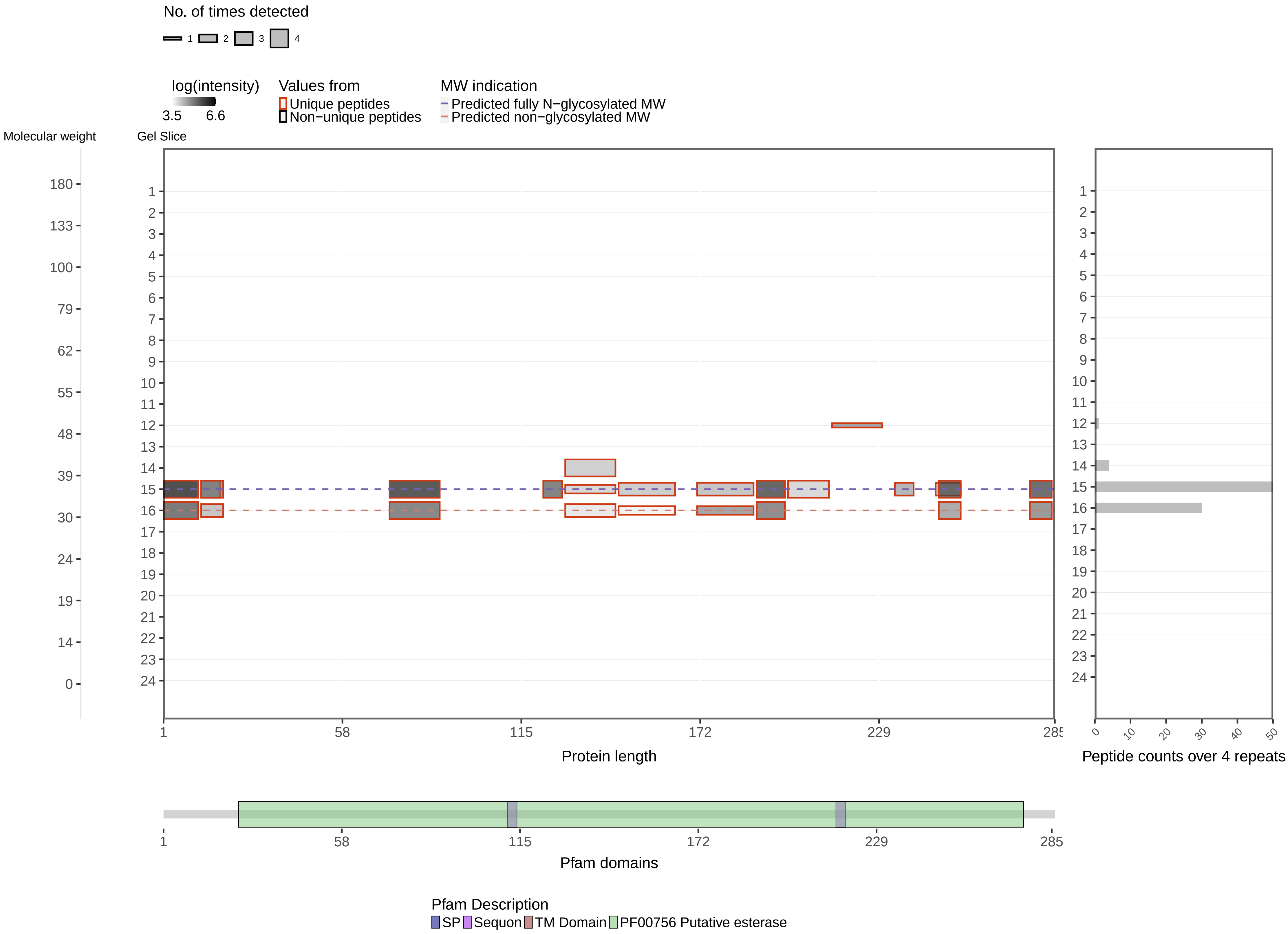


Protein sequence:

'KELPSLLLENFPELDTSRASIFGHSMGGHGALTVYLKNLDKYKSVSAFAPIANPINCPWGLKAFTNYLGENKADWEEYDATCLVSRFNNVSATILIDQGEDDKFLKDQLLPHKFEEACKKVGVPLLVRLQPGYDHSYYFIATFIDDHIRHHAQALNL*

### NbL14g15180.1 U4/U6_small_nuclear_ribonucleoprotein_PRP4−like_protein [MW = 70.1 kDa]

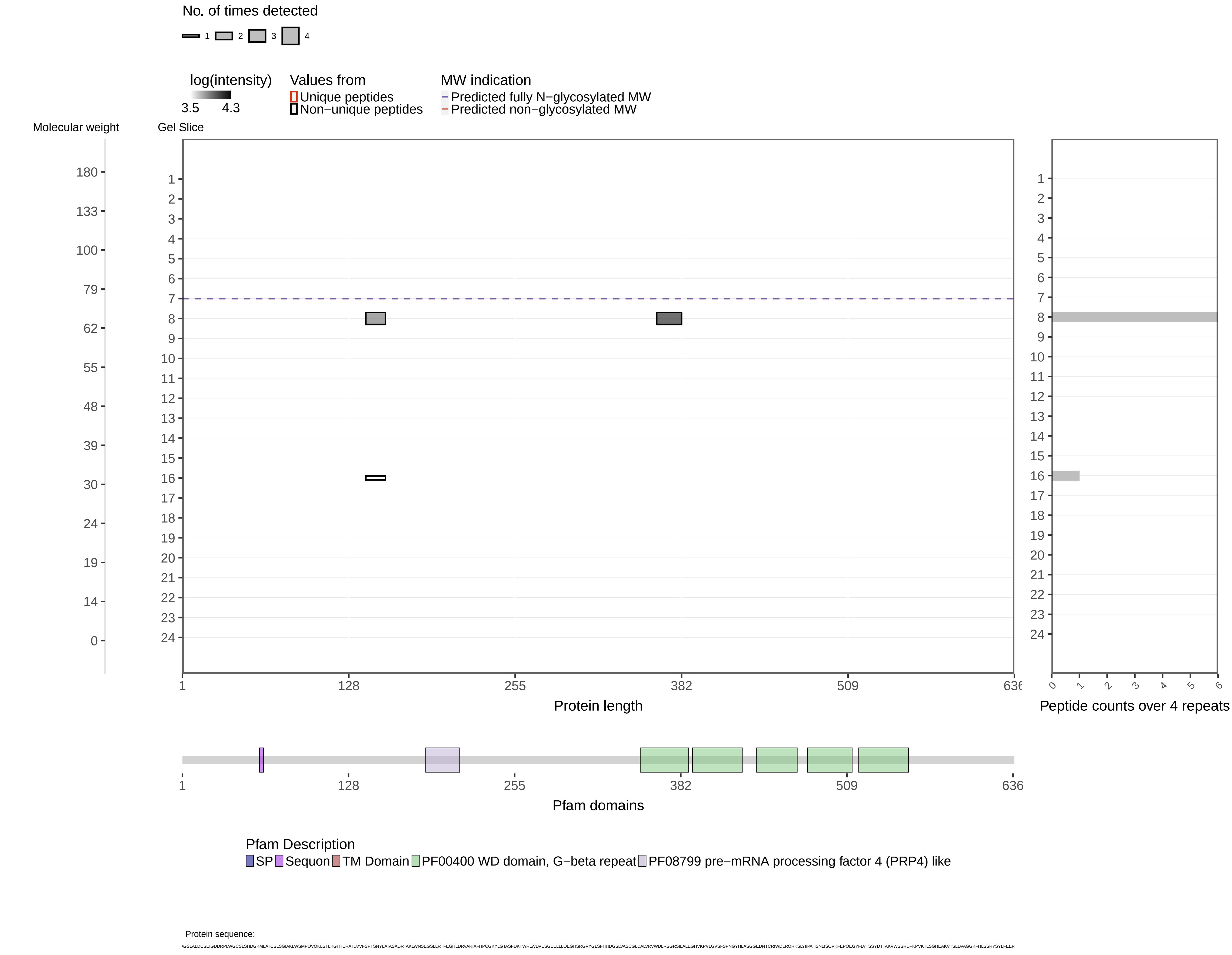

NbL14g15110.1 putative_calcium−transporting_ATPase_11_plasma_membrane−type [MW = 114.28 kDa]

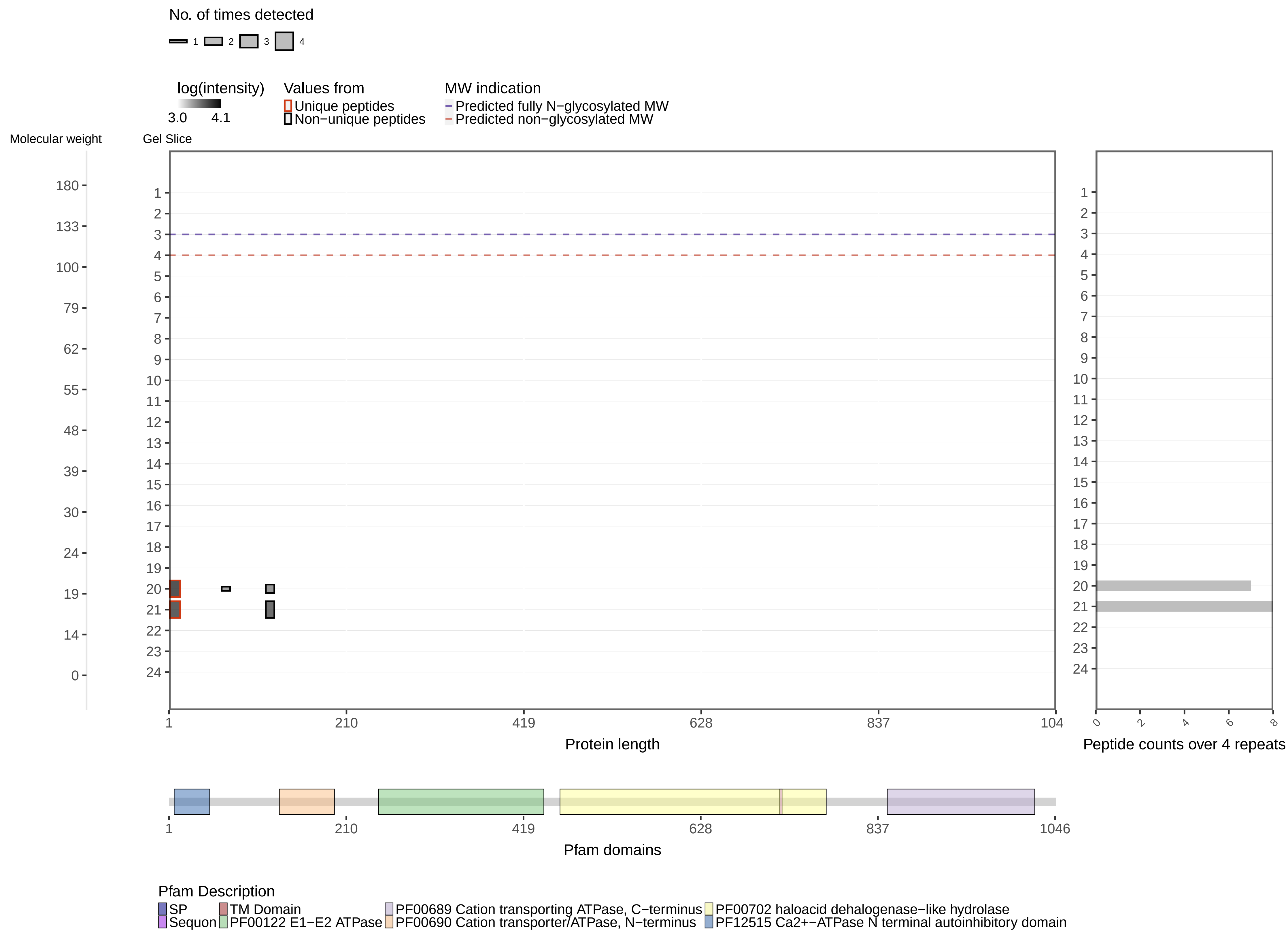


Protein sequence:

### NbL14g14920.1 villin−3−like [MW = 104.22 kDa]

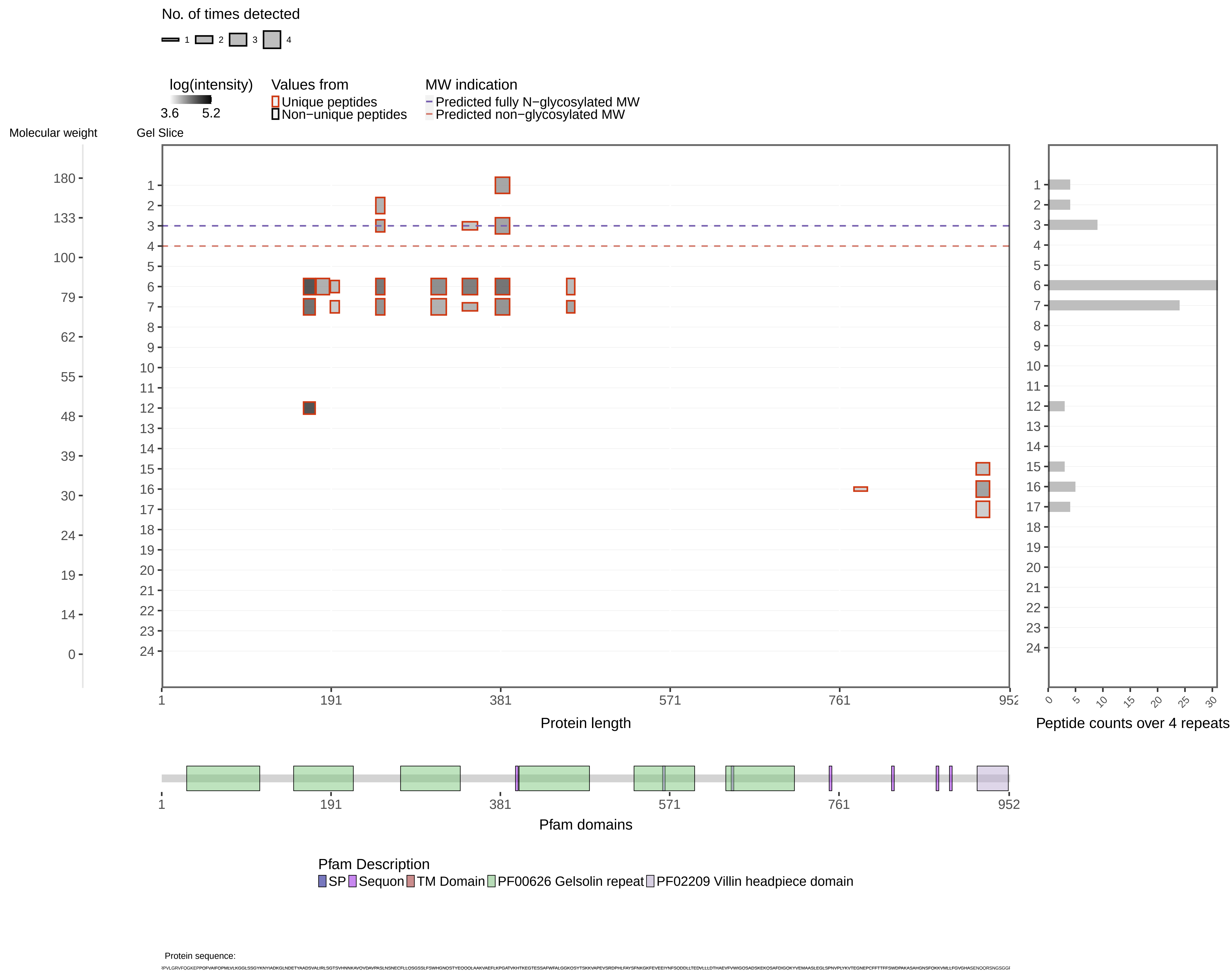

### NbL14g14900.1 magnesium−chelatase_subunit_ChlH_chloroplastic−like [MW = 153.8 kDa]

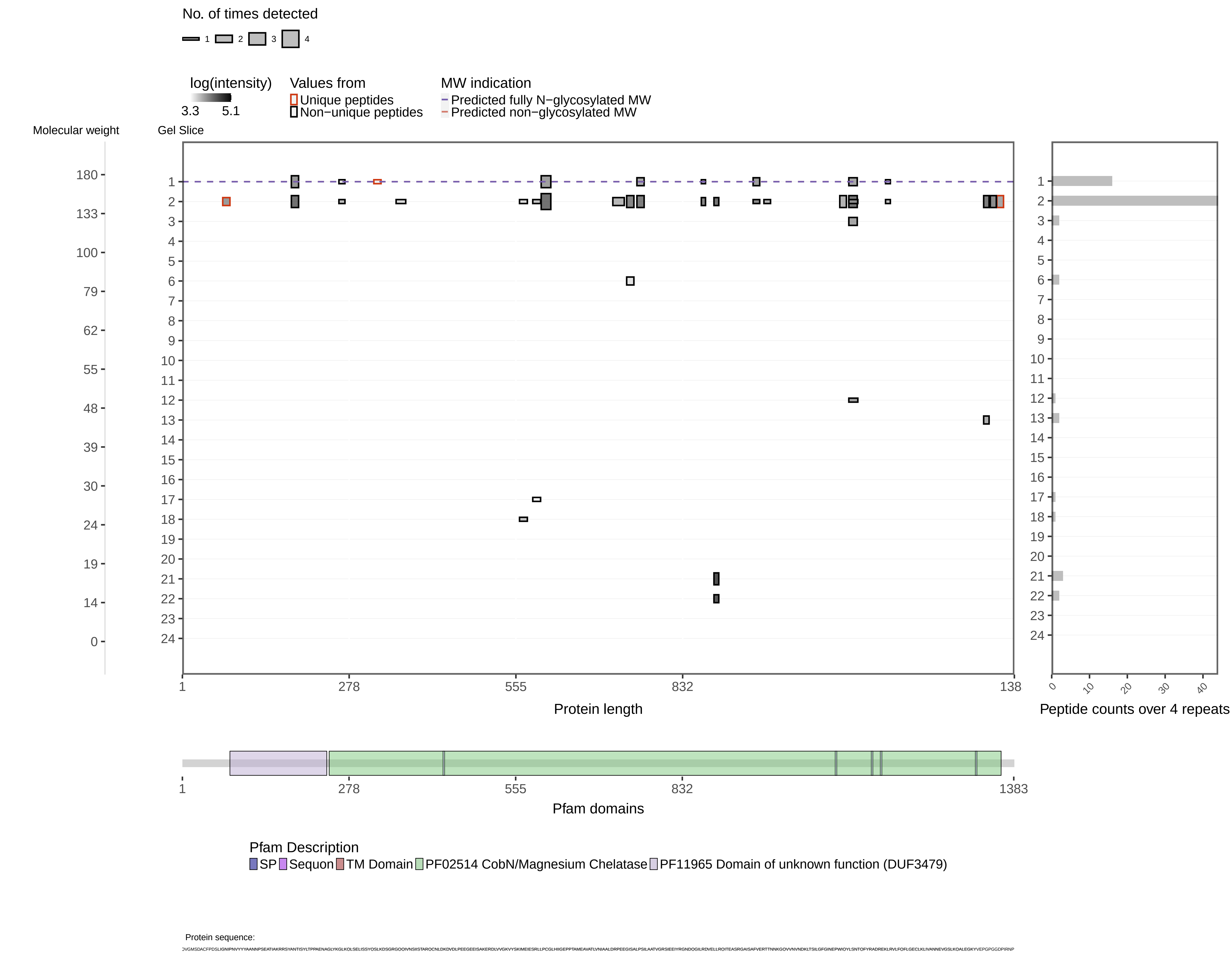

NbL14g14870.1 zinc−metallopeptidase_peroxisomal−like [MW = 112.34 kDa]

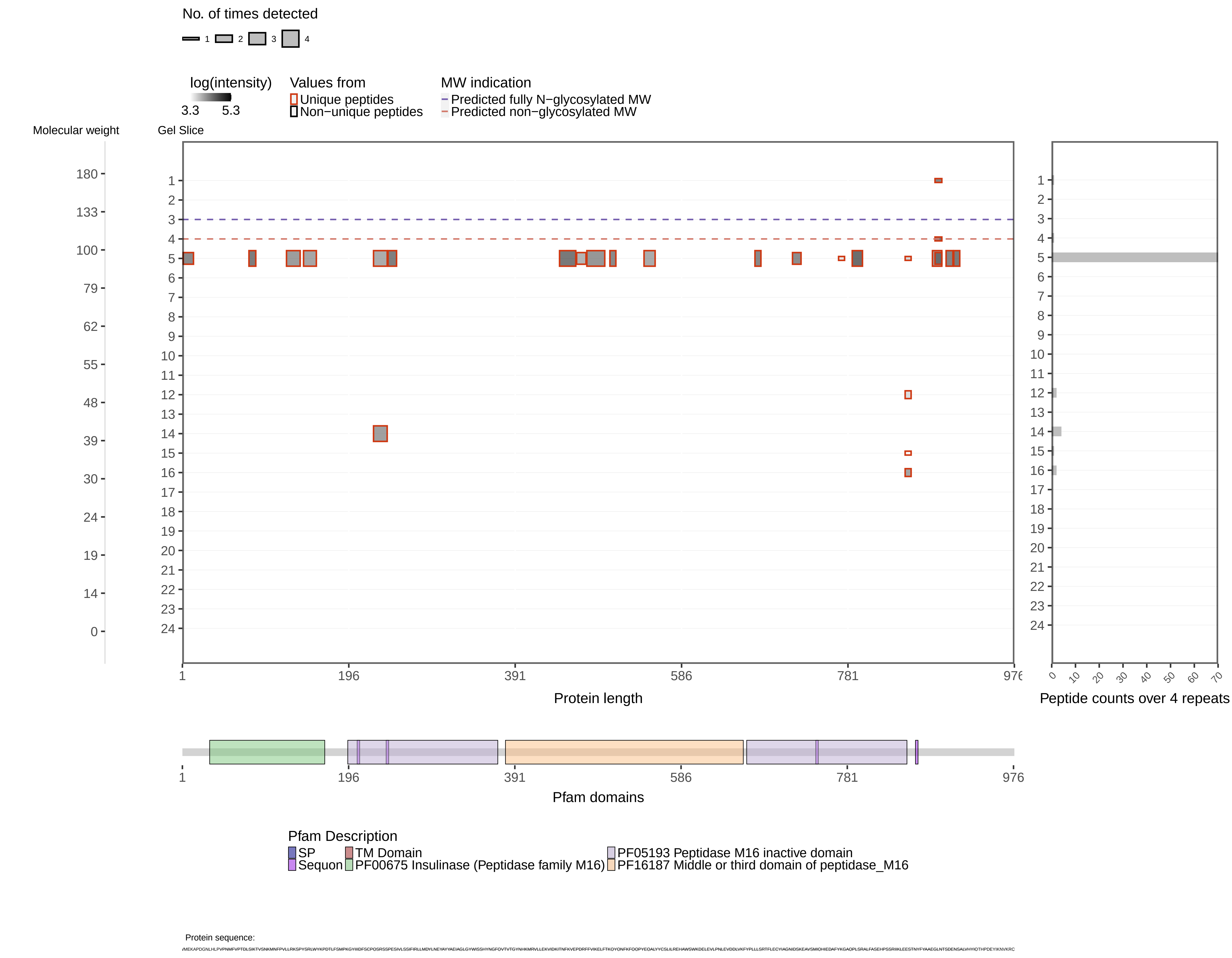

### NbL14g14850.1 insulin−degrading_enzyme−like_1_peroxisomal [MW = 92.47 kDa]

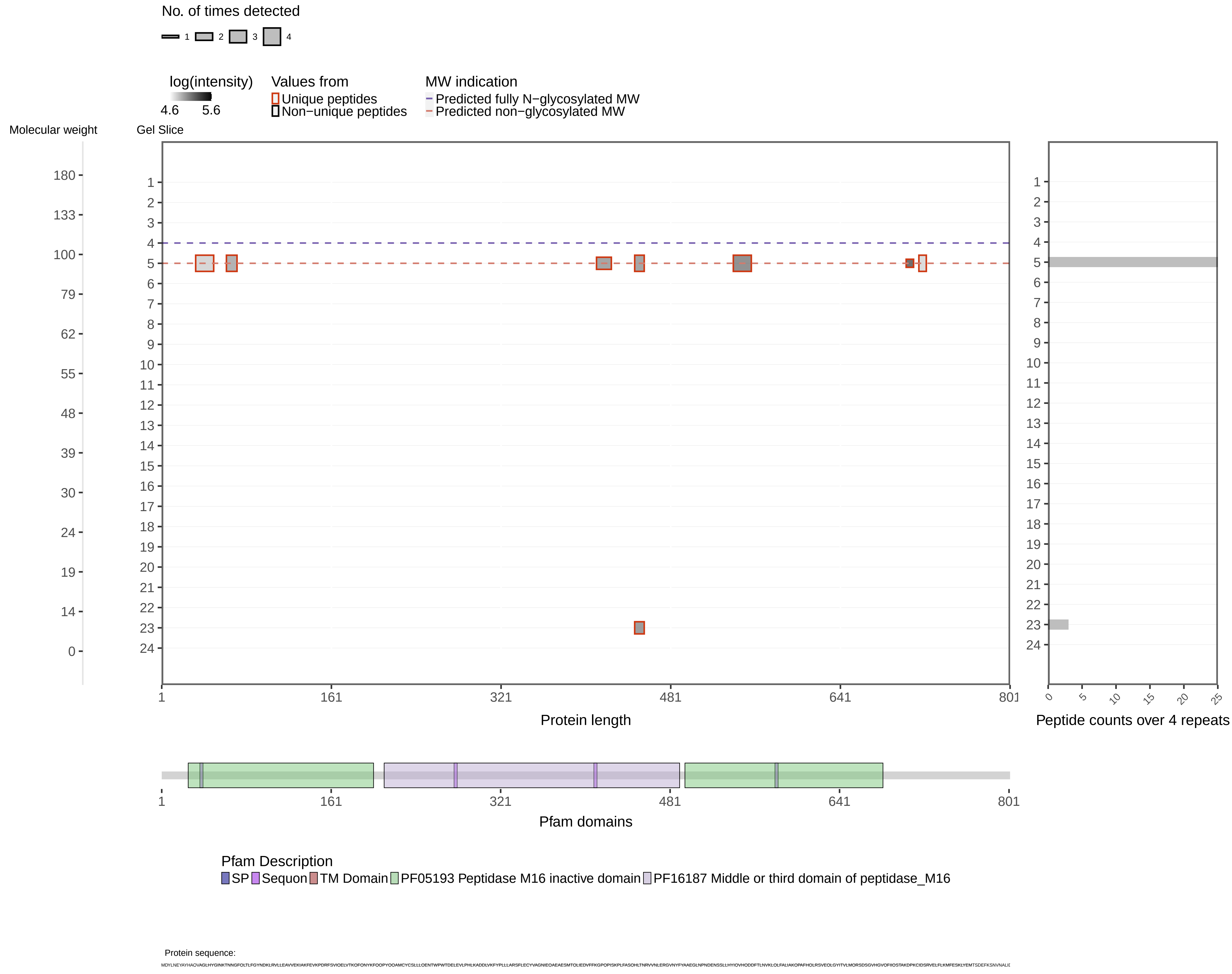

NbL14g14740.1 beta−glucosidase_BoGH3B−like [MW − SP = 66.15 kDa]

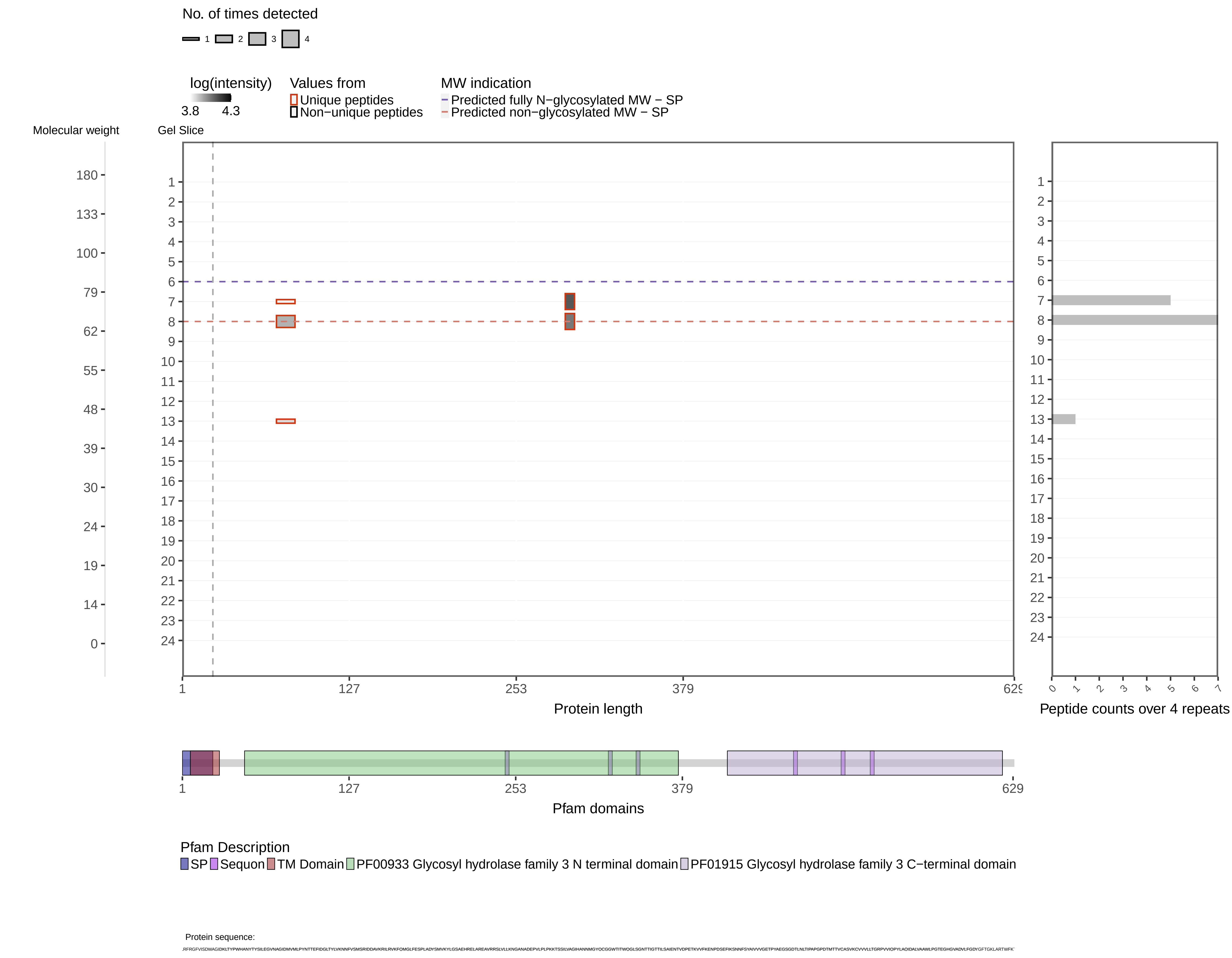

### NbL14g14640.1 magnesium−chelatase_subunit_ChlD_chloroplastic [MW = 83.19 kDa]

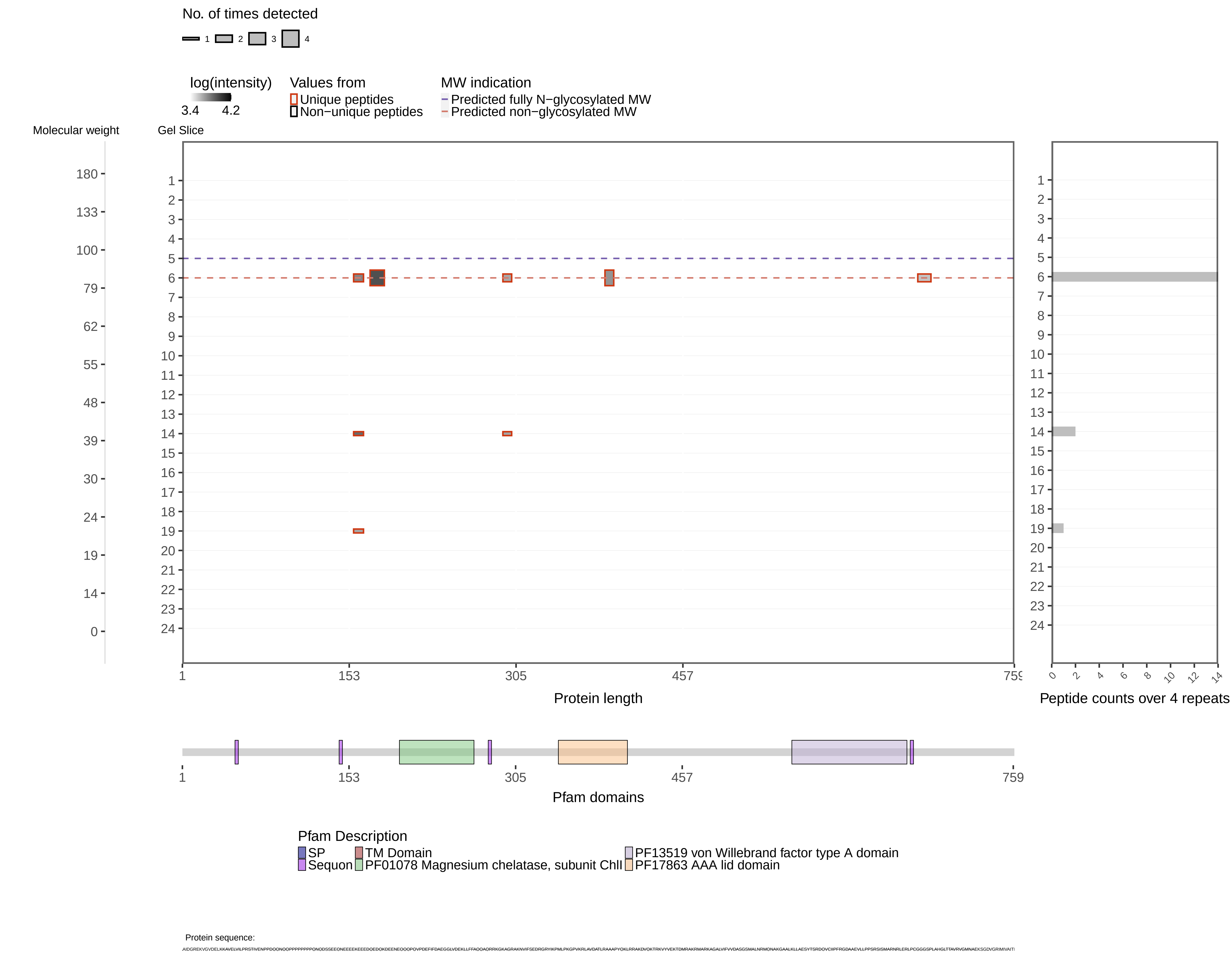

NbL14g14490.1 serine_carboxypeptidase−like_27 [MW − SP = 49.56 kDa]

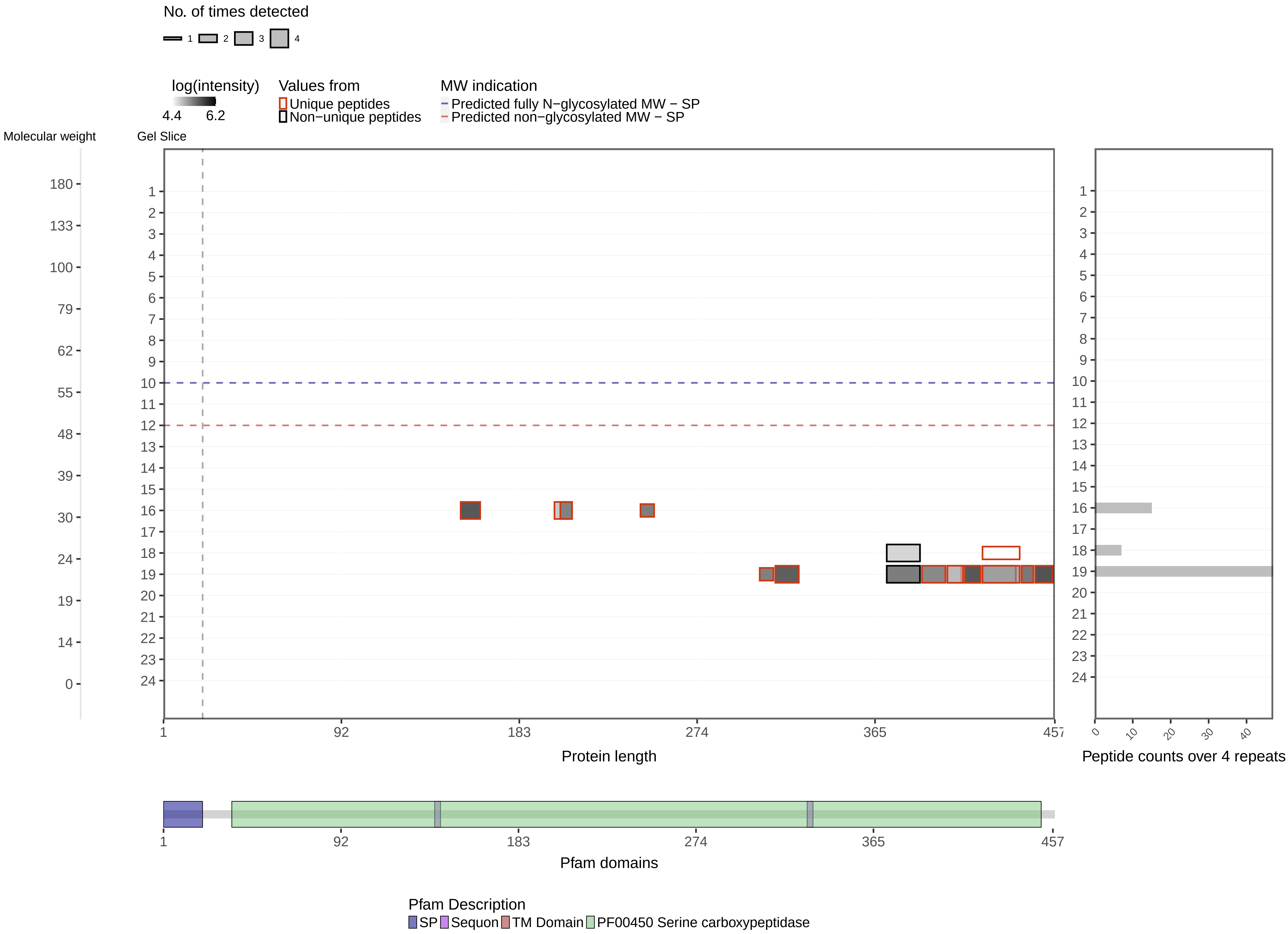


Protein sequence:

AVTDDYHDYIGTFEYWWTHGLISDSTYKLLQITCDKGSSEHPSTDCTKALLLAEQEQGNIDPYSIYTRPCASTSSLRRNLRGHYPWMSRAYDPCTARYSEVYFNLPEVQKAFHANVTKVAYPWKTCSDIVGNYWADSPLSMLPIYQELIAAGLRIWVFSGDTDSVVPVTATRYSIDALKLPTITKWYPWYDNKKVGGWSQVYKGLTLVTVTGAGHEVPLHRPRQAFILLRSFLENKPMPS*

### NbL14g14480.1 uncharacterized_LOC109211791 [MW = 28.85 kDa]

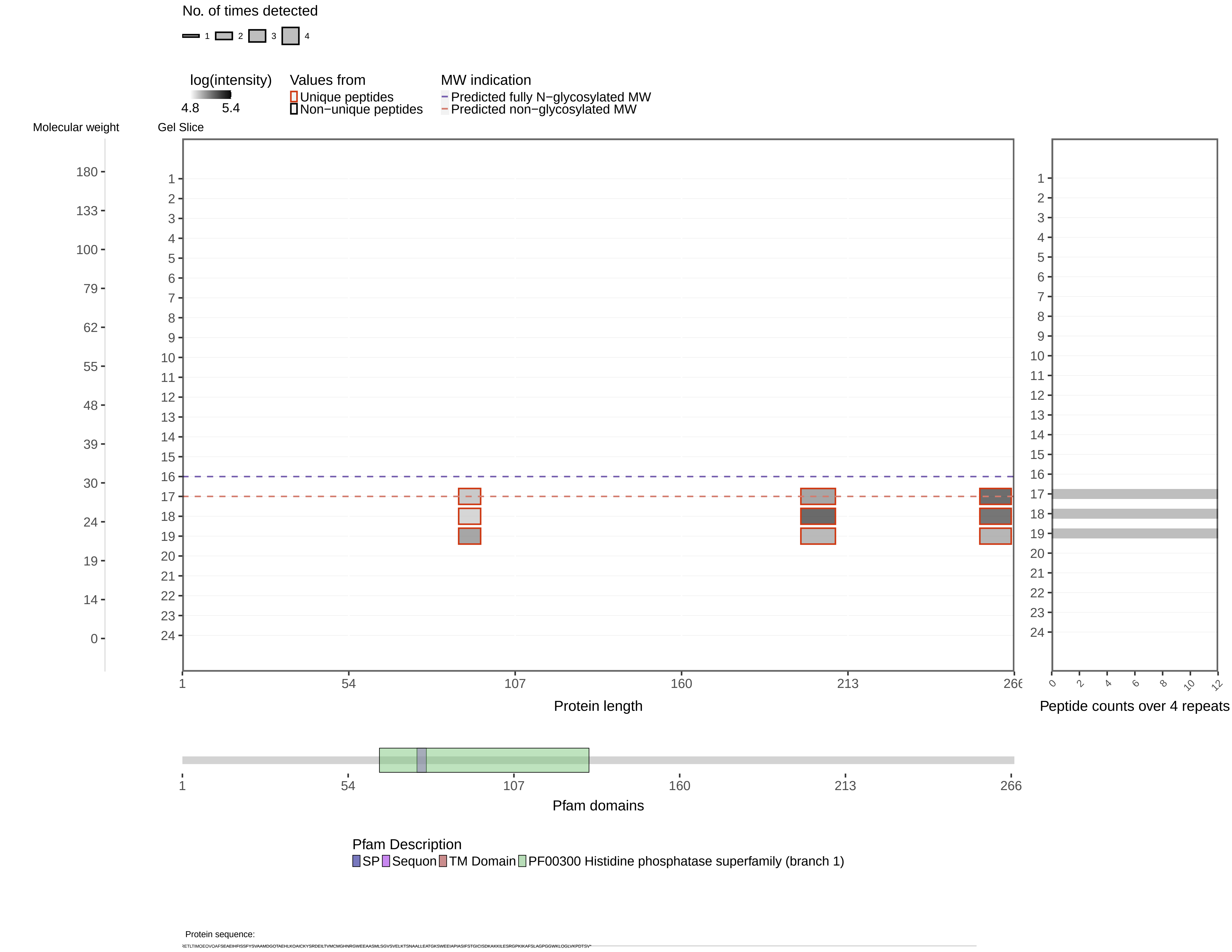

NbL14g14370.1 nuclear_inhibitor_of_protein_phosphatase_1 [MW = 45.96 kDa]

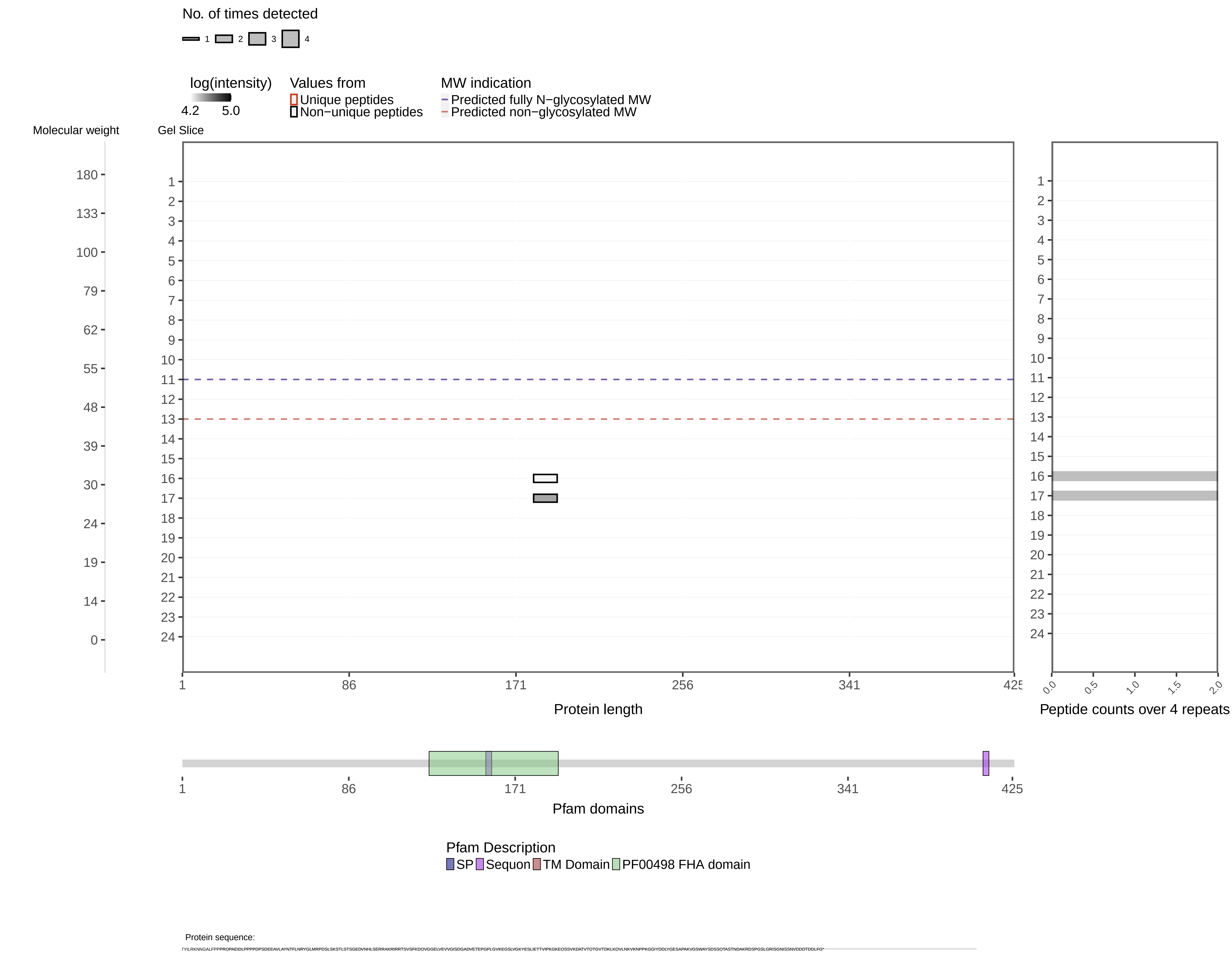

### NbL14g14310.1 trafficking_protein_particle_complex_subunit_5 [MW = 21.75 kDa]

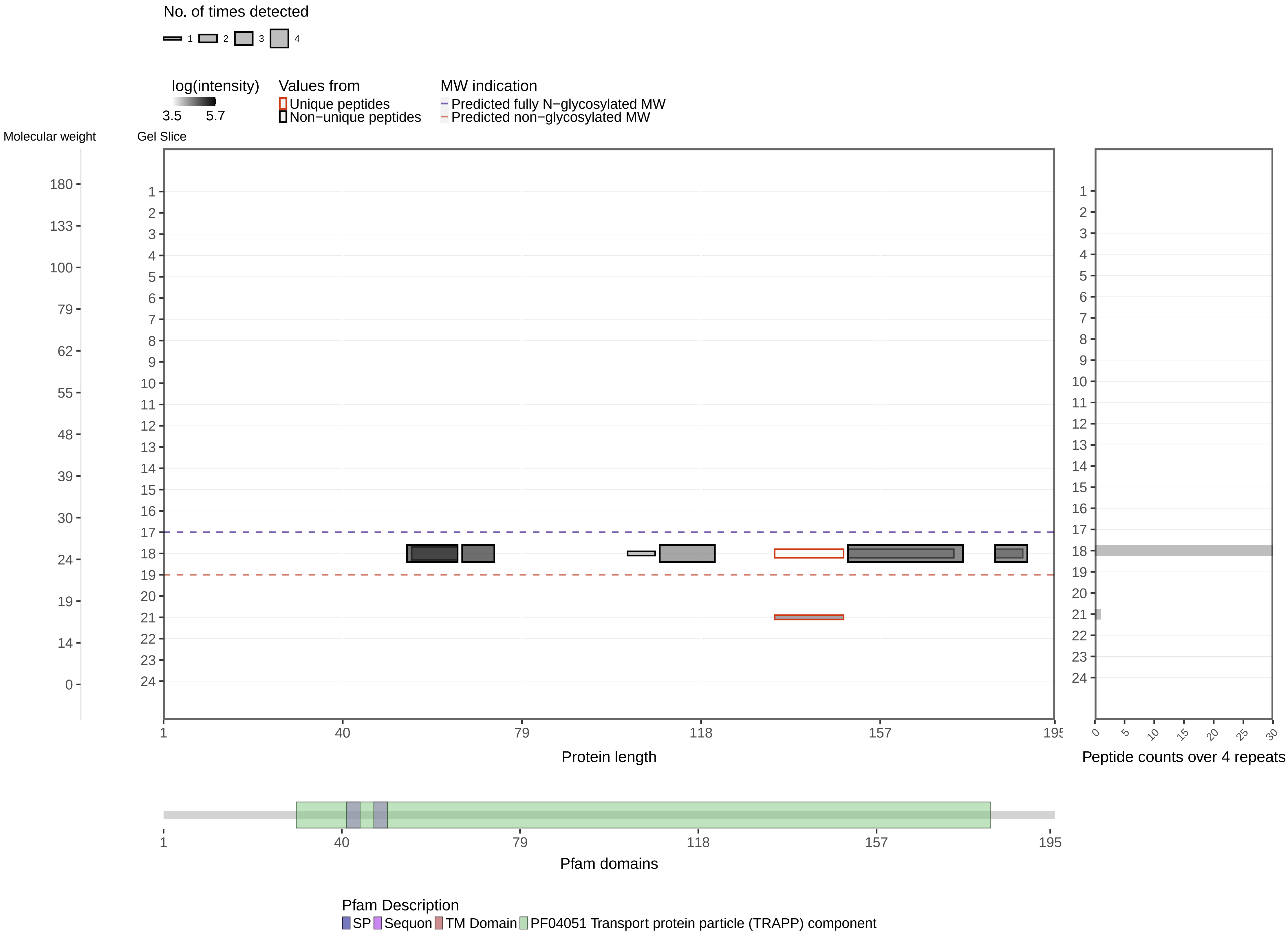


Protein sequence:

.LGILSFVHSTVWKVLFGKVADSLEKGTEHEDEYMISEKELLVNRFISIPKDMGAFNCGSFVAGIVRGVLENAGFPAVVTAHFVPVEGOORPRTTILIKFAEEVLRREARLG*

NbL14g14300.1 peptidyl−prolyl_cis−trans_isomerase_FKBP16−3_chloroplastic [MW = 27.25 kDa]

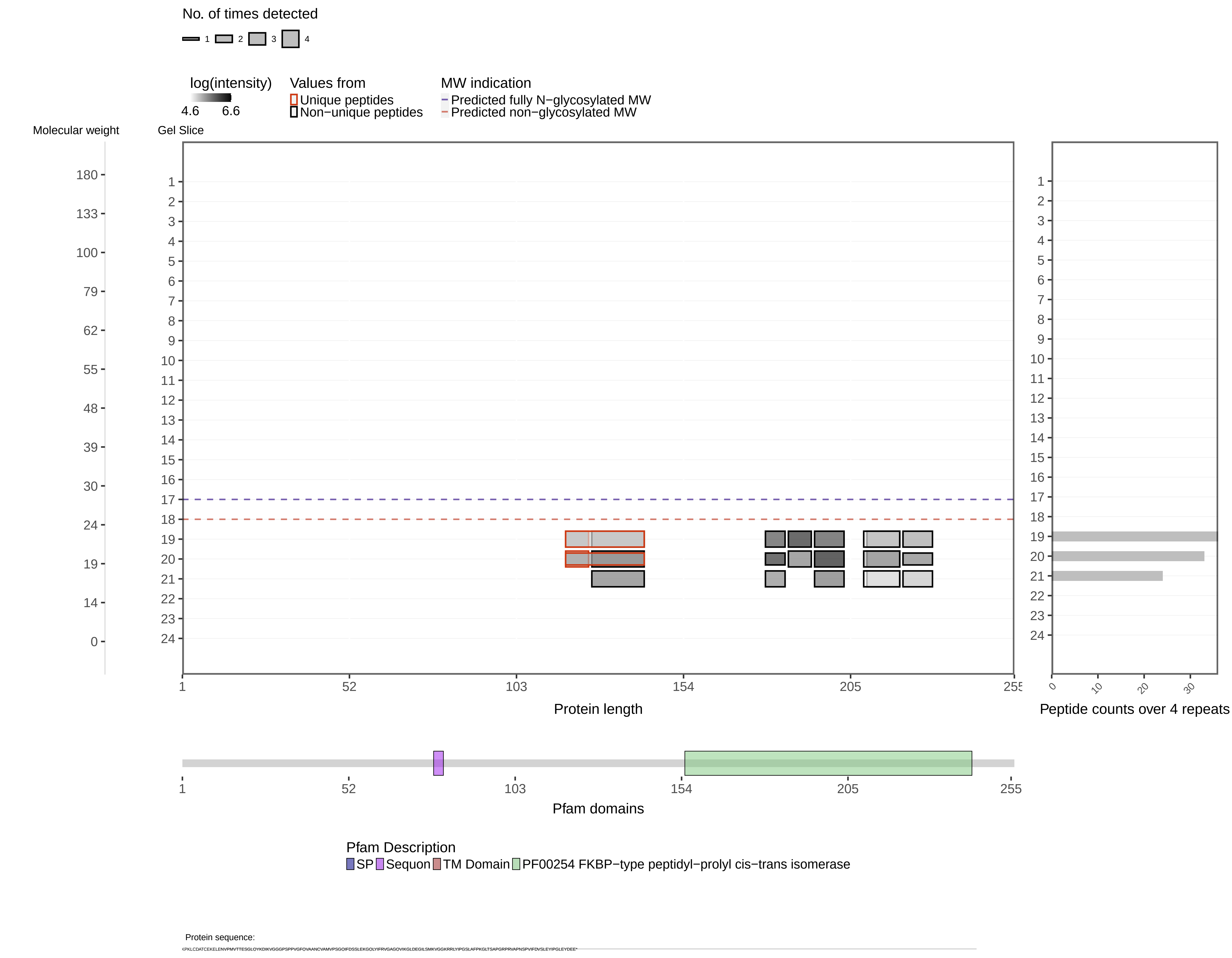

NbL14g14100.1 tropinone_reductase−like_3 [MW = 27.07 kDa]

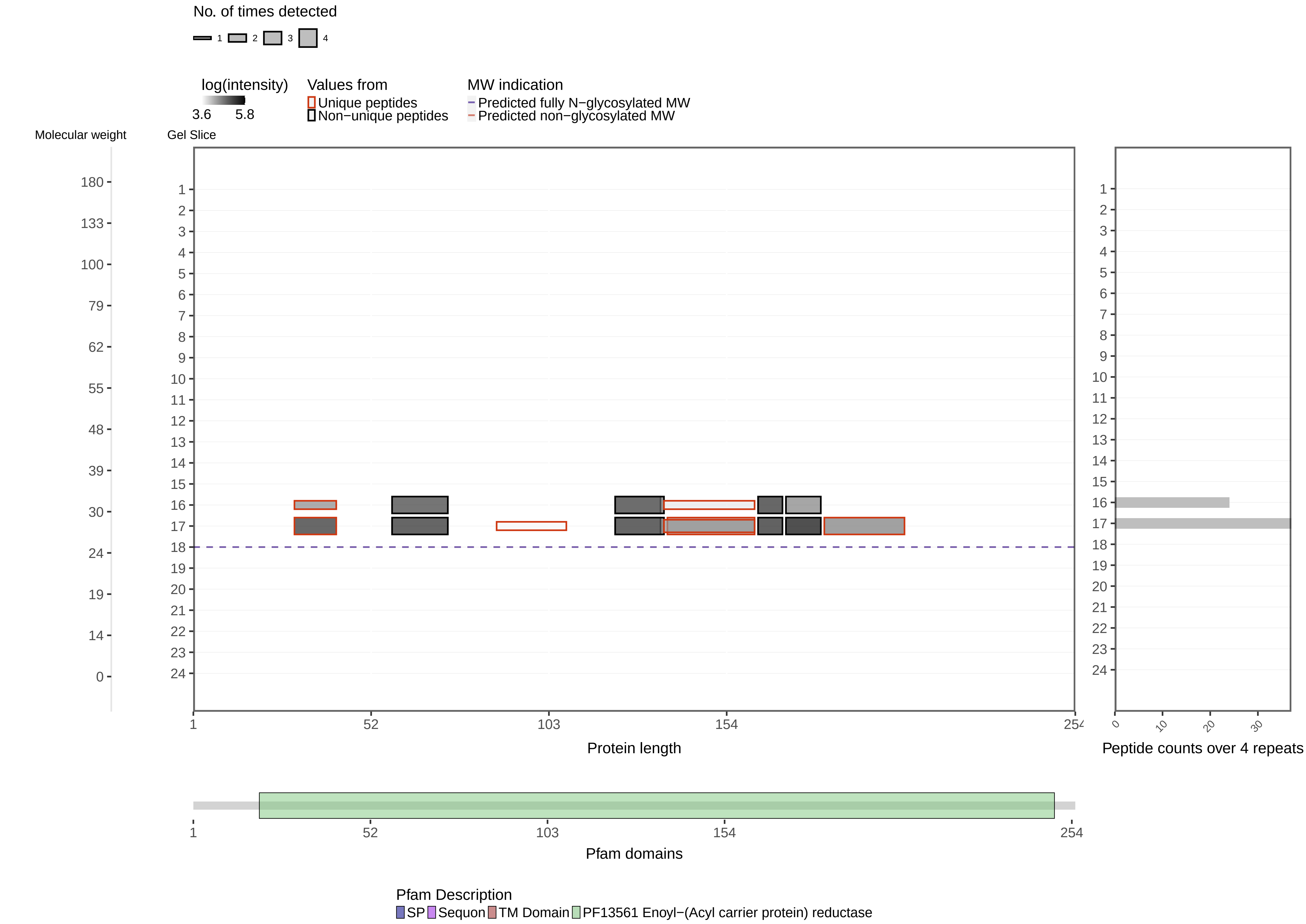


Protein sequence:

LWDINVKASILLLQDAAPYLKKGSSVVLISSISGYSPPASMGMYGVTKTALLGLTKALASEMSPDTRVNCVAPGFVPTHFADFITQNEHVRREIEGKTLLNRLGTTQDMAAAVAYLASDDAAYVTGETLVVAGGMPSRL*

### NbL14g13980.1 THO_complex_subunit_4D−like [MW = 29.2 kDa]

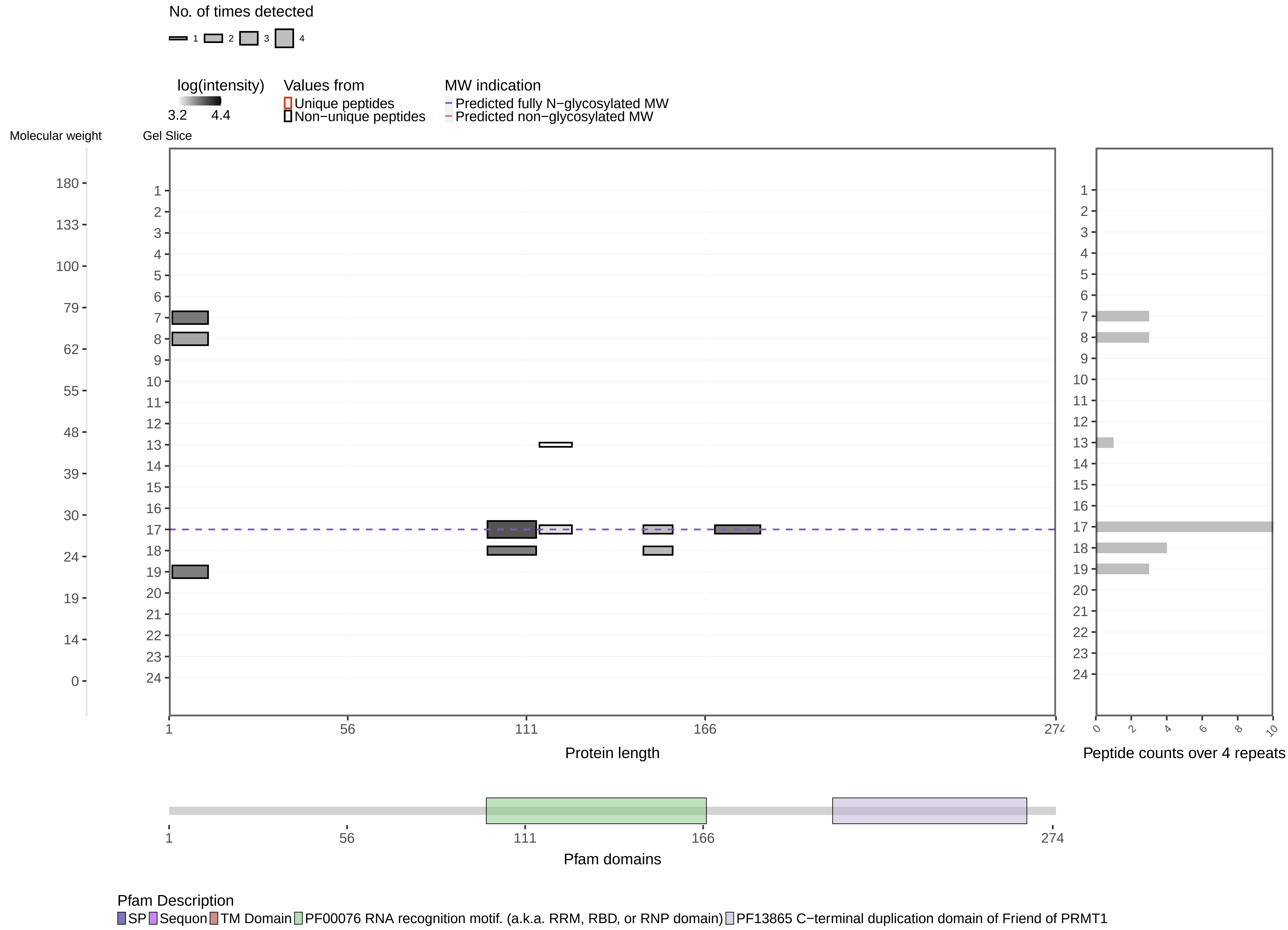


Protein sequence:

YAIHYDKNGRPSGAAEVVFARRSDAYQALKRYNNVQLDGKPMKIEVVAPPTDMPLSARVDVGRGSGRRTVVMTPGSARGRGAAPAANRGSSQRGRGGFGTASGRGRSRGHGGGRGRGGRGRKNGVEKSAEELDKELENYHATADAMQT*

NbL14g13820.1 ATP−dependent_6−phosphofructokinase_5_chloroplastic−like [MW = 59.1 kDa]

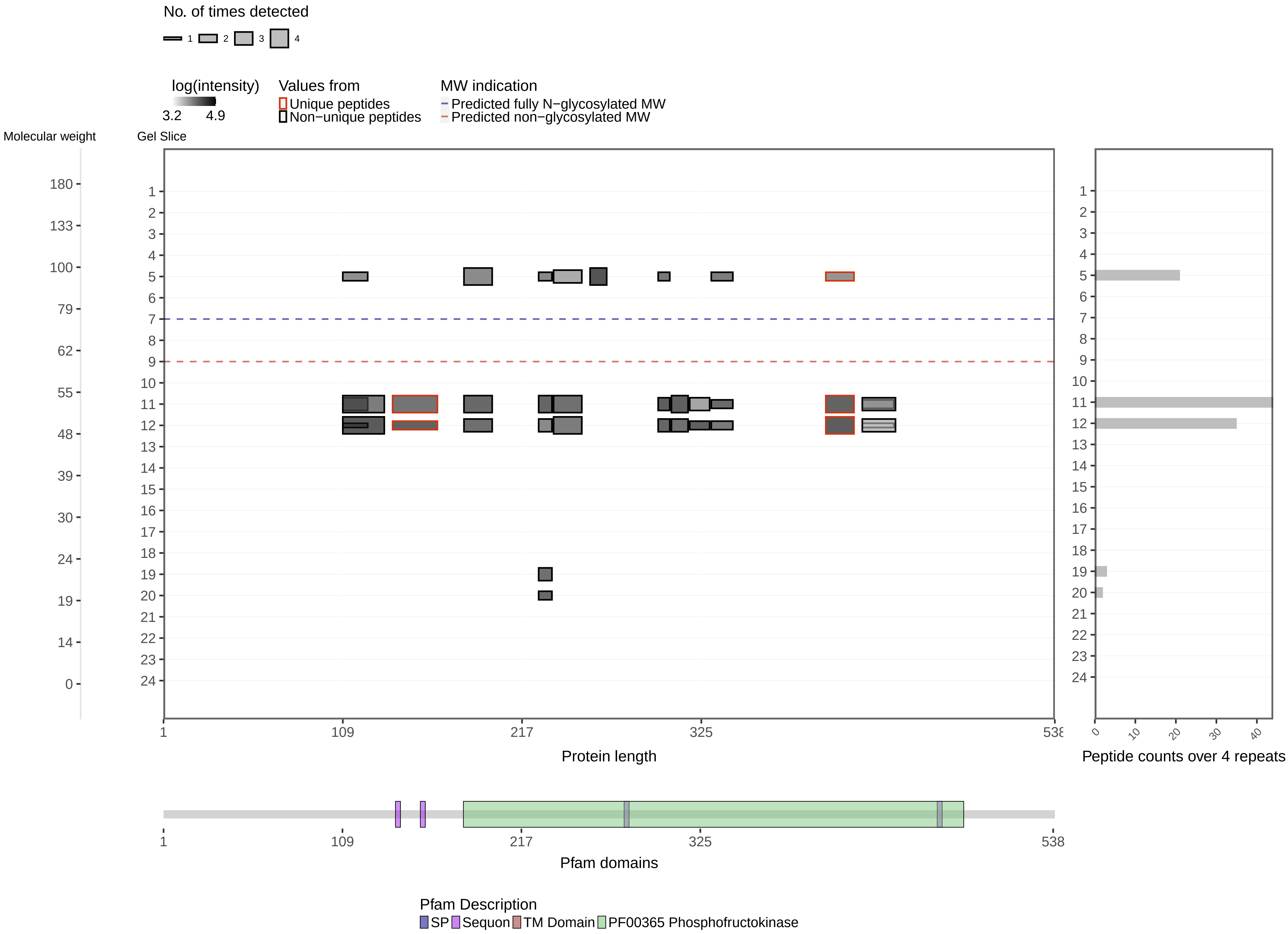


Protein sequence:
PKVSDIVDSIQERGINMLFVLGGNGTHAGANAIHDECRKRRMKVAVVGVPKTIDNDIMLMDKTFGFDTAVEEAQRAINSAYIEAHSAYRGIGIVKLMGRSSGFIAMQASLASGQIDICLIPEVPFNLHGPHGVLRHLKYLLATKGSAVVCAAEGAGQDFLEKTNAKDASGNAVLGDIGVYLQQEIKKYFKEMGETSDVKYIDPTYMIRACRANASDGILCTVLGQNAVHGAFAGYSGITVGICNTHYVYFPIPEVISKPRVVDPNSRMWHRCLTSTGQPDFL*

### NbL14g13670.1 14−3−3−like_protein_A [MW = 28.77 kDa]

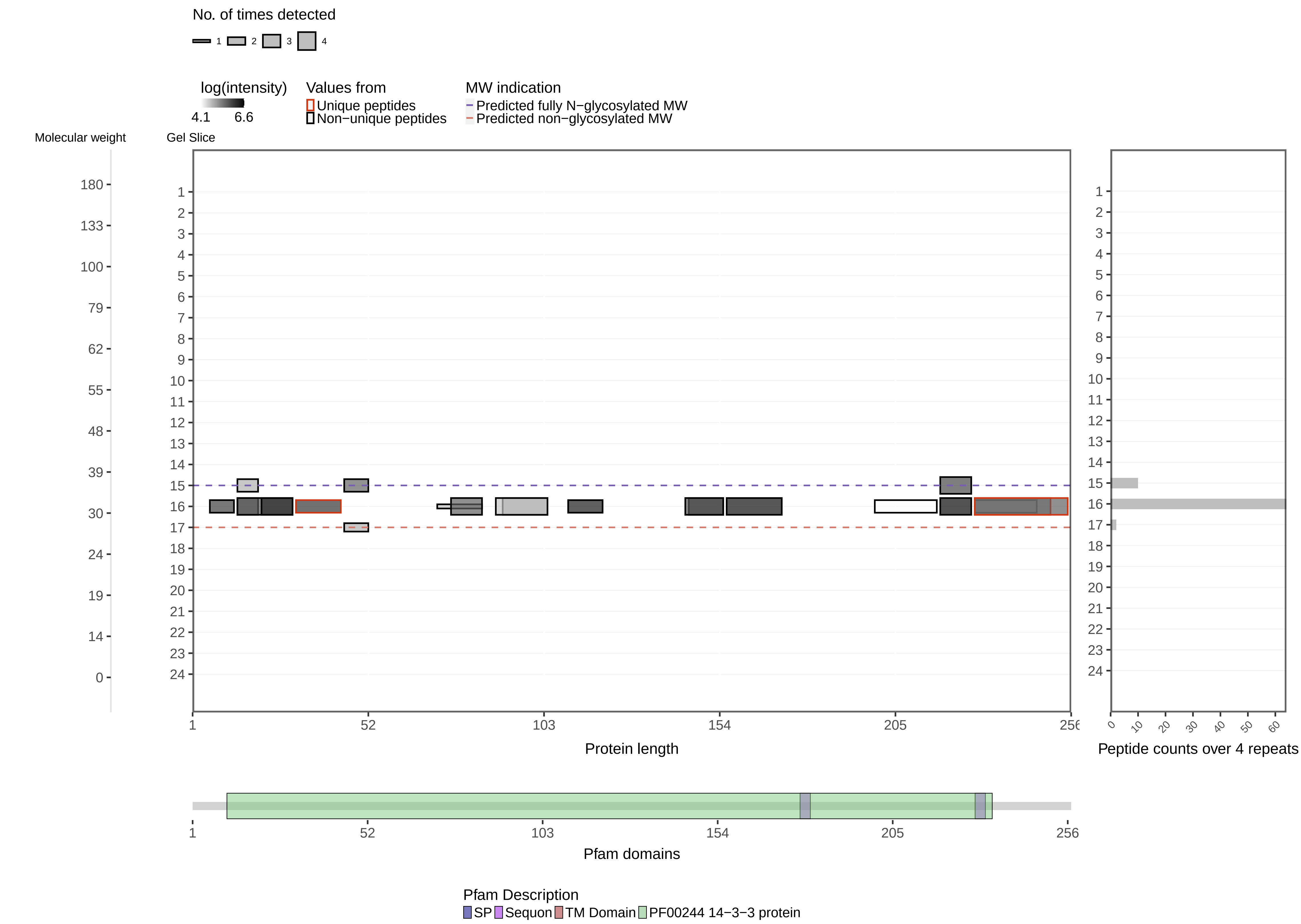


Protein sequence:

TGDSKVFYLKMKGDYYRYLAEFKTGAERKEAAENTLSAYKSAQDIANVELAPTHPIRLGLALNFSVFYYEILNSPDRACNLAKQAFDEAIAELDTLGEESYKDSTLIMQLLRDNLTLWTSDMQDDGTDEIKEPSKAEEQQ*

### NbL14g13380.1 peptidyl−prolyl_cis−trans_isomerase_FKBP20−1 [MW = 19.98 kDa]

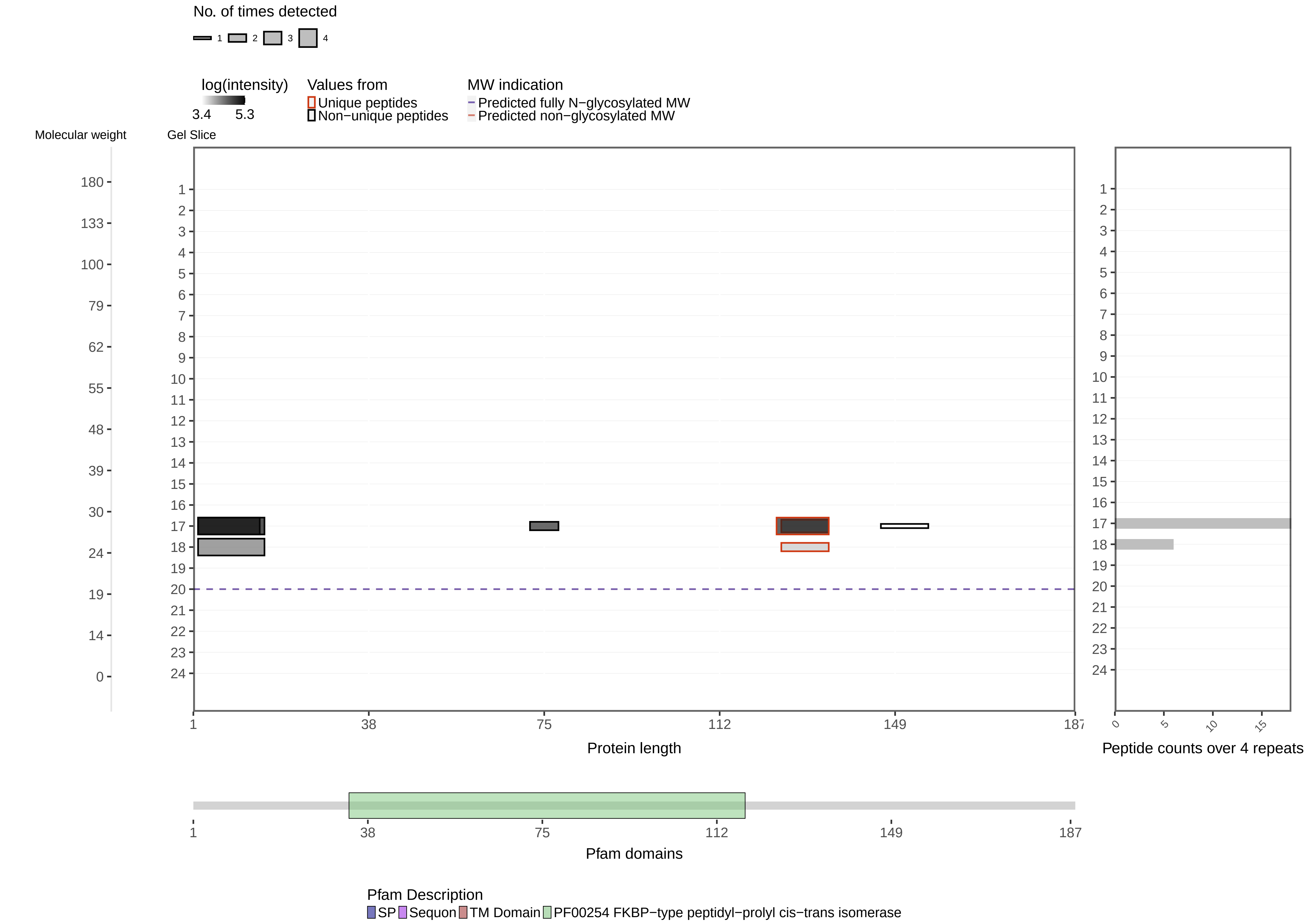


Protein sequence:

KVGEVALITCKPEYAYGSAGSPPDIPPDATLVFEVELVACRPRKGSSISSVSDERARLEELKKQREMAAAAKEEEKKKREEAKAAAAARIQAKLEAKKGKGKGKAK*

NbL14g13300.1 triosephosphate_isomerase_cytosolic [MW = 27.15 kDa]

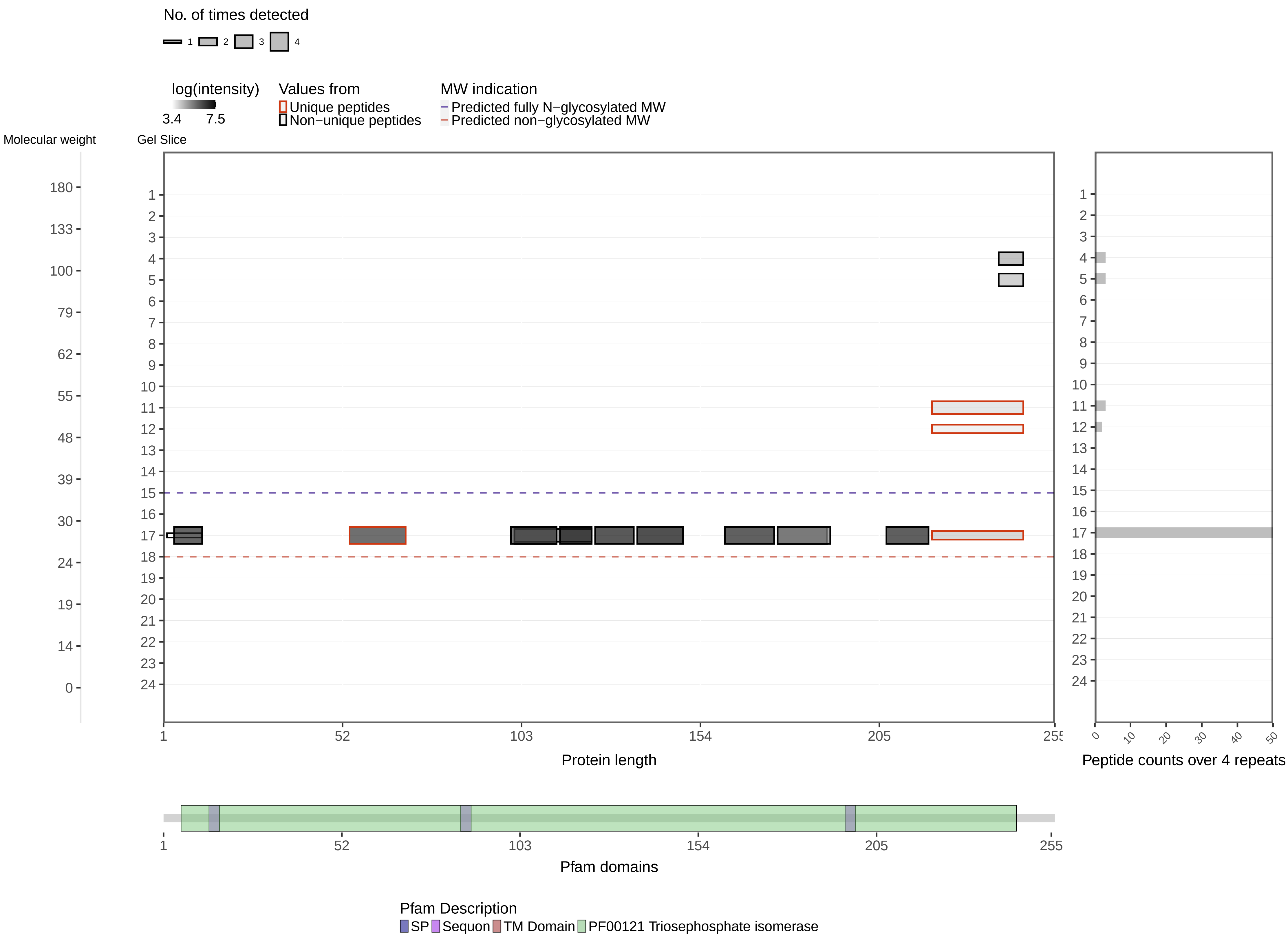


Protein sequence:

### NbL14g13290.1 actin−97 [MW = 41.7 kDa]

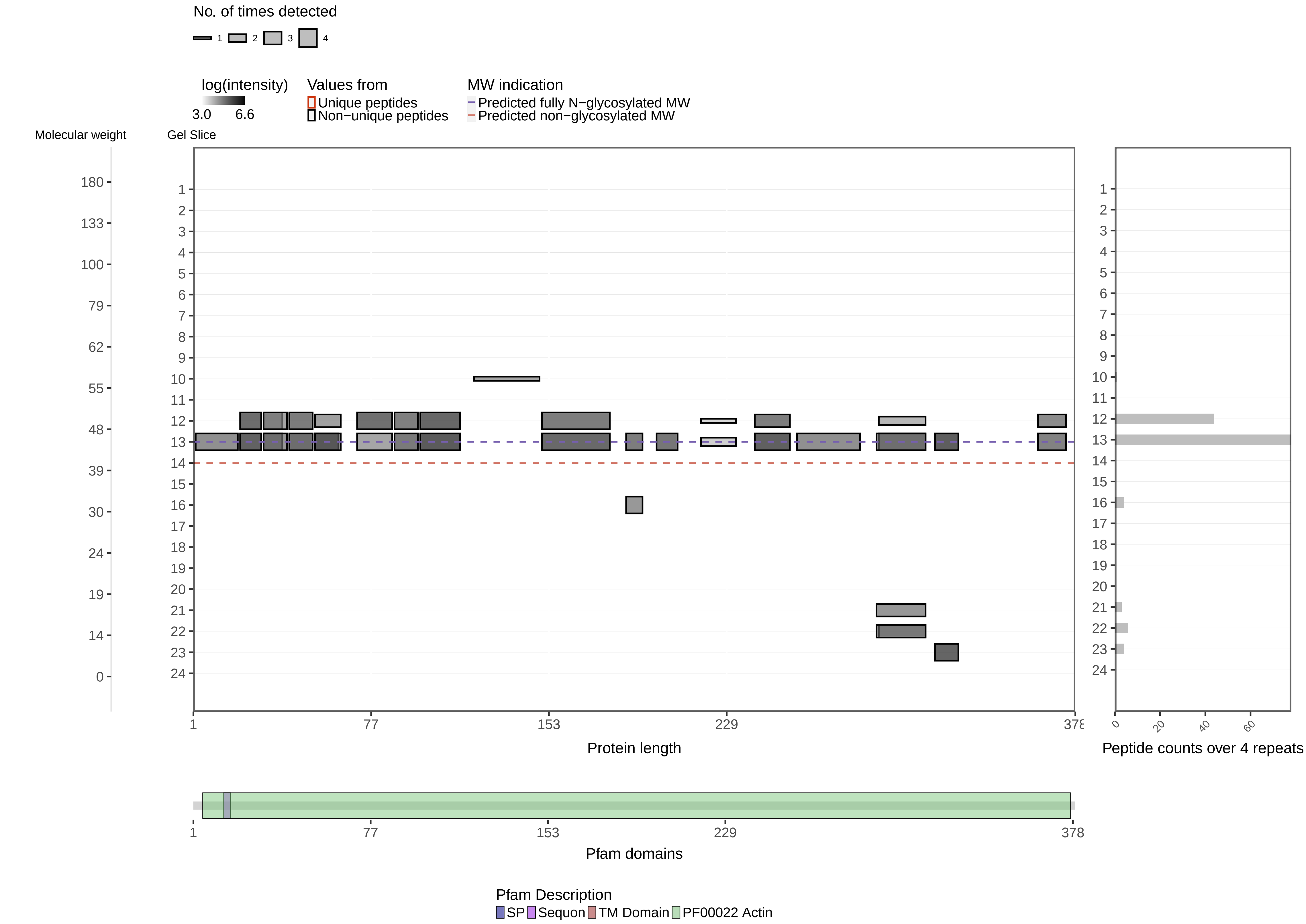


Protein sequence:

PHAILRLDLAGRDLTDSLMKILTERGYMFTTSAEREIVRDVKEKLAYIALDYEQELETAKTSSSVEKNYELPDGQVITIGAERFRCPEVLFQPSMIGMEAAGIHETTYNSIMKCDVDIRKDLYGNIVLSGGSTMFPGIADRMSKEITALAPSSMKIKVVAPPERKYSVWIGGSILASLSTFQQMWIAKAEYDESGPSIVHRKCF*

NbL14g13180.1 UDP−glucuronic_acid_decarboxylase_5−like [MW = 38.65 kDa]

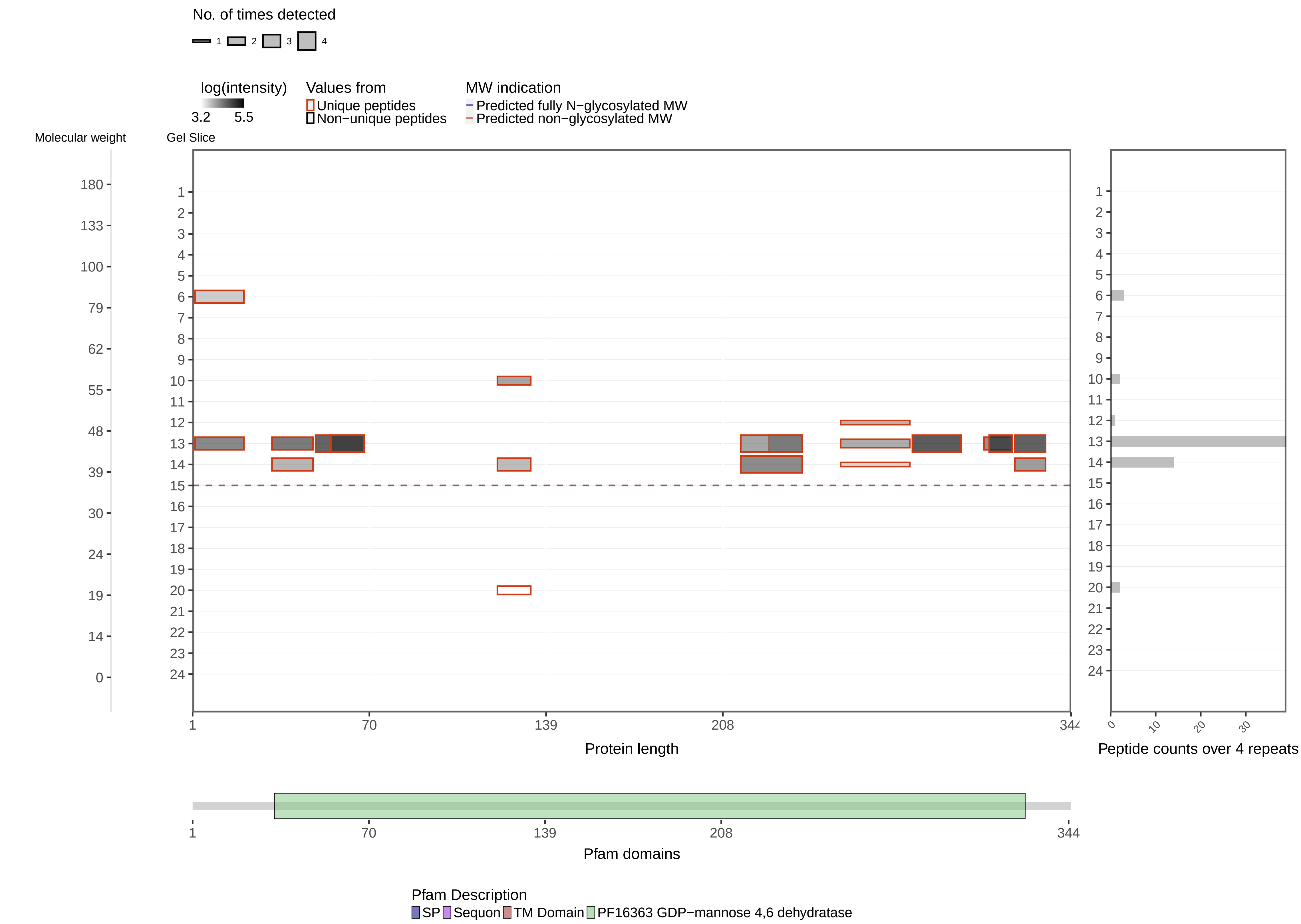


Protein sequence:

SYWGNVNPIGVRSCYDEGKRVAETLMFDYHRQHGIEIRIARIFNTYGPRMNIDDGRVVSNFIAQAIRDEPLTVQLPGTQTRSFCYVSDMVDGLIRLMEGDNTGPINIGNPGEFTMLELAETVKELINPEVQIITVENTPDDPRQRKPDITKAKELLGWEPTIKLRDGIPLMEDDFRGRLGISRKI*

### NbL14g13150.1 actin−depolymerizing_factor_2 [MW = 18.16 kDa]

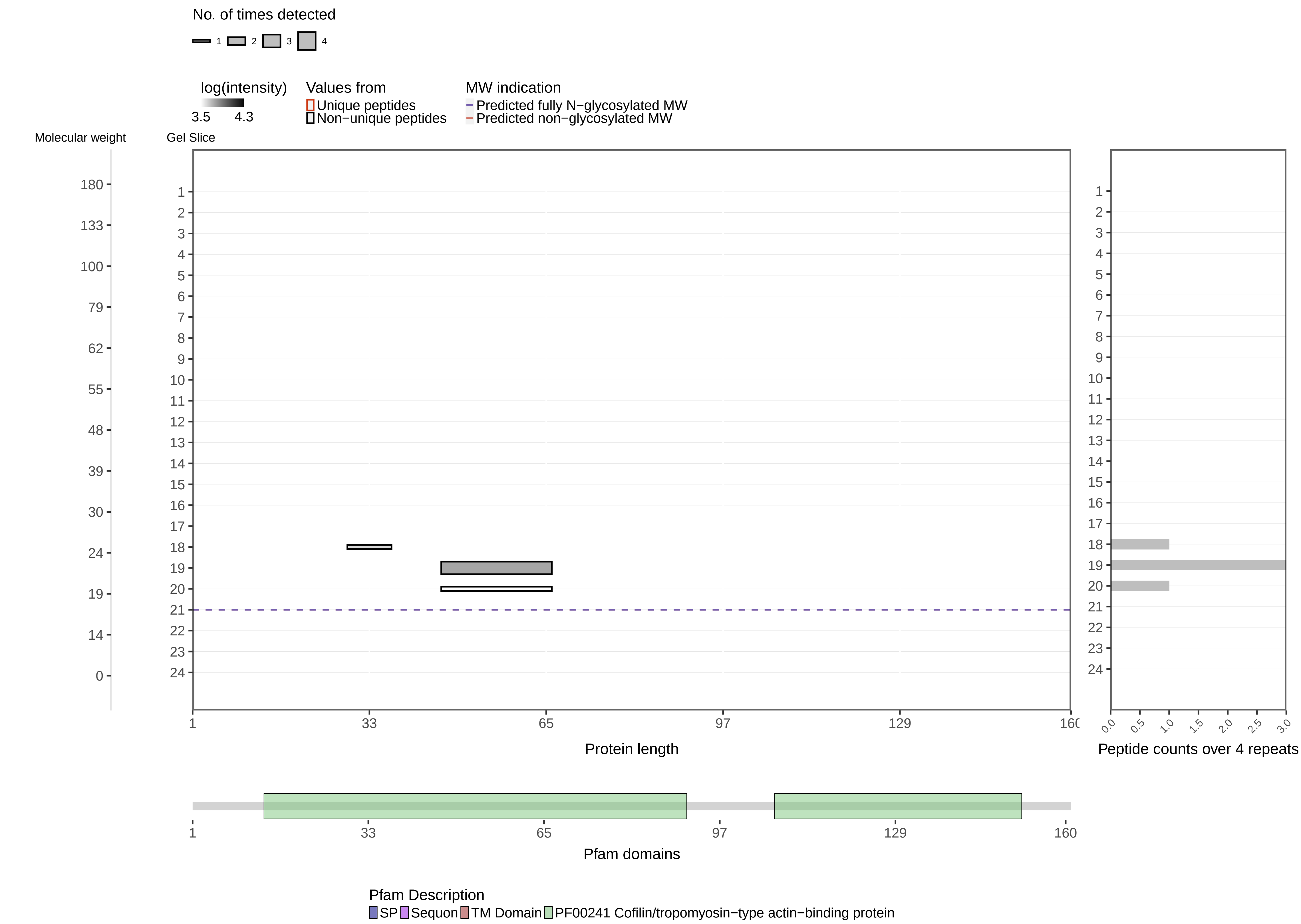


Protein sequence:

AVYDFDFTTEENVQKSRIFFIAWKSRVRPVETASCKNAGSPDTARVRSKMIYASSKDRFKRELDGFQVELQATDPTEMGLDVFRSRVGGIEC*

NbL14g13140.1 ras−related_protein_RABE1c−like [MW = 23.96 kDa]

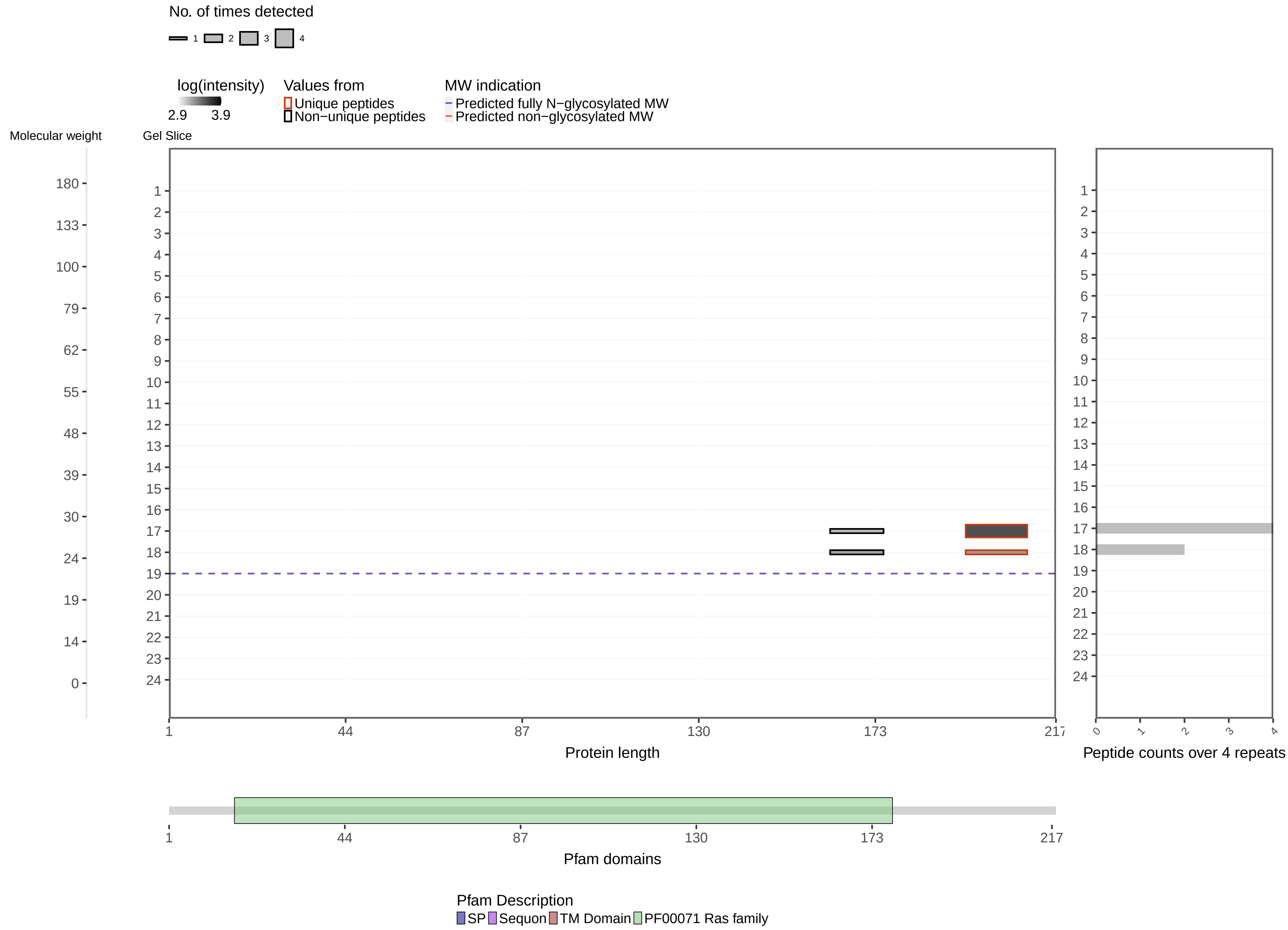


Protein sequence:

DESSFNNIRNWIRNIEQHASDNVNKILVGNKADMDESKRAVPTSKGQALADEYGIKFFETSAKTNMNVEEVFFSIARDIKQRLSESDSKTEPQAIRINQADQAGTSGQAAEKSSCCGS*

NbL14g13130.1 2−oxoglutarate_dehydrogenase_mitochondrial−like [MW = 115.58 kDa]

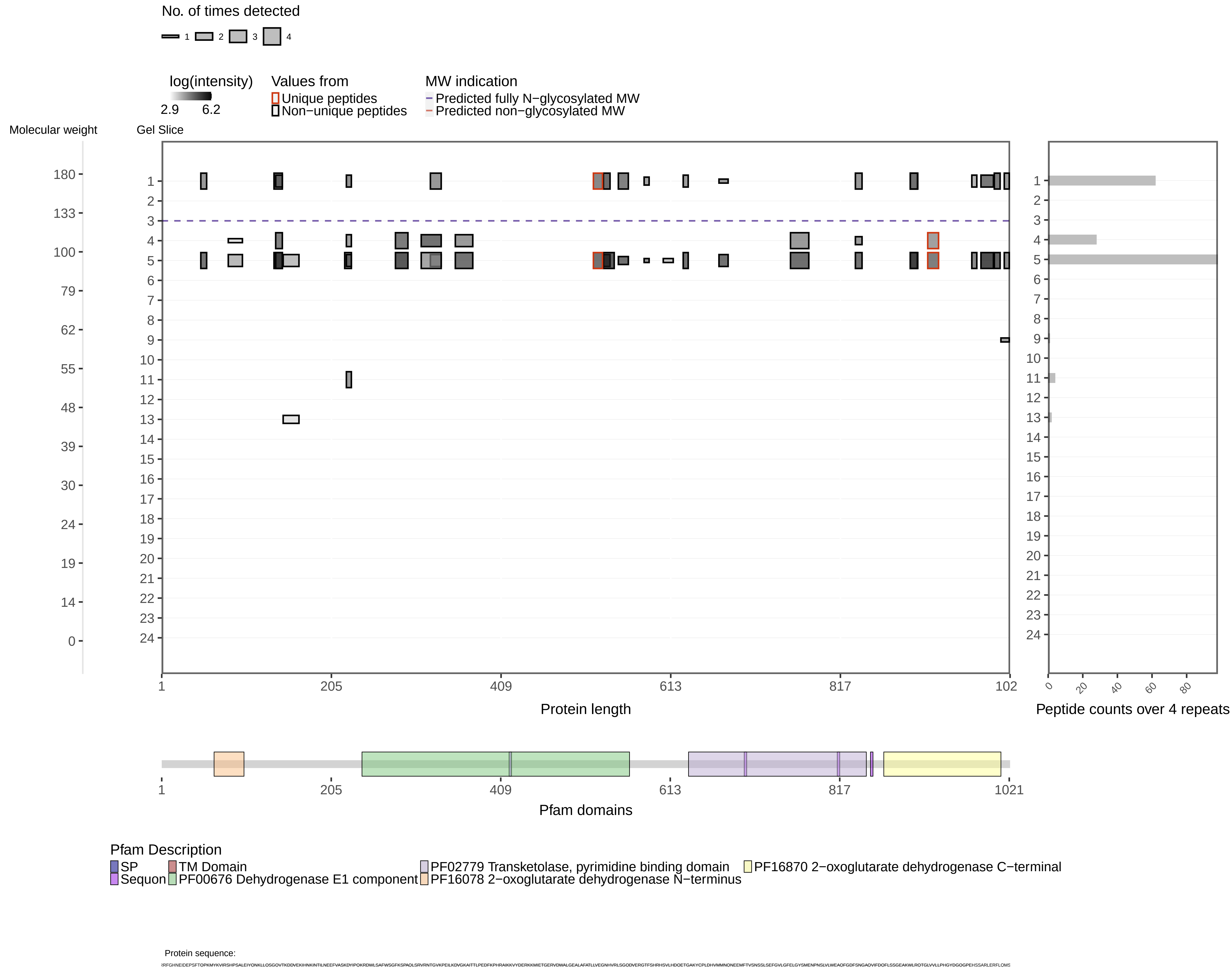

### NbL14g13000.1 histidine_biosynthesis_bifunctional_protein_hisIE_chloroplastic−like [MW = 31.54 kDa]

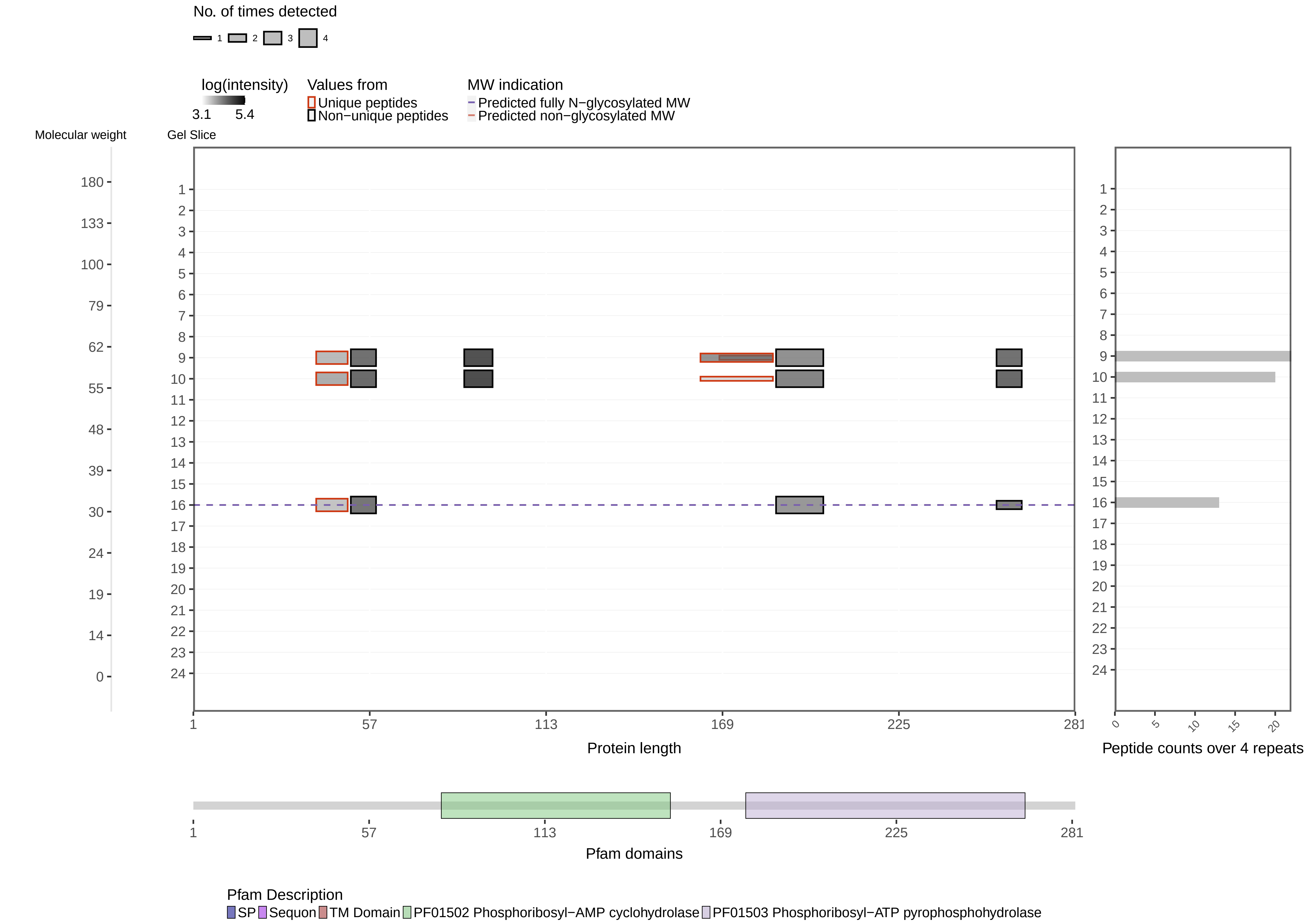


Protein sequence:

CDRDSIIYLGKPDGPTCHTGSETCYYTSVDGILKNPEVEKNELAMTTLYALESTINERKAETSSSSSGKPSWTKRLLLDDKLLCSKIREEADELCRTLEENEDKGRTASEMADVLYHAMVLLSLRGVKIEEVLQVLRQRFSKSGIEEKNSRKS*

### NbL14g12980.1 splicing_factor_3B_subunit_3−like [MW = 134.52 kDa]

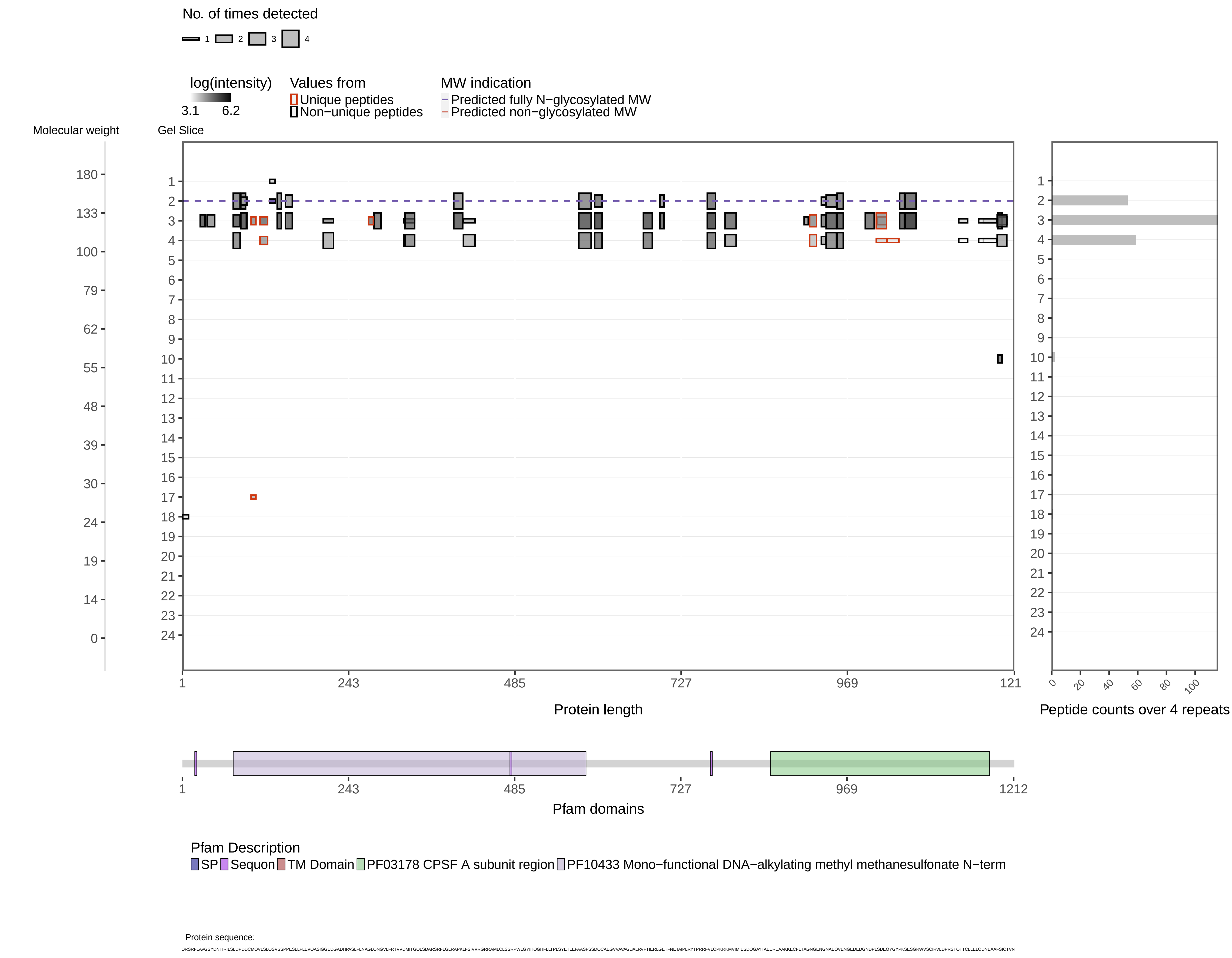

### NbL14g12870.1 diphthine−−ammonia_ligase−like [MW = 82.4 kDa]

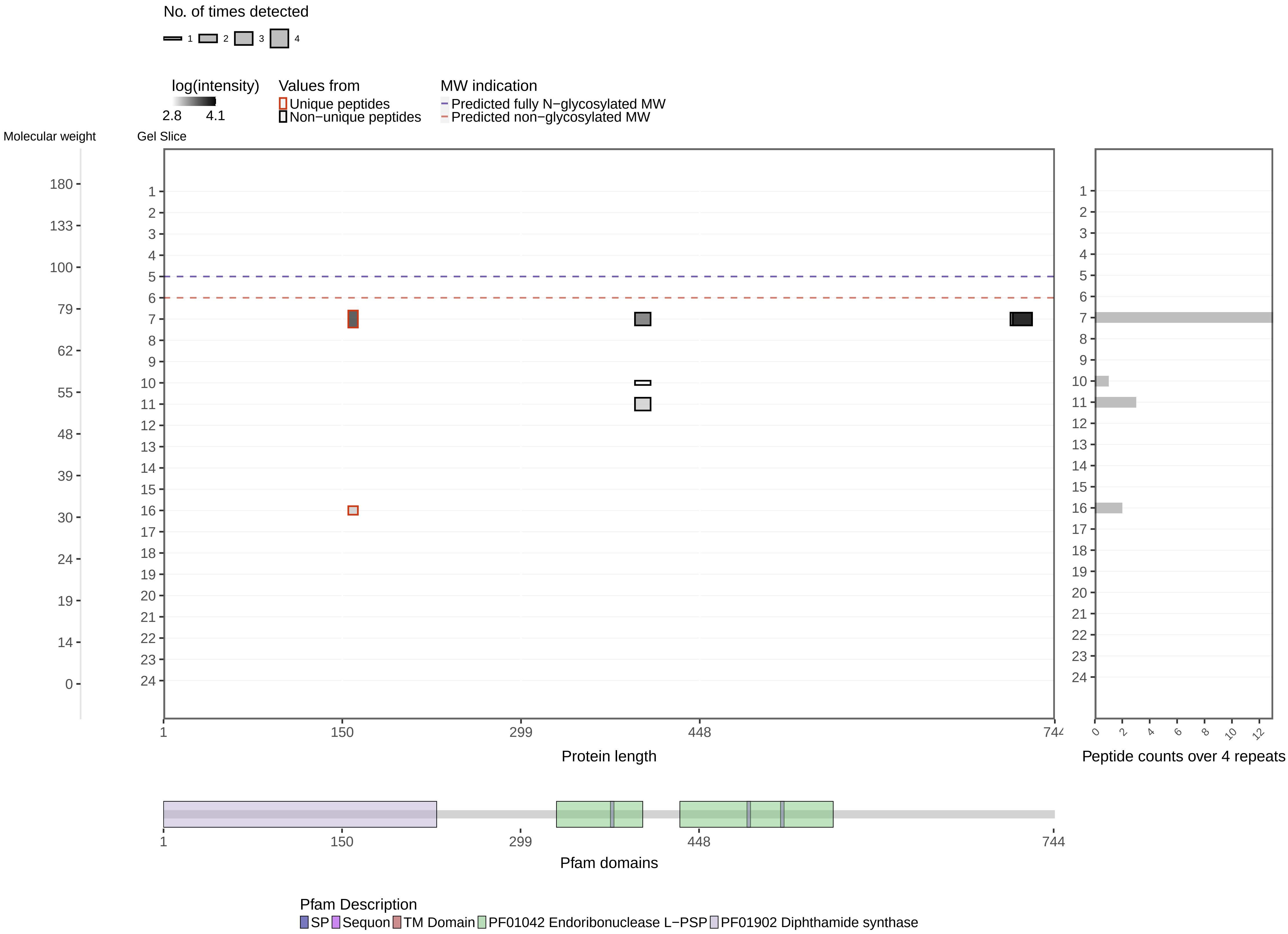


Protein sequence:

LYIHLYIADMDEFAVANETYVRYITQEKCRFGVPSRSTIELPLLLVGLGRAYIEVLVANDPTKKVLHVQSISCWAPSCIGPYSQATLHNDILHMAGQLGLDPATMVLCEGGPVAELEQALENSEAVAGSFNCSISTSAIVFVIYCSASLETSERVLVQNKTEALLNQMKSLHAEGTKKSRVLDPIFLYILVPDLPKRALVEVKPMLYTGEYMSAPSDHAMQHQSTGQGYWGFEYETWHDICLQKCVAYGKVCTAILSVTEELAAKICSPANVAGHASVKSKGPVEKEHVIMIARFCIYR

### NbL14g12820.1 RNA−binding_protein_42−like [MW = 27.09 kDa]

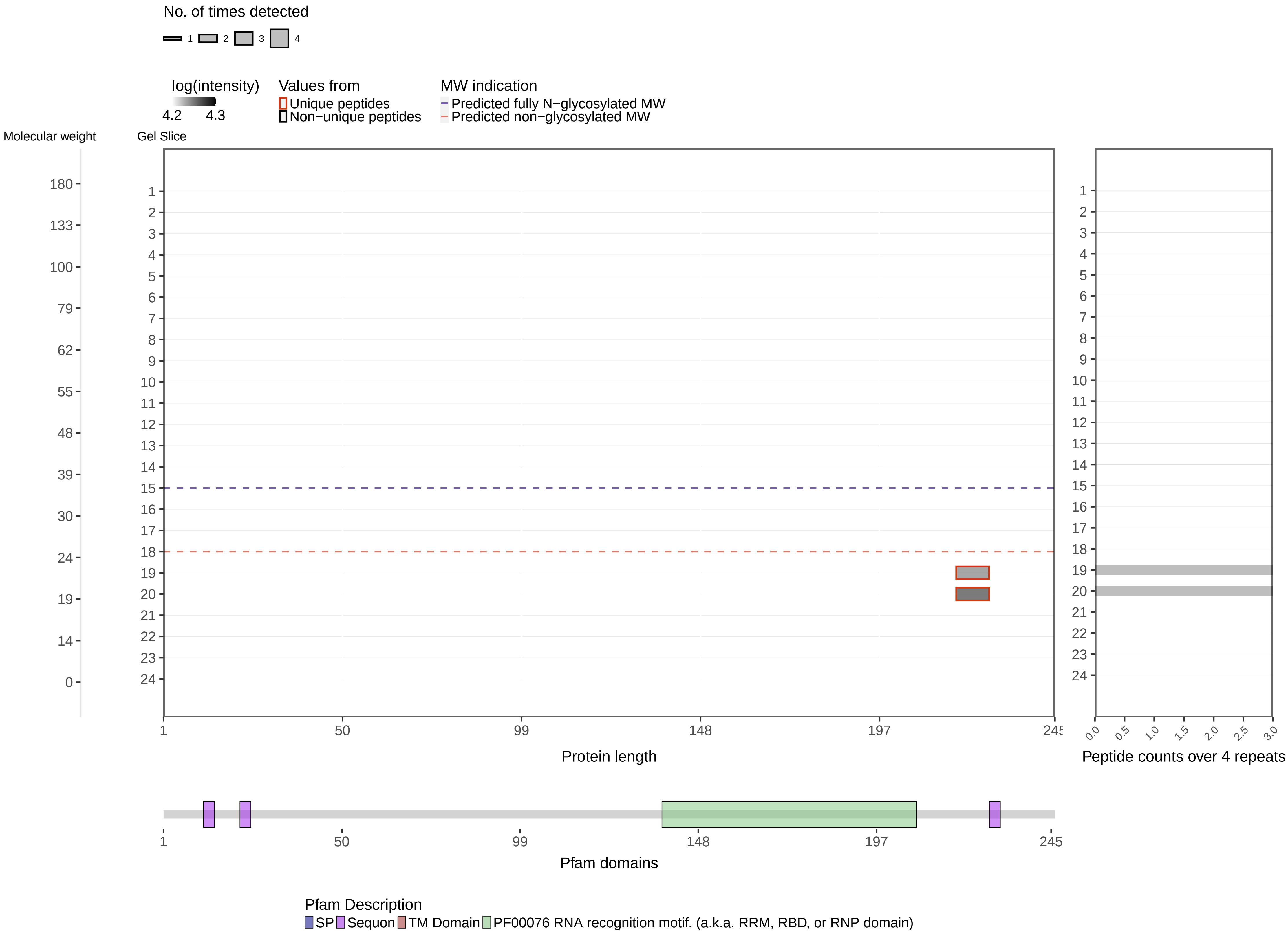


Protein sequence:

KKAVPRKAAGLSWEDPTLAEWPENDYRLFCGDLGNEVNDDVLSKAFSRFPTFNMAKVVRDKRTGKTKGYGFVSFSNLLDLAAALKEMNGKYVGNRPIKLRKSKWQERIDYEALESSKNRSHKKPKLTKKSILHK*

### NbL14g12780.1 probable_low−specificity_L−threonine_aldolase_1 [MW = 38.76 kDa]

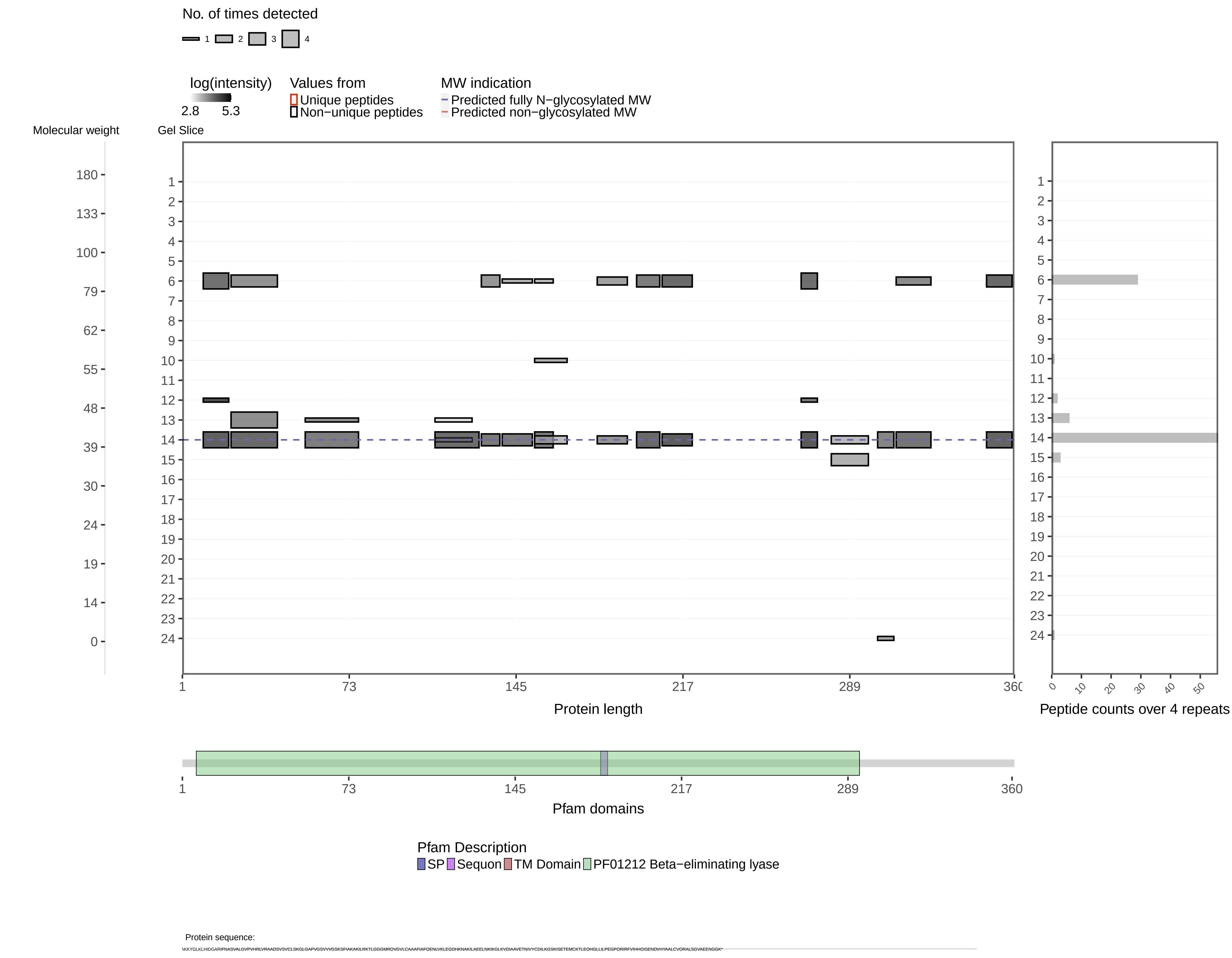

### NbL14g12710.1 1−aminocyclopropane−1−carboxylate_oxidase_homolog_1−like [MW = 46.41 kDa]

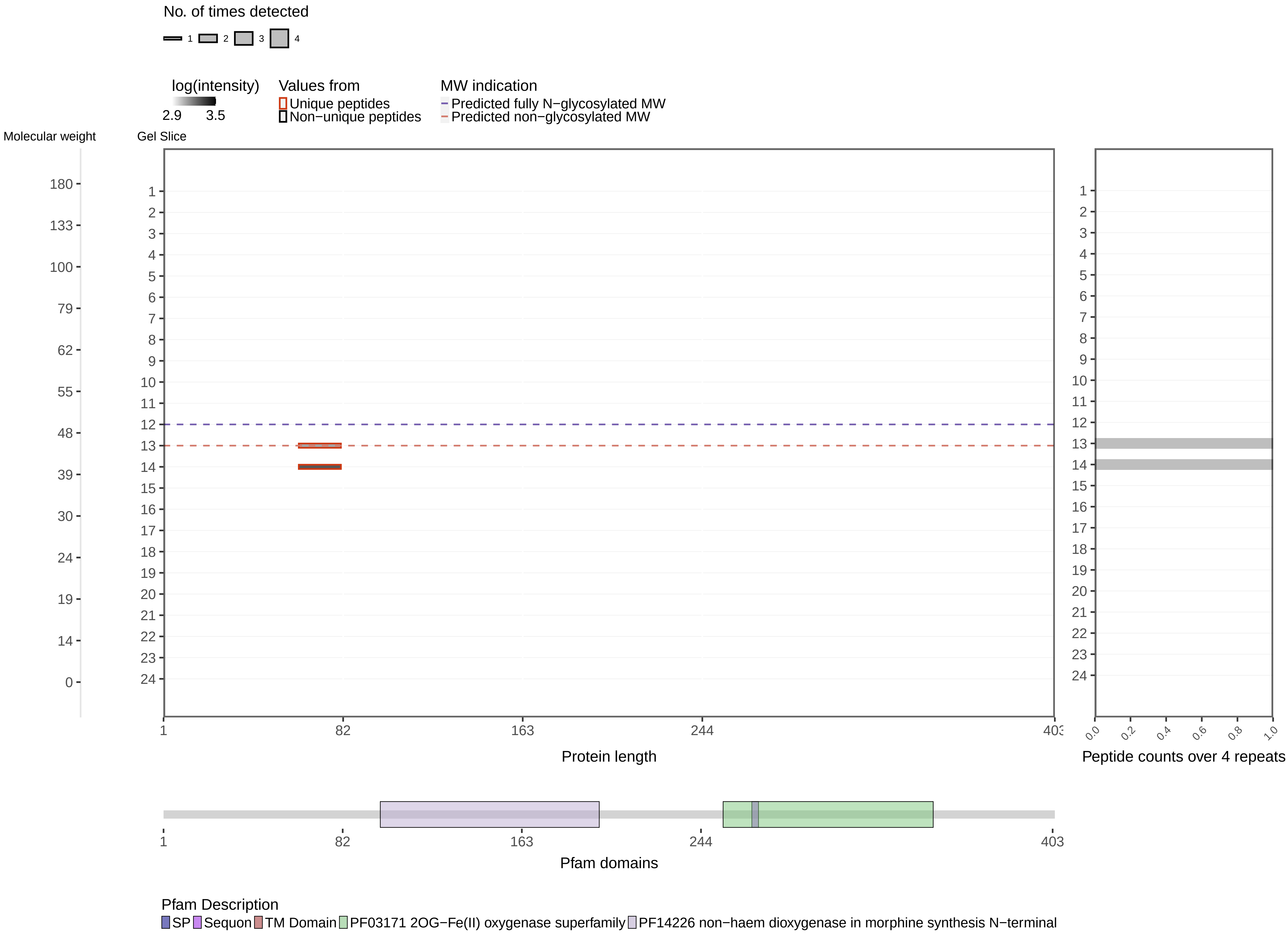


Protein sequence:

WRDTLGCIMAPNPPRSDELPETCRKELLEYSQHVRKLGITIFELLAEALGLNPNHLLEMECAKGHFILSHYYPPCPEPNKTFGITKHTDPDFFTILLQDHIGGLQIYHQNQWIDIHPVTGALVINLGDLLQLLSNDKFKSAEHRVLSKQIGPRISVACFFTTQFQPFDRLYGPIKELLSDENLPMYKETTVRDYVSYYNSKGLGARPSLLHLRL*

### NbL14g12700.1 C−Myc−binding_protein_homolog [MW = 14.44 kDa]

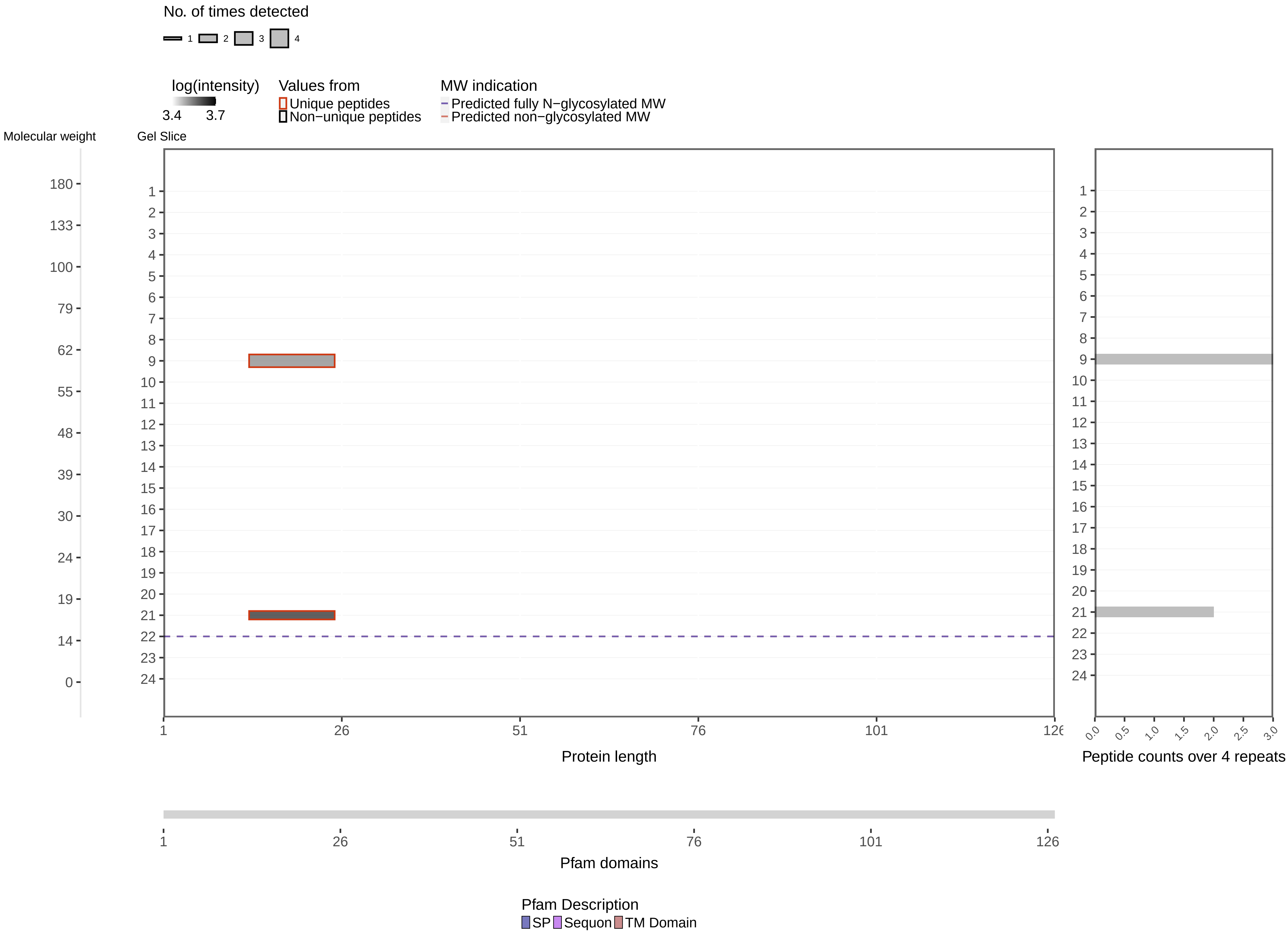


Protein sequence:

PTVSEYEKLQAELSDLQIRYDELLKEHQEKCRELEELKHTQVEASTKEPNVENAHEEELKSTVECSKEFKNAQFF*

NbL14g12630.1 alpha−glucosidase−like [MW − SP = 98.15 kDa]

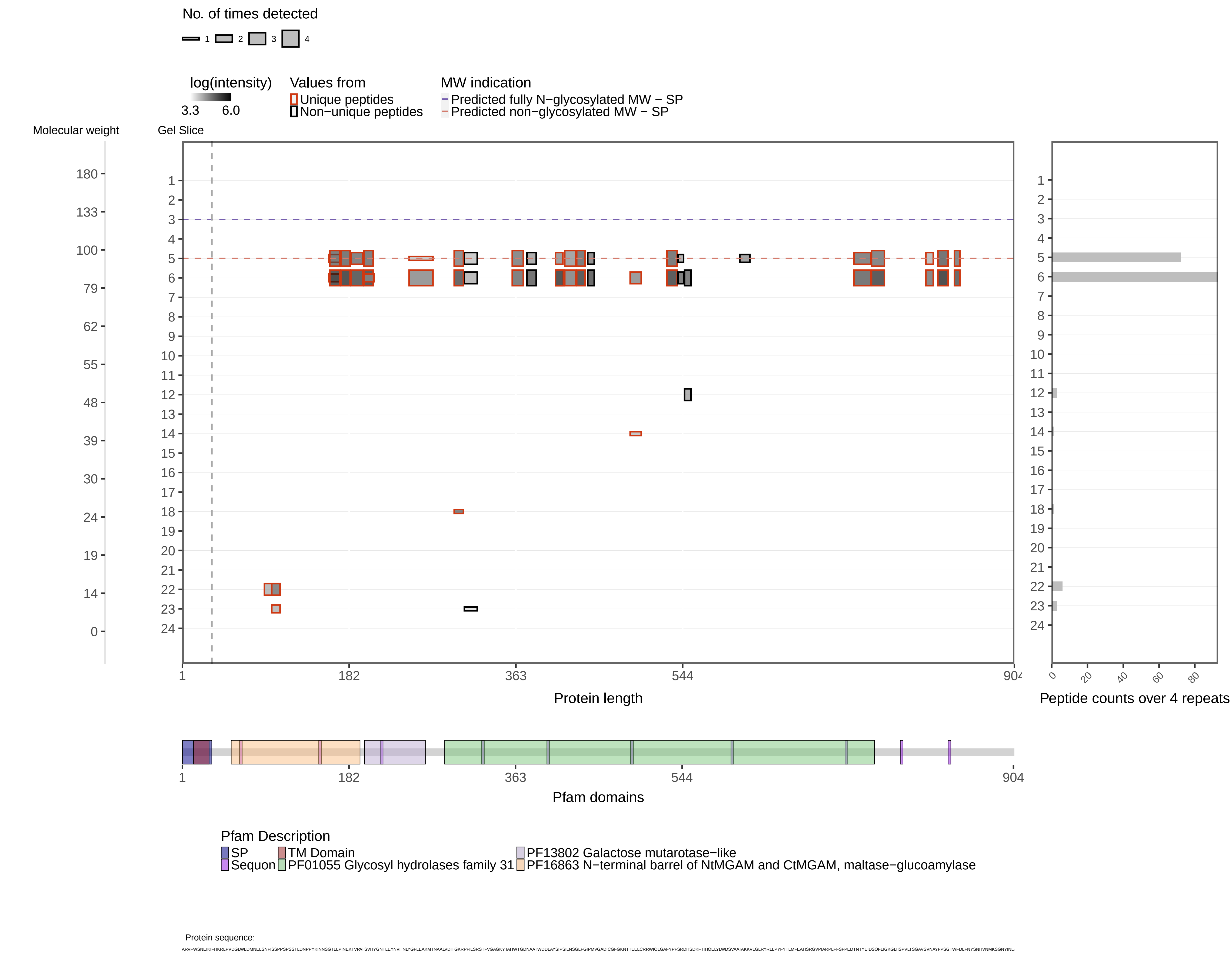

NbL14g12470.1 50S_ribosomal_protein_L10_chloroplastic [MW = 30.62 kDa]

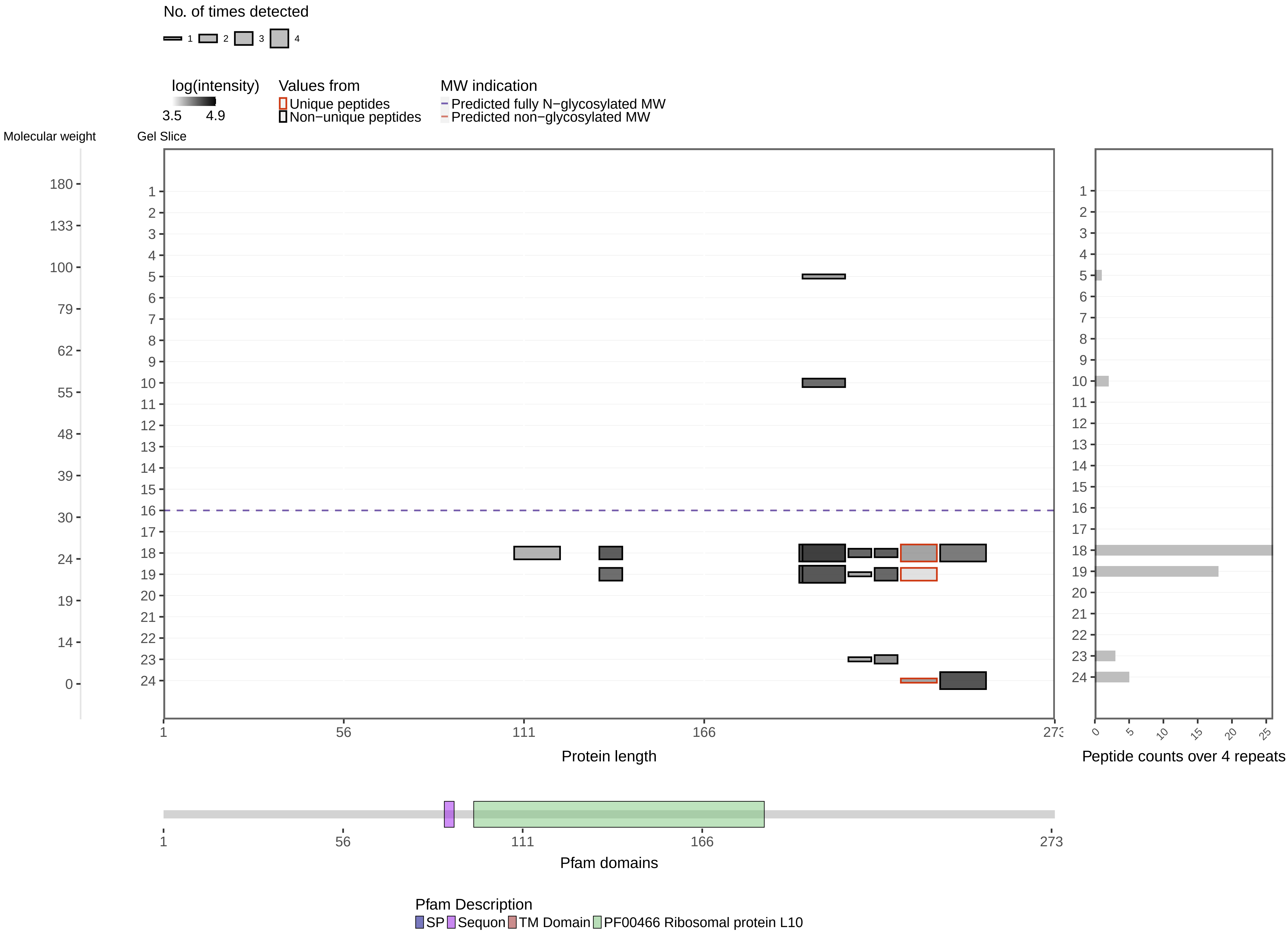


Protein sequence:

/KQFQELRSQLPDTTKLLVAKNTLVIKAIEGTKWEALKPCMKGMNAWLFVHSEEIPAALKPYRTFQKEKKLEDNDFTGAVFEGKFYGPEEFKALETLPTRAEIYSQLLGSLKGPASAVVGTIQAPARNLVMVLKAYVKKLEEEGGSQ*

NbL14g12380.1 psbP_domain−containing_protein_2_chloroplastic [MW = 28.01 kDa]

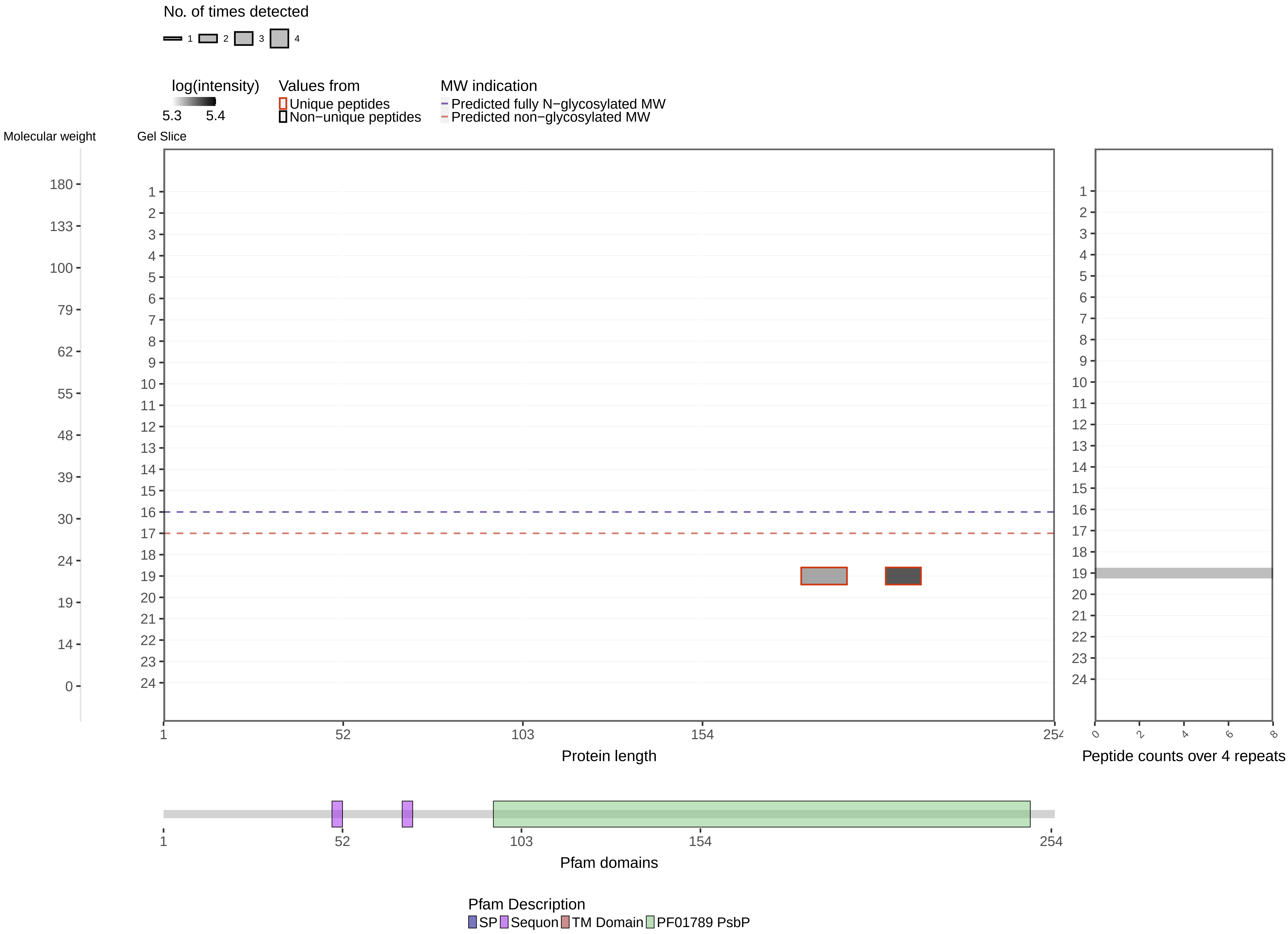


Protein sequence:

KVDKAGATVLFQDANKGSNNVGVVVIPVKITTLSQFGTPQFVADKLILAEKRKESTKEAEVVAVSERTGEGGLQVYEFEYKIDSTRGGMKRVLSAAFVASGKLYLLNIAHSDGLESPIDPVRRNTLEQILHSFDTAPST*

NbL14g12360.1 cytochrome_b−c1_complex_subunit_Rieske−3_mitochondrial [MW = 29.2 kDa]

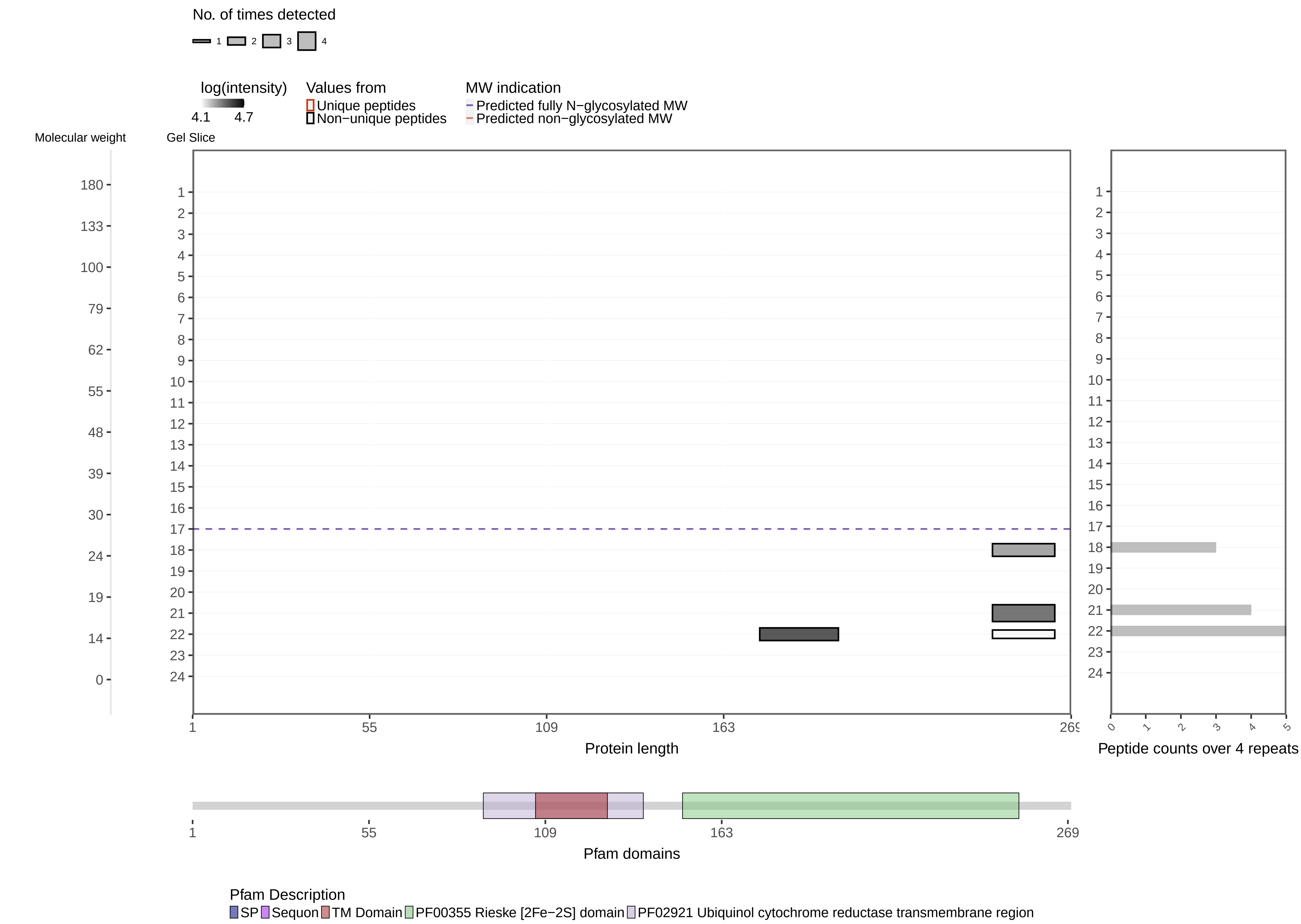


Protein sequence:

[illegible]LVRLLILKFVLSMSASKDVLALASLEVDLSSIEPGTTVTVKWRGKPVFIRRRTDEDINLANSVDLGSLRDPQQDAERVKNPEWLVVIGVCTHLGCIPLPNAGDFGGWFCPCHGSHYDISGRIRKGPAPYNLEVPTYSFMEENKLLIG*

NbL14g12350.1 glycine−−tRNA_ligase_mitochondrial_1−like [MW = 81.63 kDa]

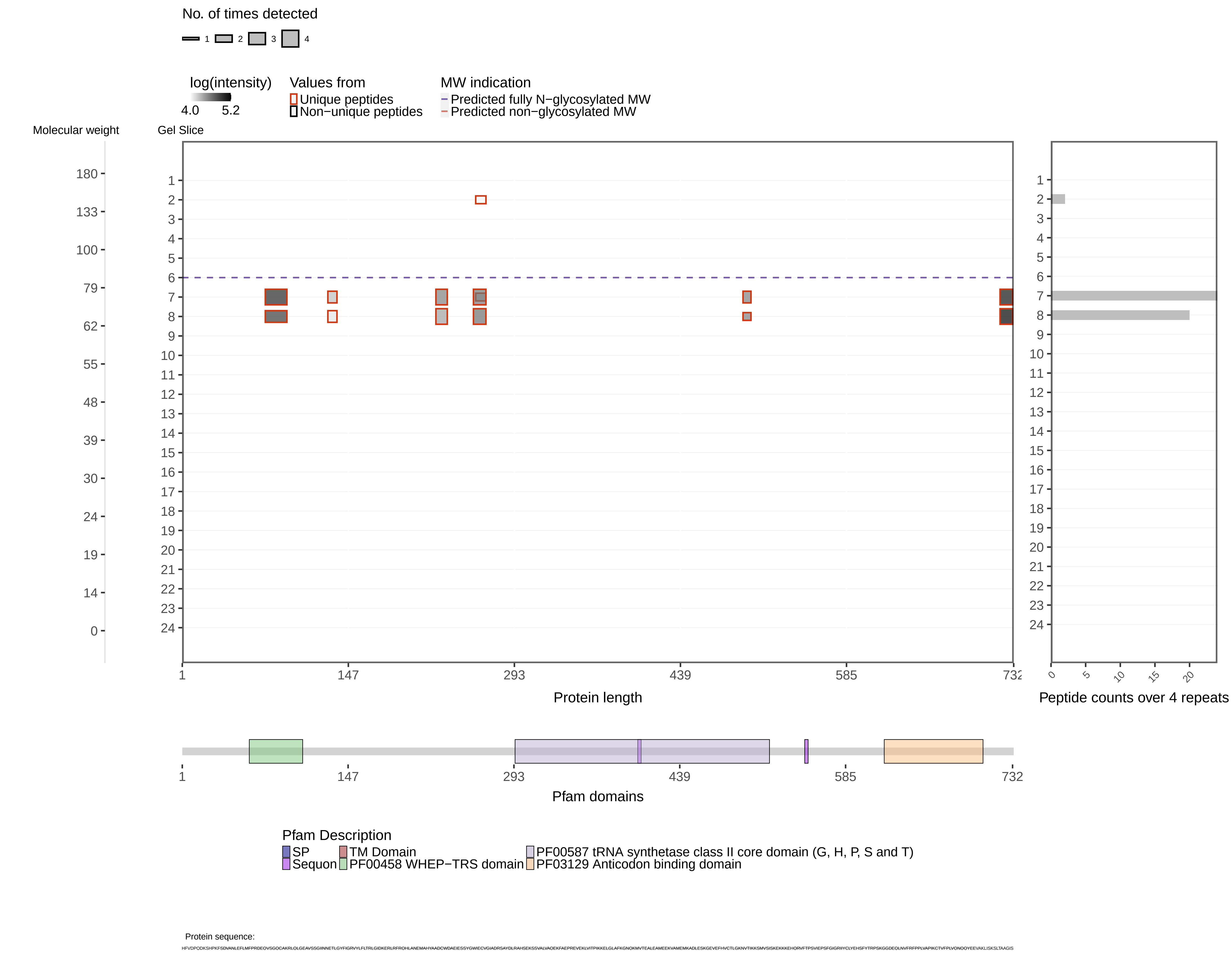

### NbL14g12330.1 transaldolase [MW = 49.64 kDa]

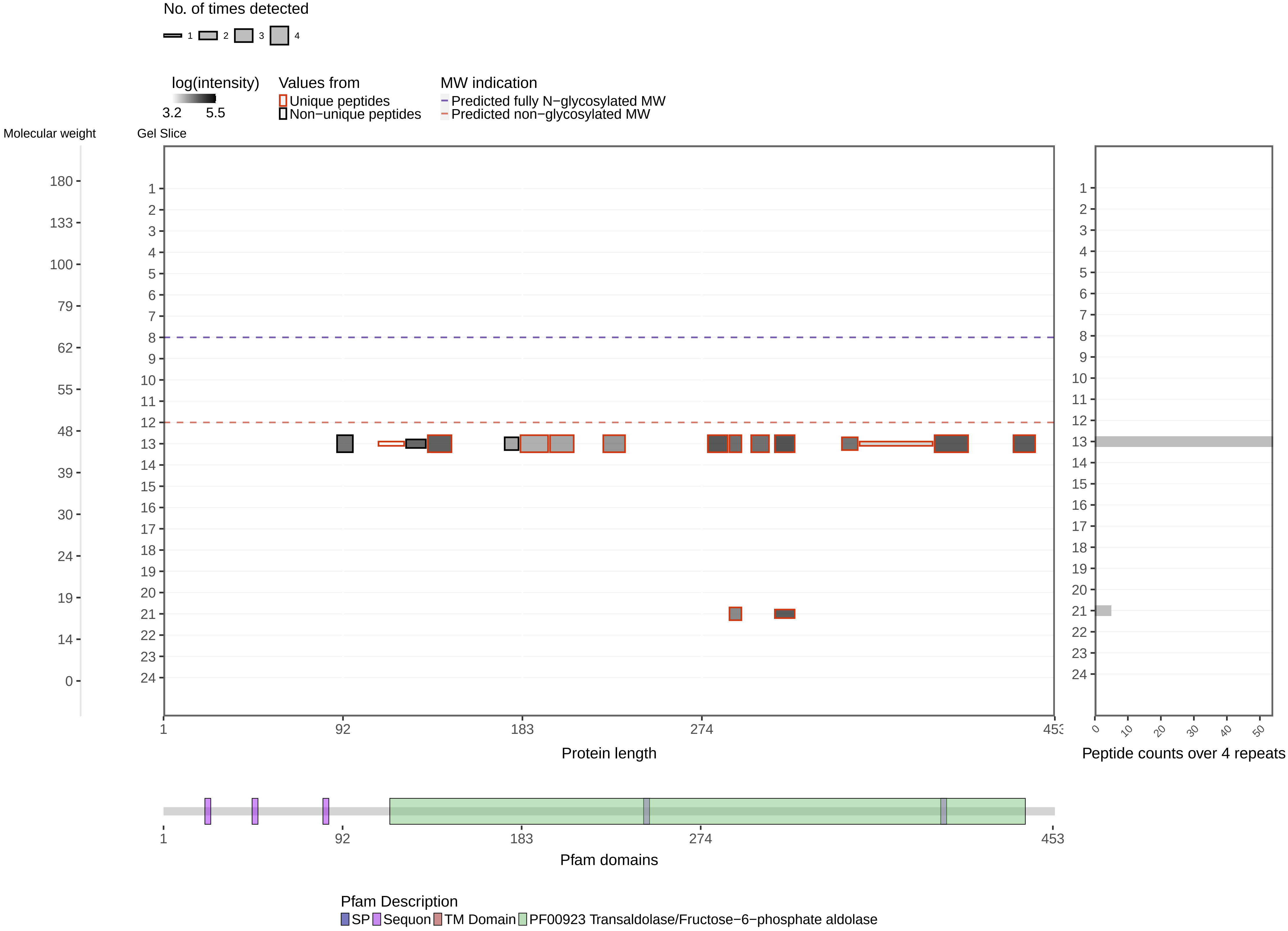


Protein sequence:

LHKKVERSNVYIKIPATAPCIPSIKEVISLGISVNVTLIFSLARYEAVIDAYLDGLEASGLSDLSRVTSVASFFVSRVDTLVDKMLEKIGTPEALDLRGKAANAQAALAYKLYQKKFSGPRWEALVKKGAKKQRLLWASTSVKNPAYPDTLYVDPLIGPDTVSTMPDQALQAFIDHGSVARTIDSNESEAEGIYSALEKLGIDWSFVGSQLELEGVDSFKKSFDSLLDSLQEKANTLKLVSL*

### NbL14g12320.1 peptidyl−prolyl_cis−trans_isomerase_FKBP19_chloroplastic−like [MW = 27.82 kDa]

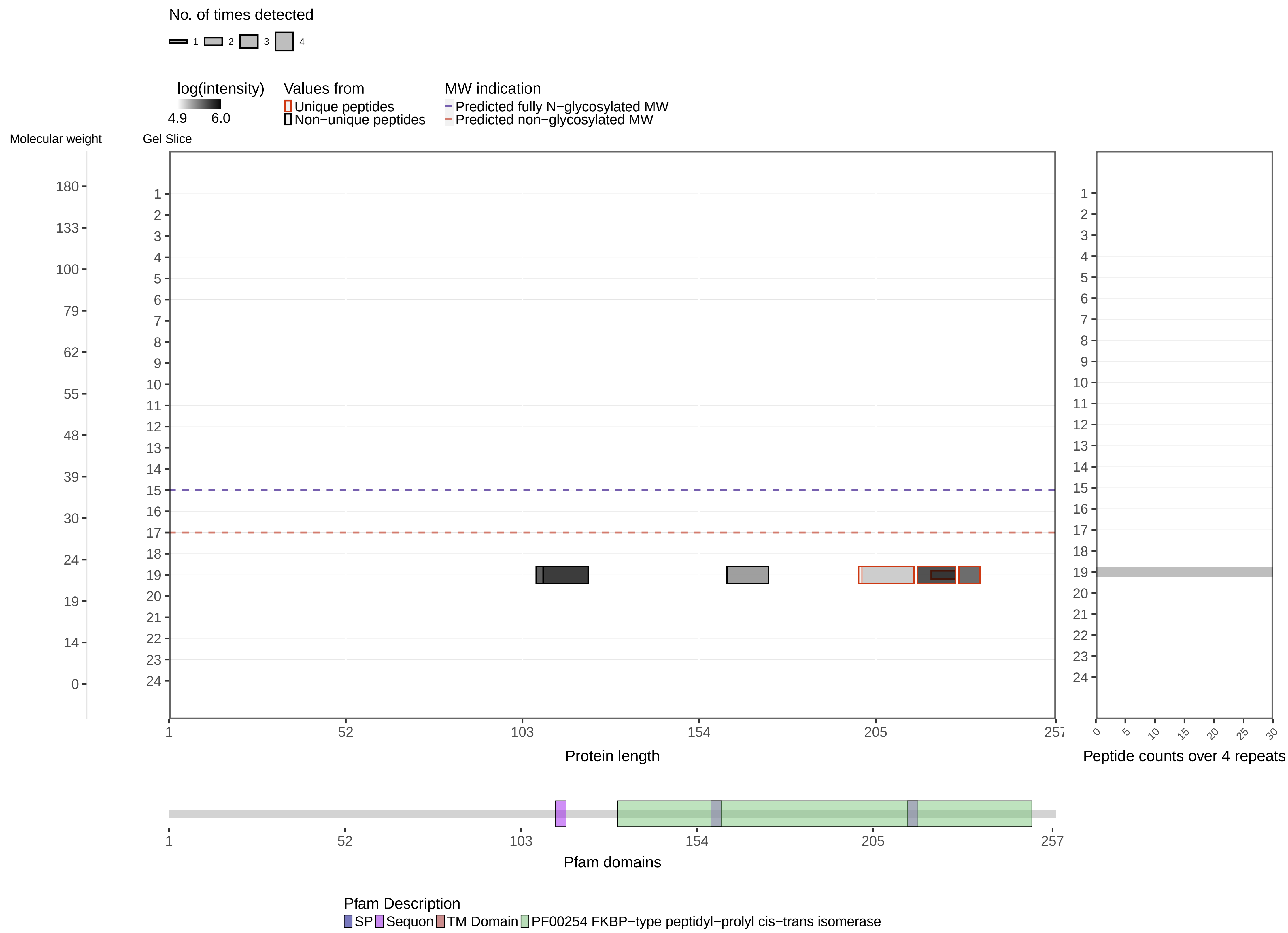


Protein sequence:

TQSGLQYKDLRVGSGPTPKIGETVVVDWDGYTIGYYGRIFEARNKTKGGSFEGDDKDFFKFRVGSQEVIPAFEEAITGMAIGGIRRIIVPPELGYPDNDYNKTGPRPTTFSGQRALDFVLRNQGLIDKTLLFDIELLKIVPK*

### NbL14g12130.1 pseudouridine−5'−phosphate_glycosidase−like [MW = 36.11 kDa]

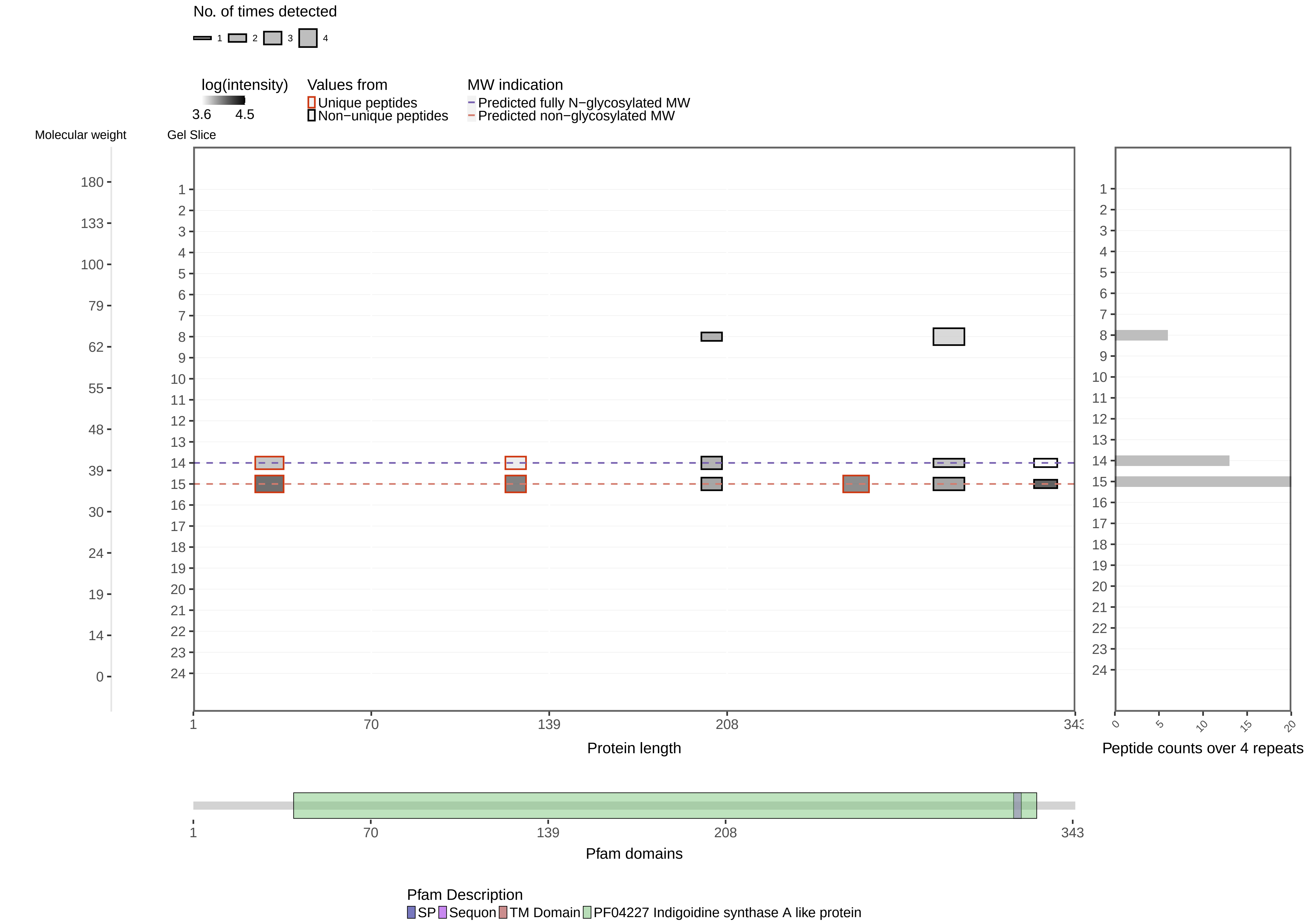


Protein sequence:

### NbL14g12100.1 eukaryotic_initiation_factor_4A−3−like [MW = 46.04 kDa]

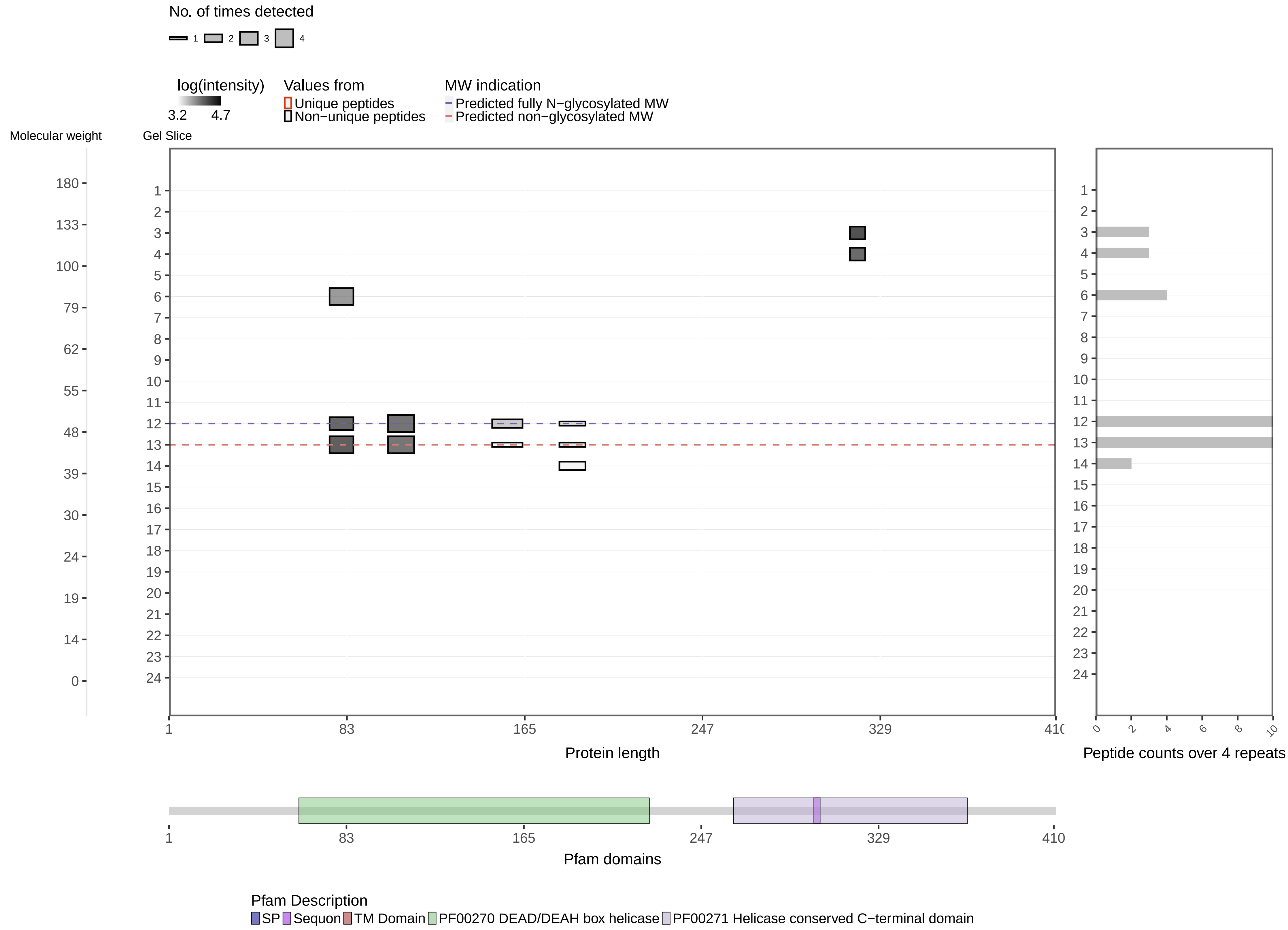


Protein sequence:

.SRGFKDQIYDVYRYLPPELQVVLISATLPNEILEITSKFMTDPVRILVKRDELTLEGIKQFFVAVEKEEWKFDTLCDLYDTLTITQAVIFCNTKRKVDWLTSKMRENNFTVSSMHGDMPQKERDAIMAEFRGGTTRVLITTDVWARGLDVQQVSLVINYDLPNNRELYIHRIGRSGRFGRKGVAINFVKSDDIKILRDIEQYYSTQIDEMPMNVADLI*

### NbL14g12070.1 uncharacterized_LOC107823533 [MW = 25.31 kDa]

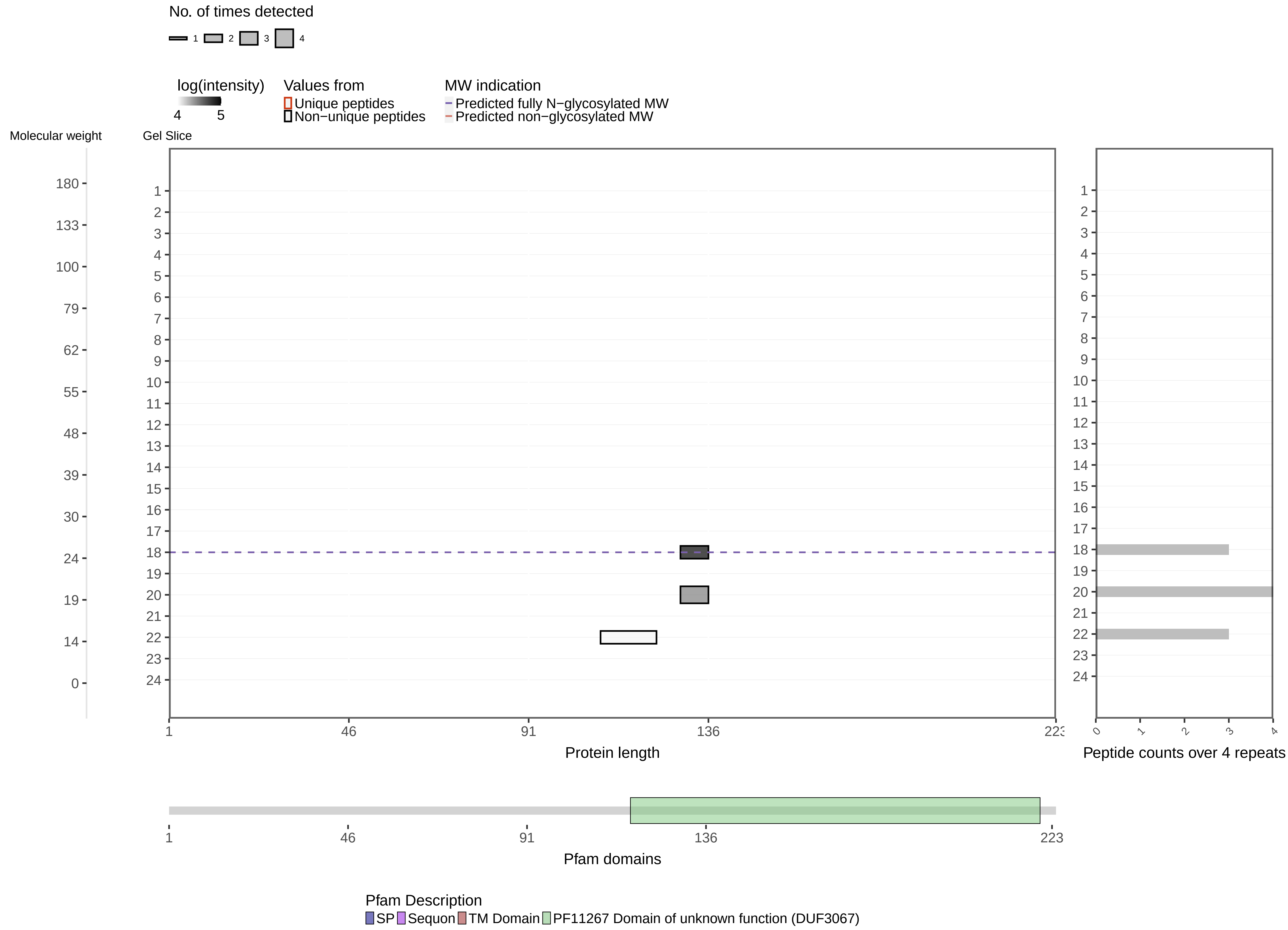


Protein sequence:

RNNLERIVGRDDSAFSGIDLAALIRNKYGRSYDVQLIKKEFMGKNLLALNVMWKYREQRSFPLTEEEYLLRLDDVANTLKCWGAVSHVRNSLEKVKERPRIGKAVSIFIDMDESGGRANEWIYK*

### NbL14g12050.1 histidinol−phosphate_aminotransferase_chloroplastic [MW = 46.28 kDa]

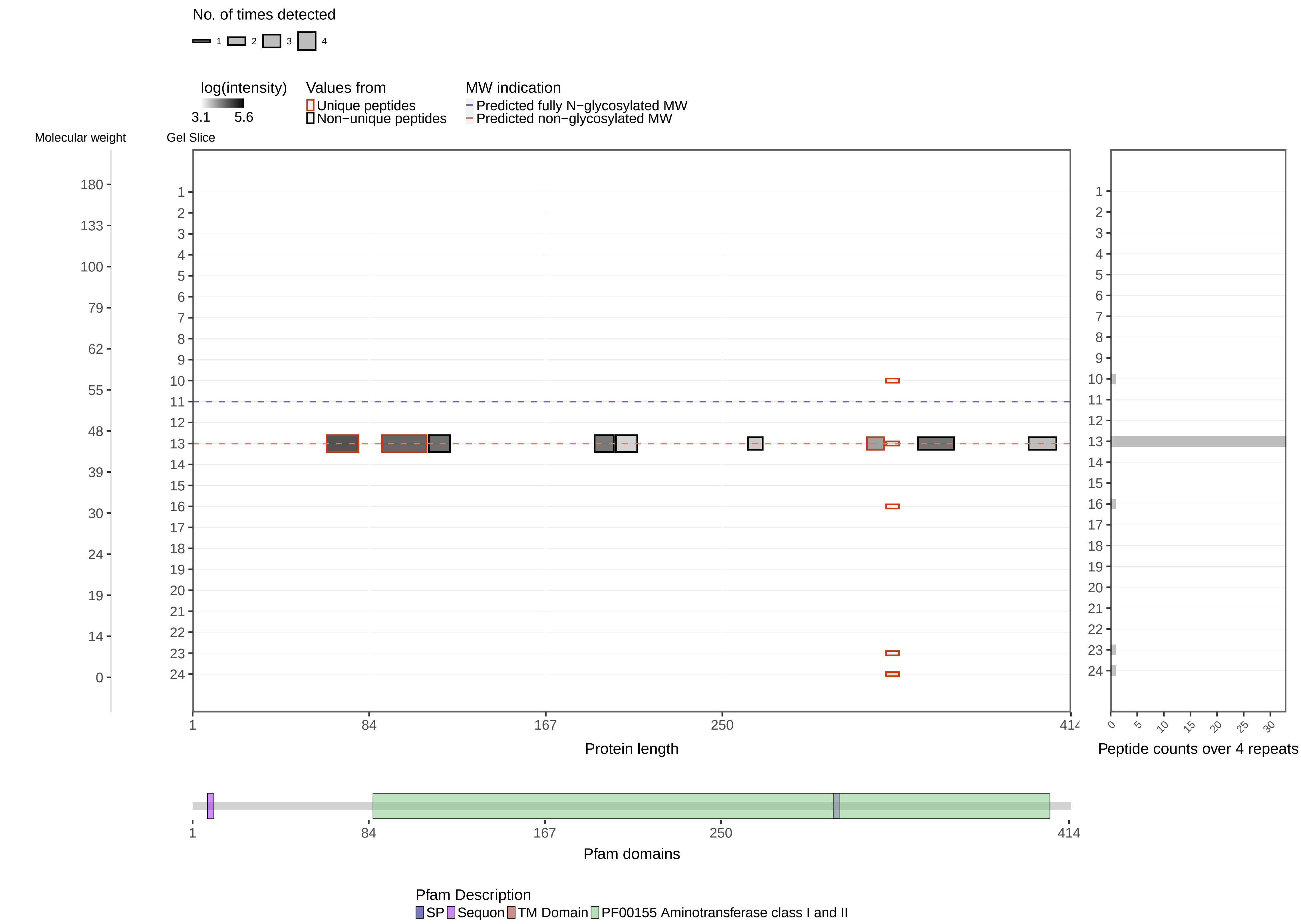


Protein sequence:

SLDVERIAEVVEHEKPKCIFLTSPNNPDGSIIDDETLLEILDLPILVILDEAYVEFSGMESKMKWVKKHENLIVLRTFSKRAGLAGLRVGYGAFPMSIIEFLWRAKQPYNVSVAAEVAACAALKNPAYLENVKMALVQERERLFNLLKEVPFLDPYPSYSNFILCKVTSGMDANKLKEDLATMGVMIRHYNSKELKGYVRVSVGKPEHTEALMKCLKHFV*

NbL14g12010.1 ATP−dependent_Clp_protease_proteolytic_subunit_2_mitochondrial−like [MW = 30.99 kDa]

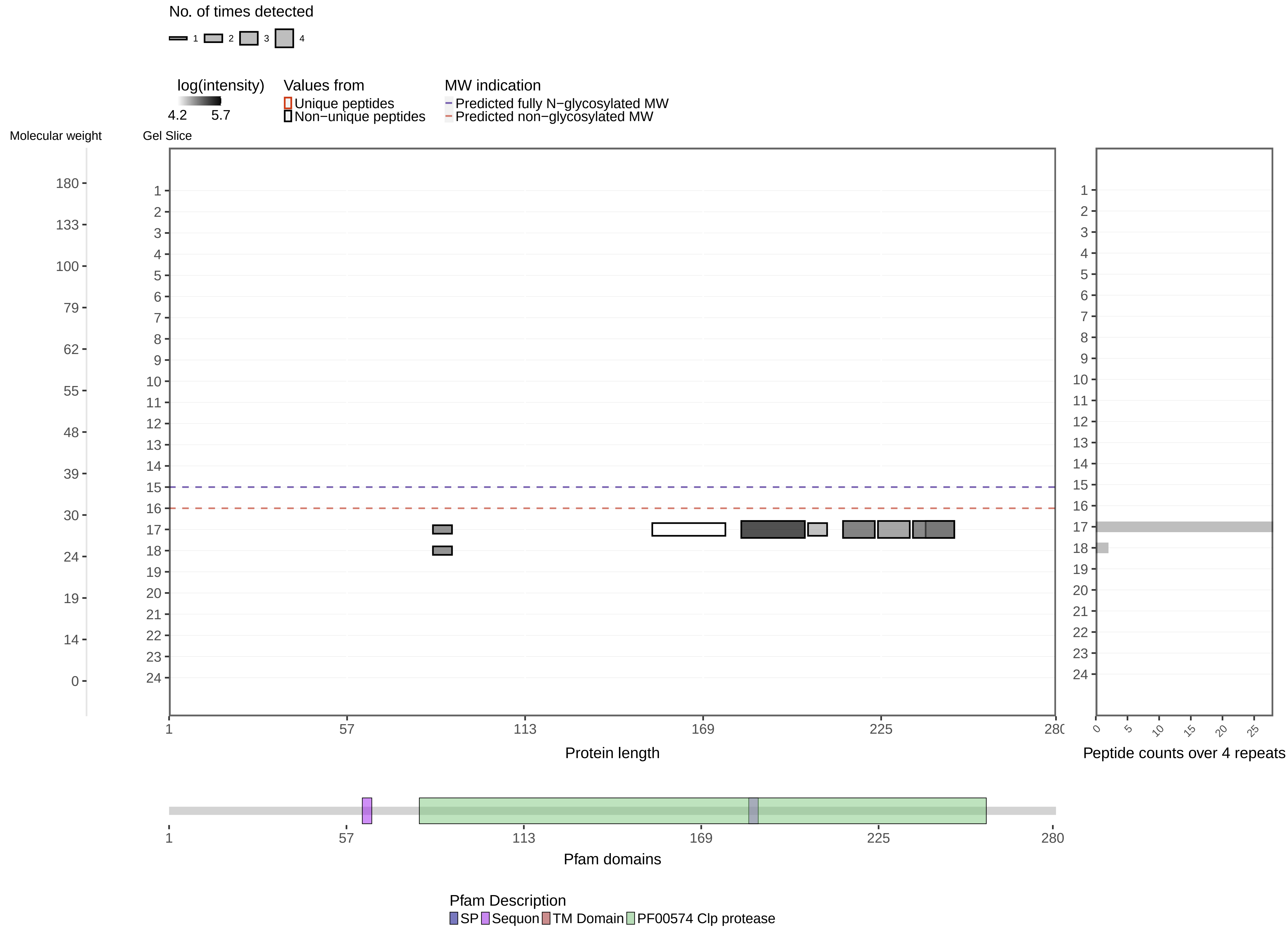


Protein sequence:

IMYLNSPGGAVTVGLAIYDTMQYIRSPIHTICLGQAASMGSLLLAAGAKGERRSLPNASVMIHQPSGGYSGQAKDLTIHTKQIVRVWDTLNDLYAKHTGQPIEIIQKNMDRDYFMTPEEAKEFGIIDEVIDERPMALVTDAVANEAKEKGSS*

NbL14g11940.1 glutamate−1−semialdehyde_21−aminomutase_chloroplastic [MW = 50.98 kDa]

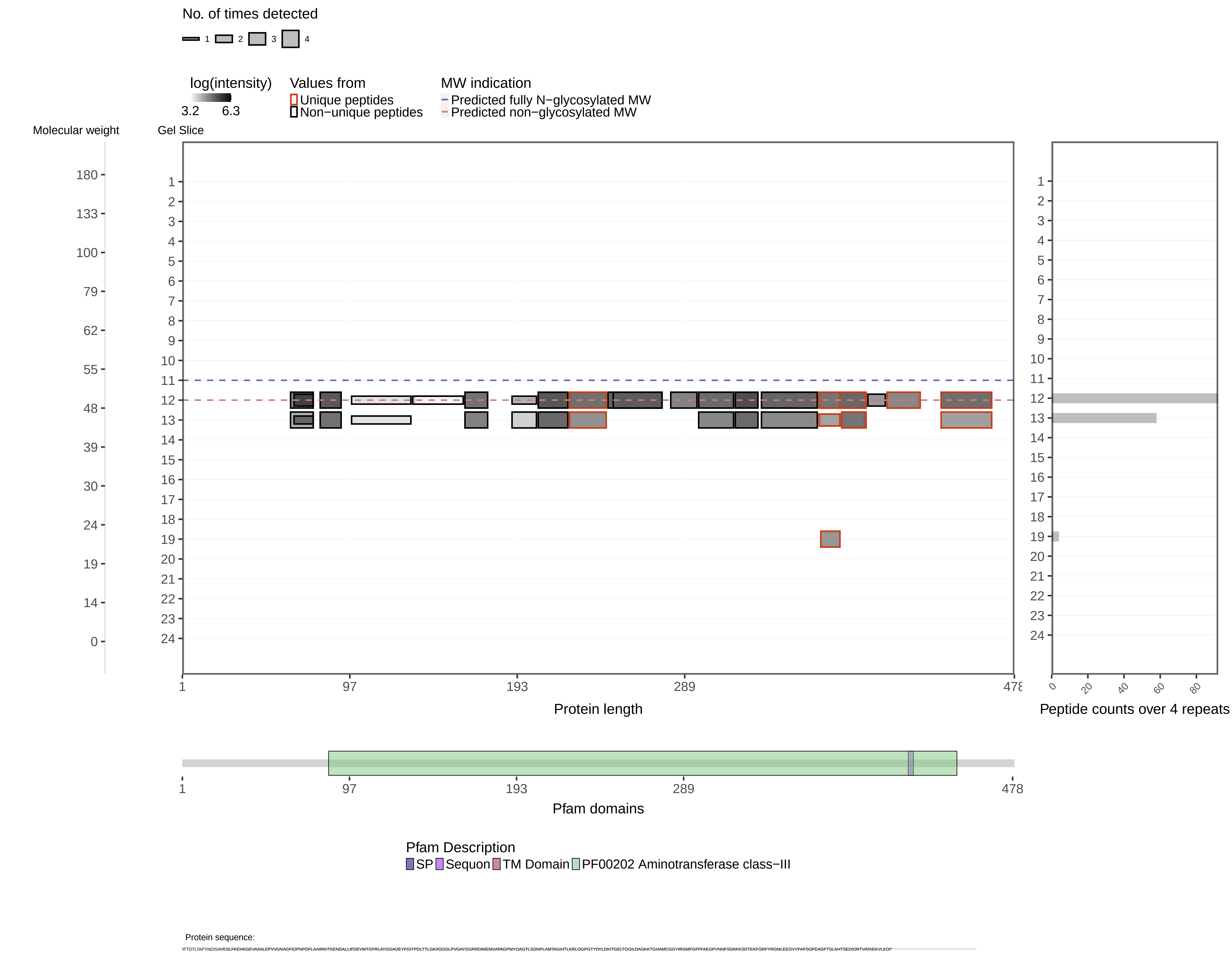

NbL14g11700.1 probable_leucine−rich_repeat_receptor−like_protein_kinase_At1g35710 [MW − SP = 62.93 kDa]

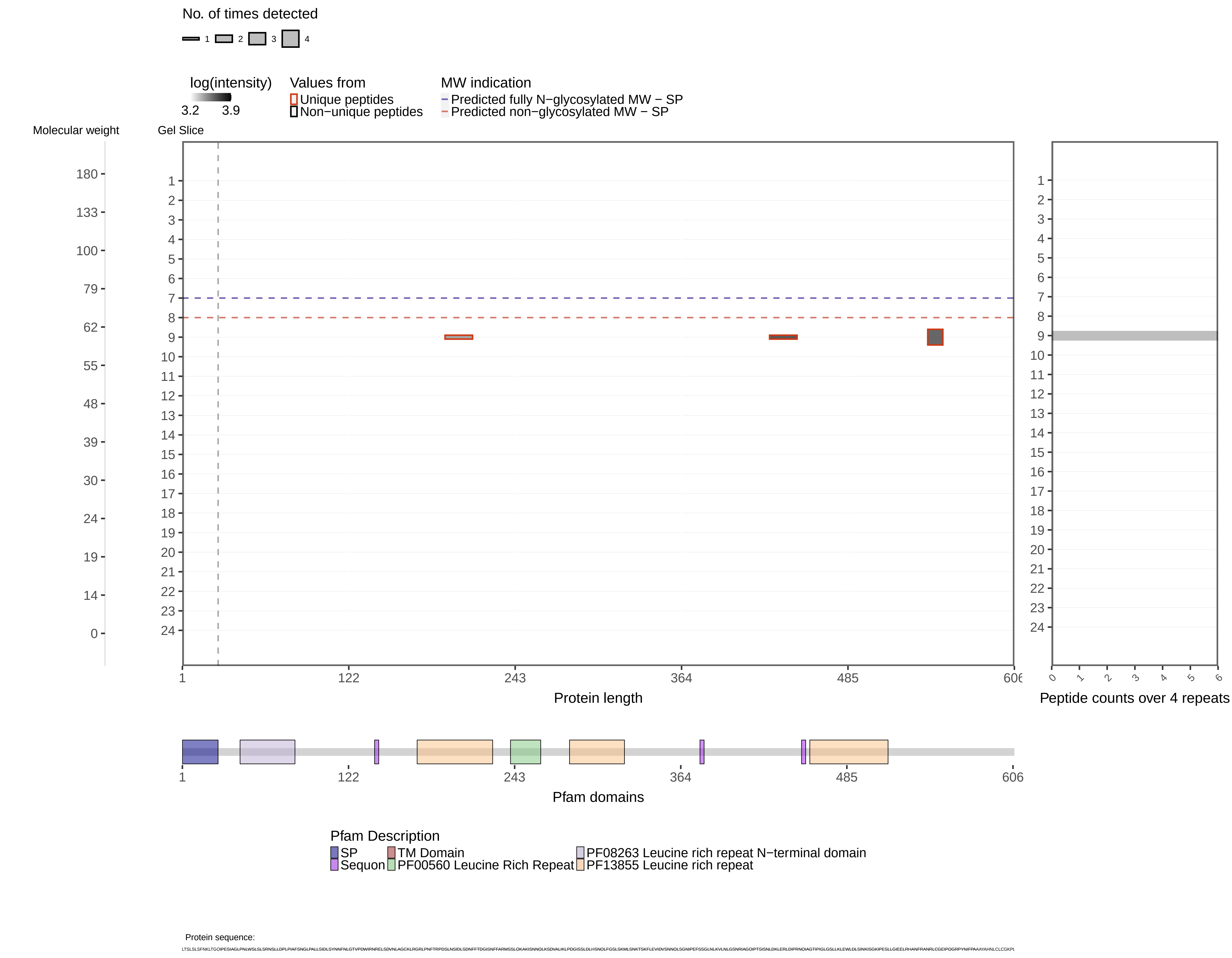

### NbL14g11690.1 40S_ribosomal_protein_S26−3−like [MW = 14.36 kDa]

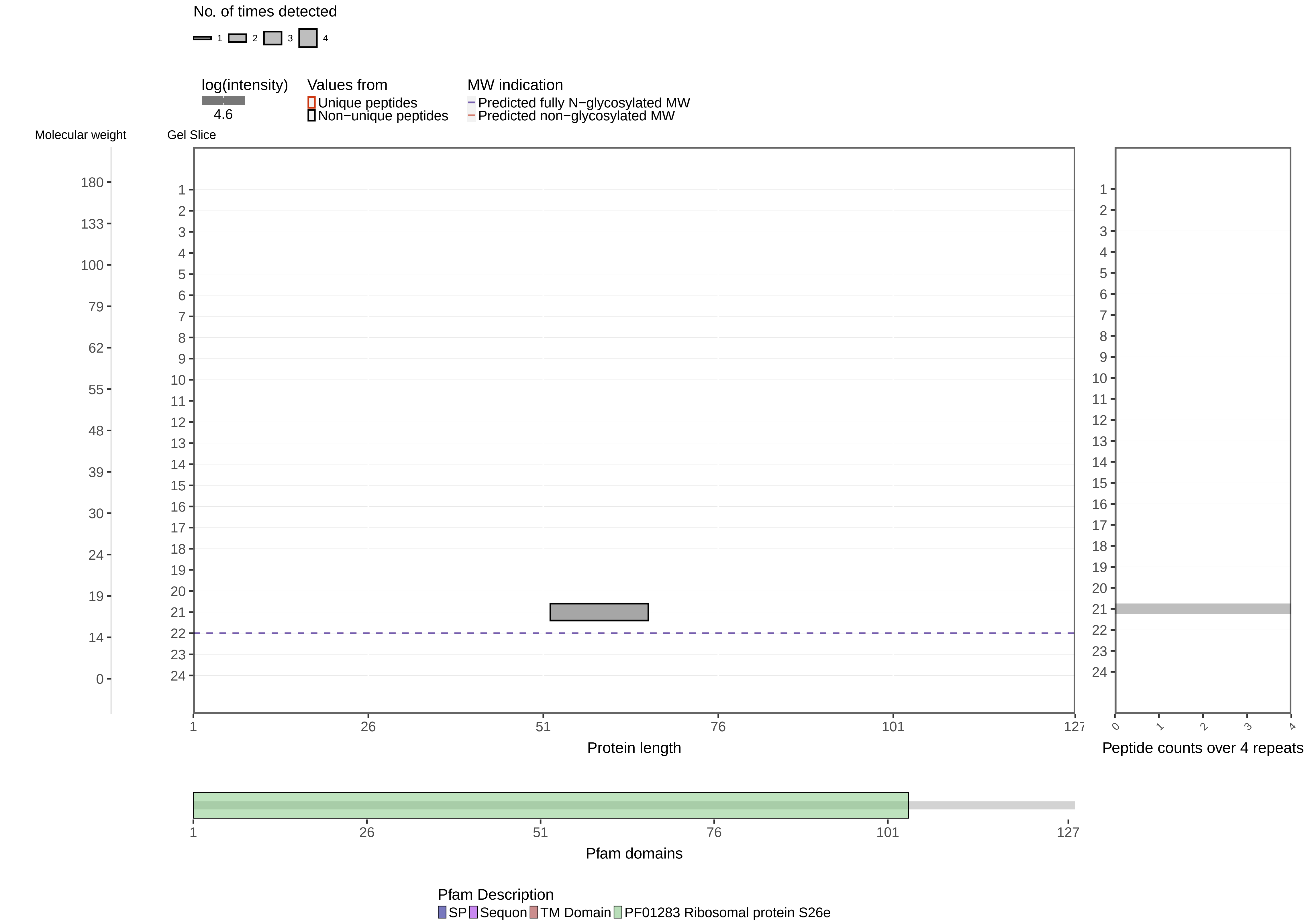


Protein sequence:

RDVQEACAFELYTLPKLYLKMQYCVSCAIHSKVVRVRSVTDRRVREPPQRFRRPRDDAPKPGQAPRPAGAPAAART*

NbL14g11680.1 uncharacterized_LOC107810722 [MW = 21.15 kDa]

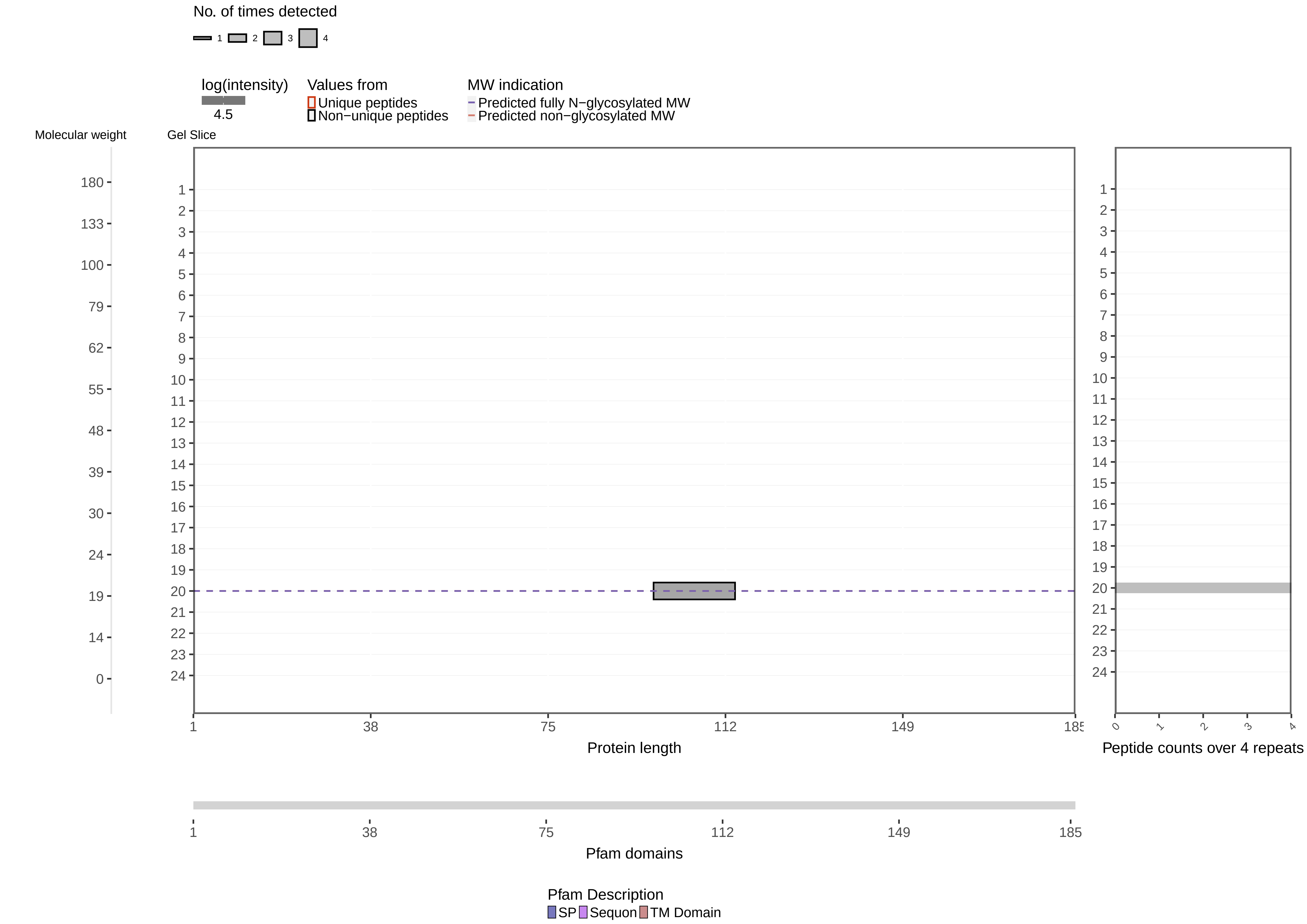


Protein sequence:
[illegible]RASLKPPGPYAWVKYVPGEPISPNNPNEGSVKRRNEKKRIGQQKAFILAEKKKRKAQLQEAHRKKMNKRVERKMAAVARDRAWTERLAELKKLEEEKKAAMA*

NbL14g11660.1 mitochondrial_intermembrane_space_import_and_assembly_protein_40−like [MW = 17.52 kDa]

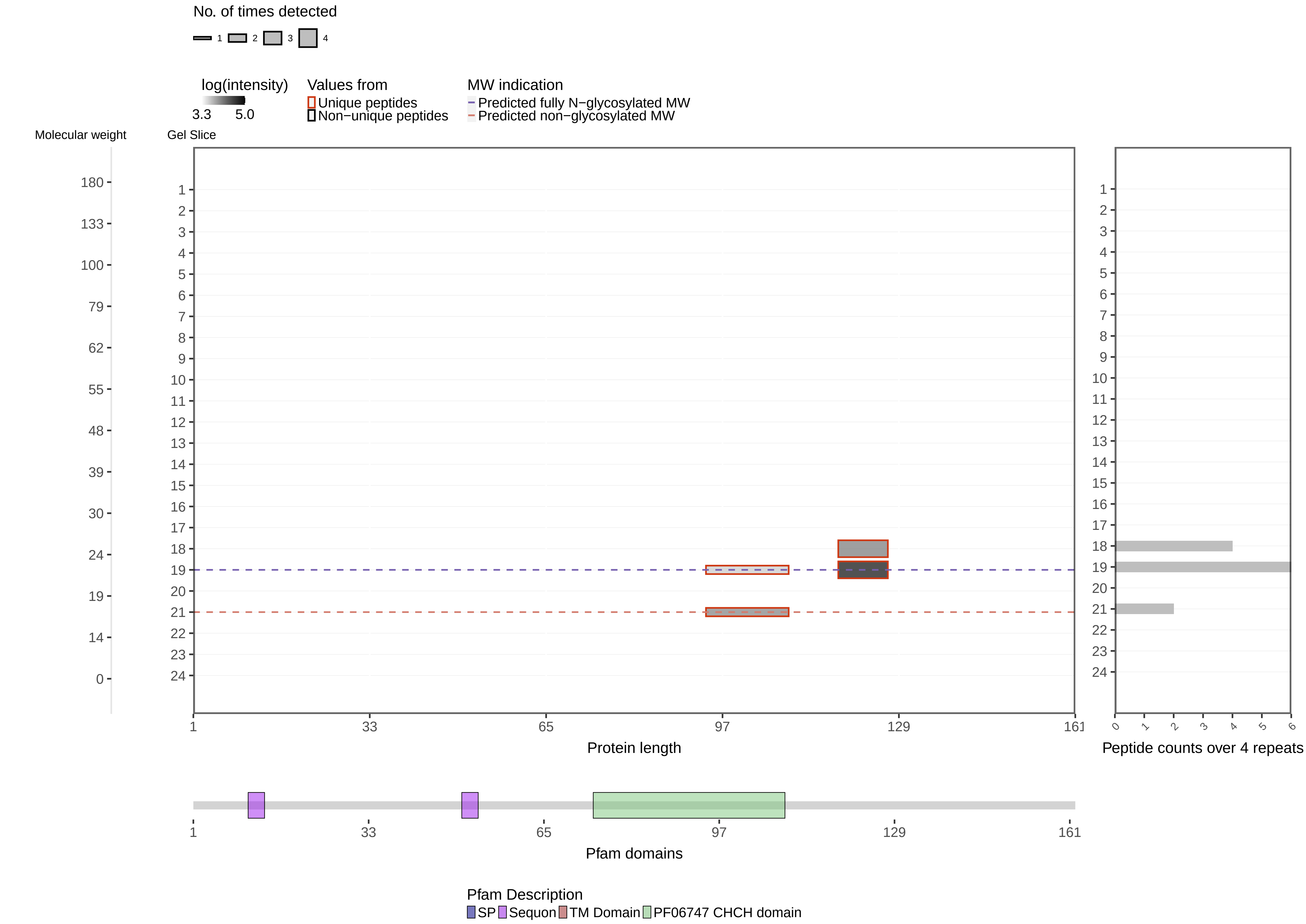


Protein sequence:

HLRSGPCGNQFSDAFLCFLKSTADEKGSDCVNPFVALQSCIKVNPNAFSKDILEDDDNTRKQDEVKKEETPKQEYRIIPPIWSVESKGSKRKA*

NbL14g11630.1 alpha−galactosidase_1−like [MW − SP = 44.49 kDa]

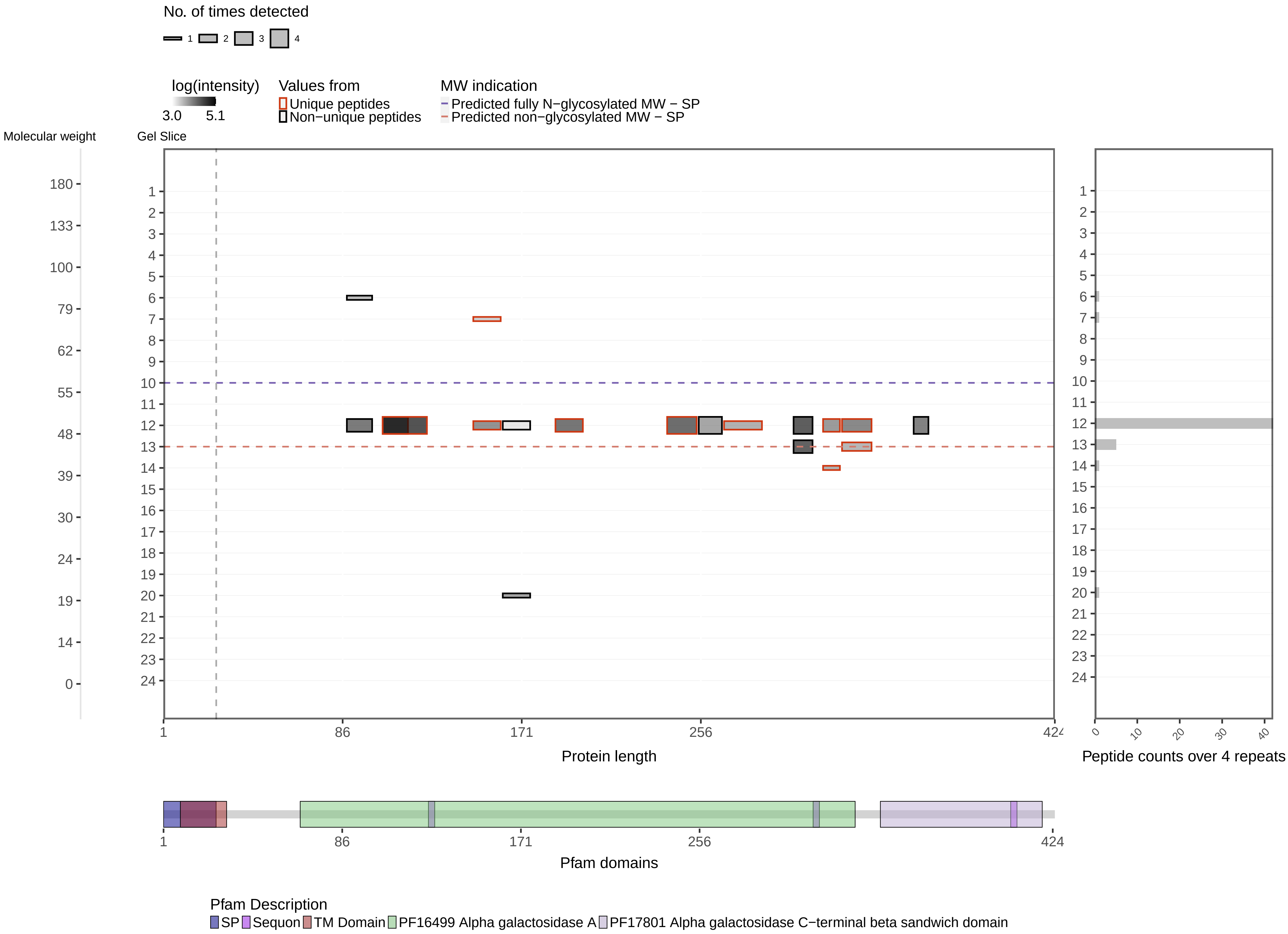


Protein sequence:

VRYPVMTRALMNAGRPIFFSLCEWGDLHPALWGGKVGNSWRNTNDITDTWESMISRADQNEVYADLARPGGWNDPDMLEVGNGGMSKDEYIVHFSLWAISKAPLIIGCDVRNITKDTMDILANKEVIAVNQDEIGVQGKKVRMEGDIEVWAGPLSGYRVALVLLNRGPQKYEITAHWDDTGIPPNSVVEARDLWEHKTLKTRFKGNLTATVNSHACKMYVLKPIA*

### NbL14g11610.1 lateral_signaling_target_protein_2_homolog [MW = 18.77 kDa]

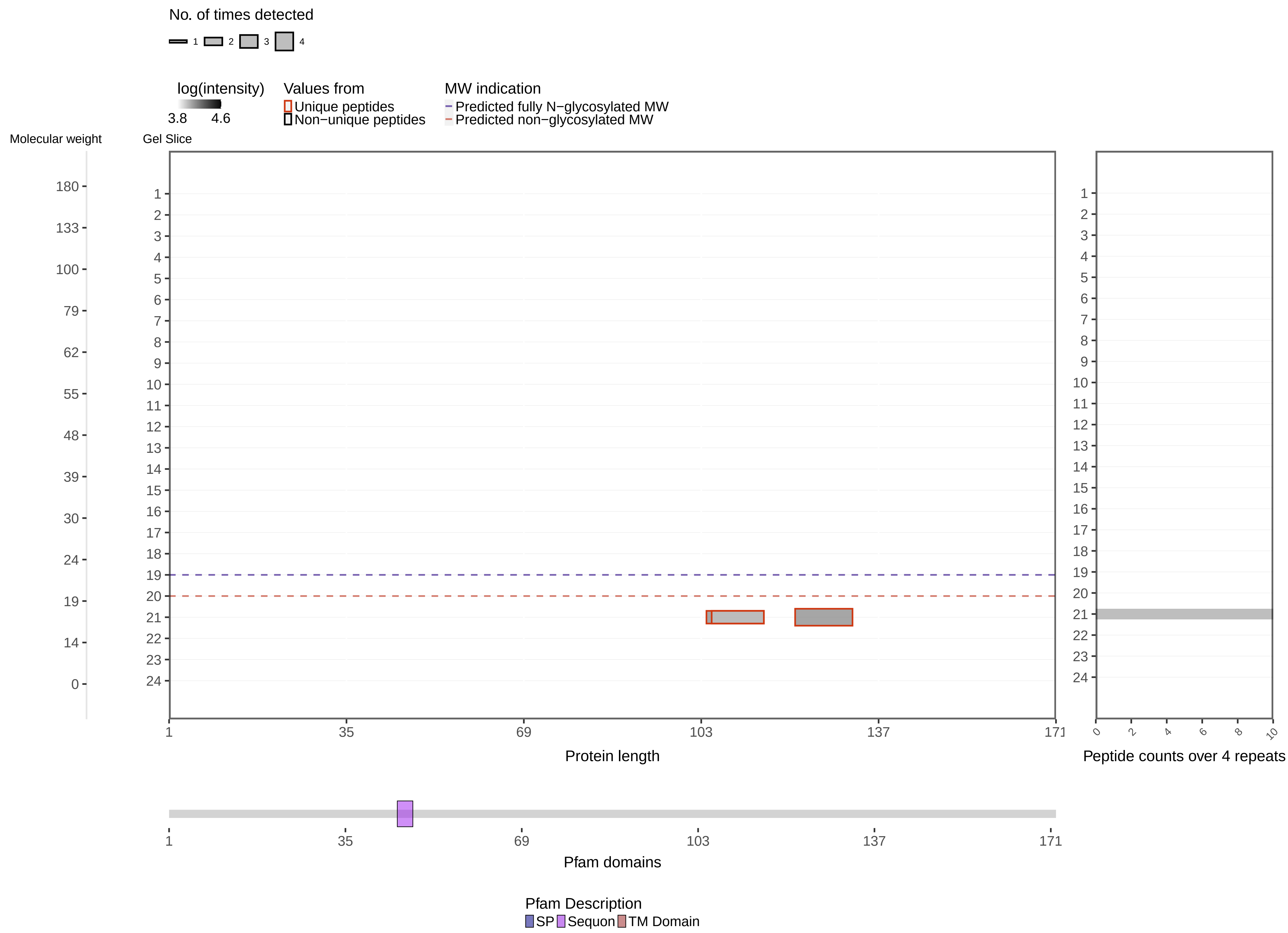


Protein sequence:

MPSVRWTSLSSSRRSSQIVRMAPEEEKMTRRSPLDFPIEWERPKPGRRPDIFPQFSPMKTPLPPPLPADPPEEDEEEEEKKEEEEDPEKEDPDKPDE*

NbL14g11580.1 vegetative_cell_wall_protein_gp1−like [MW − SP = 8.63 kDa]

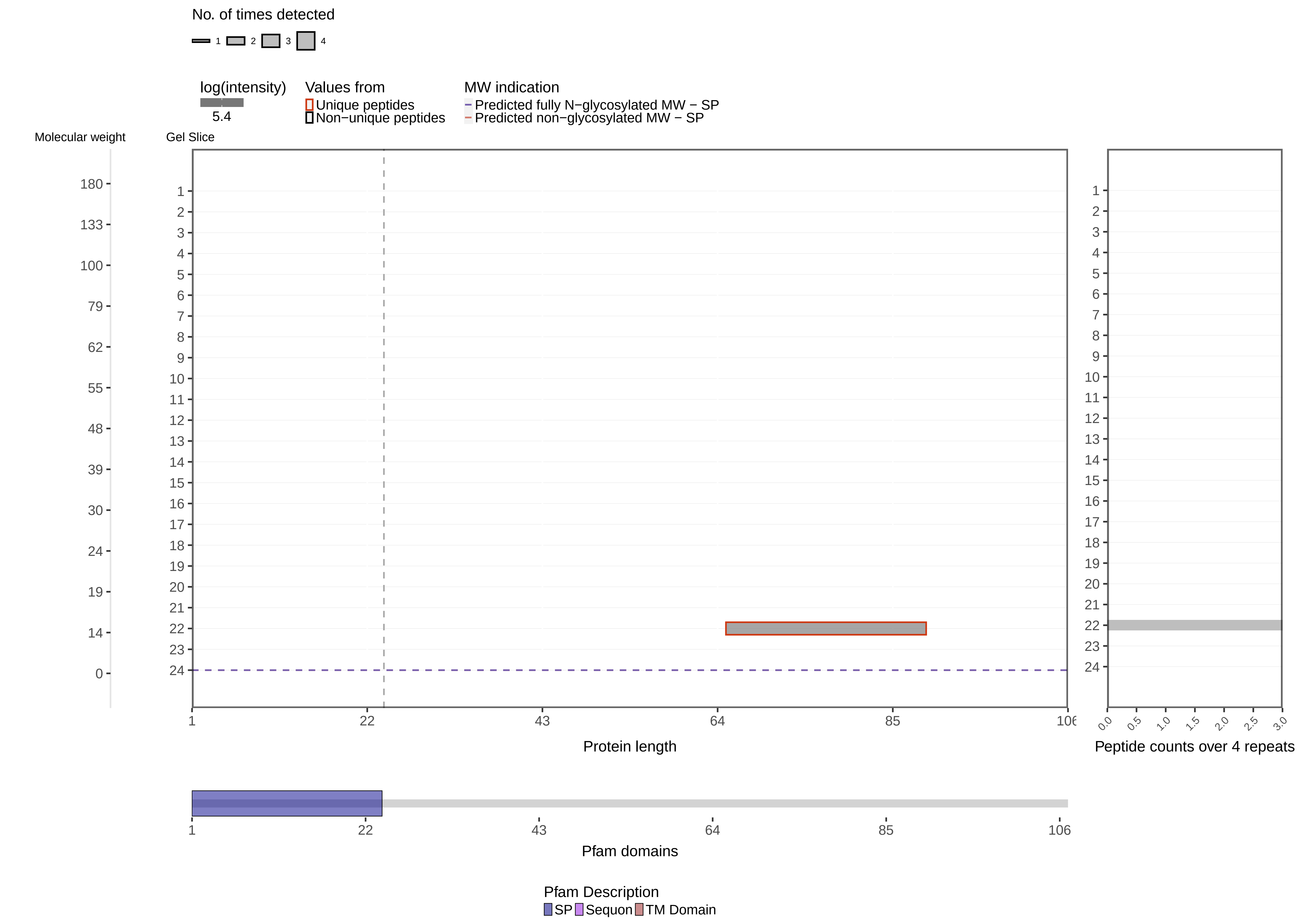


Protein sequence:

LPKPELPALPQPGLPTLPTLPMPKLPLPQLPMPQLPTLPKPDLPVPIPKLPLPELPVPLPPLPSP*

NbL14g11400.1 NADH_dehydrogenase_[ubiquinone]_1_beta_subcomplex_subunit_7−like [MW = 12.2 kDa]

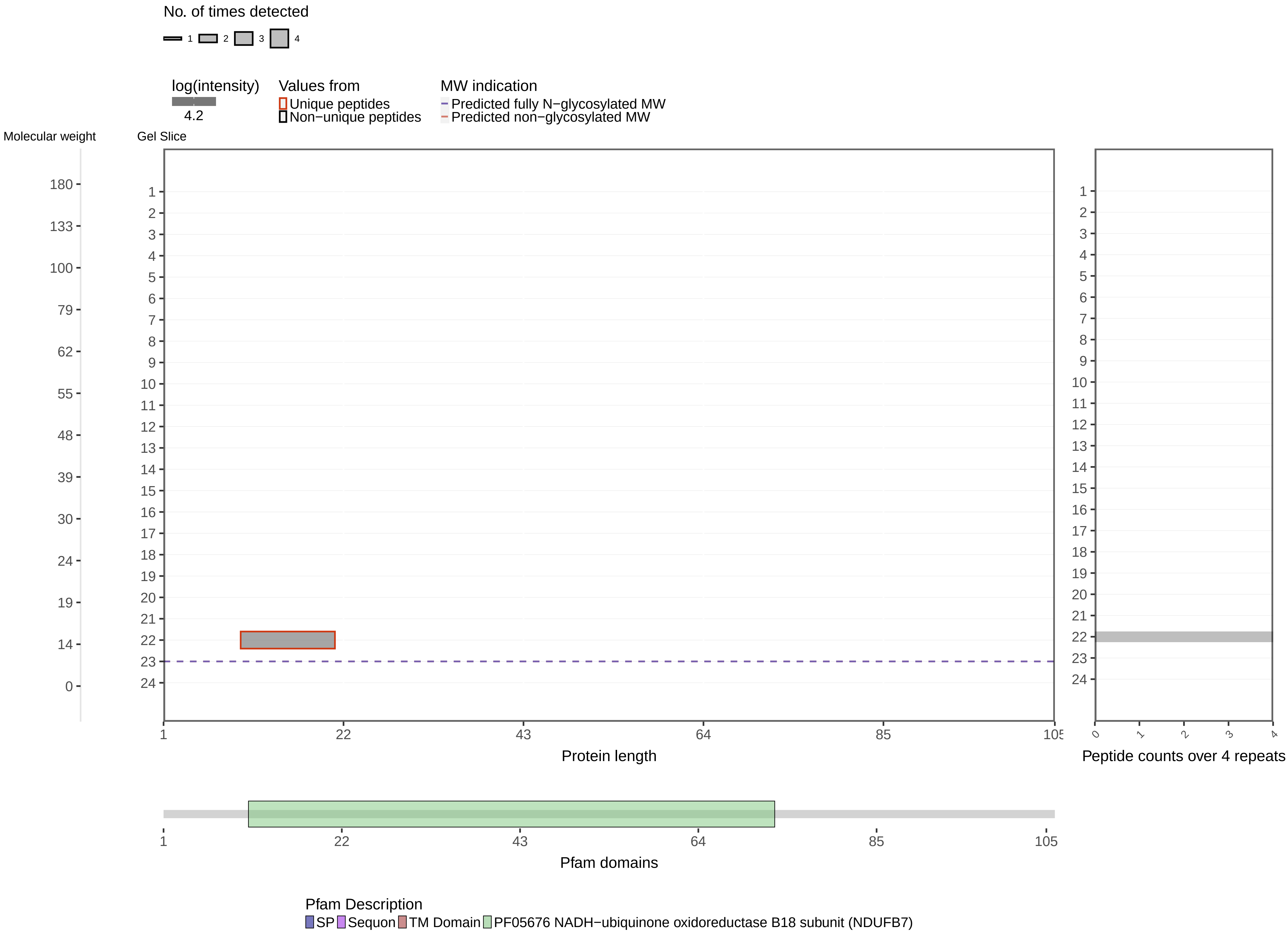


Protein sequence:

CRQSEFYLPWKCEDERHSYEKCEYELVMERMLQMQKIREEEARLKQIGKAQAQQTIPLIPKTANA*

### NbL14g11260.1 putative_4−hydroxy−4−methyl−2−oxoglutarate_aldolase_2 [MW = 17.76 kDa]

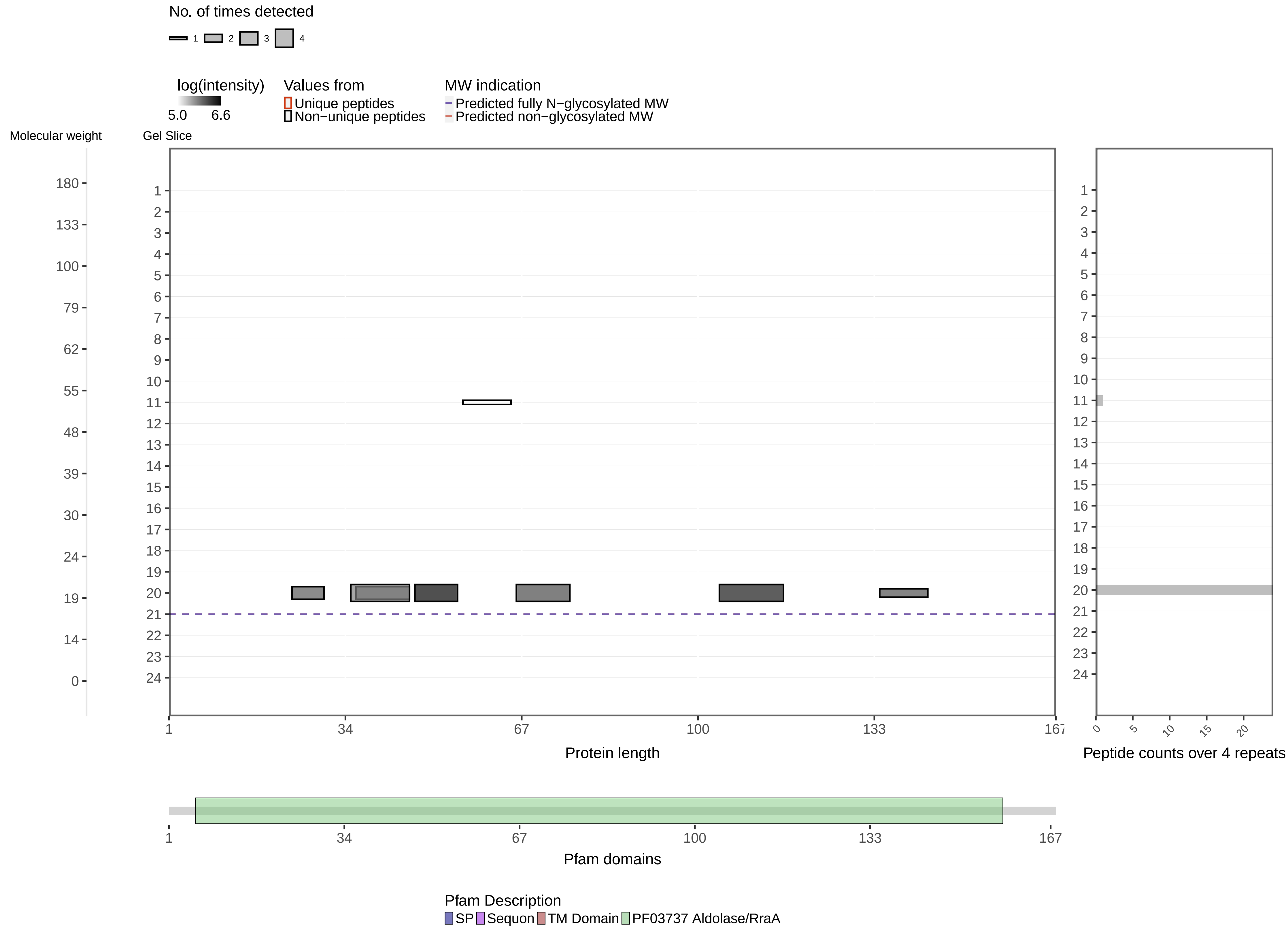


Protein sequence:

GGGSLRCAILGGNPVVQAQNNGWAGIVVNGCVRDVDEINGCDIGVRALASHPMKANKKGIGEKHVPITIAGTRICDGEWLYADTDGILISKMELSV*

NbL14g10930.1 HMG−Y−related_protein_B−like [MW = 21.03 kDa]

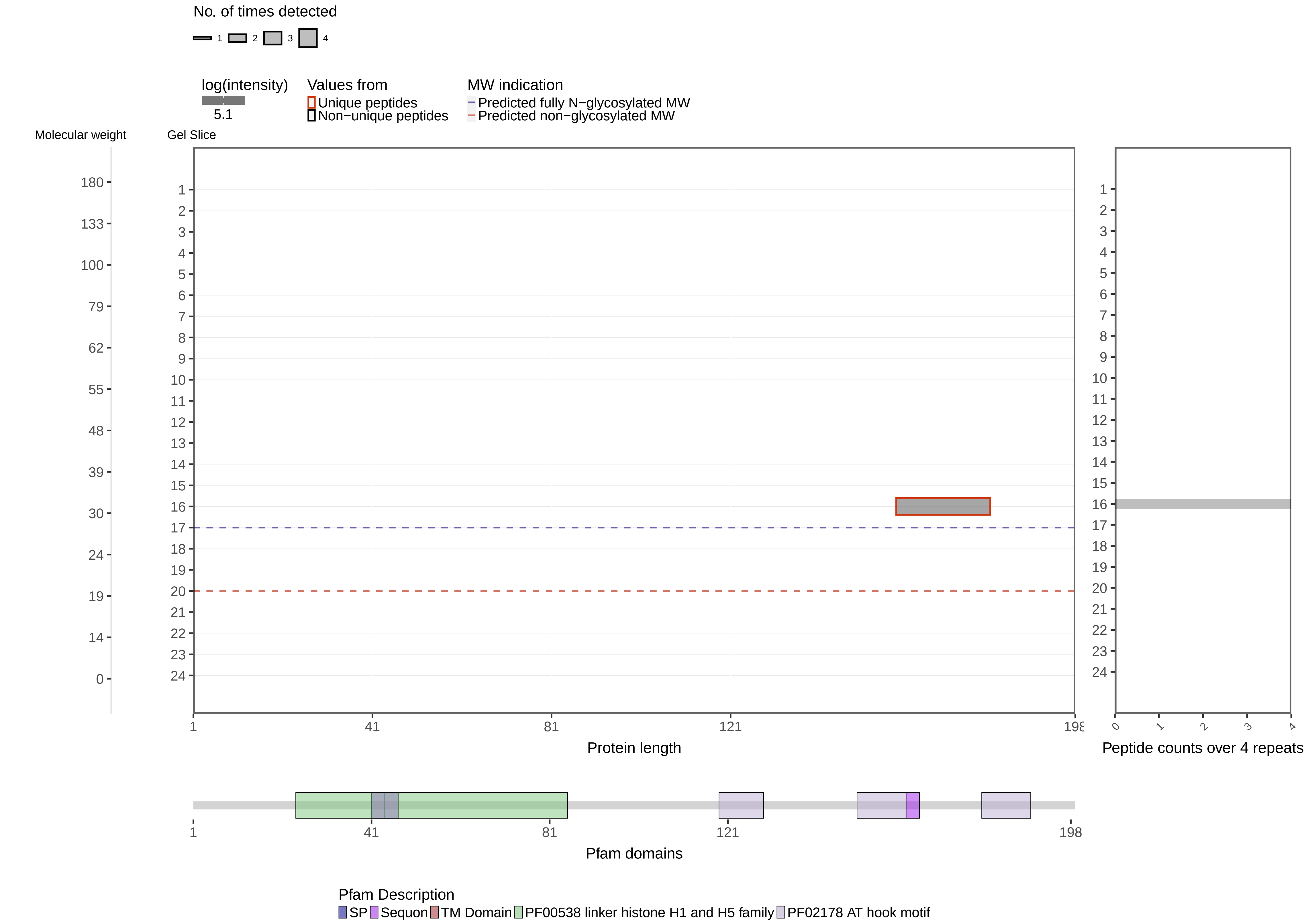


Protein sequence:
MKPDPNAPPRRGRGRPPKPKAPLPEGYVPPPPRPRGRPPKERDPHAPISVPMKVTSPSSGGTGKKRGRPRKYPNDTSSPSPAVFPPPSGTPRGRGRPPKVRNPVAAEVGA*

### NbL14g10730.1 kirola−like [MW = 19.69 kDa]

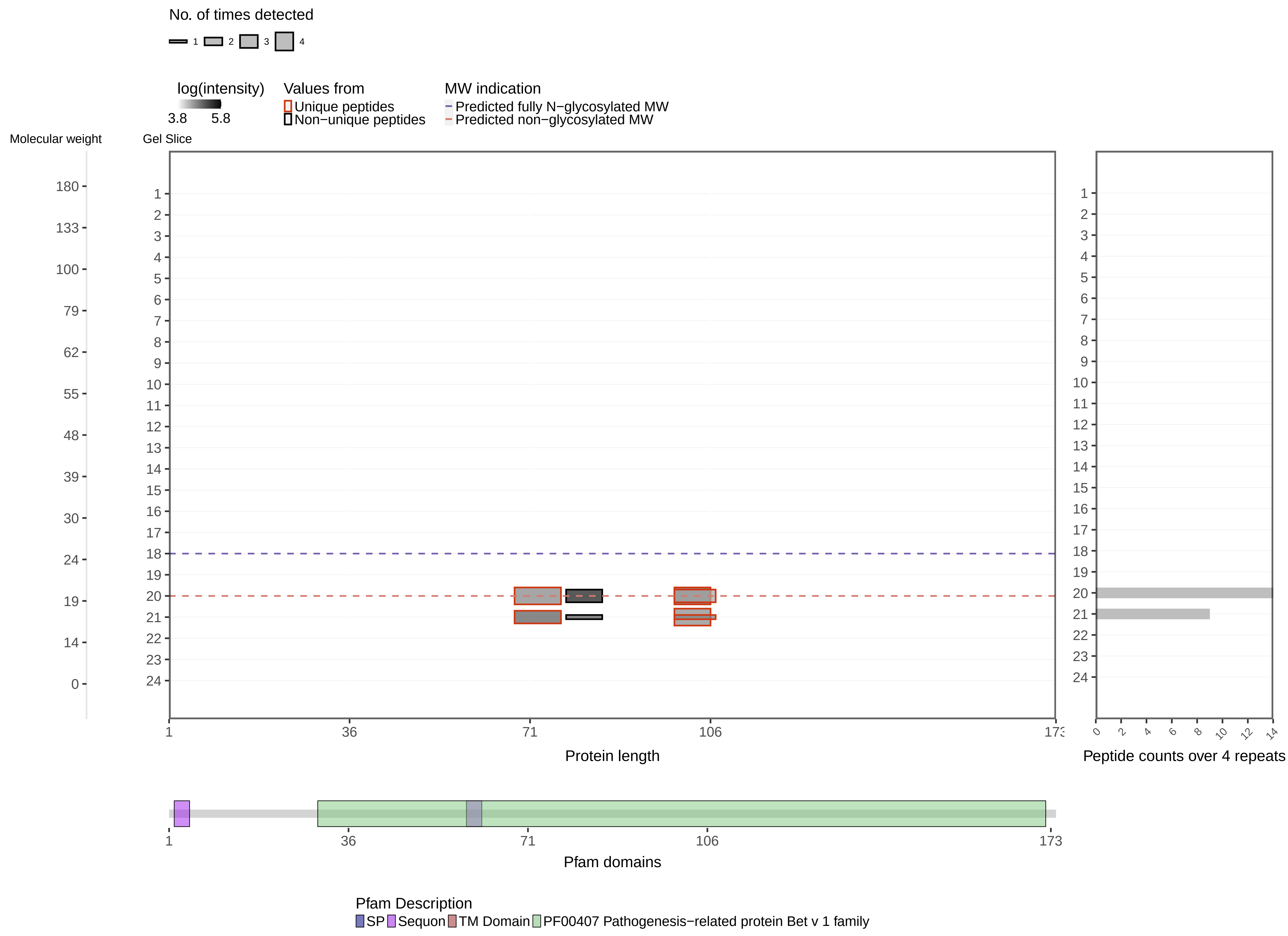


Protein sequence:

TVKVGSVVNWKYNDDGKDKICKQKIEAVDHVKKSITWKVIGGDLLELYNSFTITTSHDHQWTTWTFVYEKKTEDTPEPLVLLGFVLHVTKDIEGHLLK*

### NbL14g10560.1 probable_ribose−5−phosphate_isomerase_2 [MW = 28.85 kDa]

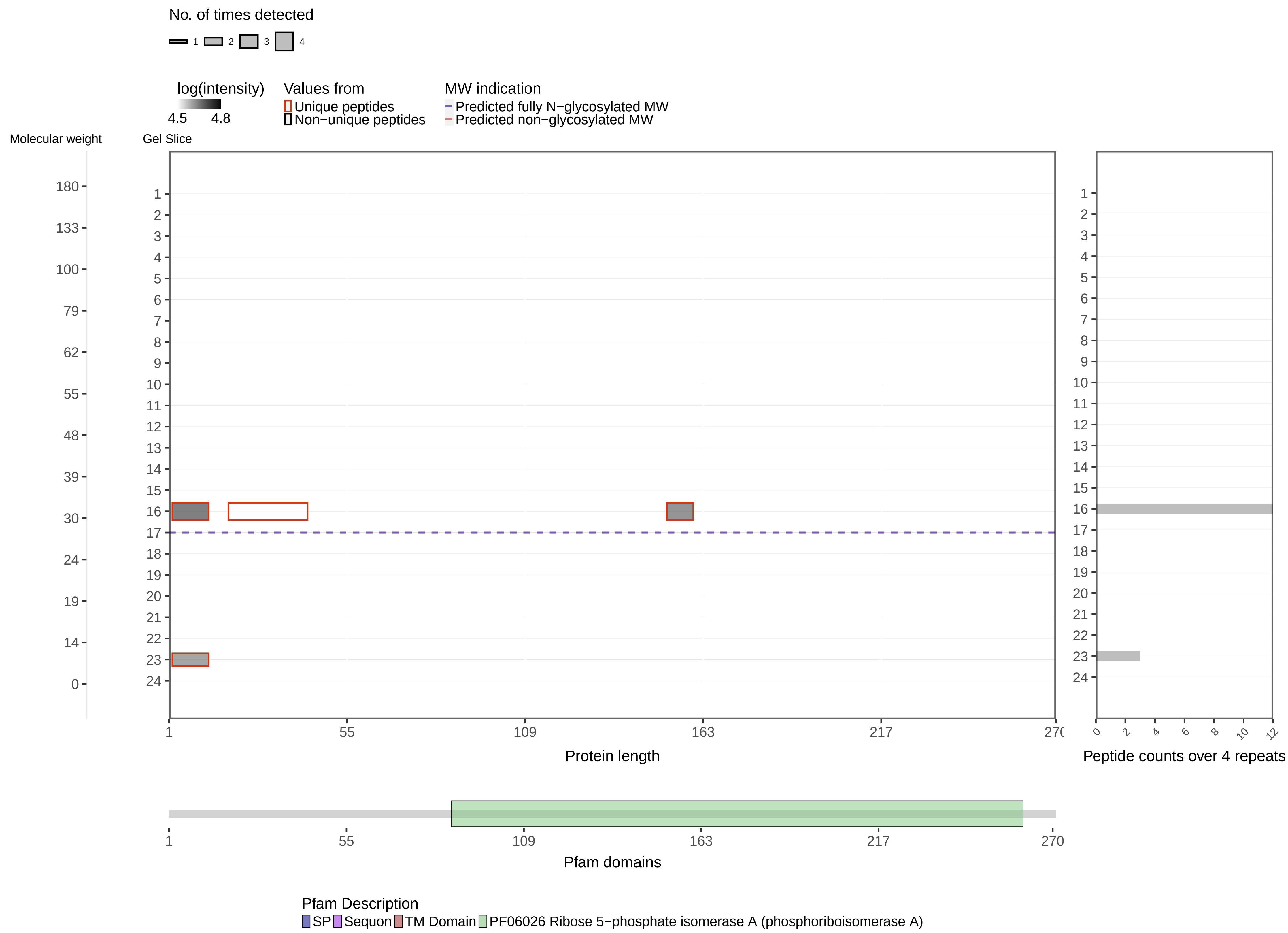


Protein sequence:

EVDREMNLVKGRGGSLLREKMVEAACKKFIVIVDESKLVSHLGGSGLAMPVEIVPFCWEFTLKRLEMLFIEAGCVGKLRRTVGGEPYVTDNGNYIIDLYFKSDMGDLKAASDAILRLAGVVEHGMFLDMATTVIVAGKLGVSVTNK*

### NbL14g10460.1 peptidyl−prolyl_cis−trans_isomerase_FKBP13_chloroplastic [MW = 23.7 kDa]

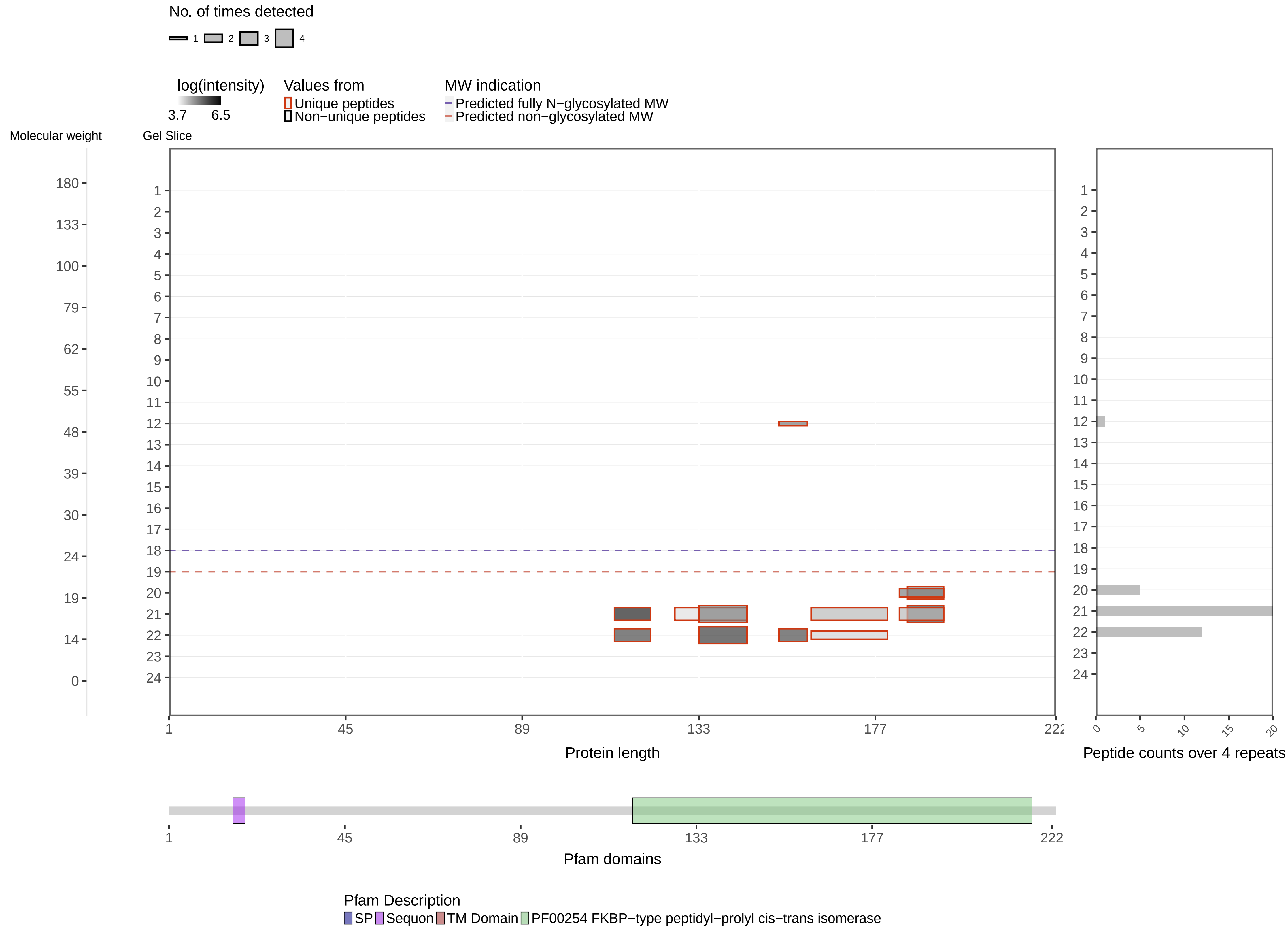


Protein sequence:

/TPSGLCYCDKVVGYGPEAVKGQLIKAHYVGKLENGQVFDSSYNRGKPLTFRVGVGEVIKGWDQGILGGDGFPPMLTGGKRKLKIPPELGYGMRGAGCRGGSCIIPPNSVLLFDVEFVGKA*

### NbL14g10350.1 60S_ribosomal_protein_L24−like [MW = 18.44 kDa]

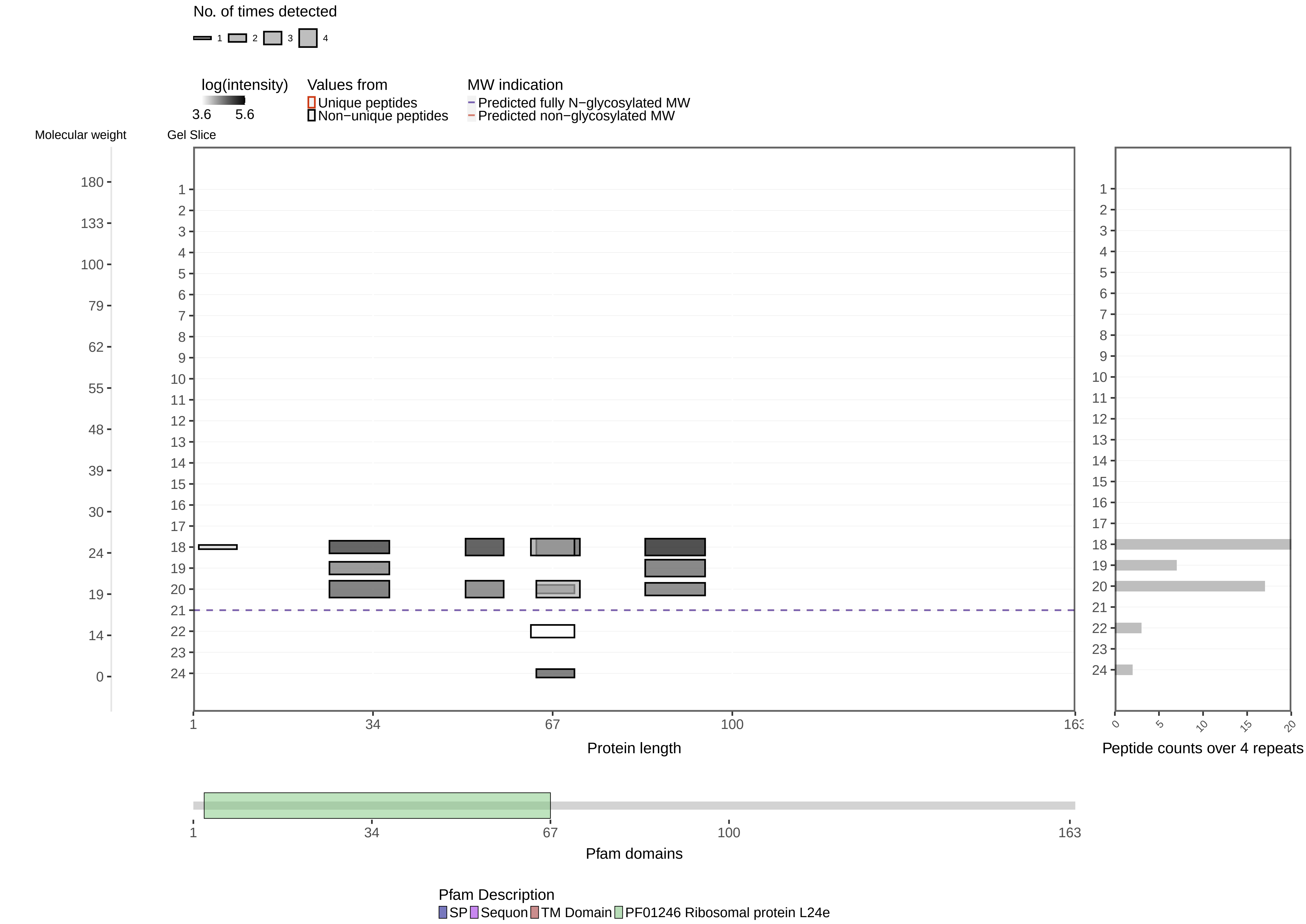


Protein sequence:

AVKKRRRATKKPYSRSIVGATLEVIQKKRTERPEVRDAAREAALREIKERIKKTKDEKKAKKAEVTAKSQKAGGKGNLPKGSKGPKLGGSGGKR*

### NbL14g10270.1 DEAD−box_ATP−dependent_RNA_helicase_38 [MW = 55.2 kDa]

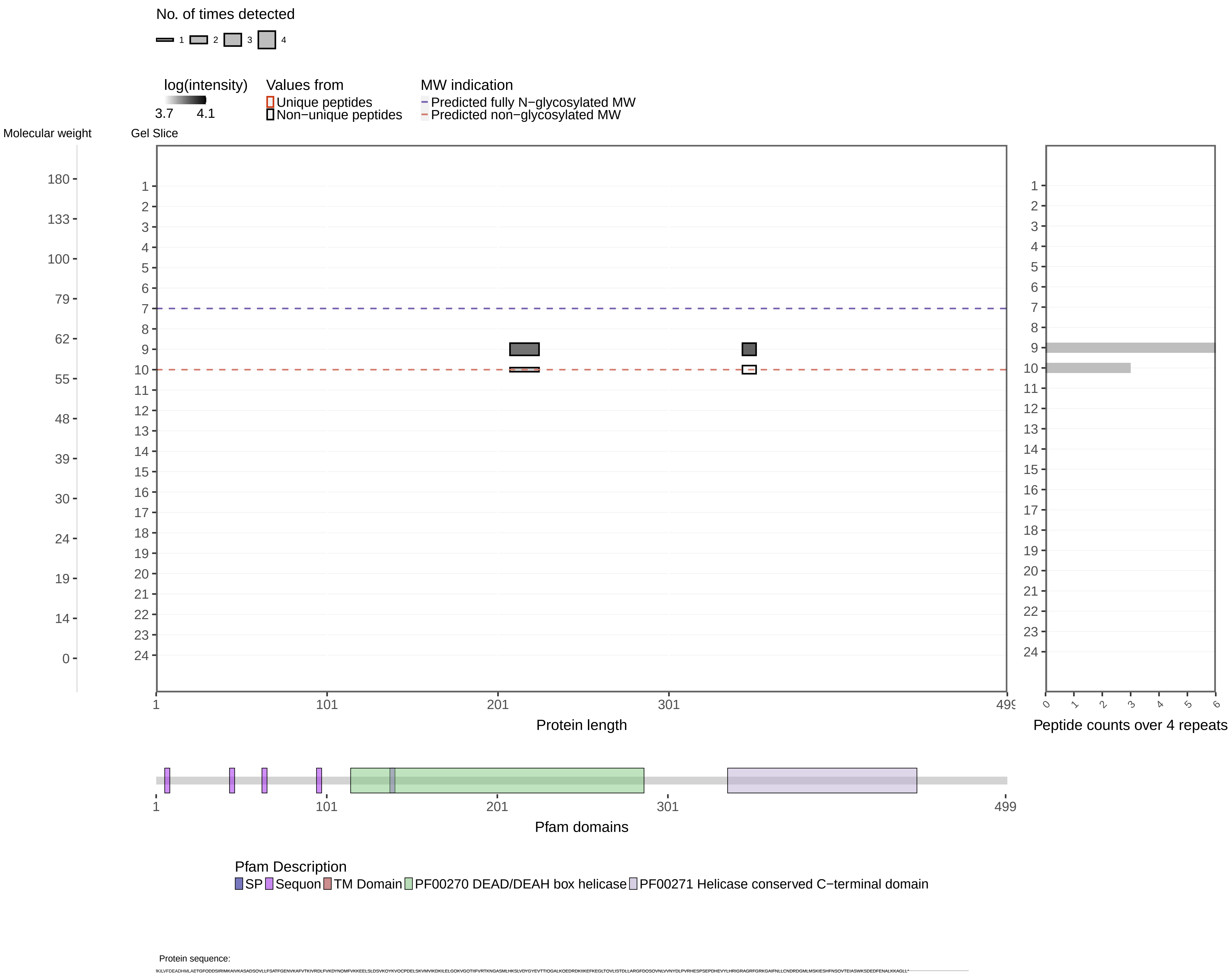

### NbL14g10190.1 protein_MODIFIER_OF_SNC1_11−like [MW = 21.18 kDa]

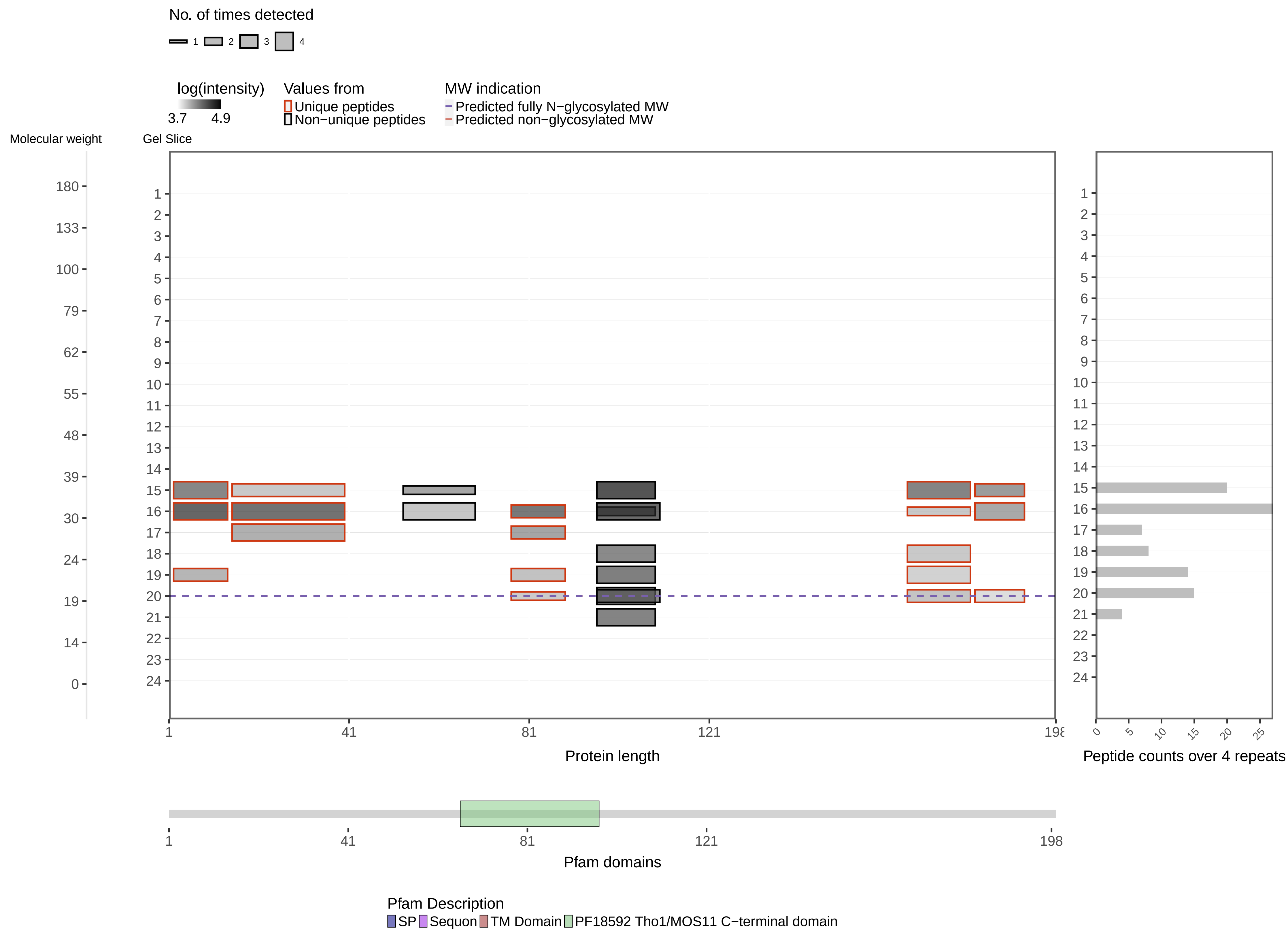


Protein sequence:

KRNSRAERFGTGSSVQGSDALKKAEEHKRQARAERFGLVKSDTTEEEAKKKARLTRFAPPVKTDPVEEDKRKARALRFSQPQSASQSQANGKGNIEQDDVAVDKAGAEPE*

NbL14g10180.1 translin [MW = 34.56 kDa]

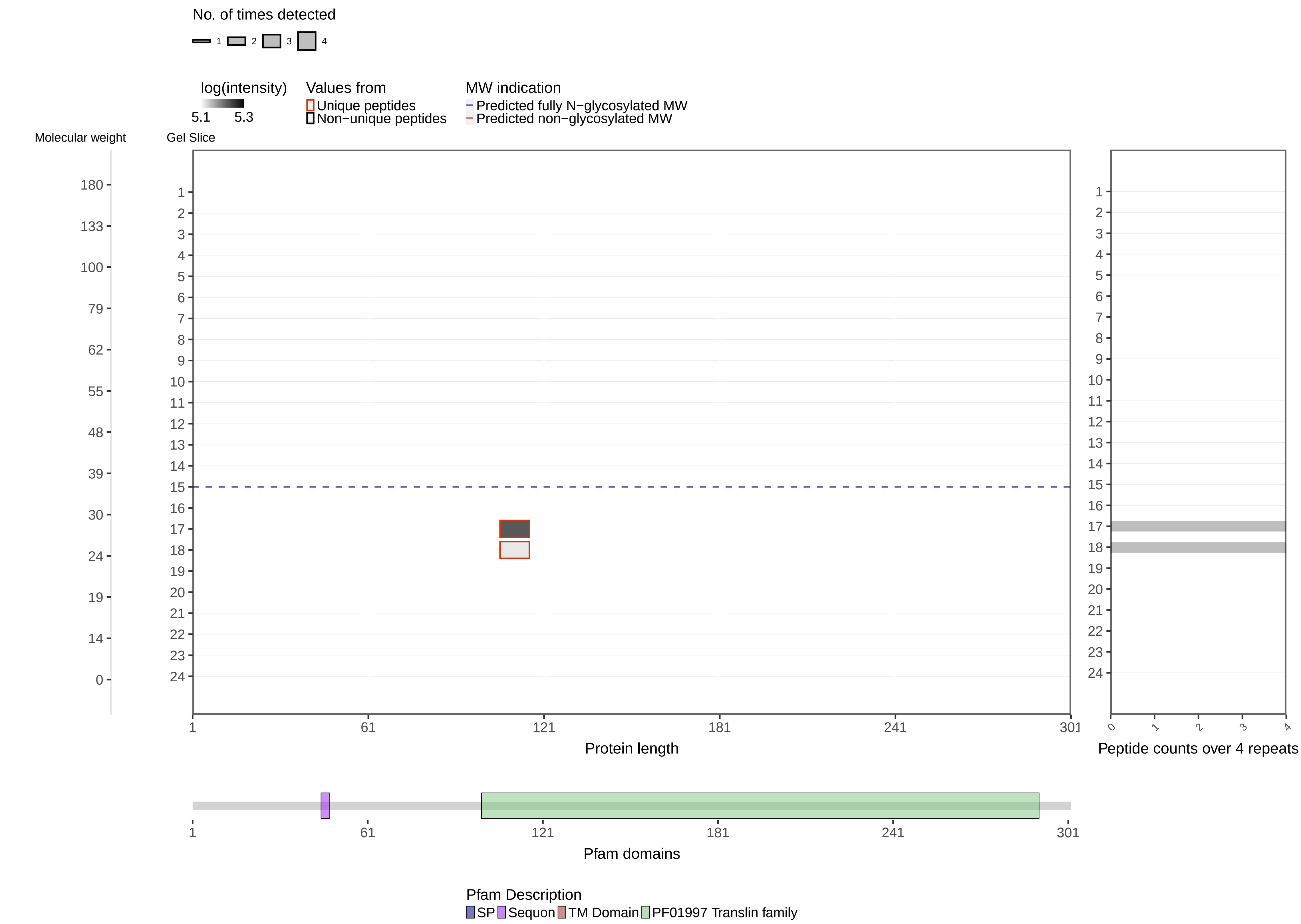


Protein sequence:

:AOIGELKELYSRLSEIVRECPGOYFRYHGDWKSETOTVVSLLAFMHWLETGTLVLHNEVEEKLGLTPTEFGLDIEDYLIGICFMSNELPRYVVNOVTAGDYDCPRKVLKFLTDLHAAFRMLNLRNDFLRKKFDGMKYDLRKVEEVFYDVKIRRLAANGESA*

NbL14g10170.1 adenosine_kinase_2 [MW = 37.46 kDa]

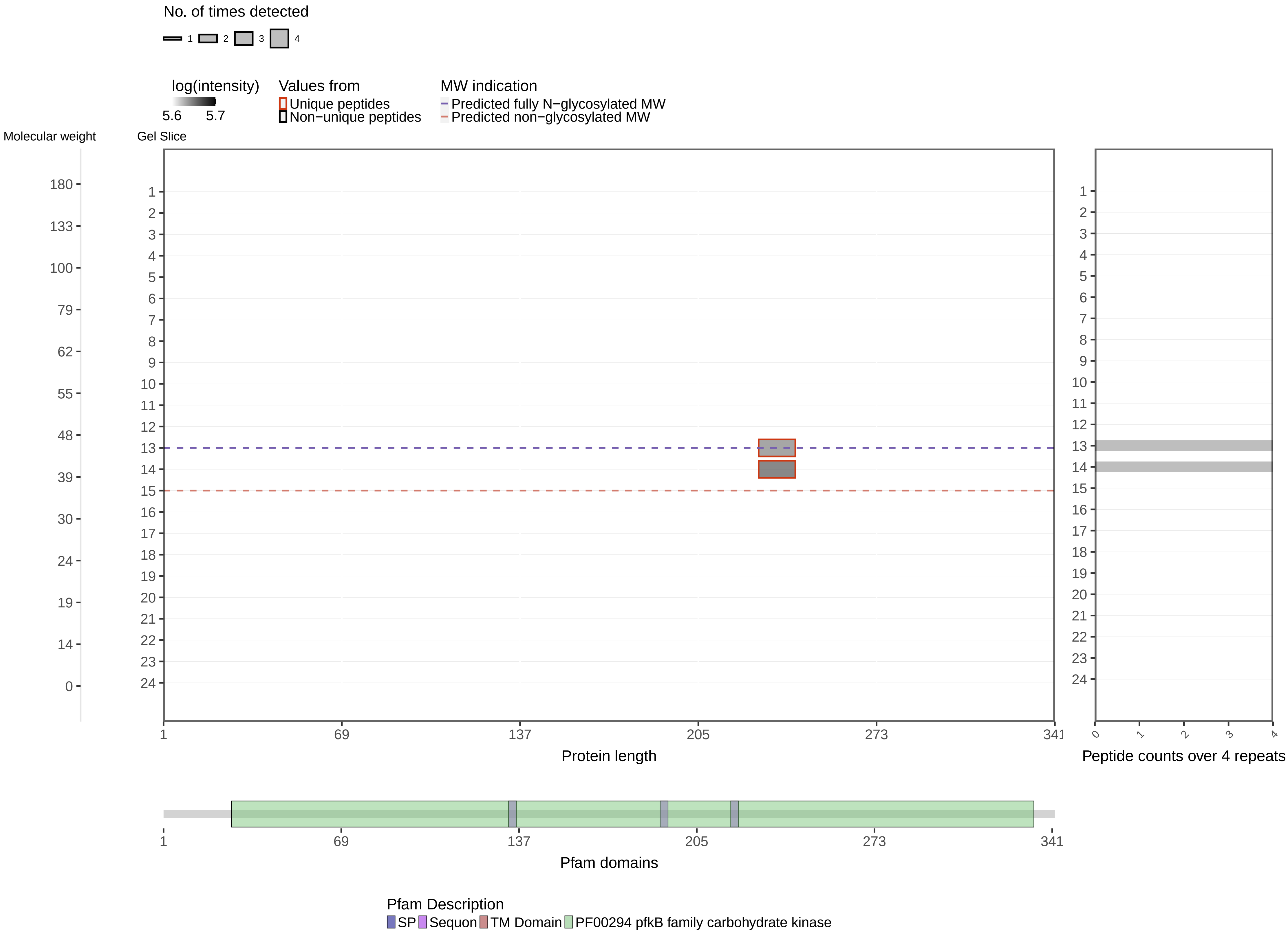


Protein sequence:

AKYYYIAGFFLTVSPESIQLVAEHAAAKHKVFSMNLSAPFICEFFKDQQEKVLPYMDFVFGNETEARTFSRVHSWETDNVEEIALKISQWAKASGTHKRITVITQGADPVVVAEDGKVKLFPVIPLPKEKLVDTNGAGDAFVGGFLAQLVQGKPIADCVKAGCFASNVIIQRSGCTYPEKPDFE*

NbL14g10160.1 phenylalanine_ammonia−lyase [MW = 77.45 kDa]

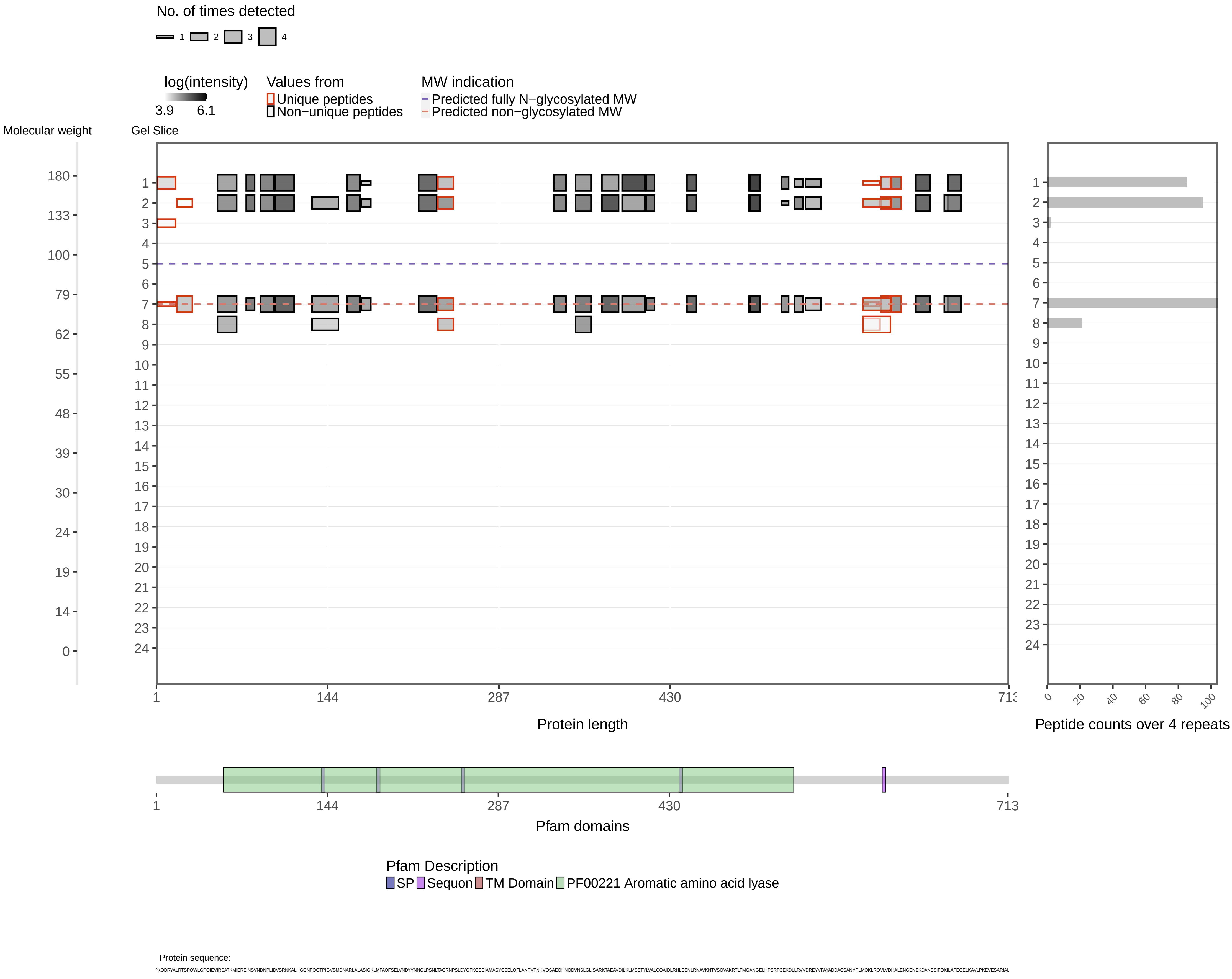

### NbL14g10150.1 29_kDa_ribonucleoprotein_B_chloroplastic [MW = 34.81 kDa]

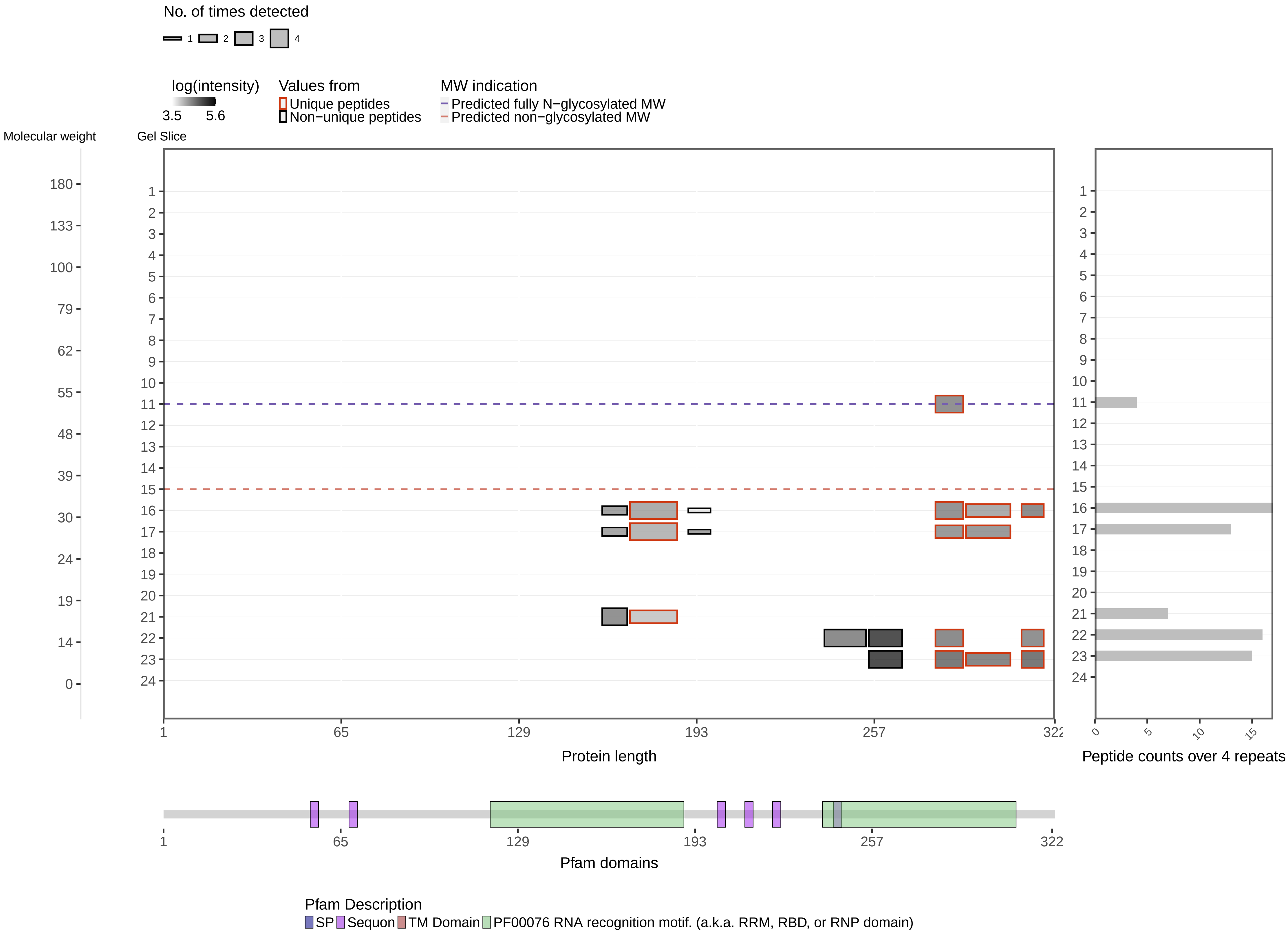


Protein sequence:

LTGRSRGFGFVTMSTKEEVEAAEOOFNGYEIDGRAIRVNAGPAPAKRENSSFGGGRGGNSSYGGGRDRNSSFGGARGGRSVDRSNRVYVGNLSWGVDDLALKELFSEOGNVVDAKVVYDRDSGRSRGFGFVTYSSSKEVNNAIDSLNGVDLDGRSIRVSAAEEOPRROF*

### NbL14g09940.1 S−adenosylmethionine_synthase_2−like [MW = 42.61 kDa]

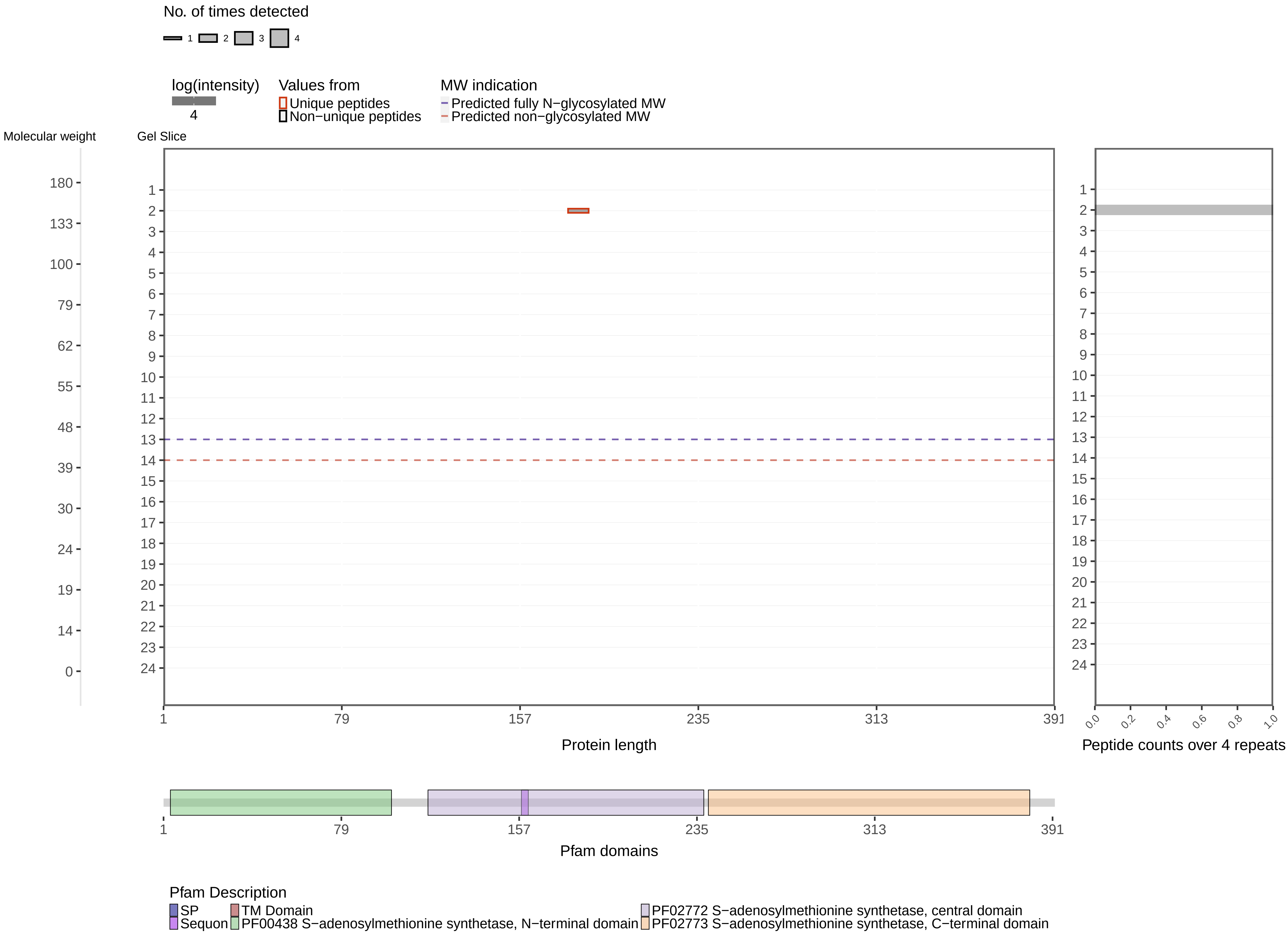


Protein sequence:

IGAMVPIRVHTVLISTQHDETVTNEQIAQDLKEHVIKPVIPAKYLDEKTIFHLNPSGRFVIGGPHGDAGLTGRKIIIDTYGGWGAHGGGAFSGKDPTKVDRSGAYIVRQAAKSVVAAGLARRCIVQVSYAIGVAEPLSVFVDTYKTGTIPDKDILTLIKENFDFRPGMIAINLDLKRGGNFRYQKTAAYGHFGRDDPDFTWETVKALNPKA*

### NbL14g09860.1 60S_ribosomal_protein_L23a−like [MW = 17.48 kDa]

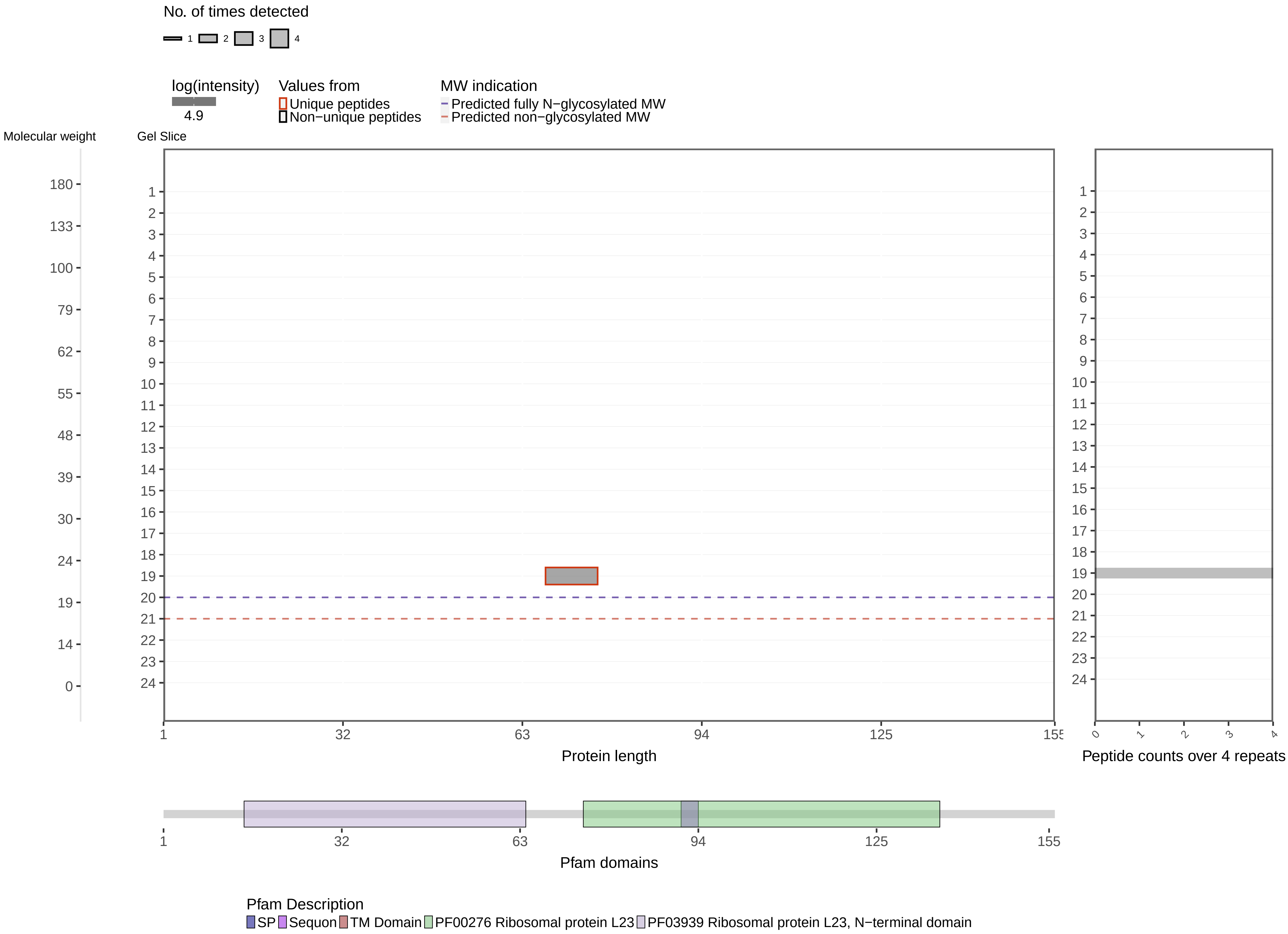


Protein sequence:

SAPGRNKLDQYEILKYPLTTESAMKKIEDNNTLVFIVDIKADKKKIKDAVKKMYDIQTKKVNTLIRPDGTKKAYVRLTPDYDALDVANKIGII*

NbL14g09790.1 THO_complex_subunit_4A−like [MW = 26.71 kDa]

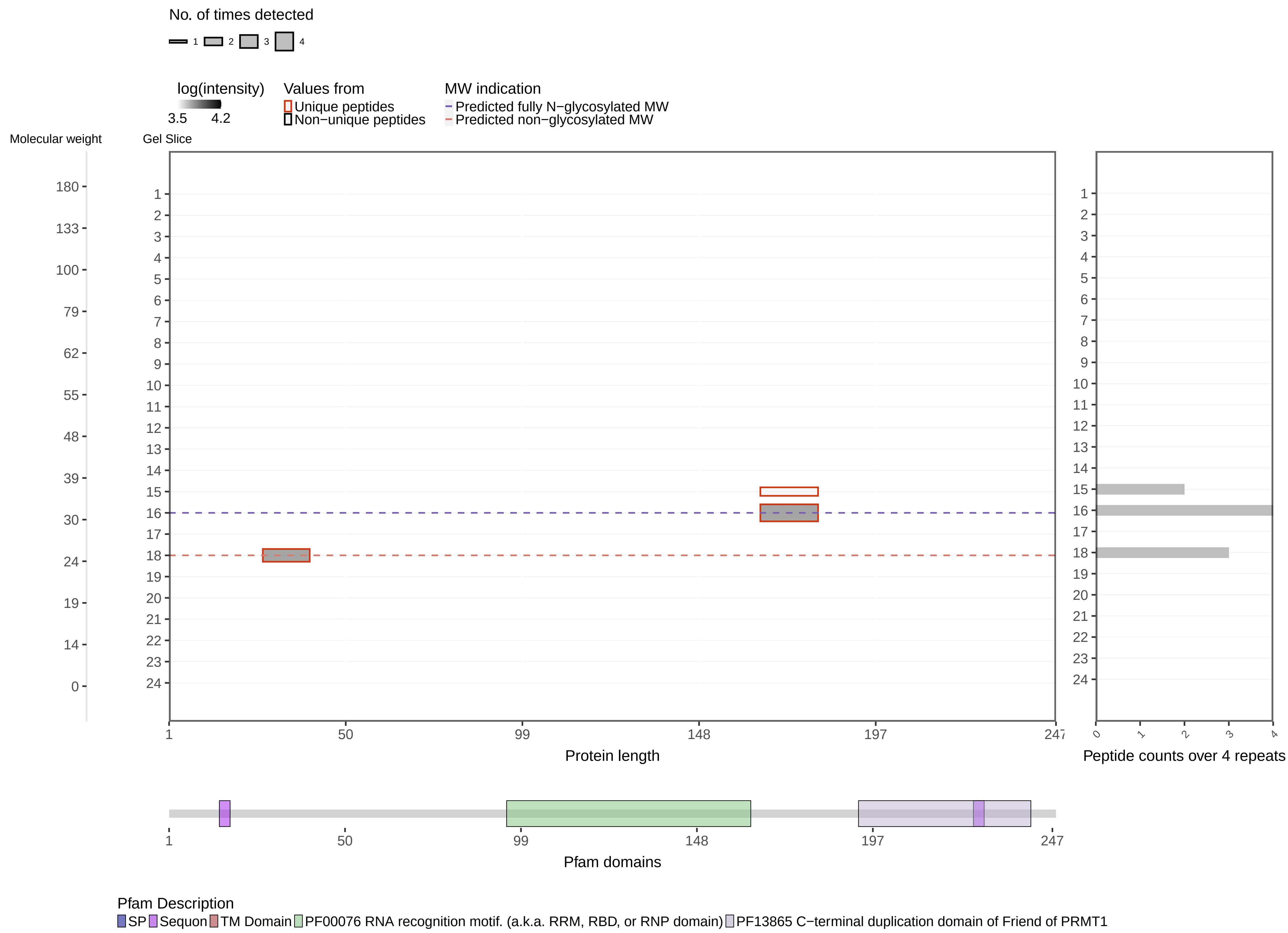


Protein sequence:

ELFSEAGDIKRYSIHYDKSGRSKGTAEVIFSRRRDAEAAIKKYNNVQLDGKPMKIEFAGPNIGAPALPPIRNRLYRNPNPAPRSQQRGGGFRRPPRGGRGSMRKEGGRGRGRGENISAEDLDADLEKYHSEAMETN*

### NbL14g09780.1 heat_shock_cognate_70_kDa_protein_2 [MW = 70.9 kDa]

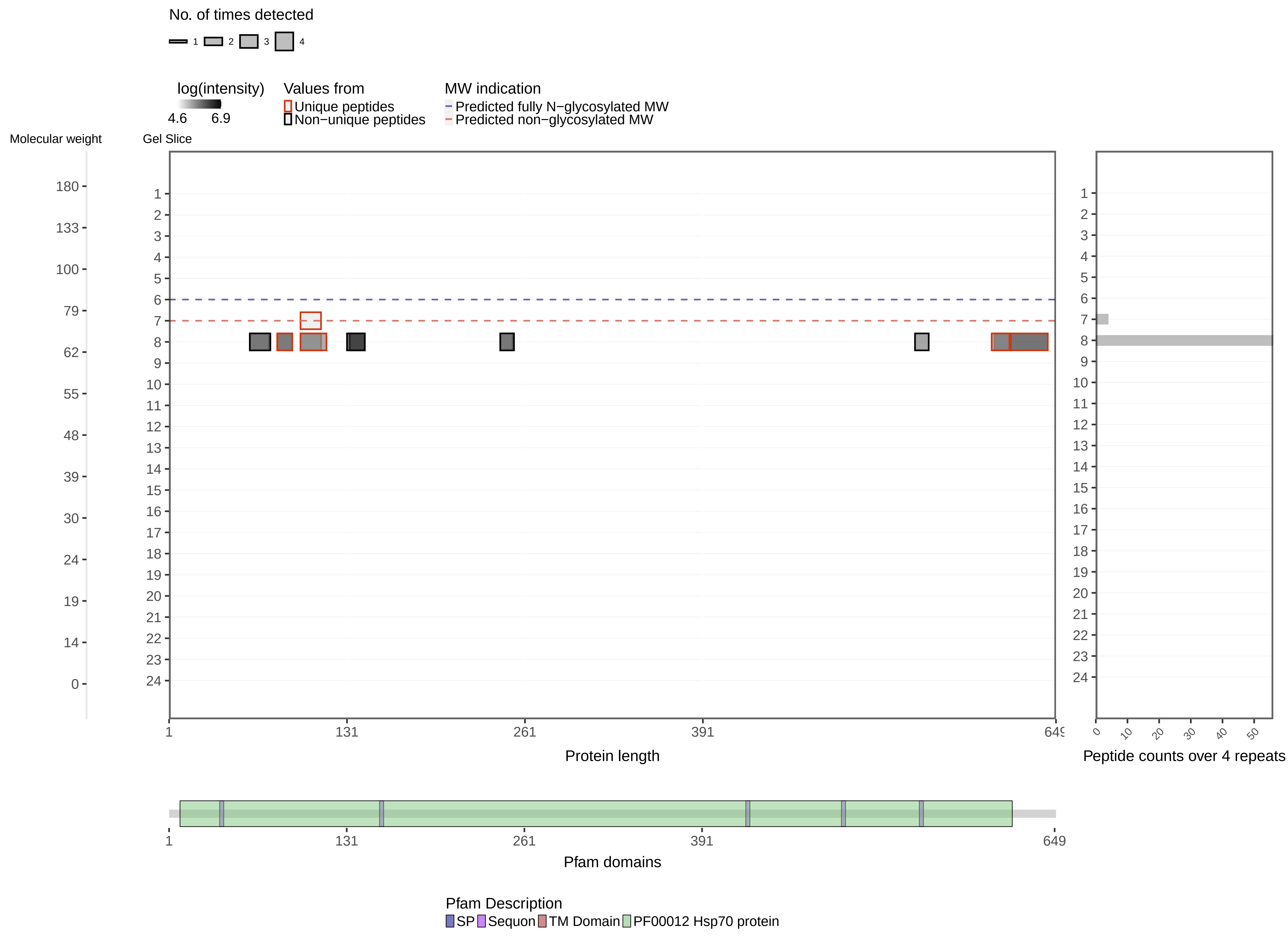


Protein sequence:

LFRKCMEPVEKCLRDAKMDKSTVHDVVLVGGSTRIPKVQQLLQDFFNGKELCKSINPDEAVAYGAAVQAAILSGEGNEKVQDLLLLDVTPLSLGLETAGGVMTVLIPRNTTIPTKKEQVFSTYSDNQPGVLIQVYEGERTRTRDNNLLGKFELSGIPPAPRGVPQITVCFDIDANGILNVSAEDKTTGQKNKITITNDKGRLSKEEIEKMVQEAEKYKSEDEEHKKKVEAKNALENYSYNMRNTVKDEKIASKLSAEDKKKIEDAIEEAIQWLDGNQLAEADEFEDKMKELEGL

### NbL14g09600.1 ribulose_bisphosphate_carboxylase/oxygenase_activase_2_chloroplastic [MW = 48.51 kDa]

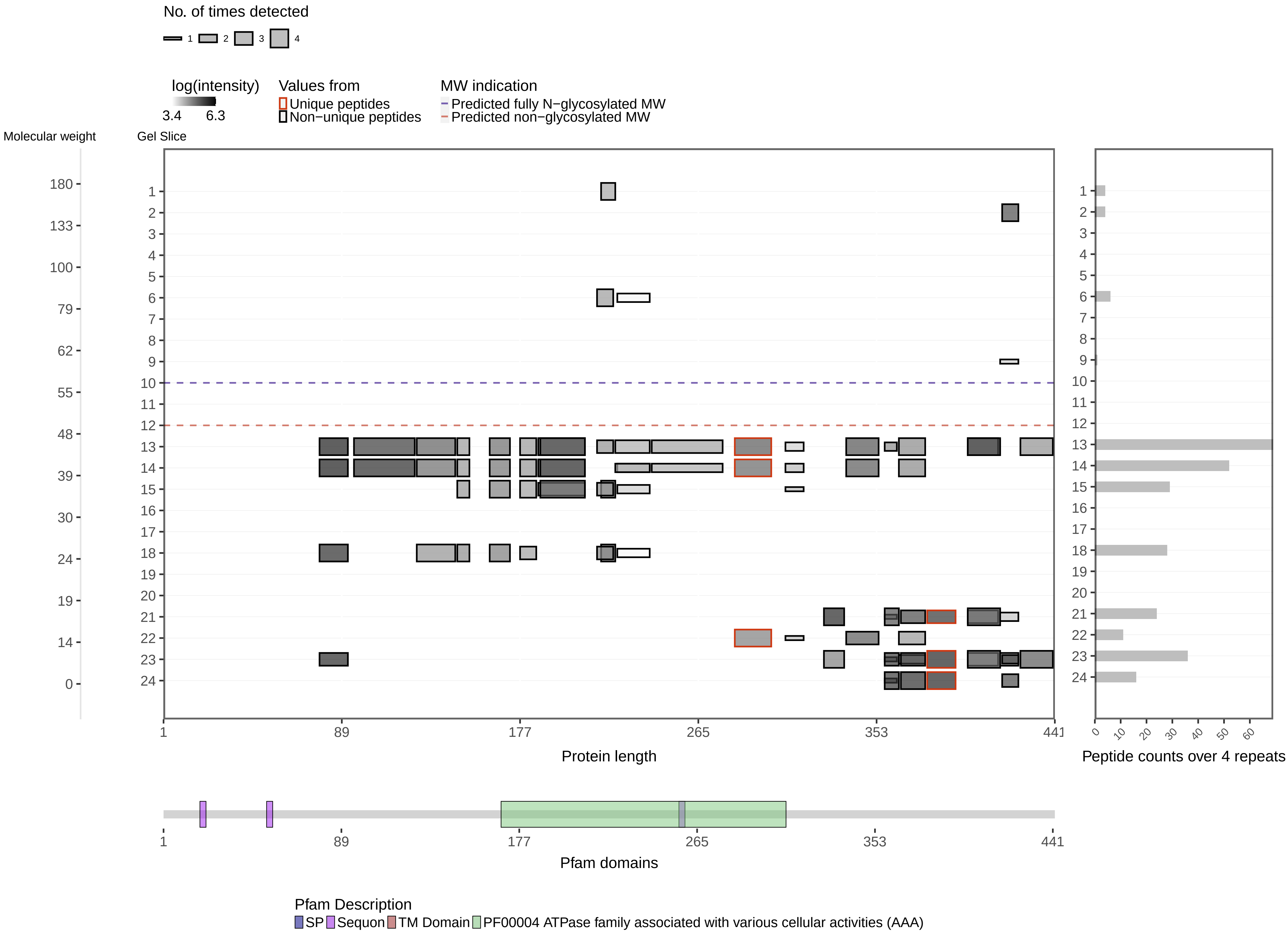


Protein sequence:

PAKLIRQRYREAAEIIRKGNMCCLFINDLDAGAGRMGGTTQYTVNNQMVNATLMNIADNPTNVQLPGMYNKQENARVPIVVTGNDFSTLYAPLIRDGRMEKFYWAPTREDRIGVCKGIFRTDNVPEEAVIKIVDTFPGQSIDFFGALRARVYDDEVRKWVSGTGIEAIGDKLLNTFDGPPTFEQPKMTVEKLLEYGNMLVQEQENVKRVQLAETYLKEAALGDANADAINTGNF*

### NbL14g09340.1 proteasome_subunit_alpha_type−4−like [MW = 30.38 kDa]

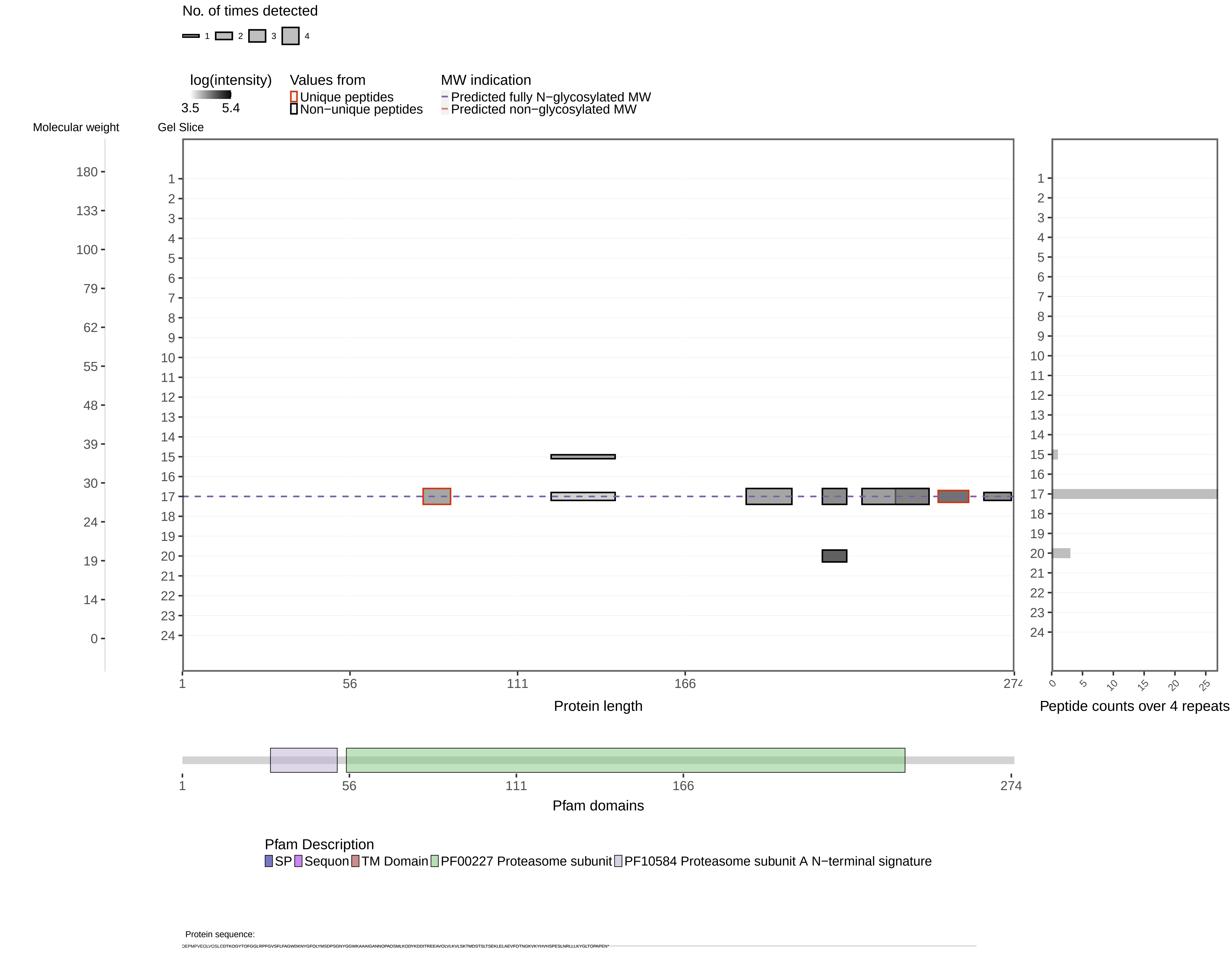

NbL14g09320.1 tubulin_beta−2_chain−like [MW = 50.28 kDa]

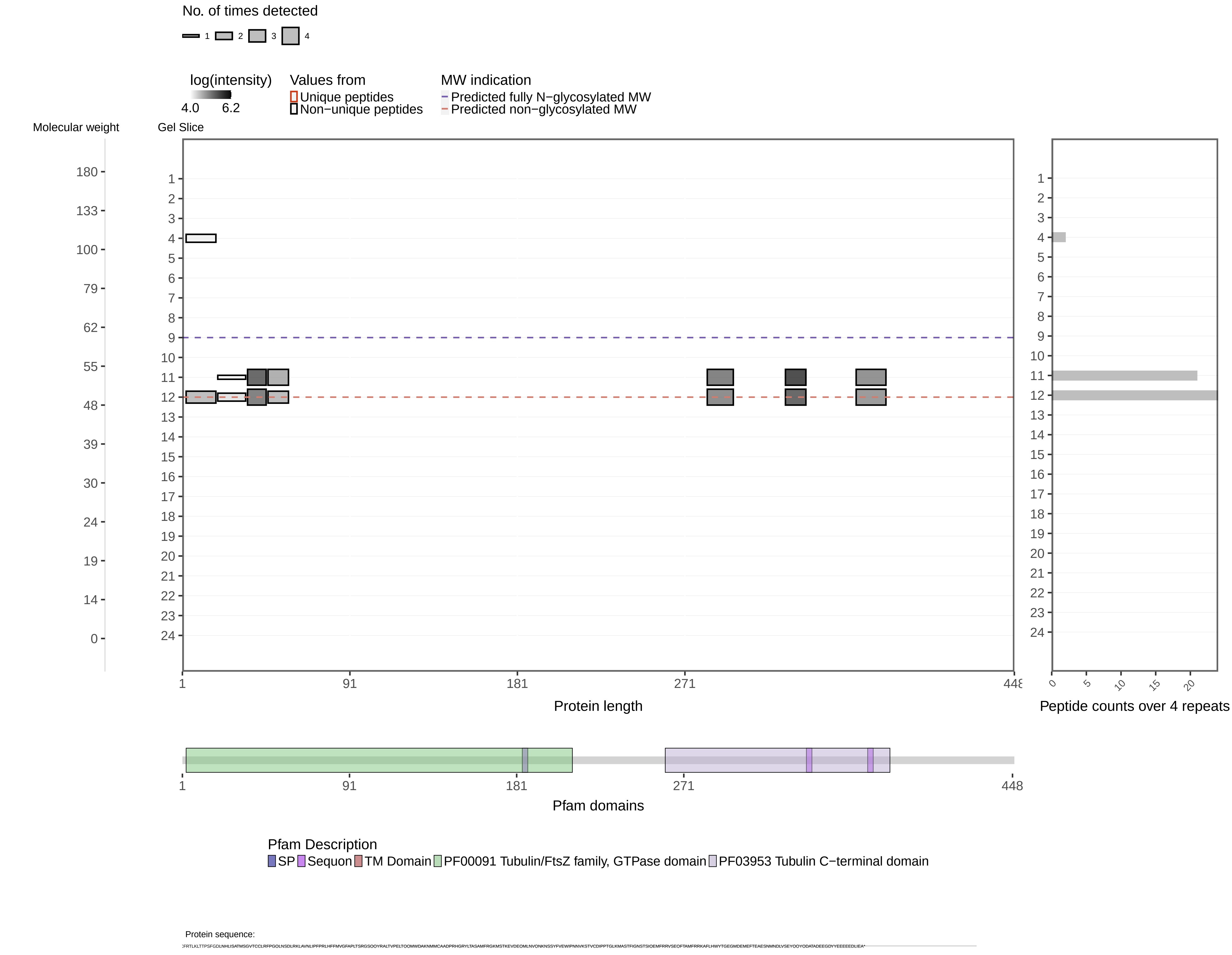

### NbL14g09270.1 probable_glutathione_S−transferase [MW = 26.54 kDa]

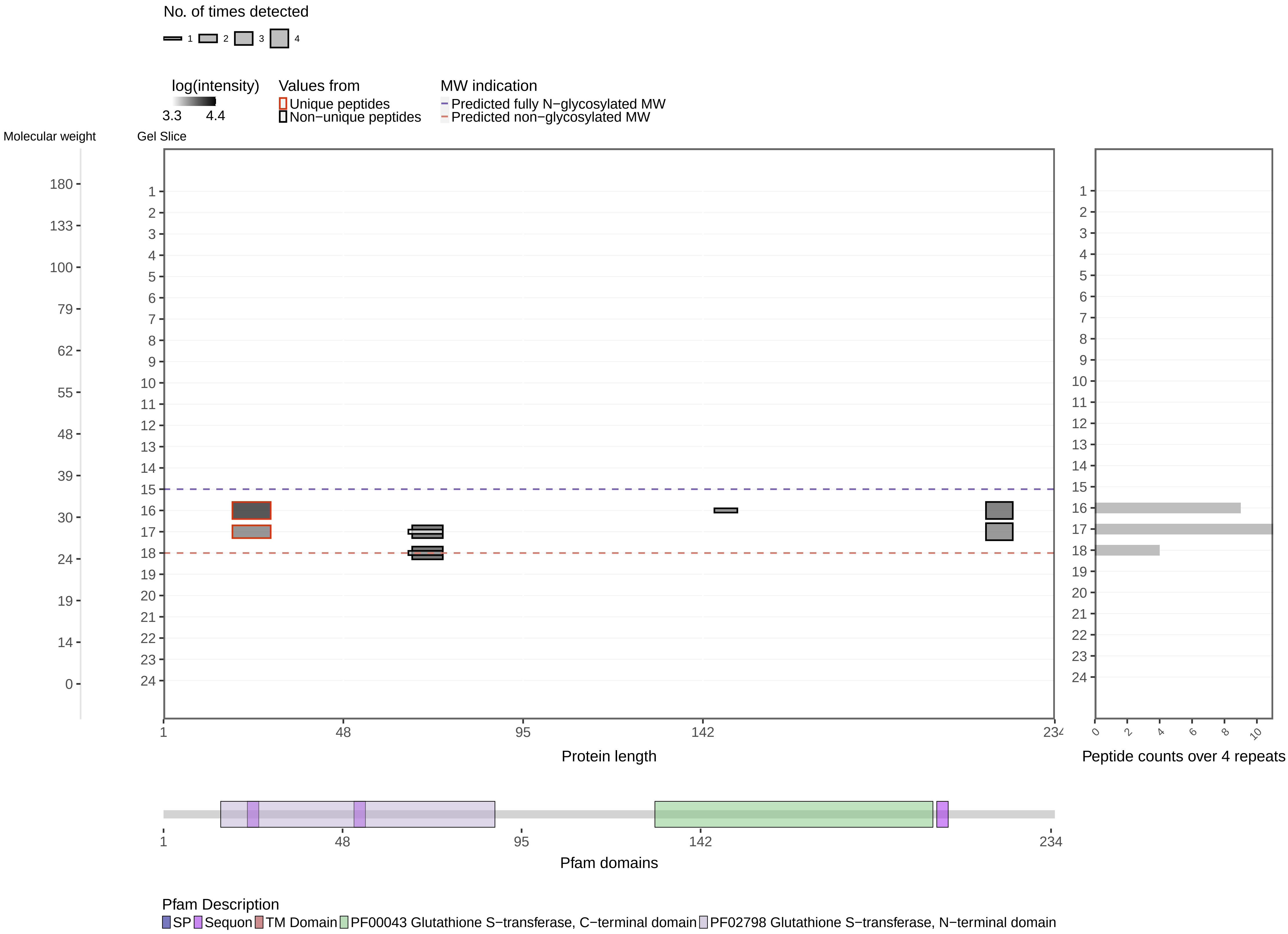


Protein sequence:

ARFLVKFLDEKCLPAVWKALWSKGEGQEKDKEEAYEVLKVLDNEVKDKKYFGGEKIGFVDIAANFVGFWIGIVEEATGVVLVTSEKFPHFCTWREEYLNCSQVKENMPNREMLLGYFKGRVRAATAPK*

### NbL14g09180.1 ribulose_bisphosphate_carboxylase/oxygenase_activase_1_chloroplastic−like [MW = 48.61 kDa]

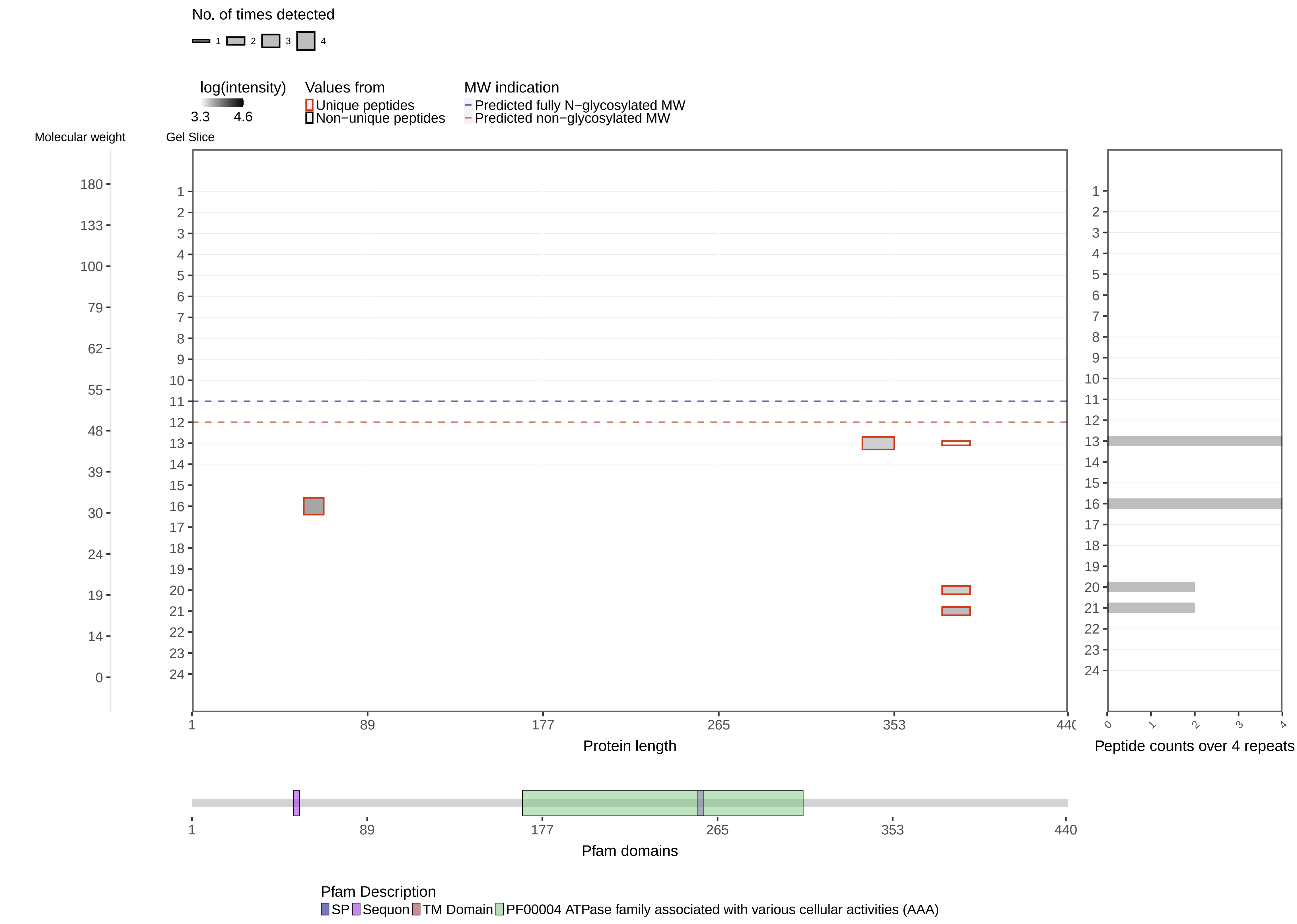


Protein sequence:

AKLIRQRYREAAEIIRKGNMCVLFINDLDAGAGRMGGTTQYTVNNQMVNATLMNIADNPTNVQLPGMYNKQENARVPIIVTGNDFSTLYAPLIRDGRMEKFYWAPTREDRIGVCKGIFRTDSVPDEHVVRLVDAFPGQSIDFFGALRARVYDDEVRKWIESTGIEQVGEKLLNSIDGPPTFEQPKMTIDKLLEYGNLLVQEQENVKRVQLADKYLKEAALGDANADAISTGNF*

### NbL14g09160.1 glucose−induced_degradation_protein_4_homolog [MW = 23.77 kDa]

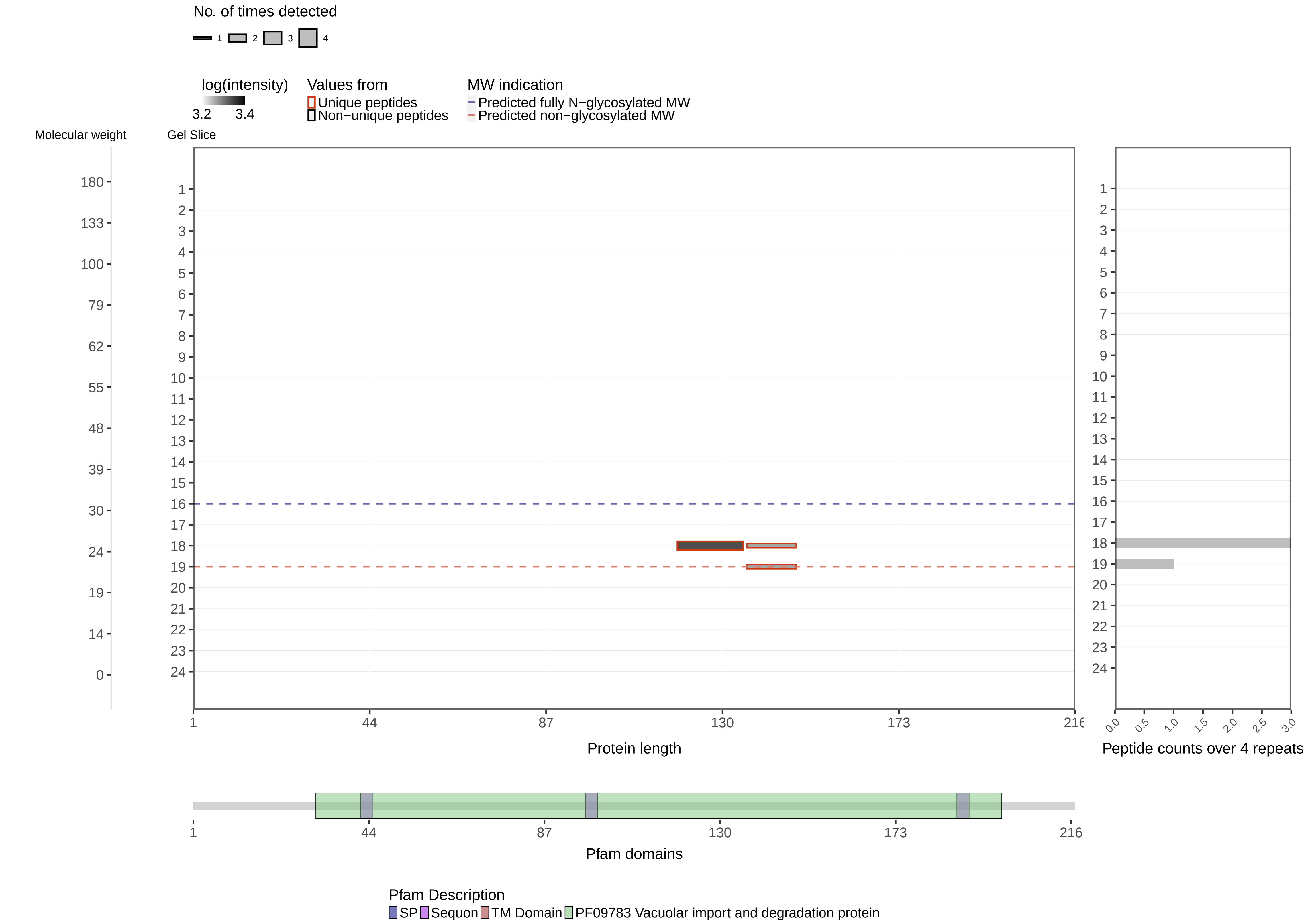


Protein sequence:

TKNYTFFTGKWGATSEDDIKHWTKFPSFSPLLSQVEVDGGKSLDLSNYPYIFMRWKEQYFVNVGTDCGLTIAGFYYVCFSCSDGSINGFYYDPNSSPFQKLELKSTNEGRSGFSFSSYELQ*

### NbL14g09090.1 acyl−coenzyme_A_oxidase_4_peroxisomal−like [MW = 47.02 kDa]

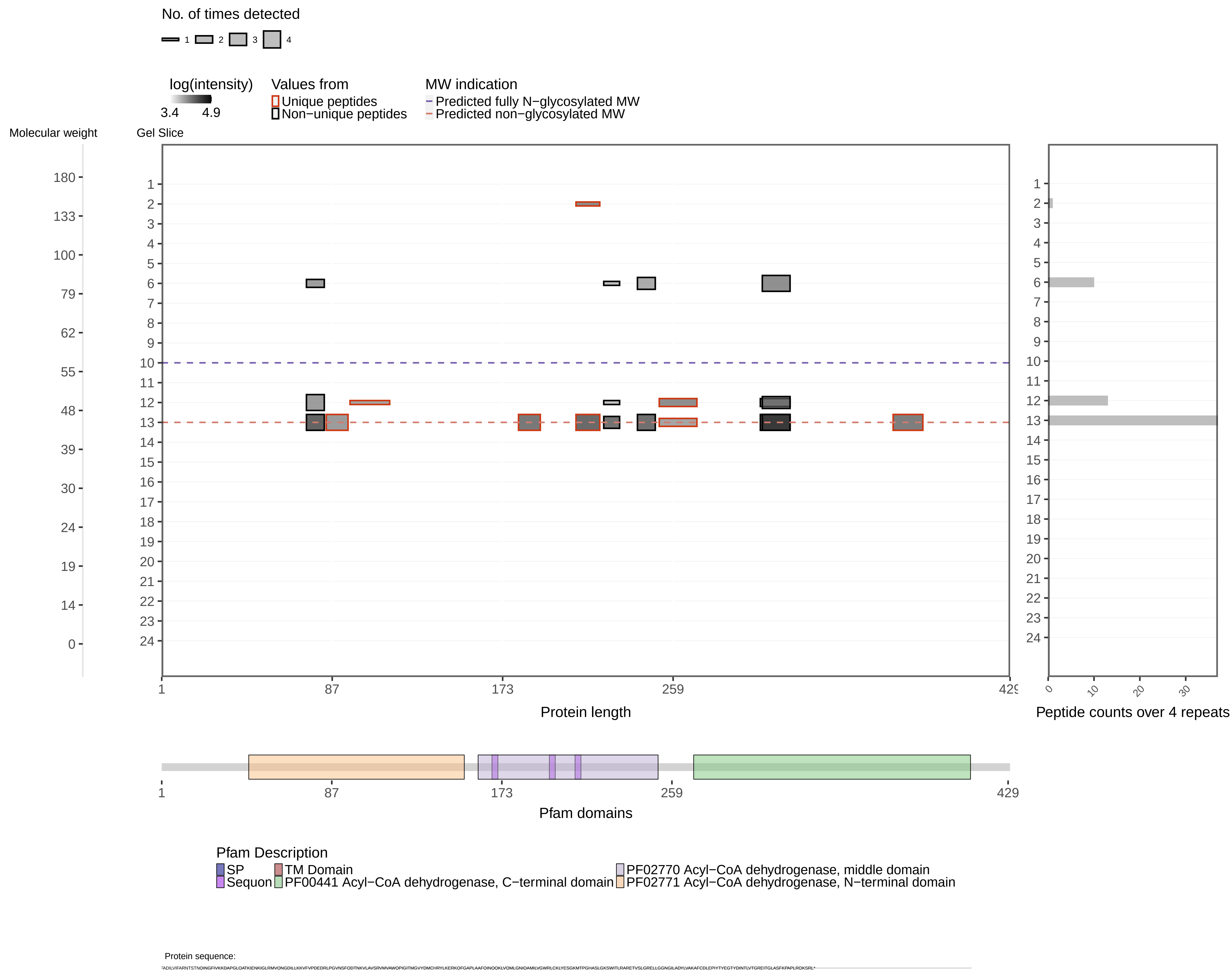

### NbL14g09010.1 uncharacterized_protein_At5g01610−like [MW = 19.21 kDa]

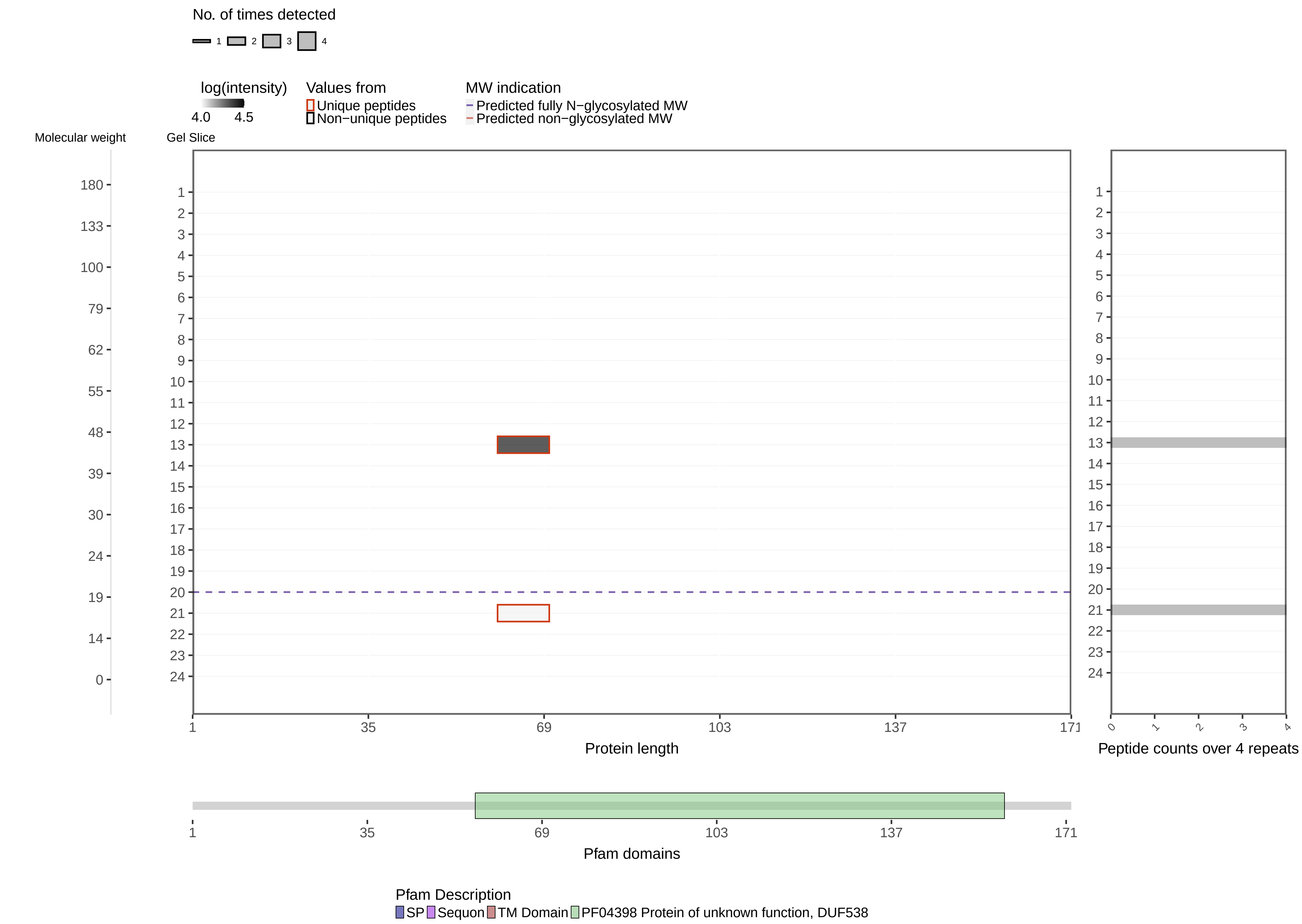


Protein sequence:

ATNYEFNEETKKFTIYIPSVCEVGYMDSSVLRFSTAVTGYLEKGKLADIEGIKTKVMMWVKVTAISSEKSKLHFTVGLKKTRSREAYEVLRDGVAVDKF*

### NbL14g08960.1 macrophage_migration_inhibitory_factor_homolog [MW = 12.07 kDa]

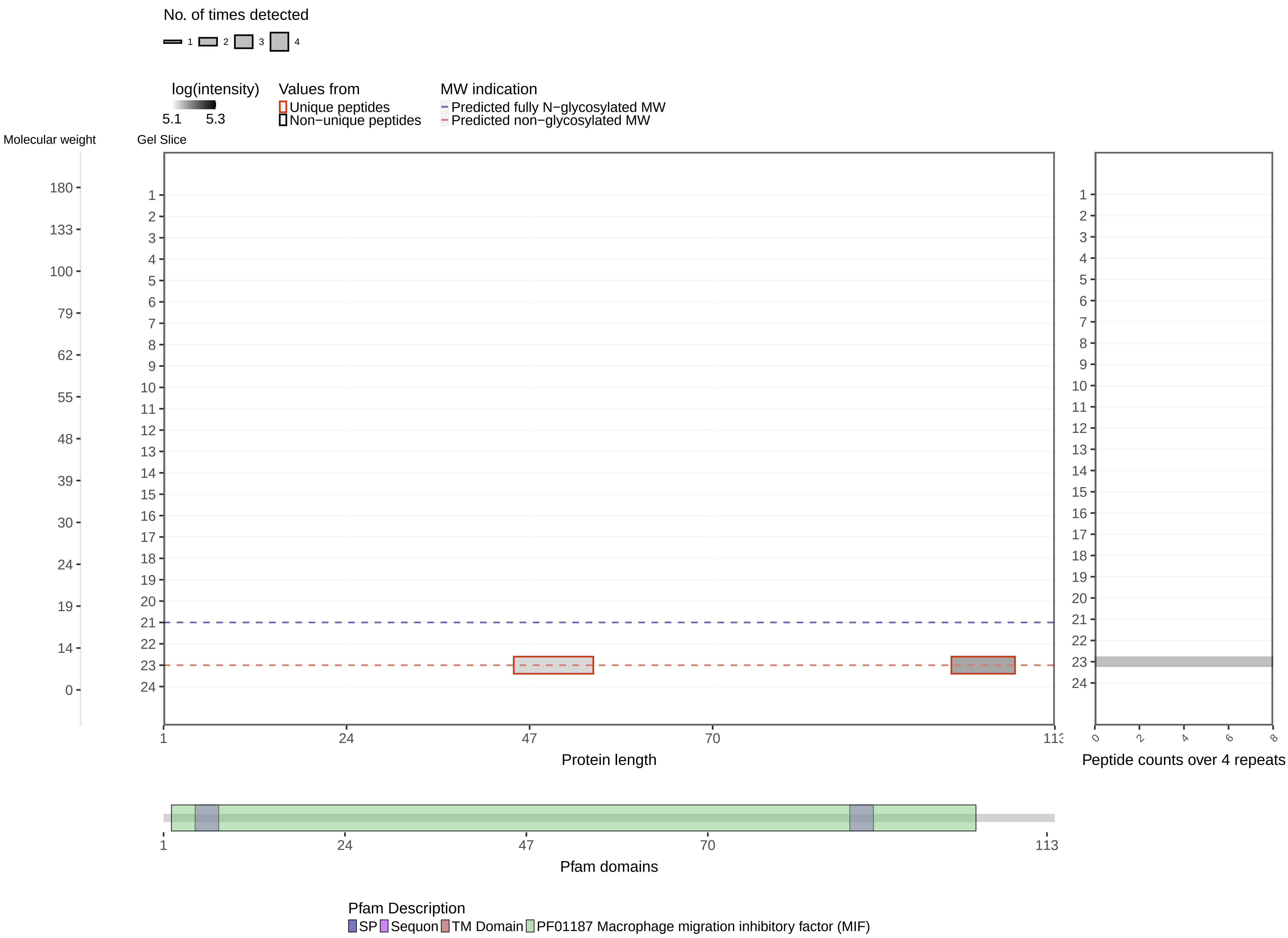


Protein sequence:

'LKGSVDISFGGNKEAAAFAEIVSMGGINSDVKRNLIATLGTILHNNFSIPRTRFFLKVYDTTMATKFSKL*

NbL14g08740.1 non−specific_lipid−transfer_protein_2−like [MW − SP = 11 kDa]

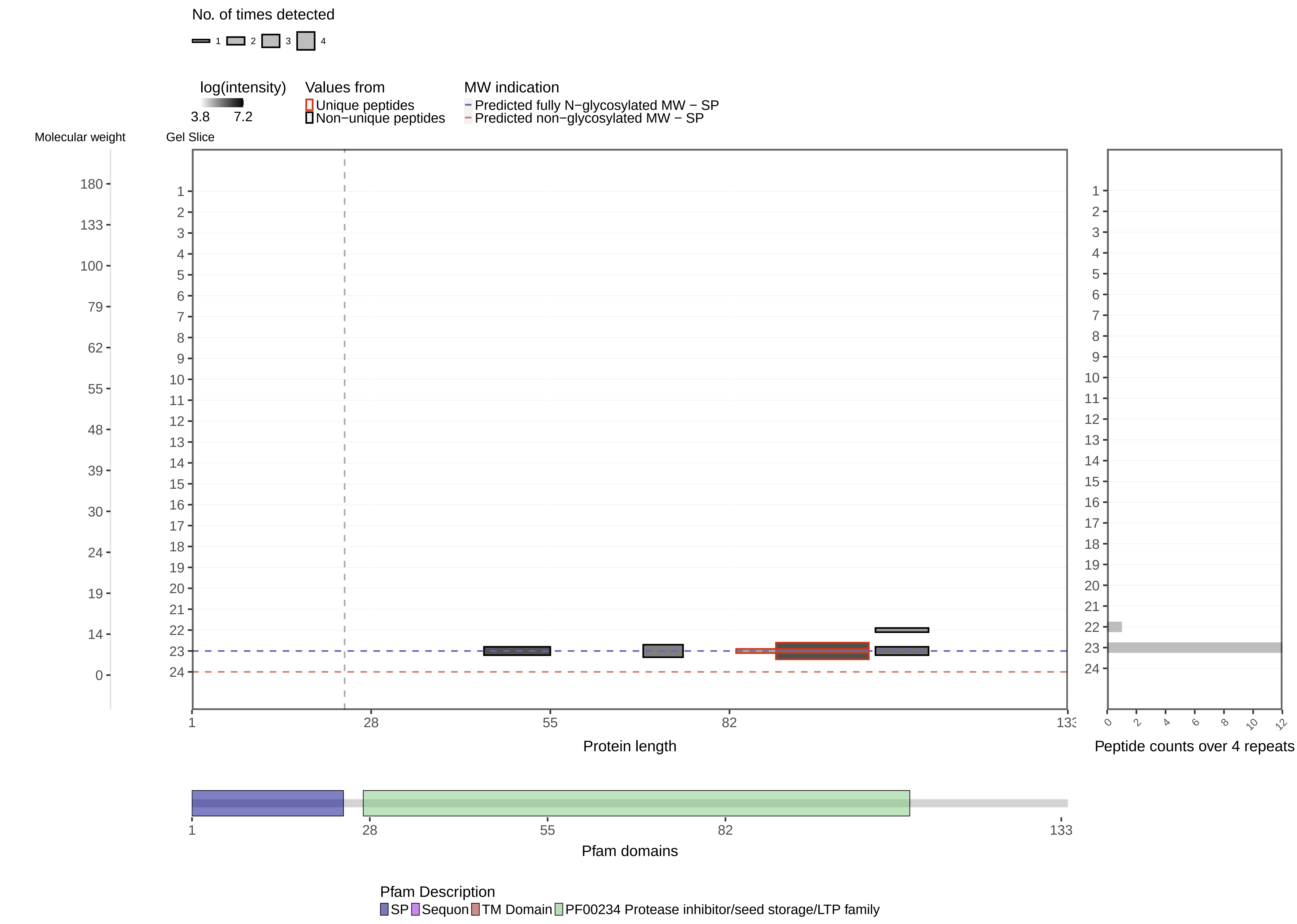


Protein sequence:

GGVKGLLGAAKSPADRKTACTCLKSAANAIKGIDMGKAAGLPAACGVNIPYKISPSTDCSKYVSPSFLSYYAFMLSSFIFY*

NbL14g08730.1 non−specific_lipid−transfer_protein_2−like [MW − SP = 10.17 kDa]

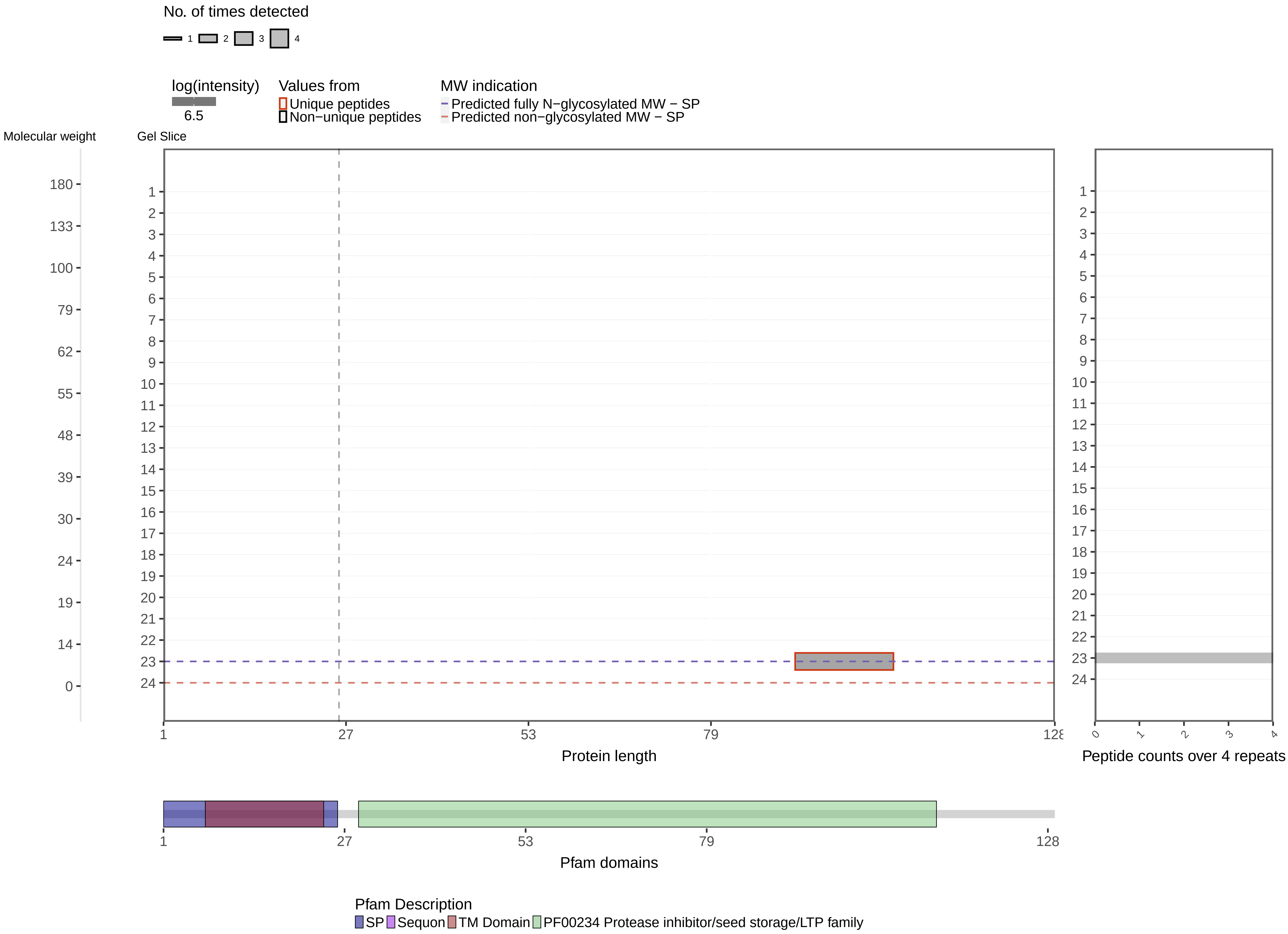


Protein sequence:

GCCGGVKGLLGAAKSSTDRKTACTCLKSAANAIKGIDMGKAAGLPGACGVNIPYKISPSTDCSKYVFQLLFFLTTSL*

### NbL14g08710.1 non−specific_lipid−transfer_protein_2−like [MW − SP = 8.56 kDa]

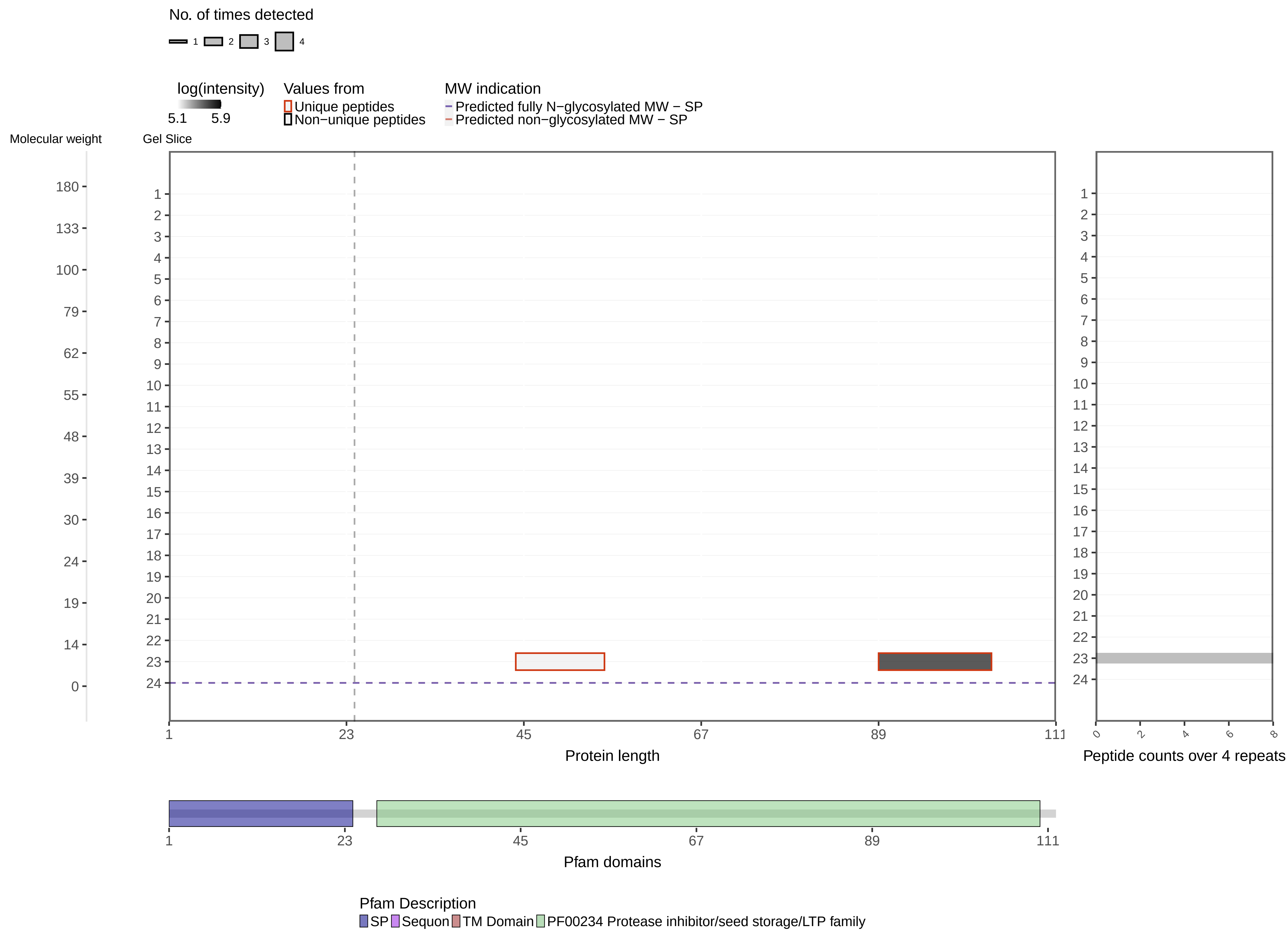


Protein sequence:

RRGPLGGCCGGVKGLLGAAKSPADRKTACSCLKSAANAIKGIDMGKAAGLPGVCGVNIPYKISPSTDCS

### NbL14g08700.1 non−specific_lipid−transfer_protein_2−like [MW − SP = 9.28 kDa]

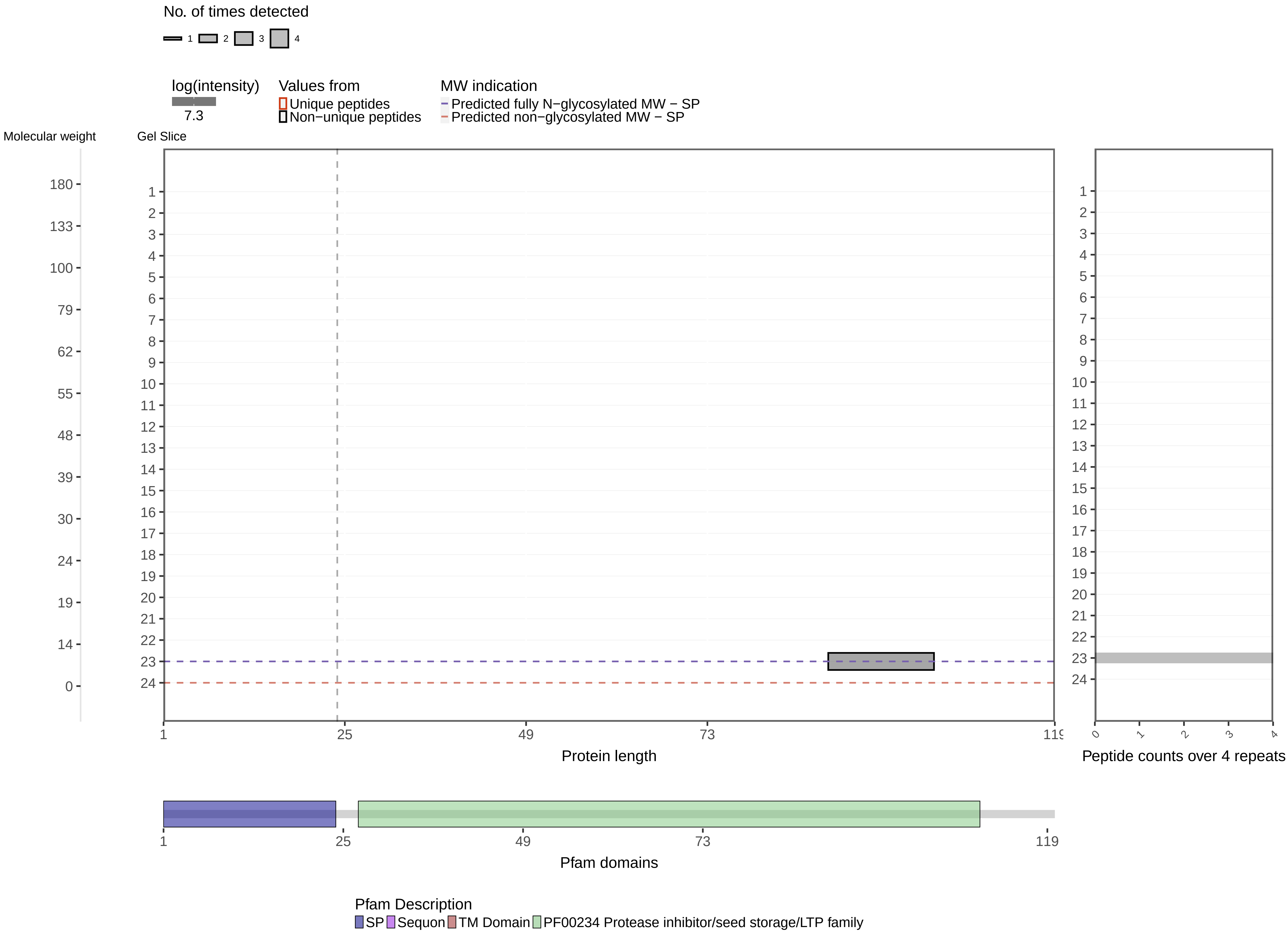


Protein sequence:

PLGGCCGGVKGLLGAAKSPADRKTACTCLKSAANAIKGIDMGKAAGLPSACGVNIPYKISPSTDCSKYVFSSS*

NbL14g08660.1 elongation_factor_P−like [MW = 25.79 kDa]

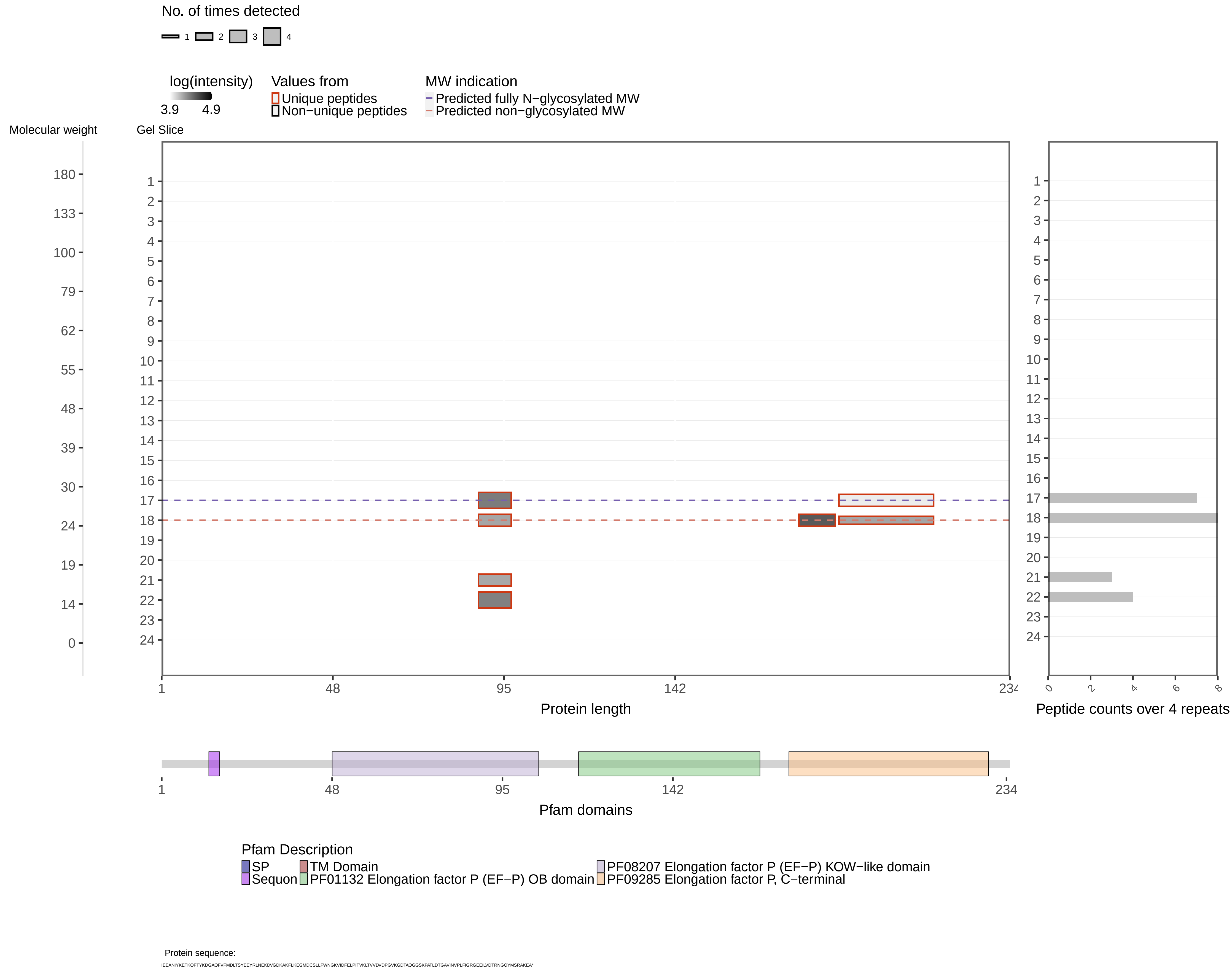

NbL14g08640.1 vacuolar−sorting_receptor_1−like [MW − SP = 67.44 kDa]

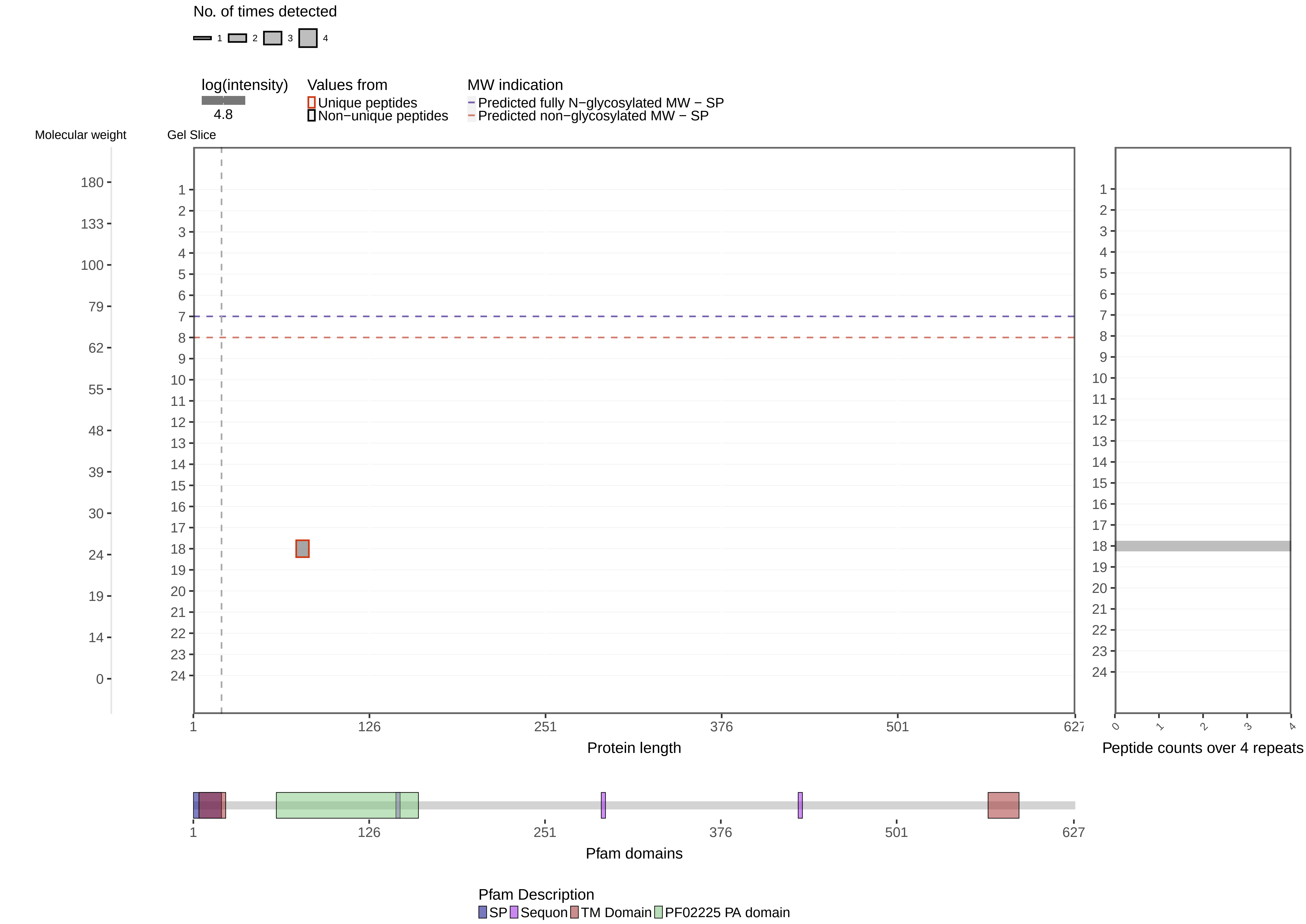


Protein sequence:

YVTDFAIRCPMKEKKYTKECADQVVKSLGFDVKQIDKCVGDPEADIDNPVLKAEQEAQIGKDFRGDVTILPTLVINNRQYRGKLDKGAVLKAICSGFEETTEPAICLTDDIQTNECLESNGGCWQDKAANITACRDTFFRGKVCECPMVQGVKFVGDGYTLCEASGALRCGINNGGCWKGTKDGRTFSACIDDHTKGCKCPPGFKGDGVNSCEDIDECKERLACQCPECKCKNTWGGYDCSCNGNLLYMHEHDTCINAGKDGKTEFSWGFVWVIILGLATAGVGAYAVYKY

### NbL14g08590.1 histone_acetyltransferase_TAP1 [MW = 28.54 kDa]

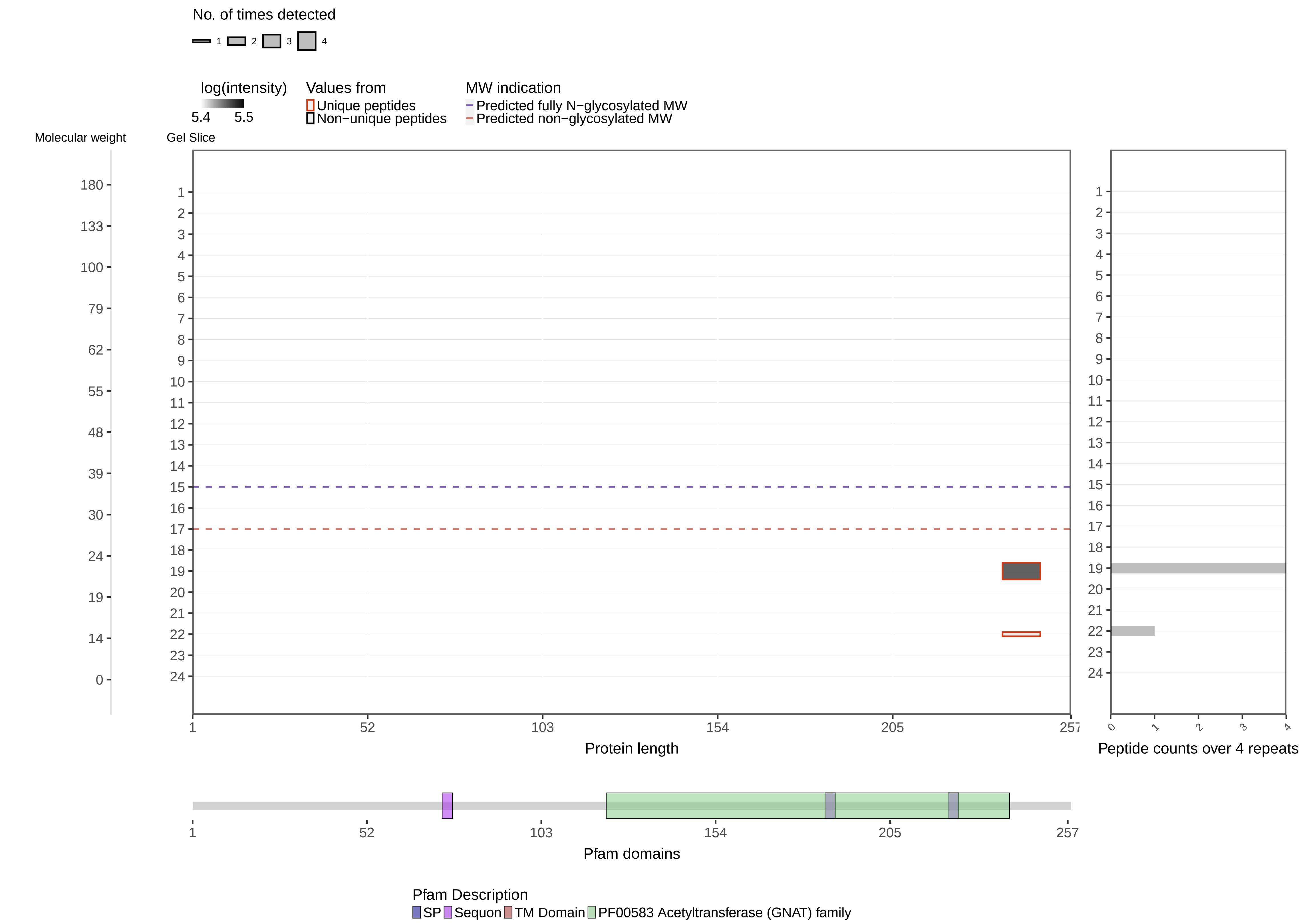


Protein sequence:

SSGGDVDVYDLQALCDKVGWPRRPLSKLAAALKNSYIVATLHSRKFSSGEEGNGEKKLIGMARATSDHAFNATIWDVLVDPSYQGQGLGKALIEKLIRTLLQRDIGNISLFADSQVVEFYKNLGFEADPEGIKGMFWYPMY*

### NbL14g08490.1 calcium−binding_protein_CAST−like [MW = 22.71 kDa]

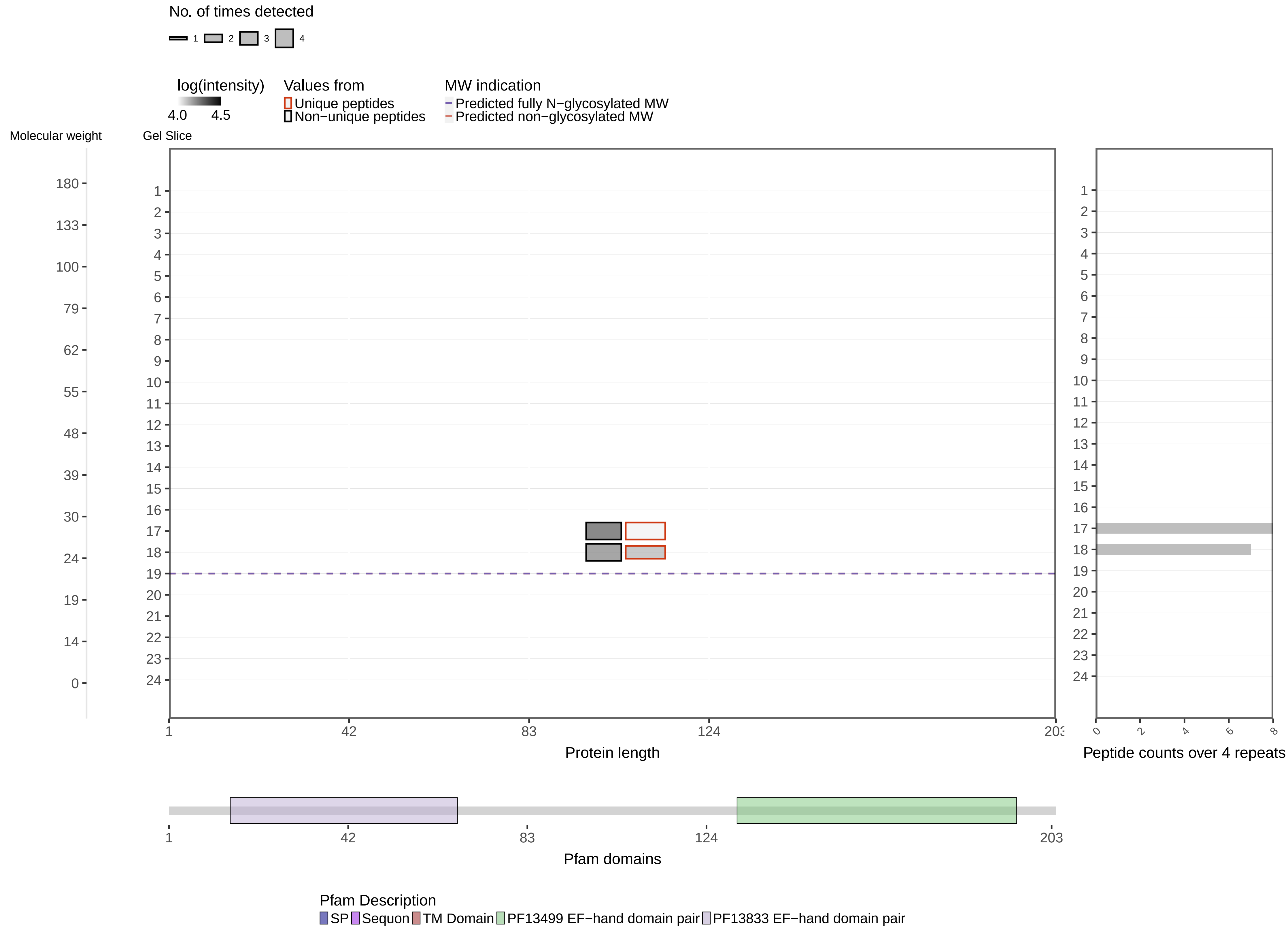


Protein sequence:

GNNGLRFEDFEALHRSLDDVFFGSKYEDKIGLDQDPDQDEVDLKDAFNVFDENGDGFISAKELQAVLEKLGLPEGNEIDRVEMMISSVDQDHDGQVDFVEFKDMMRTVIVPSS*

NbL14g08390.1 protein_transport_protein_SEC31_homolog_B [MW = 122.31 kDa]

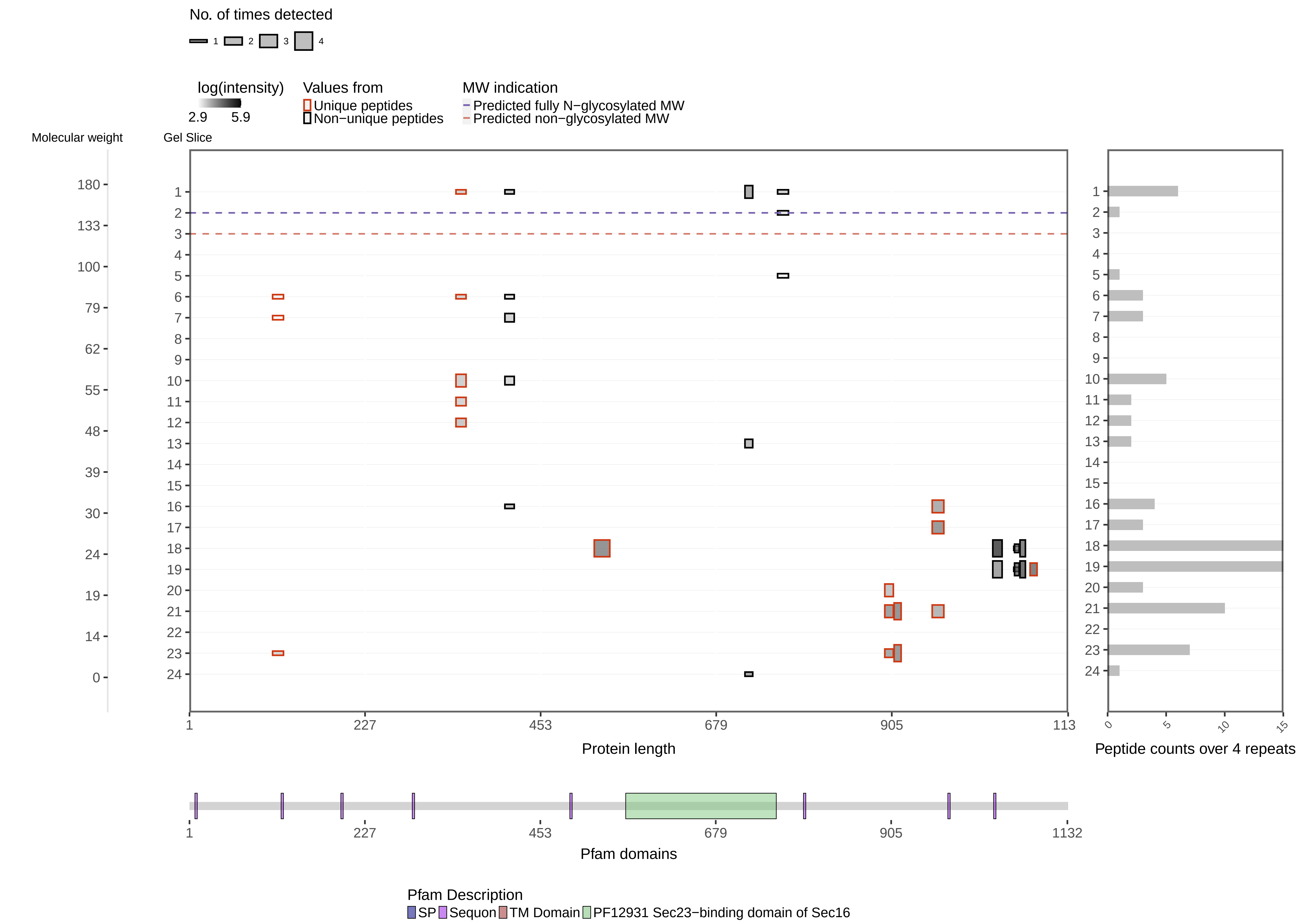


Protein sequence:

QEESADTSFDEIVQSALVVGDYKGAVAQCISANRMADALVIAHVGGASLWEHTRDQYLKTSQSSYLRVVAAMVNNDLMSLVNTRPLKSWKETLALLCTFAPQDEWTSLCDTLASRLLAAGETLPATLCYICAGNIDKTIEIWSRTLAGKRDGKSYVDLLQDLMEKTIVFALATGQKRFSASLCKLLEKYAEILASQGLLTTAMEYLKLMGSEELSPELTILRDRIALSTEPAKDASKSVAFNNSLLHAGSGYVADQSGYGMVDPSQHYYPEQPSKPQPSISNSPYAENYQQPFGS

NbL14g08290.1 50S_ribosomal_protein_L1_chloroplastic [MW = 37.05 kDa]

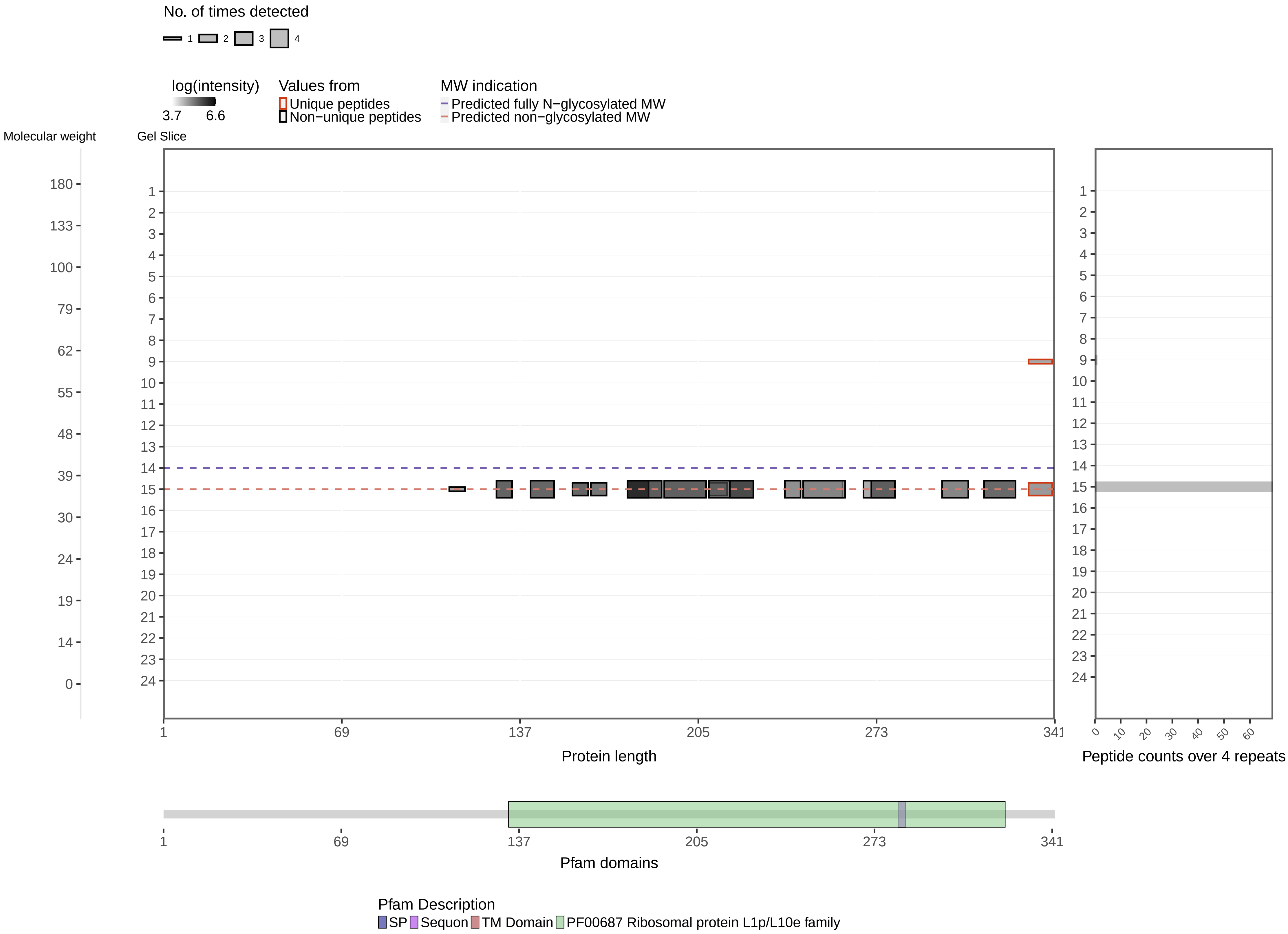


Protein sequence:

QQLRATVNLPKGTGQTVKVAVLTQGEKFDEAKNAGADLVGGEDLIEQIKGGFMDFDKLIATPDMMPKVASLGRILGPRGLMPNPKAGTVTTNIPQAIEEFKKGKVEYRADKTGIVHLPFGKANFSEEDLIINFIAAVKSIEANKPSGAKGVYWKSAHVCSSMGPSIRLNVREMLEYKLPNA*

### NbL14g08210.1 alpha−N−acetylglucosaminidase [MW − SP = 89.81 kDa]

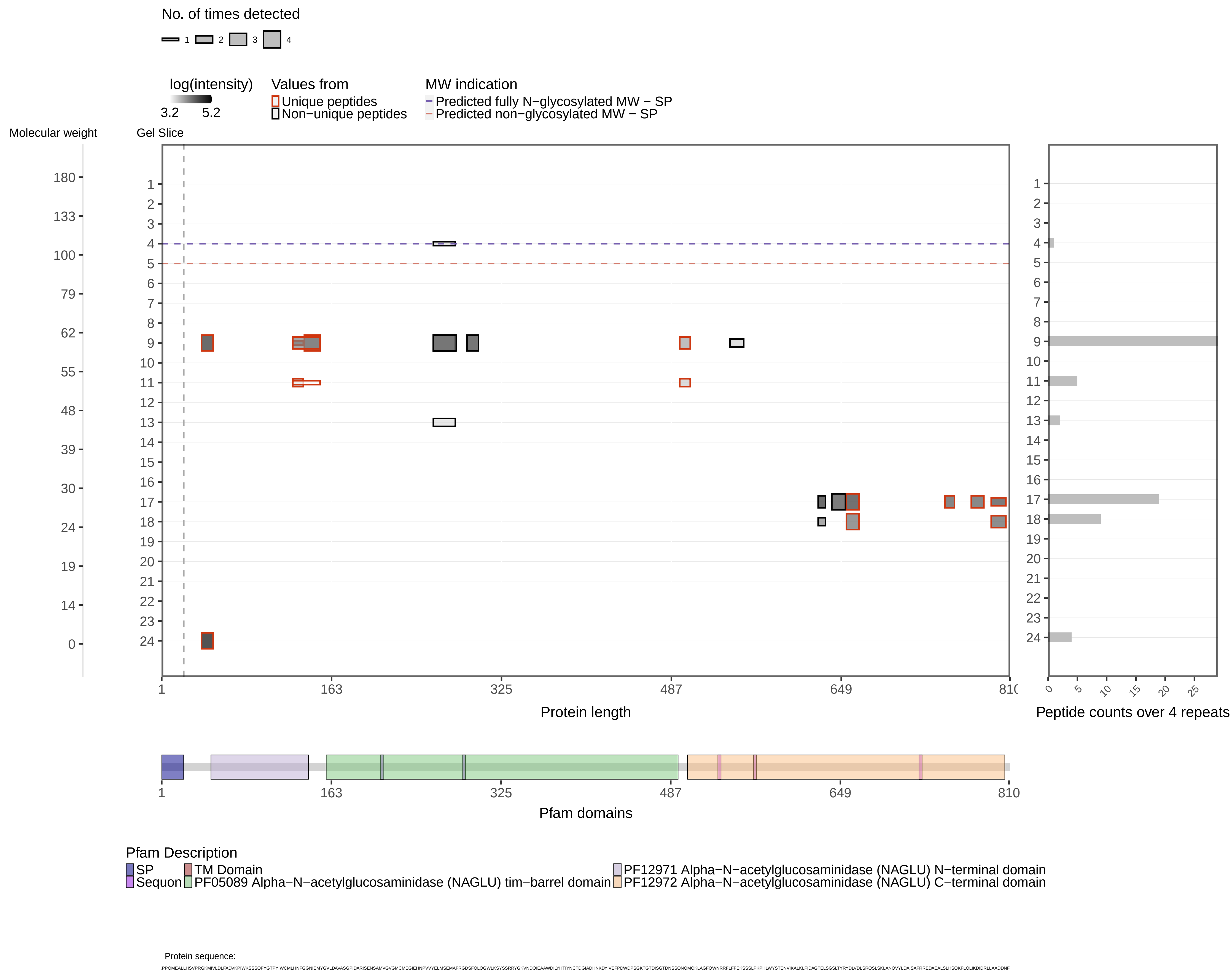

### NbL14g08120.1 ferredoxin−thioredoxin_reductase_variable_chain−like [MW = 19.24 kDa]

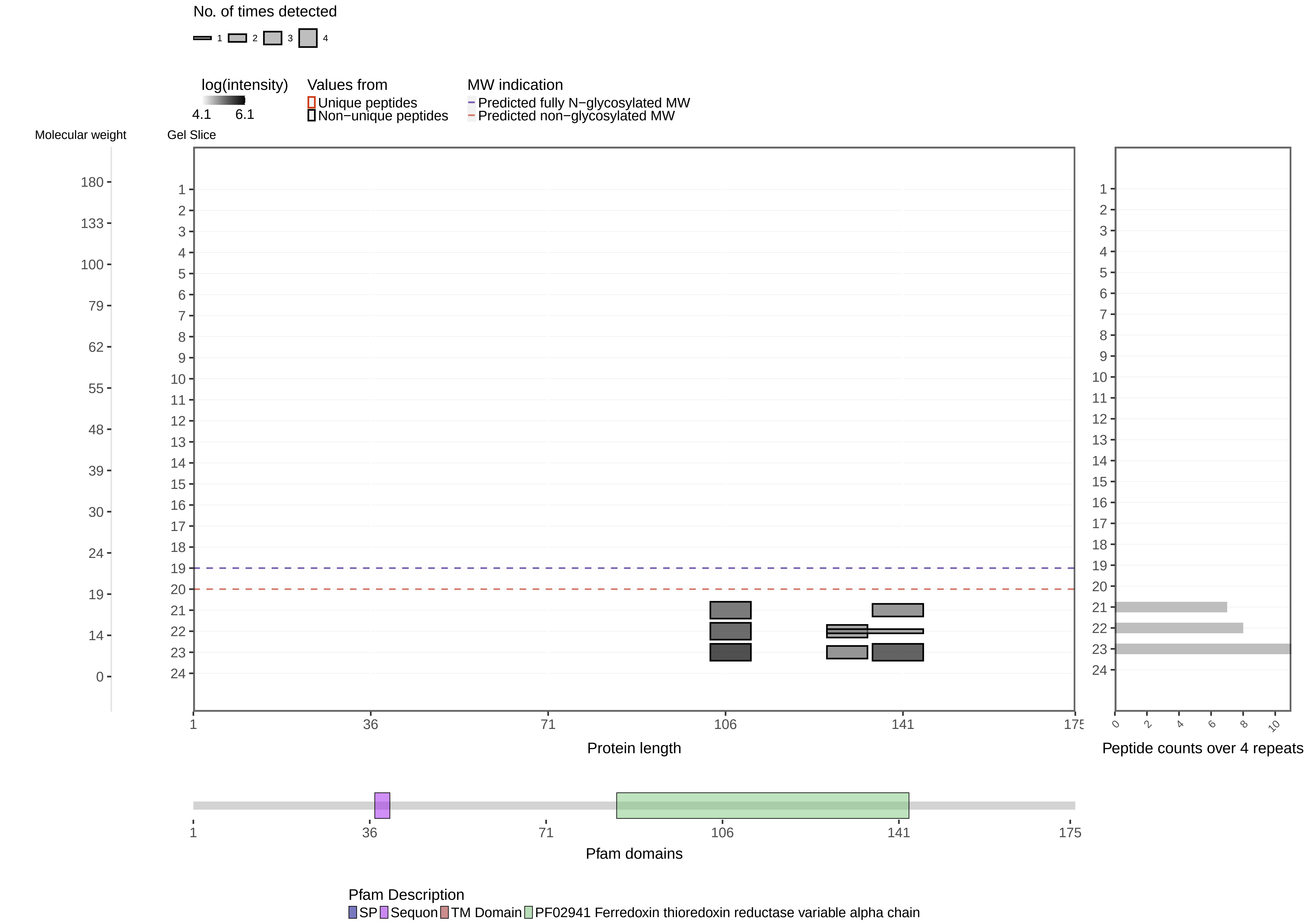


Protein sequence:

EKTAEAIGKVGSRVRVTVPLKVYHVPKVPELDLVNRIGTLKQYVGFHKGKQISANLPYKVEFVVDNLEGRDGQKVAKFMELLLCCWDHGLIINNRQMKN*

### NbL14g08060.1 tRNA−dihydrouridine(20/20a)_synthase−like [MW = 47.93 kDa]

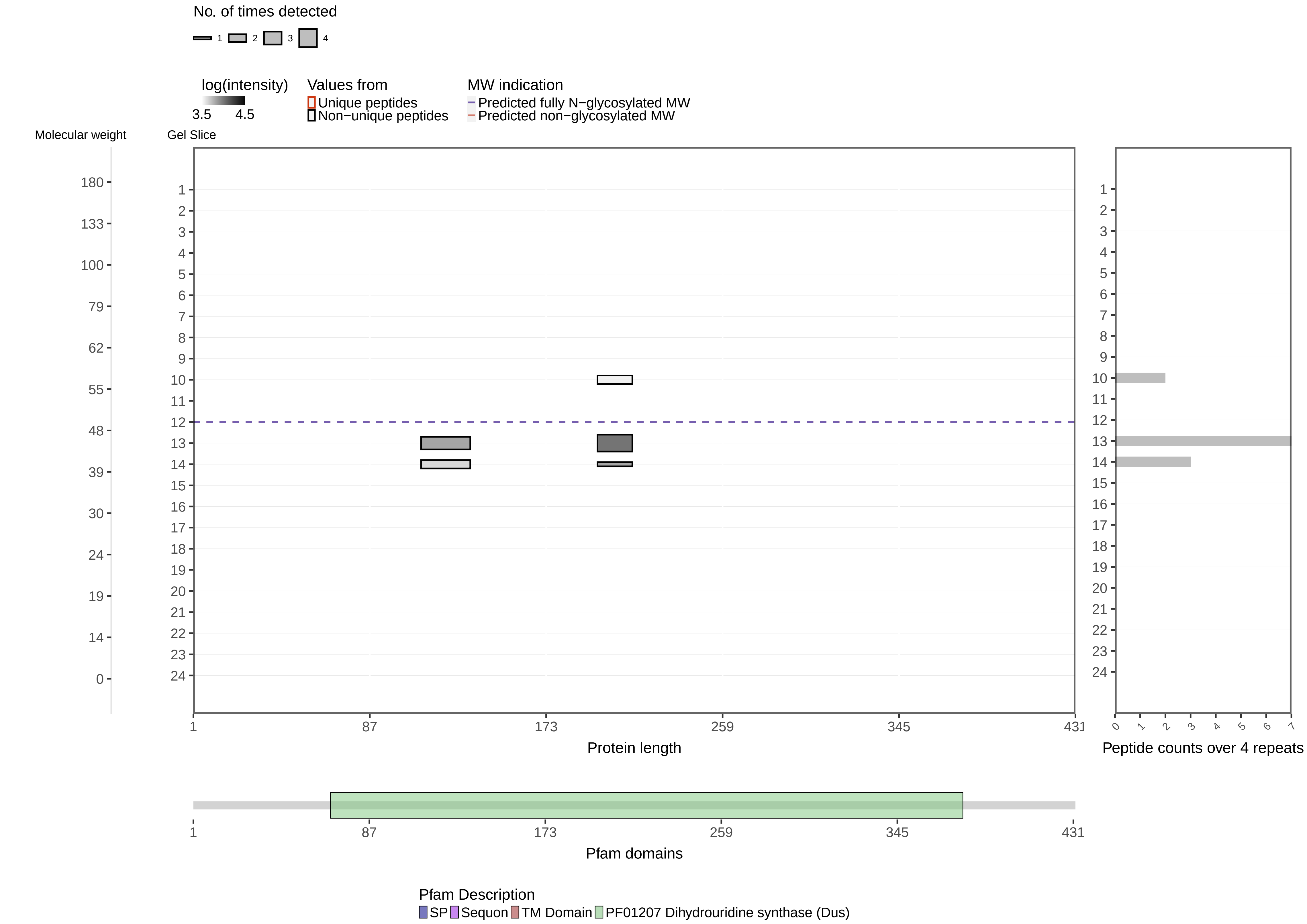


Protein sequence:

HDSYNELCDFIYKVSSQSPTRHFIIHSRKALLNGISPADNRKIPPLKYEYYYALLRDFPDLQFTINGGINSIEEVNEARTEGAHGVMLGRAAYSNPWQVLGLVDSAVYGAPLRSITRRQVLEQYQVYGDSVVGMYGGLKPSMRDVVKPLLGLFHANPGNGAWKRKVDAAFKHYTTIKSLFEETLVAIPDWVLDSPVTEVPGGCTDKFSKAKSLLPPPYTVNEEELLYA*

### NbL14g08040.1 carotenoid_910(9'10')−cleavage_dioxygenase_1−like [MW = 61.92 kDa]

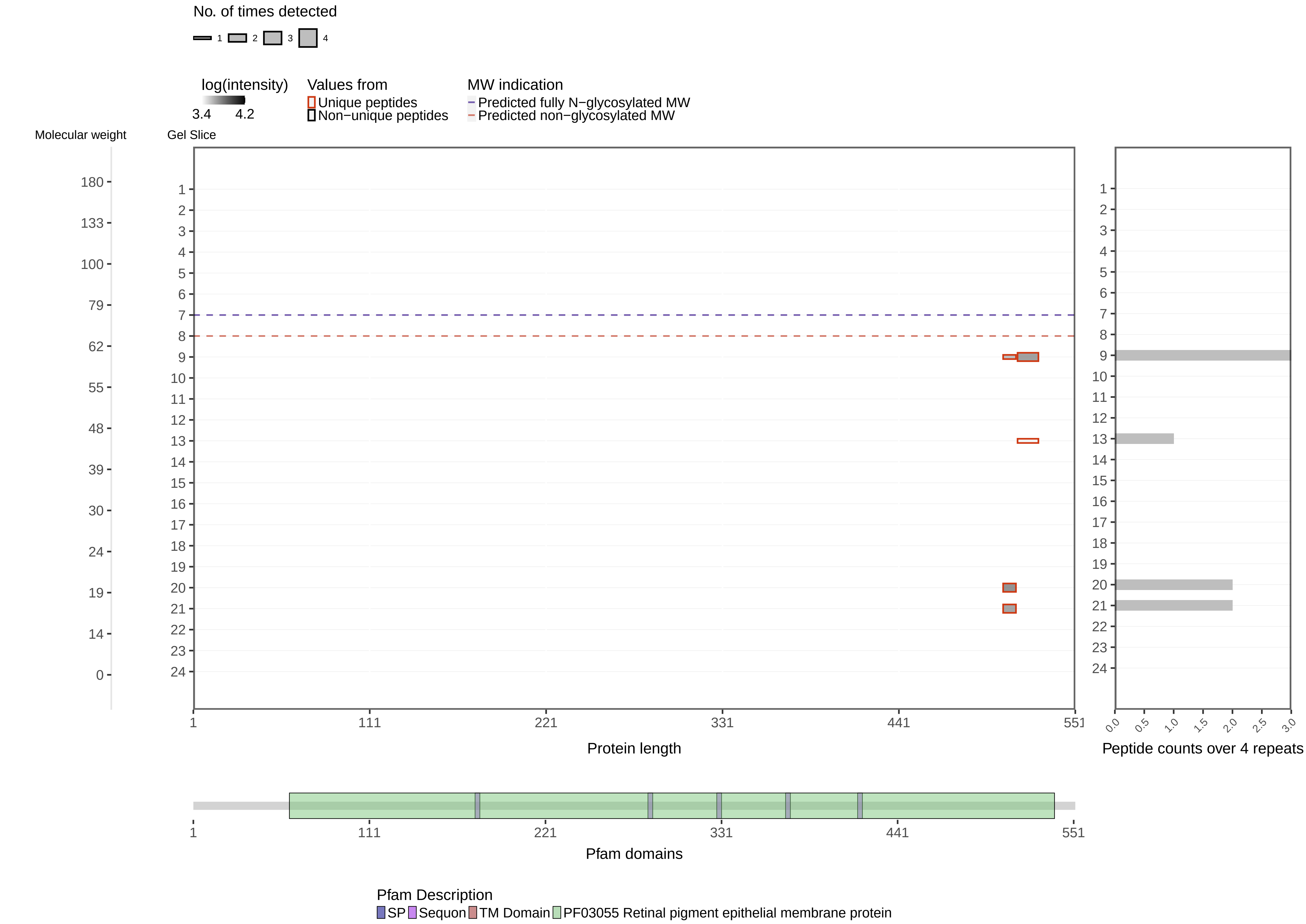


Protein sequence:

GVMQDPVPITIPASVMMHDFAITENYSIMMDLPLYFKPKEMVKNKKLAYSFDPTKKARFGILPRYAKNESLIKWFELPNCFIFHNANAWEEGDDVVLITCRLQNPDLDAINGTEKEQQRDGFTNELYEMRFNMKTGLASQKKLSESAVDFPRVNENYTGRKQRYVYGTILNHMAKITGVVKFDLHAEPDTGKTKLEVGGNVLGIFDLGPGRFGSEAIFVPCQPGTECDEDDGYLILFVHDENTGKSSVNVIDAKTMSAEPVAVVELPKRVPYGFHAFFVTEEQIQEQAKL*

### NbL14g07980.1 uncharacterized_LOC107829775 [MW = 25.35 kDa]

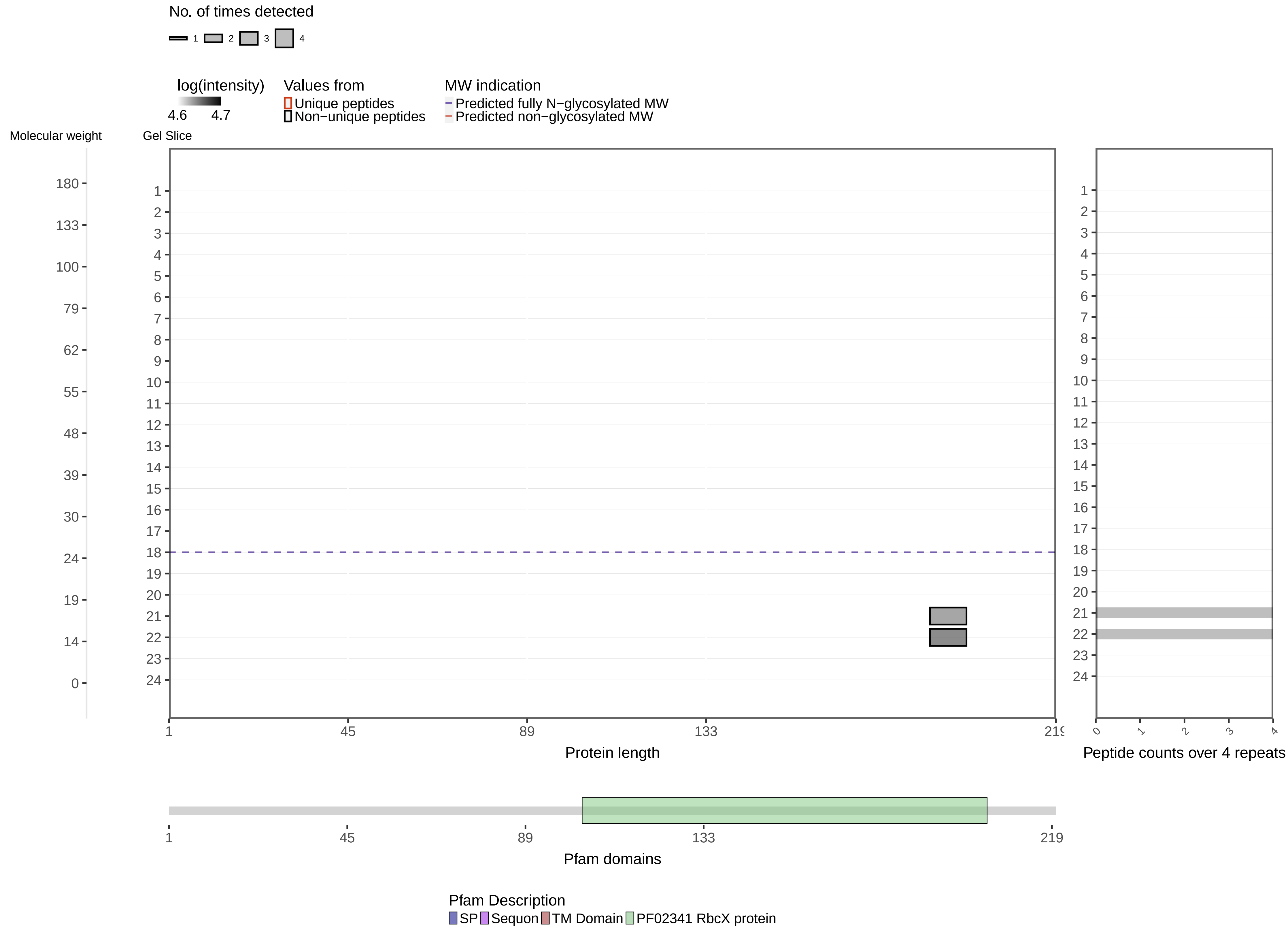


Protein sequence:

DDDSGEDVKNEIINVITYKAVRTVLQQLYEMNPPQYTWFYNFIVNHVPNTGKDFLQQLFKEKRELAERVMITRLNLYSIWMQKCDHAELYNRISDQNVELMRERLAQTVIWLSEDDEIAGSLD*

### NbL14g07950.1 uncharacterized_LOC107772121 [MW = 34.86 kDa]

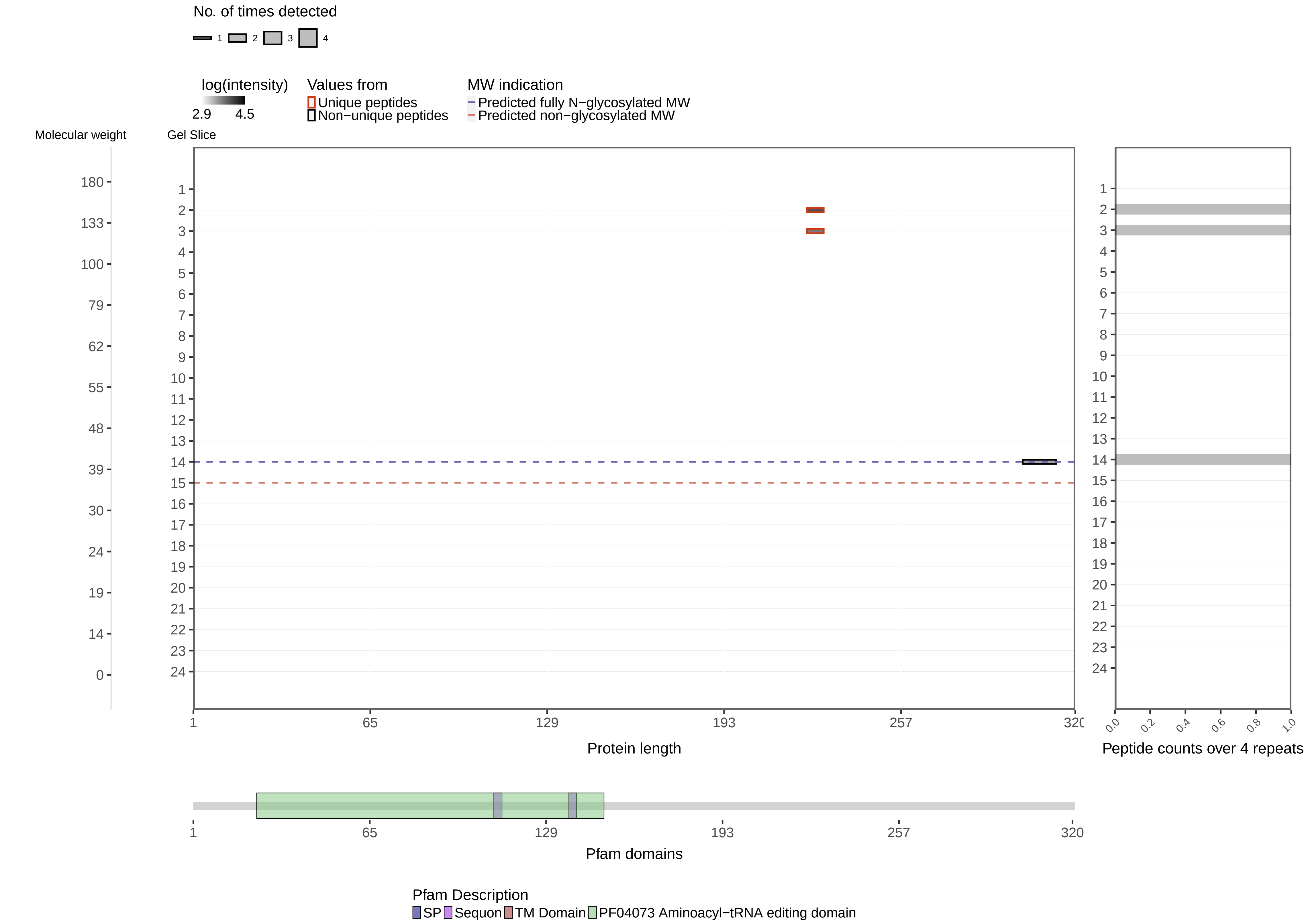


Protein sequence:

RDLDKLLSSIGKKPVYVDLEANPPVGKGQPPDLATFVPSDTIVLPGHTYKEAPLQVSDTNHAPVSNKATGVSAKAVKPSADVLKEKLSNGVSSPNSFADPEKFVEDFLEKTSSLVLSELTEENVKQDGEKLGEAISNKIRNKLATEFKSMATMFKNTAYSEGFQAGIRHQAKRP*

### NbL14g07870.1 photosystem_II_22_kDa_protein_chloroplastic [MW = 29 kDa]

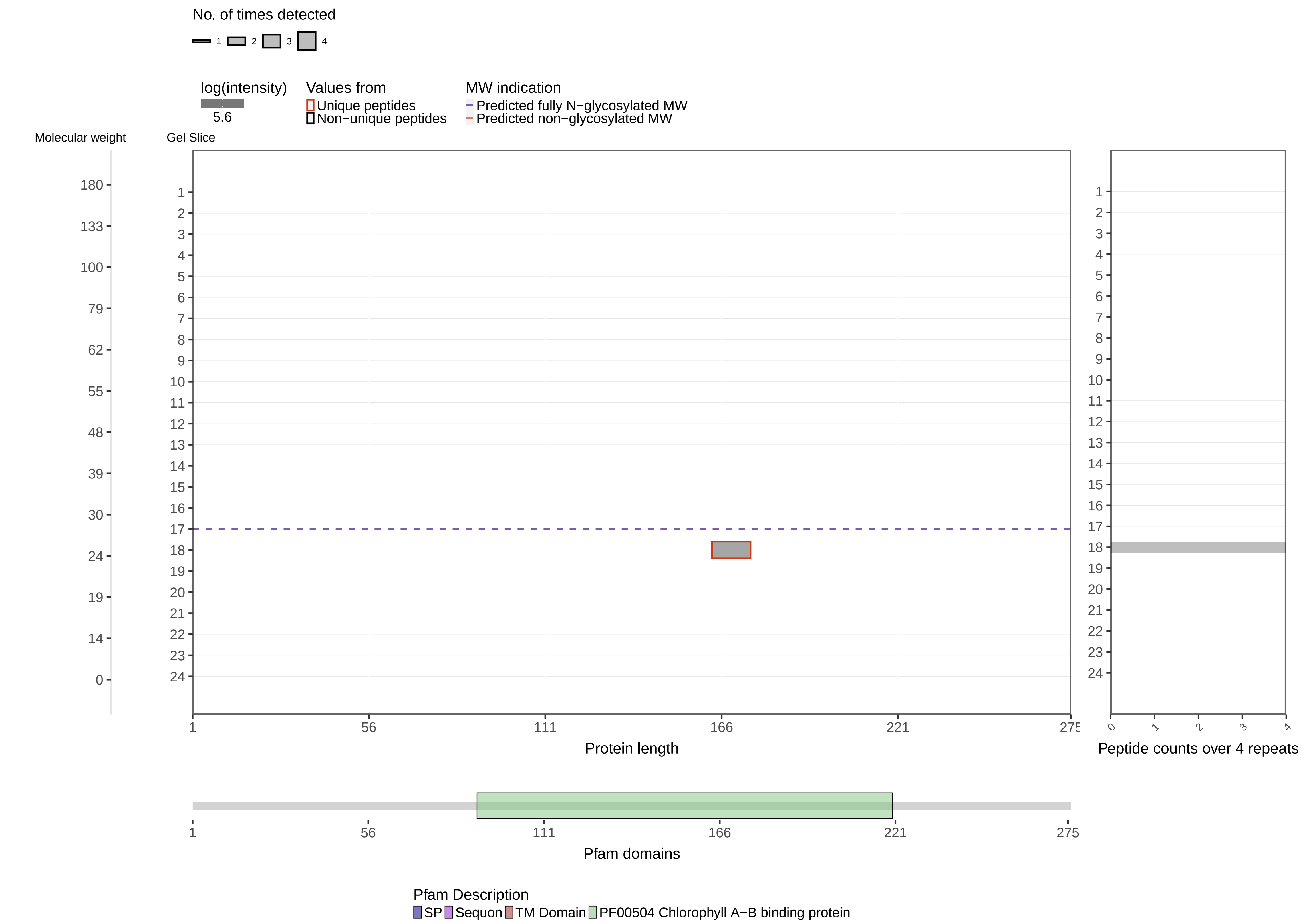


Protein sequence:

GILAQLNLETGIPIYEAEPLLLFFILFNLLGAIGALGDRGKFIDDPVPATGLDKAVIPPGKGFKSALGLSEGGPLFGFTKANELFVGRLAQLGIAFSIIGEIITGKGALAQLNFETGVPINEIEPLLLFNIVFFFVAAINPGTGKFVTDEEEE*

### NbL14g07840.1 SWI/SNF_complex_subunit_SWI3C [MW = 86.13 kDa]

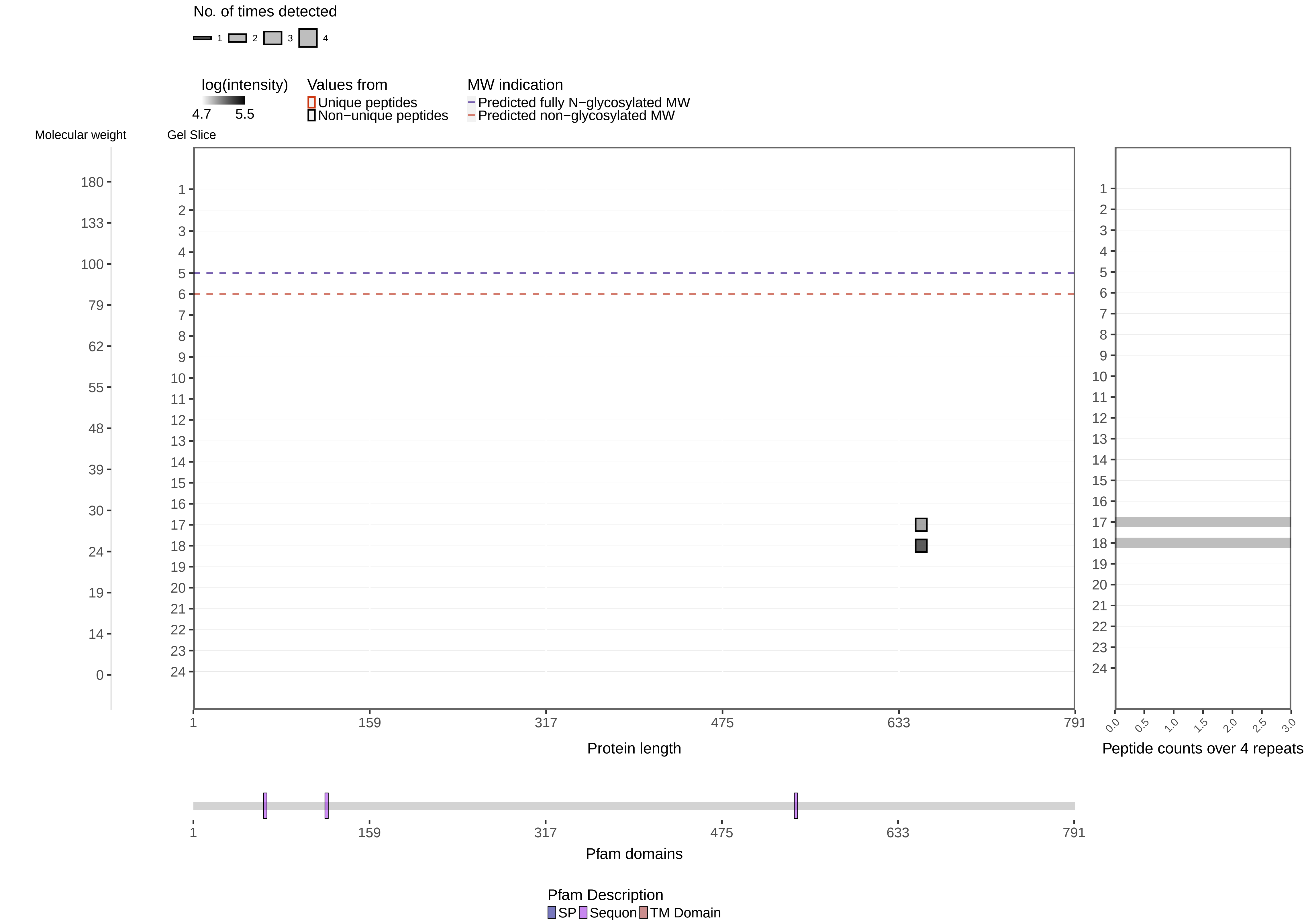


Protein sequence:

NbL14g07700.1 protein_disulfide−isomerase−like [MW − SP = 54.91 kDa]

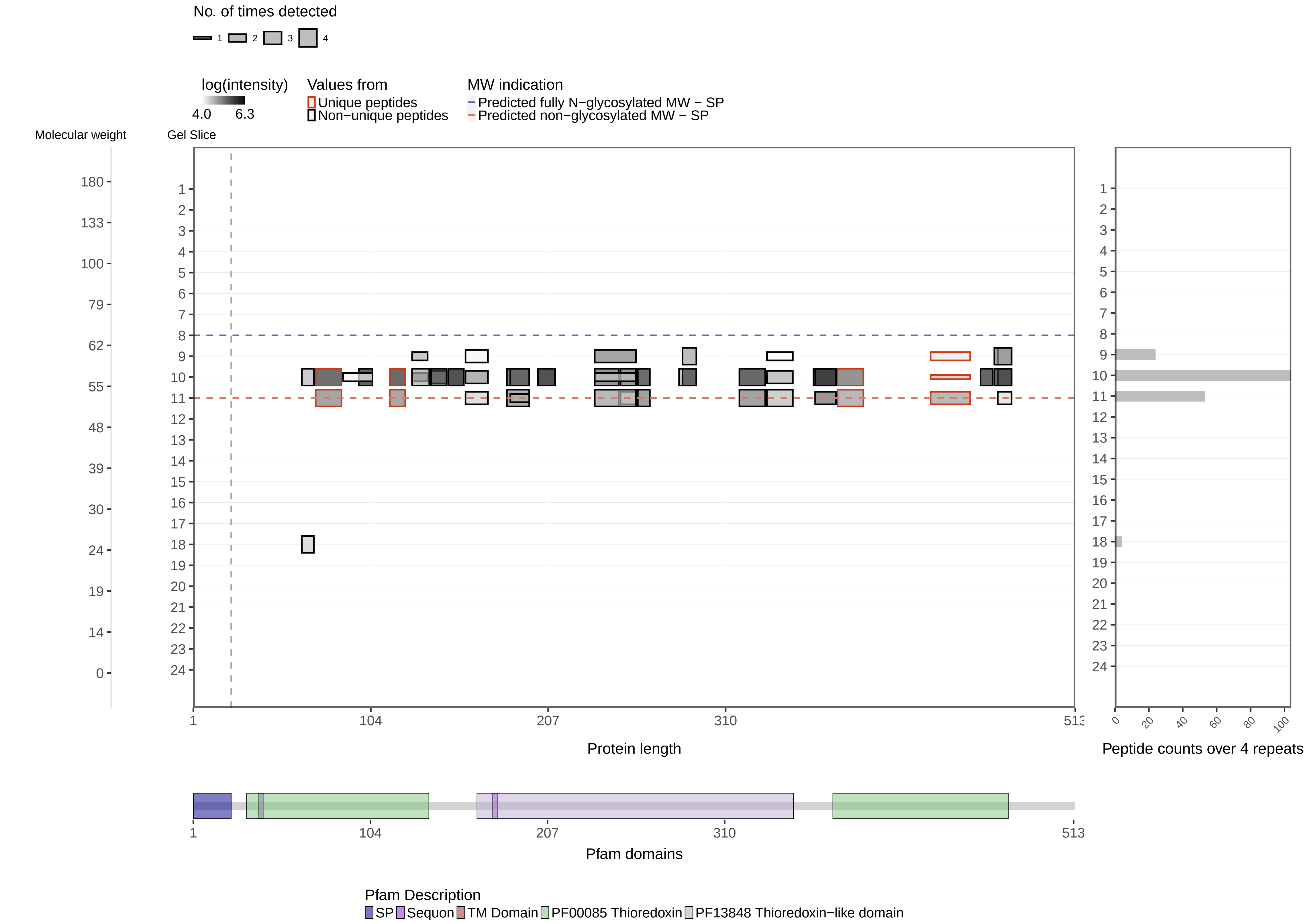


Protein sequence:

LFNKDPNNHPFVIKFFNSPNAKAMLFVNFNSDLIDTFKSKYHEVAEQYKGNDISFLIGDVEASQGAFQYFGLKEDQVPLIIIQTNDGEKYLKPNVEPDHIASWVKDFKDGKVKPFKKSEPIPEVNSEPVKVVVADTLQEMVFNSAKNVLLEFYAPWCGHCKQLAPILDEVAVSFESDADVMIAKIDATANDIPQGTFNVQGYPTLYFKTASGKISQYEGDRTKEDIIDFIHKNRDKAADQGRIATRGDFKGFQGIWHYVGCLTIYQVQ*

### NbL14g07440.1 ISWI_chromatin−remodeling_complex_ATPase_CHR11 [MW = 101.01 kDa]

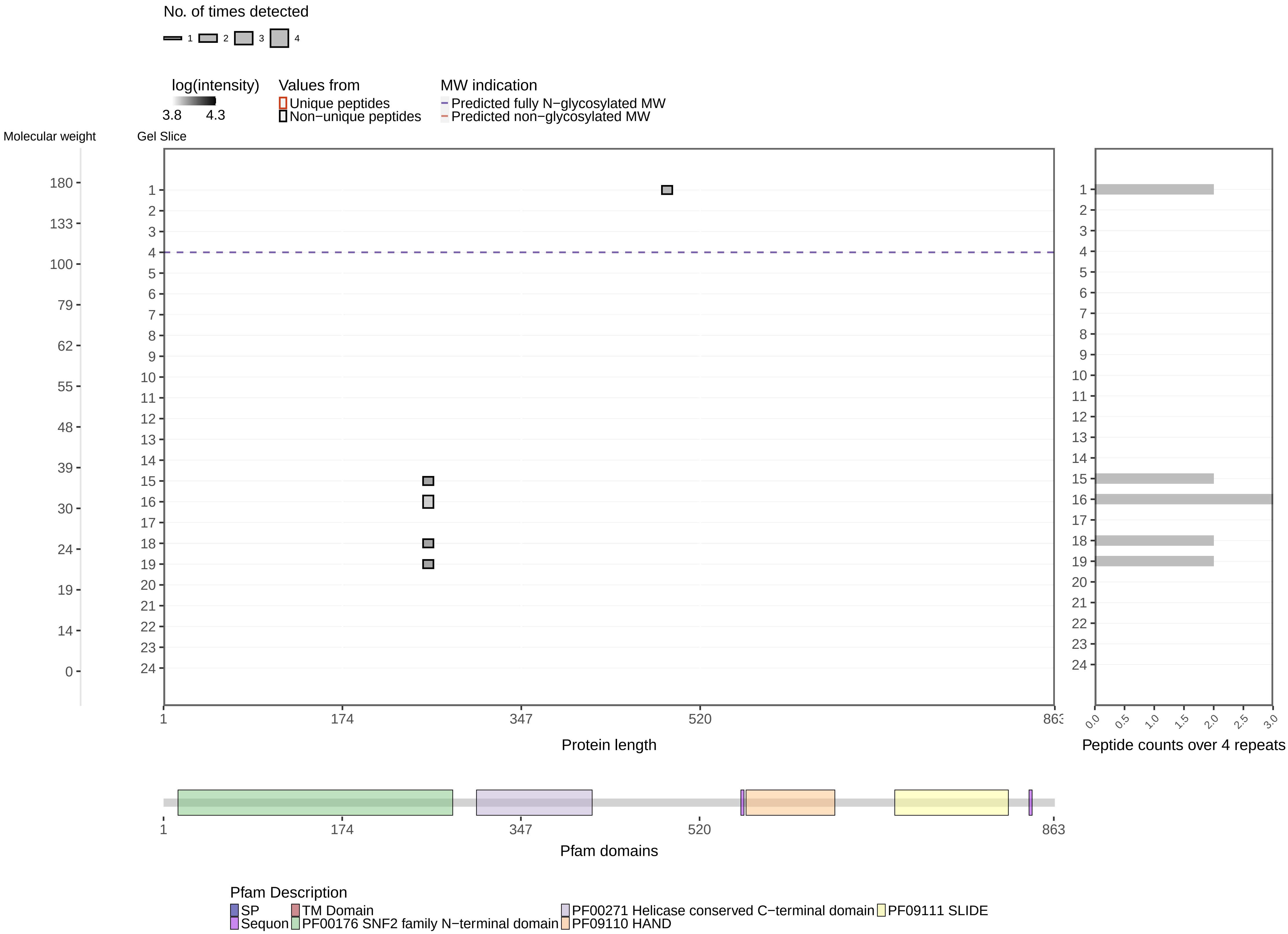


Protein sequence:

[illegible]

NbL14g07400.1 uncharacterized_LOC107810922 [MW = 37.42 kDa]

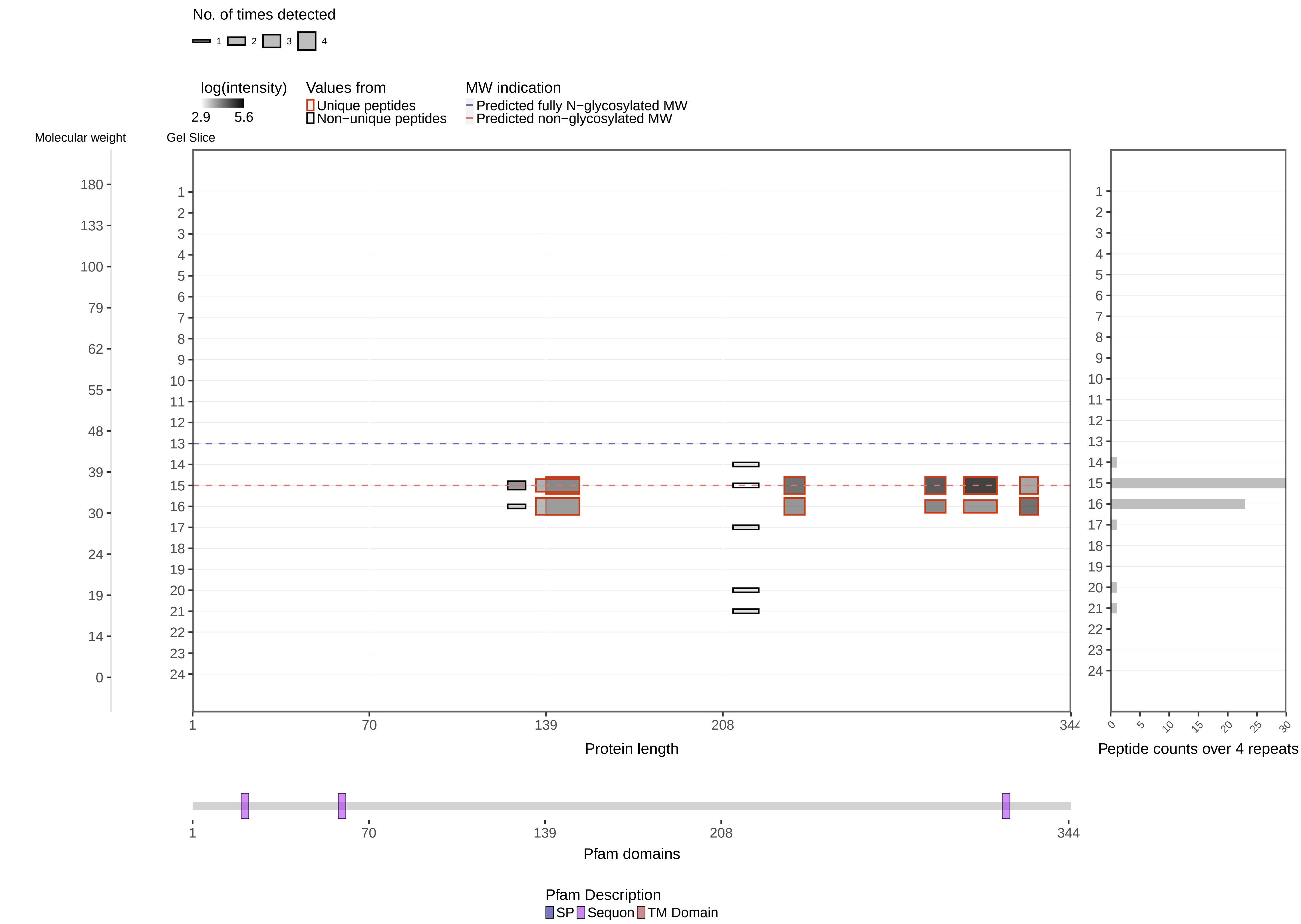


Protein sequence:
AALSWNGEGSFLFTSSSAPYDCFNNGGINEDGPLVPIGRSPRTDVLLKAEKVALDFDGCVVRLAGLYKADRGAHIYWLQKGICDIRPDHILNLIHYEDAASLSVTILKKRLRGQIFLGCDNHPLSRQEVMDLVDKSGTFDKKFEGFTGTSDPLGKKLNNSKTRAELGWEPKYPSFAQFLGVSE*

NbL14g07390.1 probable_prolyl_4−hydroxylase_4 [MW − SP = 29.94 kDa]

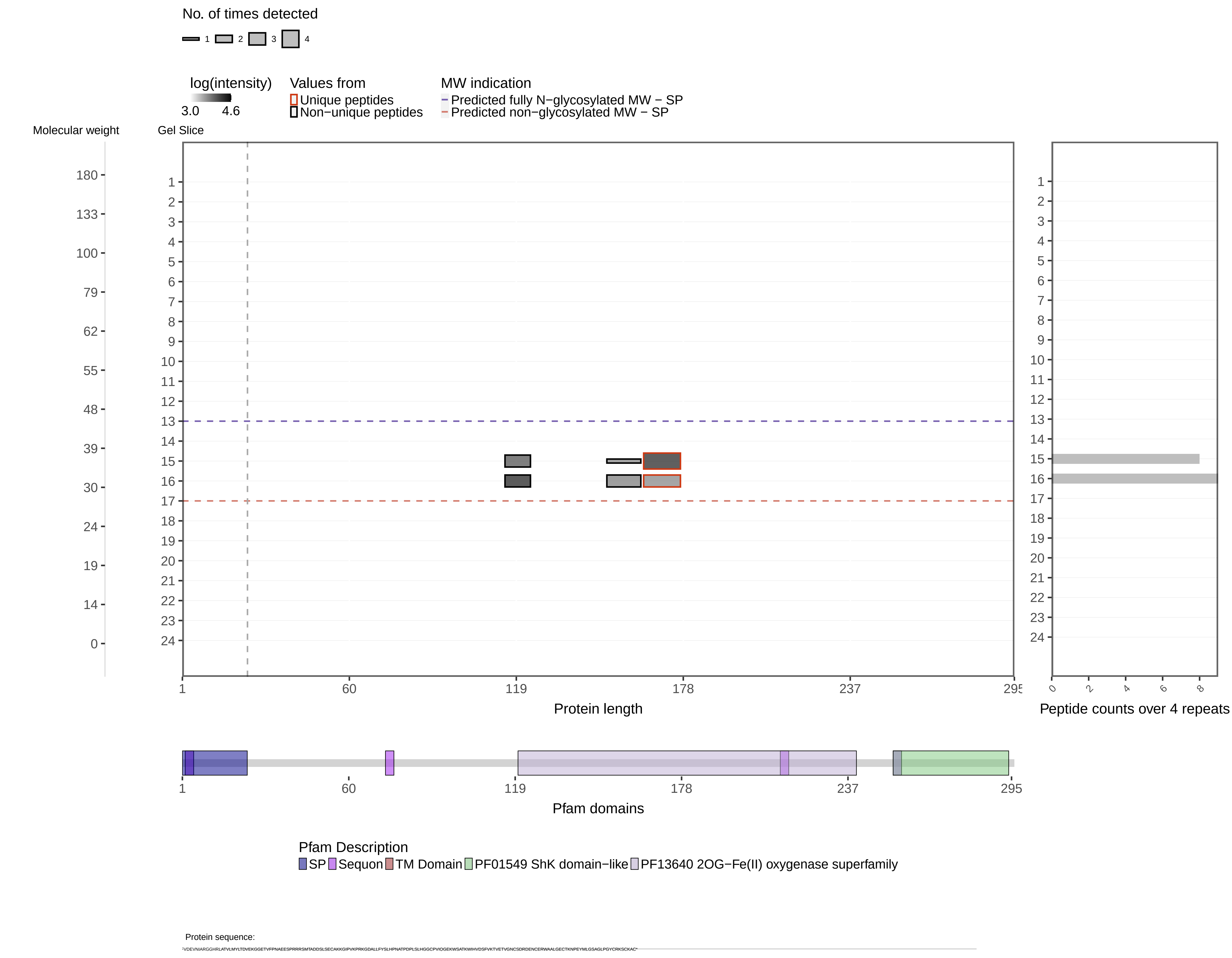

### NbL14g07260.1 photosystem_I_reaction_center_subunit_II_chloroplastic [MW = 22.63 kDa]

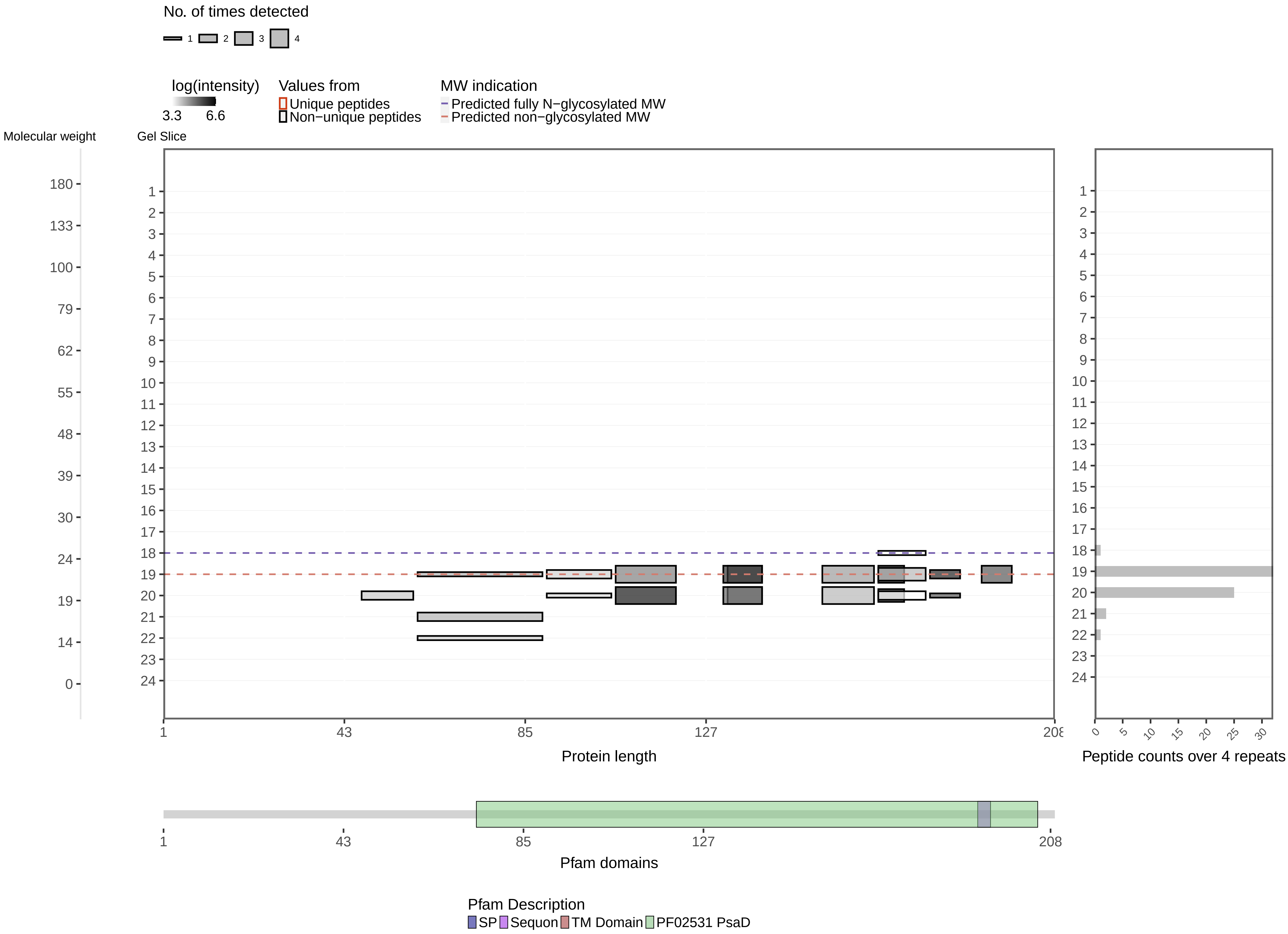


Protein sequence:

QVEEFYVITWESPKEQIFEMPTGGAAIMREGANLLKLARKEQCLALGTRLRSKYKINYRFYRVFPNGEVQYLHPKDGVYPEKVNAGREGVGQNFRSIGRNKSPIEVKFTGKQVYDL*

### NbL14g06980.1 60S_ribosomal_export_protein_NMD3_pseudogene [MW = 58.56 kDa]

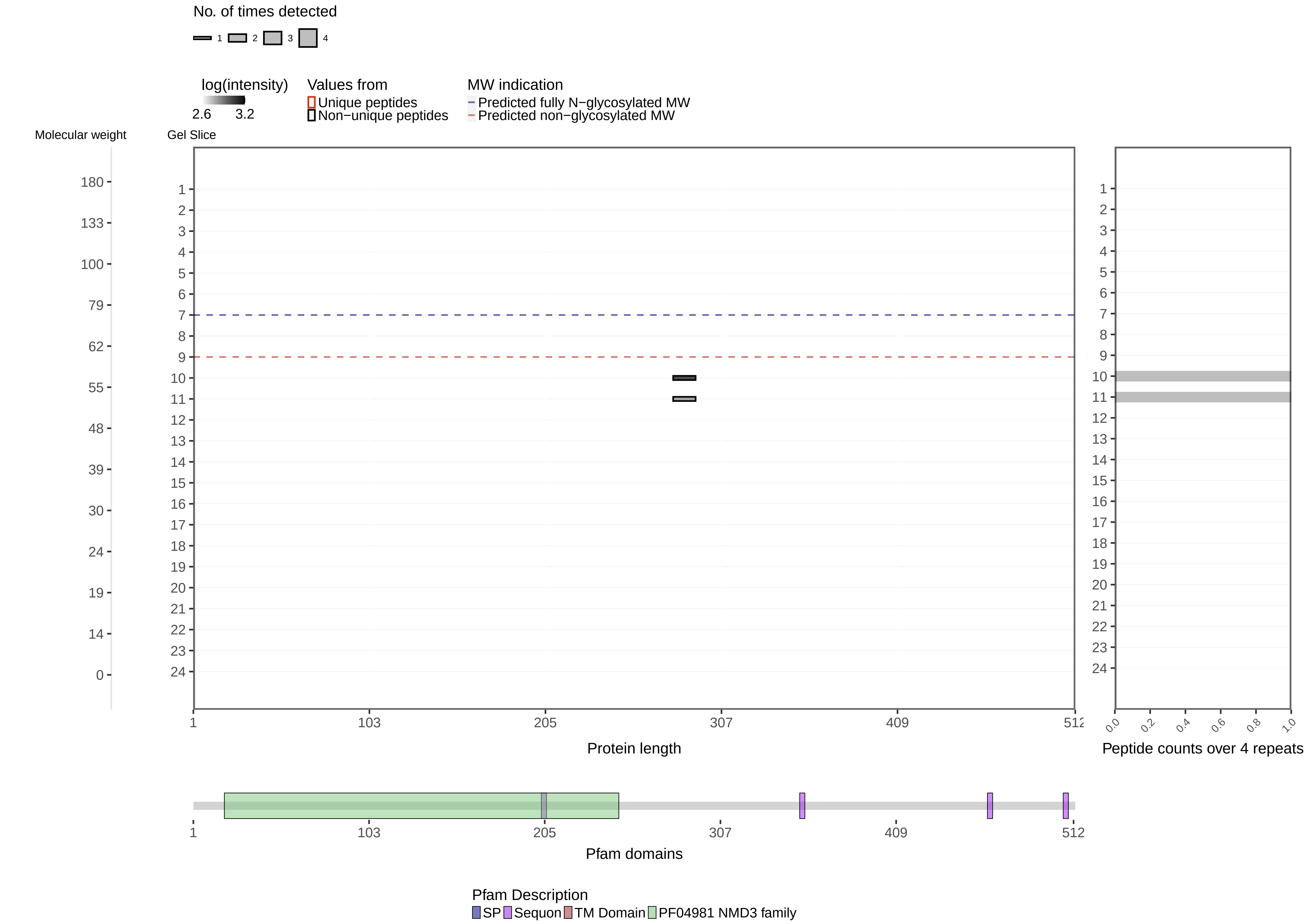


Protein sequence:

TFSVEICPVCREDLICLPPKVKDGLGNLGPLVICTKVSNNIALLDPFTLRNCFLDAEQYWRASFKSLLSSRQLVEYIVLDVESVSSEVNIGGSKYALADAQVARVSDFGKNDTIFNVRTHLGHLLNPGDYALGYDLYAANSNDNELDKYKGFVLPDVILVKKSYEEKWQKKRGKPRSWKLKSLNMEIDDNVKGRDHEEKMSSEYEQFLRDLEENPDLRFNISLYRNREYQPSEMASVTDGDDAPTVALEELLADLDLSEEEADNDSMRE*

### NbL14g06950.1 beta−glucosidase_BoGH3B−like [MW − SP = 65.7 kDa]

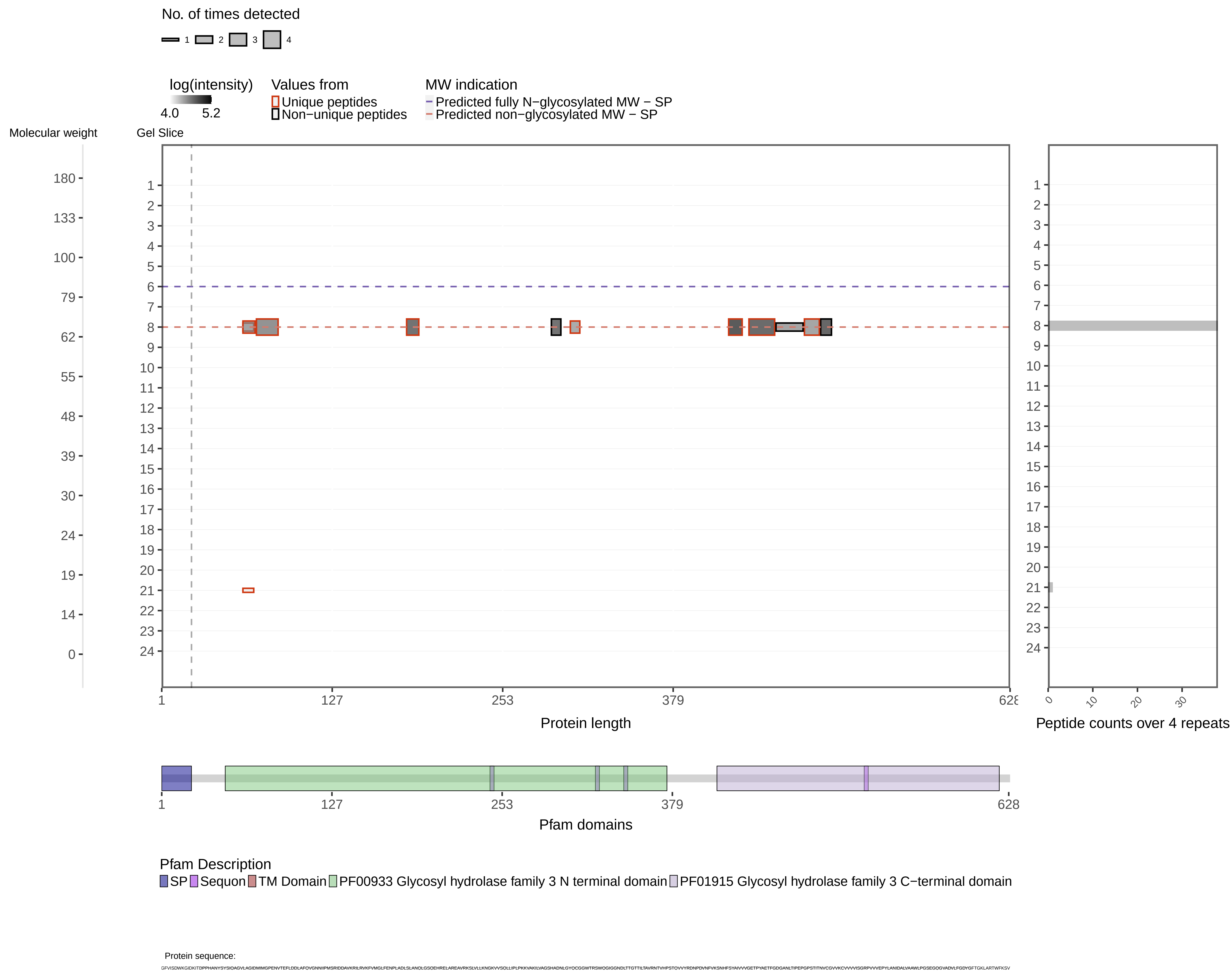

### NbL14g06940.1 lysosomal_beta_glucosidase−like [MW = 72.43 kDa]

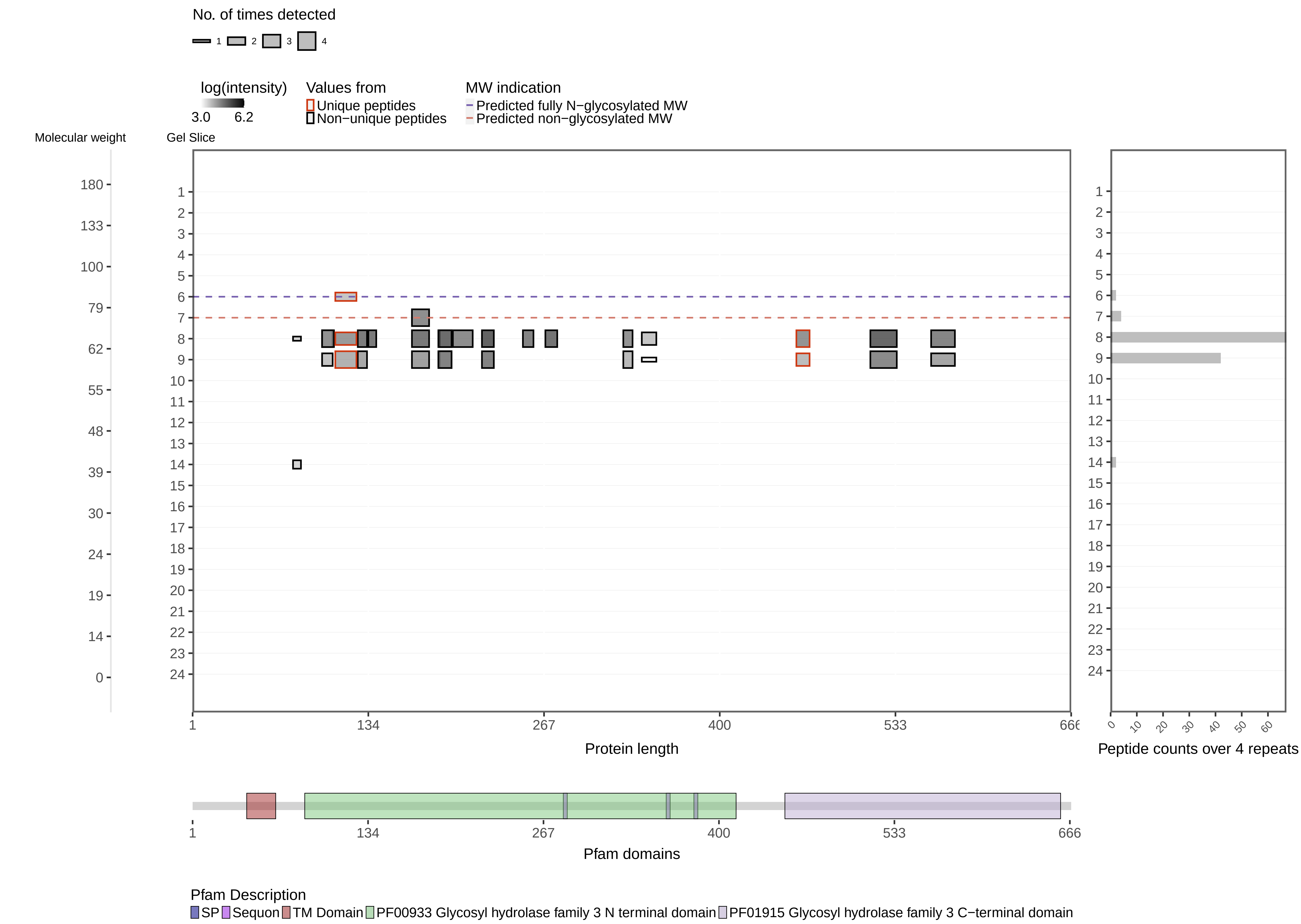


Protein sequence:

### NbL14g06720.1 2−methylacyl−CoA_dehydrogenase_mitochondrial [MW = 44.93 kDa]

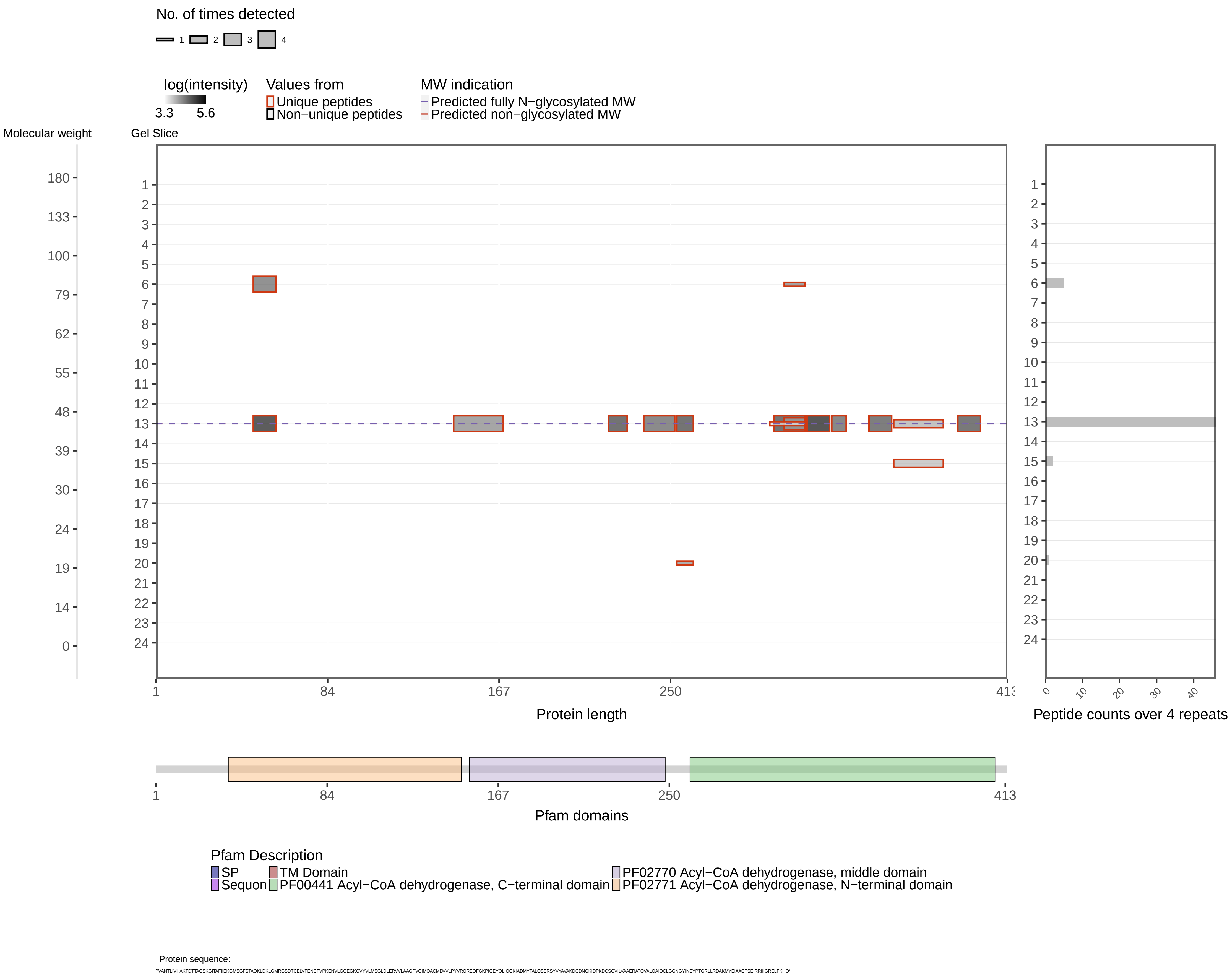

NbL14g06640.1 probable_glutathione_peroxidase_2 [MW − SP = 22.45 kDa]

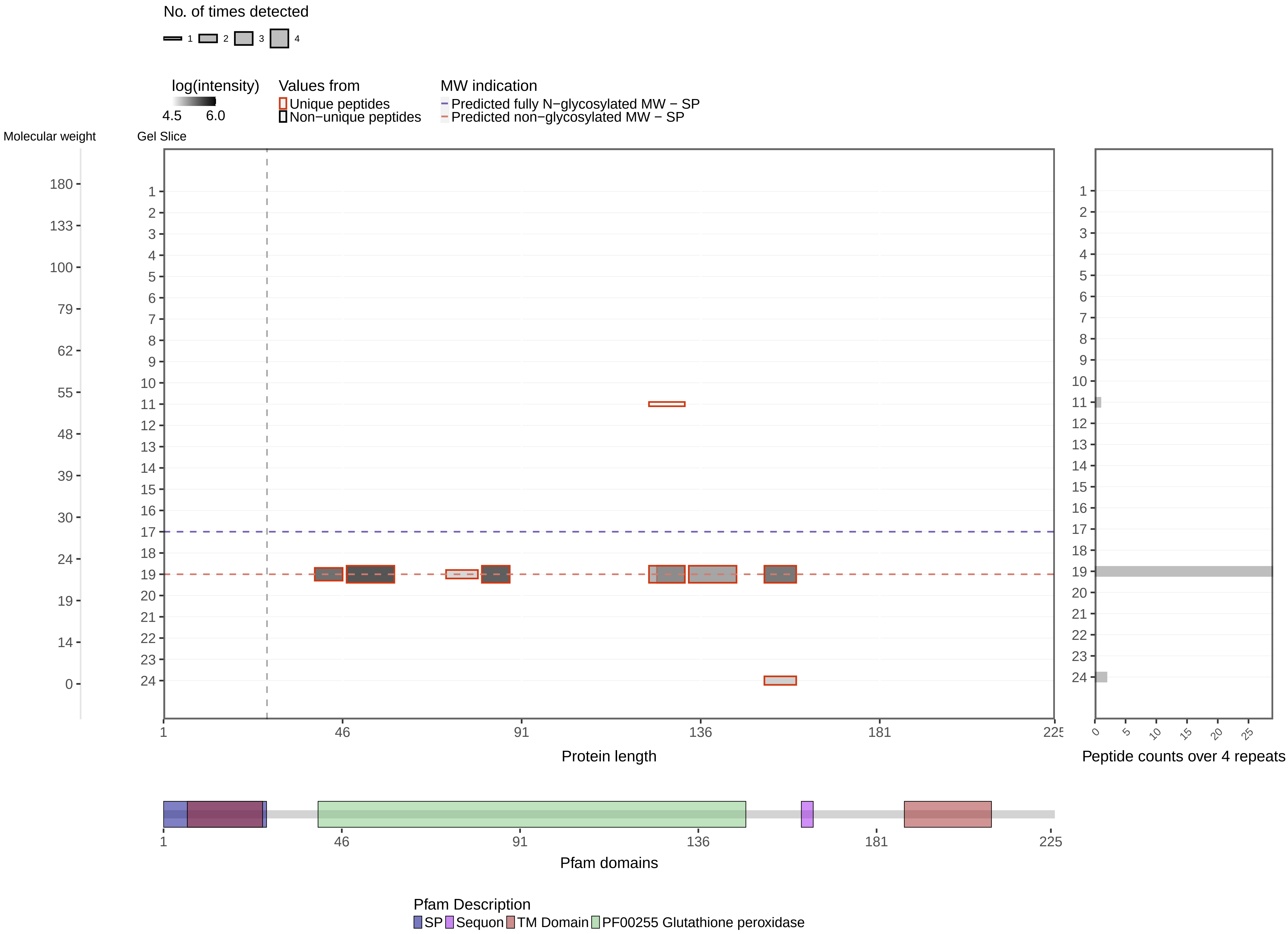


Protein sequence:
CNQFLWQEPGTNEEIQETVCTRFKAEFPVFEKVDVNGDNAAPLFKFLKSEKGGFLGNAIKWNFTKFLVNKEGKIVERYAPRTPPLQFEVTLNCYFFCSFIFYLCPSVLVIKGSGNKRKTLSYLK*

### NbL14g06100.1 heterogeneous_nuclear_ribonucleoprotein_R−like [MW = 88.15 kDa]

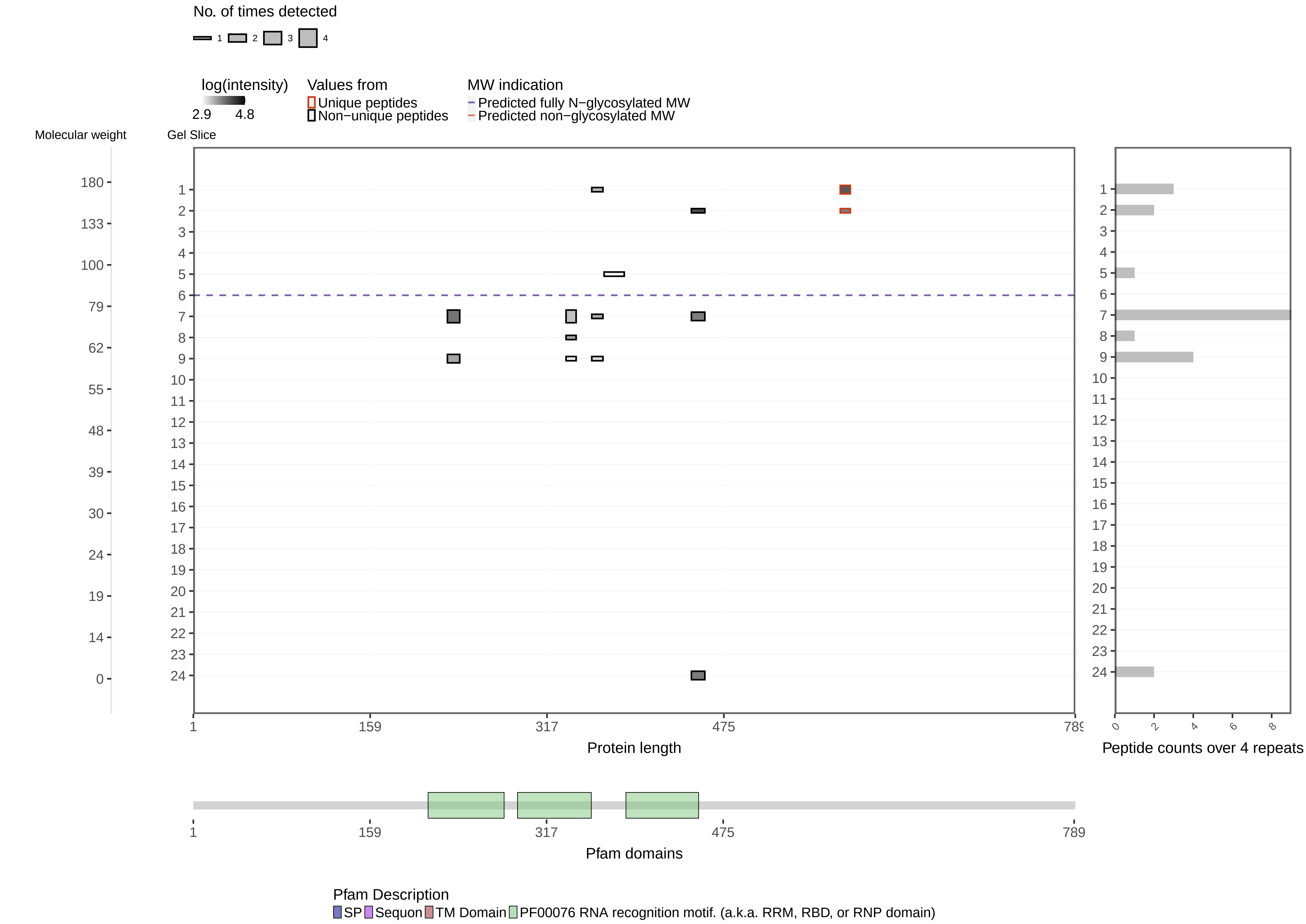


Protein sequence:

### NbL14g06040.1 dnaJ_homolog_subfamily_C_member_17−like [MW = 33.34 kDa]

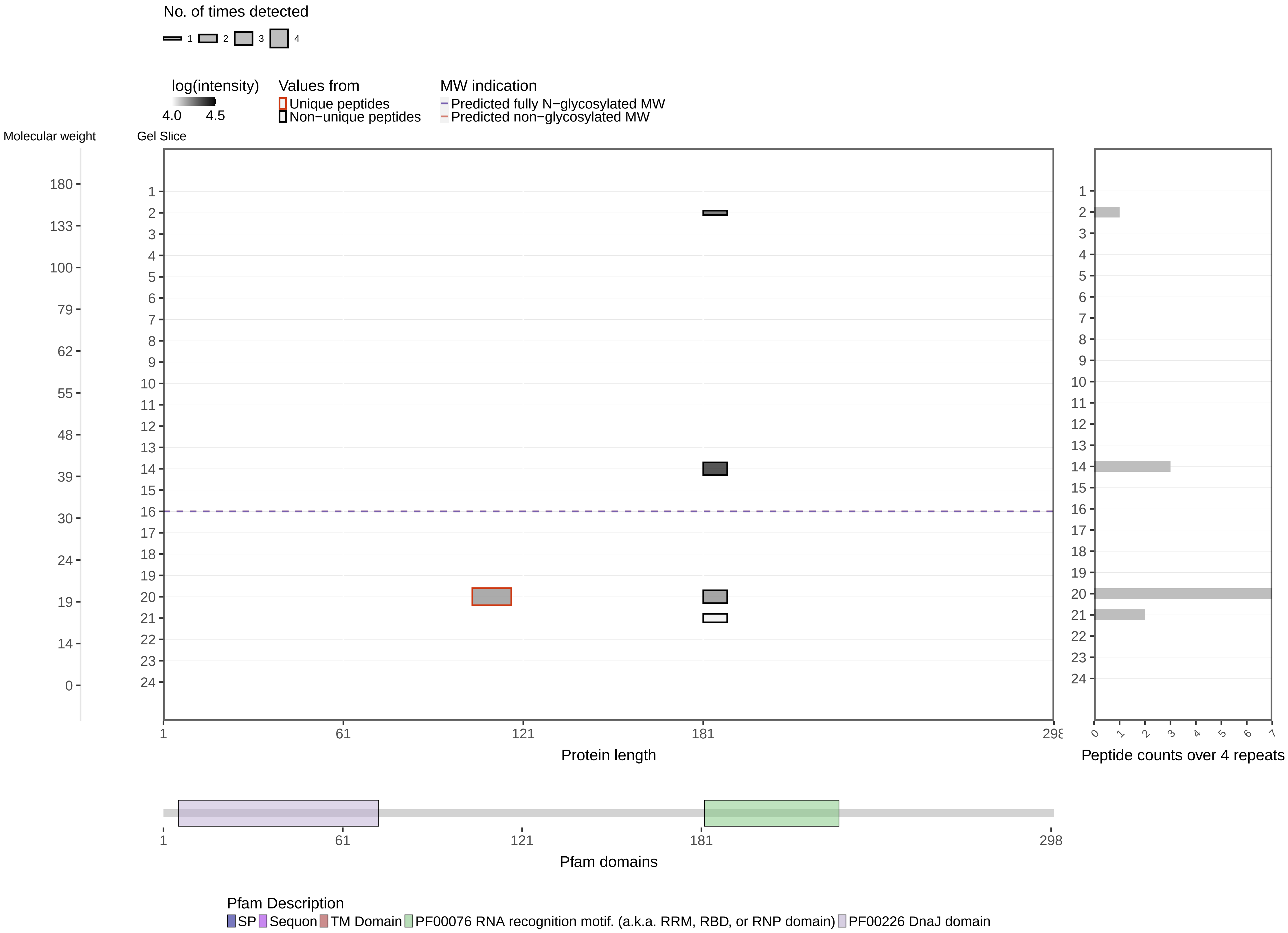


Protein sequence:

MHSNKVPTATDPSQKETHAKAKASTEGKGSSVDKEKVLKVSWEKIGEDYTAQRLRELFSEFGEVEDVVIKSSKKKGSALVVMSSKDAAKAVCGNVLGDLSNPLLVVPLQPTVQPTMPAPFFNAEKDREPEGPSLSNLVGAGYQKFEDSVLEKMRKAAQRKK*

NbL14g06000.1 eukaryotic_translation_initiation_factor_4G [MW = 194.39 kDa]

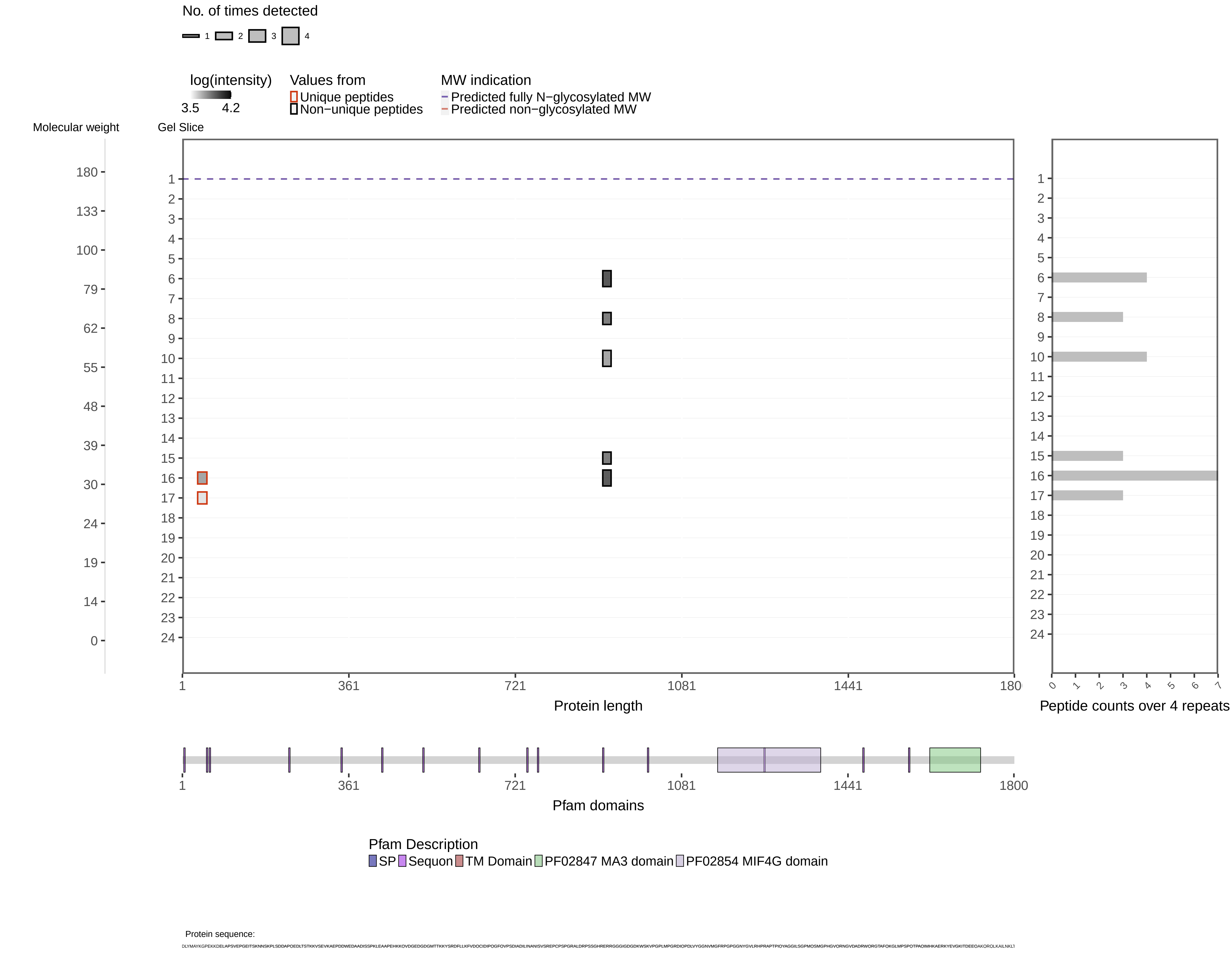

### NbL14g05930.1 10_kDa_chaperonin_1_chloroplastic [MW = 14.84 kDa]

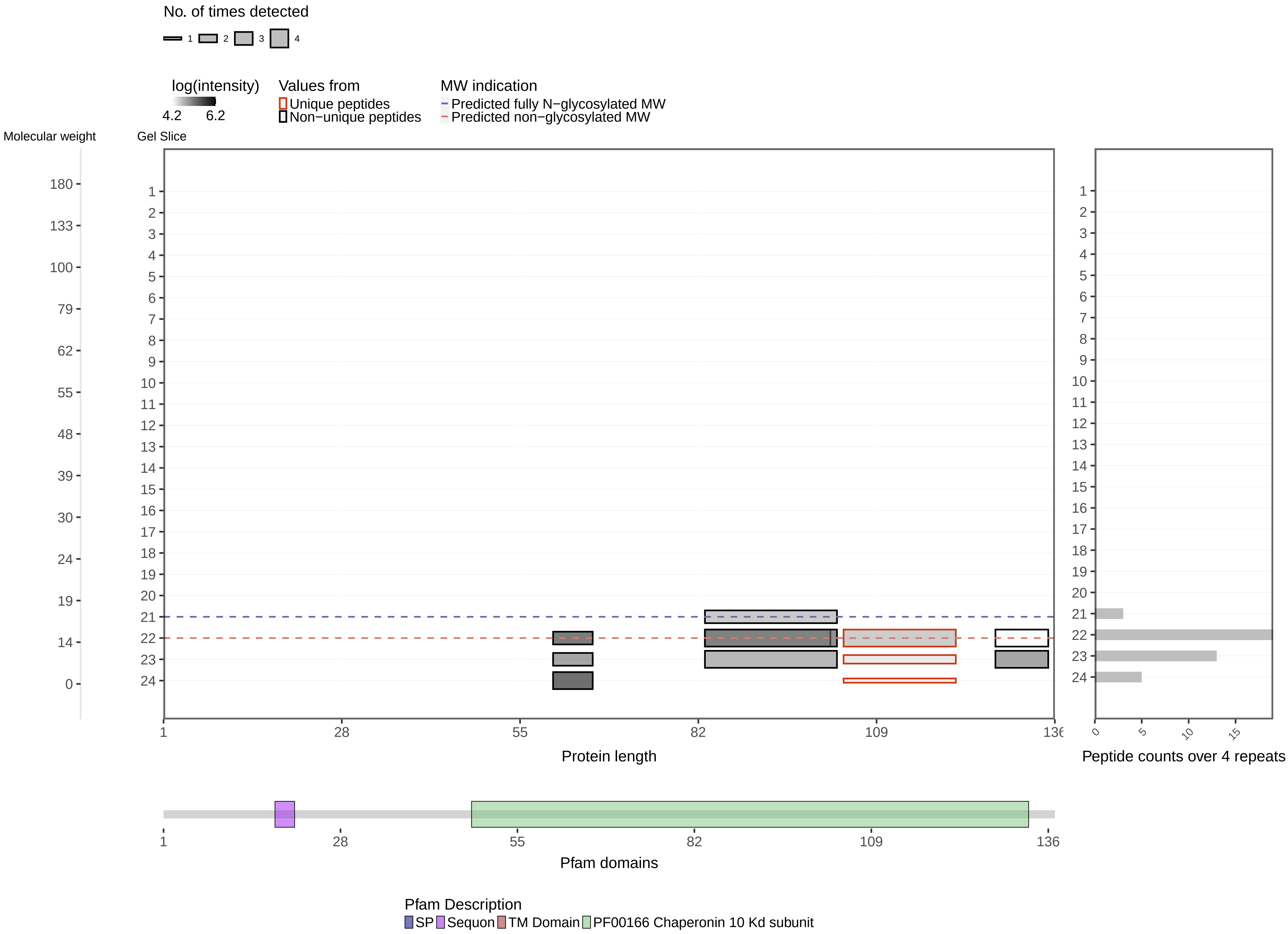


Protein sequence:

VLIRLEELSEKSAGGVLLPKSAVKFERYLMGEVLSVGSEVAQVEAGKKVLFSDINAYEVDLGTDARHCFCKESELLALVE*

### NbL14g05880.1 ras−related_protein_RABH1b−like [MW = 23.12 kDa]

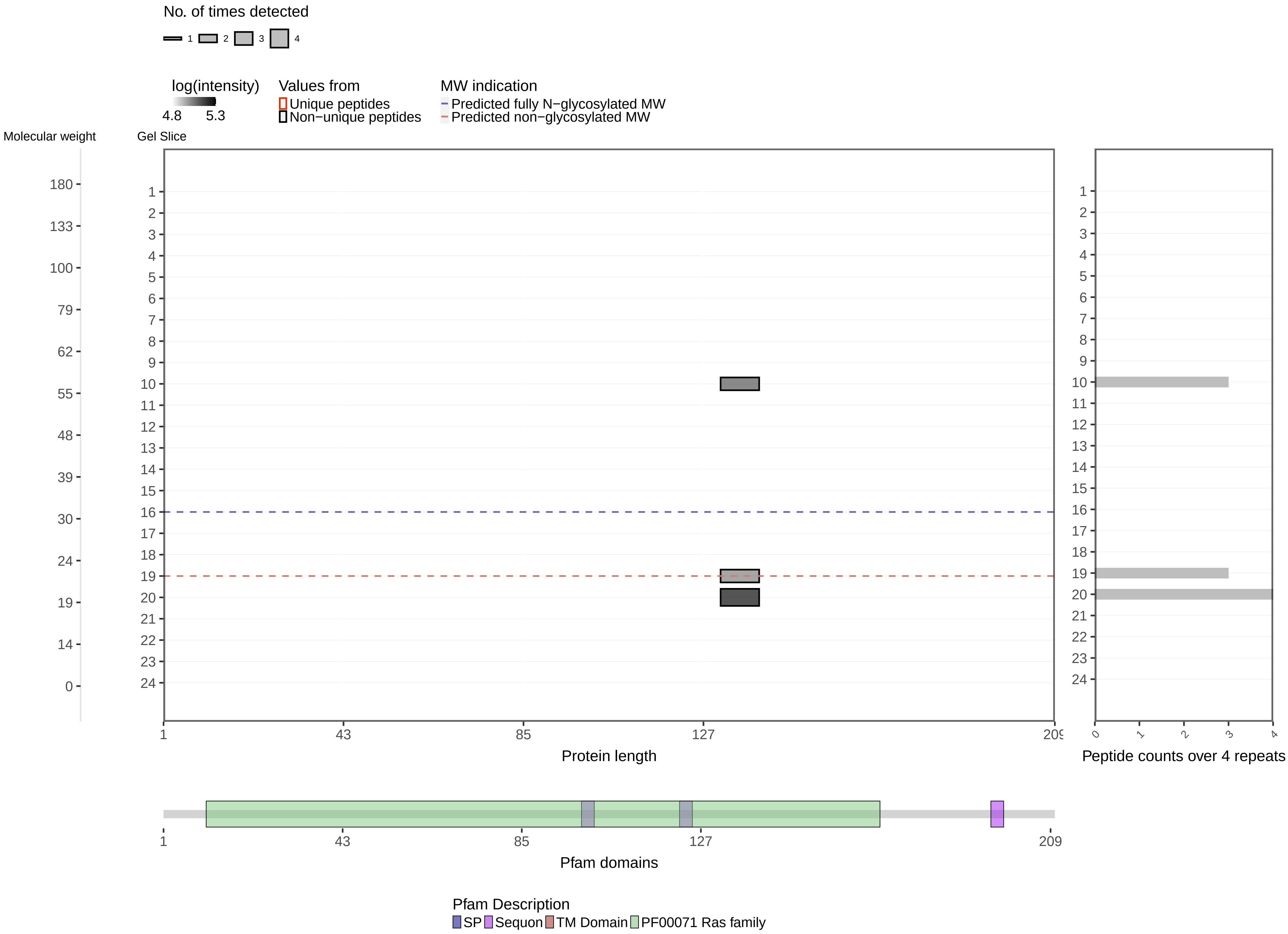


Protein sequence:

SRQSFLNTSKWIEEVRTERGSDVIIVLVGNKTDLVDKRQVSIEEGEAKARELNVMFIETSAKAGFNIKPLFRKIAAALPGMETLSSAKQEDMVDVNLKSSNPNASQSQPQSGGCAC*

### NbL14g05790.1 eukaryotic_translation_initiation_factor_3_subunit_B−like [MW = 82.58 kDa]

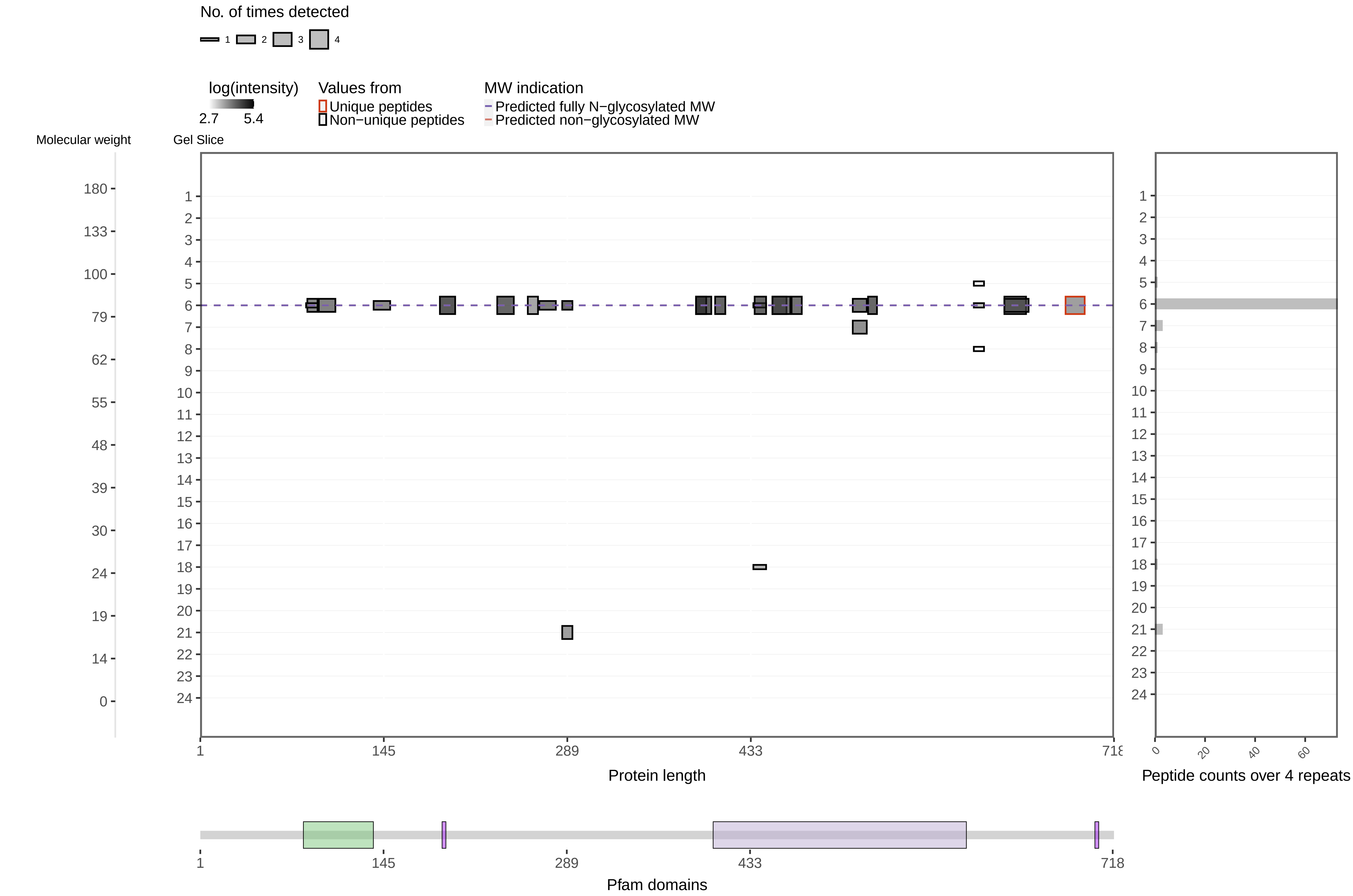


Pfam Description

SP Sequon TM Domain PF00076 RNA recognition motif. (a.k.a. RRM, RBD, or RNP domain) PF08662 Eukaryotic translation initiation factor eIF2A

Protein sequence:

VYETETFSLIDKKSIKVENVMDFSWSPADPILALFVPECGNQPARVSLVQIPSKEELRQKNLFSVSDCKMYWQSNGDYLAVKVDRYTKTKKSTYTGFELFRIKERDIPIEVLELDNKNDKIIAFAWEPKGHRFAVIHGDNPRPDISFYSVRSGTNTGRVSKLTTLKGKQANALYWSPGGRFLILAGLKGFNGQLEFFDVDELETMASAEHFMATDVEWDPTGRYVATSVTSVHEMENGFNIWSFNGKLLYRILKDHFFQYLWRPRPPSFLSKEKEEEIAKNLKRYSKKYEAEDQD

### NbL14g05640.1 protein_transport_protein_SEC23−like [MW = 84.69 kDa]

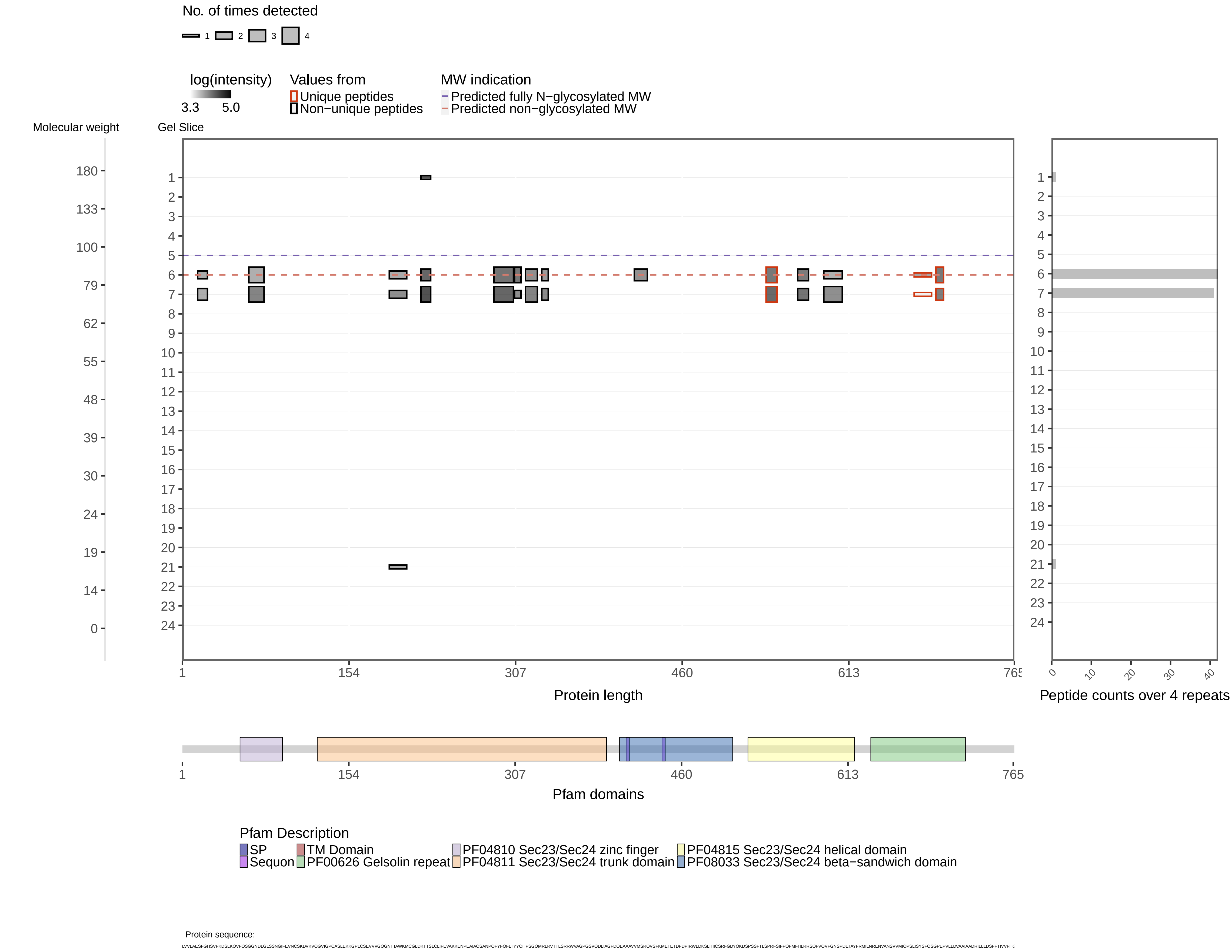

NbL14g05550.1 phosphoribosylaminoimidazole−succinocarboxamide_synthase_chloroplastic [MW = 38.64 kDa]

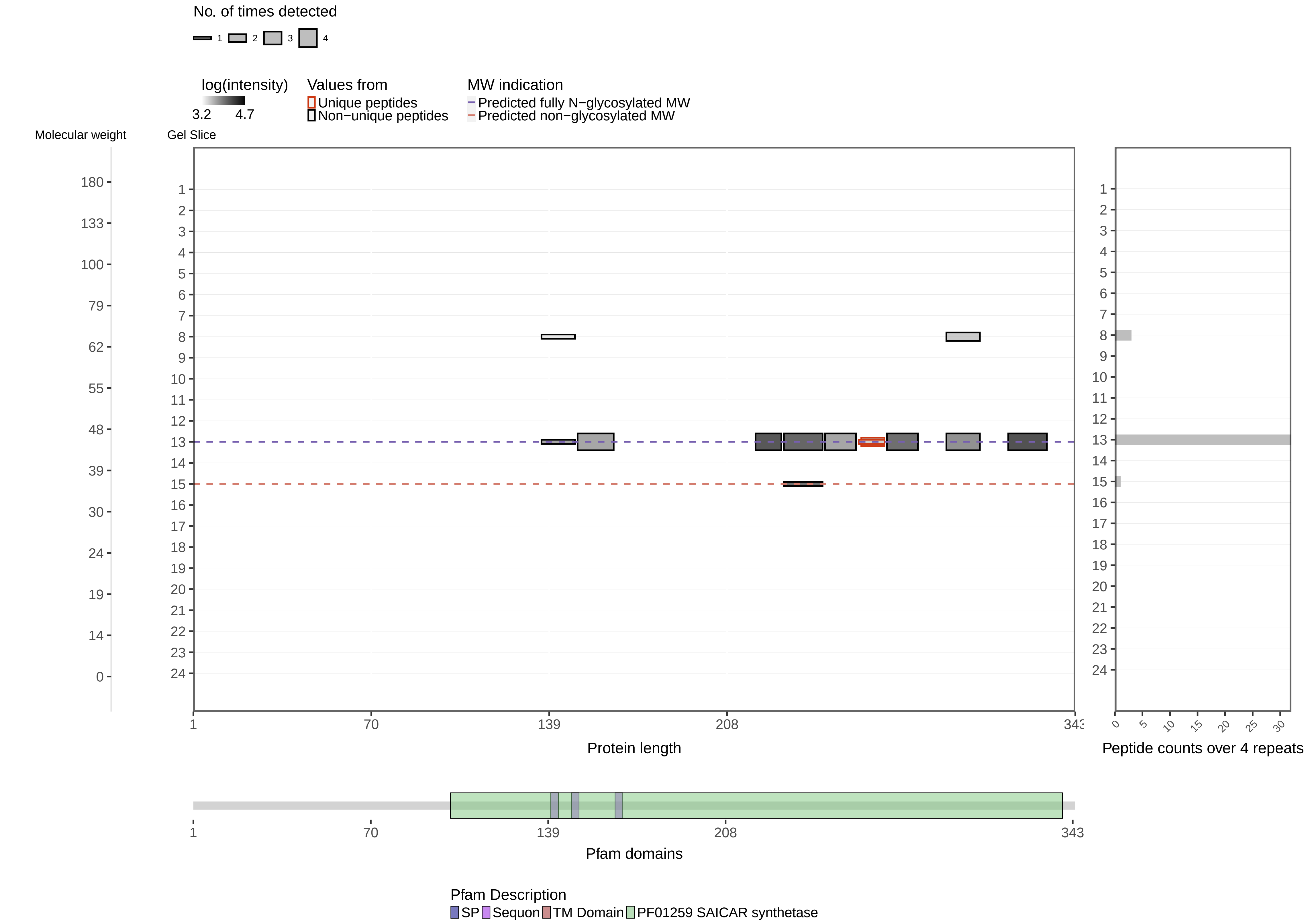


Protein sequence:

VPDRNVTIARKCSVFTVEFVVRGYVTGSTDTSLWTVYNKGVRNYCGNNLPDGLVKNQKLQENILTPTTKAADHDVPVTPDEIVQRGLMTQADYDEVSRKATSLFEFGQRVALDHGLILVDTKYEFGKGADGKIYLIDEVHTPDSSRYWIAHSYQERFQSGLEPENIDKEFLRLWFKNHCKPI*

NbL14g05380.1 uncharacterized_LOC107827413 [MW = 35.43 kDa]

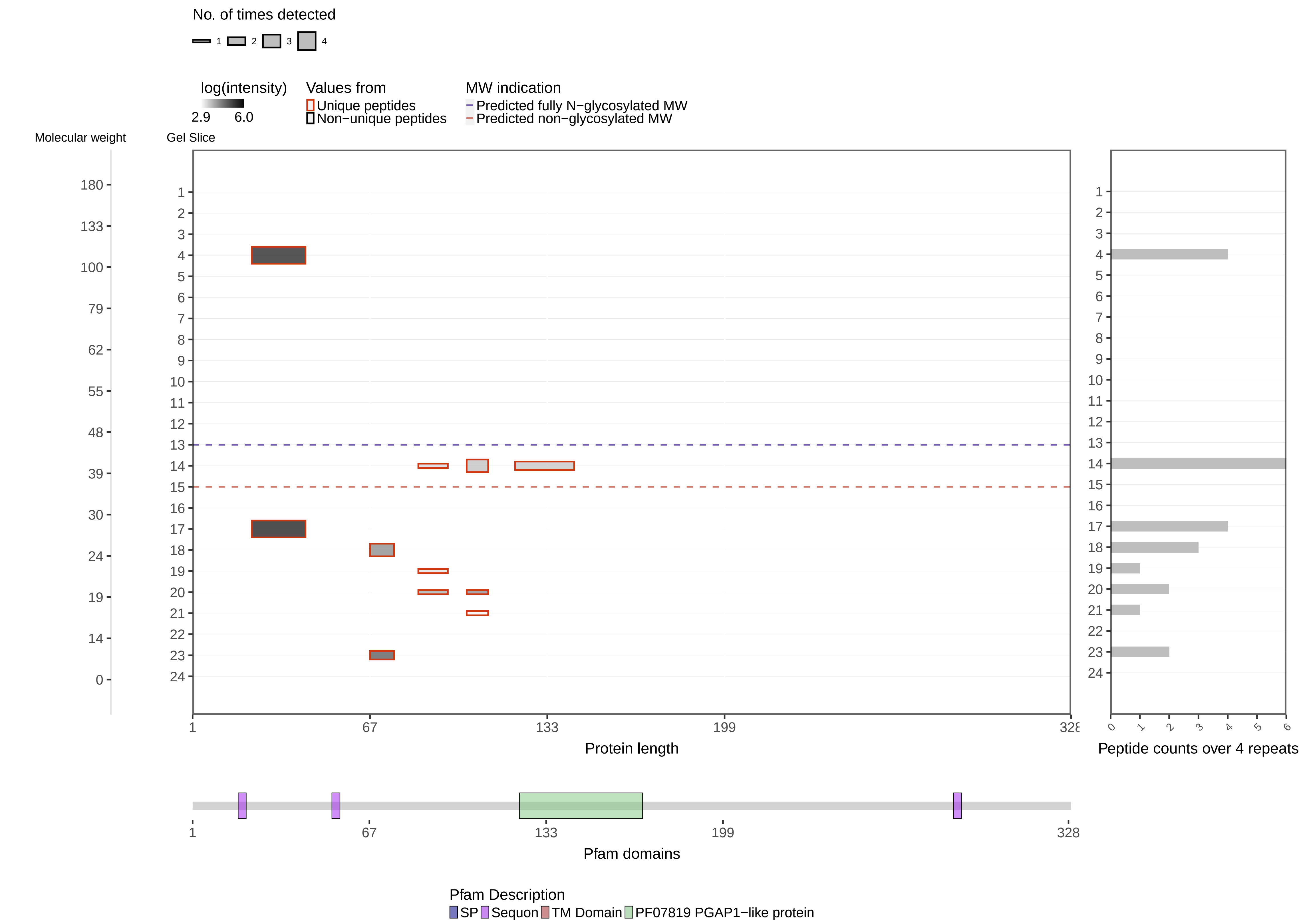


Protein sequence:

DVSLLLTLGTPHLPPPKGLPGVIDQTRGLLYYVEEHCAKAVYSPQLRYVCIAGRYIQGSRFLGSSSTYSSELEAGVDQPTSEVAAVSNTNSETASTFRARFVGQGYKQVCGQADVWGDGVVPEVSAHLKGALNITLDGVYHSPVGSDDATRPWQRKIENFREMILRVCVLCVLQL*

### NbL14g05270.1 serine/threonine−protein_phosphatase_PP2A−5_catalytic_subunit [MW = 35.77 kDa]

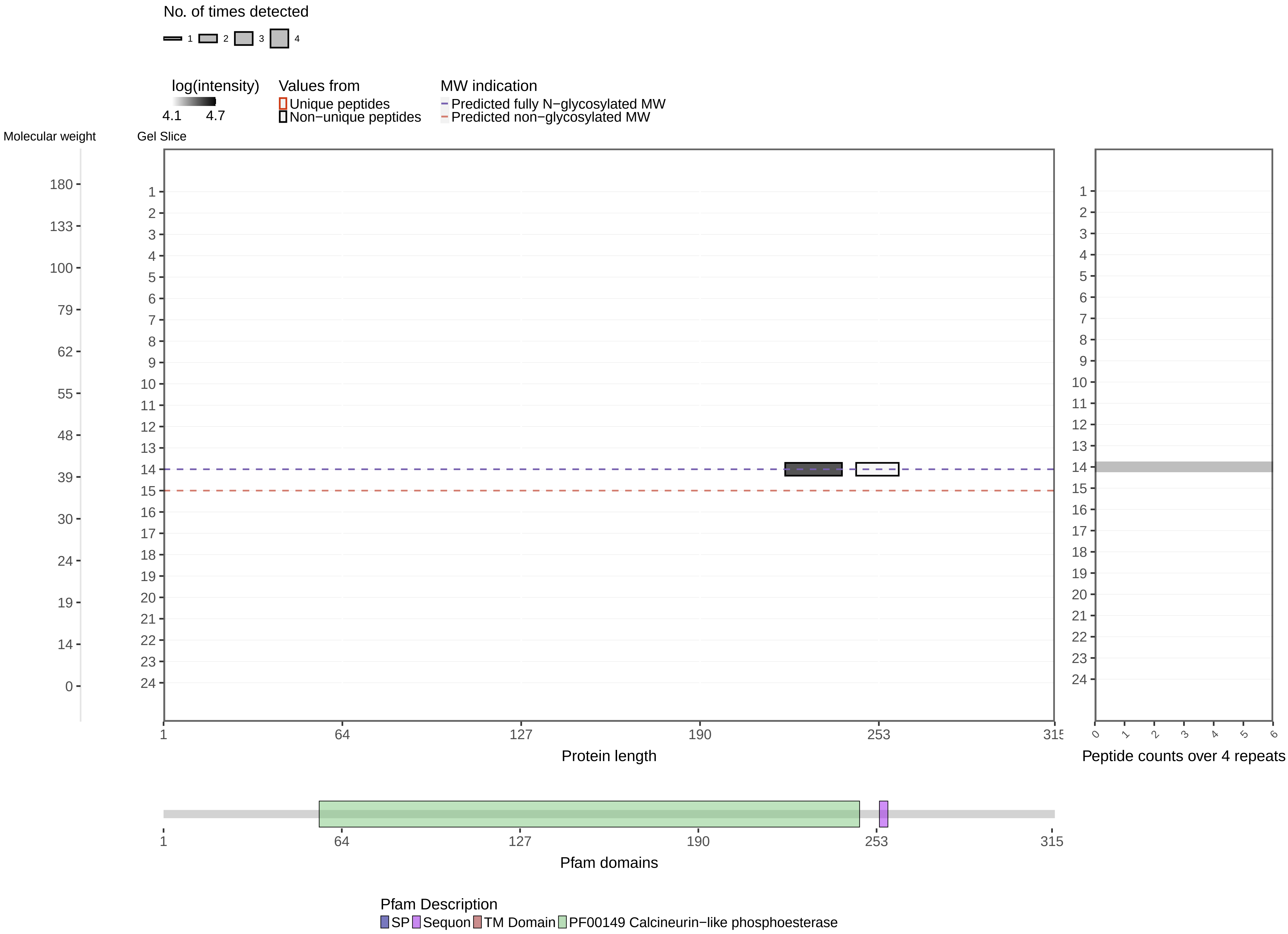


Protein sequence:

NVWKTFTDLFDYFPLTALVESEIFCLHGGLSPSIETLDNVRSFDRVQEVPHEGAMCDLLWSDPDDRCGWGMSPRGAGYTFGQDISEQFHQTNNLKLIARAHQLVMEGYNWSHEQKVVTIFSAPNYCYRCGNMASILEVDDCRGHTFIQFDPAPRRGEPDVTRRTPDYFL*

### NbL14g05230.1 V−type_proton_ATPase_subunit_D−like [MW = 29.18 kDa]

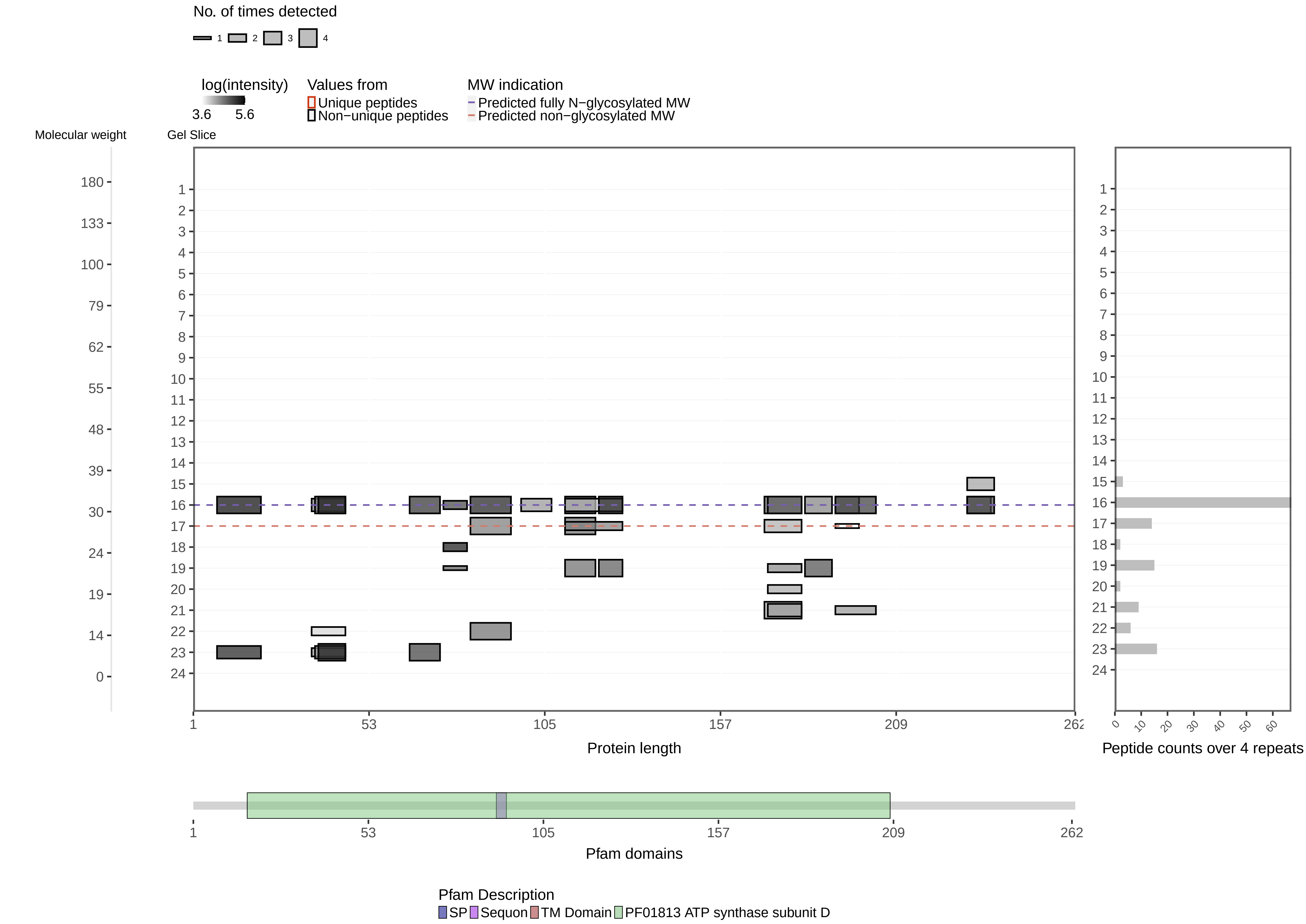


Protein sequence:

GETKNDLTGLARGGQQVQACRAAYVKSIELLVELASLQTSFLTLDEAIKTTNRRVNALENVVKPRLENTVLYIKGELDELEREDFFRLKKIQGYKKREIERQMASARQYAEERAAEELSLKKGITLGSAHNILSHASQKDDDIIF*

NbL14g05180.1 citrate_synthase_mitochondrial [MW = 52.43 kDa]

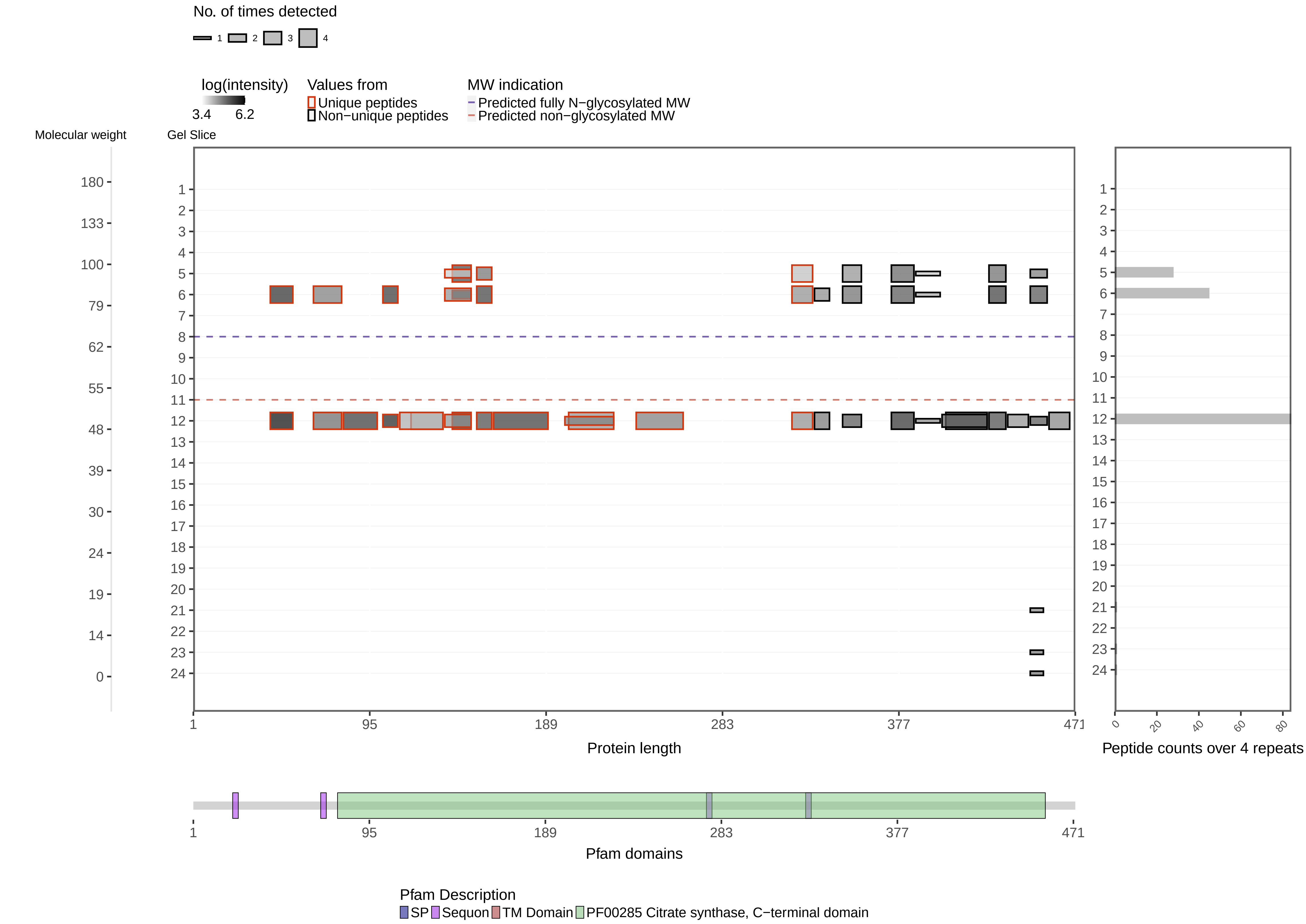


Protein sequence:

VYRRMYKNGNTIPKDDSLDYGANFAHMLGFSSSDMHELMRLYVTIHSDHEGGNVSAHTGHLVASALSDPYLSFAAALNGLAGPLHGLANQEVLLWIKSVVEECGENTSKEQLKDYVWKTLKSGKVVPGFGHGVLRKTDPRYTCQREFGLKHLPEDPLFQLVAKLYEVVPPILTELGKVKNPWPNVDAHSGVLLNYYGLTEARYYTVLFGVSRALGICSQLIWDRALGLPLERPKSVTMEWLENHCKKA*

NbL14g04980.1 clustered_mitochondria_protein [MW = 153.86 kDa]

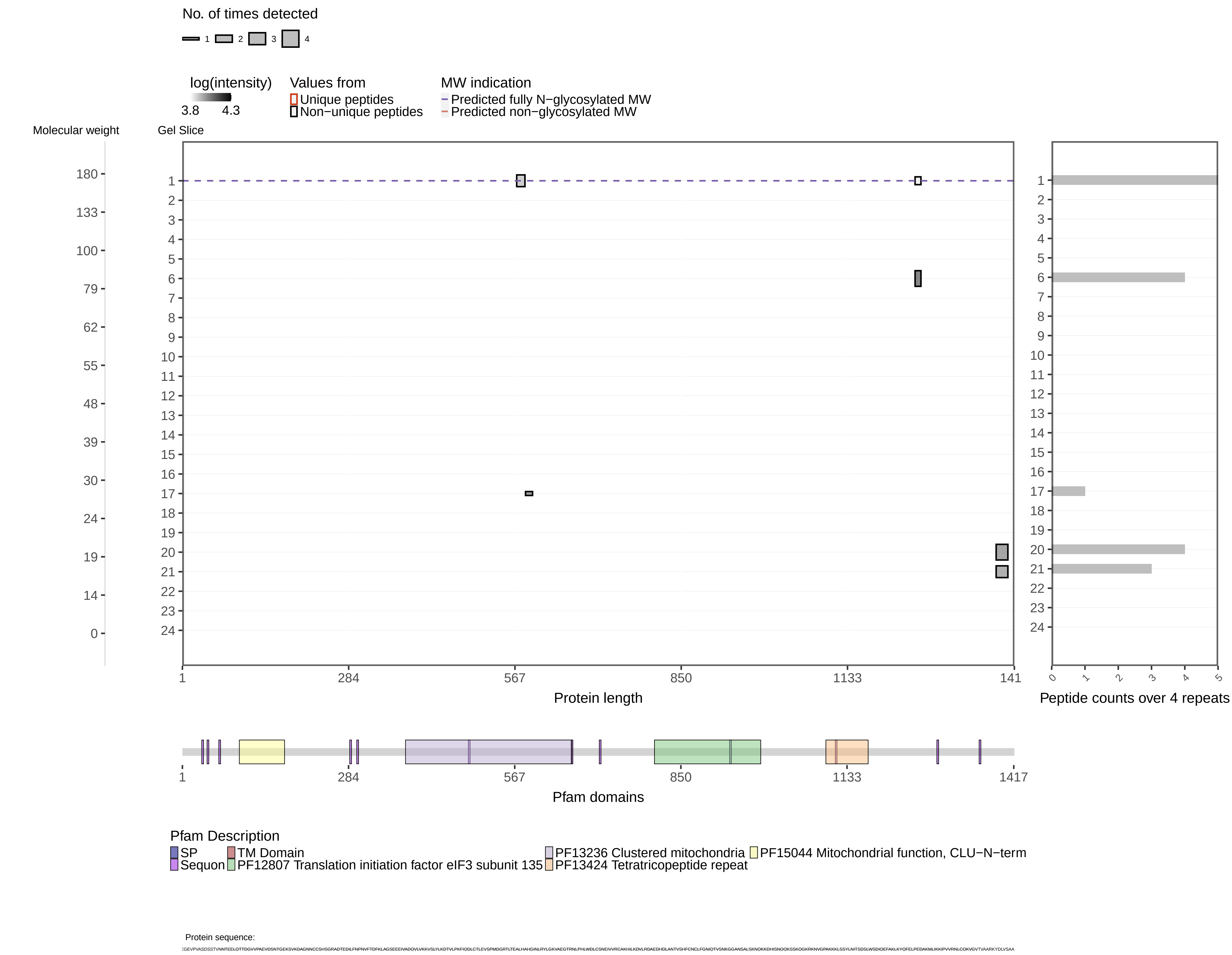

NbL14g04950.1 DNA−directed_RNA_polymerase_I_subunit_2 [MW = 133.46 kDa]

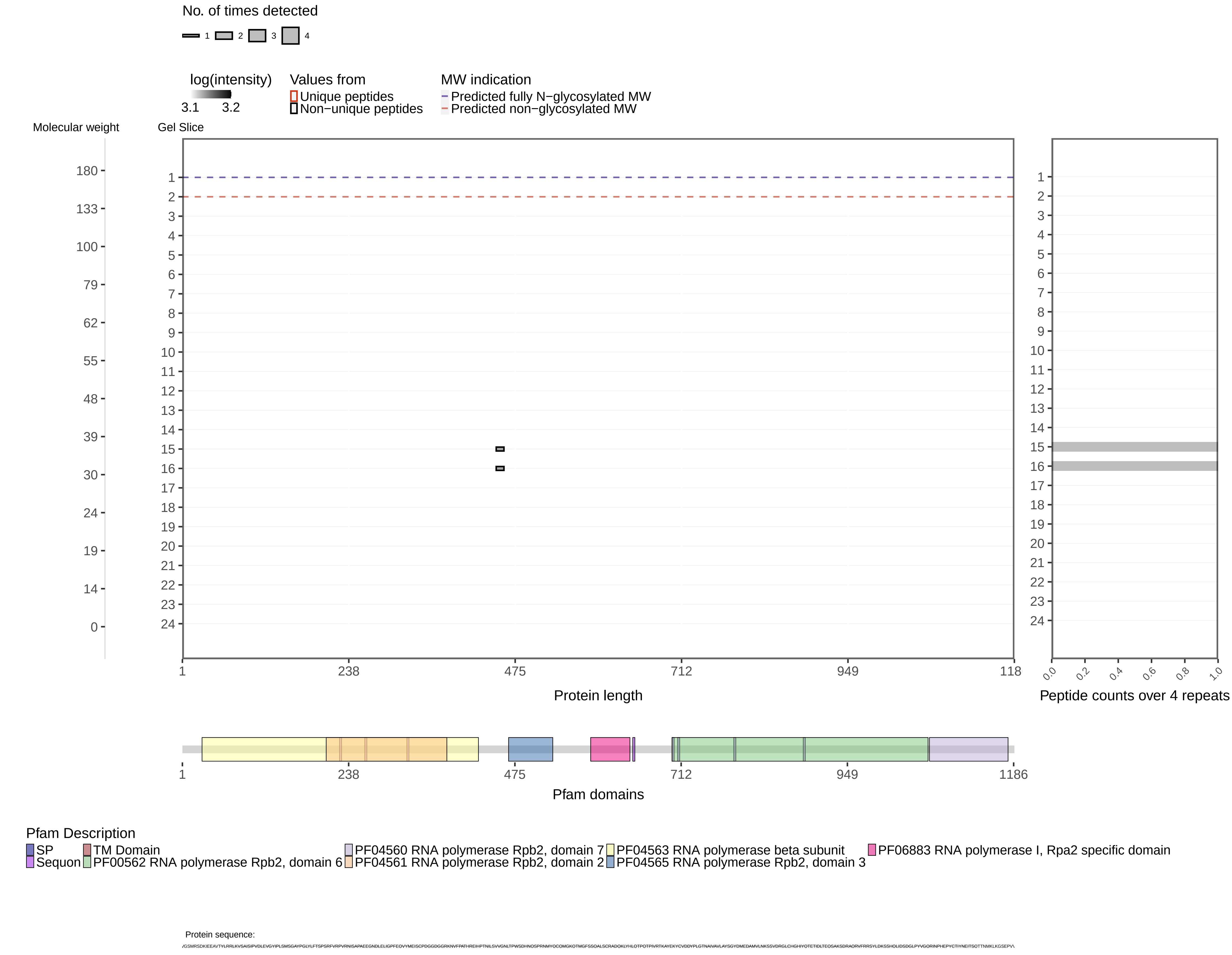

### NbL14g04880.1 putative_tRNA_(cytidine(34)−2'−O)−methyltransferase [MW = 31.36 kDa]

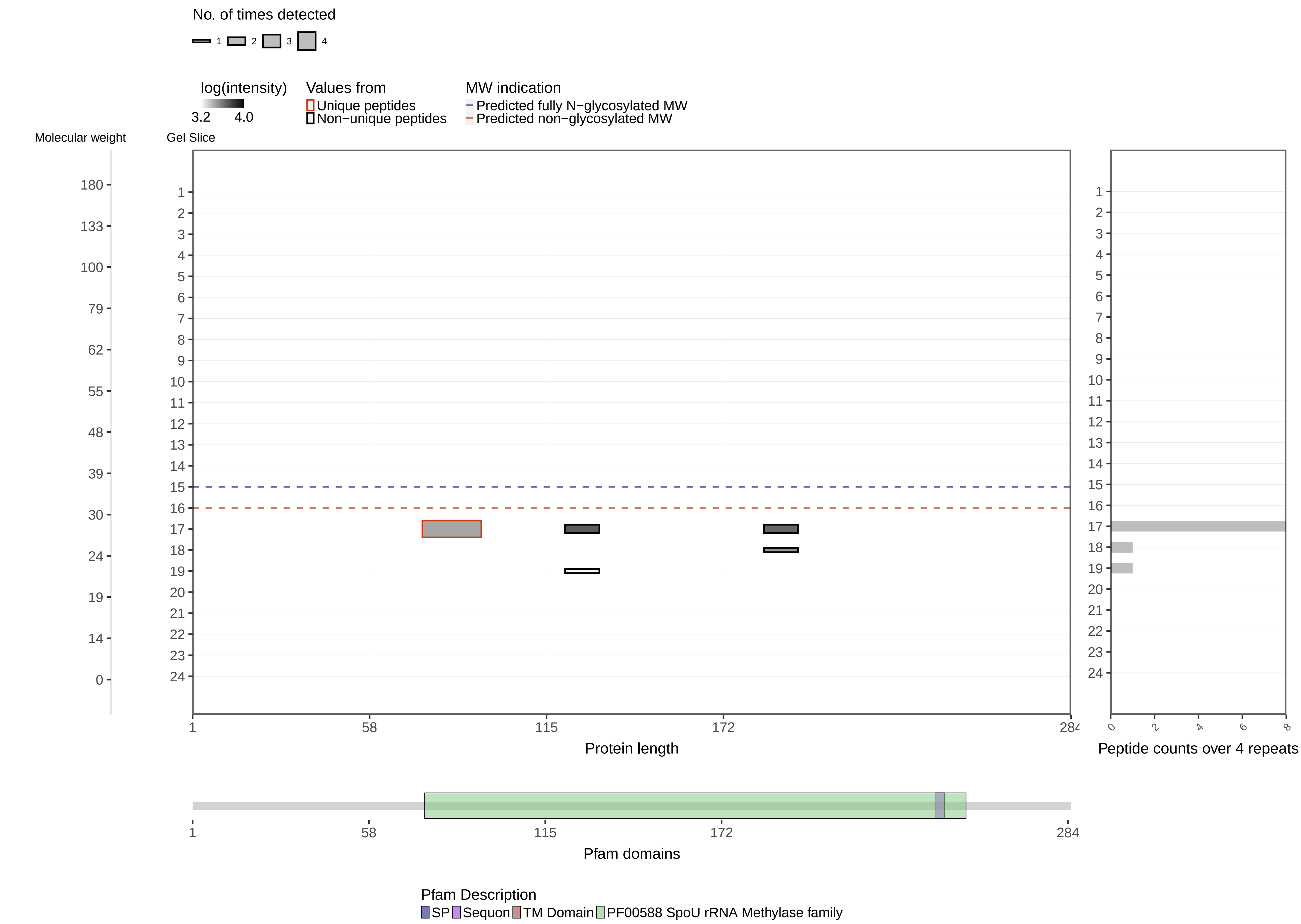


Protein sequence:

/VKVHRSWDEFREYFRGQVSFLPHSSSFKTFRVWTKIILFPSLQAGEKRLLAFTKRGTTTHSDFSYKKGDWLVFGSETKGLPPEALLDCNSESLGGGTLRIPMVETYVRCLNLSVSVGIAVYEASRQINYEQLQCPPDTCMDIETSFVTEDIYA*

NbL14g04870.1 DNA−damage−repair/toleration_protein_DRT100−like [MW − SP = 36.68 kDa]

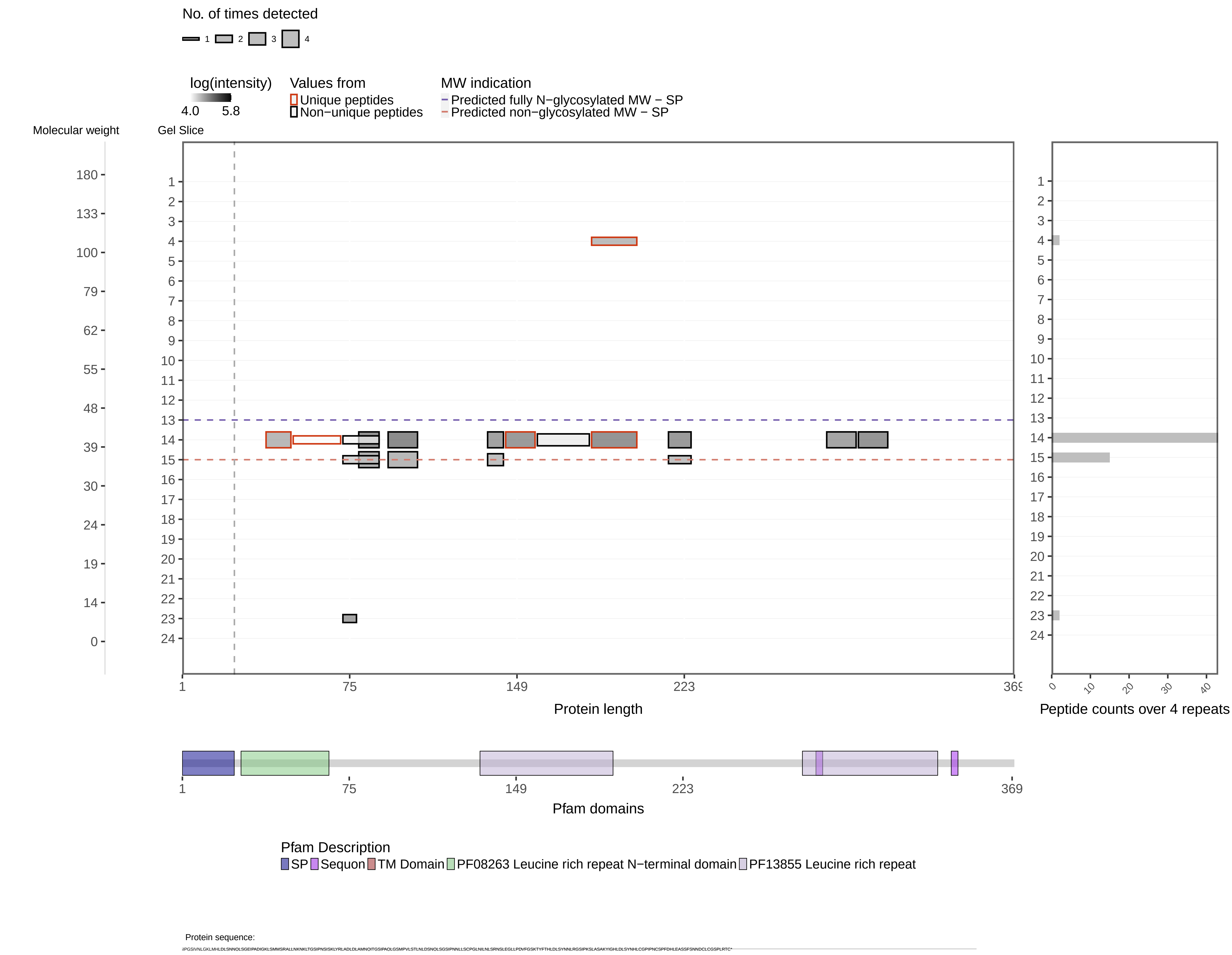

### NbL14g04820.1 pre−mRNA−splicing_factor_SYF1−like [MW = 106.71 kDa]

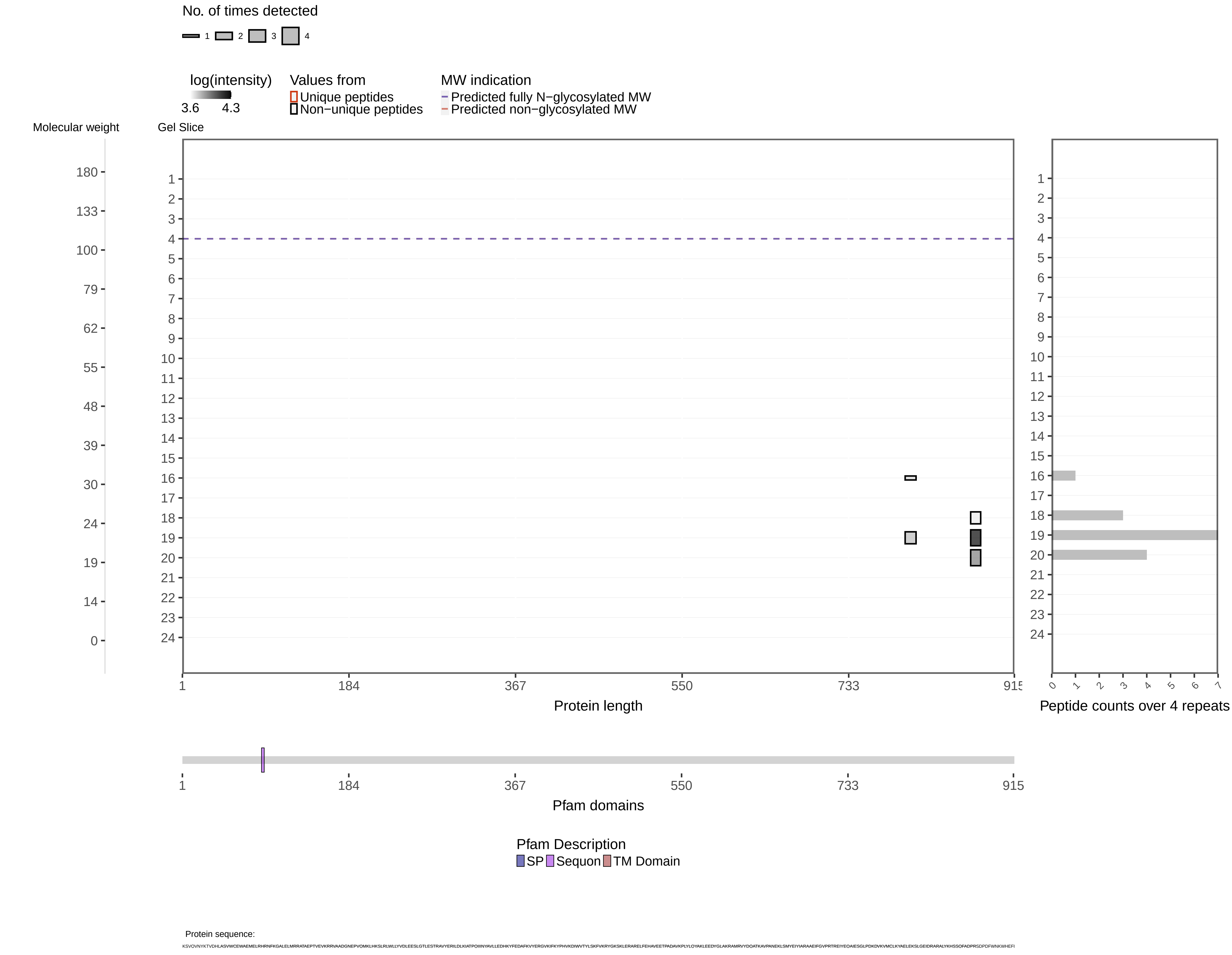

NbL14g04620.1 flavin−containing_monooxygenase_FMO_GS−OX−like_4 [MW = 59.48 kDa]

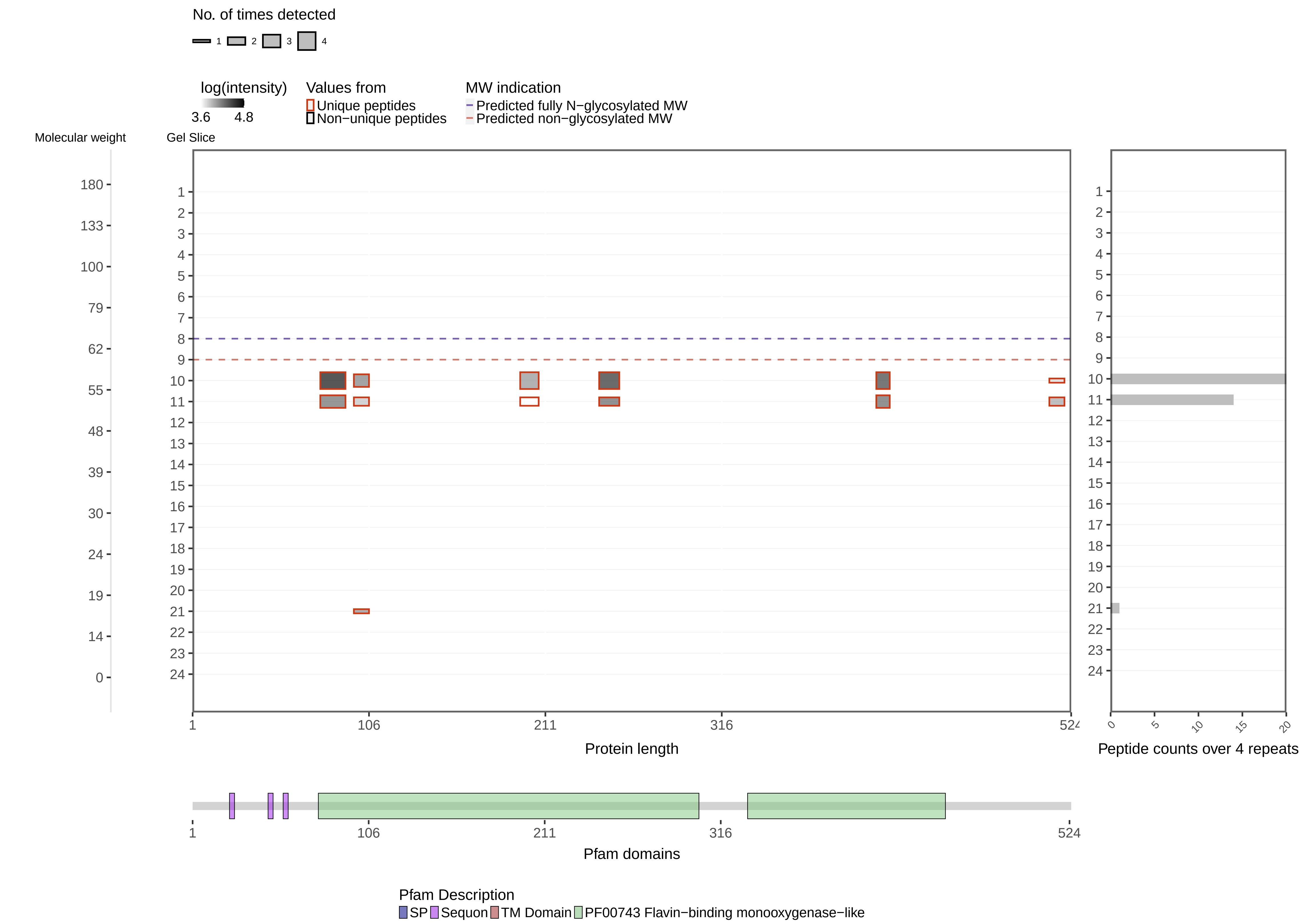


Protein sequence:

[illegible]IEVWPGKQIHSHNYRVPDPFRNQVVVLIGSAASATDISREIAEVAKEVHISSRSATSGVPLKLPGYDNIWQHSMIEAVGIDSGVNFRDGSKVIADMILHCTGYKYHFPFLETNGIVTVDDNRVGPLYKHIFPPAFAPSLSFVGLPWKVIPFPLCELQSKWIAGVLSGRISLPSKEDMNADIEALYSSMEASCIPKRYTHNMDDCQFDYDDWLAAECGSPPSEEWRKQMYFISRKNKKTQPETYRDQWDDDDLIIQAHEDFVKYIPELAQAQKLPR*

### NbL14g04590.1 14_kDa_proline−rich_protein_DC2.15−like [MW − SP = 11.5 kDa]

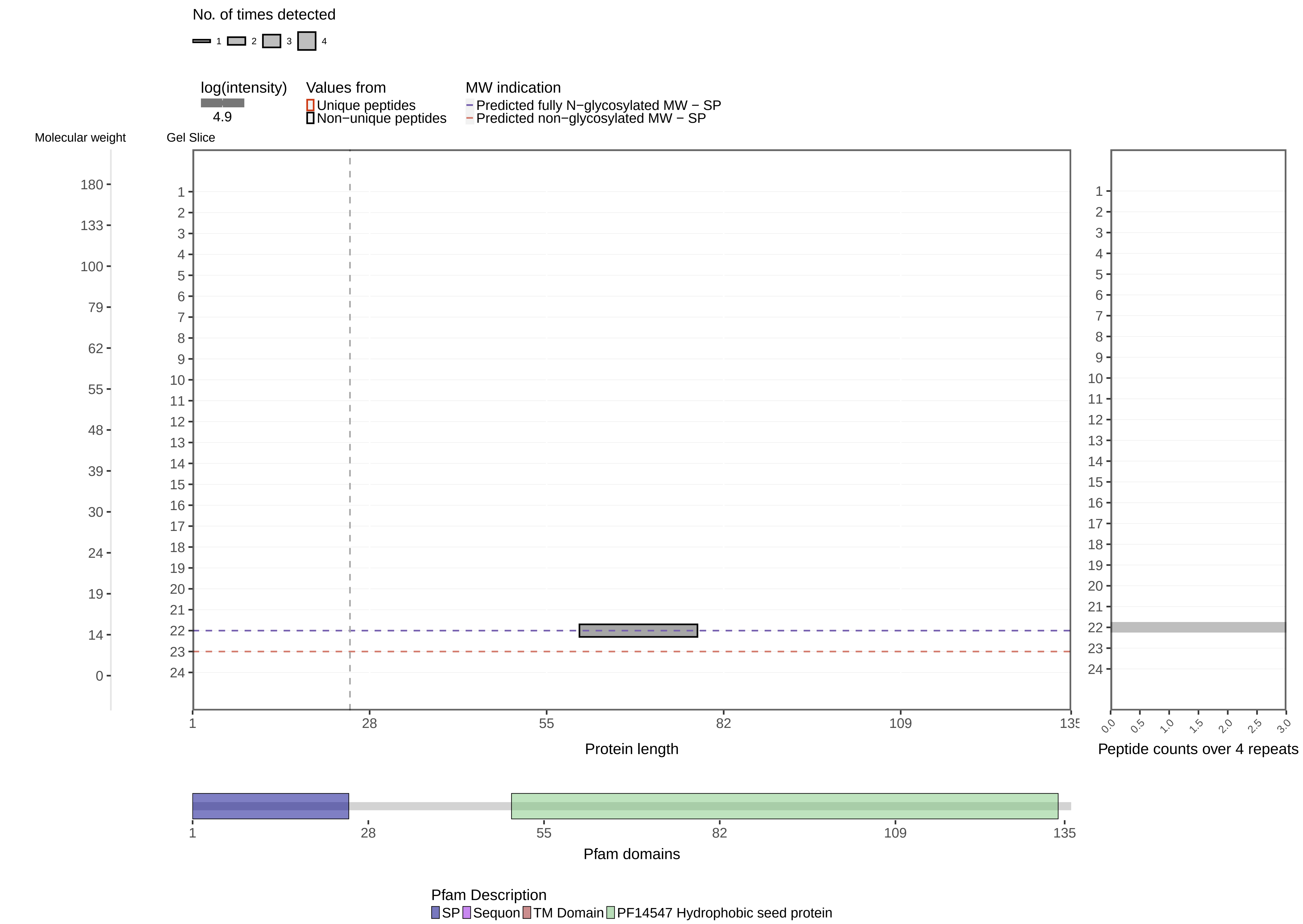


Protein sequence:

DTLKLKVCANLLNDLVHVVIGSSSKSSCCSLIENLADVDAAVCLCTAIKAKLLGTNLDTALSLSLLLNNCGKTVPKDFQCA*

NbL14g04560.1 peroxisome_biogenesis_protein_19−1−like [MW = 36.9 kDa]

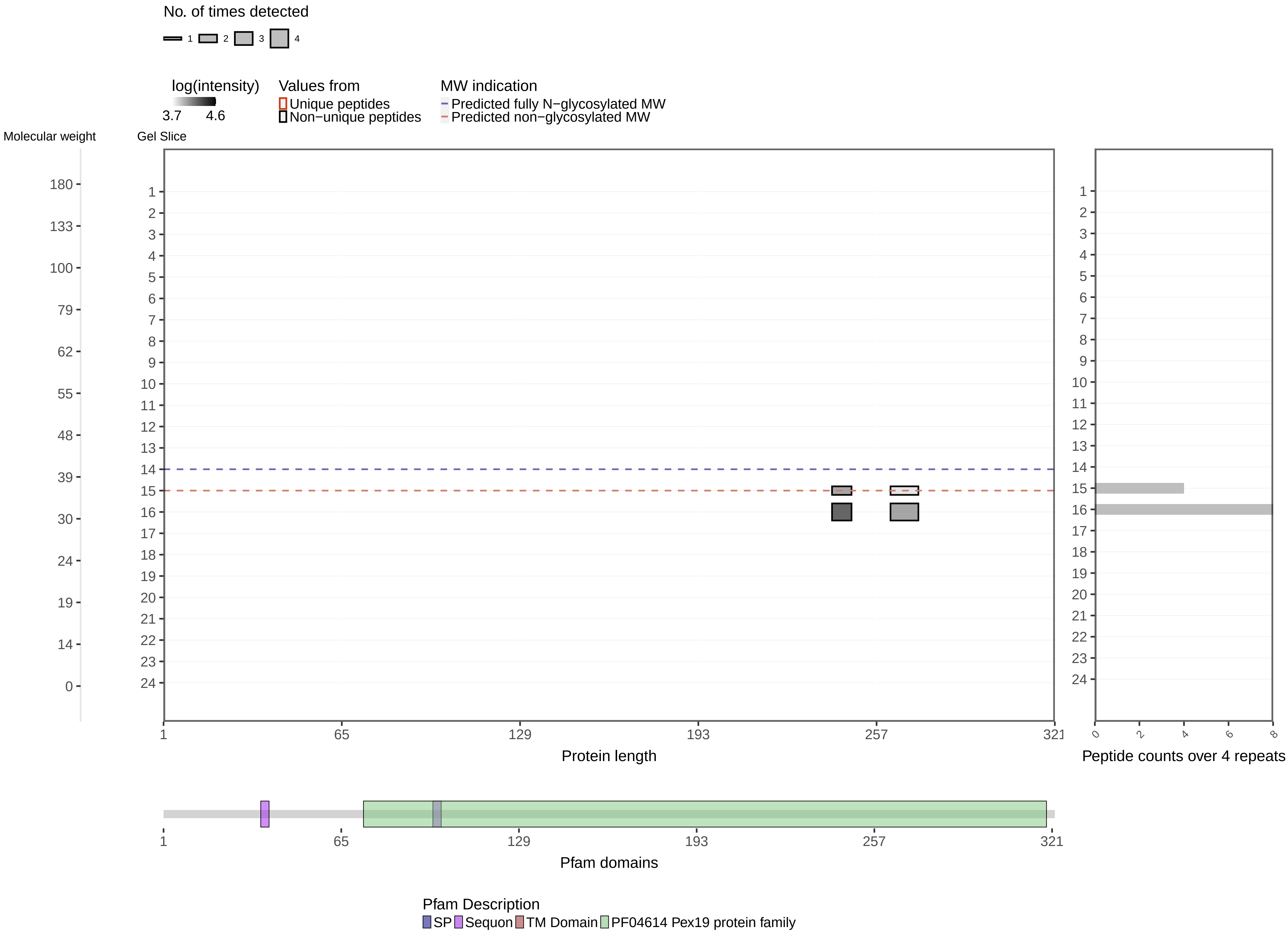


Protein sequence:

VSEALDKLREQTREALKGLESVAVPRPGLESFGSDPMMEEWAKQFEELAGSQDMESIVETMMQQLLSKEILYEPMREIGERYPKWLEDNKAKLSSEDYERYRHQYELIRDLNKVYETEPNNFNKIVELMQKMQECGQPPNDIVHELAPDFDISTLGQLSPEMLESQQNCCIM*

### NbL14g04470.1 2−oxoglutarate−Fe(II)_type_oxidoreductase_hxnY [MW = 23.65 kDa]

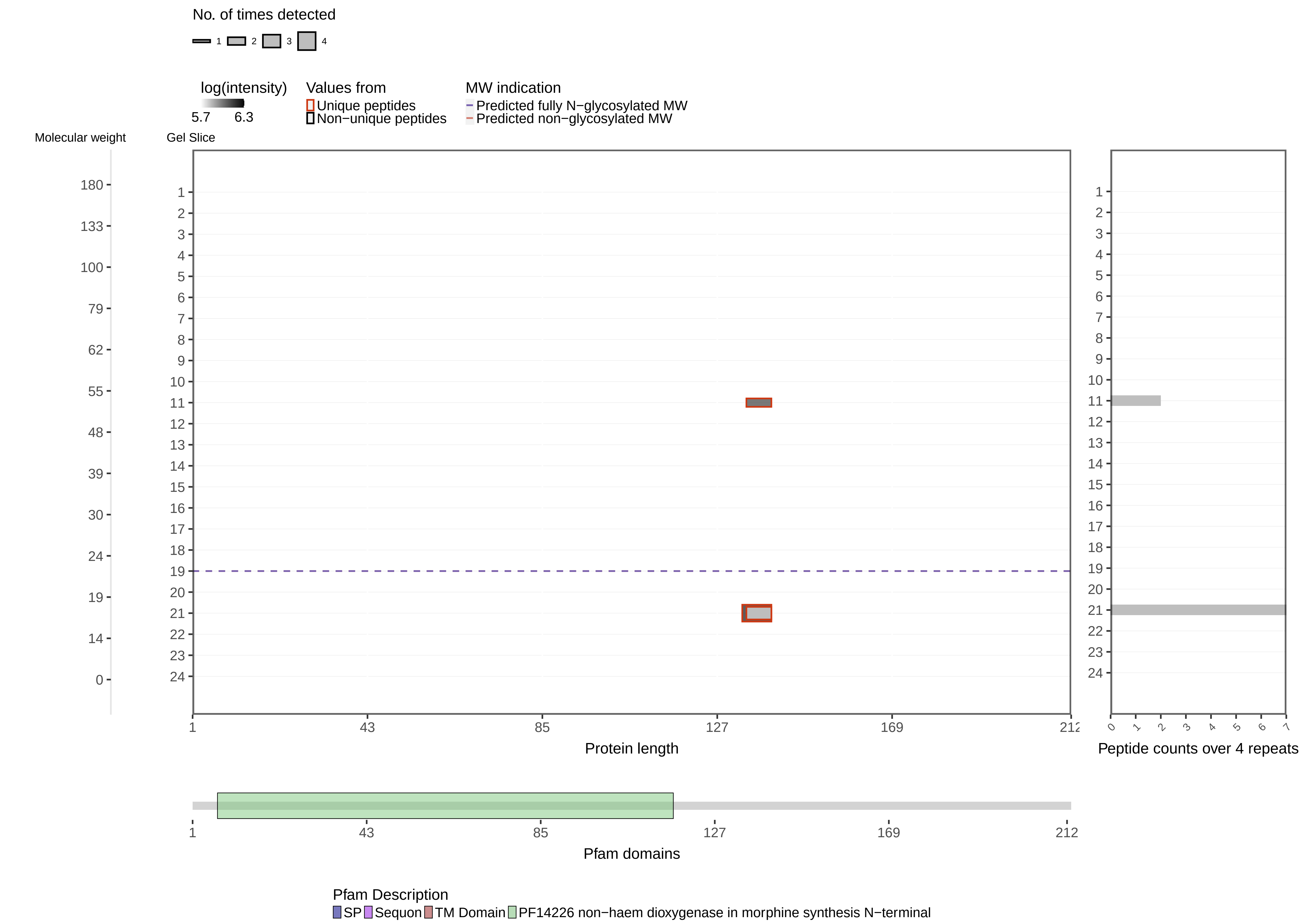


Protein sequence:

KESFYIGPLEDKRAVSNLNQWPAEEILPCWRSTMEDYYKRVLDAGIRLTSLIALALELDEDFFHKAGACDSPNAFLRLVHYPGELRLSEQVVYGASAHSDYGMLTLLATDGVGGLQHV*

NbL14g04360.1 plasminogen_activator_inhibitor_1_RNA−binding_protein−like [MW = 39.37 kDa]

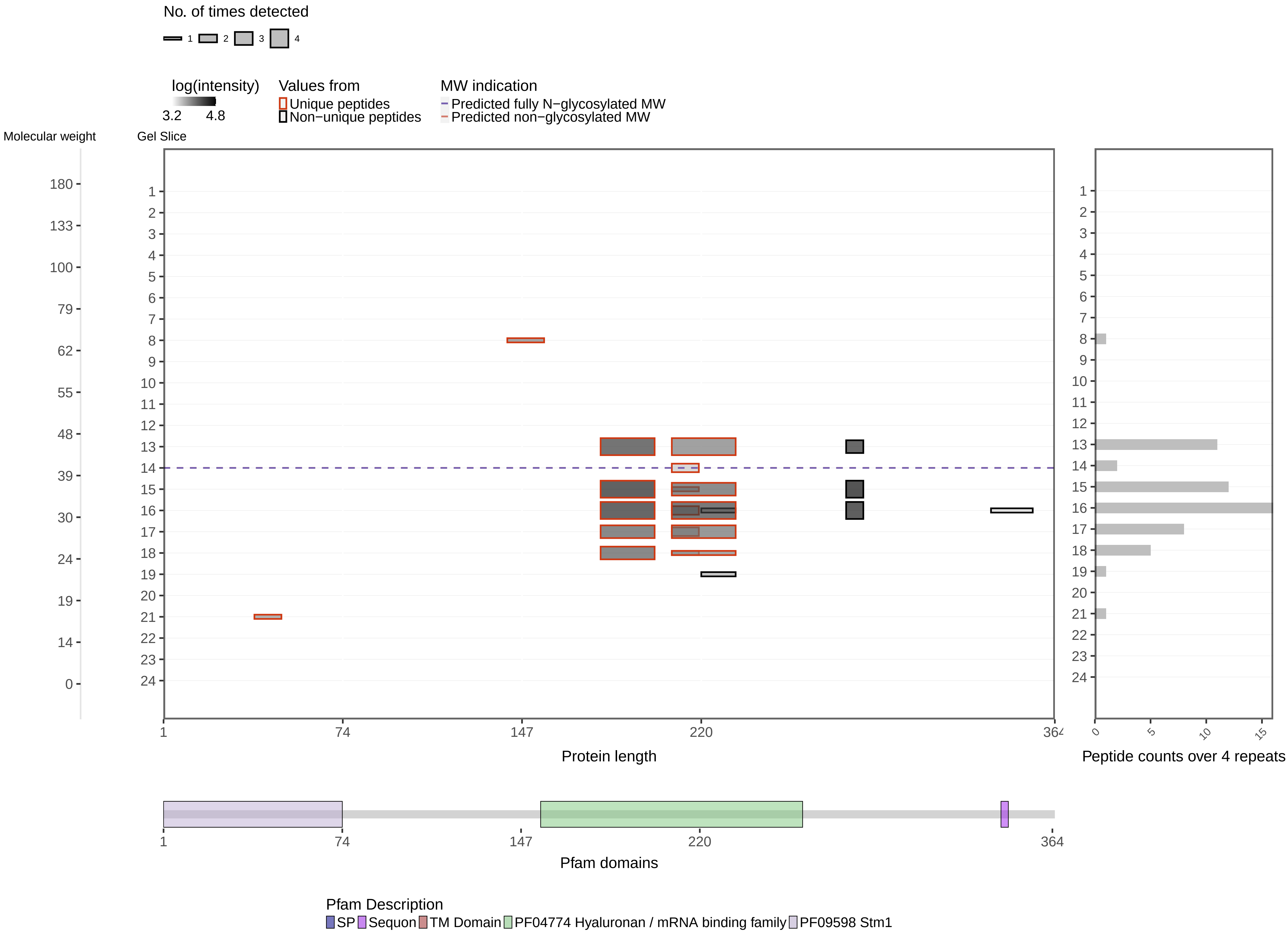


Protein sequence:

IGNEFKREGAGRGNWGTPADELAPEKEEPVNEGEKIVDTEKQAVQADAEDTSKDSPAAEPQEKEPEEKEMTLEEYEKLLEEKRKALLALKPEERKVNLDKELESMQLLSNKRSDDEVFIKLGSEKDKRKETVEKVKKTKSINEFLKPADGESYYGRGGRGGRGRGGRDRGRDGFVGNNSVSAPSIEDVRQFPSLGAK*

### NbL14g04330.1 FAD−dependent_urate_hydroxylase−like [MW = 25.55 kDa]

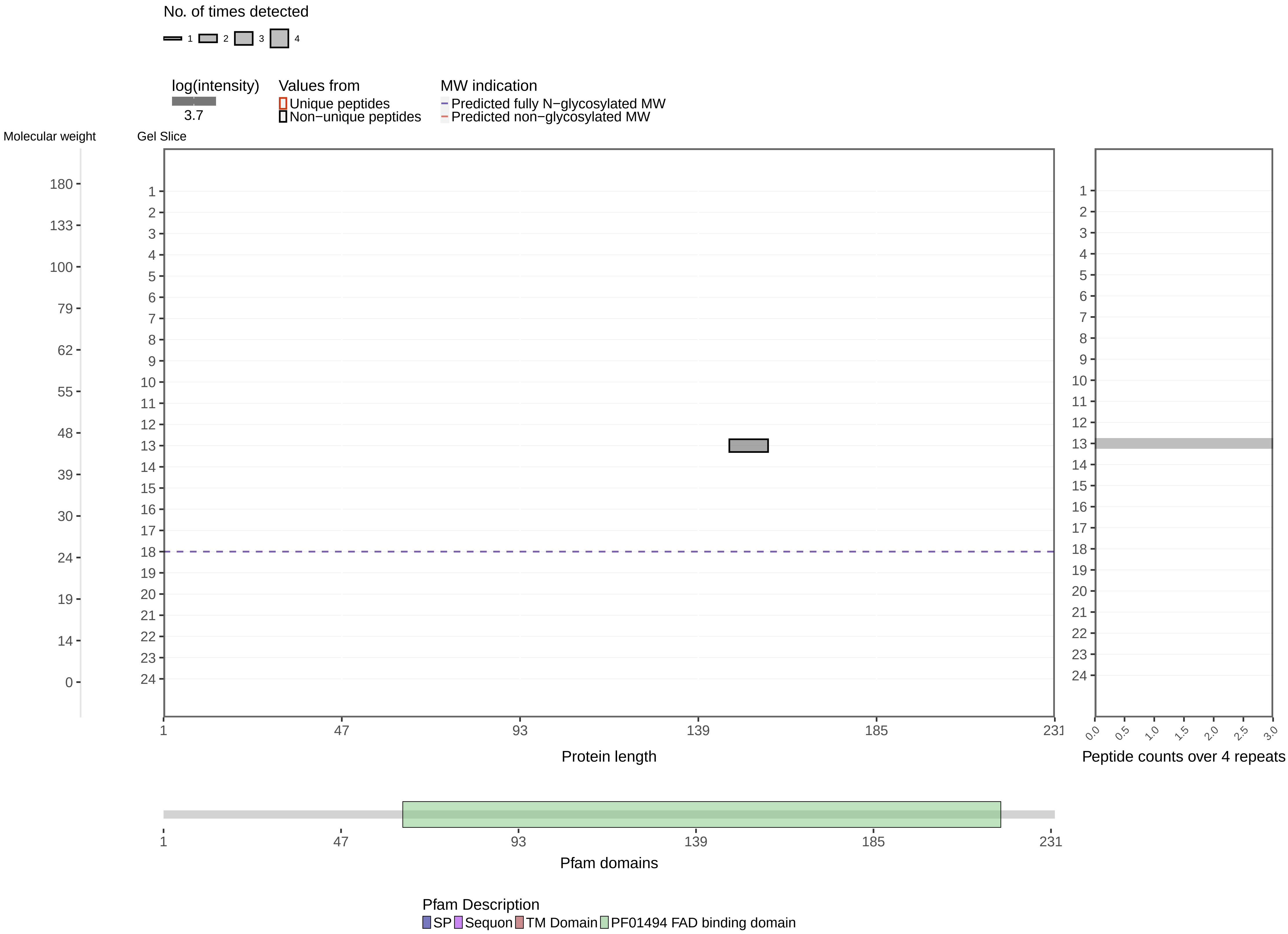


Protein sequence:

-SKNGWKALDAIGVRNDLRTQFLEIQGMVVKSEDGRELRSFRFKDEDESQEVRAVERRVLLETLASNLPPDAISFSSKLANVERSENGETMLELEDGIHISAKILIACDGIRSPWPSGWDFQSLTM*

NbL14g04140.1 uncharacterized_LOC107766858 [MW = 25.5 kDa]

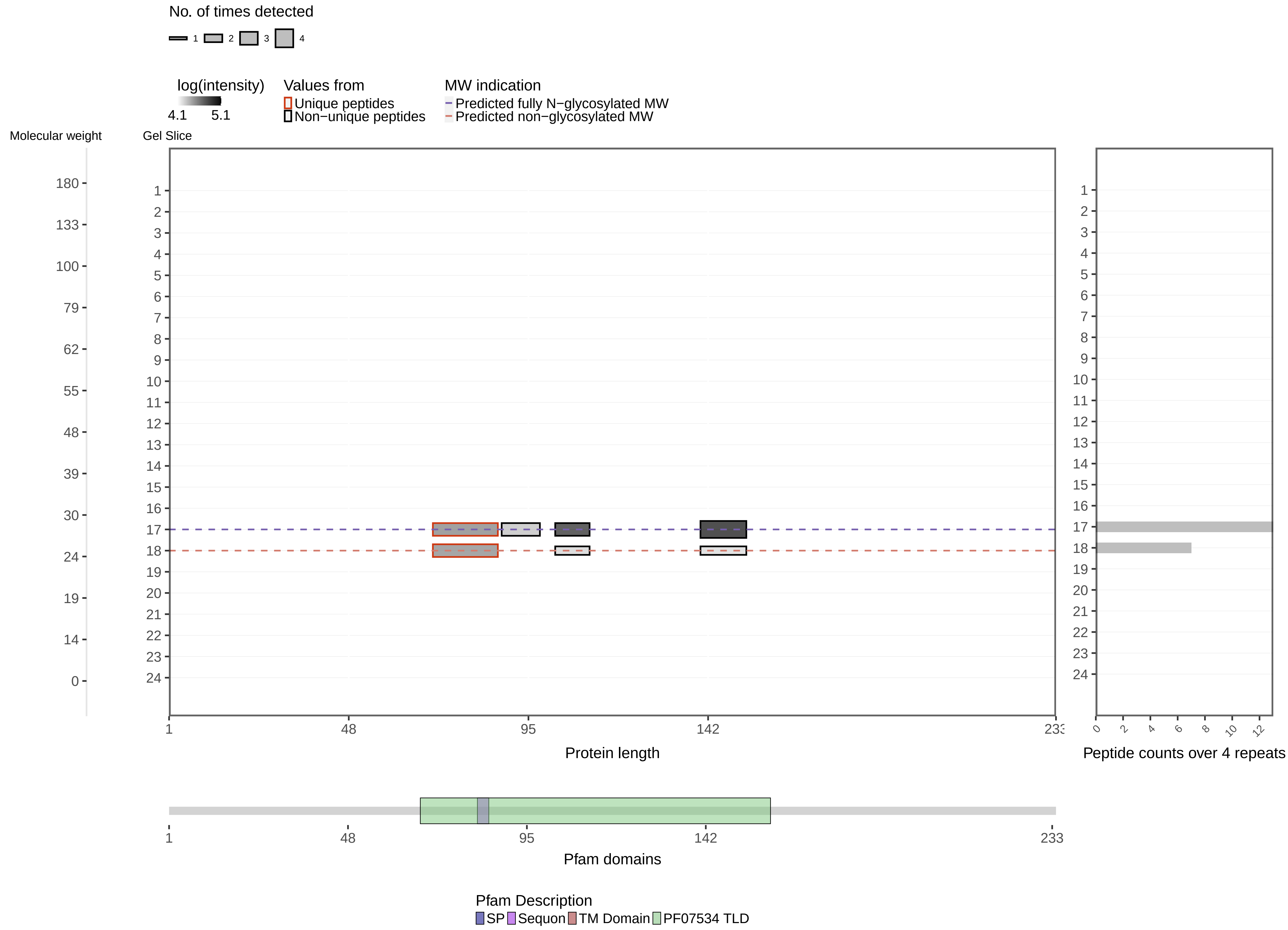


Protein sequence:

AFNPEGYRSTDDYYDTFDAFLFYWAEDEQQPIILPKVGGSGAALFDYARGGPQFGADGLLIGPPLAPVMGGFAGPDTNSGIGDLRQAKSRLGLSYAKRPDGKESLFGDESKAELDEVLVFCSPQIASLY*

### NbL14g04040.1 abscisic_acid_receptor_PYR1−like [MW = 24.98 kDa]

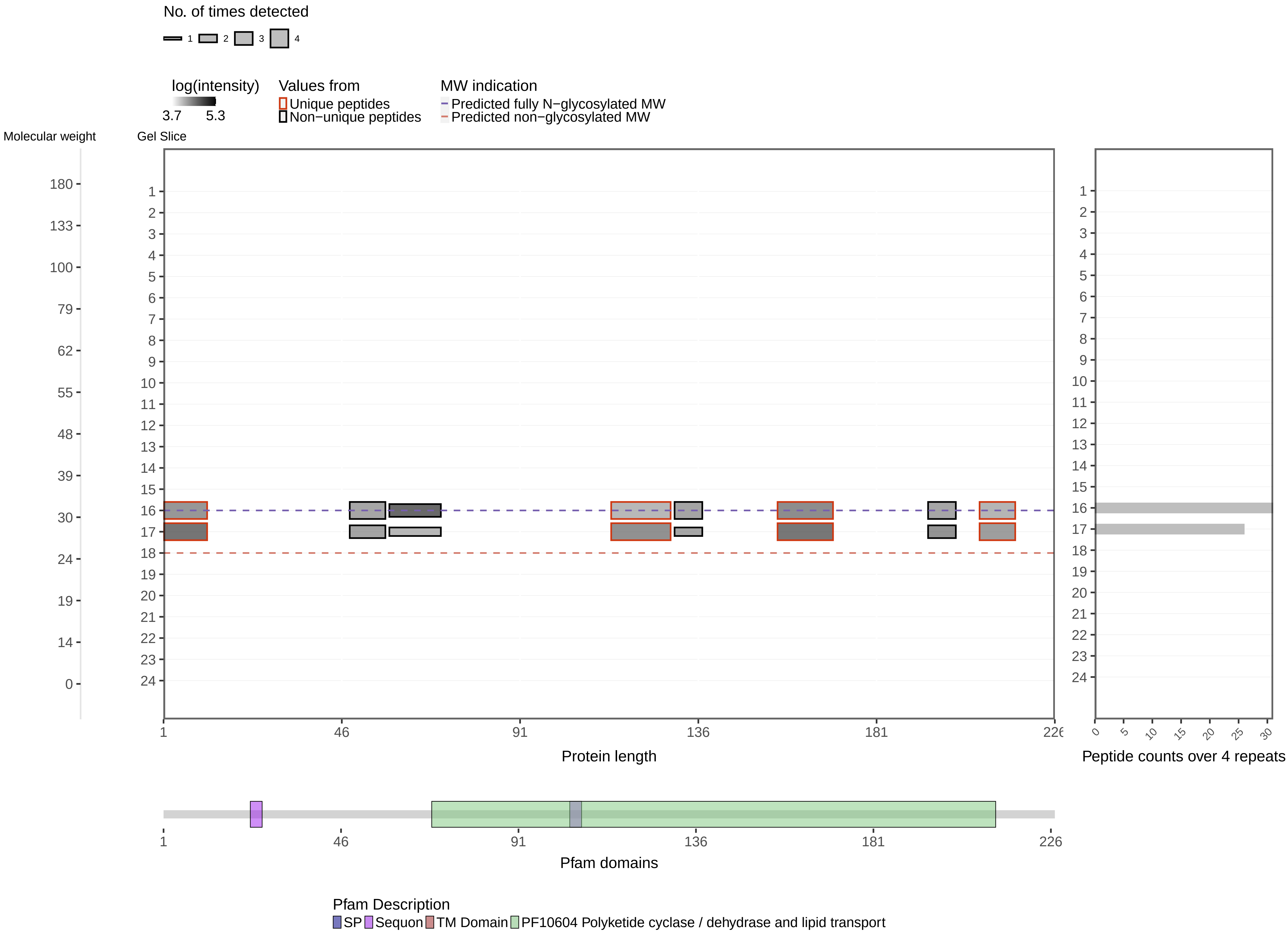


Protein sequence:

SVGENFSIVVGATRYVSVISGLPADTSTERLDILDDEKYVTGFSIIGGEHRLRNYRSVTTVHGFDQQLDGKISTVVMESYVVDVPDGNTEEDTRLFADTVVKLNLQKLGSVAEAIARGGNGVVSEN*

### NbL14g03890.1 phosphomevalonate_kinase−like [MW = 55.05 kDa]

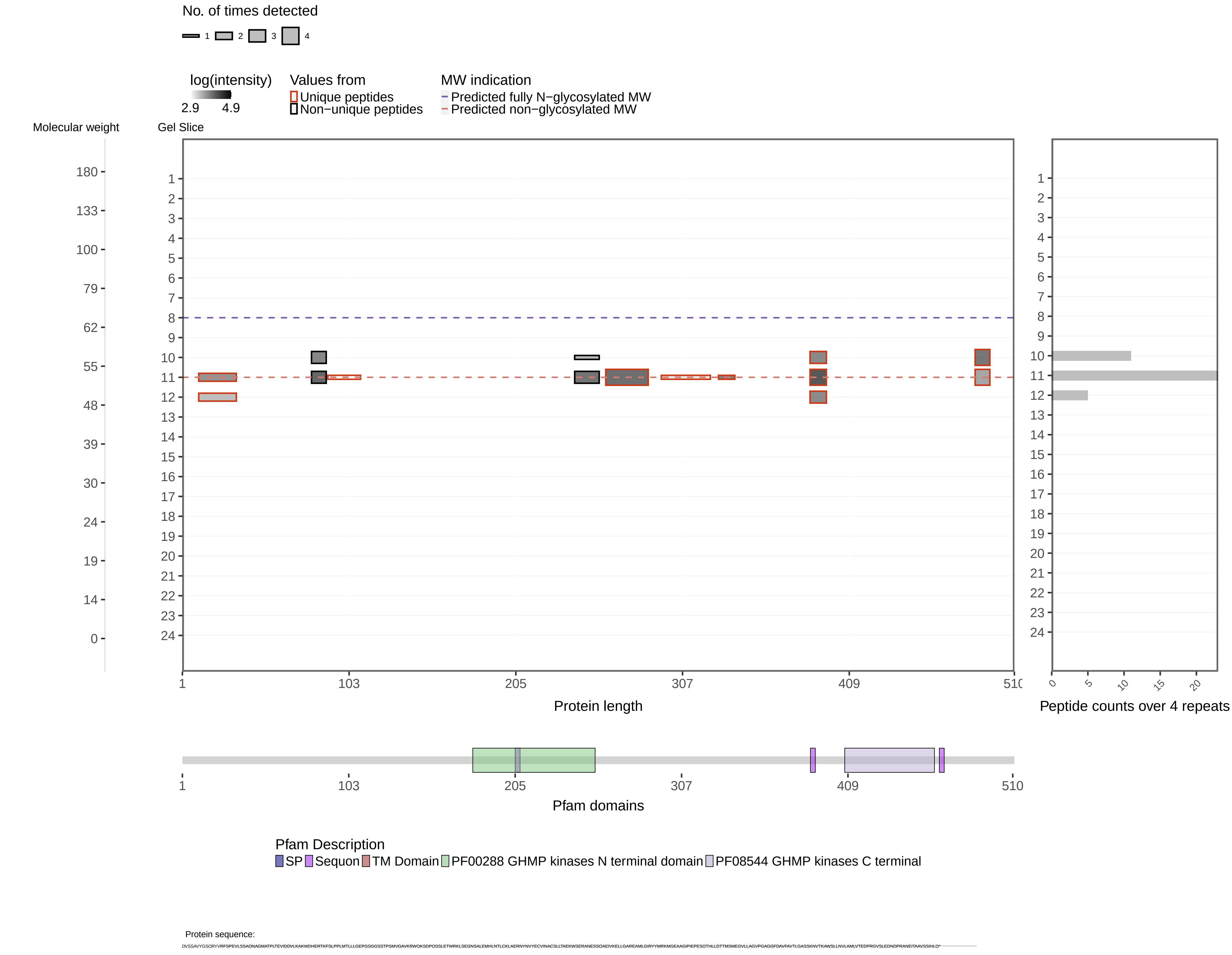

NbL14g03770.1 uncharacterized_protein_At2g34160−like [MW = 14.32 kDa]

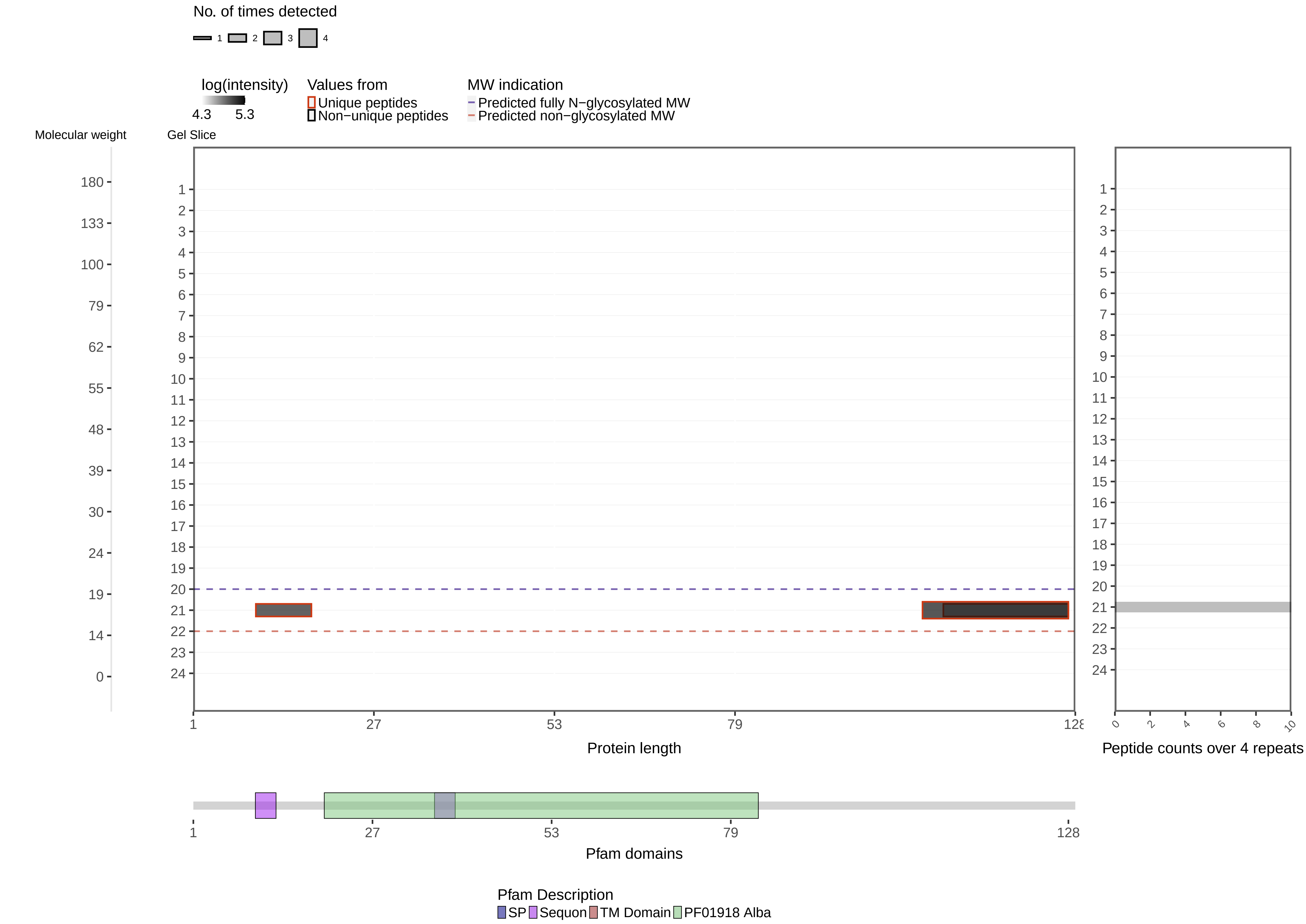


Protein sequence:

LSALGMAISTVVTVAEILKNNGFAVEKKIRTLTIDMRDEPGARPIPKAKIEIVLGKTEKFDELMAAEAEQNGDNEEQN*

### NbL14g03450.1 uncharacterized_LOC107795122 [MW = 32.75 kDa]

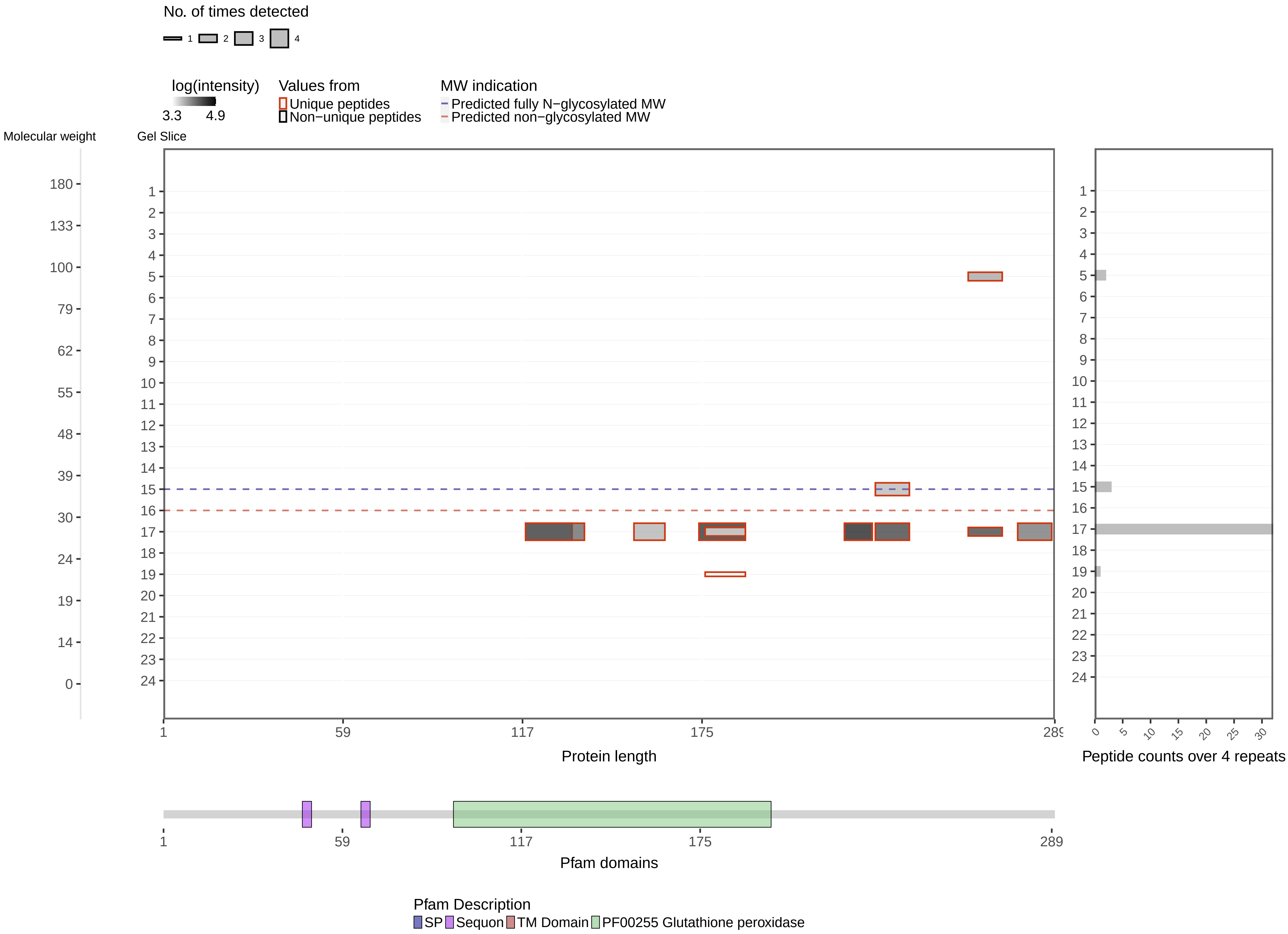


Protein sequence:

DTKSEIKLLNELHNRYKDRKPKGFAVLGFLYDKKETGKSGYGRGKQEILNTAEFPLFDKVKLNEKHEHWYGIDENKRDEDLYYFLKKETKGGDITEEFVKILVDWTGVPAGRYDPFPILHDETKKSEIAKSGEDPGTTVGLRDDPRFVLENAIATLVR*

NbL14g03440.1 probable_phospholipid_hydroperoxide_glutathione_peroxidase [MW = 25.75 kDa]

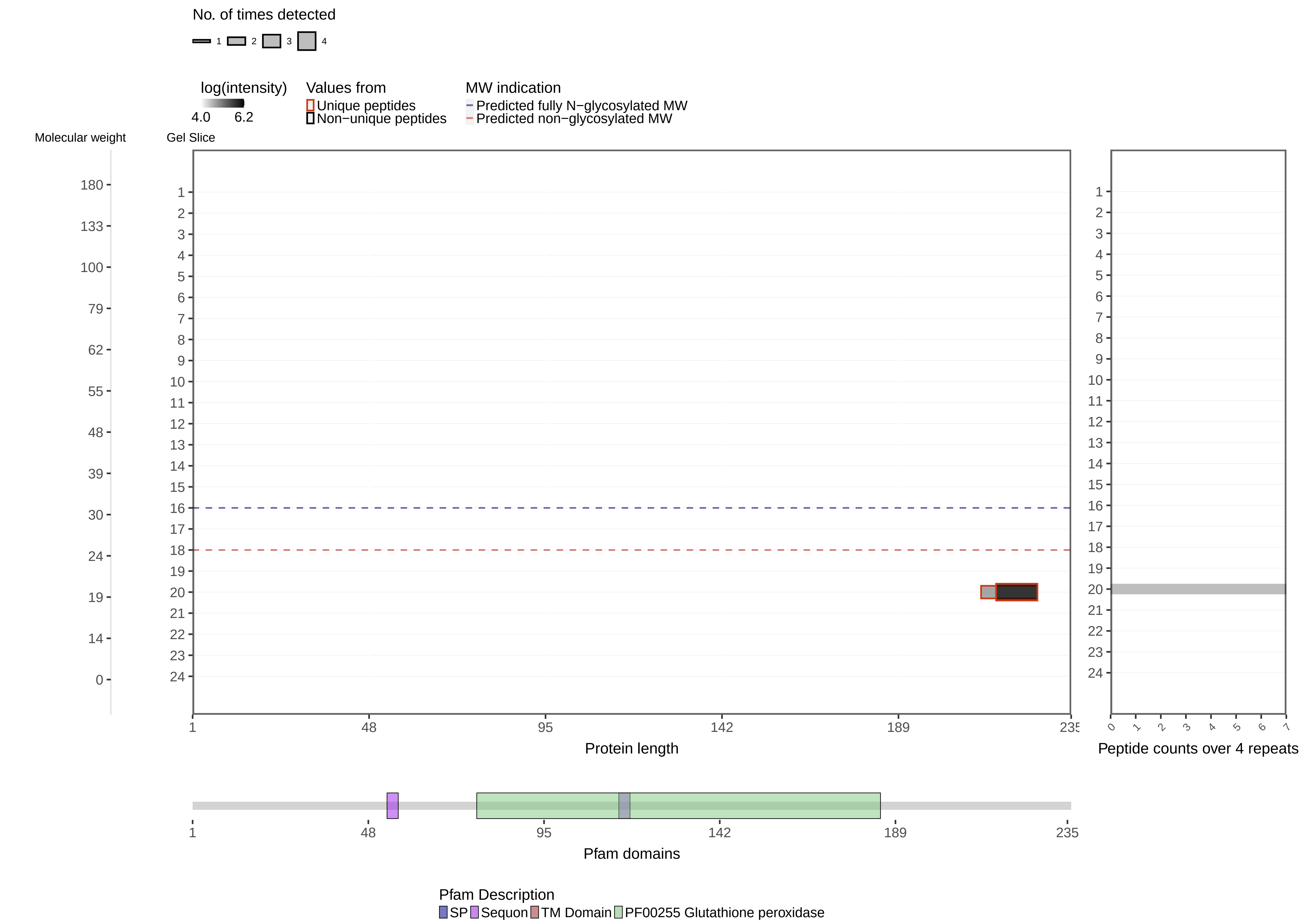


Protein sequence:
ASRCGLTTSNYTELSQVYEKYKTKGLEILAFPCNQFGAQEPGSNPEIKQFACTRFKAEFPIFDKVDVNGPNTAPVYQFLKSSAGGFLGDLVKWNFEKFLVDRNGKVVERYPPTTSPIQIEKDIQKLLAA*

### NbL14g03340.1 uncharacterized_LOC107797087 [MW = 30.98 kDa]

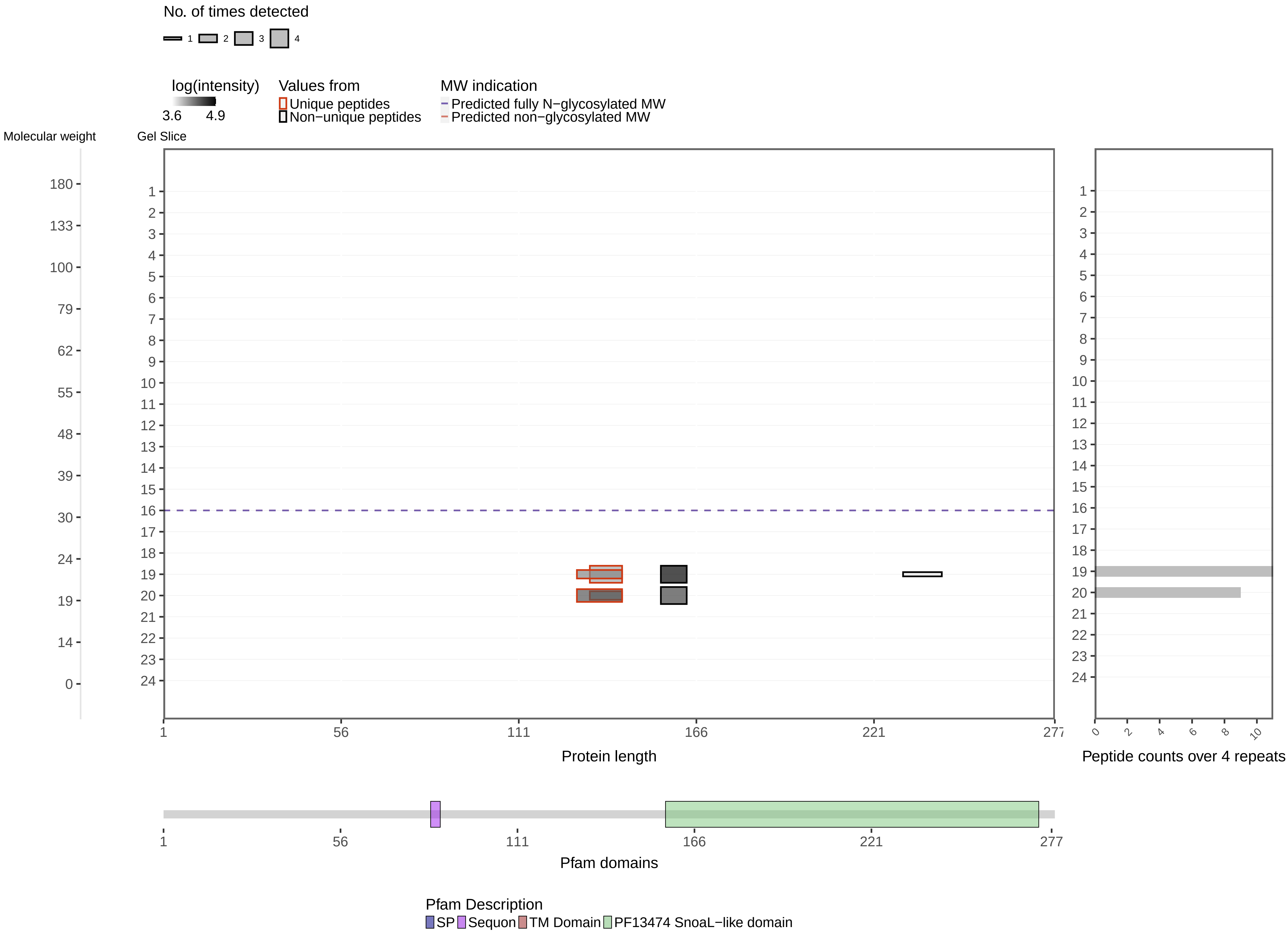


Protein sequence:

LKNAIKEENYAQAATIRDSLRLLQEDSKASVLAANARFYNSFKHGDLAAMQTLWSKGENVCVVHPGVSGITGYDLVMGSWEFVWAEYDFPLDIEIRDVQVHVRGDIGYVTCIEMVRTKGSSWGKQFATNVFEKVNGQWFMCIHHASYIDL*

### NbL14g03290.1 pectin_acetylesterase_8−like [MW − SP = 41.81 kDa]

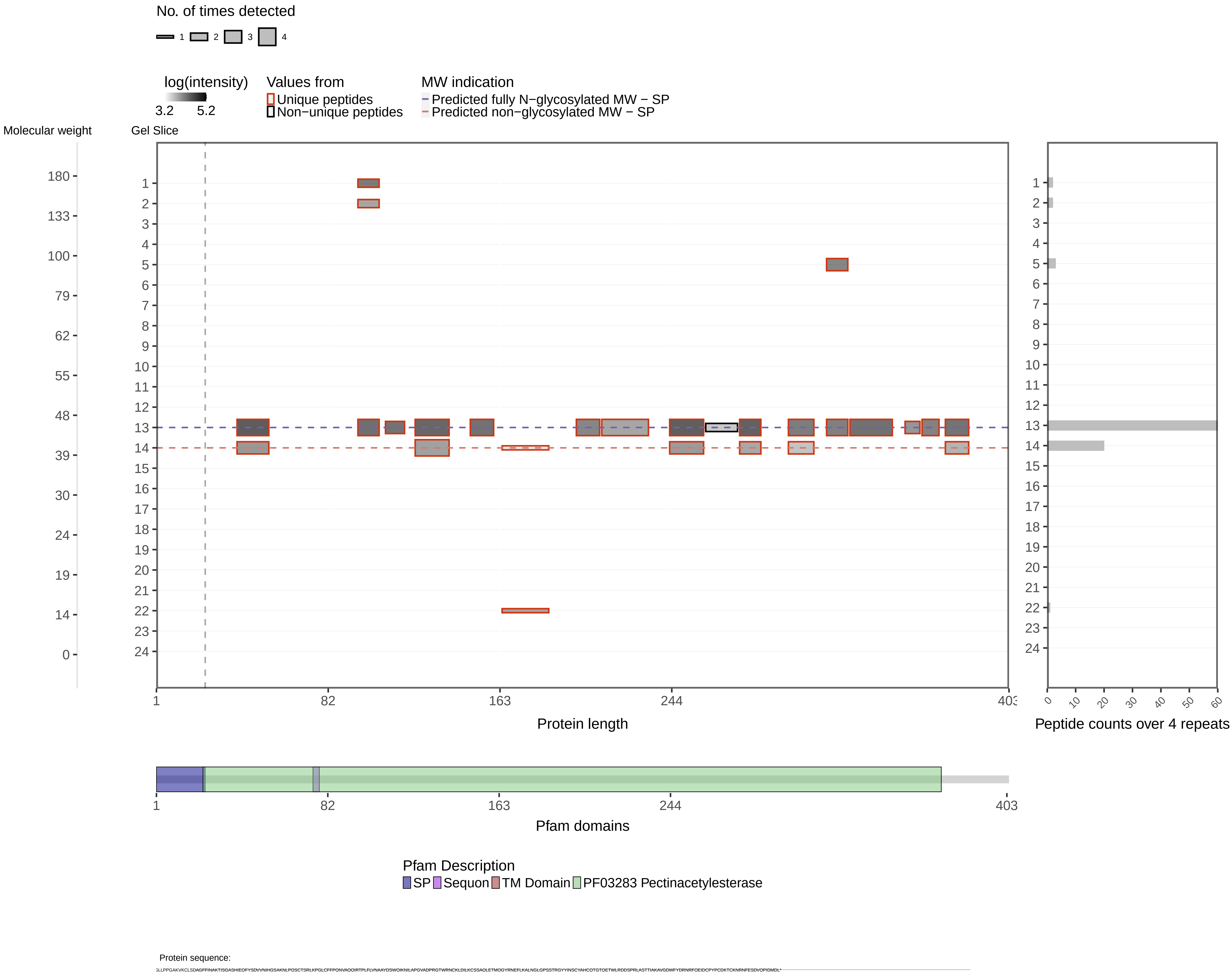

### NbL14g03150.1 protein_arginine_N−methyltransferase_1.5 [MW = 73.56 kDa]

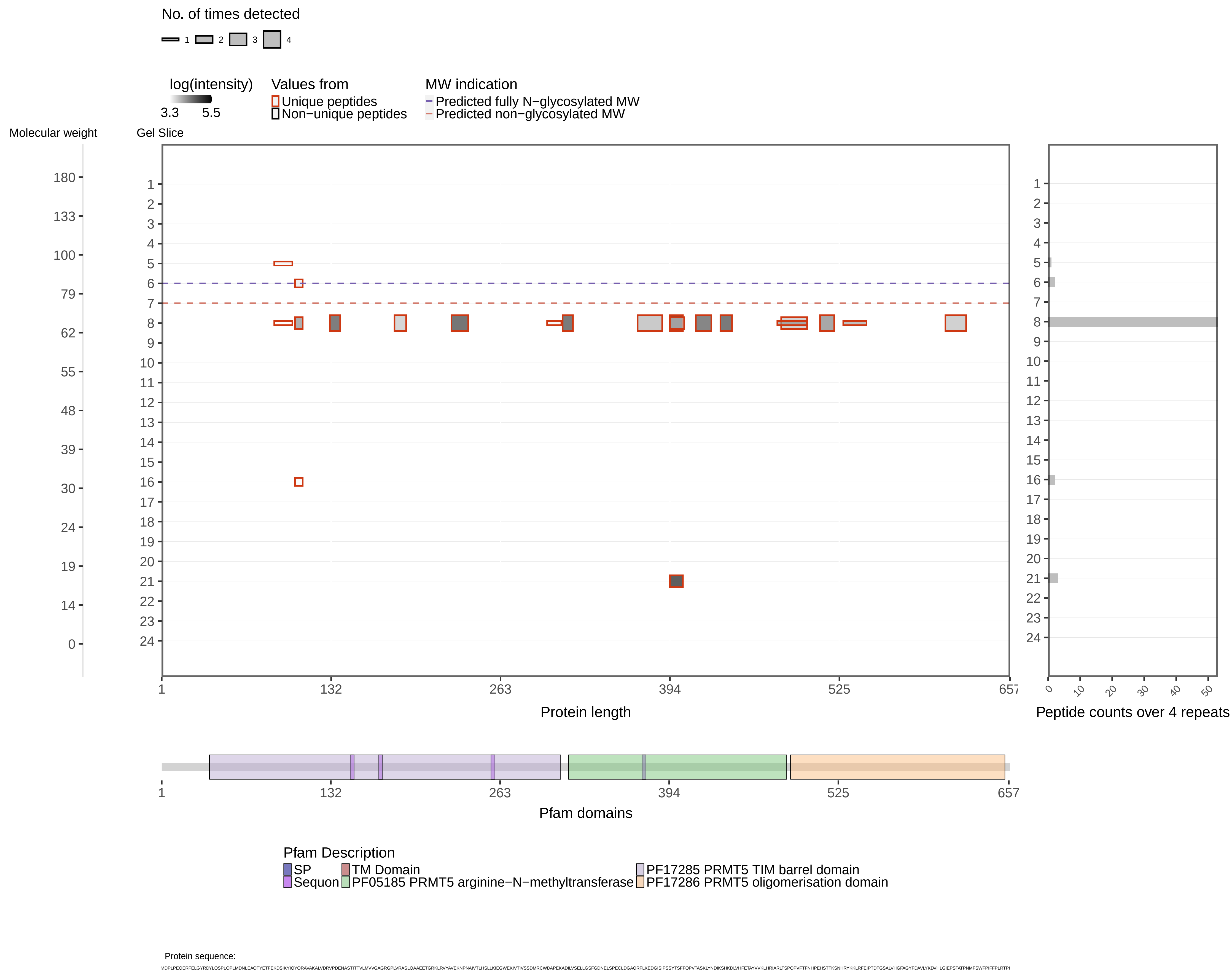

NbL14g03090.1 AIG2−like_protein_D [MW = 19.85 kDa]

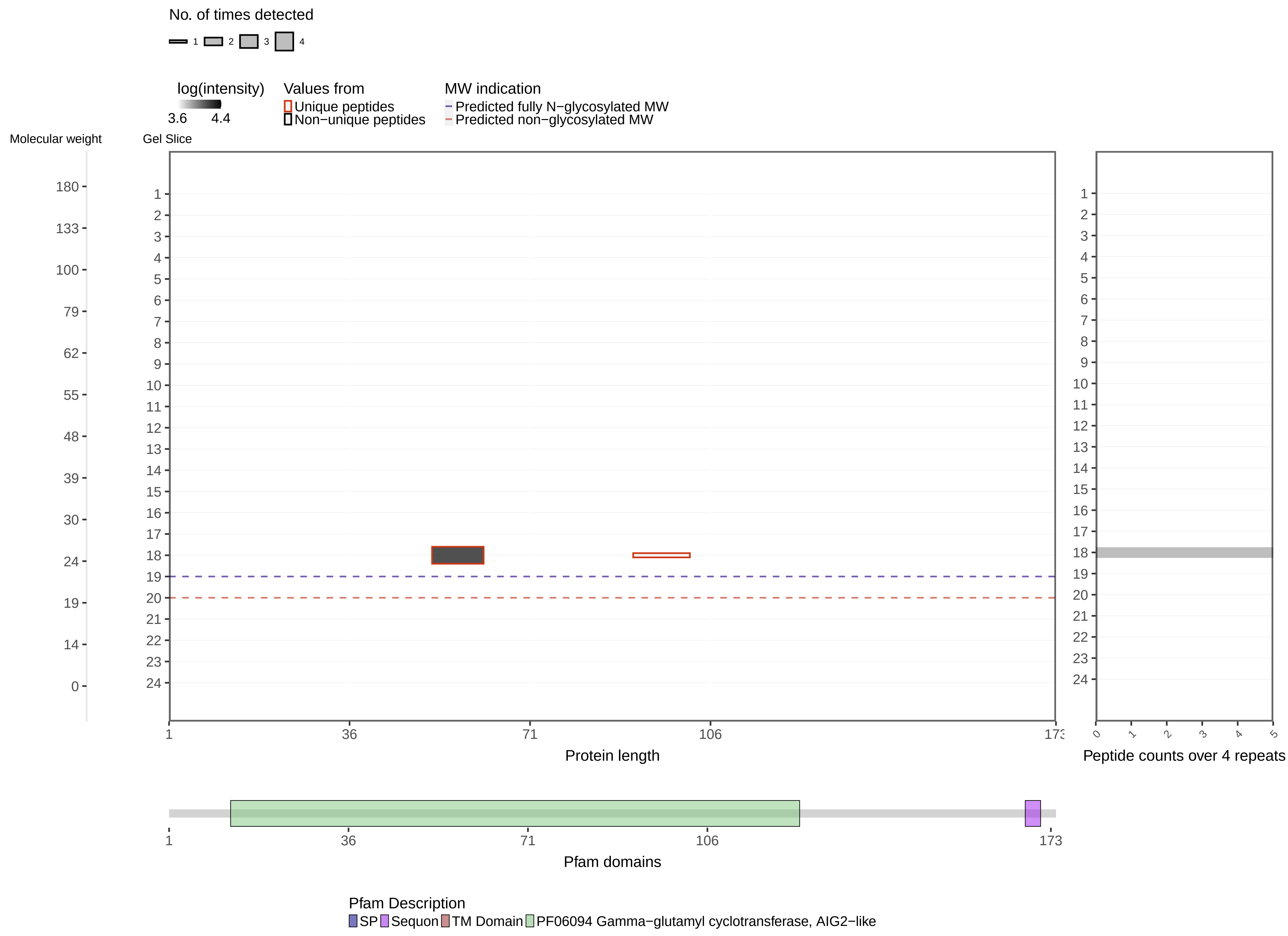


Protein sequence:

PELNILDIFEDVEYERTTVDVSLMDSSKTLLAETYVWVDKRDPNLYGEWDFEEWKQLHKGDFLKMTMGFSKELELPESKTRVATYESFYAQEDNKSKP*

### NbL14g03060.1 peptidyl−prolyl_cis−trans_isomerase_CYP18−1 [MW = 17.47 kDa]

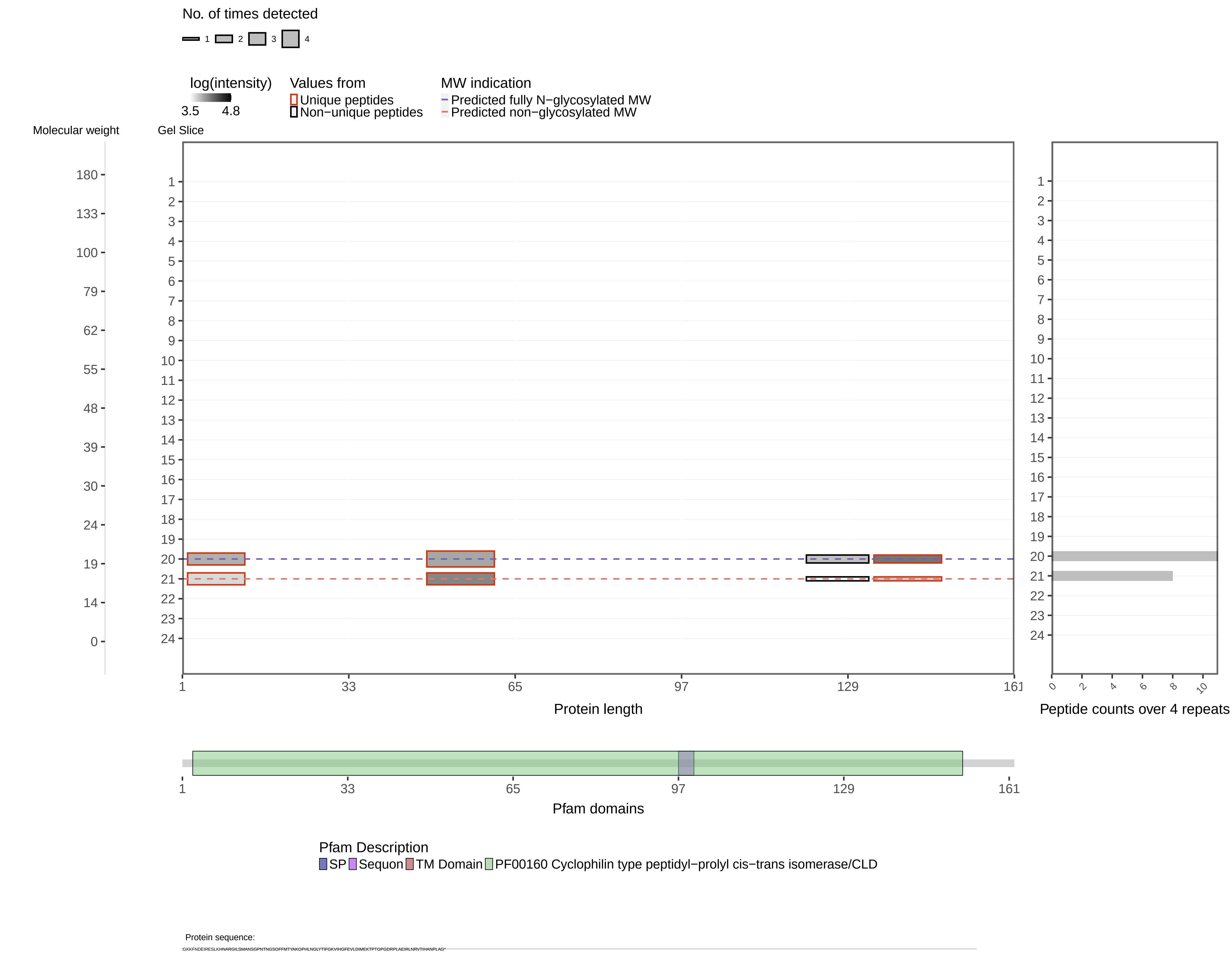

### NbL14g02490.1 3−hydroxyisobutyryl−CoA_hydrolase−like_protein_2_mitochondrial [MW = 46.56 kDa]

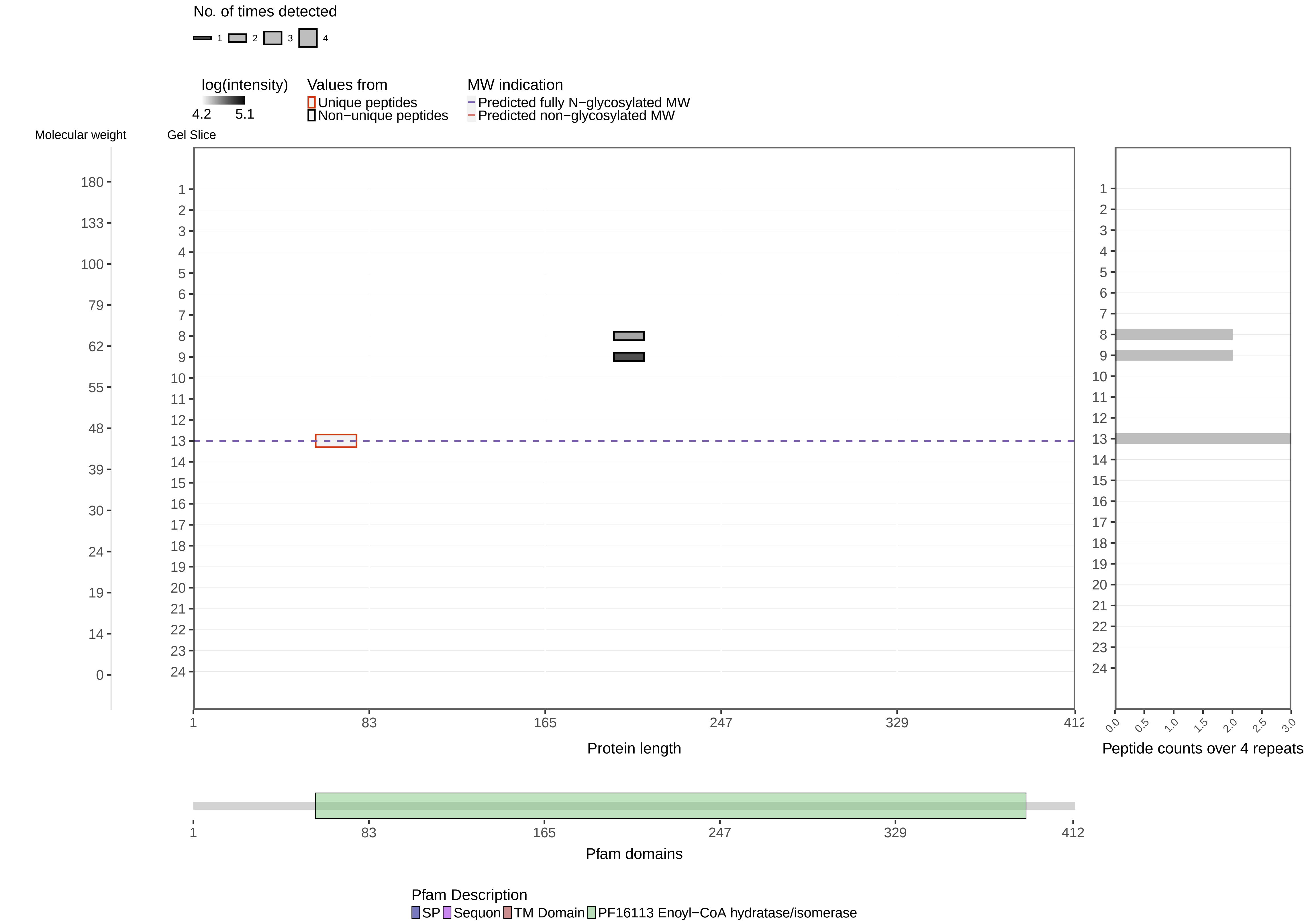


Protein sequence:

YYLSRLPGYLGEYLALTGEKLNGVEMIACGLATHYSLKERLPWMEERLGKLITDDRLVIENSLAEYGDLVYPDSKSVLHKFEKIDKFFSQEIIEEIIEVLEREAAEYHDEWCNAALNNIKVASPLSLKVALKSIREGRFQPLDQCLVREYHISVNWVSKRMSGDFCEGVRARLVDKDFAPKWGPTRLEEVTTDMVNCFFSPLGELEPELNLATALREPFM*

### NbL14g02320.1 macrophage_migration_inhibitory_factor_homolog [MW = 12.22 kDa]

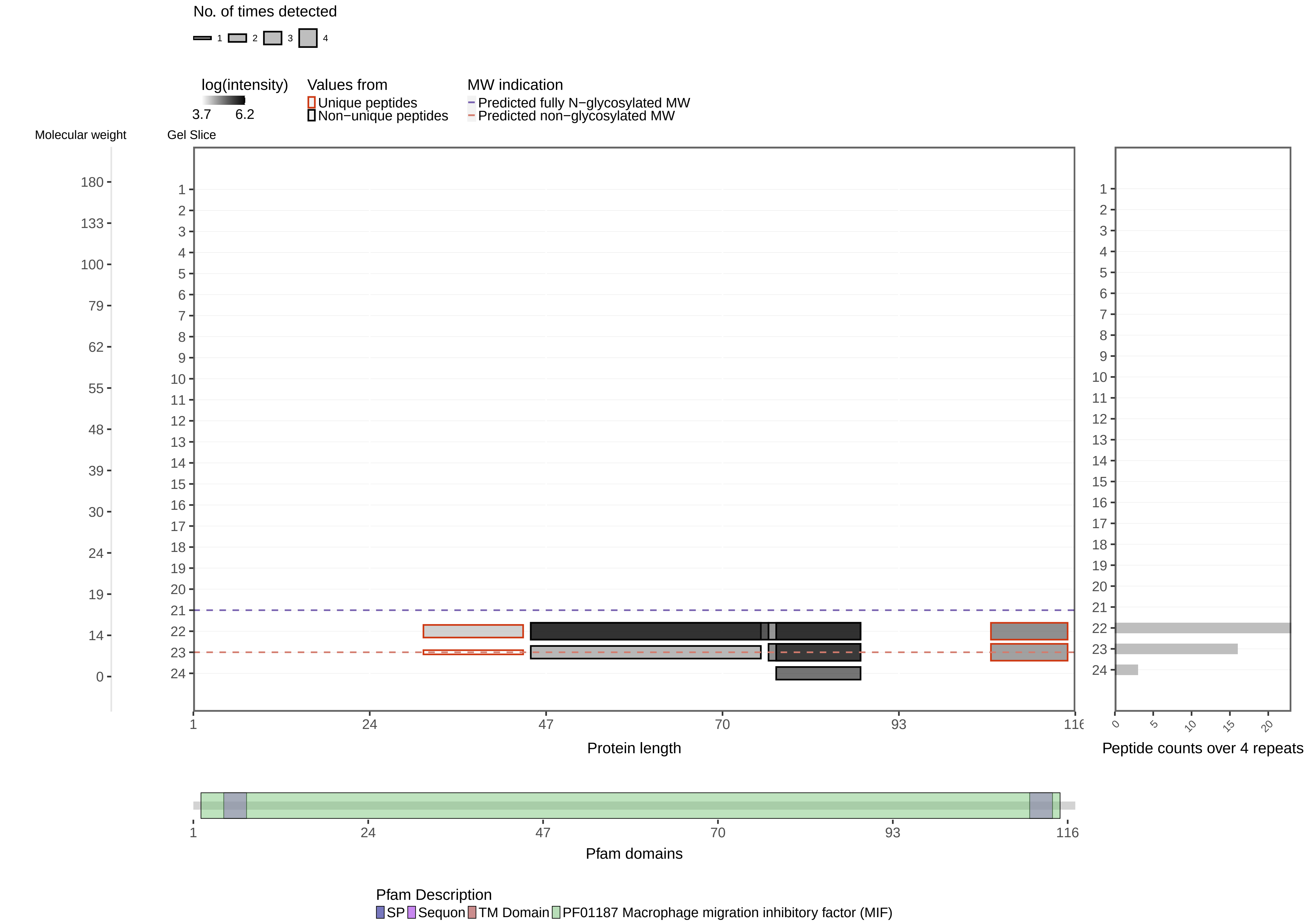


Protein sequence:

SVPMSFGGTEQPAAYGELVSIGGLNADVNKKLSAAIAGILETKLSIPKSRFFLKFYDTKGSFFGWNGSTF*

### NbL14g02040.1 upstream_activation_factor_subunit_UAF30−like [MW = 16.82 kDa]

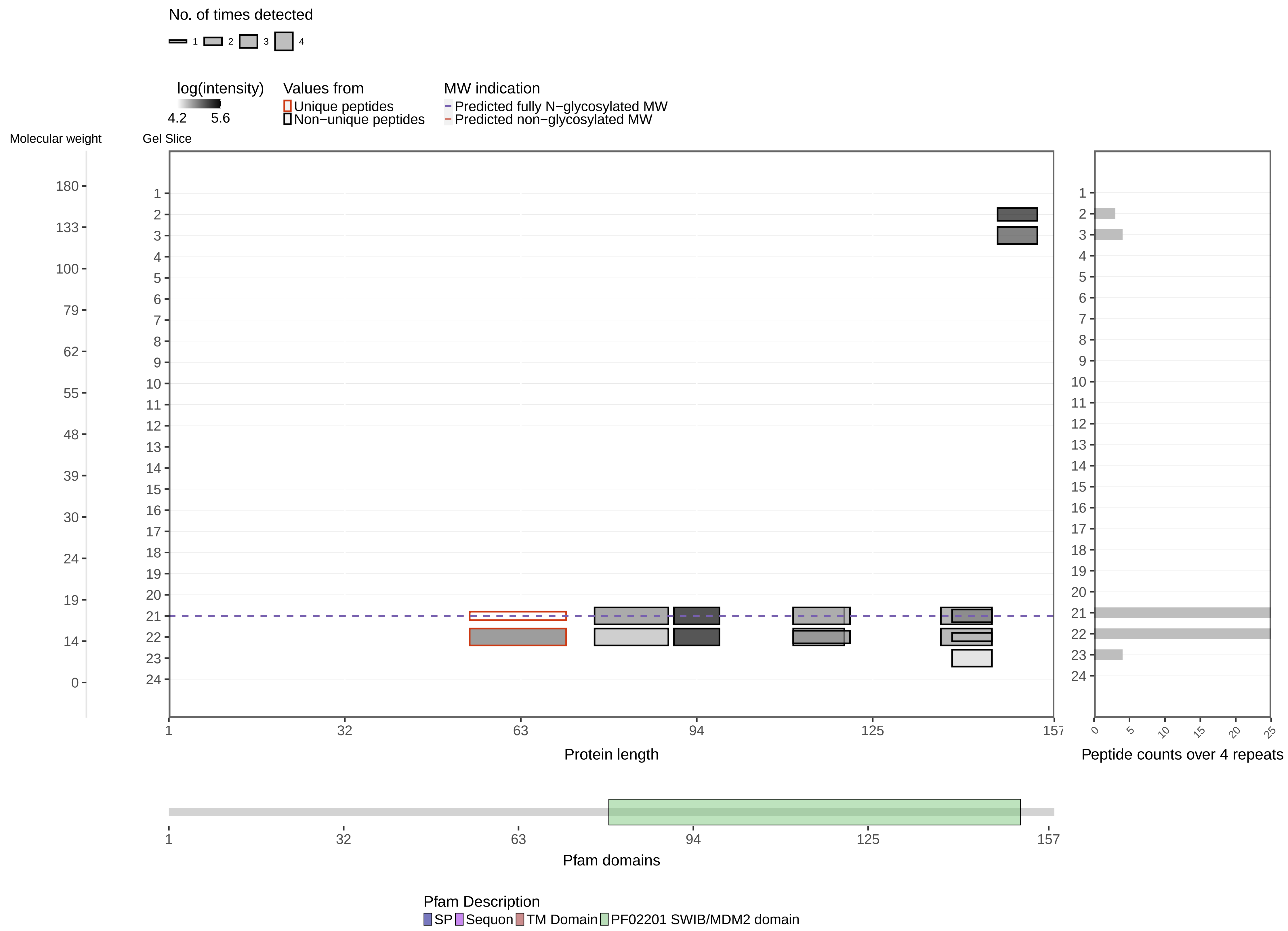


Protein sequence:

AASATGKGRHKGILKPLPVSPALGKFVGSSEISRTDAVKKVWDYIKTNNLQNPANKKEINCDEKLKTIFAAKDKVGFLEIAKLLSSHFQKAS*

NbL14g01960.1 tankyrase−like [MW = 31.96 kDa]

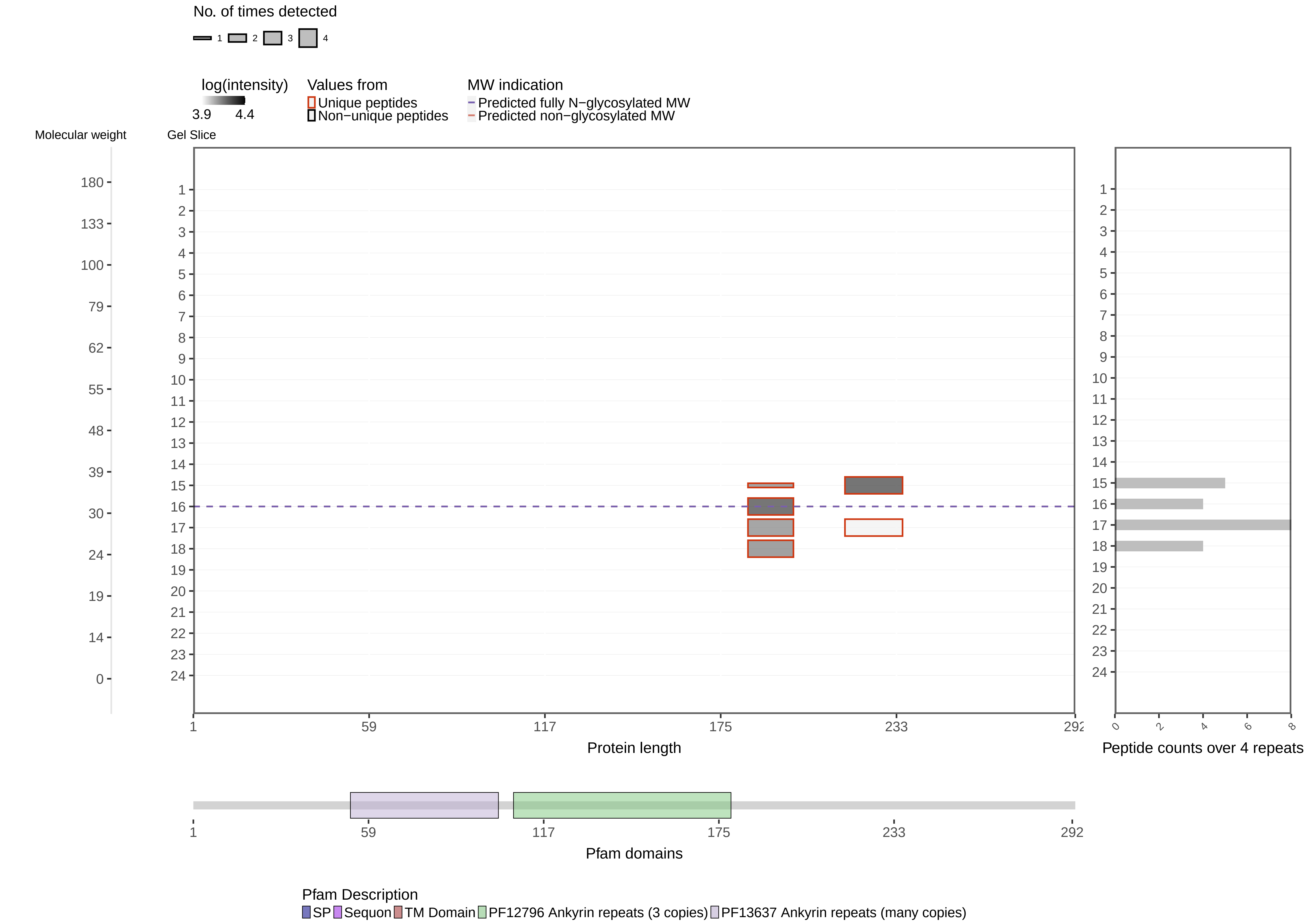


Protein sequence:

'RLLVSSGVSIKSCNRKGMTPLHYAAQGSHLELVKYLLKKGANVNTKNKAGKTPIDLASNEEVSSILRQPVTAPSKEAVNREEKRGEPESQSSSQDKIESSDGKGVATEEGEIEAEKDESLKRKGDEHETSEKSKETKKAKVSLNHLLAADDNQEDEENL*

NbL14g01640.1 plastid_lipid−associated_protein_3_chloroplastic−like [MW = 46.24 kDa]

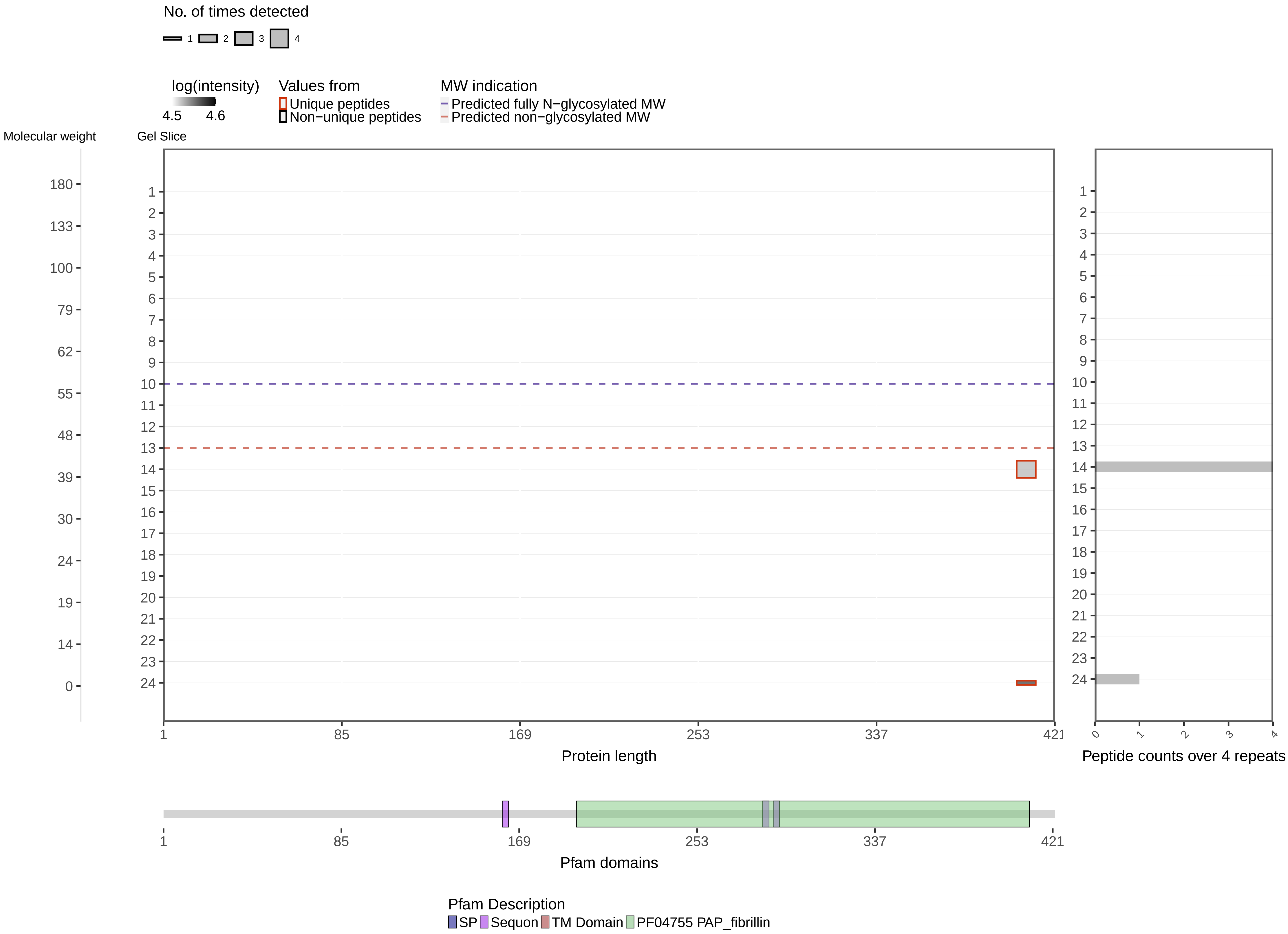


Protein sequence:

RVAELKRCLMDTVYGTNFGLRASSQIRAEALELVSQLEAANPTPSPVESPELLDGNWILVFTAFSELLPLLAVGTIPLLKVEKISQTISTSNLSIENSTTLSSPFATSSFSANAAFEVRSPSRIQVEFKEGTVNPPEVKSKIDLPENVGIFGQTISLSPIQQSLGPVENVVAGIARAISGLPSLKVTIPGERTKSWLITTYLDKDLRISRGDGGLFVLVKEESSLIDQ*

NbL14g01490.1 probable_ADP−ribosylation_factor_GTPase−activating_protein_AGD8 [MW = 43.47 kDa]

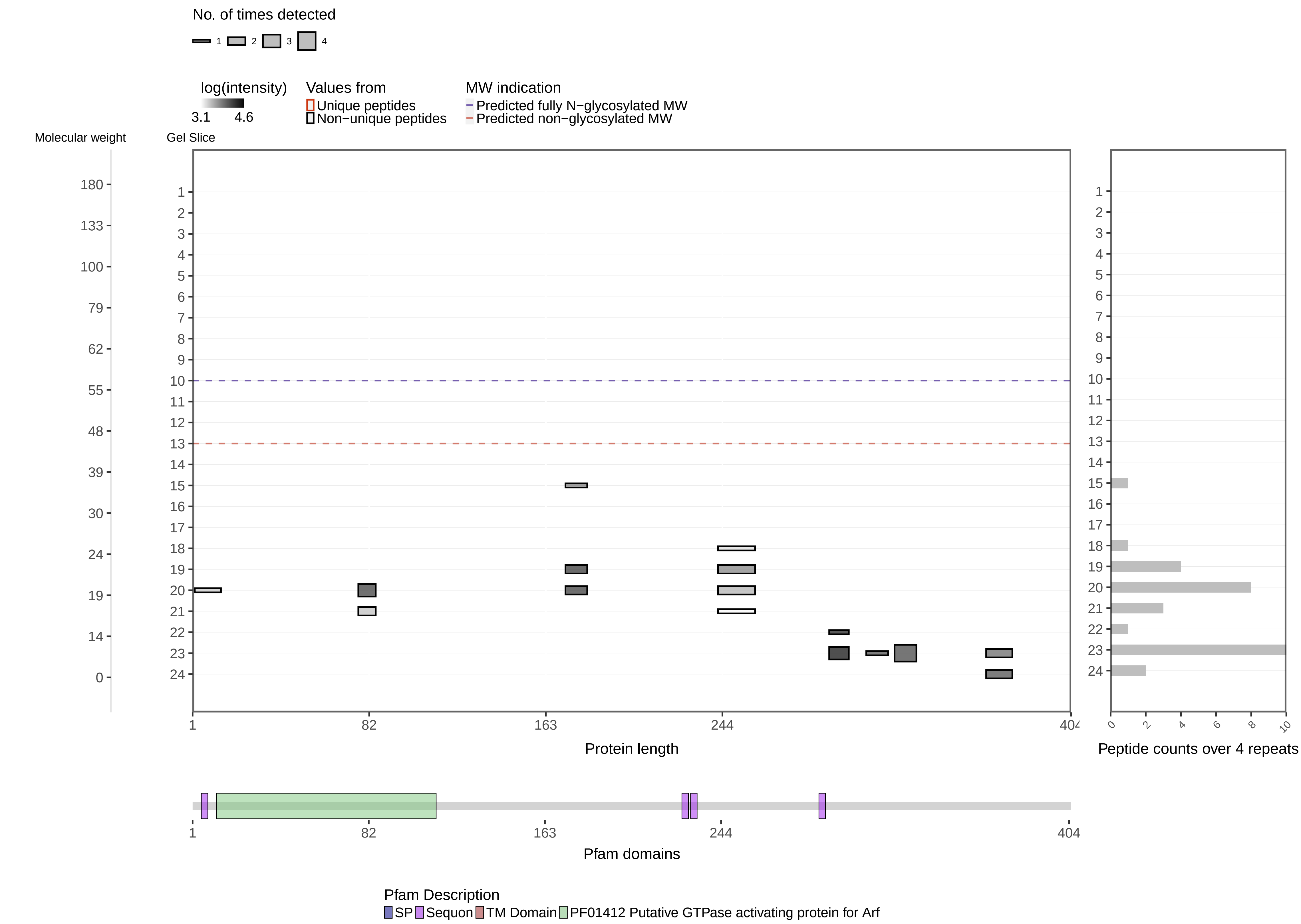


Protein sequence:

.KFGKPGGGLGARKLTKKLSESLYDQKPEEPSVQVSSSNSTSNASTVGSSFASRFEYTDNVQPAEMSSGGPRVLNHVSAPMSSSFFSDYGMDSGFTKKTSSNSSKVQIEETDEARKKFSNAKAISSAQFFGDKSKAEMEASVSLQKFSGSSAISSADLFGNDDRADLDLTAGDLINRLSFQAQQDISSLKNIAGETGKKLGSLAATFMSDFQDRIL*

### NbL14g01230.1 regulator_of_nonsense_transcripts_1_homolog [MW = 139.26 kDa]

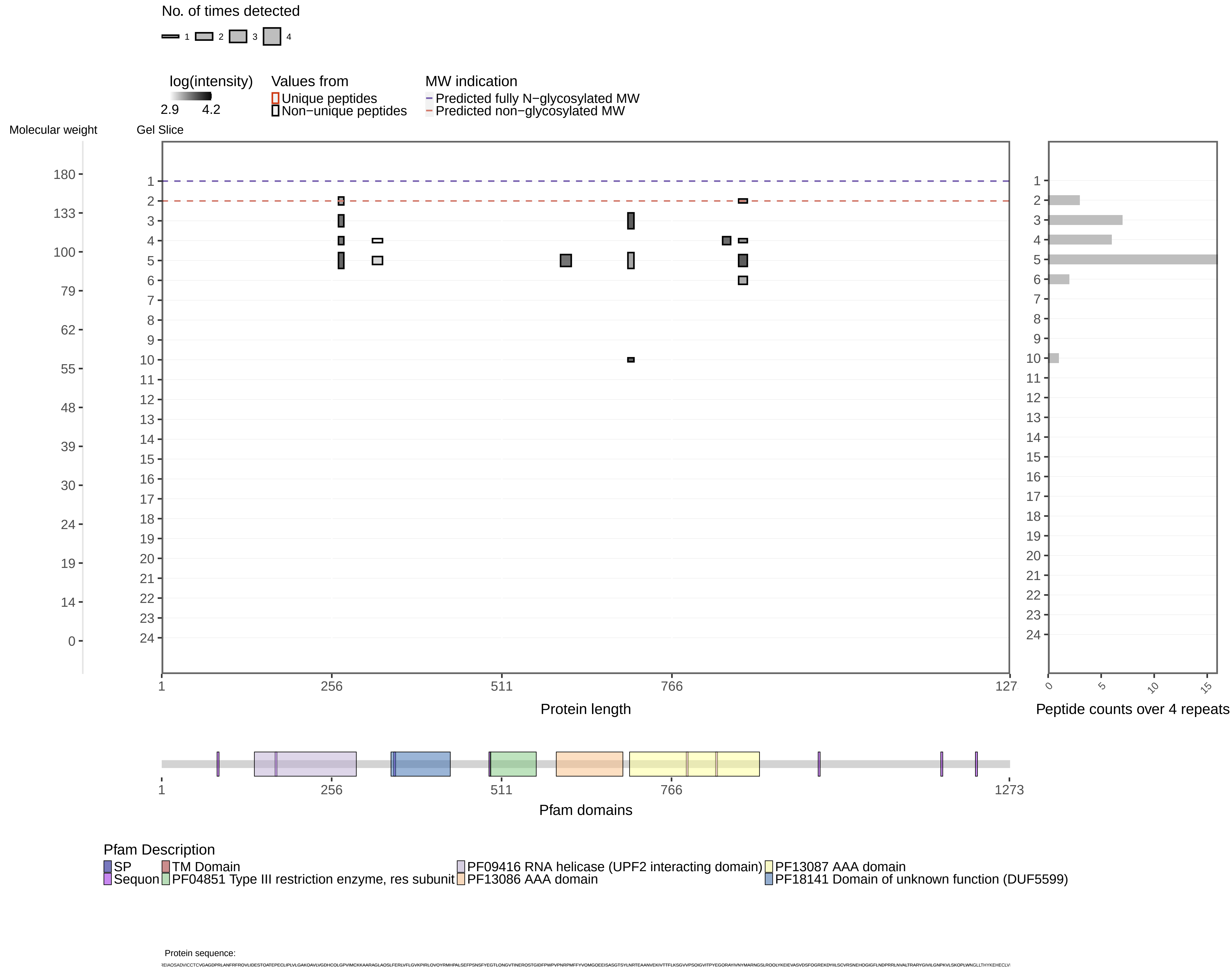

### NbL14g01190.1 60S_ribosomal_protein_L37a [MW = 7.11 kDa]

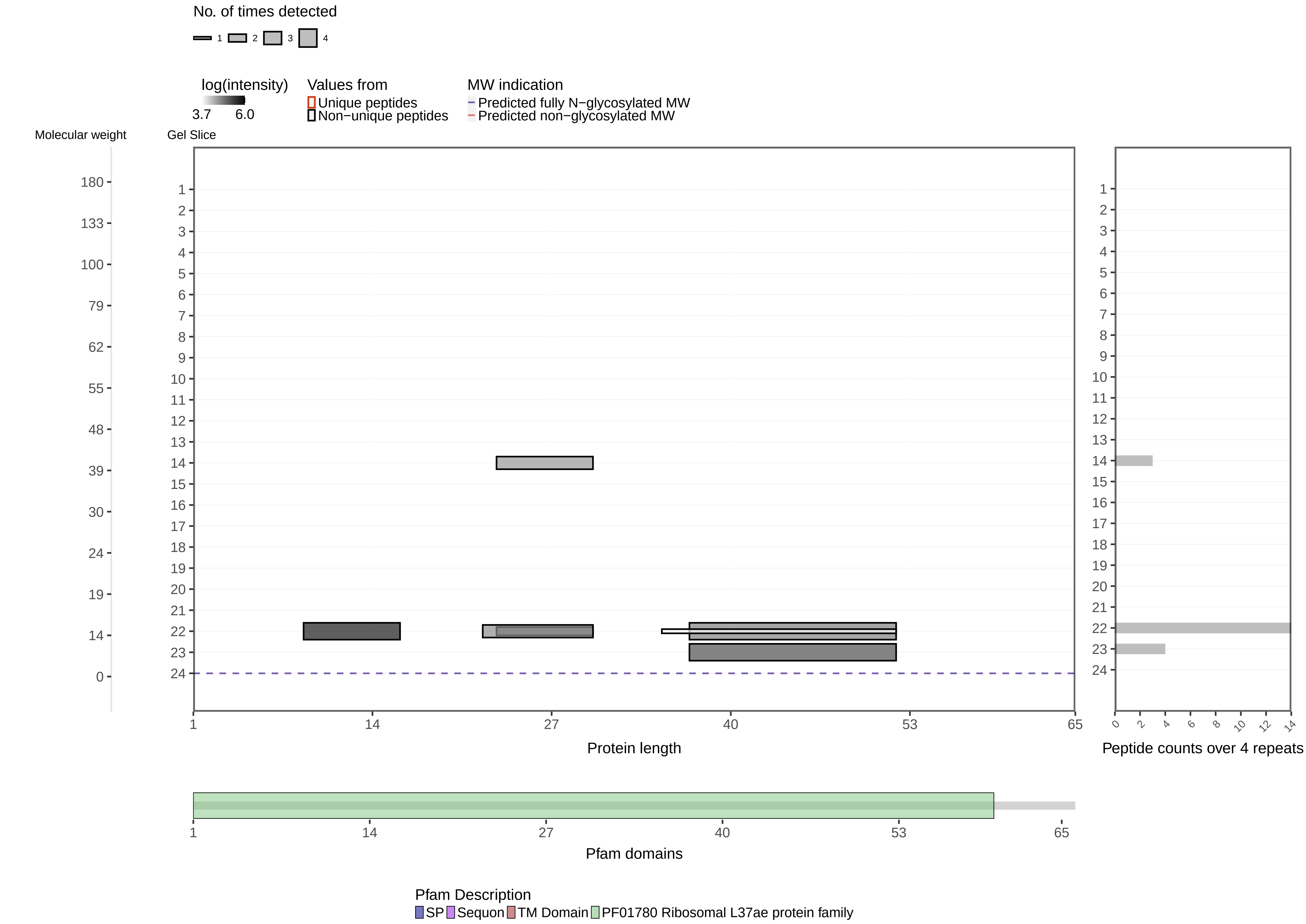


Protein sequence:

'KRKAVGIWGCKDCGKVKAGGAYTLNTASAVTVRSTIRRLREQTES*

### NbL14g01150.1 LL−diaminopimelate_aminotransferase_chloroplastic−like [MW = 50.09 kDa]

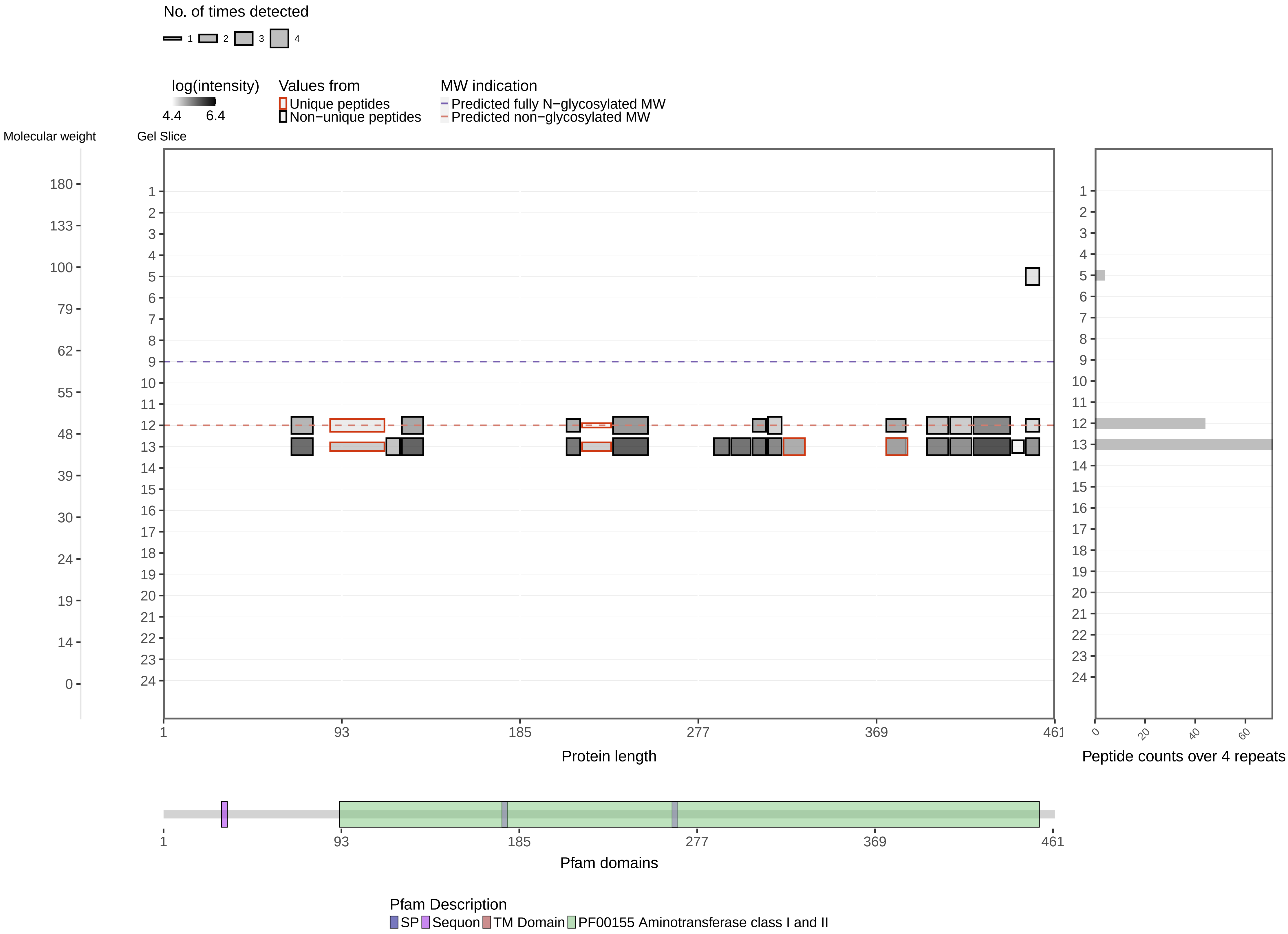


Protein sequence:

TPENGFFPDLSSVSRTDIIFFCSPNNPTGAAASREQLTKLVHFAKNNGSILVYDSAYAMYICDDSPKSIFEIPGAKEVAIEVSSFSKYAGFTGVRLGWTAIPKALLFSDGFPVAKDFNRIVCTSFNGASNIAQAGGLACLSADGFKAMKNVVSFYKENTEIIMETFKRLGYKVYGGKNAPYVWVHFPGRSSWEVFSEILEKTHVVTTPGSGFGPGGEGFVRVSAFGHRDNVVEACRRFKELYK*

NbL14g01130.1 plasminogen_activator_inhibitor_1_RNA−binding_protein−like [MW = 40.24 kDa]

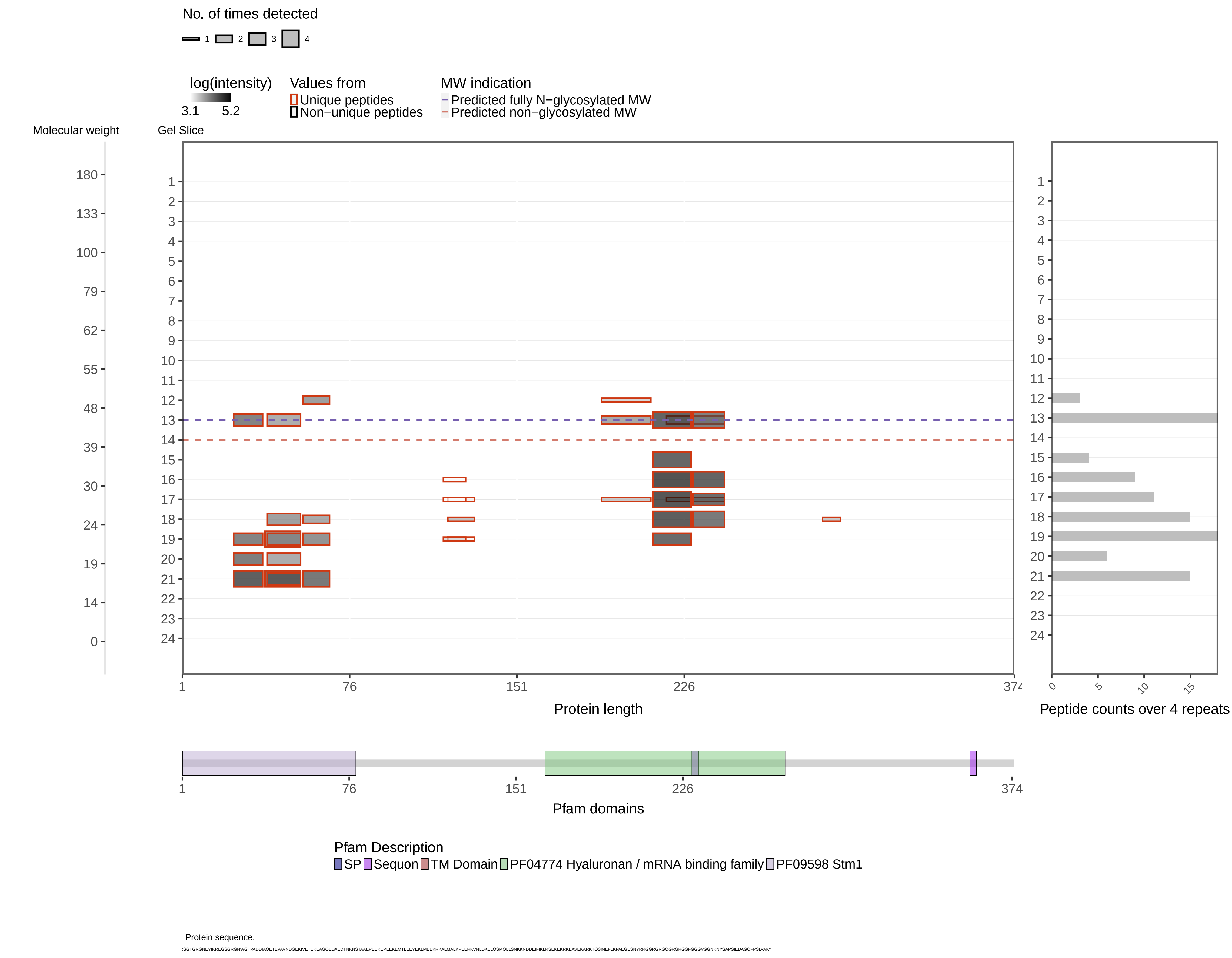

NbL14g01120.1 protein_ECERIFERUM_1−like [MW = 42.08 kDa]

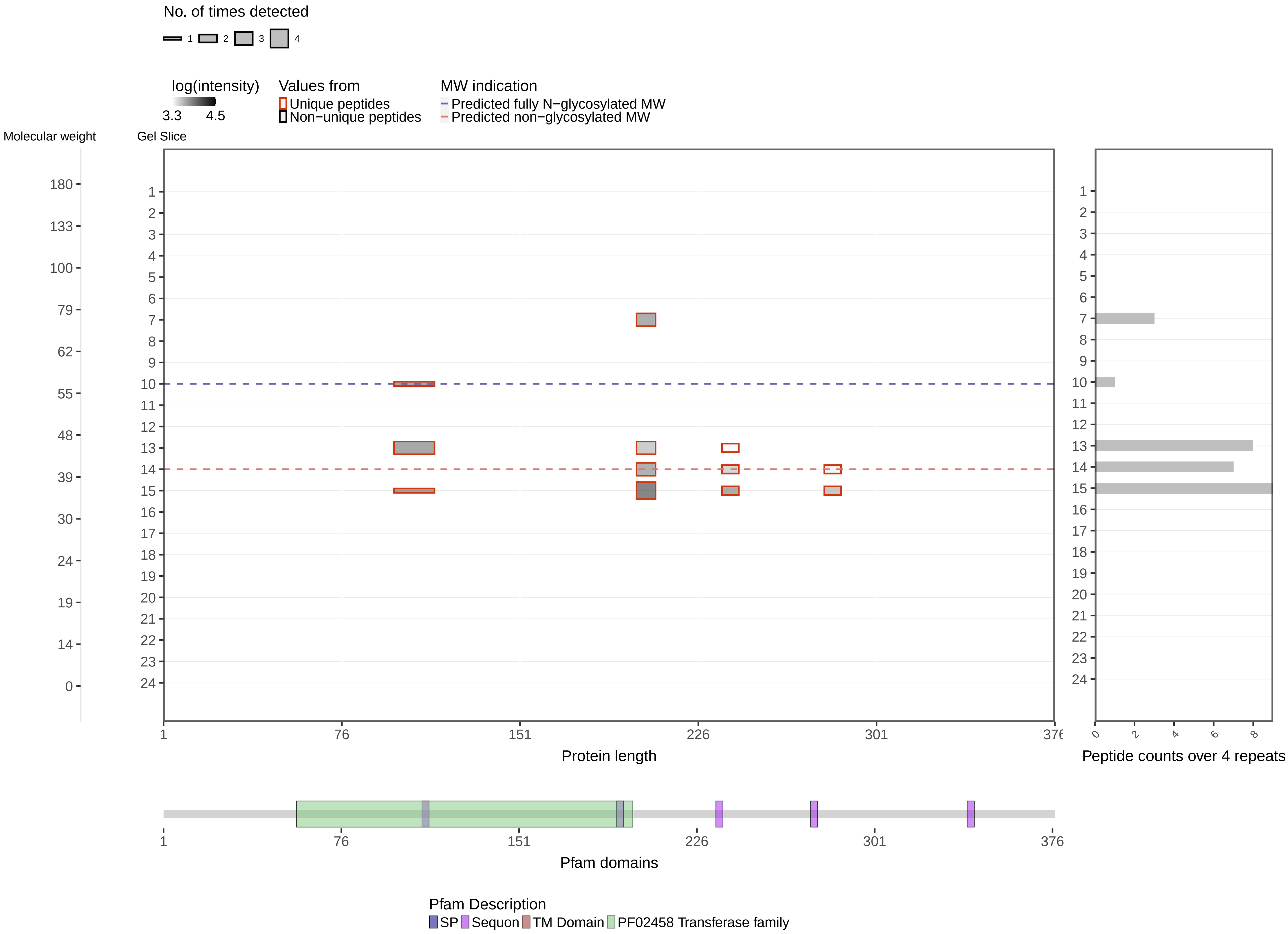


Protein sequence:

YSIGISCSLFLADPFVMTNFLKNWSKIHNKLISQIDAPKIPPFYTPNLRKIGLLPTISTSLTTKNKTTQTLIFKIPKNPLNLNNNFLKNLASKCIGEIEEKIGTNLSSKFSLFVTERANSENVIKVETCIKEEIIKDECGLIYKNGGLISVSWDDLGGDKVIFNEGNKAINISCWIINAENQDLVMITPSTDEDSSQLNIIITVTN*

### NbL14g00930.1 L−lactate_dehydrogenase_B−like [MW = 37.93 kDa]

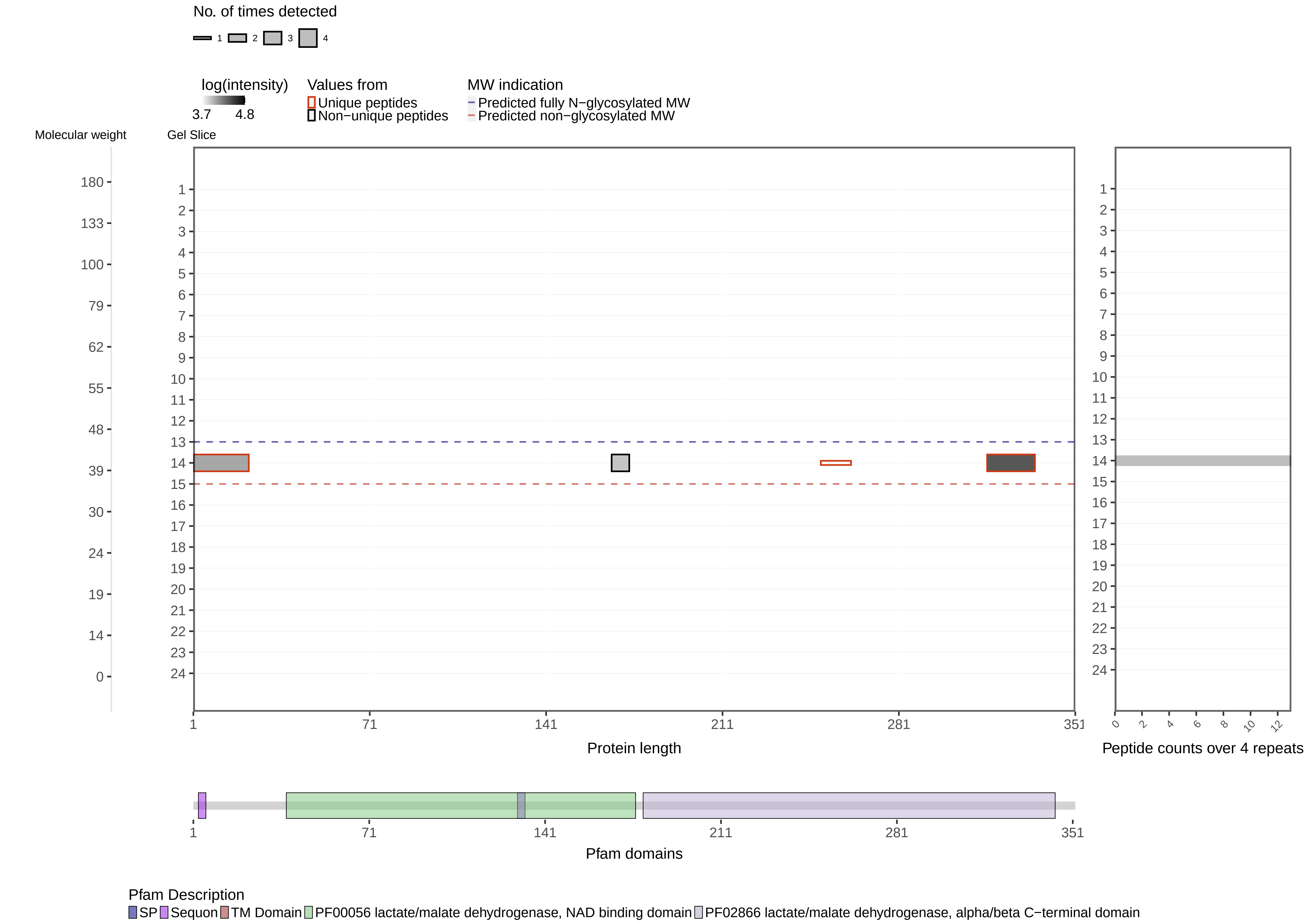


Protein sequence:

VAWKLSGFPANRVVGSGTNLDSSRFRFLIADHLDVNAQDVQAYIVGEHGDSSVALWSSISVGGVPVLSFLERQQIAFEKETLEKIHKQVVESAYEVISLKGYTSWAIGYSVANLARSIIRDQRRIHPVSVLAKGFYGIDSGDVFLSLPAQLGRSGVLGVTNVHLTDEEIQQLRNSAKTILDVQNQLGI*

### NbL14g00890.1 hsp70−Hsp90_organizing_protein_3−like [MW = 66.5 kDa]

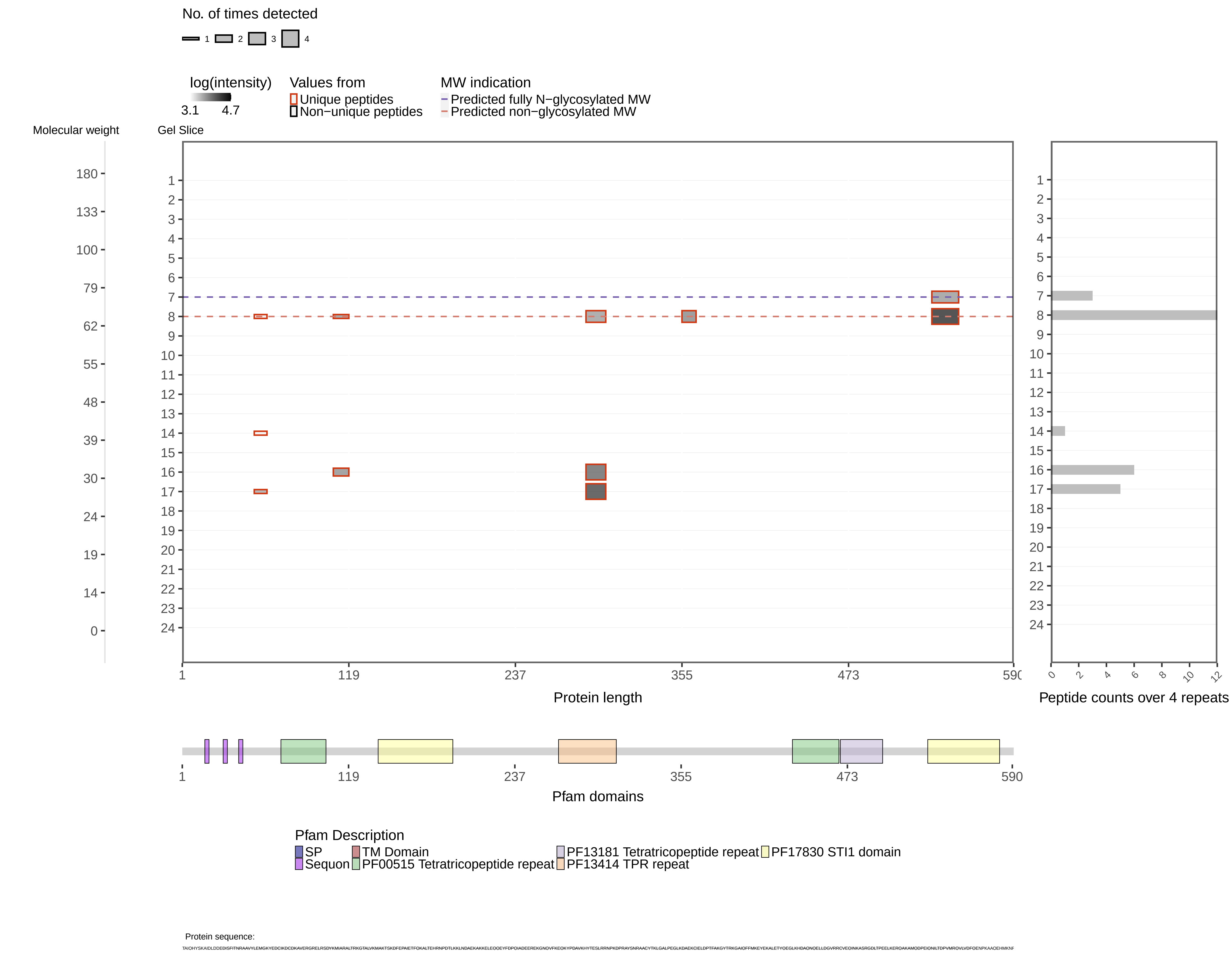

NbL14g00880.1 lipid_transfer−like_protein_VAS [MW − SP = 10.23 kDa]

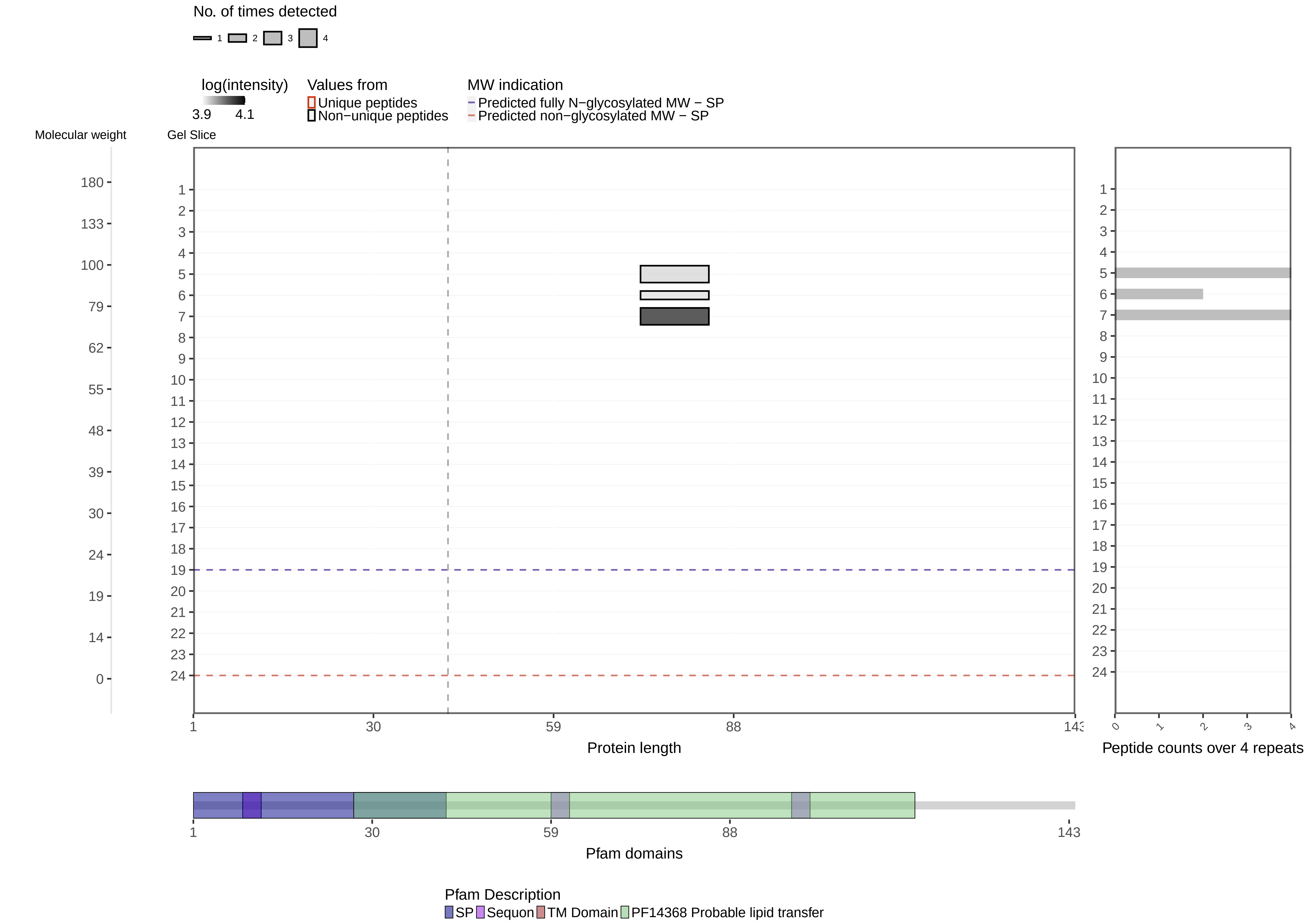


Protein sequence:

.NSTKPPAECCDPLKEAITNELDCLCKLYENPTLLPAFGINITQALALPKACNIPGDLSSCNGKAAPGPGSETLPPPGPEAAFGS*

NbL14g00720.1 serine_racemase−like [MW = 35.59 kDa]

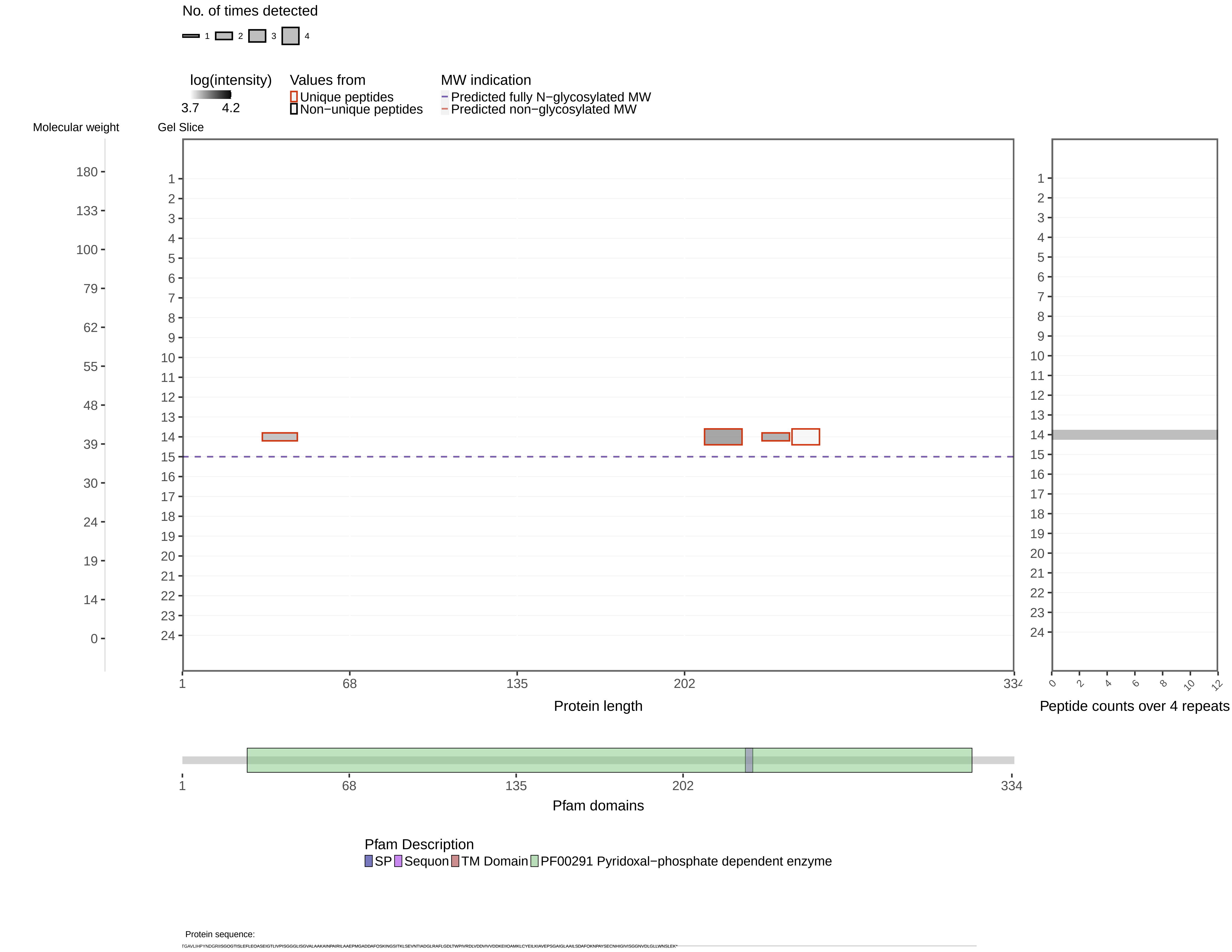

NbL14g00600.1 hydroxymethylglutaryl−CoA_synthase−like [MW = 51.04 kDa]

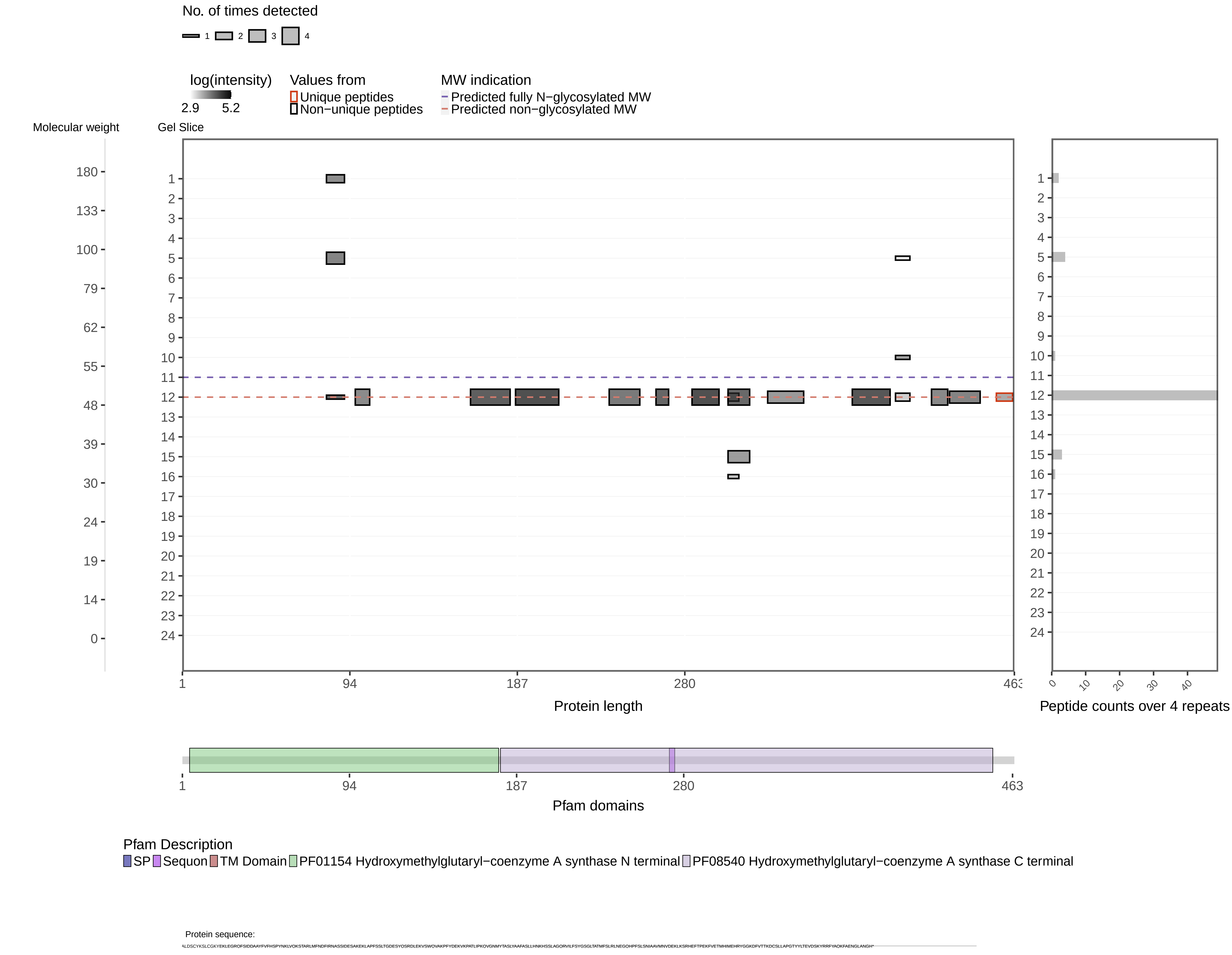

NbL14g00590.1 PGR5−like_protein_1B_chloroplastic [MW = 42.73 kDa]

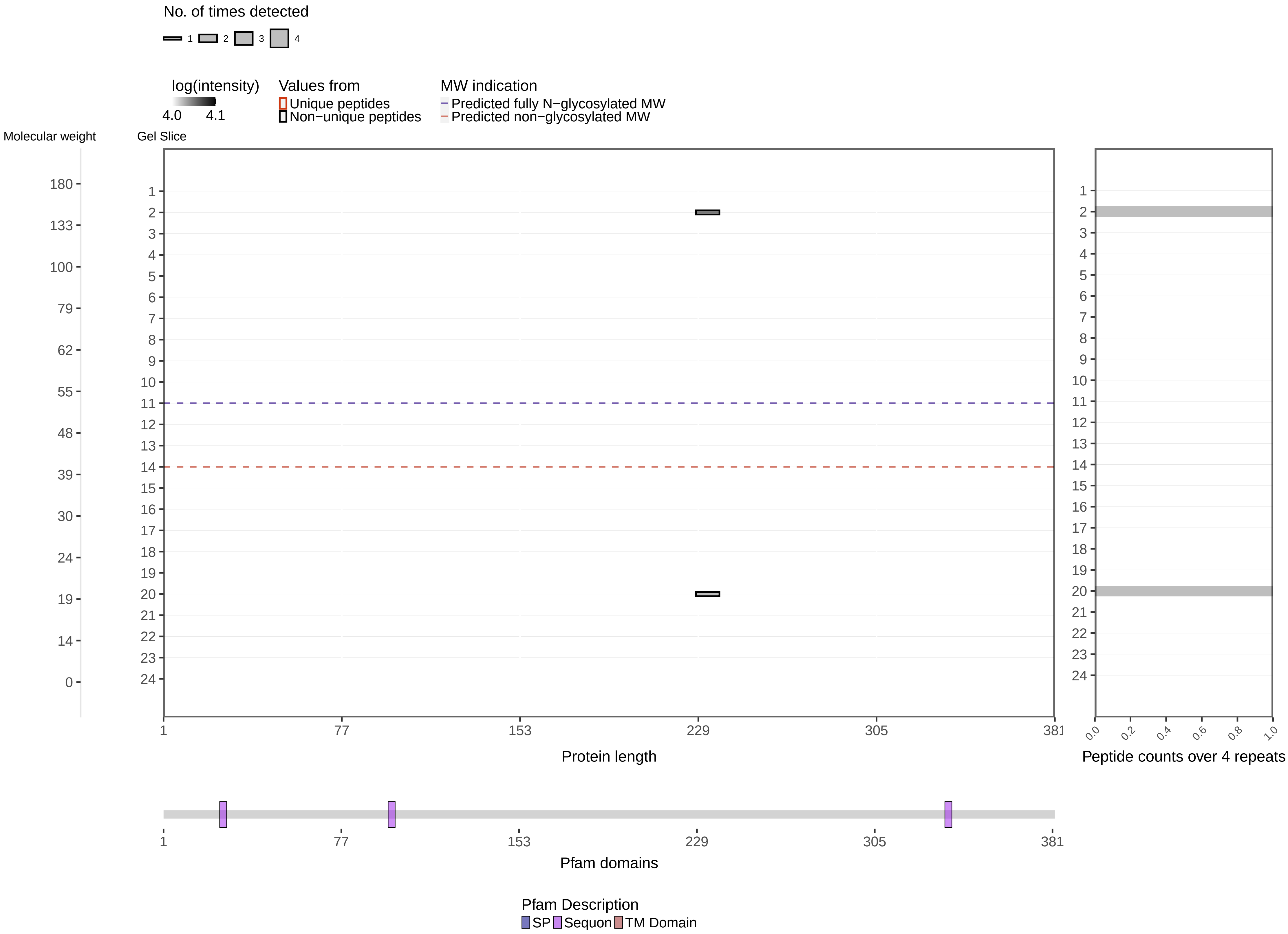


Protein sequence:

NLKEELMWEGSSVVMLSTDEQKFLEASMAYVSGNPIMTDKEYDKLKMKLKRDGSDIVVEGPRCSLRSRKVYSDLSVDYLKMFLLNVPAAVVALGLFFFLDDITGFEITYLLELPEPFSFIFTWFAALPLILYLSFTITNIVVKDFLILKGPCPDCGTENASFFGTILSISSGGSTNKIKCSGCGTDLVYDSDTRLITLPEGSSA*

NbL14g00420.1 protein_disulfide−isomerase−like [MW − SP = 56.03 kDa]

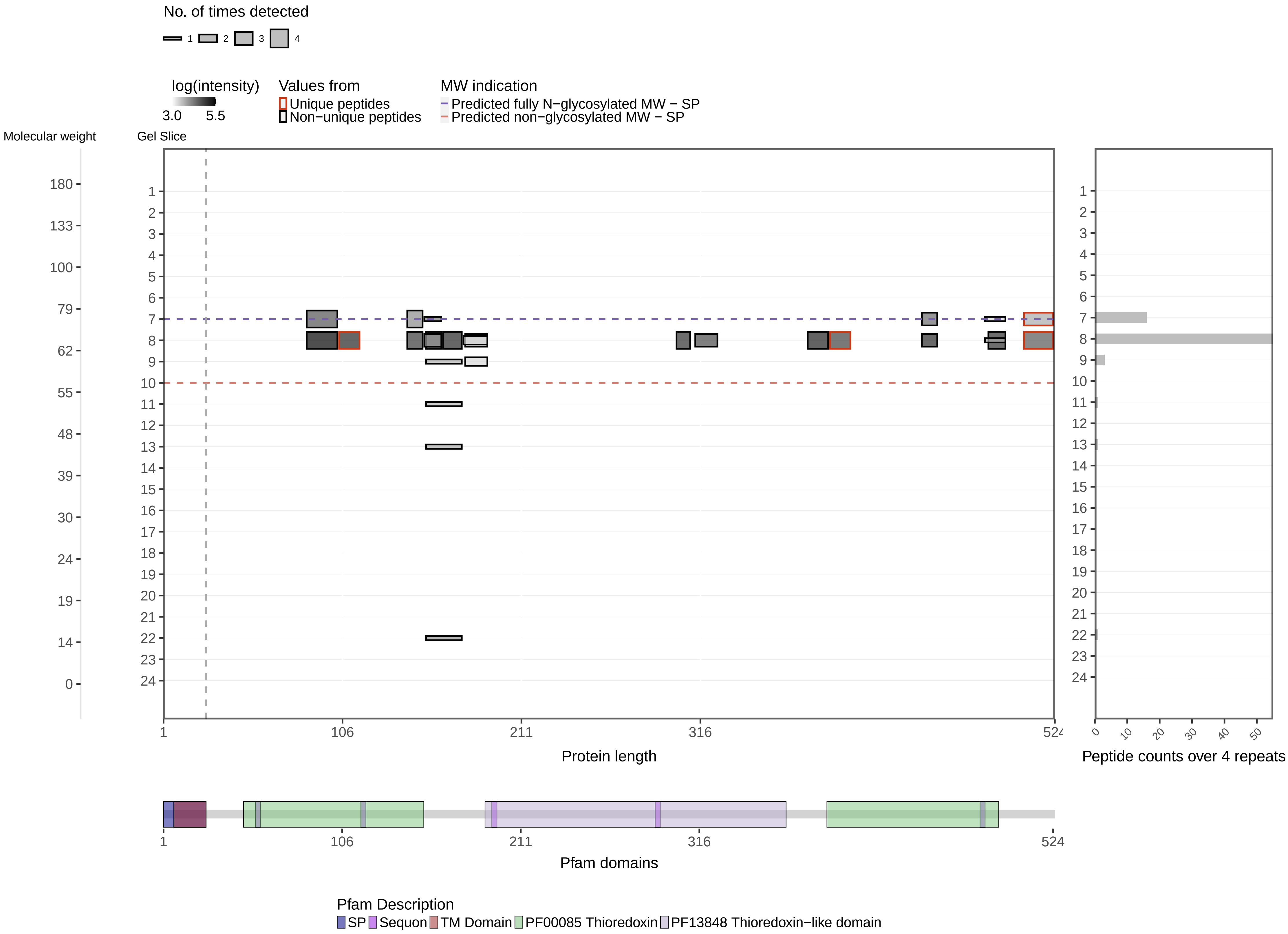


Protein sequence:

NFIEETSVPIMAILDDKPENQAFVSHFLHSPGDKVFLFLNFTMDLDHFKSKYYDLAASYKGKEANFLLGDAETGKRALEYFGLNTDQVPLIFVLAMDGTKYVQRHVQPDSLAAWLKDFNDGKLKPYLKSQPIPEVNNEPVKMVVAETLEDMVFKSGKDVLLEFYRLGCKYCEEFAPVLDEIALSFENDADVLIAKIDGTENDIPRERFEVQGFPTLYMISTAGSLSRFDGNRTKEAVIEFIQTNRGTPAPSDFAKSDLTASEHAQTDSMKDEL*

### NbL14g00410.1 calcium−dependent_protein_kinase_1−like [MW = 57.73 kDa]

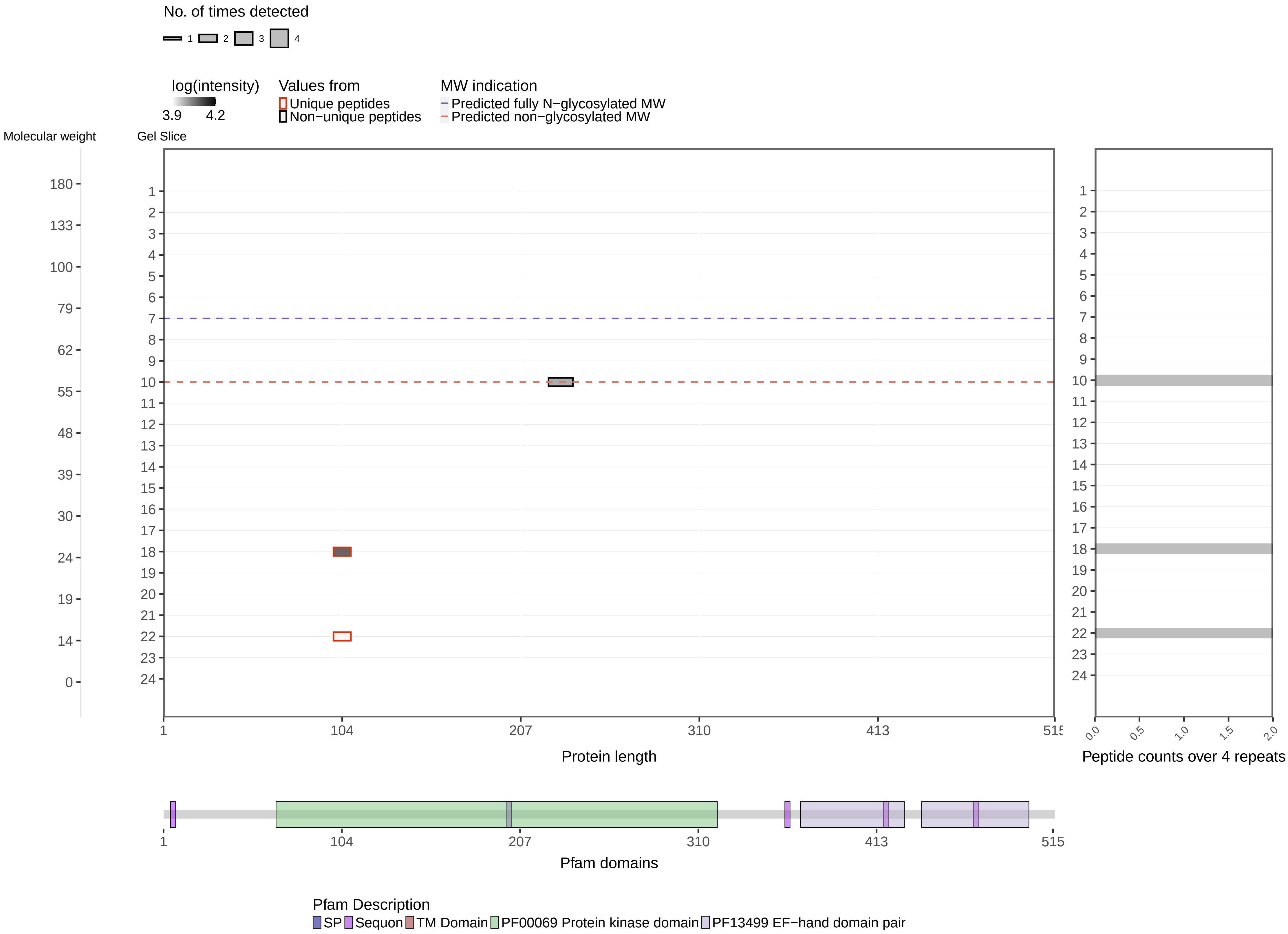


Protein sequence:

ADIWSAGVILYILLSGVPPFYGENDQSIFDAVLRGHIDFSSDPWPSVSSSAKDLVKKMLRSDPRERLSAAEVLNHPWMREDGDASDKPLDIAVLSRMKQFRAMNKLKKVALKVIAENLSEEEIIGLKEMFKSIDTDDSGTITYEELKAGLTKMGTKLSESEVRQLMEAADVDGNGTIDYLEFITATMHMNRMEREDHLYKAFEYFDKDKSGYITMDELEHALKKYNISDEKTIKEIIAEVDTDNDGRINYDEFAAMMRKGNPDFVNNRRRR*

### NbL14g00380.1 V−type_proton_ATPase_subunit_E−like [MW = 27.67 kDa]

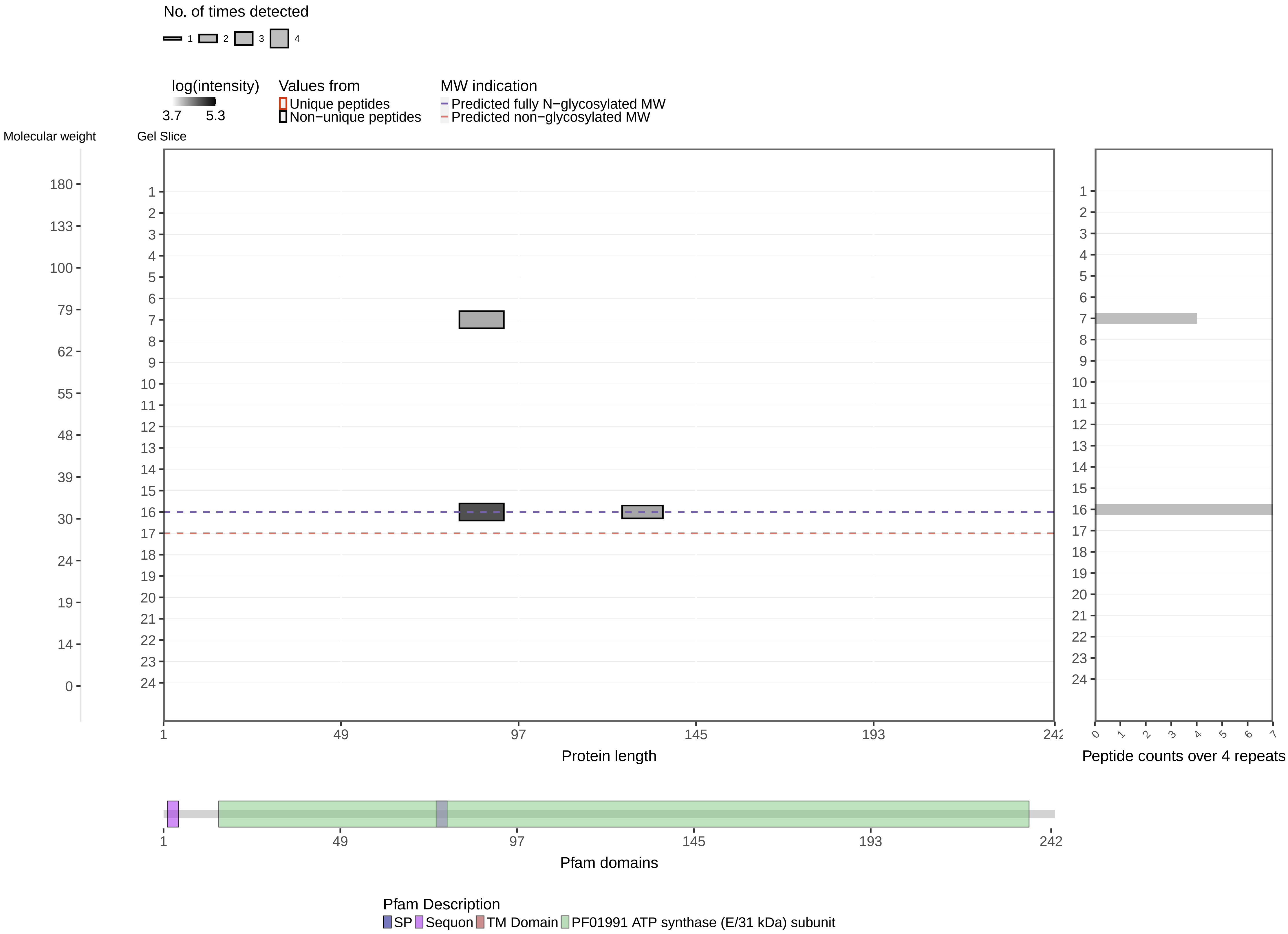


Protein sequence:

INELVEVILPHHQDSYKKLLHNLIVQSLLRLKEPSVLLRCRKHDVHLVEHVLDAAKEEYAEKANVHAPEIIVDDIHLPPAPSHHNMHGPSCSGGVVLASRDGKIVCENTLDARLEVVFRKKLPEIRKCLFGQVTA*

NbL14g00280.1 uncharacterized_LOC109214535 [MW = 25.8 kDa]

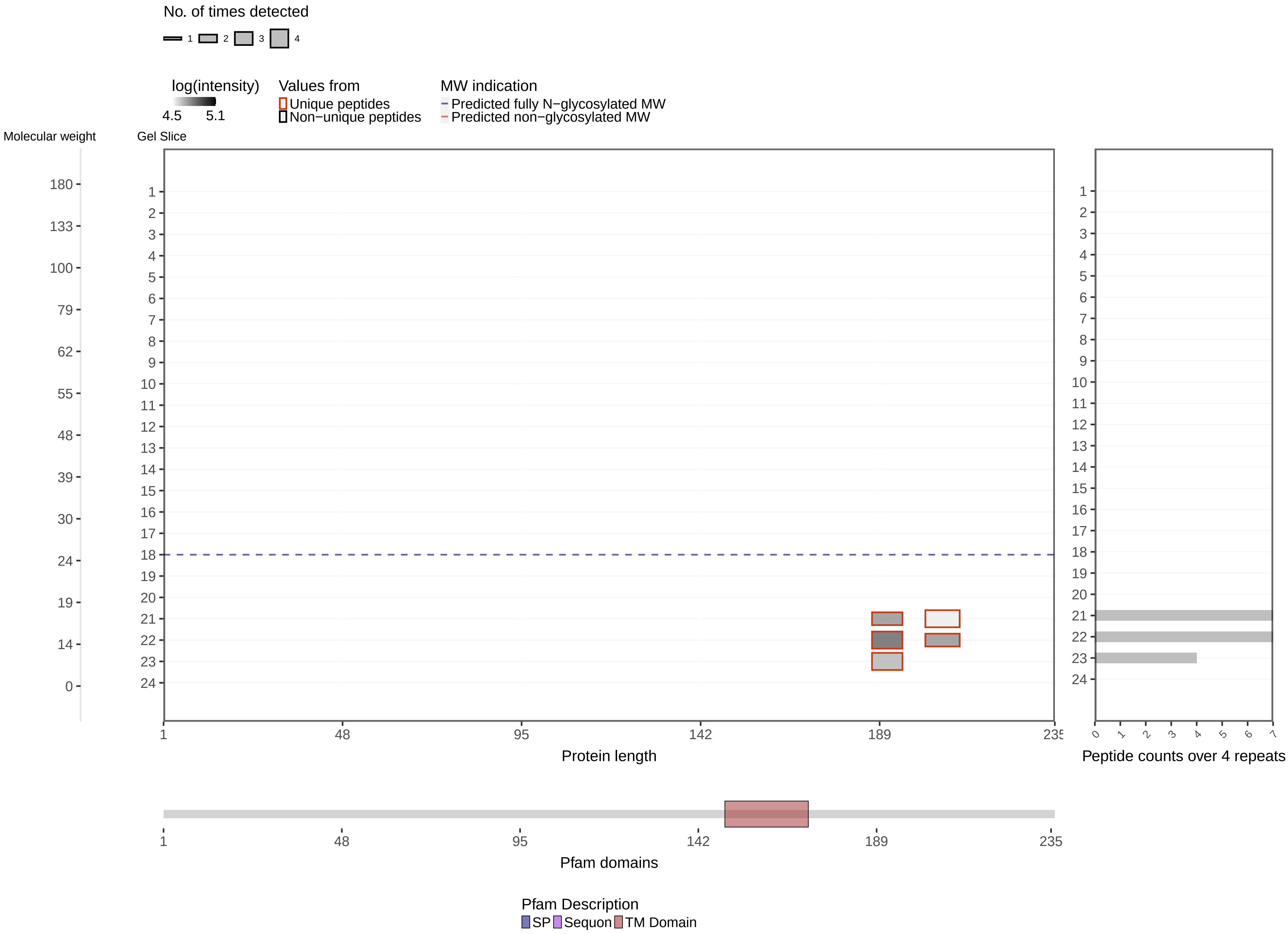


Protein sequence:

VGRGCSDCKQATPTSNREVVSSSLPKSKVGSSWREGCRESIWKQPLYPRVGIVAAIFALSLLVVHYLANRPISSTSQNSQHQKTSAAEEAPRTFFSRSPSSMSVAGKASDQPKRTPVSQEEIDSILLGGCI*

NbL14g00210.1 luminal−binding_protein_4−like [MW − SP = 70.4 kDa]

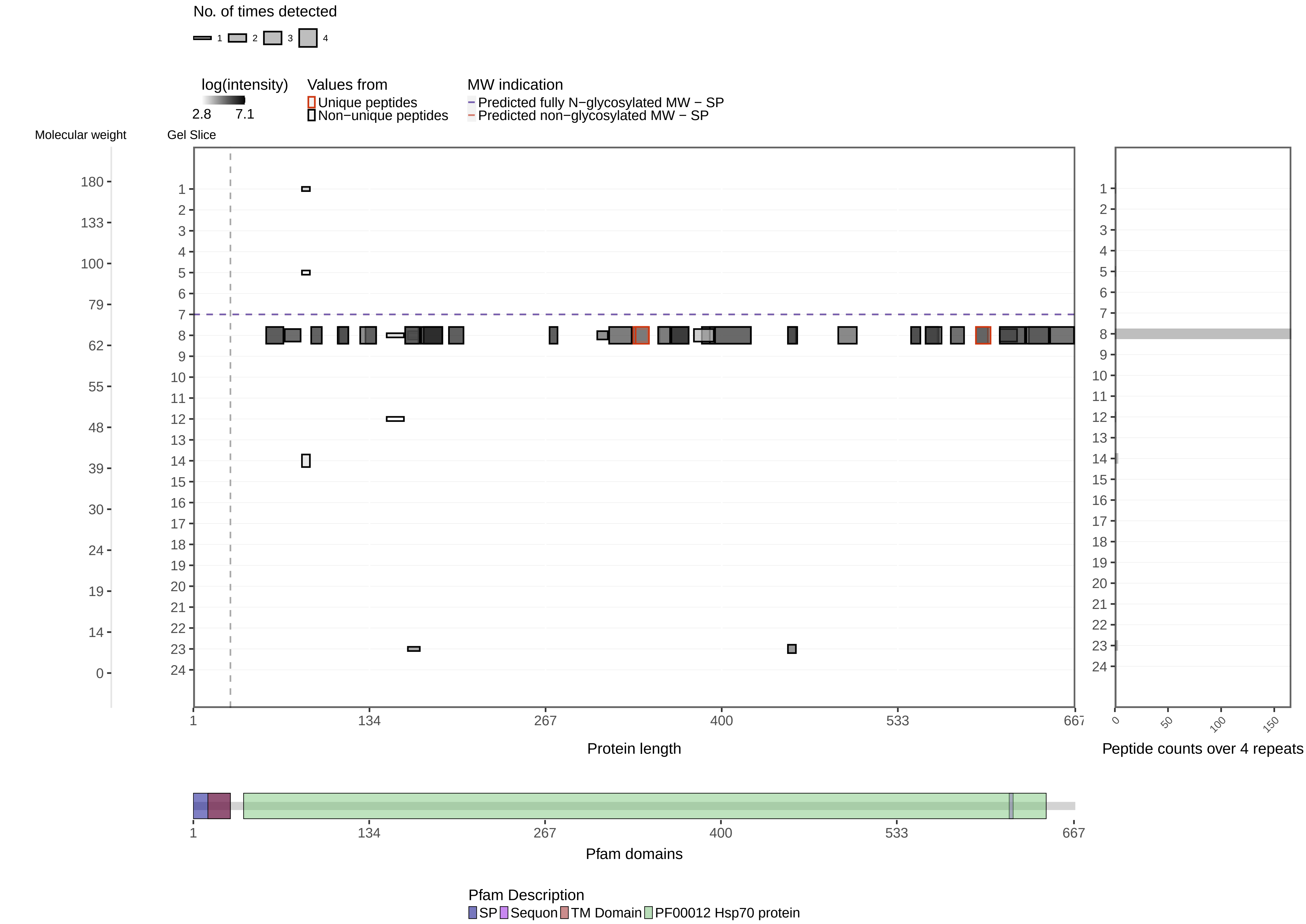


Protein sequence:

### NbL14g00110.1 beta−carotene_isomerase_D27_chloroplastic [MW = 32.98 kDa]

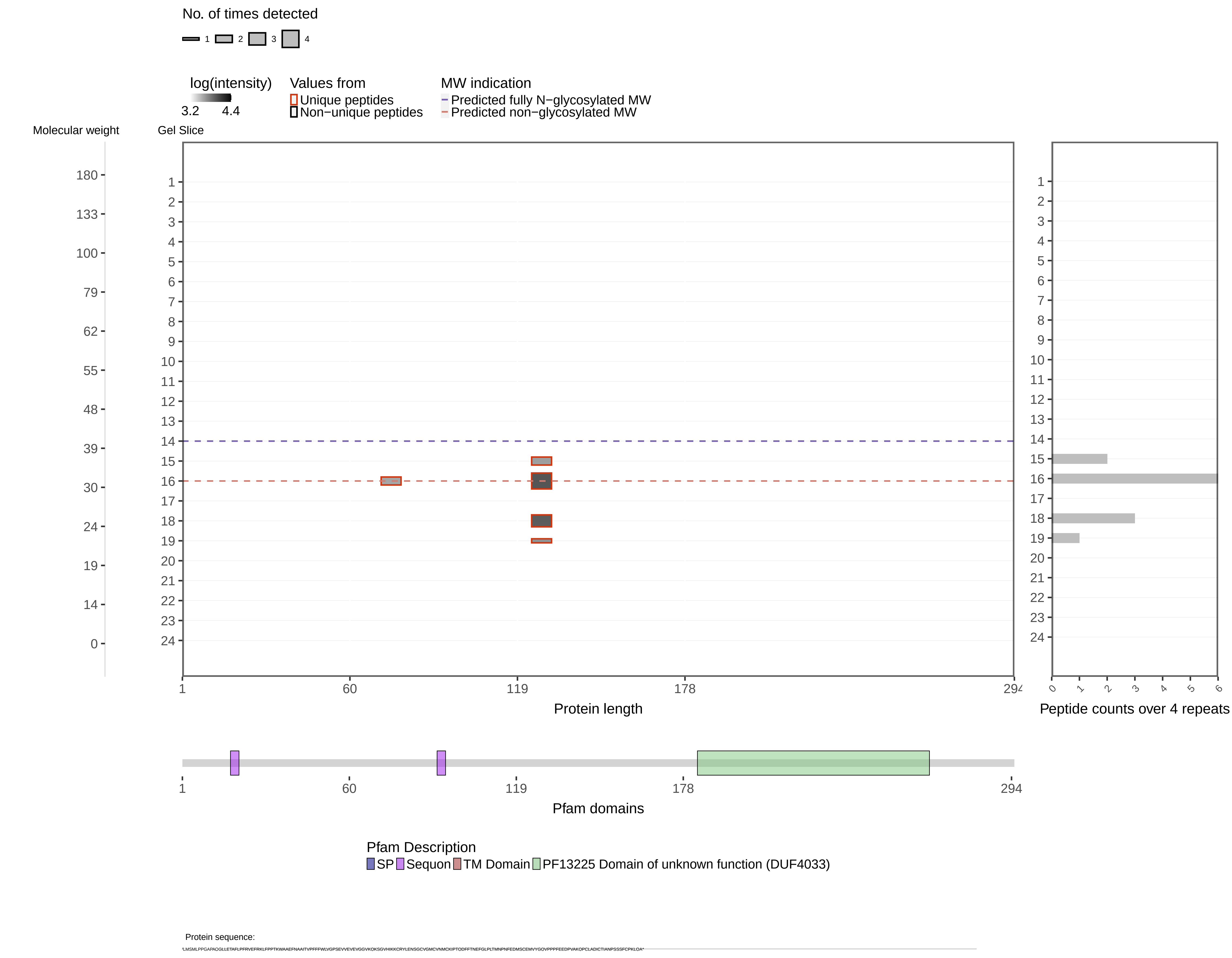

NbL14g00070.1 photosystem_I_reaction_center_subunit_N_chloroplastic [MW = 18.06 kDa]

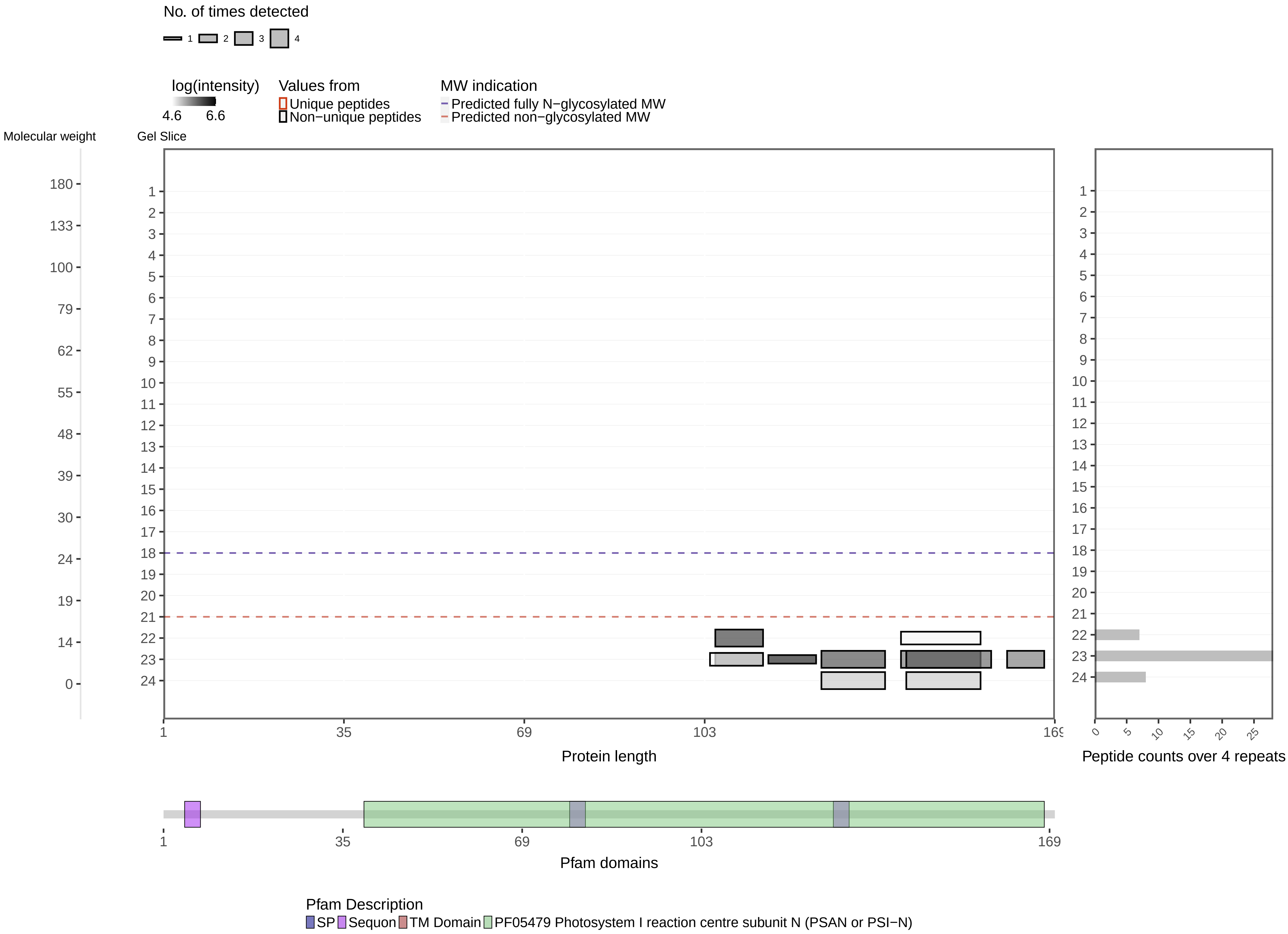


Protein sequence:

FTAASSNSSANAGVIDDYLEKSKANKELNDKKRLATSGANFARAYTVQFGTCKFPENFTGCQDLAKQKKVPFISDDLALECEGKDKYKCGSNVFWKW*

NbL13g27700.1 40S_ribosomal_protein_S9−2 [MW = 23.05 kDa]

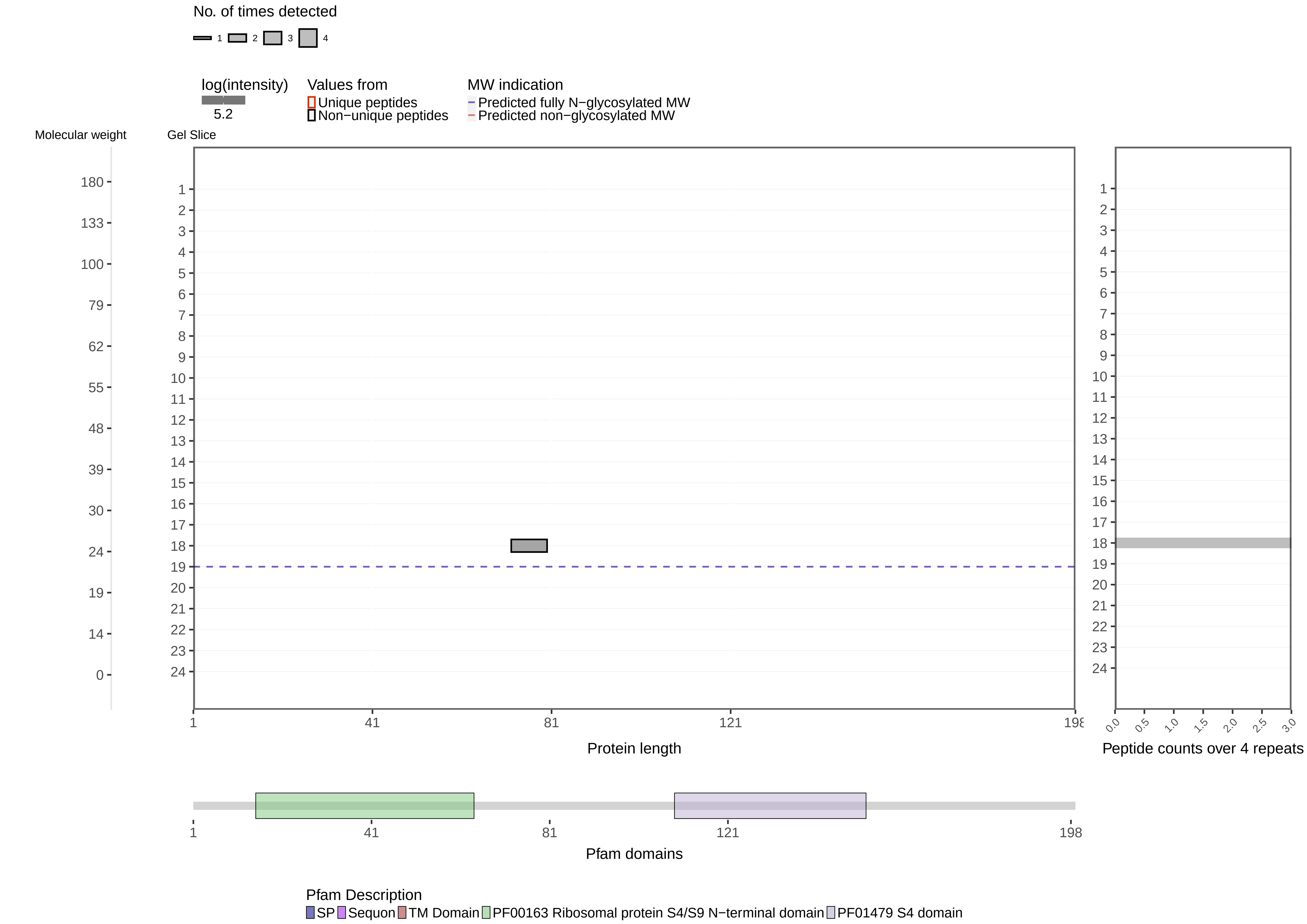


Protein sequence:

GLLDESQNKLDYVLSLTVENFLERRLQTLVFKTGMAKSIHHARVLIRQRHIRVGRQVVNVPSFMVRLDSQKHIDFSLTSPFGGGRPGRVKRKNQKAAAKKASGGDGDEDDEE*

### NbL13g27530.1 isoleucine−−tRNA_ligase_cytoplasmic [MW = 135.17 kDa]

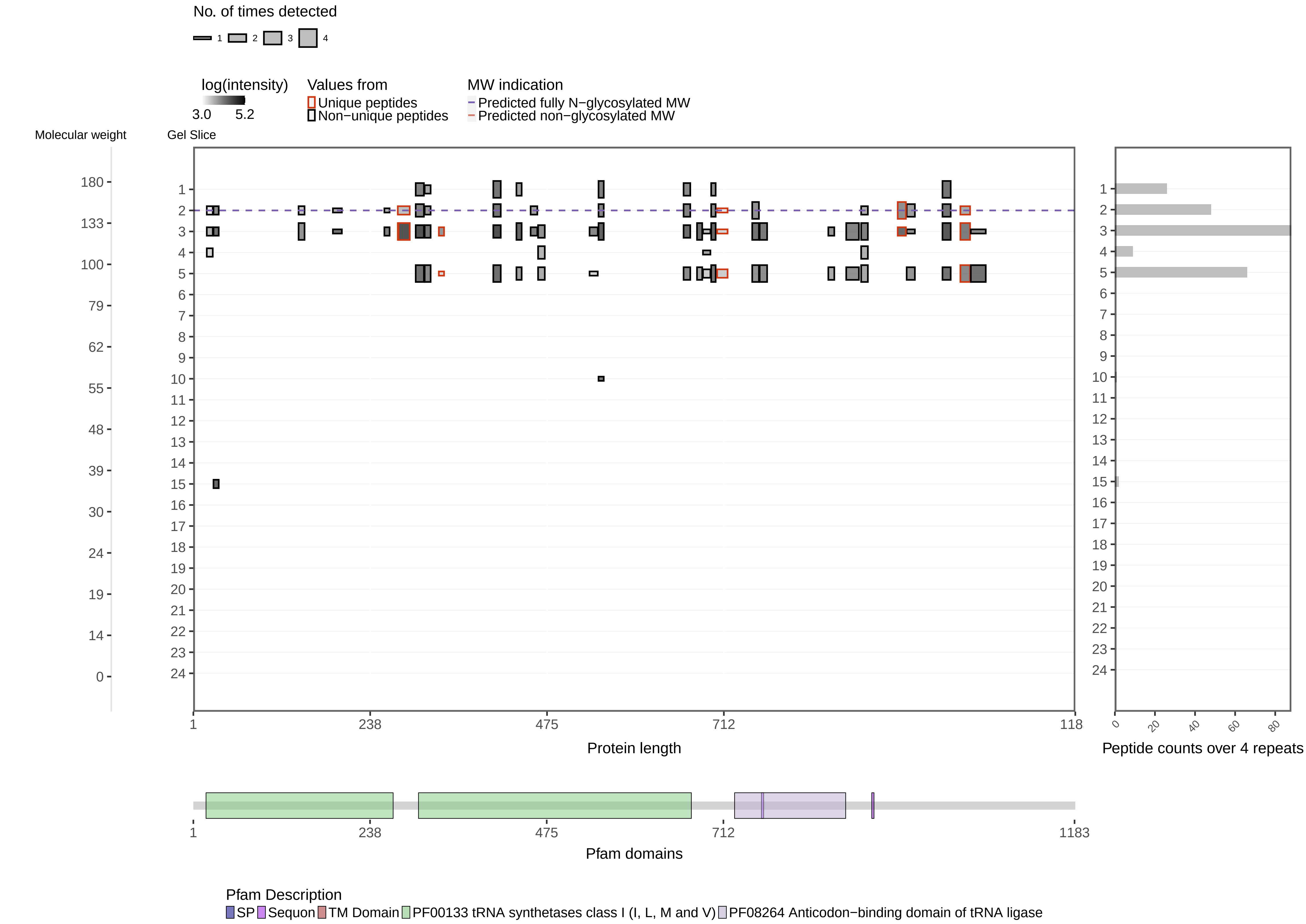


Protein sequence:

[illegible]

### NbL13g27510.1 transcription_initiation_factor_IIF_subunit_alpha [MW = 59.25 kDa]

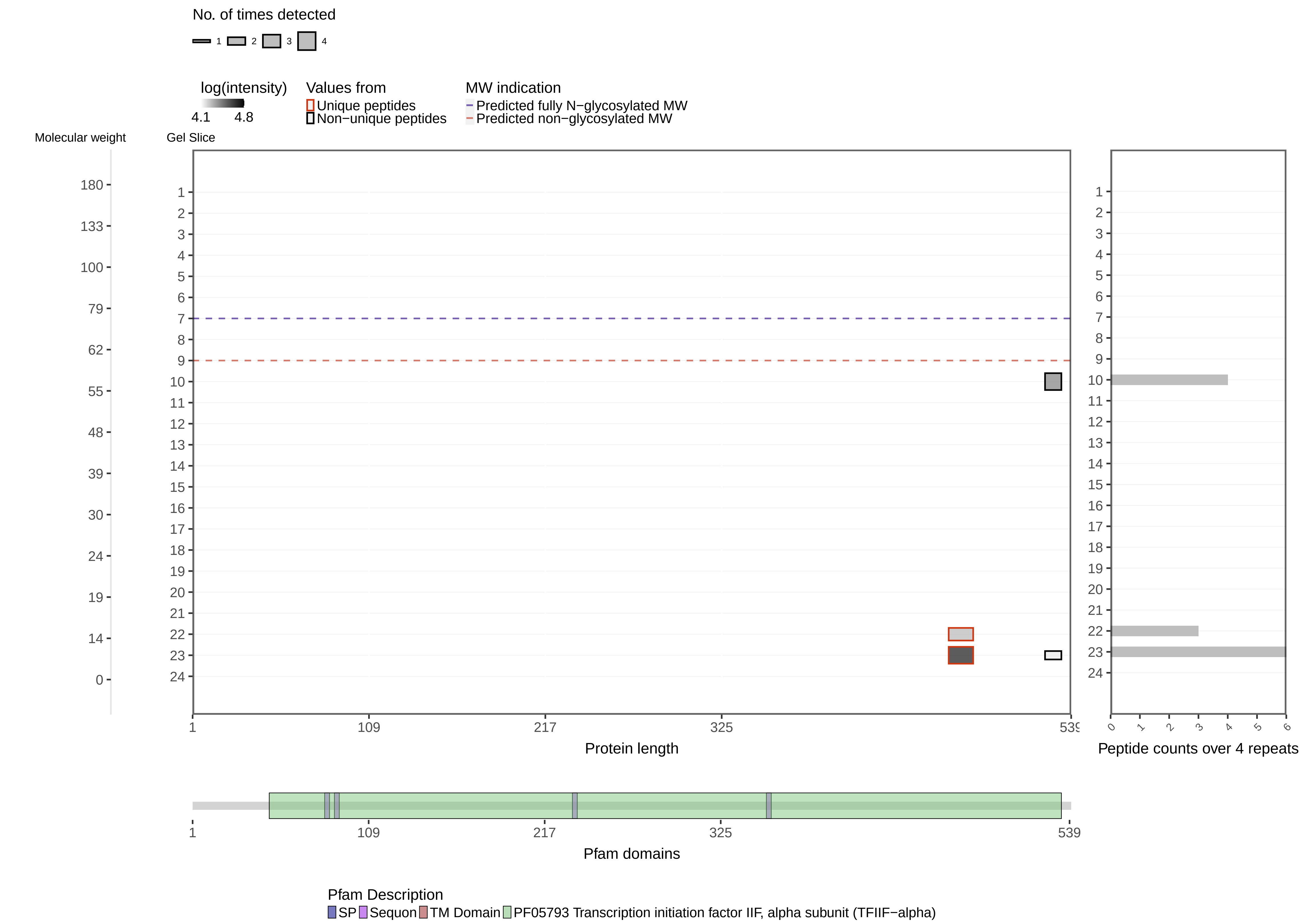


Protein sequence:

GLNKRGADDDEEGPRGGDLDLDDDDIEKGDDWEHEEIFTDDDEAVGNDPEEREDLAPEIPAPPEIKQDDEDEDEGDEEGGGLSKSGKELKKLLGKANGLNESDAEDDDDDDDTEEDISPVLAPKPKEAPKEEPIDTSSPSKPSASGPTRGTPTSKSAKTKRKAIADDSKPTNGAPLKKVKTETDAKPGKDETPSTAKNSVLPKGASPSSKTAPSPATGPVTEEEIRAVLLQRKPVTTQDLVSQFKSRLKCKEDKAAFADVLRKISKIQKTGTGTSYVVLRERTNQ*

### NbL13g27500.1 fumarylacetoacetase [MW = 49.23 kDa]

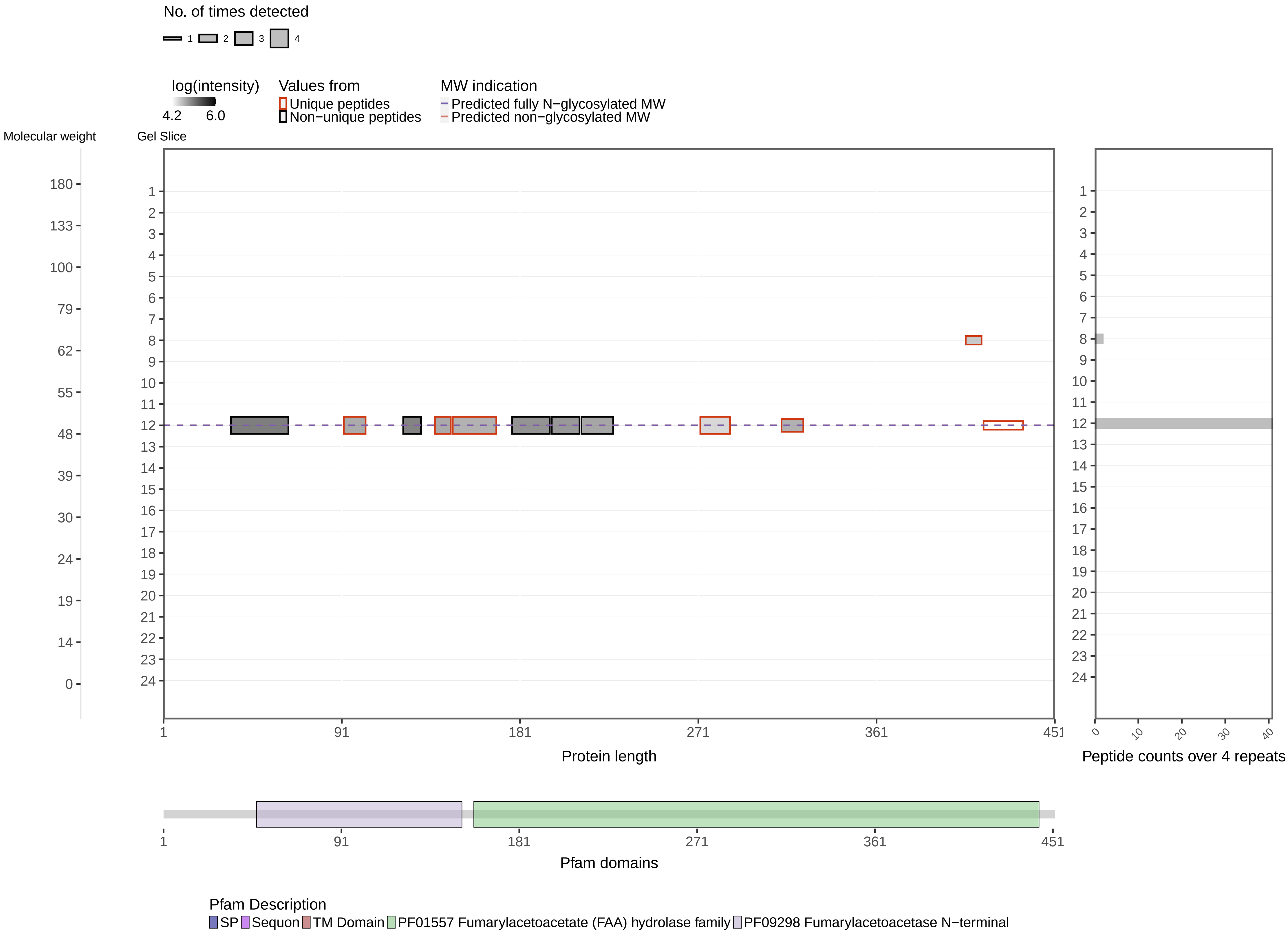


Protein sequence:

### NbL13g27390.1 cysteine_proteinase_inhibitor_1−like [MW − SP = 9.84 kDa]

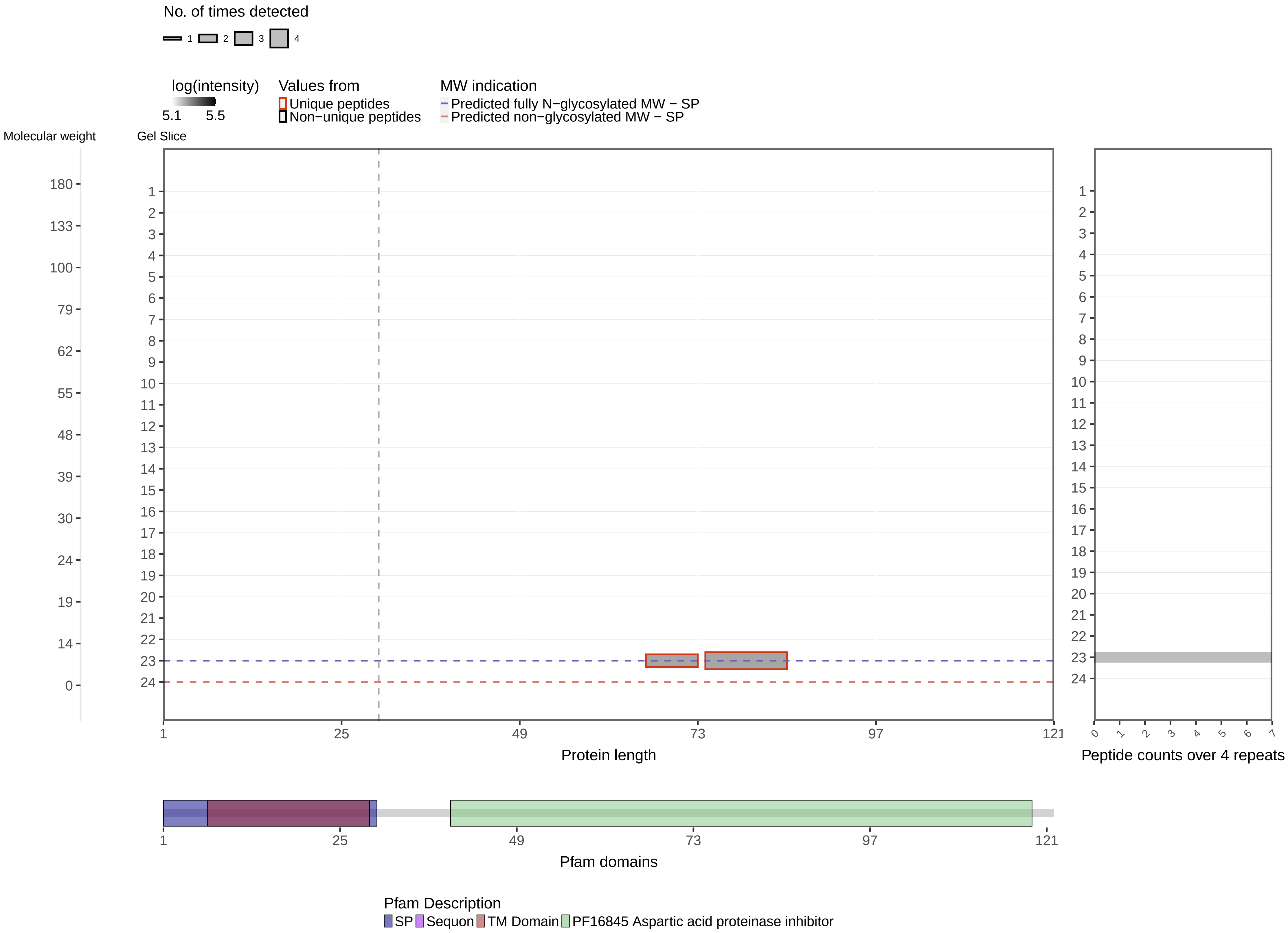


Protein sequence:

AIGQFAVNEHNKAEGTKLEFQNIAKGETQVIAGTNYRLVINAEDGGSVHKYLAVVWDKPWEKVKKLTSFKKM*

NbL13g27370.1 inositol_monophosphatase_3−like [MW = 29.49 kDa]

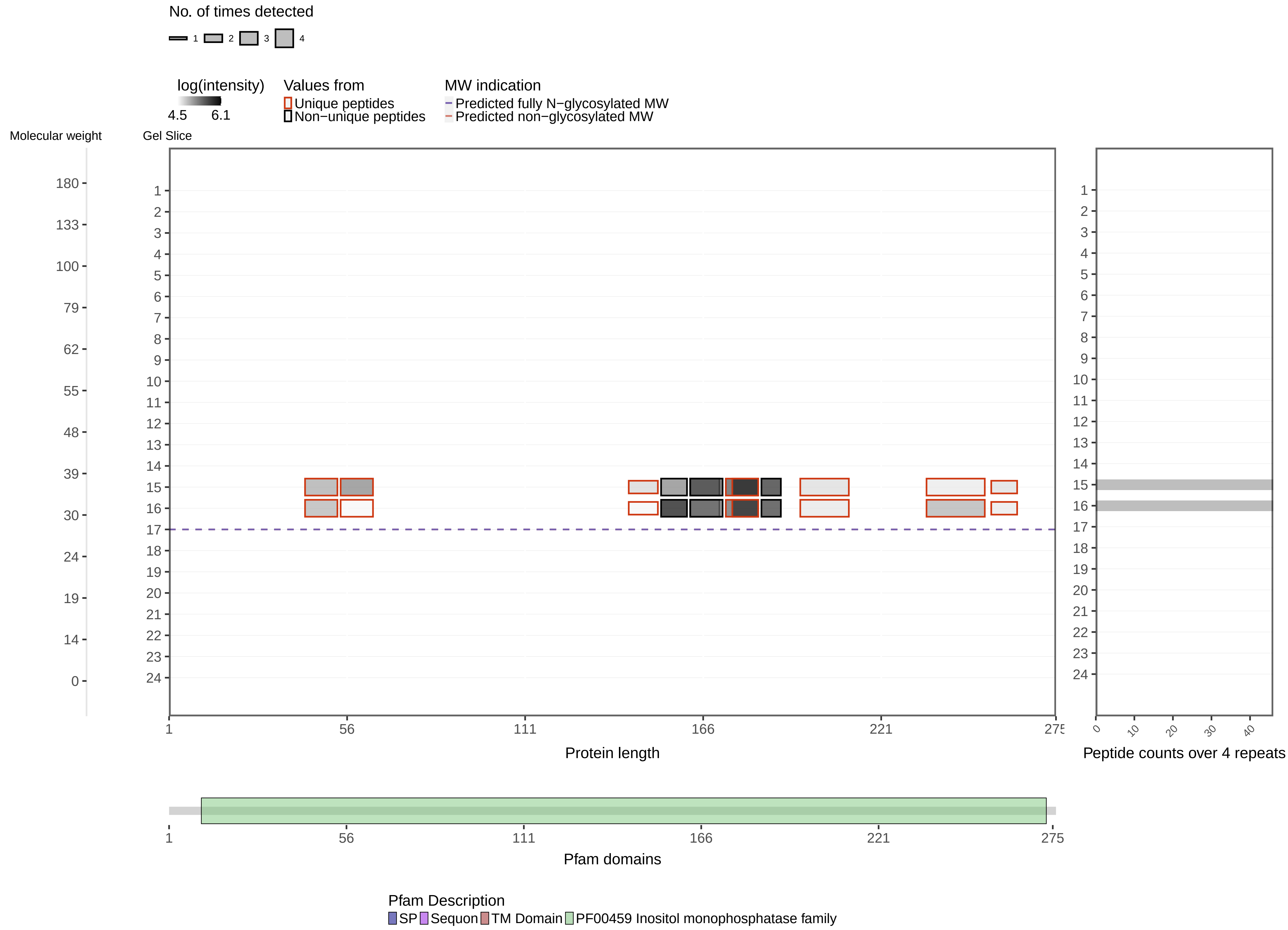


Protein sequence:

/GVVYNPIIDELFTAINGKGAFLNGKPIKVSSQSELVKALLATEAGTKRDKLTLDATTGRINSLLFKVRSLRMCGSCALNLCGVACGRLDLFYELEFGGPWDVAGGAVIVKEAGGFVFDPSGSEFDLAARRVAATNAHLKDAFIKALKESE*

NbL13g27320.1 60S_ribosomal_protein_L6−like [MW = 25.31 kDa]

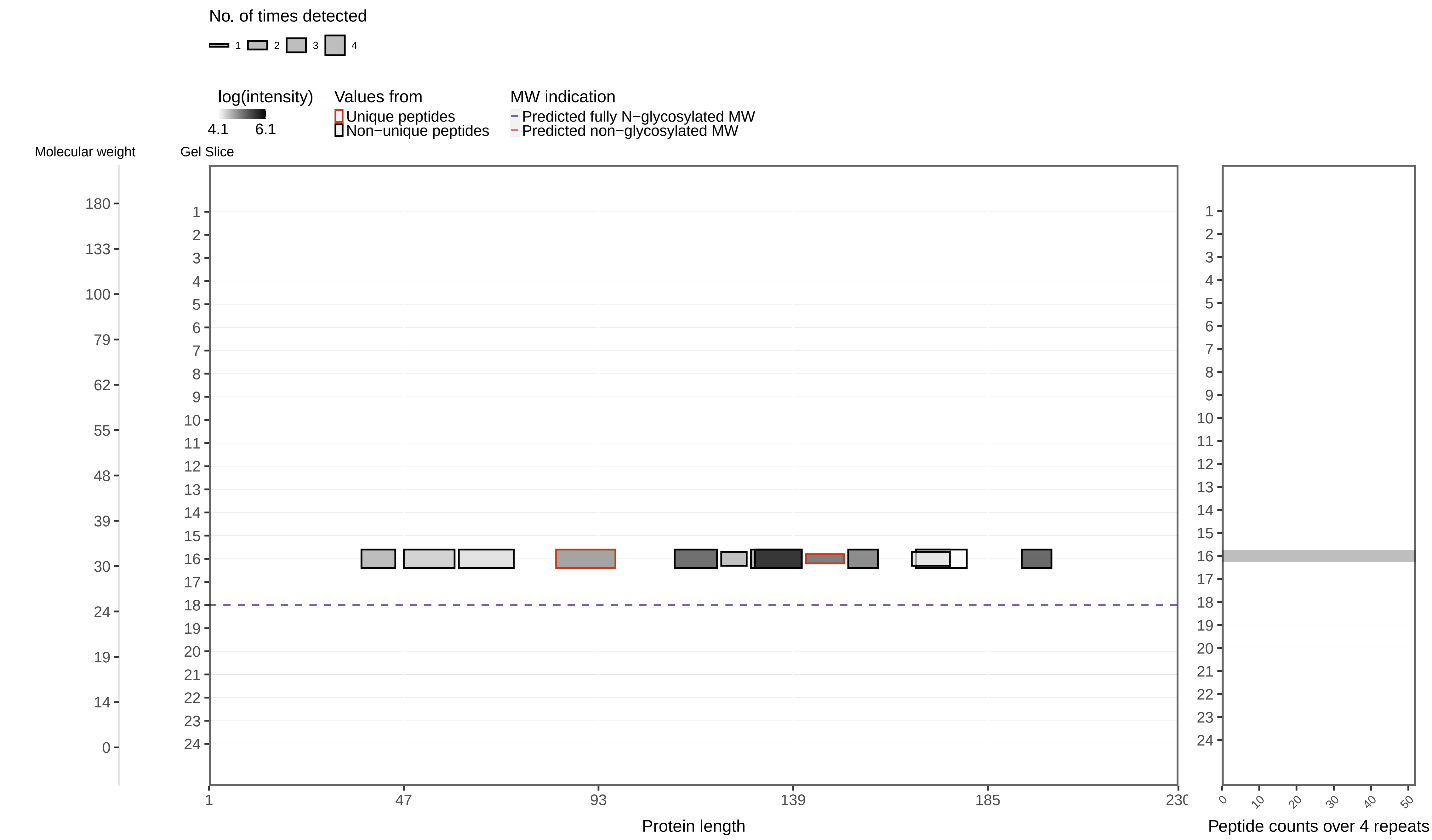


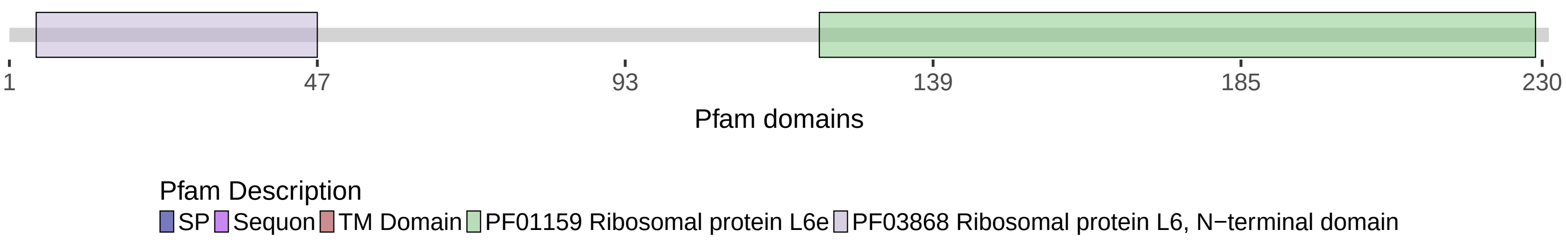


Protein sequence:

[illegible]VFLKQLKSGLLLVSGPFKLNGVPLRRVNQAYVIGTSTKVDVSGVSVEKFDDKYFAKQVEKKQKKGEGEFFEDKKEEKNVVPQGKKDDQKAVDEALIKAVECVPELKAYLSARFSLKSGMKPHELVF*

NbL13g27310.1 phosphoribosylglycinamide_formyltransferase_chloroplastic [MW = 33.19 kDa]

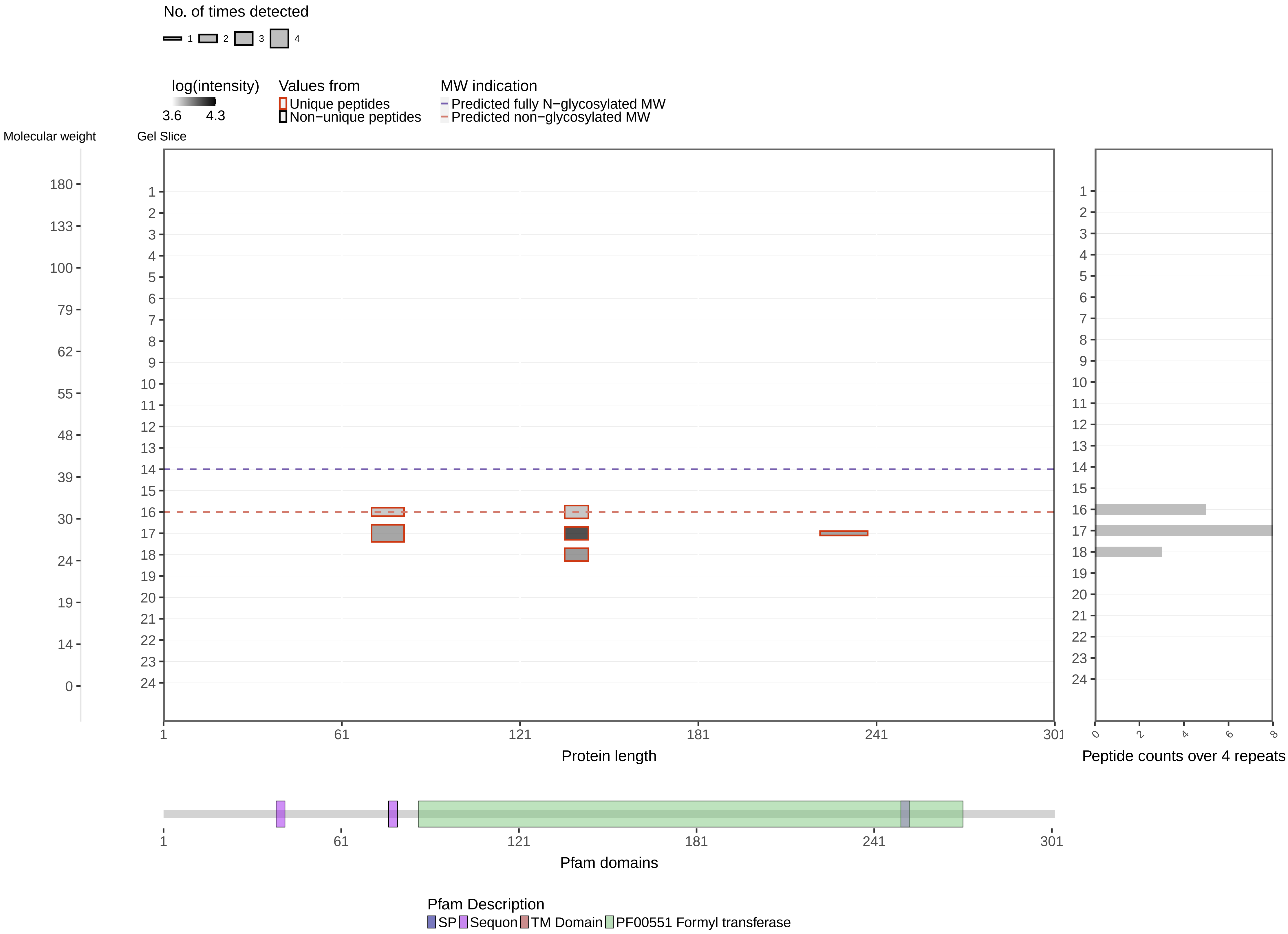


Protein sequence:

SIPVILFPKAKDTFEGLSEEDLVHALRTYNIDFILLAGYLKLIPTELVQAFPRSIFNIHPSLLPAFGGKGYYGMKVHKAVIASGARYSGPTIHFVDEHYDTGRILAQRVVPVLANDTADTLAARVLQEEHKLYVEVAAALCEERIVWREDGVPLIQSKEDPDHYK*

### NbL13g27280.1 acyl−protein_thioesterase_1−like [MW = 27.98 kDa]

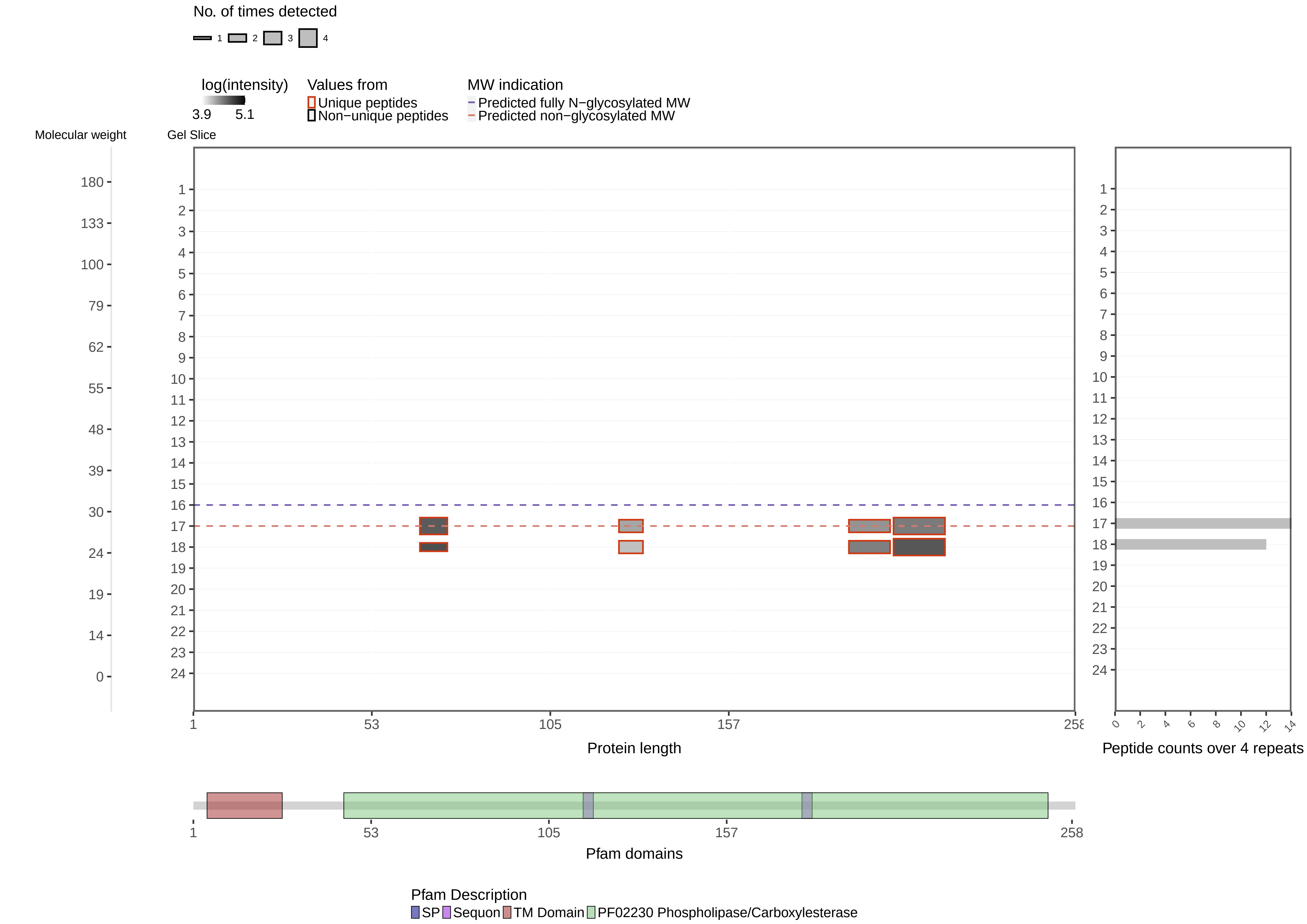


Protein sequence:

SGLLKVVKNVHAMIDKEIDAGTNPNNVFICGFSQGGALTLASVLLYPKTLGGGAIFSGWVPFNSSILEKQTPEAKKTPILWSHGIADKTVLFEAGQEGPPFLERAGMTCEFKAYPGLAHSINNEELKNLESWIKTRLQSNS*

NbL13g27150.1 UTP−−glucose−1−phosphate_uridylyltransferase [MW = 57.21 kDa]

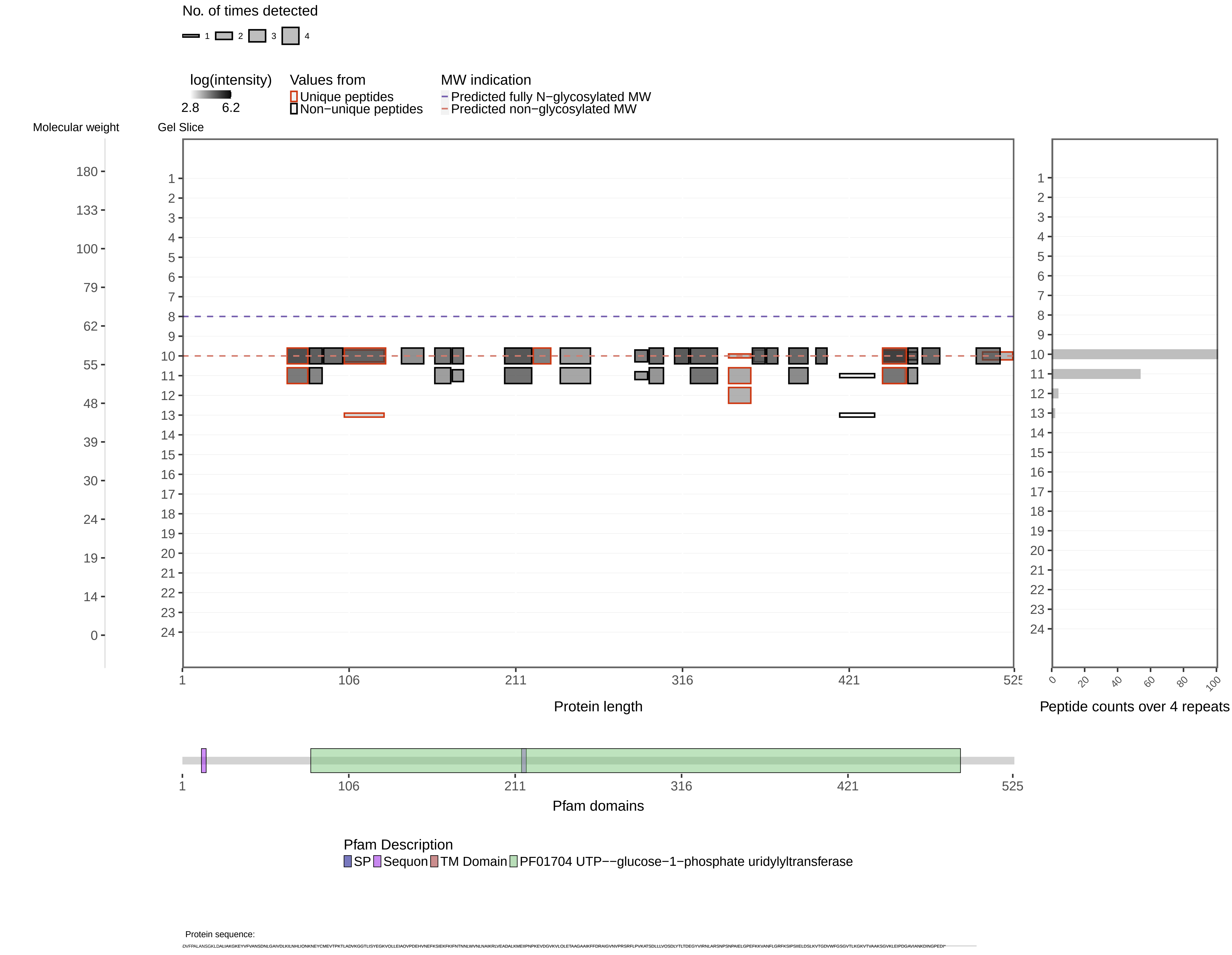

NbL13g27100.1 isocitrate_dehydrogenase_[NADP]−like [MW = 55.17 kDa]

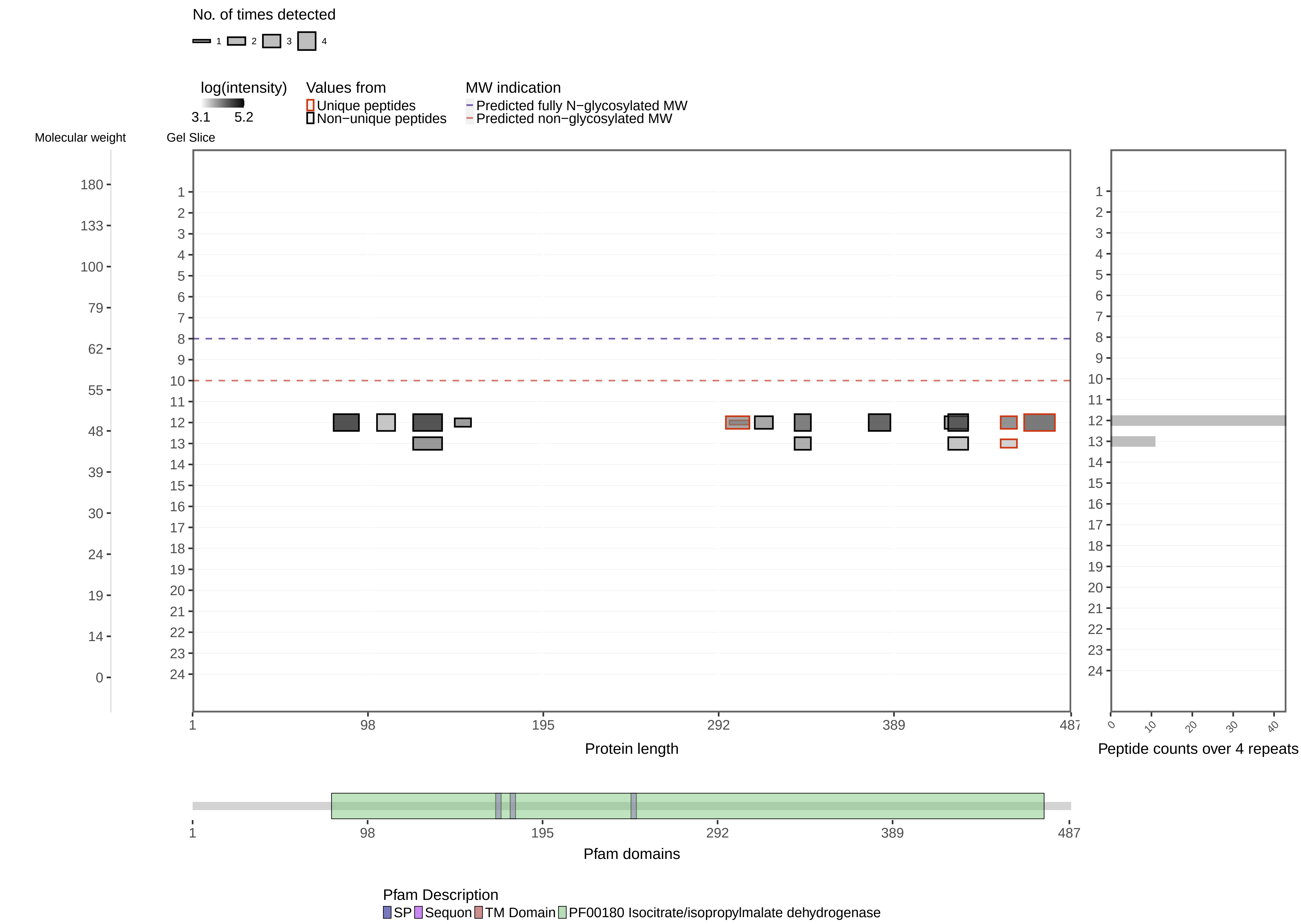


Protein sequence:

GKDEKIEFEVYNFTGAGGVALSMYNTDESIRSFAEASMNMAYQKKWPLYLSTKNTILKKYDGRFKDIFQEVYESNWKLKFEEAGIWYEHRLIDDMVAYALKSDGGYVWACKNYDGDVQSDFLAQGFGSLGLMTSILICPDGKTIESEAAHGTVTRHYRVHQKGGETSTNSIASIFAWTRGLAHRAKLDNNAALLDFTKKLEGACIGAVENGKMTKDLALIAHGSRVARHQYVNTEEFIDAVADELKARLLKAKI*

NbL13g27090.1 glutamate_decarboxylase [MW = 55.77 kDa]

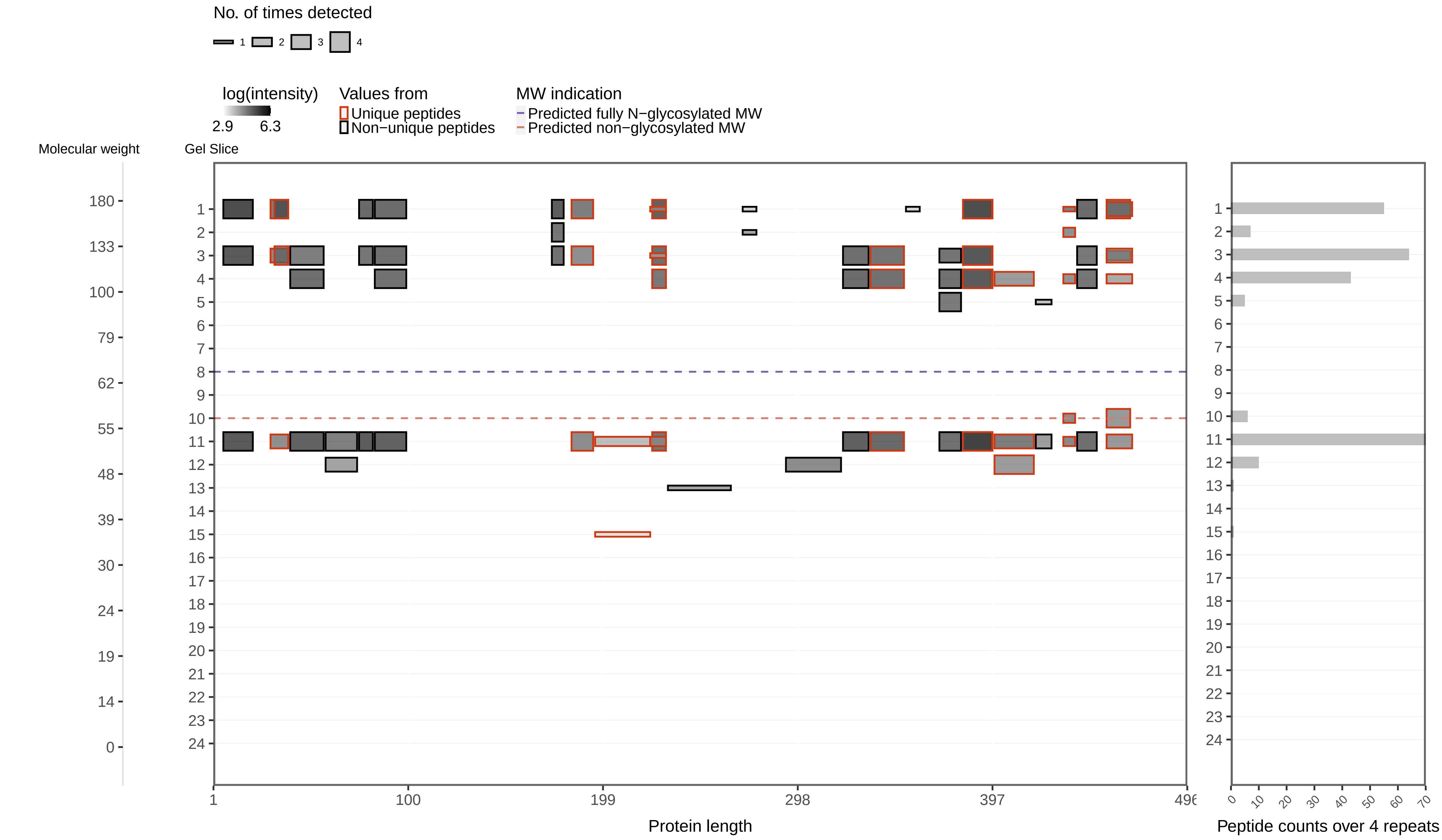


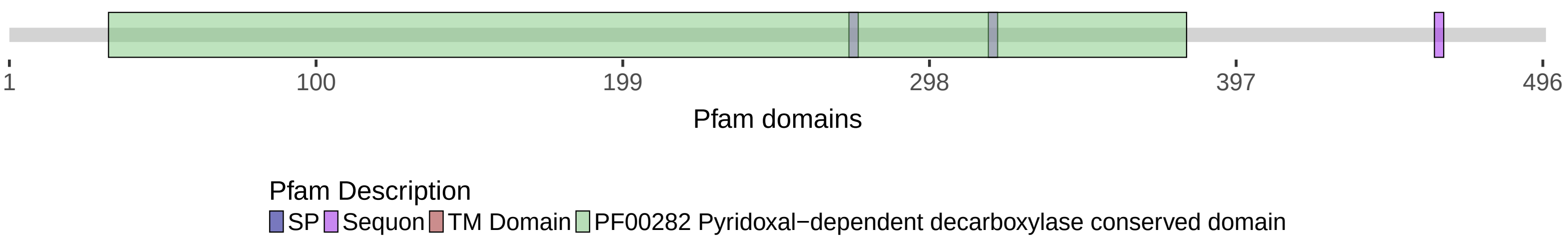


Protein sequence:

IKETGWDTPIHVDAASGGFIAPFLYPELEWDFRLPLVKSINVSGHKYGLVYAGIGWAIWRNKEDLPDELIFHINYLGADQPTFTLNFSKGSSQVIAQYYQLIRLGFEGYQNVMENCQENARVLREGLEKIGRFNIISKEIGVPLVAFSLKDNSQHNEFEISENLRRFGWIVPAYTMPPNAQHVTVLRVVIREDFSRTLAERLVIDIEKVLHELDTLPARVNAKLVVAEVNGSGVHKKTDREVQLEVTAAWKKLVADRKKTKGVC*

NbL13g27040.1 cysteine−rich_receptor−like_protein_kinase_2 [MW − SP = 67.81 kDa]

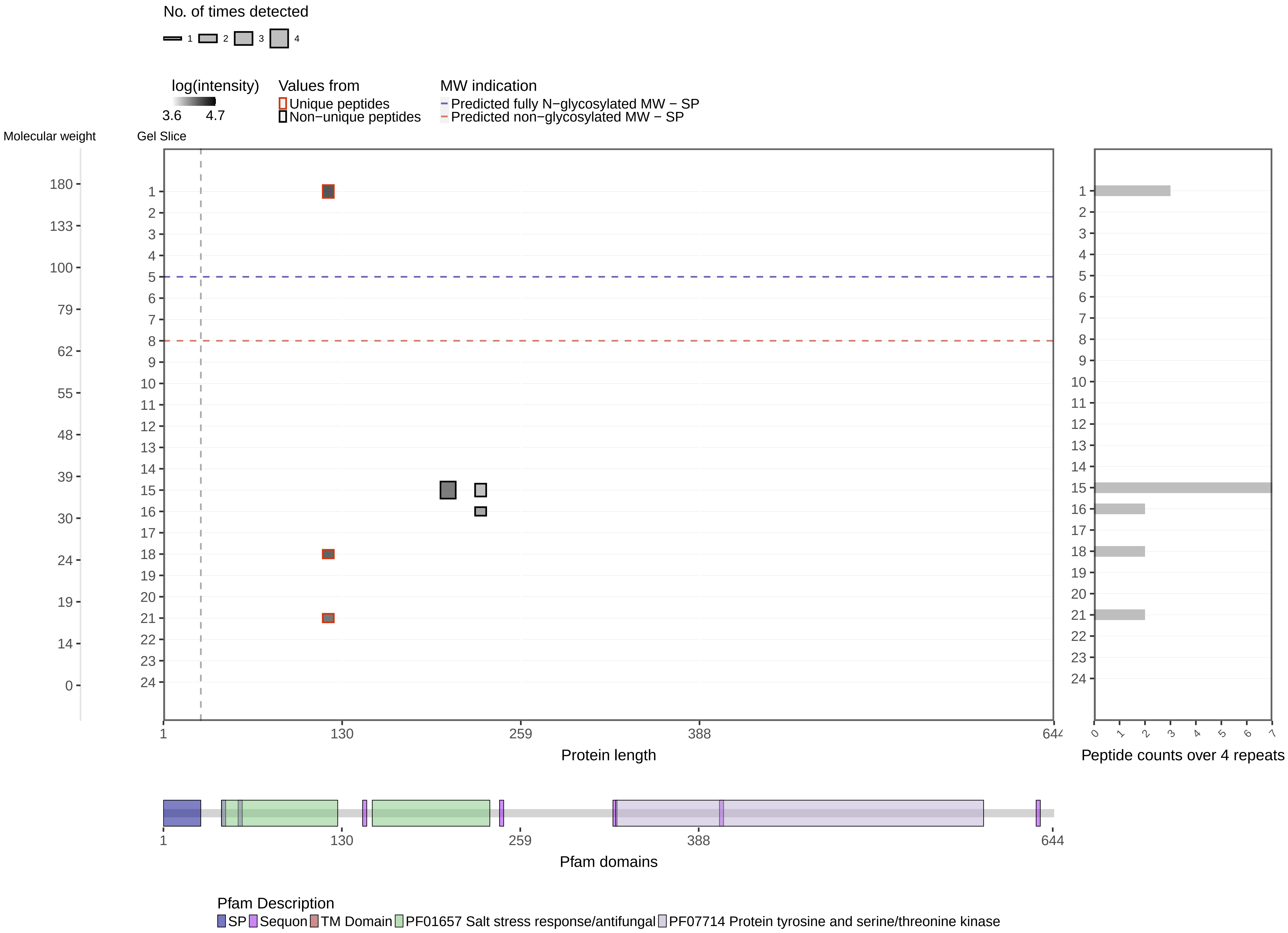


Protein sequence:

LRGPVSYRFKDLKIATQDFNESNKLGEGGFGDVYKGTLKNGHVVAVKKLAMFSSRAKAEFETEVRLISNVHHRNLIRLLGCSNKGADLLLVYEYMANGSLERYLYGDKRGMLNWKQRFDIIFGTARGLAYLHEQFHVCIIHRDIKSSNILLDGEFQPKIADFGLVRLLPEDQSHVSTKFAGTLGYTAPEYAIHGHLTEKVDVYSFGIVVLEIISGRRSNDMQIEPVTEYLLEKAWKLHETGMPEKLVDETLDPNEYNEQEAKKIIEIALMCTQSPANLRPTMSEVVVMLLSDRSTDV

NbL13g26740.1 NADH_dehydrogenase_[ubiquinone]_iron−sulfur_protein_1_mitochondrial [MW = 80.19 kDa]

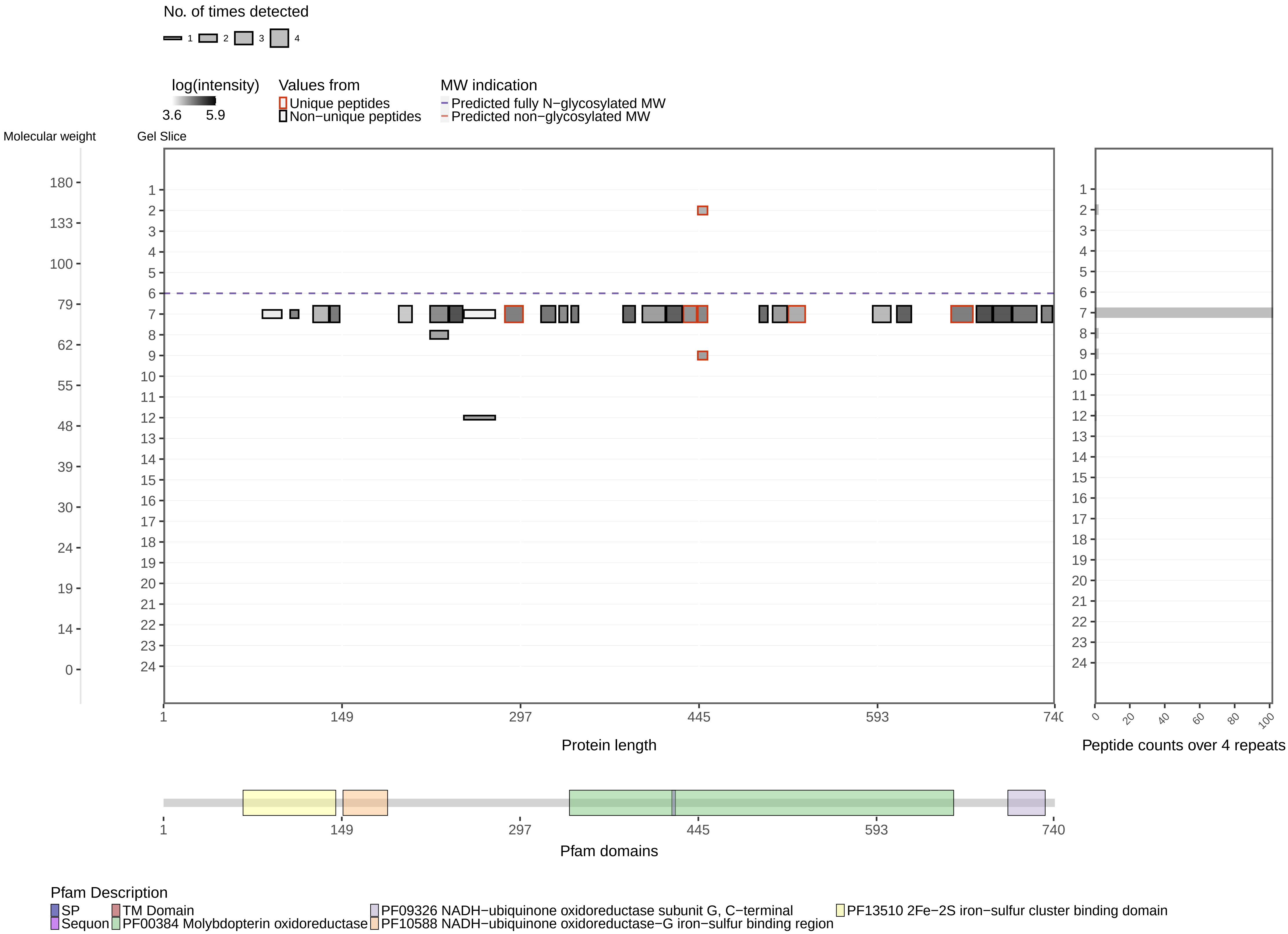


Protein sequence:

WRDALAIVAEVMHQIKPEELIGVAGKLSDAESMMALKDLLNKMGSNNIFCEGNGMQPNADLRSGYIMNTSISGLEKADVFLLVGTQPRAEAAMVNARIRKTVRANNAKVGYIGAVADFNYDHQHLGTDPQTLVEIAEGRHPFSSALKNAKNPVIIVGAGVFDREDKDAVFAAVDTIVKNNNVVRSDWNGLNVLLLNAAQAAALDLGLVPESEKCIDSAKFVYLMGADDVSLDKLPDDAFVVYQGHHGDRGVYRANVILPASAFTEKEGIYENTEGCAQITLPAVPTVGDSRDD

### NbL13g26690.1 NADH−cytochrome_b5_reductase−like_protein [MW = 36.28 kDa]

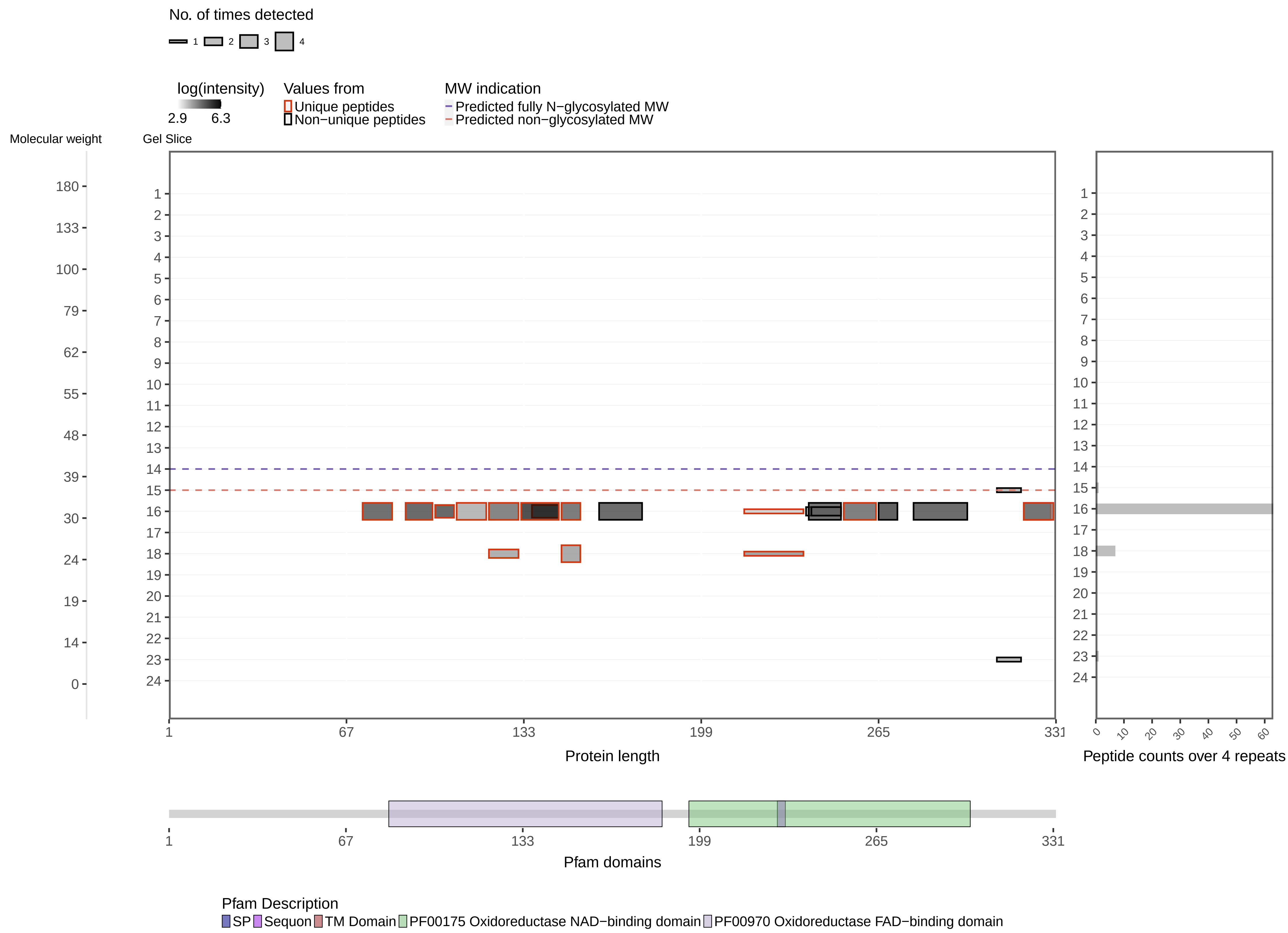


Protein sequence:

IKVYPEGKMSQHFASLKPGDVLEVKGPIEKLRYSPNMKKHIGMIAGGSGITPMLQVIEAILKNPDDHTQVSLIYANVSPDDILLKKKLDVLAATHPNLKVFYTVDSLTNDWRGGIGYVSKDMIVKGLPPPSDDTLILVCGPPGMMKHLSGEKPKPREQGELTGLLKDVGFTEHMVYKF*

### NbL13g26620.1 glutamine_synthetase [MW = 38.84 kDa]

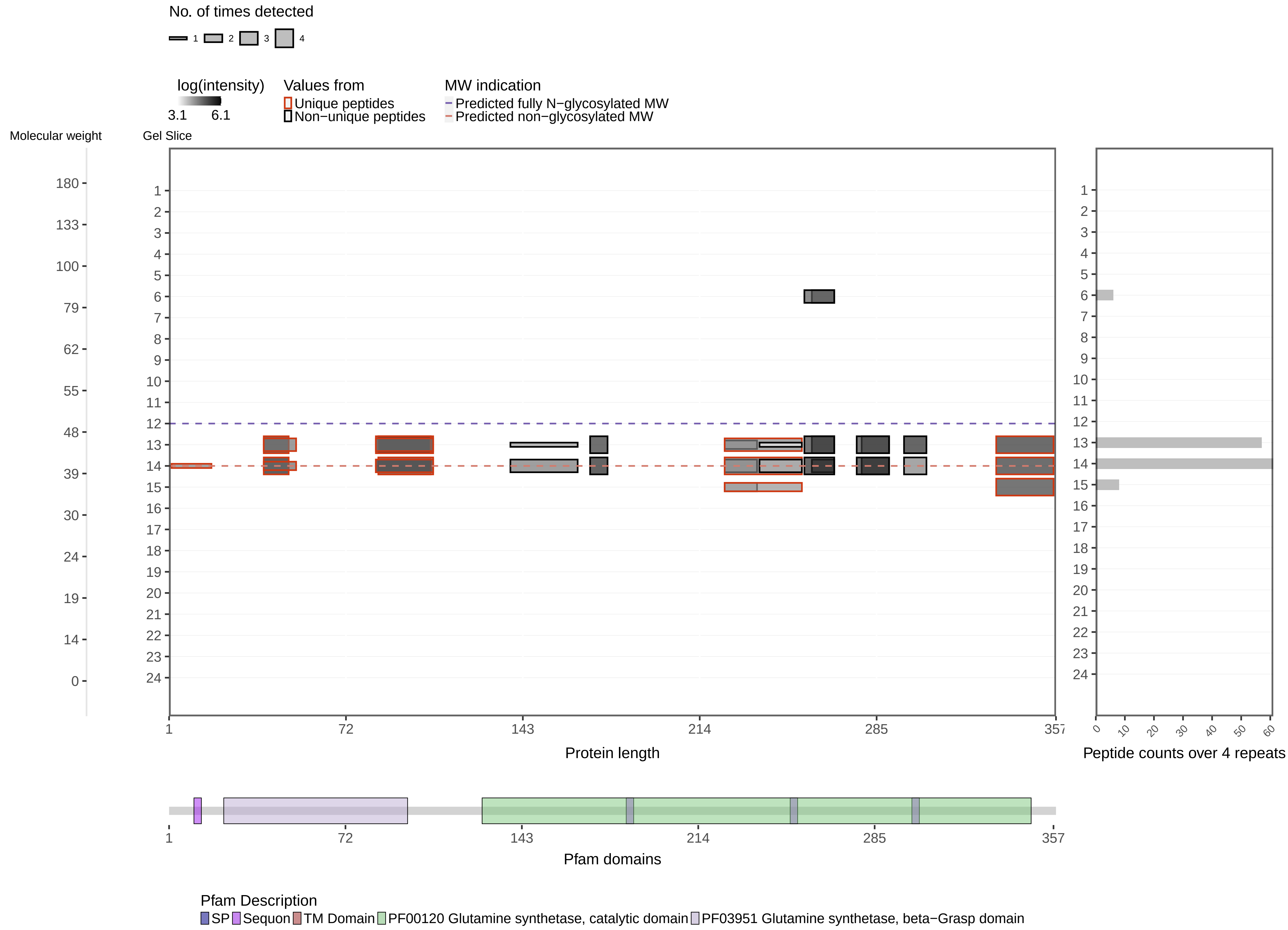


Protein sequence:

:AFGRDIVDSHYKACLYAGINISGINGEVMPGOWEFOVGPSVGISAGDEVWVARYILERIAEIAGVVVSFDPKPIPGDWNGAGAHTNYSTKSMREDGGYEVILKAIEKLGLKHKEHIAAYGEGNERRLTGKHETANISTFKWGVANRGASVRVGRDTEKAGKGYFEDRRPASNMDPYVVTAMIADTTIIGKP*

### NbL13g26560.1 probable_inactive_receptor_kinase_RLK902 [MW − SP = 67.19 kDa]

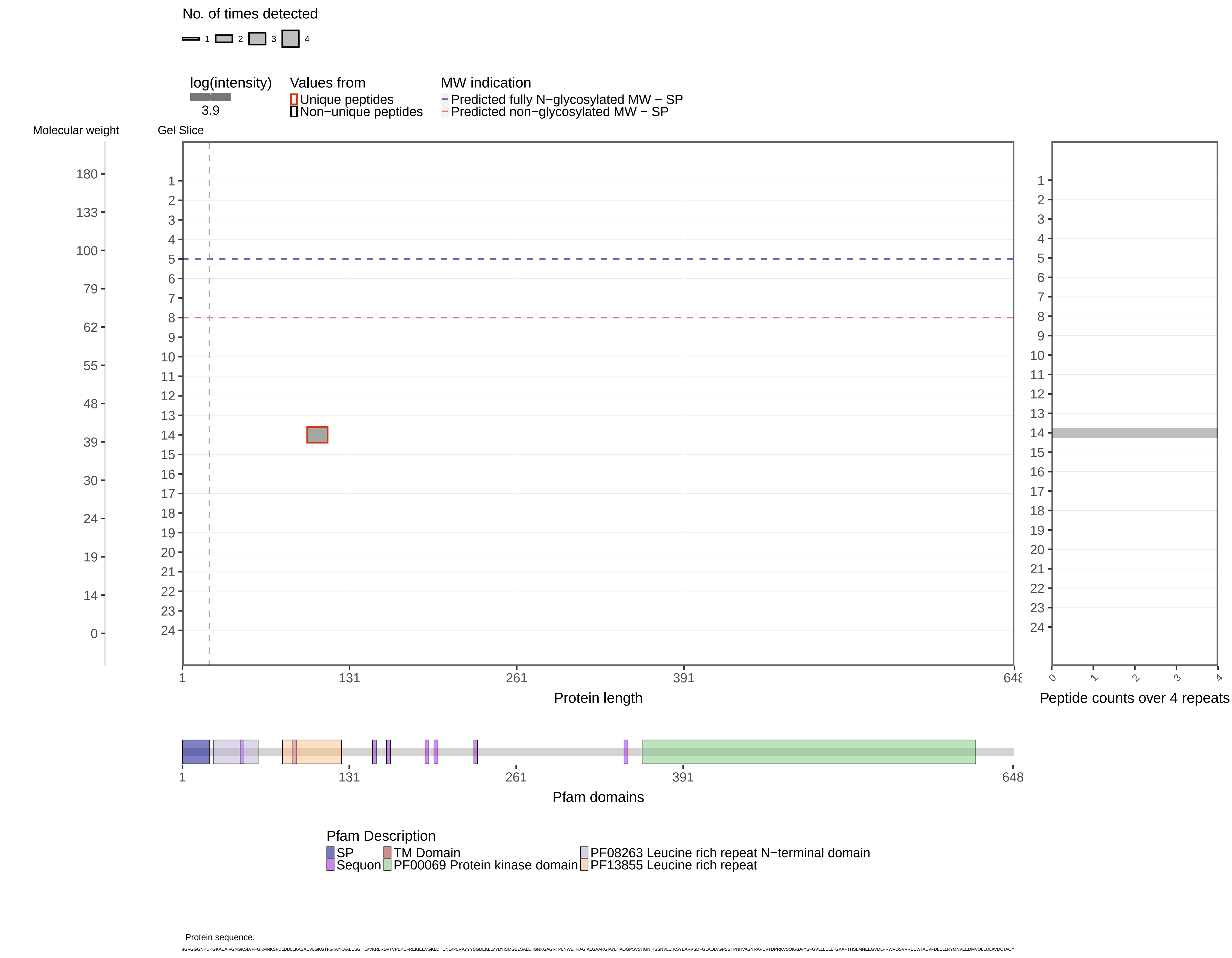

NbL13g26550.1 protein_TIFY_10B−like [MW = 26.13 kDa]

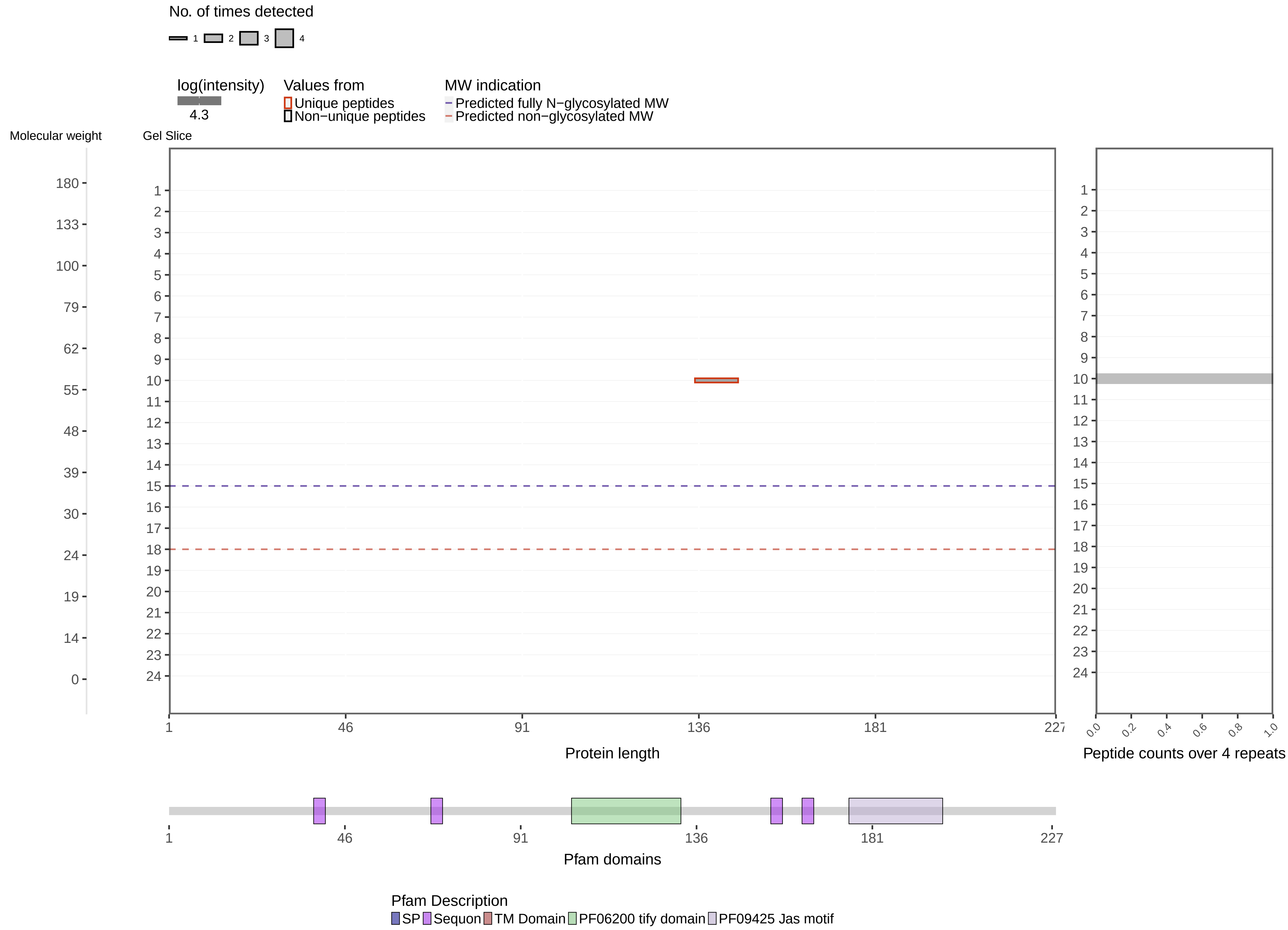


Protein sequence:

KPESTLTILYMGEVHIFQDITPEKAEVIMDLAFKSTNLHMTEILEKGNKEKSEANKSEPSTPNASTNYAKGALAMARRATLARFLEKRKHRLITARPYQYGEKTPKFPFEMHQEEETASSSFHWES*

### NbL13g26480.1 glucan_endo−13−beta−glucosidase_2−like [MW − SP = 51.79 kDa]

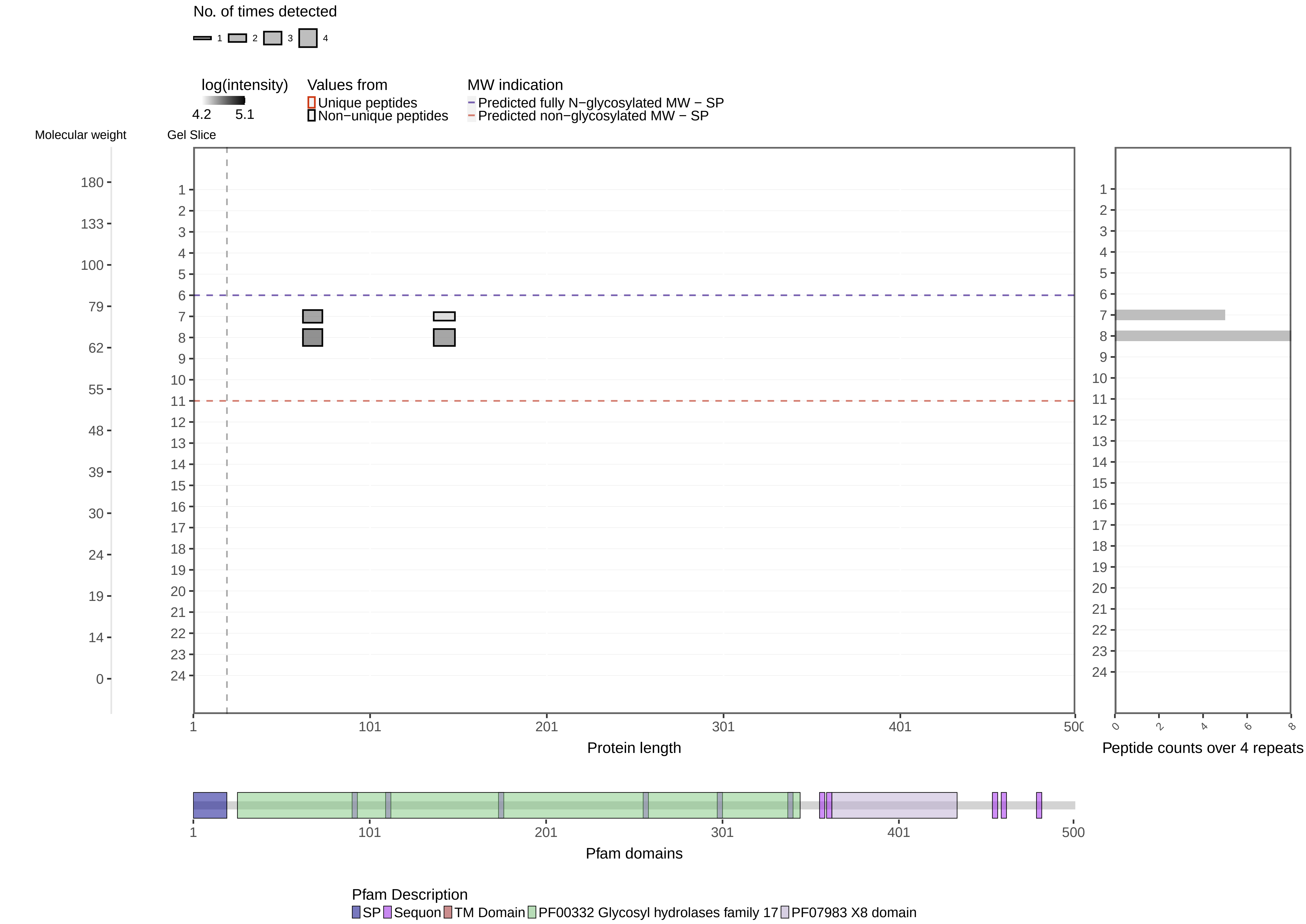


Protein sequence:

DAMVDAAYFAMLDLNFTNIPVMVTESGWPSQGDSDEPDATLDNANTYNSNLIKHVLNKTGTPKHPGIAVSTYIYELYNEDNKPGHLSEKNWGLFNANGTPVYILRLTESGSMFANDTSNQTYCAAKDGADSKMLQAALDWACGPGKVDCSALLQGEPCYEPDNVAAHATYAFDAYYHMMGRAPGTCDFNGVATITTTNPSHGSCVFSGSLGRNGTFLNGTAPAMDSTTSSSSPARYLNNTYSTILMILGILVWSKILL*

### NbL13g26420.1 HMG−Y−related_protein_A−like [MW = 18.37 kDa]

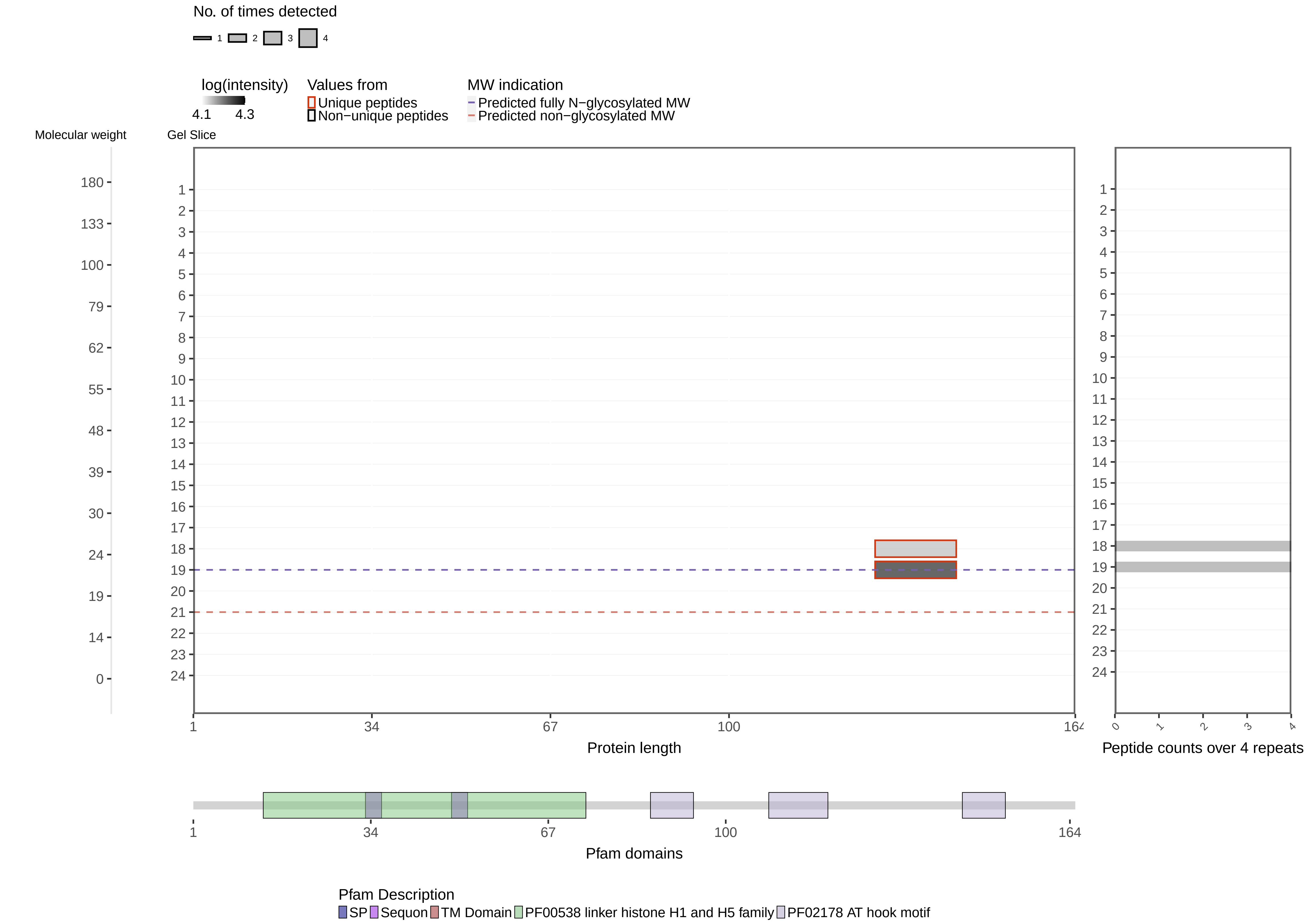


Protein sequence:

IFLKNNYIKRGPNVPPKRGRGRPPKPKTTLPQDTEIVAPKPRGRPKKDPNAPPTPKKLKPTIVPINLANPVSKTGRPRGRPRKVMPQSAQNSVE*

NbL13g26320.1 glutathione_S−transferase_DHAR3_chloroplastic [MW = 28.39 kDa]

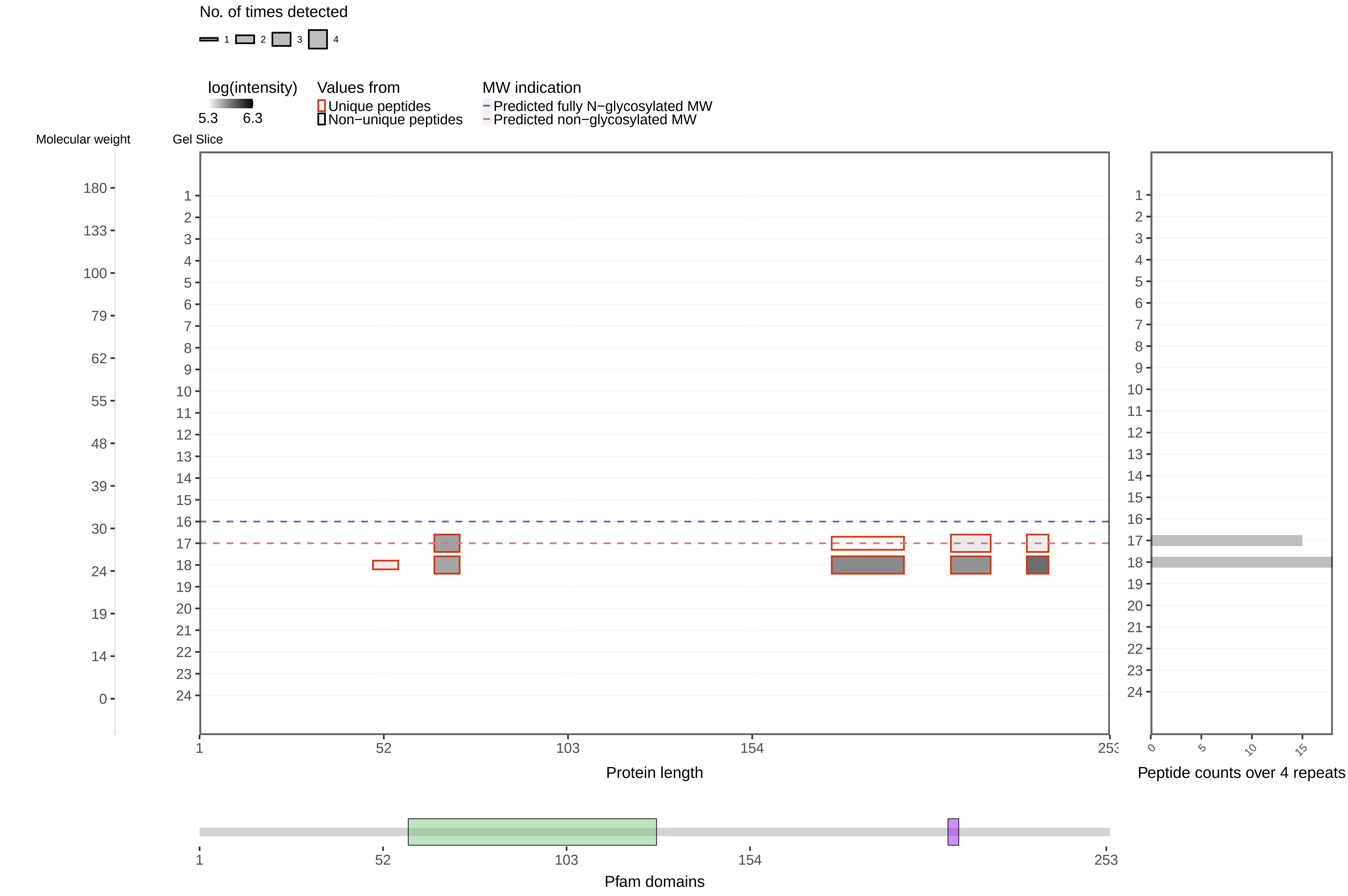


Protein sequence:

VITQALEEKFPQPPLTTPPEKASVGSKIFPKFVAFLKSKDLSDGAEQELLDELTAFNDYLKENRPFINGNEVSAADLSLGPKLYHLEIALGHYKNWSVPDSVPYVKSYMKSIFSRGSFINTRALKEDVIEGWRPKVMG*

NbL13g26210.1 8−hydroxygeraniol_dehydrogenase−like [MW = 39.37 kDa]

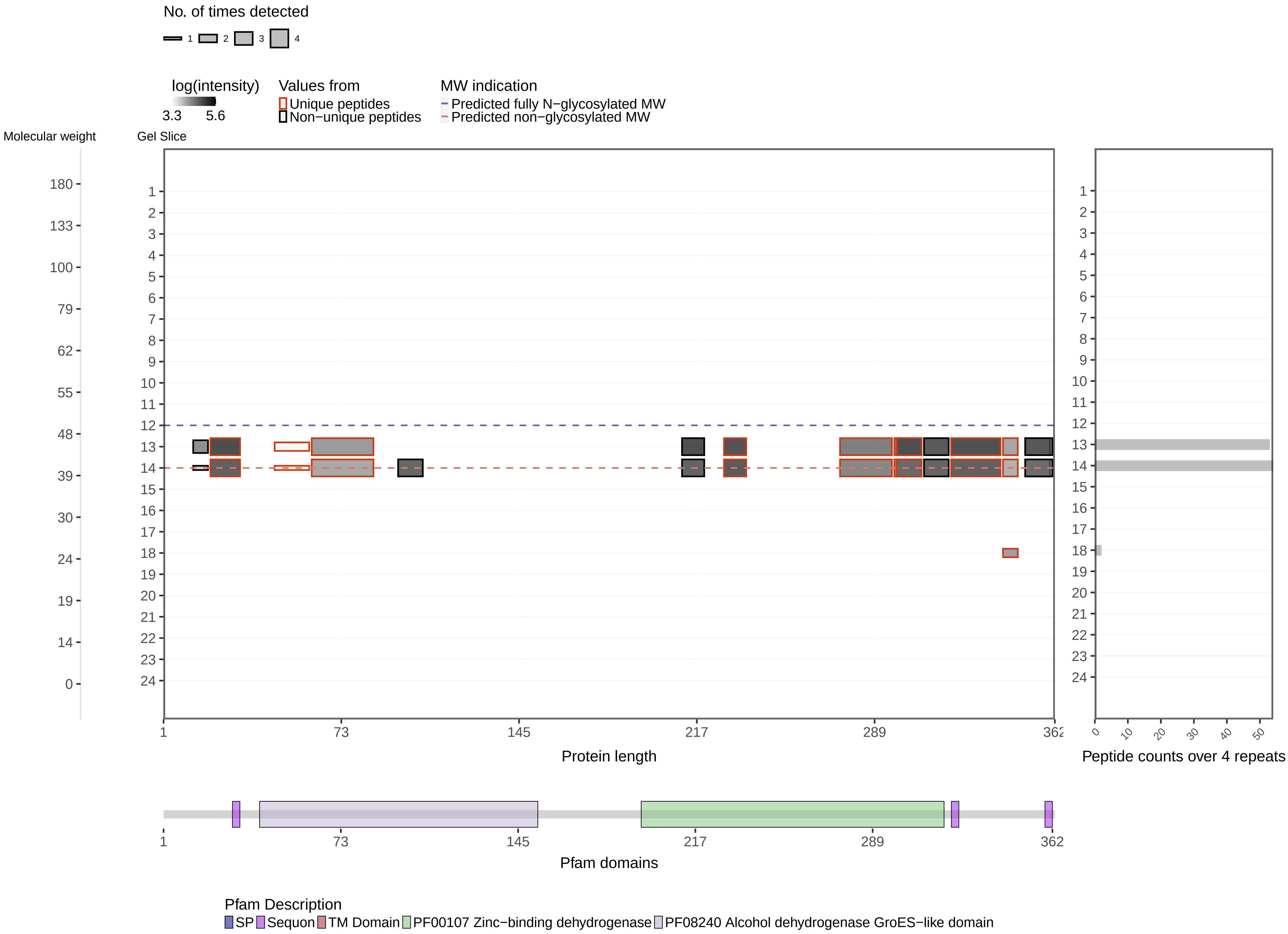


Protein sequence:

LLNAGITTYSPLKYFGLDKPGMHIGVVGLGGVGHMAVKFAKAFGTKVTVISTSASKKQEAIERLGAESFLLSRDPDQMQAATDSLDGIIDTVSAVHPLLPLINLLKTHGKLVIVGAPDKPLELPVLPLLLGRKLVAGSAIGGMKETQEMVDFAAKHNITPDVEVVPMDYVNTAMERLAESDVKYRFVLDIGNTLNKS*

NbL13g26200.1 8−hydroxygeraniol_dehydrogenase−like [MW = 39.25 kDa]

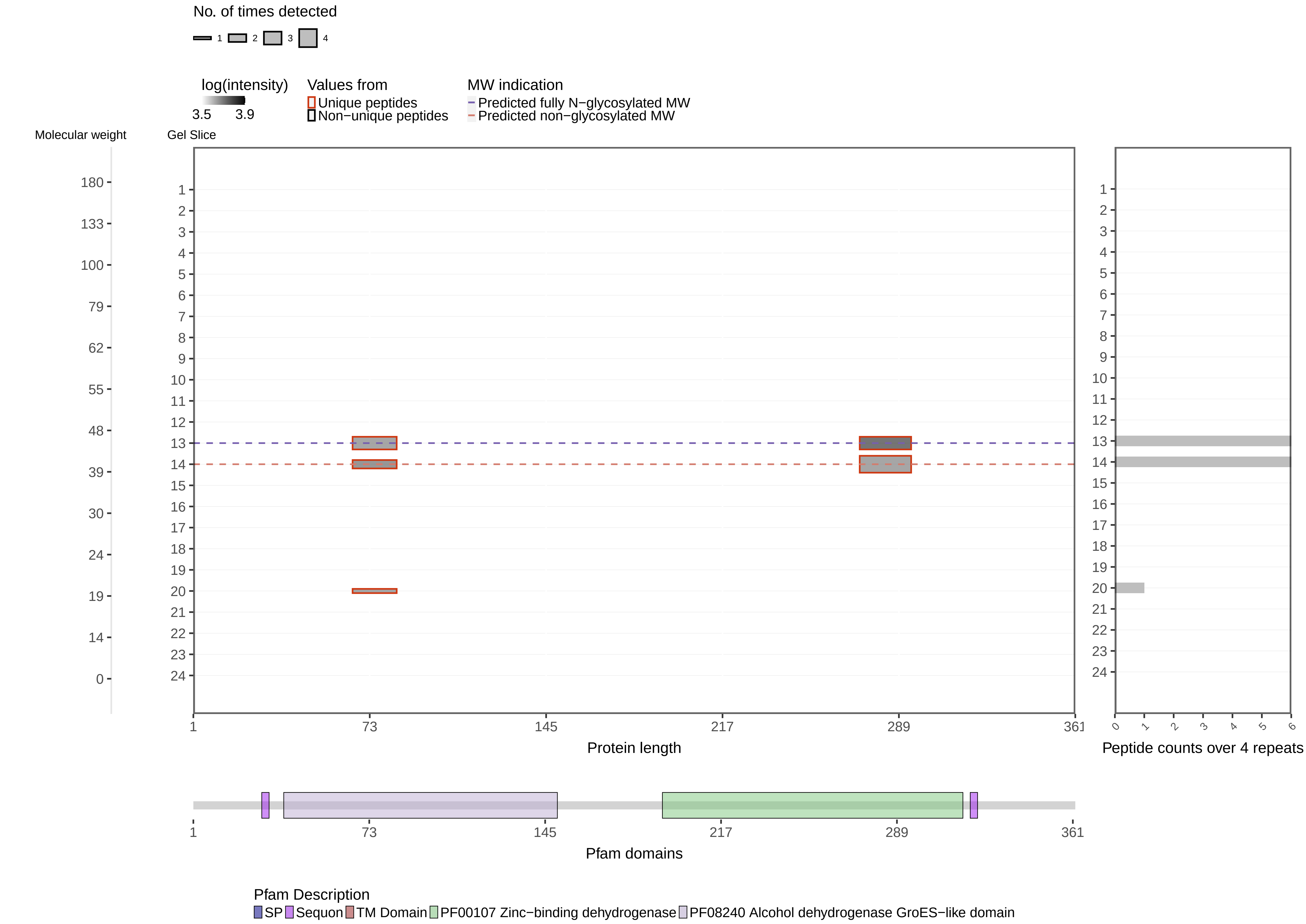


Protein sequence:

LCAGITTYSPLKYFGLDKPGMHIGVVGLGGLGHMAVKFAKAFGTKVTVISTSASKKQEAIERLGADAFLISRDPEQMQAAMSTLDGIIDTVSAVHPILPLLSLLKSHGKLVMVGAPEKPVELPVFPLLLGRKLVAGSCIGGMKETQEMVDFAAKHNITPDIEVVPMEYVNTALDRLLKSDVKYRFVLDIGNTLKAAN*

### NbL13g26170.1 probable_mannitol_dehydrogenase [MW = 38.7 kDa]

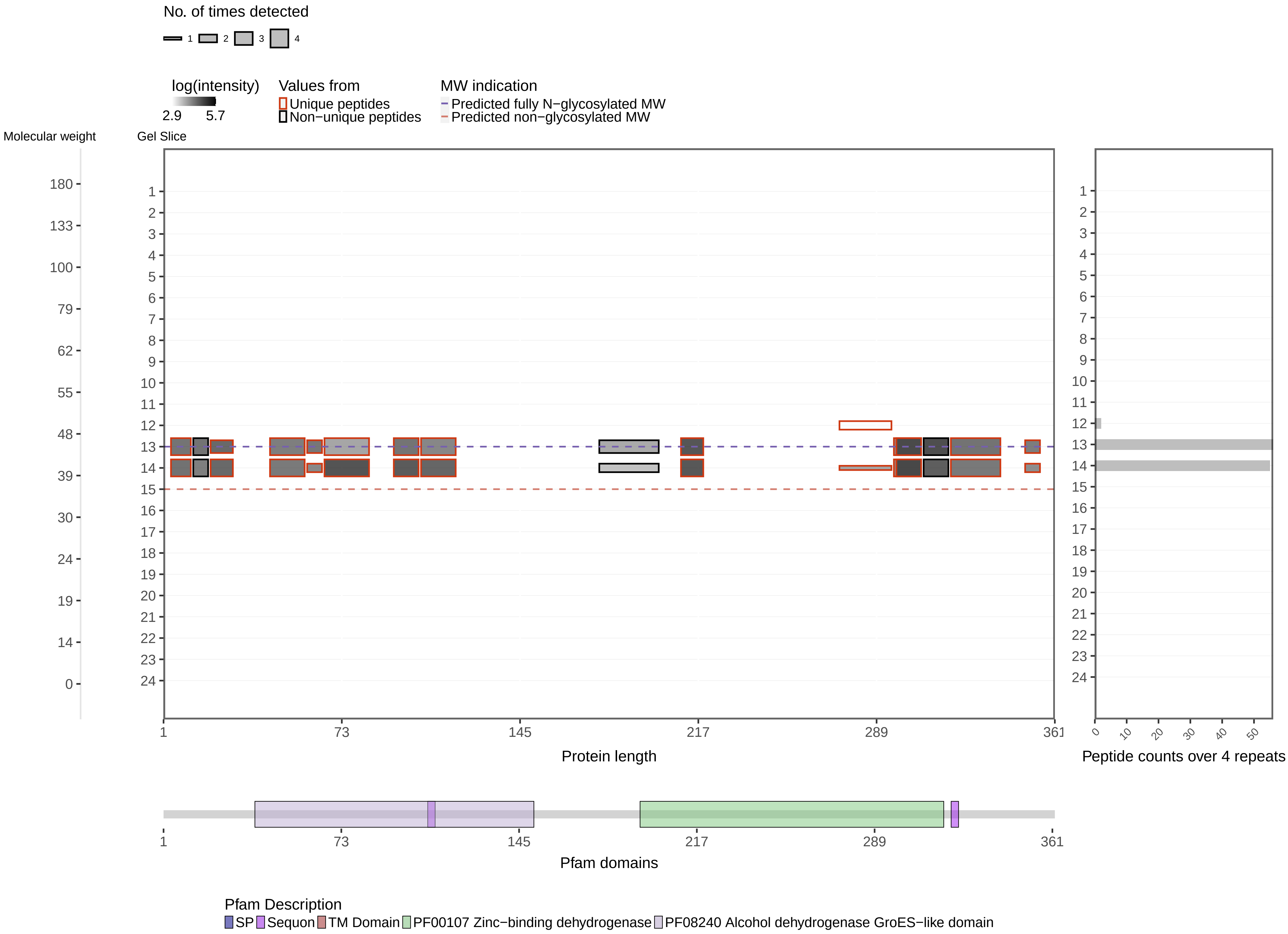


Protein sequence:

LLCAGITTYSPLRYFGLDKPGLHIGVVGLGGLGHVGVKFAKALGLKVTVISTSQSKQKEAIEHLGADSFLISTDPDQLQAAMGTMDGILDTVAATHPVLPLVGLLKTNGKLILLGALEKPLELPVFPLLMGRKLVAGSGIGGMKETQEMLDFAAKHNITADIEVIPMDYINTAMERLAKADVKYRFVIDVAKTLKAE*

### NbL13g25970.1 germin−like_protein_subfamily_3_member_2 [MW − SP = 20.88 kDa]

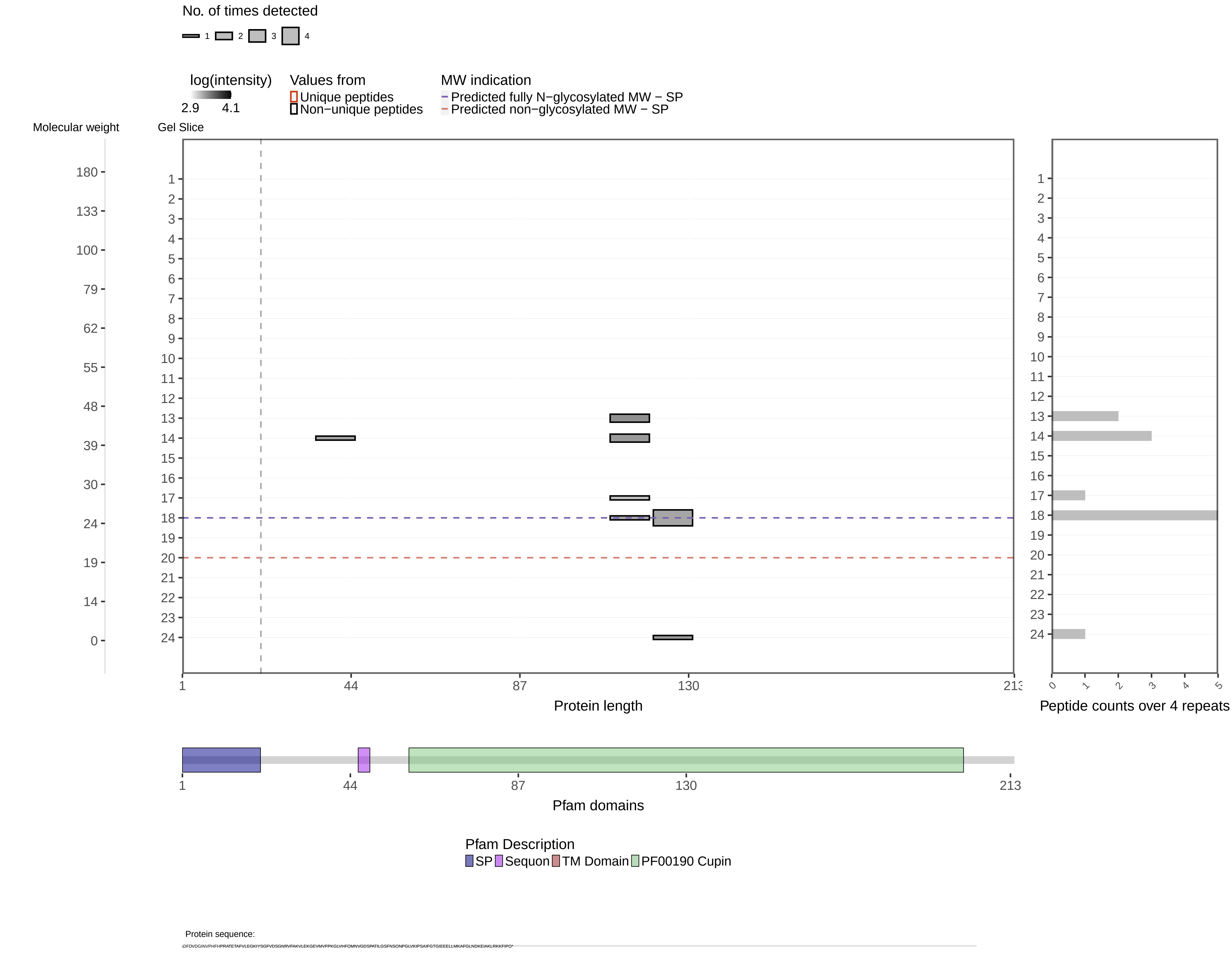

NbL13g25940.1 ACT_domain−containing_protein_ACR12−like [MW = 36.99 kDa]

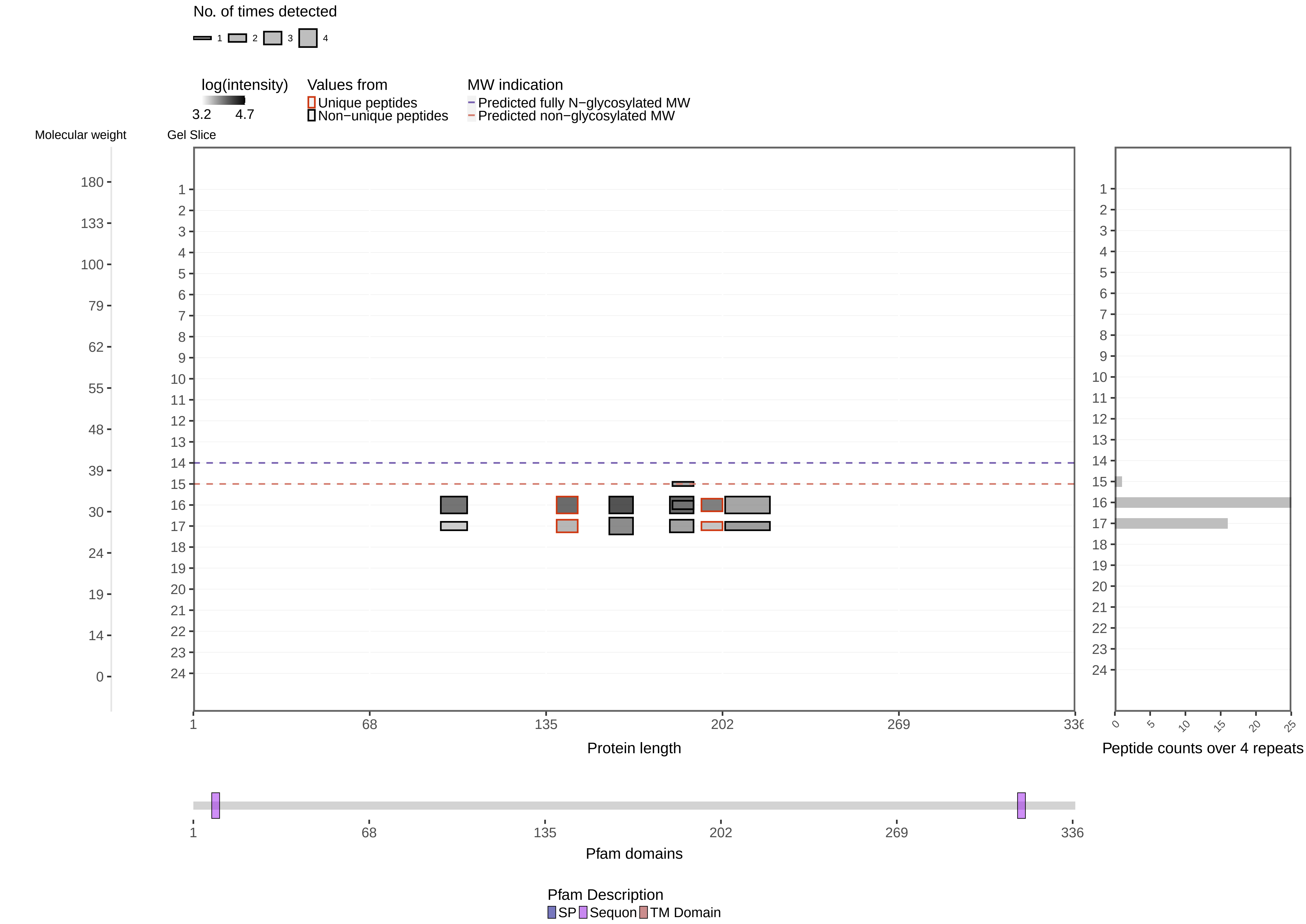


Protein sequence:

DVVKGIVTTEGSVKKTKFLITRLDTGRKVEDPDLLERIRLTIINNLIKYHPESSEQLAMGEAFGIKAPEKKPDVDIATHIHIKDDGPKRSILHVETADRSGLLVEVVKIMADINIDVESAEIDTEGLVAKDKFYVSYRGAALTSSLSQVTYCIQFSSGSGNHSHVSRLRYLFMVSNVRIFV*

### NbL13g25890.1 probable_small_nuclear_ribonucleoprotein_G [MW = 4.54 kDa]

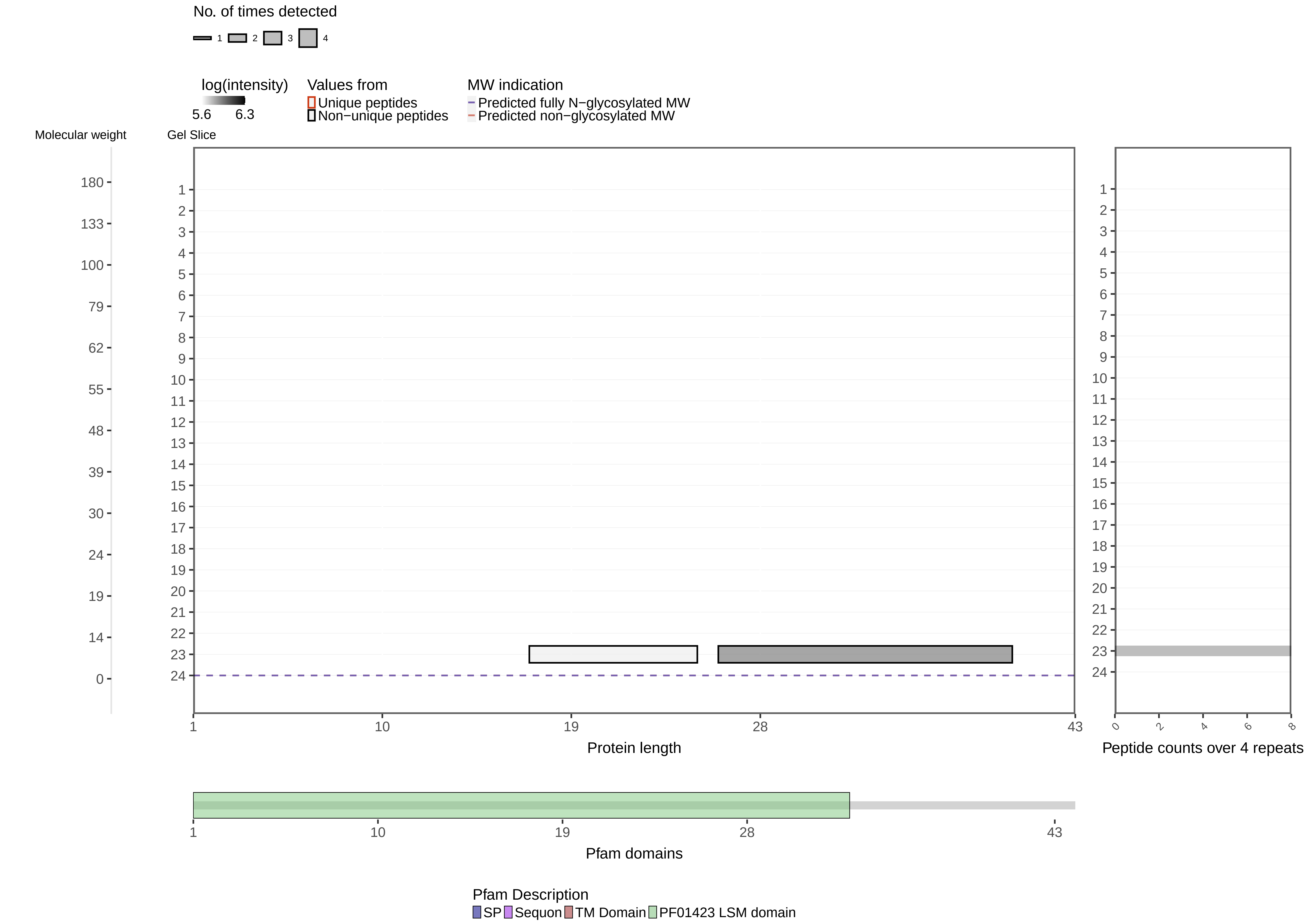


Protein sequence:

VEVNGNEKNEIGMVVIRGNSVVTIEALEPVARAQ*

### NbL13g25520.1 phosphoglycolate_phosphatase_1A_chloroplastic−like [MW = 39.79 kDa]

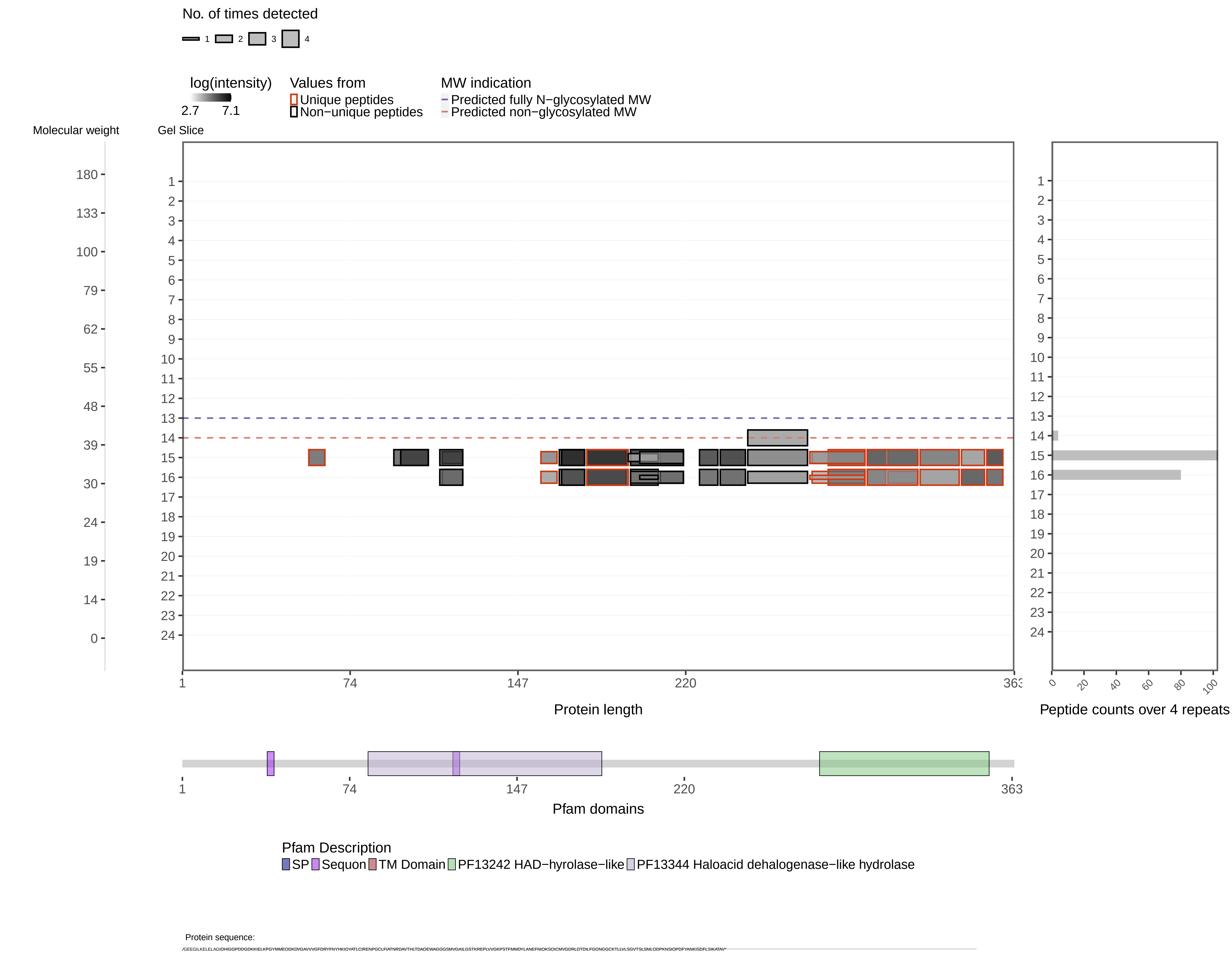

### NbL13g25510.1 omega−hydroxypalmitate_O−feruloyl_transferase−like [MW = 51.21 kDa]

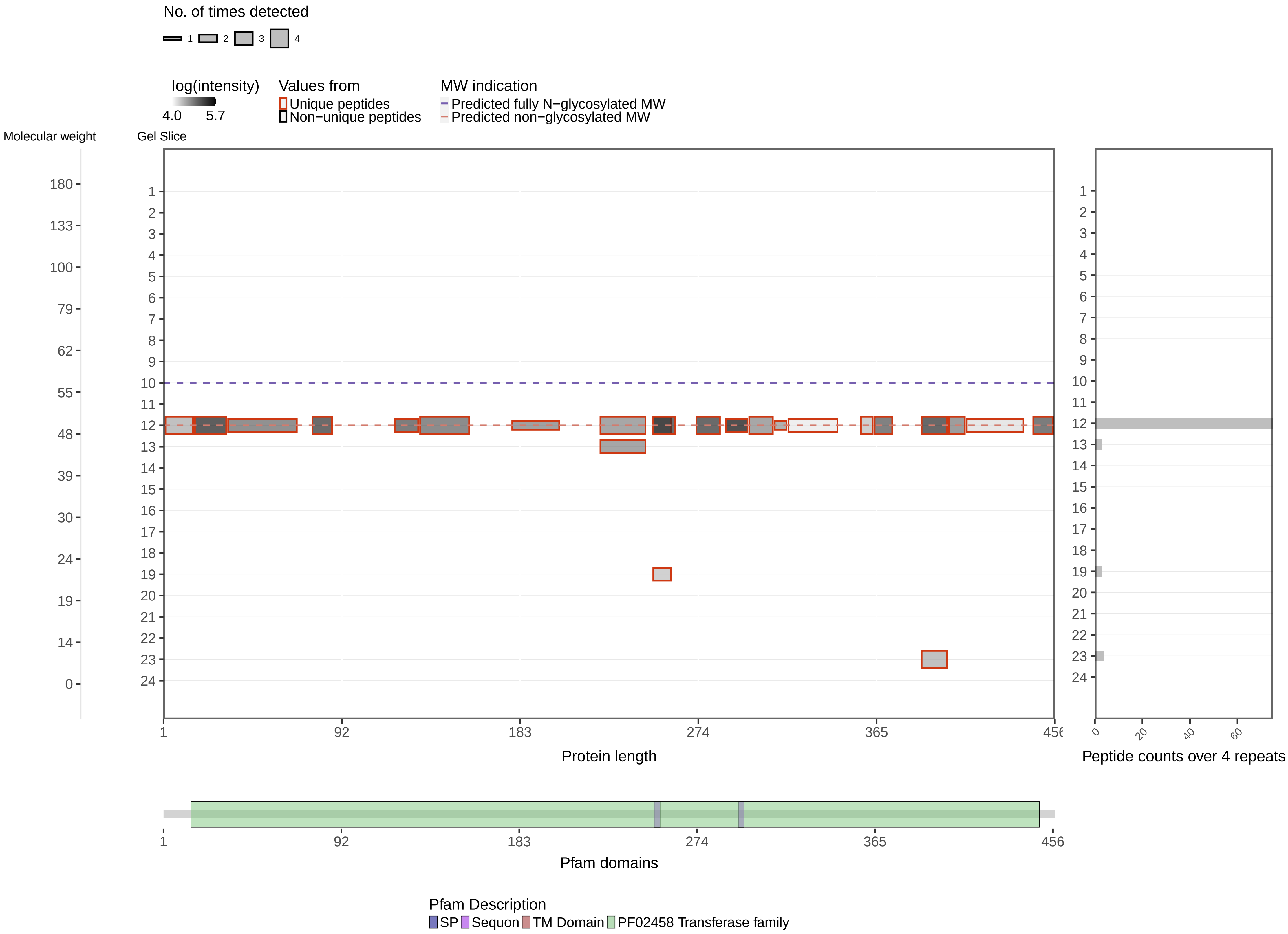


Protein sequence:

PHPELLKLEVPIGEELNPTVFEALQEELVFKIFKLNSSDISSLKDKAKDENAPHAKVTSFNVVTAHVWKCKALSFDVENNSERISTVLFAVDIRPRLNPPLPKSYTGNAVLSSYASAACHELEEWPFYKIVDMVSKGAMRMTDEYARSVIDWGEFYKGFPNGEFLISSWWRLGFSEVEYPWGKPKYSCPVVSHRKDIILFFPNIDDGISSNNDGVNILVALPAKQMEKFESYFHQFLLA*

NbL13g25460.1 uncharacterized_LOC107801208 [MW = 21.1 kDa]

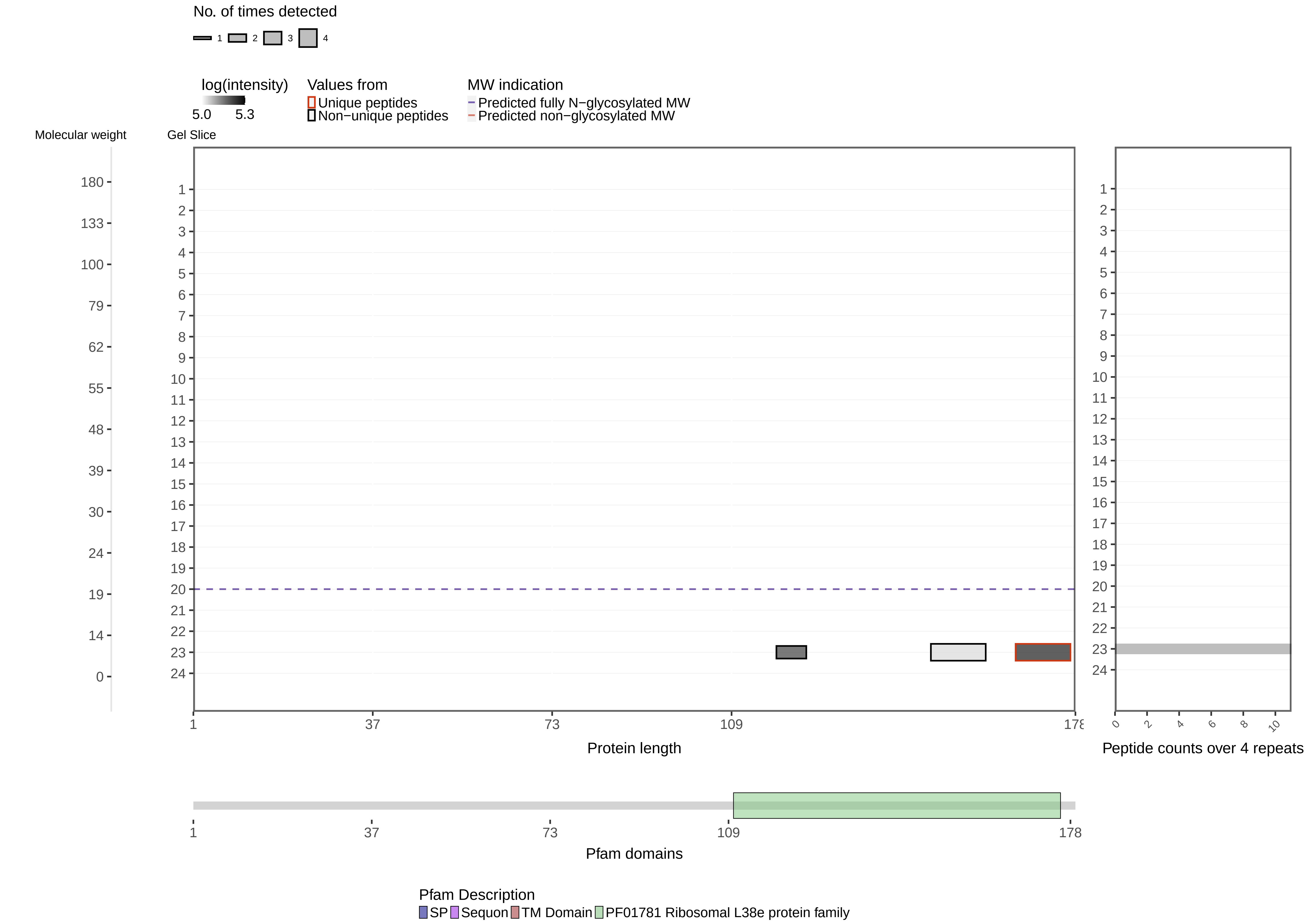


Protein sequence:

GHVRRRSPDAQQDMTRMQVSEDMTLDRKVWRSRIKPKQIHEIKDFLLTARRKDARSVKIKKNKDMVKFKVRCSKYLYTLCVSDFEKADKLKQSLPPGLSVQDL*

NbL13g25270.1 protein_transport_protein_Sec24−like_At4g32640 [MW = 117.14 kDa]

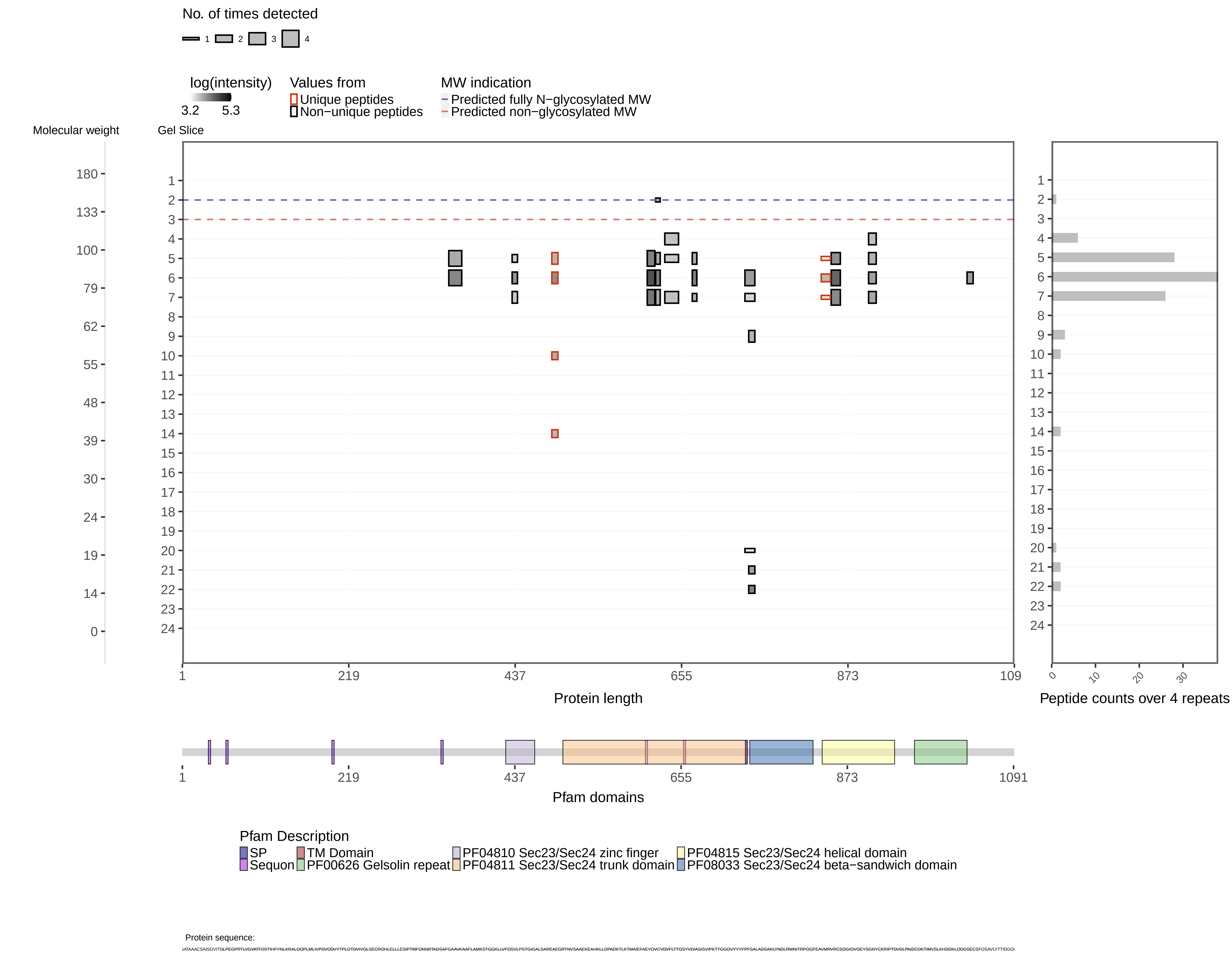

### NbL13g25260.1 uncharacterized_LOC109239406 [MW = 57.94 kDa]

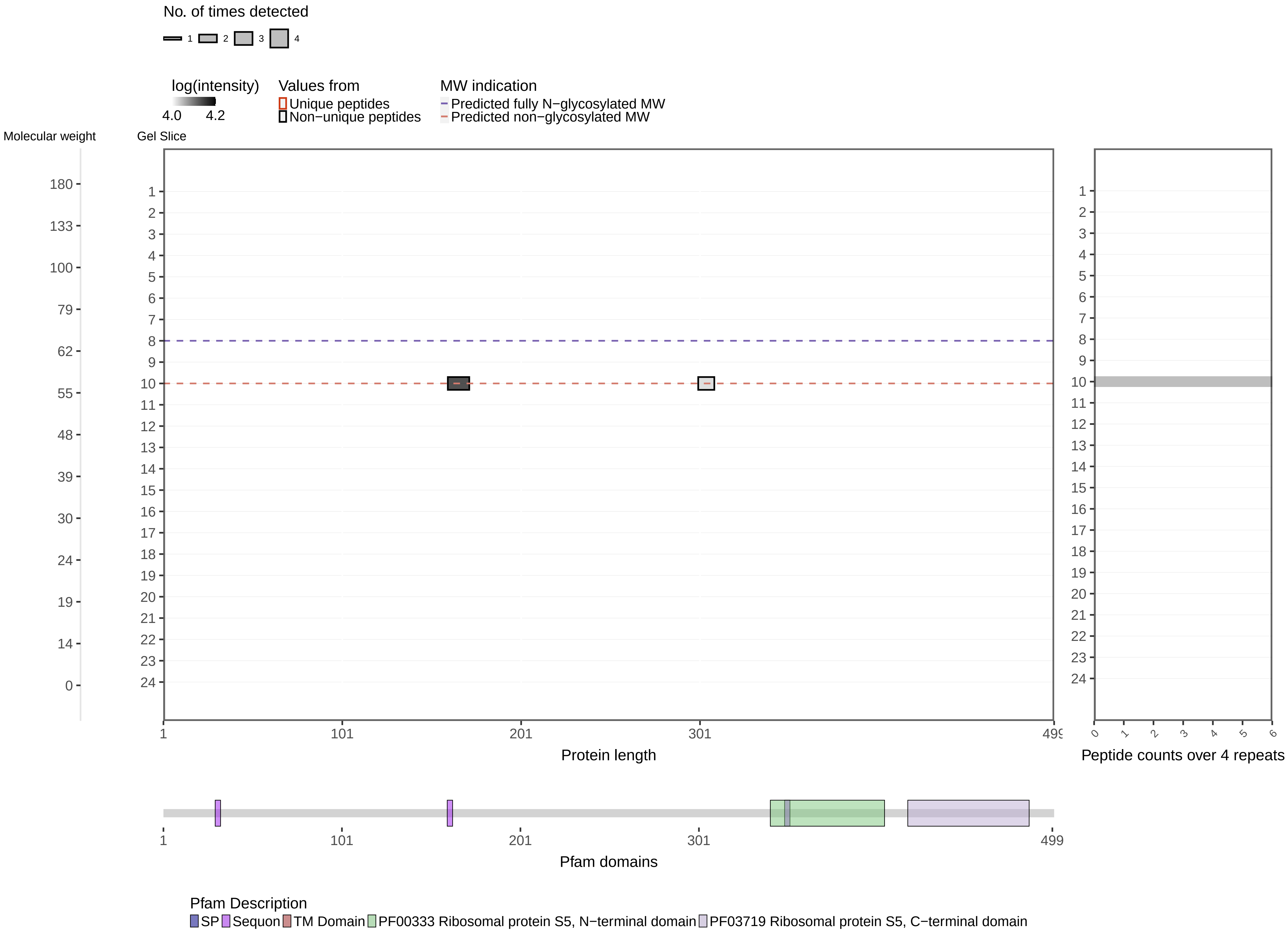


Protein sequence:

PENYGIIQHEQVGPNVDLLEHAGFEKEKEMIQGGDDDDDIEFDDMKEKDDILLEKLNAIDRKLEEKLAELDHTFGKRGKLLEEEIRDLAEERNSLTEKKRRPLYRKGFDVKLIDVNRTCKVTKGGQVVKYTALLACGNYHGVVGFAKAKGPAIPIALQKAYEKCFQNLHYVERHEEHTISHAVQTNYKKTKVYLWPAPTQTGMKAGRTVQTILHLAGFRNVKSKVVGSRNPHNTVKALFKALNSIEIPKDVEEKFGRTVVESYLL*

### NbL13g25160.1 beta−hexosaminidase_3 [MW − SP = 57.13 kDa]

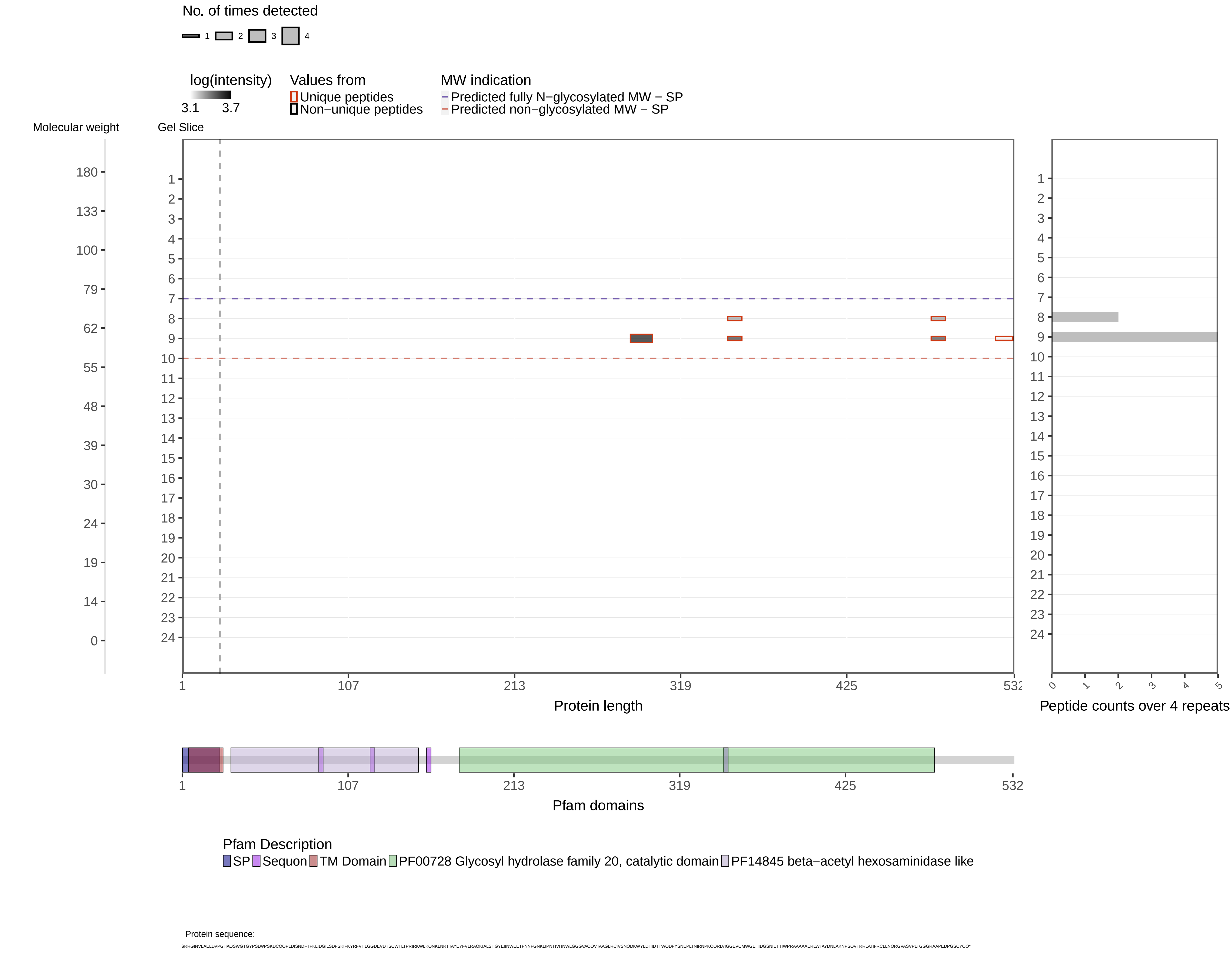

### NbL13g25120.1 protease_Do−like_10_mitochondrial [MW = 65.9 kDa]

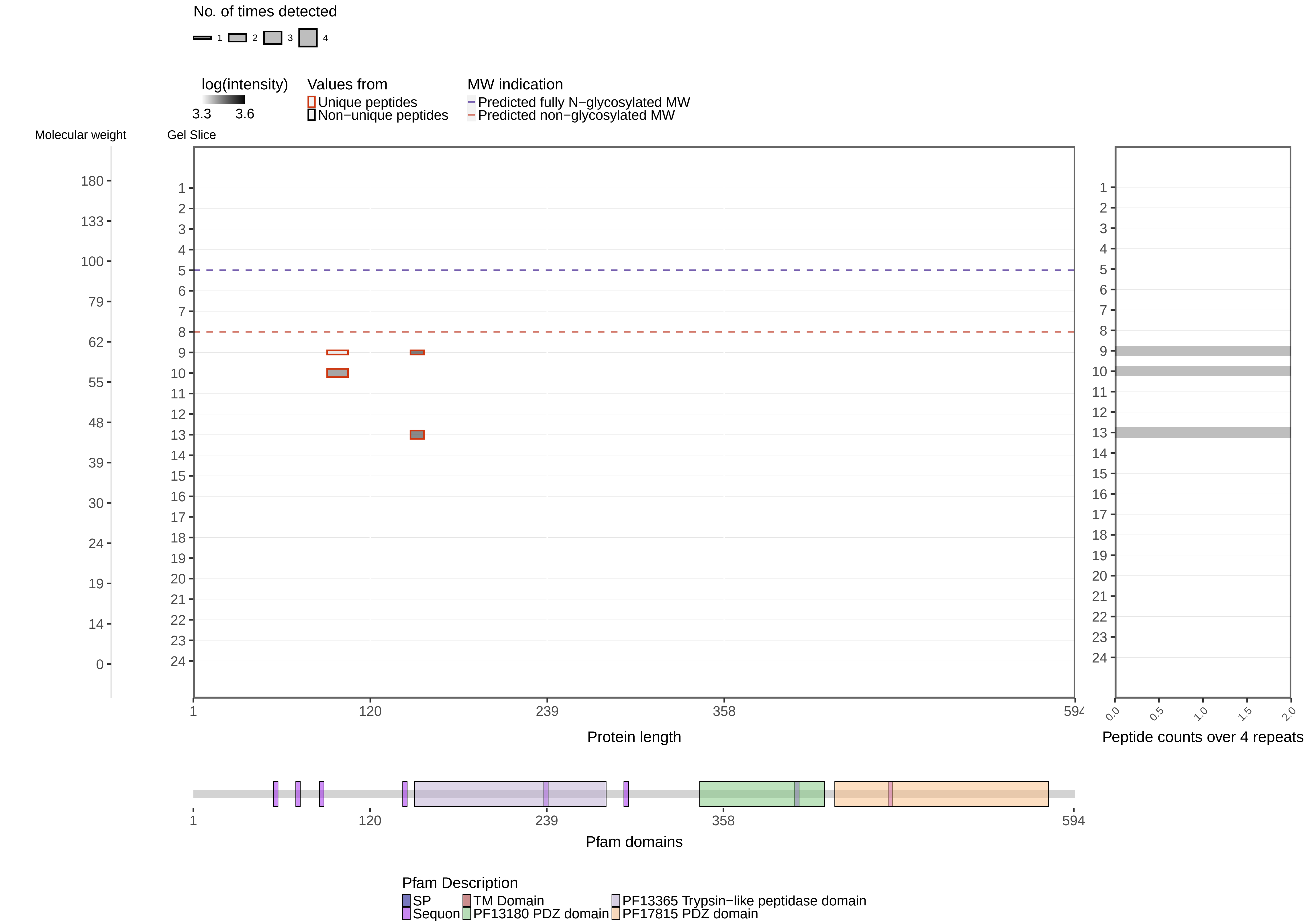


Protein sequence:

KVAGVAFQNLSGAENIGYIIPVPVIKHFIAGVEERGEYVGFCSLGLSCQPTENAQIREYFQMQSKLTGVLVSRINPLSDASRVLKKDDIVLSFDGVPIANDGTVPFRNRERITFDHLVSMKKPNETAELKVLRNGEVHDFKITLHPLQPLVPVHQFDKLPSYFIFAGLVFIPLTQPFLHEYGEDWYNASPRRLCERALRELPKKPGEQFIILSQVLMDDINAGYERLAELQVKKVNGIEVLNLKHLRQLVEDGNQKNVRFDLDDERVIVLNYETARIATSRILKRHRIPHAMSTDLTDDI

### NbL13g25080.1 reductase_2−like [MW = 67.54 kDa]

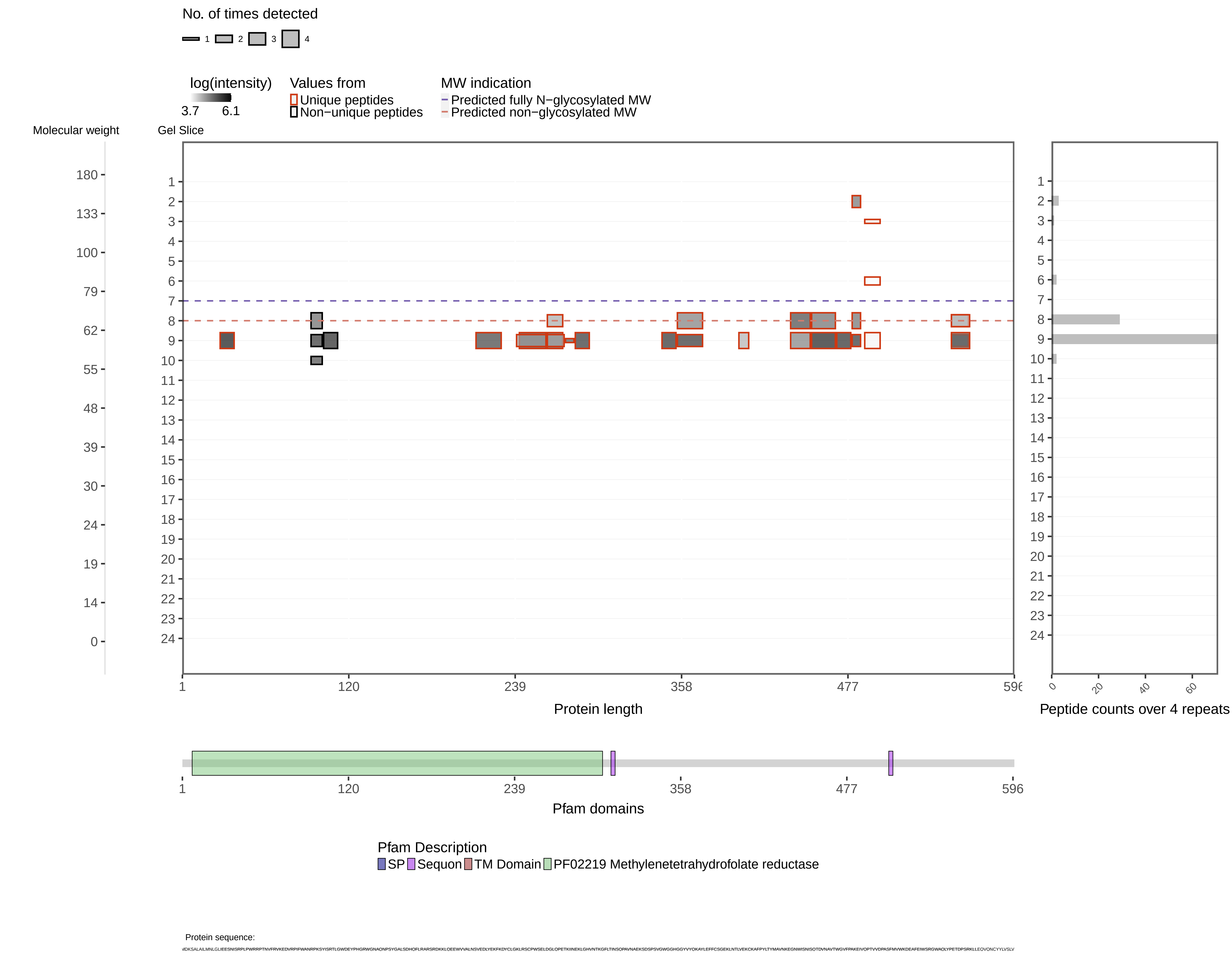

### NbL13g24990.1 membrane−associated_protein_VIPP1_chloroplastic [MW = 26.93 kDa]

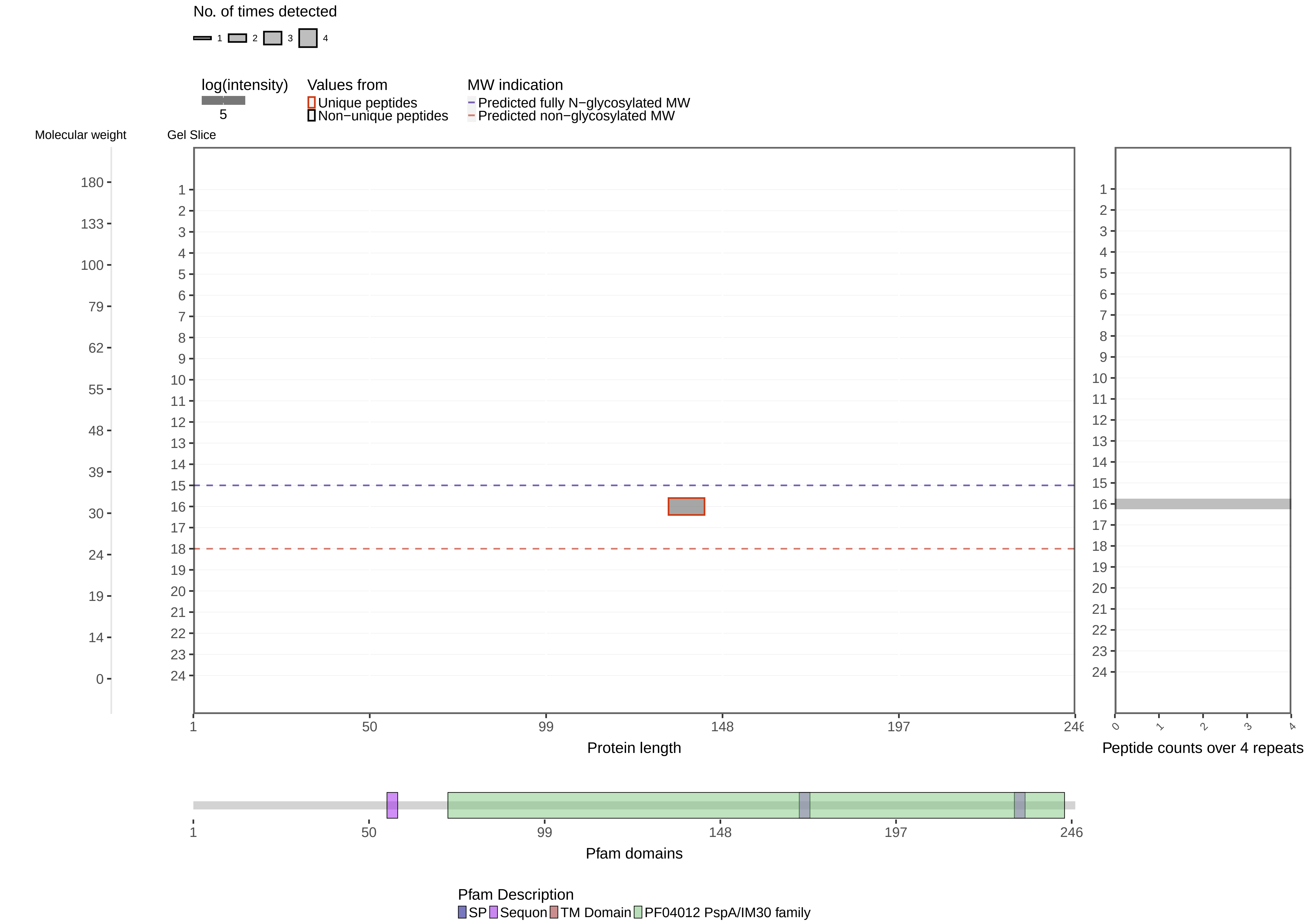


Protein sequence:

KMRQATAQVLASQKQLENKYKAAQQASEDWYRRAQLALGKGDEDLAREALKRRKAYADNASALKAQLDQQKGVVENLVSNTRLLESKIQEAKSKKDTLKARAQSAKTATKVSEMLGNVNTSSALSAFEEWKKKF*

NbL13g24760.1 14−3−3−like_protein_16R [MW = 29.1 kDa]

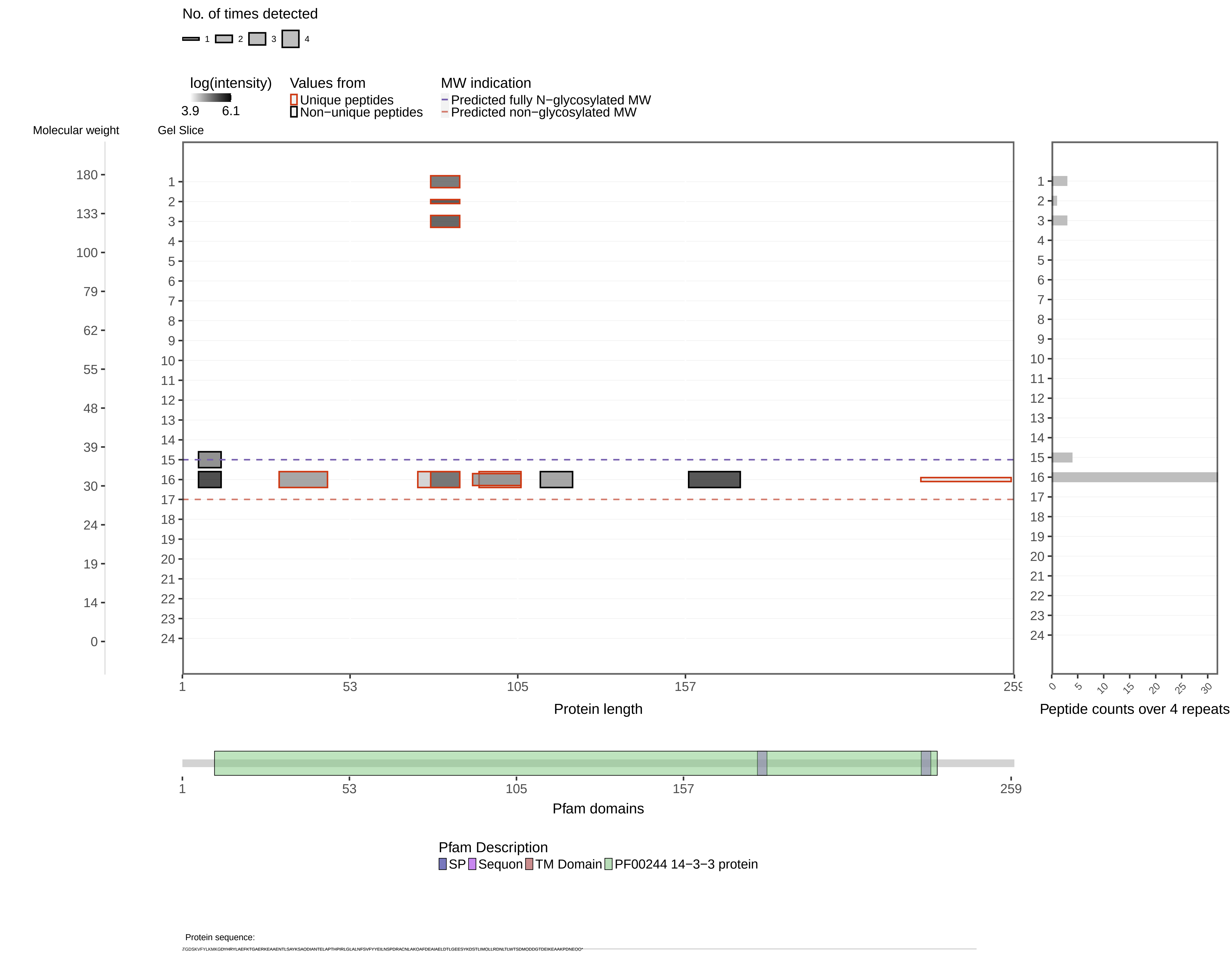

NbL13g24690.1 bifunctional_riboflavin_biosynthesis_protein_RIBA_1_chloroplastic−like [MW = 59.27 kDa]

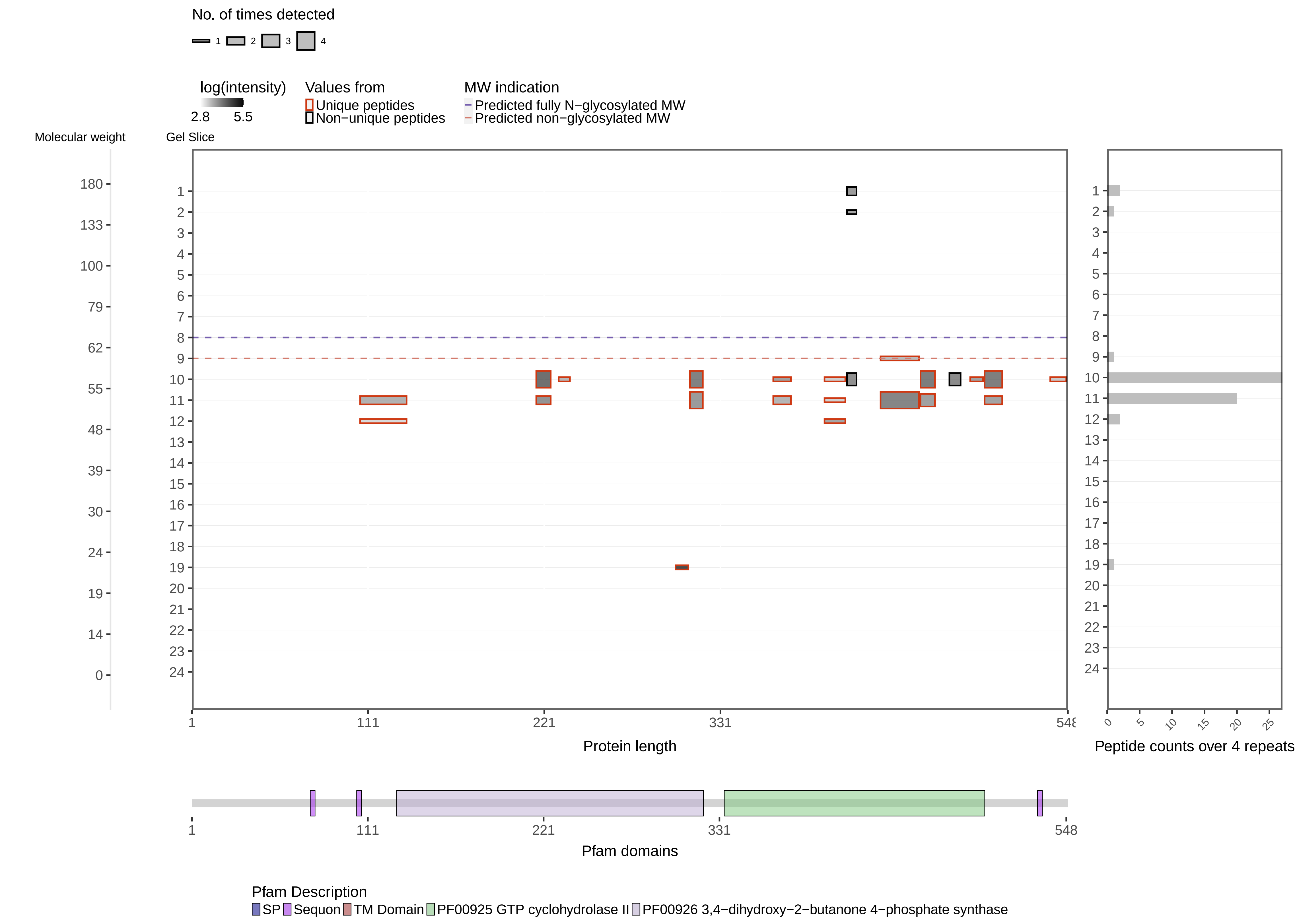


Protein sequence:

LKRAGHTEASVDLAVLAGLDPVAVLCEIVDDDGSMARLPRLRQFAEAENLKIISIADLIRYRRKREKLVELAAAAPIPTMWGPFKAYCFKSVLDGIEHIAMVKGDIGDGKDVLVRVHSECLTGDIFGSARCDCGKQLALAMEQIEQAGRGVLVYLRGHEGRGIGLGHKLRAYNLQDDGHDTVEANEELGLPVDSREYGIGAQILRDLGVRTMKLMTNNPAKYVGLKGYGLAIAGRVPLLAPITTENRRYLETKHVKLGHVYSSDANGHINGSEVGRASSDELSDNAA*

NbL13g24590.1 esterase_AGAP003155−like [MW = 24.84 kDa]

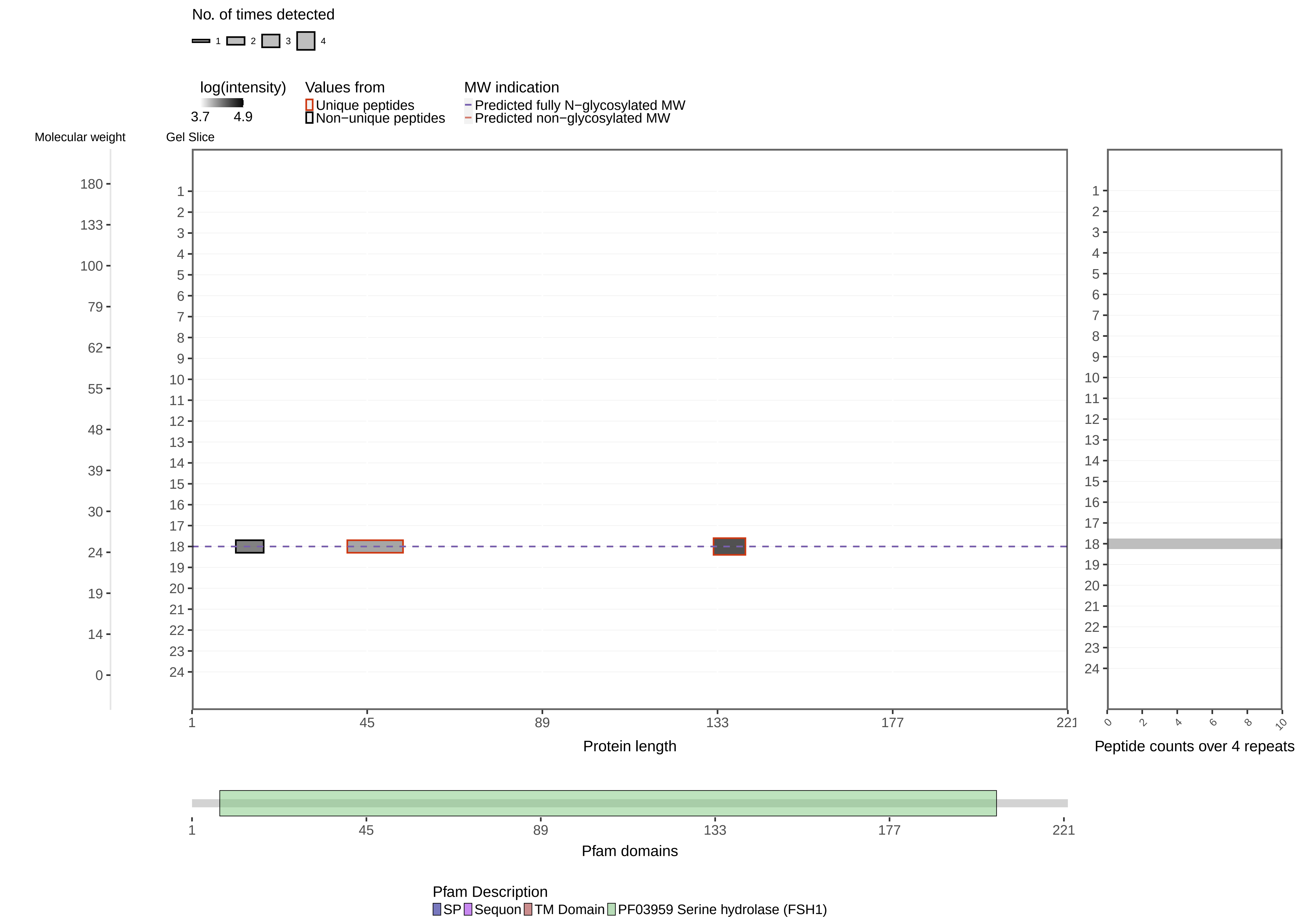


Protein sequence:
GPFDGILGFSQGAVLGAGIPGMQRERVGLTKVQEIKFVIIISGAKFGGPSFGVPQLAANVFSSPINCPSLHFLGEKDFQKKDGEILLESFVDPQVIHHPKGHTVPKLDDSSLEIMLEFIEKIQKL*

NbL13g24560.1 14−3−3−like_protein_D [MW = 28.26 kDa]

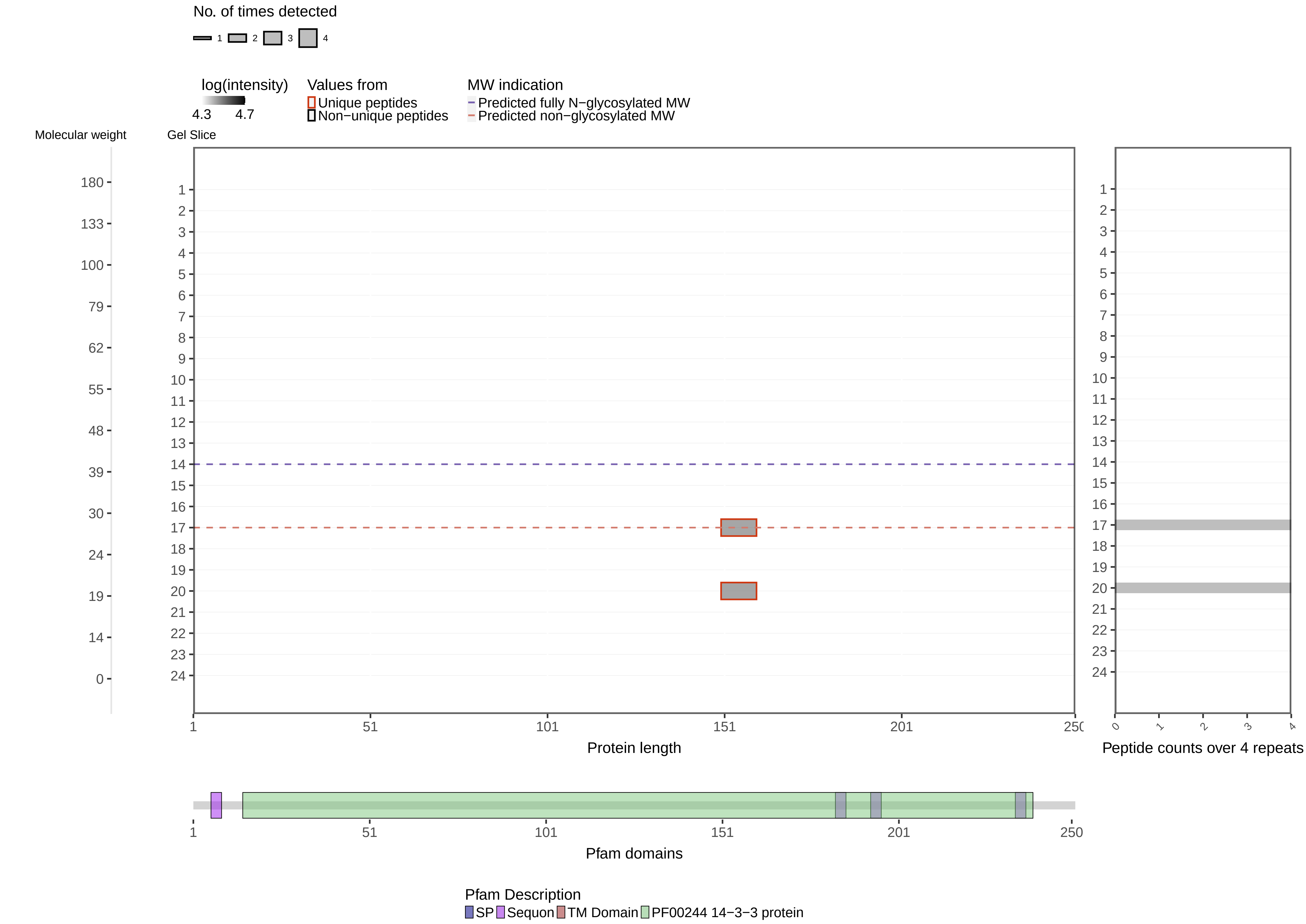


Protein sequence:

OYLIPSASAGESKVFYLKMKGDYYRYLAEFKVGNERKEAAEDTMLAYRAAQDIALAELAPTHPIRLGLALNYSVFYYEILNASEKACIMAKQAFEEAIAELDTLGEESYKDSTLIMQLLRDNLTLWTSDMQEQMDEA*

### NbL13g24550.1 protein_CURVATURE_THYLAKOID_1B_chloroplastic−like [MW = 25.65 kDa]

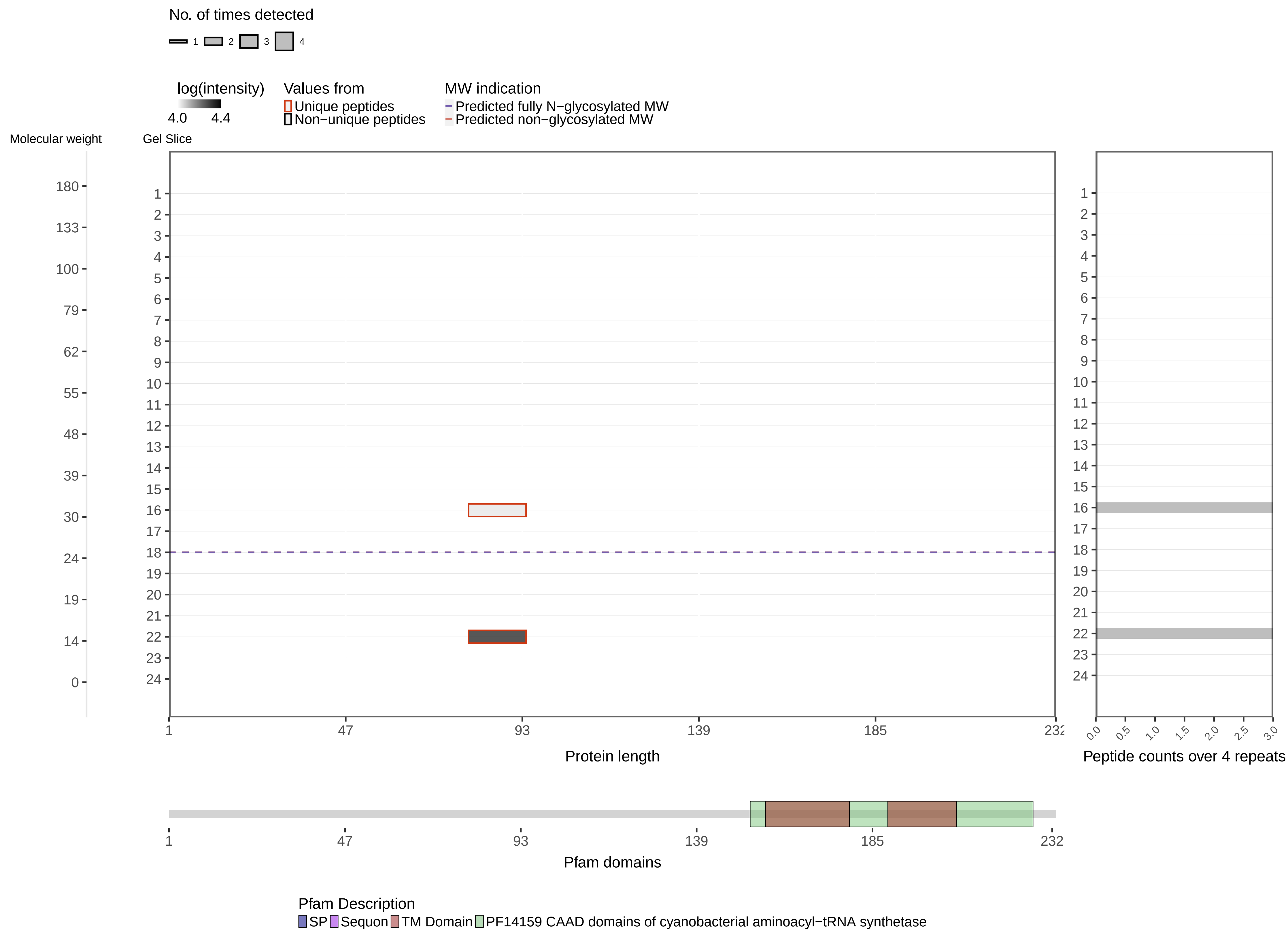


Protein sequence:

'GSLVYEPPKVEEVEEKGDGVVQAESTIEDSPLDFDSQLPKLFDKLNIKFDLEDSSSLILFGGAAVTALWLTTAVVGAIDSIPLFPKLLEVVGLGYTVWFSTRYLLFKKNRDELAAKIEELKKEVLGSNYG*

### NbL13g24320.1 diphosphomevalonate_decarboxylase_MVD2−like [MW = 22.34 kDa]

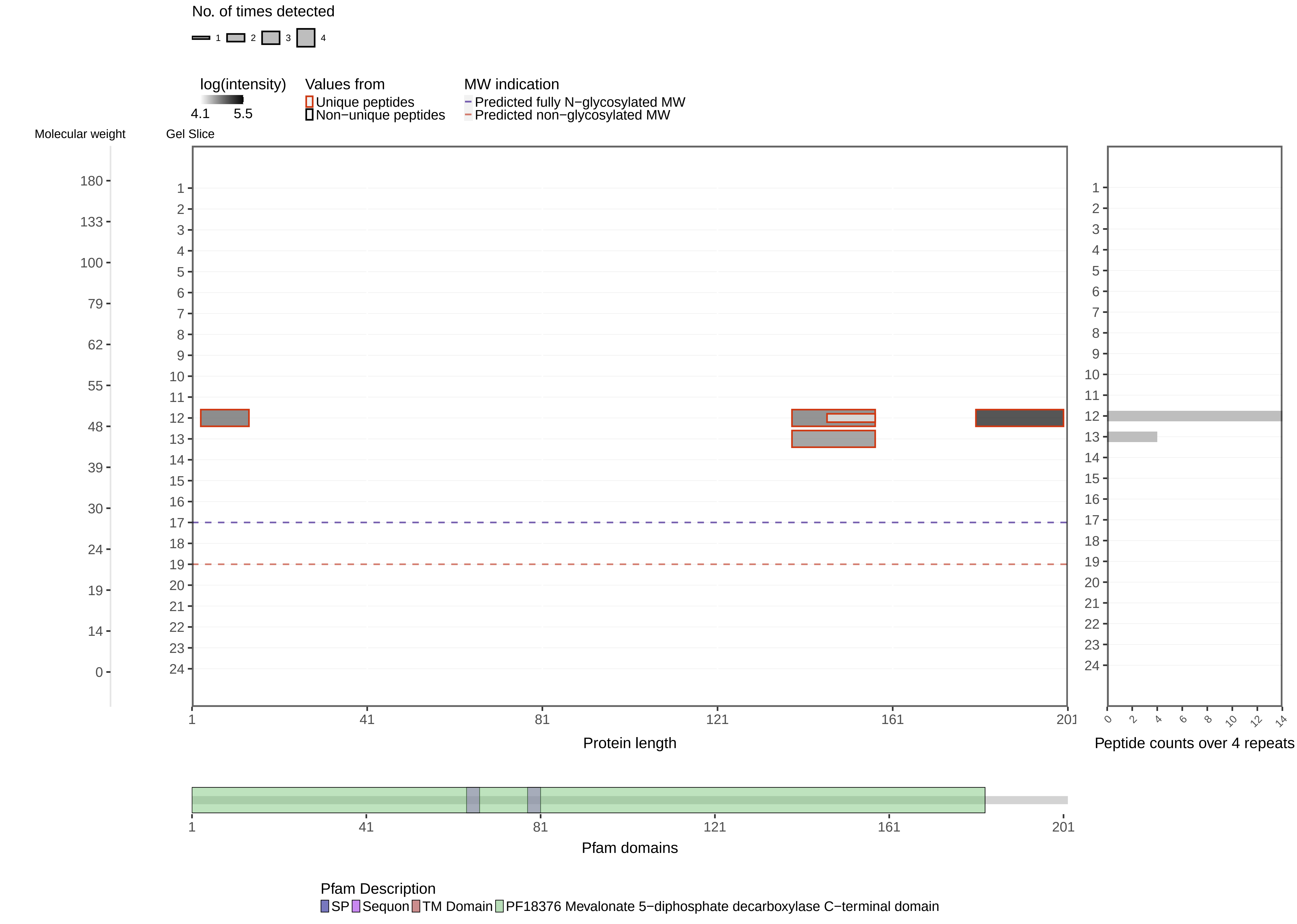


Protein sequence:

VAYTFDAGPNAVLIARNRKAATLMLQRLLFHFPPNSDTDLNSYVIGDKSILKDAGIQDIKDVEALPPPPEIKDKVPAQKYKGEISYFICTKPGRGPVLLTDDSHALLNPETGLPK*

### NbL13g24290.1 probable_aspartyl_aminopeptidase [MW = 59.88 kDa]

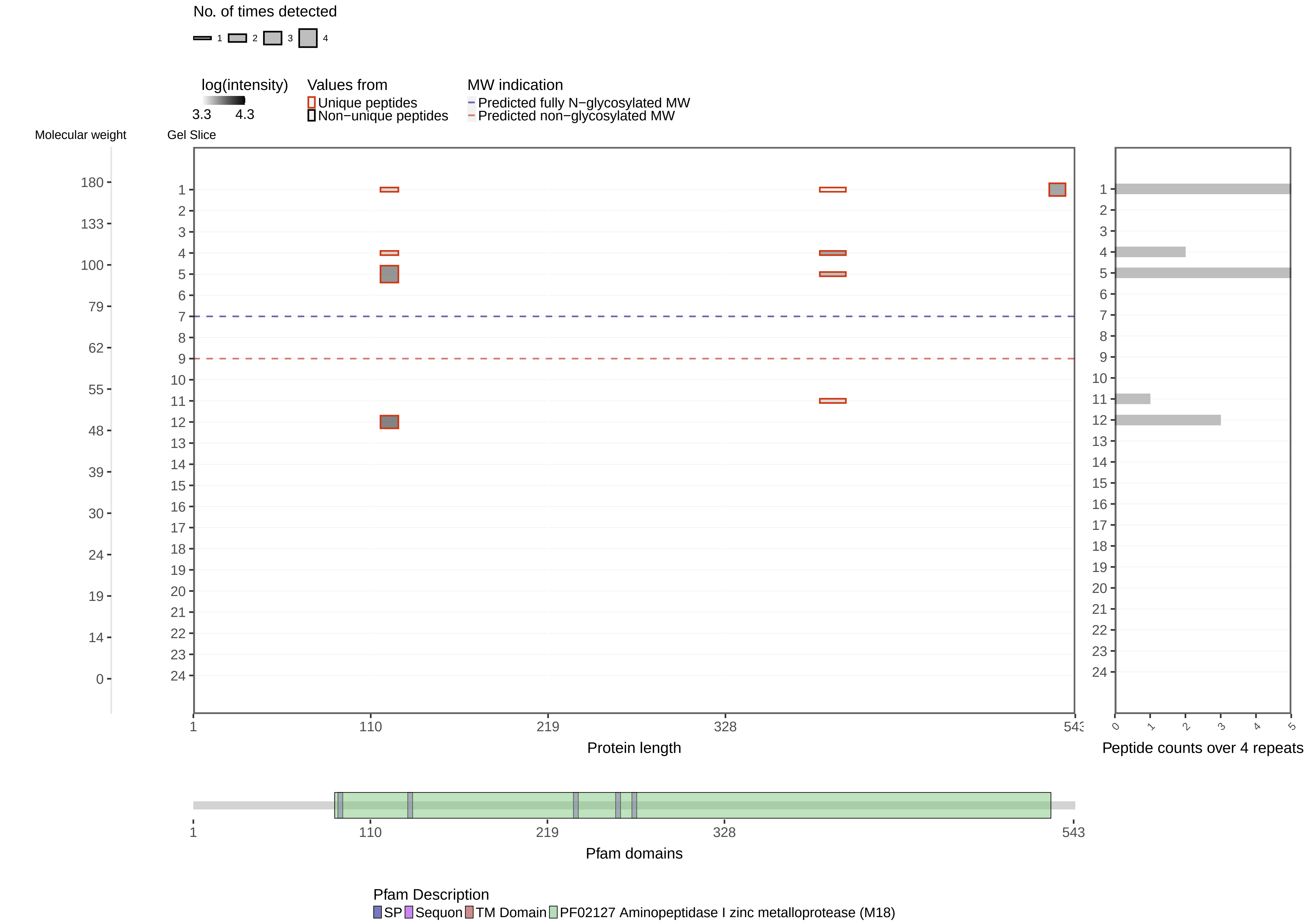


Protein sequence:

TENATADSTEKSNVSSSKDVHHQLLLQILSDELSCKIDDIVSVELNLCDTQPSCLGGANNEFIFSGRLDNLASSFCGLRALVDSCTFPEDLSNEHAIRMIALFDNEEVGSDSYQGAGAPTMFEAMRRITDCLSHQSAGESDFARAICHSYLVSADMAHGVHPNFVDKHEEHHHPELQKGLVIKHNANQRYATSGVTSFLFKEVARLHSLPTQDFVVRNDMGCGSTIGPILASGVGIRTVDCGIAQLSMHSVREICGKEDIDIAYKHFKAFYQTFSSIDRKLIVD*

### NbL13g24250.1 nifU−like_protein_4_mitochondrial [MW = 32.9 kDa]

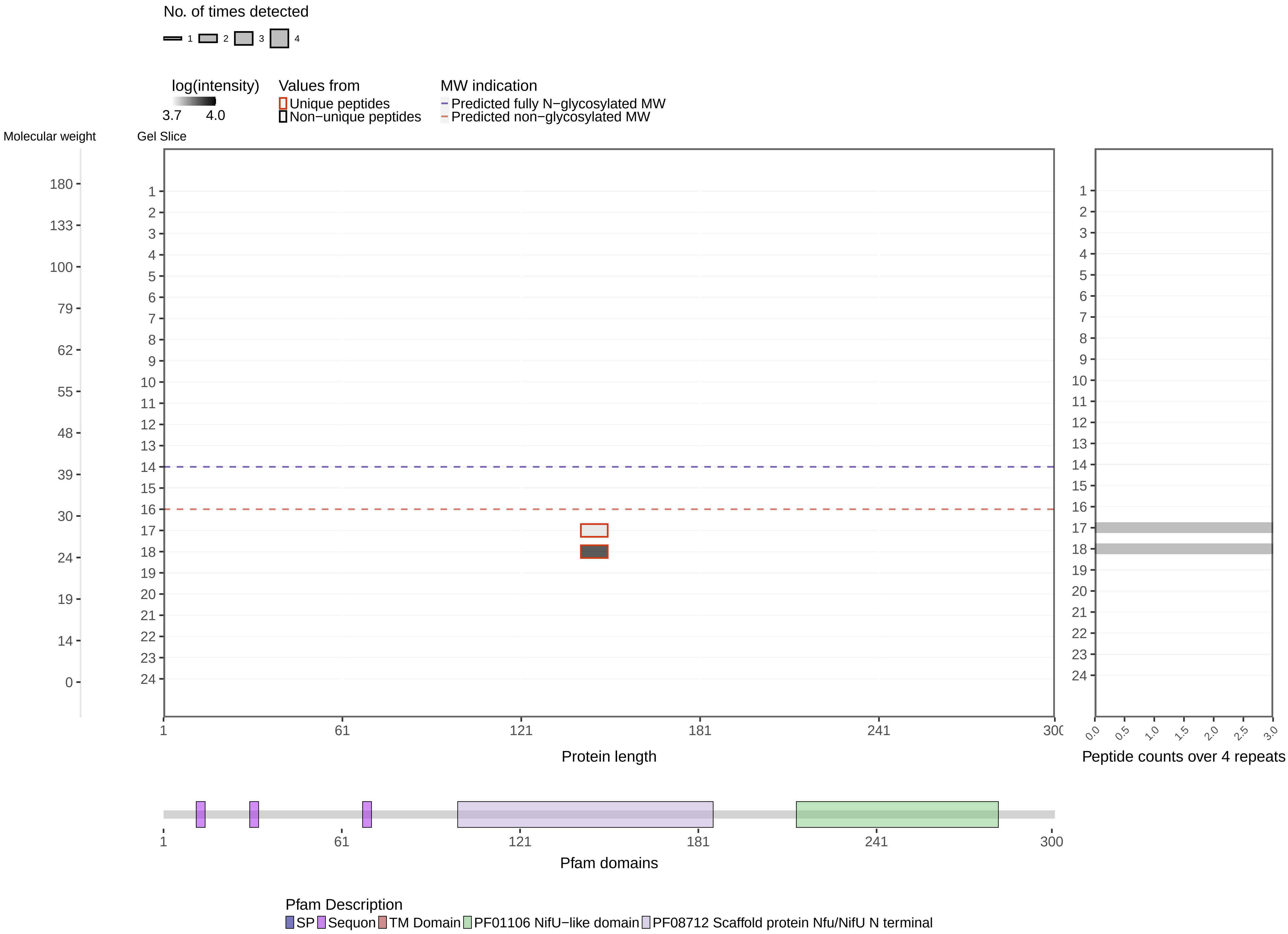


Protein sequence:

PLAKALFGIDGITRVFFGSDFVTVTKSEDASWDFLKPEIFAAIMDFYSSGKPLFLDSSTAASMDTAIHEDDSETVAMIKELLETRIRPAVQDDGGDIVYRGFDLDTGIVKLKMQGACSGCPSSSVTLKSGIENMLMHYVPEVKGVEQEMDEDEEPGLTGAPYE*

### NbL13g24160.1 protein_disulfide−isomerase−like [MW = 57.13 kDa]

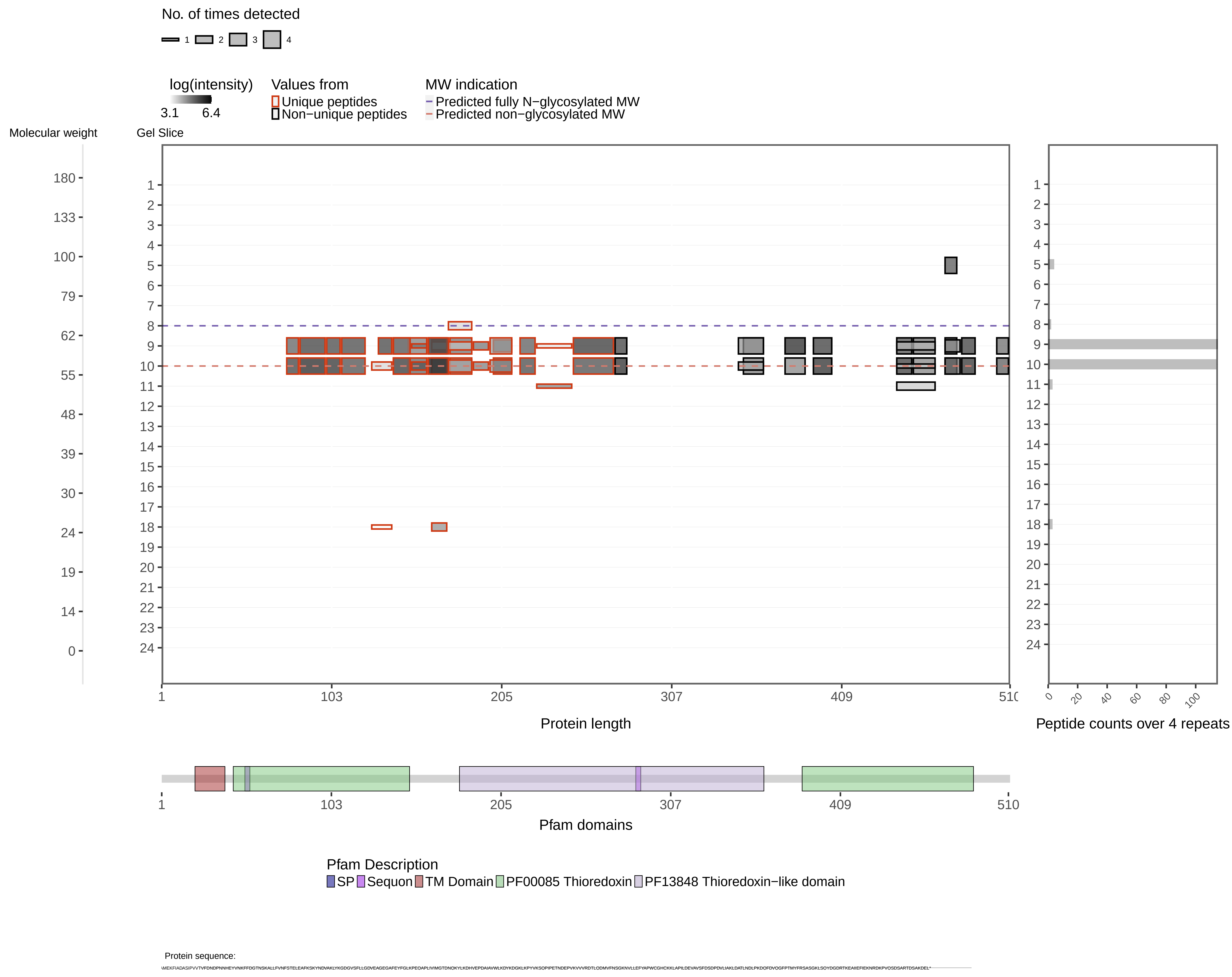

NbL13g24070.1 copper_transport_protein_ATX1−like [MW = 12.45 kDa]

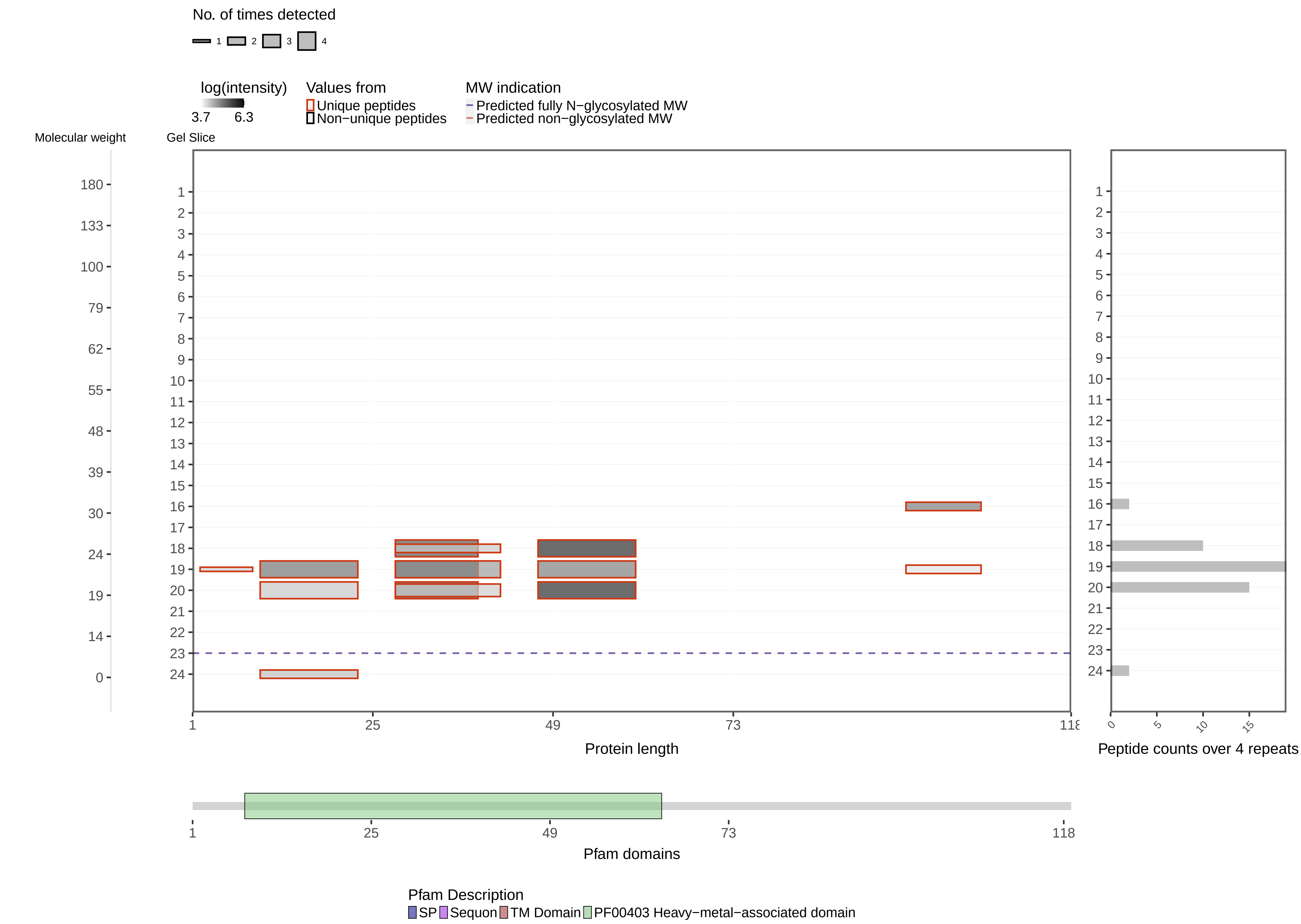


Protein sequence:

/KGNVEPEAVFQTVSKTGKKTSFWEAETPAPAPAPTEPETKPVEVKPTEEKPADTPAEPEAKPADEKPAEAVA*

NbL13g23910.1 probable_bifunctional_TENA−E_protein [MW = 26.33 kDa]

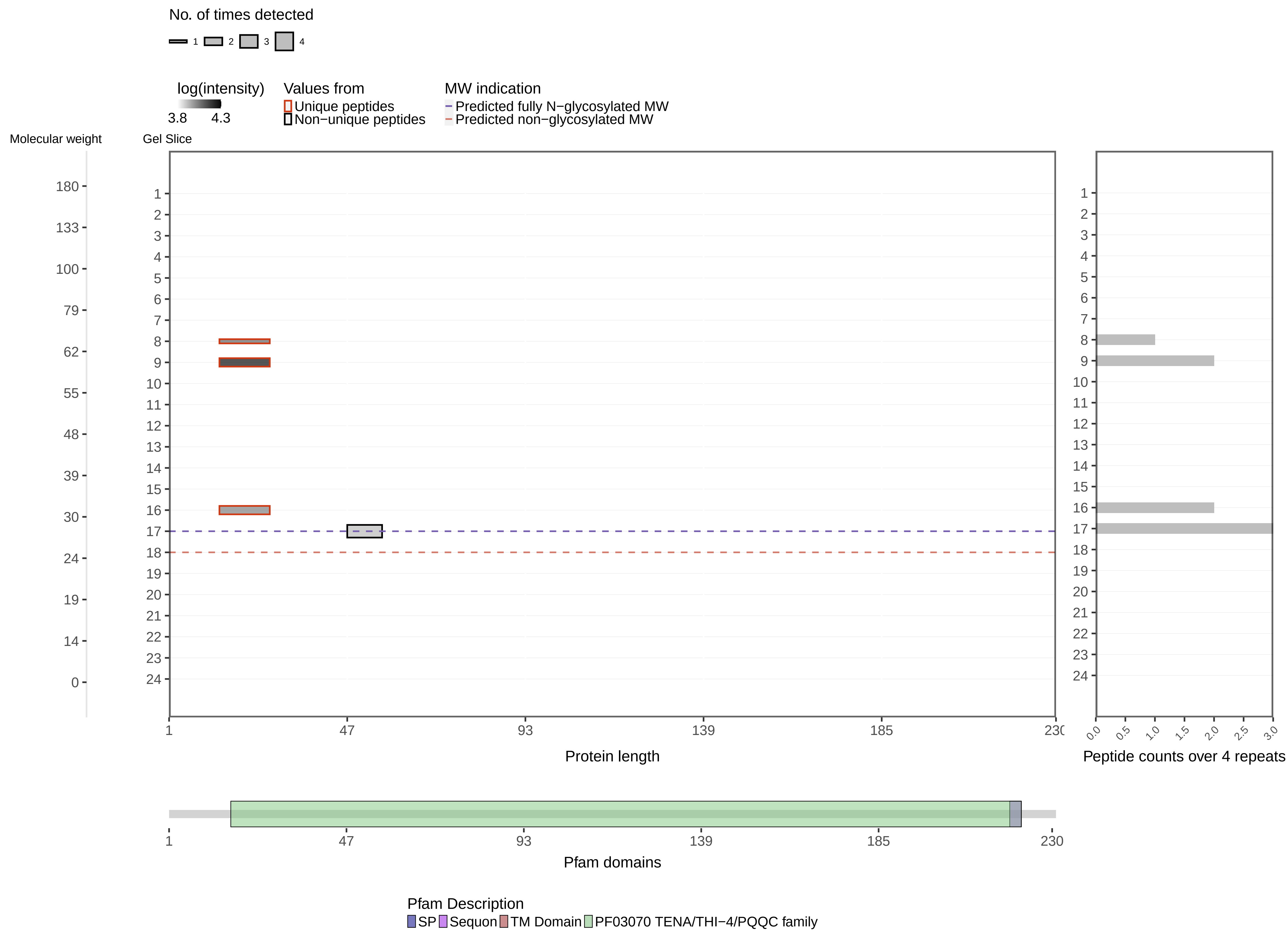


Protein sequence:

WSVSLPSIVPQKANLEYCRFLESLMSPEVEYTVAVTAFWAIEVVYQESYAHCLADGSKTLDELKETCQRWGNDDFGQYCRSLQDIANRRLEKSPEDIISKAEAIVIRVLKYEVEFWNMSRGGKLSAP*

NbL13g23800.1 pyruvate_kinase_1_cytosolic−like [MW = 57.64 kDa]

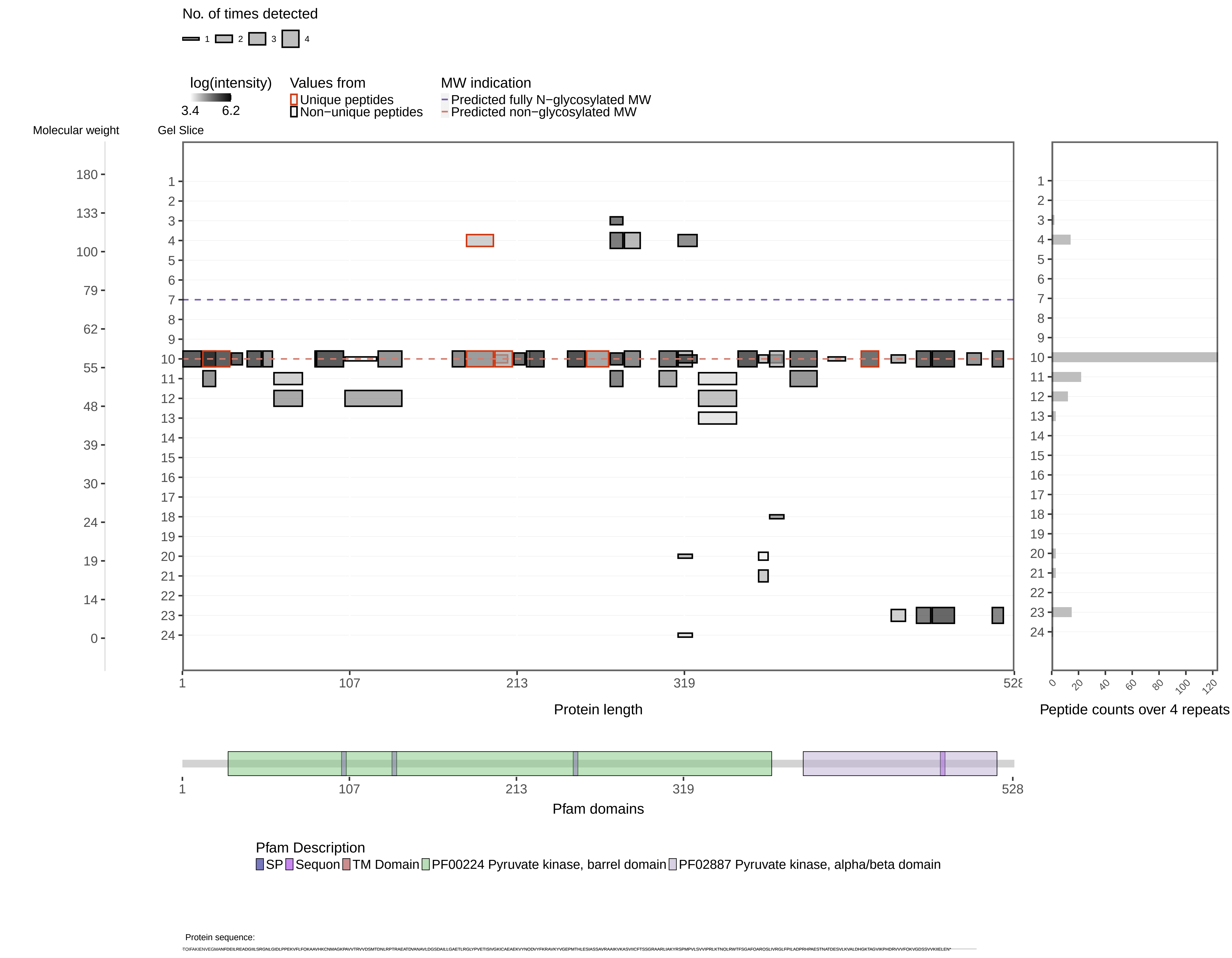

### NbL13g23780.1 uncharacterized_LOC104235088 [MW = 33.47 kDa]

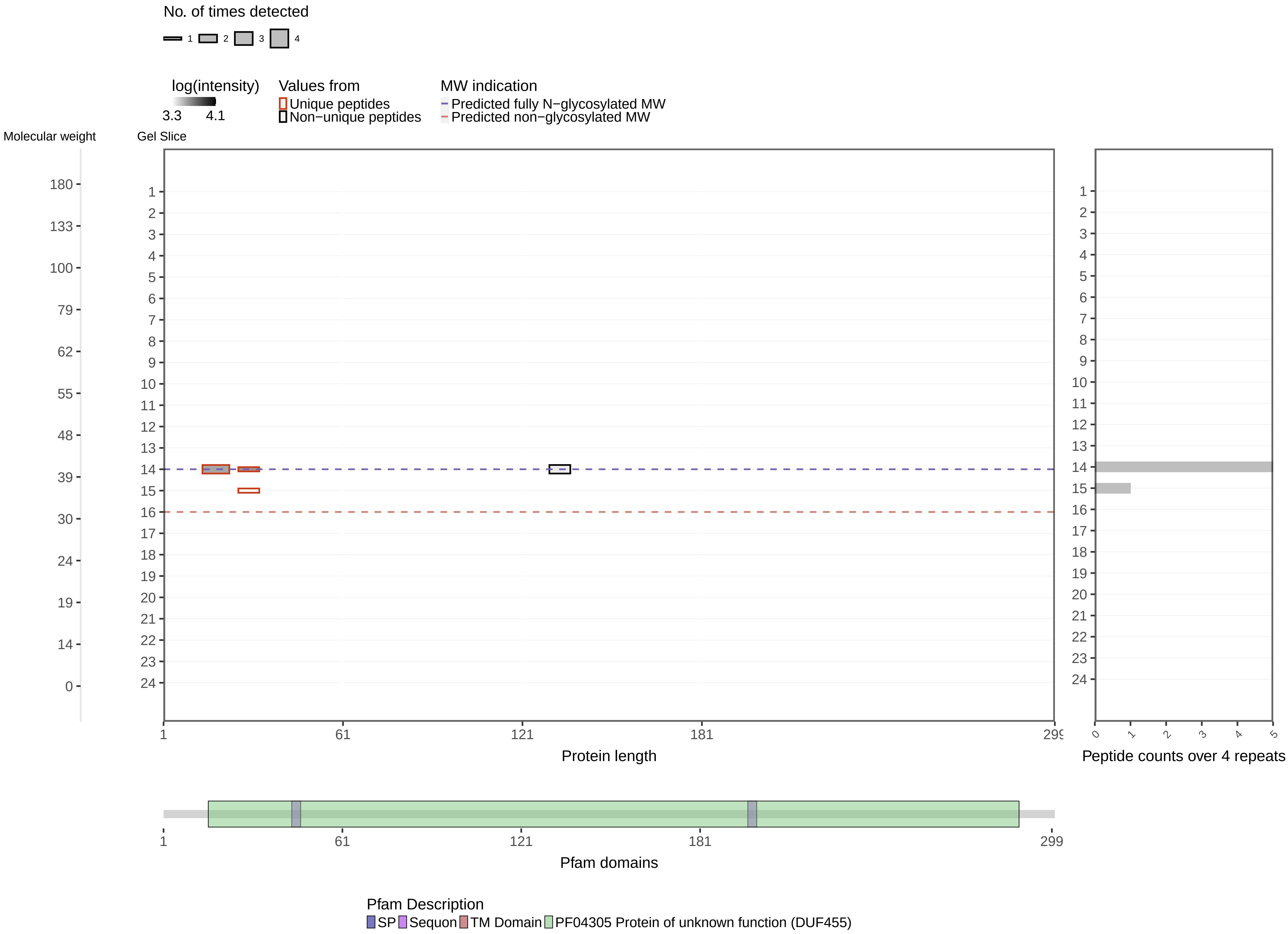


Protein sequence:

LKELGSFYGALPAHDGLWDSAIATSKDLLARLAIEHCVHEARGLDVIPTTISRFRNGGDNQTAELLENVVYPEEITHCAAGVKWFKYLCLRSRNPNPGDLPSQENSDTMDVDNEVIQRFHSTVRTHFRGPLKPPFNEQAREAAGFGPQWYEPLAVKDYTME*

### NbL13g23750.1 ribosome−recycling_factor−like [MW = 32.81 kDa]

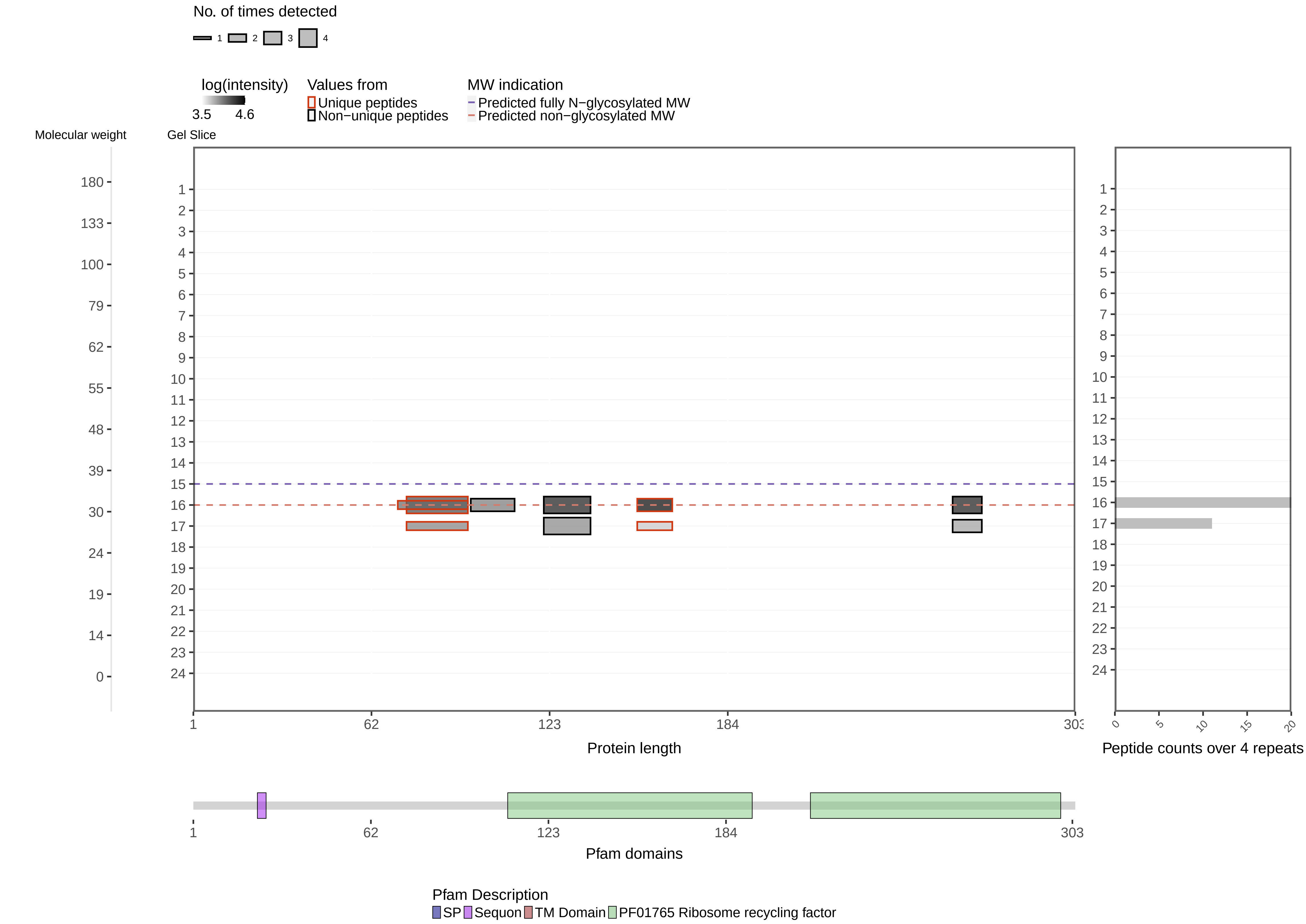


Protein sequence:

TPLCRMAVISVLDPKTLSVTPYDPNTVKELDKAIVSSPLGLNPQVDNQRLIVPIPPFPMLFDSAFLLFLAIRATSRLTKEHVQAVCKVVAKSSEDVKQSVRRARQKALDTIKKSLPRKKDKDKAASAFSEDDAKRLEKEIDDLTKKFIKSAEDMCKSKEKEITSS*

### NbL13g23650.1 5−epi−aristolochene_synthase−like [MW = 63.13 kDa]

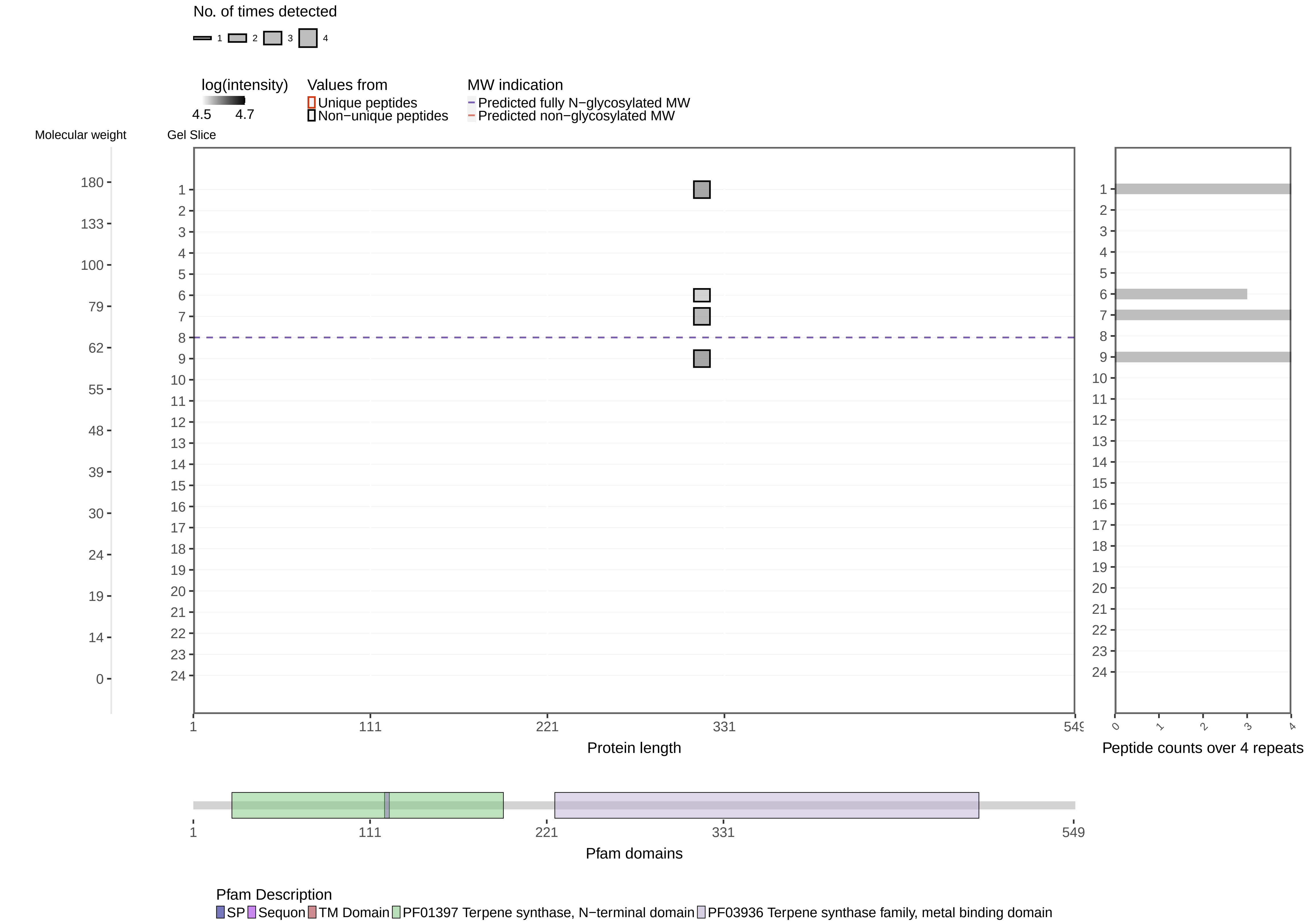


Protein sequence:

PYARDRVVECYFWALGVYFEPQYSQARVMLVKTISMISIVDDTFDAYGTVKELEAYTDAIQRWHIKEIDRLPDYMKISYKAILDLYNDYEKELSSAGRSHIVSHAIQRMKEVVRNYNVESTWFIEGYKPPVSEYLSNALATTTYYYLATTSYLGMKSVTEQDFEWLSKNPKILEASVIICRVIDDTATYEVEKSRGQIATGIECCMRDYGLSTKEAMDKFQKMAETAWKDLNEGLLRPTPVSAEFLNPILNLARIVEVTYIHNLDGYTHPEKVLKPHIVALLVDSIQI*

### NbL13g23640.1 5−epi−aristolochene_synthase−like [MW = 63.09 kDa]

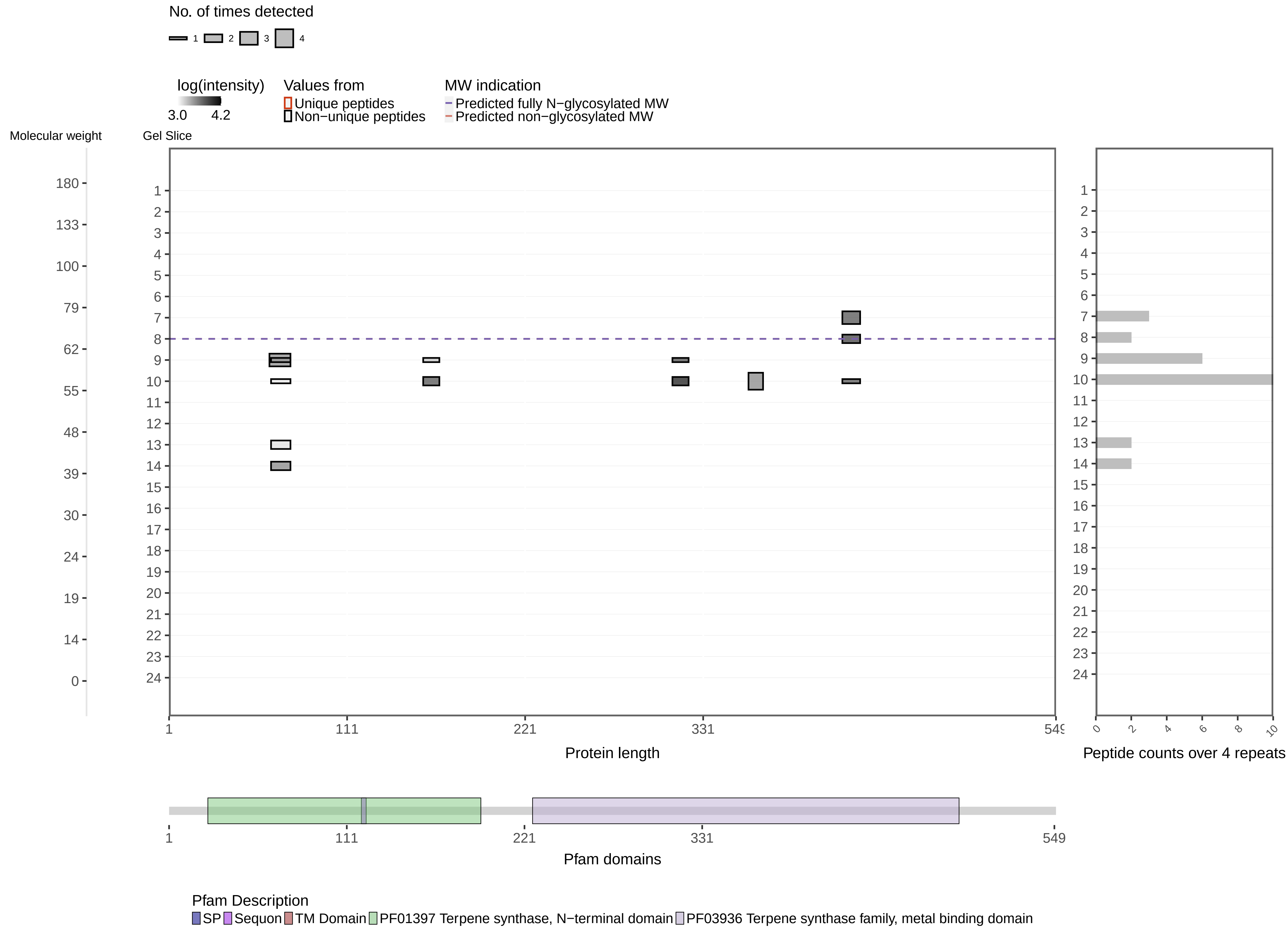


Protein sequence:

[illegible]

### NbL13g23610.1 phosphoglucan_phosphatase_DSP4_amyloplastic [MW = 46.85 kDa]

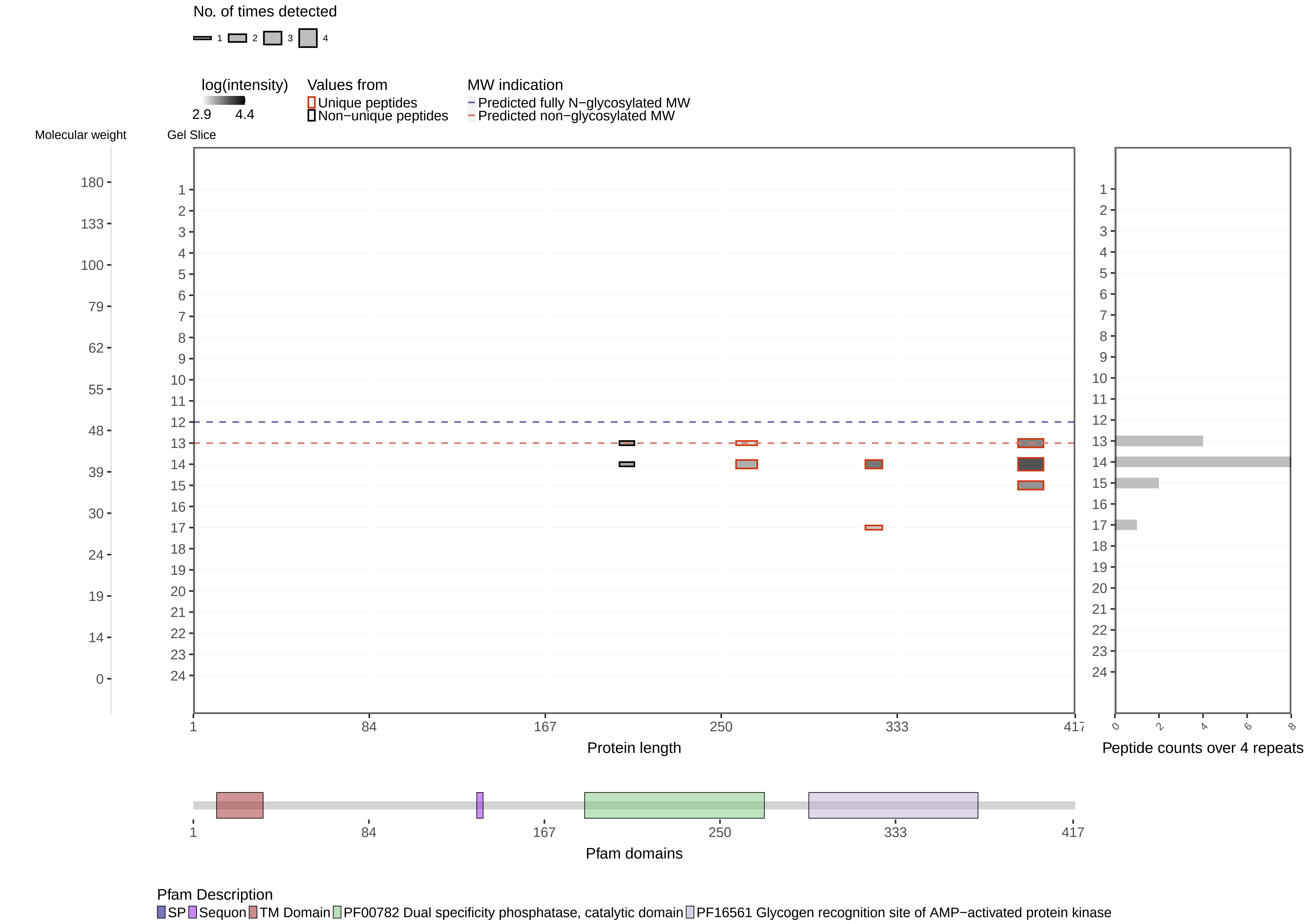


Protein sequence:

.RAEIRDFDAFDLRLRLPAVVSKLSKAVNRNGGVTYIHCTAGLGRAPAVALTYMFWVQGYKLGEAFNLLMSKRSCFPKLDAIKSATADILTGLKRKPATLTWSGDDCSTVEISGLDIGWGQRIPLEFDEERGLWTLQRDLPEGHYEYKYIVDGVWTCNEFEPITSPNKDGHVNNYVKQVLDDNPDSISAAVRNRLTSDDPDLTSDERLIIREFLEAYPDEE*

### NbL13g23340.1 malate_dehydrogenase_[NADP]_chloroplastic−like [MW = 48.31 kDa]

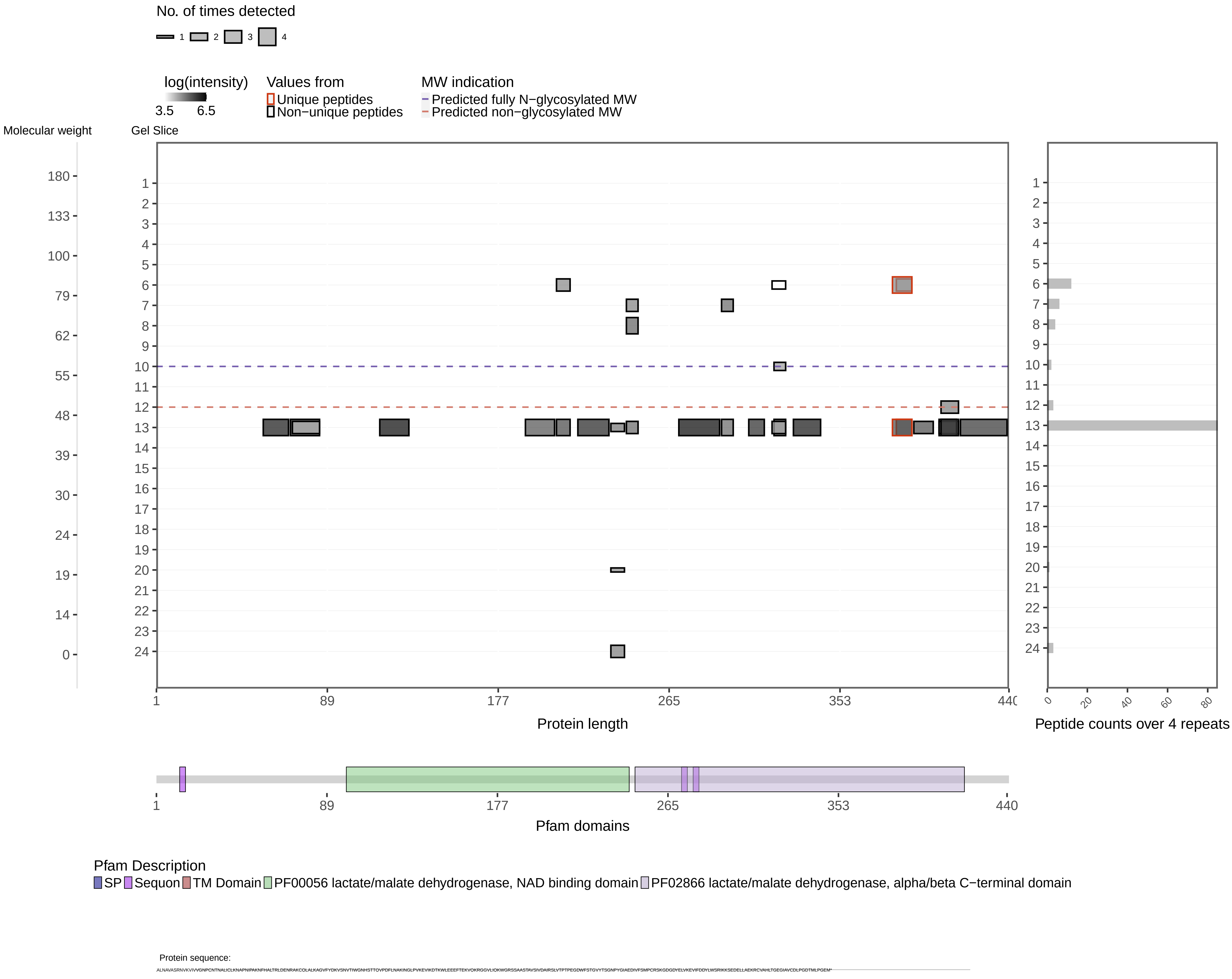

NbL13g23330.1 5−epi−aristolochene_synthase [MW = 64.08 kDa]

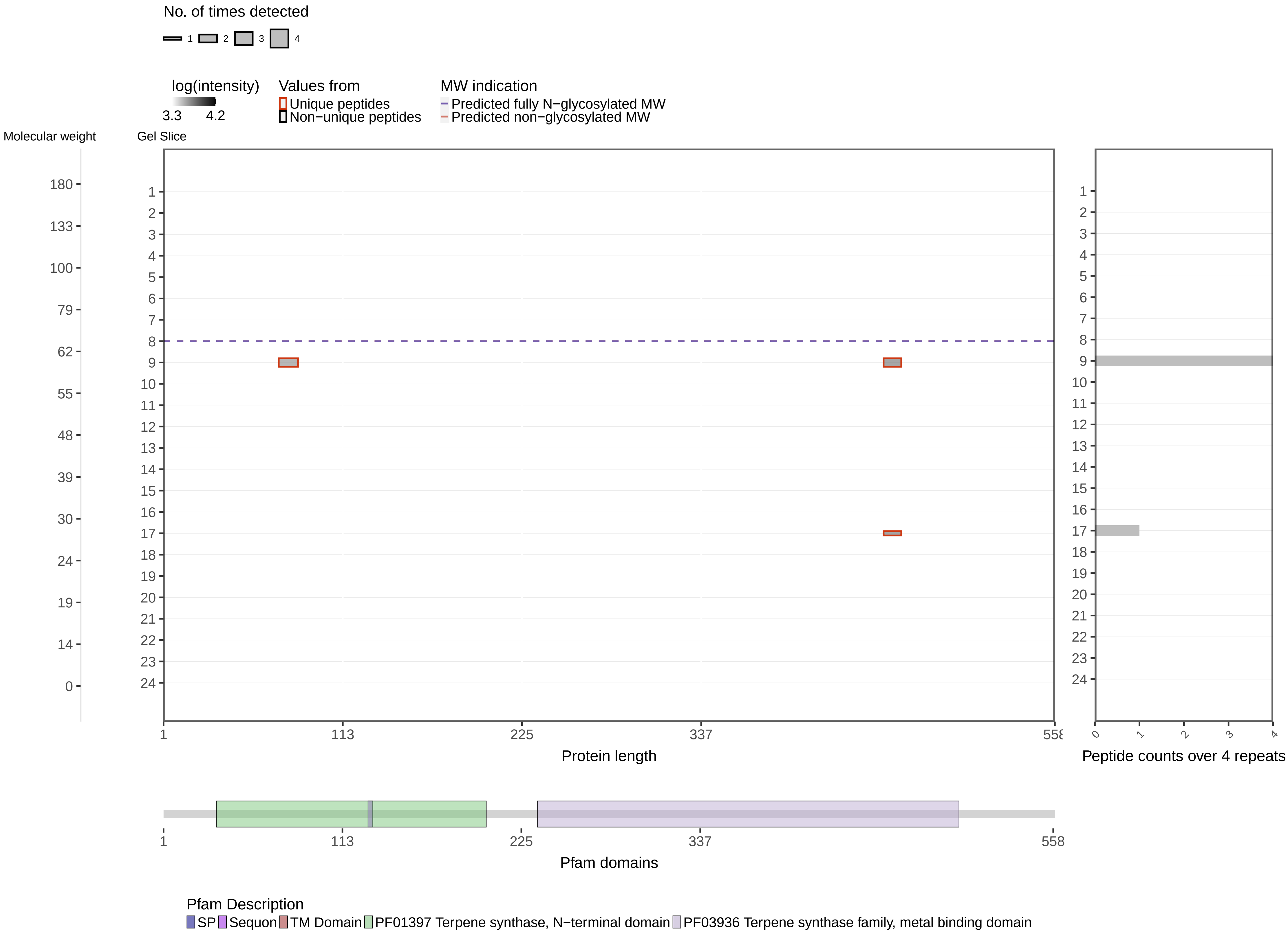


Protein sequence:

FVTTLPYARDRVVECYFWALGVYFEPQYSQARVMLVKTISMISIVDDTFDAYGTVKELEAYTNAIQRWDISEIDMLPDYMKISYKAILDLYKDYEKELSSVGRSHIVCHAIERMKEVVRNYNVESTWFIEGYKPPVSEYLSNALATTTYYYLATTSYLGMKSVTEHDFEWLSKNPKILEASVIICRAIDDTATYEVEKSRGQIATGIECCMRDYGISTEEAMTKFQEMAEAAWKDLNEGLLRPTPVSTEFLSRILNLARIVEVTYIHNLDGYTHPEKVLKPHINALVVDSIEI*

### NbL13g23310.1 protein_BTR1−like [MW = 39.37 kDa]

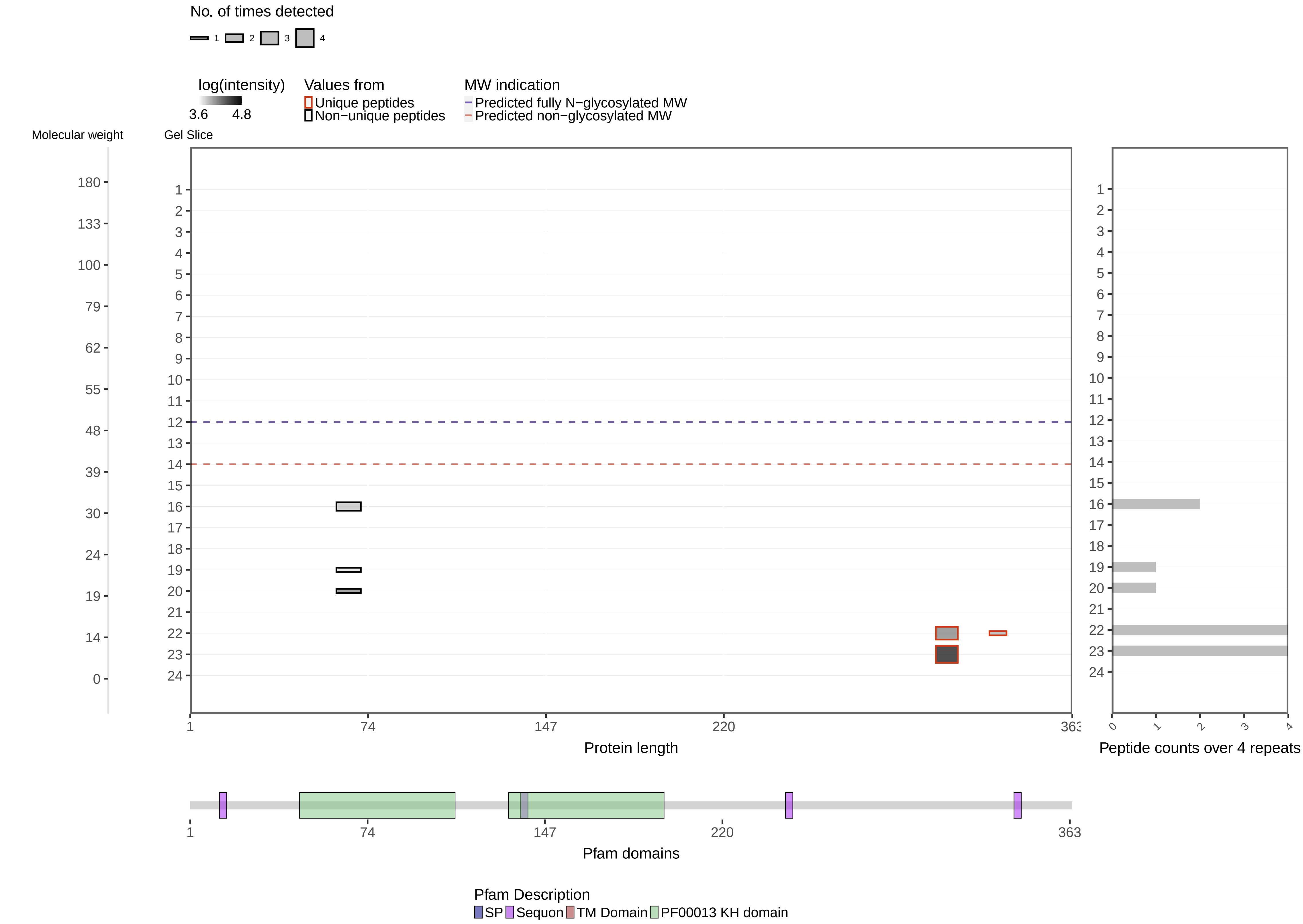


Protein sequence:

NNFPGLHDRVVTVTGTLDEQMRAIELILCKLAEDPHYMQSMNAPFQYAAVYVGMNYGPPNGVGGRYPNNRHQNKEDRNNTVTIGVADEHVGLVLGRNGRSIMEISQVCKKLLFCLVCIYAKSLLSLQLSGARIKISDRGDFLSGTSDRKVTITVSQRAIRTAESMISRKIANVSERSTVGEDLPFEELGINTVH*

### NbL13g23290.1 acyl−CoA−binding_domain−containing_protein_4 [MW = 56.75 kDa]

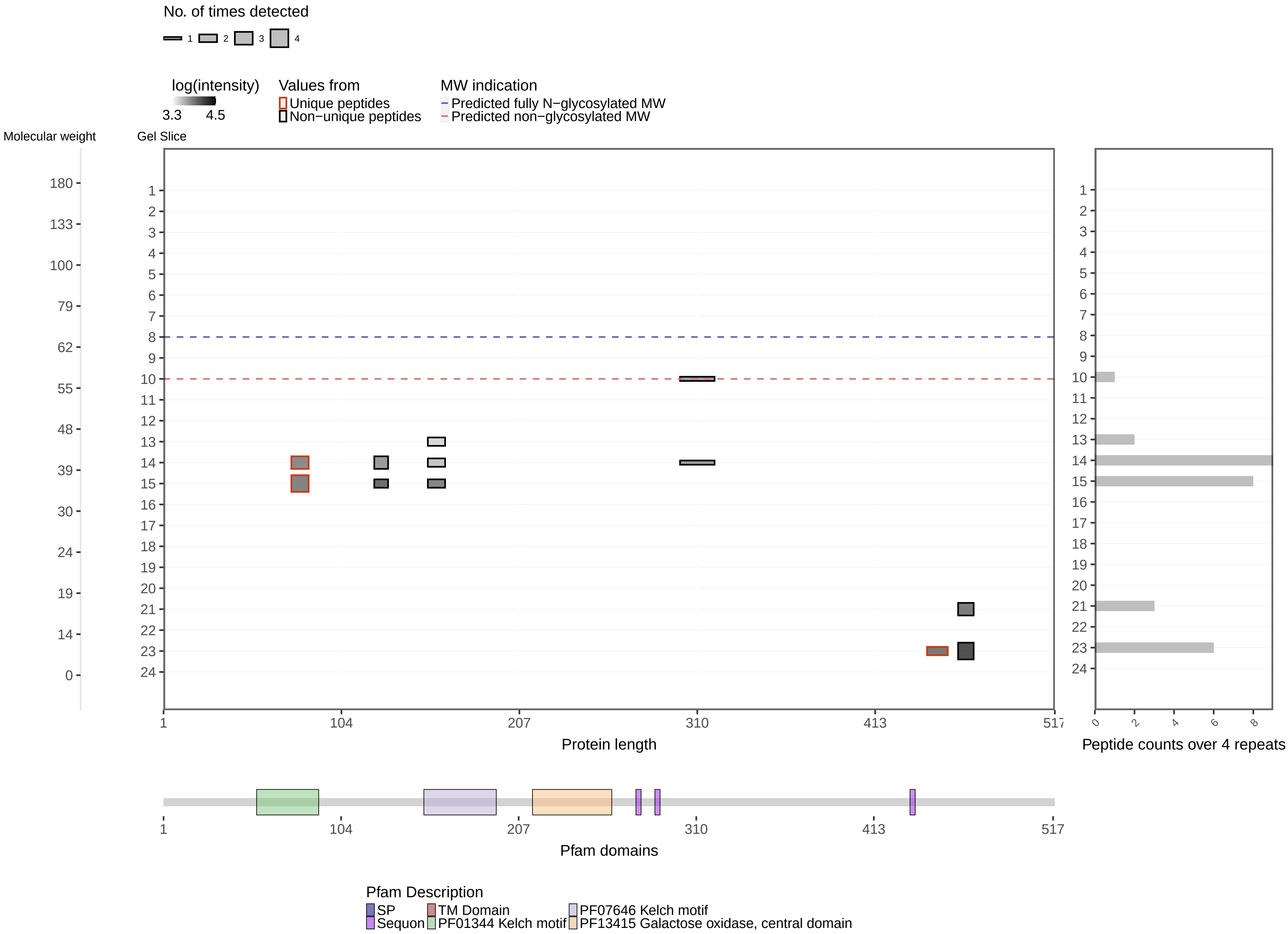


Protein sequence:

GDLVSPRAGHAGVTLGDNWYIVGGGDNRSGVPETLVLNMSKLVVSVLATVKGRDPLASEGLSVSSALLDGEKFLVAFGGYNGKYNNEVYVMRPKPRDVLHPKMLQSPAAEAAAASVRAAYALTKPEKLDLSERENSNFKEVHVDNTQQKLSAEISAHGLQKKALESSLADVQAENANLKAKIEEVNSTHTDLSKELQSVQGQLISERSTCAKLEAQIAELQKMLGSMQSIEEEVQALRKEKSELERDMERAVSVQKQGSGGAWKWITG*

### NbL13g23170.1 uricase−2_isozyme_1 [MW = 34.72 kDa]

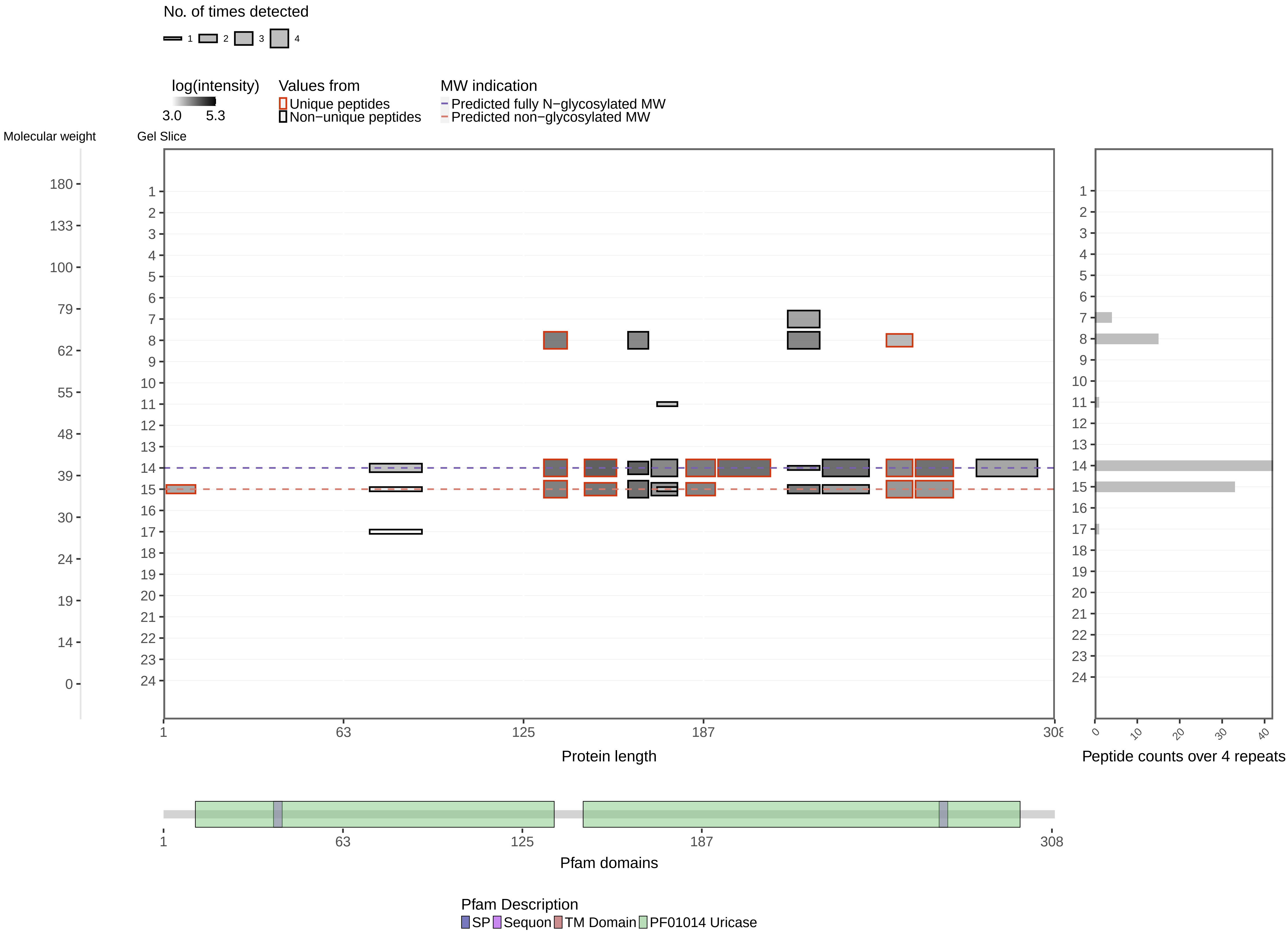


Protein sequence:

SGTLRMTSGIVGLSVLKTTKSGFEGFIRDKYTMLPETRERMMATEVTASWRYLFESLSSLPLKPLYFTERYMDVKEVLLNTFFGPSKGGVYSPSVQATLYEMAKAILGRFPDVSFIELKMPNLHFLPVNLTSKDNPVIVKFDNDVYLPTDEPHGTIQATLNRIRSKM*

### NbL13g23030.1 phosphoribosylaminoimidazole_carboxylase_chloroplastic [MW = 68.47 kDa]

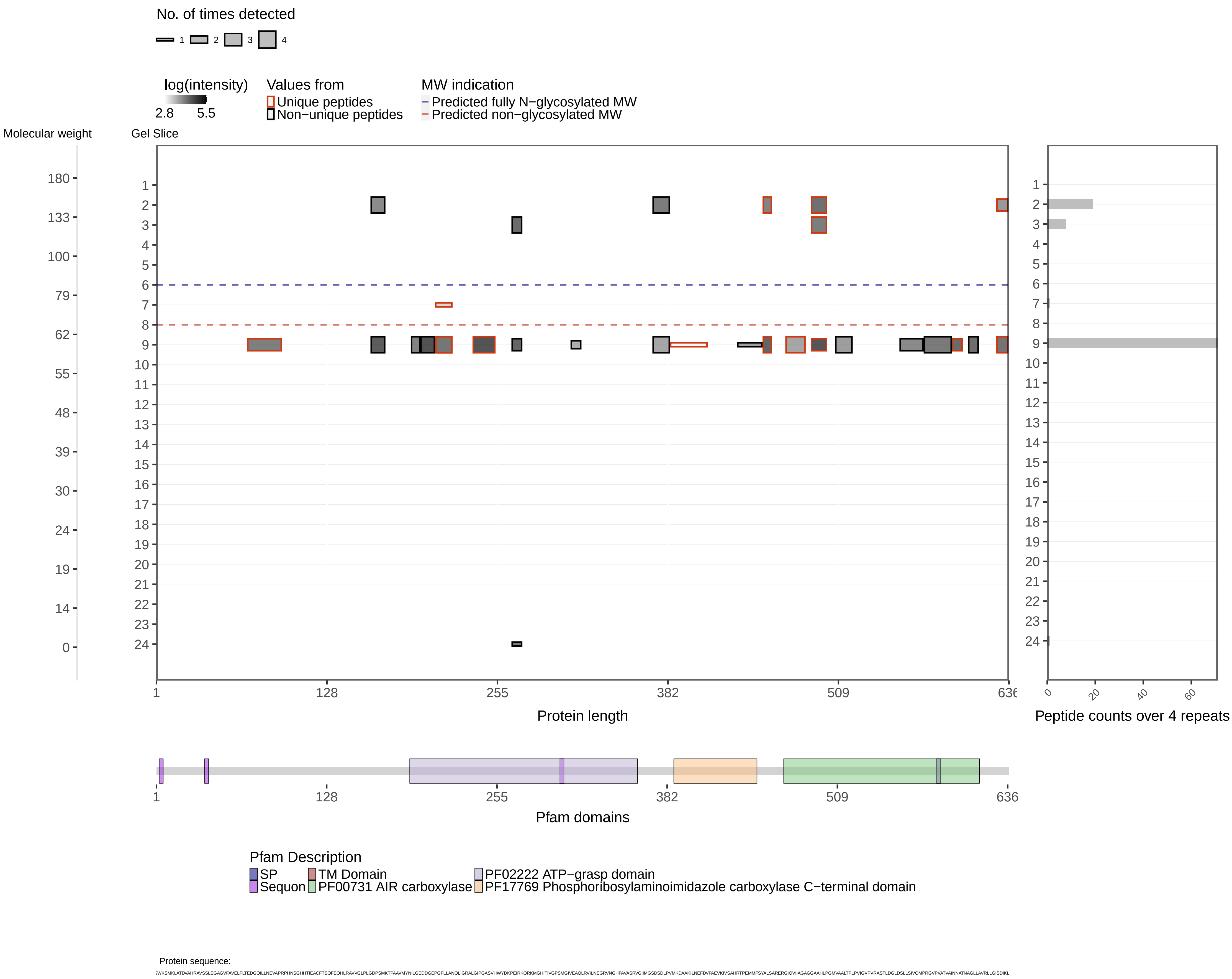

### NbL13g23020.1 ubiquitin_carboxyl−terminal_hydrolase_12−like [MW = 136.83 kDa]

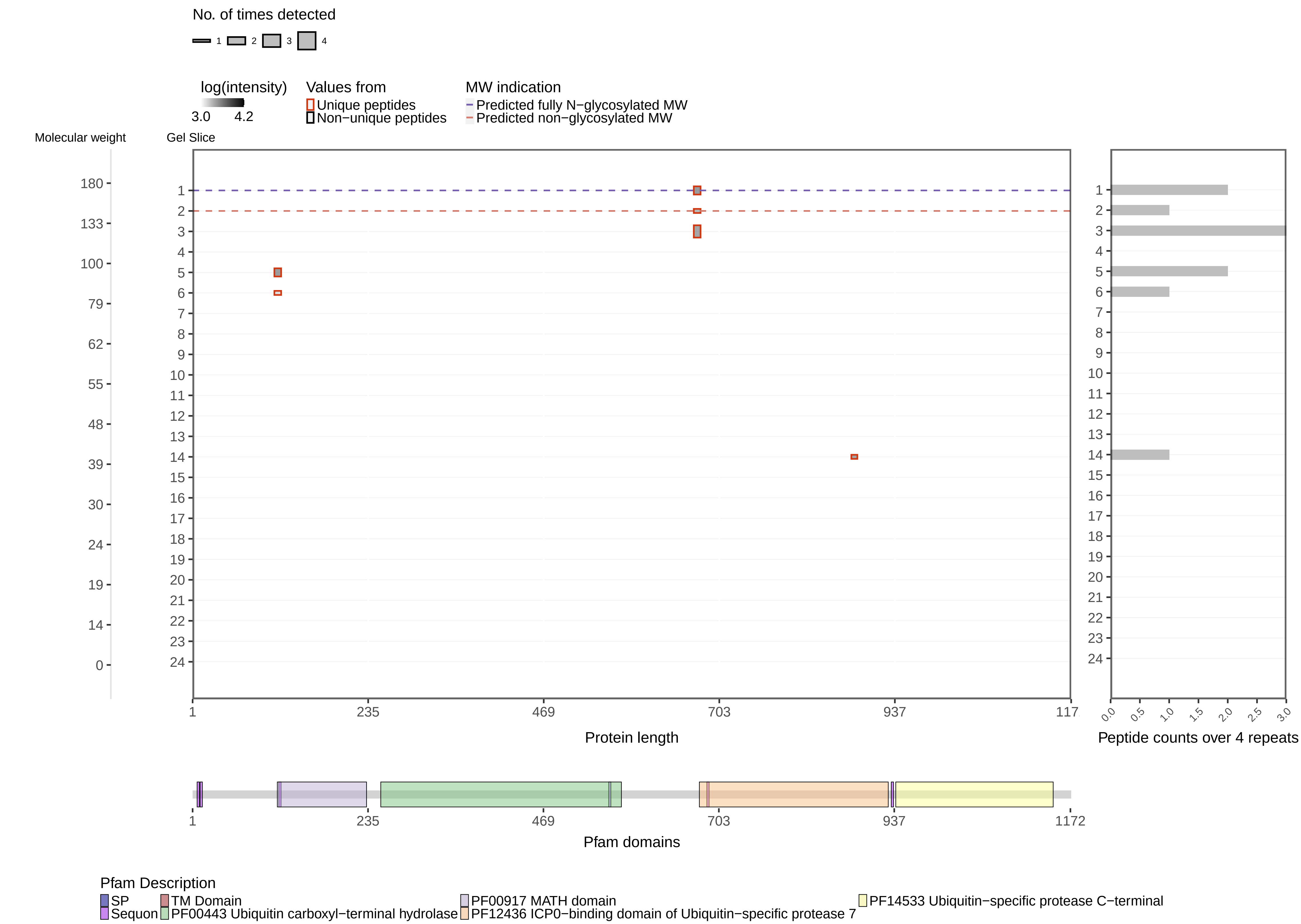


Protein sequence:

AYMLVYIRESDKEKIICNVDEKDIAEHLRVRLKKEODEKEOKRKEKAEAHLYTIIKVARDEDLGEOIGKEIHFDLVDHDKVHSFRIOKOMAFTOFKEEVAKELGIPVOLORYWLWAKRONHTYRPNRPLTPOEEAOSVGOLREVSNKANNAELKLFLEVEFDLDLRPCPPPEKTKEDILLFFKLYDPLKEEMRYIGRLFVKGSGKPLEILPKLNELAGFSPDEEIELFEEIKFDPNVMCEPIDWKLTFRGSOLEDGDIICFOKPLRSOTSEOYRFPDVPSFLEYVHNROVVRFRSLE

### NbL13g22990.1 GDSL_esterase/lipase_At5g33370−like [MW = 41.49 kDa]

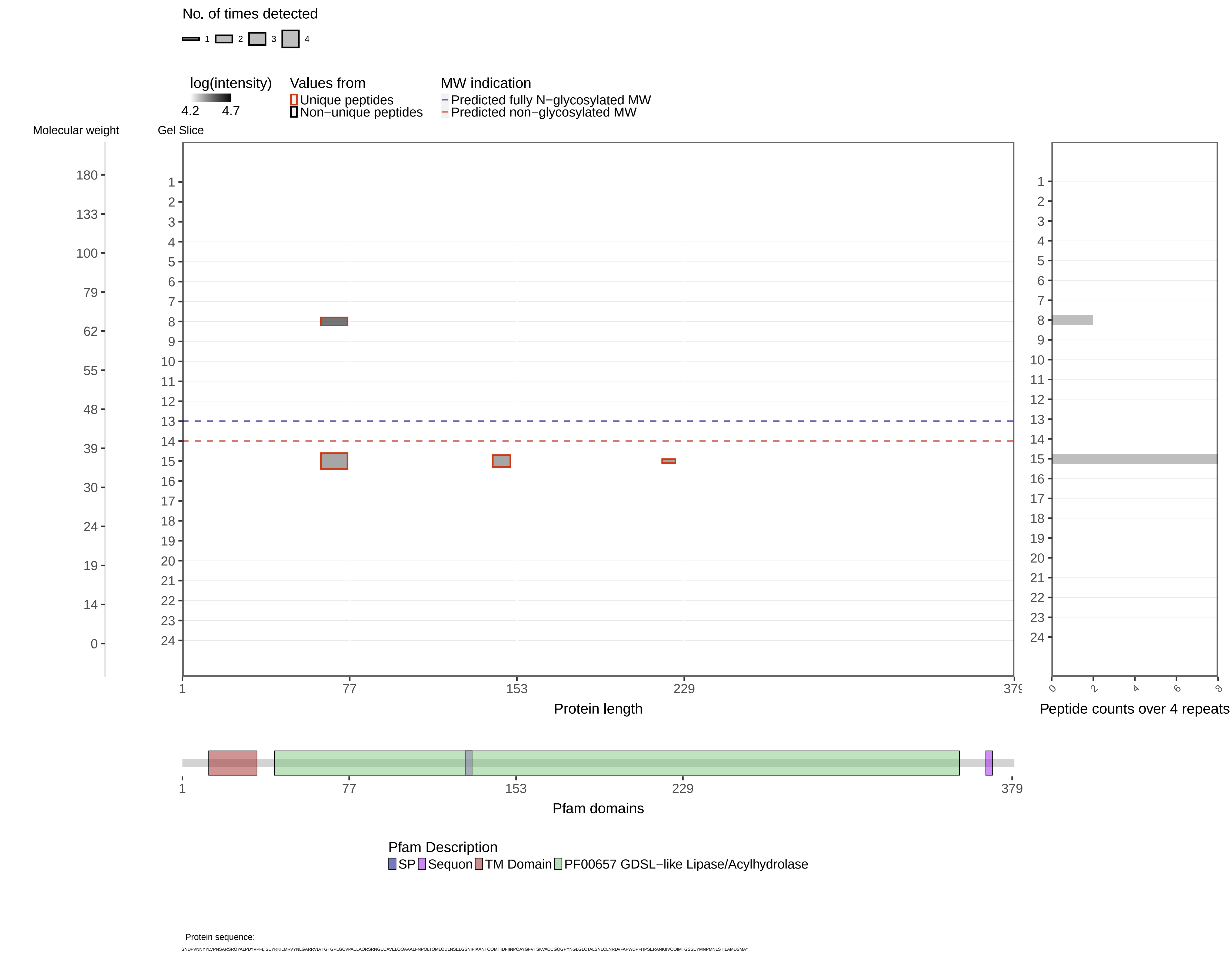

NbL13g22920.1 small_nuclear_ribonucleoprotein_Sm_D2−like [MW = 12.06 kDa]

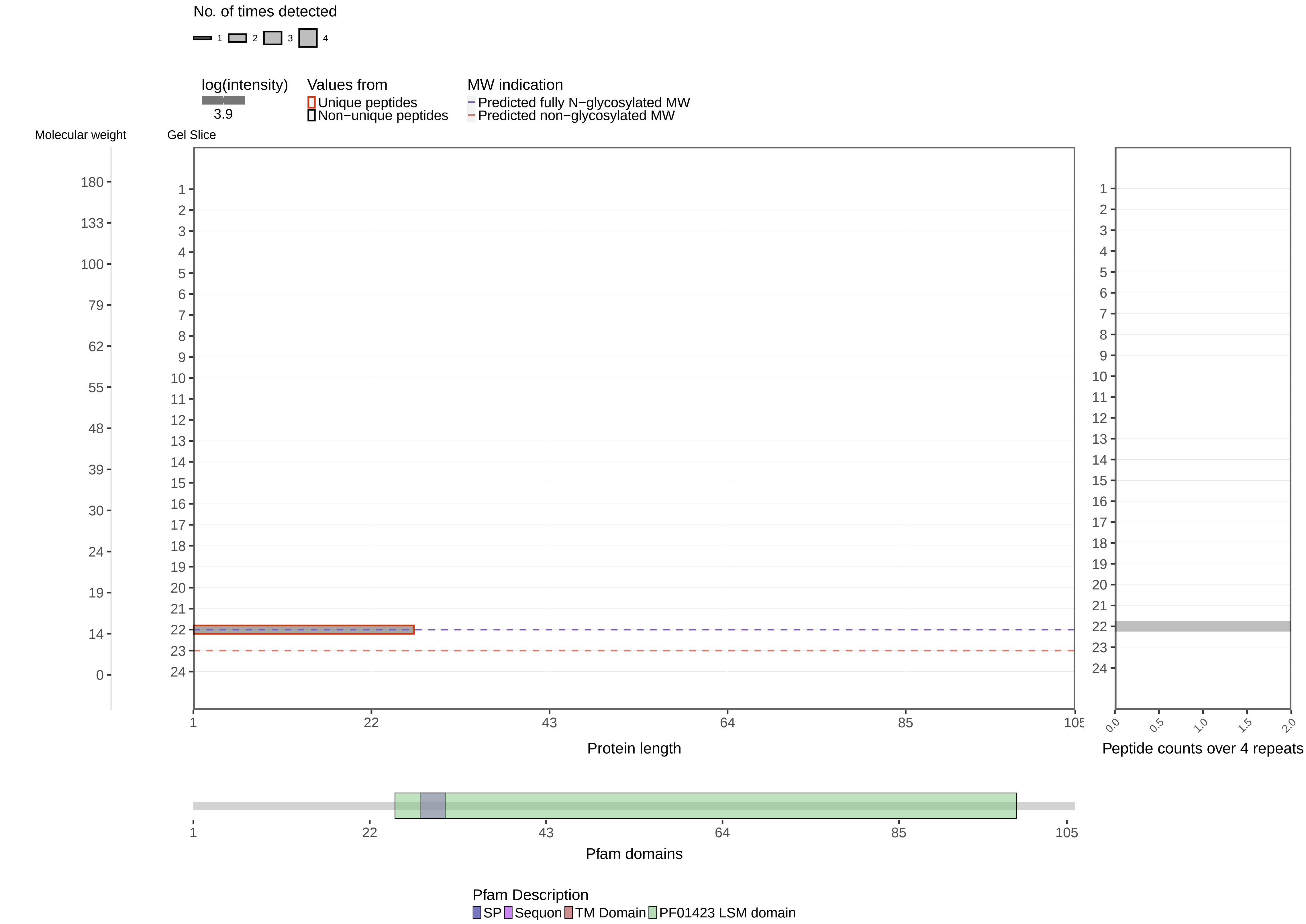


Protein sequence:

RKLLGRVRAFDRHCNMVLENVREMWTEVPKTGKGKKKAQPVNKDRFISKMFLRGDSVIIVLRNPK*

### NbL13g22910.1 NAD(P)H−quinone_oxidoreductase_subunit_O_chloroplastic [MW = 16.72 kDa]

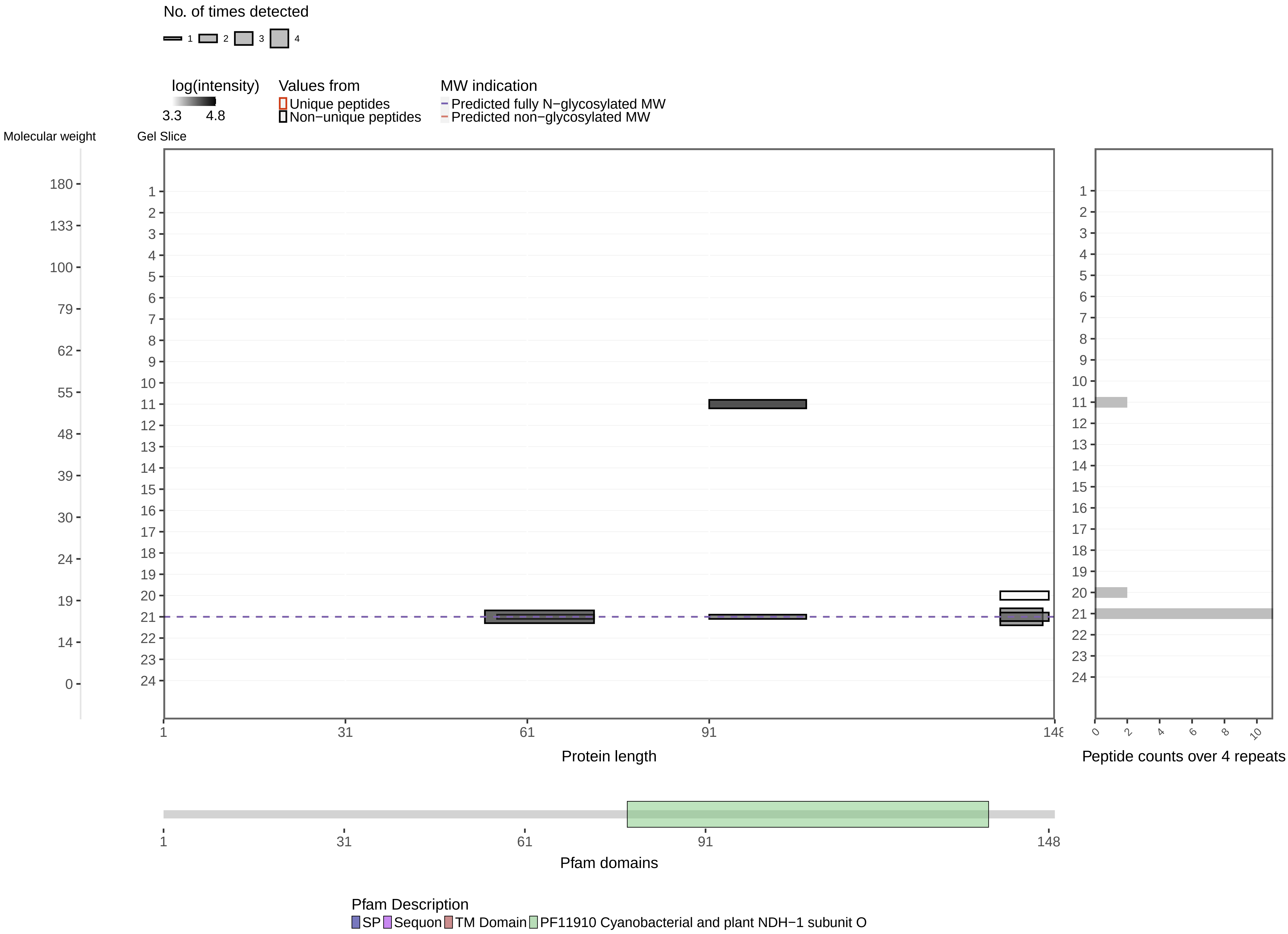


Protein sequence:

ATSSTATATKPKKPVYSMKKGQIVRVDKEKYLNSINYLSVDHPPYYKGLDYIYEDRGEIGWIGIPTAPAWLPTEMLIKSDKLDYERI*

### NbL13g22890.1 receptor_protein_kinase_TMK1−like [MW − SP = 100.84 kDa]

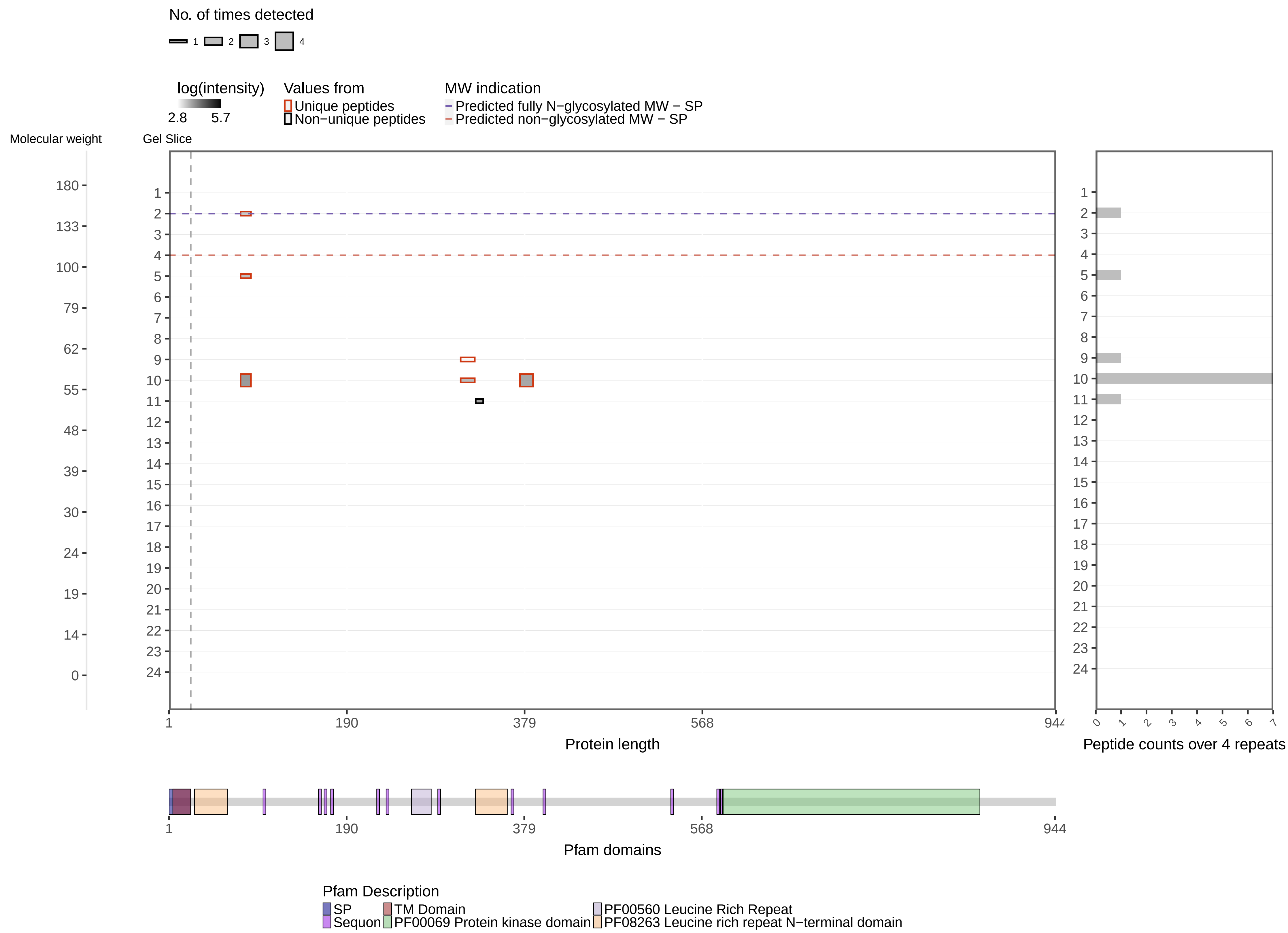


Protein sequence:

PVGSPGSGYDGDAQASRKKSNRWIGIVLFSVIGGVFVLCFIGTAAFCLYRSKQKRFSQVQSPNTVVMQPRNSGSDNDSVKFTVAGSSVVSVGAVSEVHTVSTSETGDIQMVEAGNMVISIQVLKNVTNNFSEENILGQGGFGTVYKGELHDGTKIAVKRMECGFIAGKGLTEFKSEIAVLTKVRHRHLVGLLGYCLDGKEKLLVYEYMPQGTLSSHLFNWREEALKPLEWTKRLTIALDVARGVEYLHSLAHQSFIHRDLKPSNILLGDDMRAKVADFGLVRLAPEGKGSIQT

### NbL13g22830.1 peptidyl−prolyl_cis−trans_isomerase_CYP19−3−like [MW = 19.03 kDa]

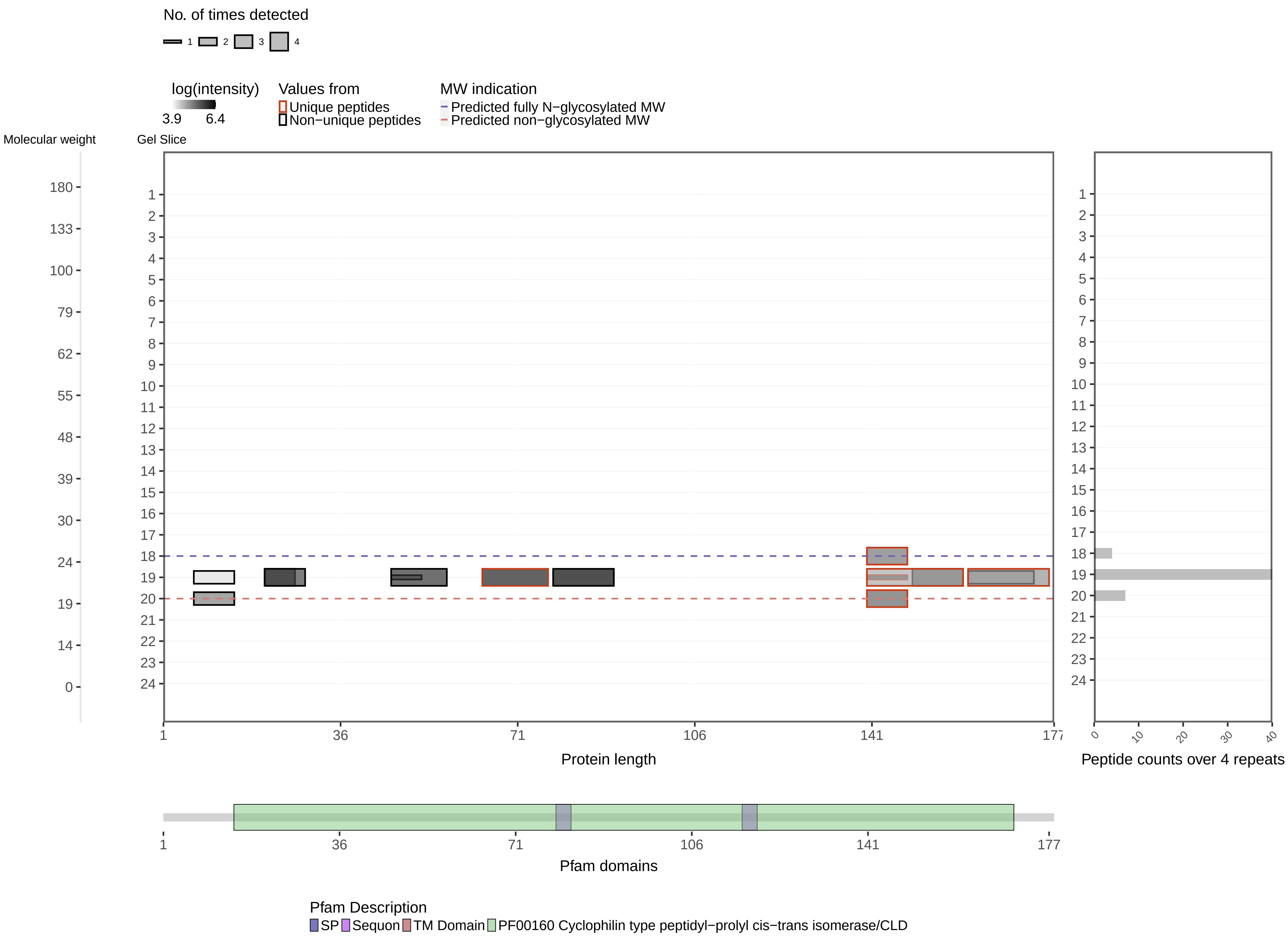


Protein sequence:

NbL13g22760.1 AT−hook_motif_nuclear−localized_protein_14−like [MW = 33.96 kDa]

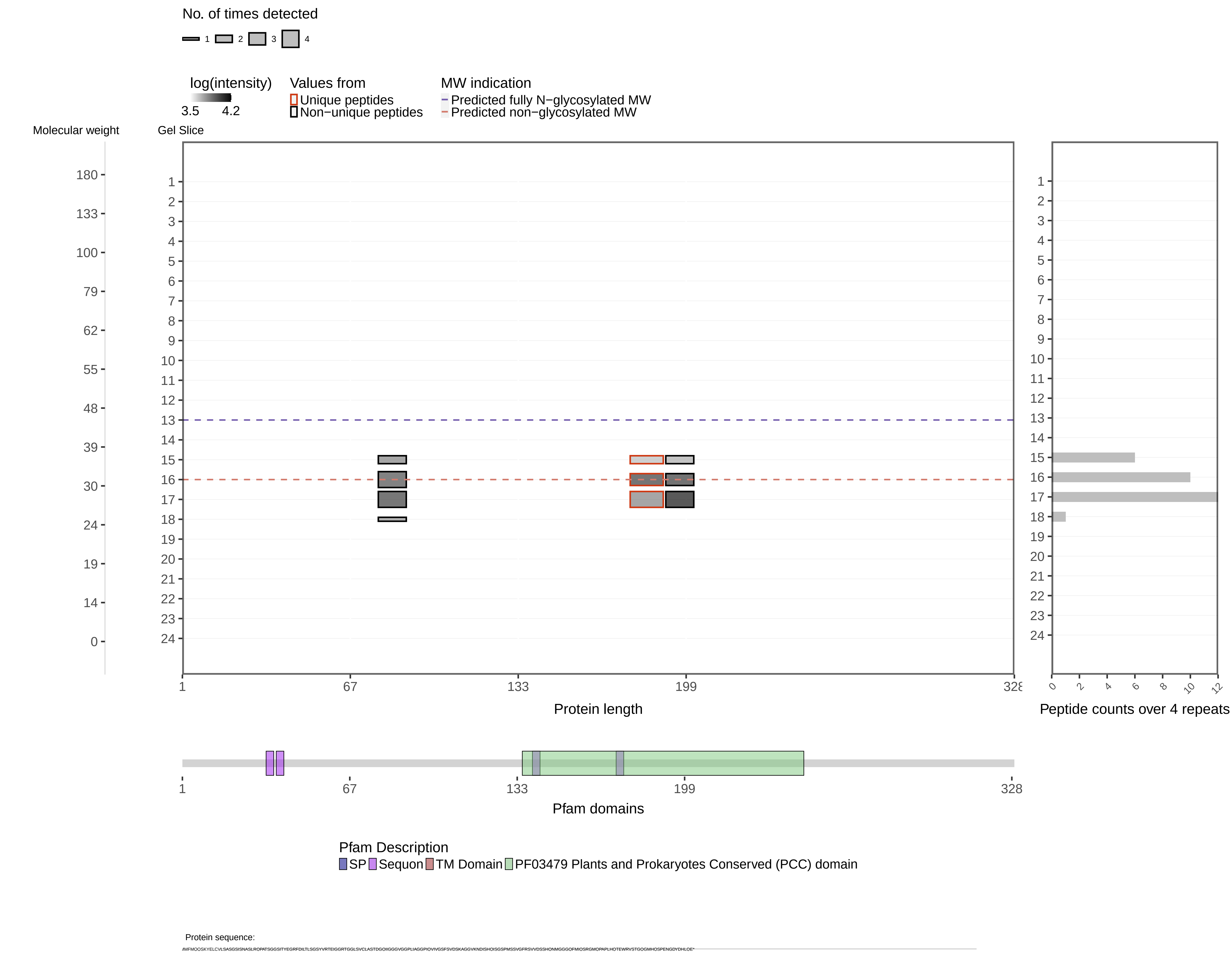

NbL13g22740.1 uncharacterized_LOC109225749 [MW = 24.42 kDa]

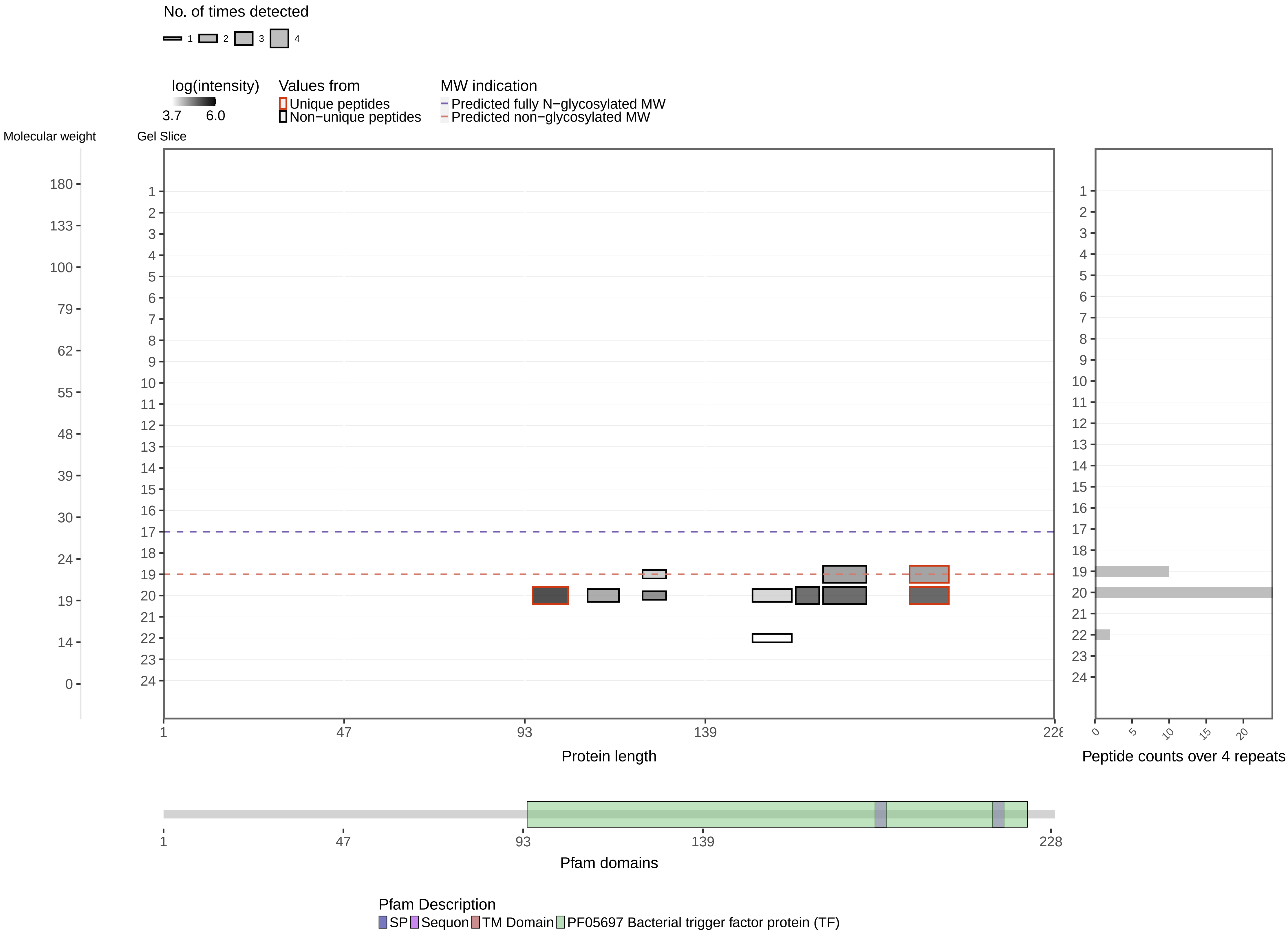


Protein sequence:

DAEKIHVRVDVDKEDTRIVFEKVLANLARSAPPVPGFRREKGGKTSKVPRDFLLQILGEDRVTNFVIREIVTSTLADYVKKENLSVKDNKISTTQTAHELKSSFTIGAEFGFNATLELEDAKTEATT*

NbL13g22560.1 5−methyltetrahydropteroyltriglutamate−−homocysteine_methyltransferase−like [MW = 84.52 kDa]

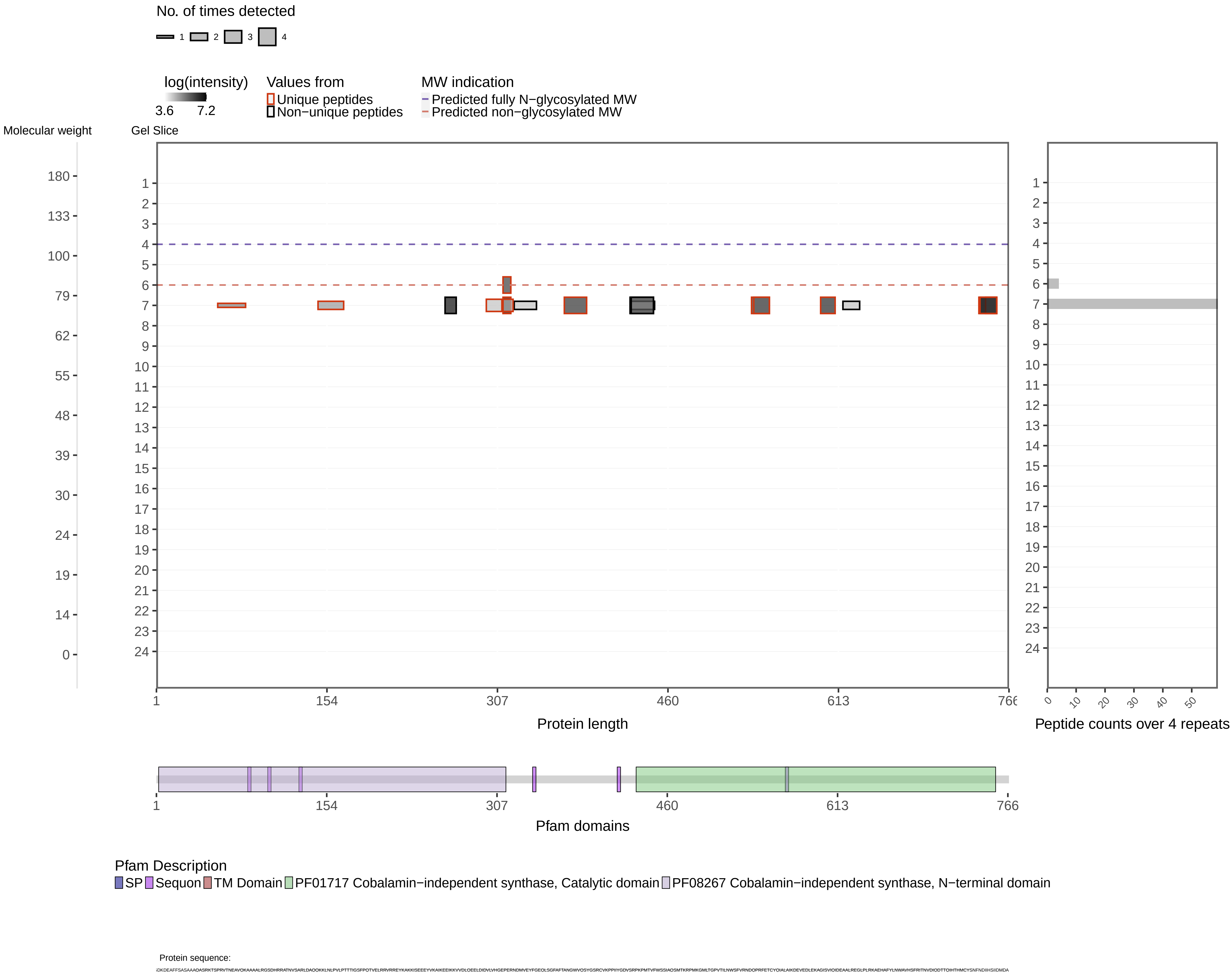

### NbL13g22460.1 calmodulin [MW = 16.83 kDa]

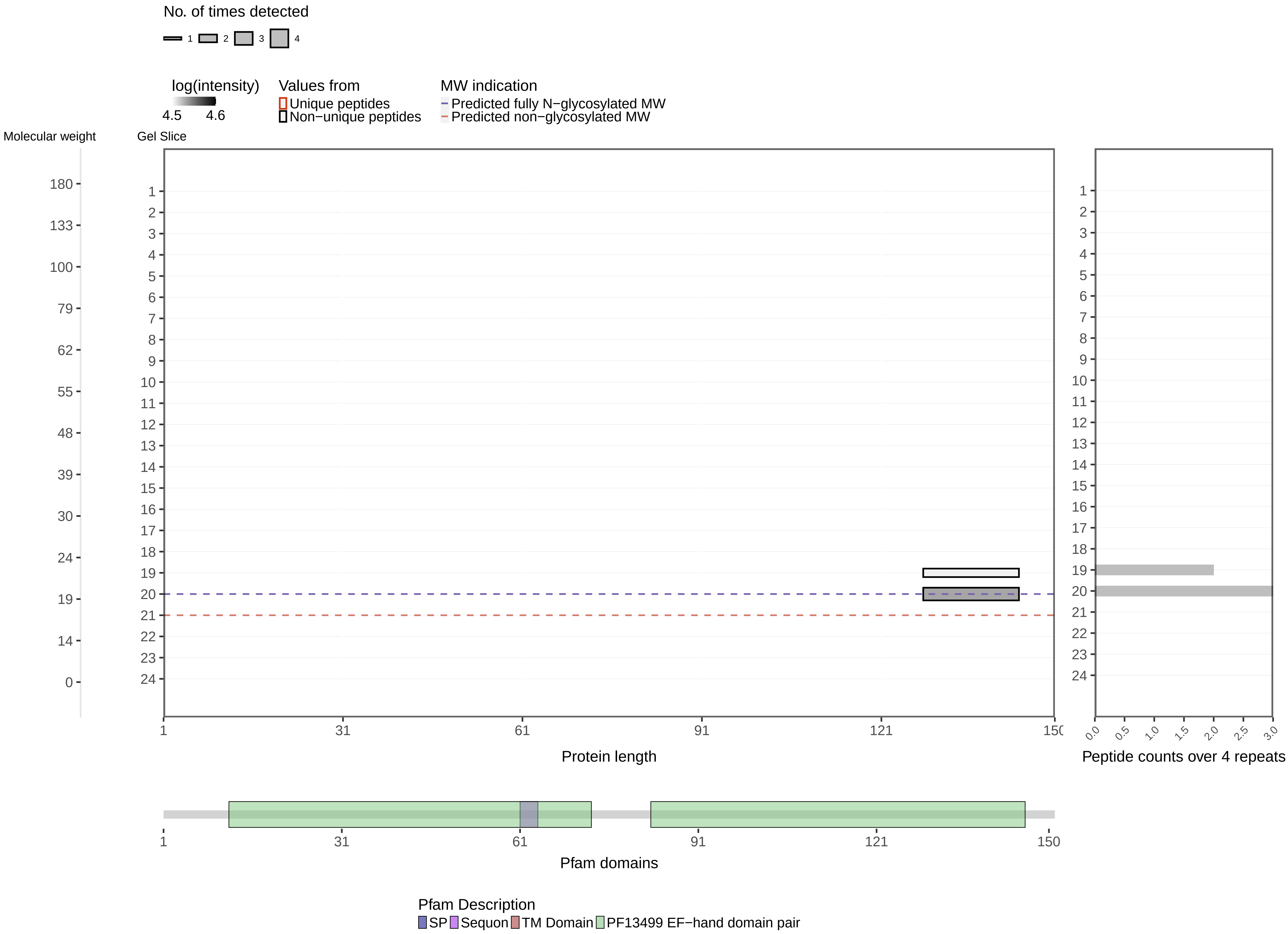


Protein sequence:

GTIDFPEFLNLMARKMKDTDSEEELKEAFRVFDKDQNGFISAAELRHVMTNLGEKLTDEEVDEMIREADVDGDGQINYDEFVKVMMAK*

### NbL13g22450.1 uncharacterized_LOC109236586 [MW = 77.89 kDa]

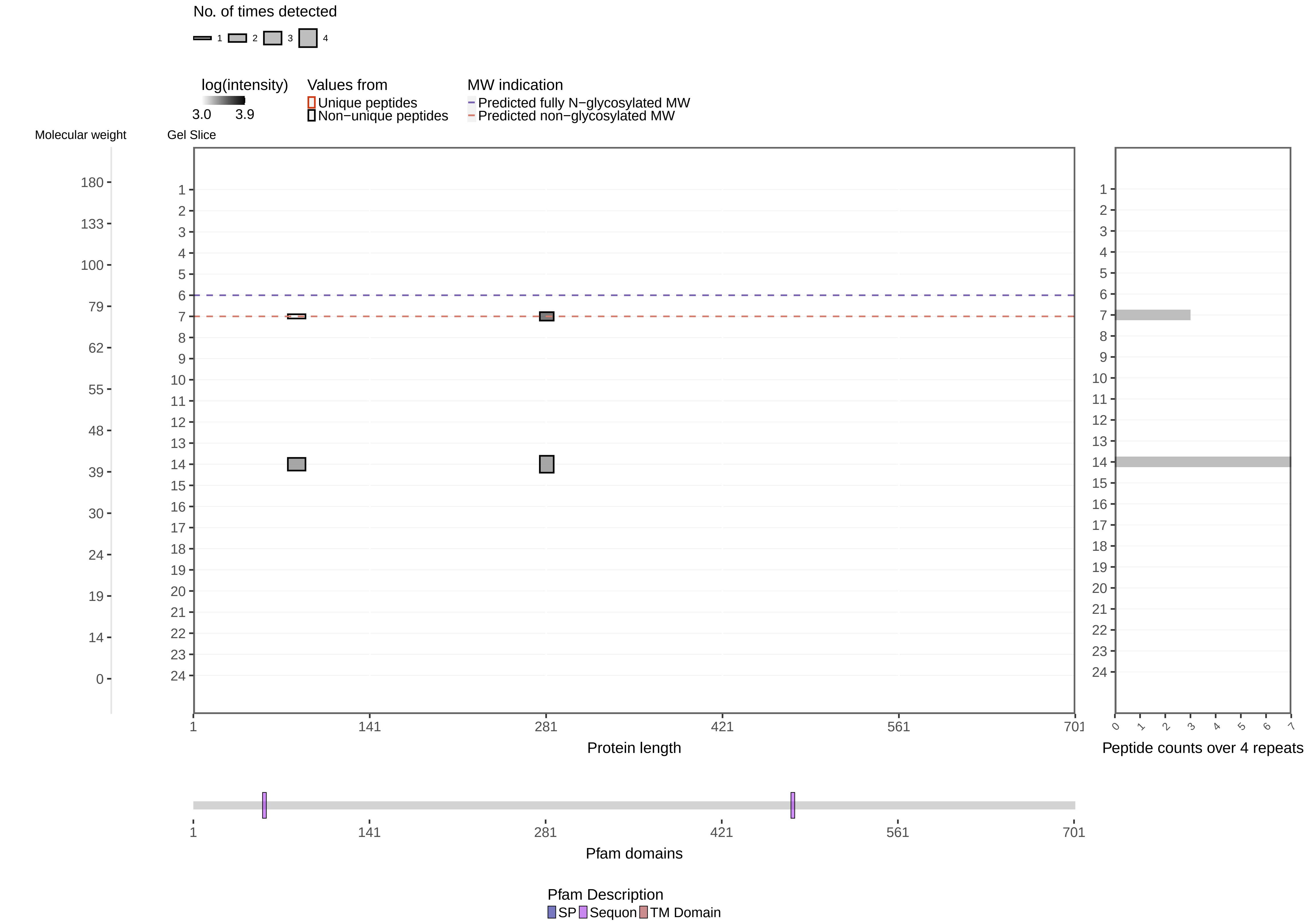


Protein sequence:

RQLGDSKANGNGLFRFQQRSASPEIVEVHEEESFGLEPDKSMKLEVGSIVLIEDSNWQKGYQINEVLTHGEIPYYIASSVEDPGTTLFLYVGSHPSQLEPAWEDMKLWYQVQRQTKVLGVMKQRGLWSKYLPQLNASGRINHPGQCRRPSSGGSCDLPWCGTPVLVTSPVGRTVADMVRLGQFGTDEAIKCCHDCLSALSAASSAGIRHGDIRPENVIYVNSGVRQPYFVLIGWGHAILEERDRPAMNLHFSSTYALQEGKLCSASDAESLVYMLYFCSGGQMPNLDSV

NbL13g22420.1 proteasome_subunit_alpha_type−3−like [MW = 27.2 kDa]

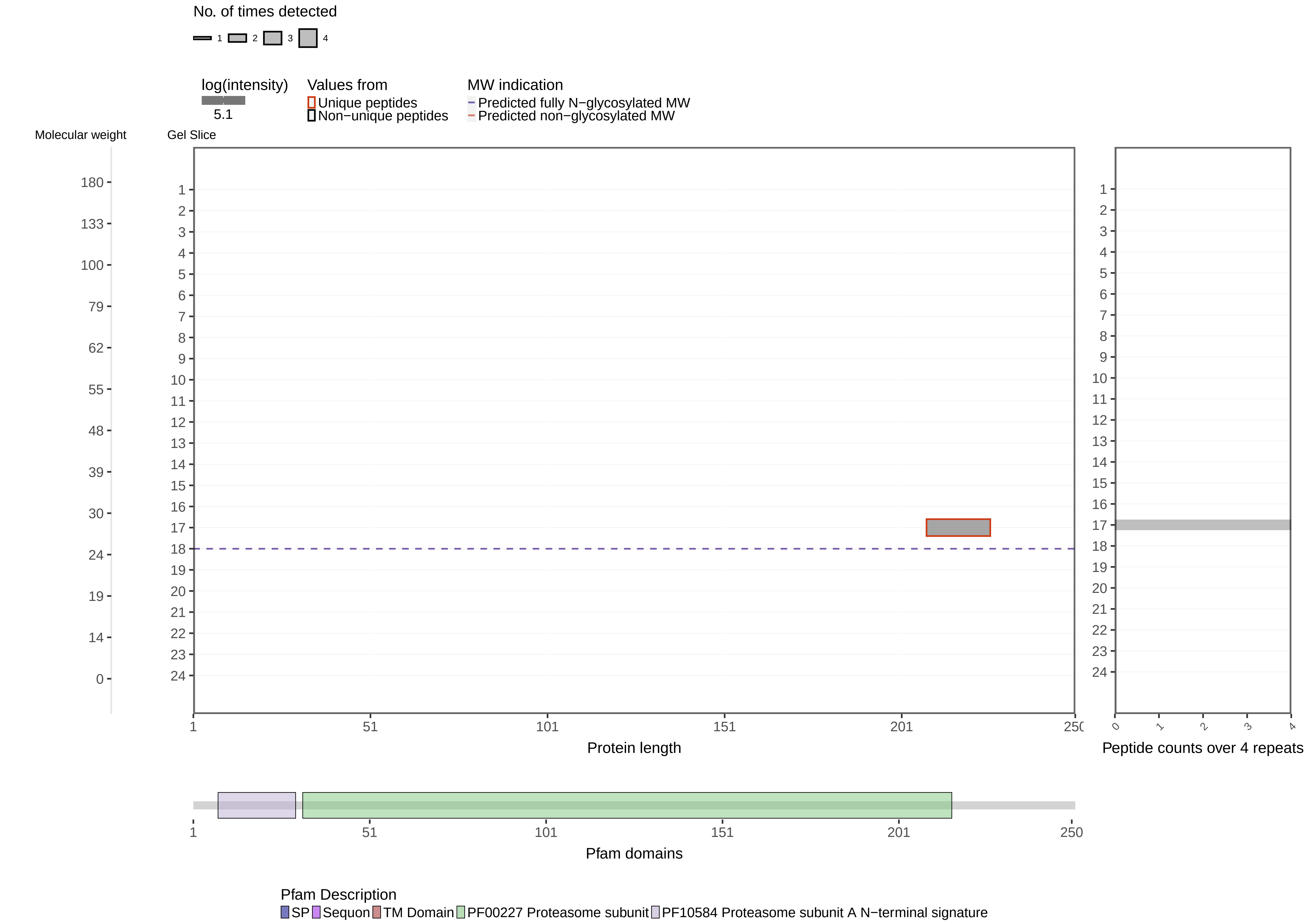


Protein sequence:

AERVASYVHLCTLYWWLRPFGCGVILGGYDRDGPQLYLVEPSGVSYRYFGAAIGKGRQAAKTEIEKLKLSEMTCRQGVIEVAKIIYGVHDEAKDKAFELEMSWICDESNHQHQKVPDDILEEAKTAAKAALEEMDAD*

### NbL13g22380.1 histone−lysine_N−methyltransferase_H3_lysine−9_specific_SUVH1−like [MW = 78.32 kDa]

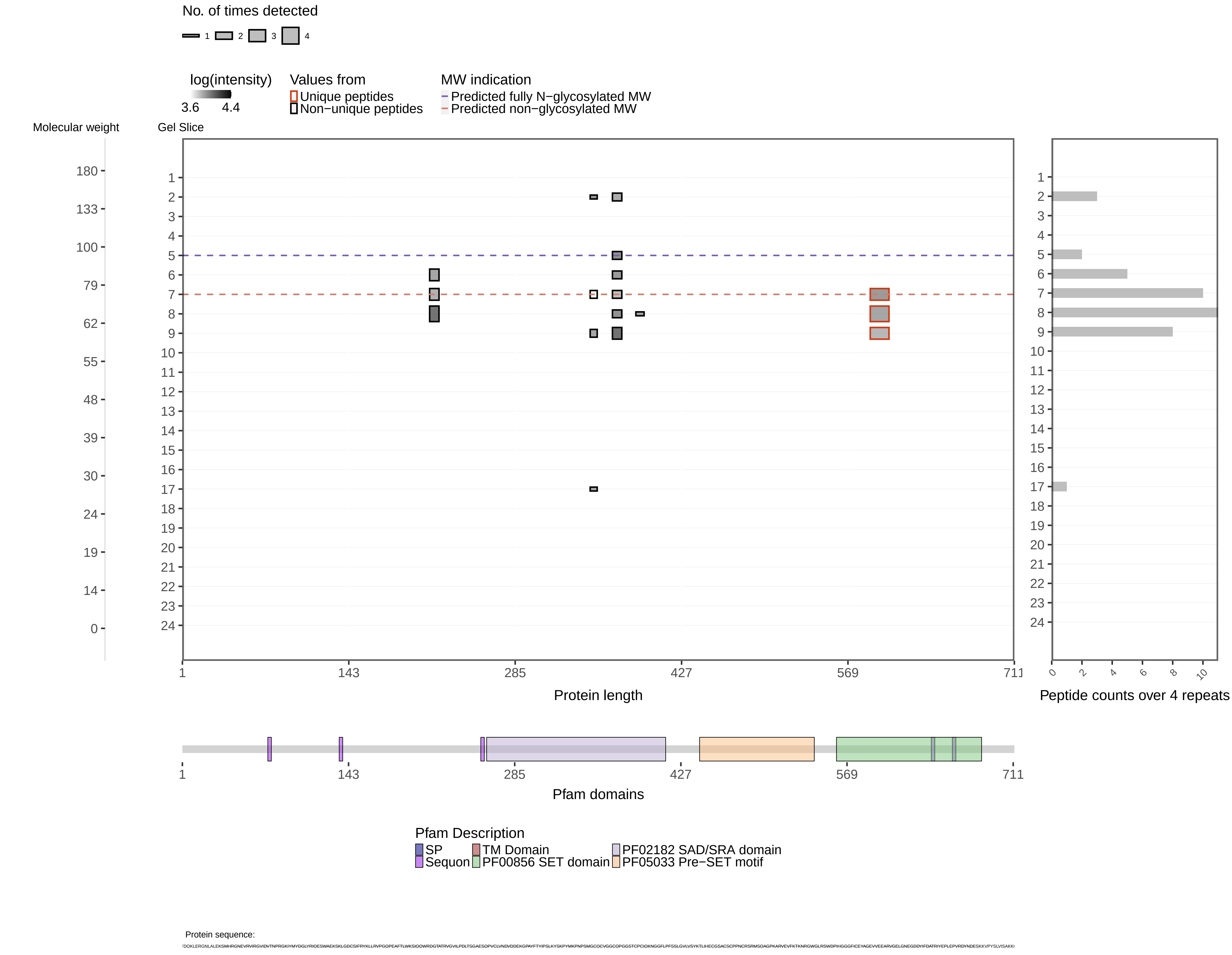

NbL13g22350.1 nascent_polypeptide−associated_complex_subunit_alpha−like_protein_1 [MW = 21.84 kDa]

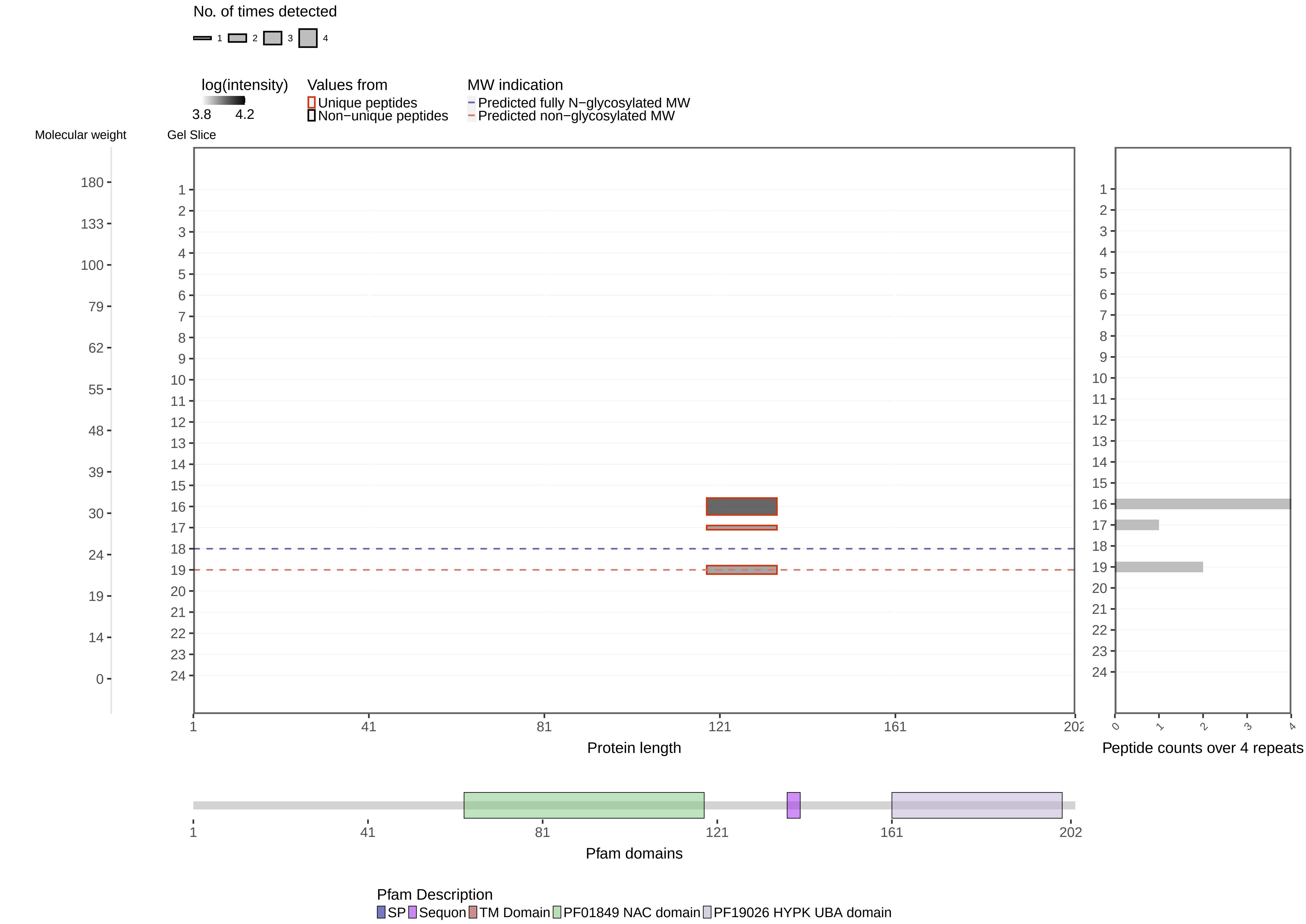


Protein sequence:

/KKSKNILFVISKPDVFKSPASDTYVIFGEAKIEDLSSQLSSQAAEQFKAPNLSNVISKPEPSTVAQDDEDVDETGVEPKDIELVMTQAGVSRAKAVKALKGADGDIVSAIMELTN*

### NbL13g22160.1 L−type_lectin−domain_containing_receptor_kinase_IX.1−like [MW − SP = 76.02 kDa]

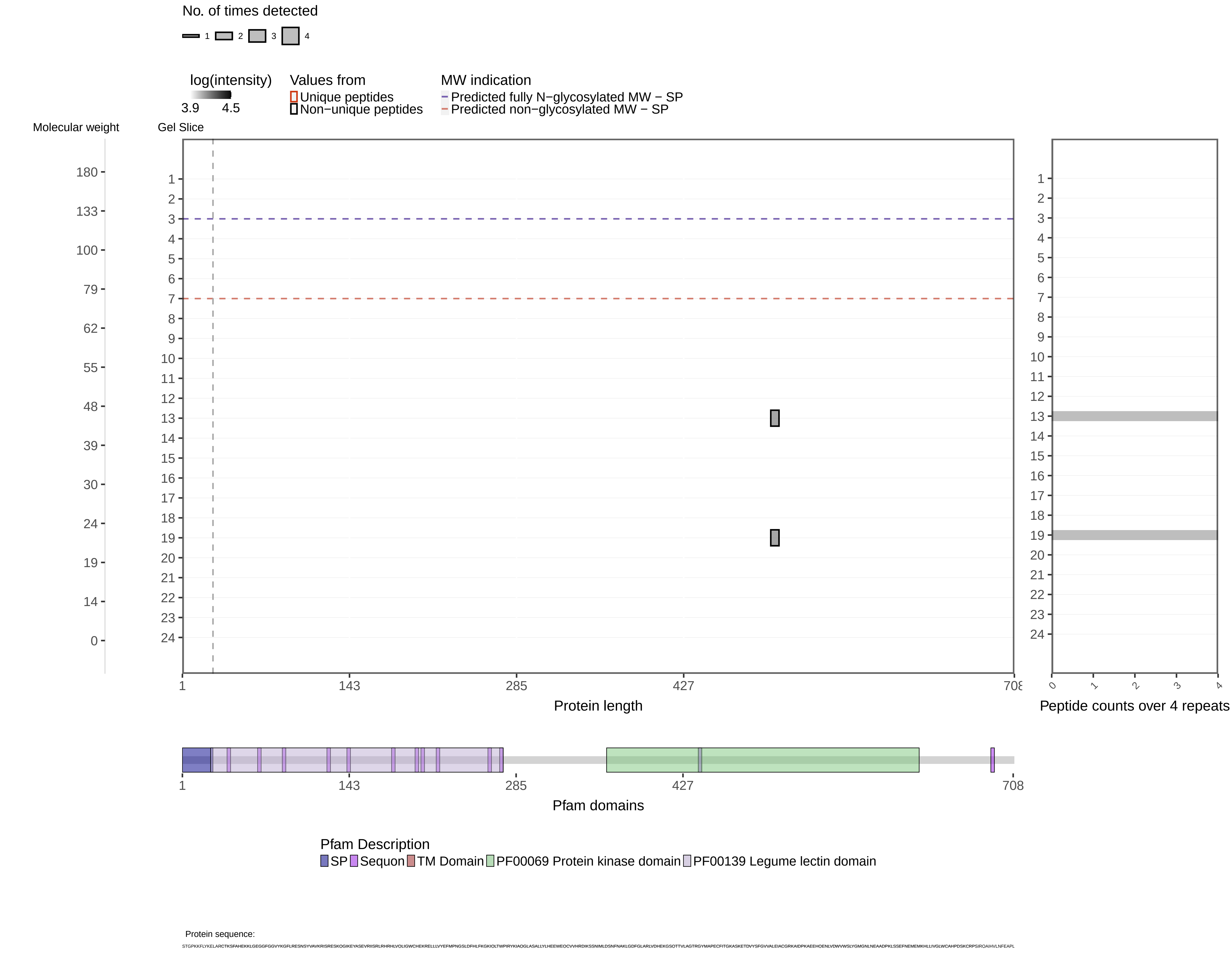

### NbL13g22150.1 heat_shock_70_kDa_protein_mitochondrial [MW = 73.22 kDa]

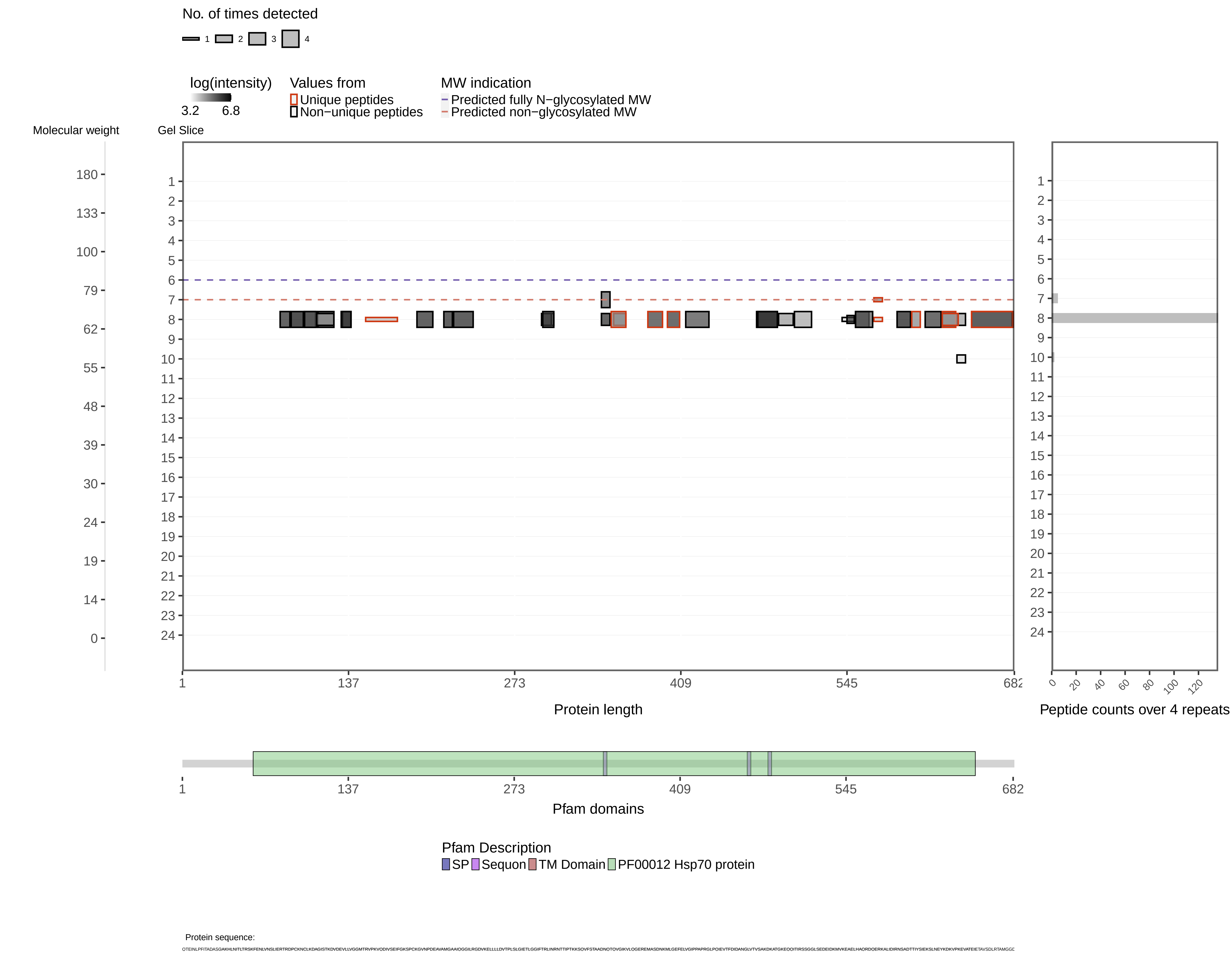

NbL13g22130.1 soluble_inorganic_pyrophosphatase_6_chloroplastic−like [MW = 32.65 kDa]

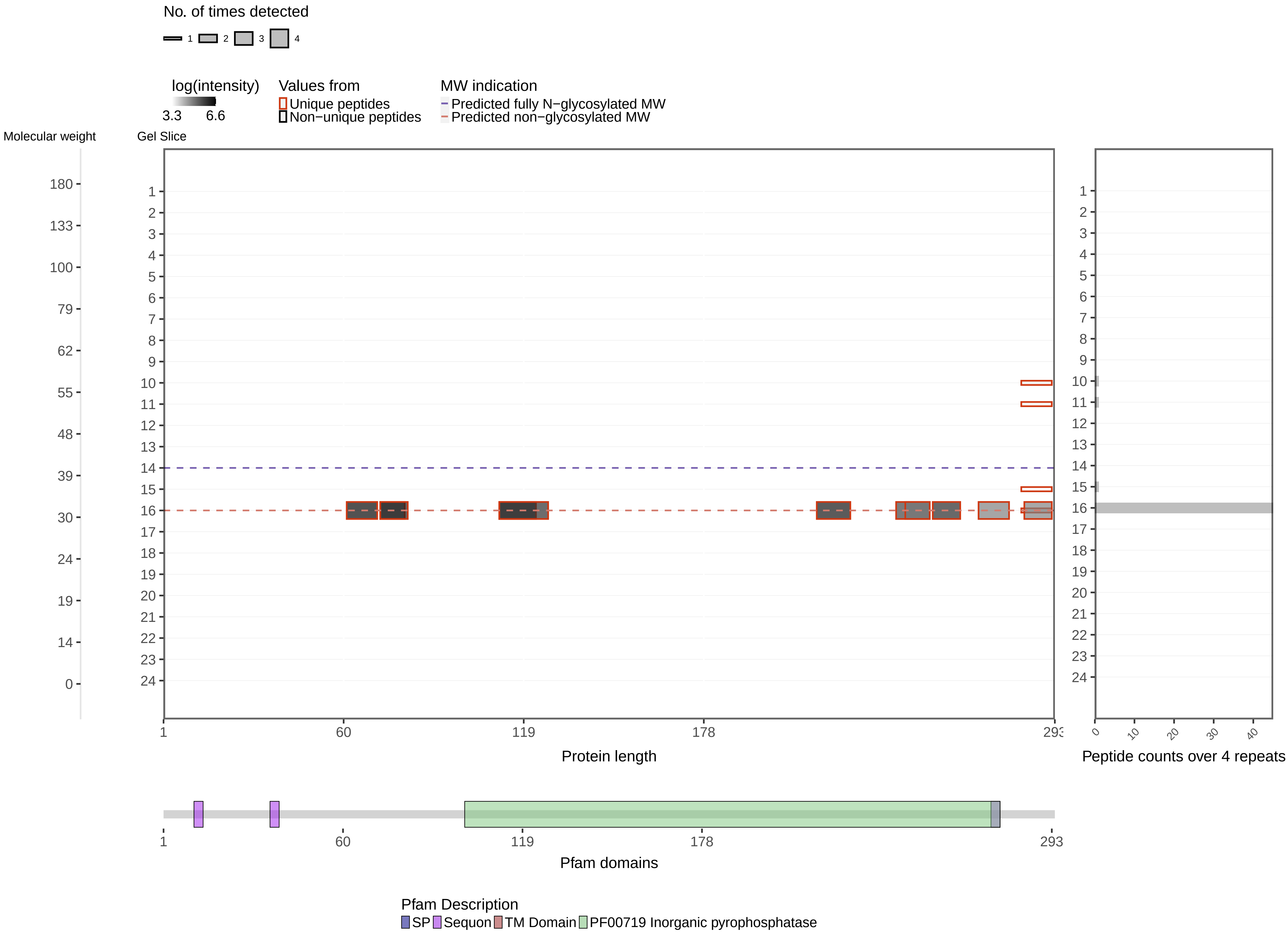


Protein sequence:

YPYNIHWNYGLLPQTWEDPSFANAEVEGAFGDNDPVDVVEIGESRAKIGQVLKVKPLAALAMIDEGELDWKIVAISLDDPRASLVNDIDDVEKHFPGTLTAIRDWFRDYKIPDGKPANKFGLGNKPANKDYALKIITETNESWAKLVKRSVPAGDLSLV*

NbL13g21930.1 protein_ABCI7_chloroplastic [MW = 53.84 kDa]

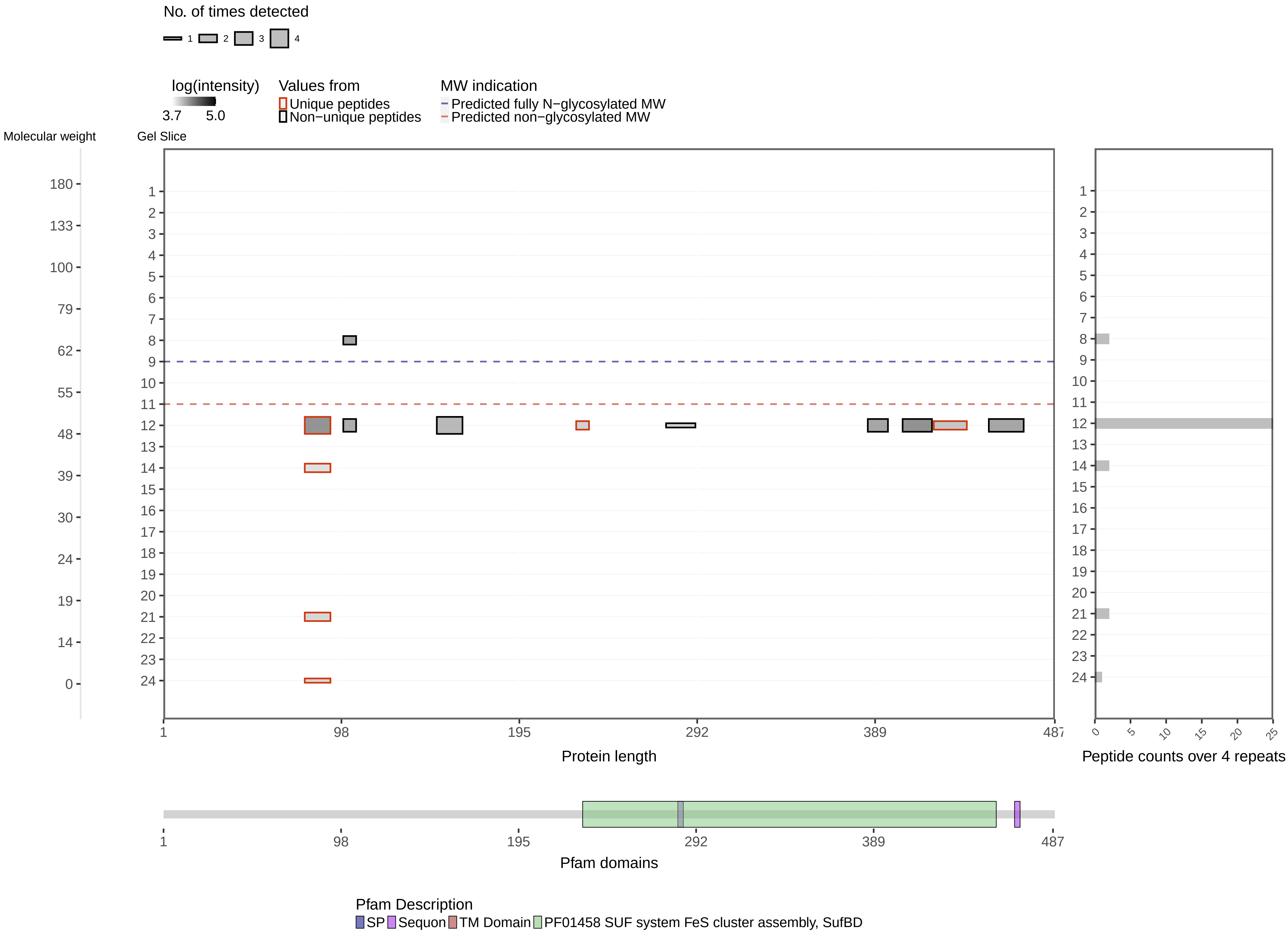


Protein sequence:

### NbL13g21860.1 glycine_cleavage_system_H_protein_mitochondrial [MW = 17.5 kDa]

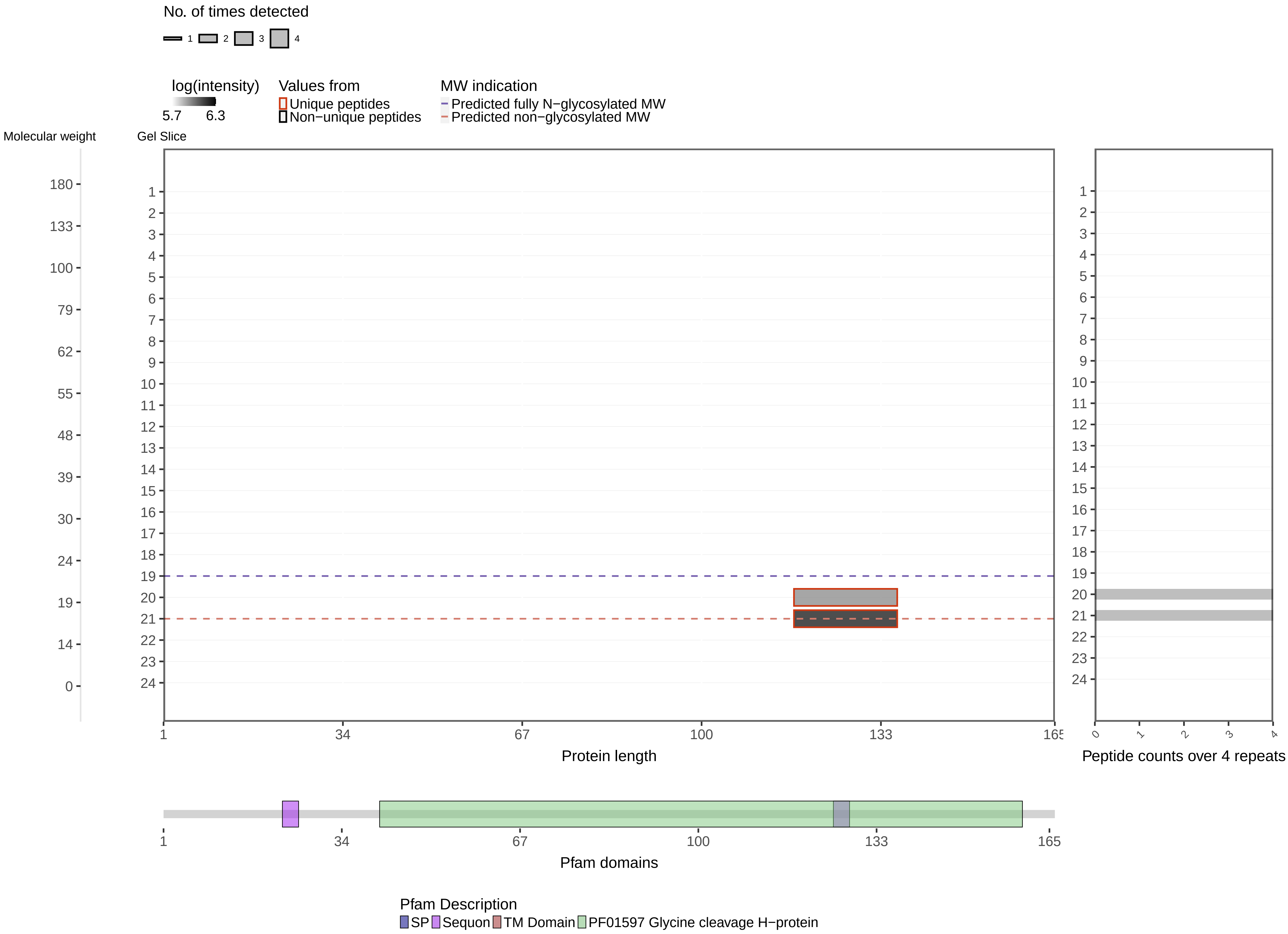


Protein sequence:

GEVVFVDLPETGAAVSQGSSFGAVESVKATSDINCPISGEIVEVNTKLTGTPGLVNSSPYEDGWMIKVKPSSPSELESLMGSKEYTKFCEEEDSH*

### NbL13g21680.1 LL−diaminopimelate_aminotransferase_chloroplastic−like [MW = 50.07 kDa]

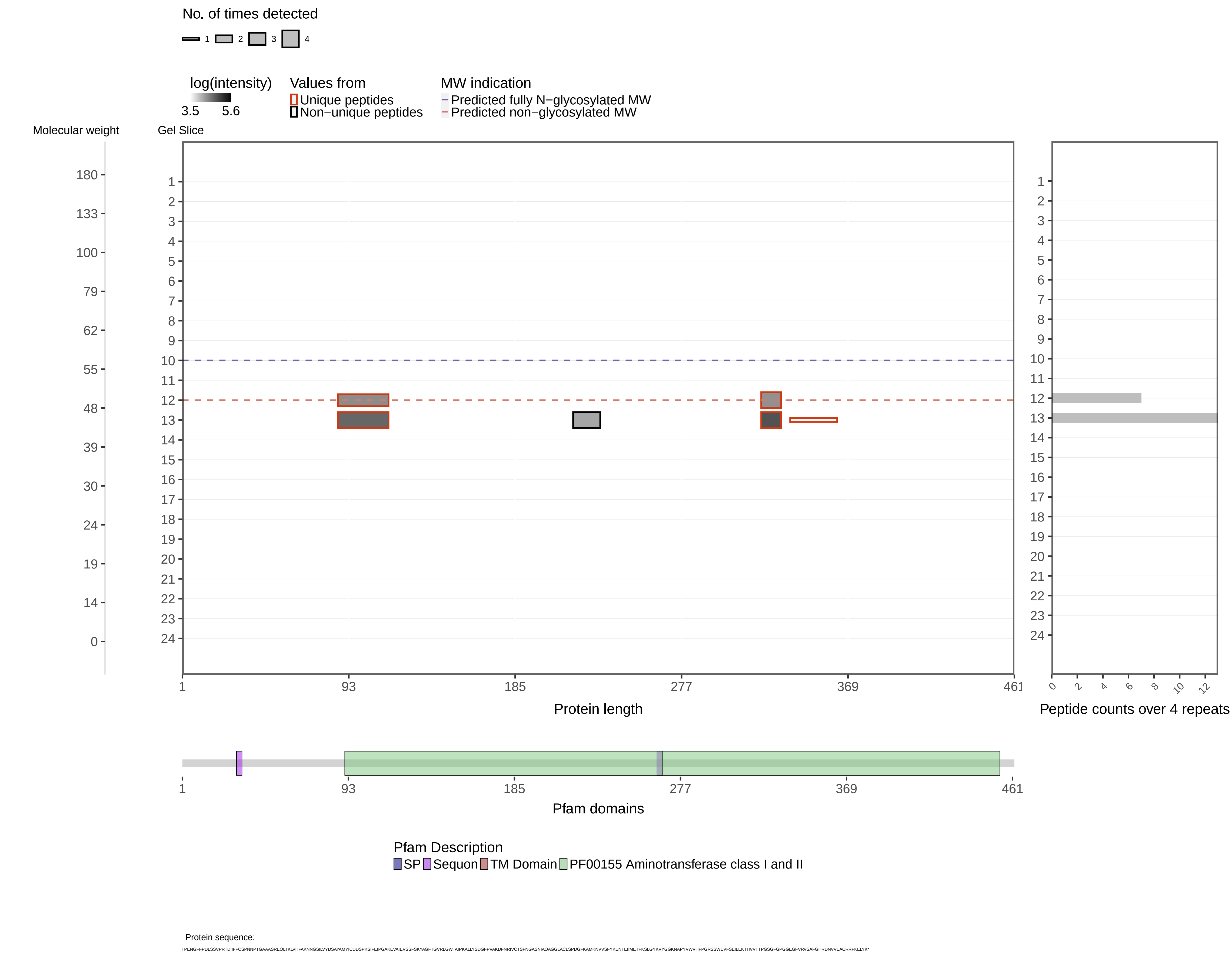

### NbL13g21520.1 FAM10_family_protein_At4g22670−like [MW = 45.01 kDa]

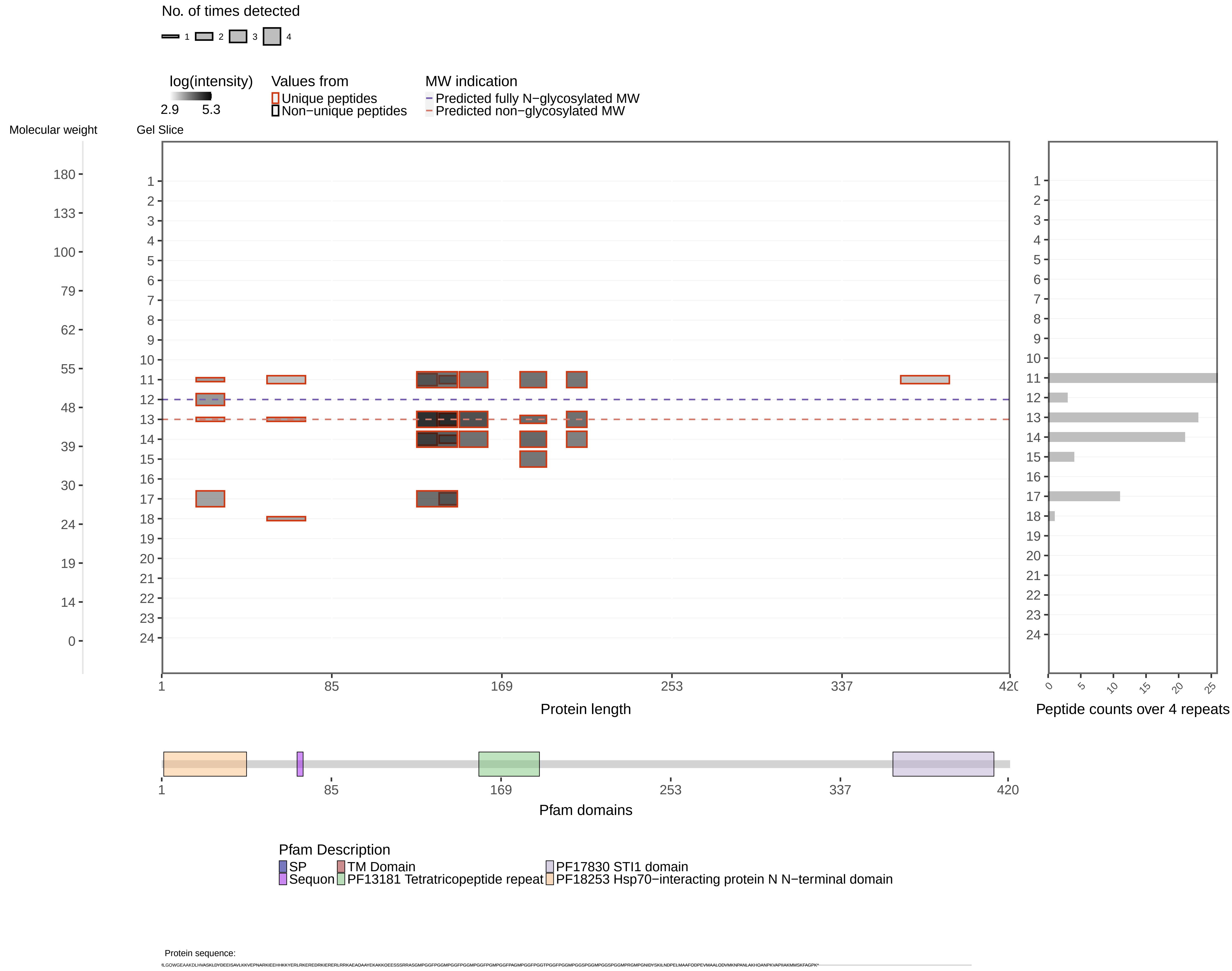

NbL13g21410.1 subtilisin−like_protease [MW = 77.95 kDa]

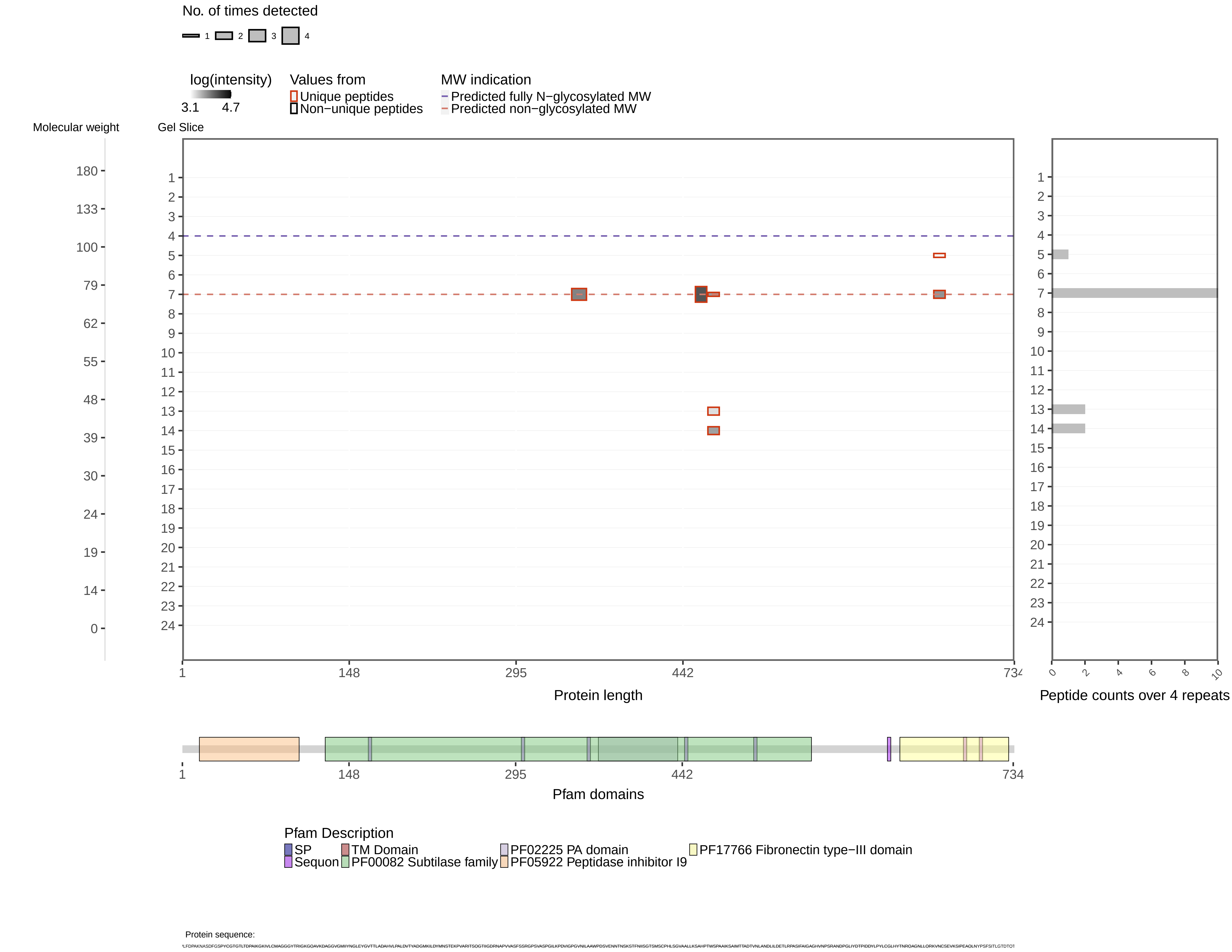

### NbL13g21150.1 protein_disulfide−isomerase−like [MW − SP = 56.02 kDa]

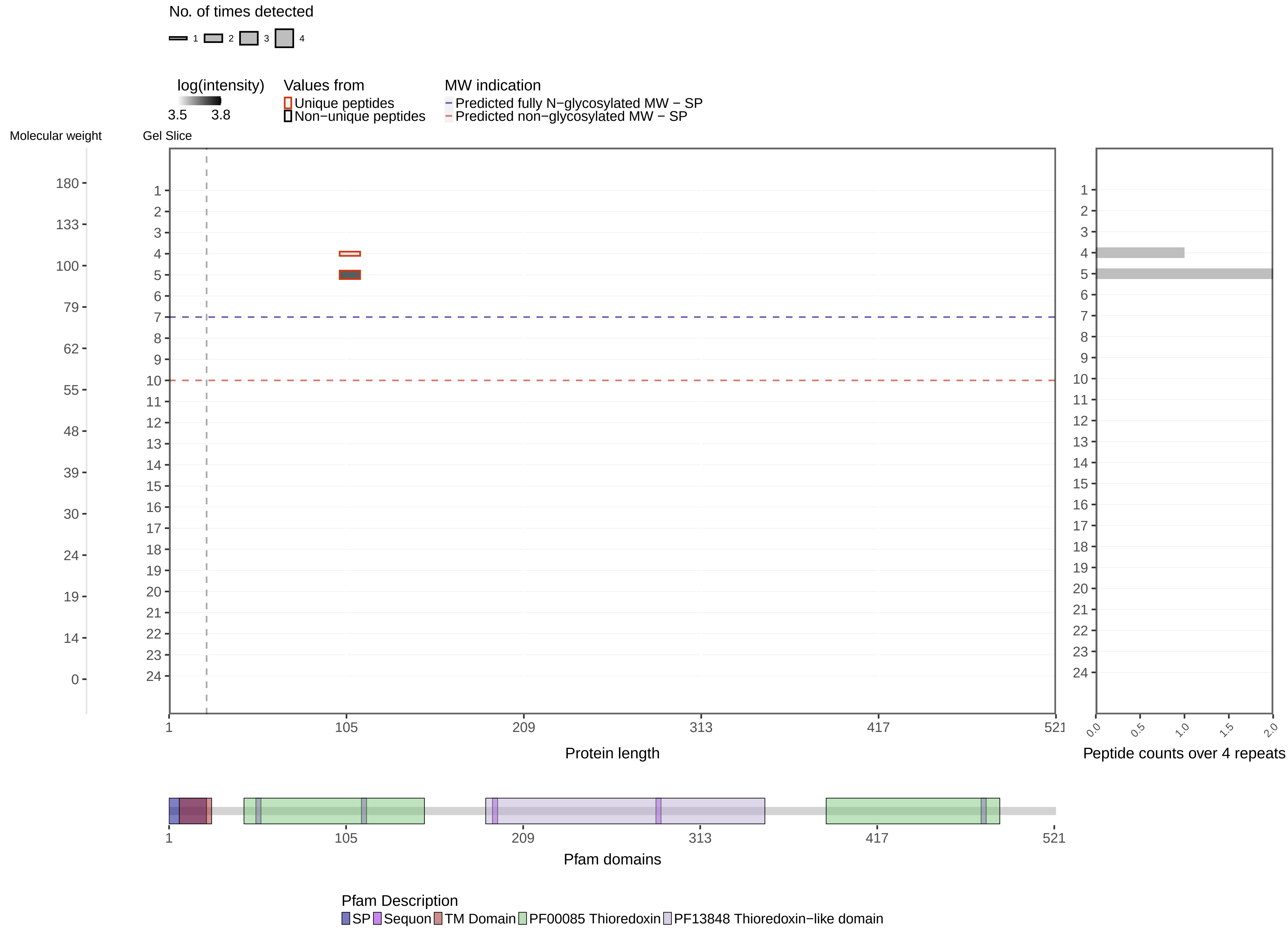


Protein sequence:

FIEETSVPIMAILDDKPENQAFVNHFLHSPGDKVFLFLNFTTDLDHFKSKYYDLAASYKGKEANFLLGDAETGKRALEYFGLNTDQVPLIFVLAMDGTKYVQRHVQPDSLGVWLKDFKDGKLKPYLKSQPIPEVNNEPVKVVVAETLEDMIFNSGKDVLLEFYRLGCKYCEEFAPVLDEIALSFENDADVLIAKIDGTENDIPRERFEVQGFPTLYMISTAGSLSRFEGNRTKEAVIEFIQTNRGTPAPSNFANSDLTASEHAQTDSMKDEL*

### NbL13g21110.1 V−type_proton_ATPase_subunit_E−like [MW = 27.71 kDa]

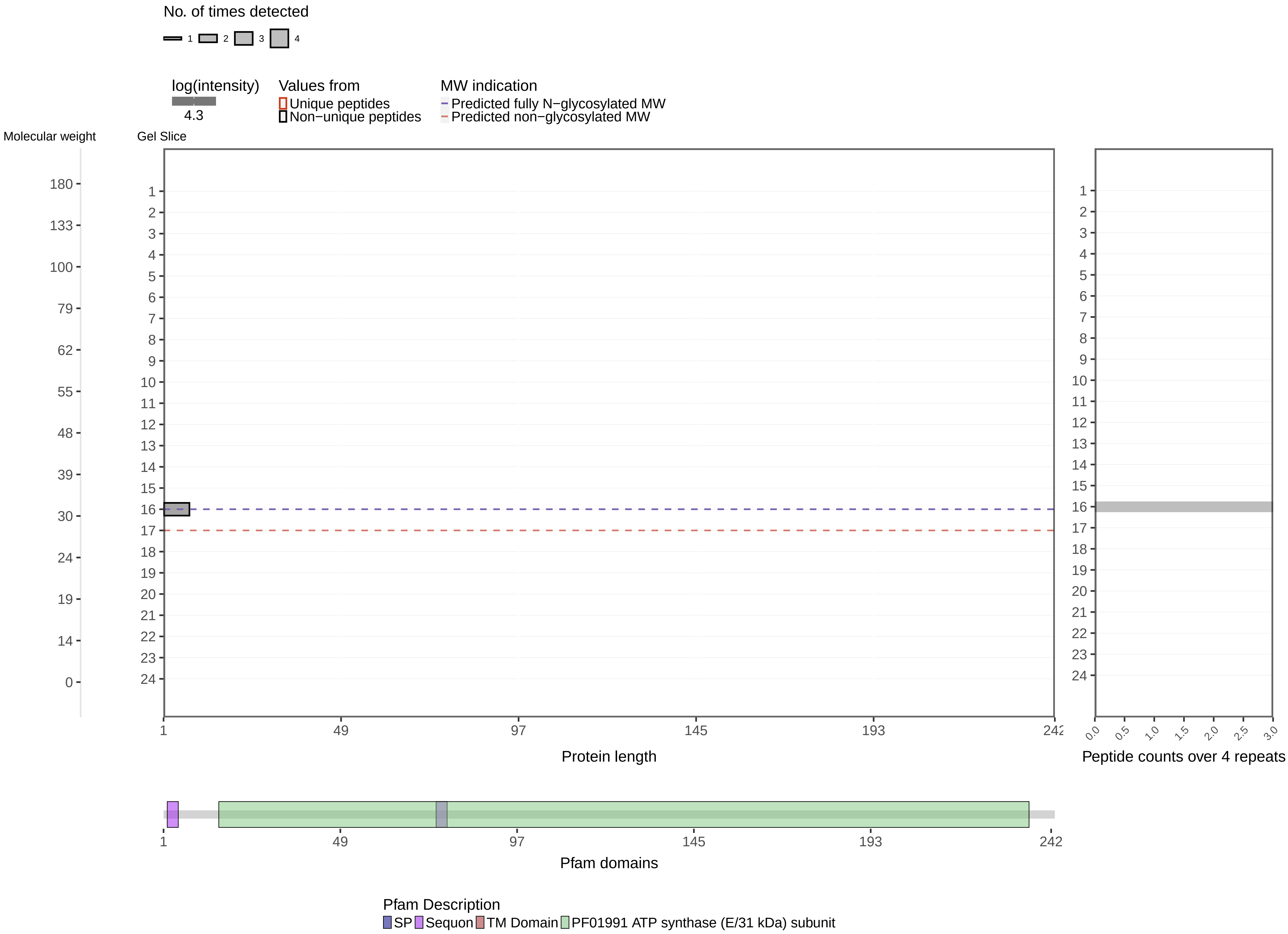


Protein sequence:

HNELVEIILPHHQDSYKKLLHNLIVQSLLRLKEPSVLLRCRKHDVHLVEHVLDAAKEEYAEKANVHAPEIIVDDIHLPPAPSHHNMHGPSCSGGVVLASRDGKIVCENTLDARLEVVFRKKLPEIRKCLFGQVTA*

### NbL13g21010.1 nucleoside_diphosphate_kinase_3−like [MW = 30.48 kDa]

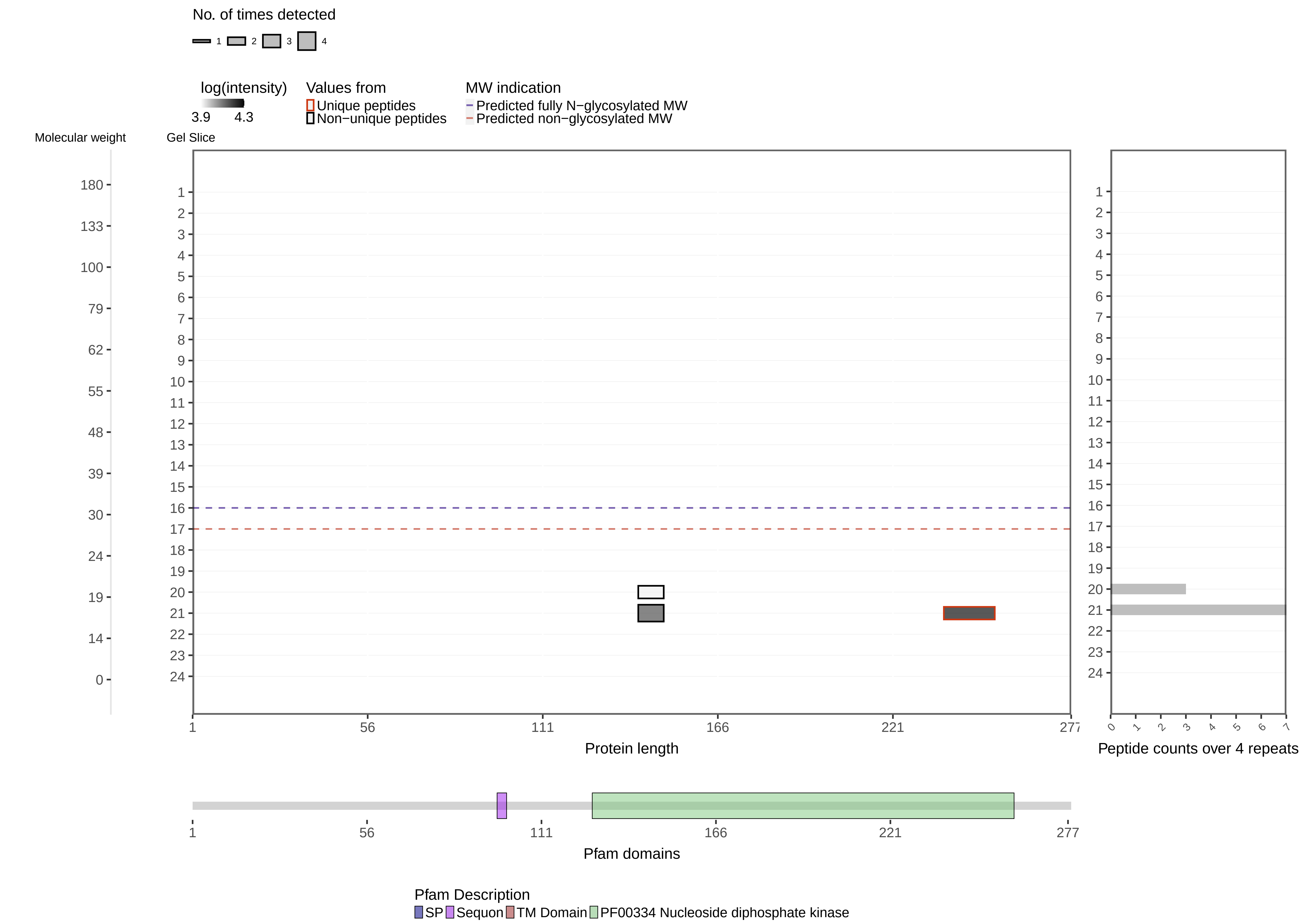


Protein sequence:

MERTFIAIKPDGVQRGLISEIISRFECKGFKLVAIKIVVPSKDFAGKHYHDLKERPFFTGLCDFLSSGPVLAMVWEGEGVIRYGRKLIGATDPQKSEPGTIRGDLAVVVGRNIIHGSDGPETAKDEIKLWFKPEELVKYTSNDEKWIYGVN*

### NbL13g20920.1 N−alpha−acetyltransferase_15_NatA_auxiliary_subunit−like [MW = 102.26 kDa]

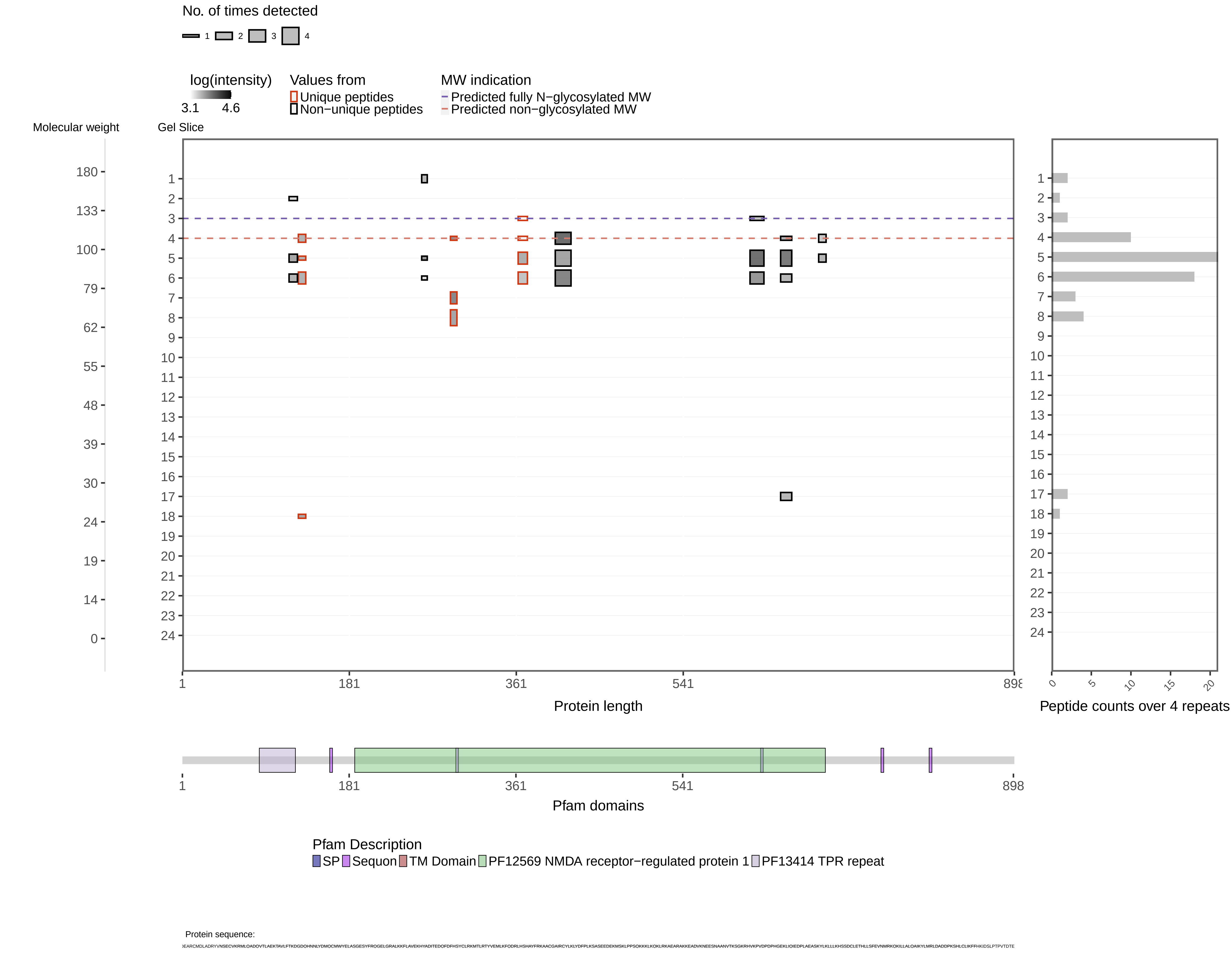

NbL13g20880.1 protein_CDI−like [MW = 32.12 kDa]

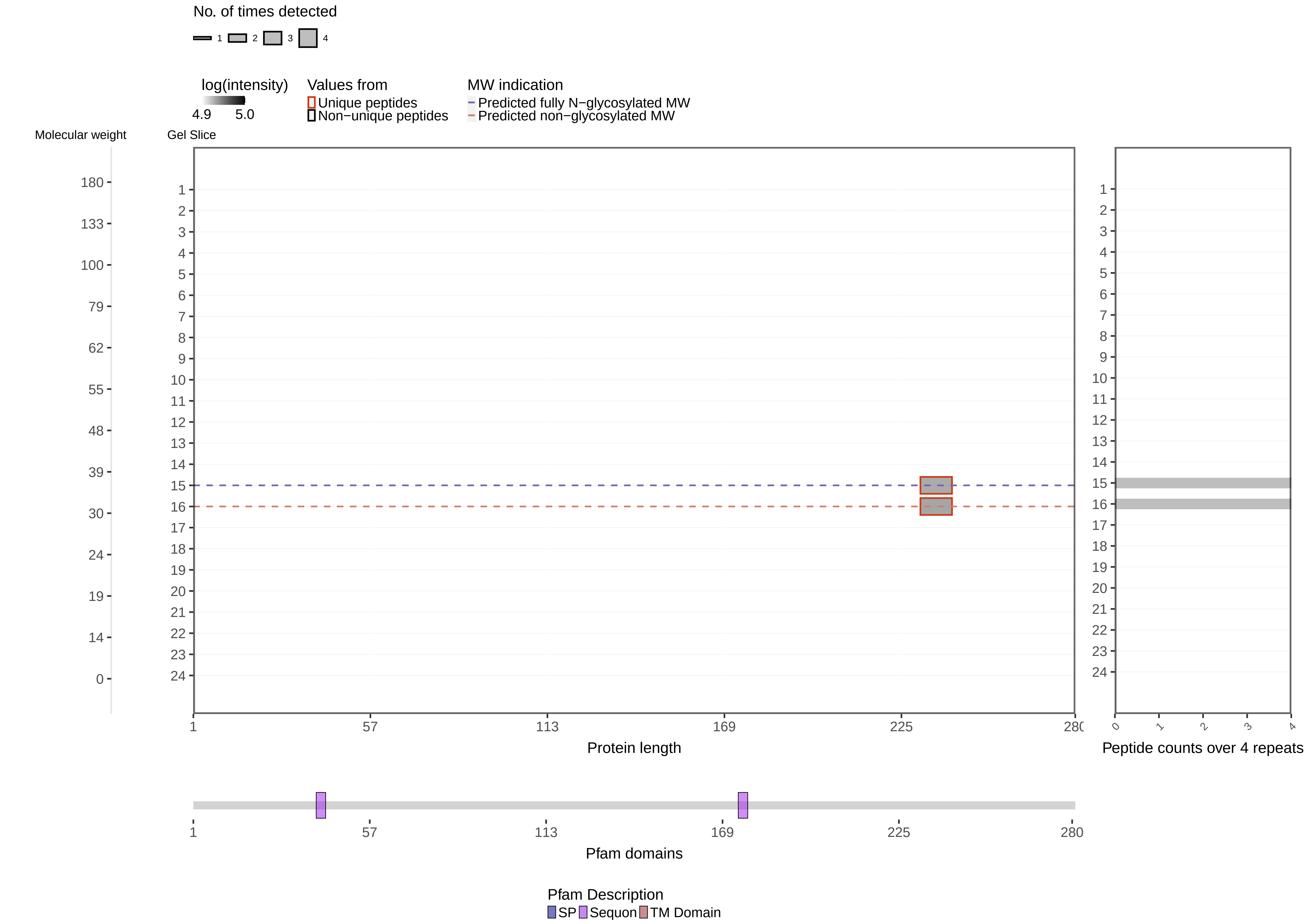


Protein sequence:

LYLEDIKELRDLVDDKYALMCVQHDYTPKETTKMDGAVQTVYPRKNWSSMVLYNCGHPKNRALTPEVVNSESGAFLHRFMWLEDKEIGEVPFVWNFLVGHNKVVEGDPATFPKAIHYTLGGPWFEAWKDCEFGDLWLKELEECEKAKEKLG*

NbL13g20870.1 D−glycerate_3−kinase_chloroplastic−like [MW = 51.26 kDa]

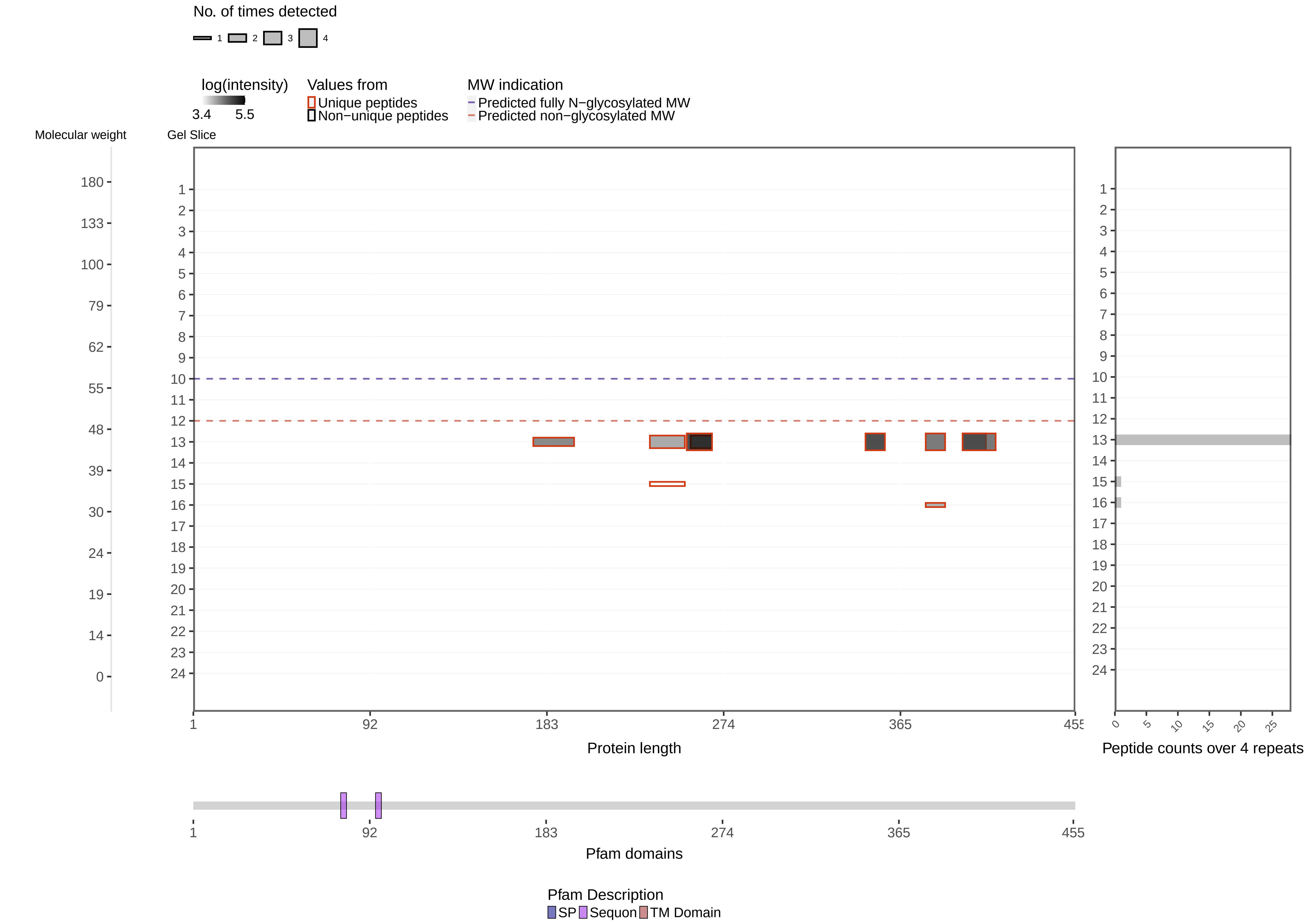


Protein sequence:

QGCGKTTLVFALEYLFKITGRKAATLSIDDFYLTAEEQARLREDNPENLLLEFRGNAGSHDLSLSVETLTELSKVTKEGVKMKLPRYDKSAYNGRGDRADPSTWPEVEGPLSVILFEGWMLGFKPLPPEVVKAVDPQLETVNKNLEAYYDAWYKYVESWIIIKIHDPSYVYQWRLQAEIAMRADGNLGMSDEEVRDFVSRYLPAYKAYLPTLYSEGPPGADPEHVLLIEIDEGRNPILGS*

NbL13g20850.1 aspartate_aminotransferase [MW = 43.28 kDa]

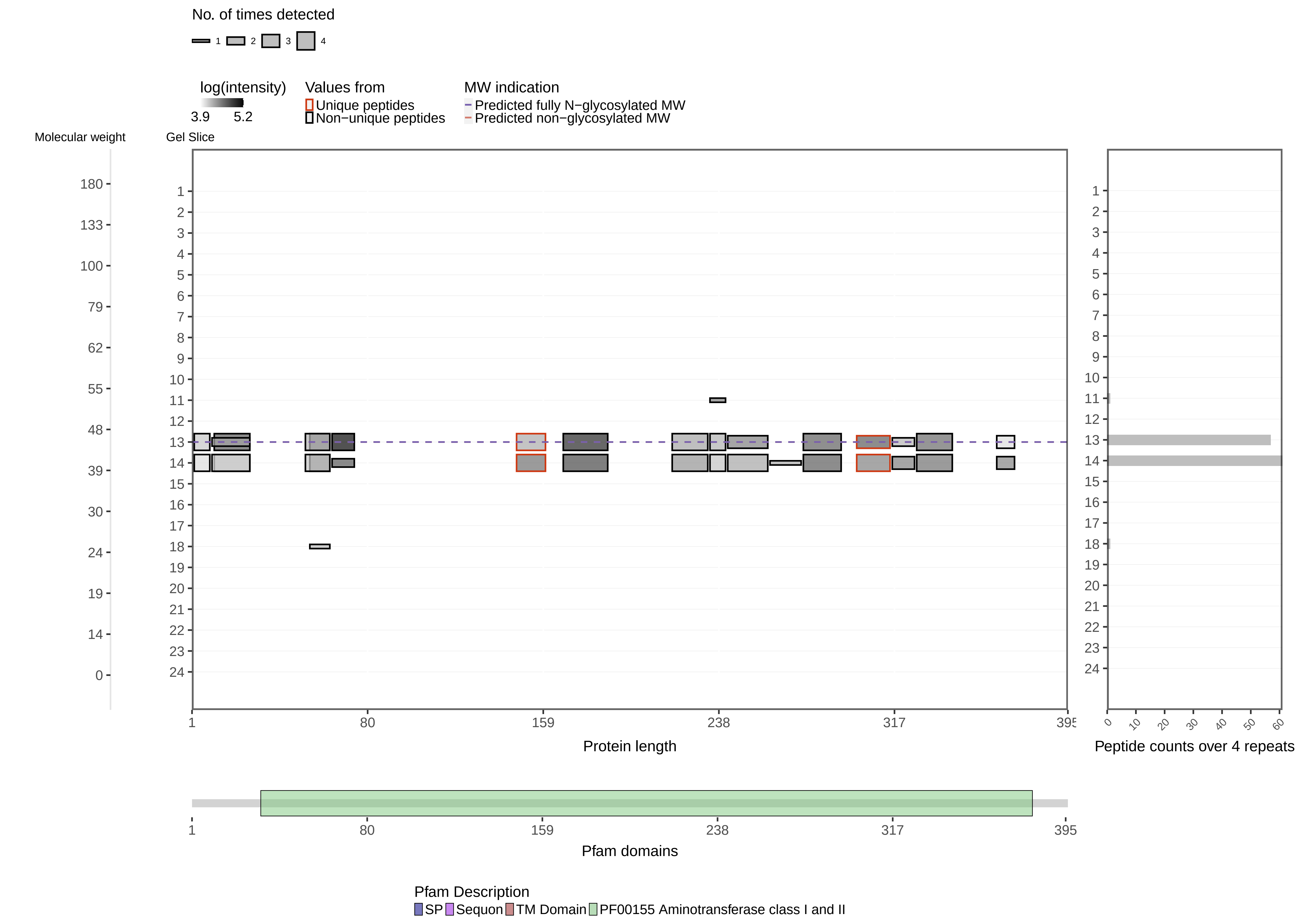


Protein sequence:

'ESLLKRISDICKEAGCWLVIDNTYEYFMYDDRKHVCIEGNHIVNIFSFSKAYGMMGWRVGYIAYPSEVEGLAAOLLKVODNIPICASIISORLALYSMEMGPEWVTDOVKVLVKNRDLLLEALSPLGEGAVKGGEGAIYLWAKLPDKYENDFEVVHWLAKRHGIVLIPGSSSGCPGHVRISFGGLIEKDCRVAAERLRKGLEELVNNGMIS*

### NbL13g20800.1 zinc_phosphodiesterase_ELAC_protein_2−like [MW = 107.82 kDa]

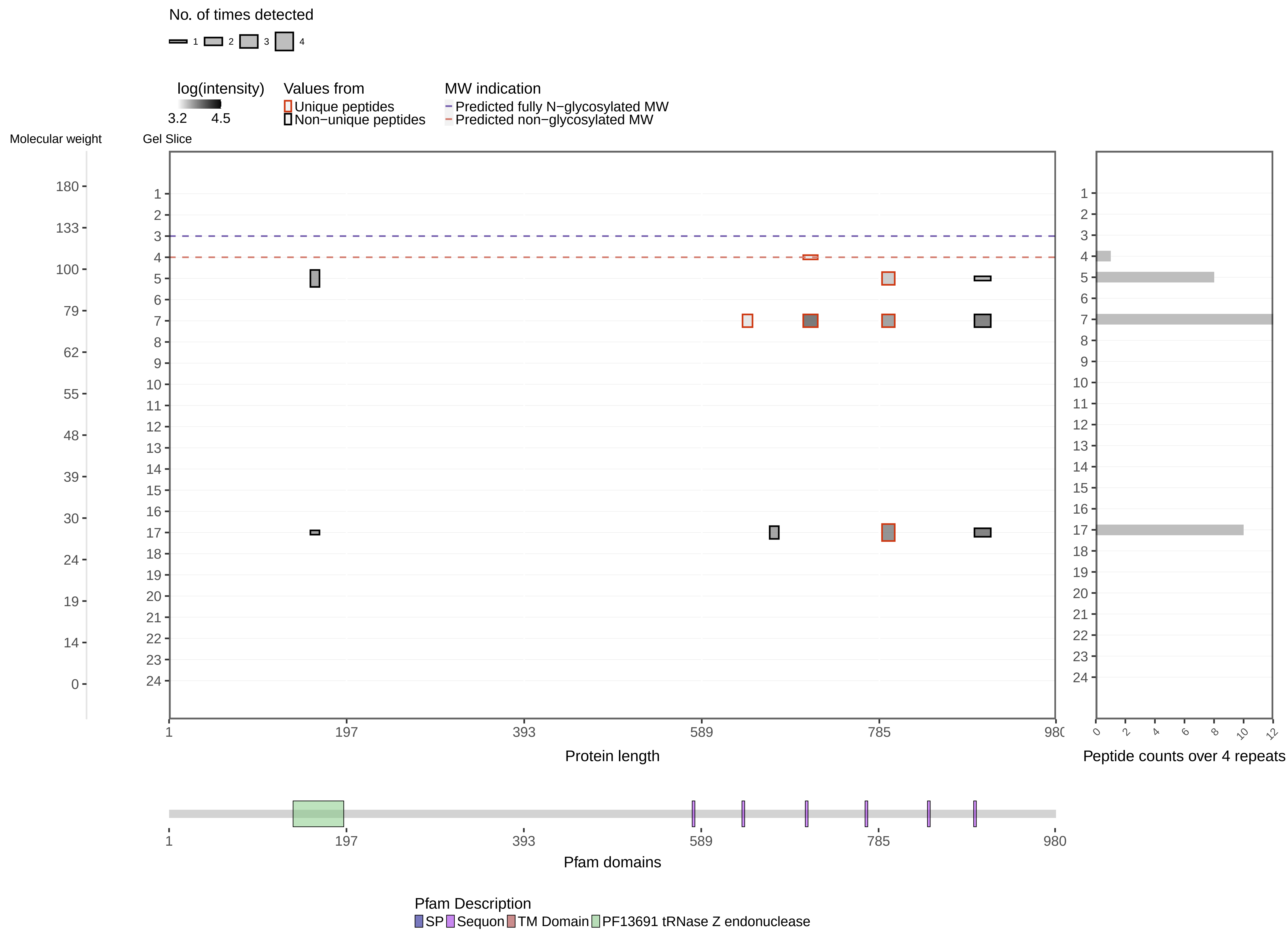


Protein sequence:

KSSARIASRLNYLCPOFFPAPGFWSLPOLKSLPSVSKTPSEFSSCOVTAENLLKFHLRPYAOLGLDRSGIPEITSRPKIIEELISEIPEILDASEHITOMLHGNNVTNGGSATMOVNKVMIEESSLHETALPSCLEGVTREDVEIVLLGTGSSOPSKYRNVSSIFVNLFSKGSVLLDCGEGTLGOLKRRFGIEGADEAVKGLRCIWISHIHADHHTGLARILALRRDLLNETPHEPLIVVGPROLKRFLDAYOKLEDLDMOFLDCRHTTEASLKTFESNGDKDVSESACVOSDOKN

NbL13g20740.1 4−hydroxy−3−methylbut−2−en−1−yl_diphosphate_synthase_(ferredoxin)_chloroplastic [MW = 82.08 kDa]

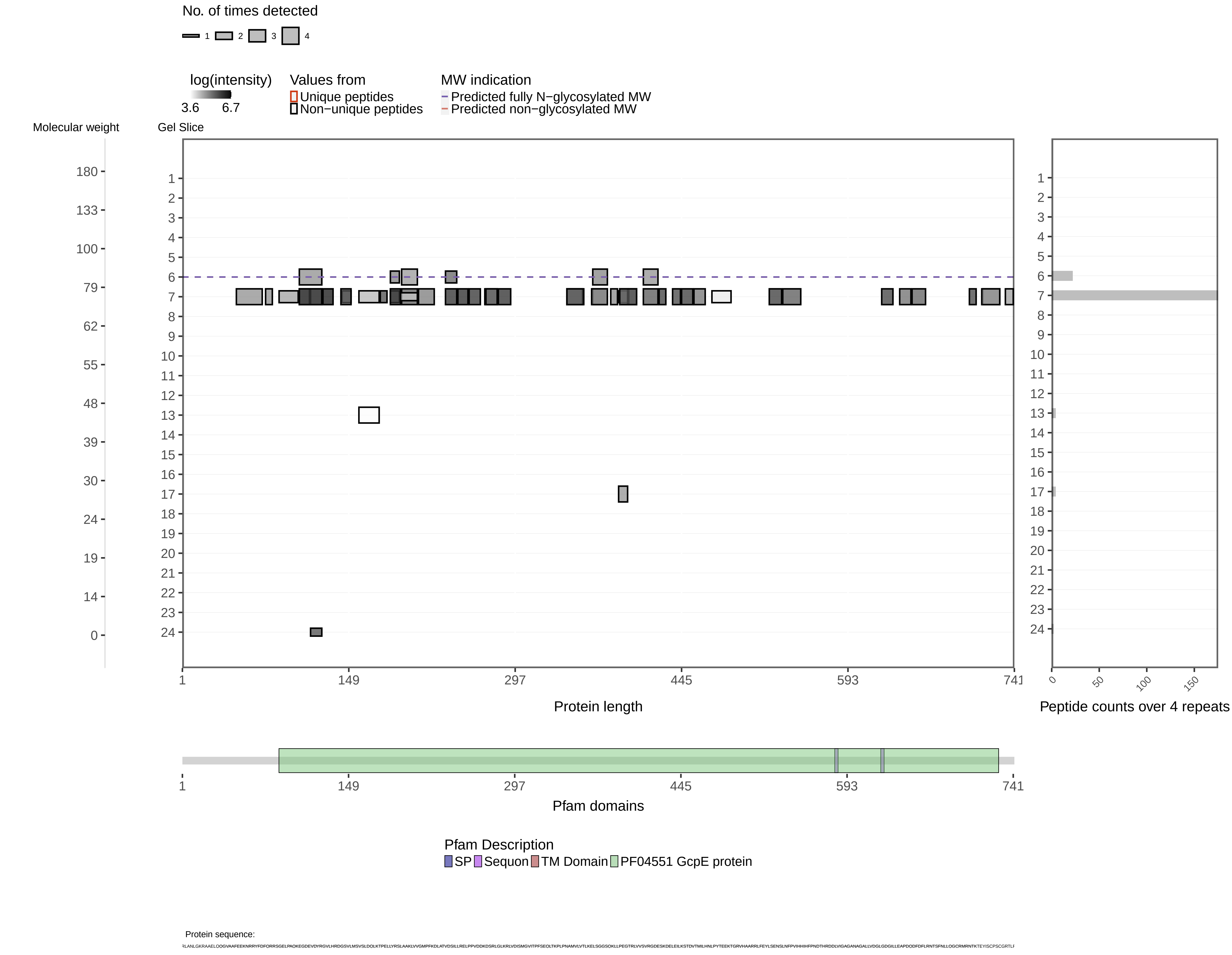

NbL13g20720.1 probable_inactive_nicotinamidase_At3g16190 [MW = 22.37 kDa]

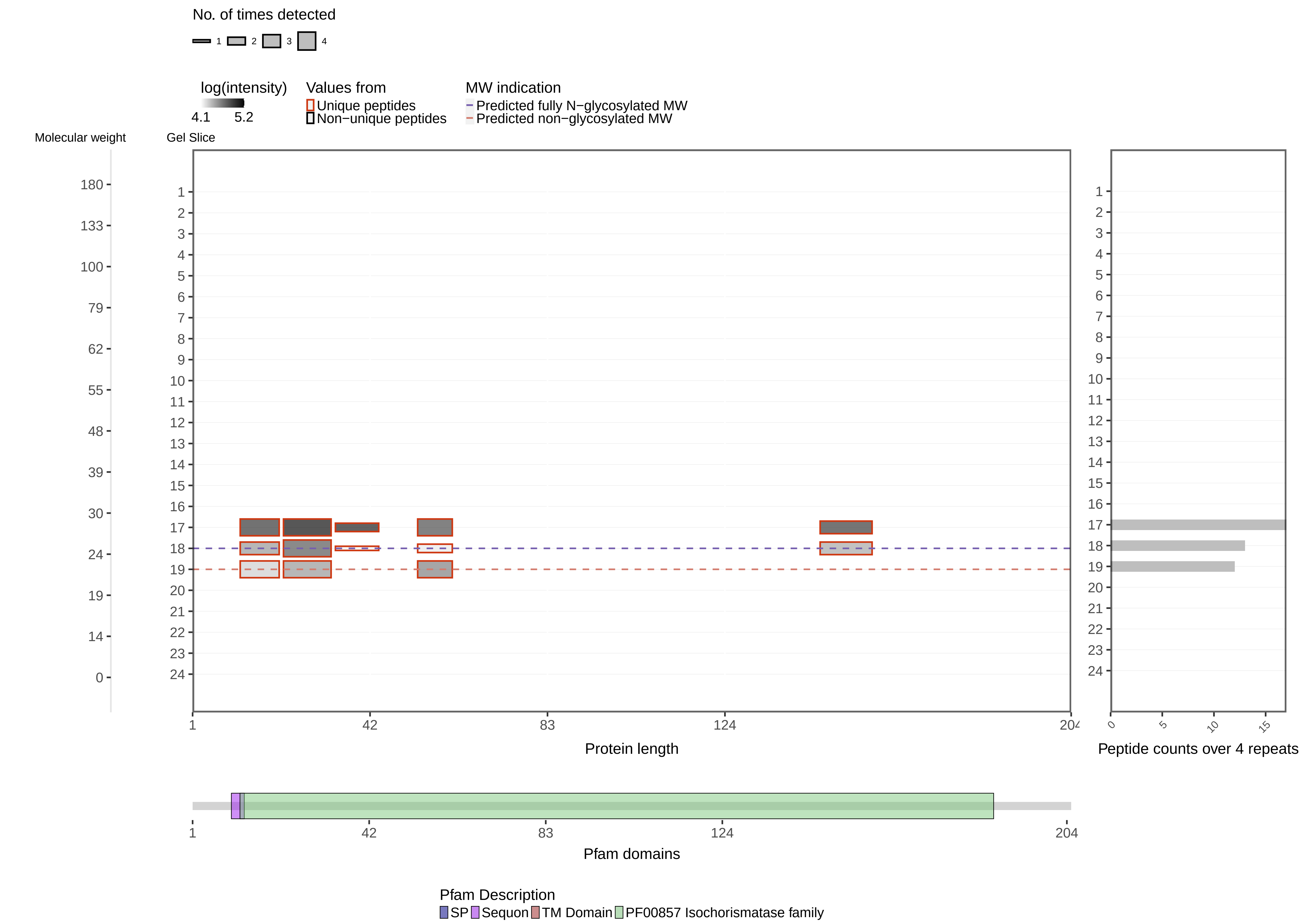


Protein sequence:

NbL13g20660.1 30S_ribosomal_protein_S17_chloroplastic−like [MW = 17.38 kDa]

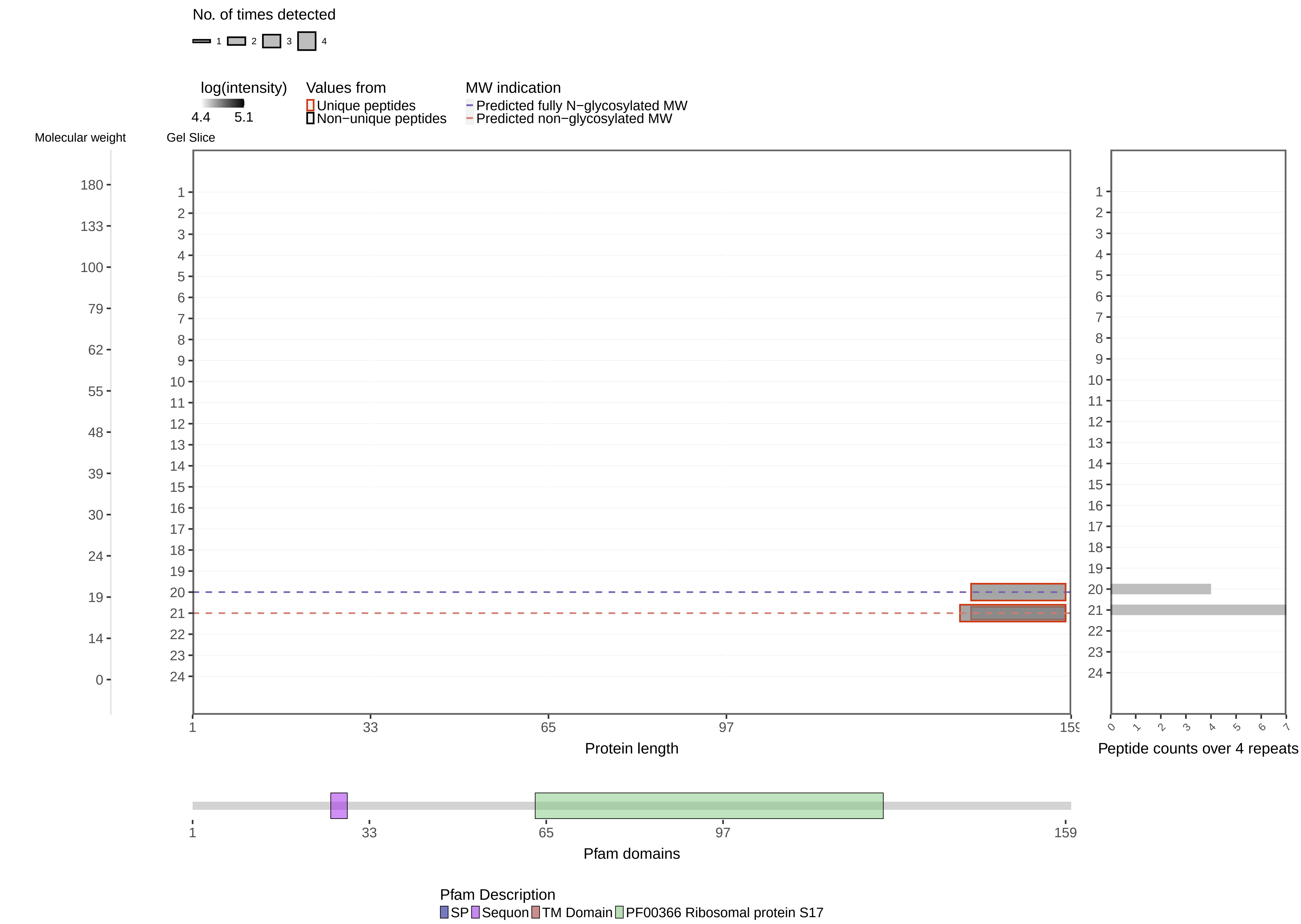


Protein sequence:

STNDKTVSVEVTRLAPHPKYKRRVRKKKKYQAHDPLNQFQVGDFVQLEKTAPISKTKTFLAVPVPPRNQPKPKGEDGQELGLPLESSLLQS*

### NbL13g20640.1 photosystem_I_reaction_center_subunit_VI−1_chloroplastic−like [MW = 15.32 kDa]

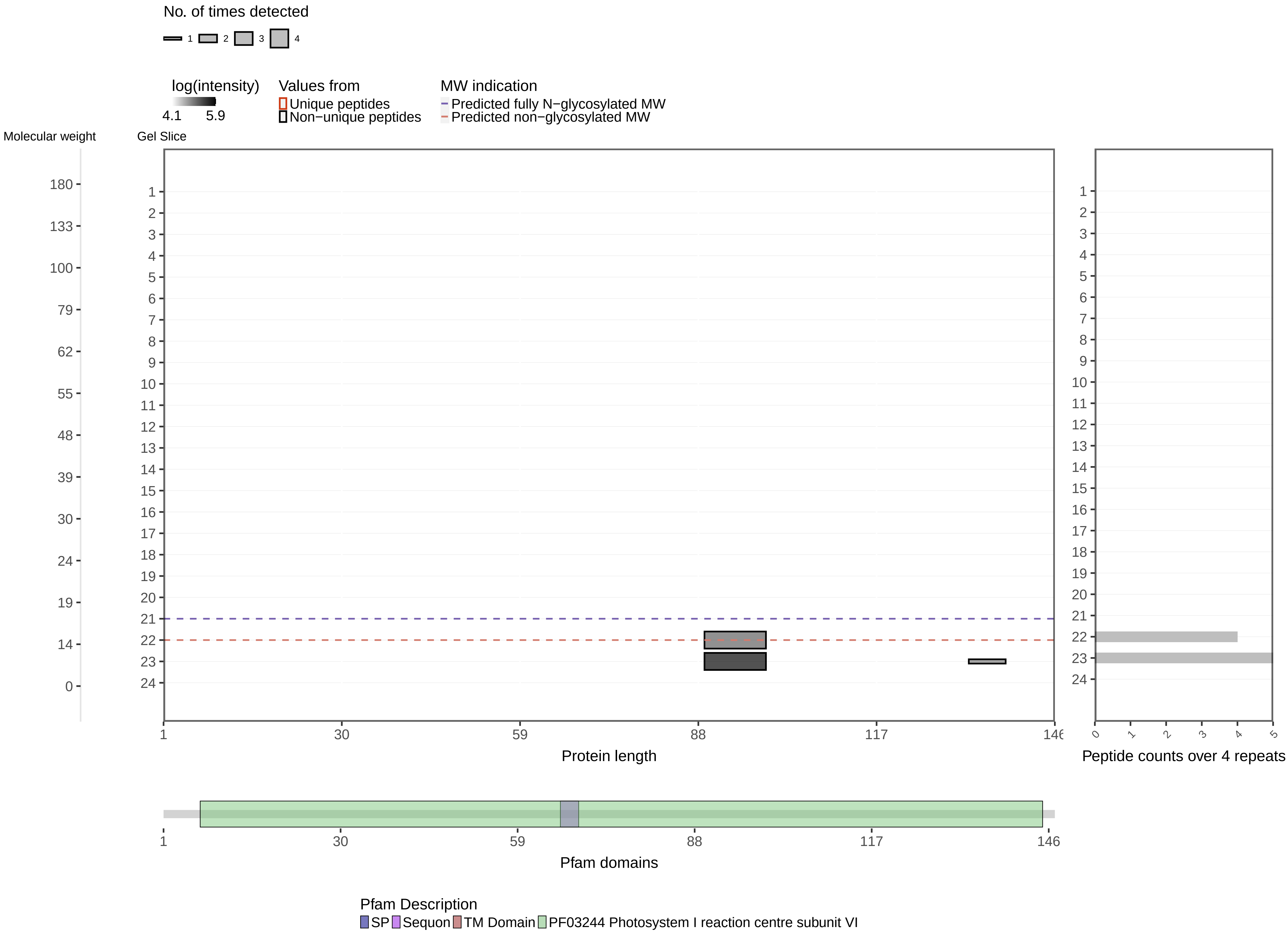


Protein sequence:

:DLEDLGNTTGQWDLYGSDAPSPYNSLQSKFFETFAAPFTKRGLLLKFLILGGGSTLAYFSSTASGDILPIKKGPQLPPKLGPRGKI*

### NbL13g20620.1 D−3−phosphoglycerate_dehydrogenase [MW = 34.47 kDa]

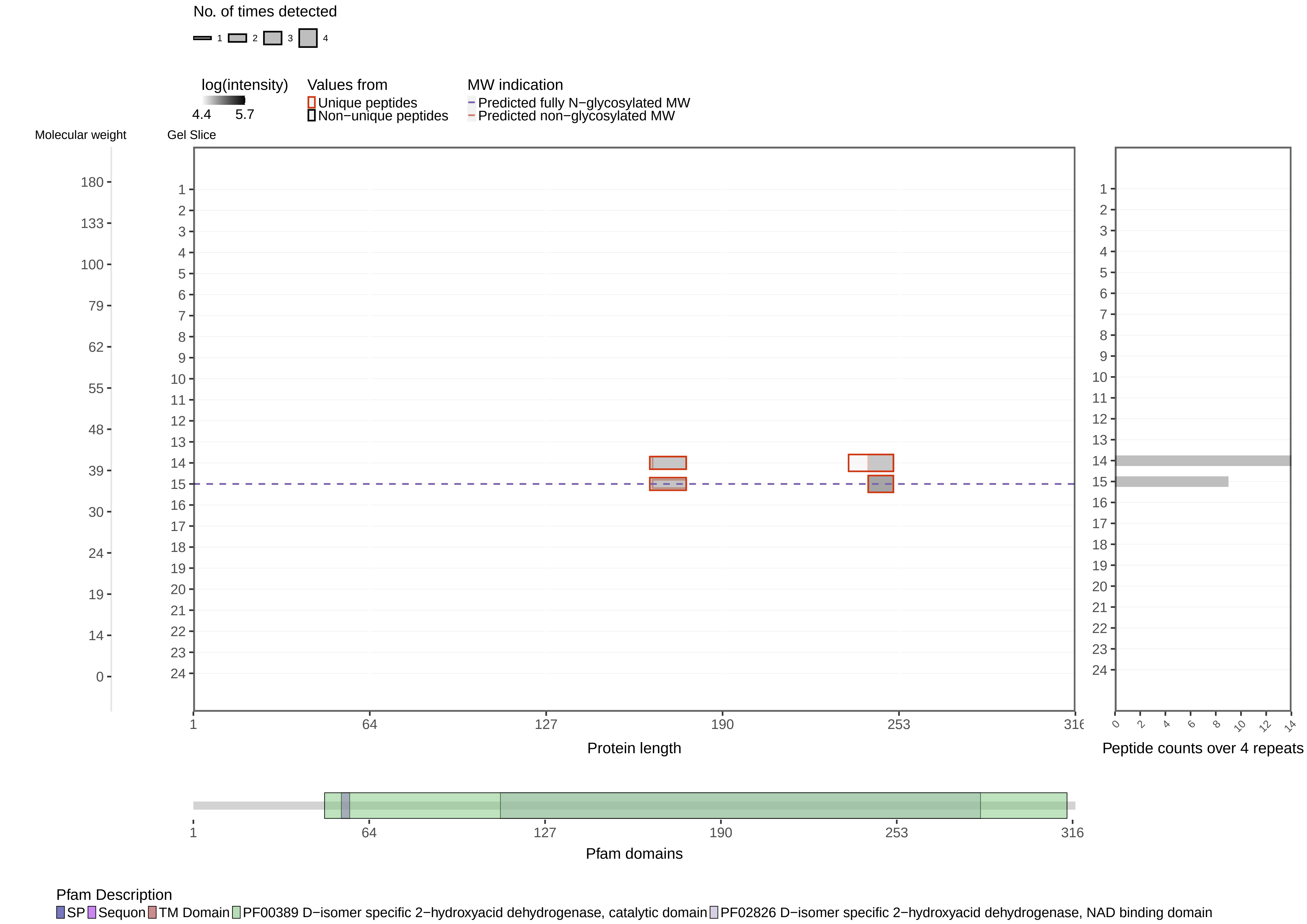


Protein sequence:

### NbL13g20570.1 ras−related_protein_Rab7 [MW = 22.98 kDa]

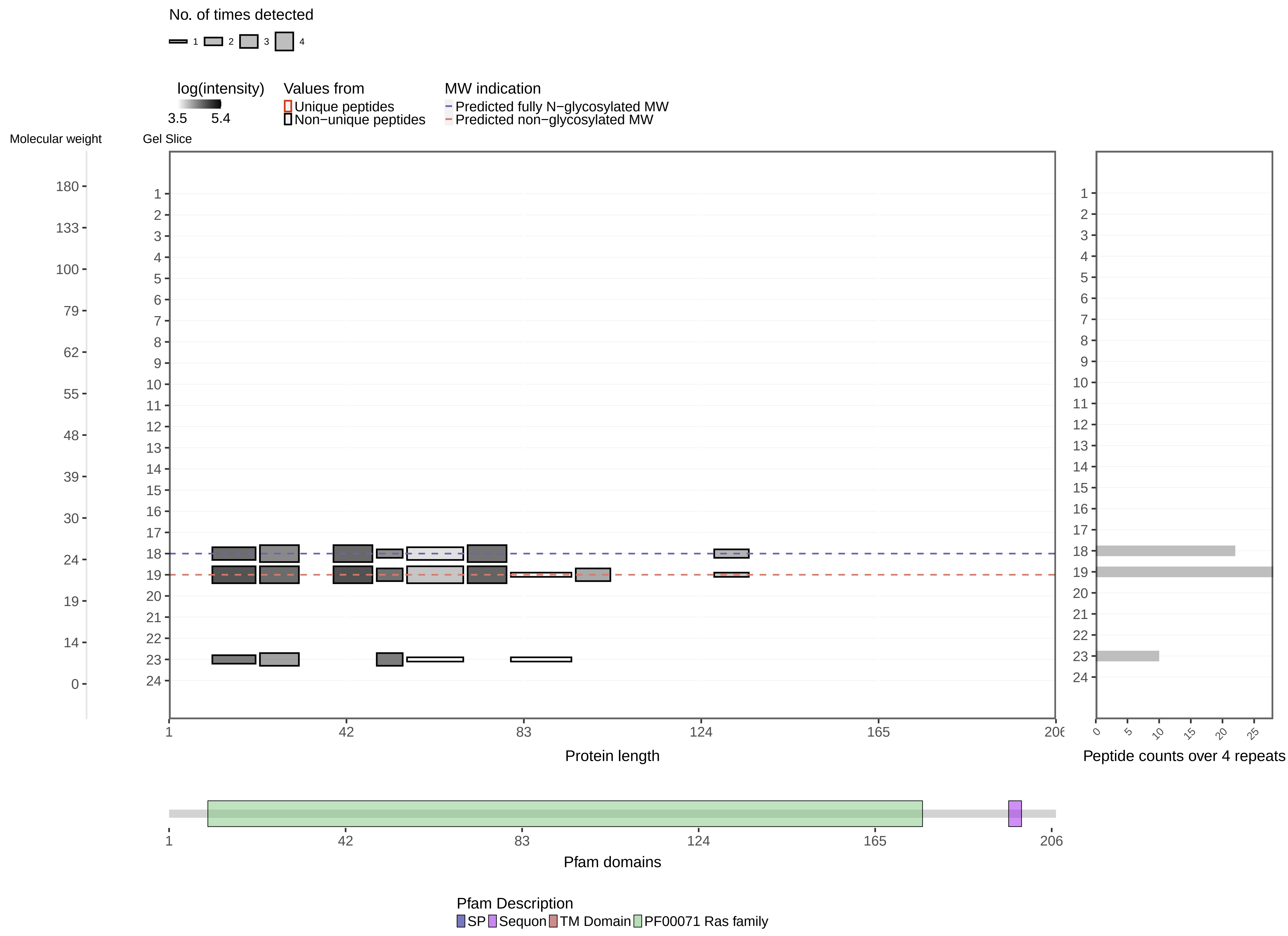


Protein sequence:

NVMKSFENLNNWREEFLIQASPSDPENFPFVVLGNKIDVDGGNSRVVSEKKAKAWCASKGIPYFETSAKEGFNVDAAFQCIAKNALKNEPEEEMYLPDTIDVAGGNQSRSTGCEC*

NbL13g20460.1 ruBisCO_large_subunit−binding_protein_subunit_beta_chloroplastic−like [MW = 64.95 kDa]

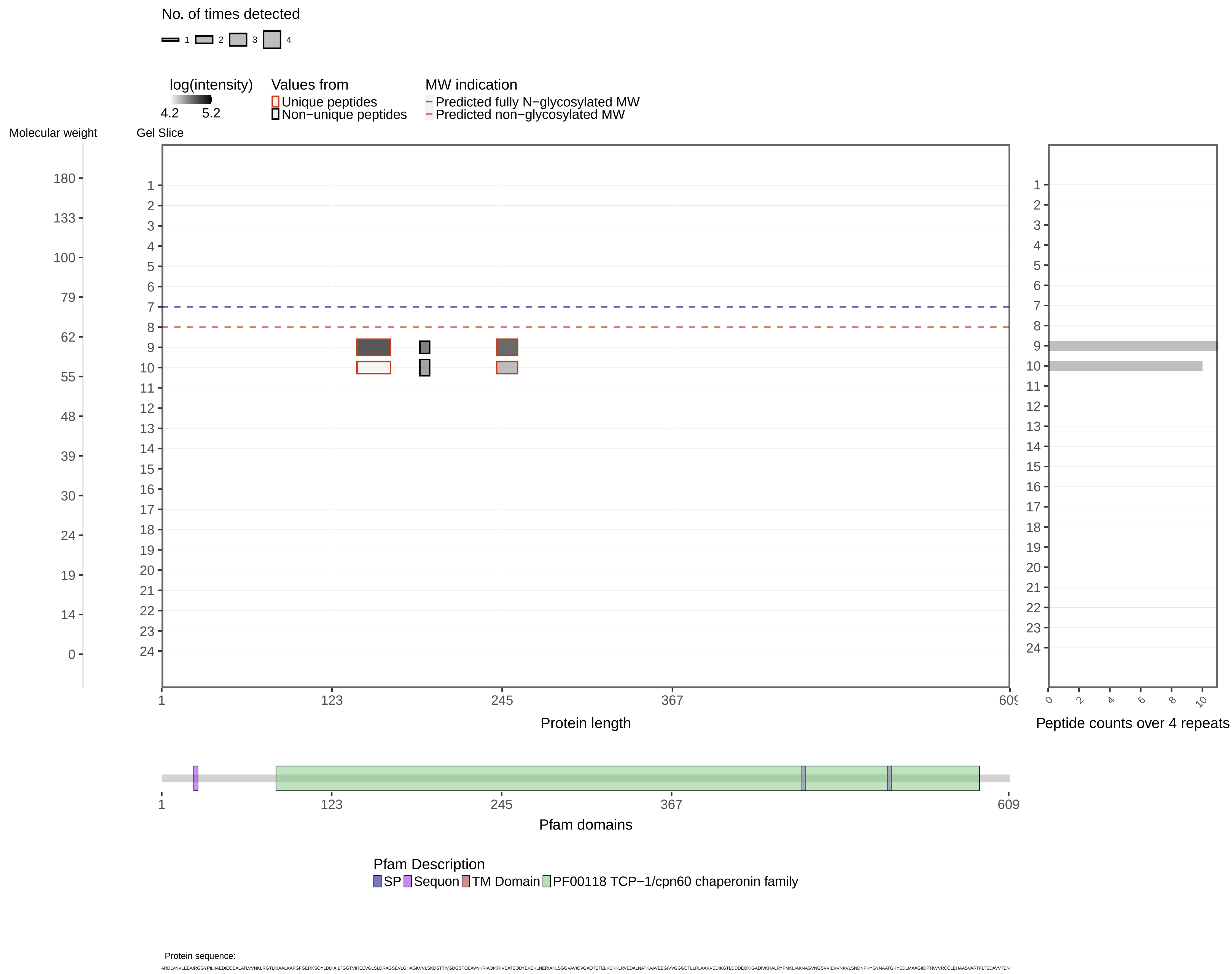

### NbL13g20300.1 clavaminate_synthase−like_protein_At3g21360 [MW = 36.24 kDa]

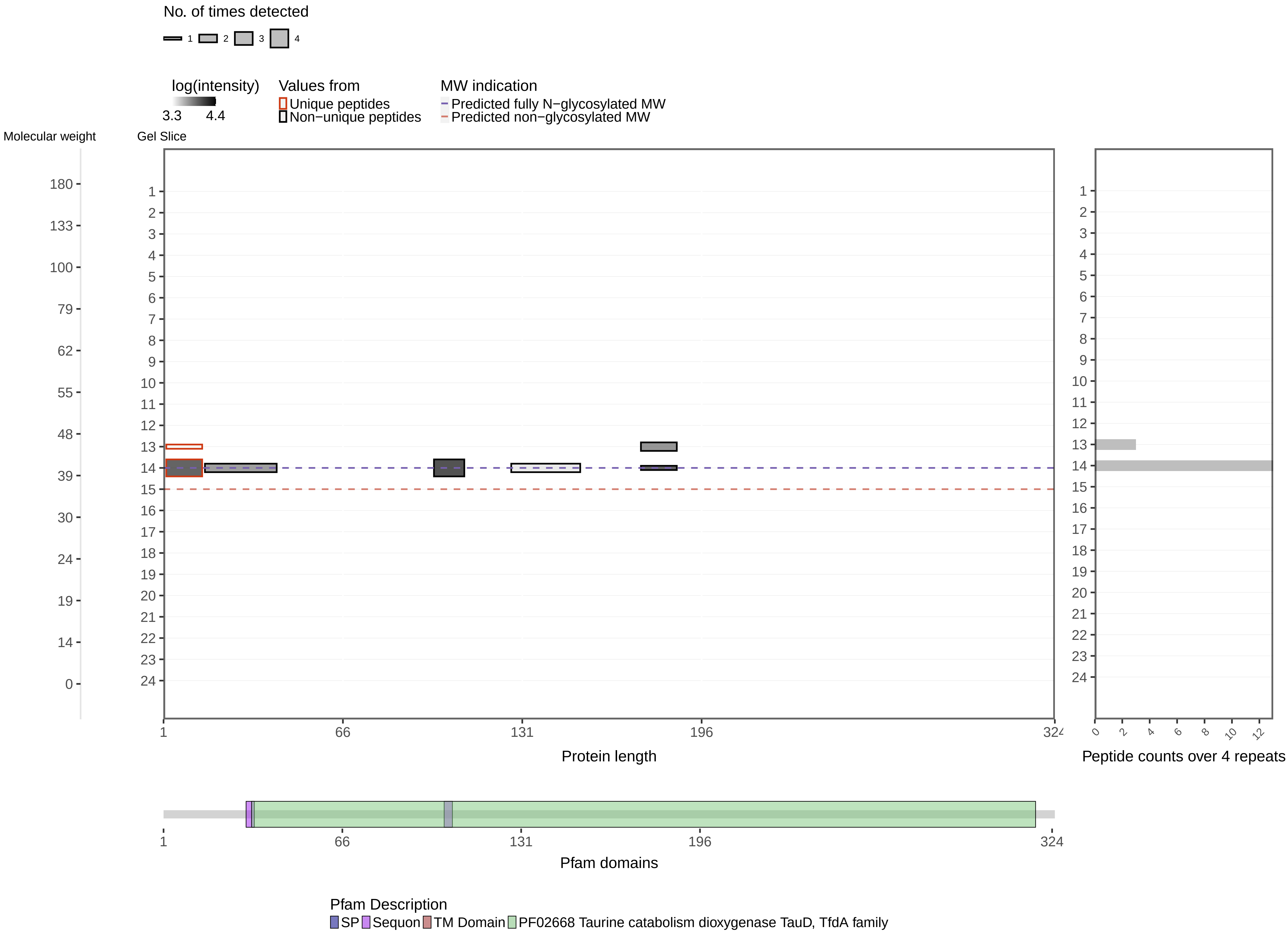


Protein sequence:

VYEKMKNKYPDFVERLEEHGLIYIRVLGEDDDPSSPIGRGWKSTFLTKDKSVAEERAAKLGMKLEWIEDGVKTVMGPIPAIKFNQERQQKIWFNSMVAAYTGWKDARNDPVKAVTFGDDQPLPAEVIHDCLRILEDESVAIPWKKGDVLLIDNLAVLHSRKPFNPPRRILASLCK*

NbL13g20290.1 lysM_domain_receptor−like_kinase_3 [MW − SP = 70.16 kDa]

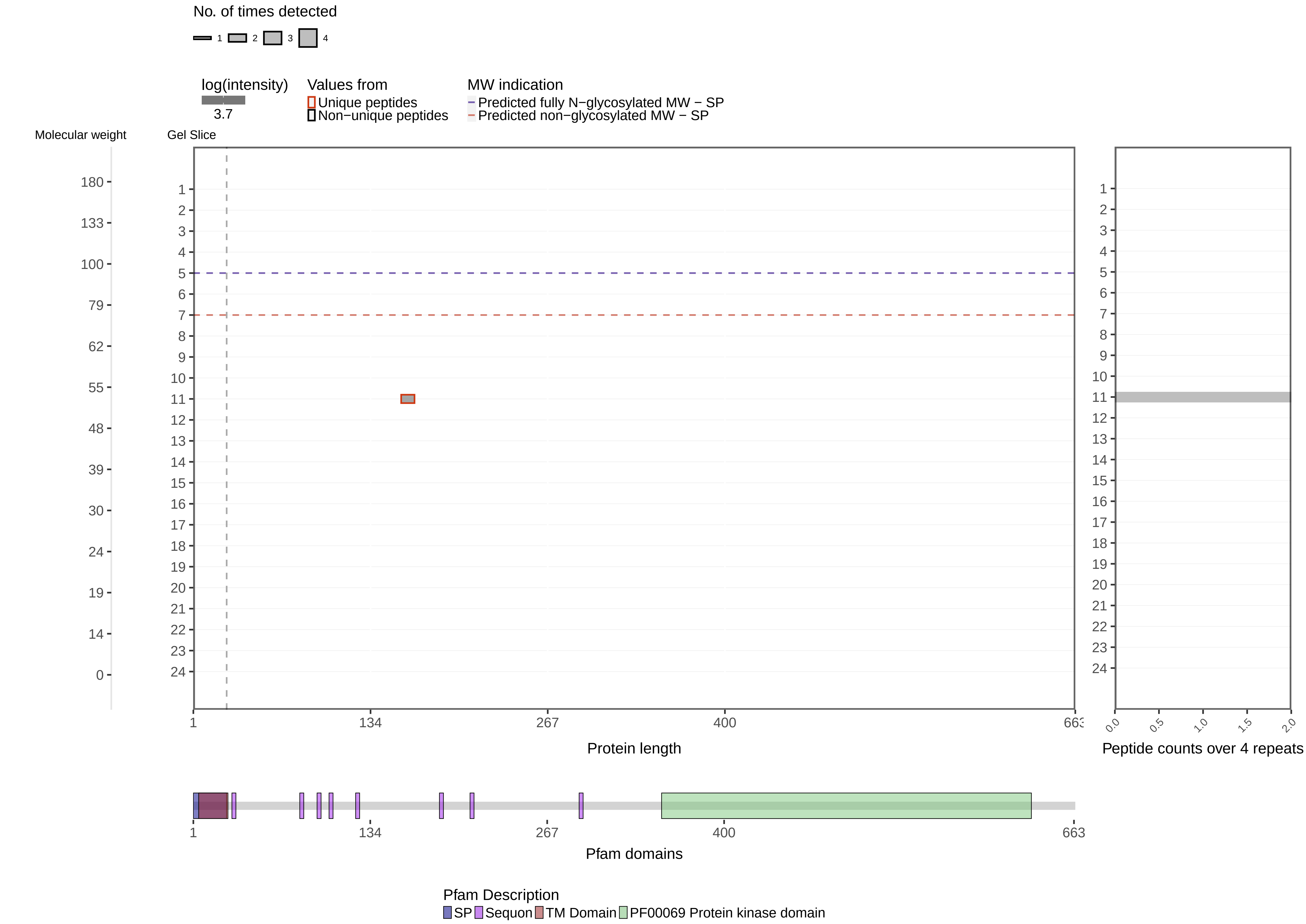


Protein sequence:

QQINIPKVIGTDVFDMEKPLVFIYEDILSSTDGFSDSNLLGHGTYGSVYYAILRNQEVAIKRMTATKTREFTAEMKVLCKVHHLNLVELIGYAVSNDELVLVYEYAQKGSLKSHLHDPQNKGHTPLSWIMRVQIALDAARGLEYIHEHTKPHYVHRDIKTSNILLDDSFRAKISDFGFSKLMGITNDGEASATRVVGTYGYVAPEYLRDGLATTKTDVYAFGVVLFEMLTGKEAVTRTEGNVMKTAERRSLVSLMLAALRNSPDSTSMTSLKDQLDPSLMDLYPSDCVFKVAMLAK

### NbL13g20270.1 hexokinase−1 [MW = 53.89 kDa]

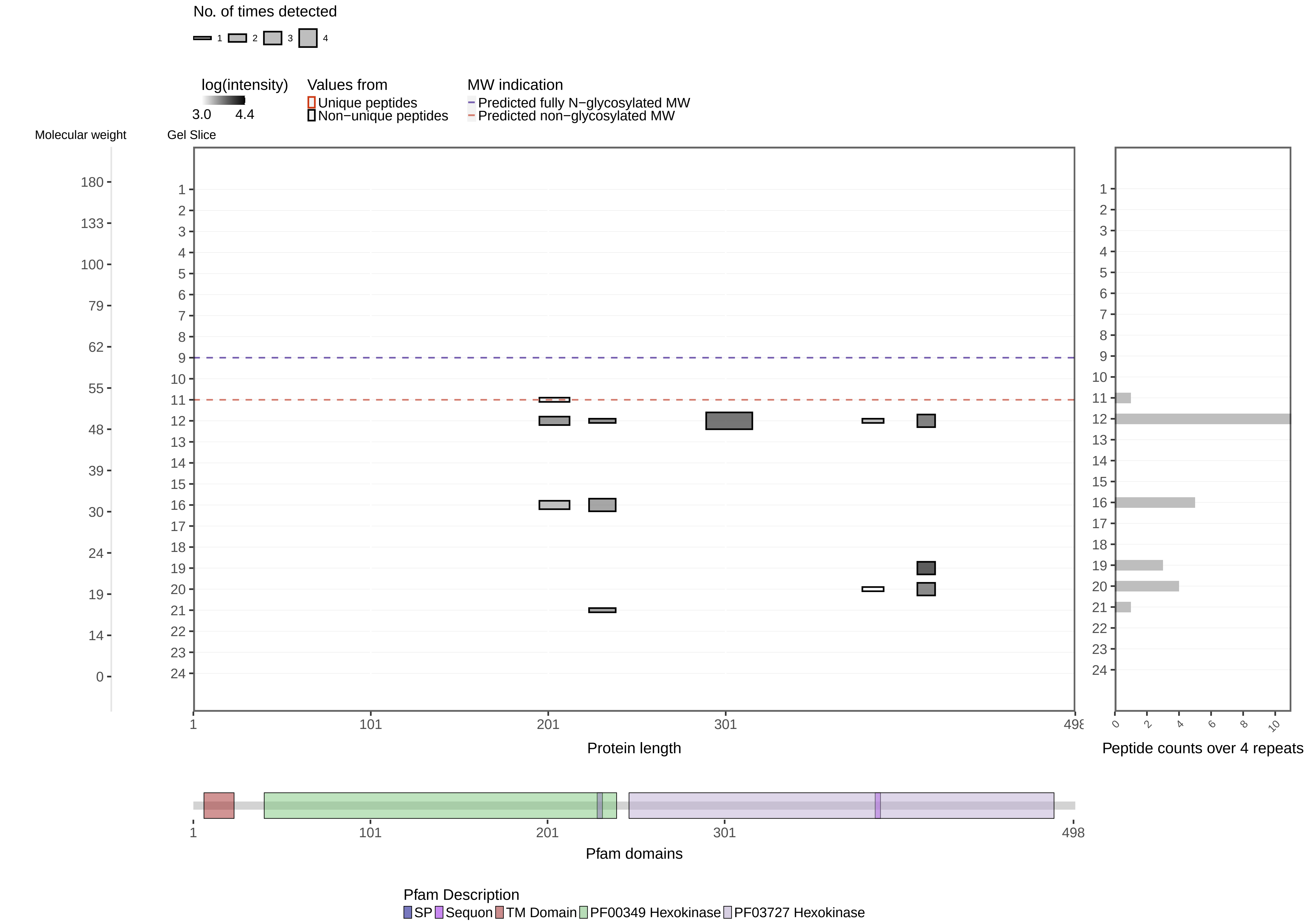


Protein sequence:

AGGKYTHNDVAVAVILGTGTNAAYVERVQAIPKWHGPMPKSGEMVINMEWGNFRSSHLPLTQYDHALDTNSLNPGDQIFEKMTSGMYLGEILRRVLLRMAEEAGIFGDEVPPKLKSPFVLRTPDMSAMHHDTSSDLRVVGDKLKDILEISNTSLKTRRLVVELCNIVATRGARLAAAGVLGILKKMGRDTPRQGGPQKMVVAMDGGLYEHYAEYRTCLENTLKELLGDELATSIVFEHSNDGSGIGAALLAASNSMYLEDKS*

### NbL13g20240.1 acid_phosphatase_1−like [MW − SP = 27.08 kDa]

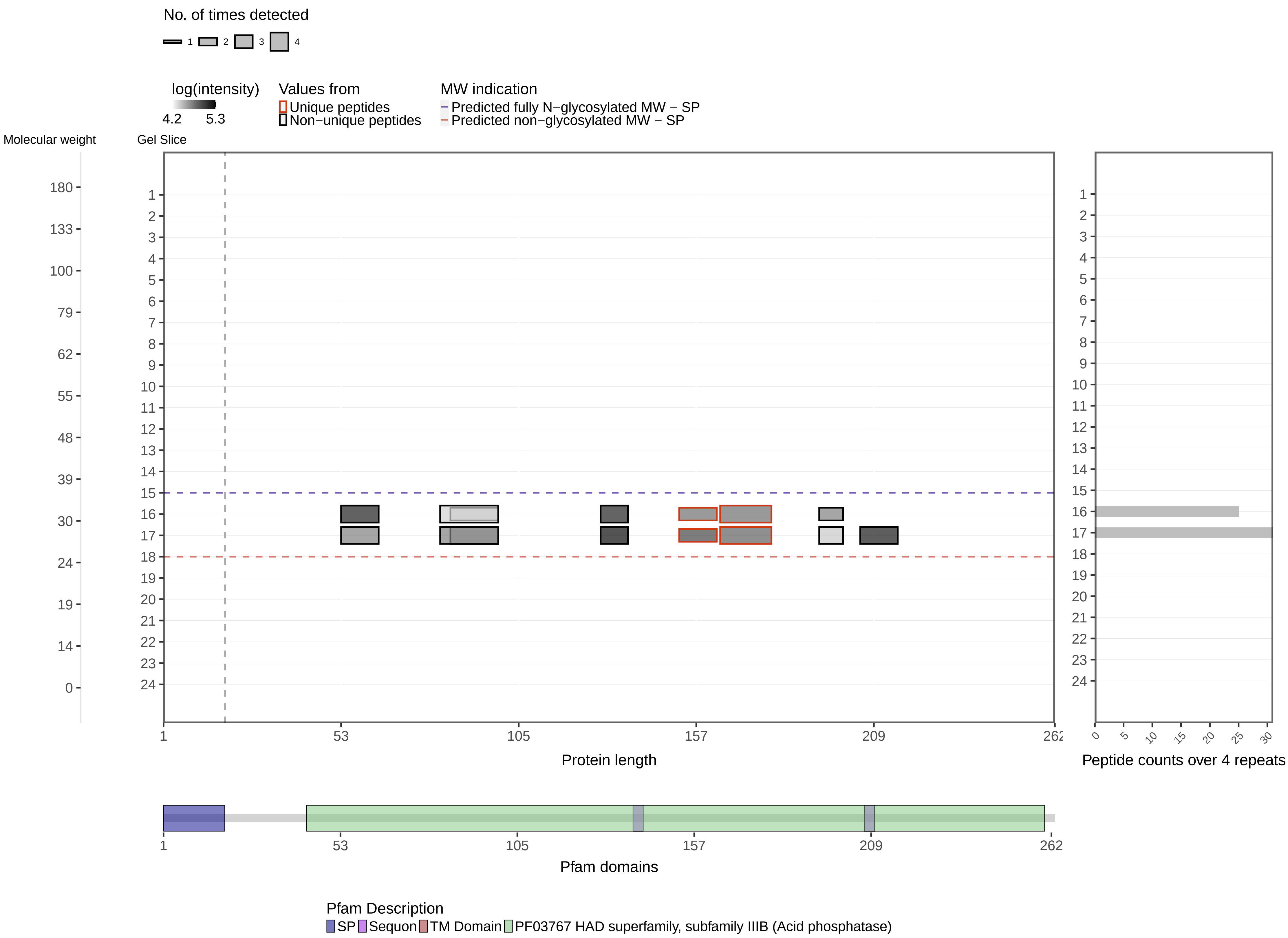


Protein sequence:

TLSNLPYYARSDVAFGATKFNGTKFDEWTREGKAPAVPGALFLYRTLLGMGIKPVFITGTKEEFRQVRIANLKKVGYHSWVKLILKGVNDTGSSVMYKSGKRAELVKAGYRIVGNIGDQWSDLLGDFVGDRTFKLPDPMYYIG*

### NbL13g20200.1 GDSL_esterase/lipase_APG−like [MW − SP = 36.29 kDa]

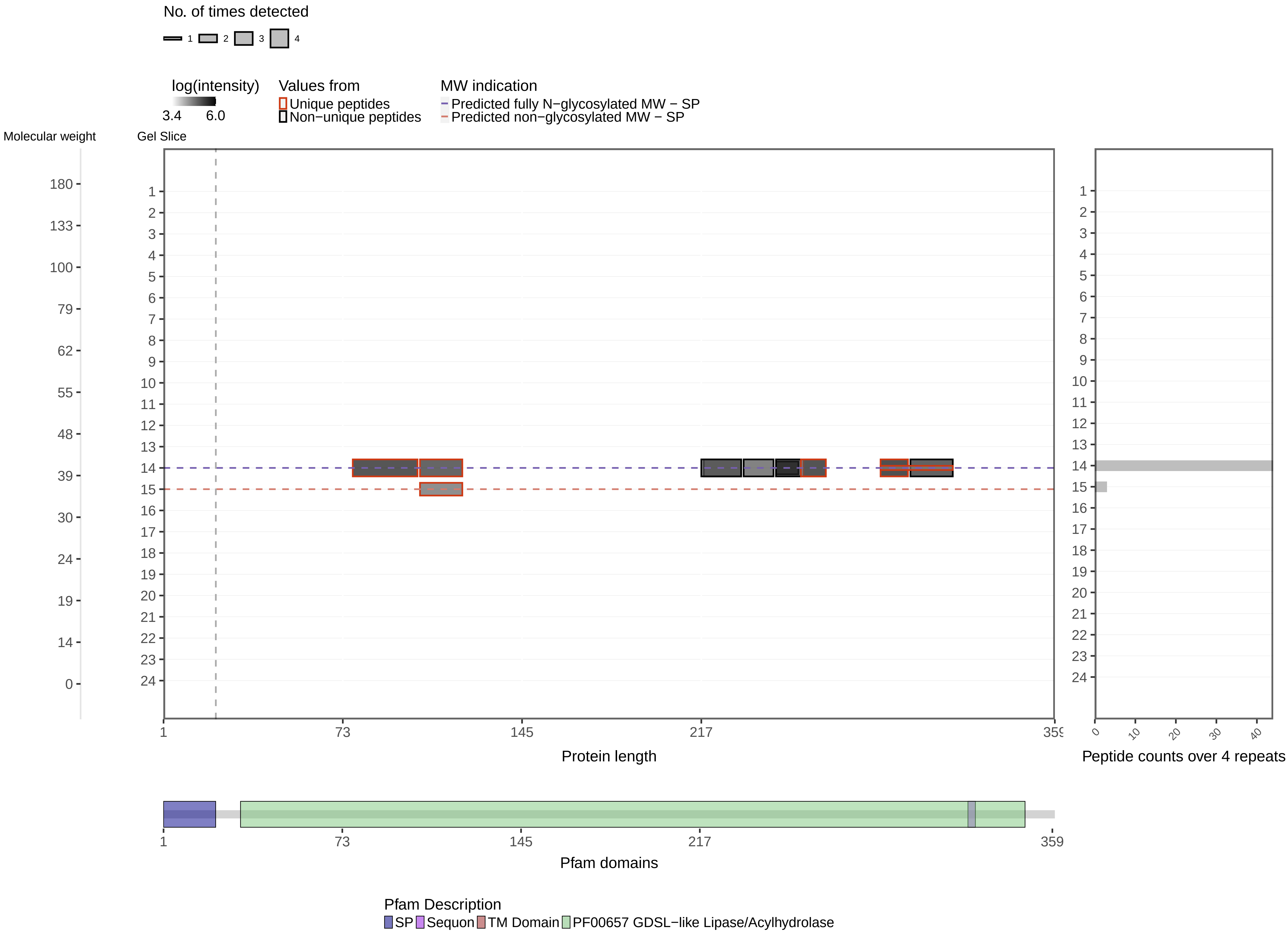


Protein sequence:

ILSAGSSDFLQNYYINPYINKLYTPDQYGSYLIGIFTSFVKDLHAQGARRIGVTSLPPLGCLPAARTLFGFHESGCVSKINTDAQQFNKKINSAATQLQKQLPGLKIAIFDIFQPLYDLVKSPSNHGFTEANKGCCGTGTVETTSLLCNPKSPGTCSNATQYVFWDSVHPSEAANQVLADSLIIQGINLIG*

NbL13g20110.1 uncharacterized_RNA_methyltransferase_BH0687 [MW = 63.51 kDa]

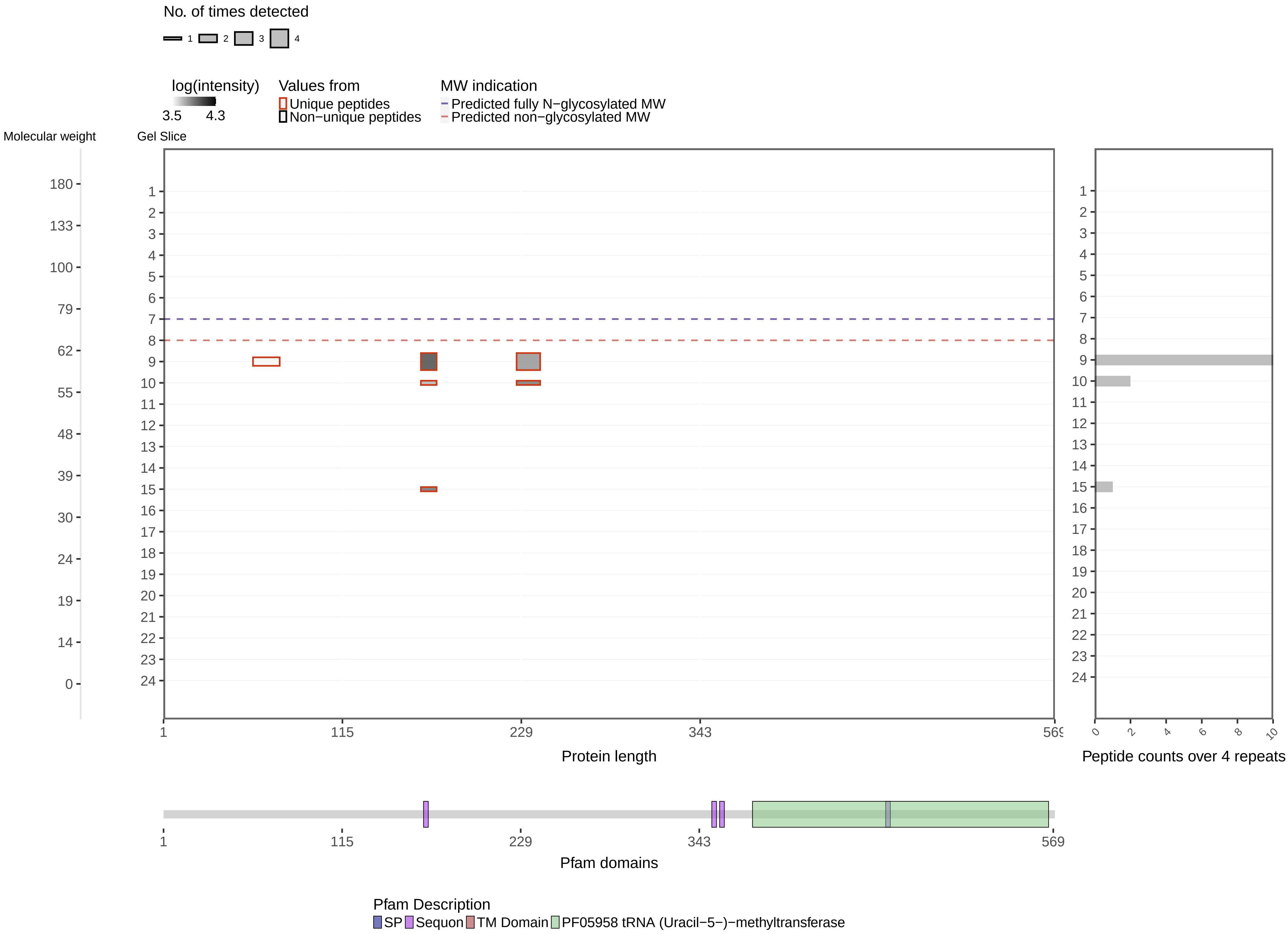


Protein sequence:

SGNEVLEVVQRCWRDPELGLSPYDVHSHSGFLKHLMLRTGRNVETGLPELMVNFVTSSYNPECLRPIVEKISAIPEVVTIVNNVNTSIGNTSIGEKEYTLYGKSTITEILRGLTFQISANSFFQTNTHQAEILYKLIEDCAFLKGDGSEIVLDLFCGTGTIGLTLAKRVKHVYGFEVVAQAVSDARQNAKLNCIDNATFIEGDLNKIDLNFASNLPKPDIVITDPNRPGMHIKLIKFLLKLKATRIVYVSCNPATCARDLNYLCYGVPEQNIEGNYRLSSLQPVDMFPHTPHIECVCLLEL

NbL13g20100.1 IAA−amino_acid_hydrolase_ILR1−like_4 [MW = 57.9 kDa]

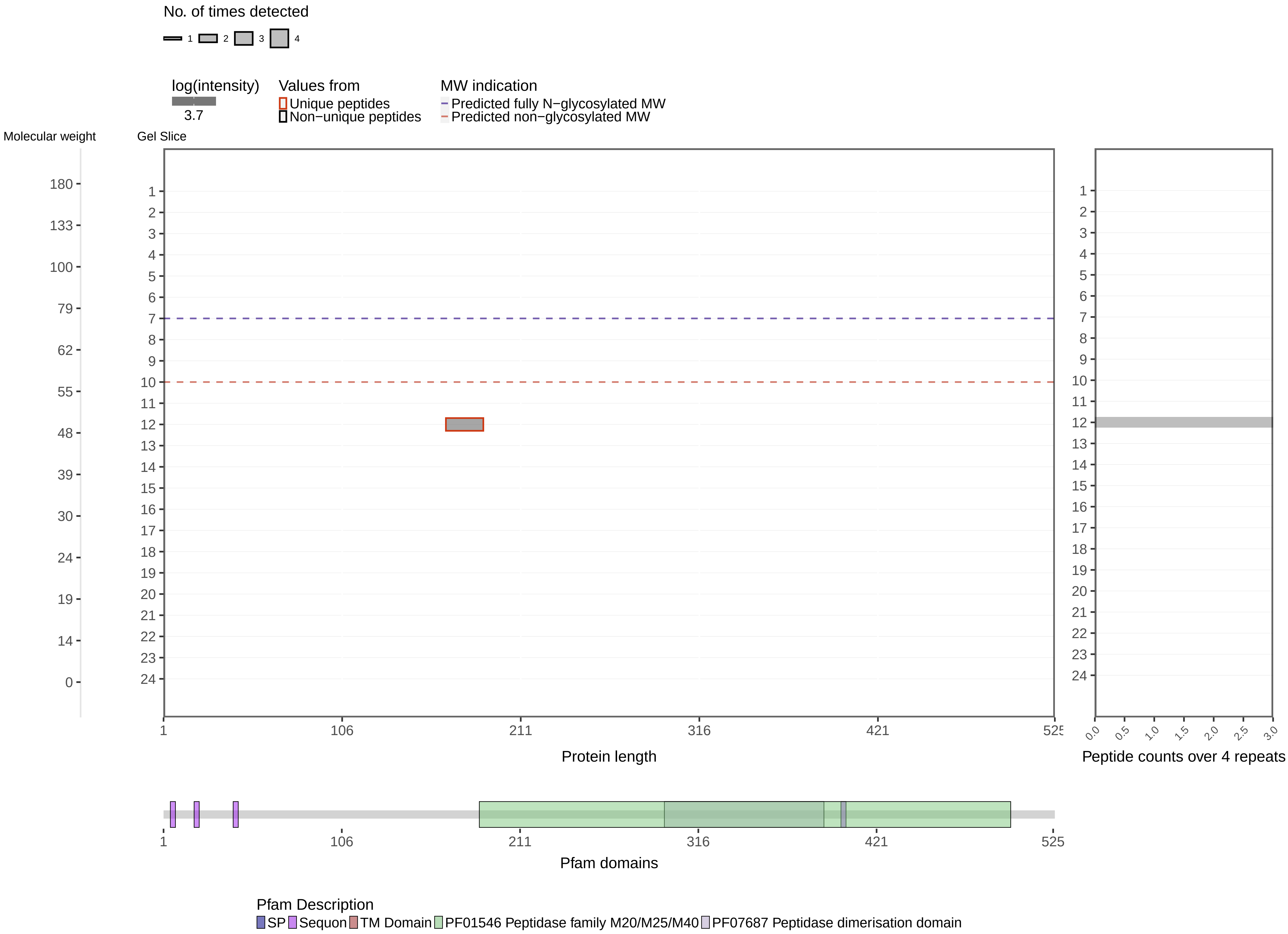


Protein sequence:

GGGGAKKMIDAGALENIEAIFGLHVNPQFPLGKVSSRPGPFLAGSGFFEAVISGKGGHAAIPQHSIDPILAASNVIVSLQHLVSREADPLDSQVVTVAKFQGGGAFNVIPDSVTIGGTFRAFSKESFQQLRQRIEEVIVGQAAVQRCNATVDFLTKEKPFFPPTVNDKNLHKHFQRVASDMLGNDHVKDMEPLMGSEDFAFYQEVIPGYFYLLGMQDETNEKLVSVHSPYFKINEEALPIGAALQASLAIRYLLEAQSLVPSSSITDHNDEL*

### NbL13g20060.1 RWD_domain−containing_protein_1 [MW = 28.06 kDa]

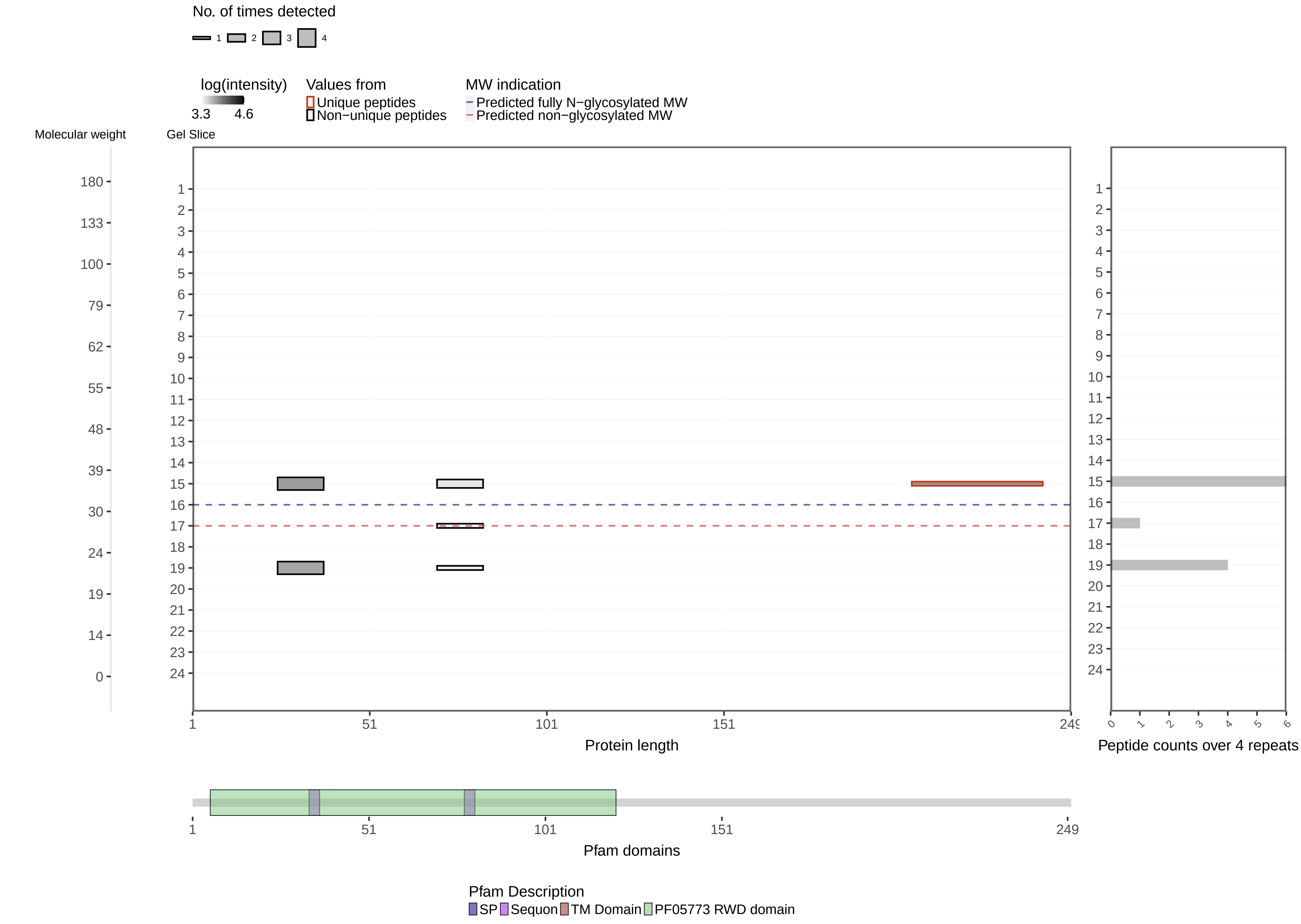


Protein sequence:

.VTSAKDWLSERFAEETGAEDAEDDEAKKEEVIIPHGEPVTIDTFLAWRERFEAELALERAKLMPDSALNAPKEKKLTGRQWFESGRASGKAAAAIAEESDEDEDDIDFDDDDFEDDEEDMLEHYLAEKTDSSSHS*

### NbL13g20040.1 60S_ribosomal_protein_L28−1−like [MW = 19.79 kDa]

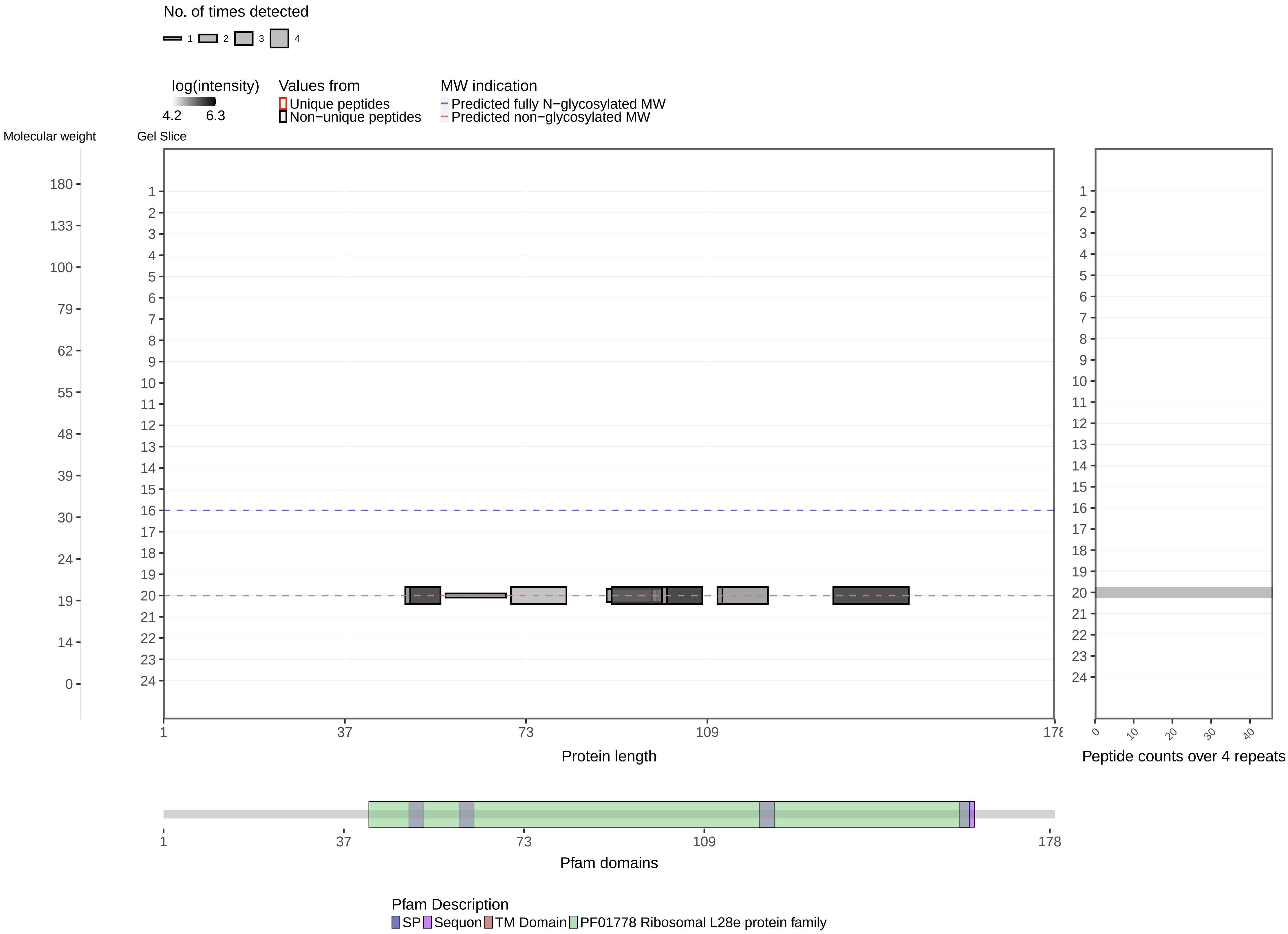


Protein sequence:

.HSYKHSGLANKKTVTIQAEGKDKSVLLATSKTKKQNKPSTLLNKSVMKKEFRRMAKAVTNQVADNYYRPDLKKAALARLSAVNRSLKVAKSGVKKKNRQA*

NbL13g20010.1 ESCRT−related_protein_CHMP1B [MW = 22.7 kDa]

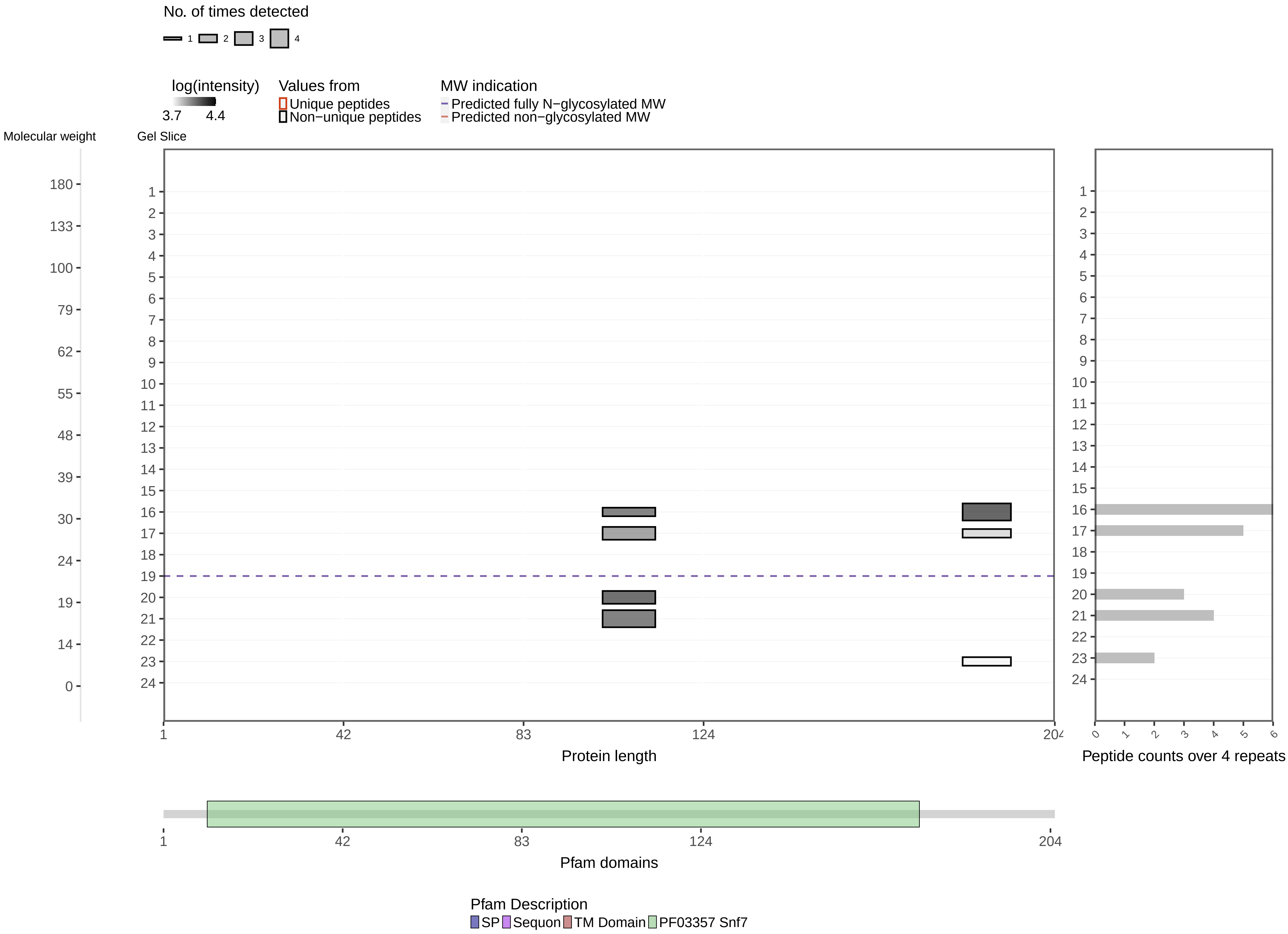


Protein sequence:

TISKSMGNIVKSLESSLATGNLQKMSETMDKFEQQFVNMEVQAEFMESSMAGSTSLSTPEDQVNSLMHQVADDYGLEVSVGLPQAAGHAIPTKDSEKVDEDDLTRRLAELKARG*

### NbL13g20000.1 40S_ribosomal_protein_S15a−2−like [MW = 14.75 kDa]

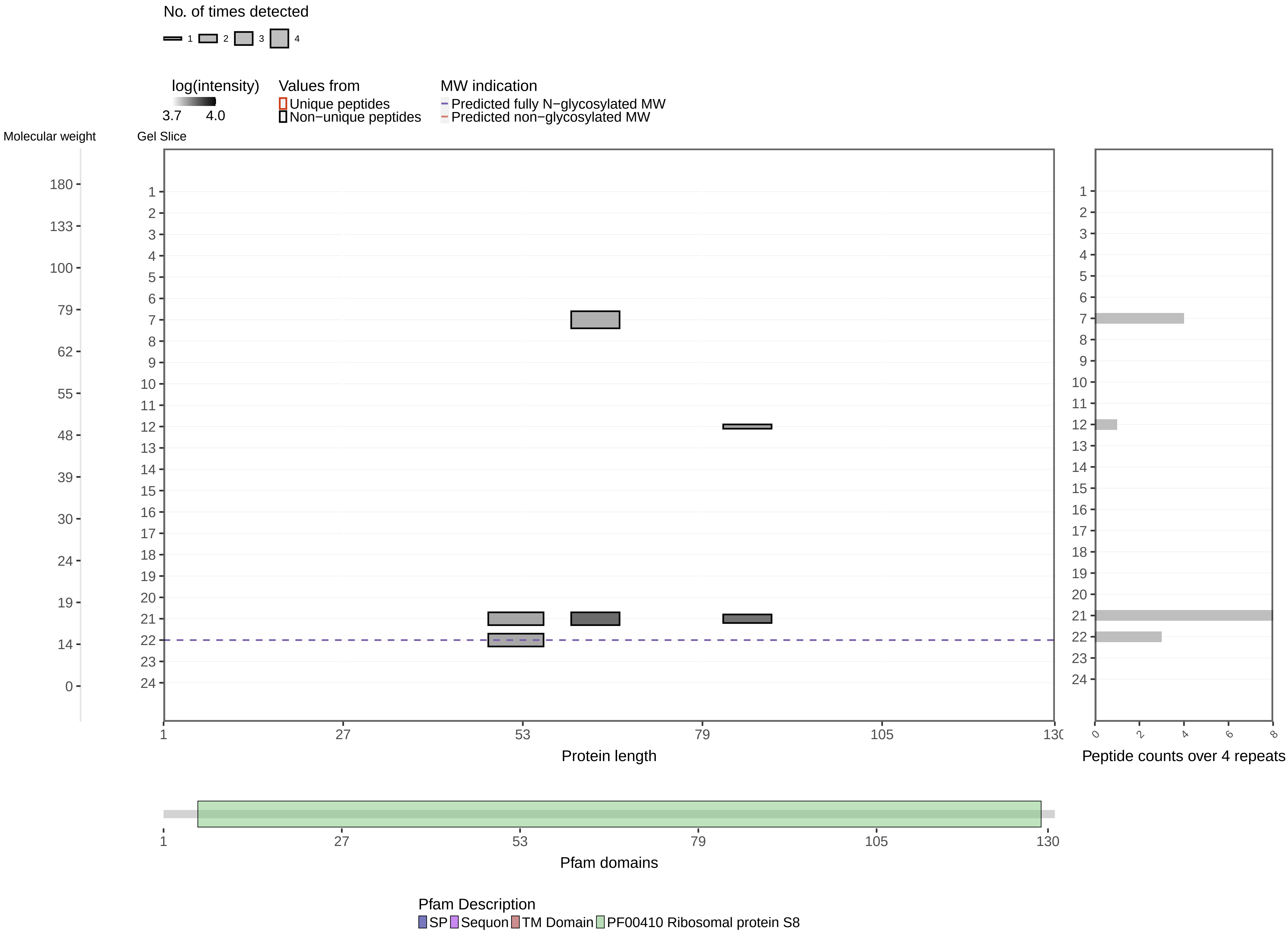


Protein sequence:

DPHRVGKITVQLLGRVNDCRALTYRQDIKAASIEQYKTRALPTRQWGFVVITTPNGVLDHEEAQRQNVGGQVIGYFY*

NbL13g19950.1 serine/threonine_protein_phosphatase_2A_55_kDa_regulatory_subunit_B_beta_isoform [MW = 57.46 kDa]

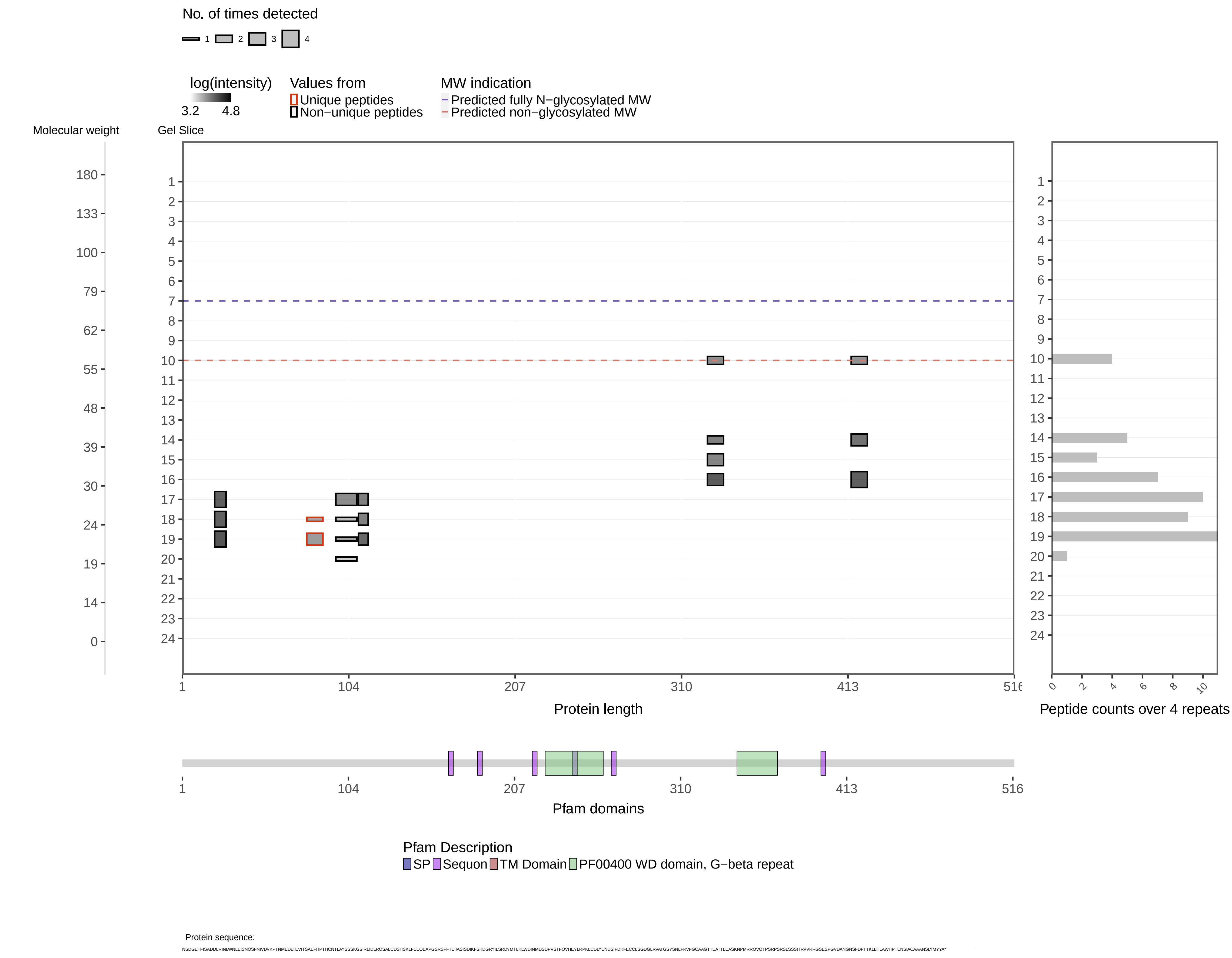

NbL13g19810.1 beta−galactosidase−like [MW = 67.48 kDa]

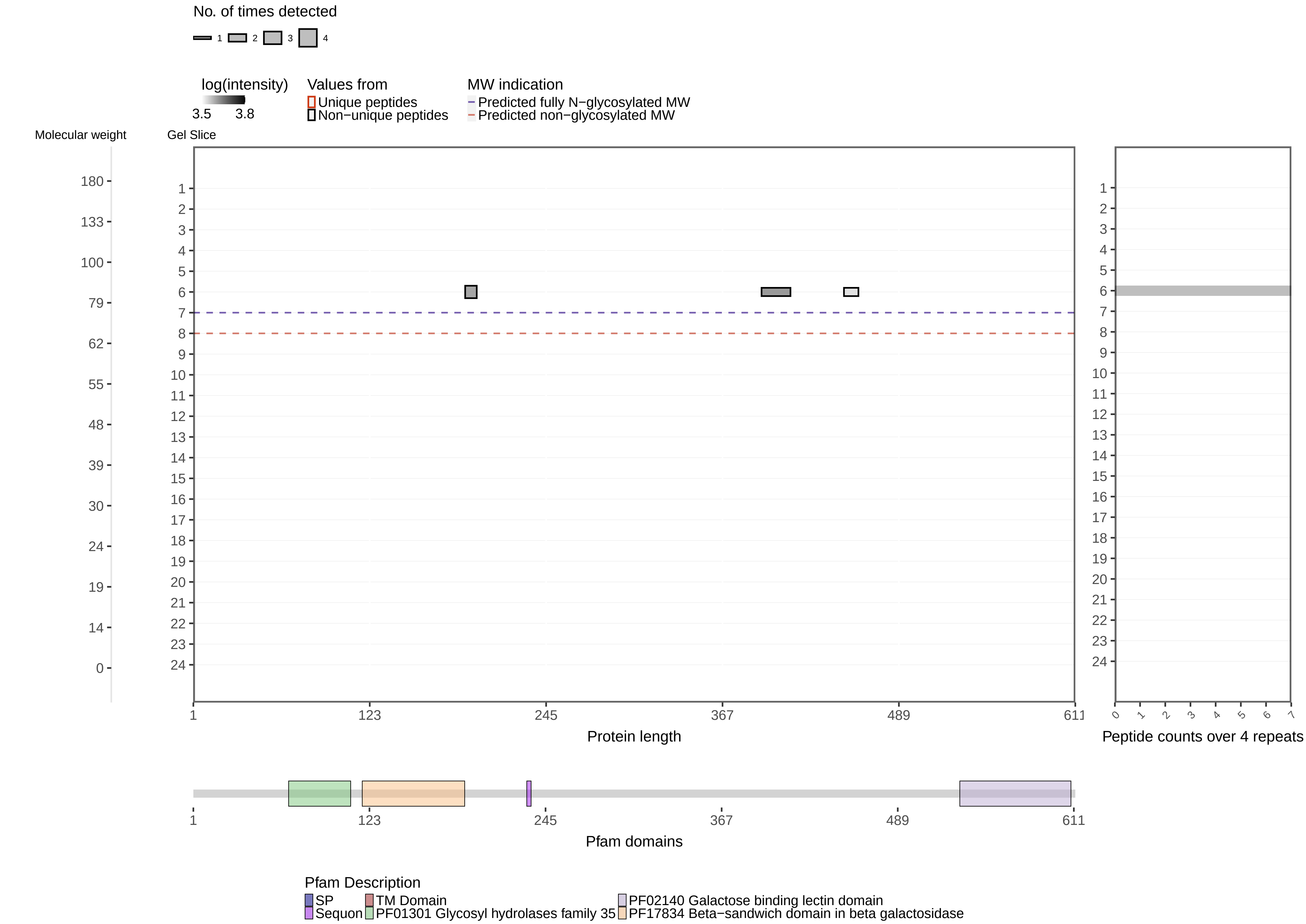


Protein sequence:

[illegible]

### NbL13g19780.1 WD−40_repeat−containing_protein_MSI4−like [MW = 47.62 kDa]

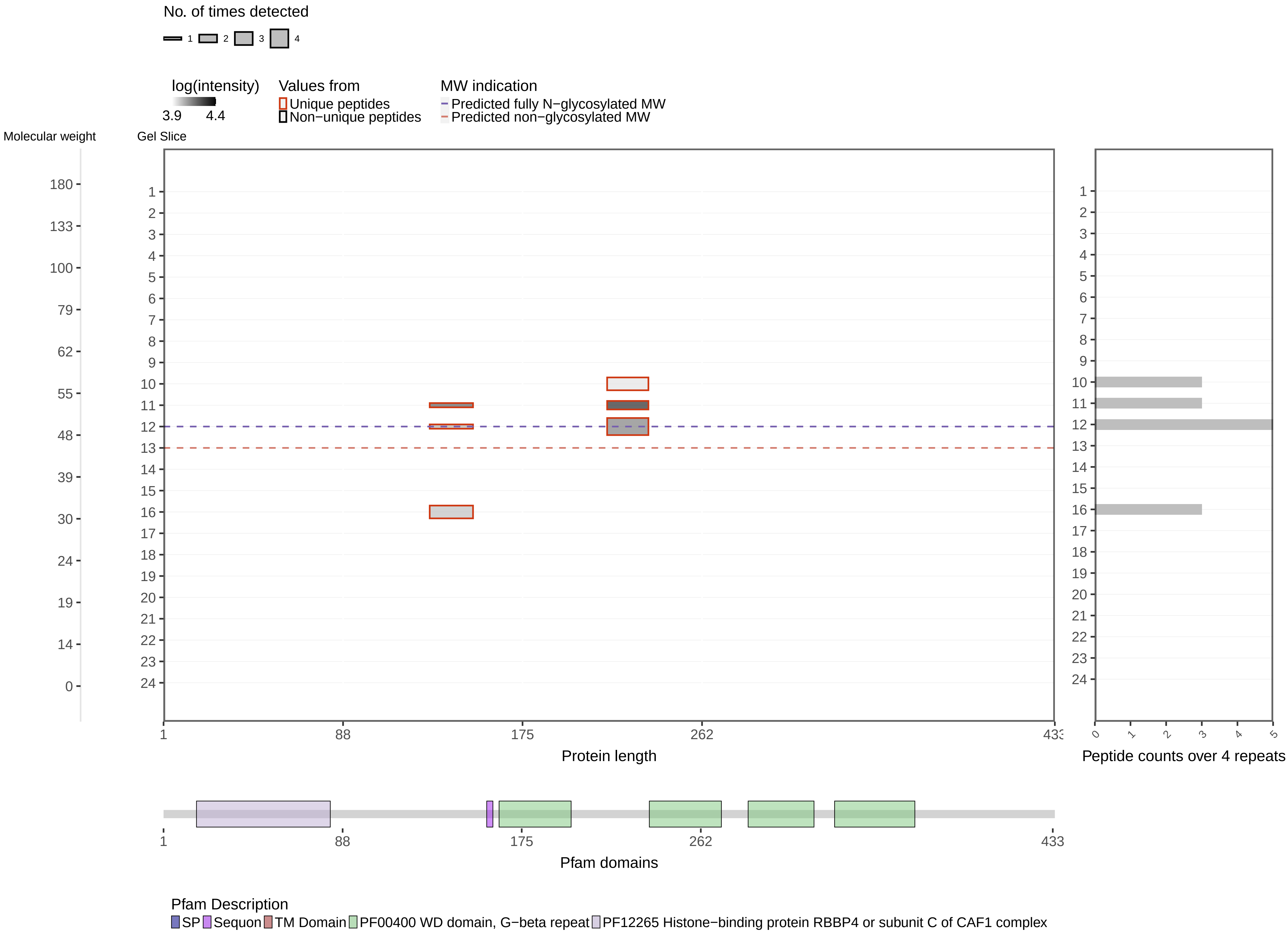


Protein sequence:

STLAADQGATKSPGSGASNSKPSAEGPTVQARGIFQGHDDTVEDVQFCPSSAQEFCSVGDDSCLILWDARTGSSPAVKVEKAHNADLHCVDWNPHDVNYILTGSADNTIRMFDRRKLTSGGVGSPIHIFEGHTAAVLCVQWSPDKSSVFGSSAEDGILNIWDHEKIGKMEDSAGRKASTSPPGLFFRHAGHRDKVVTSTGMQQIHGPLLVSQMMVKVLVEVVHYRYGE*

### NbL13g19760.1 protein_HOTHEAD−like [MW − SP = 61.68 kDa]

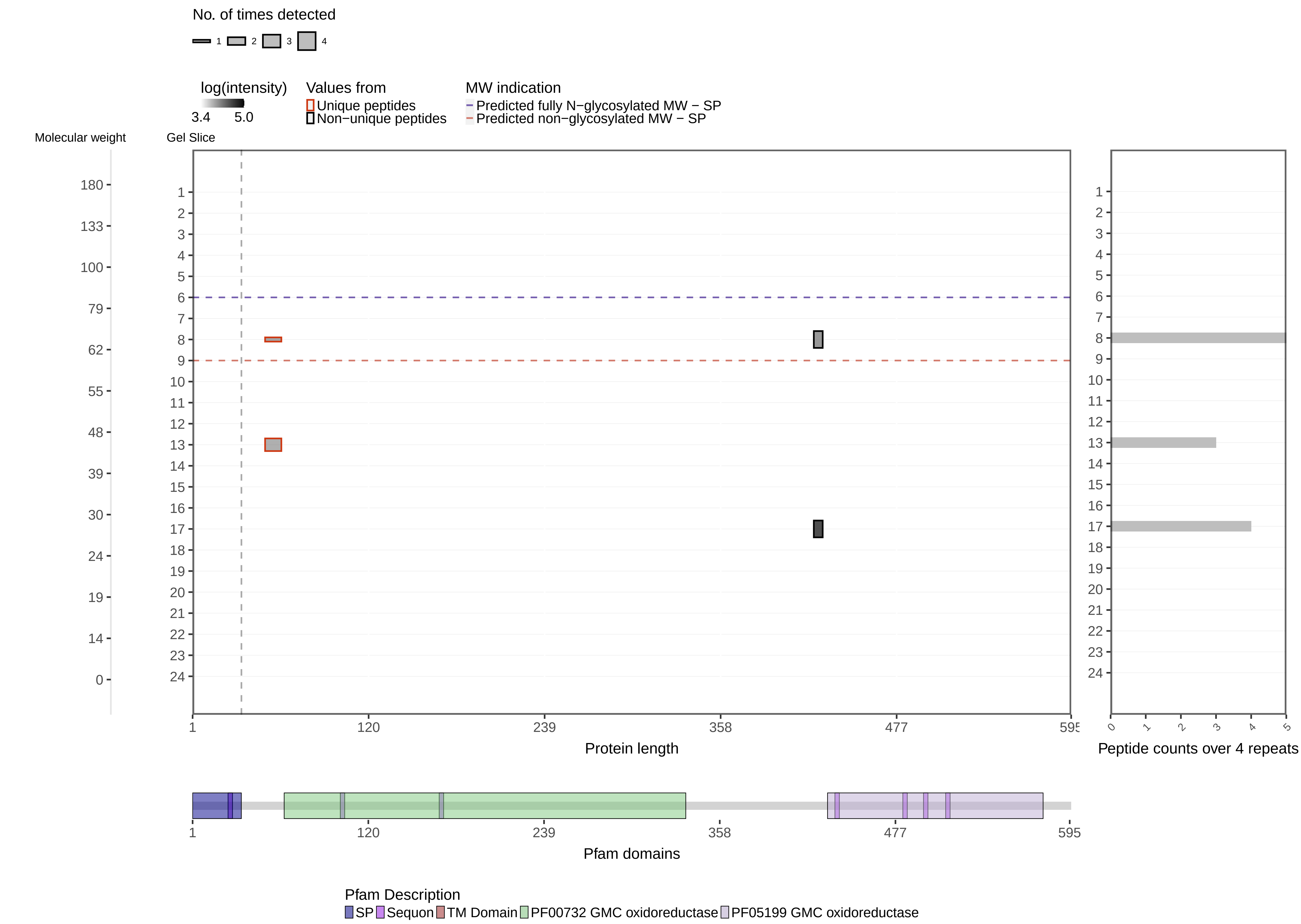


Protein sequence:

NbL13g19680.1 chaperonin_CPN60−like_2_mitochondrial [MW = 61.07 kDa]

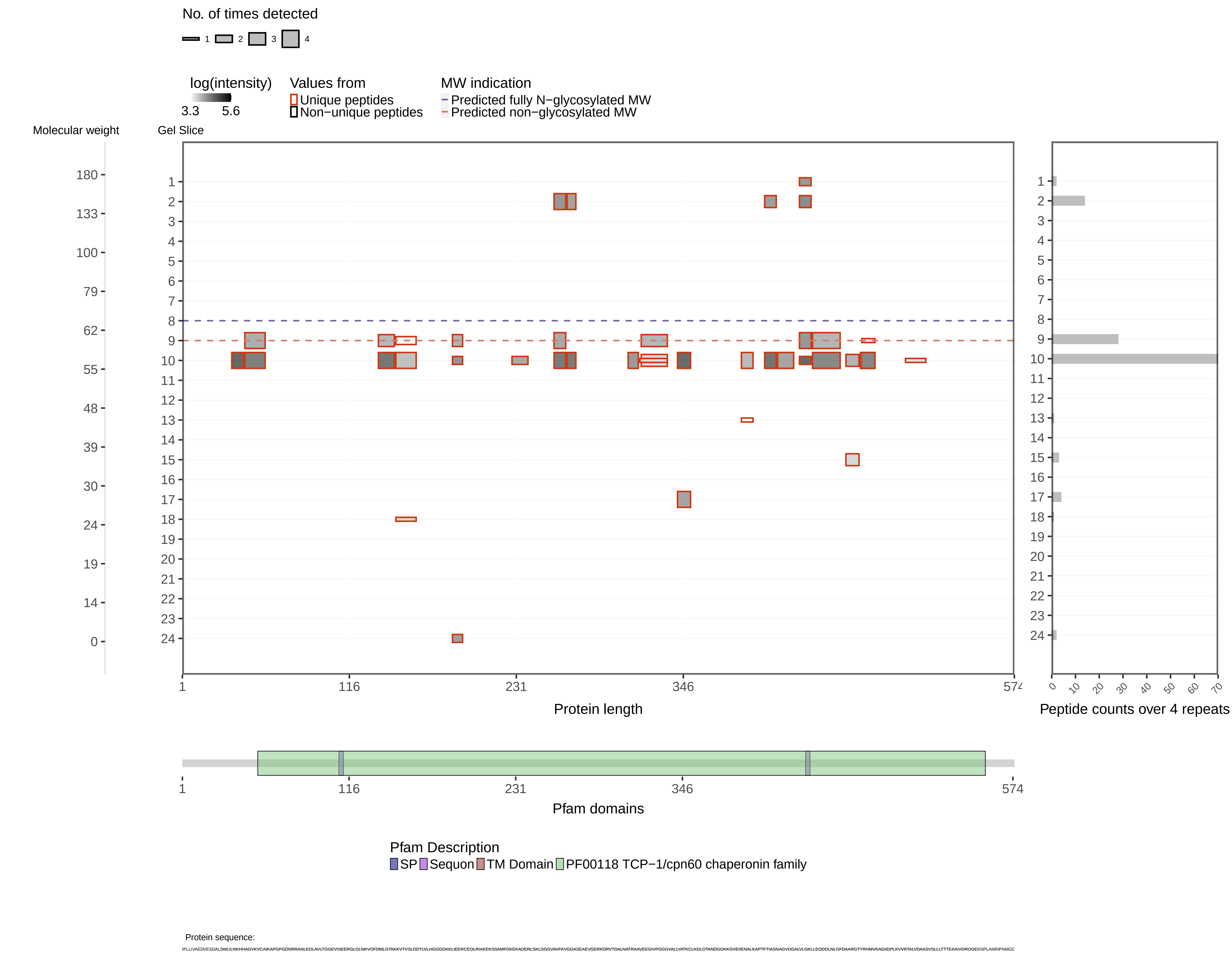

### NbL13g19600.1 glyoxylate/succinic_semialdehyde_reductase_2_chloroplastic−like [MW = 38.9 kDa]

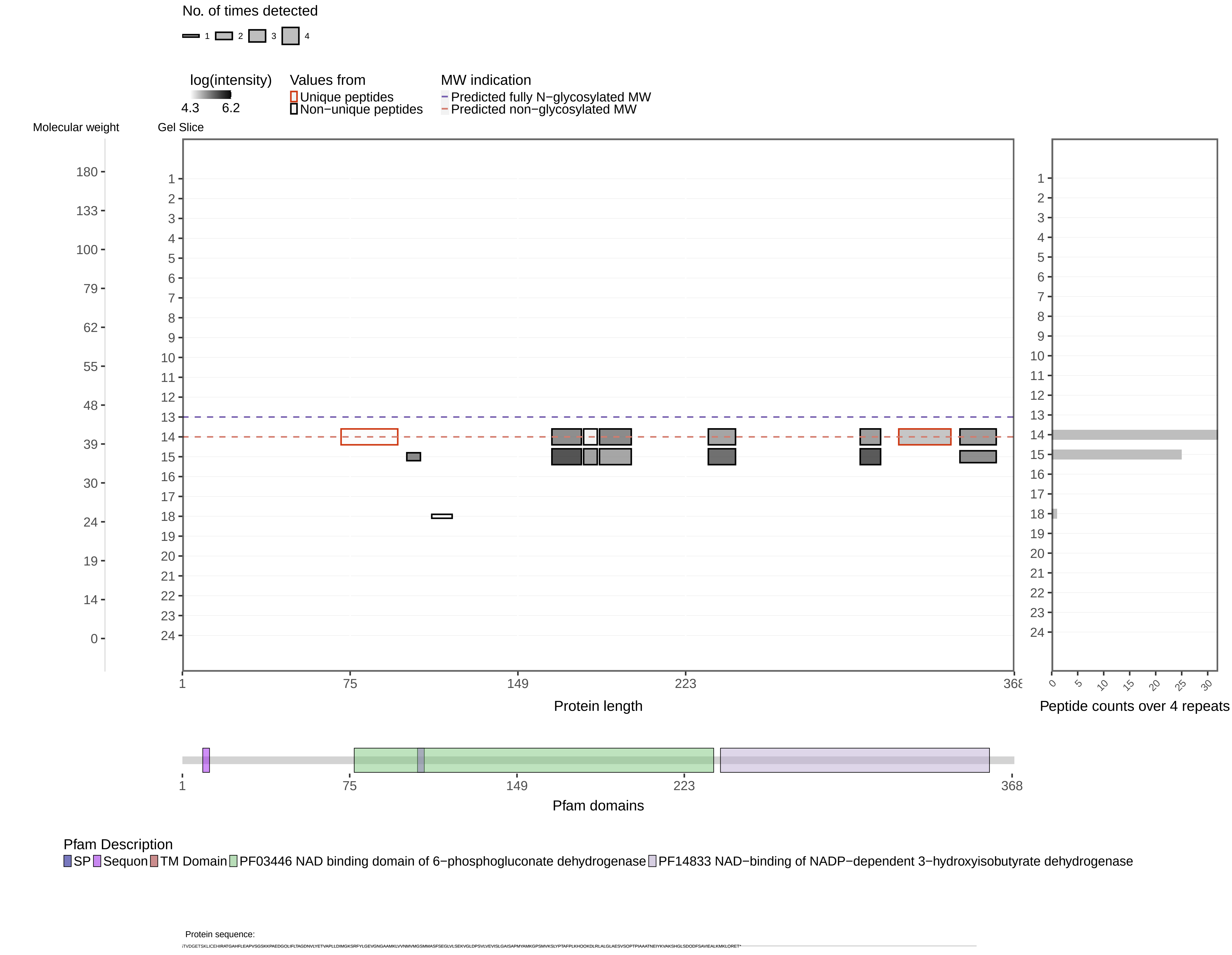

NbL13g19480.1
dihydrolipoyllysine−residue_acetyltransferase_component_2_of_pyruvate_dehydrogenase_complex_mitochondrial−like [MW = 38.23 kDa]

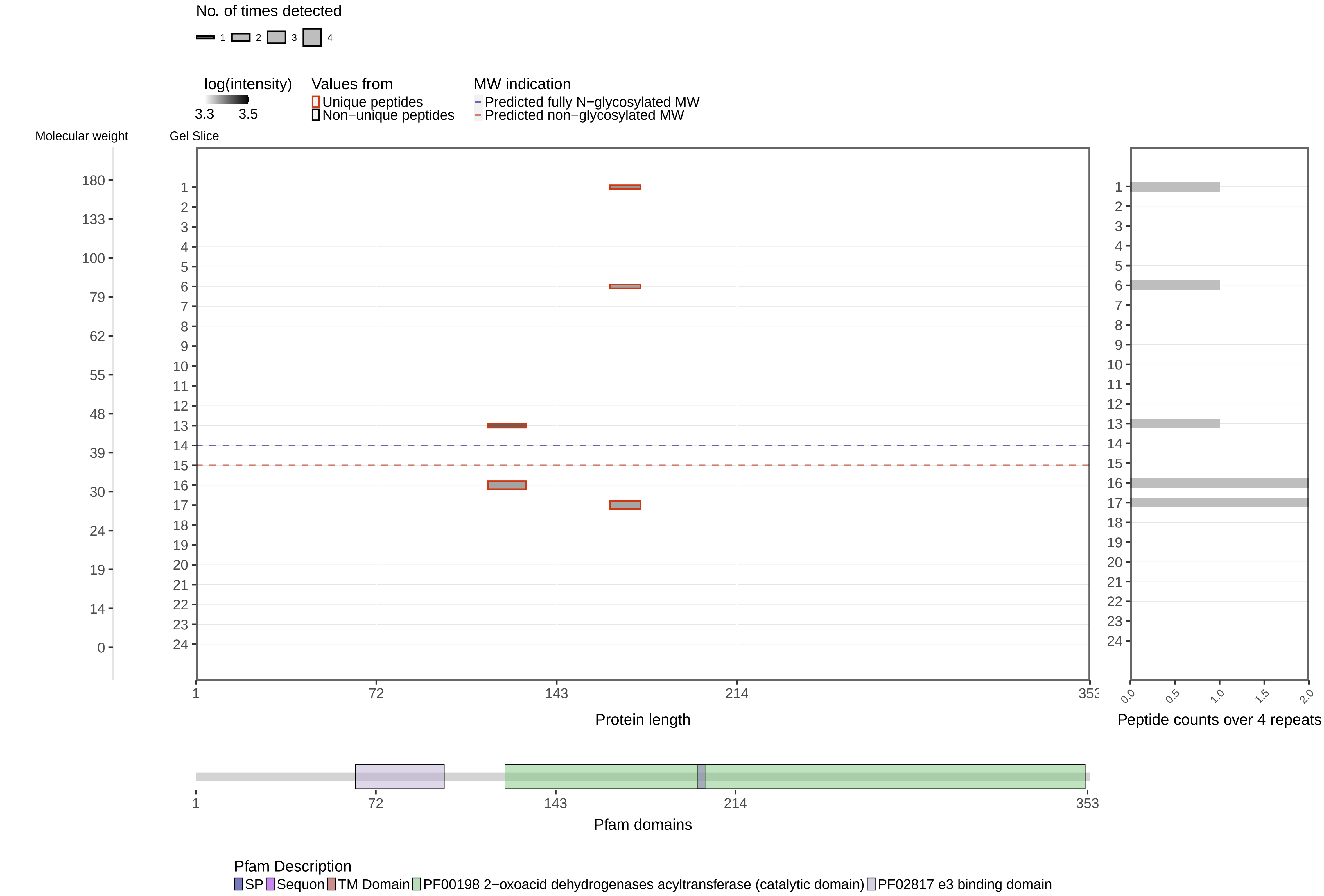


Protein sequence:

NbL13g19410.1 threonine_synthase_chloroplastic [MW = 57.7 kDa]

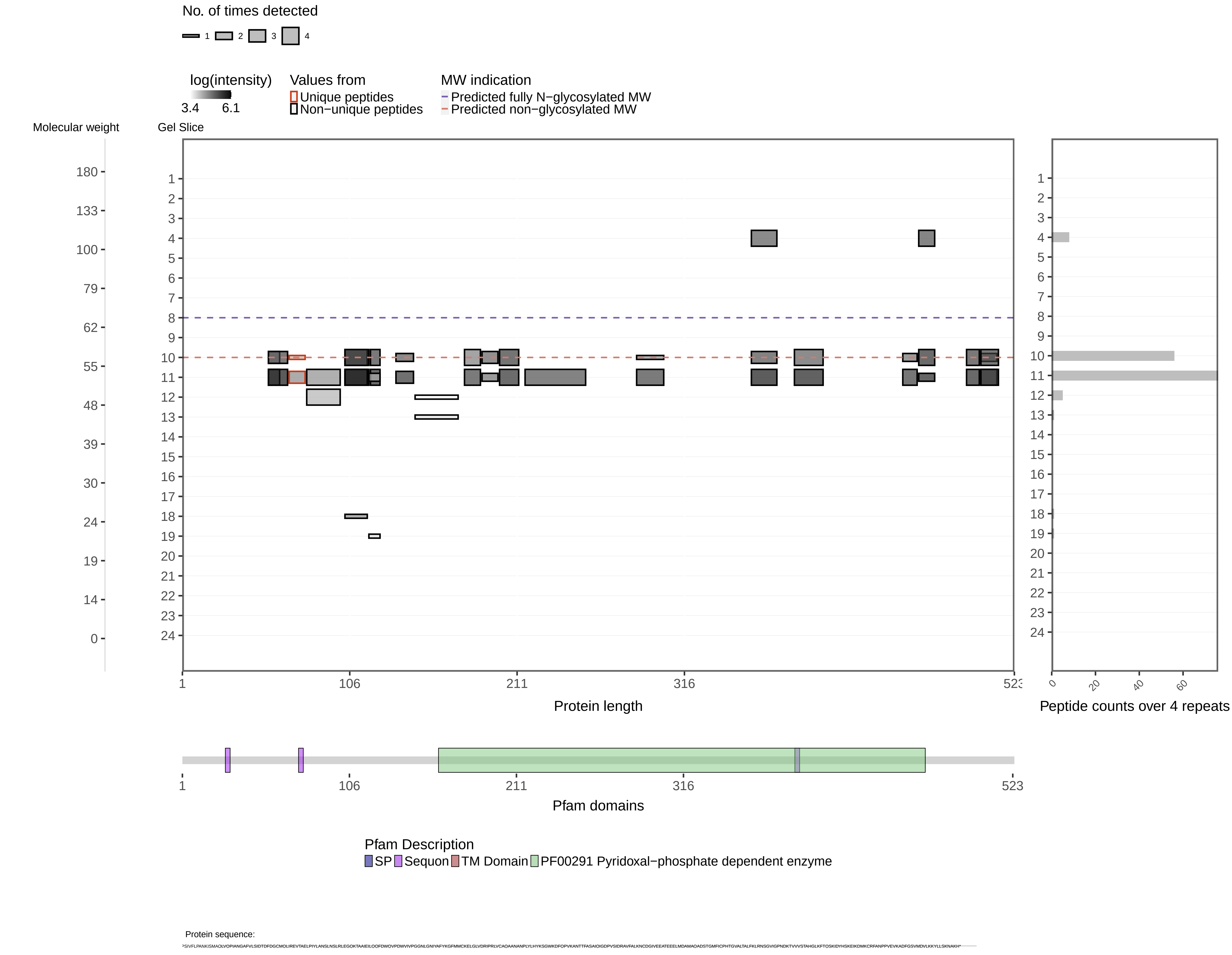

NbL13g19370.1 phospholipid:diacylglycerol_acyltransferase_1−like [MW = 74.19 kDa]

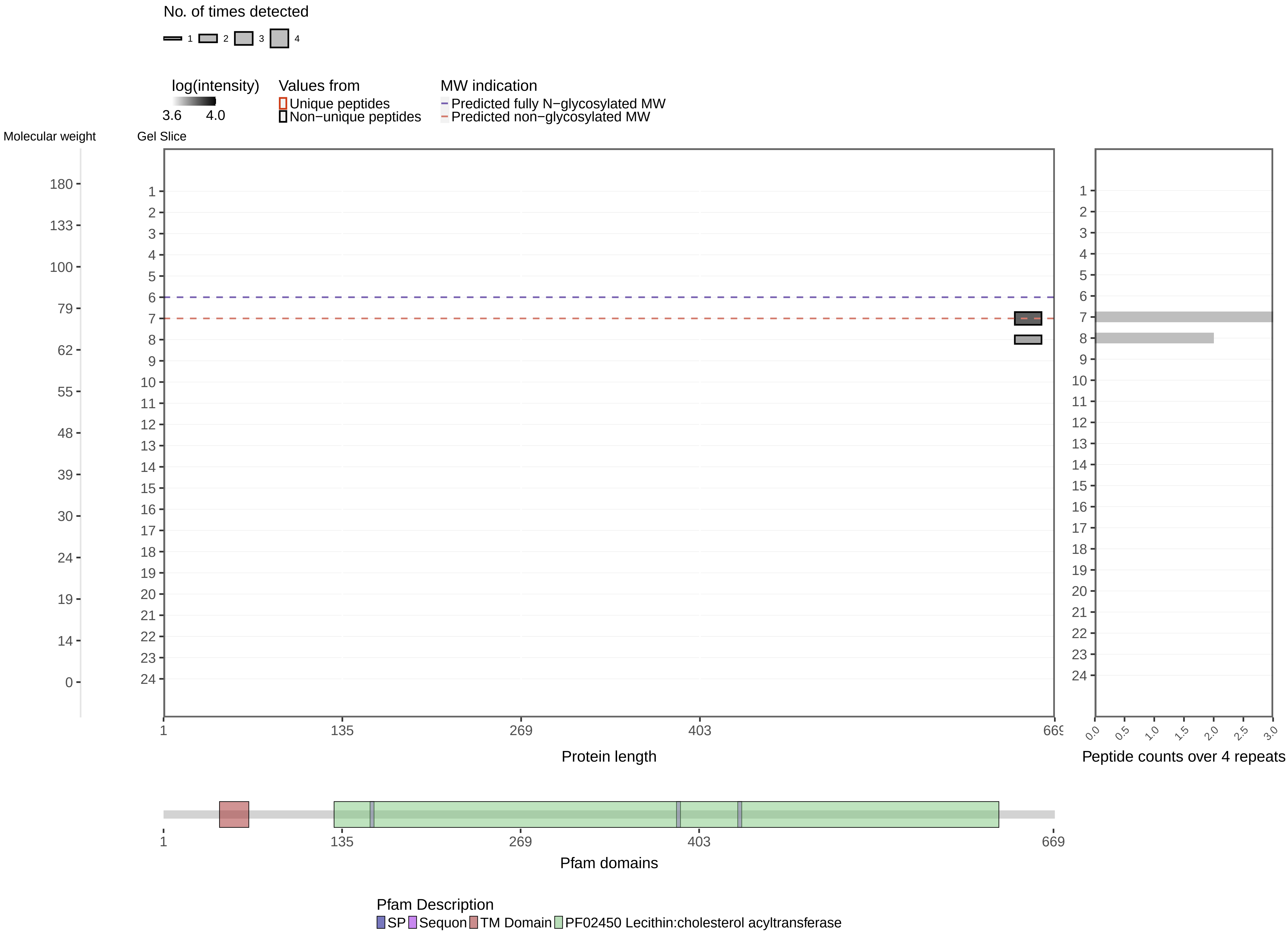


Protein sequence:

DLFHFOTLEHIMKMTRTWDSTMSMIPRGGGTIWGDLDWSPEEGYSPCKSKSKDDAAOISGSDENRTTDSNAKYYSYGRMISFGKDAAKAHSSDLKRIDFRGAVKGSNVANNSCDVWNEYHDMGVSGAKAVEEYKVYTAGEILDLLDFVAPKMMARGSAHFSYGIADDLDDLKYSHYKYWSNPLETKLPDAPDMEIYSMYGVGIOTERAYVYRRMPTAGCNIPFOIDASVDEEDEDSCLKSGVYTVDGDETVPALSAGFMCAKGWRGRTRFNPSGIKTYIREYFHAPPA

NbL13g19170.1 aldehyde_dehydrogenase_family_7_member_B4−like [MW = 54.57 kDa]

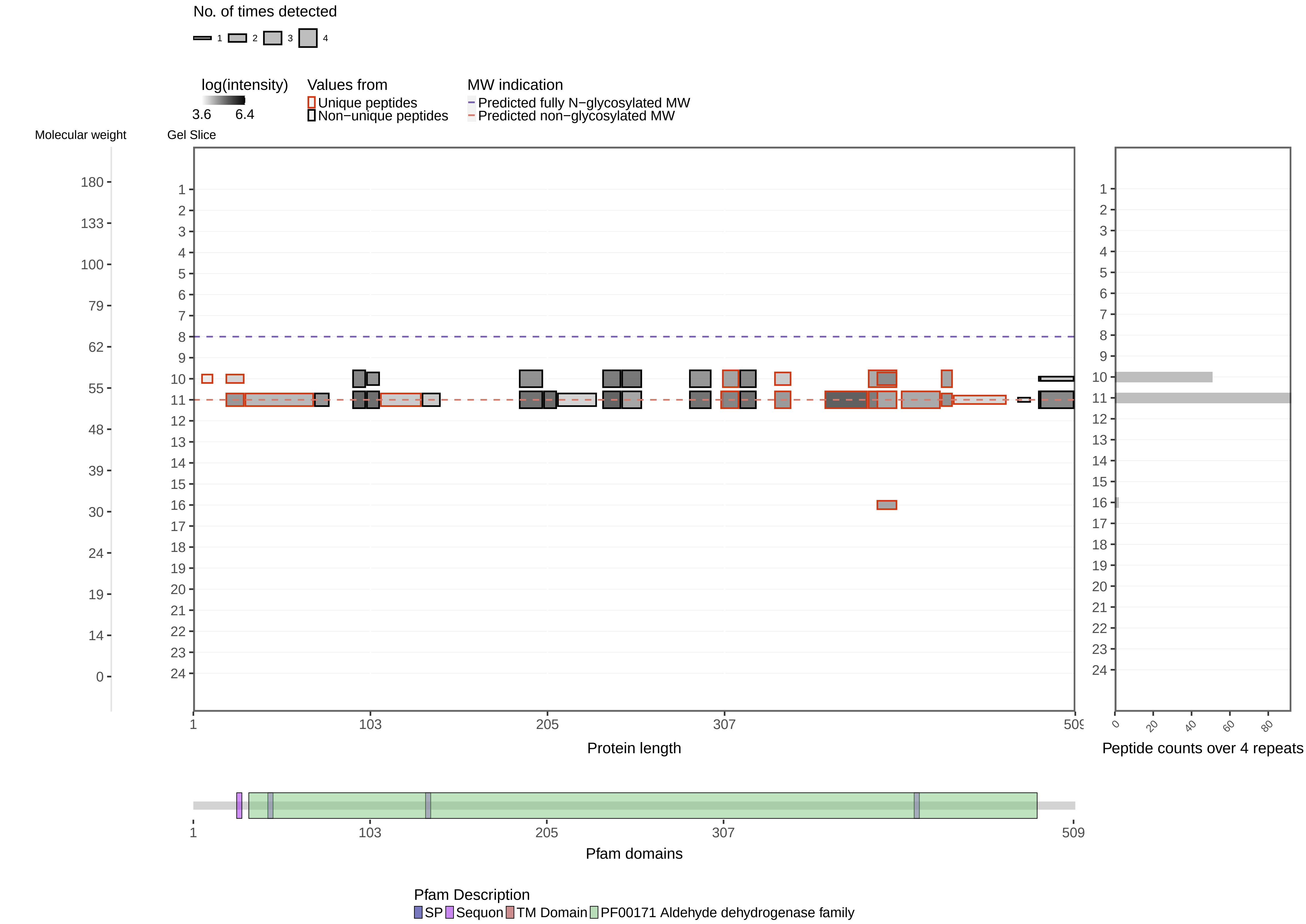


Protein sequence:

FTGSSKVGLAVQQTVNQRFGKCLLELSGNNAIIVMDDADIELAVRSVLFAAVGTAGQRCTTCRRLLVHETVYEKVLEPLVDVYKQVKIGDPLEKGILLGPLHTRTSRENFQKGIQNIKSQGGKILTGGSIIESEGNFVHPTIVEISSKAEVVKEELFAPVLYVMKFKSFEEAVEINNSVPQGLSSSIFTRNPQVMFKWIGAQGSDCGIVNVNIPTNGAEIGGAFGGEKGTGGGREAGSDSWKQYMRRSTCTINYGNELPLAQGINFG*

NbL13g19160.1 subtilisin−like_protease_SBT1.4 [MW − SP = 79.67 kDa]

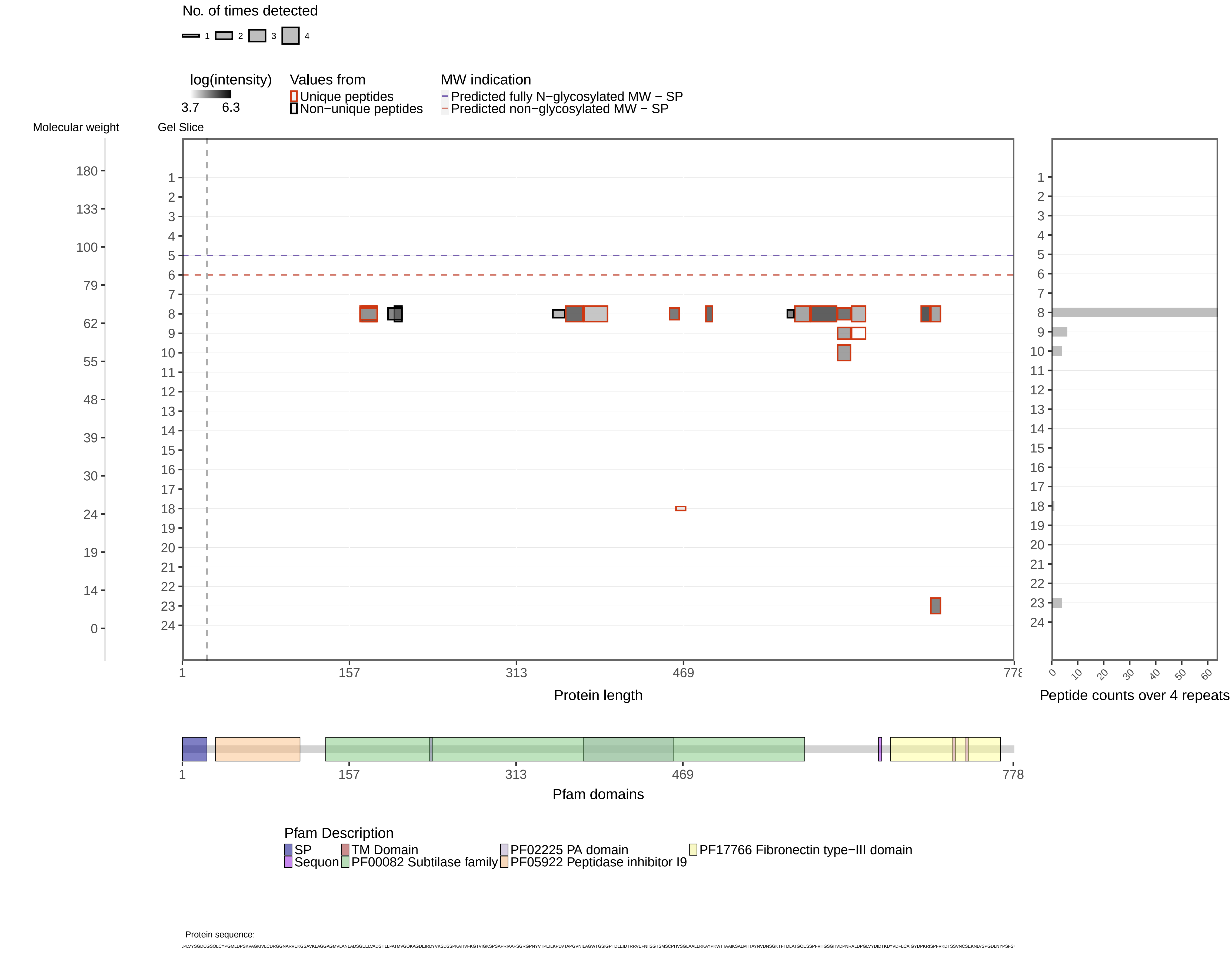

### NbL13g19070.1 auxin−binding_protein_ABP19a−like [MW − SP = 19.57 kDa]

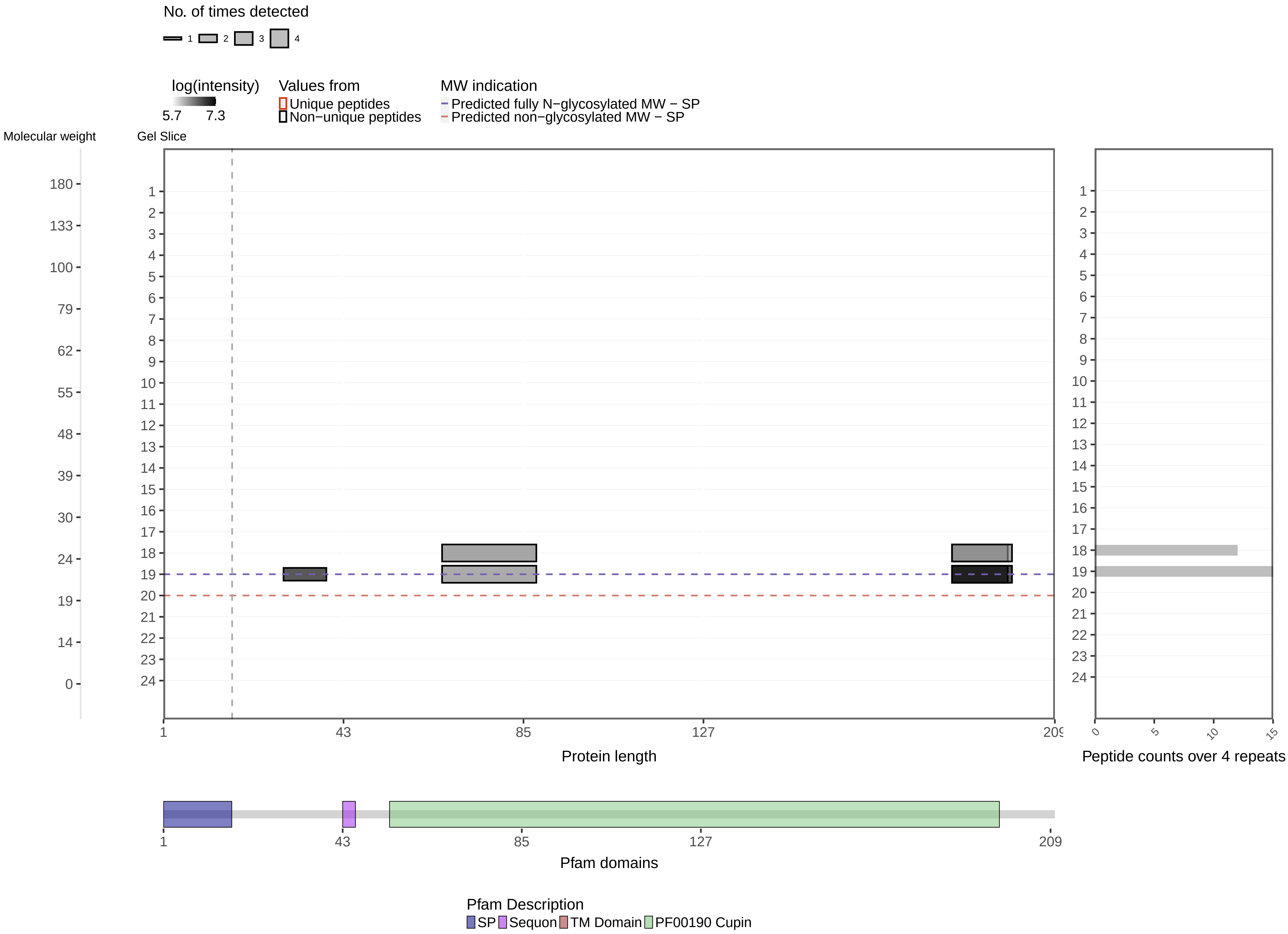


Protein sequence:

.APGGVIPFHTHPGASEVLLVVQGSITAGFVSSANAVYLKTLKKGDLMIFPQGLLHFQVSDAGYTSVGFVFFSSSSPGLQITDFALFANDLPTKLVEATTFLDEATIKKLKGVLGGTN*

NbL13g18960.1 phosphoinositide_phosphatase_SAC2−like [MW = 92.65 kDa]

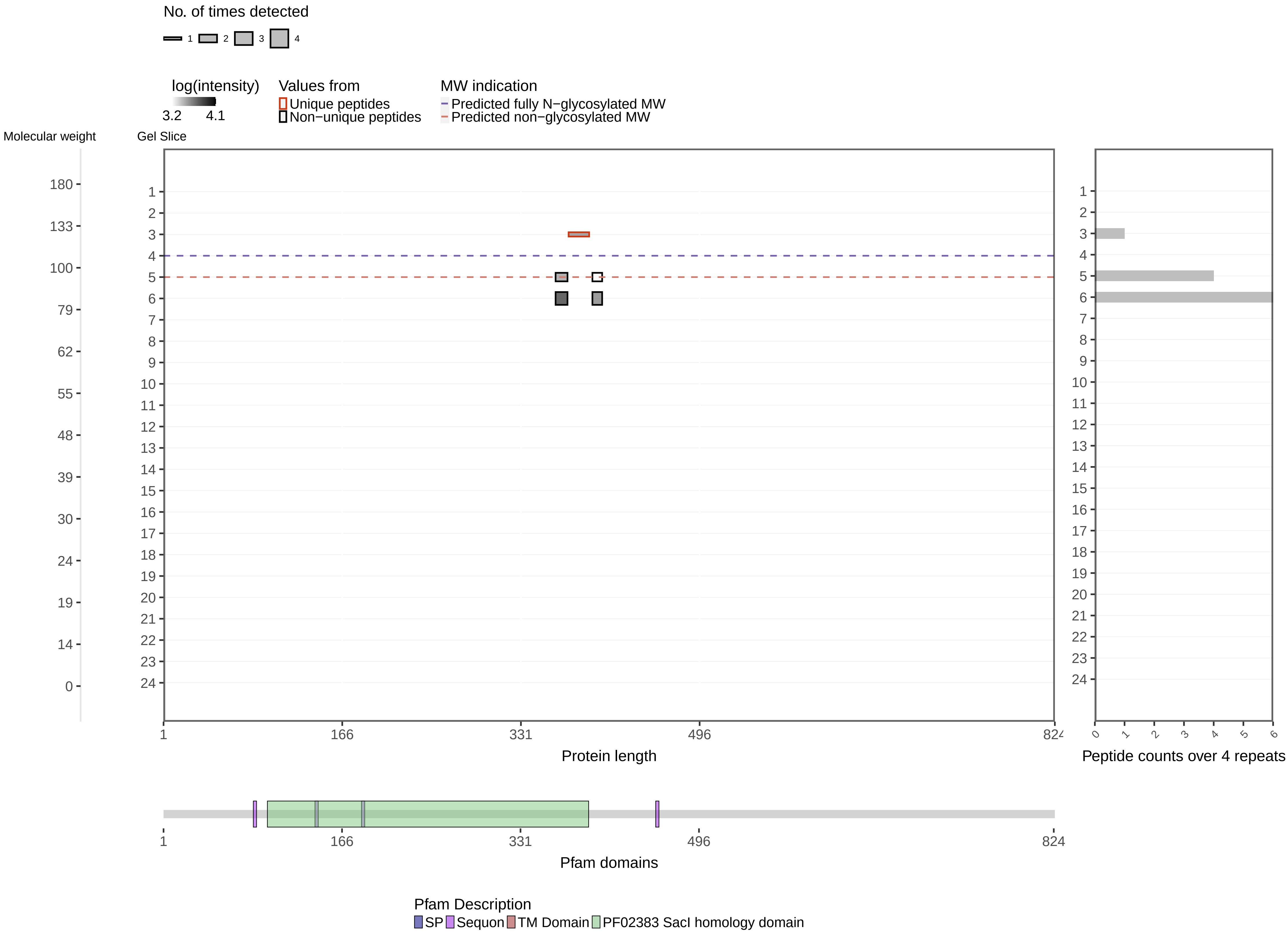


Protein sequence:

GKVAATALDLTGFLHCQLIPASKSGKLLKFSSTGNTGDQAGKDLCEVGAEPNSSSGDLCGDYYVKLSTLQRGVLRTNCIDCLDRTNVAQYAYGLVALGHQLHALGFIDVENIDLDSALADNLMKLYEAMGDTLALQYGGSAAHNKIFSERRGQWKAATQSQELFRTLQRYYSNAYMDAEKQDAINVFLGHFRPQEGKPALWELDSDQHYNVGGHGSSLAMETSRSFIKRSLSEGNLICGSSSPVKETETEQTDDSDQPLPASAKGGSKCLSESTPEISTCETDISFTRYTPS

NbL13g18780.1 protein_STRUBBELIG−RECEPTOR_FAMILY_6 [MW − SP = 44.83 kDa]

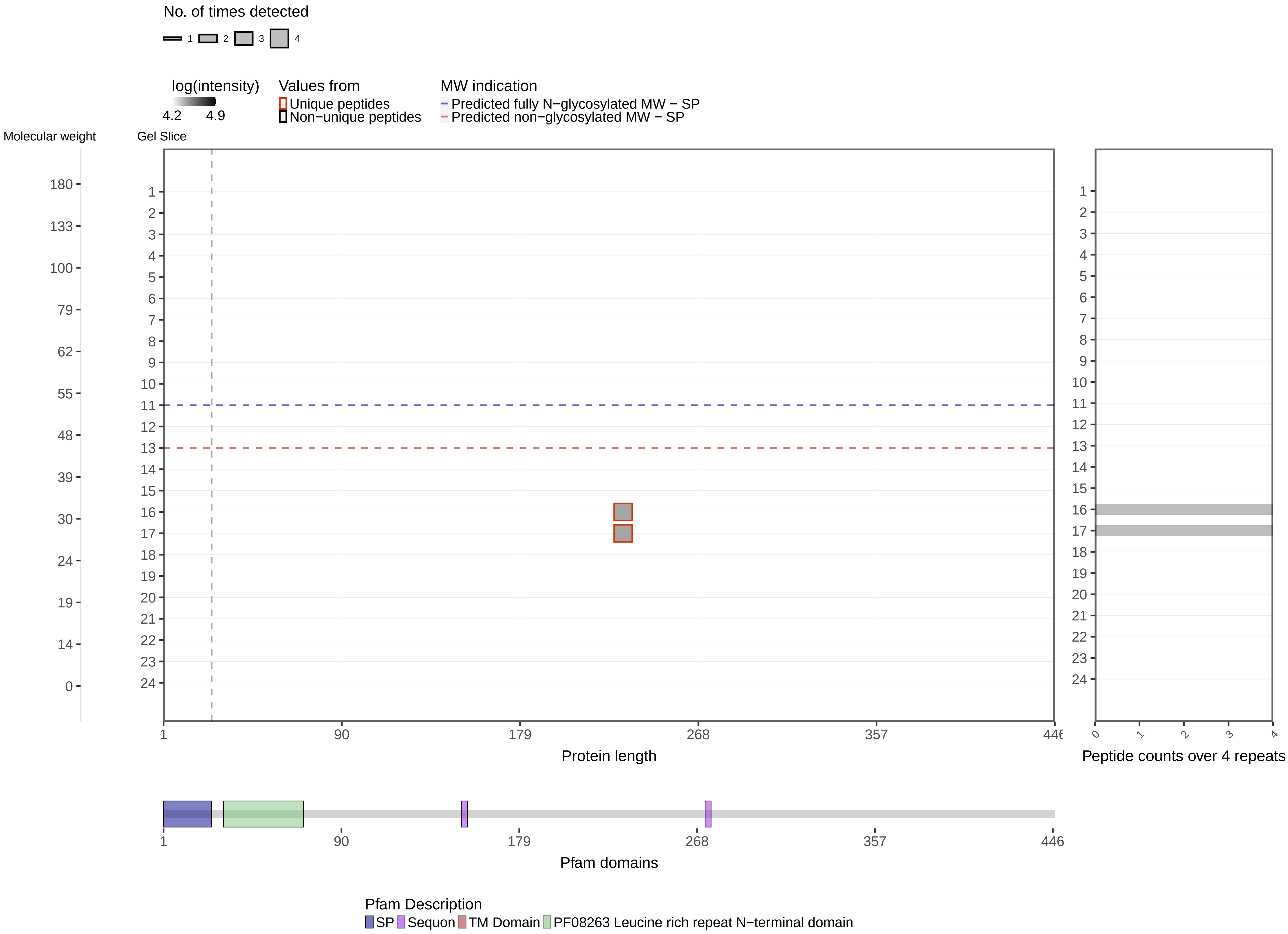


Protein sequence:

VLANLPLDDLNVENNRFTGWVPDQLKGFVKSNGNSWNSGPAPPPPPGTPPASRPDRHHKSGGNNSPSGSGGSGDGGGKSGIGGGAIAGIVISILVVGAIVAFFVIKKRSRRSSTDIEKHDNQPFAPLASEEVQEMKTNQASSAPSVKTFEAPAVVNLRPPPIERHKSFDEDDIAAKPIVPPKKVNTAKINARQYSIADLQMATDSFSIDNLIGEGSFGRVYRAQFDDGKVCFLPYD*

### NbL13g18700.1 D−3−phosphoglycerate_dehydrogenase_2_chloroplastic−like [MW = 71.54 kDa]

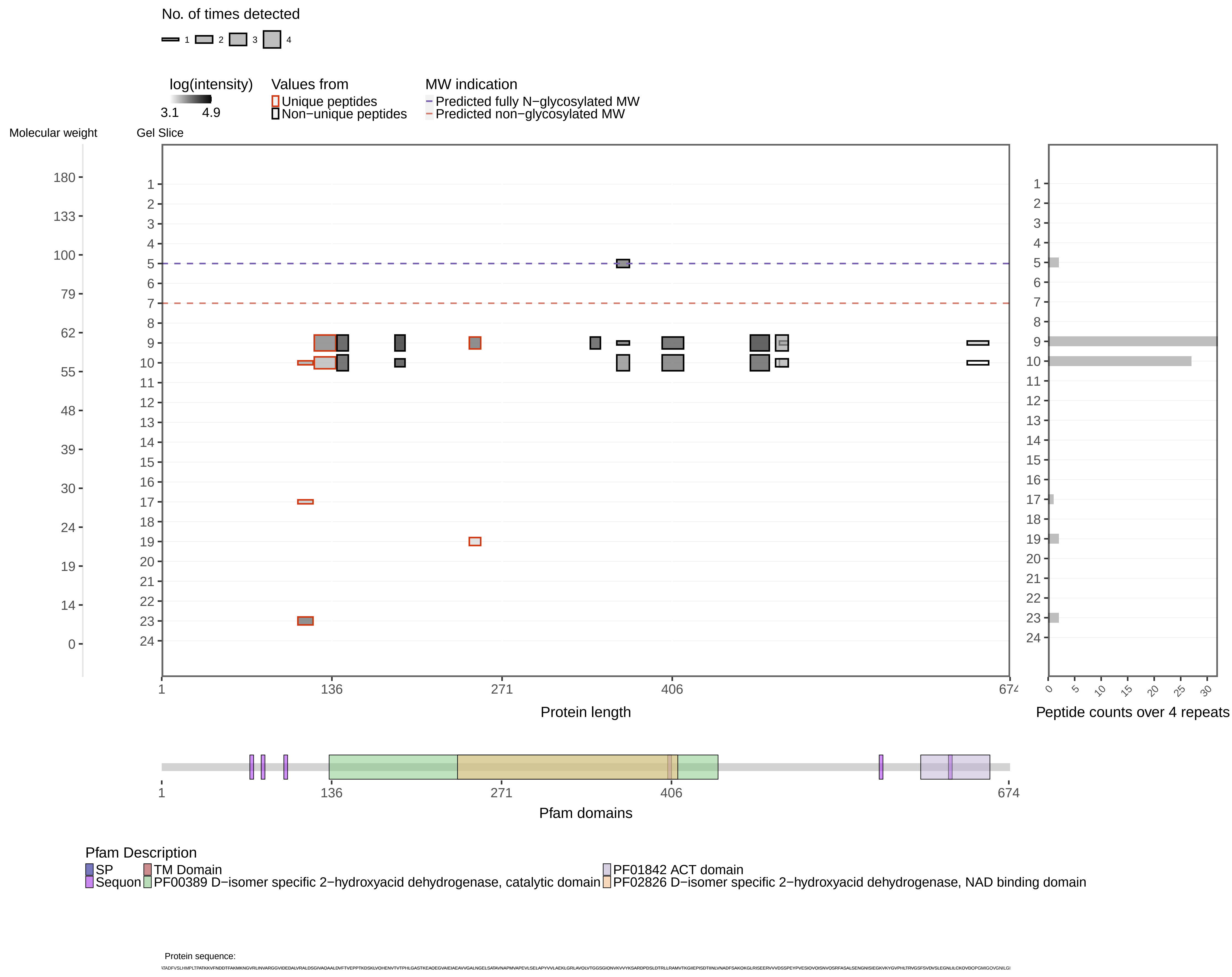

NbL13g18490.1 photosystem_II_5_kDa_protein_chloroplastic−like [MW = 11.6 kDa]

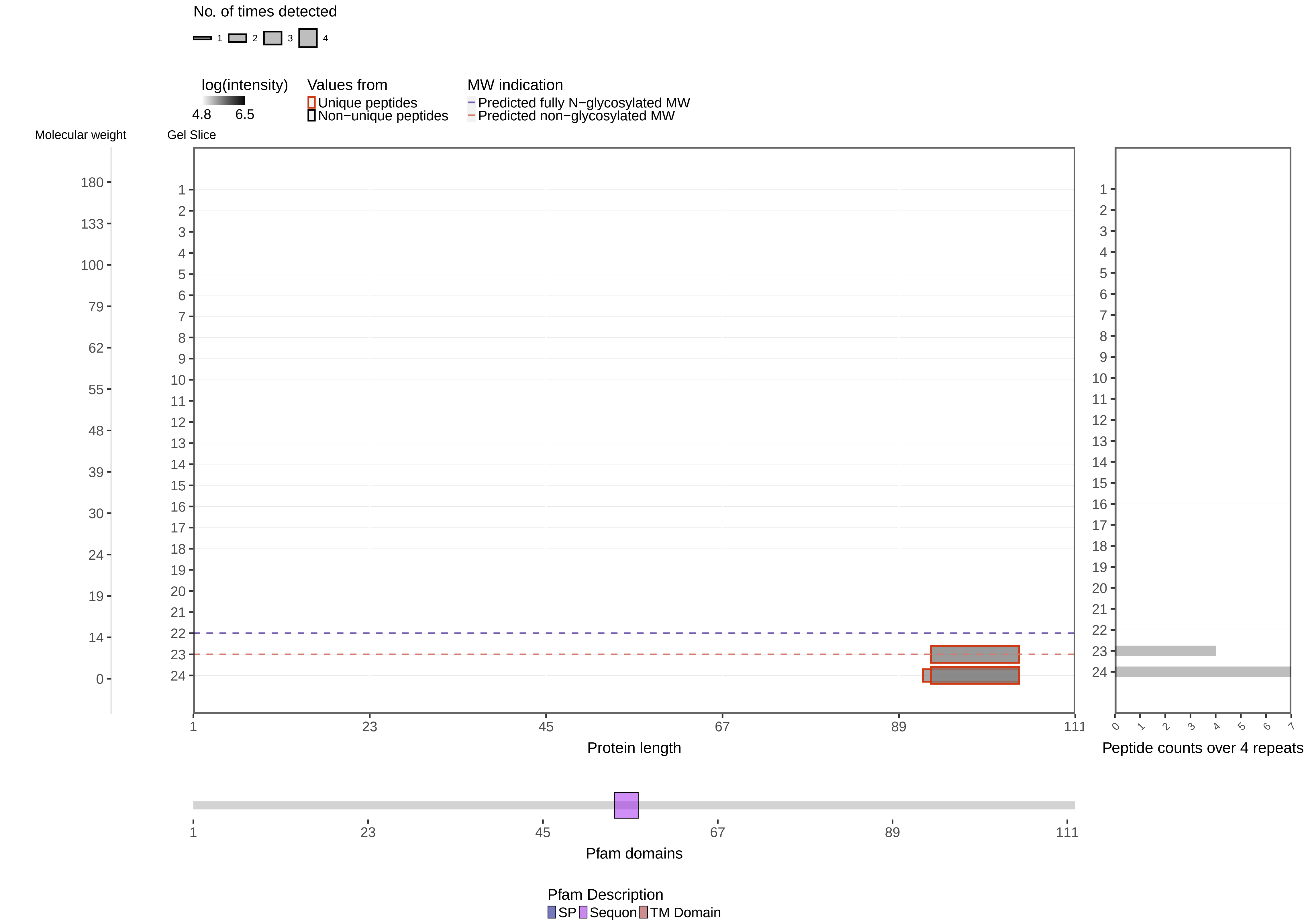


Protein sequence:

ENVVMSNKKESNNSGRRELFFAMAAVAASSVAKAAMAEEPKRGSPEAKKKYYQICVSNPTARICHNAN*

NbL13g18310.1 39S_ribosomal_protein_L47_mitochondrial−like [MW = 19.11 kDa]

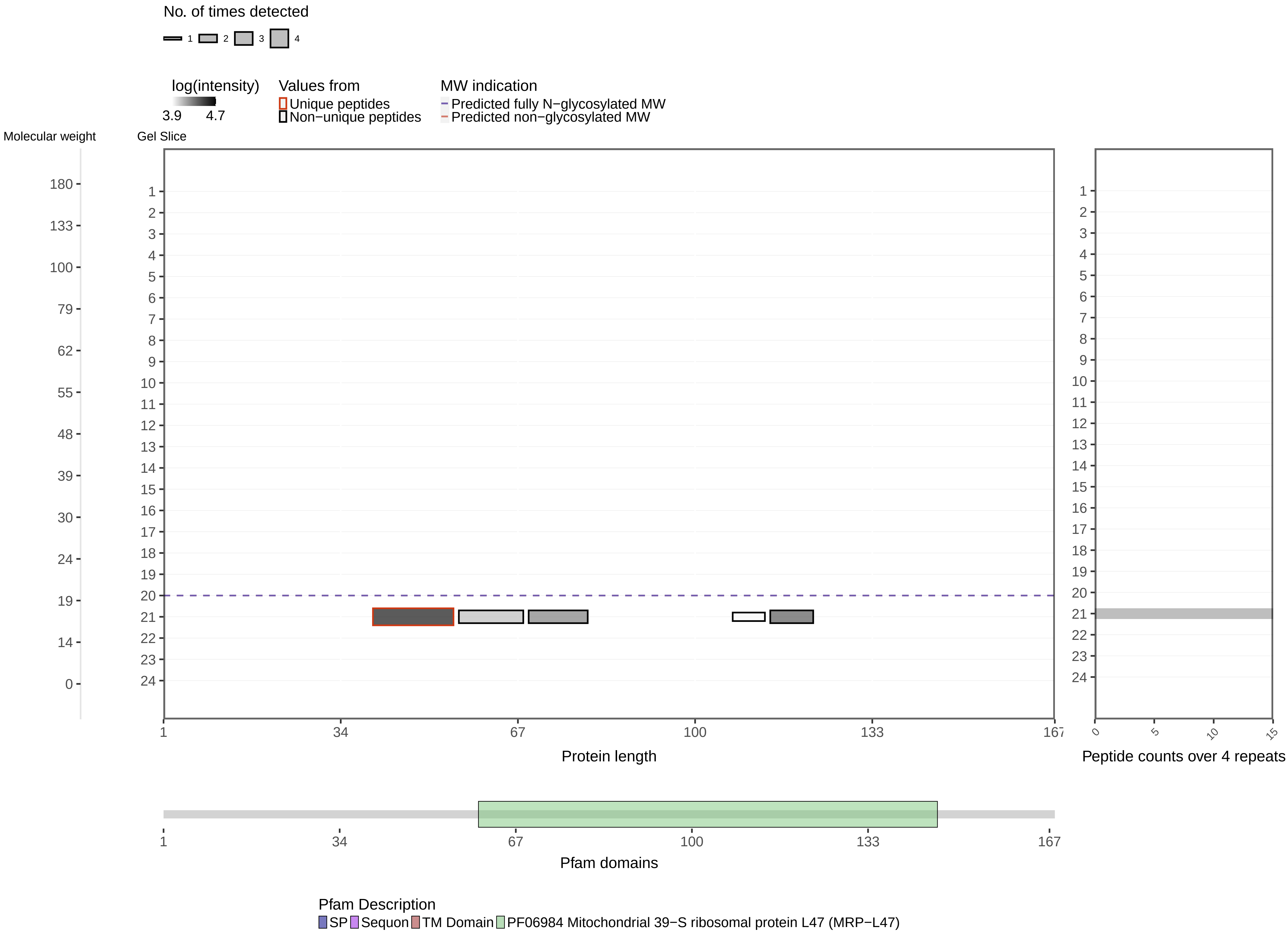


Protein sequence:

DKPVVYGRGWKASELRLKSWDDLQKLWYVLMKEKNMLMTQRQMLNAQNLRFPNPERISKVRKSMCRIKHVLTERAIDEADPRRSTEMKRMINAL*

NbL13g18160.1 betaine_aldehyde_dehydrogenase_chloroplastic−like [MW = 55.66 kDa]

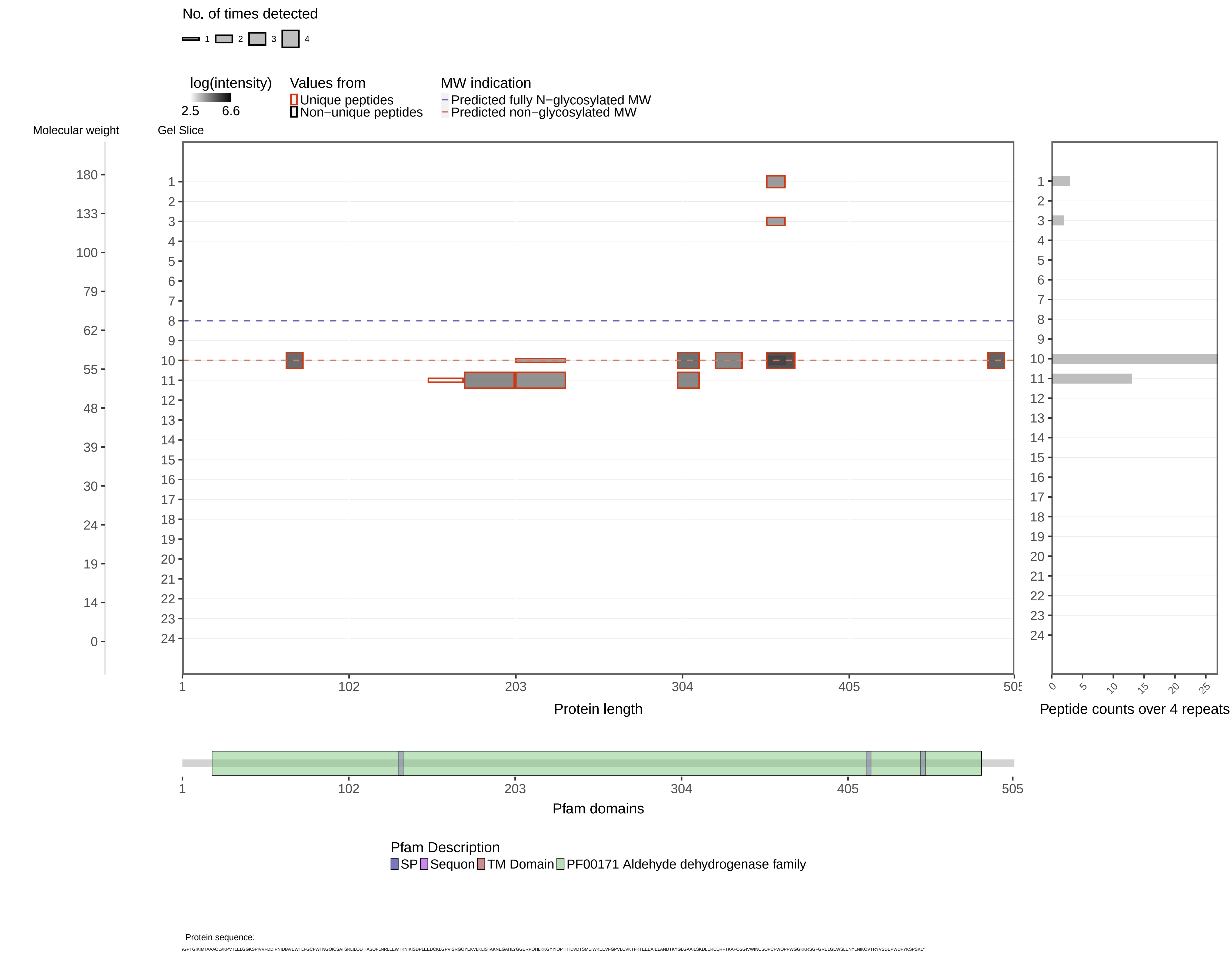

NbL13g18130.1 pyridoxine/pyridoxamine_5'−phosphate_oxidase_1_chloroplastic−like [MW = 62.08 kDa]

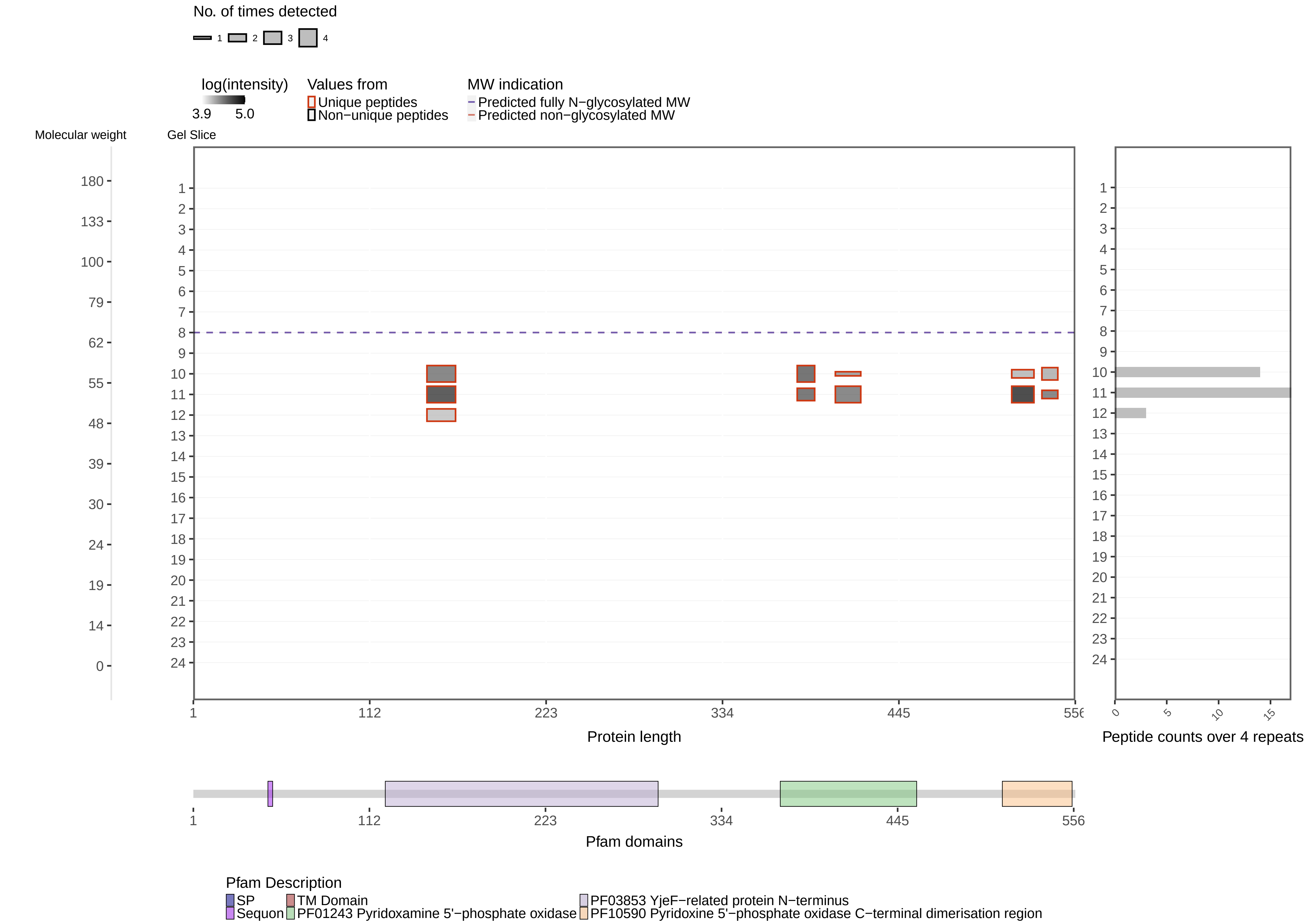


Protein sequence:

VEEGDLSGEGVKPDMLVSLTAPKLCAKSFSGTHHFLGGRFVPPSIRNKFNLQLPAYPGTSMCVRIGKPPKIDISSMRENYISPALLEEEVAADPFDQFQKWFDDAMTAGLKELNAMALSTTGKDGKPSSRIVLLKGFDNDGFVWYTNYEGSKGQQIAENPHAALLFYWDDLNRQVRIEGPVEKVSDEESEQYFHSRPRGSQLGAIASKQSTVIPGRQFLIDEYKHLEAKYADGNLIPKPKHWGGYRLRPEMFEFWQGQQSRLHDRLCYIPETVDGRREWRIERLAP*

### NbL13g17830.1 HIPL1_protein−like [MW − SP = 72.58 kDa]

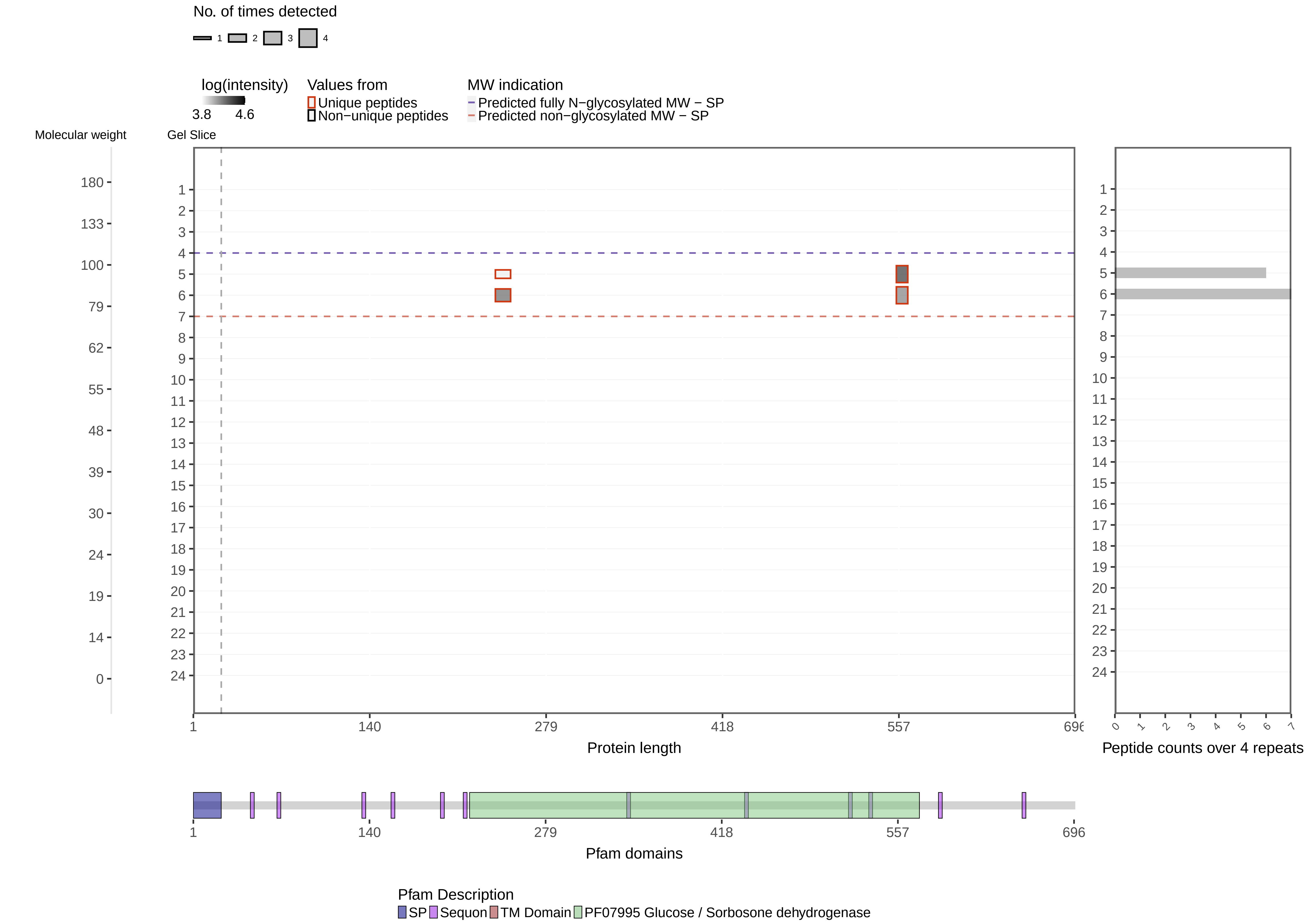


Protein sequence:

[illegible]

NbL13g17680.1 short−chain_dehydrogenase_reductase_3b−like [MW = 27.89 kDa]

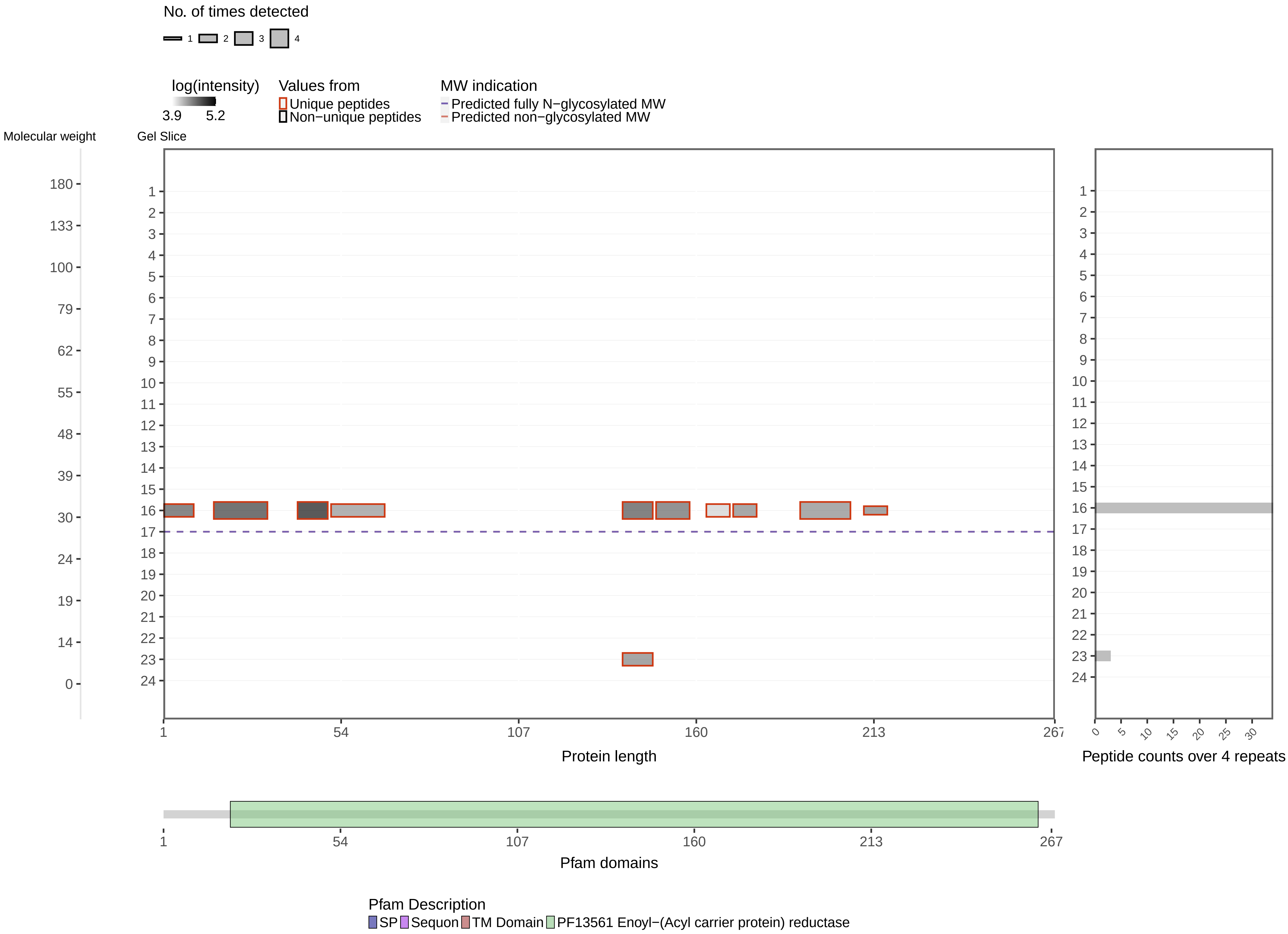


Protein sequence:

AINVRGSVACVKHVAKAMVEGGVQGKIICTGSVTATKGVTKQIDYAMSKHAILGLVKSASKGLGKYGIRVNCVSPAAVATPLLEKSMKMSVEEIEKLFNSFNCLKTGAAIKASNVADAVLFLASDDSQFVTGHNLVVDGGFHPPV*

### NbL13g17640.1 uncharacterized_LOC107771029 [MW = 19.97 kDa]

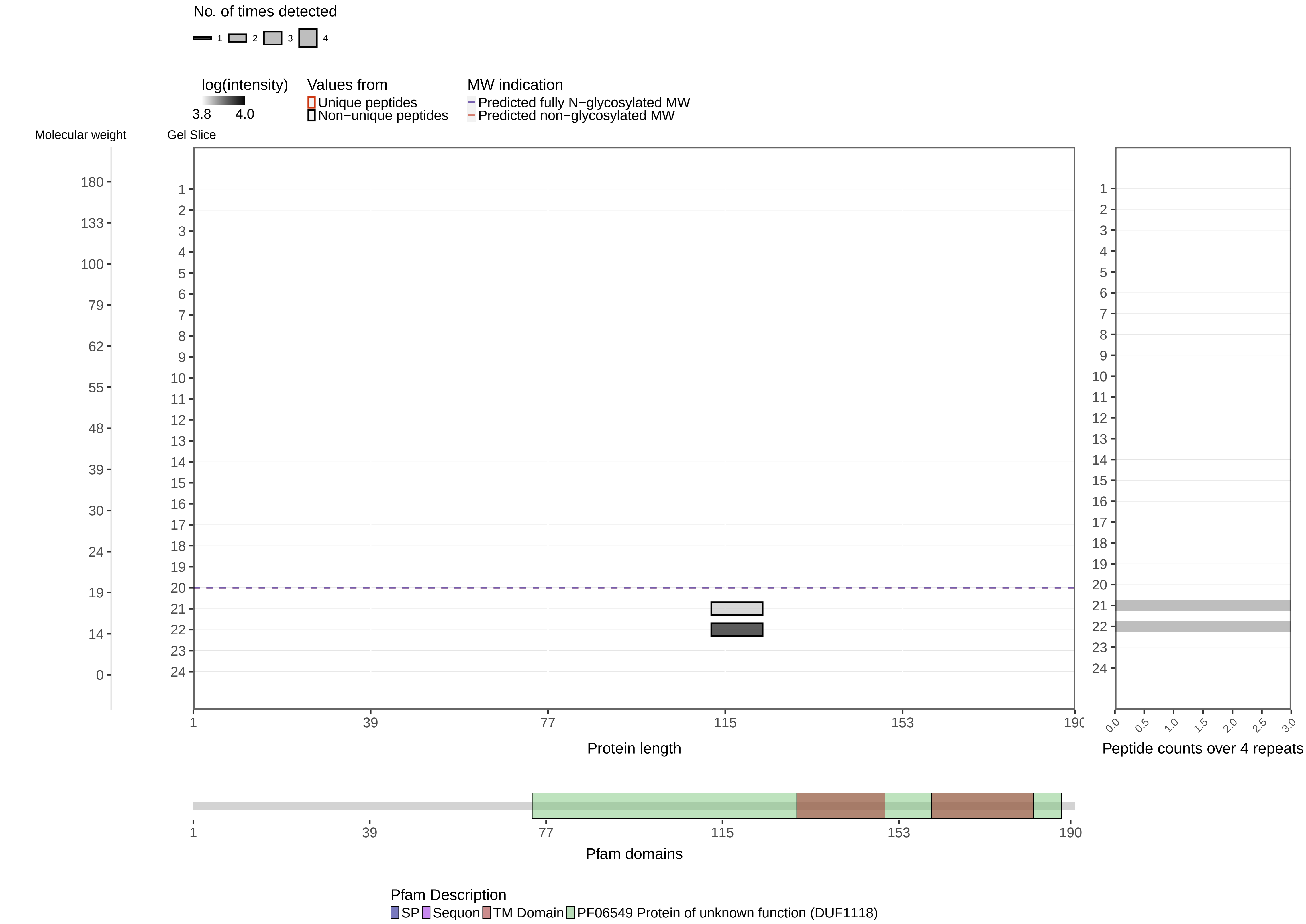


Protein sequence:

KVLSTVEKAGLLSKAEELGFTLSSIEKLGVFSKAEELGLLSLLEKAASFEPSALASAALPILVAAILTIVLIPDDSAGLVAVQAVLAGALGLGAVGLLVGSVVLDGLQEAD*

NbL13g17570.1 ubiquitin−conjugating_enzyme_E2_2−like [MW = 17.3 kDa]

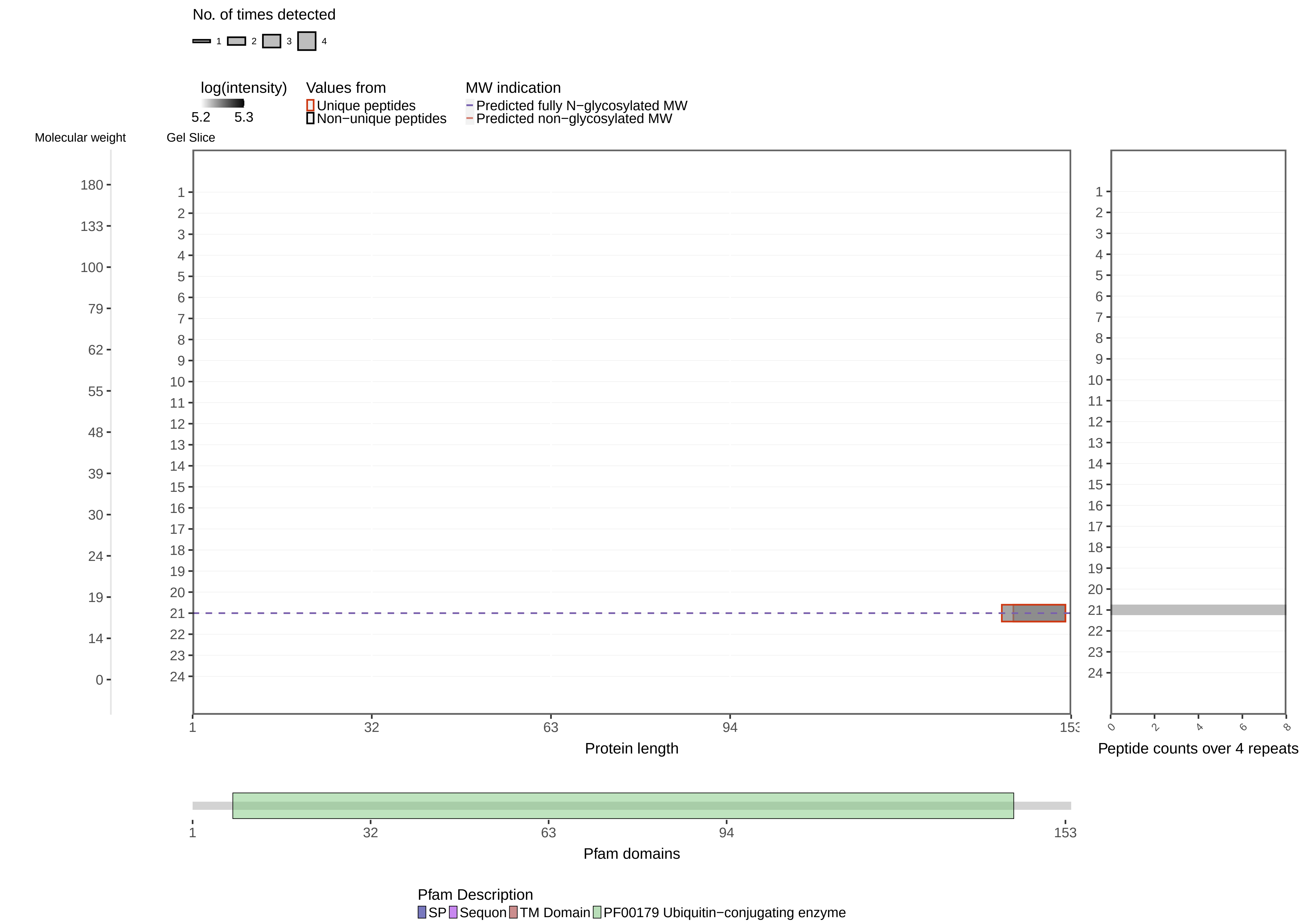


Protein sequence:

YPNKPPTVRFVSRMFHPNIYADGSICLDILQNQWSPIYDVAAILTSIQSLLCDPNPNSPANSEAARLFSENKRDYNRRVREVVEQSWTAD*

### NbL13g17510.1 aldose_1−epimerase−like [MW − SP = 36.56 kDa]

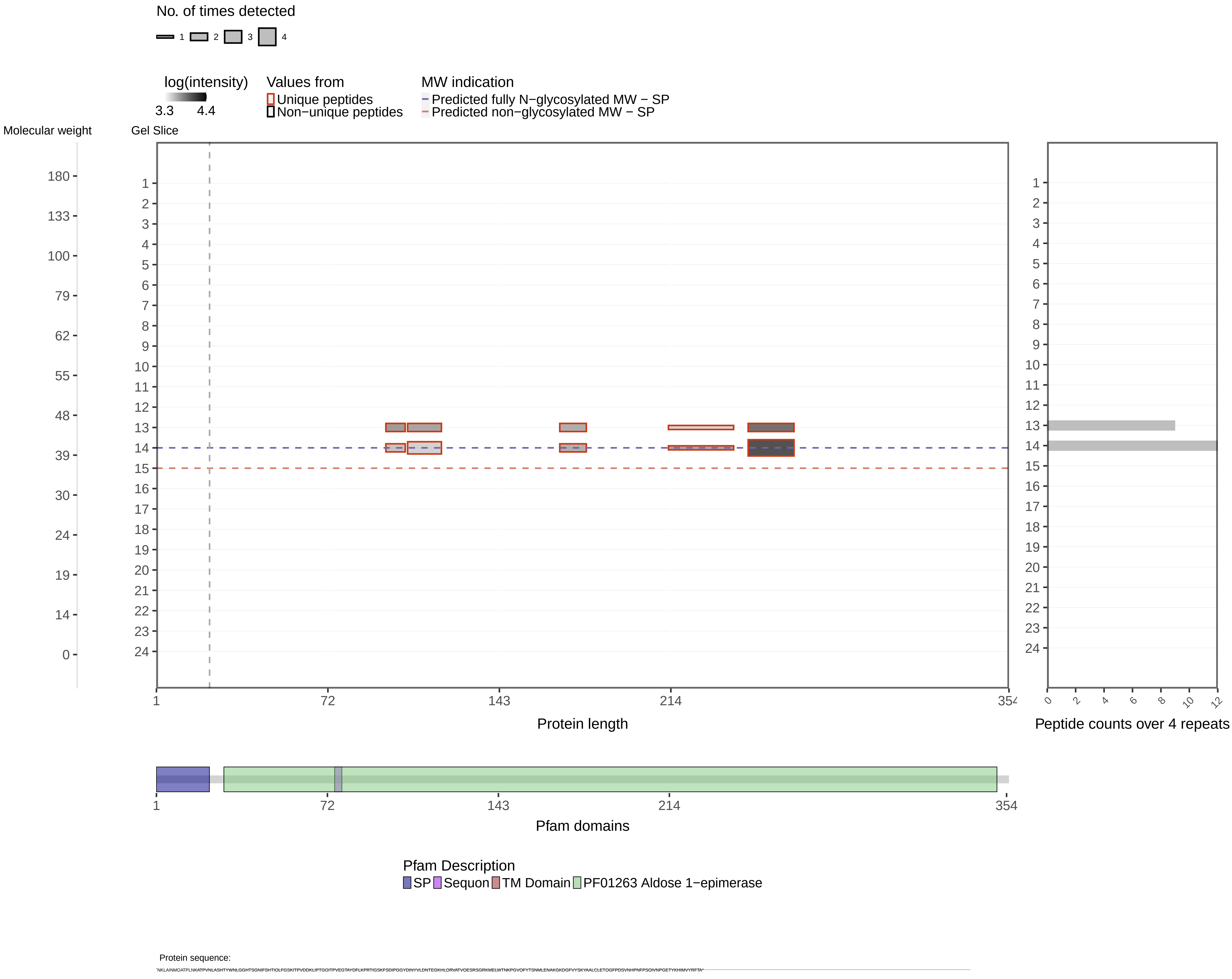

### NbL13g17500.1 vacuolar_protein_sorting−associated_protein_29 [MW = 21.06 kDa]

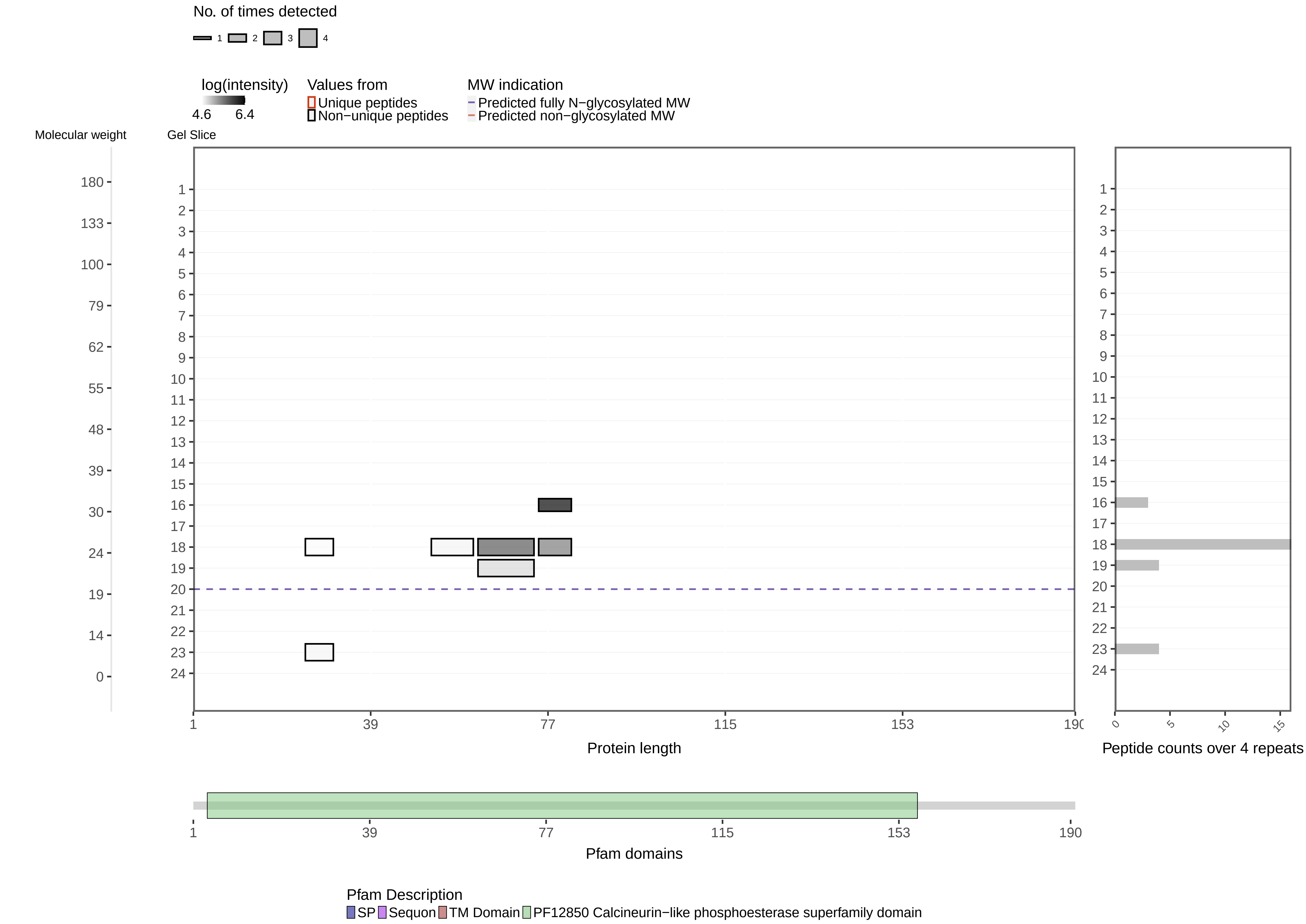


Protein sequence:

_GLCHGHQVIPWGDLDSLAMLQRQLDVDILVTGHTHQFTAYKHEAGVVINPGSATGAYSSITYDVNPSFVLMDIDALRVVVYVVYELIDGEVKVDKIDFKKTTTQNAN*

### NbL13g17400.1 pantothenate_kinase_2 [MW = 100.02 kDa]

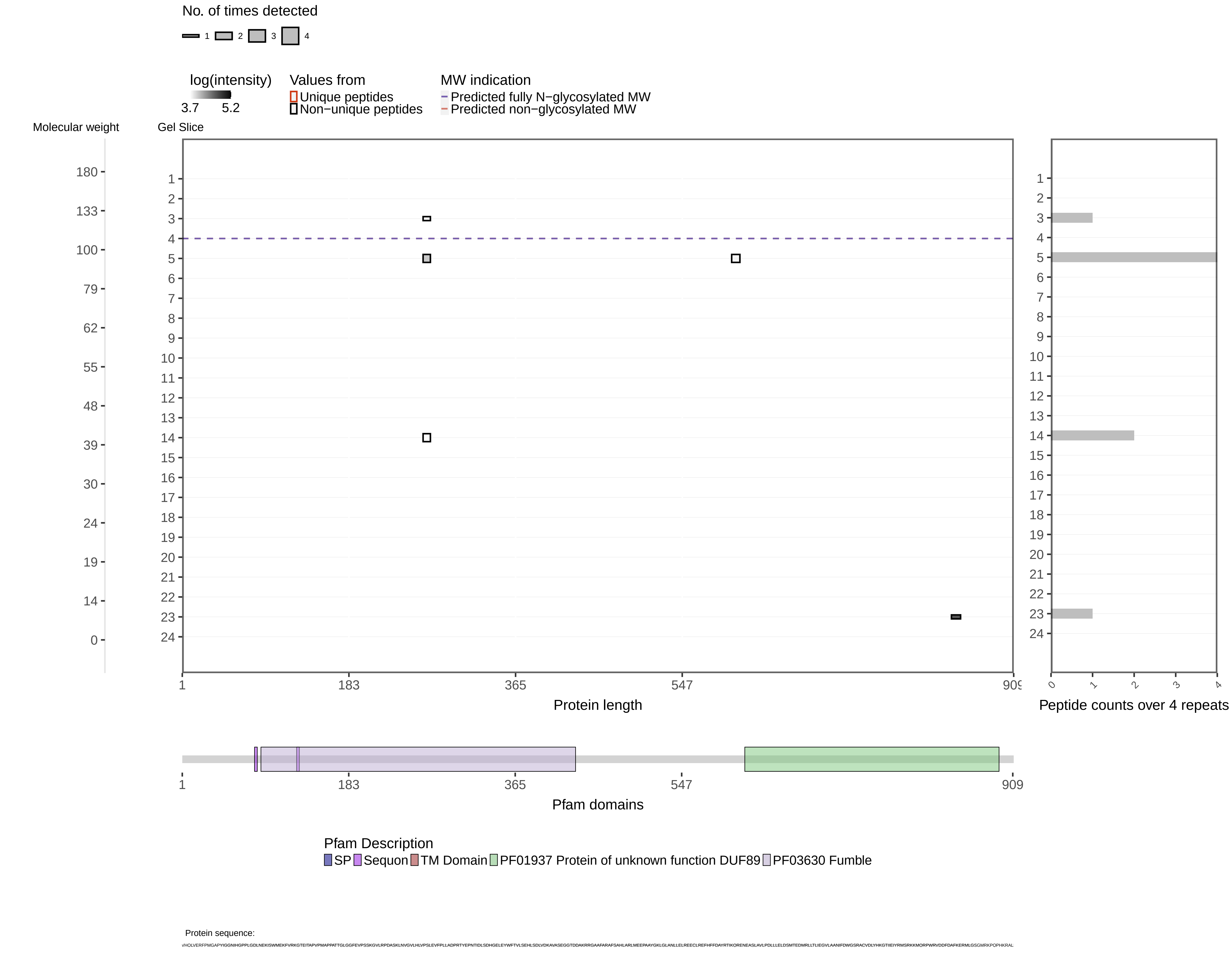

### NbL13g17280.1 glutaredoxin−like [MW = 13.57 kDa]

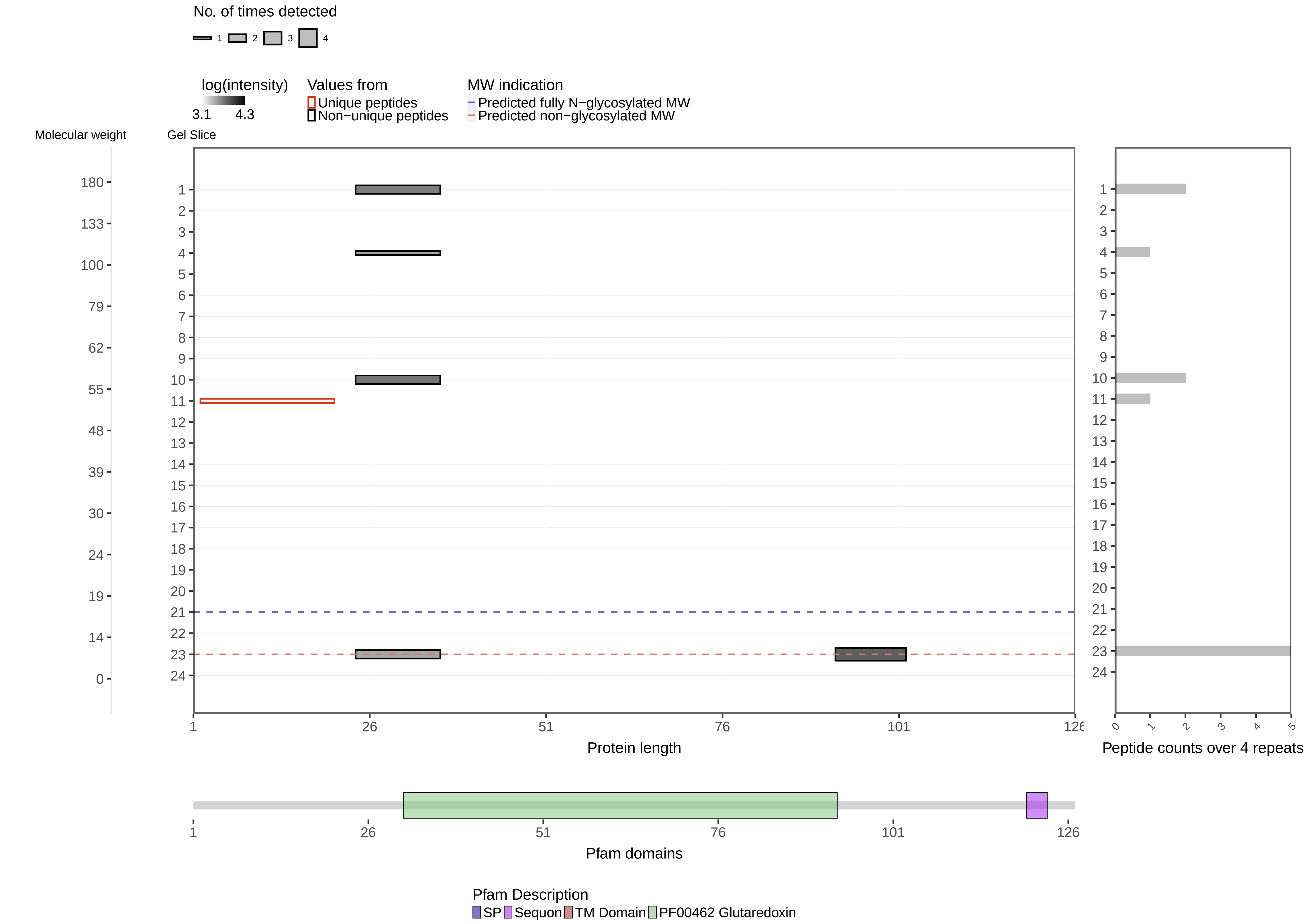


Protein sequence:

SQLGATFKVIELDQESGGNEVQQALLEWTGQRTVPNVFIGGKHVGGCDSIVEKHQQGKLLPLLKDAAAVPNTSAKV*

### NbL13g17190.1 motile_sperm_domain−containing_protein_2−like [MW = 30.51 kDa]

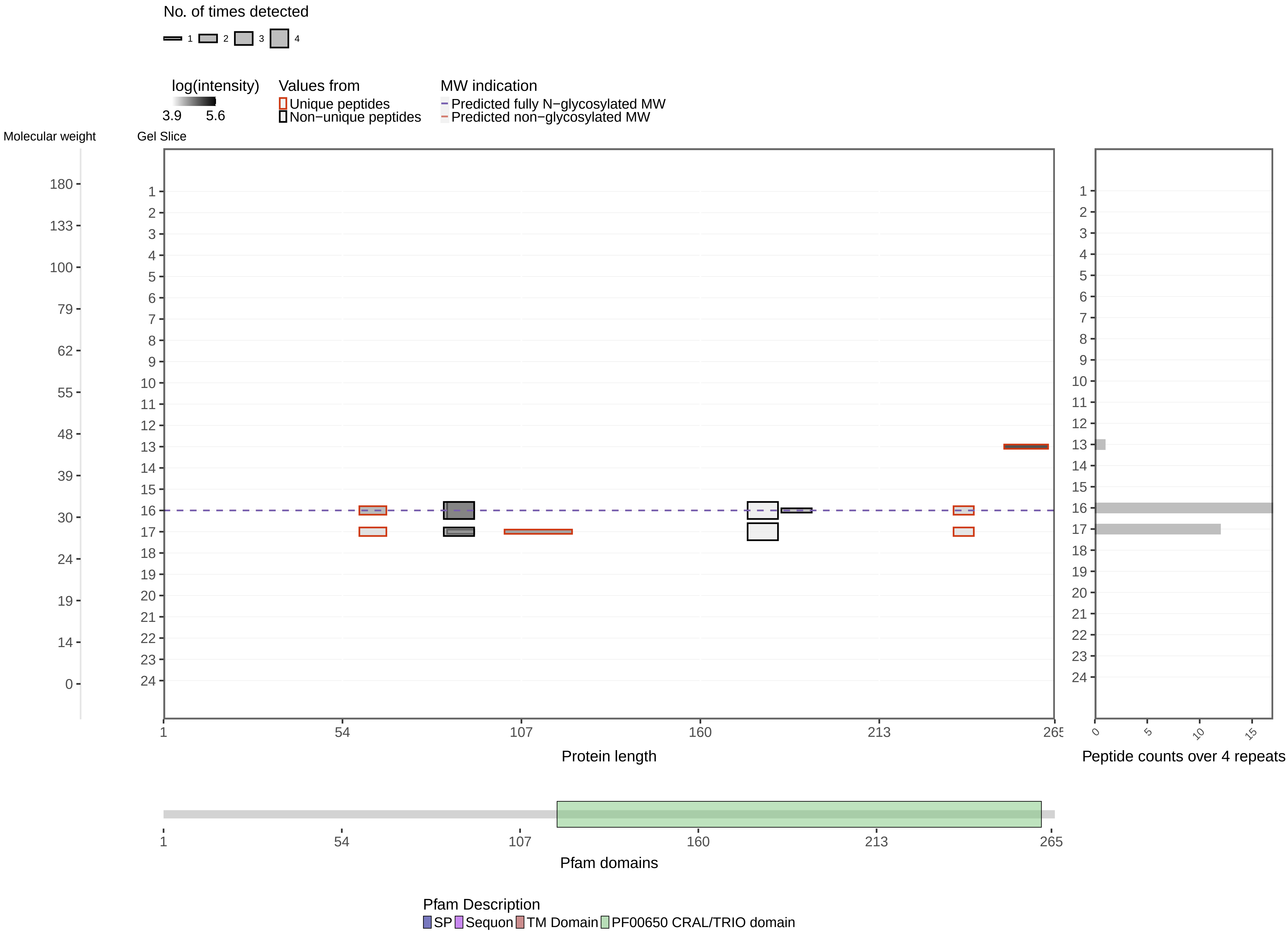


Protein sequence:

ETGKAYVHDCFDVYGRPVLIVEASKHFPQKQDPDEDEKLCVFLIEKALSRIPPGKQDILGIFDLRGFGTENADLKFLTFMFDVFYYYYPRRLGQVLFVDAPFVFKPLWQLVKPVLKSYASLVRFCSAKDVREEYFTEETVPATFRK*

NbL13g17140.1 cell_division_cycle_protein_48_homolog [MW = 90.02 kDa]

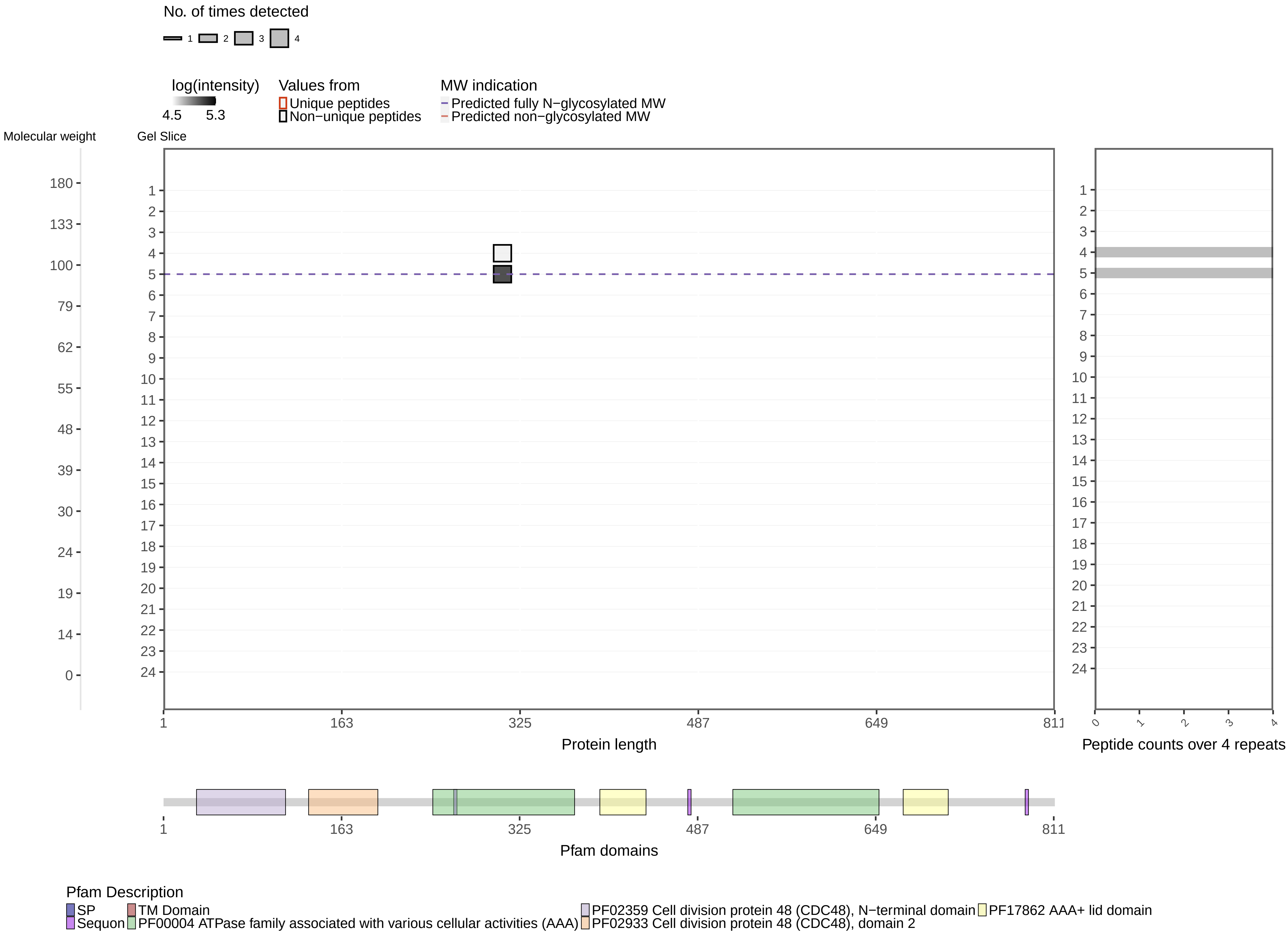


Protein sequence:

NbL13g17130.1 elongation_factor_Tu_GTP−binding_domain−containing_protein_1 [MW = 112.78 kDa]

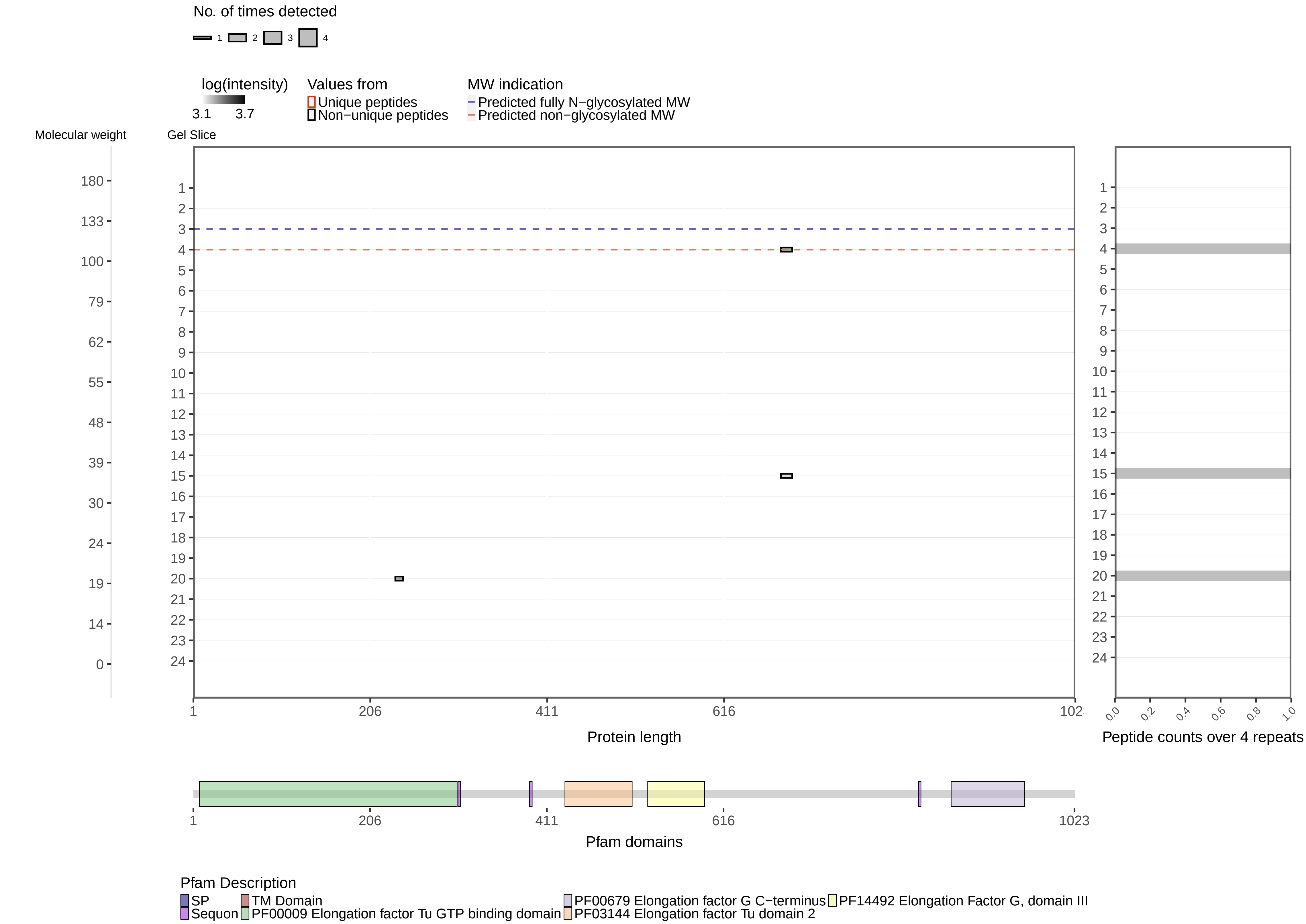


Protein sequence:

### NbL13g17050.1 notchless_protein_homolog [MW = 53.15 kDa]

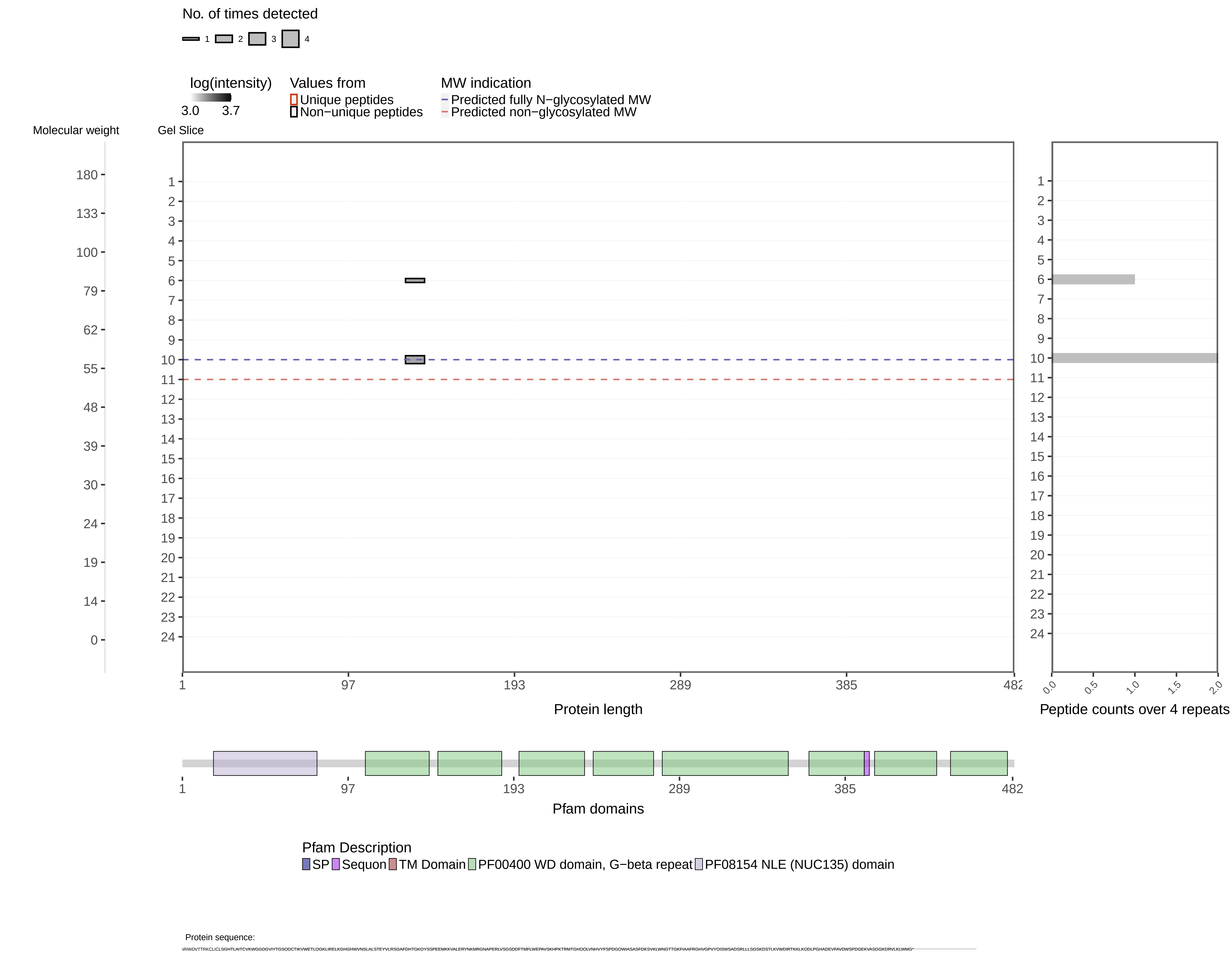

NbL13g16870.1 histidinol_dehydrogenase_chloroplastic−like [MW = 52.8 kDa]

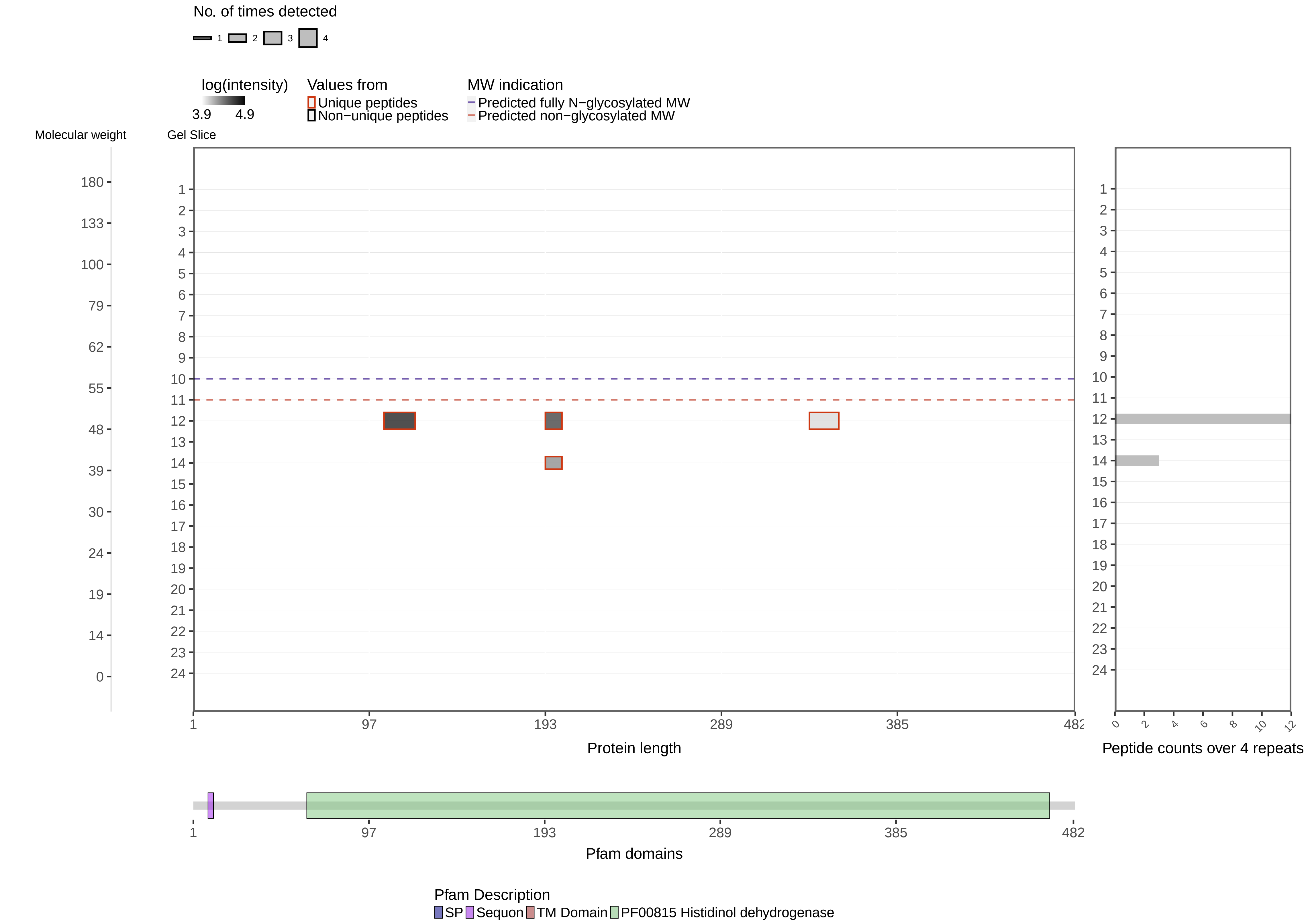


Protein sequence:

ISAMAWGTESCPKVEKIYGPGNQYVTAAKMILQNSEAMVSIDMPAGPSEVLVIADKHSSPIHVAADLLSQAEHGPDSQVVLVIAGDGVDLNAIQEEIRKQCQALPREEFALQALSHSFTVFARDMLEAISFSNMYAPEHLIINVNEAEKWESLIENAGSVFMGQWTPESVGDYASGTNHVLPTYGYARMYGGVSLDSFLKYITVQSLTEEGLRNLGPYVEKMAEVEGLDAHKRAVTLRLQDIEARQVSHSR*

### NbL13g16810.1 elongation_factor_TuA_chloroplastic [MW = 51.96 kDa]

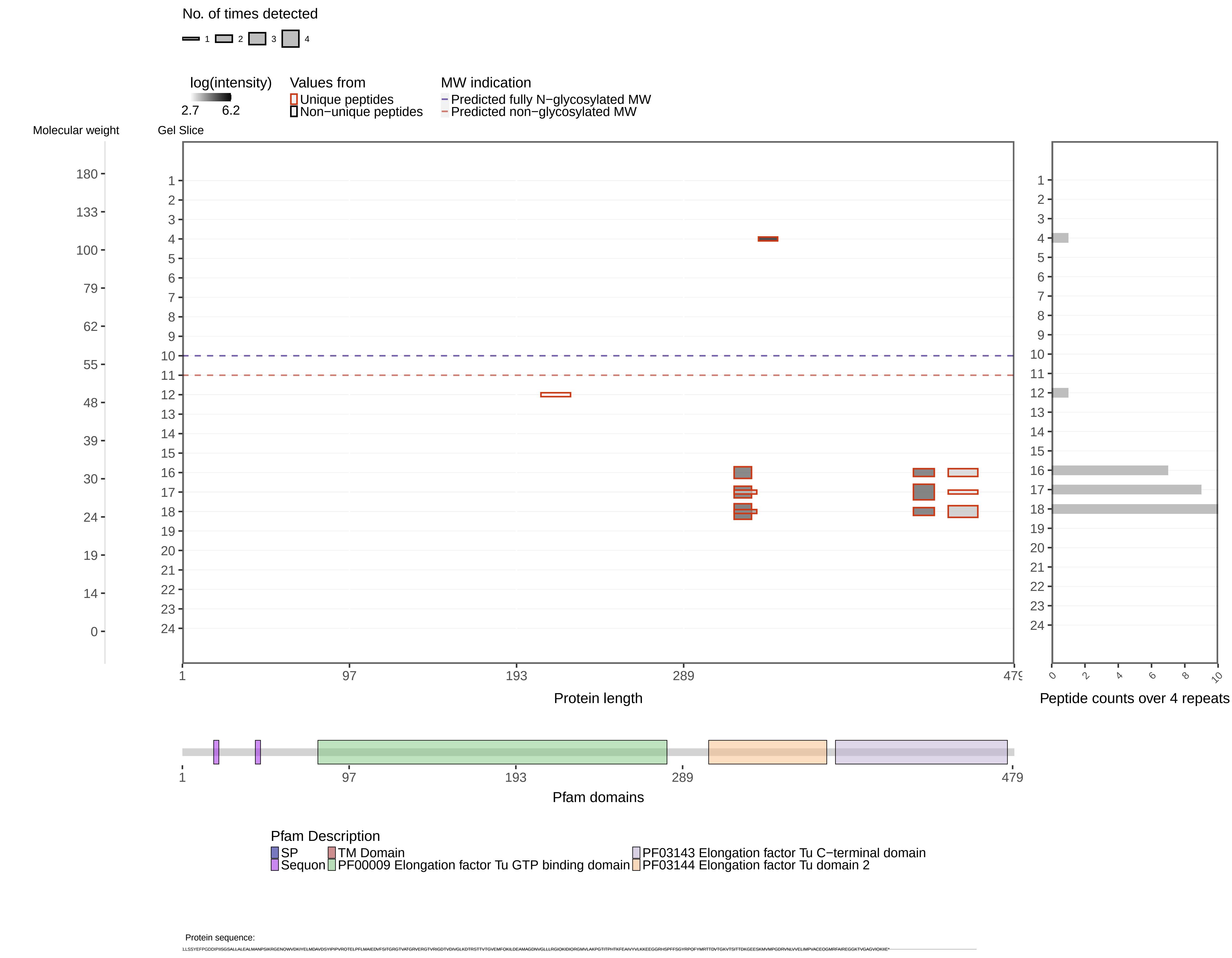

NbL13g16750.1 manganese−dependent_ADP−ribose/CDP−alcohol_diphosphatase−like [MW = 35.78 kDa]

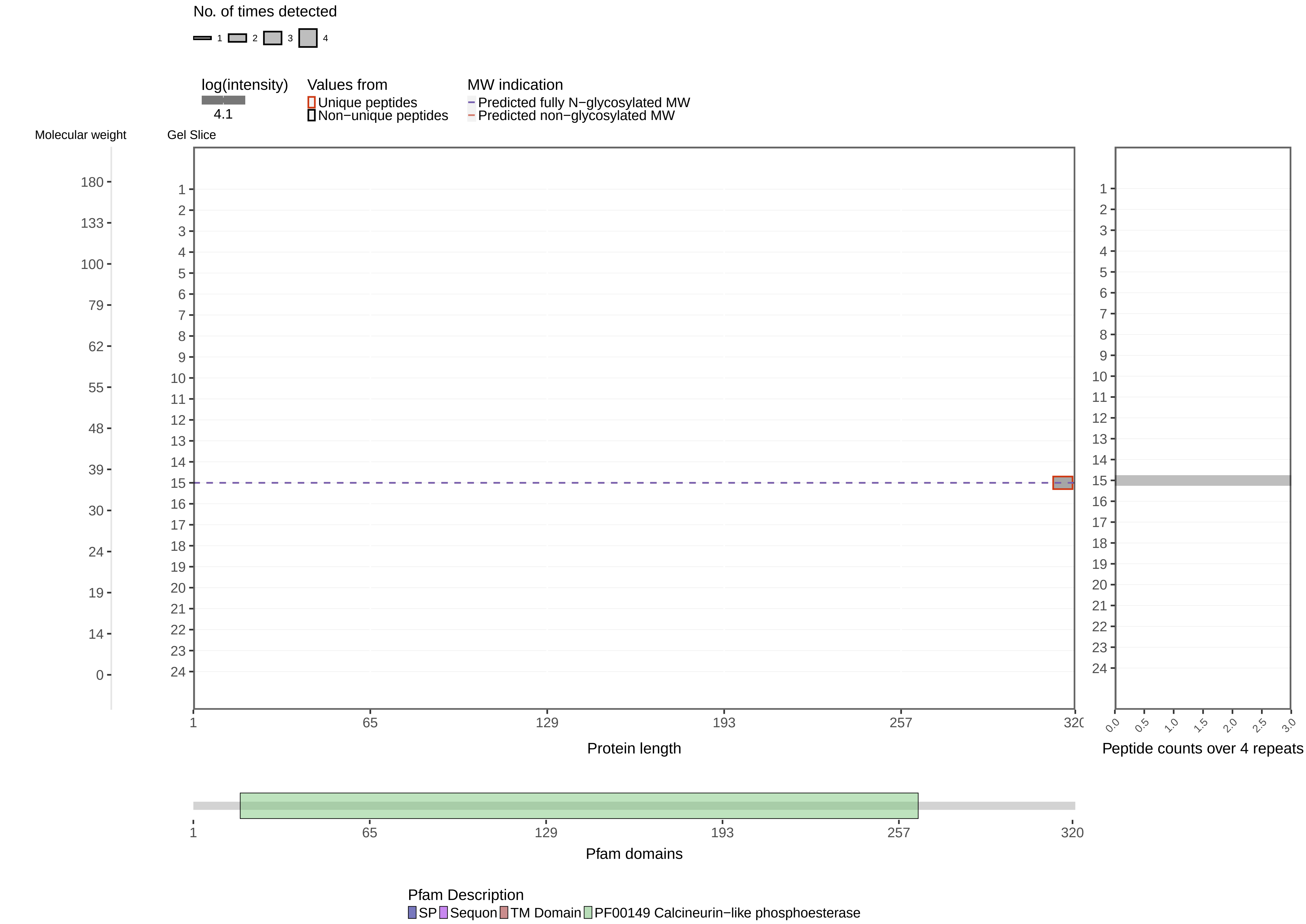


Protein sequence:

AIGWPEDHPNTLKALKVLQERNPNSDKNSPNGLVGLERRFLKFNGGVGKEQLEWLDHVLQEATELNQNVVVCCHLPLDPGASSFAALLWNYDEVMDVIHRYSCVKVCLGGHAHKGGQSVDSHGVHHRVLEAALECPPGTDAFGHIDAFNDRLSLFGNDRMKSSEMVFGH*

### NbL13g16740.1 quinolinate_synthase_chloroplastic [MW = 78.24 kDa]

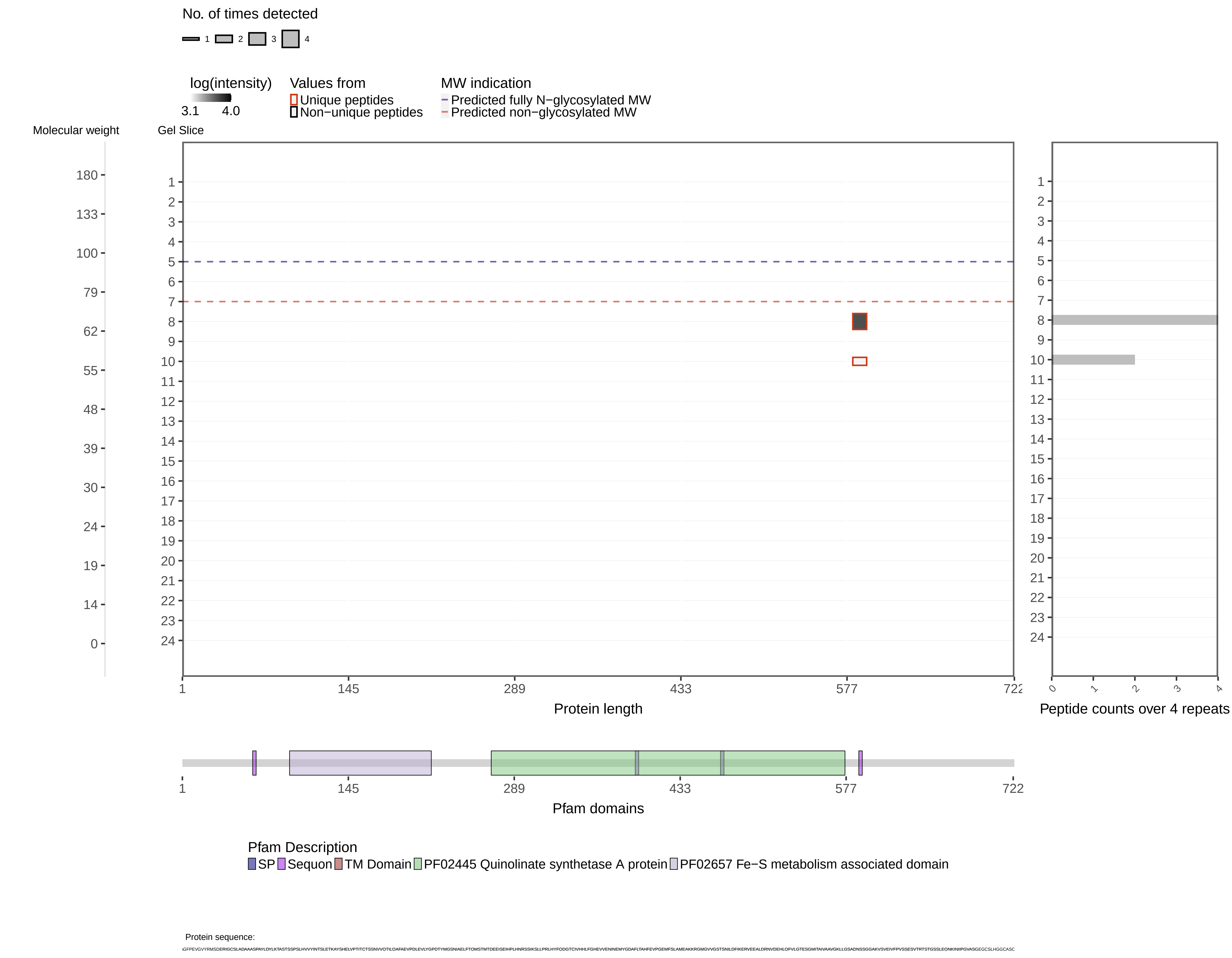

NbL13g16690.1 indole−3−glycerol_phosphate_synthase_chloroplastic−like [MW = 45.03 kDa]

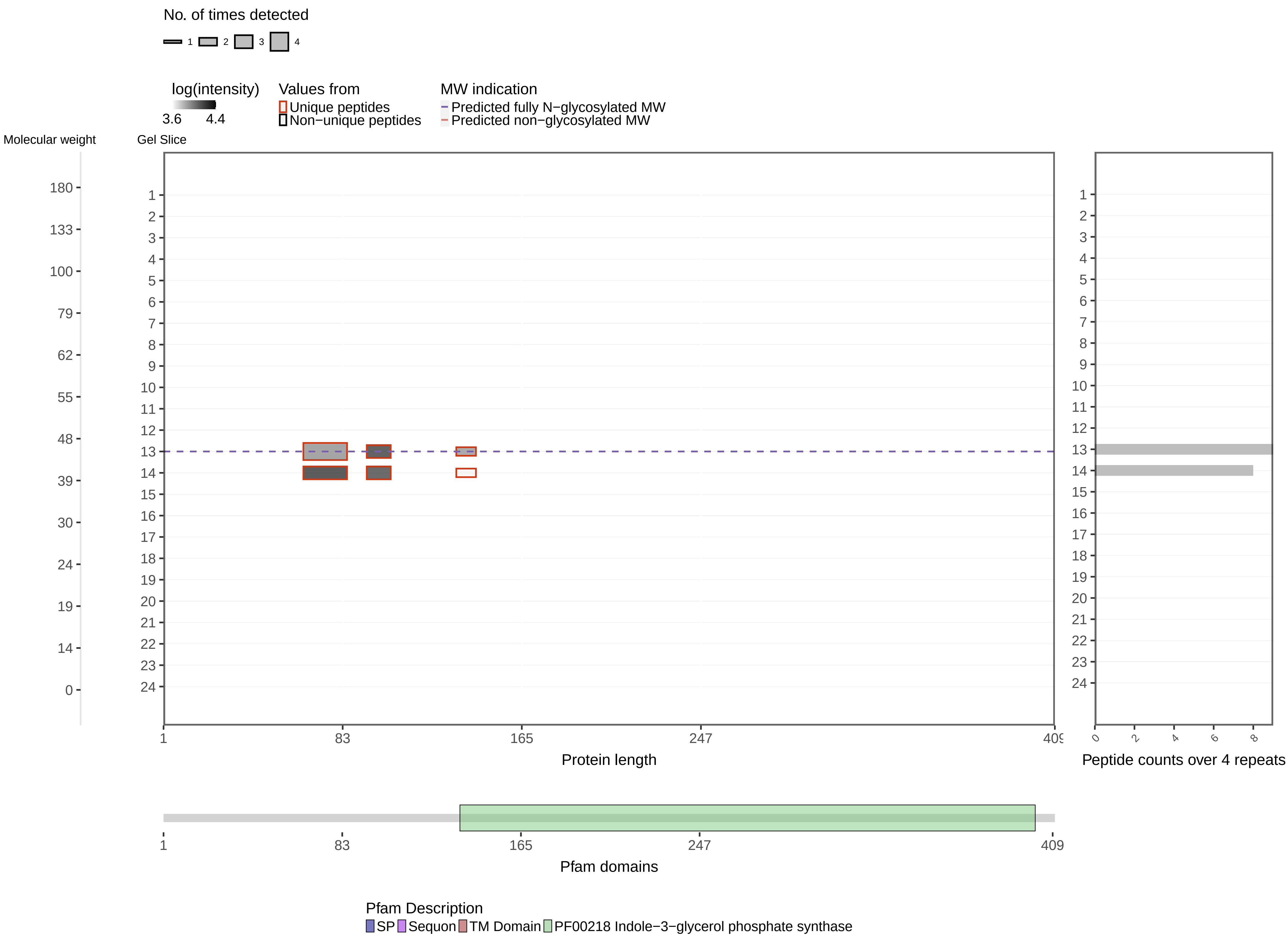


Protein sequence:

IAEVKKASPSRGVLREDFNPVEIAKAYERGGAACLSVLTDQKYFQGSFENLEAIRNAGVECPLLCKEFVIEAWQLYYARVKGADAVLLIAAVLPDLDIKYMIKICKLLGLTALVEVHDEREMDRVLGIDGVELIGINNRDLGTFKVDISNTKKLLEGERGERIREKGIIVVGESGLFTPADIAYVQEAGVKAVLVGESLVKQPDPTKGIRELFGKDISC*

NbL13g16670.1 protein−L−isoaspartate_O−methyltransferase_1−like [MW = 34.57 kDa]

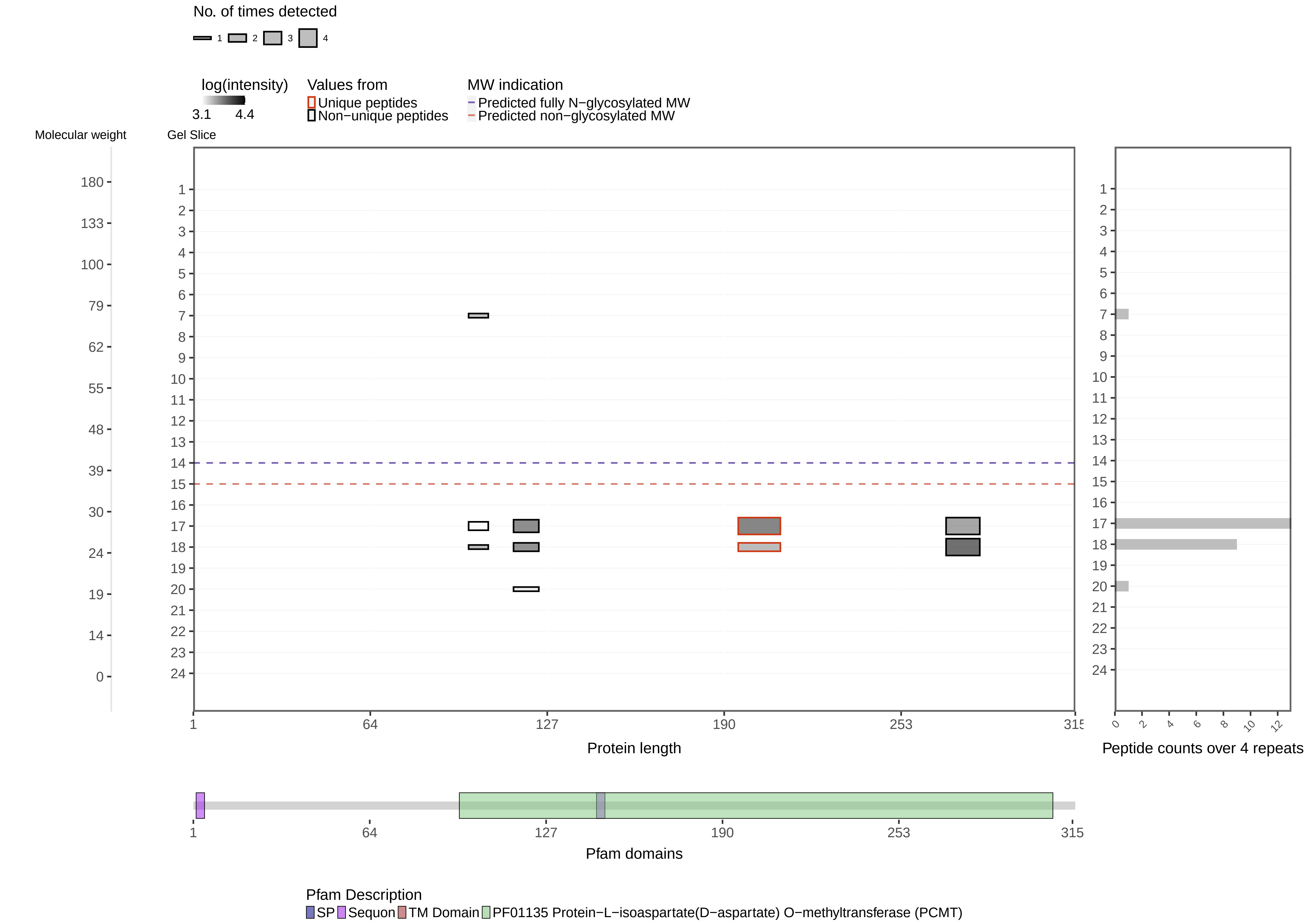


Protein sequence:

TISAPHMHAMCLELLEDRLQPGMHALDVGSGTGYLTACFALMVGPQGRAVGVEHIPELVAWSTKNVERSAAAPLLKEGSLSLHVGDGRKGWPEHAPYDSIHVGAAAADVPQDLIDQLKPGGRMVIPVGTFFQDLKVIDKNLDGSVSVRSETSVRYVPLTSREAQLKGT*

### NbL13g16650.1 protein_ABHD17C−like [MW = 42.59 kDa]

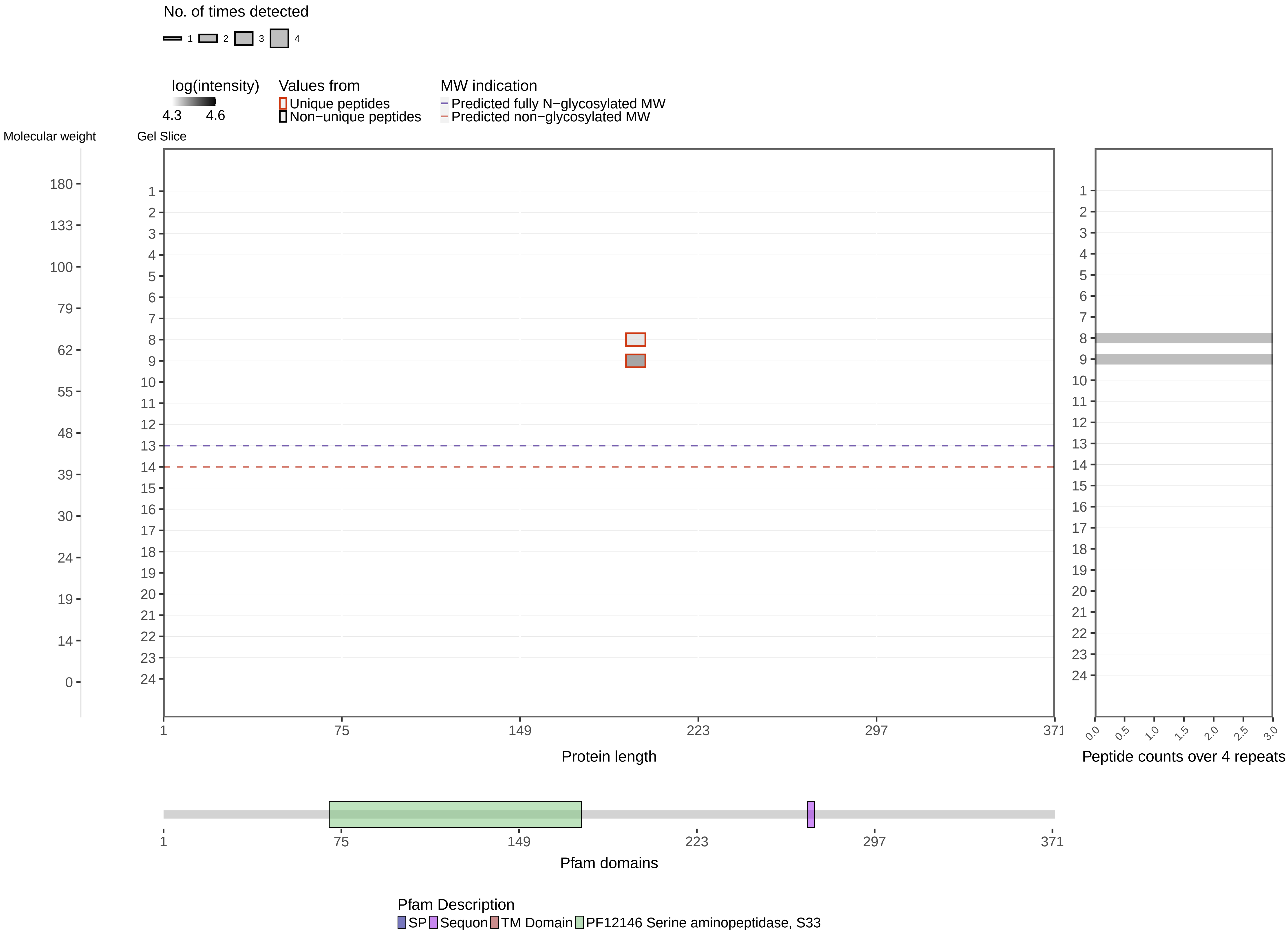


Protein sequence:

[illegible]LRVMYPVKRTYWFDIYKNIDKIPLVKCPVLVIHGTSDEVVDCSHGKOLWELCOEKYEPLWLKGGTHCDLELYPEYIRYLKKFVSTVEKPPSORNLSRKSIDRPEOSRKSTDCFEGPRKSTDWREKPRKSTDKPEKFKNYDYKFSEDKISKLRMSFDOVERSRRSVEFFEKPRRSIDOOMEKARKSVDWLDRIRAG*

### NbL13g16620.1 vignain−like [MW − SP = 38.25 kDa]

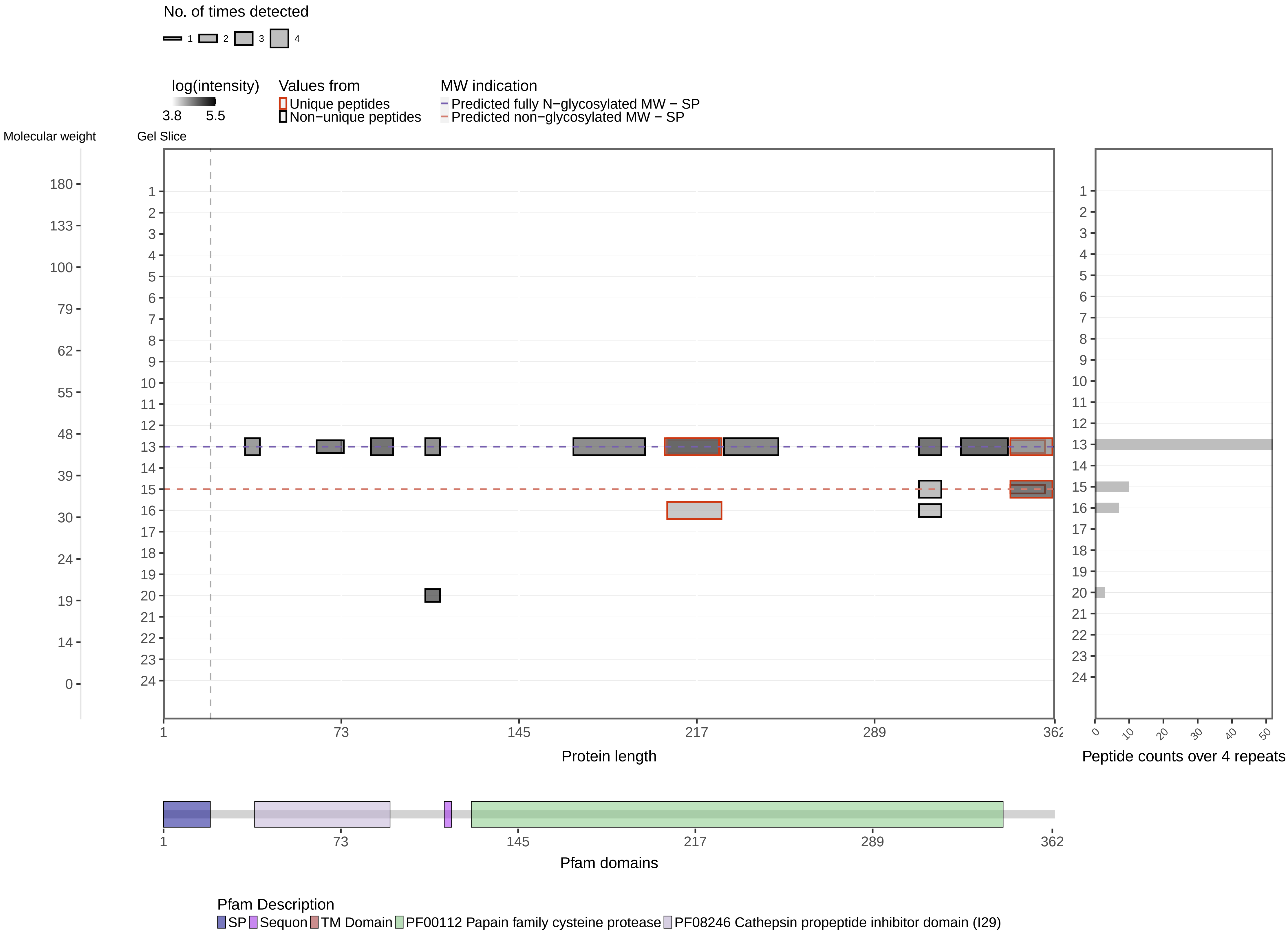


Protein sequence:

NELVSLSEQELVDCDTSQNQGCNGGLMDKAFEFIKKKGGINTEDNYPYMAEDGECDIQKRNSPVVSIDGHEDVPPNDEGSLLKAVANQPVSVAIQASGSDFQFYSEGVFTGDCGTELDHGVAIVGYGTTLDGTKYWIVKNSWGPEWGEKGYIRMQREIDAEEGLCGIAMQPSYPIKTSSSNPTGSPATTPKDEL*

### NbL13g16610.1 peptide_methionine_sulfoxide_reductase [MW = 22.1 kDa]

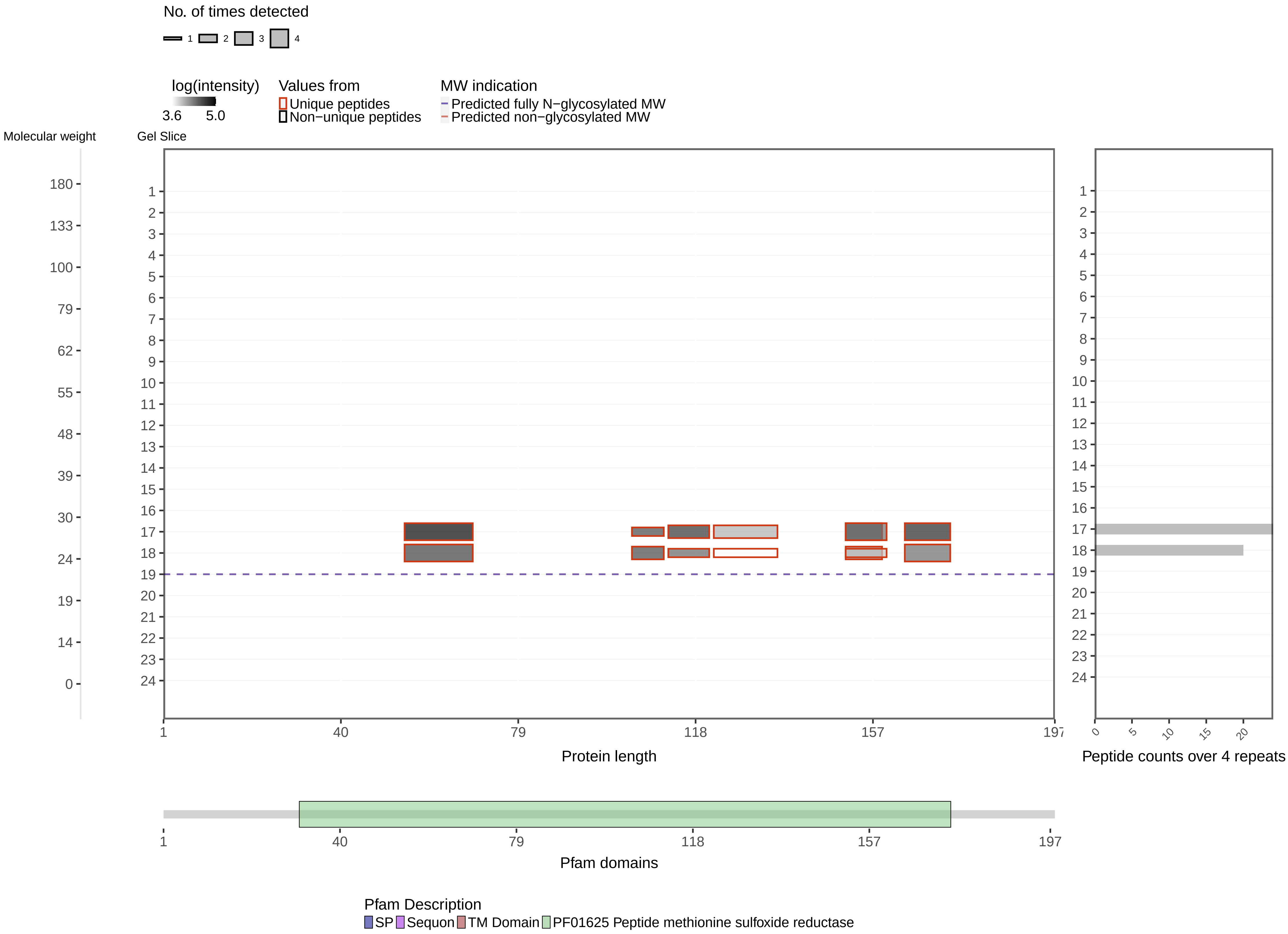


Protein sequence:

[illegible]FDPNVCPYTNLLSLFWSRIDPTTLNRQGNDVGNQYRSGIYYYNEAQAQLARESLEAKQKEFADKKIVTEILPAKRFYRAEEYHQQYLEKGGGRGAKQSAAKGCNDPIRCYG*

NbL13g16460.1 GDSL_esterase/lipase_At3g48460−like [MW − SP = 38.72 kDa]

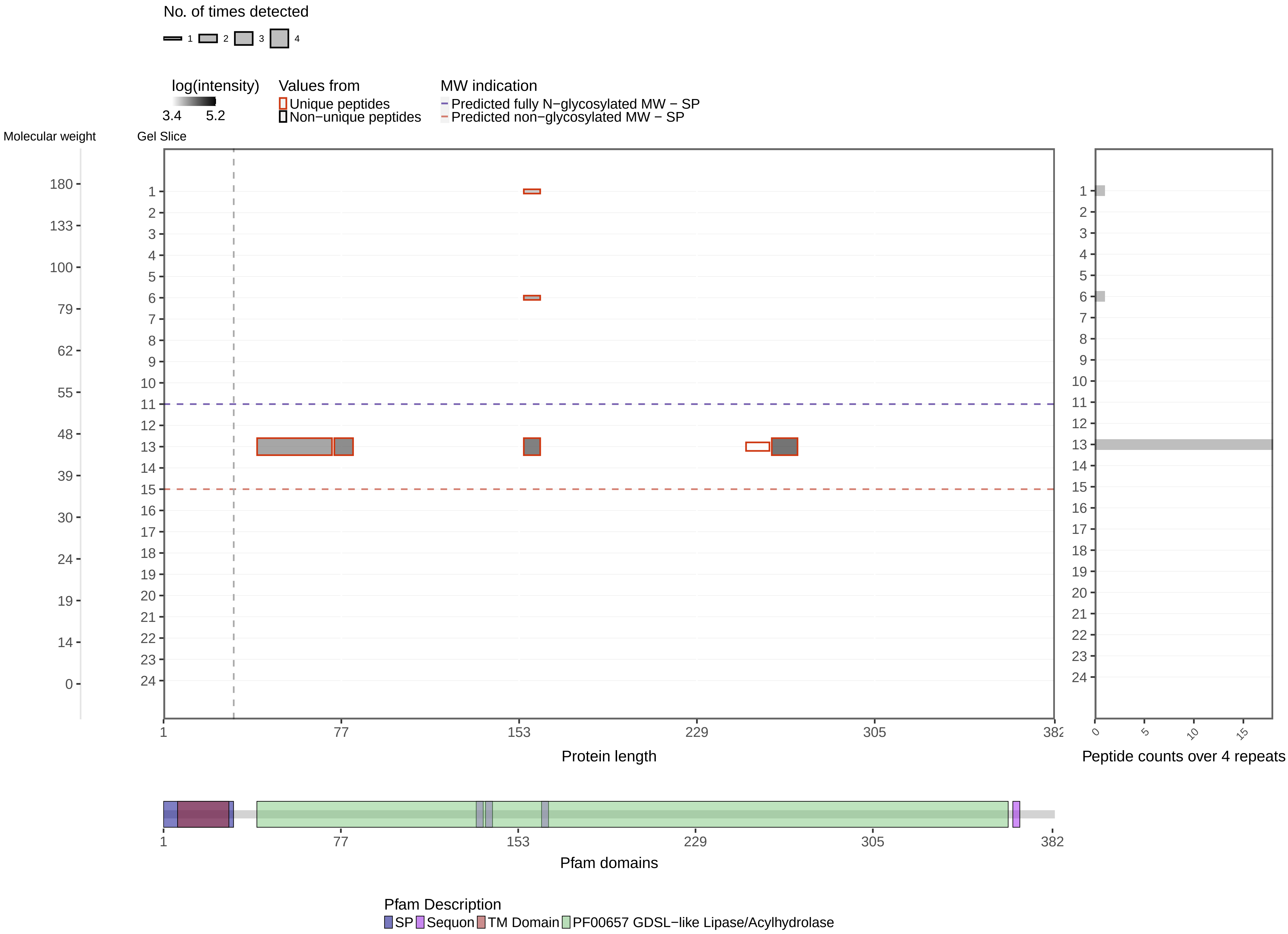


Protein sequence:

FWVGEIGANDYAYSFGSSVSPNTIQHLATTSLTGFLQVLLNKGAKYVVVQGLPPTGCLTLSMYLAPETDRDDMGCVGSVDKQSNLHNNIIQTKLDSFRKQFPQAVIIYADYWNAYSTVVKGANKYGFKEIFKSCCGSGGGNYNFDVFNTCGSPSATSCPDPSQYINWDGVHLTEAMYKAMANLFLNGTFCKPPFSYLLAKKQ*

### NbL13g16370.1 ubiquitin−like_modifier−activating_enzyme_5 [MW = 46.84 kDa]

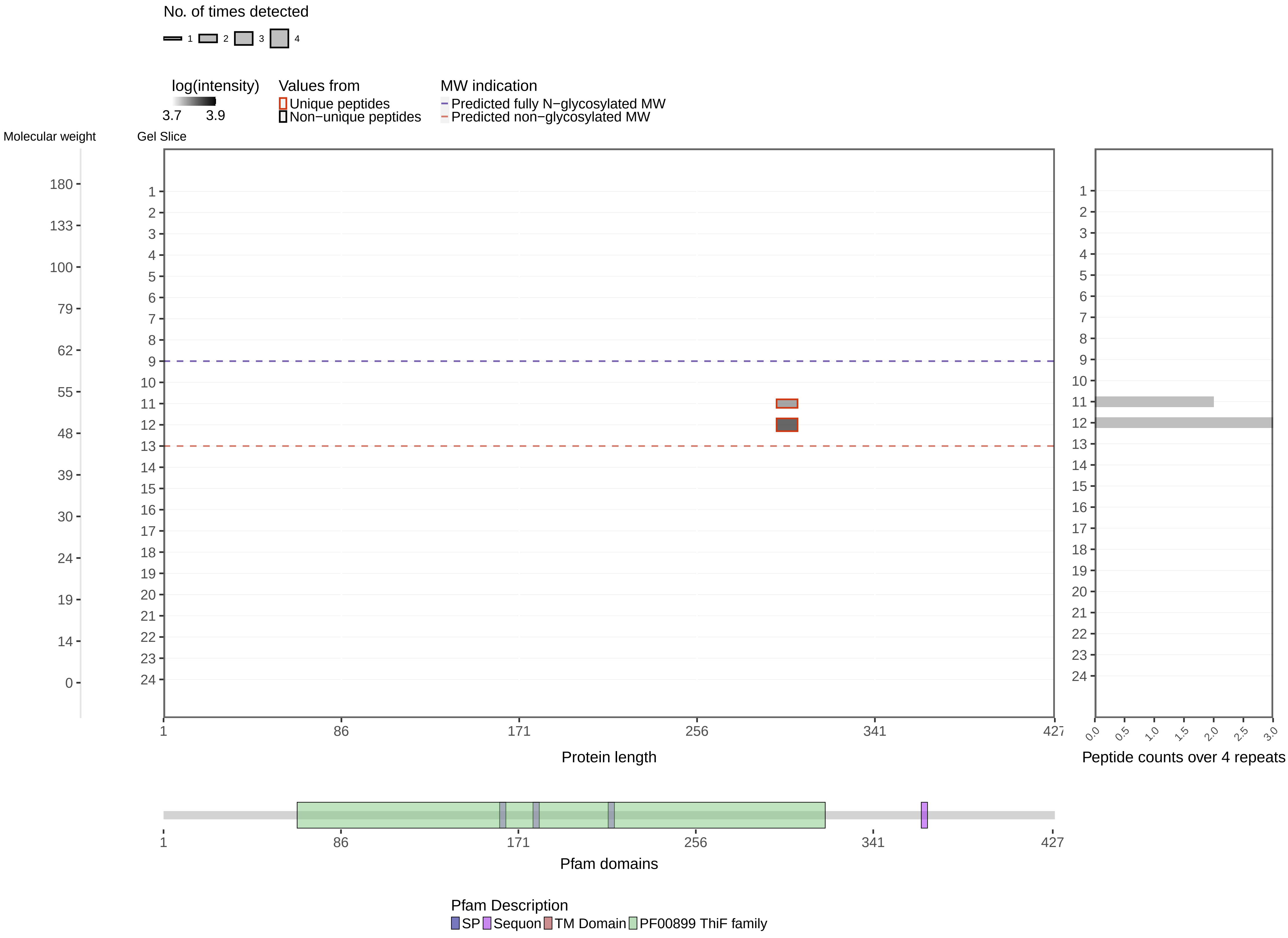


Protein sequence:

EARMVVNQACNELNQTWMESGVSEDAVSGHIQLLIPGESACFACVPPLVVASGIDERTLKREGVCAASLPTTMGVVAGLLVQNTLKYLLKFGRVSPYLGYNALKDYFPTMEMKPNTQCSNVACLEROKEYILAKPTRDAVAKAKADKEALSTPENPVHADNEWNISVLDDNEVDDAISRHSGVLPEGIVHELPVADEYPKPAITEIVGPAHDLEELRRQLEALNAD*

NbL13g16360.1 UPF0235_protein_At5g63440 [MW = 25.8 kDa]

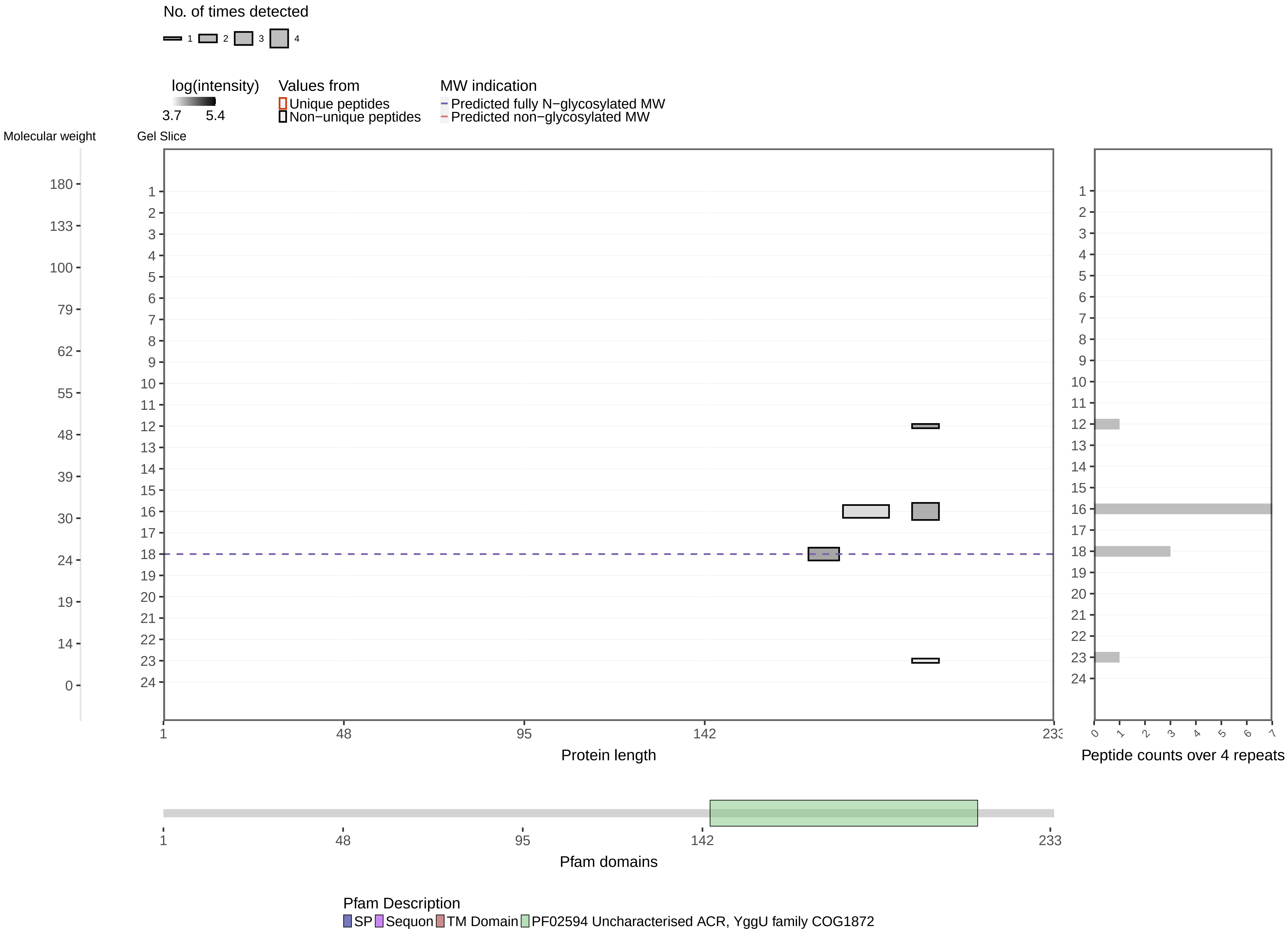


Protein sequence:

EEDLETASFIYVVDGALSTIAAETNPQDAPVPPCISQLEGGLVQVAIEVEDRAQRSAITRVNADDVRVTVSAPAARGEANNELMEFMGRVLGLKLSQMTLQRGWNSKSKLLVVEDLTARQVYEKLLEAAQP*

NbL13g16290.1 60S_ribosomal_protein_L17−1−like [MW = 19.87 kDa]

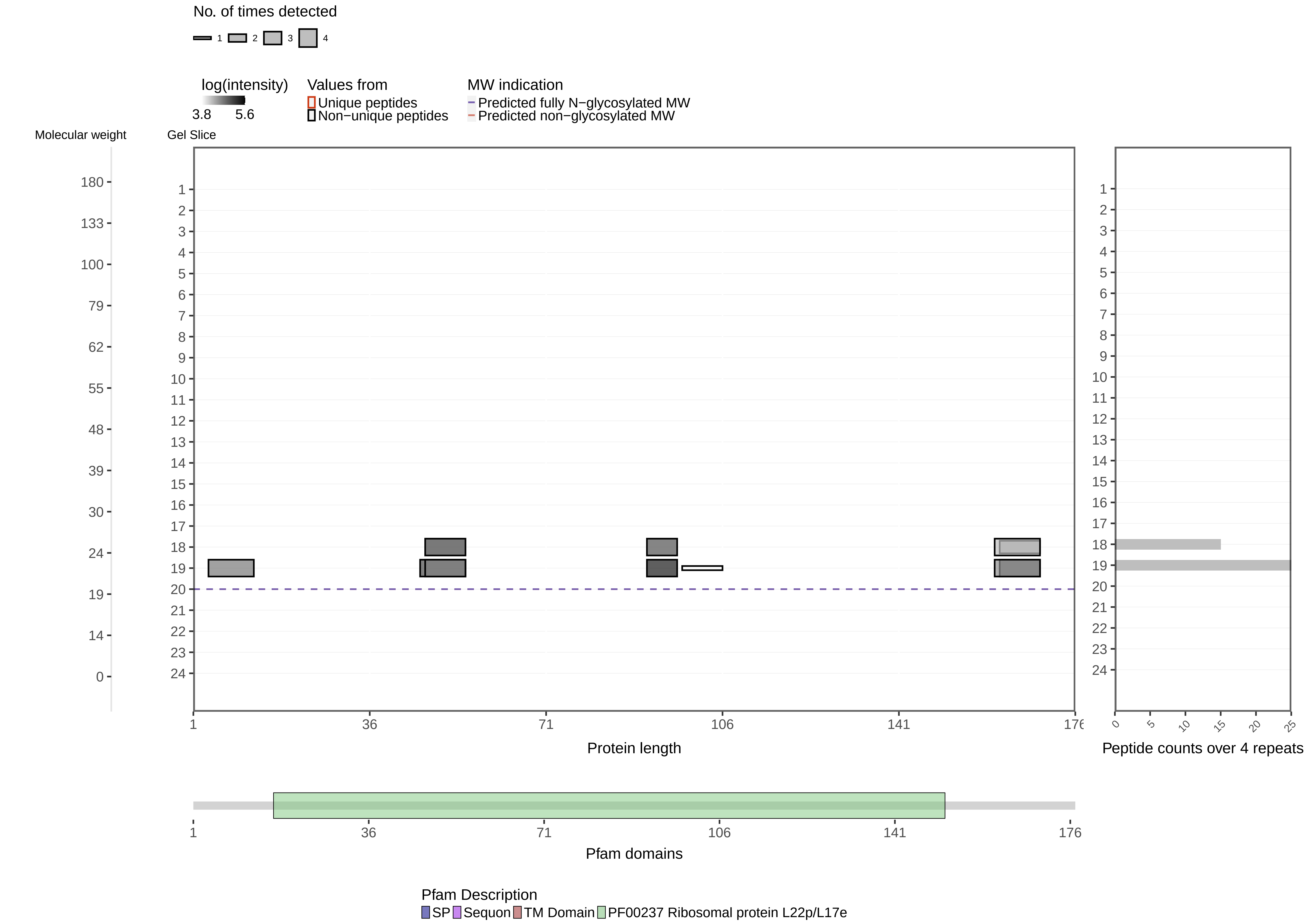


Protein sequence:

HRHSNGQGRWPVKSAKIILDLLKNAESNAEVKGLDVDSLFISHIQVNQAQKQRRRTYRAHGRINPYMSHPCHVELTLSEKEESVKKEPETQLAPRKAKASS*

### NbL13g16270.1 adenylate_kinase_4 [MW = 26.6 kDa]

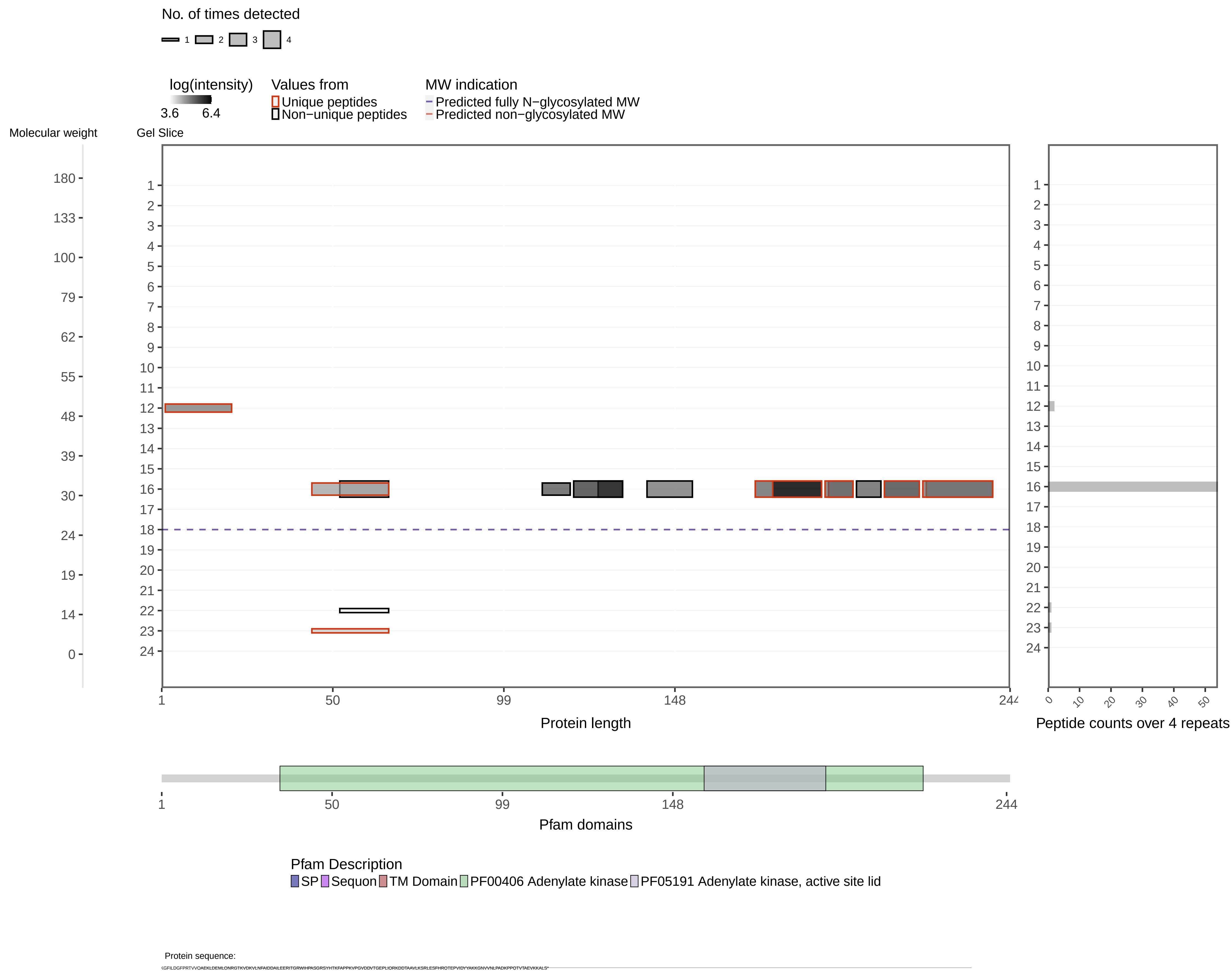

NbL13g16240.1 probable_aspartyl_aminopeptidase [MW = 52.74 kDa]

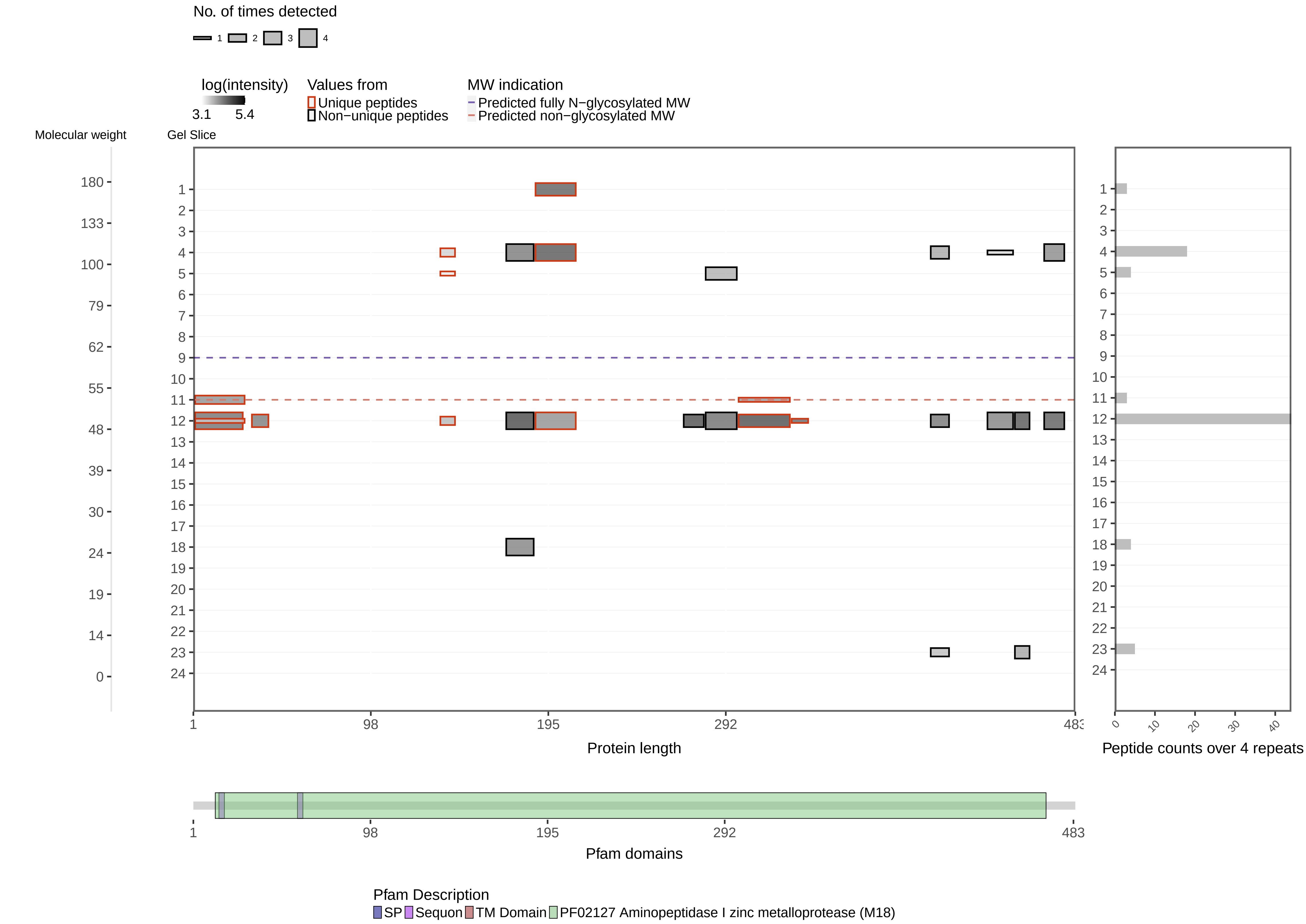


Protein sequence:

NbL13g16230.1 4−coumarate−−CoA_ligase−like_9 [MW = 59.65 kDa]

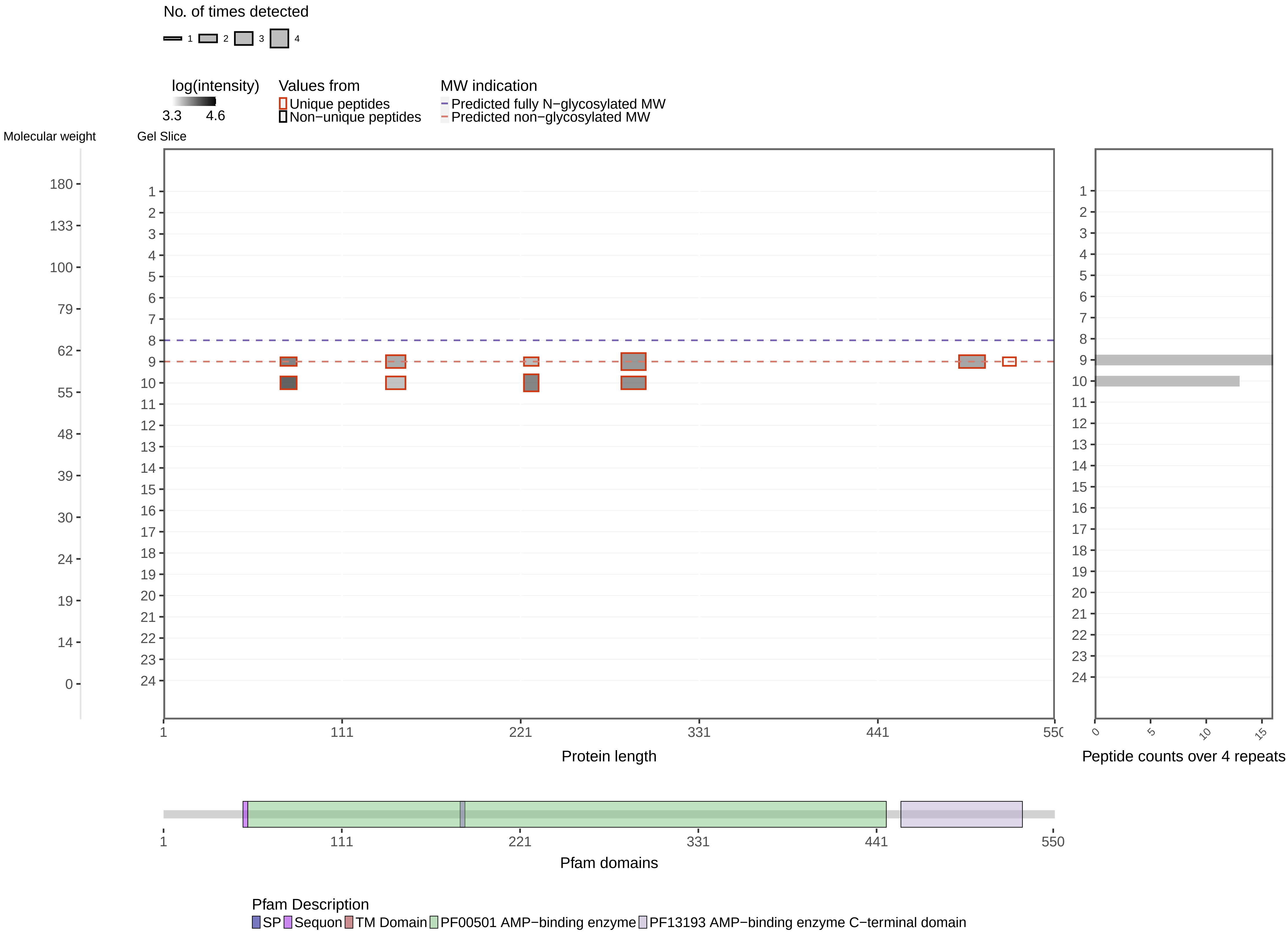


Protein sequence:

/VIIERFDFEKMLEAVEKYRVTYIPVSPPLVVAMAKSDLALKHDLYSLKLLACGGAPLGKEVAERFKSRFPNVEIIQGYGLTETTGGATGMSGPEESDRYGSAGRLSVNVEAKIVDPDNGETLPPGQRGELWIRGPTIMKGYVGDKQATSATLNSEGWLKTGDLCYFDADGFLYIVDRLKELIKYKAYQVPPAELEHLLQSIPDVADAAVIPYPDEEAGEIPMAYIVRRPGSTISESQIIDAIAKQVAPYKKIRRVAFIHAIPKSPAGKILRIELVNHASSGASARL*

NbL13g16130.1 glyceraldehyde−3−phosphate_dehydrogenase_cytosolic−like [MW = 36.73 kDa]

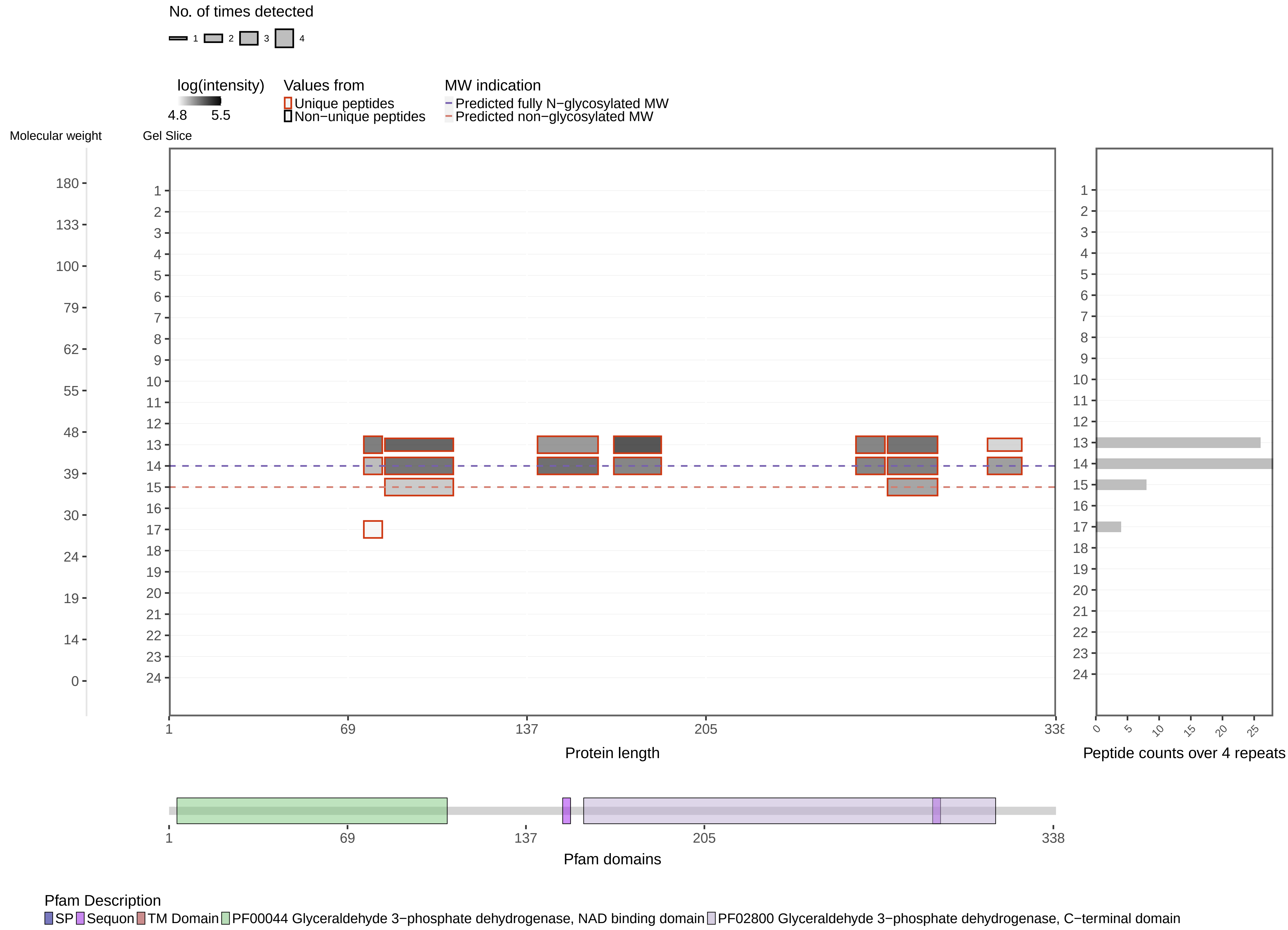


Protein sequence:

TNCLAPLAKVINDRYGIVEGLMTTVHSITATQKTVDGPSAKDWRGGRAASFNIIPSSTGAAKAVGKVLPALNGKLTGMAFRVPTVDVSVVDLTVRLEKEATYDEIKAAIKEESEGNLKGILGYTEDDVVSTDFVGDNRSSIFDAKAGIALSKNFVKLVSWYDNEWGYSTRVVDLIKHMASVQ*

NbL13g16100.1 nucleoside_diphosphate_kinase_2_chloroplastic−like [MW = 26.2 kDa]

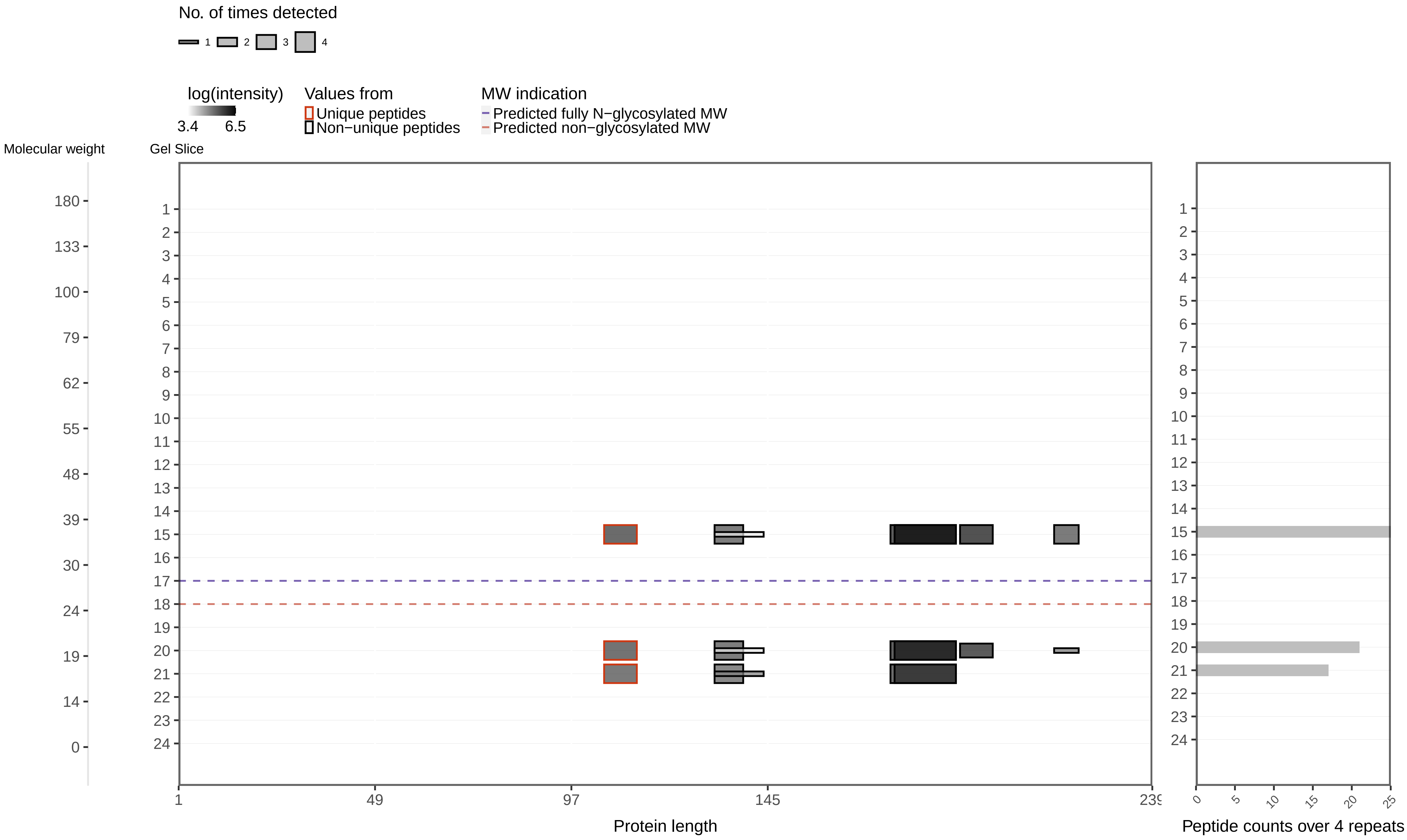


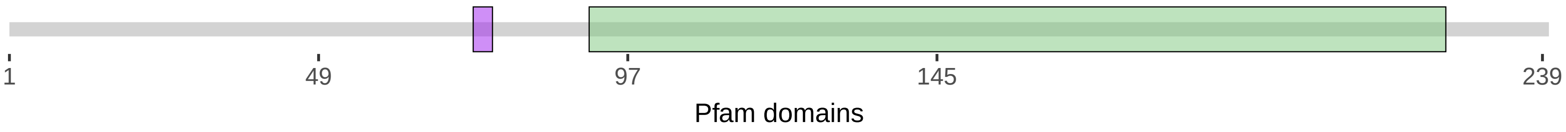


Protein sequence:

VGEIISRFEKKGFKLSGLKLFHCPKELAEEHYKDLQSKSFFPKLIDYITSGPVVCMAWEGIGVVASARKLIGATNPLNAEPGTIRGDLAVQTGRNVVHGSDSPENGKREIALWFREGELCSWTPVQEPWLIE*

### NbL13g16010.1 nuclear_transcription_factor_Y_subunit_C−1−like [MW = 25.05 kDa]

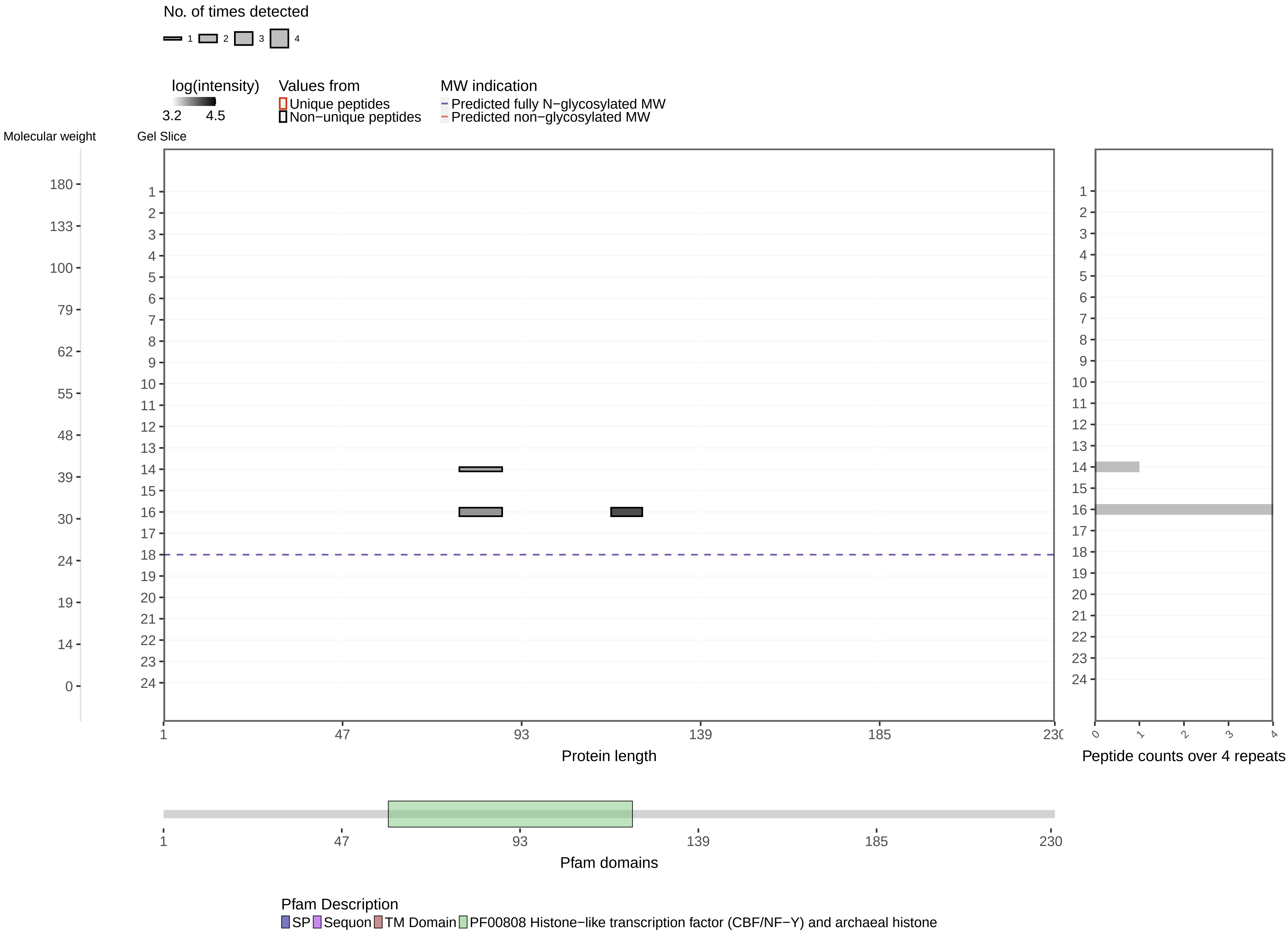


Protein sequence:

.HAEENKRRTLQKNDIAAAITRTDIFDFLVDIVPRDEIKEEGGVGLGPAGIVGSTASGVPYYYPPMGQPAPPGVMMGRPAMPGVDPSMYVQPAPPPSQAWQSAWQTAEDNSYASGGSSGQGNLDGQS*

NbL13g15930.1 thylakoid_lumenal_17.9_kDa_protein_chloroplastic [MW = 25.1 kDa]

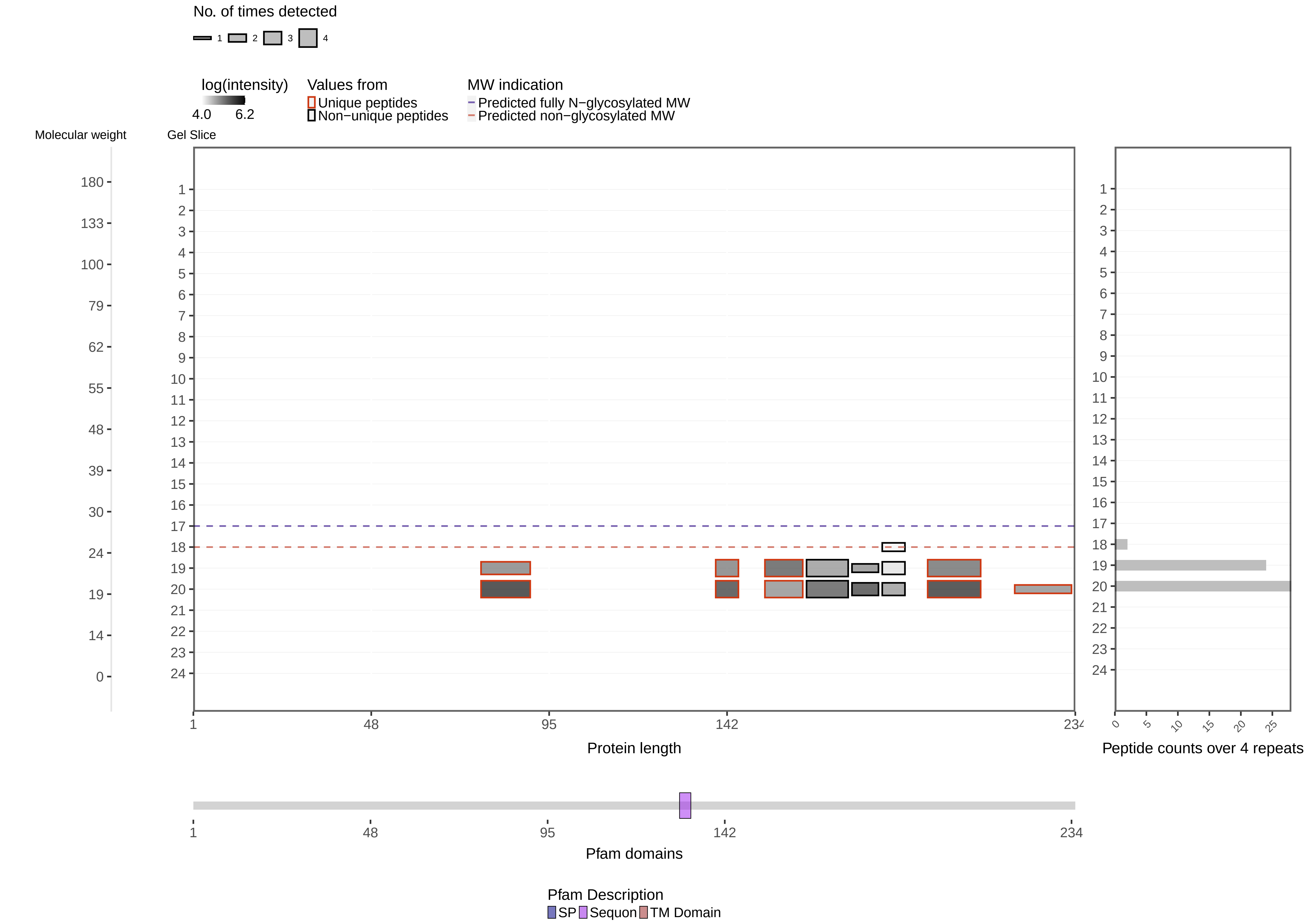


Protein sequence:

PSVNPGCVSTNPQSSSSAFPWTIPQNSTGNAIMRLQDAILKTQKNAKIQVIEDISDGKYLQAEVDGGFGRDVLEFLVKGDSVAYRAMATKVTYIYPFTTALGDSKGQEERMKKITEELGWYAPSLDSMD*

NbL13g15880.1 SUMO−activating_enzyme_subunit_1B−1−like [MW = 40.31 kDa]

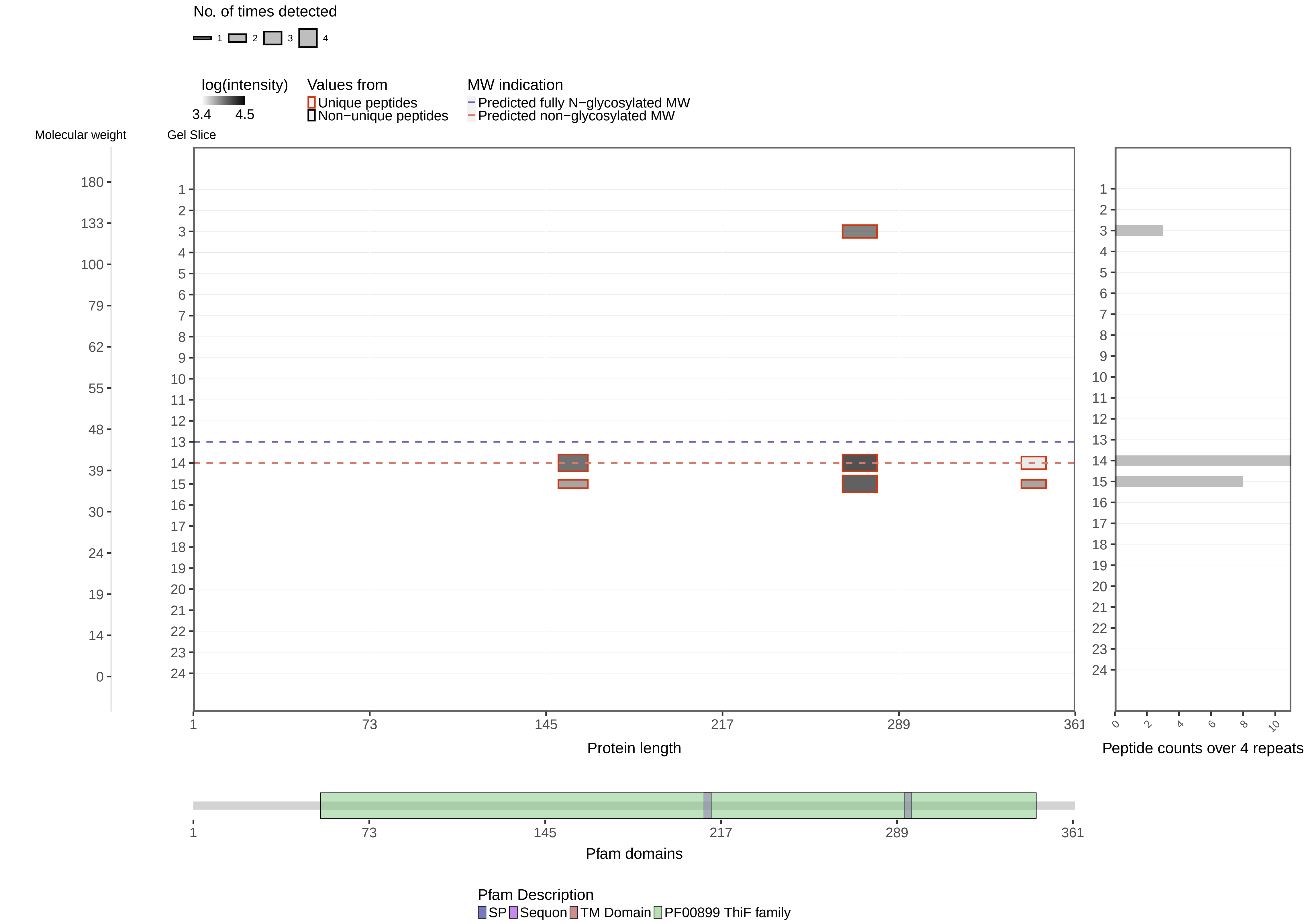


Protein sequence:
VVVSCCTLLTKKSVNAKCHKLPKRVAFYSVECRDSCGEIFVDLQNYSYCKKKNKETMECILLYPSFEEAIAVPWRSLPKRMSKLYFAMRVIERFEELEGRNPGETSADDLPNVQKLRKELCEAHCLNESQIPDSLLQRLIASRSEFPPACAIIGGILGQEVIKAISGKGDPLKNFFFFDAMDGKGIIEDISNVNS*

### NbL13g15830.1 alpha−galactosidase−like [MW = 48.41 kDa]

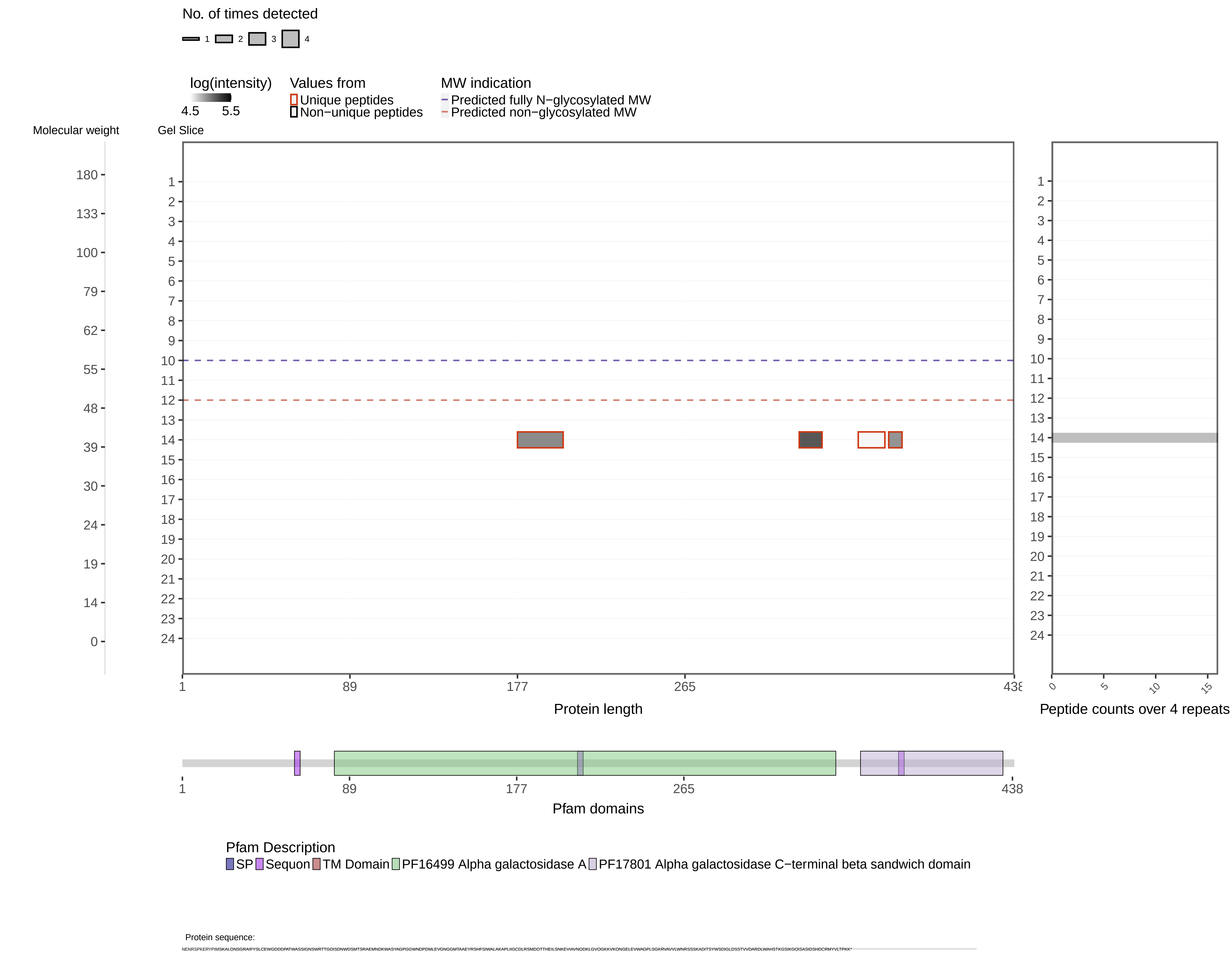

### NbL13g15700.1 asparagine−−tRNA_ligase_chloroplastic/mitochondrial [MW = 64.14 kDa]

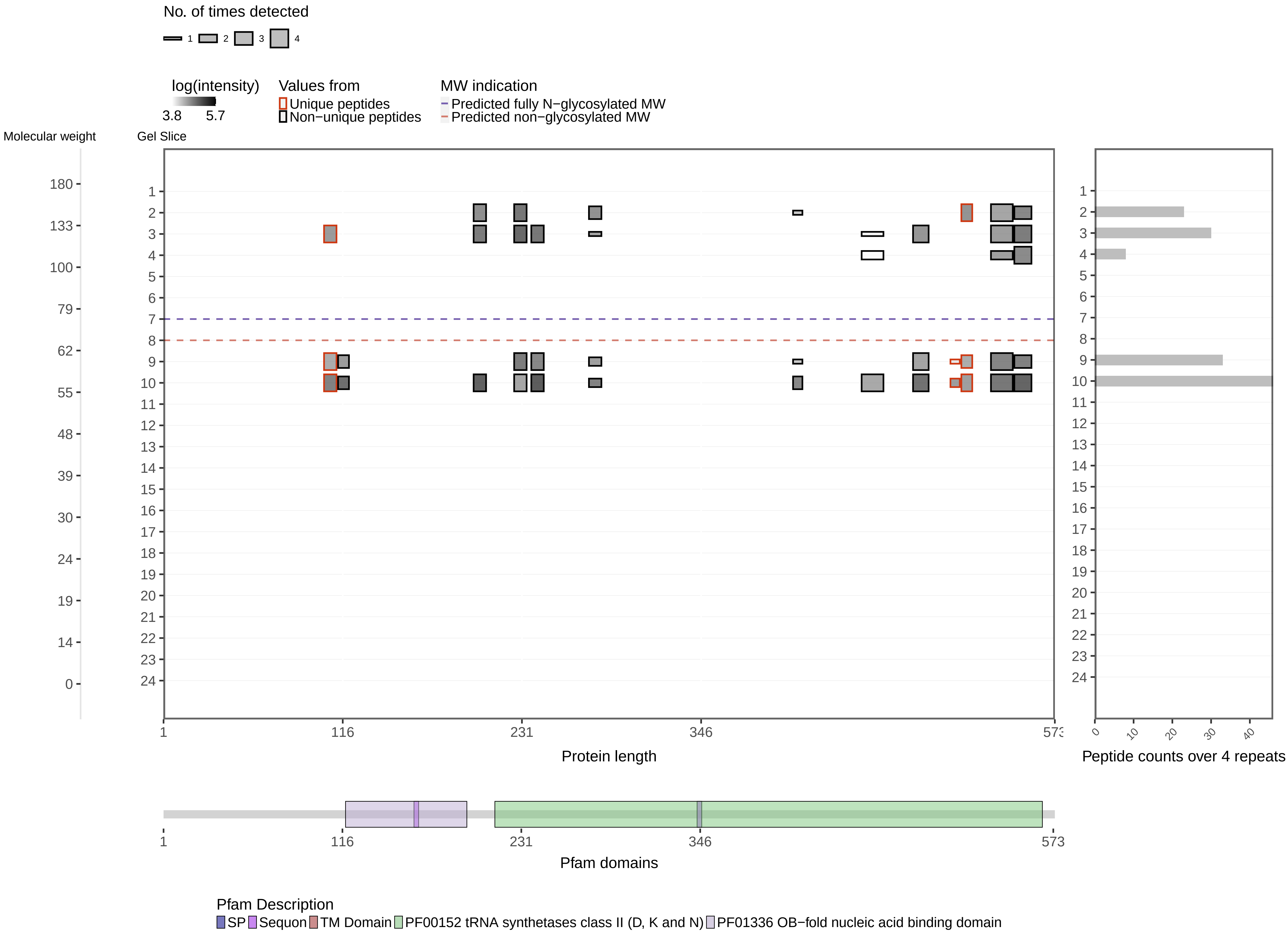


Protein sequence:

RVTTLIPTTKEGGDSPVNAIPTTESGSVDWSQDFFGKQAFLTVSGQLNGETYATAISDIYTFGPTFRAENSNTSRHLAEFWMIEPELAFADLDDDMACATAYLQYVVQYVLENCKEDMDFFNTWIEKGIISRLSDVVEKNFVQLTYTDAVELLLKAKKKFDFPVKWGCDLQSEHERYITEEAFGGCPVIIRDYPKDIKAFYMRQNDDGKTVAAMDMLVPRVGELIGGSQREERLEYLDERLDNMNLNKESFWWYLDLRRYGSVPHAGFGLGFERLVQFATGIDNIRDAIPFPRTP

### NbL13g15390.1 protein_argonaute_1−like [MW = 122.85 kDa]

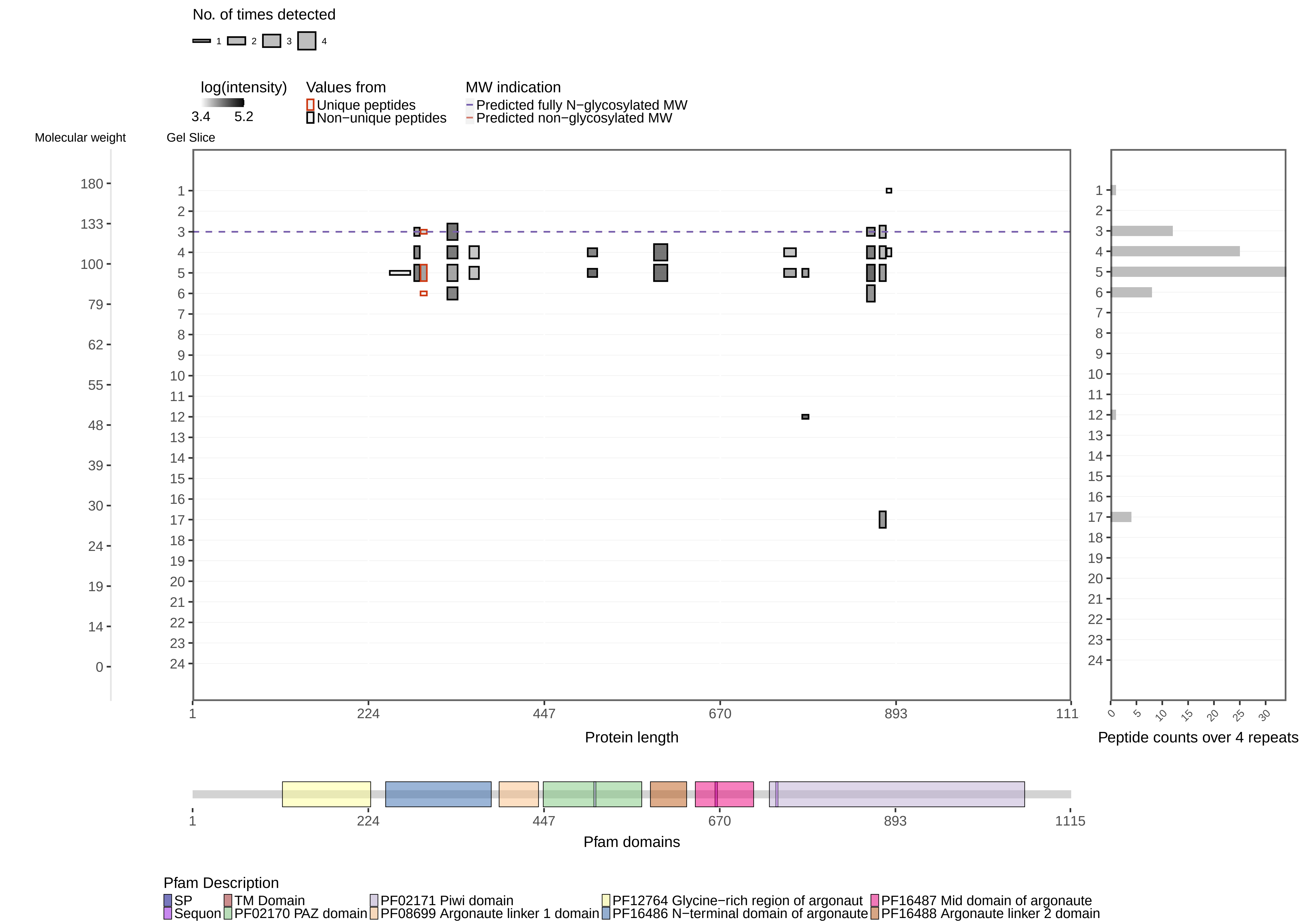


Protein sequence:

[illegible]

NbL13g15360.1 bifunctional_D−cysteine_desulfhydrase/1−aminocyclopropane−1−carboxylate_deaminase_mitochondrial [MW = 46.84 kDa]

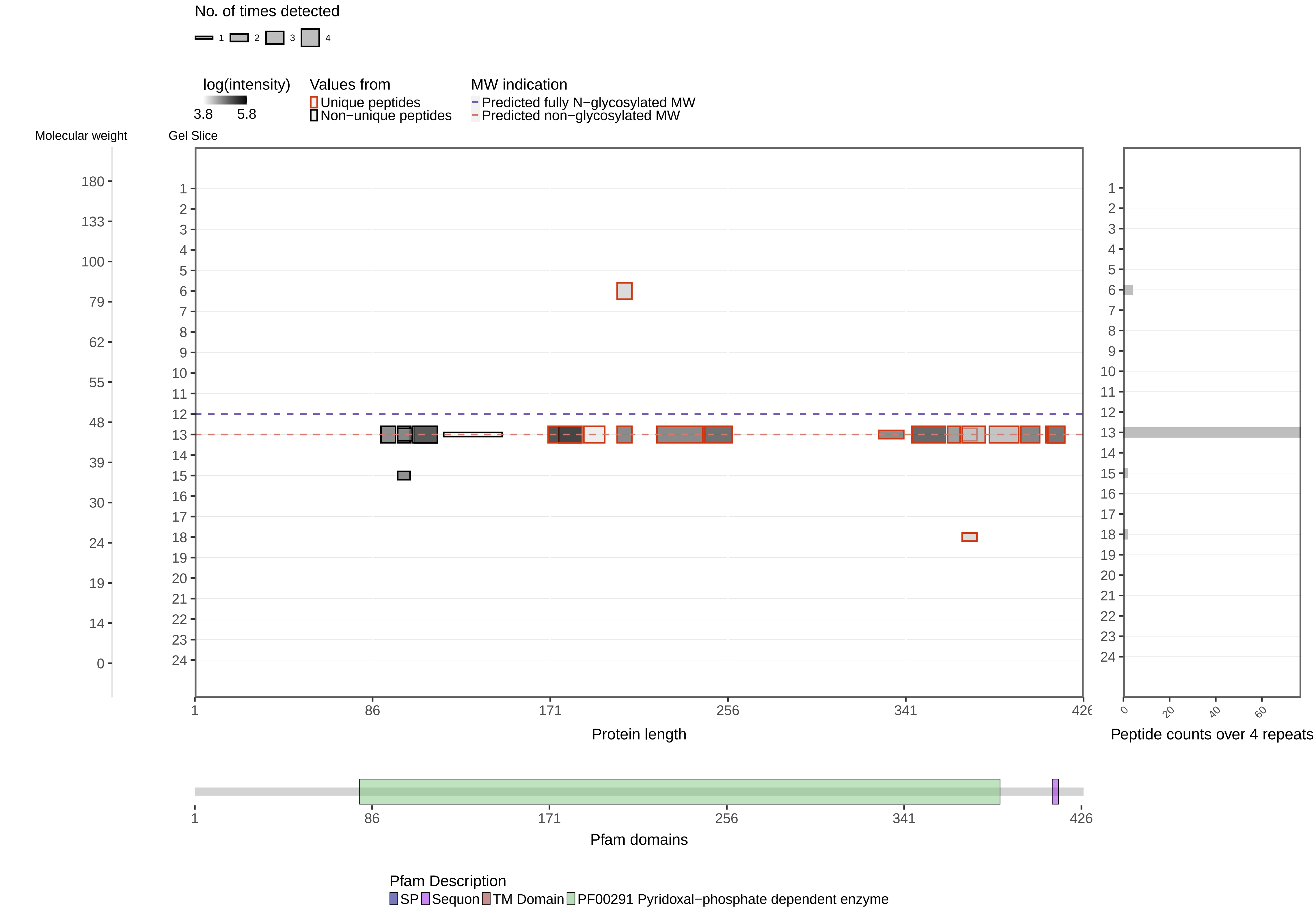


Protein sequence:

AKVGSEALTKLLKEKLLNEGRKPYVIPVGGSNSLGTWGYIEAIREVEQQLQHSSSEWKFDDIVVACGSGGTVAGLSIASRLSGLKAKVNAFCVCDDPDYFYEYVQGLLDGINAGVSSRDIVSIKNAKGLGYAMSTADELKFVKQVAEATGVILDPVYSGKAAYGMIKDMNENPTKWEGRKILFIHTGGLLGLYDKADEMGSLMGKWRKMDINESIPRQDGIGKMF*

NbL13g15240.1 calmodulin [MW = 16.93 kDa]

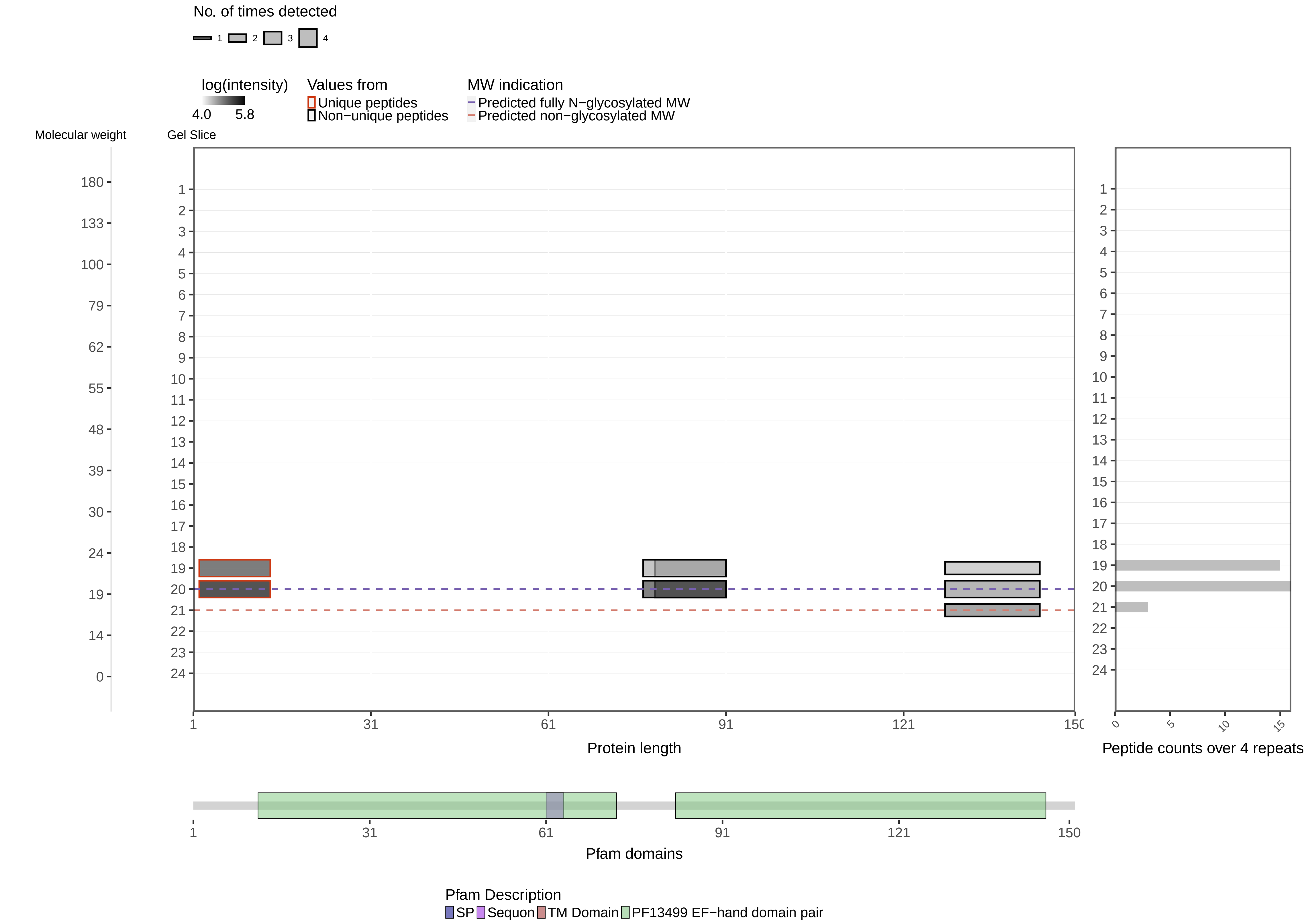


Protein sequence:

GTIDFPEFLNLMARKMKDTDSEEELKEAFKVFDKDQNGFISAAELRHVMTNLGEKLTDEEVDEMIREADIDGDGQVNYEEFVRMMLAK*

NbL13g15130.1 dihydropyrimidine_dehydrogenase_(NADP(+))_chloroplastic−like [MW = 46.79 kDa]

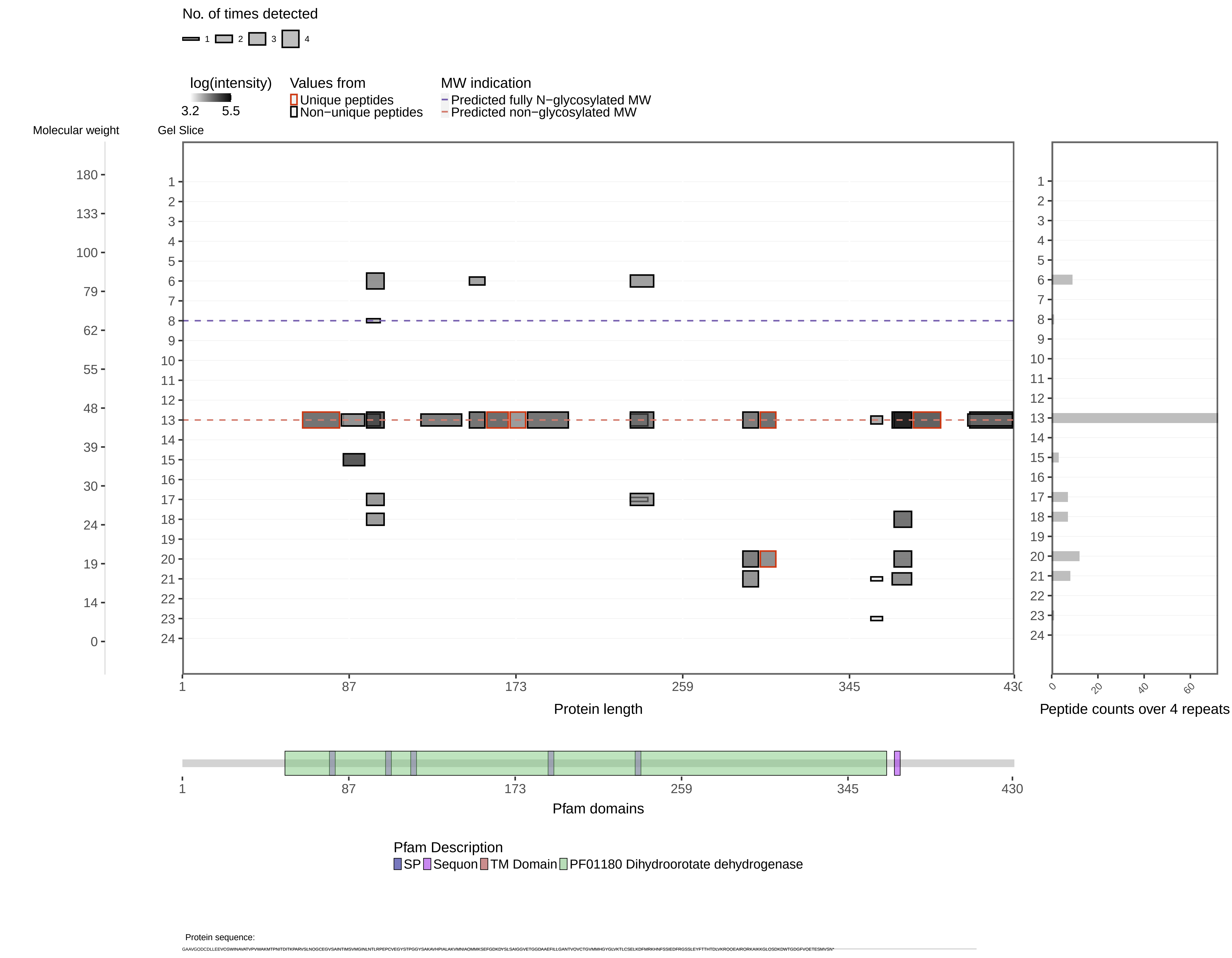

NbL13g15120.1 40S_ribosomal_protein_S3a−like [MW = 29.73 kDa]

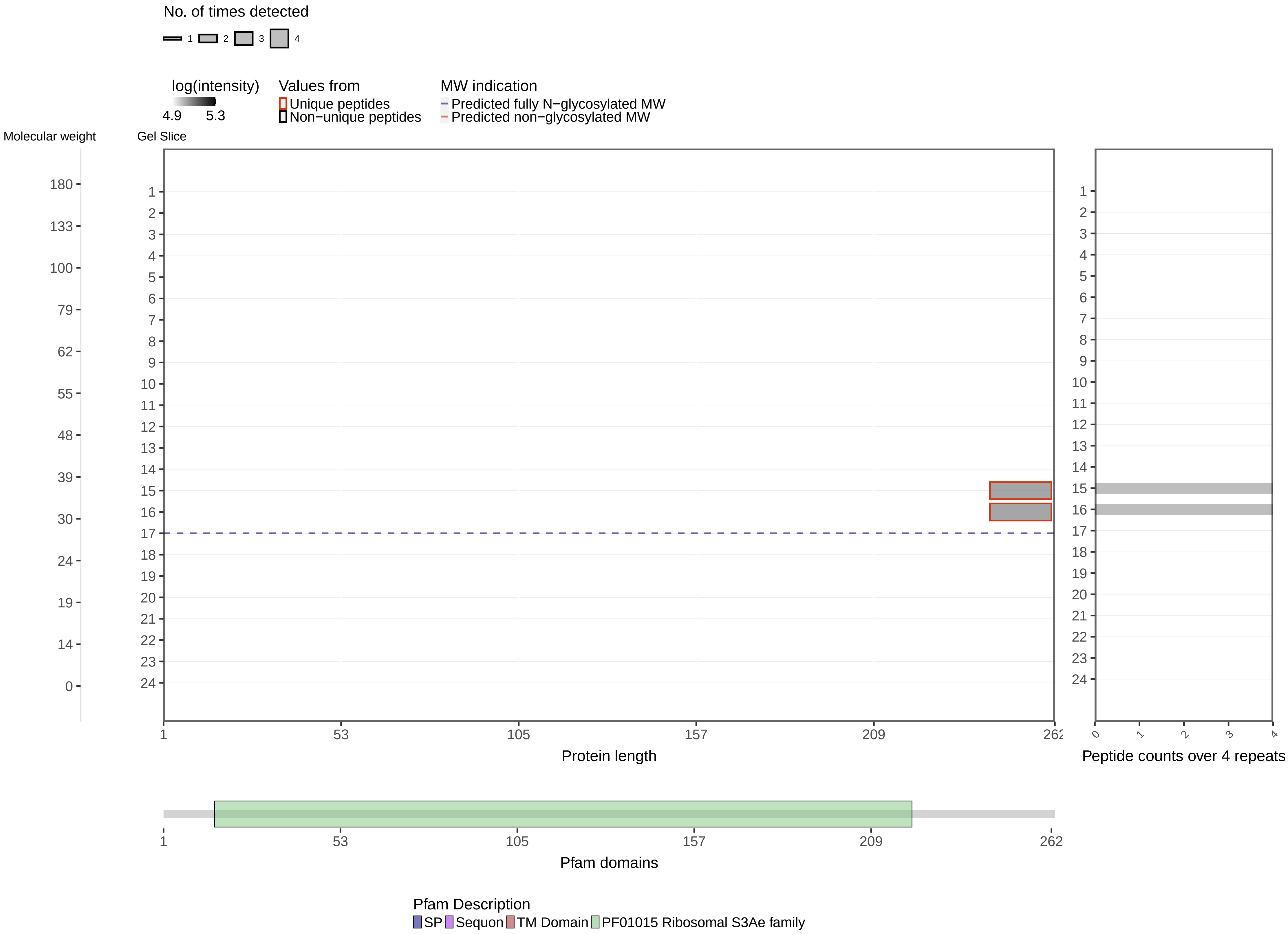


Protein sequence:

WQTLIEAHVDVKTTDSYTLRMFCIAFTKKRPNQQKRTCYAQSSQIRQIRRKMVEIMRNQASSCDLKELVAKFIPESIGREIEKATSSIFPLQNVYIRKVKILKAPKFDIGKLMEVHGDYSEDVGVKLDRPADETVAEAEPEVPGA*

NbL13g15050.1 V−type_proton_ATPase_subunit_C−like [MW = 42.86 kDa]

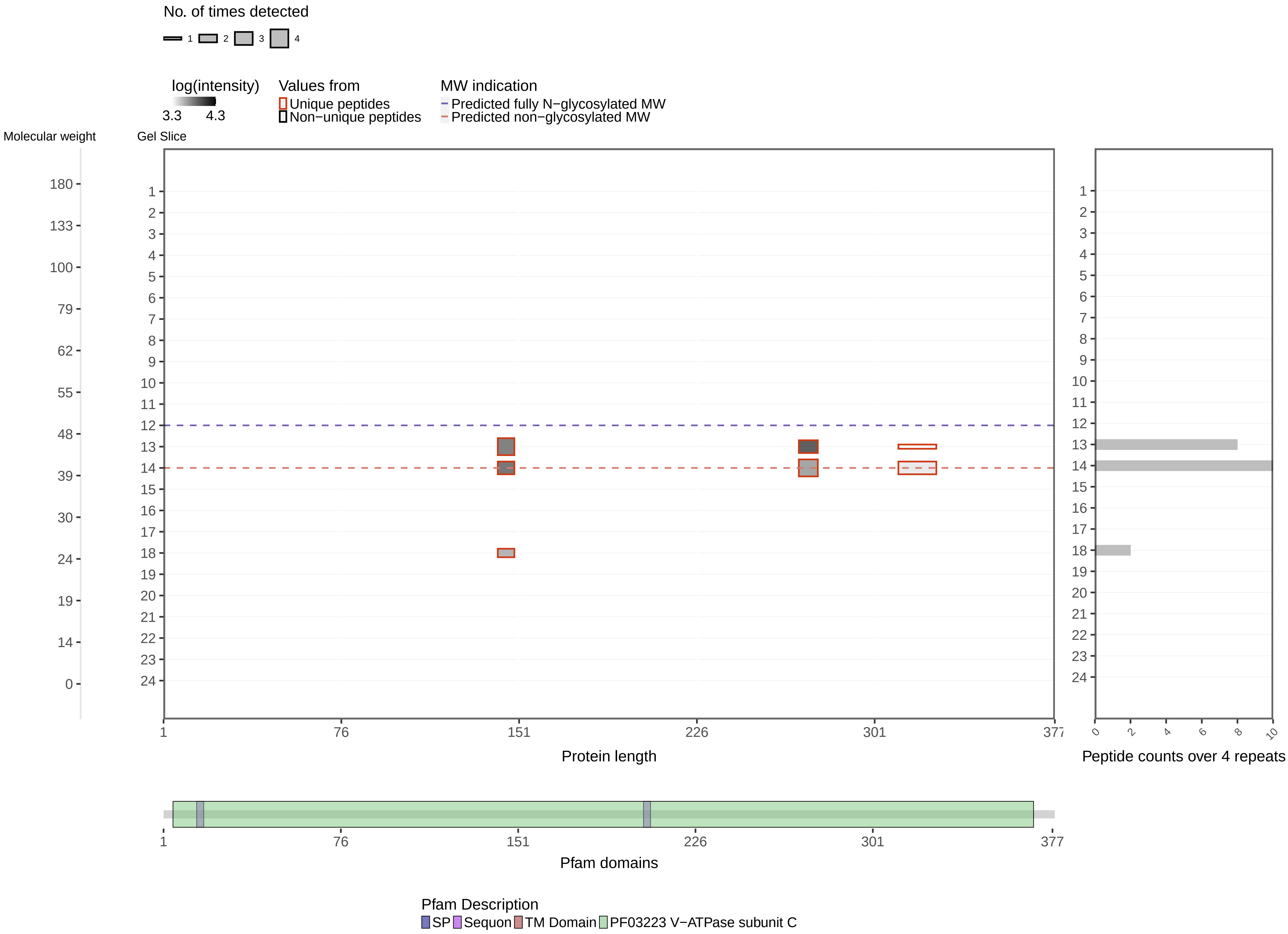


Protein sequence:

[illegible]IVTSEHLTTLLAVVSKYSQKDWLSSYENLTTYVVPRSSKMLYEDNEYALYTVTLFNRDADNFKIKARERGFQIRDFEHNPETQESRKQELEKLMQDQETLRSSLLQWCYTSYGEVFSSWMHFCAVRIFAESILRYGLPPSFLSIVLAPSTKSEKKVRSILESLCDSSNSNFWKSEDEGGMAGFGGDTEAHPYVSFTINLV*

NbL13g14950.1 pyruvate_dehydrogenase_E1_component_subunit_beta−1_mitochondrial−like [MW = 40.56 kDa]

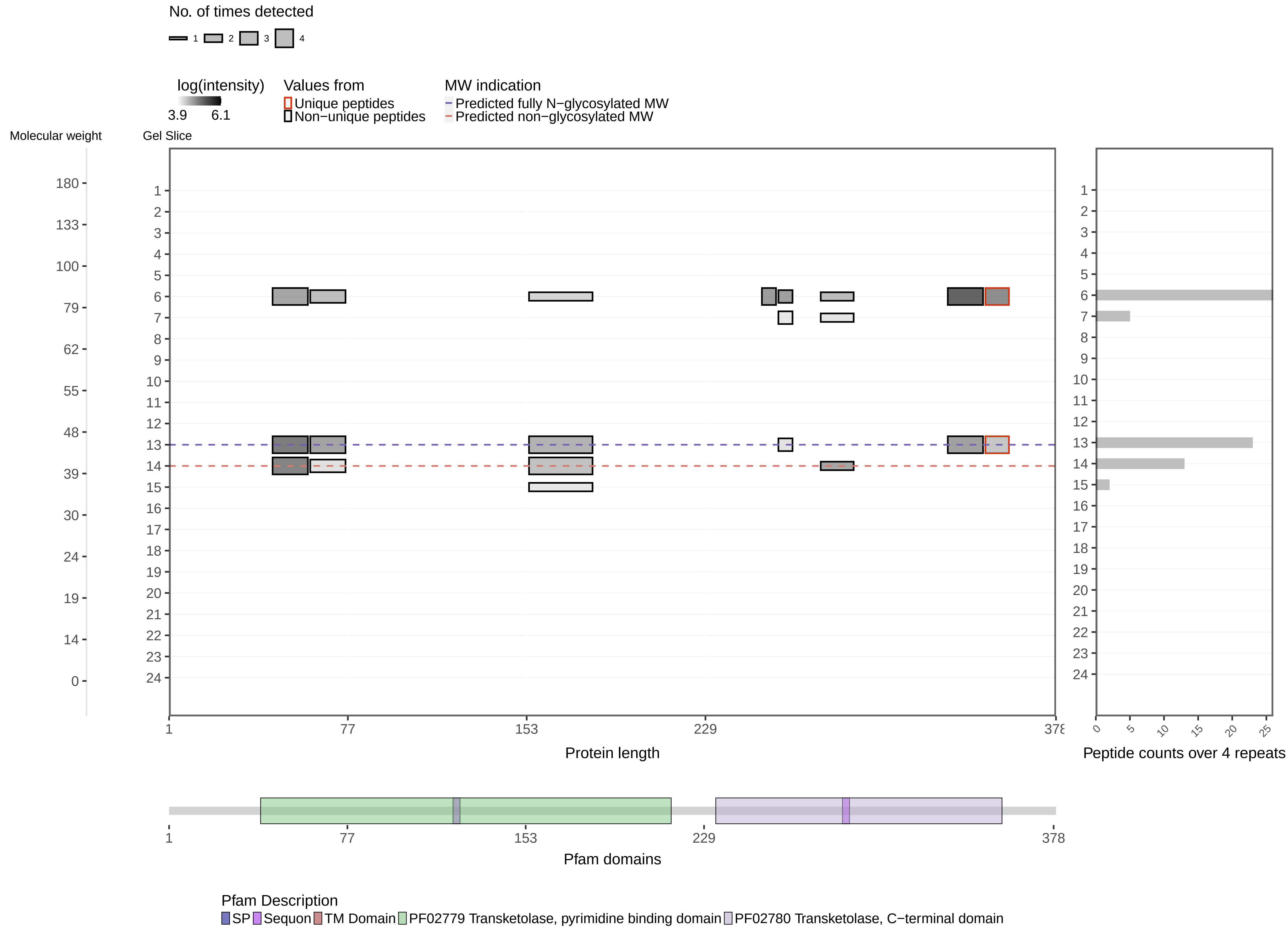


Protein sequence:

(GACPGLKVLAPYSSEDARGLLKAAIRDPDPVVFLENELLYGESFPVSAEALDSSFCLPIGKSKIEREGKDVTITAFSKMVGYALKAAEILAKEGISAEVINLRSIRPLDRPTINASVRKTNRLVTVEEGFPQHGVGAEICASVVEESFEYLDAPVERISGADVPMPYAANLERMAVPQVEGIVRAAKRACYRAAKSVPKAATA*

### NbL13g14730.1 3−oxoacyl−[acyl−carrier−protein]_synthase_3_A_chloroplastic−like [MW = 46.65 kDa]

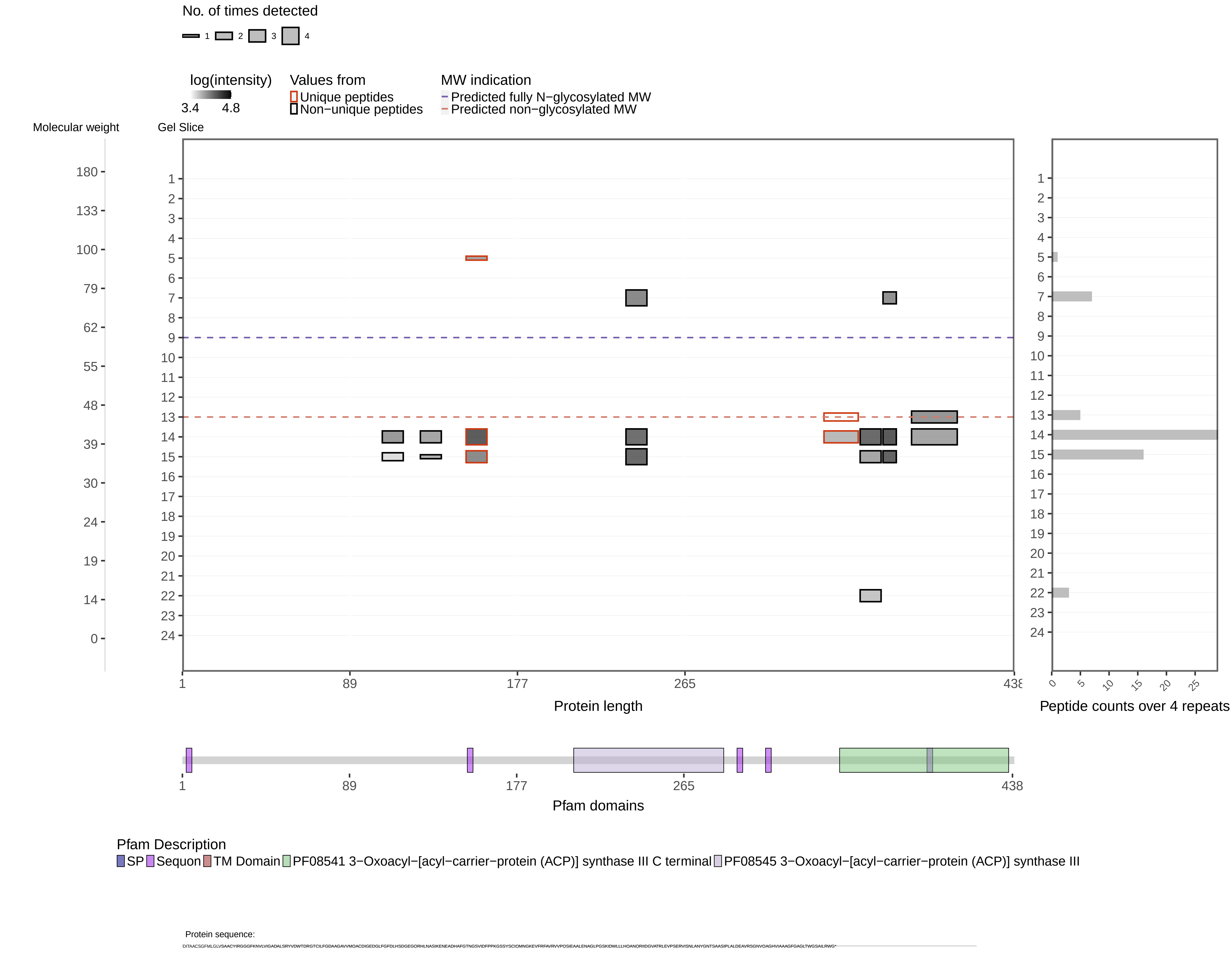

### NbL13g14600.1 DNA−binding_protein_SMUBP−2 [MW = 111.43 kDa]

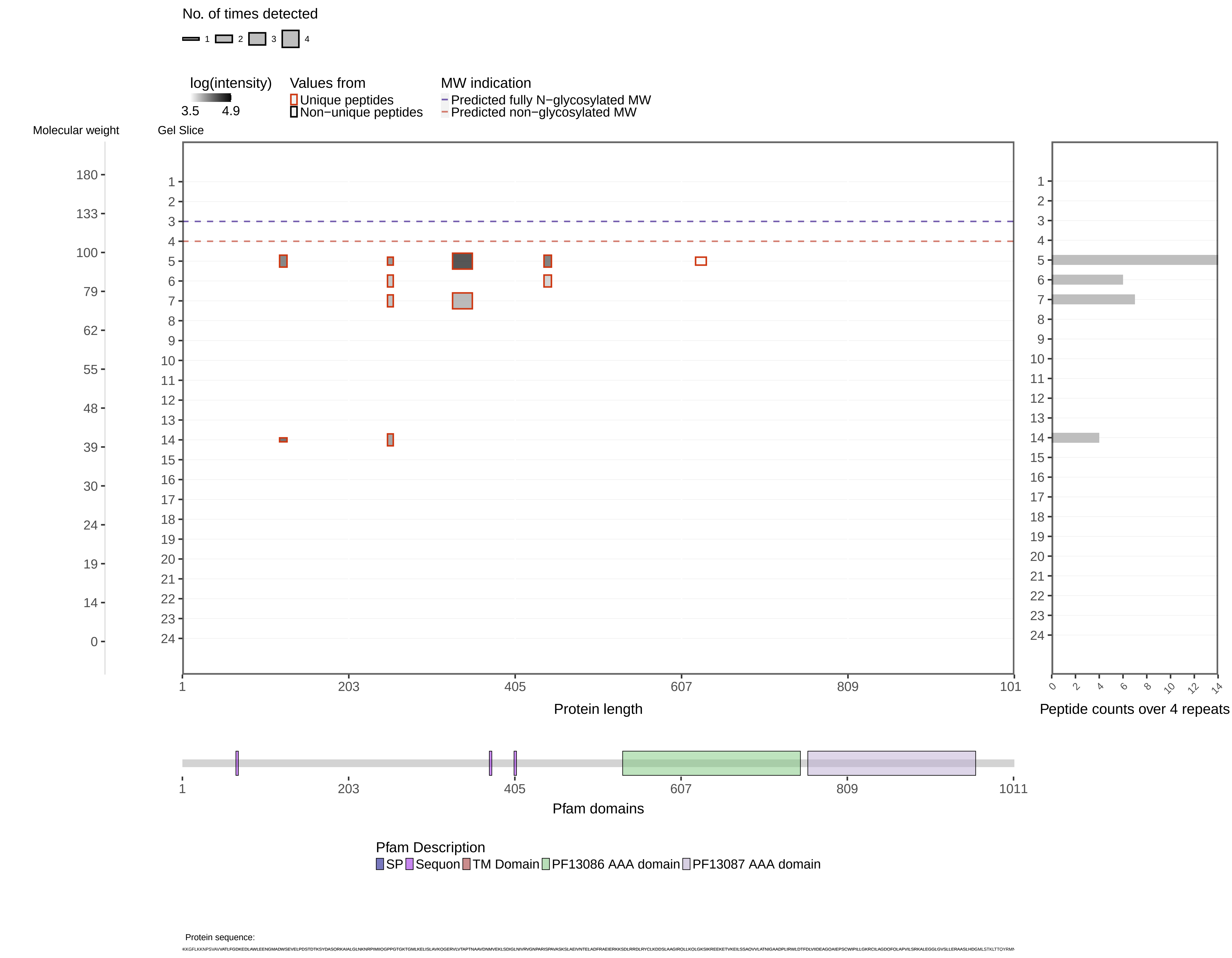

### NbL13g14590.1 uncharacterized_LOC104222403 [MW = 17.24 kDa]

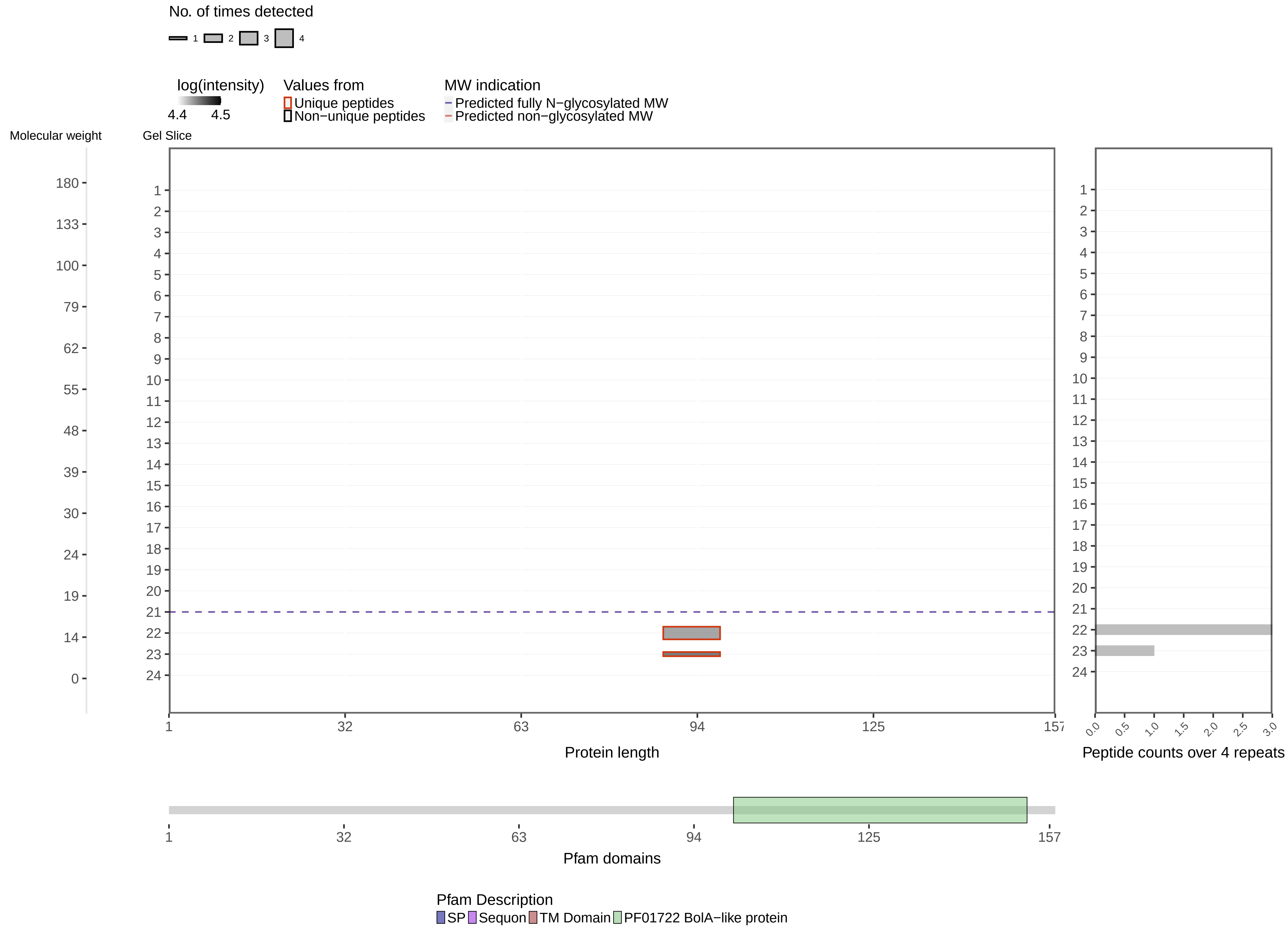


Protein sequence:

ATSDTGSIDSPLMESMQKRLKEQLNADTVVVKDAYGDGRHVSIDVVSSAFEGQSAVNRQRMVYKAIWEELQNTVHAVDQMTTKTPTEAGK*

### NbL13g14540.1 alanine−−tRNA_ligase−like [MW = 112.91 kDa]

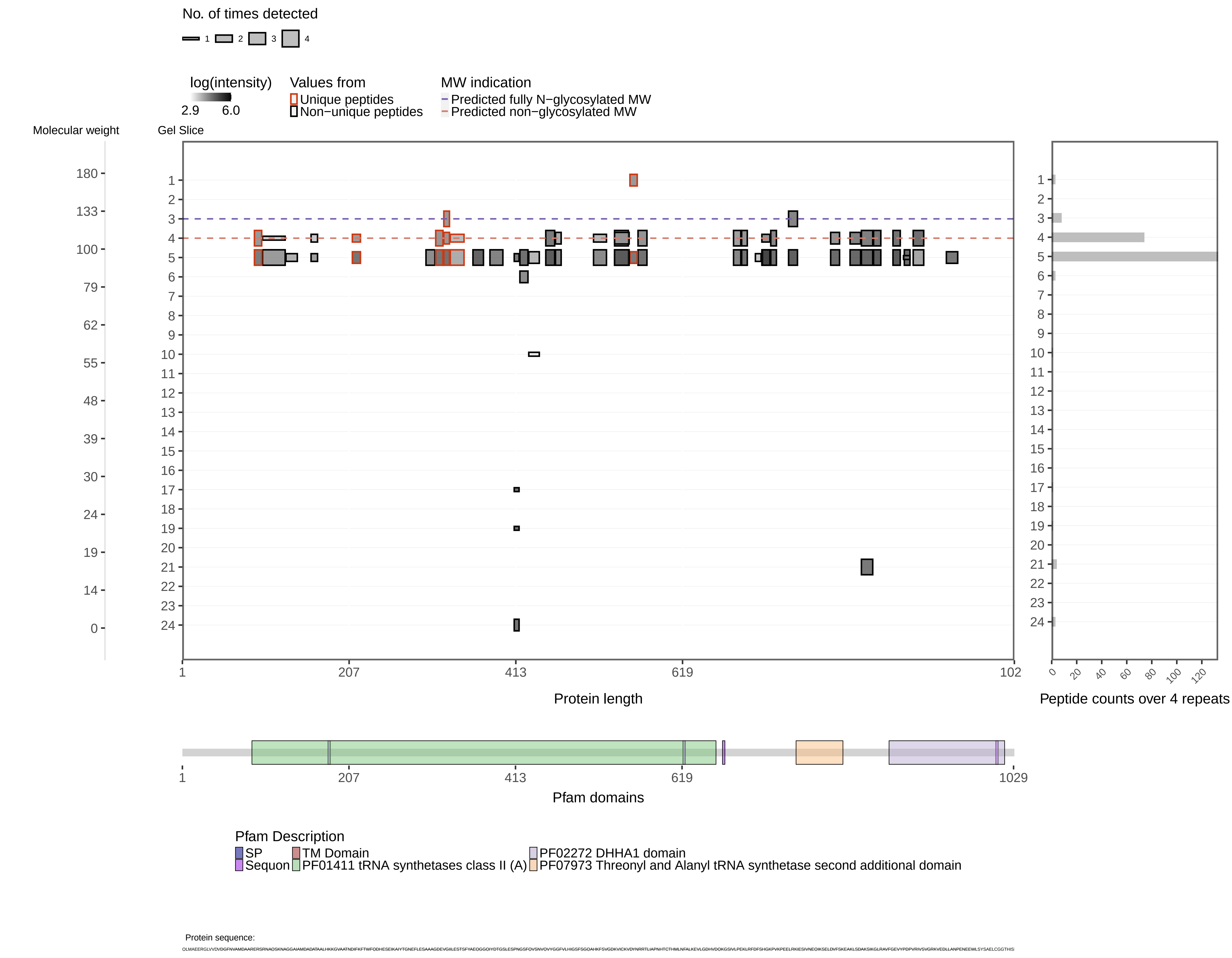

### NbL13g14530.1 cysteine_proteinase_inhibitor_6 [MW − SP = 25.01 kDa]

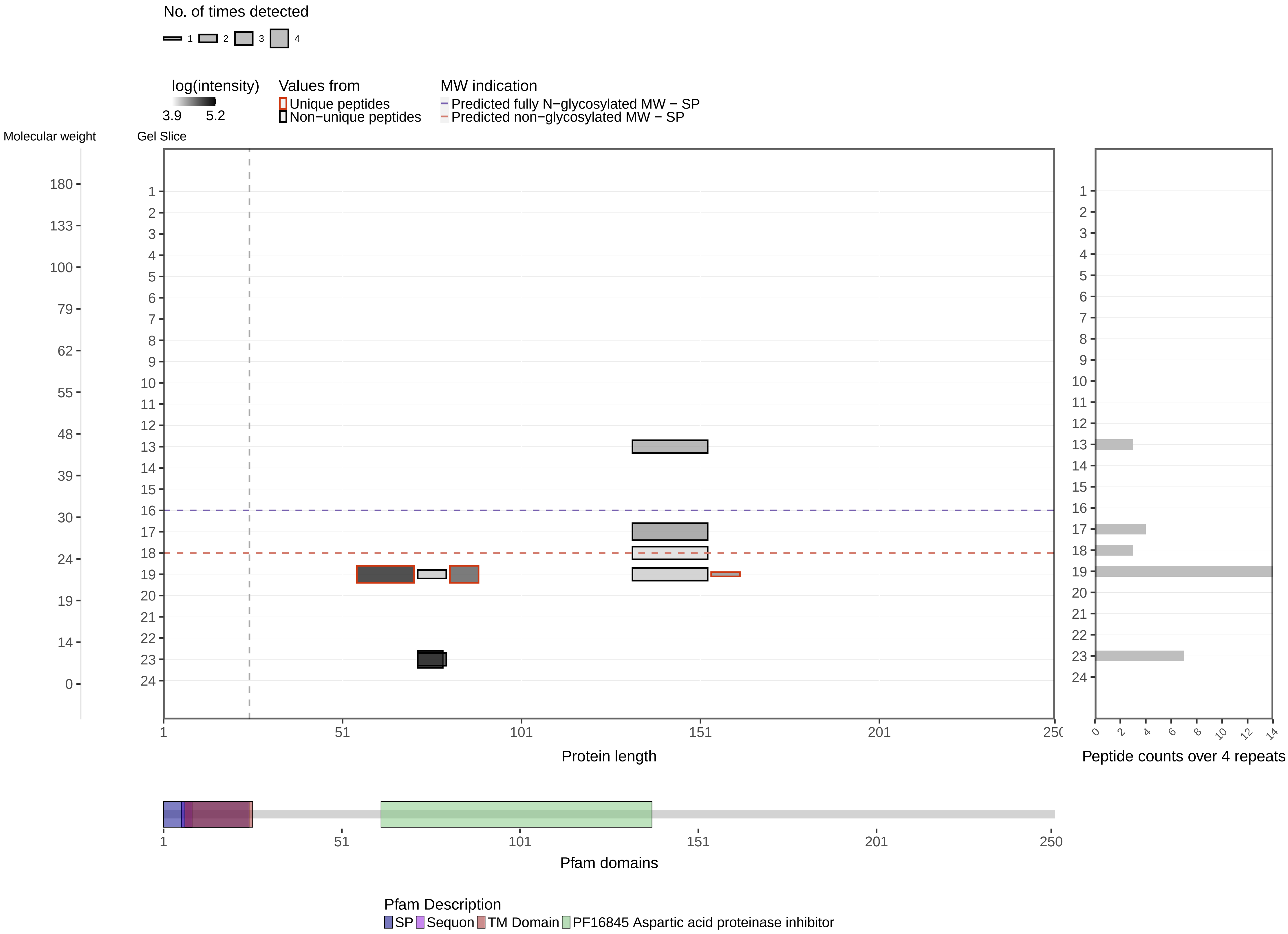


Protein sequence:

KKKLYEAKVWVKPWLNFKELQEFTHVEDVPTLTSSDLGVKQEEEGSGLKSVPVHDPVVQEAAEHAIKTIQQRSNSLLLYELQEIVHANAEVIGEDNMKLHMLIKTSRGGKREKFKVQVHHNNEGAFQLNHMEPDHS*

### NbL13g14510.1 uncharacterized_LOC104097028 [MW = 14.26 kDa]

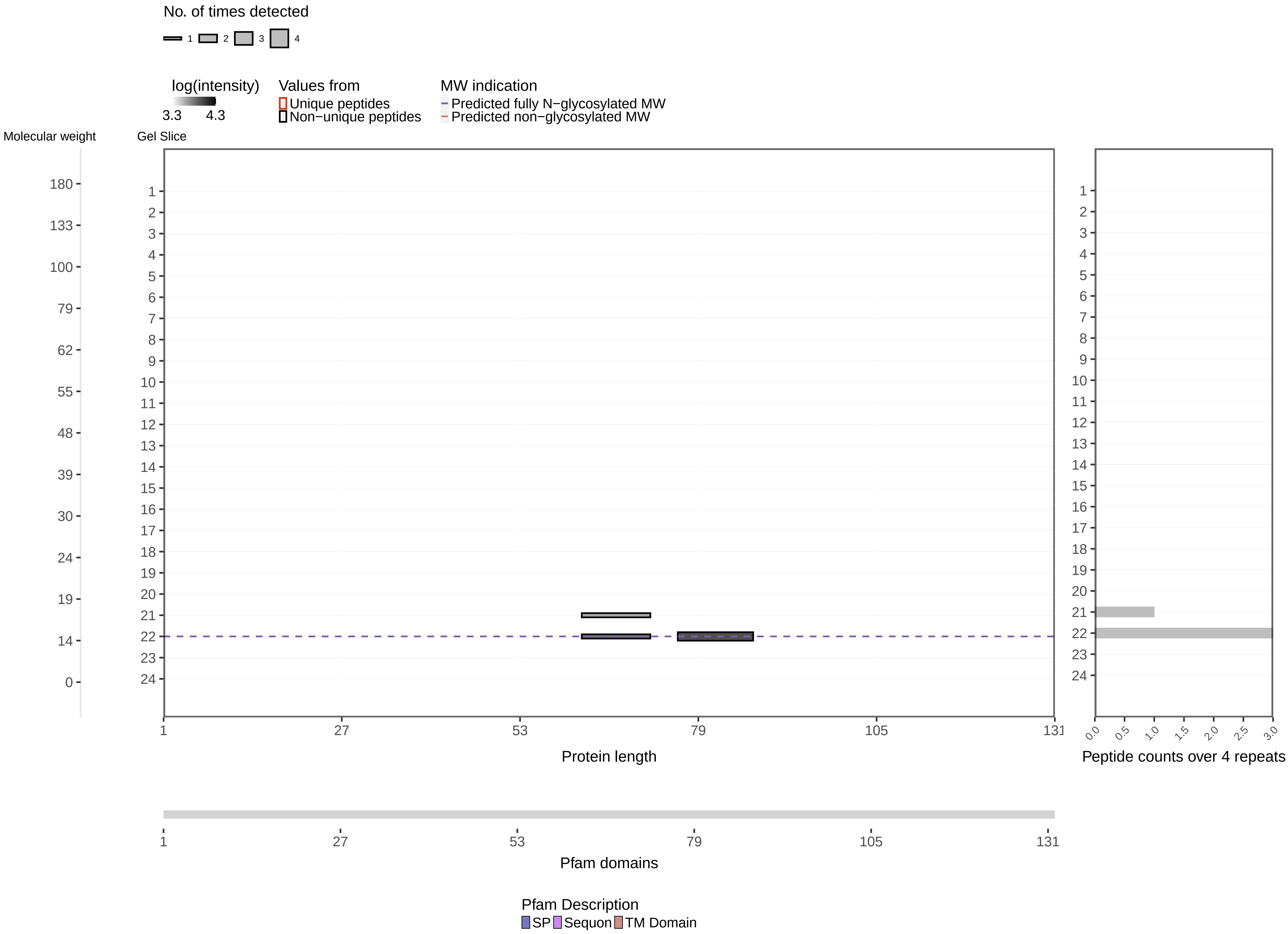


Protein sequence:

YIFAPQIRRSLLNEDEYGFRRAKRPIYHYDEGLEKTRQTLNKKLDQLNSAIDKVSSRLRGGNKMPPVPVETDPEEATM*

NbL13g14450.1 WEB_family_protein_At3g02930_chloroplastic−like [MW = 109.68 kDa]

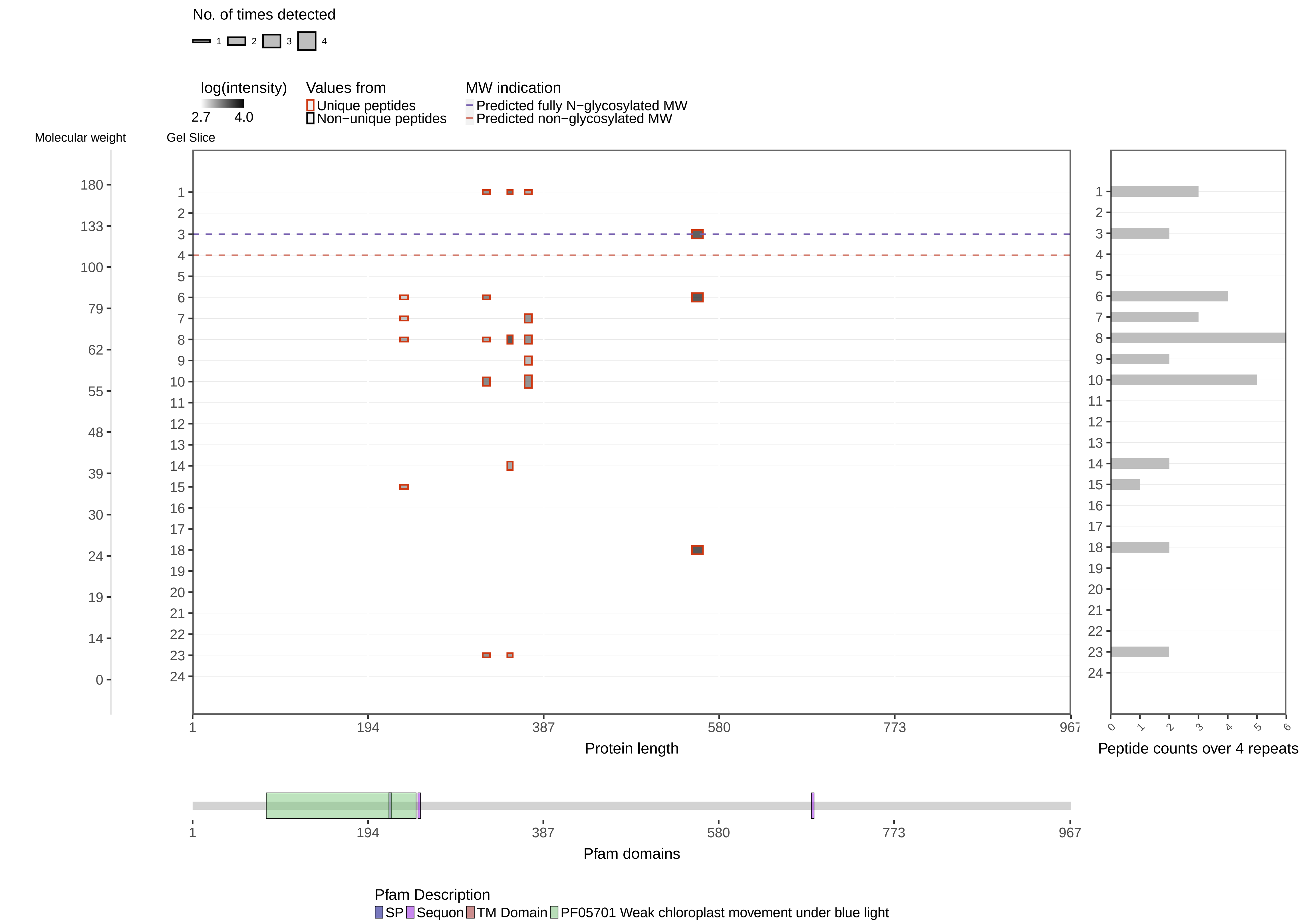


Protein sequence:

EGSNDLLHDAESEIASLKEKVGLLEMSTTRQKGDLEESERRAQIAKEEASEMAKKVESLITELETVKEEKTQAIEHEKLAAESVQSLLEEKNKLINELENSREEEEKSKKAMESLASALHEVSSEAREAKERLLSSQAEHEHYETQIEDLKLVLKATNEKYESLLDEAKEKIDDLTTSVEQSKNEHHISKAEWEDRELHLMNCVKKTEEENSSMEKEINRLVNLLKEAEQEVSFKEEAVQLKNSLHEAESEVTYLKEVLGEAKGESMKLKESLLDKENEVQNILQENEELRSREA

NbL13g14360.1 binding_partner_of_ACD11_1−like [MW = 29.39 kDa]

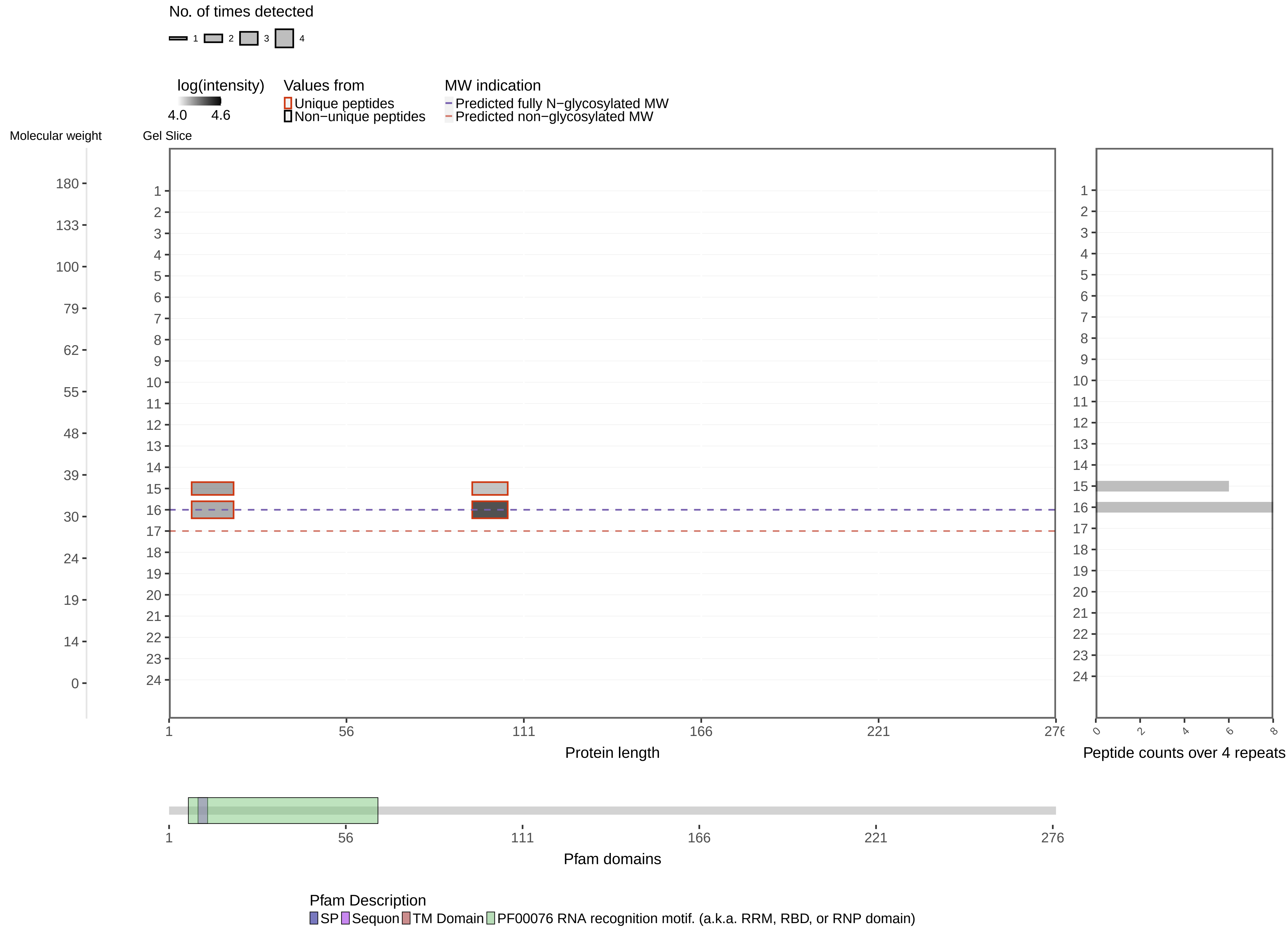


Protein sequence:

'NKAKTFDEKHQFISTASAKVASLDQKIGLSEKINMGTTIVNDKVKEMDQKFQVTEKTKSAFAAAEQTVSTAGSAIMKNRYVLTGASWVTGAFNKVTKAAGEVGQKTKEKMAEEEQGRSSAAGYVPIHAFSESPKASKTEEPANPLHLRT*

### NbL13g14210.1 ADP−ribosylation_factor_1−like_2 [MW = 21.86 kDa]

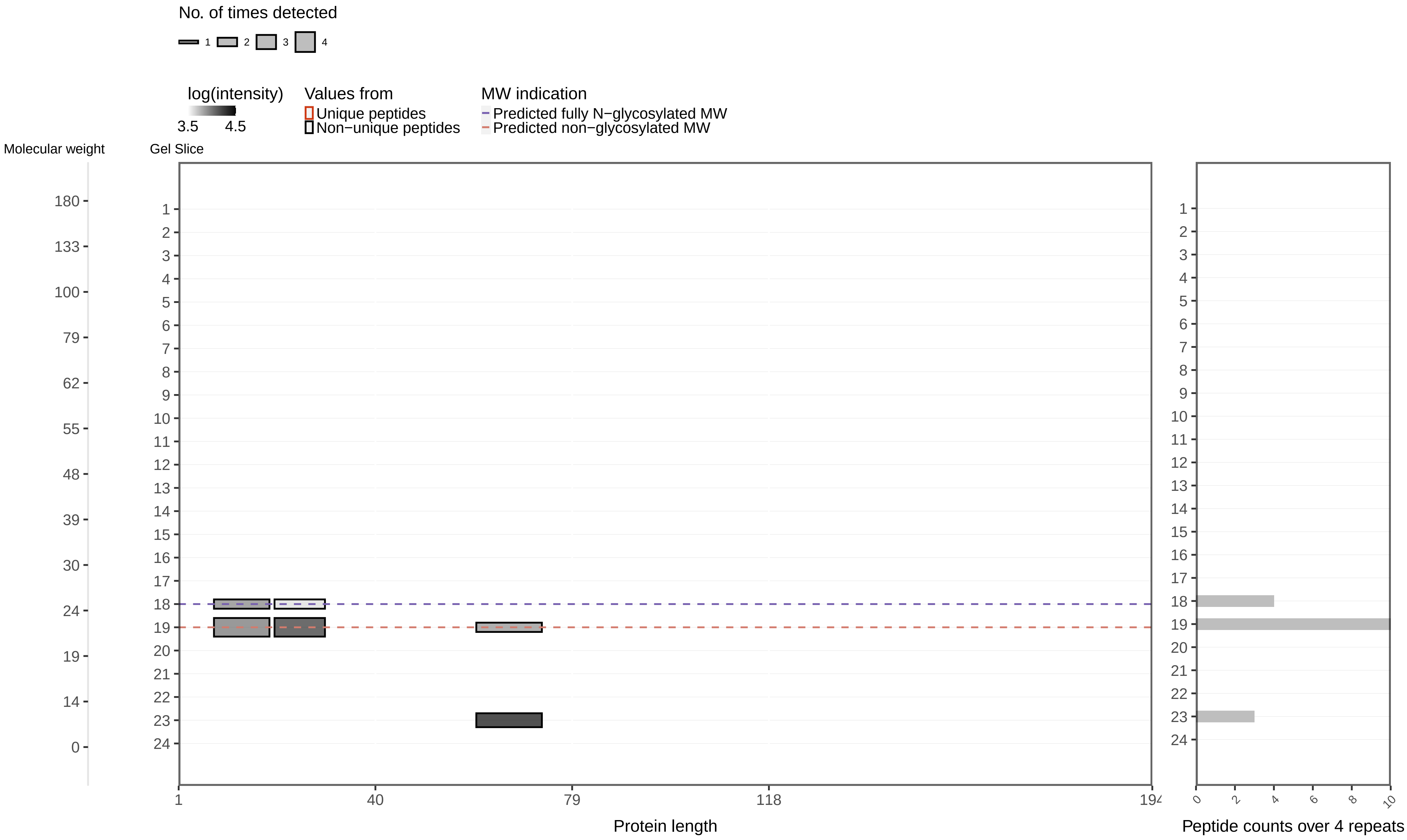


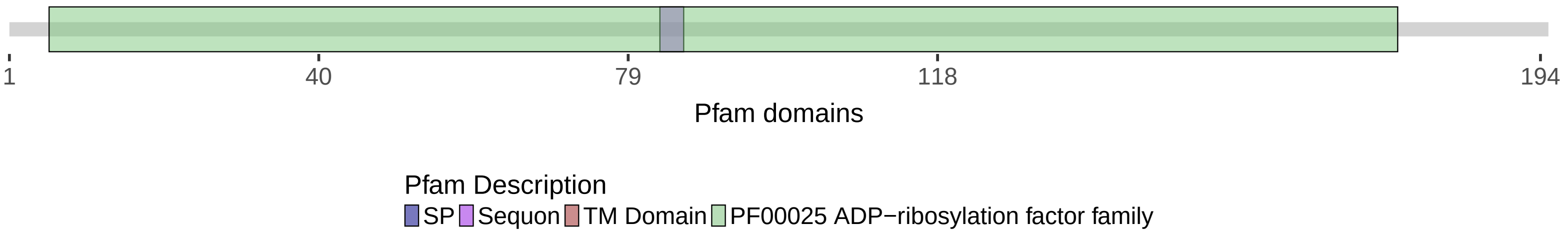


Protein sequence:

INTDGLIYVVDSLDRERIGKAKQEFQAIIRDPFMLNAIILIFANKQDMKGAMTPMEVCEGLGLYELKNRKWHIQGTCALRGDGLYEGLDWLASTLKEHKAVGYSSIGPSSF*

NbL13g14070.1 uncharacterized_LOC104210248 [MW = 13.81 kDa]

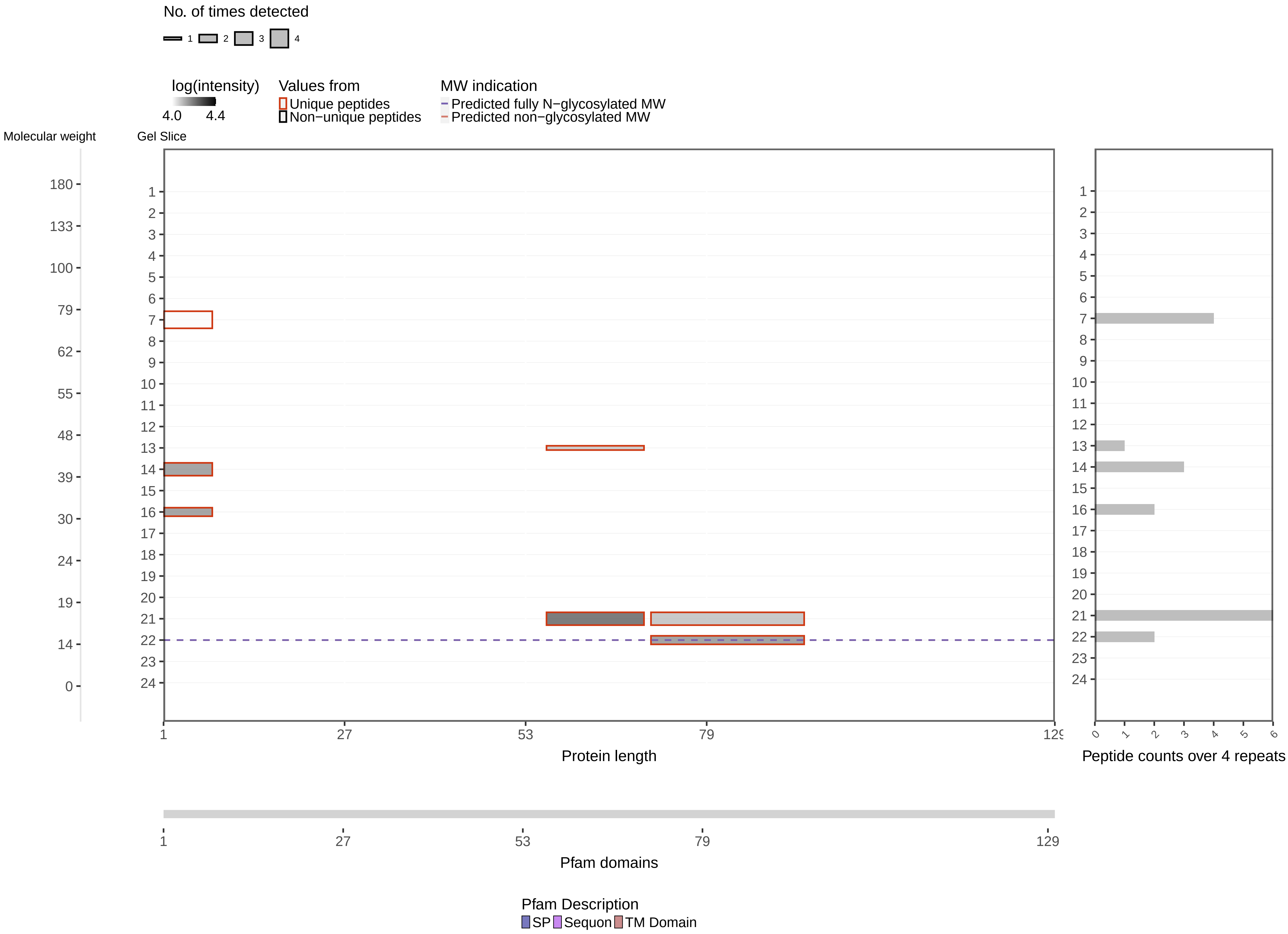


Protein sequence:

DHVRDSVVPDEVIELQSEKYWGPHPQTGVFGPAATNSAGGERGFHHSPATAAAAAESILEQKTFFRPLEDLDKPQHA*

NbL13g13970.1 DNA−binding_protein_BIN4 [MW = 44.52 kDa]

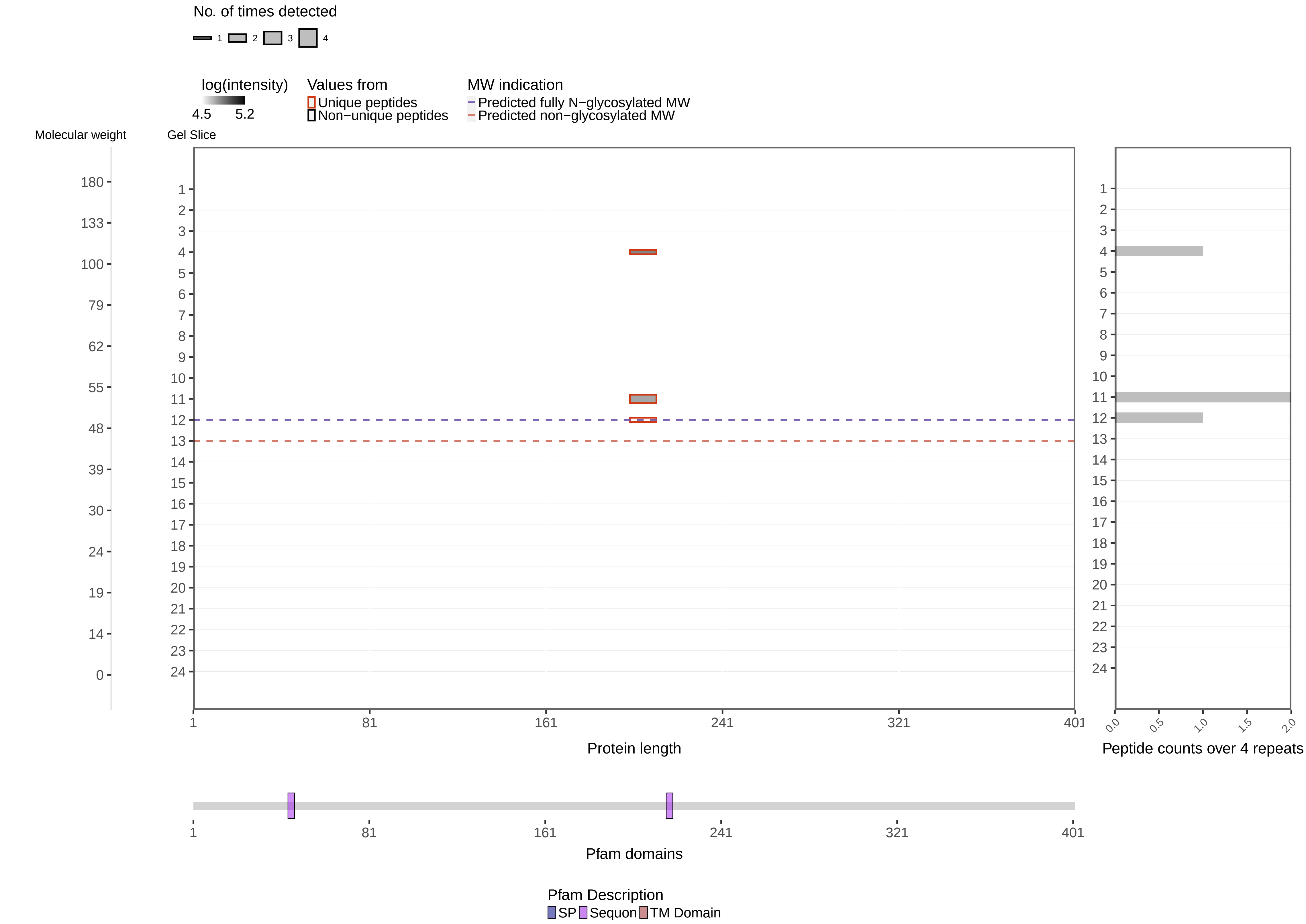


Protein sequence:

KKKLESQLKEEKEDMDAVEENIPEKQSVPNLSTSRLPLVIPEKVQRTKALVECEGDSIDLSGDMGAVGRIVISDGPSGKHEMLFDLKGTIYKTTILPSRTFCVVSFGQTEAKVEAIMNDFIQLTPQSNVYEAETMVEGTLDGFTFDSDEEAENLRRPASQGDQNENADHQINGNVKAKAKKASATEQKKGKKGVKIPNKVKKKPQAPKKSKSKK*

NbL13g13840.1 nascent_polypeptide−associated_complex_subunit_alpha−like_protein_2 [MW = 22.66 kDa]

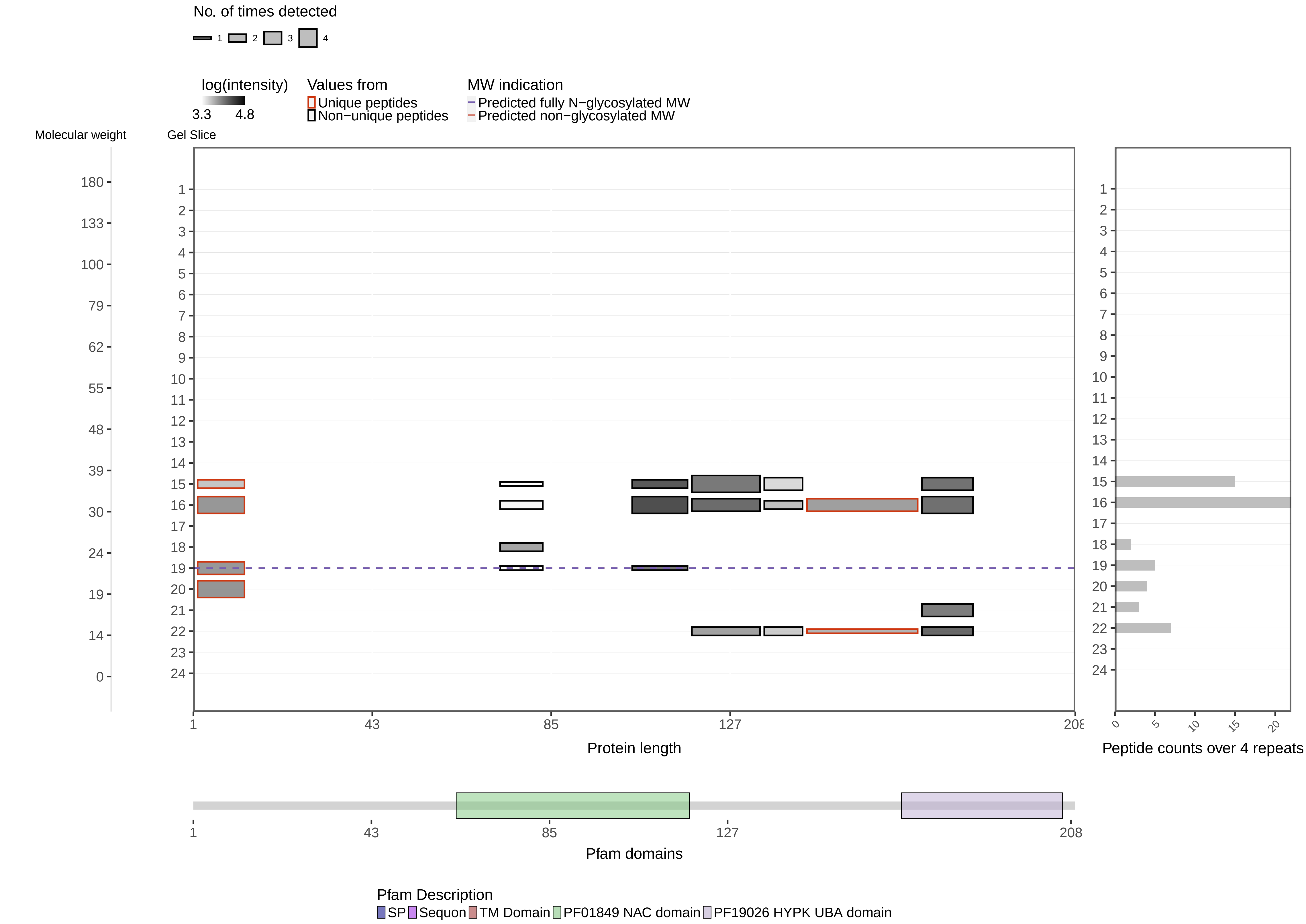


Protein sequence:

KNVLFFISKPDVFKSPNSETYVIFGEAKIEDLSSQLQTQAAQQFRMPDMGSVMAKSDVSPSTAAAEADEEEEEVDETGVEPRDIDLVMTQAGVSRSKAVKALKCHNGDIVSAIMELTT*

NbL13g13820.1 caffeic_acid_3−O−methyltransferase−like [MW = 39.74 kDa]

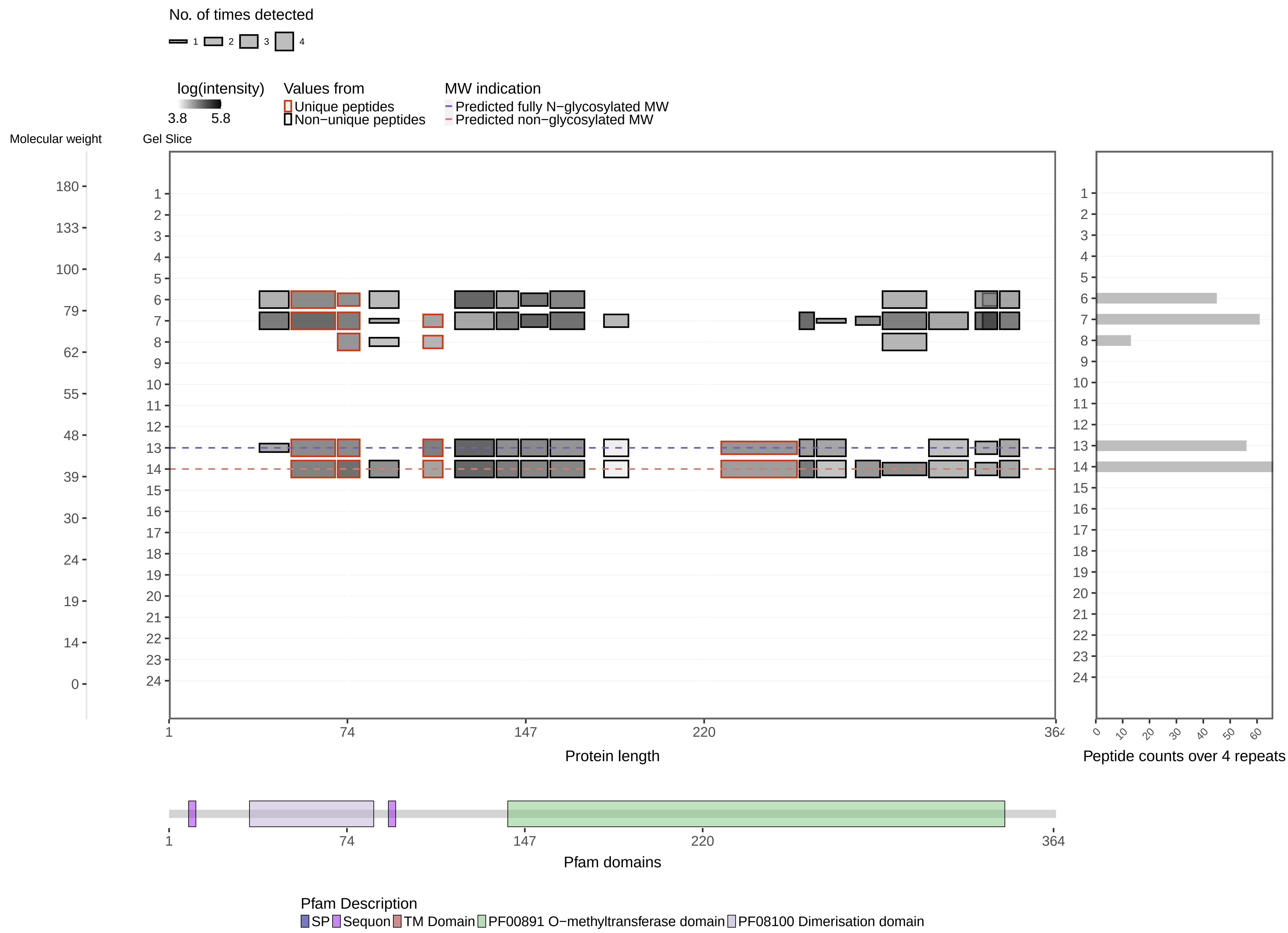


Protein sequence:

DPRFNKVFNRGMSDHSTMSMKKILEDYKGFEGLNSIVDVGGGTGATVNMIVSKYPSIKGINFDLPHVIEDAPAYPGVEHIGGDMFASVPKADAIFMKWICHDWSDEHCLKFLKNCYEALPANGKVIVAECILPEAPDTSLATKNTVHIDIVMLAHNPGGKERTEKEFQALAKGAGFTGFHKACCAYNTWVMEFNK*

### NbL13g13800.1 uncharacterized_LOC107802109 [MW = 31.37 kDa]

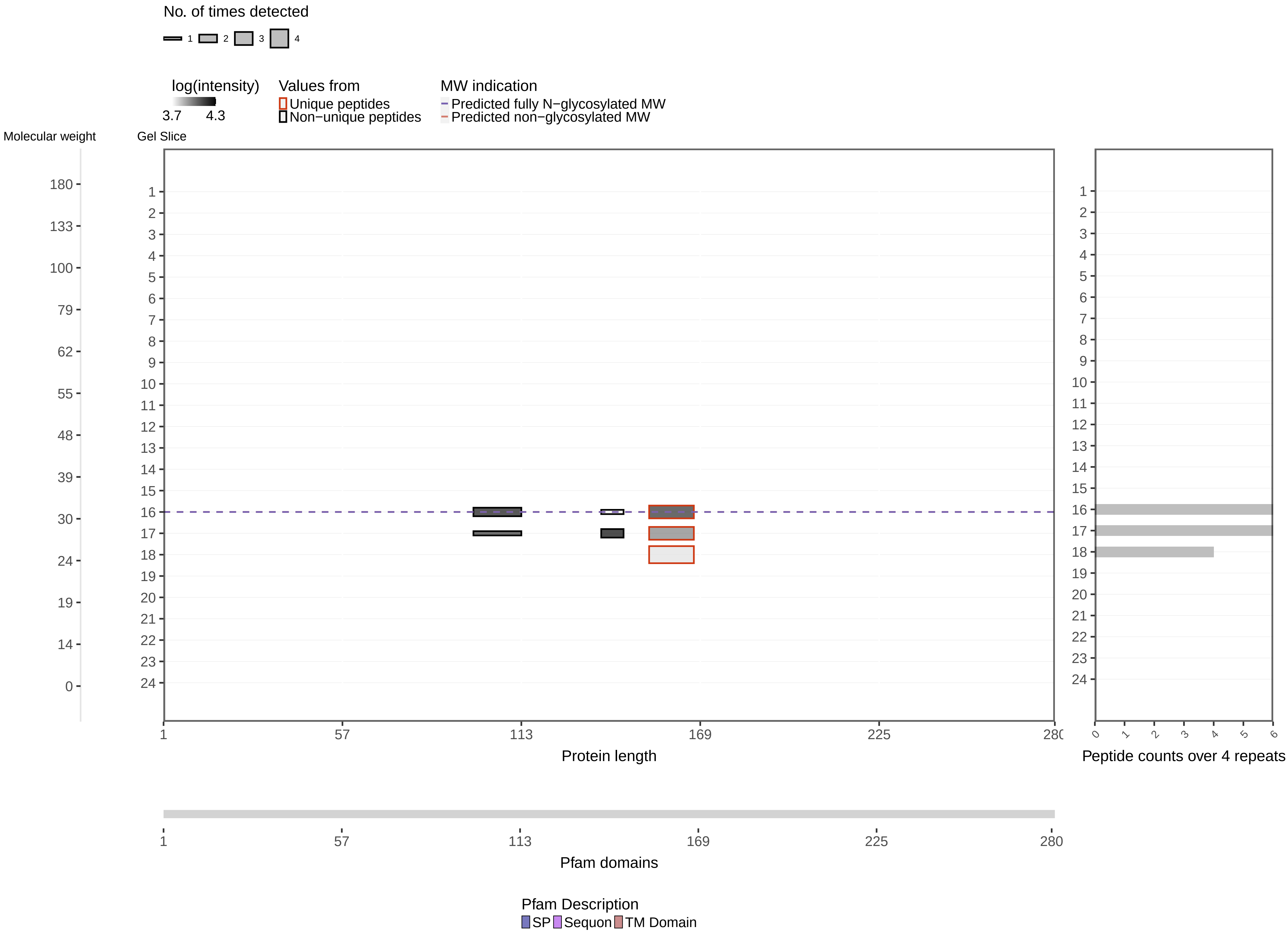


Protein sequence:

LLADSTPPVRTVEVMVVRVIGKDNKILVESHQELSNGVVRQRGRPLSEKMKPGETVDDAVFRAVKEELGSSIVKILPNSYSTKVEERVSASYPGLPACYVLHTVDVVVDGLPEEEFCTEETDEYADSSERILADGAVSCKKHYWKWVDADSL*

### NbL13g13680.1 protein_GPR107−like [MW − SP = 47.59 kDa]

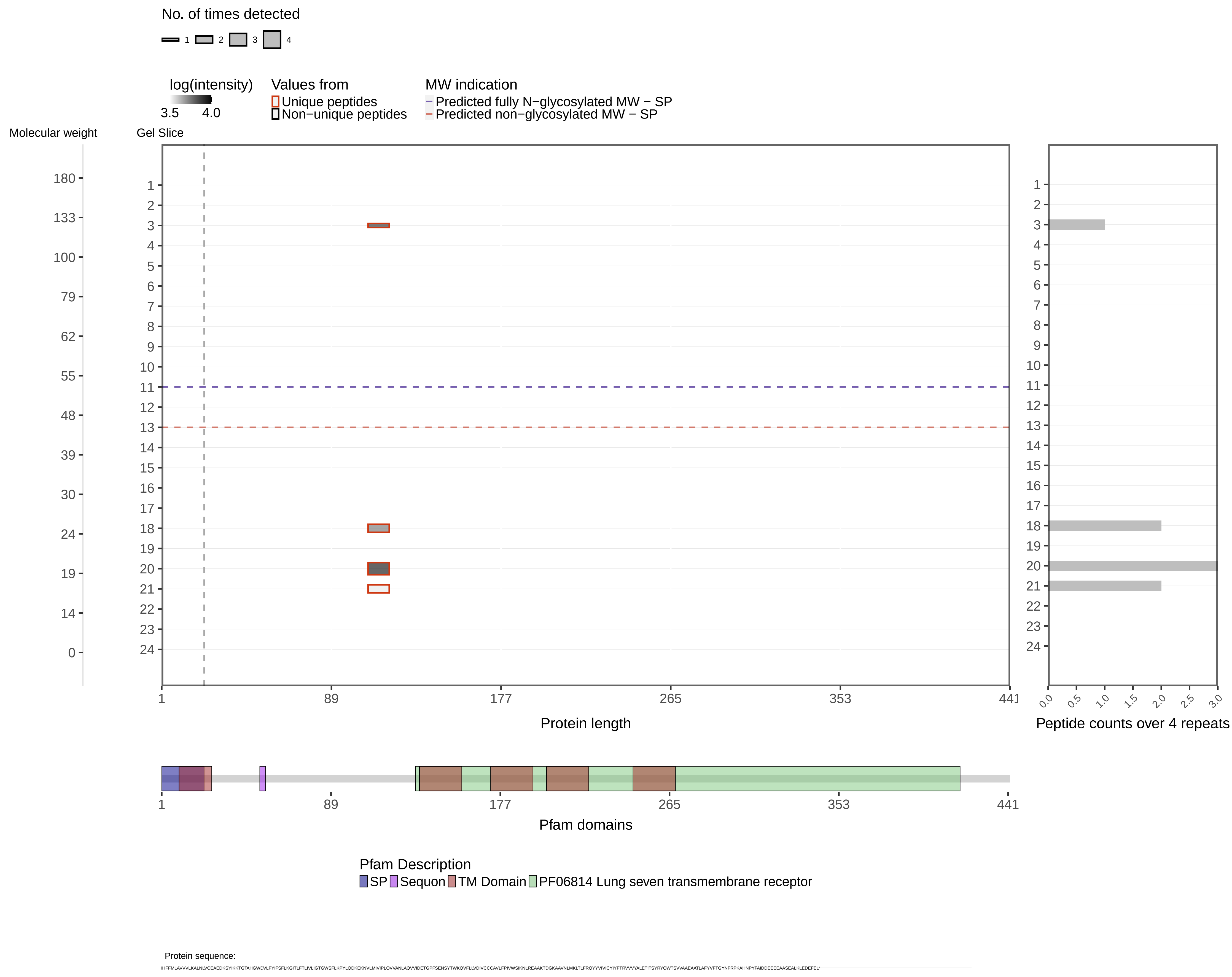

### NbL13g13660.1 GRF1−interacting_factor_3−like [MW = 21.62 kDa]

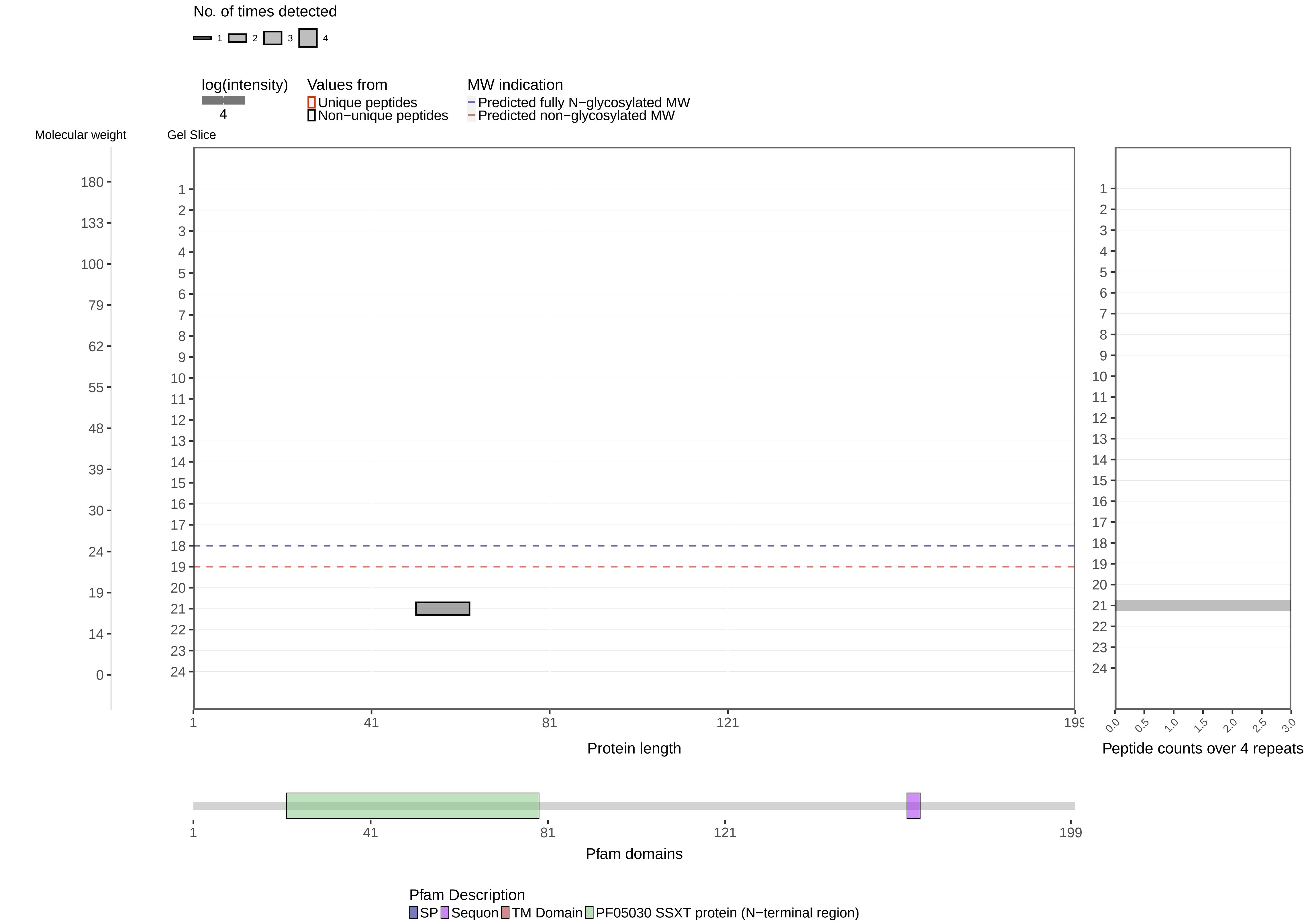


Protein sequence:

QPGSQMQQTQAALQQQPGVPISKLPFQLNALPSQEQQQQMLHFQQQQQQFQGQYGFGPTANNAMHQLMQPGLNSSGTLDARGNKQGNLEGNPSDGLGKSAARHVNSERE*

### NbL13g13530.1 cytochrome_b5−like [MW = 15.04 kDa]

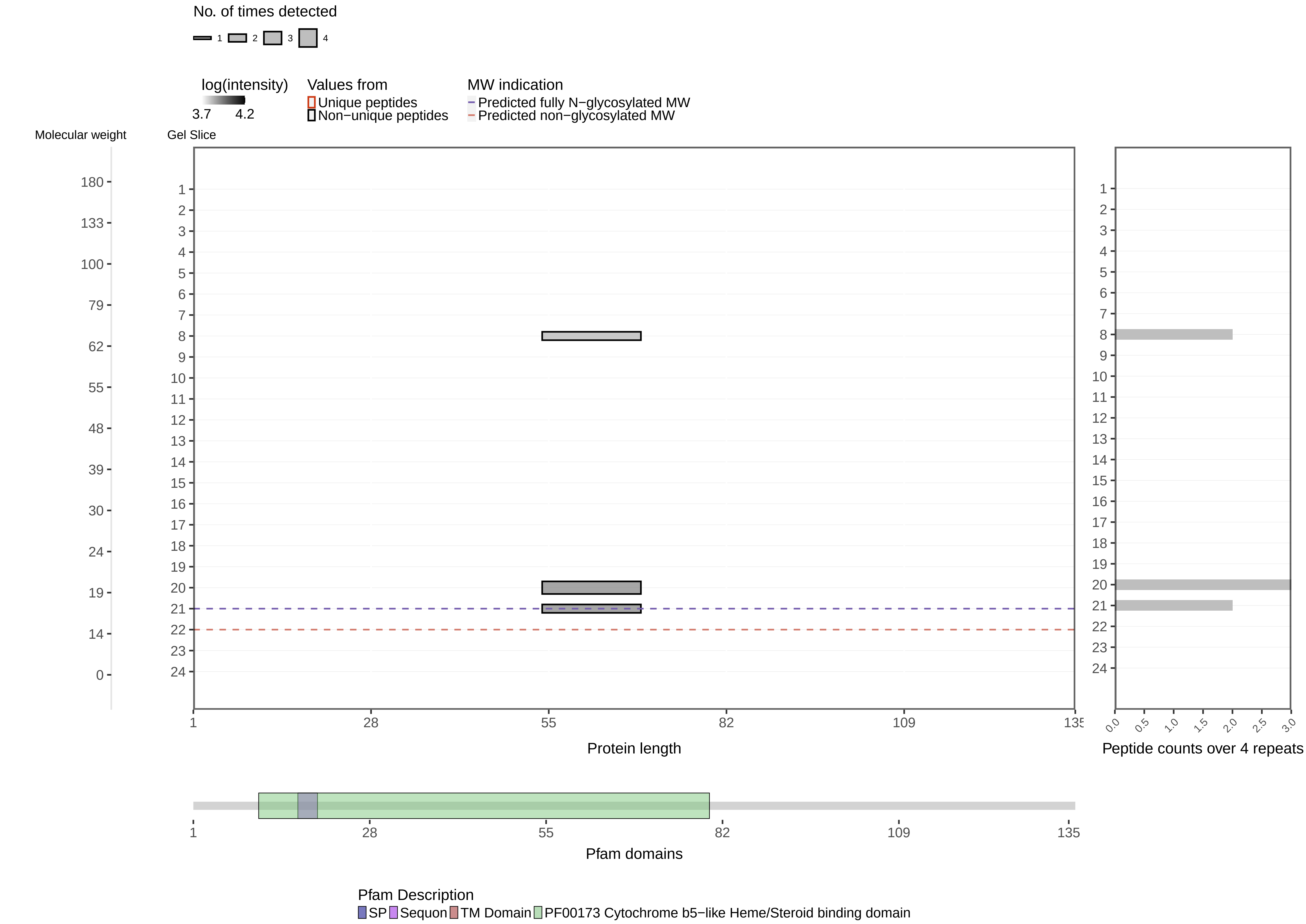


Protein sequence:

KDATNDFEDVGHSDSAREMMDKYYIGDIDVSTVPLKRSYVPPQQAPYNPDKTPEFIIKILQFLVPLLILGLAFAVRHYTKEK*

NbL13g13250.1 luminal−binding_protein_5 [MW − SP = 70.7 kDa]

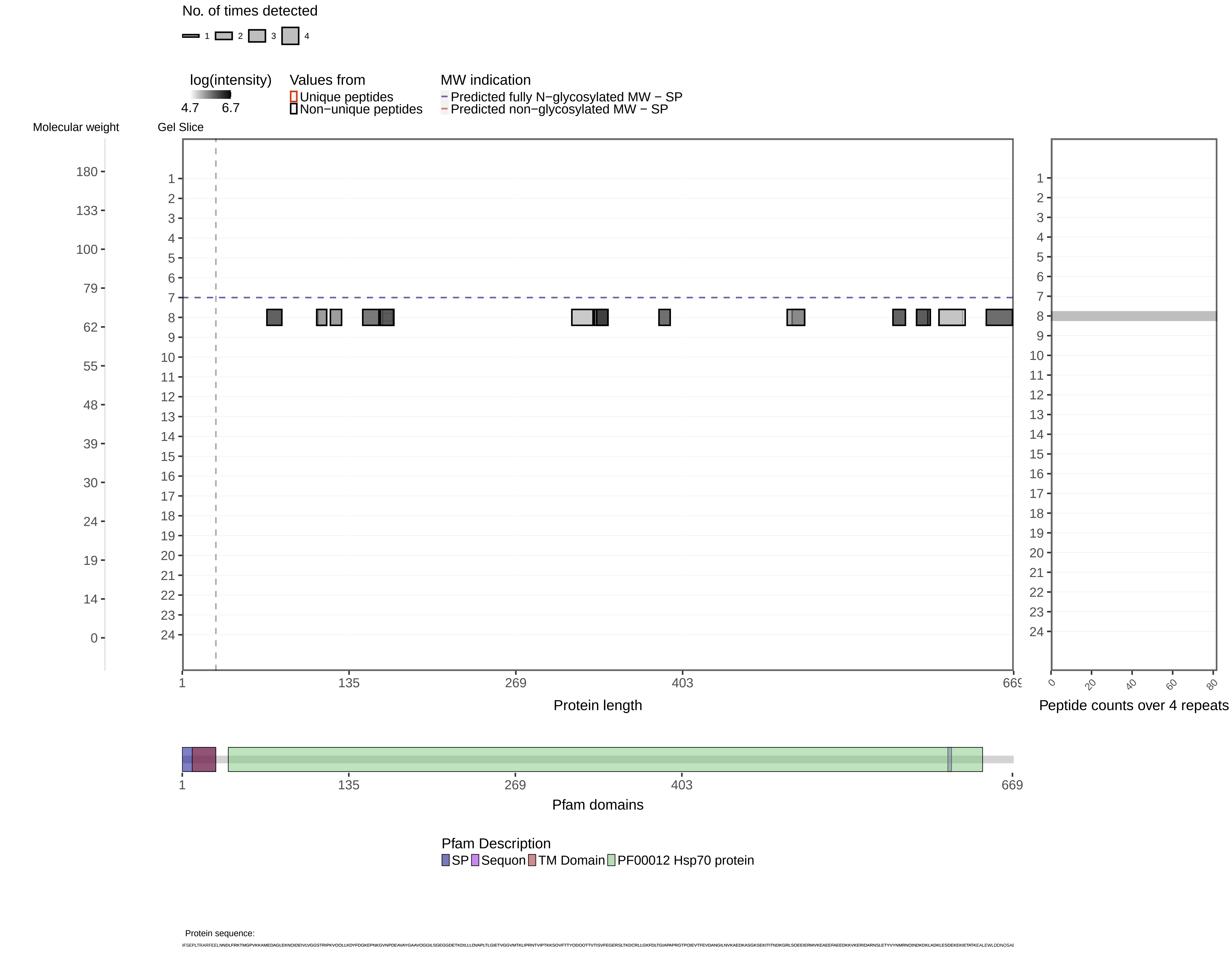

### NbL13g13030.1 CLP_protease_regulatory_subunit_CLPX1_mitochondrial [MW = 61.89 kDa]

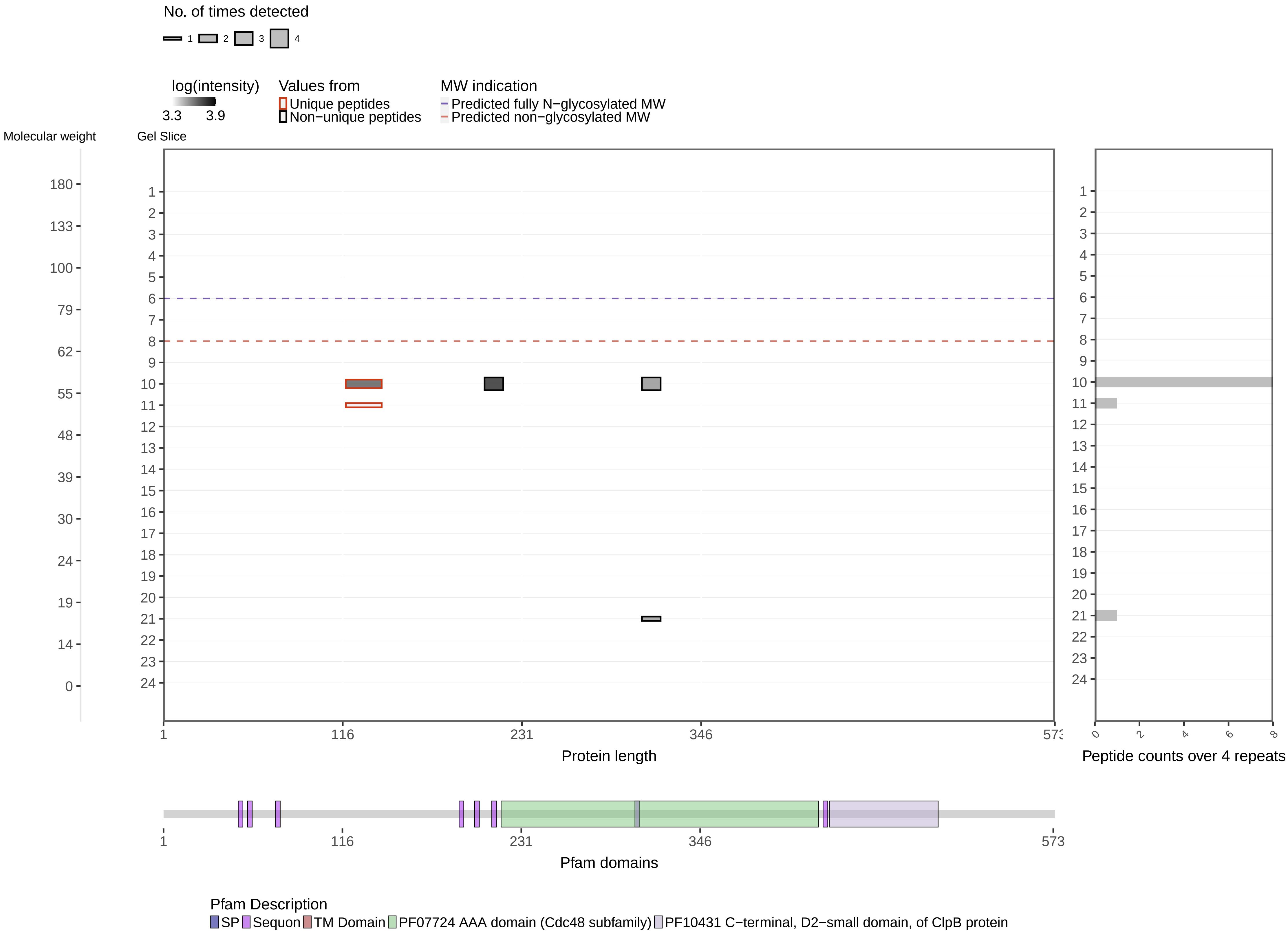


Protein sequence:

ILYKLLTVADYNVAAAQQGIVYIDEVDKITKKAESLNISRDVSGEGVQQALLKMLEGTIVNVPEKGARKHPRGDNIQIDTKDILFICGGAFIDLEKTISERRQDSSIGFGAPVRANMRTGGVTSAAVTSSLLEIVESSDLISYGLIPEFVGRFPILVNLSALTENQLVQVLTEPKNALGKQYKKMFQMNGVKLHFTEDALRLIAKKAISKNTGARGLRSILESLLMDAMYEIPDVRTGDDIIDAVVVDEESVGAEGRGCGAKILYGKGAFDRHFPQNRPKDYETNLEGSDGDHEAEQELI

NbL13g13010.1 pentatricopeptide_repeat−containing_protein_At3g49240−like [MW = 73.88 kDa]

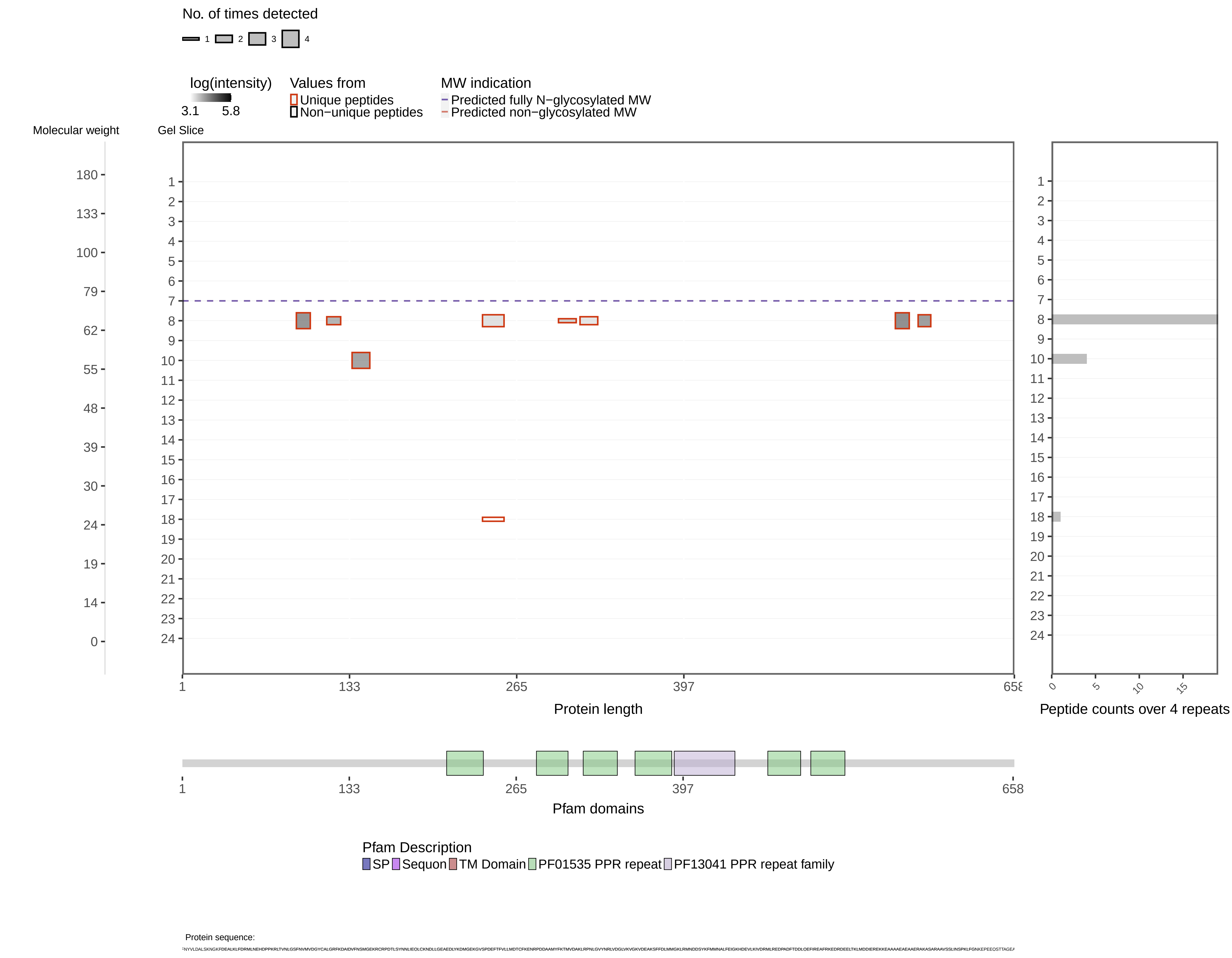

### NbL13g12940.1 dipeptidyl_peptidase_9 [MW = 87.3 kDa]

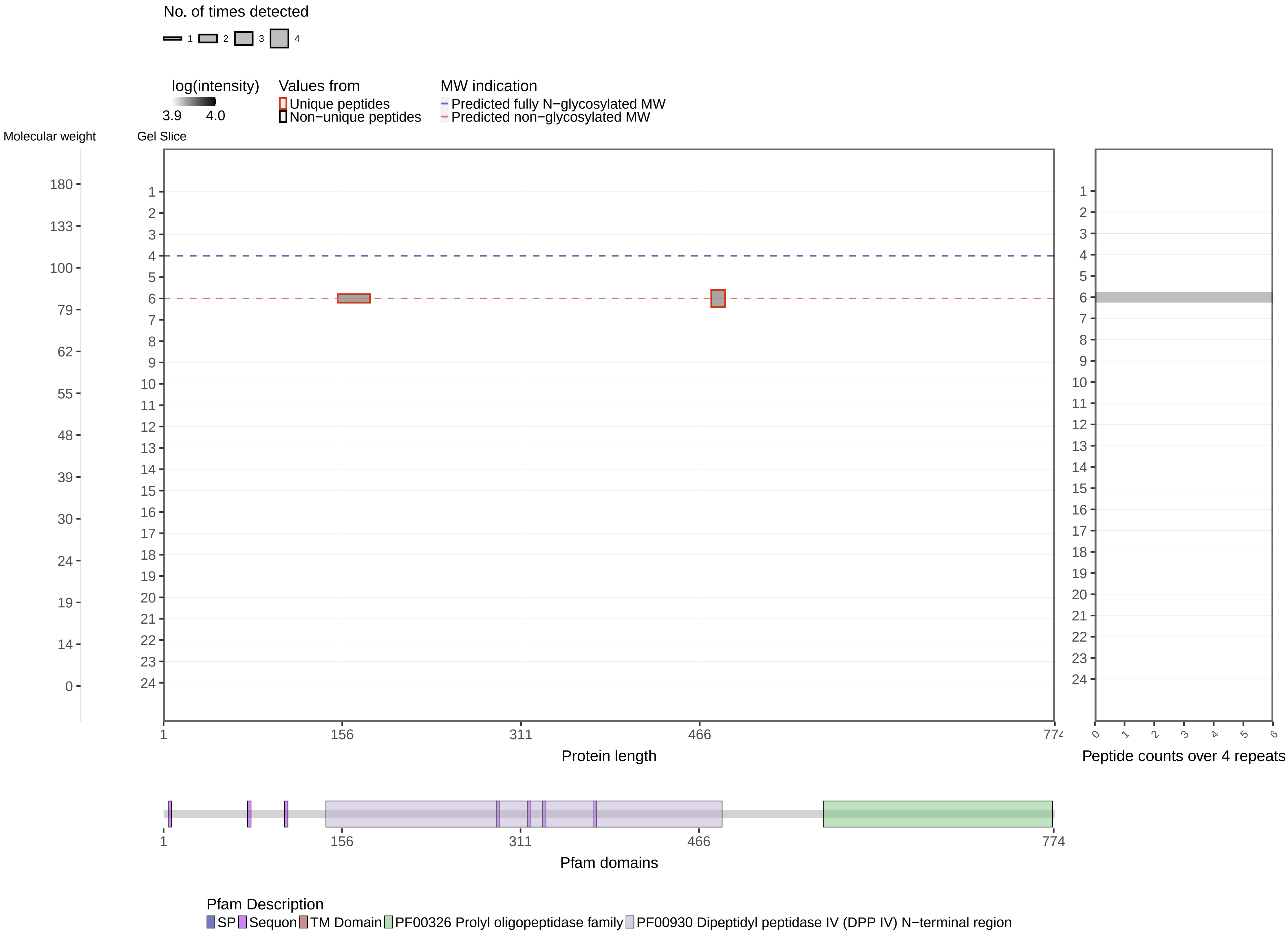


Protein sequence:

RTNGAFIWASEKTGFRHLYLHDTNGVCLGPITQGDWLVEQVAGVNEVTGLVYFTGTLDGPLESHLYCAKLFPDAHSPLQSPLRLTNGRGKHAVVLDHQMQRFIDIHDSLVSPPRISLCSLHDGSLIMHLFDQPVSVPRFRKLHLESPEIIQIQAQDGTALYGQLYKPDPVKFGPPPYRTMIEVYGGPSVQLVCDSWTNTVDMRAQYLRSKGILVWKMDNRGSARRGFVFEGALKYNCGLVDAEDQLTGAEWLLKQGLAKQGHIGLYGWSYGGYLSAMTLARFPEVFKCAVS

### NbL13g12920.1 probable_pectinesterase/pectinesterase_inhibitor_61 [MW = 65.57 kDa]

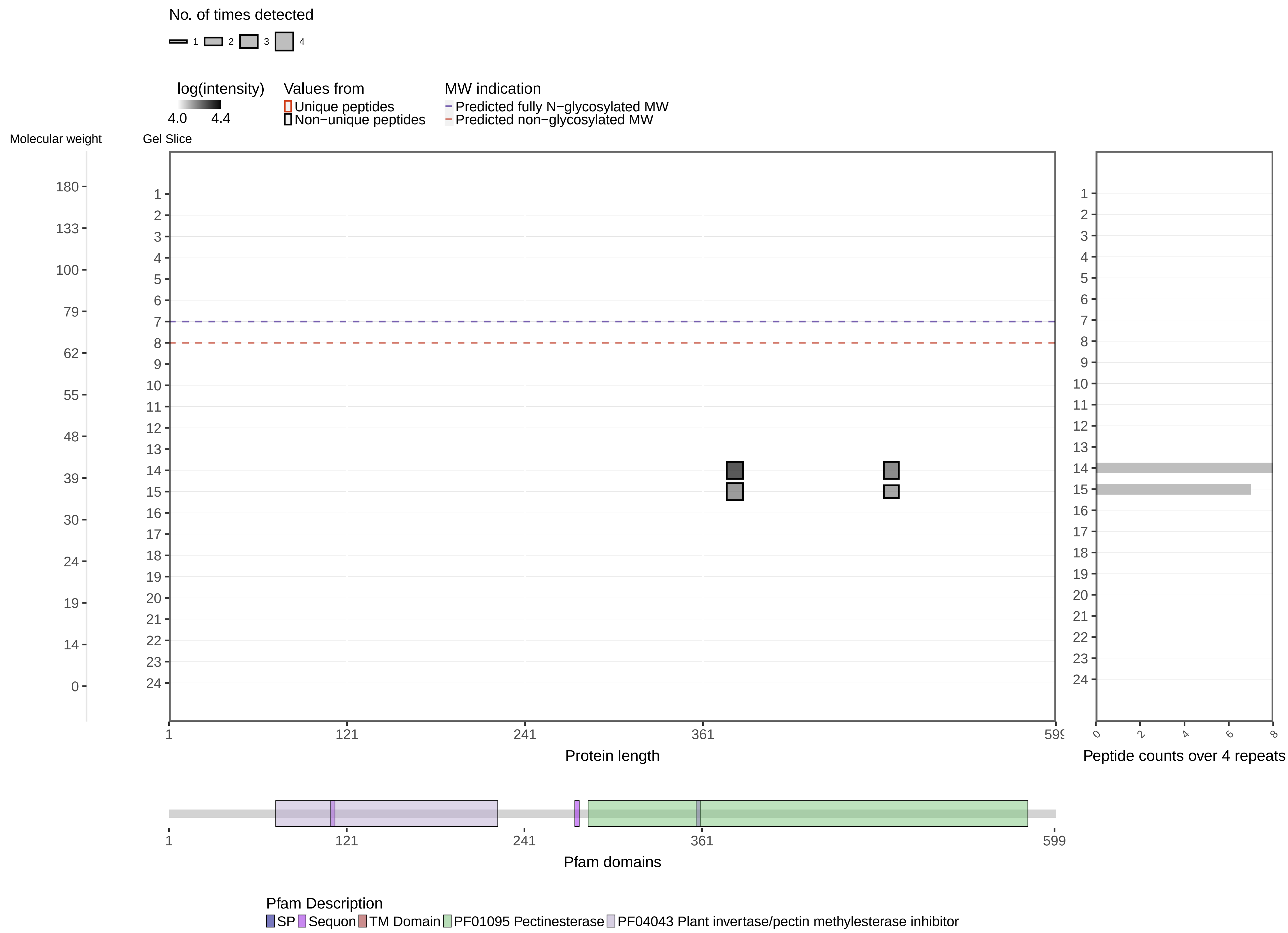


Protein sequence:

### NbL13g12900.1 protein_BOBBER_1−like [MW = 33.87 kDa]

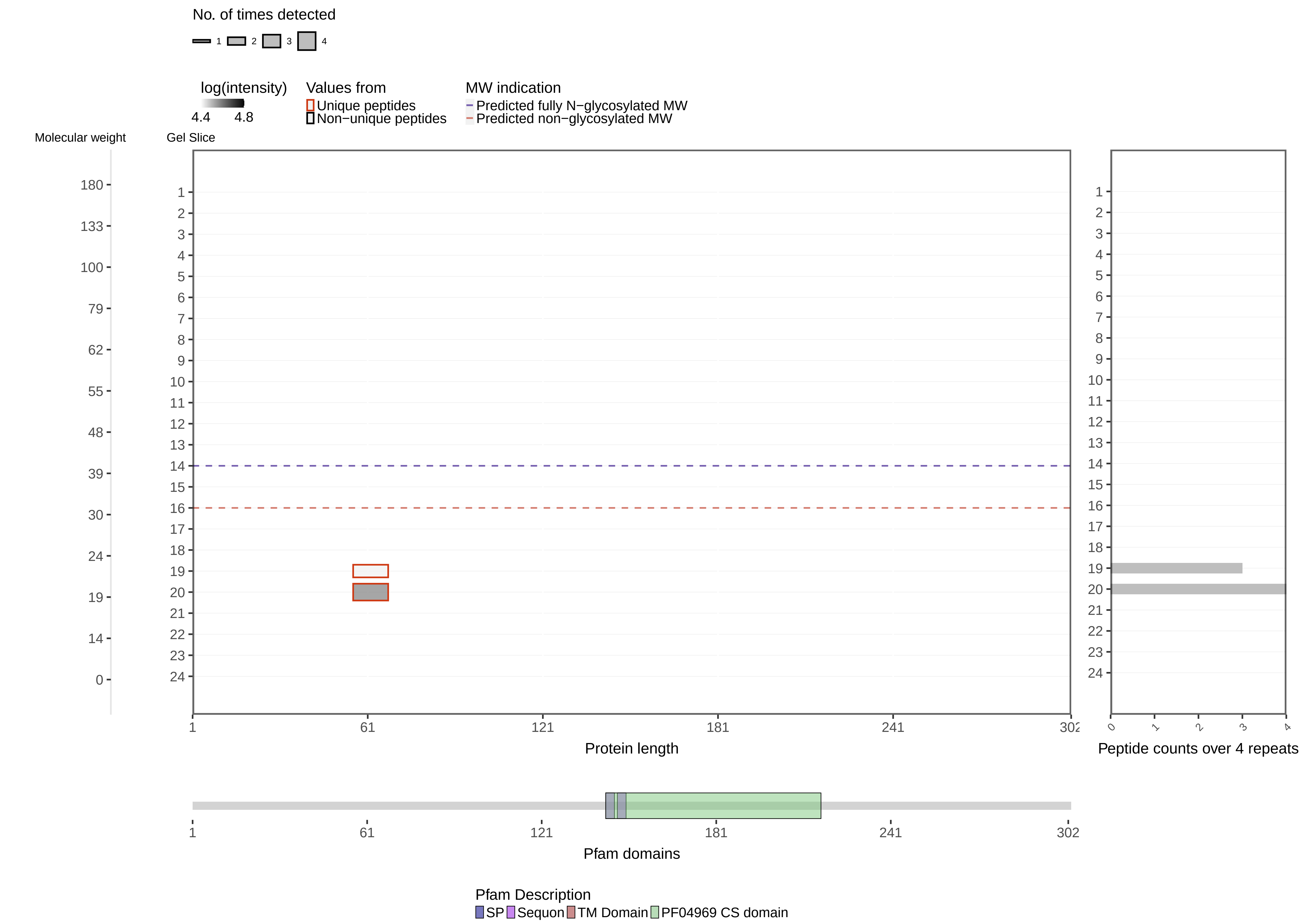


Protein sequence:

DLENYSWNQSLQEVNVNIPVPPGTKSRFIVCDISKNRLKVGLKGQPPIIDGELYRPVKVDDCFWSLEDQKSISVLLTKKDQMDWWKCCVKGEPEIDTQKAEPESSKLADLDPETRSTVEKMMFDQRQKSMGLPTSDETQKQEILKKFMAEHPEMDFSKAKIS*

NbL13g12830.1 glutamate_synthase_1_[NADH]_chloroplastic [MW = 242.8 kDa]

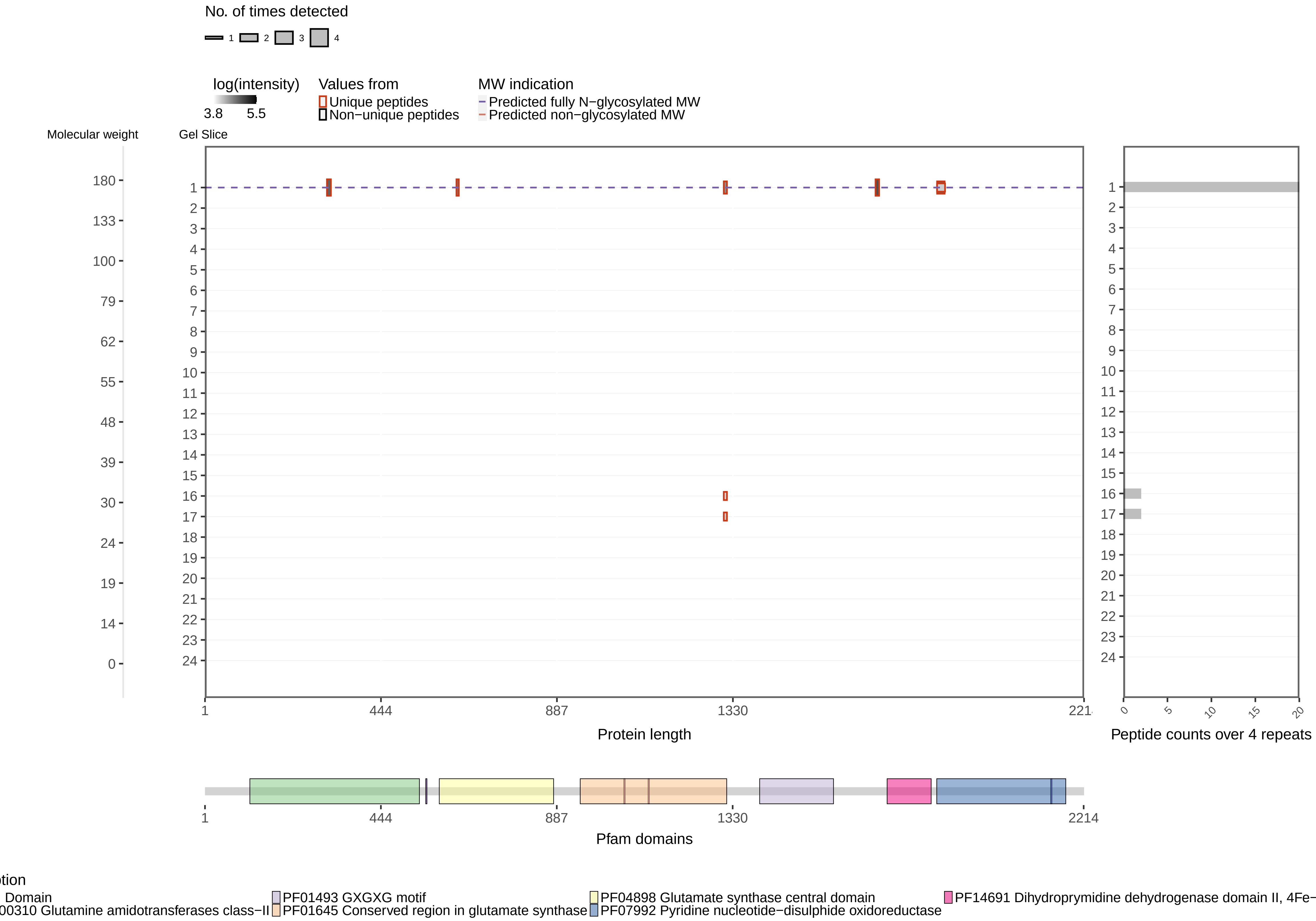


Protein sequence:

### NbL13g12770.1 probable_protein_phosphatase_2C_76 [MW = 38.79 kDa]

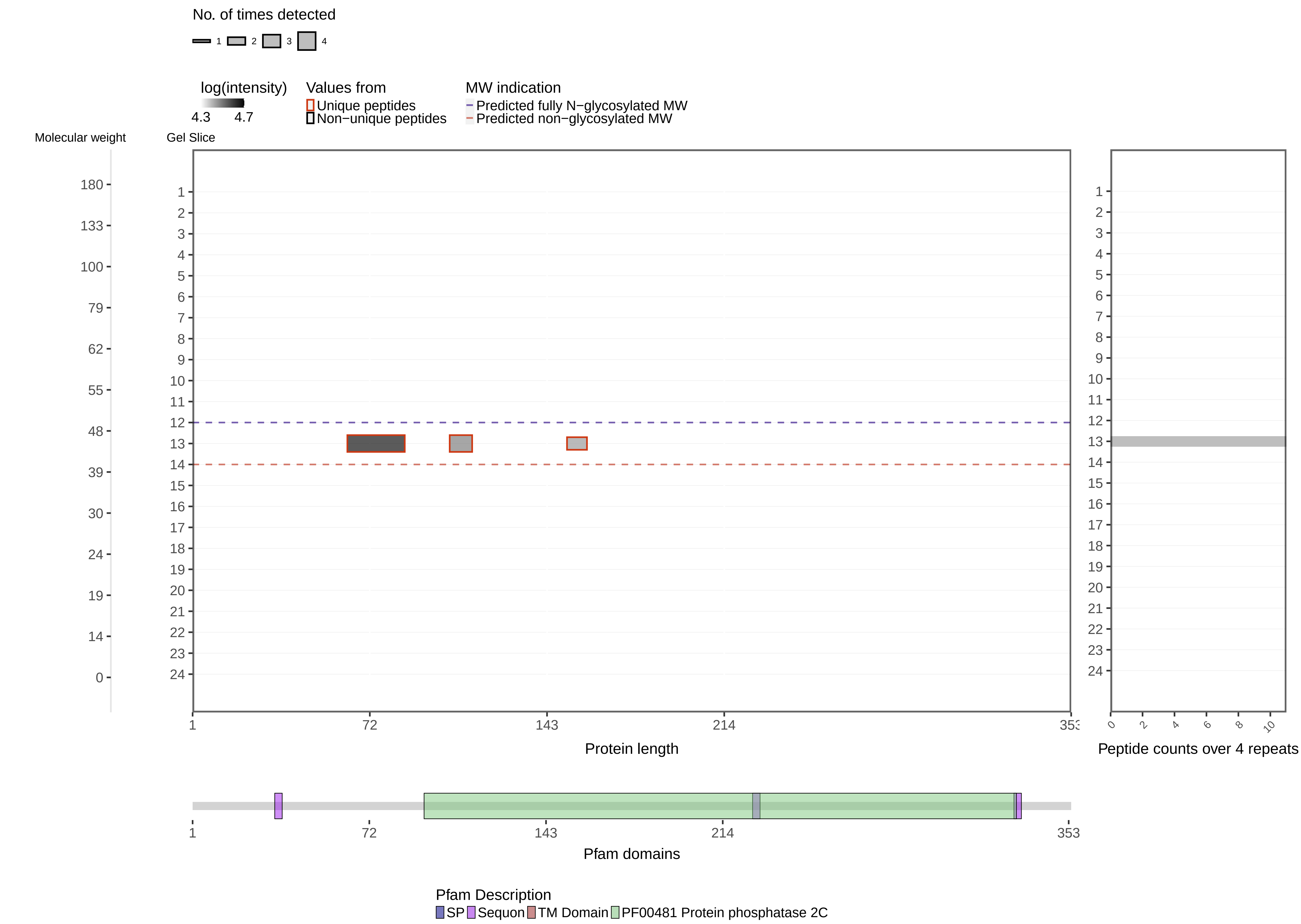


Protein sequence:

SETYQQTDMDFLDSEKDTFRDDGSTASTAVLVGNHLYVANVGDSRTIISKGGKAIALSEDHKPNRTDERKRIESAGGVVMWAGTWRVGGVLAMSRAFGNRMLKQFVVAEPEIQDQEIDEELELLVLASDGLWDVVPNEDAISLAQAEEEPEAAARKLTETAFTRGSADNITCIVVKFHHKKVEPEGSQQG*

NbL13g12740.1 probable_inactive_purple_acid_phosphatase_1 [MW − SP = 66.91 kDa]

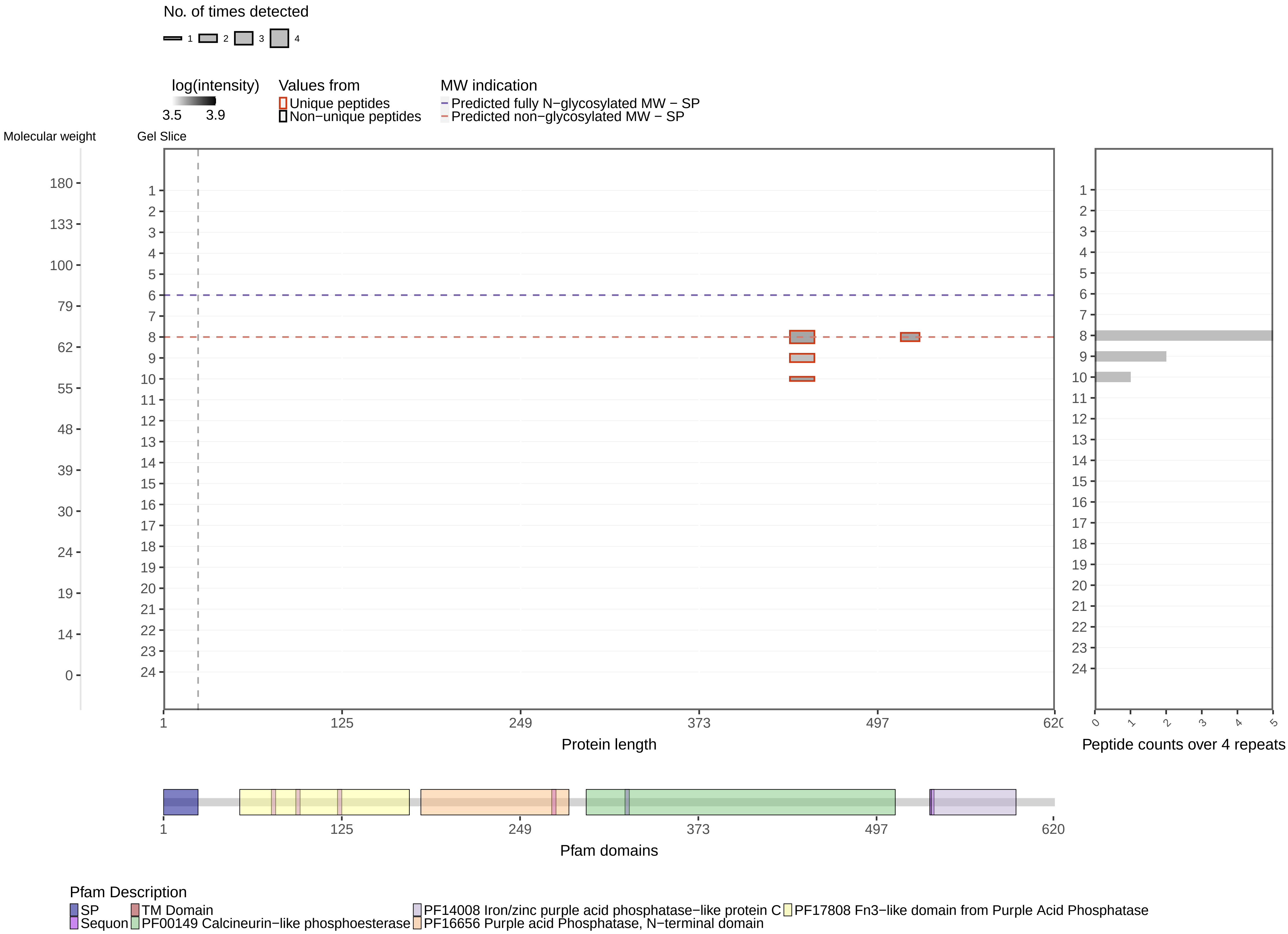


Protein sequence:

### NbL13g12590.1 probable_inactive_receptor_kinase_At4g23740 [MW − SP = 65.71 kDa]

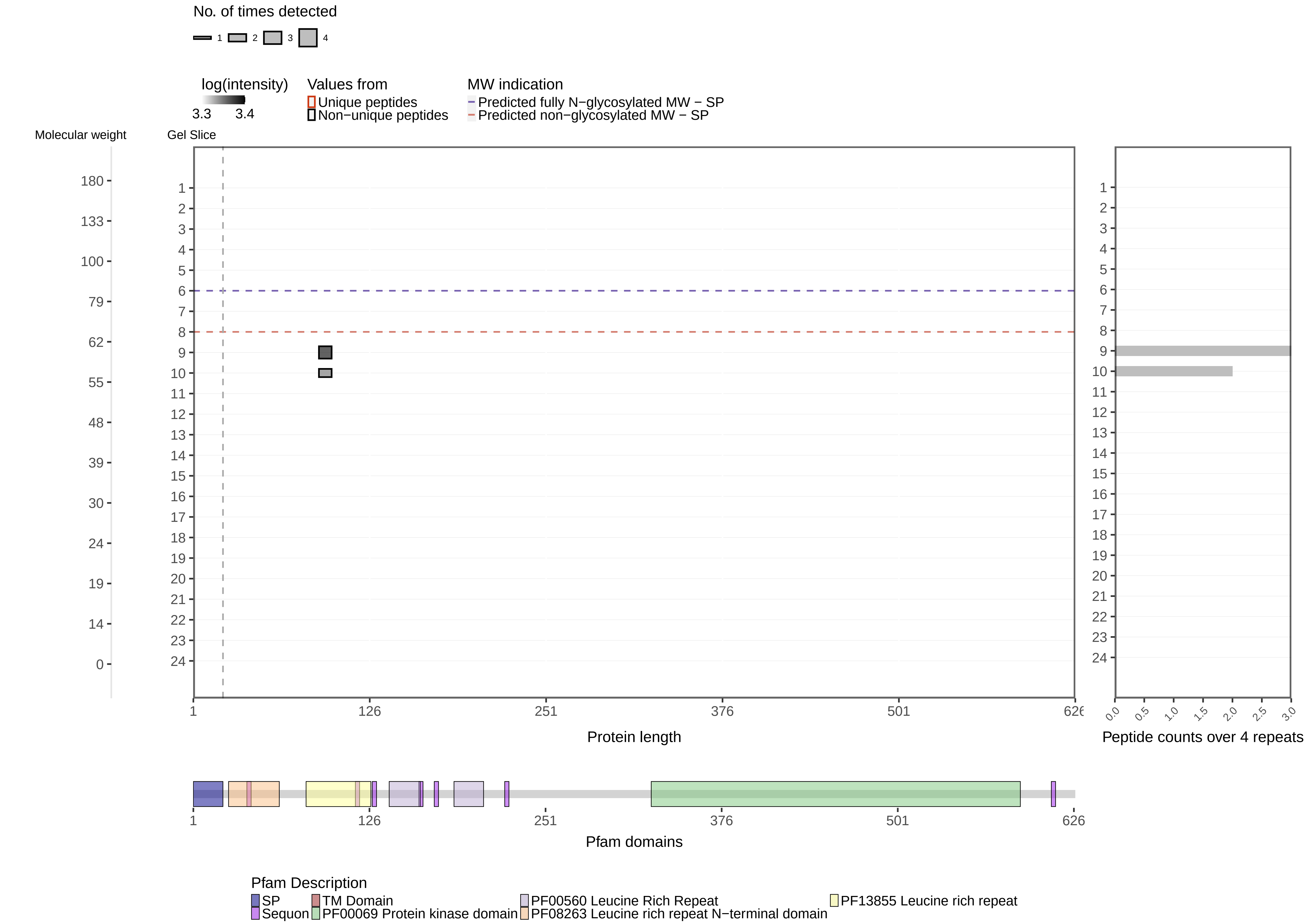


Protein sequence:

[illegible]

### NbL13g12560.1 uncharacterized_LOC107819525 [MW = 29.49 kDa]

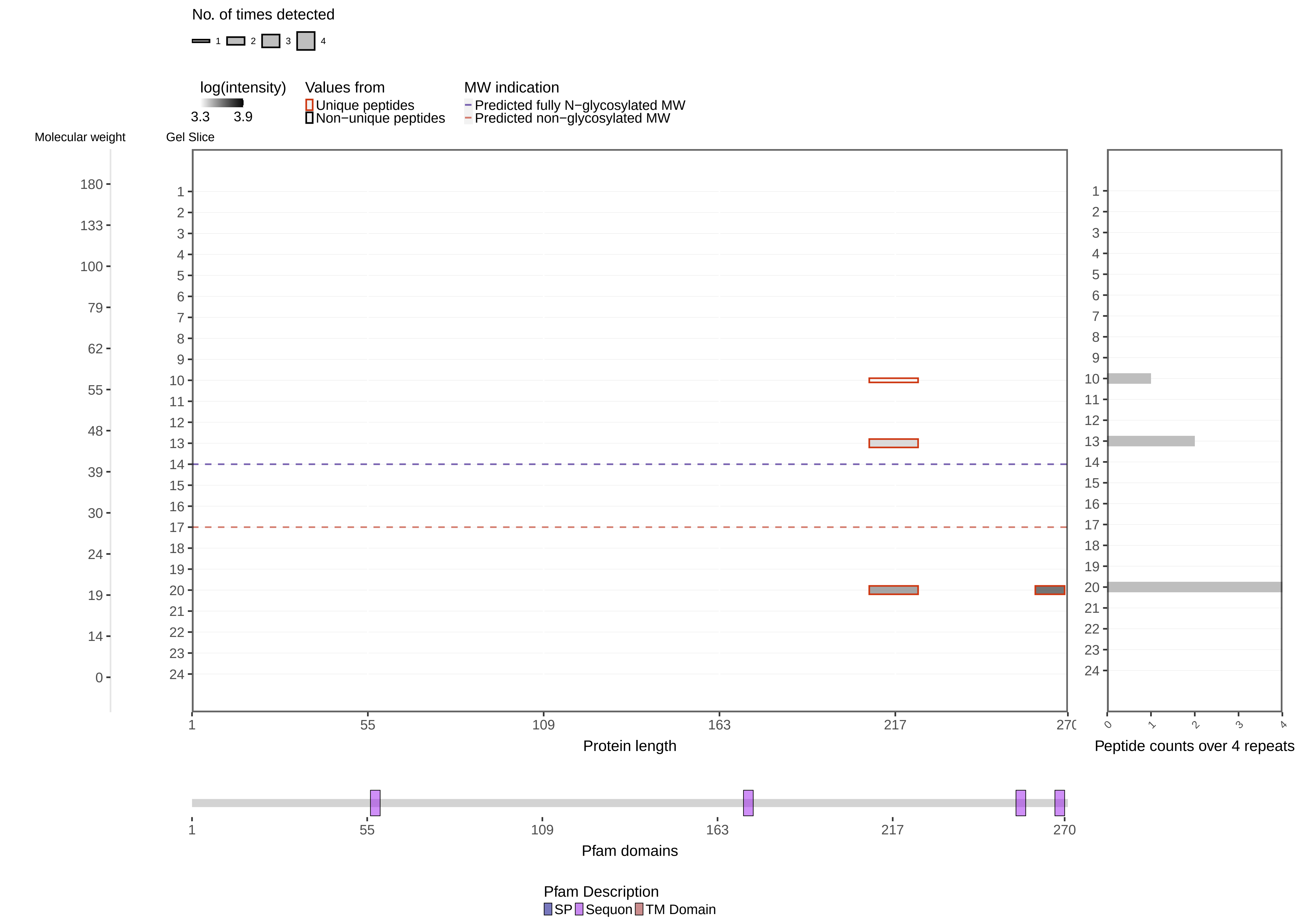


Protein sequence:

GVGIRVAIKPEYRITPPPPLSTQVGDIPRSTFNFDFDFEKKILAEAEKESQNWSRLGLENLPSRLPEQTNTGSTGDSVINKYTAFGFNREAVAIAVANYGDNPIKVKEFAERYTRLKEEMGFSSNSVADALLLNDNDTDKAISHLIGNSS*

### NbL13g12280.1 plastidial_pyruvate_kinase_2 [MW = 72.24 kDa]

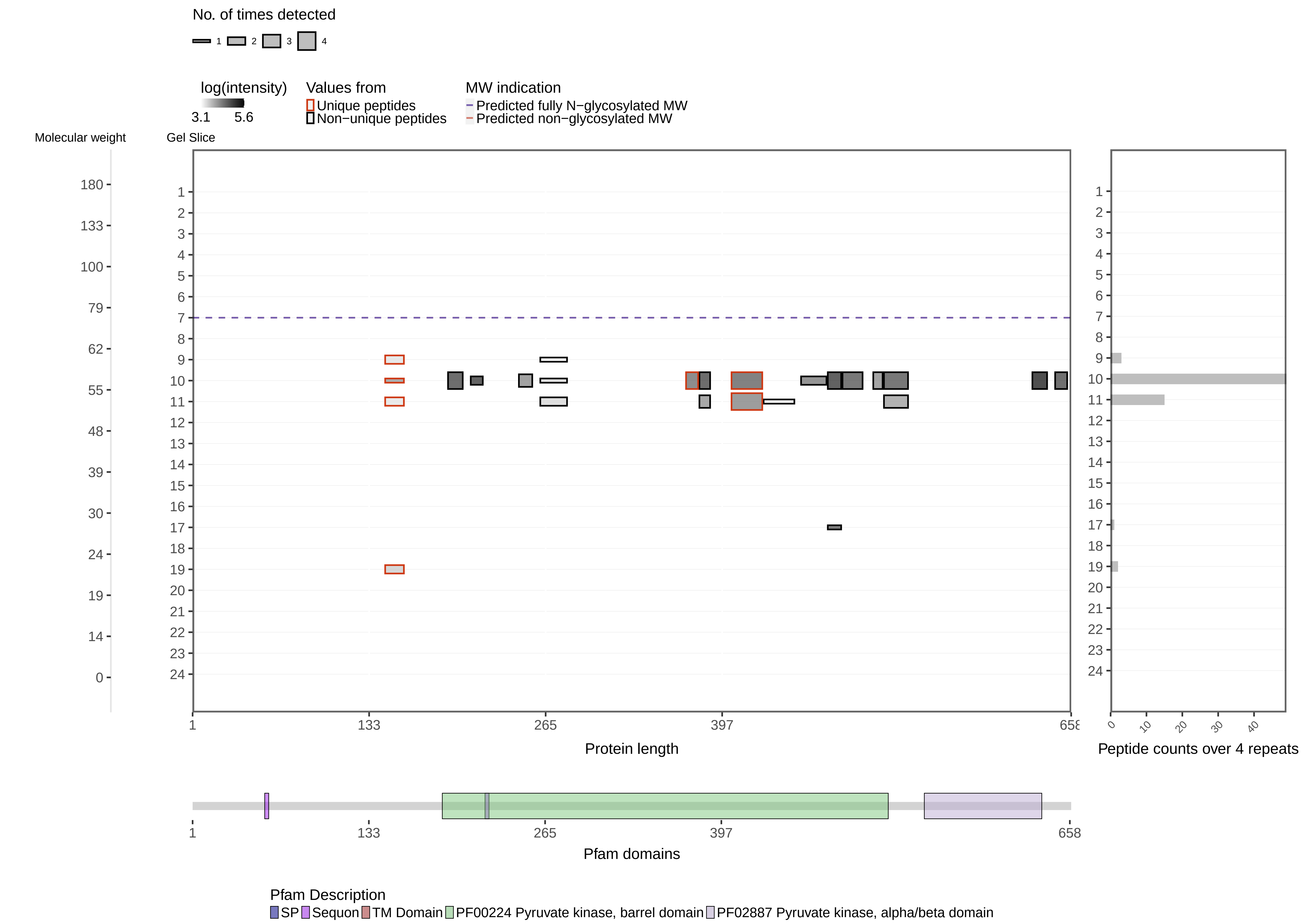


Protein sequence:

### NbL13g12170.1 glutaminyl−peptide_cyclotransferase−like [MW − SP = 29.71 kDa]

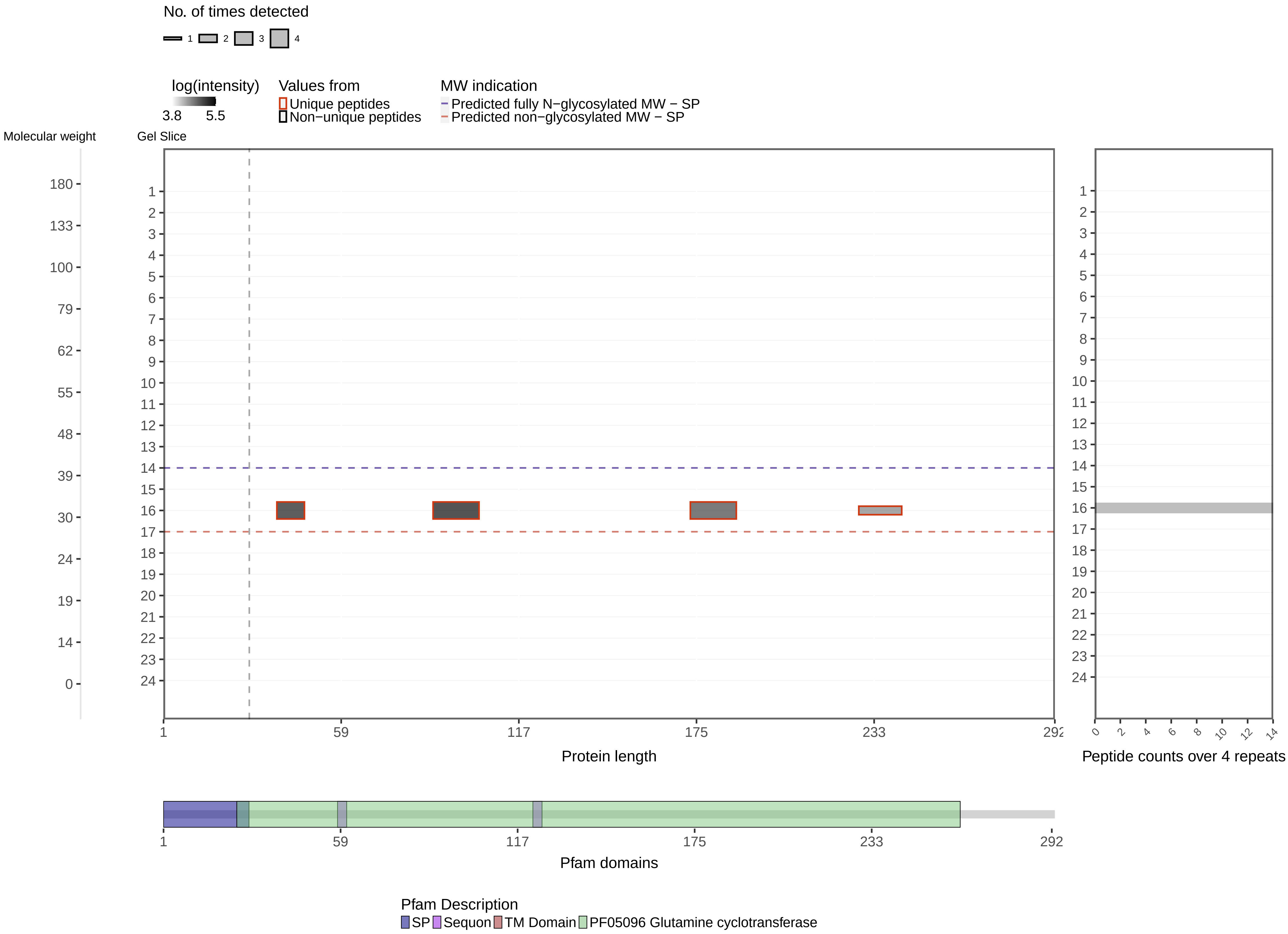


Protein sequence:

MQDGWGLATDGGVLFGSDGTSTLYEIDPQTMKVIRKRVVKSQGHEVPYLNELEYVKNEIWANVYYVTDCIARISPKDGTVIGWILLQSLREDLISRGYKDFEVLNGIAWDRDGDRILVTGKLWPKLFEIKLLPLPPNAPLIGEVNNLCIPKPFFLGELES*

NbL13g11970.1 BAHD_acyltransferase_DCR [MW = 50.16 kDa]

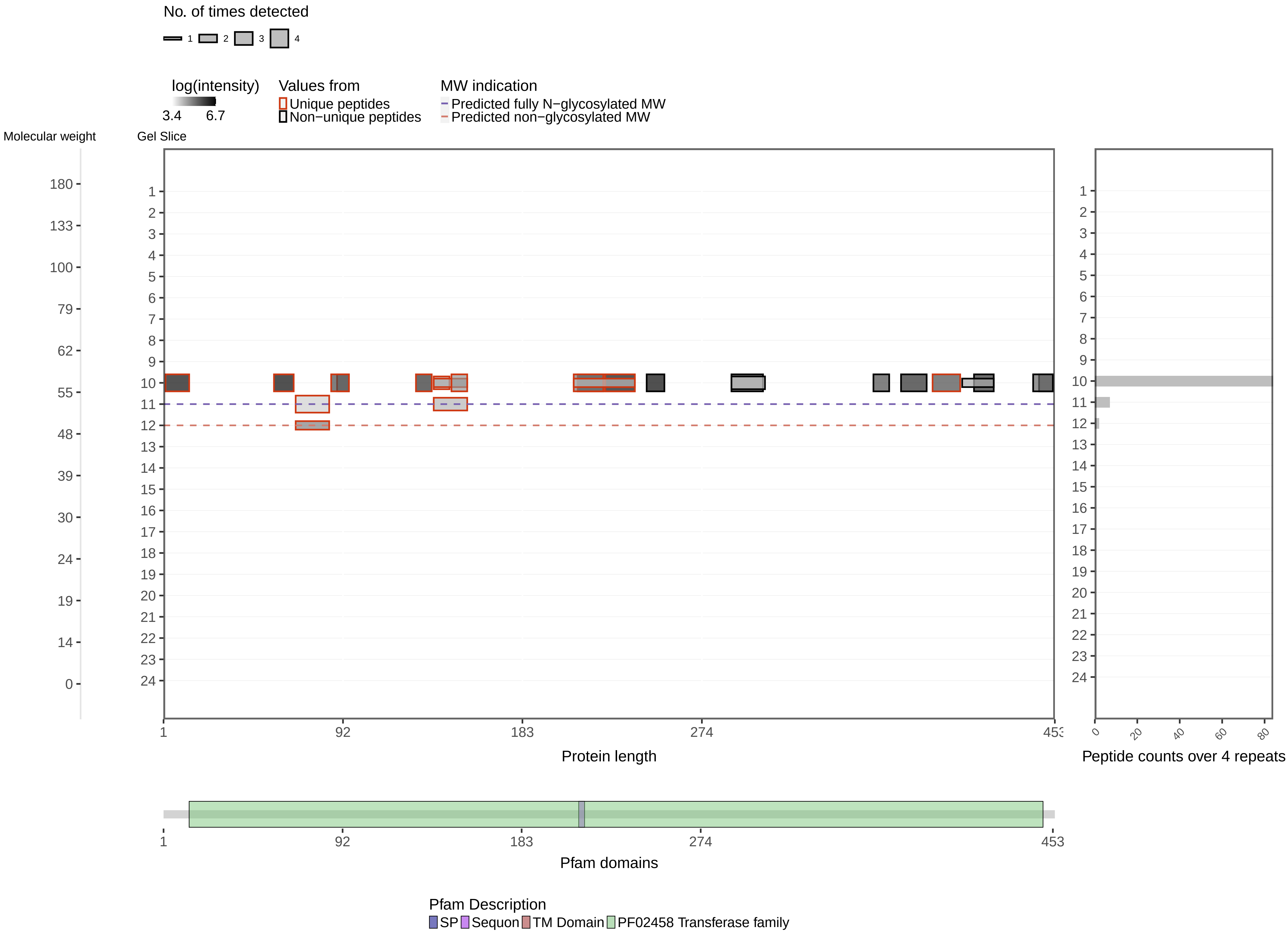


Protein sequence:

PPSDATEPDKSAPNGDVIASVDPPLRERIFKFSESAIDQIKSKVNANPPEGSSNPFSTFQSLSAHVWLAITRARQLKPEDYTVYTVFADCRKRVDPPMPESYFGNLIQAIFTVTAAGLLLSQPIEFAAGMIQQAIVKHDAKAIDERNKEWESNPKIFQYKDAGVNCVAVGSSPRFKVYDVDFGWGKPESVRSGSNNRFDGMVYLYQGKSGGRSIDVEISLEANAMERLEKDKEFLMEA*

### NbL13g11960.1 WW_domain−containing_protein_C11B10.08−like [MW = 32.91 kDa]

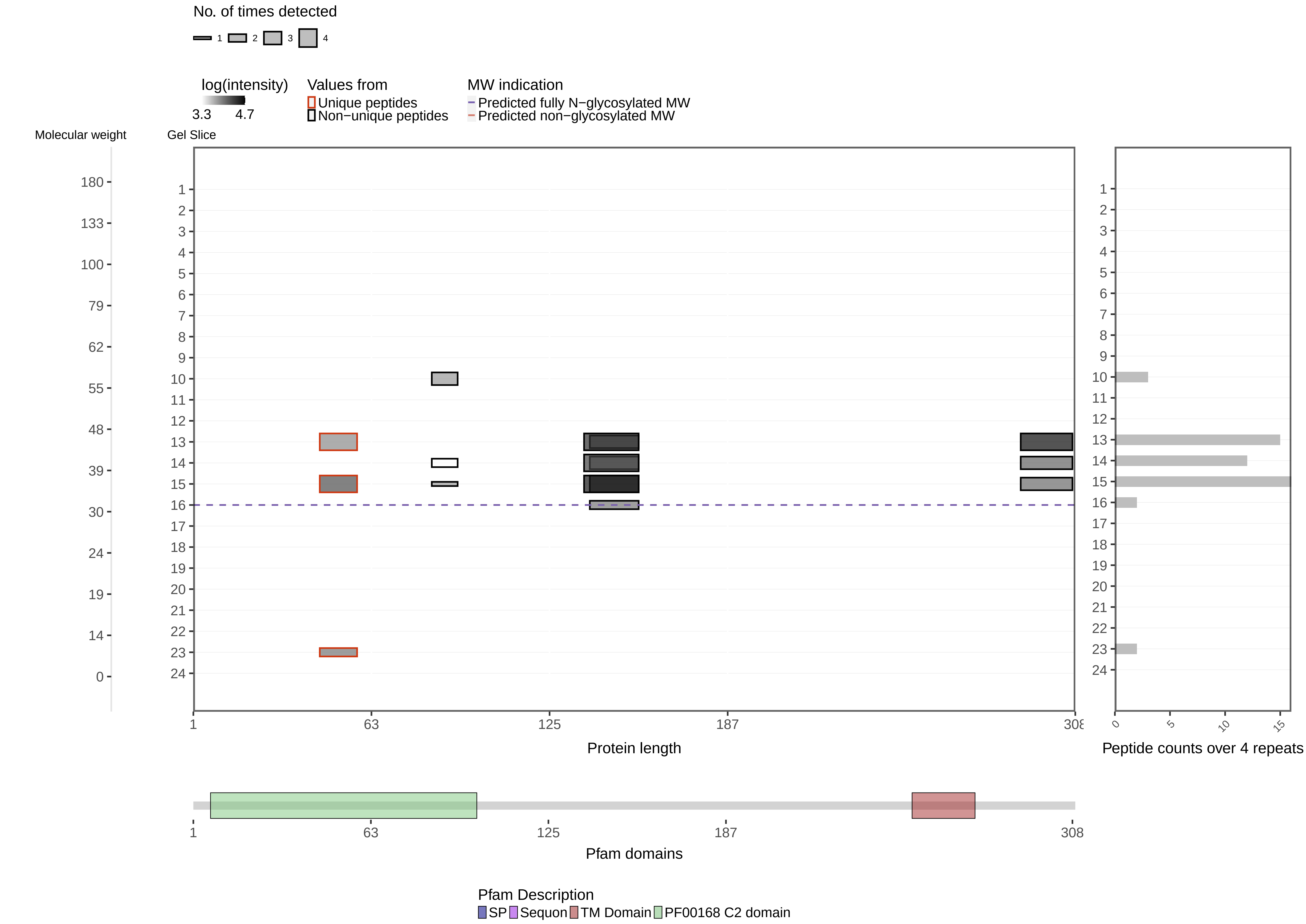


Protein sequence:

YAPPYGVPPPGSRDYPAAPPQPYGGSYGAPAPYAAPPSGYPYSAAPAPAPYGQQQPSYGAPPAPYSQGDSYGPPGYSQGSYGQQQGNYGQASYGQQSGYKYEEKKGKFGGMGLGAGLAVGAVAGALGGLAISEGIDHIEDNIADKAAEKVEDDLADDDYDGDDF*

### NbL13g11800.1 60S_ribosomal_protein_L36−1−like [MW = 12.1 kDa]

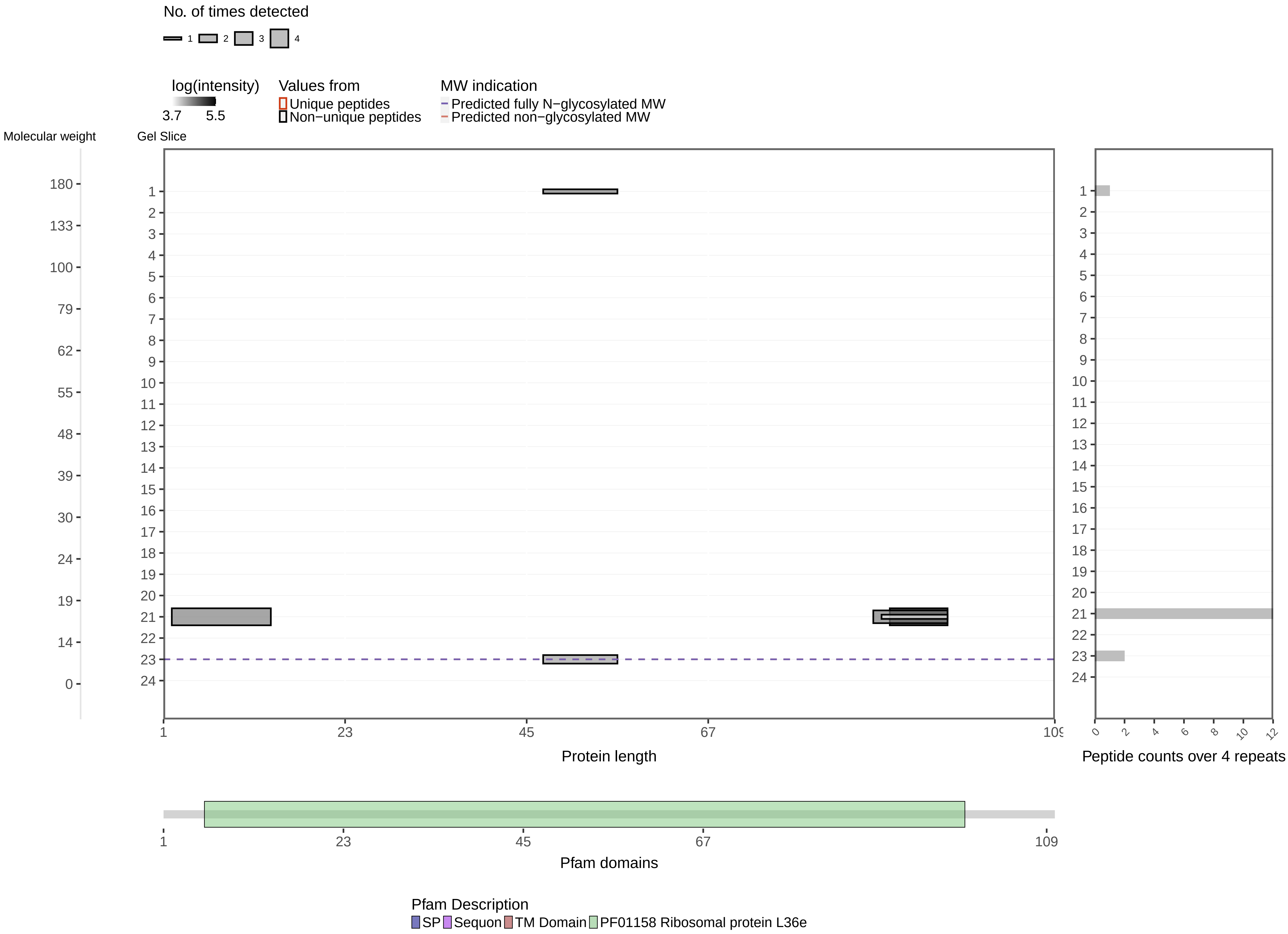


Protein sequence:

/RSLIREVAGFAPYEKRITELLKVGKDKRALKVAKRKLGTHKRAKKKREEMSSVLRKMRATGGAEKKK*

### NbL13g11780.1 cleavage_and_polyadenylation_specificity_factor_subunit_2 [MW = 81.52 kDa]

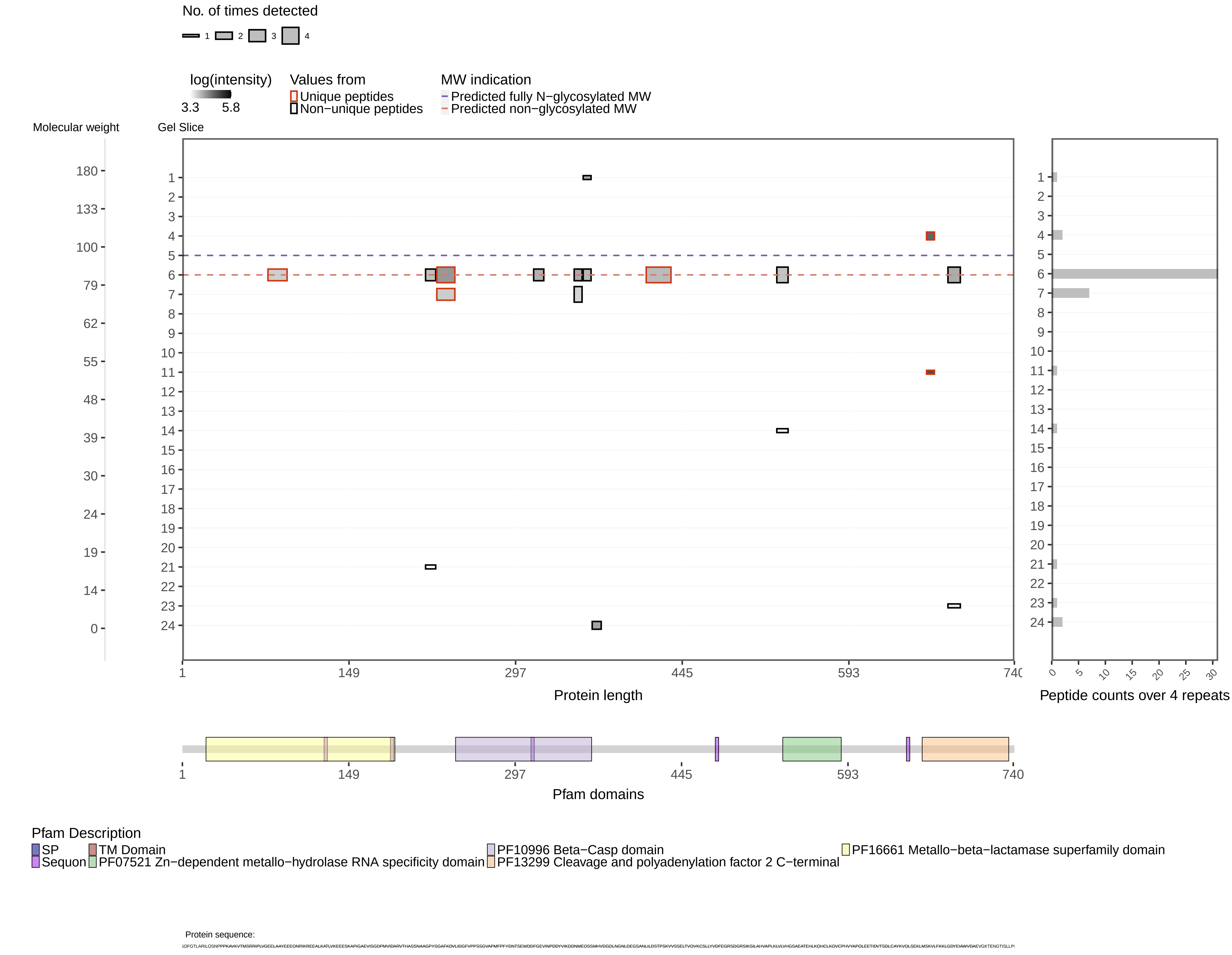

### NbL13g11710.1 oxalate−−CoA_ligase−like [MW = 56.17 kDa]

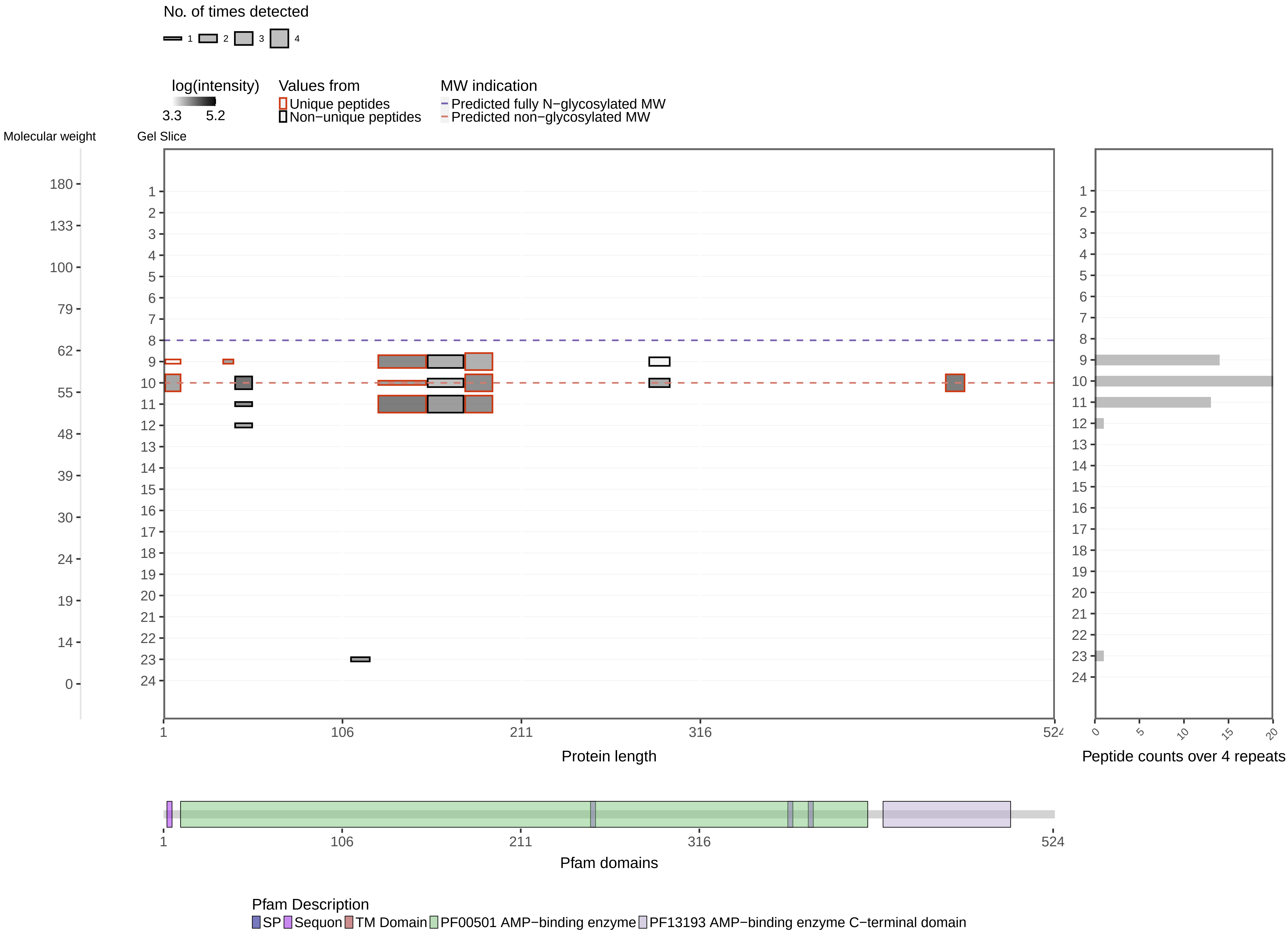


Protein sequence:

NATWYTAVPTIHQILLDCHLSKPESDYPKLRFIRSCSAALAPSVMARLEEAFGAPVLEAYAMTEATHLMASNPLPENGPHVPGSVGKPVGQEMAILDENGVVQGPNAKGEVCIRGPNVTKGYKNNPEANKSAFQFGWFHTGDVGYLDSDGYLHLVGRIKELINRGGEKISPIEVDAVLVSHAEIAQAVAFGVPDDKYGEEINCAVIPREGSNIDEAEVLRFCKKNLAAFKVPKKVFMTDSLPKTATGKIQRRLVAEHFLAQISTAKVPKFGA*

NbL13g11700.1 tubulin_beta−1_chain−like [MW = 50.45 kDa]

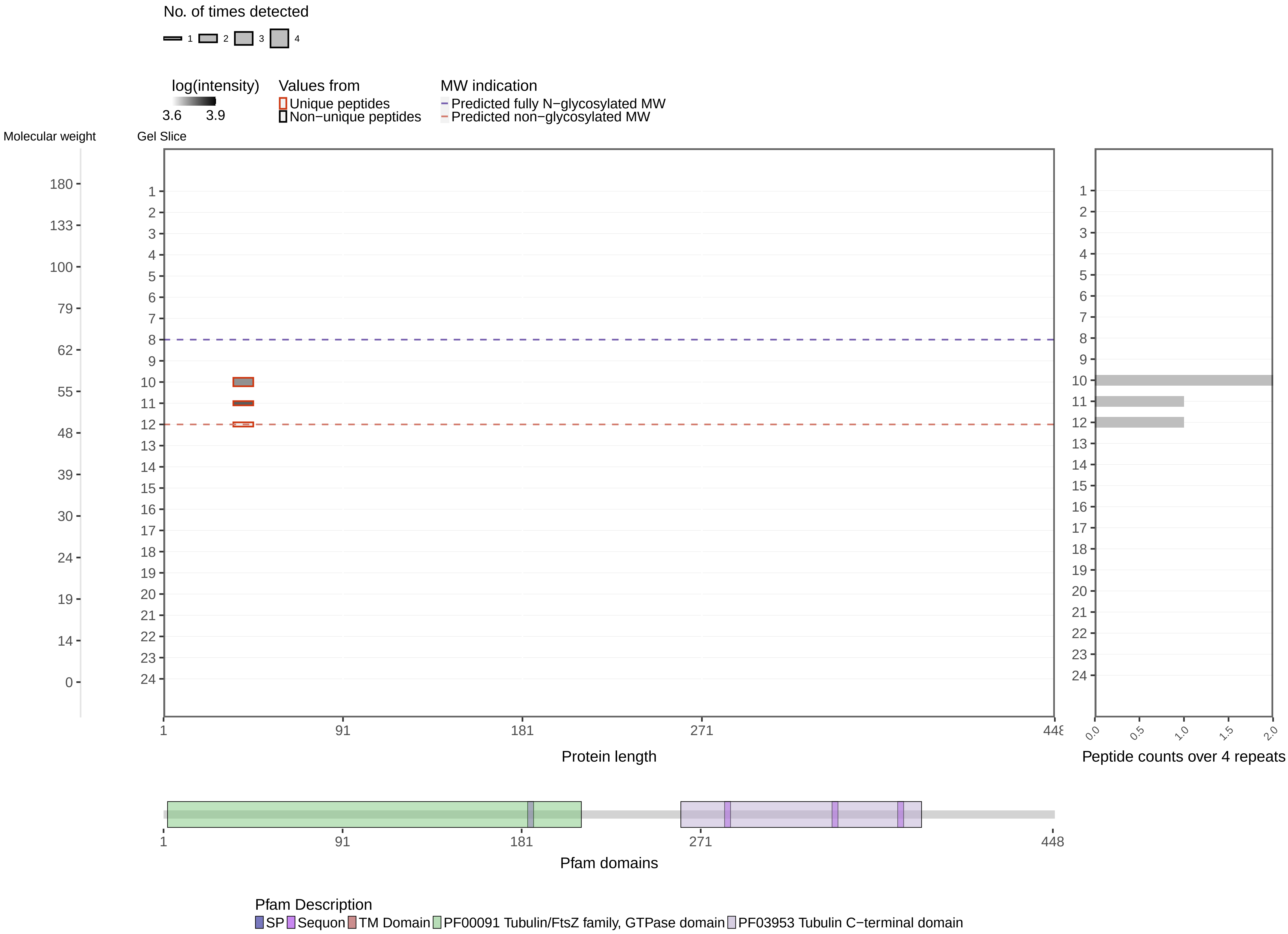


Protein sequence:

CFRTLKLTTPSFGDLNHLISATMSGVTCCLRFPGQLNSDLRKLAVNLIPFPRLHFFMVGFAPLTSRGSQQYRNLTVPELTQQMWDSKNMMCAADPRHGRYLTASAMFRGKMSTKEVDEQMINVQNKNSSYFVEWIPNNVKSTVCDIPPTGLRMASTFIGNSTSIQEMFRRVSEQFTAMFRRKAFLHWYTGEGMDEMEFTEAESNMNDLVSEYQQYQDATAEEEEYDYEDEEEEGQEA*

### NbL13g11630.1 transmembrane_ascorbate_ferrireductase_1−like [MW = 25.42 kDa]

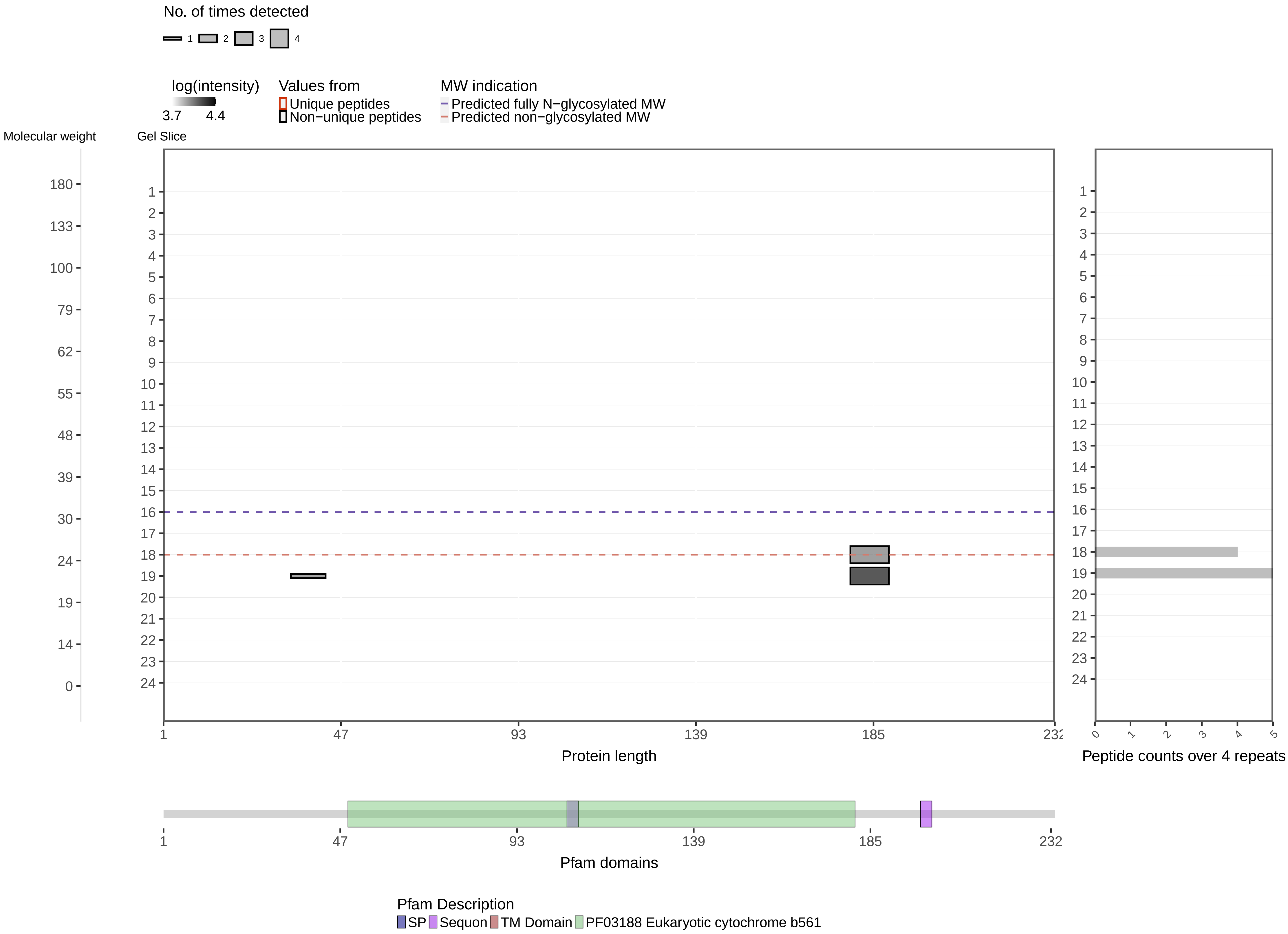


Protein sequence:

HNESNIPNMYSLHSWVGISVIVLYGIQWIYGFVVFFYPGGSAEVRRDSLPWHVLFGMFVYVMAVGNACLGLLEKLTFLEVNGLAKYGSEAFLVNFTAIATVLYGVFVFLTILSQAPTEDDYSYSAIS*

### NbL13g11620.1 remorin−like [MW = 22.85 kDa]

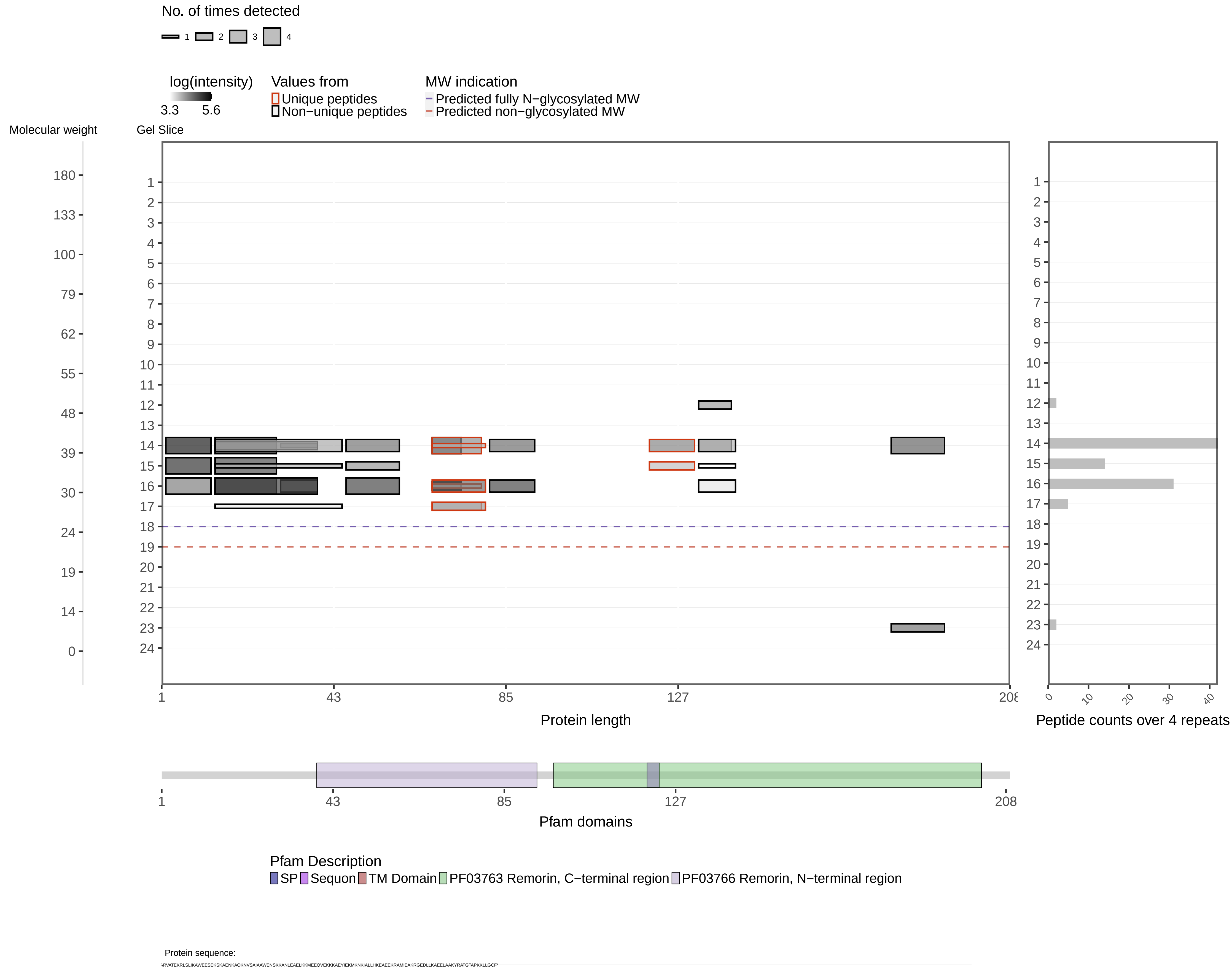

### NbL13g11590.1 probable_acetyltransferase_NATA1−like [MW = 20.81 kDa]

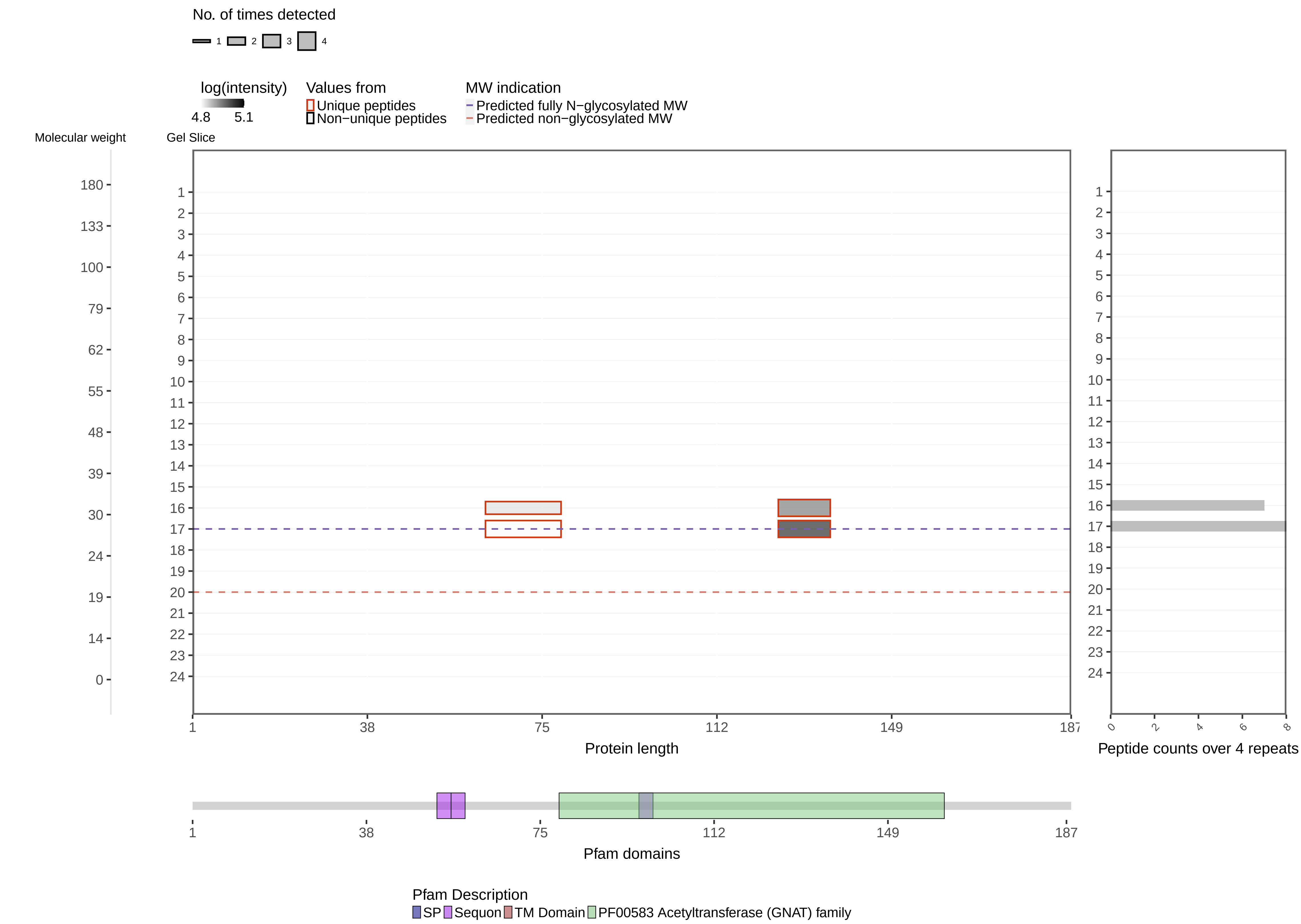


Protein sequence:

GNDAVVAGFVLFFPNYSSFLAKPGFYIEDIFVRECYRRKGFGRLLLSAVAAQAAKMGYGRVEWVVLDWNVNAIKFYEEMGAQILQEWRVCRLTGGALEAFANVDI*

### NbL13g11410.1 membrane_steroid−binding_protein_2−like [MW = 24.75 kDa]

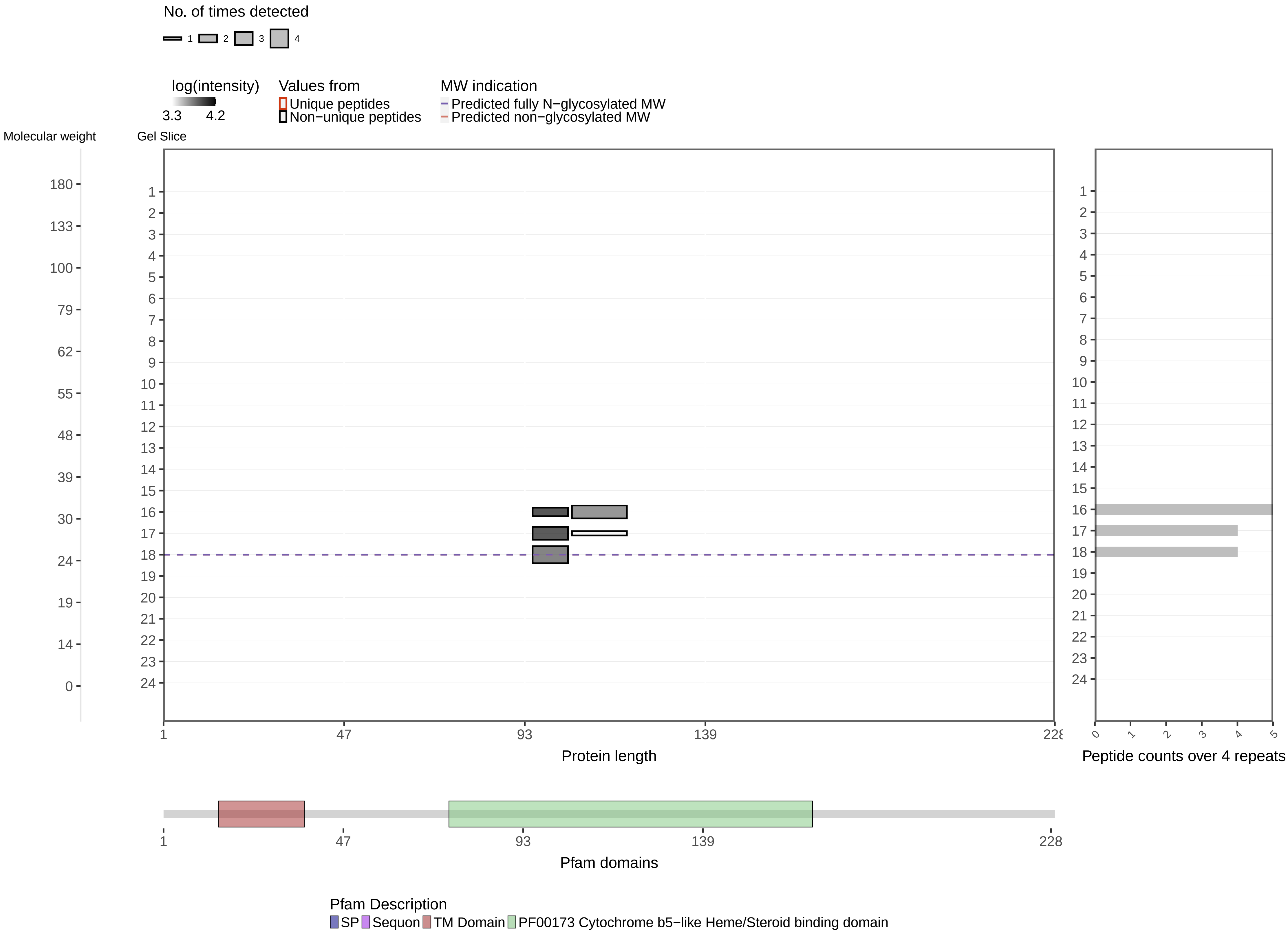


Protein sequence:

SRMFYGPGGPYALFAGKDASRALAKMSFEEKDLTGDISGLGPFELEALQDWEYKFMSKYVKVGTVKQTGPVSDGATNGESVESTDREAKPAEGFASESAKPSEDGPSESVAETVDKSDGDAEKKD*

NbL13g11340.1 superoxide_dismutase_[Fe]_3_chloroplastic [MW = 30.84 kDa]

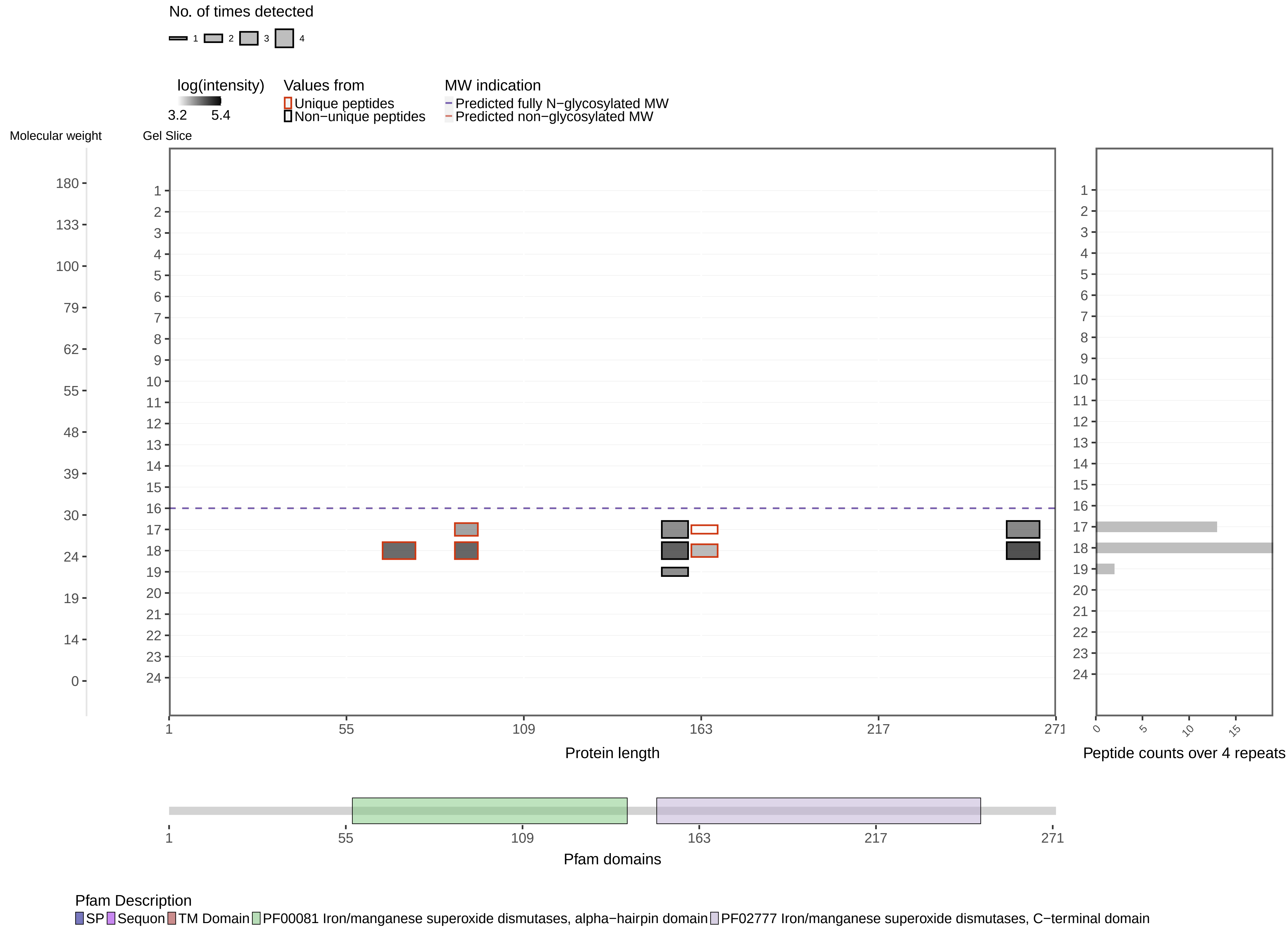


Protein sequence:

PEFSDAAQVWNHDFFWESMQPGGGDMPKLGLLQQIEKDFGSFTNFREKFTEAALTLFGSGWIWLVLKREEKRLAIVKTSNAVNPLVWNDIPLIGLDLWEHAYYLDYKNDKAKYVNVFMNHLVSWDAALGRMARAQAFVNLGEPKIPVA*

### NbL13g11330.1 ammonia−lyase [MW = 78.23 kDa]

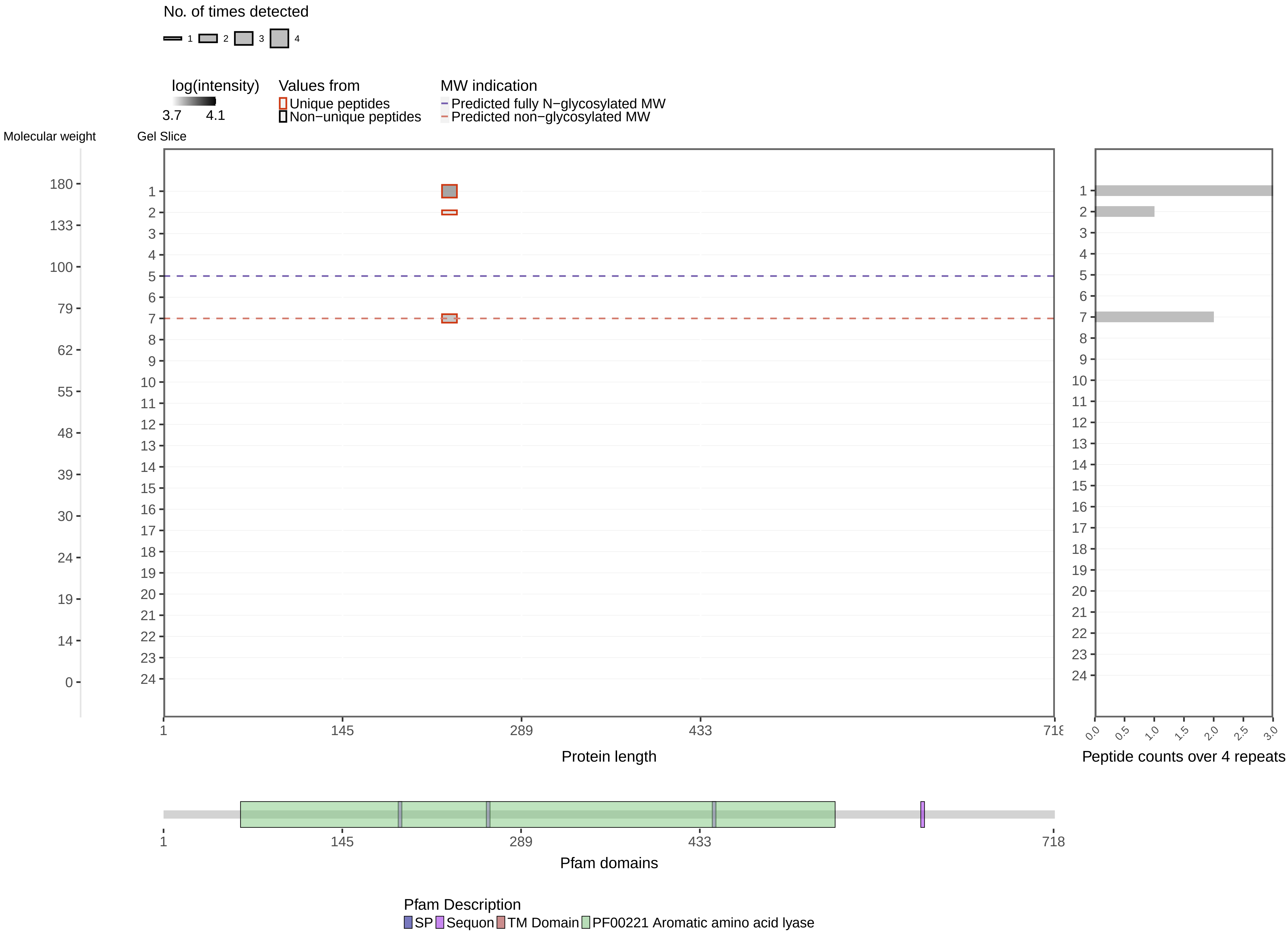


Protein sequence:

QKPKQDRYALRTSPQWLGPQIEVIRAATKMIEREINSVNDNPLIDVSRNKALHGGNFQGTPIGVSMDNTRLALASIGKLMFAQFSELVNDYYNNGLPSNLTGGRNPSLDYGFKGAEIAMASYCSELQFLANPVTNHVQSAEQHNQDVNSLGLISSRKTAEAVDILKLMSSTYLVALCQAIDLRHMEENLKASVKNVVSLVAKKVLTIGQNGELHPSRFCEKDLLKVVDREYVFAYVDDPCSCTYPLIQKLRQVLVDHALLNGEGEKNSSTSIFEKISVFEEELKAILSKEVESARC

### NbL13g11080.1 peroxidase_66 [MW − SP = 32.47 kDa]

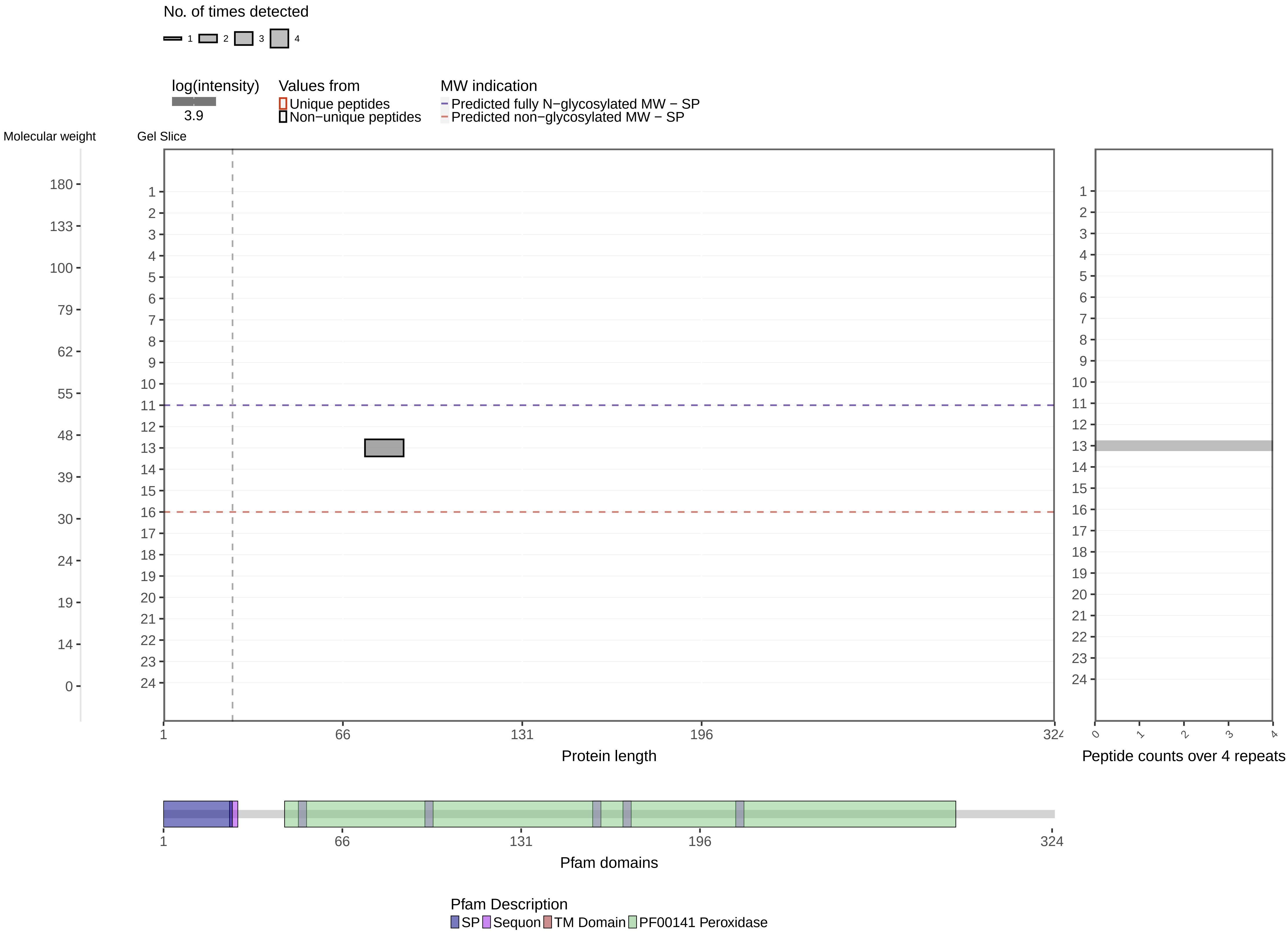


Protein sequence:

GRISRANETINLPAPSFNTSQLIQSFAKRGLGVKDLVALSGGHTLGFSHCSSFEARLRNFSSVHDTDPSLNAVFAESLKKNCPKPNRDSNAGQFLDTTSSLFDNNYYKQIVAGKGVFASDQSLLNDYRTGWIVKAFANDQASFFWEFAASMVKLGNVGVLENGEVRLNCRAVN*

### NbL13g11070.1 uncharacterized_LOC107784461 [MW = 26.57 kDa]

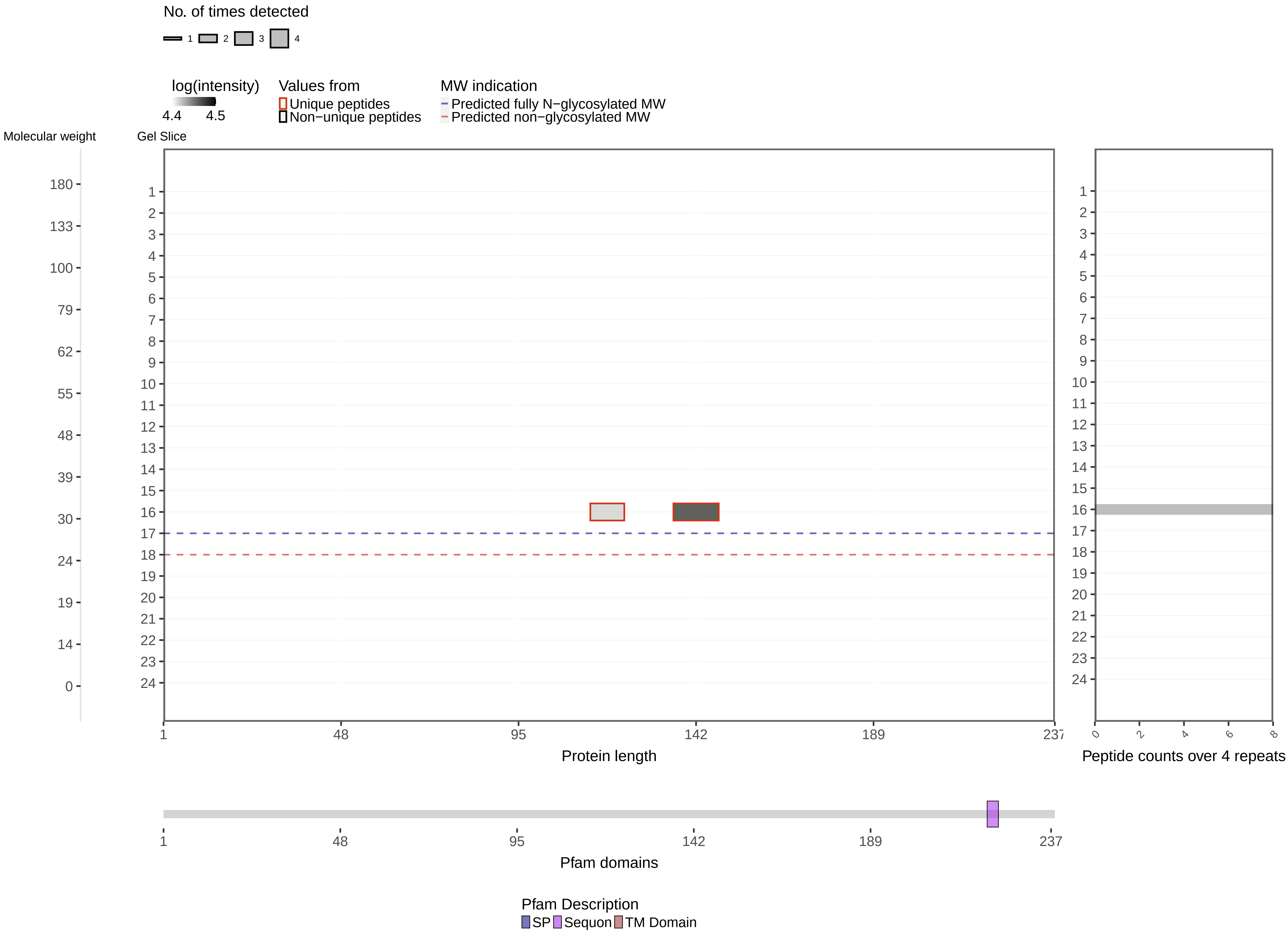


Protein sequence:

RLFSDIKIRGRVAIGLNPNIRFYRYKAGQRFGRHIDESVDIGEGKRTHYTLLIYLSGGSKSKNKDDKDSQDSVSEILVGGETVFYGPRNALVAEVPPTEGMALVHIHGDKCMLHEARNVTKGIKYVLRSDVVFA*

NbL13g11010.1 phosphoglucomutase_chloroplastic [MW = 68.69 kDa]

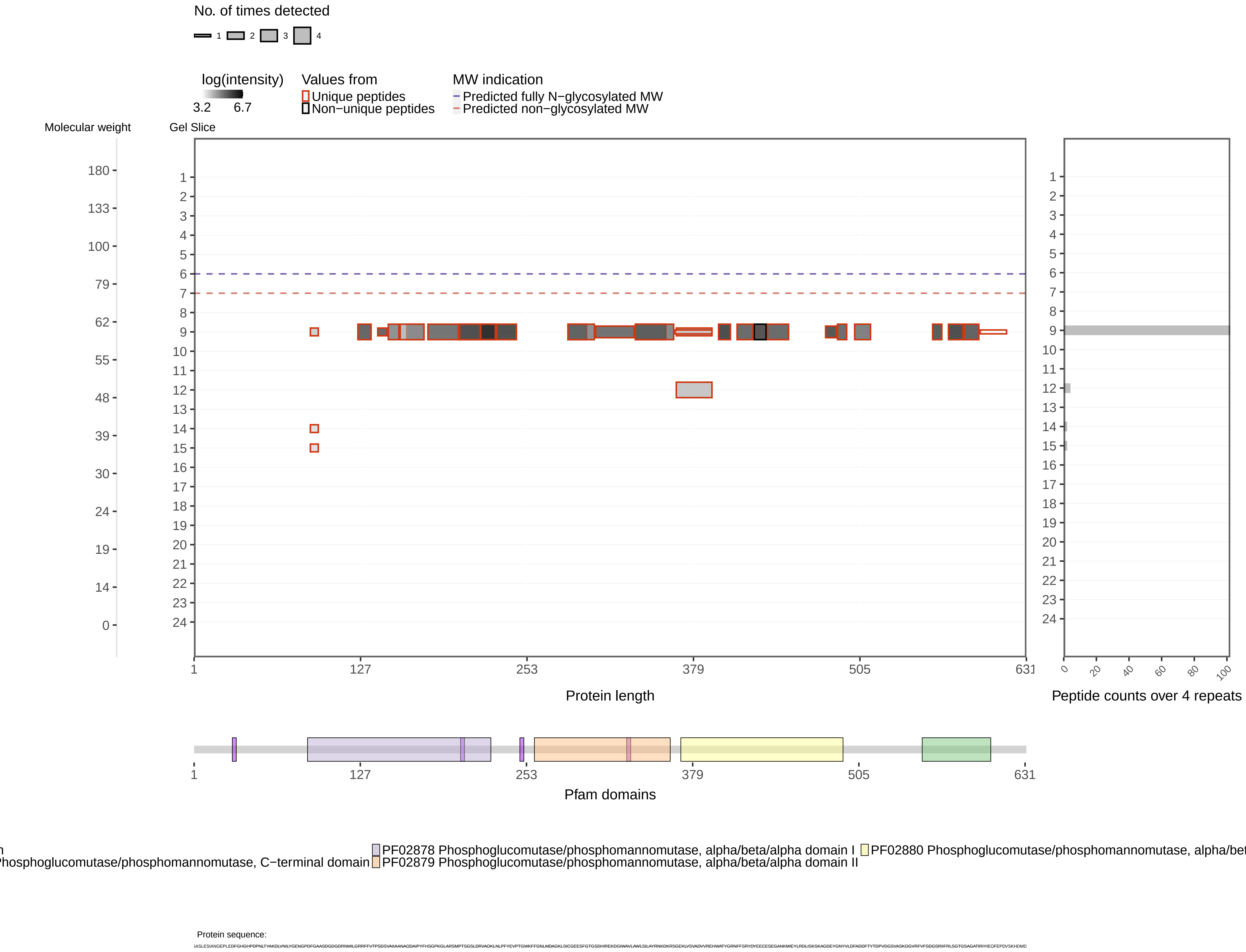

NbL13g10910.1 subtilisin−like_protease [MW − SP = 81.23 kDa]

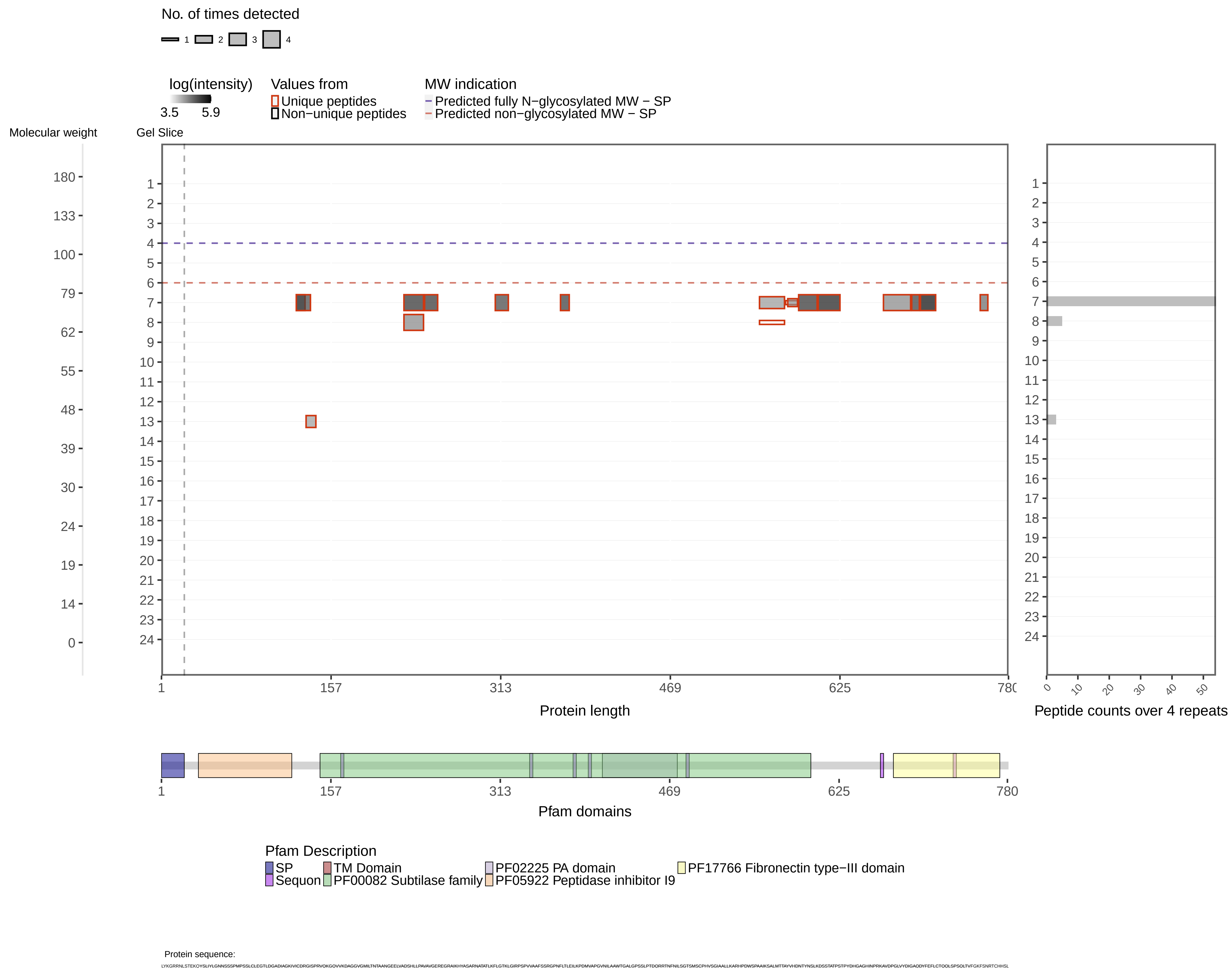

### NbL13g10850.1 uncharacterized_LOC107782590 [MW = 44.38 kDa]

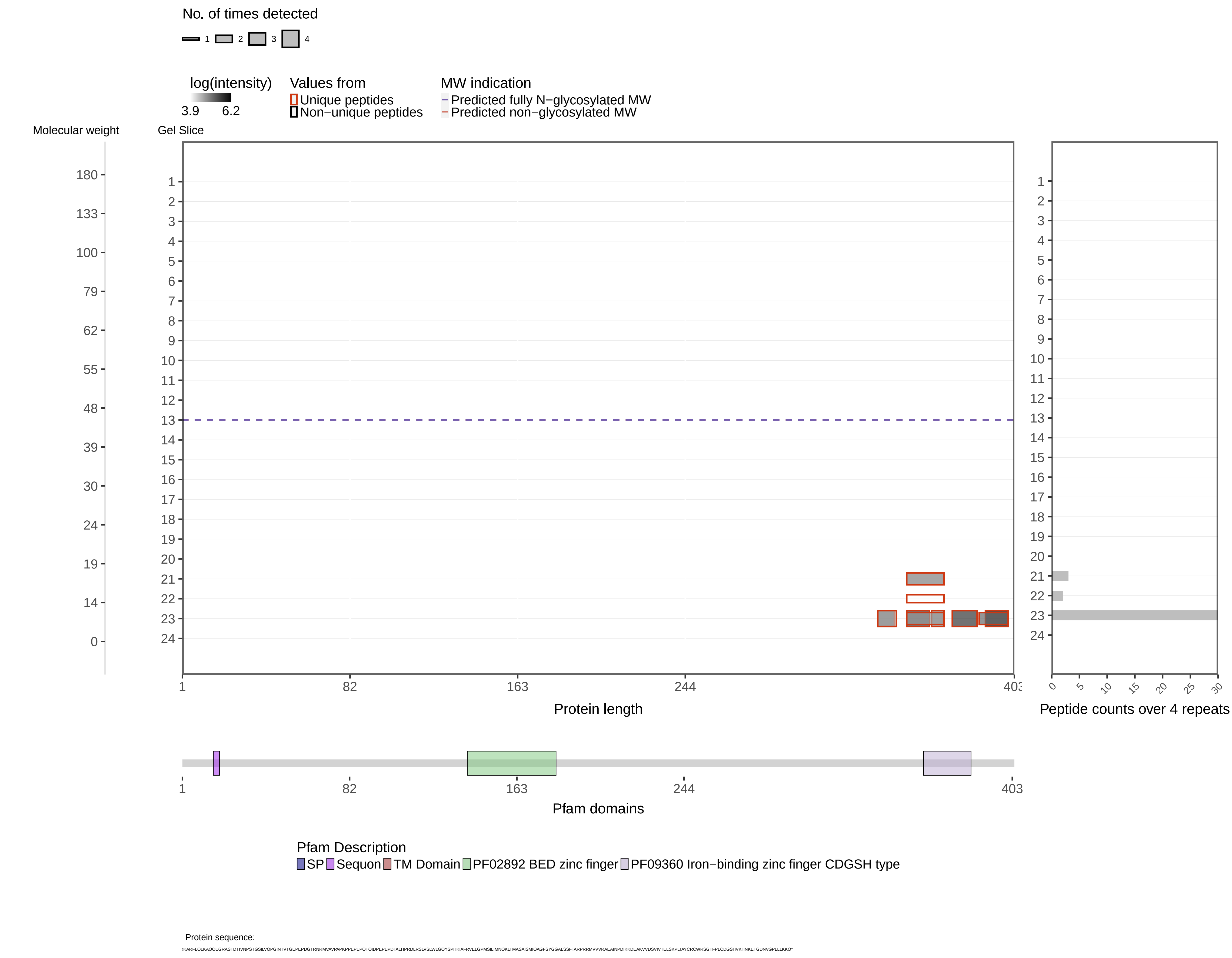

### NbL13g10810.1 cleavage_and_polyadenylation_specificity_factor_subunit_1 [MW = 158.02 kDa]

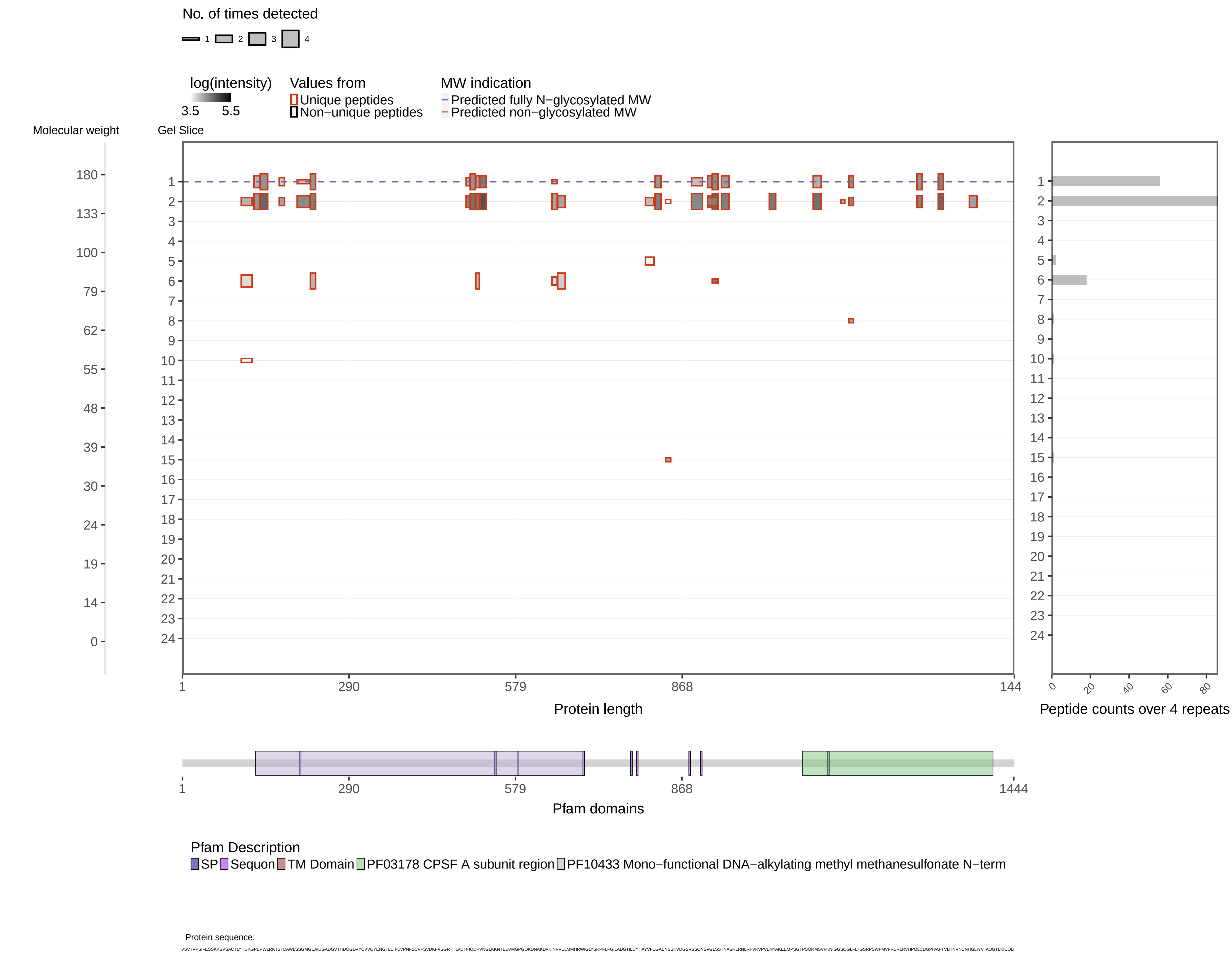

### NbL13g10800.1 ATP−dependent_Clp_protease_ATP−binding_subunit_CLPT1_chloroplastic−like [MW = 26.01 kDa]

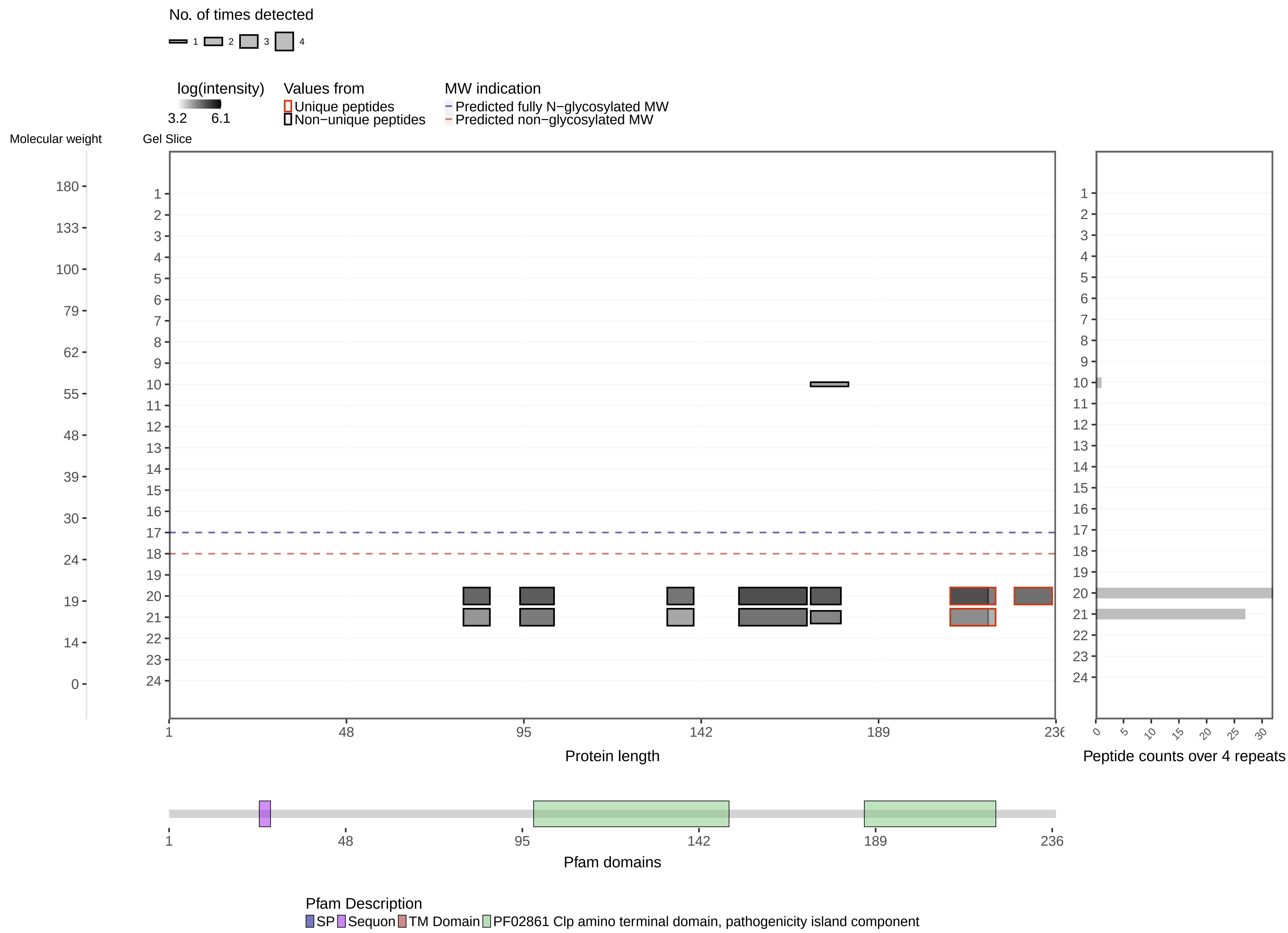


Protein sequence:

KYPNTGTEALLMGILVEGTSLAAKFLRANGVTLLKAREETVKLLGKSDMYFFSPEHPPLTEPAQRALDWAVDEKLKSGESGEITIAYLALGIWSEKESAGHKIMATLGFDDEKAKELARSMDKDIEMNYK*

NbL13g10590.1 21_kDa_protein−like [MW − SP = 18.49 kDa]

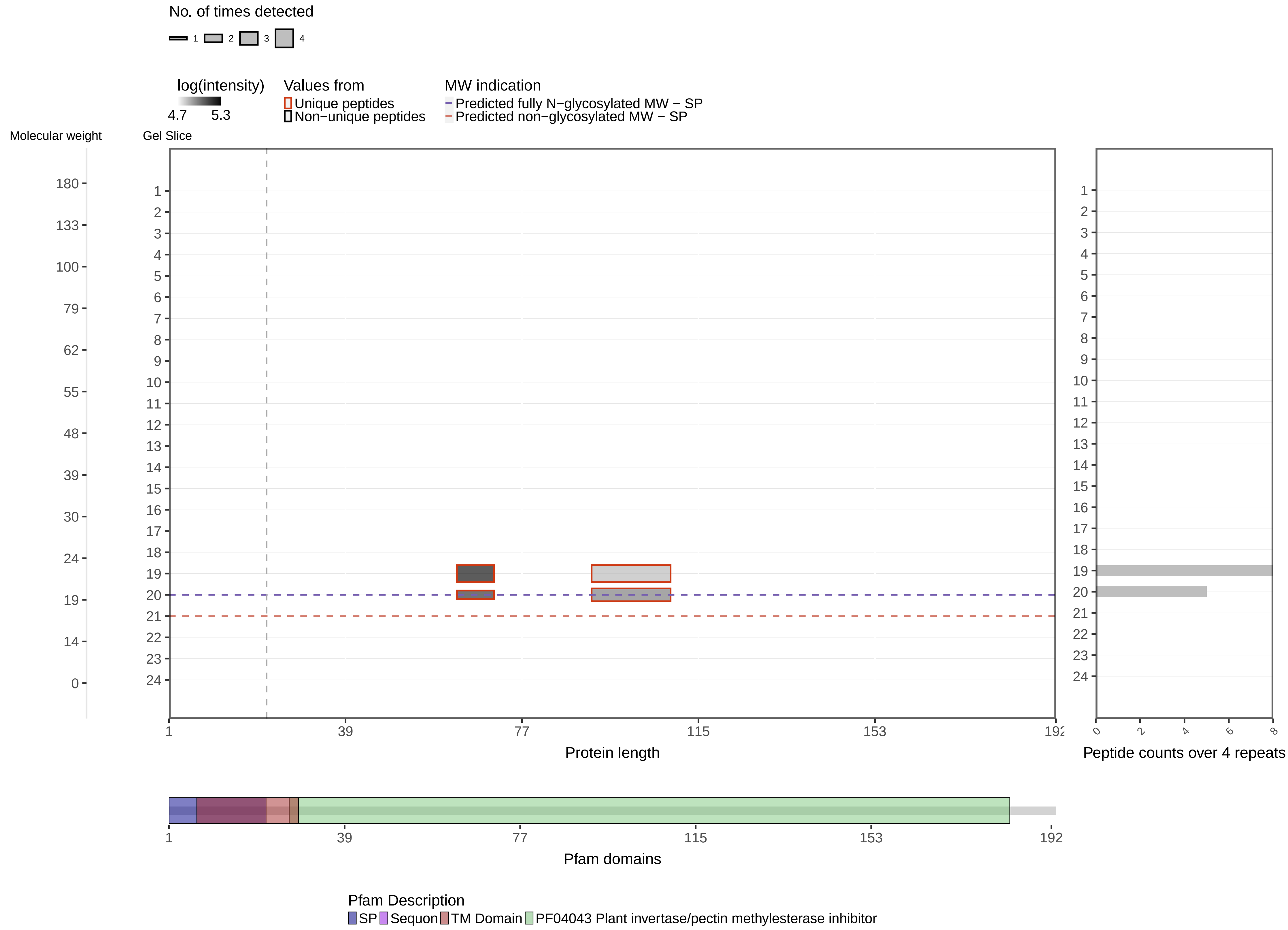


Protein sequence:

KFKGLKPREYAAIKDCIEETSDSVDRLSKSVSELKEVDHSRGKDFLWHVSNVETWVSAAITDENTCTDGFAGRALNGRIKASIRNRIANLAQVTSNALALINQYAAKH*

### NbL13g10550.1 21_kDa_protein−like [MW − SP = 18.58 kDa]

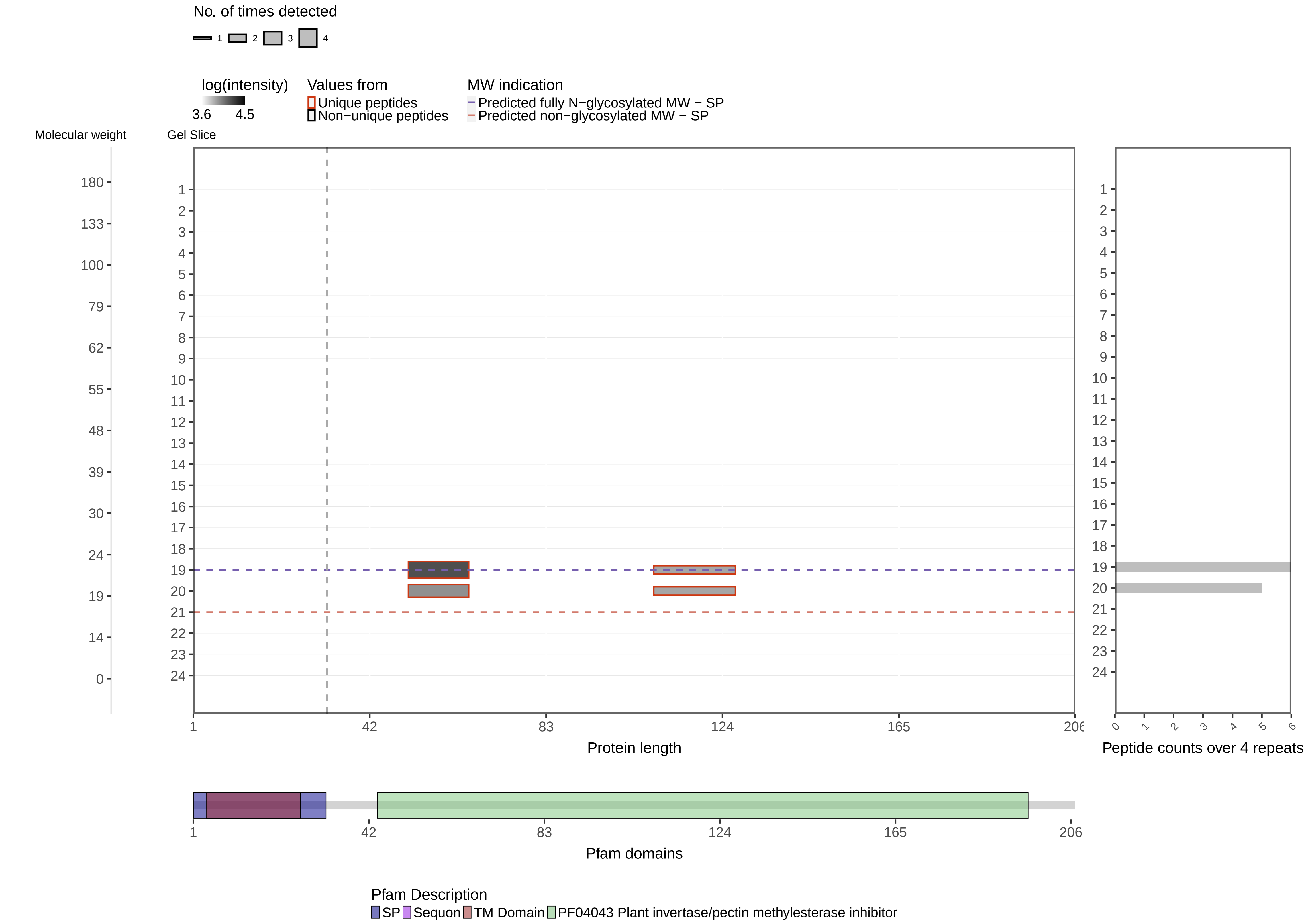


Protein sequence:

AMMLKLAHVRGLTPREVGAMHDCVEELSDTVDEMRKSLGEMKQLRGKDFDLKMNDIQTWVSAALTDEDTCTEGFAGKVMNGKVKTVVRGKILEVAHLTSNALALINSLAALHA*

NbL13g10470.1 monocopper_oxidase−like_protein_SKS1 [MW − SP = 63.57 kDa]

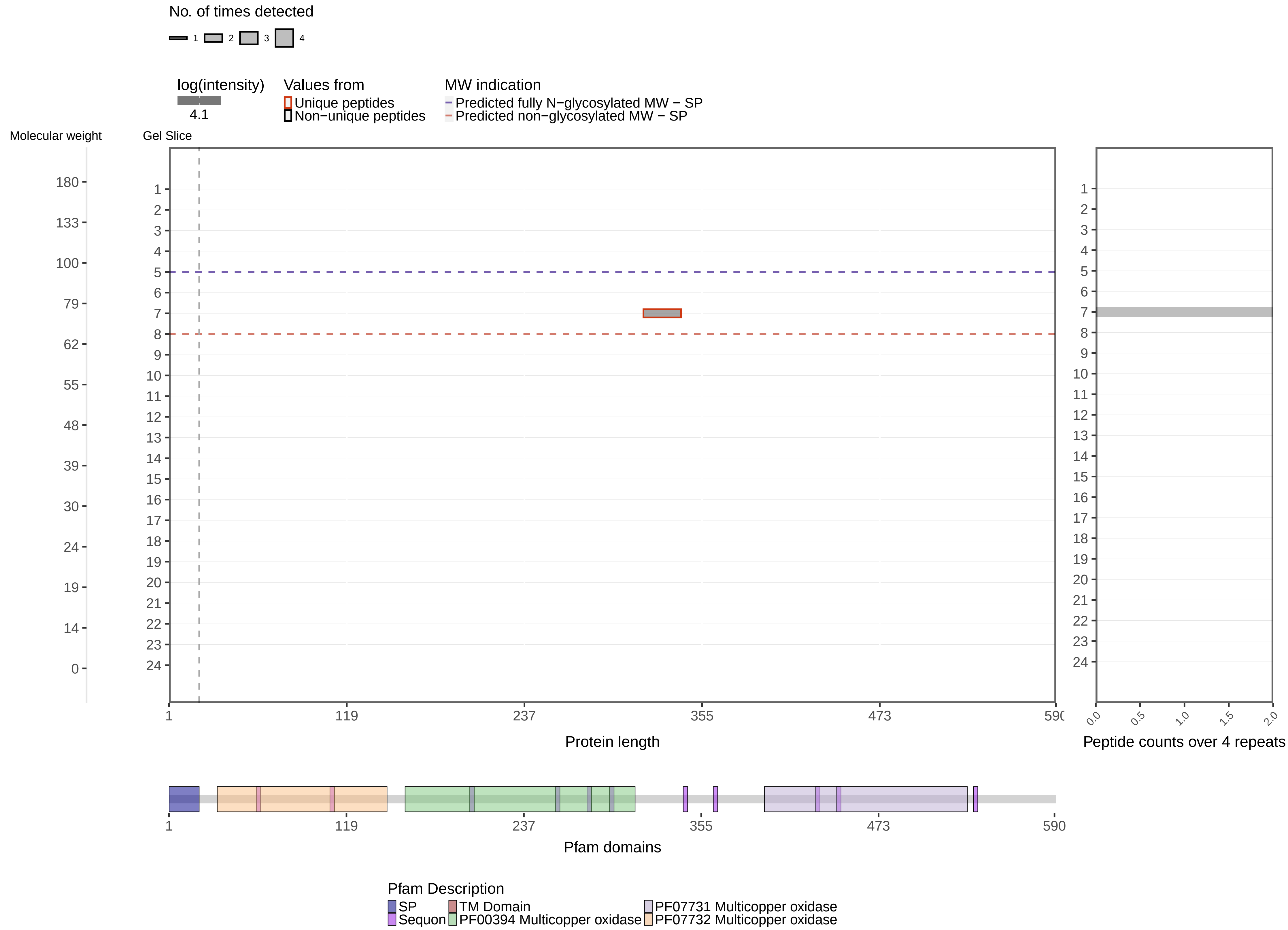


Protein sequence:

YYIVASPRFVNOSLWOKVTGVAVLHYSNSKGKAAGPLPDPPNDAYDTSYALNOAMSIRONVSASGARPNPOGSFHYGOINVTDVYLLKSVPPVVIDGKLRATYNGISFKNPETPIRLADVYHVKGDYKLDFPTKPMNRPPKVDTSLINATYKGFIEVILONNDTVAOSFHMDGYSFFVVGMGFGEWTENNRGSYNRWDAIFRSTTOVFPGGWTAIMASLDNVGVWNLRAENLDRWYLGOETYMRIINPEDHSNKTELPVPSNALYCGALASKOKPOKISAATTIFRSSEMY

NbL13g10450.1 acid_beta−fructofuranosidase_AIV−18−like [MW = 72.59 kDa]

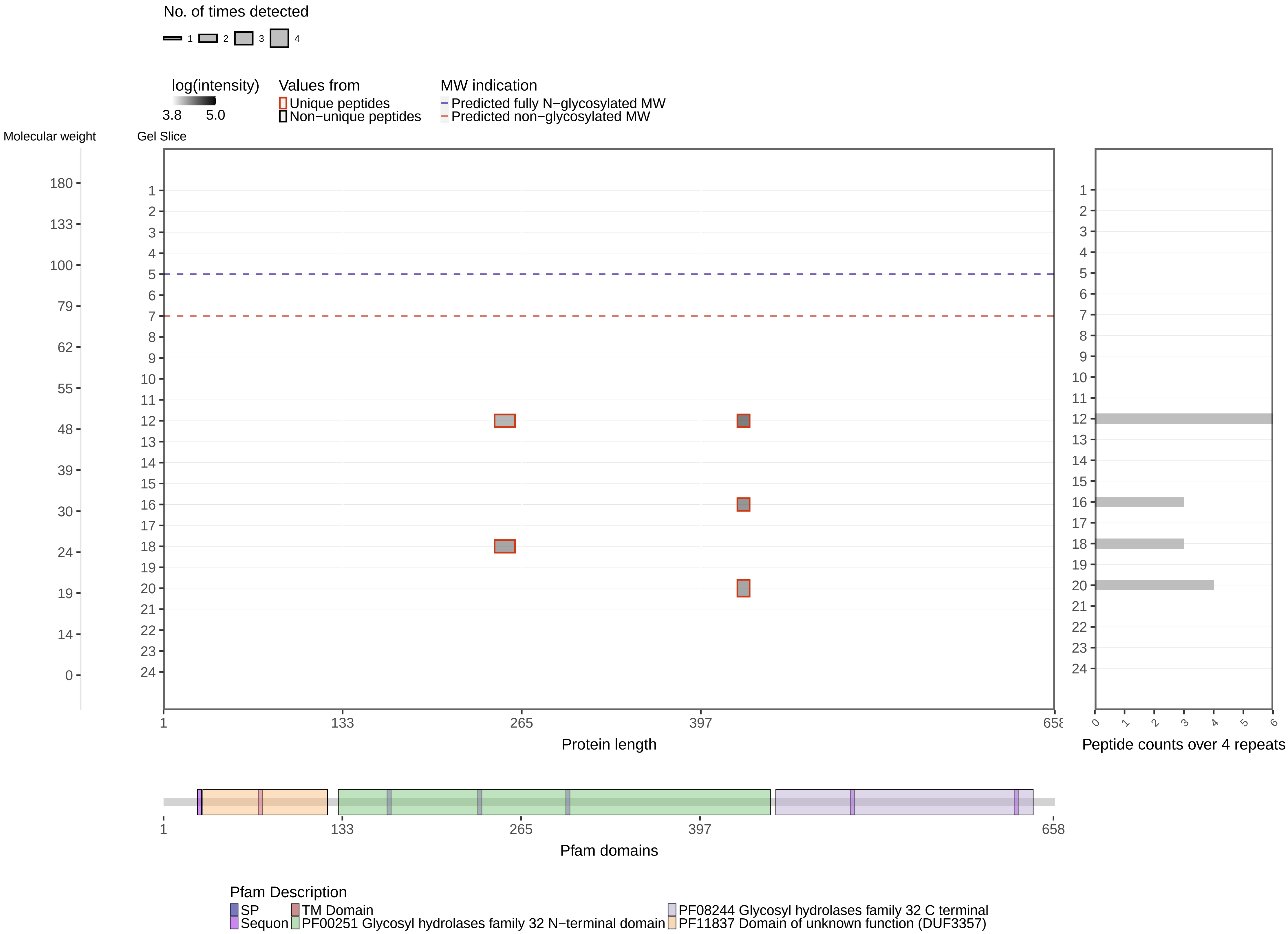


Protein sequence:

GTGMWECVDFYPVSTVEANGLDTSYNGPGIKHVLKASLDDDKHDYYAIGTYDPVKNKWTPDNPELDVGIGLRLDYGKYYASKTFYDPKEQRRILWGWIGETDSEAADLLKGWASVQSIPRTVLYDKETRTHVLQWPIKEIESLRIGDPIMKQVNLQPGSIELVHVDSAAQLDVEASFEVDKAALEGTIEADVGFNCSTSGGAAKRGILGPFGVVVIADQTLSELTLVYFYIAKGTDGRAETYFCADETRSSEAPGVAKQVYGSSVPVLDGEKHSMRILVDHSIVESFAQGGRT

NbL13g10120.1 probable_isoaspartyl_peptidase/L−asparaginase_3 [MW = 40.51 kDa]

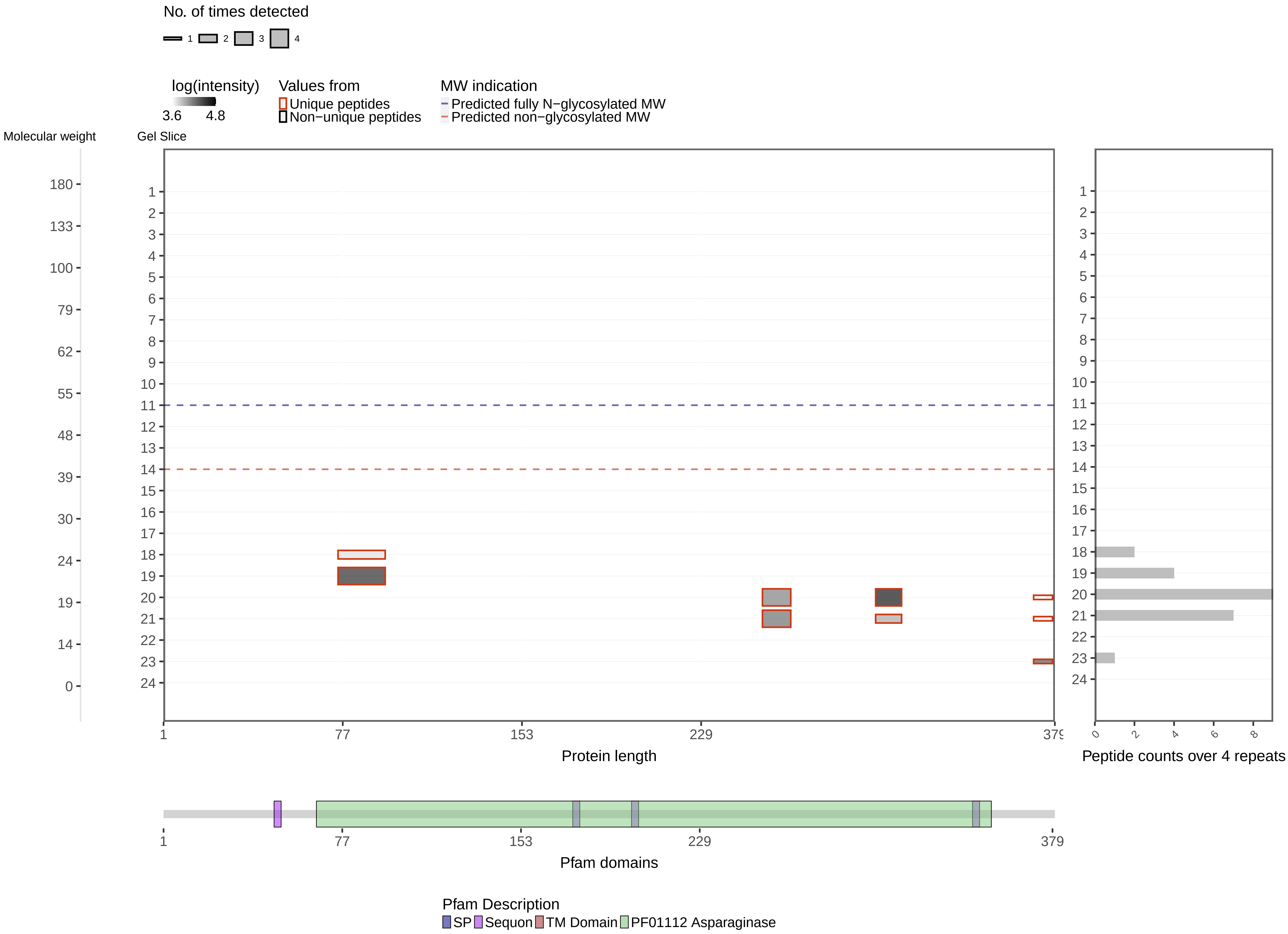


Protein sequence:

SVESTEKWMKWKENFCQPNFWKNVSPGDNCGPYHPNGPSIGRCLMANELQPNEFGSVNVGLHSHDTISMAVIDEIGRIAVGTSTNGATLKIPGRVGDGPIAGSSAYADSEVGACGATGDGDIMMRFLPCYQVVESMRLGMEPKLAAKDAISRIARKYPGFIGALFAVNRSGIHAGASHGWTFQYSVRNPGMKDVEVFTVHP*

### NbL13g10040.1 ATP−dependent_6−phosphofructokinase_4_chloroplastic−like [MW = 64.33 kDa]

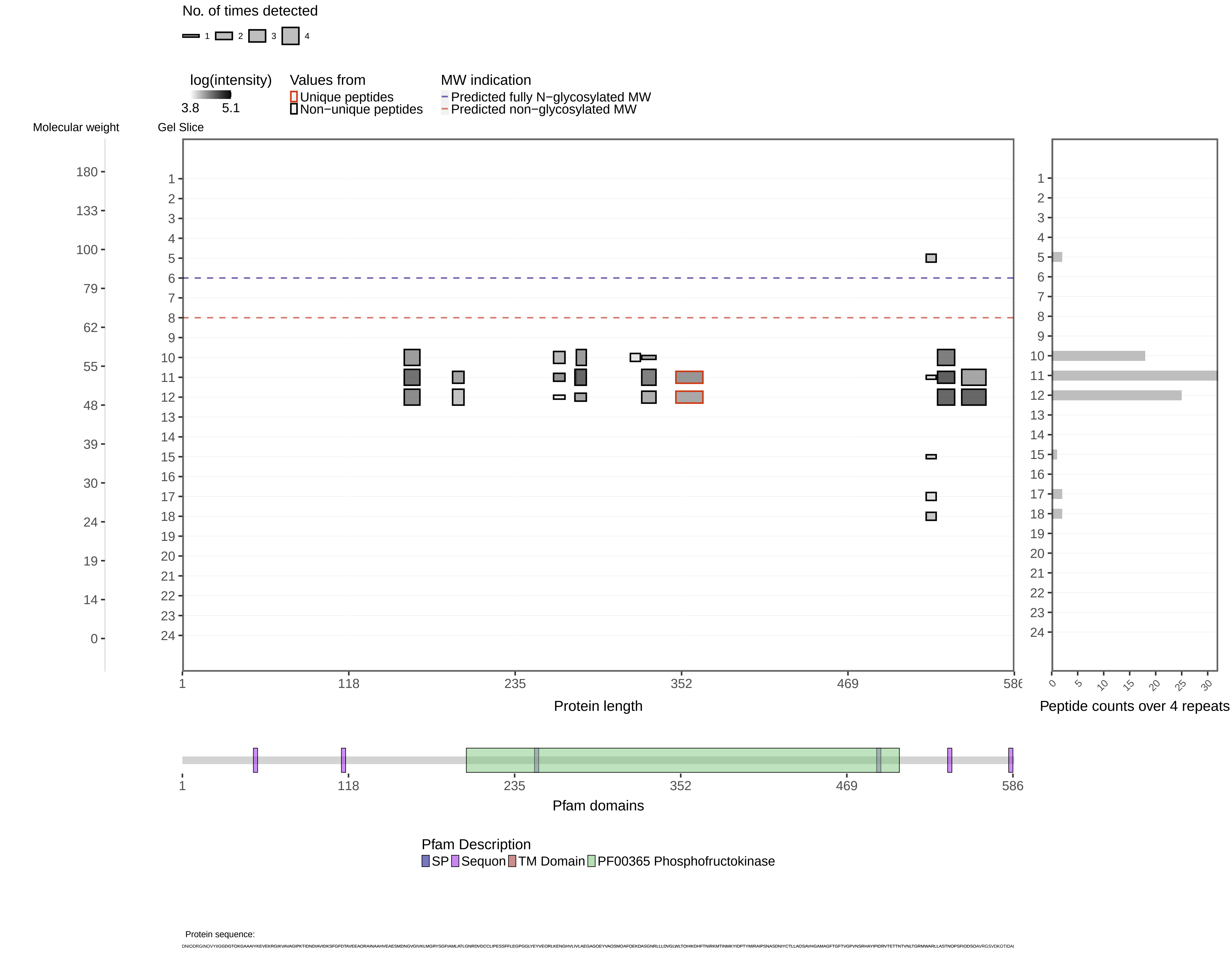

### NbL13g09900.1 protein_mago_nashi_homolog [MW = 17.79 kDa]

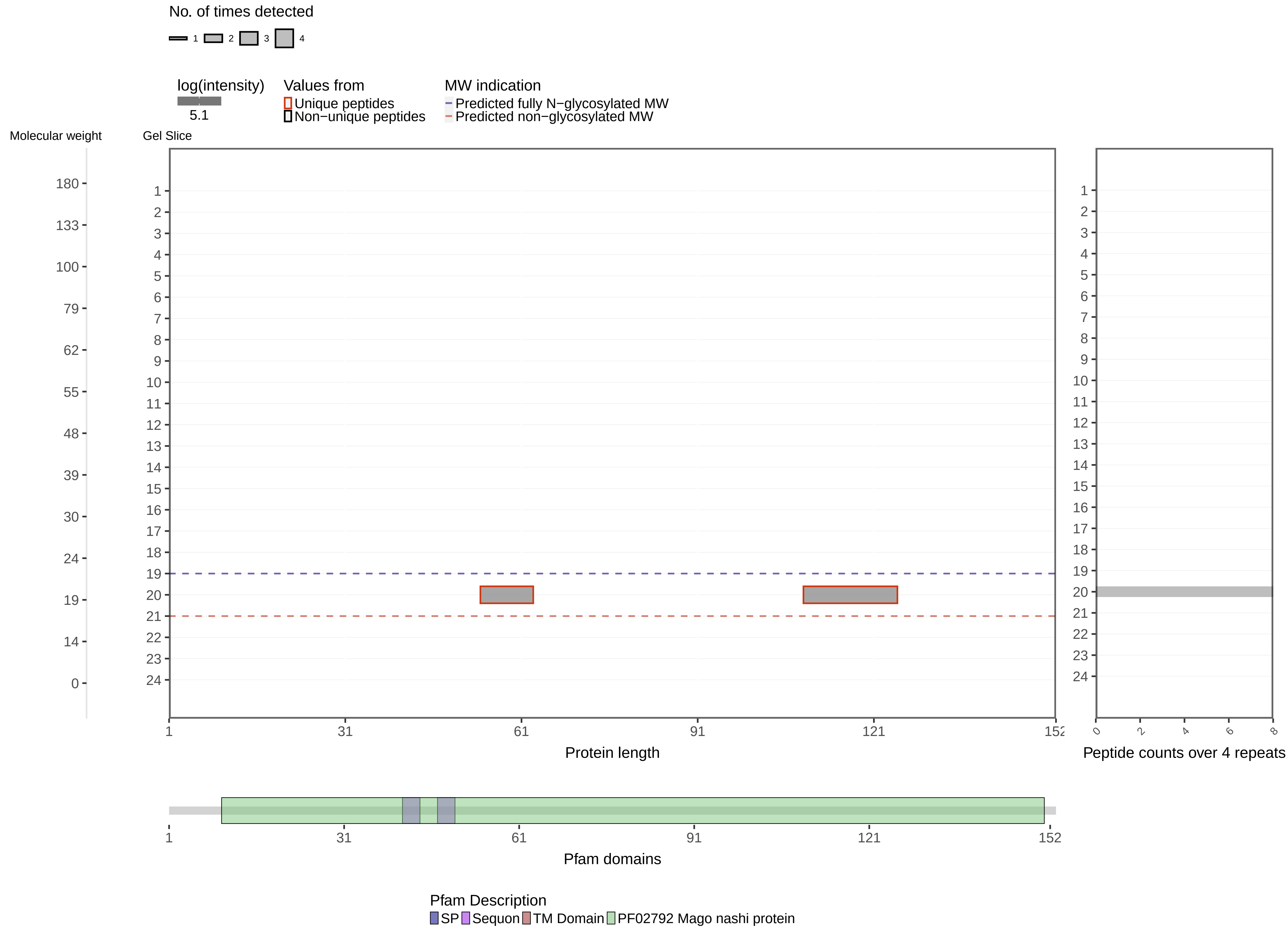


Protein sequence:

LKECRRIVADSEIMKEDDNNWPEPDRVGRQELEIVMGNEHISFTTSKIGSLMDVQTSNDPEGLRIFYYLVQDLKCFVFSLISLHFKIKPI*

### NbL13g09890.1 peptide_methionine_sulfoxide_reductase_A1−like [MW = 28.76 kDa]

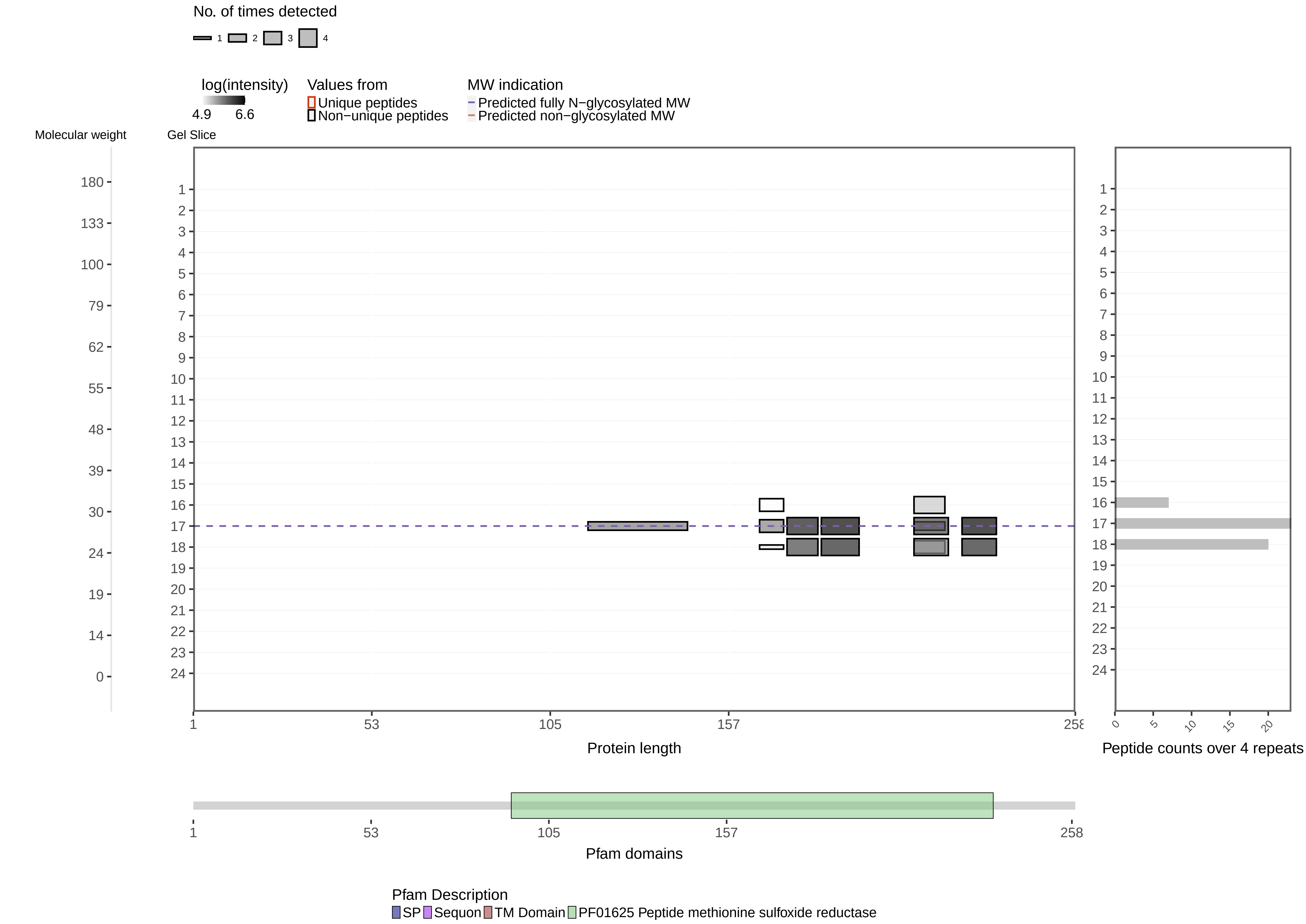


Protein sequence:

EVGYTQGYLHNPSYEDICSGTTFHSEVVRVQYDPKECNFDTLLDVFWDRHDPTTLNRQGNDVGTQYRSGIYFYTPEQEKAAIESRDKQQKILNRNIVTEILPAKKFYRAEEYHQQYLAKGGRFGFRQSTEKGCNDPIRCYG*

### NbL13g09800.1 glutamate_dehydrogenase_A−like [MW = 44.82 kDa]

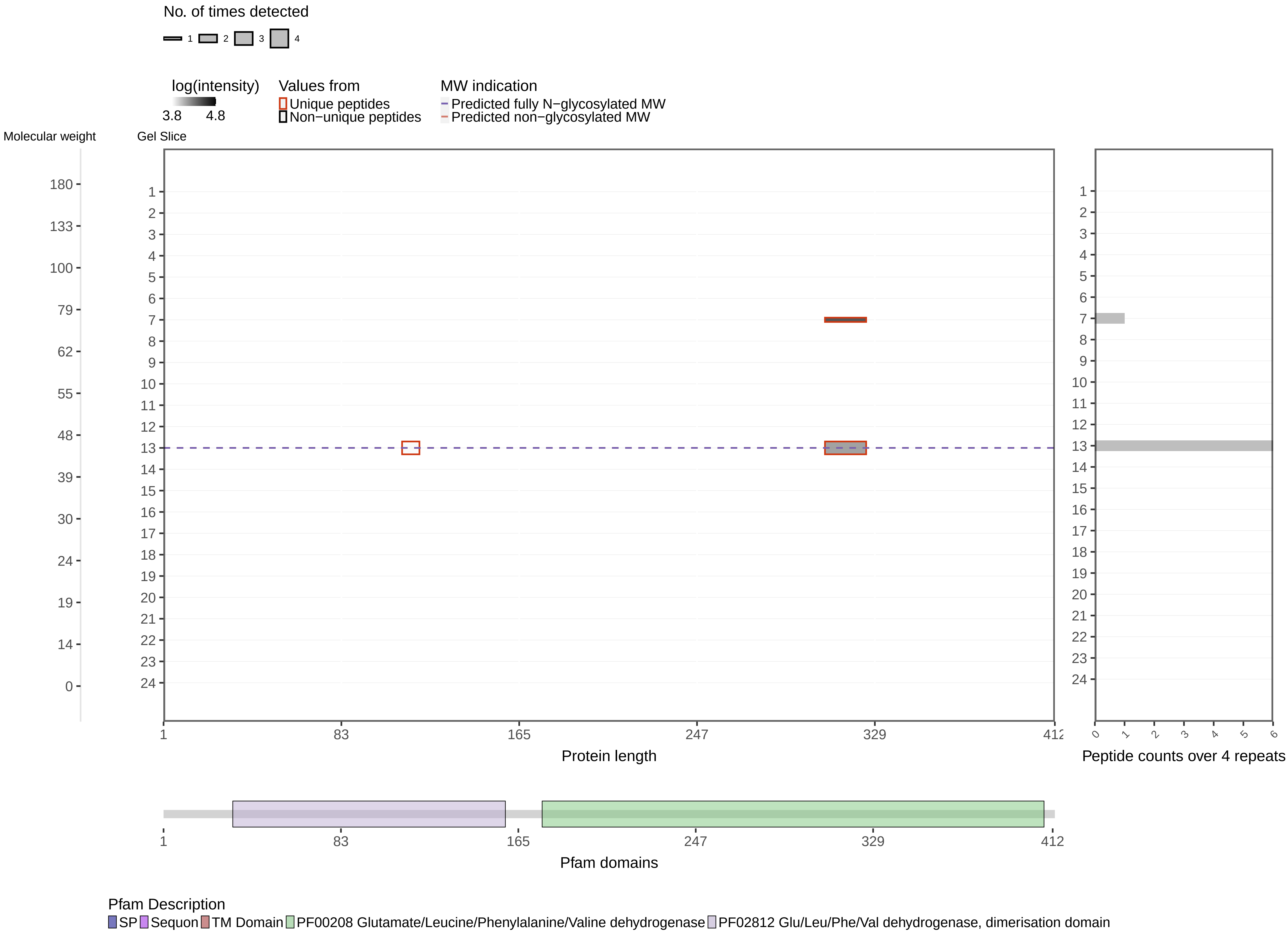


Protein sequence:

EALLAEYGKNIKDLTFAIQGFGNVGAWAGKIIHERGGKVIAVSDITGAIKNPNGLDIPALLNHREKTGKLTDFNGGDVMNSDELLTHECDVLIPCALGGVLNRENADHVKAKFIIEAANHPTDPDADEILSKKGVVILPDIYANAGGVTVSYFEWVQNIQGFMWDEEKVNAELKKYMTRAFHNLKSMCHSHNCNLRMGAFTLGVNRVARATQLRGWEA*

NbL13g09770.1 polyadenylate−binding_protein_1 [MW = 25.58 kDa]

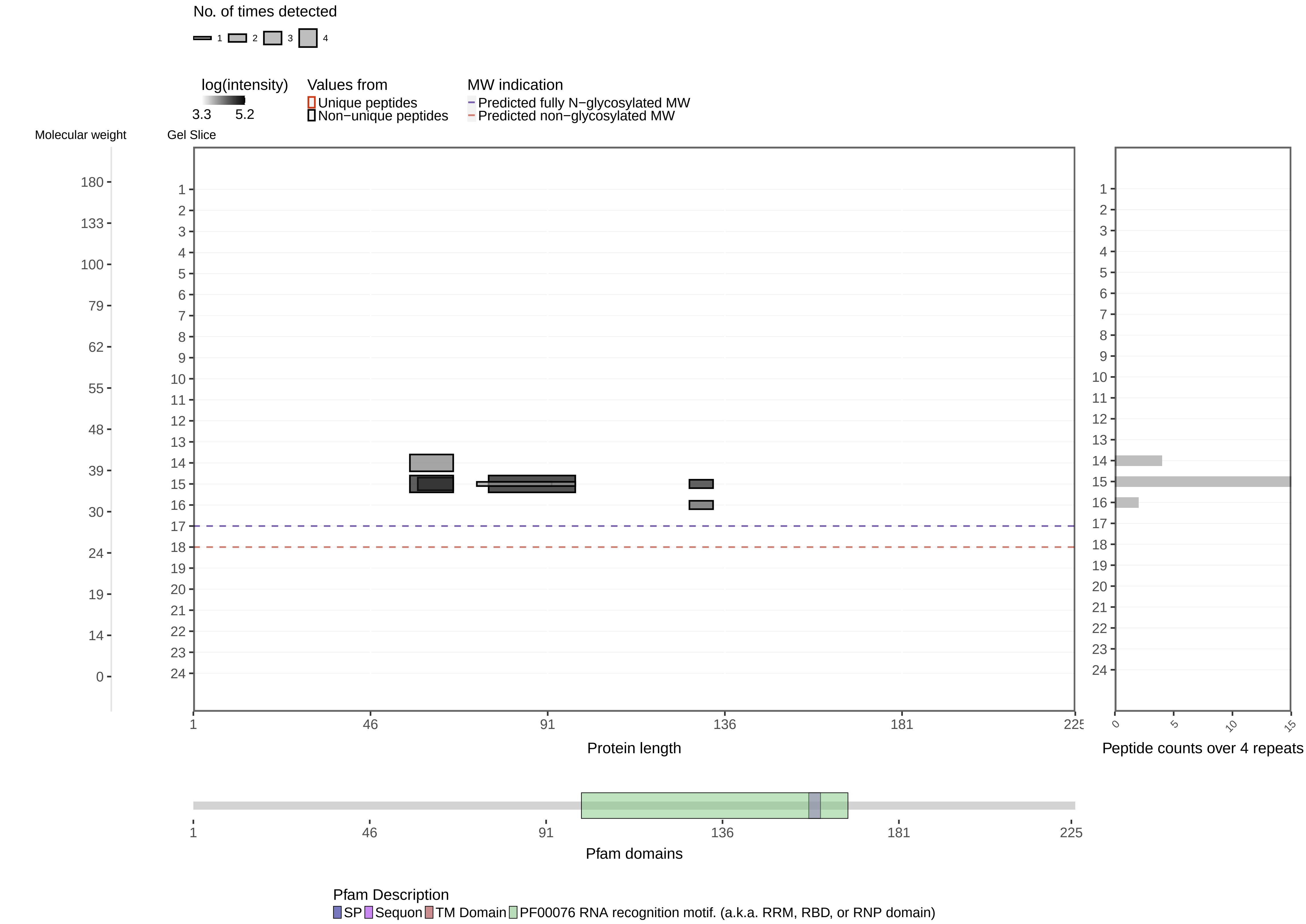


Protein sequence:

SIYVGNVDYACTPEEVQQHFQSCGTVNRVTILTDKFGQPKGYAYVEFVEVEAVQNALLLNESELHGRQLKVSAKRTNVPGLKQFRGRRFNPYAGFRPRRPFMPGAPVFPPFGRIPRFRRPTRYRPY*

### NbL13g09730.1 putative_pterin−4−alpha−carbinolamine_dehydratase [MW = 23.23 kDa]

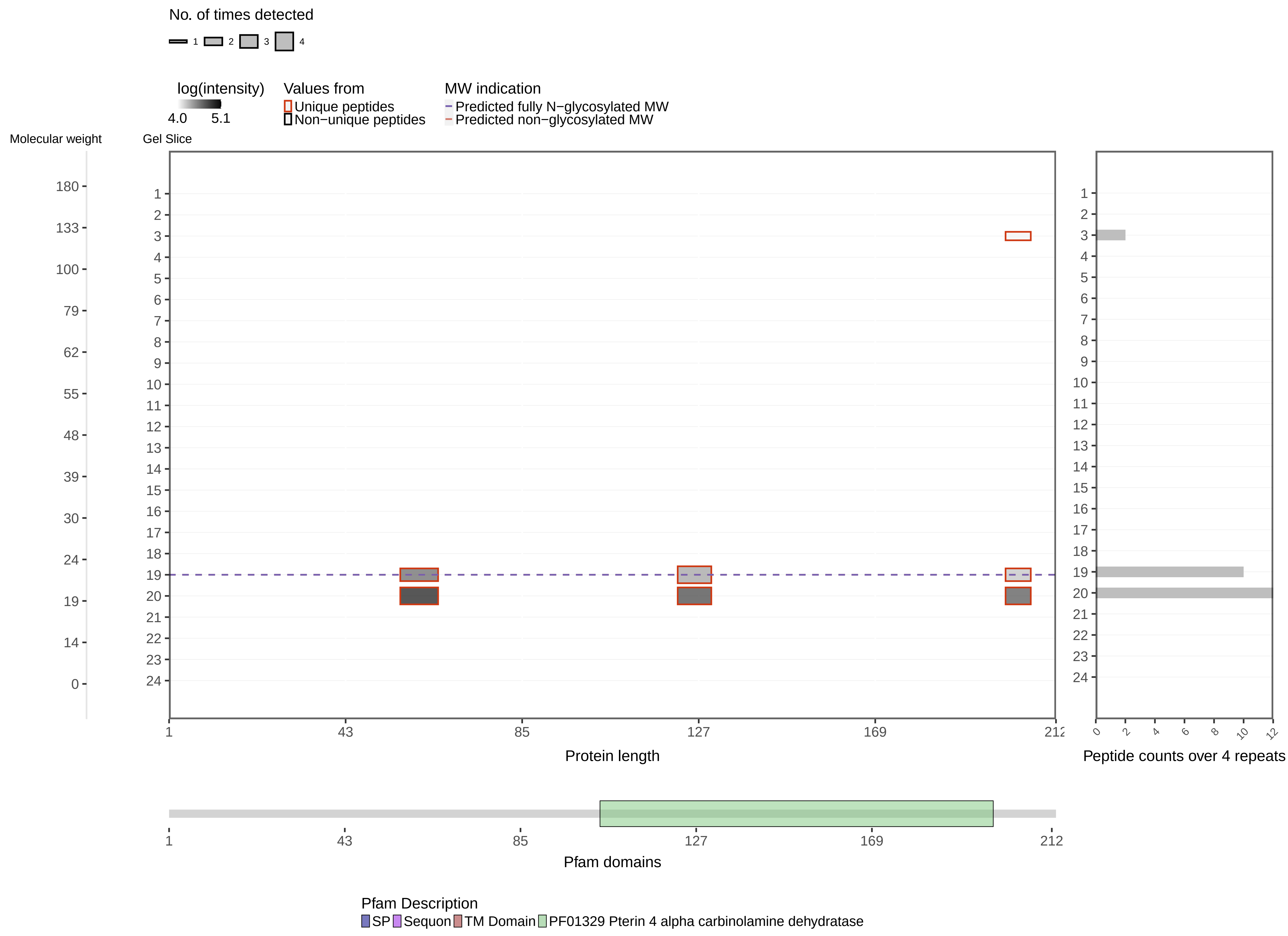


Protein sequence:

ACIVISPGQTPLSEYEARQLLFKVVGWRLAHEDGVLKLQCTWKLRDFECGVELINRIGKVVEGTGHVPTLLHLEQSNQVRAELWTASIGGLSINDFIVAAKIDQIKTSDLVPRKRVWA*

### NbL13g09710.1 superoxide_dismutase_[Fe]_chloroplastic [MW = 34.78 kDa]

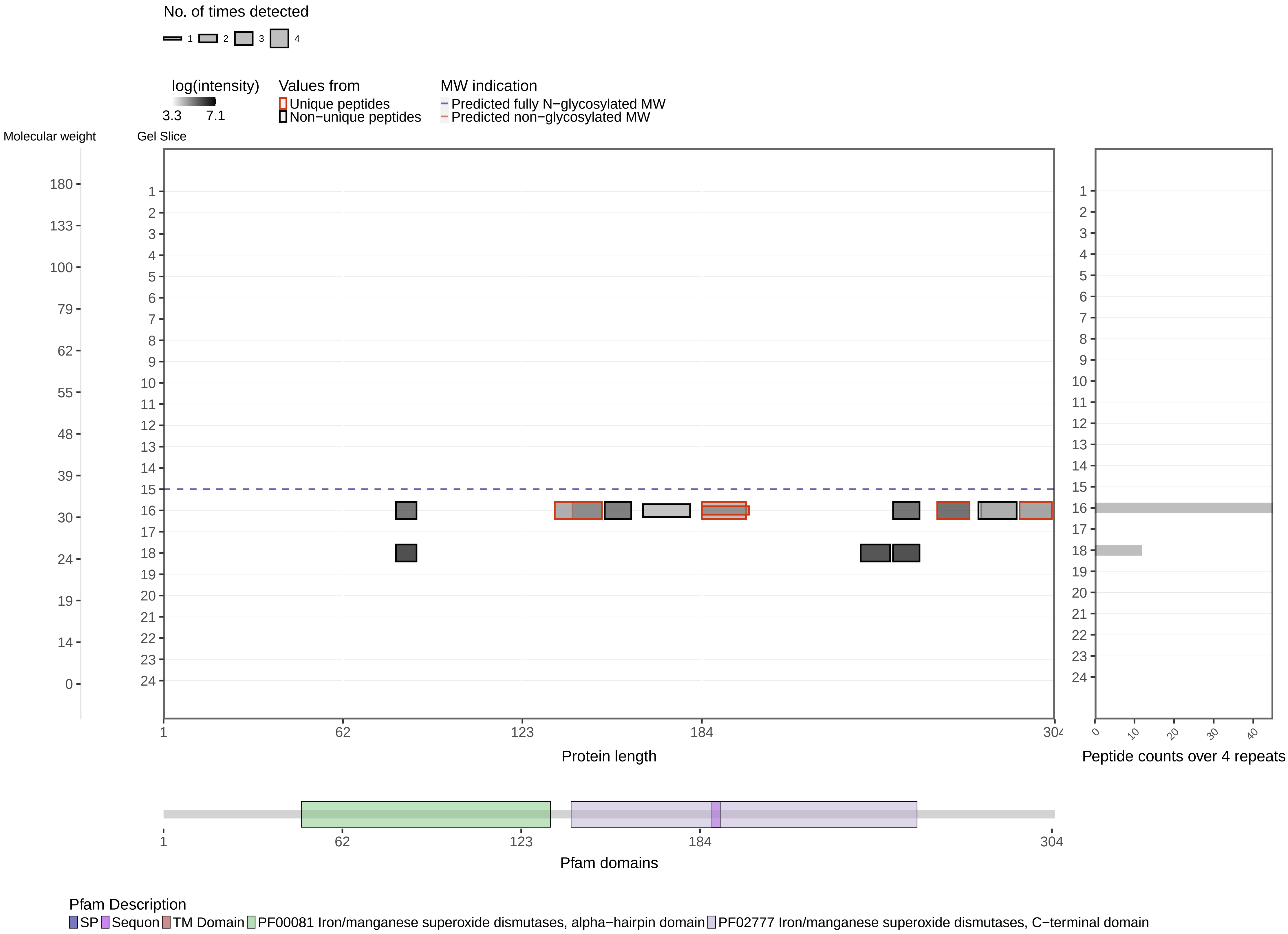


Protein sequence:

KPSGELFELINRDFGSFEAFVKEFKAAAATQFGSGWAWLAYKANRLDVGNATNPHPSNEDKKLVVVKTPNAVNPLVWDYSPLLTIDVWEHAYYLDFRNRRPDYISIFMEKLVSWEAVSSRLEVAKAQAVEREEEEERKKREEEEEYKAGGEVREMYVESTDSDAE*

NbL13g09610.1 quinone_oxidoreductase−like_protein_2_homolog [MW = 37.02 kDa]

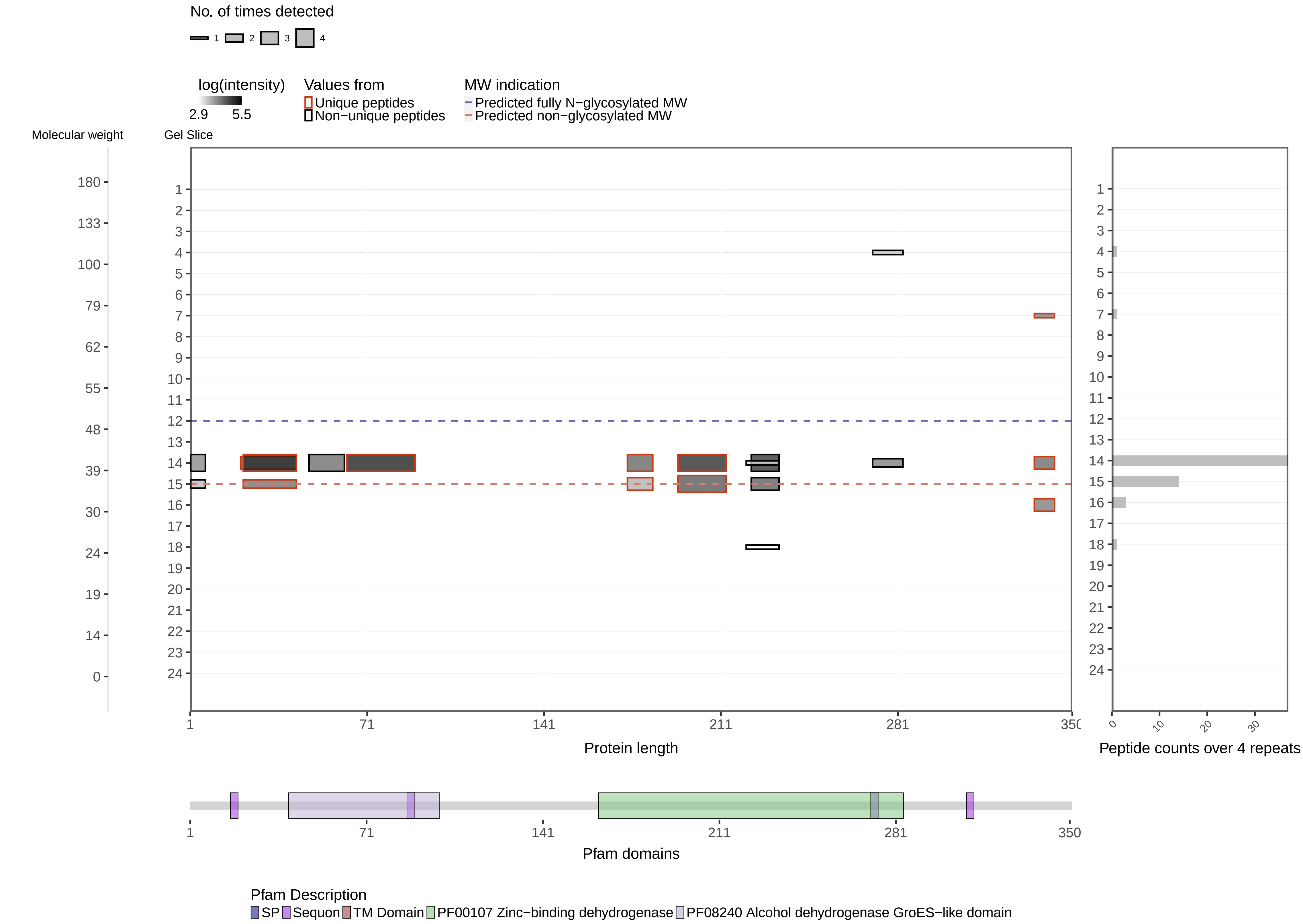


Protein sequence:

AGGVGLSAVQIGKVCGATVIAVARGNEKVQFLKSLGVDHVVDLSNANVIESAKGFLKSRKLKGVDVLYDPVGGKLTKDSMKLLNWGAQILVIGFASGDVPVIPANIALVKNWTIHGLYWGSHKIYQPNVLGDSLTELLSWLSNGLITINISHAFSLAEAHLAFTAIKDRRAIGKVMITFGDGRTLKSKL*

### NbL13g09550.1 polygalacturonase_inhibitor−like [MW − SP = 34.4 kDa]

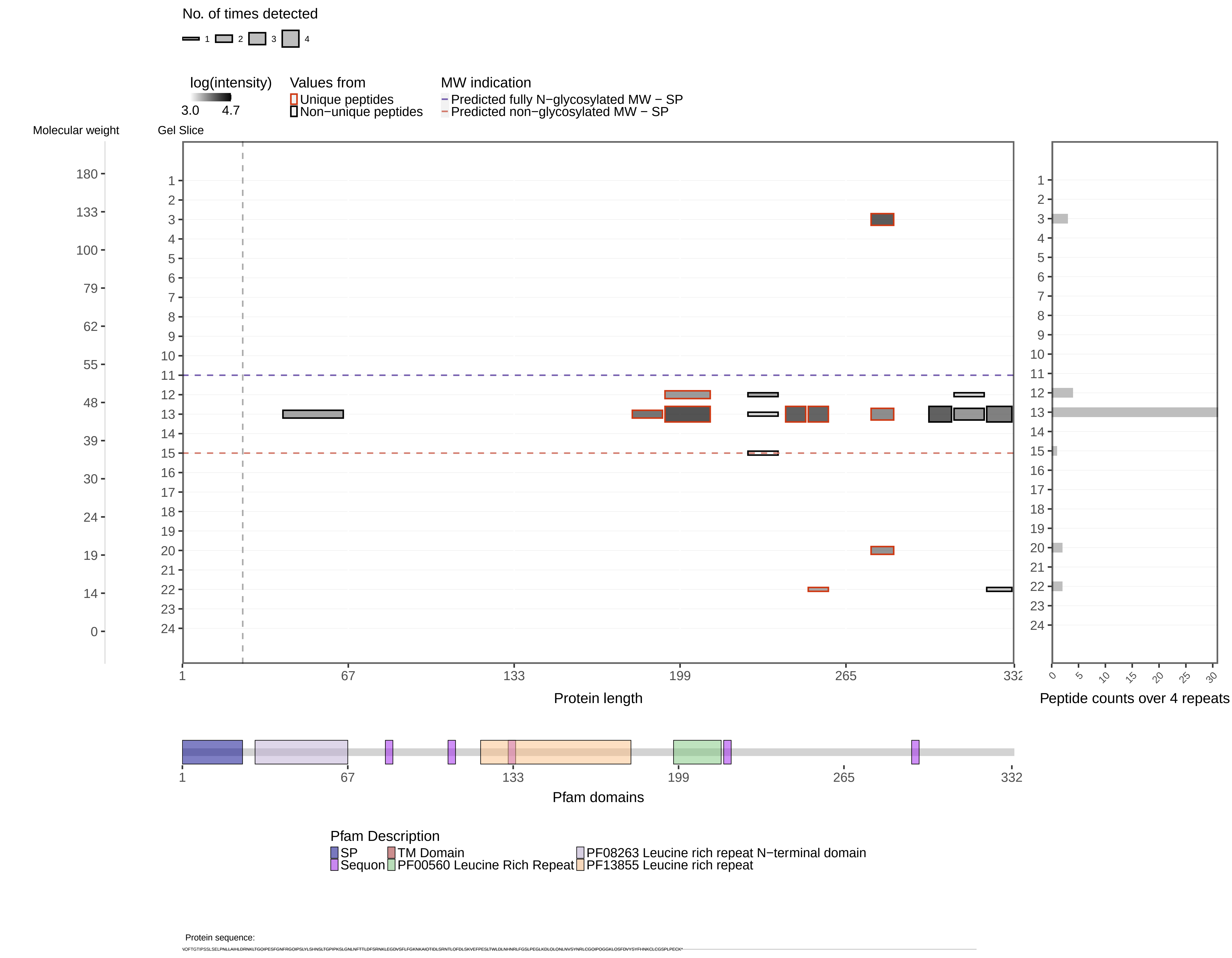

NbL13g09430.1 UBP1−associated_protein_2C−like [MW = 40.33 kDa]

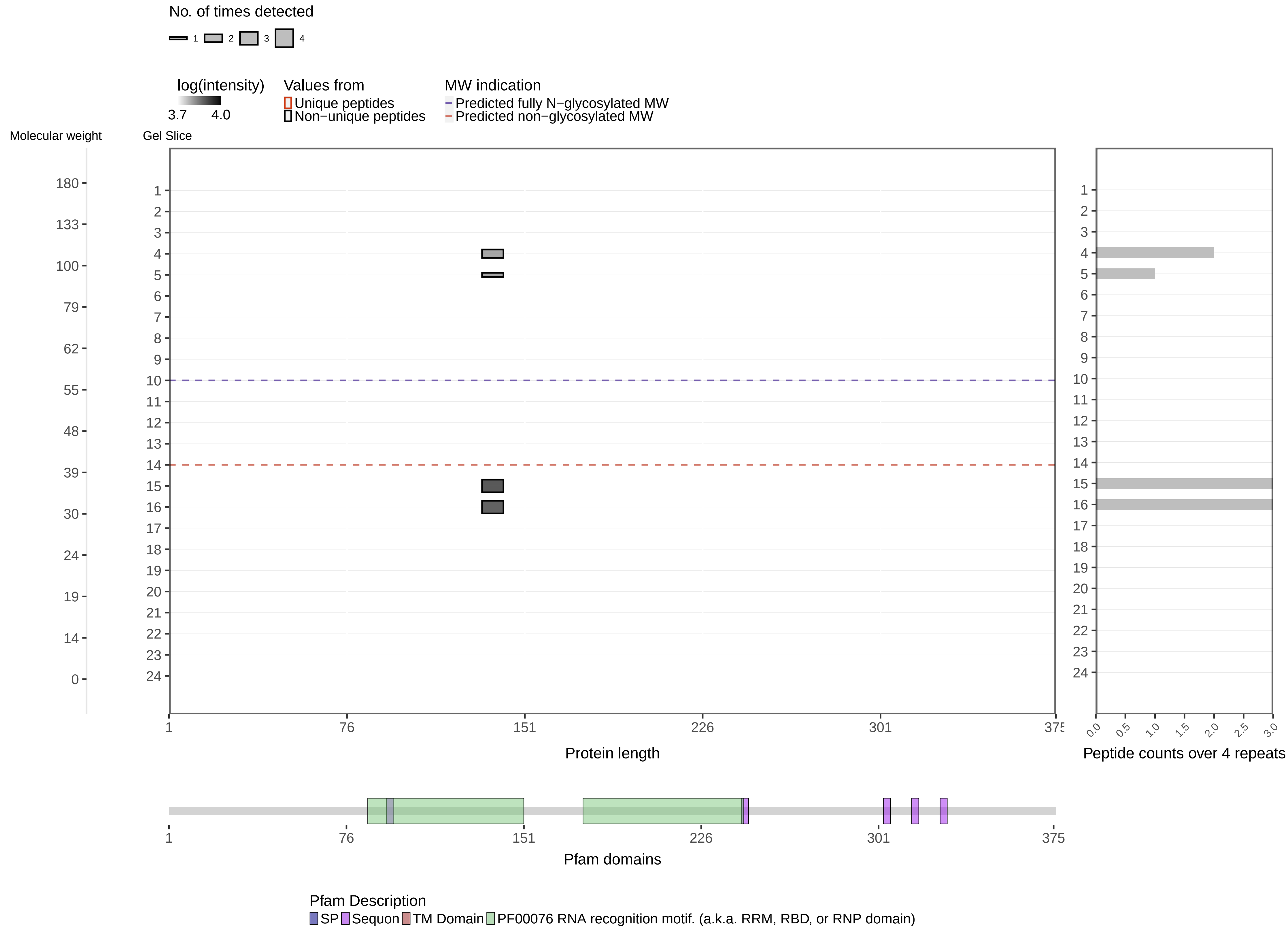


Protein sequence:

YIGGLSSDTTSEMLLSFFSRHGEIEEGSVAYNKDTKTSRGFGFVTYKTVEGAKKALNDPQRILGGRNLTVKLADNHKGKITQSQVRATMVPMSMPLAPGYPQAQKAHPGPAAPVGYGYPQTMAAYTNSTYSSQPAAYTNSTYSSQPAAYTNSTYSSLPGAAPPYSGQPQFPYPQYPVKQESQTTPASFGGYPYYMGKQ*

### NbL13g09280.1 NA [MW = 14.51 kDa]

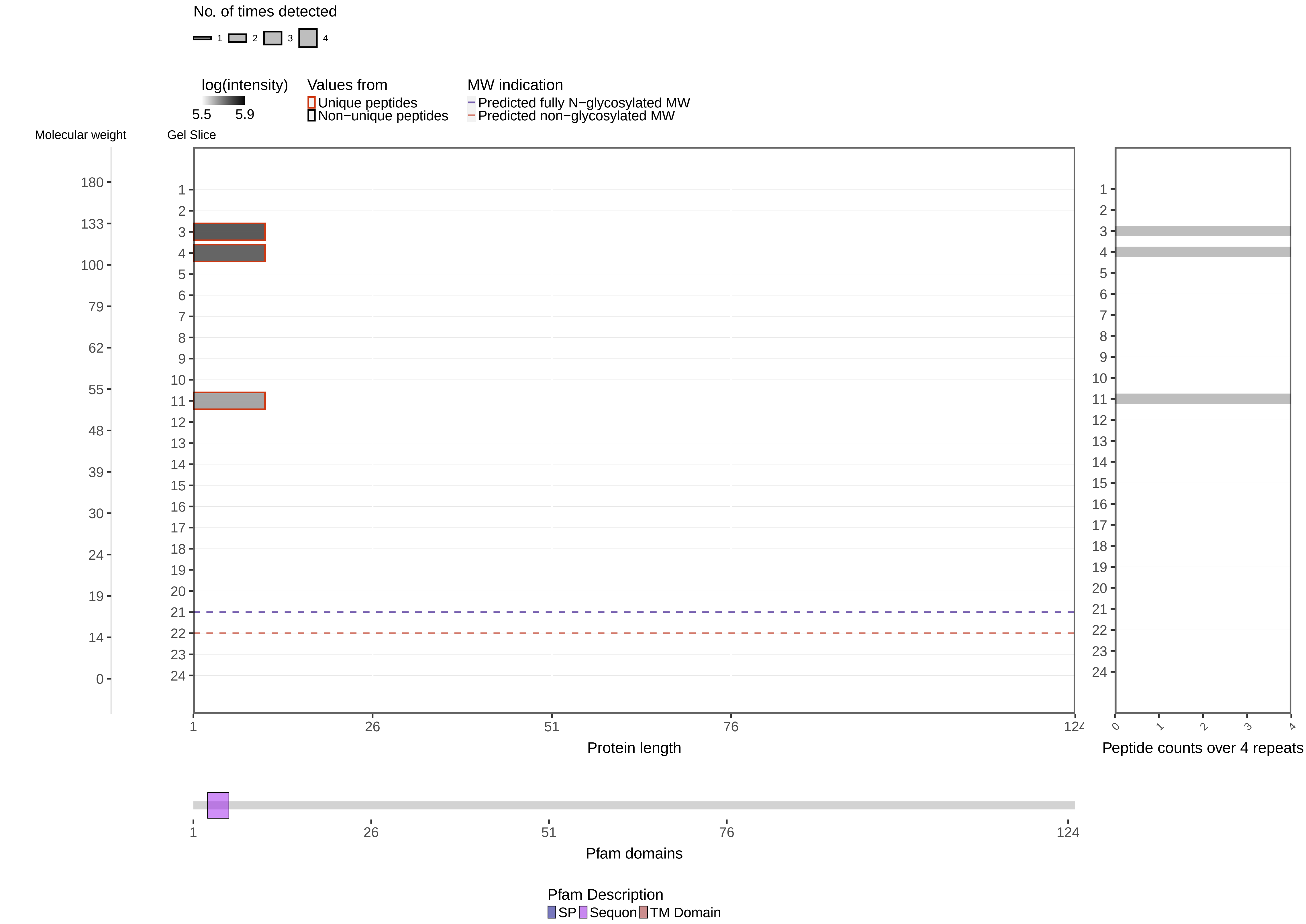


Protein sequence:

FFFSFFFFILLSSSLFSSRFSLFFFLCSSSFLSFSHEWSPAKSTNIFFFFLLFFLYPVSGVQSSPHIFVFSFVLPL

### NbL13g09190.1 serine_hydroxymethyltransferase_mitochondrial−like [MW = 49.62 kDa]

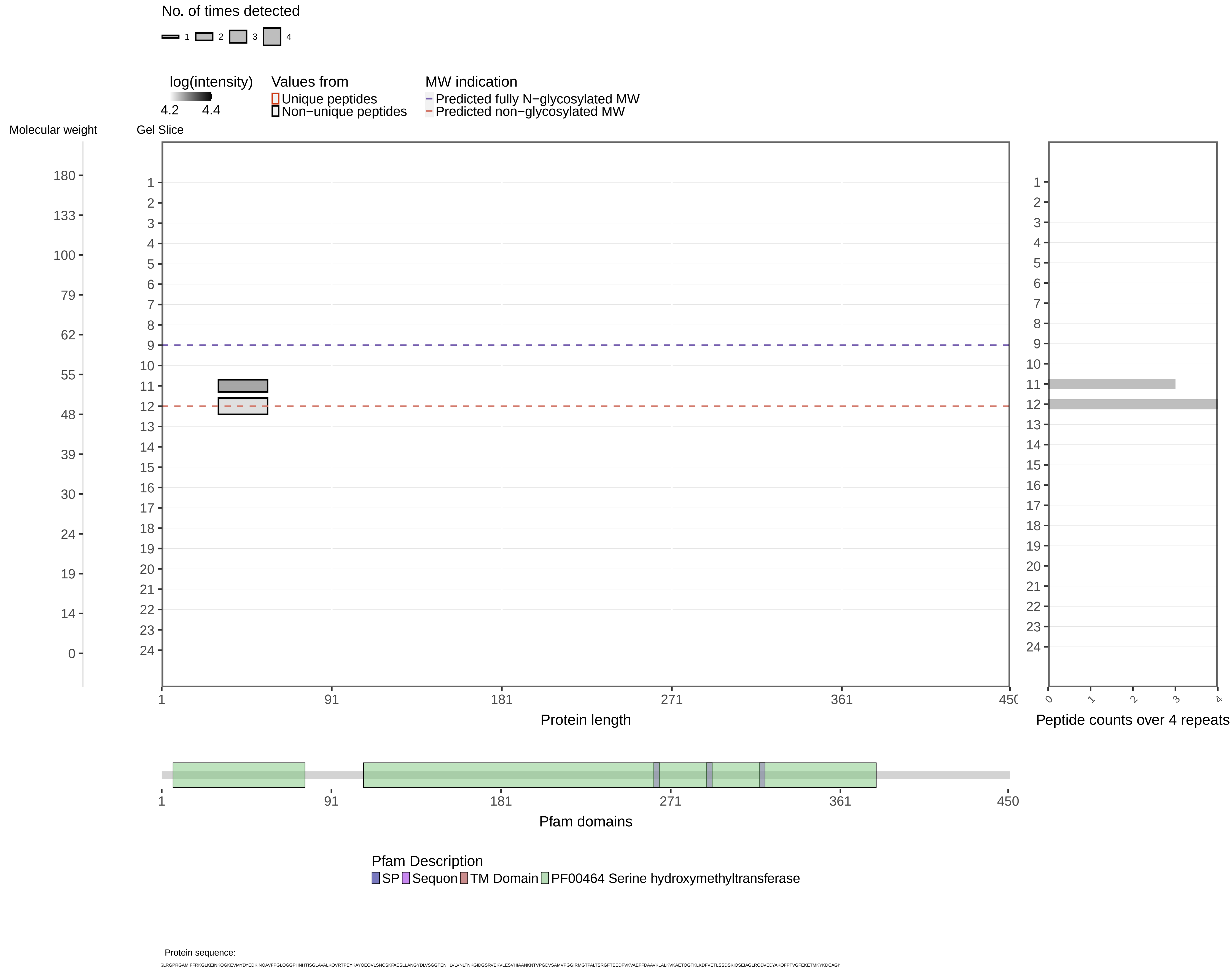

### NbL13g08950.1 annexin_D5−like [MW = 16.83 kDa]

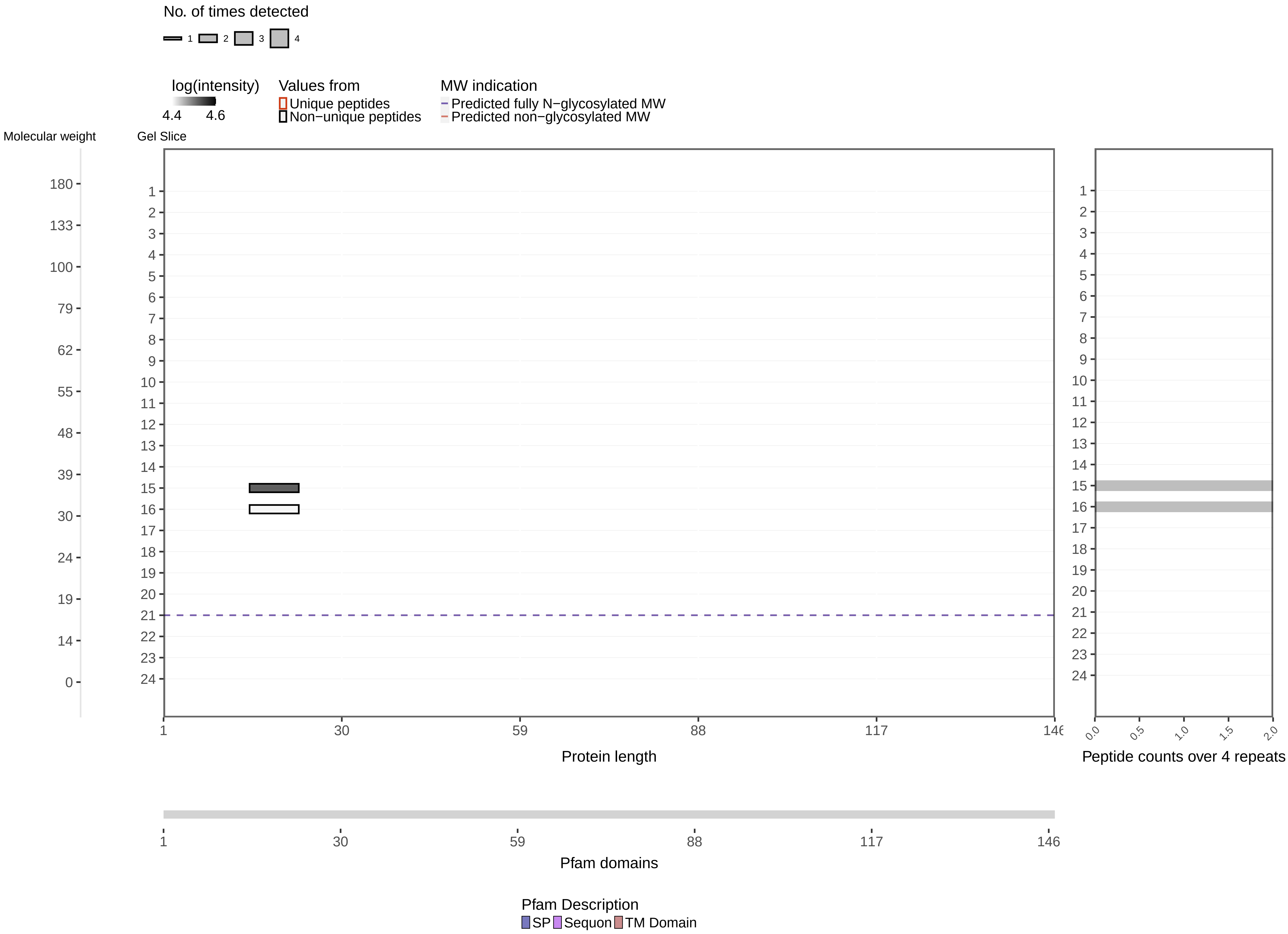


Protein sequence:

CQPRTHFQSNQEEKNICQNSFSTFSPHILYPYPHQHSYLPPNPIPTPFPATTNPPHHTHSPPLLPSTPLRLTPYPATTNPPHPTPR

NbL13g08910.1 uncharacterized_protein_At4g26485−like [MW = 21.05 kDa]

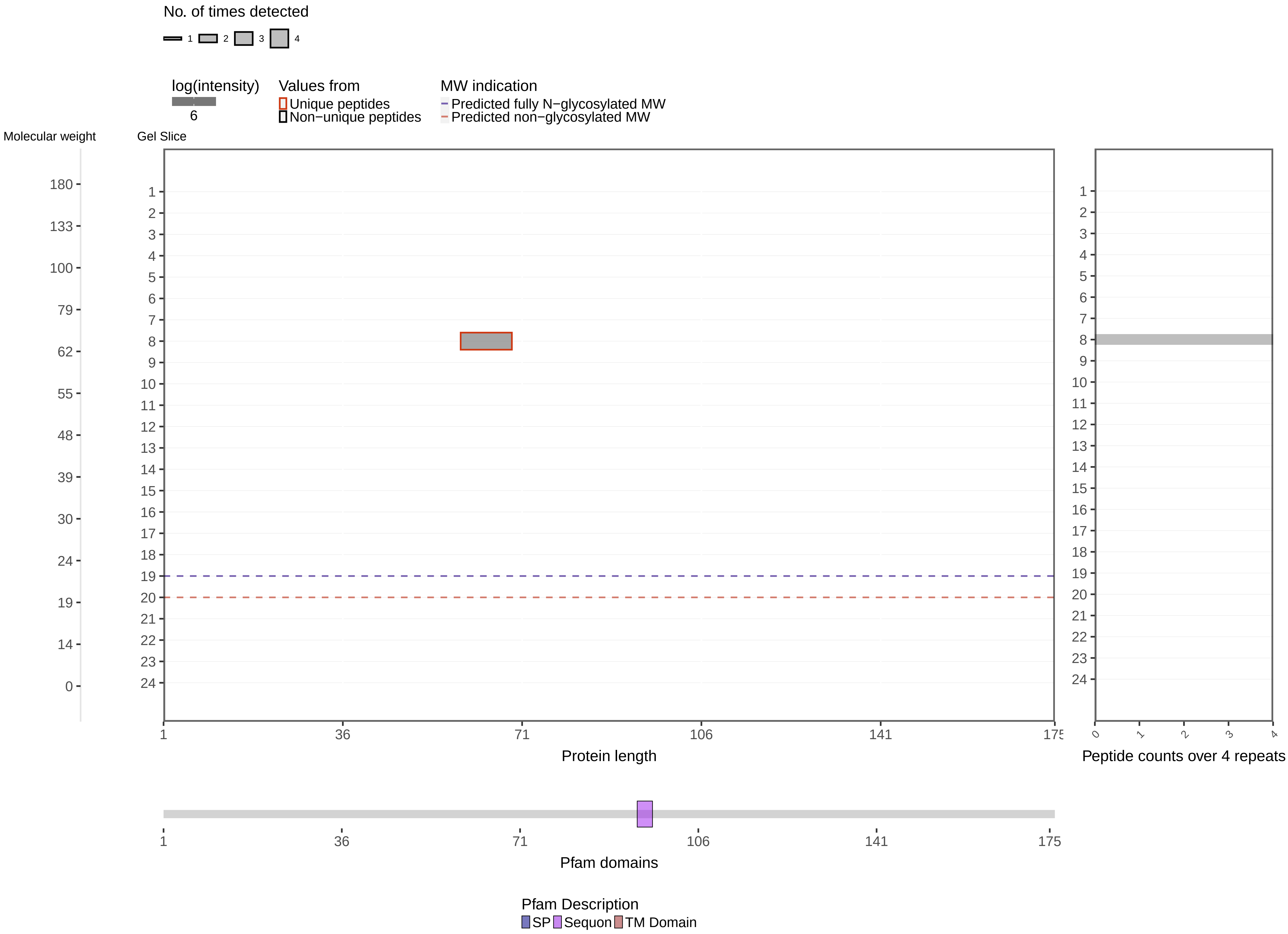


Protein sequence:

GLPARHDHRSKYFMFFKRNFSYIQQTFGRKDIDVEVEVRKAMQRASLMFRAENGWLPDPYLKILDELQFWKQSRFFRLEQMLLDIDRRLYELQGMGYYL*

NbL13g08870.1
enhancer_of_mRNA−decapping_protein_4−like_(LOC104229453)_transcript_variant_X1_mRNA_XM_009782105.1_PREDICTED:_Nicotiana_sylves
[MW = 153.02 kDa]

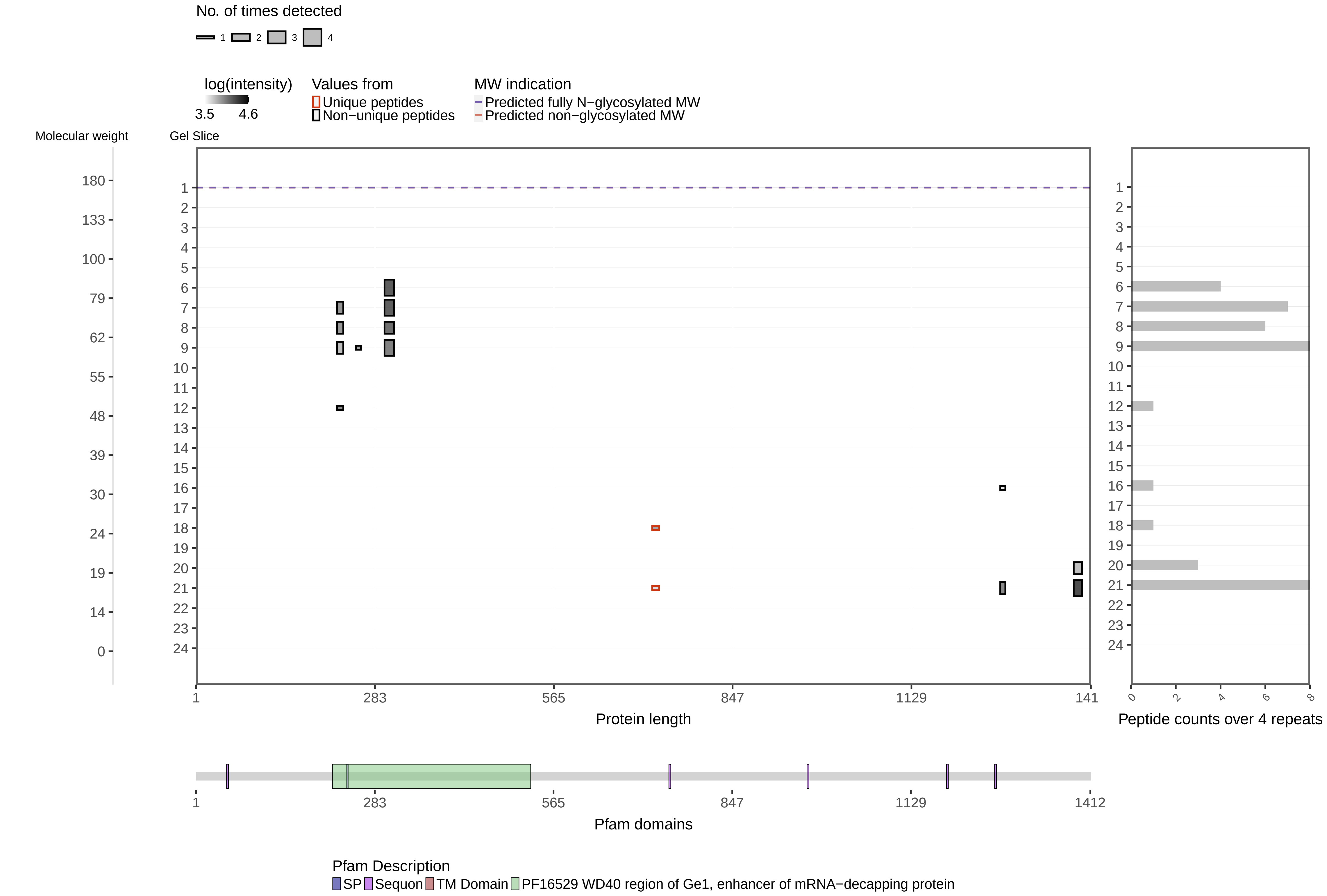


Protein sequence:
PPLSPTLSRKLSGFRGPSNSFERGTSYNEQVGDPKVVEYSVDRQKEGTTPNVSDVTSLDDEPKNDESKQSQNDVPSGISPSVKFKHPTHLVTPSEILMARSSSEVNIVNEQKSESELNIQDVVINNDARNVEVDVKVIGEAIFSQKTDVGSQEELHSFVSENKEKAFCSQASDLGIKMARECRALSPETYTVEESRQFGGAGRSEGPSQPSNTLEEDRDSAKETSEKDLDSTMSVNVHQAPAPTAKGKKQKGRNTQVSGPSSALPSFFNSTDSFNDAGLSSSTPSVEAA

NbL13g08810.1 proteasome_activator_subunit_4 [MW = 203.46 kDa]

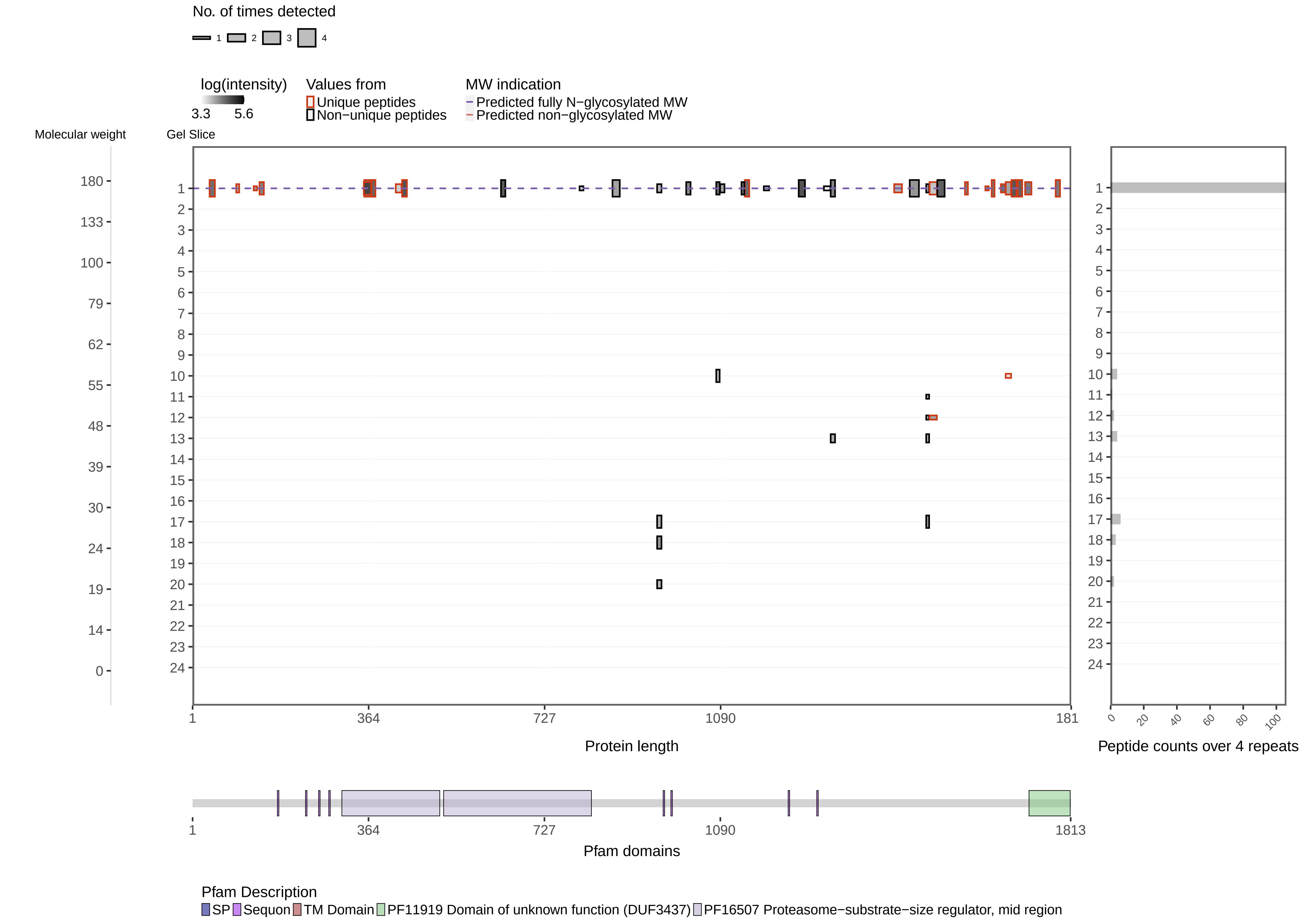


Protein sequence:

IRIIDSLGNYGSSEYDEWSNHRQSWKLESAAIIEPPVNFIVSSHSKGKRRPRWAIIDKGYMHSTWRASQSSYHVFRLSANVSPPEHIIHLMDDLLNLSLHSYETVRGLAGKSLLKMMKRWPSTISKCVLSLSQNLKNPSSPEYTVLGSCAVLATQTVLKCLTTDLKALSSFLLGILSSSHHETLKAQKAINELFVKYNIHFSGVSRNMFKVSGNSEGTDFTVLVSEIGSLSFESSNLHWRYNLMANRVLLLLSMASRNDPKSSSKILSETAGHFLQSLKSQLPQTRILAISALNTLL

NbL13g08780.1 THO_complex_subunit_3 [MW = 35.07 kDa]

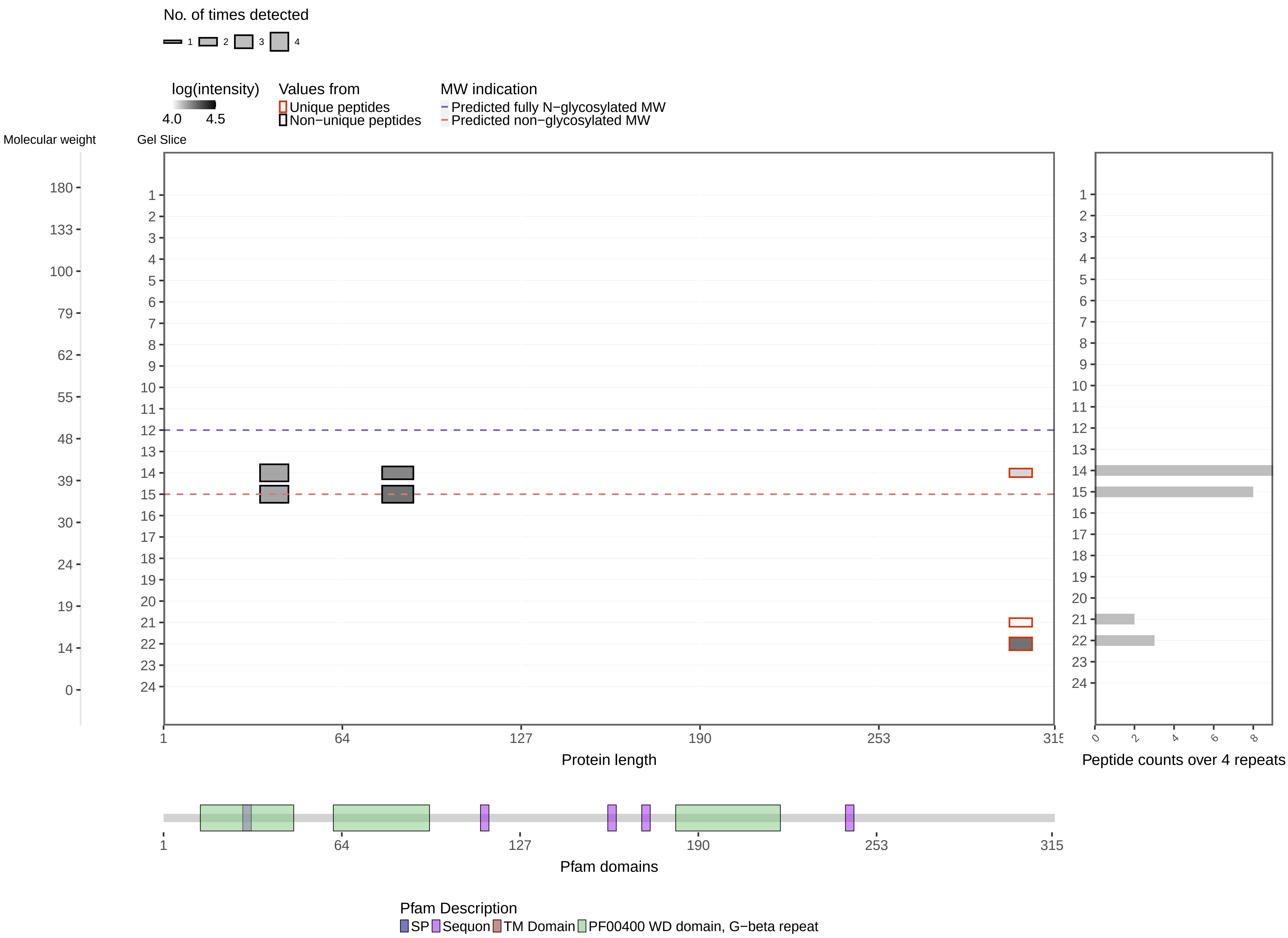


Protein sequence:

IKRKFNYEVNEIAWNMSGDMFFLTTGNGTVEVLSYPTLQPVDTLMAHTAGCYCIAIDPLGRYFAVGSADSLVSLWDIKNMLCVRTFTKLEWPVRTISFNNTGDYIASASEDLFIDIANVQTGRSVHQIPCRAAMNSVEWNPKCNLLAYAGDDKNKYQADEGVFRIFGFESA*

### NbL13g08590.1 peroxisome_biogenesis_protein_7 [MW = 35.51 kDa]

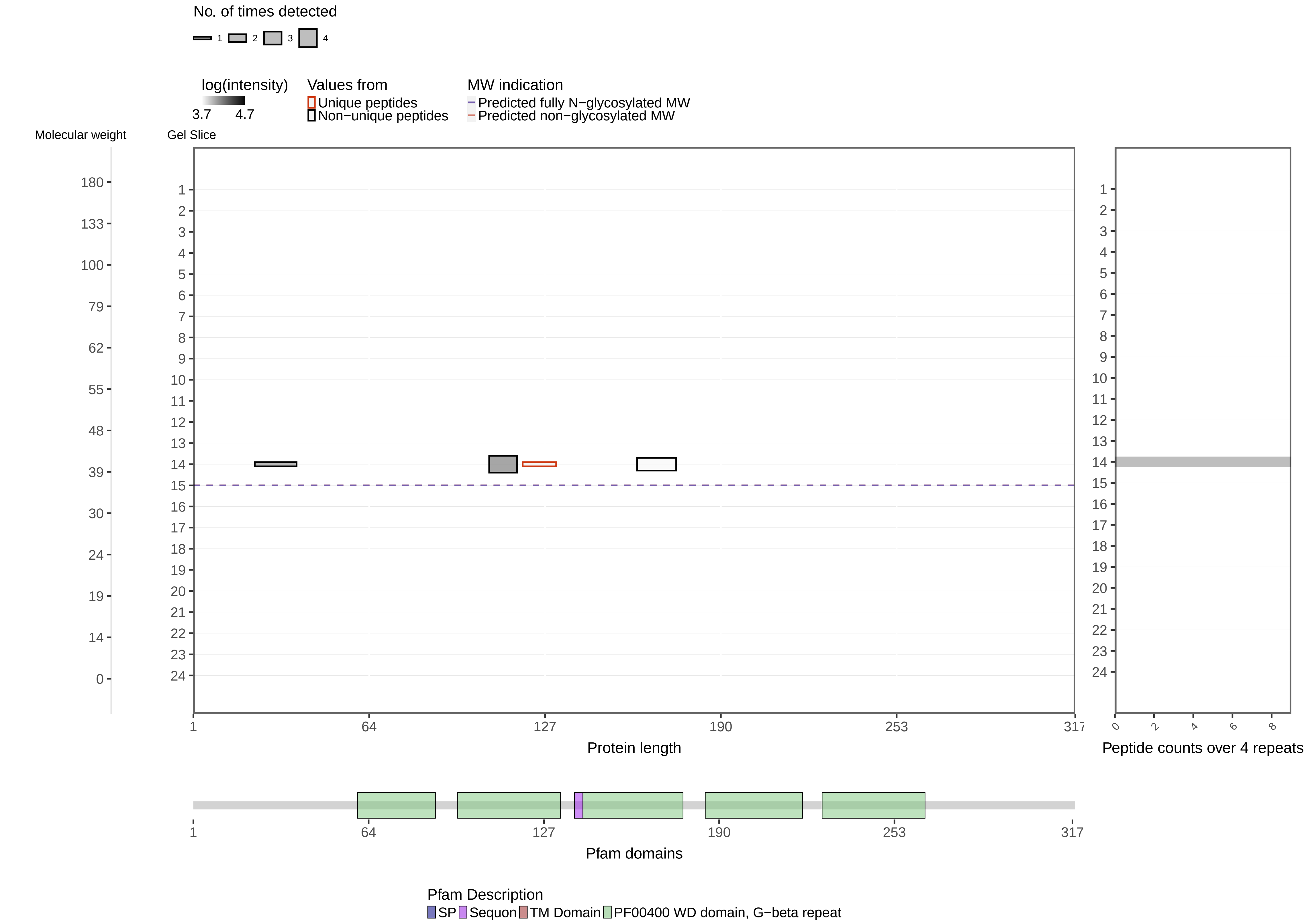


Protein sequence:

[illegible]AYCVYSTAWNPRHADIFASASGDCTTRIWDVREPGSTMILPAHEFEILSCDWNKYDDCIIATASVDKSIKVWDVRNYRVPIAVLNGHGYAVRKVRFSPHRASAMVSCSYDMTVCMWDYMVEDALIGRYDHHTEFAVGVDMSVLVEGLLASTGWDELVYVWQHGMDPRAP*

NbL13g08410.1 3−hydroxyacyl−[acyl−carrier−protein]_dehydratase_FabZ−like [MW = 24.81 kDa]

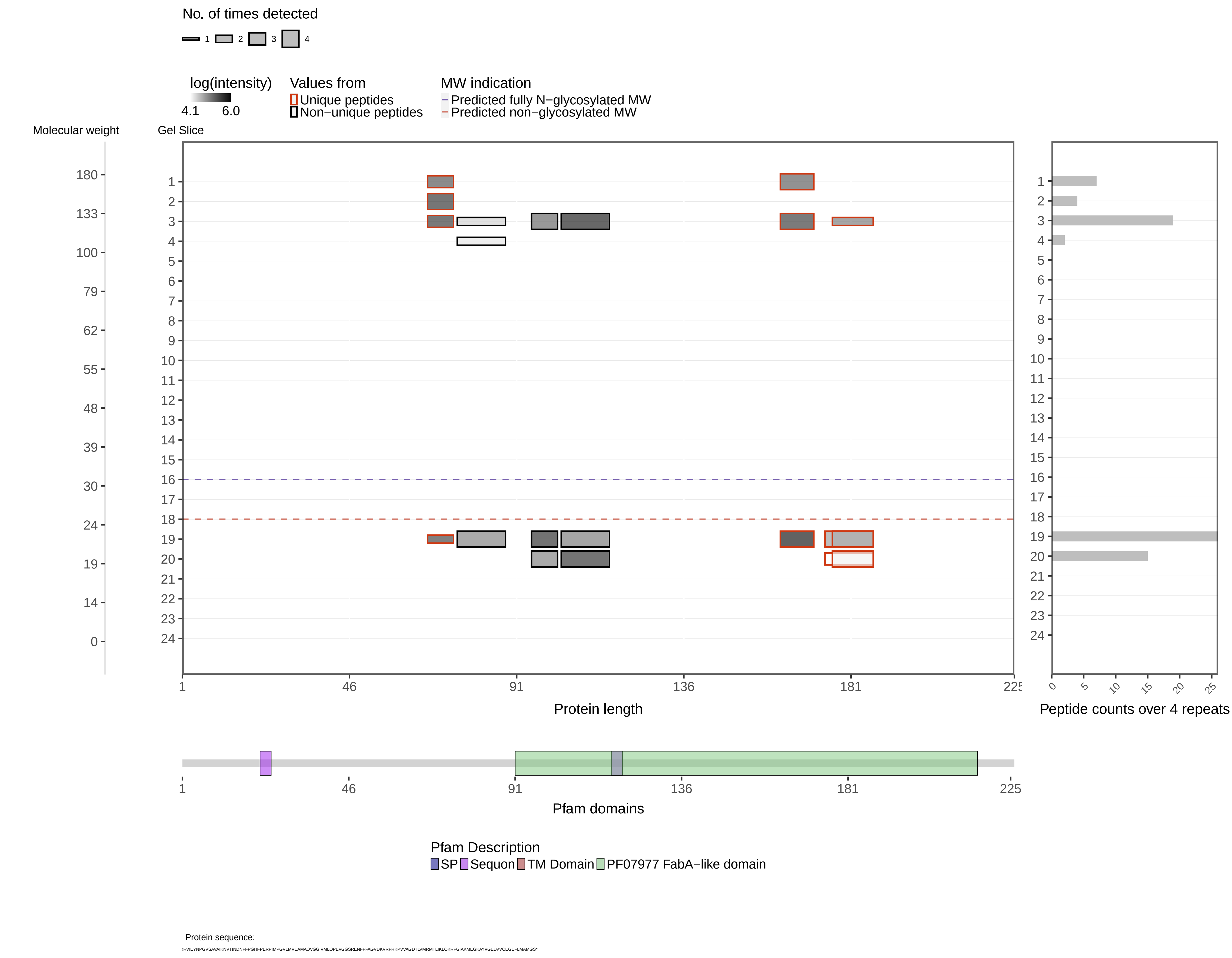

NbL13g08310.1 peroxidase_43−like [MW − SP = 32.33 kDa]

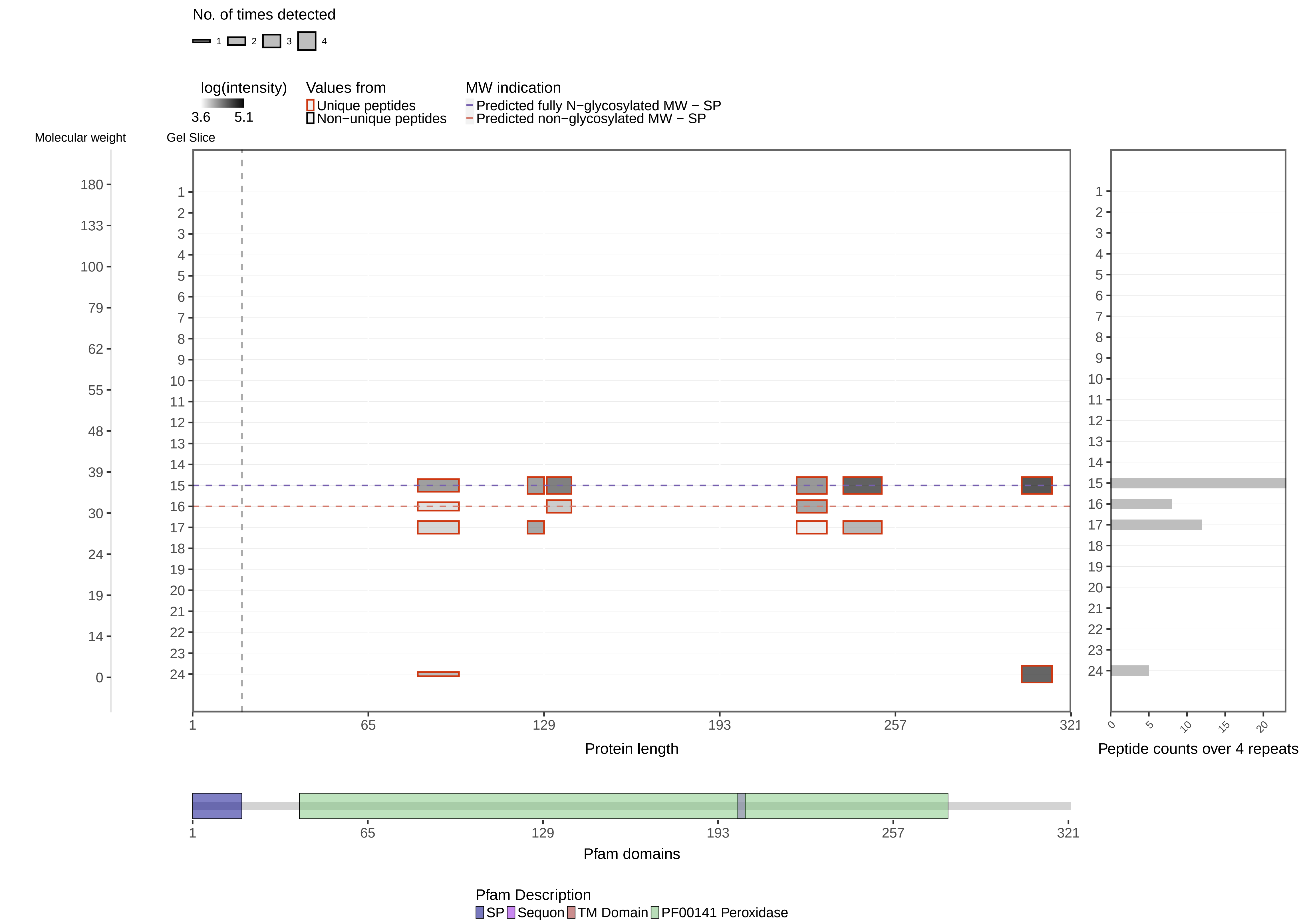


Protein sequence:

LAENMPDVEDSIQQLKAKFLEKGLSEKDLVLLSAAHTIGTTACFFMTKRLYNFSPSGGSDPSINPNFLPELKATCPQNGNVNVRLAMDRGSGEAFDSQILQNIRSGFAVLQSDANLYRDETTKRVVDSYFGILSPFLGTSFEADFAHAMLKMGRIGVLTGSQGTIRRVCASF*

### NbL13g08240.1 uncharacterized_LOC107822389 [MW = 41.63 kDa]

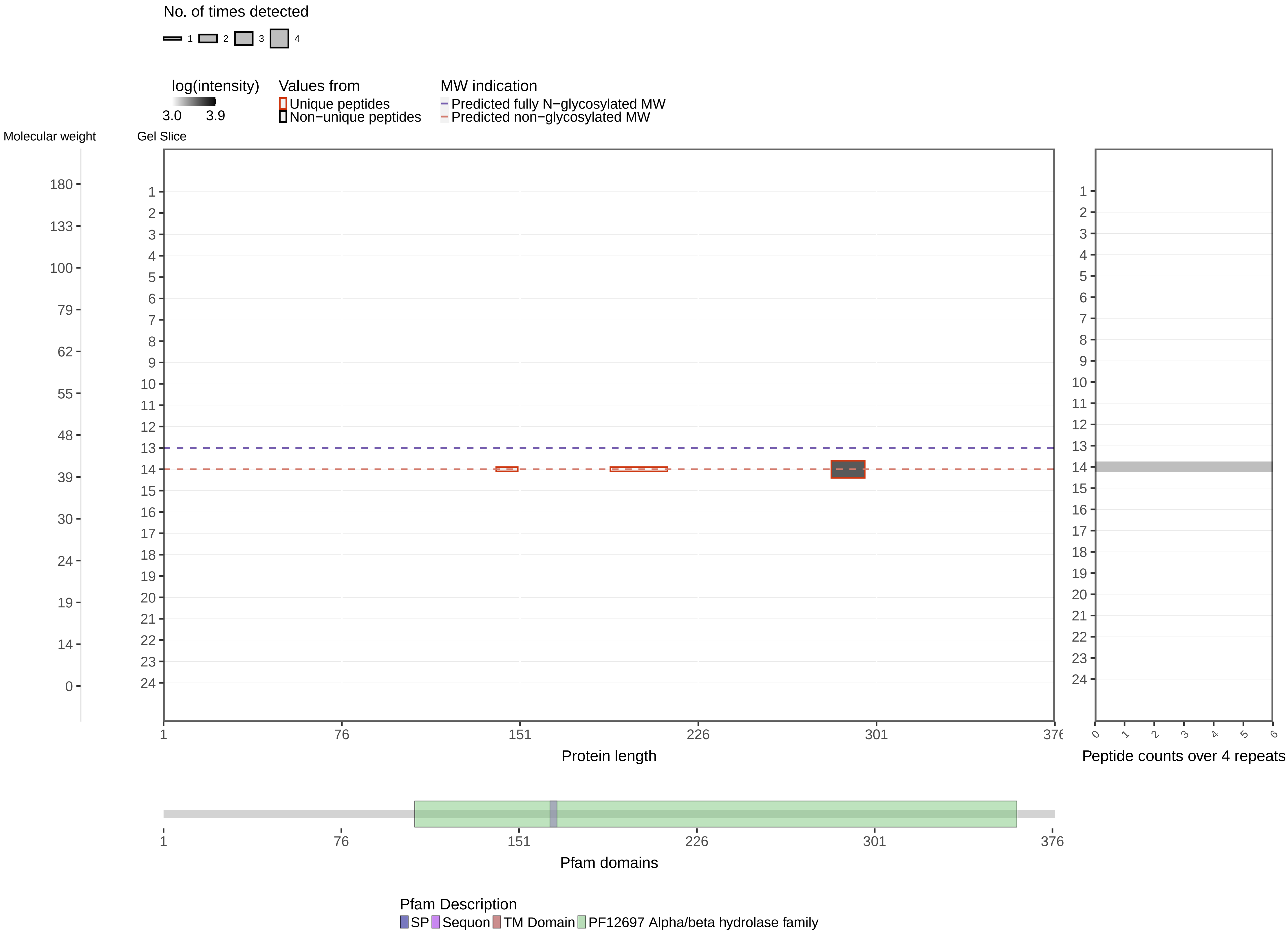


Protein sequence:

GGGHAATIVVRAAKKGLVKPTAIAAIAPTWAGPLPIVFGRDSSMETRYGLLRGTLRAPAVGWMMYNVLVSNEKSIOSOYKSHVYSDPEKVTPDIIESRYALTKROGARYVPAAFLTGLLDPVKSREEFIOLFAELEGRIPVLVLATAGSPKRSKAEMEALREAKGVSKYIEVPGALLPOEEYPEIVAEOLYRFLOEKFELEA*

### NbL13g08170.1 RNA−binding_protein_8A−like [MW = 21.84 kDa]

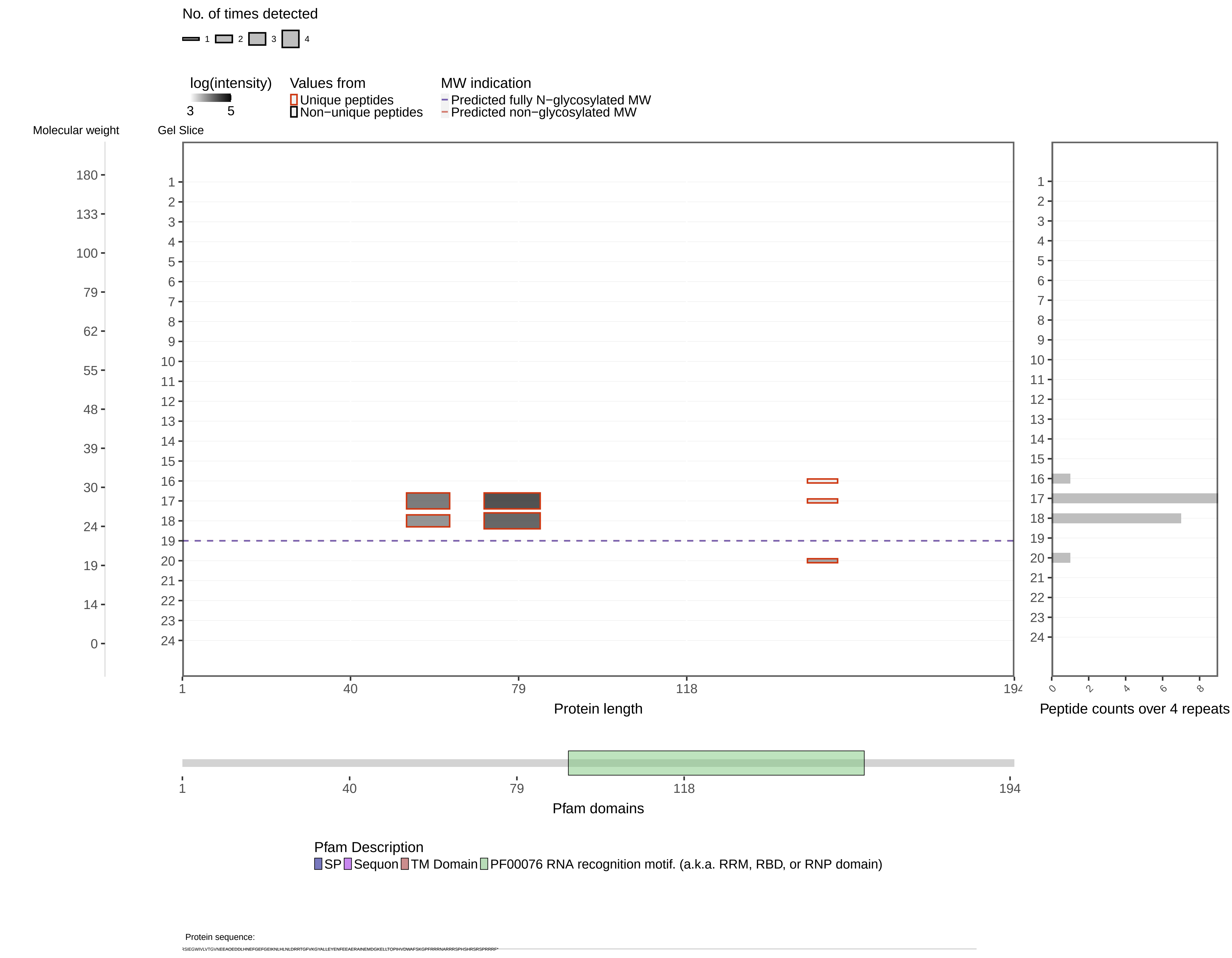

### NbL13g08120.1 malate_dehydrogenase_chloroplastic [MW = 43.23 kDa]

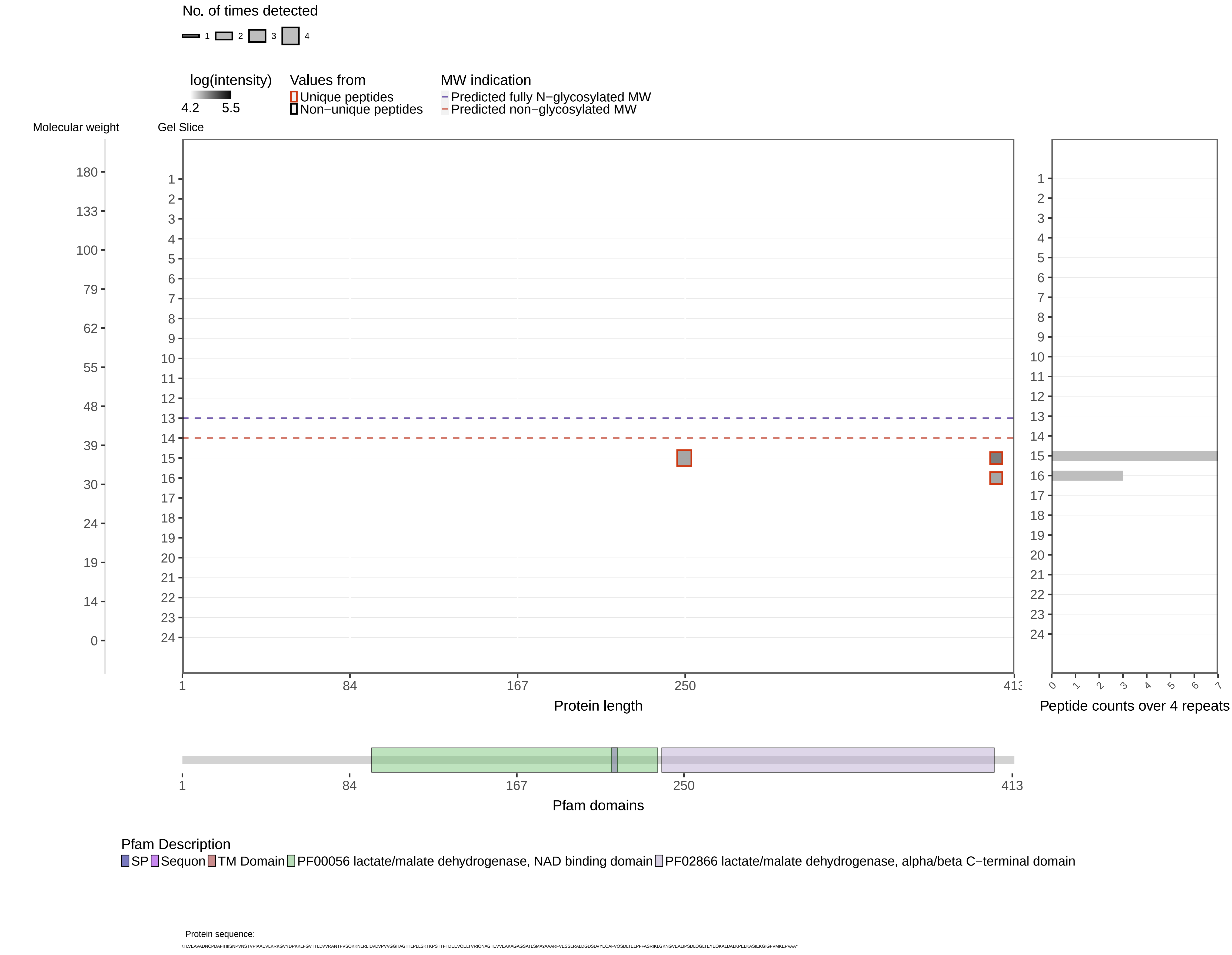

### NbL13g08110.1 diphosphate_reductase_chloroplastic−like [MW = 51.3 kDa]

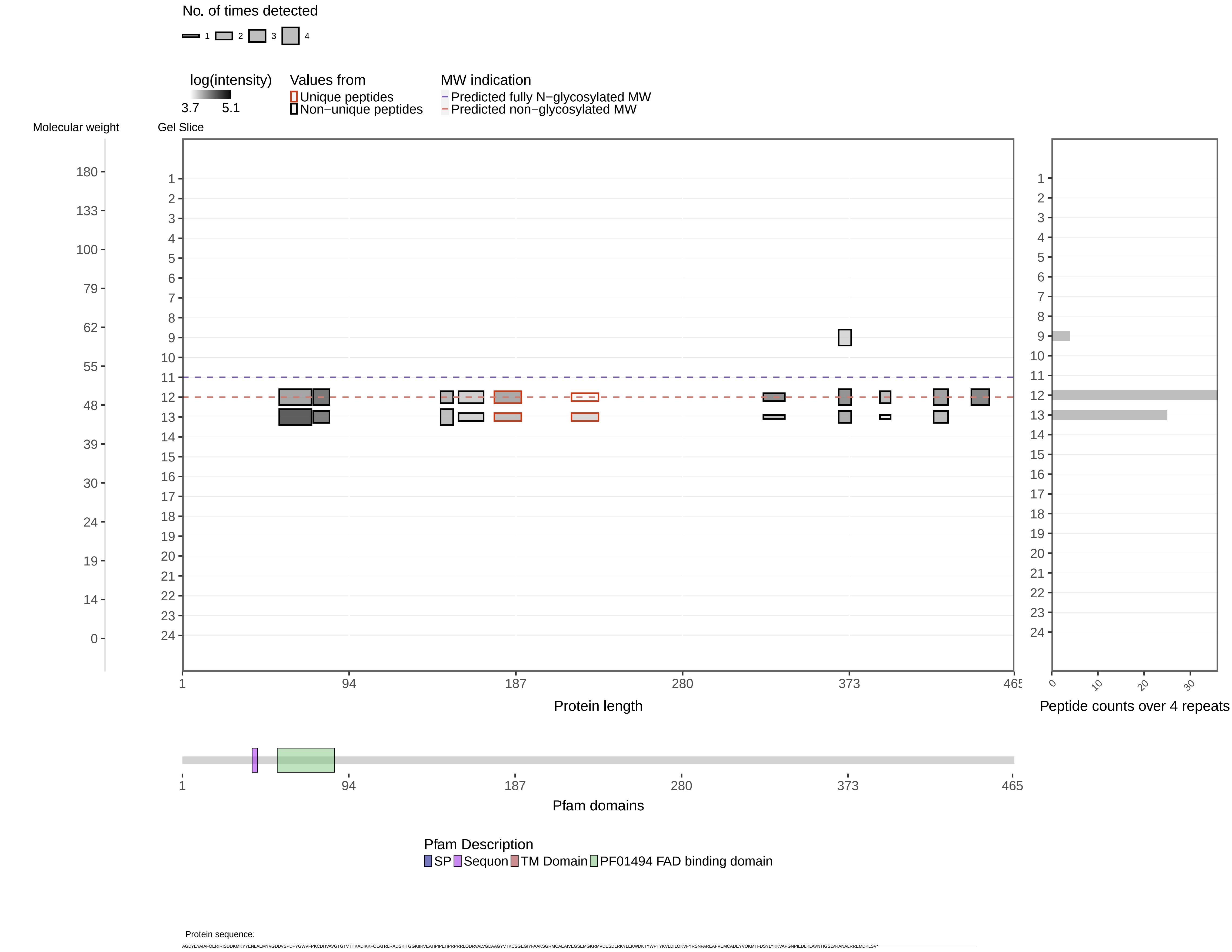

### NbL13g07930.1 vesicle_transport_v−SNARE_13−like [MW = 25.08 kDa]

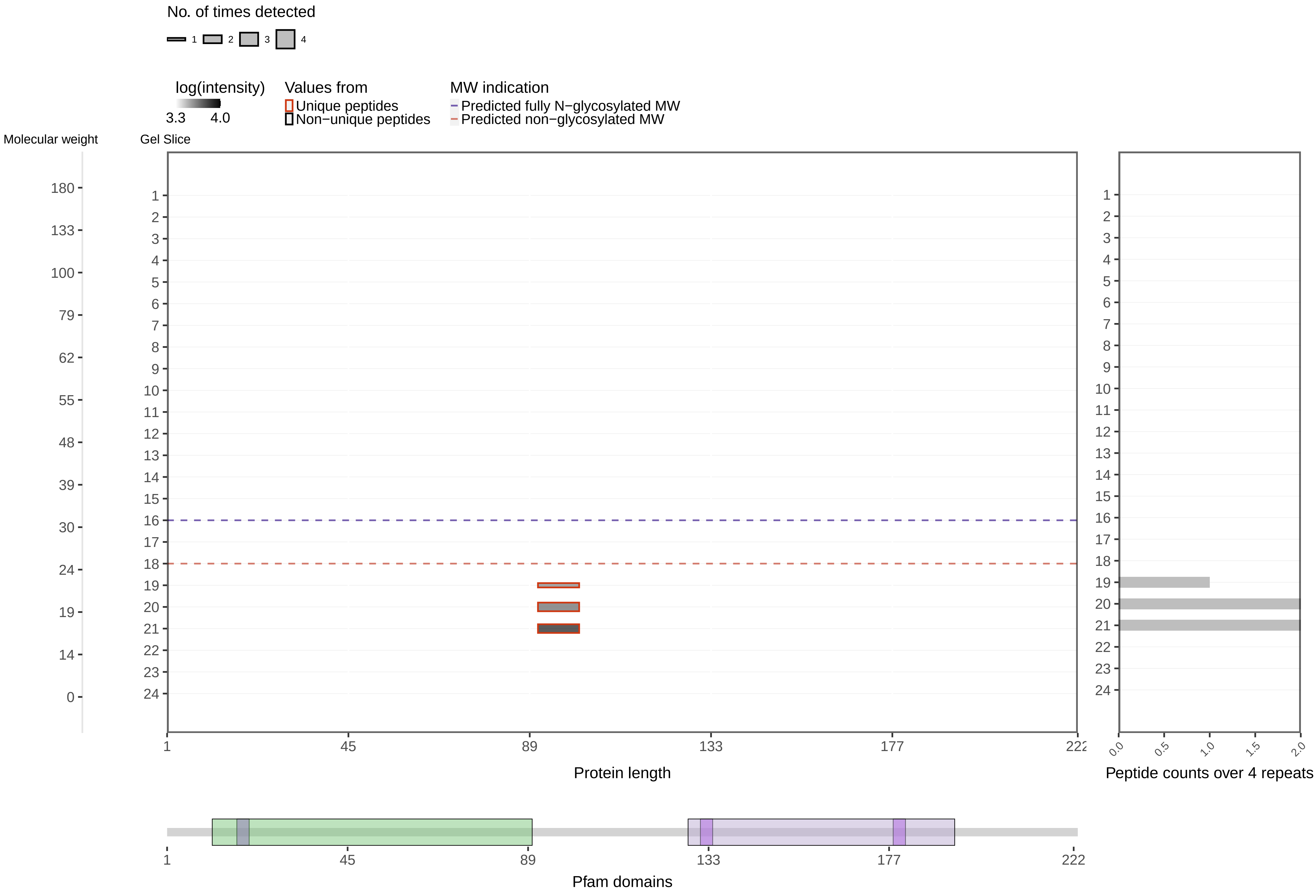


Pfam Description

SP Sequon TM Domain PF05008 Vesicle transport v−SNARE protein N−terminus PF12352 Snare region anchored in the vesicle membrane C−terminus

Protein sequence:

QAARDELLESGMADTMMVSADQRQRLMMSTERLNHSSDRIRESRKTMLETEELGVSILQDLHQQRQSLLHAHDTLHGVDDNISRSKKILTTMSRRMSRNKWIIGSIIAVLVLAILLVLYFKLAH*

NbL13g07820.1 eukaryotic_translation_initiation_factor_5A−1 [MW = 17.4 kDa]

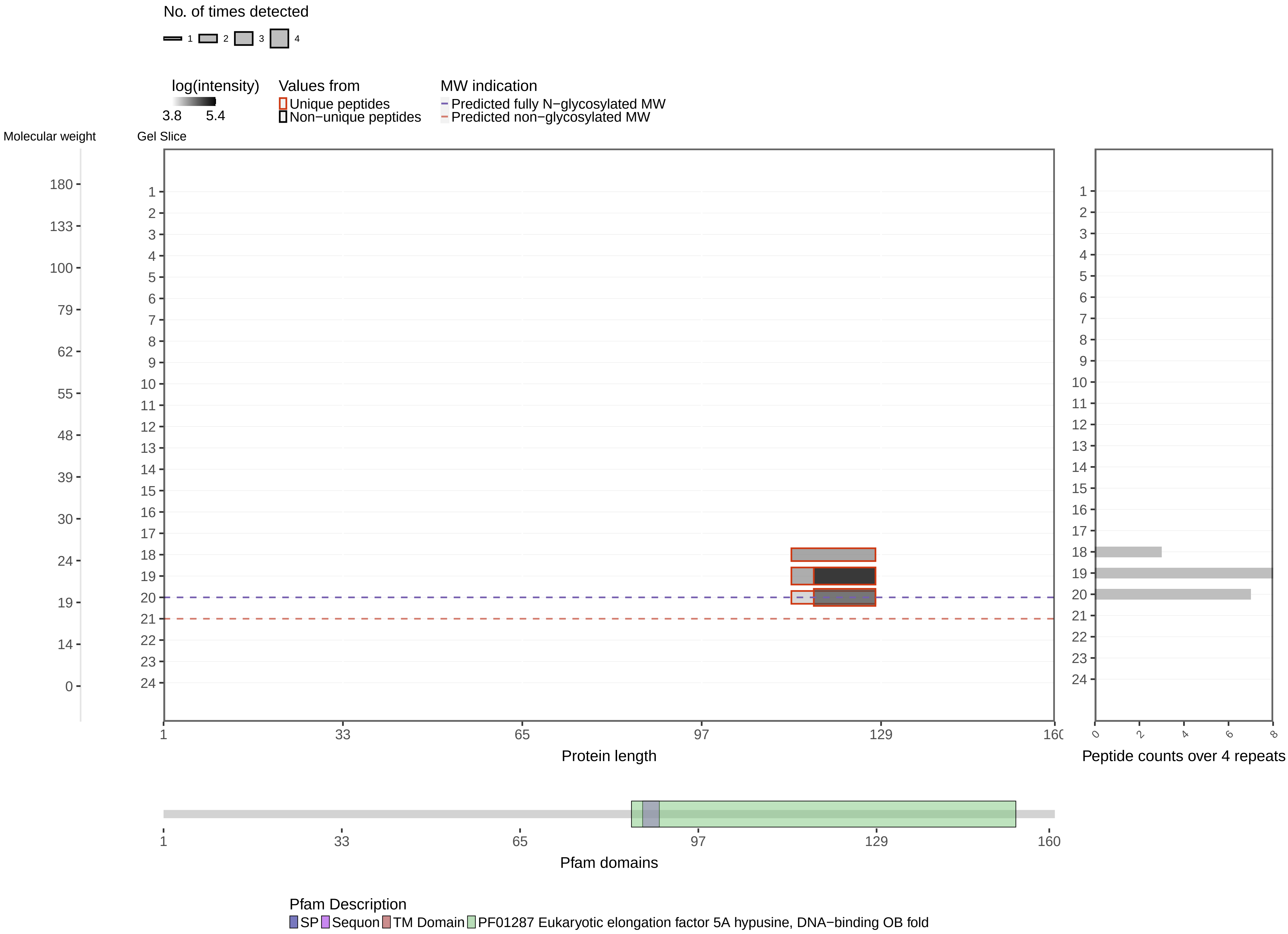


Protein sequence:

TGKKLEDIVPSSHNCDVPHVNRTDYQLIDISEDGFVSLLTENGNTKDDLRLPTDDNLLTLIKDGFAEGKDLVLSVMSAMGEEQICGIKDVGPK*

### NbL13g07660.1 heparanase−like_protein_2 [MW − SP = 58.45 kDa]

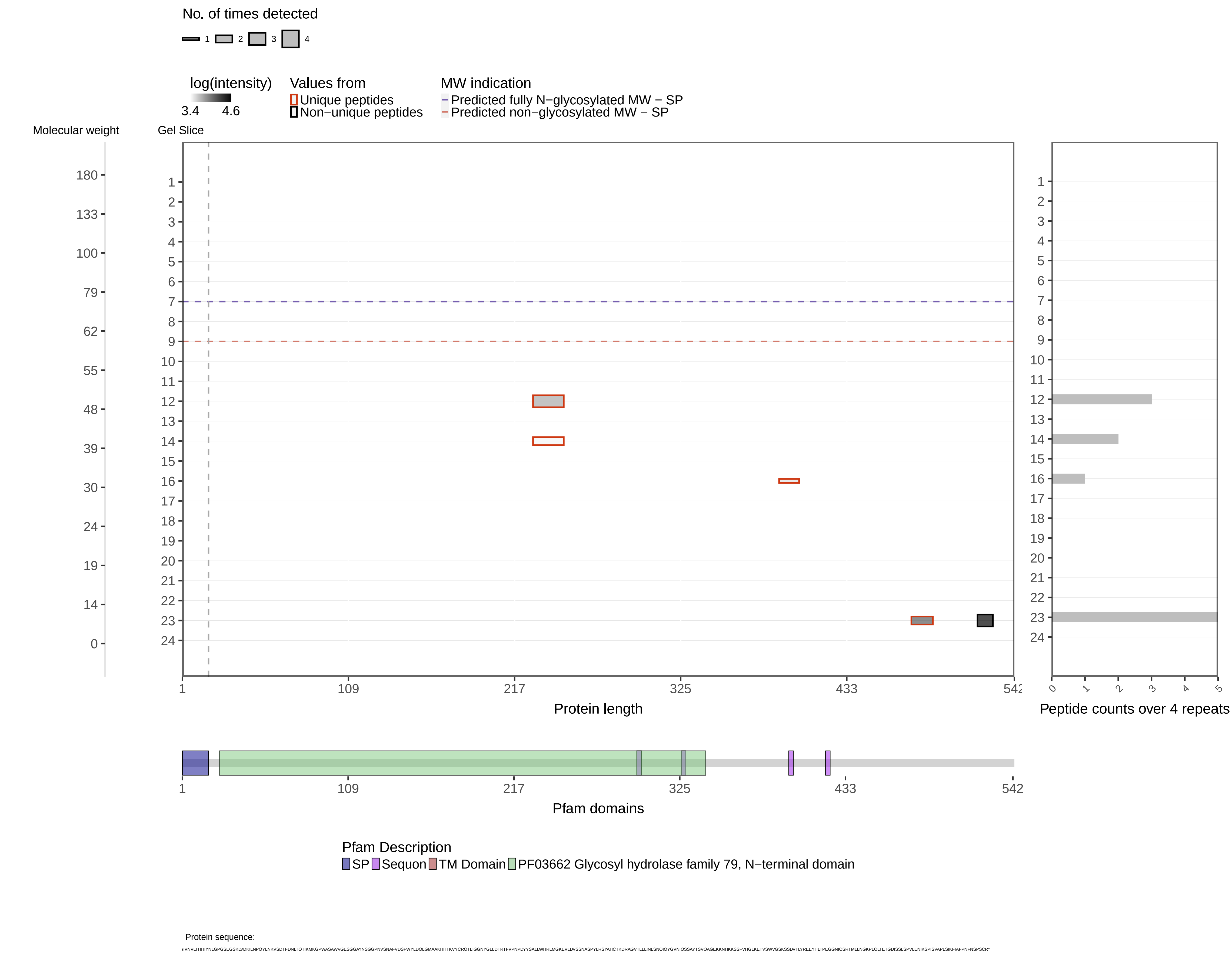

NbL13g07510.1 phenazine_biosynthesis−like_domain−containing_protein_1 [MW = 31.41 kDa]

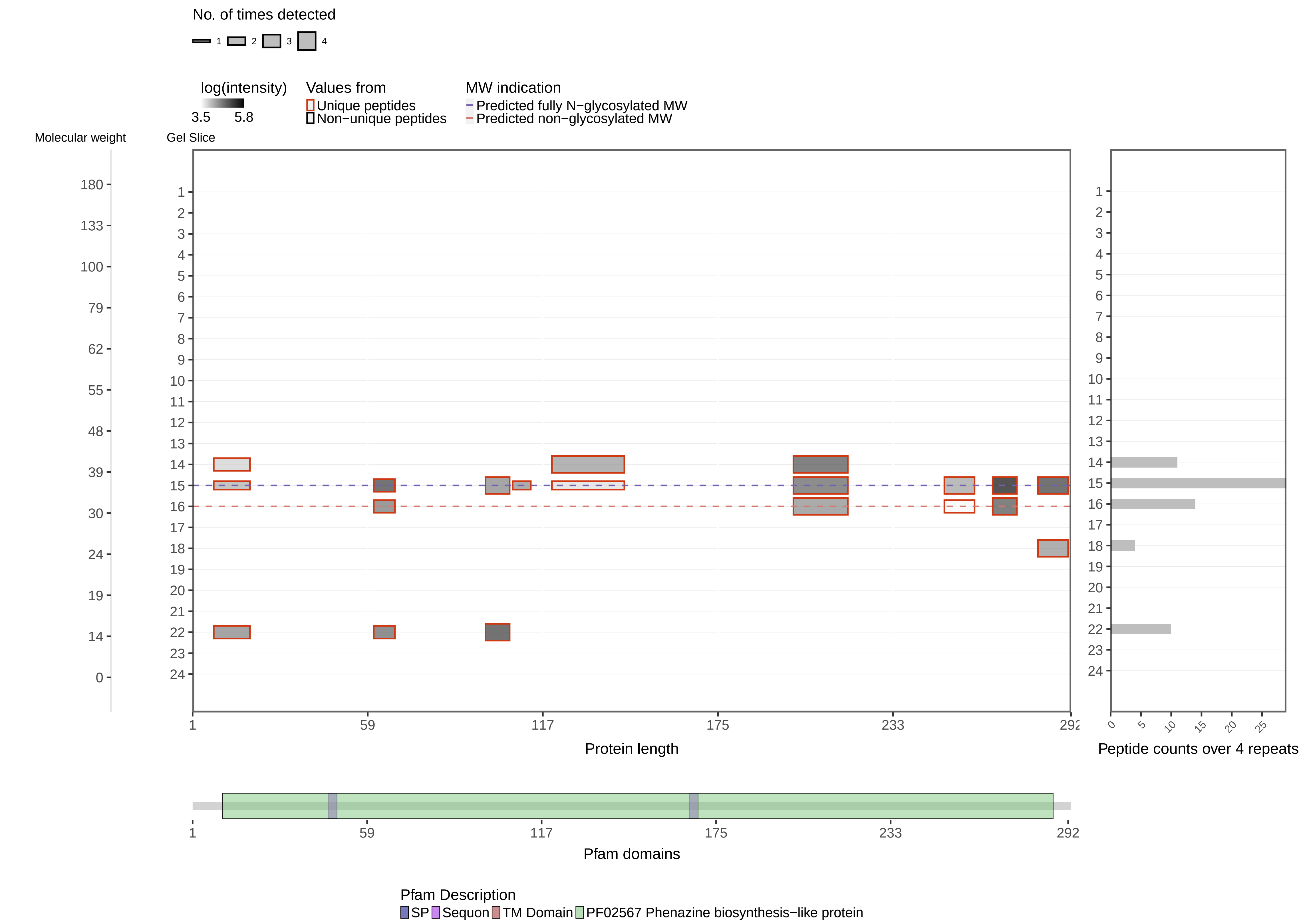


Protein sequence:

VELDFPVVQVAKTNSADVTAISESFNGASMVEINETSMSDLFILLPSGEAVVECQPQFDQIKNCPGRGIIITGPAPQGSGFDFYSRFFCPKLGINEDPICGSAHCALALYWRKKLGKCDFVALAASPRGGVVKVHLDEENQRVFLRGKAVTVMEGSLLV*

### NbL13g07360.1 ATP_synthase_subunit_gamma_mitochondrial−like [MW = 38.57 kDa]

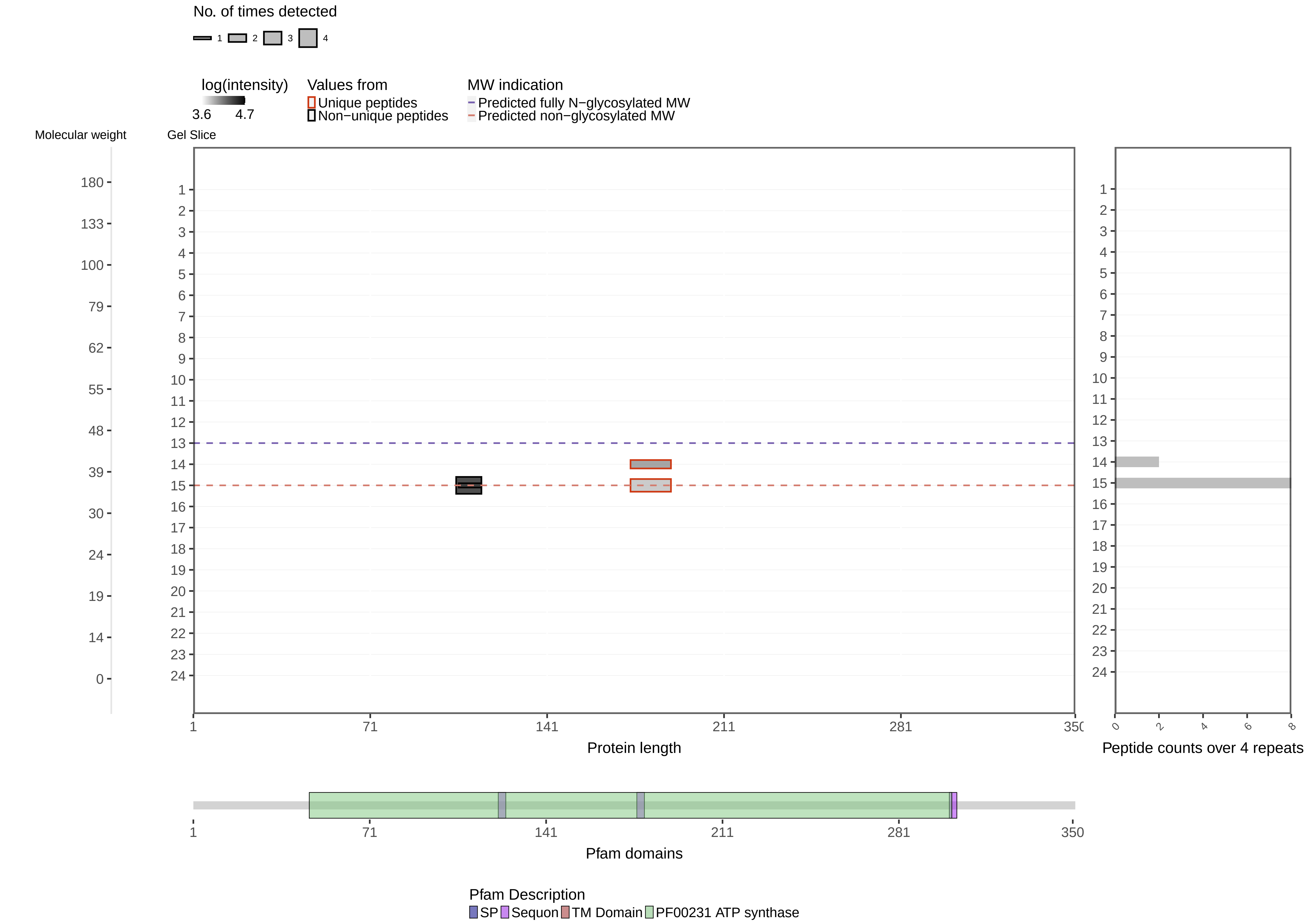


Protein sequence:

KDIELIMTELOKNPLNYTOVSVVADDILKNVEFDALRIVYNKFOSVVAFLPTMSTVLSPEVVEREAEAAGKLGELDSYEVEGAETKAEVLONLAEFOFSCVMFNAVLENACSEOGARMSAMDSSSRNAGDMLDRLTLTYNRSYSLPCLYHIHICTVKLSDMLTOSNRETOFIKLKLGLHISSDALLEK*

### NbL13g07340.1 histone_deacetylase_5 [MW = 72.43 kDa]

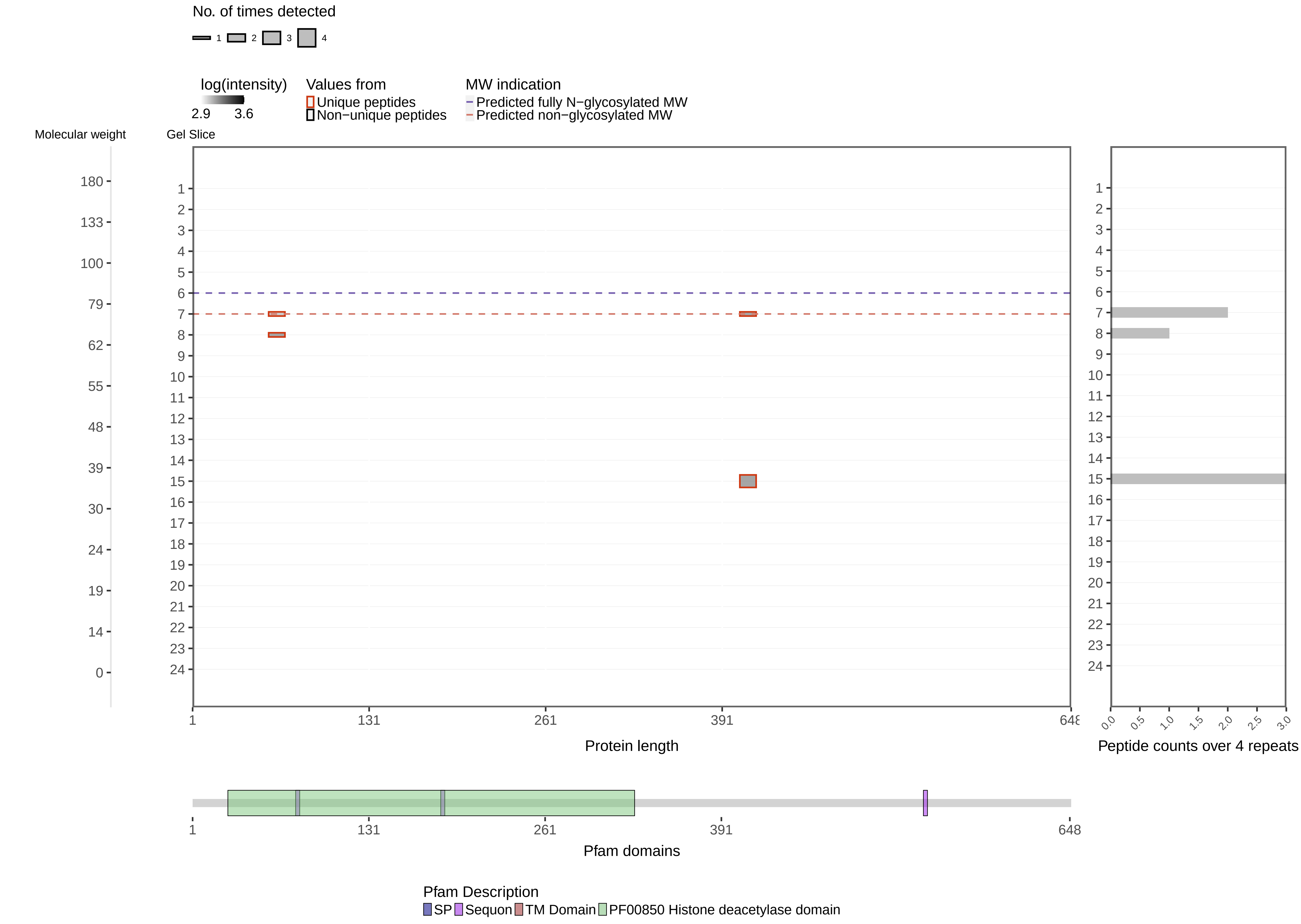


Protein sequence:

[illegible]

### NbL13g07320.1 PLASMODESMATA_CALLOSE−BINDING_PROTEIN_3−like [MW − SP = 16.62 kDa]

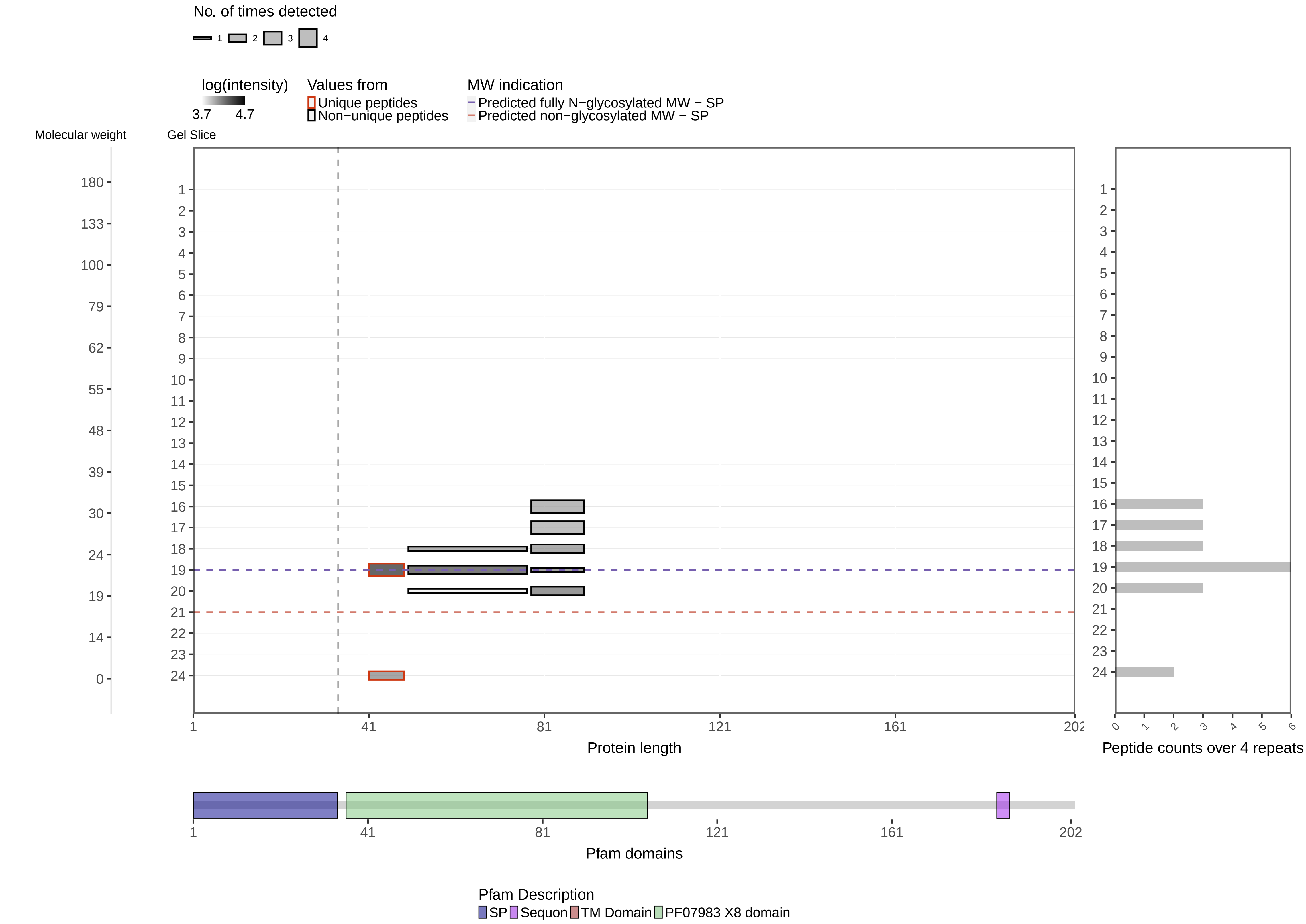


Protein sequence:

SYFQKKGQTSTACDFAGTATVTATDPSTAGCAYPVSGSATPTTPGTTSPATKTPSSSTATTGGSPYVTNTGILGGVNNGLGPSGAGMNTDISHAGISLQNASLFSFIAIISALSF*

### NbL13g07280.1 50S_ribosomal_protein_L15_chloroplastic [MW = 28.42 kDa]

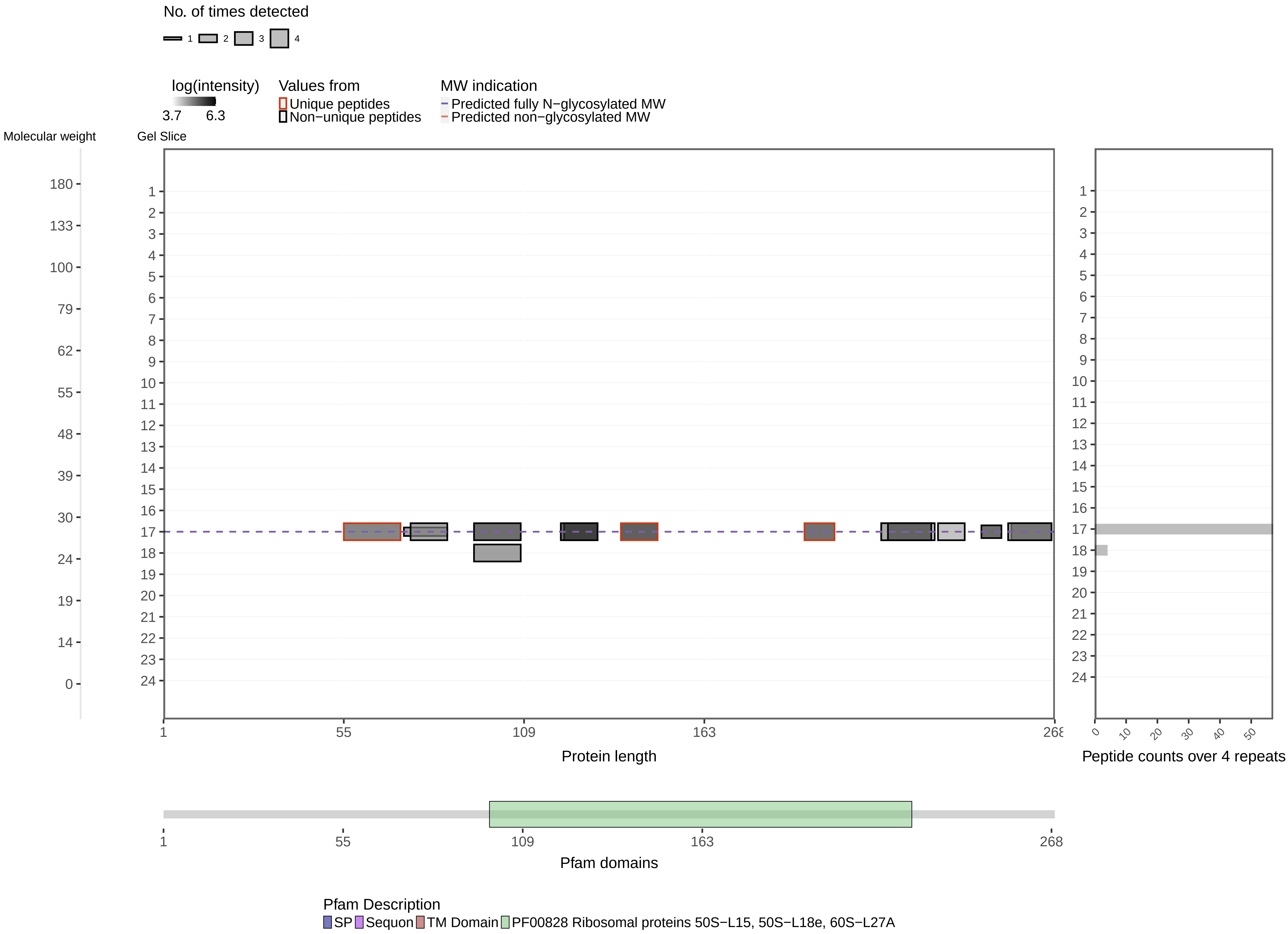


Protein sequence:

GFEGGQMPLYRRLPKLRGIAGGMSAGLPKYVPVNLKDIAEAGFQEGEEVSLESLKQKGLINPSGRERRLPLKILGDGELSLKLNFKARAFSTSAKEKLEAAGCSLTVLPGRKKWVKPSVAKNLARADEYFAKKRAAAASEAADPSSA*

### NbL13g07200.1 UDP−arabinopyranose_mutase_3 [MW = 47.86 kDa]

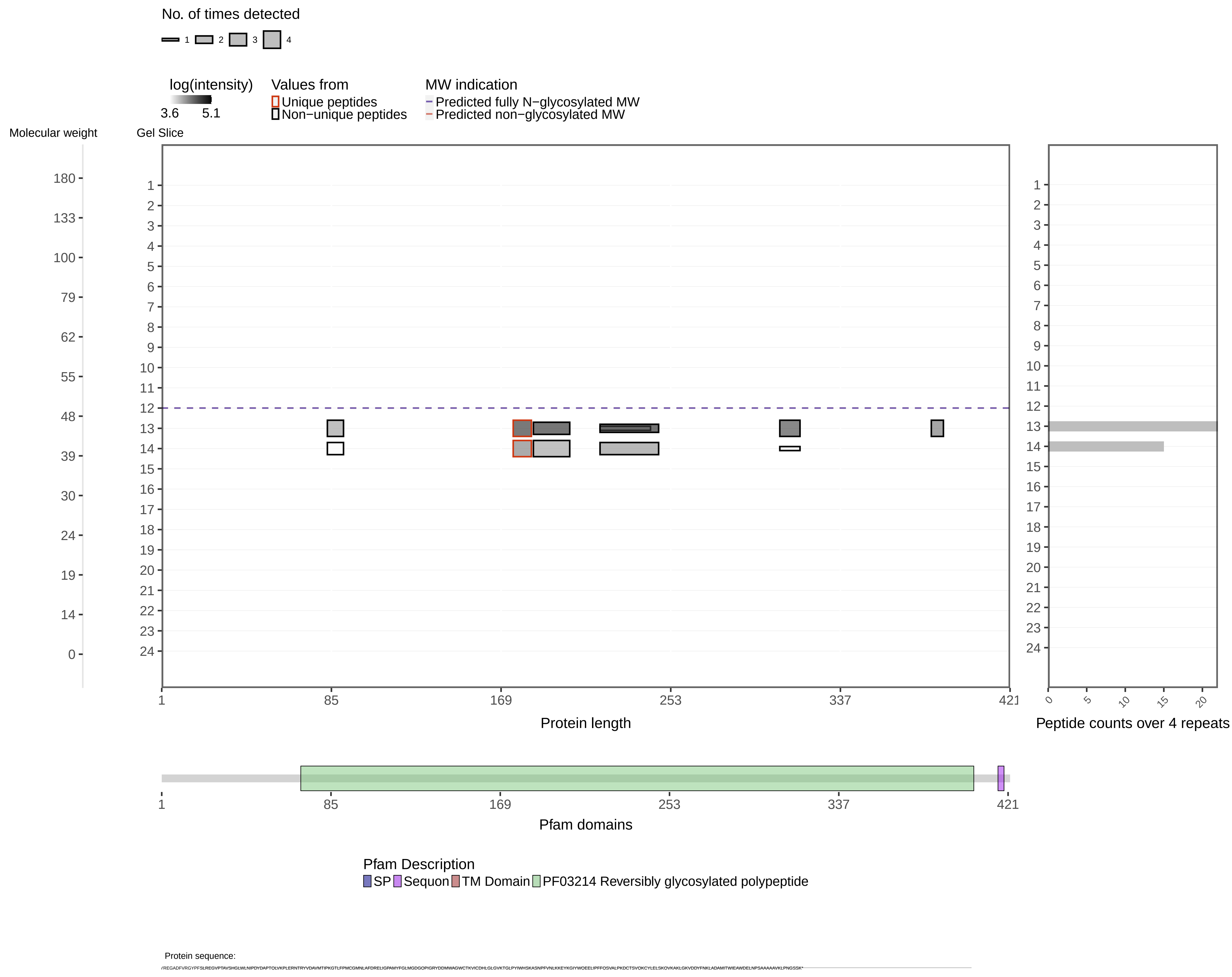

NbL13g06990.1 serine_protease_SPPA_chloroplastic−like [MW = 76.98 kDa]

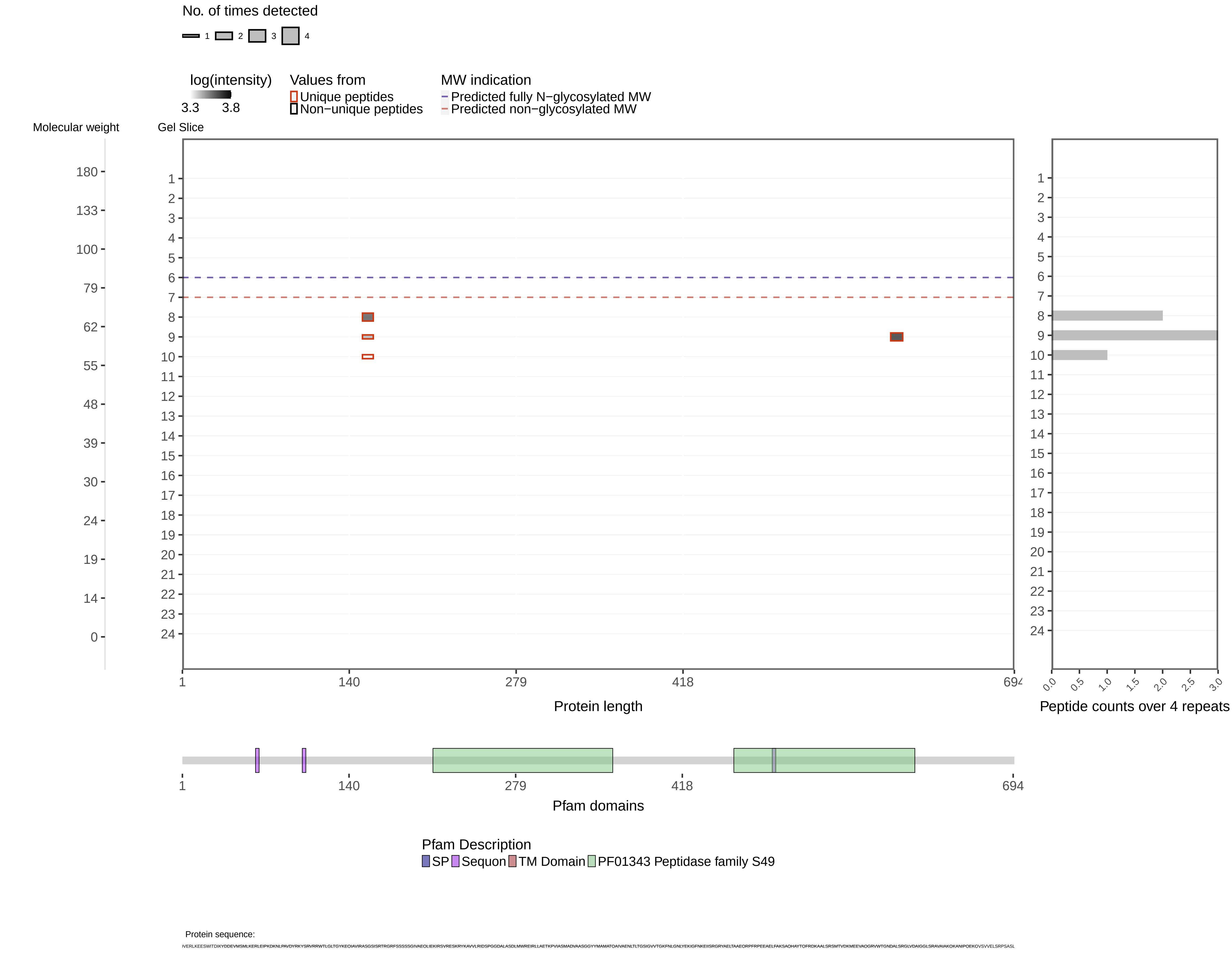

### NbL13g06970.1 L−2−hydroxyglutarate_dehydrogenase_mitochondrial [MW = 45.77 kDa]

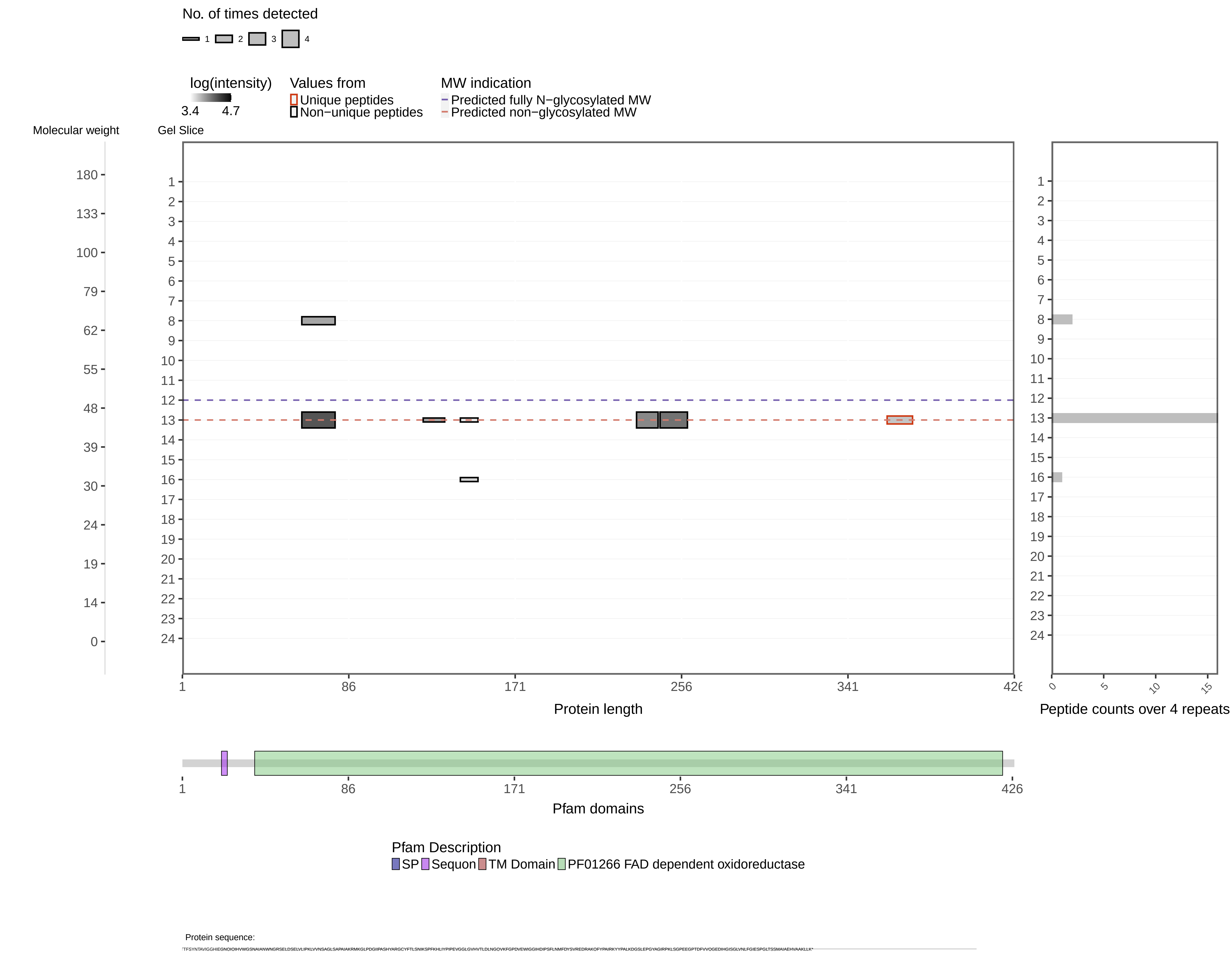

NbL13g06910.1 enolase_1_chloroplastic−like [MW = 24.4 kDa]

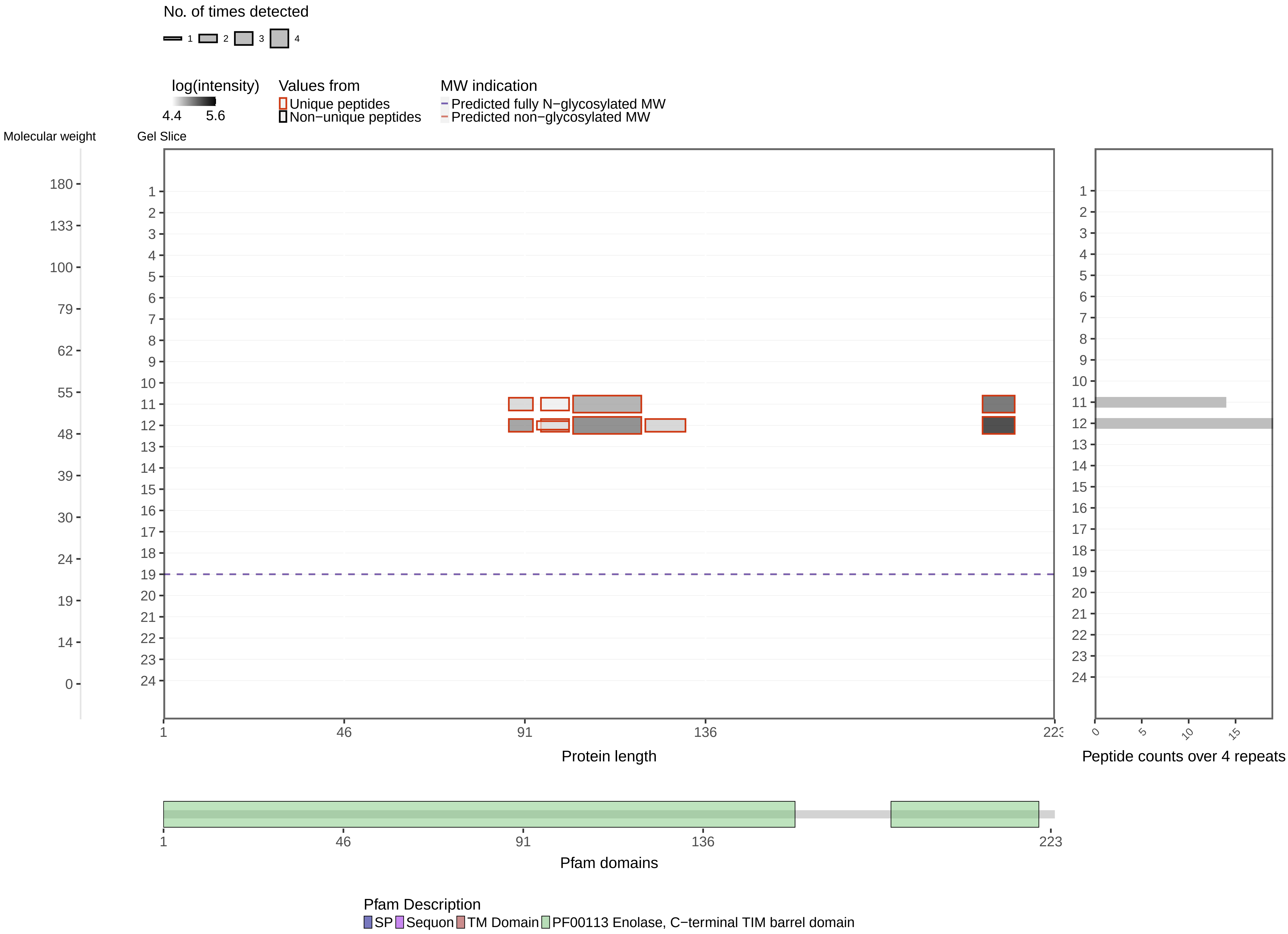


Protein sequence:

LLLKVNQIGTVTESIQAALDSKAAGWGVMVSHRSGETEDNFIADLSVGLASGQVLSTSSCSTLVNLLHSCKHQQHINILLNSPVMQIKTGAPCRSERLAKYNQLLRIEEELGDVRYAGESFRSP*

### NbL13g06900.1 enolase_1_chloroplastic [MW = 14.76 kDa]

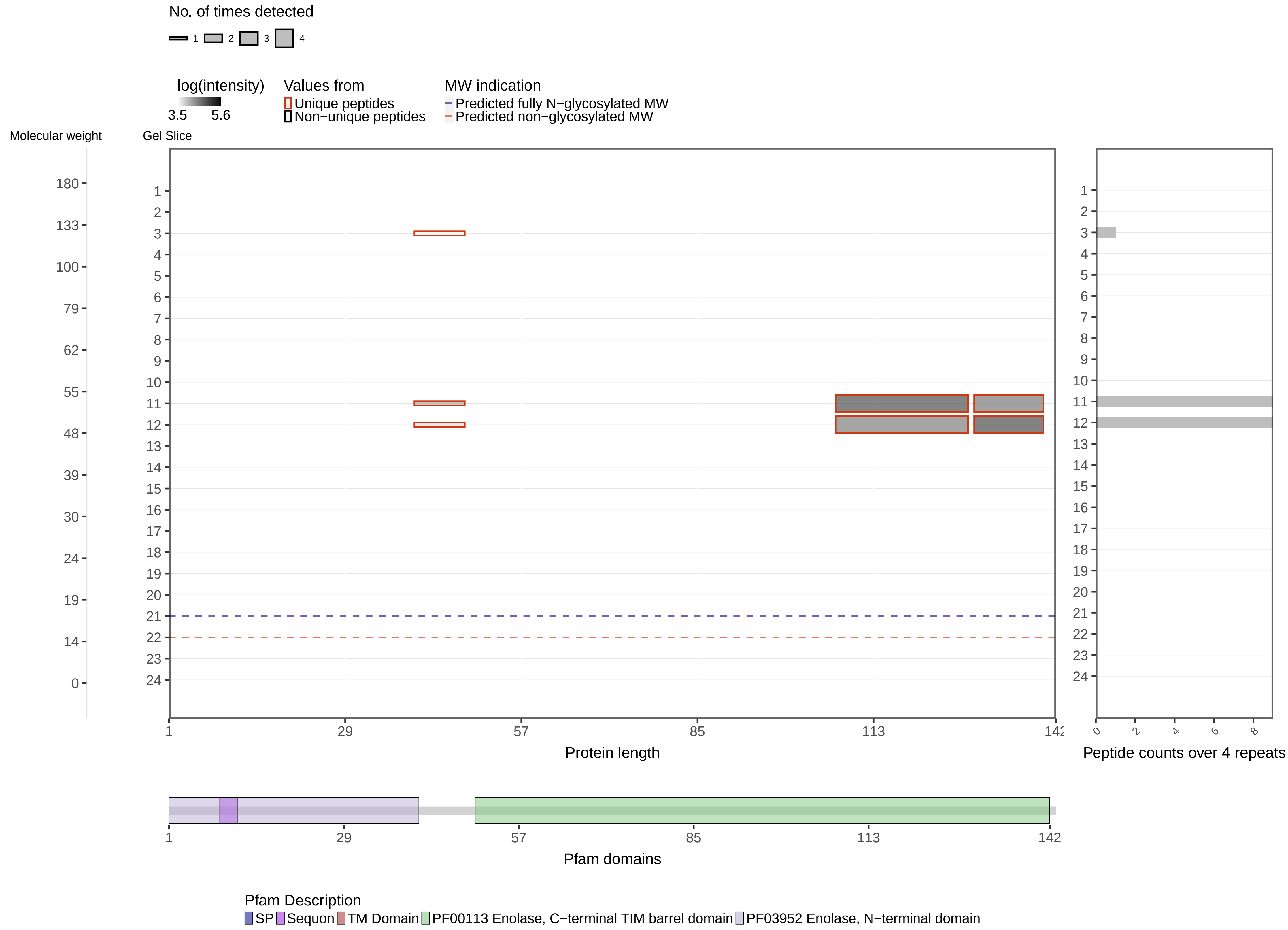


Protein sequence:

/INGGSHAGNNLAMQEFMILPVGASTFAEALCMGSEVYHTLKGIIKAKYGQDACNVGDEGGFAPNVQDNREGLVLLMDAIEKAG

### NbL13g06790.1 1−deoxy−D−xylulose_5−phosphate_reductoisomerase_chloroplastic [MW = 52.08 kDa]

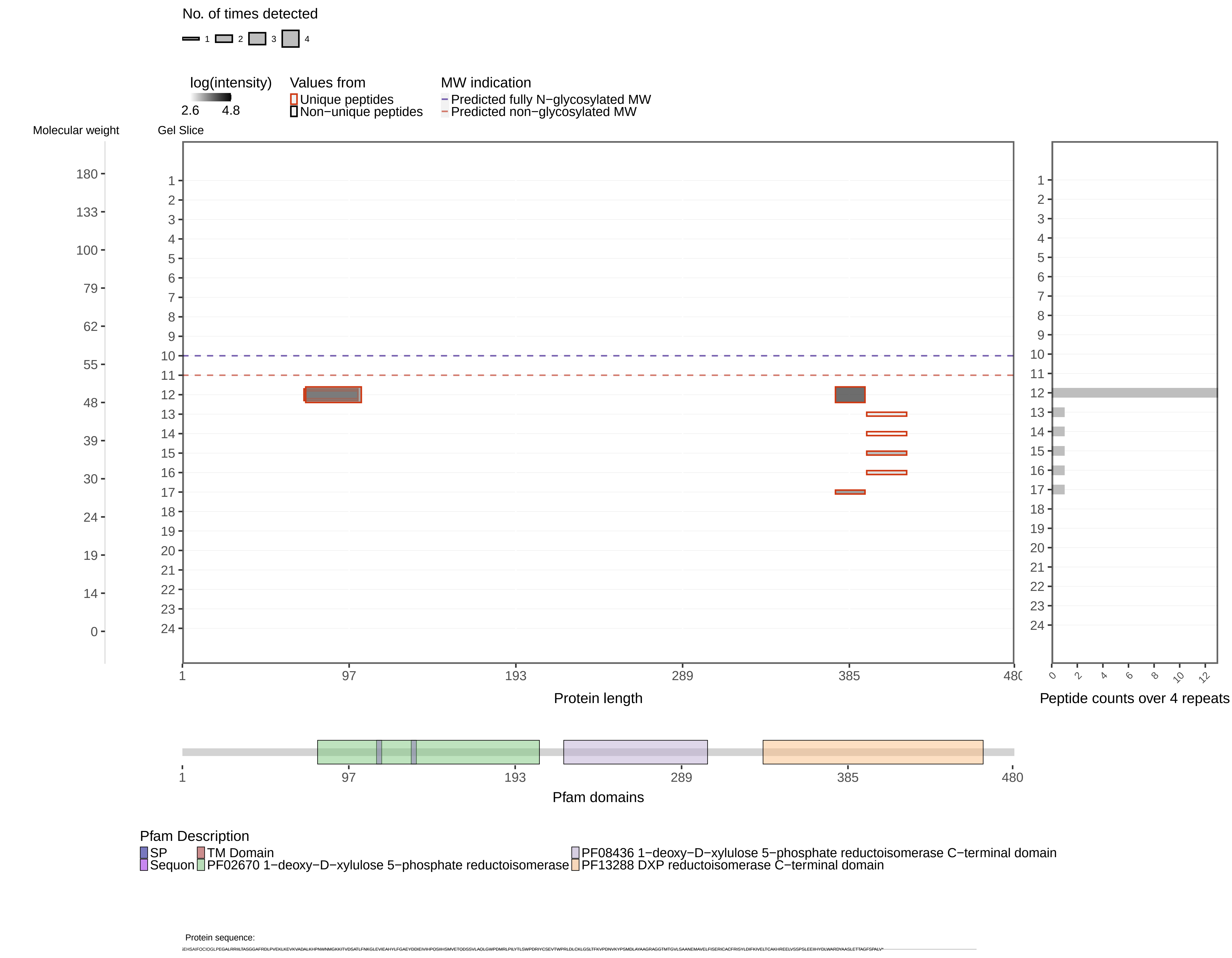

NbL13g06590.1 aldehyde_dehydrogenase_family_2_member_B4_mitochondrial−like [MW = 58.47 kDa]

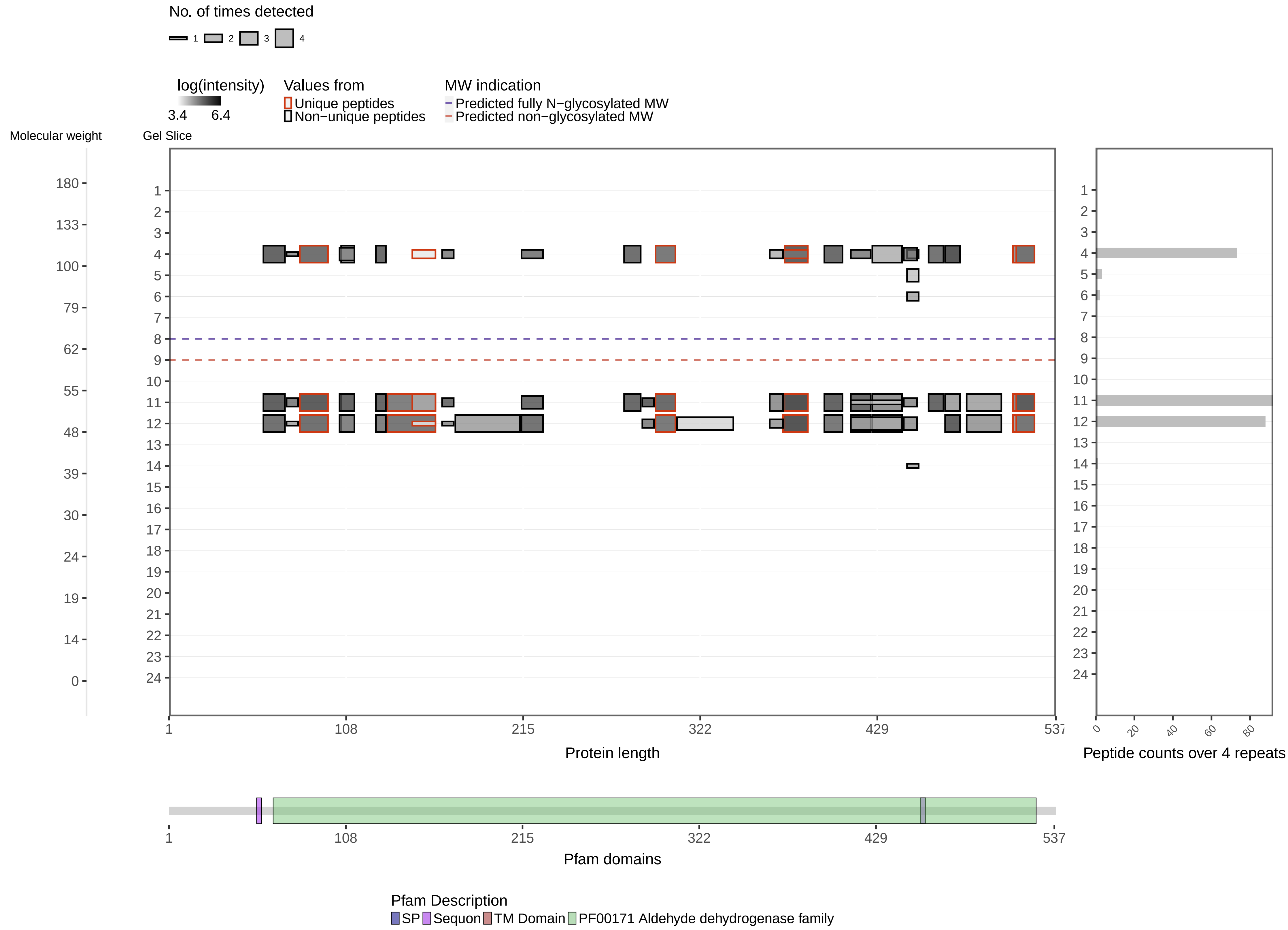


Protein sequence:

SGFGPTAGAALASHMDVDKLAFTGSTETGKTVLELAAKSNLKPFTLELGGKSPFIICEDADIDHAVELAHFALFFNQGQCCCAGSRTYVHERVYDEFIEKAKARAMRRVVGDPFKKGVEQGPQIDSEQFQKVLRYIRSGRDSSATLECGGDRFGSKGYFIQPTVFSNVKEDMLIAQDEIFGPVQCVFKFKDIGEVIKRANNTRYGLAAGVFTKNIDTANTLTRGLRAGTVWVNCYDVFDAGIPFGGYKMSGMGREKGIYSLSNYLQVKAVVTPLKNPAWI*

### NbL13g06550.1 aldose_1−epimerase [MW = 36.68 kDa]

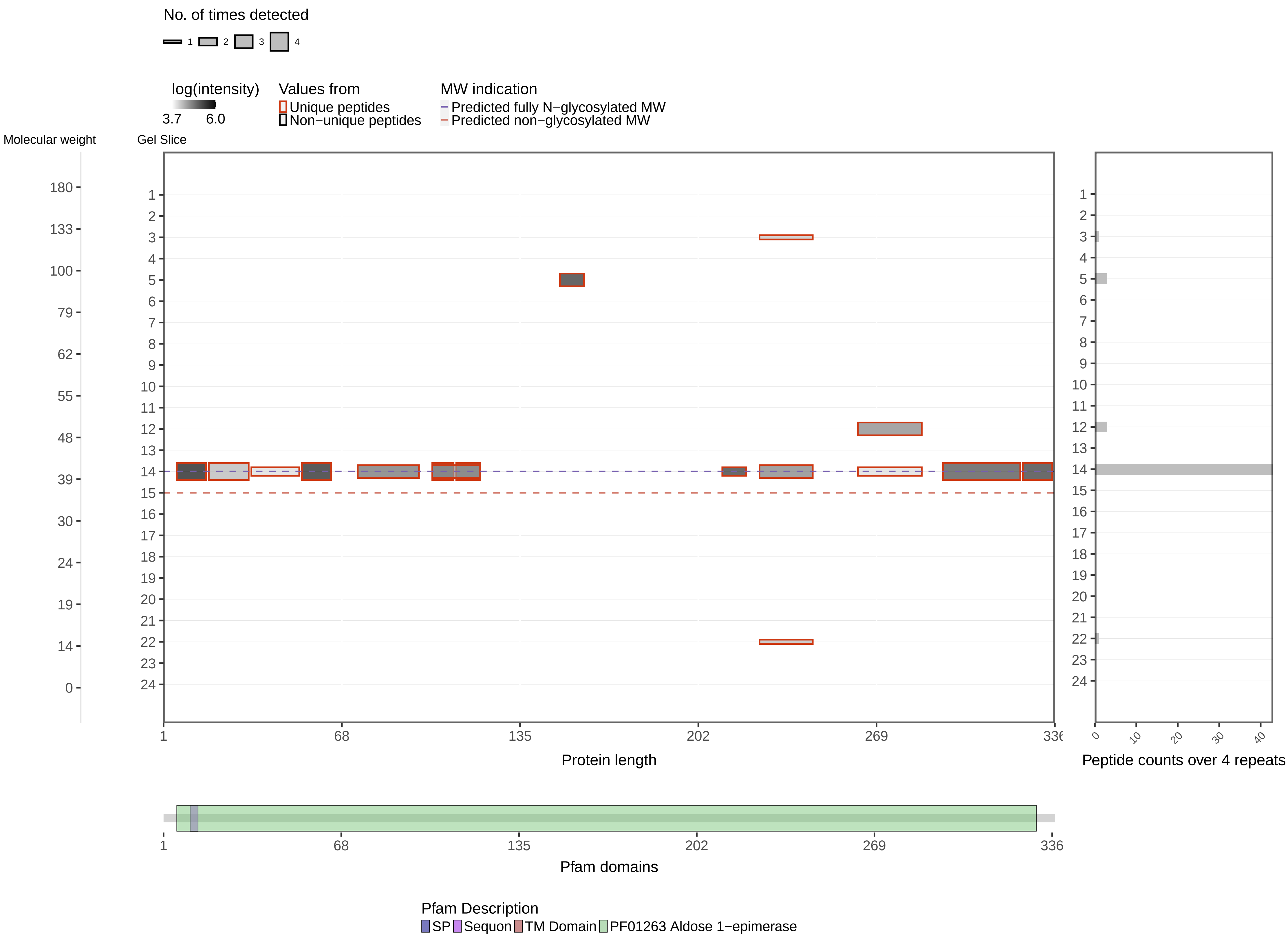


Protein sequence:
/PANKPTPVSLAQHTYWNLGGHNSGDILDNTAQIWANHITPVDQNIIPTGEIMPVKGTPFDFTAEKQIGRDIQQVGLGYDHNYVLDCGDVKDGLKHAAKIKDPKSSRVLNLWTNAPGMQFYTANYVNGVVGKGGAVYNKHAGVCLETQGFPNAVNTPNFPSVIFQPGDKYIHNMLYEFSIE*

### NbL13g06510.1 outer_envelope_protein_64_chloroplastic−like [MW = 63.68 kDa]

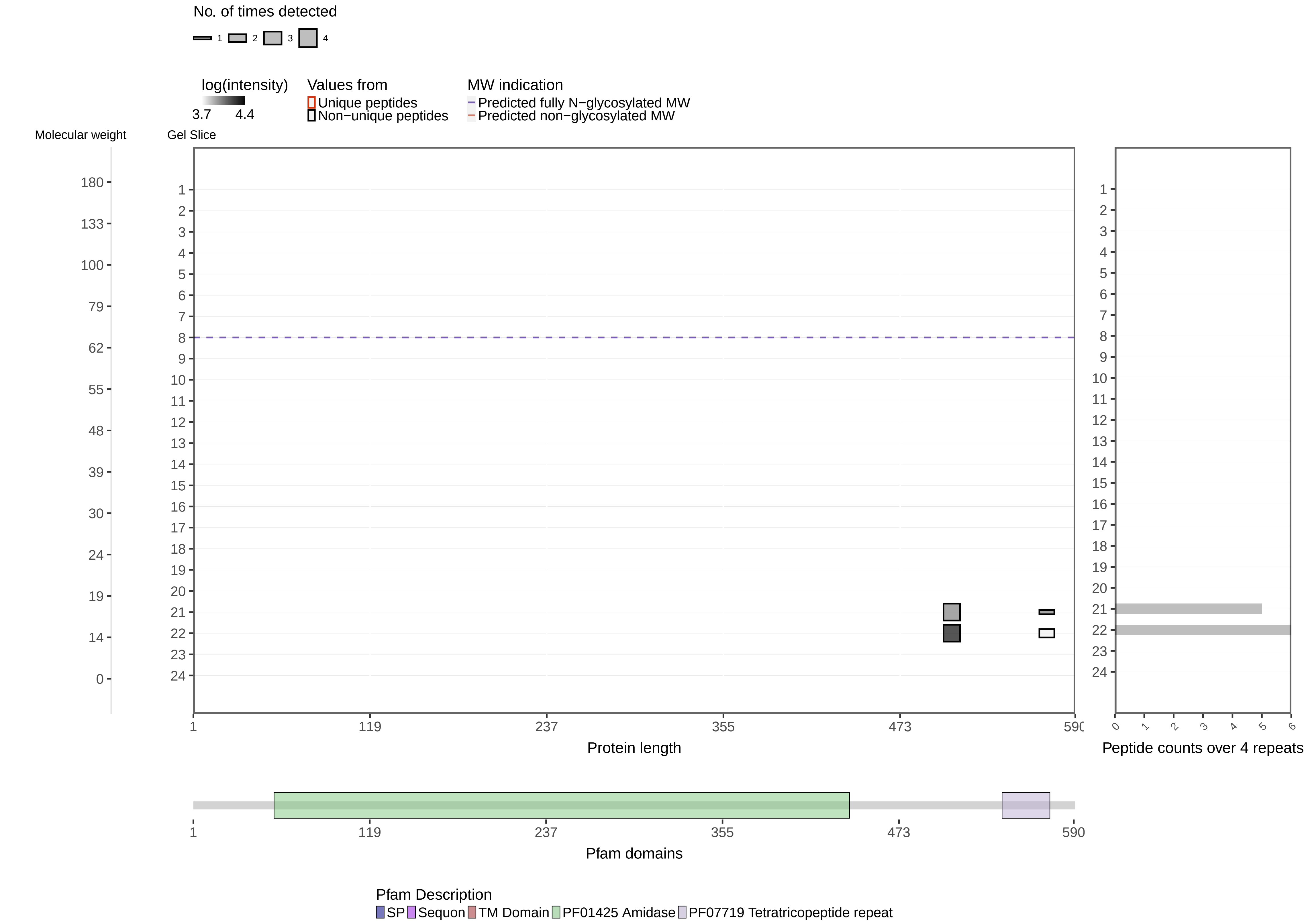


Protein sequence:

[illegible]

NbL13g06400.1 splicing_factor_3B_subunit_4 [MW = 38.89 kDa]

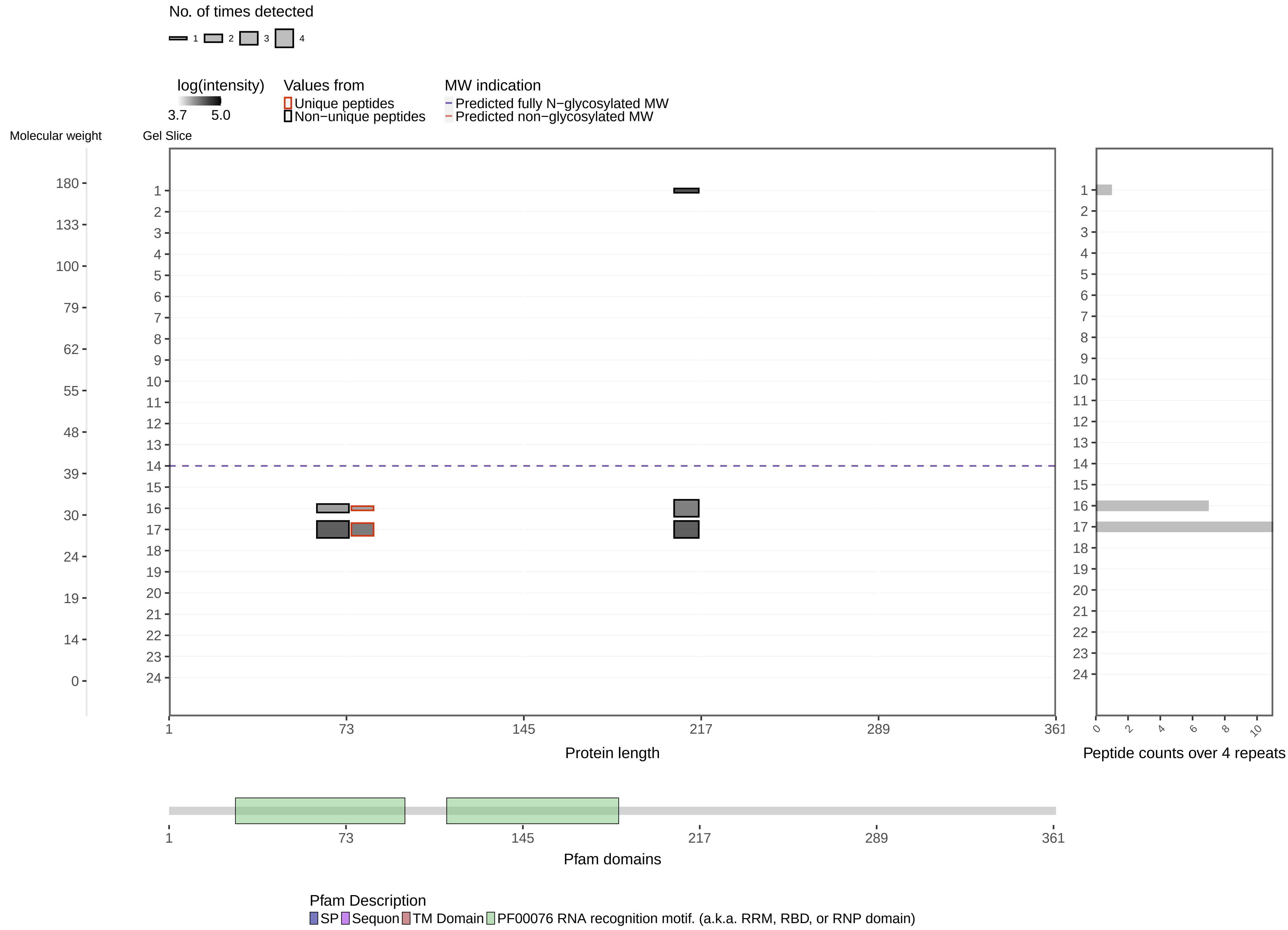


Protein sequence:

EAMNGQYLCNRQITVSYAYKKDTKGERHGTPAERVLAASNPGTQKNRPHTMFASGPPSVPQANGAVGTPGPPRPFTNGPIPPAPIPAIRPPPMQPNVYPPMQMPAPQAWQGHPMQPGPAGIPQQQMQFRGMPPPPPPQVAPPLNRPPPPPAAMAAQVWRPPPPPQHLNGGPYHSMPHMSMQPPPPPAQG*

NbL13g06230.1 T−complex_protein_1_subunit_delta−like [MW = 57.73 kDa]

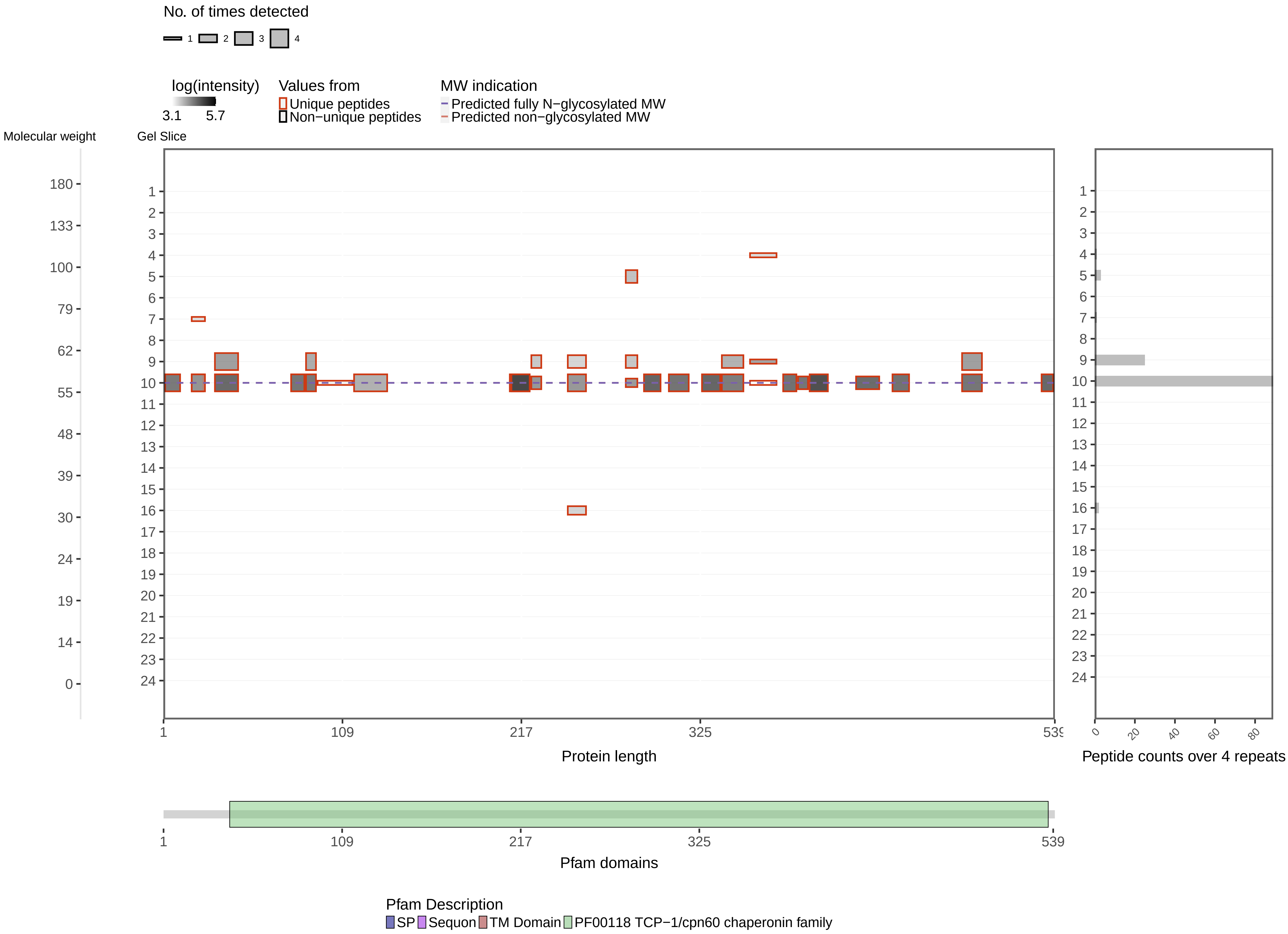


Protein sequence:

PKTDIEQSIVVSDYTQMDRILKEERNYILGMIKKIKATGCNVLLIQKSILRDAVTELSLHYLAKAKIMVIKDVERDEIEFITKTLNCLPIANIDHFRAEKLGFADLVEEISLGDGGKIVKITGIQDMGRTTSVLVRGSNQLVLDEAERSLHDALCVVRCLVSKRFLIAGGGAPEIELSRQLGAWAKVLQGMEGYCVRSFAEALEVVPYTLAENAGLNPIAIVTELRNRHAQGEINTGINVRKGQIANILEENVVQPLLVSTSAISLATECVRMILKIDDIVTVR*

### NbL13g06150.1 guanine_nucleotide−binding_protein_subunit_beta−like_protein [MW = 39.92 kDa]

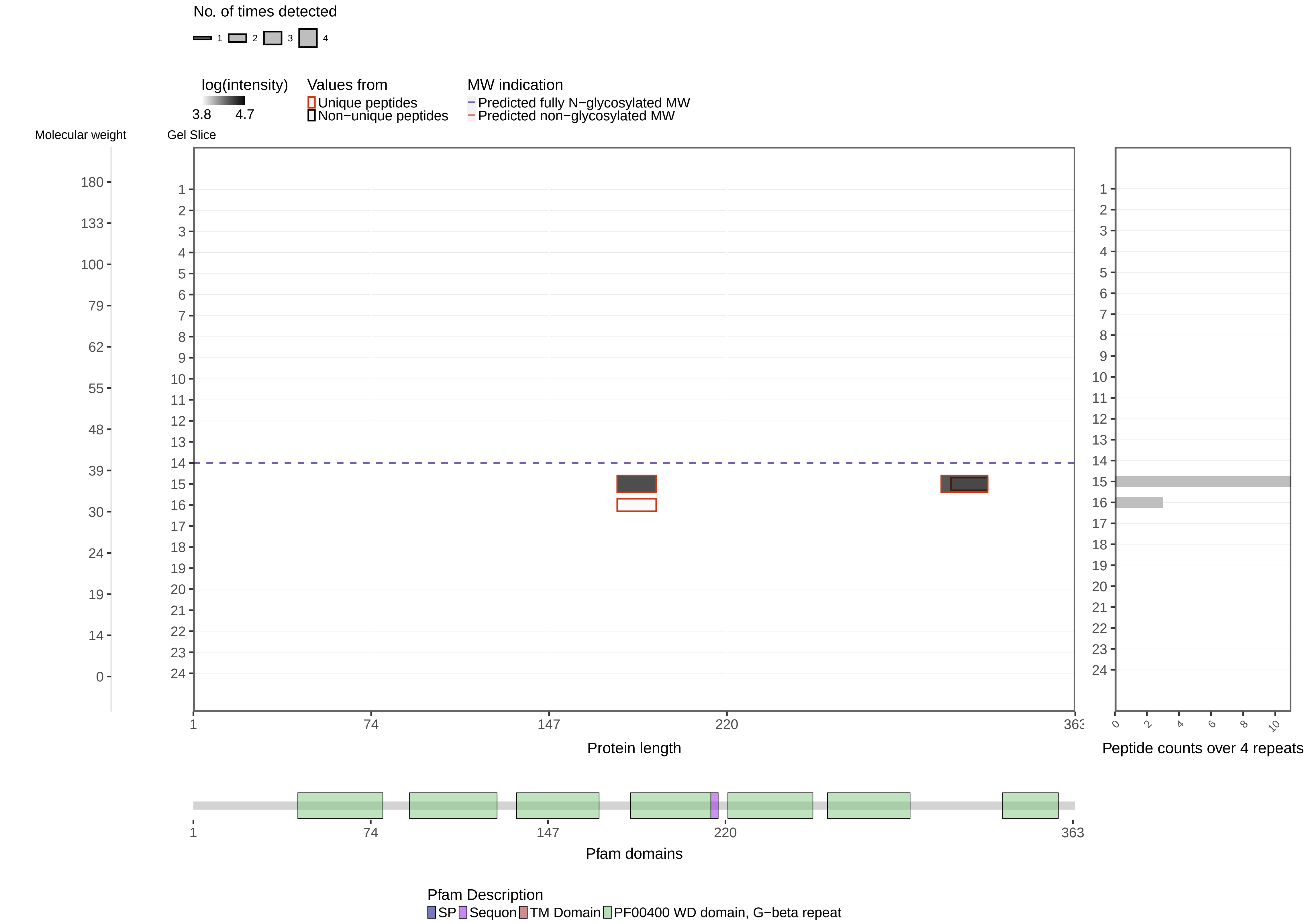


Protein sequence:

NTLGECKYTIQDGDSHSDWVSCVRFSPNTLQPTIVSGSWDRTVKIWNLTNCKLRSTLAGHAGYVNTVAVSPDGSLCASGGKDGTILLWDLAEGKKLYSLDAGSIIHALCFSPNRYWLCAATETSIKIWDLESKSIVVDLKVDLKQESEMATEGTIGSACKNKIMYCTCLSWSADGSTLFSGYTDGLIRVWGIGRY*

### NbL13g06080.1 beta−glucosidase_44−like [MW = 60.2 kDa]

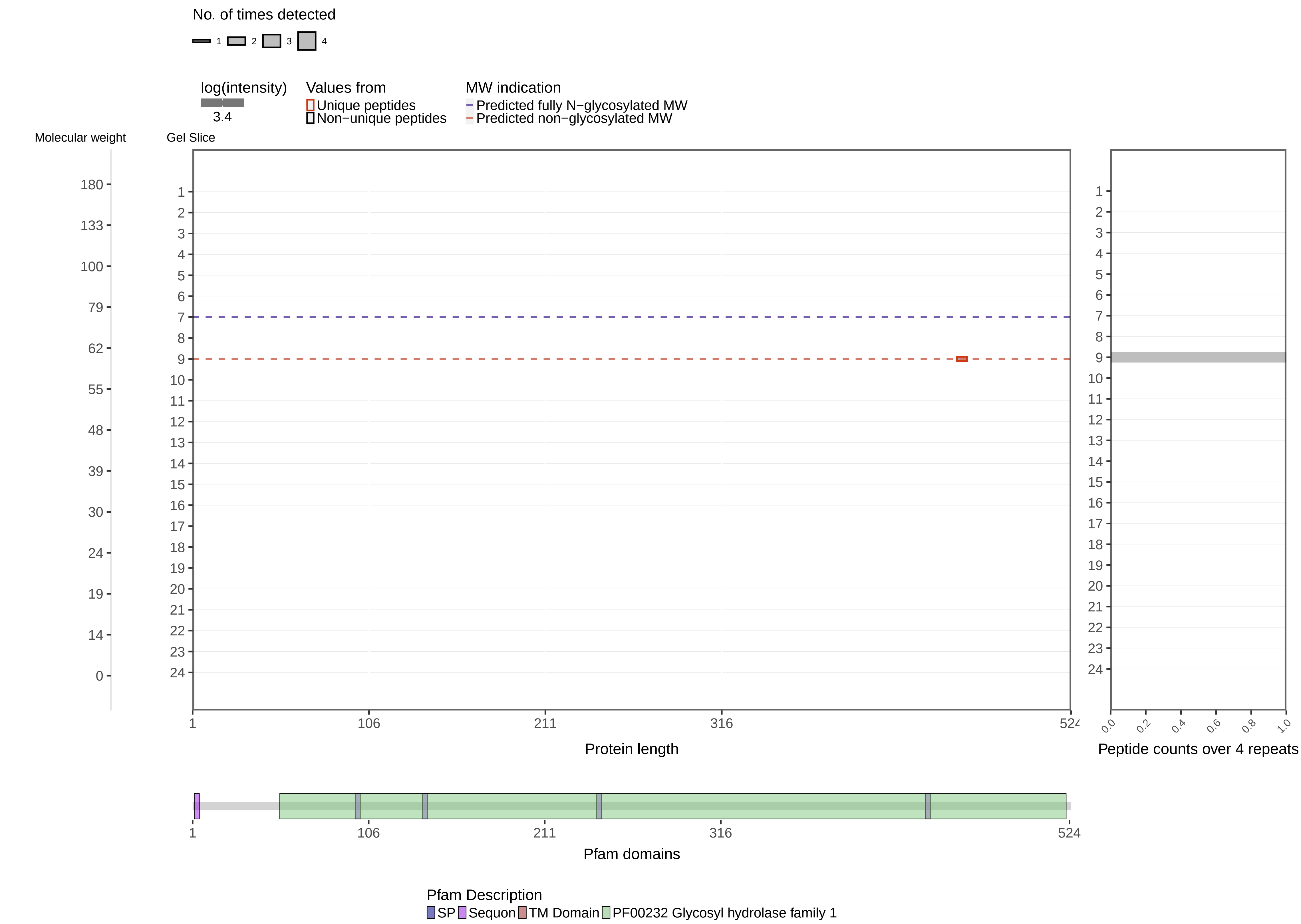


Protein sequence:

EPYIVAHNLILCHATAAQRYREKYQEKQKGKFGILLDFVWYEPLTRGKADNYAAQRARDFHLGWFLHPLVYGEYPKTMQNIVGKRLPKFTKEEVTMVKGSIDLLGINQYTAYYMYDPHYTTPQPLGYQQDWNVGFAYDRKGVPIGPRAHSDWLYIVPWGLHKAVNYVKERYGNPTMILAENGMDYAGNITLPKALYDTKRIAYYKSYLQELKKTVDEGANVIGYFAWSLVDNFEWRSGYTSRFGIVYIDFNTLKRYPKLSALWFKKLLRRHKH*

NbL13g06060.1 66_kDa_stress_protein−like [MW = 66.1 kDa]

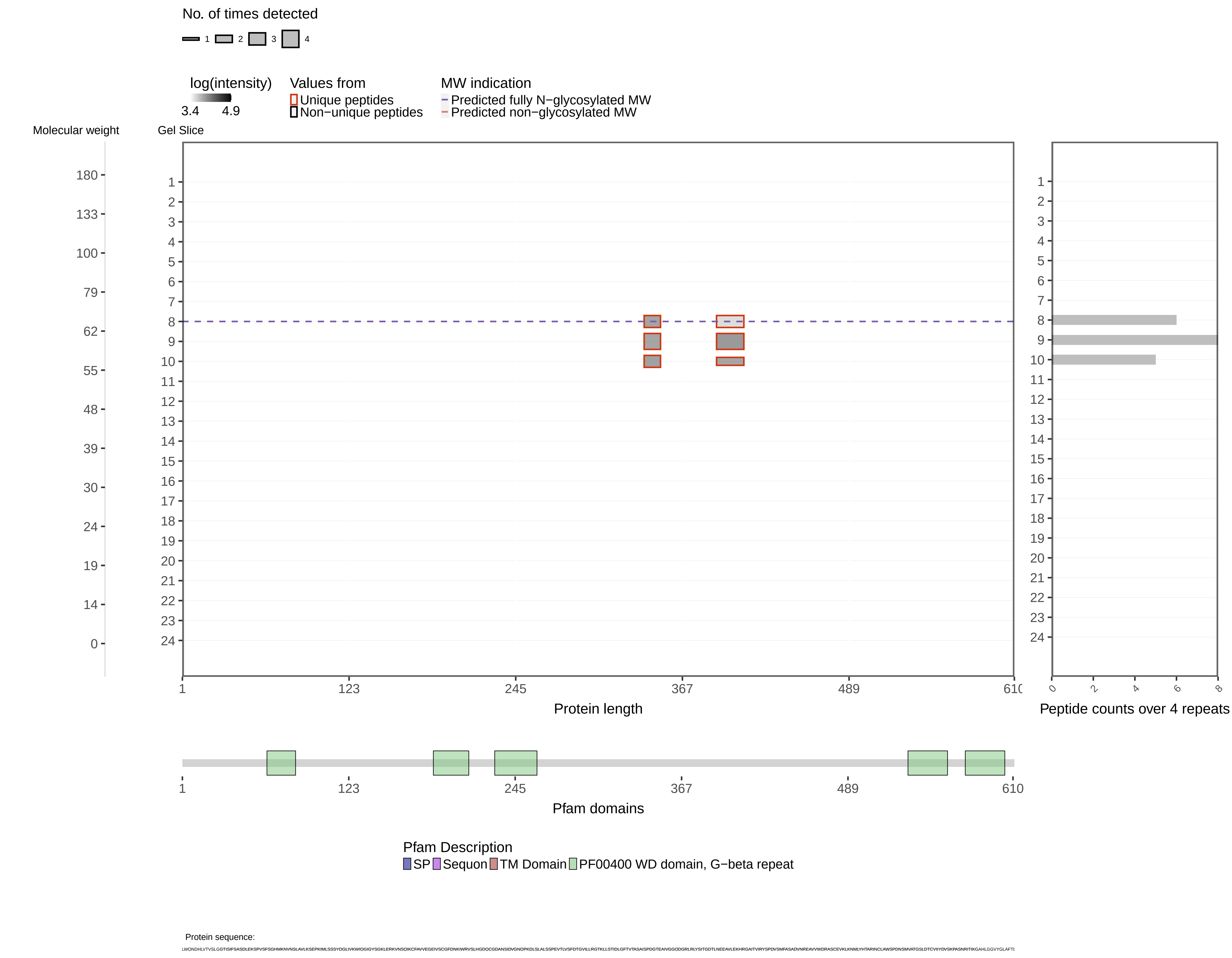

### NbL13g06020.1 histone_deacetylase_6 [MW = 16.29 kDa]

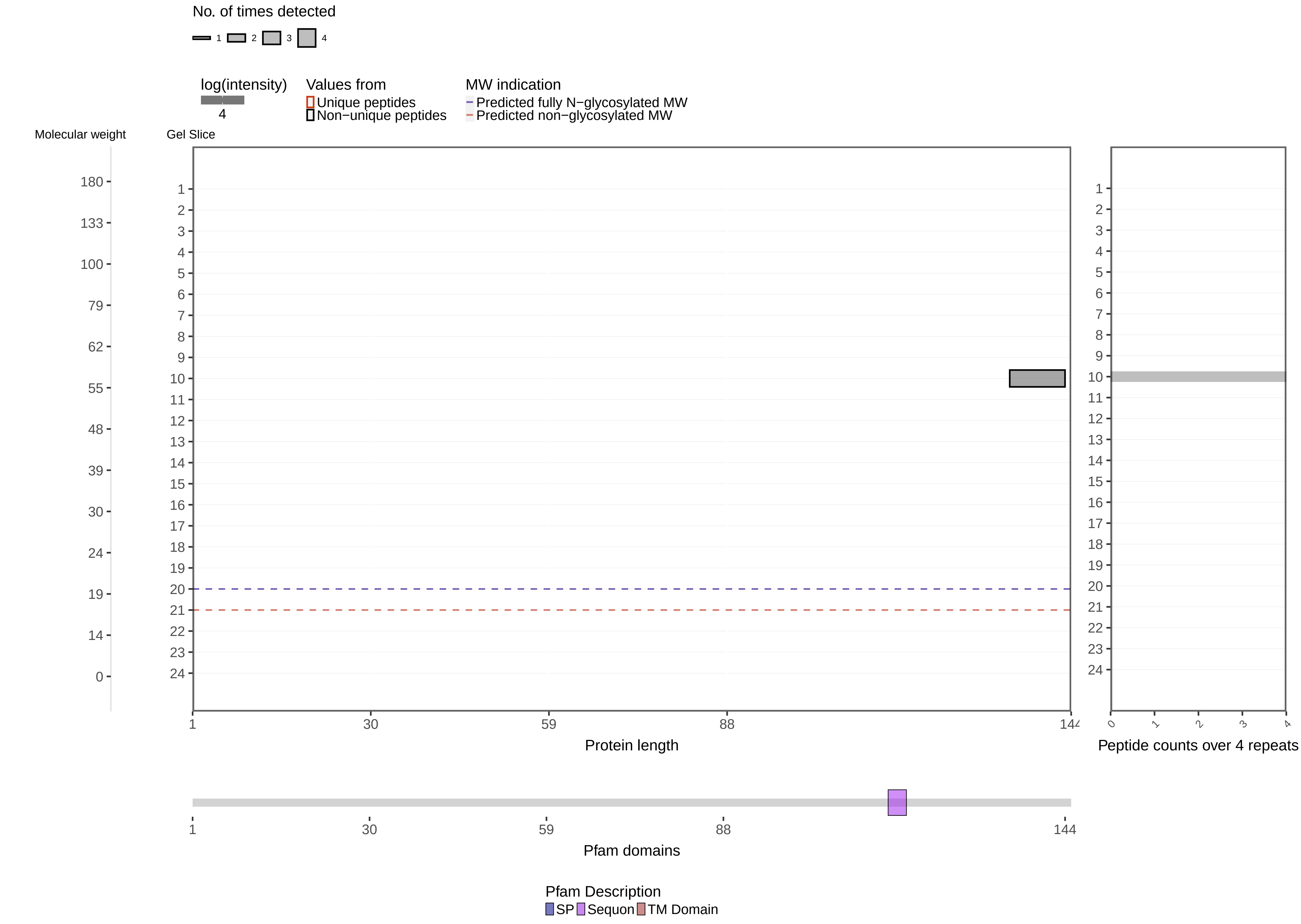


Protein sequence:

APSVPFQTTPCTTEVPEEEEEDMDQRPKPSIWNGEDYGSDADEDEHPRQRSSDSNLTPVETTTDMRDVDDQAKVDDDMVDADH*

### NbL13g05940.1 SH3_domain−containing_protein_C23A1.17−like [MW = 58.06 kDa]

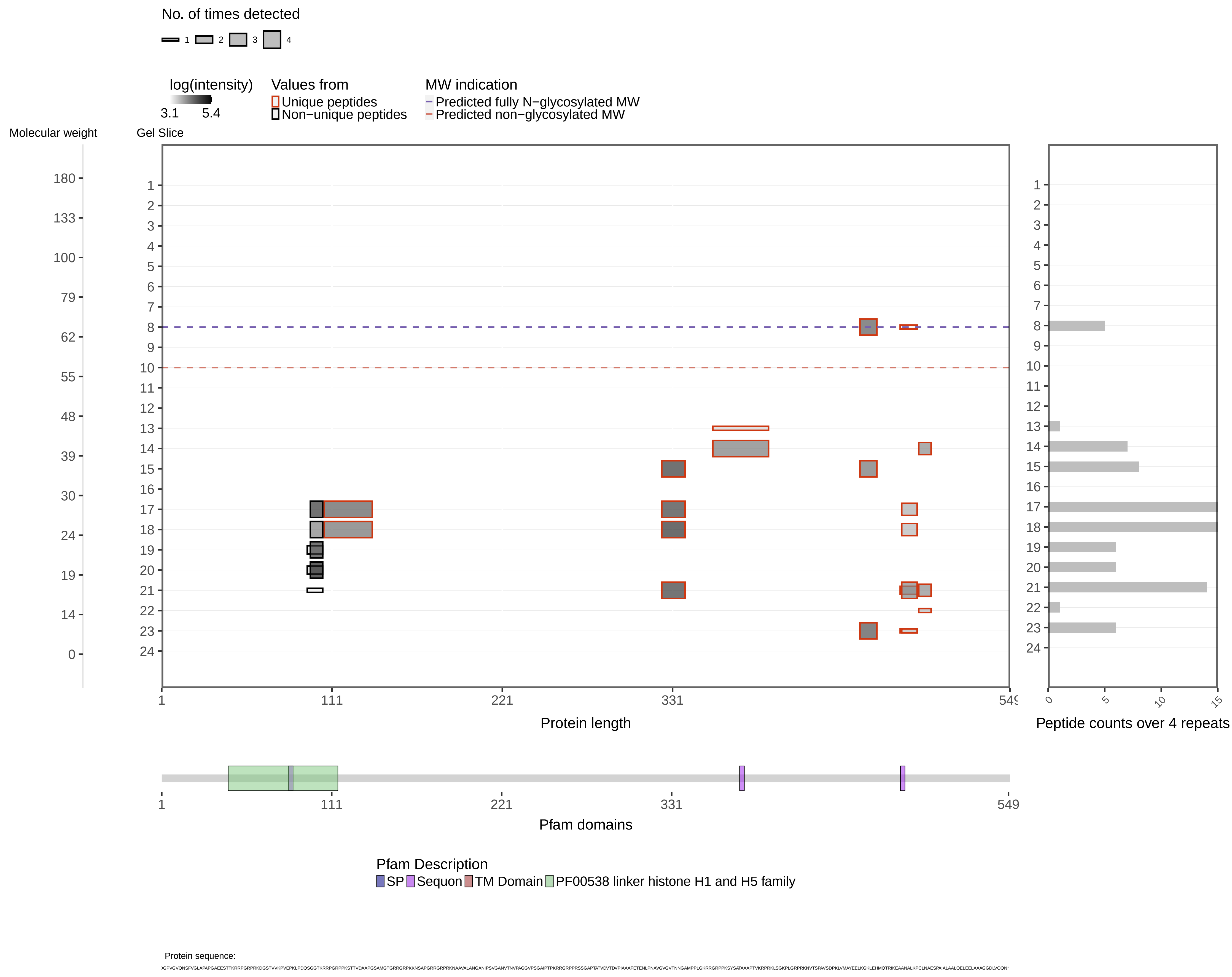

NbL13g05830.1 ubiquitin_carboxyl−terminal_hydrolase_3−like [MW = 26.01 kDa]

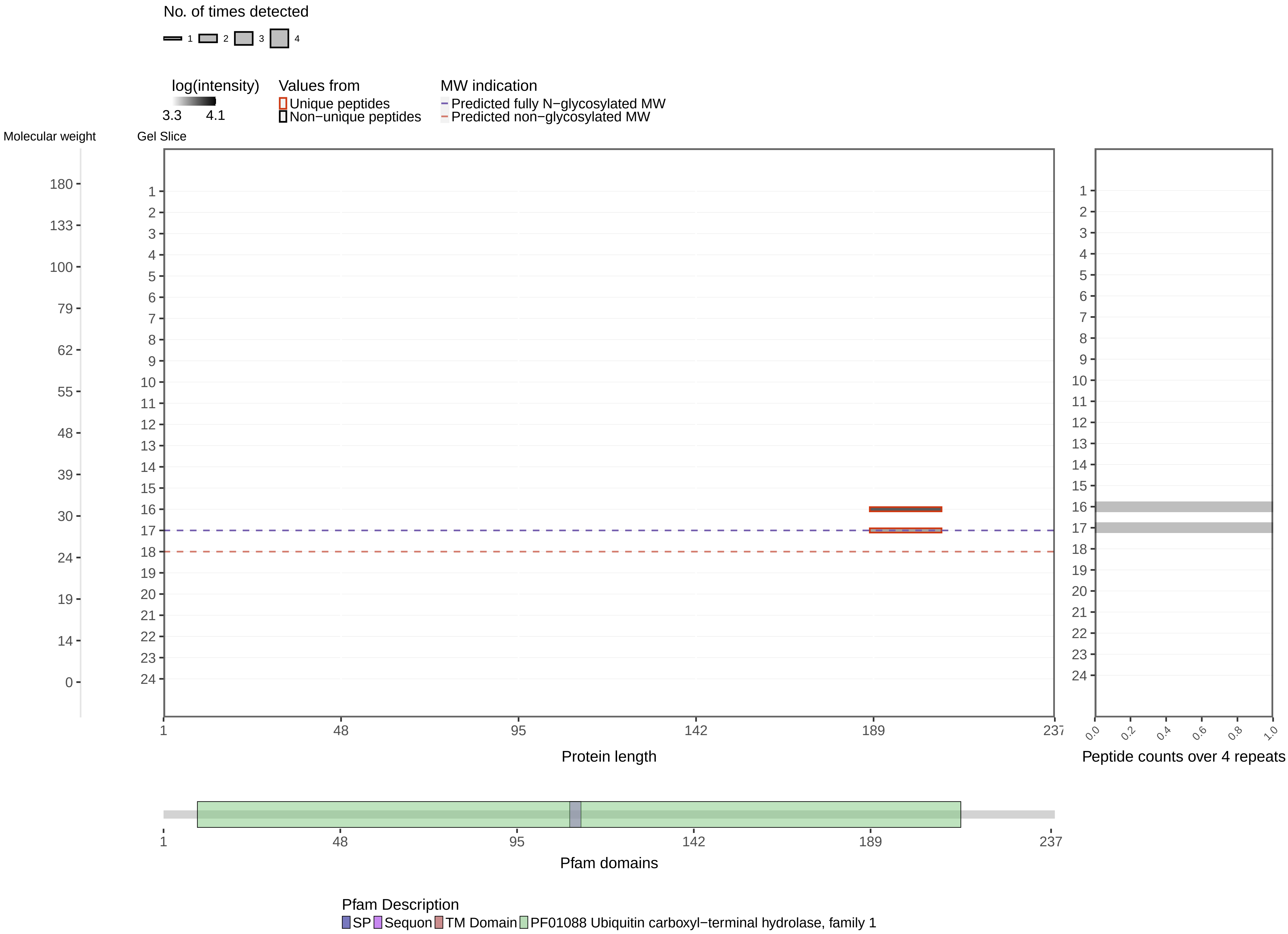


Protein sequence:

LHAIGNITSEIKLDEDSFLDKFFKSTAHMDPMERASFLENDREMEVAHSVAATAGDTEAPENVNTHFICFTCVDGQLYELDGRRSGPISHGASSPTNLLQDAAKVIEKMIQKNPESINFNVIAISKRVKGDAL*

### NbL13g05820.1 probable_enoyl−CoA_hydratase_2_mitochondrial [MW = 34.6 kDa]

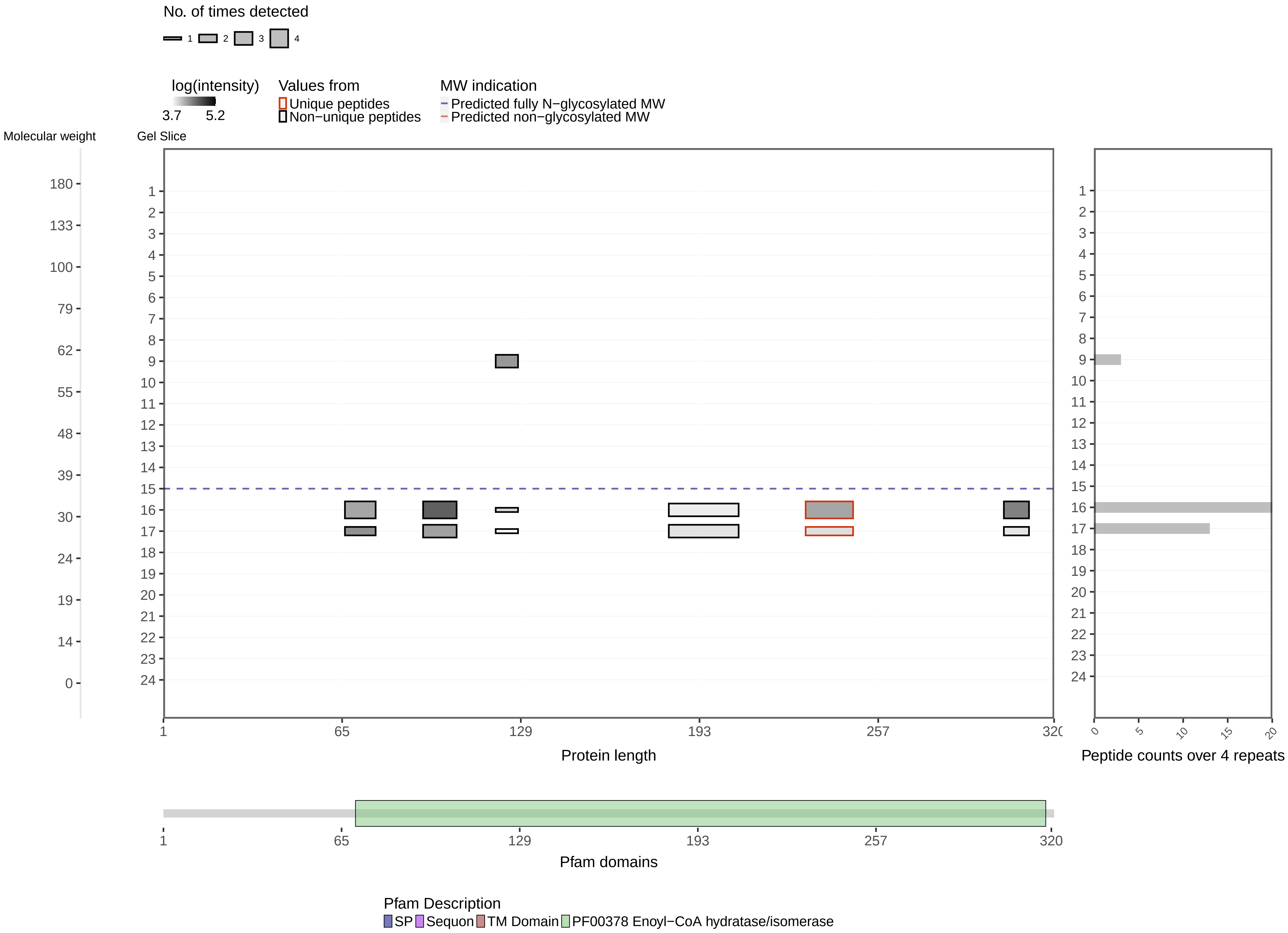


Protein sequence:

LRSTFSFLEGLHIPTIAAIEGIALGGGLEMAMSCDIRICGEDAVLGLPETGLAVIPGAGGTQRLPRLVGKSIAKDIIFTGRKINGKDAVSIGLVNYCVPAGETRLKALELARDINQKGPVALKMAKCAIDKGVELDMESALALEWDCYEQLLDTKDRLEGLAAFAERRKPRYKGE*

NbL13g05780.1 plasminogen_activator_inhibitor_1_RNA−binding_protein−like [MW = 39.51 kDa]

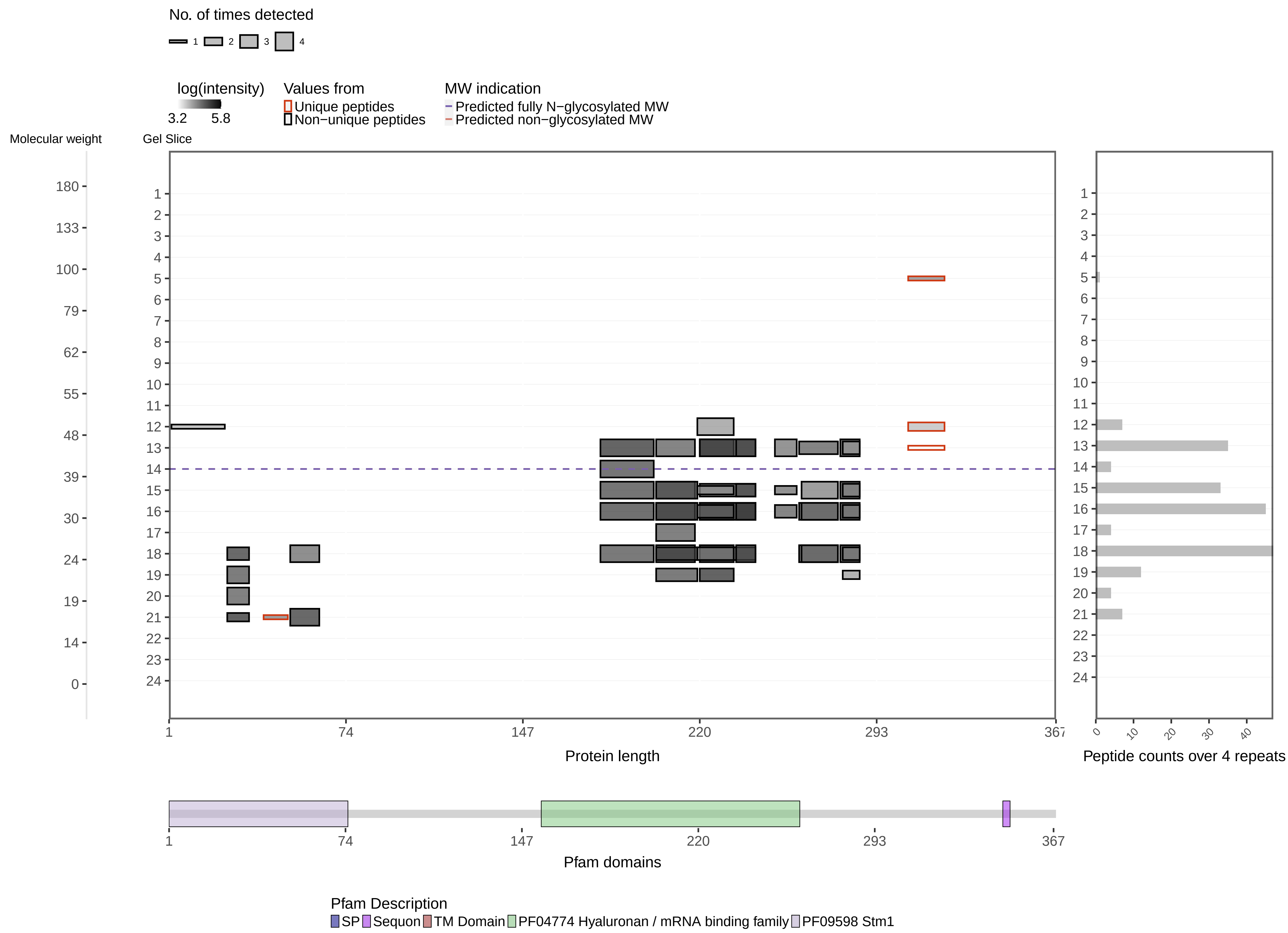


Protein sequence:

NDFKREGAGRANWGTPTDEIAPETEEPVTEGEKIVDPEKPAGEEDAGDAKKDSSAAEPEEKEPEEKEMTLEEYEKLLEEKRKALLAQKPEERKVNLDKEFESMQLLSNKKNEDDIFIKLGAEKDKRKEAVEKAKKTKSINEFLKPAEGENYYRGRGRGGRGGRGRGGYGGNGGNGGNNYSVEAPKIEDVGQFPSLGAK*

### NbL13g05760.1 GTP−binding_protein_YPTM2 [MW = 22.5 kDa]

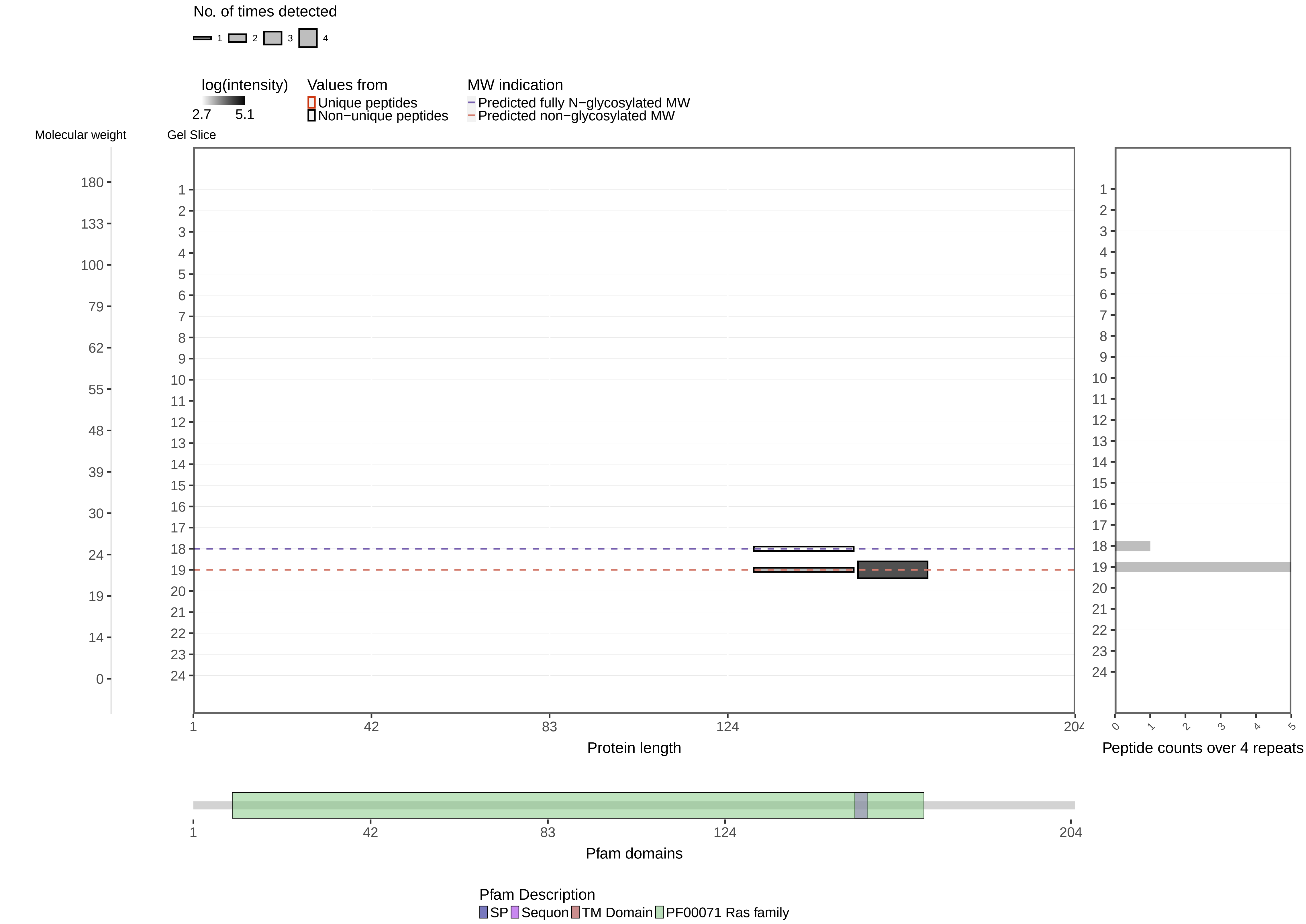


Protein sequence:

DQDSFNNVKQWLSEIDRYASDNVNKLLVGNKCDLTAQKVVSTETAQAFADEIGIPFMETSAKNATNVEQAFMAMAASIKNRMASQPASNNARPPTVQIRGQPVNQKSGCCSS*

NbL13g05730.1 50S_ribosomal_protein_L19_chloroplastic−like [MW = 24.04 kDa]

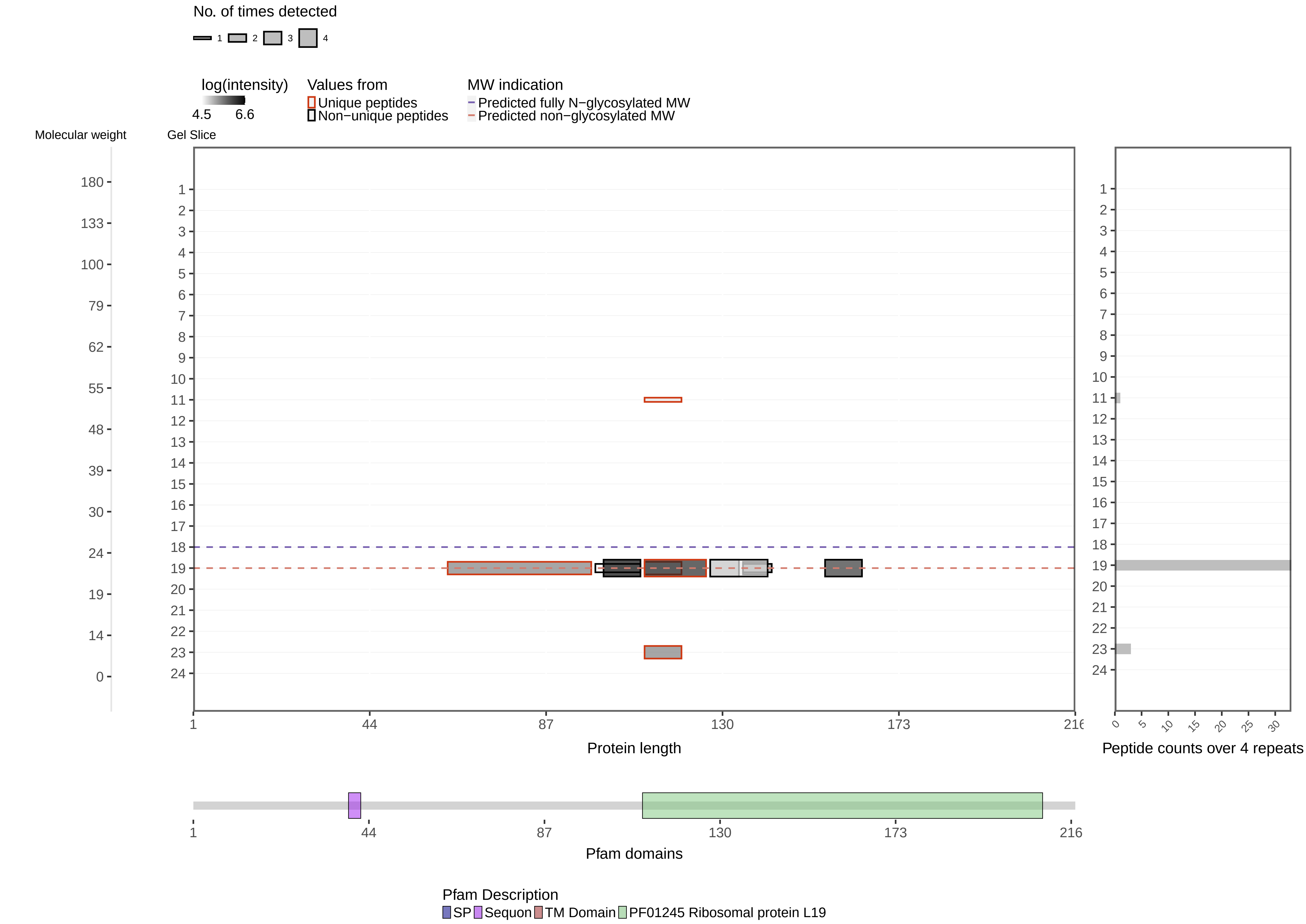


Protein sequence:

ERPPFKPRAKLGDIMGILNKQAIEASDAVRPIPDIRTGDVVEIKLEVPENRRRLSIYKGIVISKQNAGIHTTIRIRRIIAGIGVEIVFPVYSPNIKELKVVKHRKVRRARLYYLRDKLPRLSTFK*

### NbL13g05700.1 V−type_proton_ATPase_subunit_d2−like [MW = 20.92 kDa]

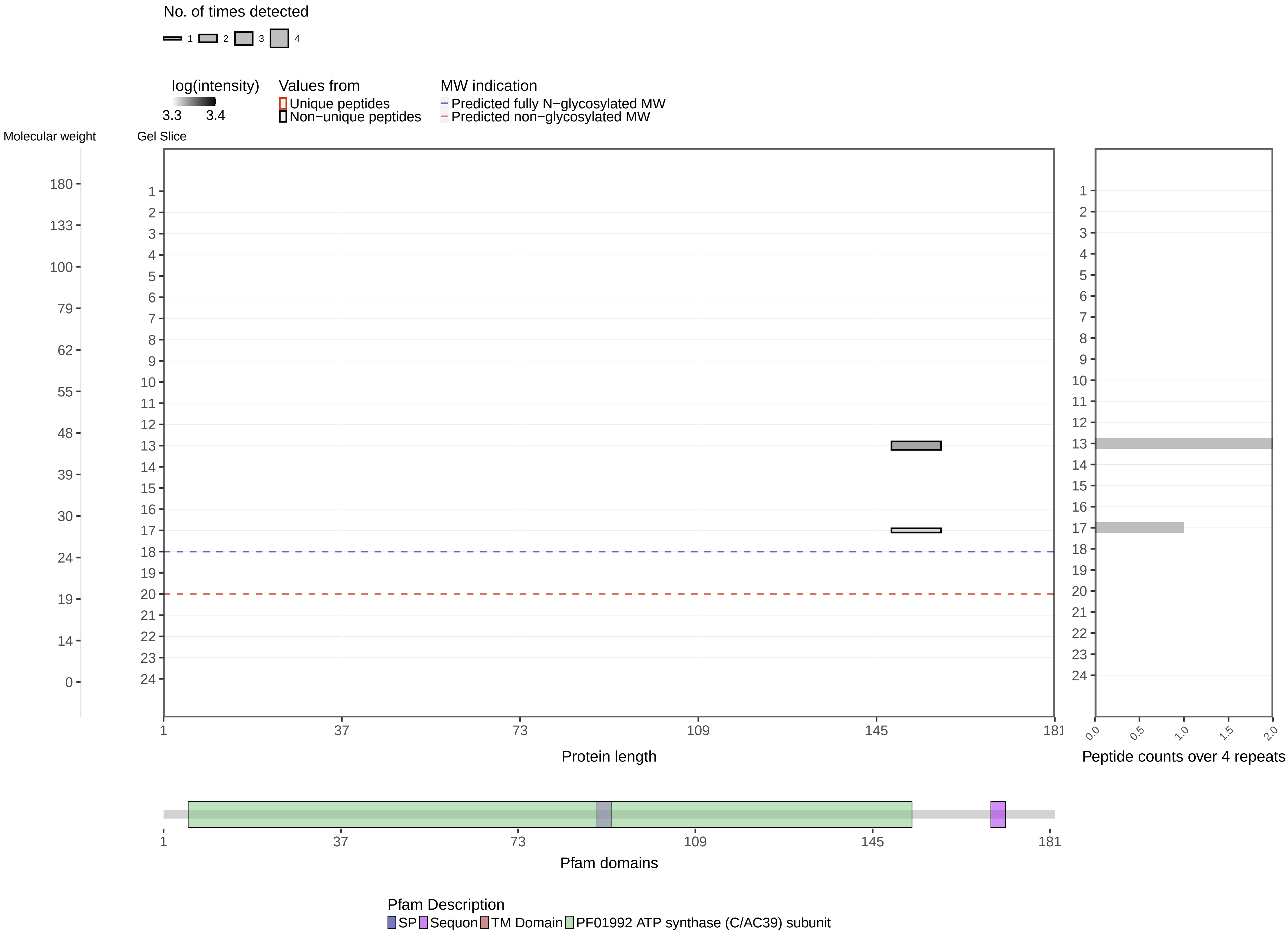


Protein sequence:

LAFEADRRAVNITINSIGTELTRDDRRKLYSNFGLLYPYGHEELAICEDIDQVRGVMEKYPPYQSIFSKLAYGESQMLDKRKSNTFAWHLNNSSTMVYFLHM*

NbL13g05620.1 ATP−dependent_Clp_protease_proteolytic_subunit−related_protein_4_chloroplastic [MW = 33.31 kDa]

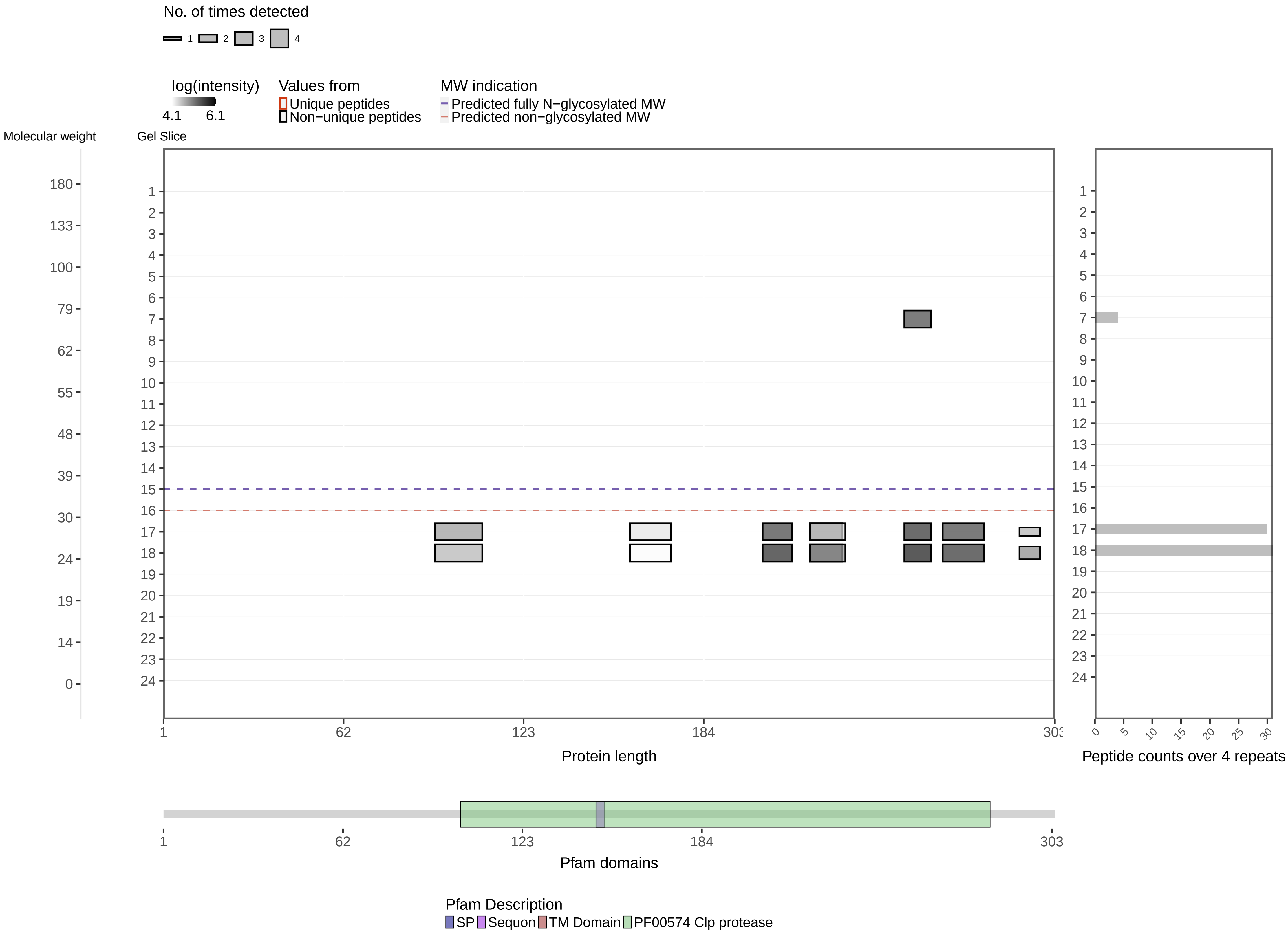


Protein sequence:
DEEKPIYLYVNSTGTTKGGEKLGYETEAFAIYDVMRYVKPPIFTLCVGNAWGEAALLLAAGAKGNRAALPSSTIMIKQPIARFQGQATDVELMRKEVKNVKAELVKLYSKHIGKSPEEIEADIRRPKYFSPSEAVEYGIIDKVLYNERGREDRGVISDLKKAQLI*

NbL13g05590.1 light−harvesting_complex−like_protein_3_isotype_1_chloroplastic [MW = 29.71 kDa]

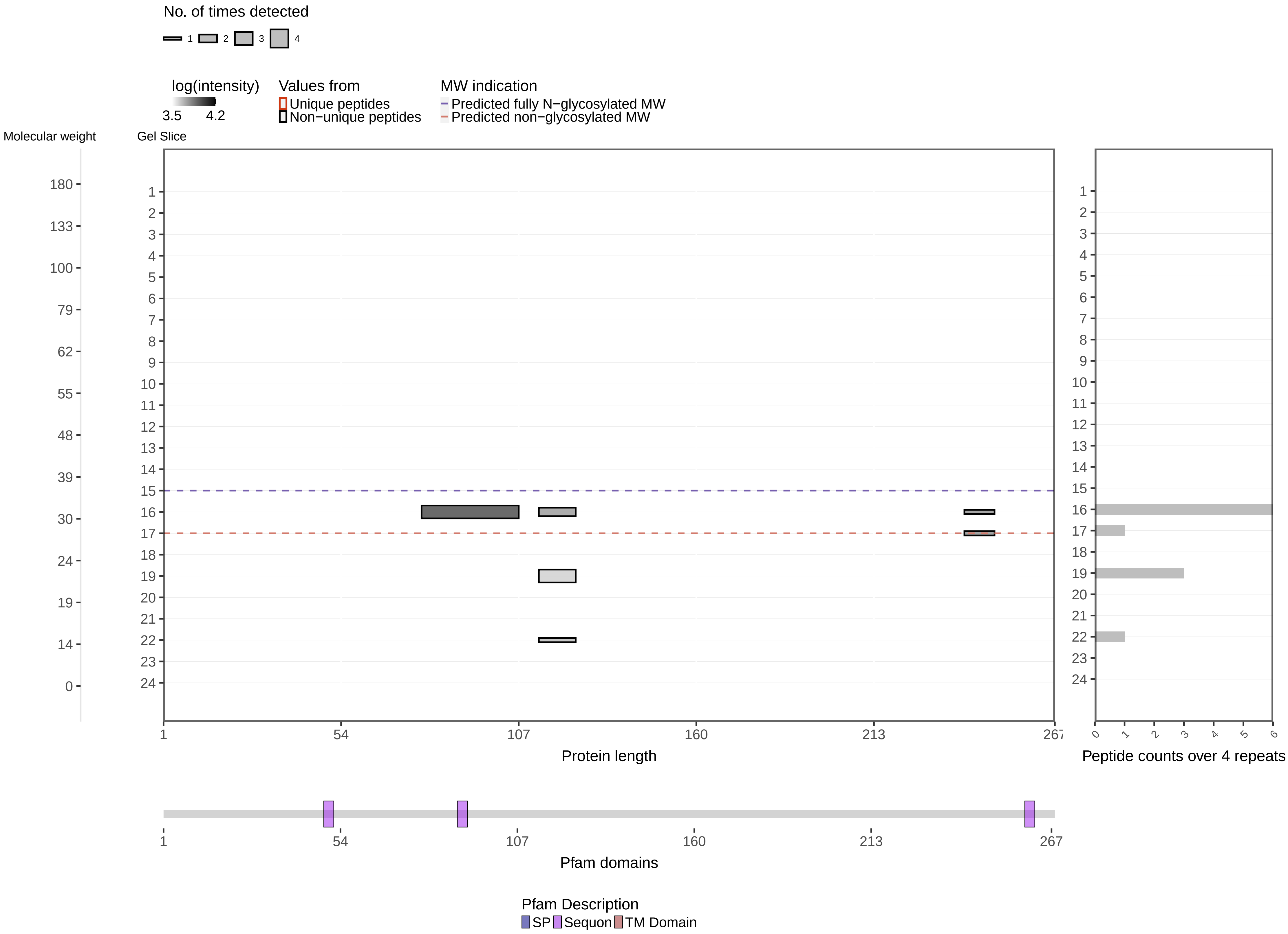


Protein sequence:

NQFKKDGETHWDSVIDAEVRRRKWLEDNPESSSNEEPVLFDTSIIPWWAWMKRFHLPEAELLNGRAAMVGFFMAYFVDSLTGVGLVDQMGNFFCKTLLFIAVAGVLLIRKNEDIETVKKLIEESTFYDKQWQASWKDENSSSSKDS*

NbL13g05560.1 (S)−ureidoglycine_aminohydrolase [MW = 36.76 kDa]

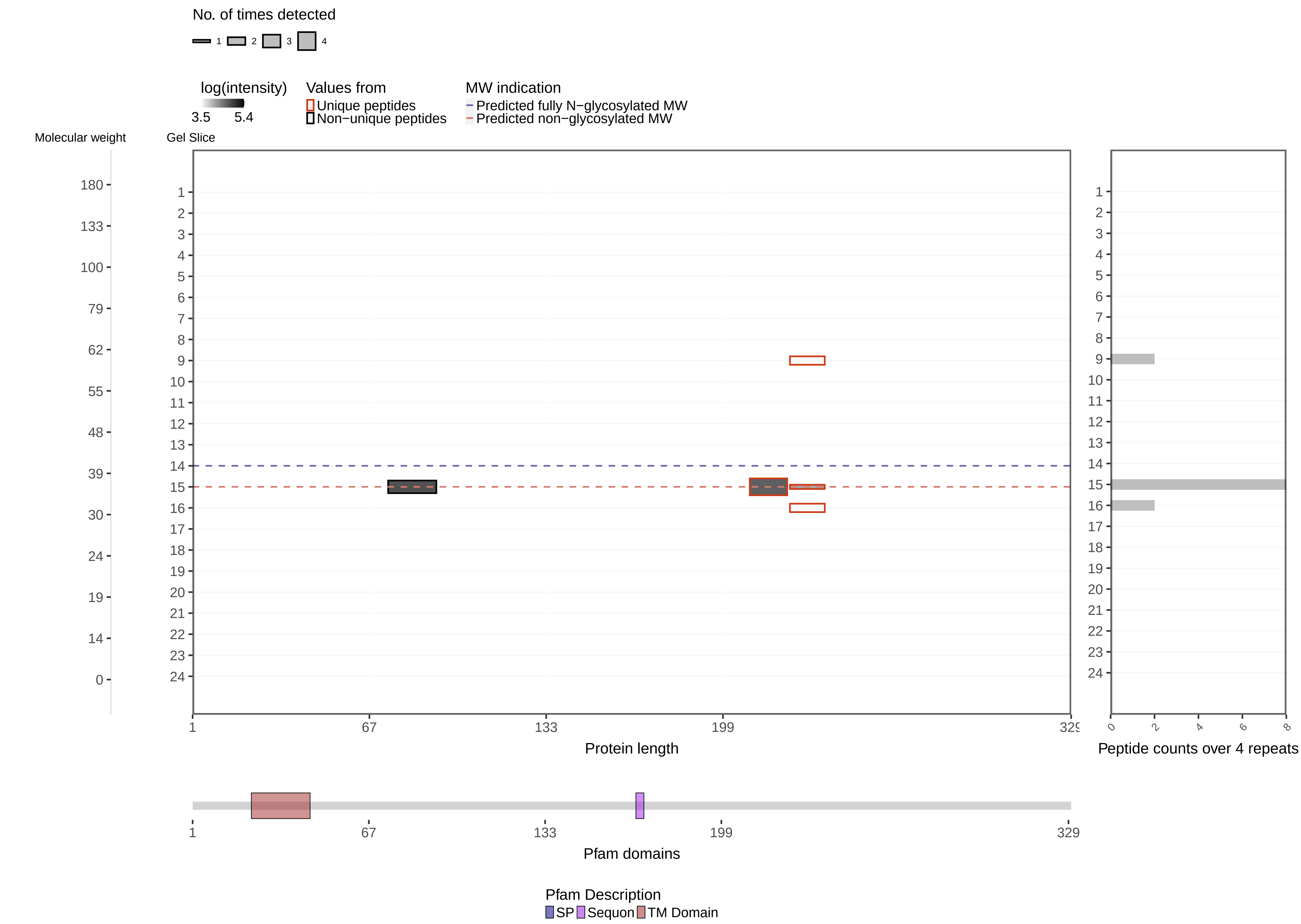


Protein sequence:
VERFIFVVQGSVVLTNISGNVHKLTVDSYAYLPPNFDHAVETDAAATLIVFERRYSRLENHVTEQIVGSTEKQPLLETPGEIFELRKLLPTSLAYDFNIHIMDFQPGEFLNVKEVHYNQHGLLLLEGQGIYRLSDSWYPIQAGDAIWMAPFVPQWYAALGKTRSRYLLYKDVNRNPL*

NbL13g05190.1 expansin−like_B1 [MW − SP = 29.79 kDa]

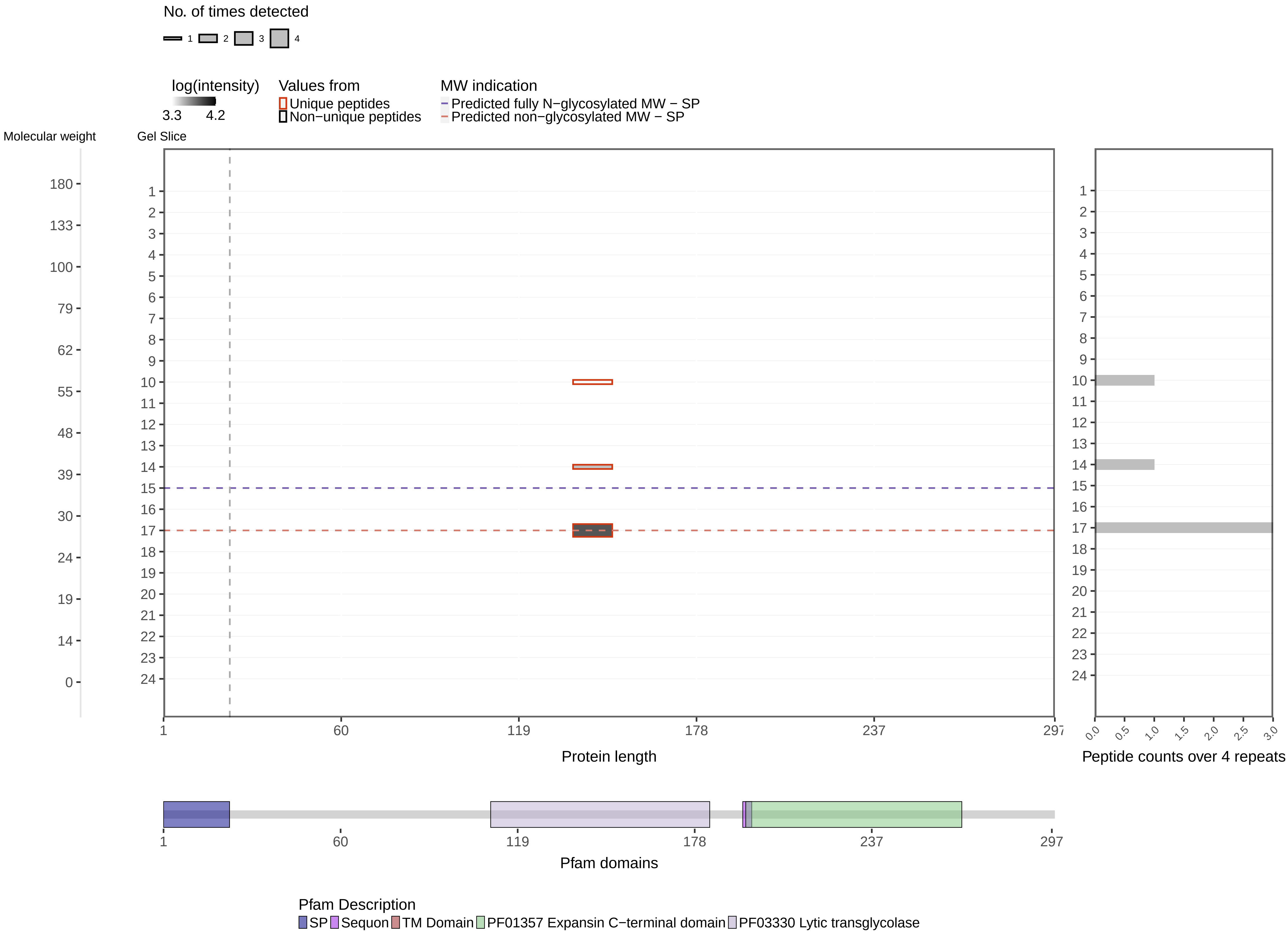


Protein sequence:

KGVTVVITDQGAGDRTDFILSQRAFARMAQTTDAAASLLSLGVVDIEYTRVSCSYPDKNITIKIEESSDNPHYLAFVIWYQQGKKDITAVQLCETQNFECKVLDRTRGAVWTATSPPRGPLQIRMLLSVDDGDEIWFVPVNNIPENWKAGDTYDSGIQVDA*

### NbL13g05120.1 phosphoacetylglucosamine_mutase [MW = 60.4 kDa]

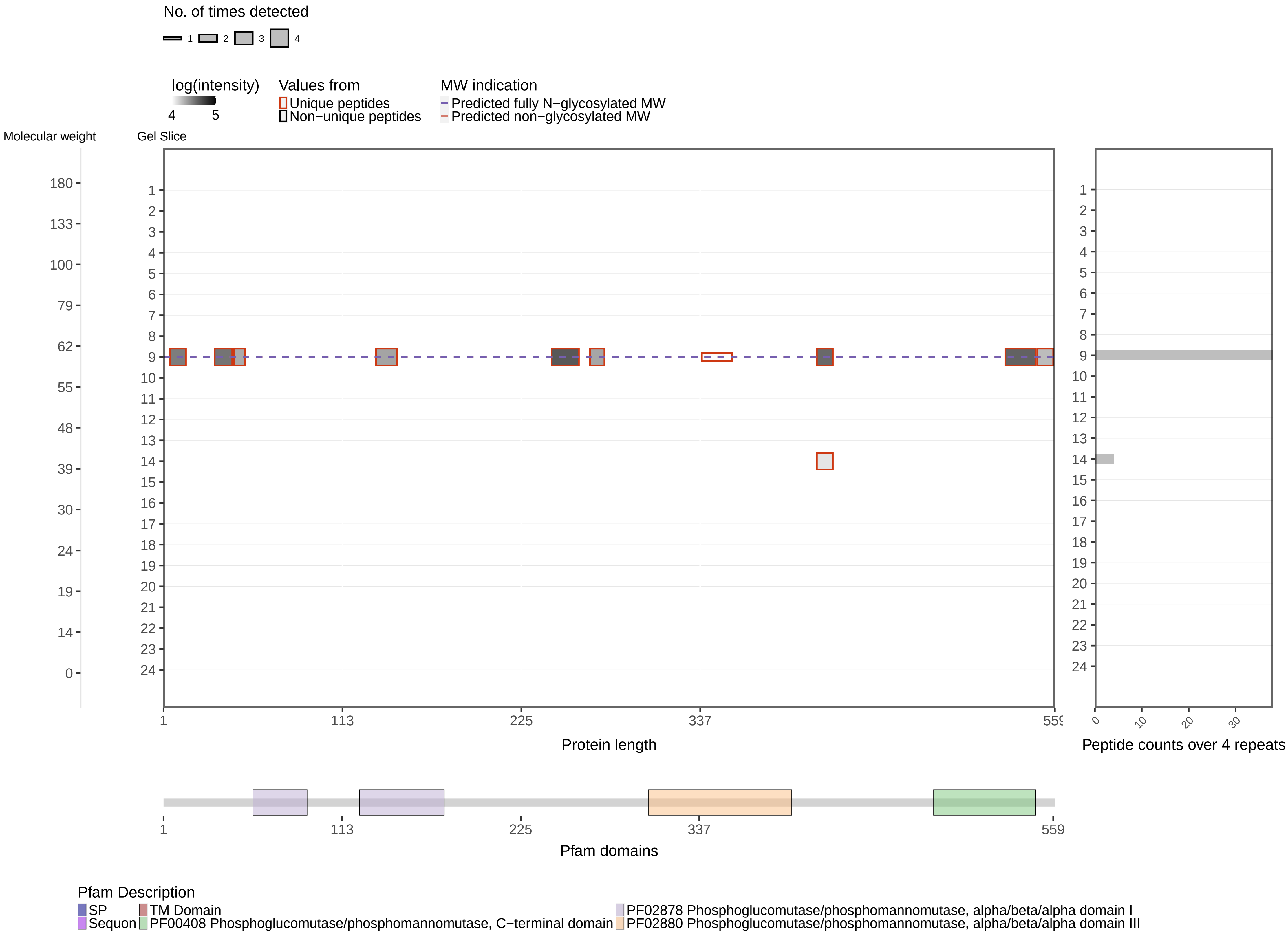


Protein sequence:

/APRSFGSNDAGLRCASLDGDADRLVYFSVLSNENNKIELVDGDKILSLFALFIKEQLSILNEGEDKKGHDSYKAHLGVVQTAYANGASTDYLKEMGLEVVLTPTGVKYLHEKASEFDIGIYFEANGHGTILFSEAFLCWLETSHKMLLSTSEGSAKQKAASRLLAVSQLINQAVGDALSGLLLVEVILKYMGWSIHRWNELYHDLPSRQLKVKVFDRTAVVTANAETVTVQPIGIQEAINAEIAKYPRGRCFIRPSGTEDVIRVYAEATTQDAADALASSVARLVDQYLGFGSS*

### NbL13g05030.1 D1−like [MW = 20.01 kDa]

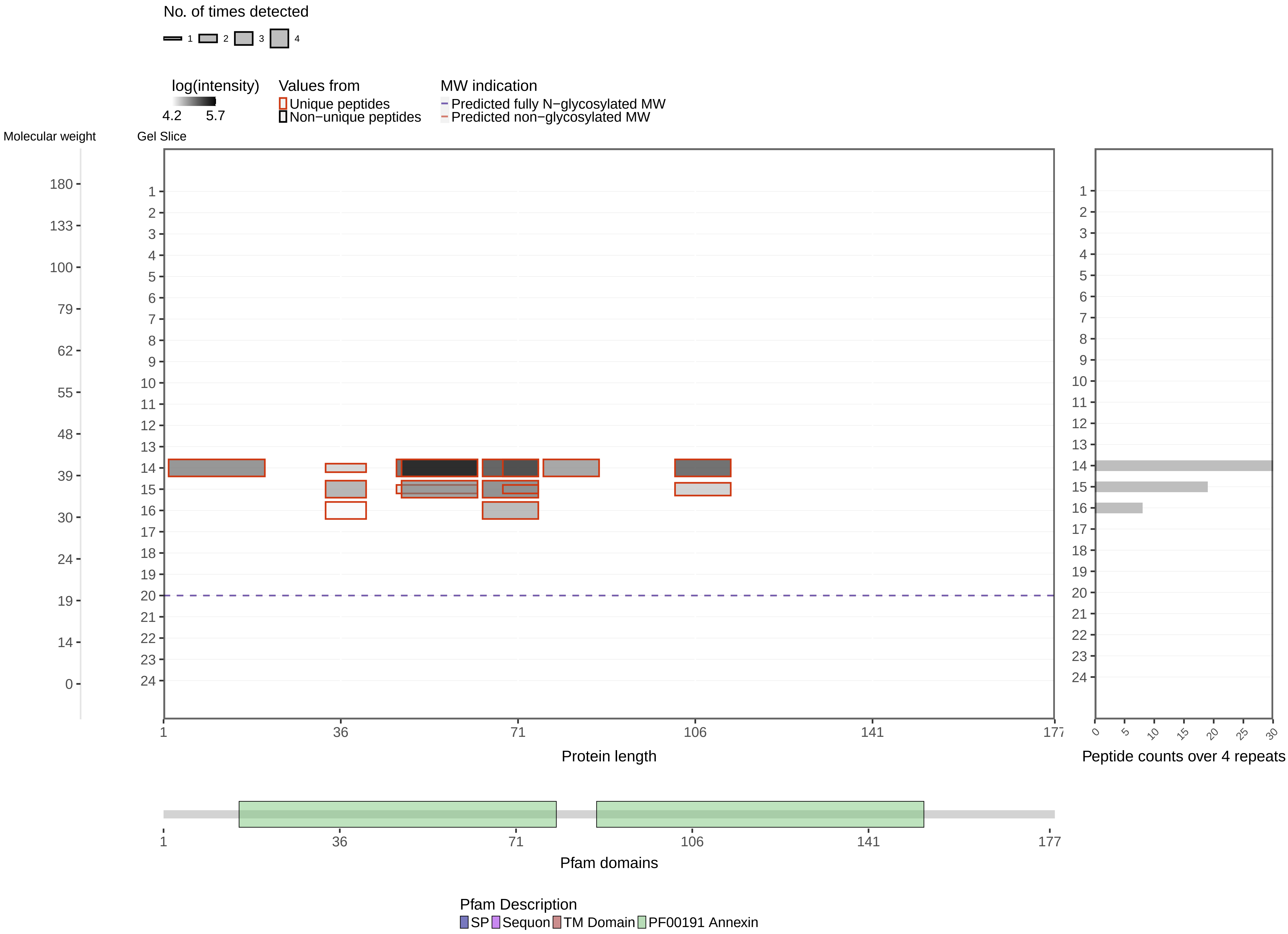


Protein sequence:

.VVVWTLDPPERDAYLAKEATKRWTKSNFVLVELACTRSPKELVLGREVYHARYKKYLEEDVAYHTTGEHRKLLVALVSSYRYGRDRAATAFGVIGGVVWT*

### NbL13g05020.1 probable_caffeoyl−CoA_O−methyltransferase_At4g26220 [MW = 32.86 kDa]

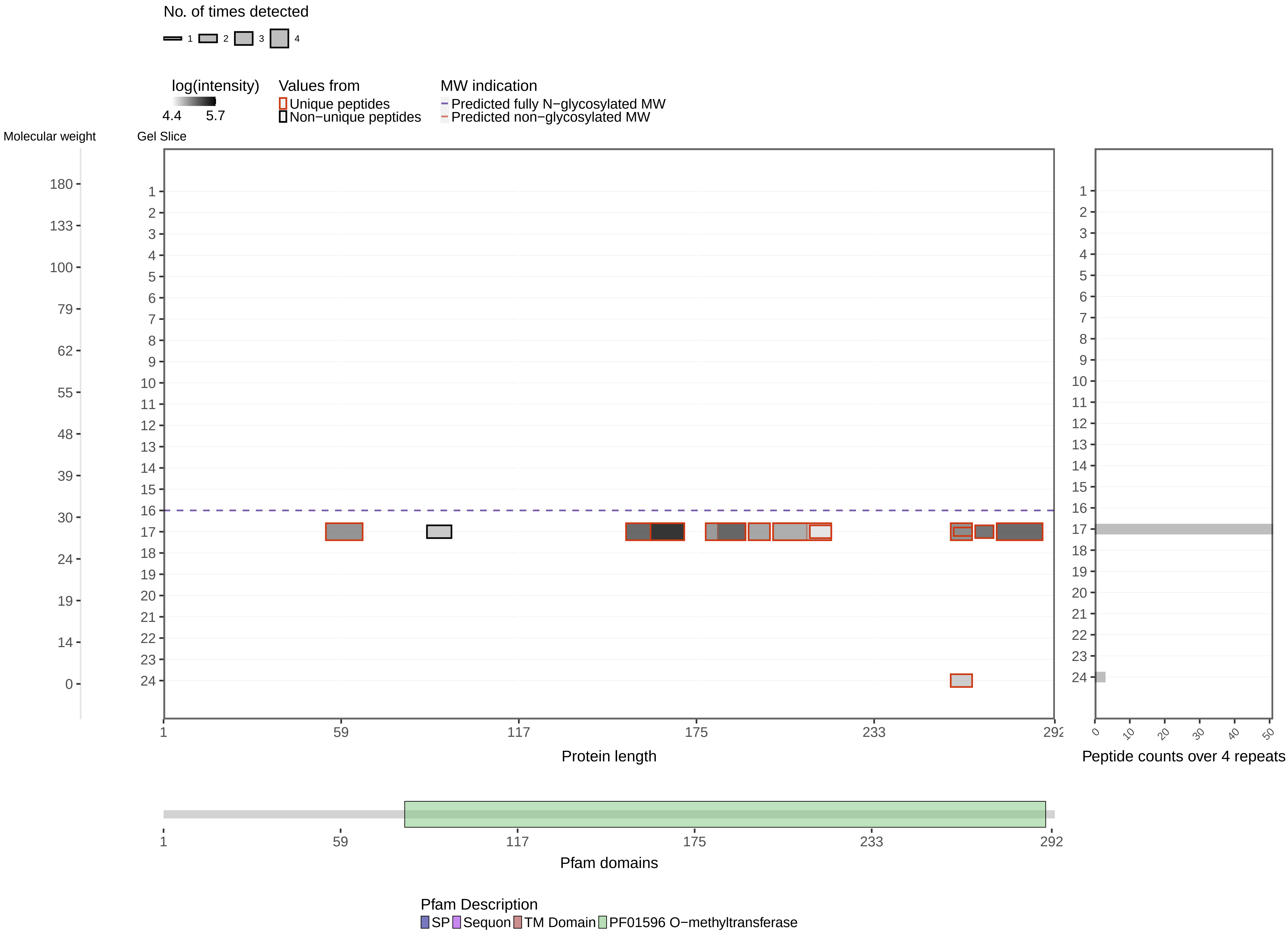


Protein sequence:

GVFTGYSVLLTALTIPDDGKIIAIDPDROSYELGLPVIOKAGVEHKIDFKDSPALPVLDKLLEDSENRDSFDFAFVDADKTNYGNYHKRLMOLVKVGGLIVYDNTLWYGTLVMPEEMVDELLRTERKYFLELNRFLASDTRIOISOVTVGDGMTICRRLF*

NbL13g05010.1 probable_caffeoyl−CoA_O−methyltransferase_At4g26220 [MW = 26.5 kDa]

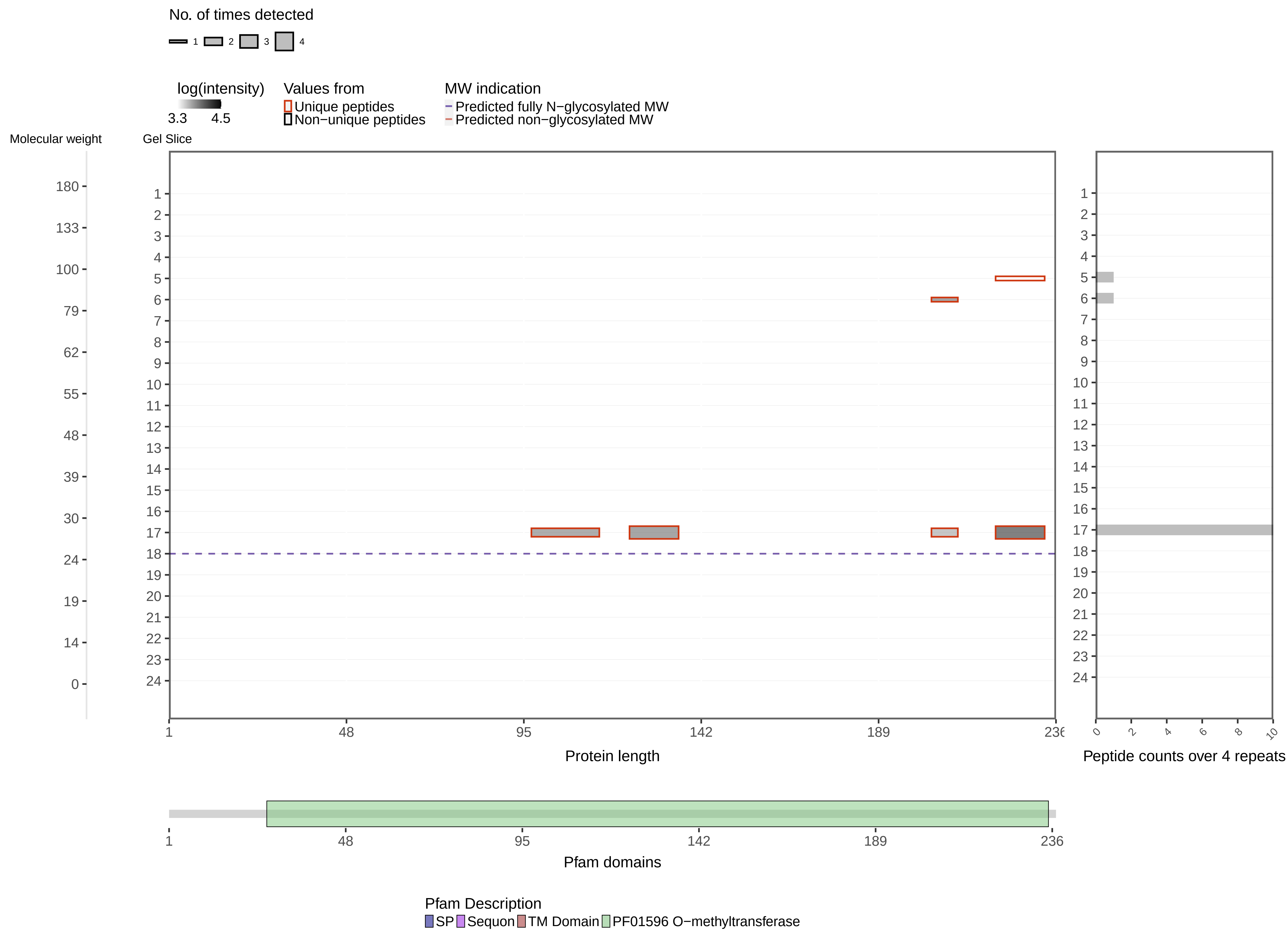


Protein sequence:

AYEMGLPVIKKAGVEHKINFIQSSALPVLDKLLEDNNNKESFDFAFIDADKVNYQKYHERLLQLVKVGGIIVYDNTLWSGTIAMPEESVPESLKAVRHHTIEFNKFLAADTRIRLSQVPLGDGITICRRL*

### NbL13g05000.1 histone_acetyltransferase_type_B_catalytic_subunit−like [MW = 53.06 kDa]

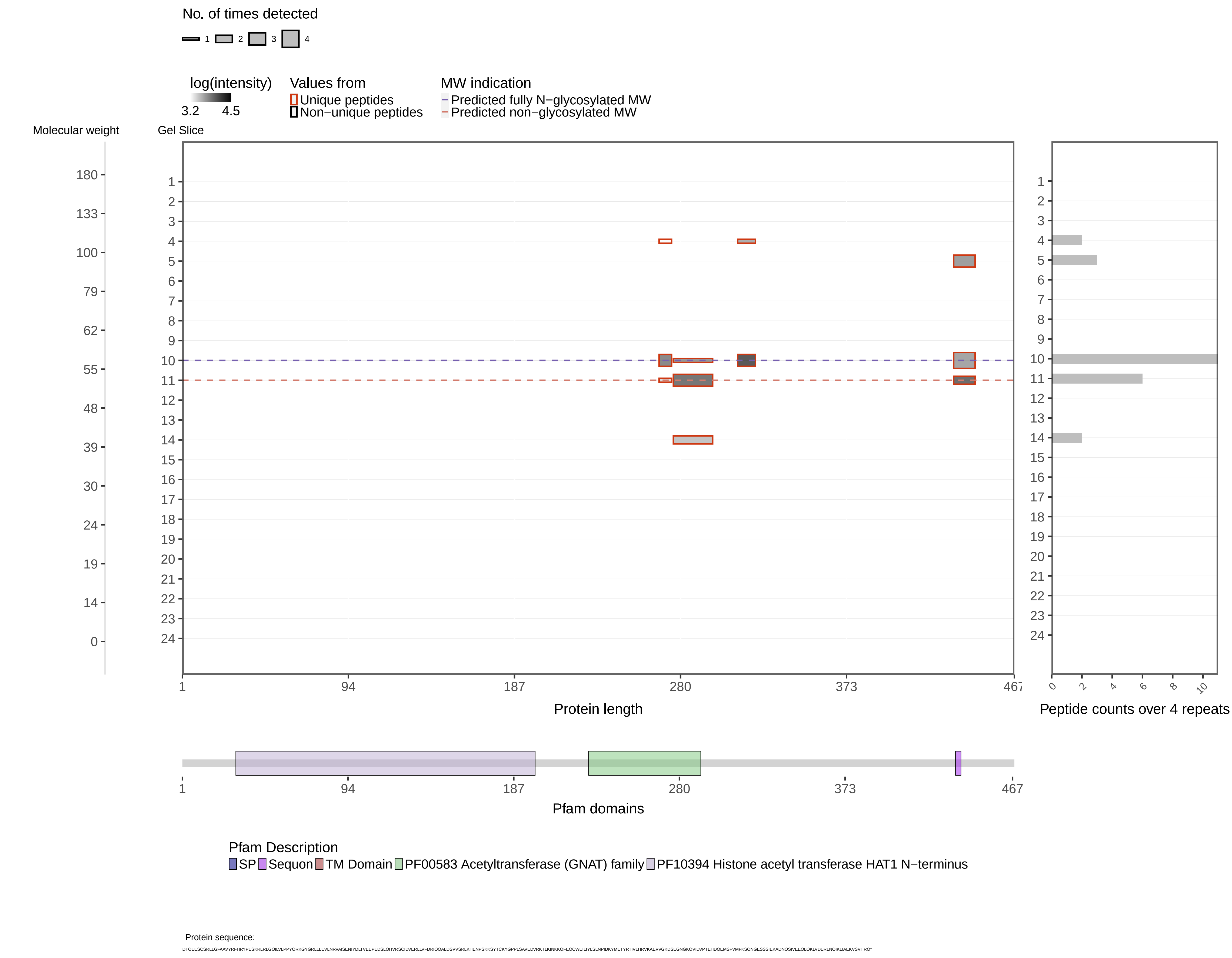

### NbL13g04930.1 ATP−dependent_6−phosphofructokinase_6−like [MW = 56.56 kDa]

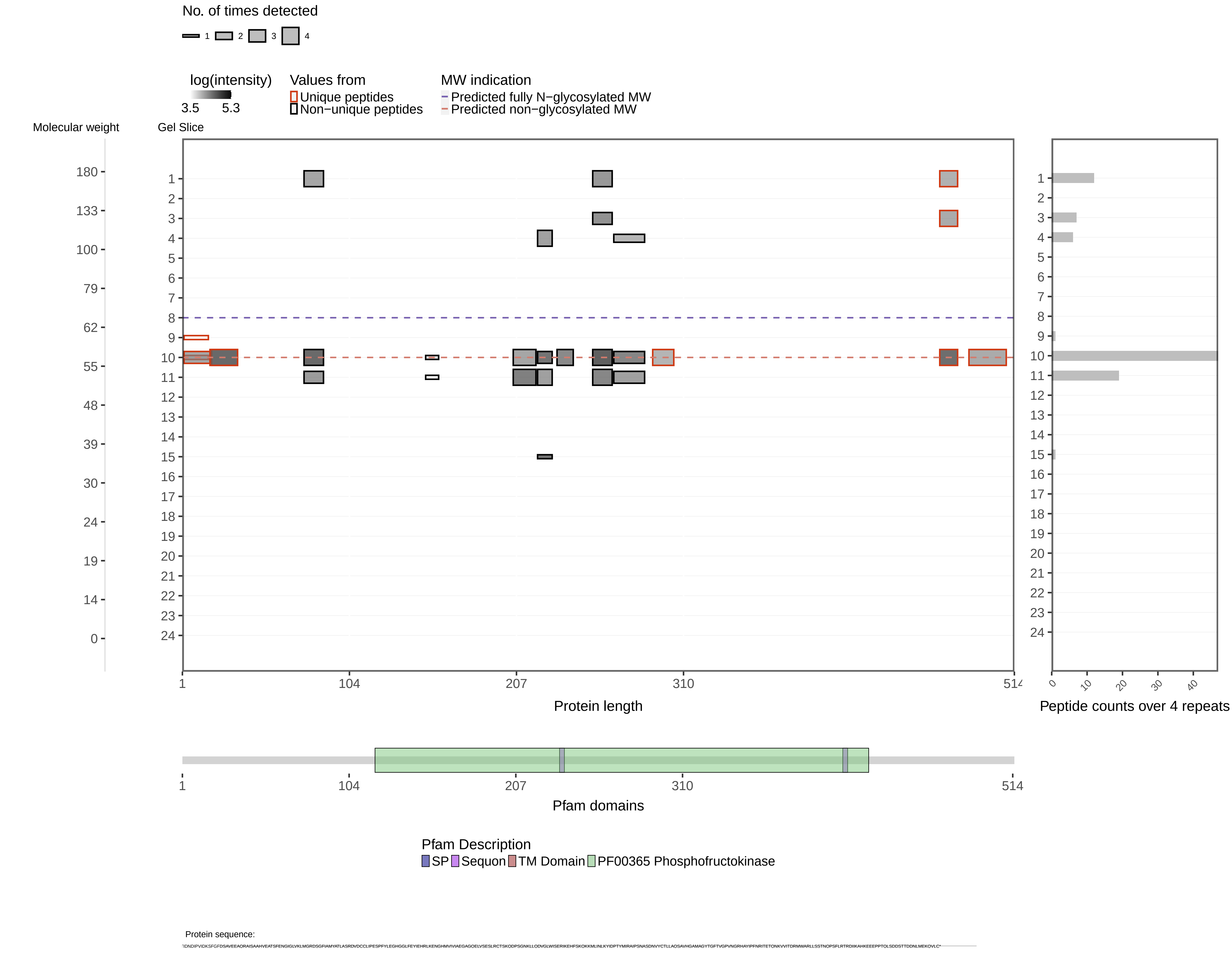

### NbL13g04910.1 leucine−rich_repeat_extensin−like_protein_4 [MW − SP = 41.64 kDa]

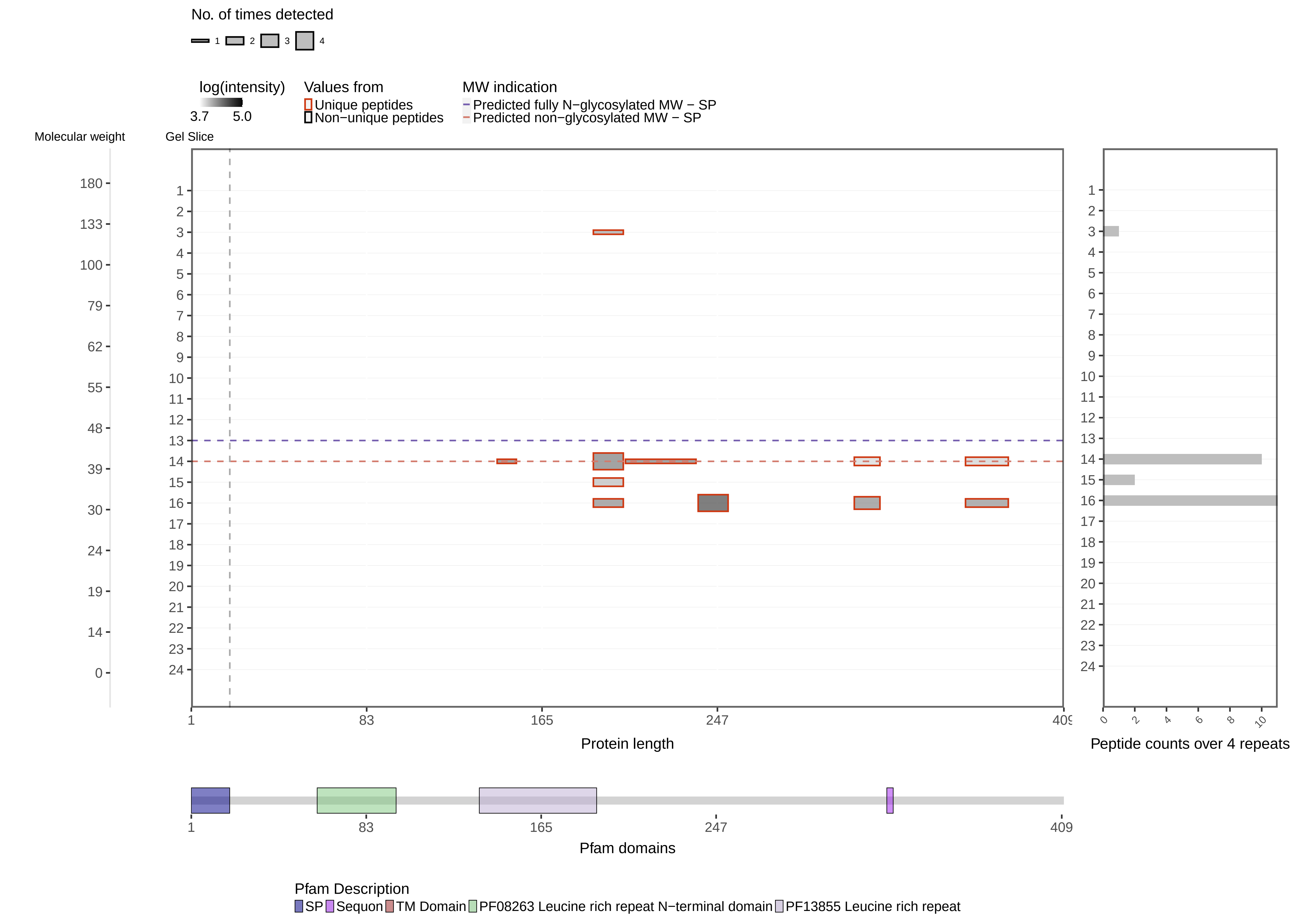


Protein sequence:

SFSGPIPEDLFNKKLDAIFLNNNQFDGELPQNLGNSPSSVINLANNKFTGNIPFSLGYMGPRIKEILFLNNQLTGCIPEGVGMWTDLQVLDVSFNSLMGHLPDTLSCLSGIEVLNLAHNKLSGDLPDLVCSLRSLINLTLTYNFFSGLSQDCDKLPYRNVGFDFSLNCIPGKQMQRPQPECSLIPGGSLSCLRIPAAKPLVCGSIISEEEIVSTPPSH*

### NbL13g04760.1 ATP_sulfurylase_2 [MW = 54.74 kDa]

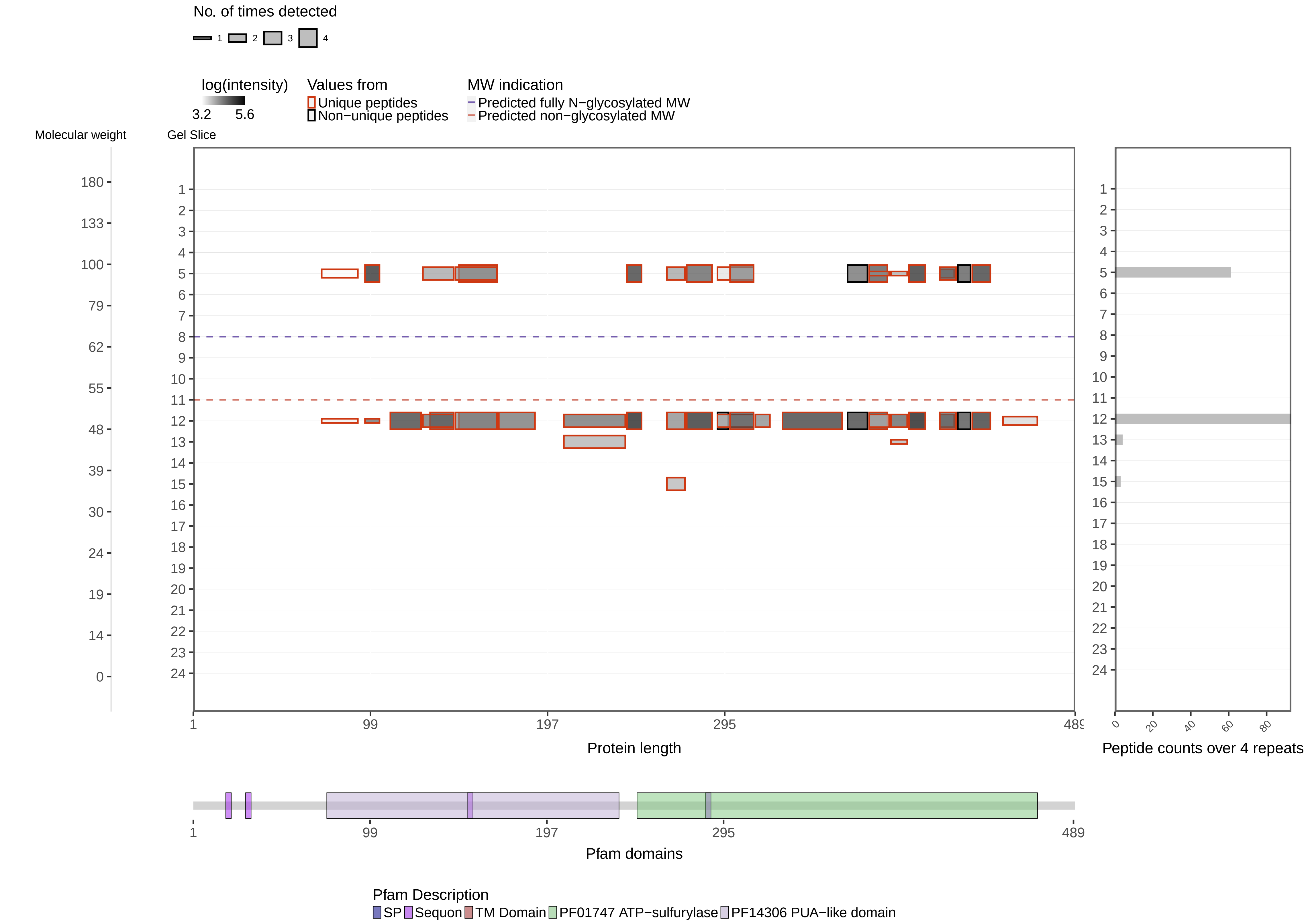


Protein sequence:

/IKPIKYNDGLDHYRLSPQQLRDEFDRREADAVFAFQLRNPVHNGHALLMNDTRRRLLEMGYKNPILLLHPLGGFTKVDDVPLDVRMEQHSKVLEDGVLDPETTIVAIFPSPMHYAGPTEVQWHAKARINAGANFYIVGRDPAGMSHPTEKRDLYDPDHGKKVLSMAPGLEKLNILPFRVAAYDTVDKKMAFFDPSRAQEFLFISGTKMRTYARTGENPPDGFMCPGGWEVLVKYYESLQAEDGVQNSAVLSAR*

NbL13g04750.1 rab_escort_protein_1 [MW = 60.71 kDa]

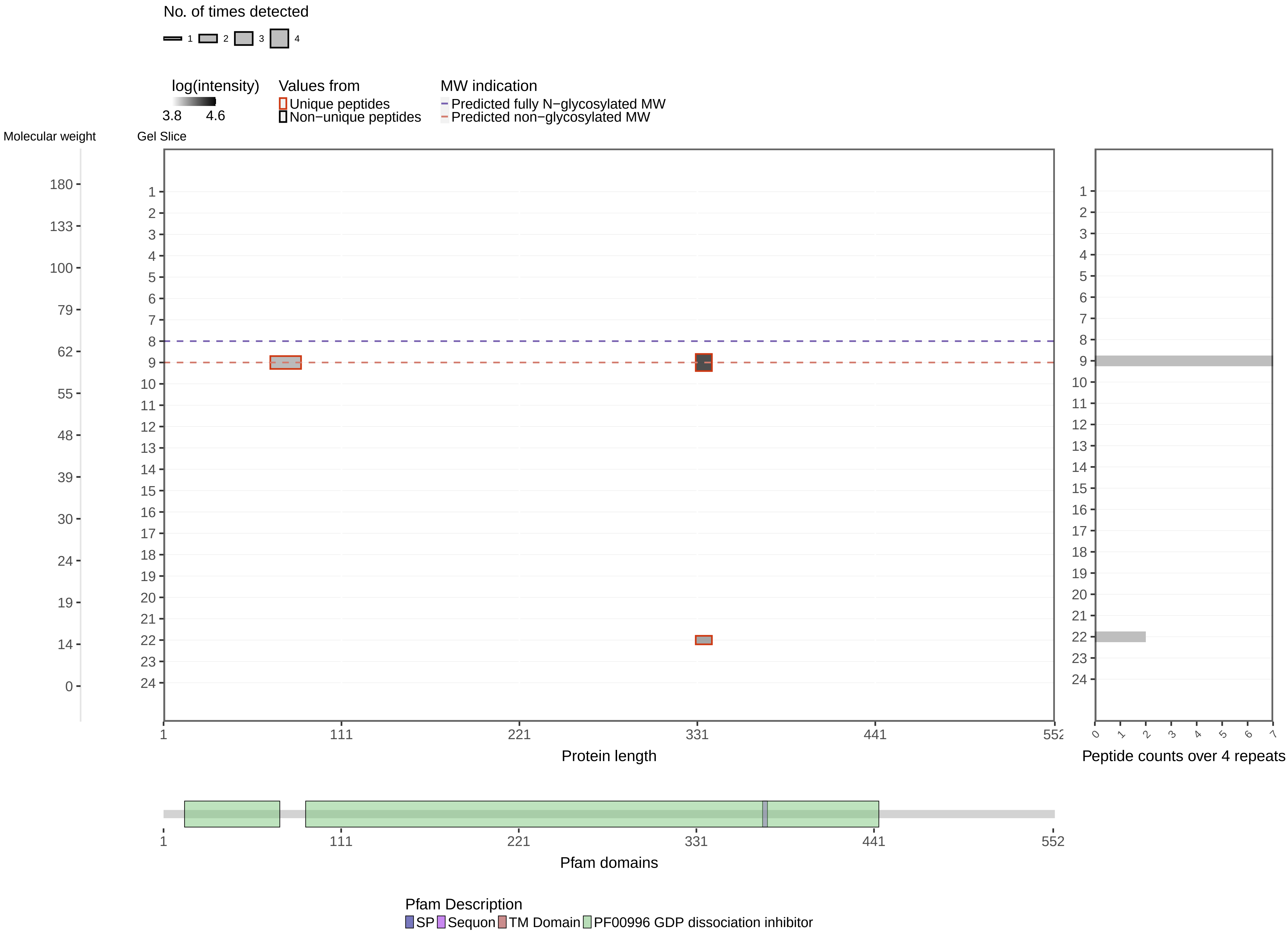


Protein sequence:

RLALYHSSVGRFPNAPGAMIYPIYGQGELPQAFCRRAAVKGCIHVLRMPVHSLLMDKATGNYKGVRLASEQELYSHKLILAPSFVFQLPPPHSSPDAQQNVYCDFGLKNTTEKLARSVCITKHSLKLDKANCLASFPPRSLFPEQVTAIHVLQLSSNVAVCPSGMFLIYLSAICEDDVQGKKLLHAAINALFCIPVSGSSENDNSAKKQSETRDAKPALLWSVLYTQELTKFQDVLDNIISASMPDGSPYYHDLLGATEKIFEELYSGEVFFPKTASSEEGAVEELED*

### NbL13g04730.1 uncharacterized_LOC109242911 [MW = 34.33 kDa]

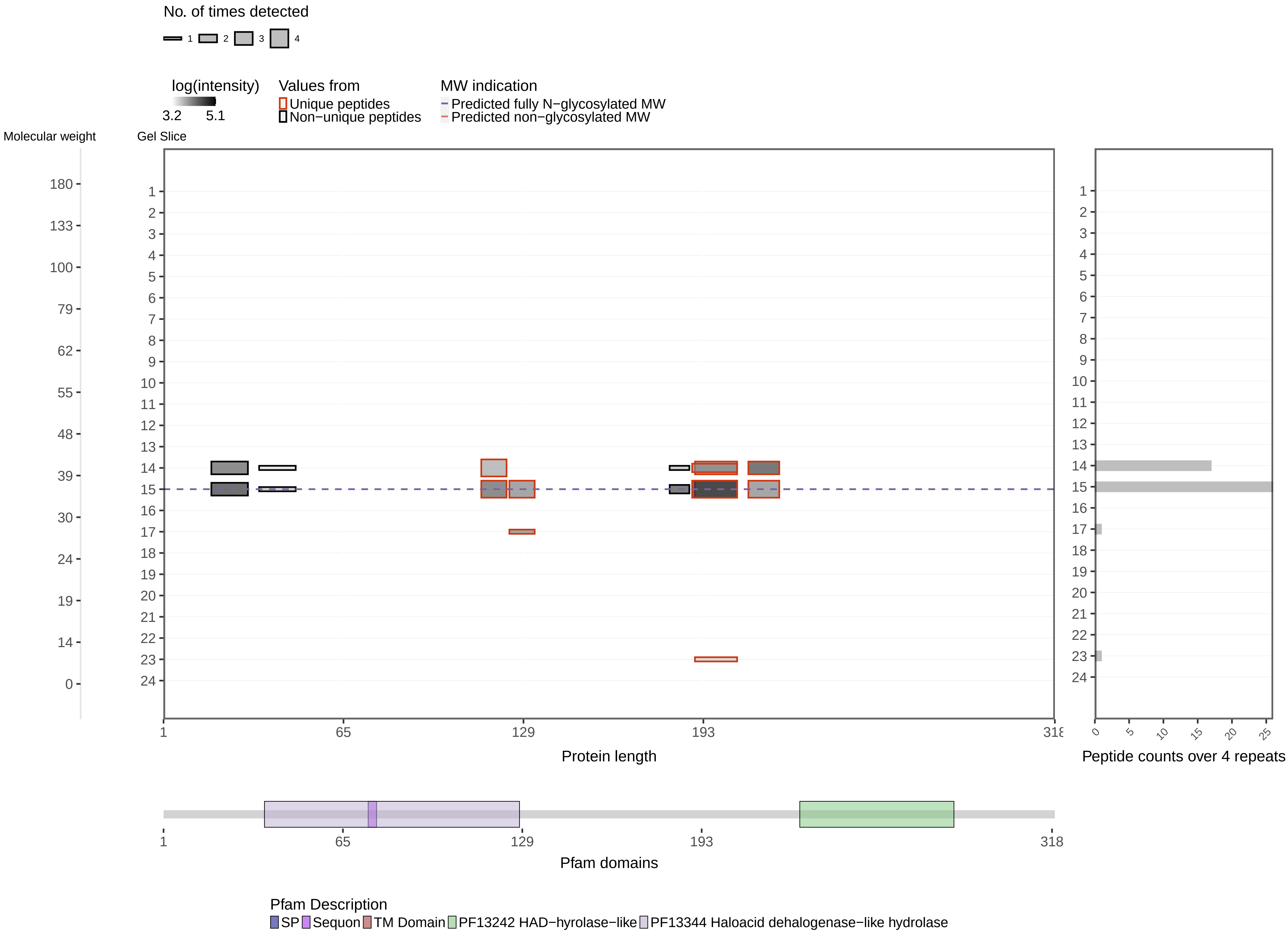


Protein sequence:

/ENAQEANFILAHGTEALGLSSGAALPMKLDDLEKILEQCAAKKIPMVVANPDFVTVEARALRVMPGTLAATYEKLGGQVKWMGKPDKIIYKSAMEMADVDVSDCVAVGDSLHHDIKGANAAGIASAFITCGIHATELGLSKFGEVADDNSIHALALQNNAYPTYVLPSFTW*

### NbL13g04670.1 uncharacterized_LOC109239937 [MW = 29.1 kDa]

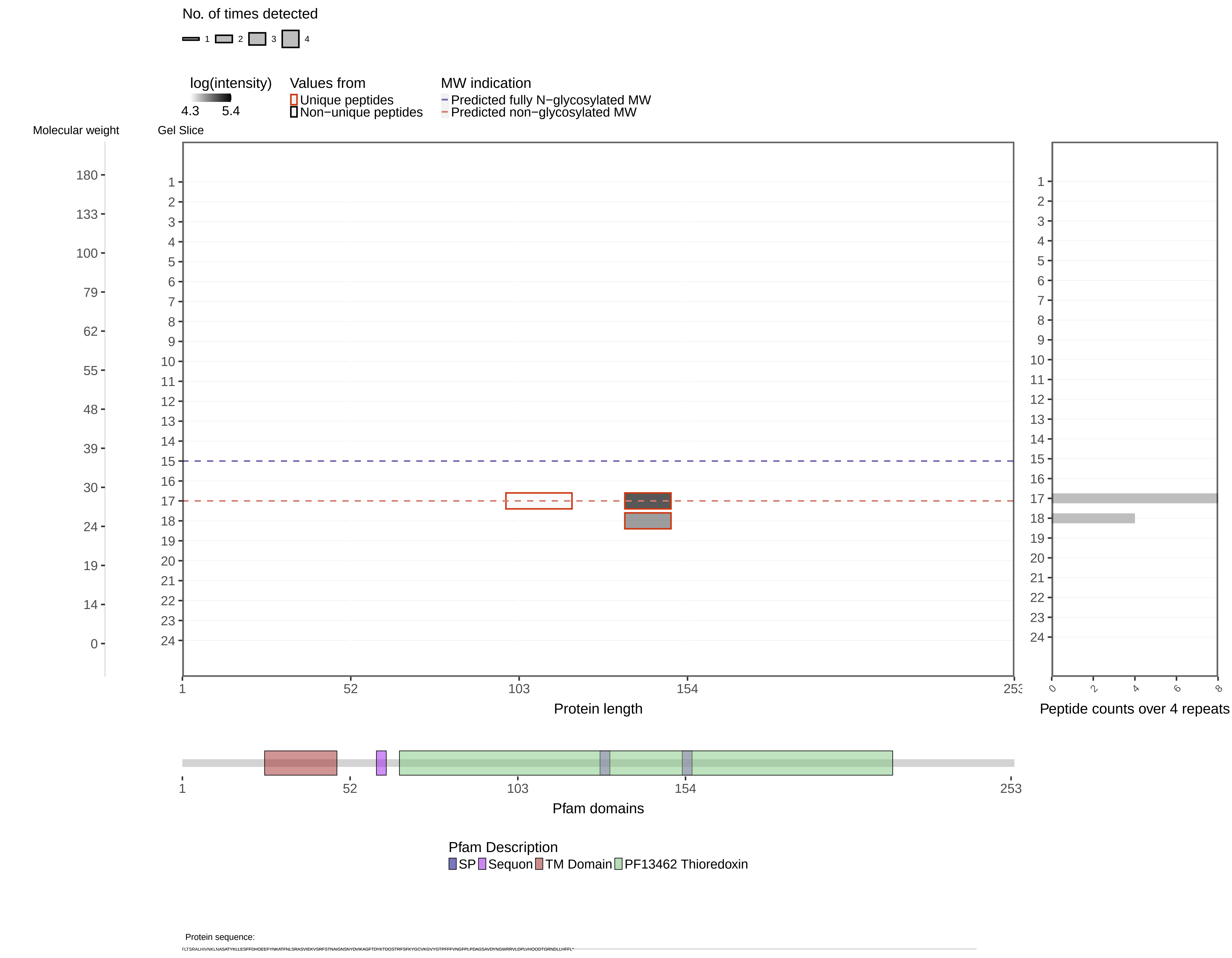

NbL13g04660.1 ribonuclease_P_protein_subunit_p25−like_protein [MW = 27.96 kDa]

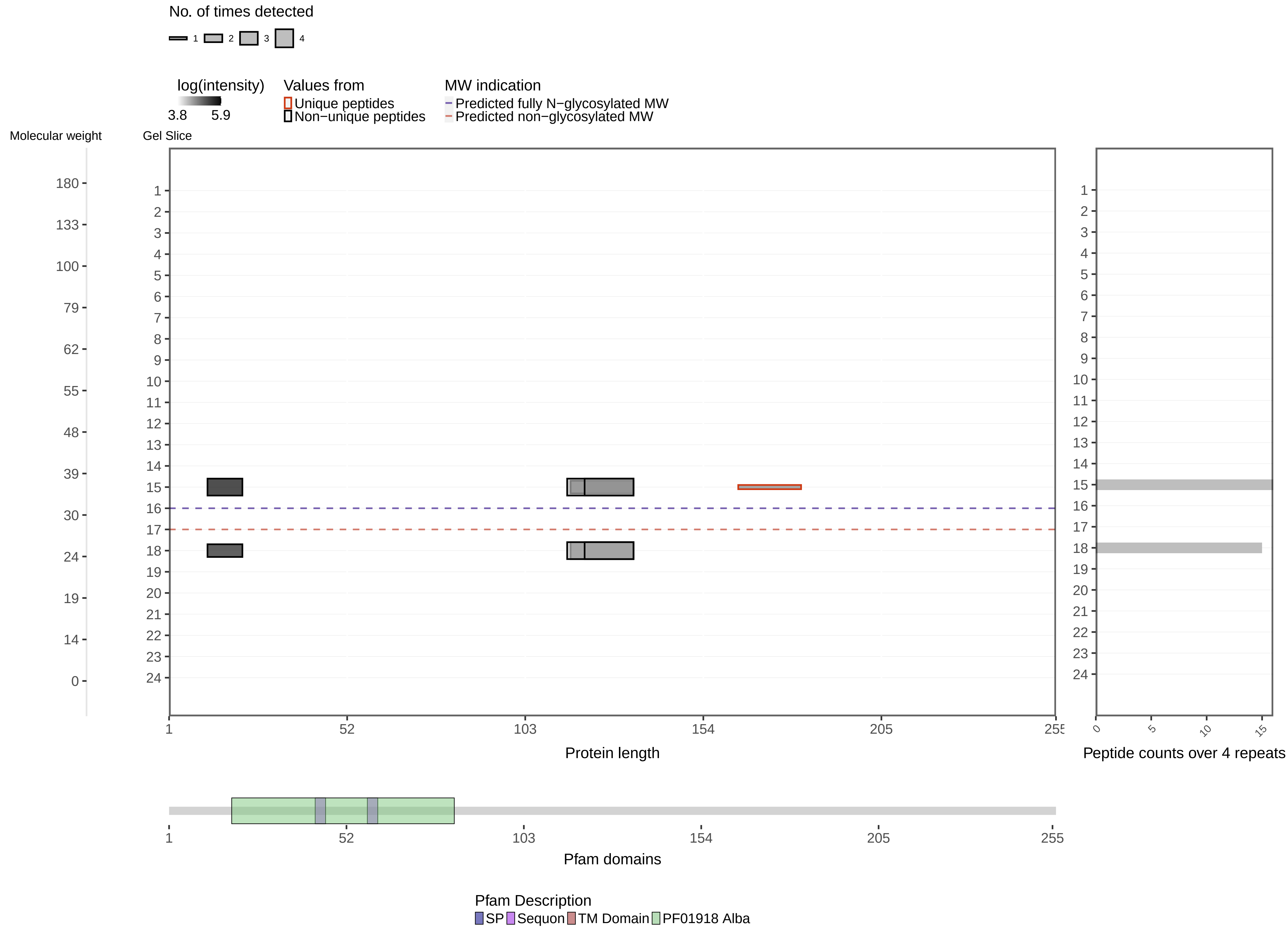


Protein sequence:

IGYQPPIPADQVKVSTDIDYEGDGSPNGRGRGRGGRGRGRSRGYSGNGVMPAEYDDGGYDRNWGYGRGRGRGRGRNFRGRGRGGFGGALDTQQDAGGYIQEAPIQGRGRGRGRGPRGRGRGFRPNGPPIHVAA*

NbL13g04590.1 subtilisin−like_protease_SBT5.3 [MW − SP = 78.45 kDa]

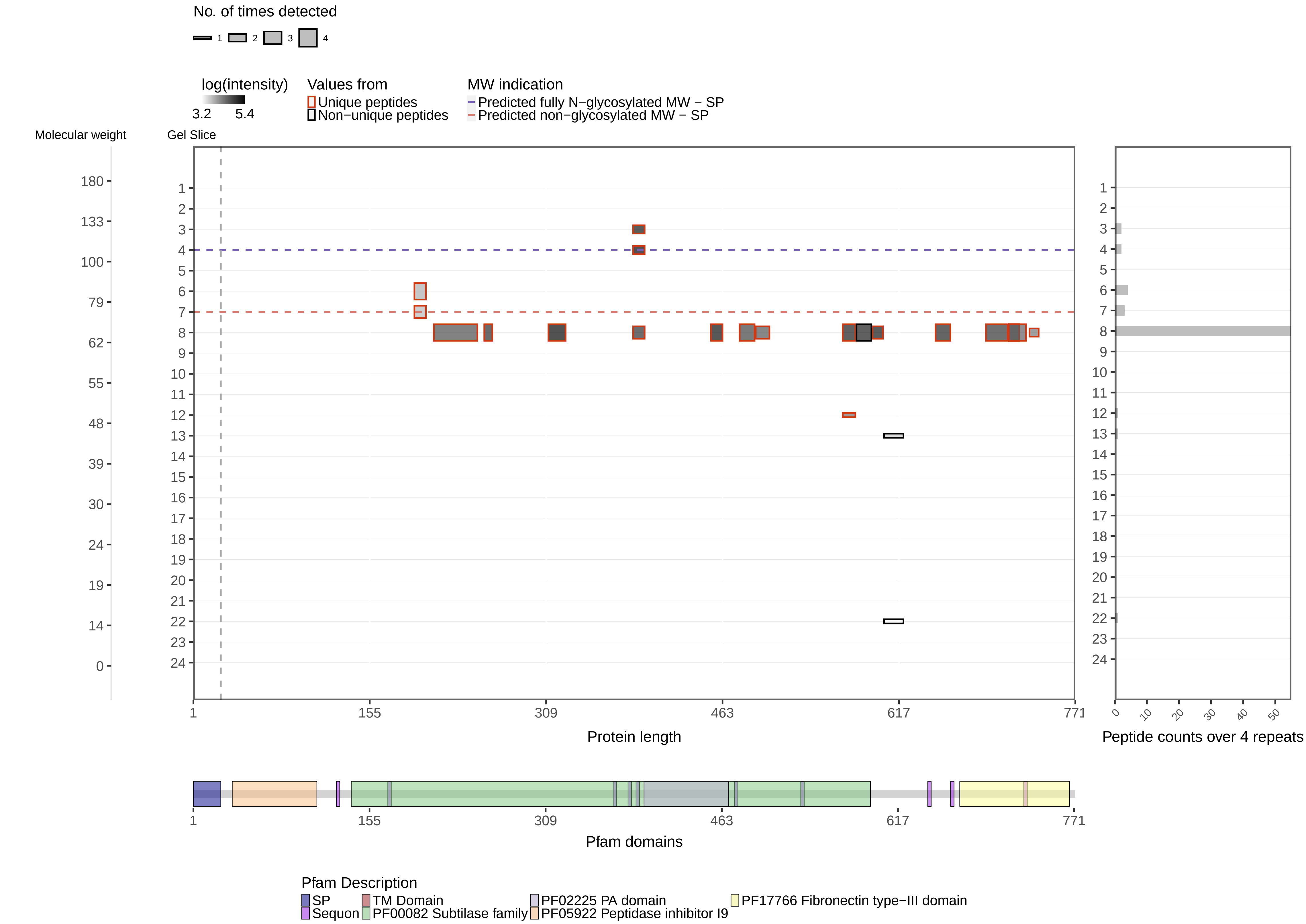


Protein sequence:

TKSSVYPLIHGNS~~TKSNDNVSEADARSCVPGSLDENKVKGKIVLCENLDDGEYFPSDKLDEVKSRGGVGFILIDDDERTVAPKFNSFSAGVVSKKDGNEILAYINSTRNPVASILPTVSITKYKPAPVVAYFSSRGPAYNTPNLLKPDITAPGVAILAAWPGNDTSEALPGQKPPIFNLLSGTSMSCPHVSGIAATVKAVNPTWSPSAVKSAIMTTAIQTNNLKAPITTVSGSKATPYDIGAGEASTSGPLKPGLVYETDVADYLQFLCSVGFNISQIKLISITVP~~EDFSCPKNSTSELV

### NbL13g04550.1 endoplasmin_homolog [MW − SP = 90.82 kDa]

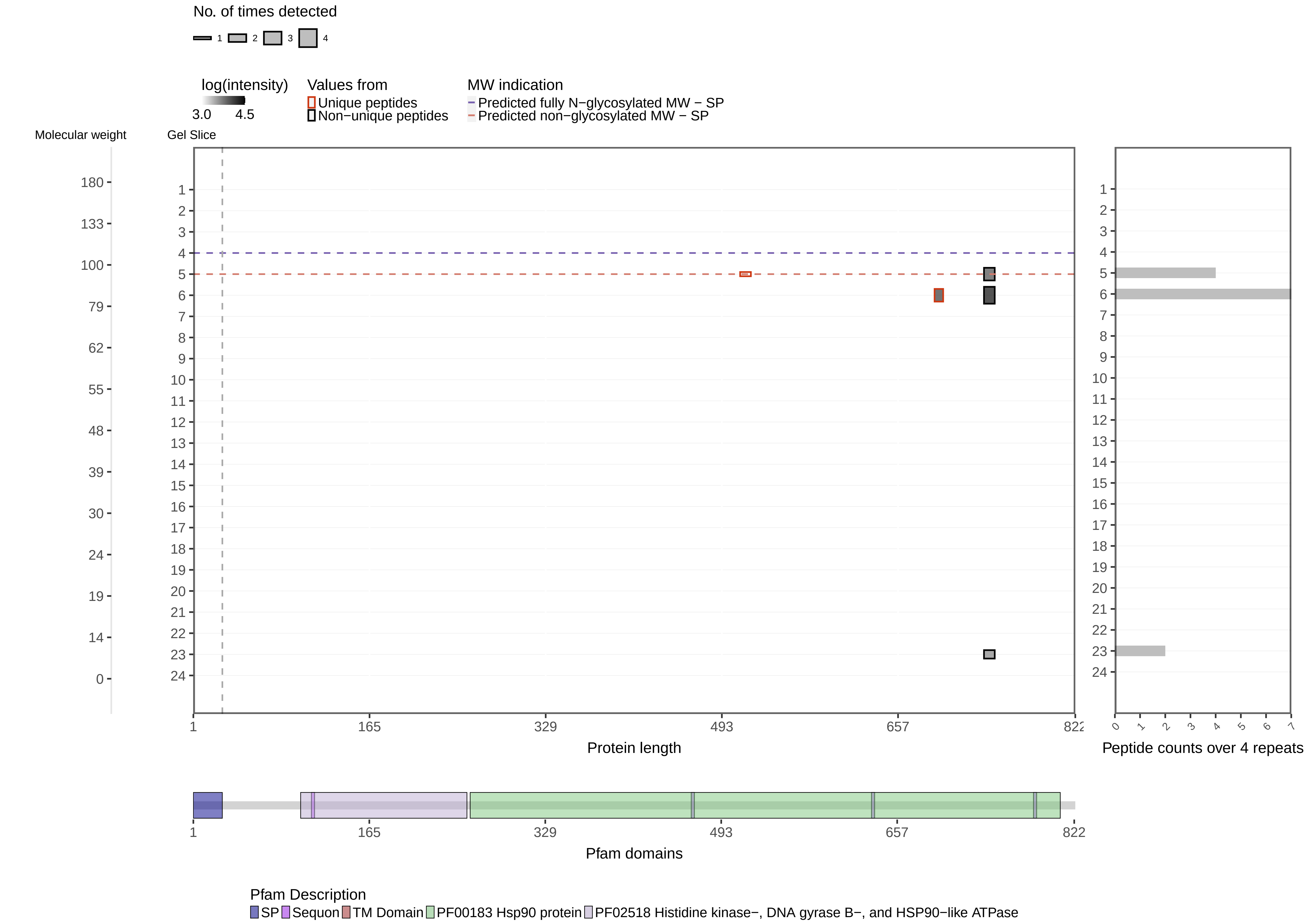


Protein sequence:

DVEFKAVLFVPPKAPHDLYESYYNSKKSNLKLYVRRVFISDEFDELLPKYLSFLMGLVDSDTLPLNVSREMLQQHSSLKTIKKKLIRKALDMIRKIADEDPDESNDKDKKDVEESGADNEKKGQYAKFWNEFGKSVKLGIIEDATNRNRLAKLLRFETSKSDGKLTSLDQYISRMKAGQKDIFYITGASKEQLEKSPFLERLTKKNYEVIFFTDPVDEYLMQYLMDYEDHKFQNVSKEGLKLGKDSKAKELKESFKELTKWWKGALASDNVDDVKISNRLADTPCVVVTSKYGW

### NbL13g04470.1 prolyl_endopeptidase−like [MW = 92.1 kDa]

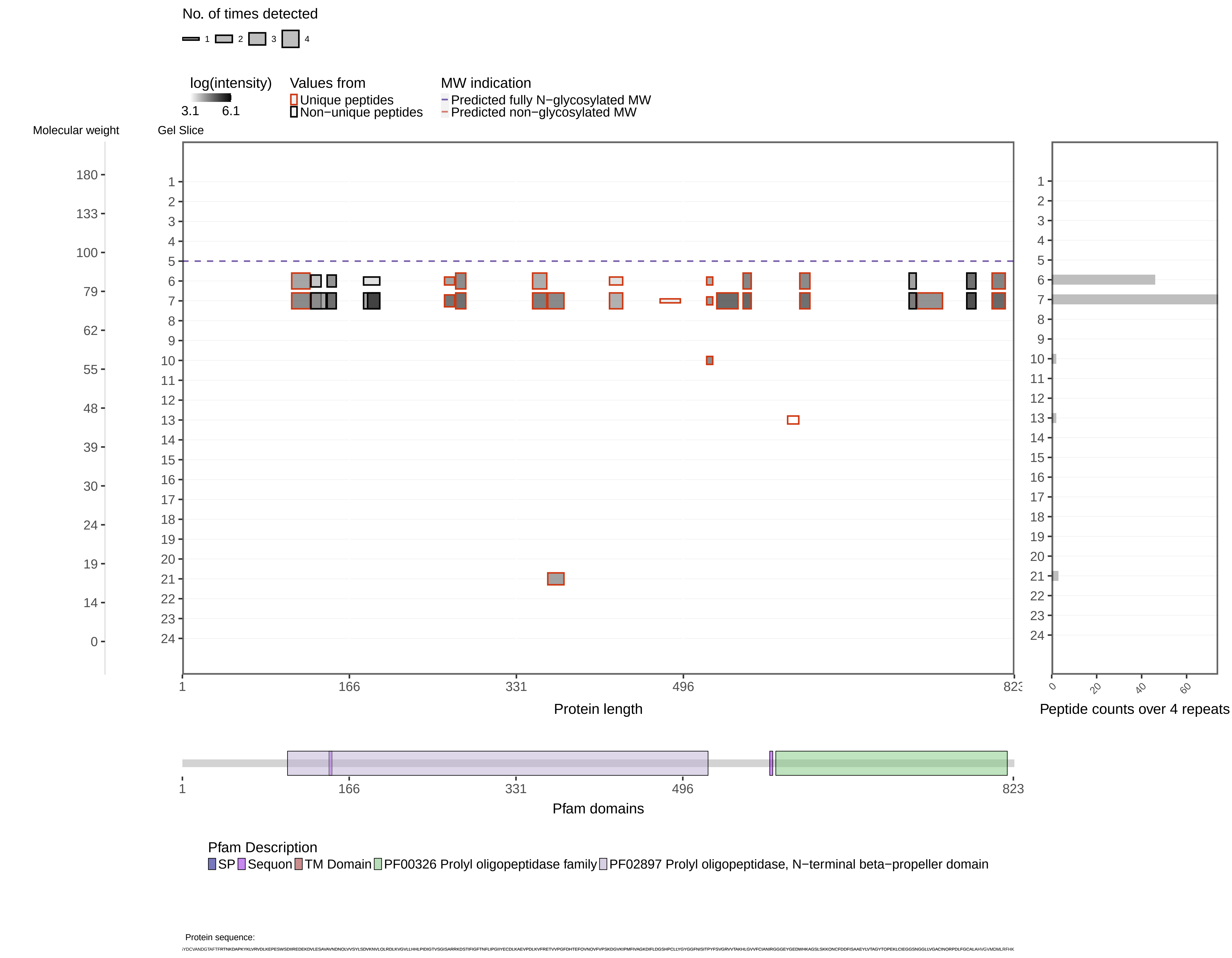

### NbL13g04400.1 probable_acyl−activating_enzyme_1_peroxisomal [MW = 60.91 kDa]

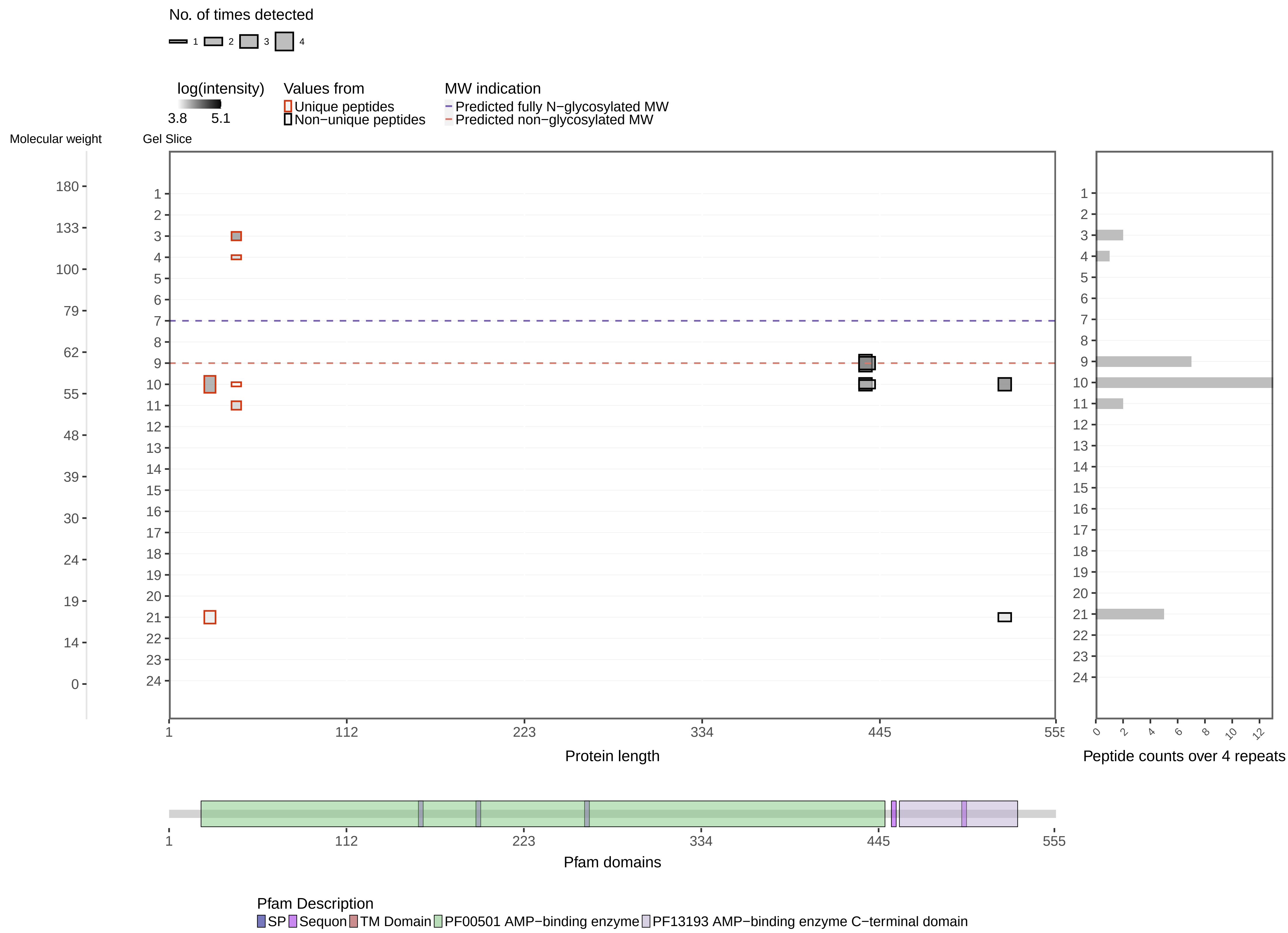


Protein sequence:

AGIFTSIAGHQVSHMGGAPTVLNMIINAPPSVRRSLPRKVAVMTGGAPPPPQVLFKMEELGFDVIHSYGLTETYGPGTVCTWKPEWSTLSPDEQAKIKSRQGLHHIGMEEVDVKDPATMKSIPPDAKTMGEVMFRGNTVMNGYLKNLKATEDAFKRGWFRSGDLAVKHPDGYIELKDRAKDIIISGGENISTIELESVIFSHPAVLEAAVVGRPDDYWGETPCAFVKLKDGNNATADEIINYCRDRLPRYMAPKTVIFDDLPKTSTGKTQKFVLRQKAKAMGSISKASKL*

NbL13g04380.1 catalase_isozyme_3−like [MW = 56.84 kDa]

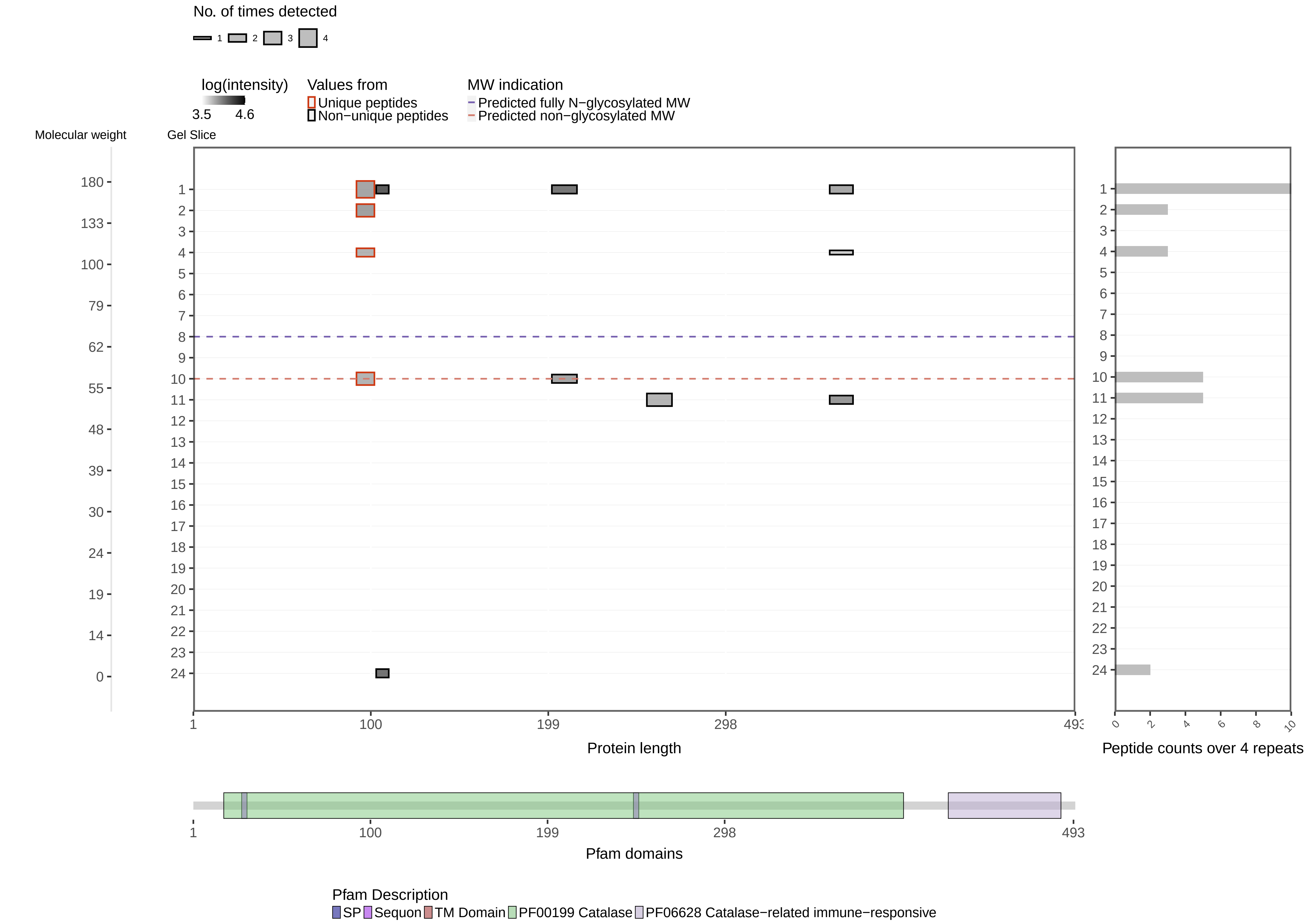


Protein sequence:

EVKCLLEEVAIKVGGANHSHATKDLYDAIAAGNYPEWKLFIQTIDPDHEDRFDFDPLDVTKTWPEDILPLQPVGRLVLNKNIDNFFAENEQLAFCPAIIVPGIHYSDDKLLQTRIFSYSDTQRHRLGPNYLQLPVNAPKCAHHNNHHEGVMNFMHRDEEVNYFPSRFDPCRHAEQYPIPPRIVTGKRDRCIIEKENNFKQPGERYRSWSPDRQERFICRWISALSDPRVTHEIRTIWISYWSQADKSLGEKIASRLNVRPTM*

### NbL13g04140.1 probable_steroid−binding_protein_3 [MW = 10.89 kDa]

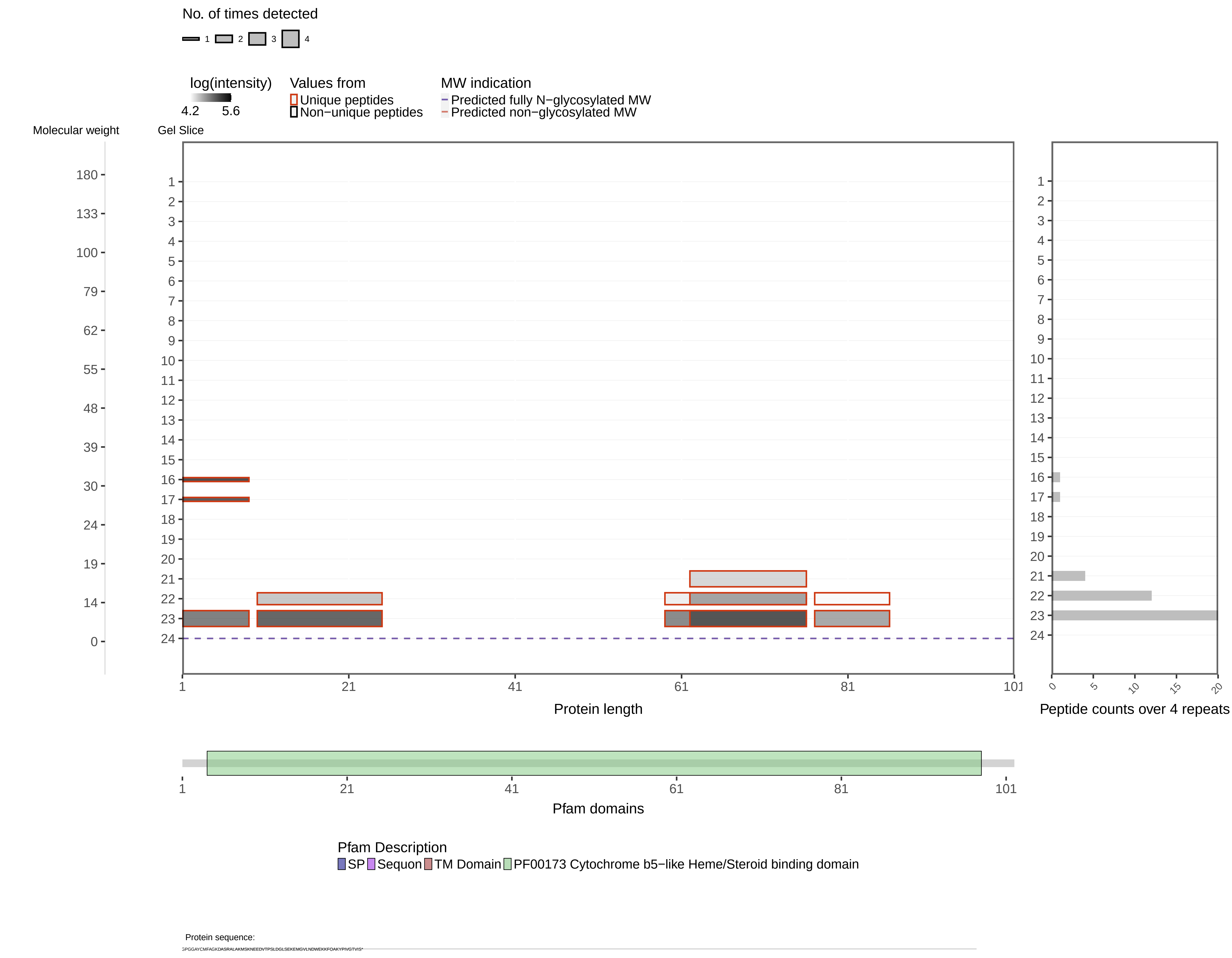

### NbL13g04130.1 cysteine−−tRNA_ligase_cytoplasmic [MW = 65.04 kDa]

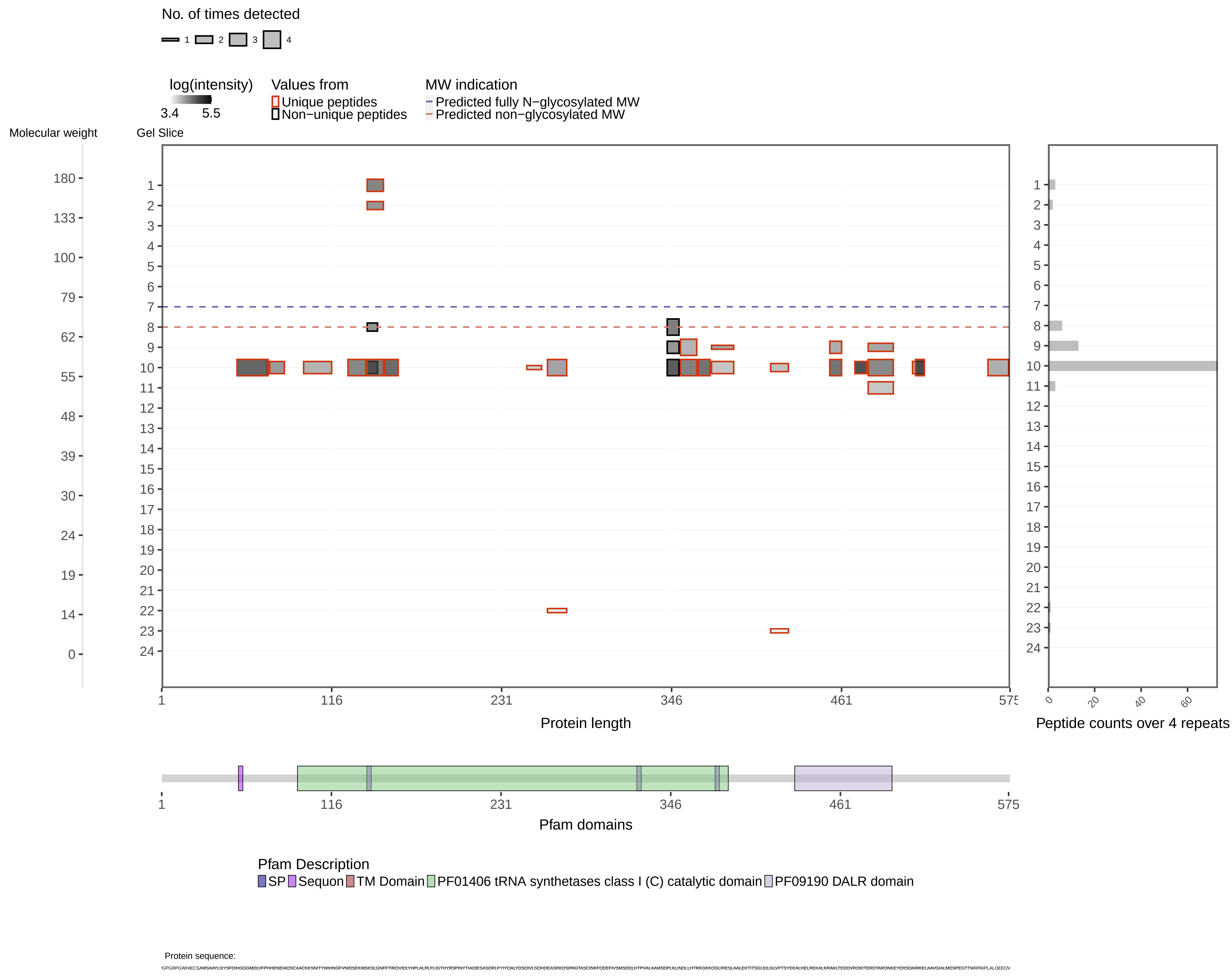

### NbL13g04050.1 signal_recognition_particle_subunit_SRP68−like [MW = 68.37 kDa]

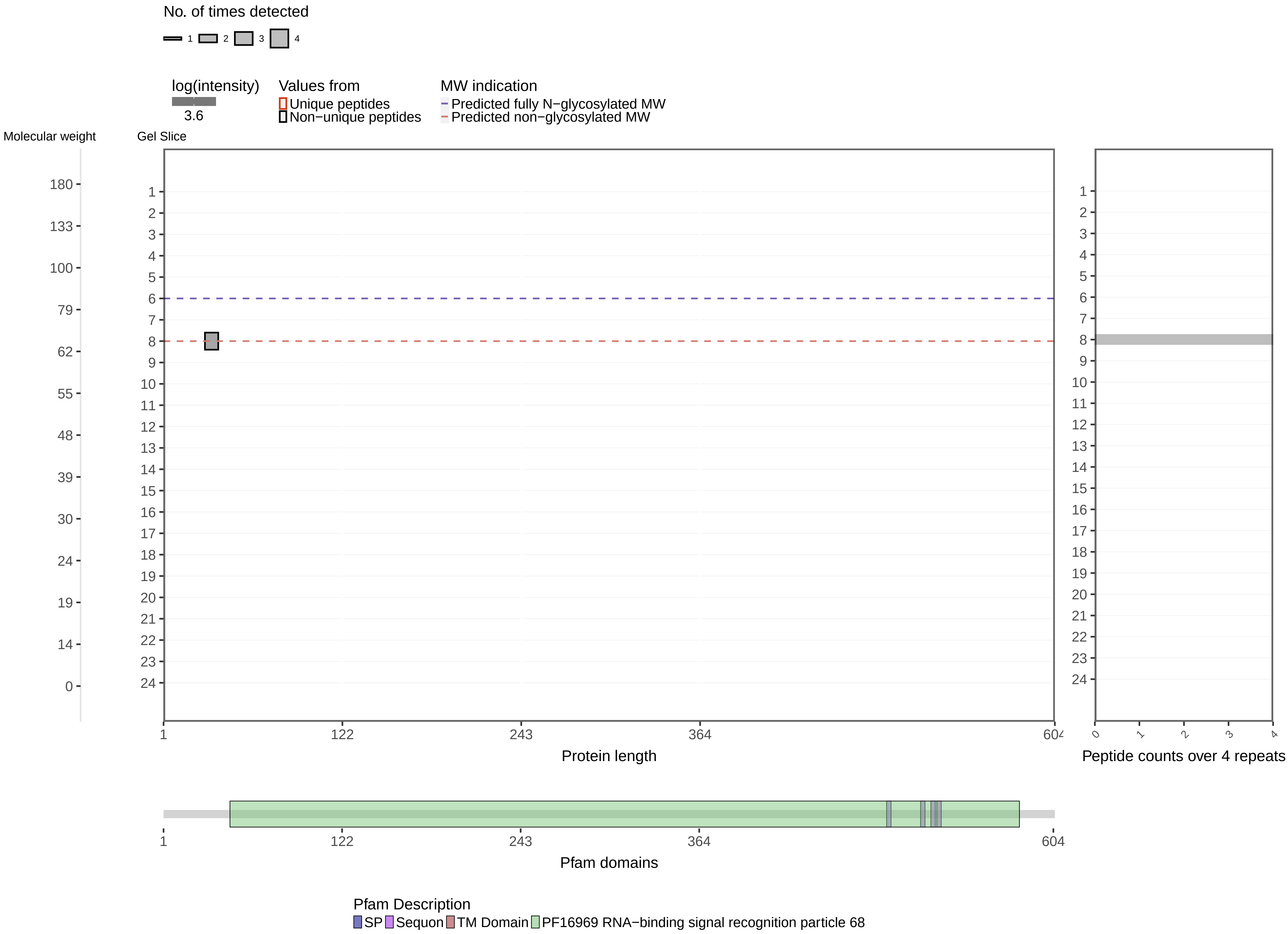


Protein sequence:

[illegible]

### NbL13g04040.1 protein_MEMO1−like [MW = 33.17 kDa]

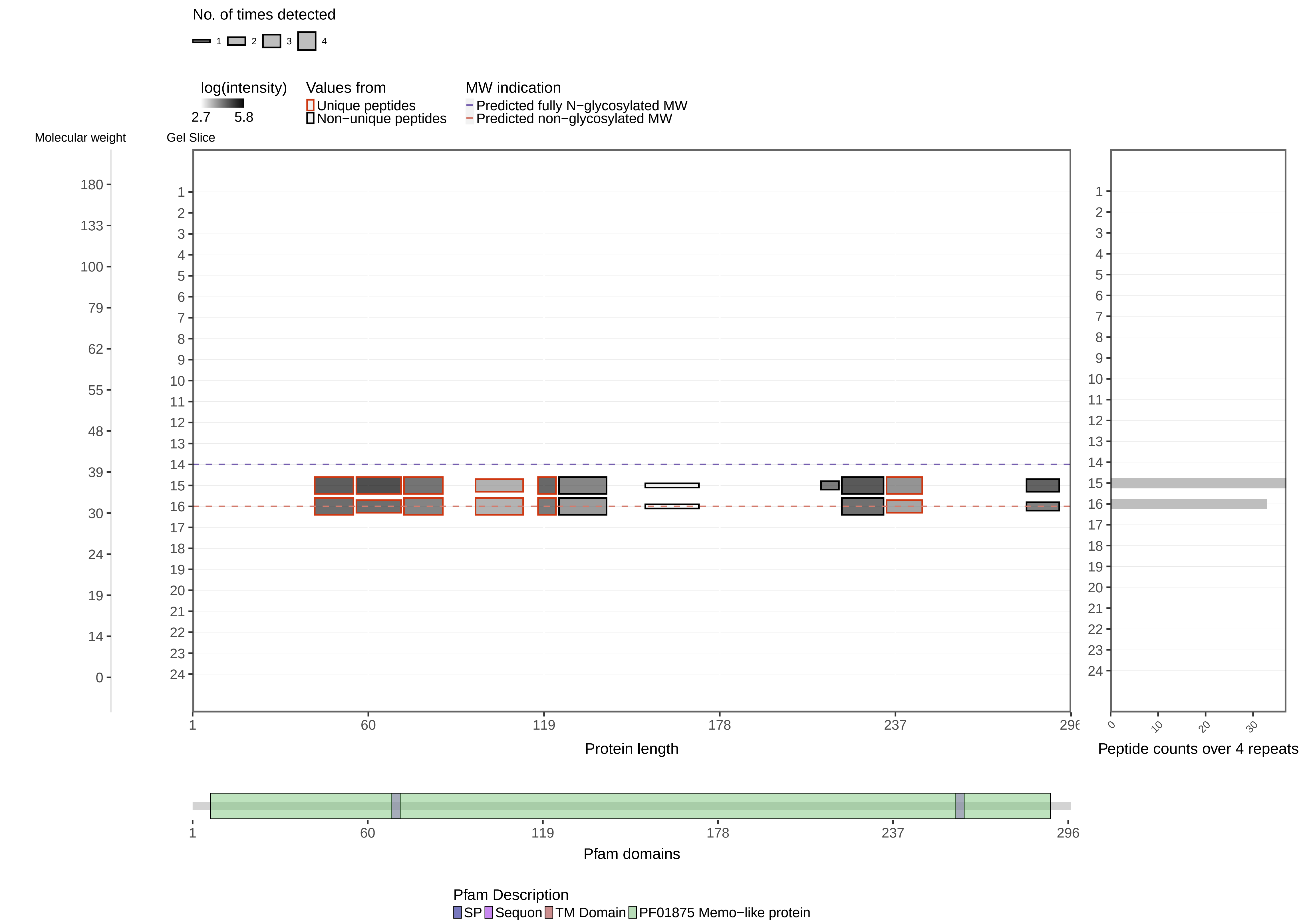


Protein sequence:

HLPYLAKVFQGYYDRYPVKIVPILVGSLSAESEALYGRLLAKYADDPKNFFSVSSDFCHWGSRFNYMHYDKSHGAIYKSIEALDKMGMDIIETGDPDAFKQYLSETDNTICGRHPISVFLHMLKNSSIKIKIRFLRYEQSSQCKSMRDSSVSYASAVGKIDA*

### NbL13g03980.1 40S_ribosomal_protein_S11−like [MW = 17.69 kDa]

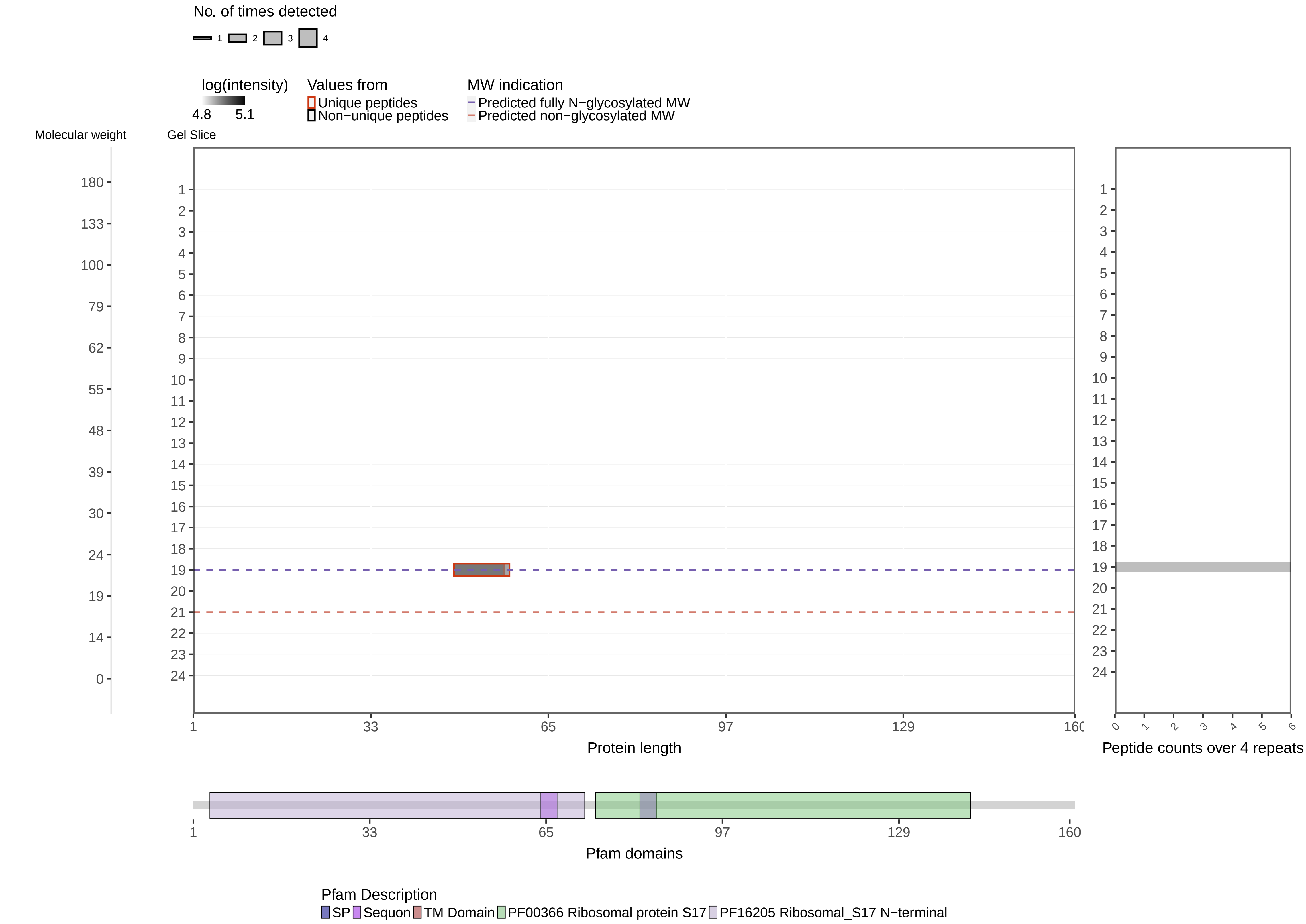


Protein sequence:

SIRGRILAGTCHSAKMNRTIIVRRNYLHYVKKYQRYEKRHSNIPAHISPCFRVKEGDHVIIGQCRPLSKTVRFNVLKVIPAGSAGGGKKAFTGM*

NbL13g03840.1 aspartate_aminotransferase_chloroplastic−like [MW = 50.77 kDa]

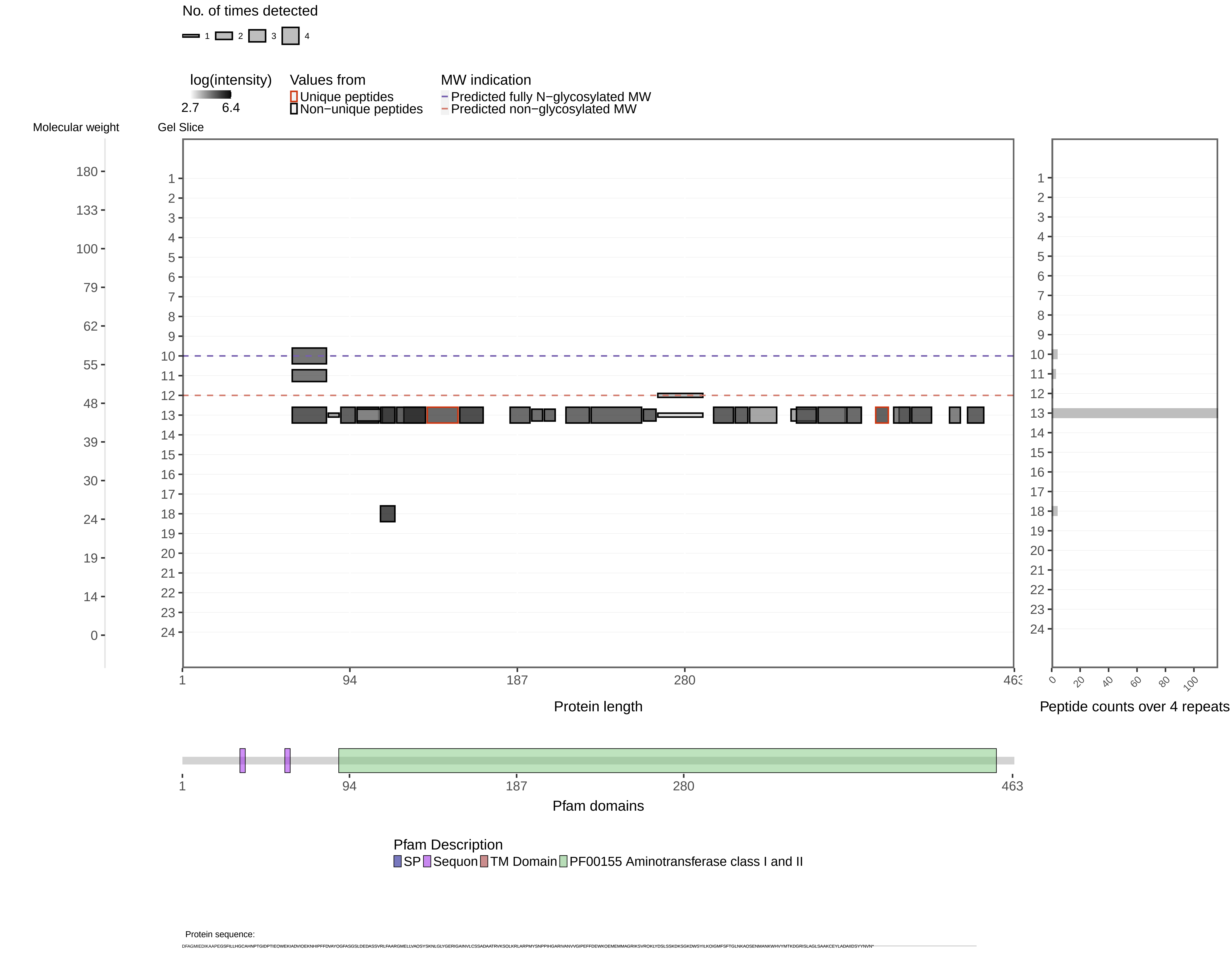

### NbL13g03780.1 glucan_endo−13−beta−glucosidase_4 [MW − SP = 52.53 kDa]

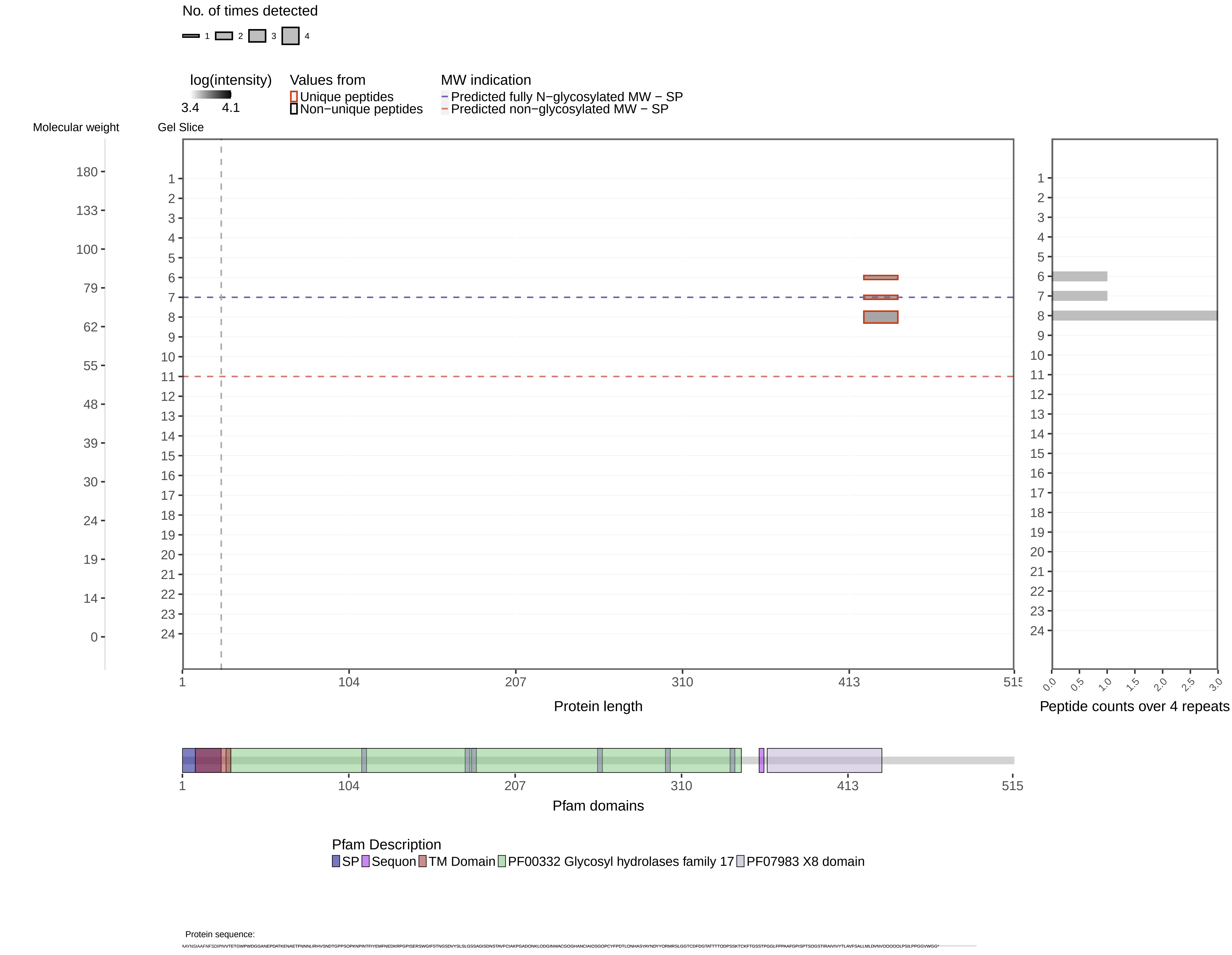

NbL13g03570.1 uridine_5'−monophosphate_synthase [MW = 51.62 kDa]

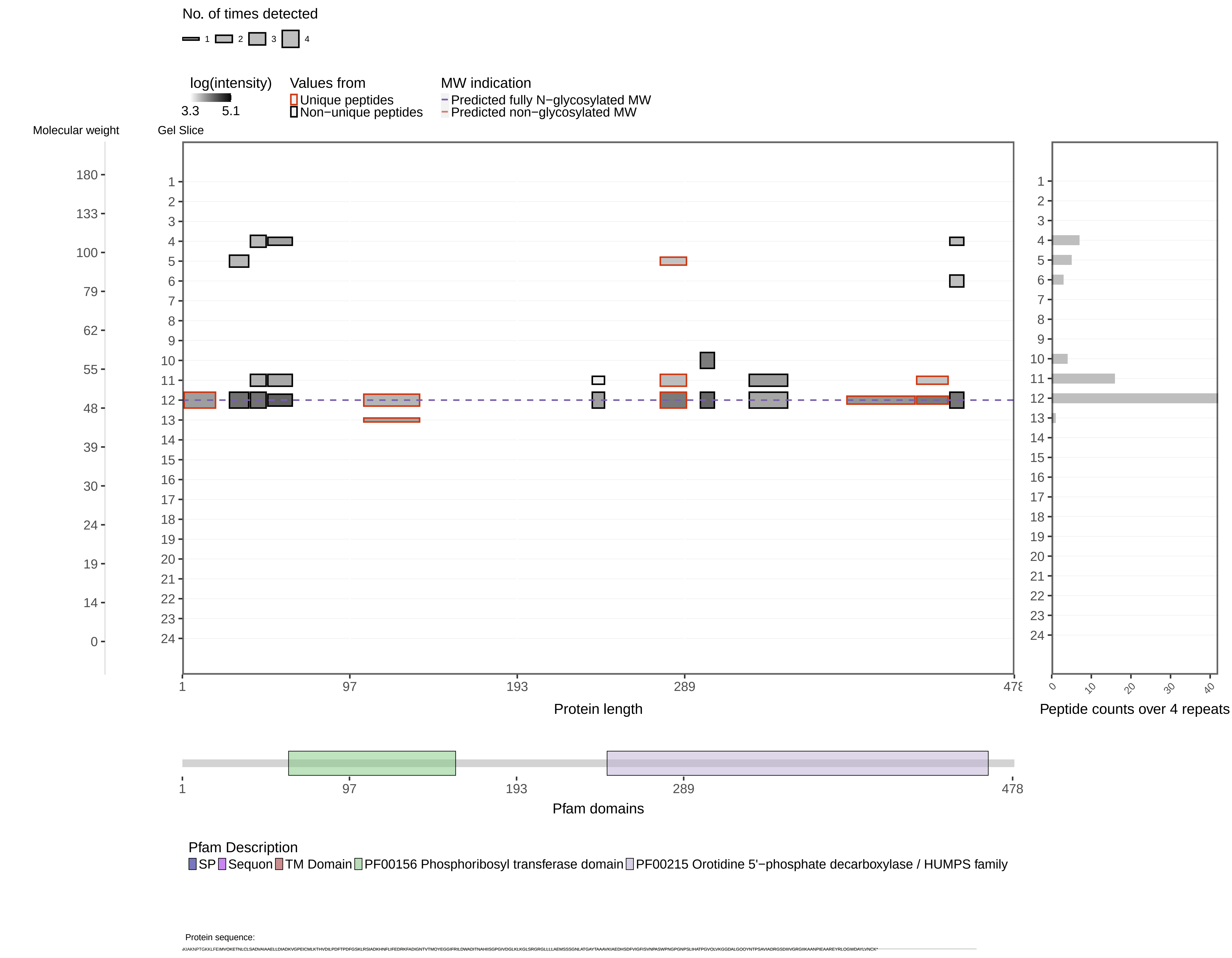

### NbL13g03490.1 methionine−−tRNA_ligase_cytoplasmic−like [MW = 91.19 kDa]

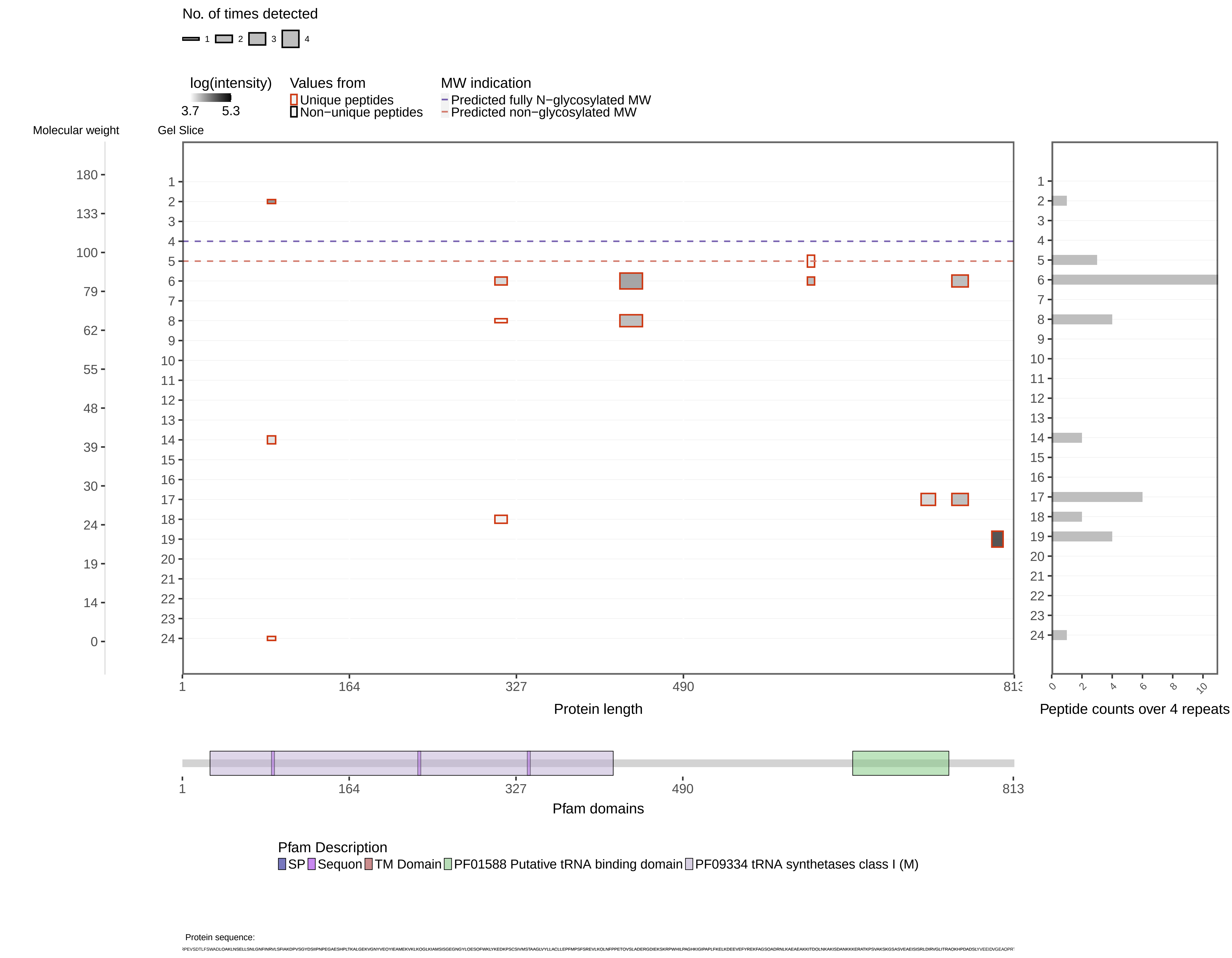

### NbL13g03440.1 subtilisin−like_protease_SBT2.5 [MW − SP = 85.1 kDa]

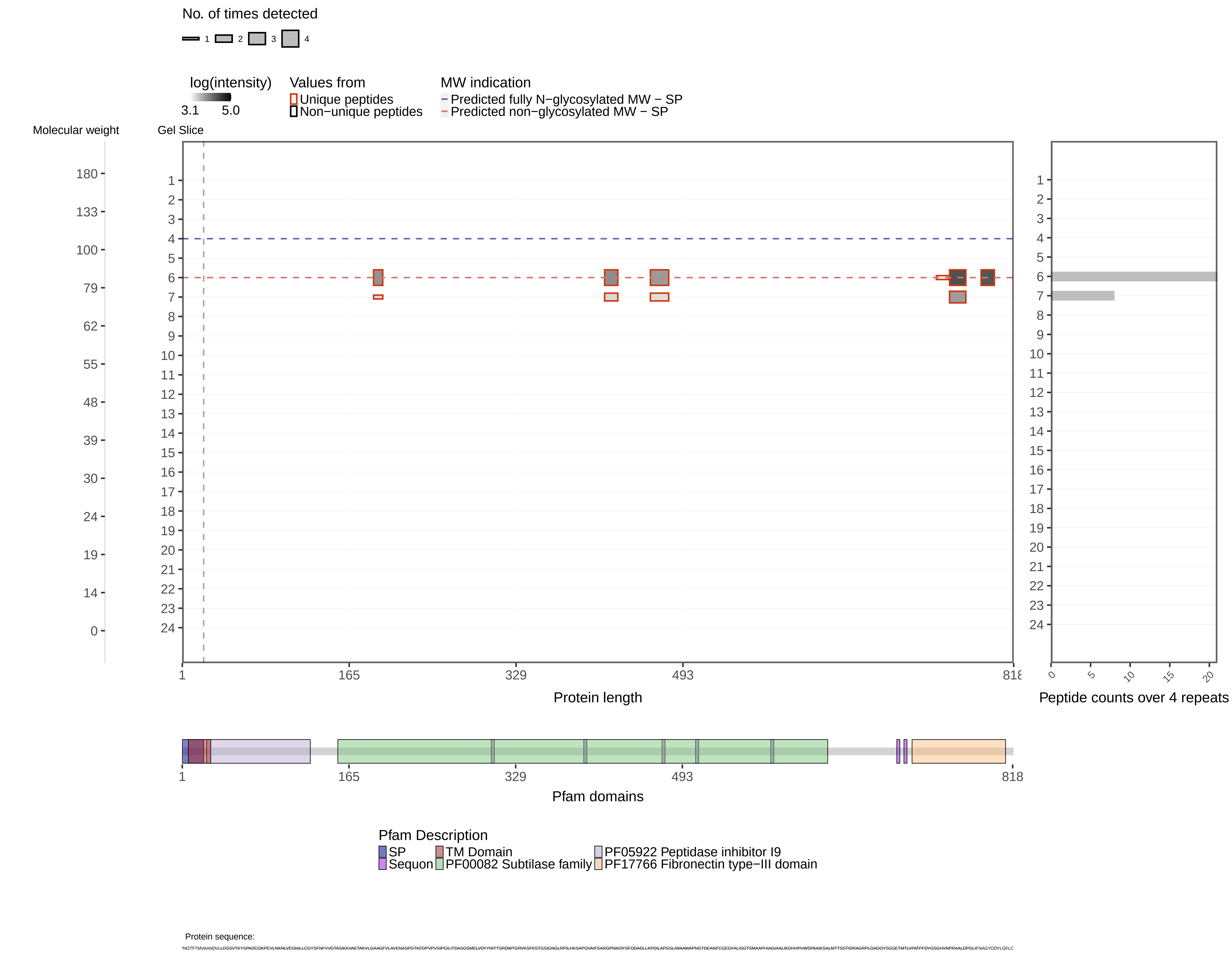

NbL13g03390.1 inosine−5'−monophosphate_dehydrogenase_2−like [MW = 53.4 kDa]

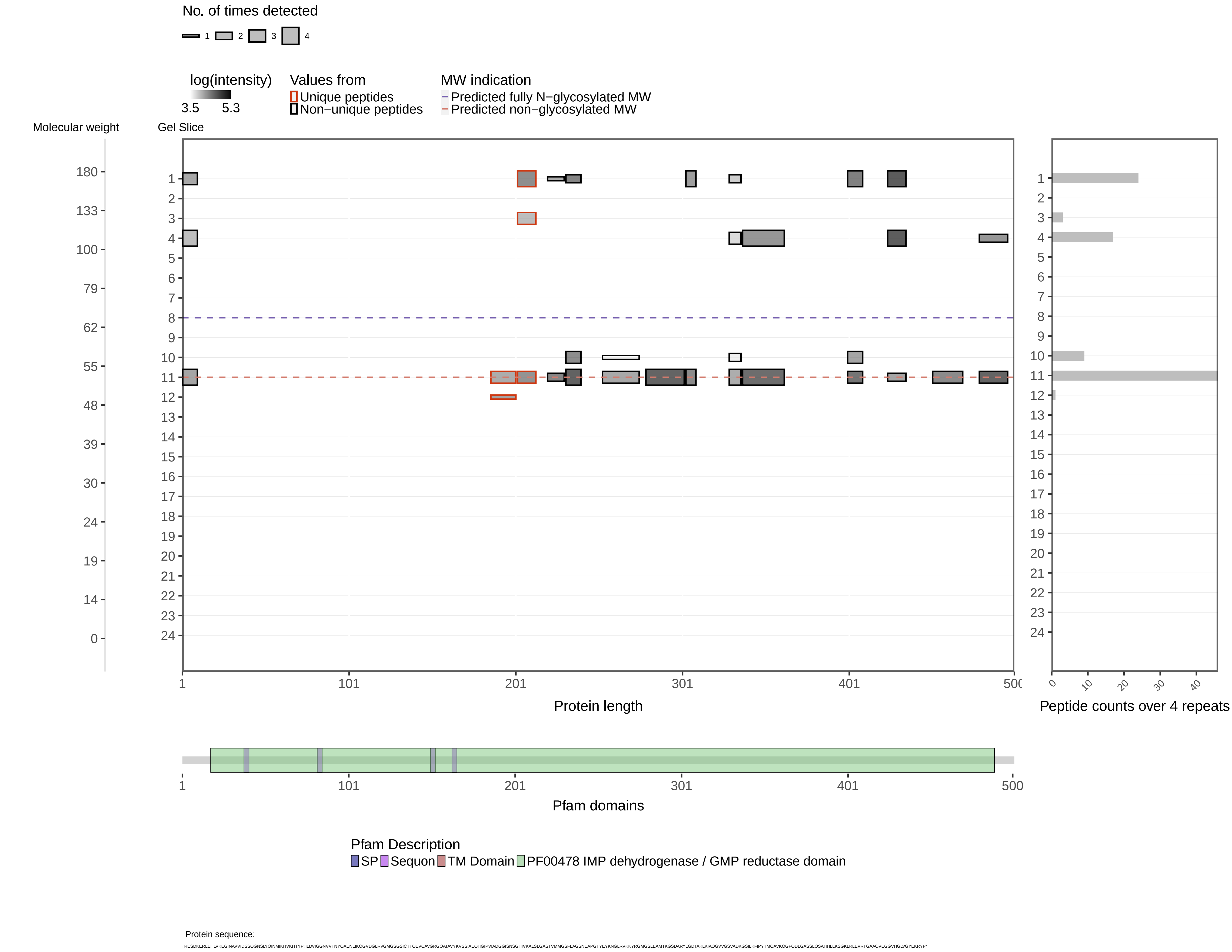

### NbL13g03170.1 nucleotide_pyrophosphatase/phosphodiesterase−like [MW = 74.73 kDa]

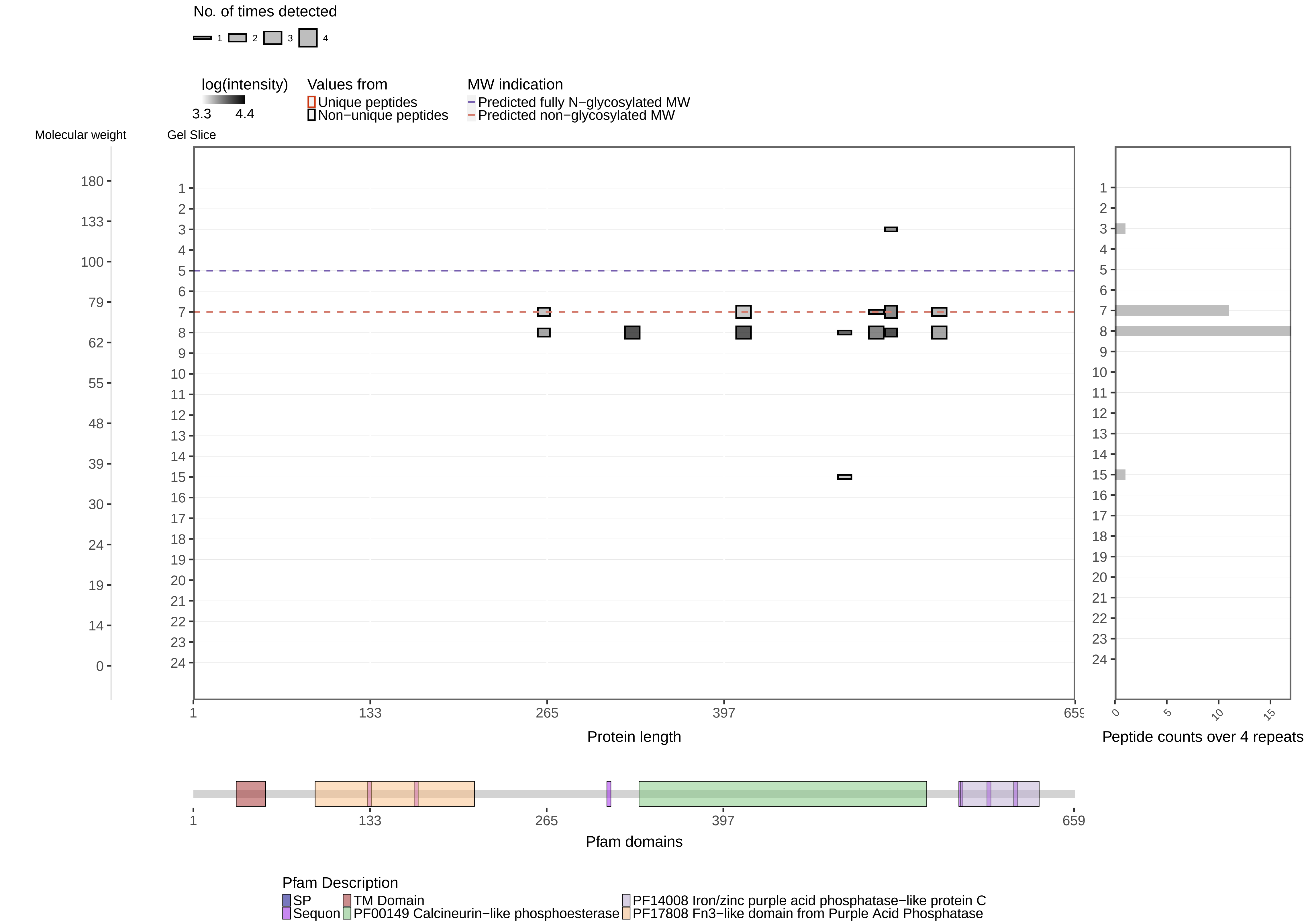


Protein sequence:

YSFKSPPFPGQESLQRVVIFGDMGKQERDGSNEYANYQPGSLMTTDTLIKDLDNIDIVFLIGDLPYANGYISQWDQFTAQVEPITSKVPFMIASGNHERTWENSGSLYTGLDSGGECGVPAETLYYVPAENRAKFWYAADYGMFHFCIGDTEHDWREGSEQYKFIEECFASANRHKQPWLIFSAHRVLGYSSNDWYAKEGSFEEPMGREHLQKLWQKYKVDMAFFGHVHNYERVCPIYQNQCVNKETSHYSGVVNGTIHVVVGGGGSHLNQFTPINTTWSVFKDYDYGF

### NbL13g03150.1 poly(rC)−binding_protein_3−like [MW = 50.02 kDa]

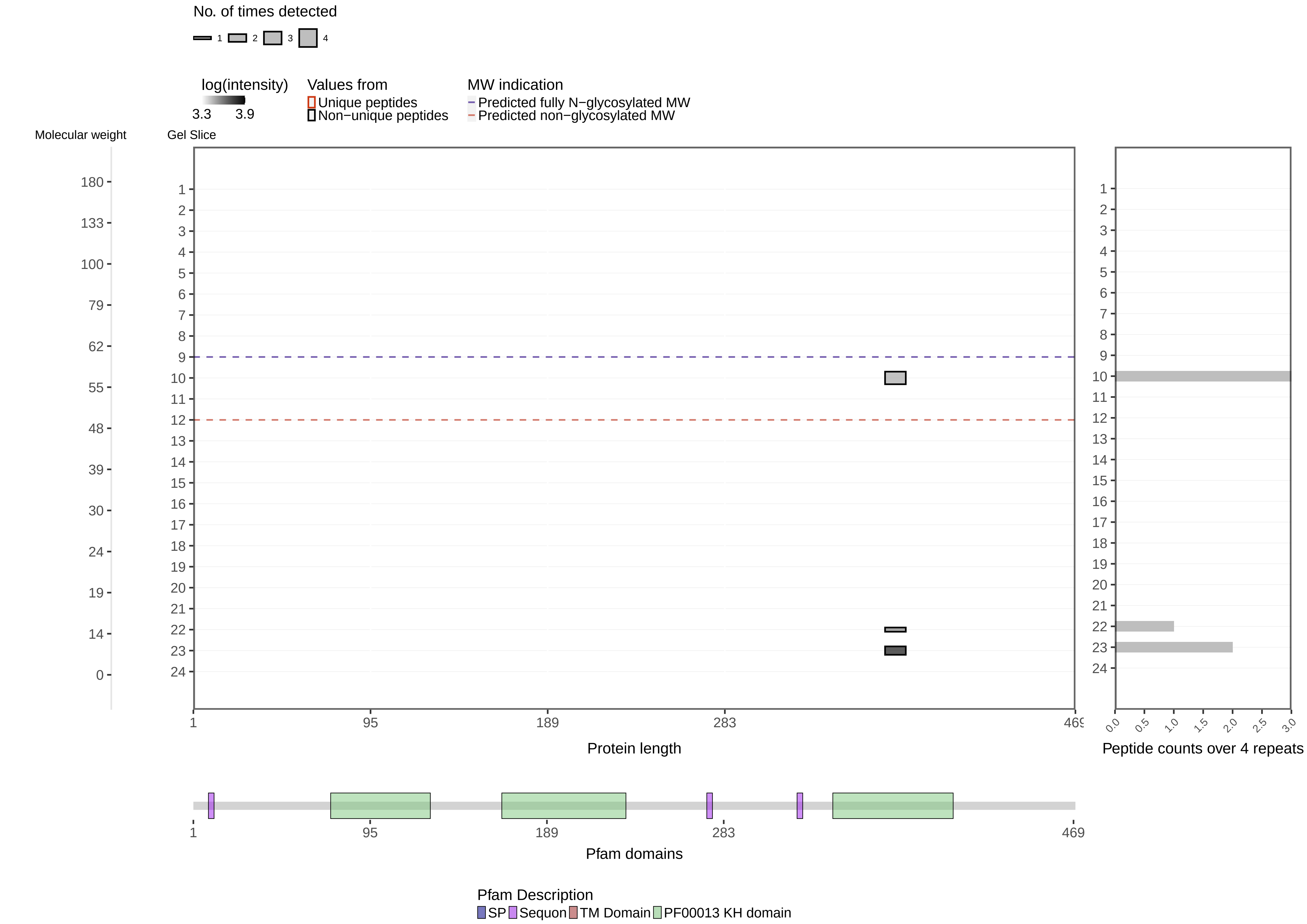


Protein sequence:

VLKALEAVIGHLRKFLVDHSILPLFEKSYNTPATQTQEREPEPWSERTMLNMSRPGLGMDYPPPLKRDSLFSERESQLDSQYHPPGLSLYGQDPGLSANRTSGLGRAGNPVITQISQTVQMQIPLAYAEDIIGIGGANIAYIRRTSGAILTVQESRGLPDEITVEIKGTSSEVQTAQQLIQEFMGSHKESLASSYGQLDTGLRQSSYSNLGSGSYSSSSYSGQQYGGGGGGYGSFGAGGYSSYRS*

### NbL13g03130.1 macrophage_migration_inhibitory_factor_homolog [MW = 12.21 kDa]

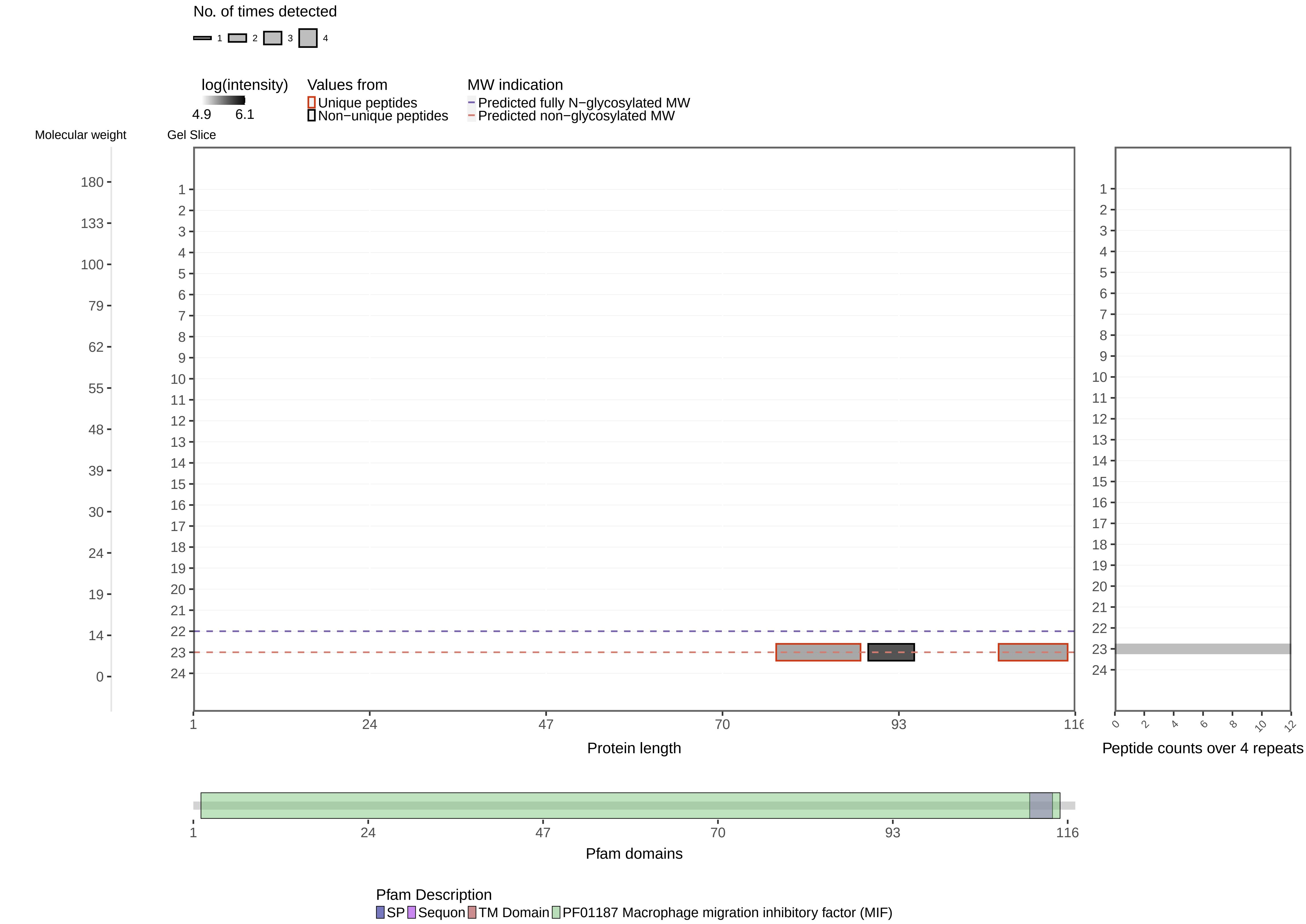


Protein sequence:

GGVPIAFAGTEAHAAYGELISIGGLGPGVNGKLSSIIAEILQTKLSIDSNRFYIKFYDSPRQFFGFNGSTF*

### NbL13g03030.1 choline_monooxygenase_chloroplastic−like [MW = 48.66 kDa]

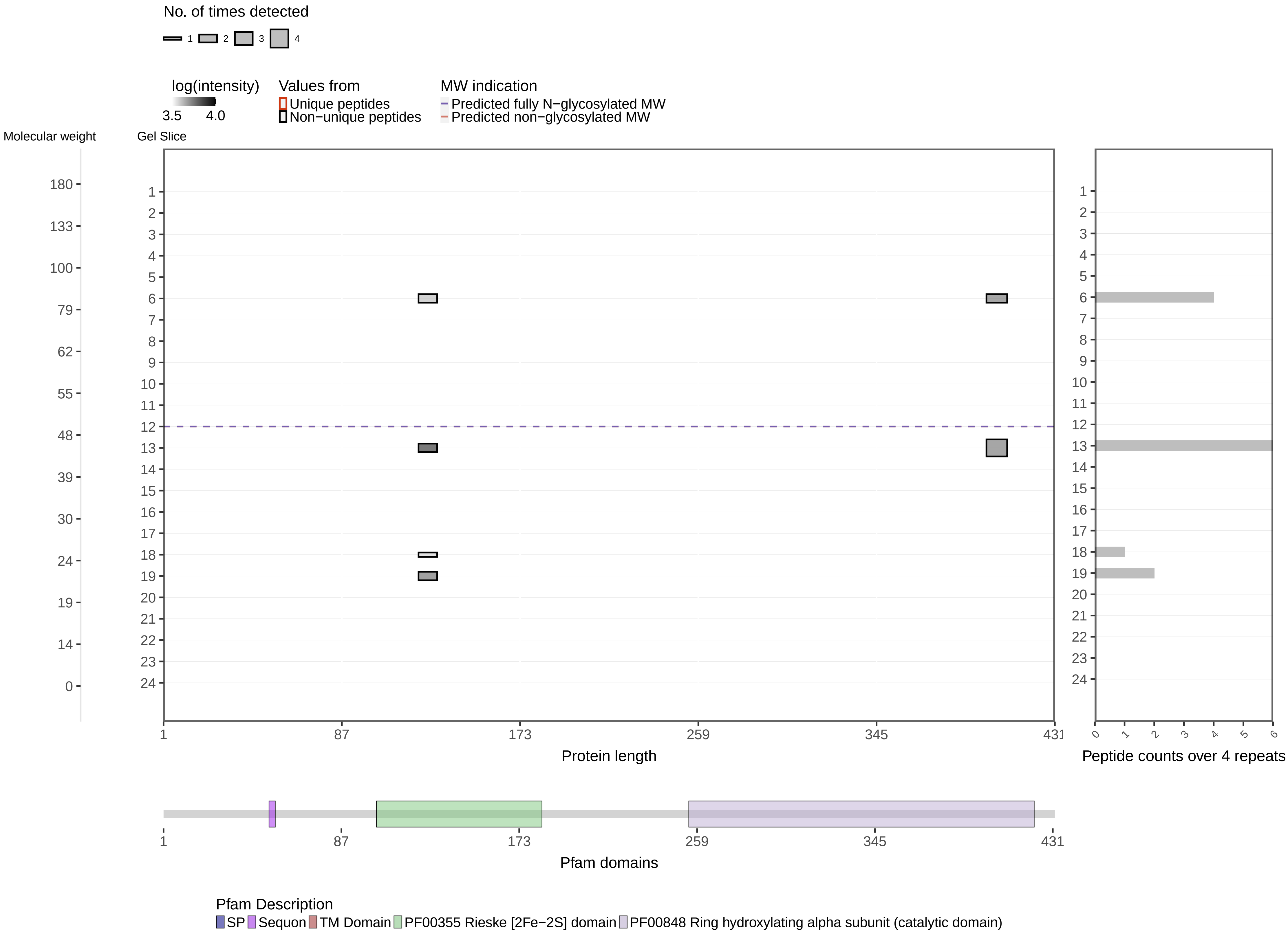


Protein sequence:

VAVWGPFILLNFENGVLSERESDFDLVGKEWLGTSSQILAEGGVDSSLSFLCRRKYTIECNWKVFCDNYLDGGYHVPYAHKGLASGLTLGSYSTTILEKVSIQRCETGSAEKGQEYDRLGSKALYAFVYPNFMINRYGPWMDTNLVLPQGPRKCLVIFDYFLDASLKGDDSFIAQSLQDSEIVQMEDIKLCEAVQRGLESPAYNSGRYAPQVEKAMHHFHCLLYENLCN*

### NbL13g02910.1 methyl−CpG−binding_domain−containing_protein_10−like [MW = 55.55 kDa]

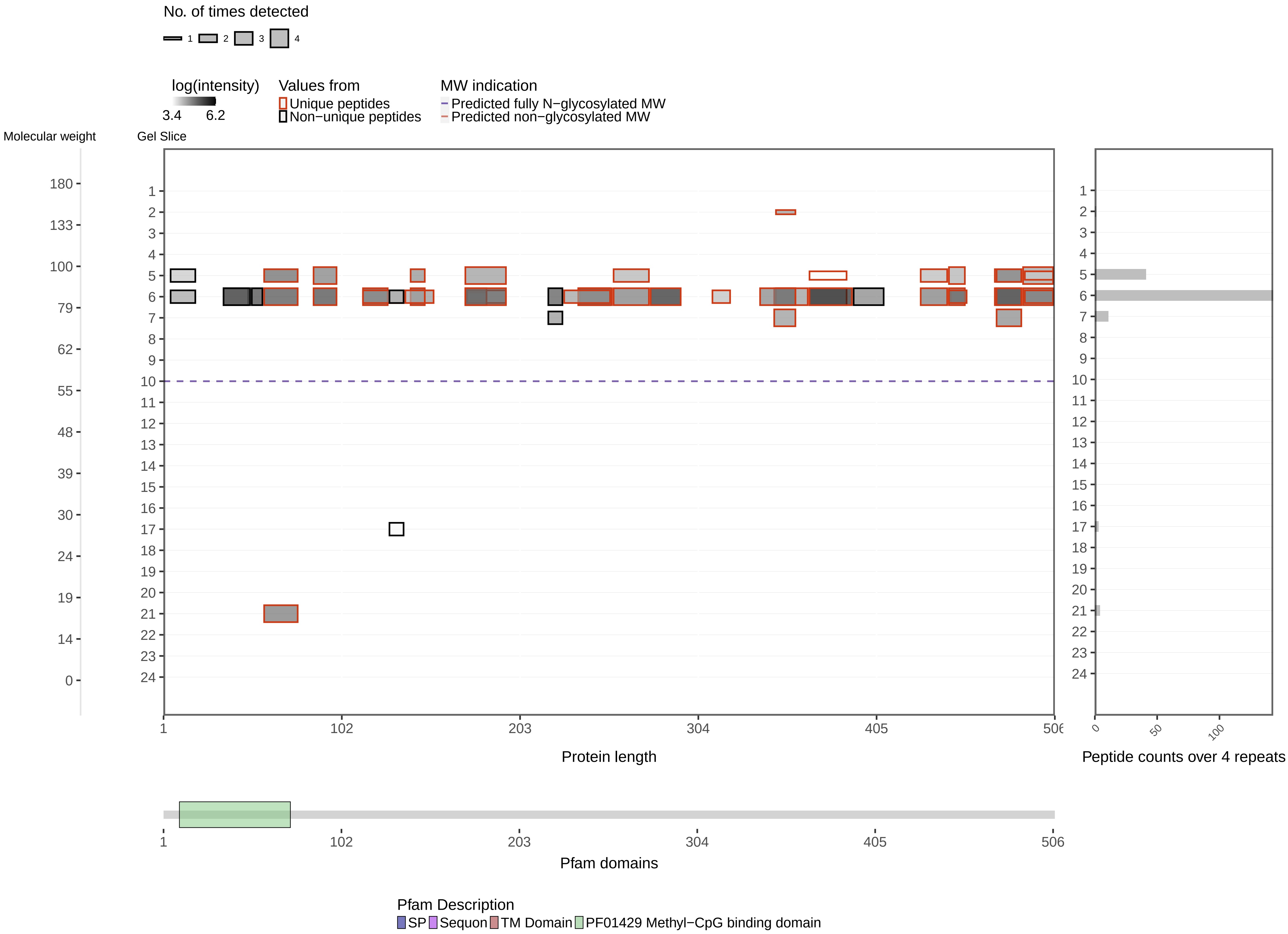


Protein sequence:

PSNVDAVEENQADRKIDGQTEDGKAEEAAEGDKDVKMADNFQHPEDAGEASVVKEAPLEKEPEATKDEEGKDVQLPESVEQKLTETKKESAAVEKDVKMADNAAEKKDEIHSSNVDAVKENQADKKSEGQTEDGKAEDAPPEAAEGDKDVKMADNFQHPEDVEEASVVKEAPIGKEADVPEATKDEEAKDVQVQESVEQKPTETKKEDFSAAEEKKHQPEQERRDENGAARKSTGEGDALMNSAISKKVQEEVTENGNEADEAYP*

NbL13g02890.1 cucumber_peeling_cupredoxin−like [MW − SP = 22.9 kDa]

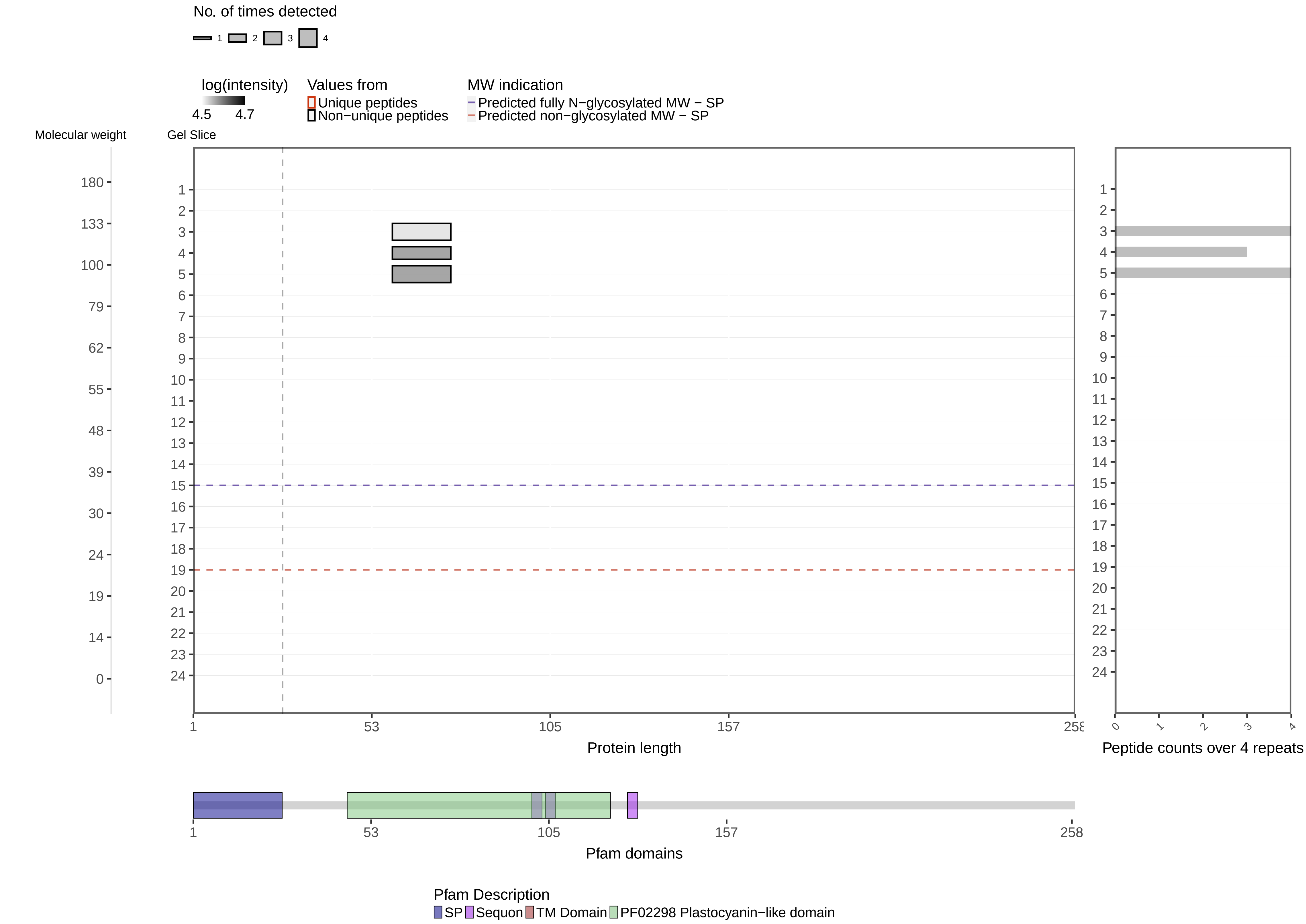


Protein sequence:

GOHCSAGOKLAINVTASSTSSPSPAPSPTTTPAPSPVPTPTPTPTPTPAPSAPSPTPMSTATPTPSPSPSMGPSAPTPTPSGGAGTSPSSSPTSSPITPGTTPSPETPPEGLIPPPAPSSAPRNVFAPALIMFMSIAISILW*

### NbL13g02870.1 blue_copper_protein−like [MW = 39 kDa]

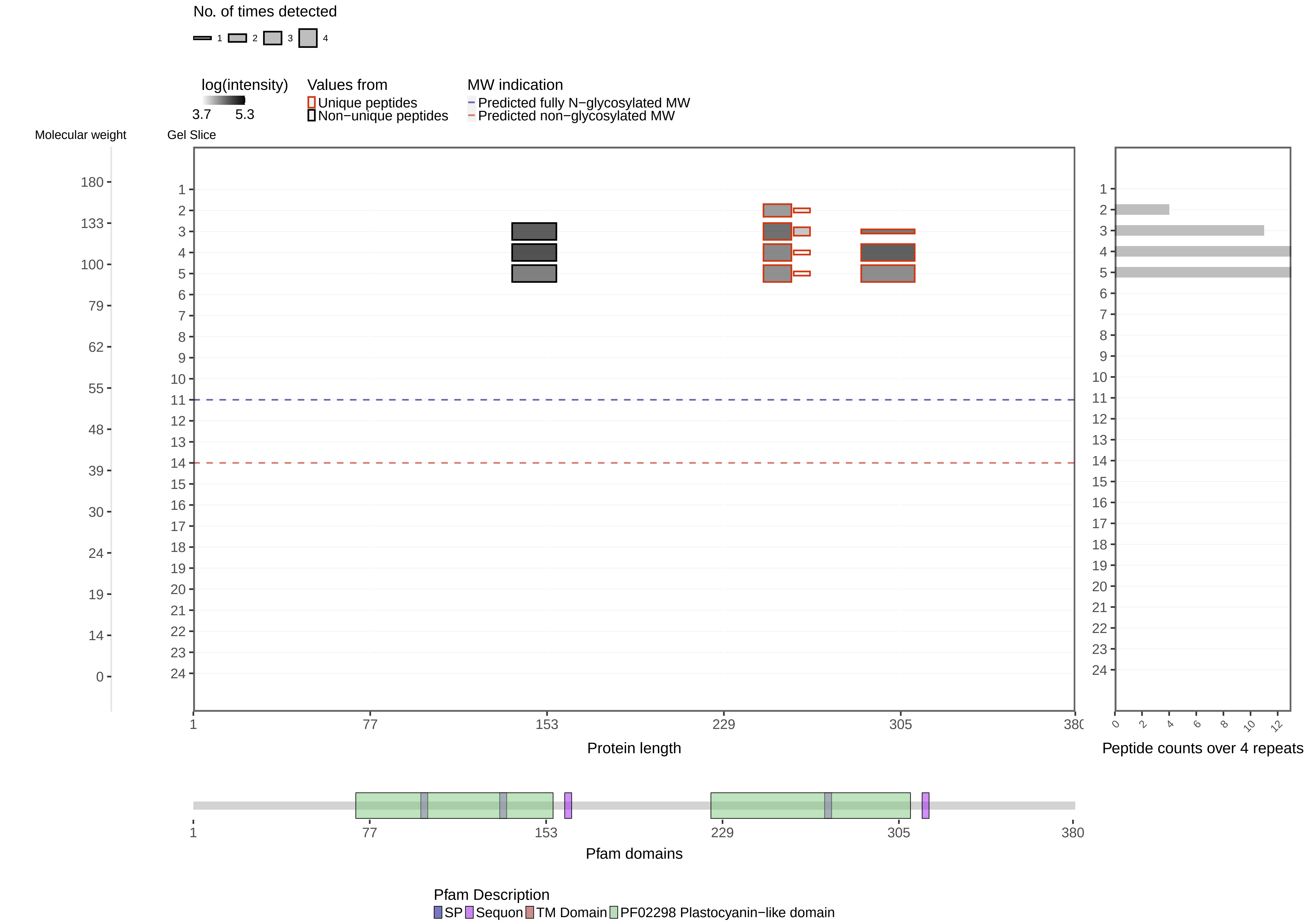


Protein sequence:

PPTAAPVTAPSASPSIAPTPSVAATPTSAPAPSVAAQTFIVGDNMGWNVPTTGGPNAYQTWANGKSFKVGDTLVFNFVNGRHNVAMVSKAAYDSCNTSSPINTISTGPARITLTNSGENYYMCTFPSHCSLGQKLAINVTGTSAAAPTPSIAATPSGSTVPSIPSGDSPVTSPPPPSASAPSLVLAALPVTFLSLAFARLLN*

### NbL13g02810.1 topless−related_protein_4−like [MW = 125.14 kDa]

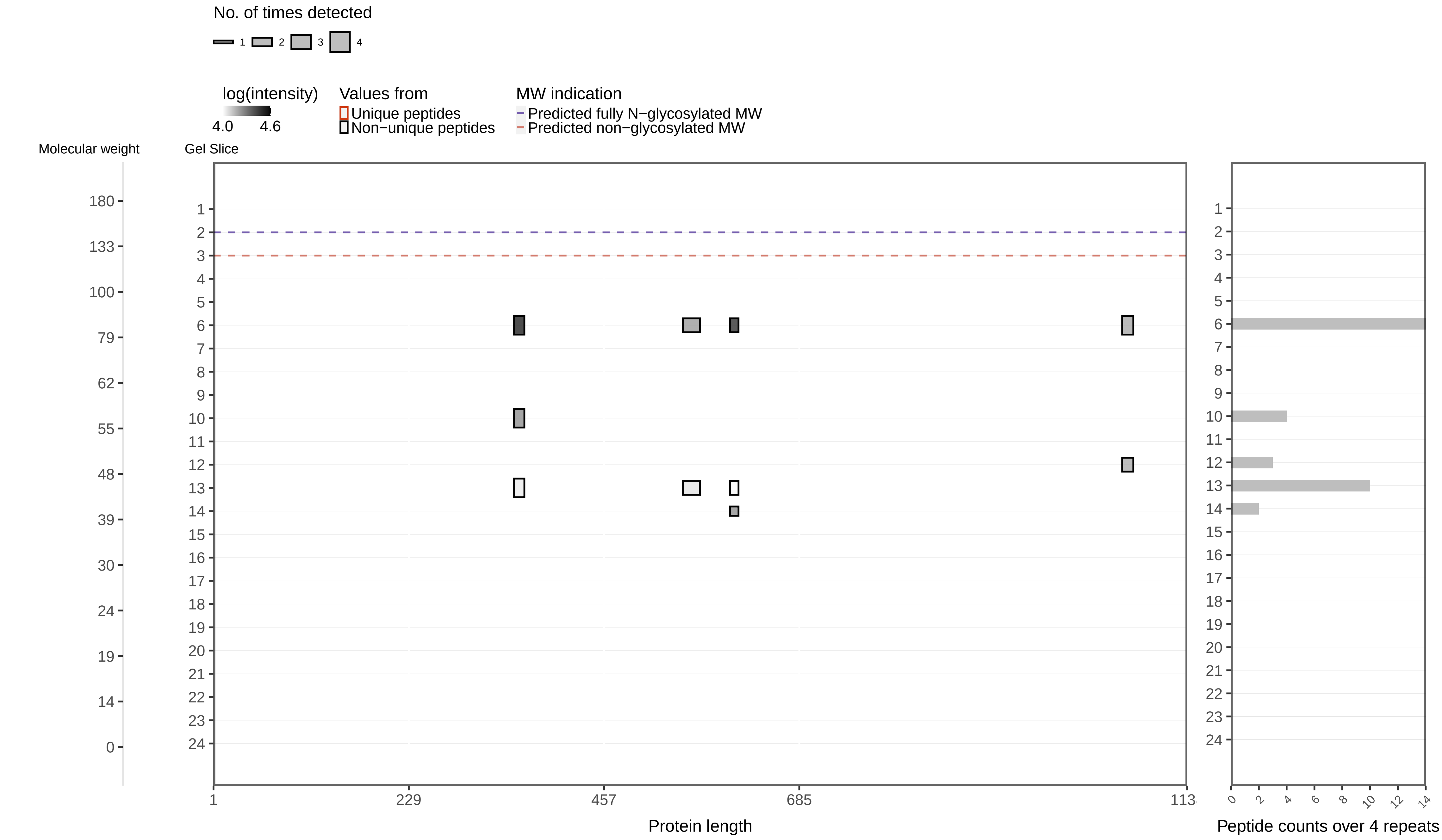


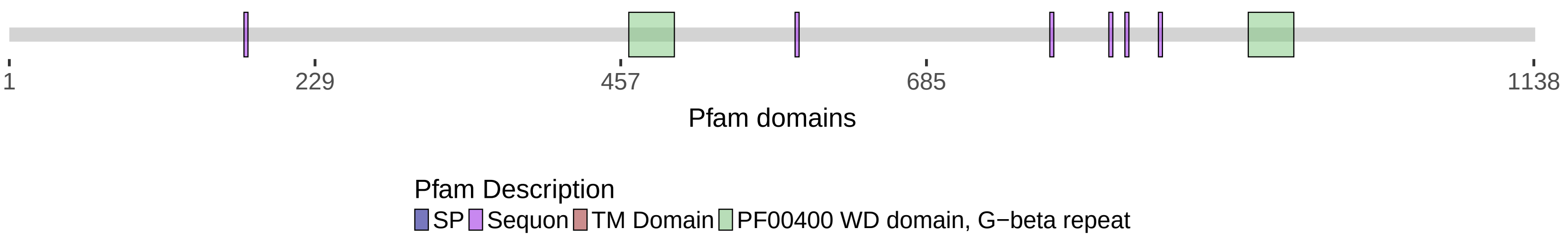


Protein sequence:

HSSTTMAYSADGTRLFSCGTNKEGESYLVEWNESEGAVKRTFIGLGKRAGGVVQFDTTKNRFLAAGDEFMIKFWDMDNINMLTSTEAEGGIPASPCLRFNKEGILLAVSTSDNGIKVLANADGLRLLRSMENRQFDASRVASASVVKPPTLGSFGPPSTSVAASFVDRGVPMTSMVSMNGENRNLGDVRPRAAEETVDKSRIWKPTEINEPSQCKSLKLPDNITATRISRLIYTNSGLAILALASNAVHKLWKWPRNERNPTGKANASIVPQLWQPSNGTLMTNDITDTNPE

NbL13g02660.1 12−oxophytodienoate_reductase_3 [MW = 40.54 kDa]

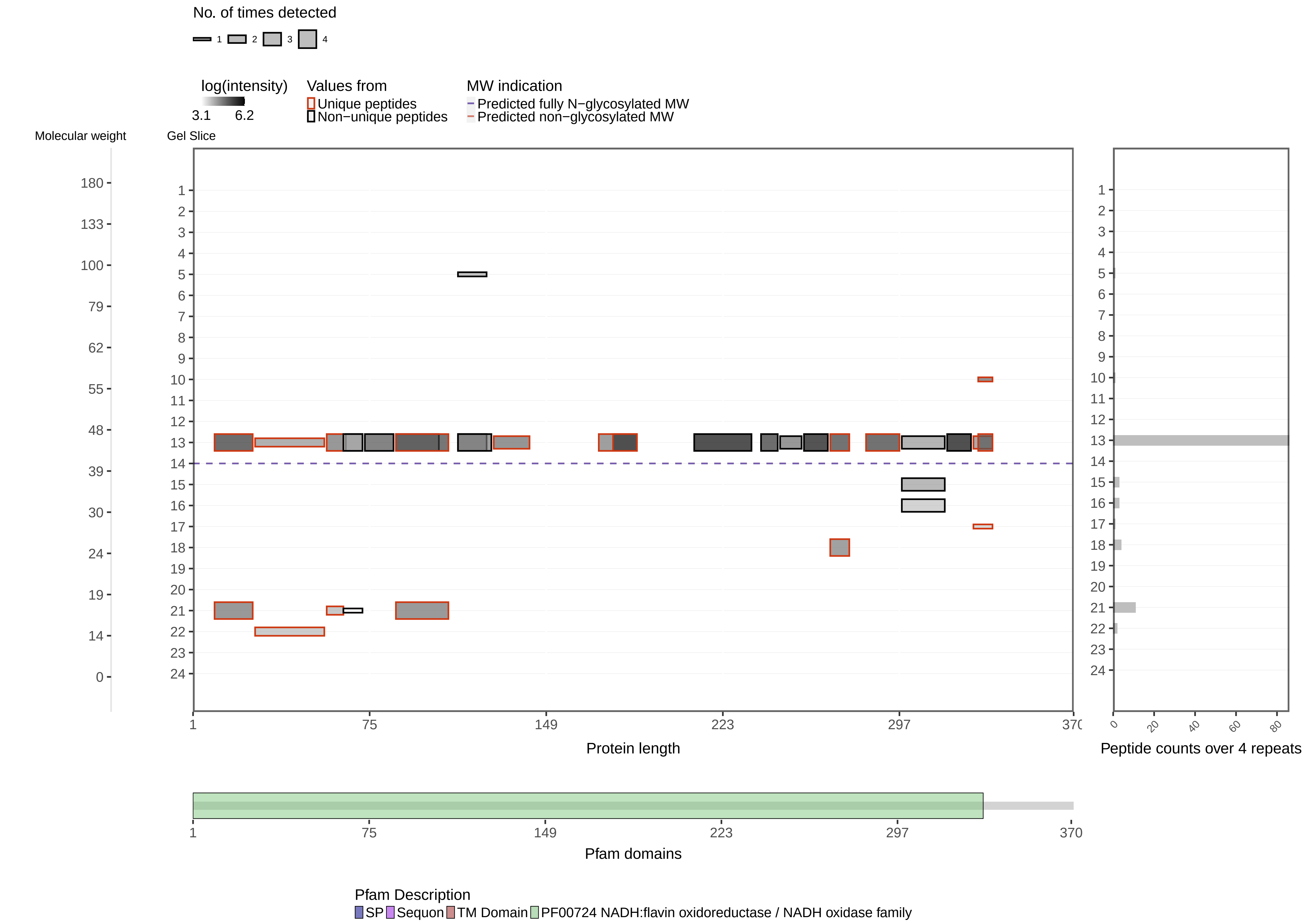


Protein sequence:
INDRTDEYGGSLGNRCKFIMQVVEAVVTAIGVDRVGVRMSPAIDHLDAMDSNPLSLGLAVVERLNKLQLHFGSKLAYLHVTQPRYVAYGQTEAGRLGSEEEEARLMRTLRNAYQGTFICSGGYTRELGIEAVAQGDADLVSYGRLFISNPDLVMRLKLNAPLTKYNRKTFYTQDPIVGYTDYPFLQGNGNSIPSSRL*

NbL13g02600.1 probable_sucrose−phosphate_synthase [MW = 118.8 kDa]

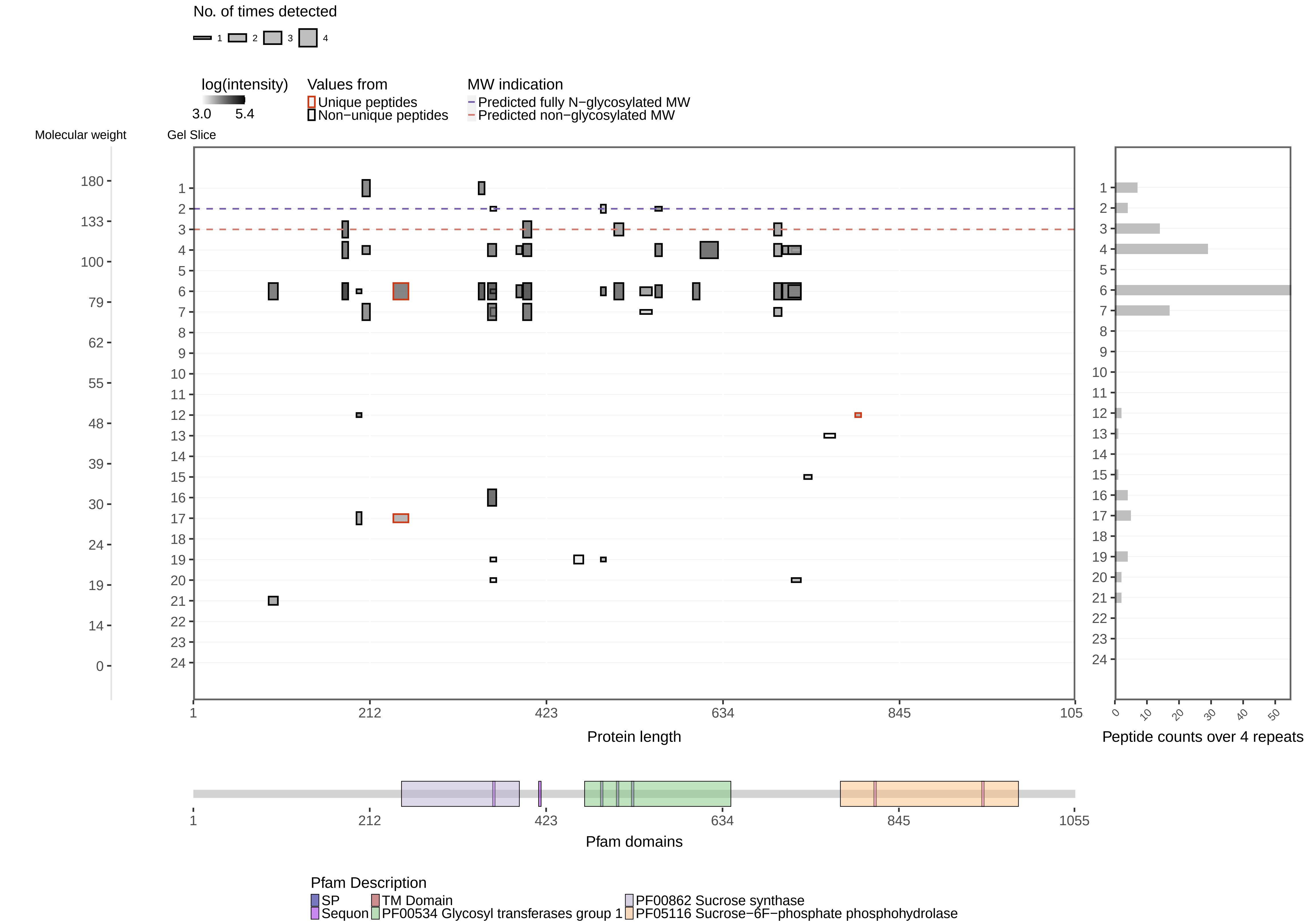


Protein sequence:

IMGNRDNIDEMSSTNSSVLLSILKMIDKYDLYGQVAYPKHHKQADVPDIYRLAAKTKGVFINPAFIEPFGLTLIEAAAYGLPMVATKNGGPVDIHRVLDNGLLVDPHDQQAIADALLKLVADKHLWAKCRANGLKNIHLFSWPEHCKTYLSRIASCKPRQPRWLRTDDDDDENSETDSPSDSLRDIHDISLNLRFSLDGEKNDNKENADNTLDPEVRKSKLENAVLSWSKGVLKSTPKAWSSDKGDQNSGAGKFPAIRRRRHIFVIAVDCDASSGLSESVRKIFEAVEKERAEGSI

NbL13g02570.1 bifunctional_purple_acid_phosphatase_26−like [MW − SP = 52.74 kDa]

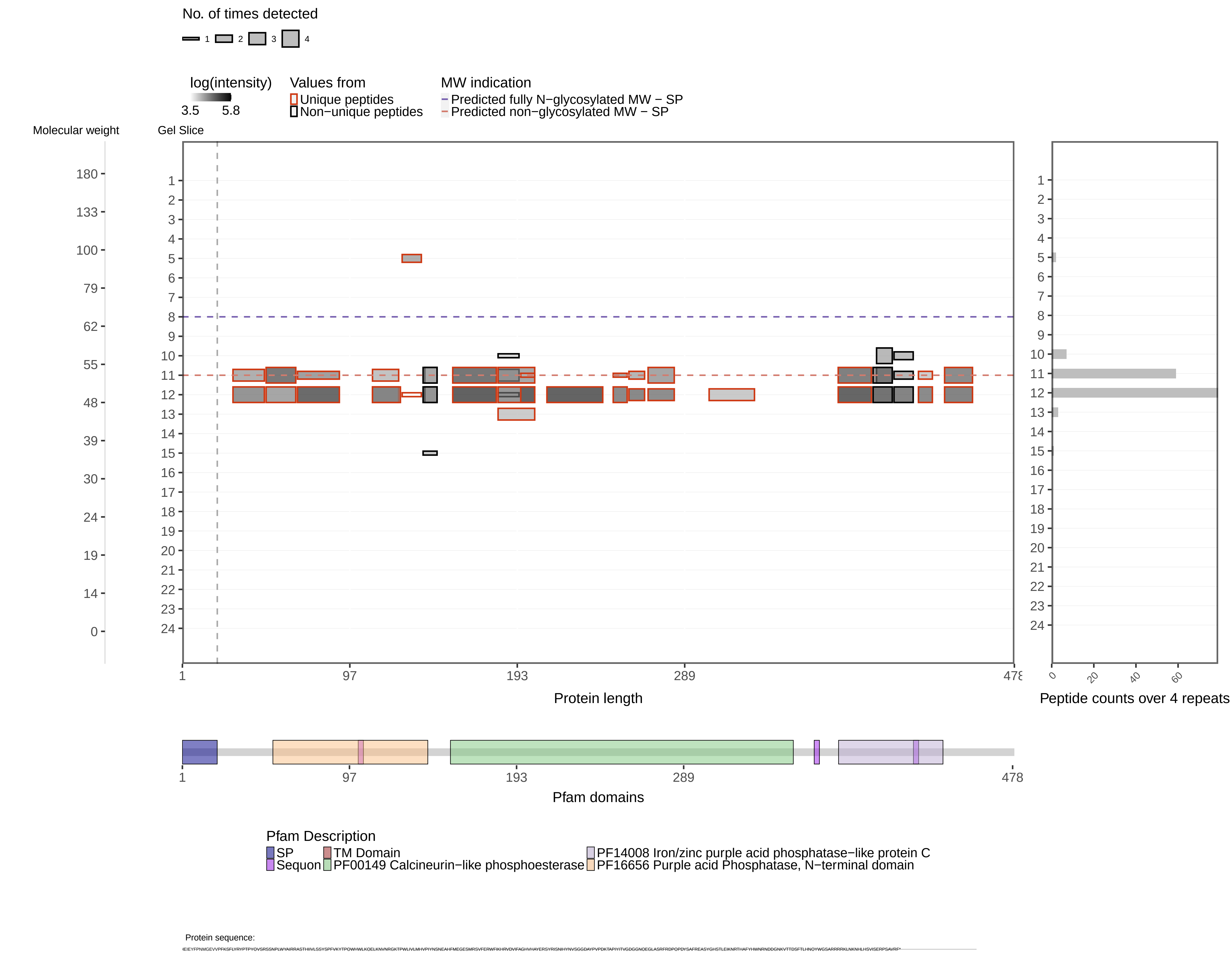

### NbL13g02470.1 pyrophosphate−energized_vacuolar_membrane_proton_pump−like [MW = 80.02 kDa]

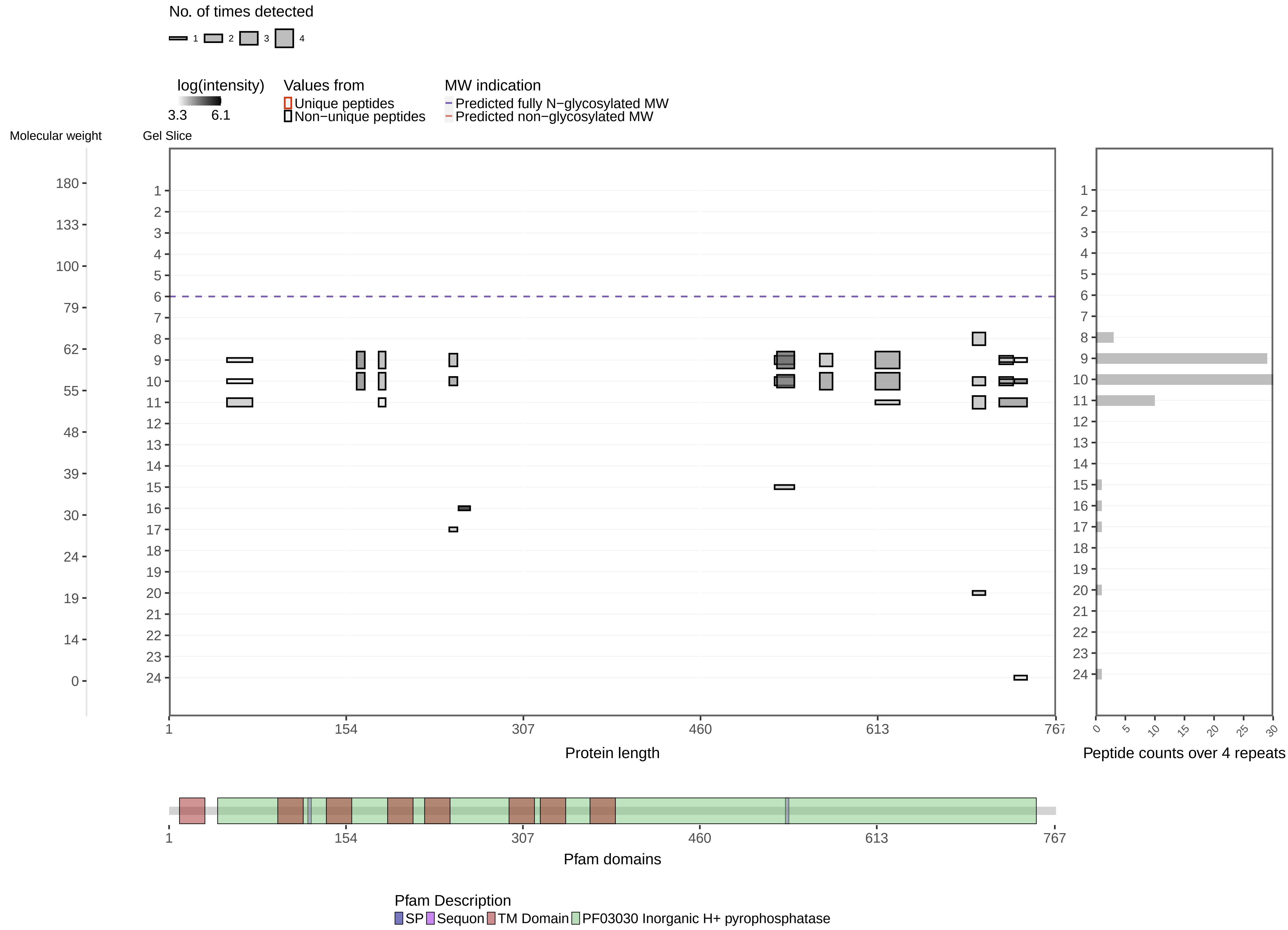


Protein sequence:

### NbL13g02460.1 arogenate_dehydrogenase_2_chloroplastic−like [MW = 42.86 kDa]

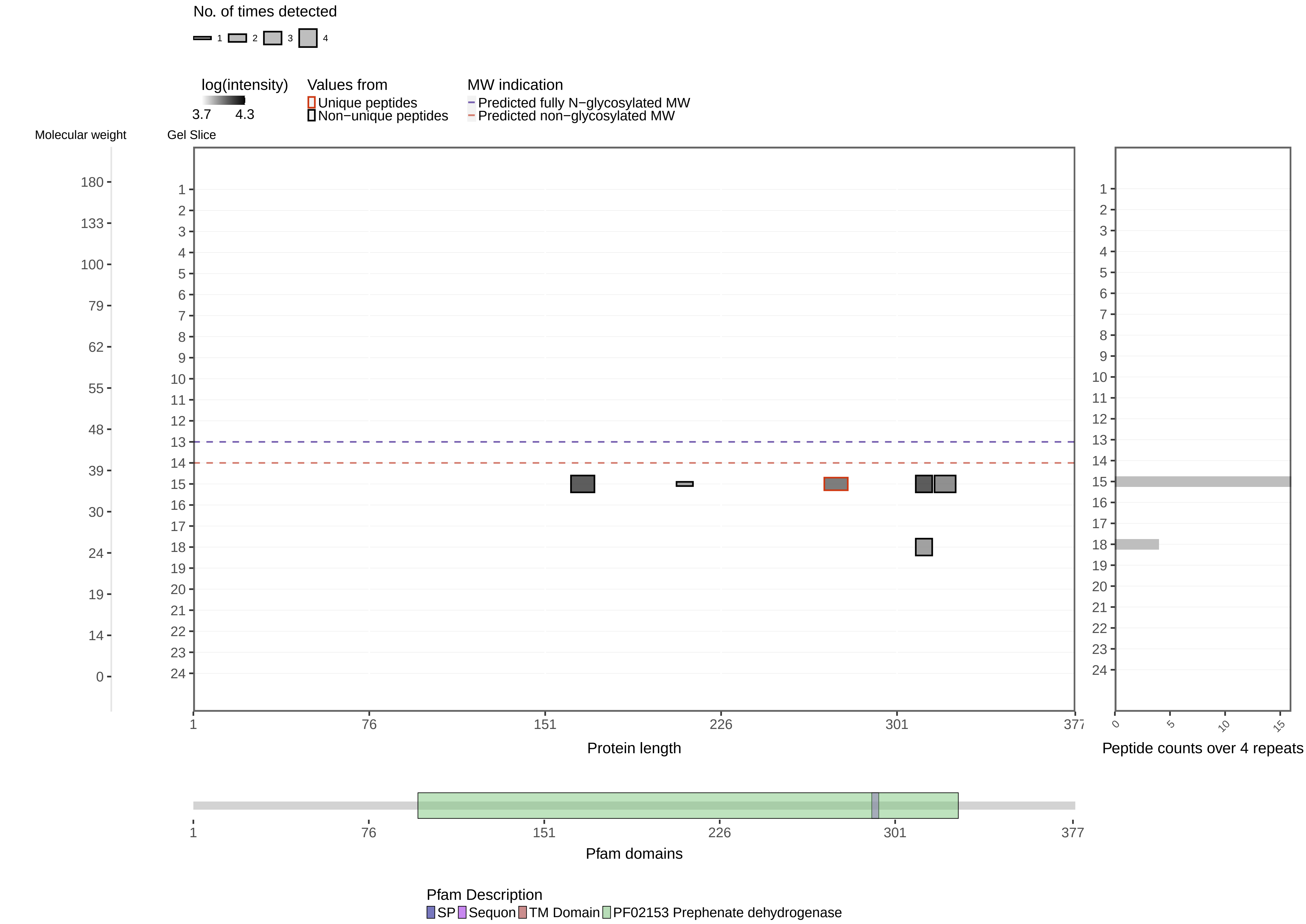


Protein sequence:

NIFLQVLPSHFDILCTHPMFGPESGKDSWKDLAFVFDKVRIGEGESRKGRVDRFLHIFEKEGCRMVQMTCAEHDRYAAGSQFITHTMGRVLEKLELETTPINTKGYETLLNLVENTSSDSFDLYYGLFMYNKNAMEQLERLDLAFEALKKELFGHLHEVLRKQLFGKAEEAGQRRILTKLPKNGYALPAPSSEAVKSENN*

### NbL13g02440.1 heparanase−like_protein_3 [MW − SP = 56.34 kDa]

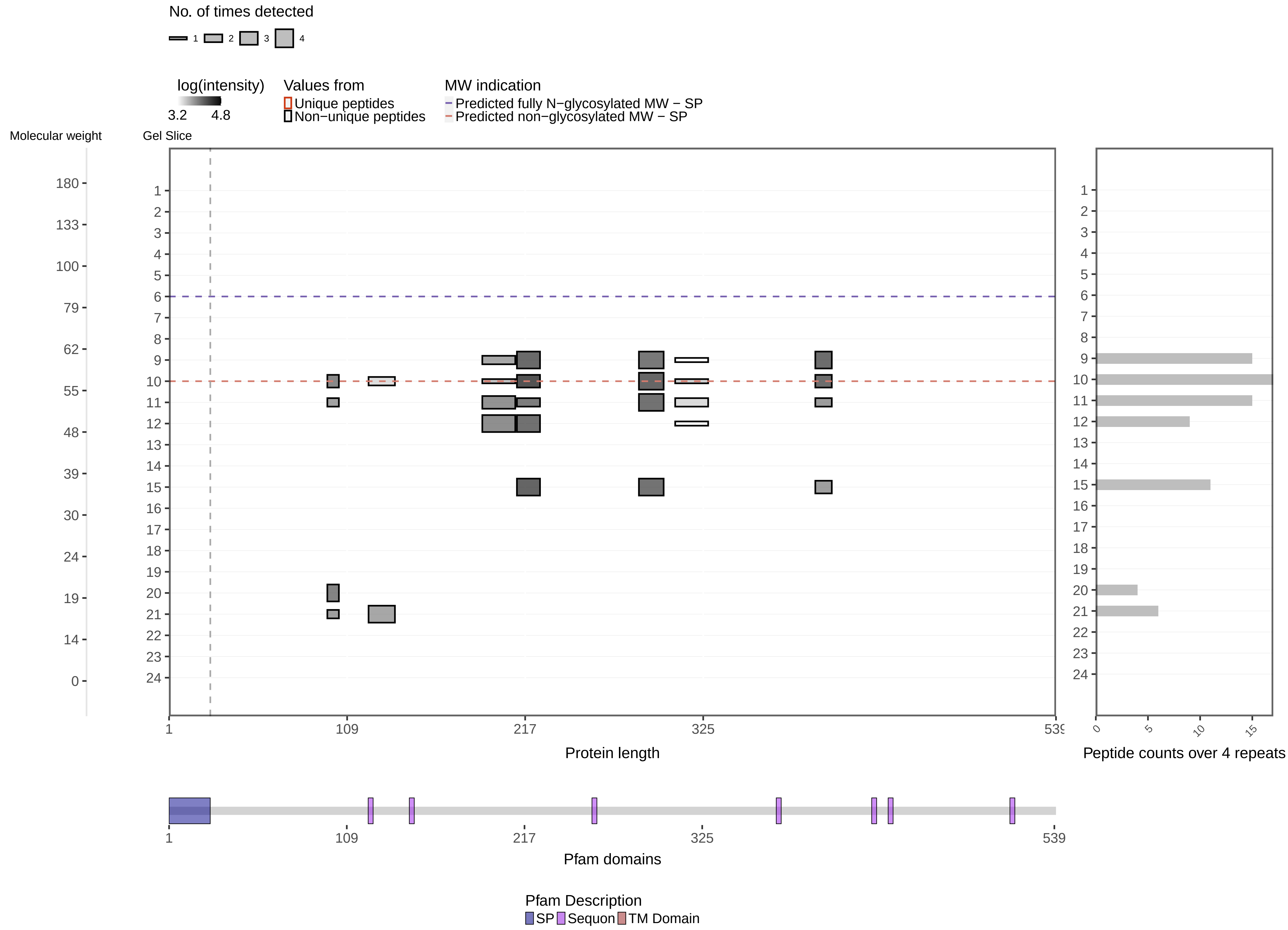


Protein sequence:

ELINKTGKSFDVATHHIYNLGPGRDEHLVDKILDPSYLDGEADTFSKLQNVLKTSGTSVVAWVGEAGGAYNSGRDHVTNAFVFSFWYLDQLGMSAAYDTNAYCRQTLIGGNYGLLNTTTFVPNPDYYSALLWHRLMGRNVLSTSFSGSKKIRAYAHCAKQSQGITLLLINLDGNTTISANVAFNGTTLRHKKHSRHSHKKSSSIRLPKSRKIASNTRQEYHLTAKDGDLQSQTMLLNGKPLTIDSSGNIPPLEPIFVNSTEPLTVAPFSIVFVHIPYVILPACS*

NbL13g02430.1 triphosphate_tunel_metalloenzyme_3−like [MW = 28.14 kDa]

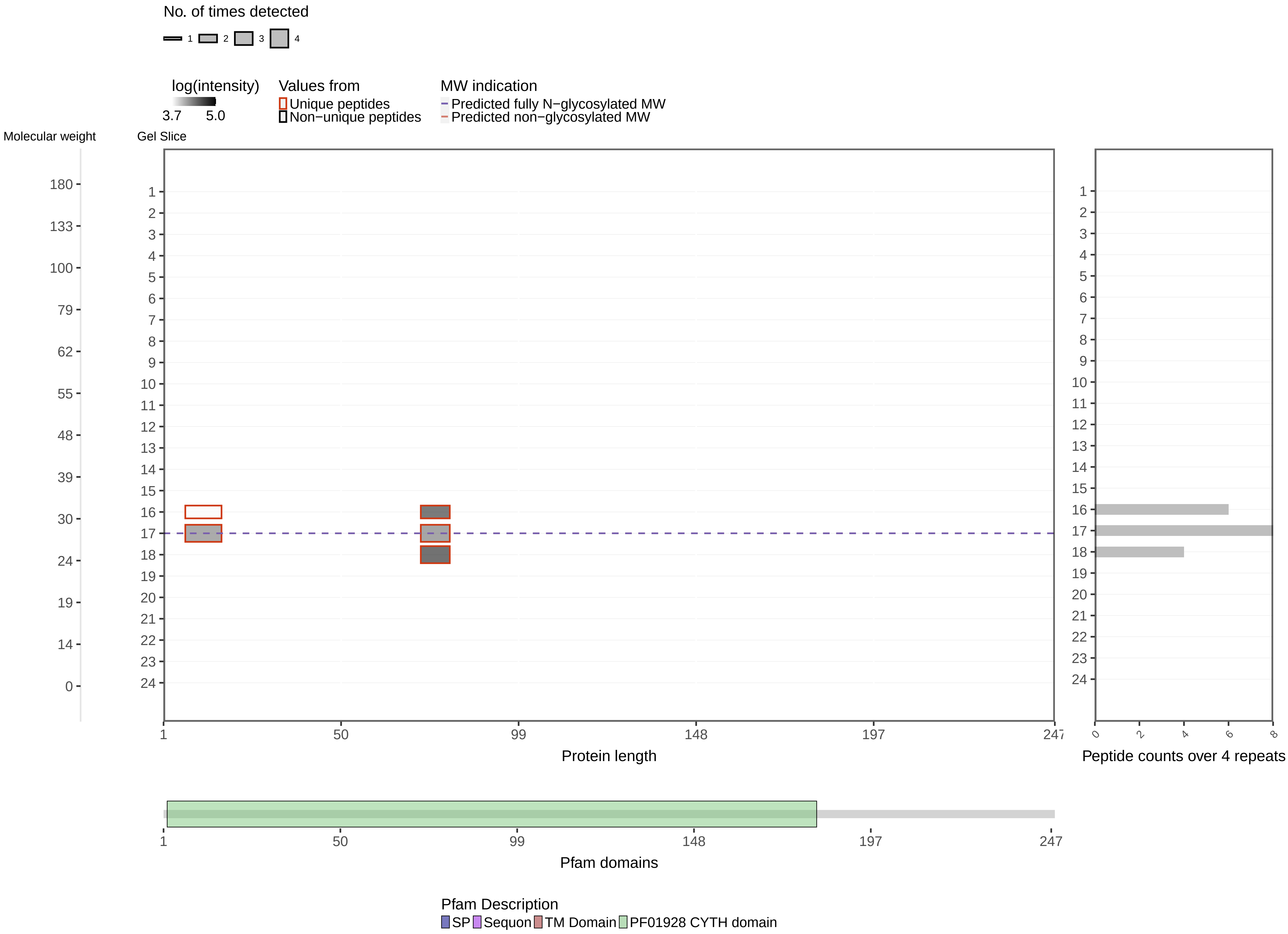


Protein sequence:

RVLKRAKEEFDVAEEGFIGLGGFKNVRNVFQWNGLELEVDETIYDFGTCYEIQCESTEPEEAKAMIEALLKENDIDYSYSRVSKFATFRAGKLPQCTSYLAPYLEGPKSSIPQYIAMKSYHLAYFRKELPCSIVFVVS*

### NbL13g02200.1 protein_CHLORORESPIRATORY_REDUCTION_6_chloroplastic [MW = 29.23 kDa]

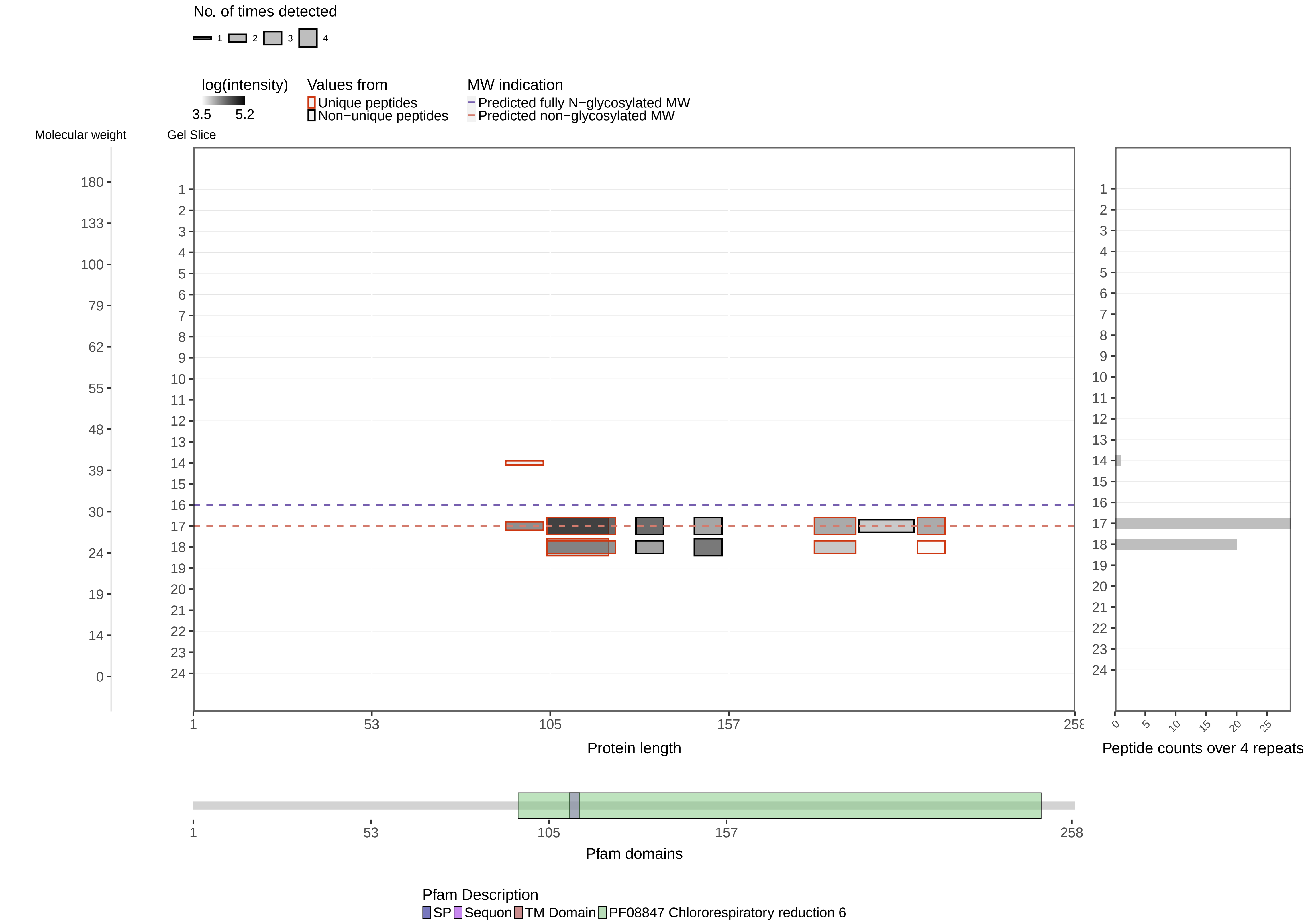


Protein sequence:

SPLSAKPKEFLERTIGFTINYKREDKYDPRELSEFPDIRLWFVRLDATYPWLPVLLDWKAGELARYAAMLVPHQMSMKMGVVFNPEALELFVMNKVFIVYSWLKQNEIPMPRLKTKDMARMLGFGIGDELFDLIDKHHIDT*

### NbL13g02130.1 110_kDa_U5_small_nuclear_ribonucleoprotein_component_CLO [MW = 110.41 kDa]

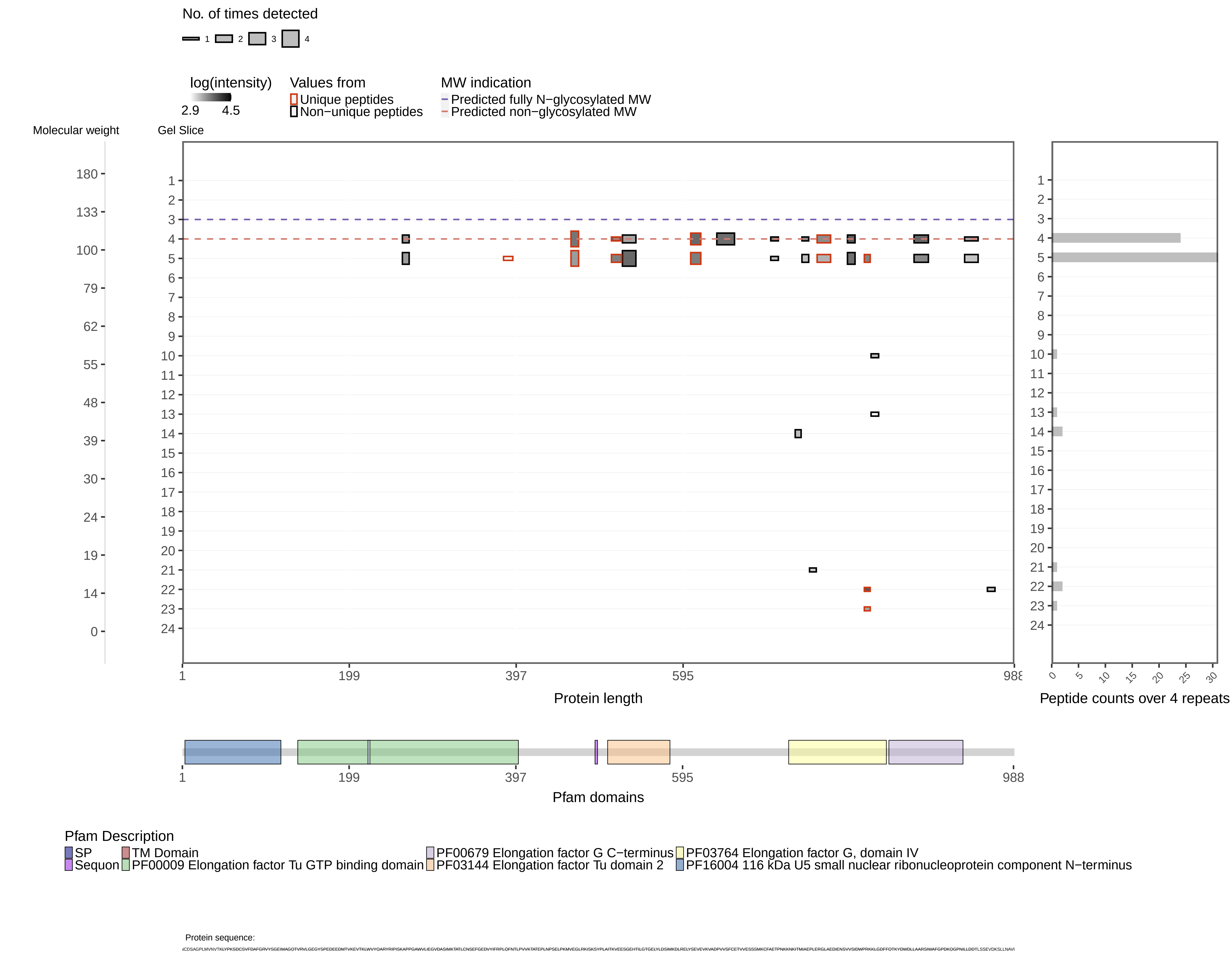

NbL13g02030.1 peptidyl−prolyl_cis−trans_isomerase_CYP28_chloroplastic−like [MW = 34.78 kDa]

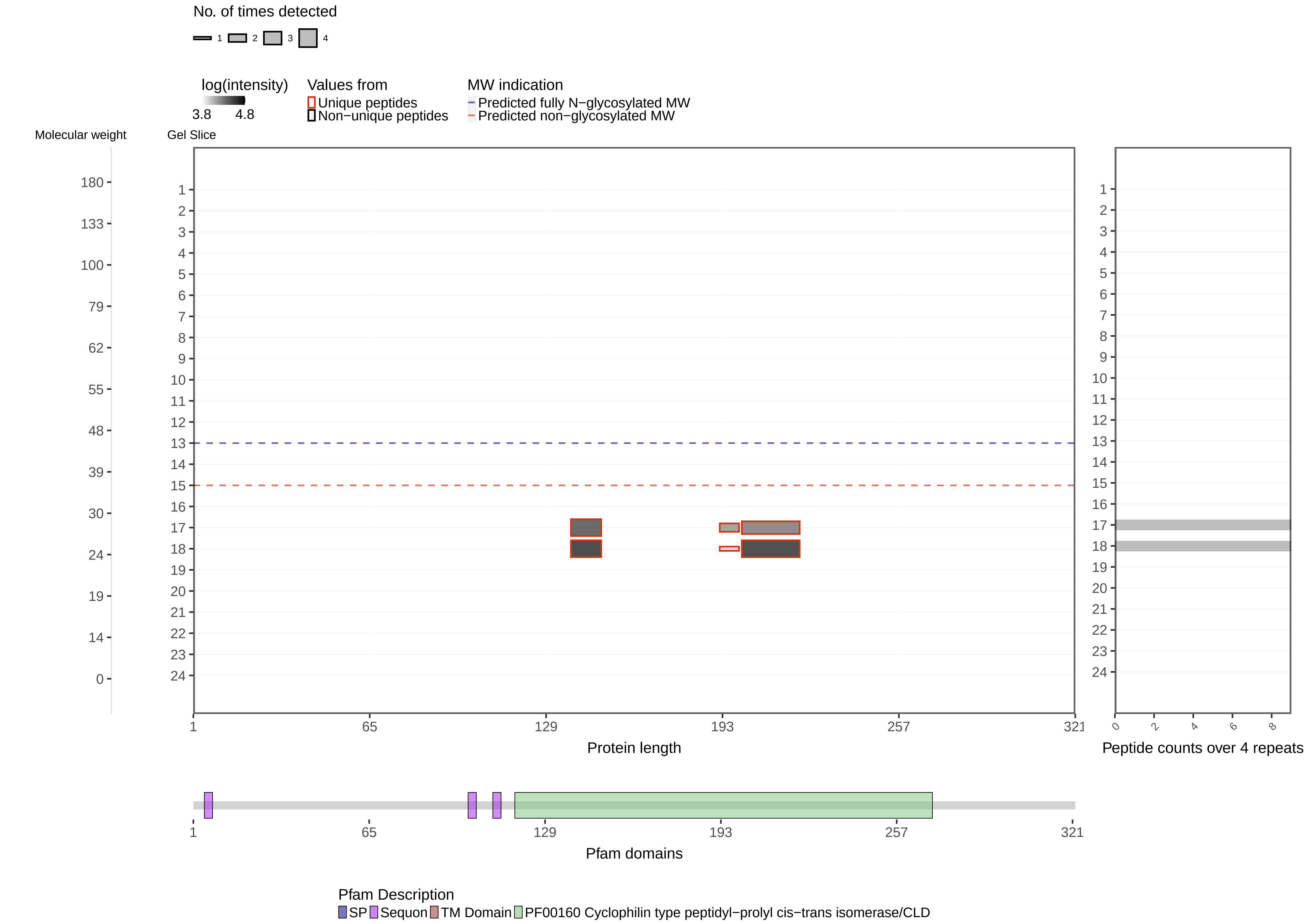


Protein sequence:

GTLVQKIFPGQFFMAGRQGRRDKGEVKPPLELIRNTETTESKAFILEHSRGGIVSLPLSENDDDDDLMLNPKYHNVEFLITTGPGPVPQLDSKNIVFGTVLEGMDIVTTIAAIPTYKPSENIRQYNGFAELLGDGRAKTARAIWNKPQKTVYISDCGELKVTKPTLSPSLP*

### NbL13g01870.1 3−isopropylmalate_dehydrogenase_chloroplastic−like [MW = 40.15 kDa]

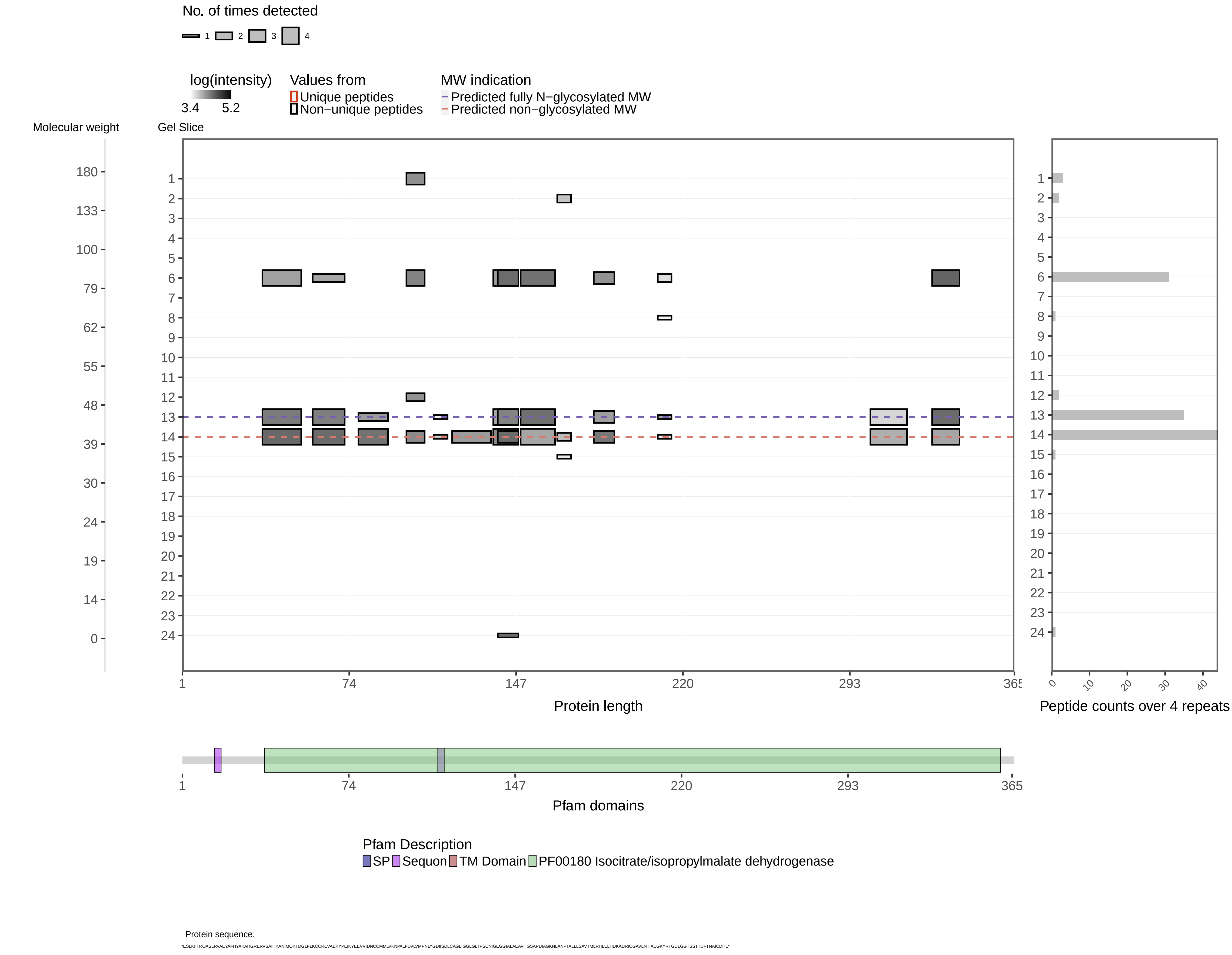

### NbL13g01840.1 chitinase−3−like_protein_1 [MW − SP = 39.78 kDa]

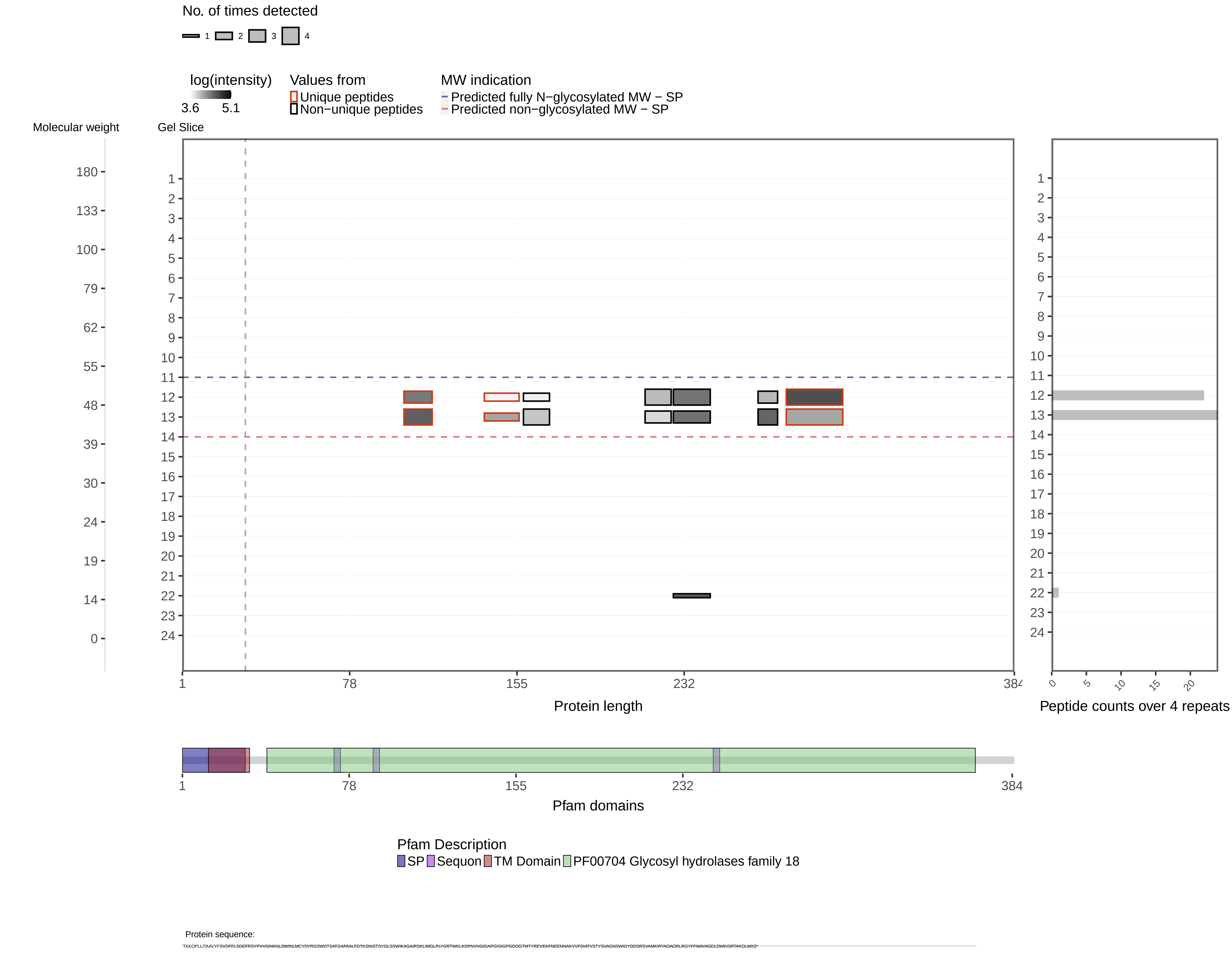

NbL13g01820.1 uncharacterized_LOC107807700 [MW = 37.58 kDa]

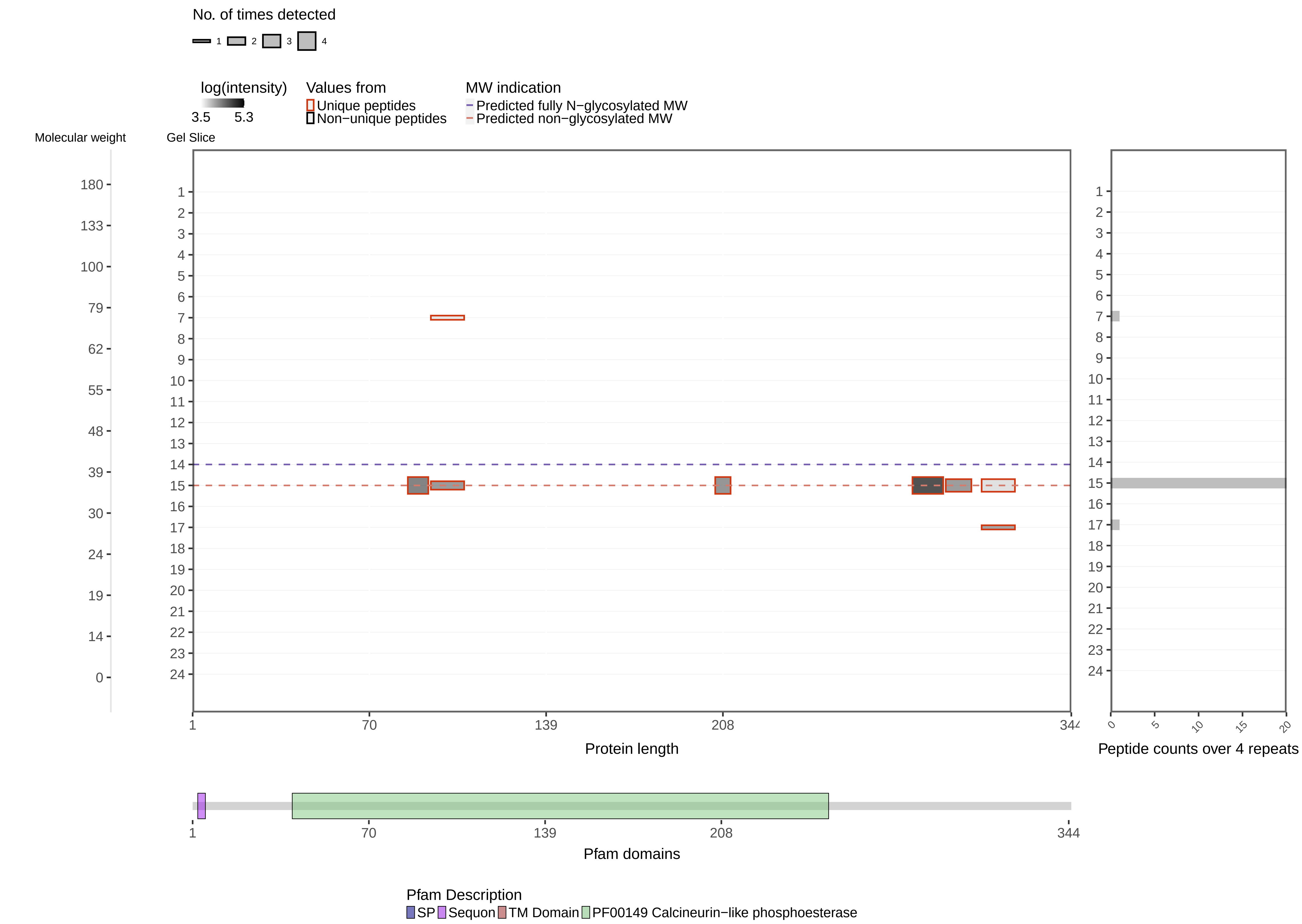


Protein sequence:

RKQLLKARYGVHDMDRSTERILQAAMGTPEDHSIIFLAHNGPTGLGAKIDDICGRDWFVGGGDHGDPDLAHALAQLKEAAPYSVPLVIFGHMHKQLALGNGFRKMIVVGNDNTIYLNGAIVPRVRTPALTAESEGTSRAFTIAEISNGRVEKIAETWISVIGDETRLEEEHILFNTPRRGSVTASLL*

### NbL13g01510.1 S−norcoclaurine_synthase_1−like [MW = 17 kDa]

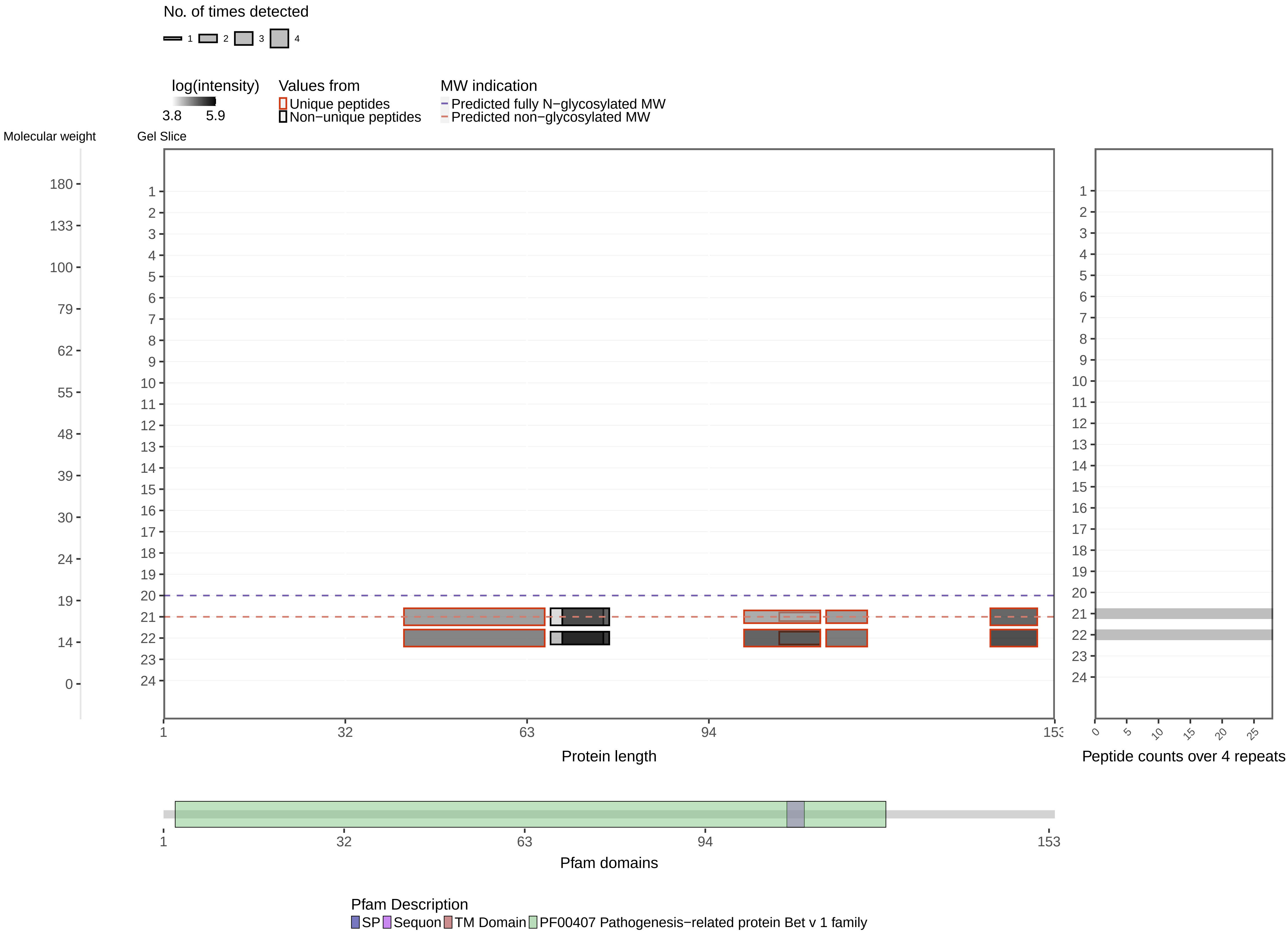


Protein sequence:

NAAYKEKFVSIDDEKRVKKVEIVEGGYLDLGFSFYGIKFEVIEKDDNLSIVKISIDFETKDAENIHLTVGNLQAFIAIMKATVEYFNQKSK*

NbL13g01500.1 S−norcoclaurine_synthase_1−like [MW = 17.21 kDa]

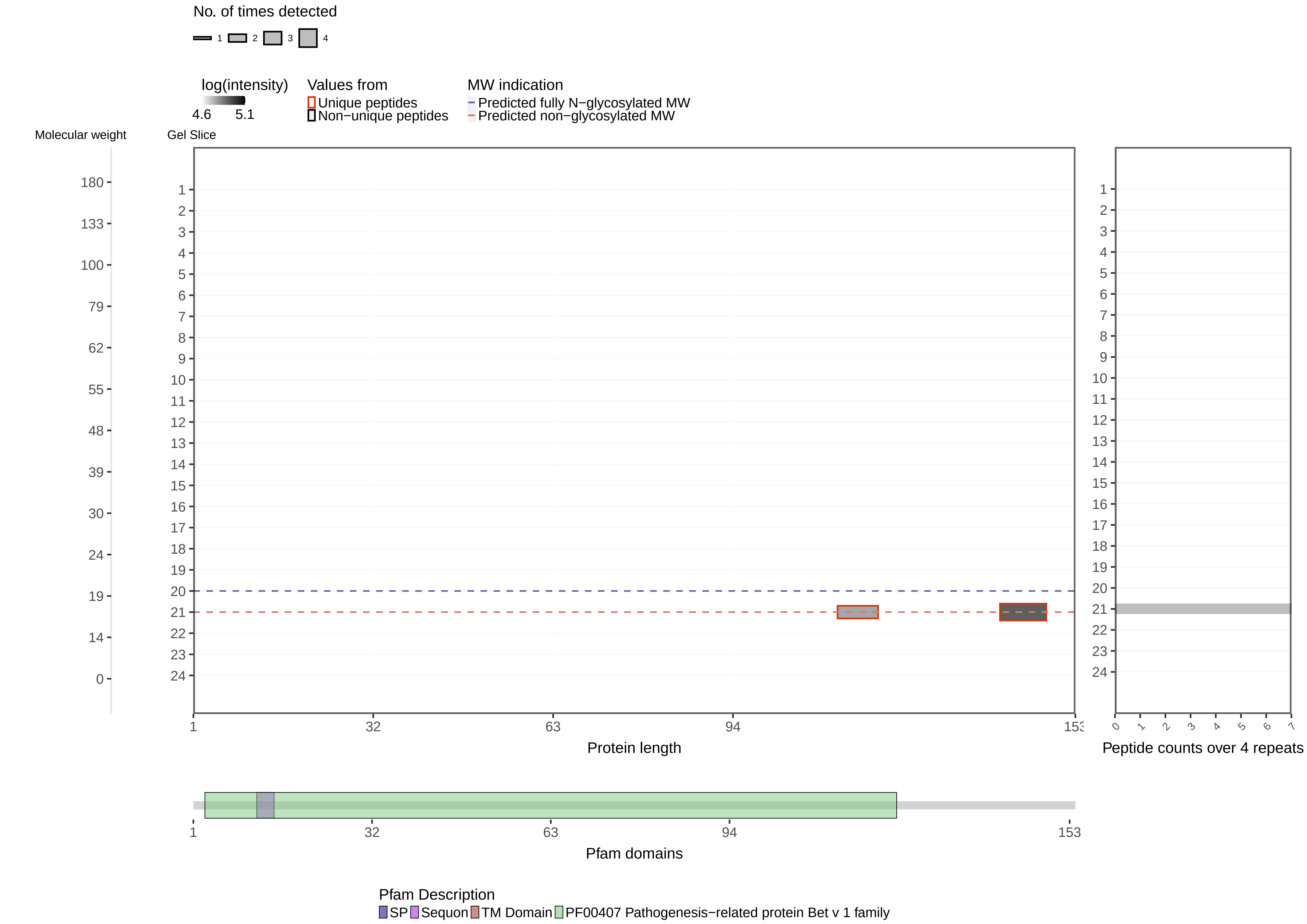


Protein sequence:

[illegible]YKEKFVSIDDEKRVKEIEMVEGGYLDLEFSFYGCKFEVIEKDENSCIVKITIGFETKDAENIHLTIGNIPTVFAILKATVEYFNEKNKQM*

NbL13g01480.1 NADP−dependent_glyceraldehyde−3−phosphate_dehydrogenase [MW = 53.27 kDa]

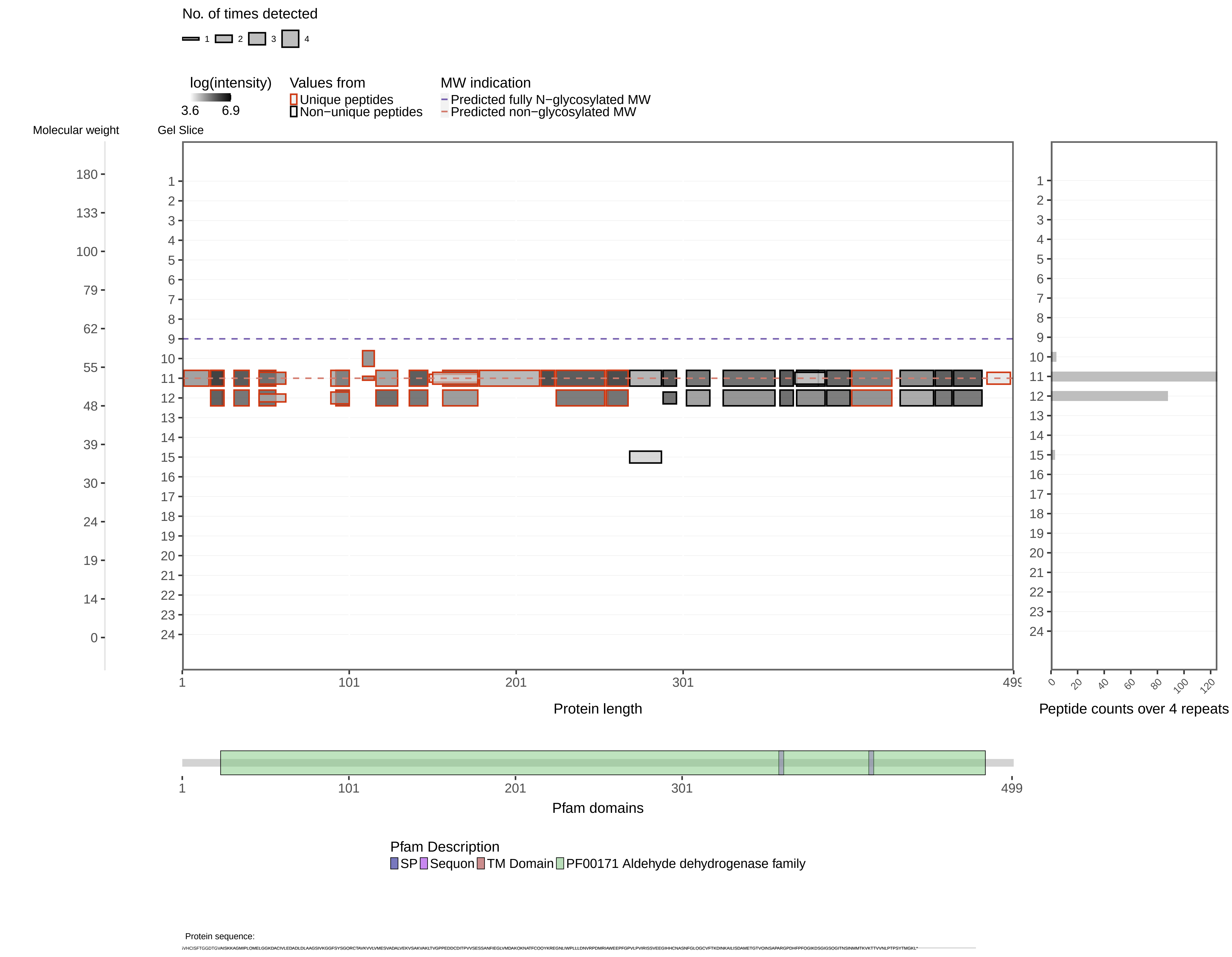

### NbL13g01230.1 serine_carboxypeptidase_24−like [MW − SP = 49.48 kDa]

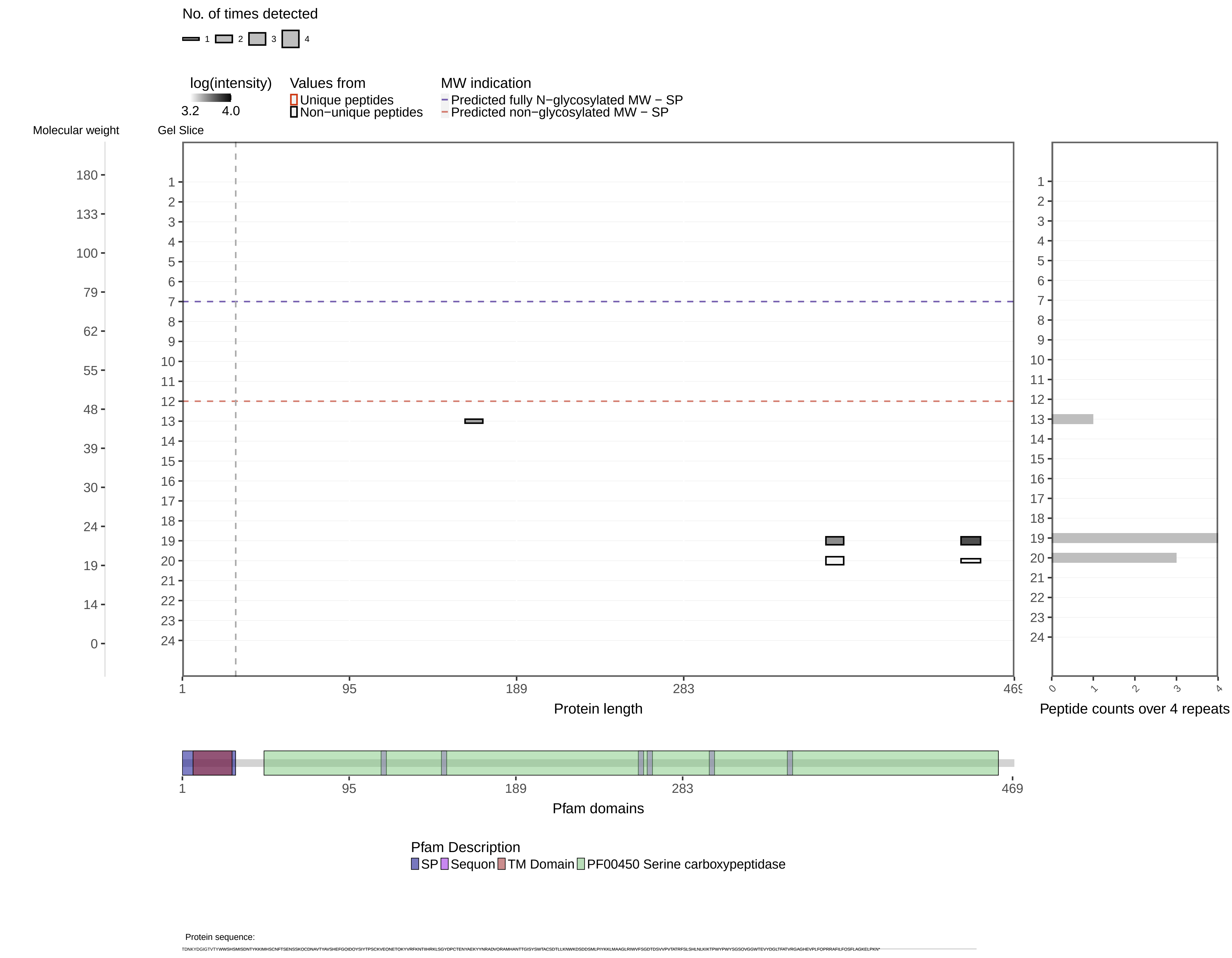

NbL13g01170.1 shikimate_O−hydroxycinnamoyltransferase−like [MW = 48.36 kDa]

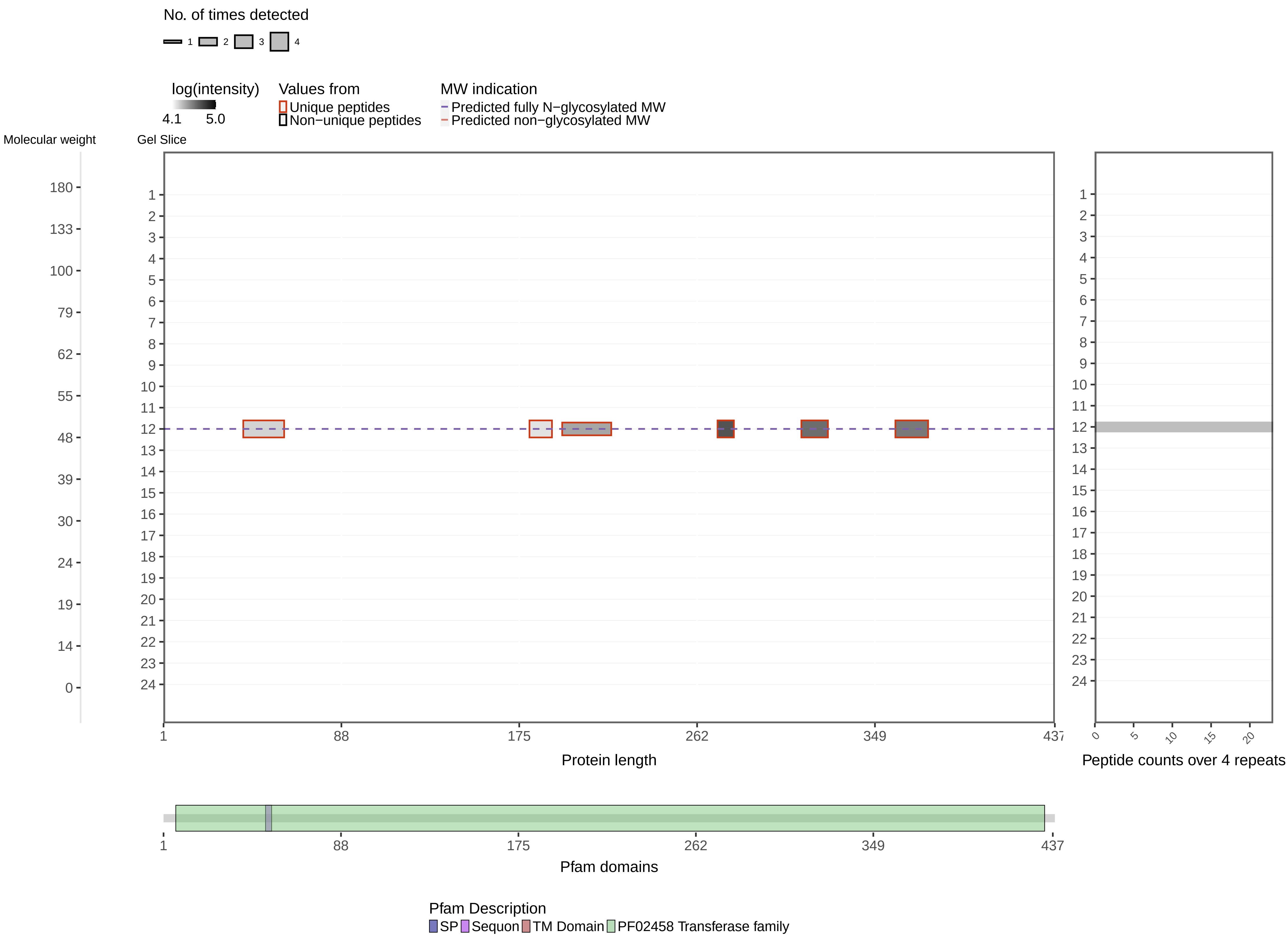


Protein sequence:

VEYHPPPSLISSSKTKESTSPKPSTTAMFKFSSDQLGLLKSKSKHEGSTYEILAAHIWRCTCKARALSDDQLTKLHVATDGRSRLCPPLPPGYLGNVVFTATPMAKSSELLEEPLTNSAKRIHSSLSKMDDNYLRSALDYLELQPDLSALIRGPAYFASPNLNINSWTRLPVHDSDFGWGRPIHMGPACILYGGTVYILPSPNNKDRNLRLAVCLDADHMPLFEKYLYEF*

### NbL13g01120.1 eukaryotic_translation_initiation_factor [MW = 85.01 kDa]

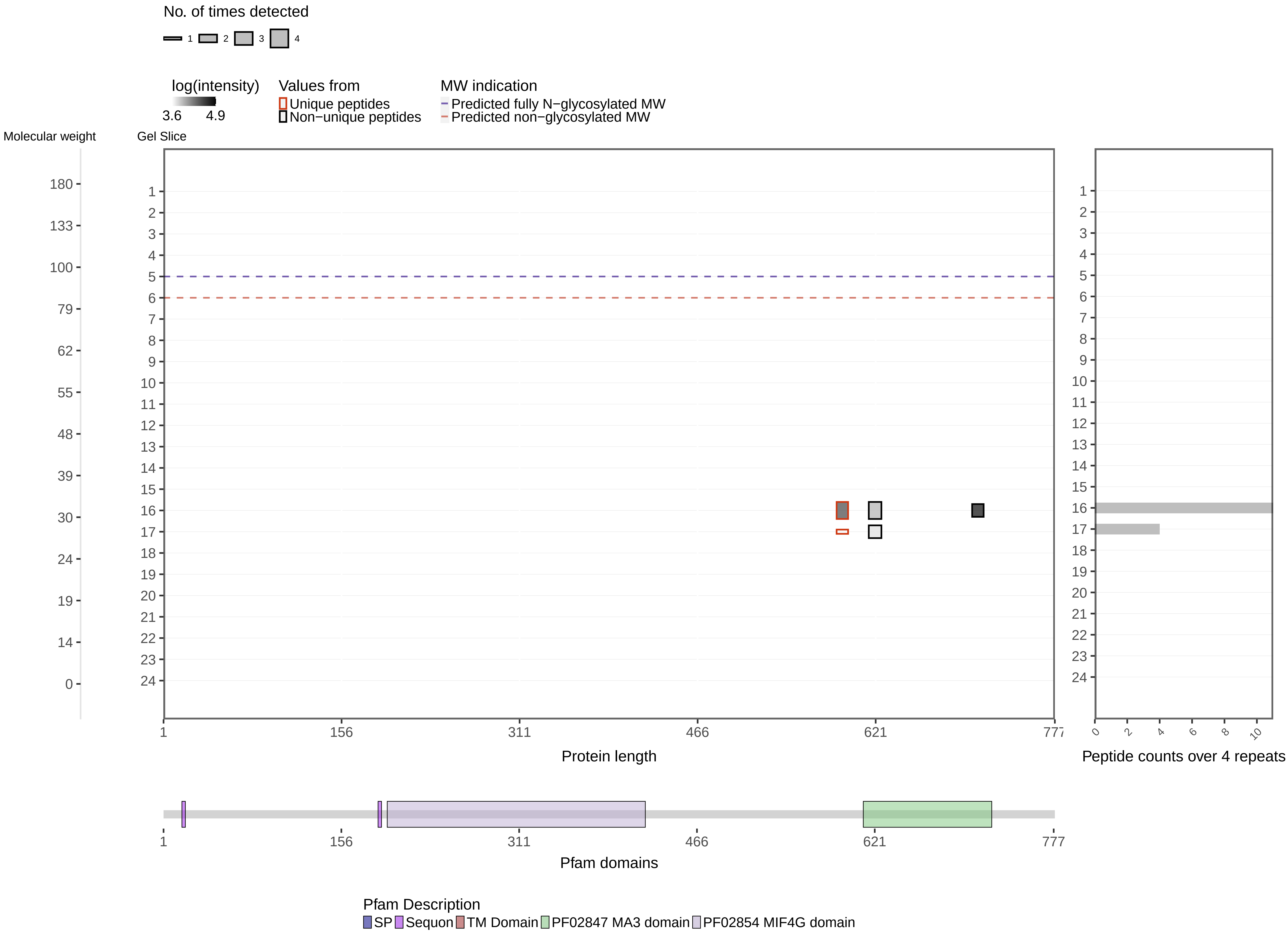


Protein sequence:

.DENQKSRRINDVYFNRLKELSTNPQLVPRLSFMVRDVLDLRSNNWVPRREEVKAKTITEIHTEAEKTLGLRPGATASMRNSRGPPSPQGSLSPGGYPMNRPGTGGMMPGMPGARKIPGMPGMDNDNWEVPRSRSMPRGNGPMVQPGGRGPPPLVGKSPSLNPRLLPQGSGGFVSGRPSALLQGSGAPPARPLGYGFGMEPASQVPGPVRSPTTVSPVAEKAPQAPAARSNPDELKRKTASLLEEYFSVRLLDEALQCVEELKSPAYHPEVVKEAISIGLDKSPPR

### NbL13g01090.1 translation_initiation_factor_IF−3_chloroplastic−like [MW = 30.09 kDa]

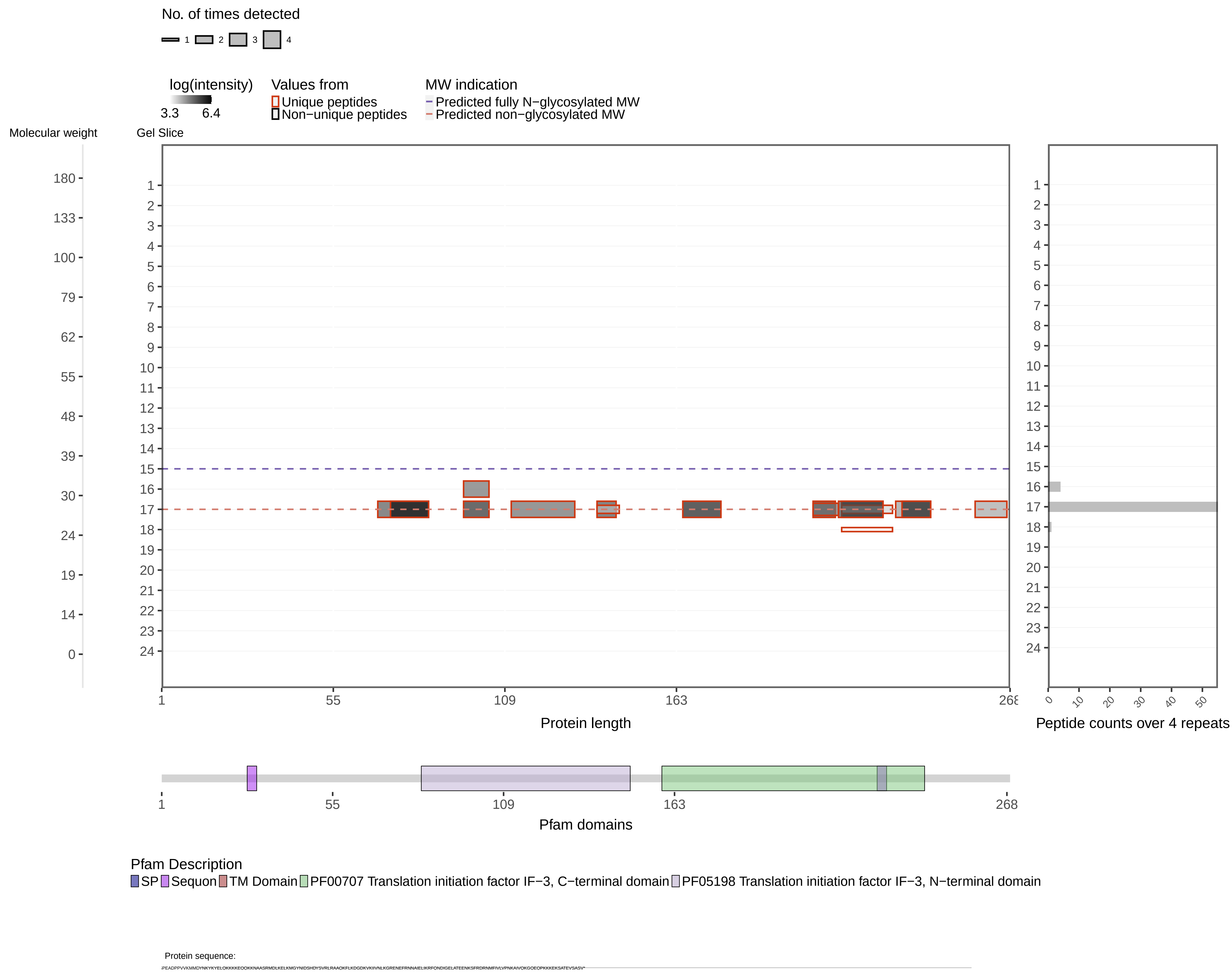

### NbL13g00950.1 serine_carboxypeptidase_II−2 [MW − SP = 50.12 kDa]

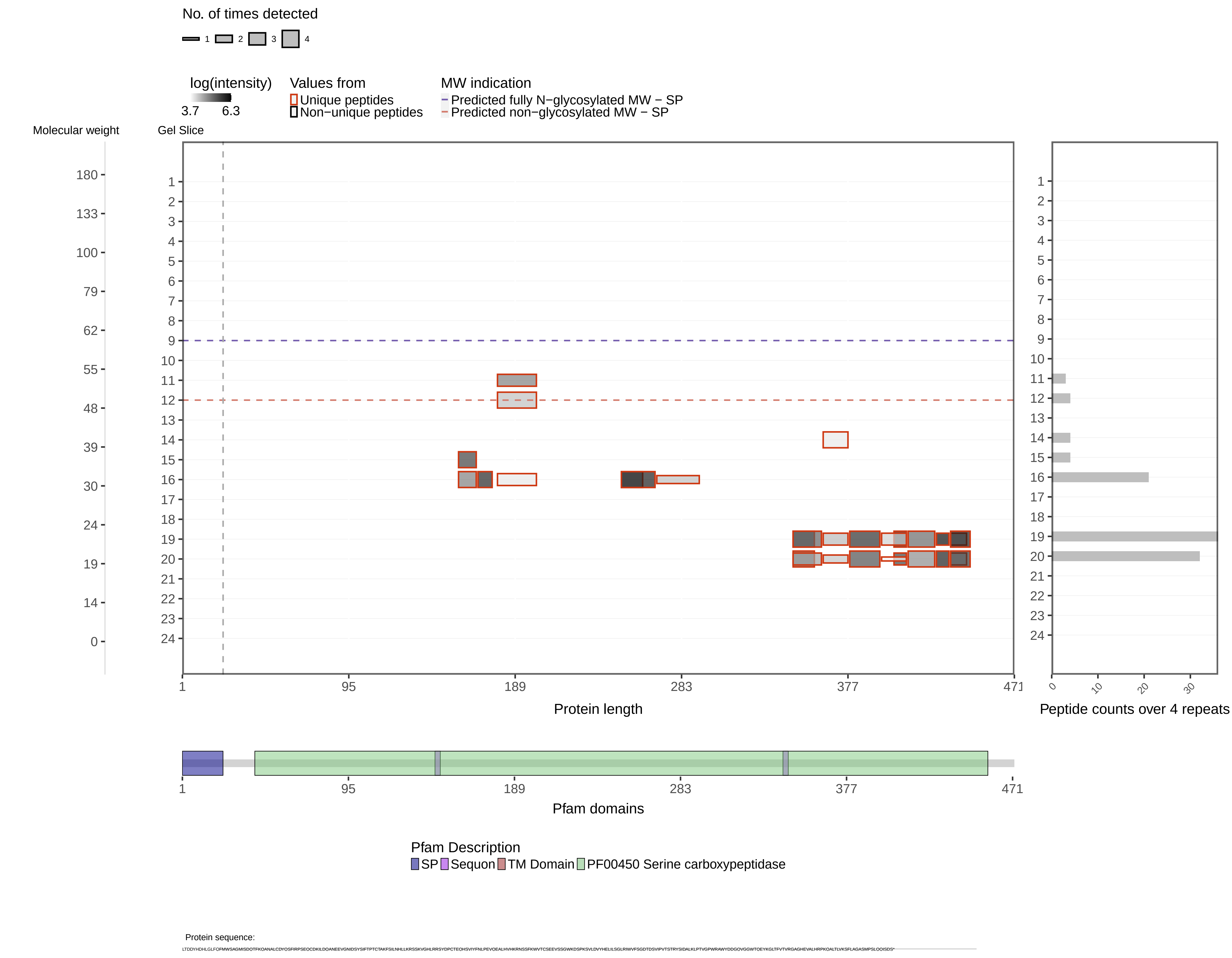

### NbL13g00890.1 50S_ribosomal_protein_L35_chloroplastic [MW = 17.15 kDa]

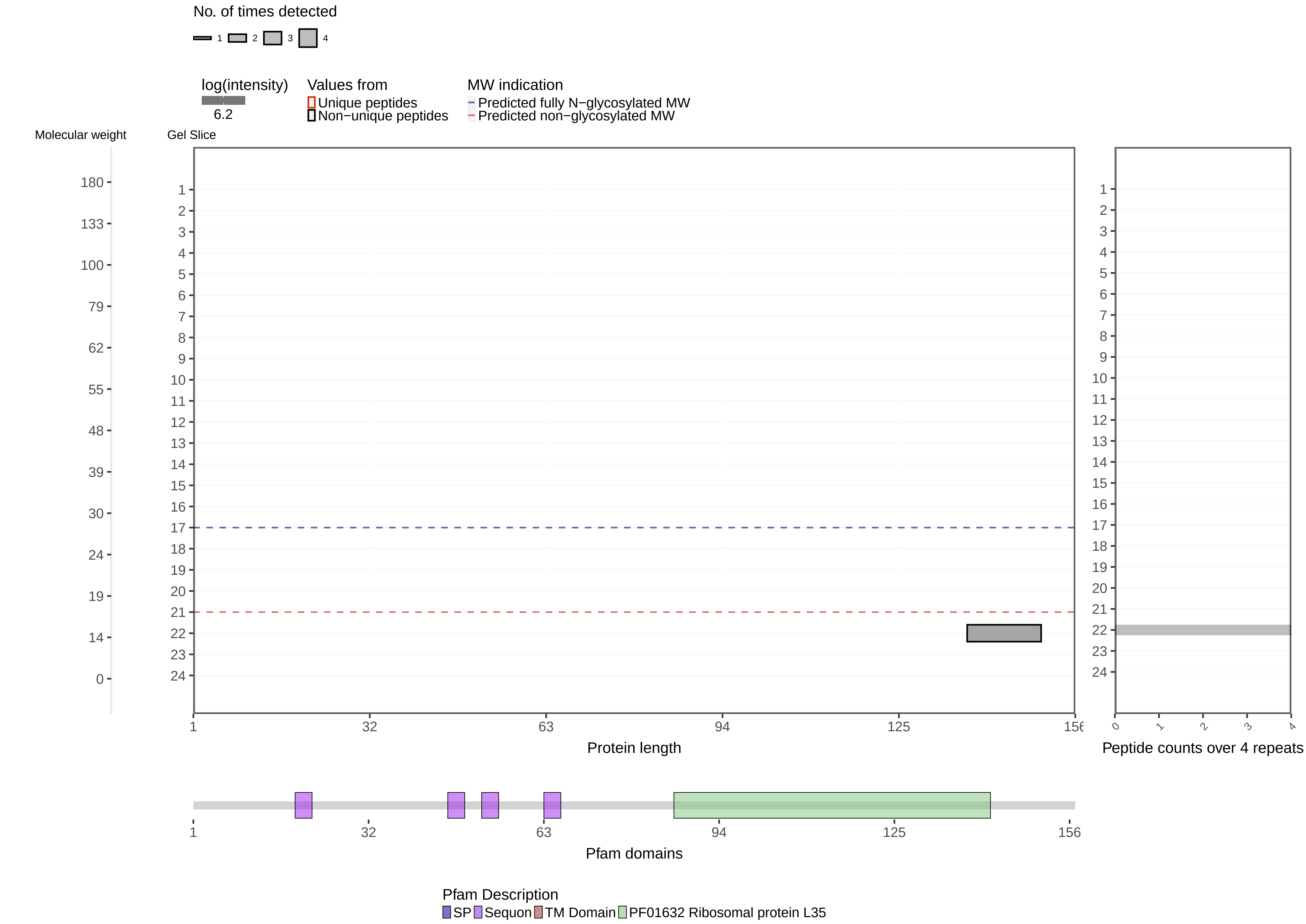


Protein sequence:

ATPTSGNPLPLTVVCAKGYKMKTHKASAKRFKVSGTGKILRRRAGKQHLLRKKNAKRRNRLSKTVQVDRSDYNNVIGALPYLKVNRAN*

### NbL13g00860.1 clustered_mitochondria_protein [MW = 202.18 kDa]

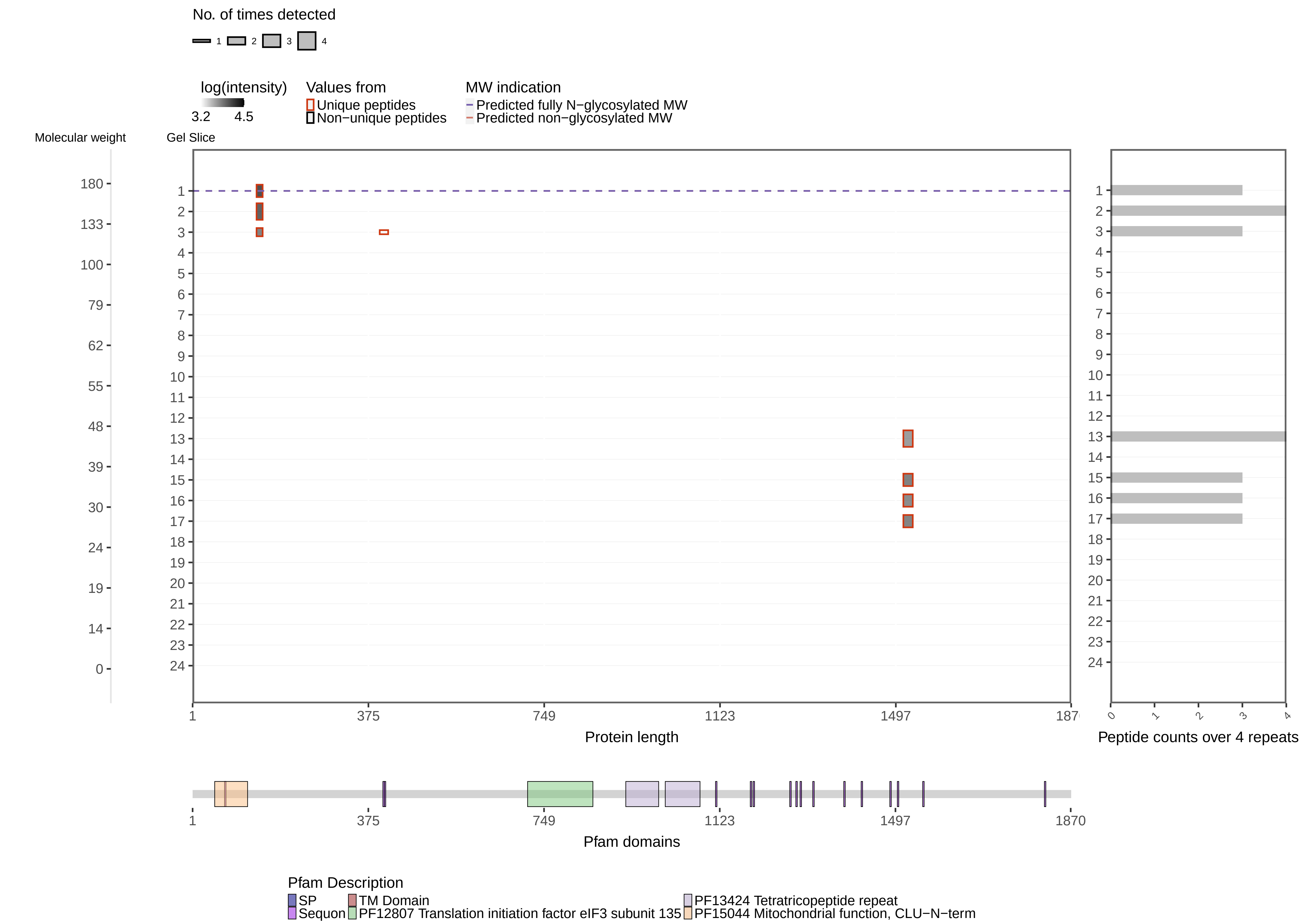


Protein sequence:

[illegible]

### NbL13g00810.1 putative_GDP−L−fucose_synthase_2 [MW = 37.46 kDa]

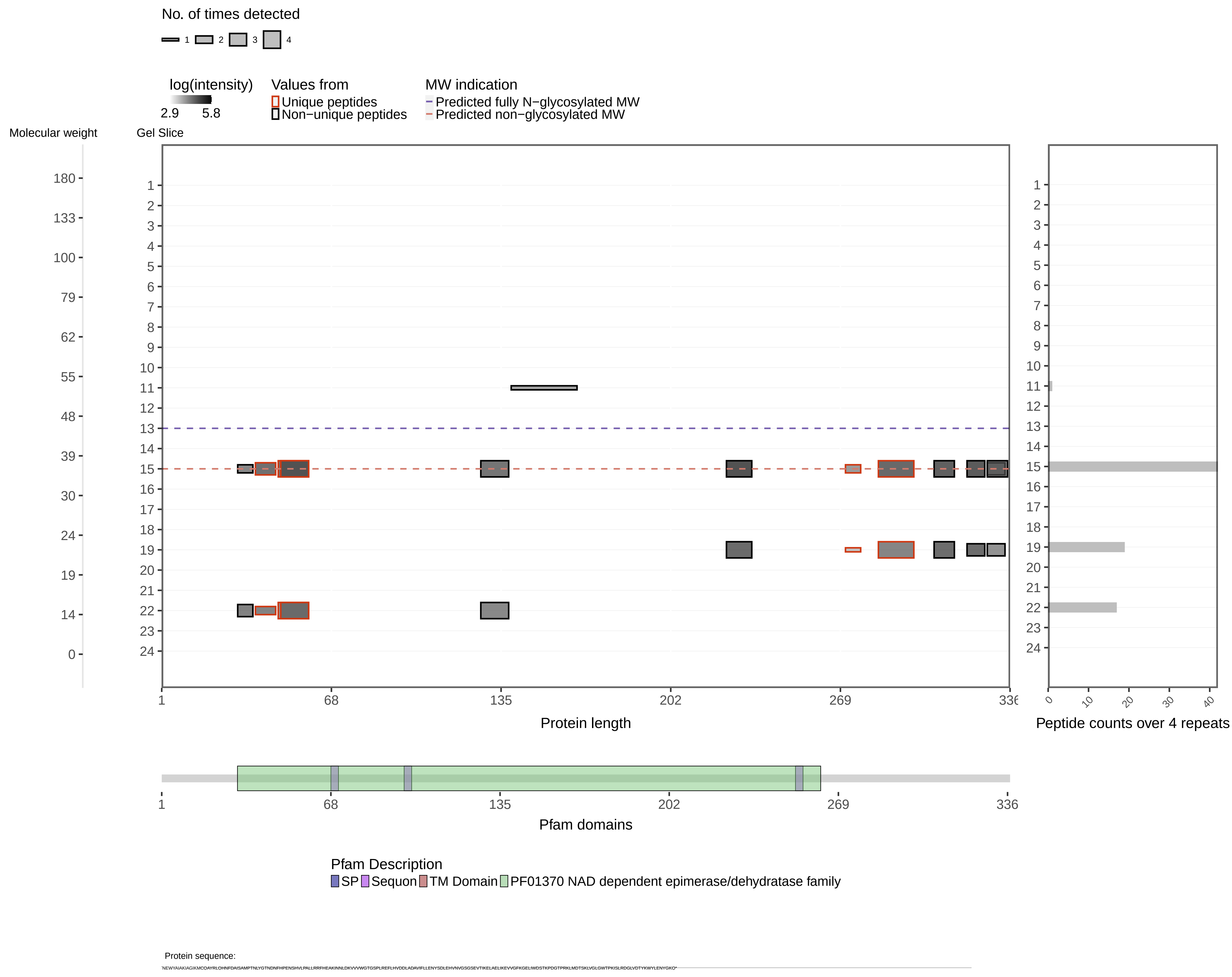

### NbL13g00690.1 tetratricopeptide_repeat_protein_1−like [MW = 30.94 kDa]

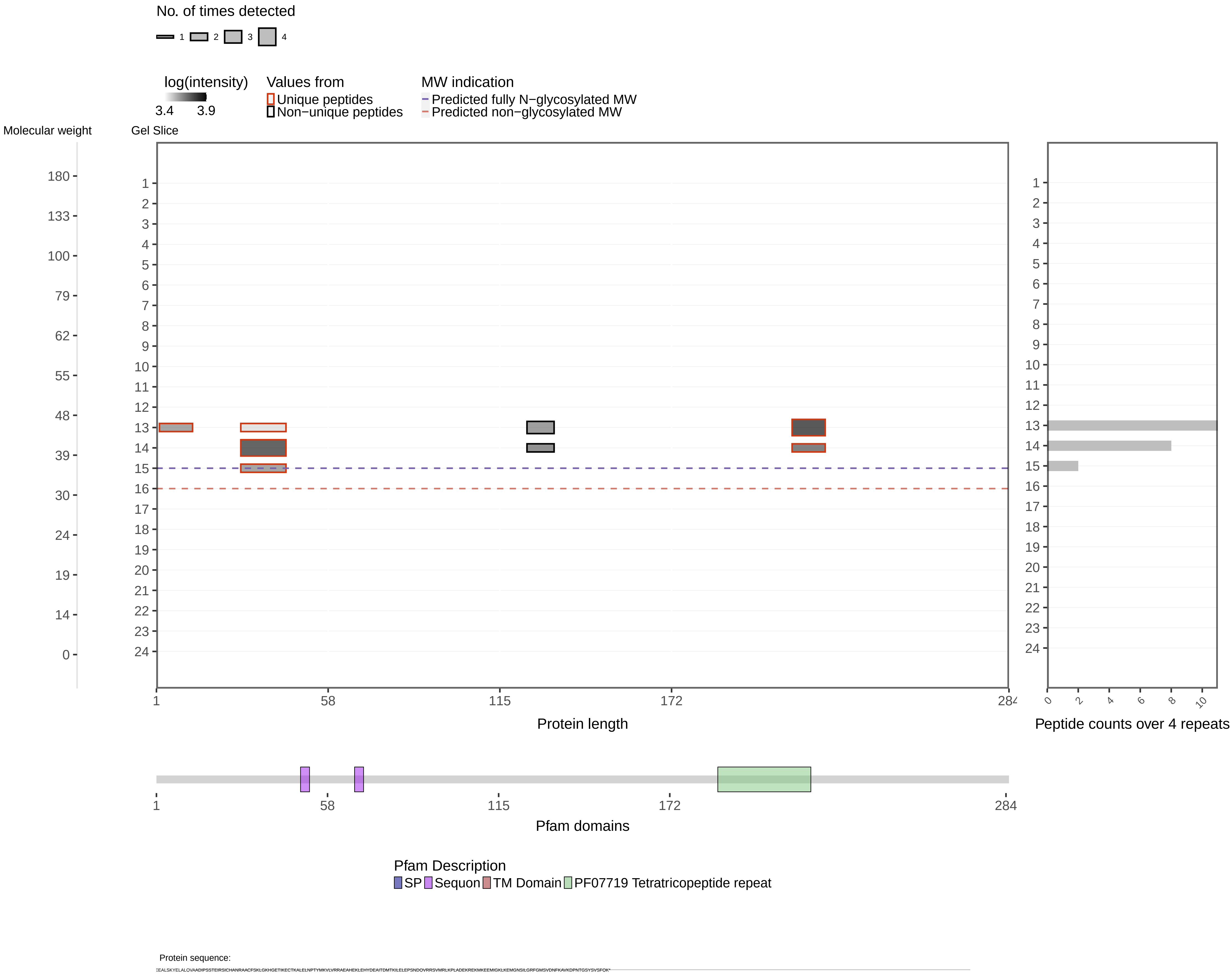

### NbL13g00670.1 HD_domain−containing_protein_2 [MW = 29.64 kDa]

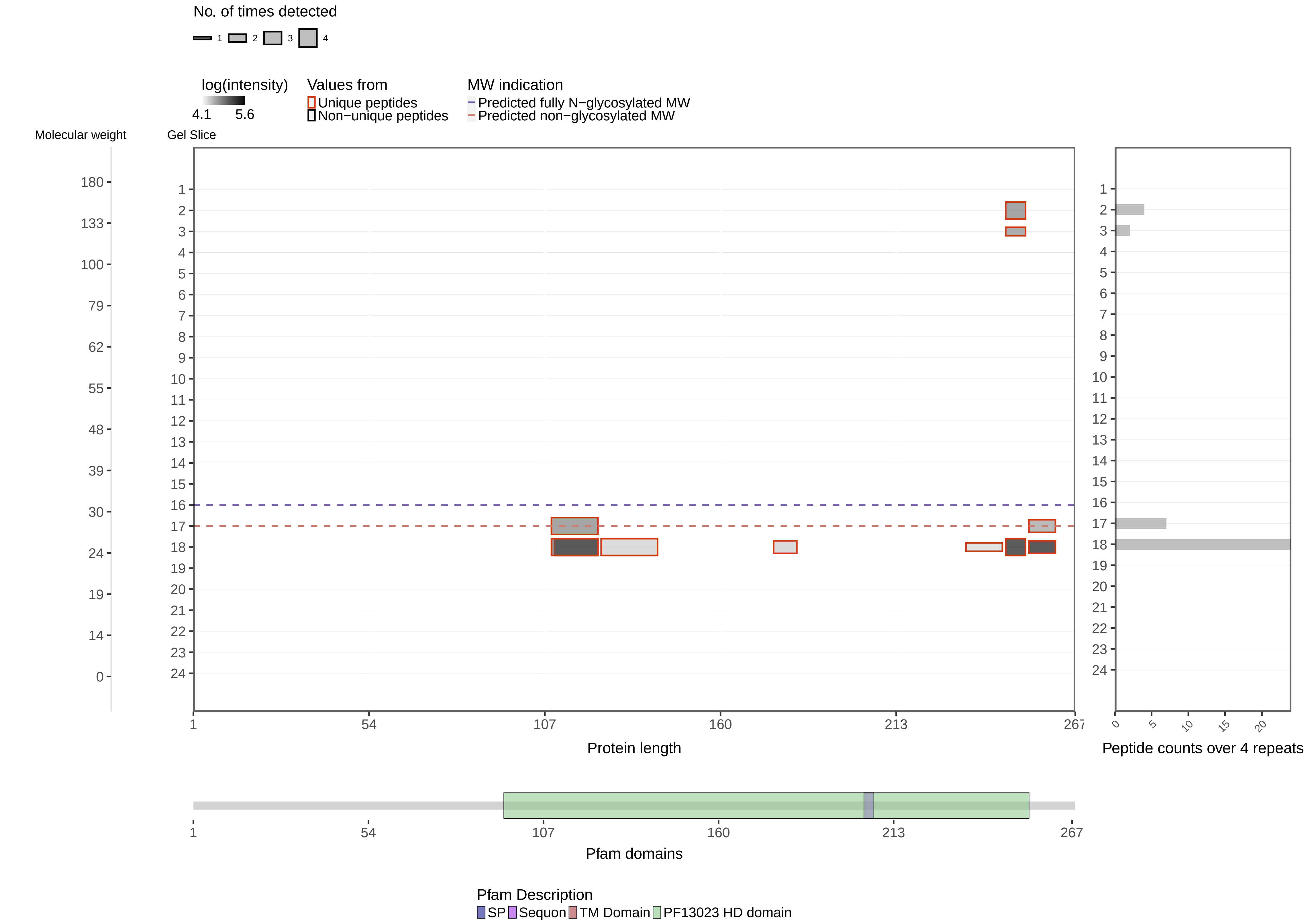


Protein sequence:

MYRMGVMALIAADLPGVDRDKCVKMAIVHDIAEAIVGDITPADGISKDEKSRRERAALEDMCKLLGGGPRAKEISDLWMEYEENSSLEAKVVKDFDKVEMILQALEYENEQGKDLEEFFQSTAGKFQTDVGKAWAAEVASRRKKSN*

### NbL13g00490.1 defensin−like_protein [MW = 18.28 kDa]

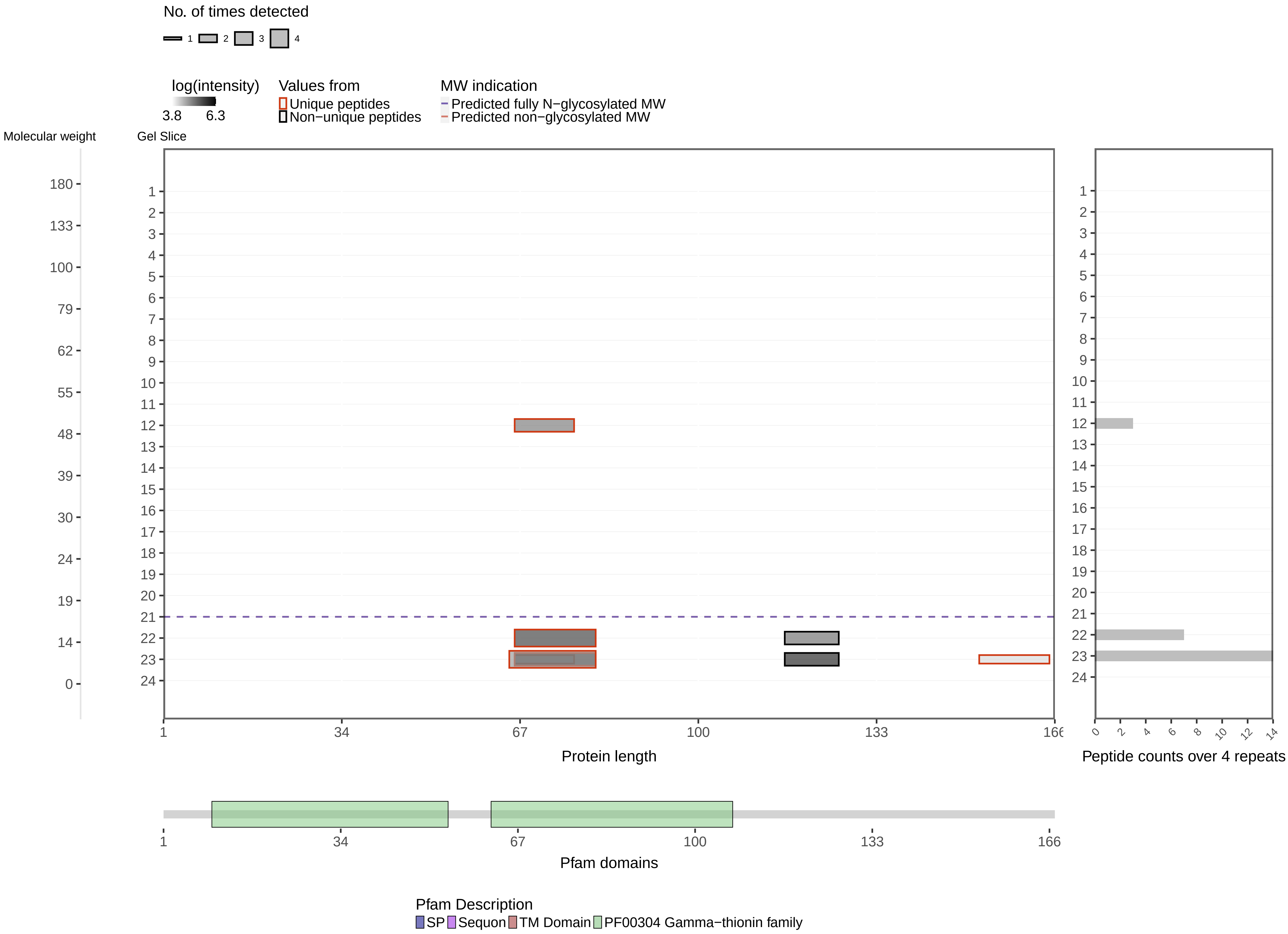


Protein sequence:

TFPGICLTKPPCRKACISEKFTDGHCSKILRRCLCTKPCVFDEKMIKTGAETLAEEAKTLAAALLEEEKMIKTGAETLAEEAKTLAAALLEEEIMDN*

### NbL13g00410.1 adenylosuccinate_lyase−like [MW = 61.29 kDa]

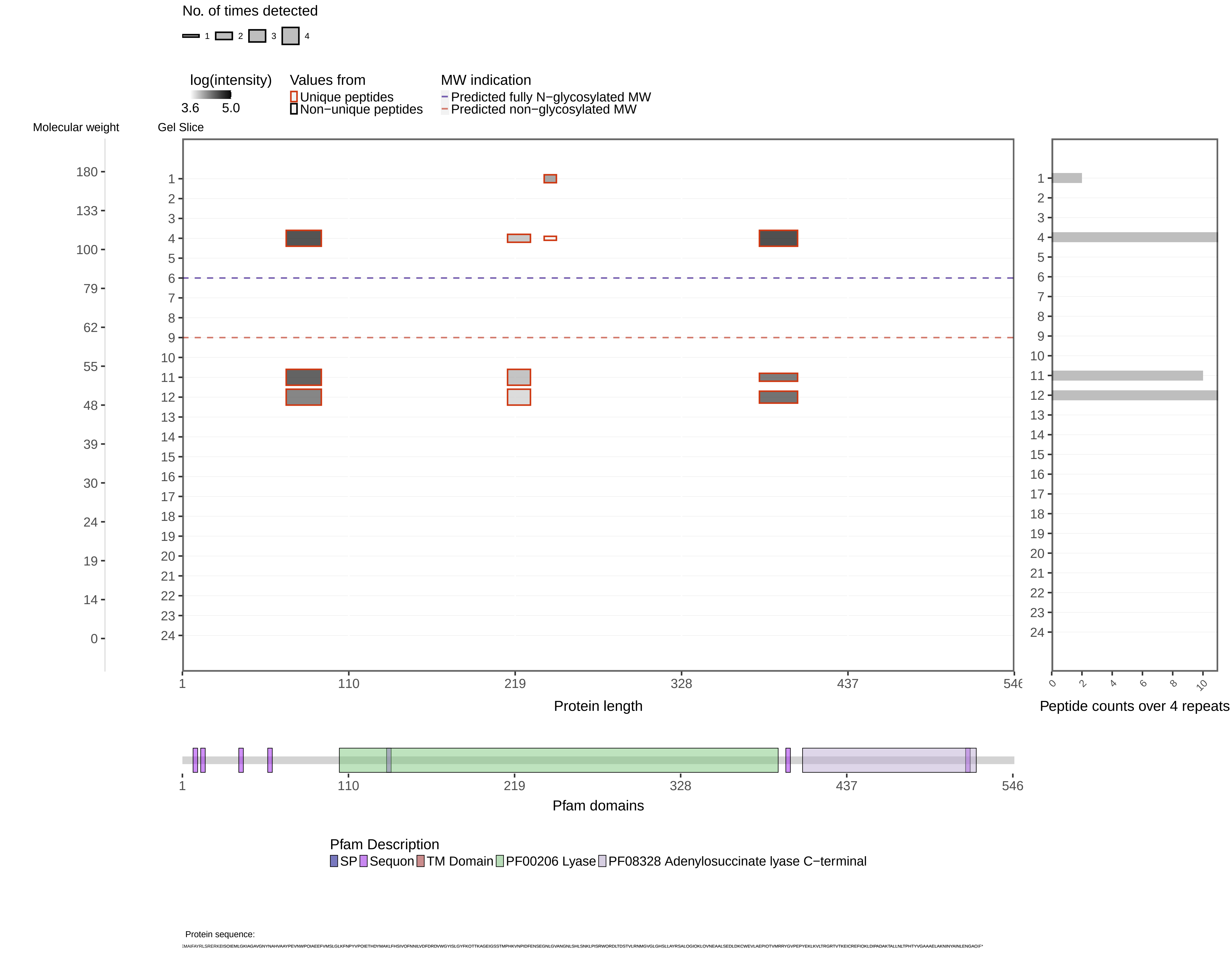

### NbL13g00400.1 50S_ribosomal_protein_L22_chloroplastic−like [MW = 30.11 kDa]

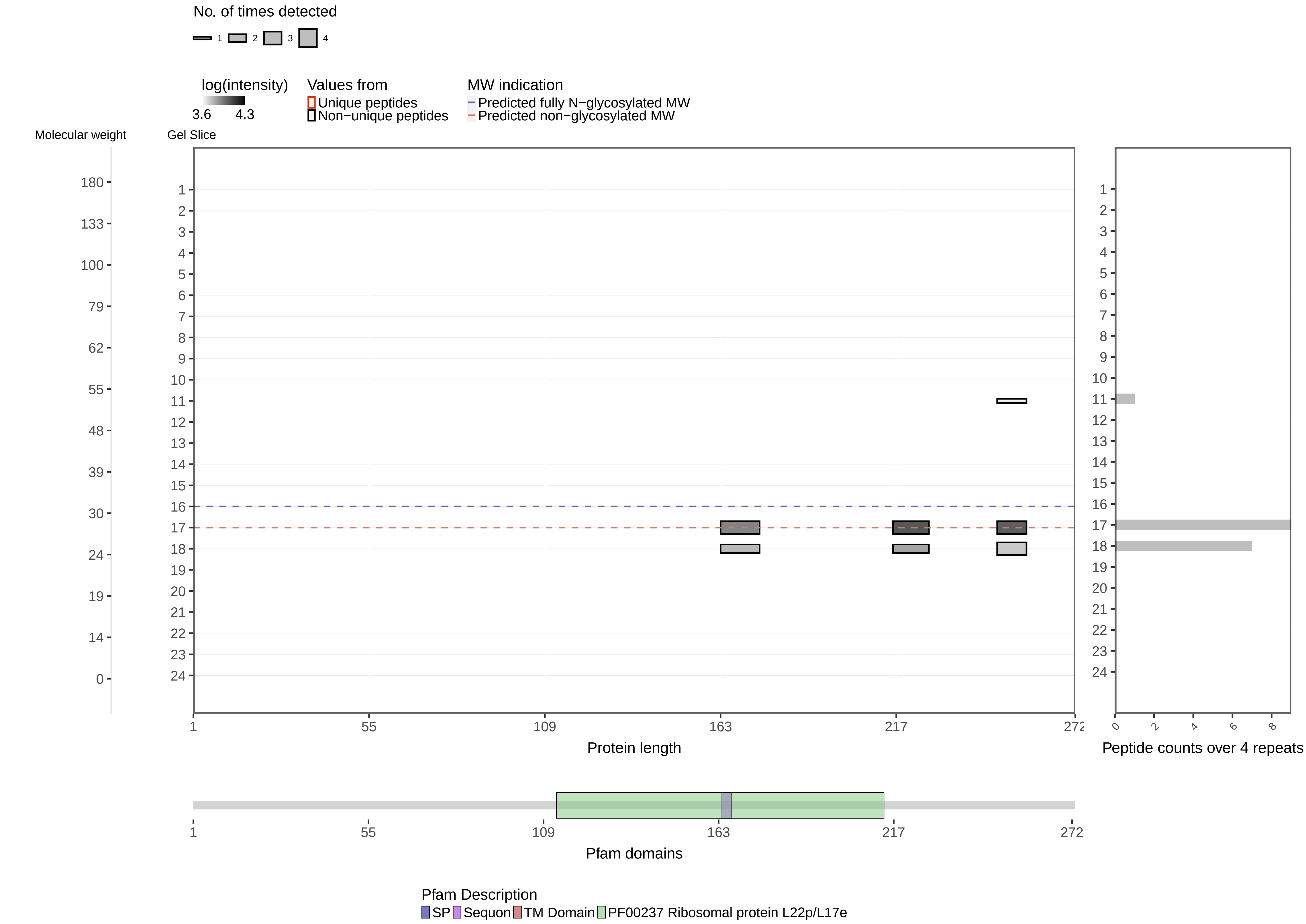


Protein sequence:

KKINLVAALVRGMRVEDALLQLQVNVKRASKTVYQVIHSARANATHNYGFDPDRLLVAEAFVGKGLFKKRVSYHAKGKCGIKVRPECRLTIVVREITPEEEAEIAKLRVHNFKKLTKRERRLVPHKLIETTPIWDRKGRAKSNGPGAAAA*

### NbL13g00280.1 xylose_isomerase−like [MW − SP = 50.72 kDa]

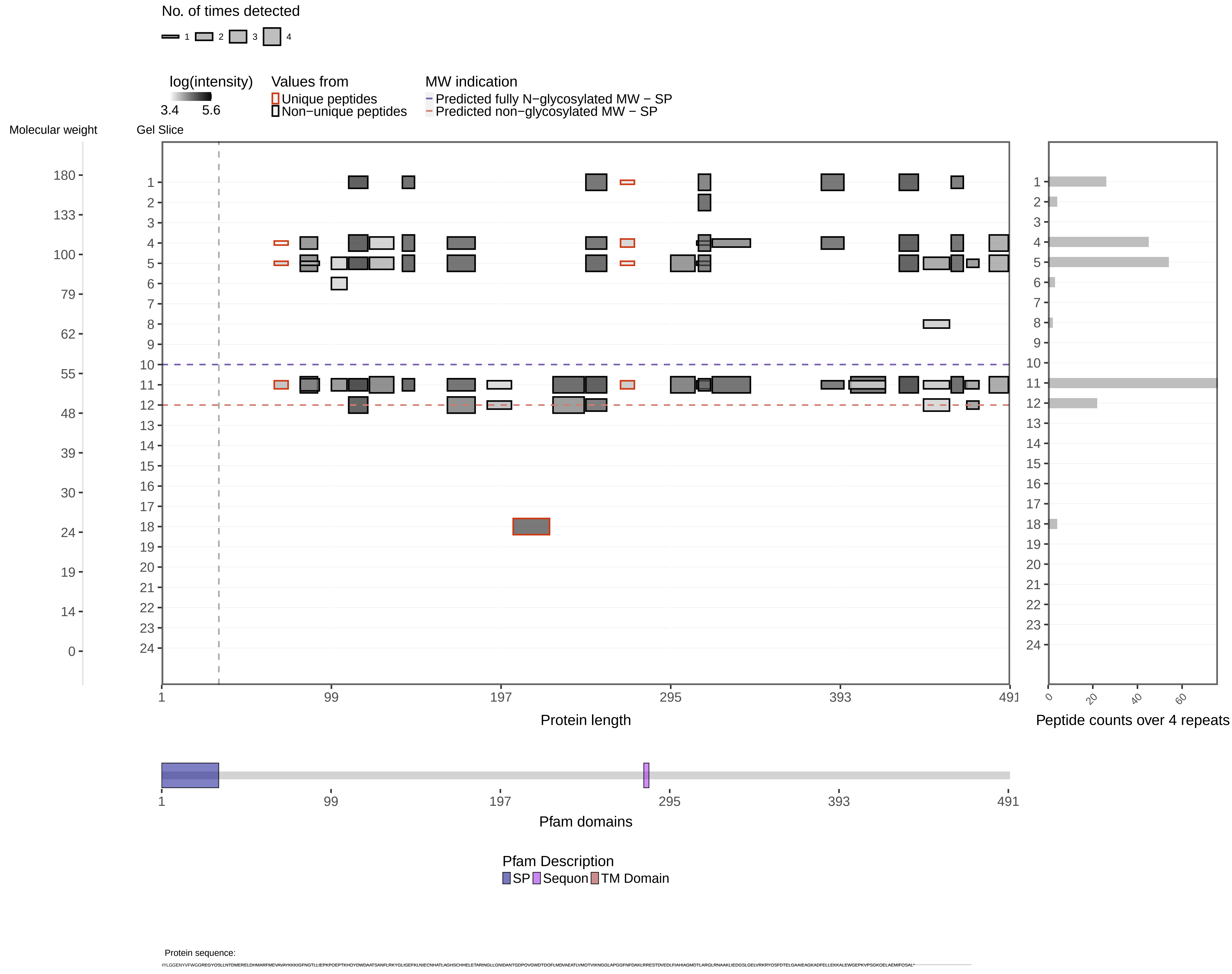

NbL13g00260.1 salutaridinol_7−O−acetyltransferase−like [MW = 49.37 kDa]

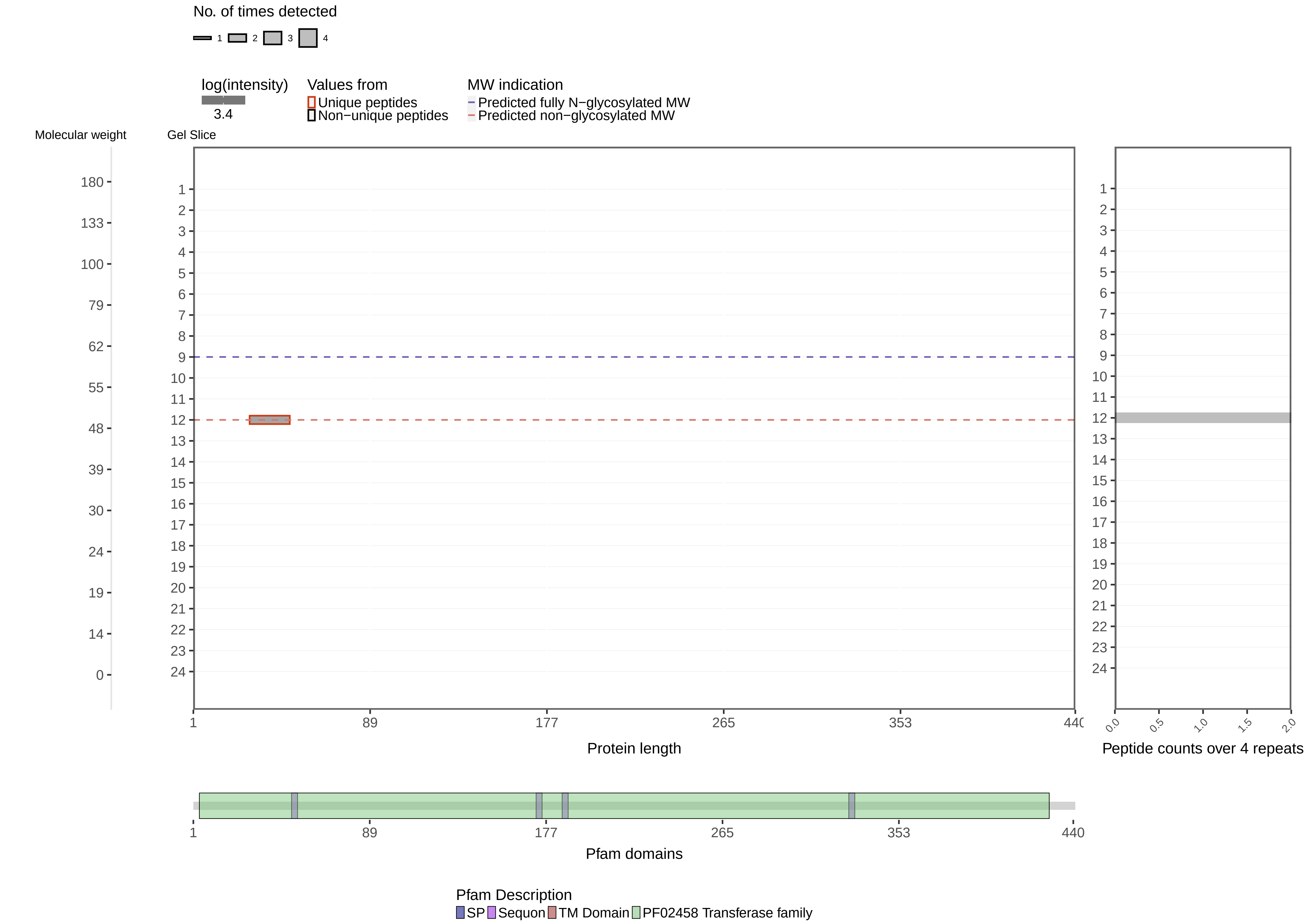


Protein sequence:

KCAAVEPVVEKLVVKRFIFTASNIVKMKSKVINSGYNNKERVTRVEVITALLWKCFMAAKGCNSIAVLPVNVRKKIVPPLPENSFGNFFLVSSAIGNVENEWFSLVGKIKNLIERIDGNYTEKIRGQNGFEFVMNNFKQVGKLLSQGDDIKVLRIGSWCNFPINGVNFGWGESILPIVAILGVKDHIVLLDSLKYSGGIEAWVVMTDQEMALFEQDKELQDFTSLDVSLL*

NbL13g00060.1 probable_xyloglucan_endotransglucosylase/hydrolase_protein_23 [MW − SP = 28.39 kDa]

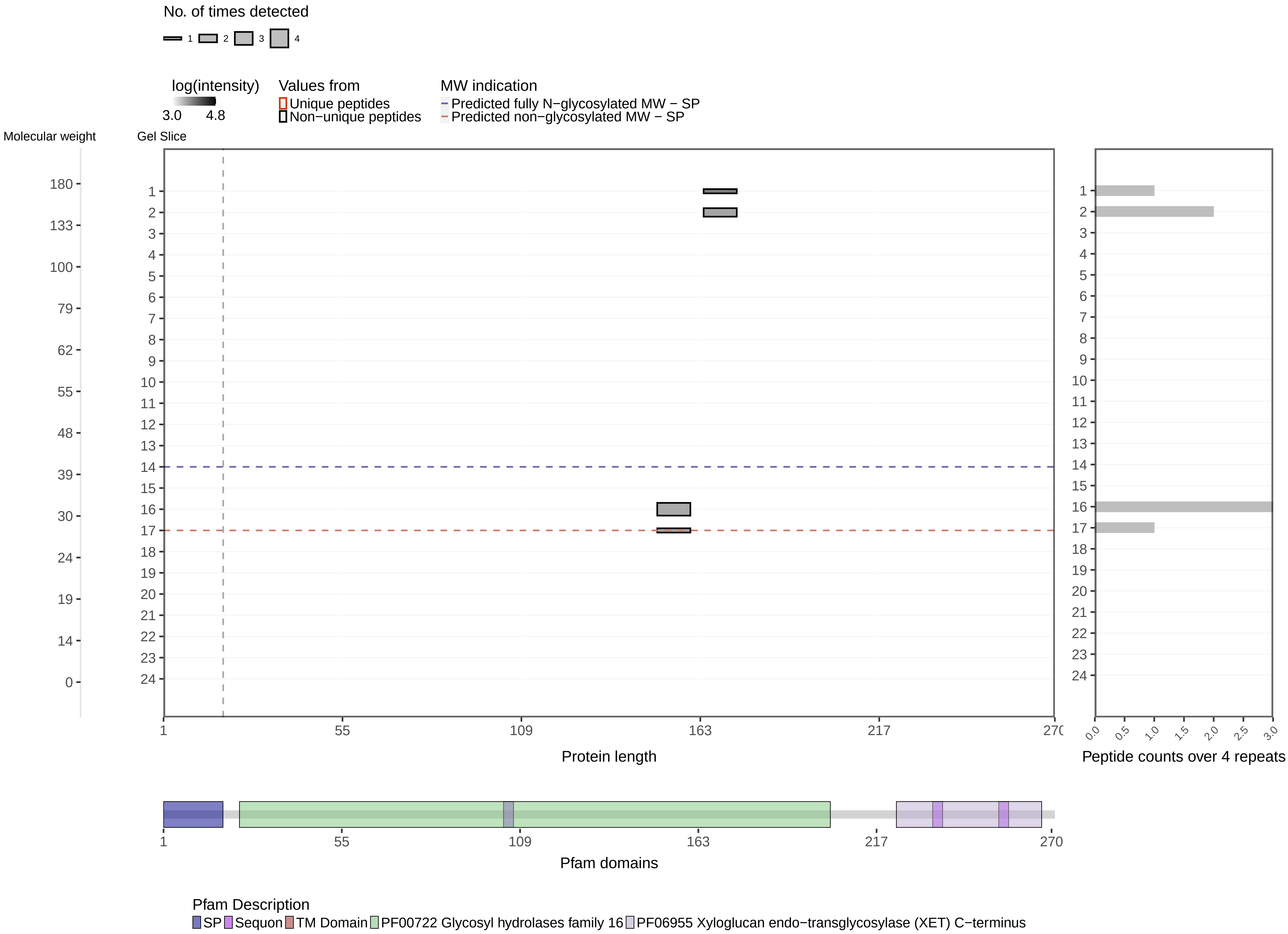


Protein sequence:

### NbL13g00040.1 calmodulin−binding_protein_60_B [MW = 71.67 kDa]

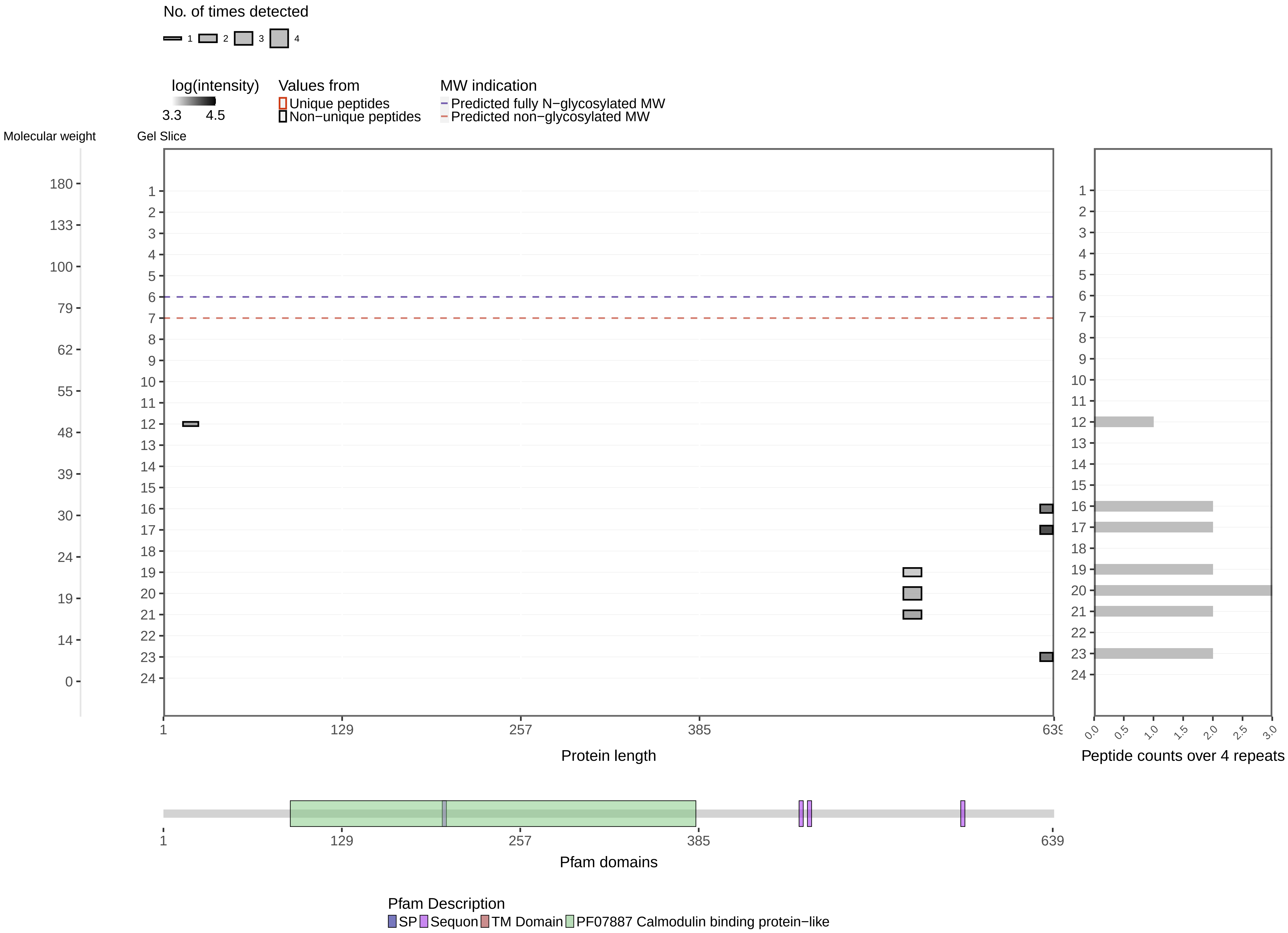


Protein sequence:

LIEHAKTCVLSGKLYVYYSDDSRNVGVVFNNIYELNGLIAGEQYYSADSLSDSQKVYVDSLVKKAYDNWNQVVEYDGKSFLNIKQNQKAGSLRNELPVGPVDYPNNMVNQLPQSRLPVSVQSEQSSMDPNFLIGGAGYNDGIVARMPNQSQMMNSSSRSQFESTPFAPQQQITNTHQLQSTSYDNNVGLALGPPQSSSYQTMTSSLPQTNLHPFEDWSHNRDKGVDEFLSEEEIRMRSHEILENDDMQHLLRLFSMGGGHGSVNVSEDGYGFPSFMPSPSPTFGYI

NbL13g00020.1 dihydrolipoyllysine−residue_acetyltransferase_component_2_of_pyruvate_dehydrogenase_complex_mitochondrial−like [MW = 58.6 kDa]

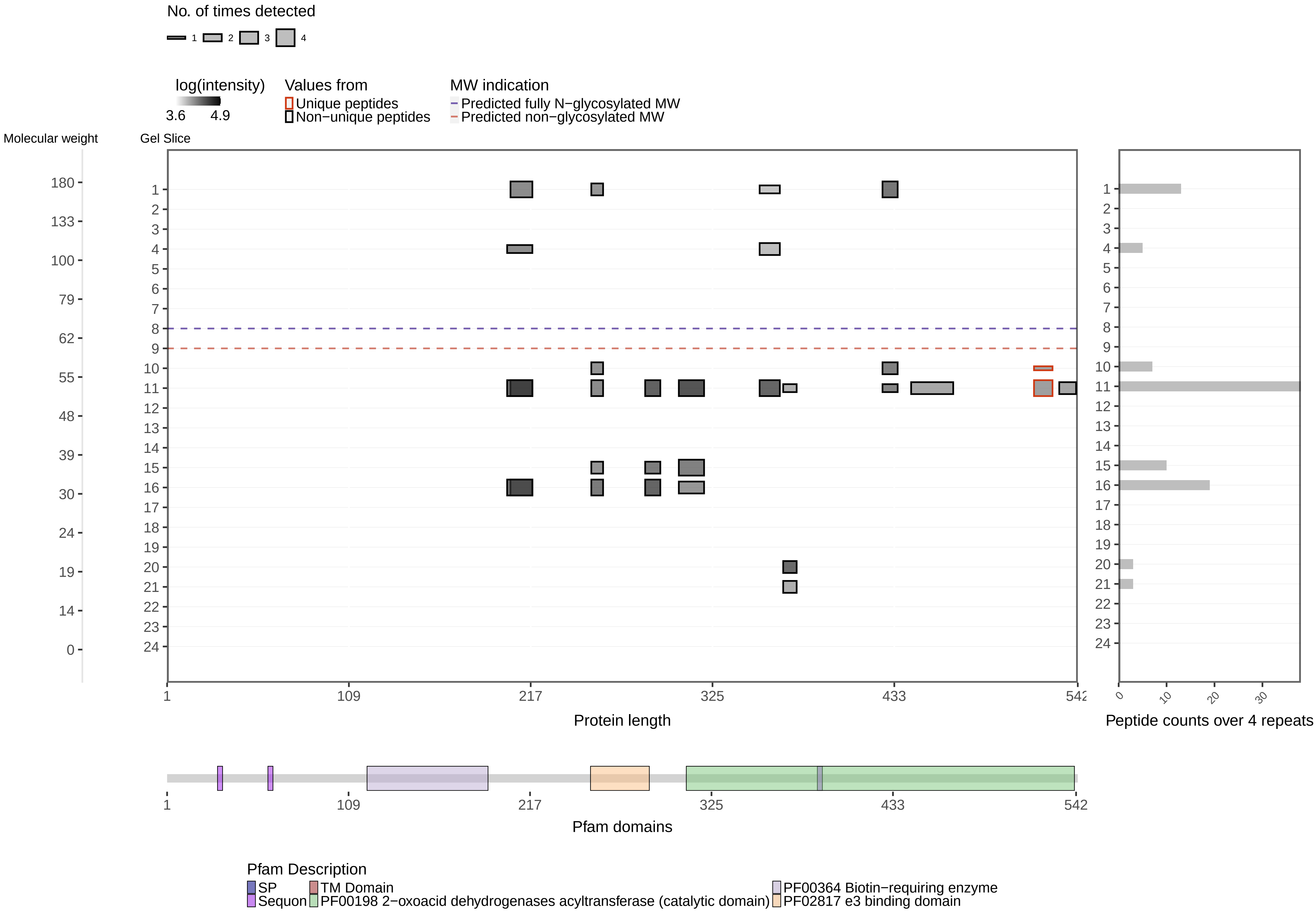


Protein sequence:

SPLARKIAEDHNIPLSSIKGTGPEGRIVKADIEDYLASRGKEALAPAPKAEASLDYTDIPVSOIRKVTASRLLLSKOTIPHYYLTVDACVDKLMELRSOLNSLOEASGGKKLSVNDLVIKAAALALRKVPOCNSSWTNDYIROYHNININVAVOTDNGLYVPVVRDADKKGLSTIAEEVKNLAOKAKENSLKPODYEGGTFTVSNLGGPFGIKOFCAIINPPOSAILAVGSAERRVIPGSGODOYTFASMMSVTLSCDHRVIDGAIGAOWLKAFKGYIESPESMLL*

NbL12g22790.1 14−3−3−like_protein_F [MW − SP = 28.98 kDa]

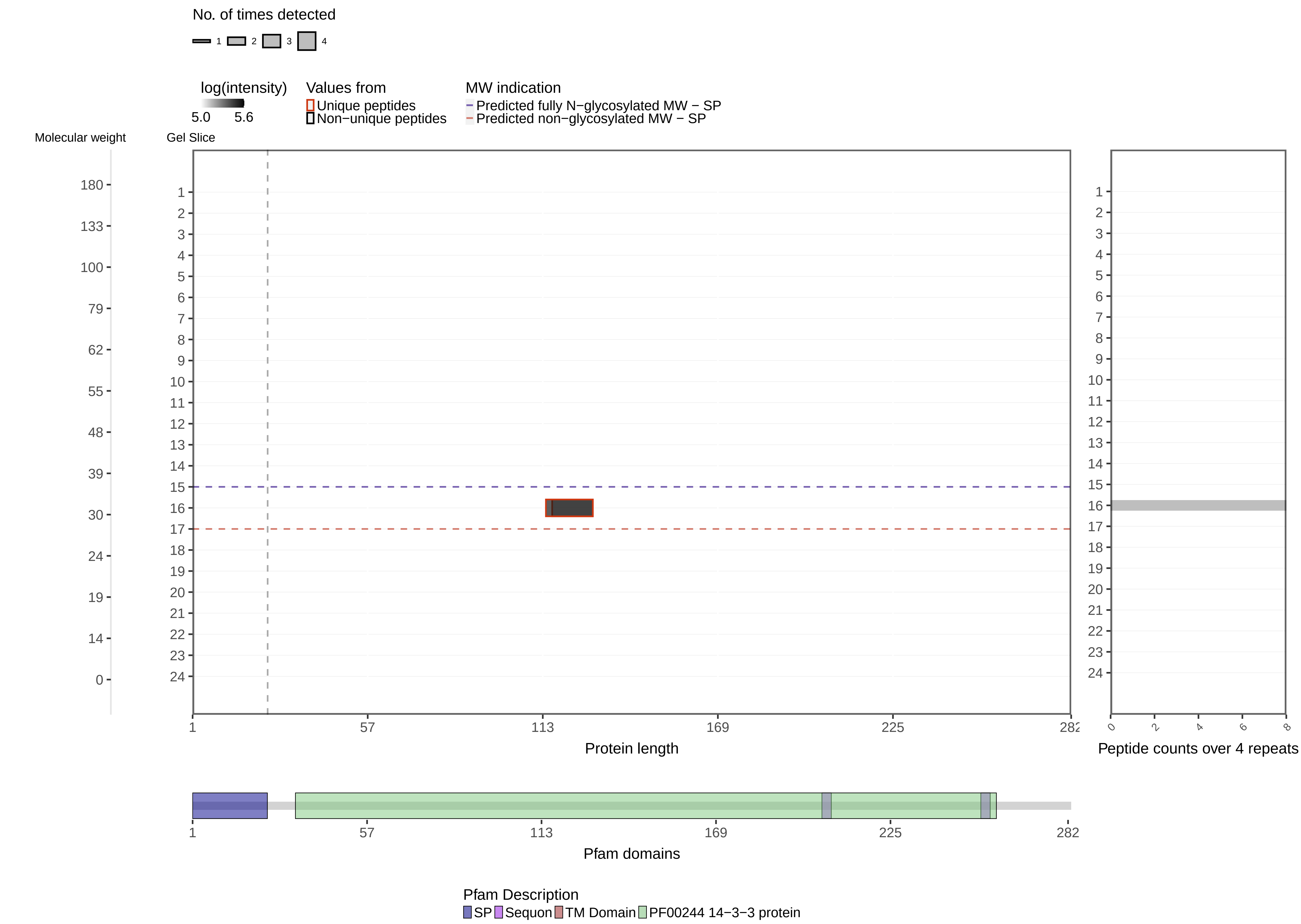


Protein sequence:

KLLDSNLIGAASIGDSKVFYLKMKGDYHRYLAEFKTGAERKEAAENTLSSYKSAQDIANAELAPTHPIRLGLVLNFSVFYYEILNSPDRACNLAKQAFDEAIAELDTLGEESYKDSTLIMQLLRDNLTLWTSDMQDEGTEDMKEVAKPENEEH*

### NbL12g22710.1 polyadenylate−binding_protein_2−like [MW = 24.84 kDa]

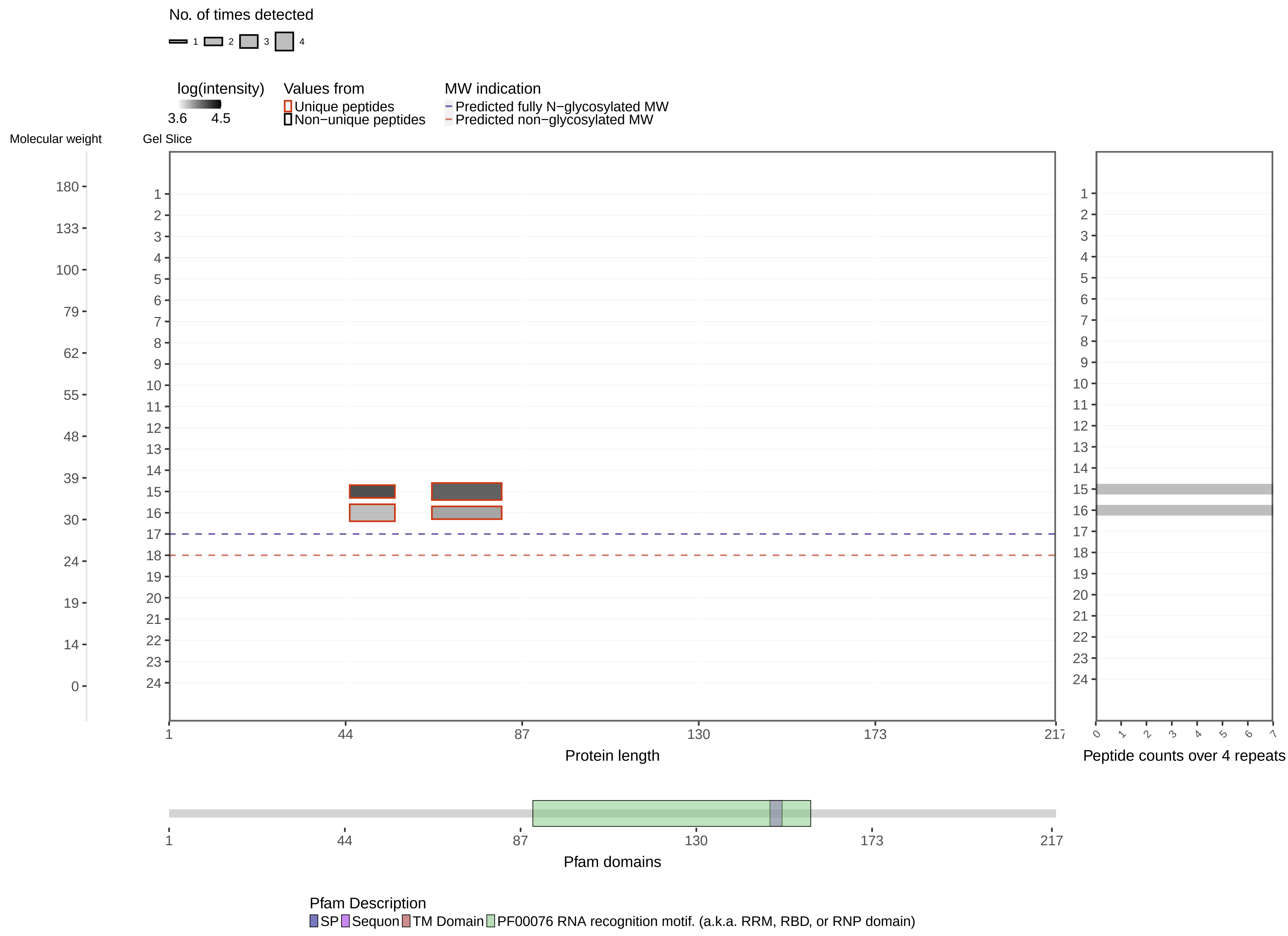


Protein sequence:

/DYACTPEEVQQHFQACGTVNRVTILTDKFGQPKGFAYVEFVEQEAIQEALQLNESELHGRQLKVMPKRTNVPGMKQFRGRRFNPYMAYGFRRPYTPPYLYSPYGFGKVPRFRRAMRFMPYY*

### NbL12g22460.1 gamma−glutamyltranspeptidase_1−like [MW − SP = 59.45 kDa]

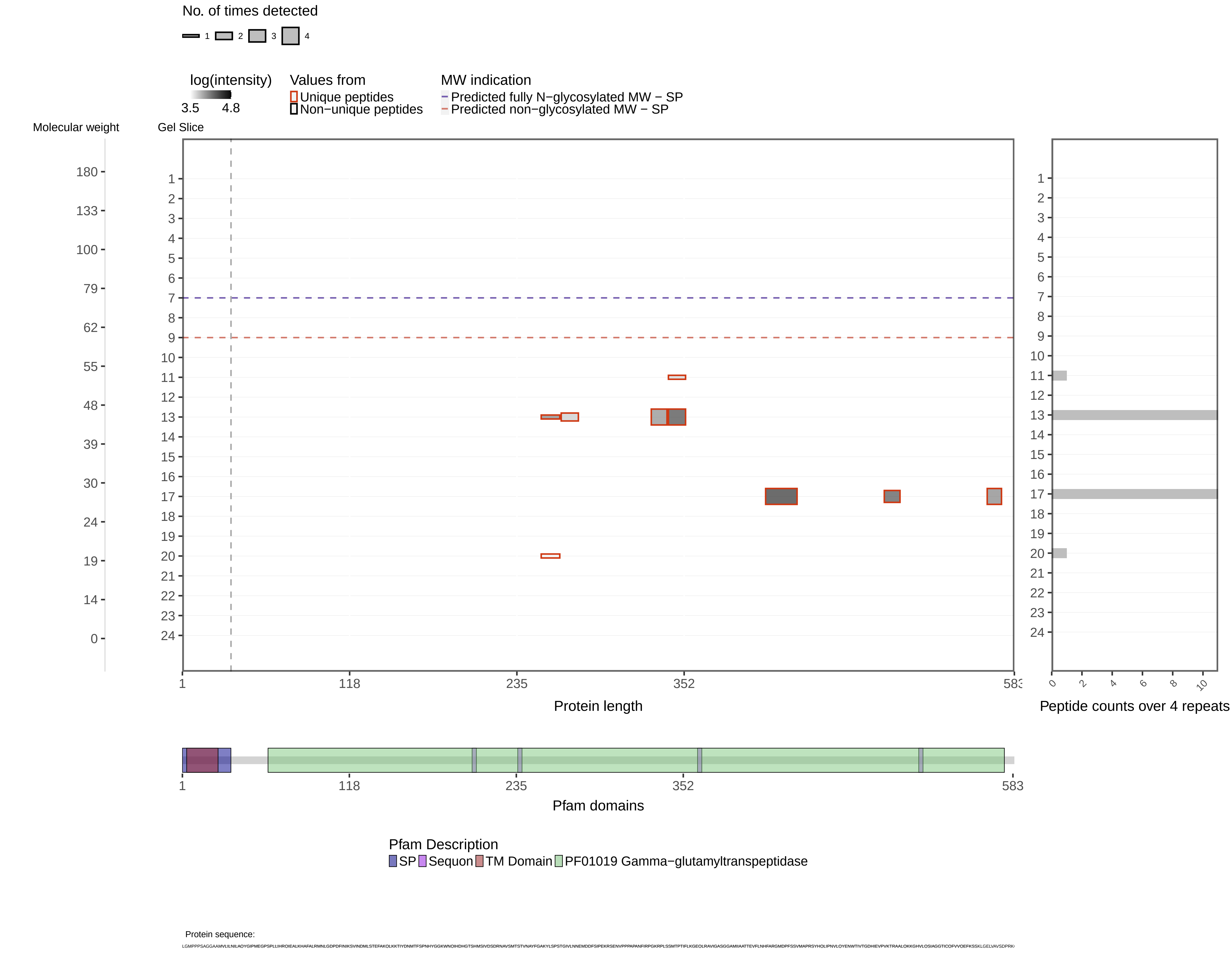

### NbL12g22410.1 RNA−binding_protein_39 [MW = 66.46 kDa]

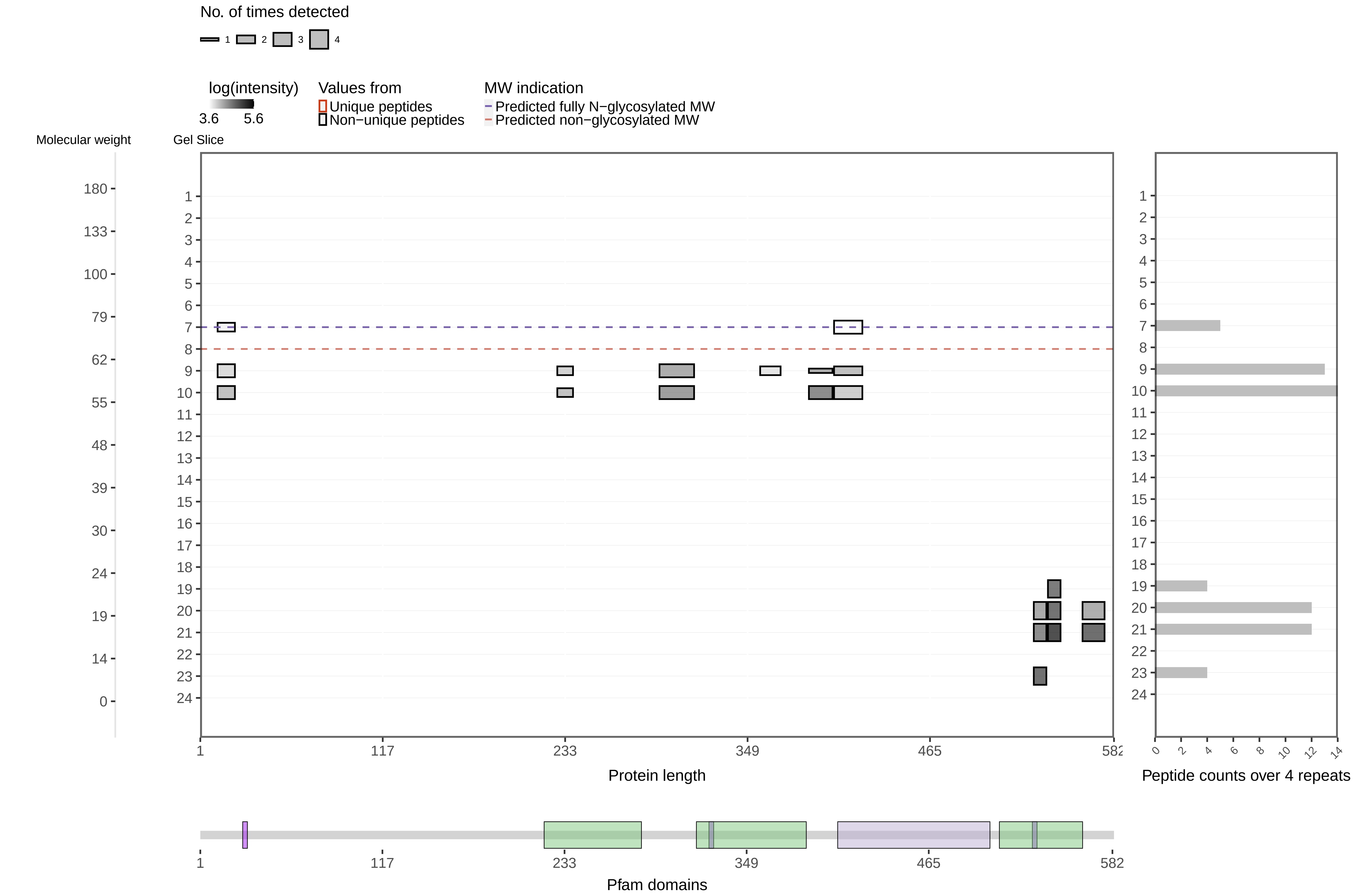


Pfam Description

SP Sequon TM Domain PF00076 RNA recognition motif. (a.k.a. RRM, RBD, or RNP domain) PF15519 linker between RRM2 and RRM3 domains in RBM39 protein

Protein sequence:

NbL12g22060.1 serine/threonine−protein_phosphatase_4_regulatory_subunit_2−like [MW = 25.37 kDa]

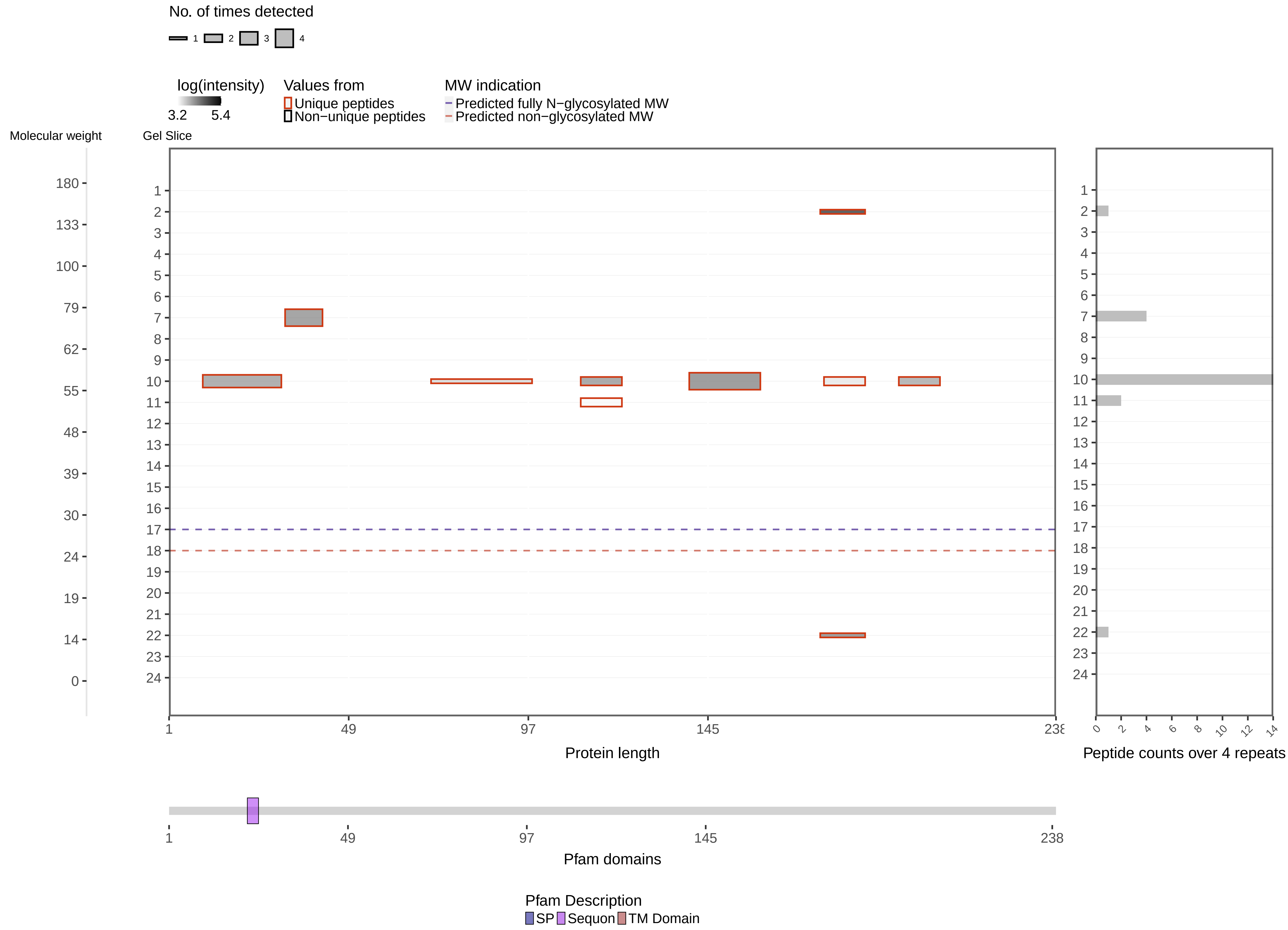


Protein sequence:

VRTETEKALQVVSTIGAPKAETSEEKKIEEVKSETKTPAENKVEEVTVAVETPAEKNIEEVTVAVETPAEKKMEEAPAATEAQKSVVPVEIKAEEKPVATEAPKSEAPVVTKAEKKPAAEAEKTVIDEKKSDKS*

### NbL12g22000.1 60S_ribosomal_protein_L13a−4−like [MW = 23.53 kDa]

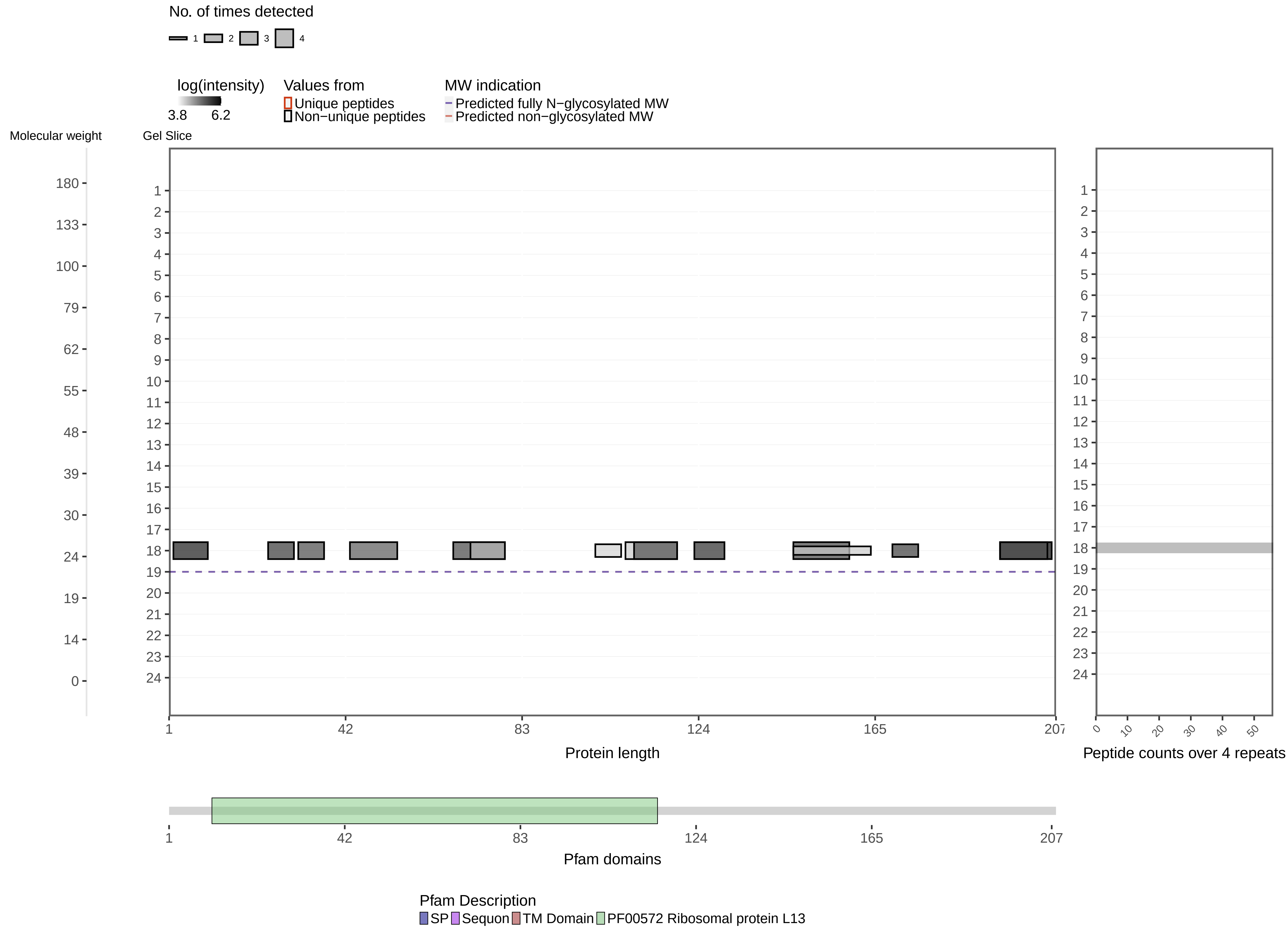


Protein sequence:

RGMIPHKTKRGAAALARLKVYEGVPPPYDKIKRMVIPDALKVLRLQAGHKYCLLGKLSSEVGWNHYDTIKELENKRKERAQVAYERRKQLAKLRVKAEKAAEEKLGPQLAVIEPIKY*

### NbL12g21990.1 U6_snRNA−associated_Sm−like_protein_LSm3 [MW = 13.91 kDa]

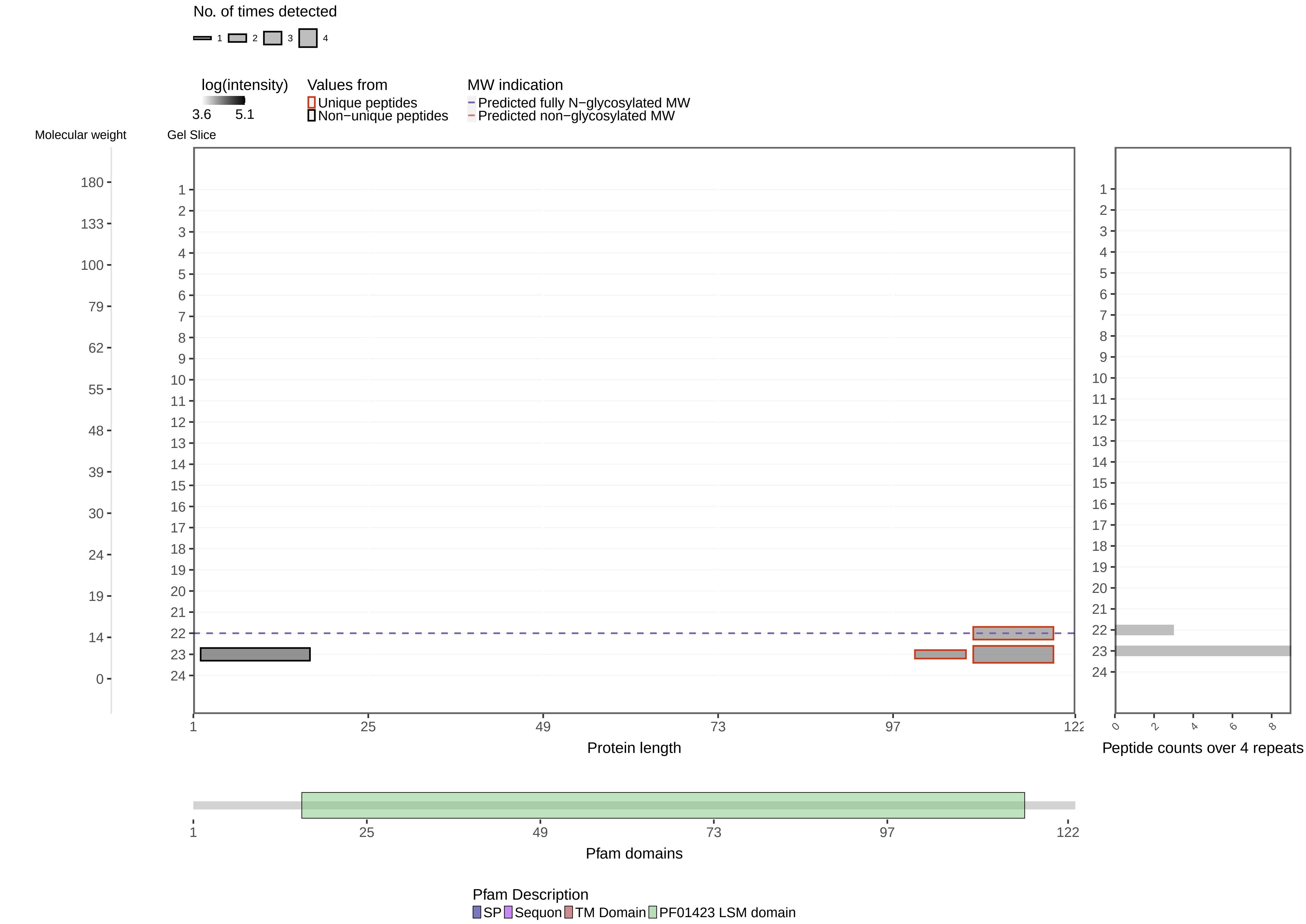


Protein sequence:

MILGDVEEIVTTVEIDDETYEEIVRTIYSCSGMCLFWIAFCKSCSLSLKTTRRTVHFLFVRGDGVILVSPPLRTV*

NbL12g21970.1 probable_plastid−lipid−associated_protein_7_chloroplastic [MW = 30.03 kDa]

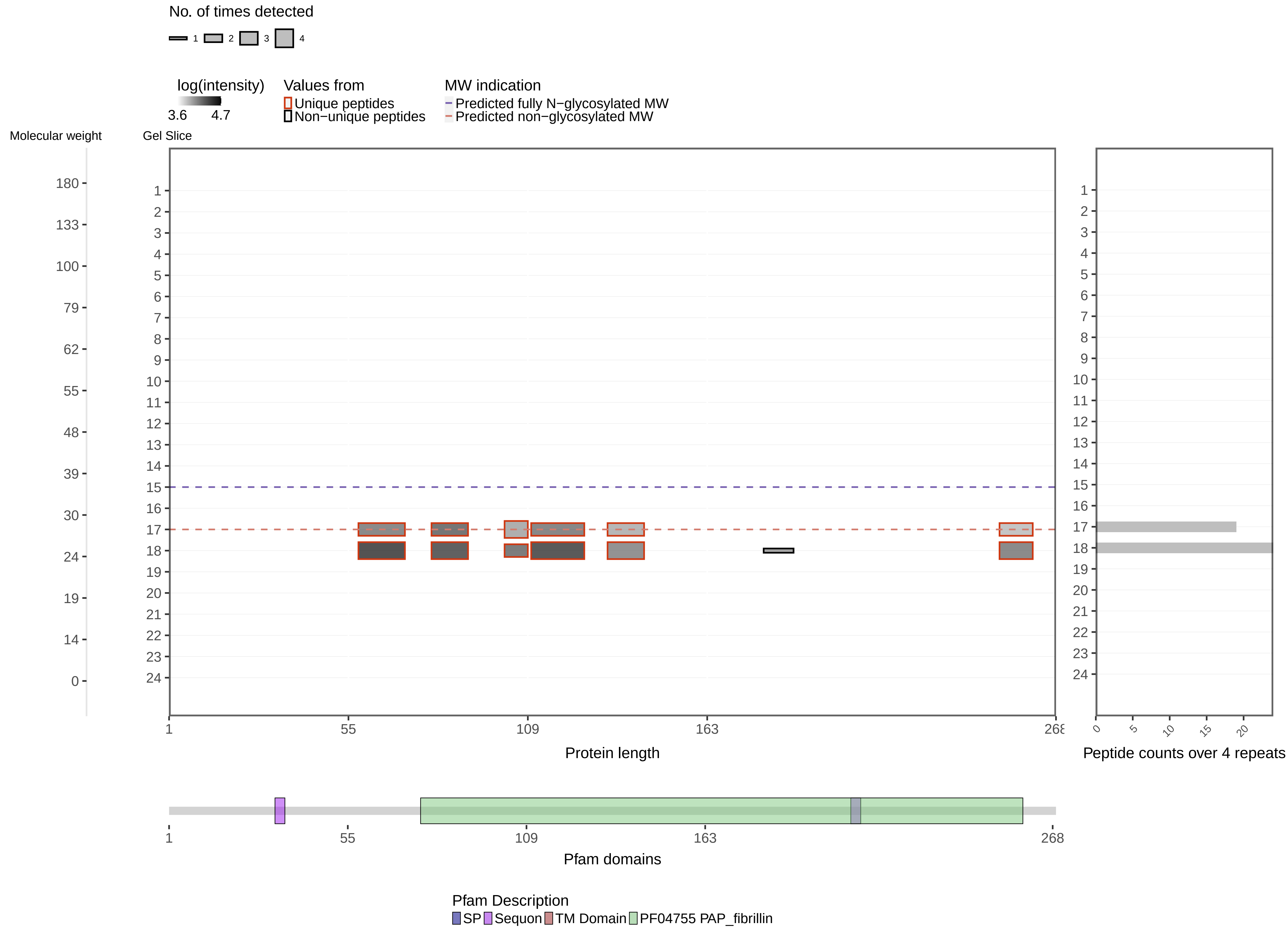


Protein sequence:

:PTLSLDKVAGSWKLIYSTISILGSRRTKLGLRDFITLGDLYQNIDISEGKAVNVIKFNARGLSLLSGELRIEATFKIASKSRVDIFYNNSAITPDQLMNVFQKNYDLLLGIFNPEGWLEITYVDENLRIGRDDKGNIFVLERSQEDKL*

### NbL12g21960.1 actin−7 [MW = 41.73 kDa]

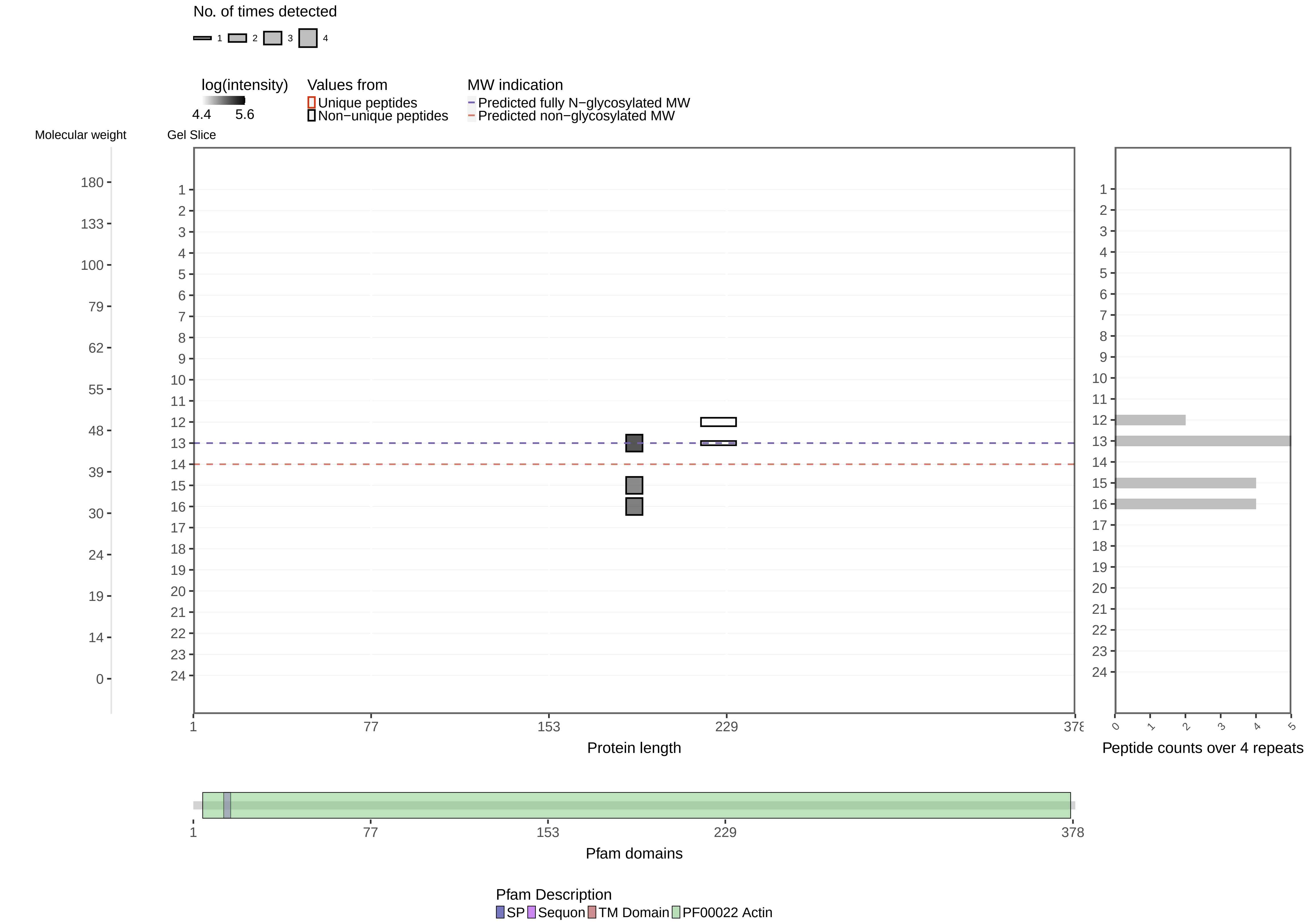


Protein sequence:

PHAILRLDLAGRDLTDNLMKILTERGYMFTTTAEREIVRDMKEKLAYVALDYEQELDTAKSSSSVEKNYELPDGQVITIGAERFRCPEVLFQPSMIGMEAAGIHETTYNSIMKCDVDIRKDLYGNIVLSGGSTMFPGIADRMSKEITALAPSSMKIKVVAPPERKYSVWIGGSILASLSTFQQMWISKGEYDESGPSIVHRKCF*

### NbL12g21880.1 acetylglutamate_kinase [MW = 36.4 kDa]

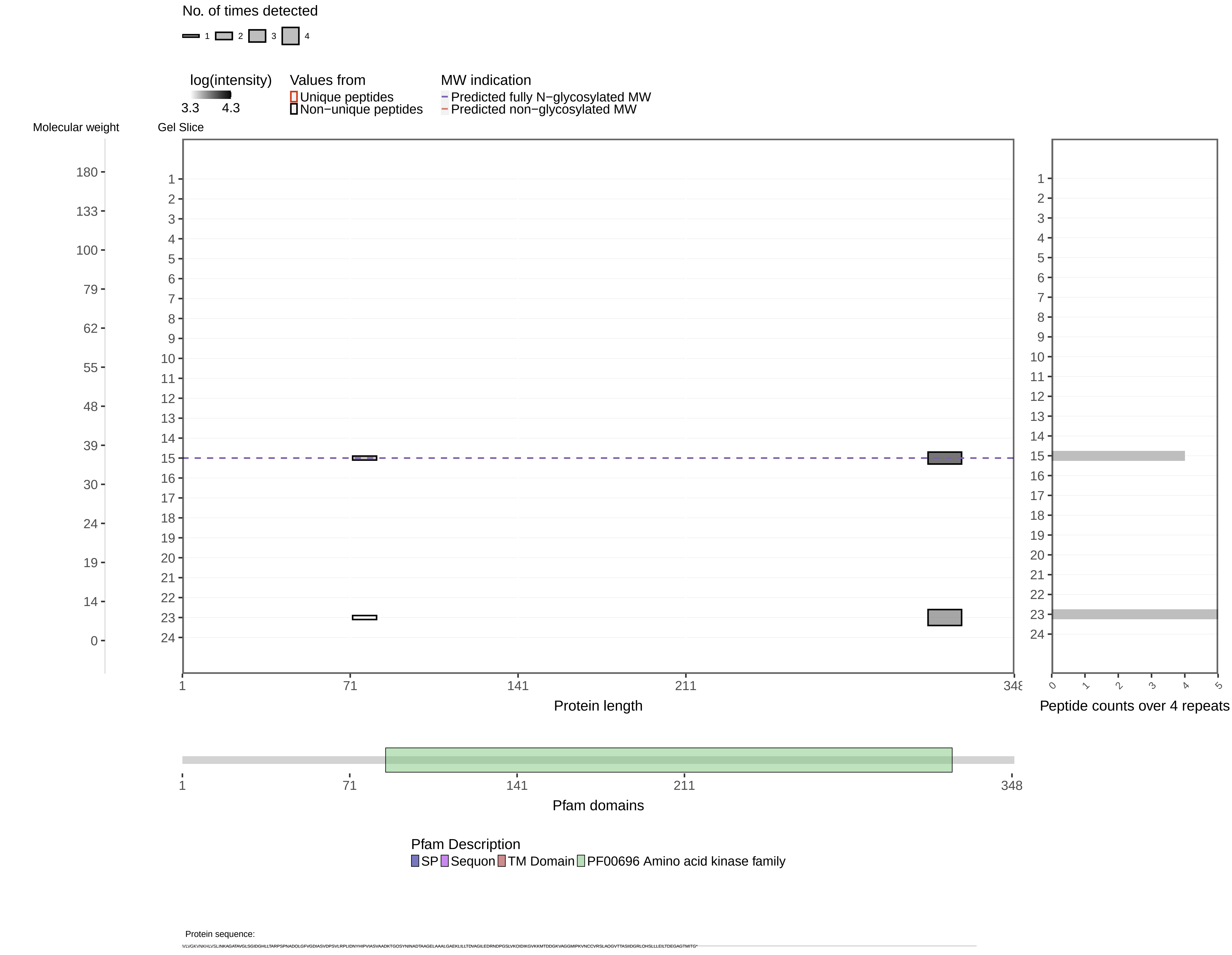

### NbL12g21860.1 40S_ribosomal_protein_S18 [MW = 17.54 kDa]

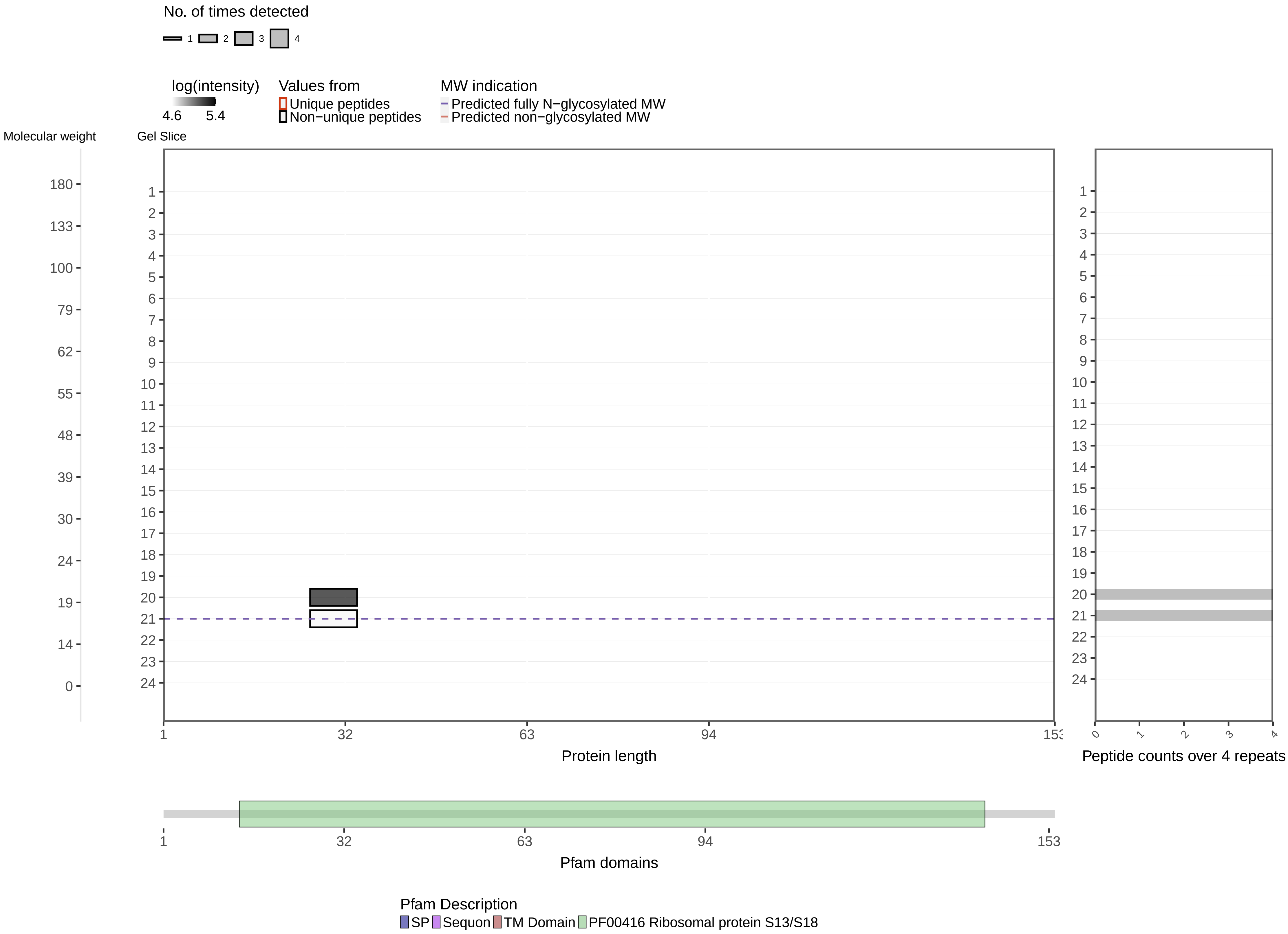


Protein sequence:

SLMVVVANPRQFKIPDWFLNRQKDYKDGKFSQVTSNALDMKLRDDLERLKKIRNHRGLRHYWGLRVRGQHTKTTGRRGKTVGVSKKR*

### NbL12g21760.1 probable_pectinesterase/pectinesterase_inhibitor_51 [MW = 61.53 kDa]

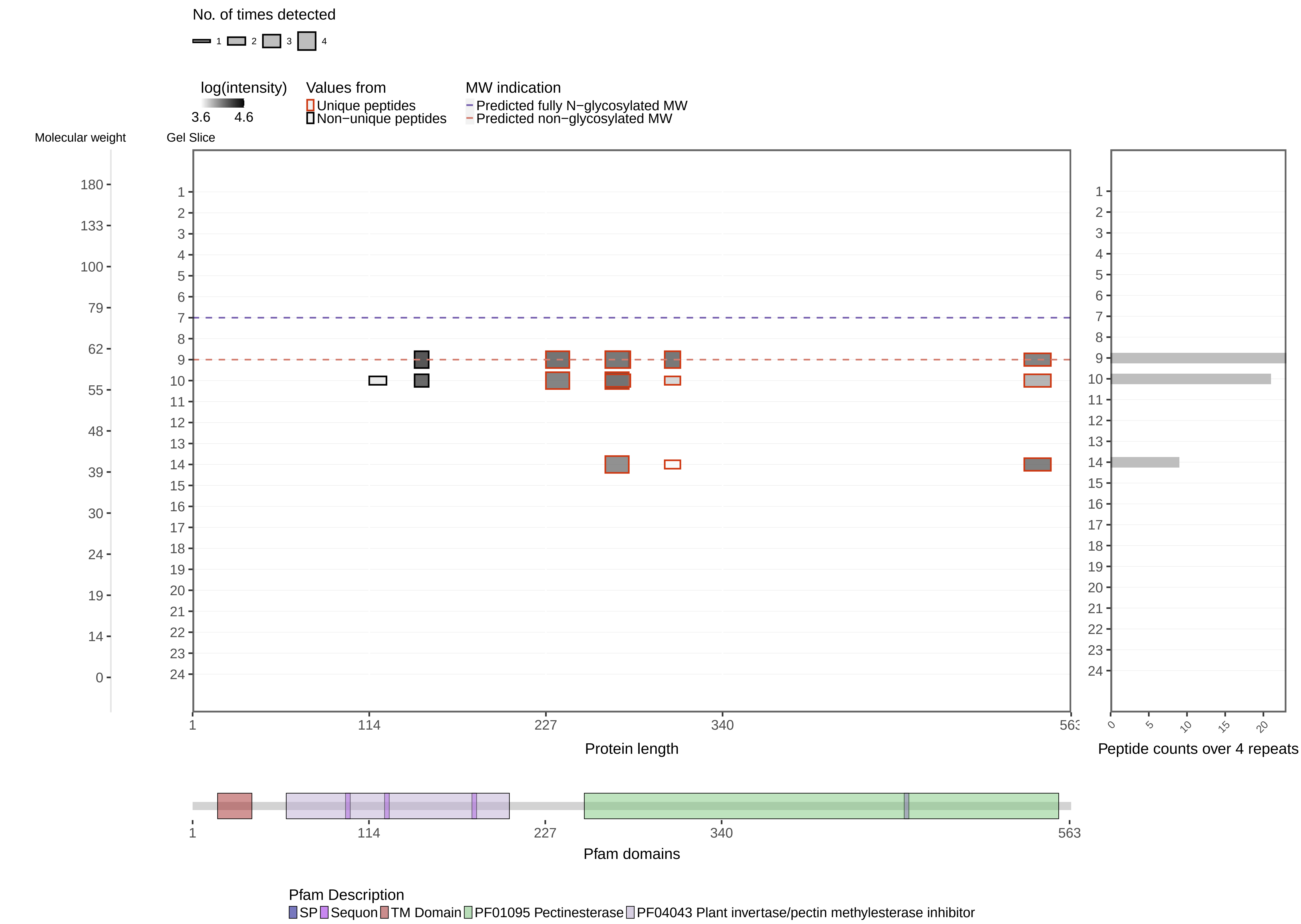


Protein sequence:

NbL12g21620.1 guanylate_kinase_2_chloroplastic/mitochondrial−like [MW = 34.17 kDa]

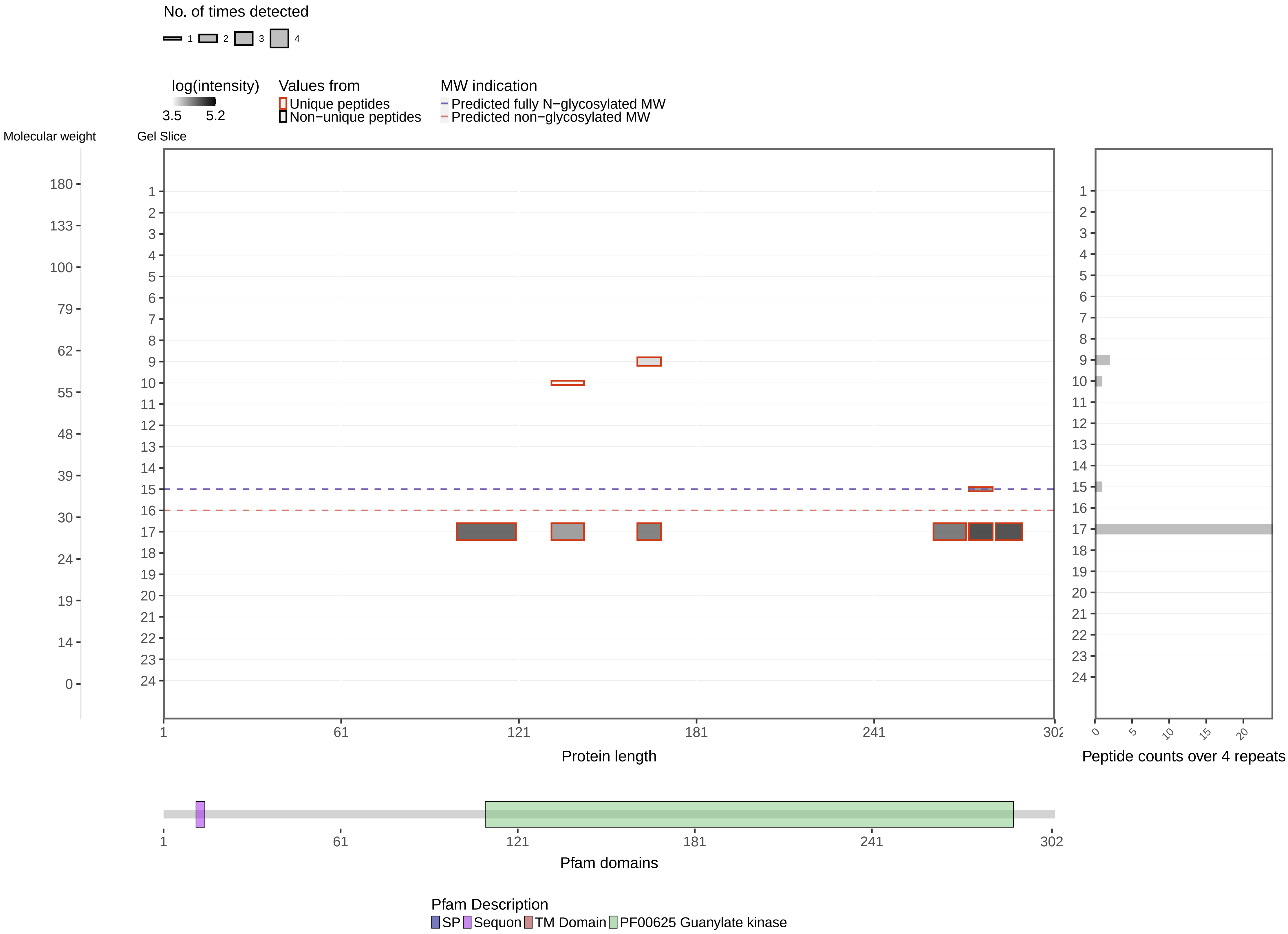


Protein sequence:

VVTATSRGKRSGEVDGKDYFFVSKEEFITMIERHELLEYALVYGDYKGIPKQQIRDQMAKGLDIVLRVDIQGAATLKRILGKSAVFVFLVAESEDALVKRLIDRKTETKETLLVRIATAREEVKHMEEFDYVVVNREGELENSVKLMESIIDAEKAKVRQRTATI*

### NbL12g21530.1 probable_polyamine_oxidase_5 [MW = 58.63 kDa]

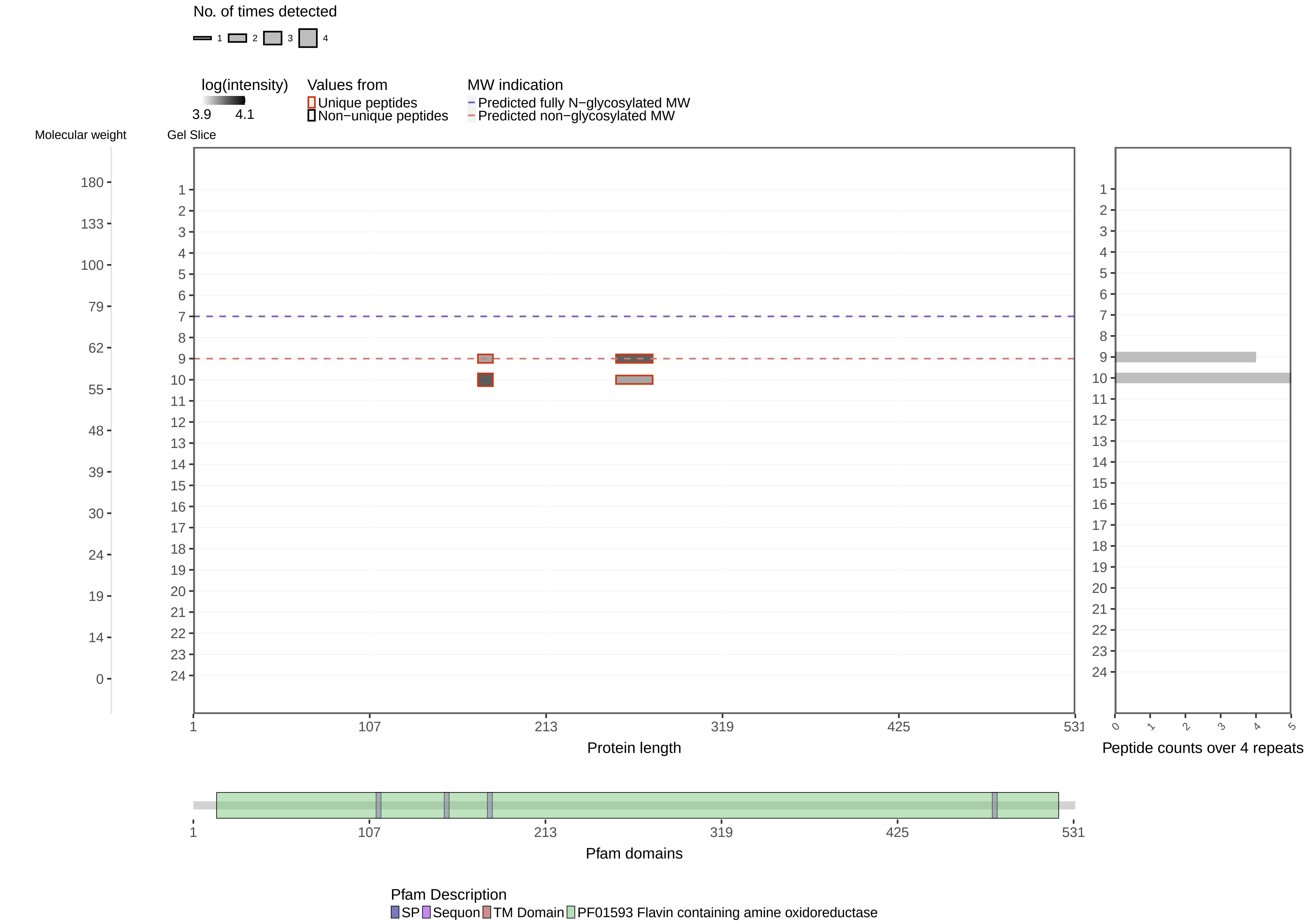


Protein sequence:

RIEWQPDDGGPLDVENGDDTKPVKLHFYDGSVMYADHVIVTVSLGVLKREIRDDSGMFSPPLPSFKTRAISRLGFGVVNKLFLQLSPTKDEDAMKFPYLQMVFHQPDSKLRHPKTPWWMRRTASLCPIYGSSNVLLSWFAGAEALELESLDNEEIIDGFSTMISDFLVNSKHLNKLCNGHAESCFKFDKVLKSQWGTDPLFLGSYSYVAVGSSGDDLDTMAEPLPKRSNYSCISPPLQVLFAGEATHRTHYSTTHGAYFSGLREANRLLQHYHRIDI*

### NbL12g21370.1 PI−PLC_X_domain−containing_protein_At5g67130 [MW = 49.01 kDa]

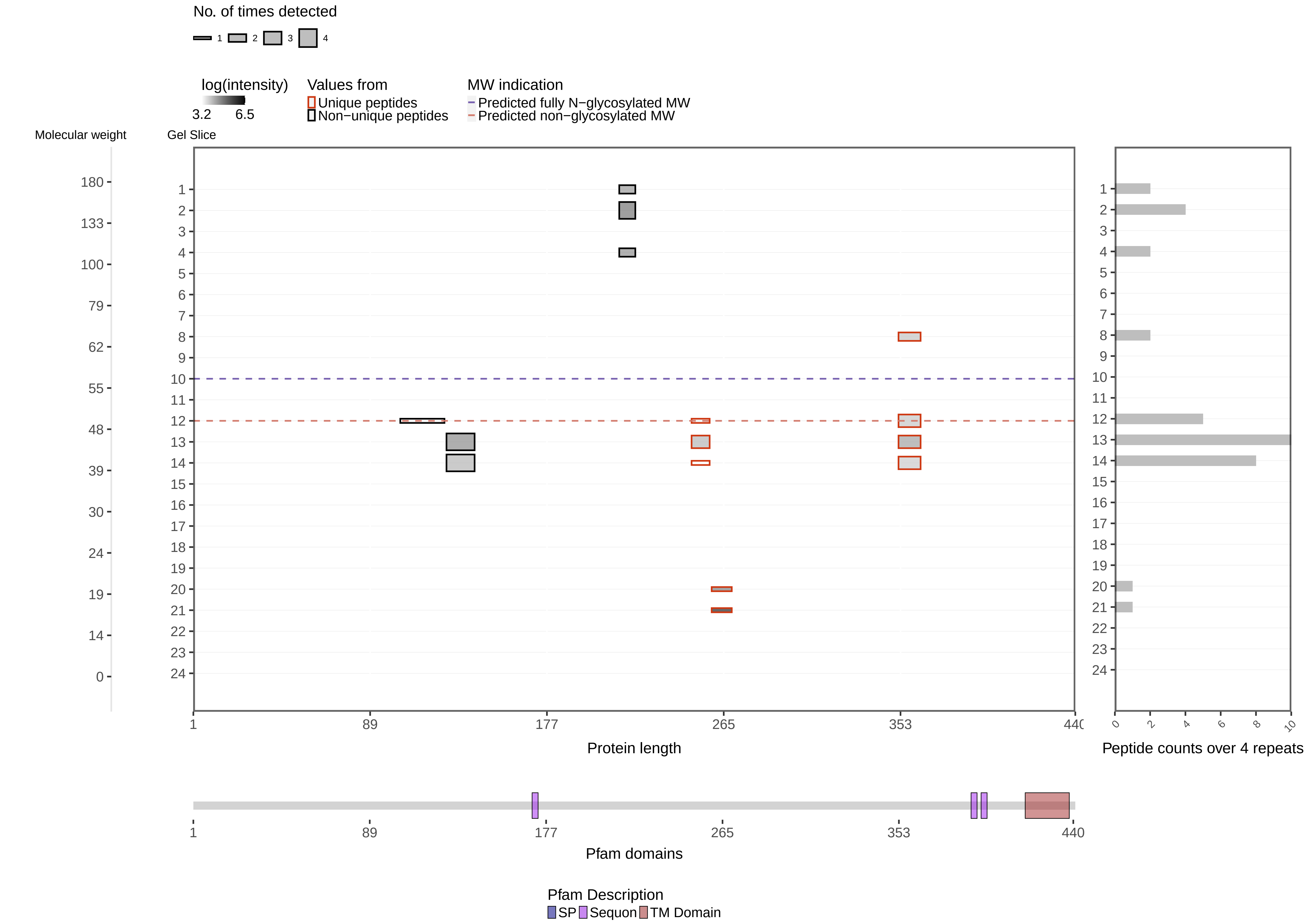


Protein sequence:

LTRVFTDAGLDKYRFFPVSKMPKKGDDWPTVNDMVKKNHRLLVFTSDSSKEAAEGIAYQWRYMVENEPGDPGVMPGACSSRKESKPLNSRSASLFLMNYFPTVAVQNGVCKEHSTQLVDTVGACFKAAGNLMPNYVAVNFYMRSDGGGVFDVLDRMNGETLCGCPTVTACQAGERFGVCRNITAANATPTATNAAGSFSGSVQLTGHASAIHFSSIIYLFLLSSVIILYLF*

### NbL12g21270.1 PI−PLC_X_domain−containing_protein_At5g67130−like [MW = 48.69 kDa]

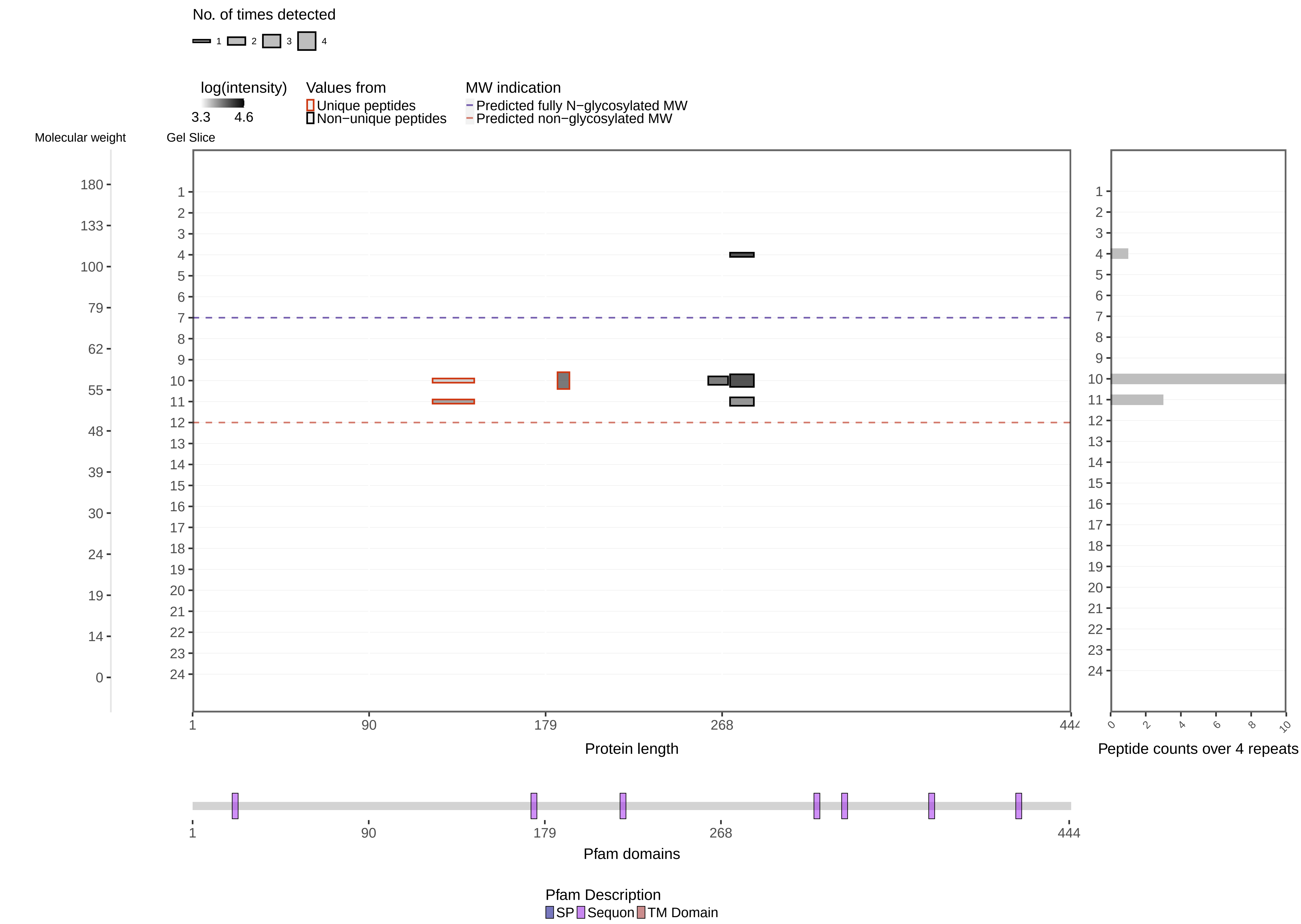


Protein sequence:

VFNASGLNKYWFPVSRMPKKGEDWPTVDDMVKQNQRLVVFTSKSSKEASEGIAYEWRYVVENQYGNGGMKVGSCPSRSESPPMSTSSISLVLQNYFPDNPNITAACVDNSAPLVNMTKTCFEAAGKRWPNFIAVDFYQKSDGGGAPEAVDEANGQLGCNCSSIAYCKASGTSHKCEAPMLSPPPPAQATTAGTSQGGNNENKNNSHSRRSKPLHWLISTIFAAILLLRA*

### NbL12g21160.1 zeaxanthin_epoxidase_chloroplastic [MW = 72.56 kDa]

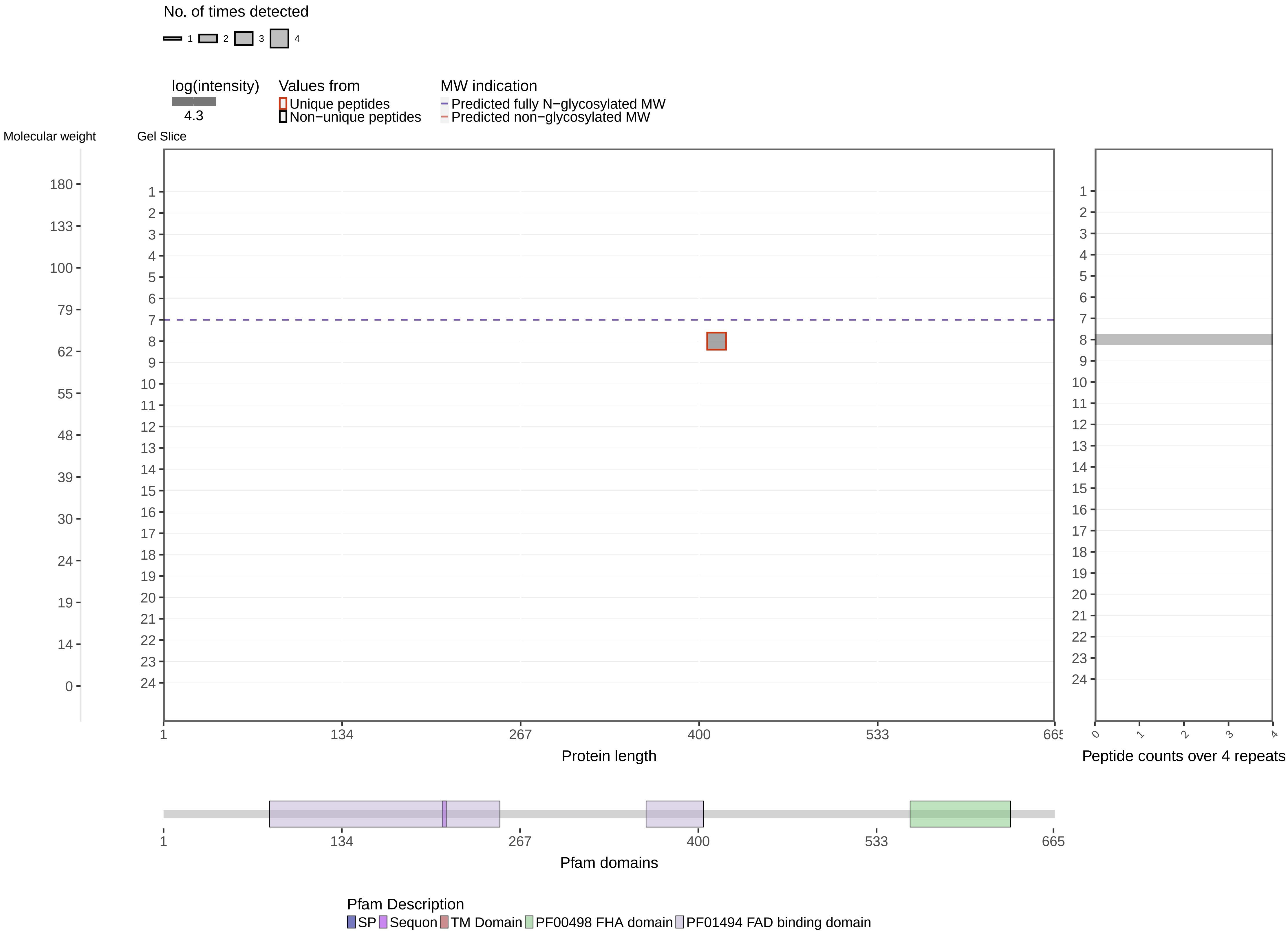


Protein sequence:

KKARLLKIFEGWCDNVIDLLVATNEDAILRRDIYDRPPTFNWGKGRVTLLGDSVHAMQPNLGQGGCMAIEDSYQLALELDKELSRSTESGTPVDIISSLRSYESSRKLRVGVIHGLARMAAIMASTYKAYLGVGLGPLSFLTKFRIPHPGRVGGRFFIDLGMPLMLSWVLGGNGEKLEGRIKHCRLSEKANDQLRNWFEDDDALERATDAEWLLLPAGNGNAASEALVLSRDEDTPCTIGSVSHTKIPGKSVVIPLPQVSEMHARISYKGGAFFVTDLRSEHGTWITDNEGRR

NbL12g21050.1 uncharacterized_LOC109225879 [MW = 178.96 kDa]

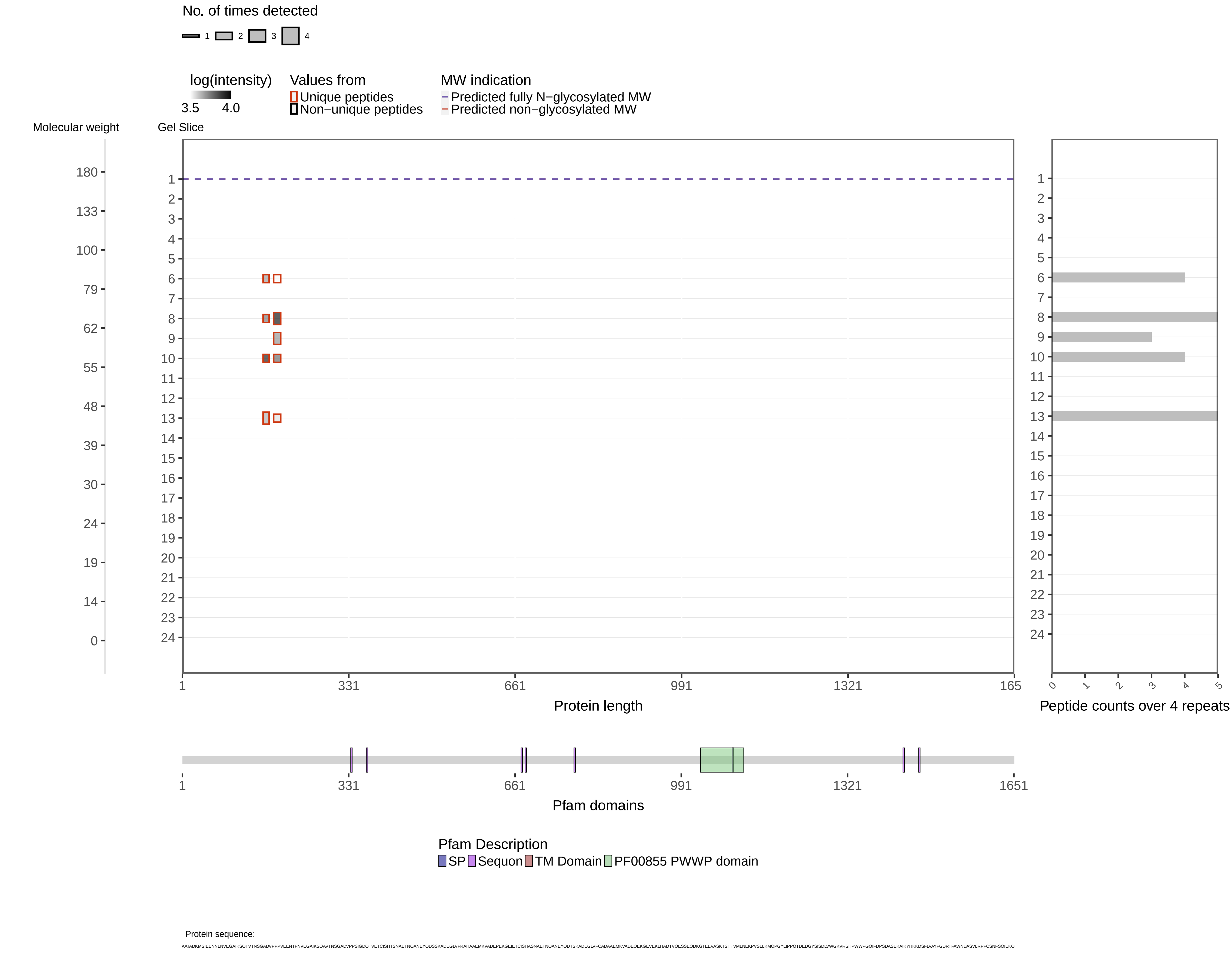

### NbL12g21020.1 uncharacterized_LOC107785953 [MW = 29.11 kDa]

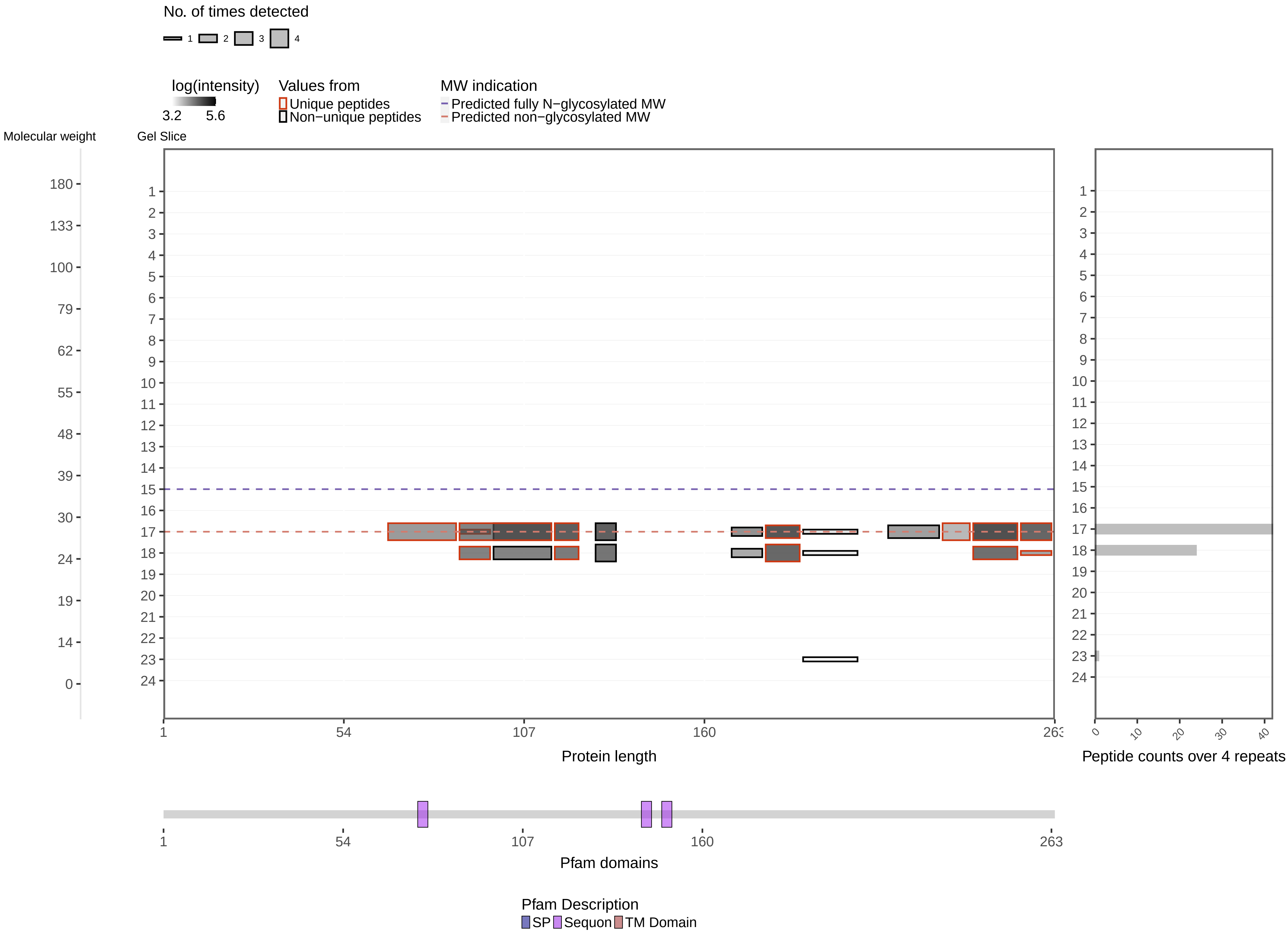


Protein sequence:

LDLSRGIPRFYIMQLKDRSLRFSNITHHANVTQCLGSIGGNVWYLGVAKPSIVDPADIKGAAVADIVQSRCGHFYVPPAVDEVQAFRISGPKFIKLNHGTWHAGPLFTDDKMDFYNLELKNTNVVDHTTHDFVKKNEVVFLLDD*

### NbL12g20890.1 vacuolar_protein_sorting−associated_protein_26A [MW = 35.66 kDa]

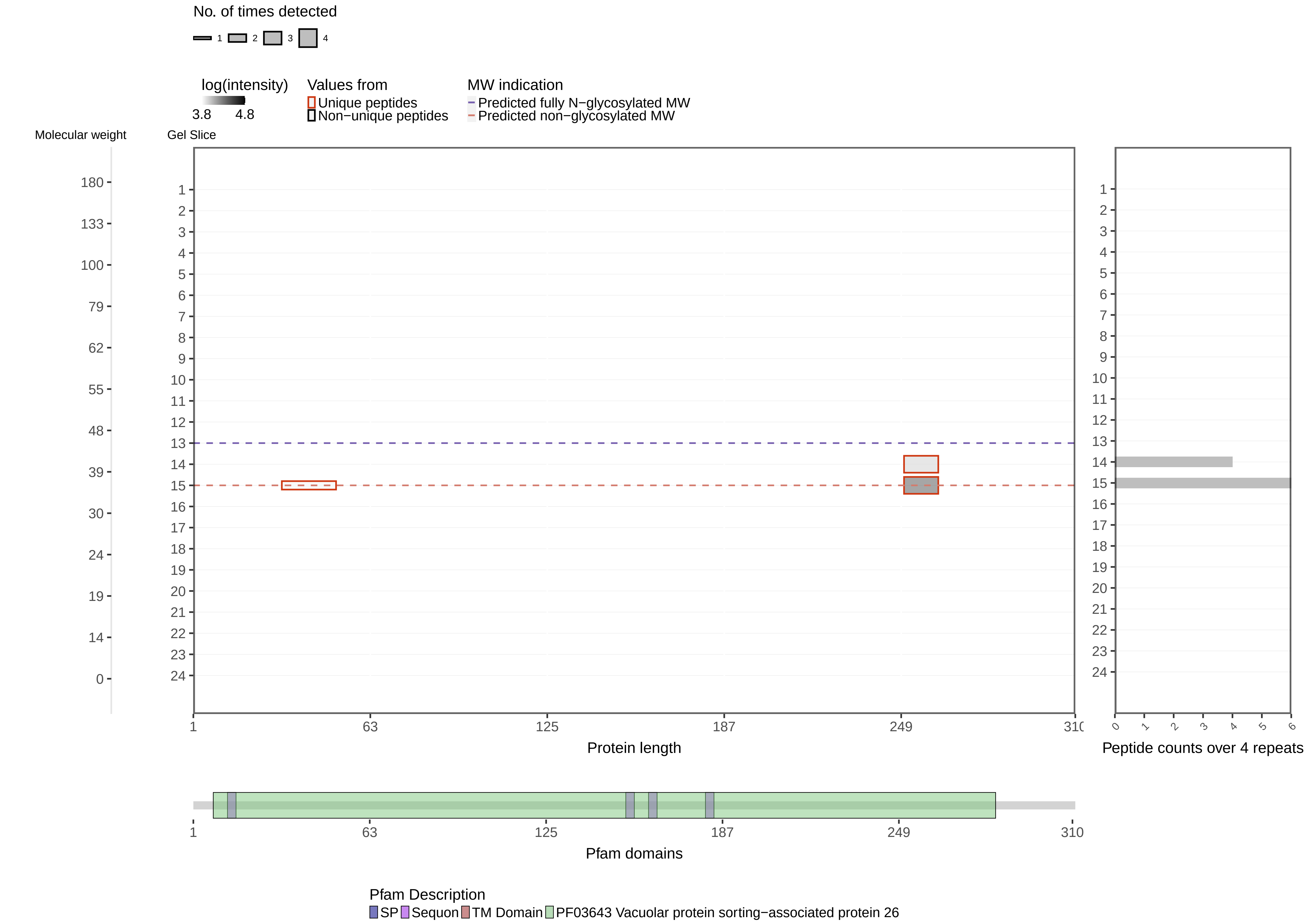


Protein sequence:

GSIVEYQYFVVRNYSPPPSINNSIKMEVGIEDCLHIEFEYNKSKYHLKDVIIGKIYFLLVRIKIKNMDLEIRRRESTGSGANTHVETETLAKFELMDGAPVRGESIPIRLFLSPYELTPTYRNINNKFSVKYYLNLVLVDEEDRRYFKQQEITMFRLAETSVINQAVGA*

### NbL12g20750.1 GDSL_esterase/lipase_At5g33370−like [MW − SP = 37.25 kDa]

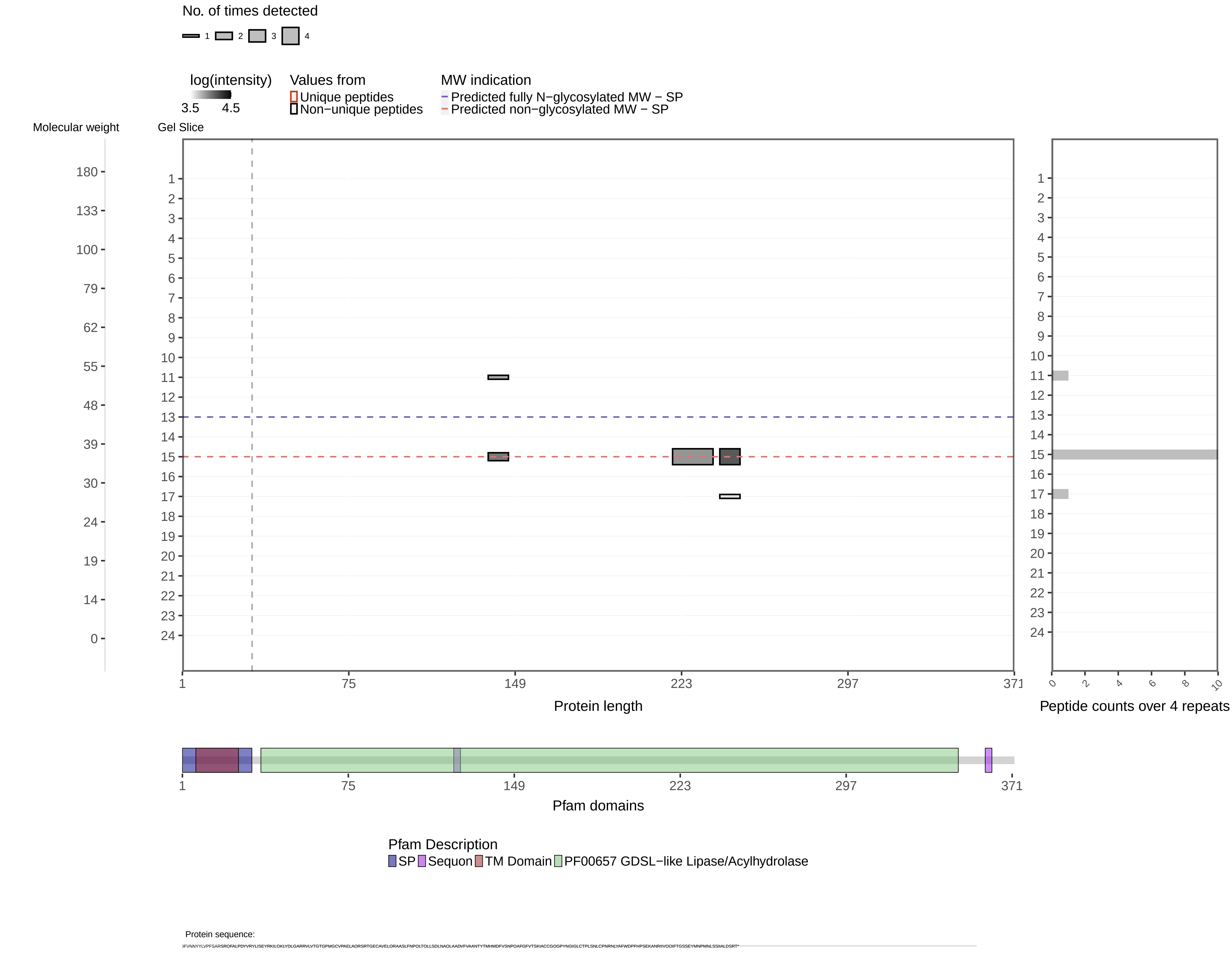

NbL12g20700.1 uncharacterized_LOC104249736 [MW = 25.63 kDa]

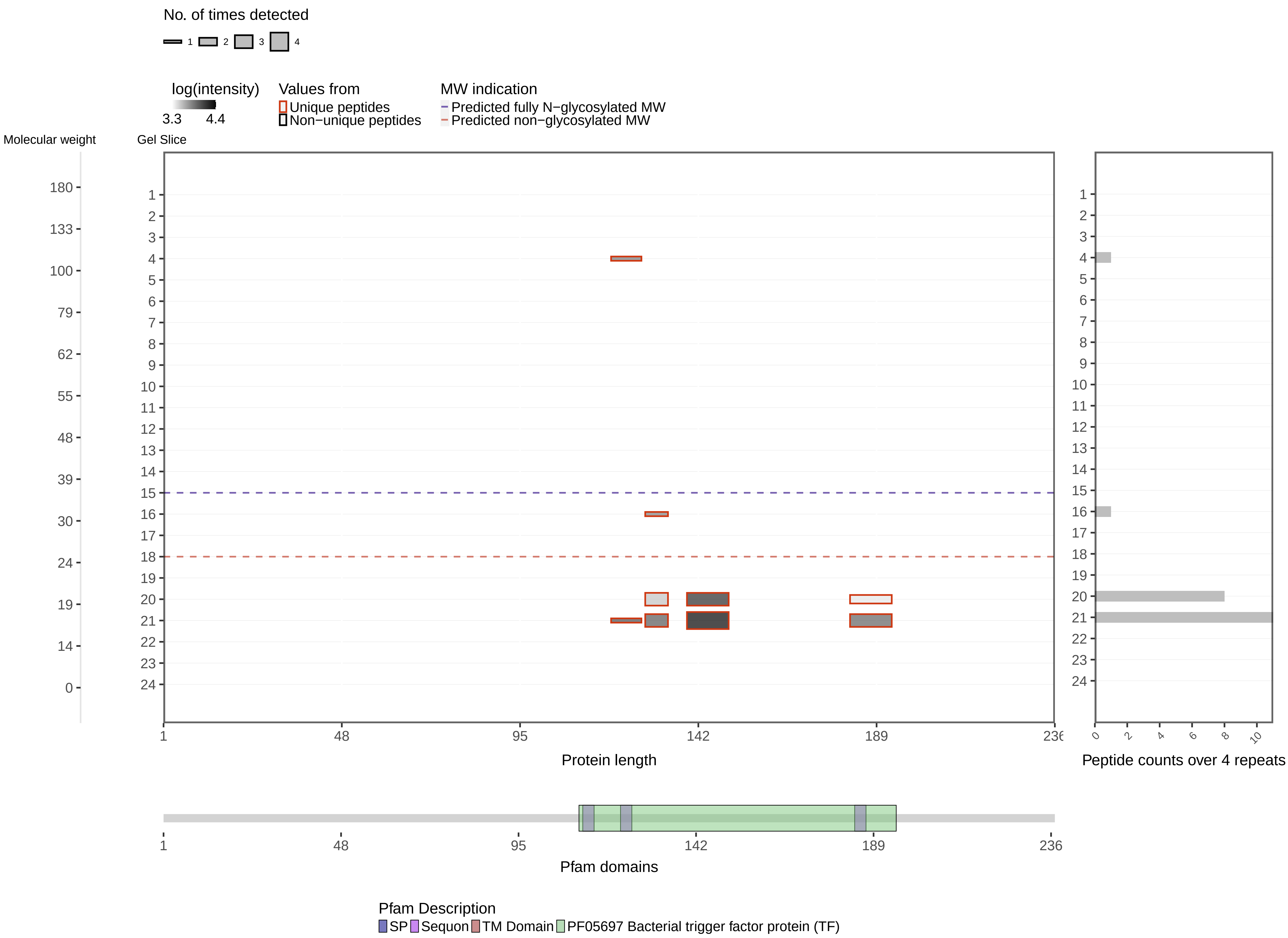


Protein sequence:

:FEDFSVTTSGVNGSNKLKISVNVSGAKTEEIFGKVFSRMVEDAQPIPGFRRVKGGKTPNIPRDILIEILGYSKVYKQVIKKIINSTISEYVEKVDPPLPSPSLCISVILSLHIHTLLLYLKDLDRYRIIKCYGCI*

### NbL12g20610.1 PHD_finger_protein_ALFIN−LIKE_4−like [MW = 27.77 kDa]

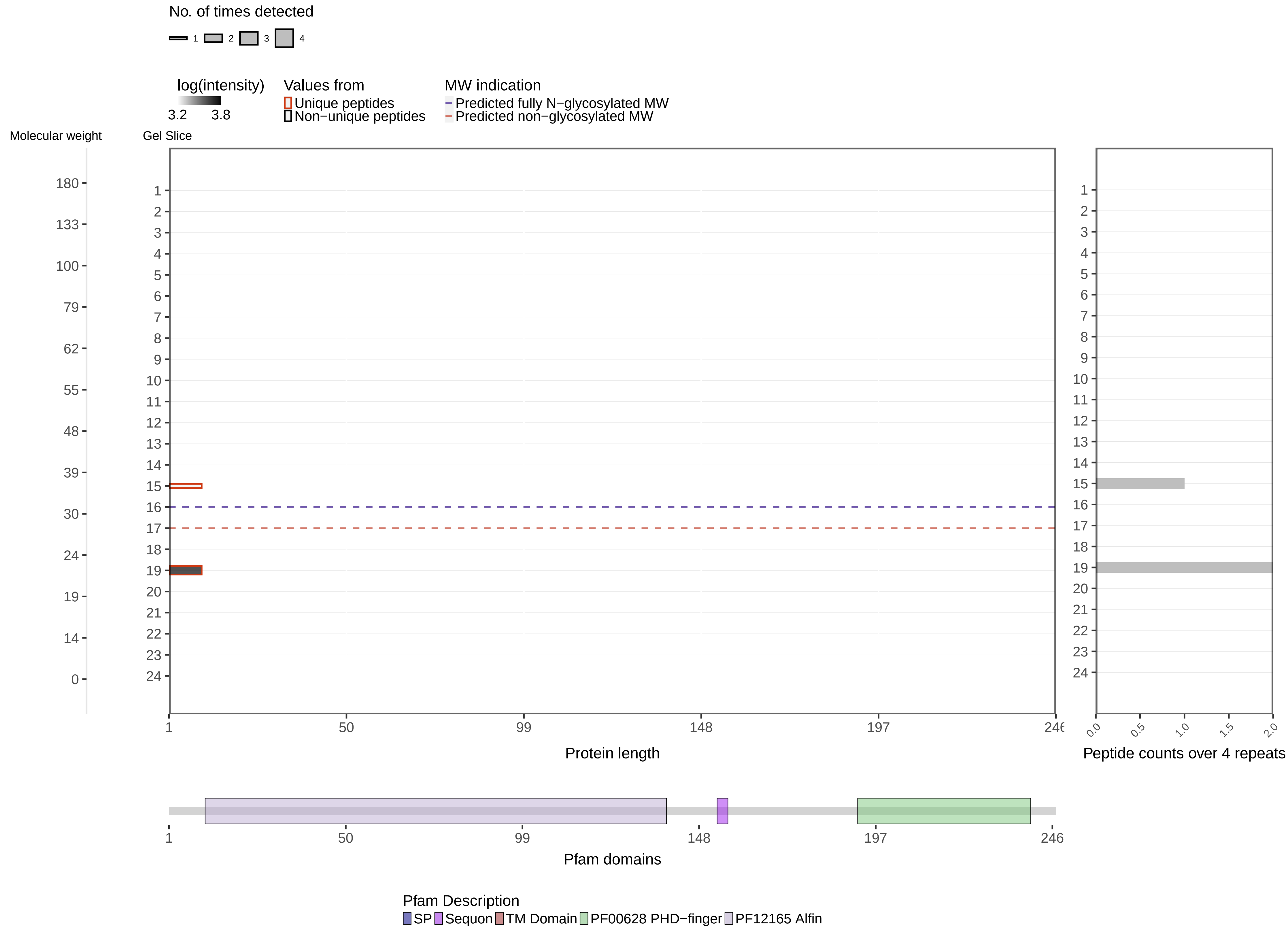


Protein sequence:

ARFGFDKADRKRLFNMINDLPTIYEVVTGAAKKQVKERSSVSNHSSTKSKSSSKVGKYSKVEVKDEDDGLDEEEEEHGDTLCGACGENYASDEFWICCDICERWFHGKCVKITPAKAEHIKQYKCPTCSNKRPRP*

NbL12g20600.1 threonine_synthase_chloroplastic−like [MW = 54.34 kDa]

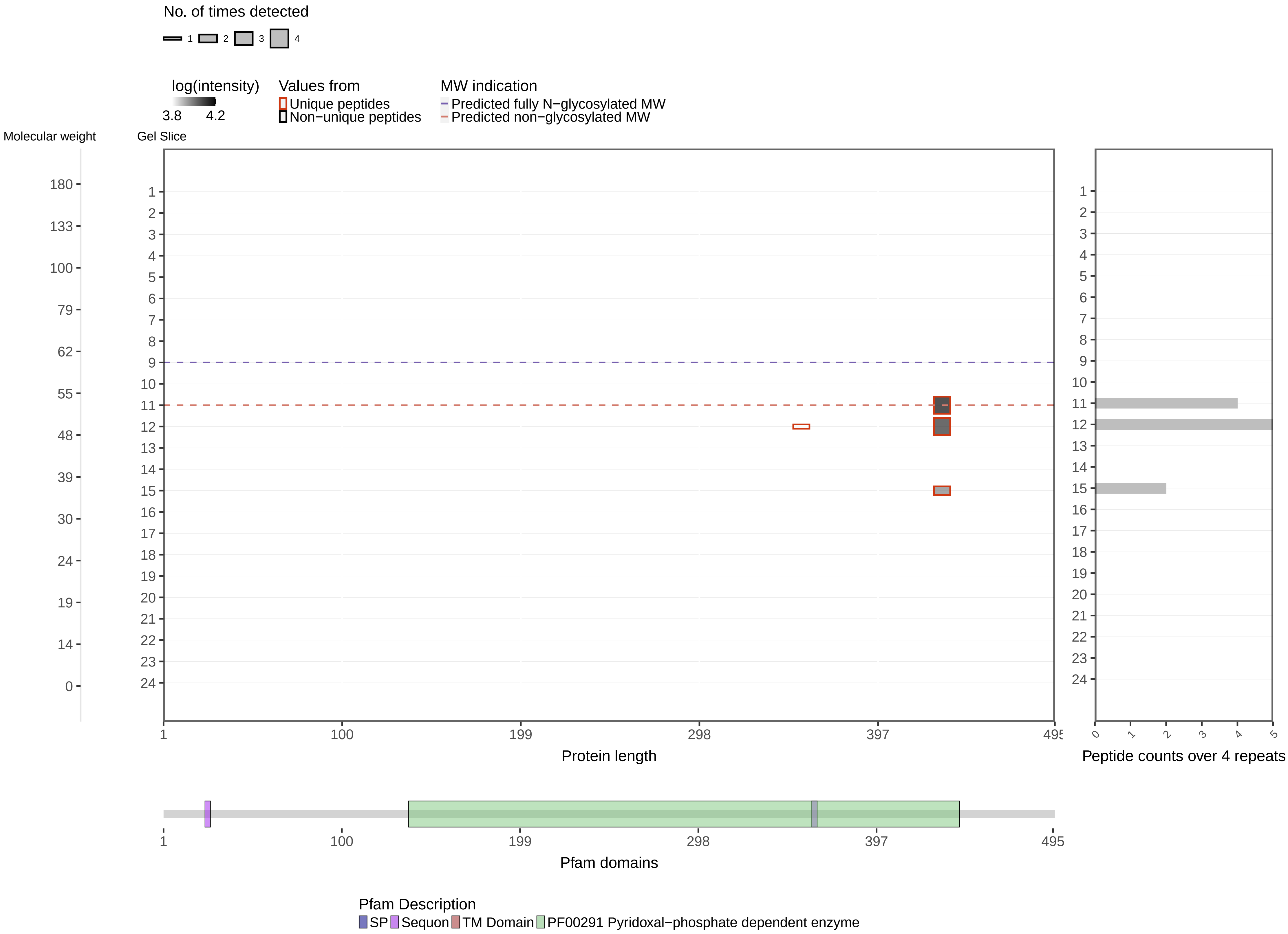


Protein sequence:

MAQLVQPIANGAFVISIDTDFDGCMKLIREVTSELPIYLANSLNSLRLEGQKTAAIEILQQFDWEVPDWVIIPGGNLGNIYAFYKGFKMCQEMGLVDRIPRLVCAQAANANPLYMYYKSGWTNFQPVKANTTFASAIQIGDPVSIDRAVYALKNSNGIVEEATEEELMDATALADSTGMFICPHTGVALTALMKLRNSGVIAPTERTVVVSTAHGLKFTQSKIEYHSKEIKDMASRFANPPVQVKADFGSVMDVLNNYLKSKS*

NbL12g20510.1 CBS_domain−containing_protein_CBSX1_chloroplastic−like [MW = 26.28 kDa]

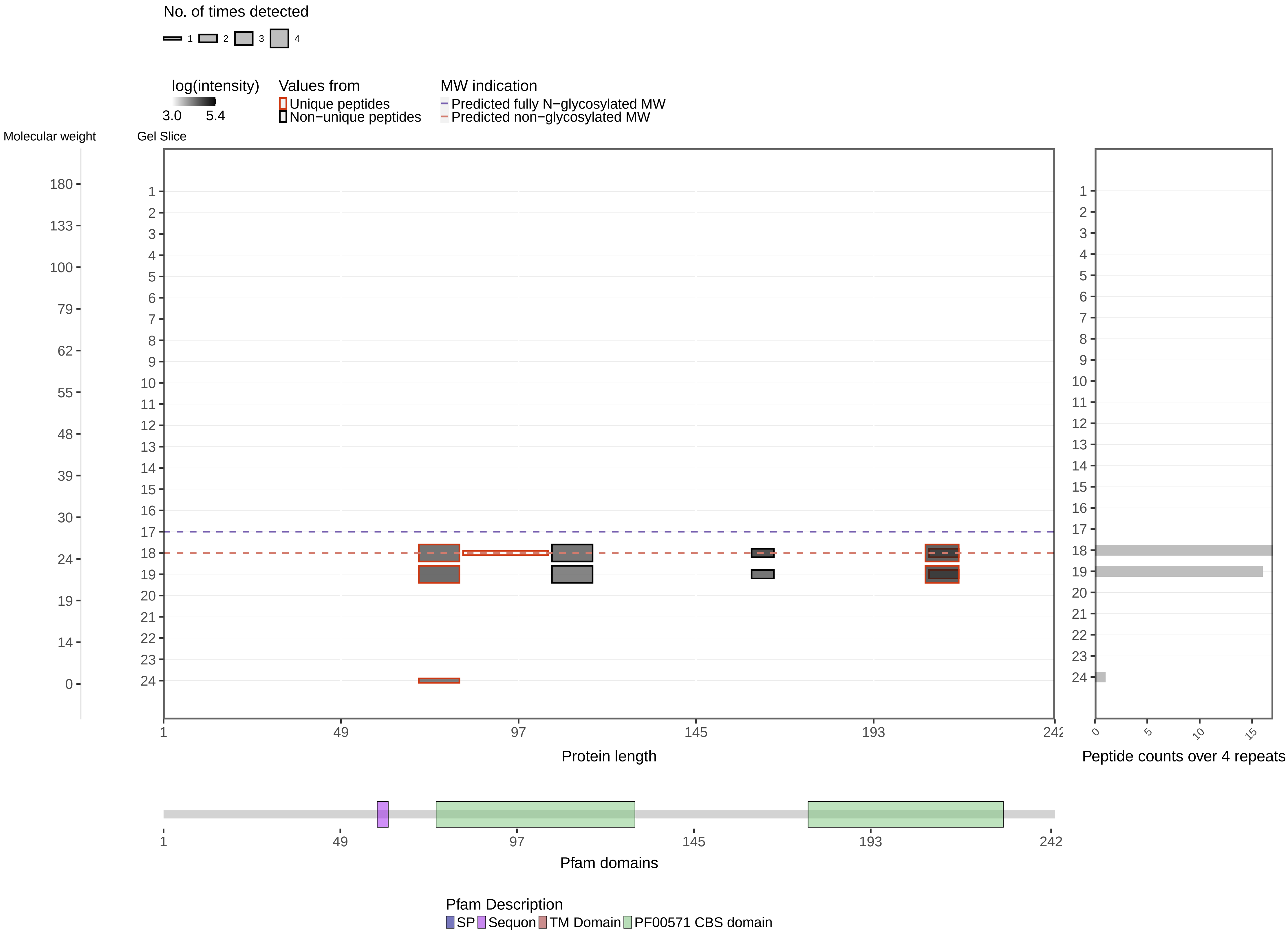


Protein sequence:
GFPVIDDNWKLVGLVSDYDLLALDSISVVQILLLGTGTTDADMFPEVDSNWKTFNEIQKLLGKTKGKVVGDLMTPAPLVVRESTNLEDAARLLLKTKYRRLPVVDGDGKLVGIITRGNVVRAALQVKRANEMEG*

### NbL12g20150.1 kinase_II_subunit_alpha−like [MW = 39.33 kDa]

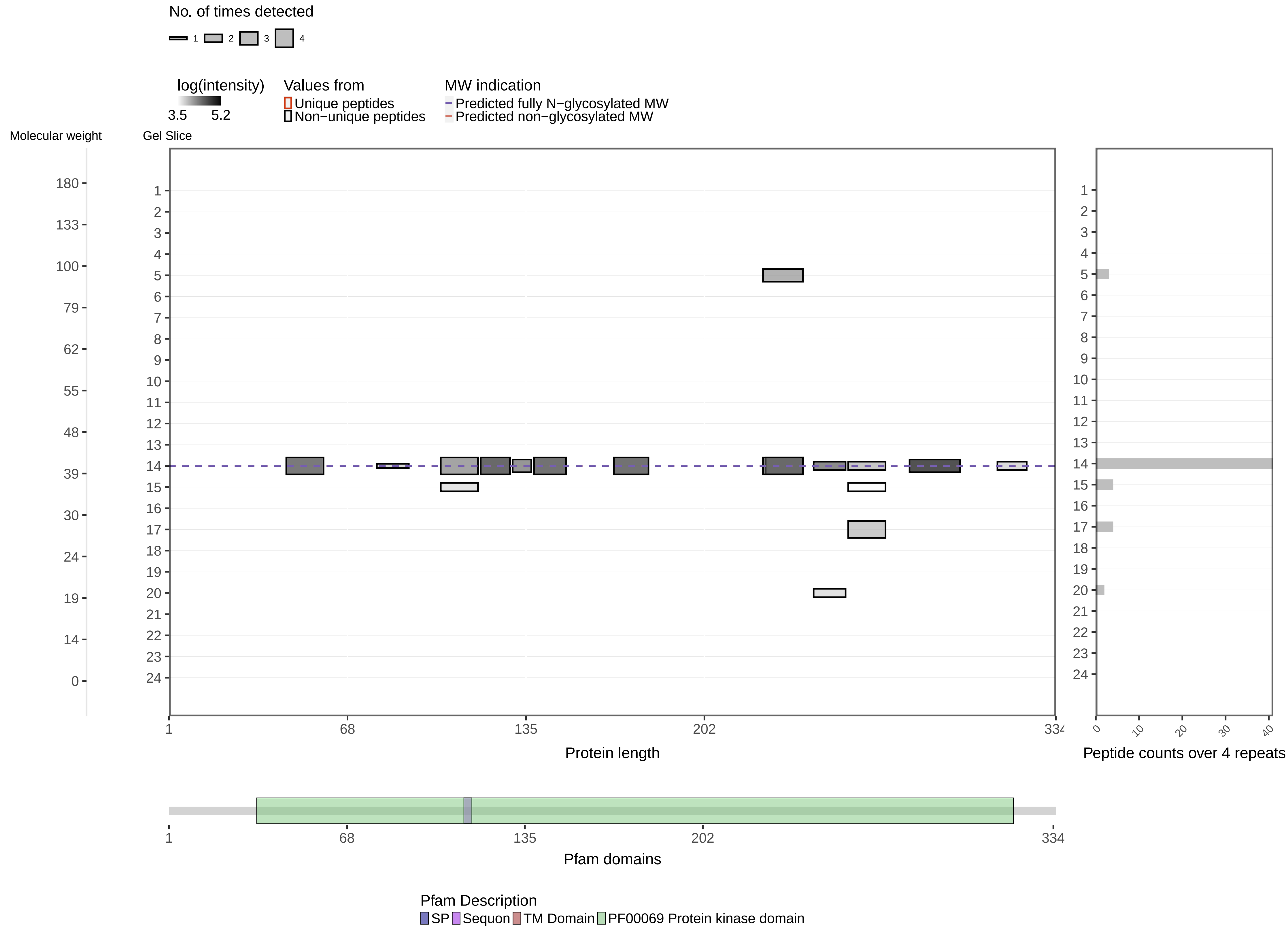


Protein sequence:

VMIDHELRKLRLIDWGLAEFYHPGKEYNVRVASRYFKGPELLVDLODYDYSLDMWSLGCMFAGMIFRKEPFFYGHDNODOLVKIAKVLGTDELNAYLHKYOLELDPOLEAMVGRHSRKPWSKFINADNOHLVSPEAIDFLDKLLRYDHOSRLTAKEAMAHPYFLOVRAAETSRMRAO*

### NbL12g19960.1 60S_ribosomal_protein_L26−1−like [MW = 16.71 kDa]

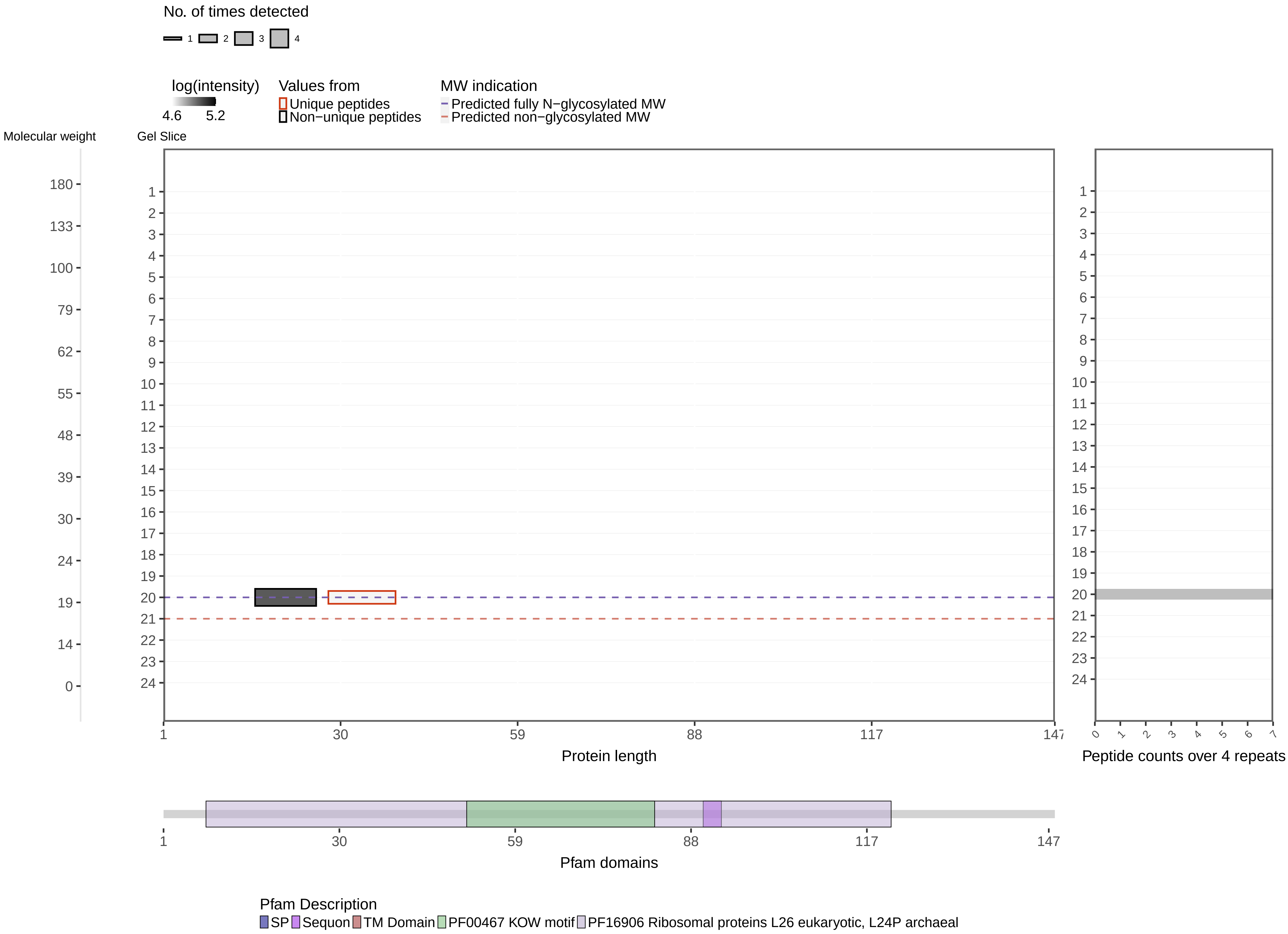


Protein sequence:

GTYKGREGKVVQVYRKKWVIHIERITREKVNGSTVNVGIHPSKVVVTKLRLDKDRKSLLDRKAKGRAAADKDKGTKFTAEDIMQTVD*

### NbL12g19640.1 superoxide_dismutase_[Fe]_3_chloroplastic [MW = 22.96 kDa]

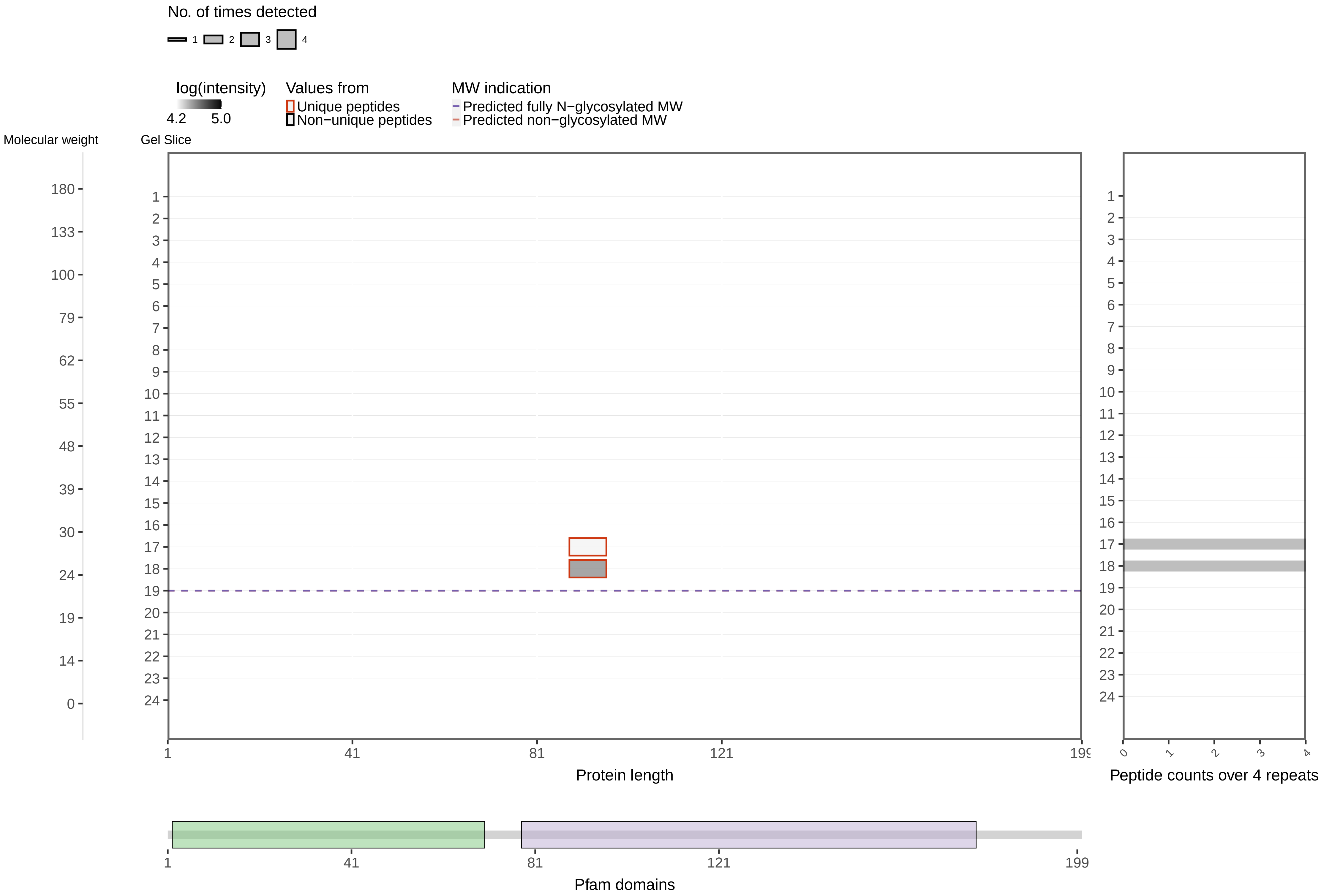


Pfam Description

SP Sequon TM Domain PF00081 Iron/manganese superoxide dismutases, alpha−hairpin domain PF02777 Iron/manganese superoxide dismutases, C−terminal domain

Protein sequence:

)IEKDFGSFTNYREKFTEAALTLFGSGWIWLVLKREEKRLAIVKTSNAVNPLVWNDIPLIGLDLWERAYYLDYKNDKAKYVNVFMNHLVSWDAALGRMARAQAFVNLGEPKIPVA*

### NbL12g19580.1 proteasome_subunit_beta_type−5−like [MW = 31.11 kDa]

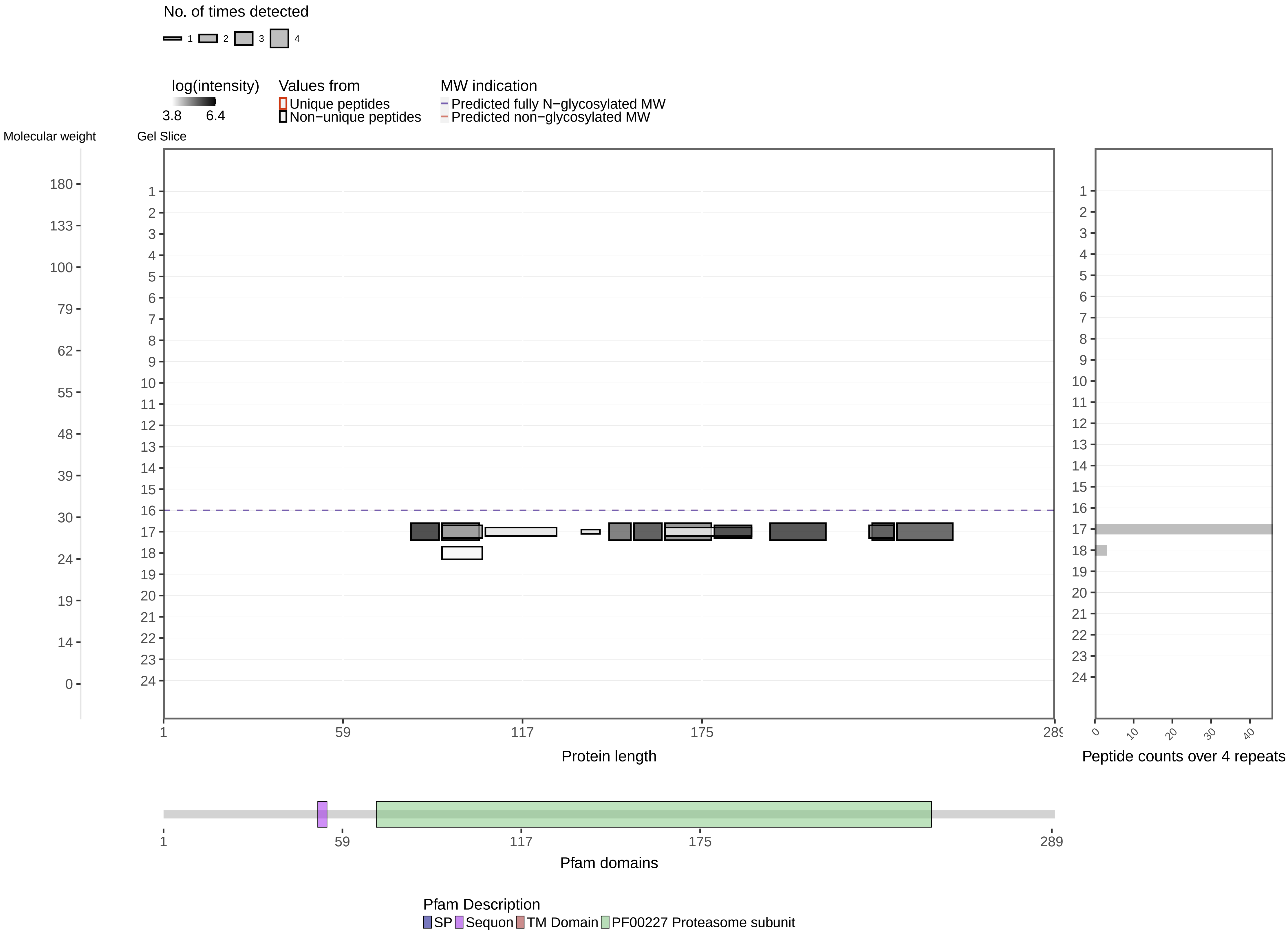


Protein sequence:

CRLHELANKRRISVAGASKLLANILYSYRGMGLSVGTMIAGWDEKGPGLYYVDSEGGRLKGNRFSVGSGSPYAYGVLDSGYRFDLSVEEAAELARRAIYHATFRDGASGGVASVYHVGPNGWKKLSGDDVGELHYNYYPVELESVEQEMAEVPVA*

NbL12g19410.1 phosphomannomutase/phosphoglucomutase [MW = 65.64 kDa]

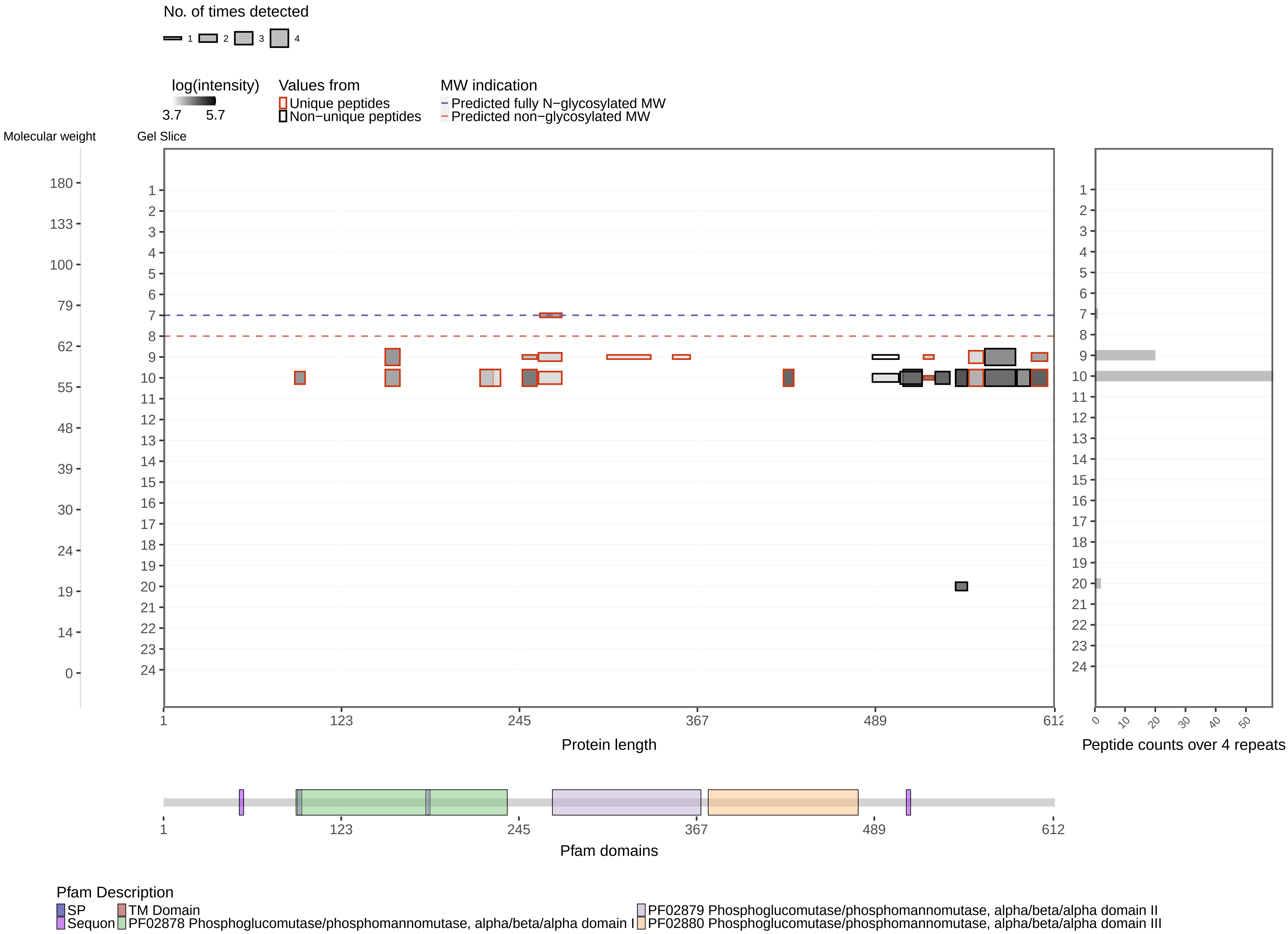


Protein sequence:

NbL12g19340.1 acetylornithine_aminotransferase_mitochondrial [MW = 49.27 kDa]

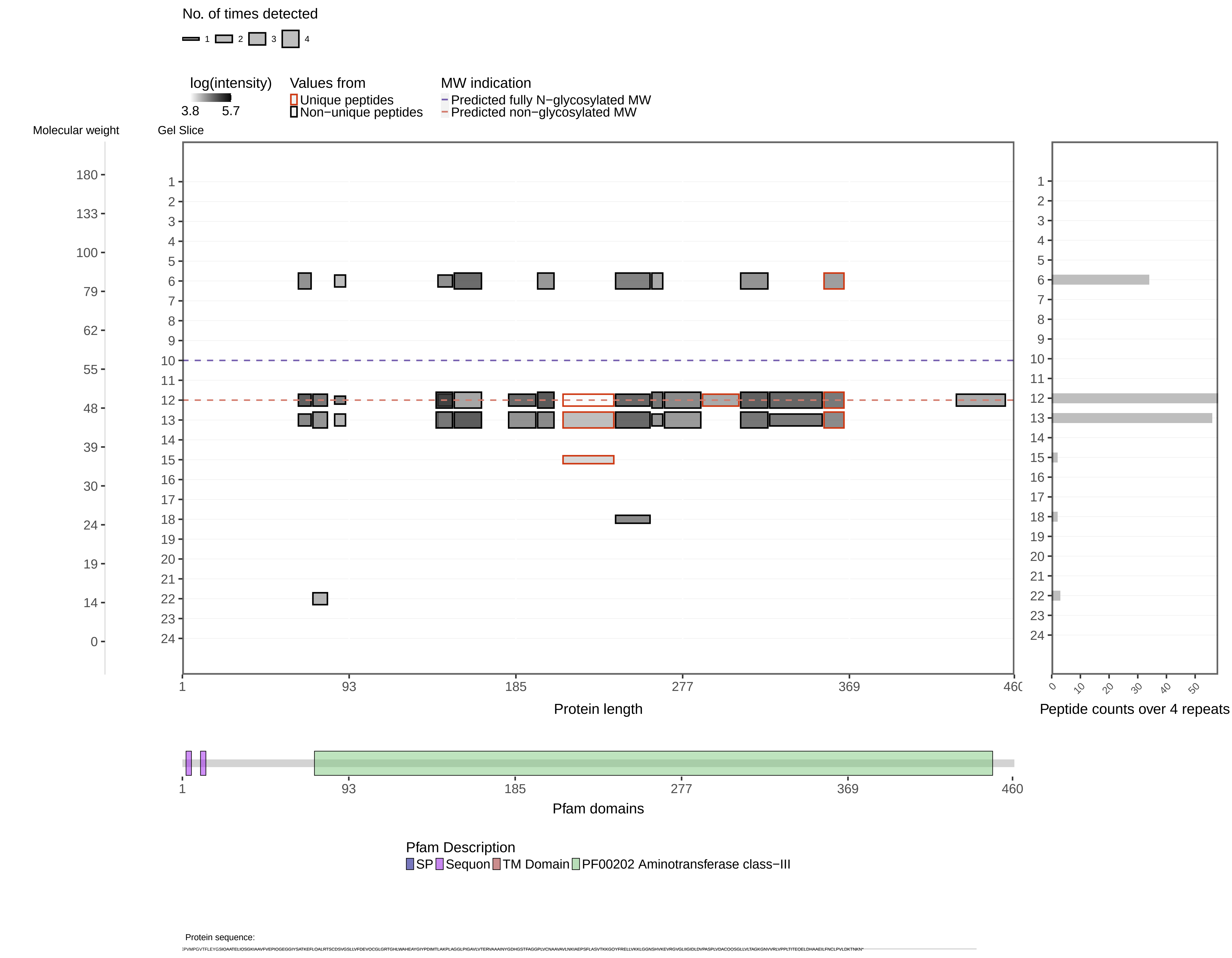

NbL12g19140.1 uncharacterized_LOC109205533 [MW = 11.93 kDa]

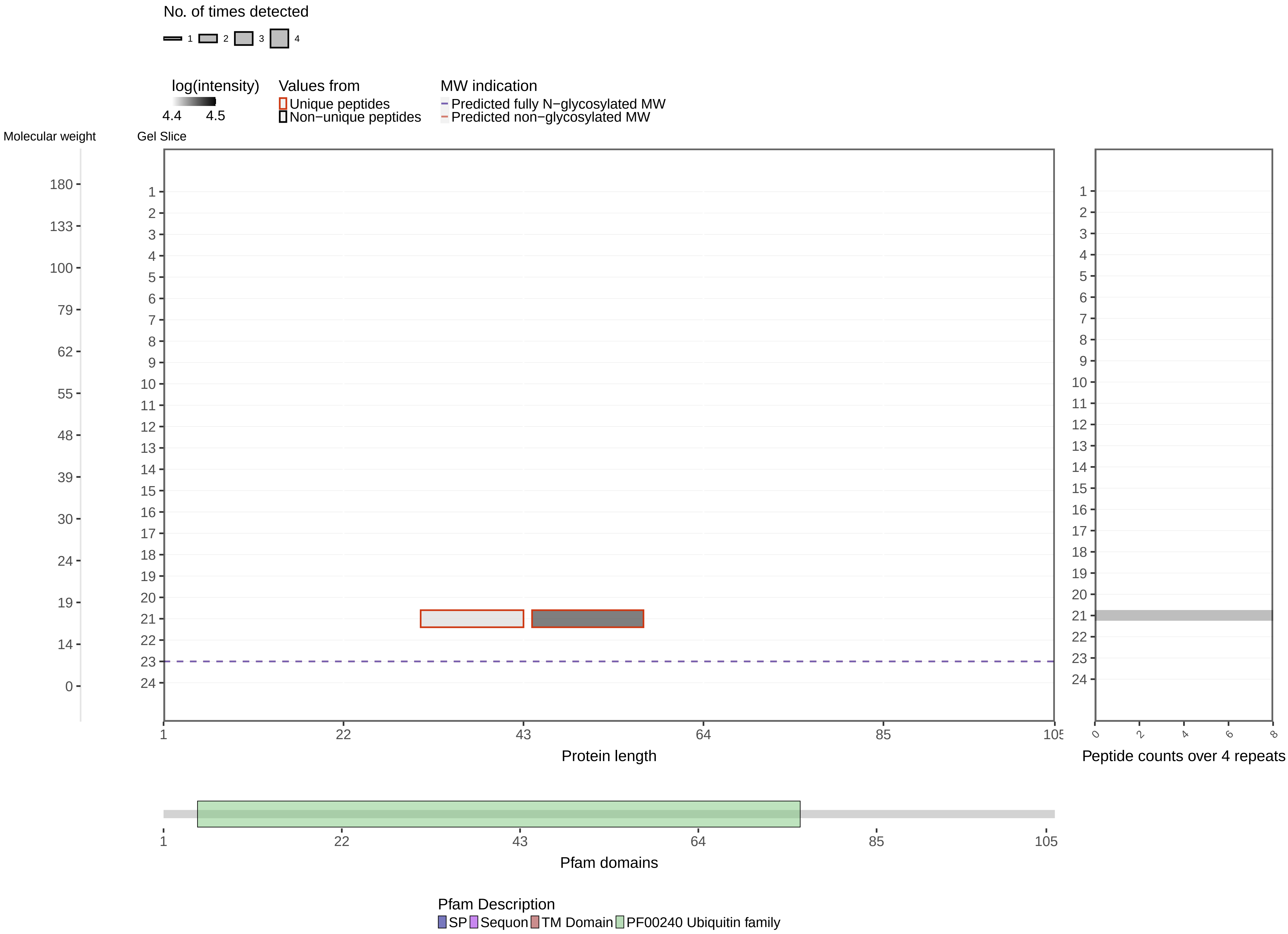


Protein sequence:

DQRLILVATGDVLDDSKSLADQKVENDAVVALTLRKDDNEFEDANIVRPNDFYQSRDAESGASW*

NbL12g19130.1 chlorophyll_a−b_binding_protein_6A_chloroplastic−like [MW = 26.24 kDa]

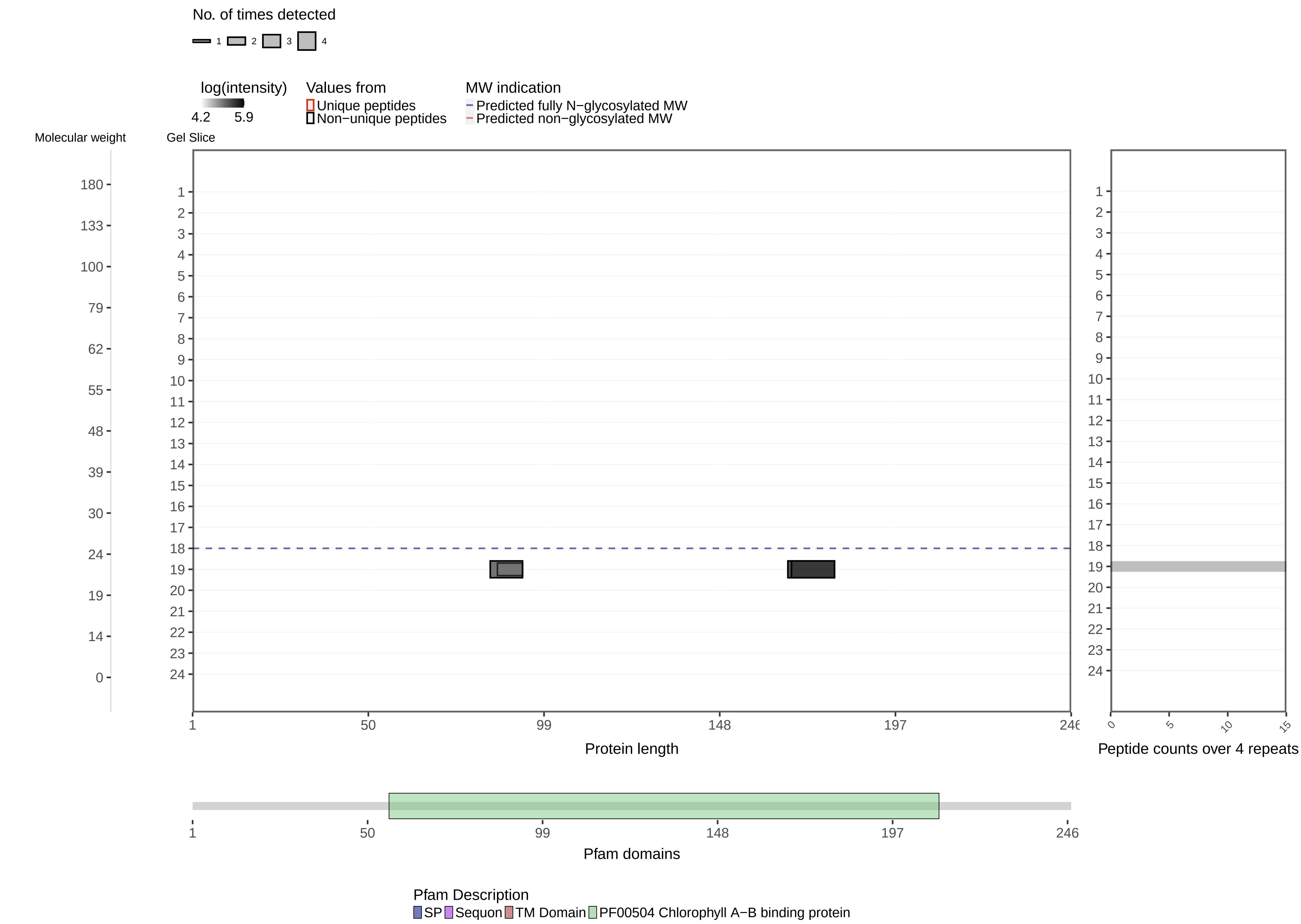


Protein sequence:

GLGNWVKAQEWAAIPGGQATYLGQPVPWGTLPTILAIEFLAIAFVEHQRSMEKDPEKKKYPGGAFDPLGYSKDPKKFEELKVKEIKNGRLALLAFVGFCVQQSAYPGTGPLENLATHLADPWHNNIGDVIIPRGILP*

### NbL12g19100.1 chorismate_synthase_1_chloroplastic [MW = 20.84 kDa]

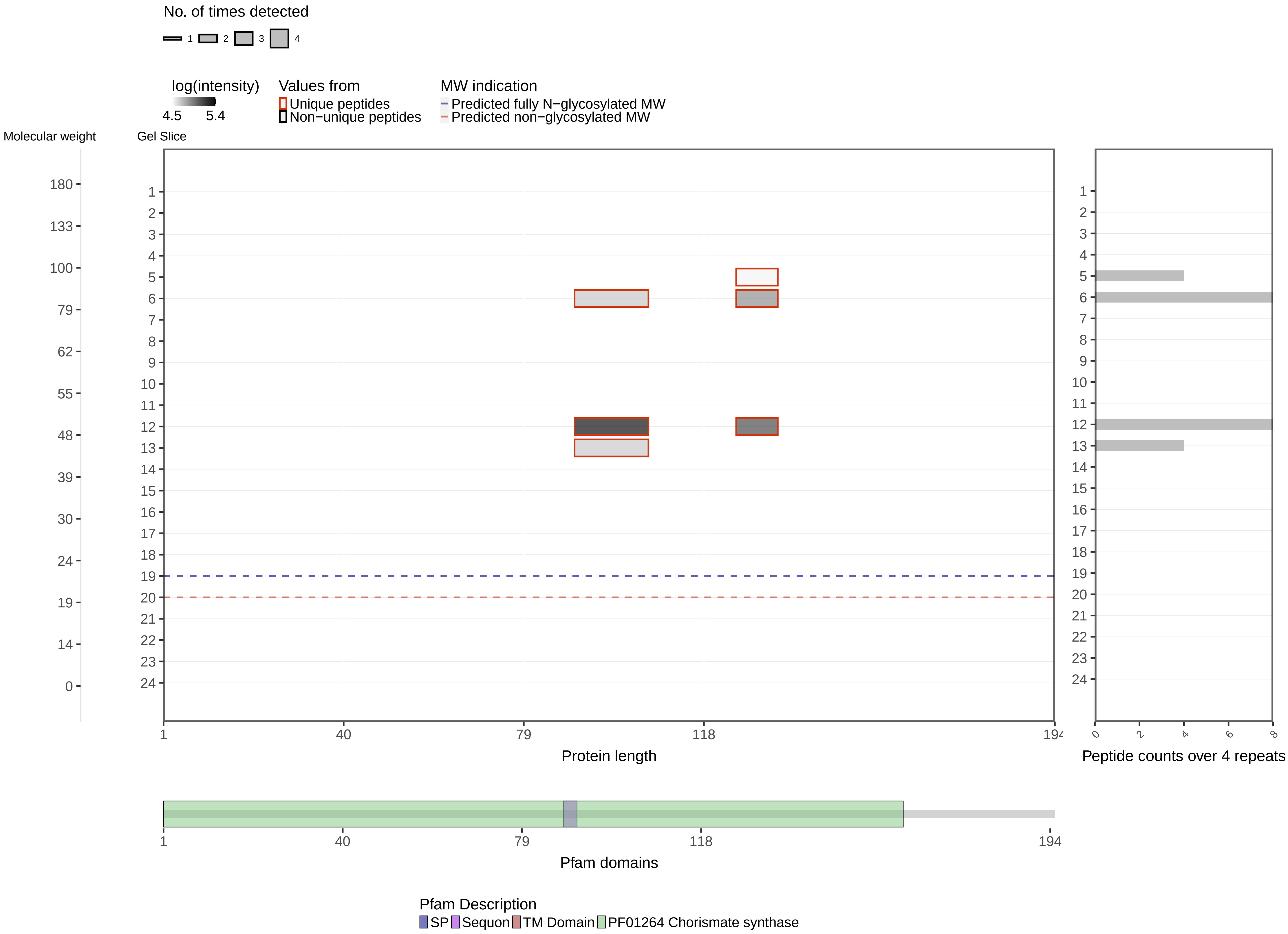


Protein sequence:

IRTRTNRSGGIQGGISNGEVINMKIAFKPTSTIARKQQTVTRDKHDTELIARGRHDPCVVPRAVPMVEAMVALALVDQLMAHYAQCMLFPINPALQEPLQSSTPESAEVTL*

### NbL12g19050.1 V−type_proton_ATPase_subunit_G_1 [MW = 12.33 kDa]

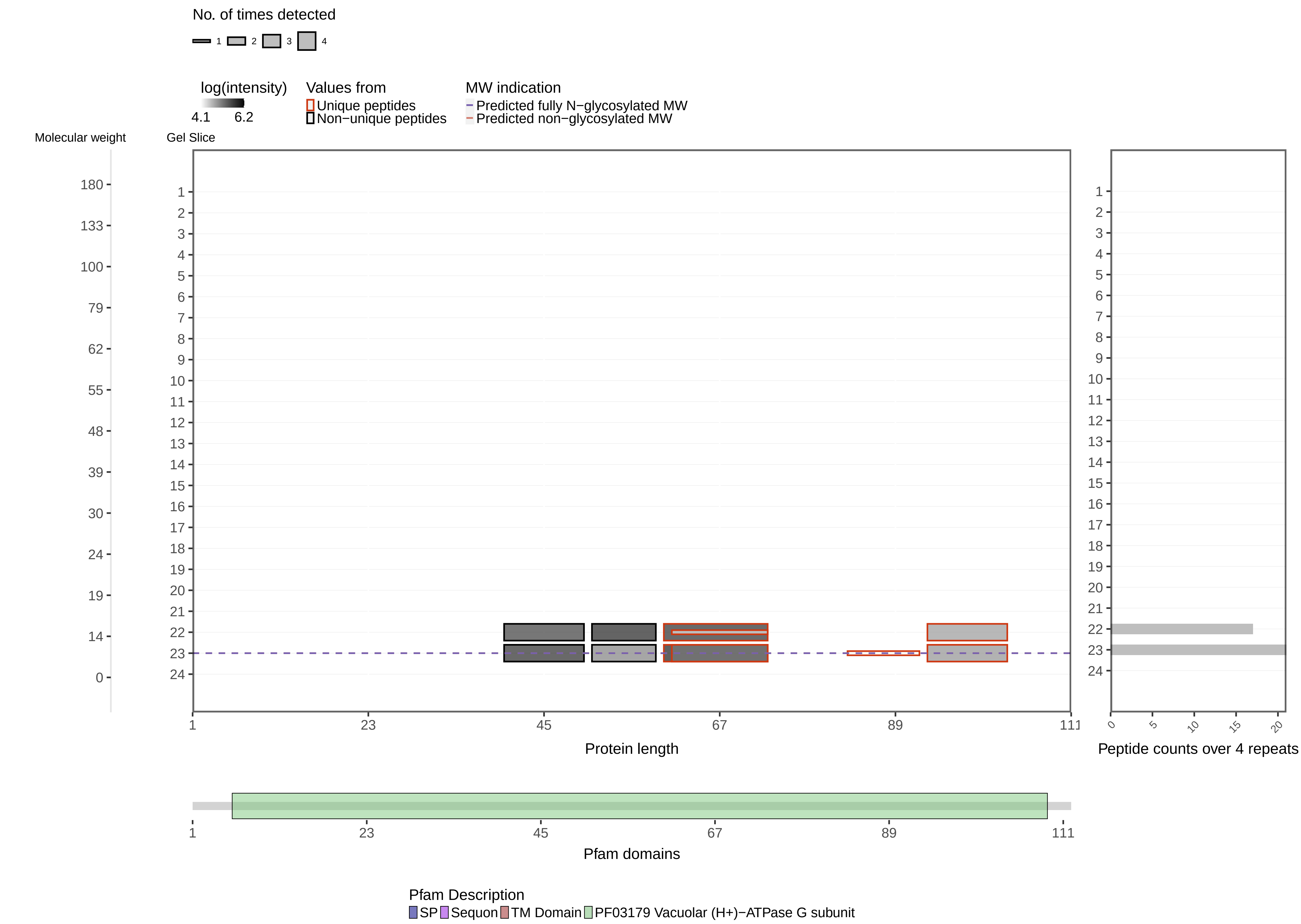


Protein sequence:

EKEIAEFRAYMEAEFQRKLEQTSGDSGANVKRLEQETDAKIEHLKTEAERVSPDVVQMLLRHVTTVKN*

NbL12g19020.1 protein_disulfide_isomerase−like_1−3 [MW − SP = 61.17 kDa]

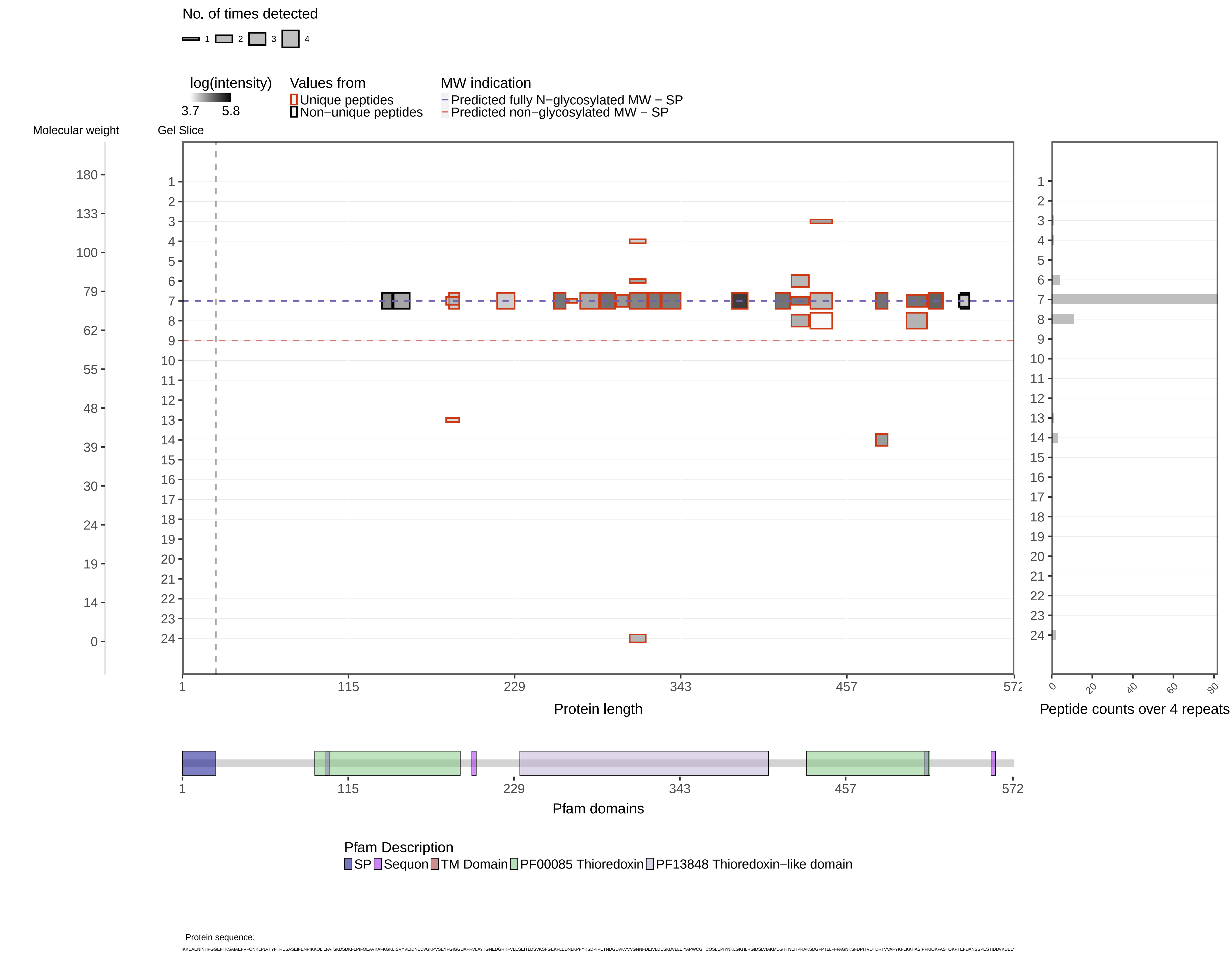

### NbL12g18780.1 leukotriene_A−4_hydrolase_homolog [MW = 68.54 kDa]

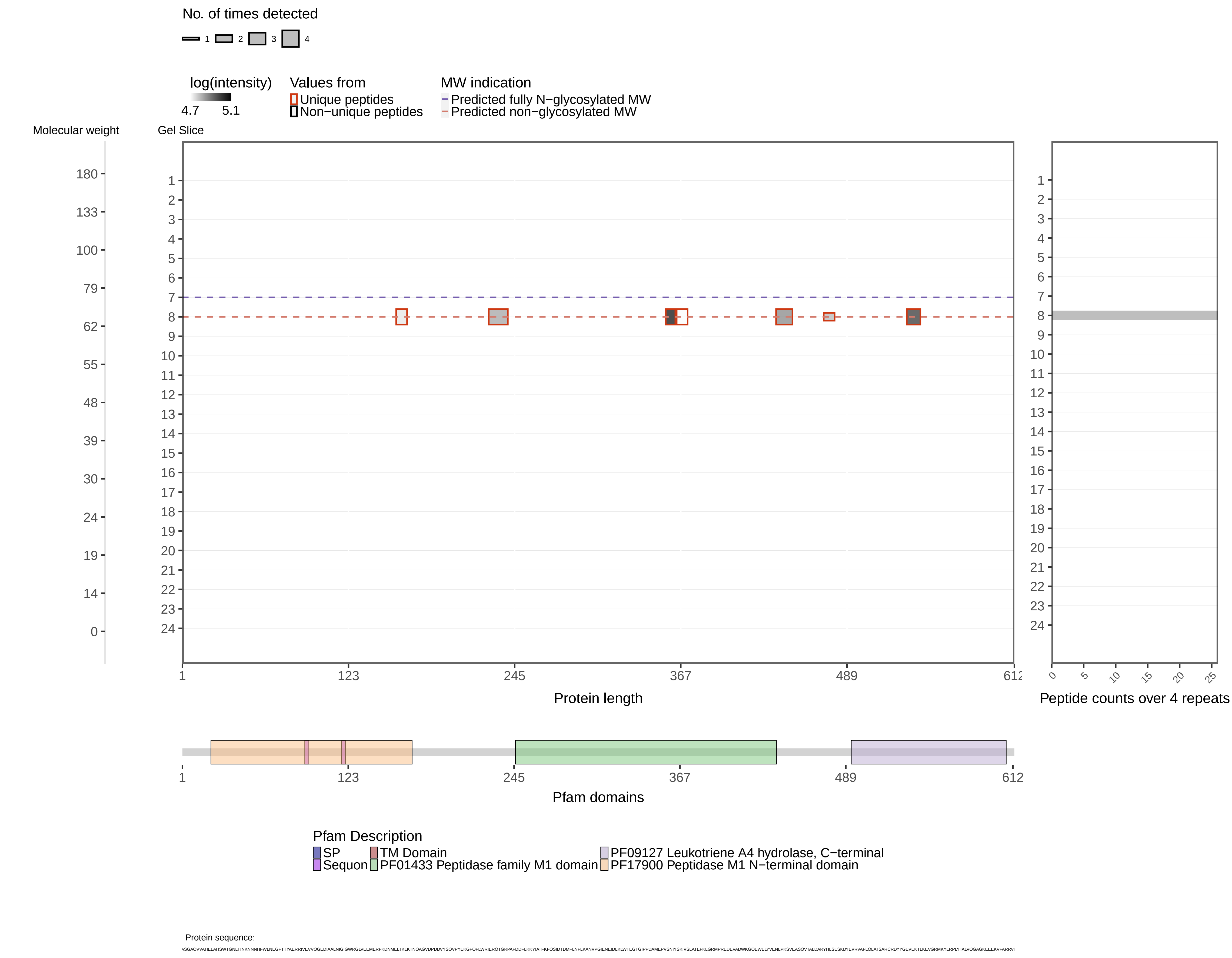

### NbL12g18630.1 uncharacterized_LOC107816357 [MW = 265.67 kDa]

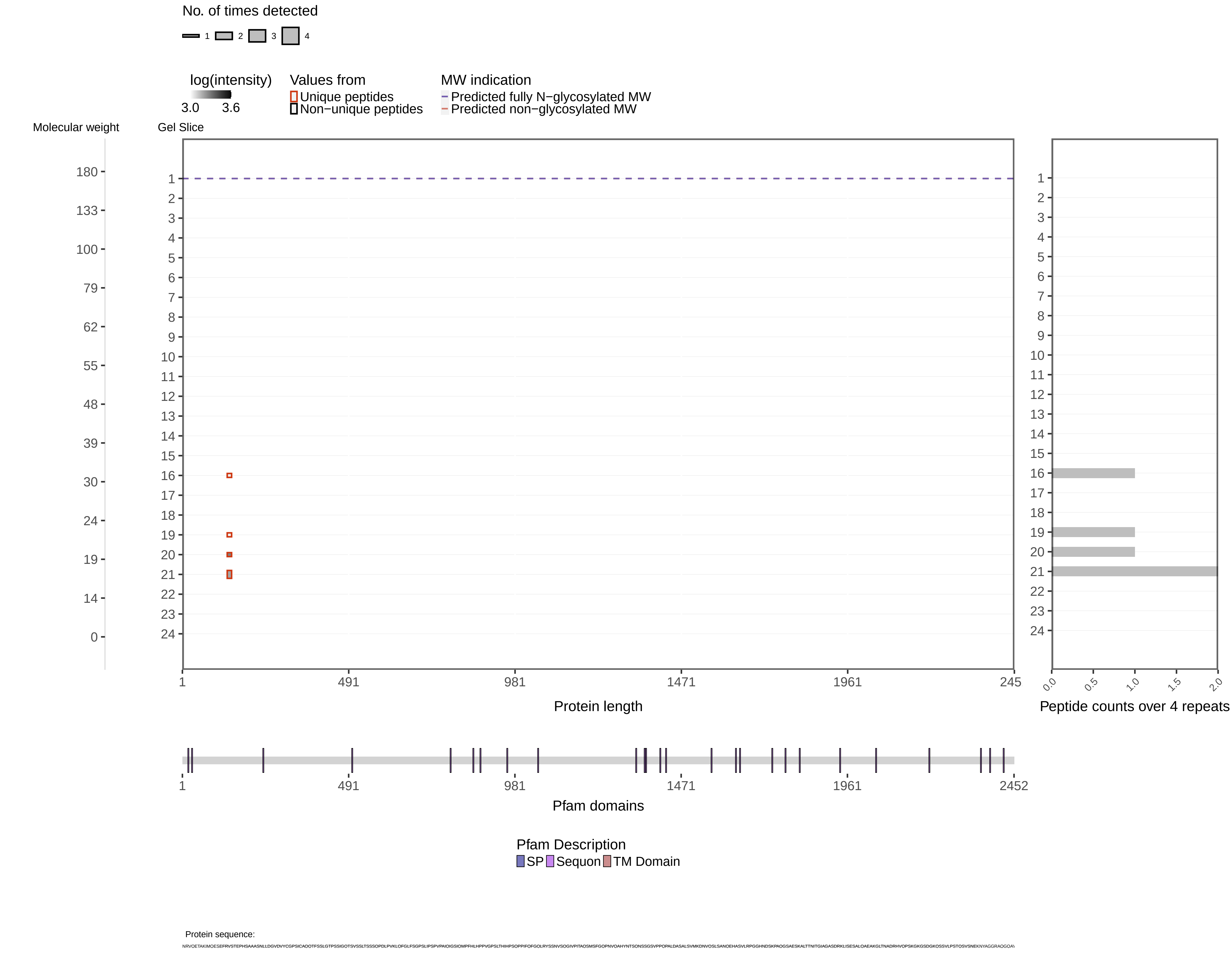

### NbL12g18540.1 methionine_aminopeptidase_1D_chloroplastic/mitochondrial [MW = 33.77 kDa]

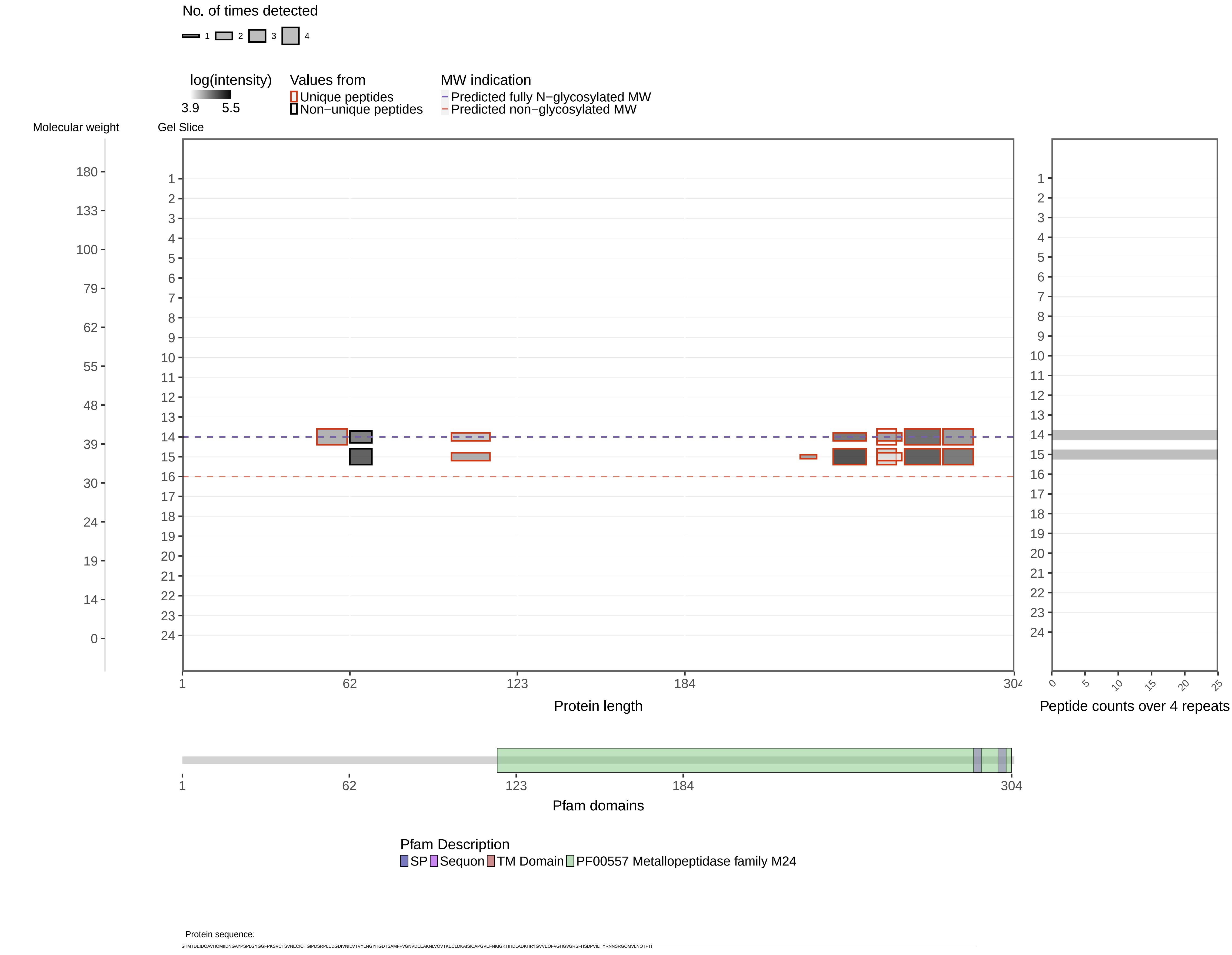

NbL12g18430.1 3−ketodihydrosphingosine_reductase−like [MW = 28.79 kDa]

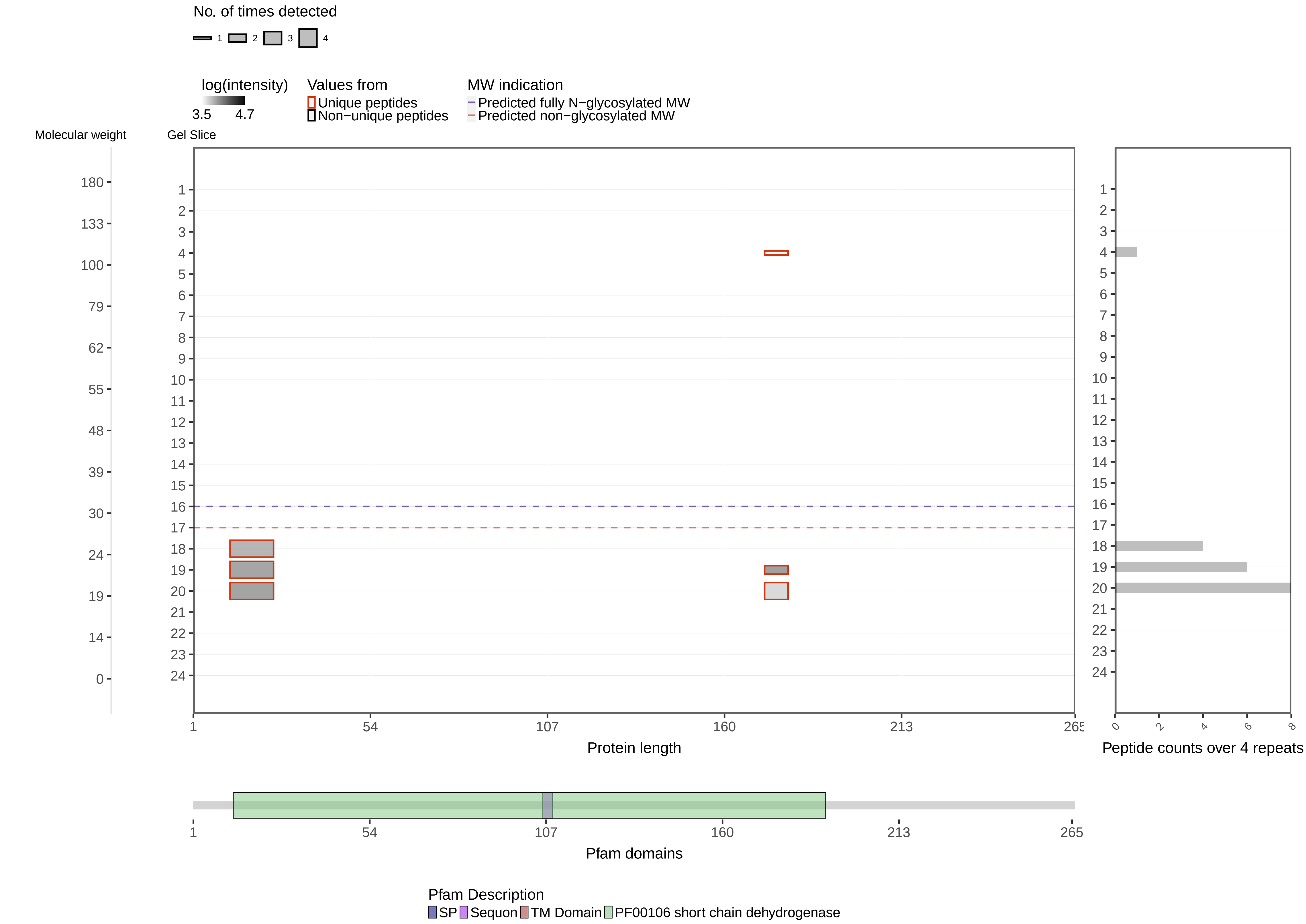


Protein sequence:

SVGAFHCAOOVLPGMVERGRGTVLFSGCSASLNGIAGYSELCCGKFAMRALSOCVAREFOPLGIHVAHVIIHGNVGAPRGSMVSTSONRLLVGEOOOOOOSGGRDGLMDPDALAOSYWHLHIODRSAWTOEIDLHPSNPRYM*

NbL12g18370.1 peroxidase_10−like [MW − SP = 32.89 kDa]

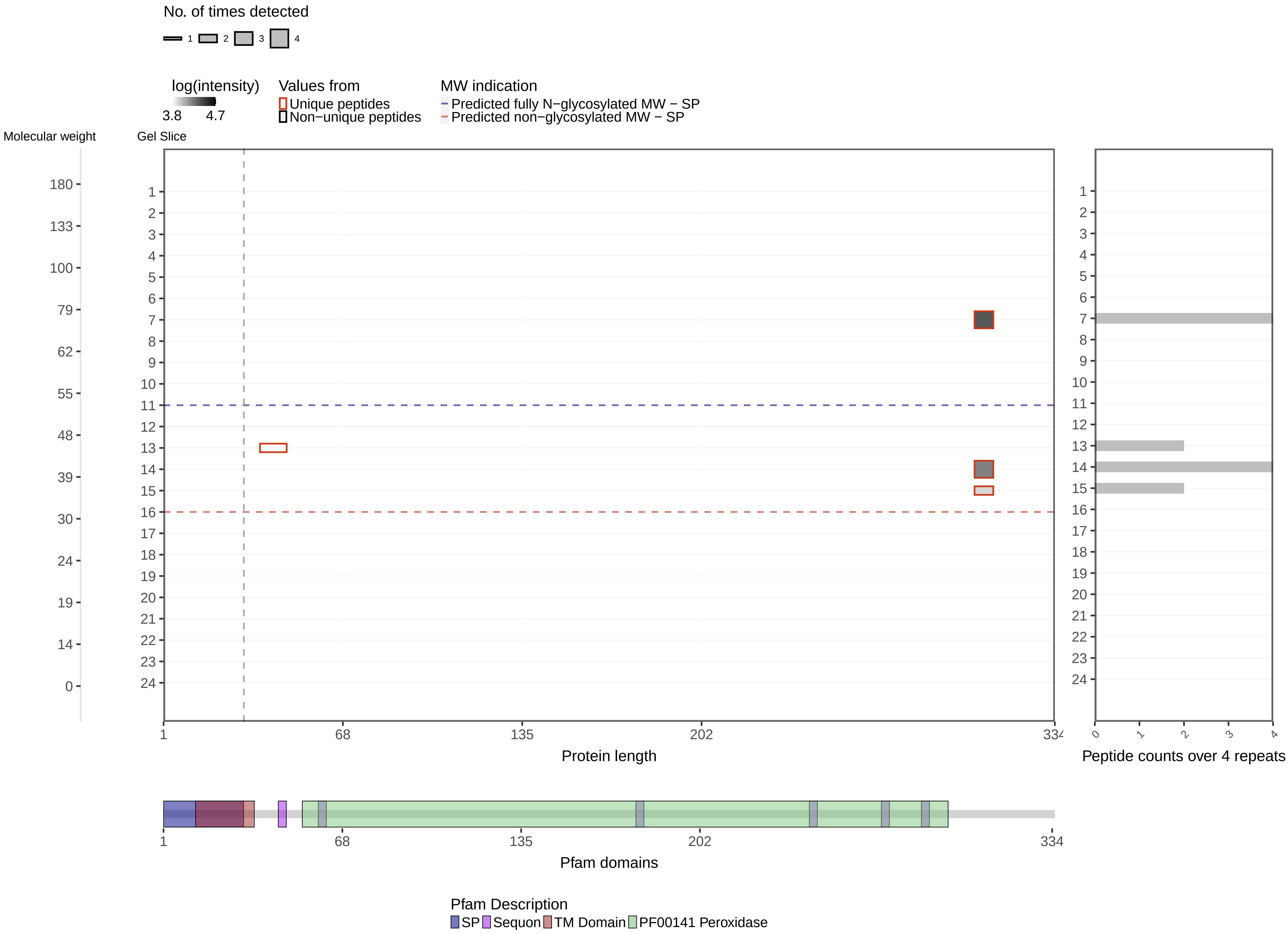


Protein sequence:

RDSLIASQSAANAQIPSPFEPLQNITAKFVSKGLDVKDVVVLSGAHTIGFAQCFTFKQRLFDFDGSGNPDPTLDSSLLGNLRSVCPNQNDSDSNLAPLDAVTINKFDNVYFKNLMNNSGLLQSDQALMNDNNTASMVSTYSKYPYLFSKAFAASMVKLTNLGVLTGQDGEIRKNCRLMN*

### NbL12g18290.1 proteasome_subunit_beta_type−6−like [MW = 25.17 kDa]

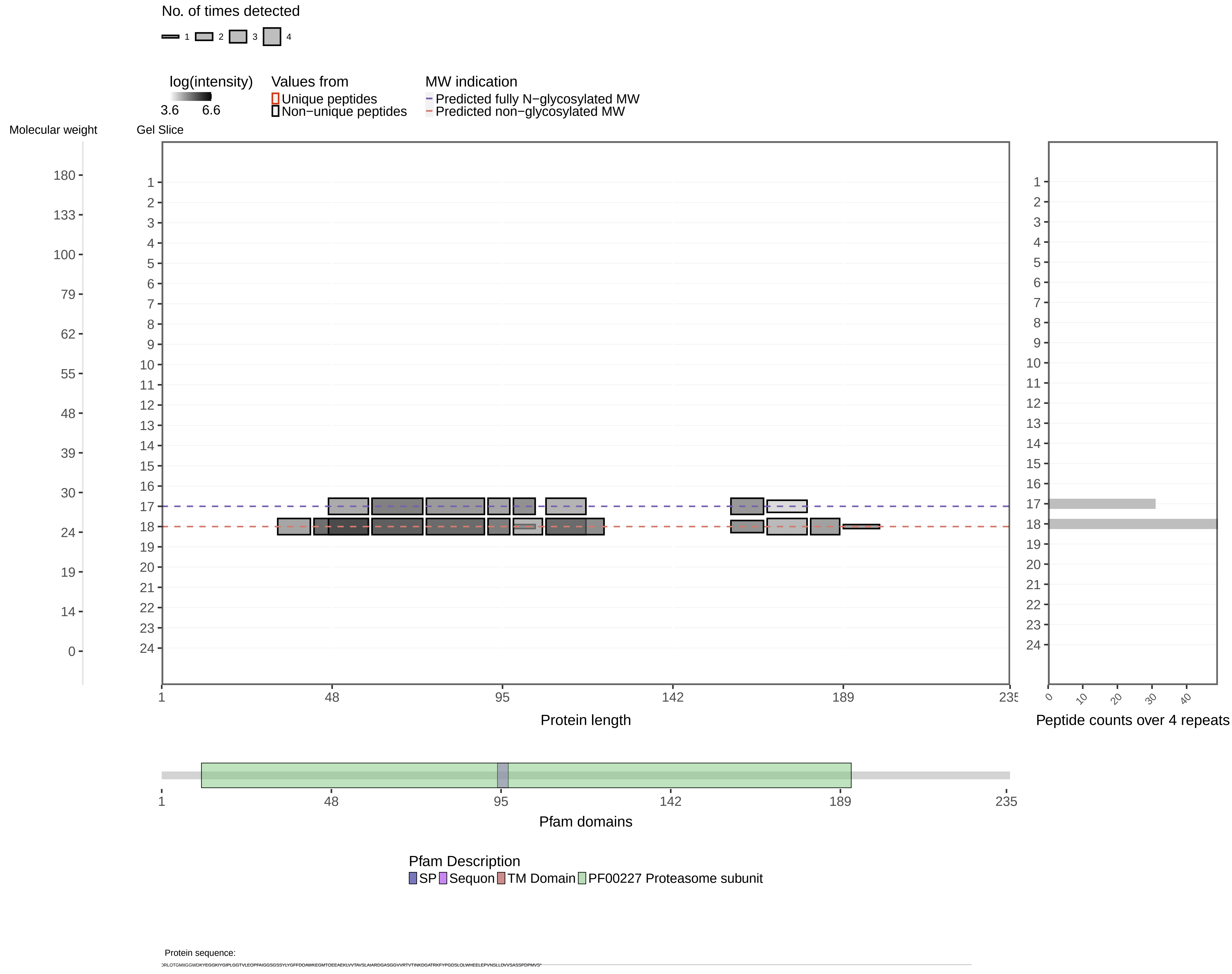

### NbL12g17850.1 alanine−−glyoxylate_aminotransferase_2_homolog_1_mitochondrial−like [MW = 42.22 kDa]

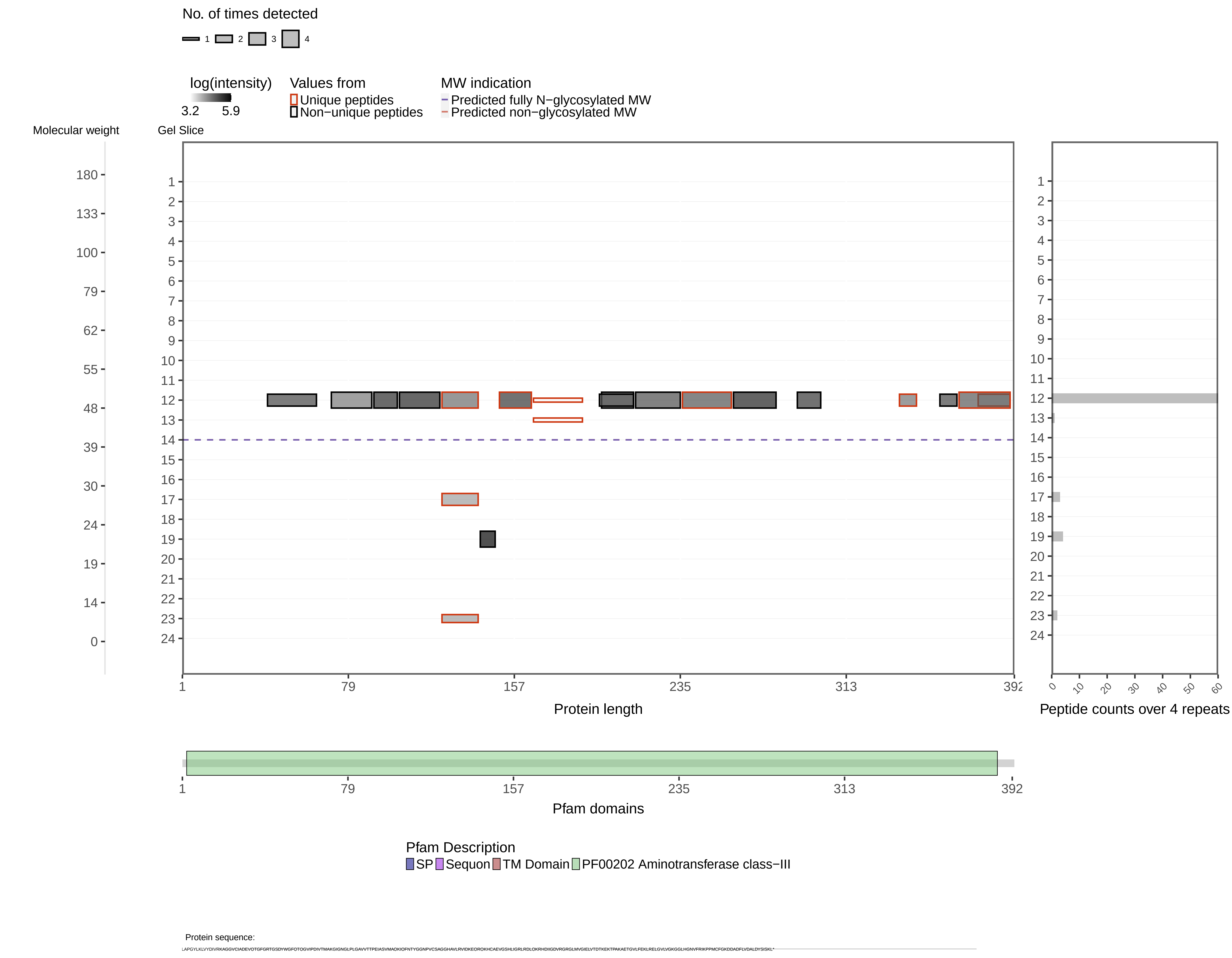

### NbL12g17760.1 uncharacterized_LOC107825767 [MW = 65.74 kDa]

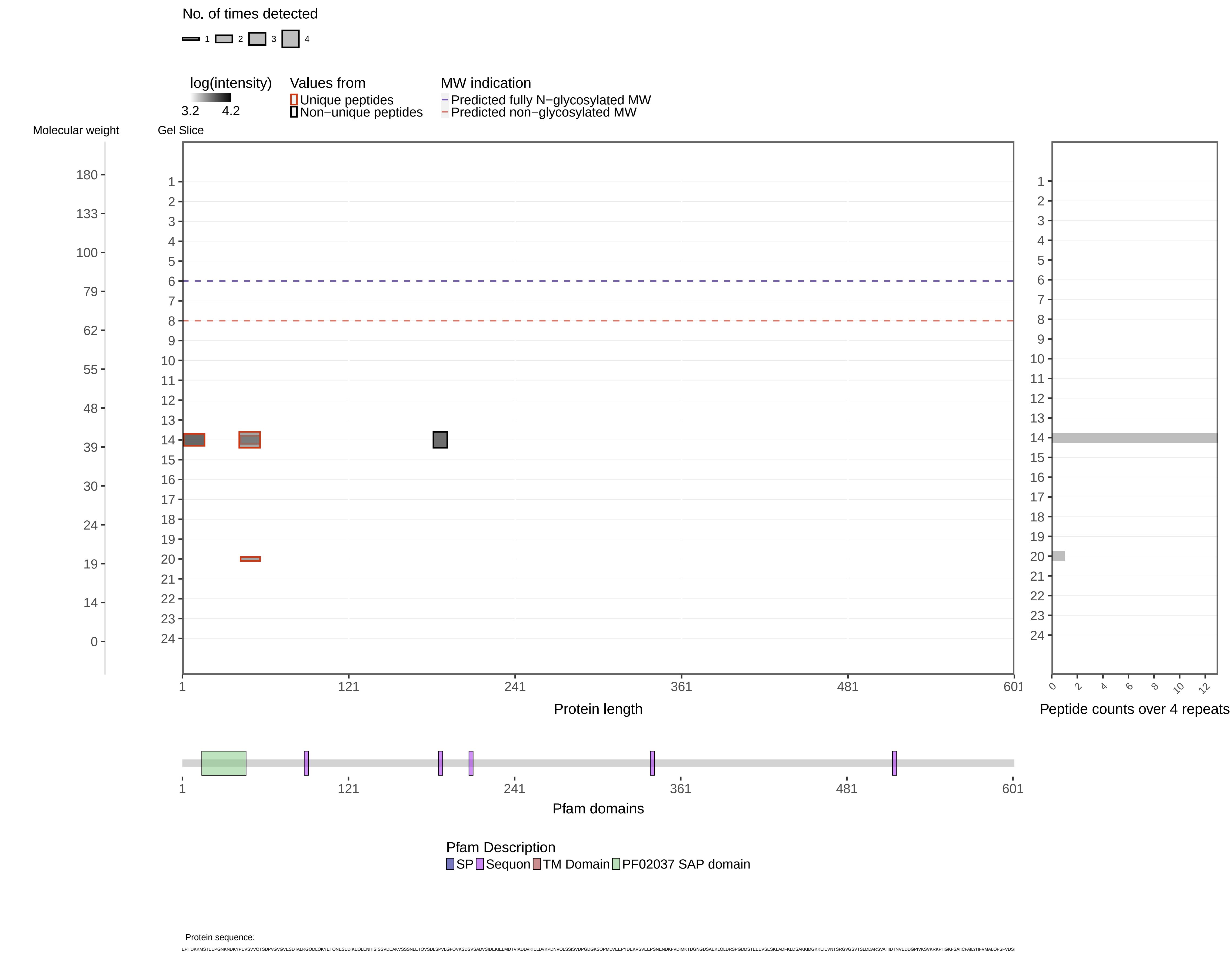

### NbL12g17750.1 apoptotic_chromatin_condensation_inducer_in_the_nucleus−like [MW = 110.72 kDa]

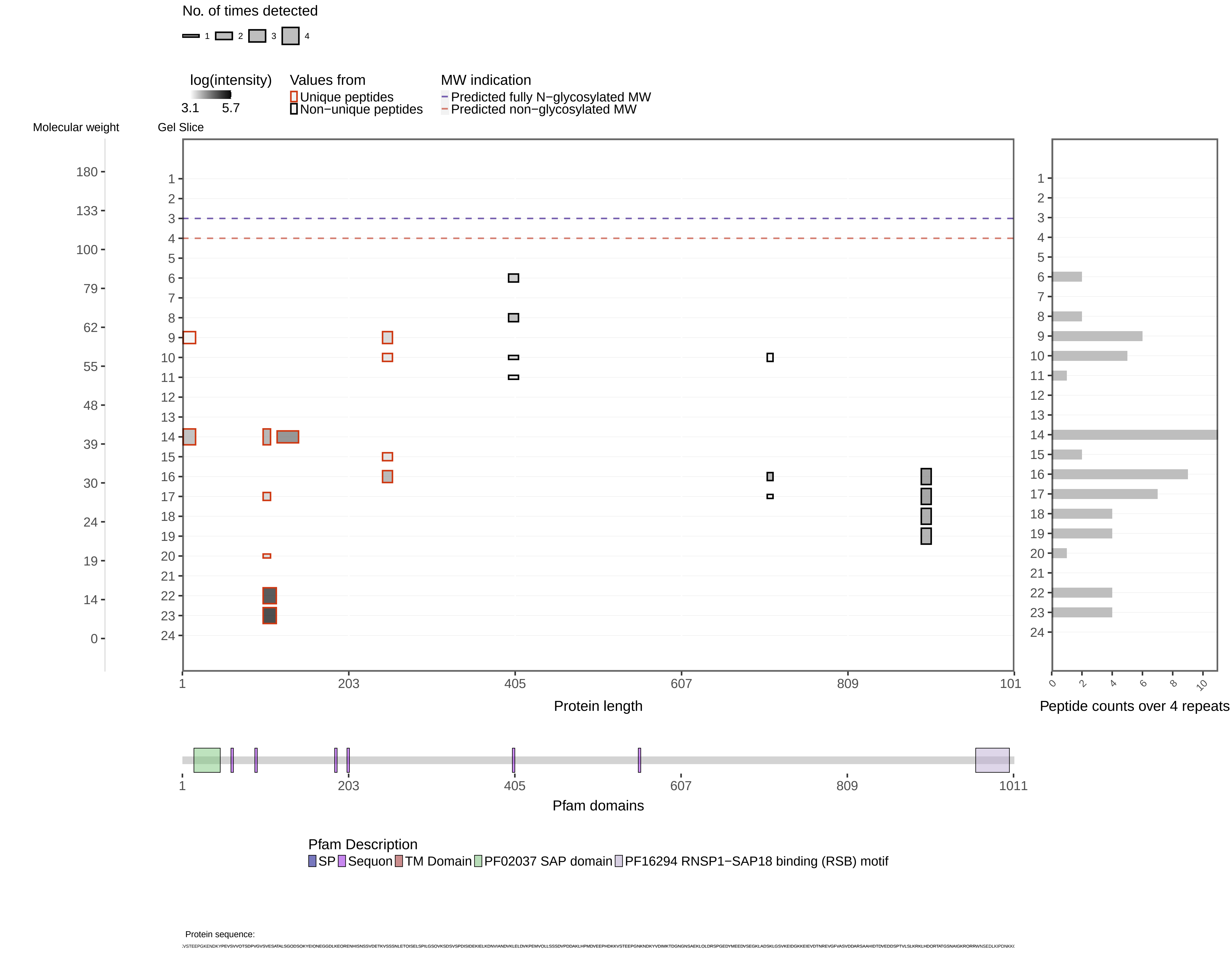

NbL12g17690.1 photosynthetic_NDH_subunit_of_lumenal_location_4_chloroplastic [MW = 22.09 kDa]

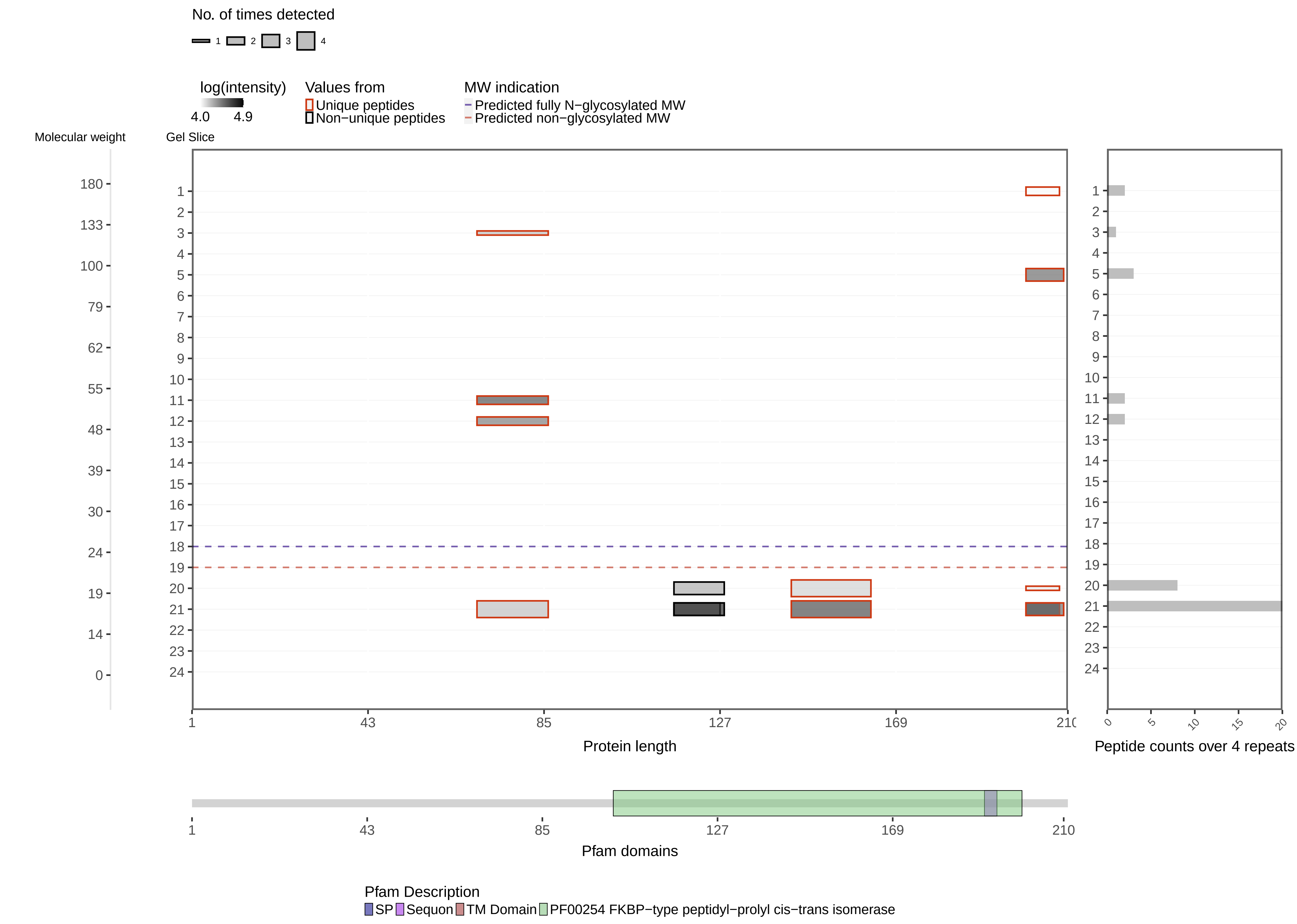


Protein sequence:

IAVGSGQEAPYNQLINIHYTARFGDGIVFDSSYKRGRPLTMRLGMGKVIKGLDQGILGGEGVPPMHIGGKRKLQIPPHLAYGPEPAGCFSGDCNIPGNATLVYDIKFVDIYSGNRK*

NbL12g17670.1 L−ascorbate_oxidase [MW − SP = 61.41 kDa]

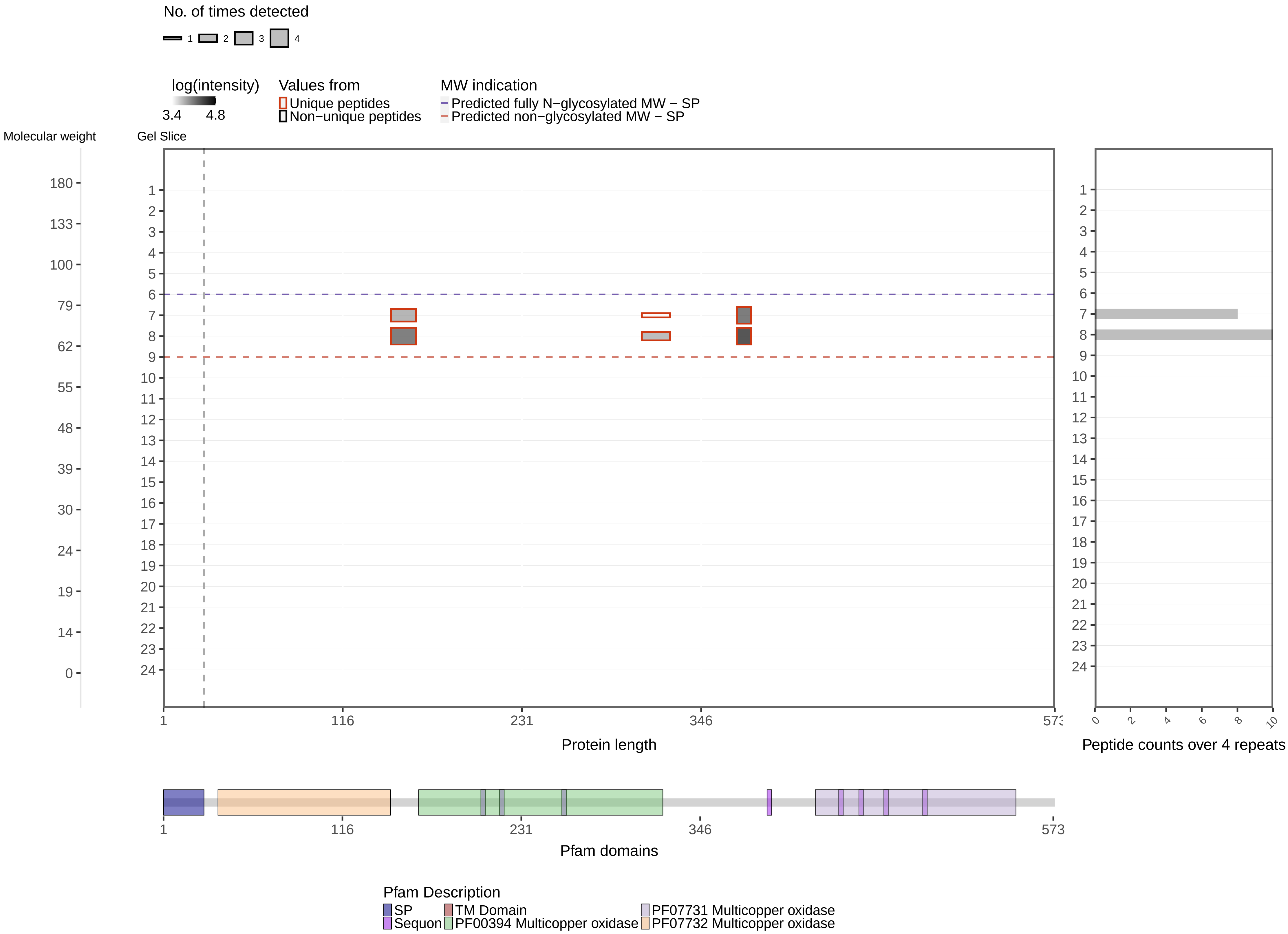


Protein sequence:

NLFLYSGETYSVLIKADQDPSRNYWASTKVVSRPSATPNGLGIISYYPNHPRRAPPTIPPAGPHWNDVAPRVAQSVAIKSHKDFIHAPPKTSDRIIVMLNTQNRINGFVRWSINNVSFNMPHTPYLIALKHNLLHTFEQTPPPENYNQENYDIFSVQKNVNATTSNSIYRLKFNTTVDIILQNANTMNPNNSETHPWHLHGHDFWVMGYGNGKFNQSIDPKNYNLVNPIMKNTIPVHPYGWTALRFRADNPGVWAFHCHIESHFFMGMGVVFEEGIDKVGKLPTSIMGCGDSK

NbL12g17630.1 bifunctional_aspartate_aminotransferase_and_glutamate/aspartate−prephenate_aminotransferase−like [MW = 52.49 kDa]

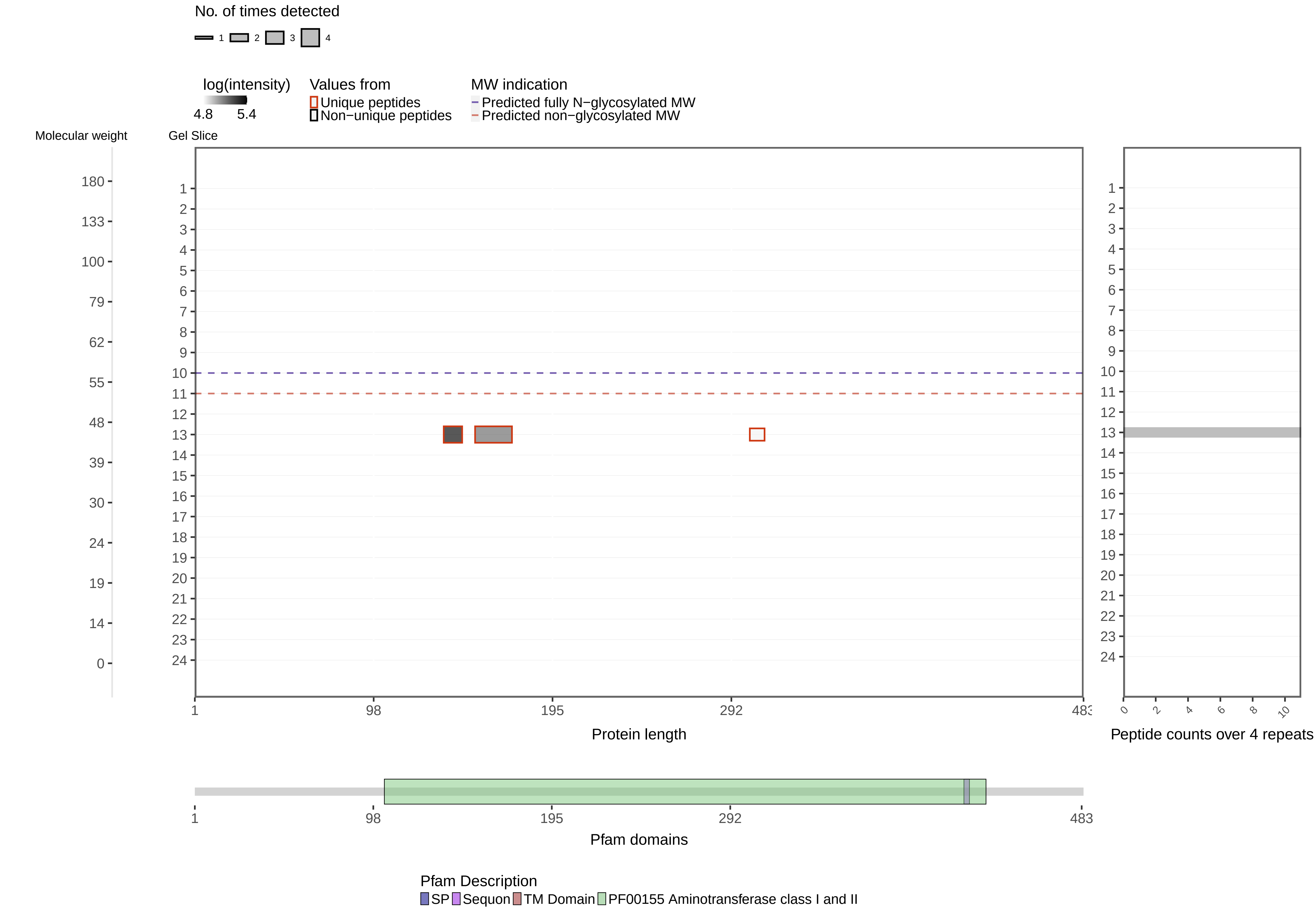


Protein sequence:
ESKLTEKSRLLILCSPSNPTGSVYPRKLLEEIAEIVARHPRLLVISDEIYEHIIYAPATHISFASLPGMWDRTLTINGFSKAFAMTGWRLGYIAGPKHFVSACNKIQSQFTSGASSISQKAAVAALGLGYAGGEAVATMVKAFRERRDYLVKSFGEIDGVKISEPRGAFYLFIDLSSYYGVEVDGFGSINDSESLCRYLLDKAQEMHLGMTHASVSPMQHPSRPCRQQWRESRKHWLLLDFMSQFKTSINYAC*

### NbL12g17590.1 inositol−3−phosphate_synthase [MW = 56.46 kDa]

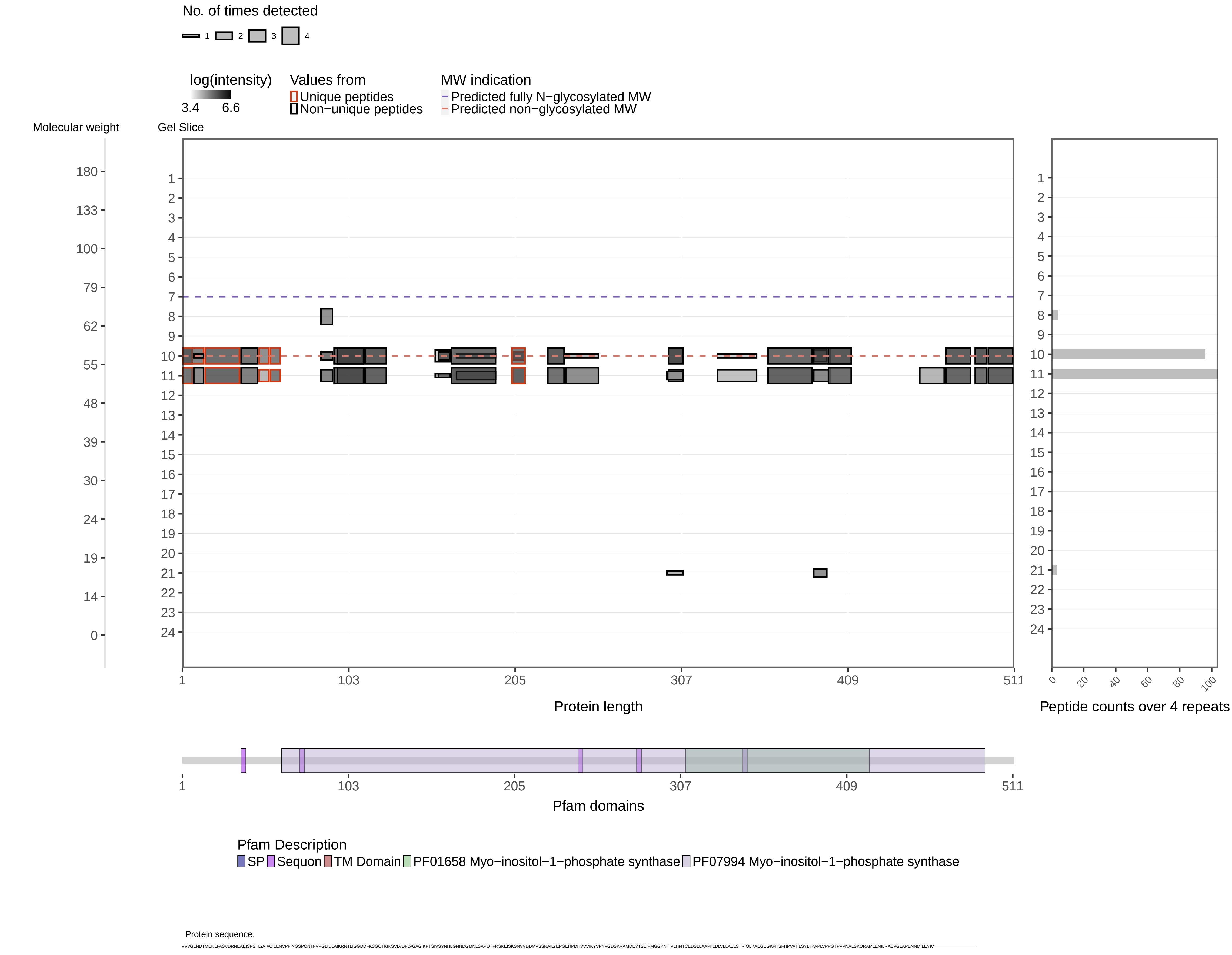

### NbL12g17270.1 annexin_D2−like [MW = 35.91 kDa]

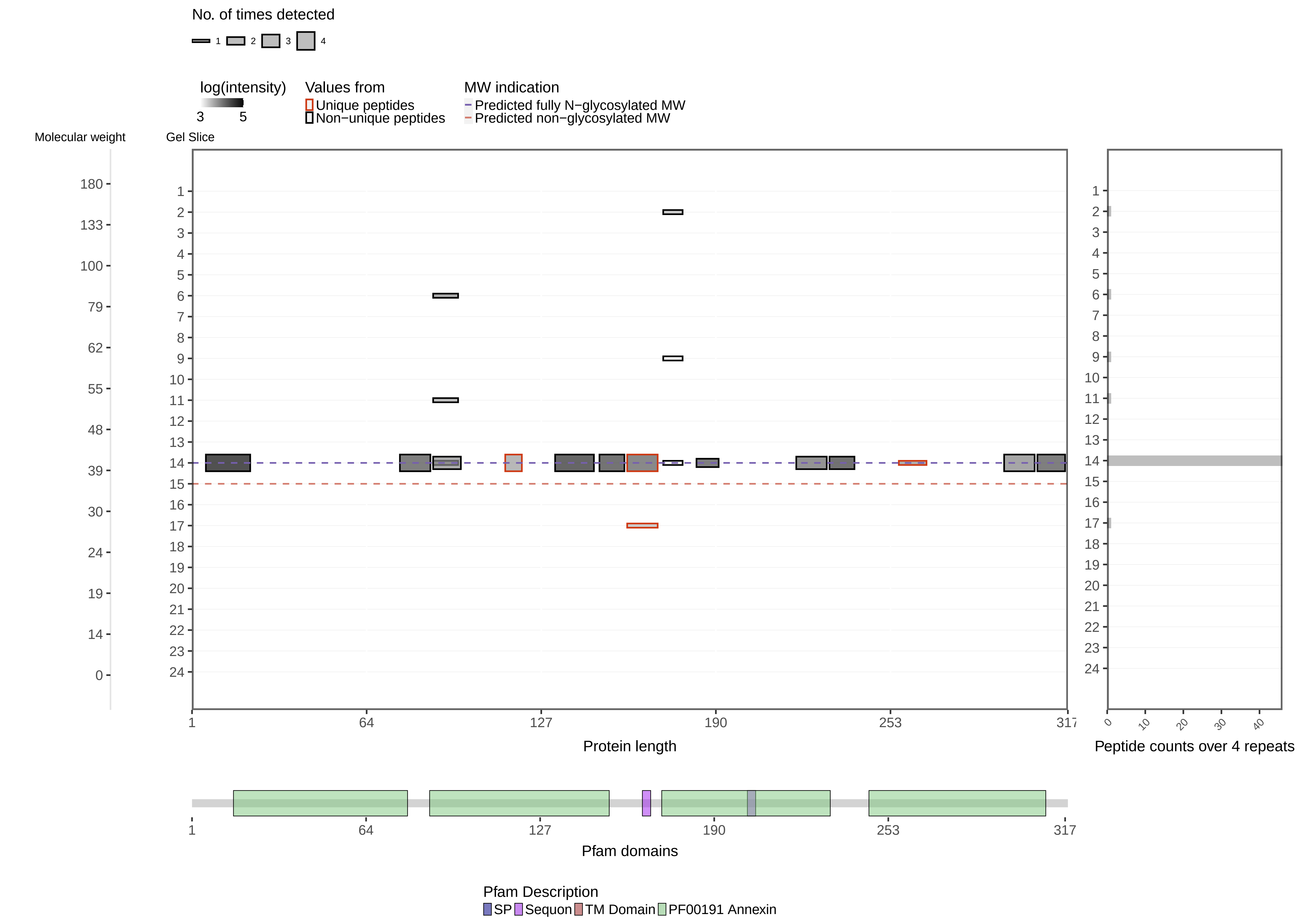


Protein sequence:

-RKLLVPLVTAFRYEGAEVNMTLARKEANILHEKISDKAYNDEELIRILSTRSKAOLNATFNHYLDOHGSEINKDLETDSDDEYLKILSAAIECLKTPEKYFEKVLRLGIKGTGTDEWDLTRVVTTRAEVDMERIKEEYOKRNSVPLDRAIAGDTSGDYERMLLALIGHGDA*

### NbL12g17250.1 asparagine_synthetase_[glutamine−hydrolyzing]_2 [MW = 67.91 kDa]

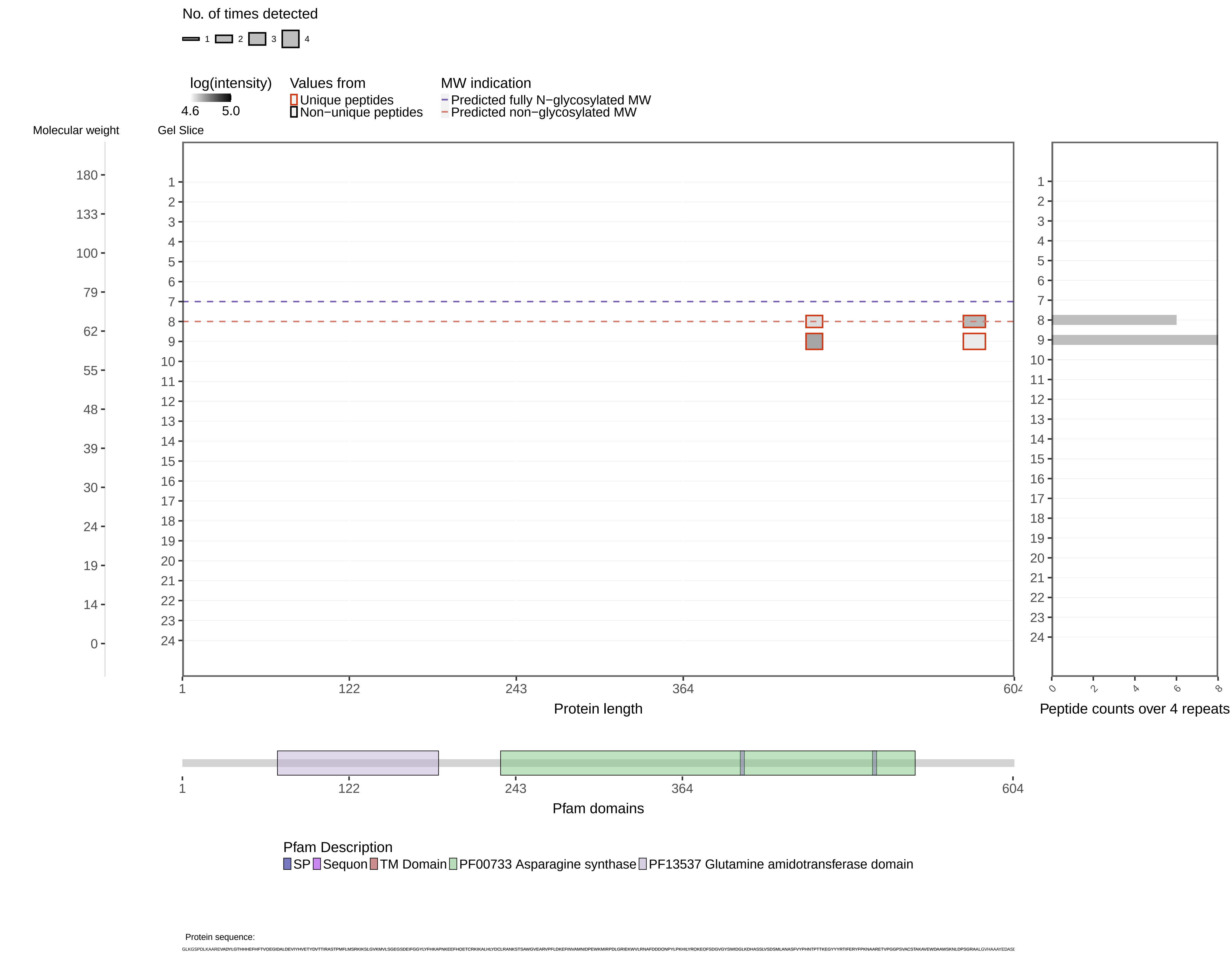

NbL12g17060.1 aspartic_proteinase−like_protein_2 [MW − SP = 49.84 kDa]

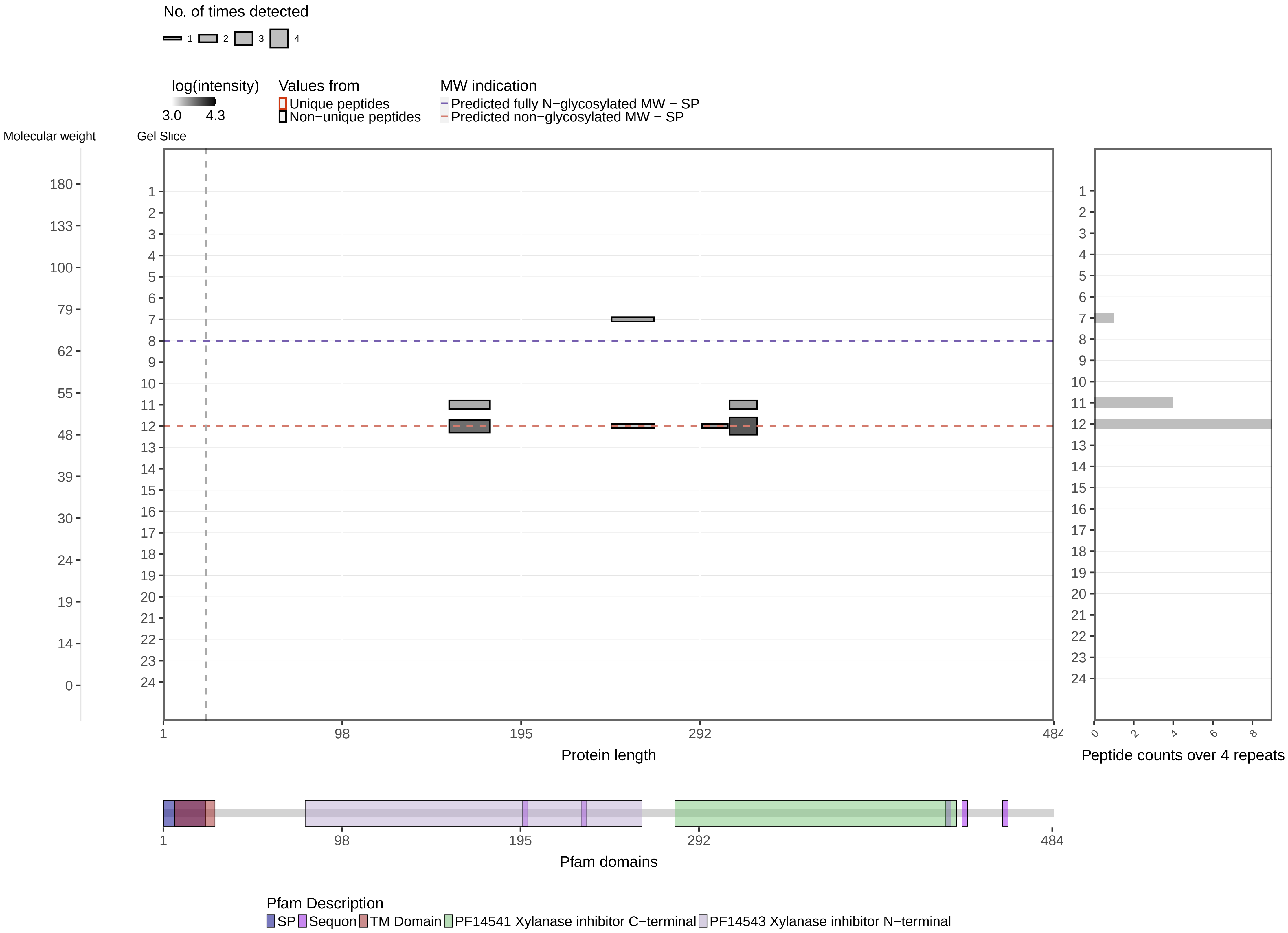


Protein sequence:

ANSSLISQLAISGKVKKVFAHCLDGNNGGGIFAIGQVVQPKVKSTPLVPNEPHYNVVMKGIEVNGKALDIPTTIFDAGSNRGTIIDSGTTLAYLPNKVYNALMSKLMVKQPDLPTHLVEGSFHCFYYSGKVDDGFPVVTFQFANSLYLTVYPHDYLFQVNDDEWCIGWQDSGMQTKDGKEITLLGDLVLSNKLVLYDLENQSIGWTEYNCSSSIKVRDETSGKDYTIAAHNISSAWNCNSRMVFAFFVLIISILCNV*

NbL12g16930.1 uncharacterized_LOC104214608 [MW = 75.67 kDa]

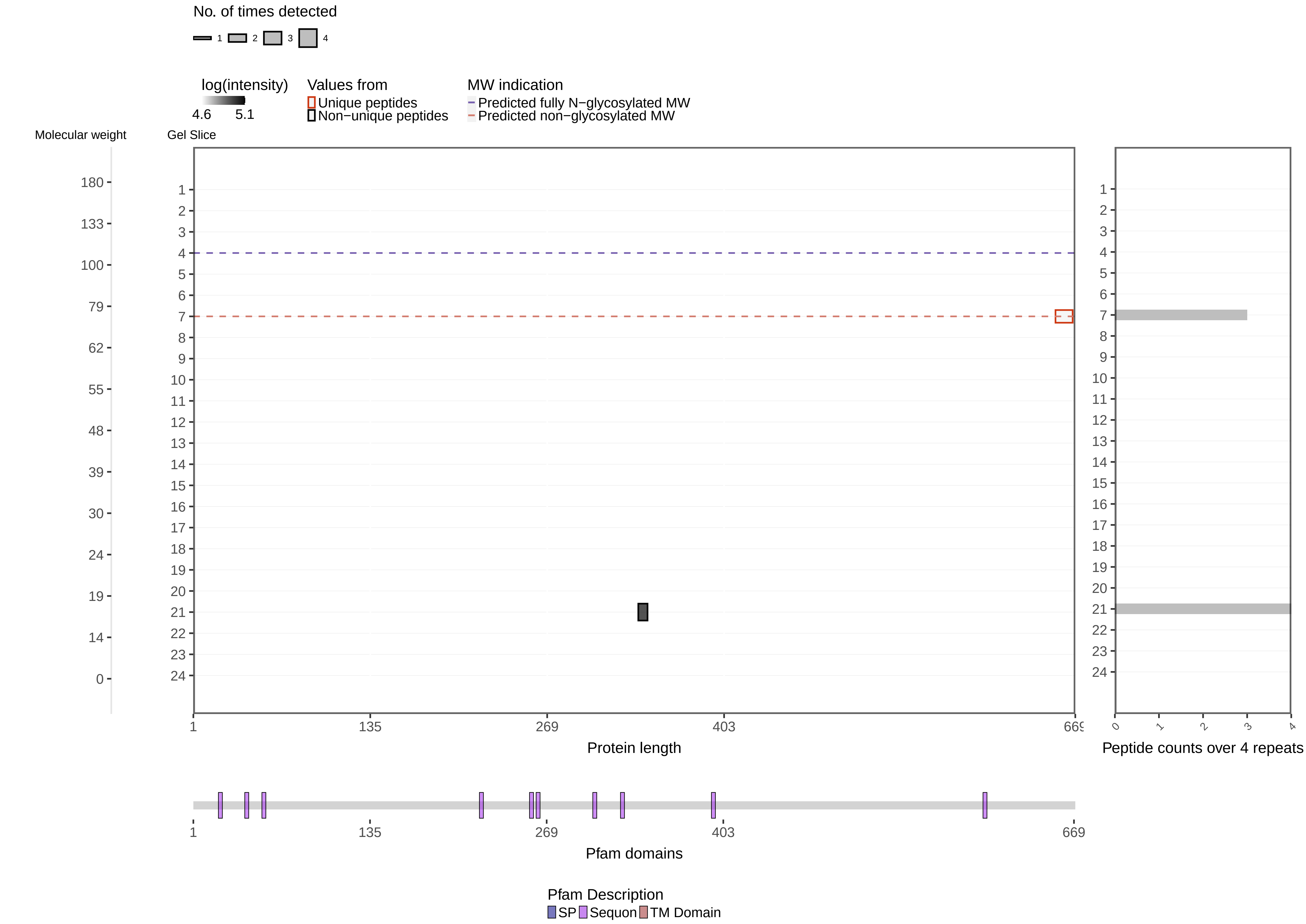


Protein sequence:

LRNGSNFDNKPVTFRGLTPNILKFNFFPRMYHNLHDLRLIQPLLHEFLPGSSFNEISQLQASVQRPNDGLINTTLYVNFLSSYIVEIKNQNVLGPVSFLADLGGLYCISIGLFFYILVQCEYRIKKLRNEDCVMRKVRSRRKAQDRWDKLRKYVMYTYAPNQLEEVYDMRNDGCCTGVKMESLNRKSSSVRGRGSTRLDTISFSRKVSLPSEKRAISEQIDTQGATHFLPESAADRRKSTRLKEERFQENAVPPSTENSKPEDLDIVIQSPVFSAADNLTLPPPPPLESQAADHI

### NbL12g16900.1 probable_UDP−arabinopyranose_mutase_5 [MW = 39.11 kDa]

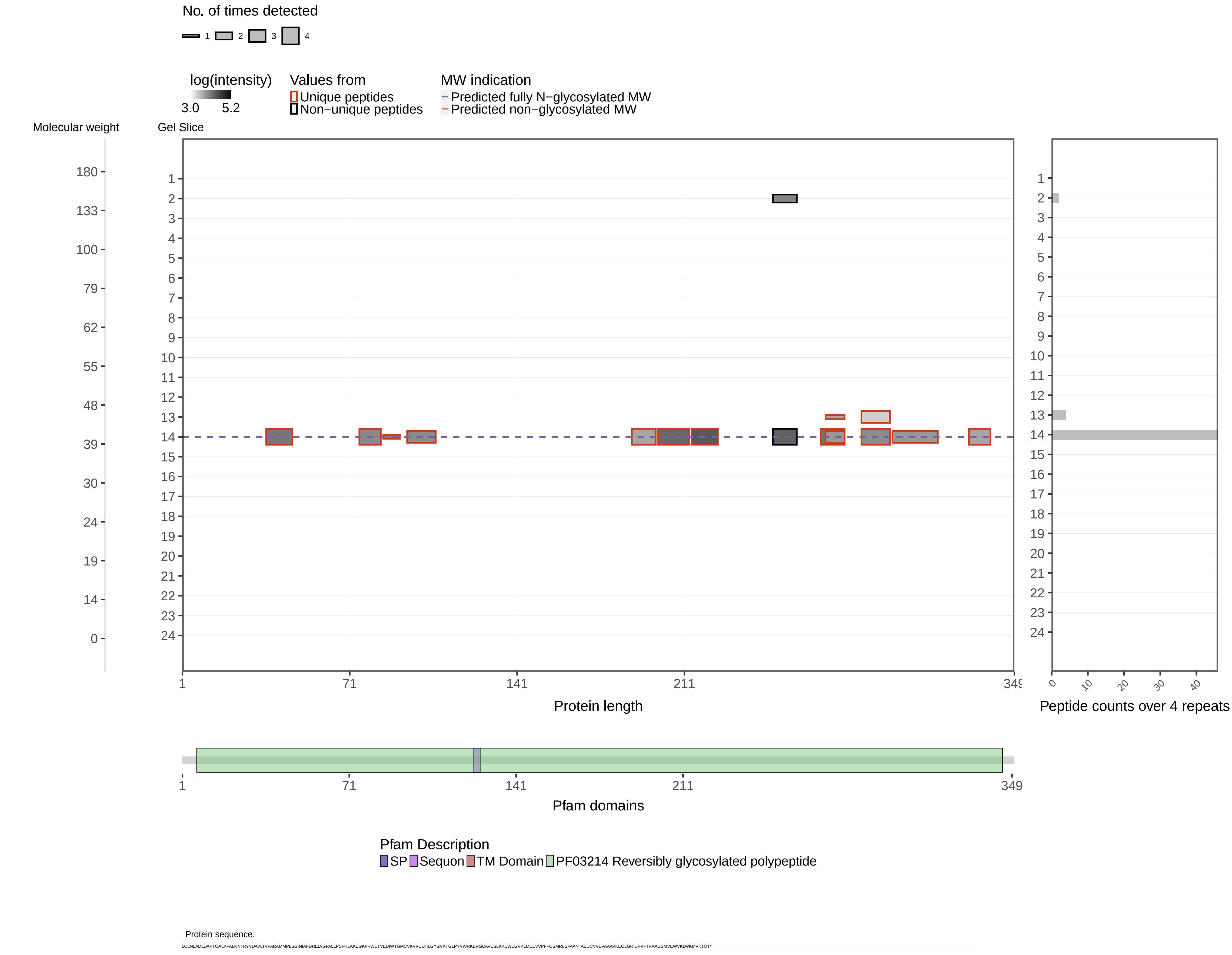

### NbL12g16800.1 violaxanthin_de−epoxidase_chloroplastic [MW = 54.82 kDa]

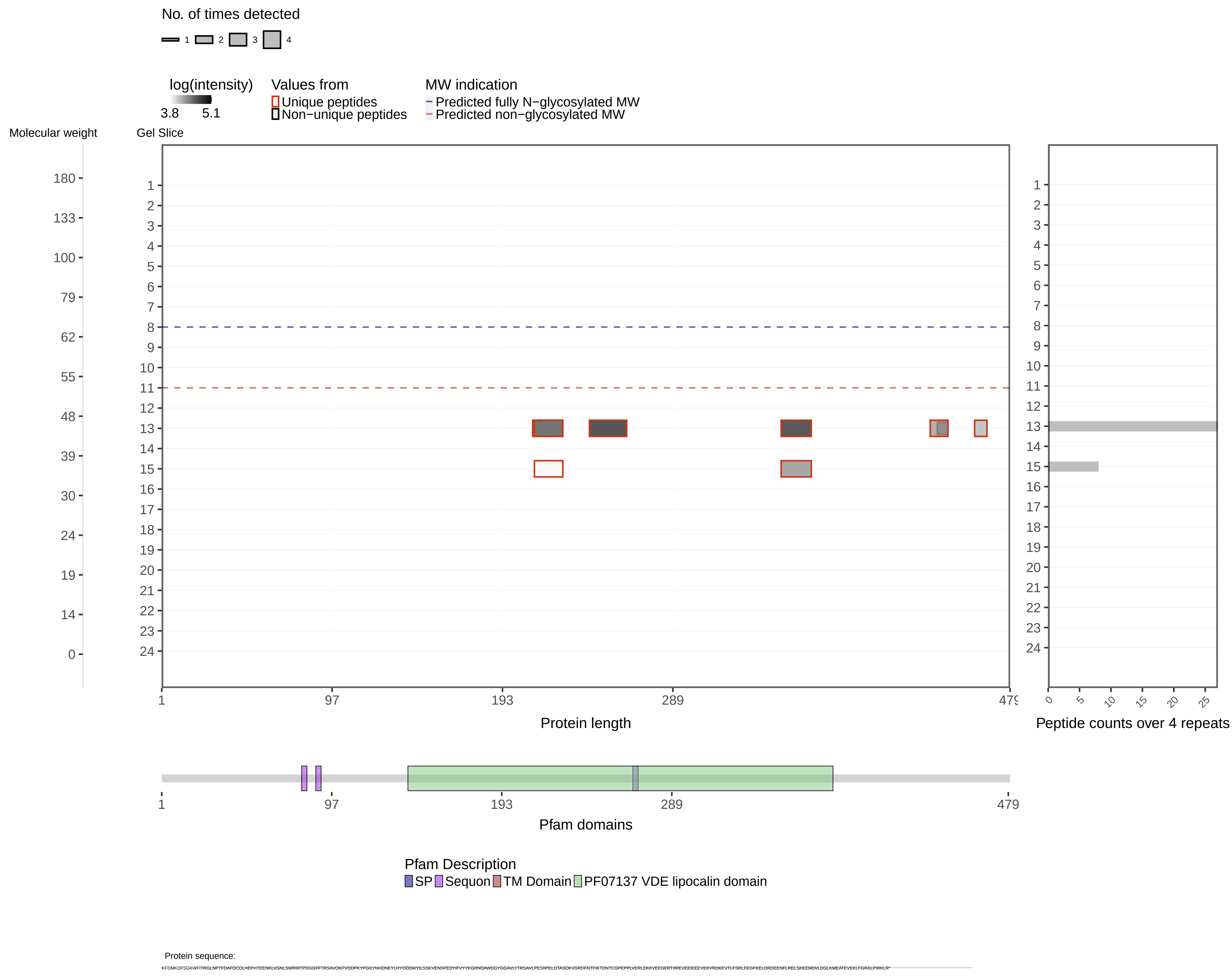

NbL12g16720.1 peroxiredoxin_Q_chloroplastic−like [MW = 24.04 kDa]

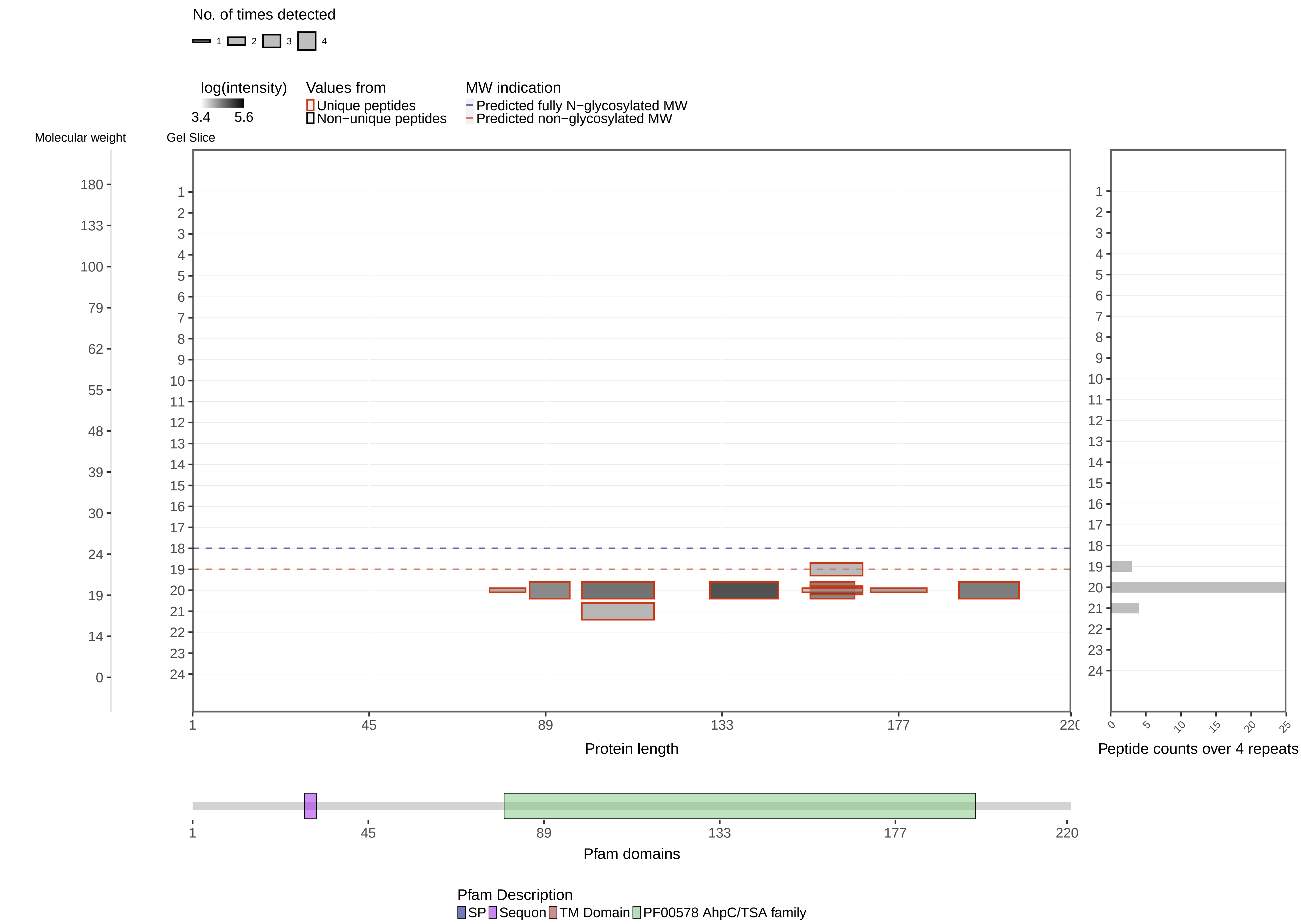


Protein sequence:

PVVVYFYPADESPGCTKQACSFRDSYEKFKQAGAEVVGISGDDPESHKAFAKKYRLPYTLLSDEGDKVRKEWGVPADLFGTLPGRQTYVLDKNGVVQLVYNNQFQPEKHIDETLKLLQSL*

### NbL12g16640.1 uncharacterized_LOC104113886 [MW = 107.6 kDa]

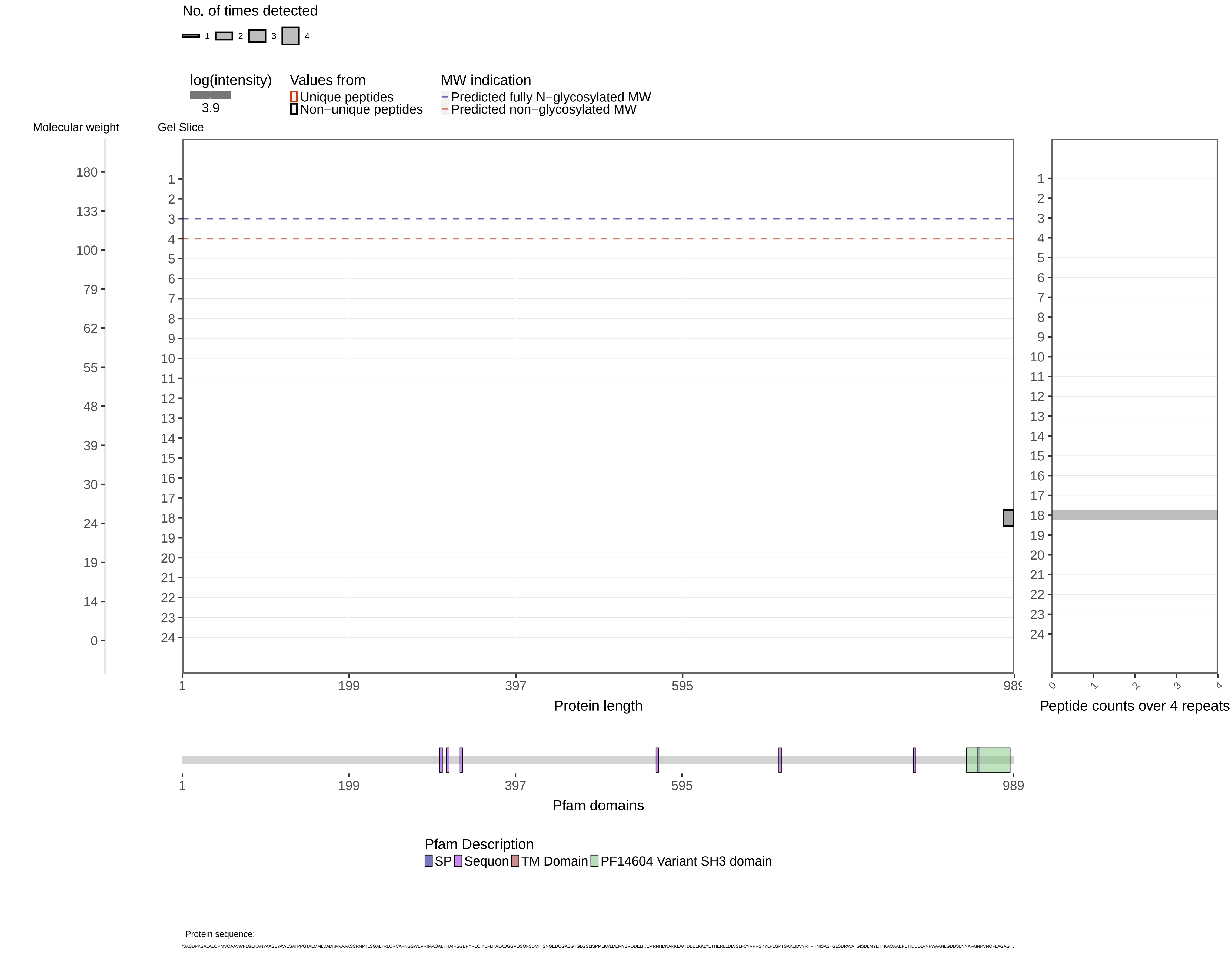

NbL12g16630.1 glucan_endo−13−beta−glucosidase_6−like [MW − SP = 50.11 kDa]

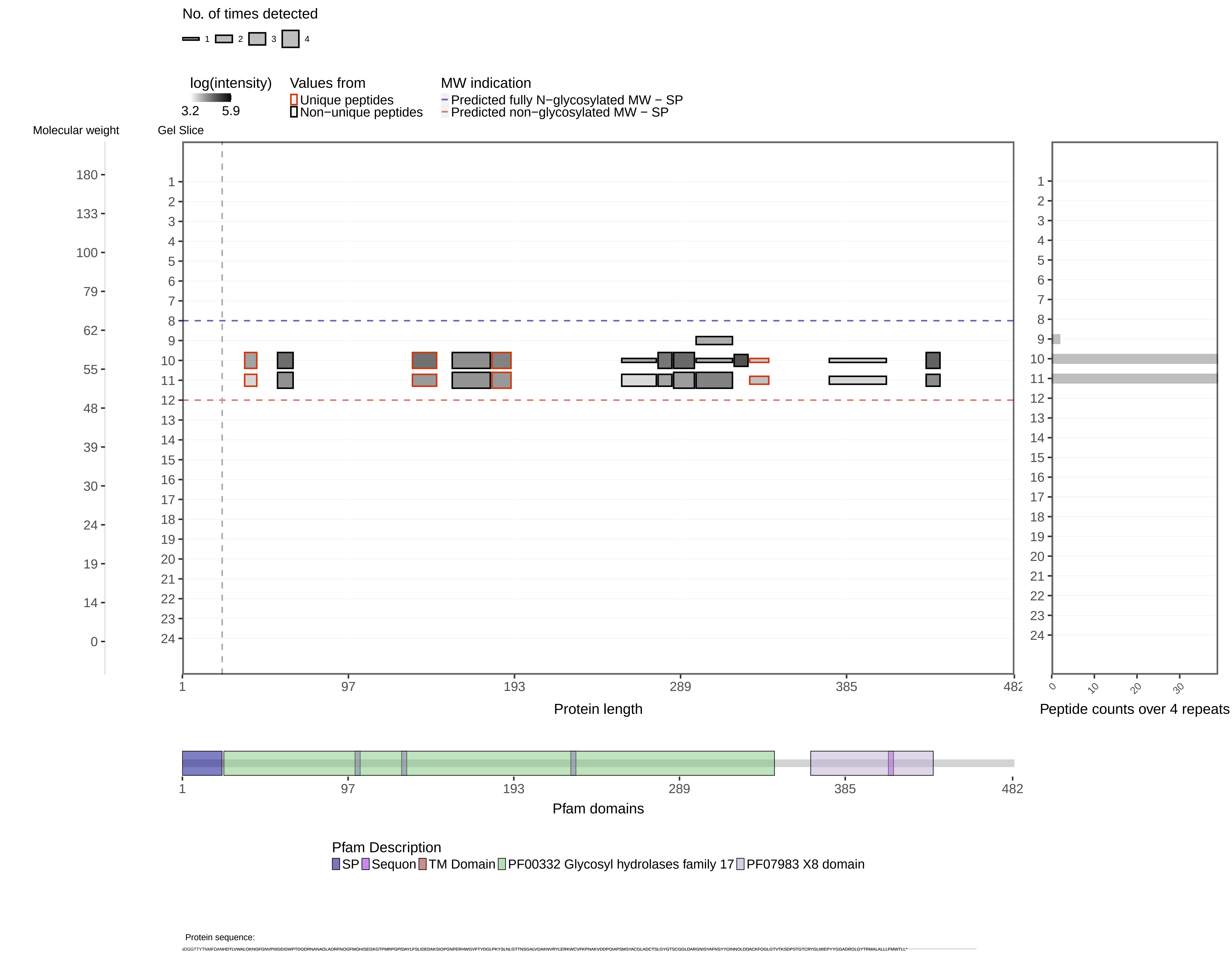

NbL12g16420.1 uncharacterized_LOC109231956 [MW = 37.8 kDa]

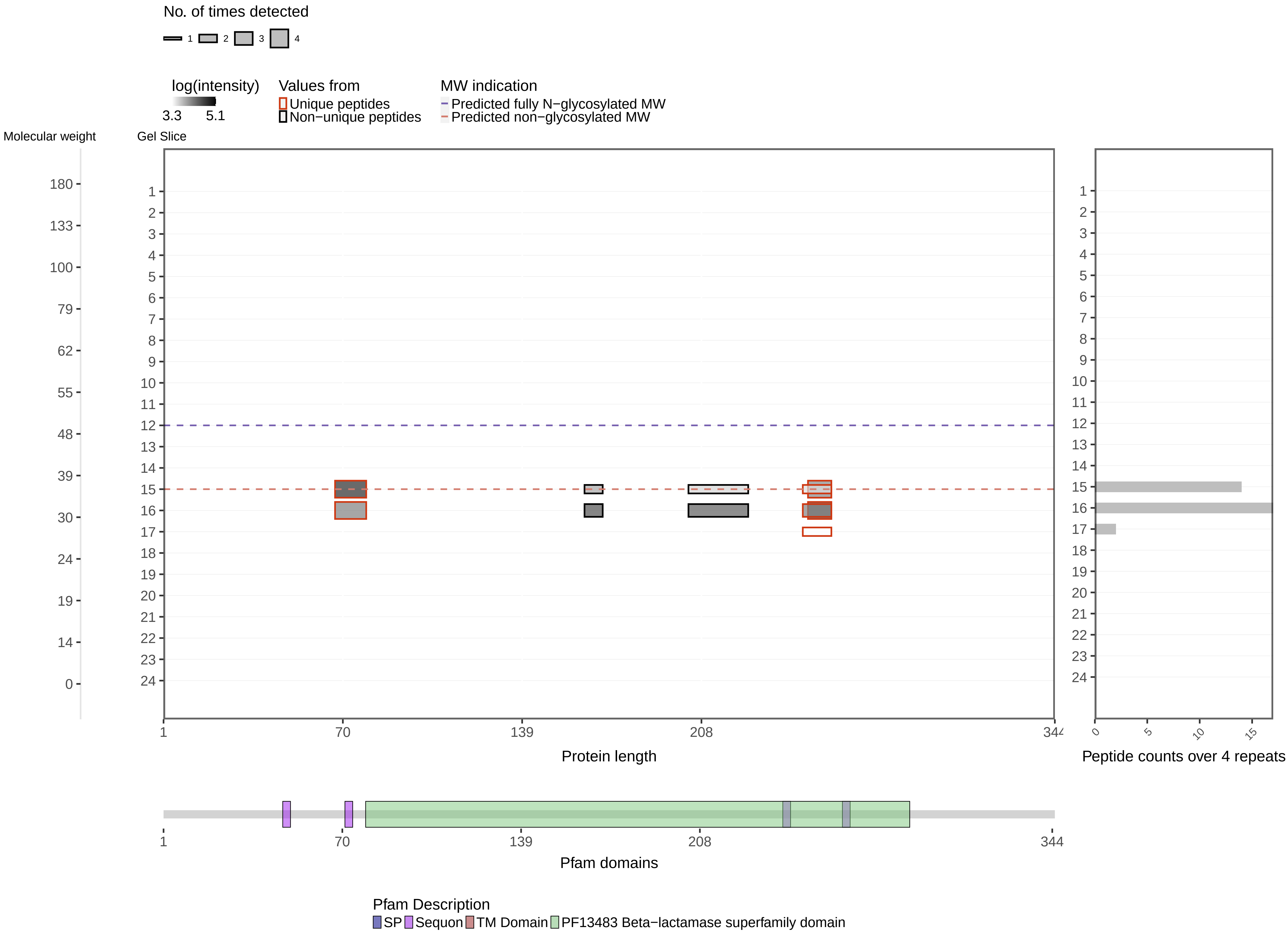


Protein sequence:

PNLRVIATPNAKTLLDPLFSNVIYLEPGODSEIEVSNGLOVNIKATAGPVLGPPWORPENGYLVTSPKGALTLYYEPHCVYNKTFLEKYKADIVITPVIKOLLPNFTLVSGOEDAVOLAKLLSARFVVPMKNGDLDSKGFLASIIESEGTMESFKELLSKELPDAKVLEPVPGEPLDISIAVOGO*

### NbL12g16410.1 phenylalanine−−tRNA_ligase_chloroplastic/mitochondrial−like [MW = 54.19 kDa]

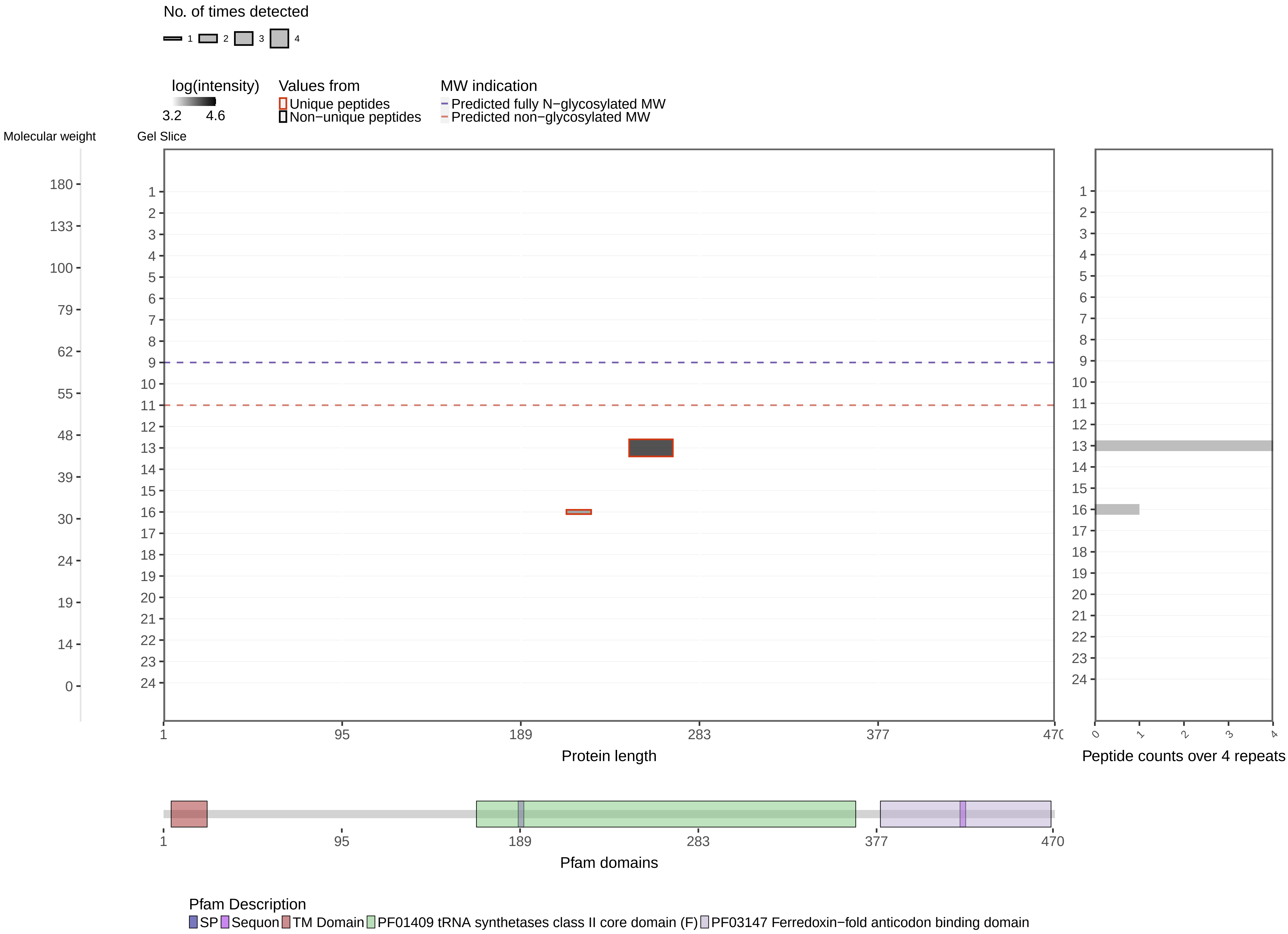


Protein sequence:

RRDSIDATHYPVFHQMEGVRVFAPNDWEGSGADATSHAADDLKKCLEGLARHLFGGVEMRWVDTYFPFTNPSFELEIYFQNEWMEVLGCGVTEQEILKRSGRTDNVAWAFGLGLERLAMVLFDIPDIRLFWSTDERFTSQFSSGRLGLKFKPFSKYPPCYKDMSFWISDSFTENNLCEVVRGIAGDLAEEVKLIDNFTNKKGMTSHCYRIAYRSMERSLTDEEINDLQWKVREQVESKLKVVLR*

### NbL12g16130.1 SAL1_phosphatase−like [MW = 44.32 kDa]

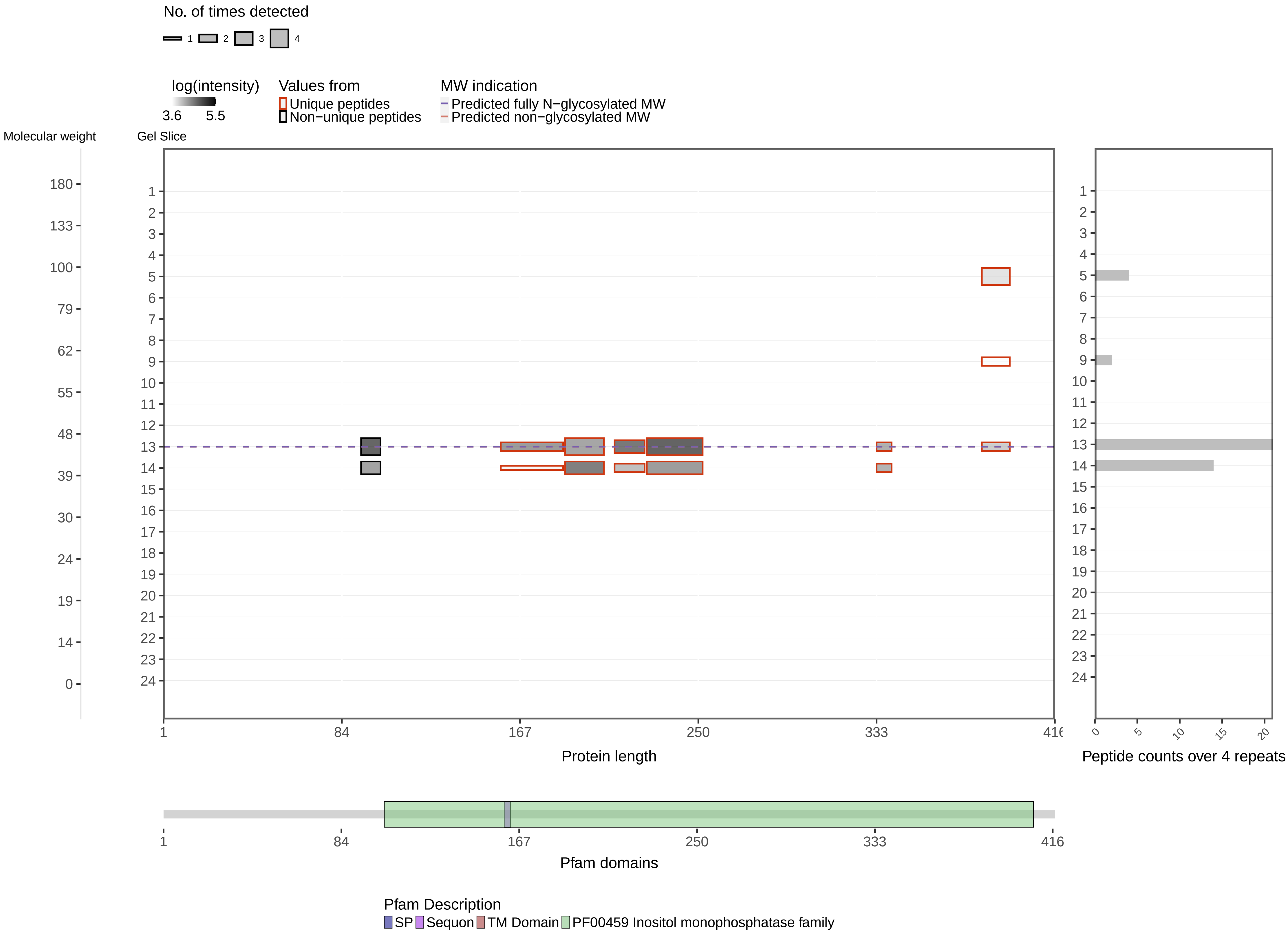


Protein sequence:

HWVLDPIDGTKGFVRGDQYAIALALLDEGKAVLGVLACPNLPLSSVASHNPQDSQEKVGCLFYAQVGSGTYMQSLDGSTLIKVHVTDLENPEEASFFESFEAAHSLHDLSSSIAKKLGVRAPPVRIDSQAKYGALSRGDGAIYLRFPHKGYREKIWDHAAGYLVVAEAGGVVSDAAGNPLDFSRGRYLDLHEGIIVTNQKLMPALLKAVKESLNEKSSSL*

### NbL12g16060.1 alcohol_dehydrogenase_1B [MW = 36.82 kDa]

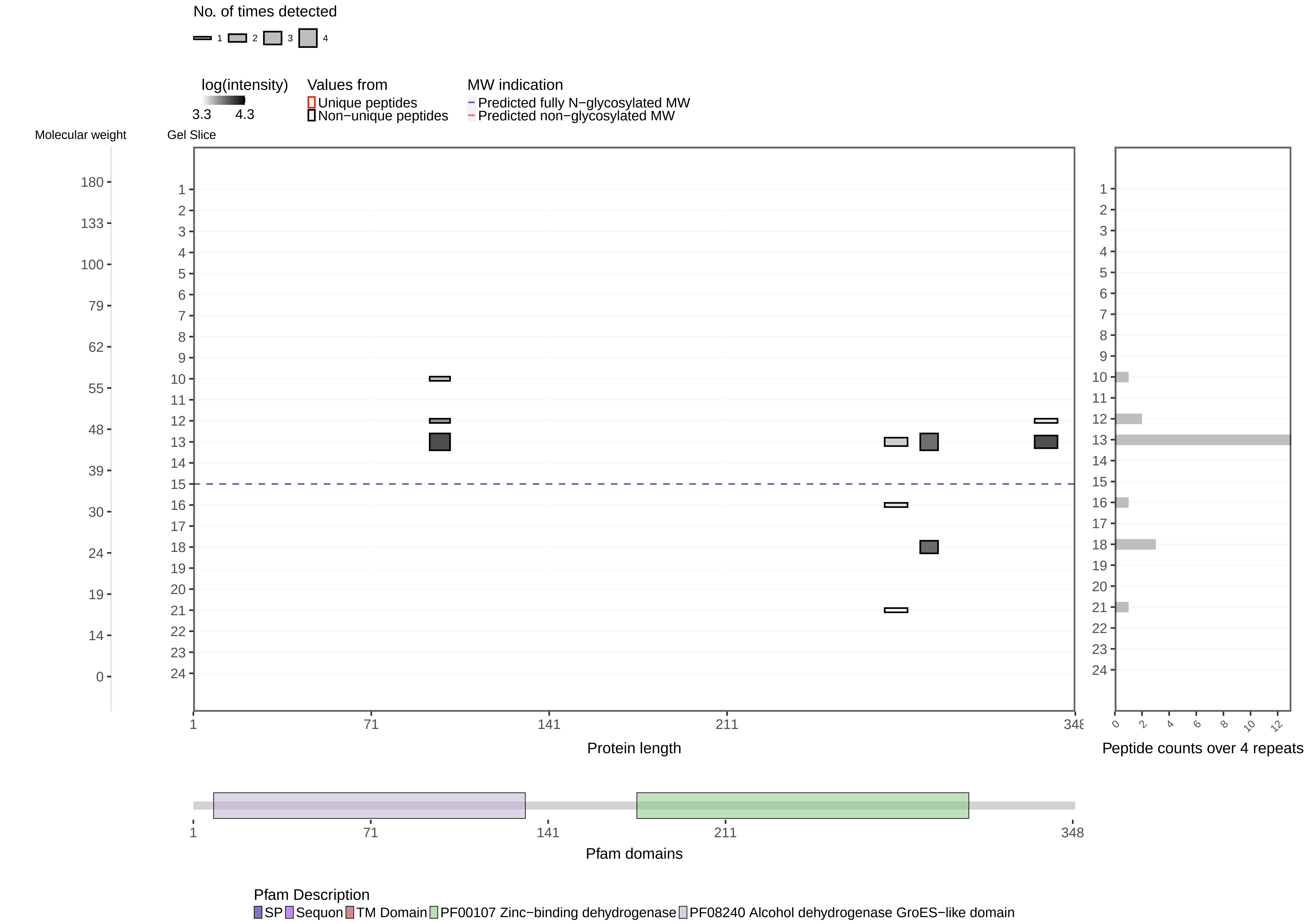


Protein sequence:

AAEVRPGAAIAVIGIGGVGSSCLQIARAFGASEIIAVDVQDEKLQKAKMFGATHCVNALKEDAVKRIKDITGGMGVDVAVEALGKPQTFLQCVQSVRDGGKAVMIGLTLSGAKGEVDINHLVRRQIKVIGSYGARARQDLPKLIKLAESGIFDLSAAVSRTCKFEEAGQAYKDLDQGAISGRCVVEIM*

### NbL12g16050.1 eukaryotic_translation_initiation_factor [MW = 22.85 kDa]

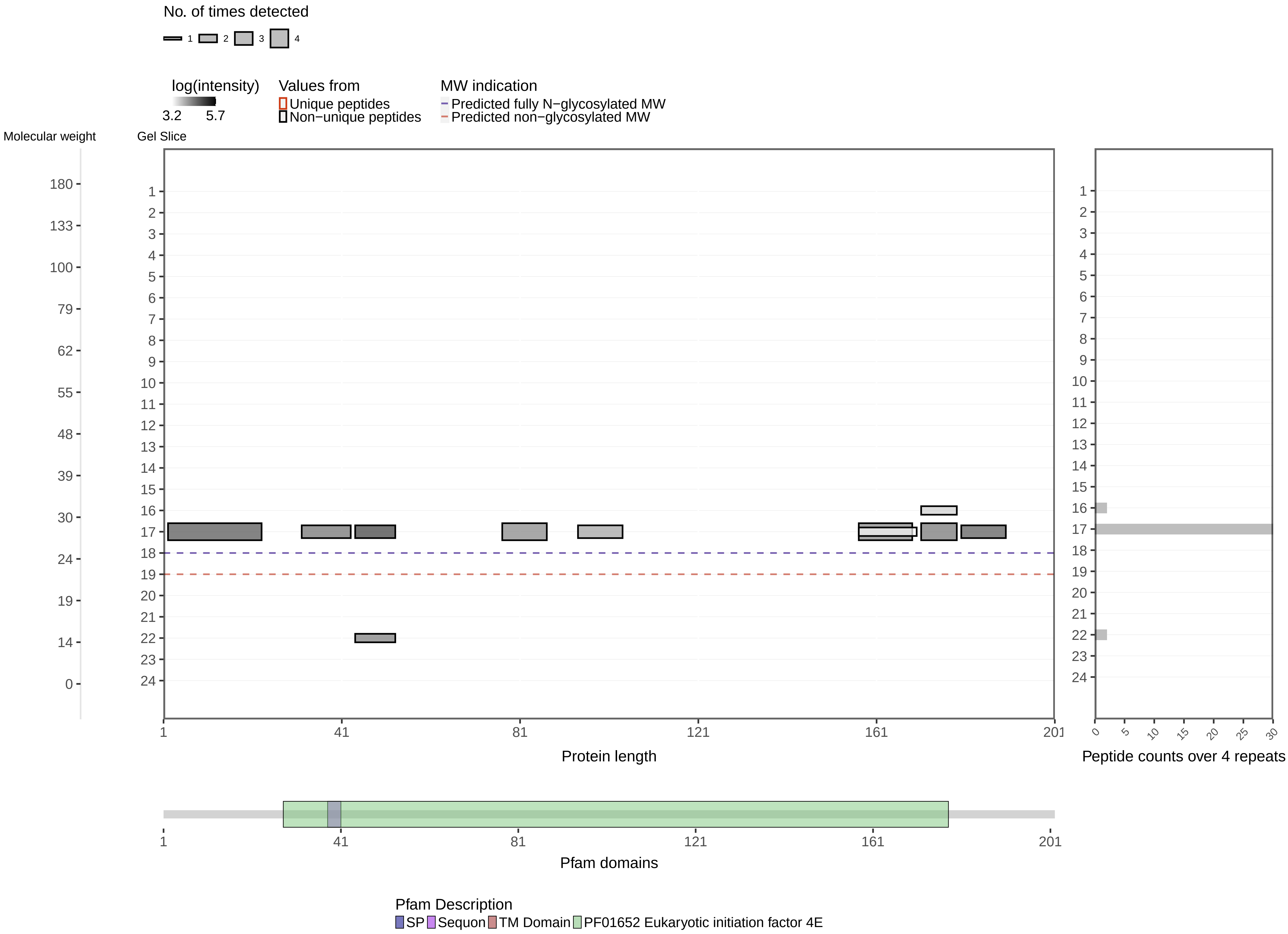


Protein sequence:

AGIEPKWEDPECASGGKWTVTSSRKANLETMWLETLMALVGEQFDESEEICGVVASVRRSQDKLSLWTKTASNEAIQMSIGRKWKEIIDAEKISYSFHDDSKRERSAKSRYTV*

NbL12g16020.1 calcium−dependent_protein_kinase_SK5−like [MW = 62.55 kDa]

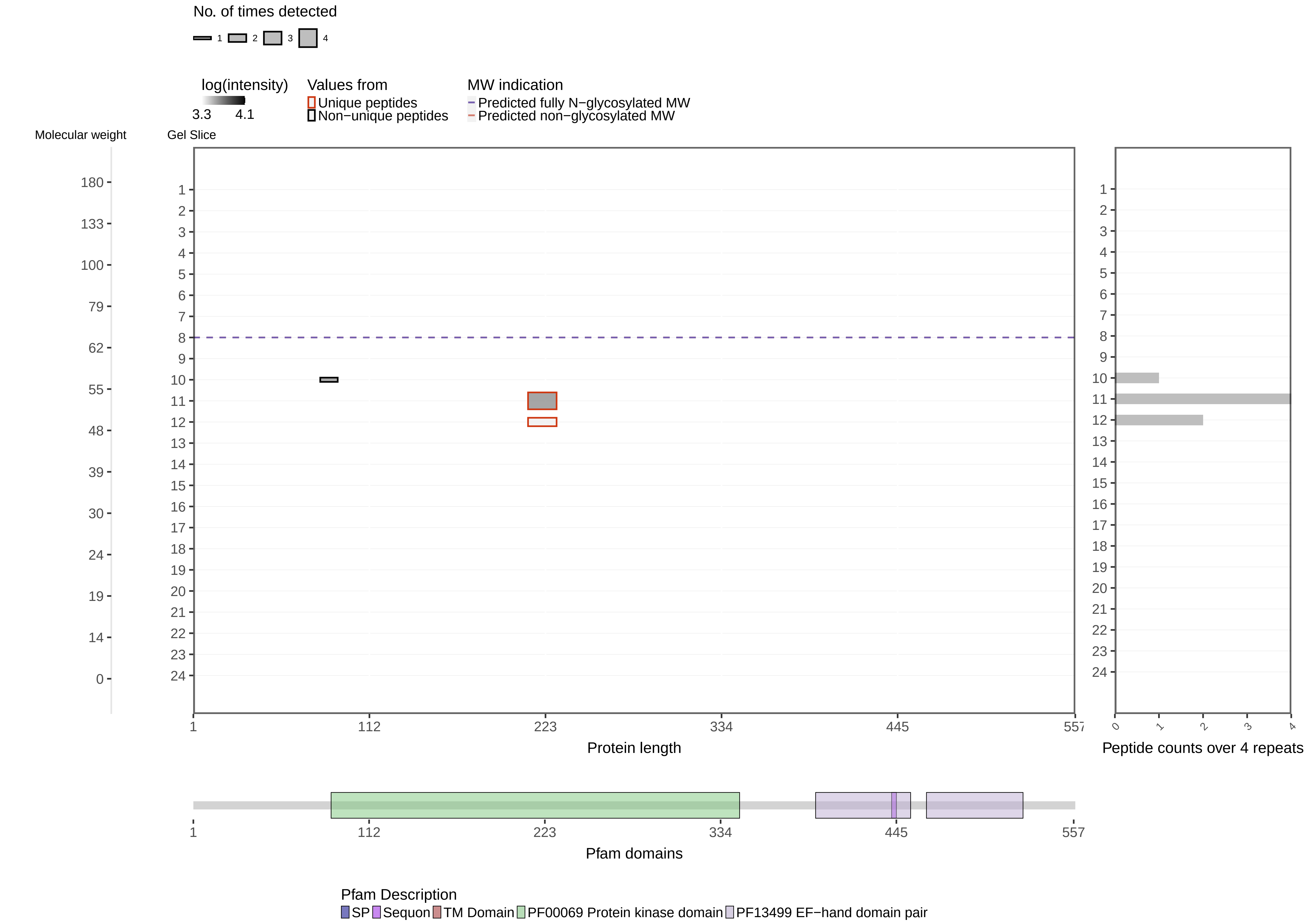


Protein sequence:

YGHESDVWSAGVILYILLSGVPPFWAETDMGIFRQILRGKLDLESEPWPGISDSAKDLIRKILDRNPKRRLTAHEVLCHPWIVDDTMAPDKPLDSAVLSRLKQFSAMNKLKKMALRVIAERLSEEEIGGLKELFKMLDTDNSGTITFEELKEGLRRVGSELMESEIKDLMDAADIDNNGTIDYGEFIAATVHLNKLEREENLLSAFSYFDKDGSGYITIEELQQACKEFGLSELNLDEIIKDIDQDNDGQIDYKEFSAMMRKGTGGGVGRRTIRNNLNLGEALGLVQSEENL*

### NbL12g16010.1 acyl−coenzyme_A_thioesterase_13−like [MW = 18.35 kDa]

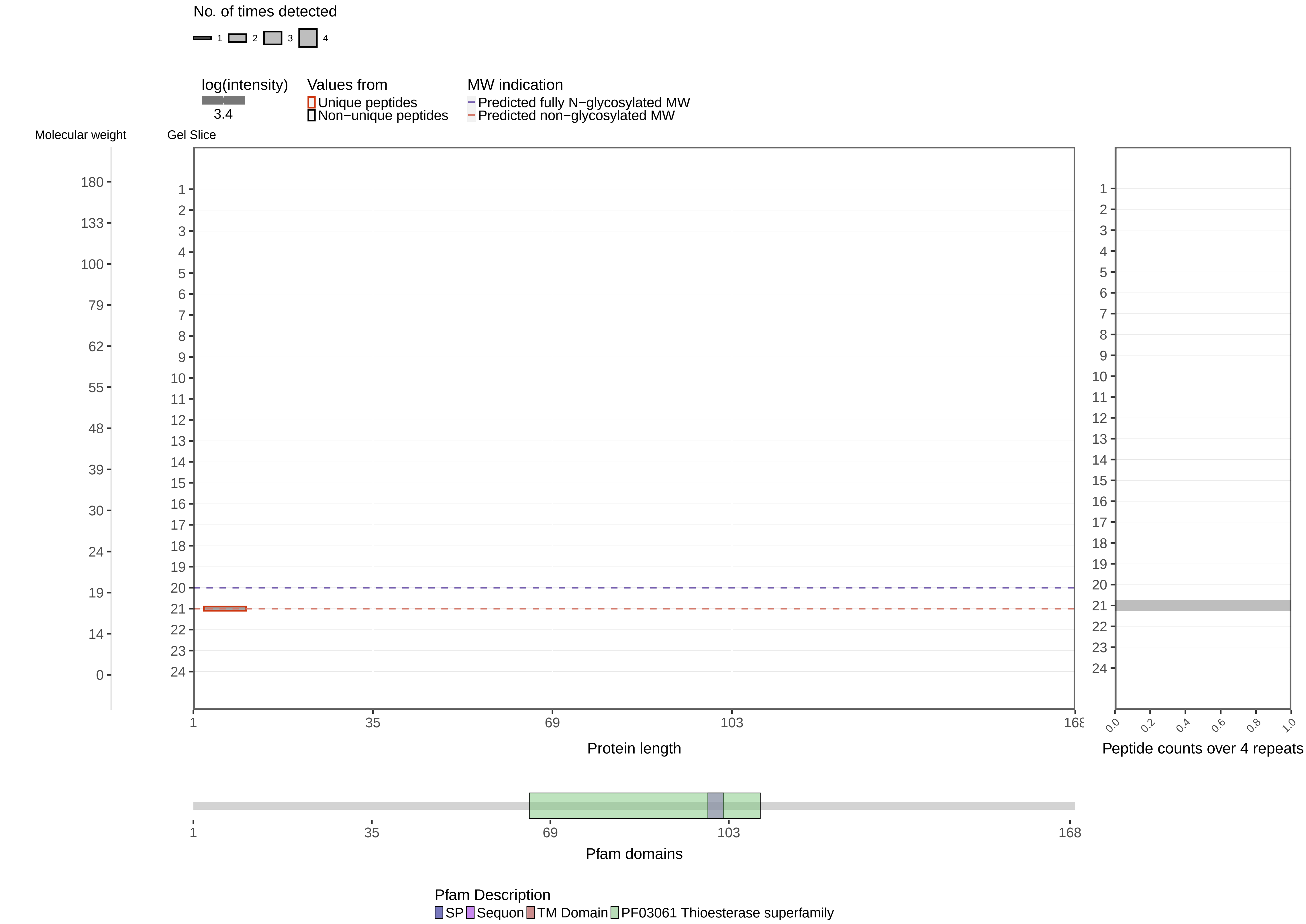


Protein sequence:

GATATLVDILGSAVIHTVGAPFTGVSVEINVSYLDAAFIGVSCPLSCFSDLNLVHFFLNTELLNTGMRIYLADPQLVWDCIFVVKVKLGCELALFFFS*

NbL12g15990.1 probable_methyltransferase_PMT26 [MW = 89.94 kDa]

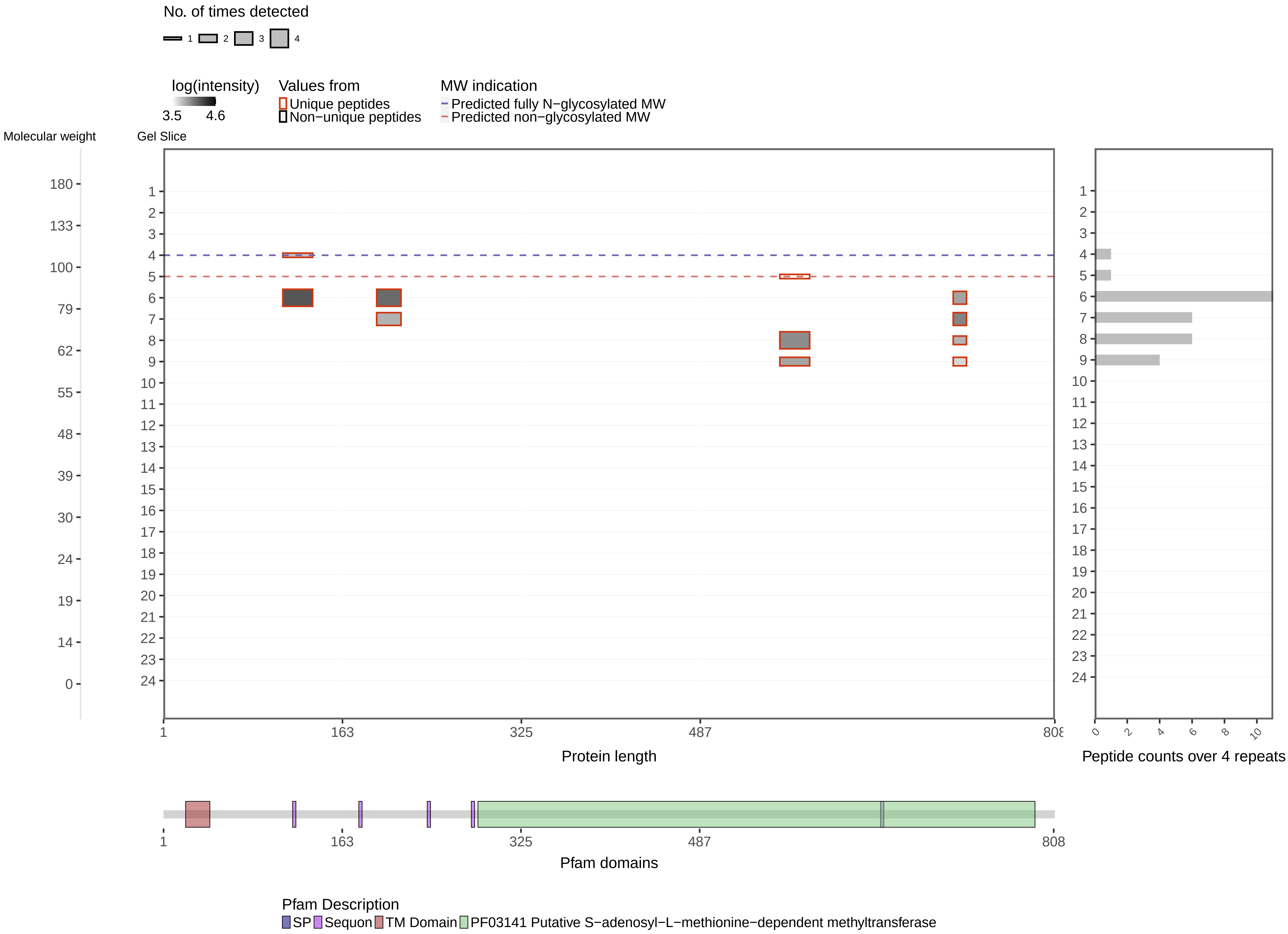


Protein sequence:

FIQQSFPDIAWGKRTRVILDVGCGVASFGGYLFDRDVLAMSLAPKDEHEAQVQFALERGIPAISAVMGTKRLPFPSRVFDVVHCARCRVPWHIEGGKLLLELNRLLRPGGLFVWSATPVYQKLPEDVEIWEAMKKLTKAMCWELISKTKDRVNGVGVAVYRKPMDNECYEQRSEDAPPLCQESDDPNAAWNVPLQACLHKAPAATSERGSQWPESWPARLSKSPYWLLSSQVGVYGKPAPEDFTADYEHWKHVVKNSYINGMGINWSTVRNVMDMRAIYGGFAAALRD

### NbL12g15920.1 4−hydroxyphenylpyruvate_dioxygenase−like [MW = 49.26 kDa]

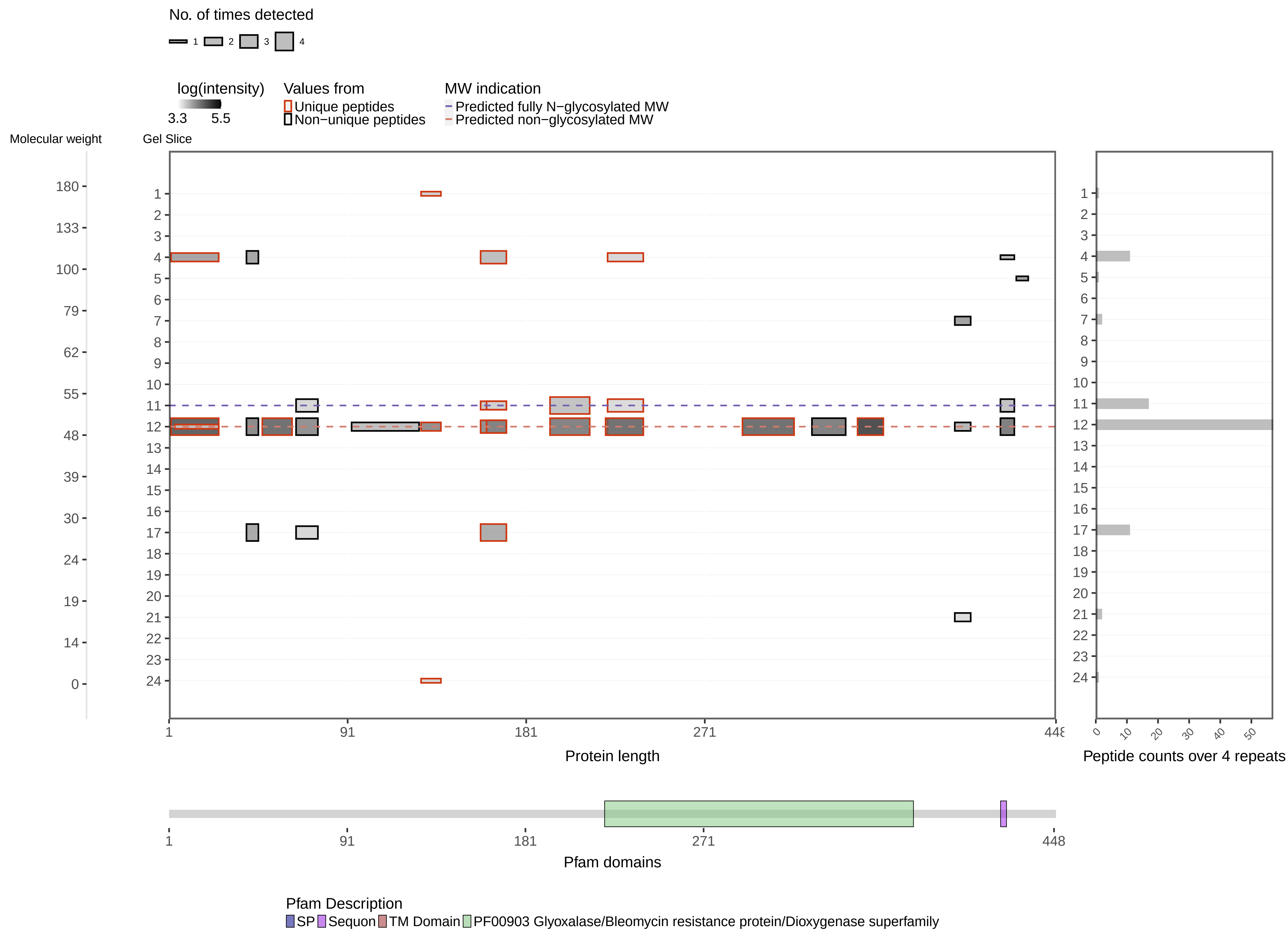


Protein sequence:

YKELDYGIRRLDHAVGNVPELGPVVDYIKRFTGFHEFAEFTAEDVGTAESGLNSMVLANNDETVLLPLNEPVYGTKRKSQIQTYLEHNEGAGMQHLALVTEDIFKTLTEMRKRSGVGGFEFMPSPPPTYYKNLKNRAGDVLTDEQIQACKELGILVDRDDQGTLLQIFTKPVGDRPTIFIEIIQRIGCMLKDEKGQVYQKGGCGGFGKGNFSELFRSIEEYEKTLEAKQANPVTAA*

NbL12g15830.1 WD_repeat−containing_protein_26−like [MW = 27.03 kDa]

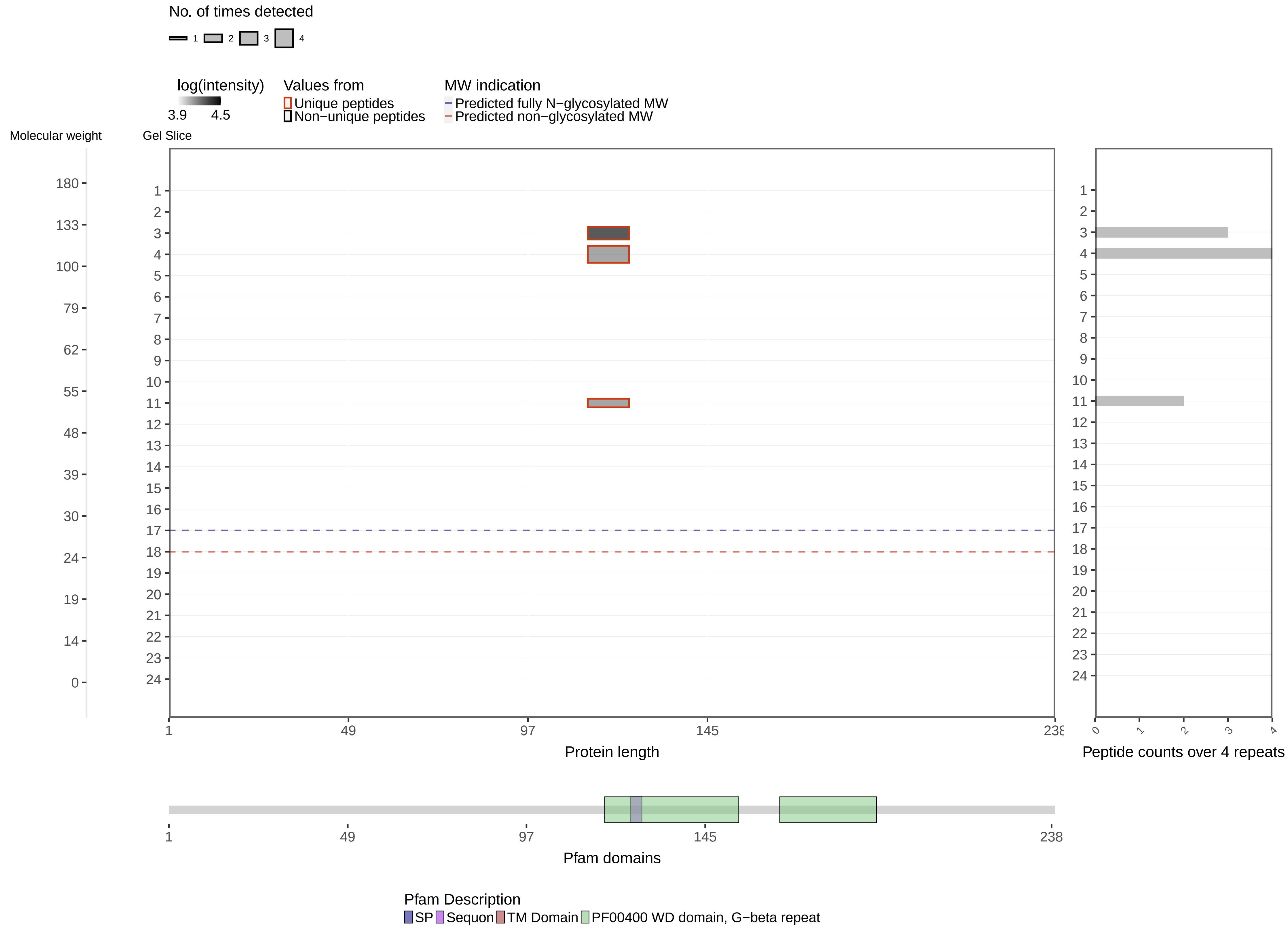


Protein sequence:

QCGRDQIPSQTLQILRNHSDEVWFLQFSHDGKYLASSAADCSVIIWEVKLDGLFCLKHQFSGHQKPVSYMSWSPDDHQLLTCGVEVVRRWDVESGECVHIFEKNGLGLISCGWAPDGKRILWRCYGQEH*

NbL12g15820.1 oxygen−evolving_enhancer_protein_2−2_chloroplastic [MW = 37.57 kDa]

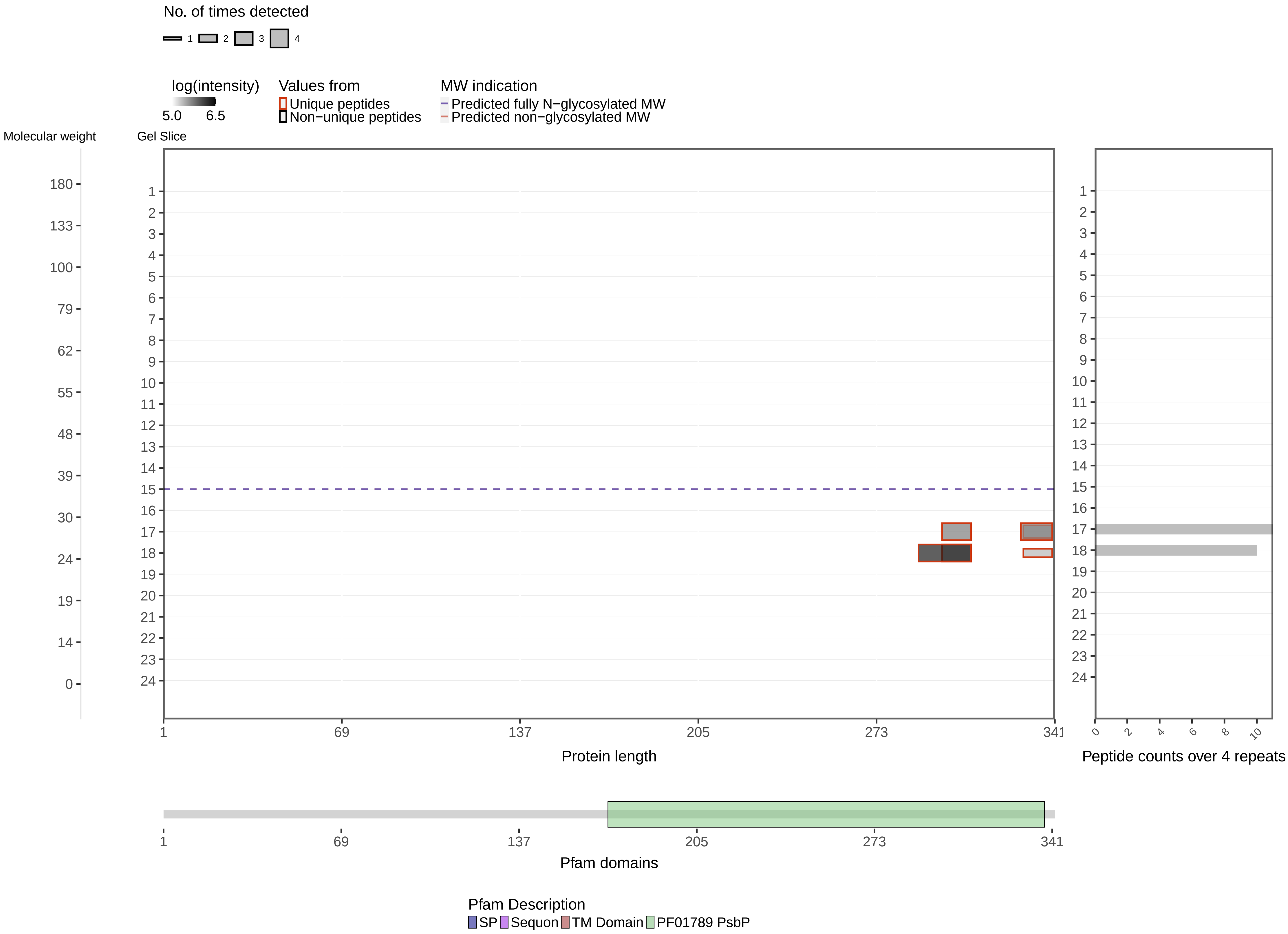


Protein sequence:

EAANVFGKPKTDTDFQTYNGDGFKLQIPSKWNPNKEVEYPGQVLRFEDNFDATSNVIVAITPTDKKSITDFGSPEQFLSQVDYLLGRQAYSGKTDSEGGFESDAVAIANVLETSSAEVGGKPYYYLSVLTRTADGNEGGKHQLITATVNNGKLYICKAQAGDKRWFKGAKKFVENTATSFSLA*

### NbL12g15730.1 V−type_proton_ATPase_subunit_G_2 [MW = 12.5 kDa]

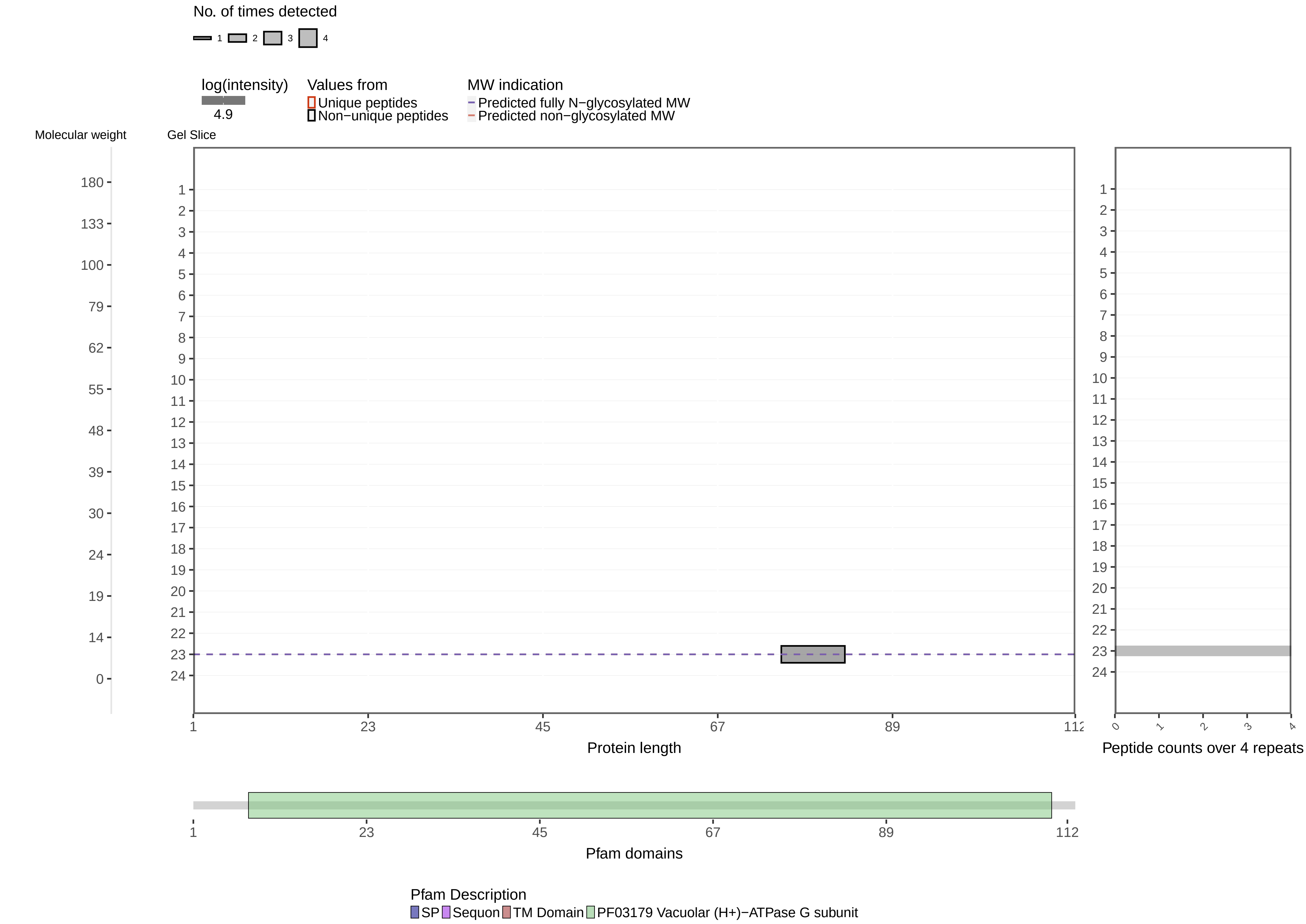


Protein sequence:

AEKEIAEFRAYMEAEFQRKVEQTSGDSGANVKRLEQETFAKIQHLKTEAESISHDVVQMLLRQVTTVKN*

### NbL12g15550.1 proteasome_subunit_beta_type−3−A−like [MW = 22.67 kDa]

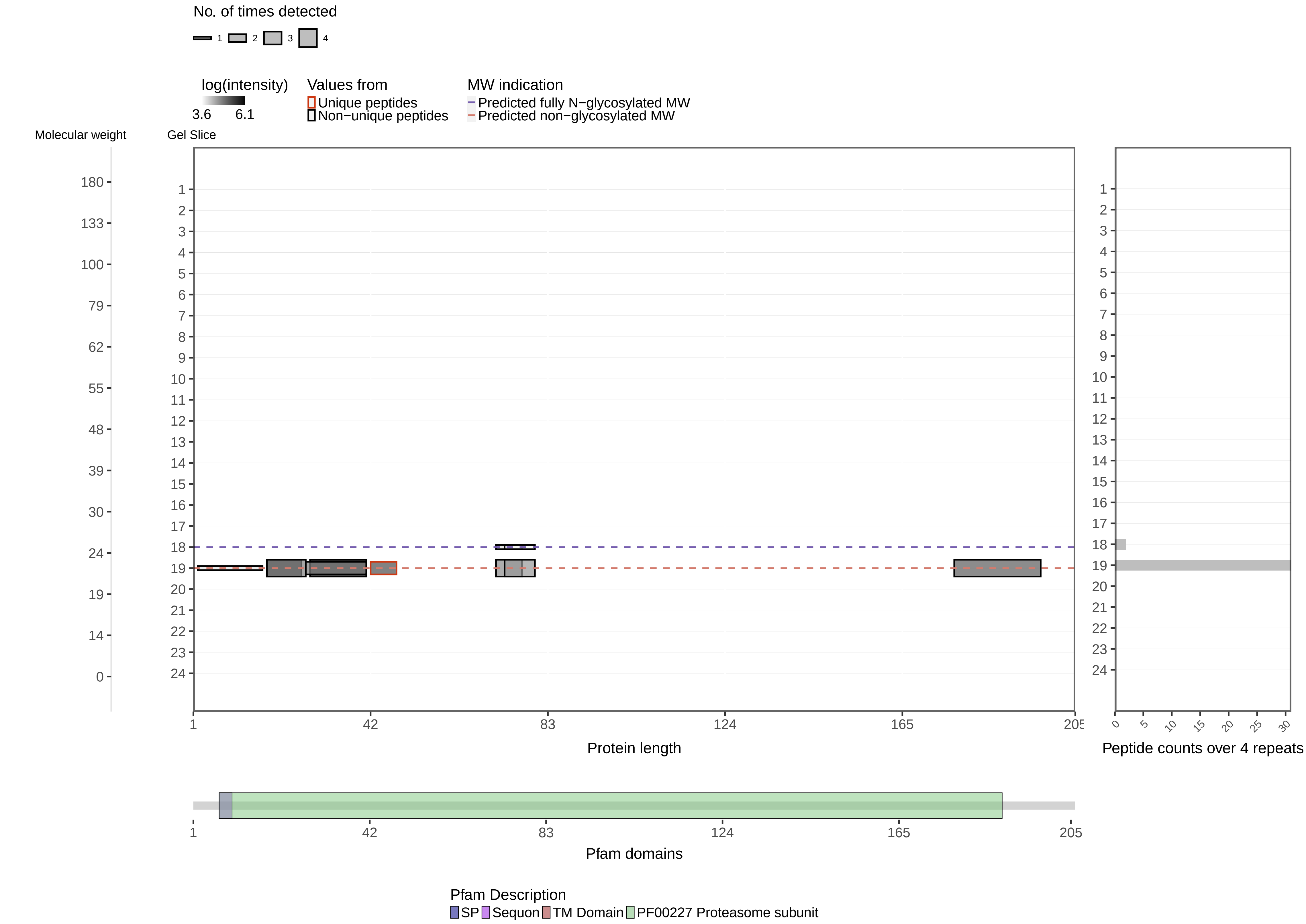


Protein sequence:

VSALLYEKRFGPYFCQPVIAGLGDDDKPFICTMDSIGAKELAKDFVVAGTASESLYGACEAMFKPDMGPEELFETISQALIASVDRDCLSGWGGHVYVVTPTEVTEKILKGRMD*

NbL12g15520.1 uncharacterized_LOC109225451 [MW = 29.8 kDa]

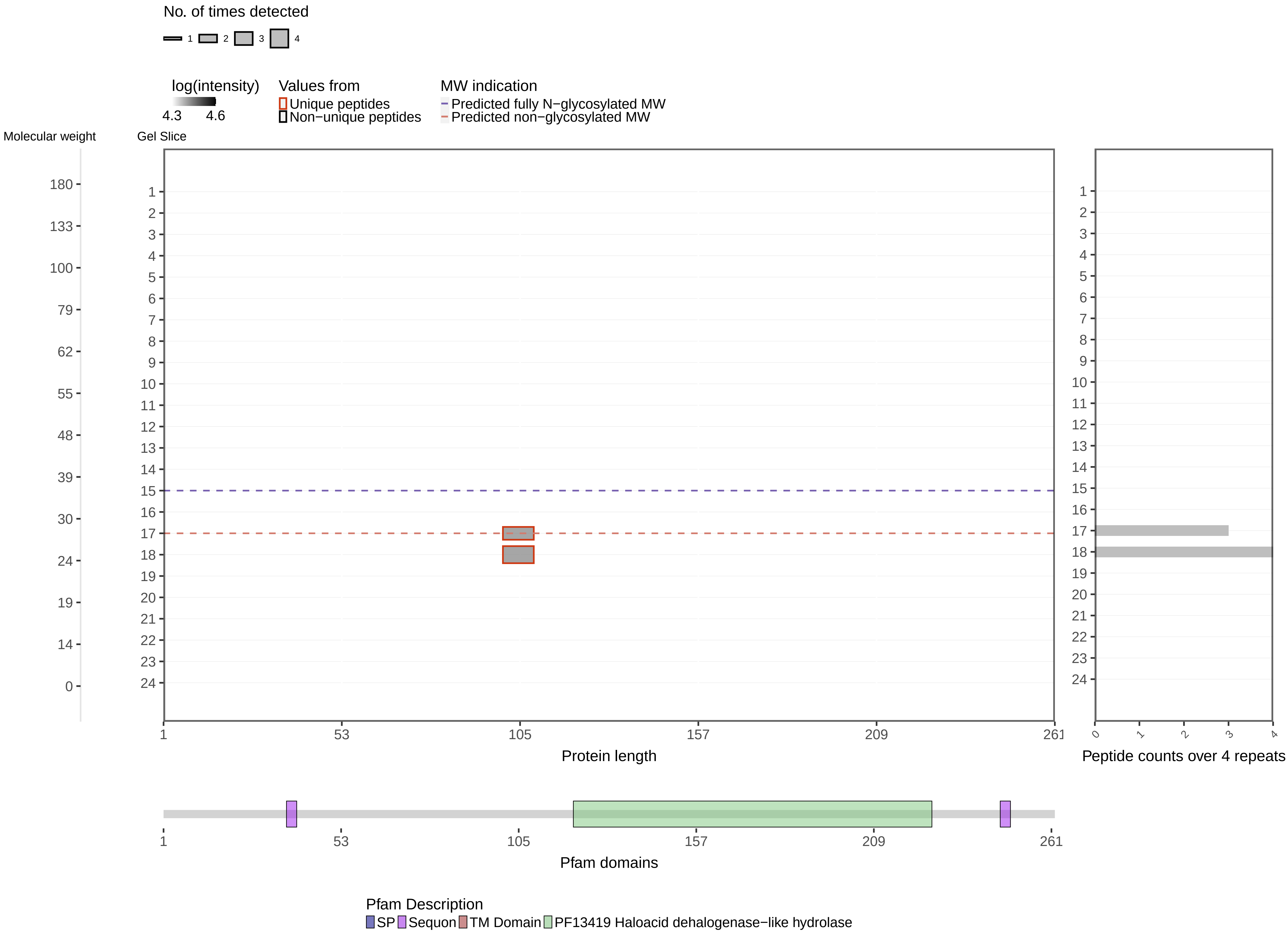


Protein sequence:

FDLEGLKNCMQRAYSYIEGVEGLLNSLKENGYEIHAFTNYPIWYQMIEDKLKLSNYLSWTFCSCIFGKRKPDPDFYLEVVKHLNVEASSCIFVDDRMRNVEAAVEVGLNGLQFKNADLLRKDLSLLGVNISTNENPDLAECSI*

NbL12g15370.1 2−alkenal_reductase_(NADP(+)−dependent)−like [MW = 39.52 kDa]

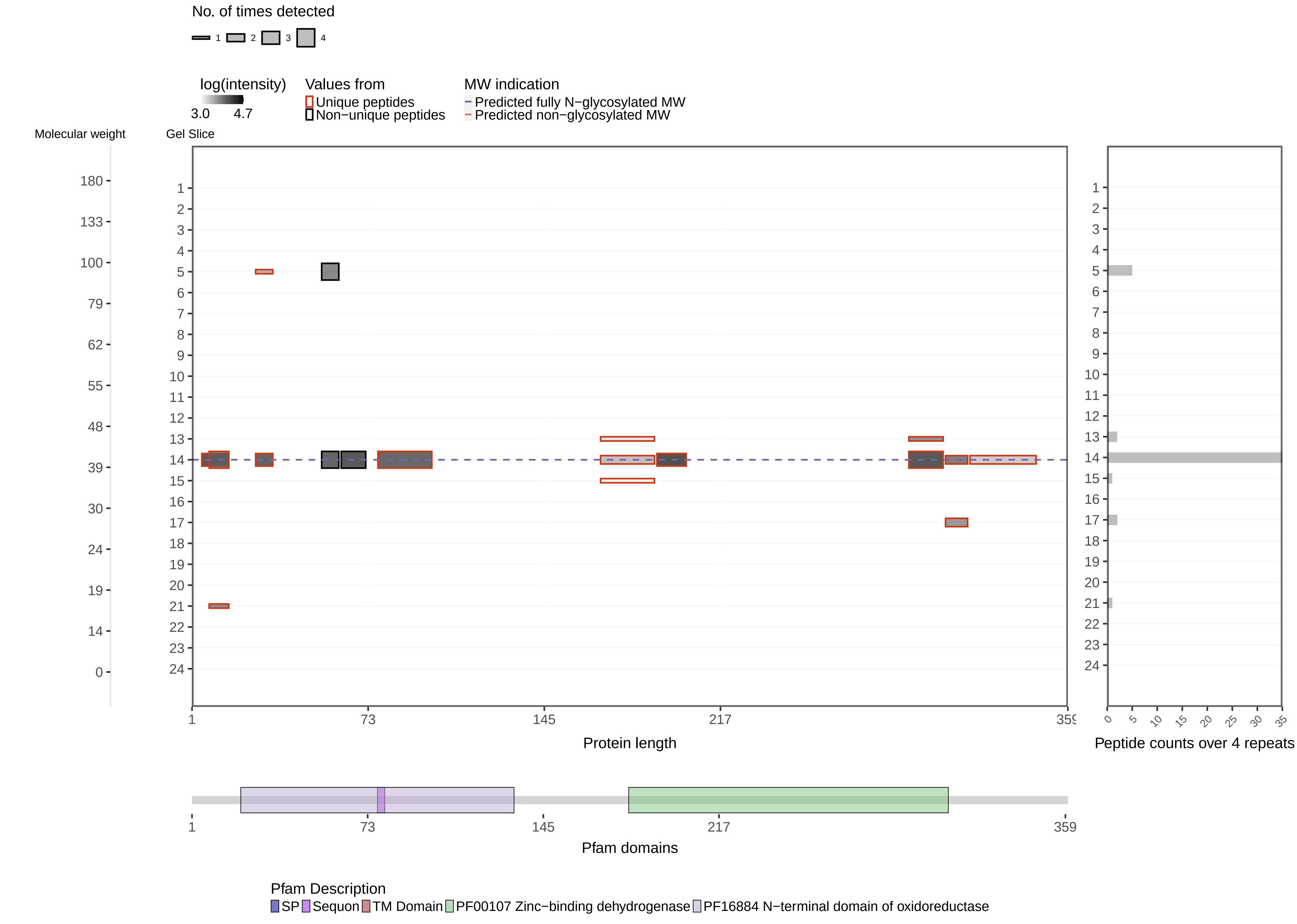


Protein sequence:

RGEYVFVSAASGAVGQLVGQLAKLHGCYVVGSAGSKQKVDILMDKLGFDGAFNYKEEADLDKALKRHFPKGIDIYFDNIGGSMLDAALINMRINGRIAICGMVSQQSLSDPQGIHNLIQLITKRVKMQGFLQSDYLHLFPRFLEDVTSLCKQGEITYLEDMNEGLESGPSAFVGLFSGKNIGKQVIRVAHD*

NbL12g15350.1 probable_methylenetetrahydrofolate_reductase [MW = 67.13 kDa]

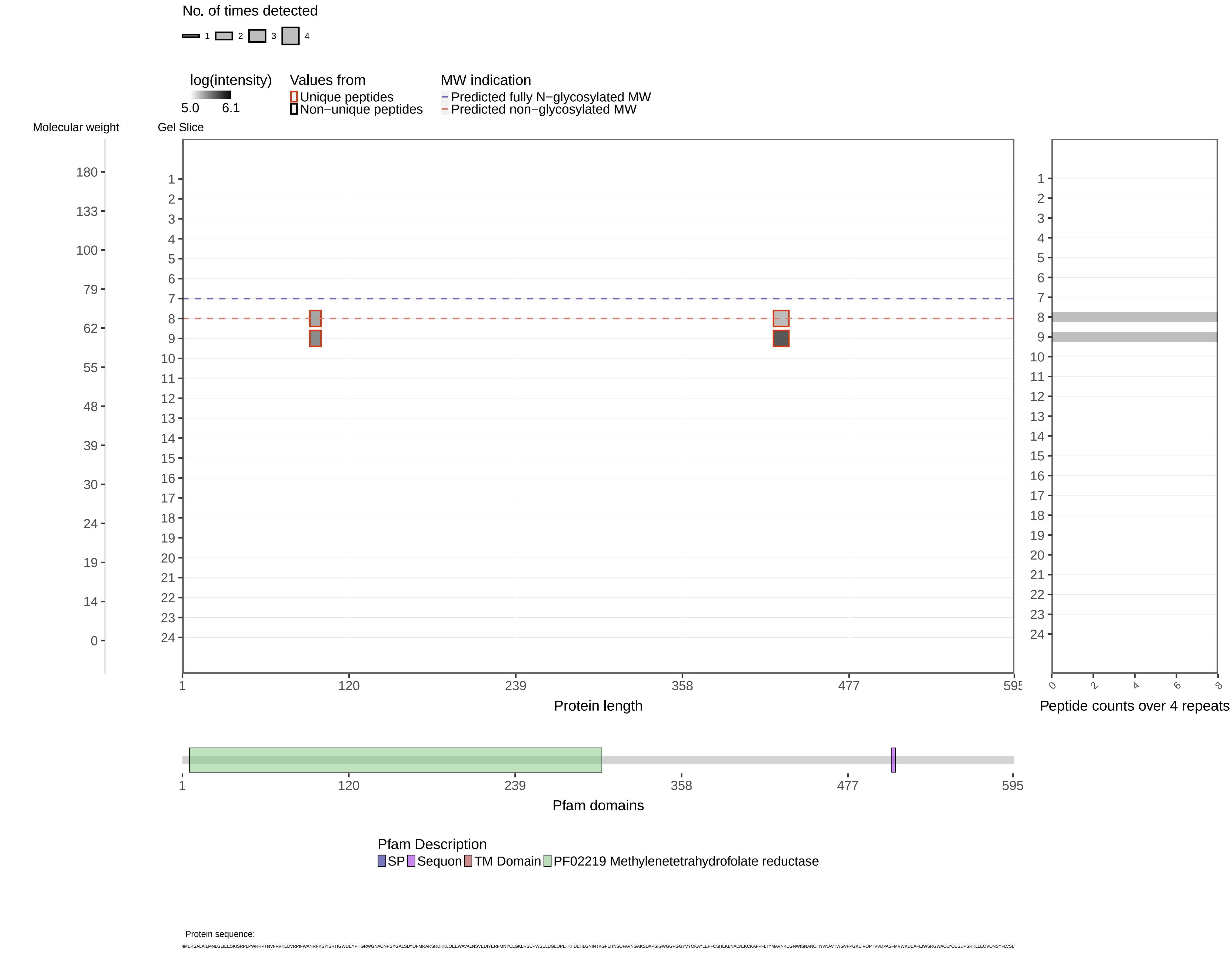

### NbL12g15300.1 isopentenyl−diphosphate_Delta−isomerase_I [MW = 27.12 kDa]

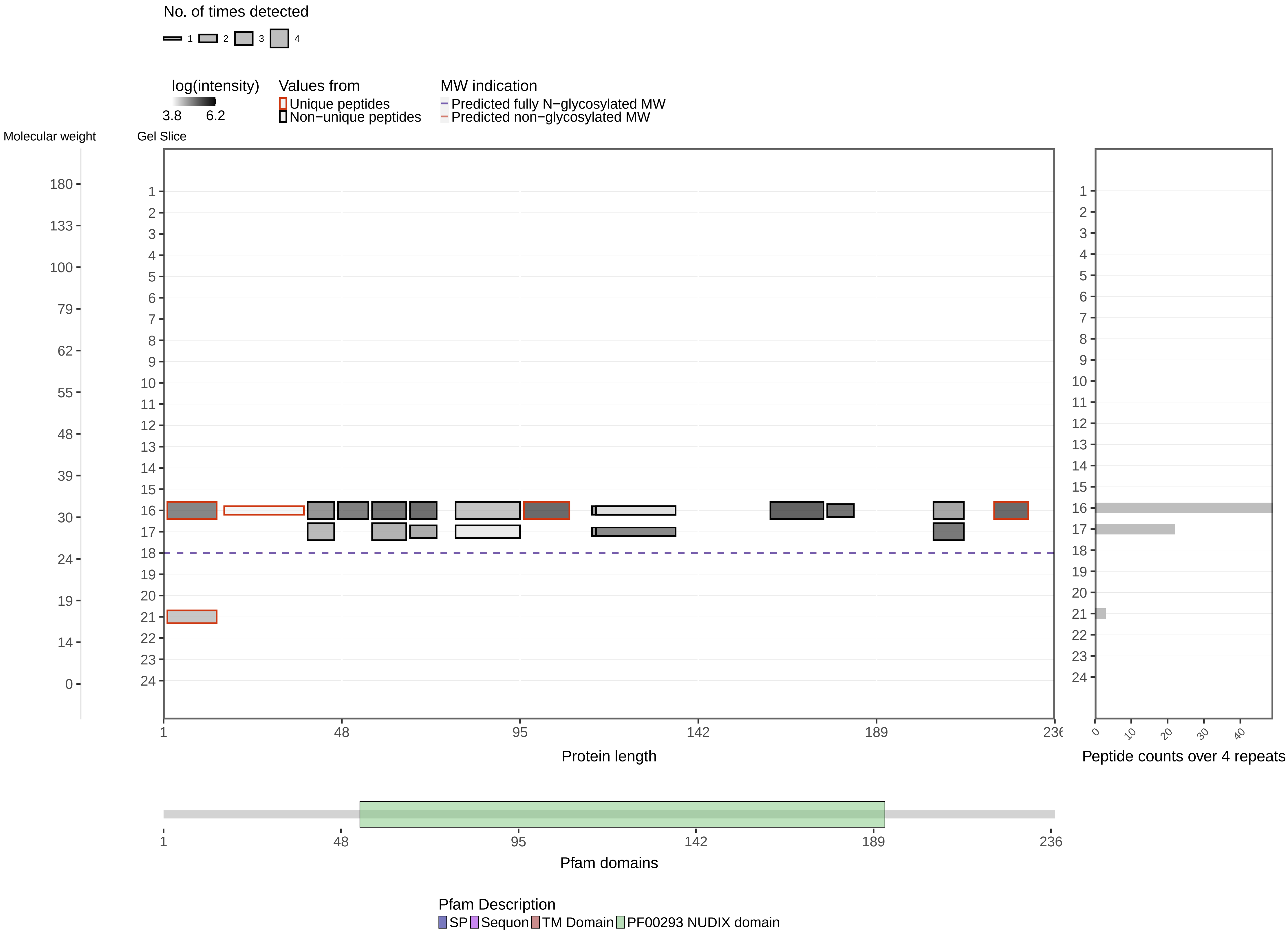


Protein sequence:
GVRNAAQRKLLDELGIPAEDVPVDQFTPLGRMLYKAPSDGKWGEHELDYLLFIVRDVNVNPNPDEVADIKYVNQEQLKDLLRKADAGEEGLKLSPWFRLVVDNFLFKWWDHVEKGTIQEAADMKTIHKLT*

NbL12g15270.1 uncharacterized_LOC107805381 [MW = 23.48 kDa]

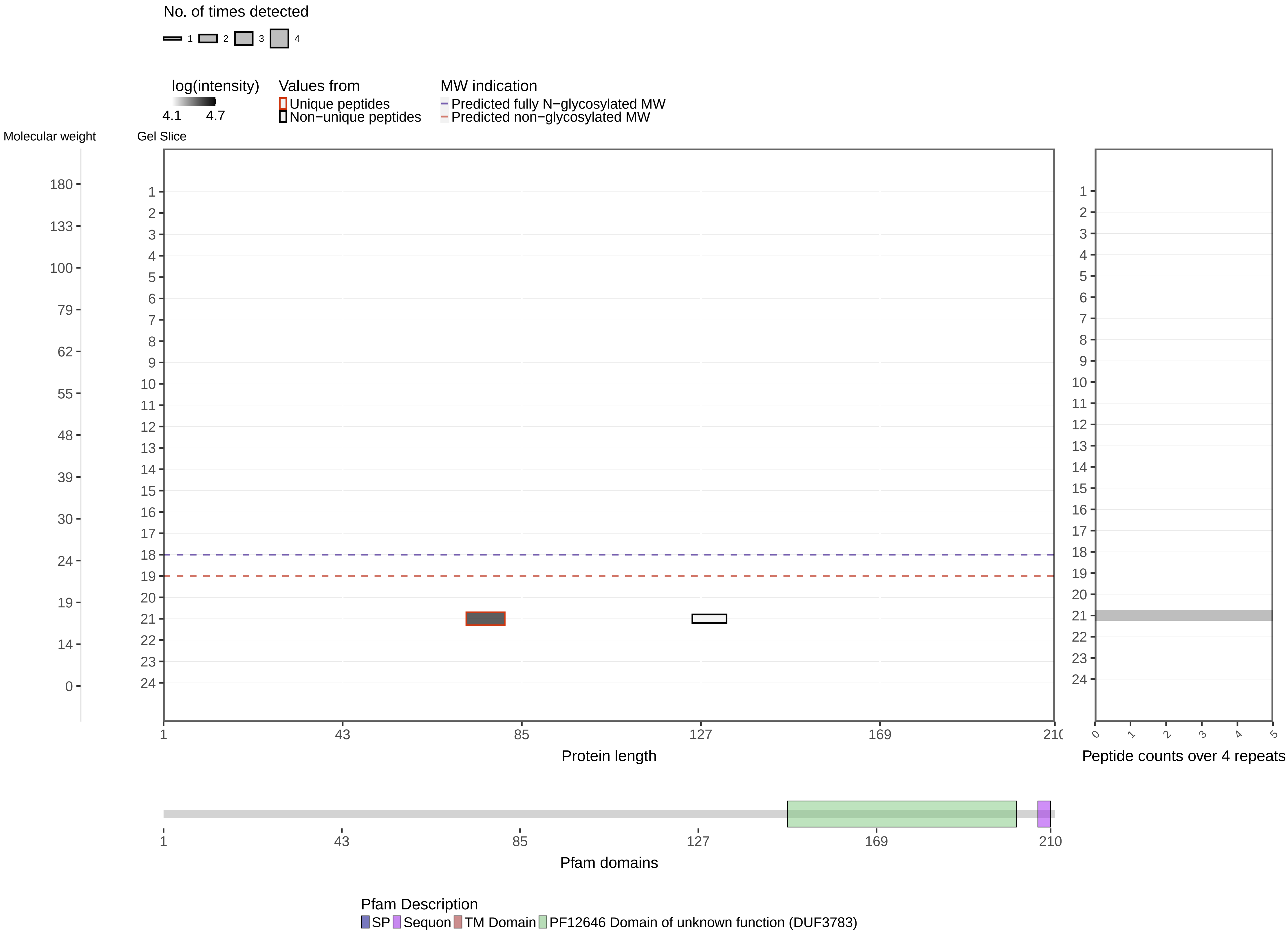


Protein sequence:

EVEEAAKIQQLLKEMDGEFLQVIFCTEDMISRSLWEAMNTKQPNLDASKIAKQLPRICFLSGLTGEEMMMFIDAFEESELEDPVFAALVENSADKPLQELIDEIMGDHELLSARNSS*

NbL12g15170.1 protein_BTR1−like [MW = 35.87 kDa]

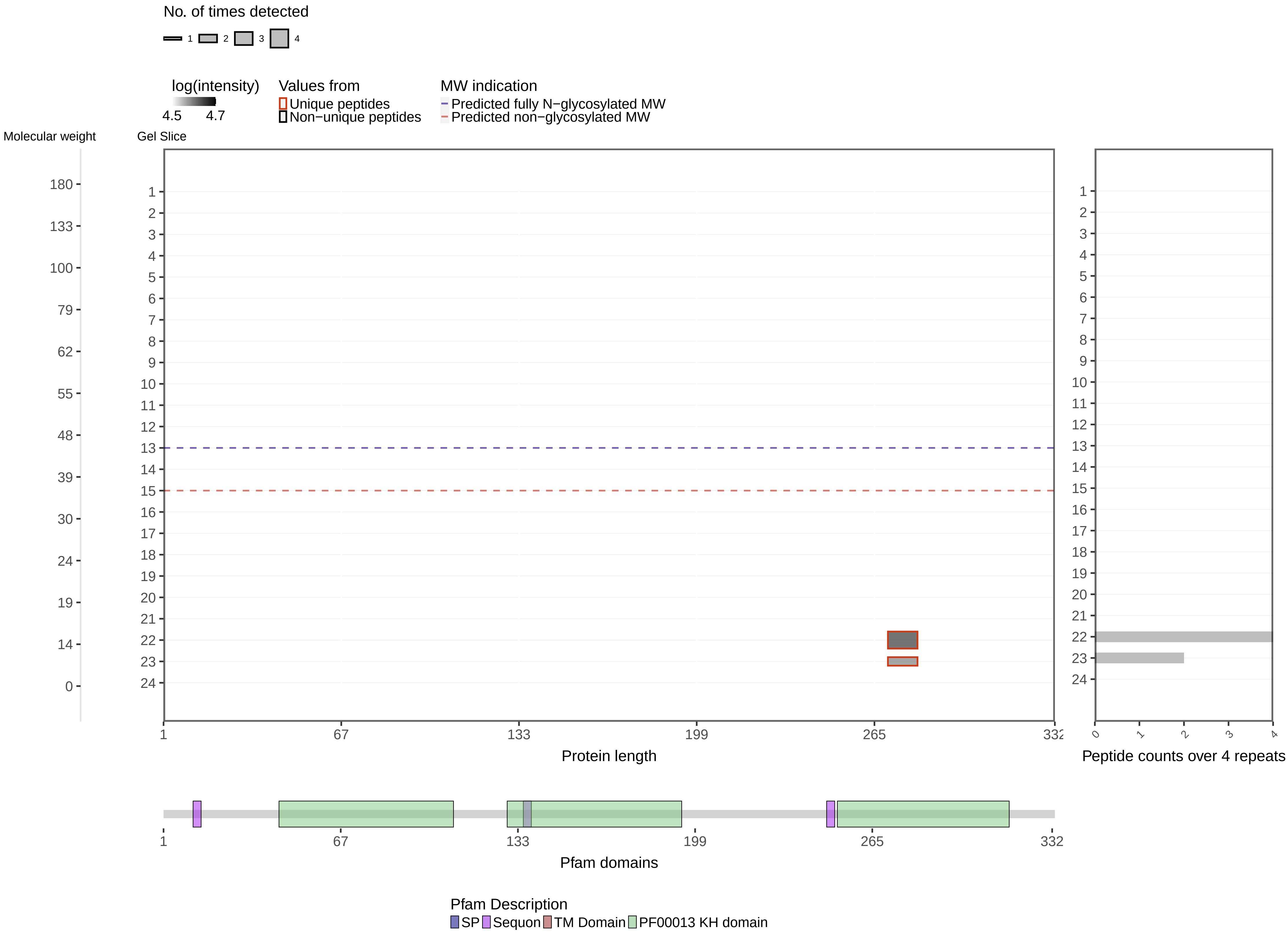


Protein sequence:

FIEDSHAGIKISSQDDNFPGLHDRLVIVTGTLEEQMRAIELILHKLAEDSHYIQNMNAPFPYAVYVGMNYGPRNGVGGRYPNNRYQNNVQSNSEDRNNSVTIGVADEHVGLVLGRNGRSIIEISQQSGARIKISDRGDFISGTSDRKVTITGSQRAISMAESMISQKLARVSERDEFVKG*

NbL12g15020.1 copper_transport_protein_ATX1−like [MW = 12.87 kDa]

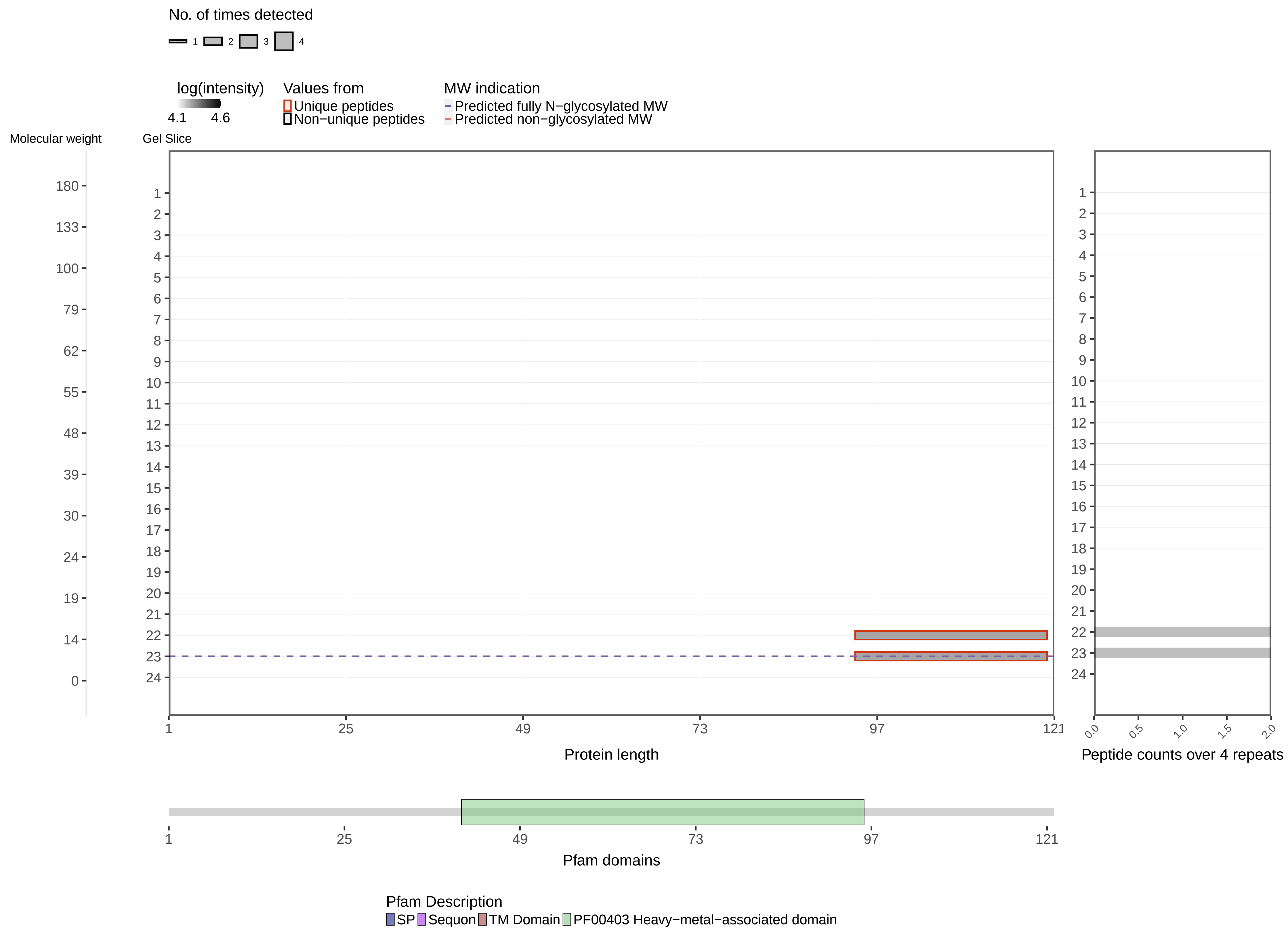


Protein sequence:

[illegible]GCVGAVKRVLGKMEGVENFDIDLKEQKVTVKGNVQPEAVLQTVSKTGKPISFWEAGPAQTEEKPAEAVGTA*

NbL12g14900.1 dnaJ_protein_homolog [MW = 46.66 kDa]

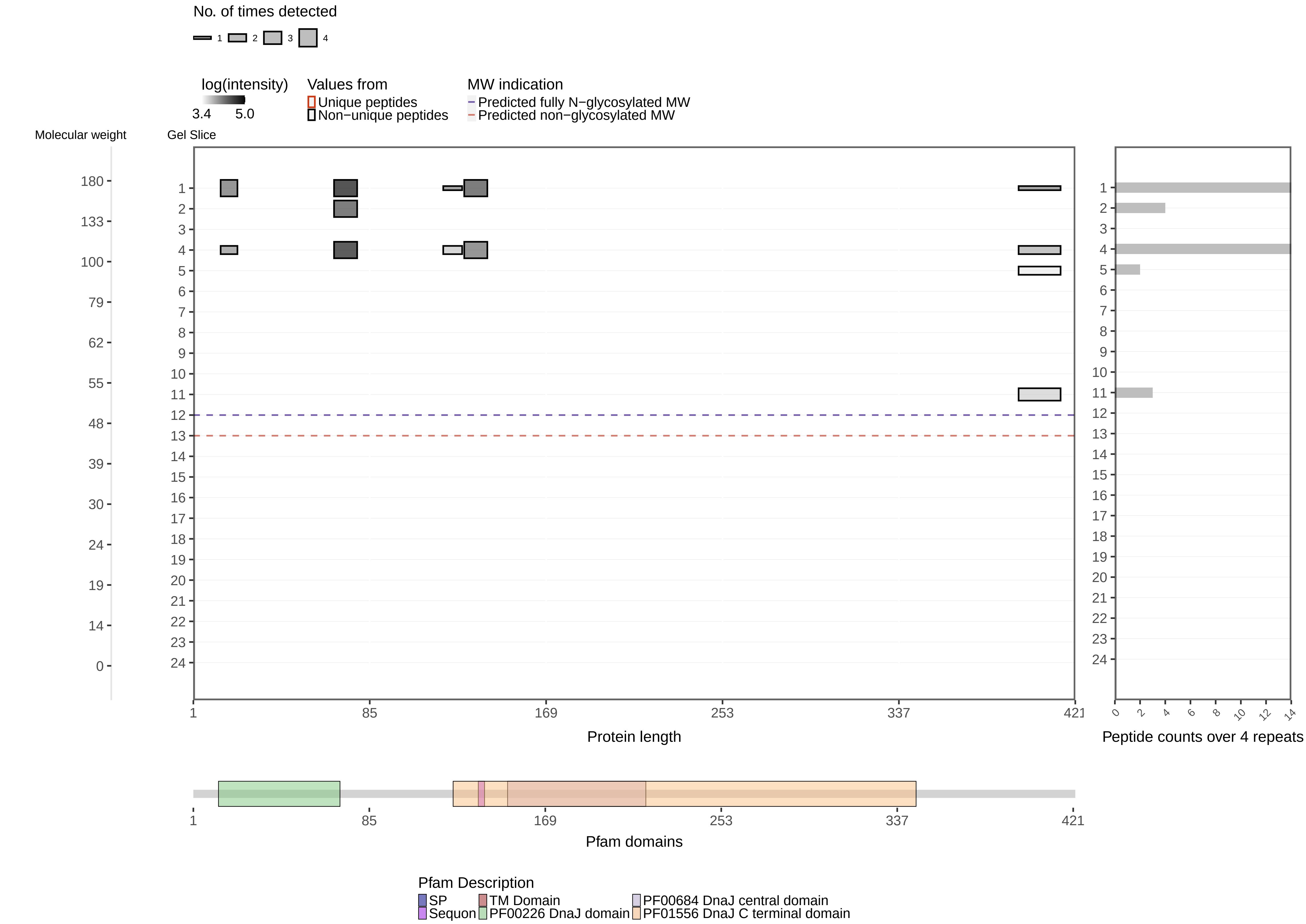


Protein sequence:

ECKGTGETISDKDRCGQCKGDKVVQEKKVLEVVVEKGMQNGQKITFPGEADEAPDTITGDIVFVLQQKEHPKFKRKGDDLFVEHTLSLTEALCGFQFILTHLDNRQLLIKSQPGEVVKPDQFKAINDEGMPMYQRPFMRGKLYIHFTVDFPETLSLEQCKNLEAVLPPKPKTQMTDMELDECEETTLHDVNIEEEMRRKQQQAQEAYDEDEDMHGGAQRVQCAQQ*

### NbL12g14860.1 ubiquitin_carboxyl−terminal_hydrolase_12−like [MW = 139.89 kDa]

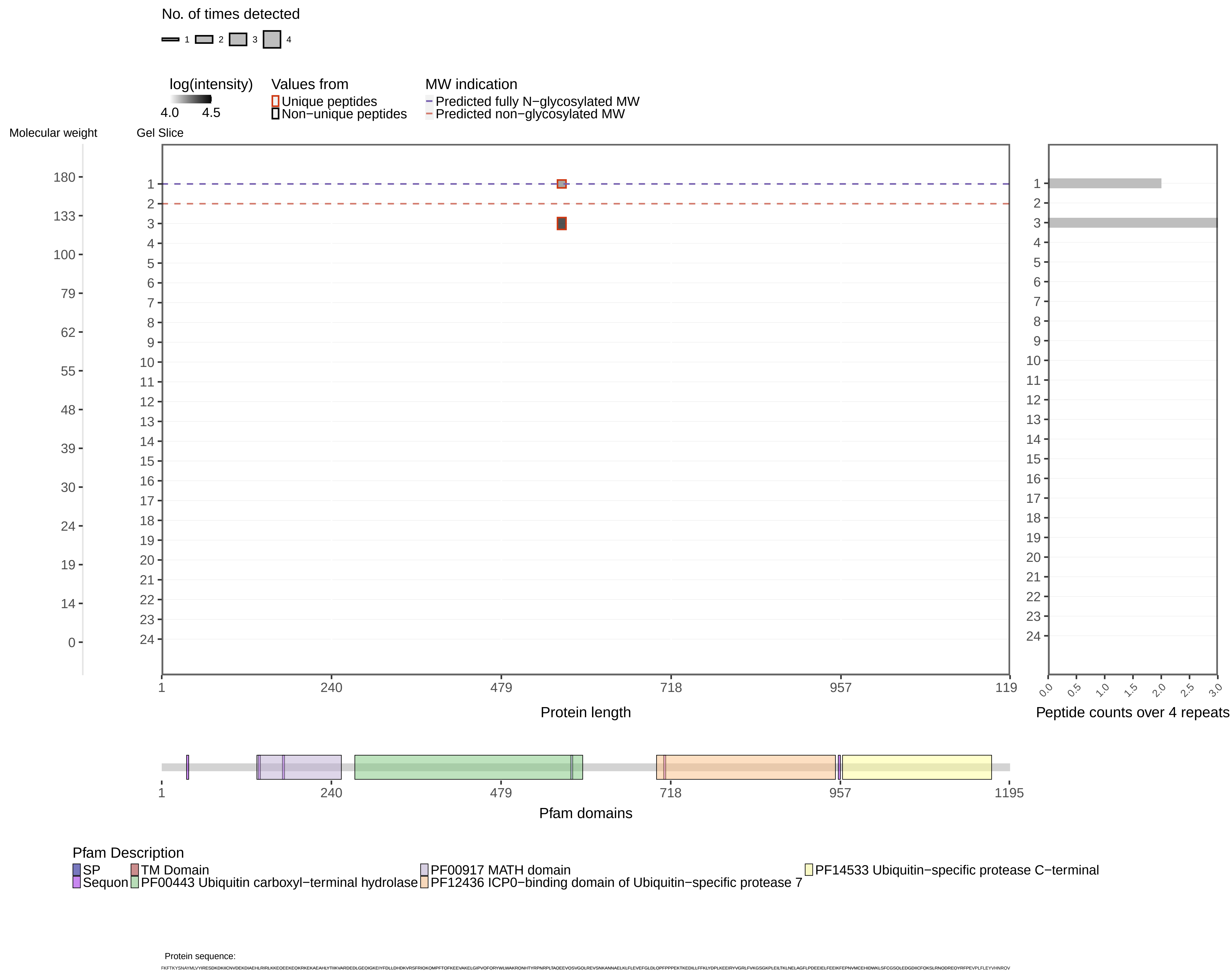

NbL12g14810.1 serine/threonine−protein_phosphatase_PP1−like [MW = 37.95 kDa]

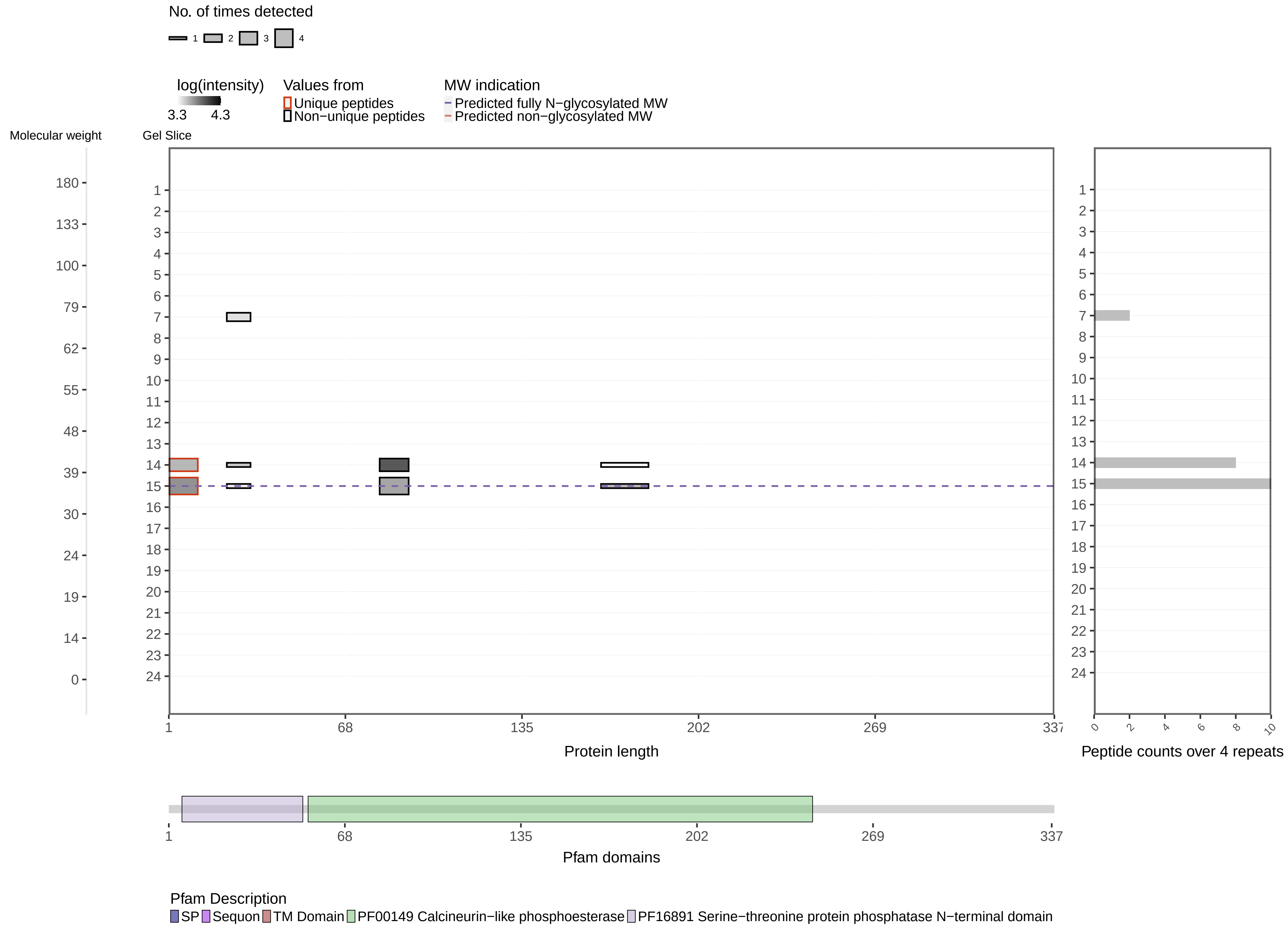


Protein sequence:

LIDEKILCMHGGLSPDLNNLDQIRNLQRPTDVPDAGLLCDMLWSDPSKDVQGWGMNDRGVSYTFGADKVTEFLEKQDLDLICRAHQVVEDGYEFFANRQLVTIFSAPNYCGEFDNAGAMMSVDETLMCSFQILKPAEKKSKFNFGSTTTAKPGSPTGMKSFLYVKVCEPASCGSQNC*

NbL12g14740.1 cysteine_desulfurase_1_chloroplastic−like [MW = 52.13 kDa]

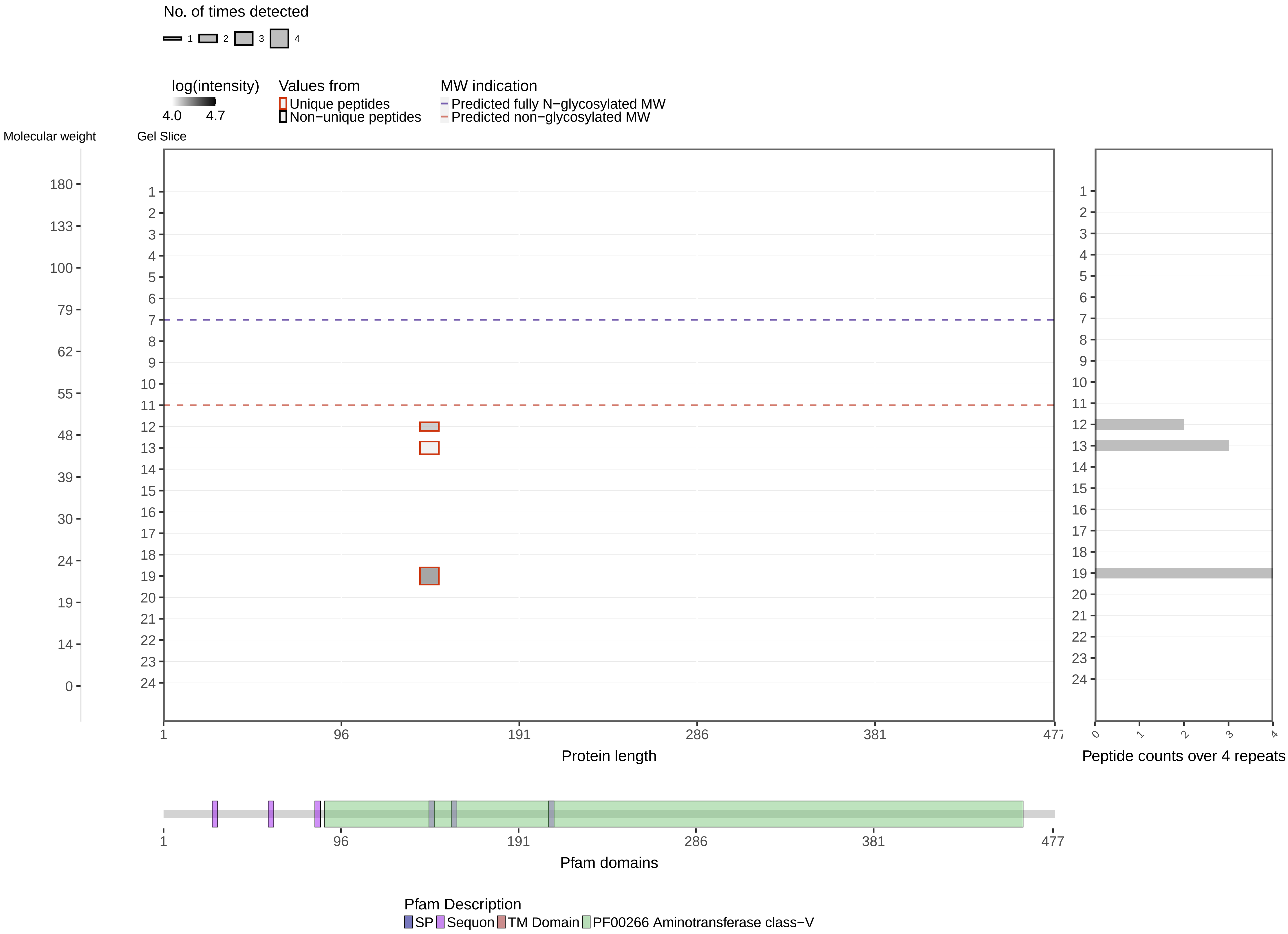


Protein sequence:

TKLLVIHHVSNMLASVLPMDEVISWAHDVGAKVLVDACQSVPHMVVDVKTLDADFLVASSHKMCGPTGVGFLYGKSEILSAMPPFLGGGEMIADVYLDHSTYAEPPSRFEAGTPAIGEAIGLGAAIDYLSEIGMQQIHDYEMELANYLYDSLSSVSDVRIYGPAPSRAVKRAALCSFNVKDVHPTDIATFLDQQHGVAIRSGHHCAQPLHRHLGISSSARASLHFYNTKDDVDRFIEALKDTISFFTSFK*

### NbL12g14680.1 nitrilase−like_protein_2 [MW = 35.68 kDa]

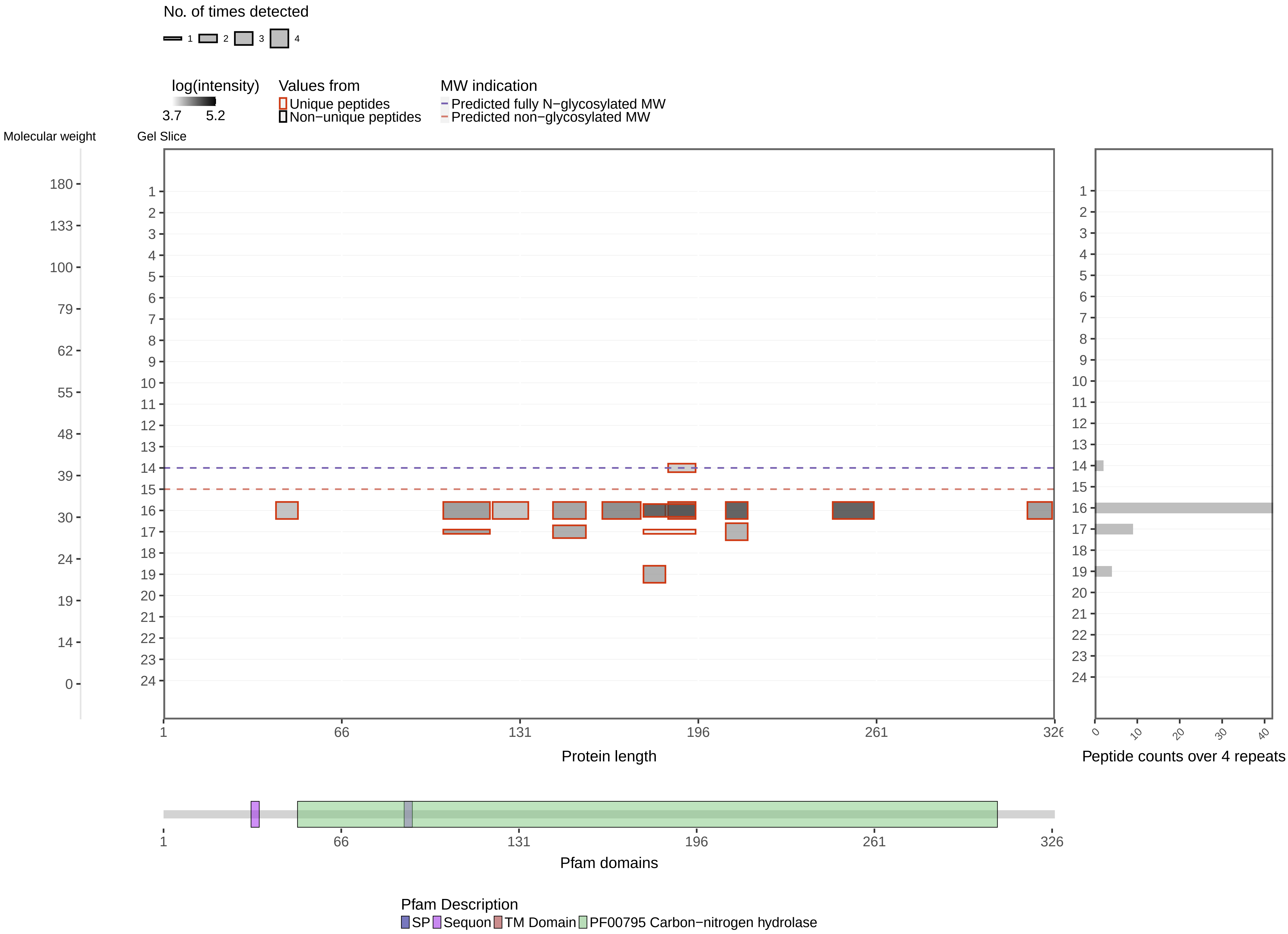


Protein sequence:

NGIIRSTYSKMHLFDVDVPGGAVYKESSFTEAGKDIVVADSPCGRLGLTVCYDLRFPELYQQLRFNHDAQILLVPAAFTMVTGQAHWEILLRARAIETQCYVIAAAQAGKHNEKRESYGDTLIIDPWGTIVGRLSDRSSTGITVADIDFSLIDSVRAKMPVSKHRKPAEFWNQHP*

NbL12g14610.1 putative_uncharacterized_protein_DDB_G0270496 [MW = 48.61 kDa]

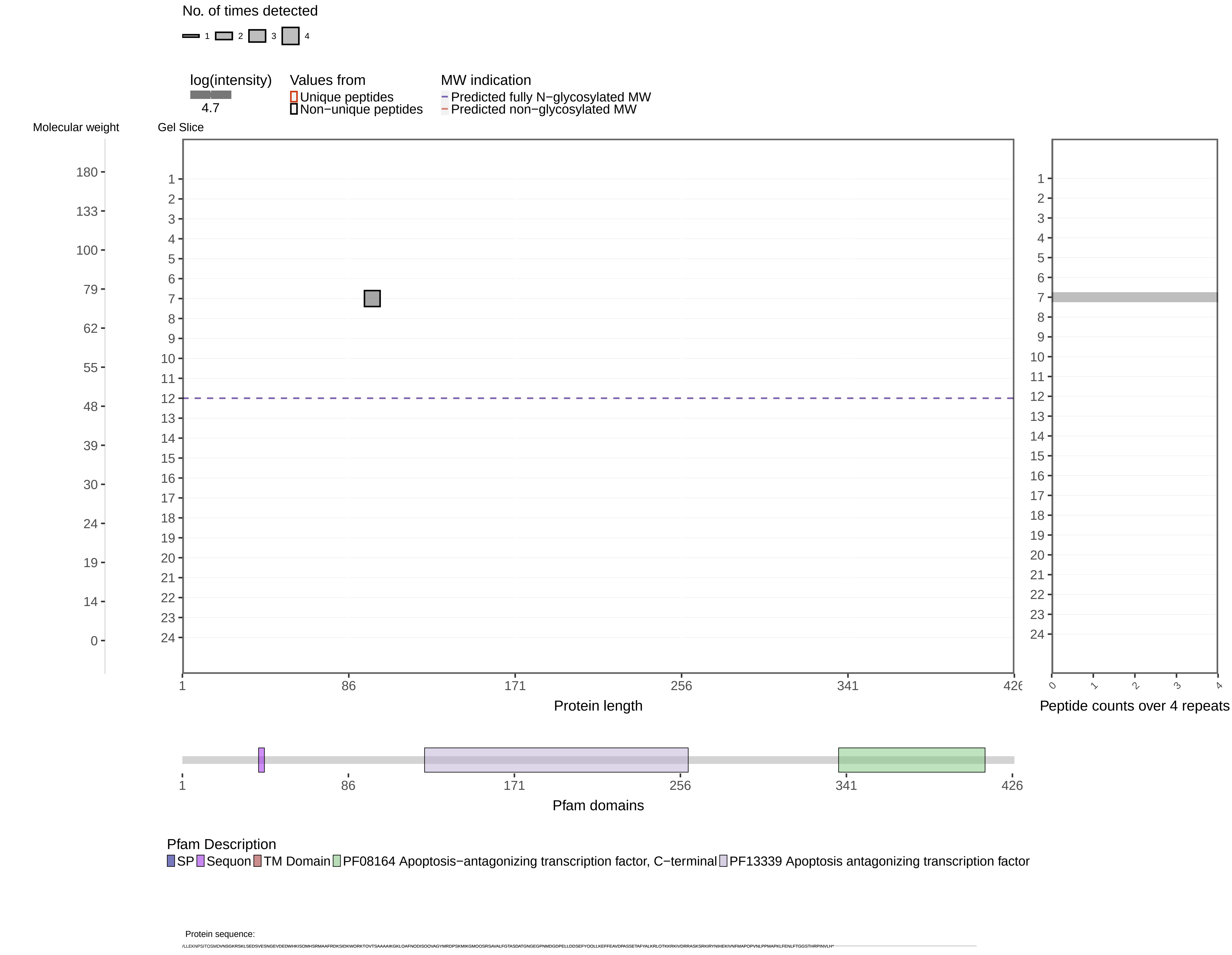

NbL12g14540.1 60S_ribosomal_protein_L35 [MW = 14.32 kDa]

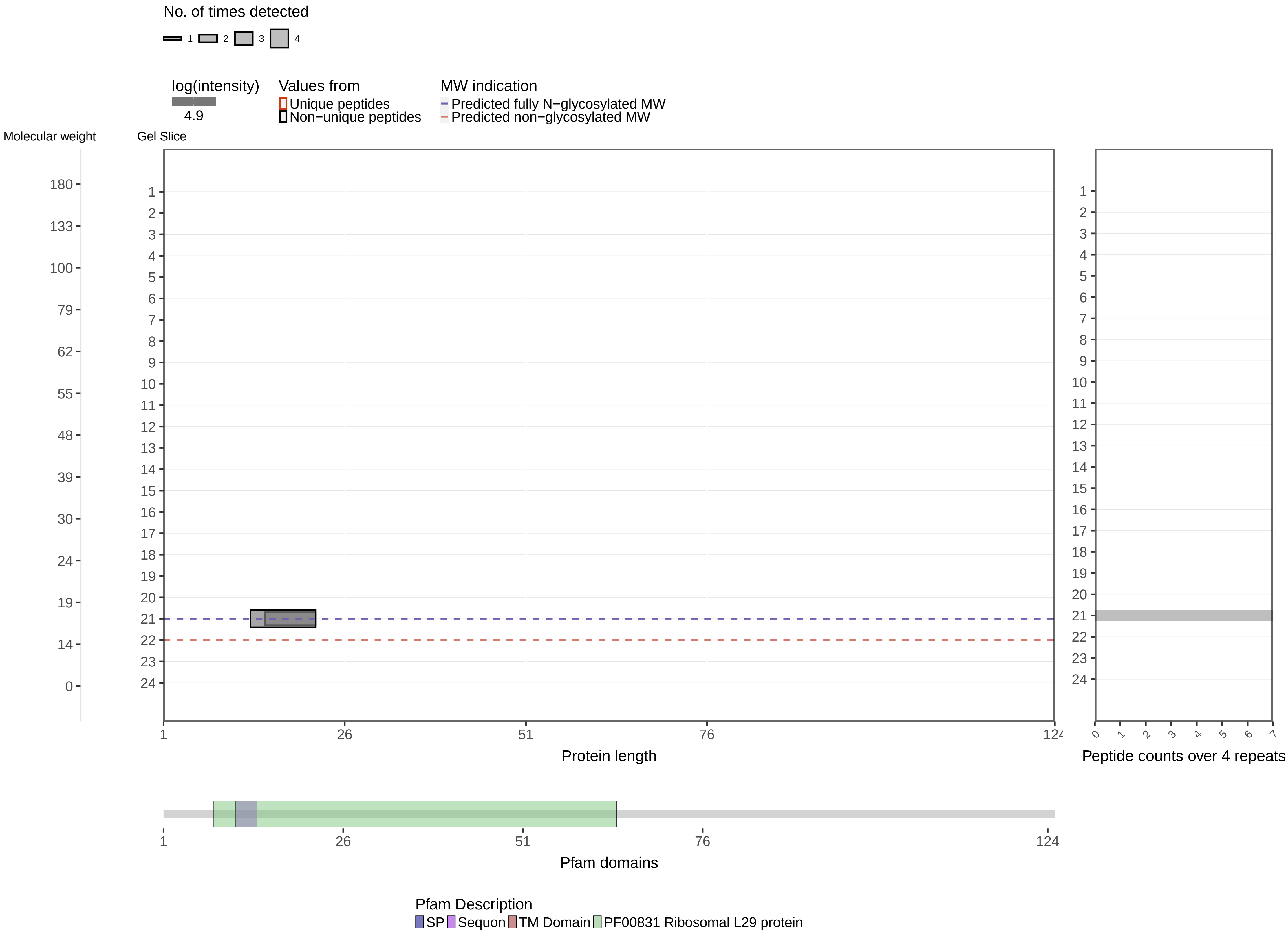


Protein sequence:

[illegible]VVRLSIAQVLTVISQKQKTALREAYKNKKYLPLDLRPKKTRAIRKRLTKHQVSLKTEREKKKEMYFPIRKYAIKV*

### NbL12g14460.1 glutathione_S−transferase_DHAR2 [MW = 23.79 kDa]

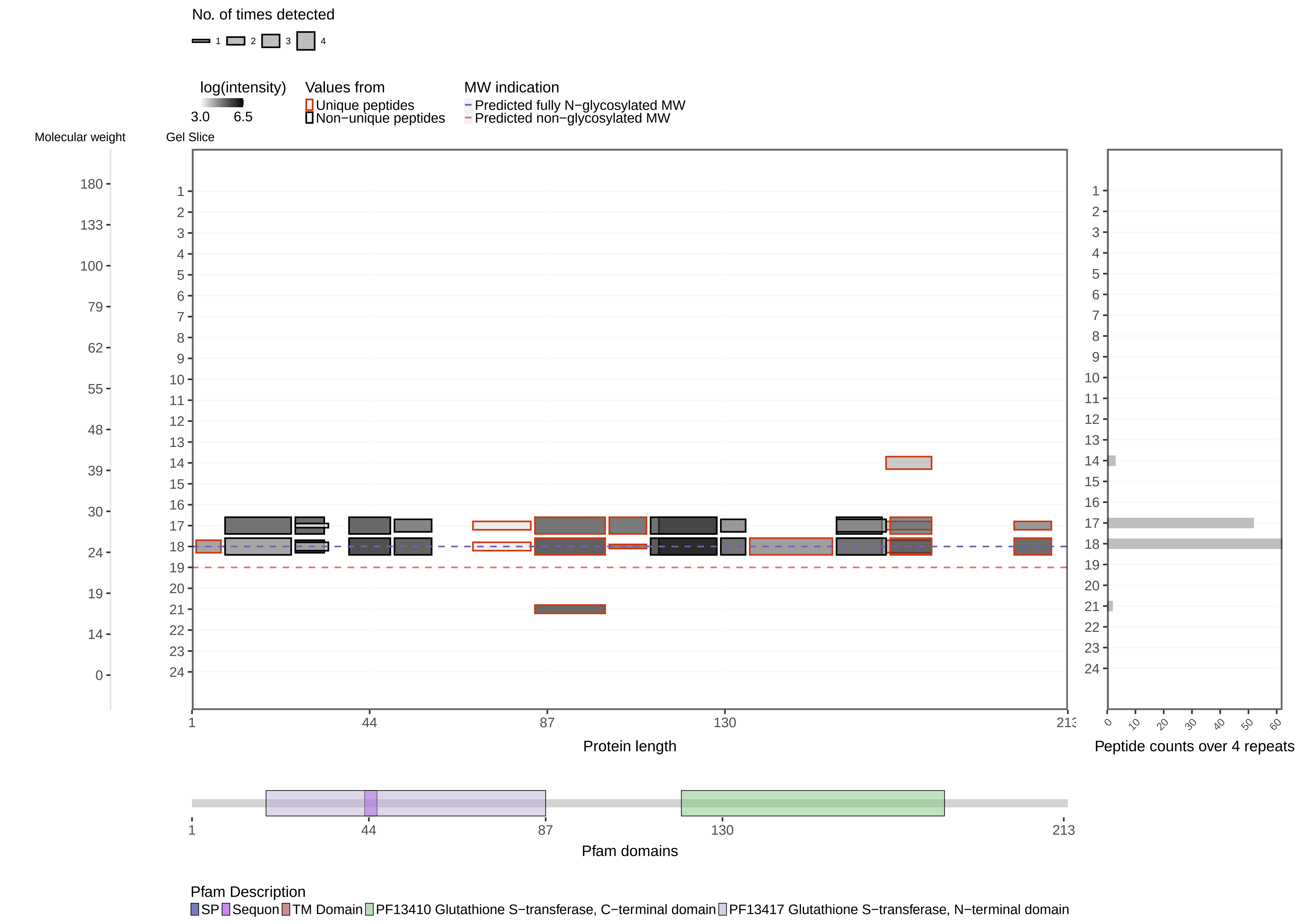


Protein sequence:

EFASVGSKIFPTFVSFLKSKDASDGTEQALLDELKALEEHLKAHVPYVNGENICSVDLSLAPKLYHLEVALGHFKKWSVPESFSHVRNYMKLLFERESFQKTKCAKEYVIAGWAPKVNP*

### NbL12g14340.1 2−oxoglutarate_dehydrogenase_mitochondrial−like [MW = 115.63 kDa]

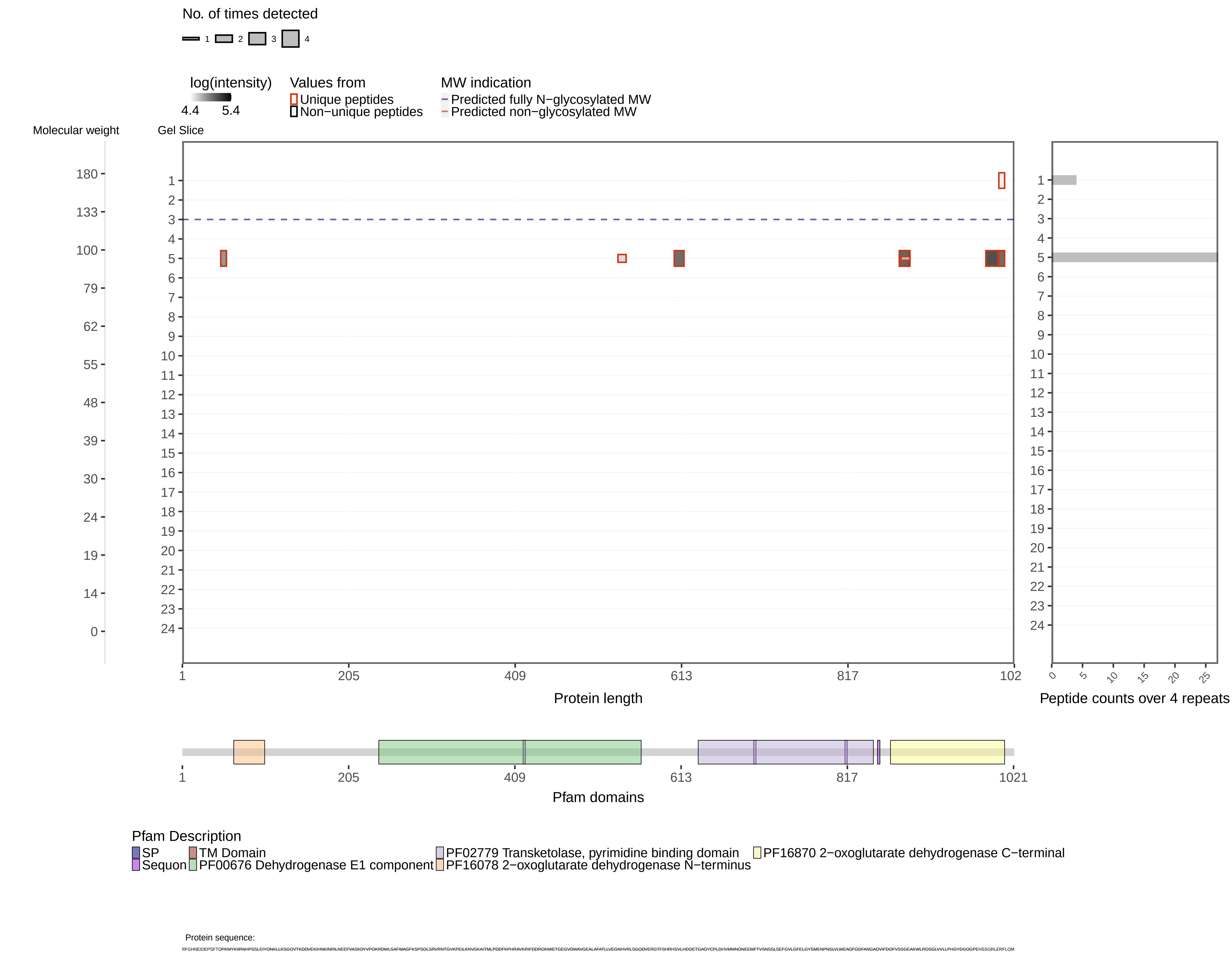

### NbL12g14310.1 histone_H4−like [MW = 11.43 kDa]

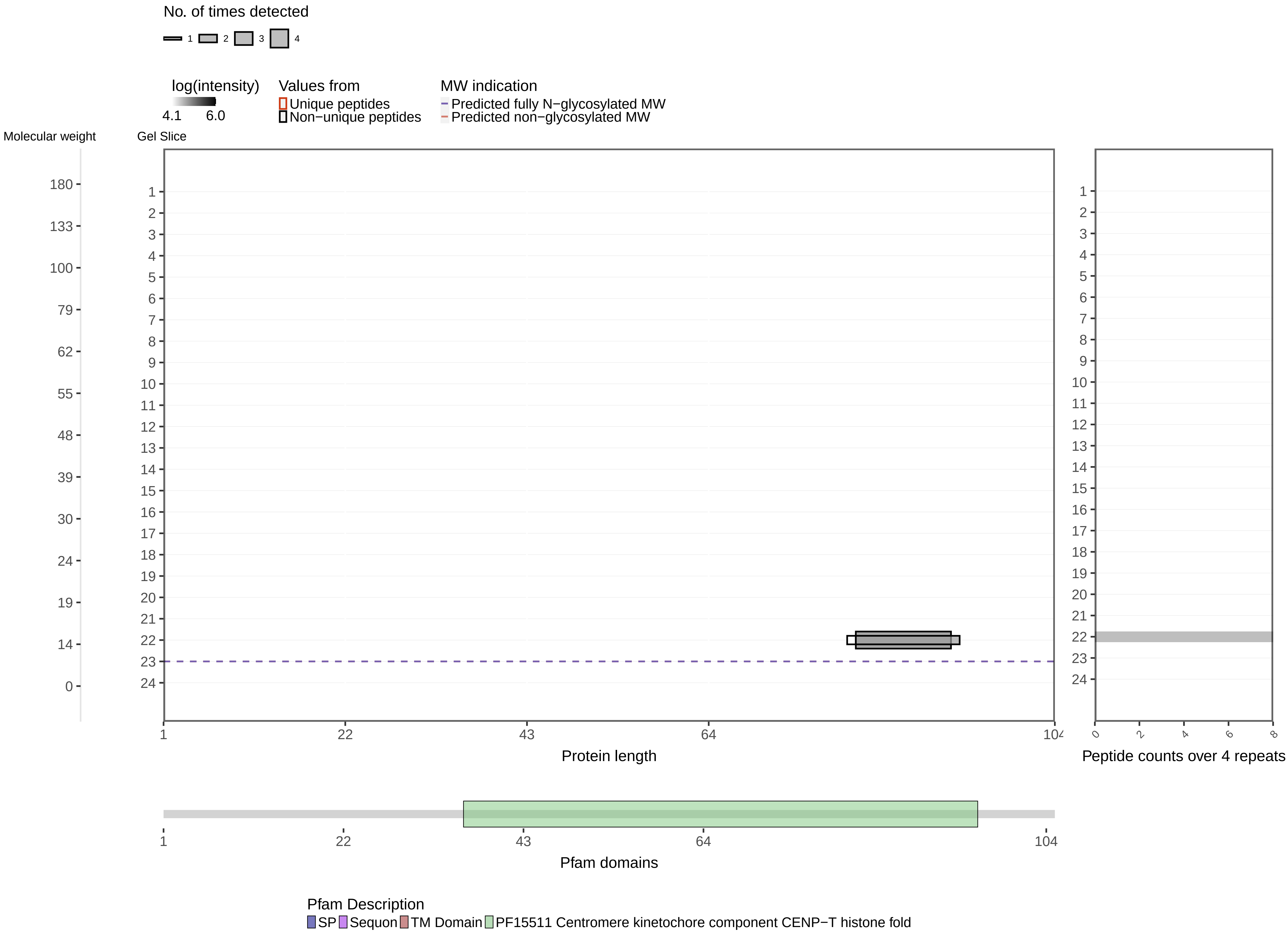


Protein sequence:

.ARRGGVKRISGLIYEETRGVLKIFLENVIRDAVTYTEHARRKTVTAMDVVYSLKRQGRTLYGFGG*

NbL12g14280.1 UDP−glucuronic_acid_decarboxylase_6−like [MW = 38.65 kDa]

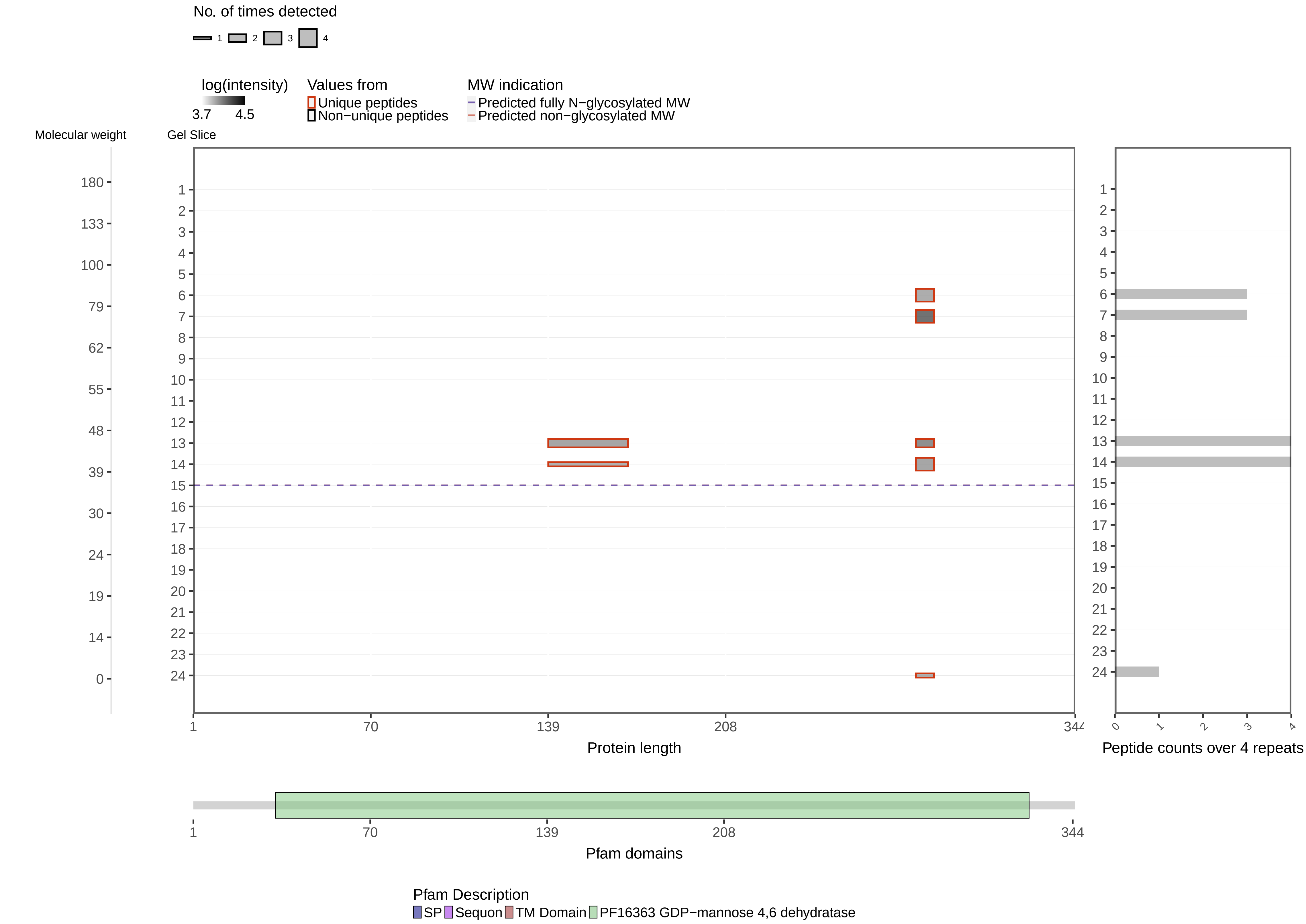


Protein sequence:

ETYWGNVNPIGVRSCYDEGKRVAETLMFDYHRQHGIEIRIARIFNTYGPRMNIDDGRVVSNFIAQAIRDEALTVQLPGTQTRSFCYVSDMVDGLIRLMEGDNTGPINIGNPGEFTMIELAENVKELINPGVKIITVENTPDDPRQRKPDITKAKELLGWEPKIKLRDGIPLMEEDFRGRLGIARKK*

NbL12g14240.1 ubiquitin−conjugating_enzyme_E2_7 [MW = 18.81 kDa]

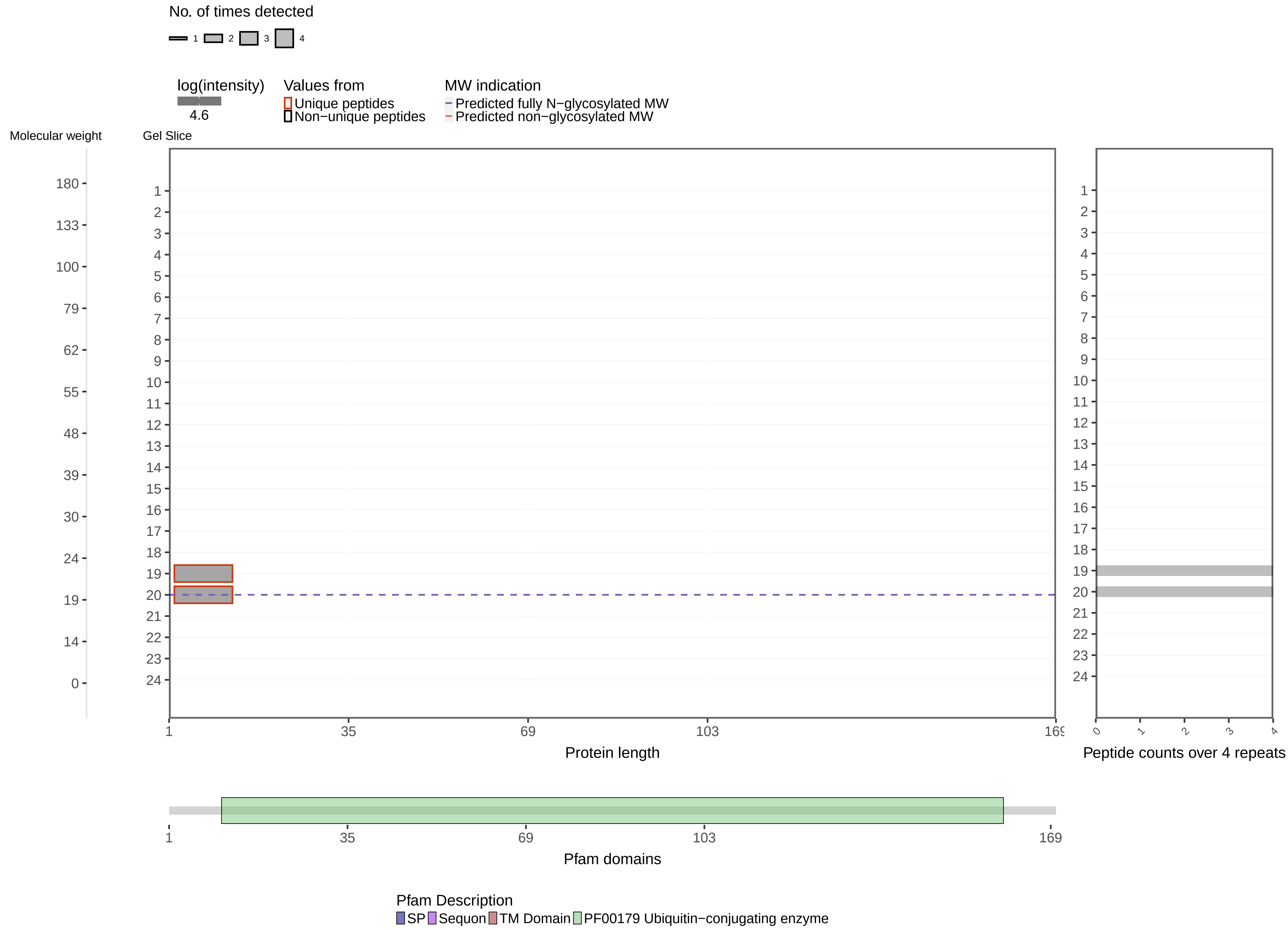


Protein sequence:

TIRFTSEVWHPNVYSDGKVCISILHPPGDDPNGYELASERWSPVHTVESIILSIISMLSSPNDESPANVEAAKEWRDNRDEFKKKVSRCVRRSQEMT*

NbL12g14020.1 pathogenesis−related_protein_STH−2−like [MW = 17.72 kDa]

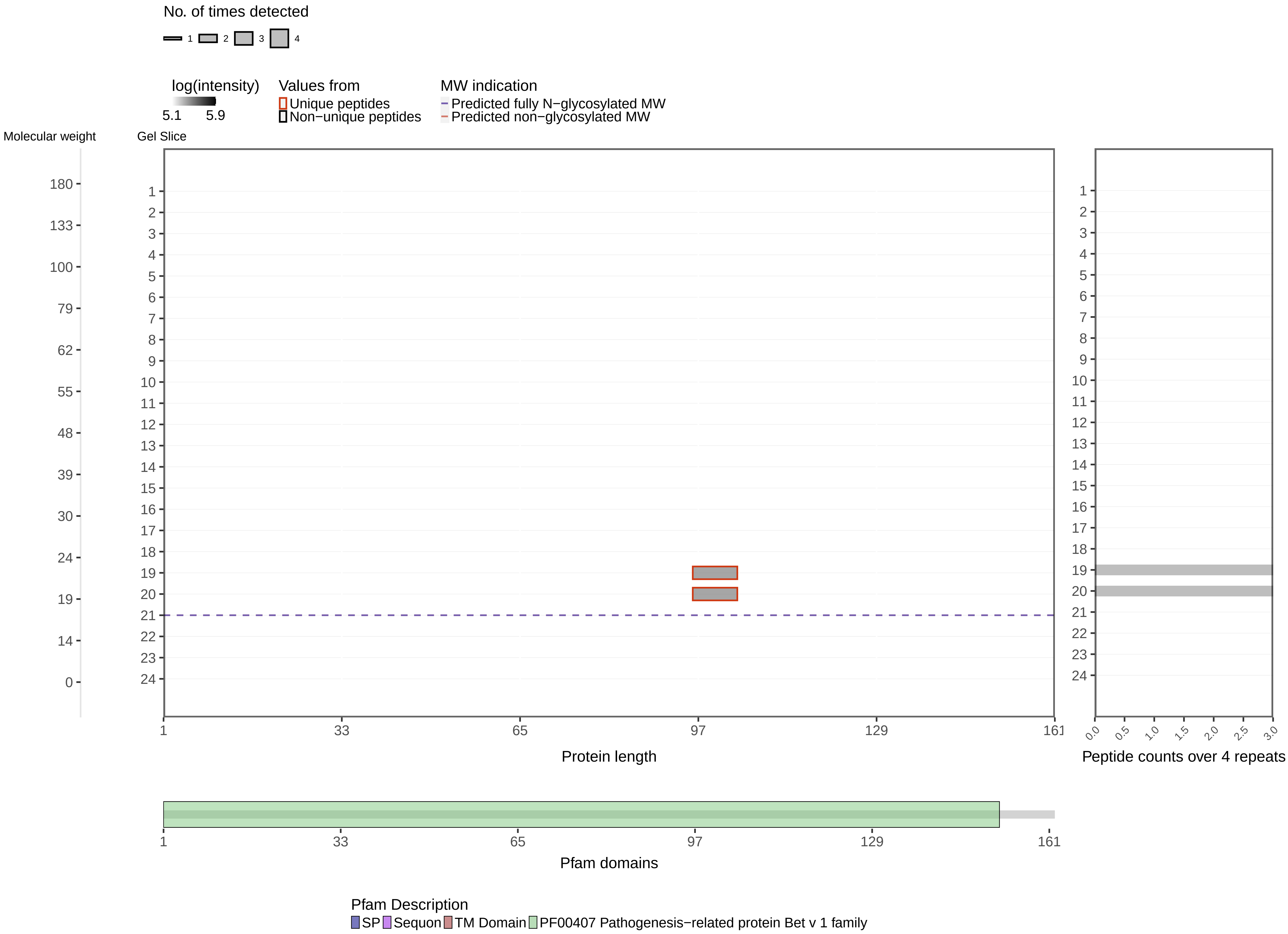


Protein sequence:

VKHRIDEVDKEKMVCKYTMIEGNALRDKLDSISYEVKFEESNNGGCICKMTTKYIGIGDFIVKEEEIKAGKDGATGIYKAVESHLPQNPSLYA*

### NbL12g13940.1 50S_ribosomal_protein_L7/L12 [MW = 23.86 kDa]

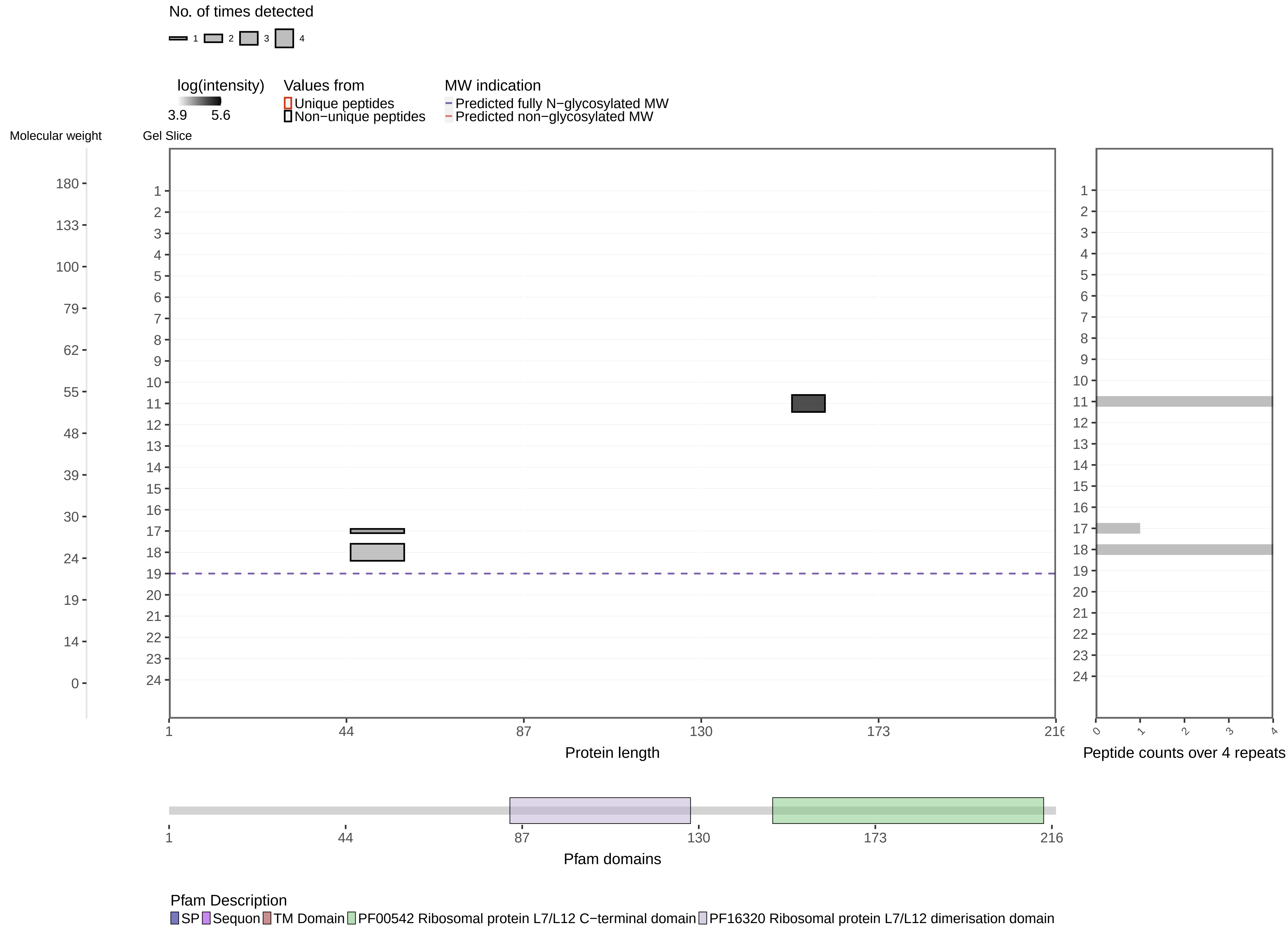


Protein sequence:

LVEVAELSSIIMKRLGMTEQPMVGVMKAGAAGLAAAAIKGSEAAKEDKKPEKTVFELKLESFDAAQKIKIIKEVRSCTELGLKEAKELVEKTPAIFKKGVSKEEGEQIIEKMKAVGAKVVLE*

NbL12g13890.1 probable_histone−arginine_methyltransferase_1.3 [MW = 59.13 kDa]

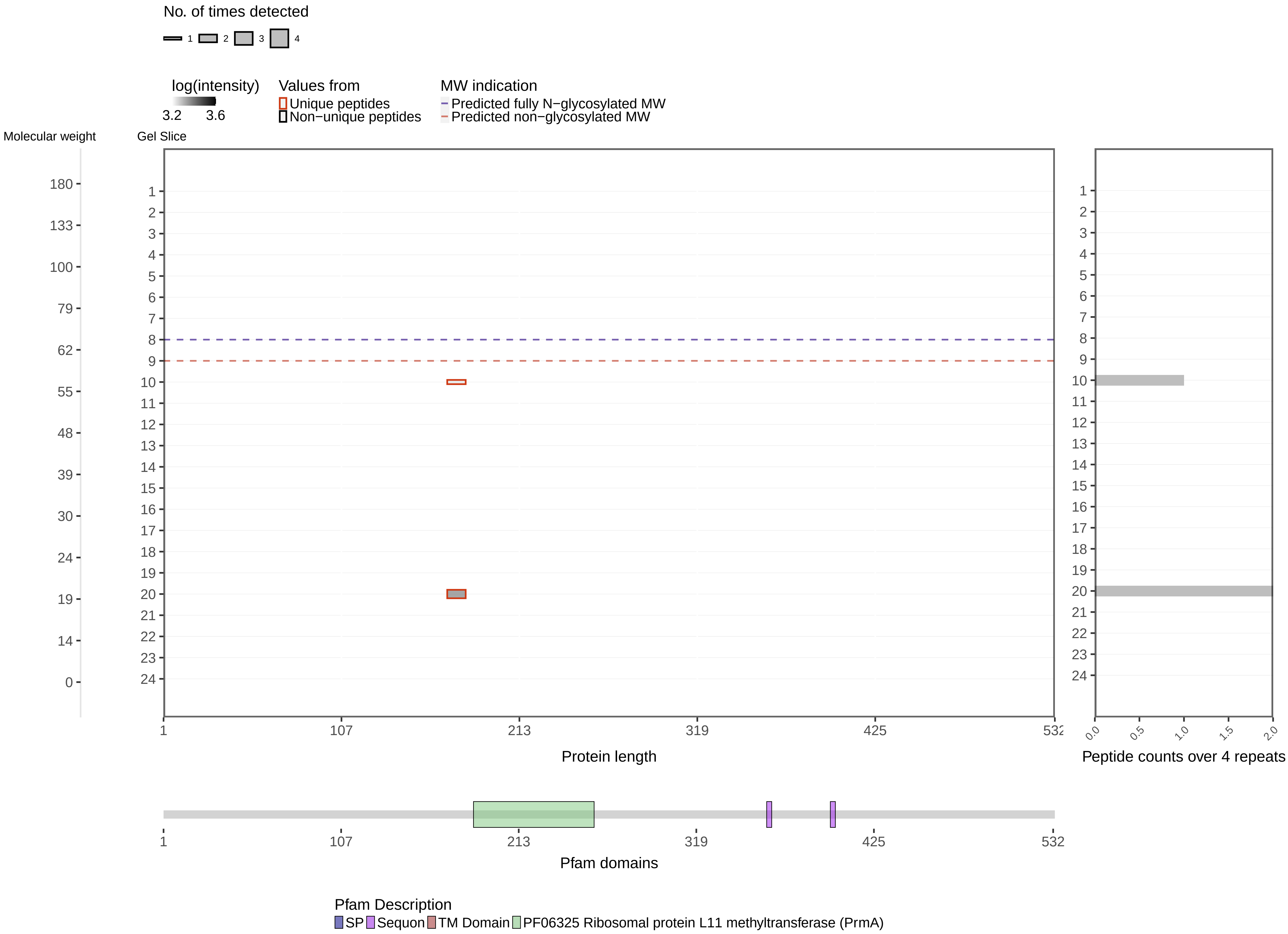


Protein sequence:

KADILISEPMGTLLVNERMLESYVIARDRFLVPNGKMFPTVGRIHVAPFSDEYLYMEIANKAIFWQQQGYFGVDLTPLQRSAYEGYFSQPVVDAFDPRLLVAPAISHVINFTSIKEEDLYEIDIPLRFTSSASTRIHGLACWFDVLFNGSTVPRLLTTAPGAPTTHWYQLRCVLSQPLYVMPGQEITGCLRLVAHKAQSYTIYLTLSATVGDVLQTSSGKLDLKEPYYRMSQPQSYSVAQDQQPNQLLQAQDILTQSRDEDGSVPVQQPSPNSGAVLHSL*

### NbL12g13710.1 induced_stolen_tip_protein_TUB8−like [MW = 20.91 kDa]

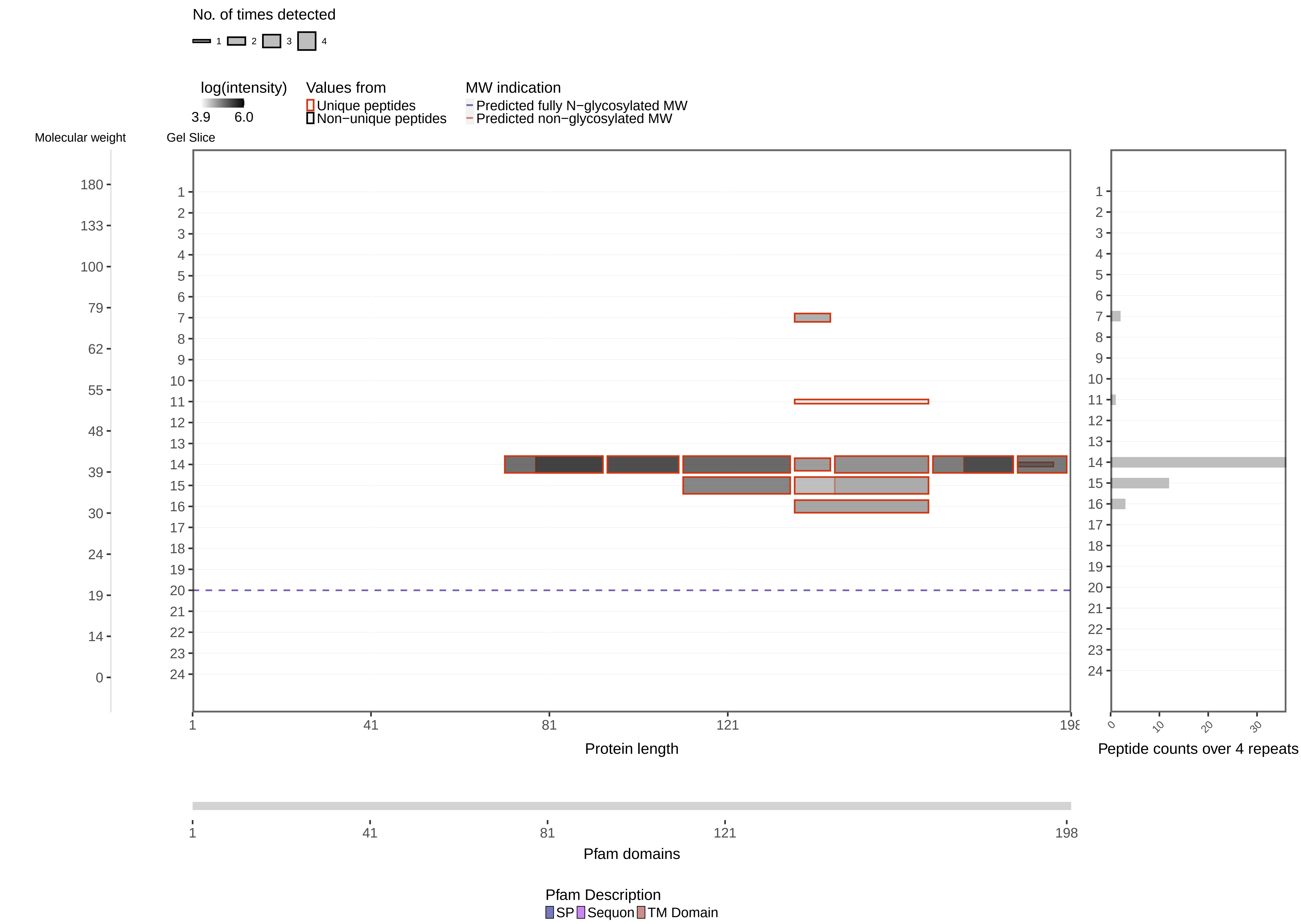


Protein sequence:

AEENVAEKNVAETTPAEETAPVEEKAAPAAEYTVAEEATTEVETEAPVAKETVAEEAAKAETVTVAAAAVETPAVELAEVKTEAEEGKAVETTEAPAAAKATTEVPVEKIEE*

### NbL12g13680.1 60S_ribosomal_protein_L6−like [MW = 27.78 kDa]

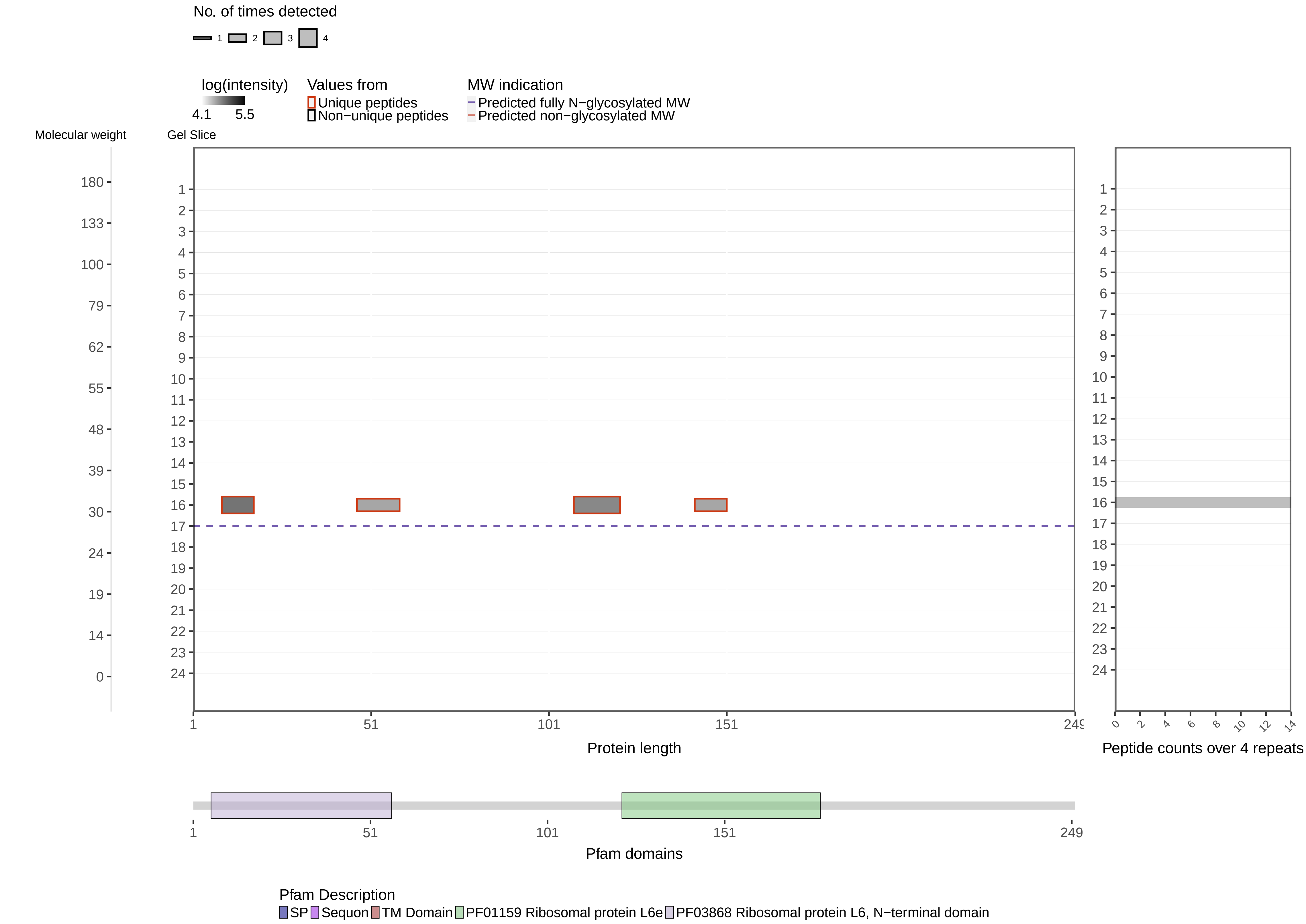


Protein sequence:

GLLLVTGPFKLNGVPLRRVNQAYVIGTSTKVDVSGVNVEKFDDKYFAKQVEKKQKKGEEEFFEDKKEVWIFLHYVICDFYIKSKAMGHLYNICARGHGAQCRRRPFGNYLICFLTSYANSLSIELVRLCLCMCACAC*

### NbL12g13660.1 UTP−−glucose−1−phosphate_uridylyltransferase−like [MW = 51.91 kDa]

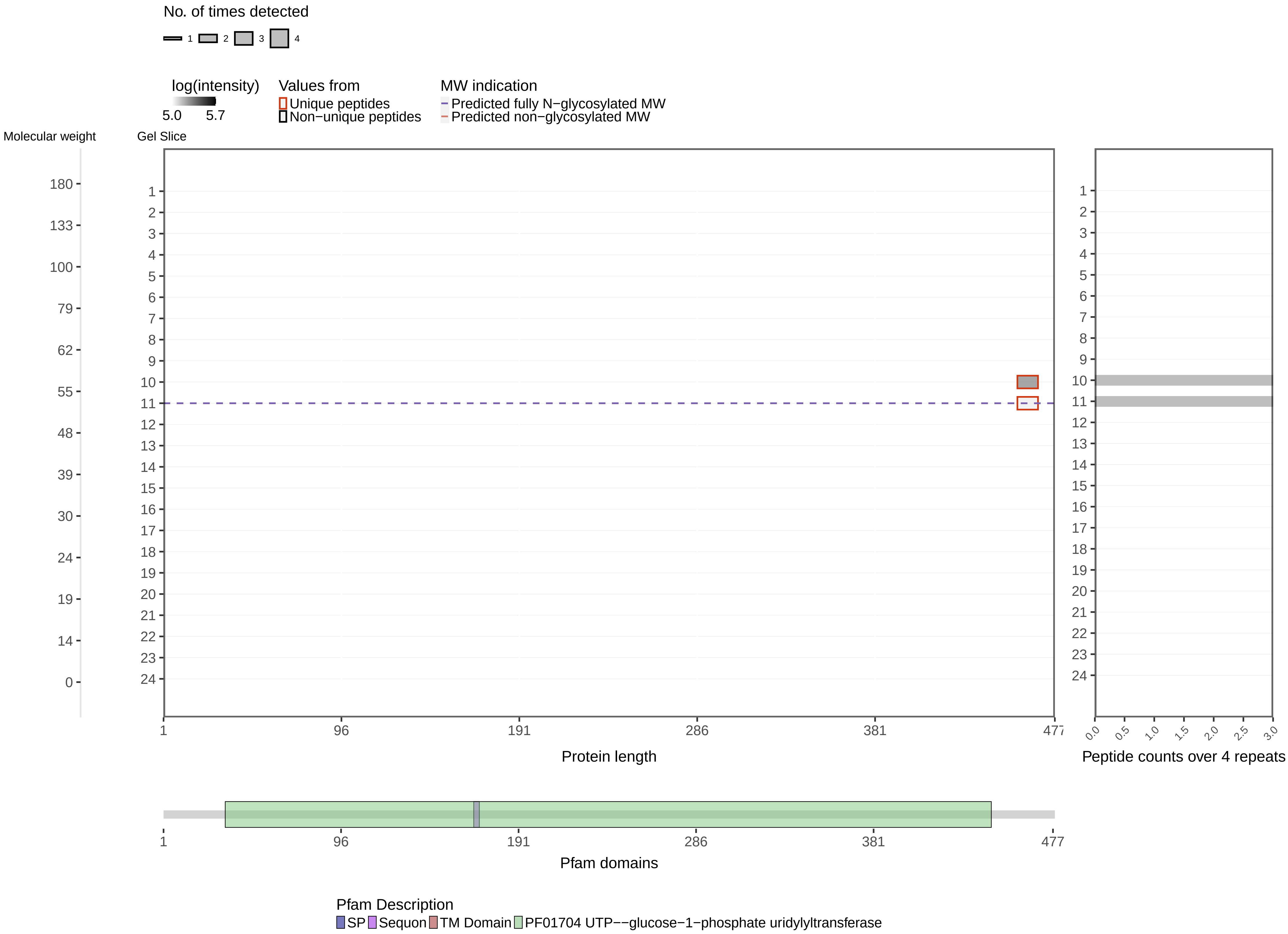


Protein sequence:

FVANSDNLGAVVDLKILNHLIQNKNEYCMEVTPKTLADVKGGTLISYEGKVQLLEIAQVPDQHVNEFKSIEKFKIFNTNNLWVNLKAIKRLVEADALKMEIIPNPKEVDGIKVLQLETAAGAAIRFFDRAIGANVPRSRFLPVKASSDLLLVQSDLYTLTDDGYVIRNPARTNPSNPSIELGPEFKKVANFLSRFKSIPSIIELDSLKVTGDVYFGAGITLKGKVNVAAKAGVKLEIPDGAVLANKDVNGPEDI*

### NbL12g13650.1 glutamate_decarboxylase−like [MW = 55.96 kDa]

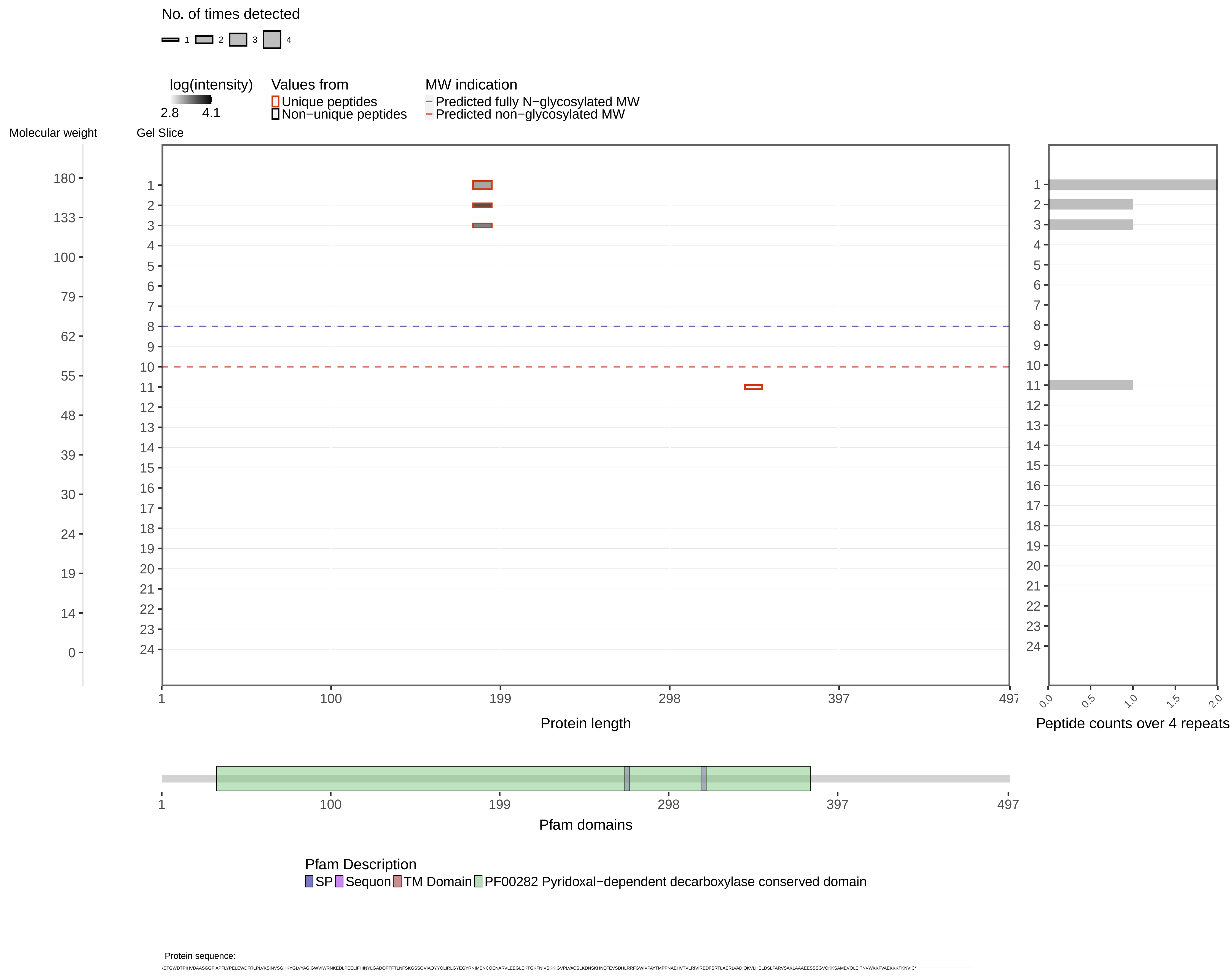

NbL12g13600.1 pre−mRNA−splicing_factor_ISY1_homolog [MW = 35.21 kDa]

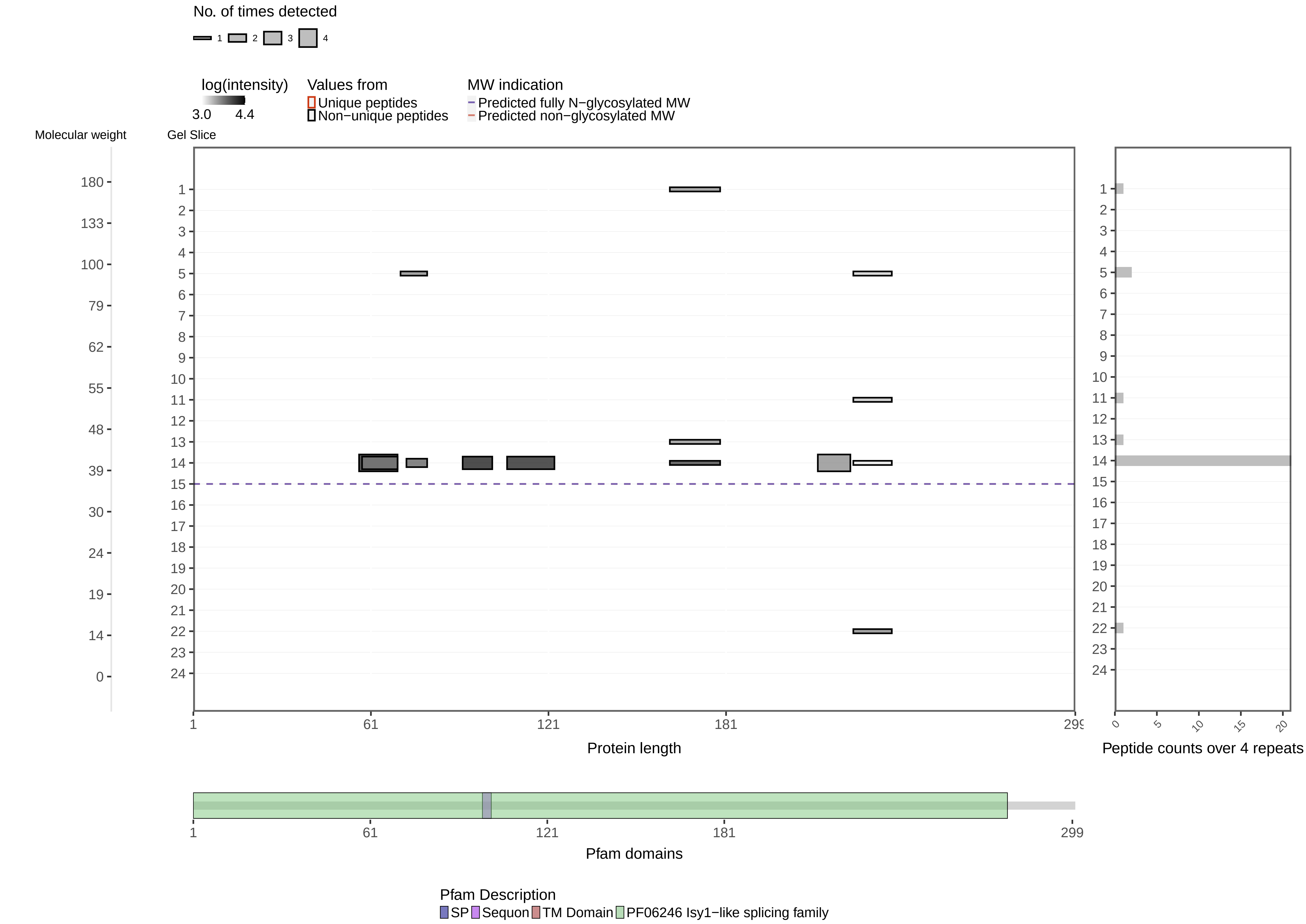


Protein sequence:

LPGVRELFEKPPELRKRRTRYDIYKRIDASYYGYRDEEDGILEKLEGPAEKKMOAAAVKEWMEMEEIKREARKAVKSGEVVEVGLGSKILFEEEEDVIEEERROKEREEEKEKEFVVHVPLPDEKEIERMIVEQKKMELLSKYASEDLMEEOMEAKAMLNIOR*

### NbL12g13520.1 HMG−Y−related_protein_A−like [MW = 21.68 kDa]

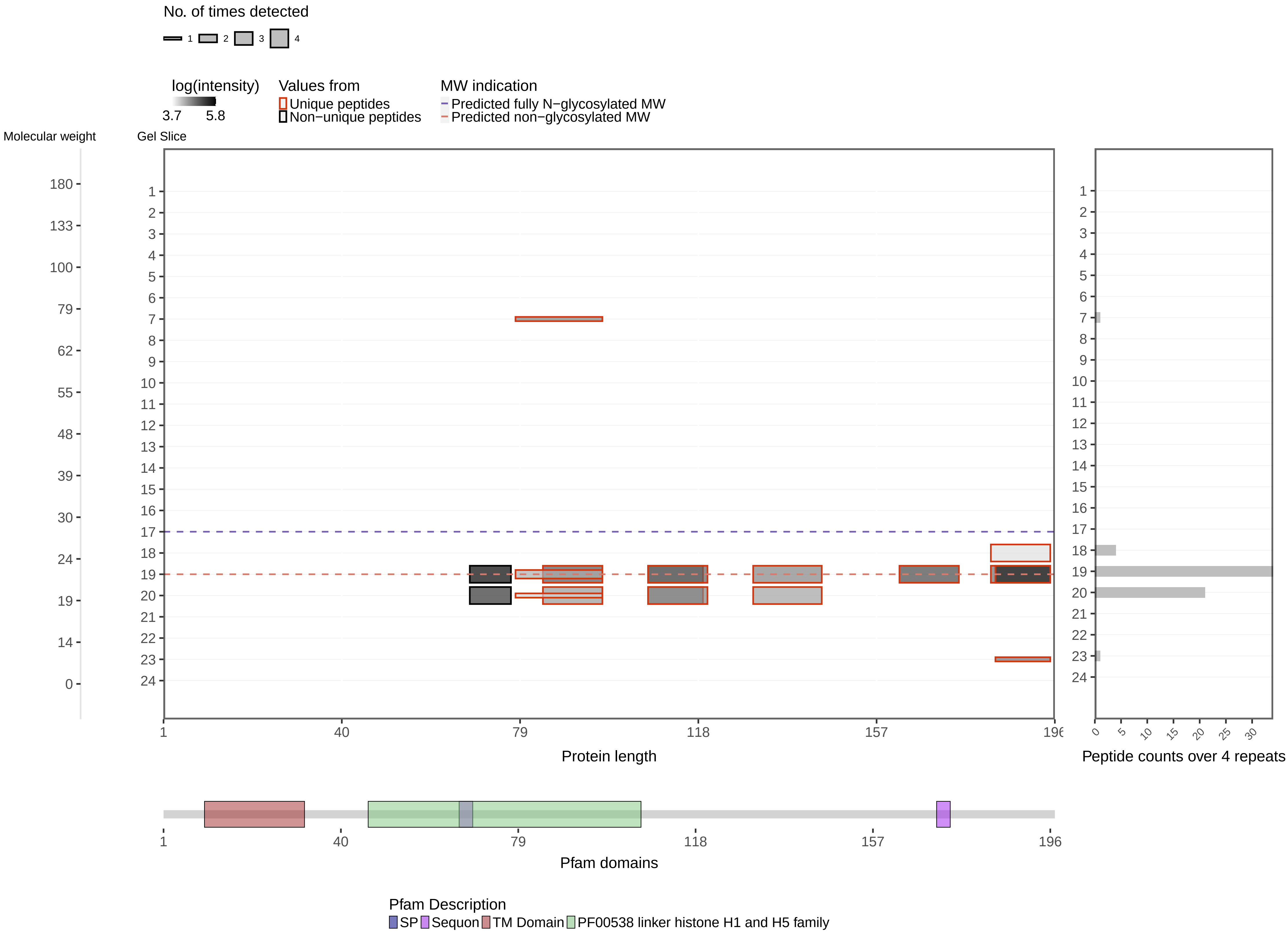


Protein sequence:

EFLTLNLDNMKQTGELIFLKNNYIKPGTDVPQKRGRGRPPKPKTPLPPGAESAEIASPRPRGRPRKDPNAPPTSKKPKPTPAAPNSTGRPRGRPRKVRPQPAQDDVEES*

### NbL12g13510.1 uncharacterized_LOC104103198 [MW = 49.69 kDa]

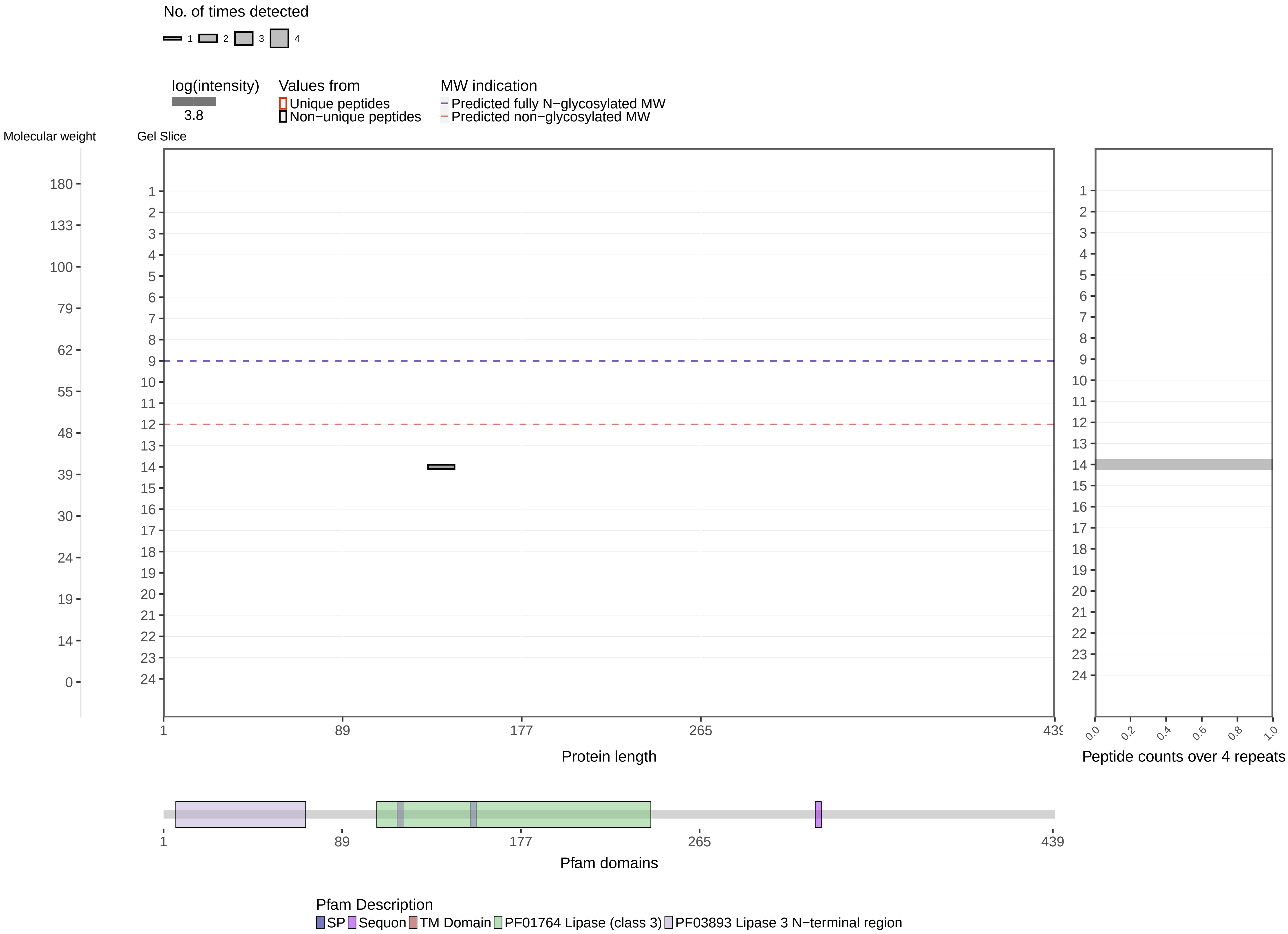


Protein sequence:

RCYAVAPARCMSLNLAVKYADIIHSVVLQDDFLPRTATPLEDIFKSIFCLPCLIFLVCLRDTFIPEGRKLQDPRRLYAPGRMYHIVERRFCRCGRYPPDVRTAIPVDGRFEHIVLSHNATADHGIIWIERESEKALARLKEASAETTTTPPKVPKIERLKTLEKEHKDALERAVSLNIPYAVEADEEESAESKTEESSQKQEEDAMTNKAQCSDARTNWDEVVEKLFNRDESGK*

### NbL12g13340.1 NA [MW = 28.57 kDa]

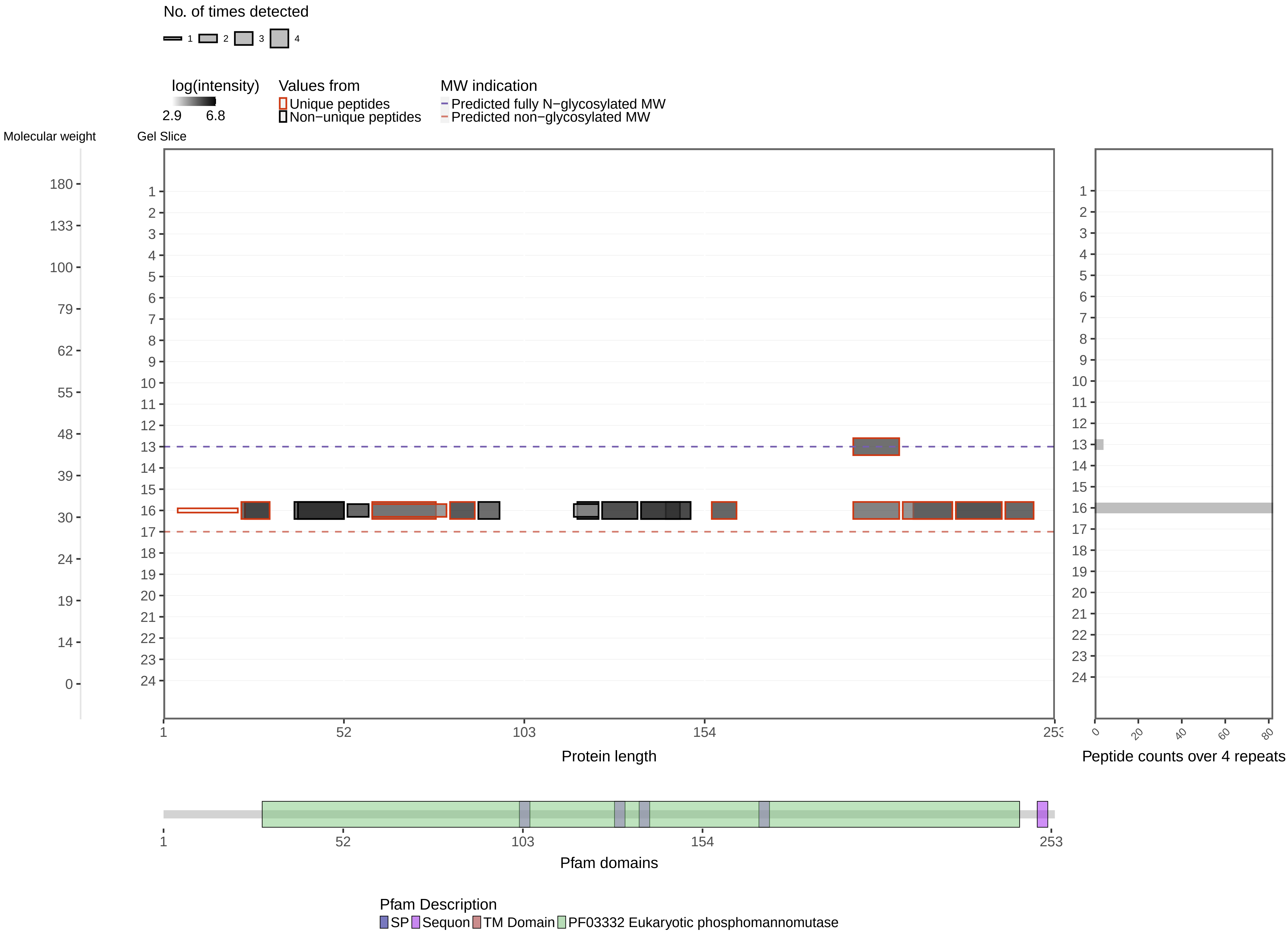


Protein sequence:

KRGTFIEFRSGMLNVSPIGRNCSQEERDEFEKYDKVHKIRQTMVSVLREKFAHLNLTFSIGGQISFDVFPQGWDKTYCLRYLEEFNEIHFFGDKTYKGGNDHEIYESERTVGHTVTSPEDTVKQCSEQFLGKDNGSS*

NbL12g13270.1 probable_polygalacturonase [MW − SP = 49.72 kDa]

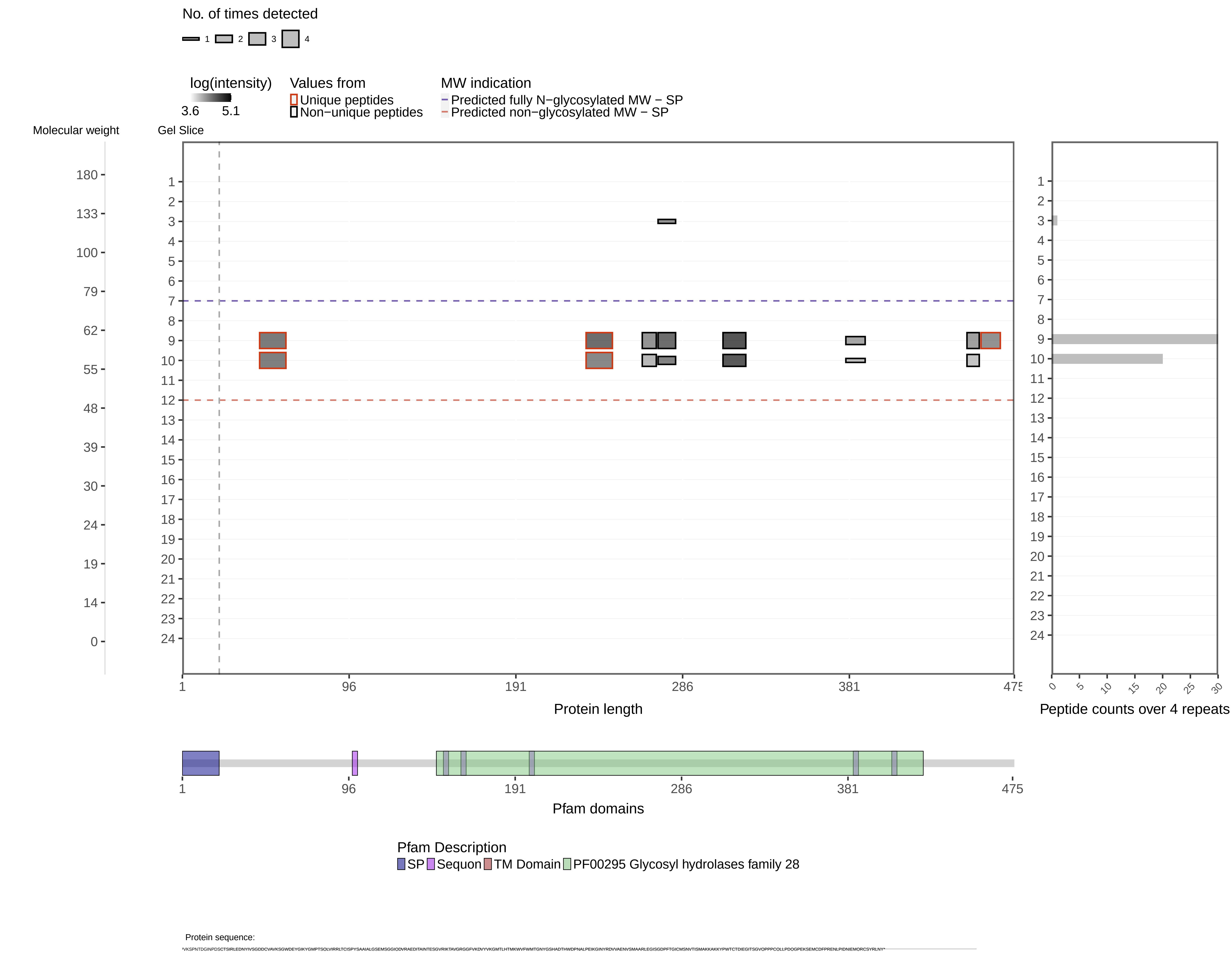

NbL12g13260.1 mitogen−activated_protein_kinase_homolog_MMK2−like [MW = 42.94 kDa]

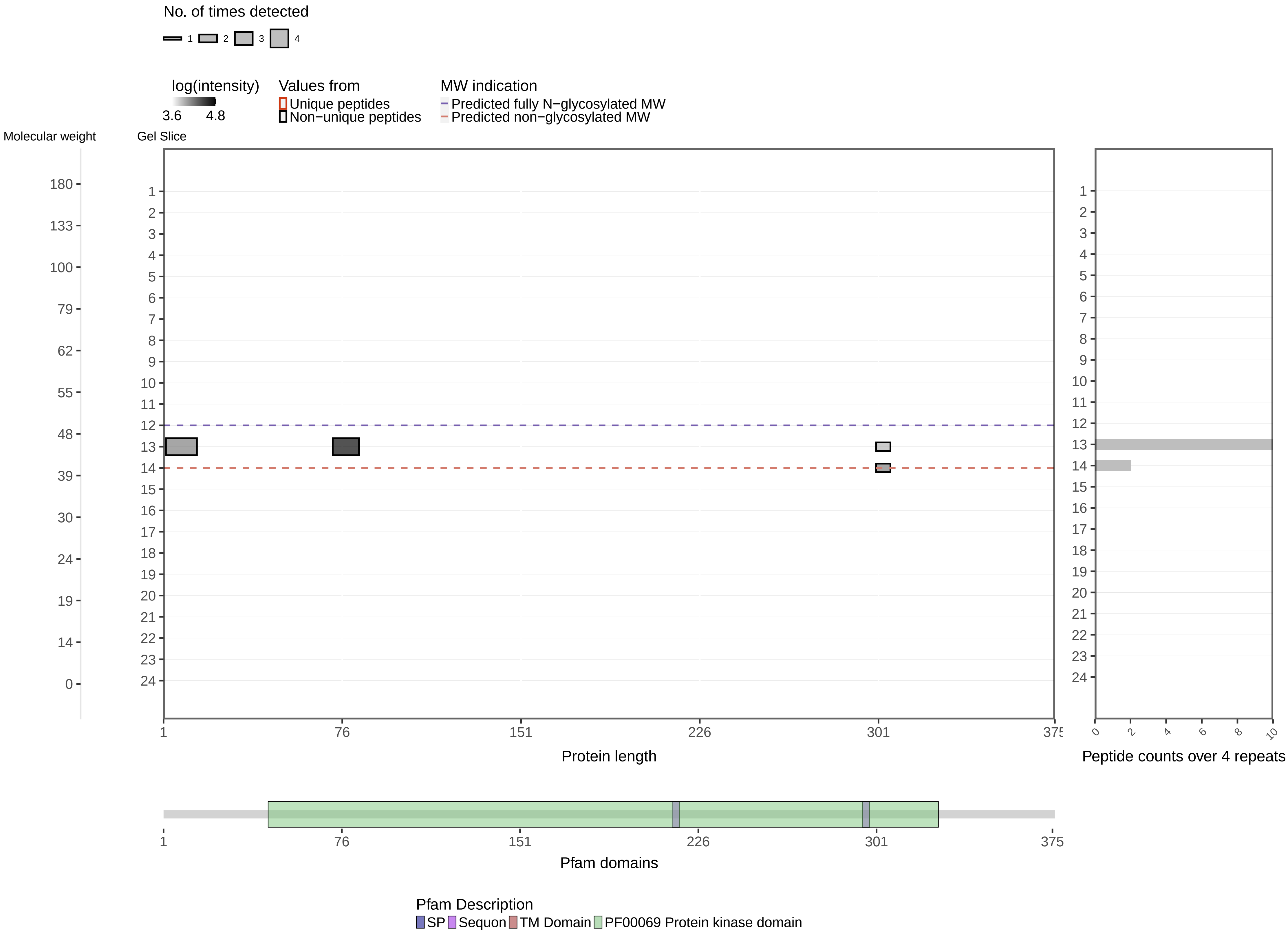


Protein sequence:

/NAKCDLKIGDFGLARTTTETDFMMEYCVTRWYRAPELLLNCSEYTSAIDVWSVGCILGEILTRQPLFPGRDYVHQLRLITELIGSPDDASLGFLRSNNARRYVRQLPRYPRQQFSARFPNSSPRAVDLLEKMLIFDPSRRITVDEALSHPYLAPLHDSNEEPVCPRPFSLDFEQPSFTEDNIKELIWREAVKFNPDPTH*

### NbL12g12910.1 cyclic_pyranopterin_monophosphate_synthase_mitochondrial [MW = 25.05 kDa]

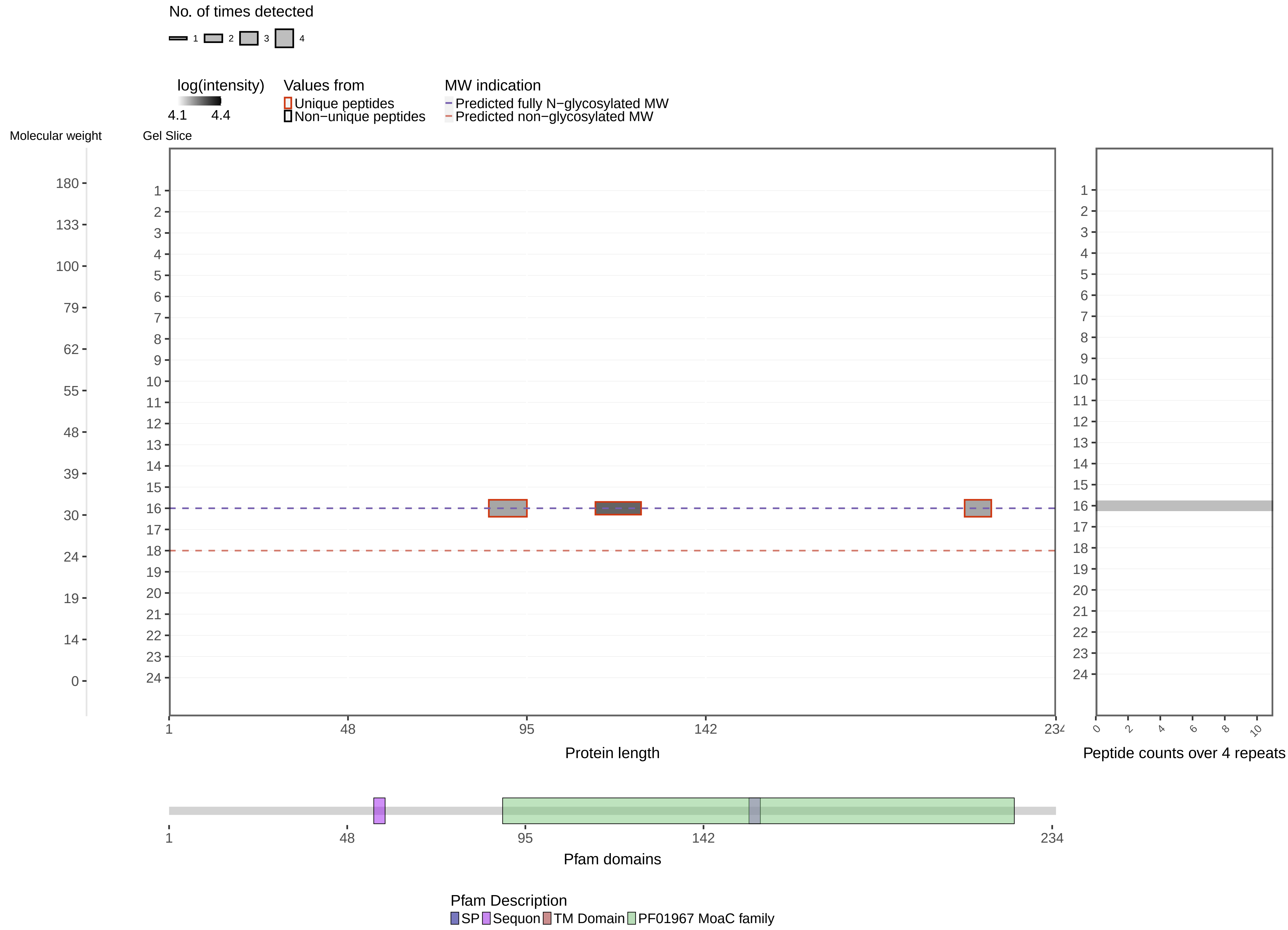


Protein sequence:

AHGKVILGREVFDLVSANQMGKGDVLSVAKLAGICGAKQTSNLIPLCHNINLTHVRVDLALNPHDFSVEIEGEAASDGKTGVEMEALTAVTVAGLTVYDMCKAASKYIQITDIRLERKTGGKSGDWSRDK*

### NbL12g12750.1 transketolase_chloroplastic−like [MW = 79.79 kDa]

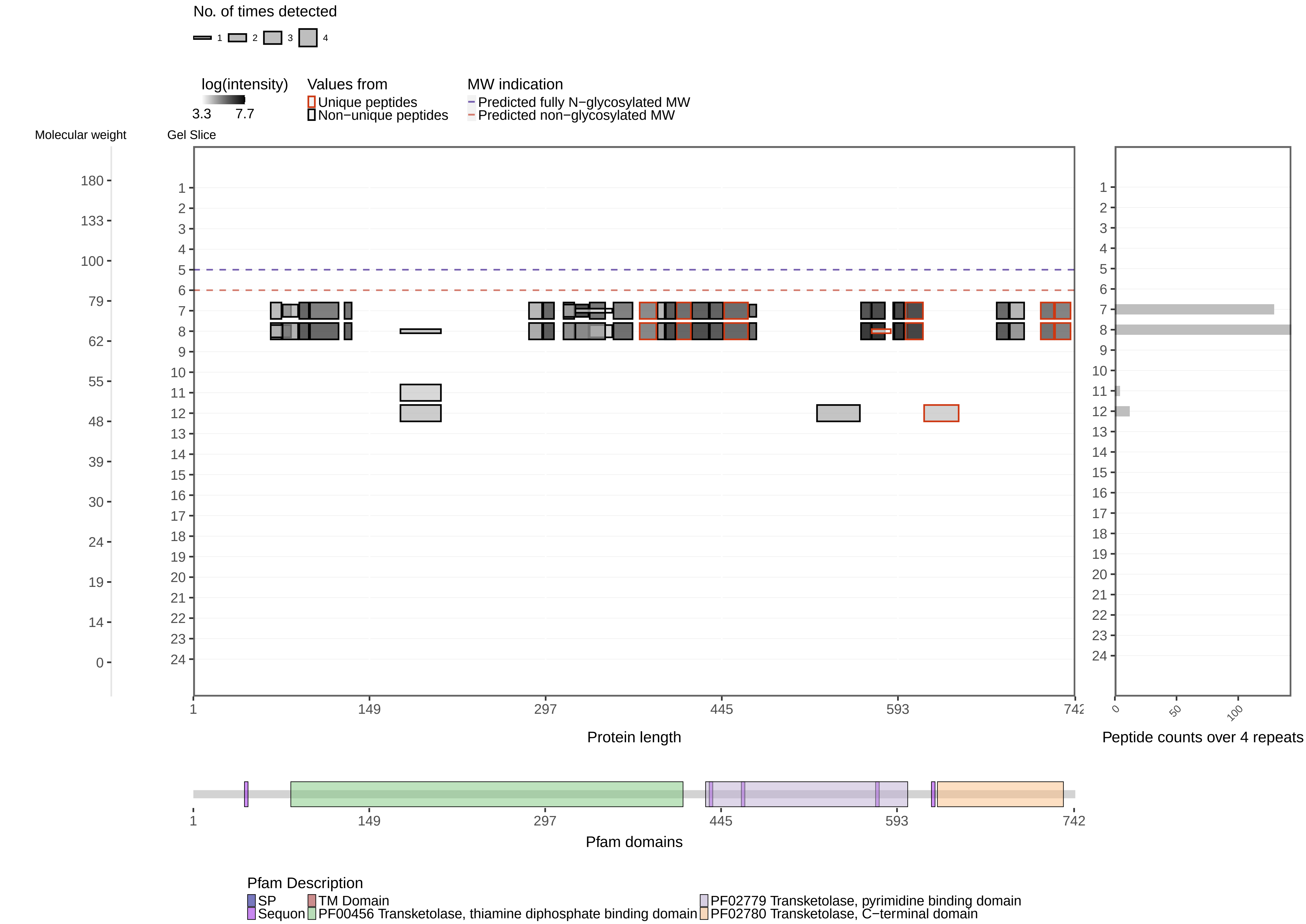


Protein sequence:

### NbL12g12650.1 protein_transport_protein_Sec61_subunit_beta−like [MW = 12.77 kDa]

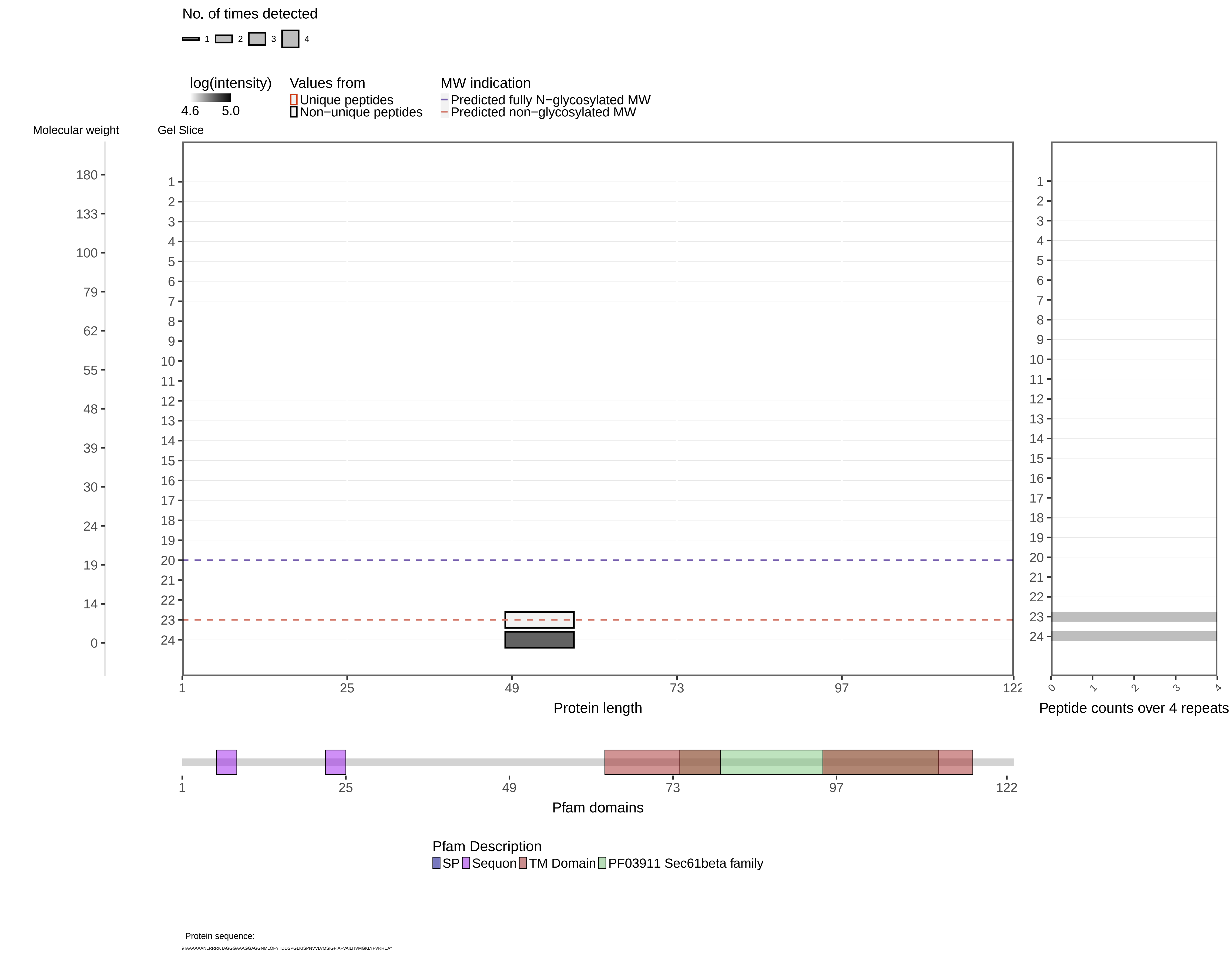

#### NbL12g12610.1 uncharacterized_LOC109212372 [MW = 28.38 kDa]

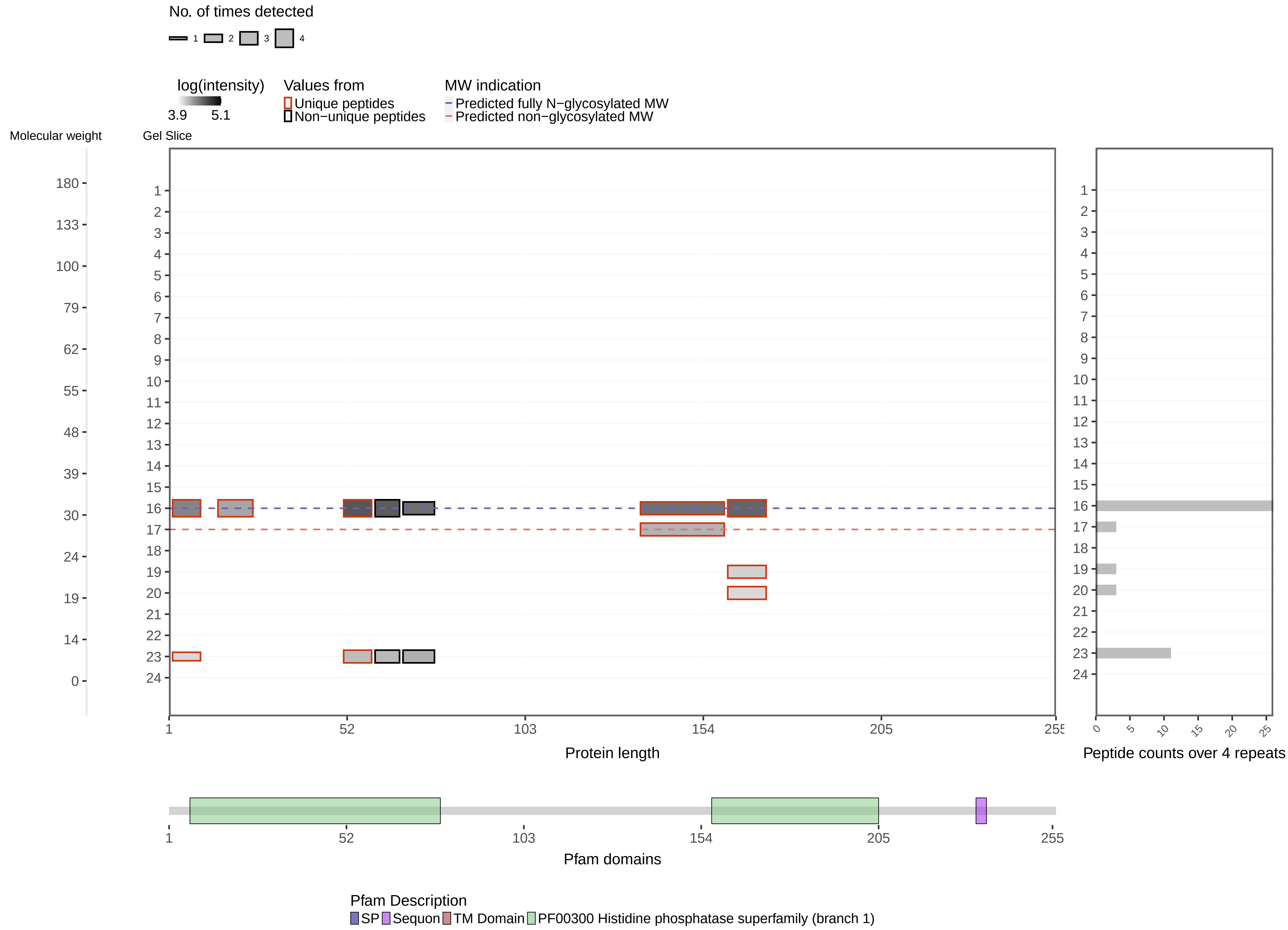


Protein sequence:

LCEMLNLVAIRSNAAPKDGDFKFSISQYEAELPAGTVDHTVEPVYQKLPEWQETLDSARARYVEVVKALADKYPSENLLLVTHGEGVGSIFTELNKDATVLEVEYCGHVLAKRSIQFGENQSFTAGEFVYEKQTGILSAAK*

### NbL12g12590.1 60S_ribosomal_protein_L27−like [MW = 17.9 kDa]

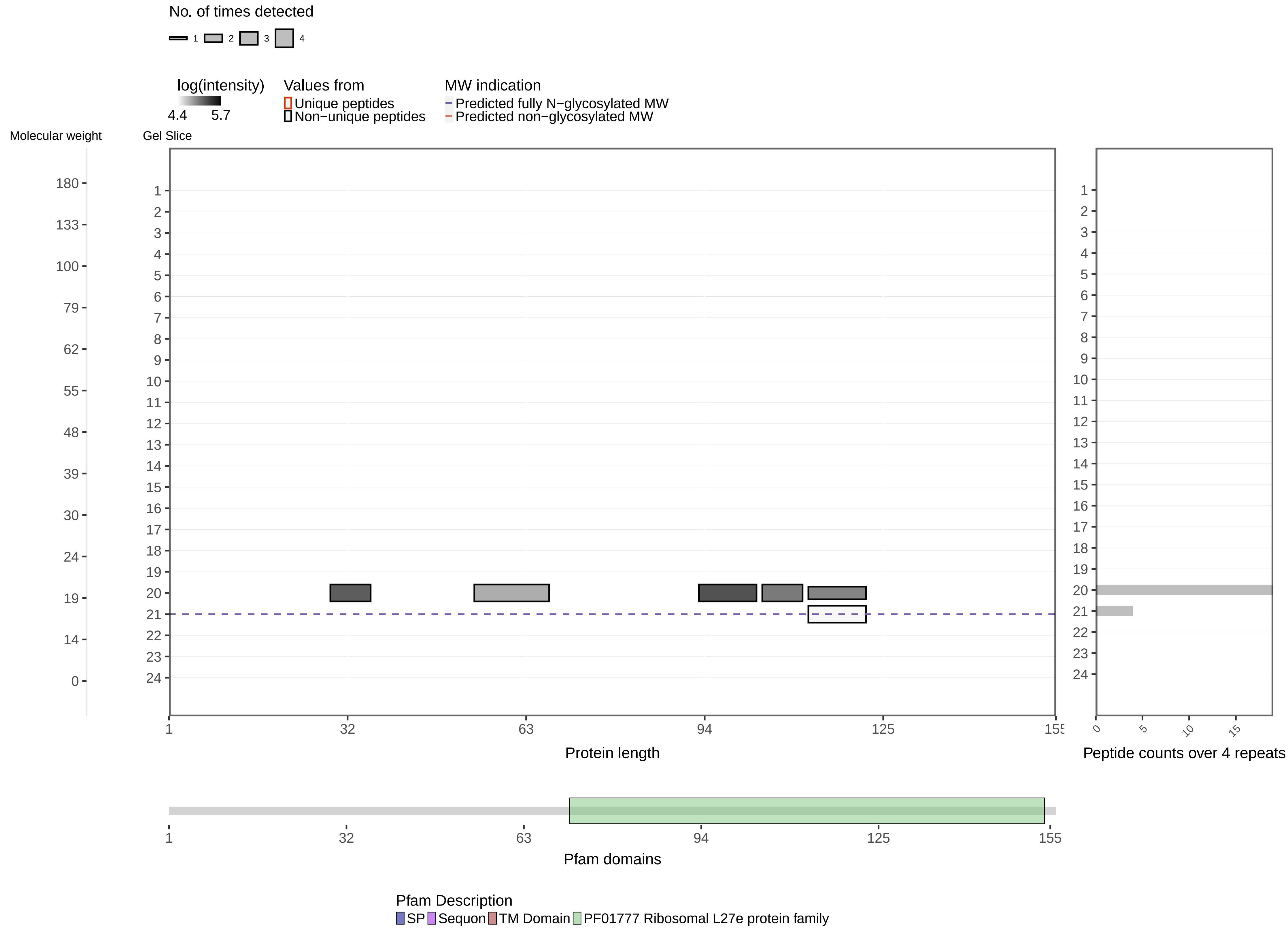


Protein sequence:

ISKYPKKVIRKDSAKKQAKKSRVKAFIKLVNYNHIMPTRYTLDVDLKDVVNVDVLQSRDKKVTAAKEAKARLEERFKTGKNRWFFTKLRF*

### NbL12g12520.1 nucleoside_diphosphate_kinase_1−like [MW = 15.71 kDa]

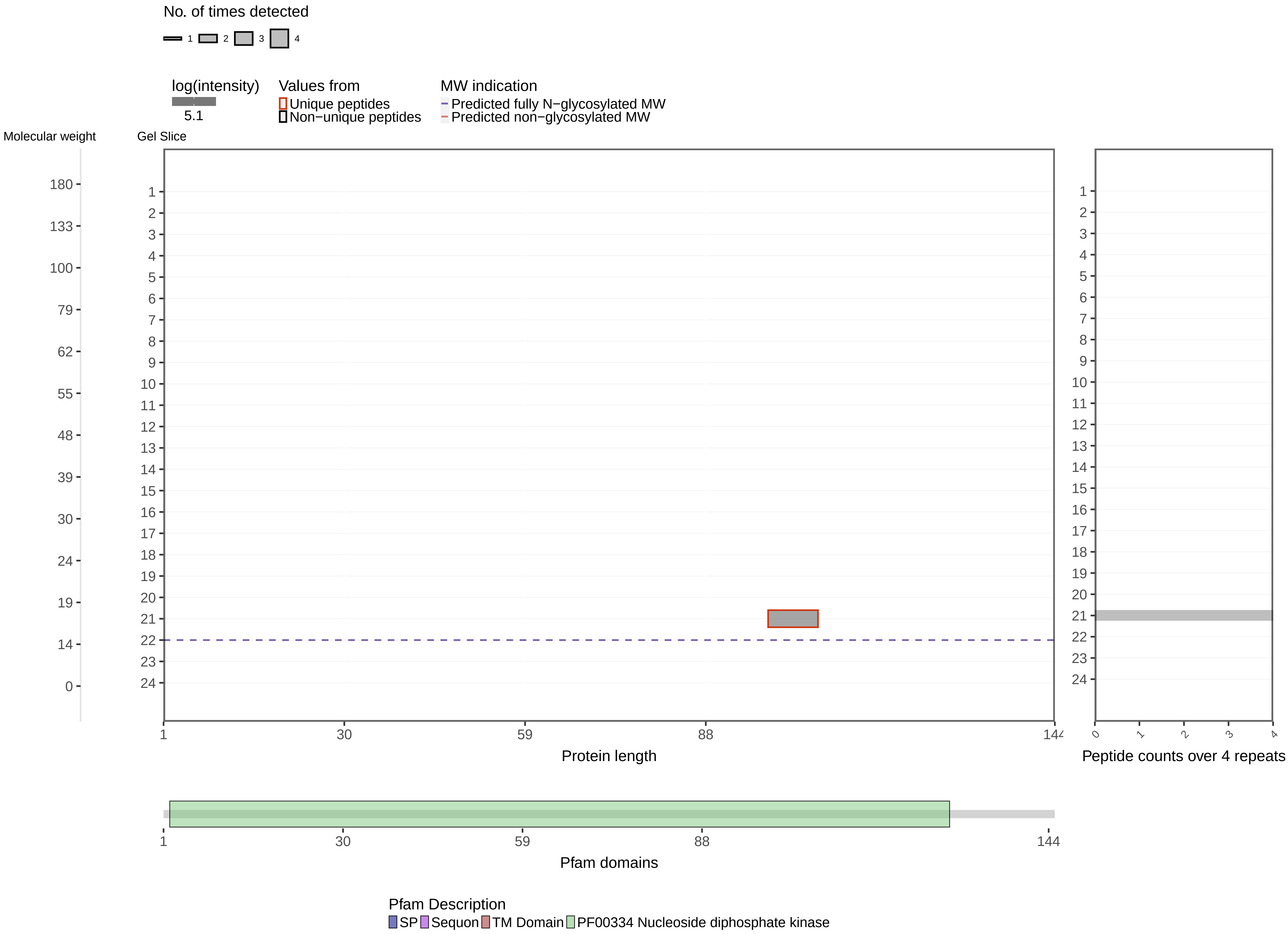


Protein sequence:

IVSGPVVAMVRAGKGVVSTGRKIIGATNPLESAAGTIRGDYSIDIGRNVIHGSDSVESAKKEIALWFPEGVVQWQSSLHSWIYE*

NbL12g12390.1 protein_transport_protein_SEC23−like [MW = 85.26 kDa]

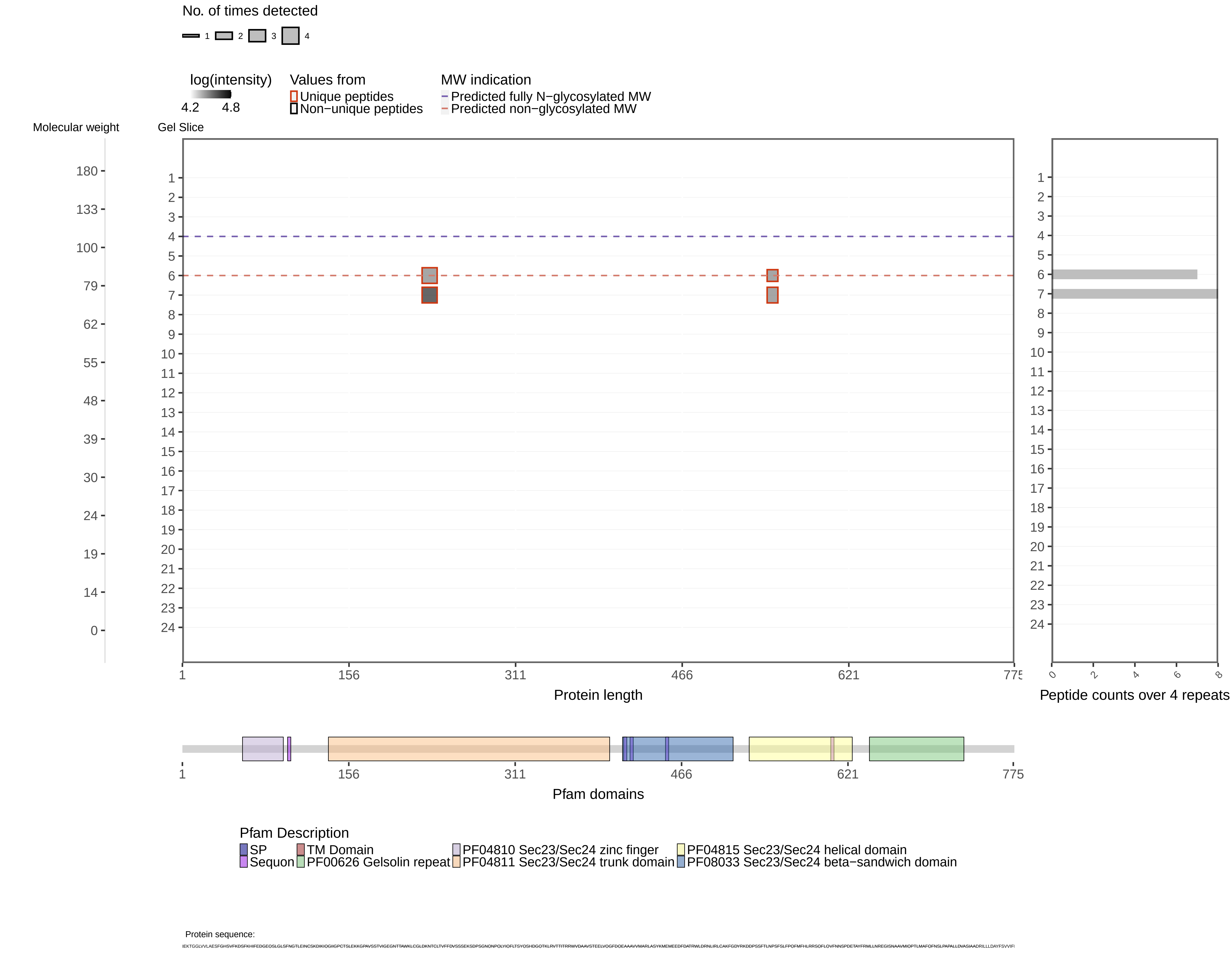

### NbL12g12320.1 serine_hydroxymethyltransferase_4−like [MW = 54.44 kDa]

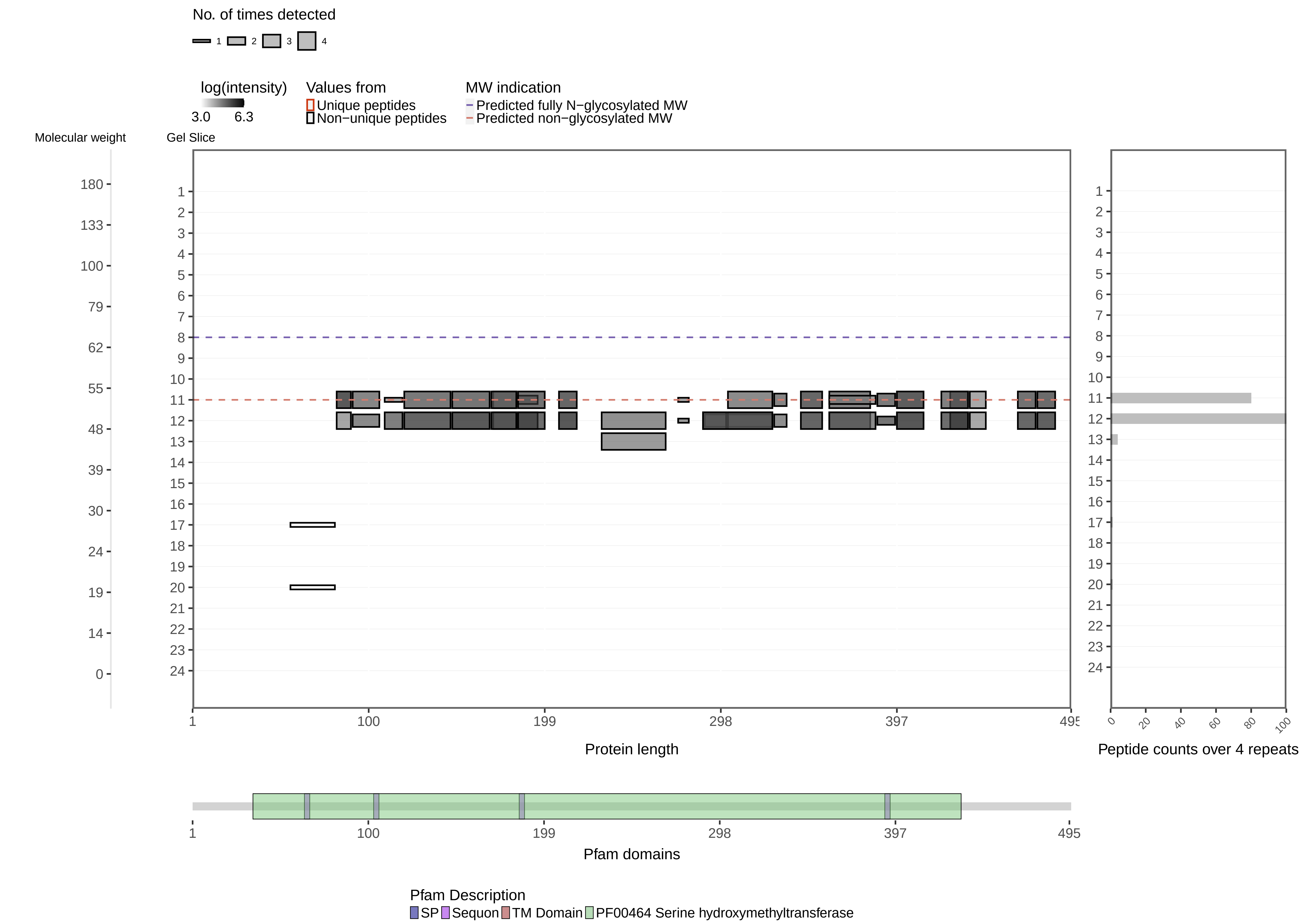


Protein sequence:

CDMAHISGLVAAQEAADPFEYCDLVTTTTHKSLRGPRAGMIFYRKGPKPPKKGQPEDAVYDFEDKINFAVFPSLQGGPHNHQIGALAVALKQAATPGFKAYAKQVKANAVALGNYLMSKGYKLVTGGTENHLVLWDLRPLGLTGNKVEKLCDLANITVNKNAVFGDSSALAPGGVRIGTPAMTSRGLVEKDFEQIAEFLHRAVTITLNIQKEYGKLLKDFNKGLVNNKEIEELKADVEKFSSSFDMPGFKLSEMKYKD*

### NbL12g12300.1 glycine−rich_RNA−binding_protein_2_mitochondrial−like [MW = 21.12 kDa]

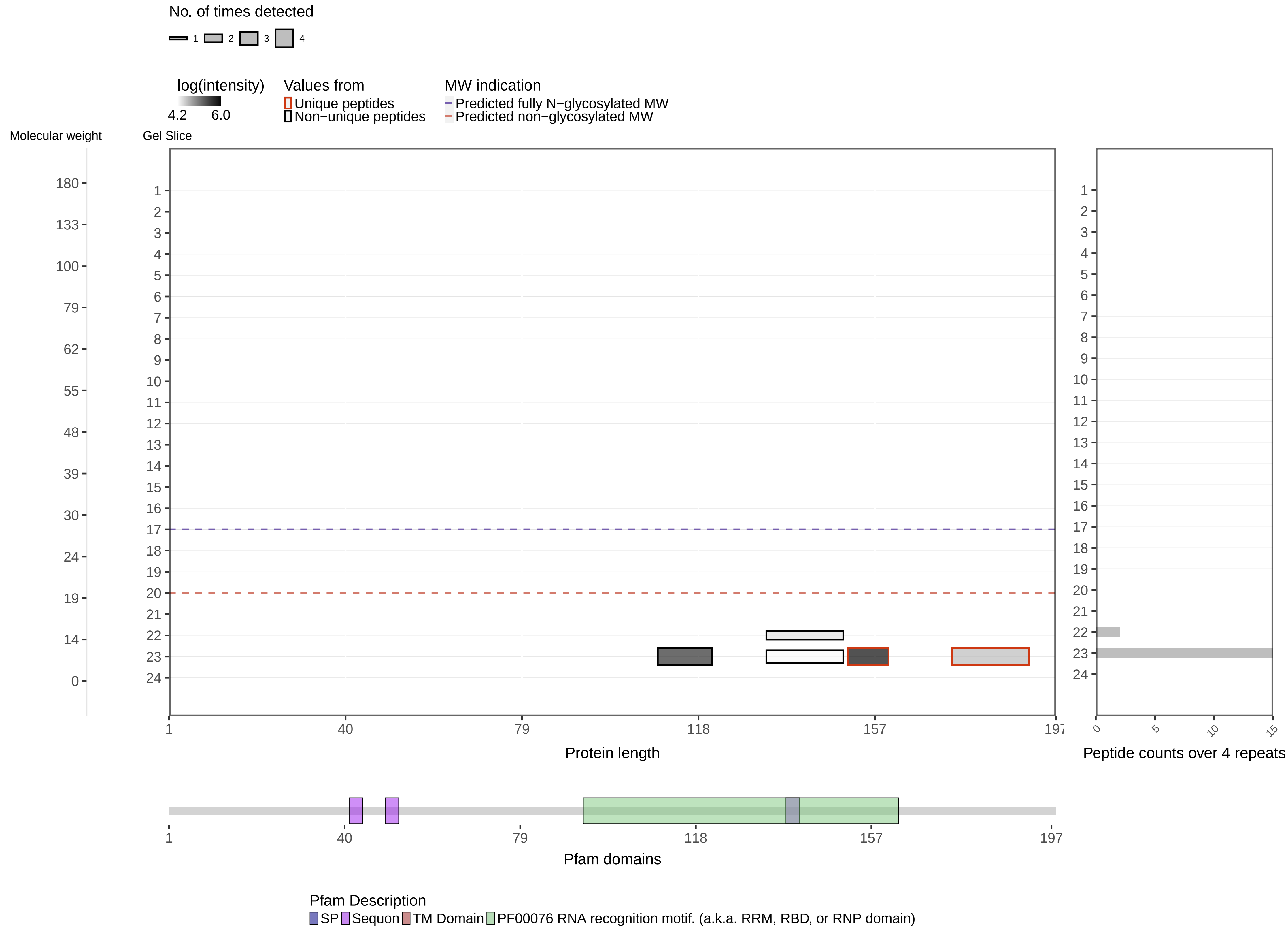


Protein sequence:

CMSTKLFIGGLSWGTDDQSLRDAFATFGDVVDARVIVDRDSGRSRGFGFVNFSDDESANEAIKAMDGQELQGRNIRVSIAQERAPRSGGFGGSGGFGGGYGQARDNDGY*

NbL12g12250.1 50S_ribosomal_protein_L9 [MW = 24.56 kDa]

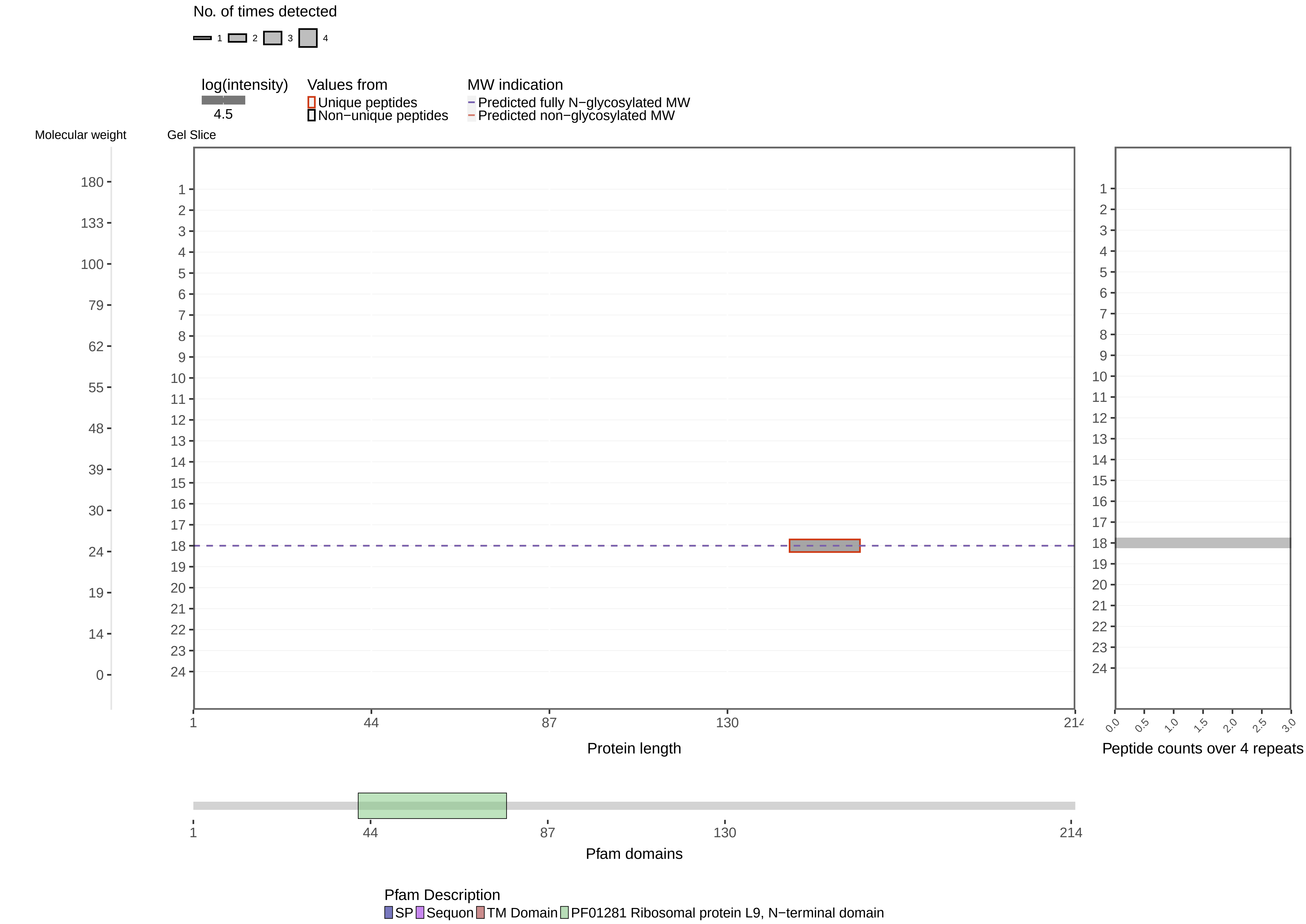


Protein sequence:

YQREEVEEVKIVPTTKEDKMKEYQMAANRLDRTKLVLRKLAKKPLTKDDSGRTSIQEPIGKDEIVAEVARQLSVHIEPEHLVLDTPLHVMGEFDIPLRMPRSIPLPEGKVQWILQVKIRGR*

NbL12g12130.1 cytochrome_b561_and_DOMON_domain−containing_protein_At5g47530−like [MW − SP = 41.15 kDa]

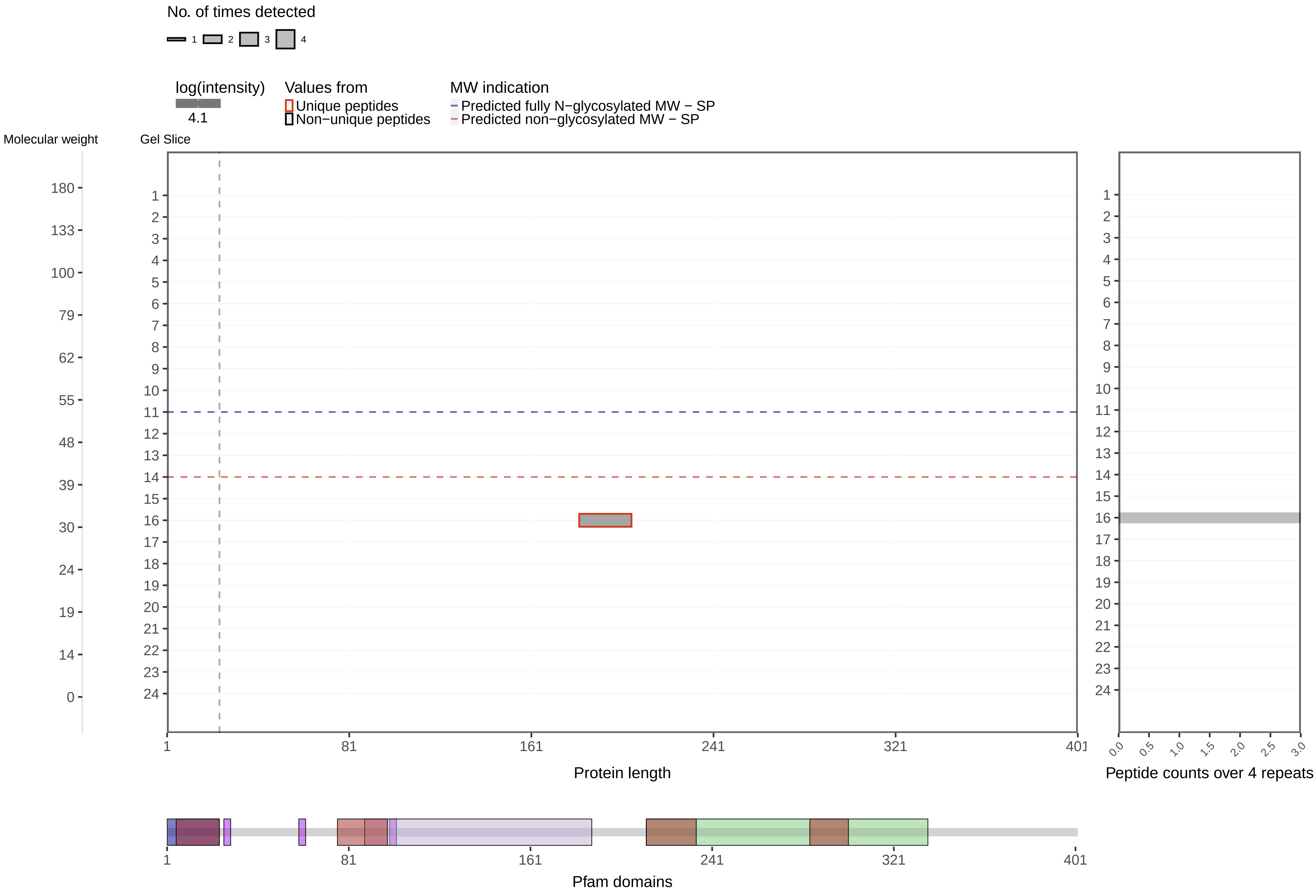


Protein sequence:

IDFASGQTSAAAGGASASSRQRRRNTHGVLNAISWGVMMPMGAVFARYLKVFKSANPAWFYLHVACQTSAYAVGVAGWGTGLKLGSDSVGIKFNTHRNIGITLFCLGTLQVFALLLRPKPDHKYRLYWNIYHHTIGYTVIILSIINVFEGFDALNGQKYWKRAYIGVIIALGAIAVLLEAFTWFIVIKRKKTSVSDKYPNGNGANGVNAYANREV*

### NbL12g11970.1 probable_inactive_receptor_kinase_At5g10020 [MW − SP = 112.45 kDa]

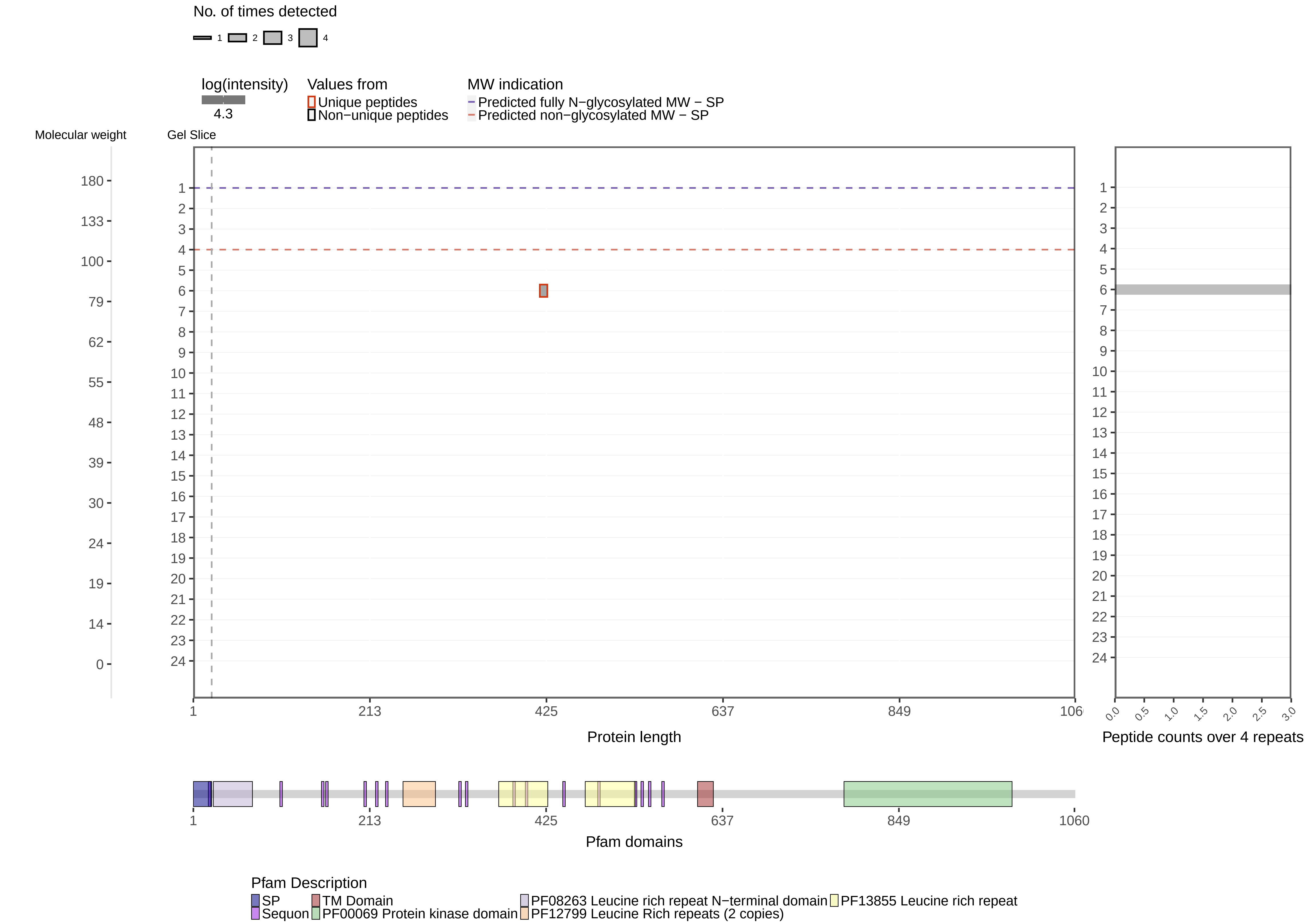


Protein sequence:

[illegible]

NbL12g11950.1 protein_HEAT_INTOLERANT_4 [MW = 33.76 kDa]

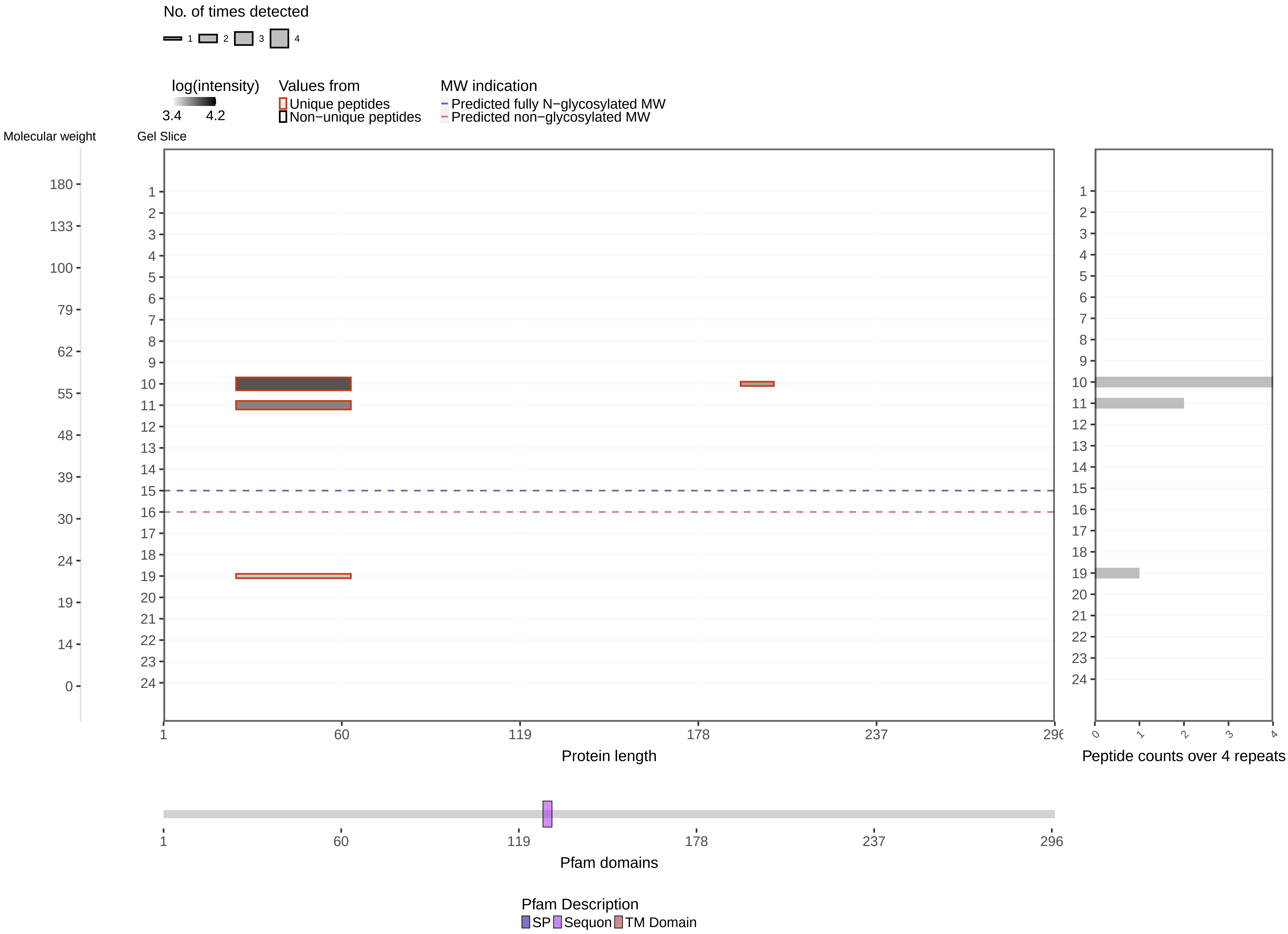


Protein sequence:

LEDAFEEGGELHGKKVYLFGCTEPOLVYFOGOSKVTCIPVVVAVVSPFPPSDKIGINSVOREAEEILPMNOMKMDWVPYIPLENRESOVERLKSOIFILRCTORRAGLKHLKLERVKKFEYCLPYFYOPFAEDELEOSTIVOILFPTEPPVSFOILVDYTIAIPO*

### NbL12g11890.1 60S_ribosomal_protein_L35a−3−like [MW = 12.92 kDa]

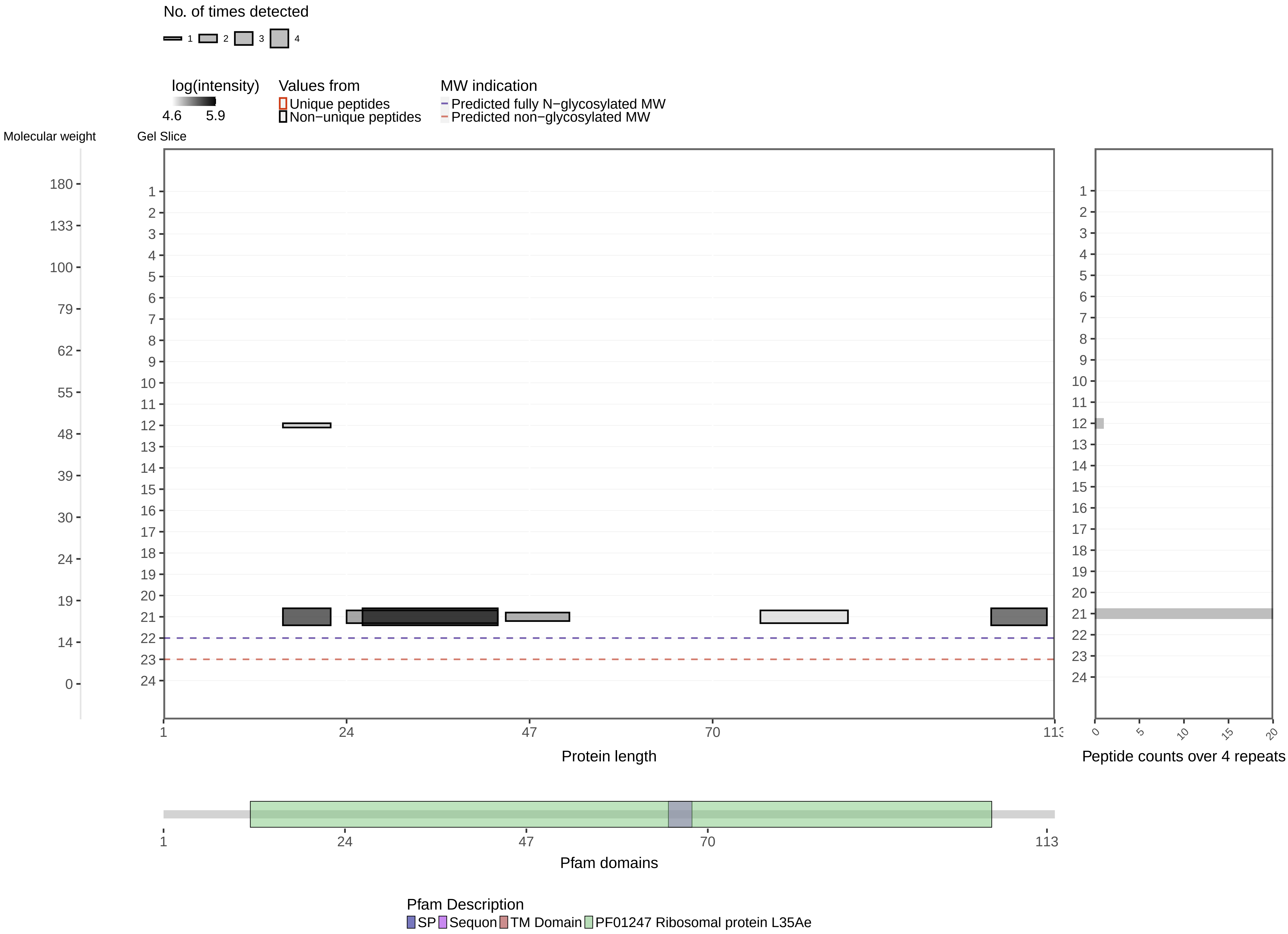


Protein sequence:

EVDWYLGKRMAYVYKAKTKKNNSHYRCIWGKVCRPHGNSGVVRAKFKSNLPPKSMGAKVRVFMYPSNI*

### NbL12g11650.1 uroporphyrinogen_decarboxylase_chloroplastic−like [MW = 43.46 kDa]

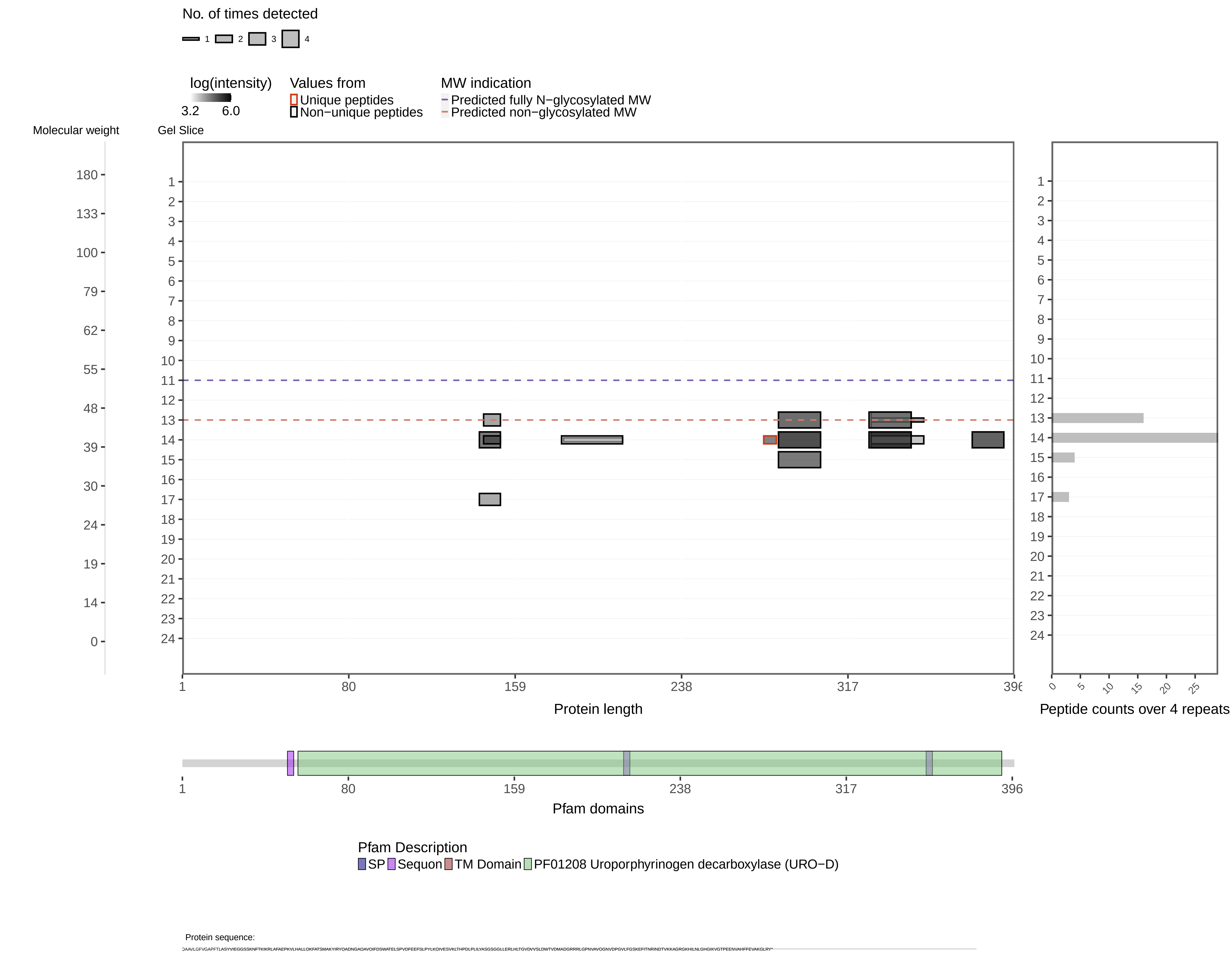

NbL12g11550.1 ABC_transporter_I_family_member_6_chloroplastic−like [MW = 34.54 kDa]

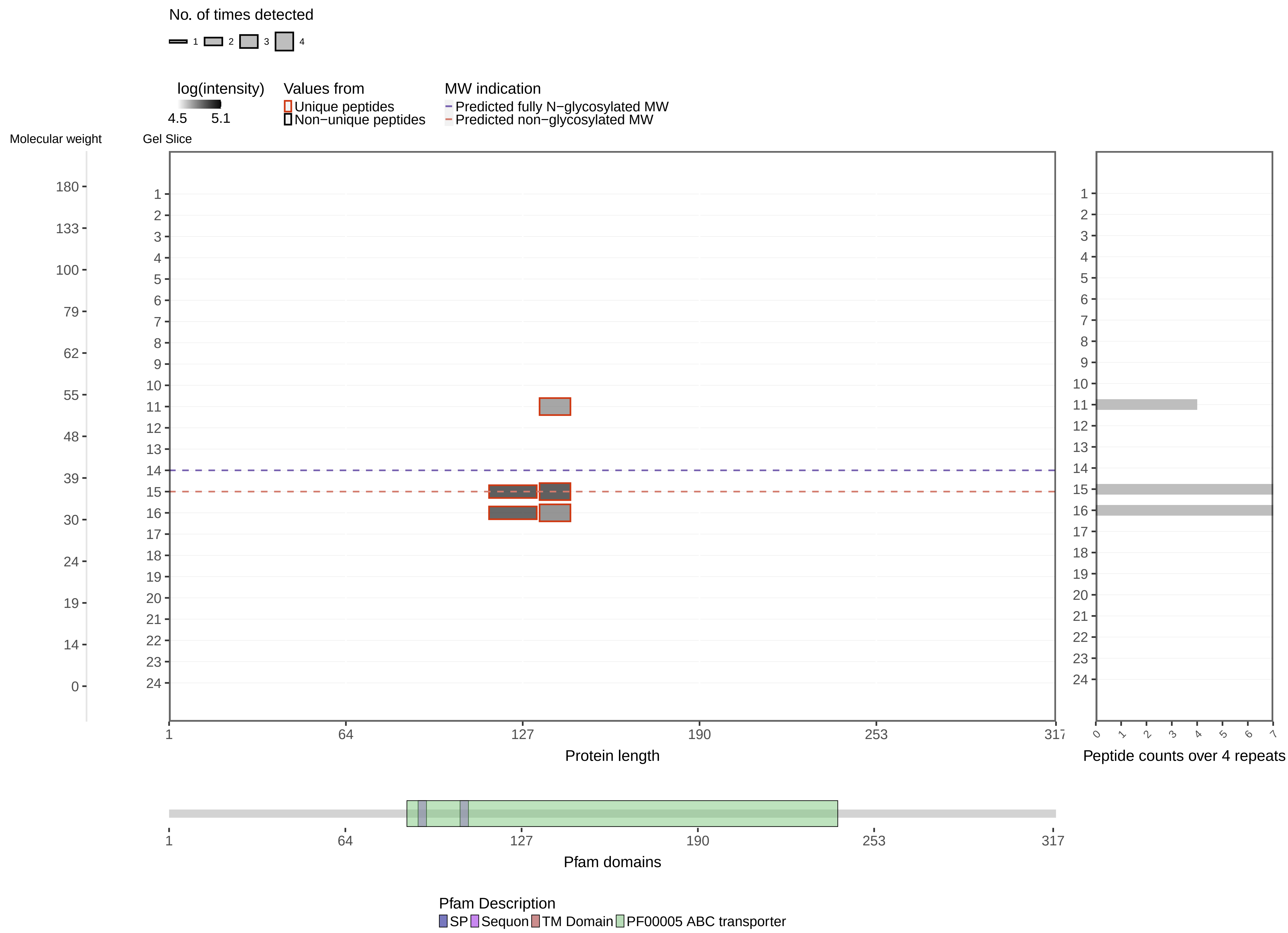


Protein sequence:

ERSLAGLFMSFOSPVAIPGVSNIDFLNMAYNAORRKLGLPELGPIEFYGYIVPKLELVNMKIDFLNRNVNEGFSGGERKRNEILOLAVLGADLAILDEIDSGLDVDALRDVAKAVNGLLSPKNSVLMITHYLRLLEFIKPTYIHIMENGRIVKTGDISIAKVLEKEGYKAISGA*

### NbL12g11540.1 hydroxyacylglutathione_hydrolase_cytoplasmic [MW = 34.13 kDa]

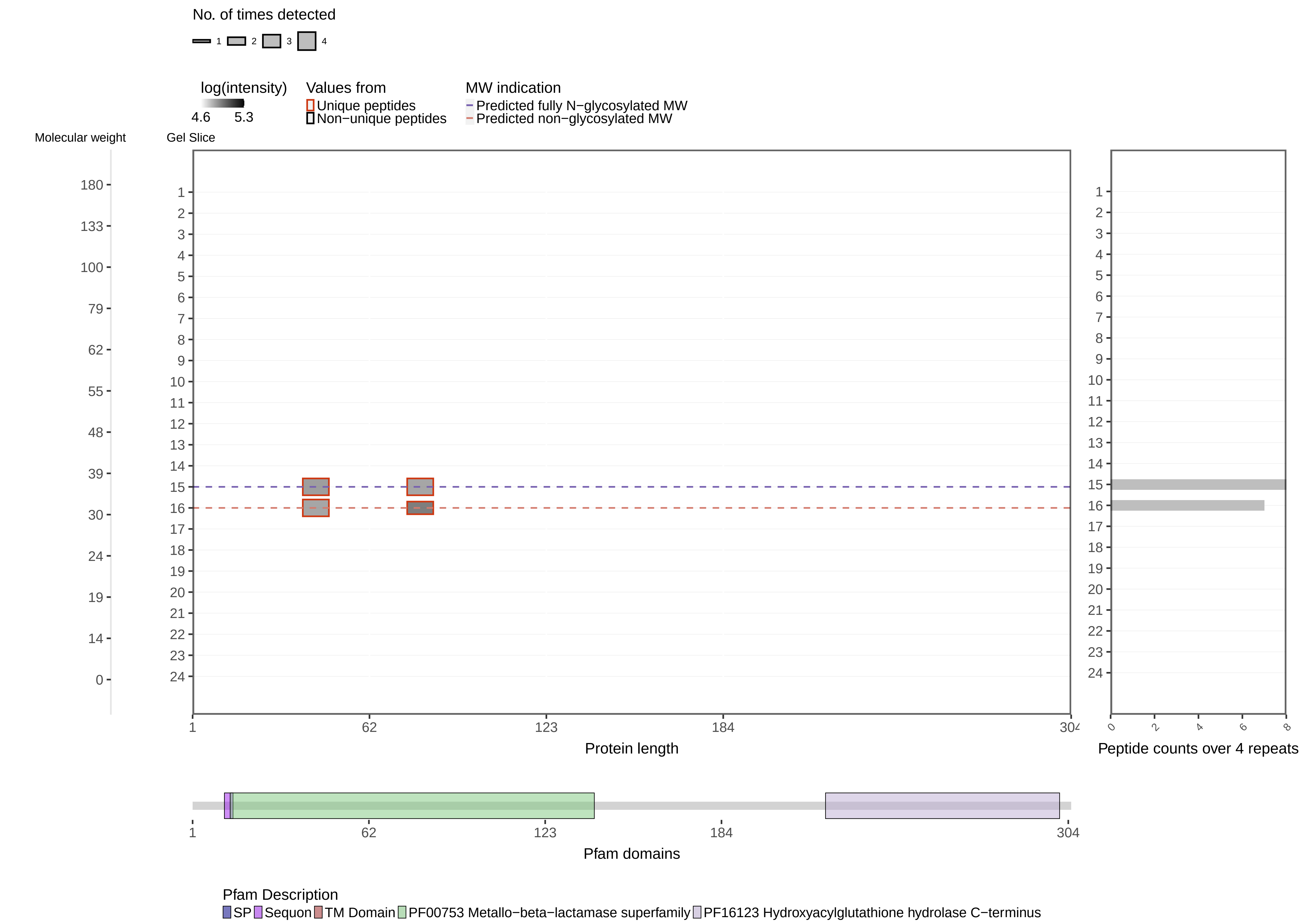


Protein sequence:

ITLSSVFPDLGLSWCTFLHFHLHFVFSVSRFRPFMVHLSSFSPFIAGCGKFFEGTAEQMYQSLCVTLASLPKPTRVYCGHEYTVKNLQFALTVEPDNAKIAQKLTWAEHQRRVGLPTIPSTIEEEMETNPFMRADLPEVQEKVGFKSPVEALREIRQRKDNWKG*

NbL12g11490.1 superoxide_dismutase_[Fe]_chloroplastic [MW = 27.94 kDa]

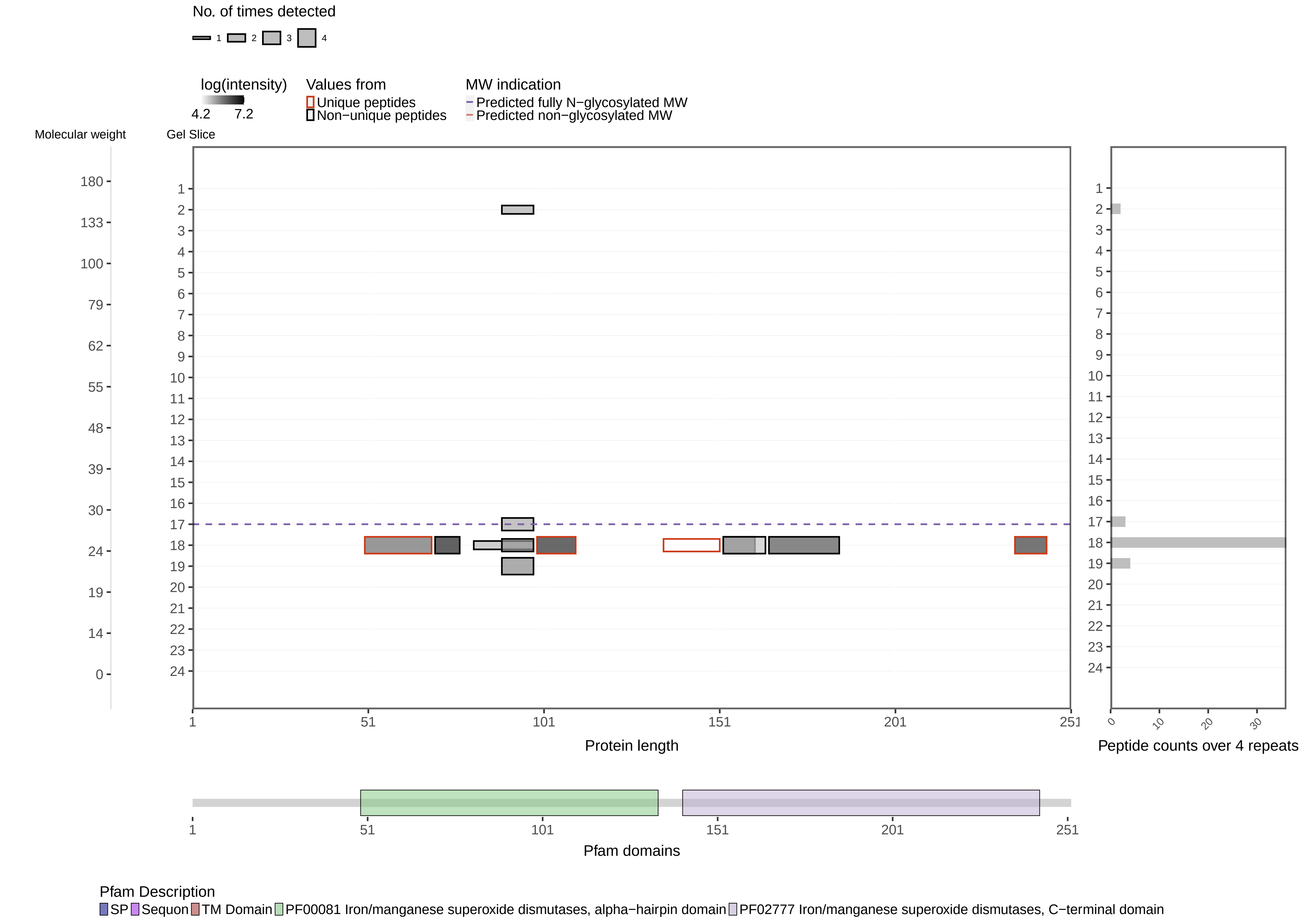


Protein sequence:

[illegible]PLPAFNNAAQAWNHQFFWESMKPNGGGEPSGDLLELINRDFGSYDAFVKEFKAAAATQFGSGWAWLAYKPEEKKLALVKTPNAENPLVLGYTALLTIDVWEHAYYLDFQNRRPDYISIFMEKLVSWEAVSCRLKAATA*

NbL12g11310.1 probable_carboxylesterase_120 [MW = 36.65 kDa]

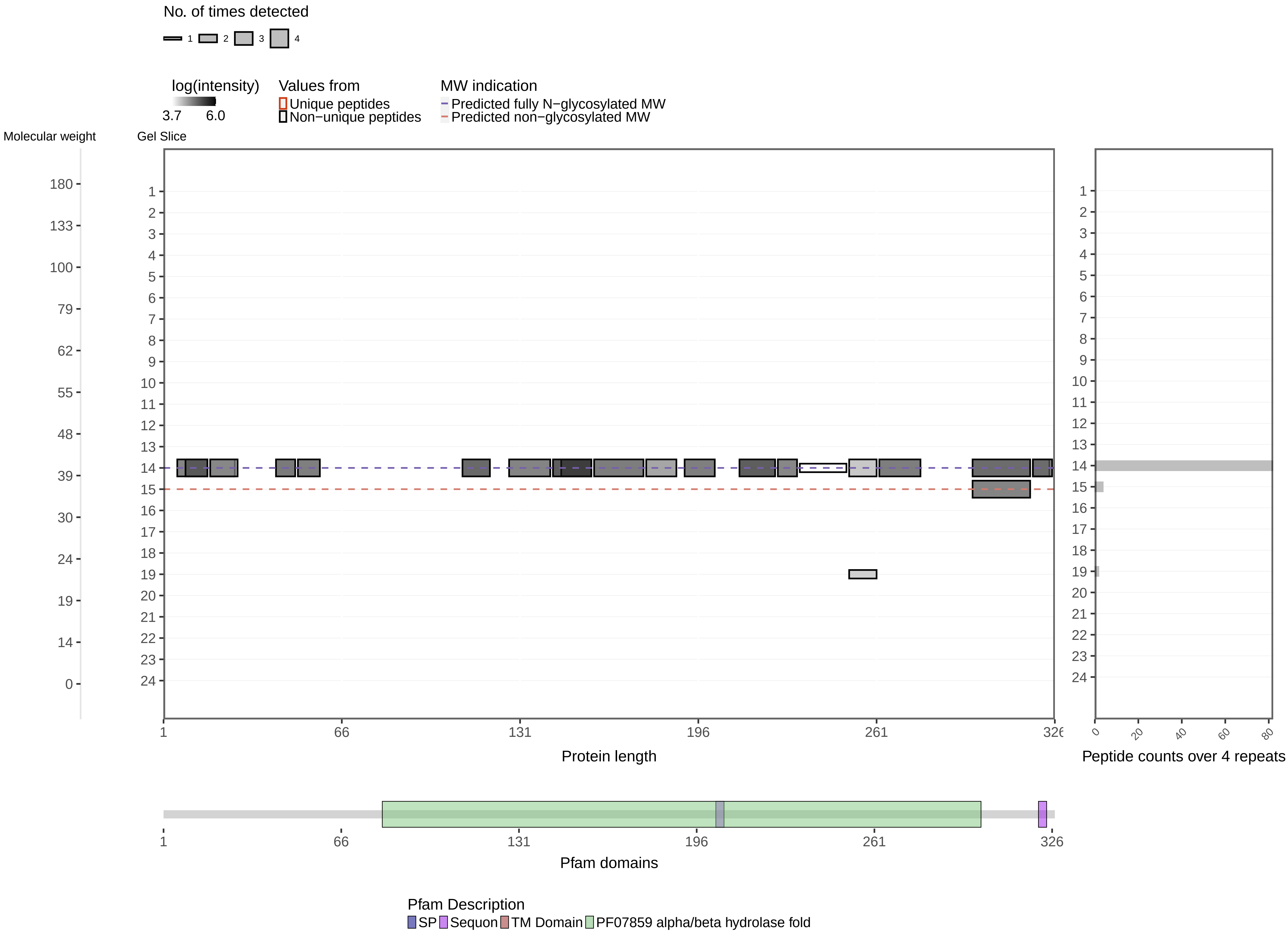


Protein sequence:

NYADLSRVYLYGTSAGGNLAYHAGLRAAAAANELEPIRIKGLILHHPYFSGKNRTESEEKLKDDOLLPLHAIDKMFDLCLPKGVDHDHEYCNPCANGGSKHLDDMKELGWKVLVTGVCEDPLVDAGRNCVKLMEEKGIKVYKFFRDGYHAMEVFDOSTAAALYDATRDFLNASKN*

NbL12g11010.1 5'−3'_exoribonuclease_4 [MW = 111.7 kDa]

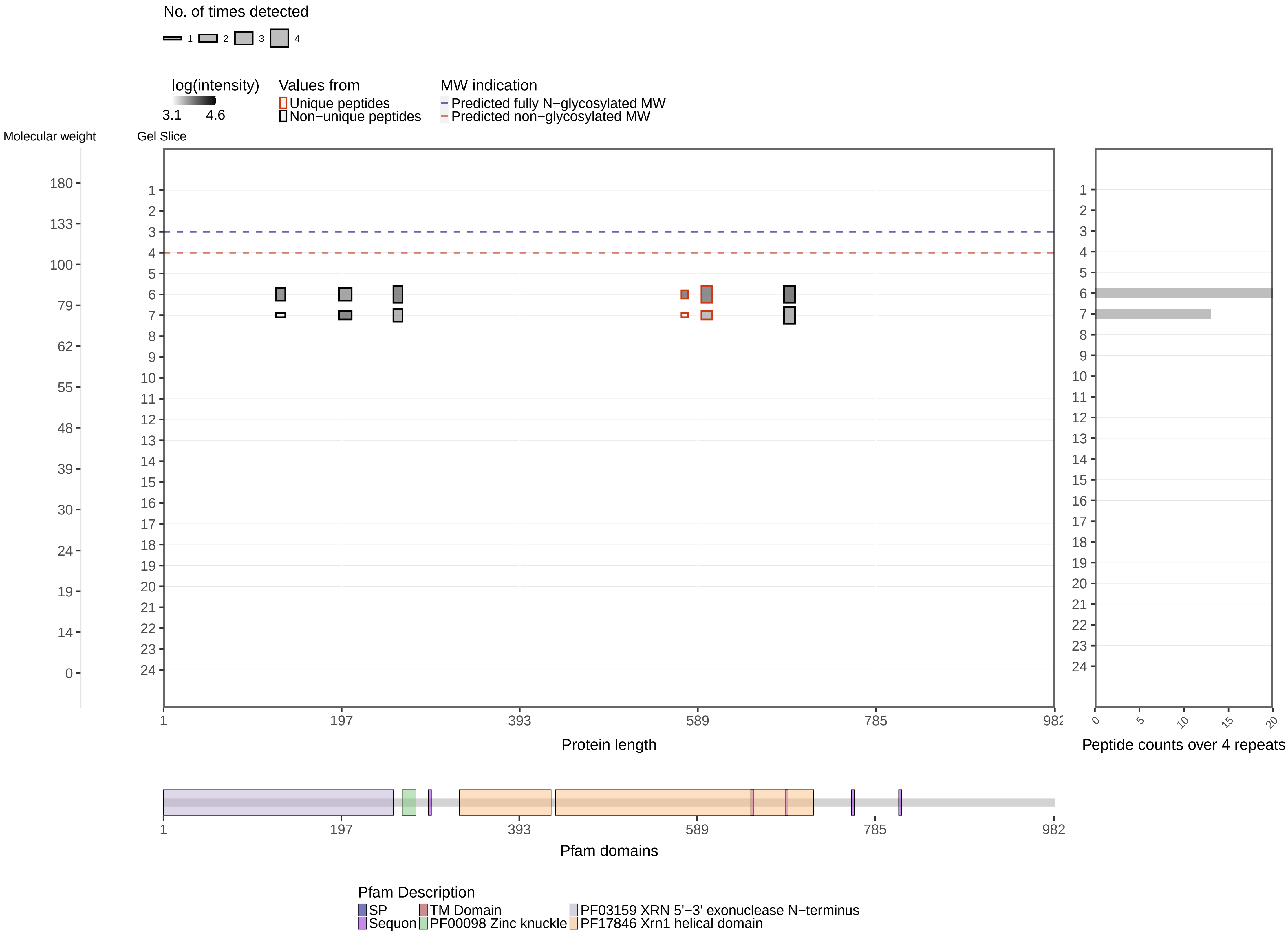


Protein sequence:

YYQGVCSWQWFYPYHYAPFASDLKGLADLEITFFPGEPFKPFDQLMGVLPAASANALPEKYRMLMMDSSSPISDFYPTDFELDMNGKRFAWQAVVKLPFIDEKKLLAETKKLEDTLSEDEQLRNSVLFDLLYVHHHHHTLAPYIISYYHHNNKVPPGEQRPWLIDANLSGGMNGFIWLSERNGLRDGVCSPLNGLEDISNNKILNITFLNPVPHKHIPKPPEGVYMPKKILRAVDIKPFPVLWHEDHGTRRQLGKDRGPVPGAIAGPTLGEAAHRLLNNTLNIKPGGTNSGVL[illegible]

NbL12g11000.1 probable_glucan_13−alpha−glucosidase [MW − SP = 102.83 kDa]

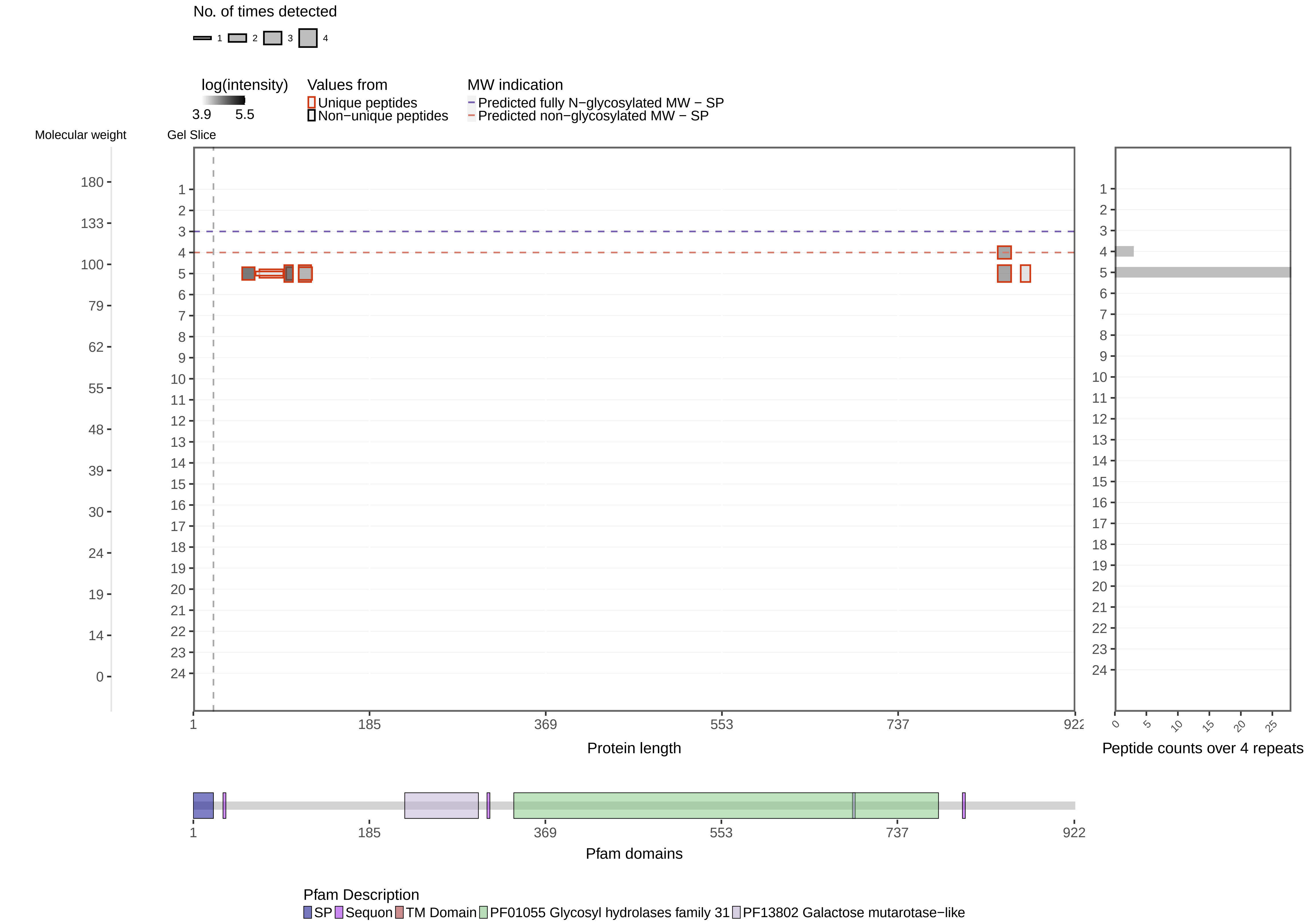


Protein sequence:

HKEASEKGYYVKDATGKDYDGWCWPGSSSYVDLLNPEIRSWWGDKFSLDCYTGSTQYLHIWNDMNEPSVFNGPEVTMPRDALHHGGVEHRELHNAYGYYFQMATSNGLLKRGDGKDRPFVLGRAFFAGSQRYGAIWTGDNTAEWEHLRVSVPMVLTLSISGIVFSGADVGGFFGNPDTELLVRWYQLGAYYPFFRGHAHHDTKRREPWLFGERNTQLMREAIHVRYMYLPYFYTLFREANSSGTPVARPLWMEFPGDEKSFSNDEAFMVGNGLLVQGIYTERAK

### NbL12g10960.1 aspartate−−tRNA_ligase_2_cytoplasmic−like [MW = 60.28 kDa]

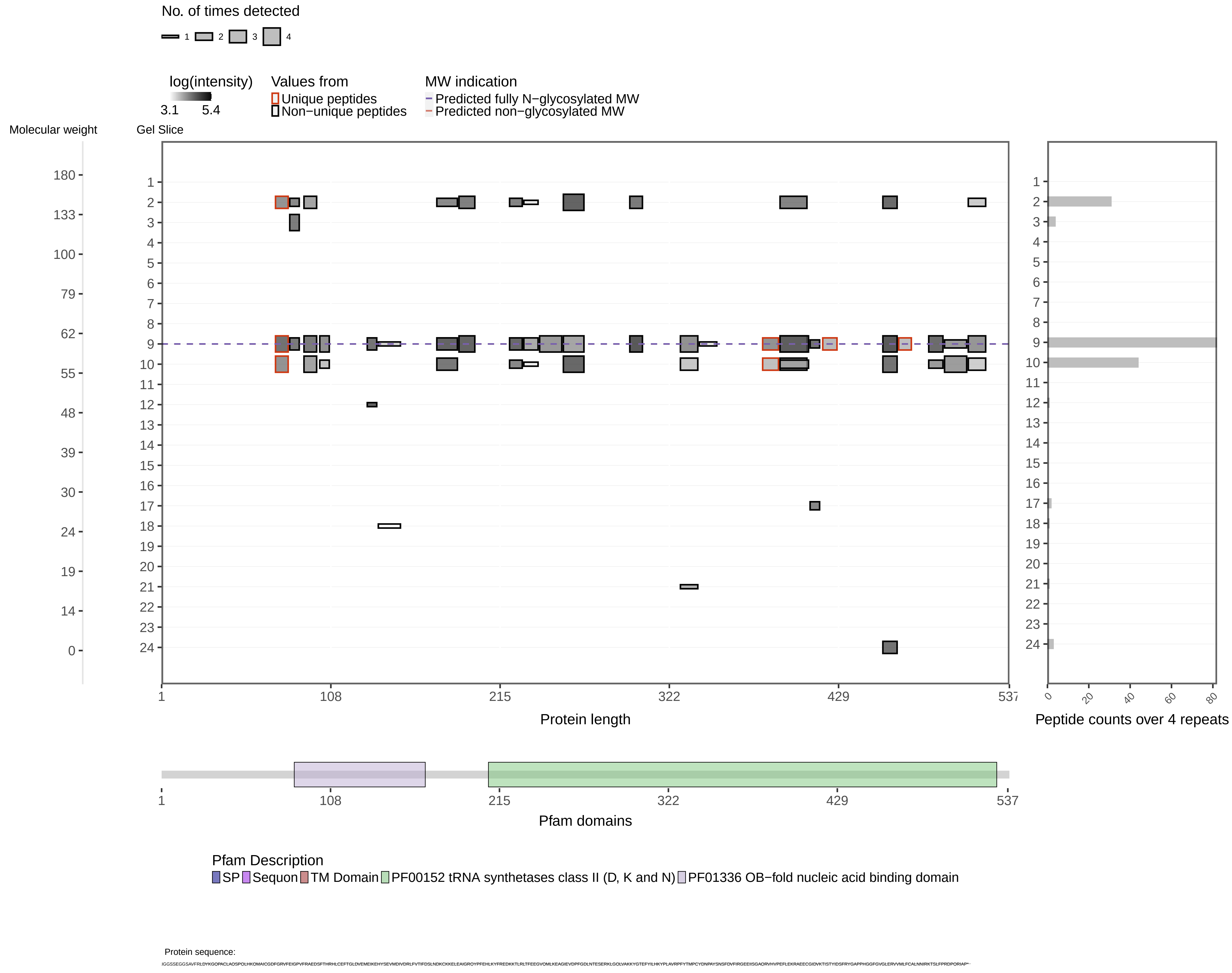

### NbL12g10940.1 peptide_chain_release_factor_PrfB1_chloroplastic [MW = 51.2 kDa]

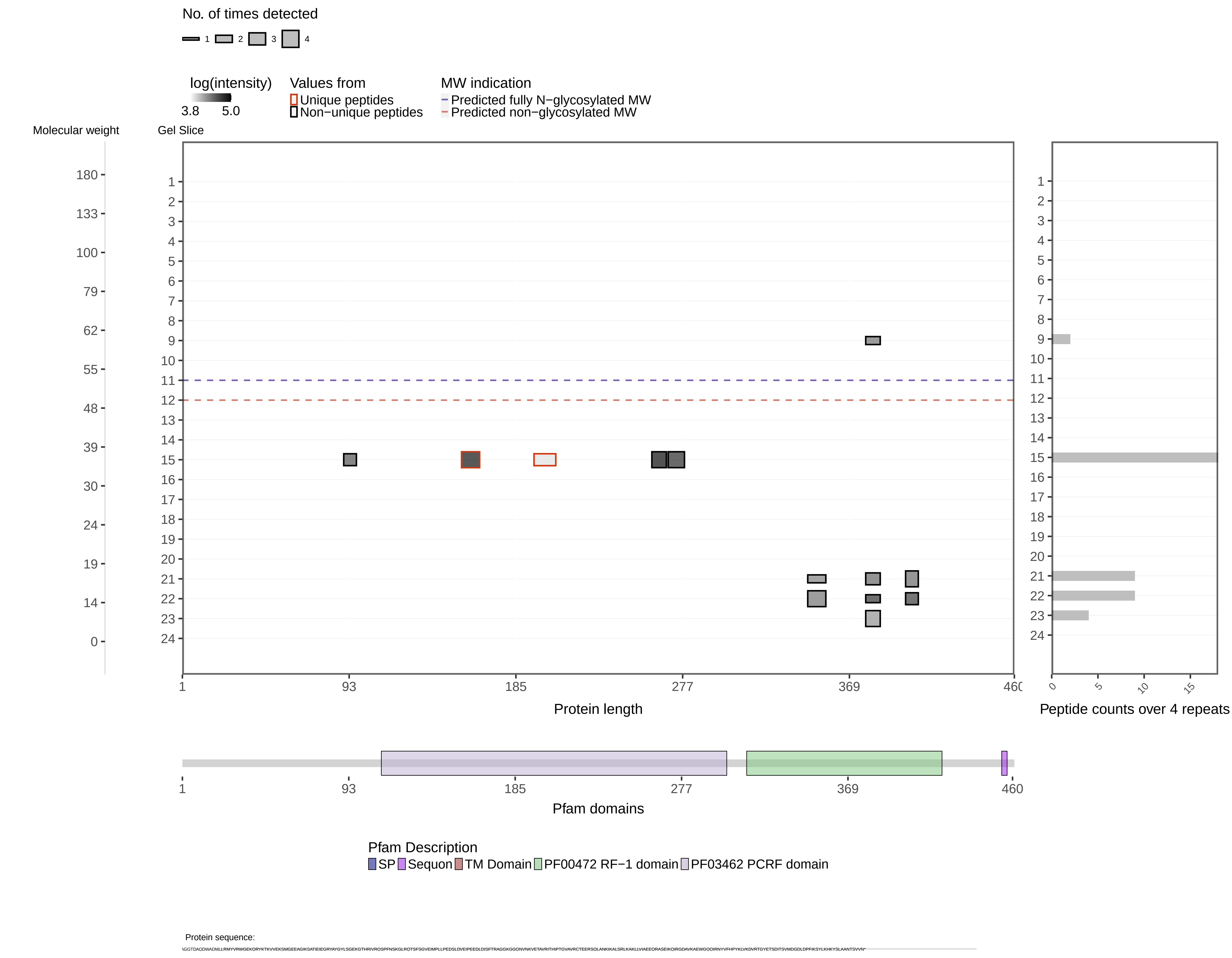

### NbL12g10910.1 calreticulin−3−like [MW − SP = 46.66 kDa]

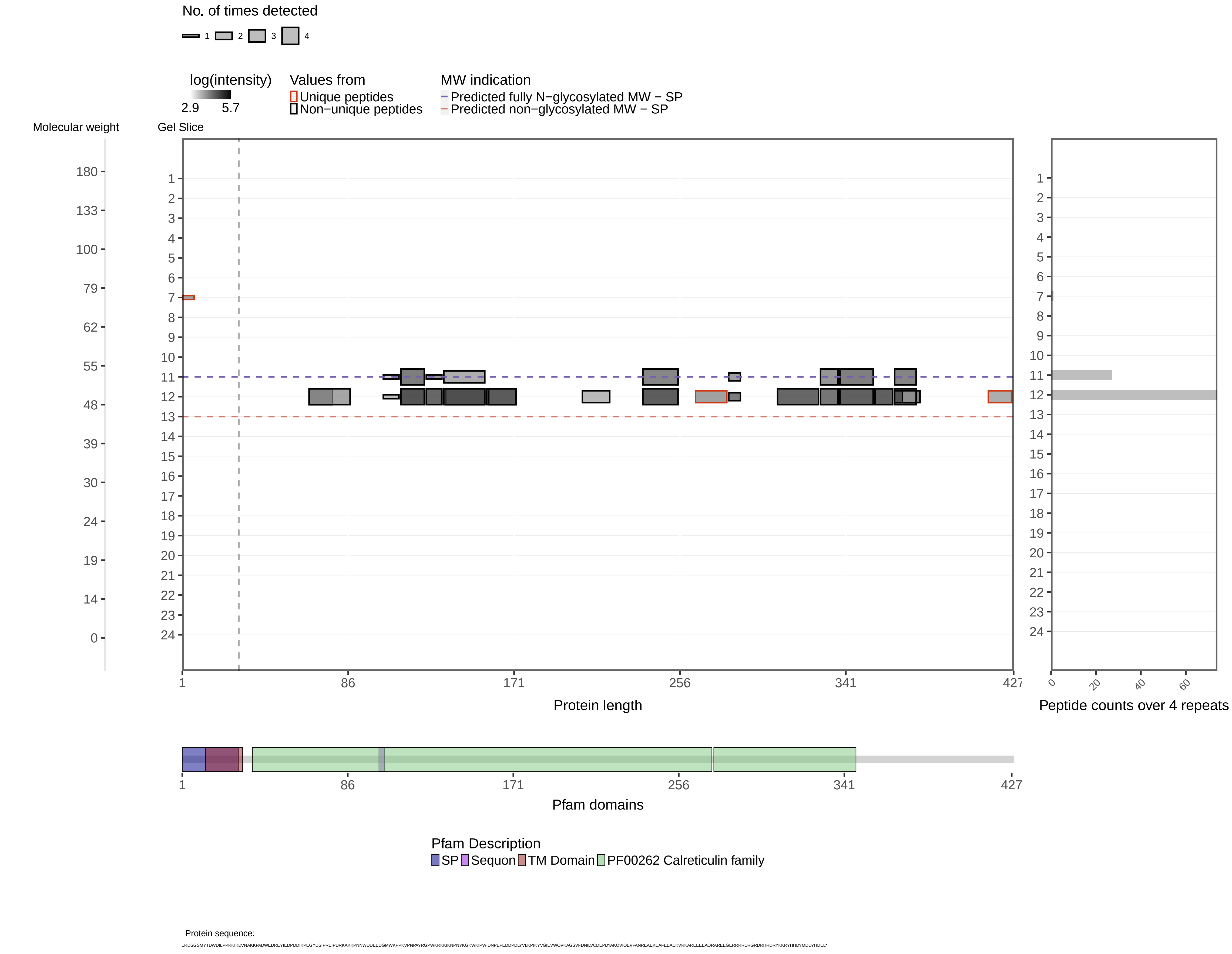

### NbL12g10780.1 branched−chain−amino−acid_aminotransferase_3_chloroplastic−like [MW = 45.35 kDa]

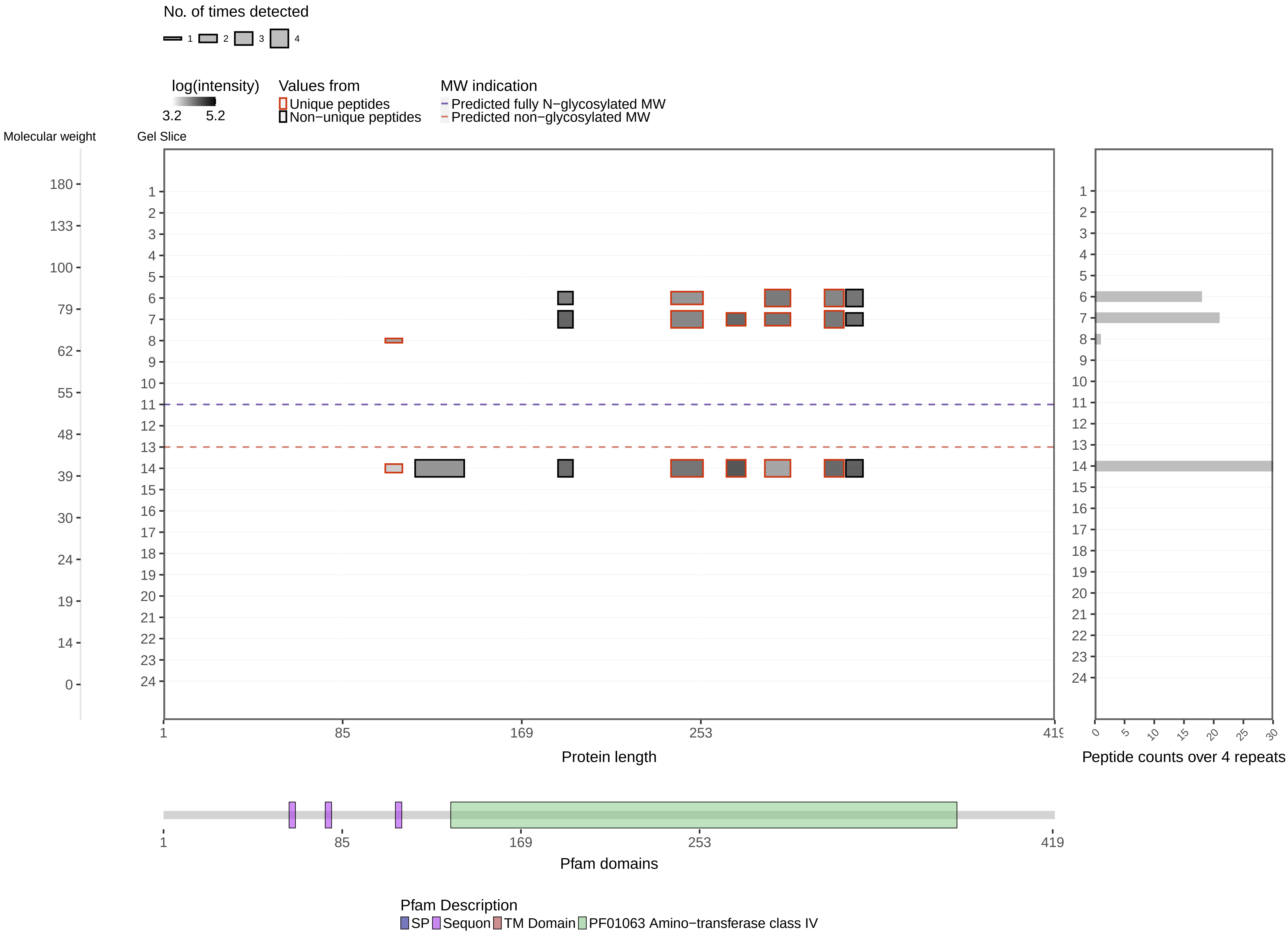


Protein sequence:

[illegible]IPPSGKGSLYIRPLLMGSGAVLGLAPAPEYTFLIYVSPVGNYFKEGLAPINLVVETEMHRATPGGTGGVKTVGNYAAVLKAQSAAKAKGYSDVLYLDSVQKRYLEEVSSCNVFIVKGNLIVTPAIKGTILPGITRKSIIDVALSQGFEVEERLVSVDELLDADEVFCTGTAVVVSPVGSITHQGKRVTYGNNGVGIVSQQLYSALTSLQMGLAKDKMGWIVELK*

### NbL12g10770.1 protein_CROWDED_NUCLEI_4−like [MW = 109.76 kDa]

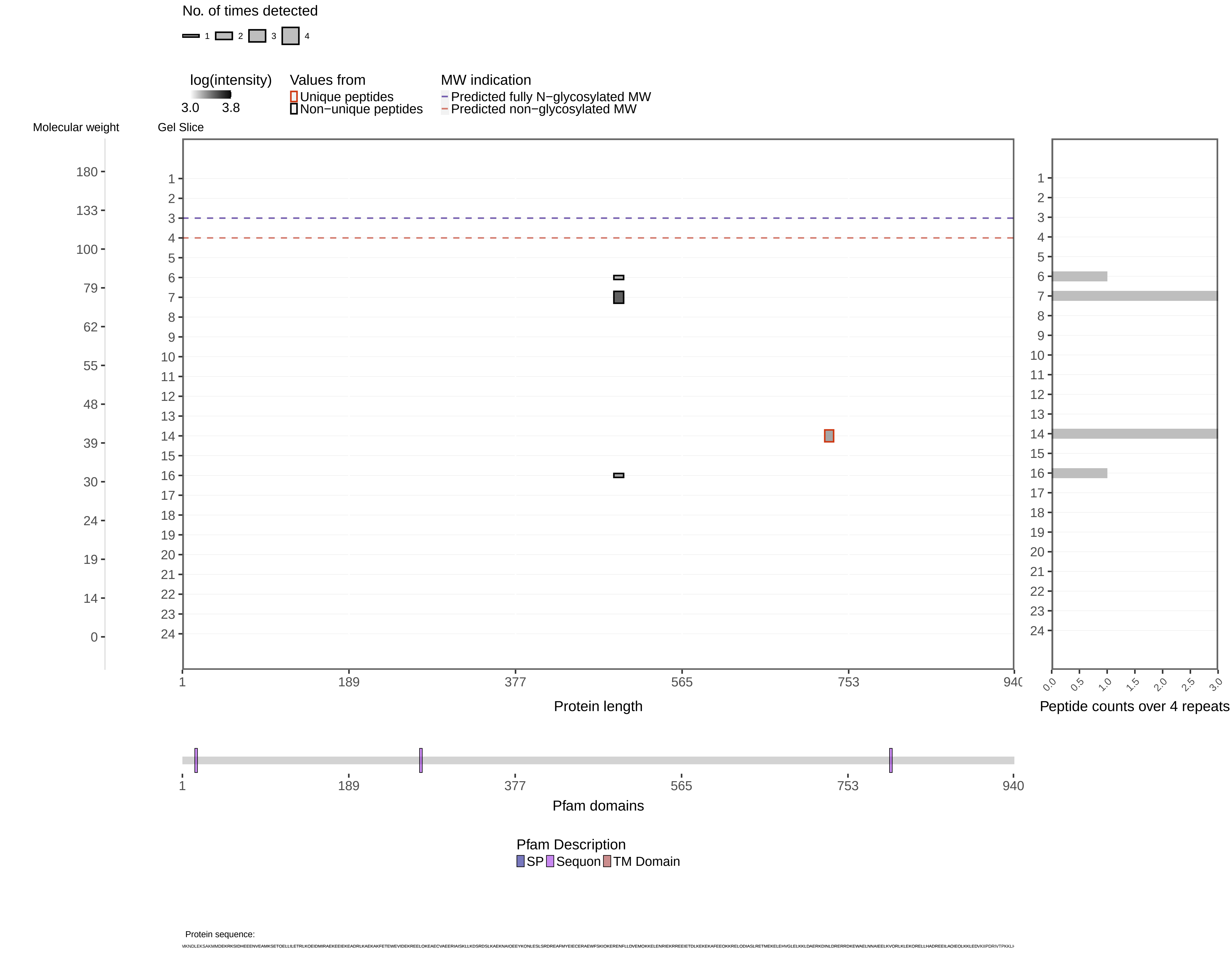

NbL12g10760.1 lysosomal_Pro−X_carboxypeptidase−like [MW − SP = 55.04 kDa]

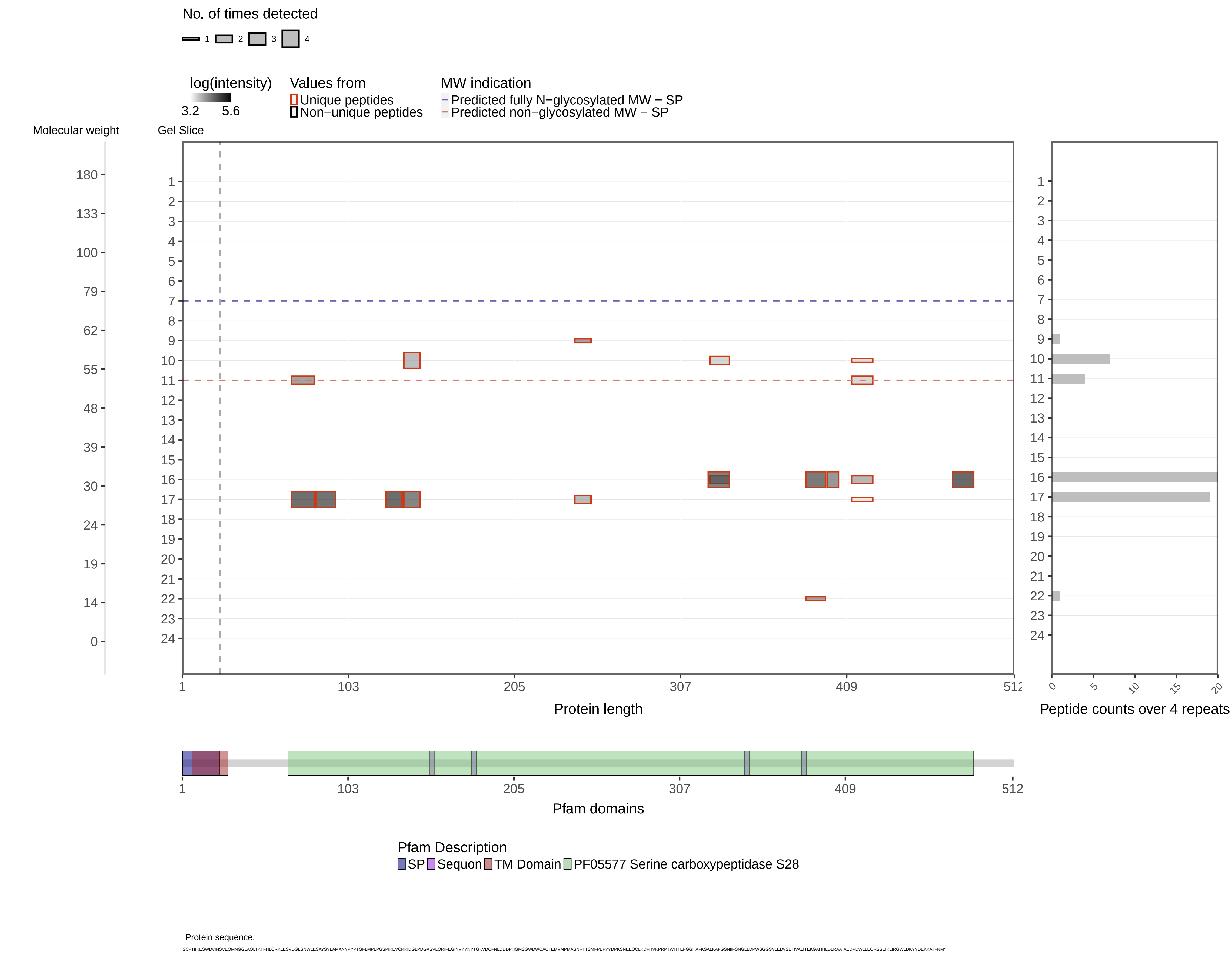

### NbL12g10700.1 probable_xyloglucan_endotransglucosylase/hydrolase_protein_7 [MW − SP = 29.89 kDa]

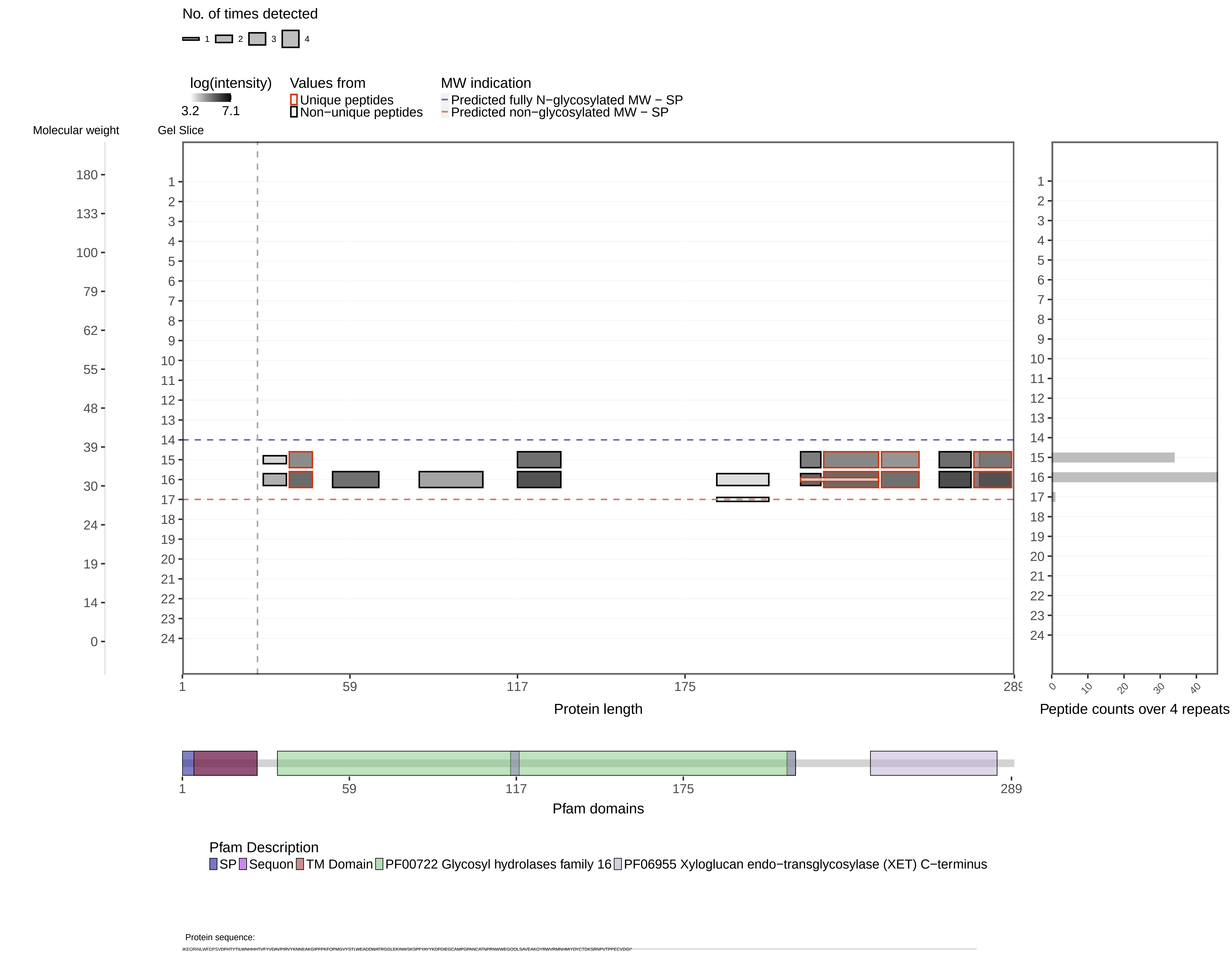

### NbL12g10680.1 cysteine_desulfurase_mitochondrial [MW = 50.39 kDa]

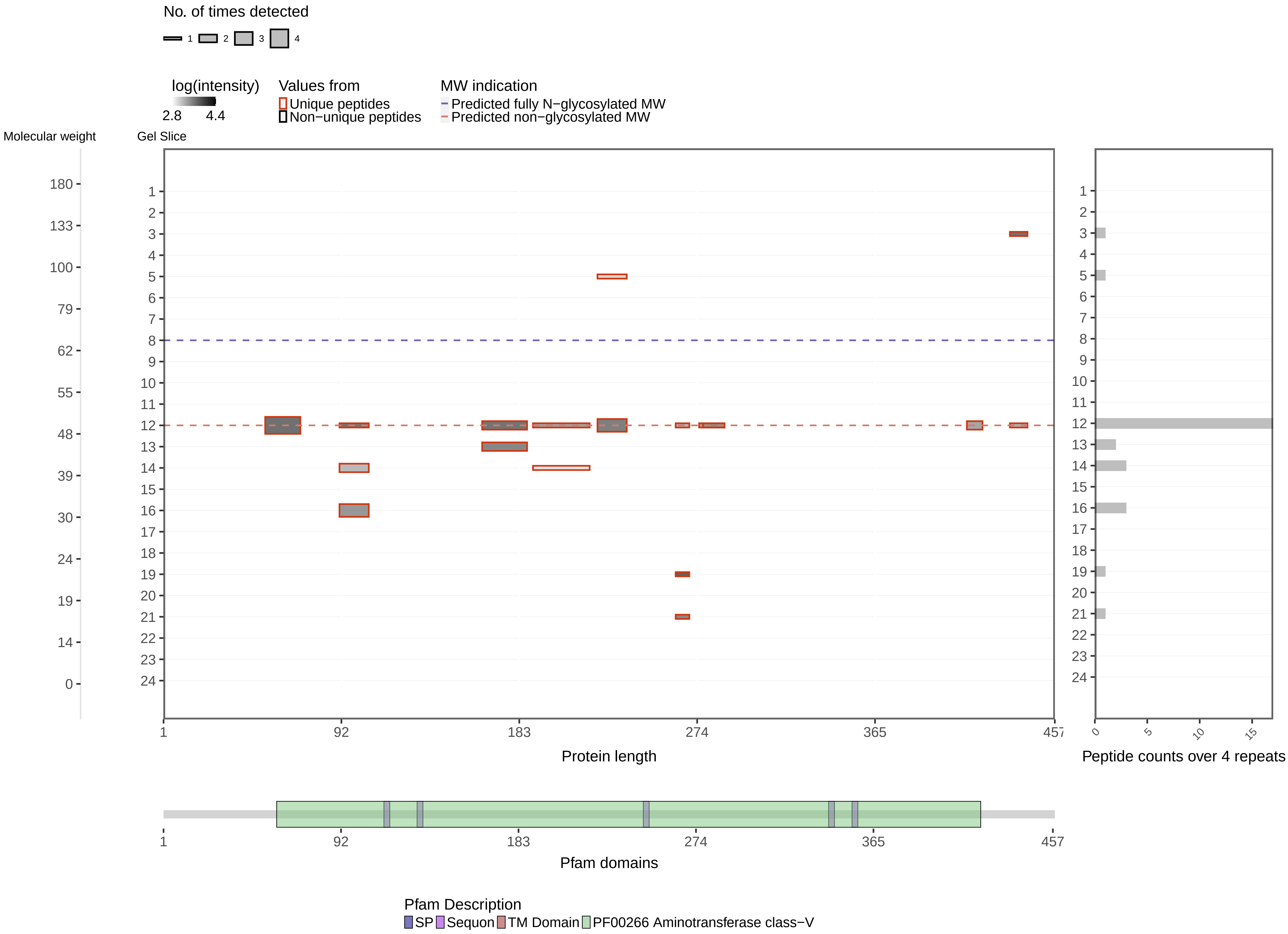


Protein sequence:

EIGKICKEFNVPFHTDAAQALGKIPIDVEKMNISLMSLSGHKIYGPKGVGALYIRRRPRIRVEPQMNGGGQERGIRSGTVPTPLVVGFGAACELAMKEMEYDAQRIKGLQERLLNGIRSKIDGVVVNGSFERRYAGNLNLSFAYVEGESLLMGLKEVAVSSGSACTSASLEPSYVLRALGVEEDMAHTSIRYGIGRFTTEQEIDRAMDLTVKQVEKLREMSPLYEMVKEGIDIKSIQWAQH*

NbL12g10670.1 dentin_sialophosphoprotein [MW = 79.26 kDa]

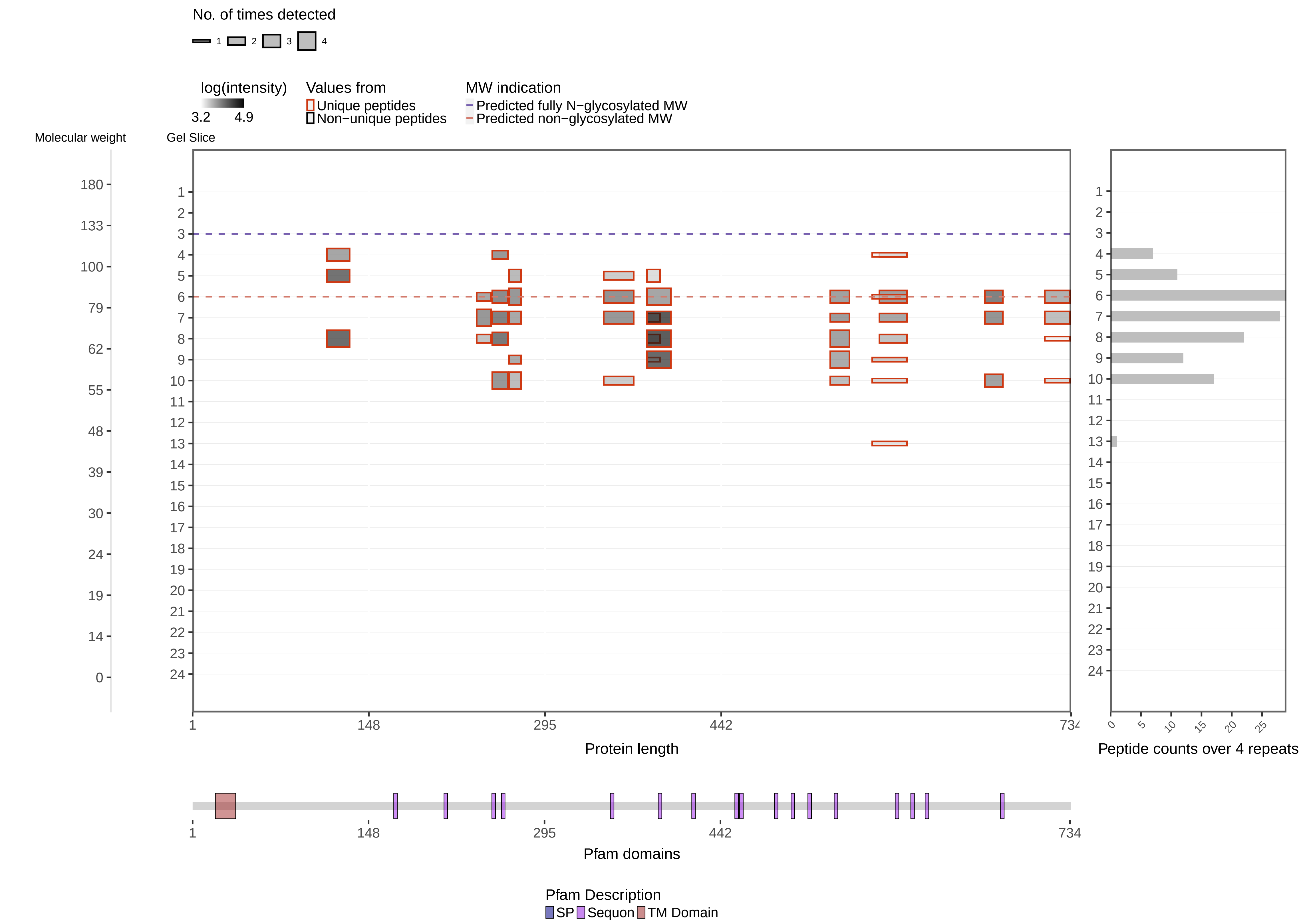


Protein sequence:

NLAMNNEEQSKKGEDELQDTKNNNEASDESLNKSEADAGENKMVEQSSSSDANAAGEASNGSNSEDNSDPAAGNNEATTQQTETGNAASETNRNENNSNNNSDAAAGTNEANEHQQGNENSASHTQNENNSSDNSNPAAGNENNSSDNSNPAAGNENNSSDNSNPAAGNENNSSDDSNSAAETNEANKQQQENDSSVSGTTQNEKNSGEDSNSAEGTNKANKKQQEDEKNSANEAASVGDDTQNISNEQQSEKSDSNSSSNEESVTSSNSSESVVSVDQNGNSVTAGATEKST

### NbL12g10620.1 40S_ribosomal_protein_S14−2 [MW = 16.34 kDa]

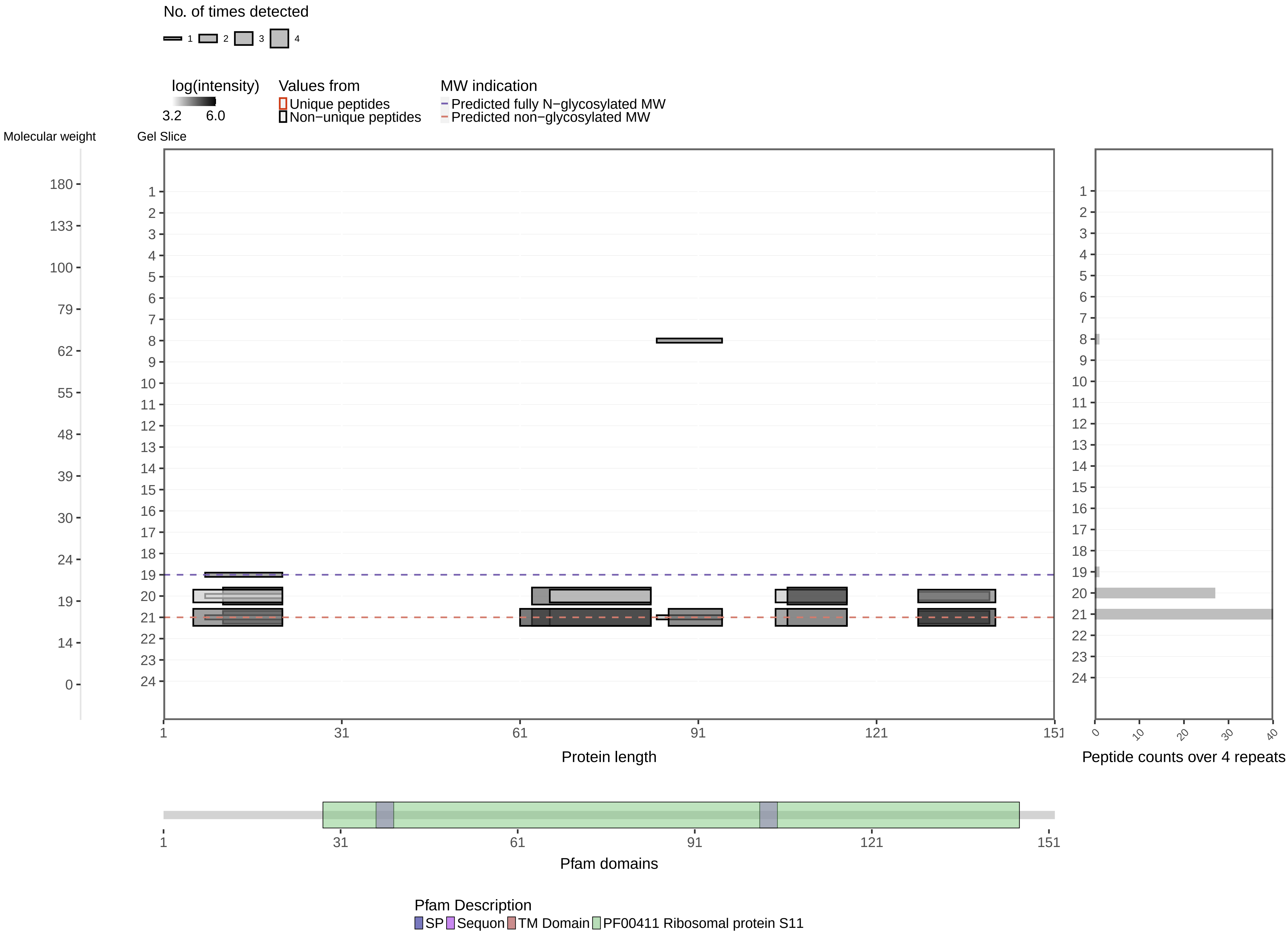


Protein sequence:

ADRDESSPYAAMLAAQDVSQRCKELGINALHIKLRATGGNKTKTPGPGAQSALRALARSGMKIGRIEDVTPIPTDSTRRKGGRRGRRL*

### NbL12g10600.1 la−related_protein_6A−like [MW = 46.4 kDa]

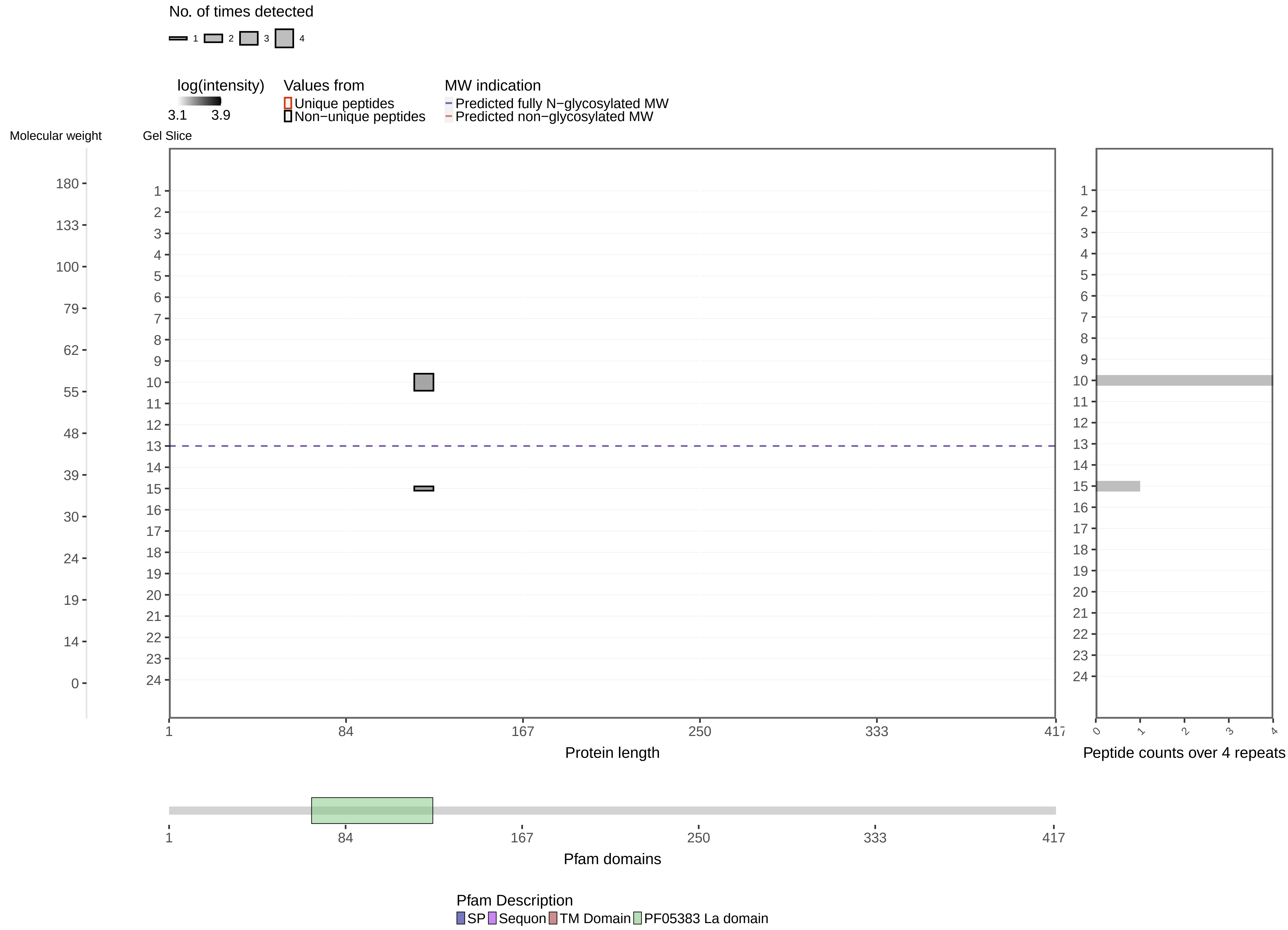


Protein sequence:

AEKLLSGKLHALVEYNTVEAAEKAVAILNDEQDWRFGLRVKLLKKTNKPGQNKKGWRDPDSDRNMNIQASGQAVNEEYHSSEHRVLQKKPGQSKKGRSDPDSDRNTNVQASDLAVNEEHHSNEHHVDSPDEEEGDHRSKETVGEHAQKEKNVHRVQTRNRGRGRRNKRGTNGHGHGTTSYNHAVESSKPPPGPRMPDGTRGFAMGRGRPLSSSPS*

### NbL12g10590.1 uncharacterized_LOC109208446 [MW − SP = 39.86 kDa]

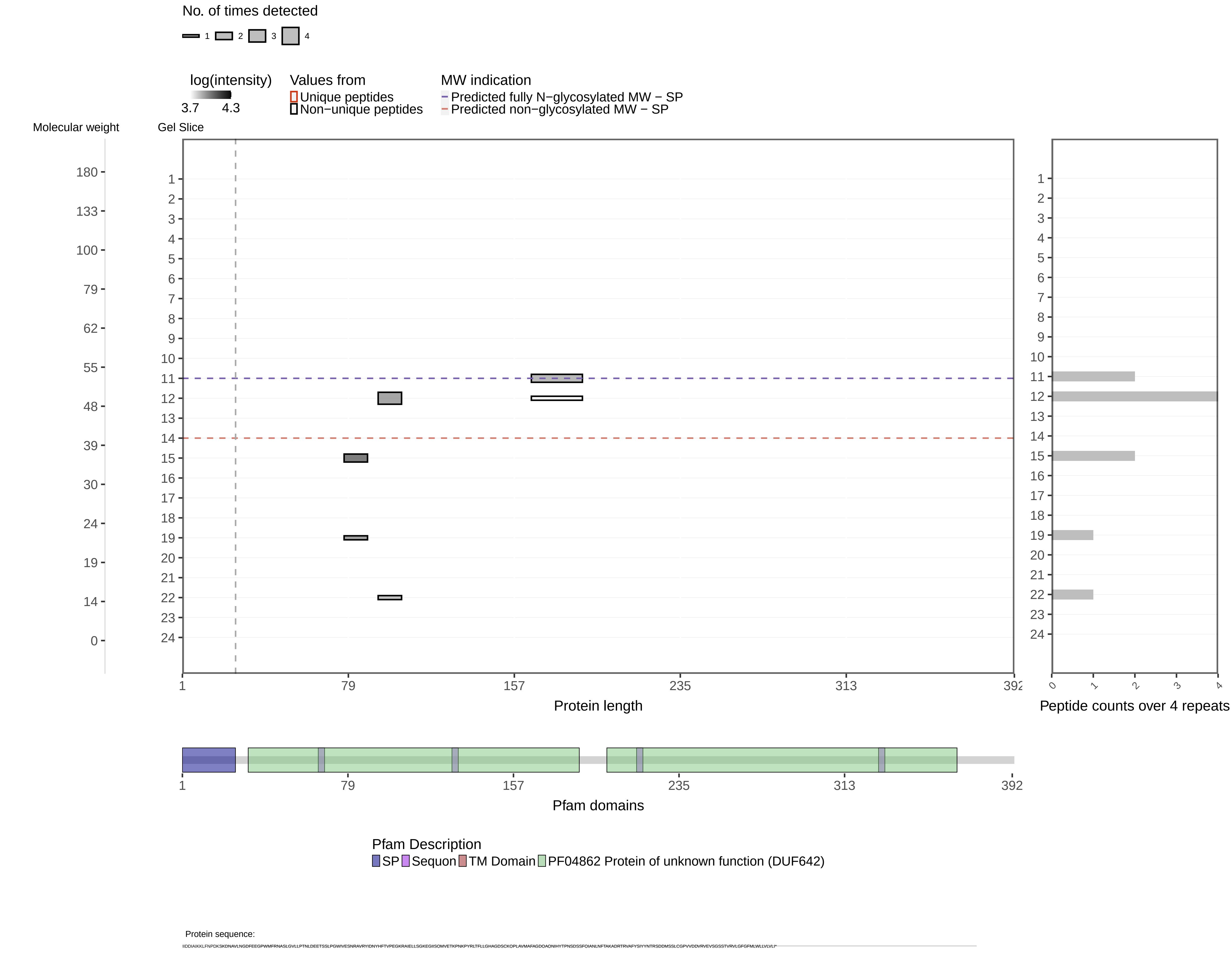

### NbL12g10450.1 40S_ribosomal_protein_S4 [MW = 29.85 kDa]

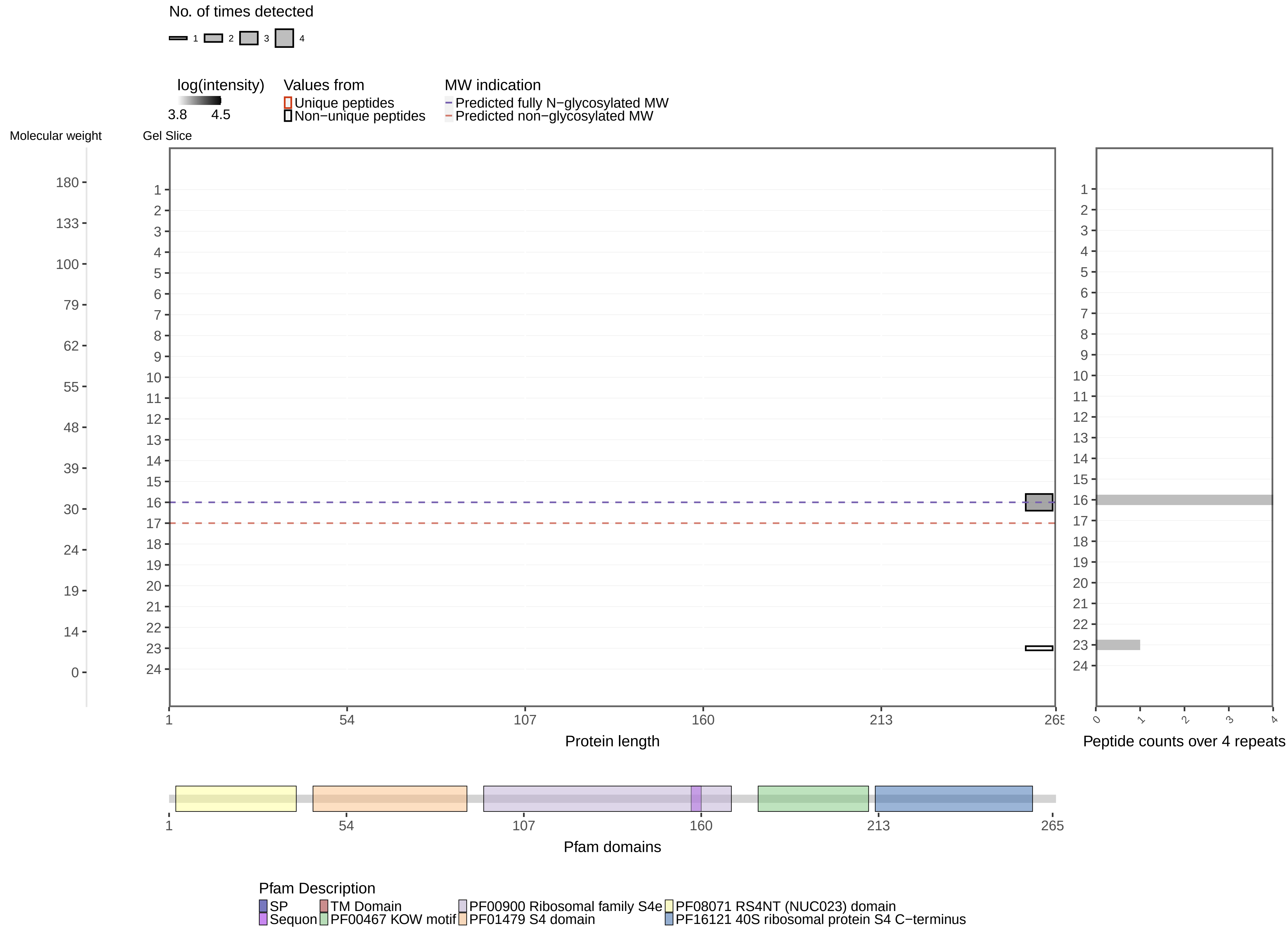


Protein sequence:

AKFKLCKVRSVQFGQKGIPYLNTYDGRTVRYPDPLIKANDTIKLDLESNKIVDFIKFDVGNVVMVTGGRNRGRVGVIKNREKHKGSFETVHIQDALGHEFATRLGNVFTLGKGTKPWVSLPKGKGIKLSIIEEARKRLAAQSATAA*

### NbL12g10430.1 60S_ribosomal_protein_L32−1 [MW = 15.63 kDa]

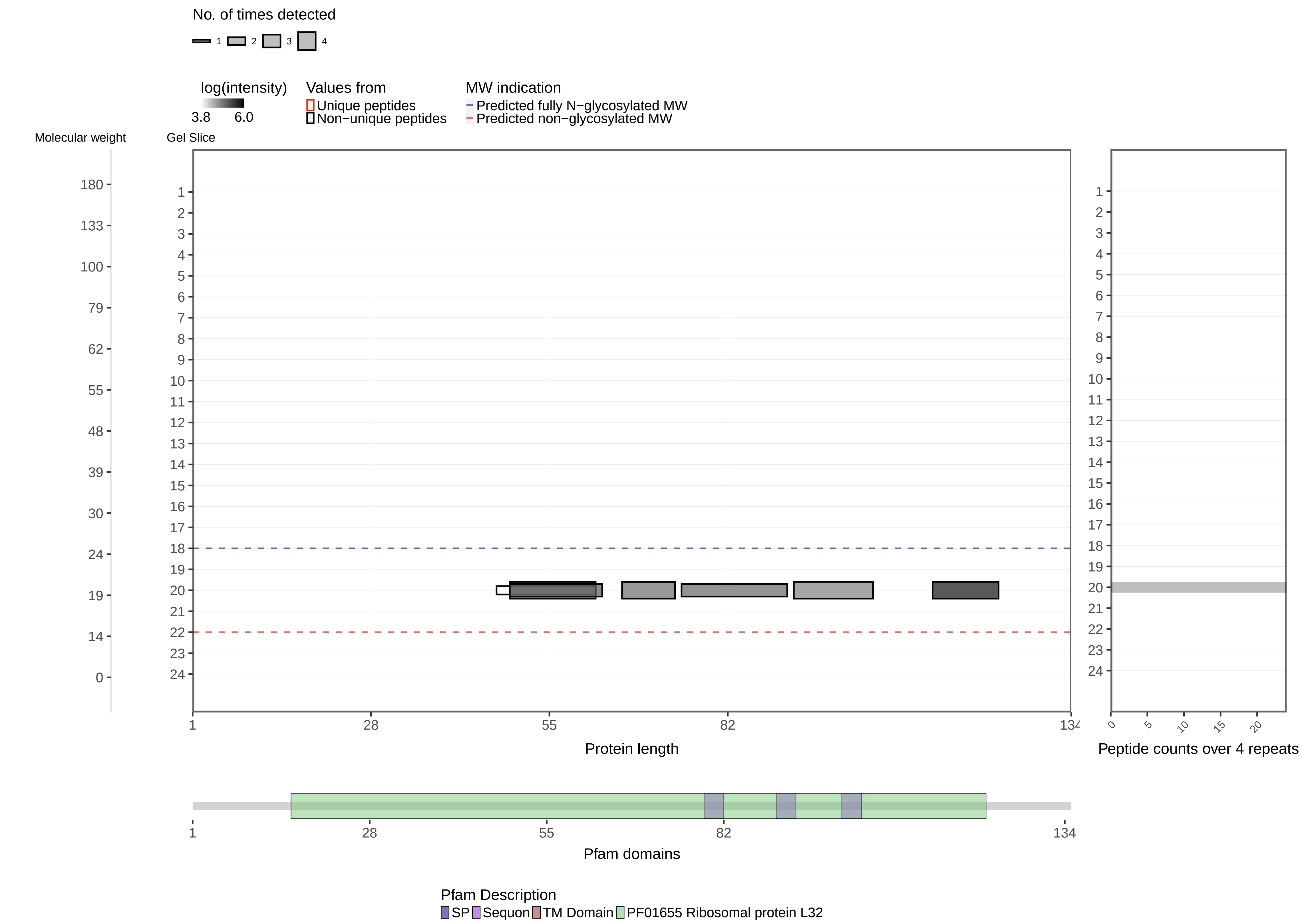


Protein sequence:

PNIGYGSDKKTRHYLPNGFKKFVVHNASELEVLMMHNRTYCAEIAHNVSTRKRKEIVERAAQLDVVITNKLARLRSQEDE*

### NbL12g10370.1 accelerated_cell_death_11 [MW = 23.42 kDa]

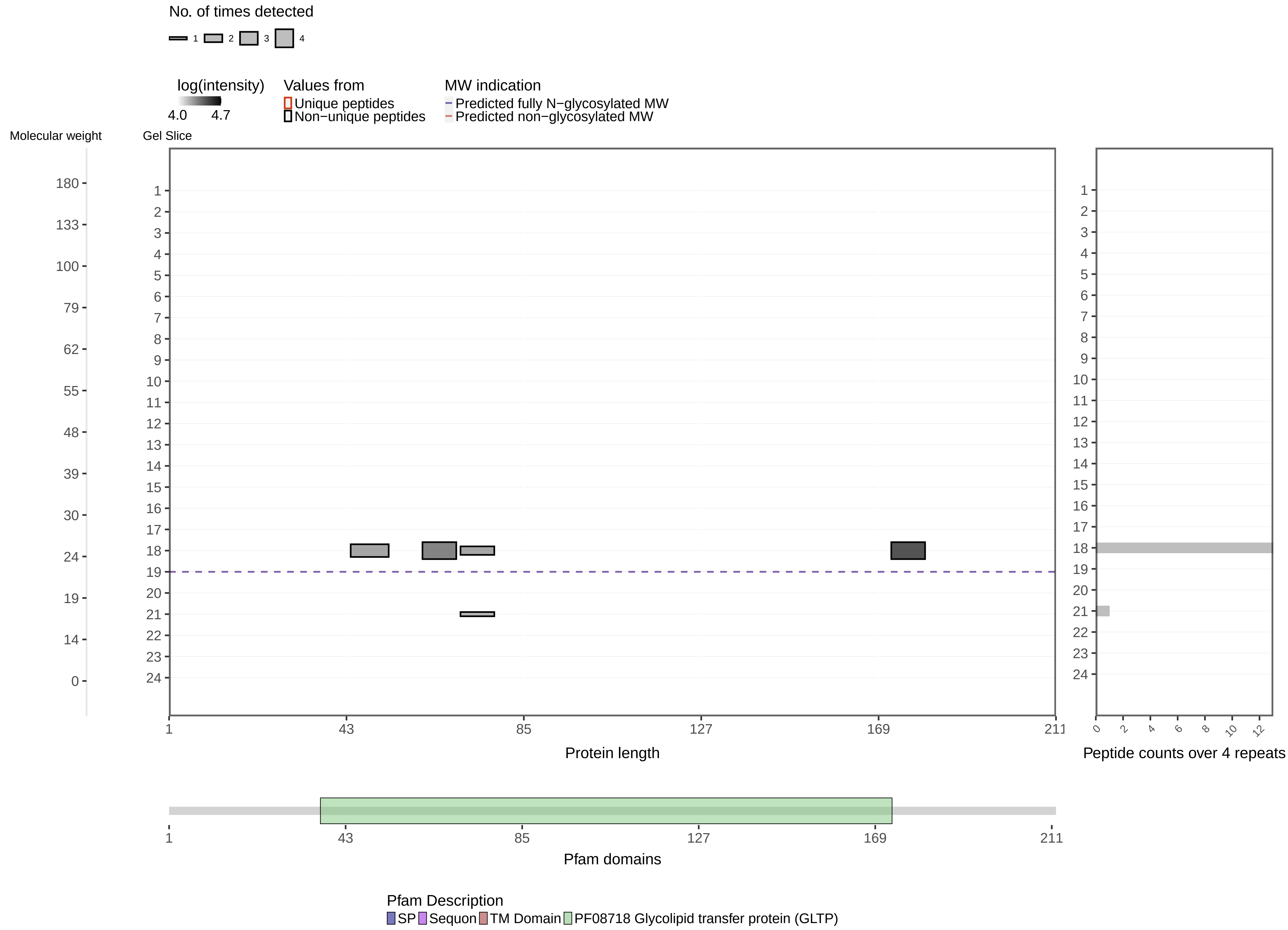


Protein sequence:

ADCVRRAGSHTRNLLRVKRGLDMVKVLFEQIIASEGNSLKDPASKAYQQVFAPYHGWAIRKAVSAGMYALPTRQQLMNKLNEDEASARTQMQNYVASSGTVILYIDKLFISRDLGIDW*

### NbL12g10110.1 40S_ribosomal_protein_S25 [MW = 11.77 kDa]

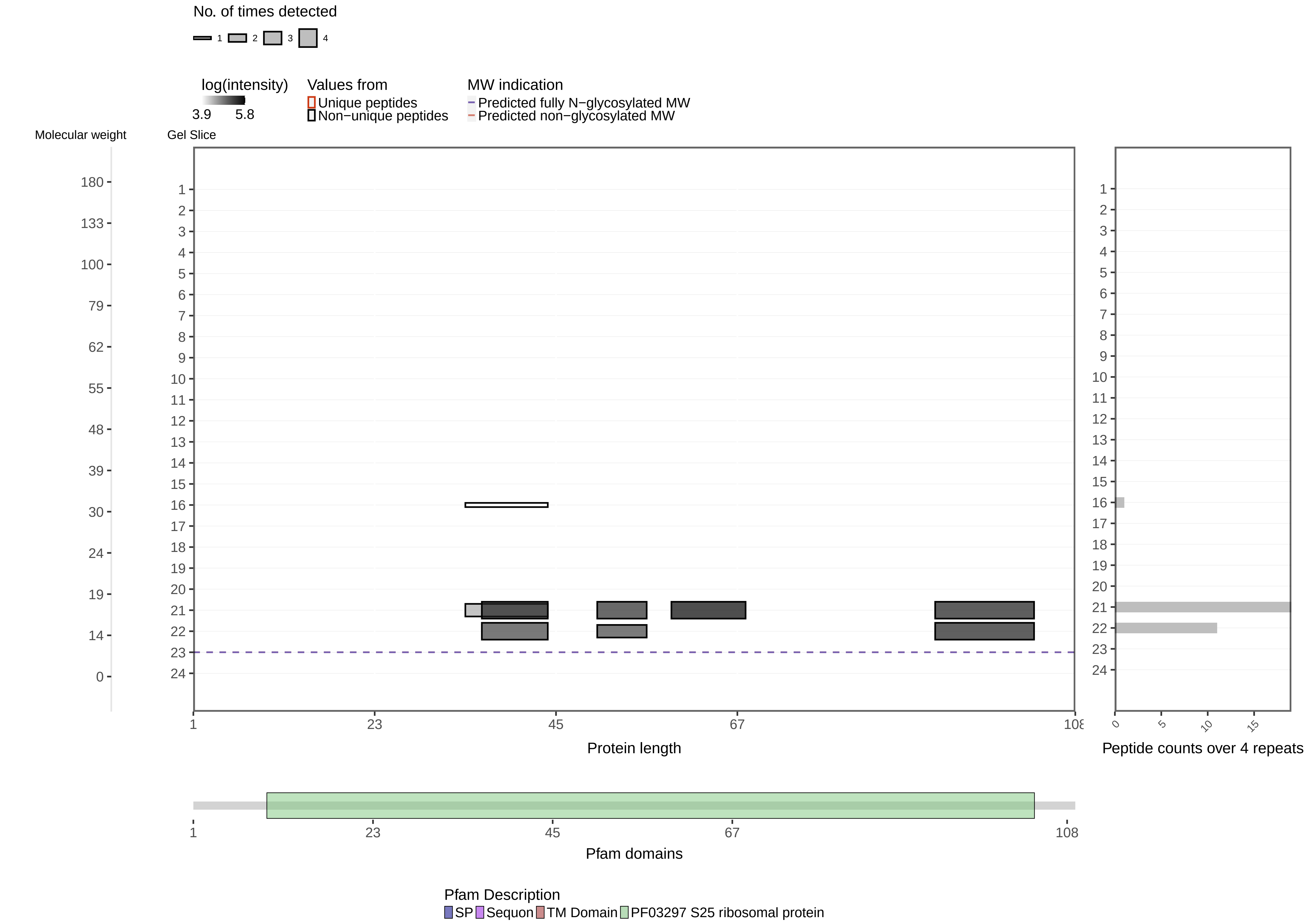


Protein sequence:

MVLFDKATYDKLLSEAPKYKLITPSVLSDRLRISGSLARKAIRDLMARGSIRMVSAHASQQIYTRATNT*

### NbL12g10100.1 50S_ribosomal_protein_L20−like [MW = 14.52 kDa]

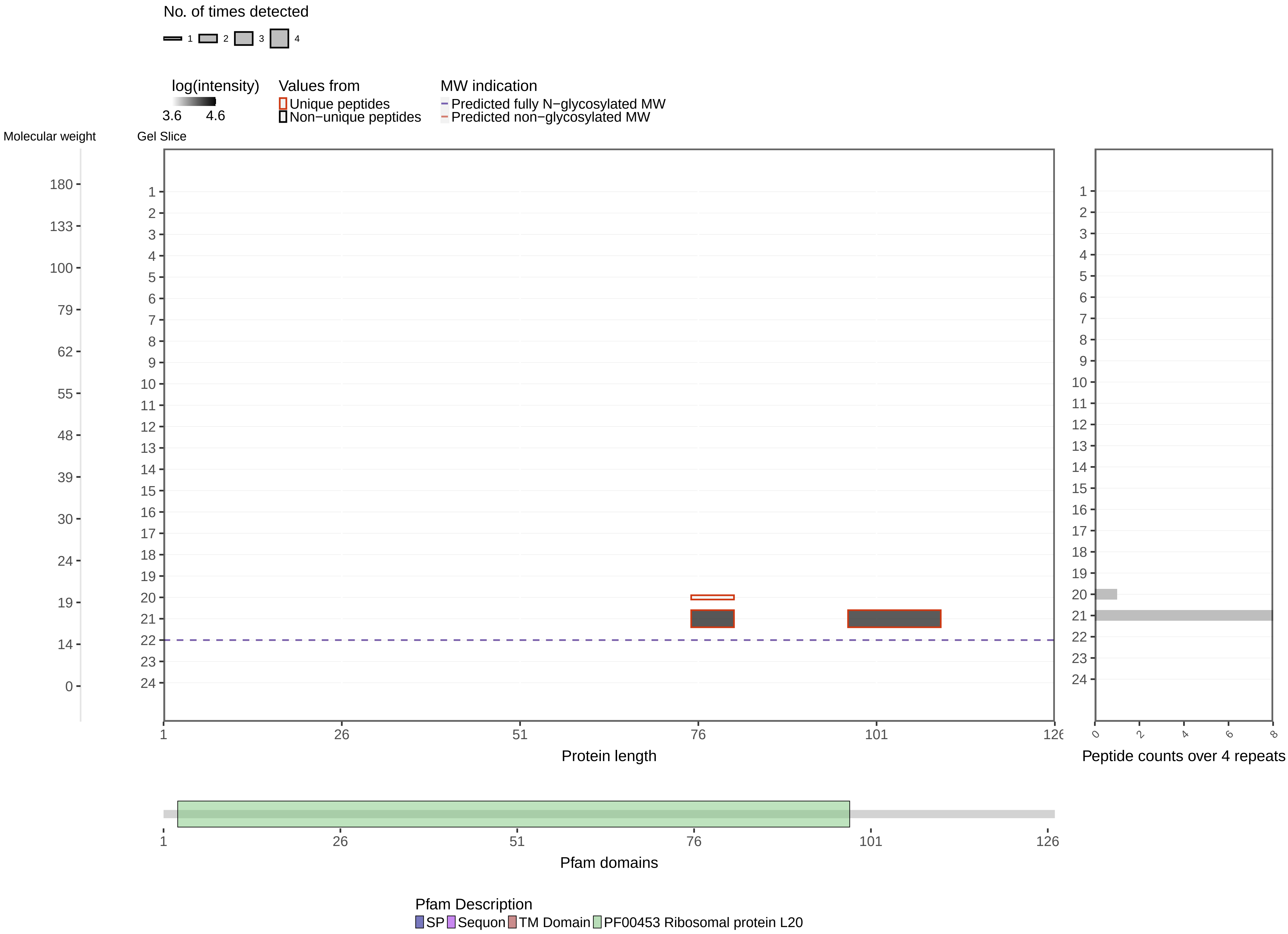


Protein sequence:

WIQRINAGTRQHGVNYGNFMHGLMKENVQLNRKVLSELSMHEPYSFKALVDISSSAFPGNKKAIVPPKKEGLAIFL*

### NbL12g10070.1 probable_carboxylesterase_17 [MW = 40.96 kDa]

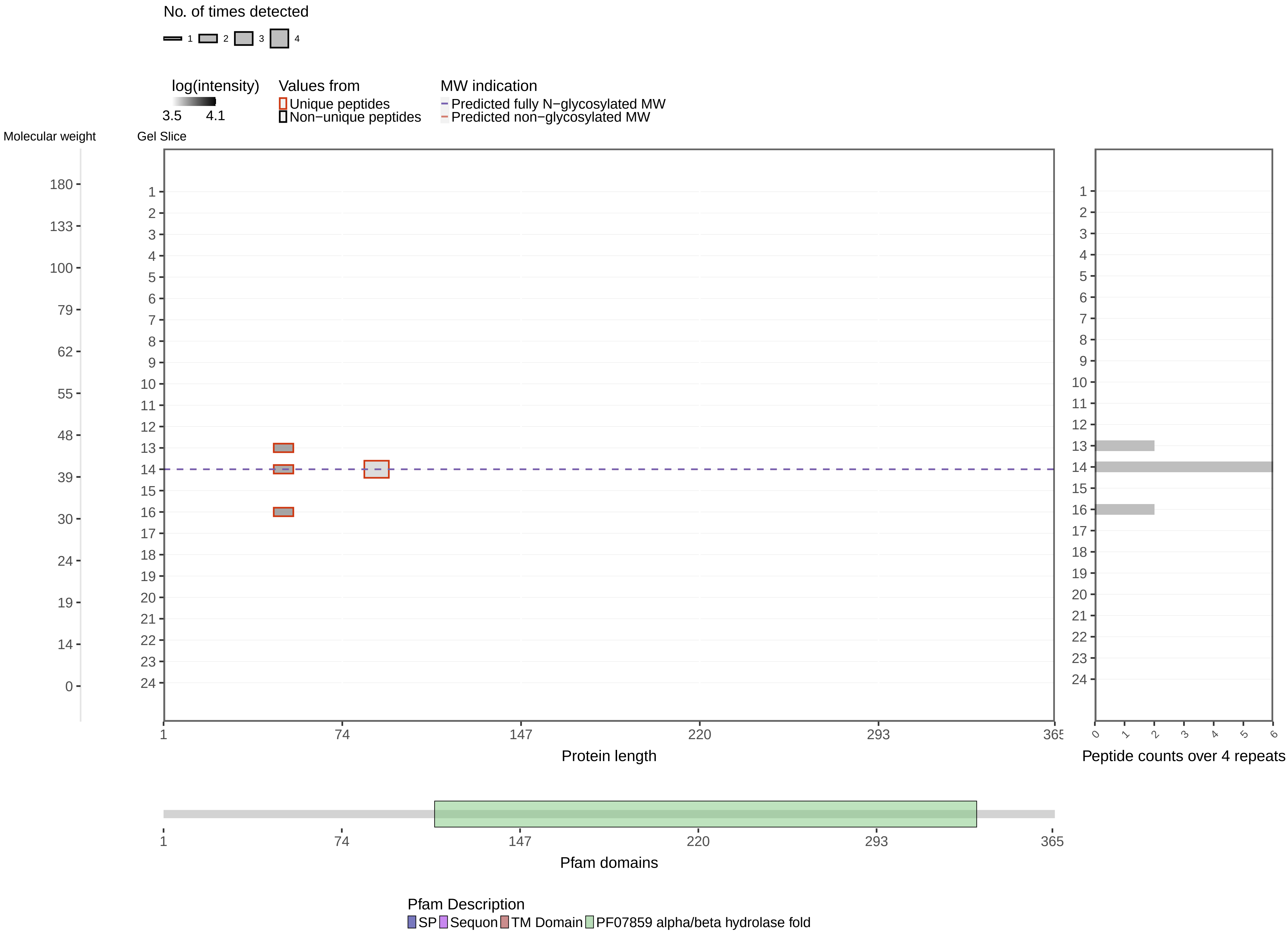


Protein sequence:

ALLWLRDLSROOGYEPWLKDYADFNRVFLIGDSSGGNIVHOVAVRAGEETLSPMRLAGAIPIHPGFVRSYRSKSELEOEOTPFLTIDMVDKFLALALPIGSNKDHPITCPMGDAAPAVEELKLPPYLYCIAEKDLIKDTEMEFYEAMKKGEKDVELFINNGVGHSFYLNKIAVKMDPVTGSETEKLYEAVAEFISKH*

NbL12g10060.1 DNA−damage−repair/toleration_protein_DRT111_chloroplastic−like [MW = 42.02 kDa]

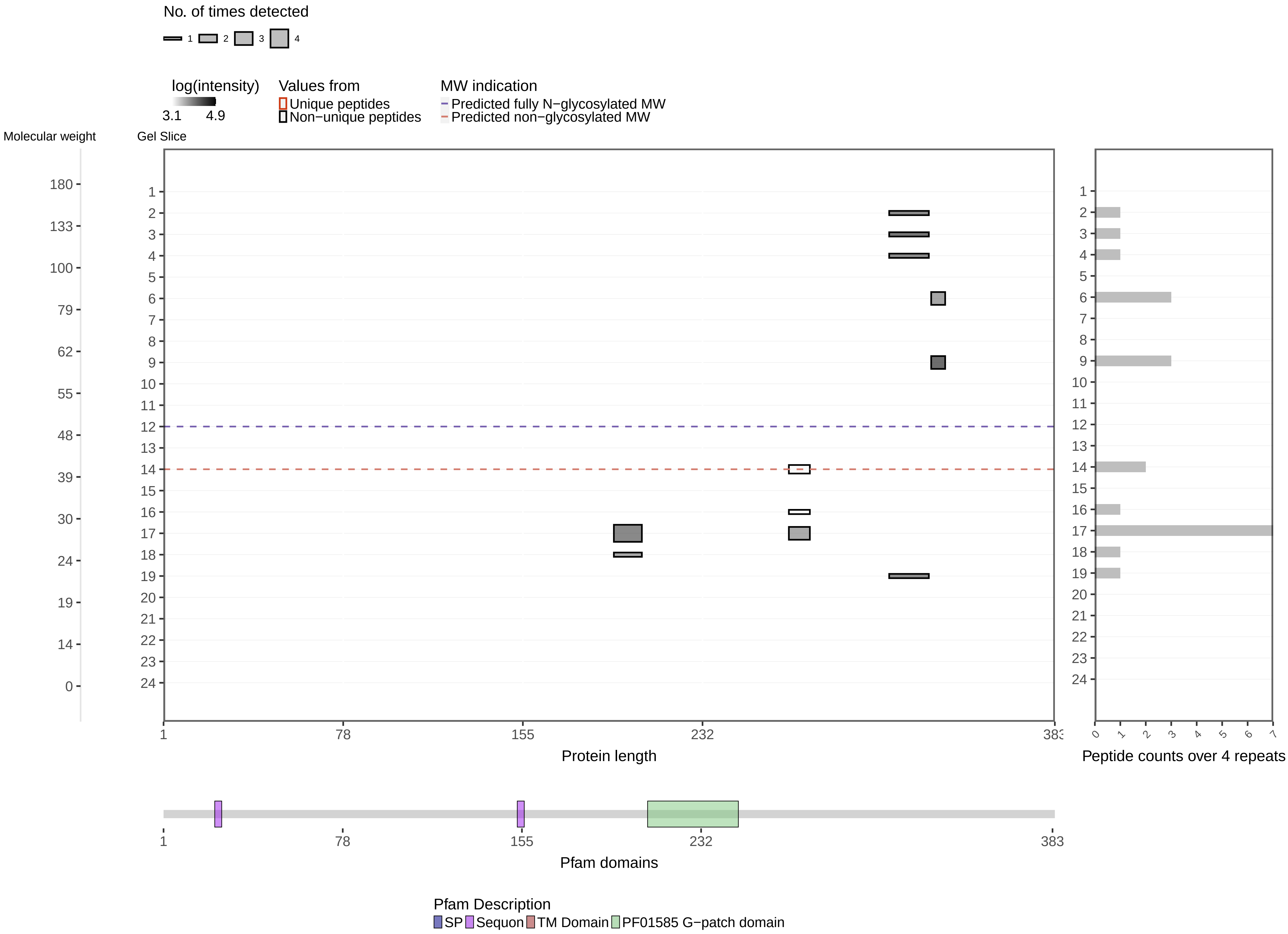


Protein sequence:

GSPSPPSGNEFSIGRSESGGLGLGAEGKMTAAQRMMAKMGWKEGQGLGKLEQGITTPLMAKKTDKRGGVIVASEAKQQQQQQQHPEKKVKSVNFNMPPTRVVLLKNMVGPGEVDDDLEGEVAEECTKFGTVTRVLIFEITEANFPHEEAVRIFVQFERAEQATKALIELEGRFFGGRIVHACFYDEERFSNNELAPMPGEIPGF*

NbL12g10030.1 probable_choline_kinase_2 [MW = 40.48 kDa]

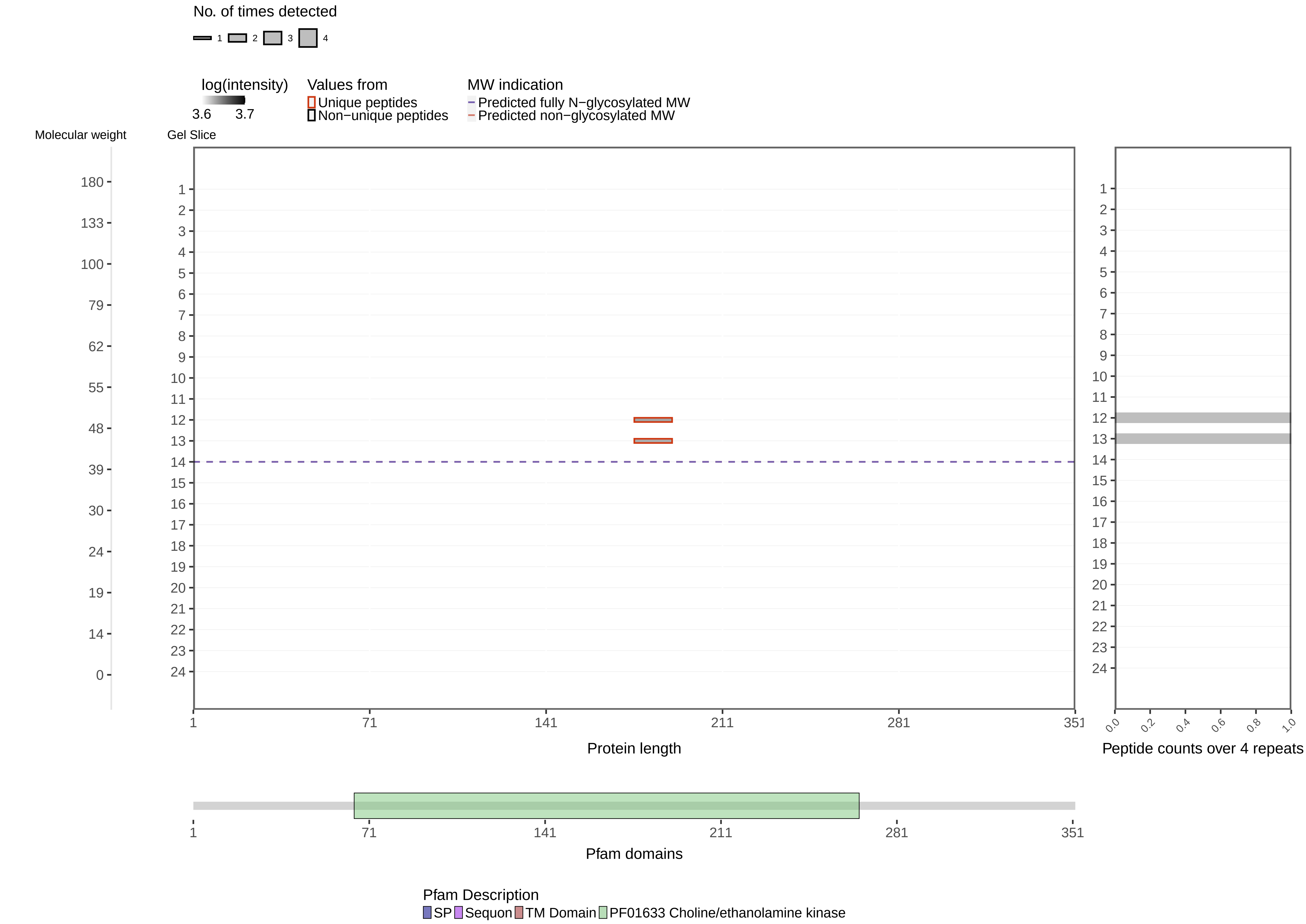


Protein sequence:

.NVAKRLASAEEAKDFQLDHLQGEISALEKSLAGNHLSMGFCHNDLQYGNIMMDEEIRSITFIDYEYAGYNHVAFDIANHFCEMVADYHTETPHIMDFSKYPGLEERKRFLSAYLSSSGGSPSEQELEKLAQEVEKYTLASHLFWGLWGIISHHVNKIDFNYLEYARQRFQPYWSKKPELLGSSGSKKG*

### NbL12g09980.1 flavin−dependent_oxidoreductase_FOX2−like [MW − SP = 58.95 kDa]

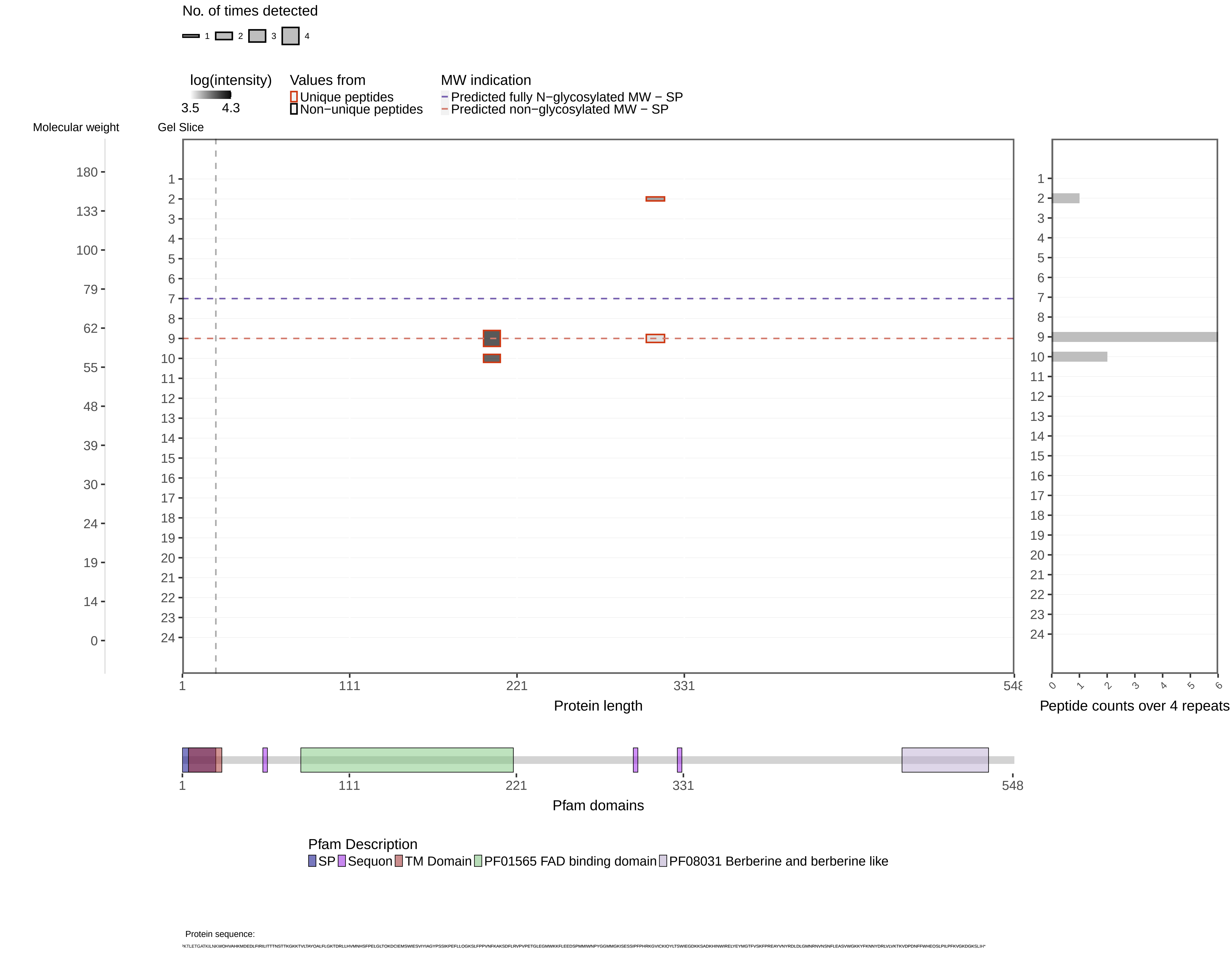

NbL12g09940.1 ATP_synthase_subunit_alpha_chloroplastic [MW − SP = 57.08 kDa]

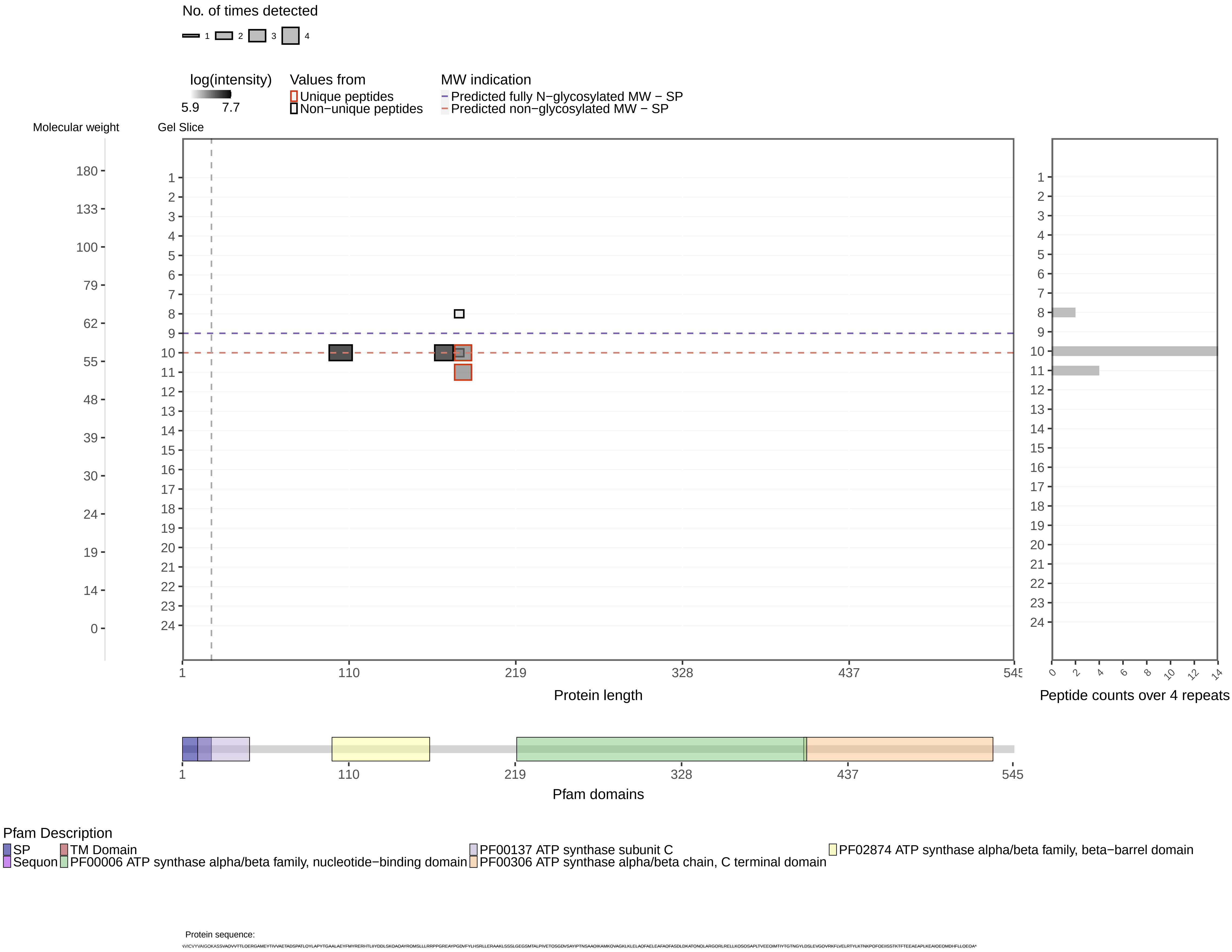

NbL12g09770.1 haloacid_dehalogenase−like_hydrolase_domain−containing_protein_At3g48420 [MW = 41.56 kDa]

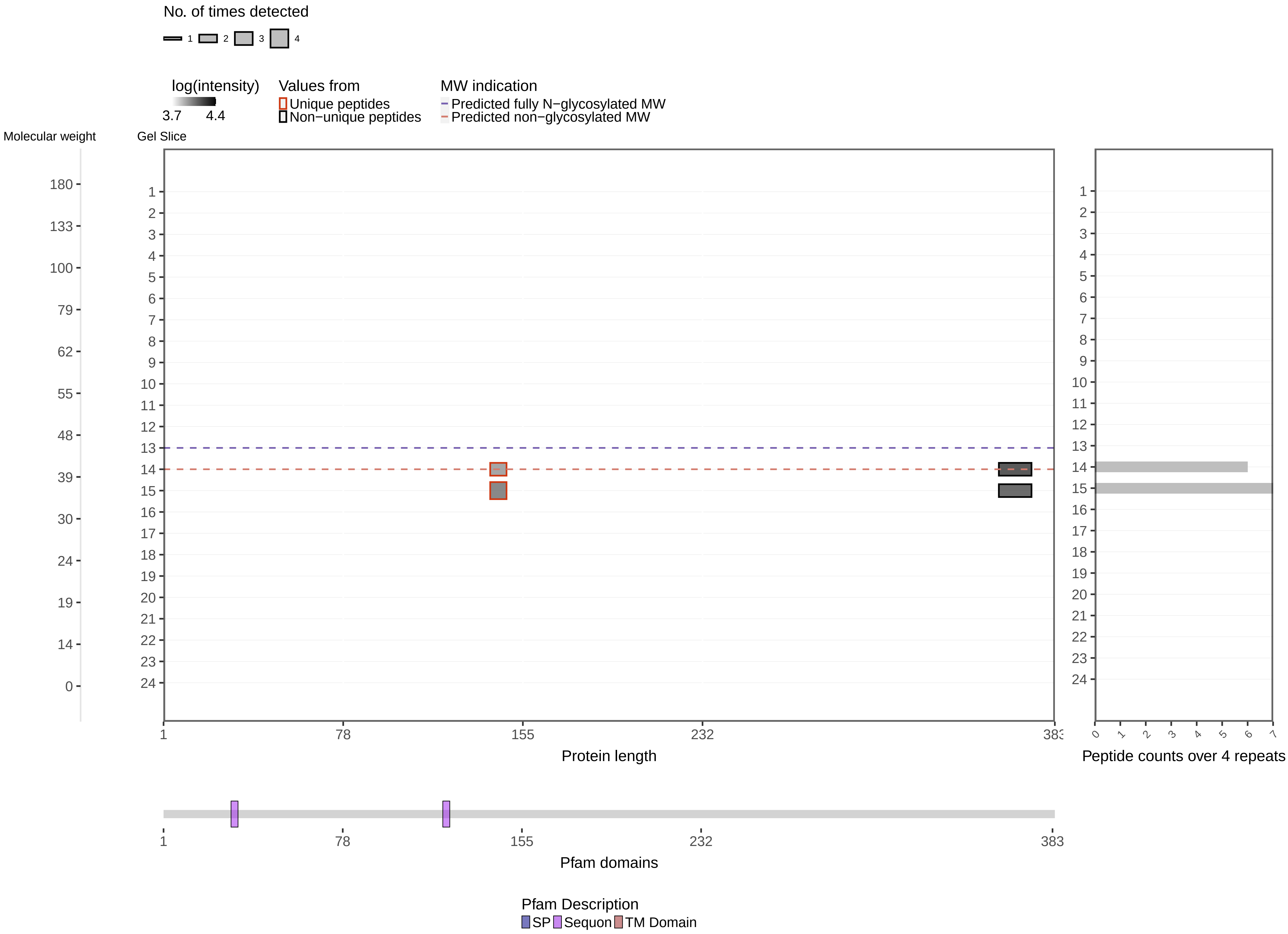


Protein sequence:

VMSKSLPLRPGVEEFMDEAYEEGVPVVMLTSYCKCGEKPARSIIEKLGNDKMARIKILGMEEVKQSFYGQLVFGEGVASDLGEQLAKEARKAASAEEQRIAKEVASMLKLSVDIDTTSSEEMKNVVAALRAGAEYANVPLDNCVLVAGSQSGVAGAERIGMPCVVVRSSSTARAEFPGAKAIMDGFGGGDLSISRLRQIQGLDV*

NbL12g09750.1 non−specific_lipid−transfer_protein_1−like [MW = 11.18 kDa]

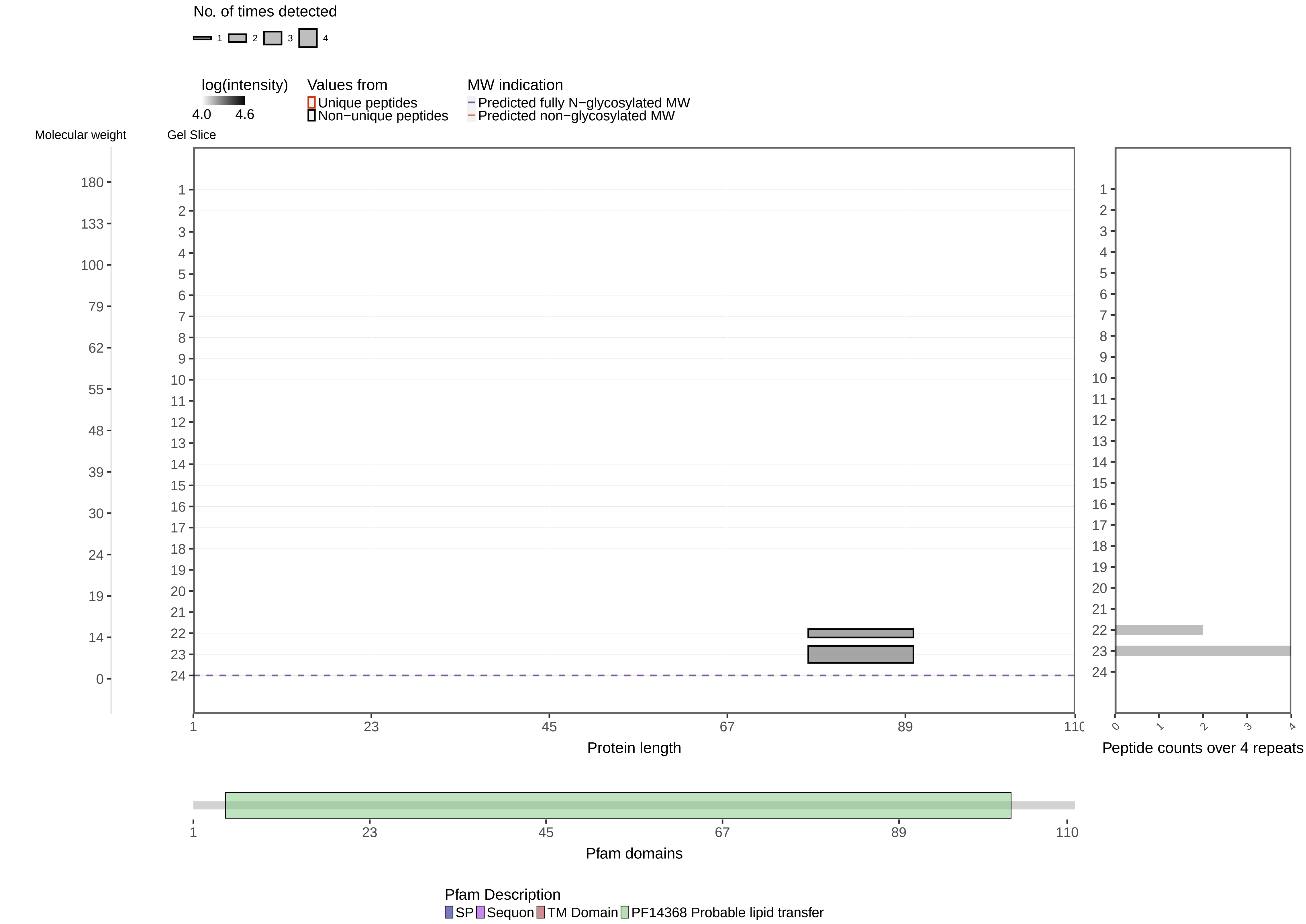


Protein sequence:

ISRIKVPPACCGKIRALLKTSPKCLCAALLSPAAKNAGVNPAIAVTIPKRCNIKNRPVGKRCGRYVVP*

NbL12g09720.1 peptidyl−prolyl_cis−trans_isomerase_FKBP17−1_chloroplastic−like [MW = 23.88 kDa]

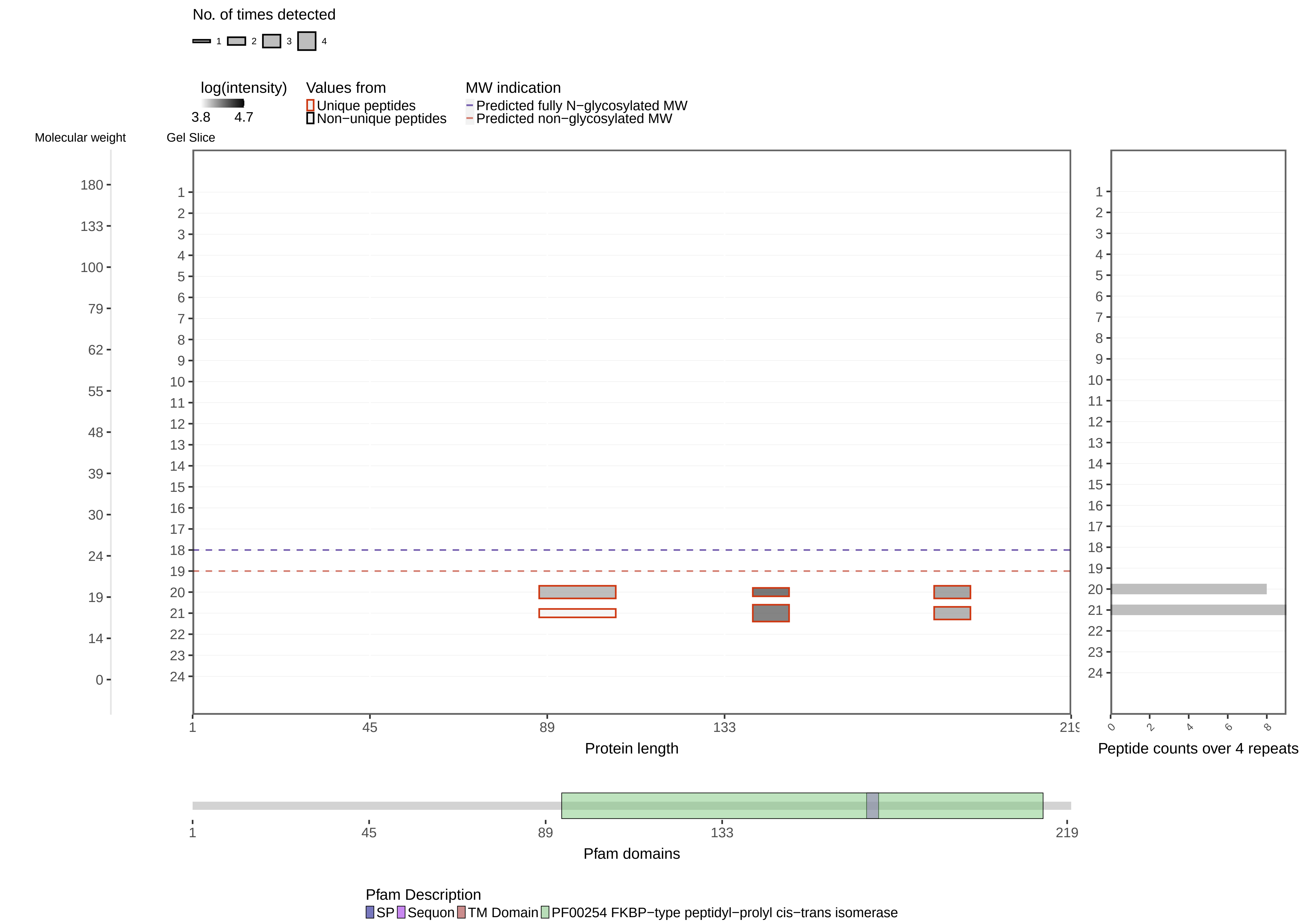


Protein sequence:

GDVVAIHYYGRLAAKQGWRFDSTYDHKDETGEPIPFSFILGSGKVISGIESAVRSMKVGGLRRVIIPPSKGYQNTSQEPLPPNYFDRQRLFTTIFNPTRLANGEGATLGTVIFDIELVSLRHL*

NbL12g09700.1 leucine−−tRNA_ligase_cytoplasmic−like [MW = 122.94 kDa]

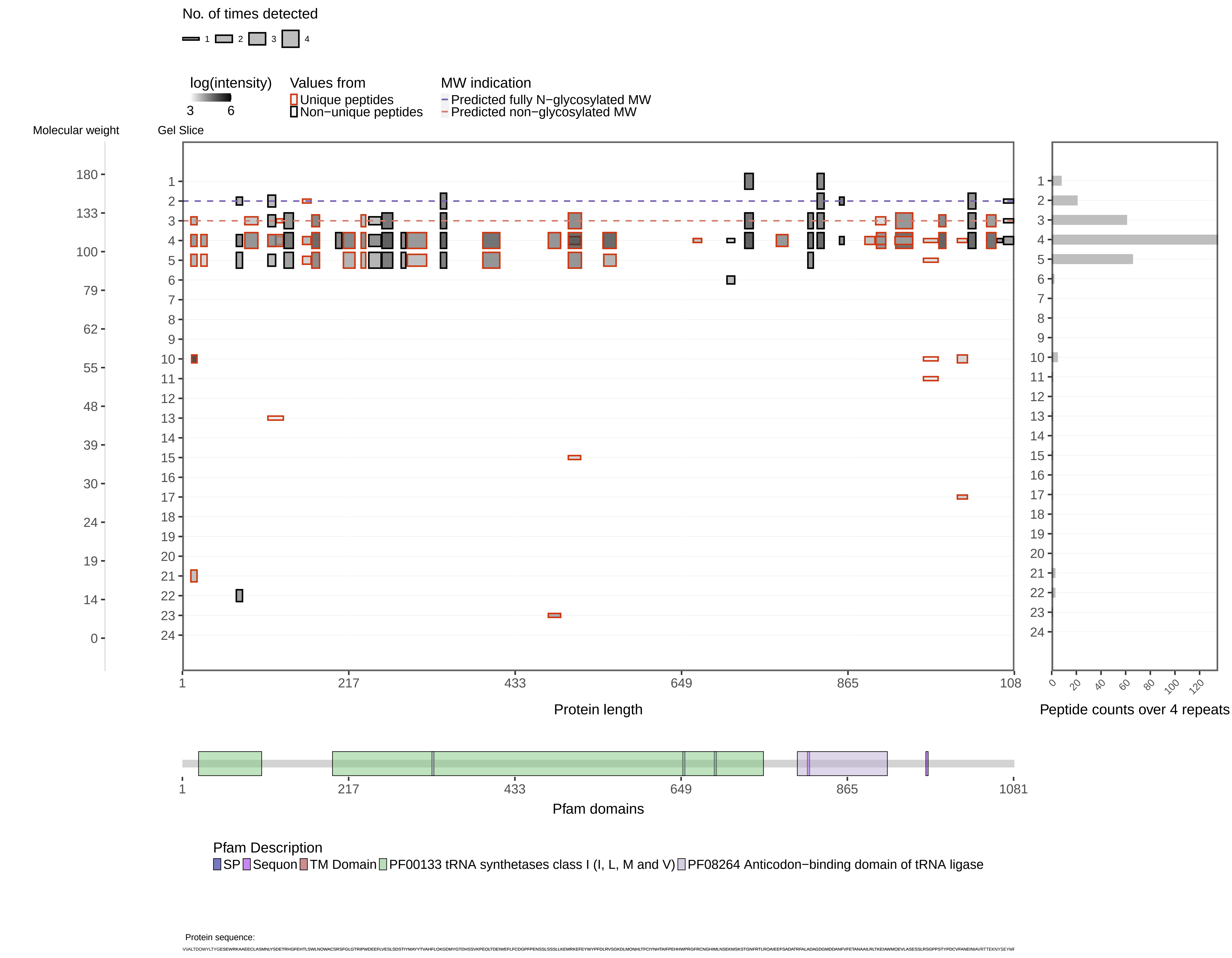

### NbL12g09640.1 protein_argonaute_2−like [MW = 120.9 kDa]

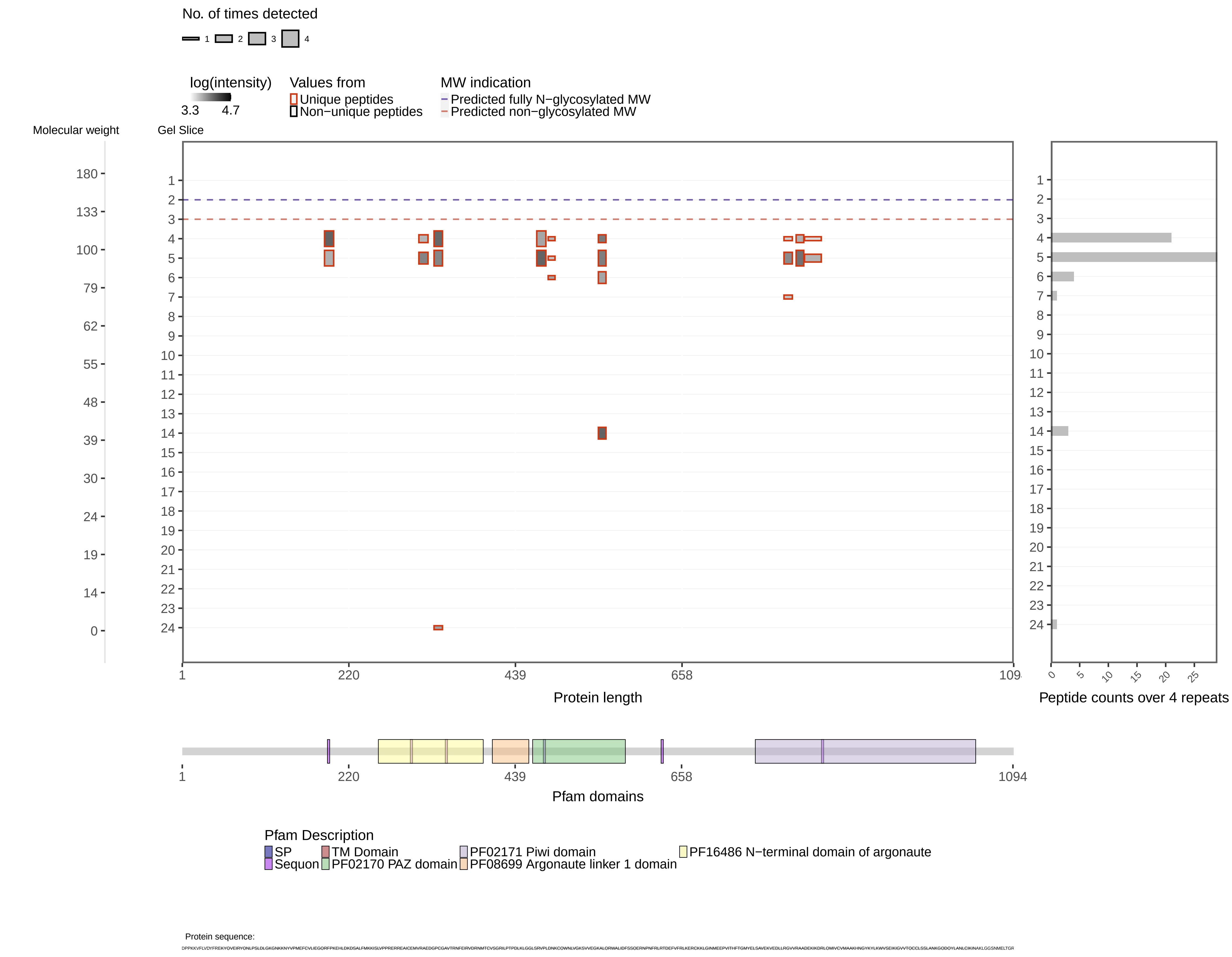

### NbL12g09550.1 protein_RABF2a [MW = 21.73 kDa]

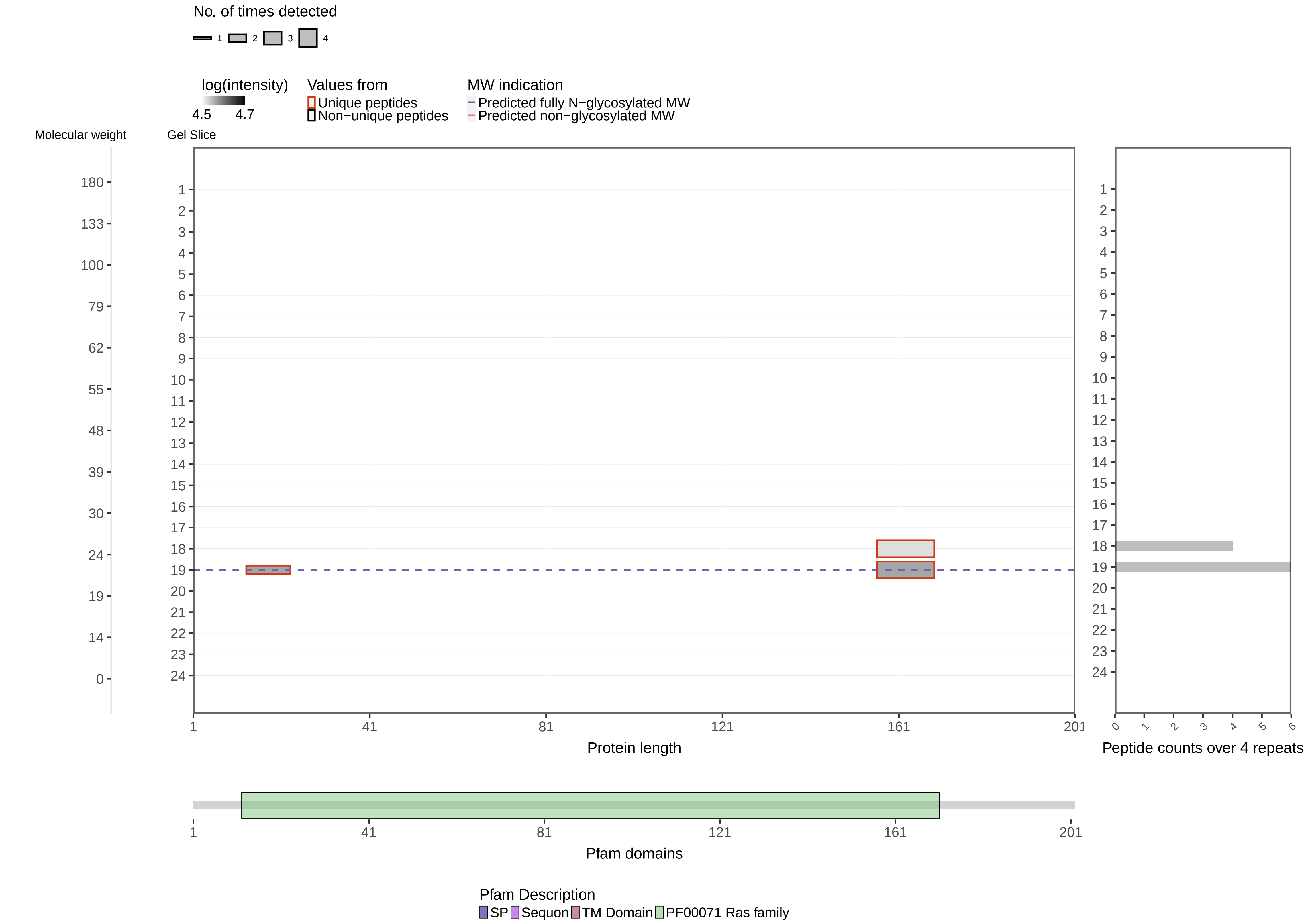


Protein sequence:

/FDITNQASFDRAKKWVQELQAQGNPNMVMALAGNKADLLDARKVAAEEAQTYAQENGLFFMETSAKTASNVNDIFYEIAKRLPRLQPAQNPSGMVLMDRPAQTPASASCCS*

### NbL12g09520.1 cathepsin_B−like [MW − SP = 36.51 kDa]

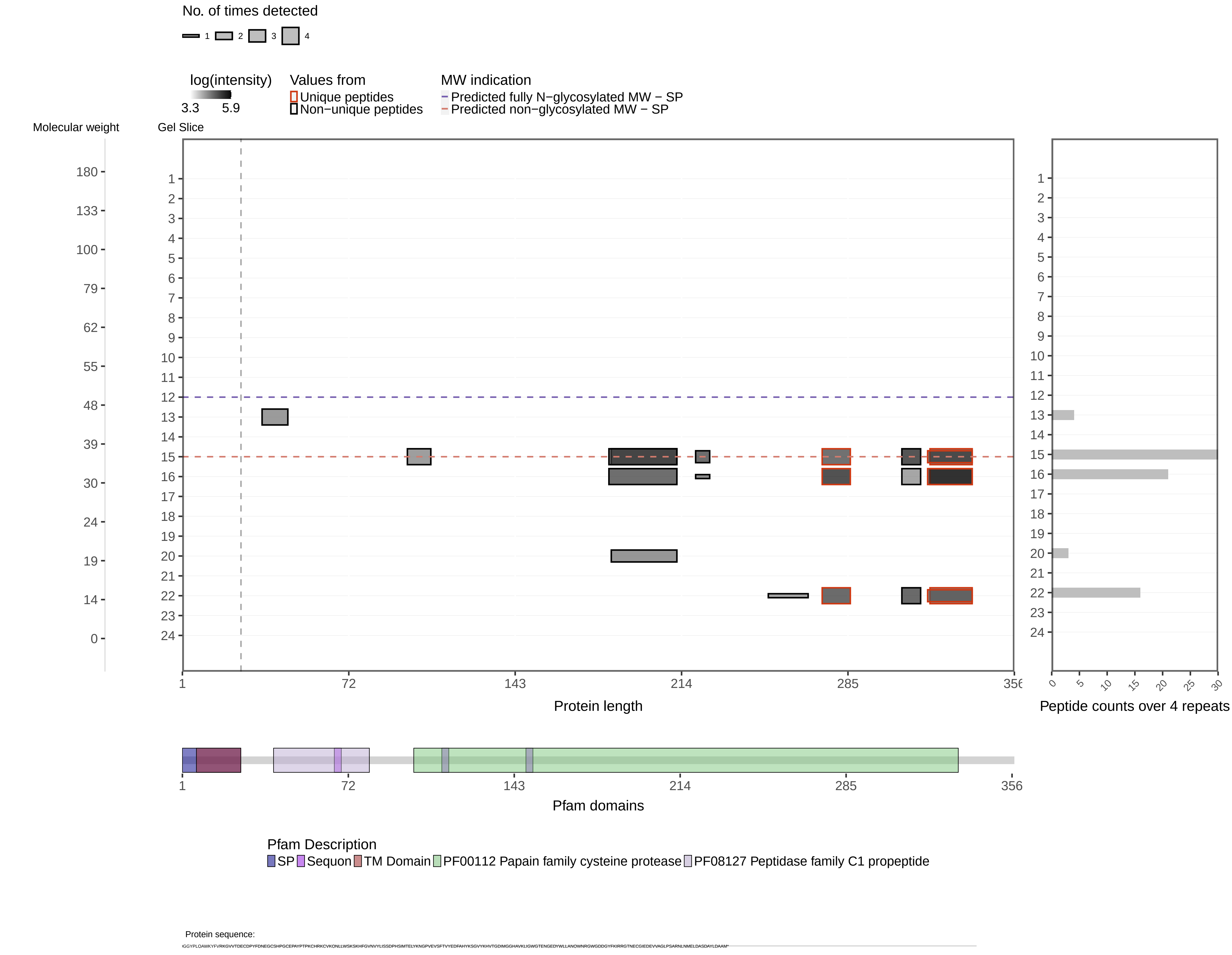

### NbL12g09480.1 phosphatase_IMPL1_chloroplastic−like [MW = 38.44 kDa]

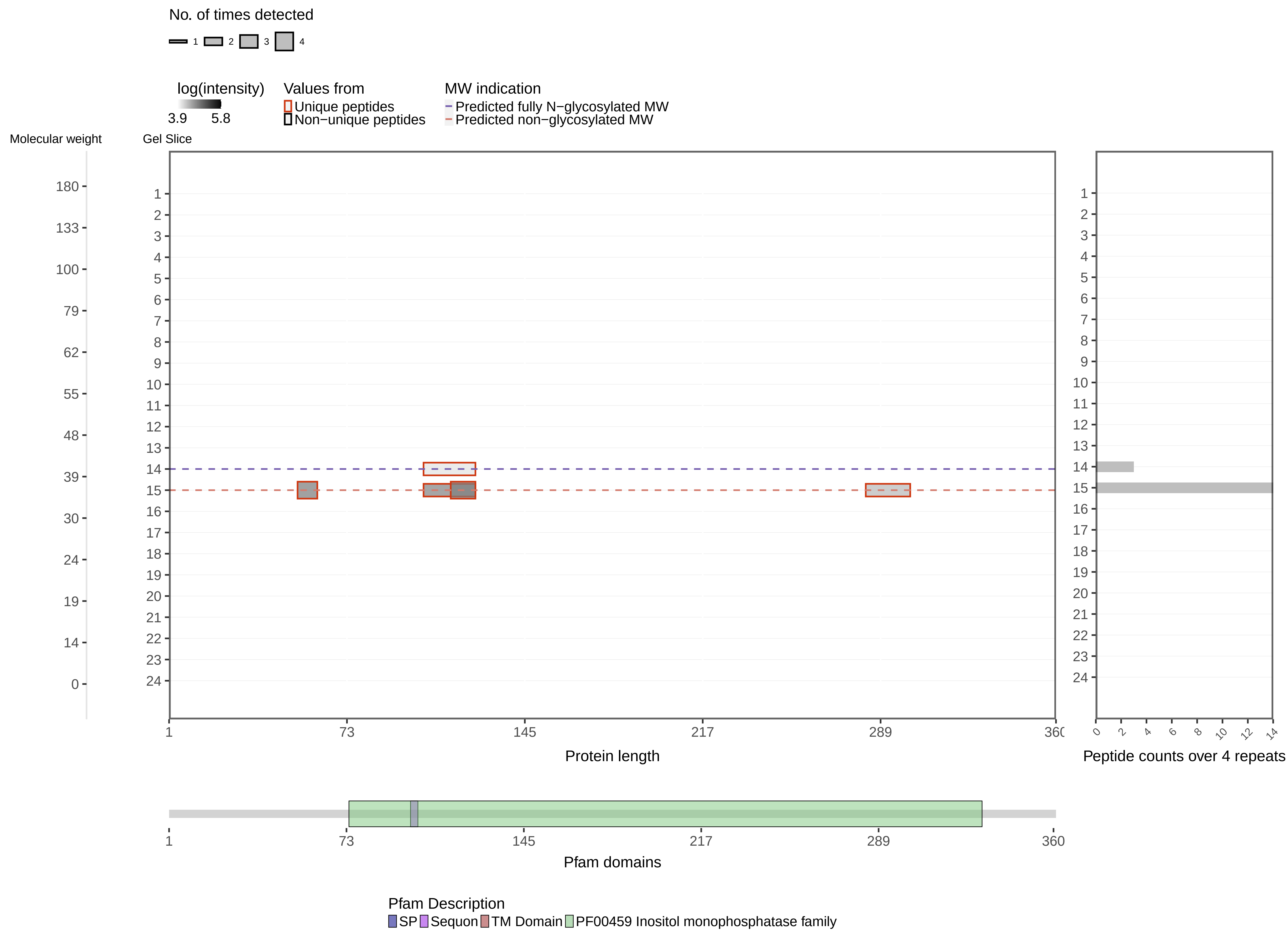


Protein sequence:

\VSVGVLFRGKPAAATVVEFVGGPMCWNTRIFSAAAGKGAFANGEKIHVSGTDKVEOSLLVTGFGYEHDDPWATNMELFKDFTDVSRGVRRLGAAAVDMCHVALGIAEAYWEYRLKPWDMAAGVLIVEEAGGKVSCMDGGKFSVFDRSVLVSNGLLHAKLLEKIGPATENLKGKGIDFSVWFKPENYHTEV*

NbL12g09330.1 glycine_cleavage_system_H_protein_2_mitochondrial [MW = 16.78 kDa]

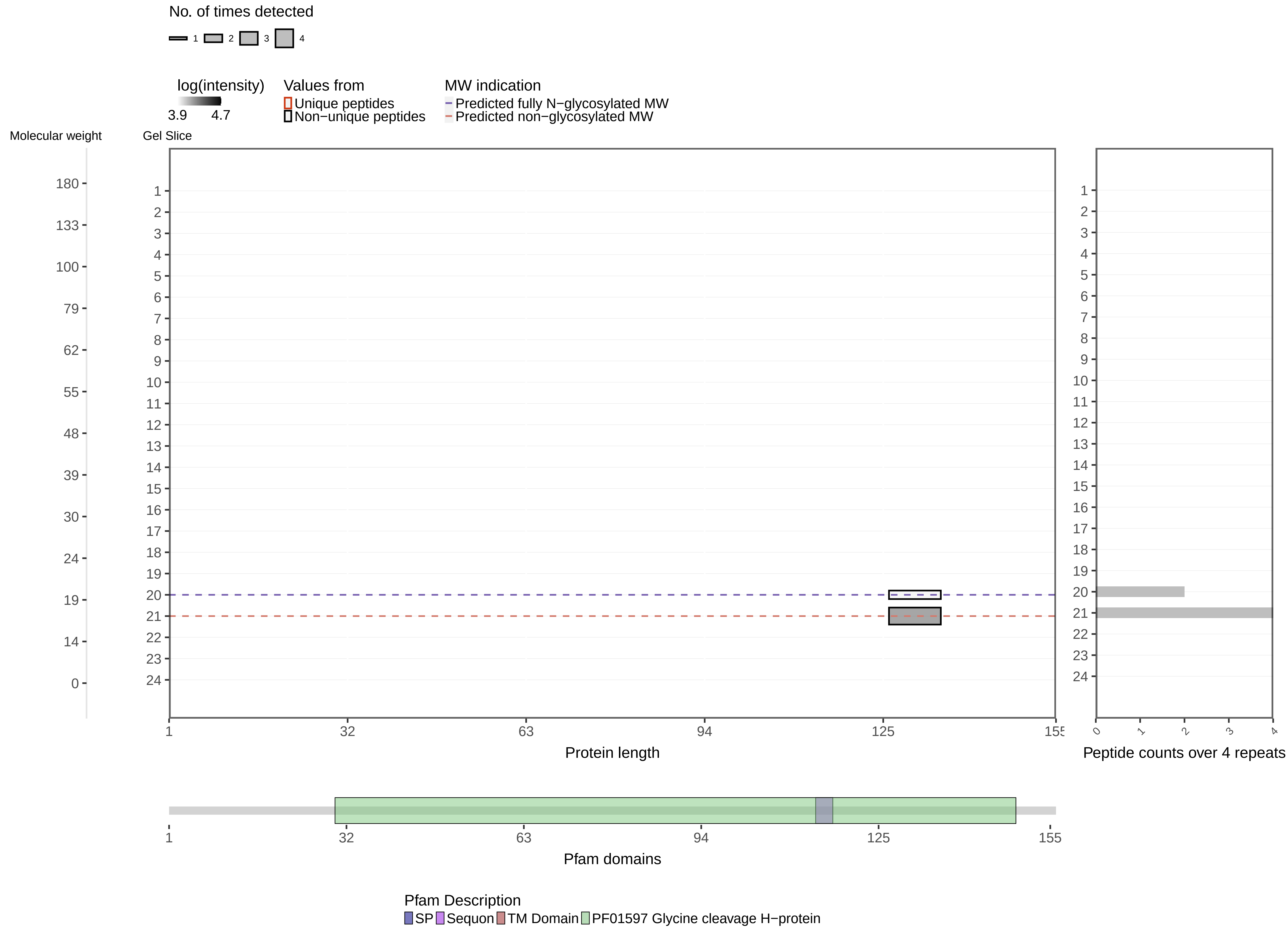


Protein sequence:

ELPEVGASVNQFDSFGAVESVKATSDINSPVSGKVVEVNEKLDSSPALINGSPYQEGWIIKVEMNNPDELKSLMDPDKYSSFCDEEDAKH*

### NbL12g09290.1 14_kDa_zinc−binding_protein [MW = 19.67 kDa]

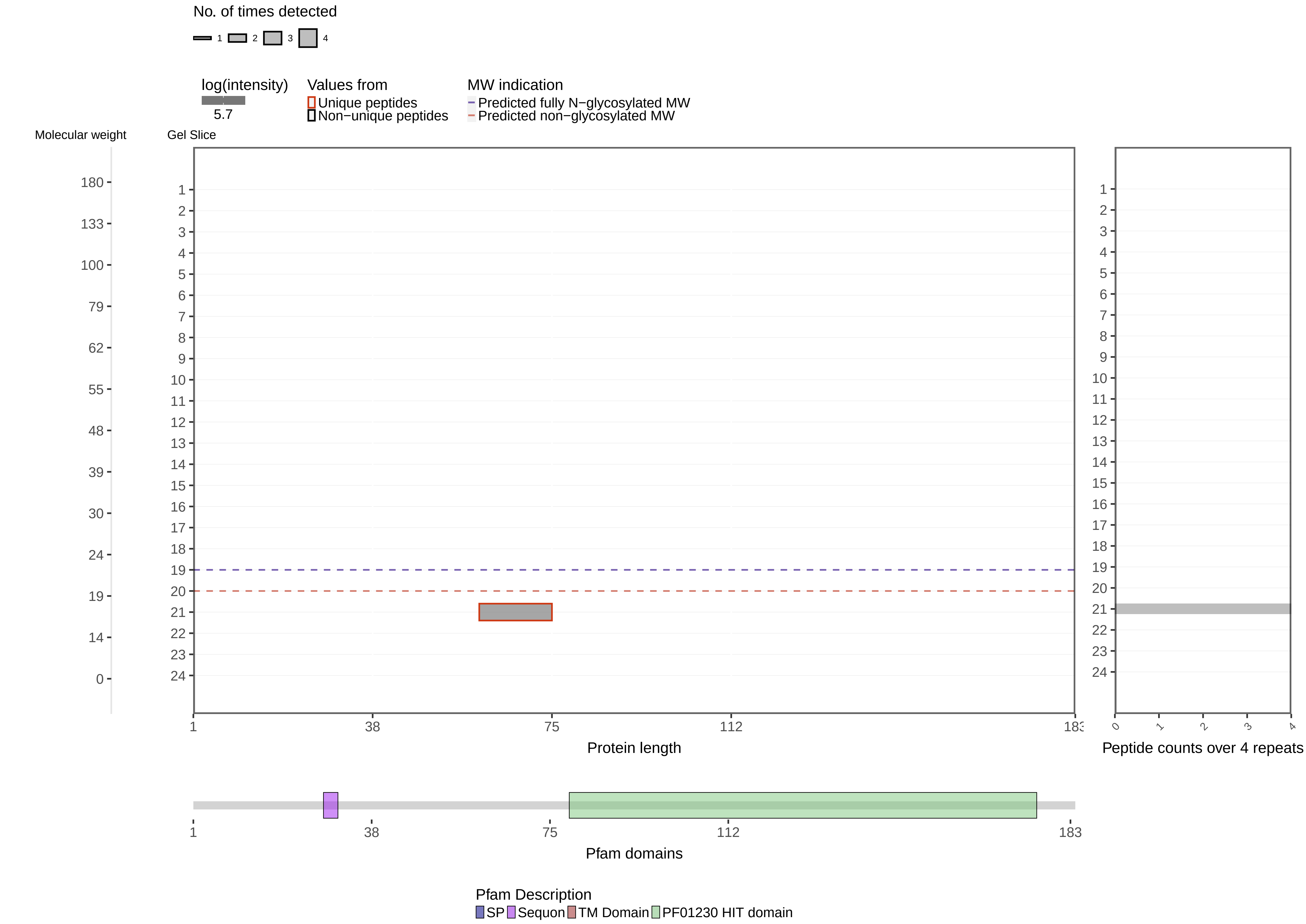


Protein sequence:

KEIPASVVYEDEKVLGFRDINPQAPVHVLVIPKLRDGLTELGKAEQRHEDILGNLLYAAKIVAEKEGIVDGFRVVINSGSSACQSVYHLHLHVLGGRQLKWPPG*

### NbL12g09210.1 pyrroline−5−carboxylate_reductase [MW = 28.97 kDa]

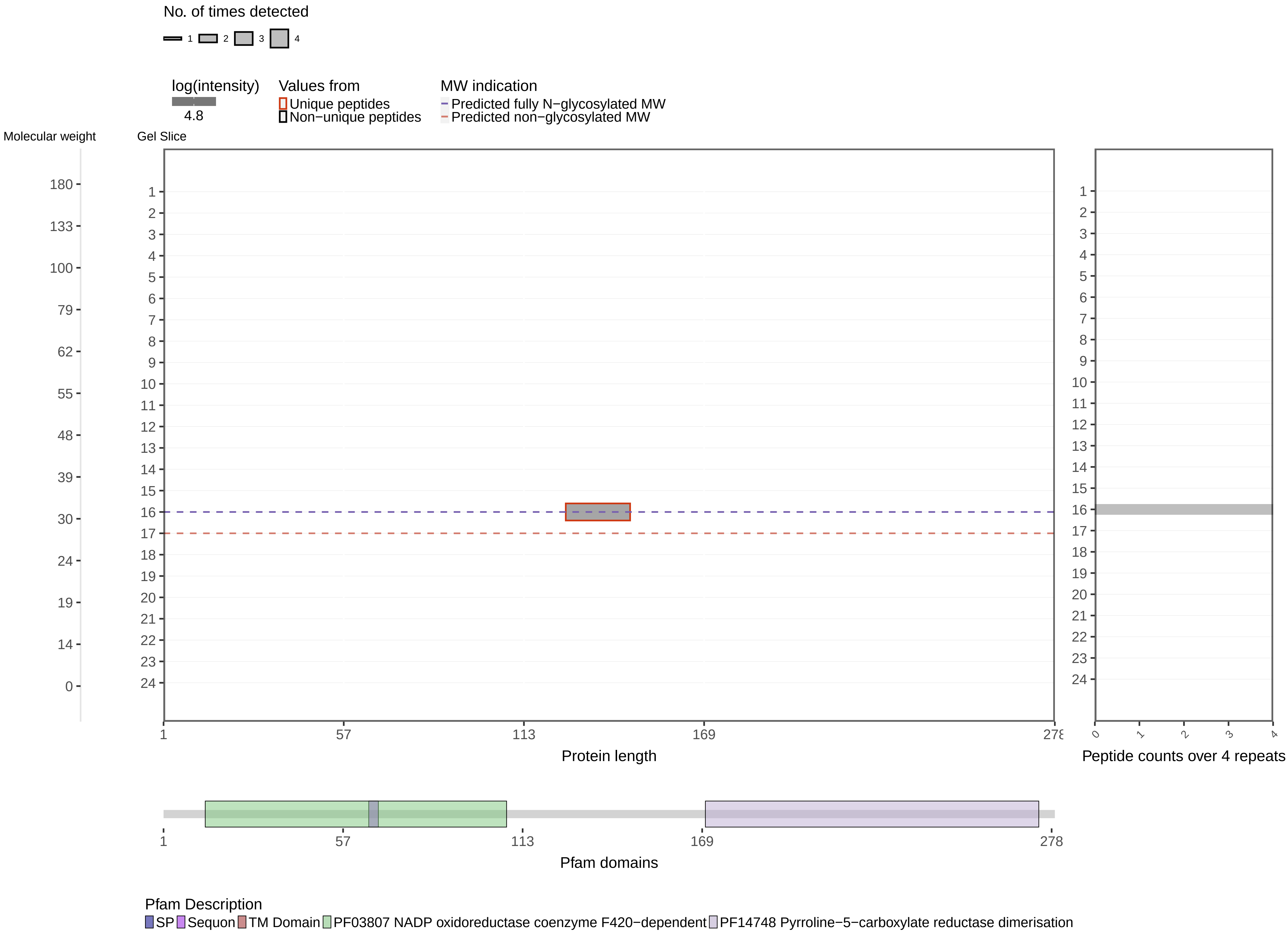


Protein sequence:

MPNTPSAVGEAATVFTLGEKATTEDGELISQLFGAIGKVWRADEKLFDAVTGLSGSGPAYIFLAIEALADGGVAAGLPRELALGLASQTVLGAASMVKGMGKHPGQLKDDVASPGGTTIAGIHELEKAGFRGILMNAVVAAAKRSRELSQN*

### NbL12g09040.1 V−type_proton_ATPase_subunit_F−like [MW = 14.5 kDa]

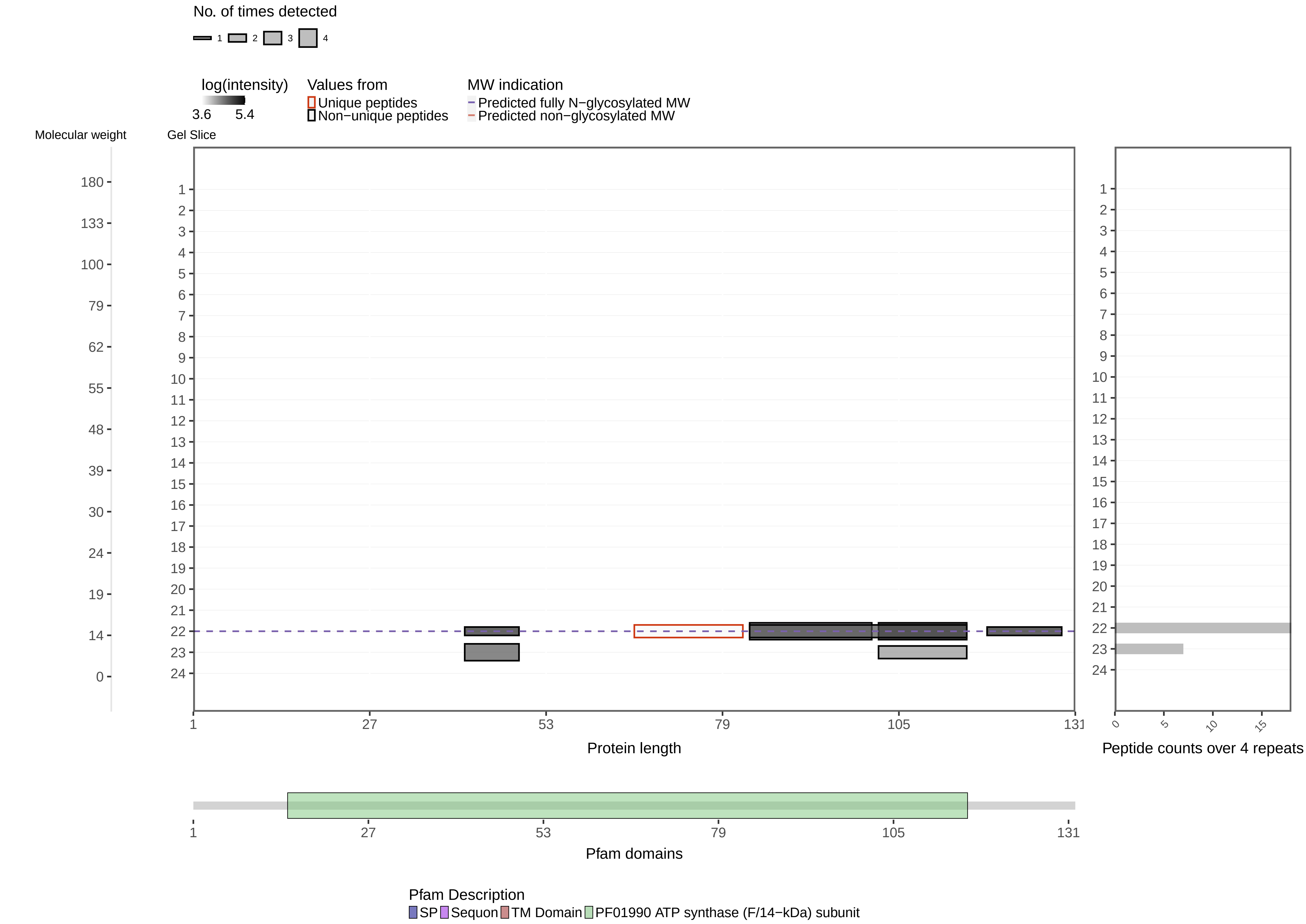


Protein sequence:

VKQIEDAFKEFTTREDVAIVLISQYIANMIRFLVDSYNKPIPAILEIPSKDHPYDPAHDSVLSRVKYLFSTESVASGRR*

### NbL12g08980.1 allantoate_deiminase [MW = 53.89 kDa]

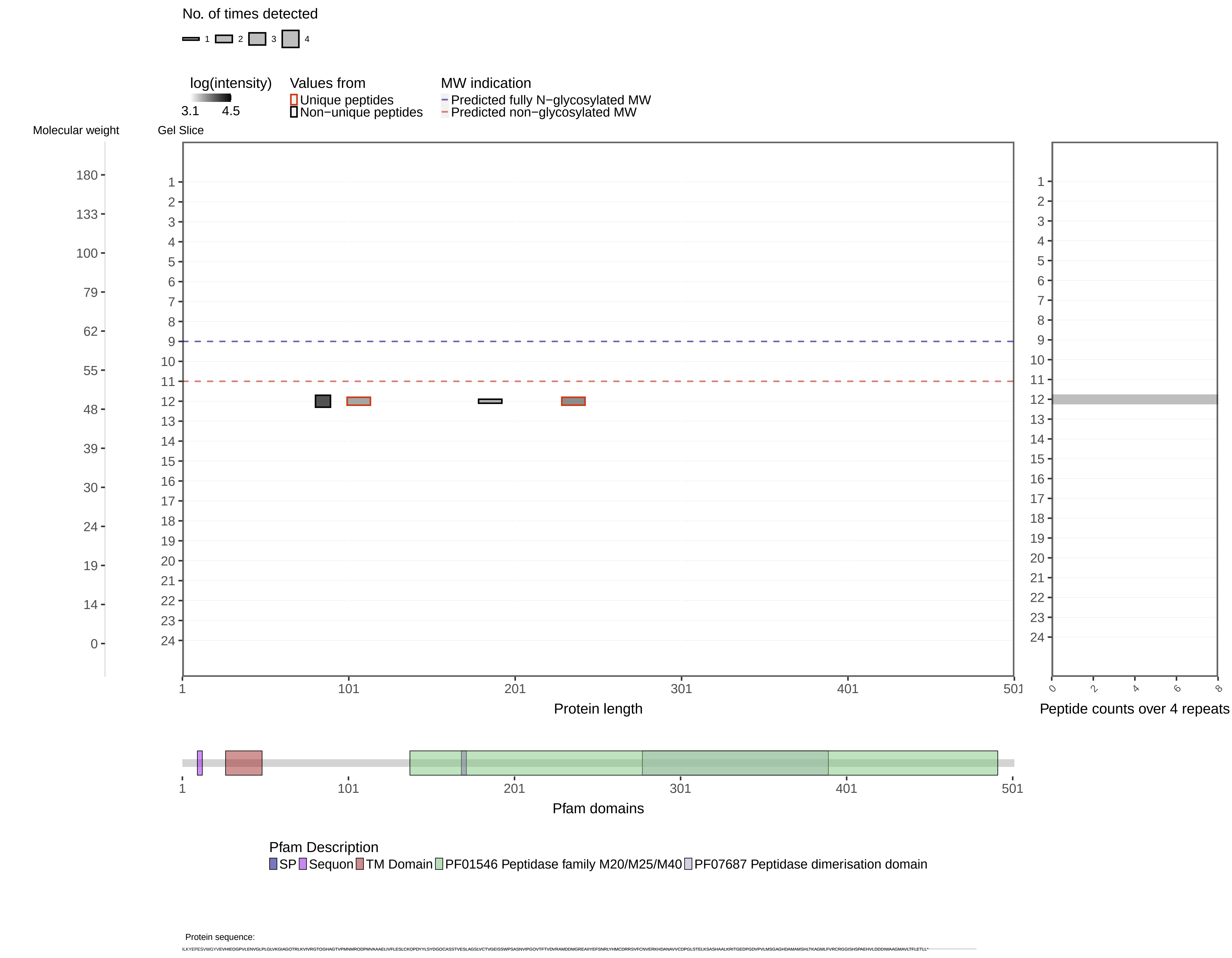

NbL12g08840.1 protein_transport_protein_Sec24−like_At3g07100 [MW = 115.04 kDa]

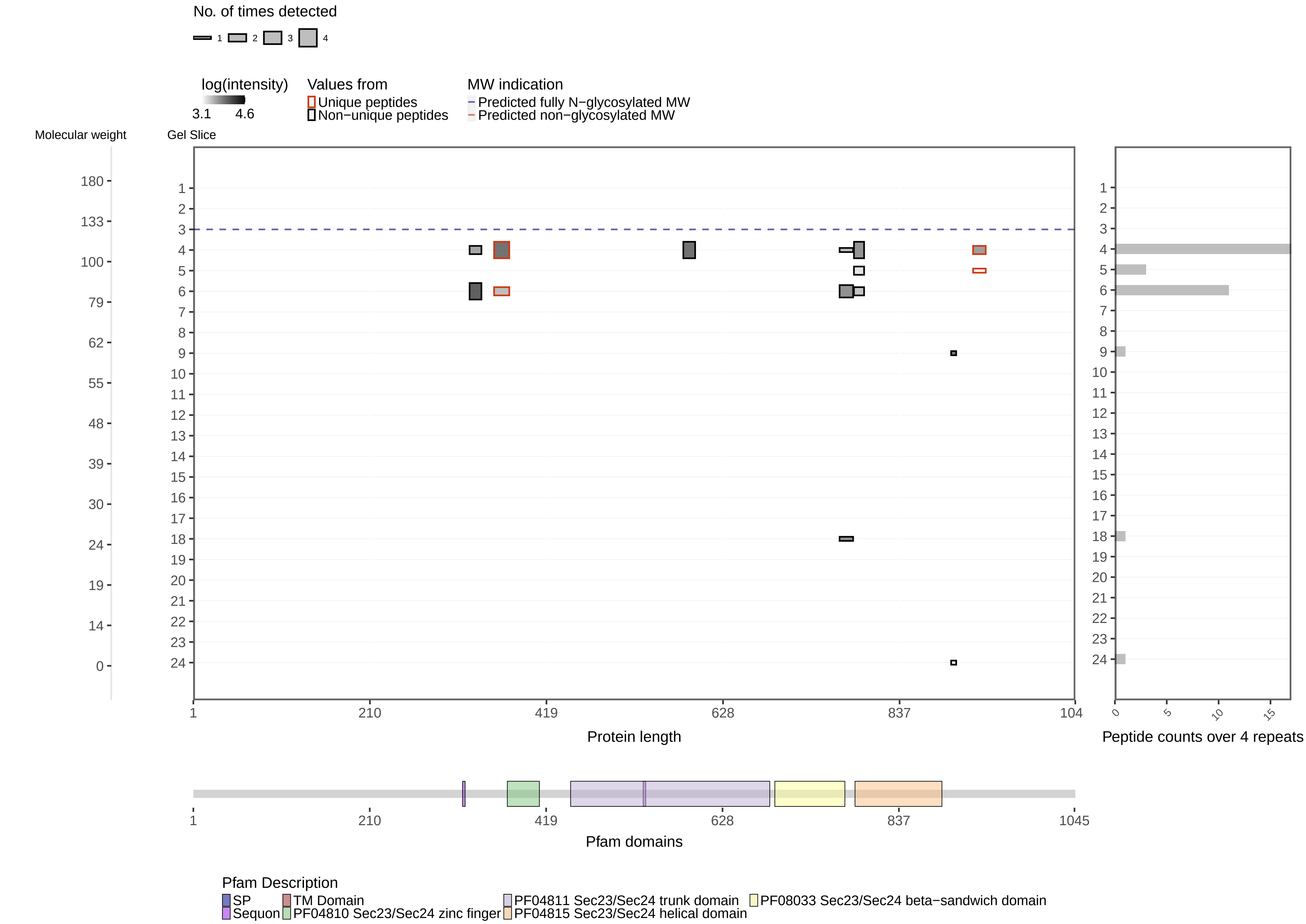


Protein sequence:

NMKSSFTQPQMMVMSDLEDVFVPLPDDLLVNLSESRTVVDAFLDSLPSMFQDNVNVESAFGPALKAAFMVMSQLGGKLLIFQSSLPSLGVGRLRLRGDDLRIYGTDKEHTLRVPEDPFYKQMAADFTKYQIAINIYAFSEKYTDLATLGTLAKYTGGQVYYYPSFQASIHKDRFYHELTRDLTRETAWESVMRIRCGKGVRFTTYHGNFMLRSTDLIALPAVDCDKAYAMQLSLEETLLTFQTVFFQIALLYTSSSGERRIRVHTAAAPVVSDLGEMYRLADTGAIISLFTRLAIEI

NbL12g08820.1 thioredoxin_reductase_NTRB−like [MW = 39.13 kDa]

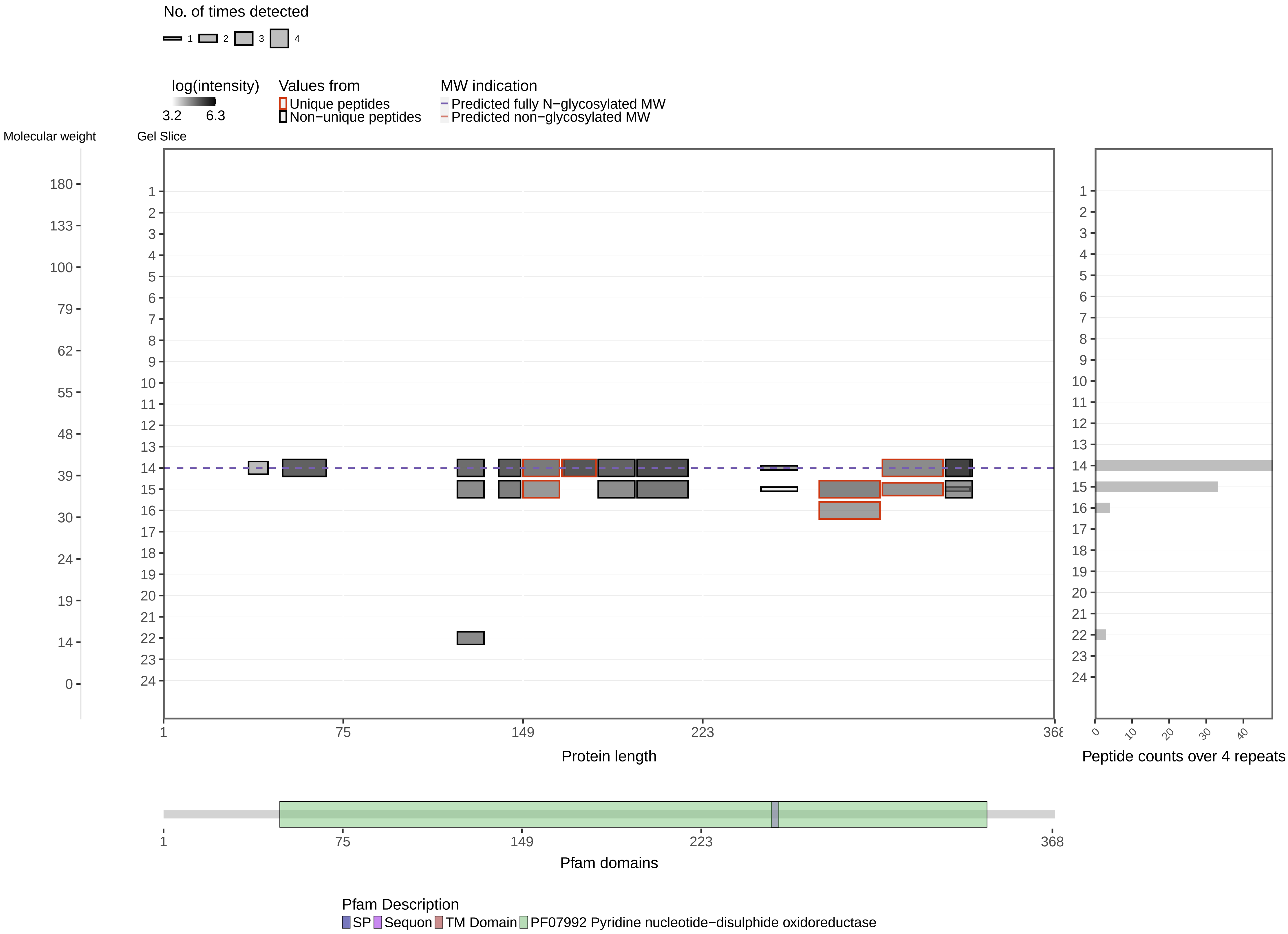


Protein sequence:

NDGFWNRGISACAVCDGAAPIFRNKPLAVIGGGDSAMEEANFLTKYGSKVYIIHRRDEFRASKIMQSRALSNPKIEVIWNSSVVEAYGEKLLGGLKVKNVVTGEVSDLQVSGLFFAIGHEPATKFLDGQLELDSDGYVVTKPGTTLTSVKGVFAAGDVQDKKYRQAITAAGCRCMAALDAEHYLQEIGAQEGKSD*

### NbL12g08770.1 MORC_family_CW−type_zinc_finger_protein_2A−like [MW = 83.88 kDa]

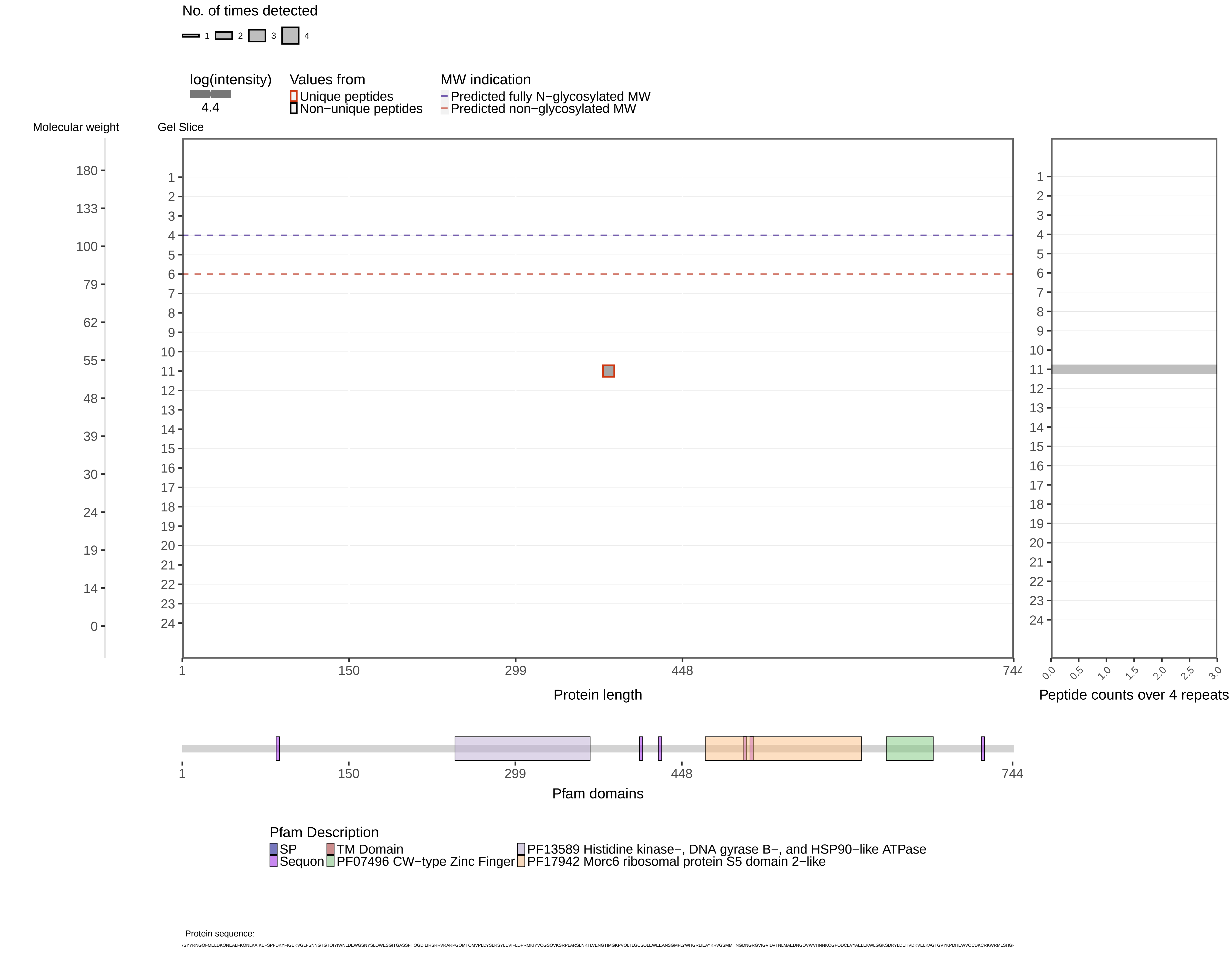

NbL12g08650.1 exosome_complex_exonuclease_RRP44_homolog_A [MW = 104.73 kDa]

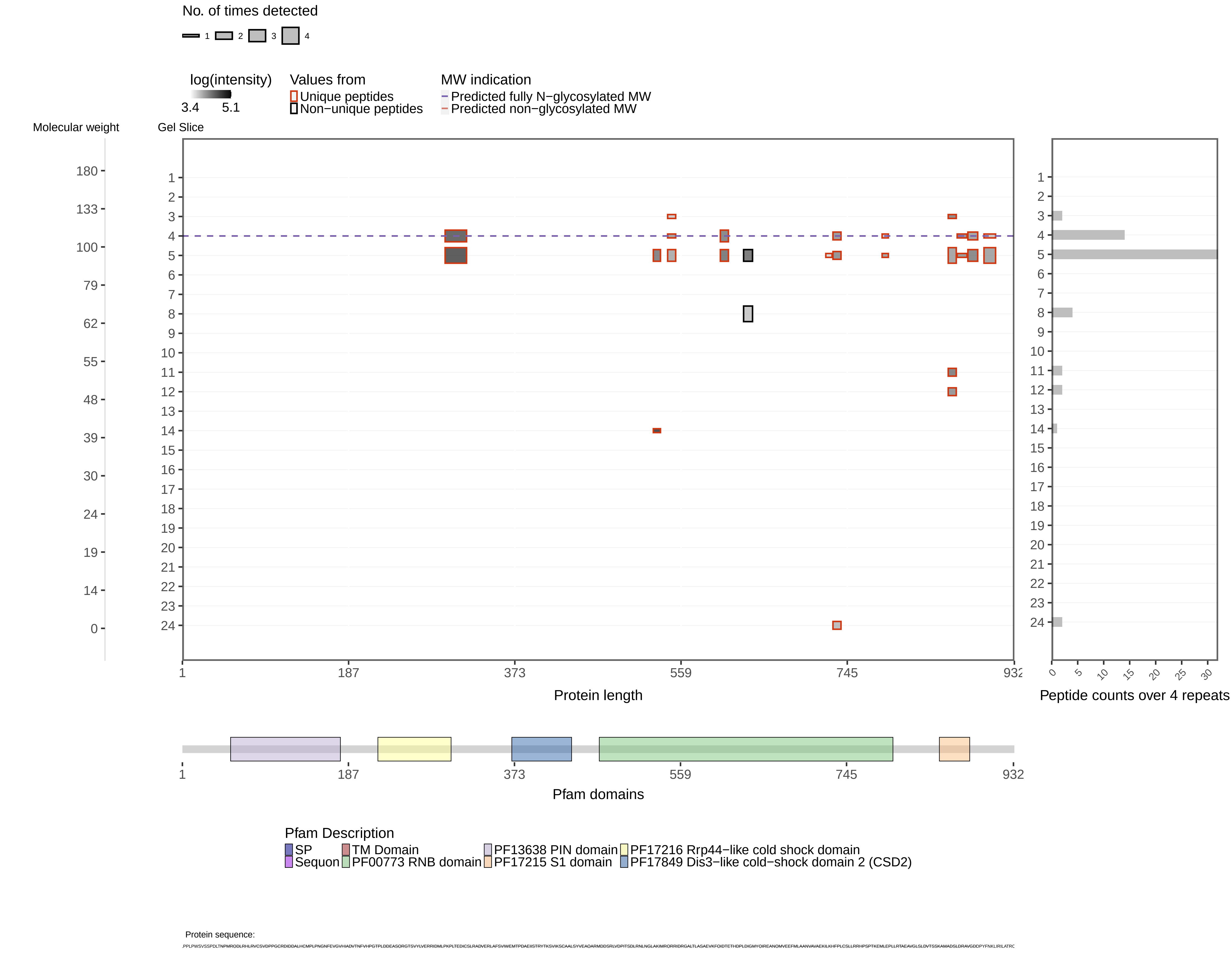

NbL12g08330.1 isocitrate_dehydrogenase_[NAD]_regulatory_subunit_1_mitochondrial−like [MW = 38.87 kDa]

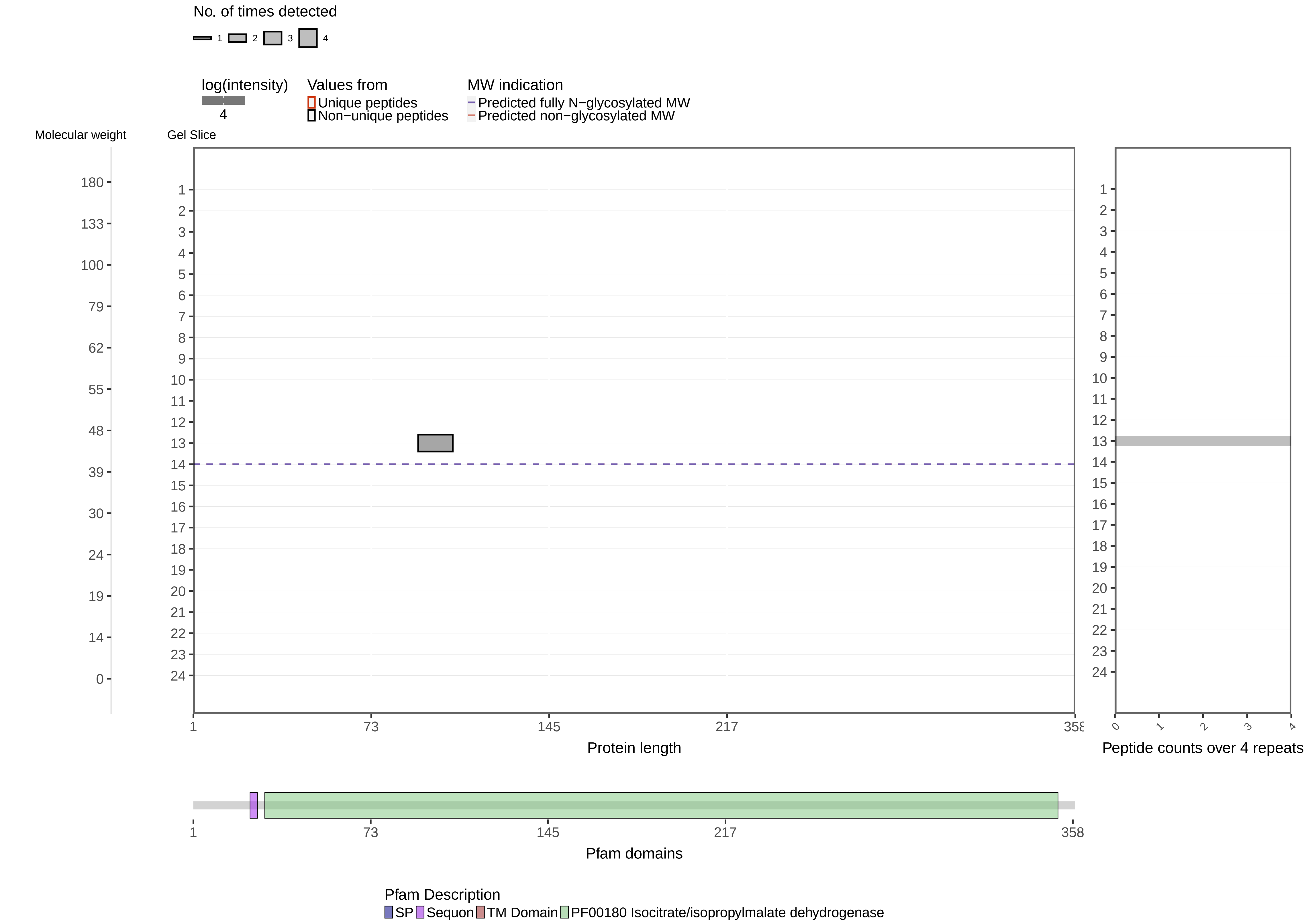


Protein sequence:

CSERIAKFAFEYADVNKRKKVTAVHKANIMKLADGLFLESCREVAAKYPEIQYEEIIVDNCCMQLVSRPEQFDVMVTPNLYGNLIANTAAGLAGGTGVMPGGNVGTDQAVFEQGASAGNVGNEKILKLKKANPVALLLSSAMMLRHLEFPSFADRLETAVKRVIAEGKVRTKDLGGDCTTQEVVDAIIALD*

NbL12g08300.1 phosphoserine_aminotransferase_2_chloroplastic−like [MW = 47.45 kDa]

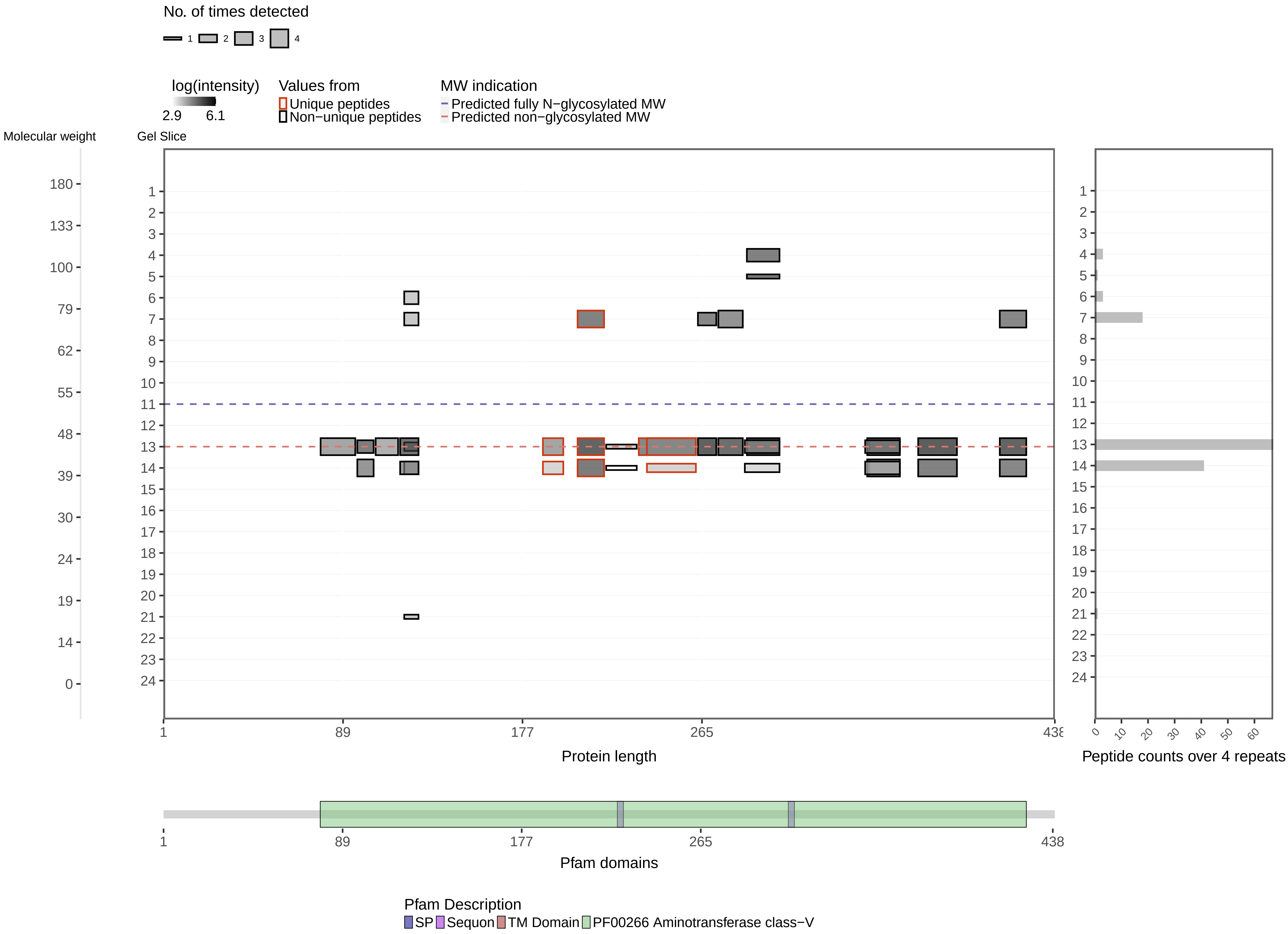


Protein sequence:

AFDSLQQTPGAKYLHICANETIHGVEFKDYPKPSNANTILVADMSSNFCSKPVDVSKFGVIYAGAQKNVGPSGVTIVIIRKDLIGNAQESTPVMLDYKIHAENNSLYNTPPCYGIYMCGLVFEDLLAQGGLVEVEKKNKKKAEILYNAIDSSNGFFRCPVEKSVRSLMNVPFTLAKPELEAEFVKAAAAEKMVQLKGHRSVGGMRASIYNAMPLAGVEKLVAFMKDFQAKHA*

NbL12g08260.1 isozyme_1_(LOC107799583)_mRNA_HF564631.1_Nicotiana_tabacum_mRNA_for_catalase_1 [MW = 57.89 kDa]

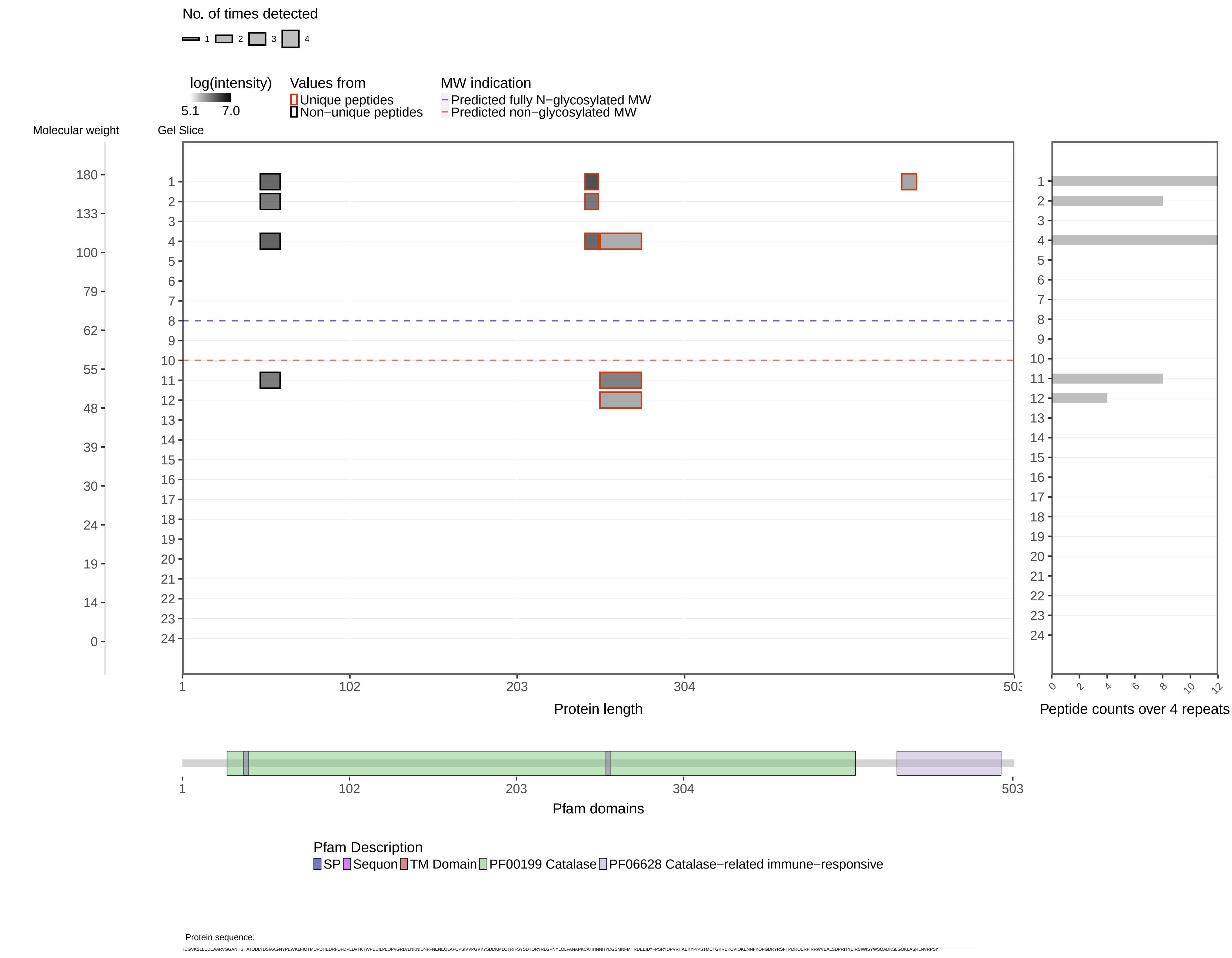

### NbL12g08140.1 thiol−disulfide_oxidoreductase_LTO1 [MW = 40.07 kDa]

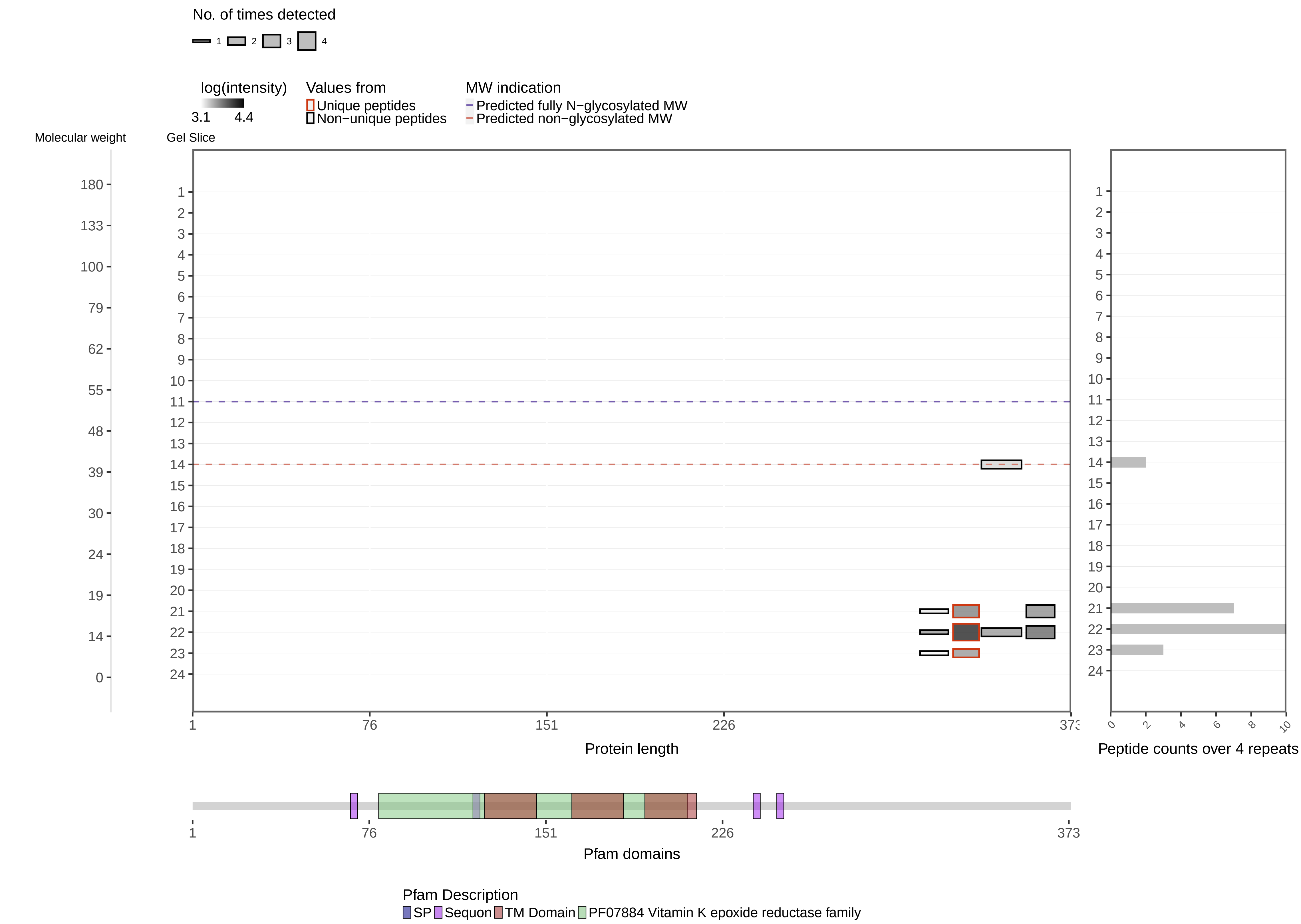


Protein sequence:

MAAASAYFLYILSTQFSGEFCLYCLASALLSFGLFISSIMGLGSQEVQKEVGLQLLIVLLVVSTLNNSYSASQPTNTSSAMTELEYFTTEITSPSTPFAISLAKHLRSIGAKMYGAFWCSHCQEQKEMFGREASKLLDYVECFPDGVKKGIYMANACQEAKLEGFPTWVINGEVLSGEKKLSELAELSGFTMKELTEAK*

NbL12g08050.1 DNA−directed_RNA_polymerase_II_subunit_1 [MW = 205.42 kDa]

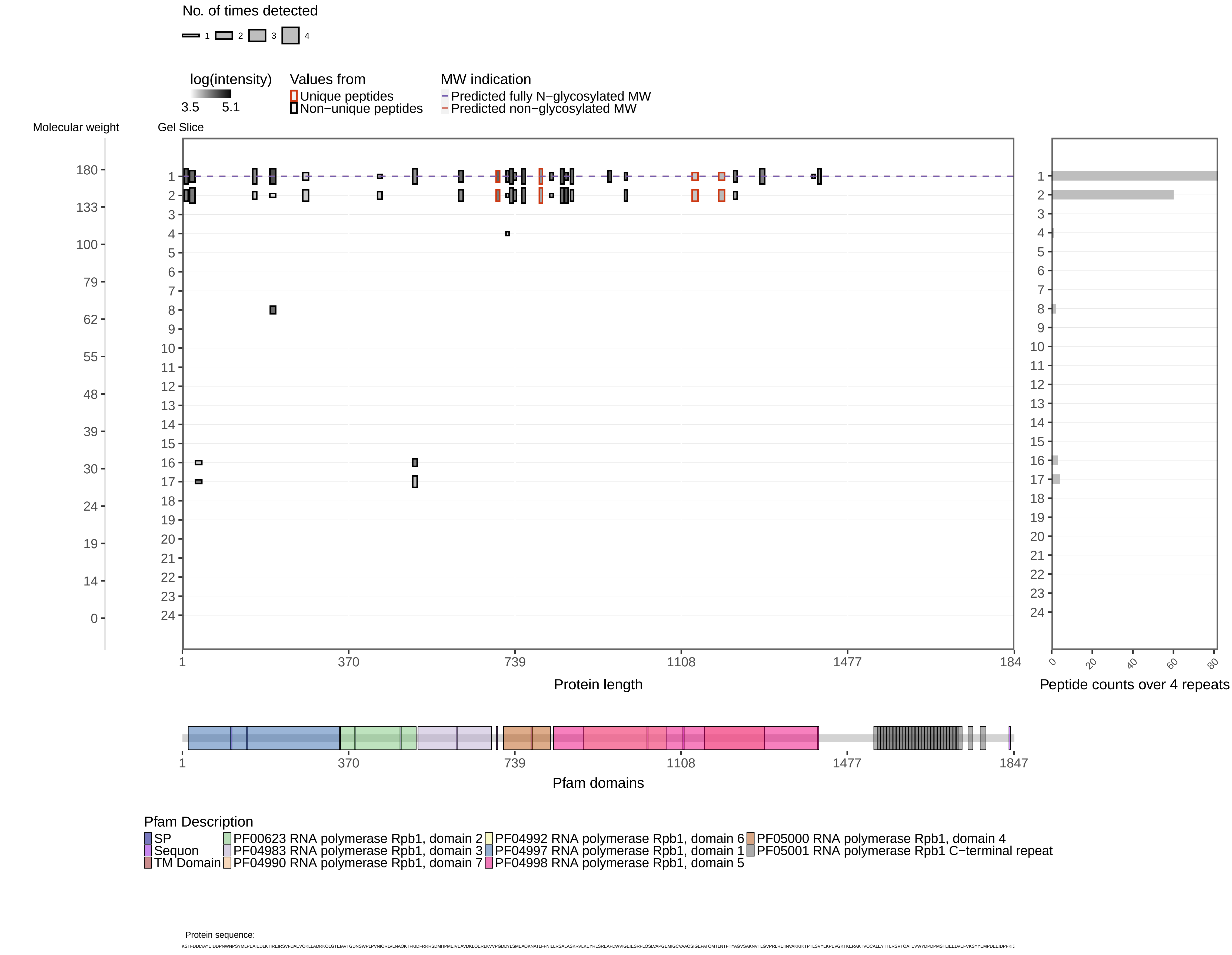

NbL12g07870.1 uncharacterized_LOC107826763 [MW = 23.25 kDa]

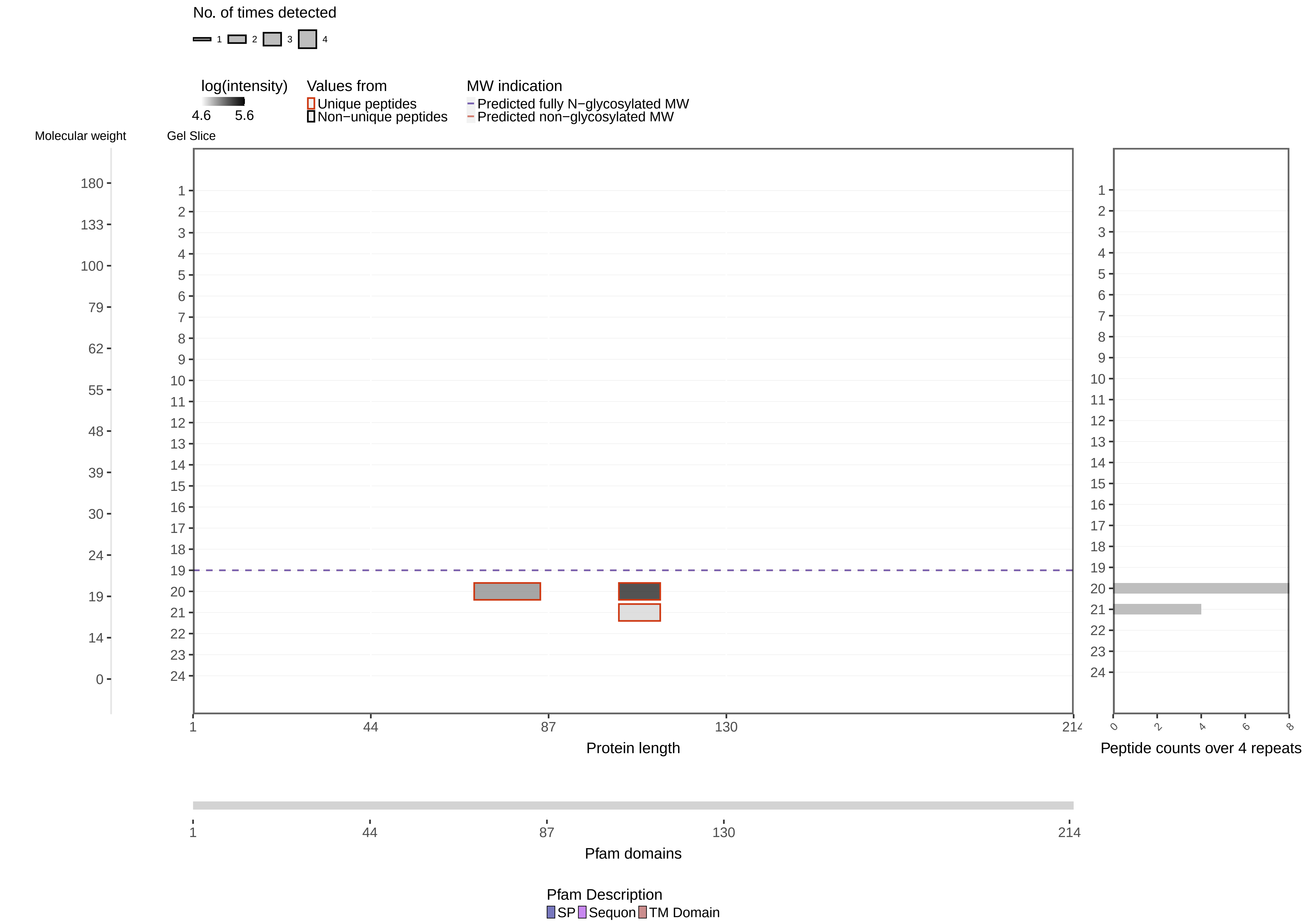


Protein sequence:

ORNPYLSEGSRQAAATAAALAKKYGADITVVVIDEKEKEAVPEHETQLASIRWHLSEGGYQEFKLLERLGEGSKPTAIIGEIADEMNLDVVIMSMEAIHSKHVDANLLAEFIPCPVLLLPL*

### NbL12g07820.1 26S_proteasome_non−ATPase_regulatory_subunit_4_homolog [MW = 42.25 kDa]

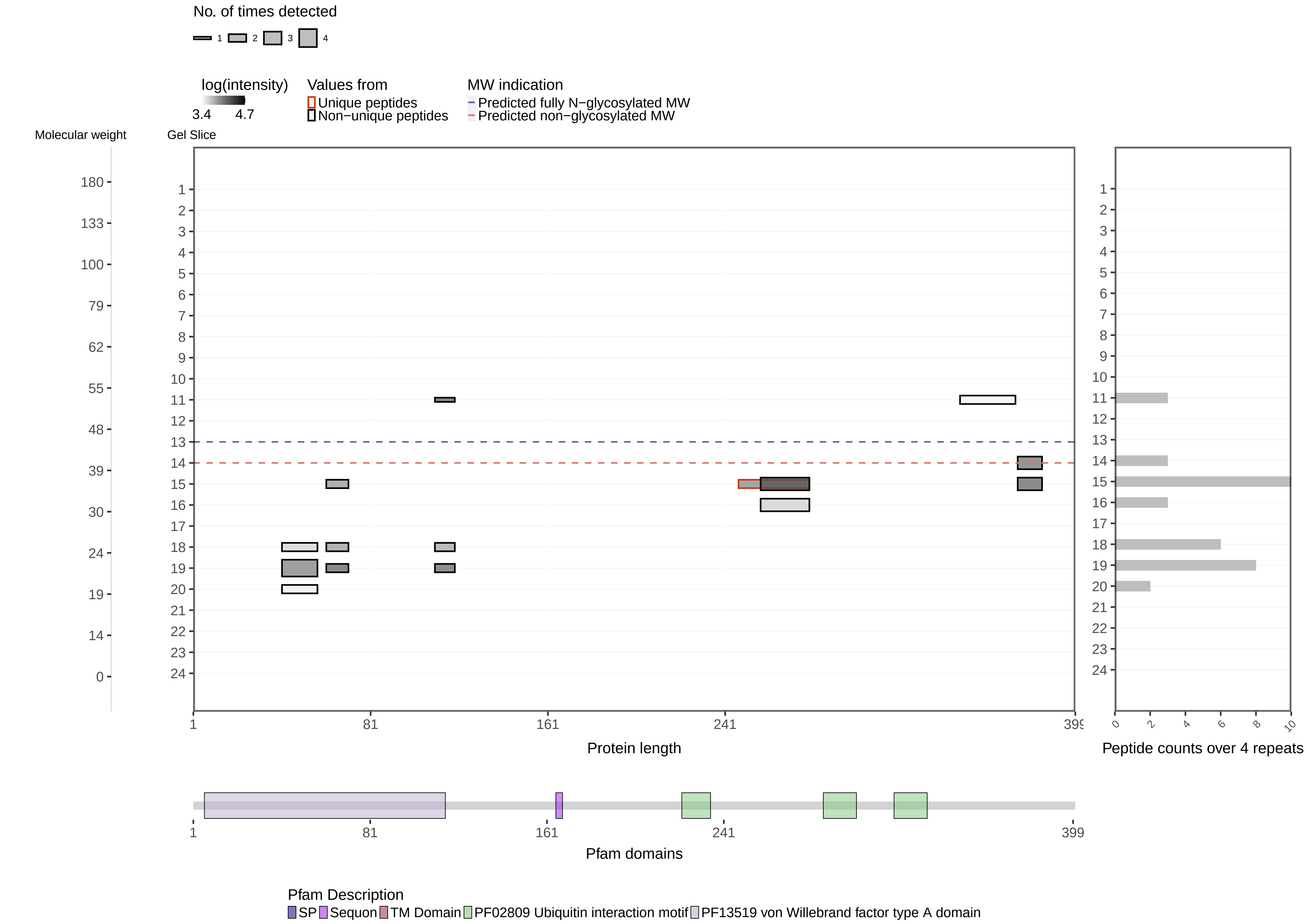


Protein sequence:

TGDGEGGSGFAAAAAAAGGVSGFDFGVDPNLDPELALALRVSMEEERARQEAAAKKAADETATQEKGETSQDVTMAENVNAGTPEPENKATDLMEDENALLQQALAMSMDDASSNVATRDTDMSEAASEDQDLALALQLSVQDSTNDQSSQTDMSKLLADQSFVSSILASLPGVDPNDPSVKDLLASMQGQSEKKDEDTDKEQKEEKK*

### NbL12g07810.1 alpha−mannosidase [MW = 34.18 kDa]

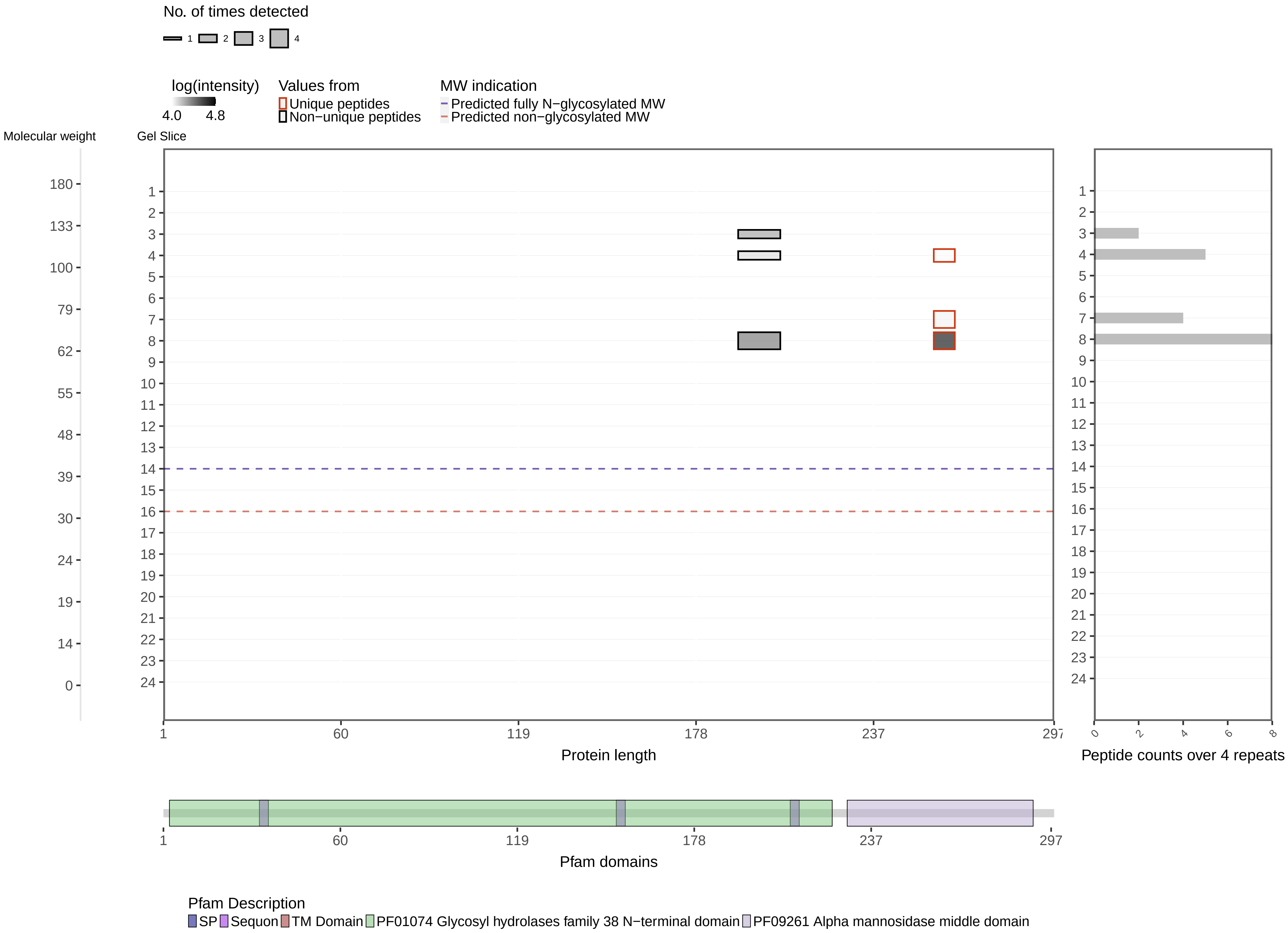


Protein sequence:

'DIRVNDFINSAITQANVTRTNHIMWTMGDDFQYQYAESWFKEMDKLIHYVNKDGRVNALYSTPSIYVDAKHATNQSWPLKTDDYFPYADGANSYWTGFFTSRPAFKRYVRMLSGYYMAARQLEFLTGRKSEGFNTFSLGDALGVTQHHDAVTGTQNSIQLM

### NbL12g07800.1 alpha−mannosidase [MW = 46.45 kDa]

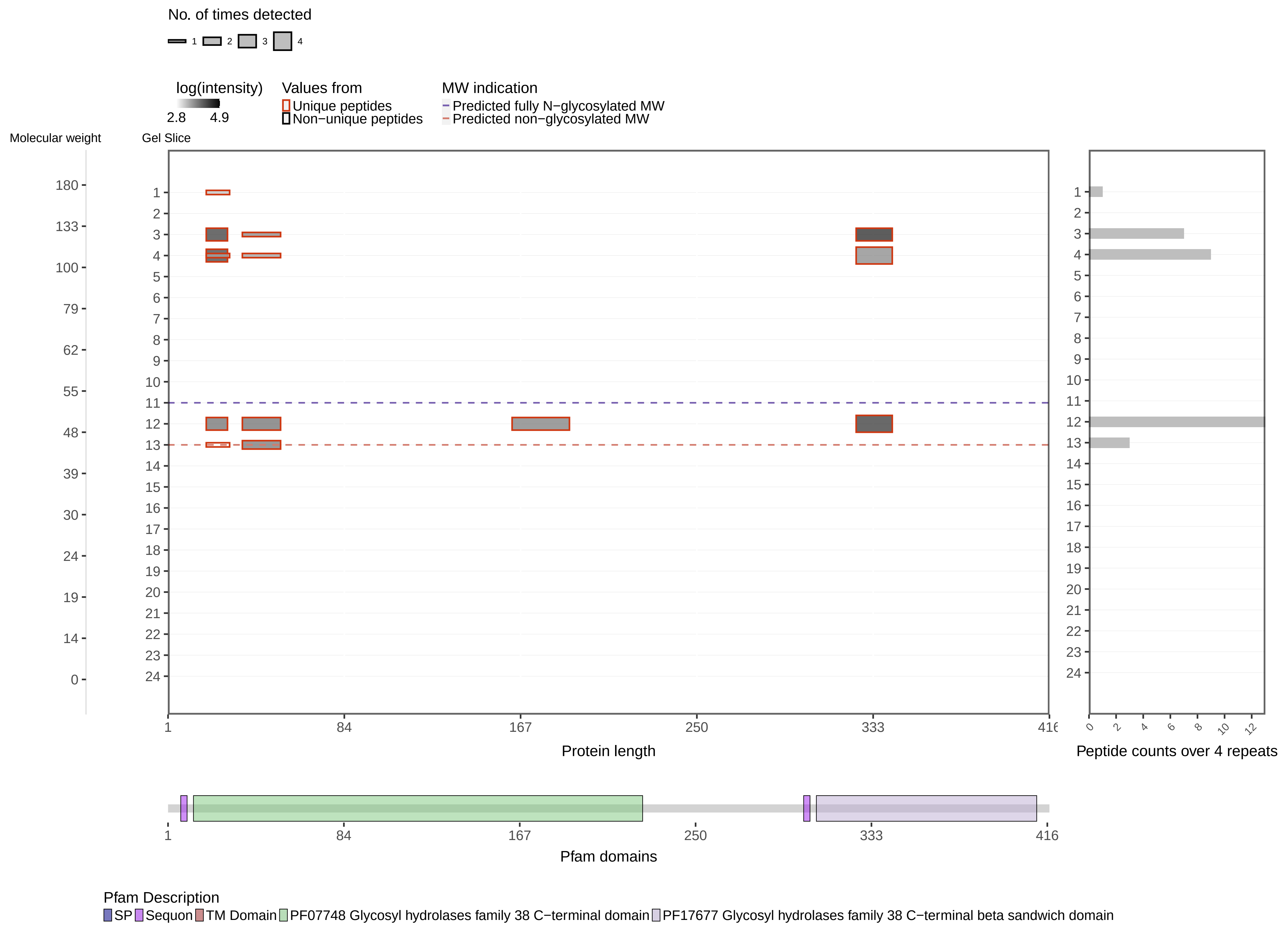


Protein sequence:

/LVDRATGGASINDGEIELMLHRRLLYDDARGVGEALDETVCVGTACEGLRVRGNYYLGIHKNSDGSRWRRTTGQEIYSPLVLAFGHENQEEWRASHLTKATIMNPNYSLPPNVALITLQELDNGGVLLRLAHLYEAGEDTEYSTITKVQLKDMFAGKTIKEIKETSLSANQGKNEMKKMSWNIEHDTGSEPAPIRGVPVDISSLVVELGPMEIRTFVVEF*

NbL12g07720.1 3−dehydroquinate_synthase_chloroplastic [MW = 47.87 kDa]

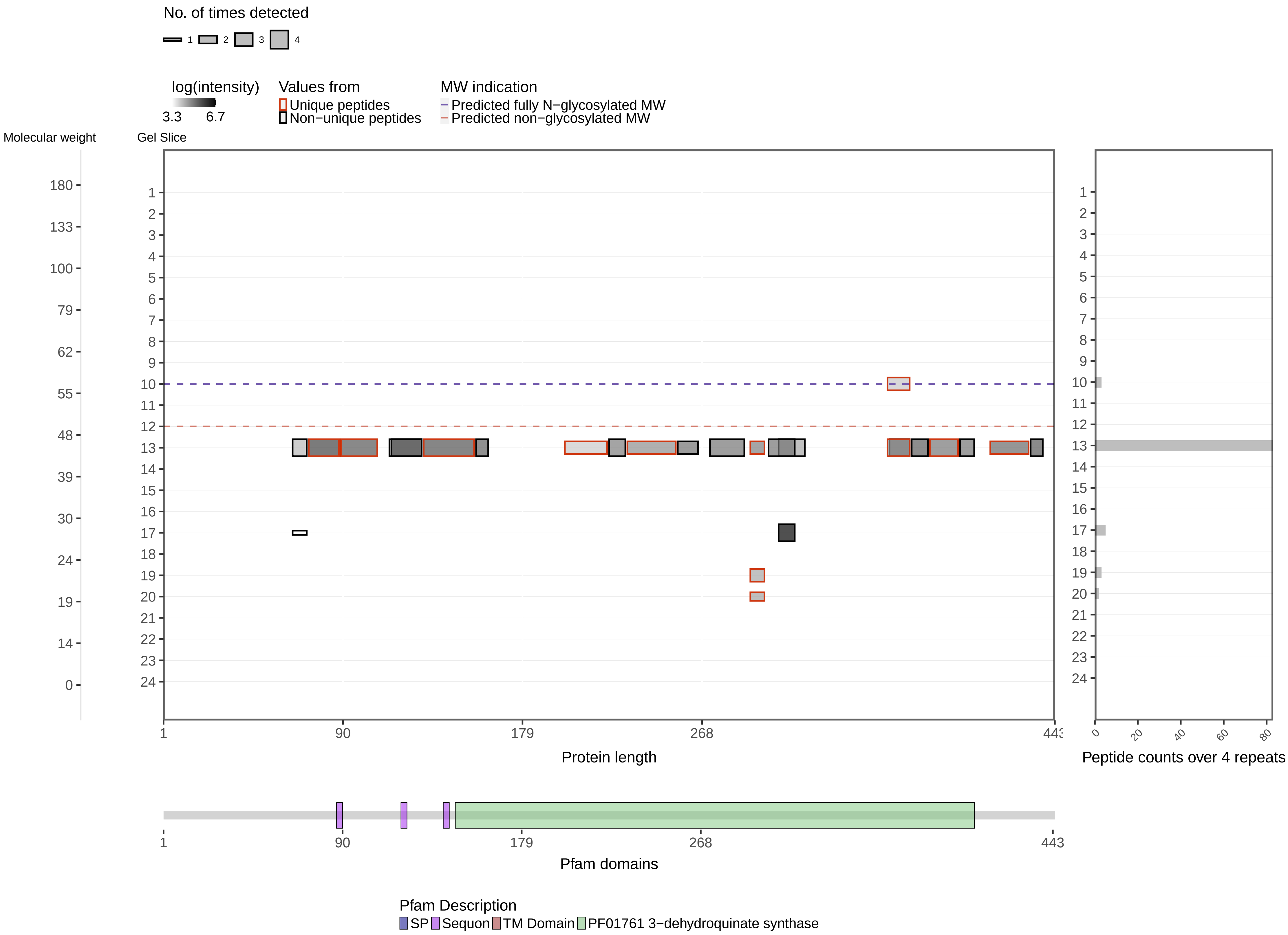


Protein sequence:

MAQVDSSVGGKTGINHPLGKNMIGTFYQPQCVLVDTDTLDTLPDRELASGLAEVIKYGLVRDAEFFEWQEQNMPLLLARDRTAFAYAIKRSCENKAEVVSLDEKESGLRATLNLGHTFGHAVETGFGYGQWLHGEAVAAGTVMAVDMSRRLGWIDDSLVQRVHNILQQAKLPTAPPGTMTVEMFKSIMAVDKKVADGKLRLILLEGPLGNCVFTGDYDQKALDETLRAFSKS*

### NbL12g07650.1 extradiol_ring−cleavage_dioxygenase−like [MW = 30.35 kDa]

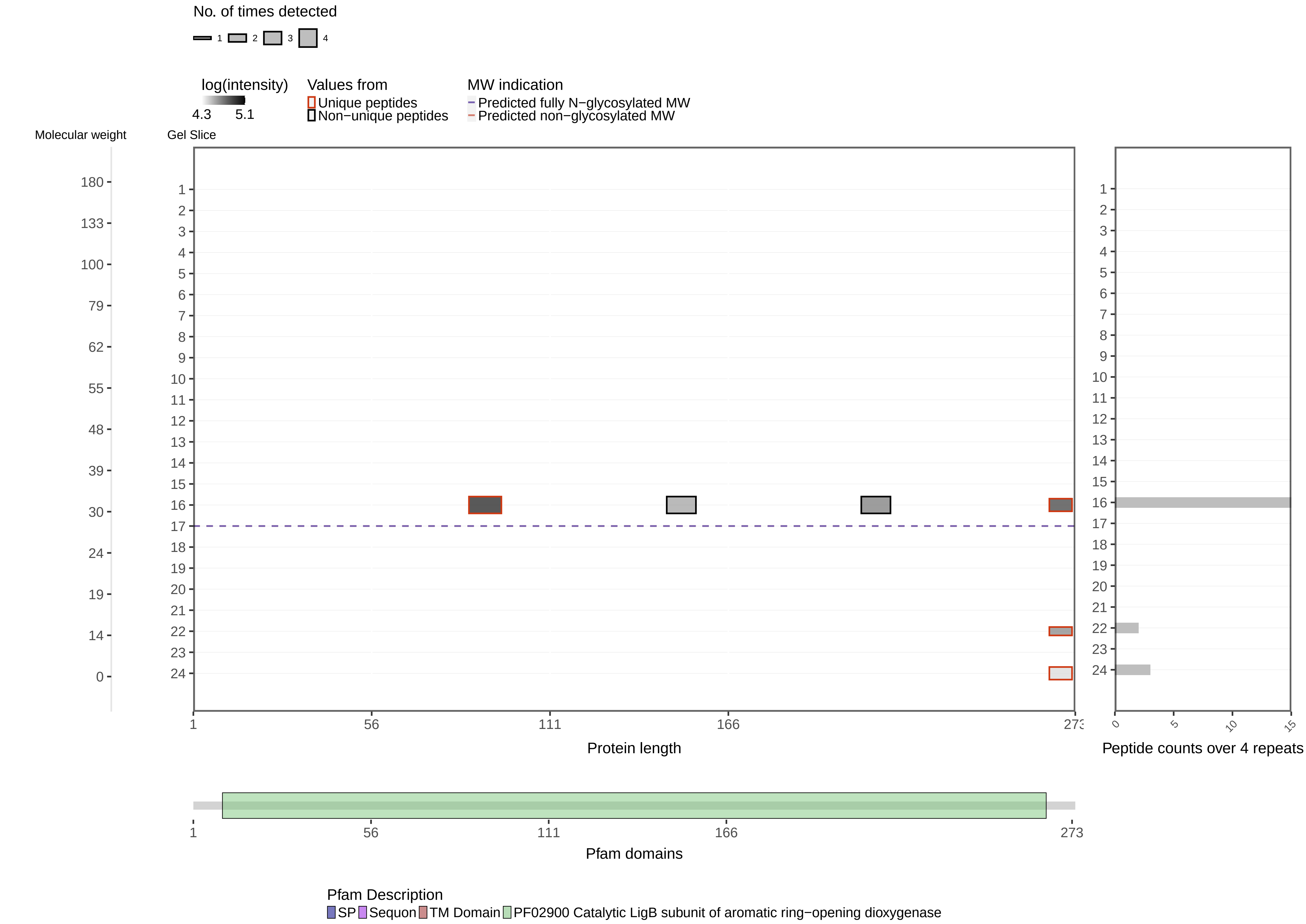


Protein sequence:

PLMLMYPEADIPVCQLSVQPNRDGTYHYNMGRALASLKDEGVLIIGSGSATHNLRALGSSKSVVSWALEFDNWLKDALLSGRHQDVNNYDMKAPHAKVAHPWPEHIYPLHVALGASGEGTNGELIHHSWDLGALSYASYKFSSLNRPS*

### NbL12g07490.1 ferredoxin−−NADP_reductase_leaf−type_isozyme_chloroplastic [MW = 40.32 kDa]

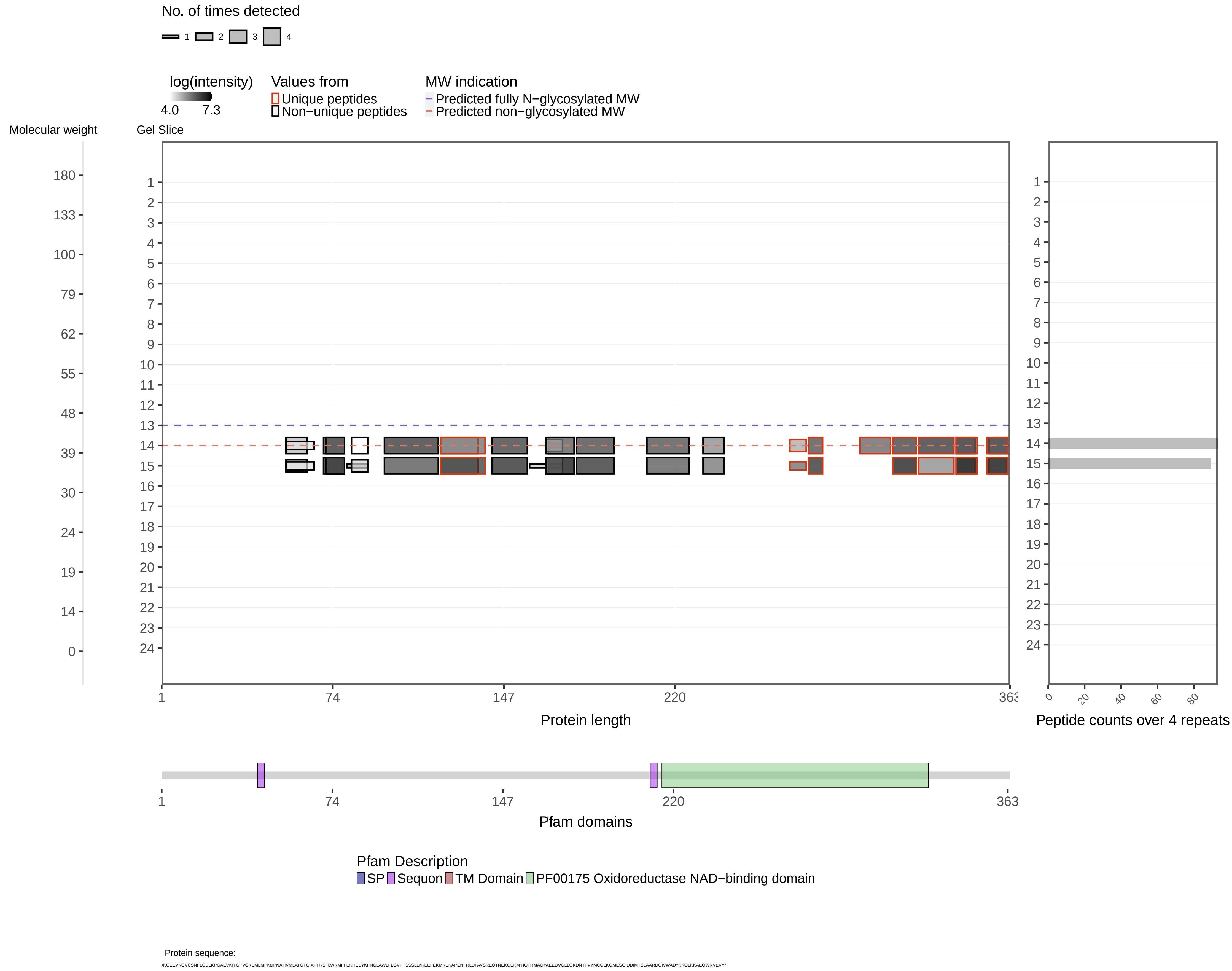

### NbL12g07300.1 DNA_repair_protein_UVH3−like [MW = 54.81 kDa]

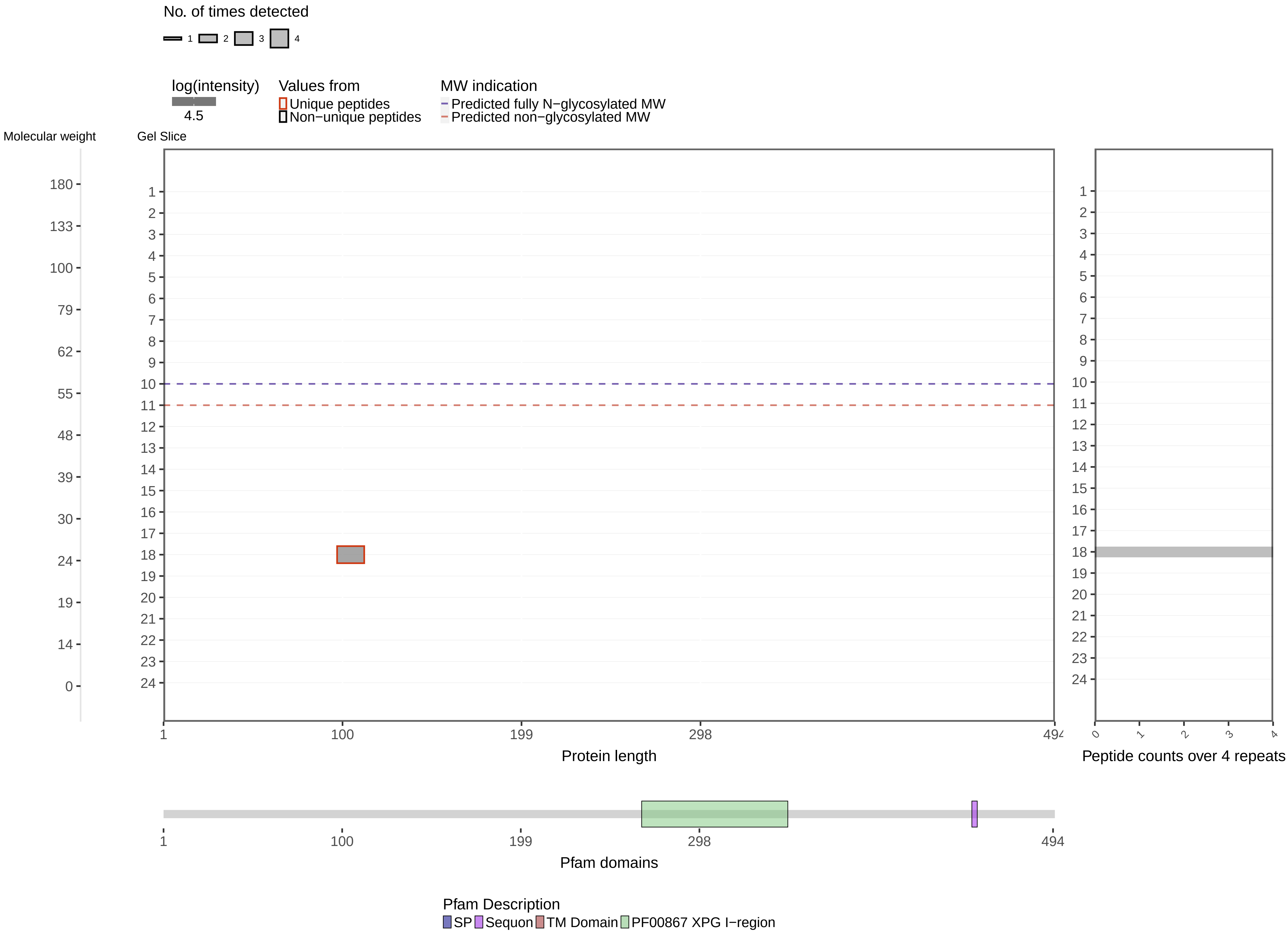


Protein sequence:

EREKLGDEQRKLERNAESVSSEMFAECQELLQMFGLPYIIAPMEAEAQCAYMELTNLVDGVVTDDSDAFLFGARNVYKNIFDDRKYMETYFMKDVENELGLDREKLIRMALFLRSYYTEGVSGIGIVNAIEVLNAFPEEDGLQKFREWVESPDPSILGGLDAQVGSSTRKRGSEVGDPDMSCSSSNVEGNSESDENVKSEDRVQKLKQIFMNKHRNISKNWHIPSSFPSDAVISAYASPRVDKSTEPFAWGKPDVSVLRK*

NbL12g07170.1 mitochondrial_outer_membrane_protein_porin_of_34_kDa [MW = 29.61 kDa]

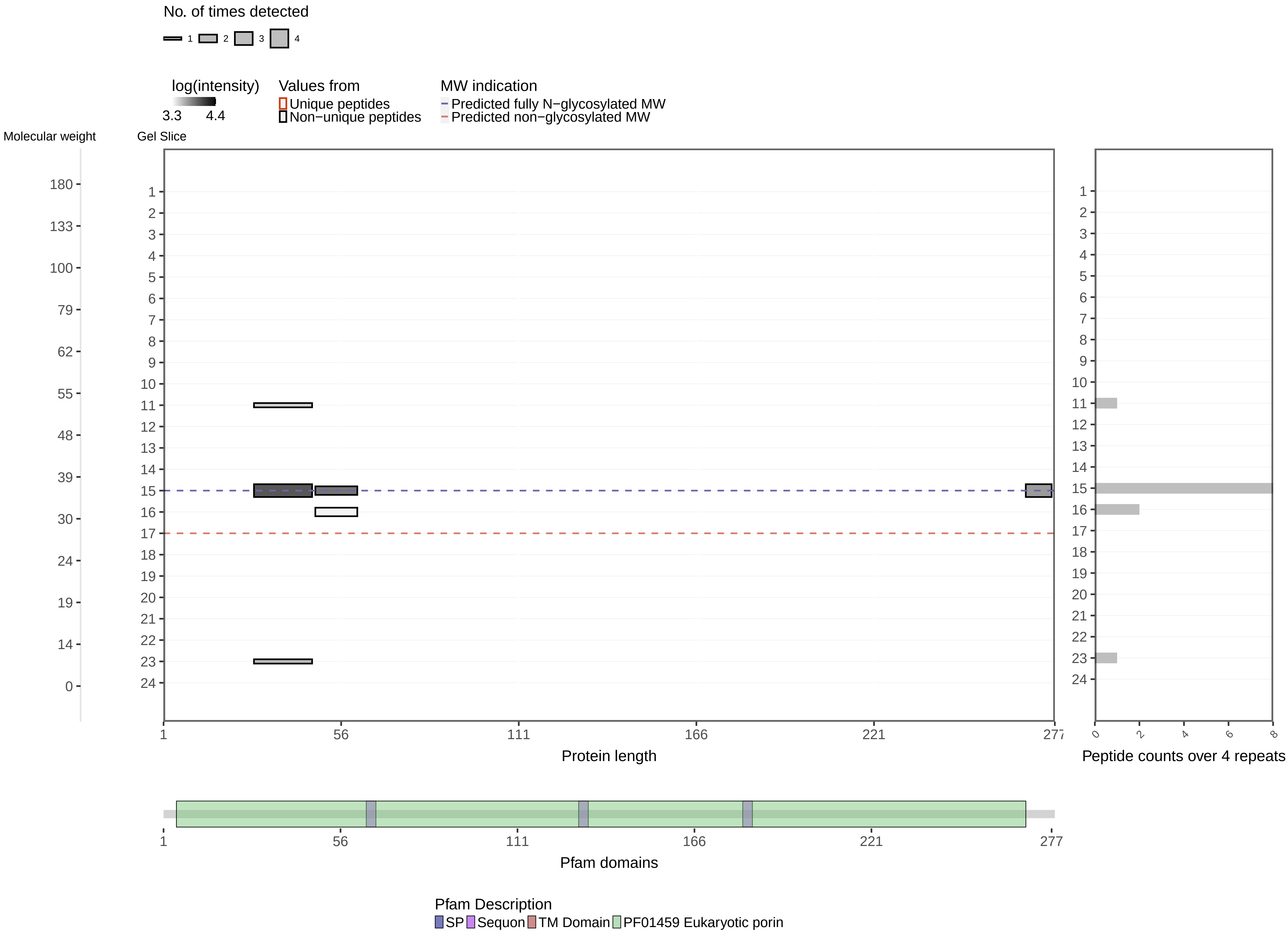


Protein sequence:

PIVNFSGVVGTNILALGTDVSFDTKTGDFTKCNAGLSFTNADLVASLNLNDKGDNLSASYYHTVSPFTSTAVGAEVTHCFSTNENTITVGTQHRLDPLTSVKARVNNFGKASALLQHEWRPKSLFTISGEVDTKSVDKGAKFGLALALKP*

### NbL12g07070.1 NAP1−related_protein_2−like [MW = 28.27 kDa]

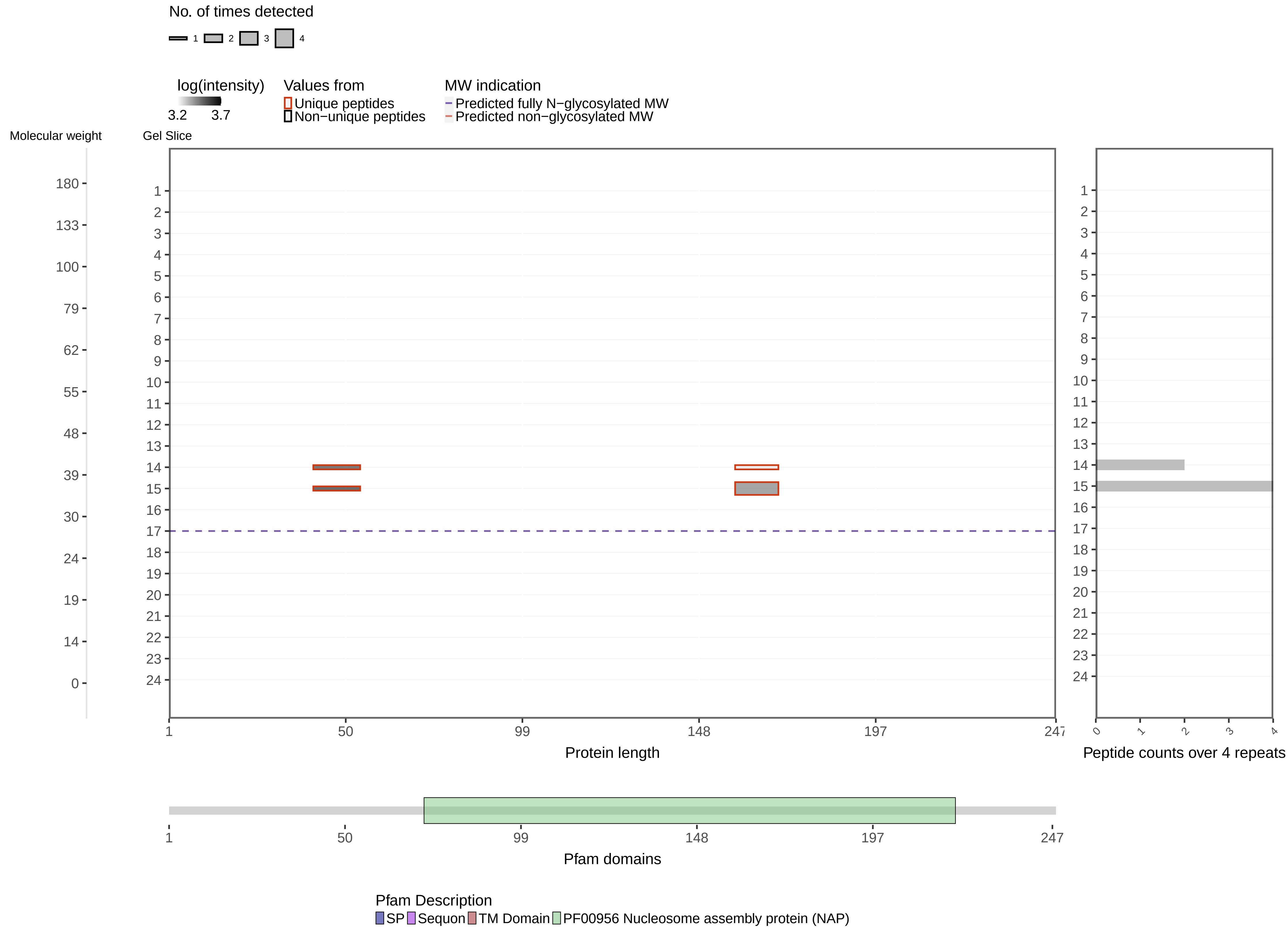


Protein sequence:

MTSGYSLTFKFKPNPYFEDTKLVKTYTFLDEGTTTVTGMPIKWKEGMDAAIGGTPEKKGNKRPPSVESFFSWFSETQLKDIADGLSDEVGEIIKDDLWPNPLKYFNNEADEEESDGDEDDEEGDDDEEDEDDC*

### NbL12g07060.1 protease_Do−like_8_chloroplastic [MW = 54.41 kDa]

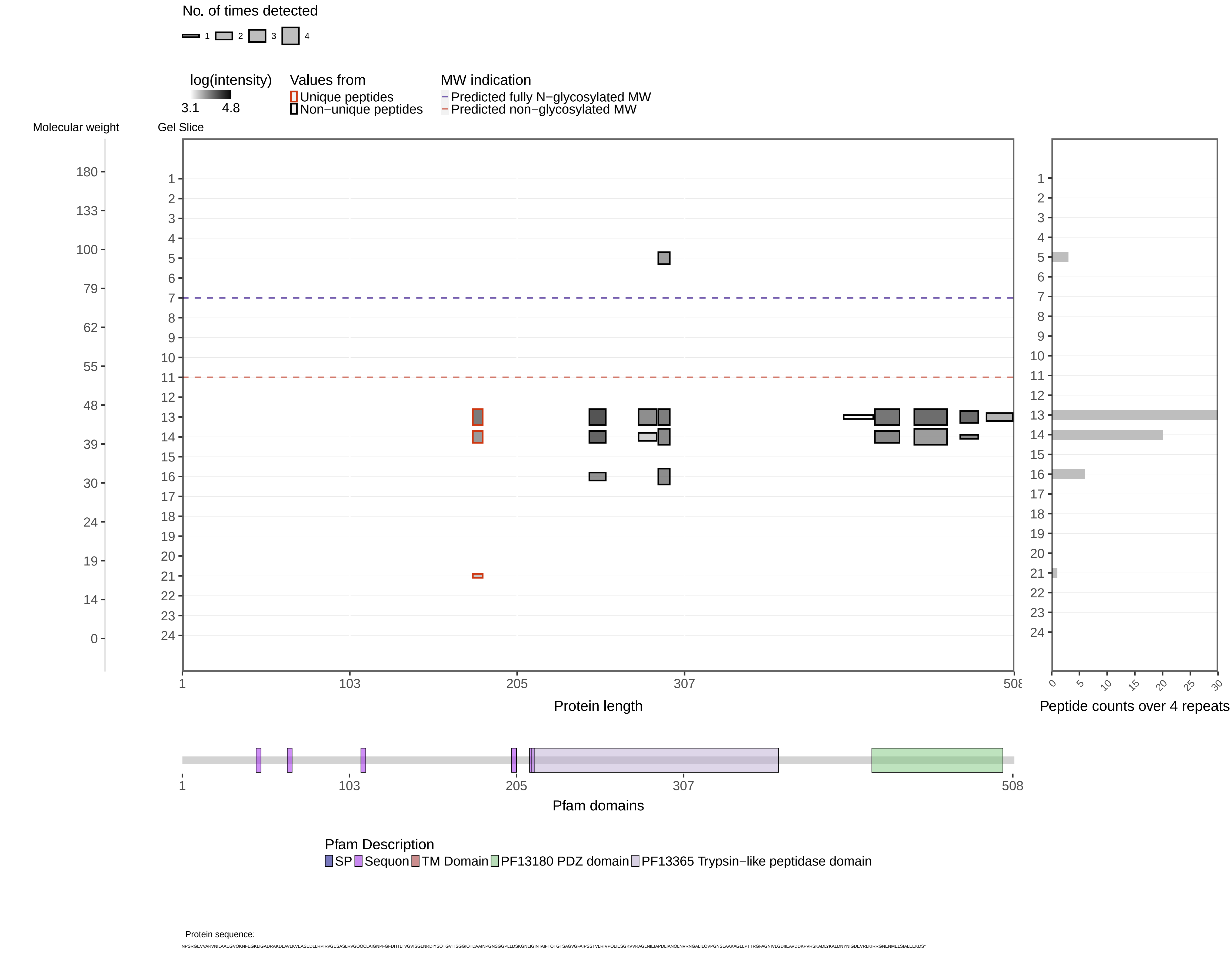

NbL12g06860.1 UDP−glucose_6−dehydrogenase_3 [MW = 52.9 kDa]

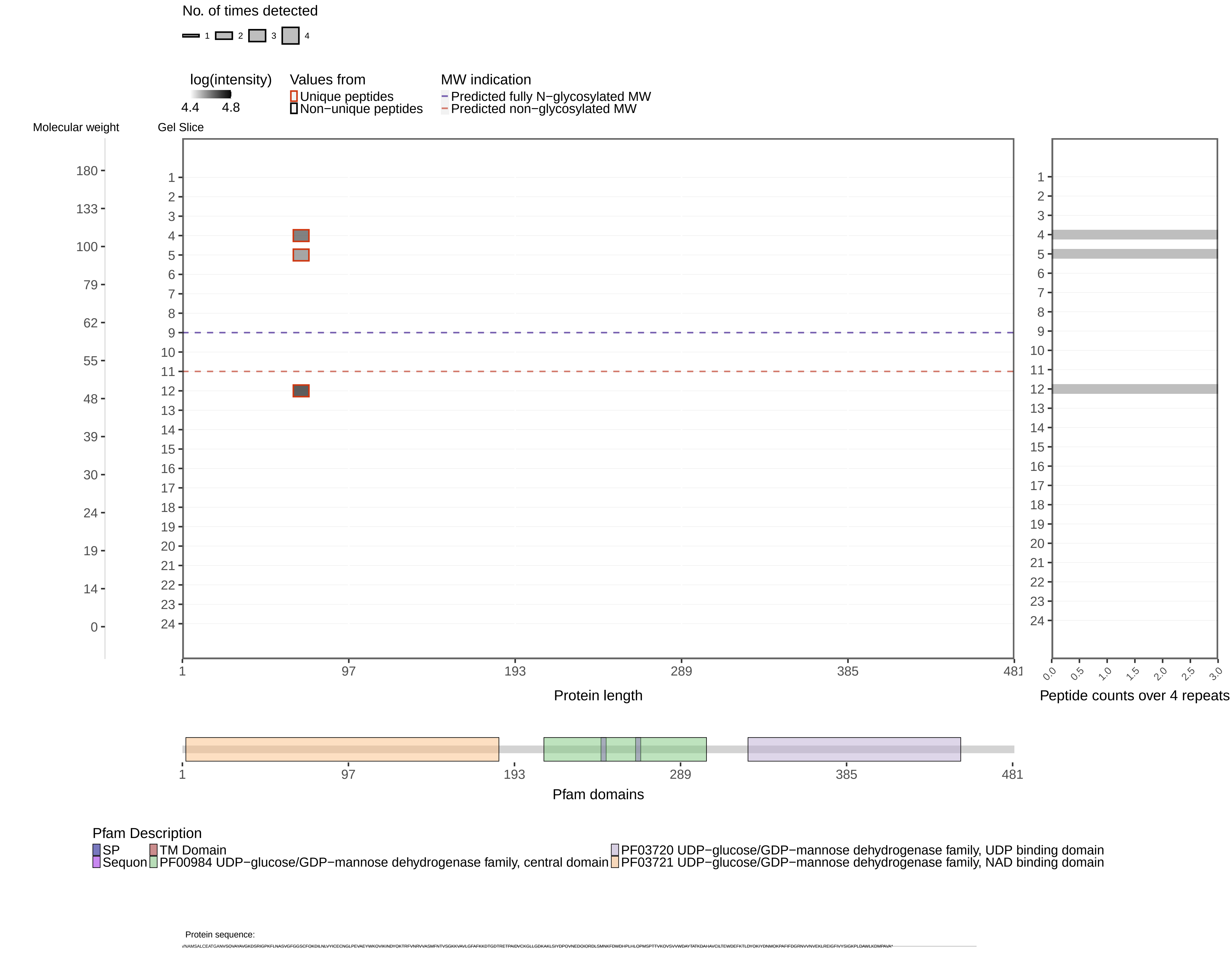

### NbL12g06790.1 protein_CHLORORESPIRATORY_REDUCTION_7_chloroplastic [MW = 24.17 kDa]

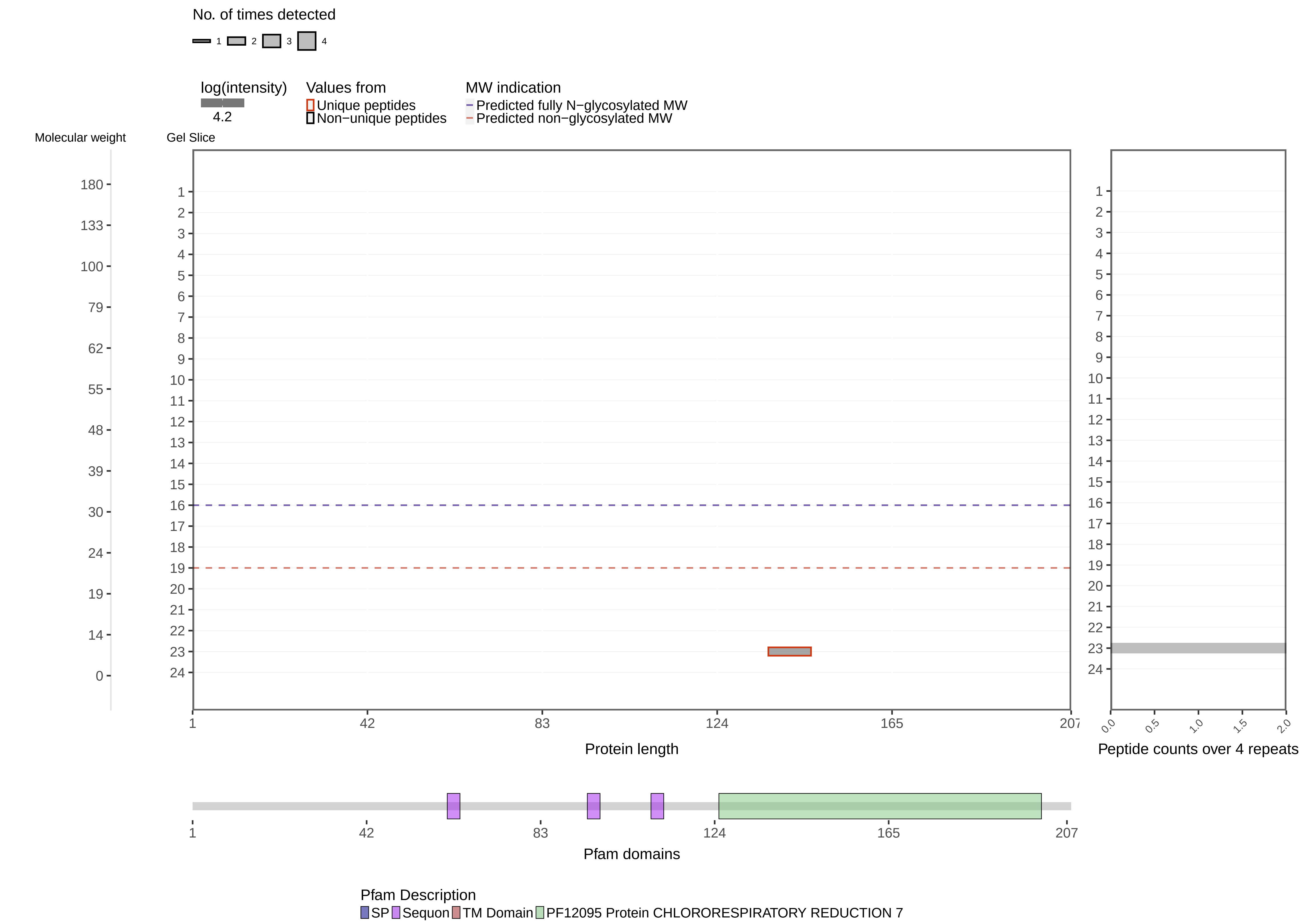


Protein sequence:

FQPFNSSFFHSVTCRCIRLNGTKTLAIRRRRANIGTDTYVLMEPGKSEEFVSEEELRERLKGWLENWPGNELPKDLARFEEIDDAVEYLVKSVCELEIDGDVGSIQWYEVRLQQDME*

### NbL12g06680.1 protein_COBRA−like [MW − SP = 47.37 kDa]

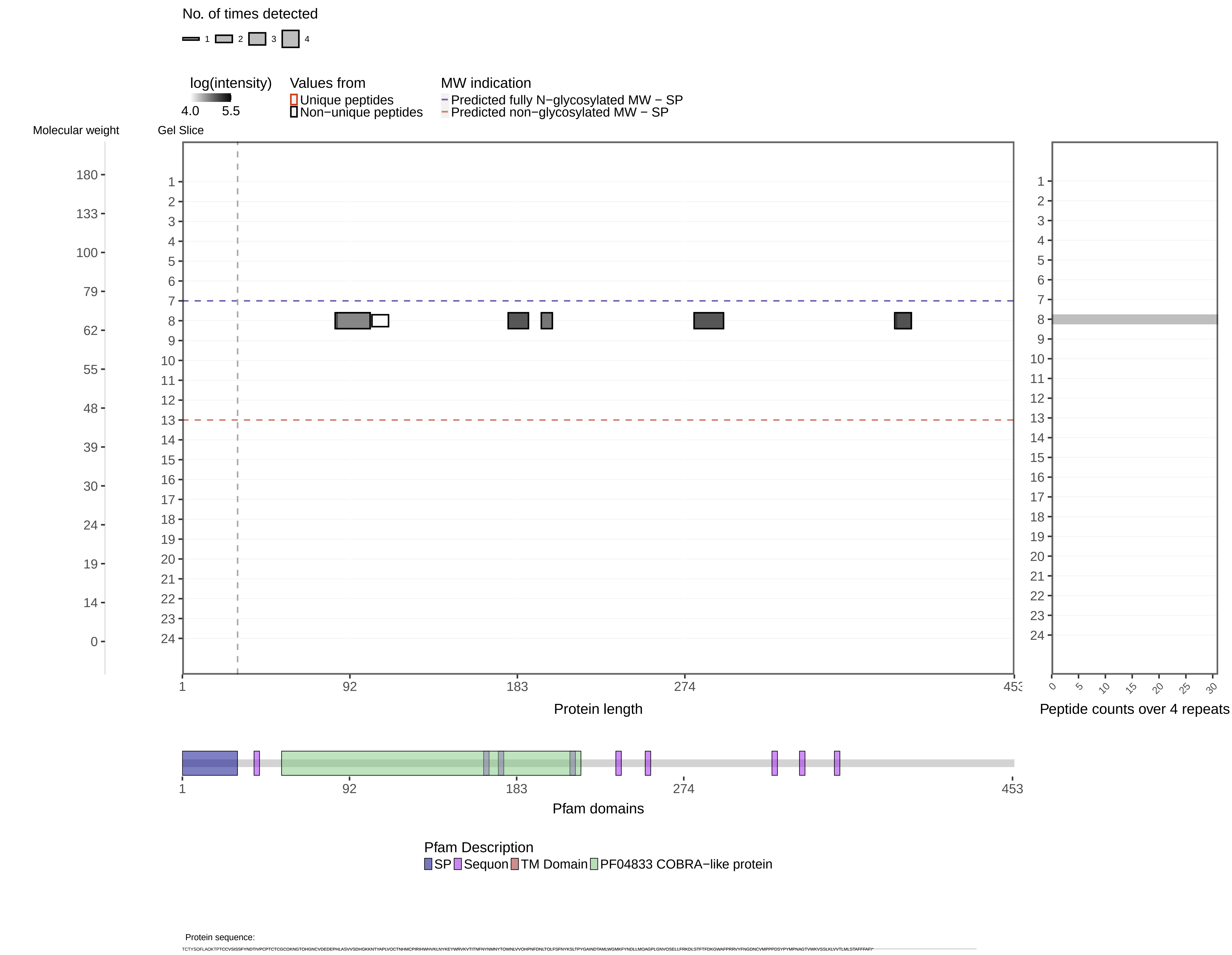

NbL12g06640.1 alpha−14−glucan−protein_synthase_[UDP−forming]_2−like [MW = 40.62 kDa]

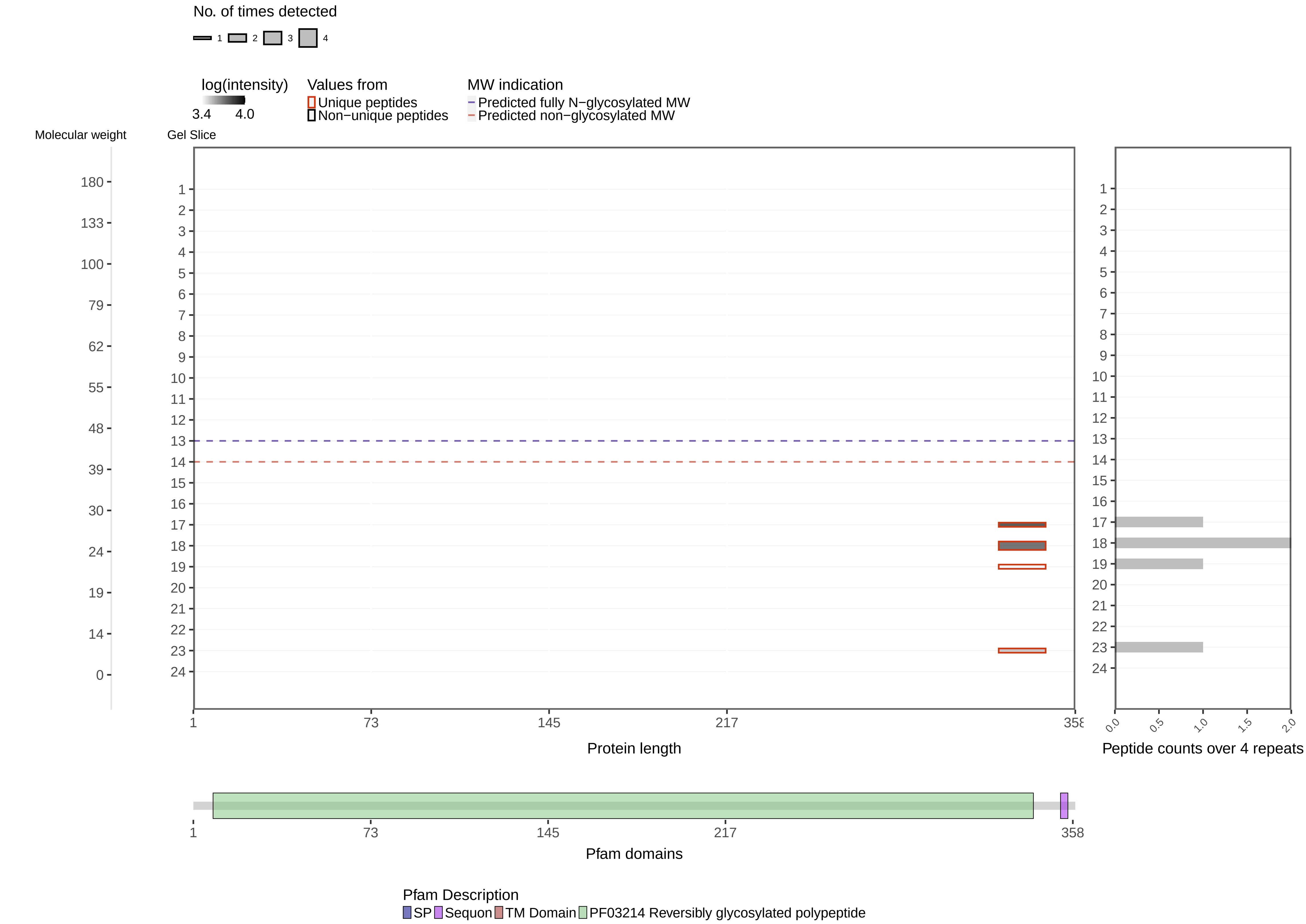


Protein sequence:

LNIPDYDAPTQLVKPLERNTRYVDTVLTIPKGTLFPMCGMNLAFDRELIGPAMYFGLMGDGQPIGRYDDMWAGWCCKVICDHLGLGIKTGLPYIYHSKASNPFVNLKKEYLGIFWQEEIIPFFQSHTFSKDCTSVQKCYVELSKQVKEKLGKVDPYFVKLADAMITWIEAWDMLNPASKDSSKLPNGSSK*

### NbL12g06470.1 STS14_protein−like [MW − SP = 17.37 kDa]

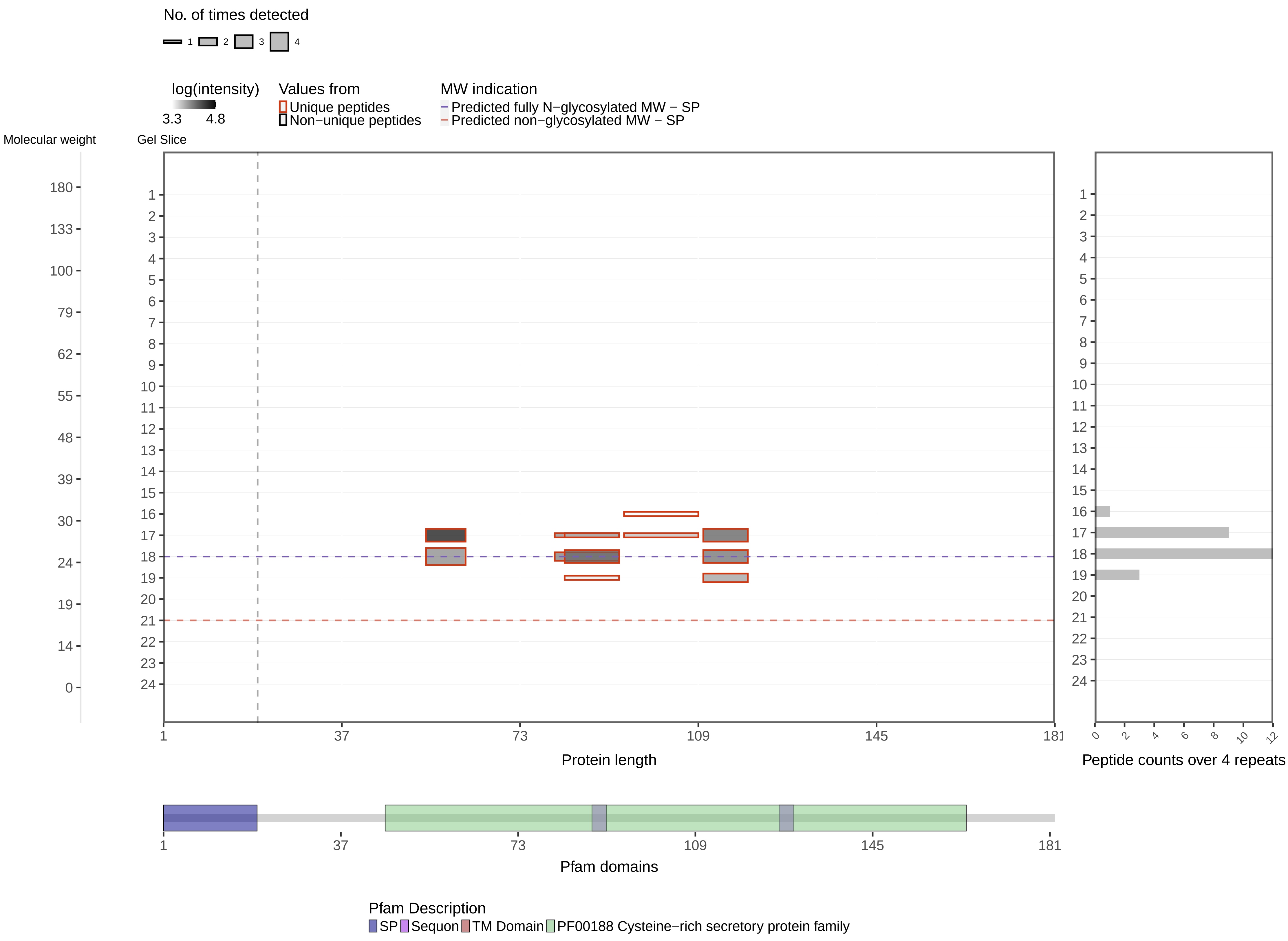


Protein sequence:

IKQNCNFANLTSGKYGGNQLWASGMAVTPRMAVDNWVAEKKFYNYANNSCVGDHKCGVYTQVVWKQSLELGCAQSTCSNRPSTLTICFYNPPGNVIGEKPY*

NbL12g06360.1 acid−binding_protein_2 [MW = 29.21 kDa]

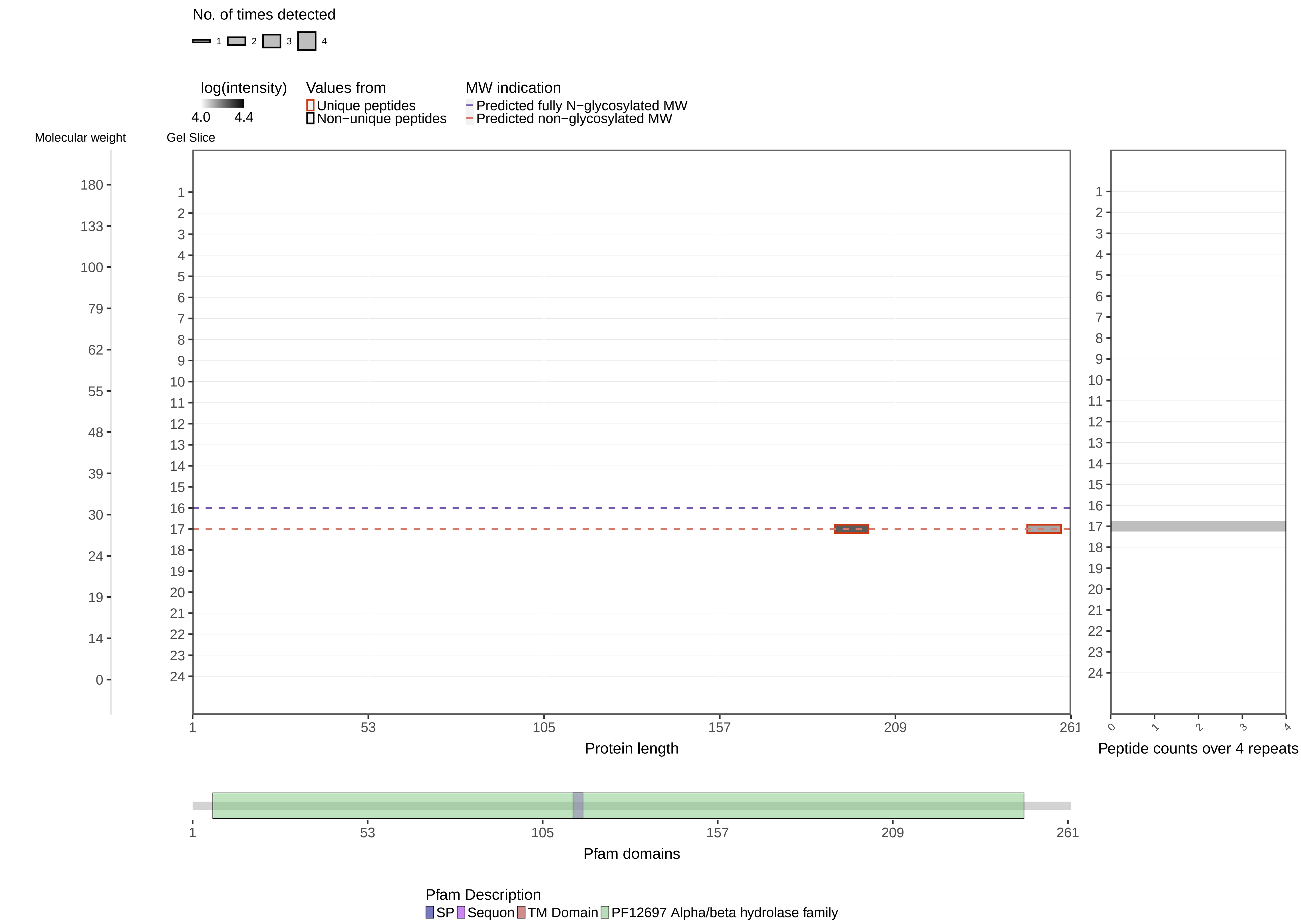


Protein sequence:

FVLEKYYERTPAESWLDTQFLPYGSPEEPLTSMFFGPKFLAHKLYQLCSLEDLALASSLVRPSSLFMEDLAKAKYFTDEGFGSVKRVYIVCTEDKAIPEEFQRWQIDNIGVTEAIEIKGADHMAMLCEPQKLCAALLEIAHKYN*

NbL12g06270.1 patatin−like_protein_2 [MW = 35.72 kDa]

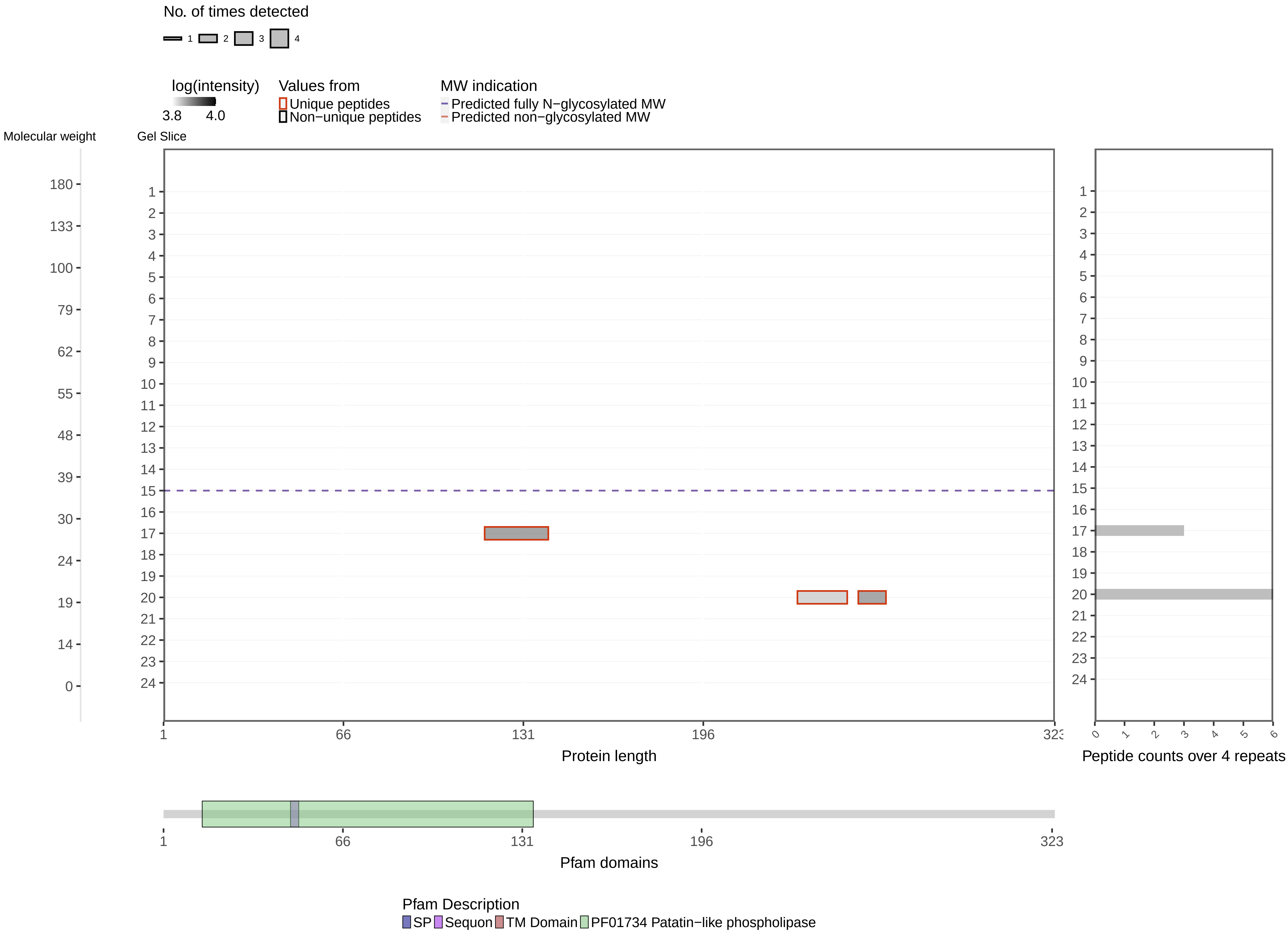


Protein sequence:

GMEPMDCKKMLVLSLGTGIGKEEKKYSAAVASTWGVLGWLYNNGASPLLDVFGDASADIVDIHLSTMFQSLQNEKNYLRIQDDSLTGEAASMDVSTTKNMKALVQIGNDLLKKPVSRVNLDTGRYEEVSGEGTNEEALIRFAKLLSEQRKLRQPVEEFTSTRKIFQESVKRQ*

### NbL12g06200.1 calmodulin−like_protein_1 [MW = 21.21 kDa]

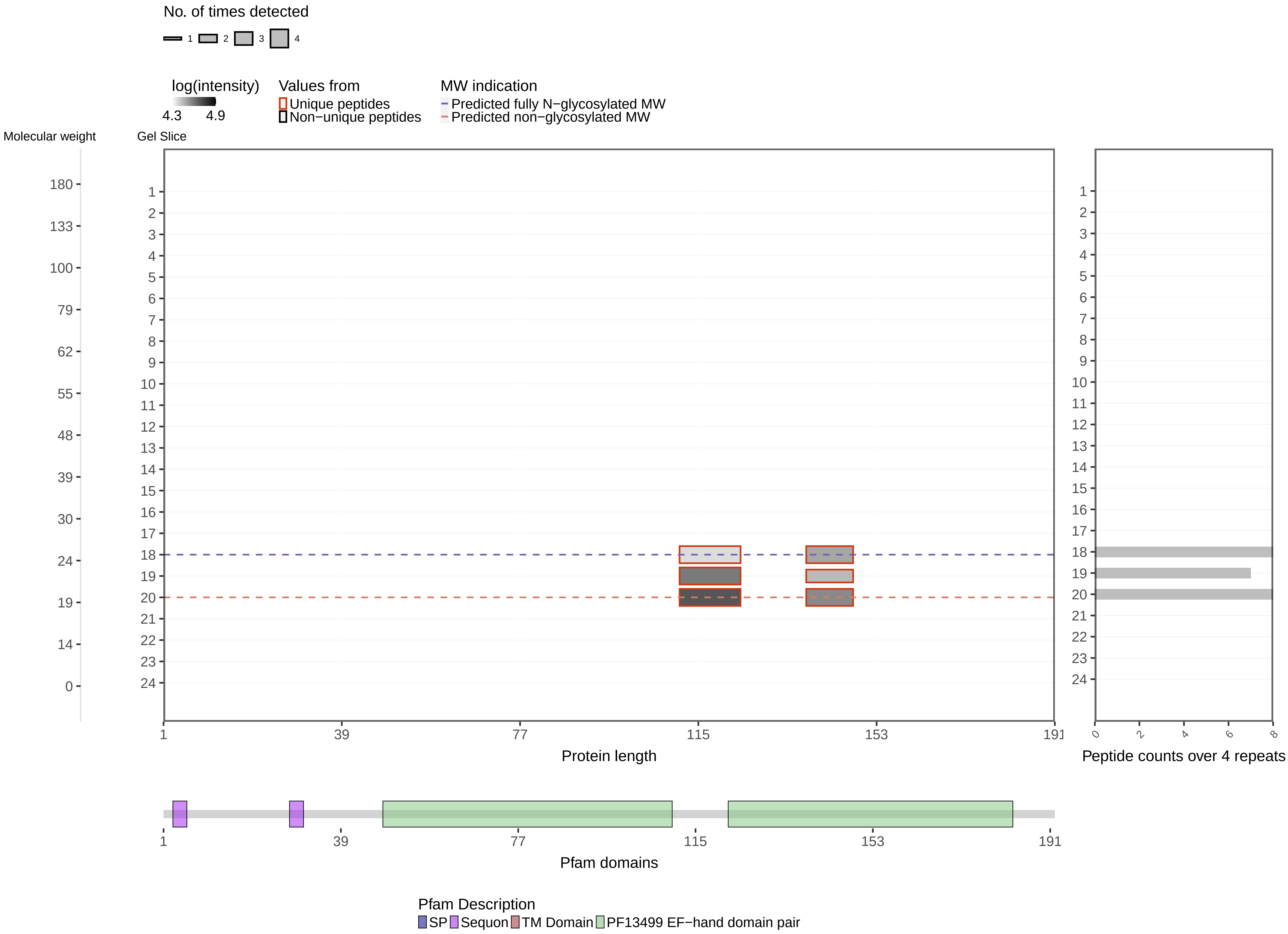


Protein sequence:

TKTYDVGDAFQAADTDGDGFIKFEEFMRVQNLEGGVNPTDIKSAFKAFDLDGDGKISADELLQVQRMLGEKCSLENCKKMVRGVDANGDGLIDMDEFVTMMTRTMKLV*

NbL12g06150.1 CBS_domain−containing_protein_CBSX1_chloroplastic−like [MW = 25.36 kDa]

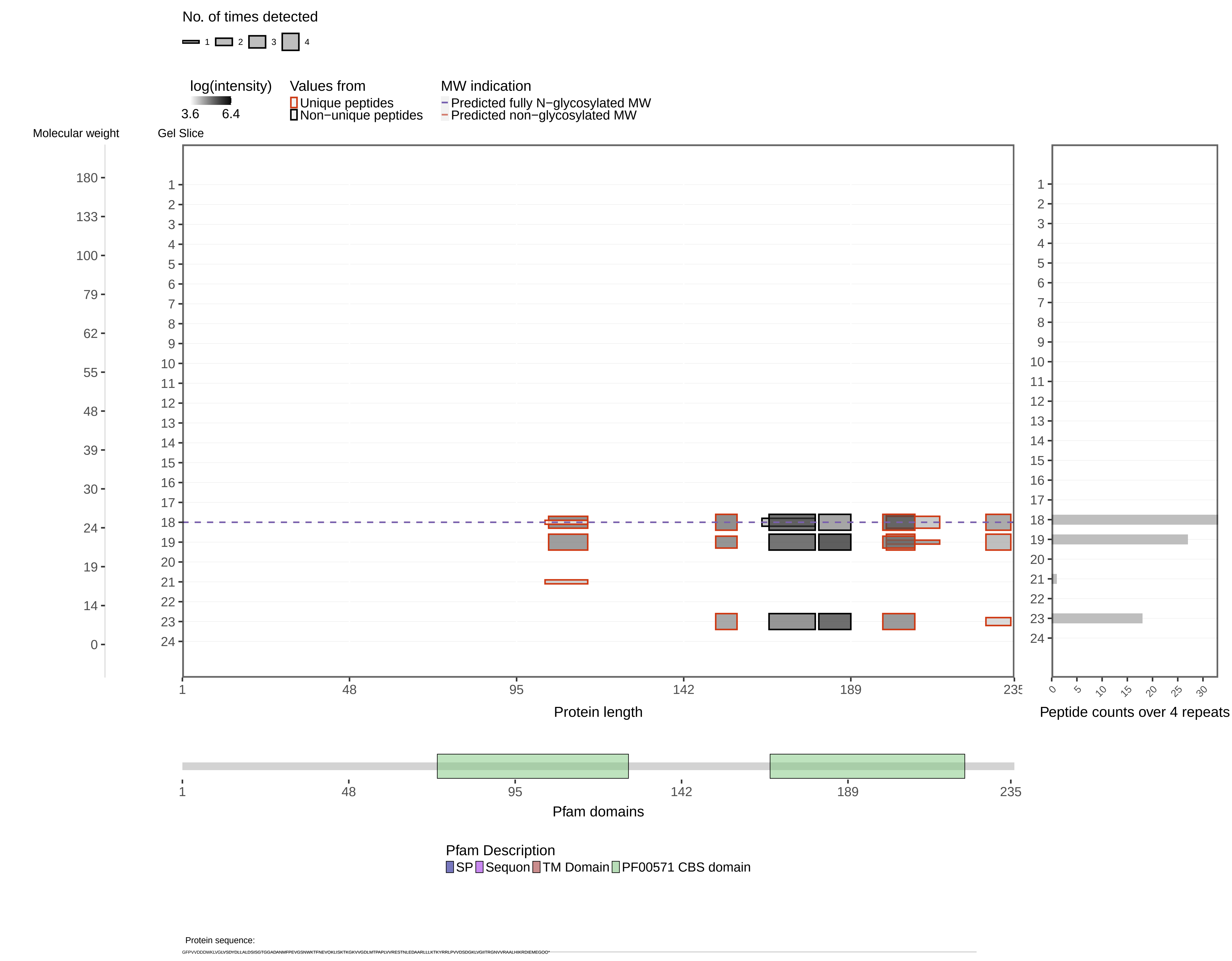

NbL12g06070.1 probable_esterase_KAI2 [MW = 29.97 kDa]

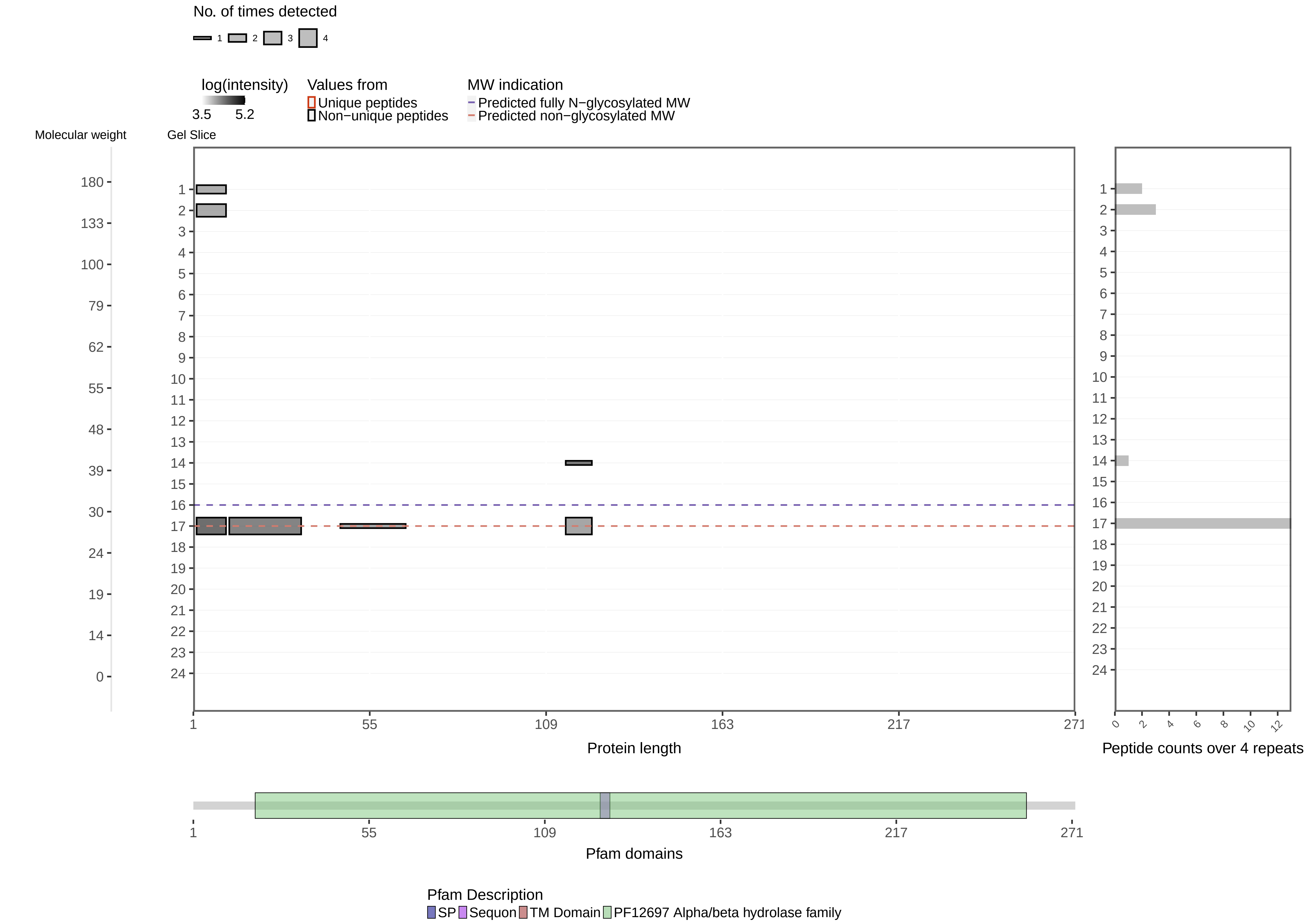


Protein sequence:

YLNDSDYYGGFEQEDLDQLFEAMRSNYKAWCSGFAPLVVGGDMDSVAIQEFSRTLFNMRPDIALSVLQIIFLSDLRHLLAHVTVPCHLIQSMKDLAVPVVVSEYLHQHLGGESIVEVMSTEGHLPQLSSPDVVIPVLLRHIRHDIVV*

### NbL12g05960.1 60S_ribosomal_protein_L26−1 [MW = 16.75 kDa]

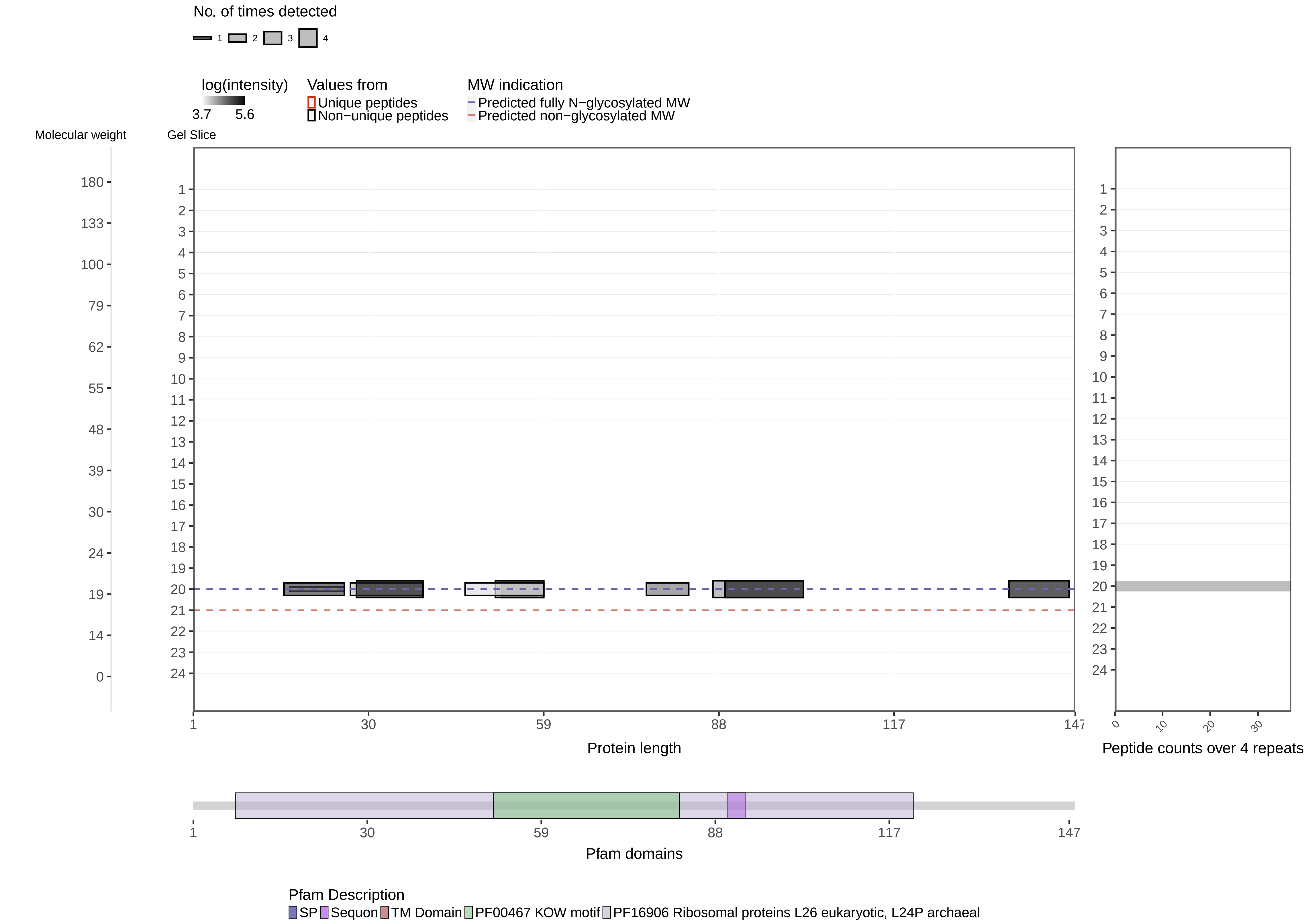


Protein sequence:

GTYKGREGKVVQVYRKKWVIHIERITREKVNGSTVNVGIHPSKVVVTKLRLDKDRKSLLDRKAKGRAAADKDKGTKFTAEDIMQTVD*

NbL12g05890.1 prostamide/prostaglandin_F_synthase−like [MW = 31.47 kDa]

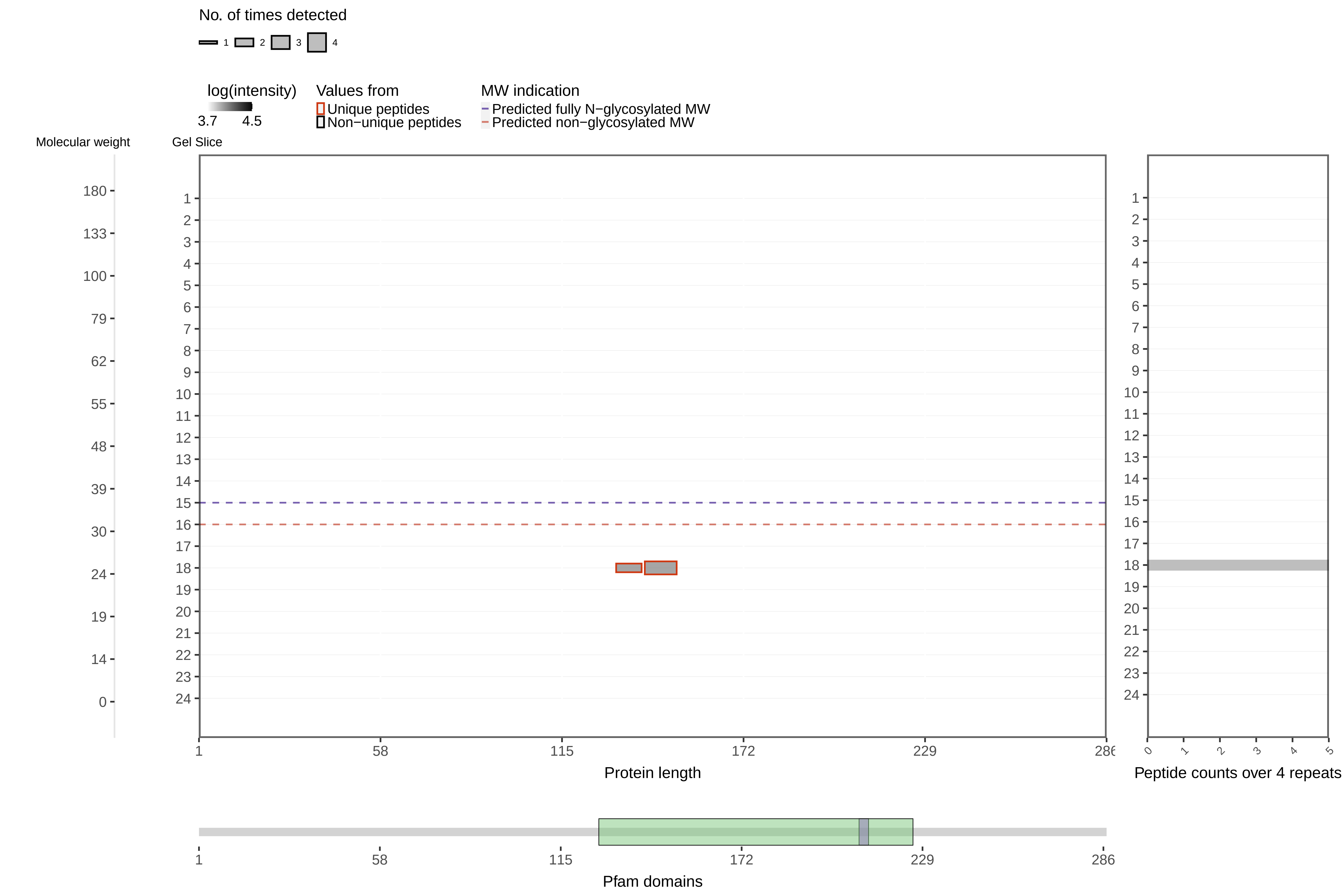


Protein sequence:

RERFESAGVKLIAVGIGAPSKARILAERLPFPLDCLYADPDRRAYDVLGLYYGFGRTFFNPASAKVLSRFKELQEATKNYTIEATPDDISGVLQQGSNCCMHGRMKEPVTTLPWKISLMFAAKFLLYKAICIPKEVDEYRIKVSPCLCCKSGCRY*

NbL12g05850.1 malate_dehydrogenase_glyoxysomal−like [MW = 37.66 kDa]

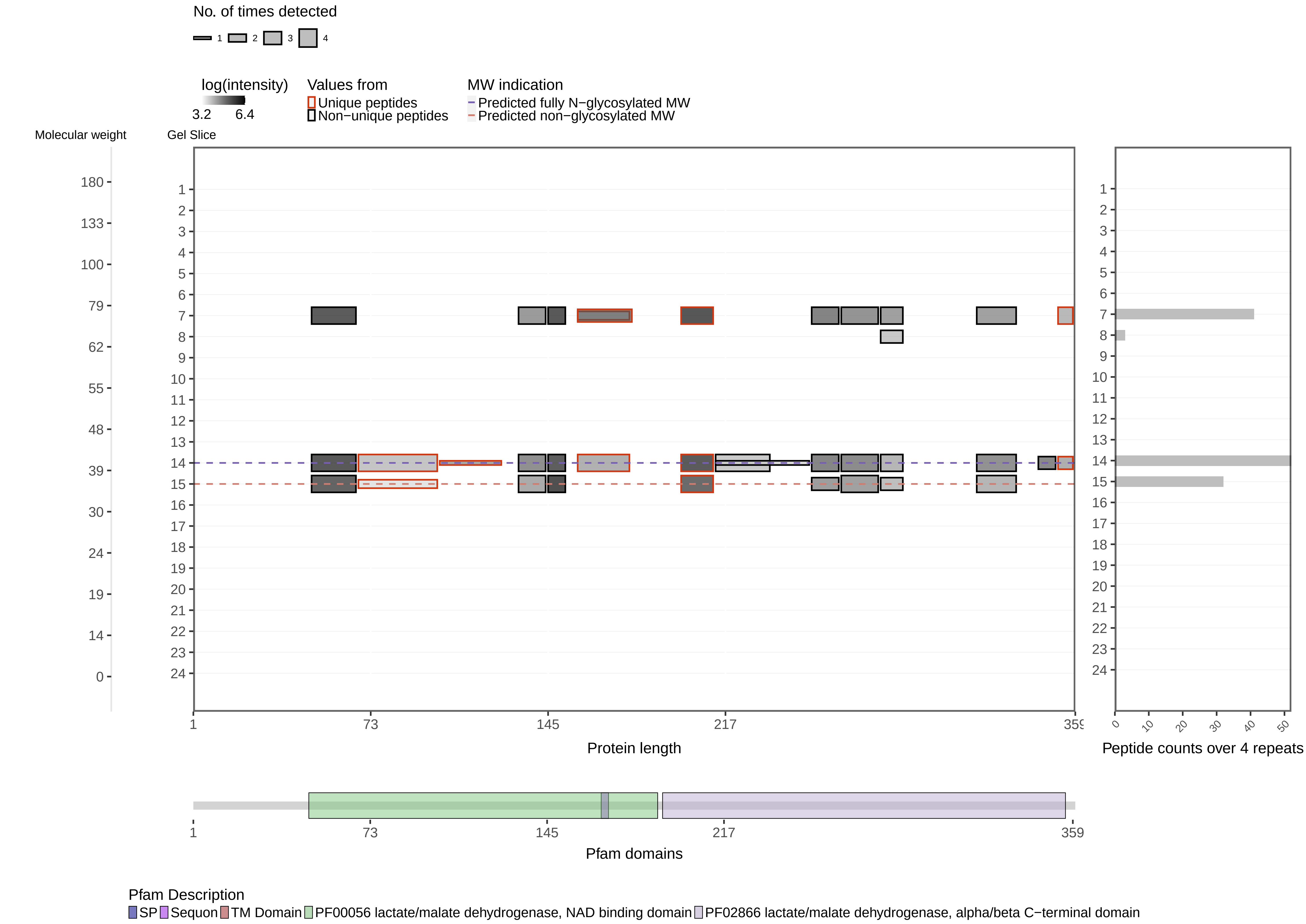


Protein sequence:

ISTVPIAAEVFKKAGTFDPRRLLGVTKLDIVRANTFVAEVLGLDPREVDVPVVGGHAGVTILPLLSQVKPPCSFTPEETEYLTSRIQNGGTEVVEAKAGAGSATLSMAYAAVKFADACLHGLRGDAGIVECAFVSSQVTELPFFASKVRLGRNGVEEIYPLGPLNEYERSGLEKAKKELATSIQKGVNFVMK*

### NbL12g05790.1 serine_hydroxymethyltransferase_mitochondrial−like [MW = 57.15 kDa]

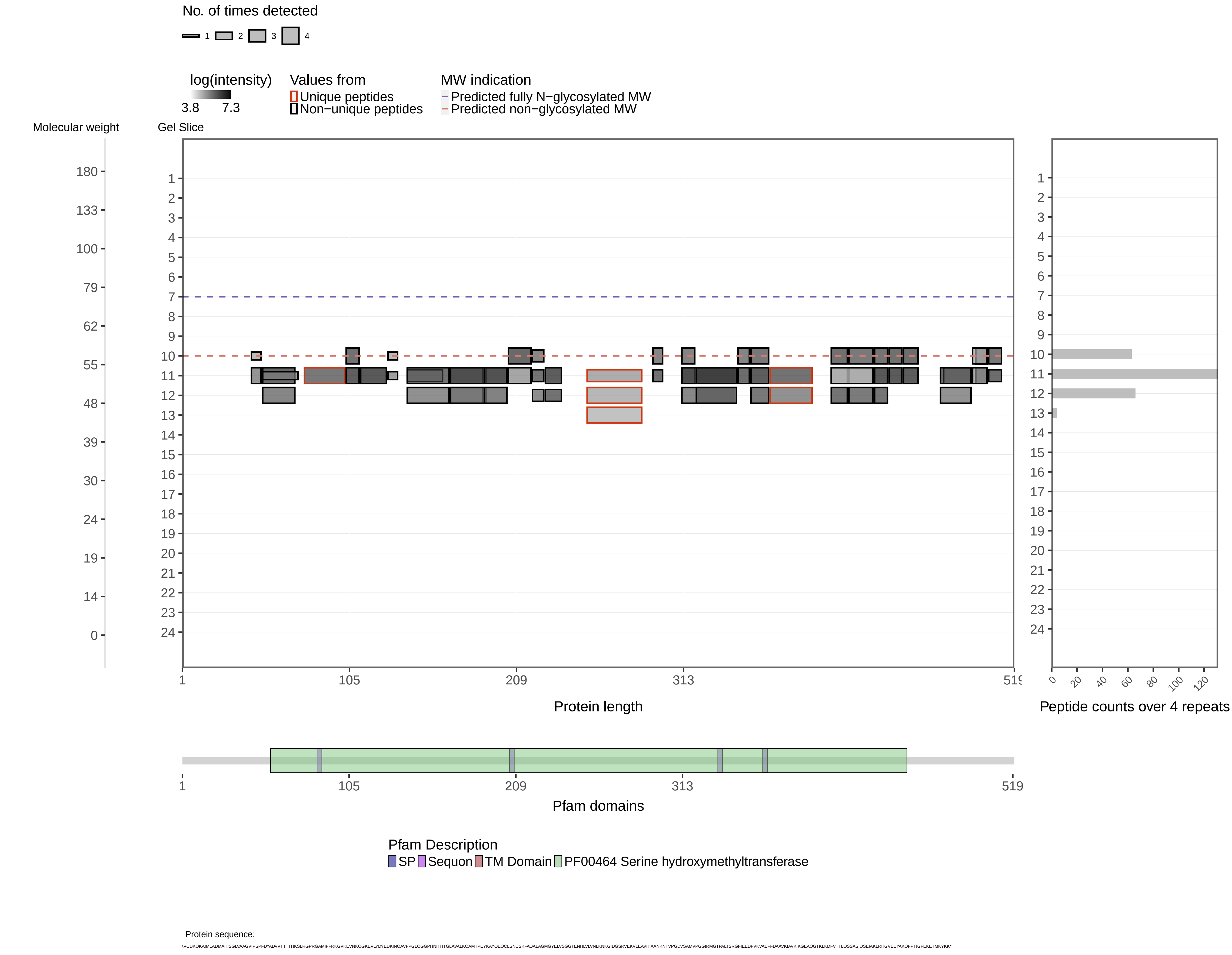

### NbL12g05570.1 ubiquitin_receptor_RAD23d−like [MW = 40.88 kDa]

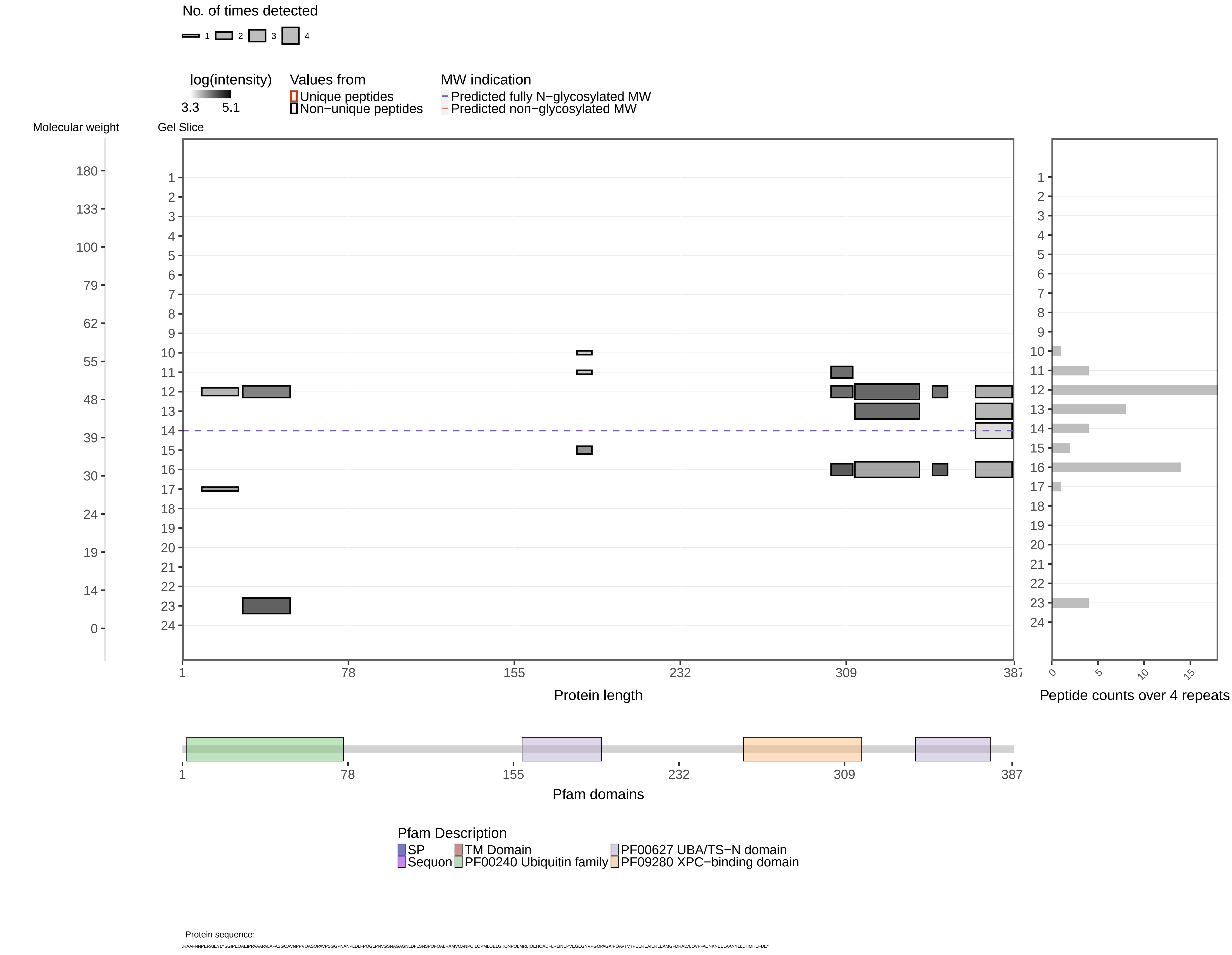

### NbL12g05560.1 T−complex_protein_1_subunit_zeta_1−like [MW = 58.91 kDa]

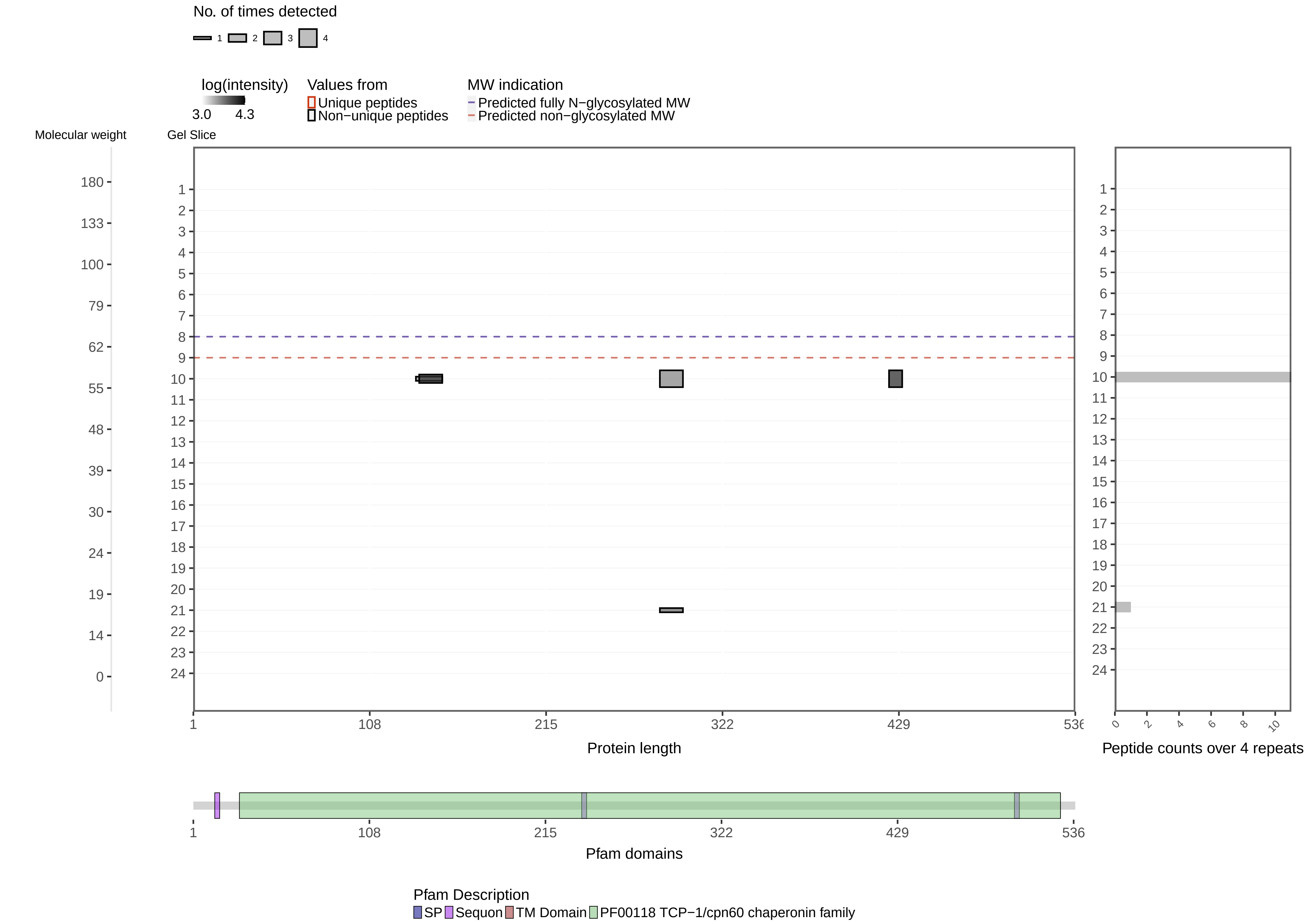


Protein sequence:

[illegible]NAEQREAMVAAERRSVDERVQKIIDLKNKVCAGNDNNFVIINQKGIDPPSLDILARAGIIALRRAKRRNMERLVLACGGEAVNSVDDLTPDCLGWAGLVYEHVLGEEKYTFVENVKNPHSCTILIKGPNDHTIAQIKDAVRDGLRAVKNTIEDEAVVLGAGTFEVAARQHLINEVKKTVKGRAQLGVEAFANALLVIPKTLAENSGLDTQDVIIALTGEHDKGNVVGLNHHTGEPIDPQMEGIFDNYSVKRQIVNSGPVIASQLLLVDEVIRAGRNMRKPN*

### NbL12g05480.1 probable_Xaa−Pro_aminopeptidase_P [MW = 72.9 kDa]

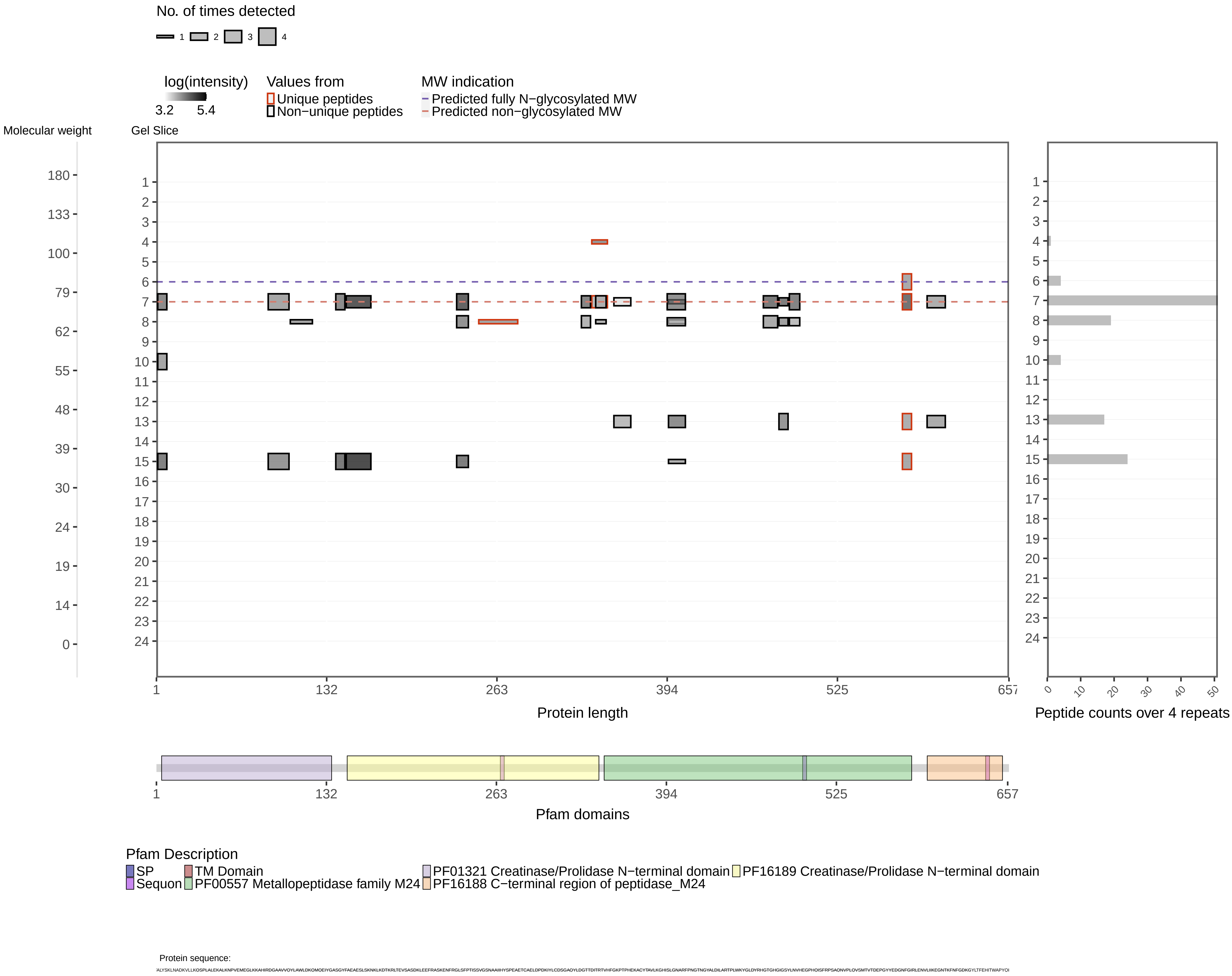

### NbL12g05450.1 G−box−binding_factor_1−like [MW = 36.55 kDa]

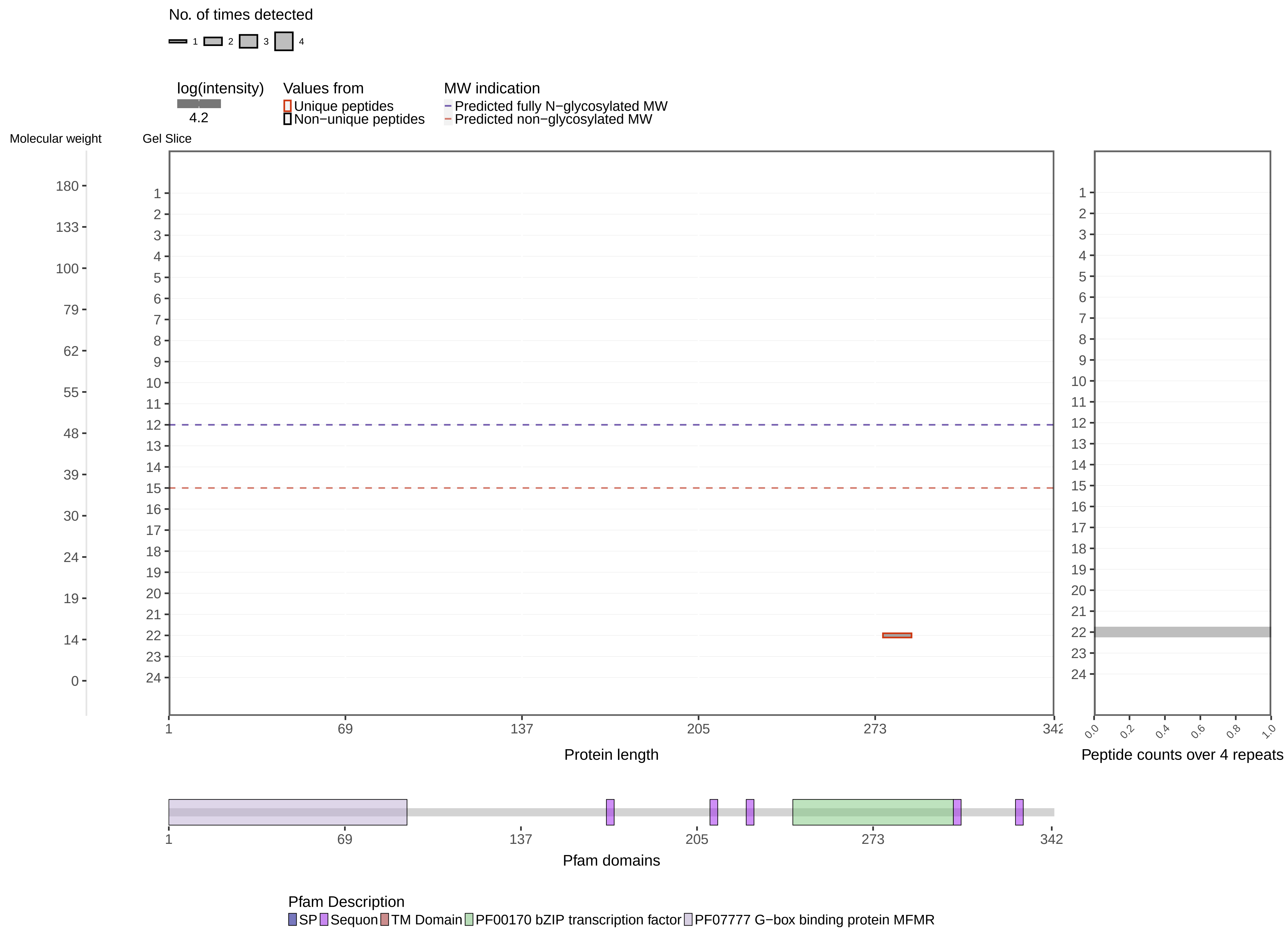


Protein sequence:

ANDENDNHEFSANKSRNFDLMLANGASSQNNPATGNPVAMPATNLNIGMDLWNASSAGPGMIKMRSNQSGVAPAPGIGREWIQDERELKRQKRKQSNRESARRSRLRKQAECEELQQRVETLSNENHALKEEMRKLSEECEKLTSENNSIKEELTRLCGPEAVSKLESNANATHLQSNLDEANS*

### NbL12g05210.1 proteasome_subunit_beta_type−1 [MW = 29.12 kDa]

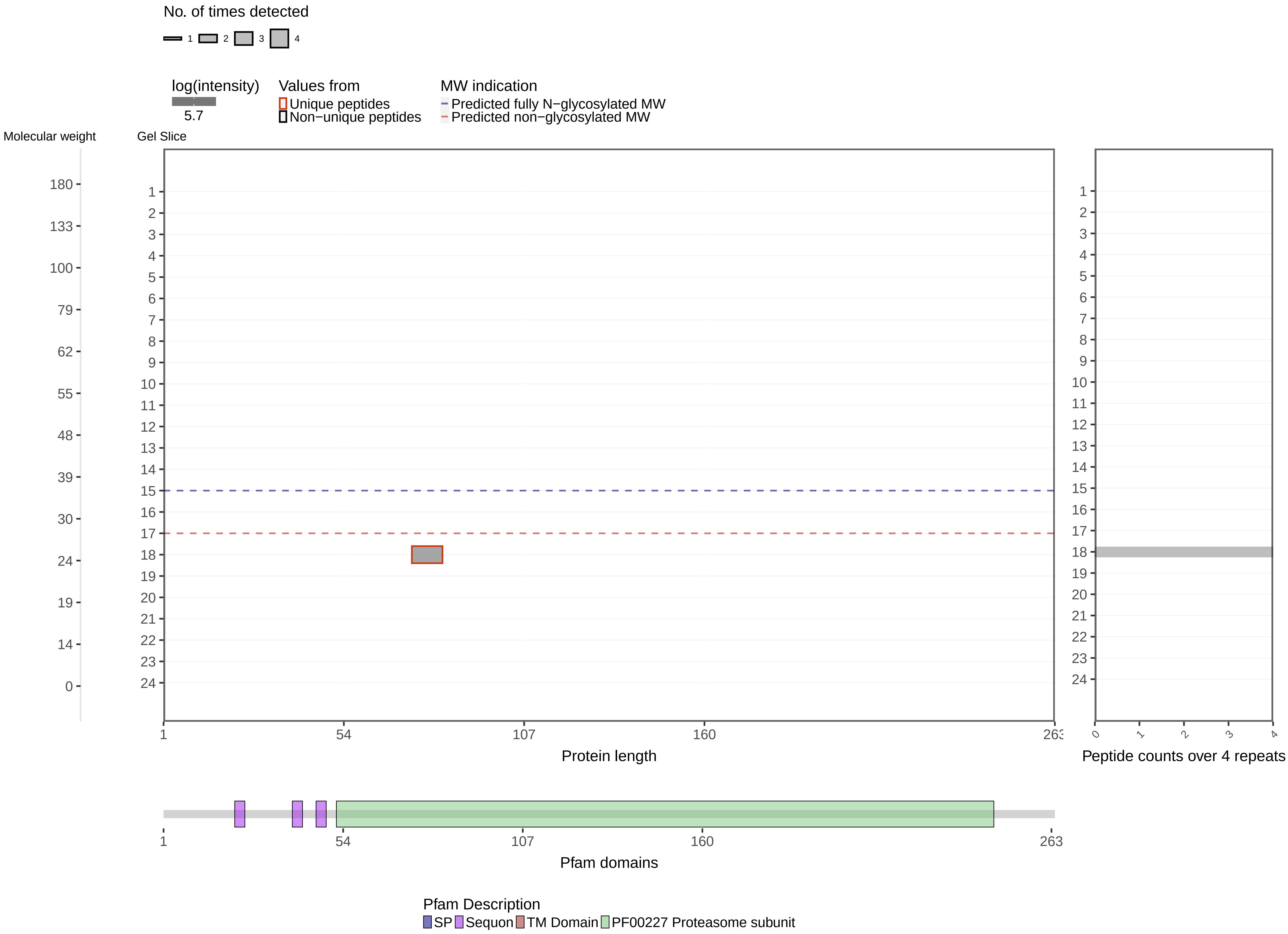


Protein sequence:

LIYOHOHNKOMSCPAMGOLLSNTLYYKRFFPYYSFNVLGGLDSEGKGCVFTYDAVGSYERVGYSSOGSGSTLIMPFLDNOLKSPSPLLLPAKDAVTPLSESEAIDLVKTCFASATERDIYTGDRLEIVILNANGIRREEMELRKD*

NbL12g05140.1 D−2−hydroxyglutarate_dehydrogenase_mitochondrial [MW = 63.49 kDa]

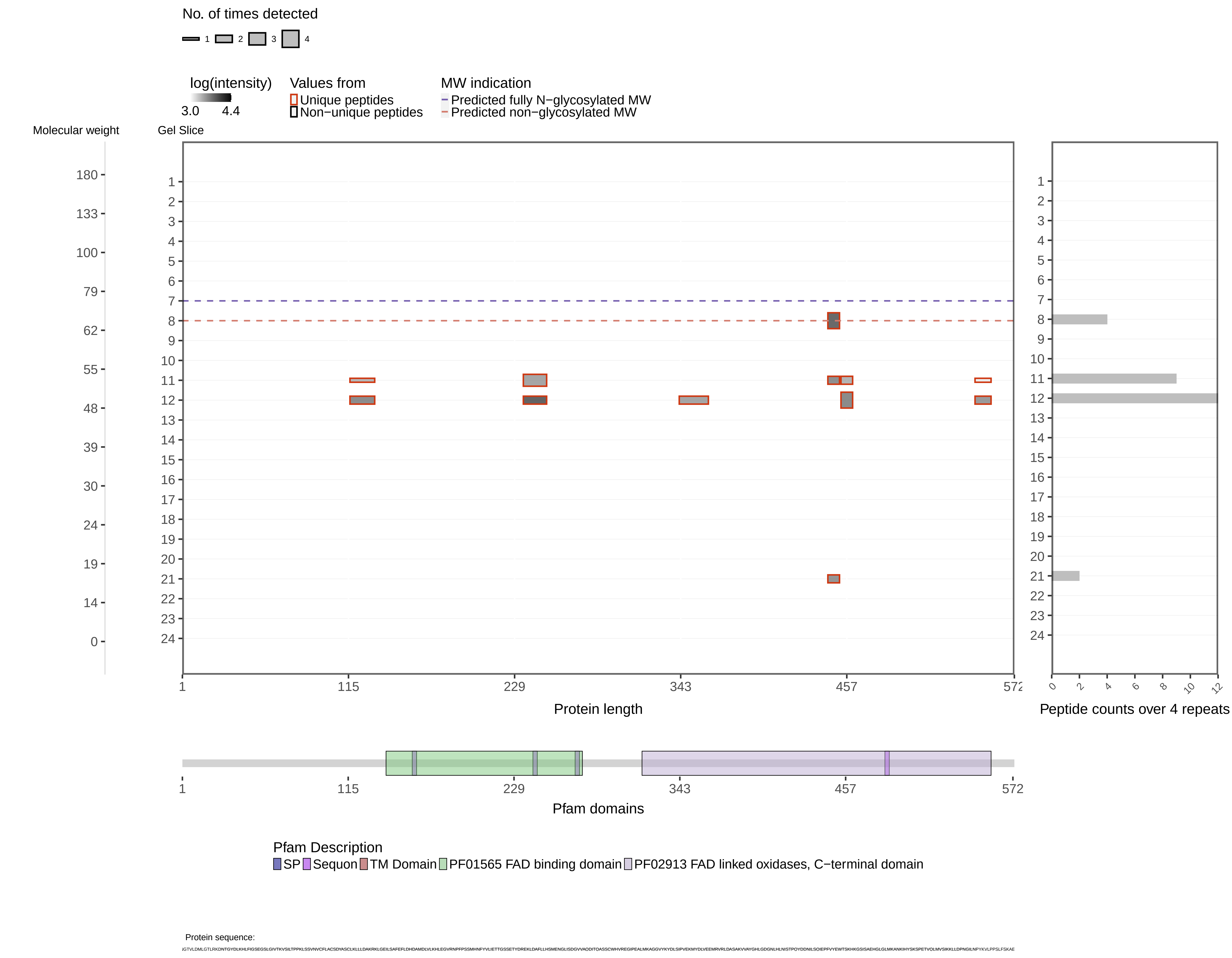

### NbL12g05080.1 fructose−bisphosphate_aldolase_1_chloroplastic [MW = 42.59 kDa]

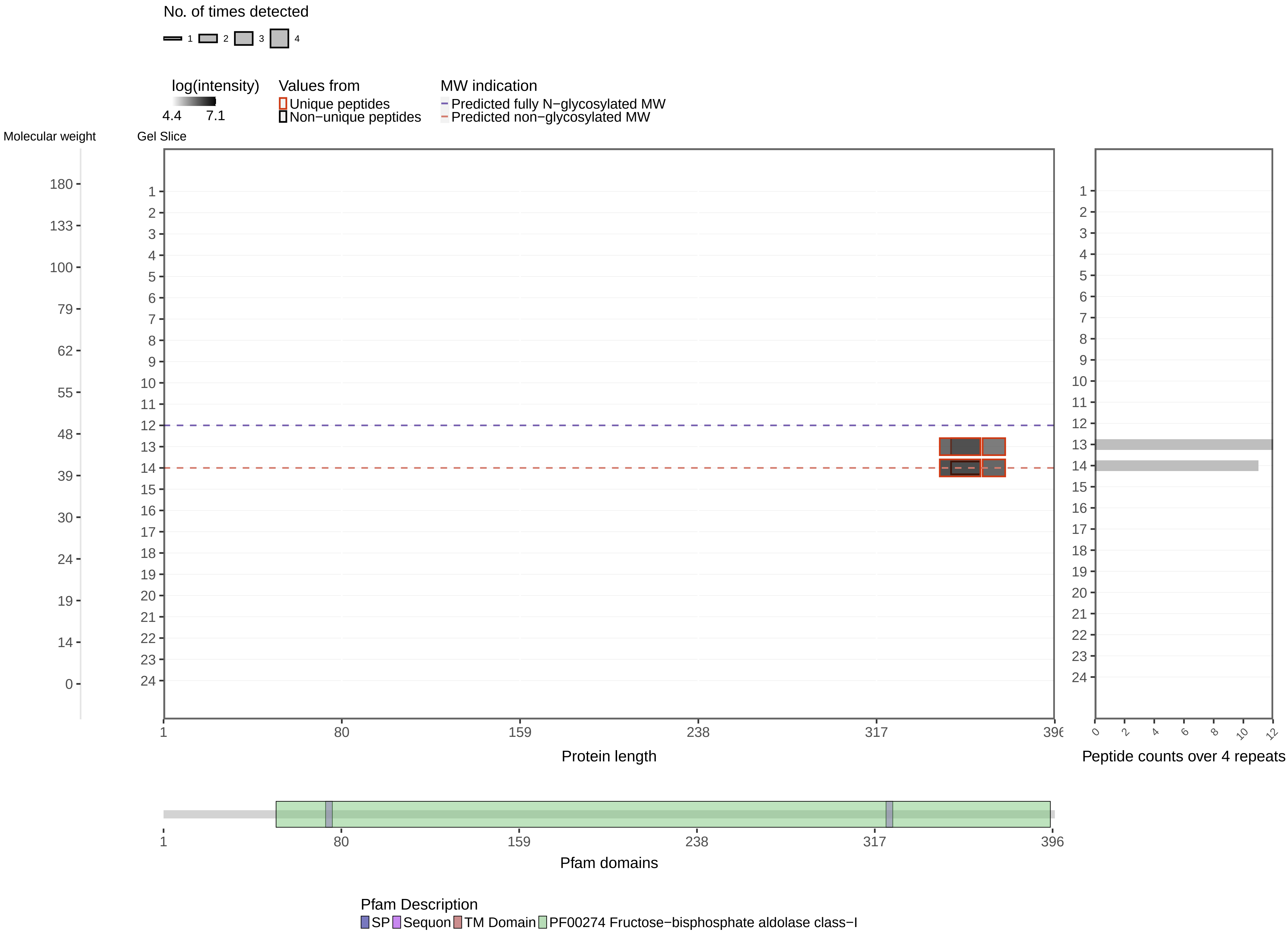


Protein sequence:

VSIPNGPSALAVKEAAWGLARYAAISQDNGLVPIVEPEILLDGEHNIDRTFEVAQKVWAEVFFYLAENNVMFEGILLKPSMVTPGAECKERATPDQVADYTLKLLQRRIPPAVPGIMFLSGGQSEVEATLNLNAMNQSPNPWHVSFSYARALQNTCLKTWGGRPENVEAAQEALLIRANANSLAQLGKYTGEGESEEAKKGMFVKGYVY*

NbL12g05030.1 nucleolin−like [MW = 128.69 kDa]

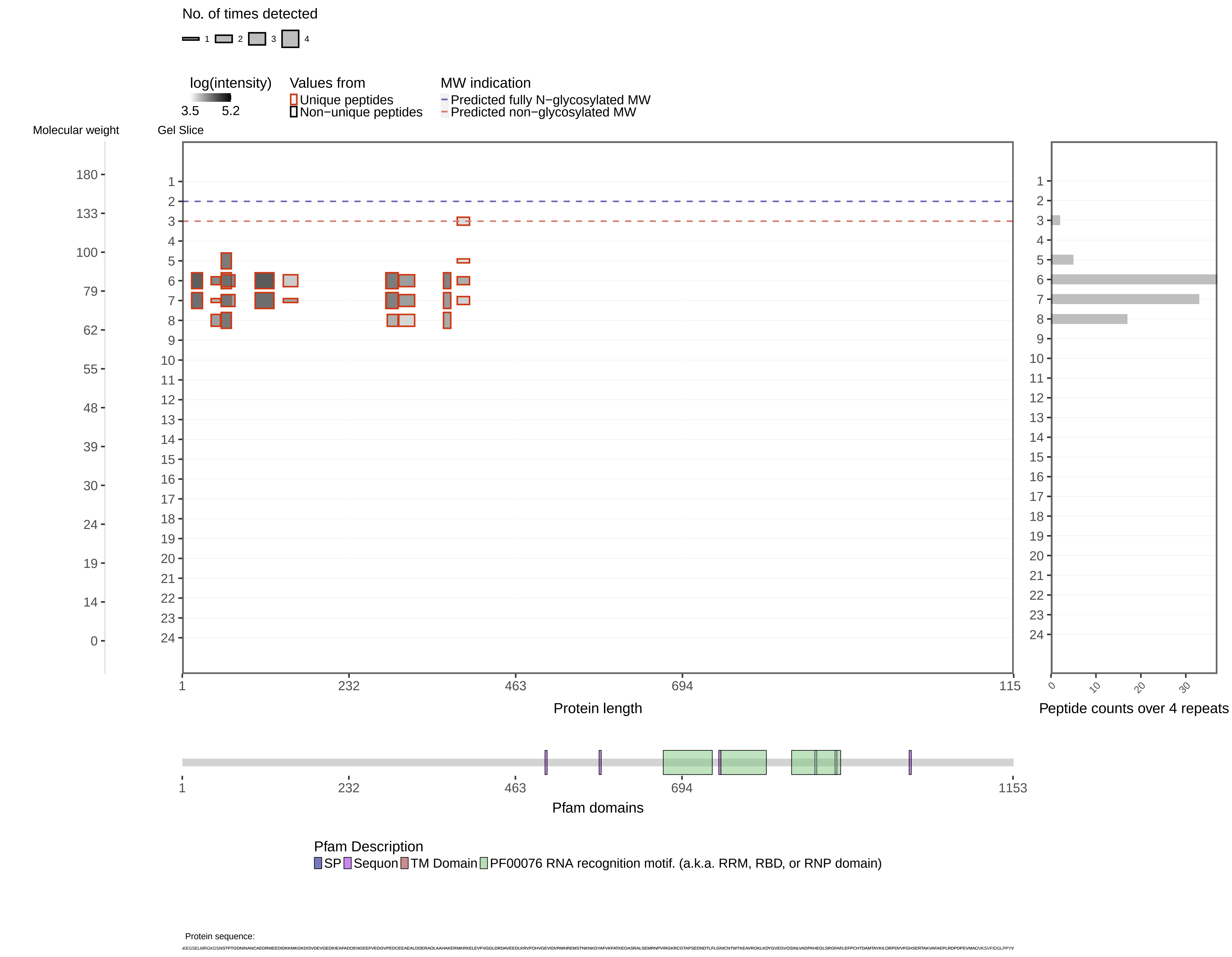

### NbL12g05020.1 peptidyl−prolyl_cis−trans_isomerase_Pin1−like [MW = 13.53 kDa]

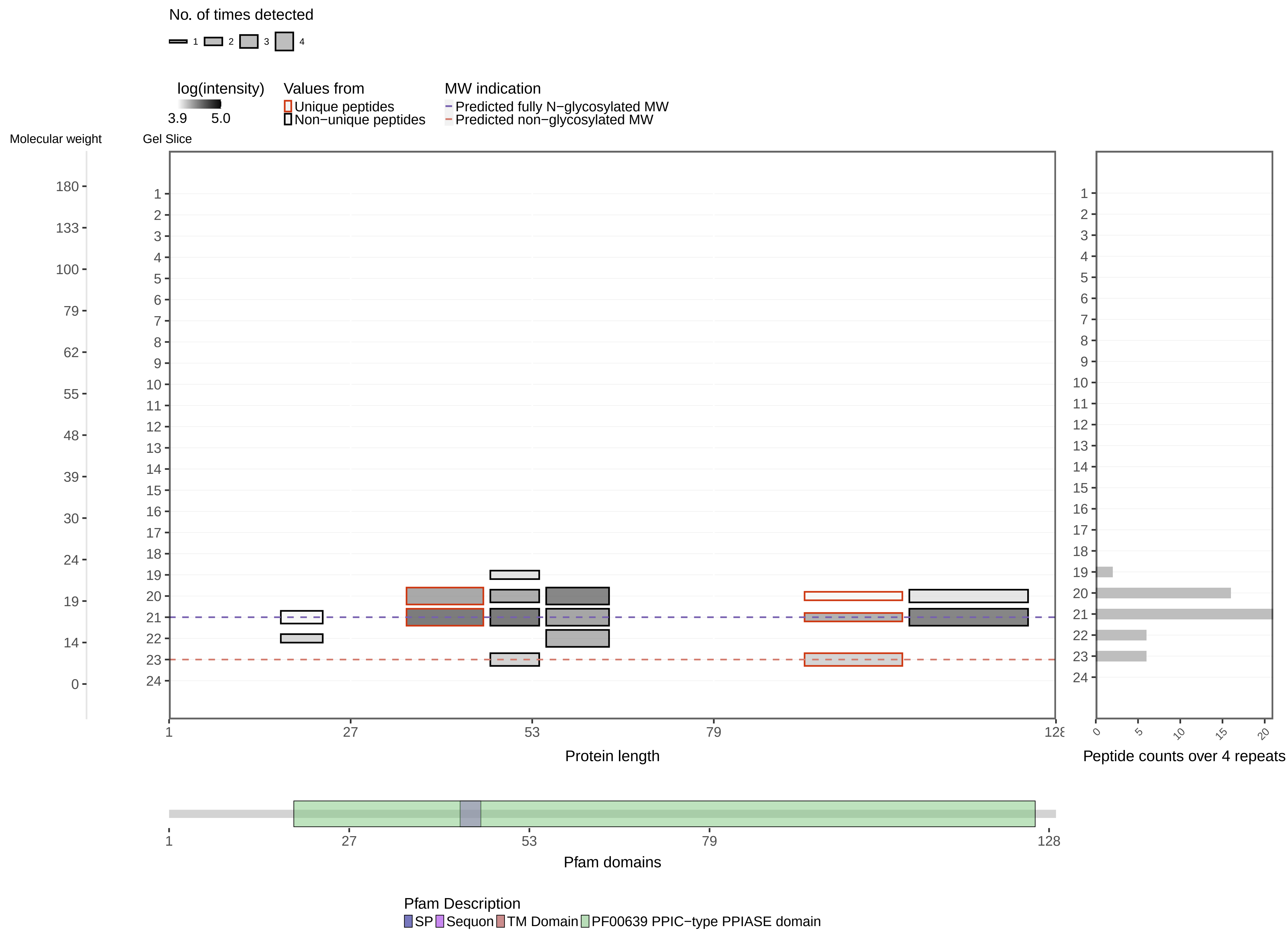


Protein sequence:

/SQLKSLREDIVSGKAKFDDVAATHSDCSSAKRGGDLGPFGKGQMQKPFEVATYALKVGEISDIVETDSGVHIIKRTA*

### NbL12g05010.1 small_nuclear_ribonucleoprotein_Sm_D3−like [MW = 14.45 kDa]

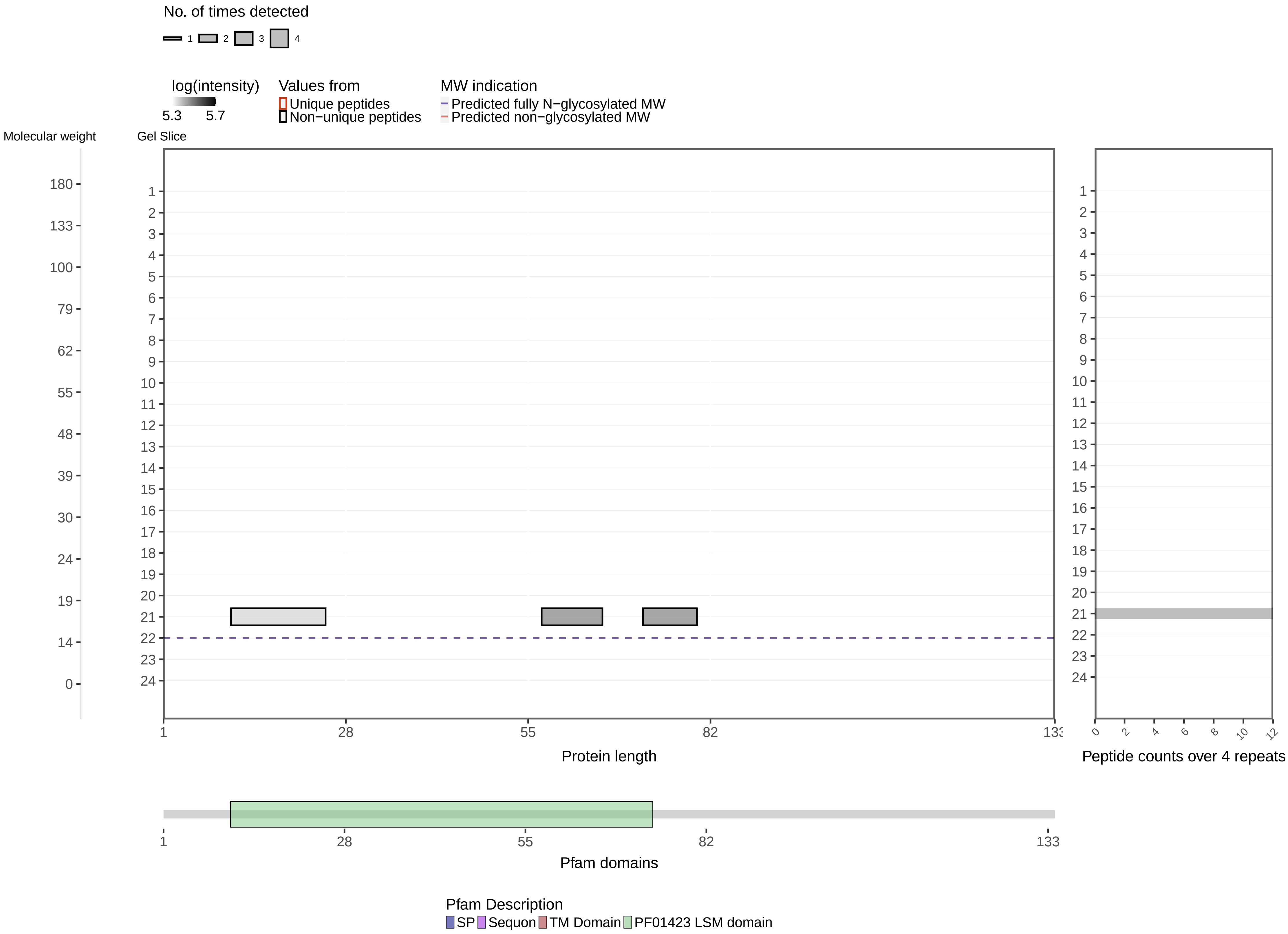


Protein sequence:

OGRVSQLEHVFIRGSKVRFMIIPDMLKNAPMFKRLEARIKGKGTSLGVGRGRAIAMRAKAQAAGRGAAPGRGVVPPVRR*

NbL12g04810.1 basic_endochitinase−like [MW − SP = 26.81 kDa]

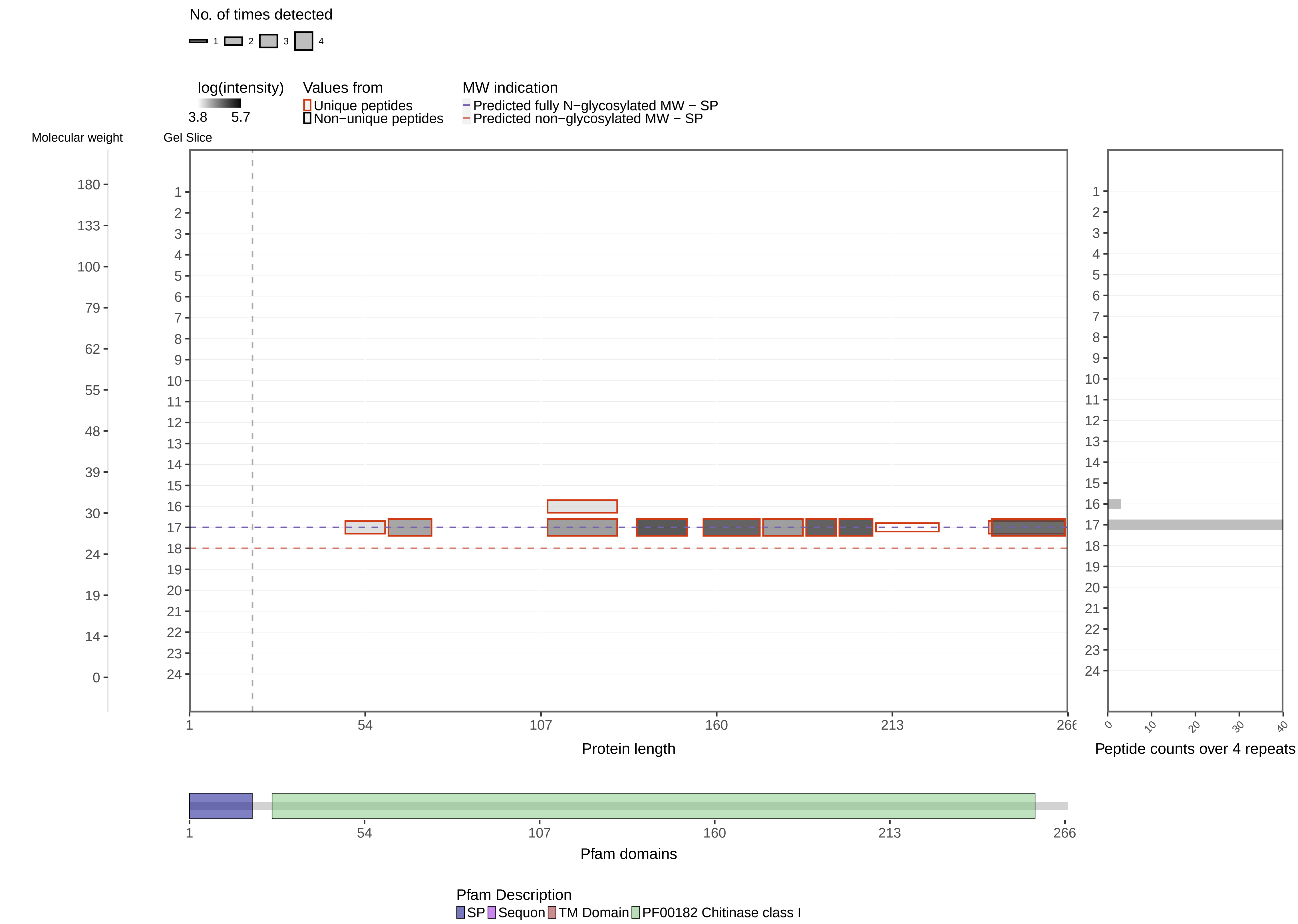


Protein sequence:
SQQWPCAPGKSYFGRGPIQISYNYNYGPAGRAIGRDLLNNPDLVANDPVVSFKTALWFWMTPQQPKPSAHDVITGRWTPSAADLAARRVPGFGVITNIINGGIECSKGSNAQMQSRIGFYRRYCQILGVDPGNNLDCASQRPFGQ*

NbL12g04760.1 ubiquitin_domain−containing_protein_DSK2b−like [MW = 58.35 kDa]

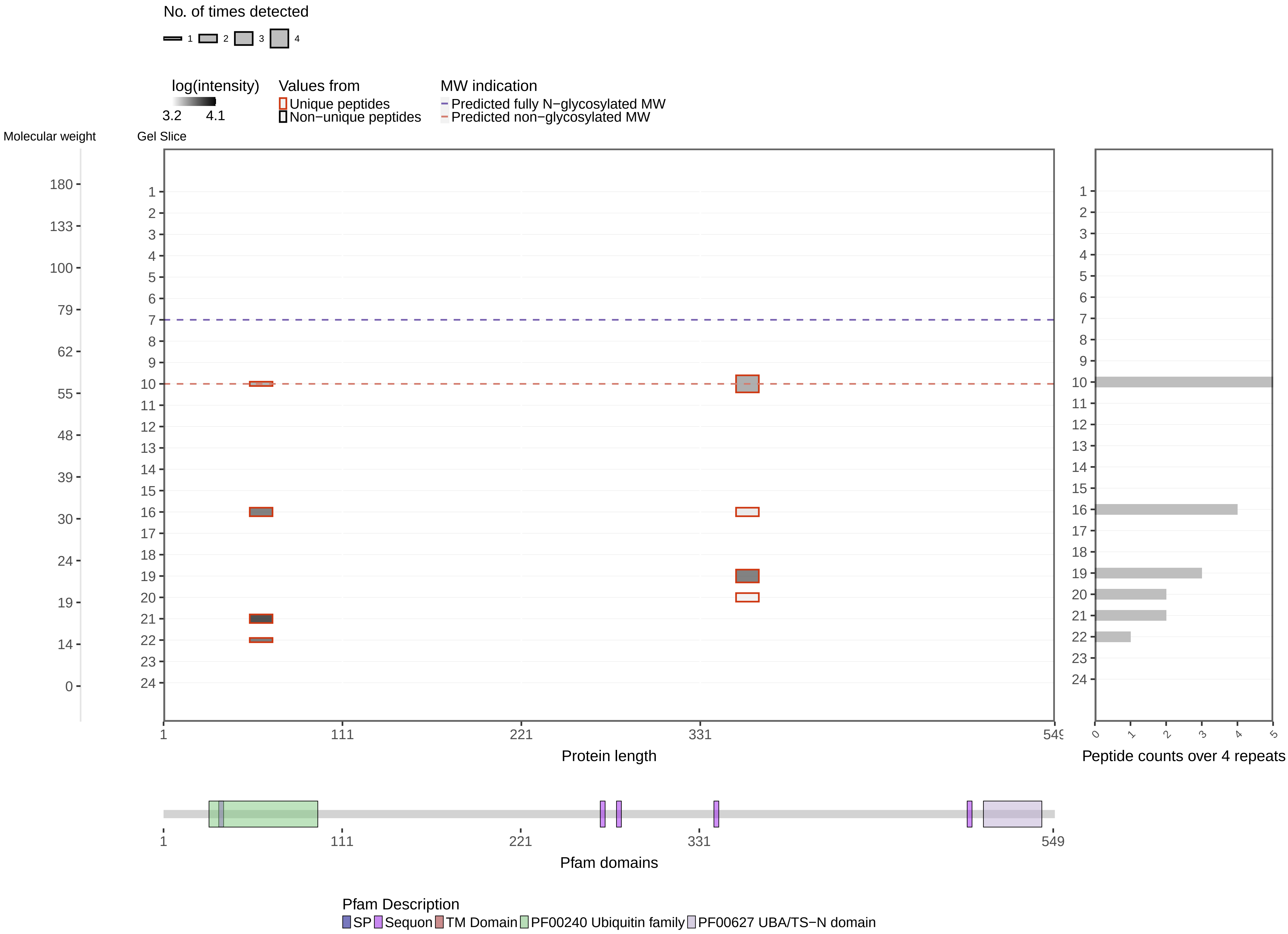


Protein sequence:

IVQEPFLNATTMAGDTRNDSGANPFAALLGAQGGGQAGNQATSLPVTGSETTANPPAPNTNPLPNPWASAGTGAAQTNTSARSNAAADTRALPLSGLGALPDLQRMLGGMPDASSLNQLMQNPAISQMMQSLLSNPQYMNQISGLNPQLRSMLDSNPHLREMMQNPEFIRQLTSPETMQQVMSLQQGLLSQLGRQQTNRGQGQDAGRTAFDNMGMEMLMNMFGGLGTGGLGVSNRSNVPPEELYATQLAQLQEMGFFDTQENIRALIATAGNVHAAVERLLGNT[illegible]

NbL12g04600.1 eukaryotic_translation_initiation_factor_6−2 [MW = 26.42 kDa]

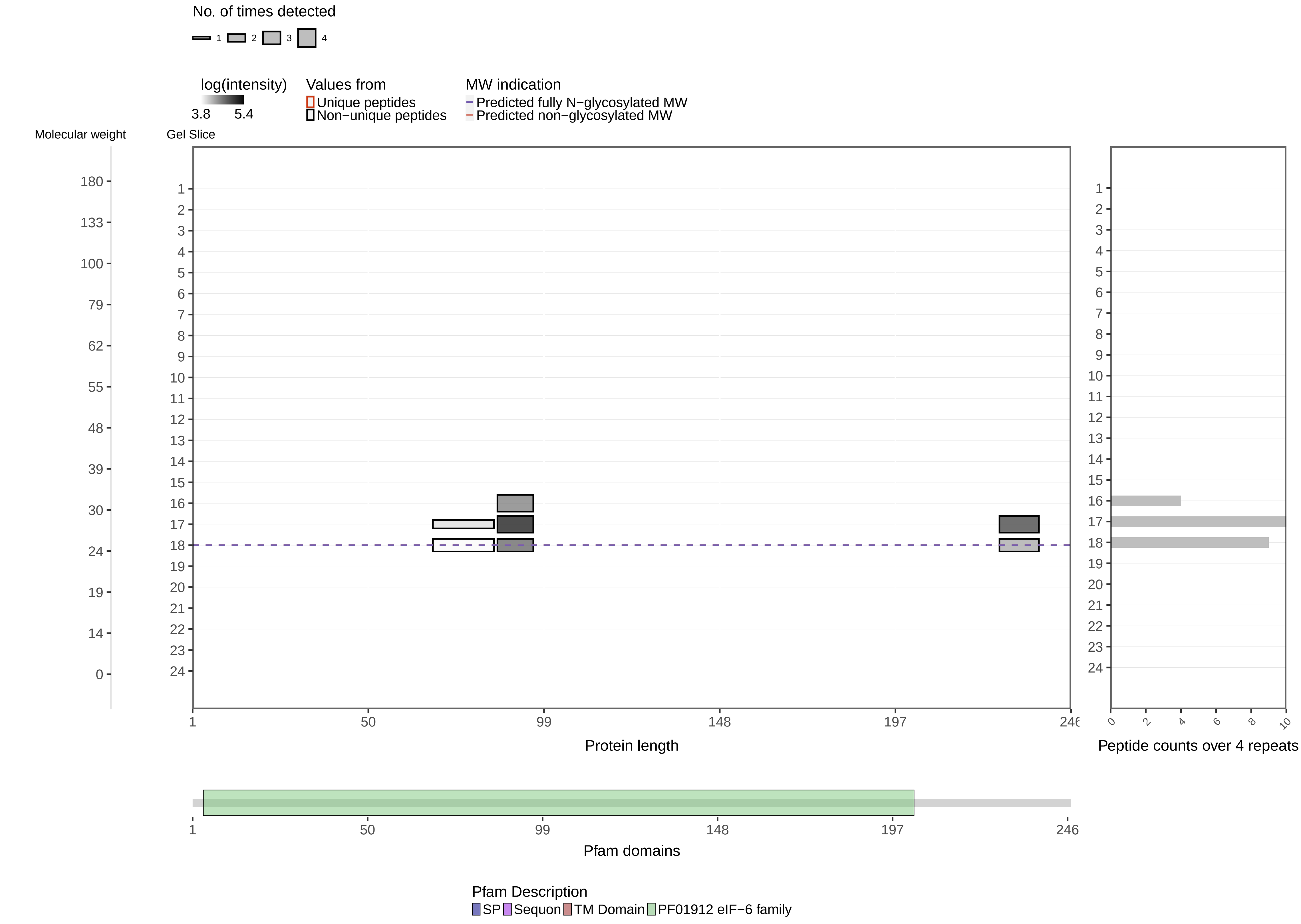


Protein sequence:

NDHVALTHTDLDKETEEMIADVLGVEVFRQTIAGNILVGSYCSFSNRGGLVHPHTSIEDLDELSTLLQVPLVAGTVNRGSEVIAAGLTVNDWTAFCGSDTTATELSVIESVFKLREAQPSAIVDEMRKSLIDSYV*

### NbL12g04470.1 probable_acyl−activating_enzyme_6 [MW = 60.98 kDa]

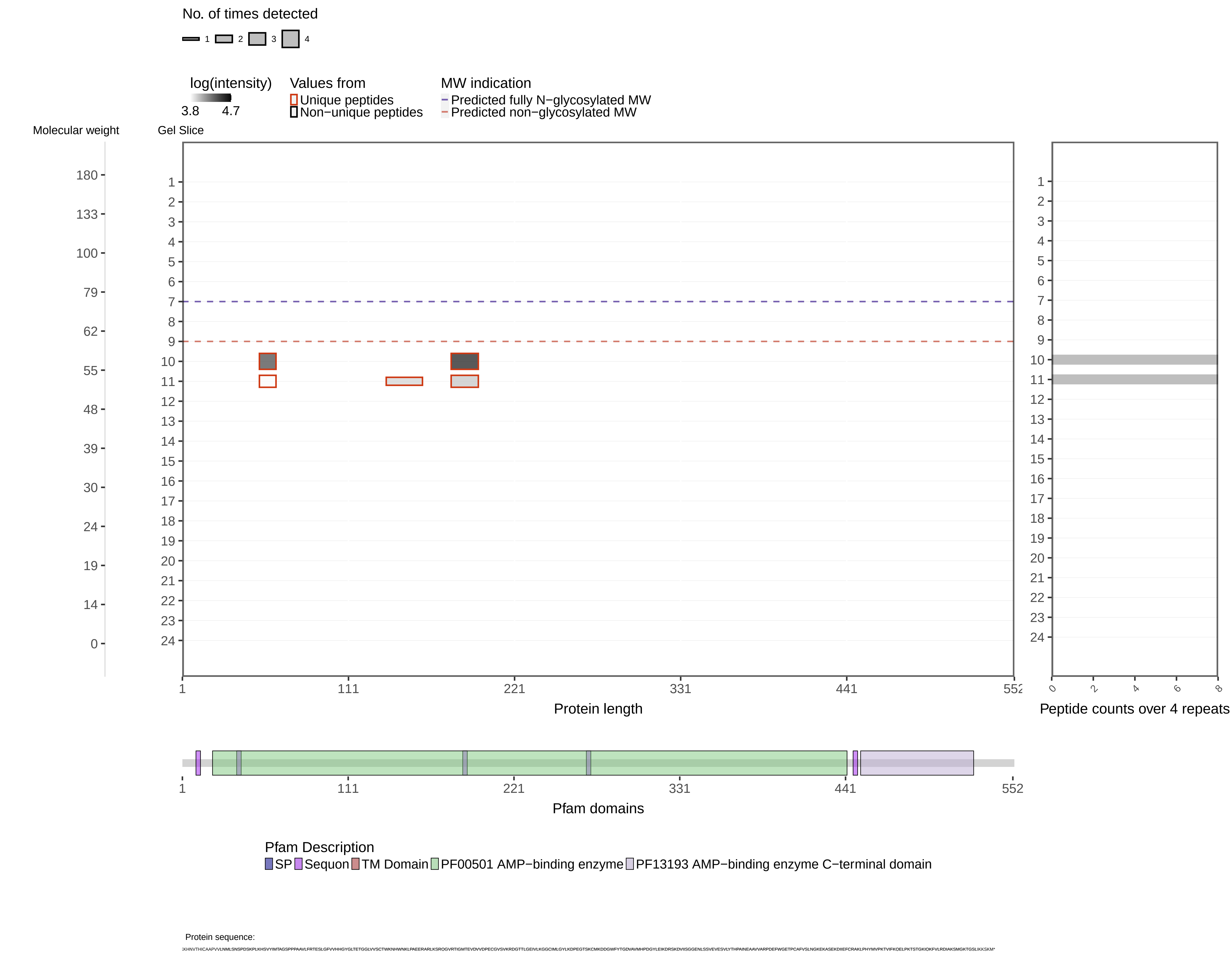

### NbL12g04450.1 protein_YLS9−like [MW = 167.14 kDa]

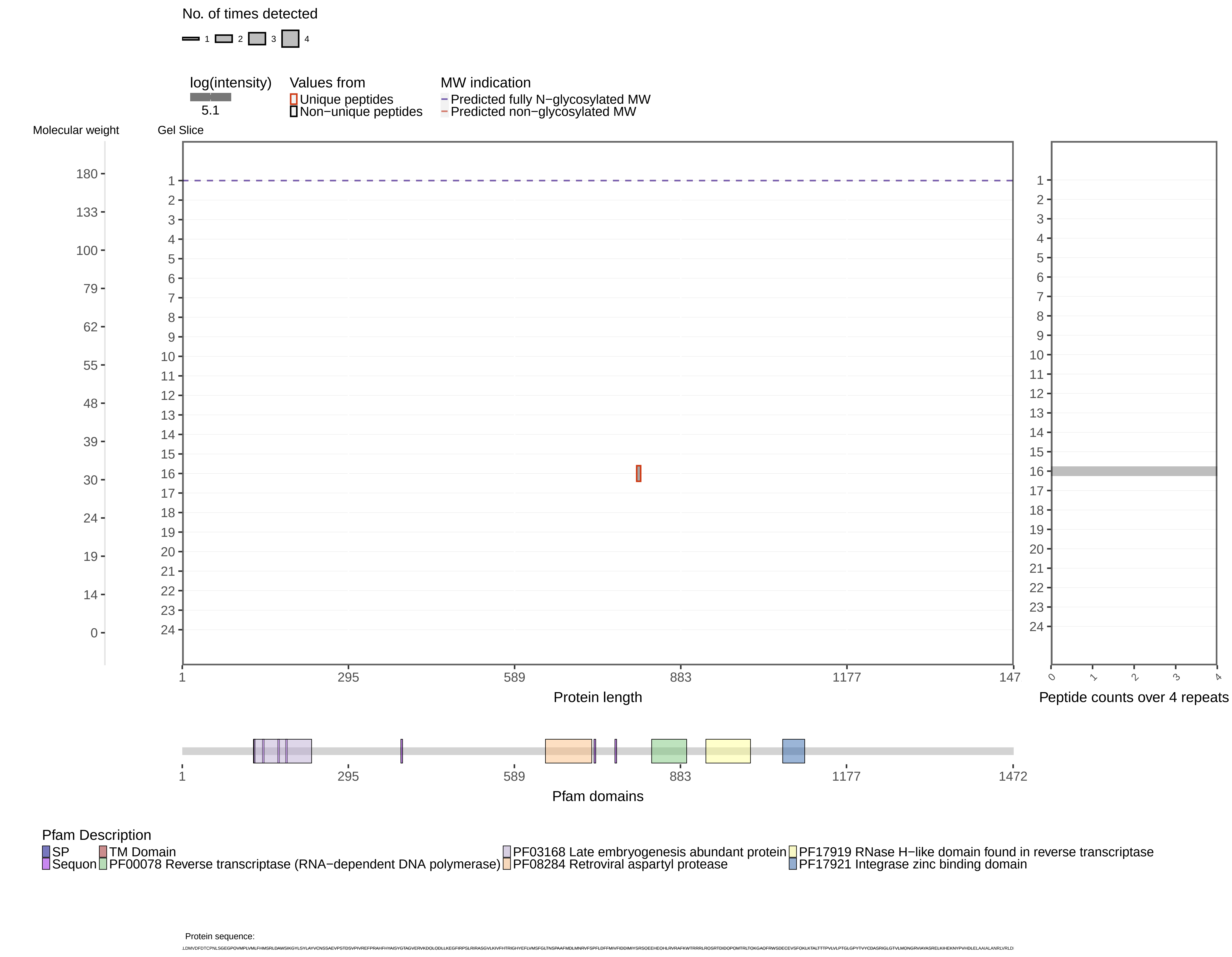

NbL12g04430.1 ras−related_protein_Rab5 [MW = 21.87 kDa]

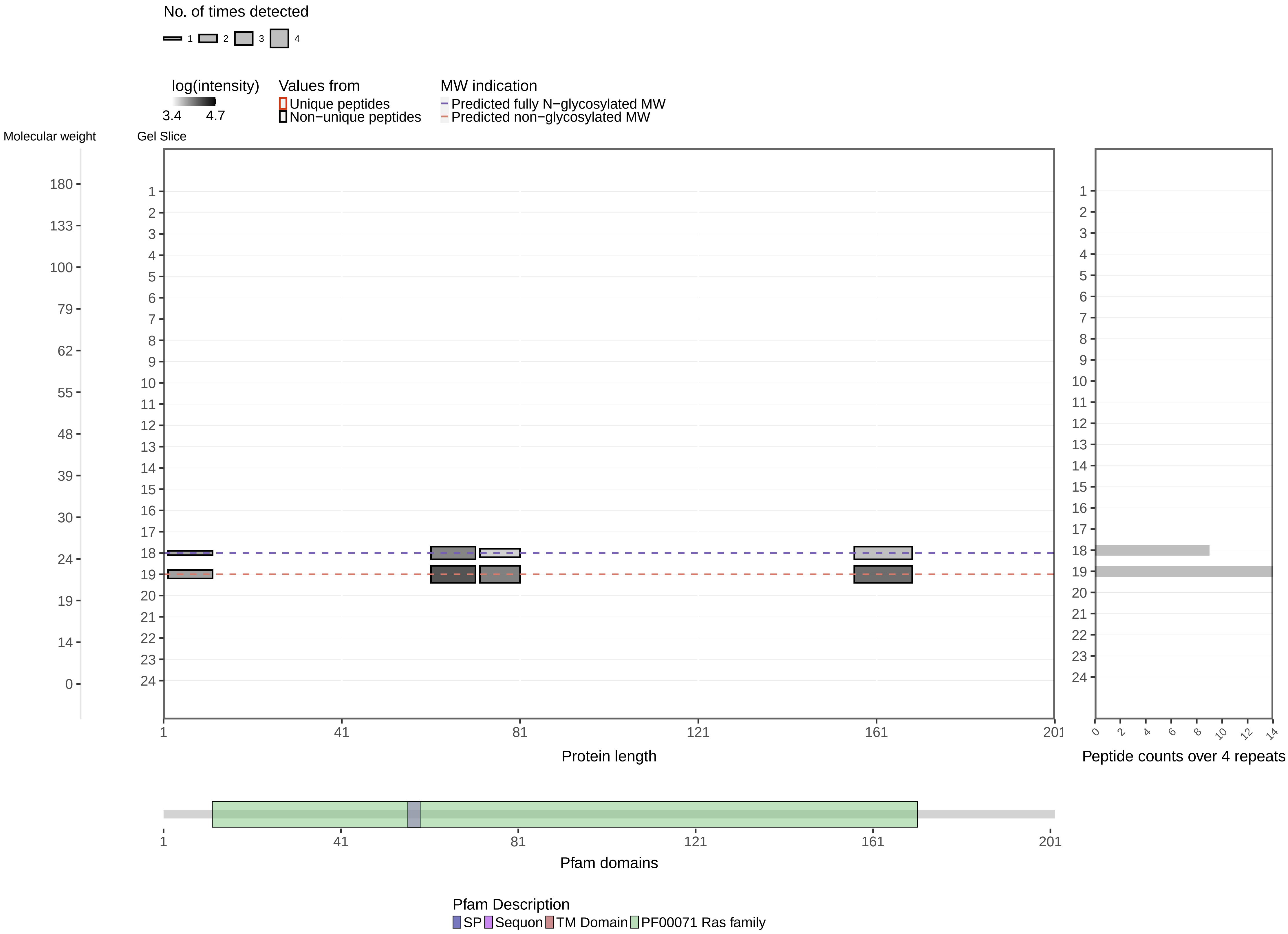


Protein sequence:

VYDITSTESFARAKNWVQELQKQGNPNMVMALAGNKADLEDKRKVTAEEARLYAEENGLFFMETSAKTATNVNDIFYEIAKRLPRAQPAQNPAGMVLVDRPEQGSQAASCCT*

### NbL12g03940.1 protein_PHLOEM_PROTEIN_2−LIKE_A9−like [MW = 19.3 kDa]

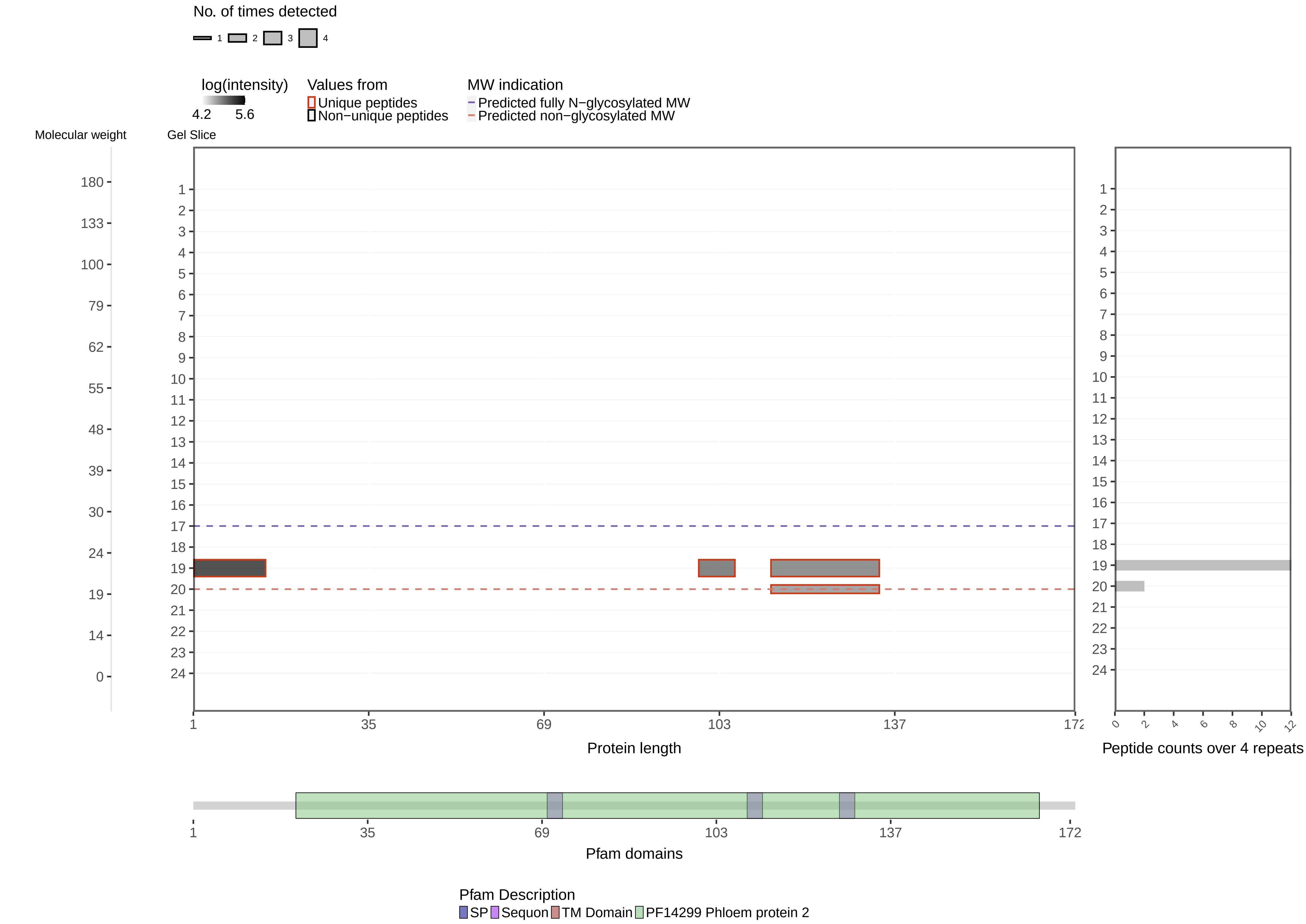


Protein sequence:

'DVKFHLKLTADAFGWNDLPIYFMVKSGSGSLWKKVNLTKDGPTNPDGTYVVPNNLTVTVGKGGNDENKLYFGLYEVWSGKWKGGLRIQKVVITKSVEG*

### NbL12g03820.1 delta(24)−sterol_reductase−like [MW = 66.11 kDa]

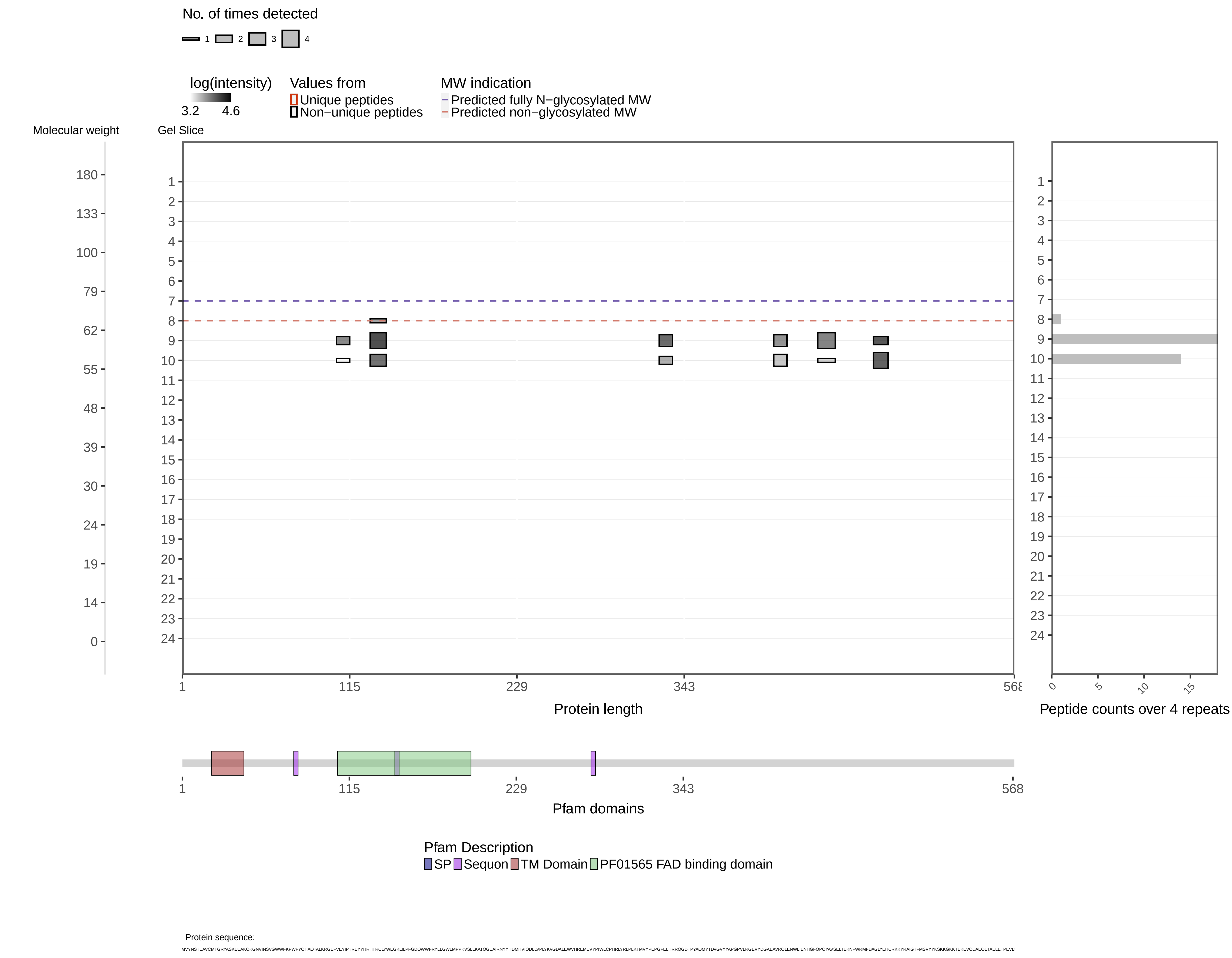

NbL12g03730.1 photosystem_I_P700_chlorophyll_a_apoprotein_A1 [MW = 141.14 kDa]

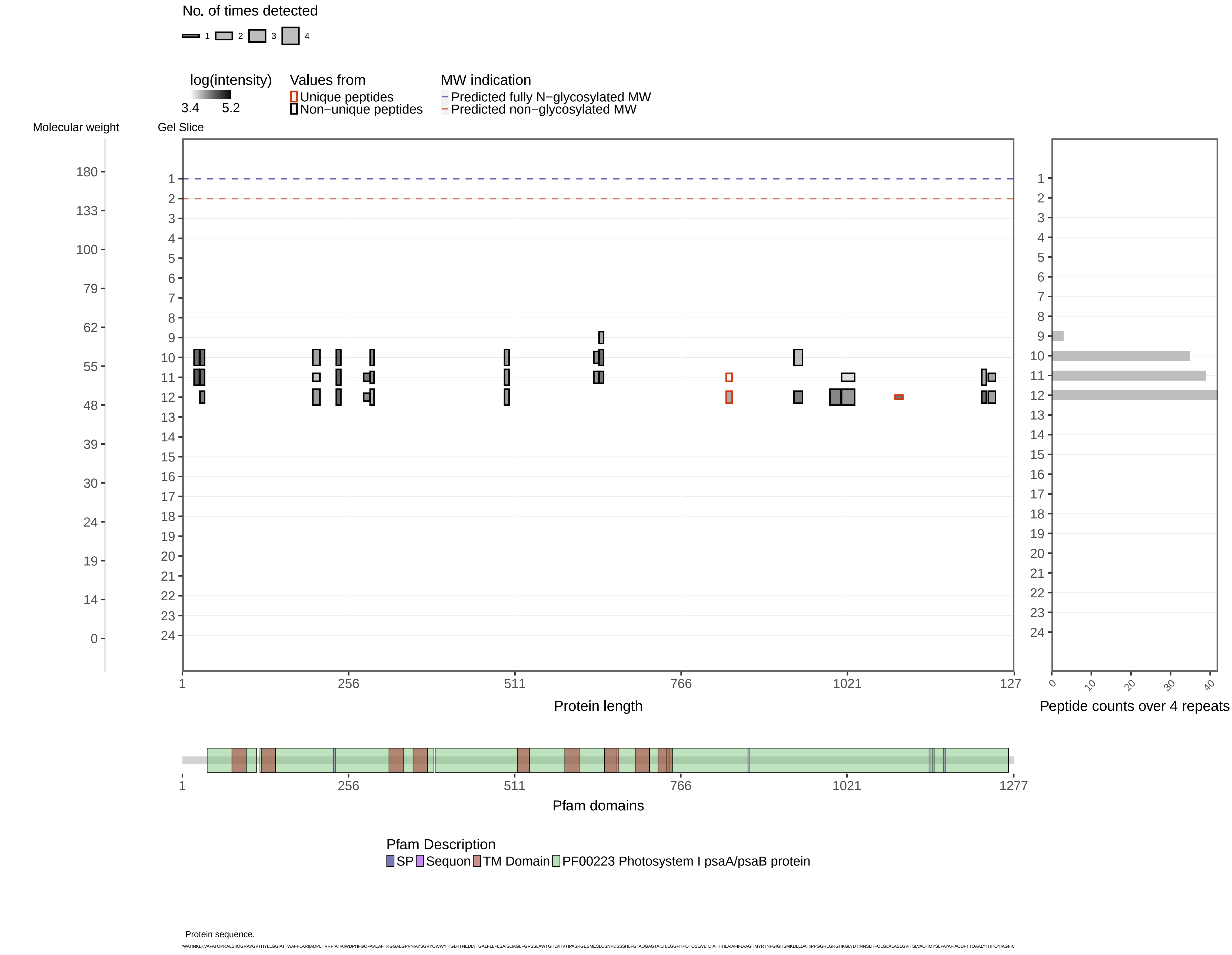

### NbL12g03720.1 photosystem_II_CP43_reaction_center_protein [MW = 51.98 kDa]

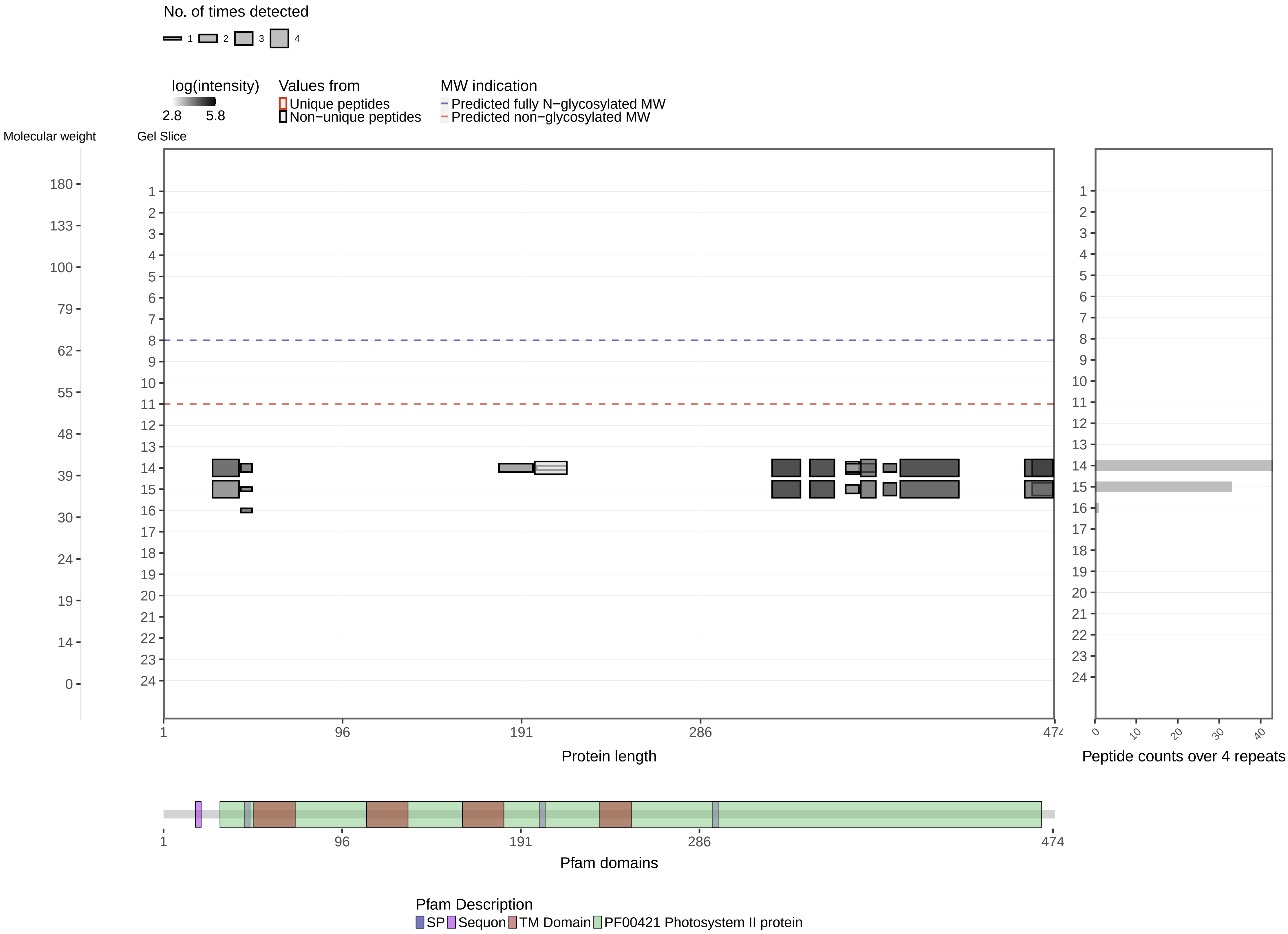


Protein sequence:

VDDLEDIIGGHVWLGSICILGGIWHILTKPFAWARRALVWSGEAYLSYSLGALSVFGFIACCFVWFNNTAYPSEFYGPTGPEASQAQAFTFLVRDQRLGANVGSAQGPTGLGKYLMRSPTGEVIFGGETMRFWDLRAPWLEPLRGPNGLDLSRLKKDIQPWQERRSAEYMTHAPLGSLNSVGGVATEINAVNYVSPRSWLATSHFVLGFFLFVGHLWHAGRARAAAAGFEKGIDRDFEPVLFMTPLN*

NbL12g03690.1 ferredoxin−−NADP_reductase_root−type_isozyme_chloroplastic−like [MW = 41.95 kDa]

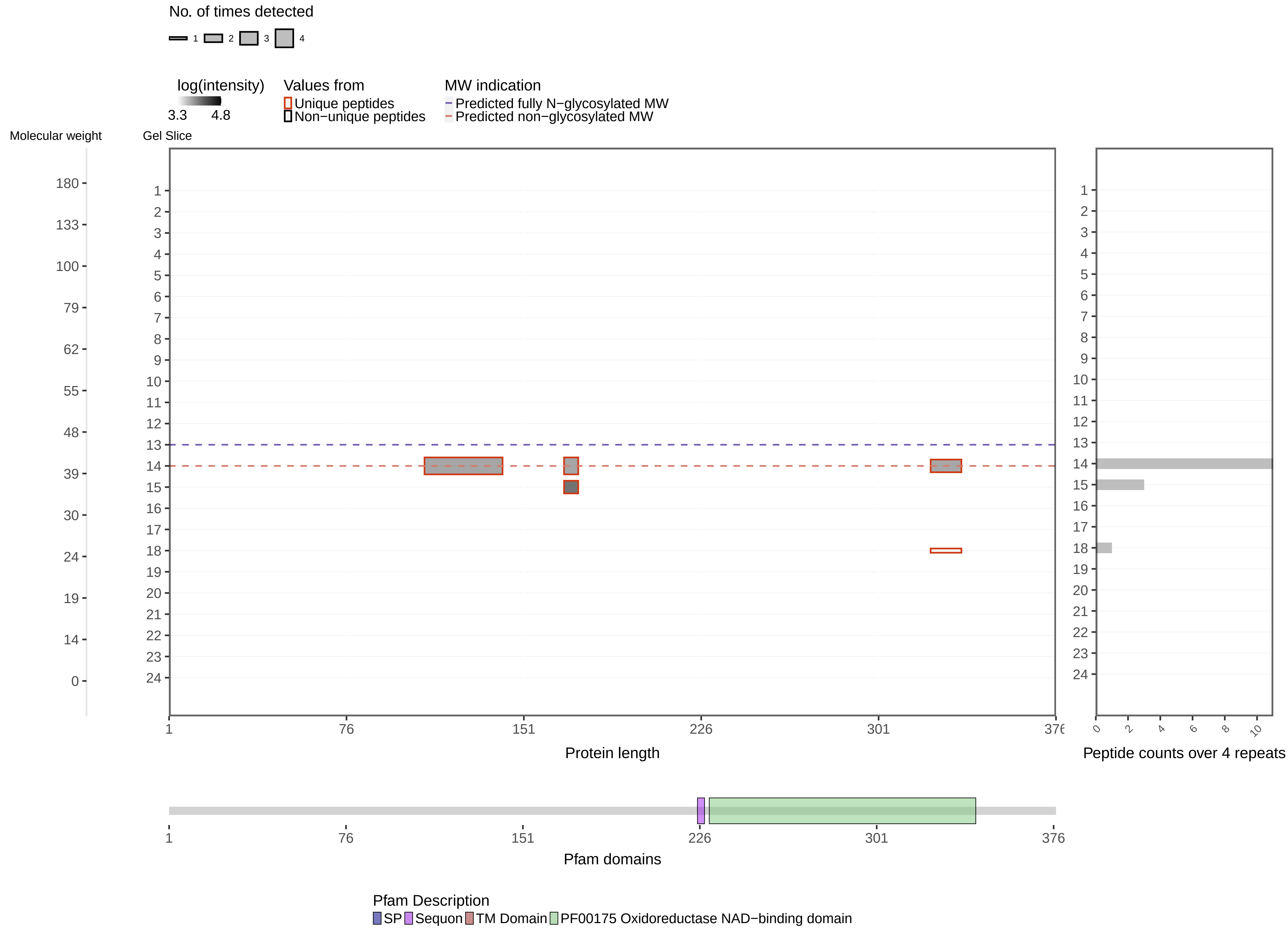


Protein sequence:

YYDPETGKEDPSKNGVCSNFLCDSKPGDKVKITGPSGKIMLLPEDNPNATHIMIGTGTGVAPFRGYLRRMFMESVPTKFNGLAWLFLGVANTDSLLYDDEFTKYLNDYPGNFRYDRALSREOKNNKGGKMYVODKIEEYSDEIFKLLDEGAHIYFCGLKGMMPGIODTLKRVAEGRGESWEOKLSOLKKKKOWHVEVY*

NbL12g03650.1 C−factor−like [MW = 32.61 kDa]

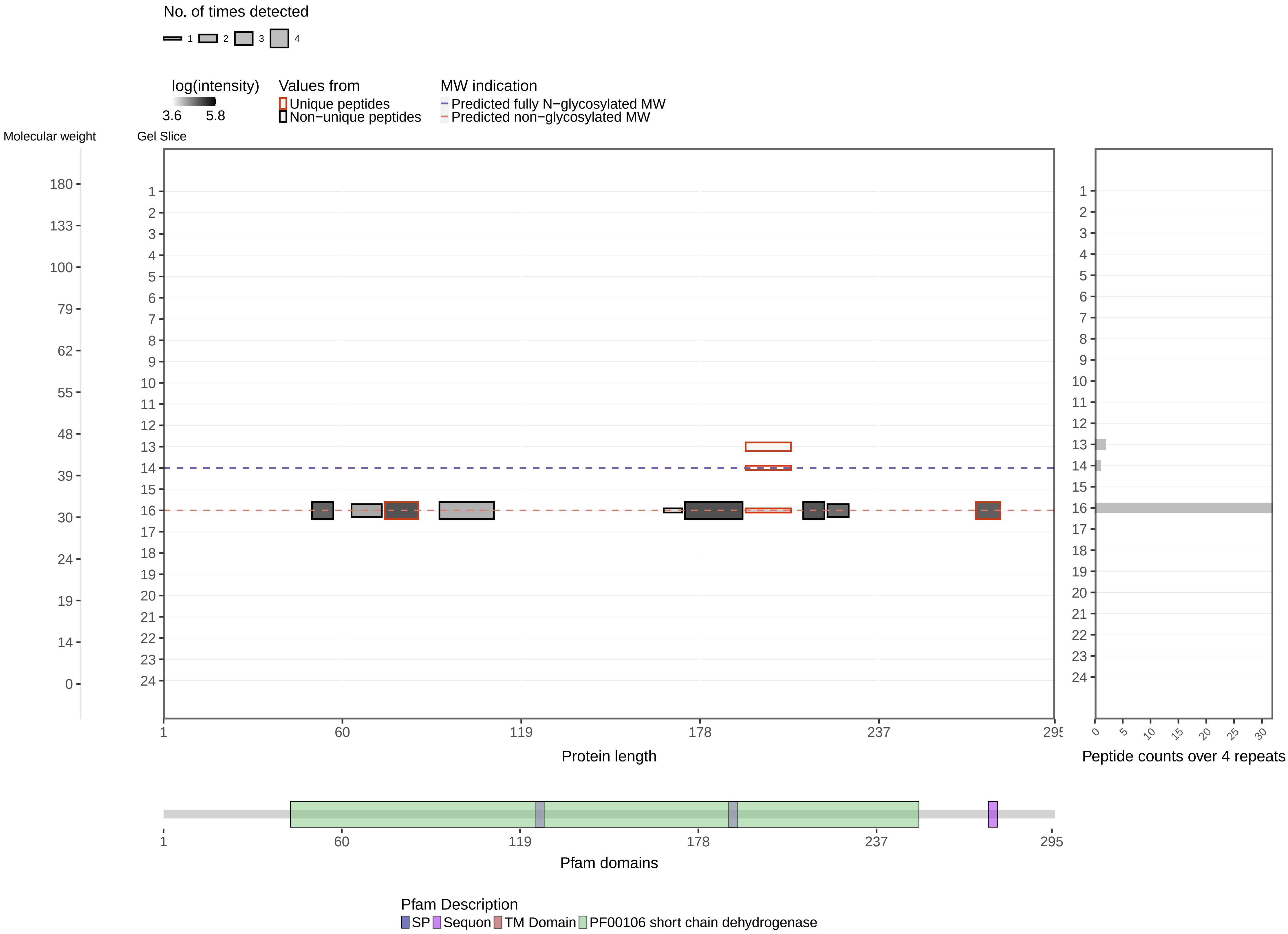


Protein sequence:

VMQPETTLSKVQRSSLLLAYDINAVGPILVIKHMWPLLKAGGGSGTDKDFAVVANLSARVGSIGDNSLGGWHSYRASKTALNQLTKNVAVEFARKKDPIICILLHPGTVDTDLSKPFQRNVPKEKLFTKEYSVQKLLSIINNTKRSDNGKFFAWDAQQIPW*

### NbL12g03630.1 obg−like_ATPase_1 [MW = 44.37 kDa]

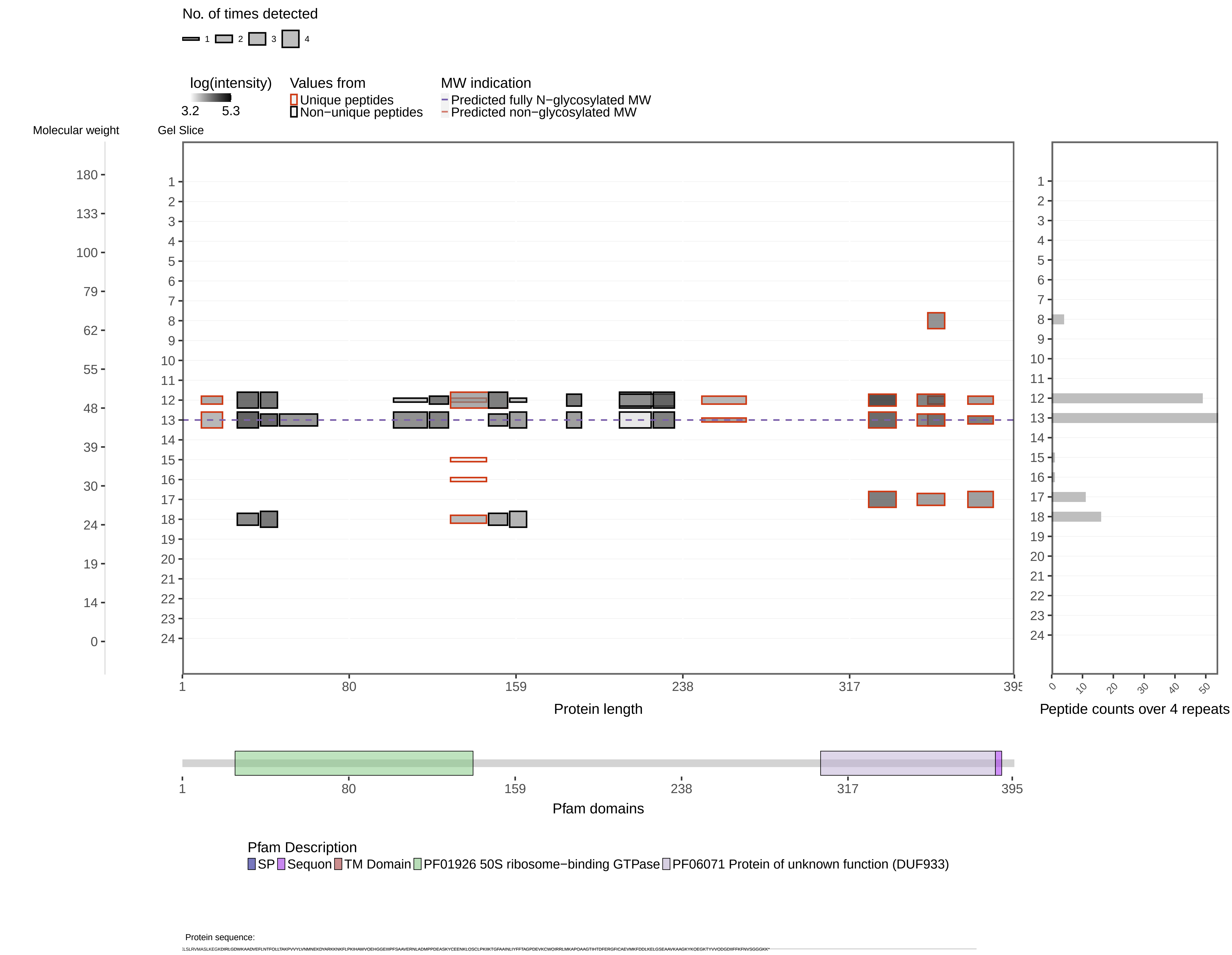

NbL12g03540.1 reticuline_oxidase−like_protein [MW − SP = 58.63 kDa]

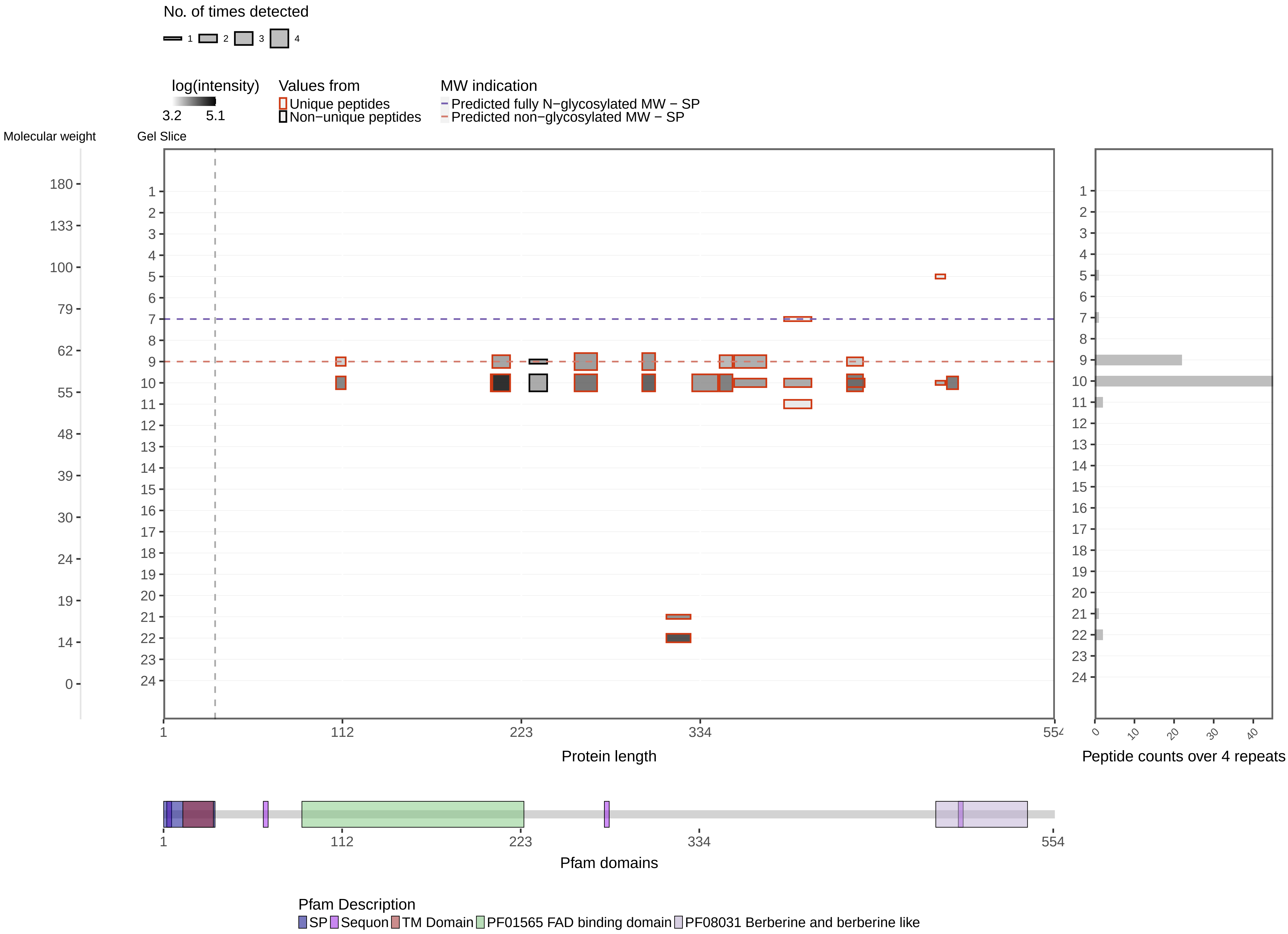


Protein sequence:

VFTVPKTLEQNGTKIIYKWQQVADKIDEDLFIRVIMNVVDKKDKKGEKTIQMAYNSLFLGTSDRLLEIMNENFPELGLTQKDCTEMSWIKSILYIAGYPSNTPPEVLLQGKSLFKNYFKAKSDFVKEPIPEDGLEGLWKRLMEEDSPLMIWNPYGGMMAKISESETPFPHRKGVIYMVQYLTLWNDPDKELATKHFDWIRRLYNYMTPYASMFPRQAYVNYRDLDLGINKNGSSSFIQASIWGNKYFKDNFNRLVKVKTKVDPENYFLHEQSIPTLPISRNGKGKSMNQ*

NbL12g03520.1 reticuline_oxidase−like_protein [MW − SP = 58.91 kDa]

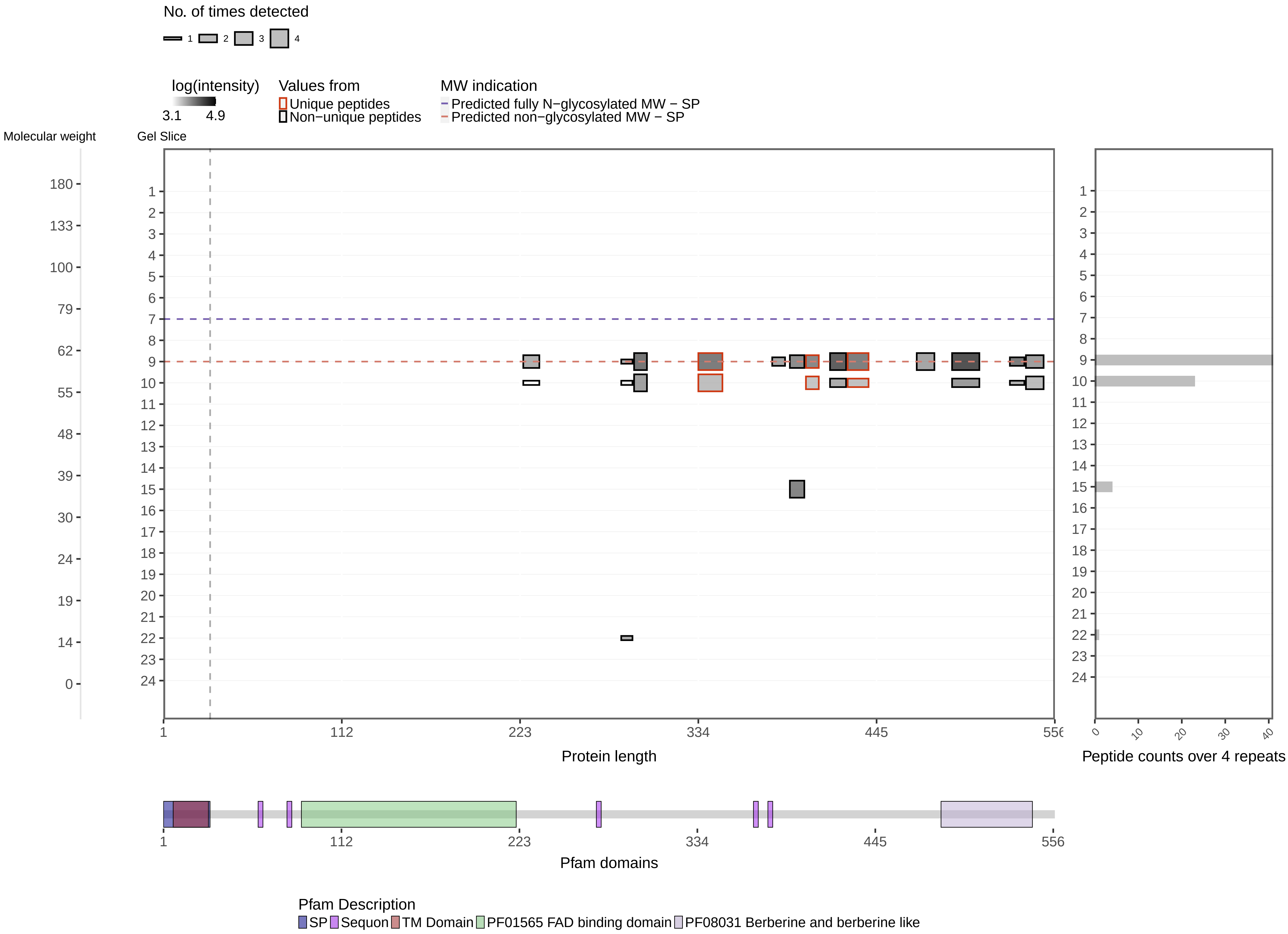


Protein sequence:

FLDQNATDAVLQWQNVVNKIDNDLFIRLLVQPITVKNKEKKVKGKKNAQQKSIRVTFIALFLGDSSRLMSLVSNELPALGLTKQDCFEMSWIDSVLQWANFDNKTKPIALLNRTGDPLNFLKRKSDYVQEPIPRDVLESIFQKMISLGKAGLVFNPYGGRMTEIPEGETPFPHRAGILFKIQYSVNWNEEGAAAEKEYLTQIRDLYSFMTPFVSKNPRQAYLNYRDLDIGTNDQGPHSLEEGRVYGTMYFKNNFDRMVKVKSAVDPTNFFRNEQSIPTQGTTKPRKIGE*

NbL12g03370.1 DNA_damage−binding_protein_1 [MW = 122.23 kDa]

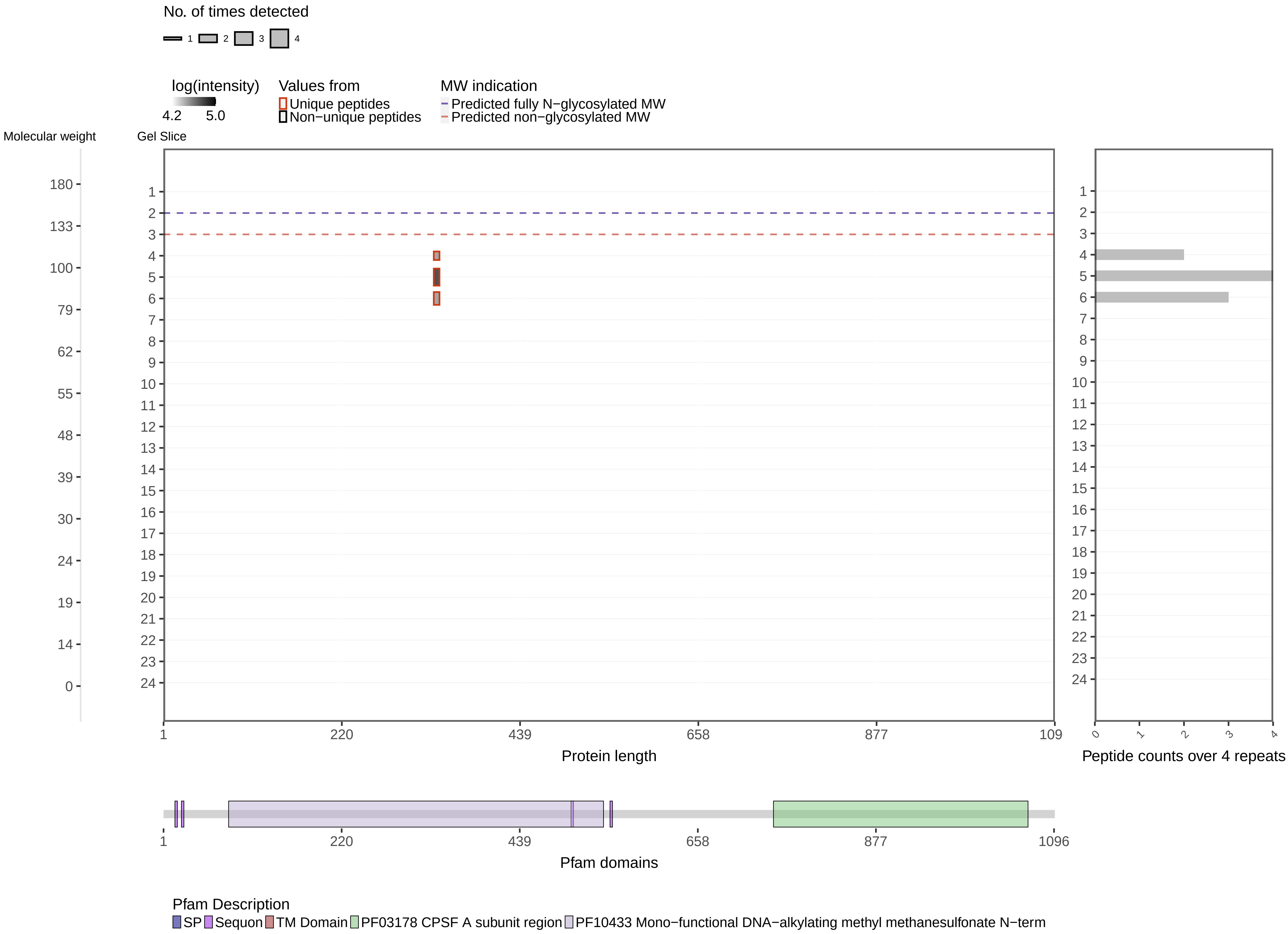


Protein sequence:

ISCLDINPIGENPNFSQIAAVGMWTDISVRIYSLPDLNLITKEQLGGEIIPRSVLMCSFEGISYLLCALGDGHLLNYVLSMSSGELTDRKKVSLGTQPITLRTFSSKDATHVFAASDRPTVIYSSNKKLLYSNVNLKEVSHMCPFNVAAFPDSLAIAKEGELTIGTIDEIQKLHIRSIPLGEHARRISHQEQTRTFALCSVKYTQSNADDPEVHFVRLLDDQTFEFISTYPLDQFEYGCSILSCSFSDDSTVYYCVGTAYVMPEENEPTKGRILVFIVEDGKLQLIAEKETKGAVYSLNAFN

### NbL12g03330.1 40S_ribosomal_protein_S4−like [MW = 29.86 kDa]

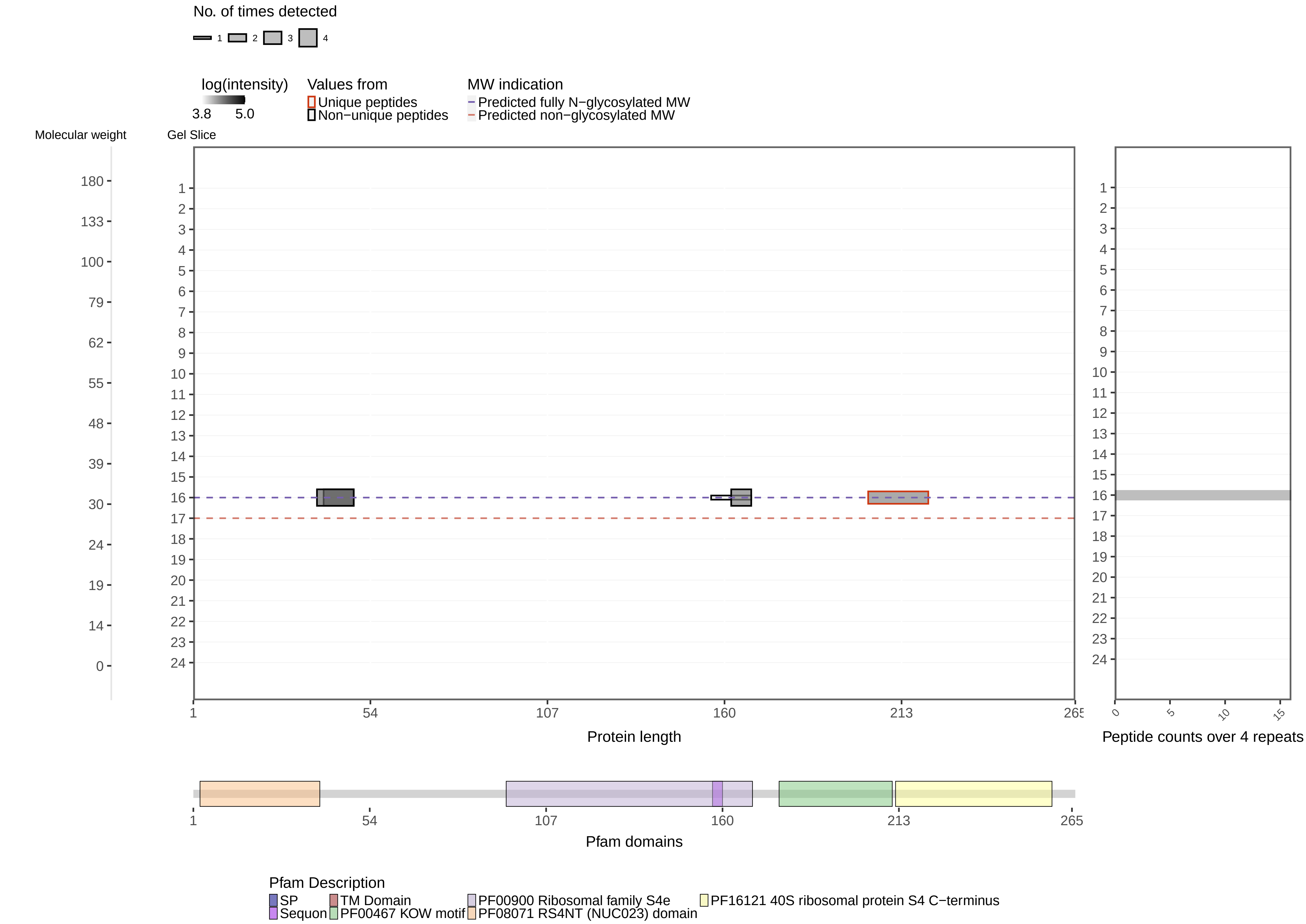


Protein sequence:

:AKFKLCKVRSVQFGQKGIPYLNTYDGRTIRYPDPLIKANDTIKLDLENNKIVDFIKFDVGNVVMVTGGRNRGRVGVIKNREKHKGSFETVHIHDAQGHEFATRLGNVFTLGKGTKPWVSLPKGKGIKLSIIEEARKRLAAQSATTA*

NbL12g03250.1 UBP1−associated_protein_2C−like [MW = 43.84 kDa]

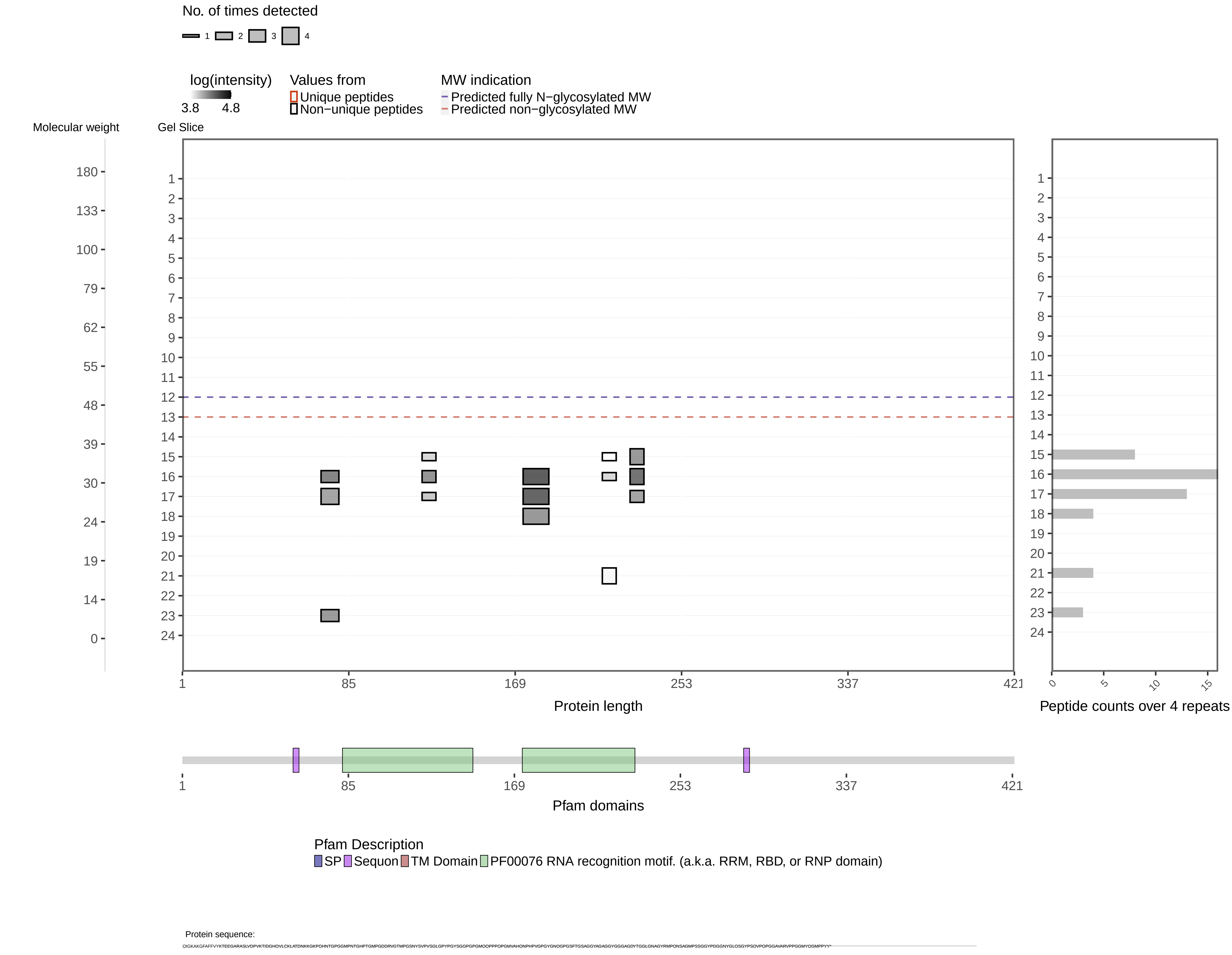

NbL12g03230.1 p21−activated_protein_kinase−interacting_protein_1 [MW = 36.72 kDa]

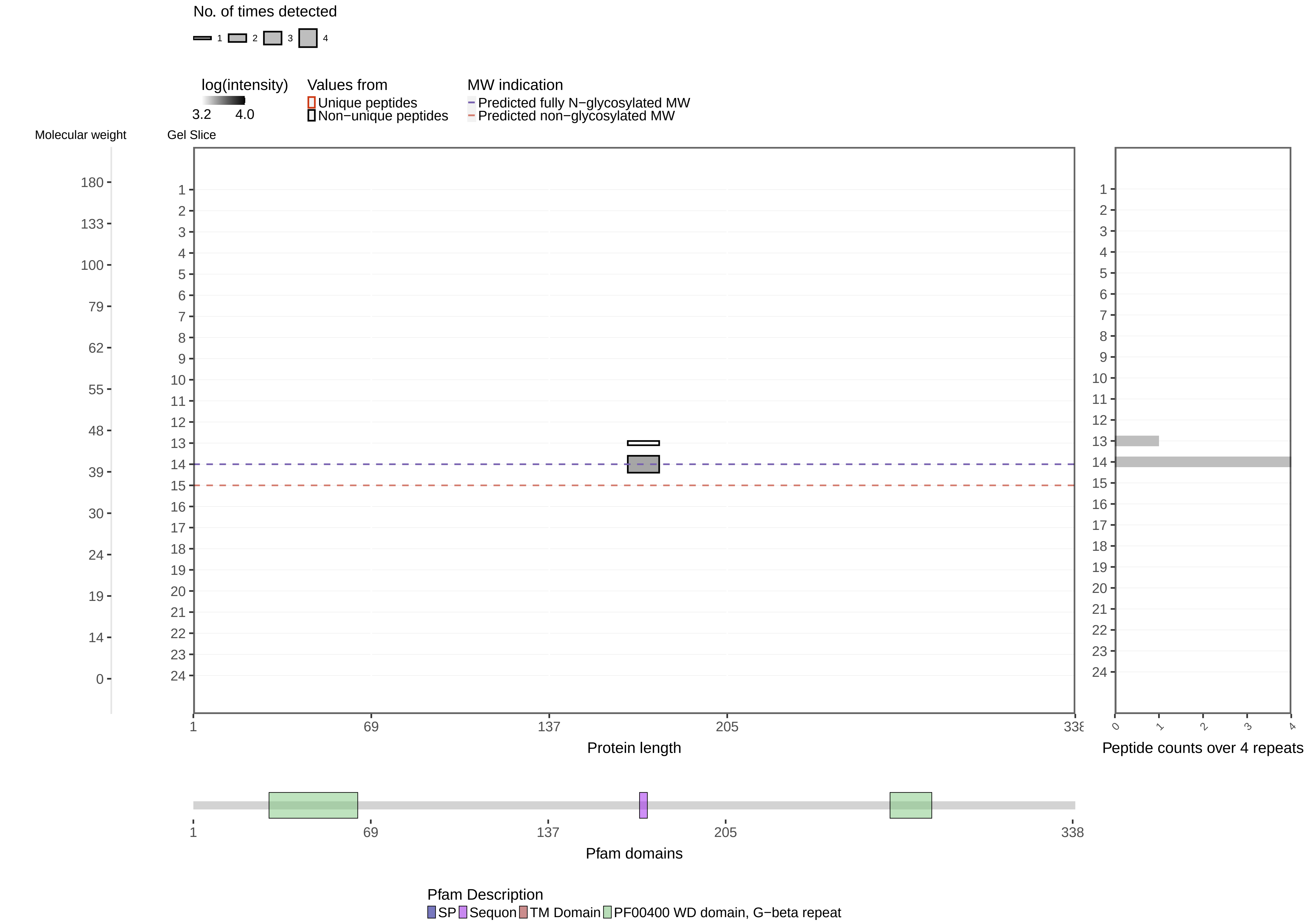


Protein sequence:

RRSFYCRLGKEASLVNYSDTGEKFYMVMDDKISVHESEDAKIILELDNKKKVLCATPGTHGILFTGGEERNIKAWDITSGKVAYSIEDAHSTRVKGIVVLYKNNDGDAEENQHIVASASSDGIIRVWDVRMTRKEKPIPLAEANTKSRLTCLAGSSIKSVKRPLVGNTASNGQQLDAGEES*

NbL12g03200.1 uncharacterized_LOC107786033 [MW = 24.77 kDa]

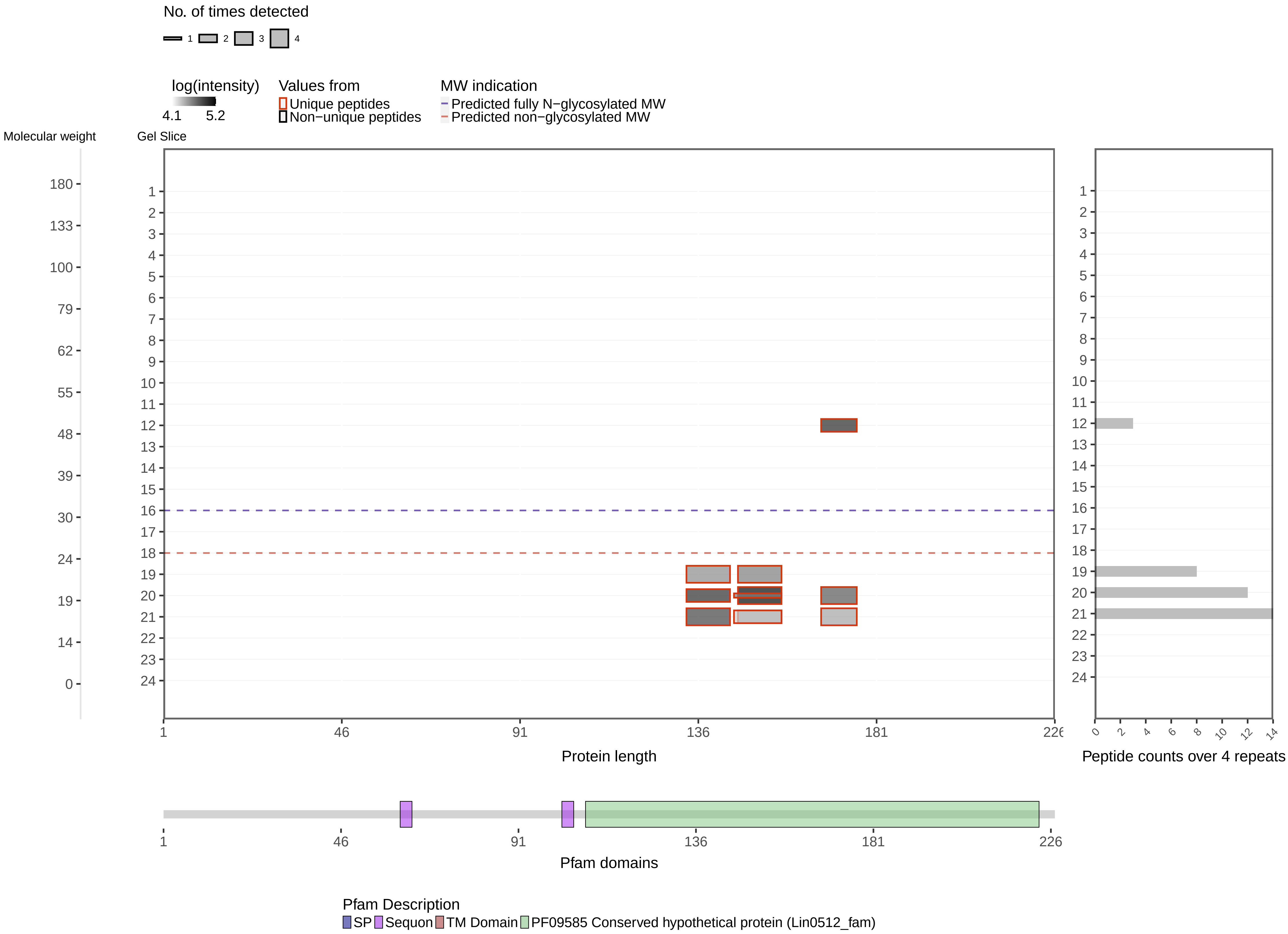


Protein sequence:

NSSMTLLFVEMGVGYDQHGQDITSAAMRACKNAISSNSIPAFRRGSIPGVSFEQMKLQIKLGVPRPLQQSLDLEKVKSVFPYGRILIIEVVDGGLICSSGVQVEDMGDKNDDCYIVNAAIYVGY*

NbL12g03170.1 proteasome_subunit_alpha_type−5−like [MW = 26.08 kDa]

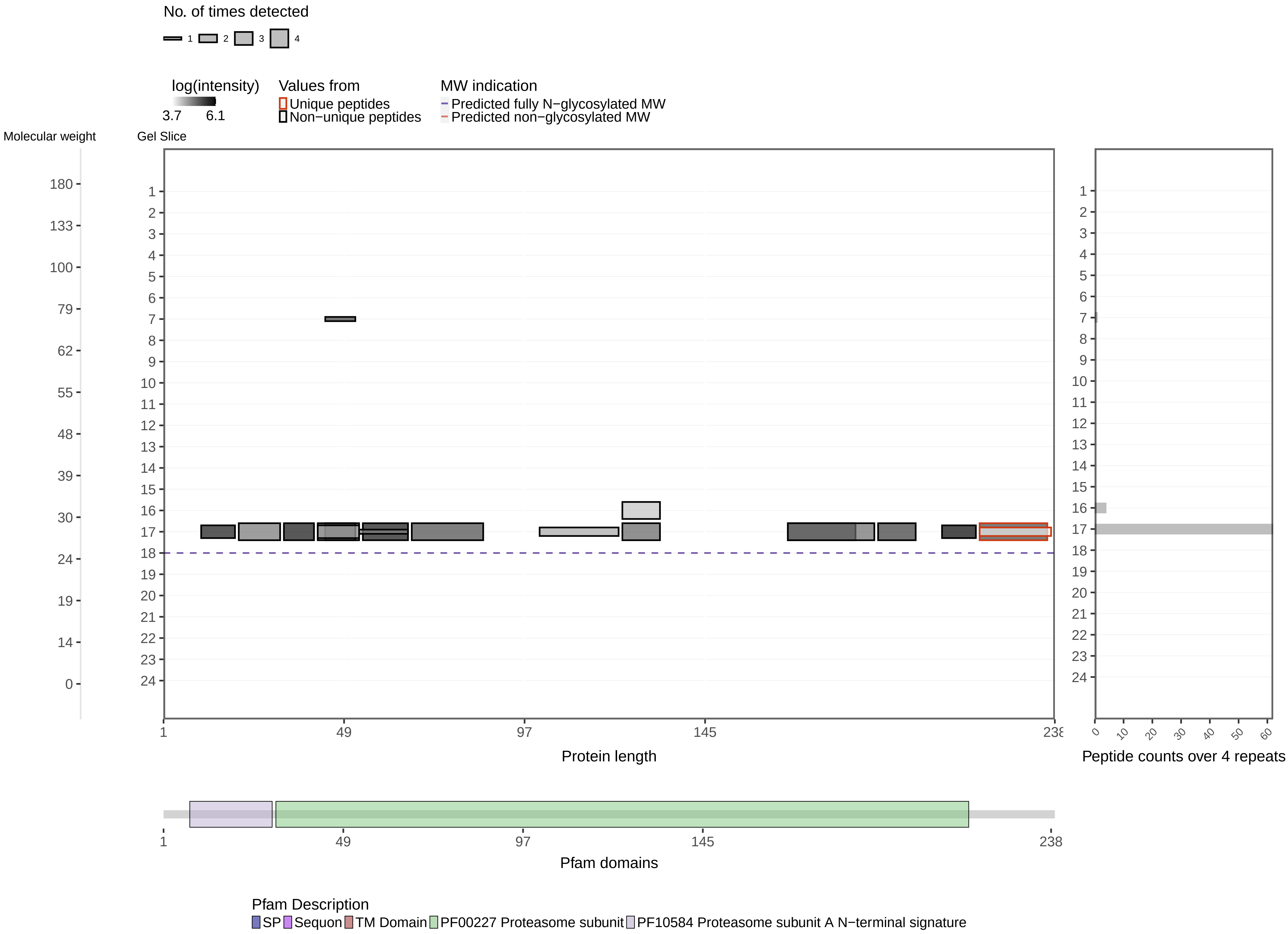


Protein sequence:

TVESTTQALCDLALRFGEGDEESMSRPFGVSLLIAGHDENGPNLYYTDPSGTFWQCNAKAIGSGSEGADSSLQEQYNKDLTLKEAEAIALSILKQVMEEKVTPNNVDIARVSPTYHLYSPSEVEDVISRL*

### NbL12g03070.1 40S_ribosomal_protein_S11−like [MW = 17.66 kDa]

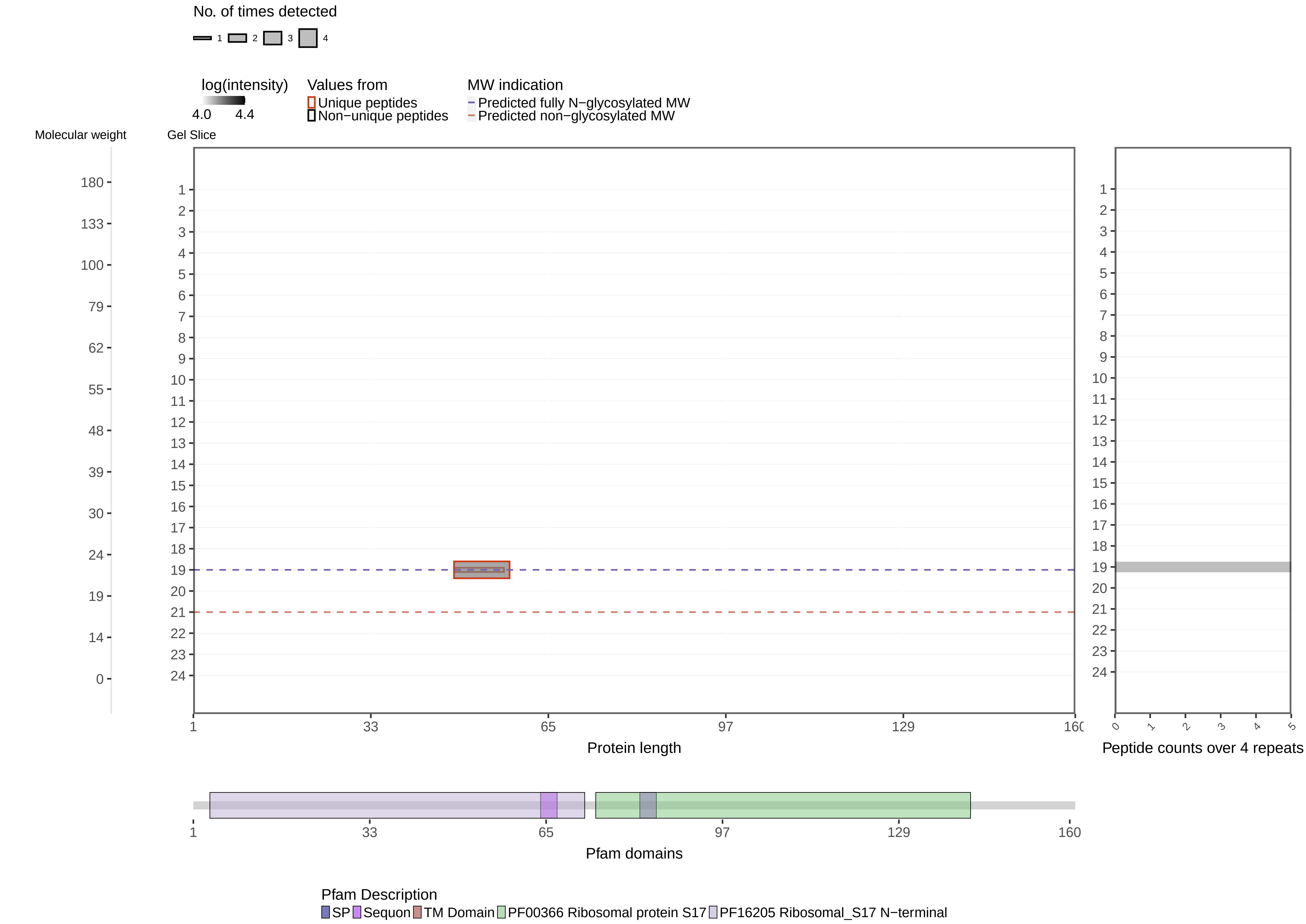


Protein sequence:

'SIRGRILAGTCHSAKMNRTIIVRRNYLHYVKKYQRYEKRHSNIPAHISPCFRVKEGDHVIIGQCRPLSKTVRFNVLKVIPAGSAGGGKKAFTGM*

### NbL12g03020.1 uncharacterized_LOC109211931 [MW = 33.69 kDa]

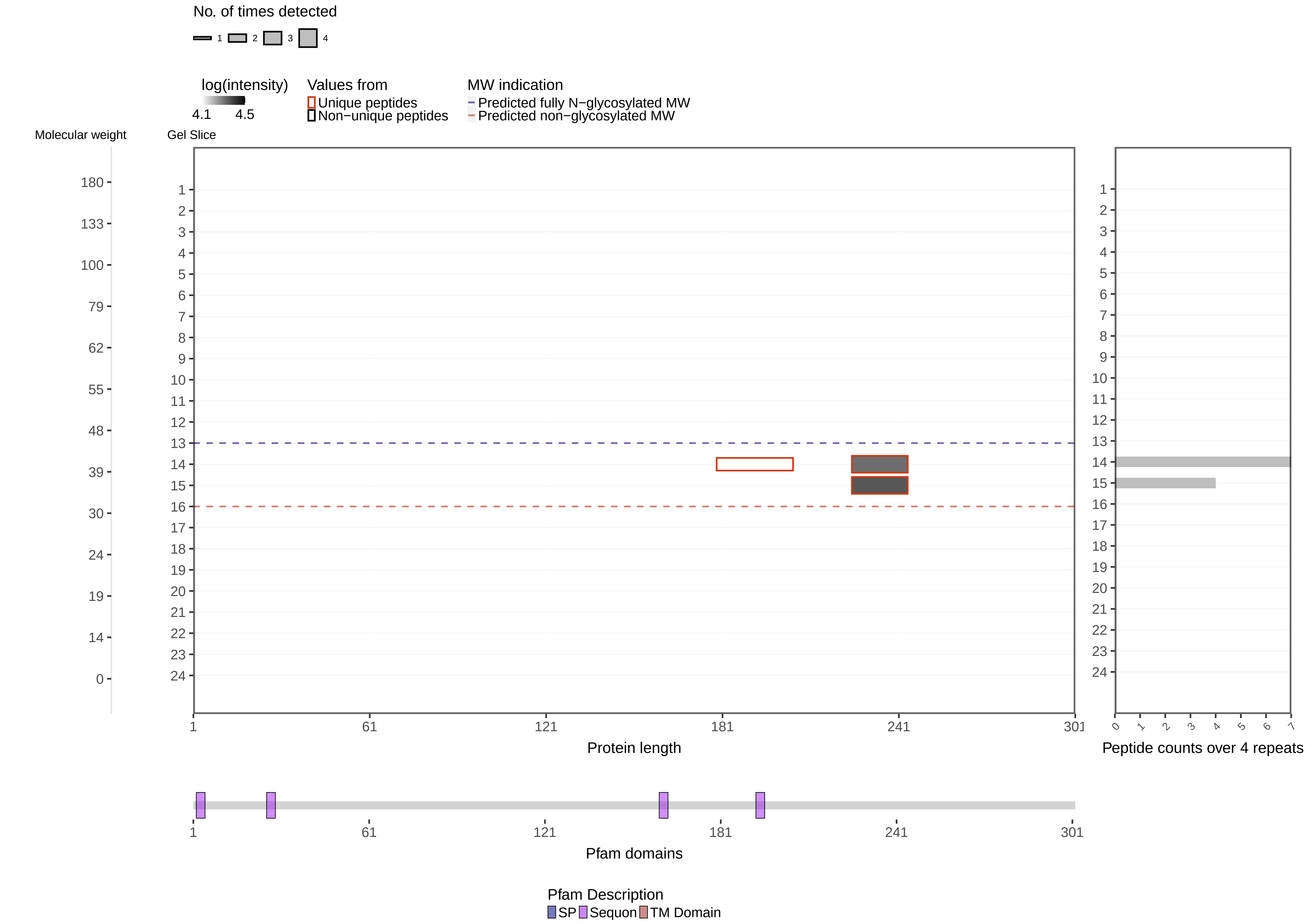


Protein sequence:
GELGIYAFKTLKNANIQVRVERLTNKSGTPNMMDIEAFSLRYREKLDEDEVAGSIPNNVSLEVSSPGVERVIRIPQDLERFKDRPMYVKYVSDEVAEGGSSSEHDGVLRLVSFNLETNACIWGIADVRVNREKAGKGKPLSKKQRDWRLETSFTSLRLVRLYSEF*

### NbL12g02940.1 uncharacterized_protein_At2g34160−like [MW = 18.29 kDa]

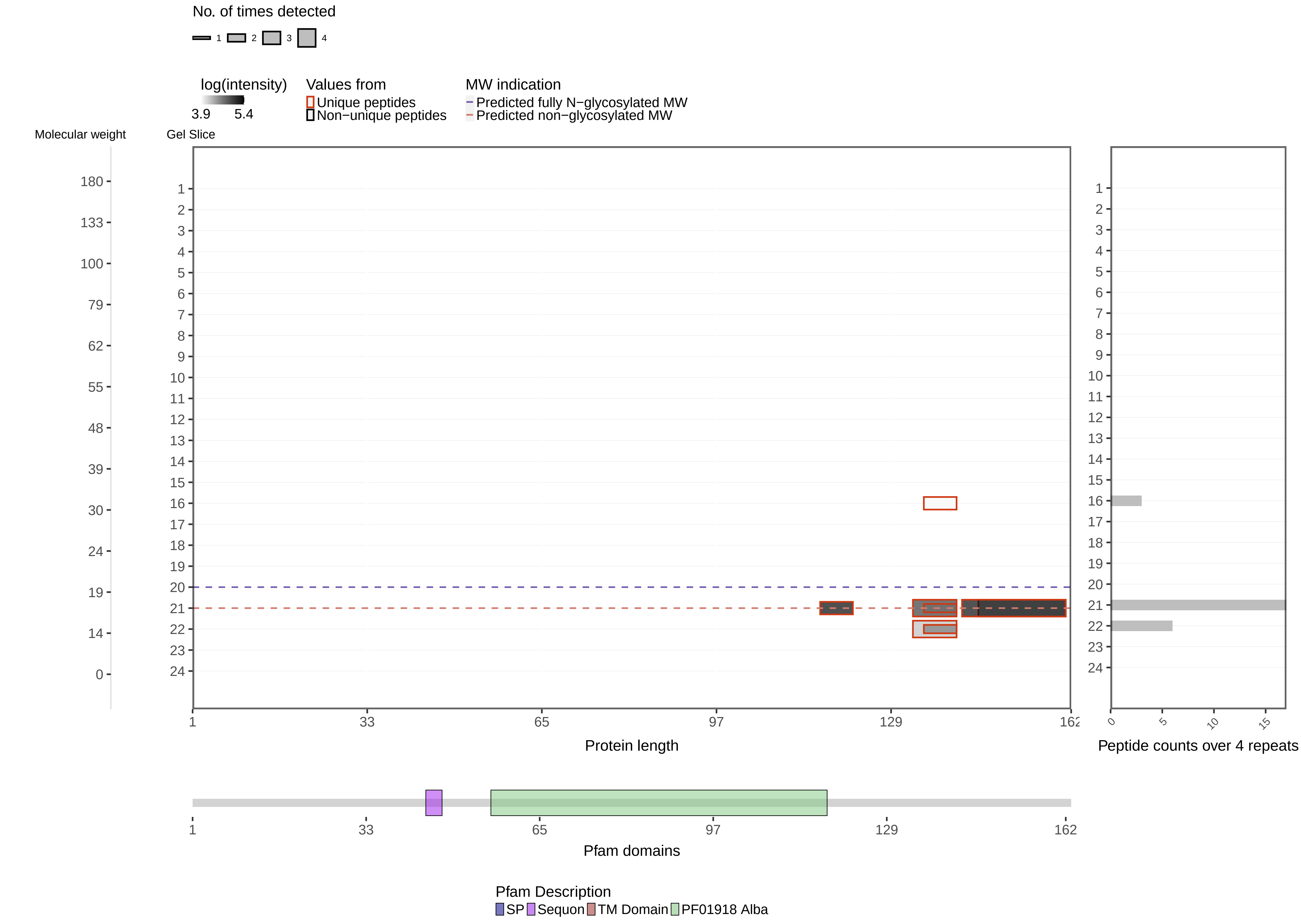


Protein sequence:

FFYVNLAKRYMQQYNEVELSALGMAISTVVTVAEILKNNGFAVEKKIRTLTVDMRDEPGARPIPKAKIEVVMGKTEKFDELMAAEAEQNGDNEE*

NbL12g02780.1 serine_carboxypeptidase−like [MW − SP = 53.98 kDa]

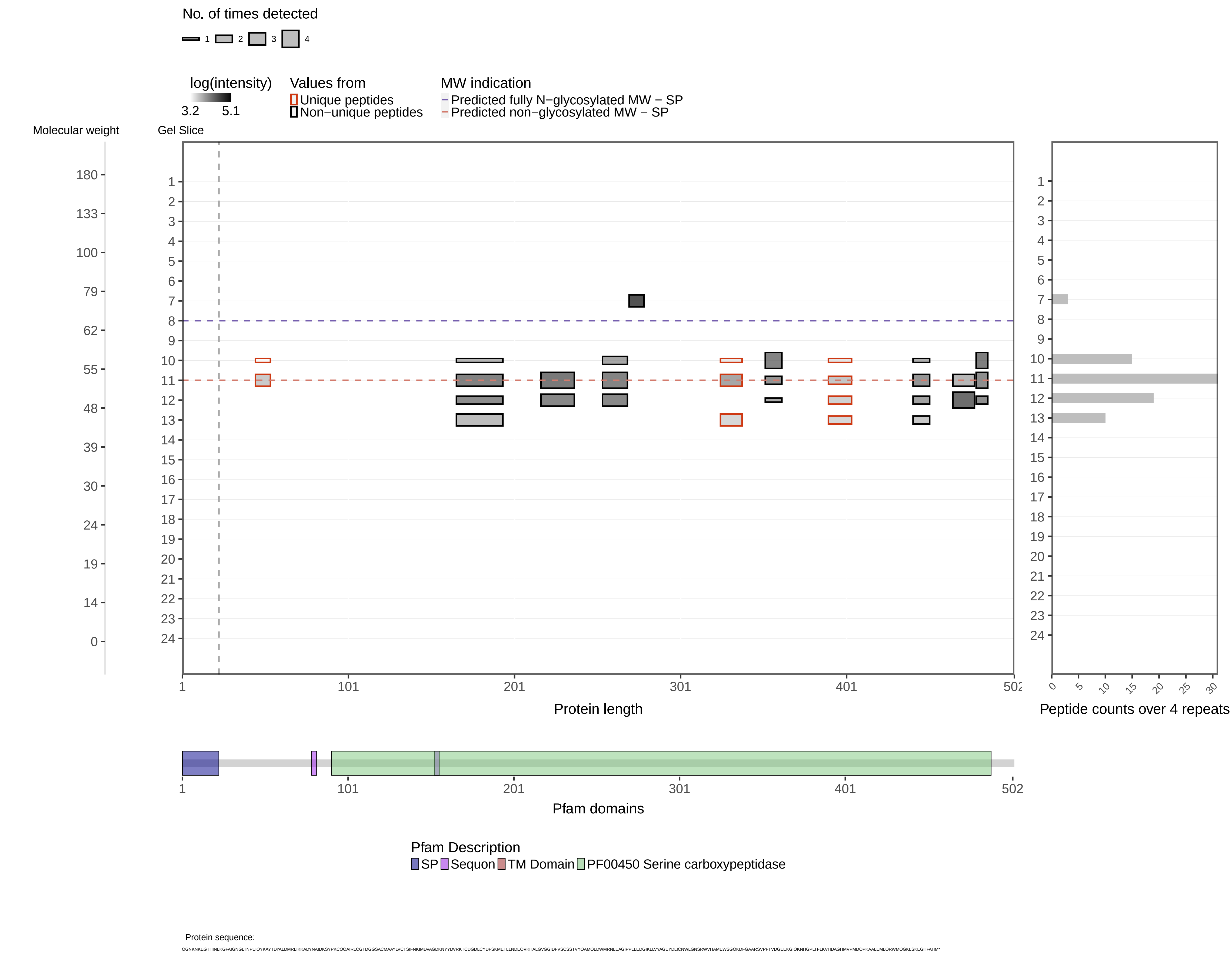

NbL12g02670.1 protein_DETOXIFICATION_21−like [MW = 53.08 kDa]

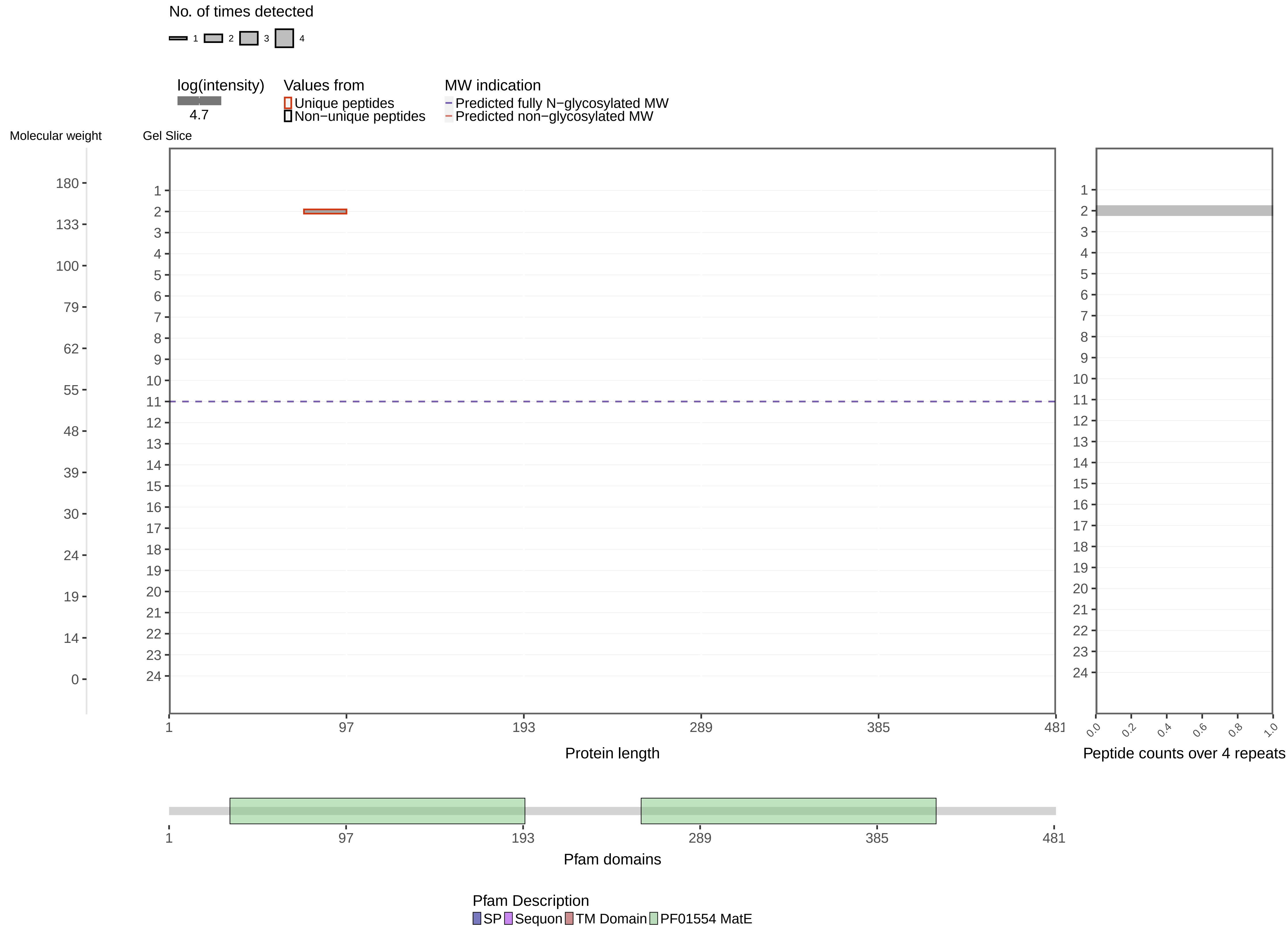


Protein sequence:

### NbL12g02560.1 60S_ribosomal_protein_L12−like [MW = 17.81 kDa]

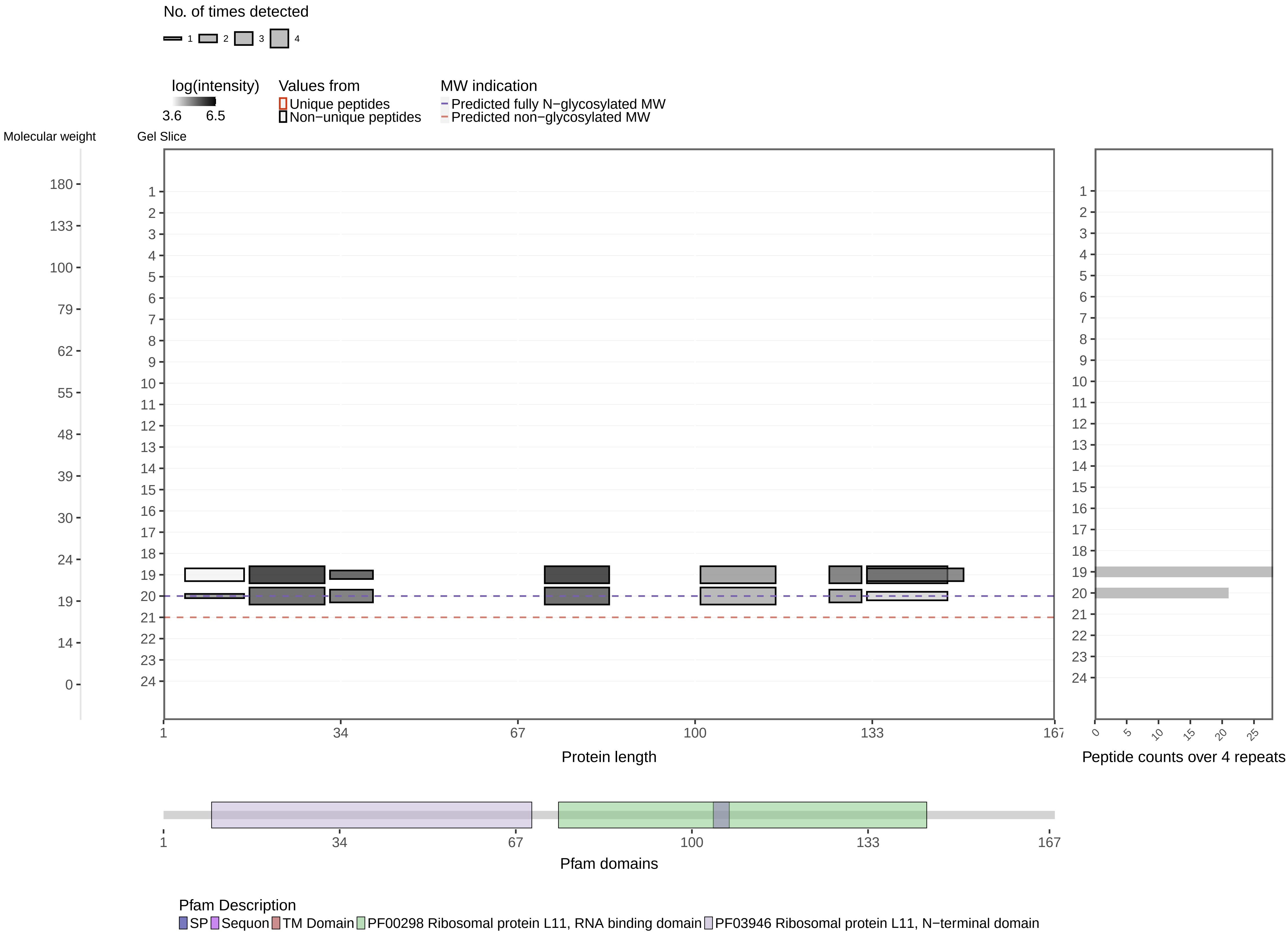


Protein sequence:

AKVSVVPSAAALVIKALKEPERDRKKTKNIKHNGNISLDDVIEIAKVMKPRSMAKDLTGTVKEILGTCVSVGCTVDGKDPKDLQQEIDDGDVEIPLD*

### NbL12g02440.1 ATP_synthase_subunit_d_mitochondrial−like [MW = 19.76 kDa]

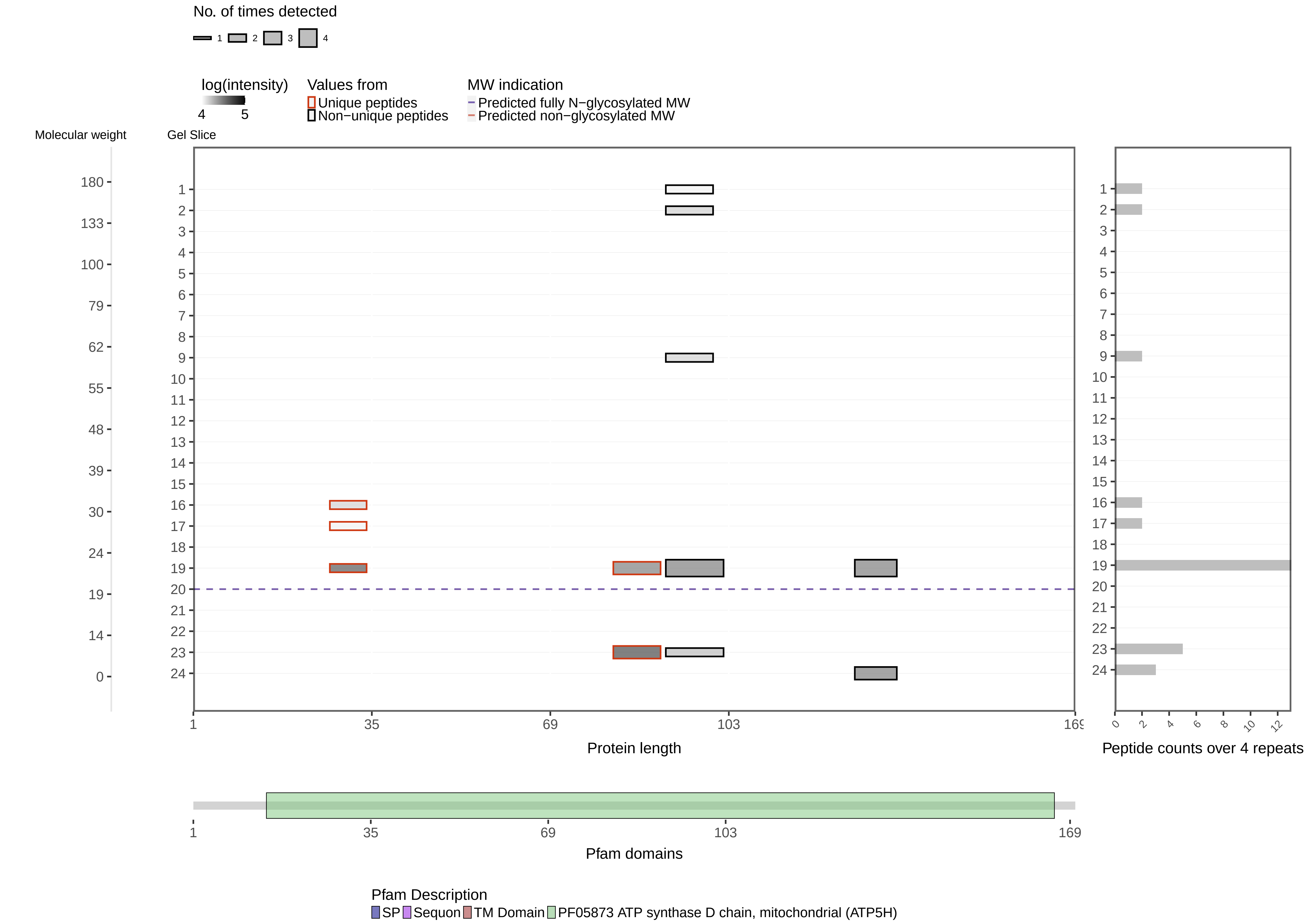


Protein sequence:

IGSRLVDMYKEAYDSIEIPKFVDTVTPQYKPKFDALLVELKEAEQKSLKESERLEKEIADVQELKKKLSTMTAEEYFEKHPELKKKFDDEIRNDYWGY*

### NbL12g02420.1 elongation_factor_1−alpha−like [MW = 49.26 kDa]

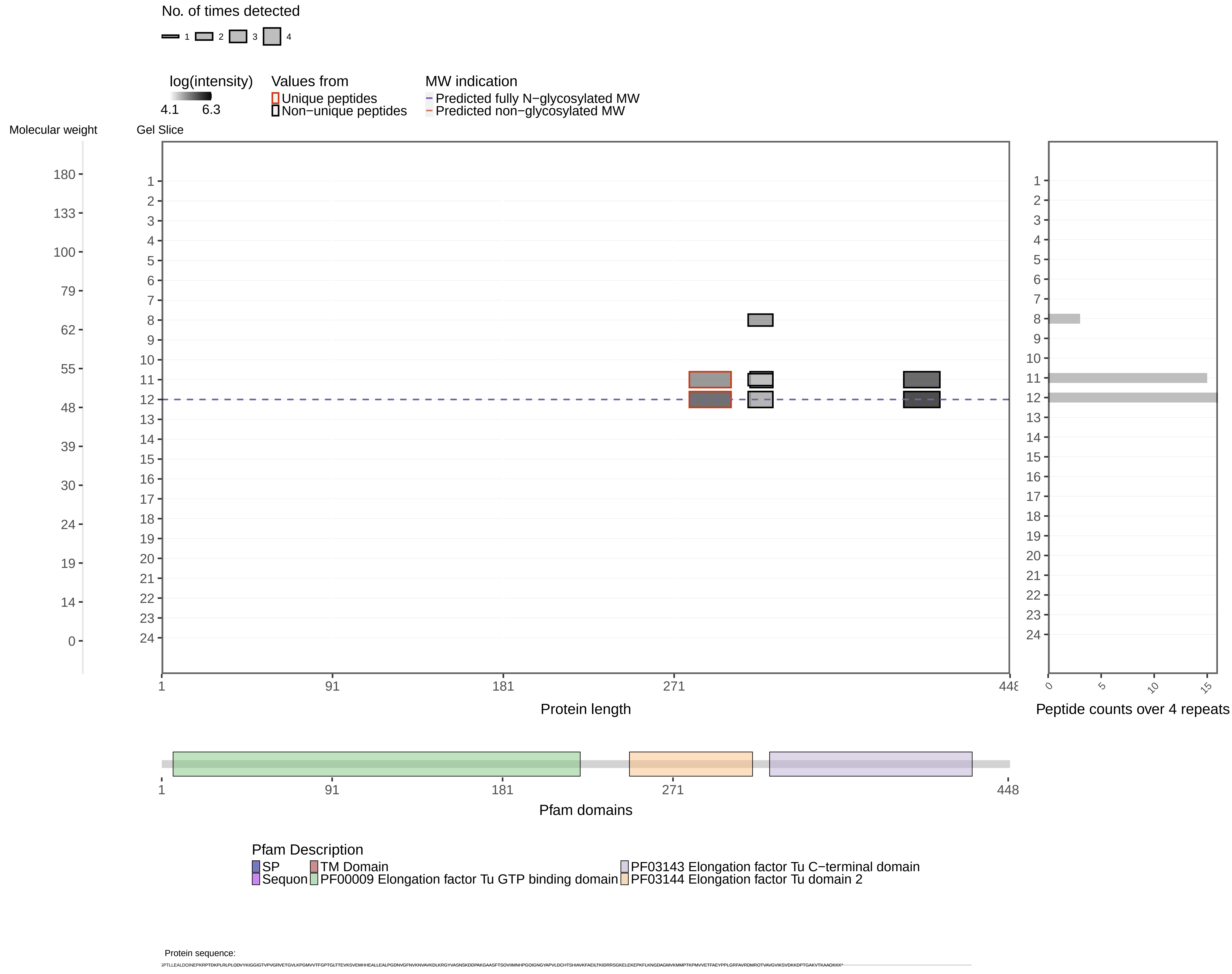

### NbL12g02390.1 endochitinase_EP3−like [MW − SP = 26.75 kDa]

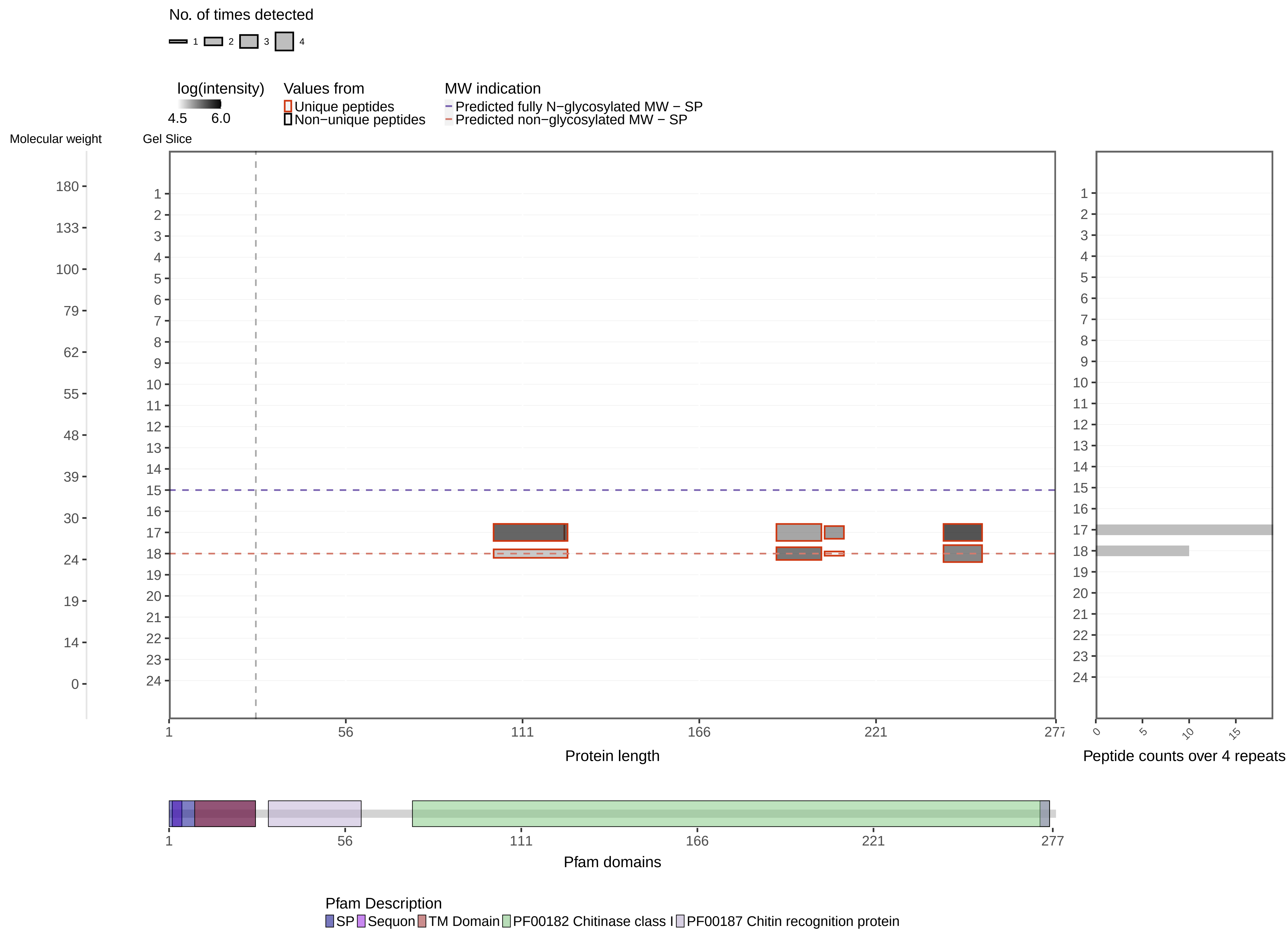


Protein sequence:

REIAAFFAHVTHETGHMCFINEINGPSGDYCDEENTEYPCVSGKKYYGRGPIQLSWNFNYGPAGKSIGFDGLNDPDIVARDAVISFKTALWYWMNNCHSLITSGQGFGPTIRAINGRIECDNGNPQTVARRVEYYTEYCQQLGVETGDNLTC*

### NbL12g02360.1 mitogen−activated_protein_kinase_3 [MW = 42.89 kDa]

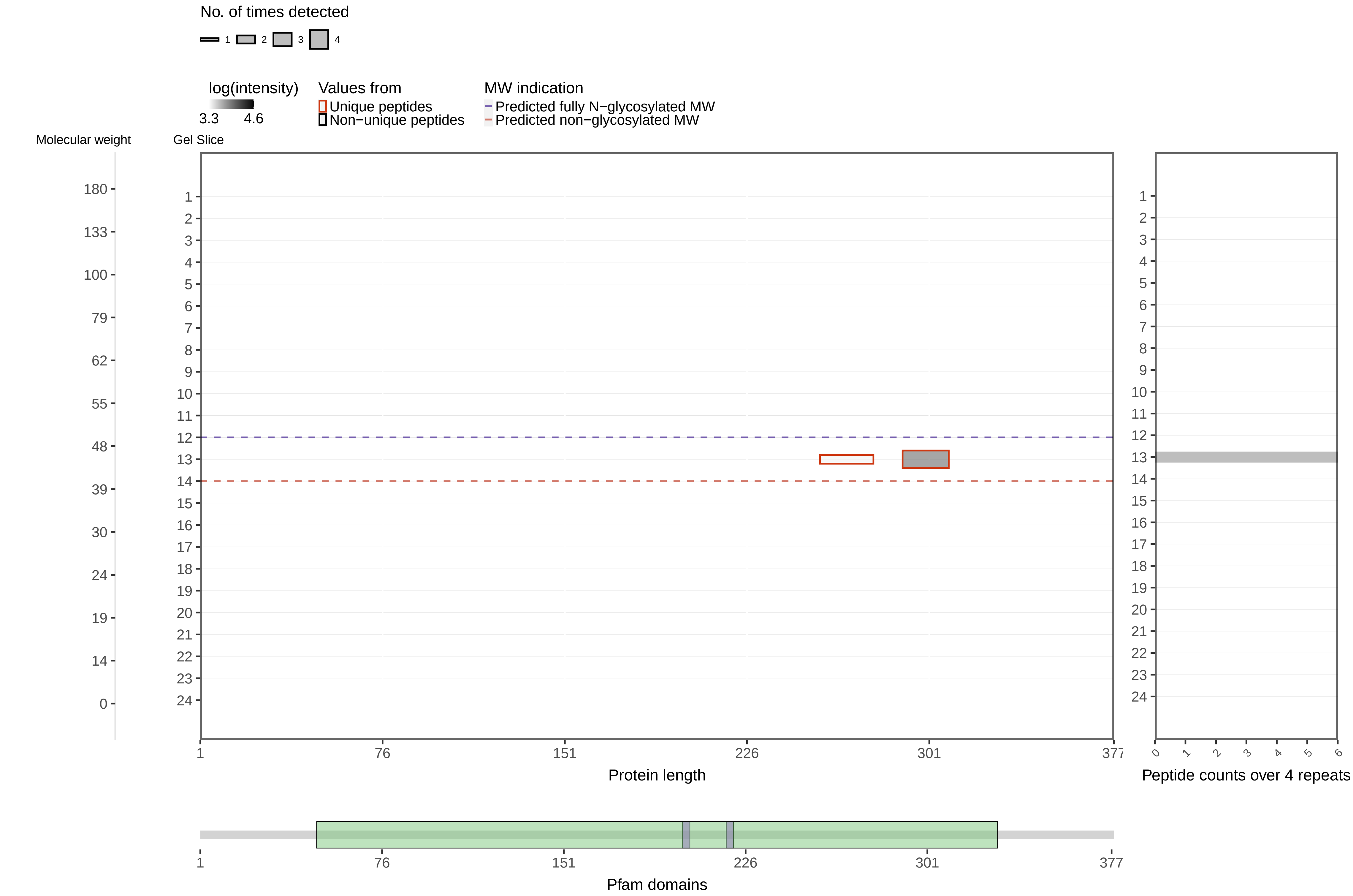


Pfam Description

SP Sequon TM Domain PF00069 Protein kinase domain

Protein sequence:

NLLLNANCDLKICDFGLARPNIENENMTEYVVTRWYRAPELLLNSSDYTAAIDVWSVGCIFMELMNRKPLFAGKDHVHOIRLLTELLGTPTEADLGFLONEDAKRYIROLPOHPROOLANVFPHVNPLAIDLVDKMLTLDPTRRITVEEALDHPYLAKLHDAGDEPICPVPFSFDFEOOGIGEEOIKDMIYOEALSLNPEYA*

NbL12g02350.1 peroxidase_2_cytosolic−like [MW = 27.41 kDa]

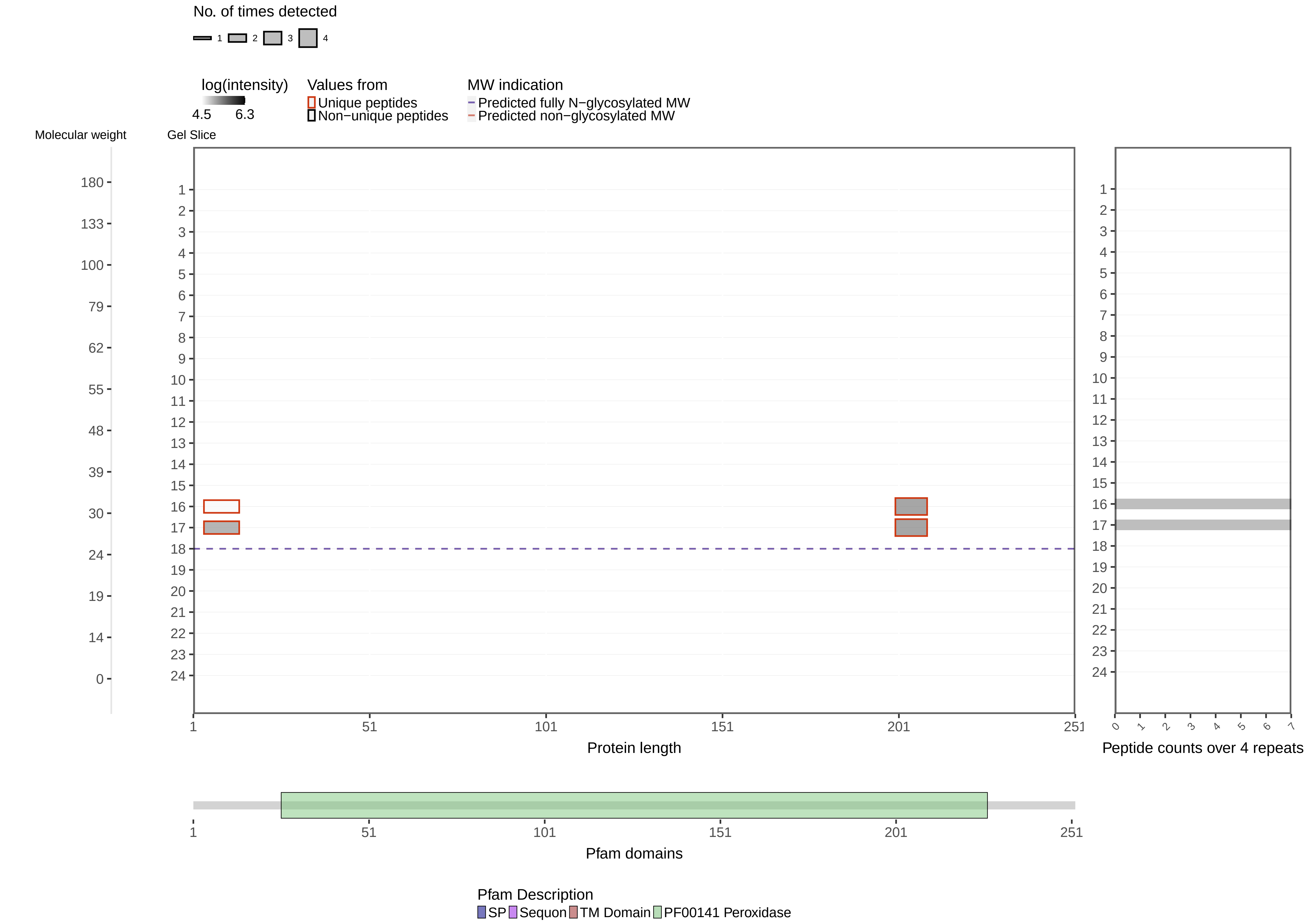


Protein sequence:

VPFHPGREDKTEPPVEGRLPDATKGSDHLRDVFVKQMGLSDKDIVALSGGHTLGRCHKERSGFEGPWTTNPLIFDNSYFTELLSGEKEGLLLLPSDKALLSDPAFRPLVEKYAADEDAFFADYAEAHLKLSELGFAEA*

### NbL12g02210.1 histone_H2A [MW = 15.78 kDa]

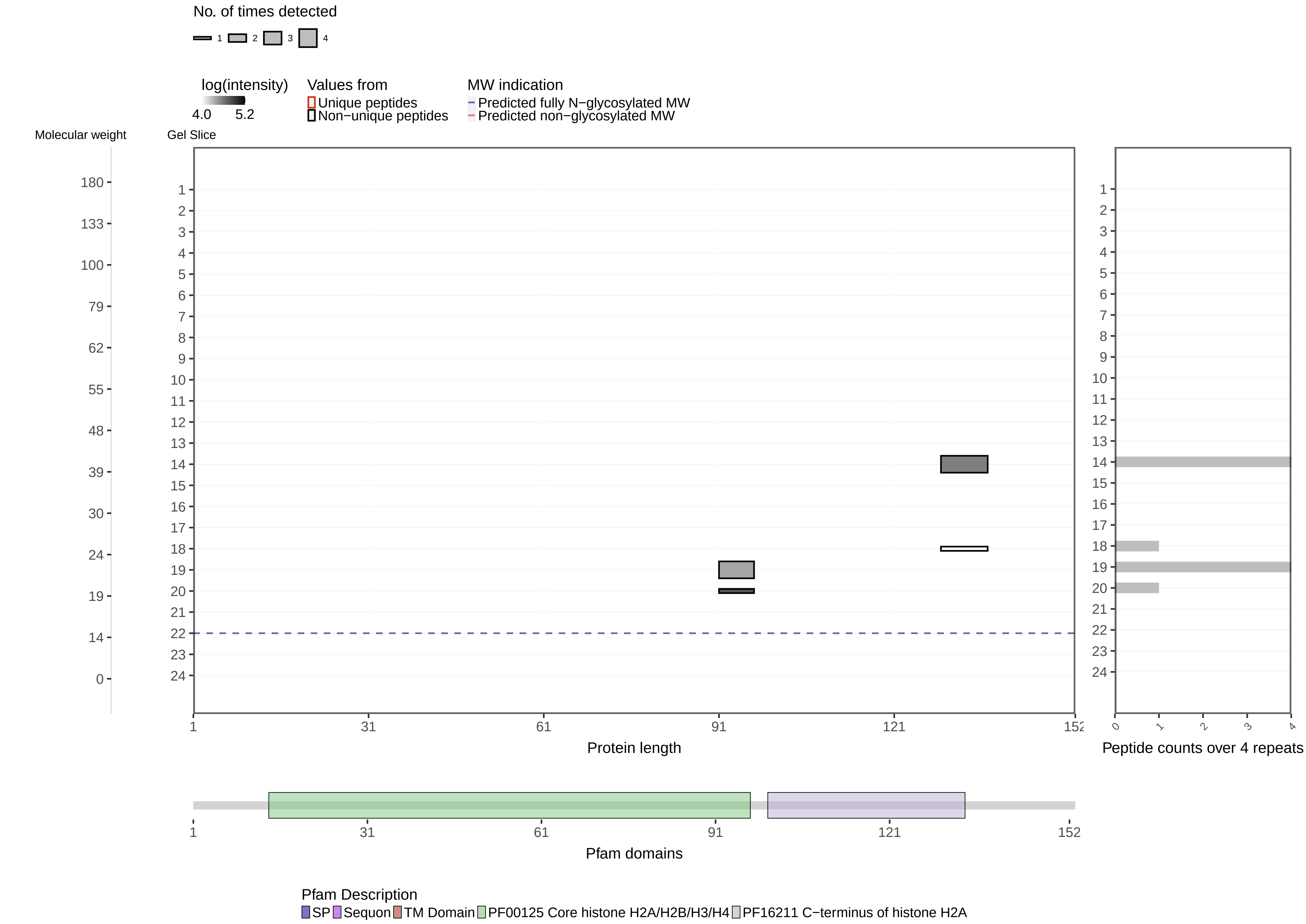


Protein sequence:

AAVLEYLAAEVLELAGNAARDNKKNRIIPRHVLLAVRNDEELGKLLAGVTIAHGGVLPNINPILLPKKTGGAEKEPKSPAKATKSPRKAAA*

### NbL12g02110.1 protein_plastid_transcriptionally_active_16_chloroplastic [MW = 54.52 kDa]

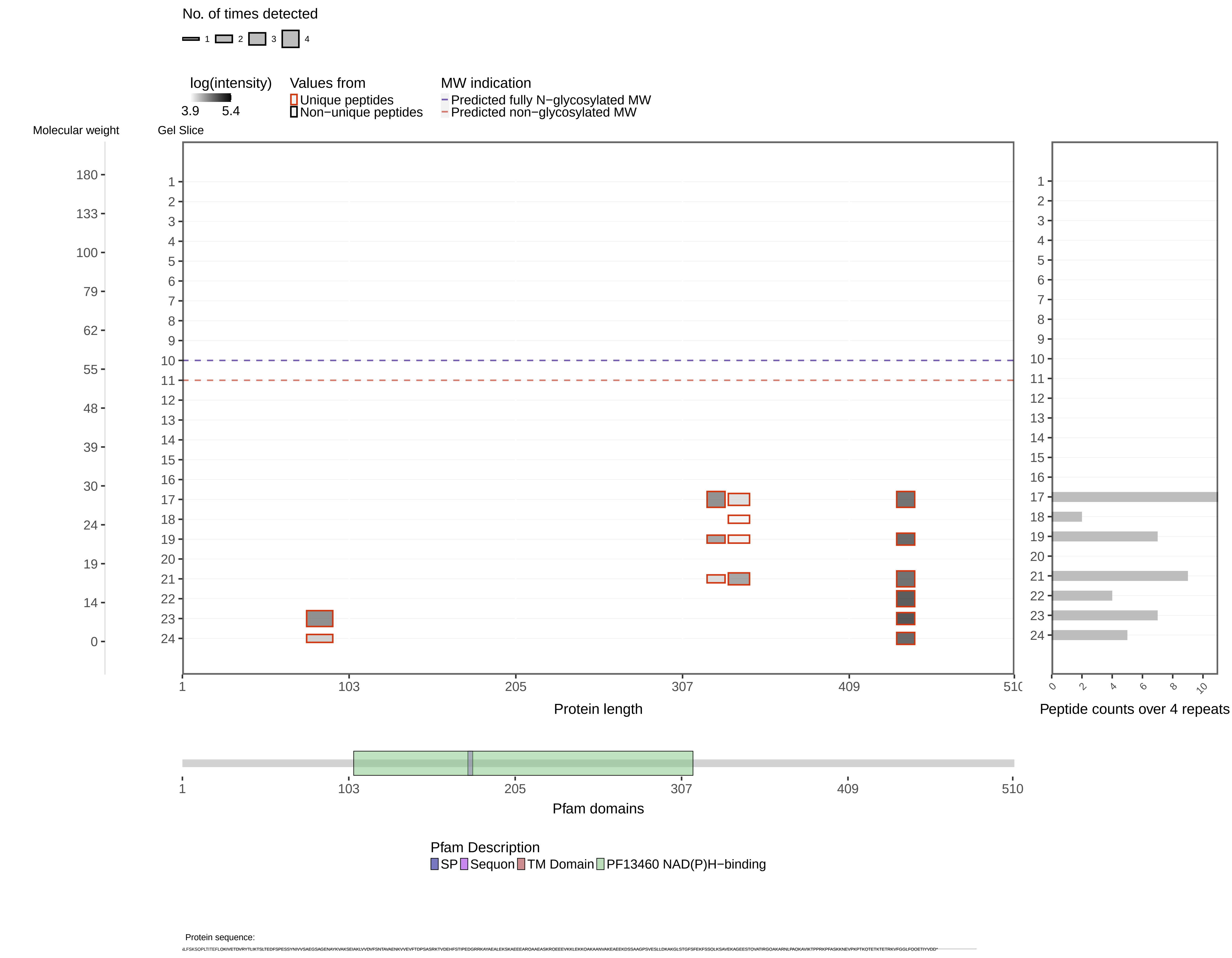

### NbL12g01970.1 serine/threonine−protein_phosphatase_PP1_isozyme_1−like [MW = 36.28 kDa]

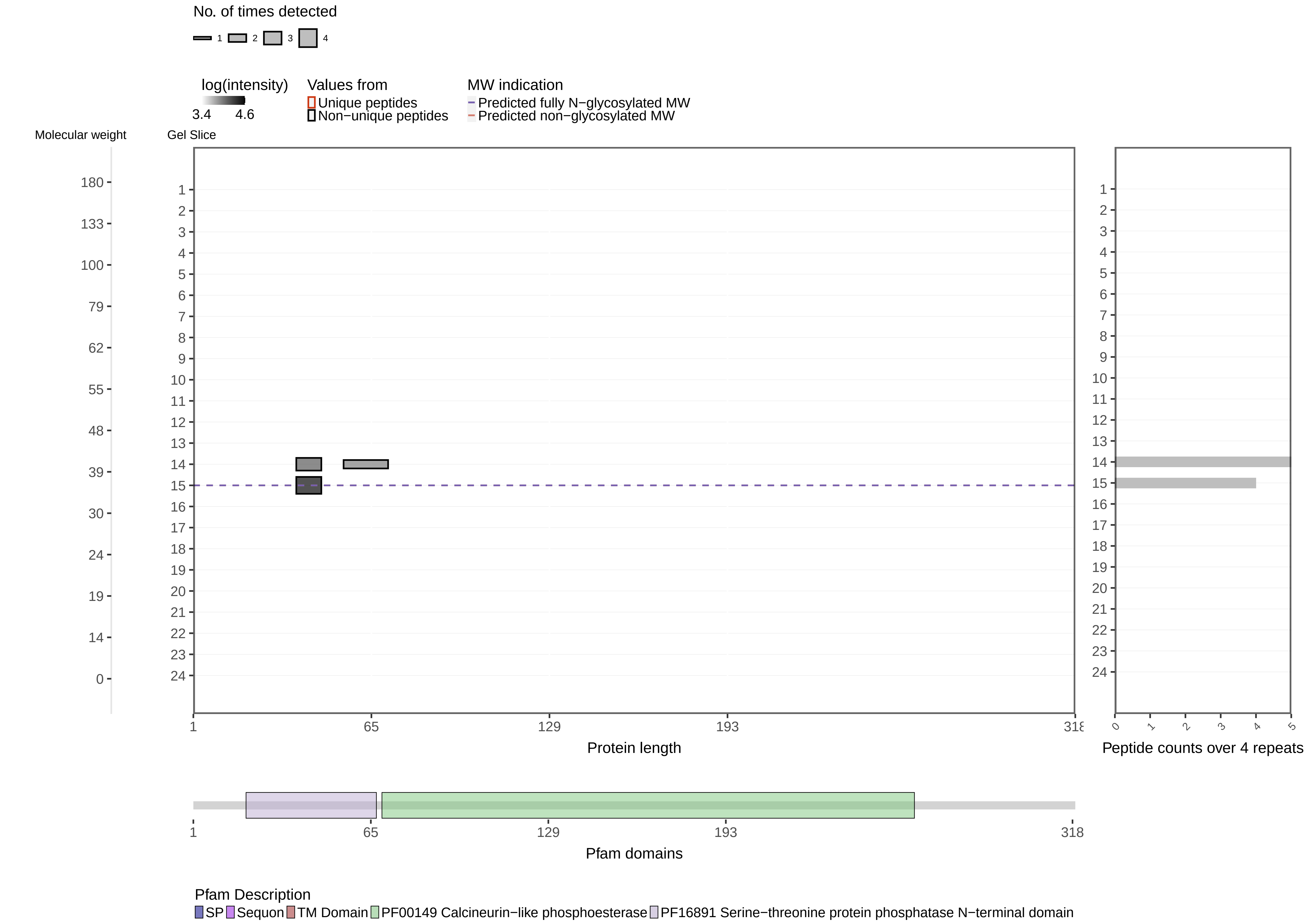


Protein sequence:

DECKRRFNVKLWKCFTECFNCLPVAALIDEKILCMHGGLSPDLTNLDQIRNLPRPTDVPDSGLLCDLLWSDPSREVKGWGMNDRGVSYTFGPDKVAEFLMQHDMDLVCRAHQVVEDGYEFFAERQLVTIFSAPNYCGEFDNAGAMMSVDESLMCSFQILKPTDRKPRFL*

NbL12g01920.1 M−phase_phosphoprotein_6−like [MW = 20.71 kDa]

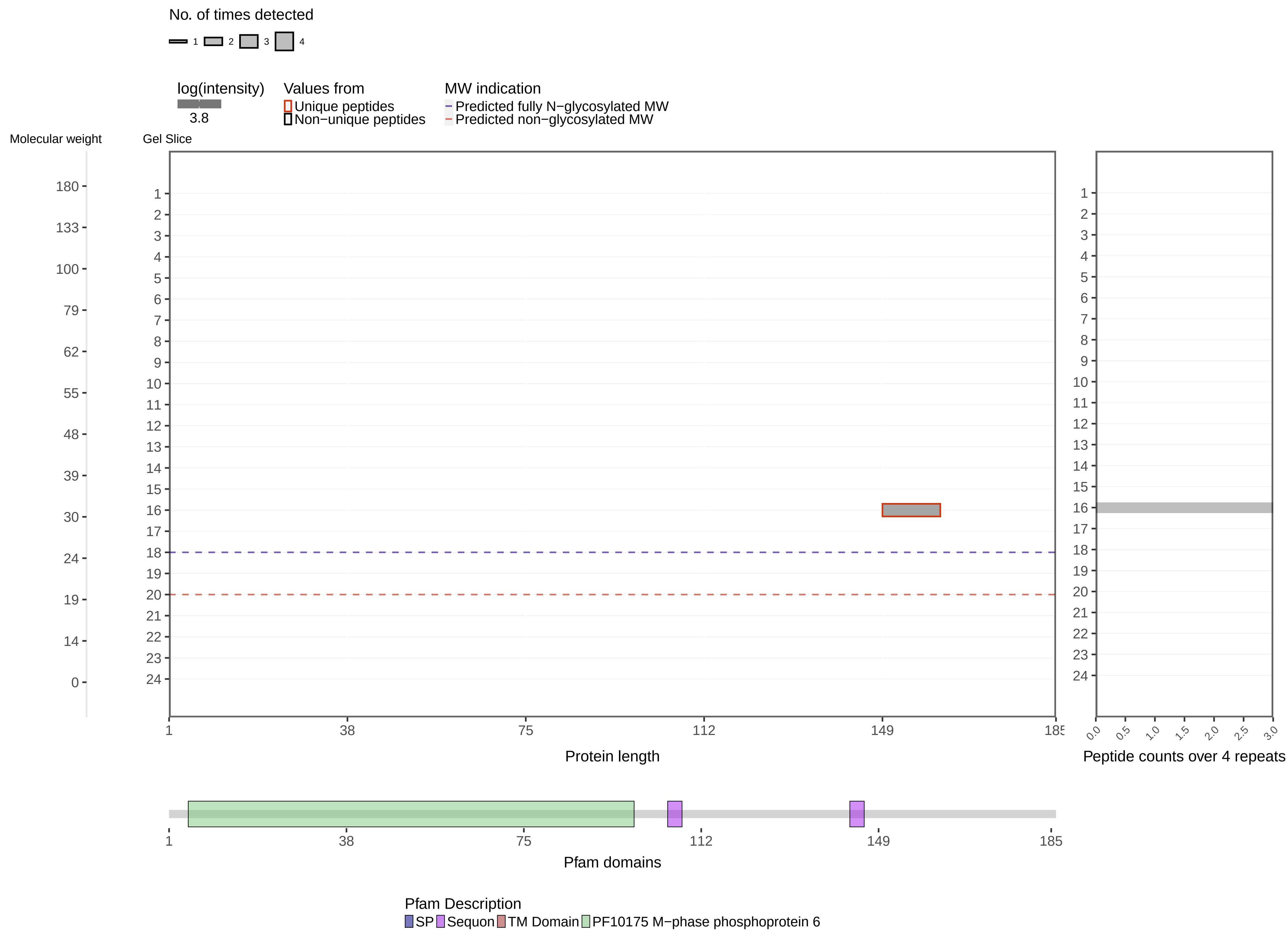


Protein sequence:

[illegible]SKPHSEGSAACSSETSEINSKRENGTSQCGLENSNMDDSSLSPNEDLKRKQDDTSSEVQYPNKSHKRVLGNAESSPSSSRSSQKPHRLDWSVLKPPKSQKKKR*

NbL12g01790.1 beta−glucosidase_BoGH3B−like [MW = 66.12 kDa]

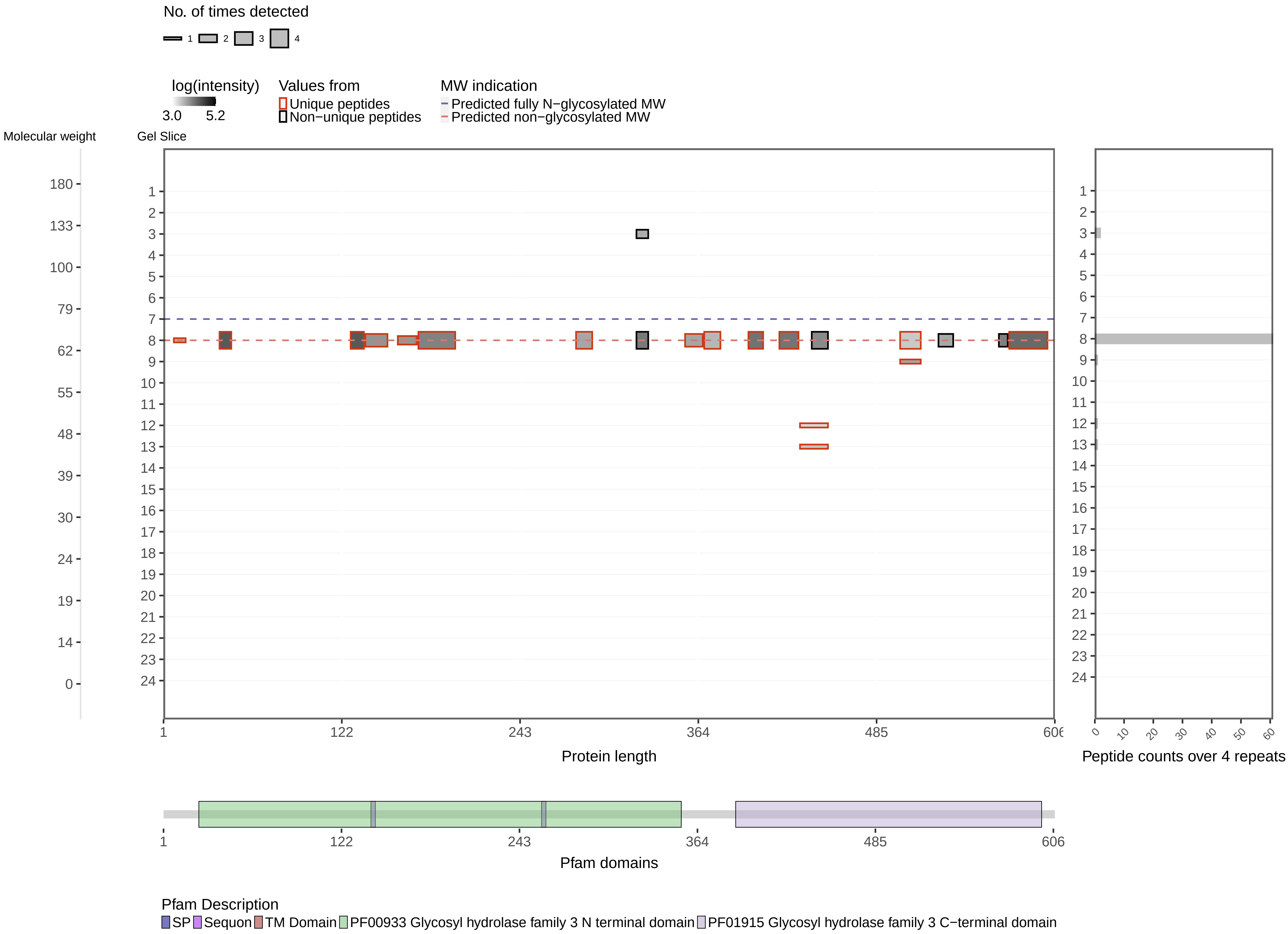


Protein sequence:

LDRLSHPHGSNYDQCILTAINAGIDMVMVPFRYQLFLDHLRCLVESGKIPMTRIDDAVERILRVKFVAGAFEYPLSDRSLLDTVGCNQHRELAREAVRKSLVLLKNGKDVTKPFLPLERNAKKILVAGKHADDLGFQCGGWTKTWDGMSGRITIGTTILEAIKDAVGGETEVVYEENPSPETFASQDFSYSIVVVGEPPYCESGGDSQDLKIPLGGEELISLVADRVPTLVILISGRPLYIEPSLLEKMDAFVAAWLPGTEGAGITGVVFGDYEFQGKLPMTWFKSVDQLPLHQEQ

### NbL12g01760.1 phosphomethylpyrimidine_synthase_chloroplastic [MW = 72.08 kDa]

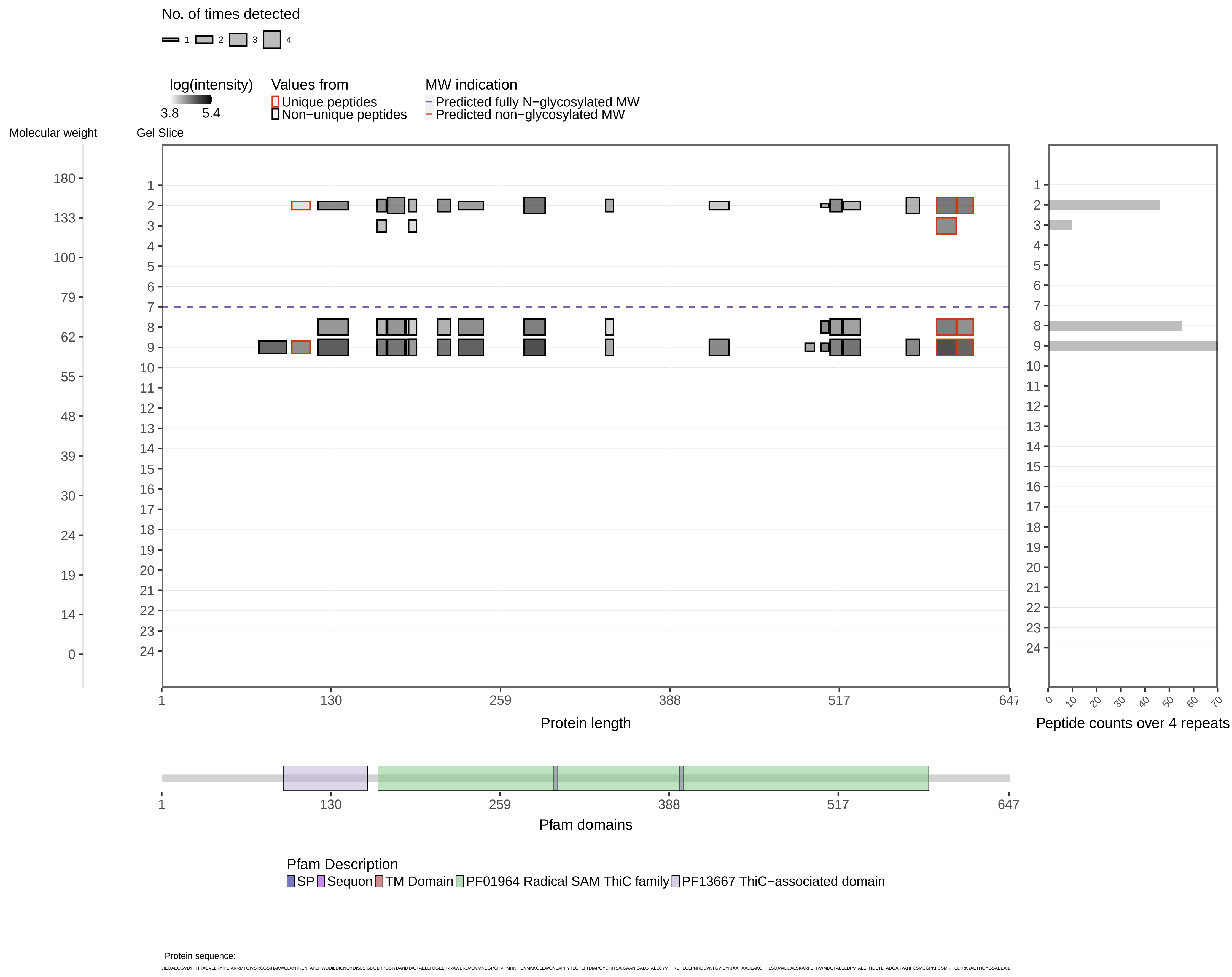

NbL12g01750.1 uncharacterized_LOC107777172 [MW = 10.76 kDa]

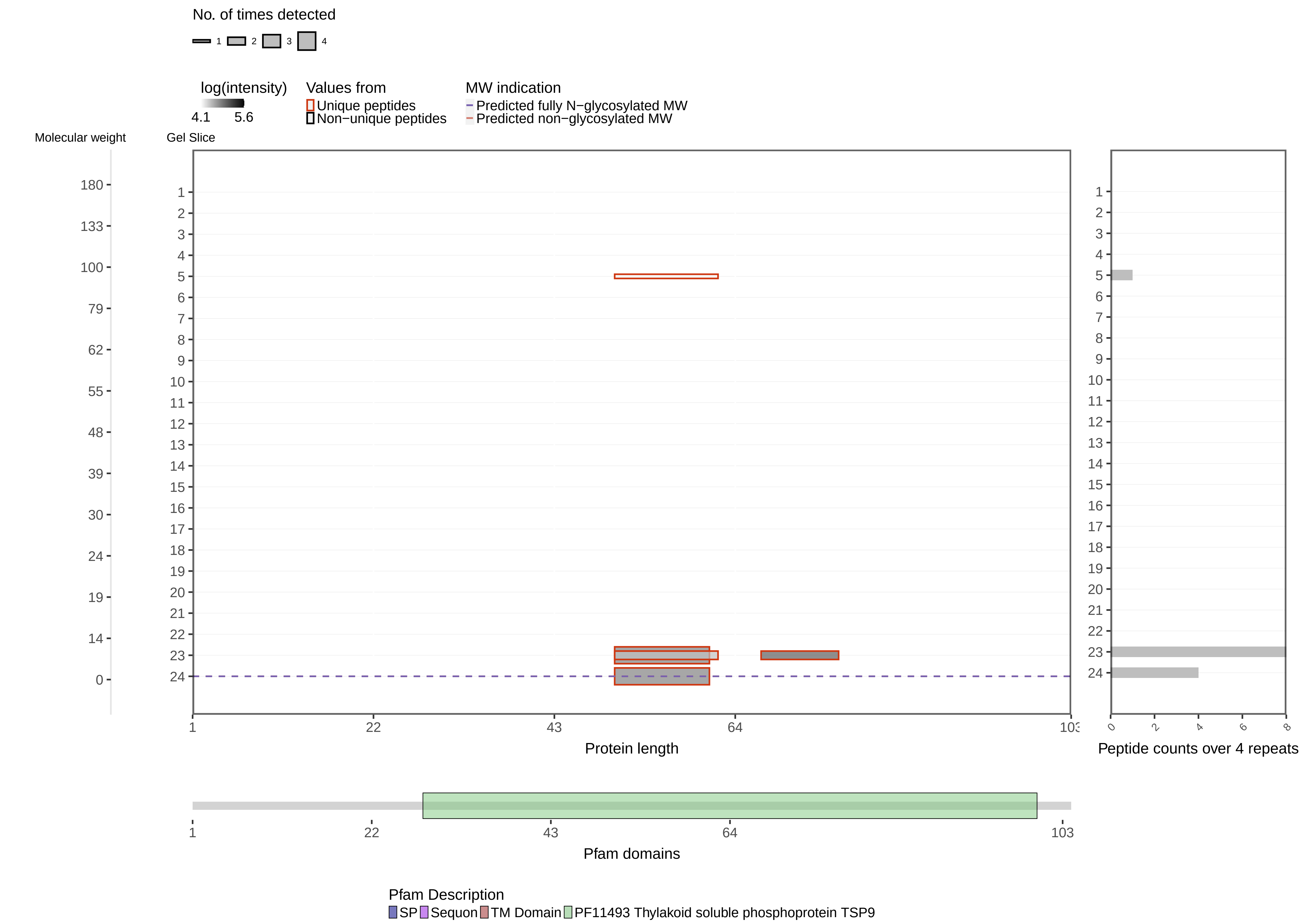


Protein sequence:

[illegible]ILDFVLGAITKEDQLLETDPILKKVEGKSTGTTTISSKKSVSVPPKKNSDGGFGGFGGLFAKKE*

### NbL12g01730.1 anthranilate_synthase_alpha_subunit_2_chloroplastic−like [MW = 66.49 kDa]

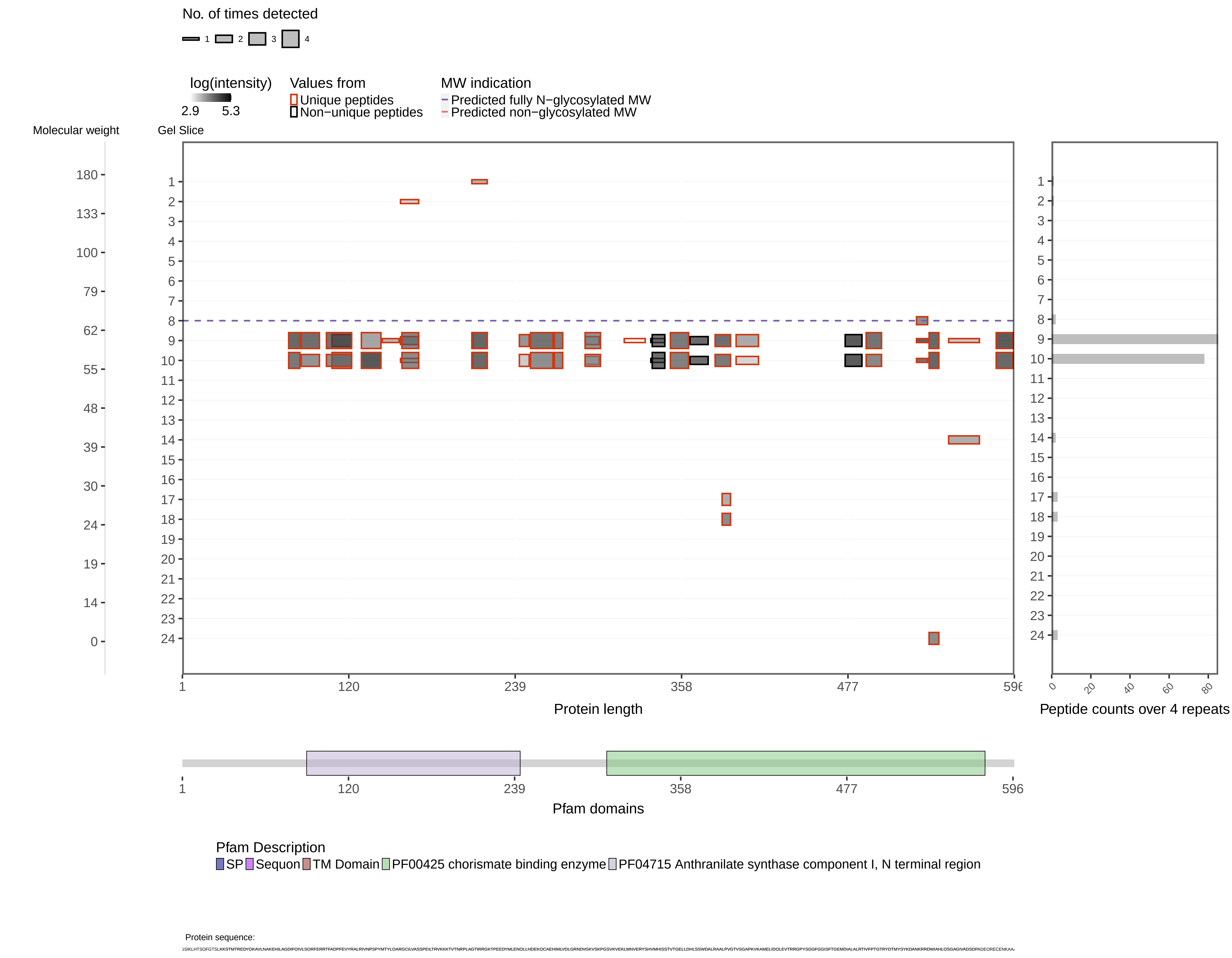

NbL12g01670.1 protein_PMR5−like [MW − SP = 40.5 kDa]

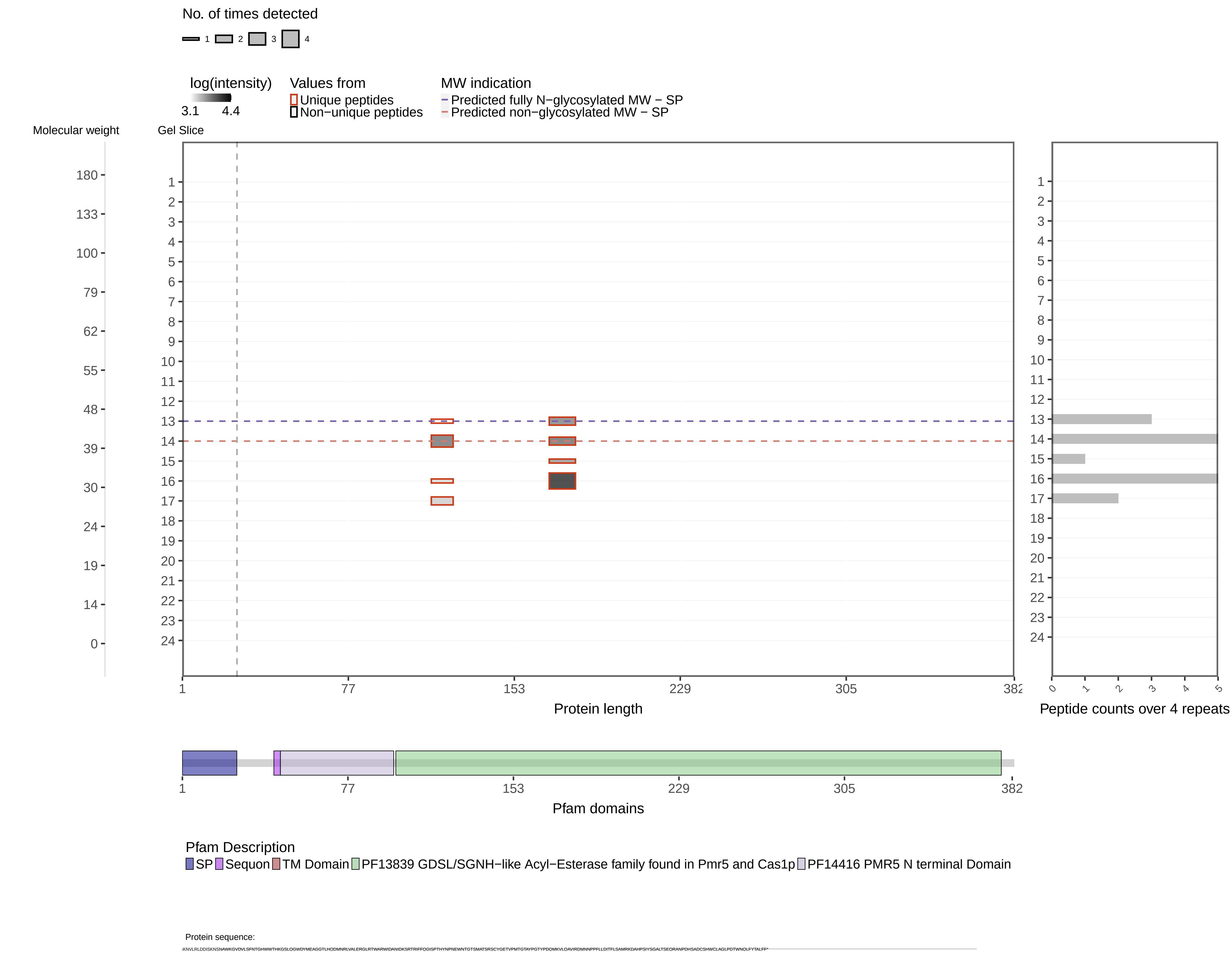

### NbL12g01630.1 ran−binding_protein_1_homolog_c−like [MW = 24.79 kDa]

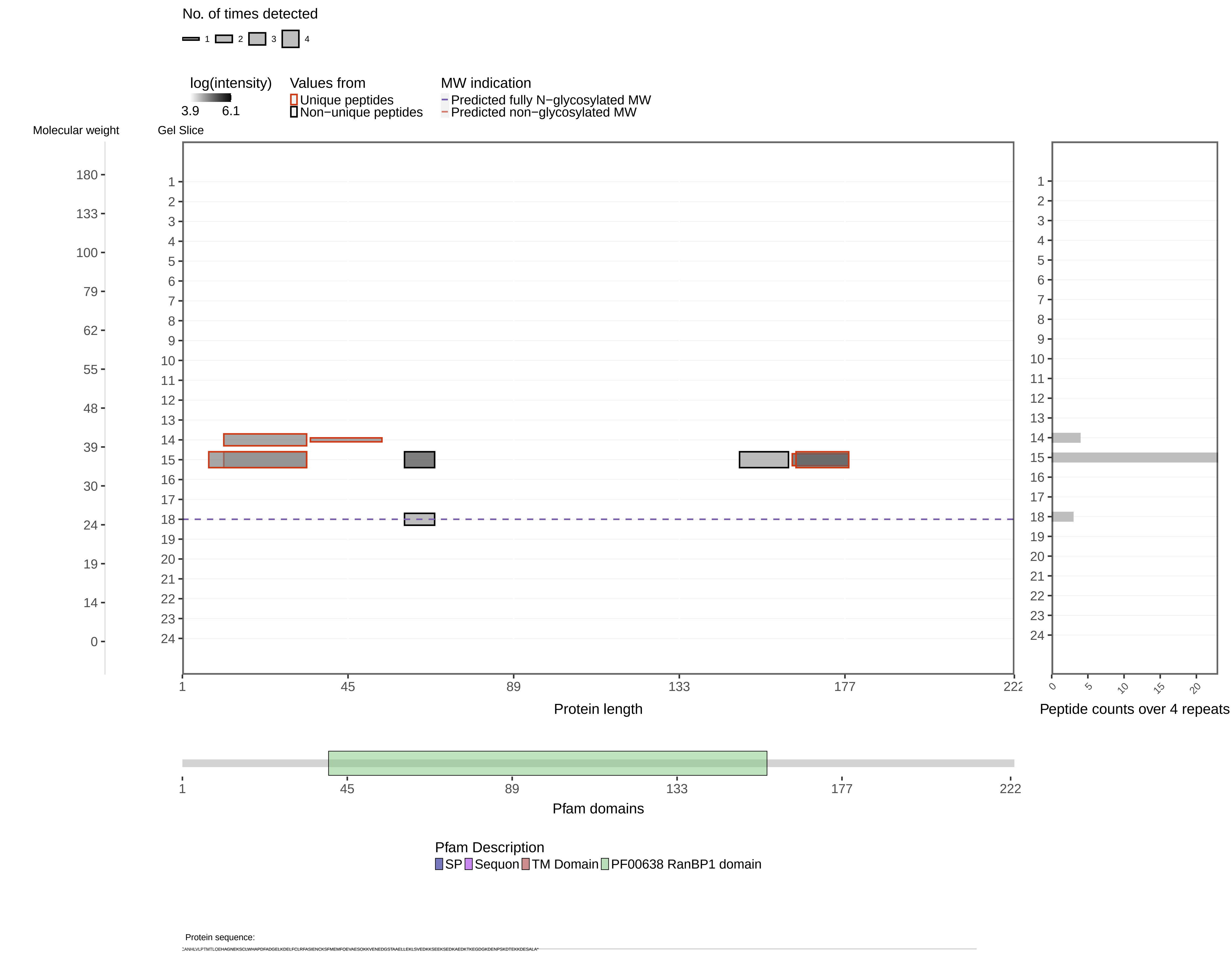

### NbL12g01620.1 ERBB−3_BINDING_PROTEIN_1−like [MW = 43.52 kDa]

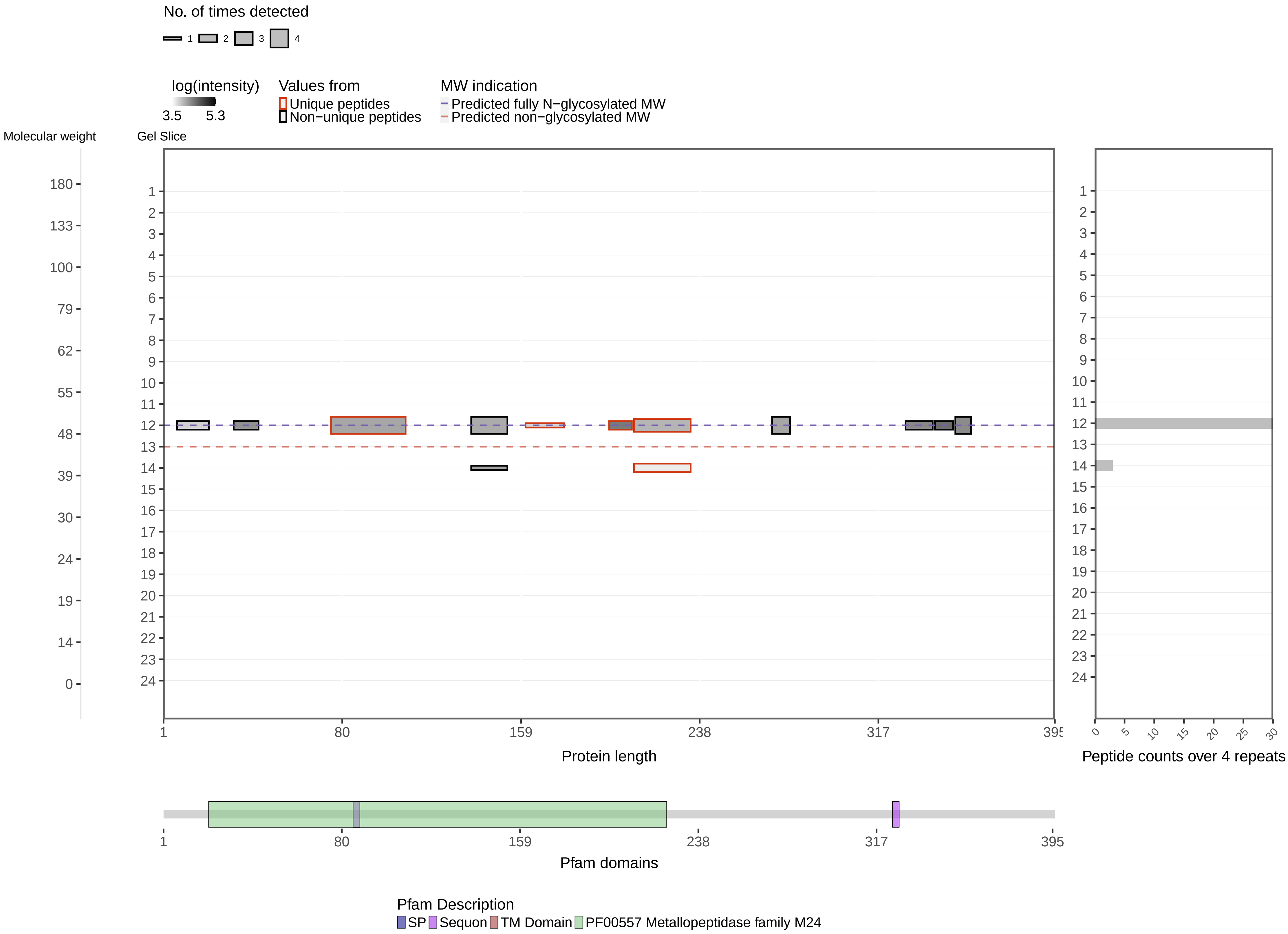


Protein sequence:

SHQMKQFVIDGNKVVLSVSNPETRVEDAEFEENEVYSIDIVTSTGEGKPKLLDEKQTTIYKRAVDKSYNLKMKASRFIFSEISQKFPVMPFTARDLEEKRARLGLVECVNHELLQPYPVLHEKPGDLVAHIKFTVLLMPNGSDRITTHTLQELKPAKTIDDEPEIKTWLALPVKSKKKGGGKKKKAKKGEKTEELSQAEPMEGESNGAES*

### NbL12g01610.1 BAG_family_molecular_chaperone_regulator_1−like [MW = 22.68 kDa]

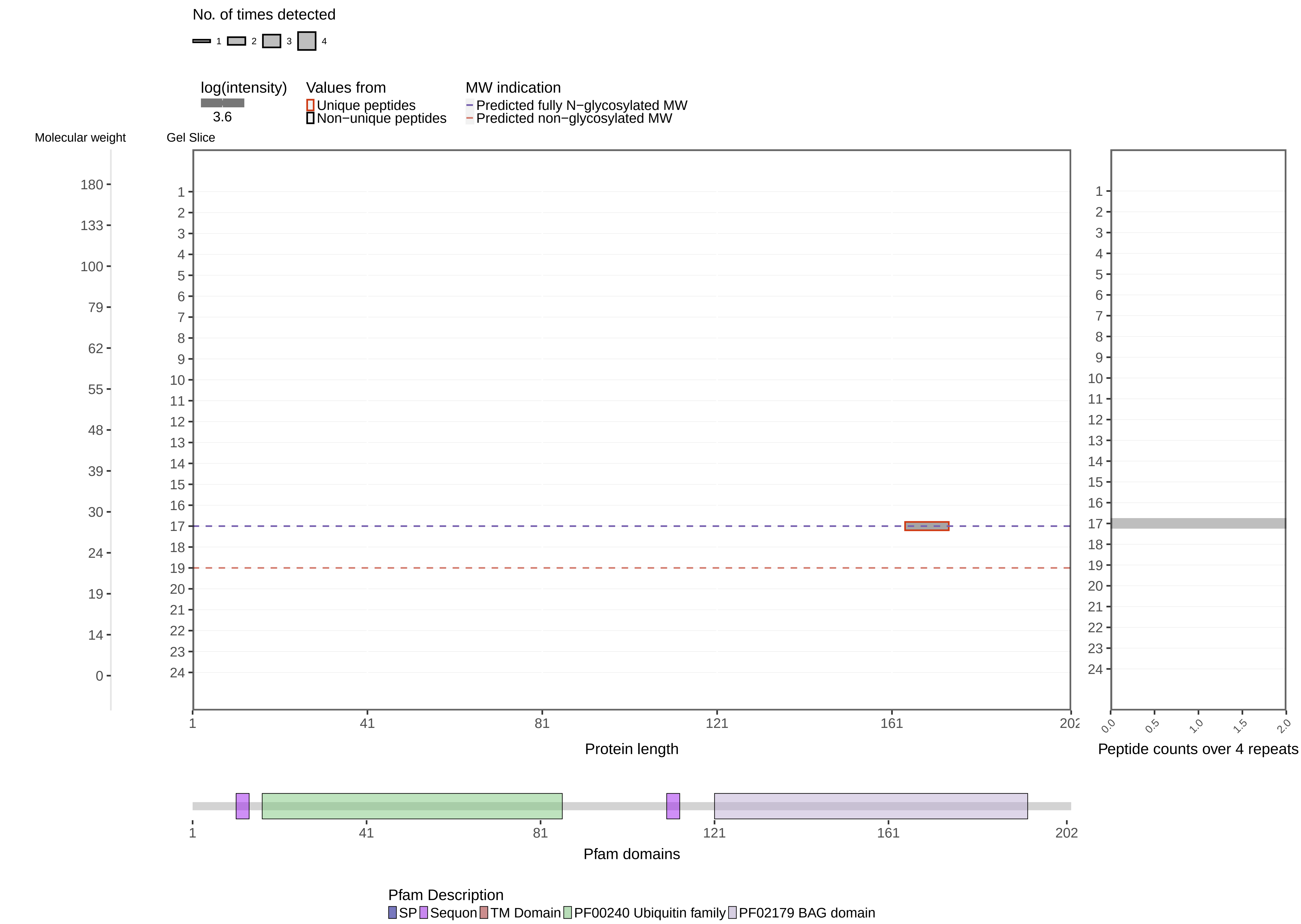


Protein sequence:

LHQEDMPNLLEDPMSKERKNIQNTSVSCEAVAGVRVEVDKLSHRILAIEEAMQRGTKVEDREFVVLTELFMIQLLKLDSIEADGEARAERRKEVHRIQSFVDMLDNFKARNSKL*

### NbL12g01580.1 ubiquitin−activating_enzyme_E1_1−like [MW = 128.33 kDa]

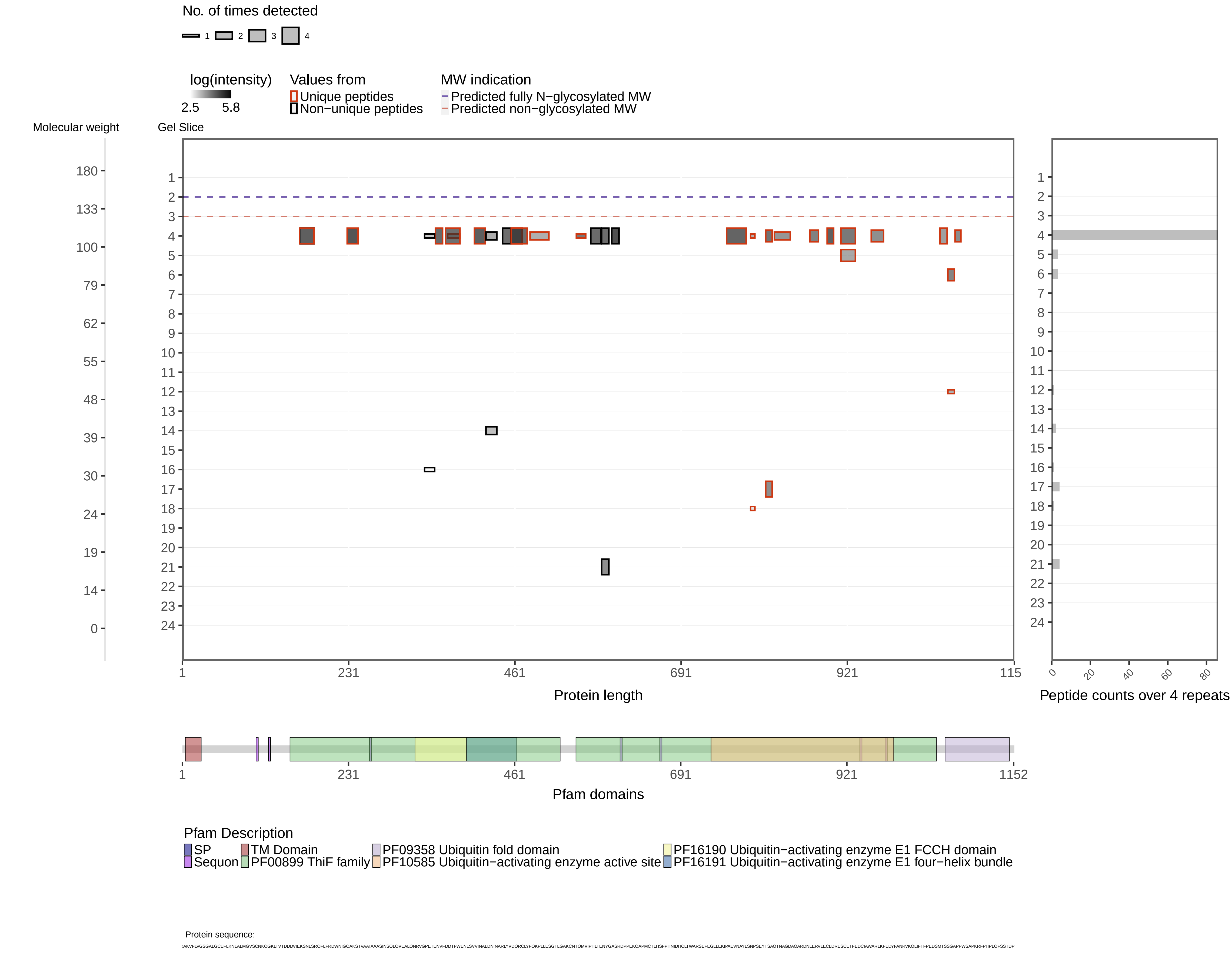

### NbL12g01570.1 gamma−interferon−inducible_lysosomal_thiol_reductase−like [MW − SP = 25.32 kDa]

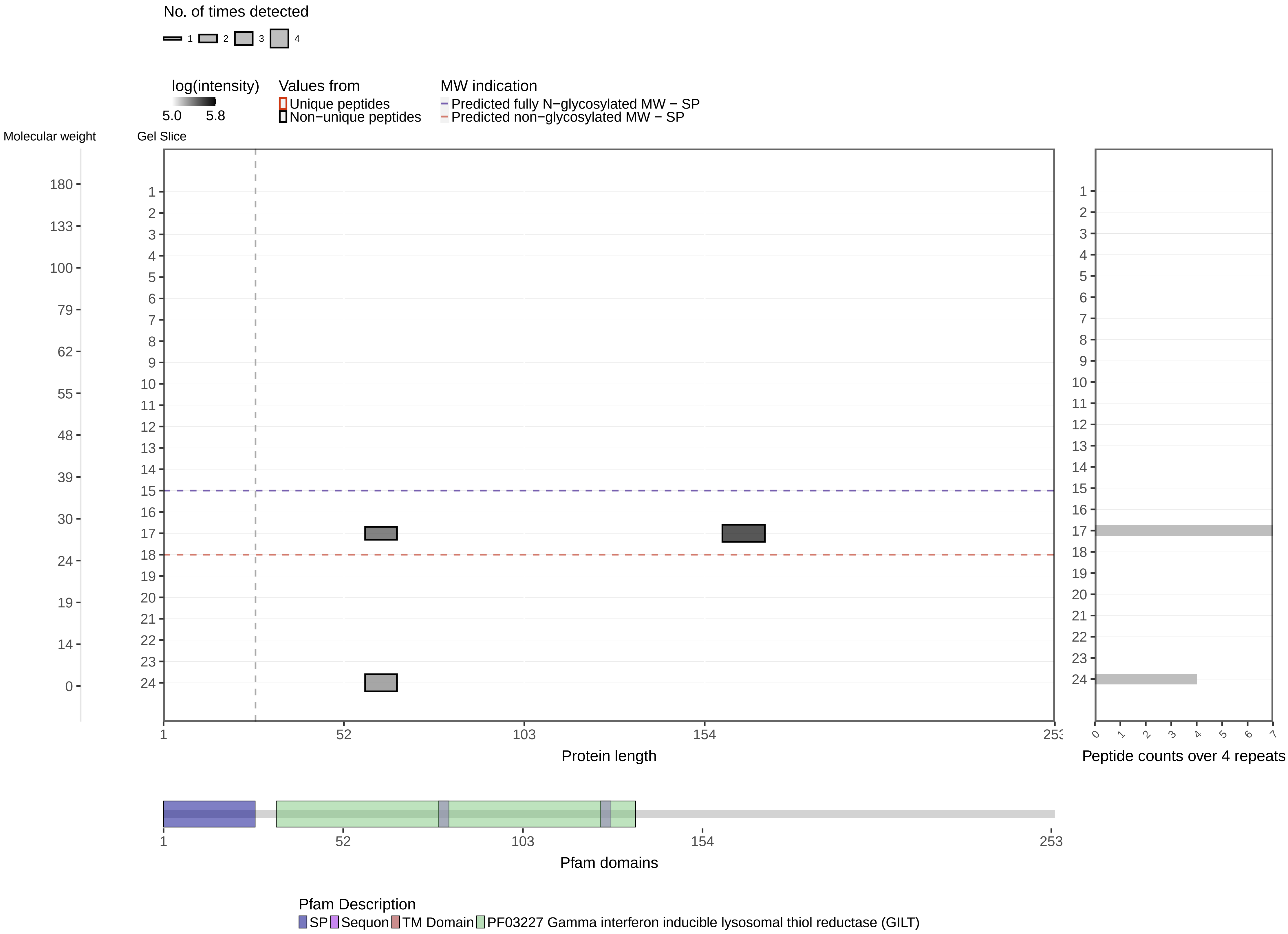


Protein sequence:

YCVESLVYHKNYTQWETCFEKLNLKAKPVKDCVGSGRGKELELRYAAETNALQPPHKYVPWVVVDGQPLYDEYMDFISYICKAYKGAPVPGCSSTVNVIKTGRKFTPFCLMHTASSKLSTIGSAITSWMNRVNPAASA*

### NbL12g01560.1 40S_ribosomal_protein_S4−like [MW = 32.02 kDa]

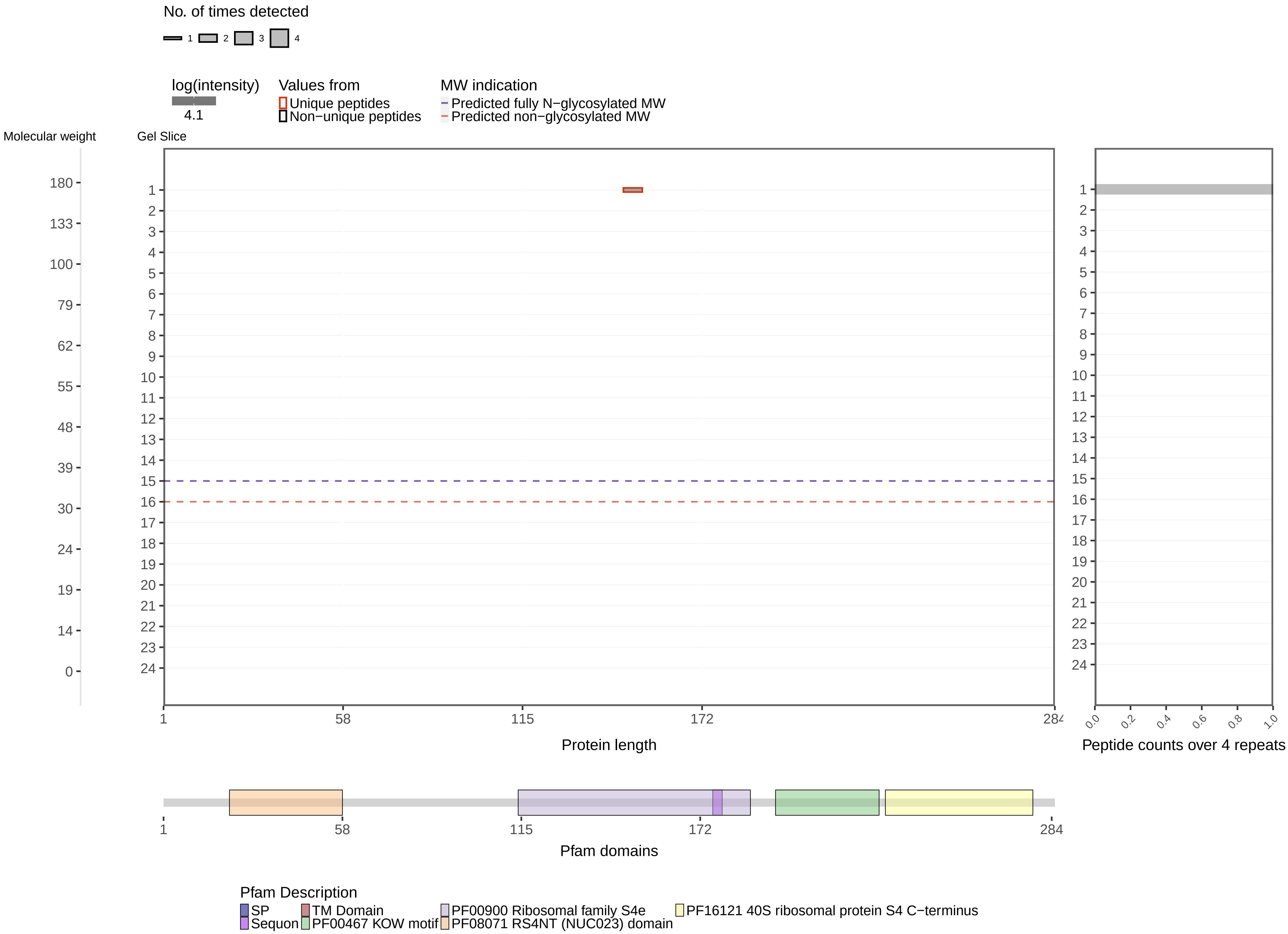


Protein sequence:

RLHSVRDEEAKFKLCKVRSIQFGQKGIPYLNTYDGRTIRYPDPLIKANDTVKLDLESNKIVDFIKFDVGNVVMVTGGRNRGRVGVIKNREKHKGSFETVHIQDALGHEFATRLGNVFTLGKGTKPWVSLPKGKGIKLSIIEEARKRLAAQSATTA*

### NbL12g01480.1 40S_ribosomal_protein_S8−like [MW = 25.12 kDa]

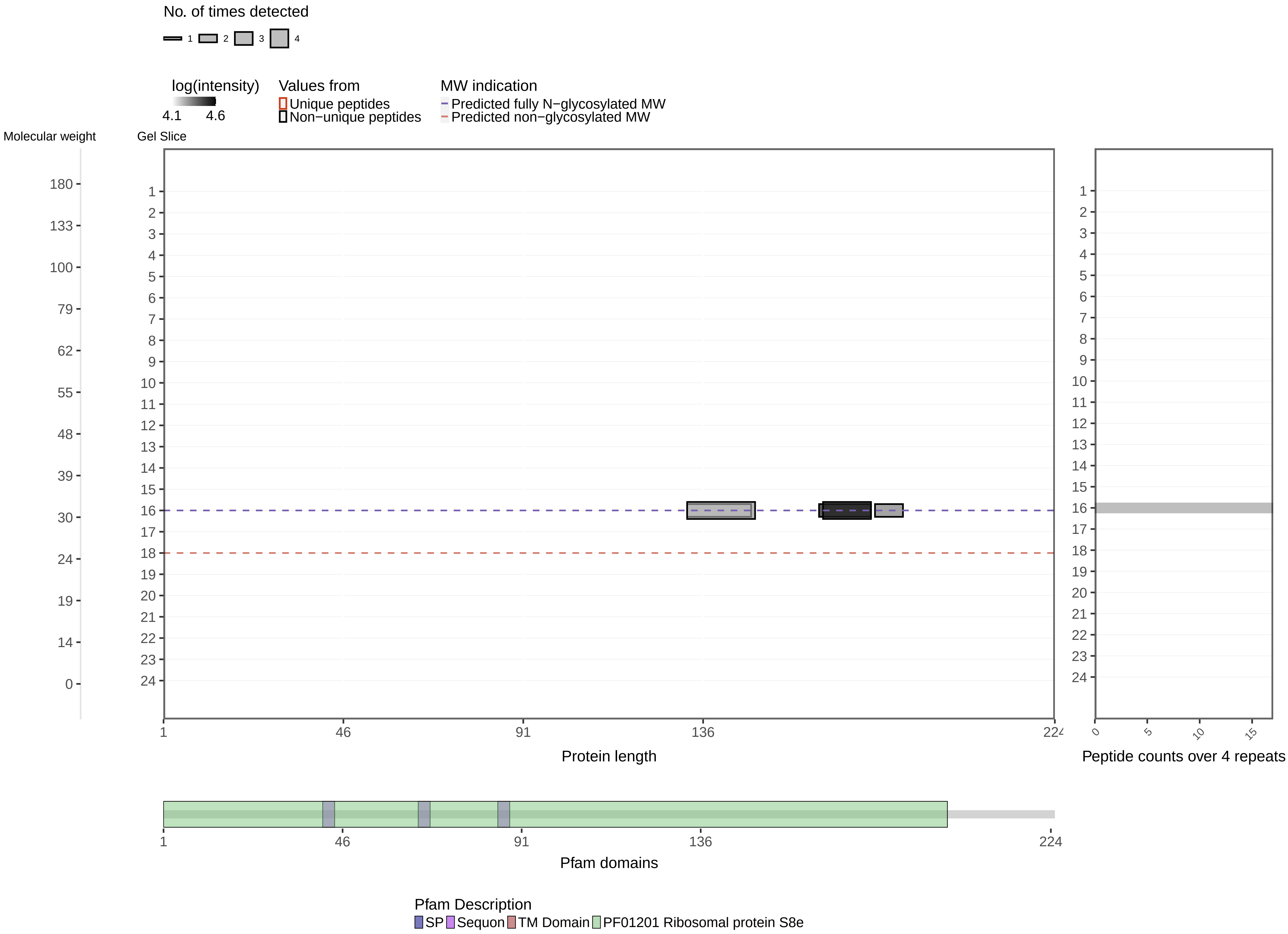


Protein sequence:

[illegible]SAIVQVDAAPFKQWYLQHYGVDIGRKKKGAAKKETTEEGEAAAPAEEAKKSNHVARKLEKRQKDRKLDPHLEEQFGSGRLYACISSRPGQCGRADGYILEGKELEFYMKKLQKKKGKAGSGAAA*

NbL12g01410.1 26S_protease_regulatory_subunit_6B_homolog [MW = 46.34 kDa]

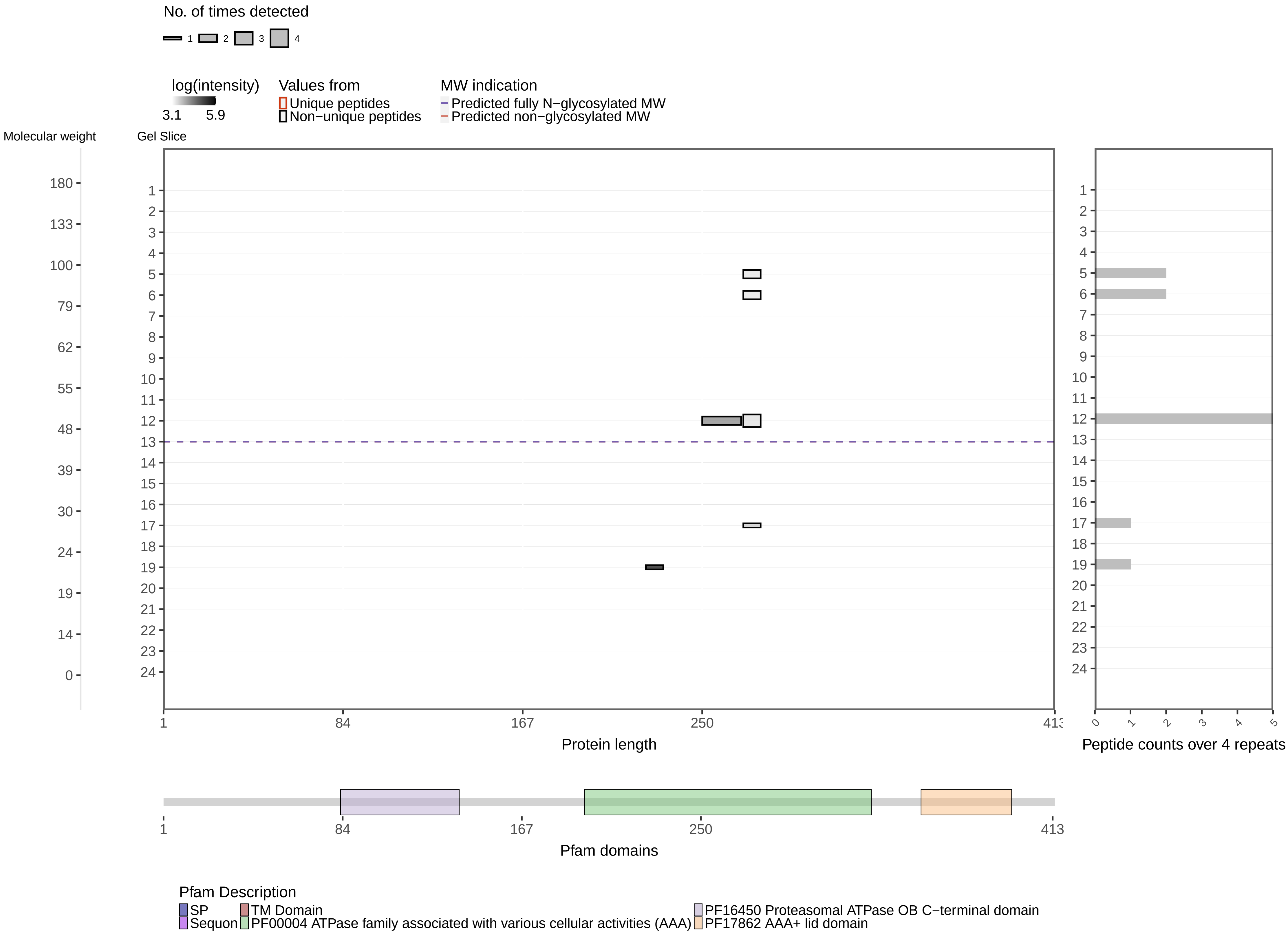


Protein sequence:

VLLYGPPGTGKTMLAKAVAHHTTAAFIRVVGSEFVQKYLGEGPRMVRDVFRLAKENAPAIIFIDEVDAIATARFDAQTGADREVQRILMELLNQMDGFDQTVNVKVIMATNRADTLDPALLRPGRLDRKIEFPLPDRRQKRLVFQVCTAKMNLGDEVDLEDYVSRPDKISAAEISAICQEAGMHAVRKNRYVILPKDFEKGYRTNVKKPDTDFEFYK*

NbL12g01390.1 40S_ribosomal_protein_S26−3−like [MW = 14.48 kDa]

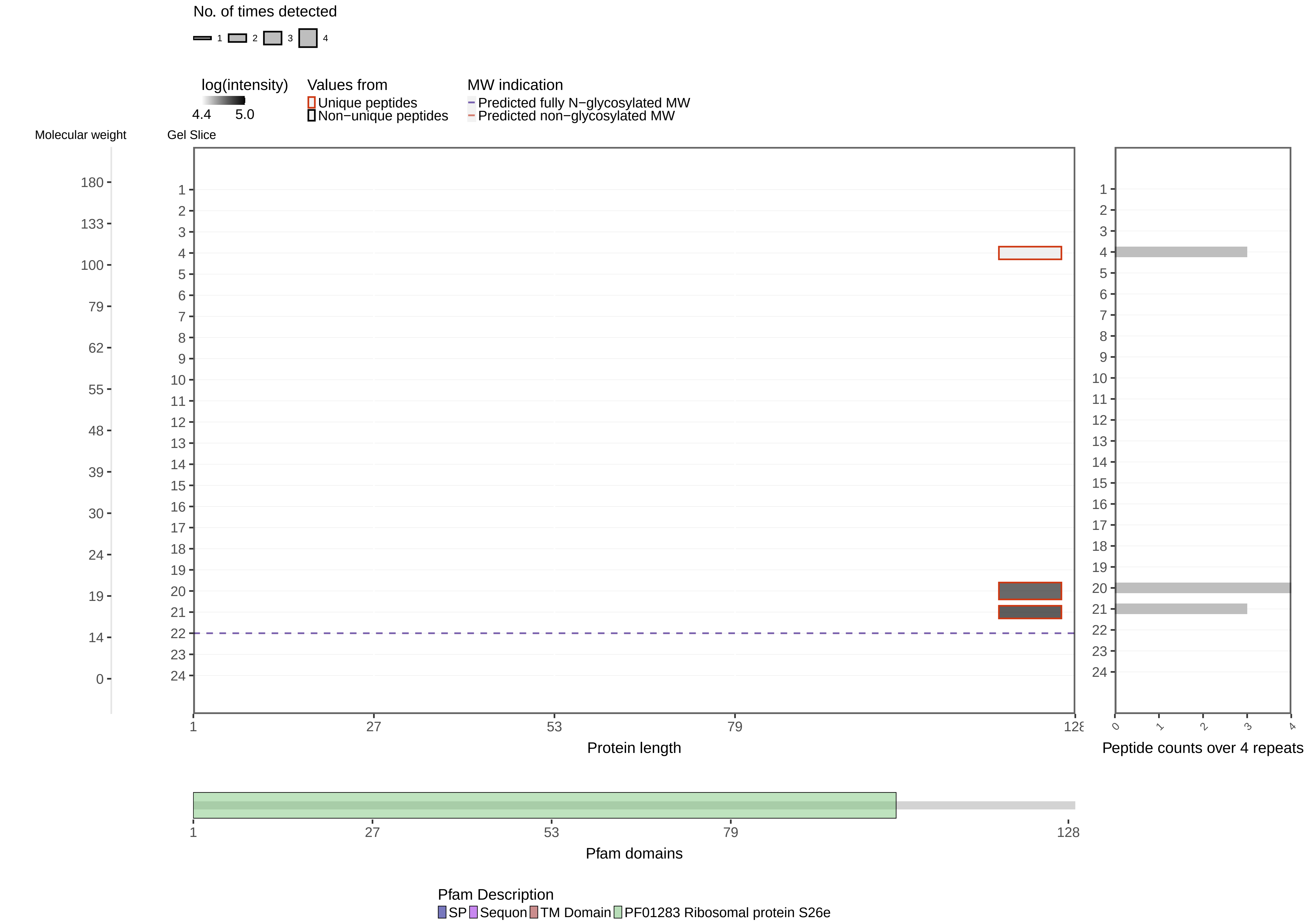


Protein sequence:

RDVQEACAFETYTLPKLYLKMQYCVSCAIHSKVVRVRSVTDRRVREPPQRFRRPRDDLPKPGQAPRPAGGAPAAPRT*

NbL12g01350.1 uncharacterized_protein_At5g48480−like [MW = 17.7 kDa]

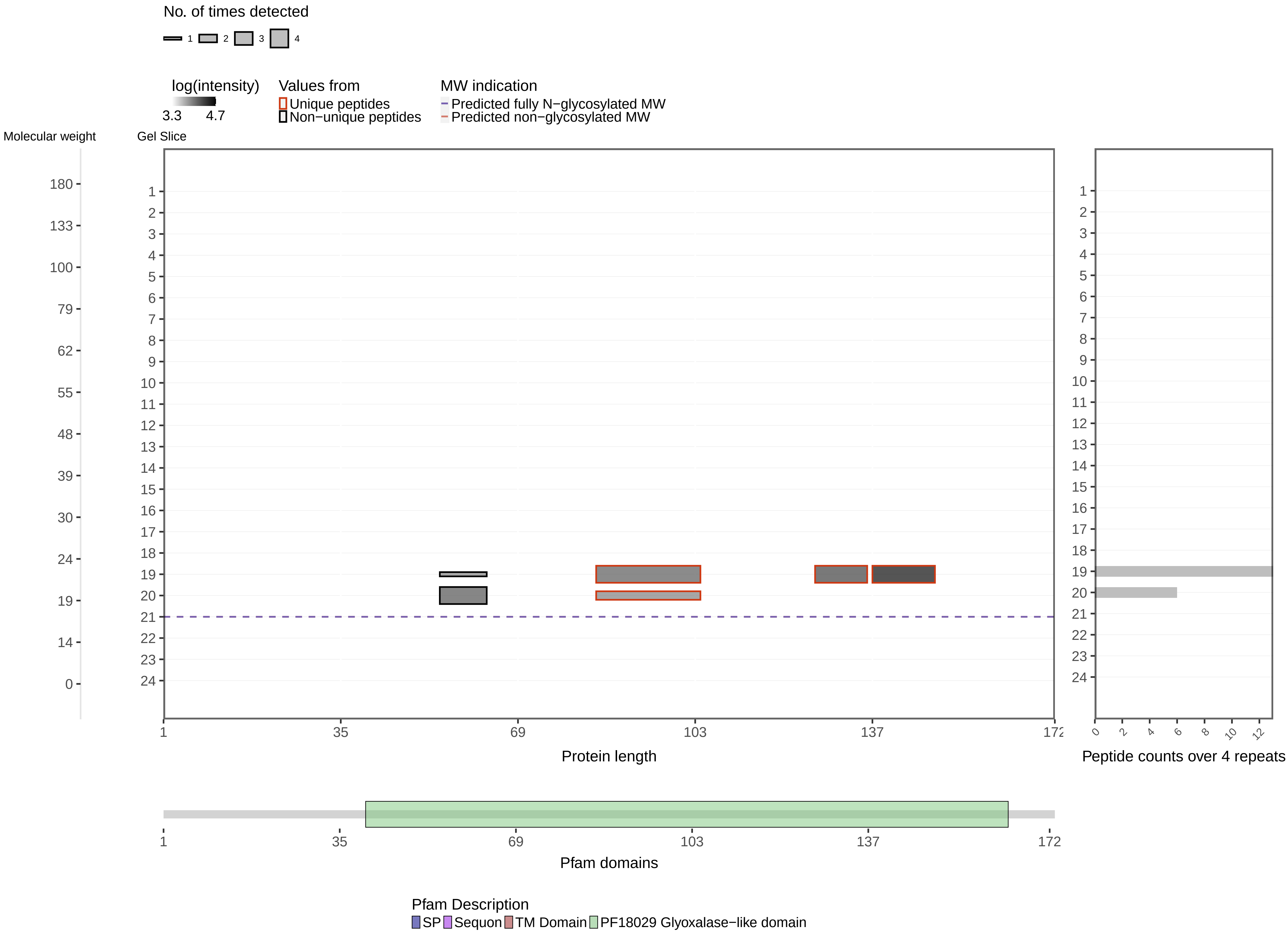


Protein sequence:

DEIPLISSVELKLGSSSFLVSDLTDDDSSAPVKVATTGCVFCLETEDVEAAVAKAVSAGAVSEGKIAEGDGAAGFGGRVGKLKDPYGNVWMICSPVKKSQ*

### NbL12g01340.1 uncharacterized_LOC104248456 [MW − SP = 18.71 kDa]

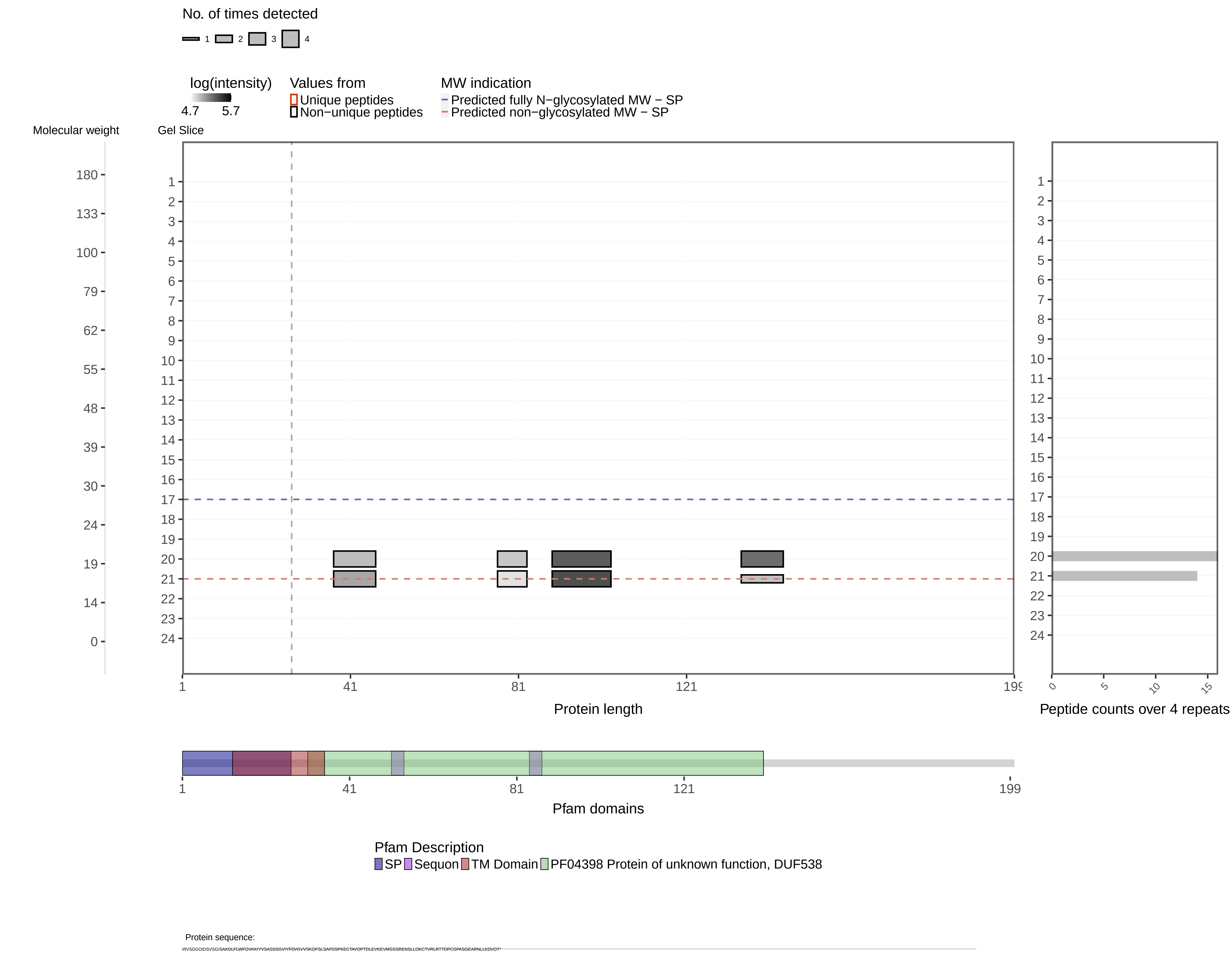

### NbL12g01330.1 adrenodoxin−like_protein_mitochondrial [MW = 17.89 kDa]

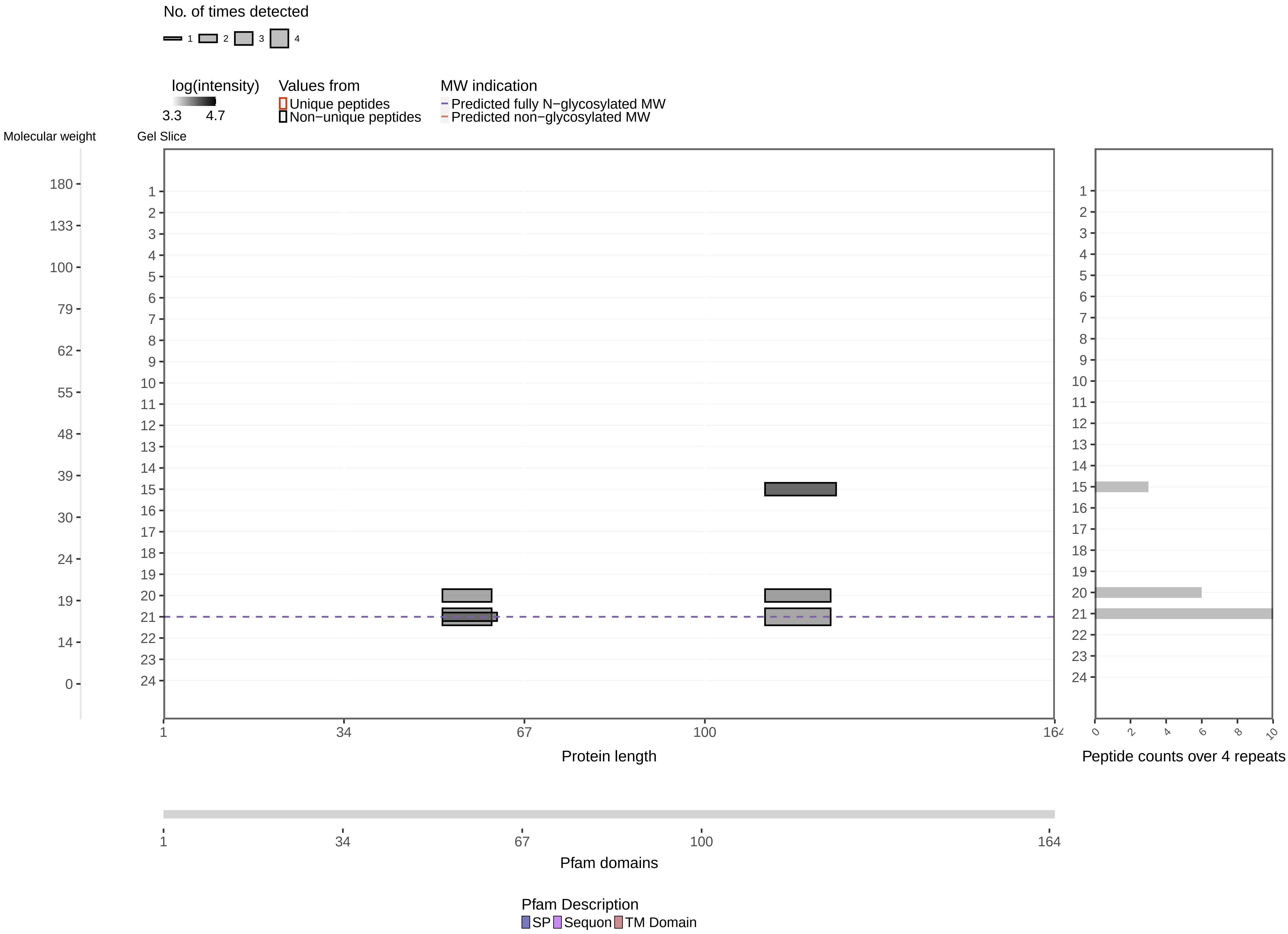


Protein sequence:

QTLLKALTNHGLIDPDSHRLEDIDACSAECEVHIAQEWLEKLPPATYDEQYVLKRNSRARVLNKHSRLGCQVVLSPELQGMVVAIPEPKPWDIP*

### NbL12g01280.1 60S_ribosomal_protein_L5−like [MW = 37.81 kDa]

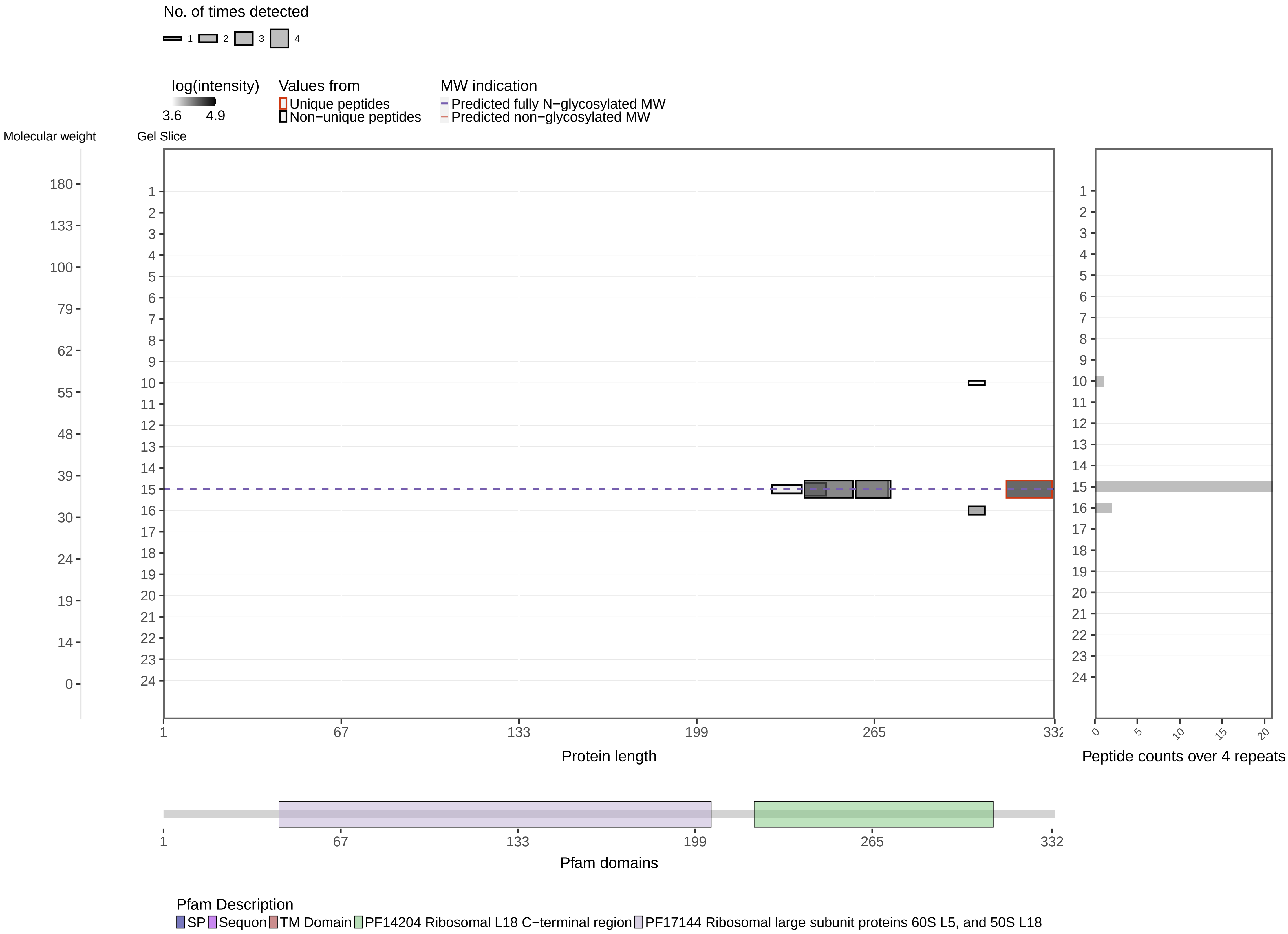


Protein sequence:

VNGEDYSVEPAESRRPFRALLDVGLIRTTTGNRVFGALKGALDGGIDIPHSEKRFAGFGKDSKQLDADVHRKYIYGGHVSAYMKTLMEDEPEKYQSHFSEYIKTGLEADDLEEMYKKVHAAIRADPSSKKSDKPPPKQHKRYNLKKLTYDERKARLIERLNALNAAAGNDDDEEDDN*

NbL12g01190.1 uncharacterized_LOC104090208 [MW = 22.02 kDa]

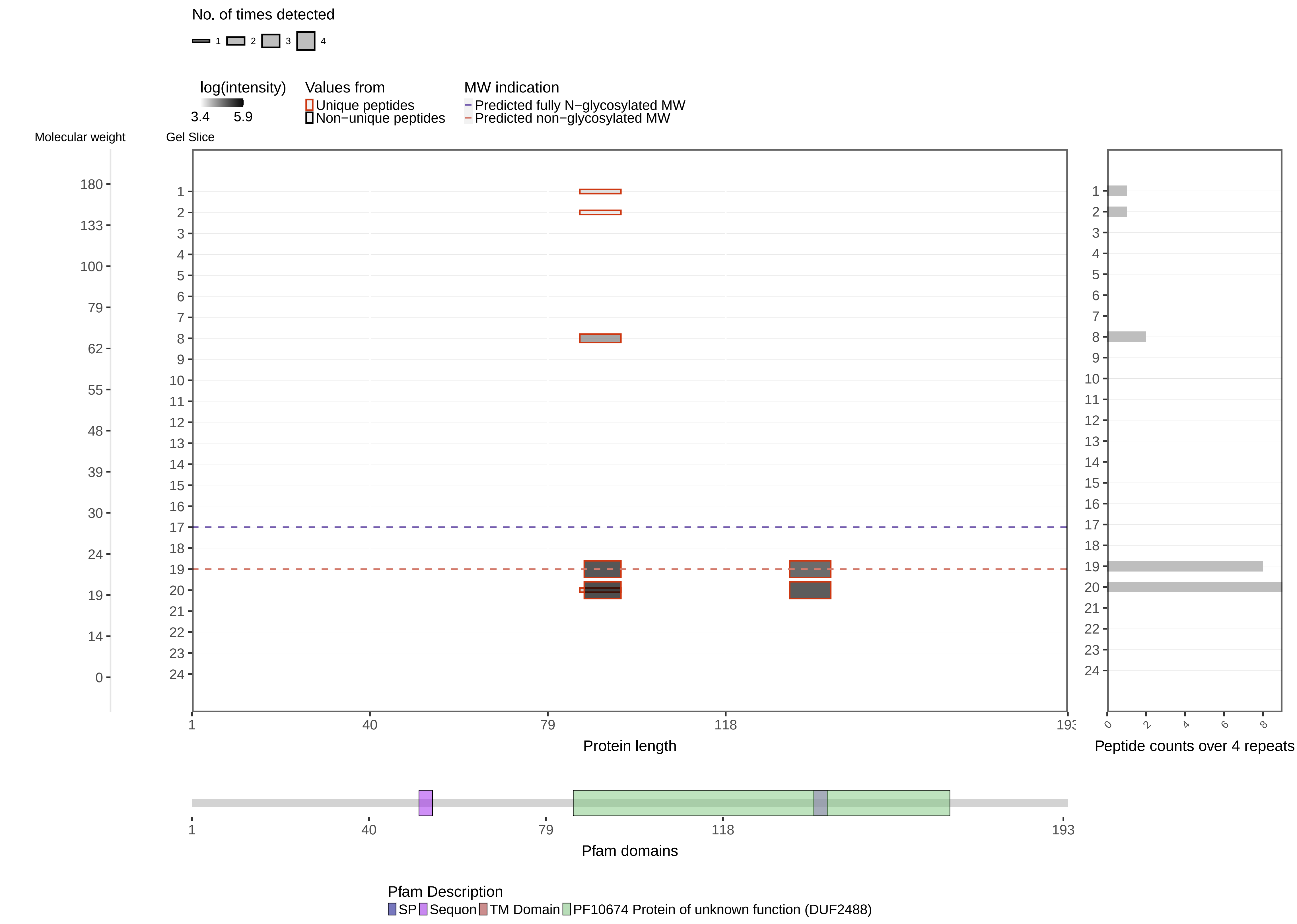


Protein sequence:

TERKKYYFLVANAKFMLDEEEHFQEQLFERLRLFGERDKGQDFWLVIEPKFLDKFPNITKRLKRPAVALVSTNGPWITVTNFRLSYLFFKPSPKIGGSLTVIYSIFKPSSR*

NbL12g00990.1 uncharacterized_LOC104215819 [MW − SP = 11.14 kDa]

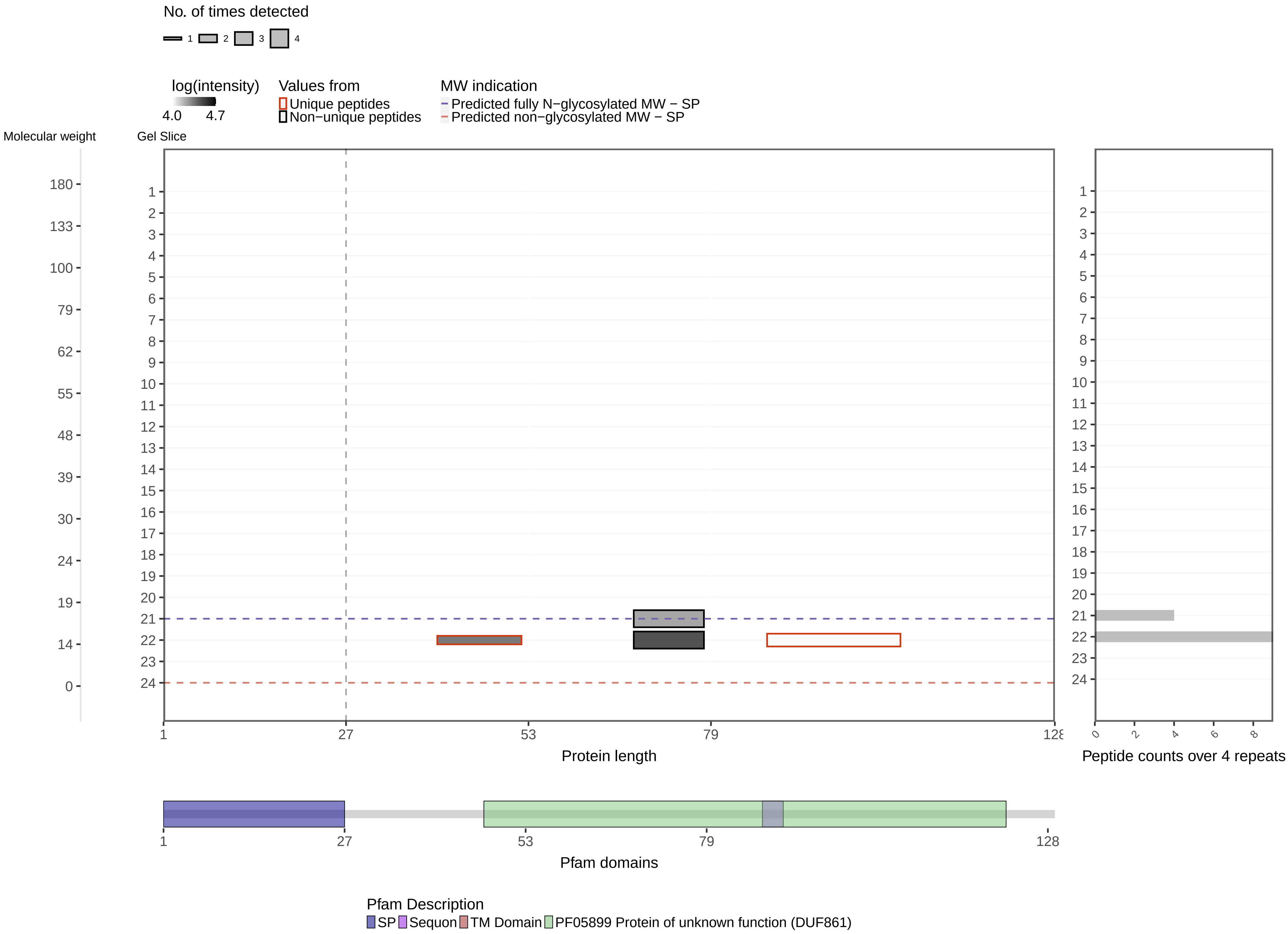


Protein sequence:

SWPKWGCSPGKYQLKFDAEETCYLLRGKVKVYPKNNSSDEIVEFGAGDLVIIPKGLSCTWDVSVAVDKHYKFDSS*

NbL12g00980.1 WD_repeat−containing_protein_70 [MW = 71.56 kDa]

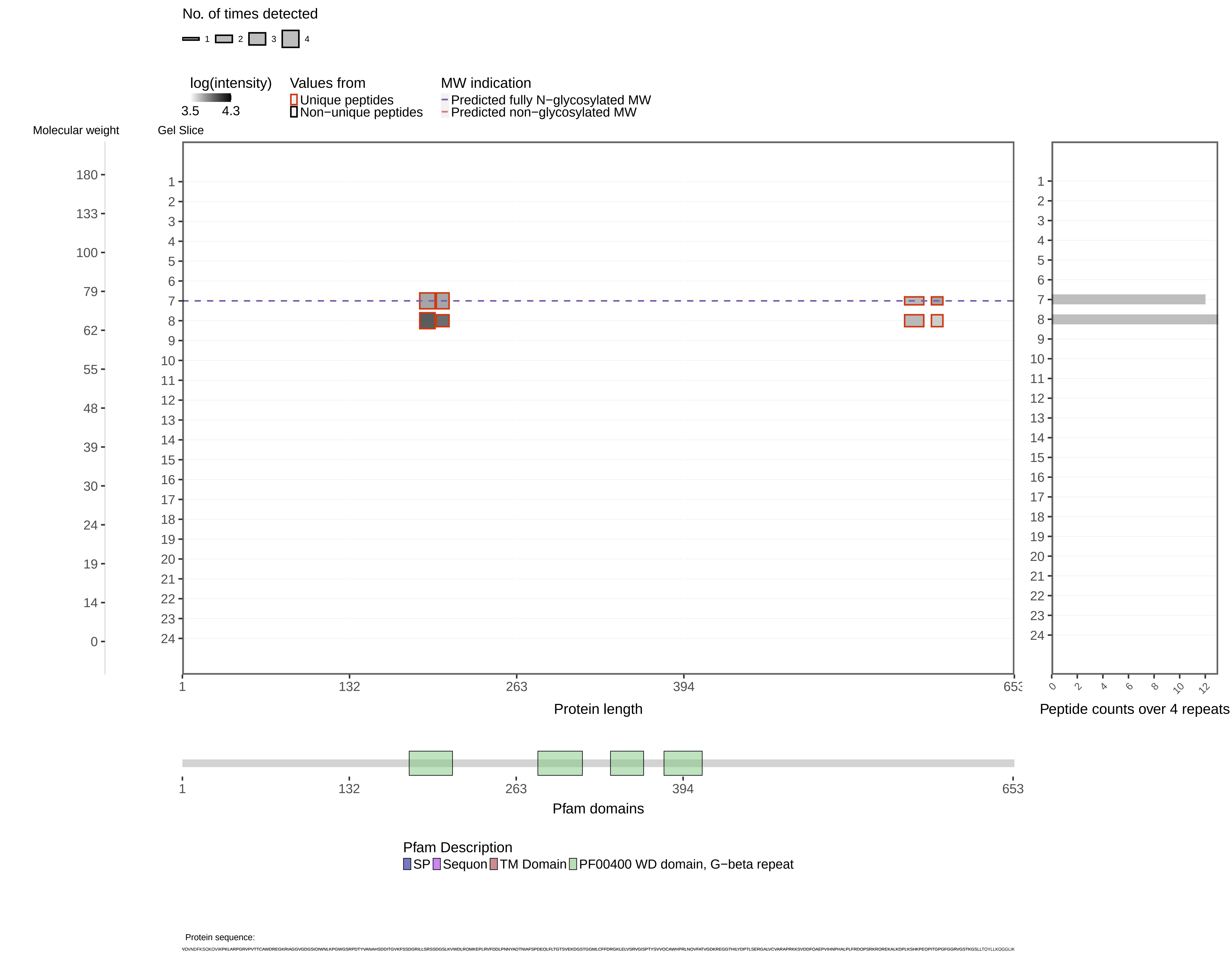

### NbL12g00930.1 protein_FLUORESCENT_IN_BLUE_LIGHT_chloroplastic−like [MW = 31.99 kDa]

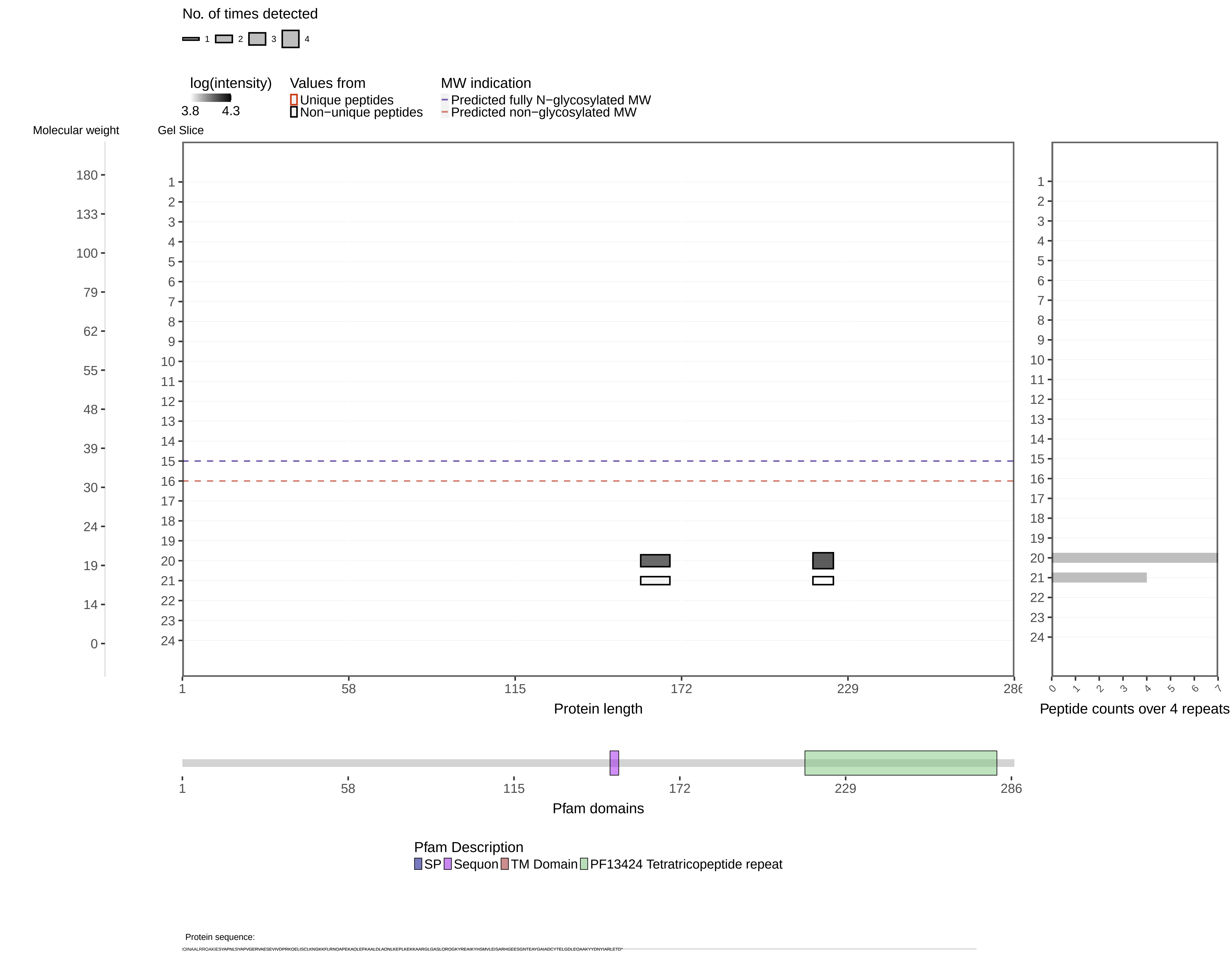

### NbL12g00850.1 60S_ribosomal_protein_L14−1 [MW = 15.4 kDa]

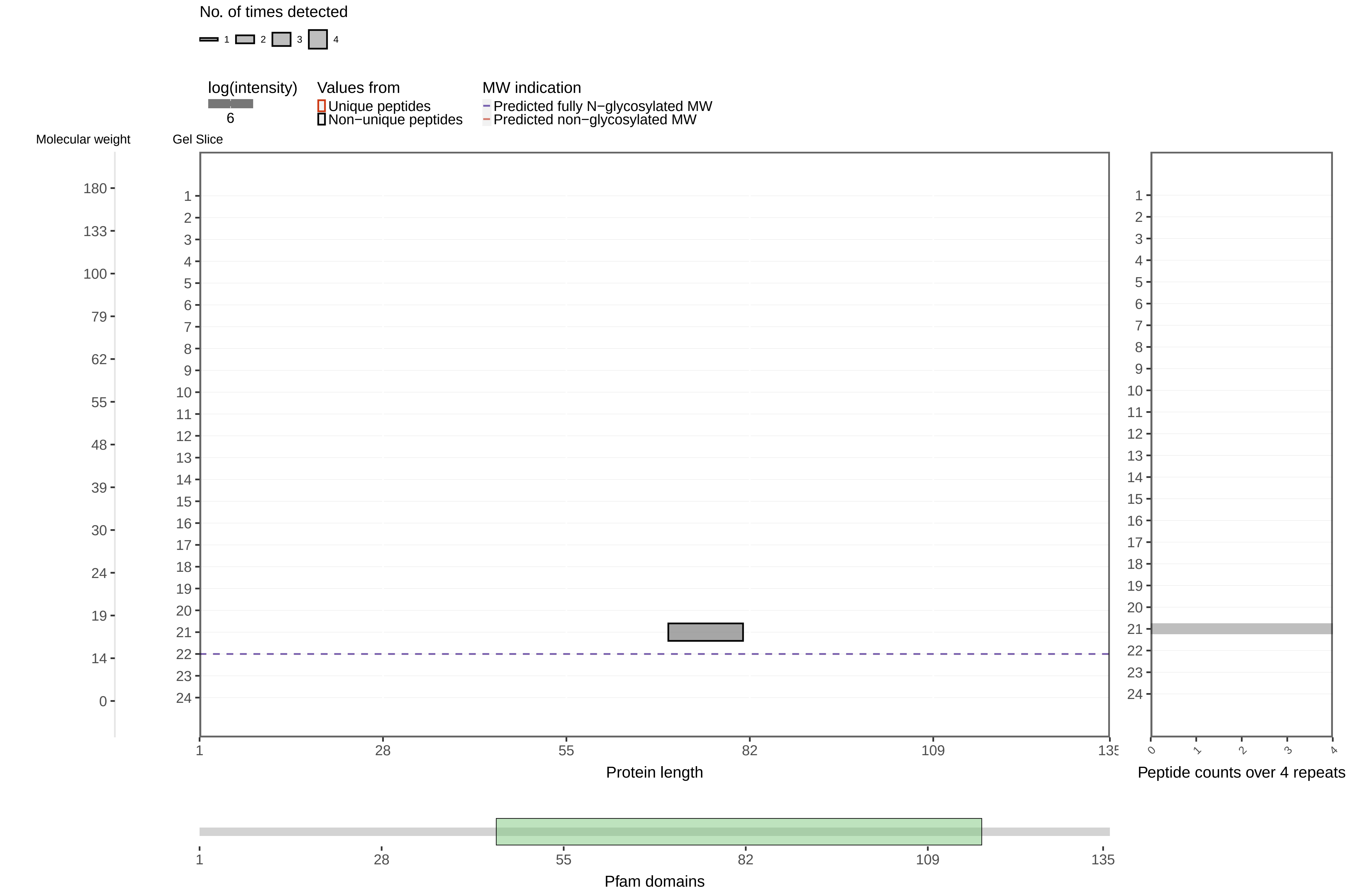


Protein sequence:

SLTDIKIDIKRIPKKKTLVAAMEAADVKNKWESSSWGRKLIVQKRRASLNDFDRFKLMLAKIKRAGVVRQELAKLKKETVA*

NbL12g00680.1 F−box/kelch−repeat_protein_SKIP30−like [MW = 39.34 kDa]

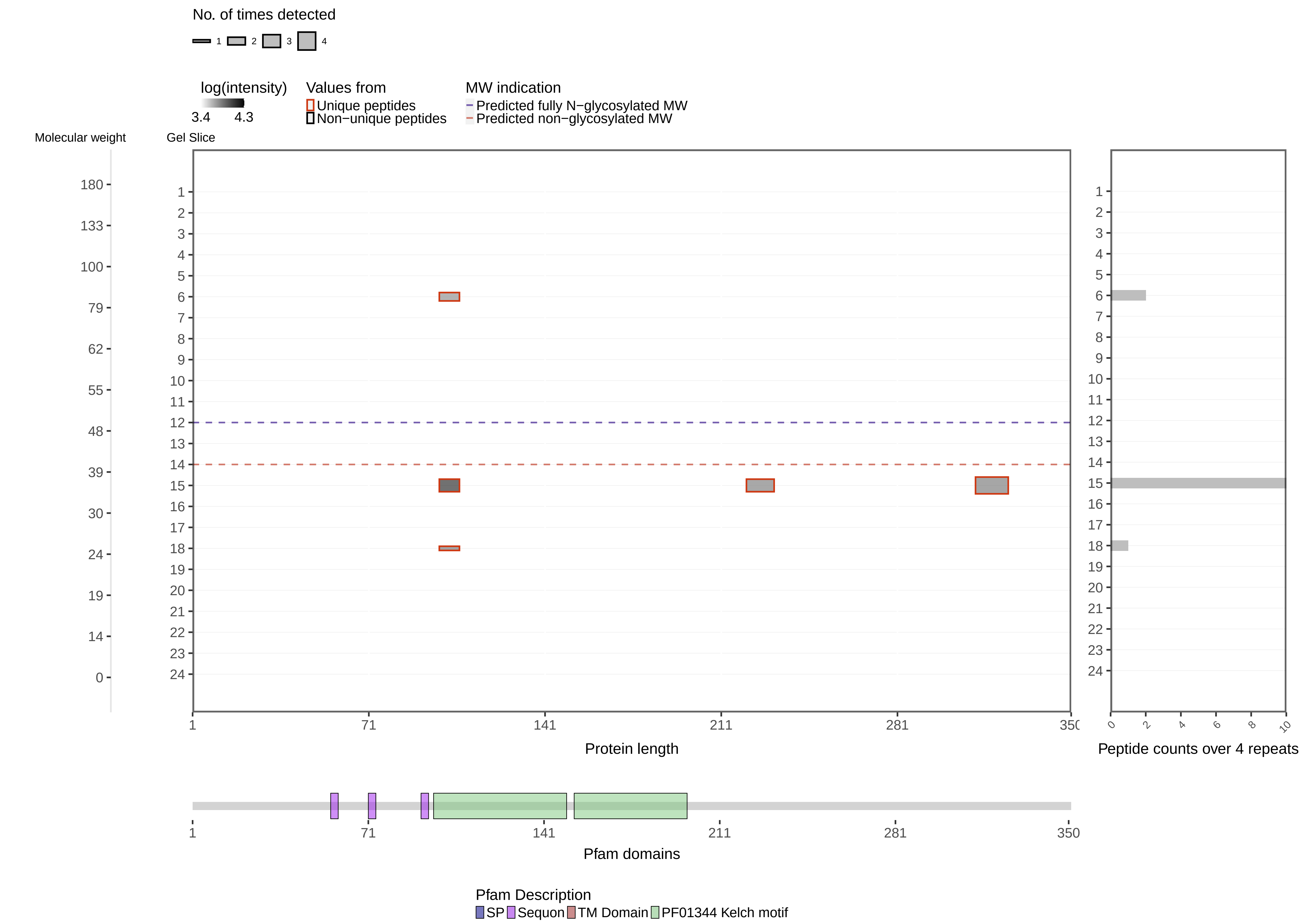


Protein sequence:
EGKIIVAGGFTNYRKSICNAEIYDPEKDVWVQLPDLHHAHNSACSGVVVSGKVHVLHKGLSTIQVLEIVKQGWTVHEYSWLQGPMTVVREKLYVMSNWFIYKQEKEVREMVVSASEFRRRIGFAMIGLGDDIYIVGGVNGPEHWNCNIKVLSDVDVLTLGNERLVWRKVAPMTRCRGTILACTQLRI*

### NbL12g00670.1 tRNA_(guanine−N(7)−)−methyltransferase_non−catalytic_subunit_WDR4 [MW = 47.11 kDa]

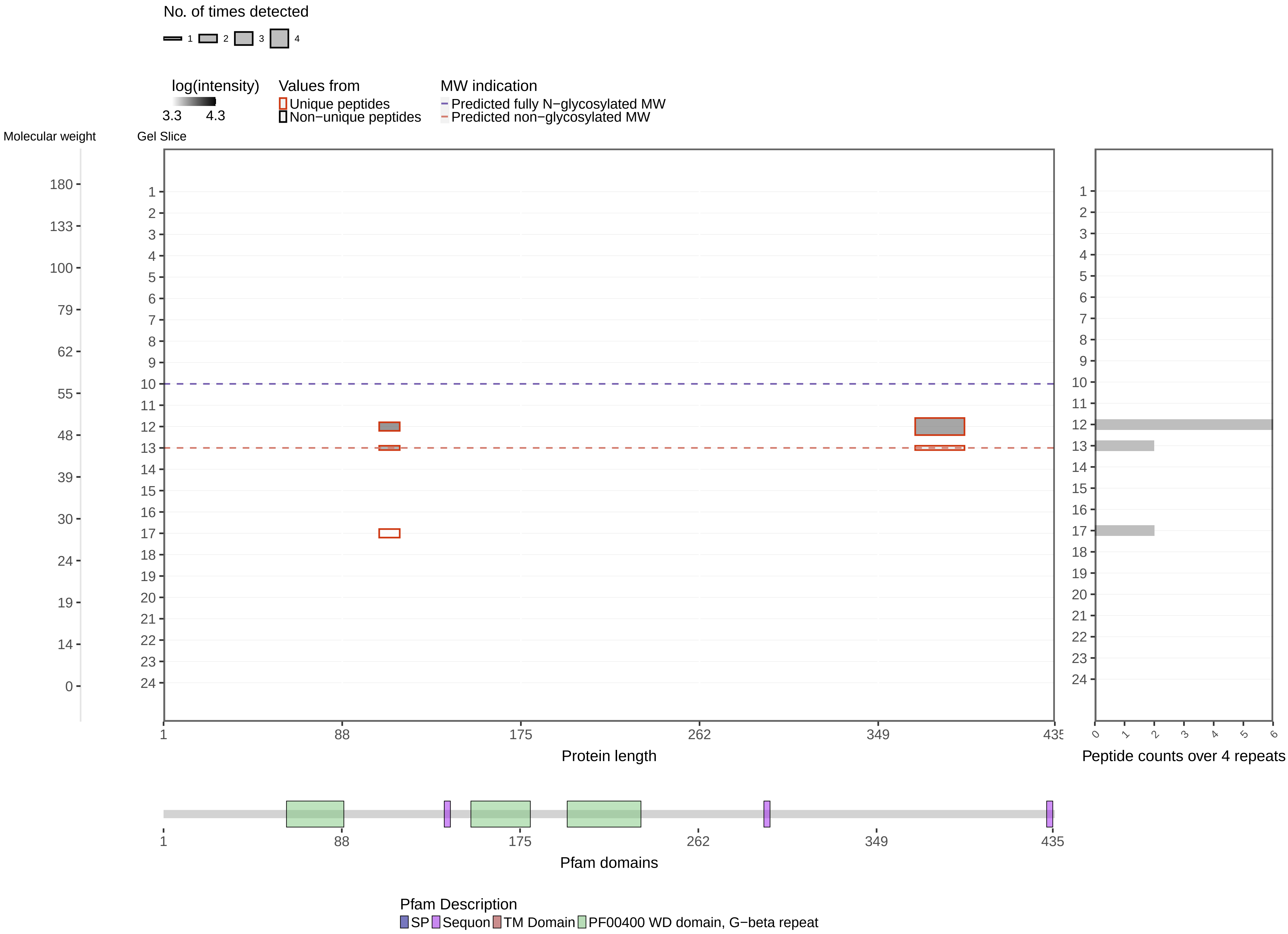


Protein sequence:

SCLAFICSKDSQQRYLLSGGGDSTVRLWDFTCGSLLDTCHIGPETGLSQSKGVDDSLLAVTDLCATRGGSLIAVAIQSSLAGVVLLSCNLSARSLHFARVVPIPGETFIPTSIEAAYSSNQLWMVMGASTLCASDSAPLARVKVVDGFCESNQDSVEHEMRVLEDKDIQGGEQLLQTLQGSSSIEKDALATAAEAVKTAMCNLLIKKQYPSENRDFRKRGRNDKKVNKS*

### NbL12g00550.1 elongation_factor_Tu_mitochondrial−like [MW = 49.06 kDa]

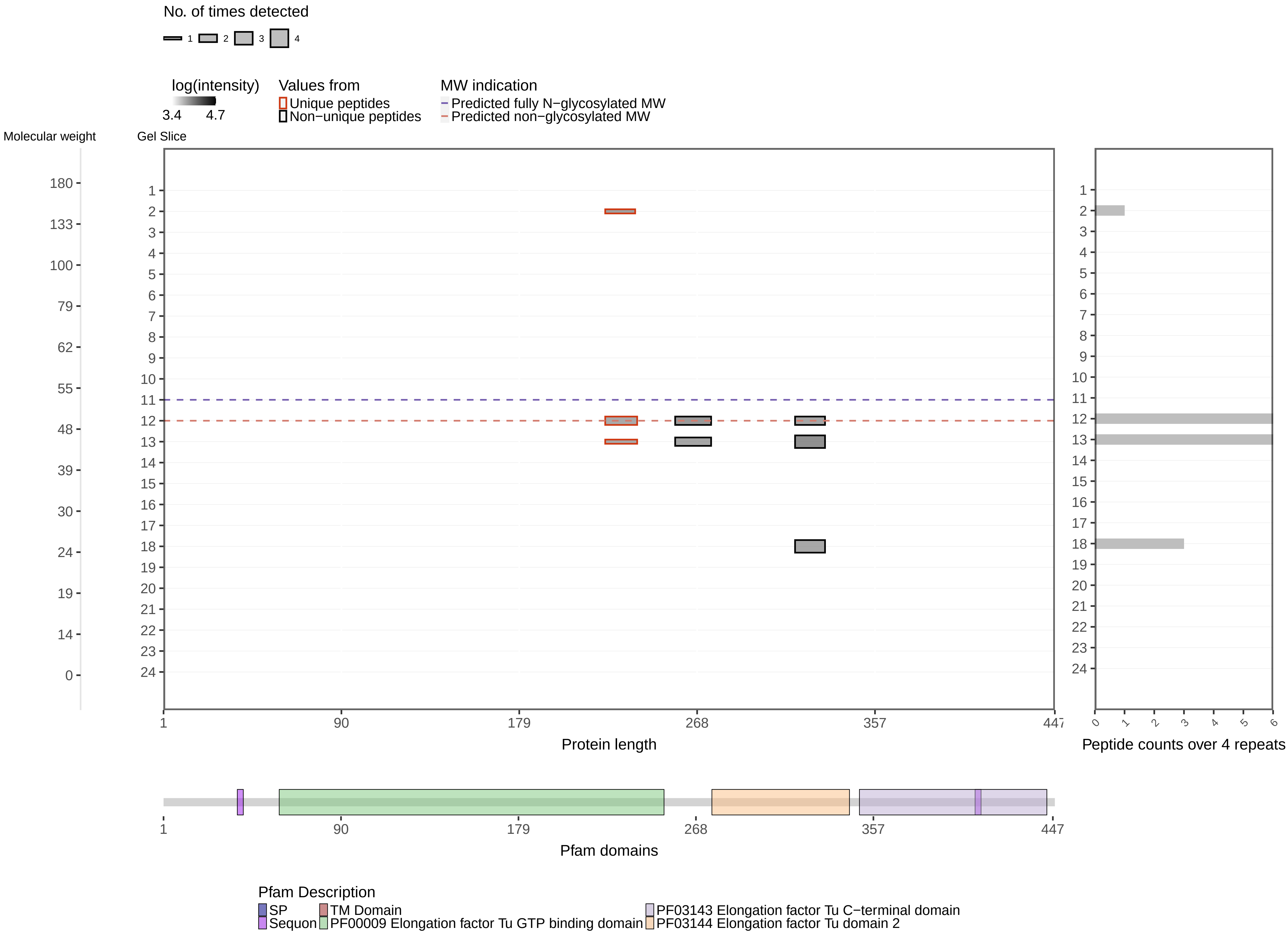


Protein sequence:

KFPGDEIPIIRGSALSALQSTNEEIGKKAILKLMEAVDEYIPDPVRQLDKPFLMPIEDVFSIQGRGTVATGRVEQGTIKVGEDVEILGLMQGNLKSTVTGVEMFKKSLDYGQAGDNVGLLLRGLKRDDIMRGMVIAKPGSVKTYKKFEAEIYVLTKDEGGRHTAFFSNYMPQFYMRTADITGKVELPENVKMVMPGDNVTATFELMSPVPLDAGQRFALREGGRTVGAGVVSKVIS*

### NbL12g00450.1 glutathione_S−transferase [MW = 18.69 kDa]

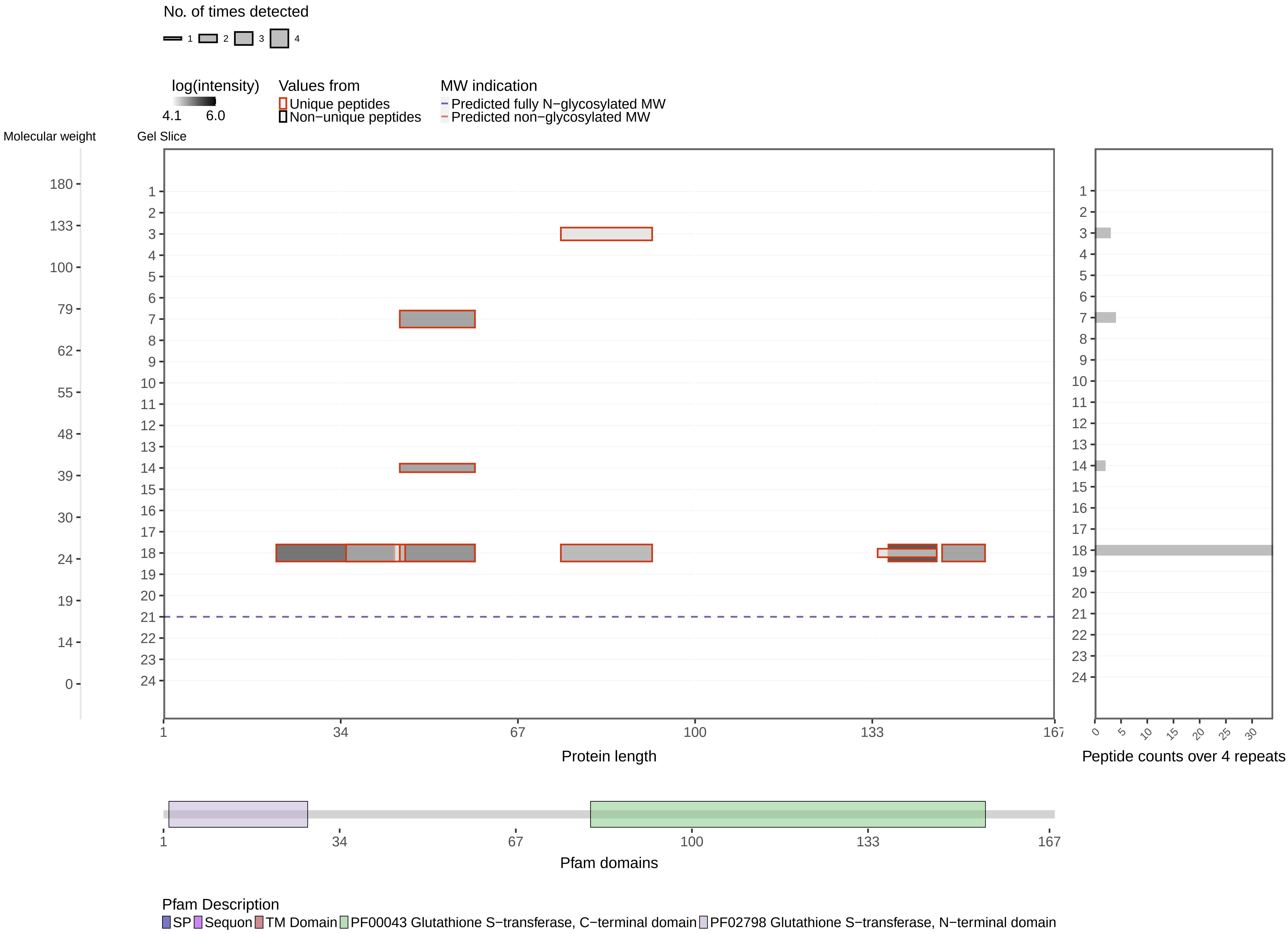


Protein sequence:

LVVKPMLGMVTDDVAVAENEEKLGKILDVYESRLKESKYLGGESFTLADLHHAPVVHYLMGTRVKNLFDARPHVKAWCADILARPAWSKALELHKQ*

NbL12g00430.1 universal_stress_protein_A−like_protein [MW − SP = 19.55 kDa]

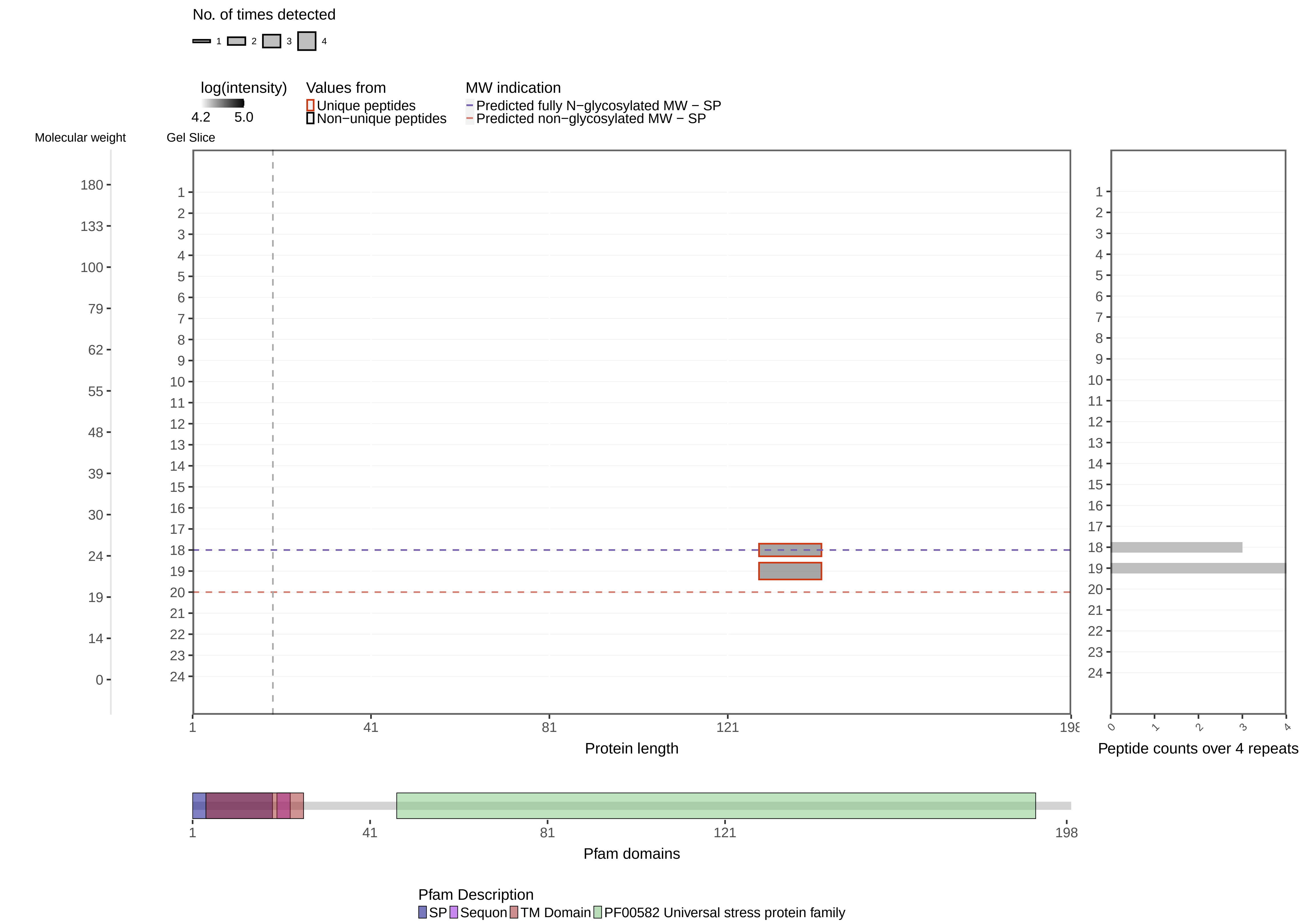


Protein sequence:

PASVIGLTGLGSVEILPSIETDLKKIAARVVEKAKEICTAKSVNDVIVEVEEGDARNVLCEAVEKHHASILVVGSHGYGVLRRTVLGSVSDYCAHHAHCSVMIVKKPKHKS*

NbL12g00400.1 bifunctional_epoxide_hydrolase_2−like [MW = 36.42 kDa]

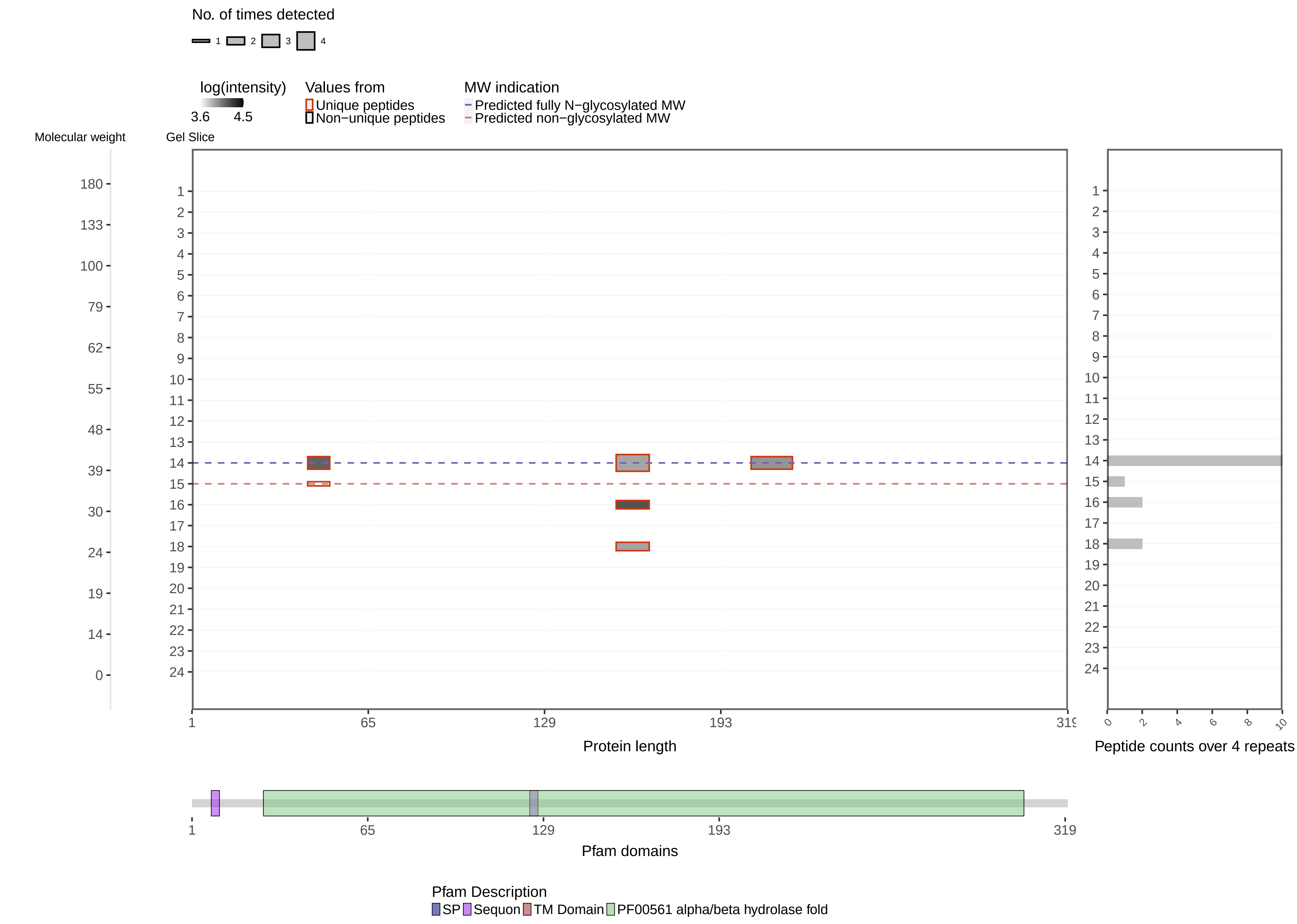


Protein sequence:

GDDYYICRFQEYGQVEEEFARVDTARLIAKFLISRNPAPLRVSKEKGFGGSSHTPIKLPSWLSEEDINYYANKFSQKGFTGGLNYYRAMDLNWELTAPWTGQQIKVPVKFIVGDLDLTYNTPGVKEYIHKGGFQKMVPFLQEVVIMEGVGHFINEEKPEEINDHIYDFIQKF*

### NbL12g00390.1 pectinesterase−like [MW = 64.09 kDa]

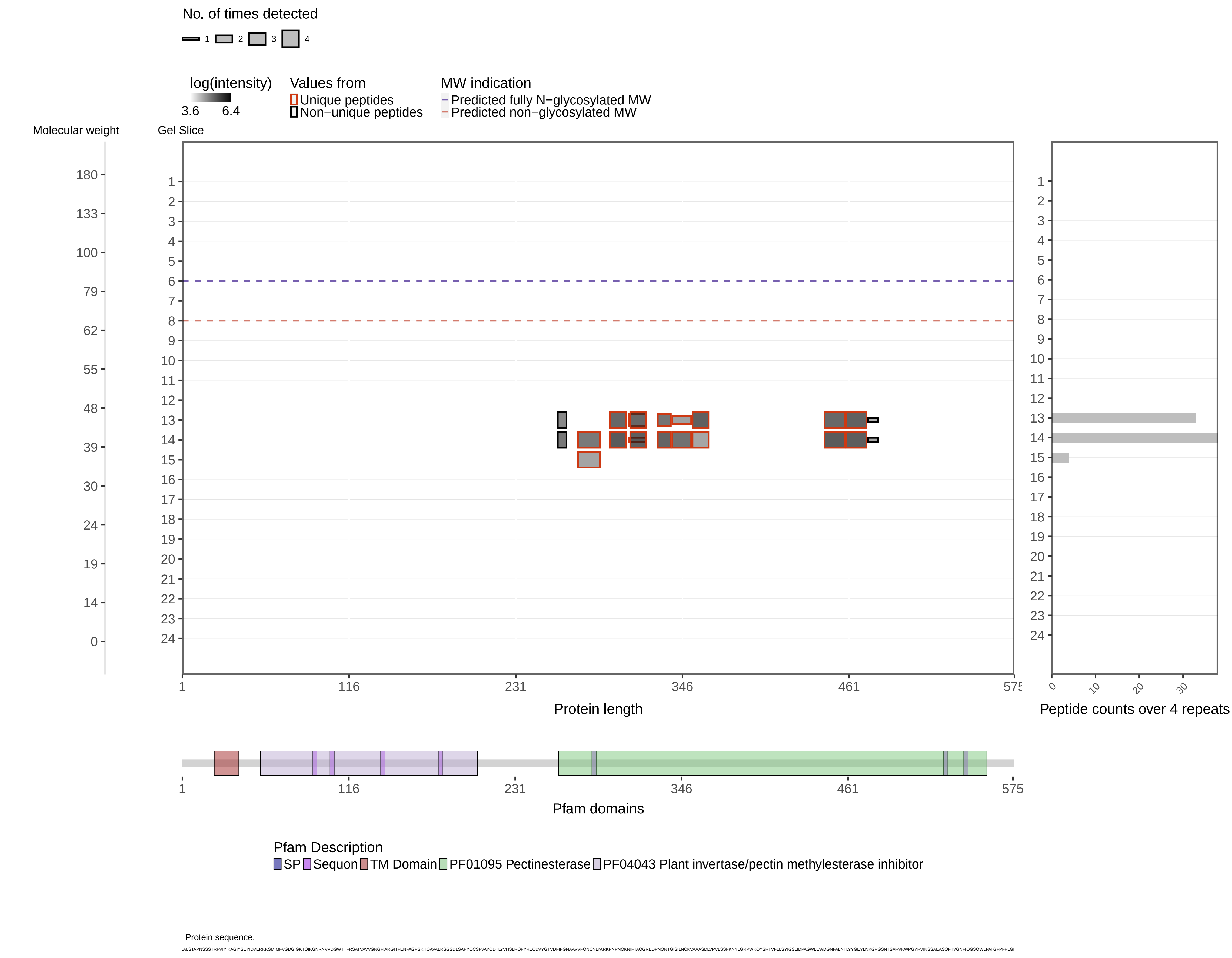

### NbL12g00370.1 60S_ribosomal_protein_L19−3 [MW = 24.79 kDa]

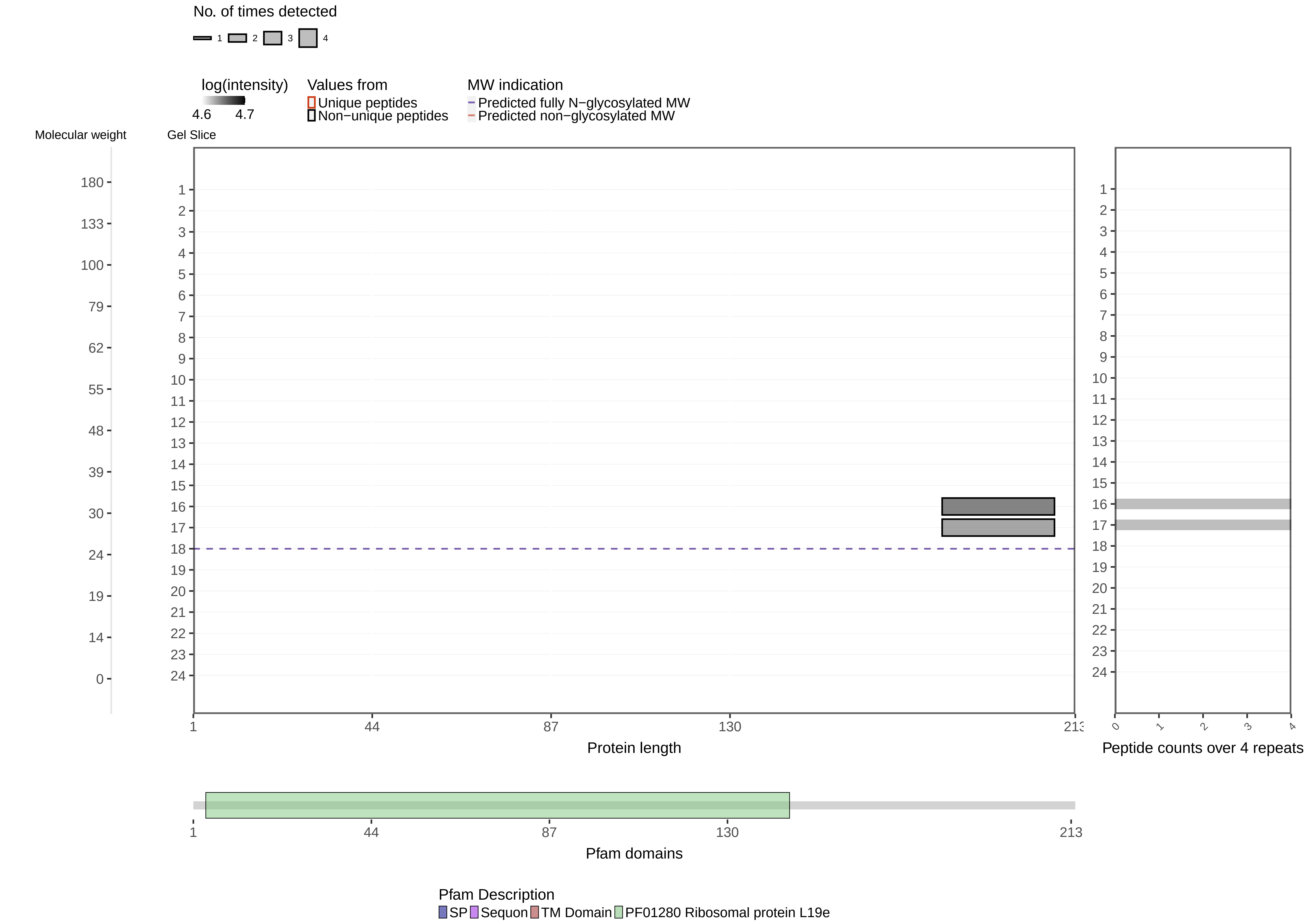


Protein sequence:

WMRRMRVLRRLLRKYRESKKIDRHMYHDMYMKVKGNVFKNKRVLMENIHKTKAEKAREKTLSDQFEARRAKNKASRERKFARREERLAQGPGERPVQPAAPAAVAPTQPAQGSKKSKK*

### NbL12g00350.1 probable_polygalacturonase [MW − SP = 46.36 kDa]

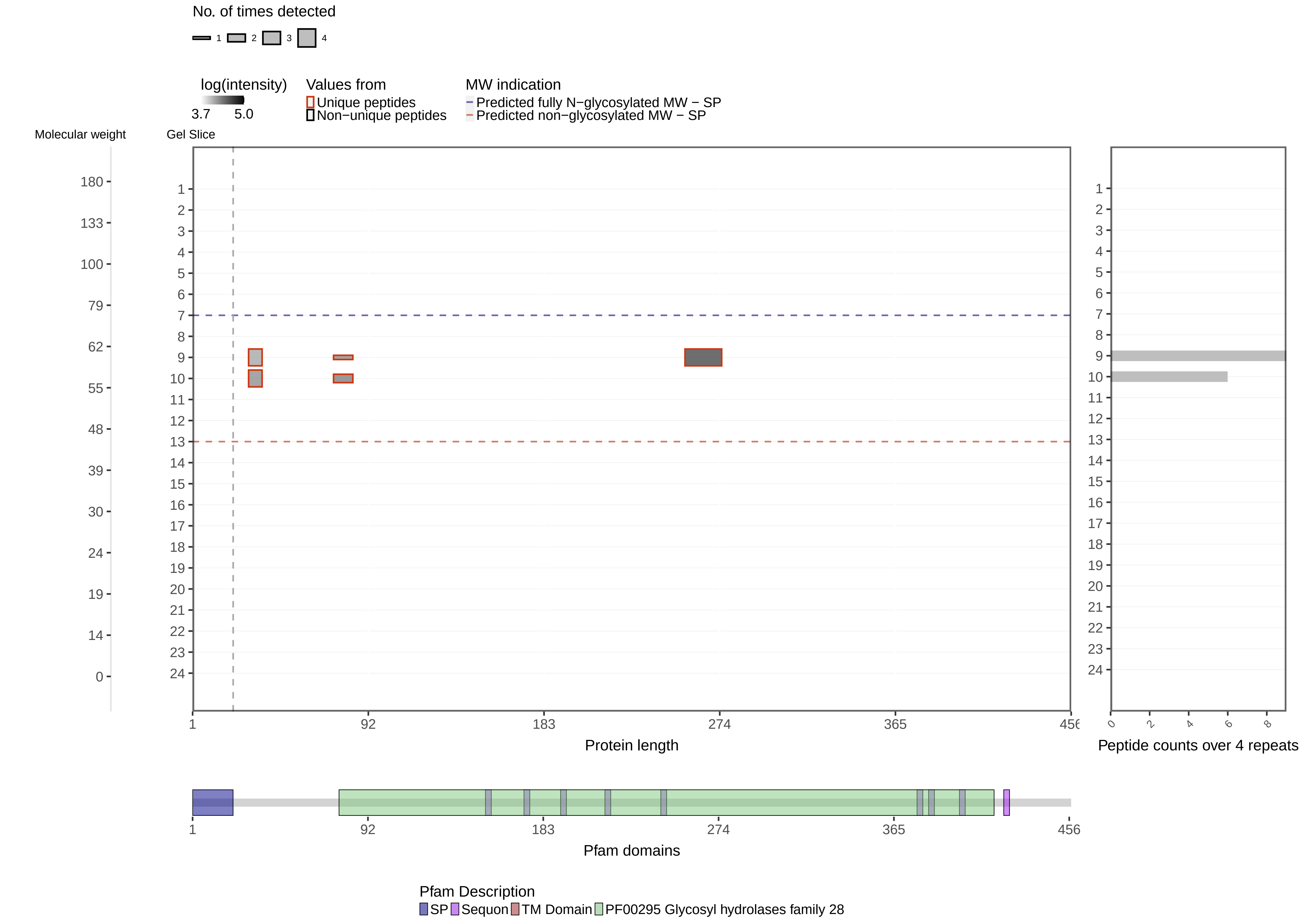


Protein sequence:

ITVHSPANSPYTYGIVPDSSEHVCIENSNISMGHDAVVLKSGWDEYGISYGKPTTNVHVRGVRLQSCAGAGLAFGSEMSGGISNVLAEHVYLHDSLFGIELKTARGRGGYIKDIIVTDVVMANLQVGIEATGHCDSHPDEKFDPSALPVVSGITFEDIVGTNISTAGNFSGLAESPFTSICLSNVTFSVSSNPTPWLCSDISGSSQNVSPDPCPELQNSFSGTTSTCFALLHPYSQVAFS*

NbL12g00220.1 calvin_cycle_protein_CP12−1_chloroplastic−like [MW = 14.3 kDa]

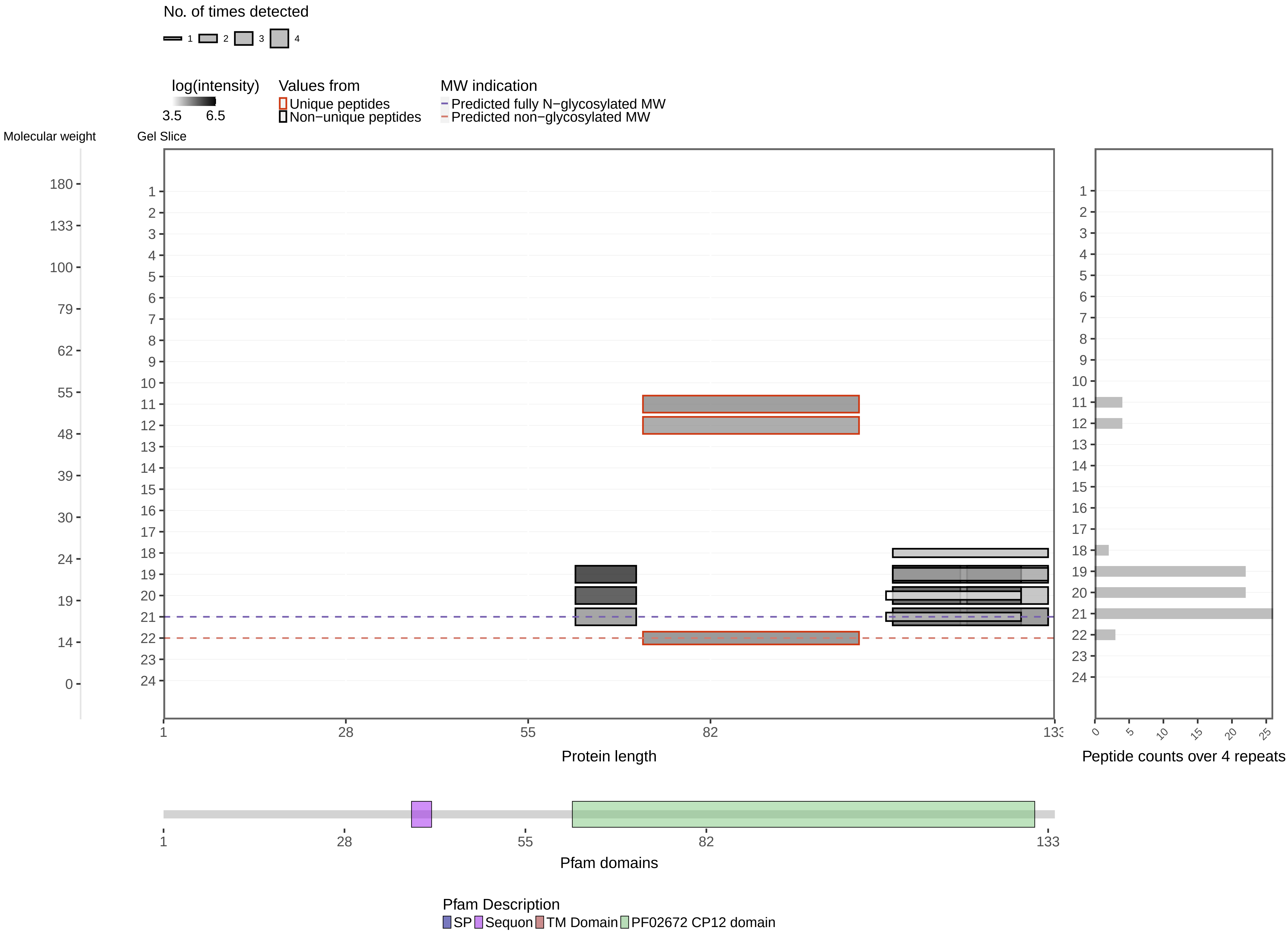


Protein sequence:

SATPDNKLSDLVAESVKEAEEACAENPVSGECAAAWDVVEEASAAASHARDKKKESSDPLENYCKDNPETDECRTYDN*

### NbL12g00170.1 nodal_modulator_3 [MW − SP = 129.1 kDa]

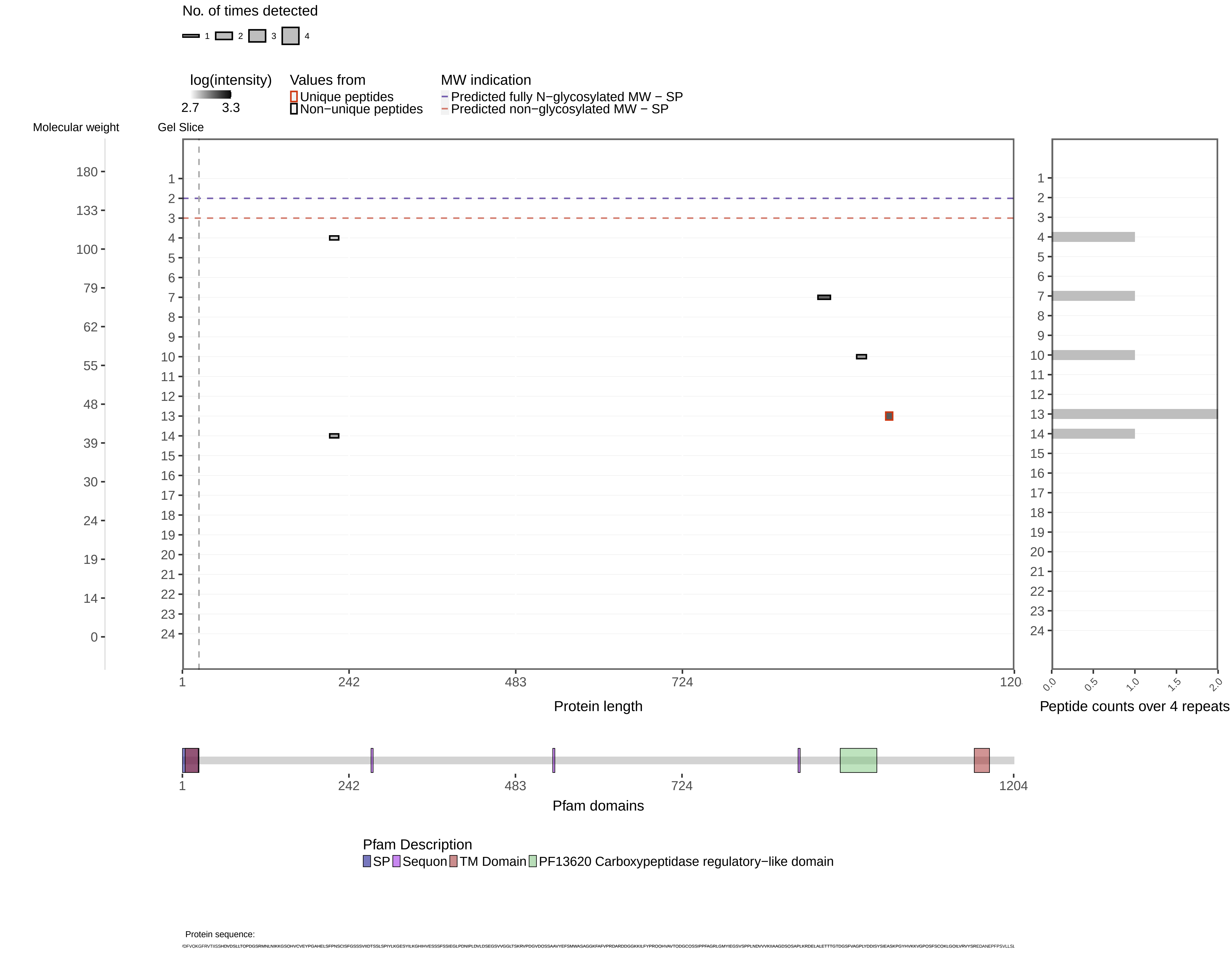

### NbL12g00140.1 ABC_transporter_B_family_member_21−like [MW = 139.82 kDa]

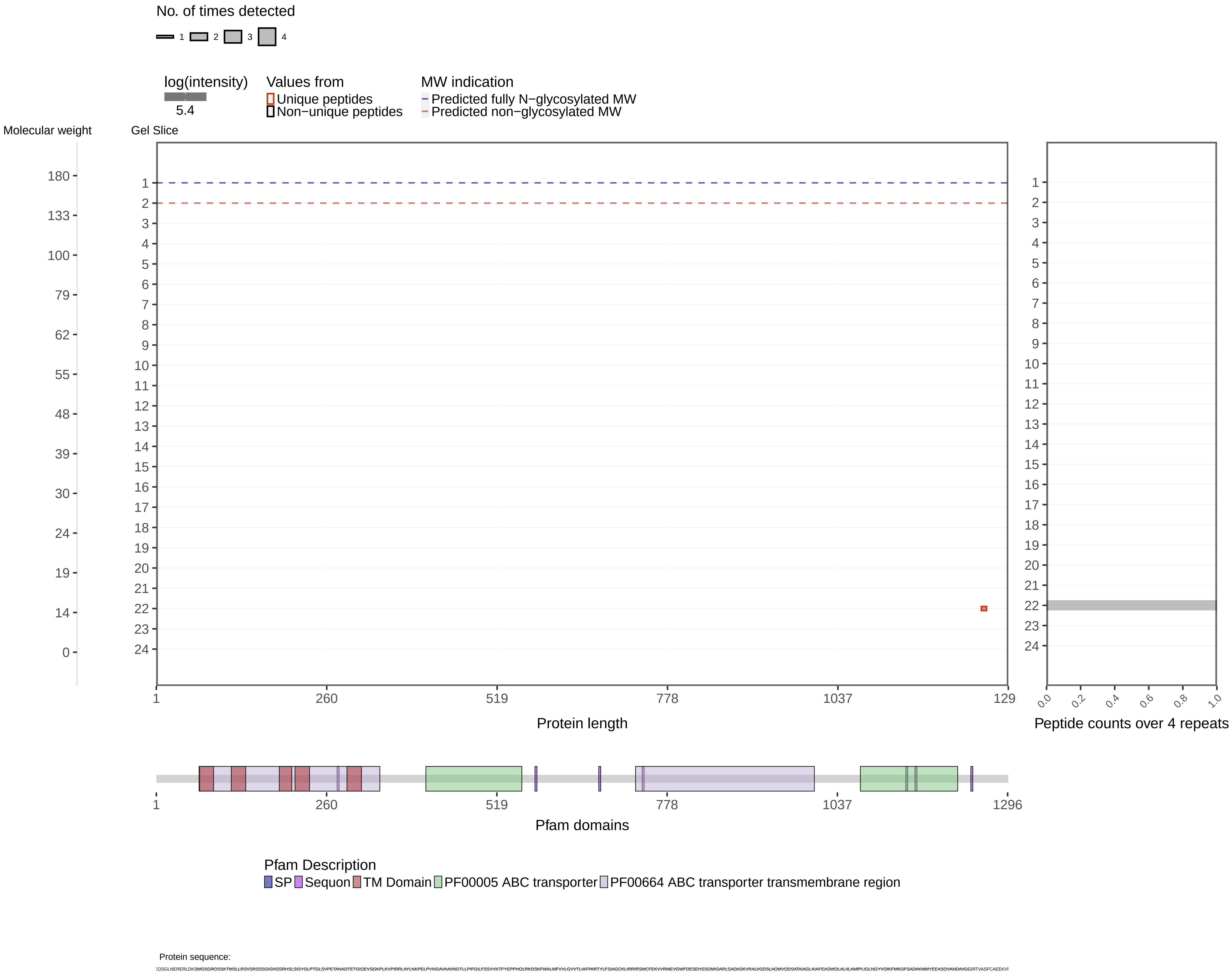

NbL11g24050.1 translocase_of_chloroplast_159_chloroplastic [MW = 158.33 kDa]

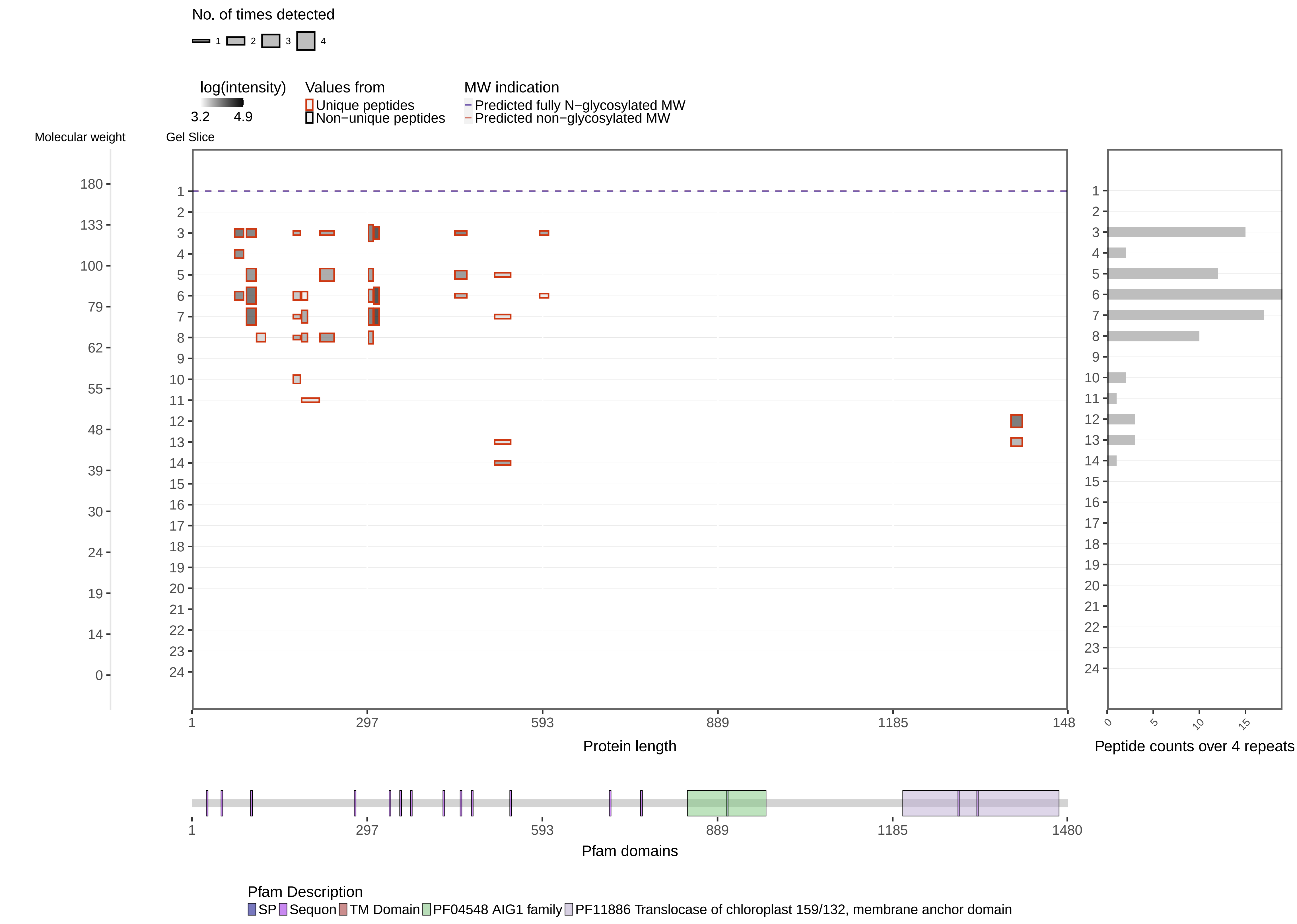


Protein sequence:

GLGSSLRSLRPAPQPNRPNLFTSSSLQNSGESENNLSEEEKKKLEKLQQIRVKFLRLIHRLGLSSDESIAAQVLYRLALIARRQNSPLFSLEAAKMRARQLEAEGEDDLDFSVNIQVIGKSGVGKSATINSIFGEEKTPINAFGPATTSVKEISGVVEGVKIRVFDTPGLKSSVMEQGFNRSVLSSAKKFTKKNPPDIFLYVDRLDAQTRDLNDLPMLKTITSCLGPSIWRSAIVTLTHGASAPPDGPSGSPLSYEVFVTQRSHVVQQSIGQAVGDLRMMSPSLMNPVSLVENHPS

NbL11g24030.1 WD_repeat−containing_protein_89_homolog [MW = 42.43 kDa]

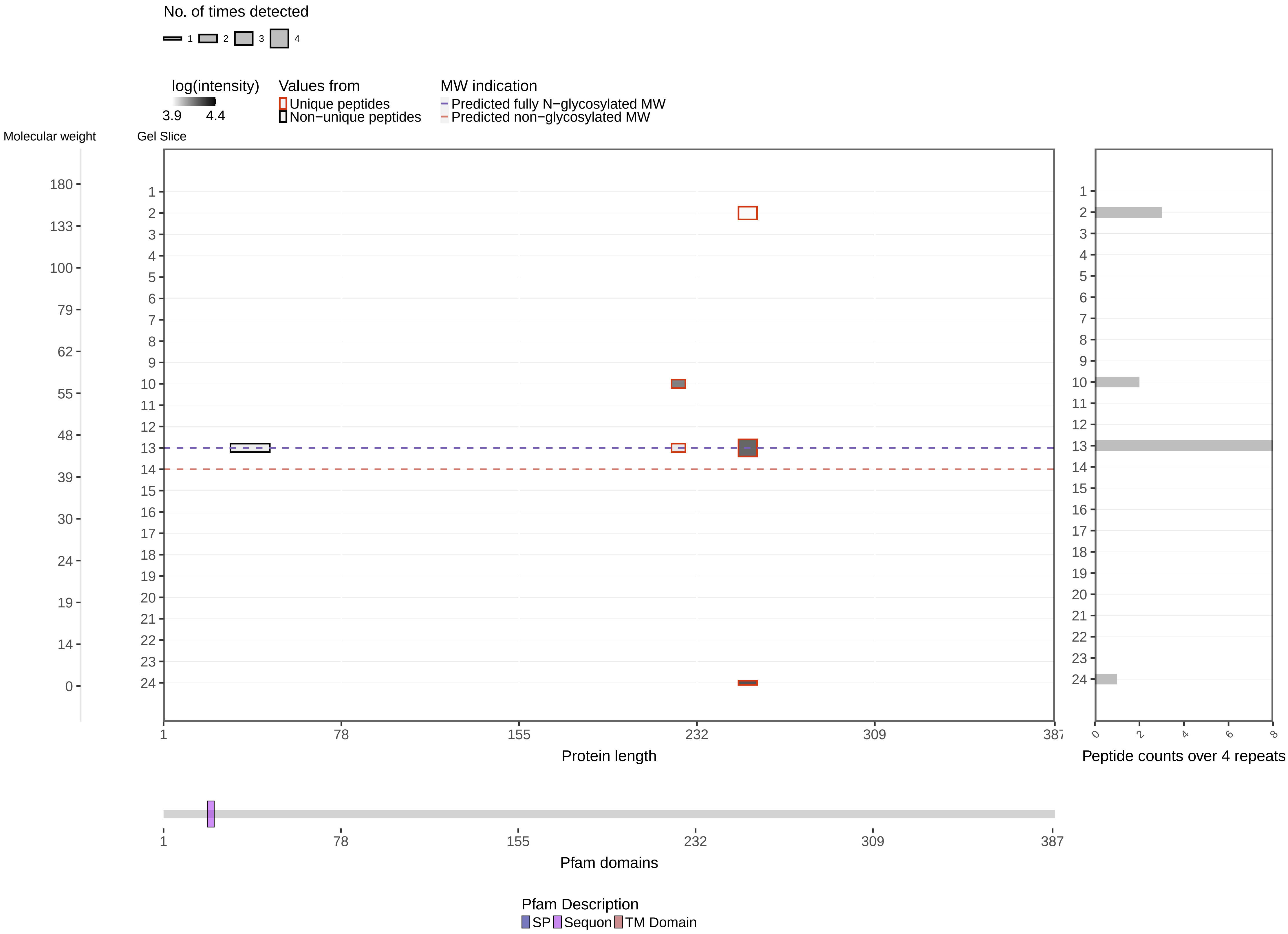


Protein sequence:

DDRLISASVDGLICLFDTGGDINDDDHLVSVLNVGTSVGKVGFLGEKNQKLWCLTHIETLSFWDWKDERIEASFEDARALASKSWITNDVDYFVDCHYSVEEDRFWVIGGTNGGSLAYCPVNYKGTRVIGPPEAILSGGHEGIIRSVLSGVPGETMDNHGIFGWTGGEDGRLCCWLSDGSDNSQSWISTLALKSEKTRTKSRHSPY*

### NbL11g24020.1 uncharacterized_protein_OsI_027940 [MW = 23.3 kDa]

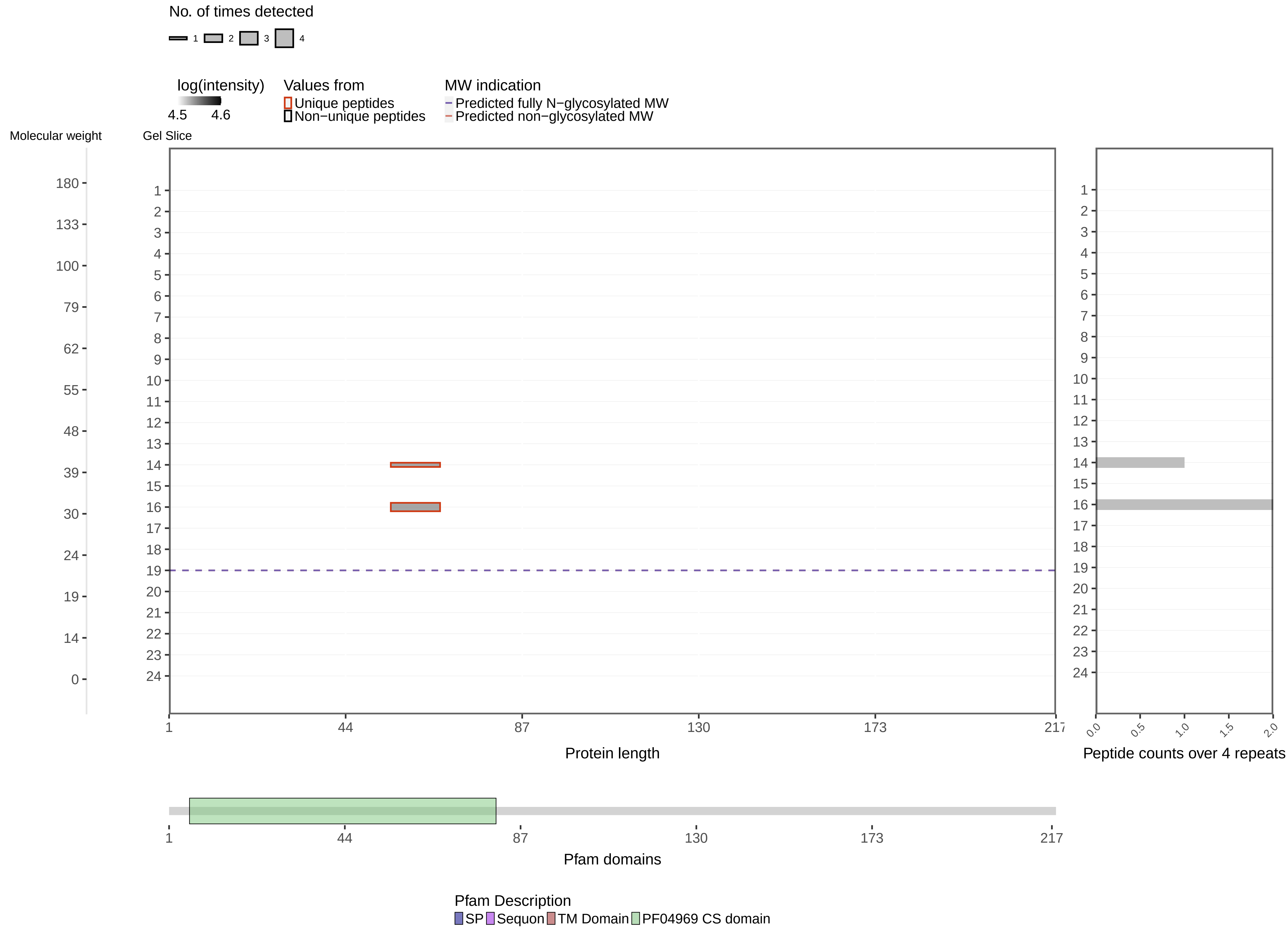


Protein sequence:

KVDWDKWVDEDDEADAGPGDMDMNSMDFSKFGDMGGMMGGMGGDMAGMMGGMGGMMGGMGGDMAGMMGGMGGDMGDDLDDSDDEEQEVSKPSAKADEKVDGEPKEEGSKVVDEAK*

NbL11g24010.1 ABC_transporter_C_family_member_14−like [MW = 168.99 kDa]

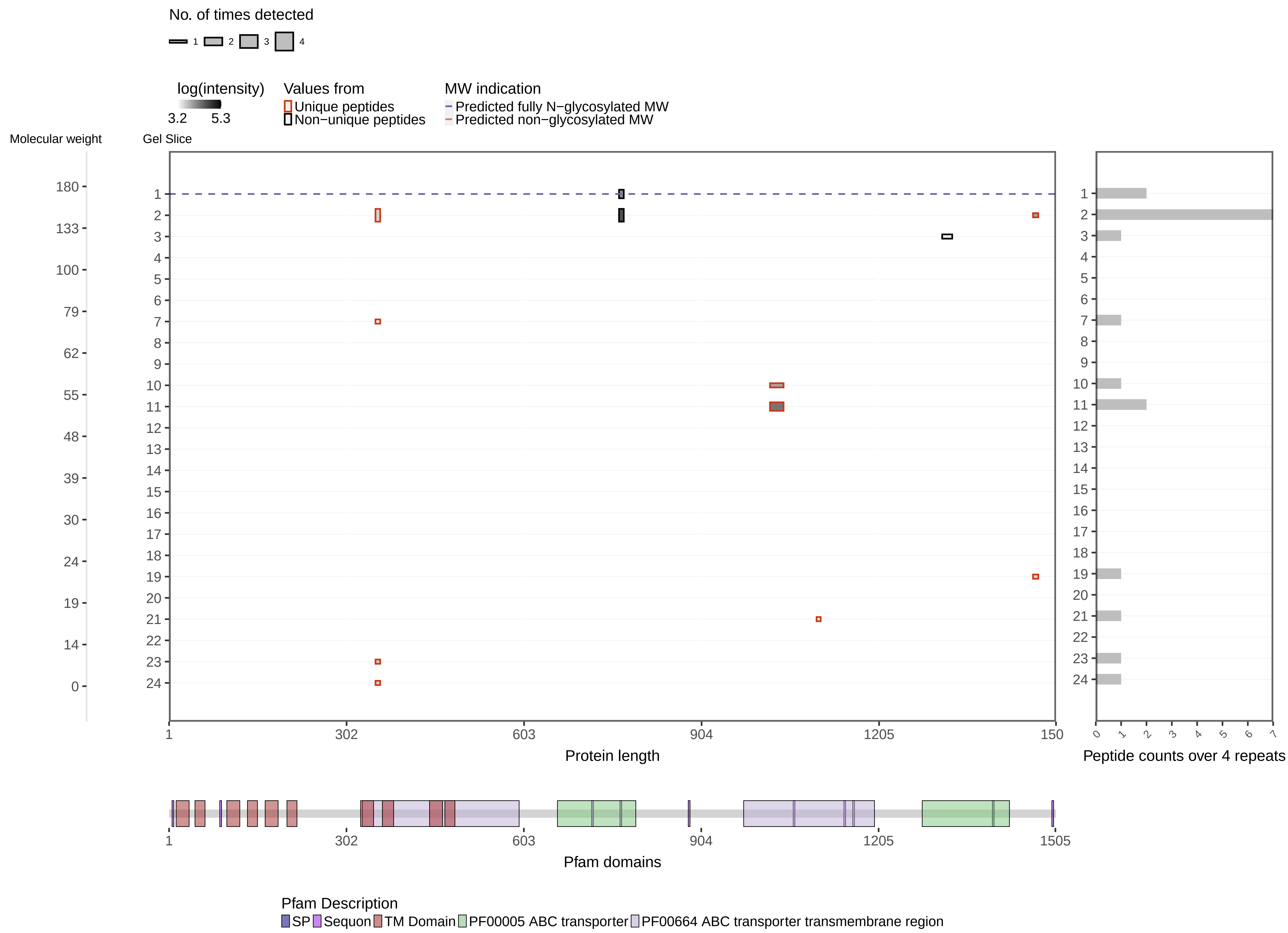


Protein sequence:

[illegible]

### NbL11g23970.1 mitochondrial_zinc_maintenance_protein_1_mitochondrial [MW = 11.85 kDa]

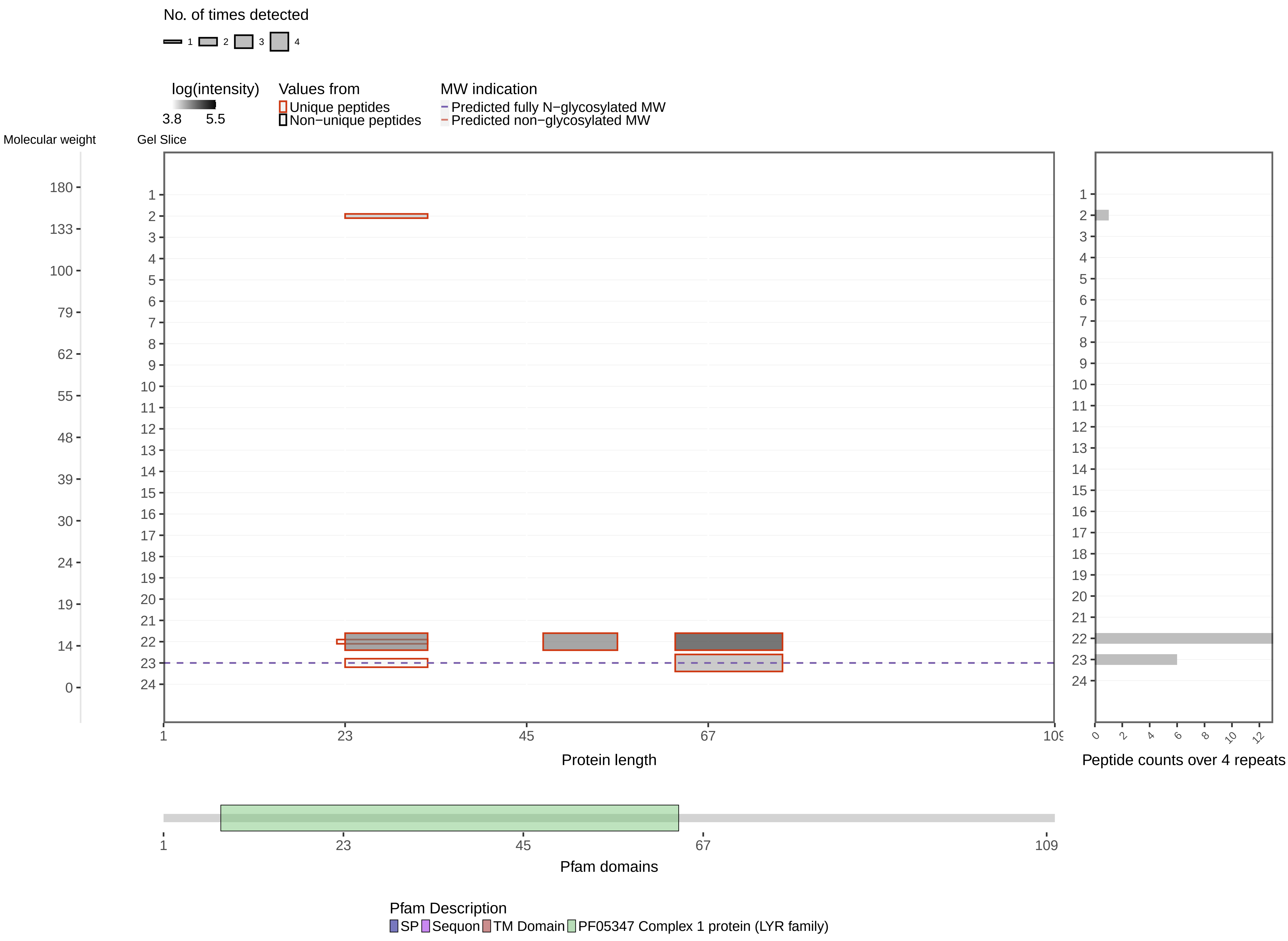


Protein sequence:

IKFEENRHVTSEADIQRLLDDARDASEFISTMIVQAKSSPSGAFVVKPEKEHAGATLEIPSEEILKKSV*

### NbL11g23950.1 small_nuclear_ribonucleoprotein_Sm_D2−like [MW = 12.53 kDa]

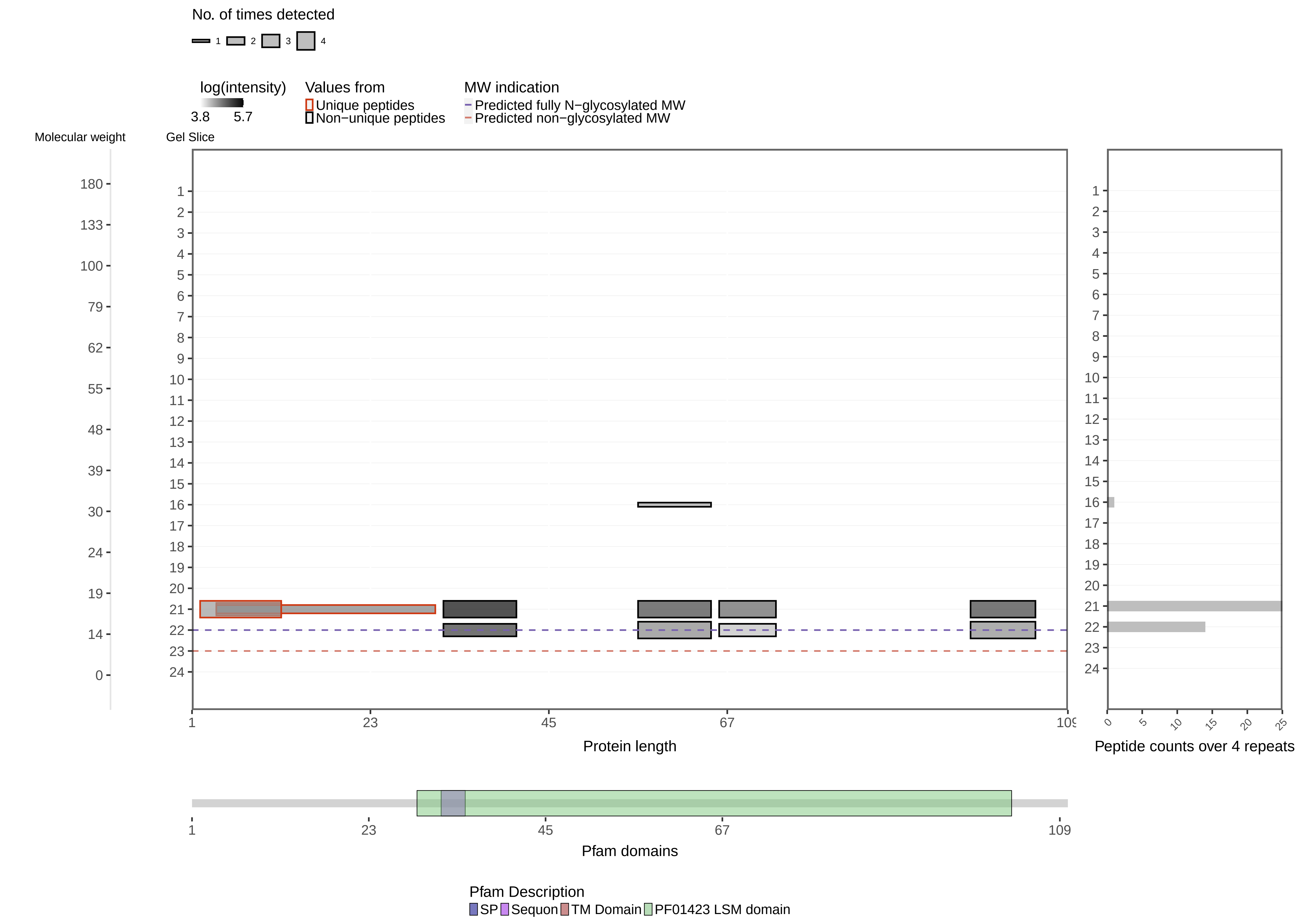


Protein sequence:

NNRKLLGRVRAFDRHCNMVLENVREMWTEVPKTGKGKKKAQPVNKDRFISKMFLRGDSVIIVLRNPK*

### NbL11g23890.1 26S_proteasome_non−ATPase_regulatory_subunit_1_homolog_A−like [MW = 107.94 kDa]

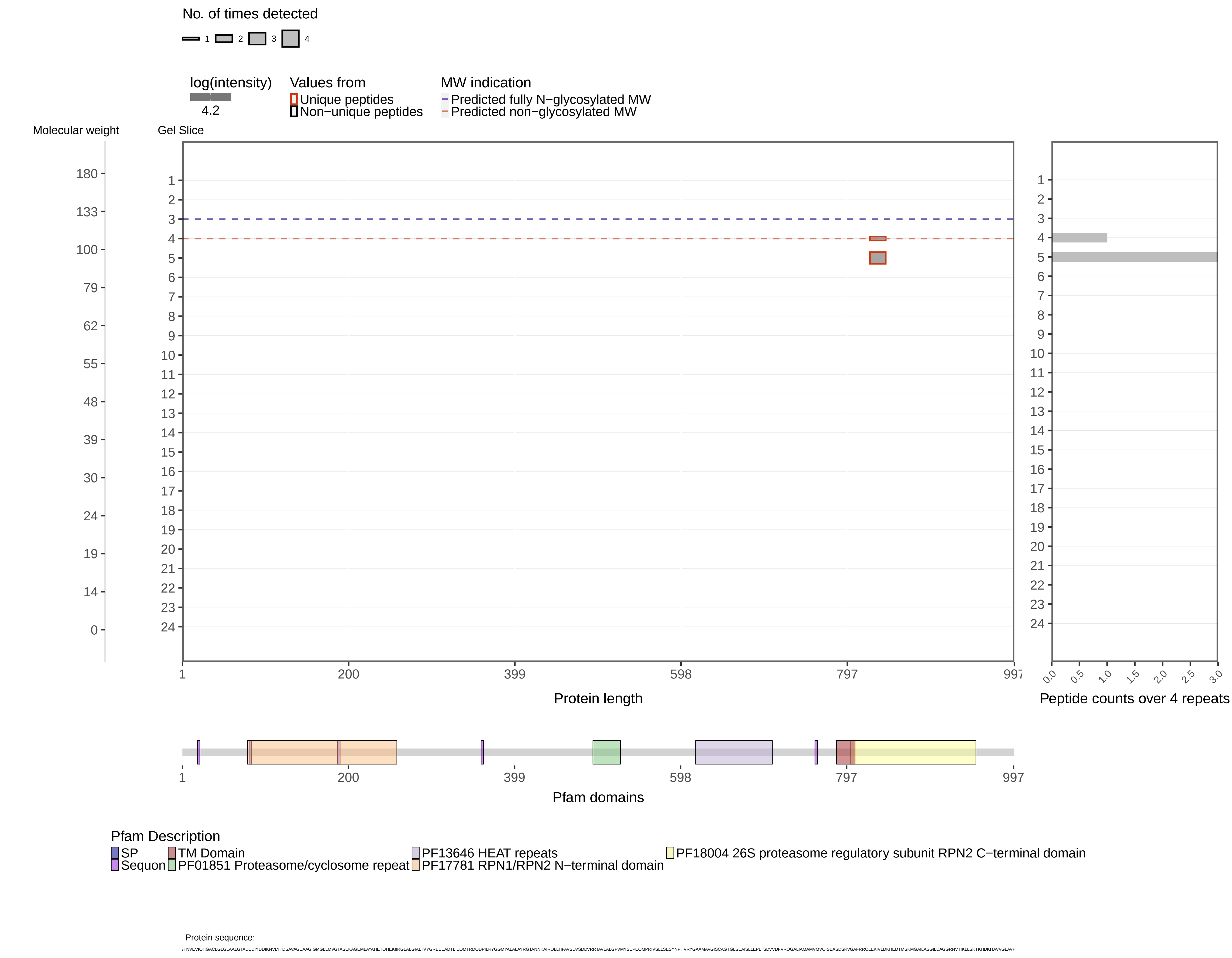

### NbL11g23870.1 60S_ribosomal_protein_L13a−4−like [MW = 23.48 kDa]

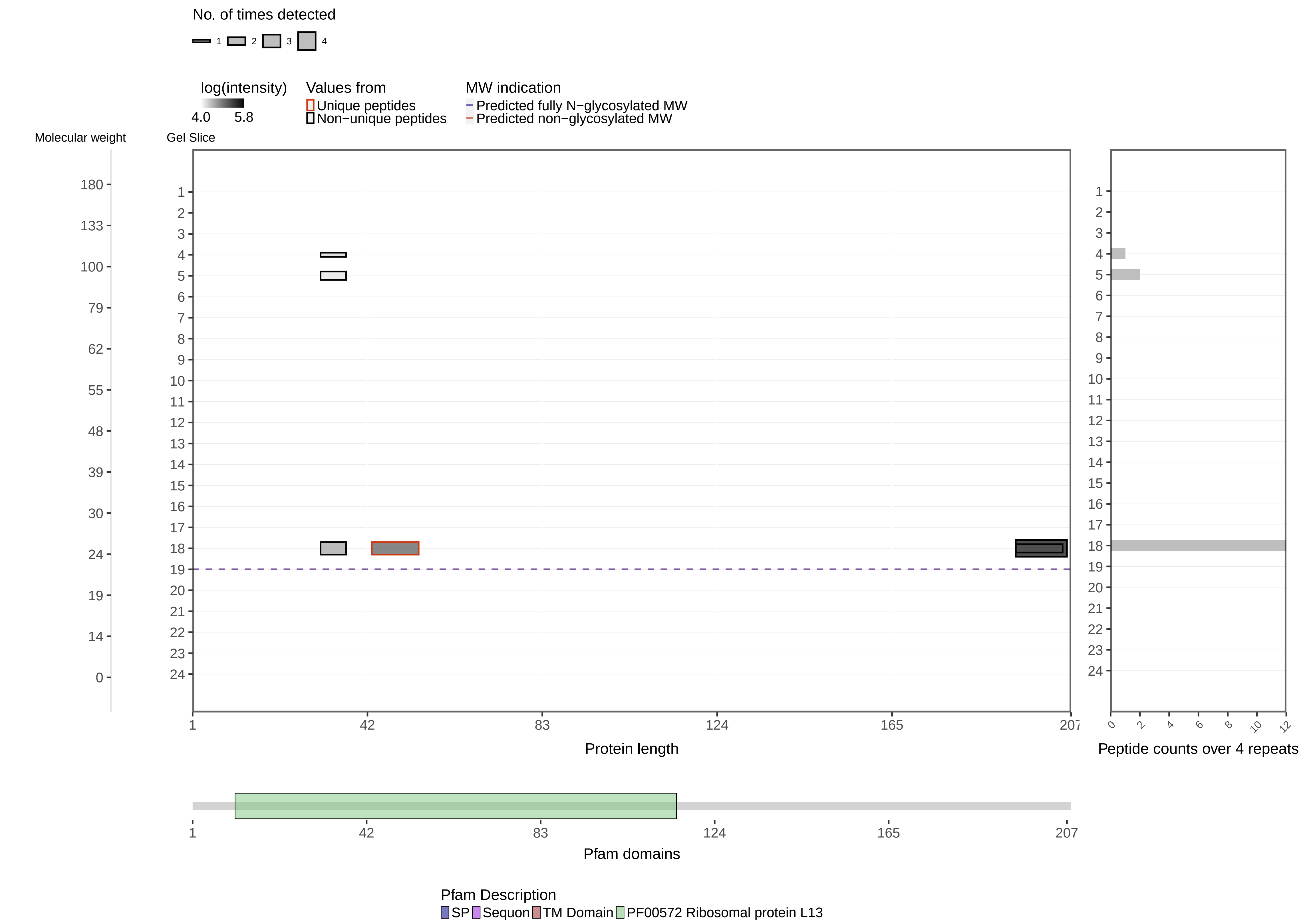


Protein sequence:

[illegible]GMIPHKTKRGAAALARLKVYEGVPPPYDKIKRMVIPDALKVLRLQAGHKYCLLGKLSSEVGWNYYDTIKELENKRKERAQVAYERRKQLAKLRVKAEKAAEEKLGPQLAIIAPIKY*

### NbL11g23830.1 epoxide_hydrolase_A−like [MW = 36.33 kDa]

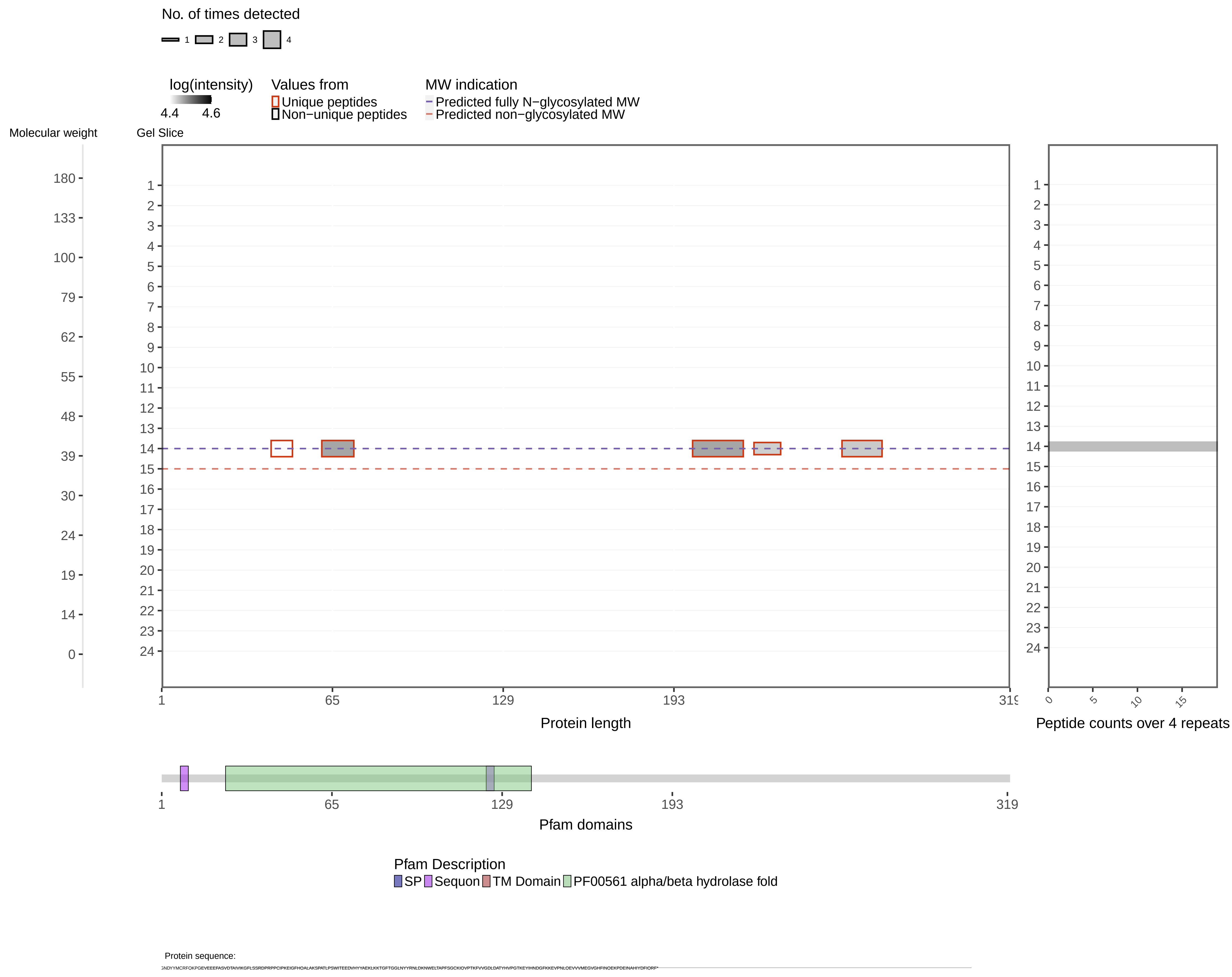

NbL11g23820.1 60S_ribosomal_protein_L18−2 [MW = 20.99 kDa]

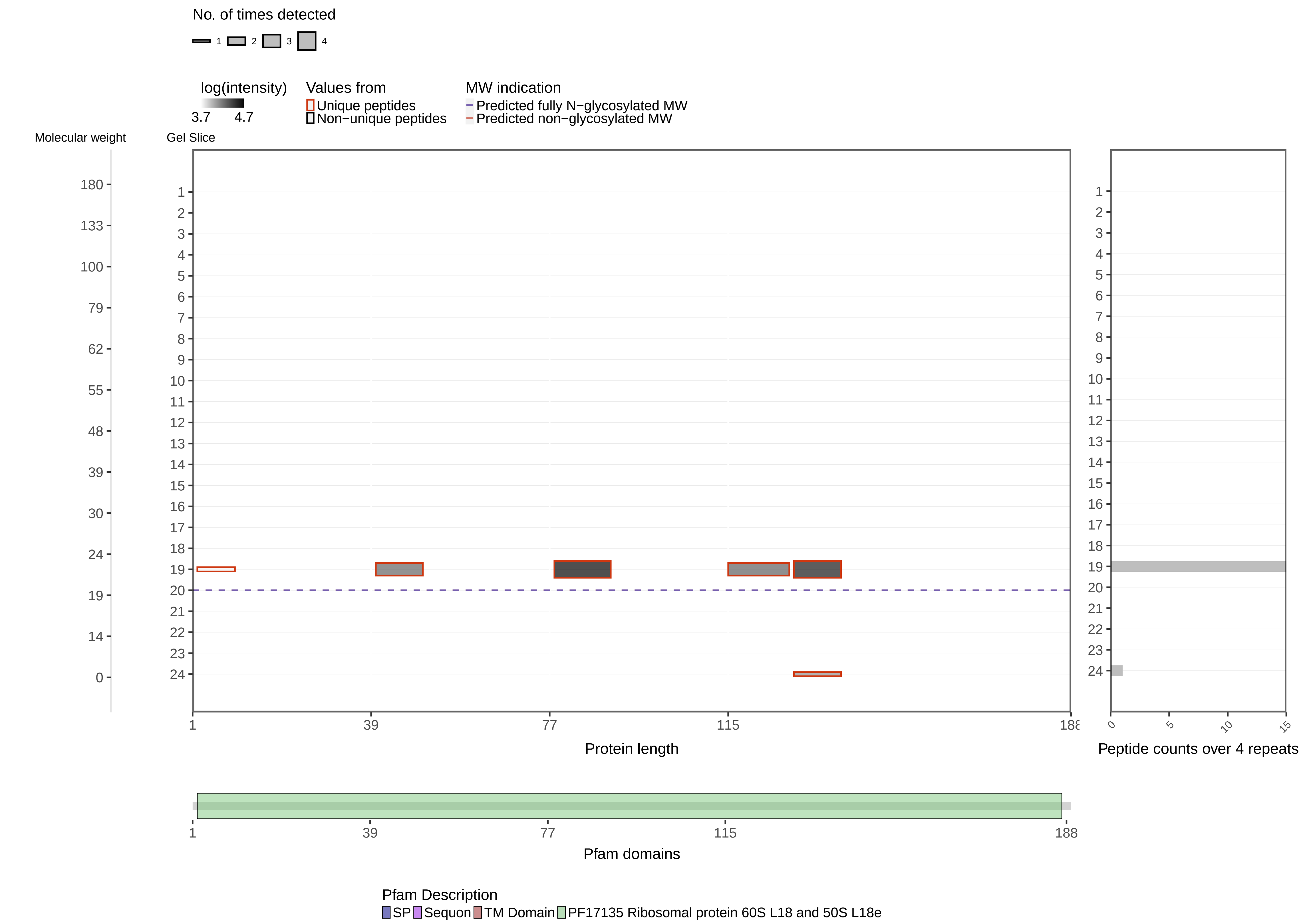


Protein sequence:

GTVTDDVRAYEVPSLKVTALRFTETARARIEKAGGECFTFDQLALRAPLGQNTVLLRGPRNSREAVKHFGRAPGVPHSHTKPYVRAKGRKFERARGKRKSRGFKV*

NbL11g23770.1 U1_small_nuclear_ribonucleoprotein_A [MW = 28.35 kDa]

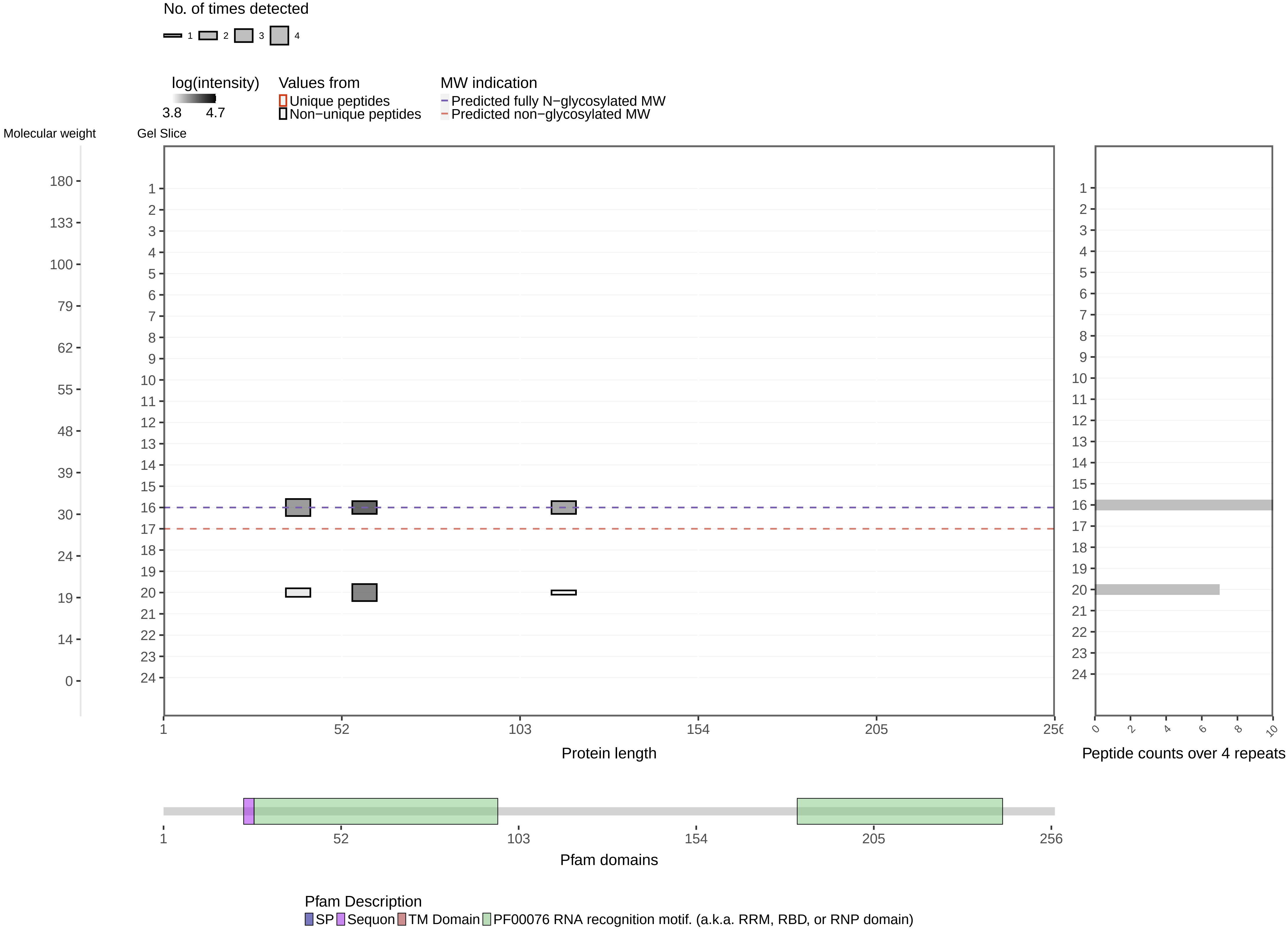


Protein sequence:
FVPREKRKRHEDKGRKKKDOODANOAGMALNLAYAGAYGAAPPFSOIPYMGGAKAAVPEAPAPPNSILFVONLPHOSTPMMLOMLFFOYPGFKEVRMVDTKPGIAFVEYGDEMOSTVAMOALOGFKITPENPMLITYAKK*

### NbL11g23700.1 glutathione_reductase_chloroplastic−like [MW = 57.73 kDa]

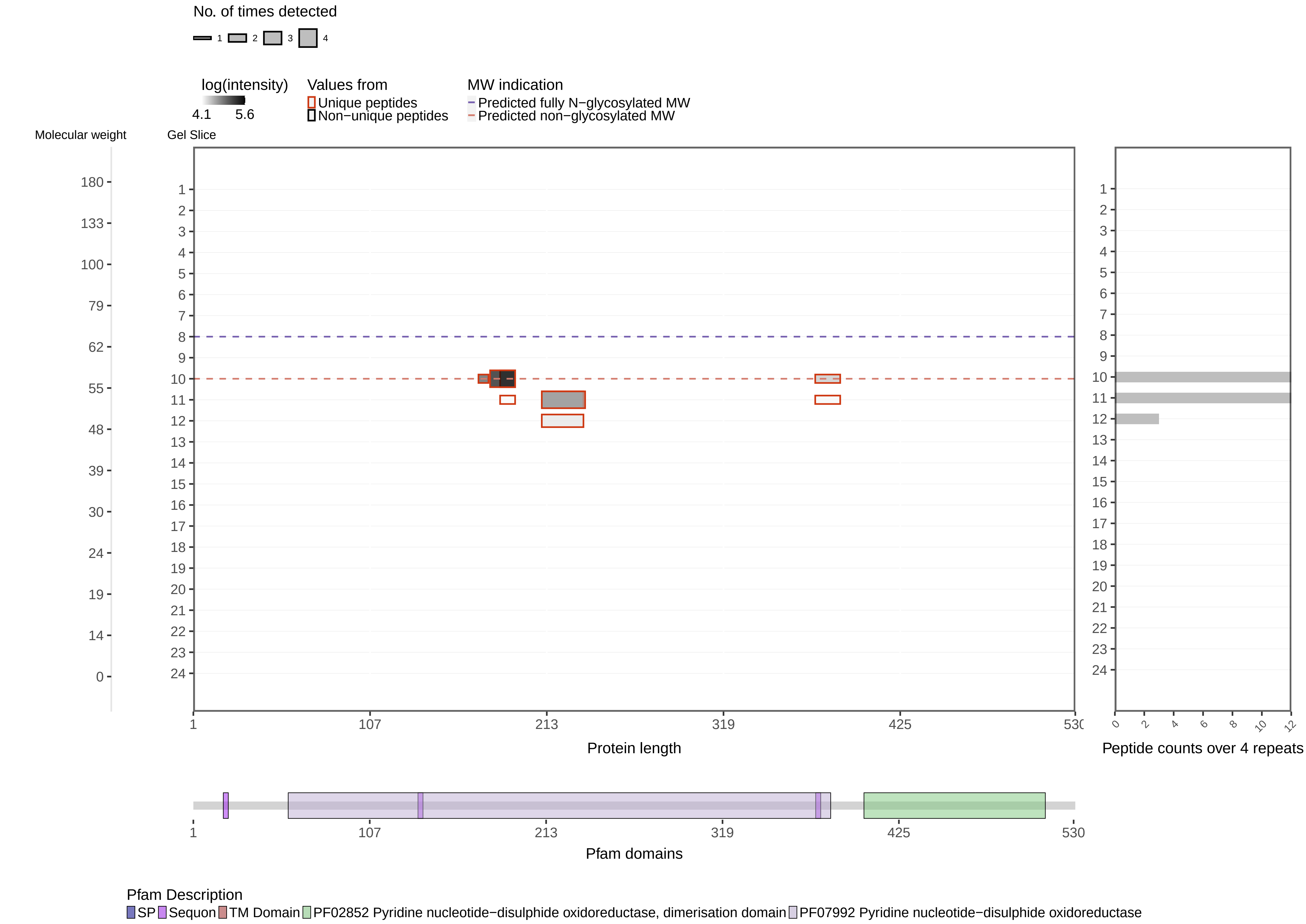


Protein sequence:

GMGATVDLCFRKELPLRGFDDEMRAVVARNLEGRGITMHPCTTLTKLEKTEDGIKVHTDHGEVIVADVVLFATGRRPNTKRLNLDAVGVELDKMQAVKVDEYSRTNVPSIWAIGDVTNRINLTPVALMEGTCFAKTVFGGKPTKPDYSHIPCAVFCIPPLSVVGLGEEQAIEQVNGEVSVYTSTFNPMKNTISGRQEKSVMKLLVEAETDKVLGASMCGPDAPEIMQGIAVALKCGATKEQFDSTVGIHPSAAEEFVTMRSETRRVSSNKPKTNL*

### NbL11g23550.1 TBC1_domain_family_member_13−like [MW = 54.99 kDa]

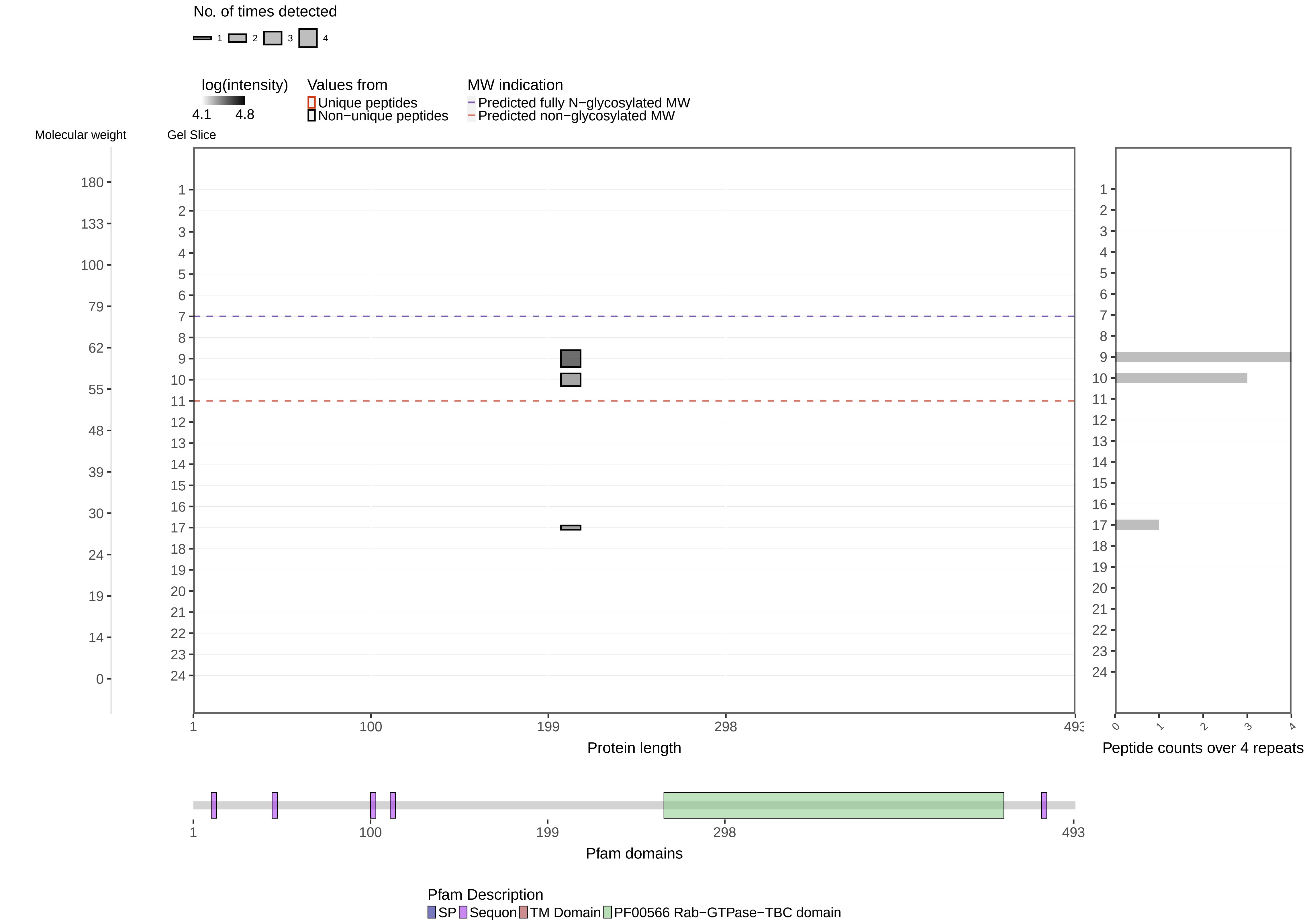


Protein sequence:

VEDKGVLPRSEISHGEHPLSLGQNSIWNQFFQDTEIIEQIDRDVKRTHPDLHFFSGDTPFAKSNQDALRNILIVFAKLNPGIRYVQGMNELLAPLFYVFRNDPNEENTASAEADTFFCFVELLSGFRDHFCQQLDNSVVGIRSTITKLSQLLKEHDEELWRHLEMTTKVNPQFYAFRWITLLLTQEFNFVDSLLIWDTLLSDPEGPQETLLRVCCAMLIIVRRRLLAGDFTSNLKLLQNYPSSNISHLLYVANKLRVKSAG*

### NbL11g23520.1 50S_ribosomal_protein_L13−like [MW = 23.63 kDa]

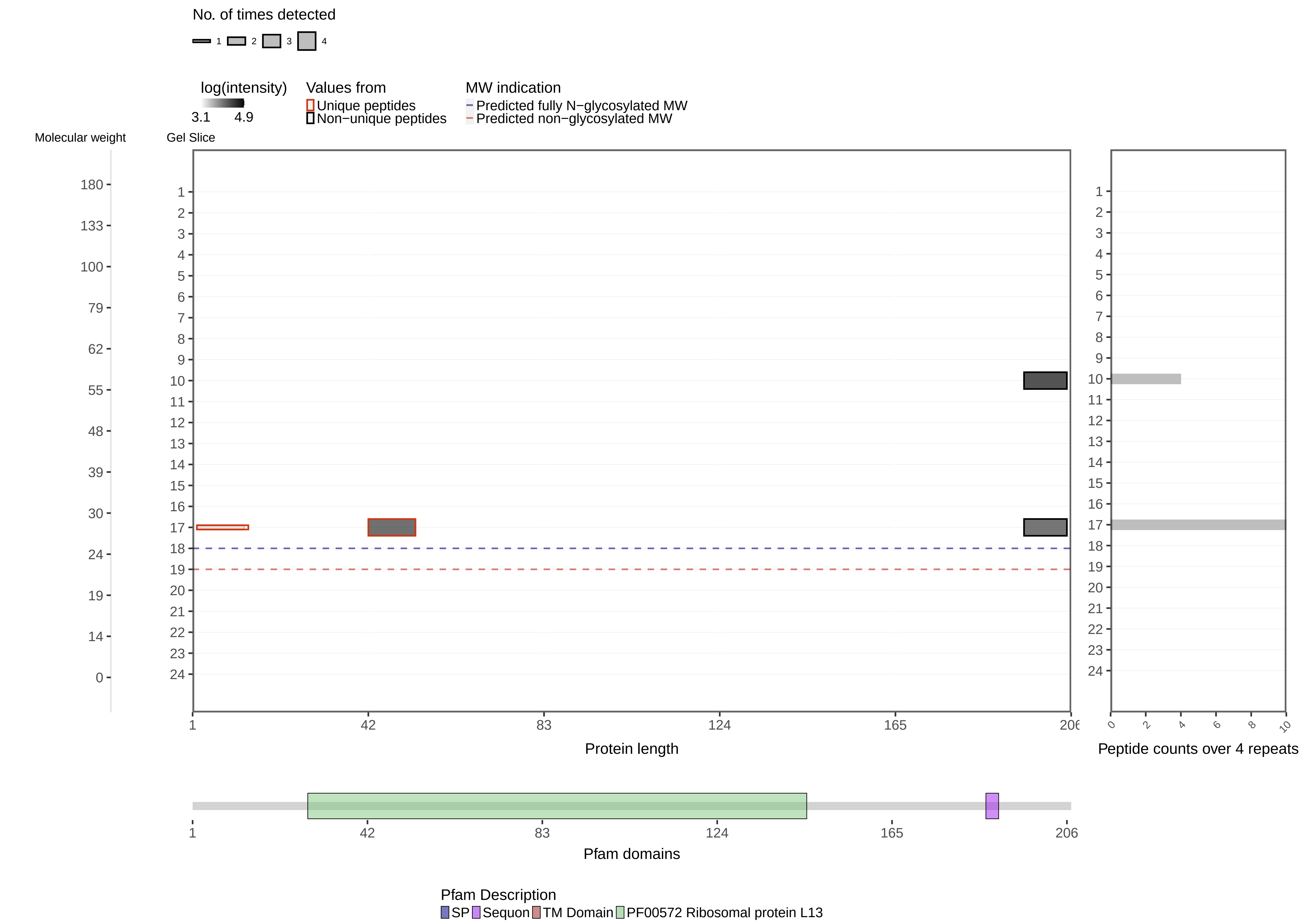


Protein sequence:

HTGYIGHLKERSLKDQMAKDPTEVIRKAVLRMLPRNKLRDDRDRKLRIFAGDEHPFSDRPLEPYVMPPRQVRELRPRTRRAMIRAQKKAEQQQQGTNNTSKRKKNKEVEEEAVSA*

### NbL11g23480.1 protein_BOBBER_1−like [MW = 29.77 kDa]

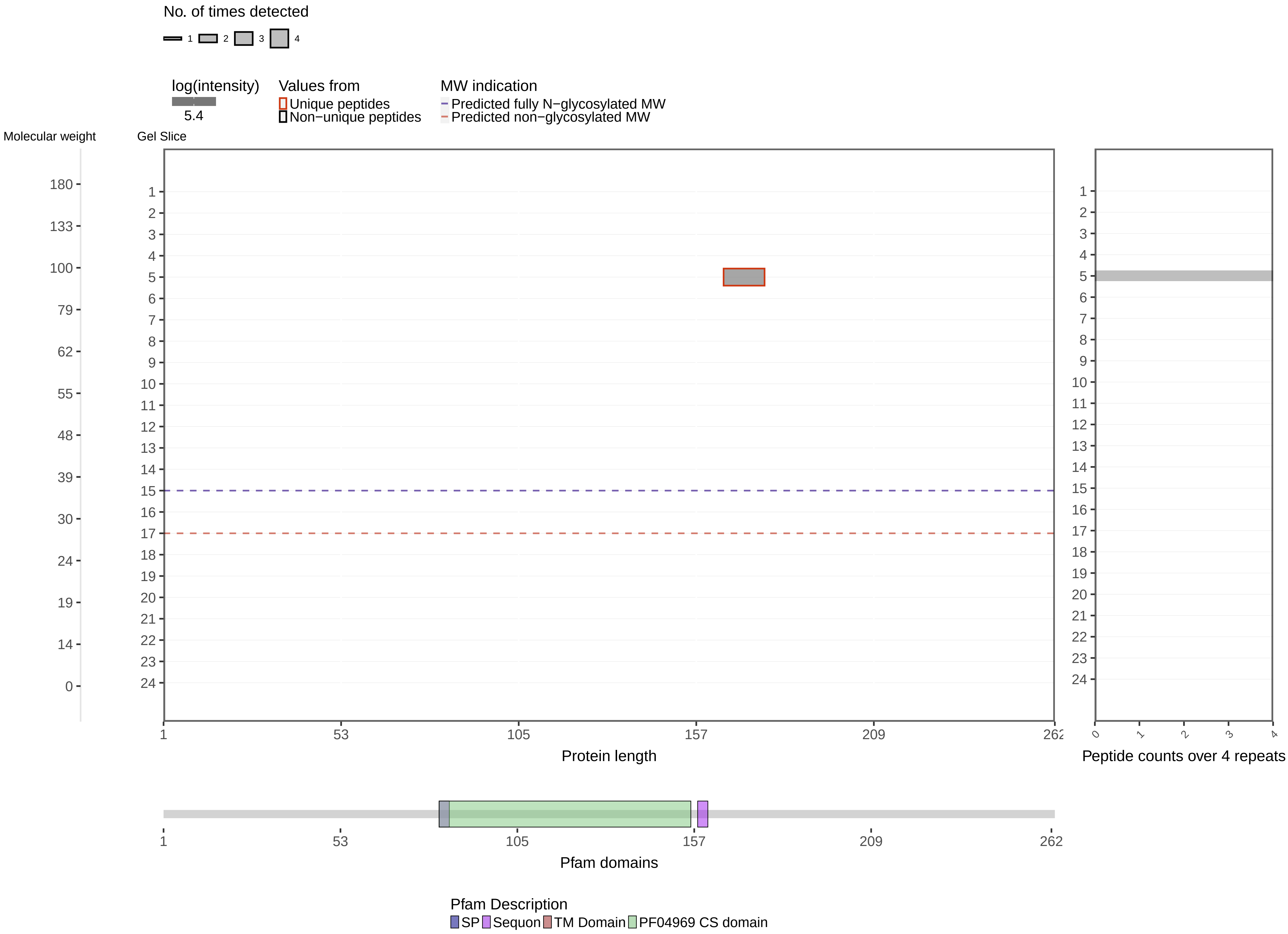


Protein sequence:

KNQPPIVEGELLEPVKADECFWSLEDQKQVTILMTKRNQSDWWKSLFKGSPEIDTQKAEPEPSRLSELDLETRSAVEKMMFDQKQKTLGKPTSDEIQNQDMVKKFMEENPEIAKHFTNSPANANGKGGKPNVRMMPSGGMP*

### NbL11g23400.1 adenosylhomocysteinase [MW = 60.9 kDa]

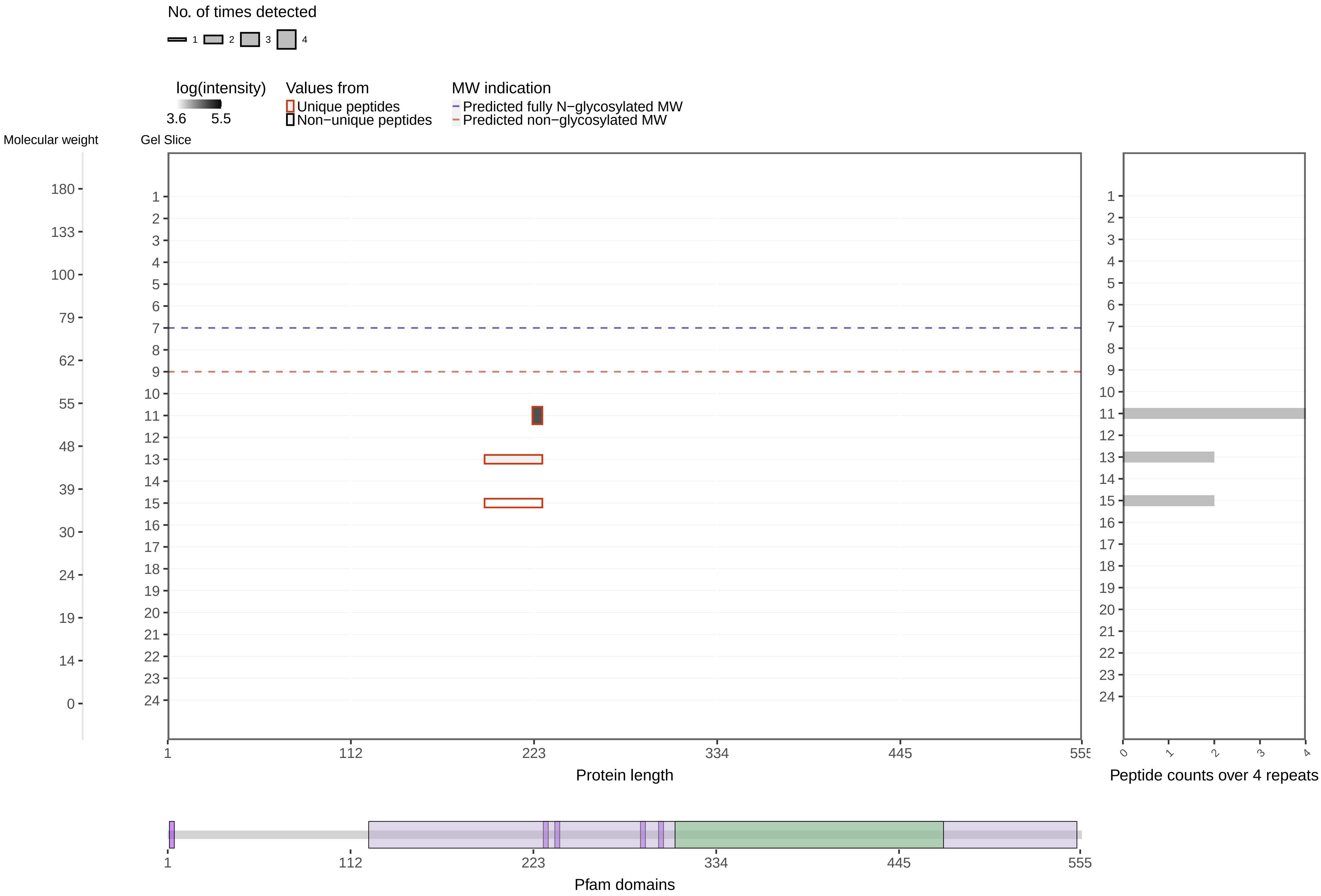


Pfam Description

SP | Sequon | TM Domain | PF00670 S−adenosyl−L−homocysteine hydrolase, NAD binding domain | PF05221 S−adenosyl−L−homocysteine hydrolase

Protein sequence:

RLVGVSEETTTGVKRLYQMQANGTLLFPAINVNDSVTKSKFDNLYGCRHSLPDGLMRATDVMIAGKVALVAGYGDVGKGCAAALKQAGARVIVTEIDPICALQATMEGLQVLTLEDVVSDVDIFVTTTGNKDIIMVDHMRKMKNNAIVCNIGHFDNEIDMLGLETYPGVKRITIKPQTDRWVFPDTNSGIIVLAEGRLMNLGCATGHPSFVMSCSFTNQVIAQLELWNEKSSGKYEKKVYVLPKHLDEKVAALHLGKLGAKLTKLSKDQADYISVPVEGPYKPAHYRY*

### NbL11g23380.1 selenium−binding_protein_1−like [MW = 53.68 kDa]

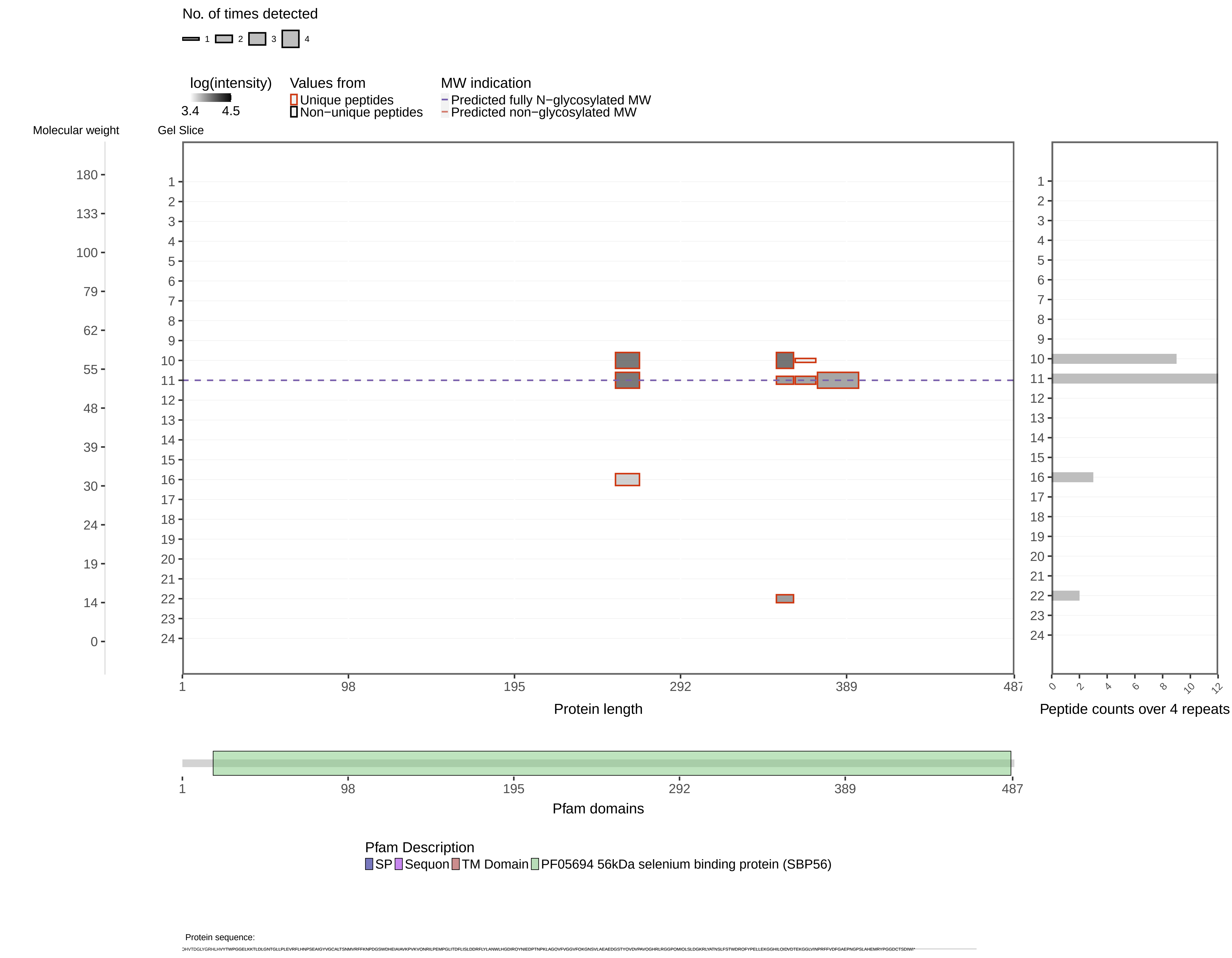

### NbL11g23360.1 receptor−like_kinase_TMK4 [MW − SP = 98.48 kDa]

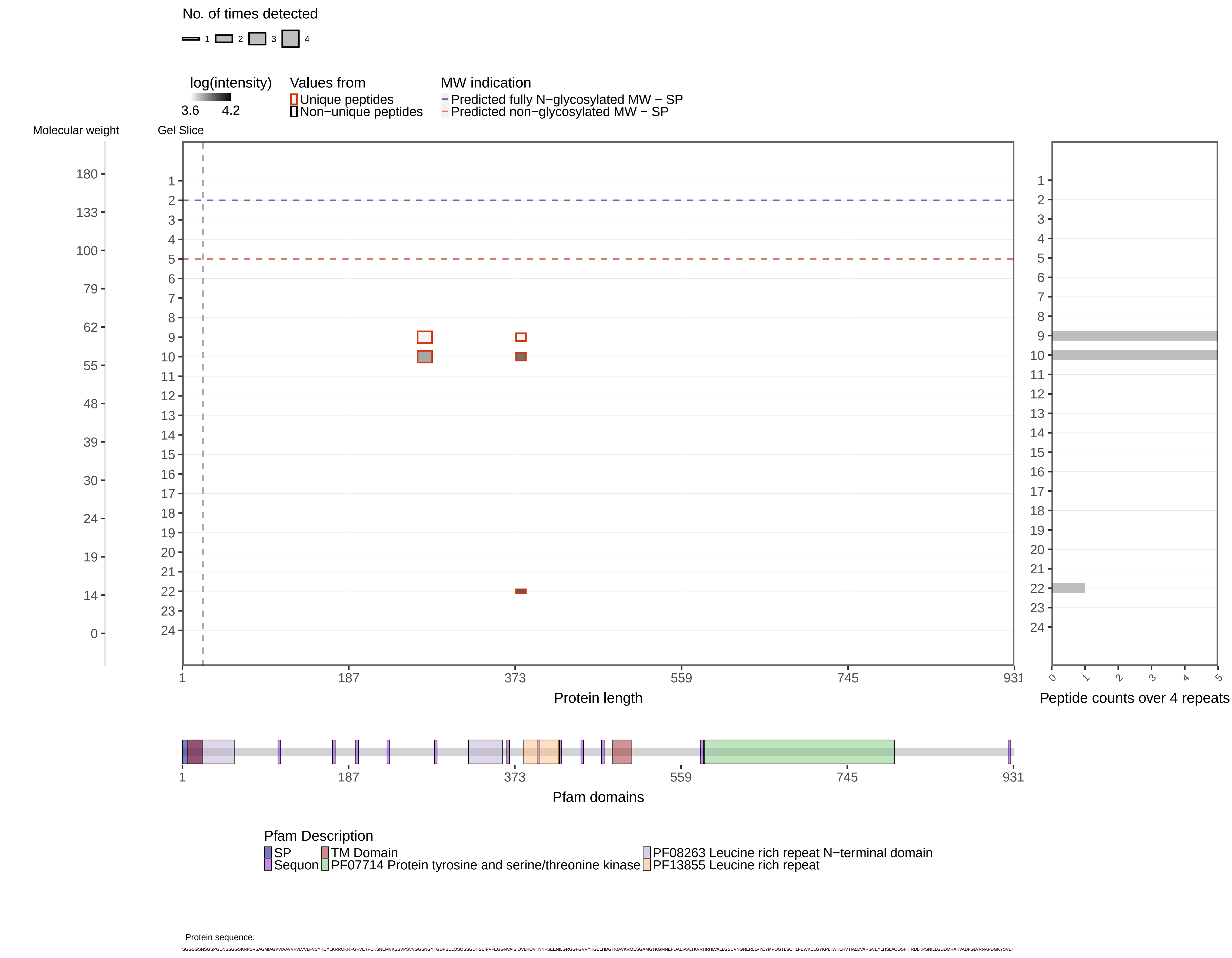

### NbL11g23220.1 cysteine_proteinase_inhibitor_B−like [MW = 19.34 kDa]

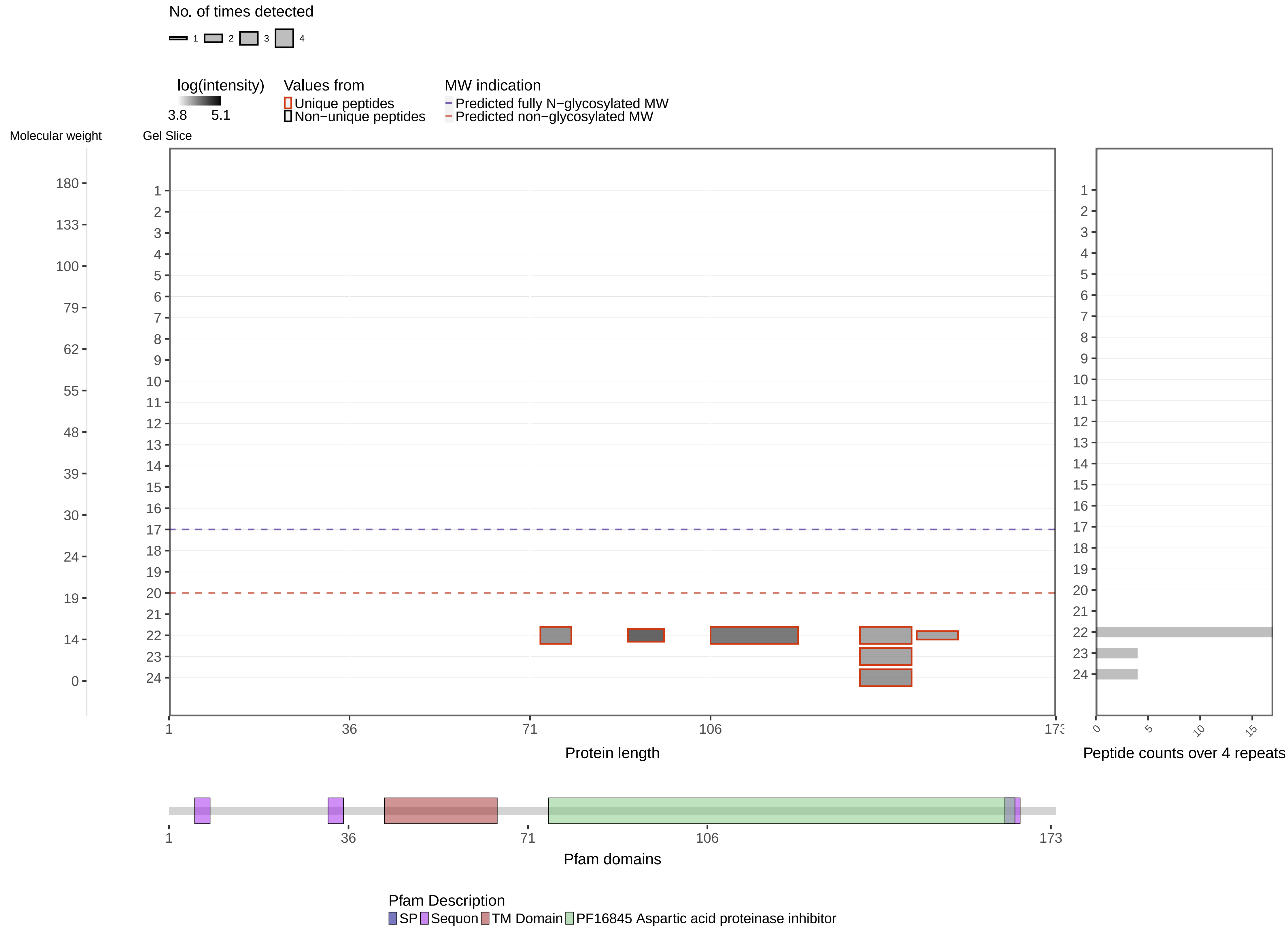


Protein sequence:

DVKTNKEIQDLGKFCVEEYNRNLQKQKQKVNNNNGLLSFSEVVEAEKQVVSGIKYYLKISATTSLGSPKMFNAVLVVKAWEKKKELLNFSPSPATK*

### NbL11g23210.1 50S_ribosomal_protein_L24−like [MW = 17.6 kDa]

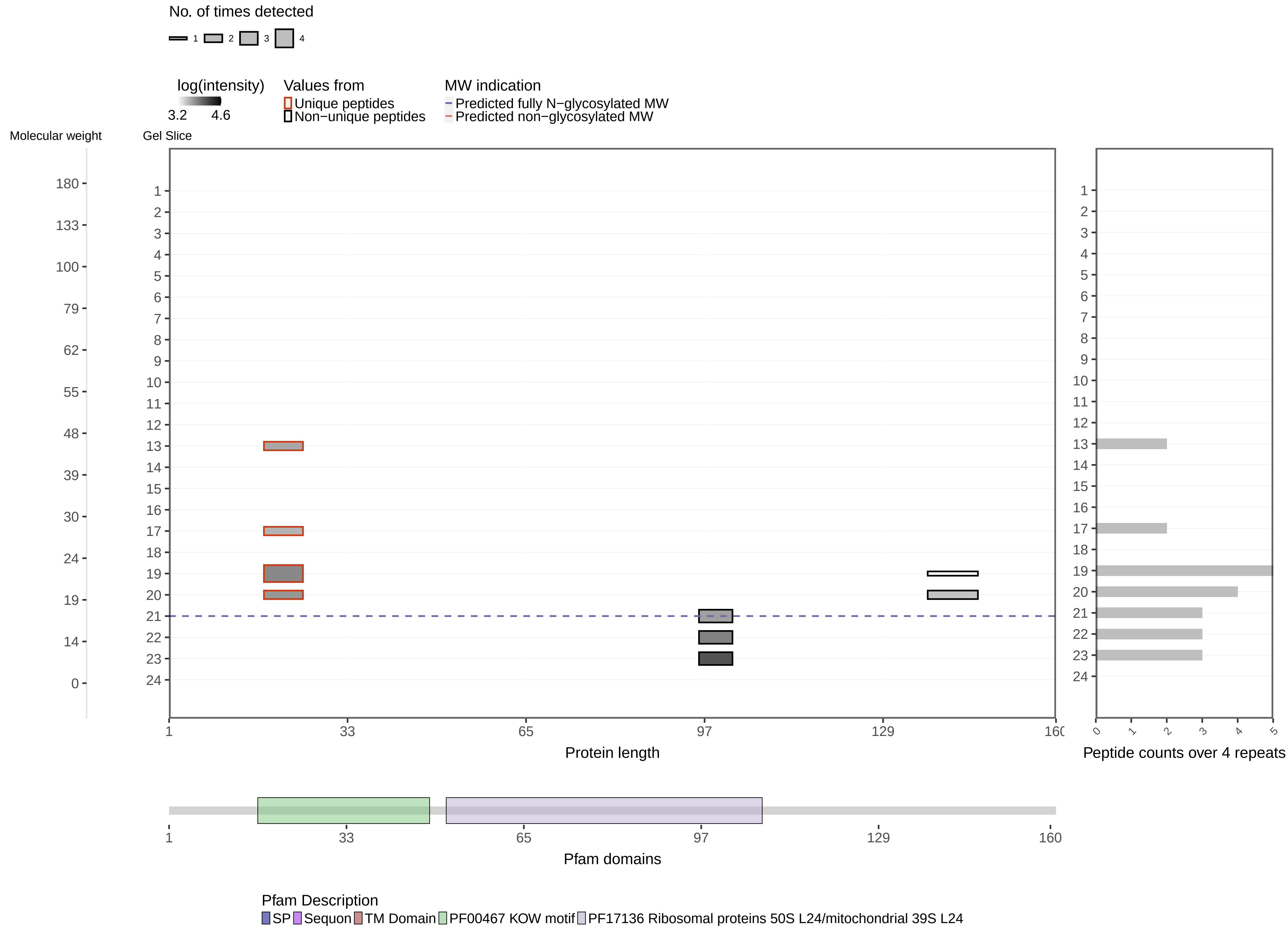


Protein sequence:

GIFSVEAPLHVSNVQVVDPVTGKPTKVGIRFLEDGSKVRVSRGIGASGSIIPRPEILKIRTTPRPTVAGPKDTPMEVVMERTYDPKTGKGMPDL*

### NbL11g23010.1 phosphoglucan_water_dikinase_chloroplastic [MW = 132.06 kDa]

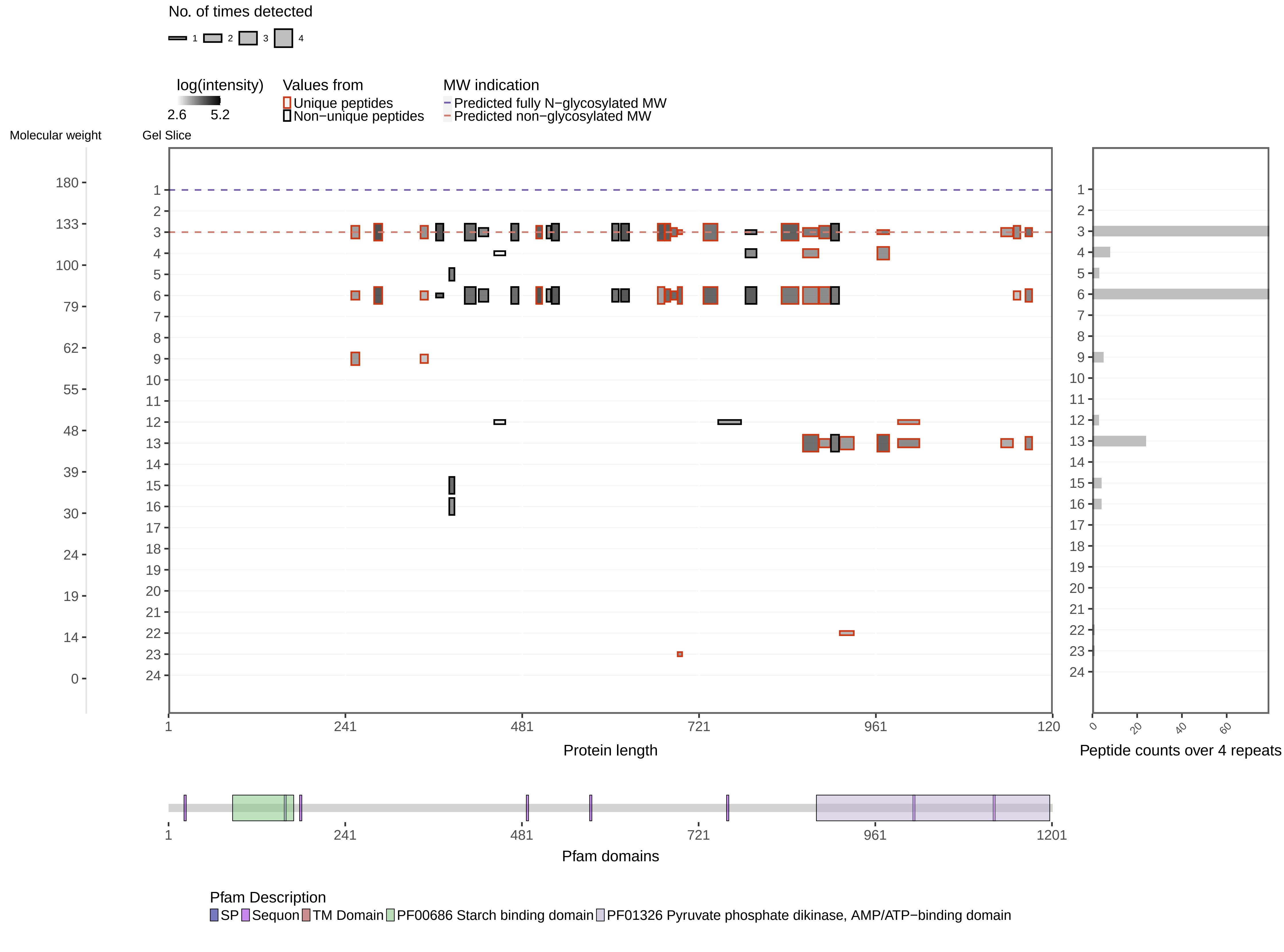


Protein sequence:

### NbL11g22950.1 probable_aldo−keto_reductase_2 [MW = 38.58 kDa]

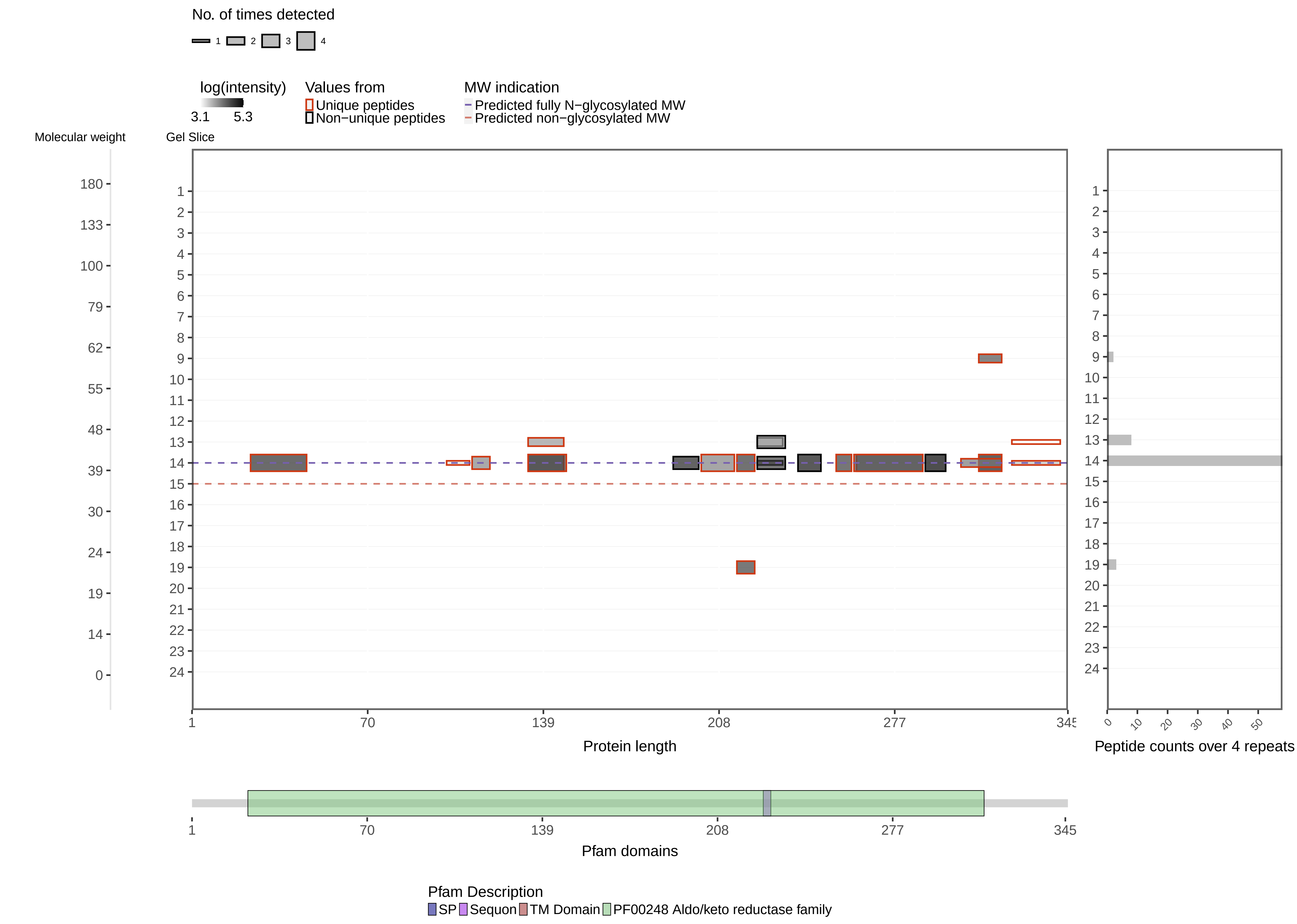


Protein sequence:

ACASTIRRAHAVHPITAVQMEWSLWTRDLEEEVVPTCRELGIGVVPYSPLGRGFFSAGTKLIENLSEDDFRKNFPRFRPENFELNKQVFEKLNVMAARKGCTPSQLALAWVLHQGDDVCPIPGTTKIENFNENVRALSVKLTTEDMKELESYASVVKGDRHLYMSSTWINSETPPLSSWKTE*

NbL11g22910.1 metacaspase−4−like [MW = 45.58 kDa]

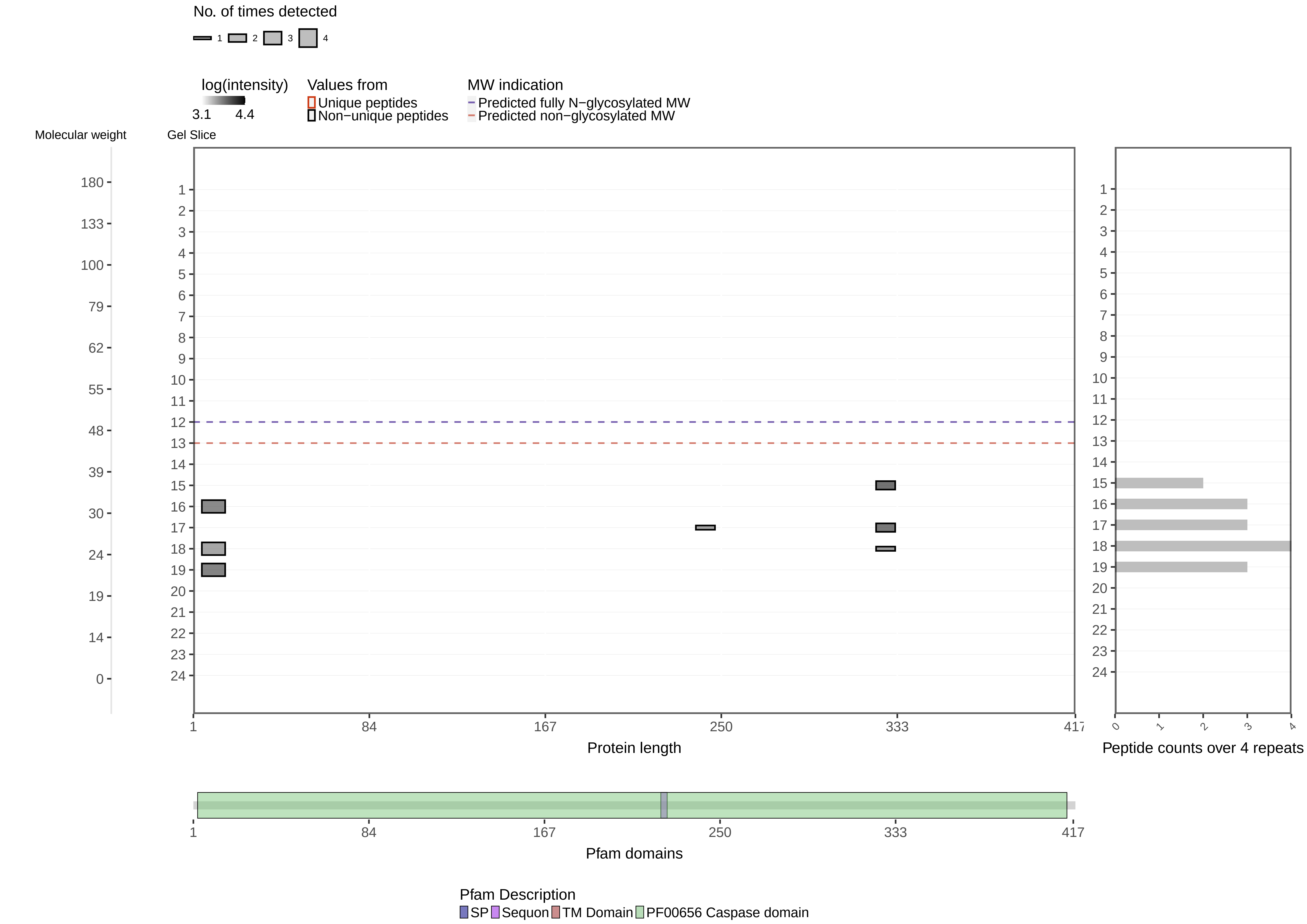


Protein sequence:

RAEEEESFAESSVIETEDGDQVHVKNKSLPLSTLIEILKQKTGKDDIDVGKLRPTLFDVFGEDASPKVKKFMKVIFNKLQHGNGEGESGSFLGMVGNLAQEFLKQKLDENDESYAKPAMETHVEGKQEVYAGSGSRGLPDSGILVSGCQTDQTSADATPAGGDSYGALSNAIQEILAESDGPITNEEVVNKARKKLQKQGYTQRPGLYCSDHHVDAPFVC*

NbL11g22890.1 pirin−like_protein [MW = 36.81 kDa]

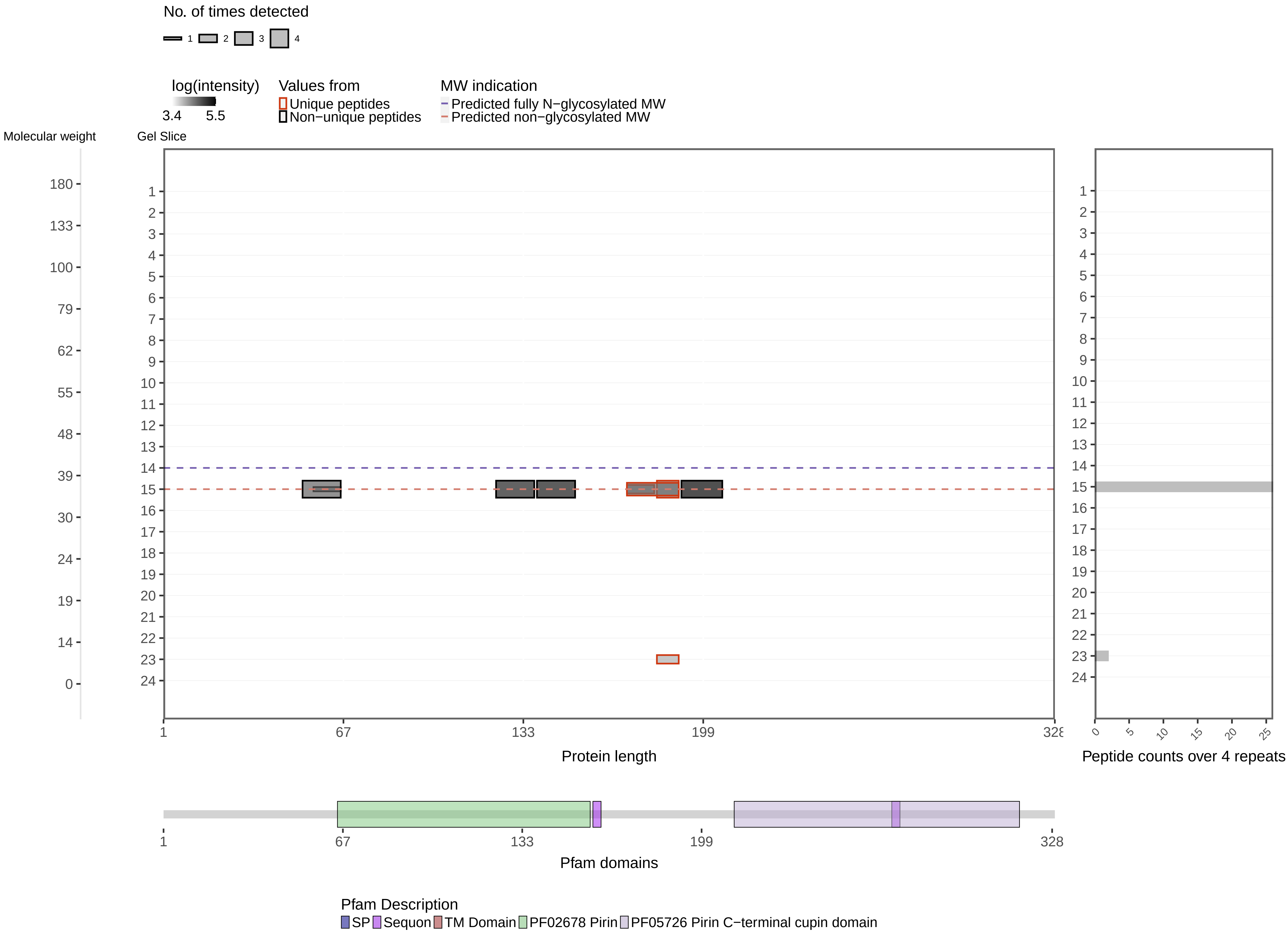


Protein sequence:

[illegible]GLQLWINLSSKDKMIEPRYQELLNEDIPRAENEGVEVKIIAGEAMGVQSPVYTRTPTMYLDFTLQPTSYYHQATPESWNAFVYIVEGEGVFGIPNSGPVSALHCLVLGPGEGLSVWNKSSKPLRFVLLGGQPINEPVVQHGPFVMNSQAEIDQTFEDYQYCKNGFENARYWRSGH*

### NbL11g22880.1 signal_peptide_peptidase−like_5 [MW − SP = 53.48 kDa]

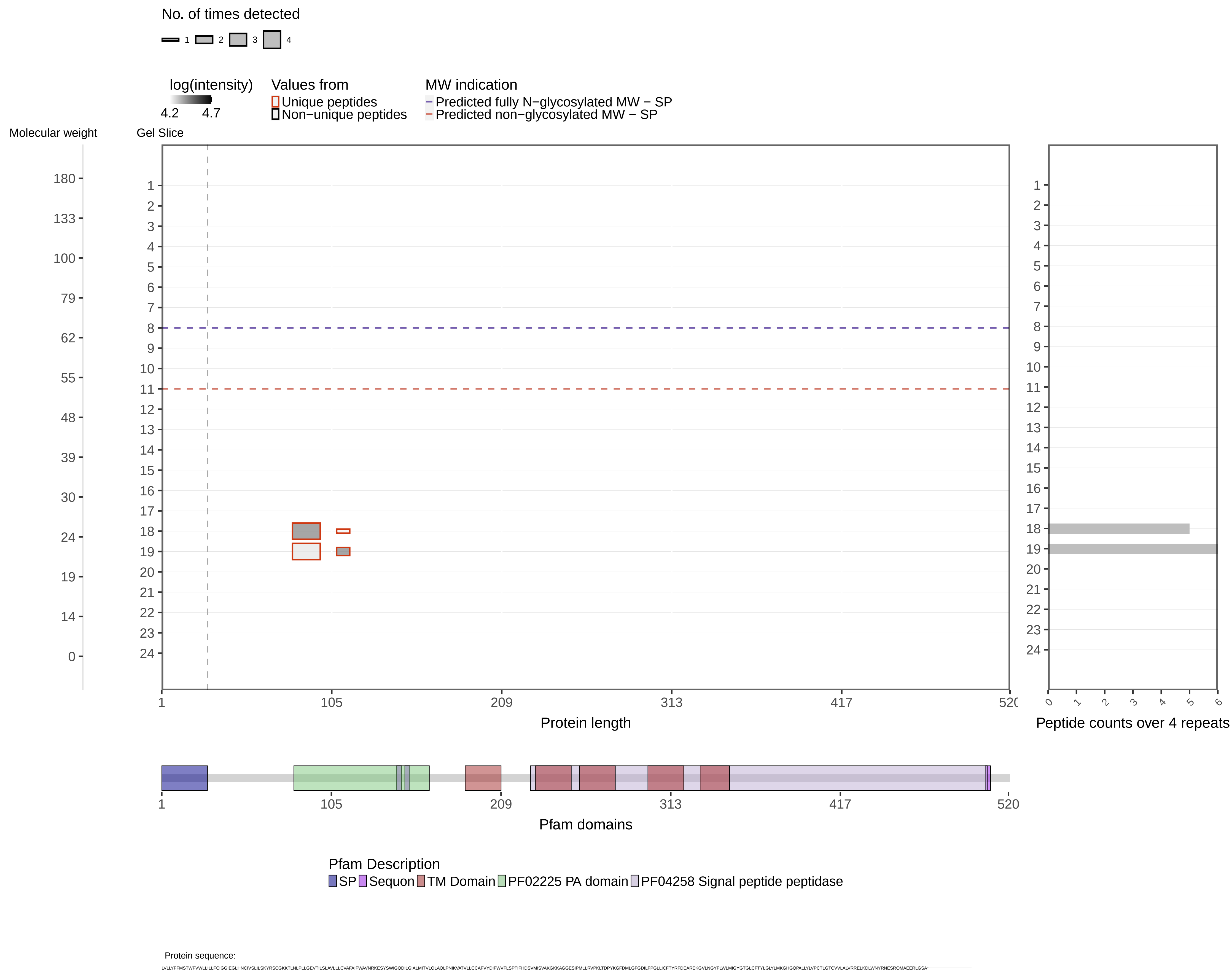

### NbL11g22840.1 importin_beta−like_SAD2 [MW = 118.76 kDa]

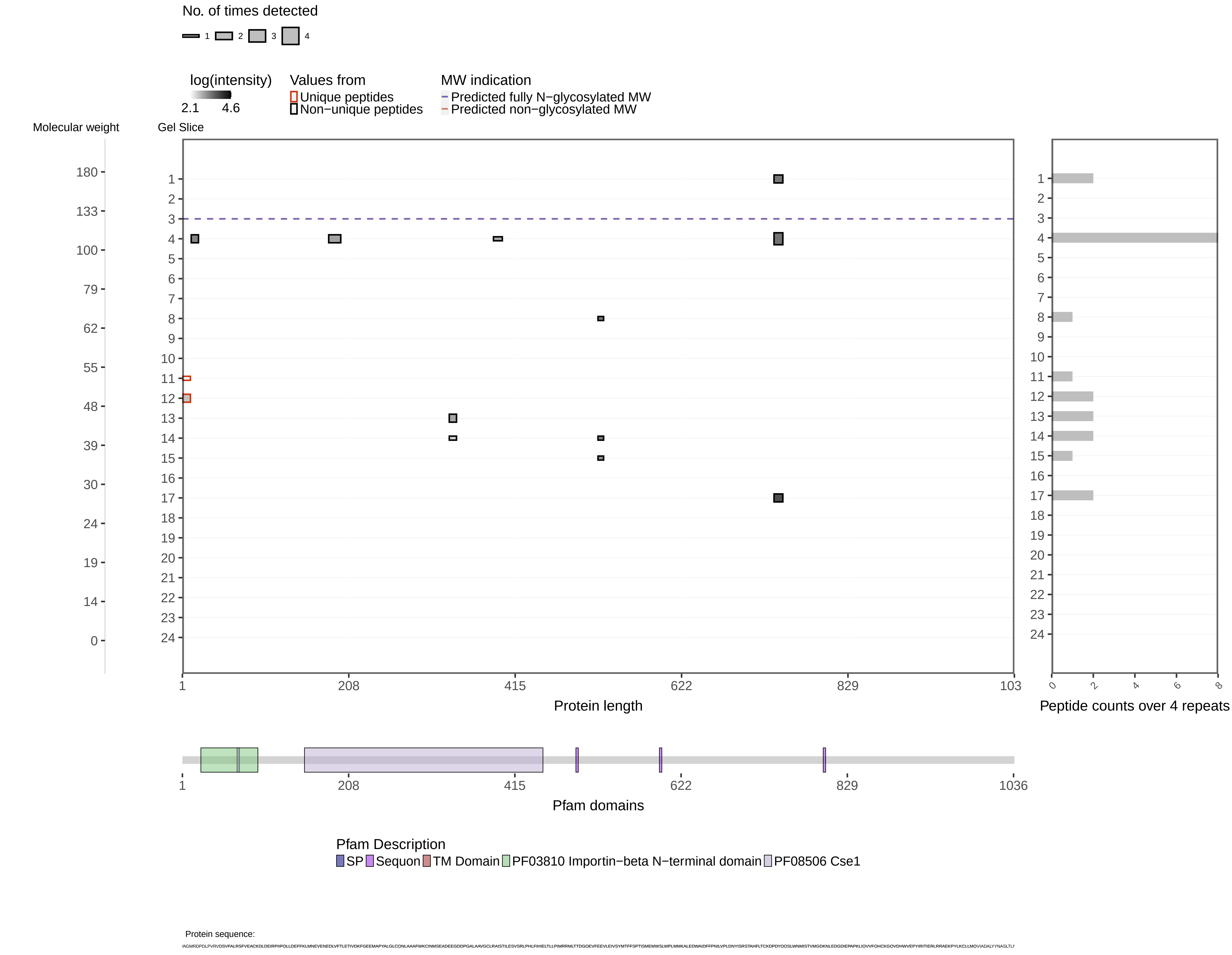

### NbL11g22640.1 chitinase−like_protein_1 [MW = 51.38 kDa]

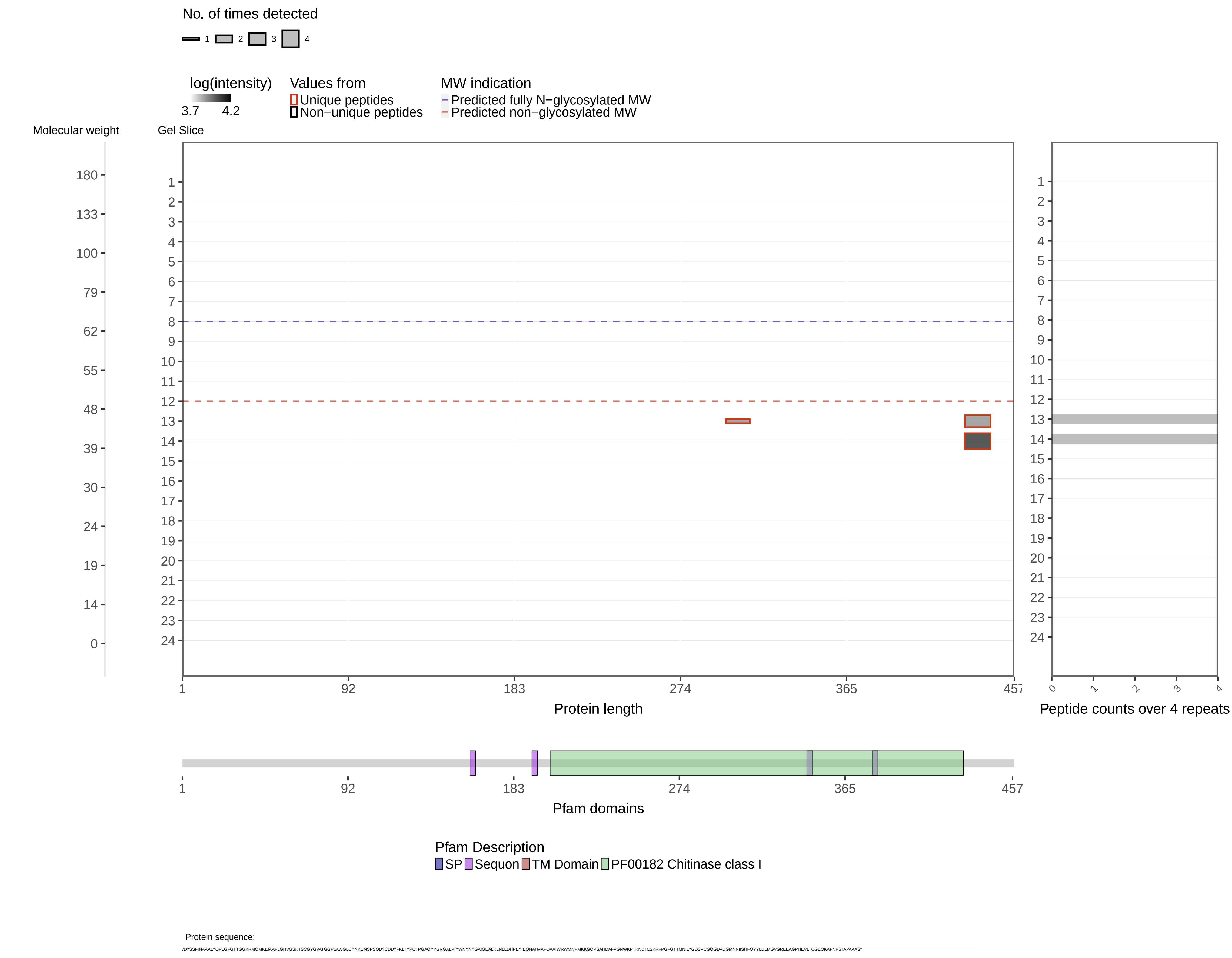

### NbL11g22510.1 spermatogenesis−associated_protein_20−like [MW = 96.58 kDa]

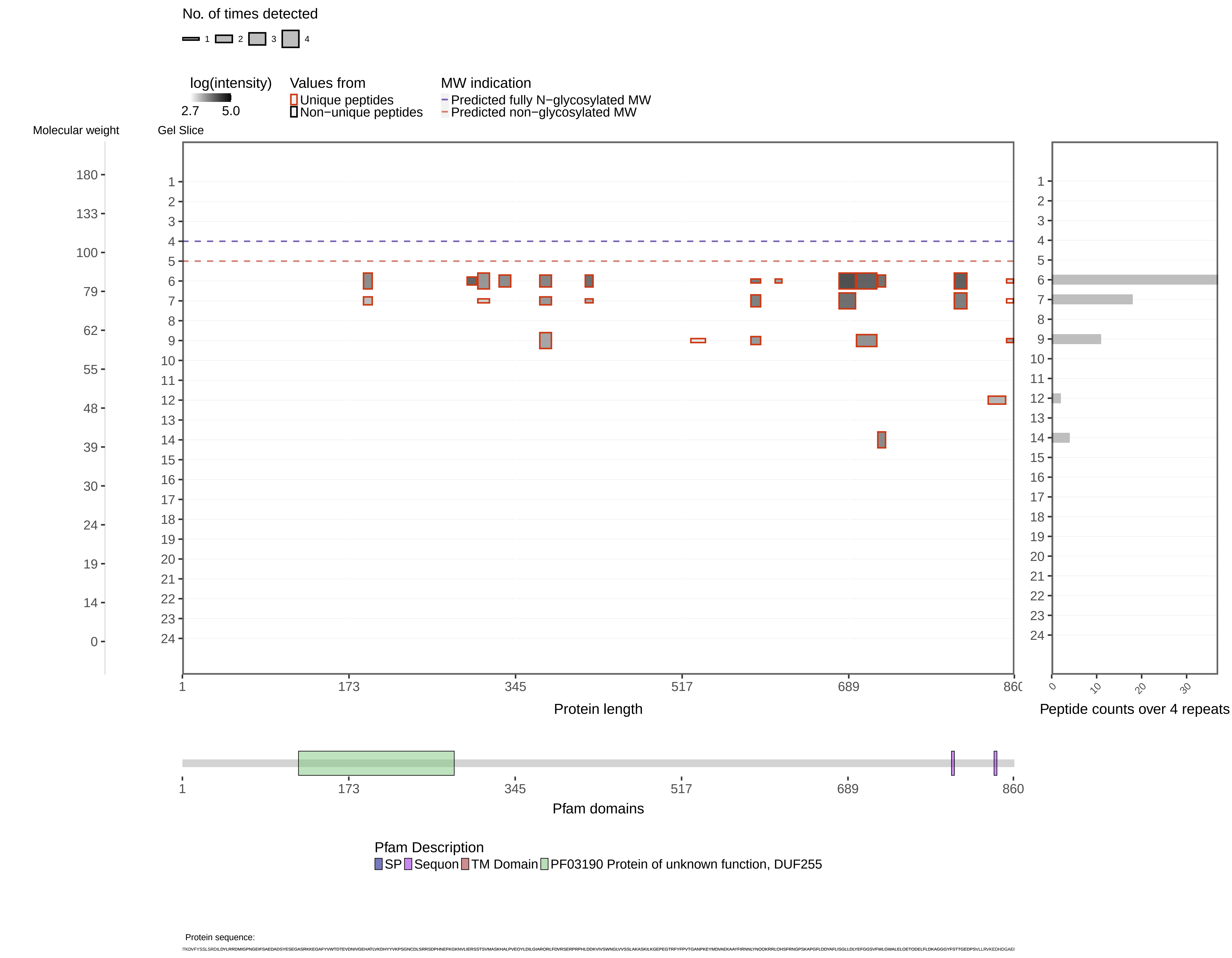

NbL11g22500.1 peptide_chain_release_factor_APG3_chloroplastic [MW = 47.12 kDa]

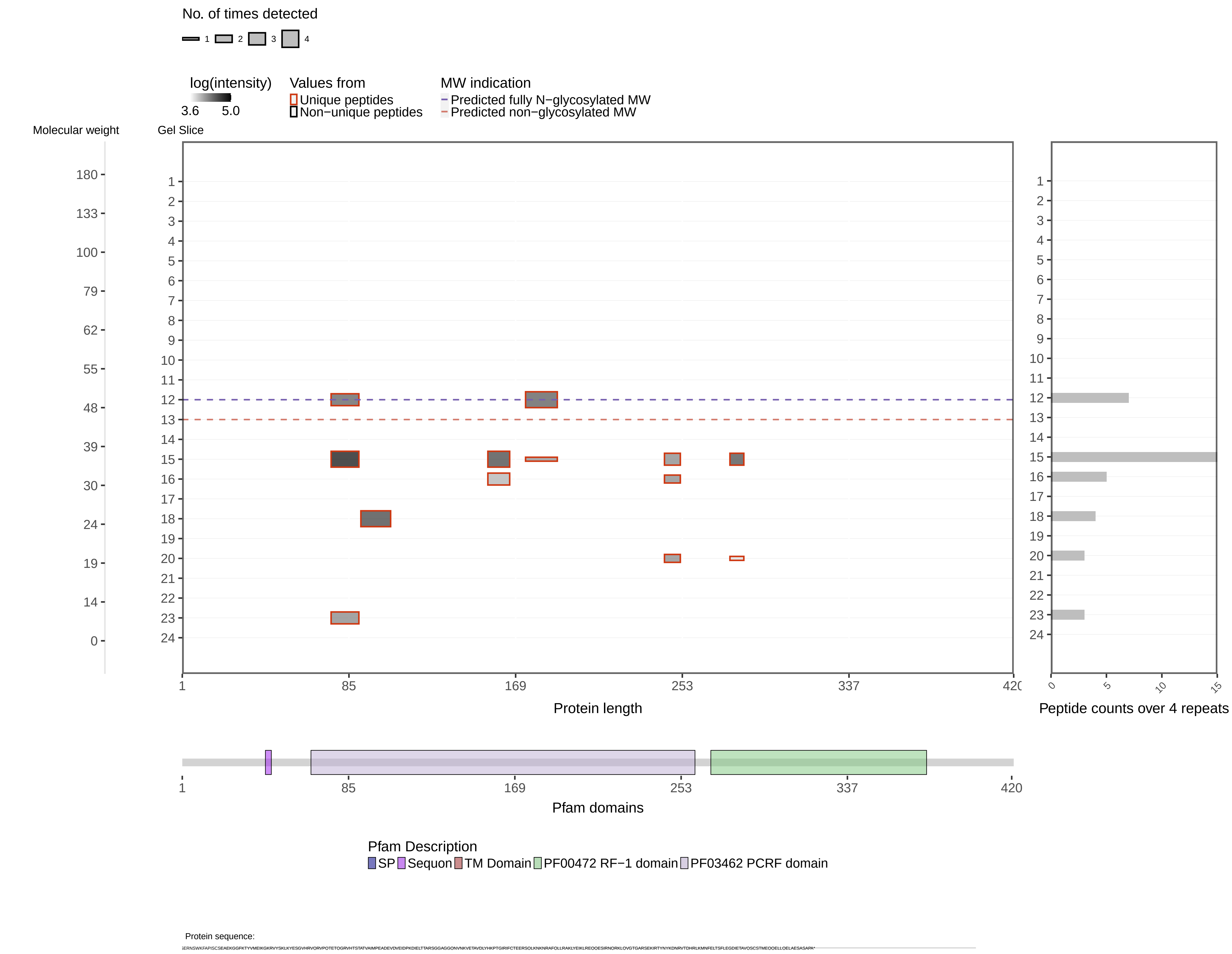

NbL11g22410.1 protein_TRANSPORT_INHIBITOR_RESPONSE_1−like [MW = 65.19 kDa]

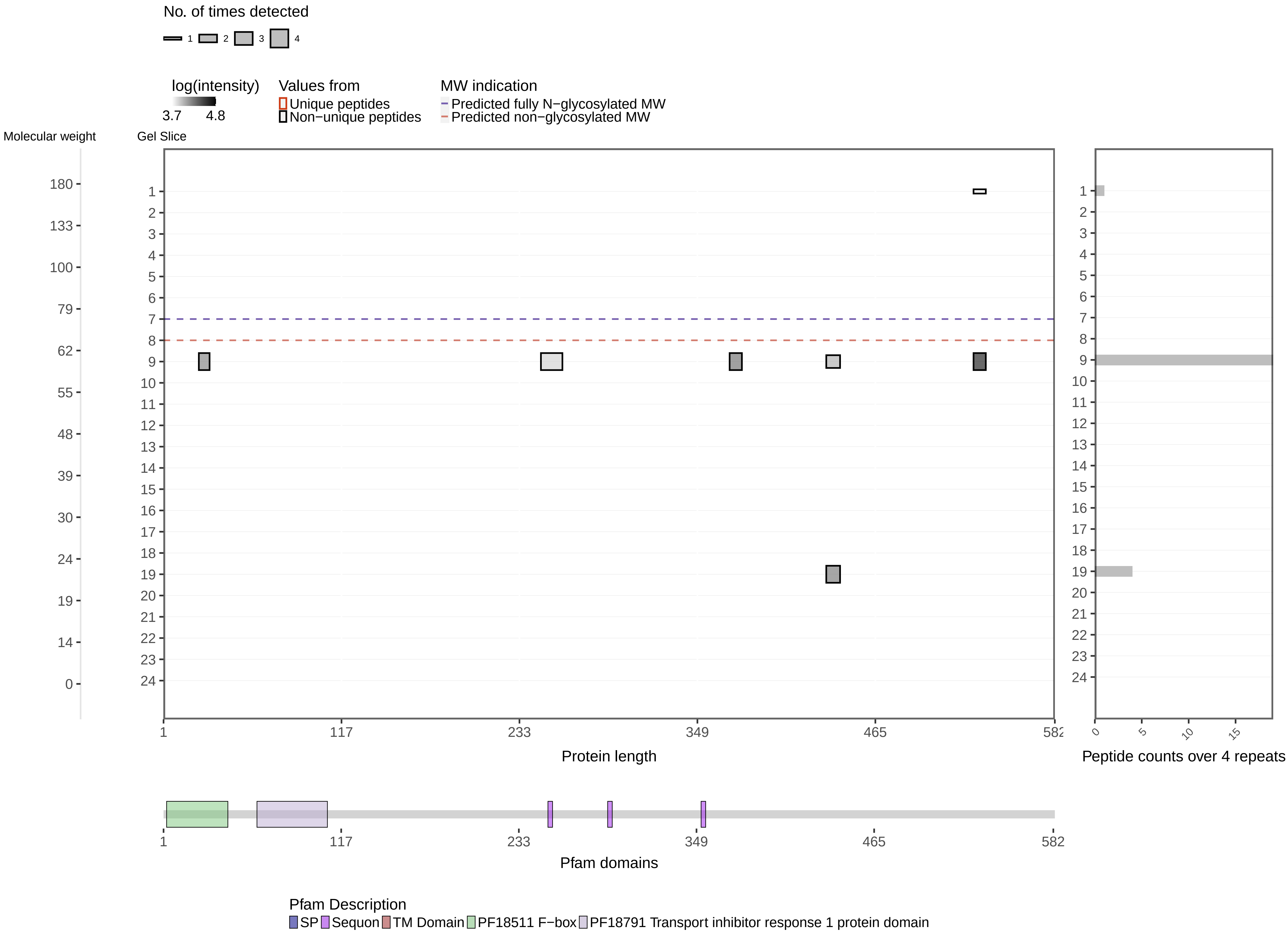


Protein sequence:

.PAIYPVCSRLTSLNLSYATSQNPDLGKLISQCHNLQRLWVLDYIEDIGLEELAANCKDLQELRVFPSDPFAPEPNATLTEQGLVAVSDGCPKLQSVLYFCRQMTNAALVTIAKNRPNMIRFRLCIIEPRSPDYLTLGPLDAGFGAIVENCKELRRLSLSGLLTDRVFEYIGTHAQKLEMLSIAFAGDSDLGLHHVLSGCESLRKLEIRDCPFGDKALLANAAKLETMRSLWMSSCSVSFGACKLLAQKIPRLNVEVIDERGPPDTRPESCPVEKLYIYRTVAGPRFDTPGYVWTMDEI

### NbL11g22370.1 protease_Do−like_2_chloroplastic [MW = 68.51 kDa]

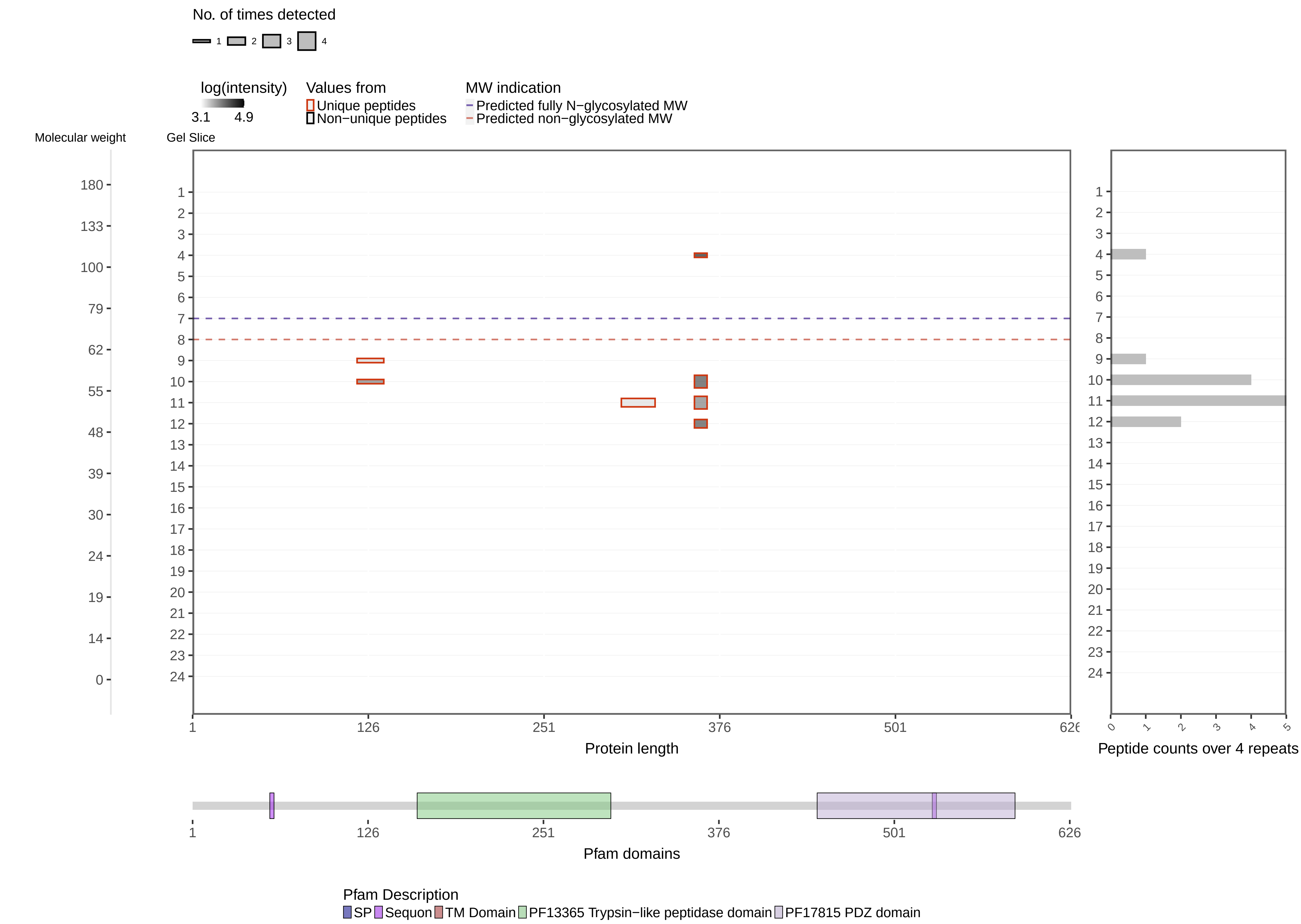


Protein sequence:

### NbL11g22260.1 serine/threonine−protein_phosphatase_BSL1 [MW = 96.04 kDa]

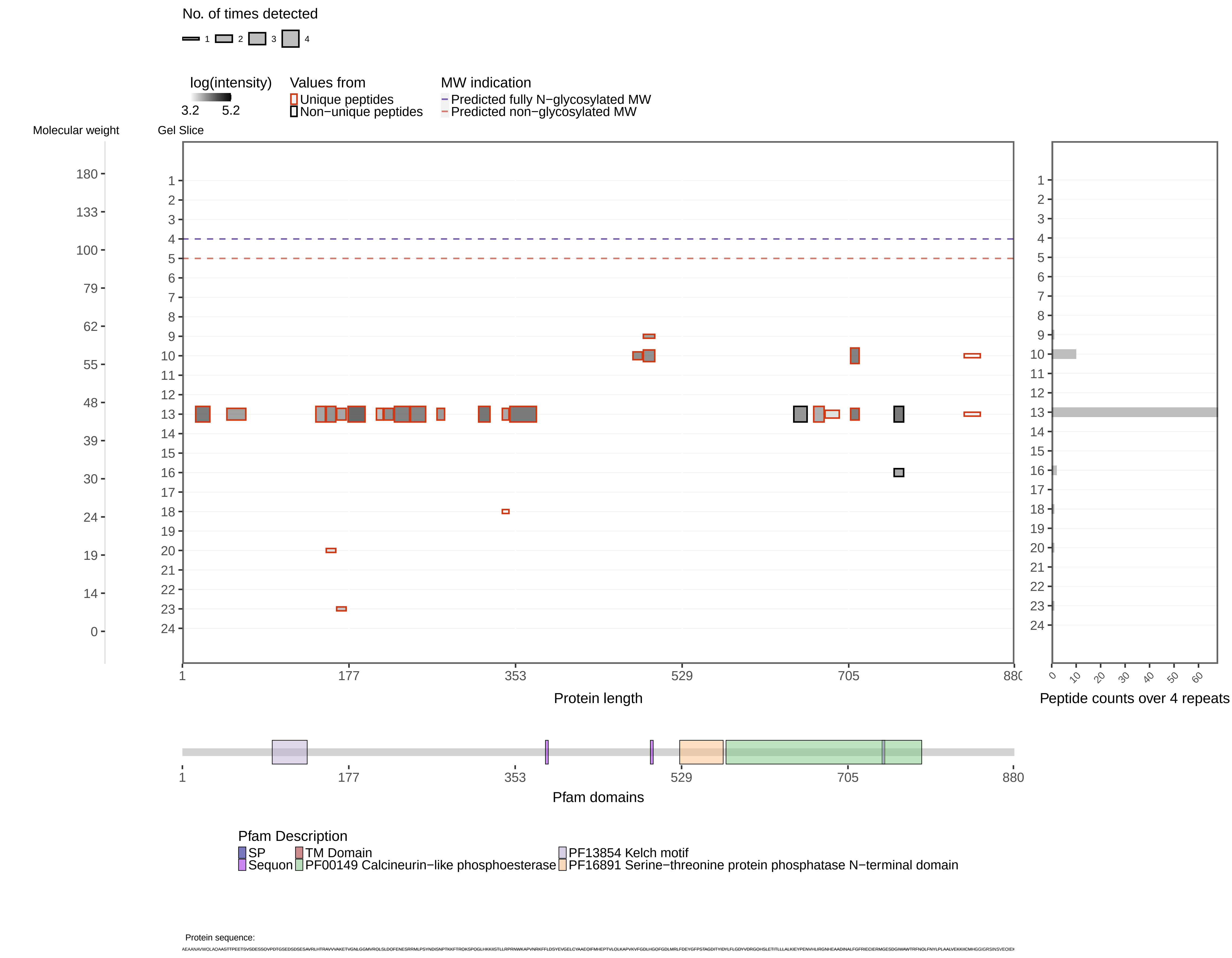

NbL11g22100.1 exosome_complex_component_RRP4_homolog [MW = 36.53 kDa]

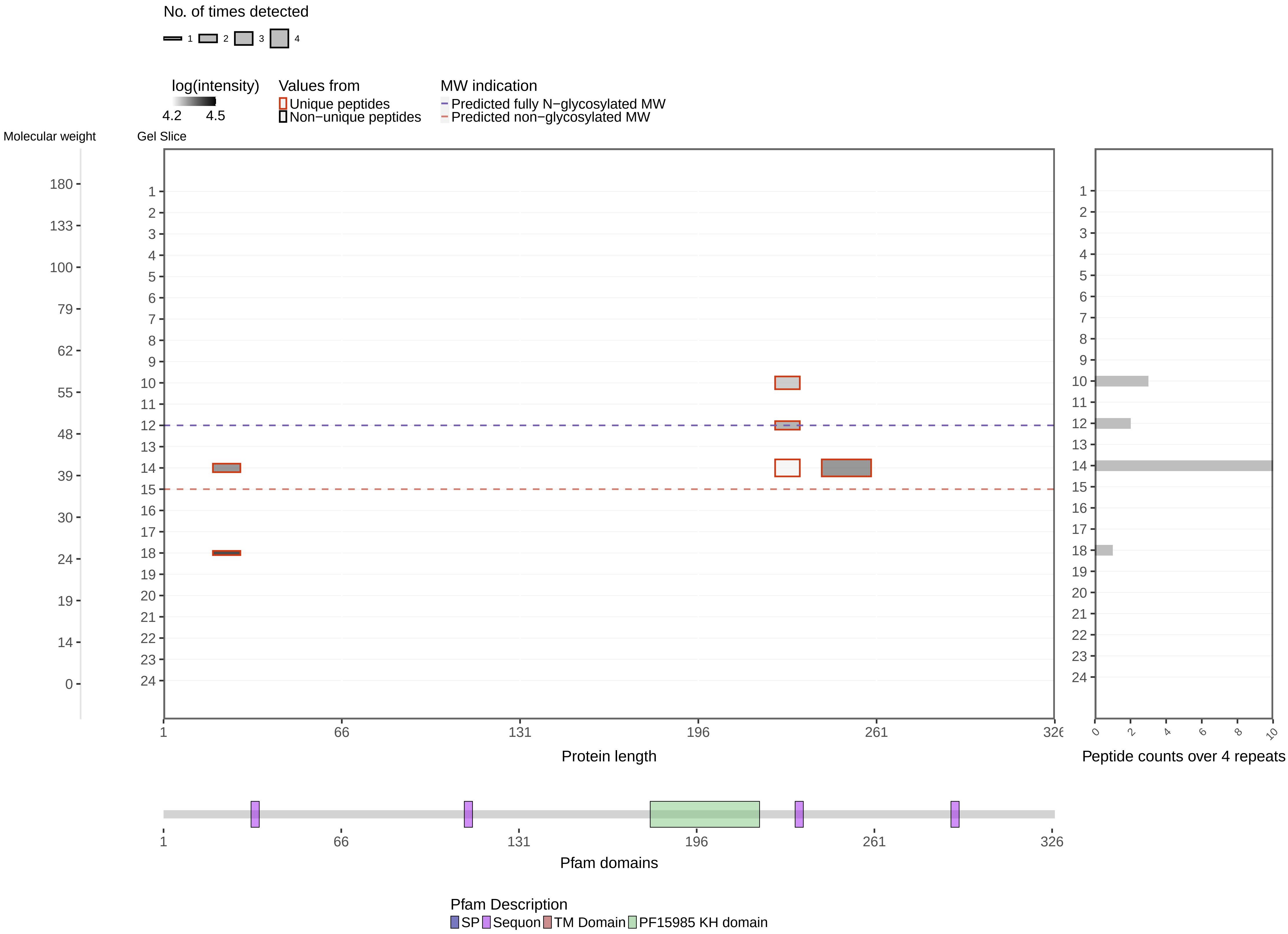


Protein sequence:

CAEVRGFQHDGSLHLQARSQKYGKLERGRLLTIPPYLVKRRKQHFHHLDQYGVDLILGSNGFIWIGEHVEVKDAMVEDQVNKSDPQNTRSGSLLGSSEEPTSTPPETRQYICRIANAITVFAALDFMVTLESVIEMVNLSLSLNLGVDEMLKAEFHVQVAEKEAERRSTLTKRR*

### NbL11g22020.1 NA [MW = 25.99 kDa]

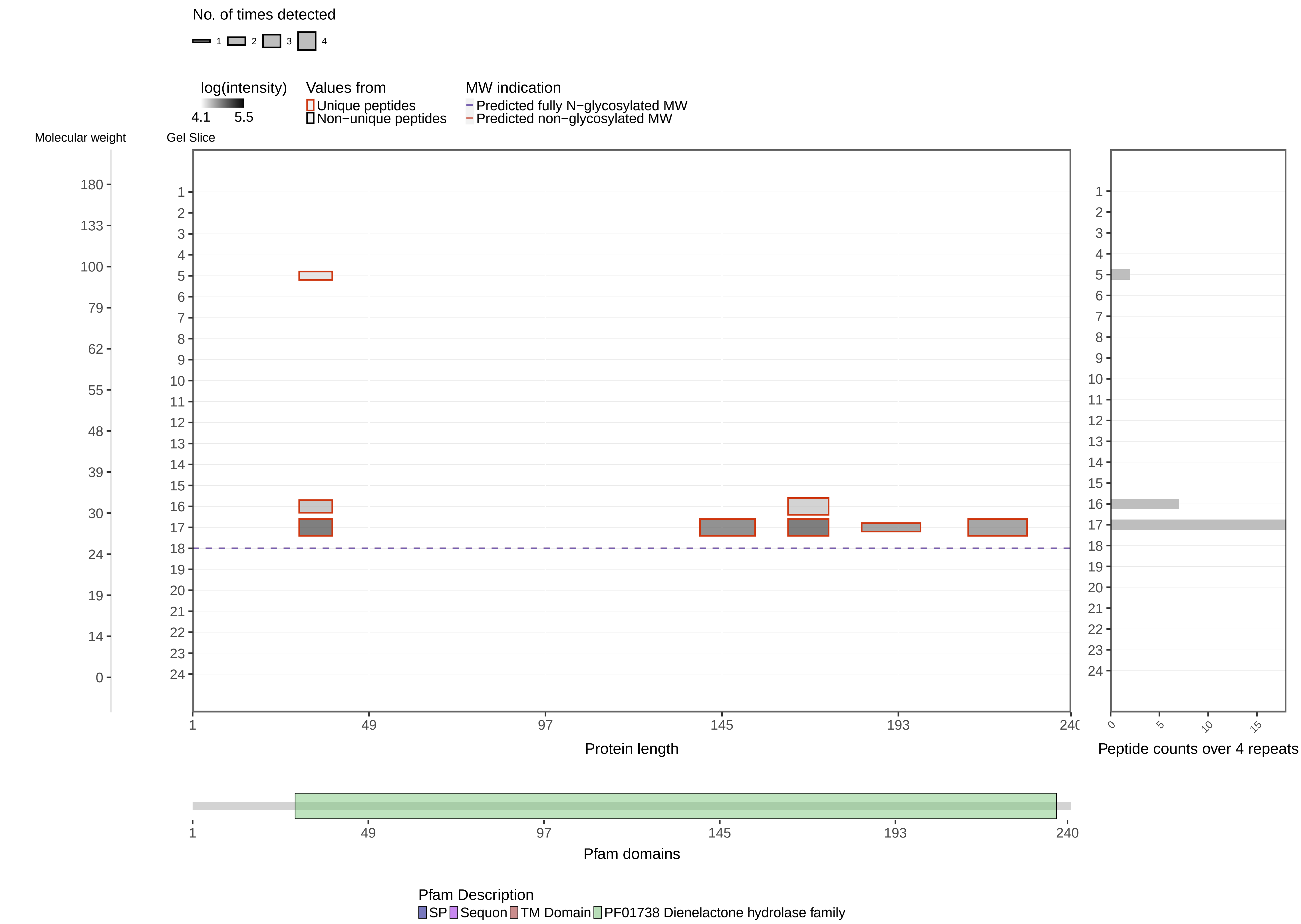


Protein sequence:

AKKVVAALKDKGISAIGAAGFCWGAKVVTDLAKSDYIQAAVQLHPSLVKVDDIKEVKAPIAILAAEIDRISPPELIKQFEEILSSKPEVDSFVKIFPSVKHGWTVRYNDEDKEAVQSAEEAHRHMLDWFTKYVK*

NbL11g22010.1 NA [MW = 26.03 kDa]

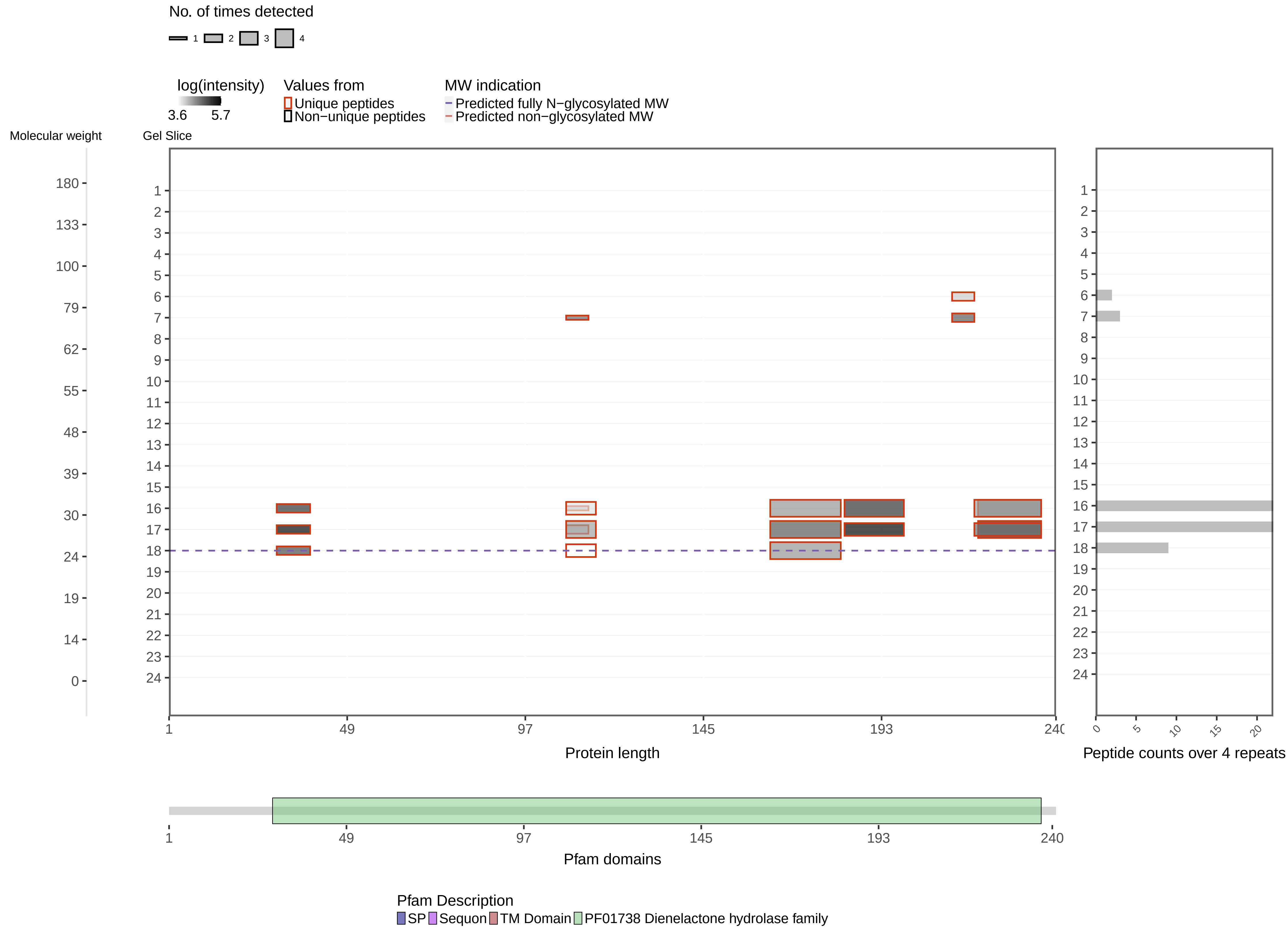


Protein sequence:

KQVVAALKDKGISAIGAAGFCWGAKVVVQLAKSDDIQAAVLLHPSFVTVDDIKEVKAPIAILGAEIDKMSPPELVKQFEEILSSKPEVDNFEKIYPGVAHGWTVRYDVEDKKAVESAEEAHQDMLEWFTKHVK*

### NbL11g21960.1 isoamylase_2_chloroplastic [MW = 95.96 kDa]

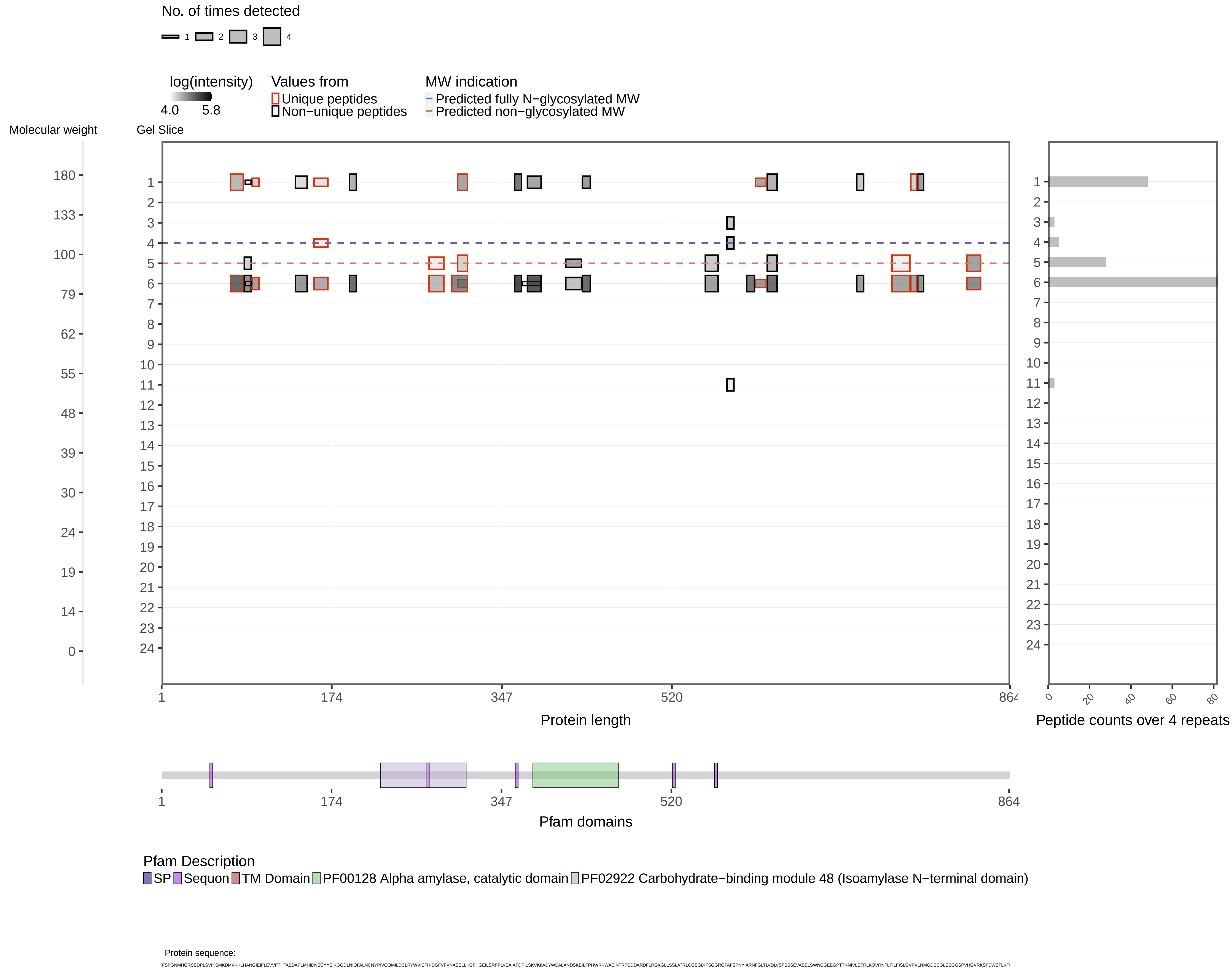

NbL11g21890.1 DEAD−box_ATP−dependent_RNA_helicase_24 [MW = 80.67 kDa]

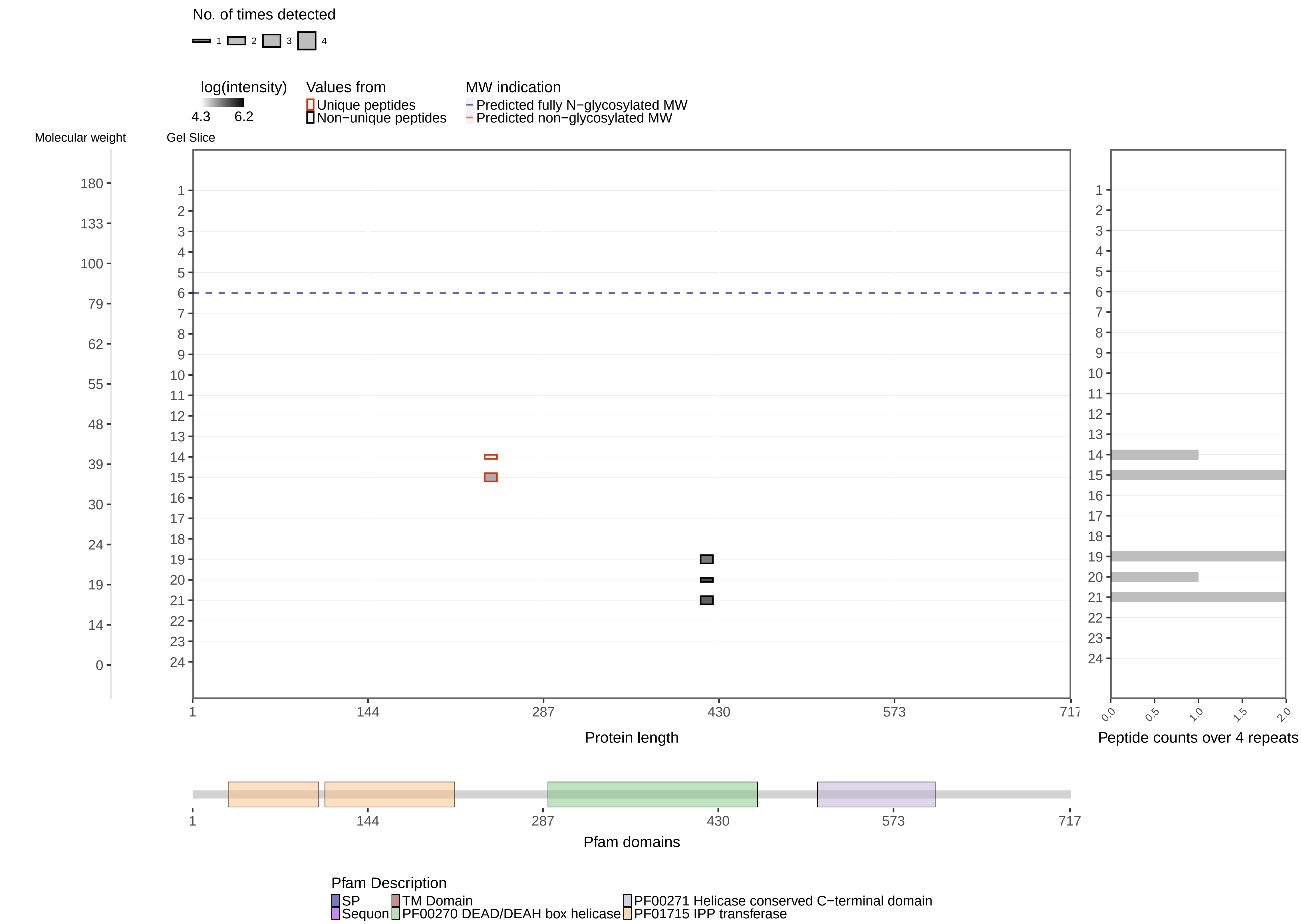


Protein sequence:

[illegible]

NbL11g21760.1 chloroplast_stem−loop_binding_protein_of_41_kDa_a_chloroplastic−like [MW = 43.51 kDa]

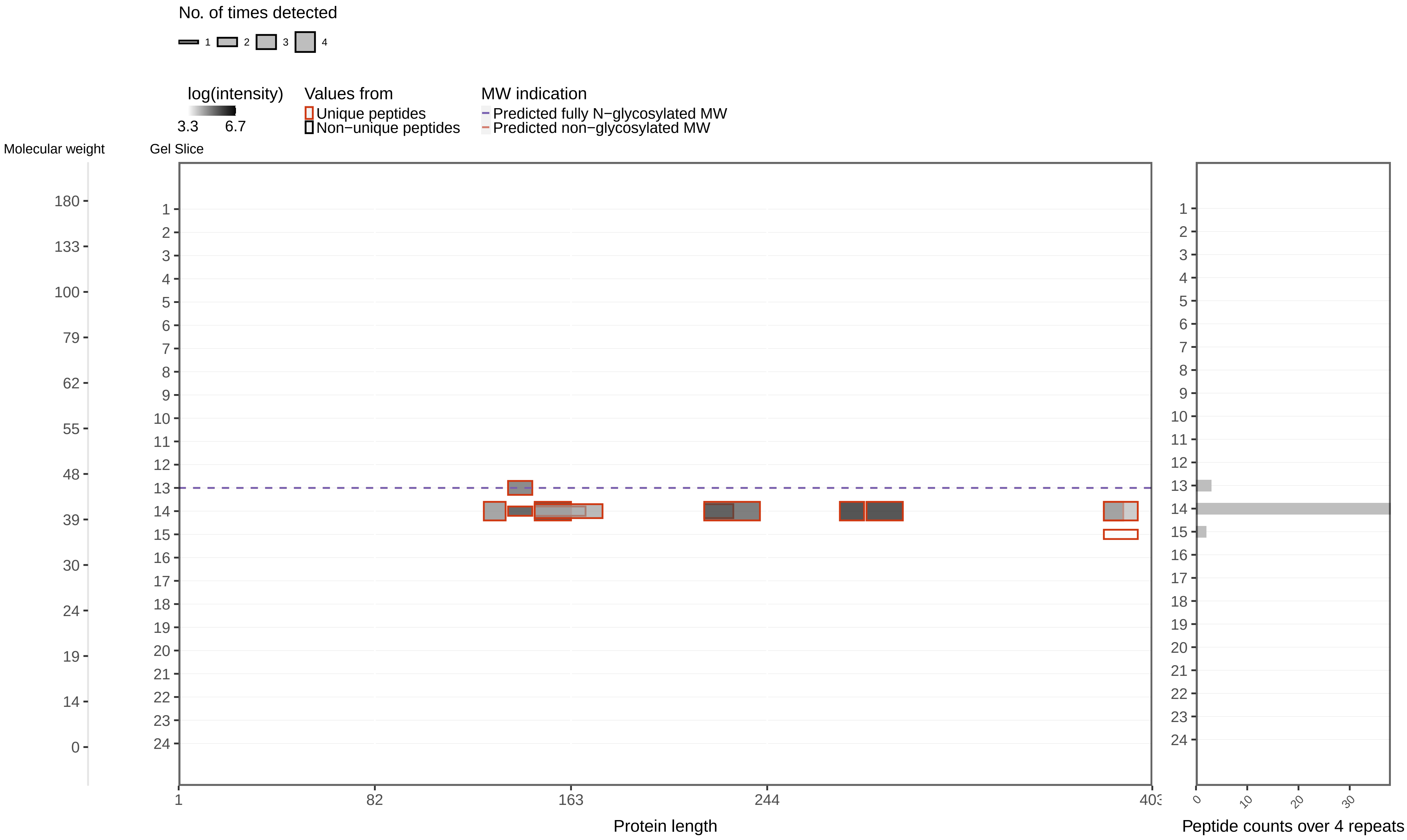


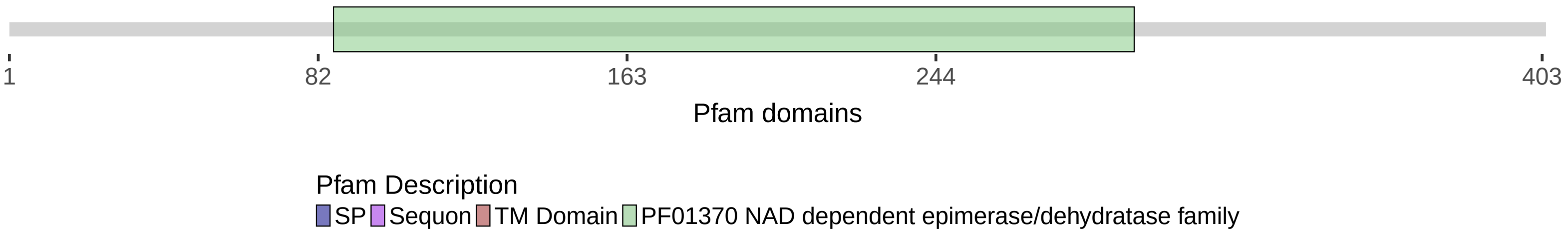


Protein sequence:

GIYKSTDEPPHVEGDAVKADAGHVGVEKYISEIFDSWSSFRPOYMIGSGNNKDCEEWFFDRIVRGRPVPIPGSGMOLTNIAHVRDLSSMLTAAVKNPAAASGHIFNCVSDRAVTLDGMAKLCAKAAGLSVEIVHYDPKAVGVDAKKAFPFRNMHFYSEPRAAKEILGWSATTNLPEDLKERFDEYVKIGRDKKEIKFELDDKILEALKVPVAA*

### NbL11g21680.1 ribosome−recycling_factor_chloroplastic [MW = 35.41 kDa]

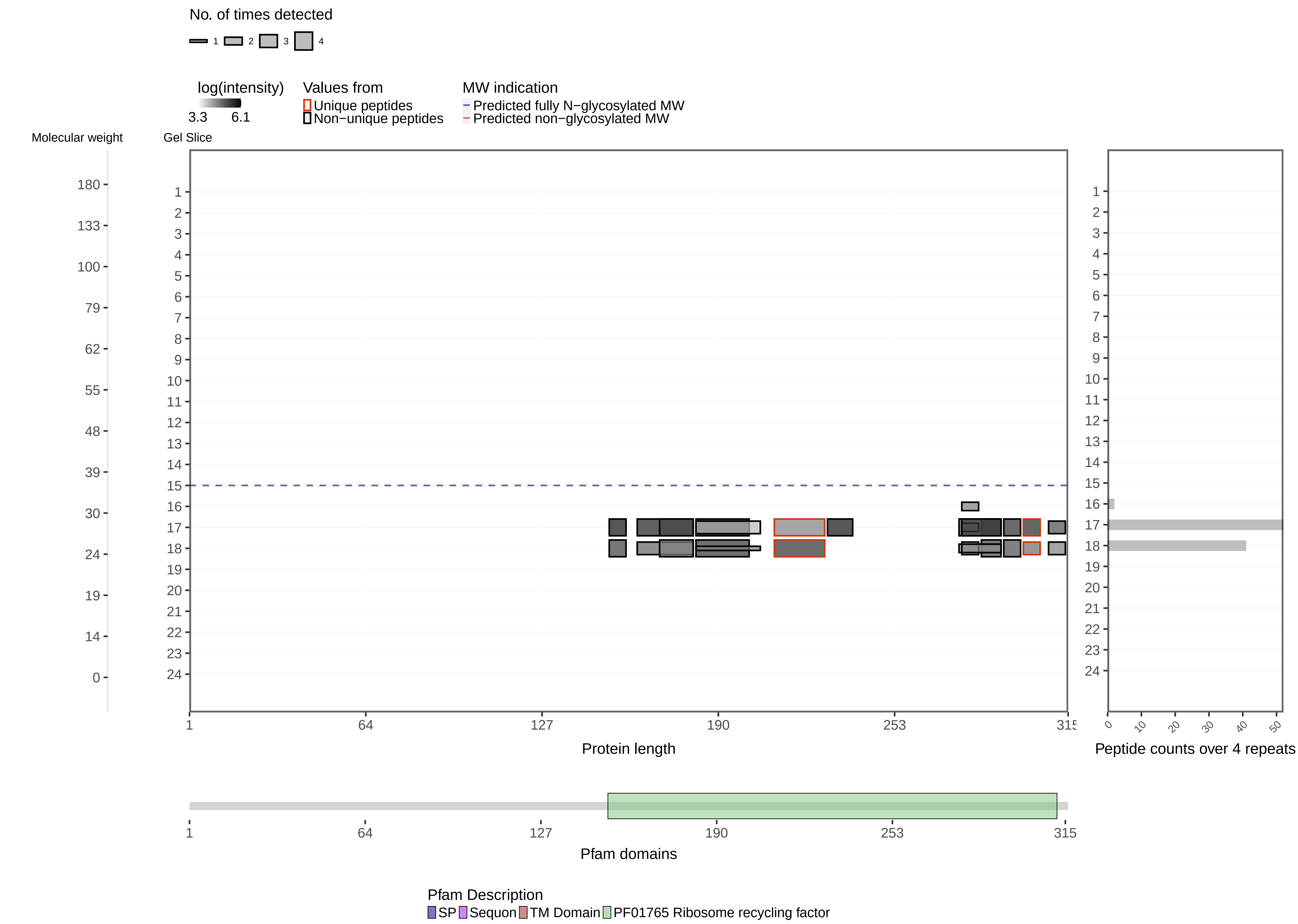


Protein sequence:

IEKTIENVRSNFNSIRTGRASPAMLDKIEVDYYGTPVSLKSIAQISTPDSSSILVQPYDKSSLKAIEKAIVGSDLGMTPNNDGELIRMSVPQLTSDRRKELSKIVSKQAEEGKVALRNIRRDALKAYEKLEKEKKLSEDNVKDLSGDLQKVTDEYMKKIDTVFKQKEKELLTV*

NbL11g21530.1 dehydrogenase_(LOC107772922)_mRNA_AJ299256.1_Nicotiana_tabacum_mRNA_for_malate_dehydrogenase [MW = 39.85 kDa]

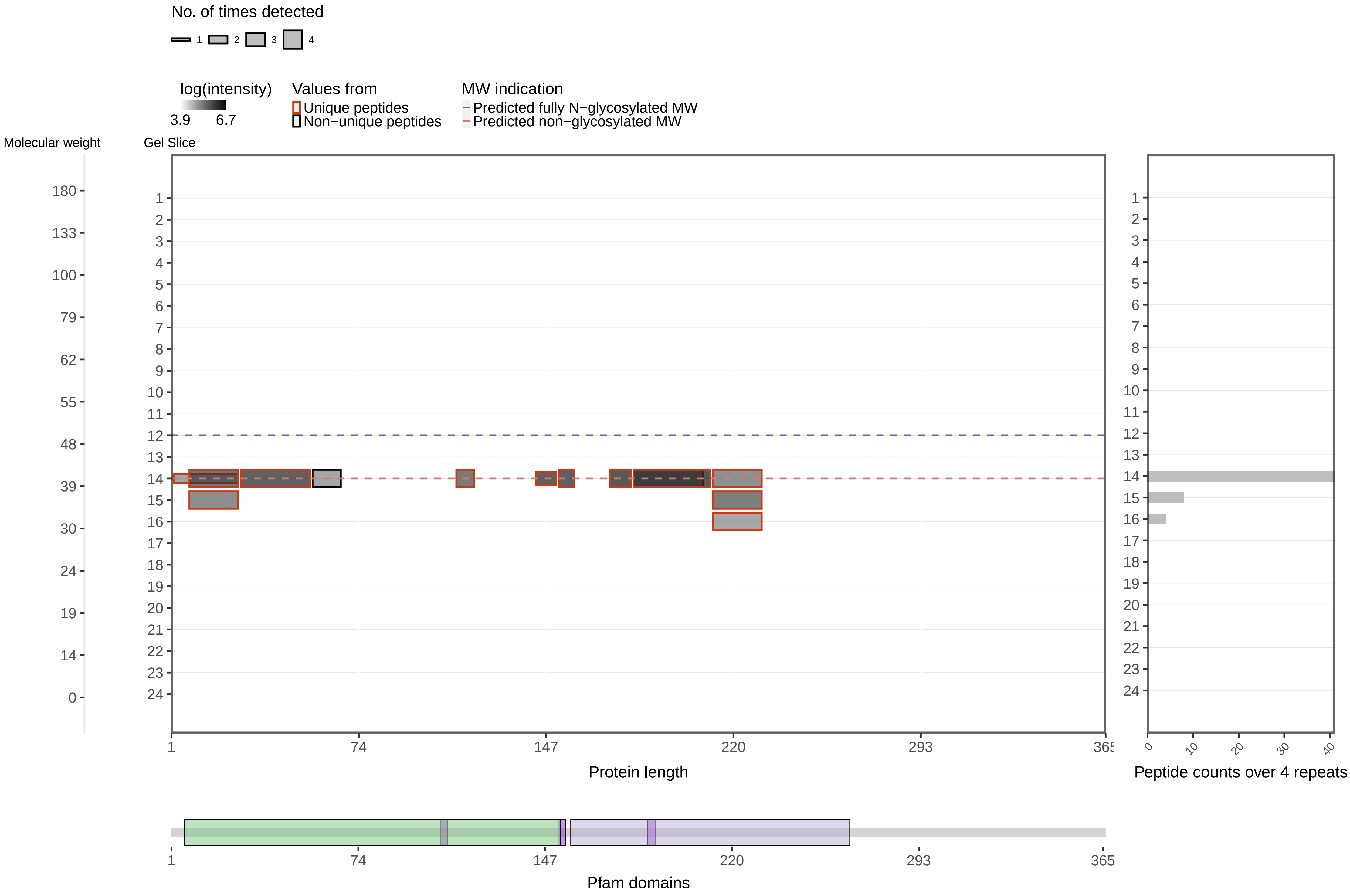


Pfam Description
SP Sequon TM Domain PF00056 lactate/malate dehydrogenase, NAD binding domain PF02866 lactate/malate dehydrogenase, alpha/beta C−terminal domain

Protein sequence:
ERLNVQVSDVKNVIIWGNHSSSQYPDVNHATVATPAGDKPVRELVADDAWLNGEFISTVQQRGAAIIKARKLSSALSAASSACDHIRDWVLELQRELGFLWVFTLMGHTVSQLVLFIPSQLHAKMENGLLFKVFQLMSSQERSWTPQLRNYLRKKHWHIPALPKLQHFLNCLFNIWGVYVILEVASVLNKAIVDS*

### NbL11g21460.1 activating_signal_cointegrator_1−like [MW = 31.61 kDa]

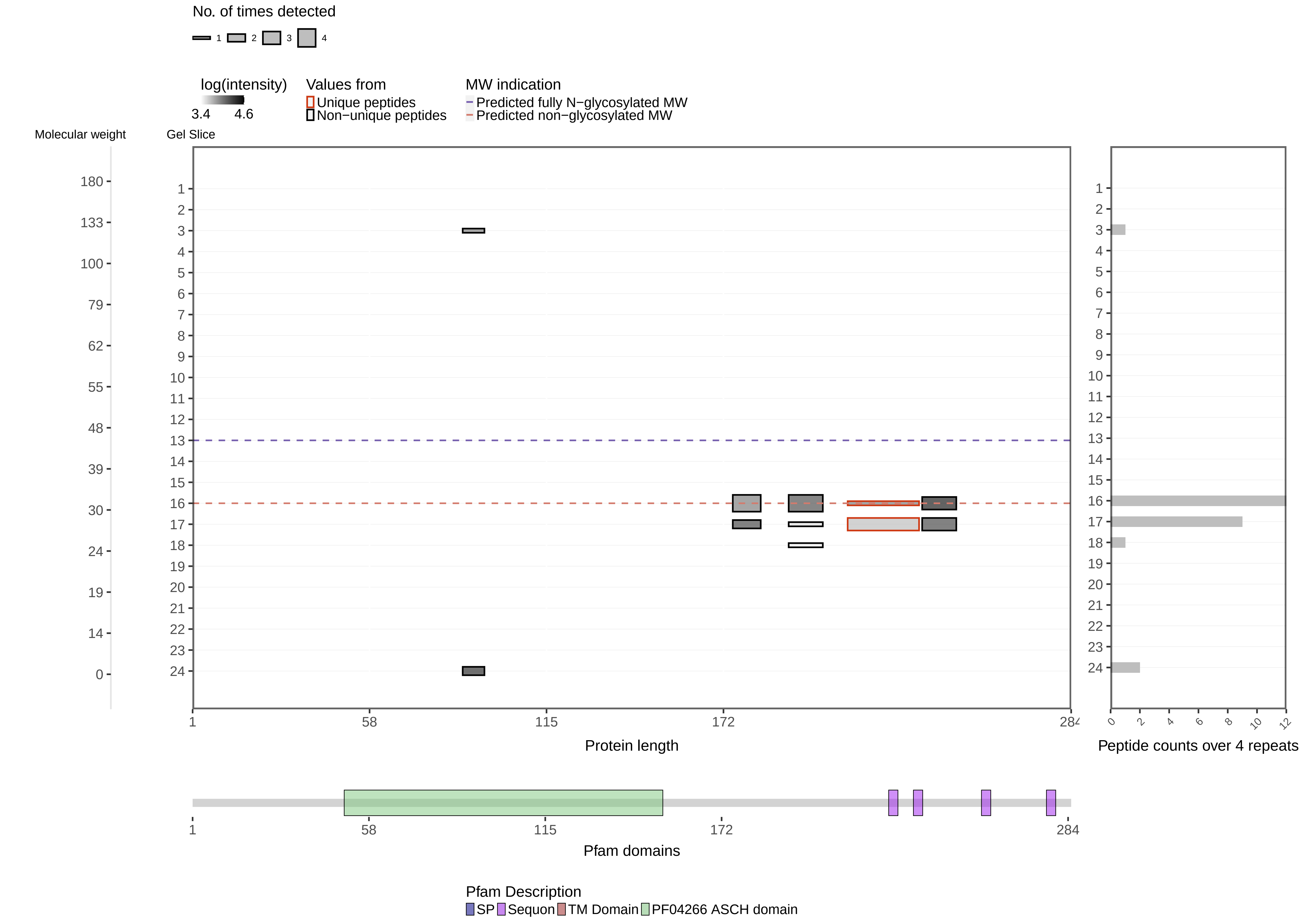


Protein sequence:

VVGCVTCEELVGWDQIPEGVRLEGQTKFCWLCEQPQKLIVPFEMRGFQGVYNLERKIYEAAIRGISPVAPPLPVKFPLPNPRDPFSLKPGSLASILNSSSTSEVNRSENLAAAIAGARAAATQFSKNSSSQSSTVEVGDEDVSSNTRNKSRRIQ*

NbL11g21410.1 probable_plastid−lipid−associated_protein_4_chloroplastic [MW = 28.63 kDa]

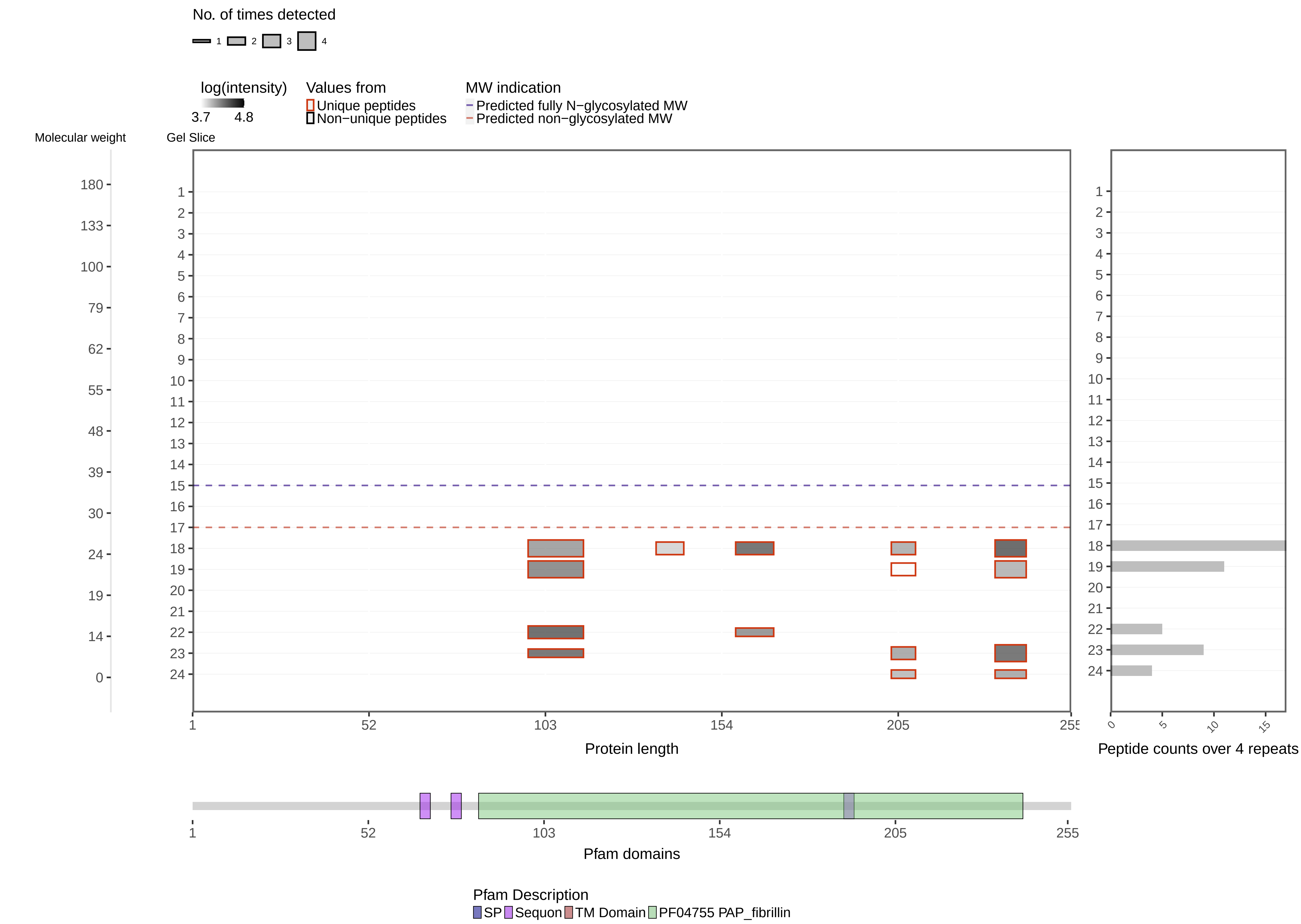


Protein sequence:

KLEACNPTKEPLKSNLLNGKWELIYTTSKSILQTQRPKILRSRVNYQAINGDTLRAQNMEGLPFFNQVTADLTPLNSSKVAVKFDYFKIGGIIPVKAPGRARGELEITYLDEELRVSRGDLGNLFILKMVDPSYRVPTRD*

NbL11g21400.1 uncharacterized_LOC109207172 [MW = 72 kDa]

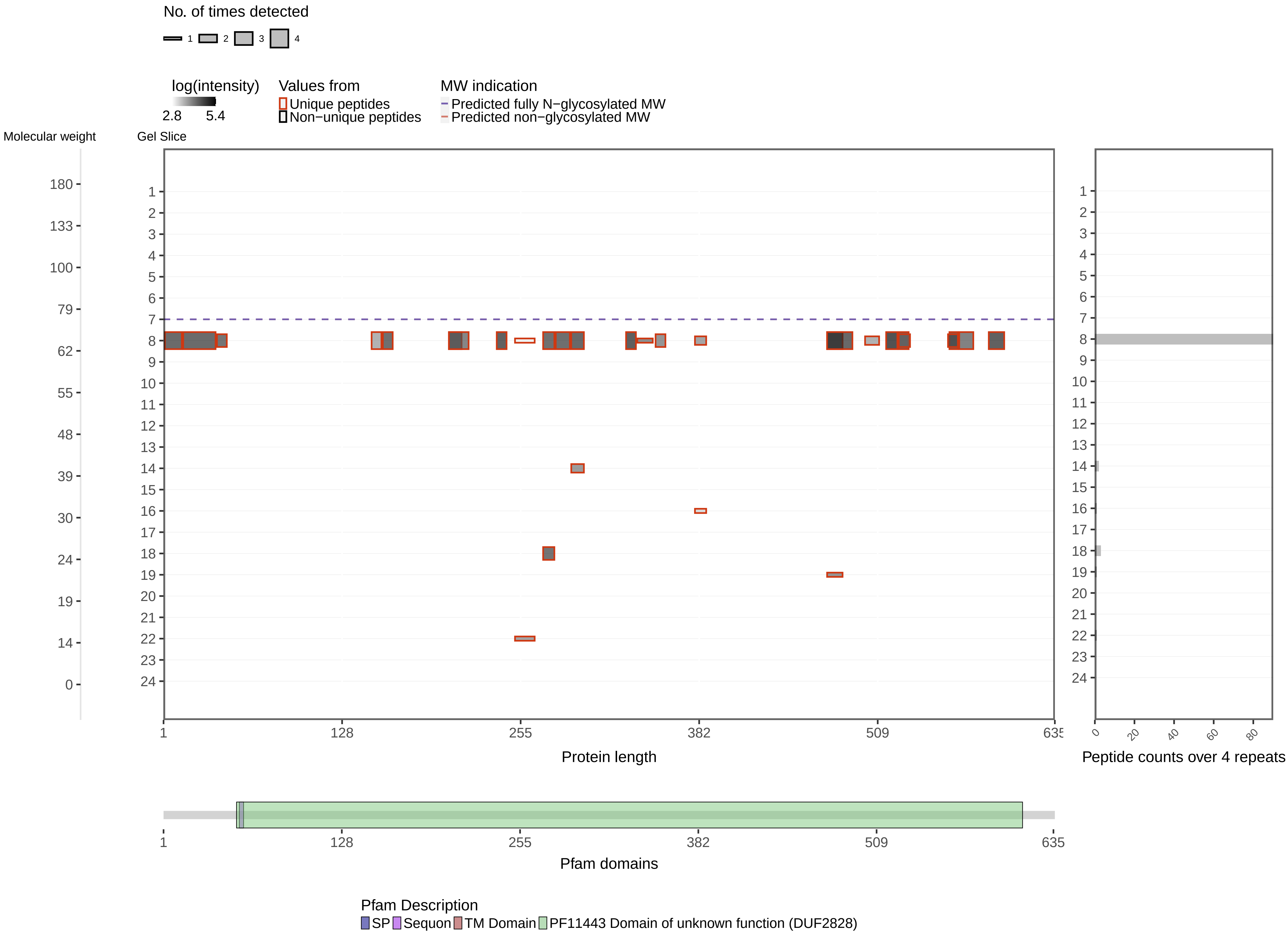


Protein sequence:

YGEYNGVEEGHYAYRVRDRLRKDVLVPLHKALELPEVYMSAKKWNSLPYNRVASVAMKNYKELFLKHDKERFDKYLEDVKSGKAKIAAGALLPHEIIGALSDGDGGEVAELQWKRMVDDLCKKGKLSNCIAVCDVSGSMSGIPMEVSVALGVLVSELSEEPWKGKLITFSADPEMQKVEGDTLKEKTEFVRRMDWGMNTDFQKVFYRILEVAVQGKLSEDQMIKRVFVFSDMEFDQASENAWETDYQAIQRKFVETGYKNVPEIVFWNLRDSRSTPVLEKQSGVALVSG

NbL11g21390.1 electron_transfer_flavoprotein_subunit_beta_mitochondrial−like [MW = 27.6 kDa]

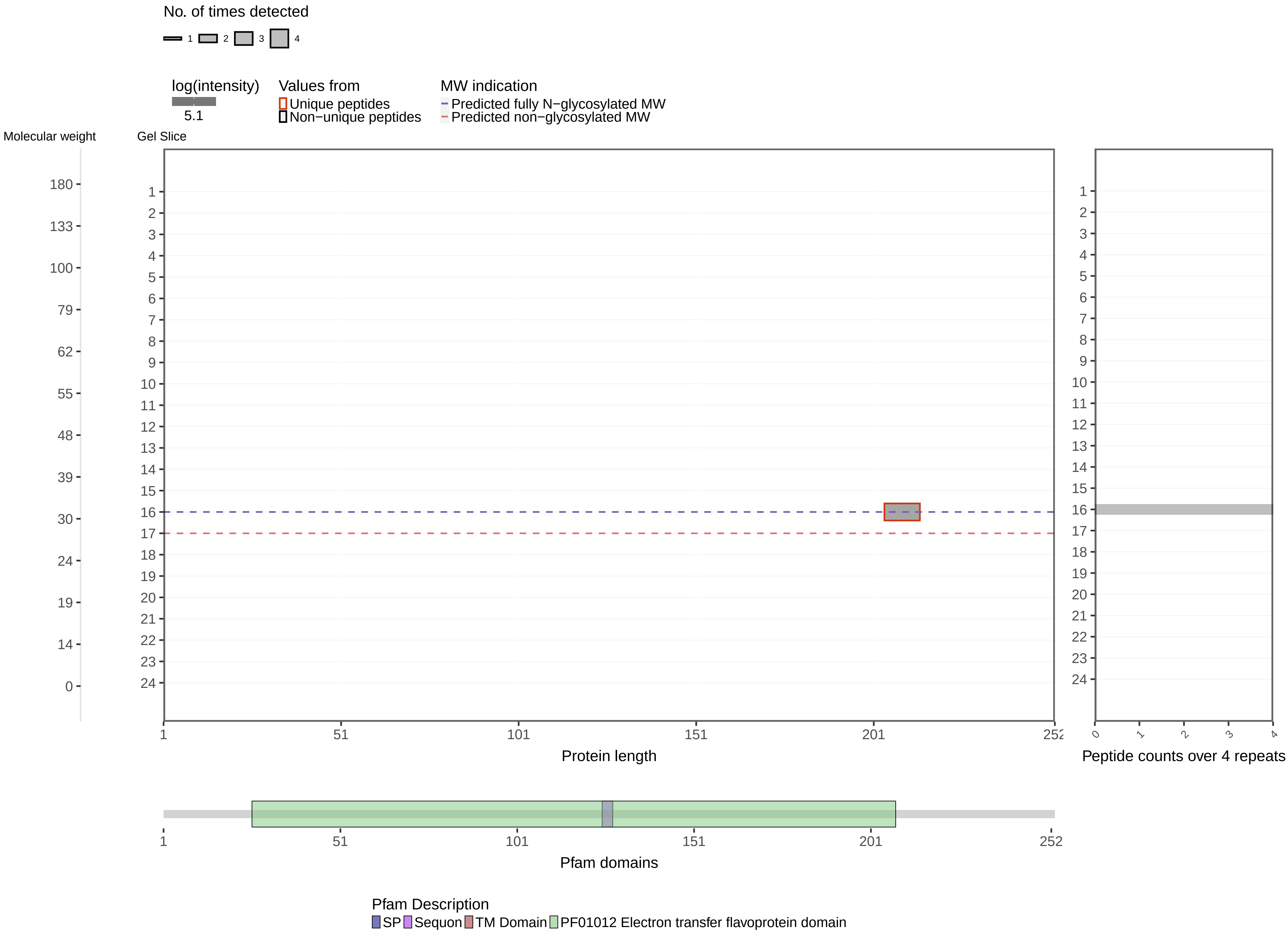


Protein sequence:

LLGKQAIDNDQNQTGQMVAGLLKWPQGTFASKVVLDKEKQVATVDREVDGGIETLSLDLPAVITTDLRLNQPRYATLPNIMKAKSKPIKKFTPQDLNMEIKSDFEVVEVTEPPKRKAGLILSSVDELIDKLKNEARVI*

NbL11g21240.1 proteinase_inhibitor_I−B−like [MW = 16.68 kDa]

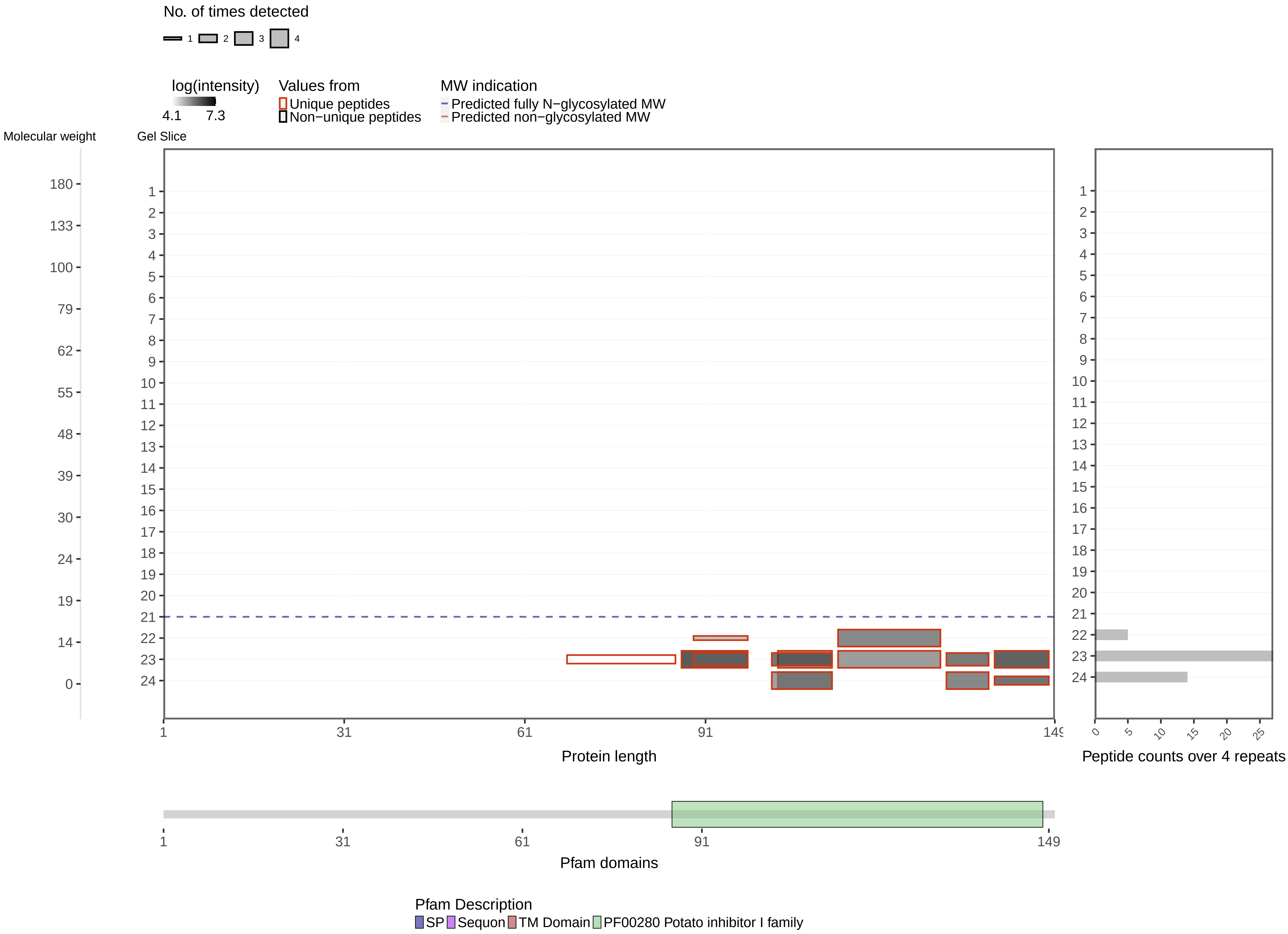


Protein sequence:

KFPMAKENQVETLDDASLSLICQGKQKWPEVVGMPGRTAKKIIEKENPLVKVHFLFPDMLQPLDLDCSRVFVLVNWKFIVQITPSVG*

NbL11g21220.1 proteinase_inhibitor_I−B−like [MW = 16.65 kDa]

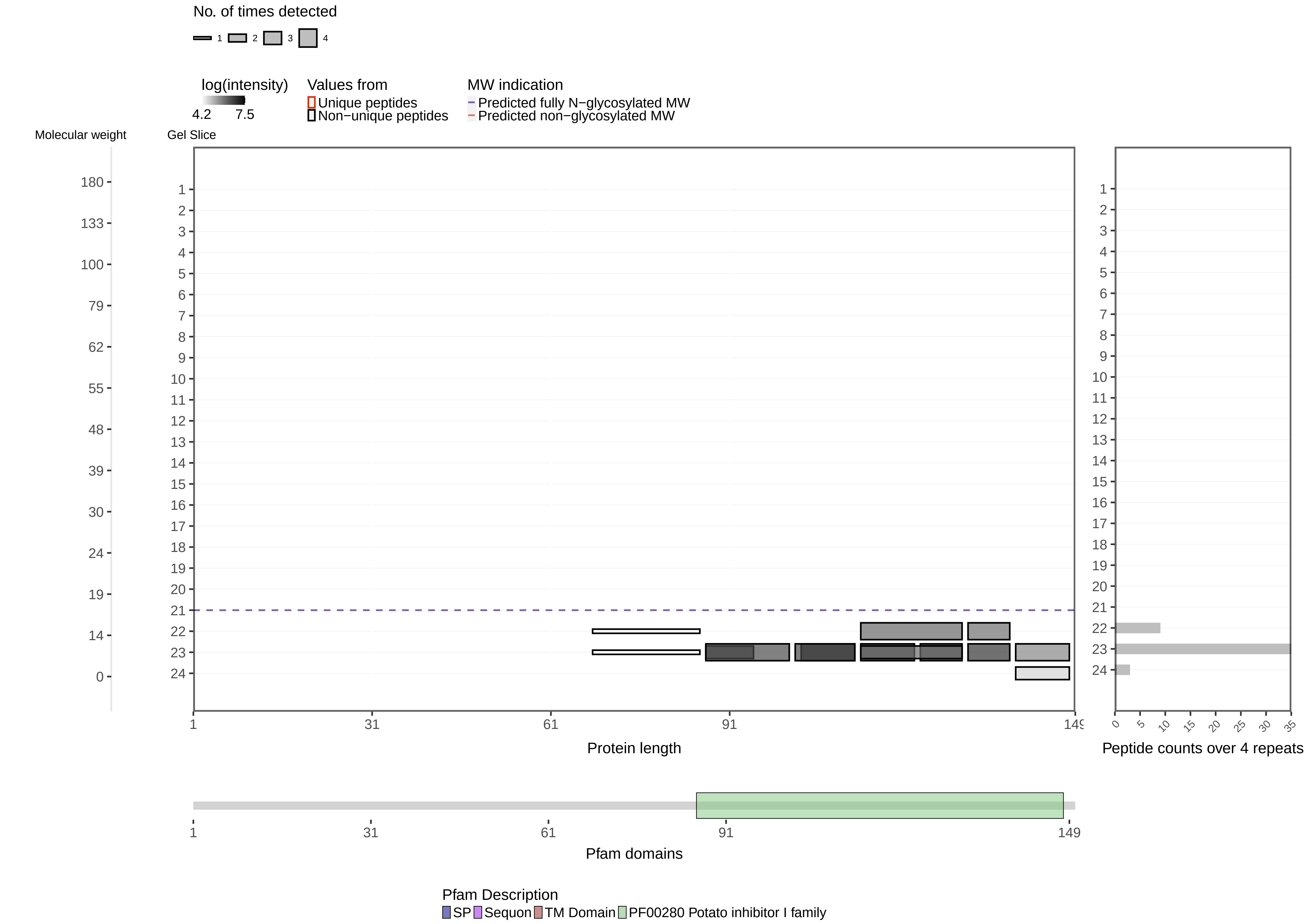


Protein sequence:

FPMAKENQVETLDDPSLSIICPGKQSWPELVGKPAATAKRIIEKENPIAKVQFLFPGMVRPLNYVCGRVFVVVNWKLIVRDTPSMG*

### NbL11g21120.1 40S_ribosomal_protein_S15 [MW = 17.02 kDa]

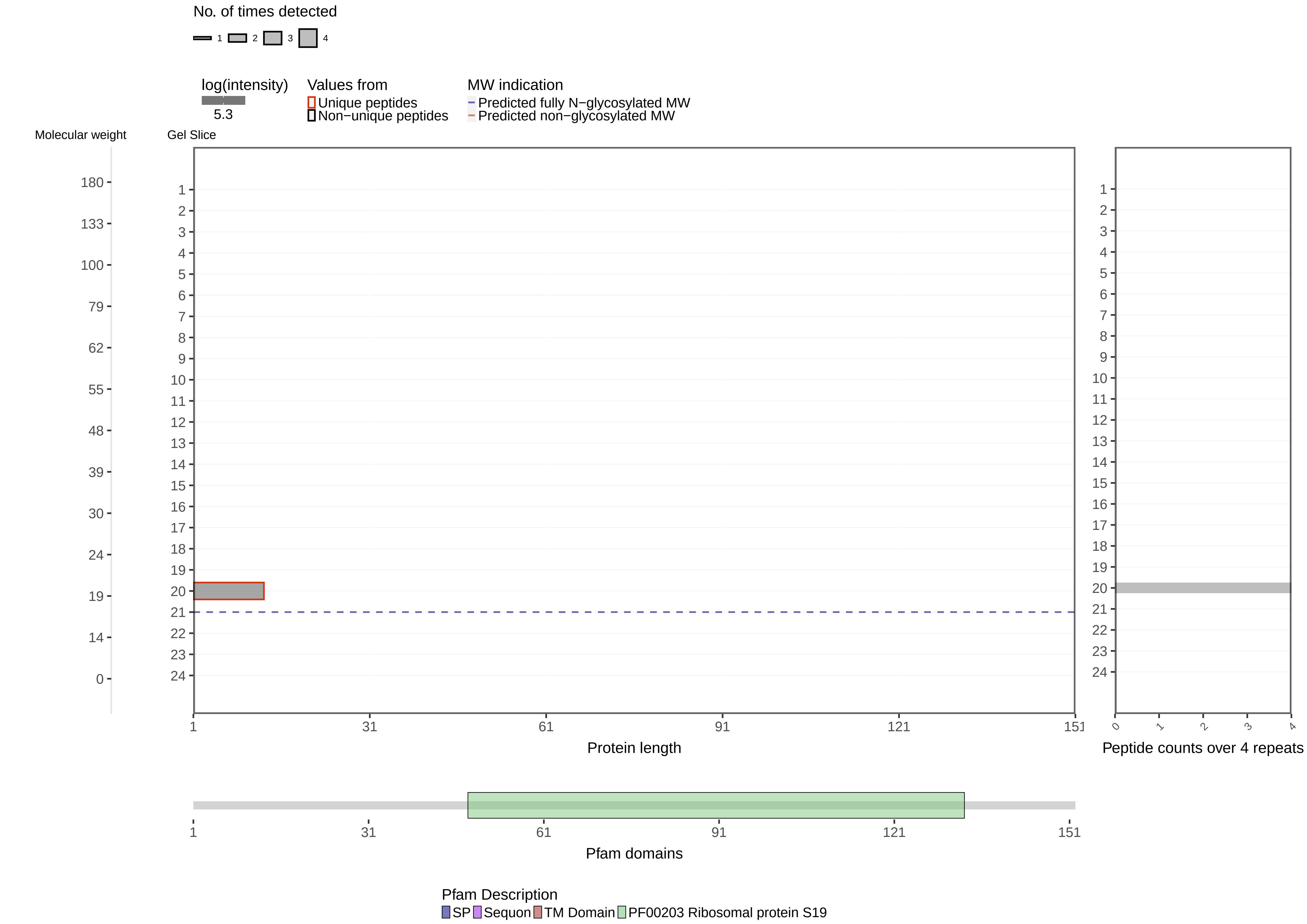


Protein sequence:

LIKKLRKAKREAPPGEKPEPVKTHLRNMIIVPEMIGSVIGIYNGKTFNQIEVKPEMIGHYLAEFSISYKPVKHGRPGIGATHSSRSIQLK*

### NbL11g21000.1 protein_LYK5−like [MW − SP = 67.5 kDa]

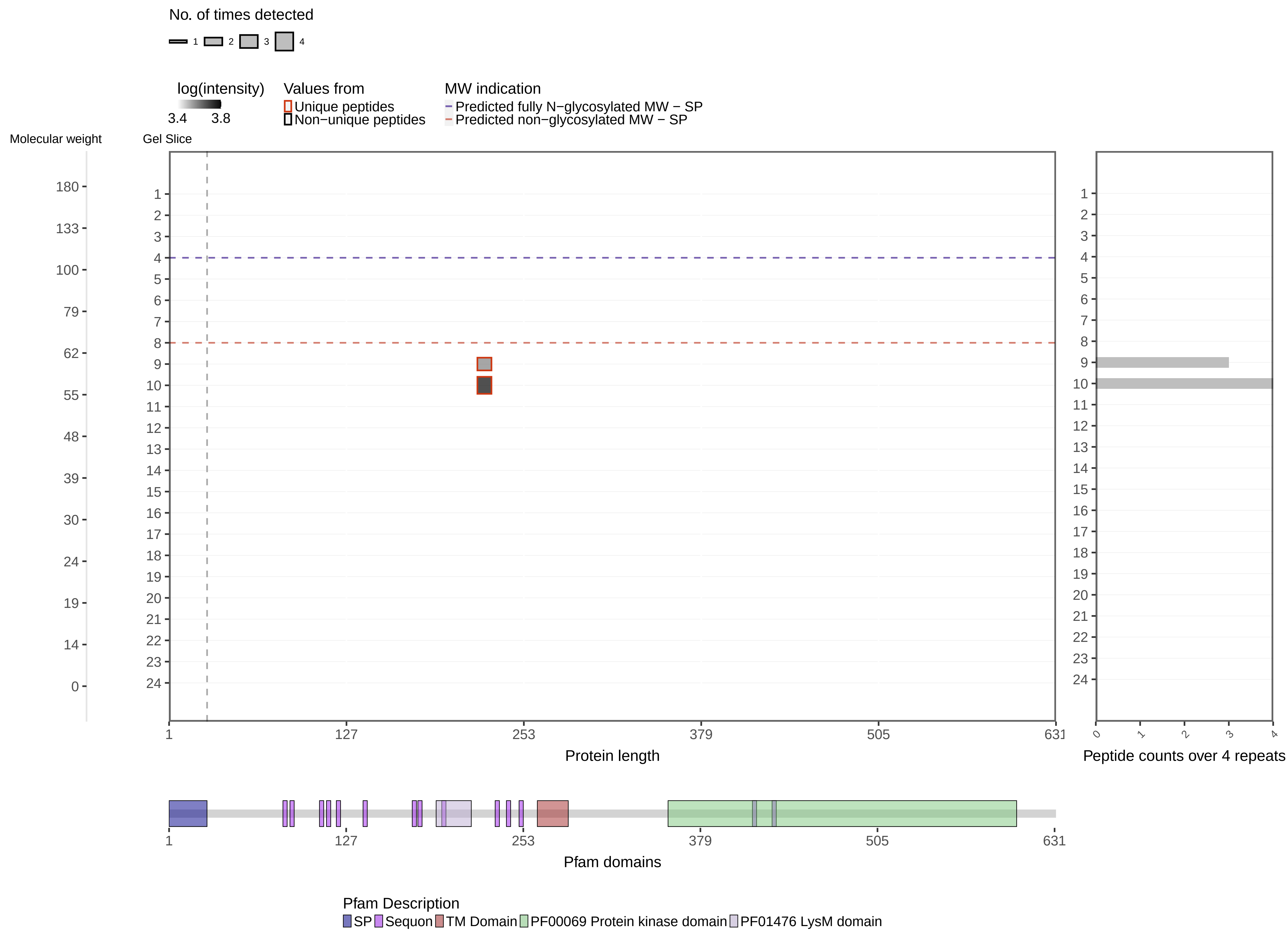


Protein sequence:

LGSKKESLNKDVEVNLMADVSDCLDKYKMYTIEQLWEATDGFDEGCLIQGSVYKGIIDGEVFAIKKMKWNAREELKILQKVNHGNLVKLEGFCIDPKEANCYLVYEYVENGSLHSWLHEDKKENLSWKTRLKIAIDVANGLLYIHEHTRPRVVHKDIKSSNILLDSNMRAKVANFGLAKSGCNAITMHIVGTQGYIAPEYLTDGIVSTKMDVFSFGVVLLELMSGREAIDNEGRVLWAKVGEILEGSEERKEKKLQEWMDDSLLREECTIESVMNVMSVAISSLNKDPSKRPSMV

NbL11g20980.1 acylamino−acid−releasing_enzyme−like [MW = 91.1 kDa]

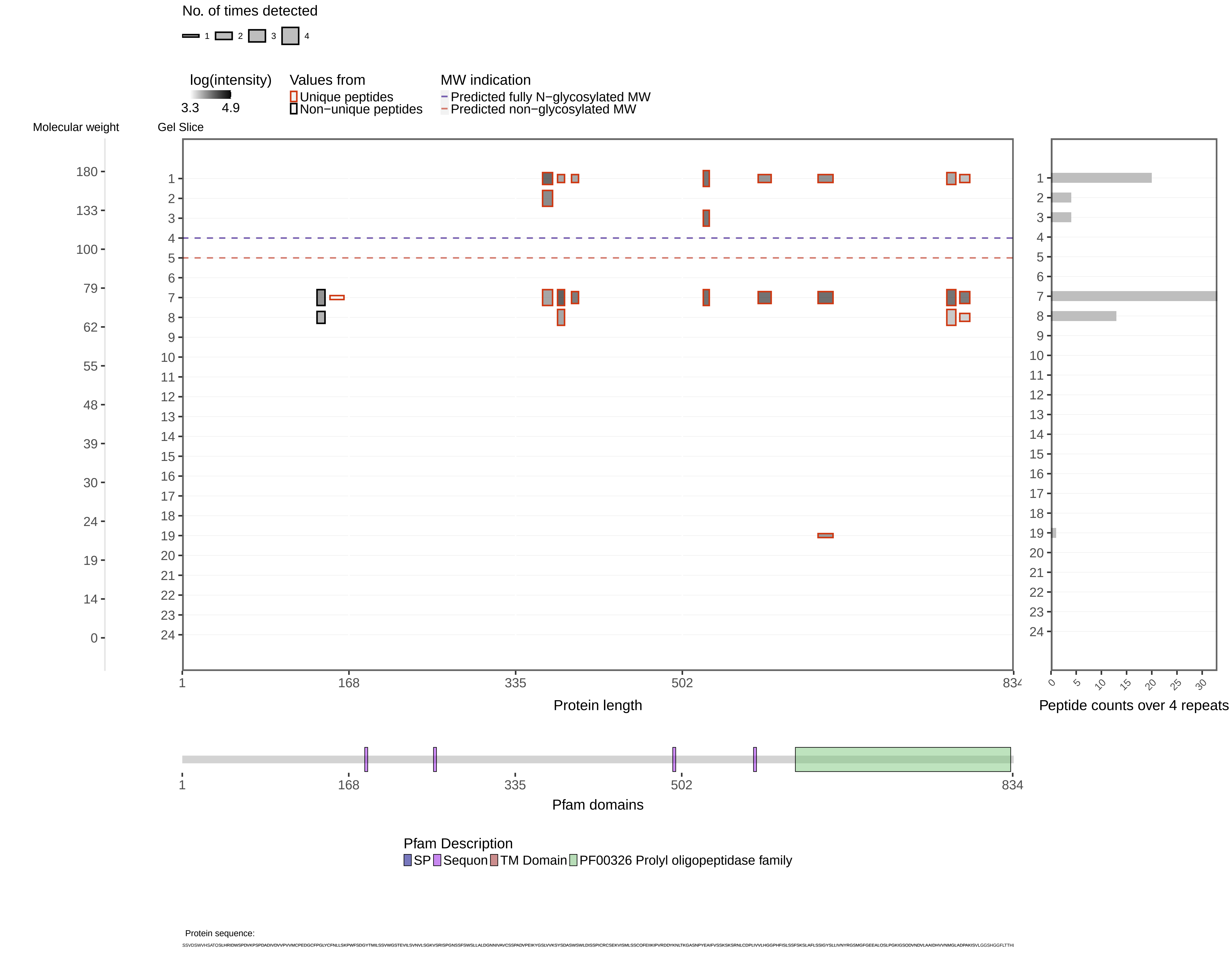

NbL11g20910.1 GDP−mannose_35−epimerase_1 [MW = 42.98 kDa]

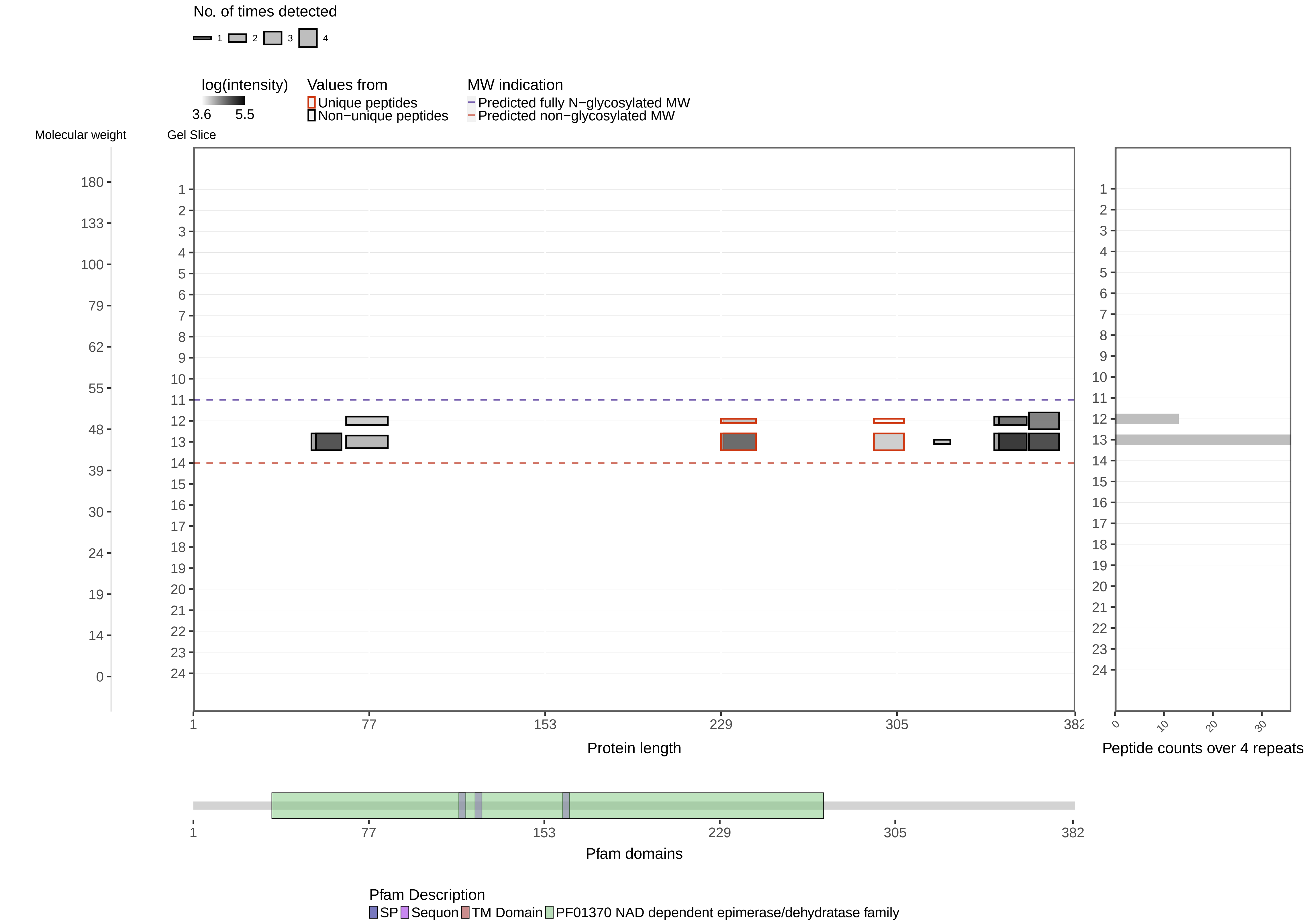


Protein sequence:

LEKLATEELCKHYNKDFGIECRIGRFHNIYGPFGTWKGGREKAPAAFCRKAQTSADKFEMWGDGKQTRSFTFIDECVEGVLRLTKSDFREPVNIGSDEMVSMNEMAEMVLSFEDKKLPVHHIPGPEGVRGRNSDNTLIKEKLGWAPSMKLKDGLRITYFWIKEQIEKEKVQGADVSVYGSSKVVGTQAPVQLGSLRAADGKE*

### NbL11g20840.1 sucrose_synthase_2−like [MW = 92.72 kDa]

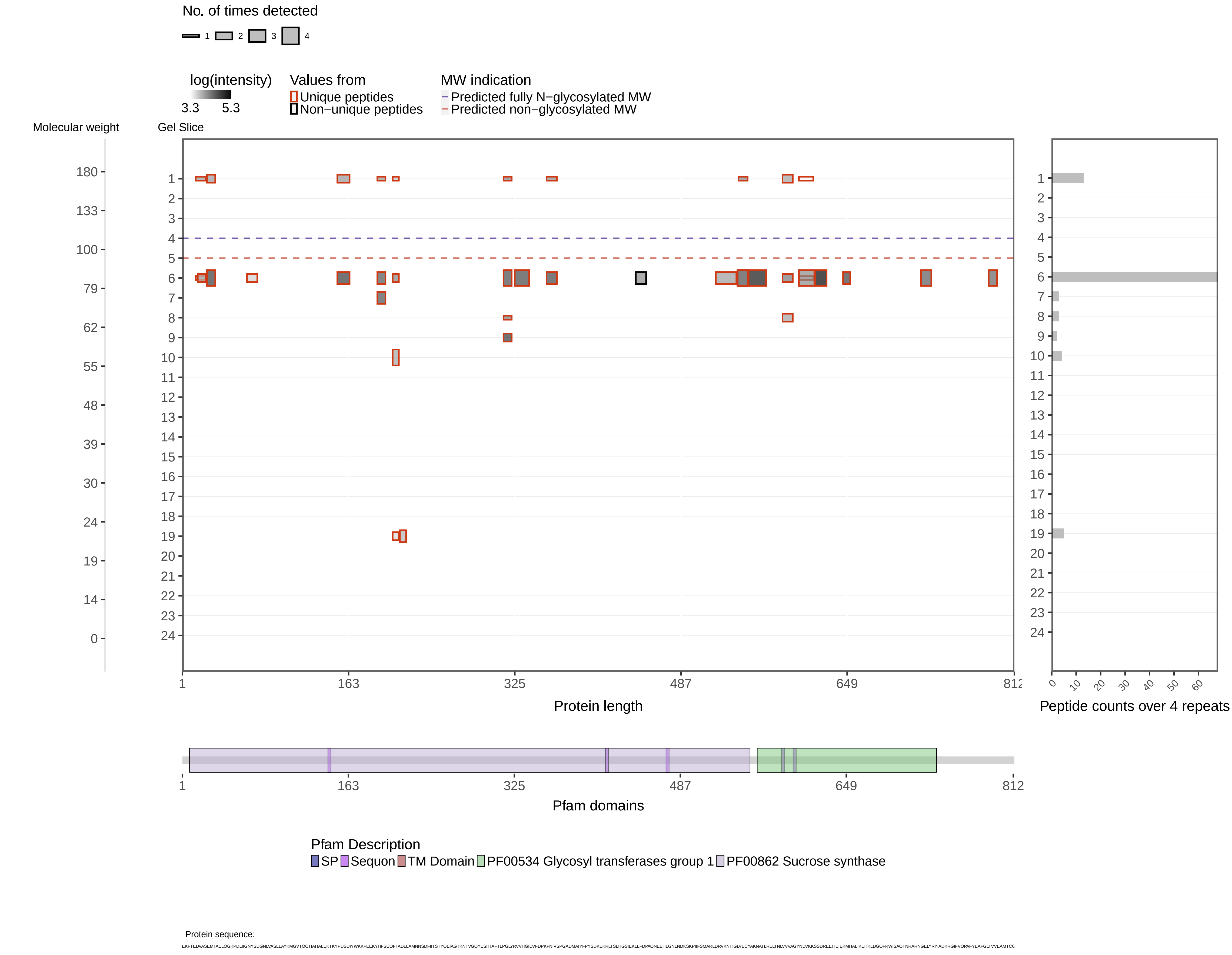

NbL11g20780.1 60S_ribosomal_protein_L19−1 [MW = 24.67 kDa]

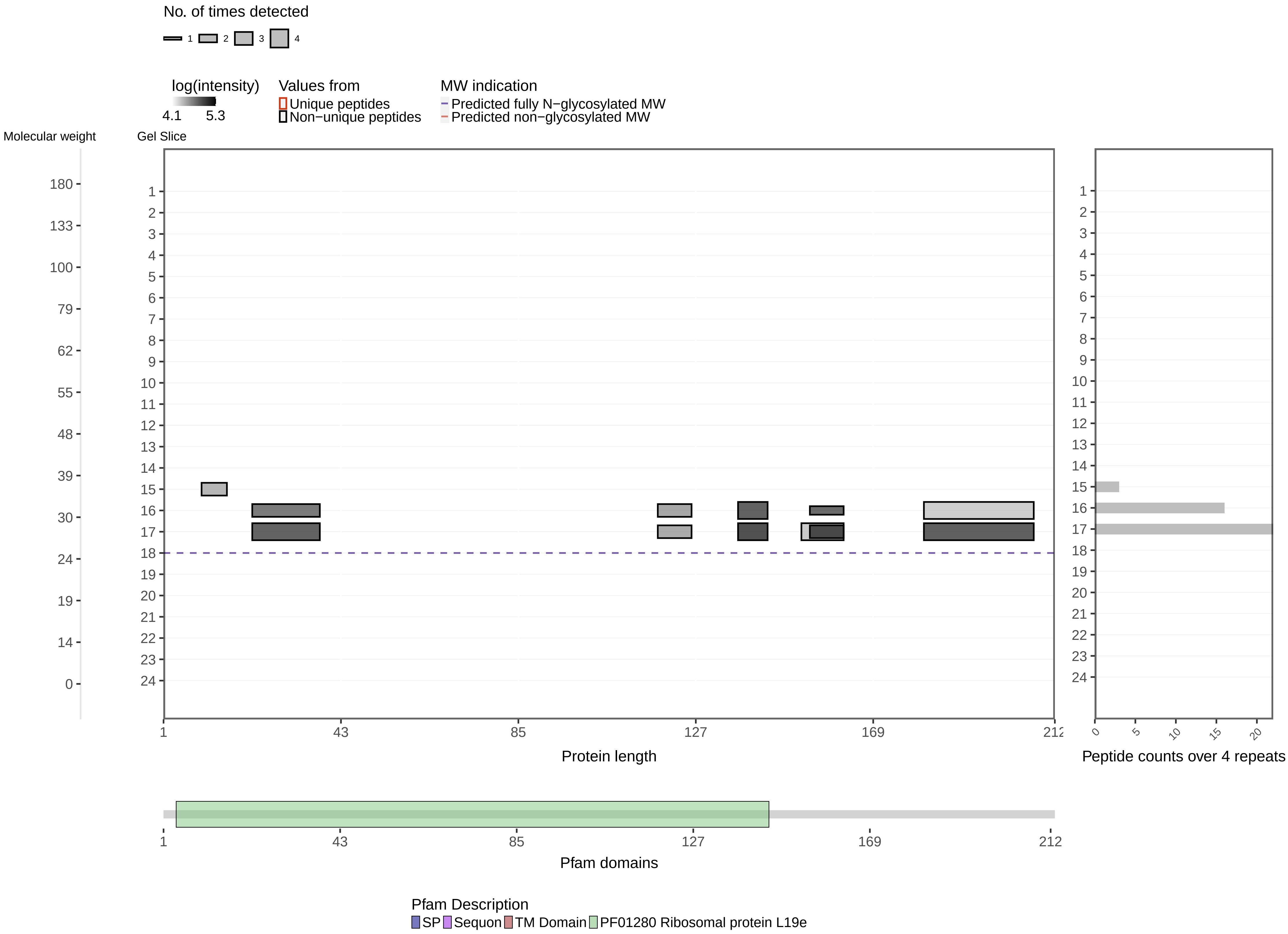


Protein sequence:
WMRRMRVLRRLLRKYRESKKIDRHMYHDMYMKVKGNVFKNKRVLMENIHKTKAEKAREKTLSDOFEARRAKNKASRERKFARREERLAOGPGERPVOPAASATPAOTAOGGSKKSKK*

NbL11g20760.1 probable_polygalacturonase [MW − SP = 46.86 kDa]

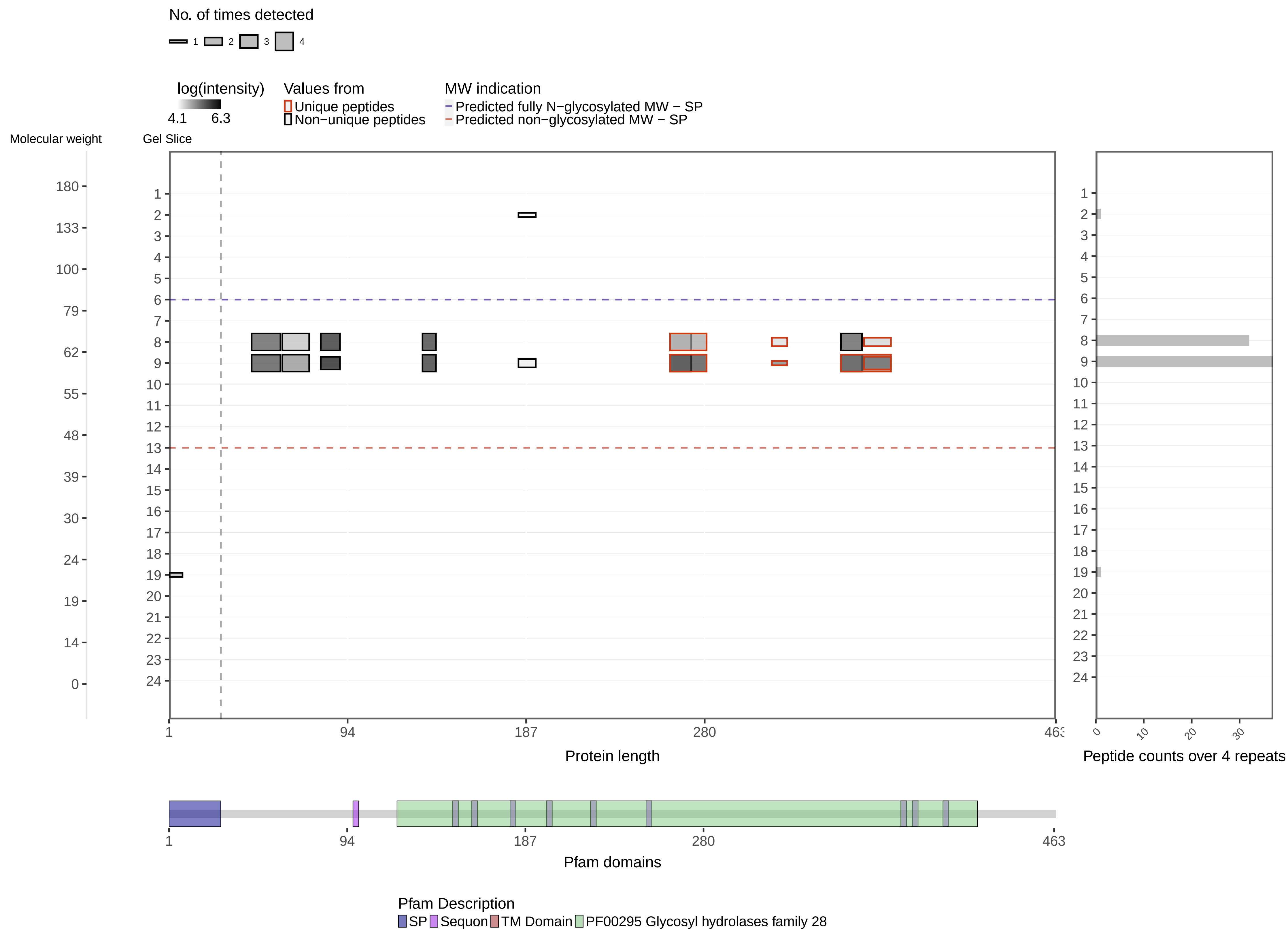


Protein sequence:

VIQNLTVHSPQGSPYTNGIVPDSSEHVCIENSNITIGYDAVVLKSGWDEYGISYGKPTSNVHVRGVRLQSSAGSAVAFGSEMSGGISNVLVEHISLRNSLFGIELKTARGRGGYIKDILITNVVMENVQVGIKATGYYDSHPDEKYDPSSLPIVSGITFKDIVGTNISIAGNFTGLSESPFTSLCLSNISFSISSDPSTPWLCSDVSGSSENVFPEPCPELQSSLSSYSSACFALPNPYTEVAVL*

### NbL11g20620.1 CTP_synthase−like [MW = 61.81 kDa]

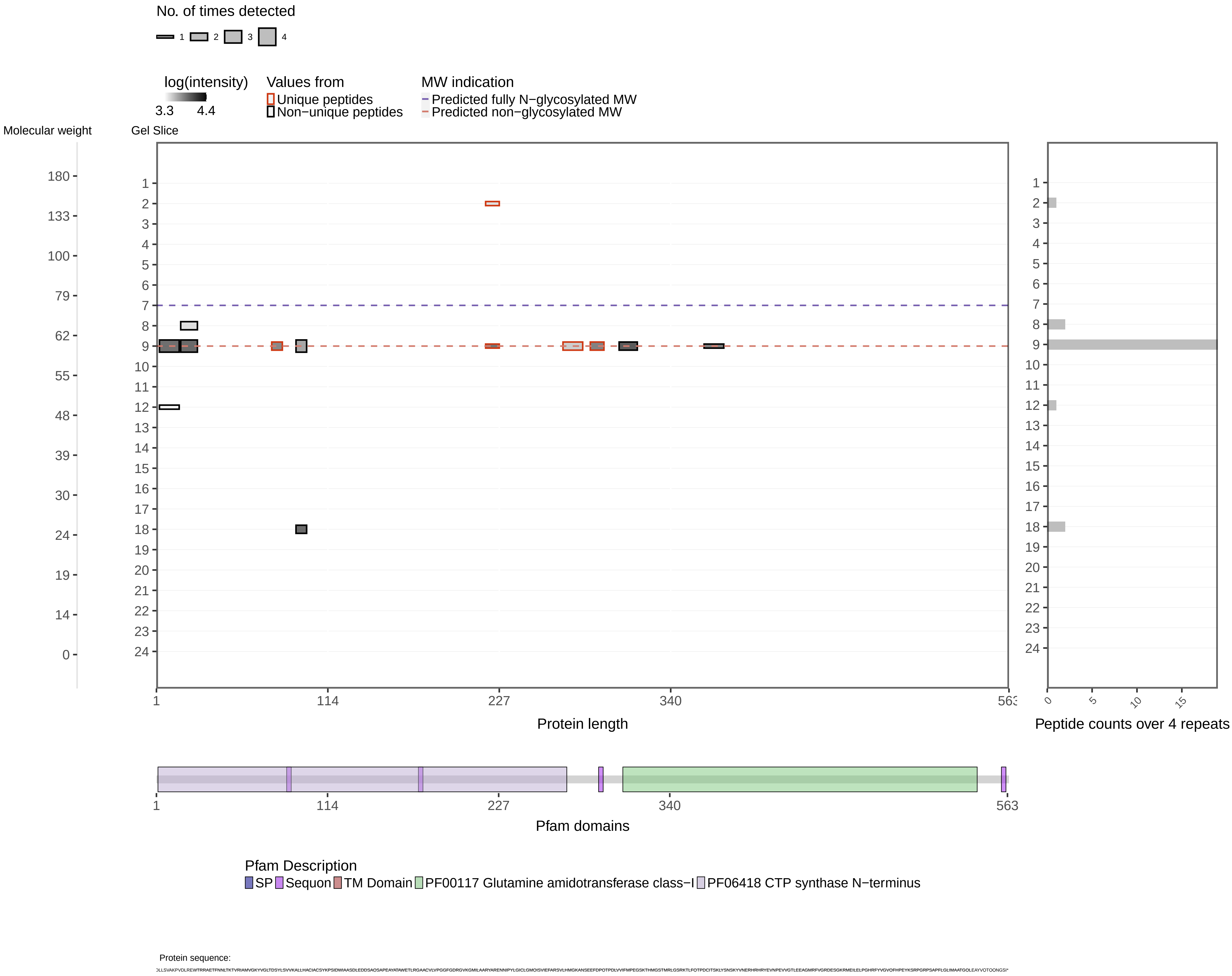

NbL11g20540.1 probable_beta−D−xylosidase_2 [MW = 87.92 kDa]

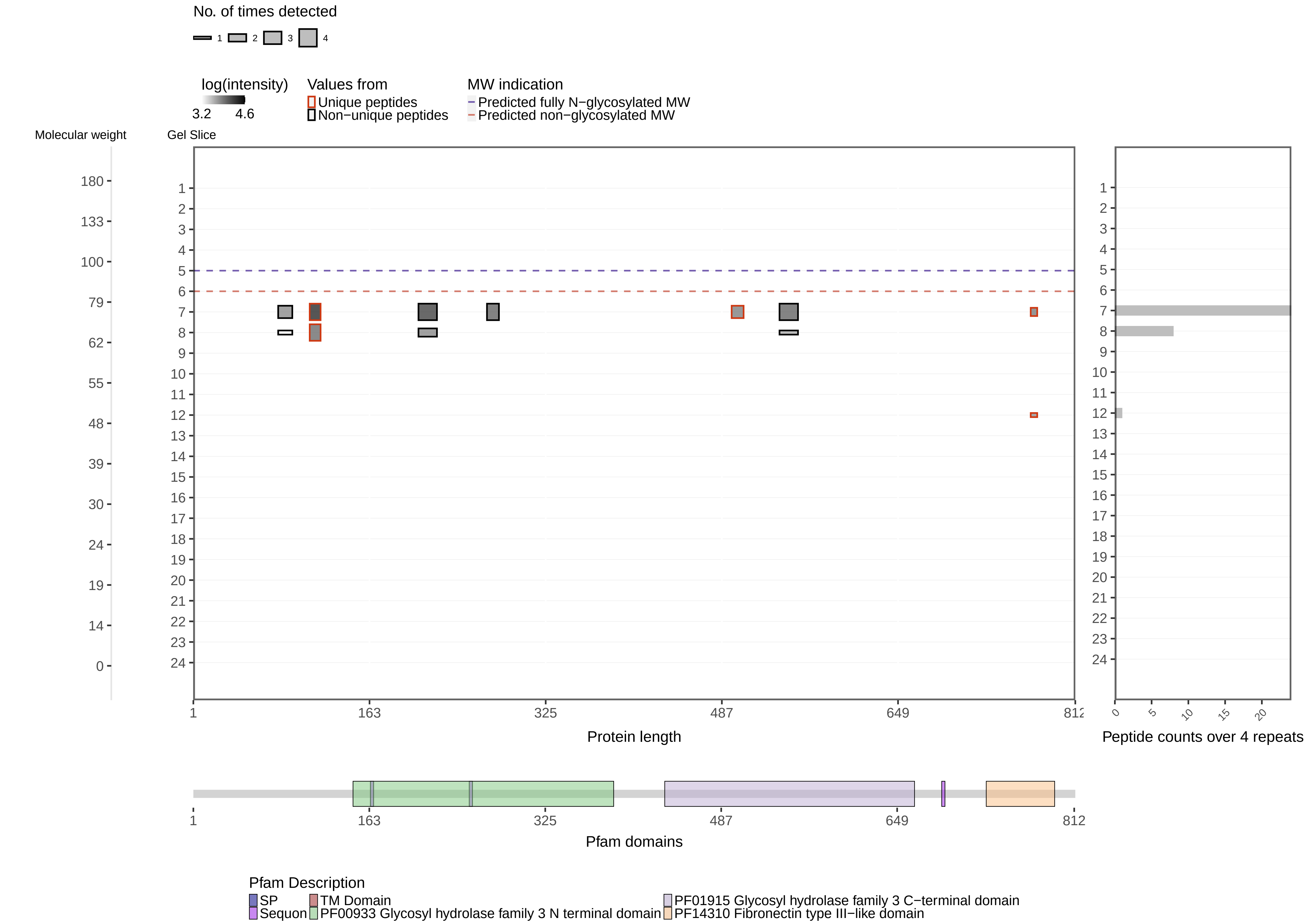


Protein sequence:

NTVAVQMRLGMFDGEPSAQPYGHLGPKDVCSPAHQELALEAARQGIVLLKNHGPALPLSPRSHRSVAVIGPNSDVTLTMIGNYAGVACGYTSPLQGIAKYAKTIHRQGCGDVACSDDRLFPGAVDAARQADATVLVMGLDQSIEAEFRDRTGLLLPGFQQDLISQVSKASRGPVILVLMSGGPVDVTFAEKDPRIGGIVWVGYPGQAGGAAIADVLFGAHNPGGKLPMTWYPQEYLNNLPMTTMDMRANLAKGYPGRTYRFYKGPLVYPFGHGLSYTKFINTIAEAPIMLAI

NbL11g20300.1 probable_pre−mRNA−splicing_factor_ATP−dependent_RNA_helicase_DEAH2 [MW = 82.14 kDa]

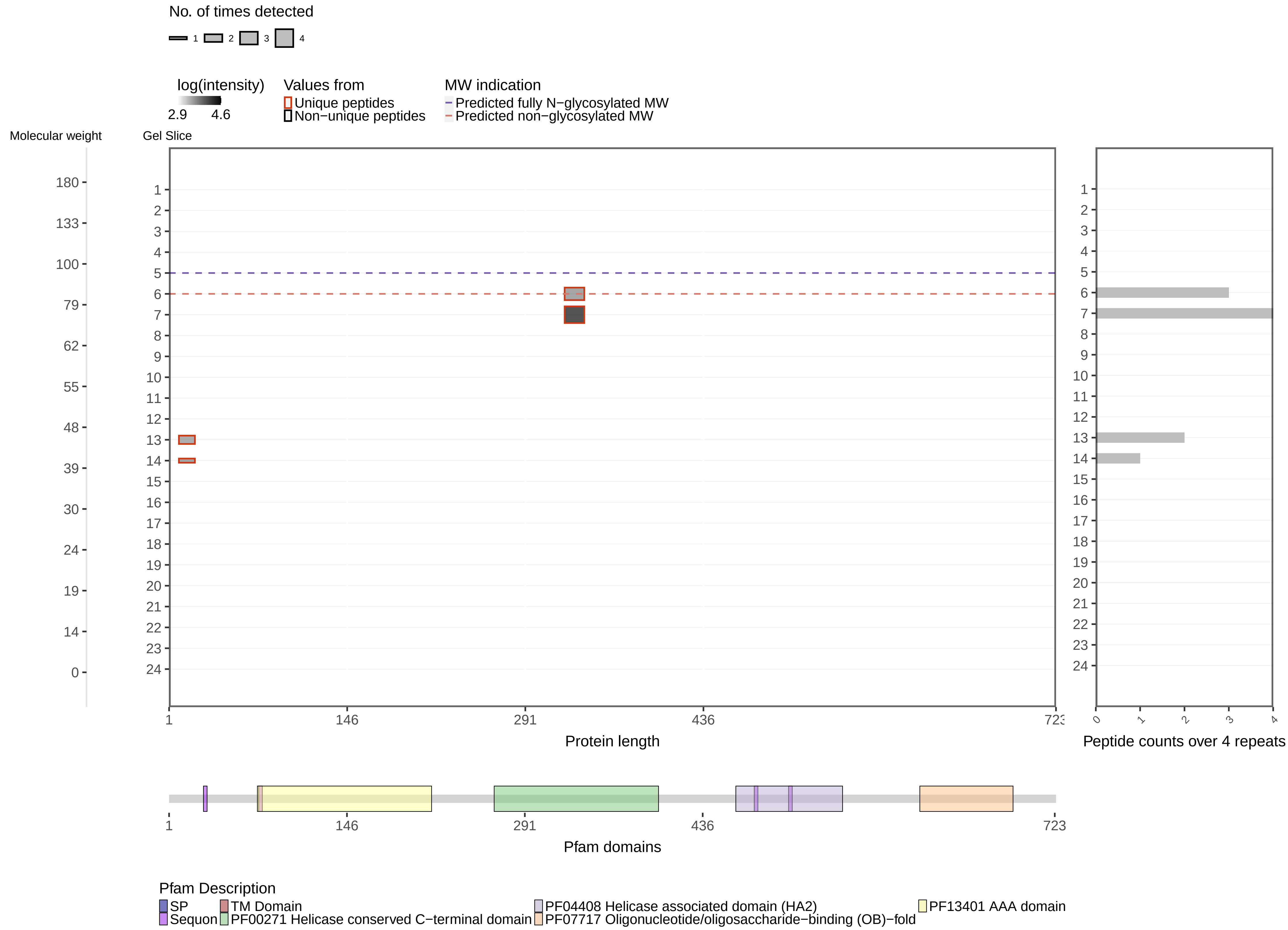


Protein sequence:

TSLTIDGIVYVIDPGFAKQKVYNPRVRVESLLVSPISKASAHQRSGRAGRTQPGKCFRLYTEKSFHNDLQPQTYPEILRSNLANTVLTLKKLGIDDLVHFDFMDPPAPETLMRALEVLNYLGALDDEGNLTRLGEIMSEFPLDPQMAKMLVVSSEFNCSNEILSISAMLSVPNCFVRPREAQKAADEAKNRFGHIDGDHLTMLNVYHAYKQNTEDPQWCYENFINQRALKSADNVRQQLSRIMARFNLKLCSTDFNSRNYYVNIRKAMLAGYFMQVAHLERTGHYLTVKDNQV

### NbL11g20280.1 nifU−like_protein_1_chloroplastic [MW = 23.39 kDa]

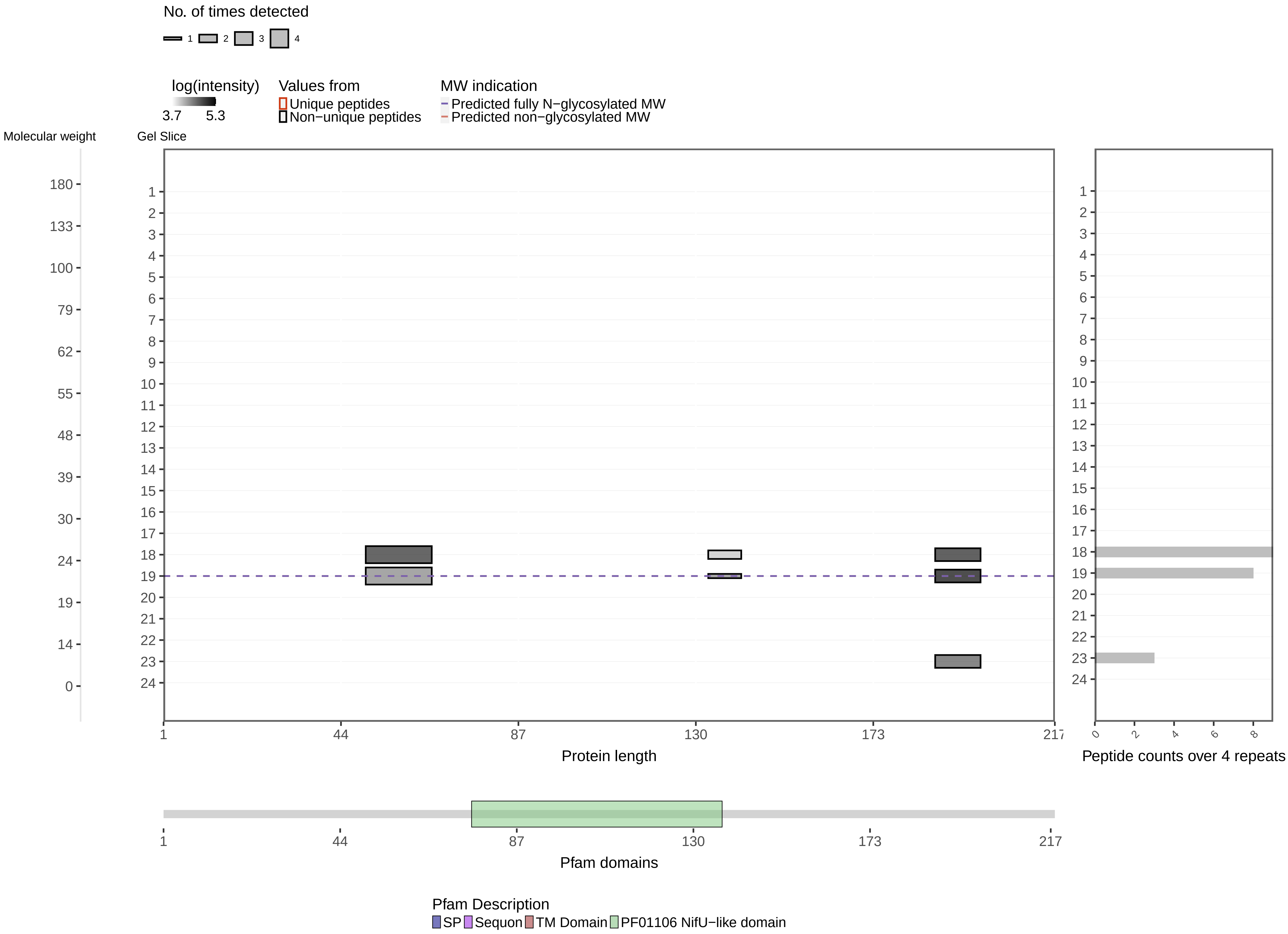


Protein sequence:

VSVEDGVVSLQLQGACESCPSSTTTMKMGIERVLKEKFGEAIKDIRQVYDEGVVETTVEAVNAHLDILRPAIKNFGGSVEVLSVEEGNCNVKYVGPDSIGSGIKAAIKEKFPDIVNVVFTA*

### NbL11g20170.1 uncharacterized_LOC104217660 [MW = 27.78 kDa]

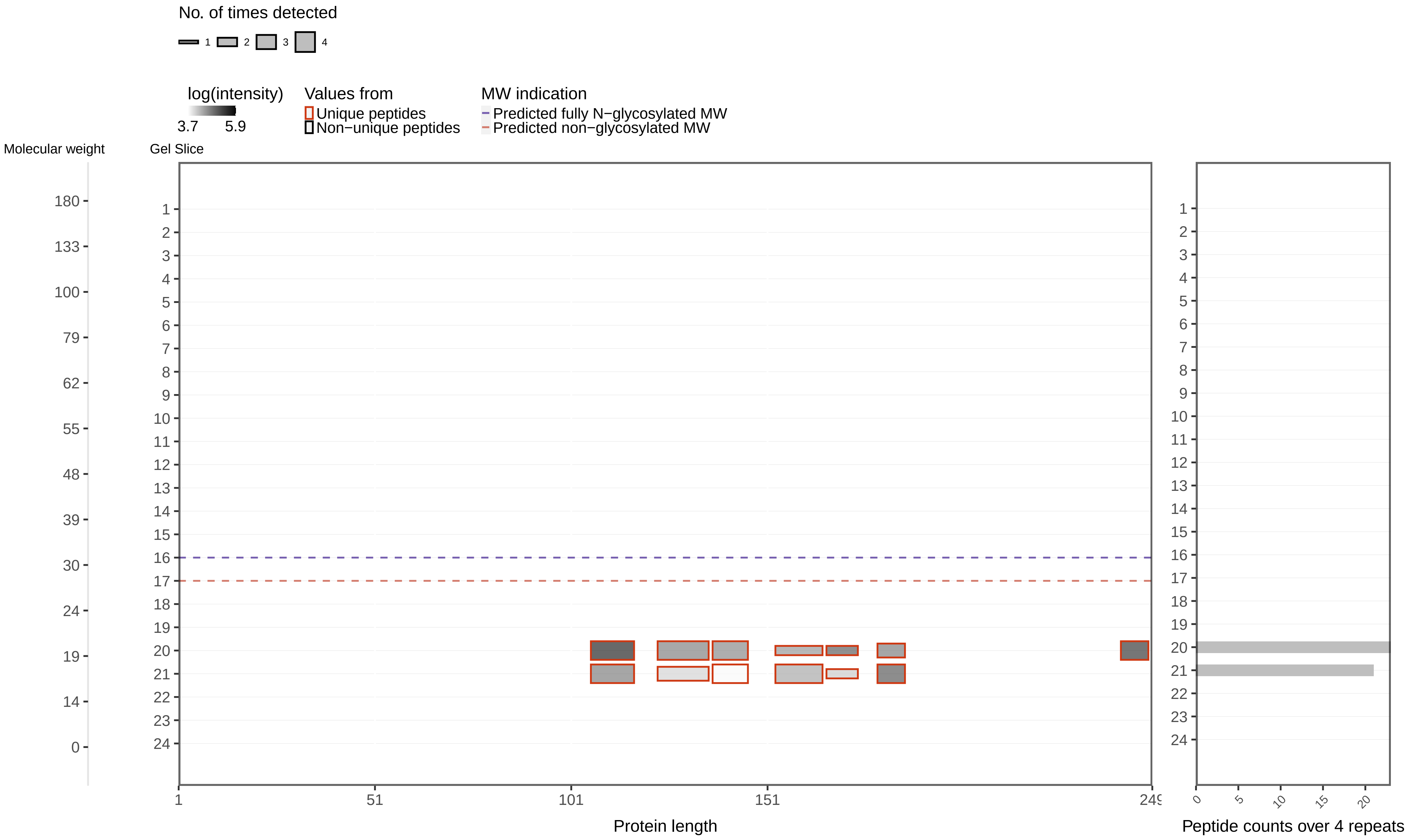


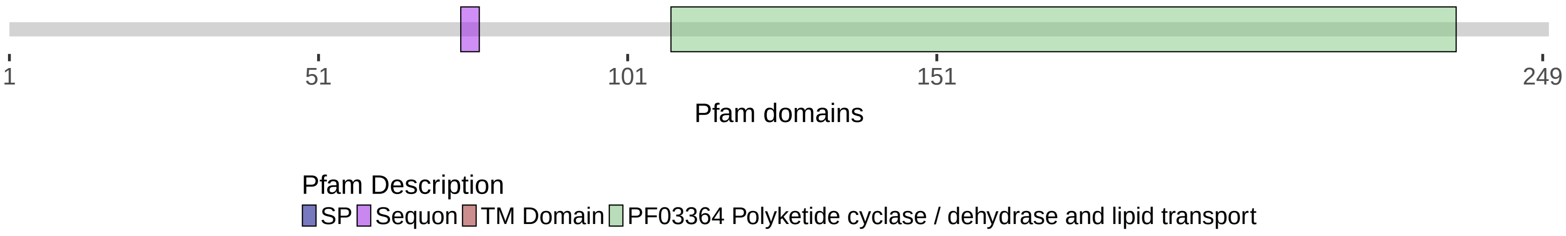


Protein sequence:

SVAYKCYSDREAIPQWMPFISSVKILEDKPDLSRWSLKYKAFGQDIEYSWLARNMQPTPNQKIHWRSLEGLPNRGAVRFFPKGPSSCTIELTVSYEVPQLLVPVASALQPFLENLLGRGLERFATFAKSYSADTPK*

NbL11g20090.1 uncharacterized_LOC109224790 [MW − SP = 50.26 kDa]

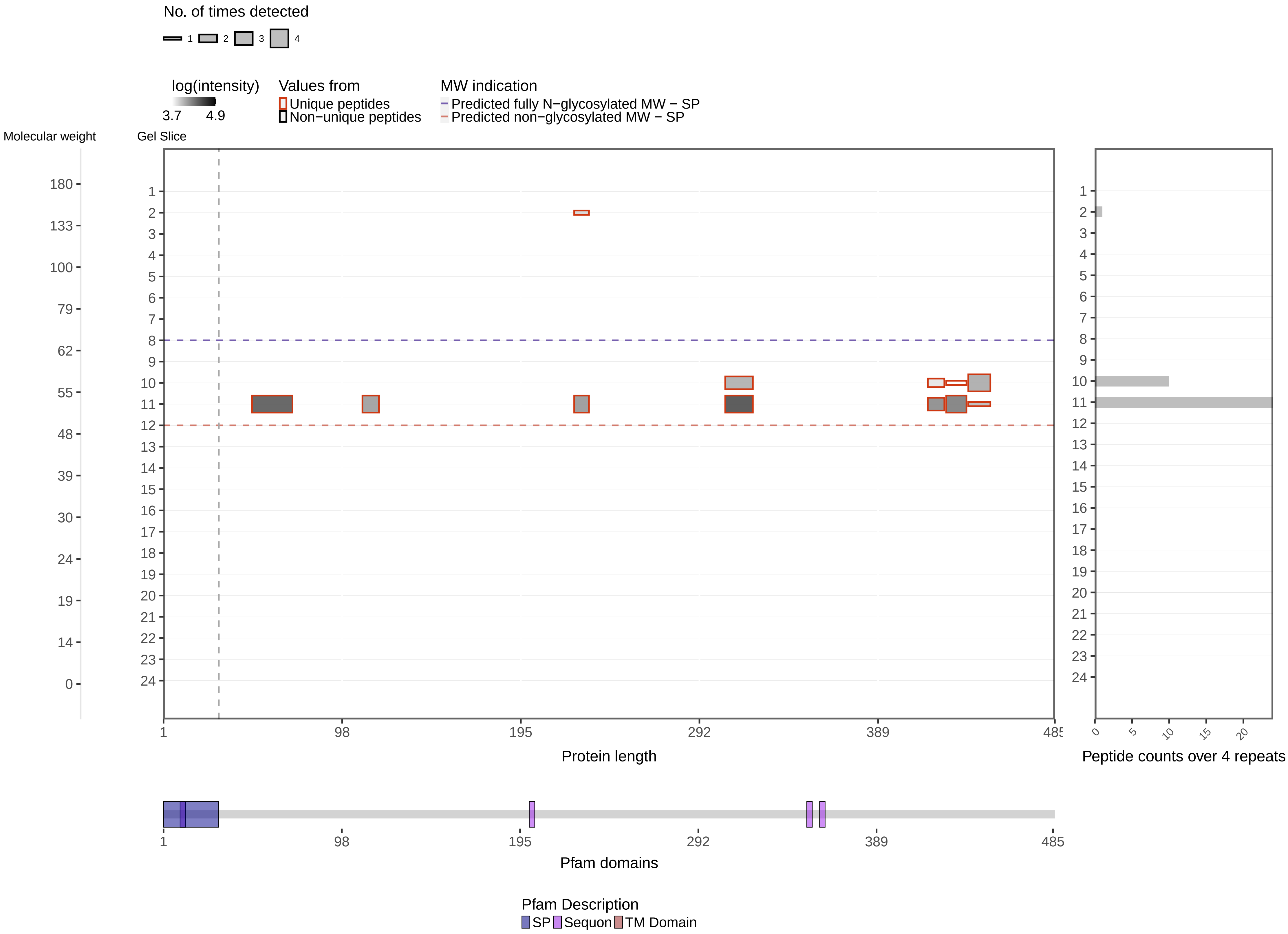


Protein sequence:

GWVFKHKLRTSKLYPWLMLYFRADATKGFSGGYHYDTRGMLKTLPESPNFKVKLTLDVKQGGGPKSQFYLIDIGSCWKNNGAPCDGDVLTDITRYSEMIINPETPAWCSPTNIGNCPPFHITPNNTKIYRNDTSHFPYSAYHYYCAPGNAEHLEKPYSTCDPYSNPQAQELVQLLPHPIWADYGYPSKQGDGWVGDGRTWELDVGGLSSRLYFYQDPGTPPARRIWTSLDVGTEIFVSNKDEVAEWTLSDFDVLLTS*

### NbL11g20060.1 L−cysteine_desulfhydrase [MW = 50.29 kDa]

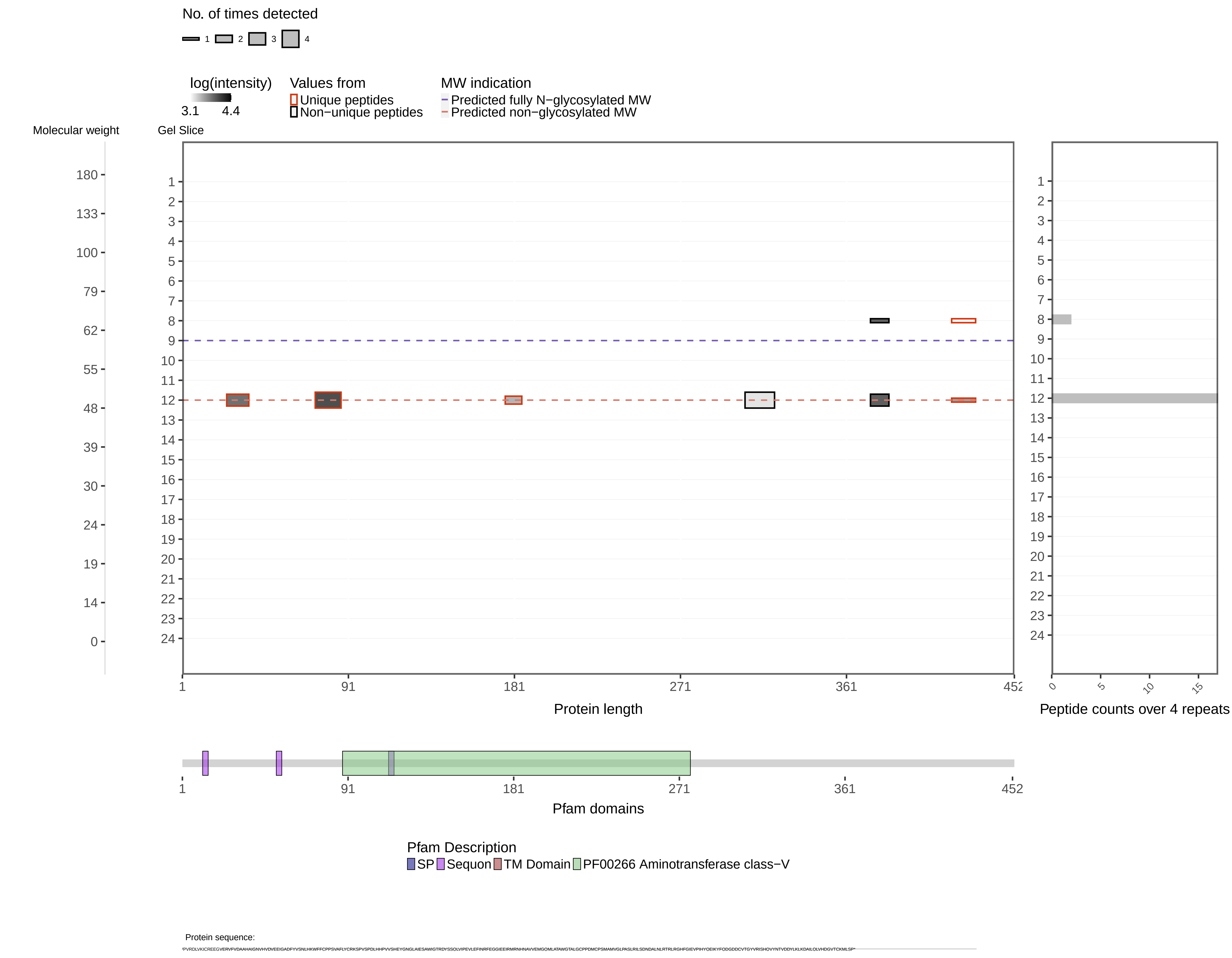

### NbL11g20050.1 protein_transport_protein_SEC23−like [MW = 95.02 kDa]

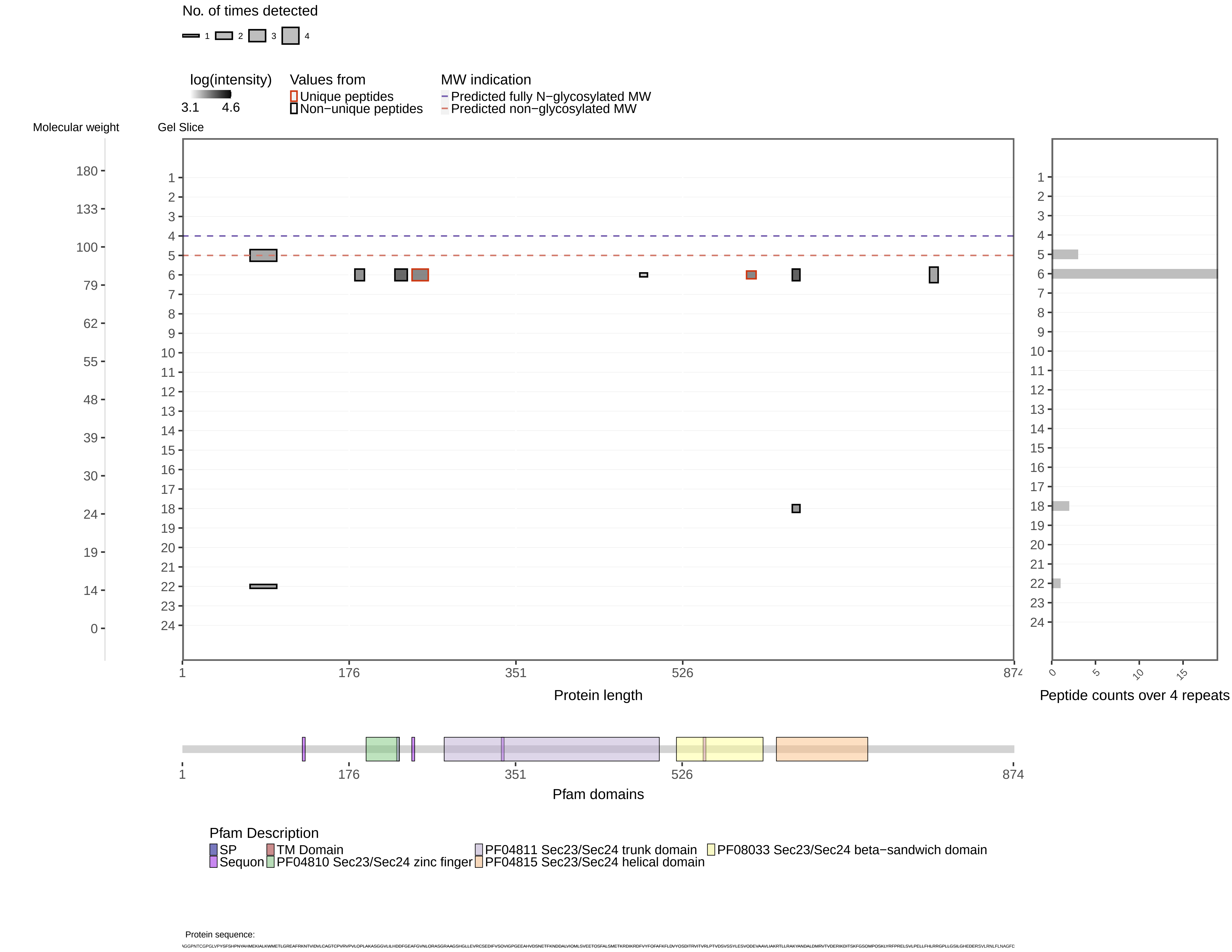

NbL11g19920.1 ADPATP_carrier_protein_1_mitochondrial−like [MW = 42.33 kDa]

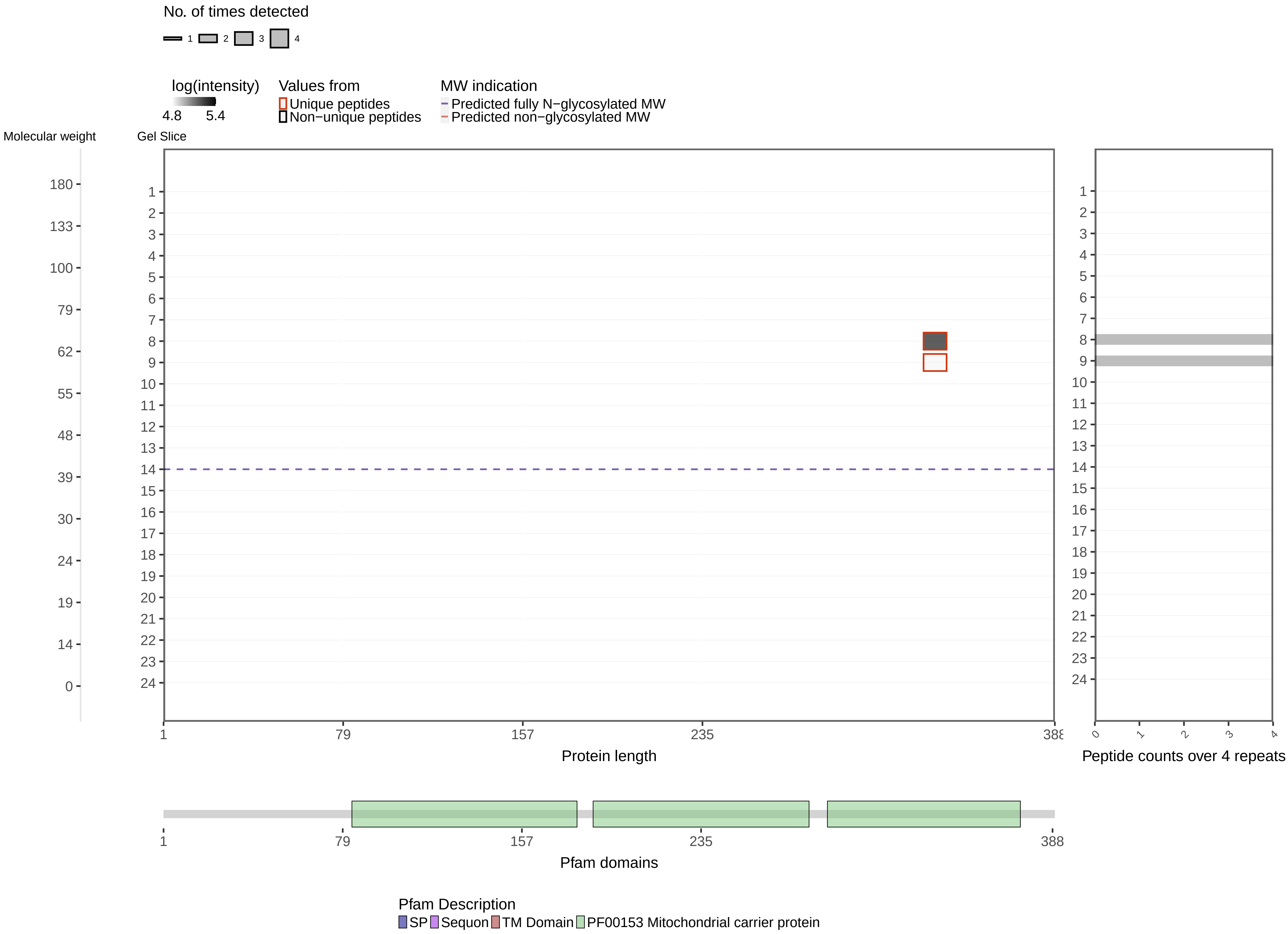


Protein sequence:

ERDGYWKWFAGNLASGGAAGASSLLFVYSLDYARTRLANDAKAAKKGGERQFNGLPDVYRKTIKSDGIVGLYRGFTISCVGIIVYRGLYFGIYDSLKPVVLVGNLQDSFVASFLLGWGITIGAGLASYPIDTVRRRMMMTSGEAVKYKGSLDAFSQIMRKEGAKSLFKGAGANILRAVAGAGVLAGYDKLQLVVLGKKYGSGGGS*

NbL11g19860.1 uncharacterized_protein_At4g28440−like [MW = 16.17 kDa]

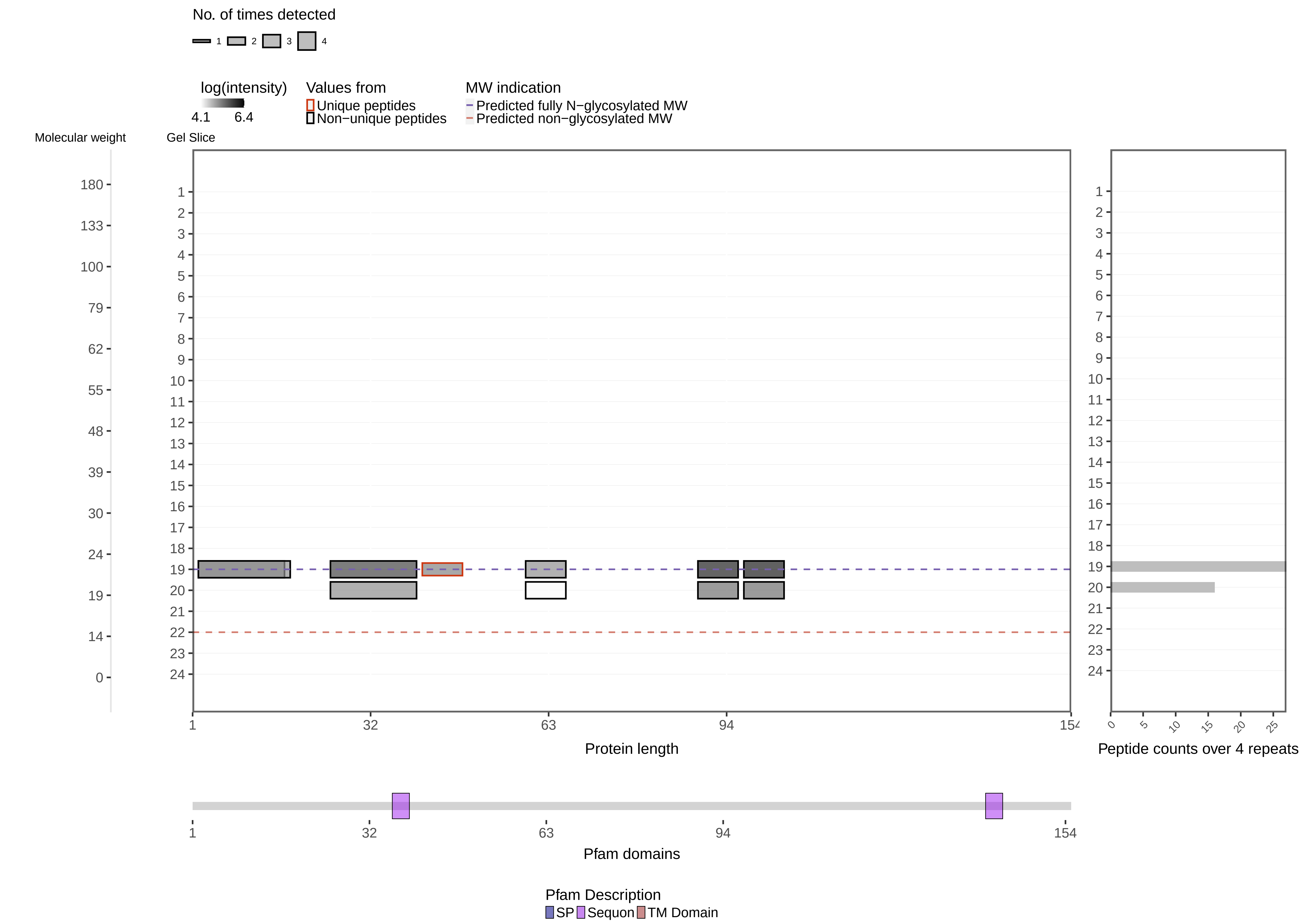


Protein sequence:

NPRAPPRISECLVGDDTGSILFTARNEQVDLLKAGATVILRNAKIDMFKGSMRLAVDKWGRVEVTEPADFAVNEQNNLSLVEYELVNVEE*

NbL11g19850.1 26S_proteasome_non−ATPase_regulatory_subunit_2_homolog_A−like [MW = 98.35 kDa]

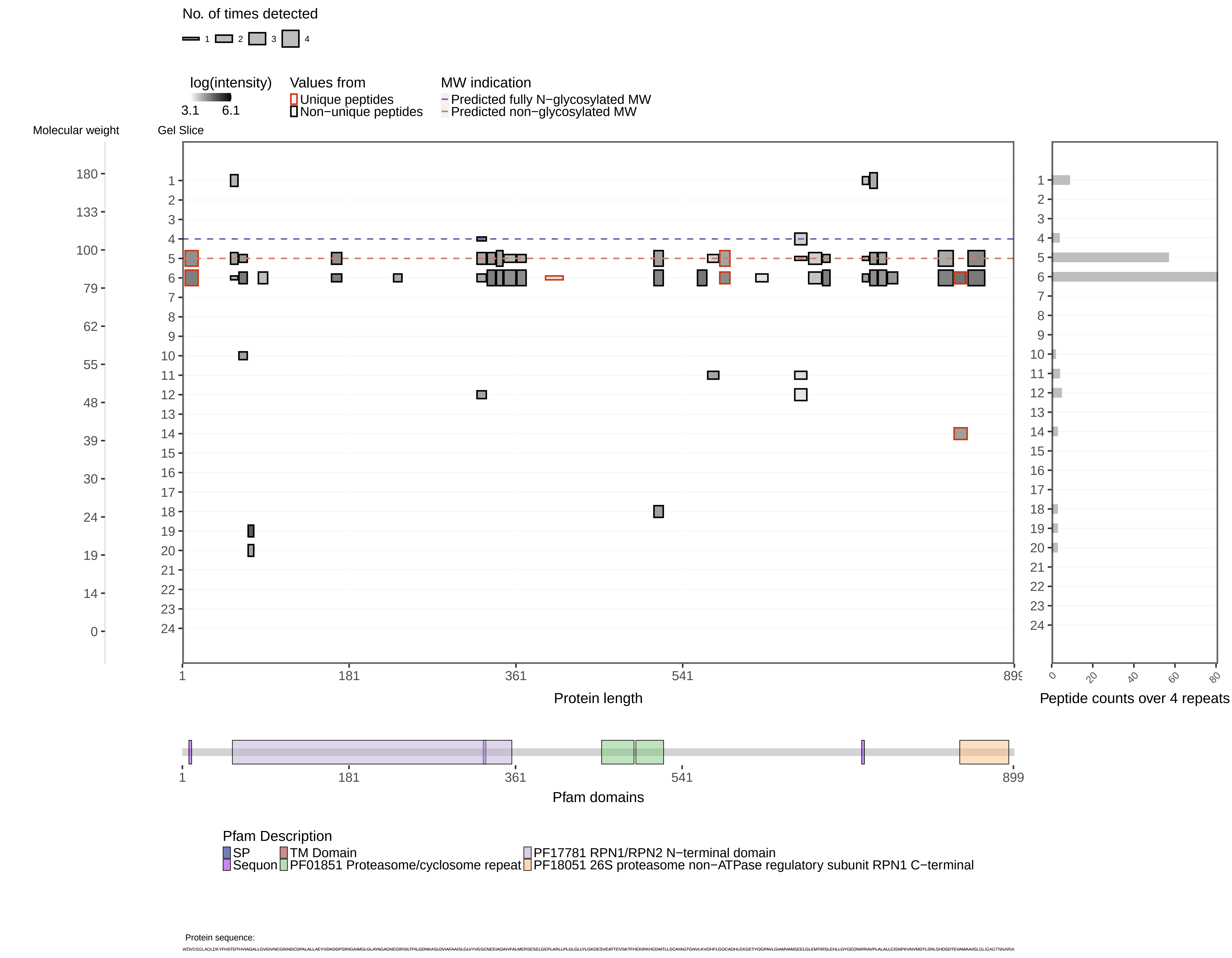

### NbL11g19780.1 acyl−coenzyme_A_oxidase_3_peroxisomal−like [MW = 82.77 kDa]

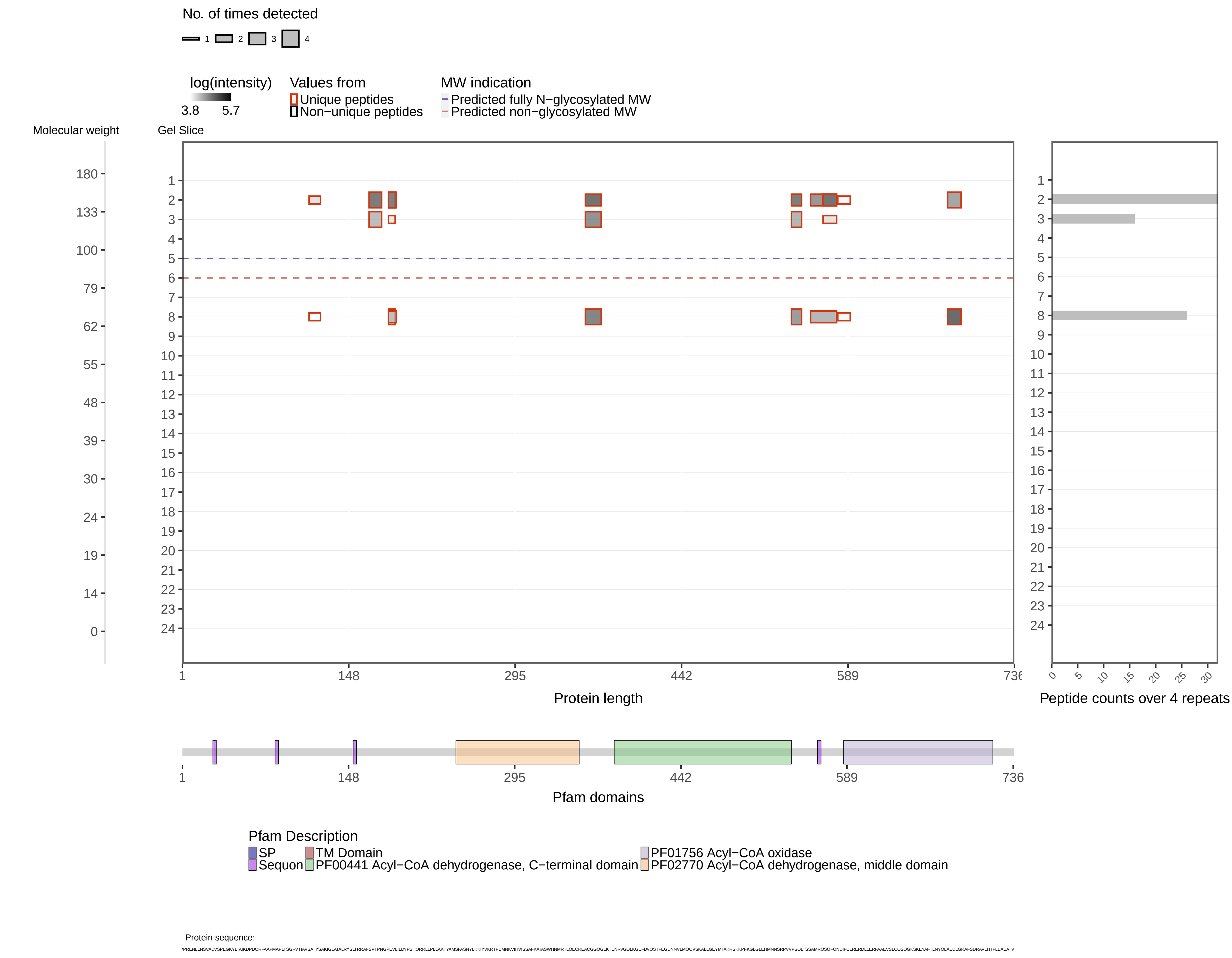

### NbL11g19670.1 5−methyltetrahydropteroyltriglutamate−−homocysteine_methyltransferase−like [MW = 84.57 kDa]

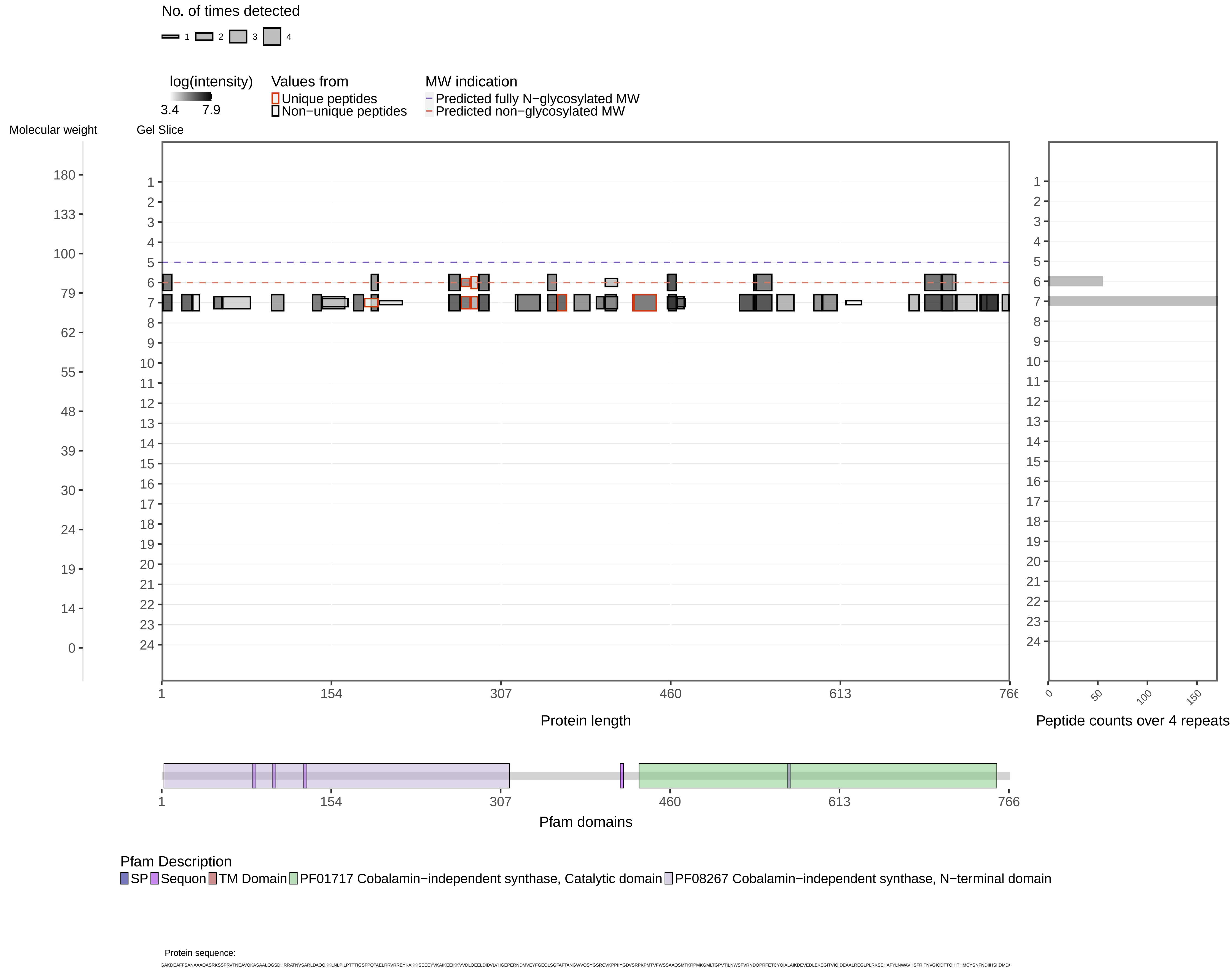

### NbL11g19630.1 glyceraldehyde−3−phosphate_dehydrogenase_GAPCP2_chloroplastic−like [MW = 53.34 kDa]

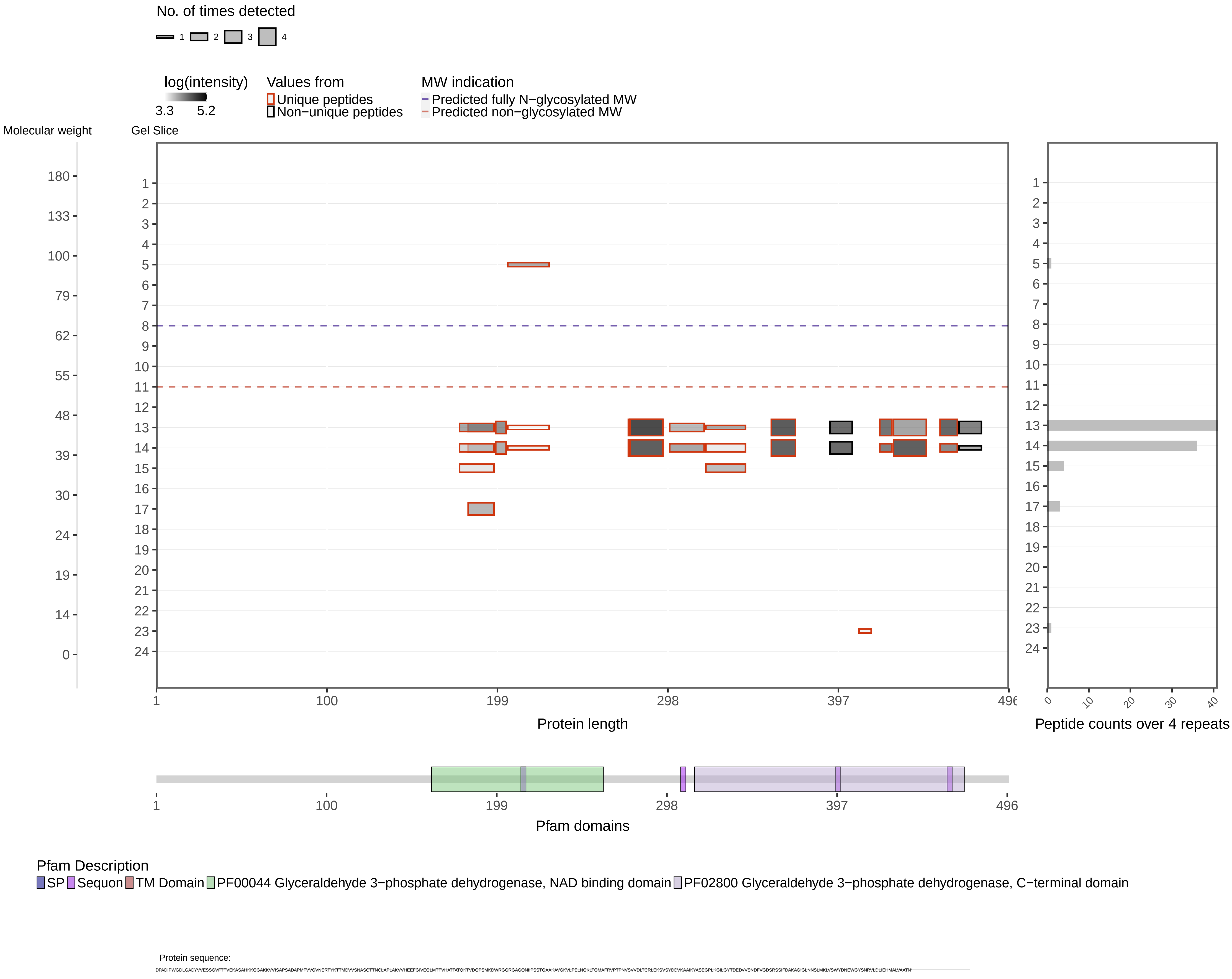

NbL11g19610.1 nuclear_cap−binding_protein_subunit_1 [MW = 98.05 kDa]

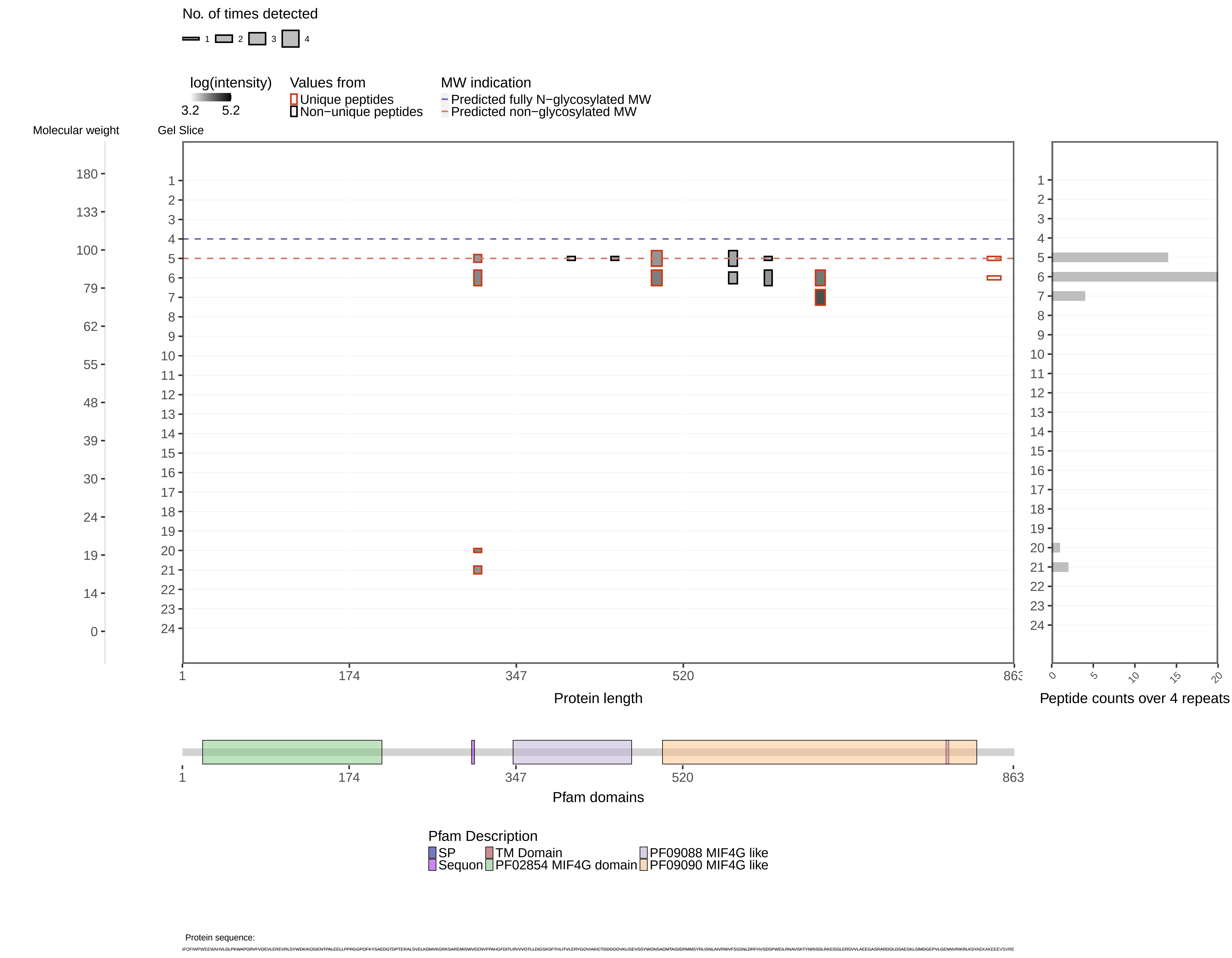

### NbL11g19510.1 renalase [MW = 40.5 kDa]

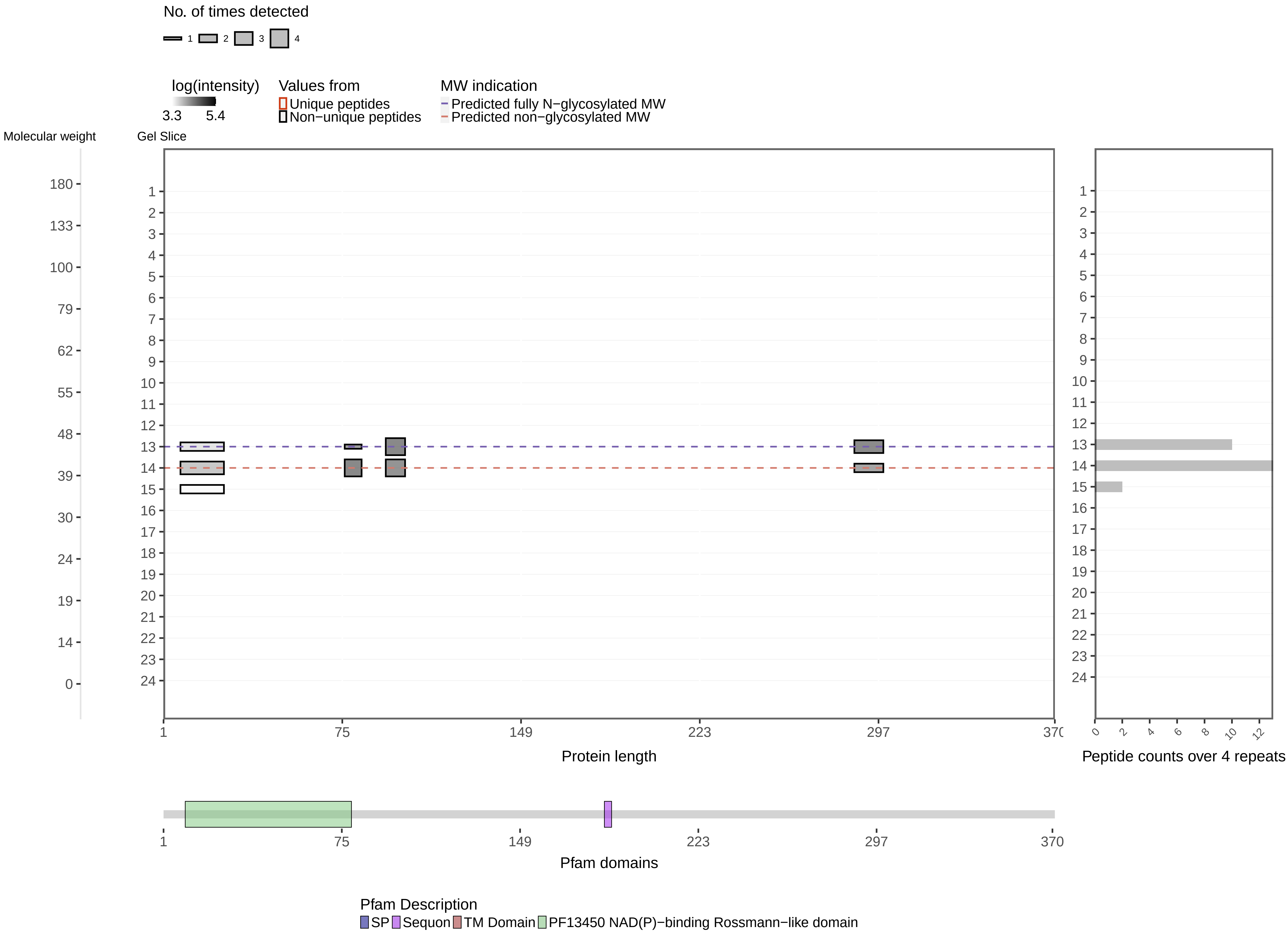


Protein sequence:

VVTSDKSTFSPKFTNVTGKPMPIDMGKFPEIPLKMTEIPVSPRFALMLAFEELLSEIPIRGFSFKNSEVLRWAHCDSSKPRRSHNSECWVLHSTPEYARDVIAKTGLQKPSSATLTKVAEELFQEFQSTKLNIPQAFFNKAHRWGSAFPATSIAGDEKCLWDAKKRIAICGDFCVSPNVEGAIVSGLAAAAKCSEVFSRL*

### NbL11g19400.1 probable_calcium−binding_protein_CML21 [MW = 25.89 kDa]

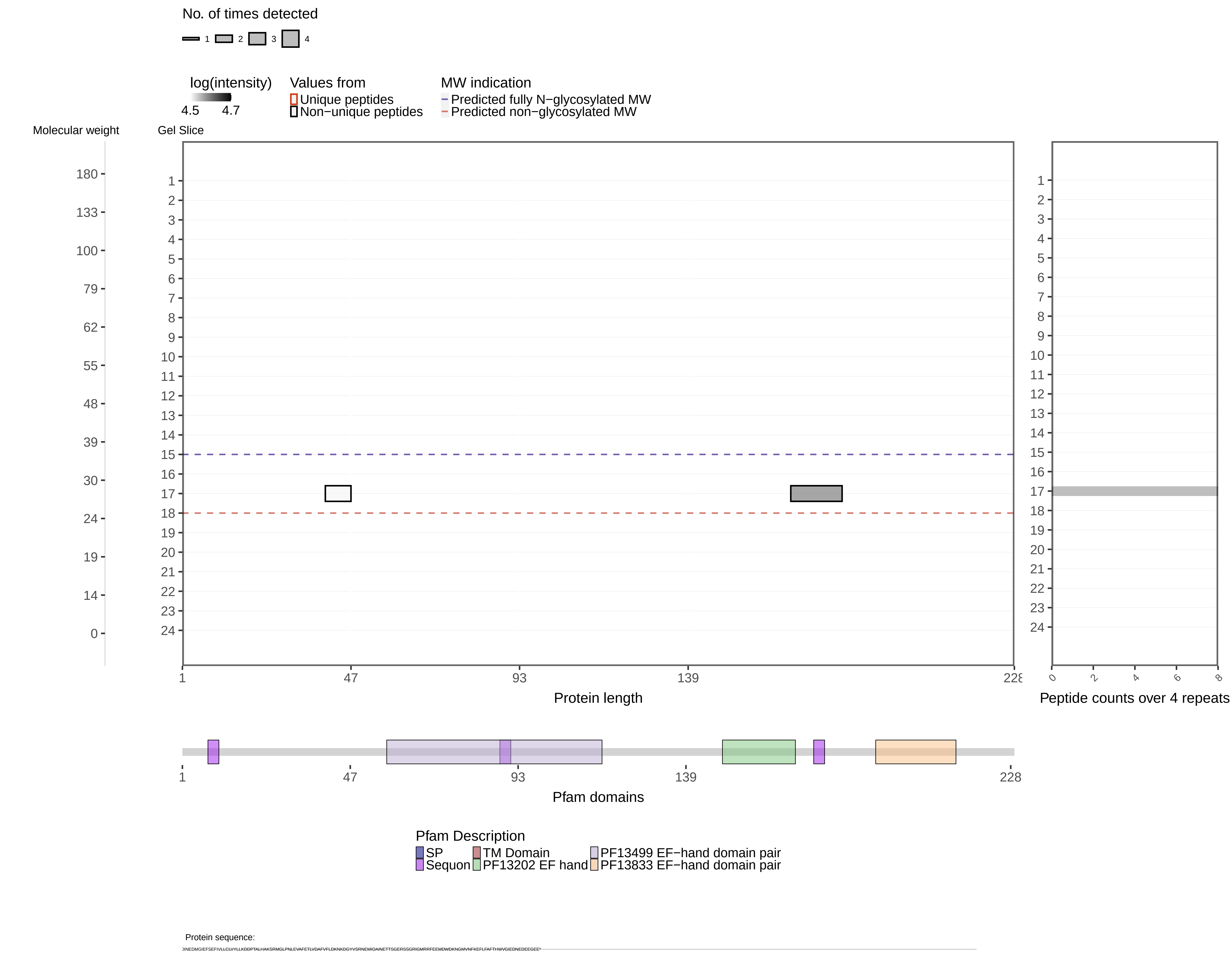

NbL11g19350.1 peptidyl−prolyl_cis−trans_isomerase_FKBP16−1_chloroplastic [MW = 27.94 kDa]

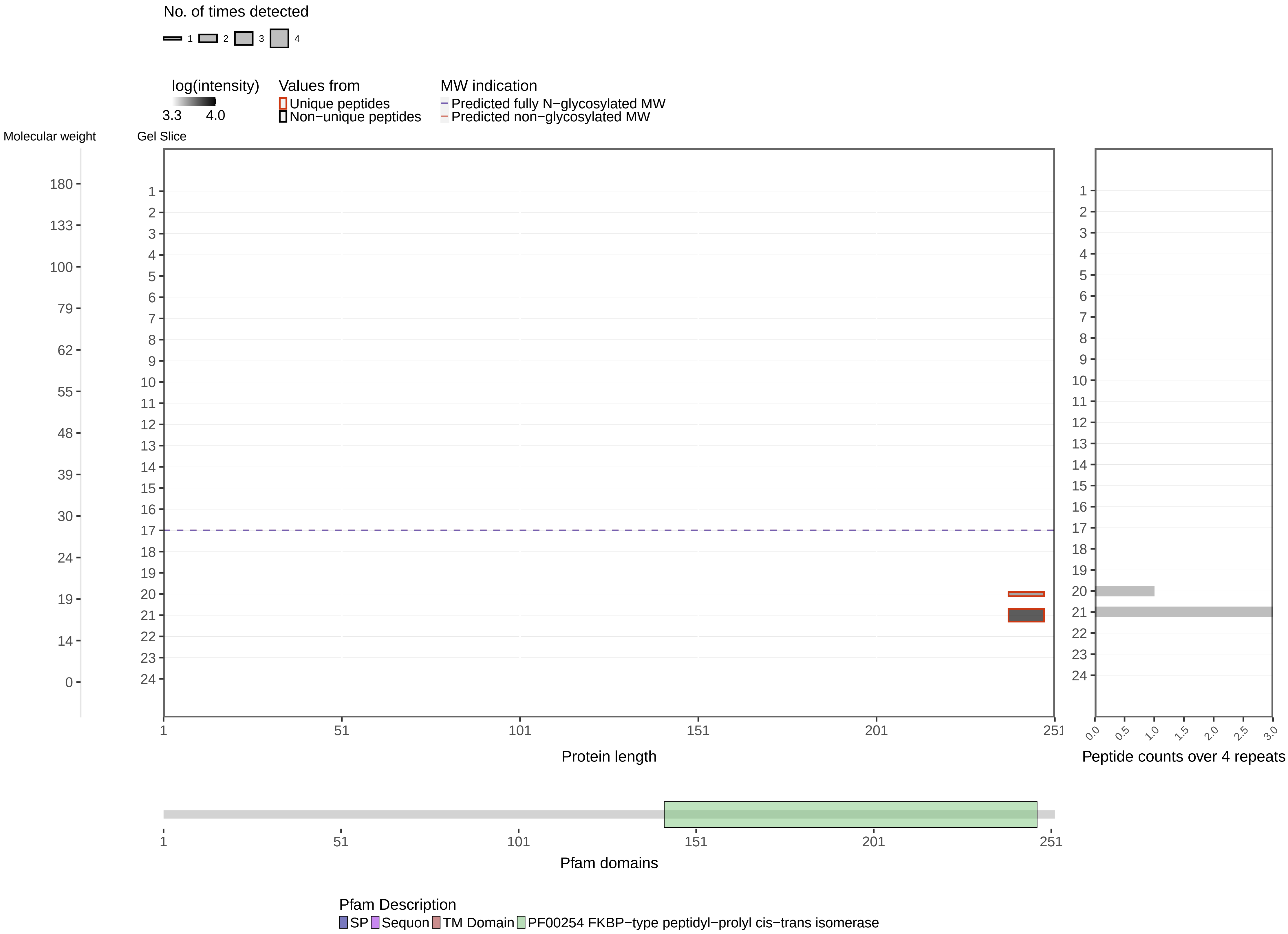


Protein sequence:

KEPDVIRFEAFRTLKLASGVRIQDVVEGEGPEAHEGDIVEINYVCRRSNGYFVHSTVDQFSGESSPVILPLDDTQIIEGLKEVLIGMKPGGKRRALIPPSLGYINENLKPVPEEFGPRRSLMSHVKEPLIFEVQLLKVL*

### NbL11g19340.1 heat_shock_cognate_protein_80−like [MW = 80.17 kDa]

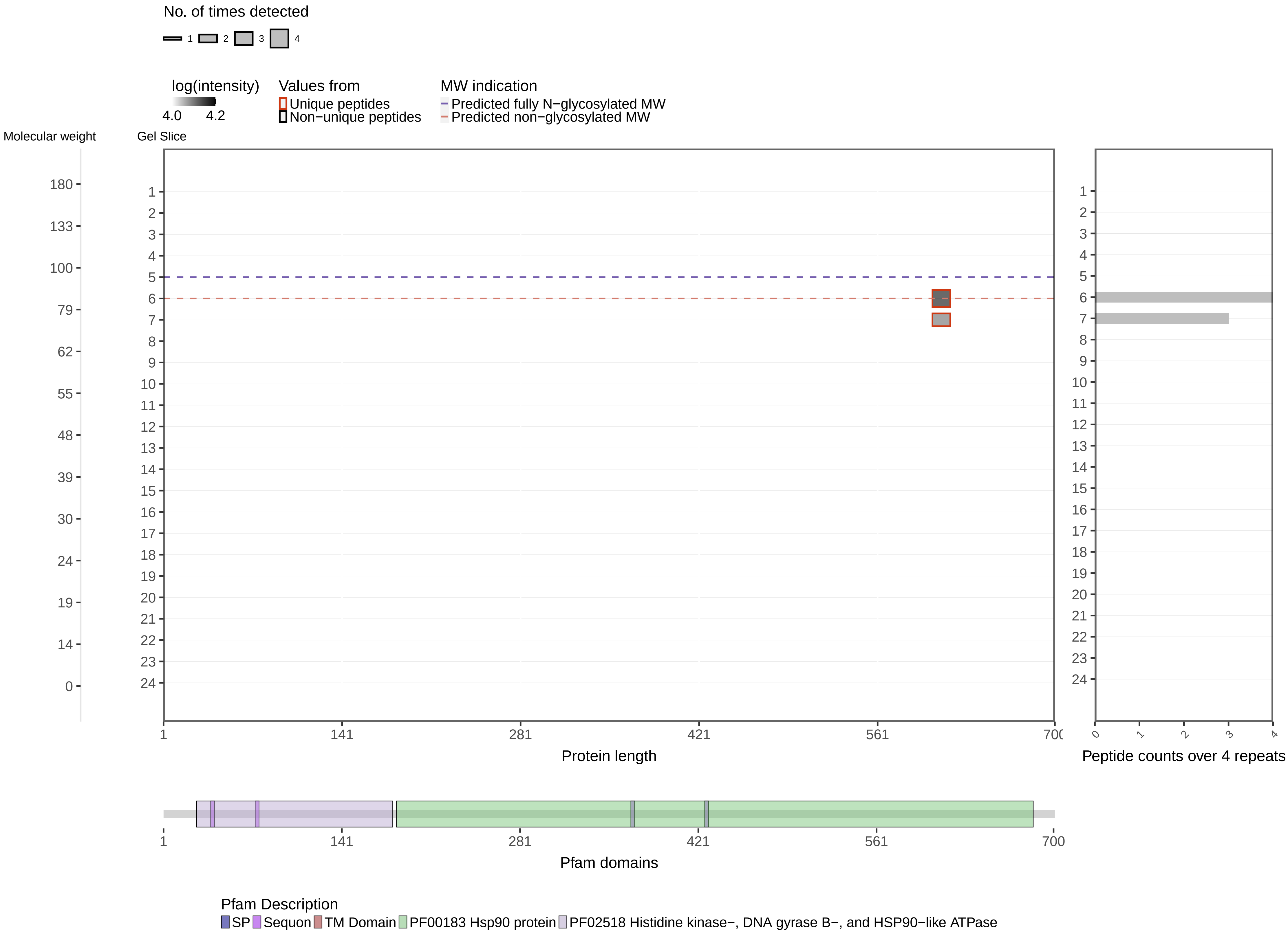


Protein sequence:

FIMDNCEELIPEYLSFVKGIVDSEDLPLNISREMLQQNKILKVIRKNLVKKCIELFFEIAENKEDFDKFYEAFAKNIKLGIHEDSQNRSKFAELLRYHSTKSGDDMTSLKDYVTRMKEGQNDIYYITGESKKAVENSPFLEKLKKKGYEVLYMVDAIDEYCIGQLKEFEGKKLVSATKEGLKLDESEDEKKKKEELKEKFEGLCKVIKDVLGDKVEKVVVSDRVVDSPCCLVTGEYGWTANMERIMKAQALRDSSMAGYMSSKKTMEINPENAIMDELRKRADADKNDKSVKDLVL

NbL11g19310.1 farnesyl_pyrophosphate_synthase_1−like [MW = 48.87 kDa]

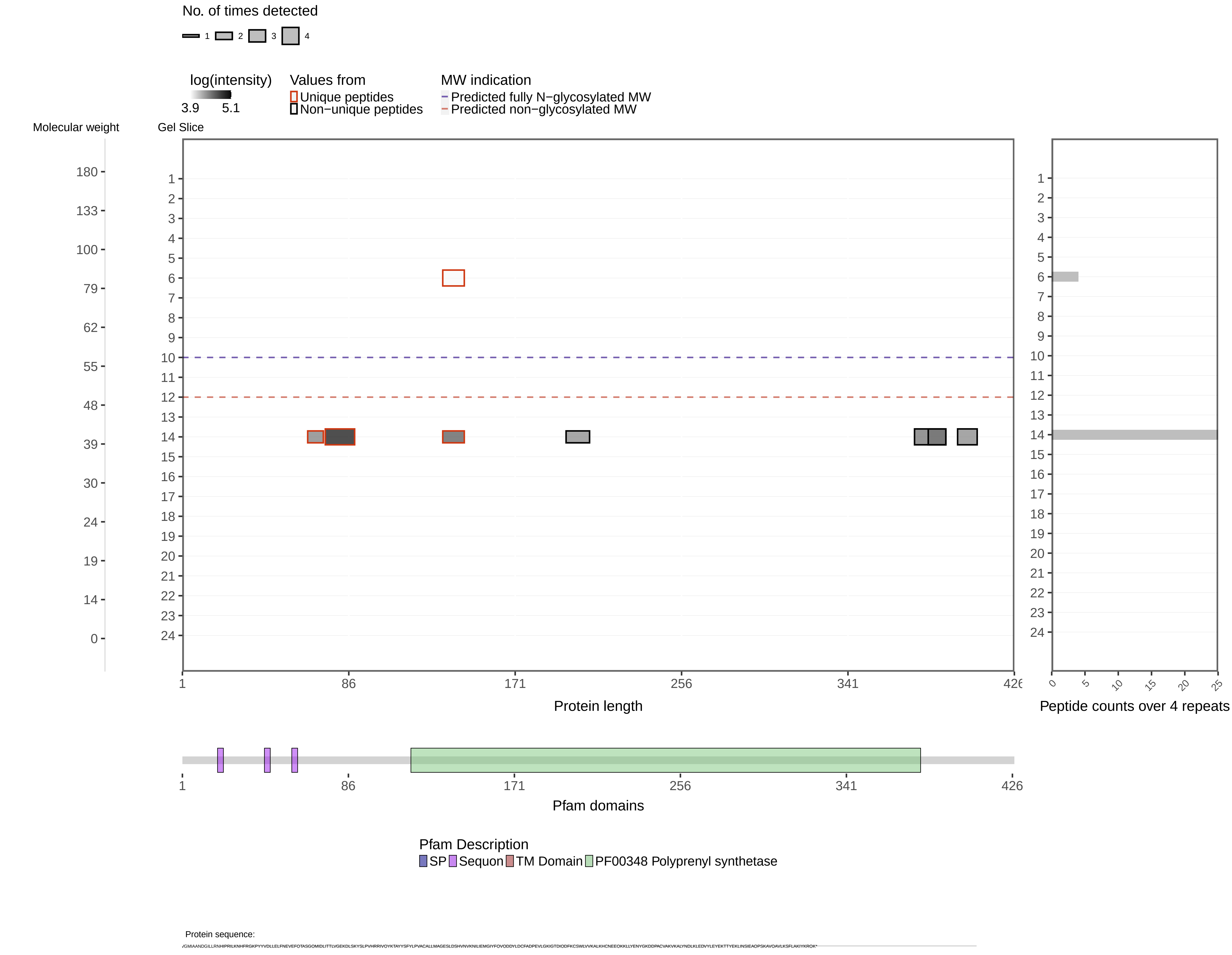

### NbL11g19250.1 fasciclin−like_arabinogalactan_protein_1 [MW − SP = 41.12 kDa]

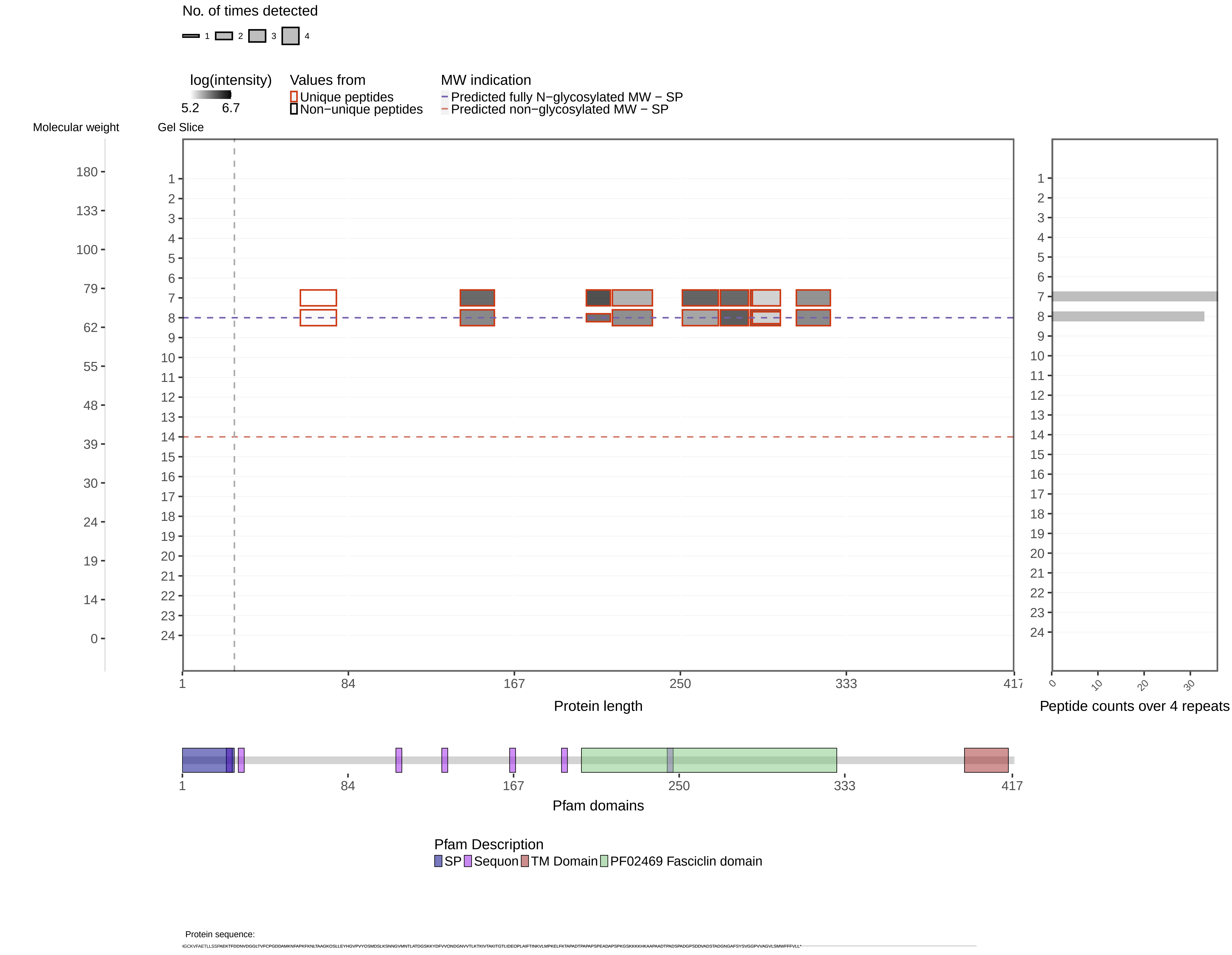

### NbL11g19180.1 30S_ribosomal_protein_S10_chloroplastic−like [MW = 20.71 kDa]

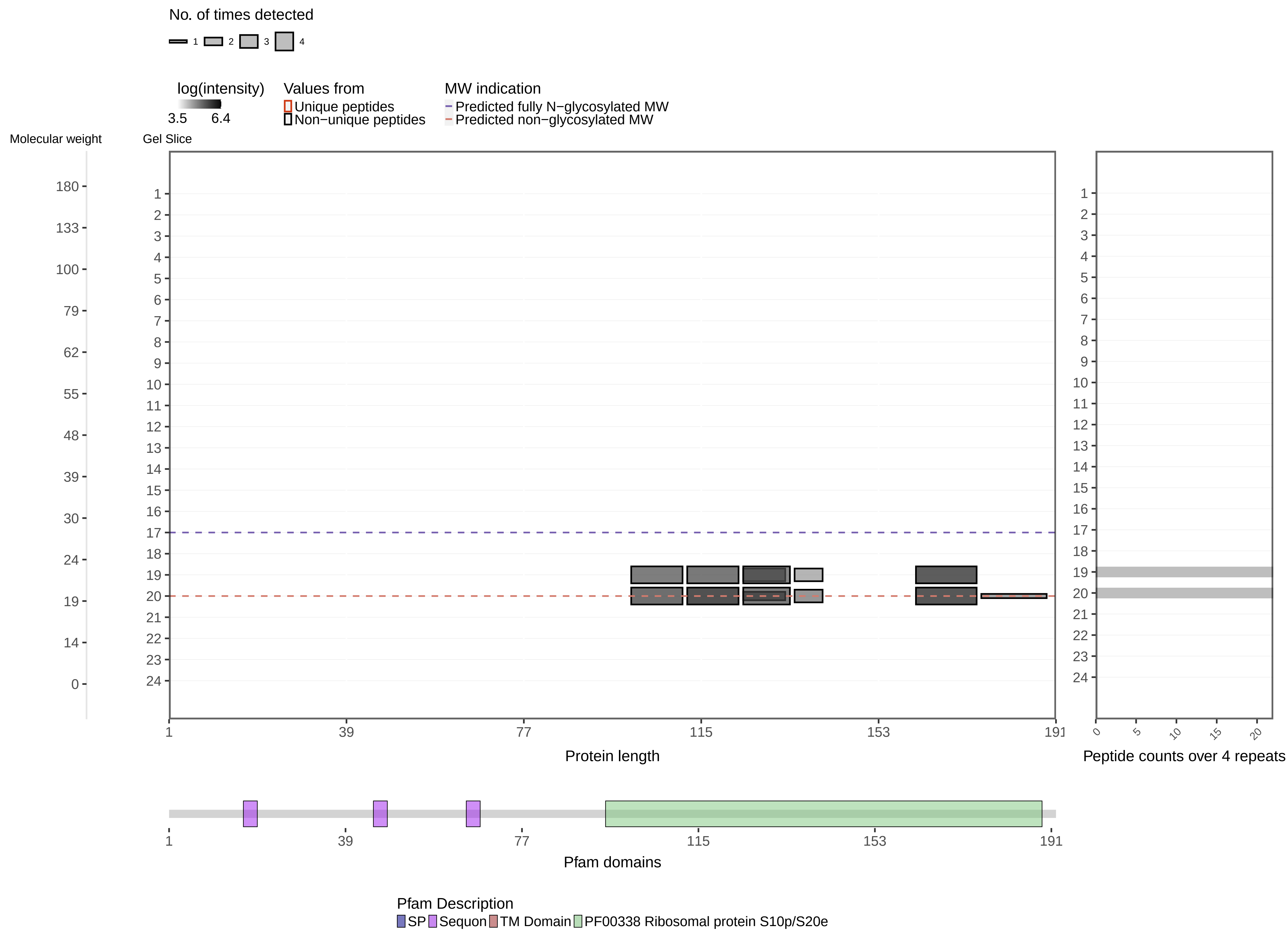


Protein sequence:

GVDADKAAPKQKIRIKLRSYWVPLIEDSCKQIMDAATTTNAKTVGPVPLPTKKRIYCVLKSPHVHKDARFHFEIRTHQRLIDILYPTAQTIDKLMLLDLPAGVDVEVKL*

NbL11g19150.1 40S_ribosomal_protein_S8−like [MW = 25 kDa]

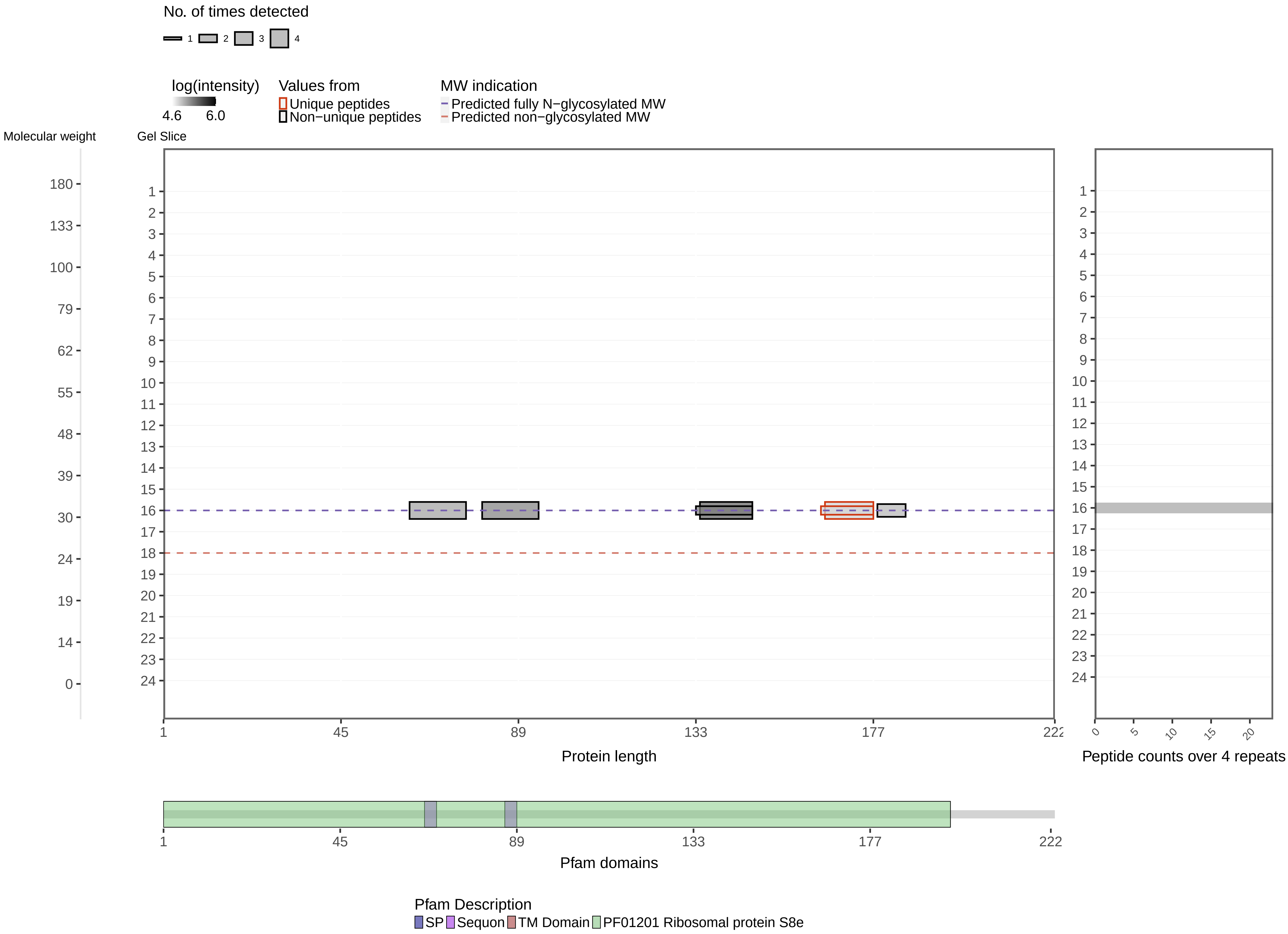


Protein sequence:

LVKSAIVQVDAAPFKQWYLQHYGVDIGRKKKSAVKKEGEEAETAPAAEEKKSNHVQRKLQKRQQDRKIDPHVEEQFASGRLLAAISSRPGQCGRADGYILEGKELEFYMKKLQKKKGKGASGAA*

NbL11g19000.1 proline_synthase_co−transcribed_bacterial_homolog_protein−like [MW = 26.8 kDa]

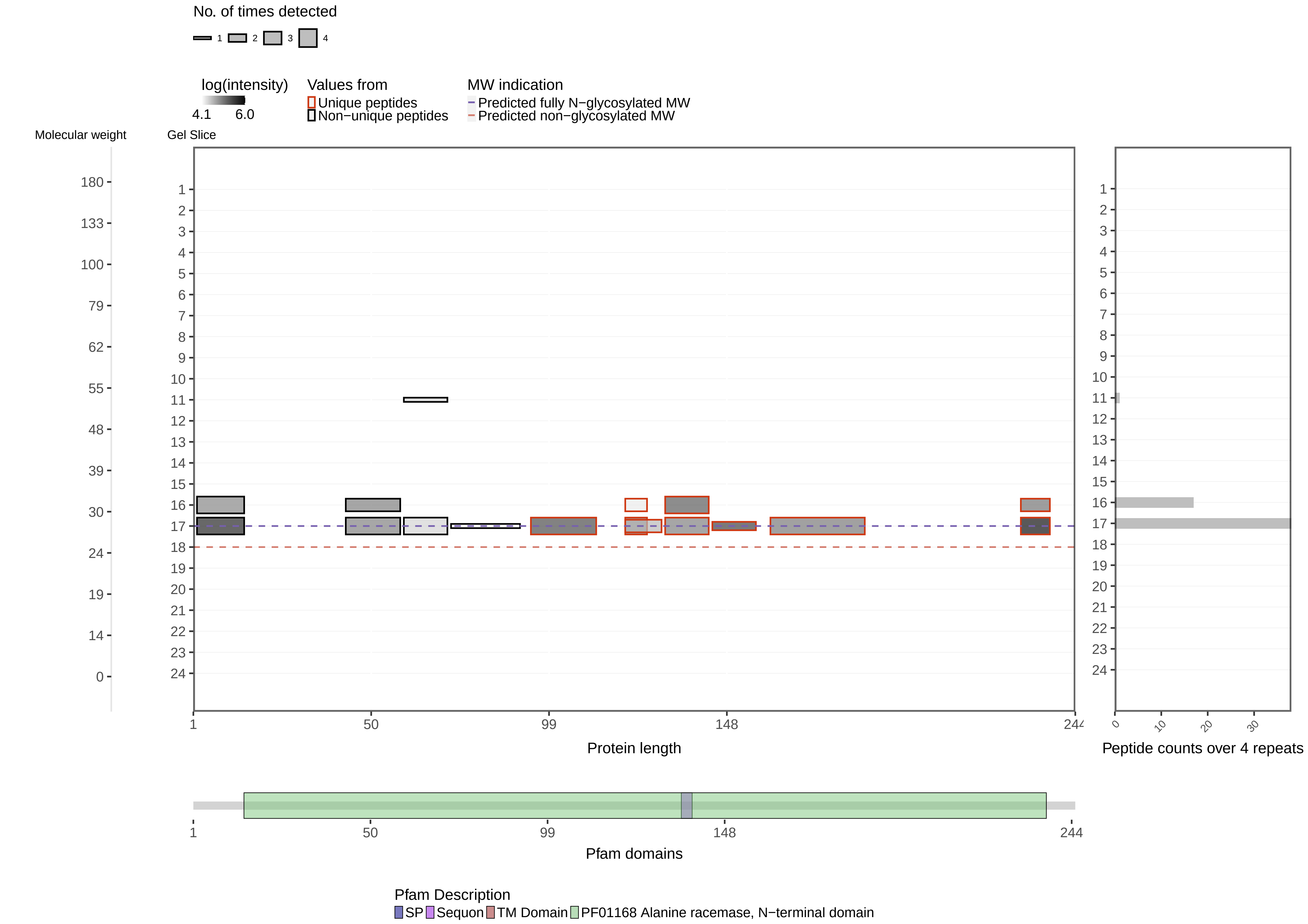


Protein sequence:

NQKLANHLDRMVSSIDRQPLKVLVQVNTSGEESKSGIDPSNCVELAKHVKLDCPNLEFSGLMTIGRPDYTSTPENFKTLLNCRTEVCKVLGMGESQCELSMGMSSDFELAIEMGSTTVRIGSTIFGPREYPKRQ*

NbL11g18960.1 reactive_Intermediate_Deaminase_A_chloroplastic−like [MW = 20.65 kDa]

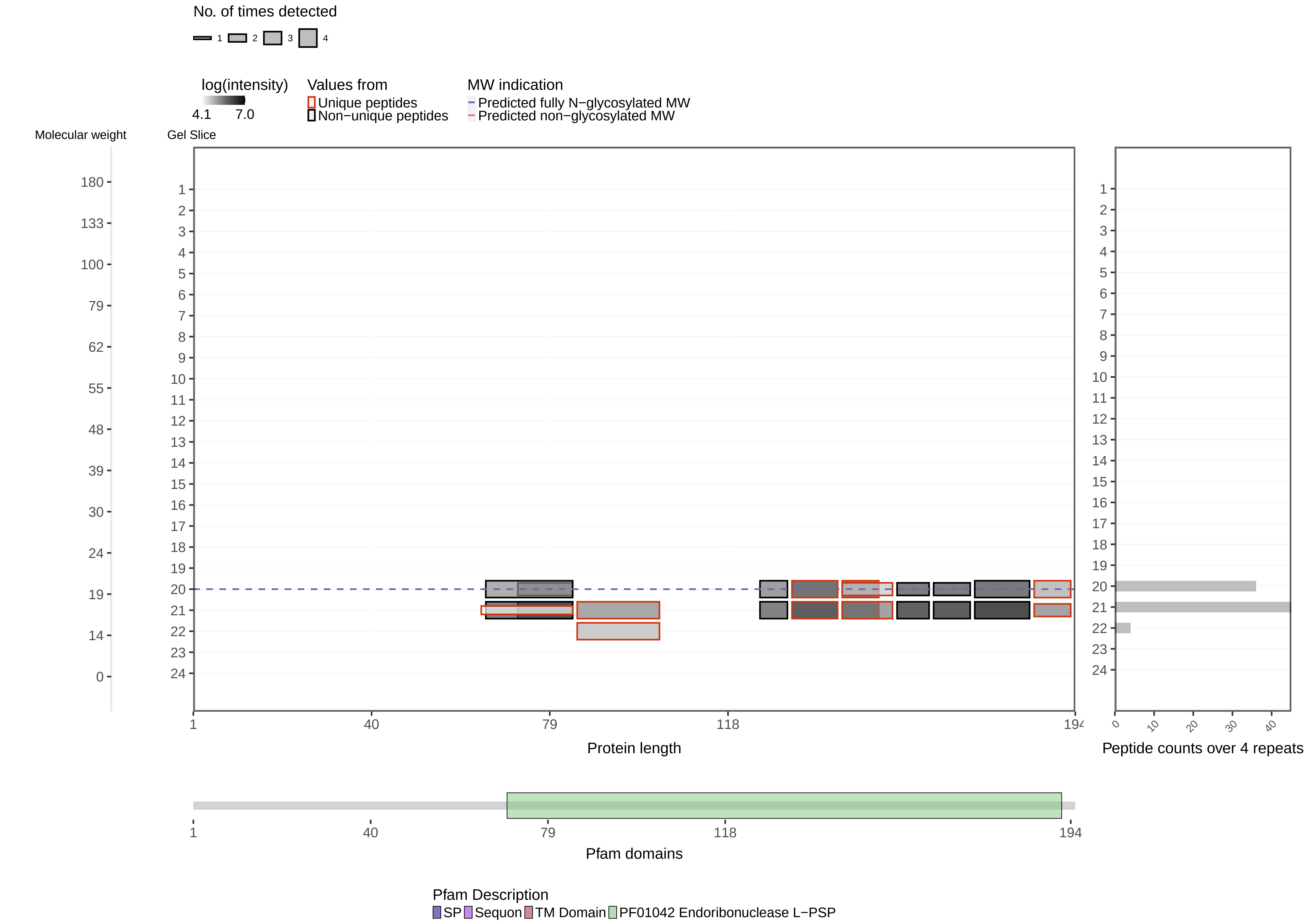


Protein sequence:

AIKANNFVFVSGVLGLVPETGKFVSDSVEDQTEQLNIDFQVLKNMGEILKASGVSYSAVVKTTILLADLKDFKTVNEIYAKYFQSPAPARATYQVAALPMDARIEIECIAAL*

### NbL11g18890.1 protein_LHCP_TRANSLOCATION_DEFECT [MW = 19.64 kDa]

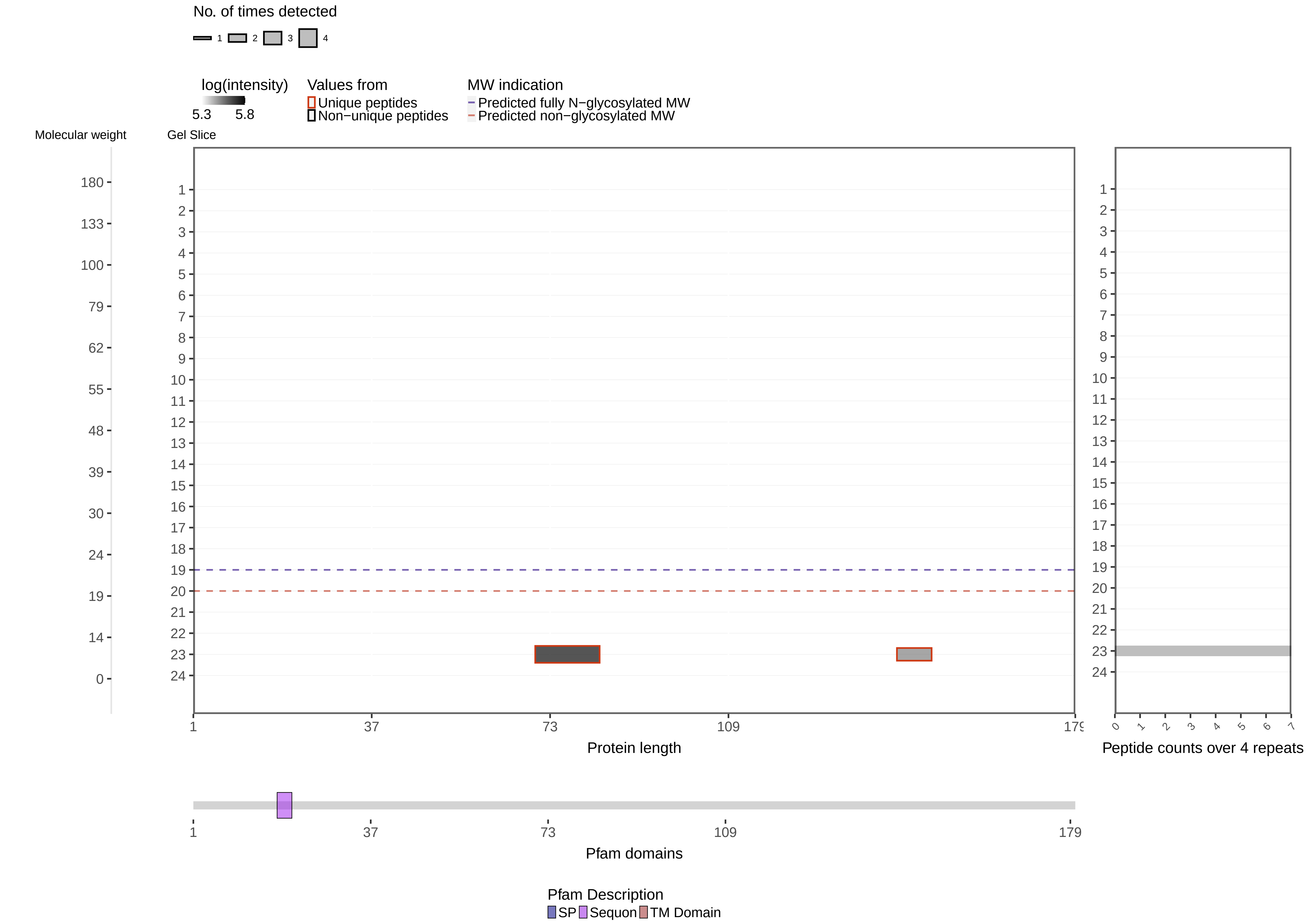


Protein sequence:

GVYGSQSRDDFDRDDVEQYFNYMGMLAVEGTYDKMEALLSQNIHPVDILLLMAATEGDLPKIEELLRAGADYTVKDADGRTALDRAASDEIKDFILNYKVQKA*

NbL11g18850.1 tryptophan_synthase_beta_chain_1 [MW = 55.22 kDa]

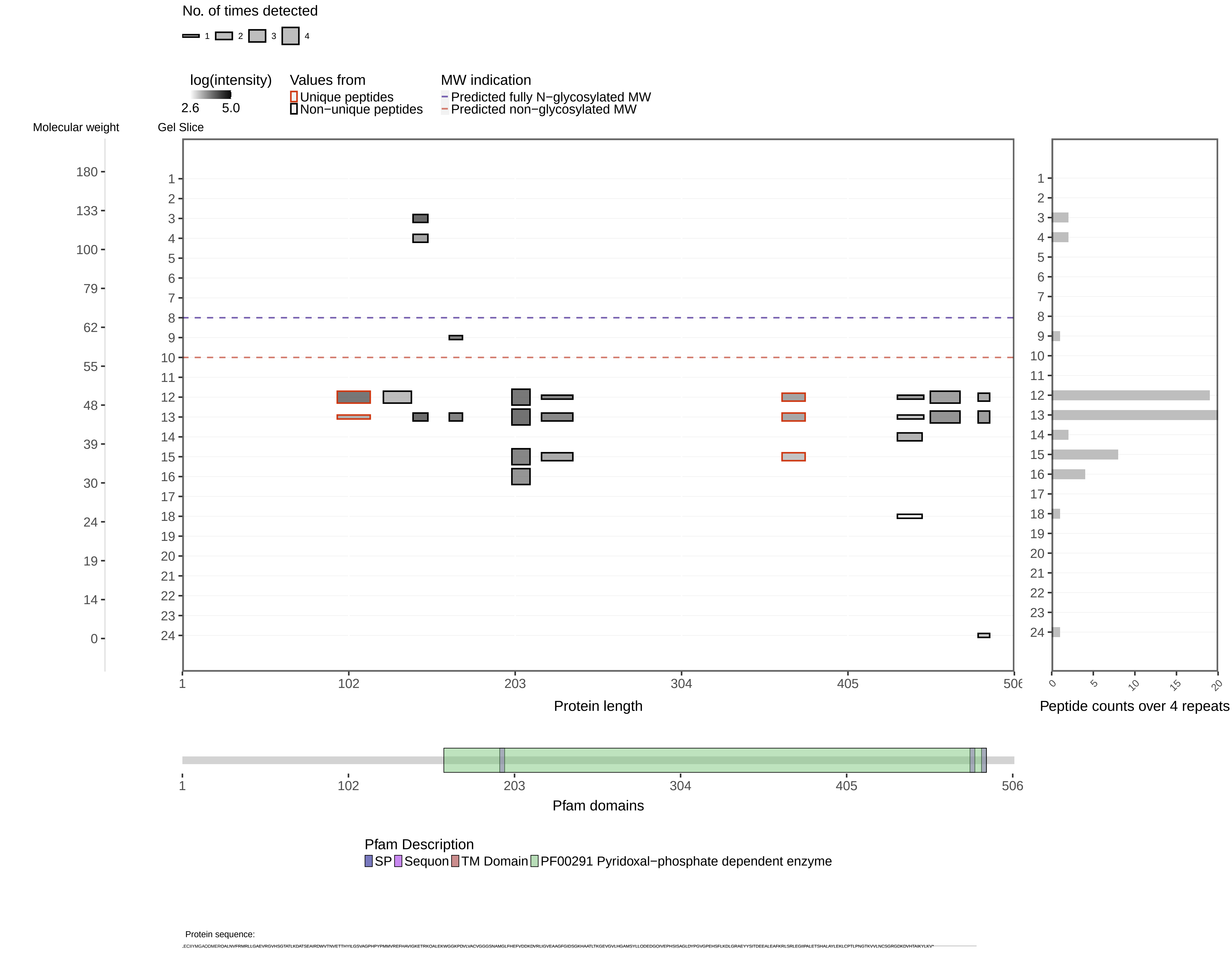

### NbL11g18750.1 trafficking_protein_particle_complex_subunit_3−like [MW = 20.83 kDa]

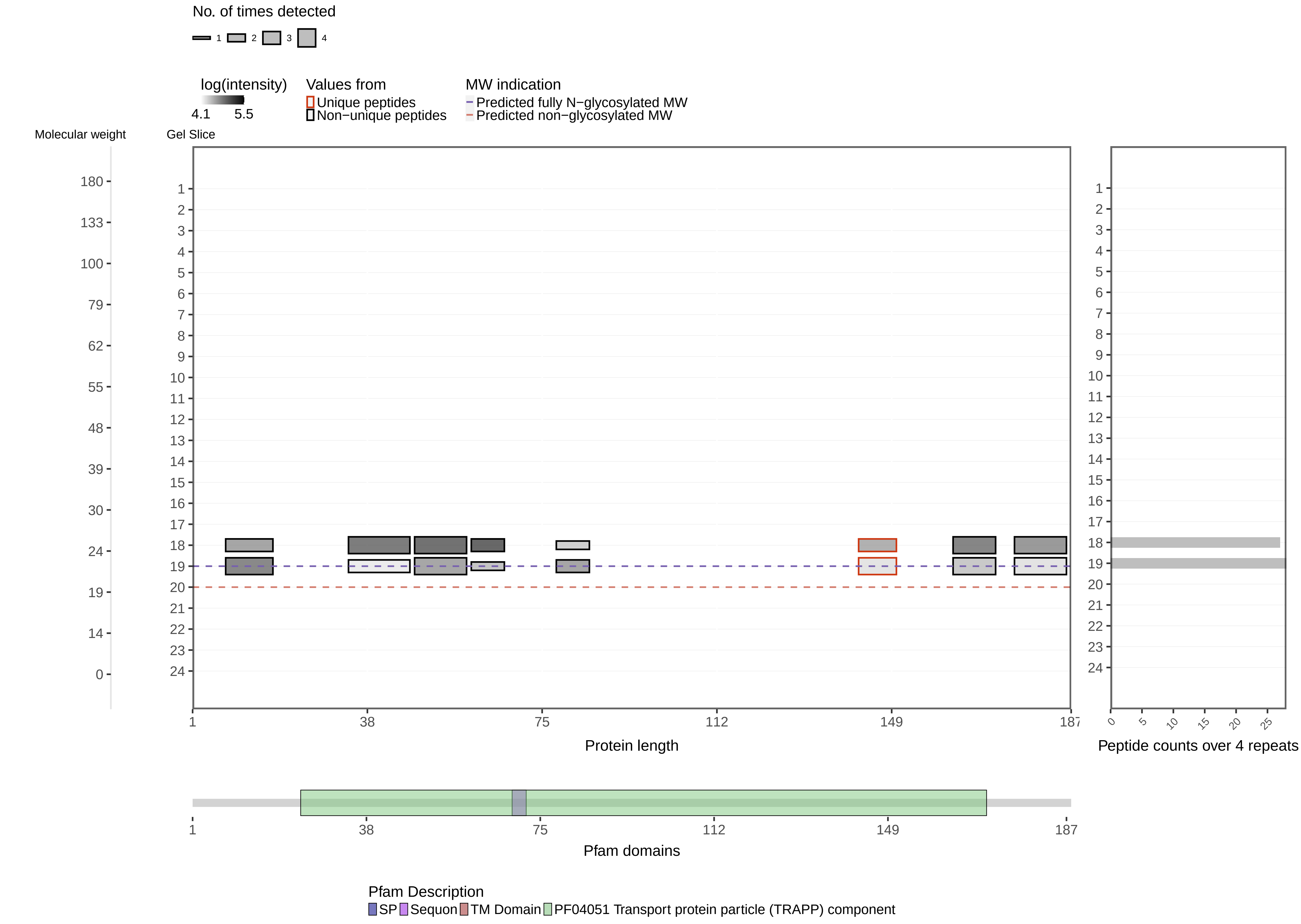


Protein sequence:

:VIAKVGLKMFLGVTATVGNWDAEGTTCSLILEDNPLVDFVELPDTCQGLYYCNILSGVIRGALETVSMKTEVTWIRDMLRGDDAFELQLKLLKQVPEEYPYKDDE*

NbL11g18680.1 60S_ribosomal_protein_L8 [MW = 28.21 kDa]

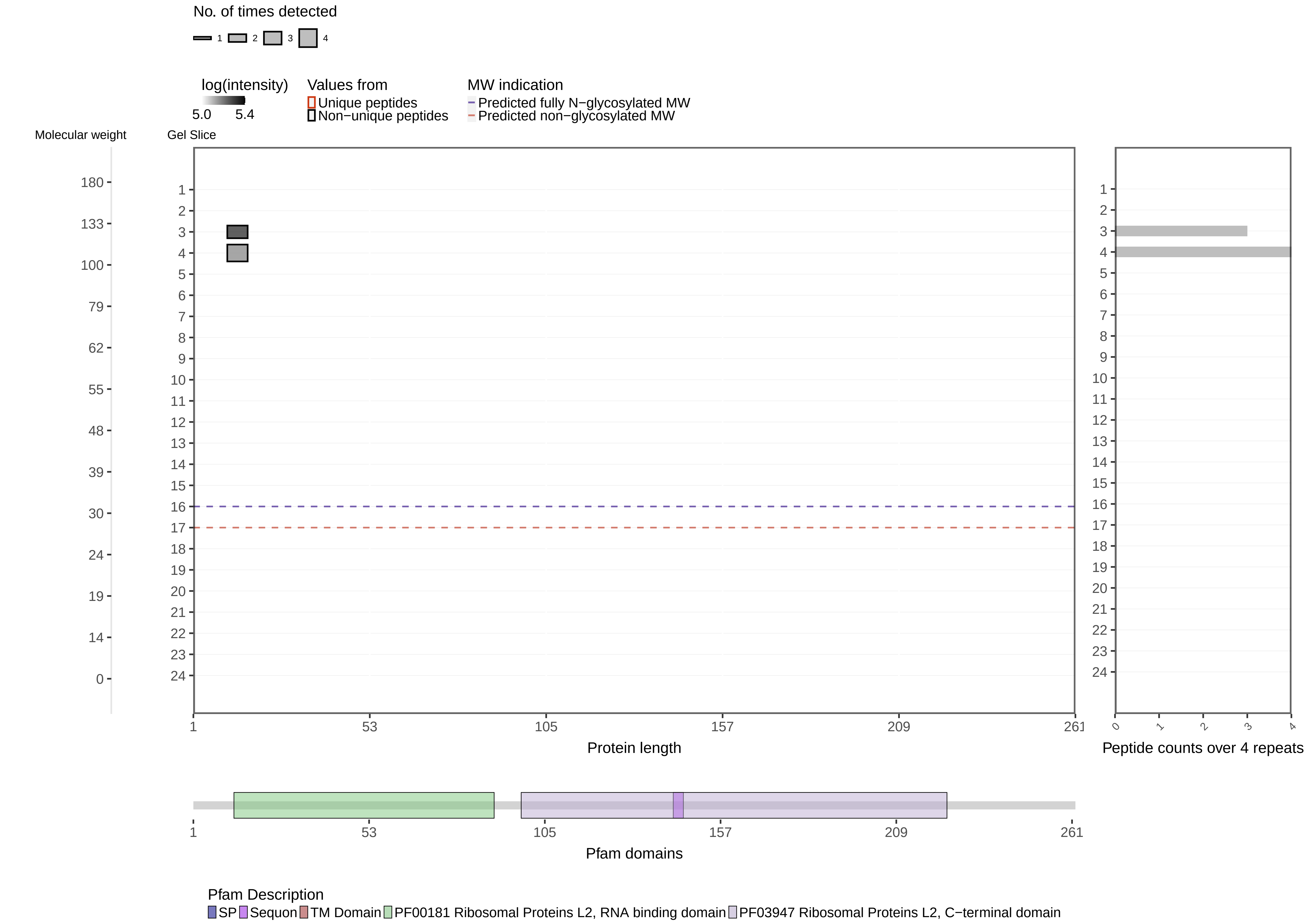


Protein sequence:

IKVGDRGVFARCSGDYAIVISHNPDNGTTRVKLPSGAKKIVPSGCRAMIGQVAGGGRTEKPMLKAGNAYHKYRVKRNCWPKVRGVAMNPVEHPHGGGNHQHIGHASTVRRDAPPGQKVGLIAARRTGRLRGQAAATAAKADKA*

NbL11g18650.1 indole−3−acetic_acid−amido_synthetase_GH3.6−like [MW = 68.64 kDa]

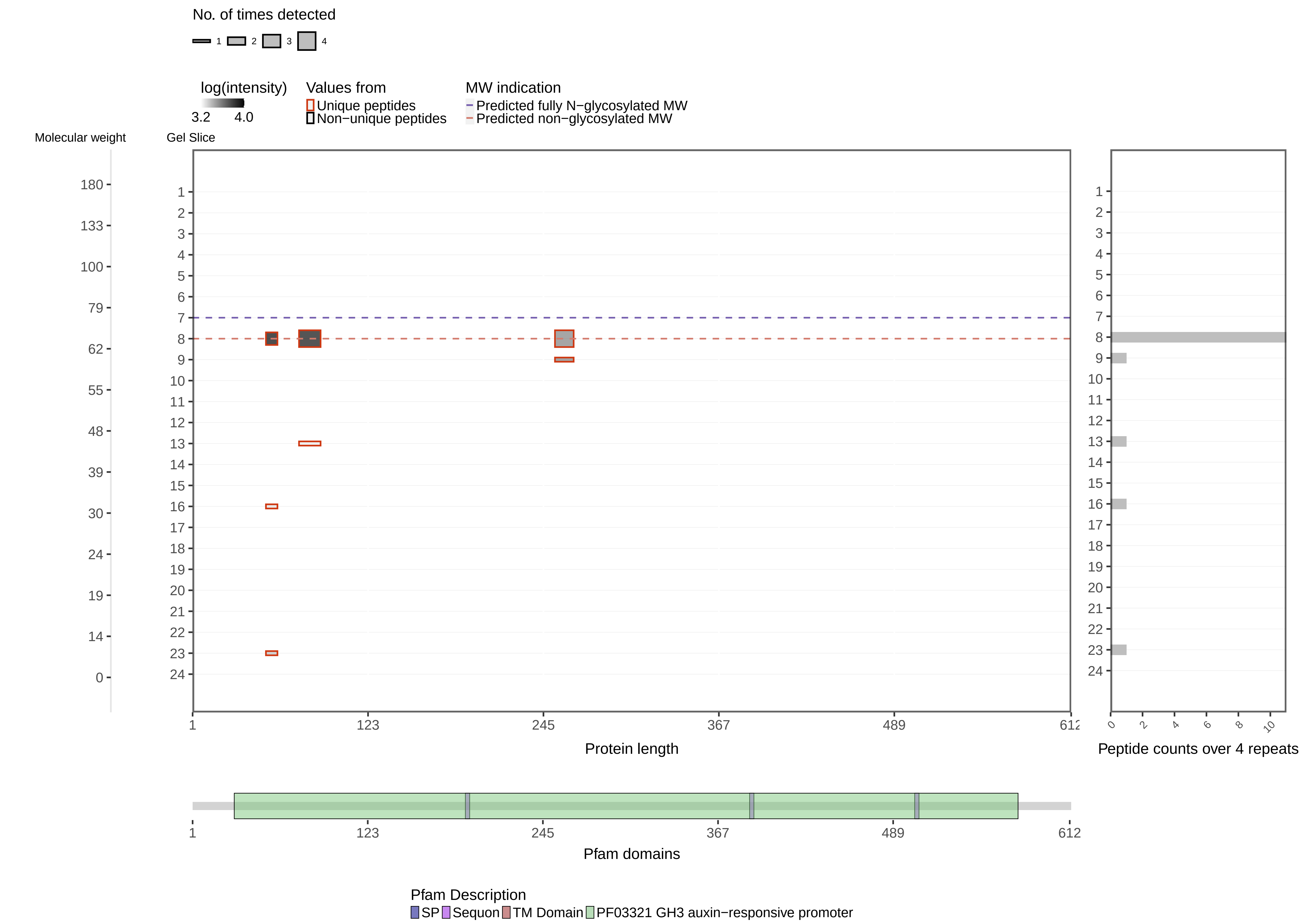


Protein sequence:

NbL11g18630.1 probable_NAD(P)H_dehydrogenase_(quinone)_FQR1−like_1 [MW = 21.59 kDa]

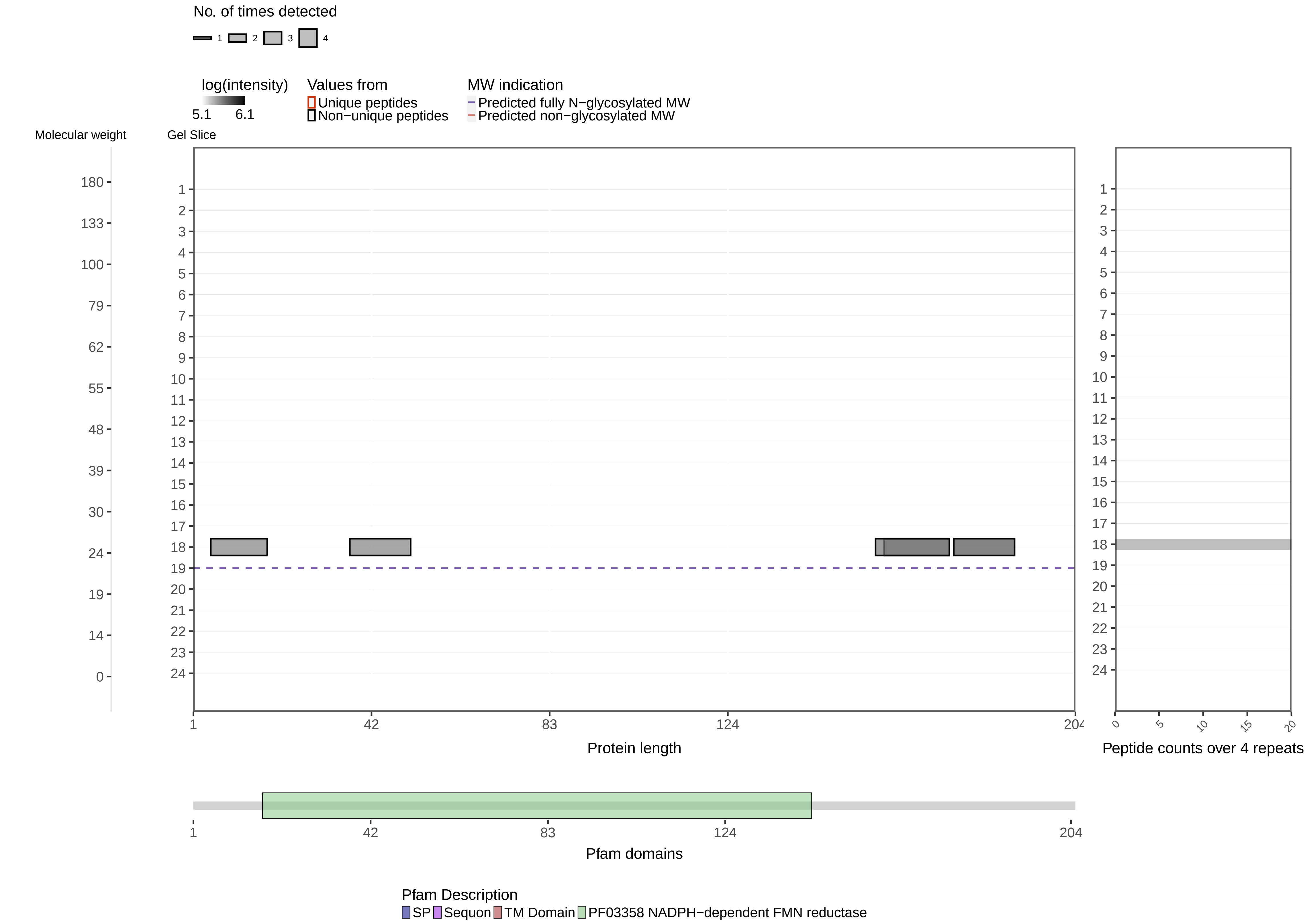


Protein sequence:

KAFLDATGGLWRTQQLAGKPAGIFYSTGSQGGGQETTPLTAITQLVHHGMIFVPIGYTFGAGMFEMEKVKGGSPYGAGTFAGDGSRQPSDLELQQAFHQGKYIAGIAKKLKGAA*

### NbL11g18610.1 bifunctional_L−3−cyanoalanine_synthase/cysteine_synthase_2_mitochondrial [MW = 39.33 kDa]

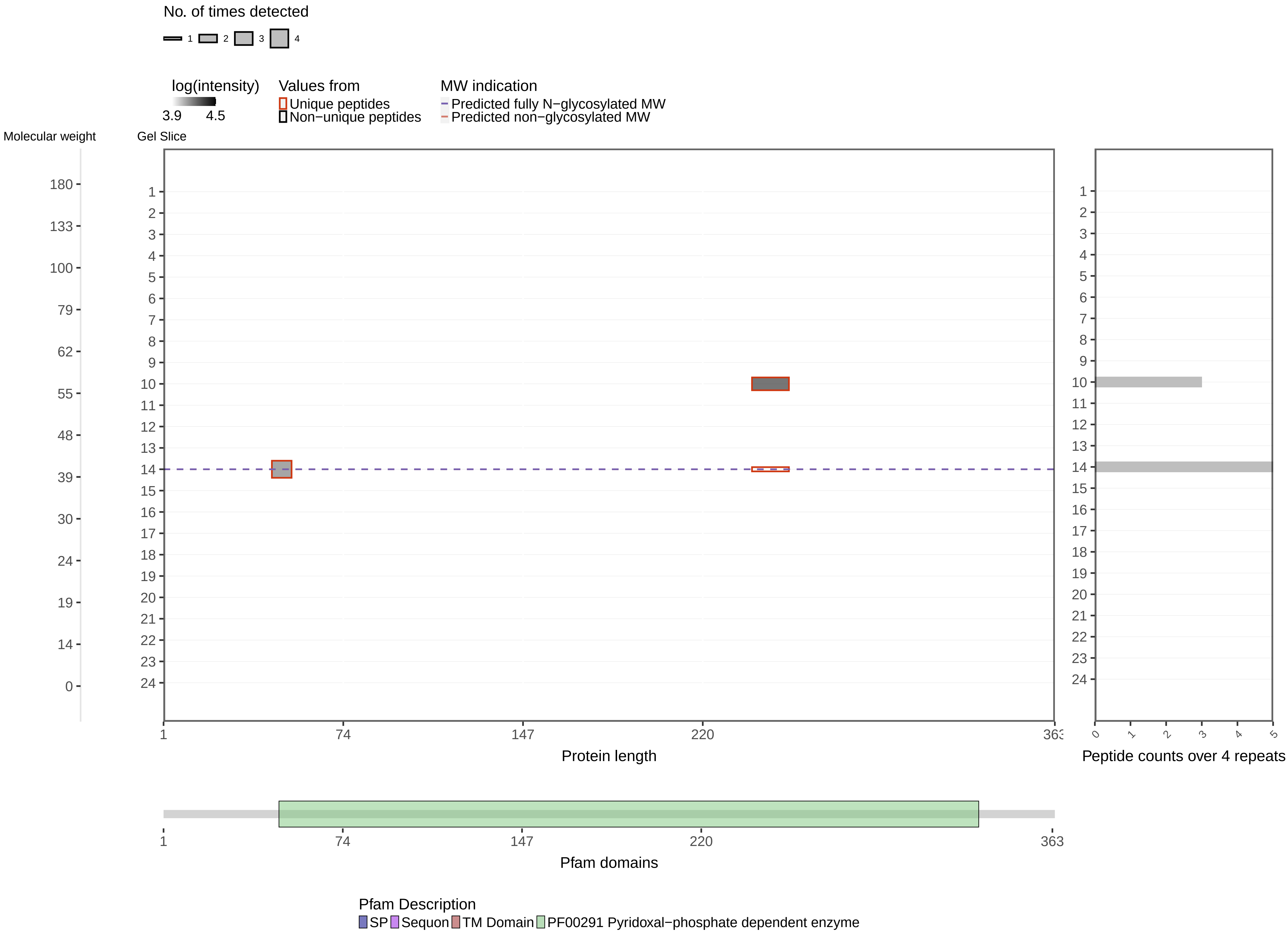


Protein sequence:

KAYDLLESTPNAFMLQQFSNPANTQIHFETTGPEIWEETQGNVDIFVMGIGSGGTVSGVGQYLKSKNPNVKIYGVEPTESNILNGGKPGPHHITGNGVGFKPDILDMDIMEEVLMVSSEDAVNMARELAVKEGLMVGISSGANTVAALQLAQKPENKGKLIVTIHASFGERYLSSVLYQDLRKEAENMQPVSVD*

NbL11g18420.1 mitogen−activated_protein_kinase_homolog_MMK2 [MW = 43.13 kDa]

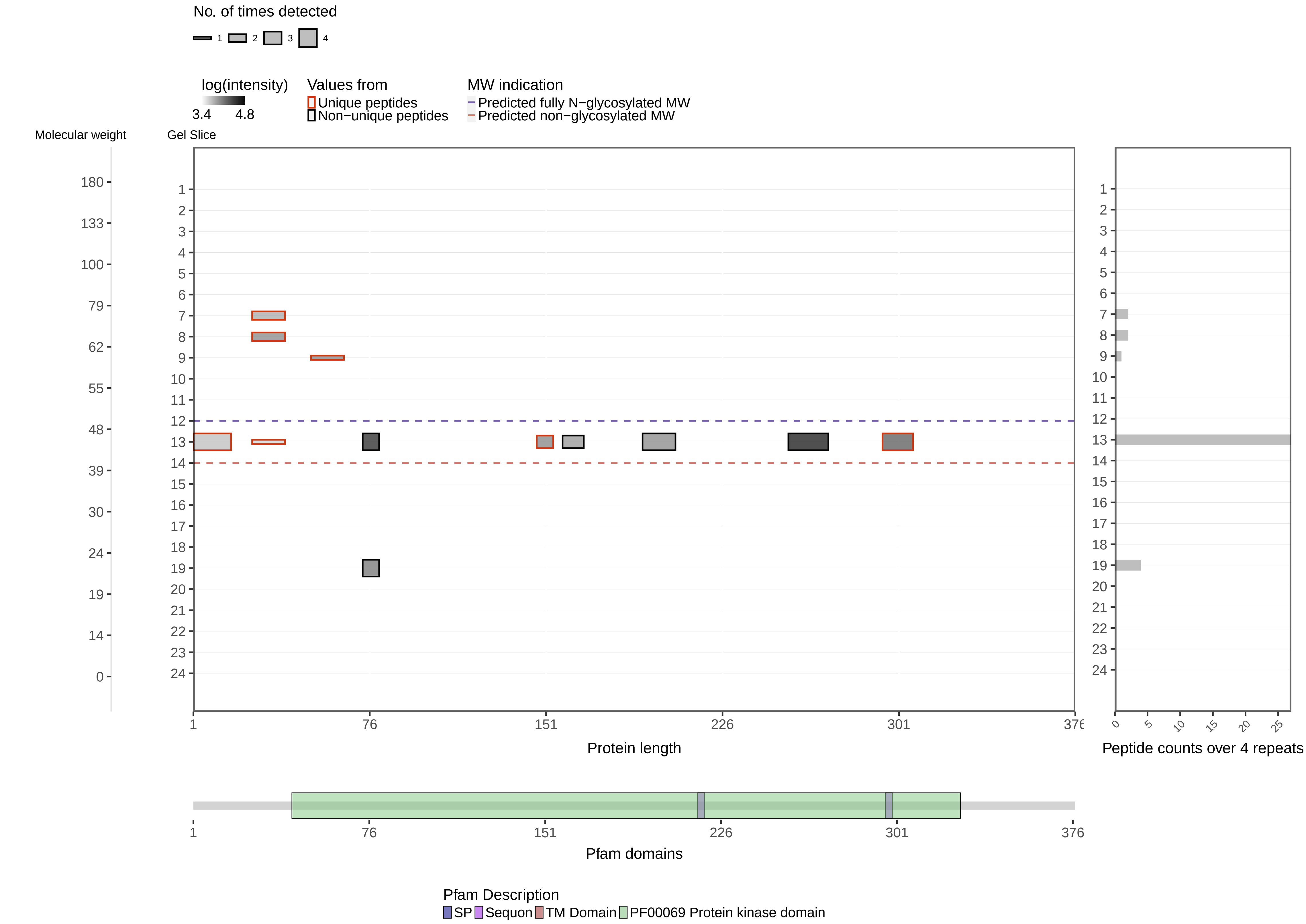


Protein sequence:

NANCDLKVGDFGLARTTSETDFMTEYVVTRWYRAPELLLNCSEYTAAIDIWSVGCILGEMMTRQPLFPGRDYVHQLRLITELIGSPDDASLGFLRSDNARRYVRQLPQYPRQQFAARFPNSSPGAVDLLEKMLVFDPSRRVTVDQALCHPYLAPLHDINEEPICPKPFSFDFEQPSFTEENIKELIWRESVKFNPDPTH*

NbL11g18370.1 desiccation_protectant_protein_Lea14_homolog [MW = 18.37 kDa]

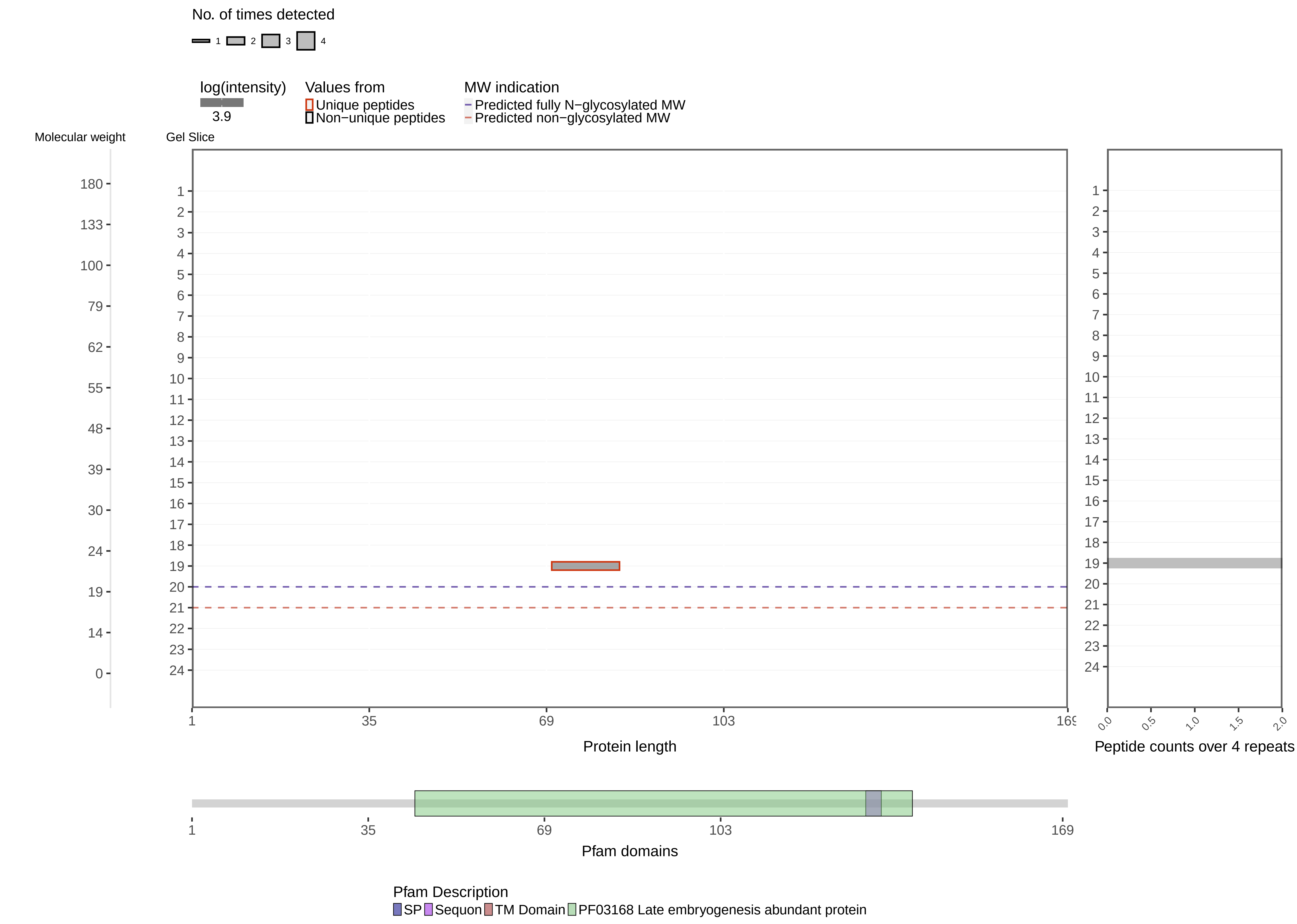


Protein sequence:

VILSGTIPDPGSIKADDTTMLDVPVKVPHSVLVSLAKDIGADWDIDYTLELGLTIDLPVIGNFTIPLSHSGEIKLPTLSDLWKGGEEEEKEDTEKVKEI*

NbL11g18190.1 eukaryotic_translation_initiation_factor_1A−like [MW = 16.35 kDa]

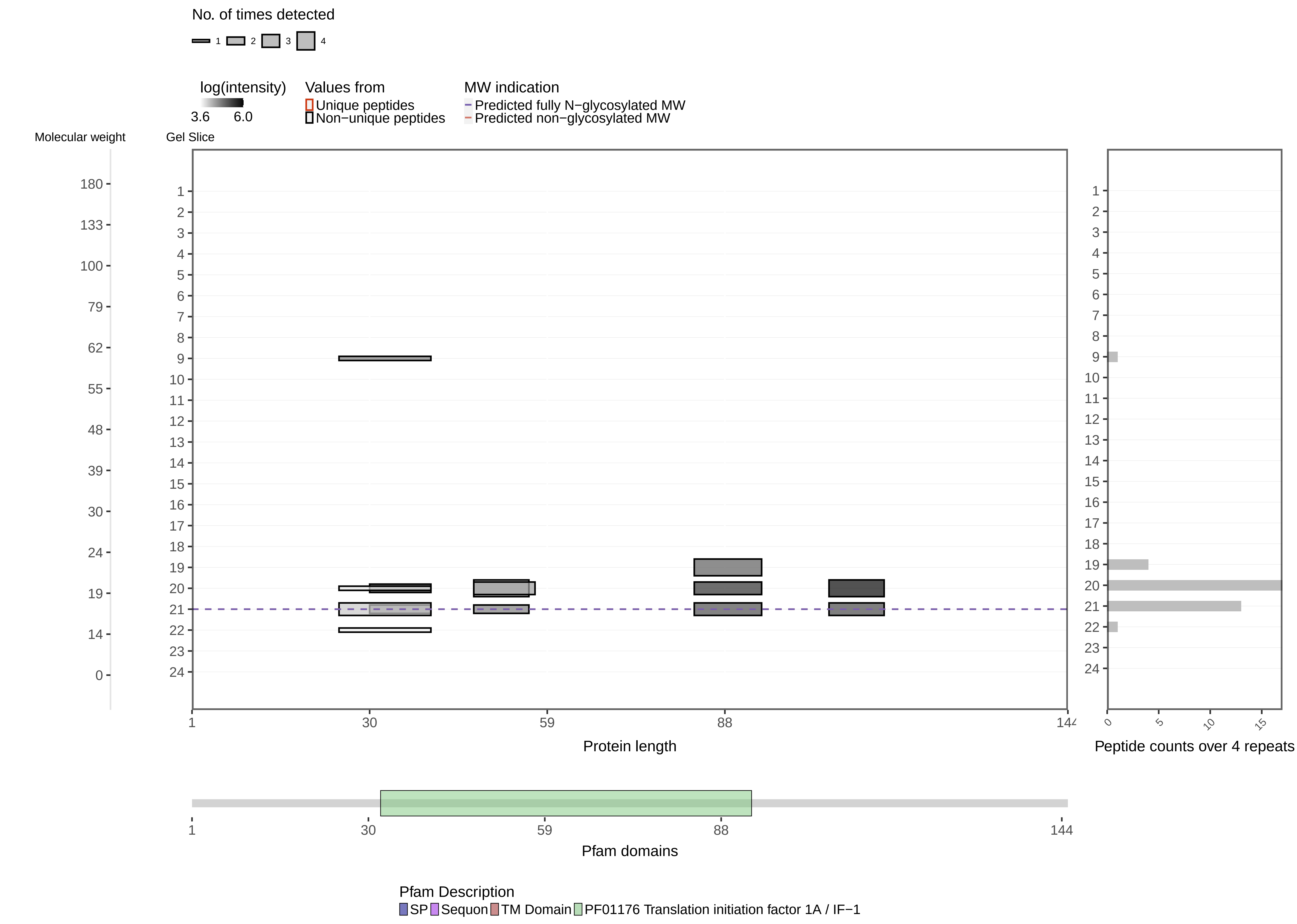


Protein sequence:

RLCHIRGKMHKKVWIAAGDIILVGLRDYQDDKADVILKFMPDEARLLKAYGELPENTRLNEVNGLDEEDDGDGDDYIEFEDEDIDKI*

NbL11g18180.1 protein_CURVATURE_THYLAKOID_1A_chloroplastic [MW = 17.94 kDa]

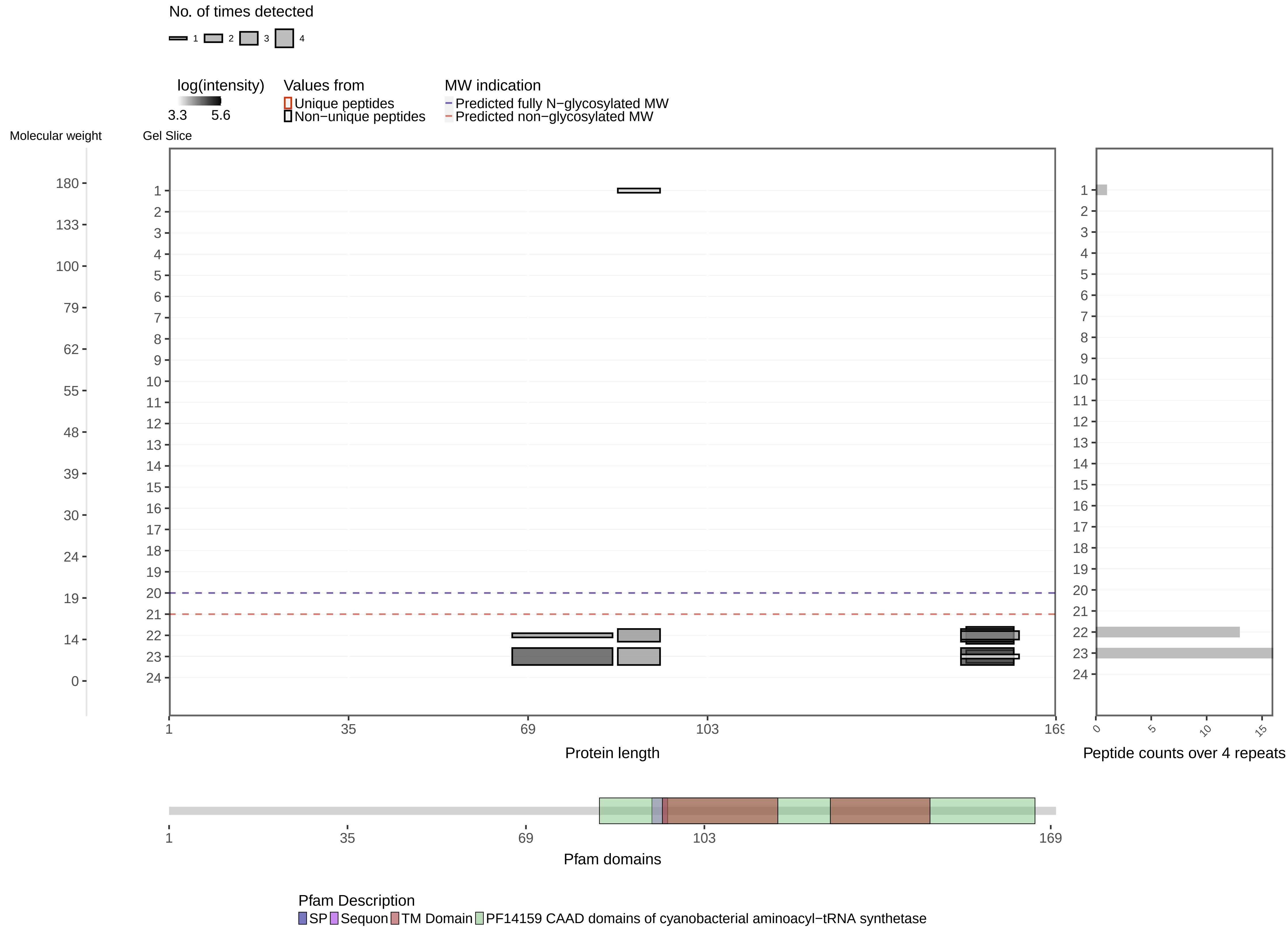


Protein sequence:

ESGAVDTSELLTDLKEKWDAVENKSTVIVYGGGAIVAVWLSSIVVGAINSVPLLPKIMELVGLGYTGWFVYRYLLFKSSRKELAEDIEQLKKKIAGTE*

NbL11g18160.1 GDSL_esterase/lipase_At4g01130−like [MW = 19.55 kDa]

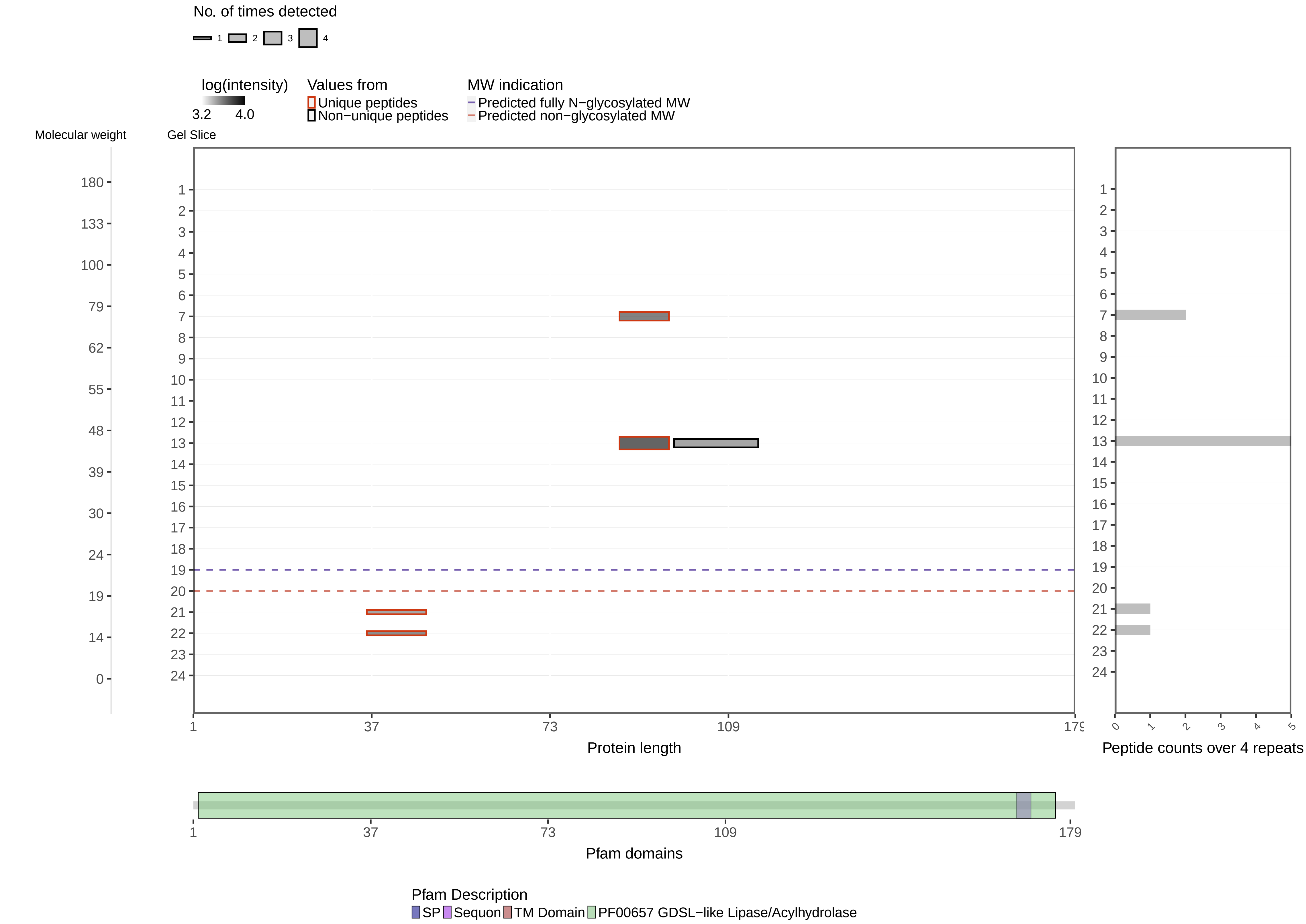


Protein sequence:

VKVDEFQHKGKTNLPSPDIFGKSLYTFYIGQNDFTGNLARIGISGVKEYLPQVVSQIASTIKEIYALGGRTFWVLNLAPIGCNPAFLVELPHNTSDIDQFGHE*

### NbL11g18140.1 homoserine_kinase−like [MW = 17.33 kDa]

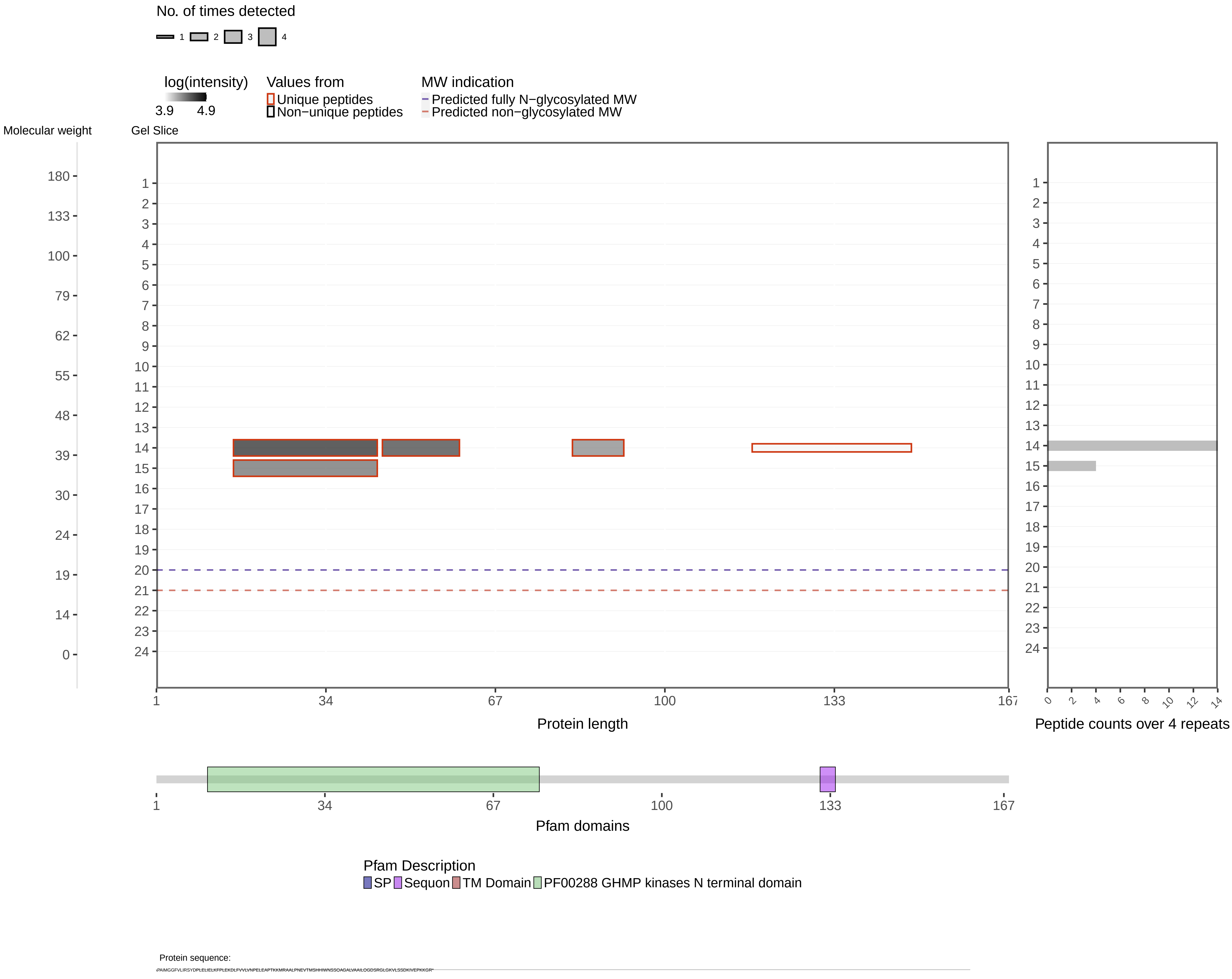

### NbL11g17970.1 rhodanese−like_domain−containing_protein_4_chloroplastic [MW = 46.61 kDa]

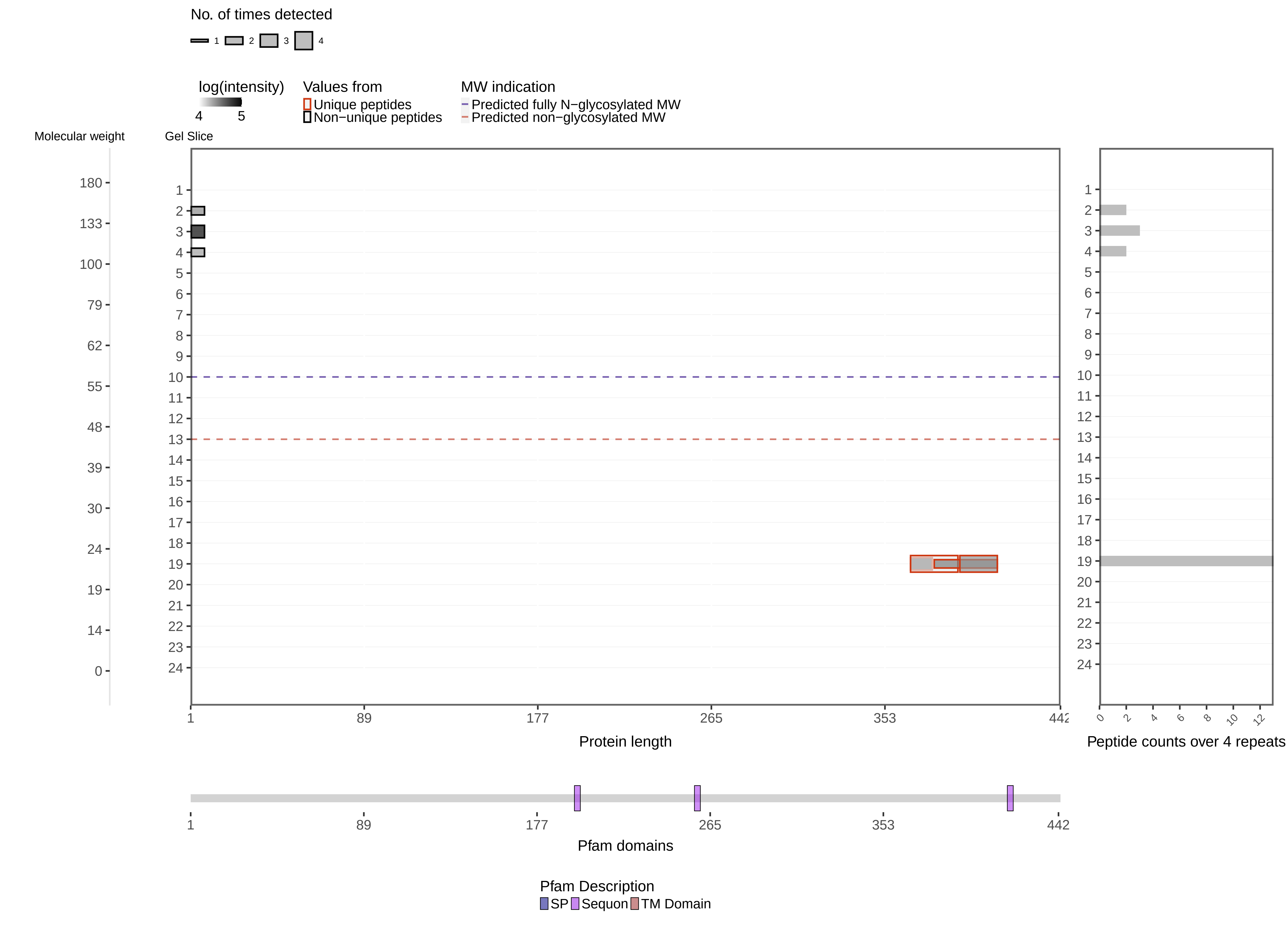


Protein sequence:

### NbL11g17840.1 clustered_mitochondria_protein_homolog [MW = 89.68 kDa]

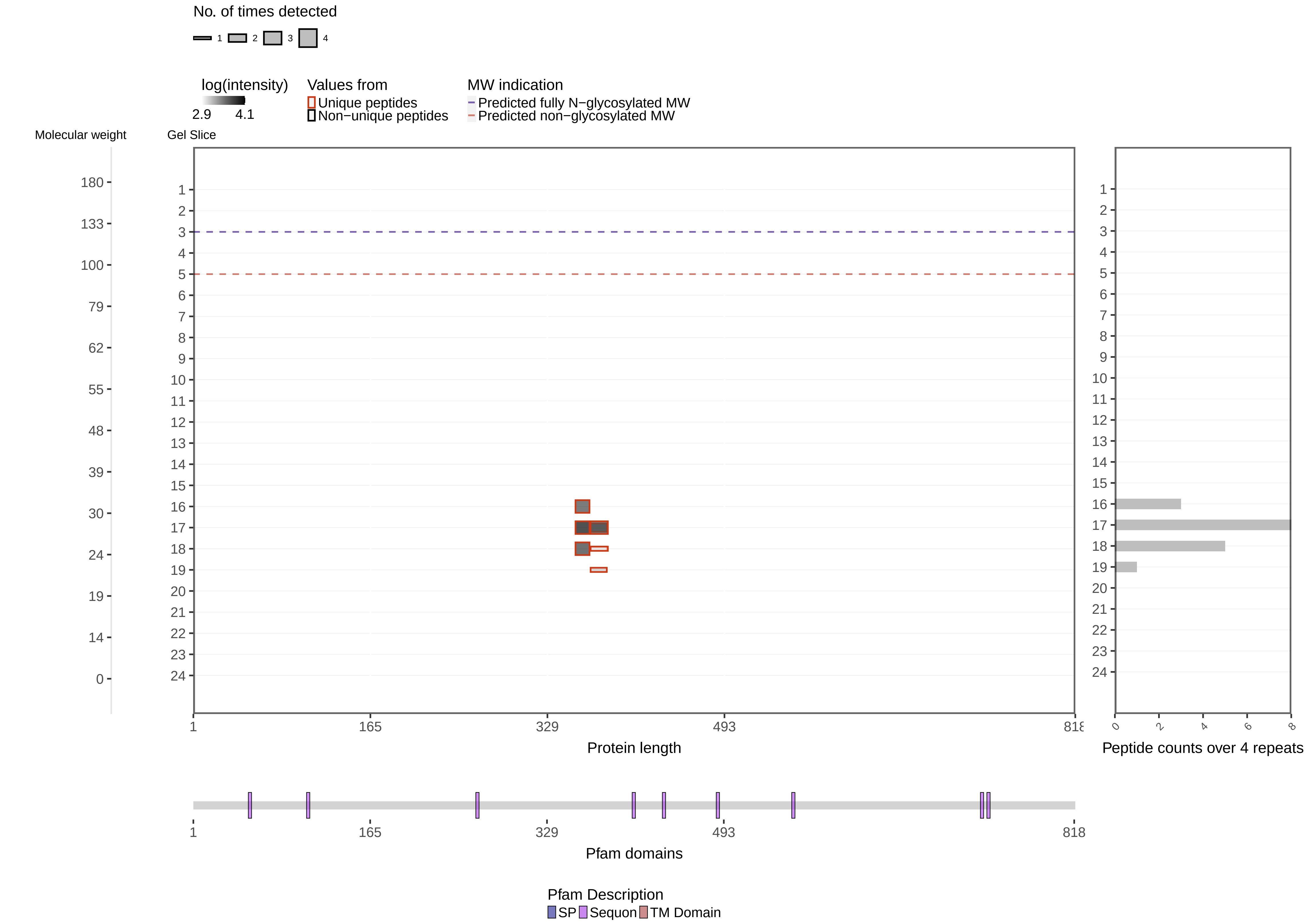


Protein sequence:

NIQDLVTDSADQVQNETEDTDNKEEIQLSDLKGGEISDVISANASIQPGHVDVSPMEQGSVQAHNVPTSDNSPKVDLCEKDSSRNLDPSYISNLTLQDMDHLKVKAASSLTCDASRELSRKLSASAAPFSPSAGIARVAPLPMNINLPSPPGTLPPVGPWPMNMSLHQGPPTMLPNPMCSSPHHLYPSPPHTPNMMHPIRFMYPPYSQPQTLPPSTFPISSSNFHPNHYAWQCNIAPTASEYVPATVWPGCHPVEFSVSPPVIEPLTDSISAAKEPSDSPESIILATSLPVDLN

### NbL11g17710.1 exportin−2 [MW = 109.56 kDa]

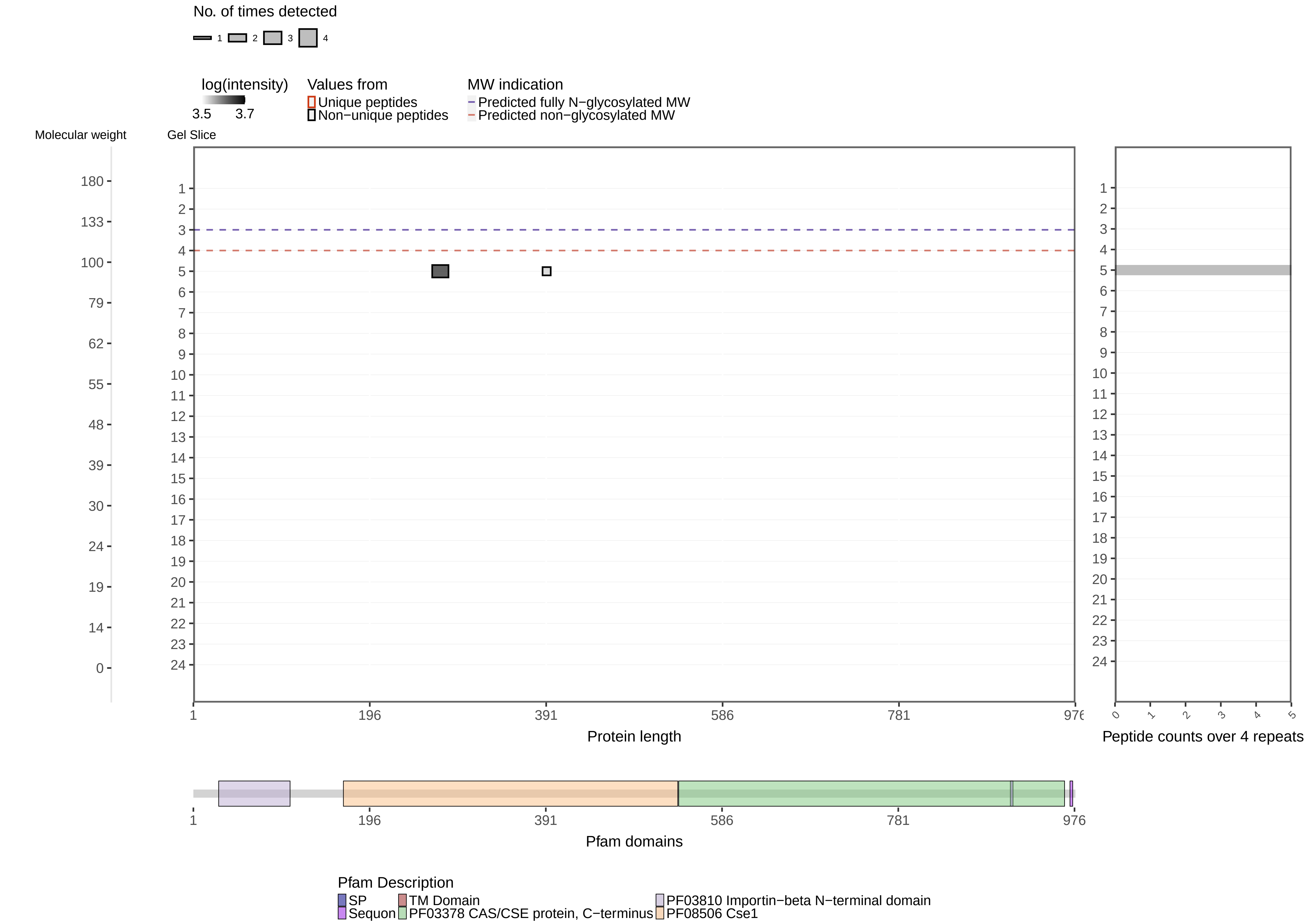


Protein sequence:

SVIVPELQSRDVNAFPMLKAGALKFFTMFRNQLPKAVAMALLADVVRFLAAESNVVHSYAASCIEKLLLVKDEGTRARYTATDISPFLLVLMSNLFSALEKPESEENQYIMKCIMRVLGVAEISRDVASACITGLTNVLNRVCQNPKNPIFNHYLFESVAVLIRRACEKNPTLISAFEGSLFPSLQMILANDVSEFFPYAFQLLSQLVELNRPPVPQHYVQIFEILLLPESWKKSANVPALVRLLQAFLRKAPHELNQQGRLSNVLGIFNTLISSPSTDEQGFYVLNTVIENLGYDVISPF

NbL11g17620.1 bifunctional_fucokinase/fucose_pyrophosphorylase [MW = 118.2 kDa]

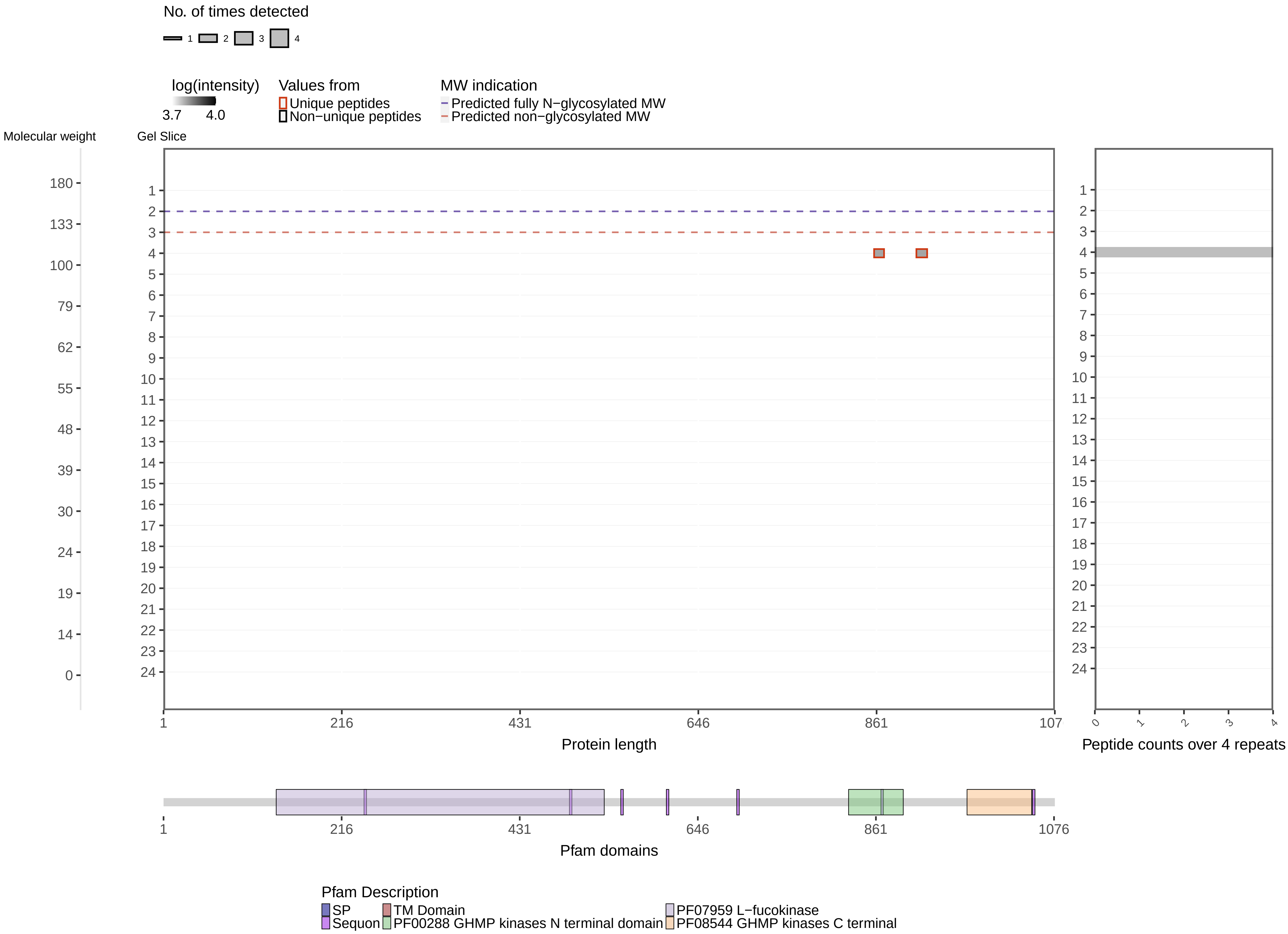


Protein sequence:

[illegible]

### NbL11g17570.1 ras−related_protein_RABA3−like [MW = 27.71 kDa]

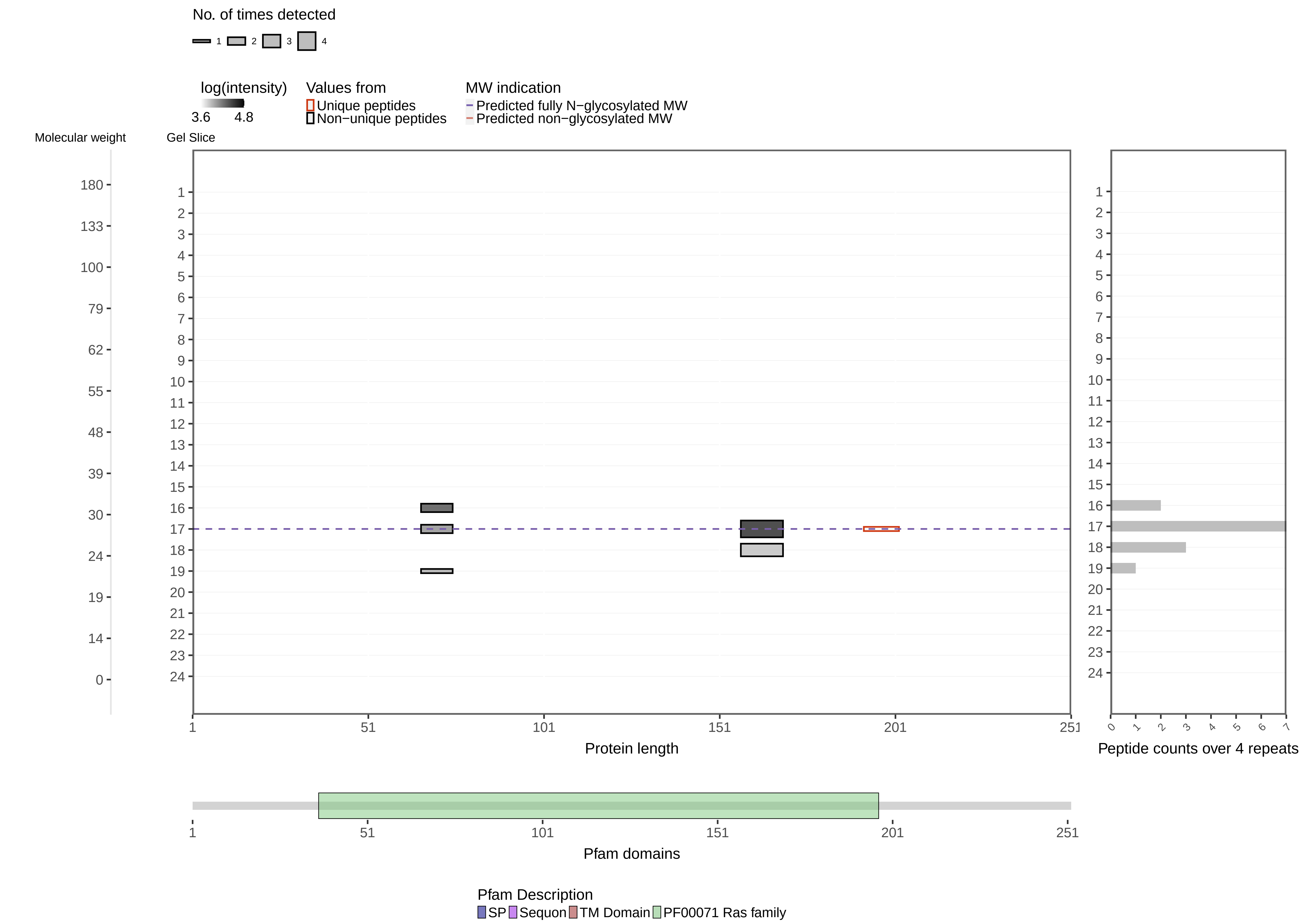


Protein sequence:

MLVYDITKRQSFDNVARWVDELRVHADSSIVITLIGNKADLVDLRAVPTEDAVEFAERQGLFFSETSALSGHNVESAFLKLLEEIYNVVSRKTLLIGSGANGNHIKETNNGLLKGLKIDVISGSDFEVSEMKKLSSCSC*

NbL11g17540.1 uncharacterized_LOC104241754 [MW = 29.54 kDa]

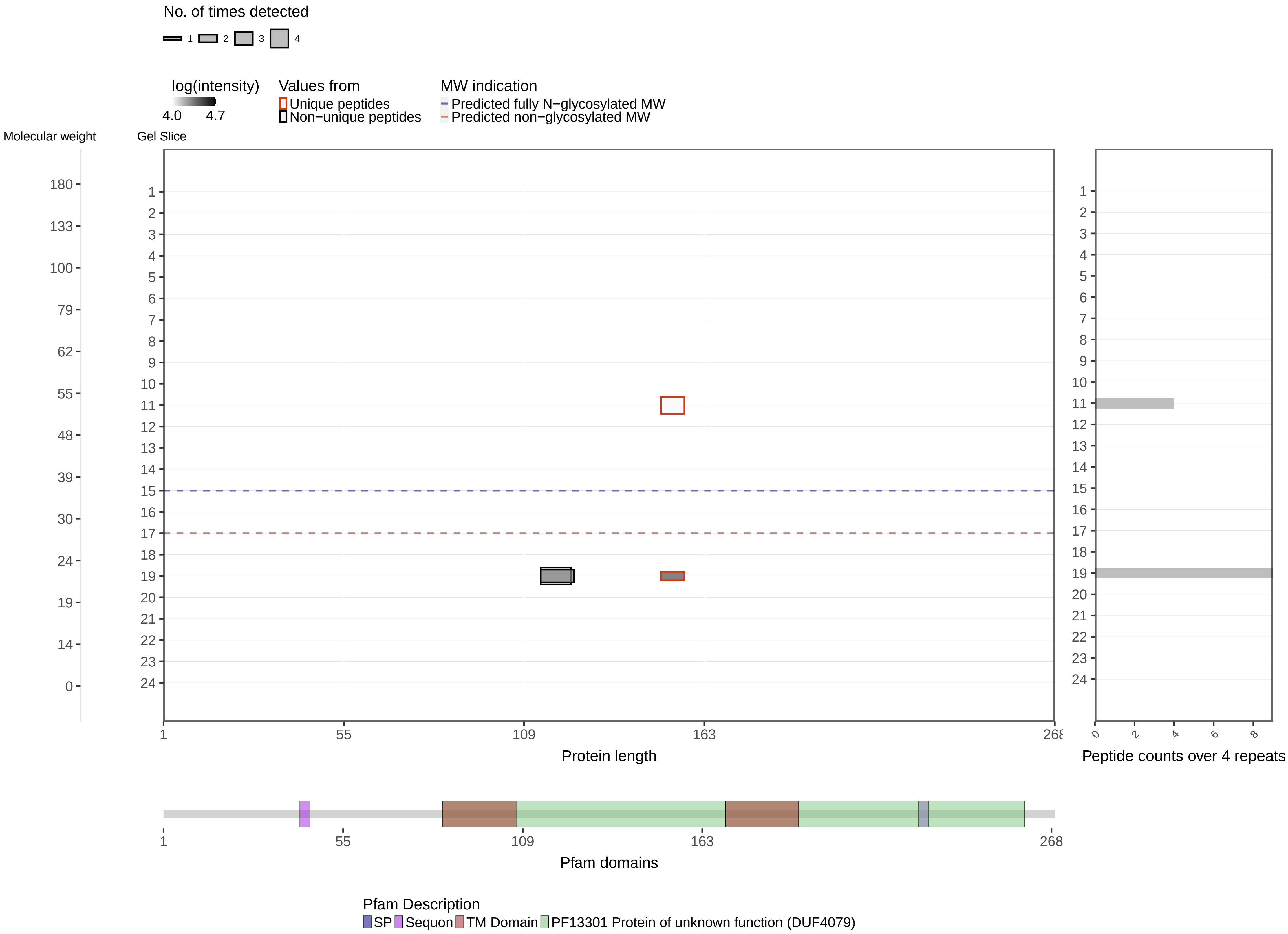


Protein sequence:

ELKKQVKPLPVTPEGTPIQPPKPSPVEARIQQLTEQRKELIKGSYKDKHFNAGSILLAWGVFESIAGGVNTWFRTGKLFPGPHLFAGAGITILWAAAAALVPAMQRGNETARNLHIALNALNVILFIWQIPTGIDIVFKVFEFTSWP*

### NbL11g17480.1 homeobox−leucine_zipper_protein_ATHB−12−like [MW = 28.95 kDa]

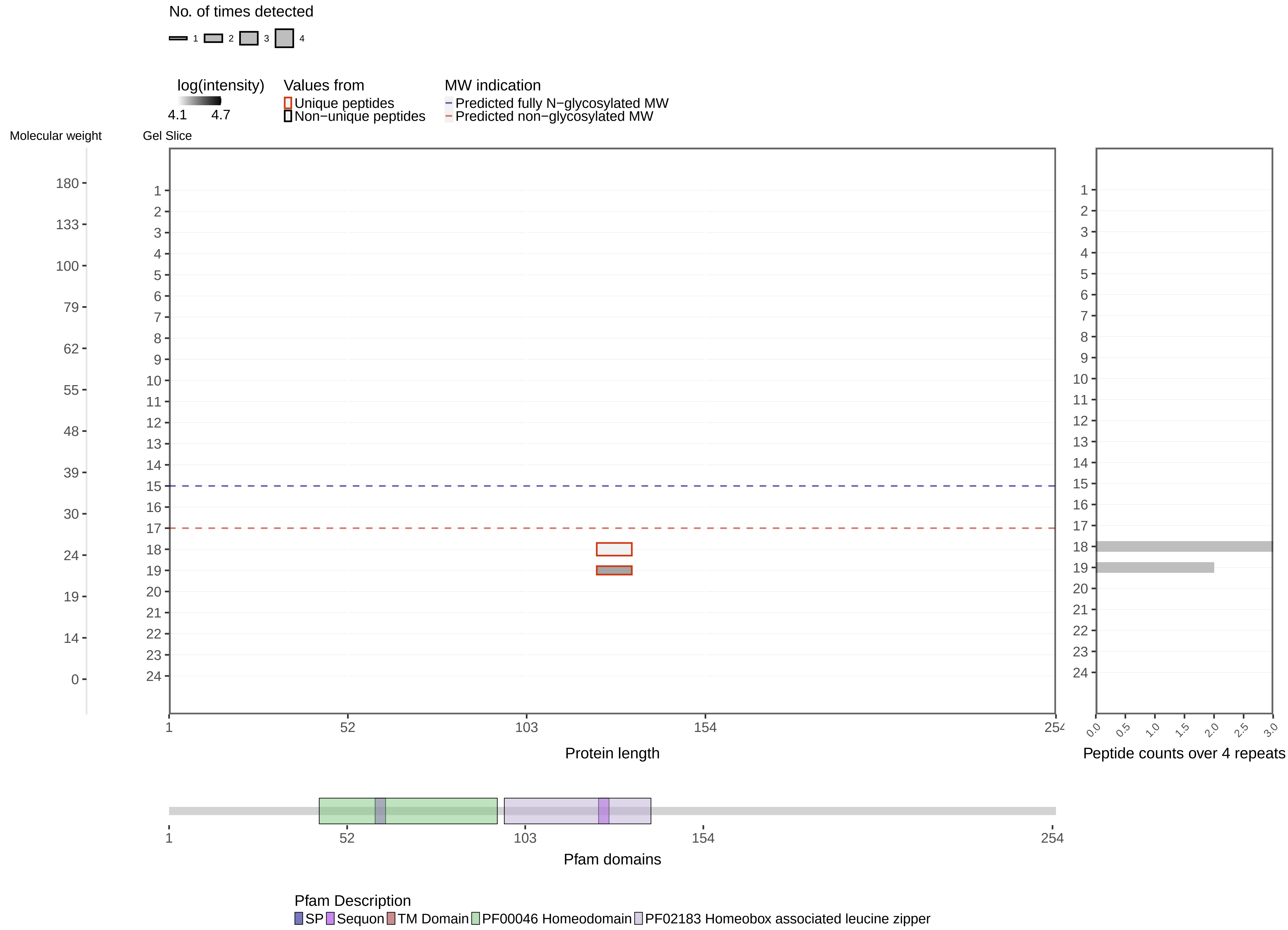


Protein sequence:

QYNSLKKENQSLLFQLQKLNDLMQKSEKEGGQYCSIGVDQESYNRHDNTIKSKEMDGKPSLSFDLSEHGVNGVISDDDSSIKADYFGLDEESDHLLKMVEAGDSSLTSPENWGTLEDDGLLDQQPNSCNYDQWWDFWS*

### NbL11g17270.1 aspartyl_protease_AED3−like [MW − SP = 44.78 kDa]

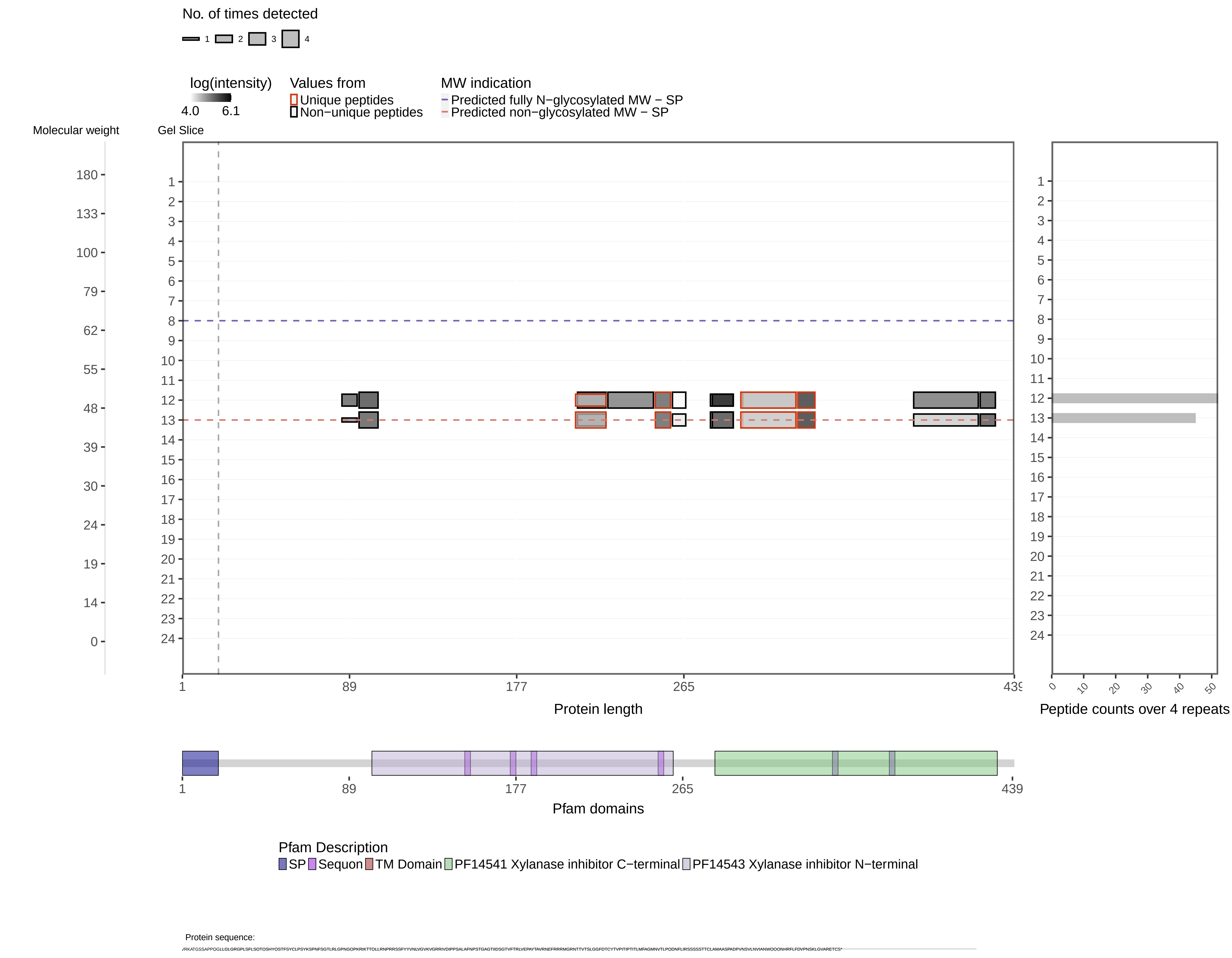

NbL11g17240.1 thioredoxin−like_protein_CITRX1_chloroplastic [MW = 20.56 kDa]

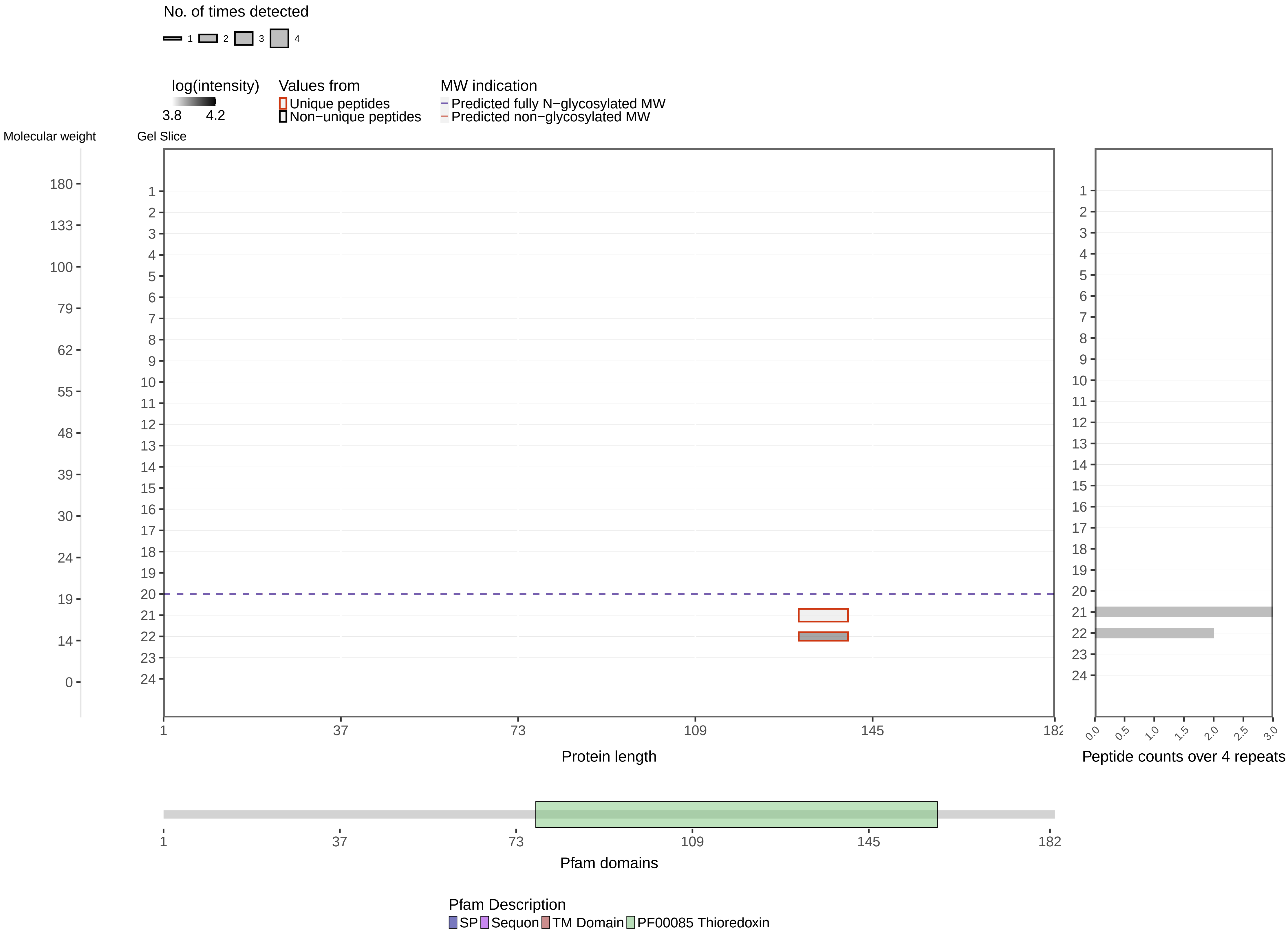


Protein sequence:

KVSAKDIQELIKGERNVPLIIDFYATWCGPCILMAQELEMLAVEYESNALIVKVDTDDEYEFARDMQVRGLPTLYFISPDPNKDAIRTEGLIPIQMMRDIINNDL*

### NbL11g17210.1 peptidyl−prolyl_cis−trans_isomerase_CYP22−like [MW = 20.61 kDa]

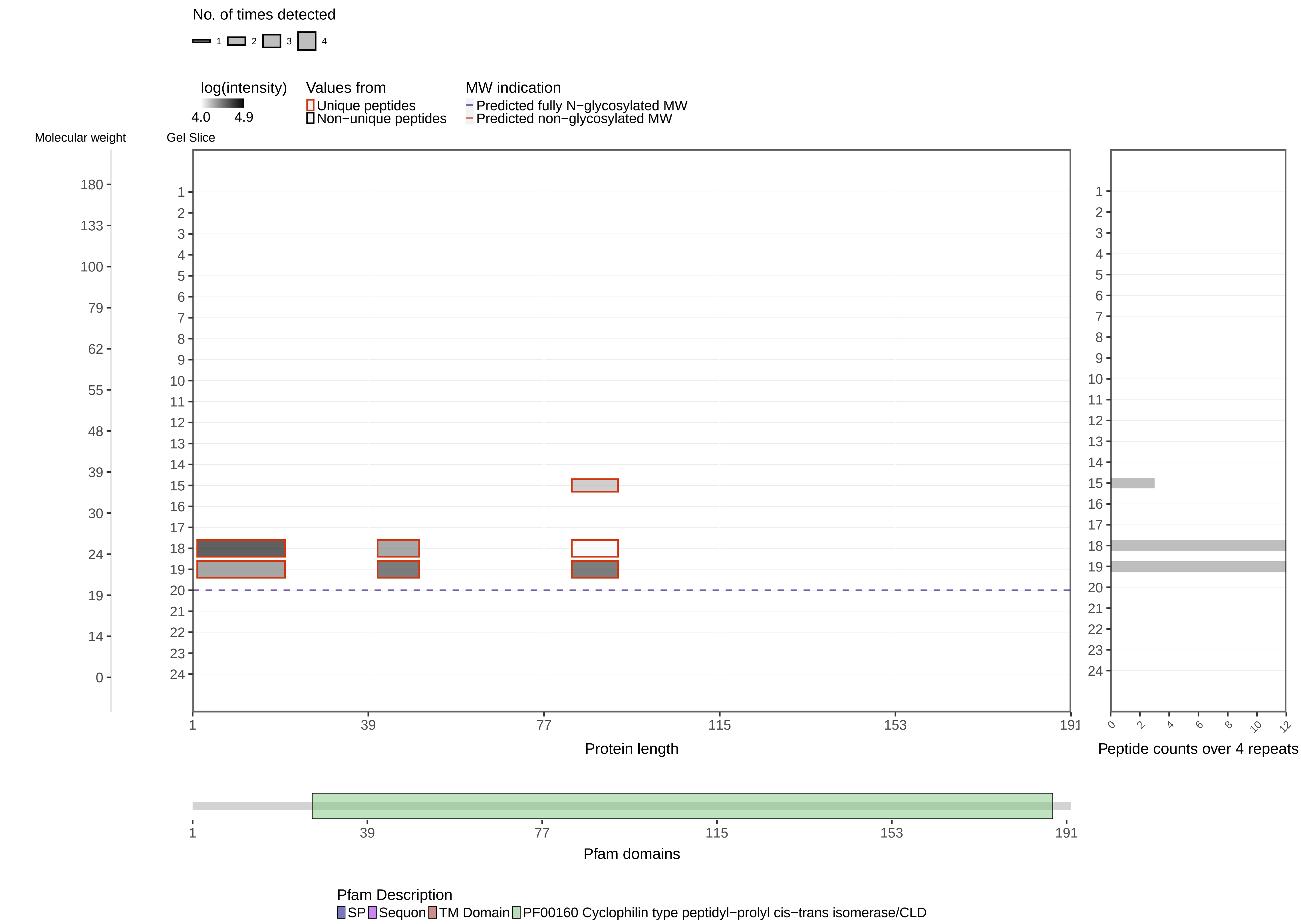


Protein sequence:

)FMVQGGDFLKGDGSGCVSIYGSKFEDENFIAKHTGPGLLSMANSGPNTNGCQFFITCAKCDWLDNKHVVLGRVLGDGLLVVRKIENVAVGANNKPKLACMIAECGEM*

### NbL11g17200.1 uncharacterized_LOC109235373 [MW = 121.78 kDa]

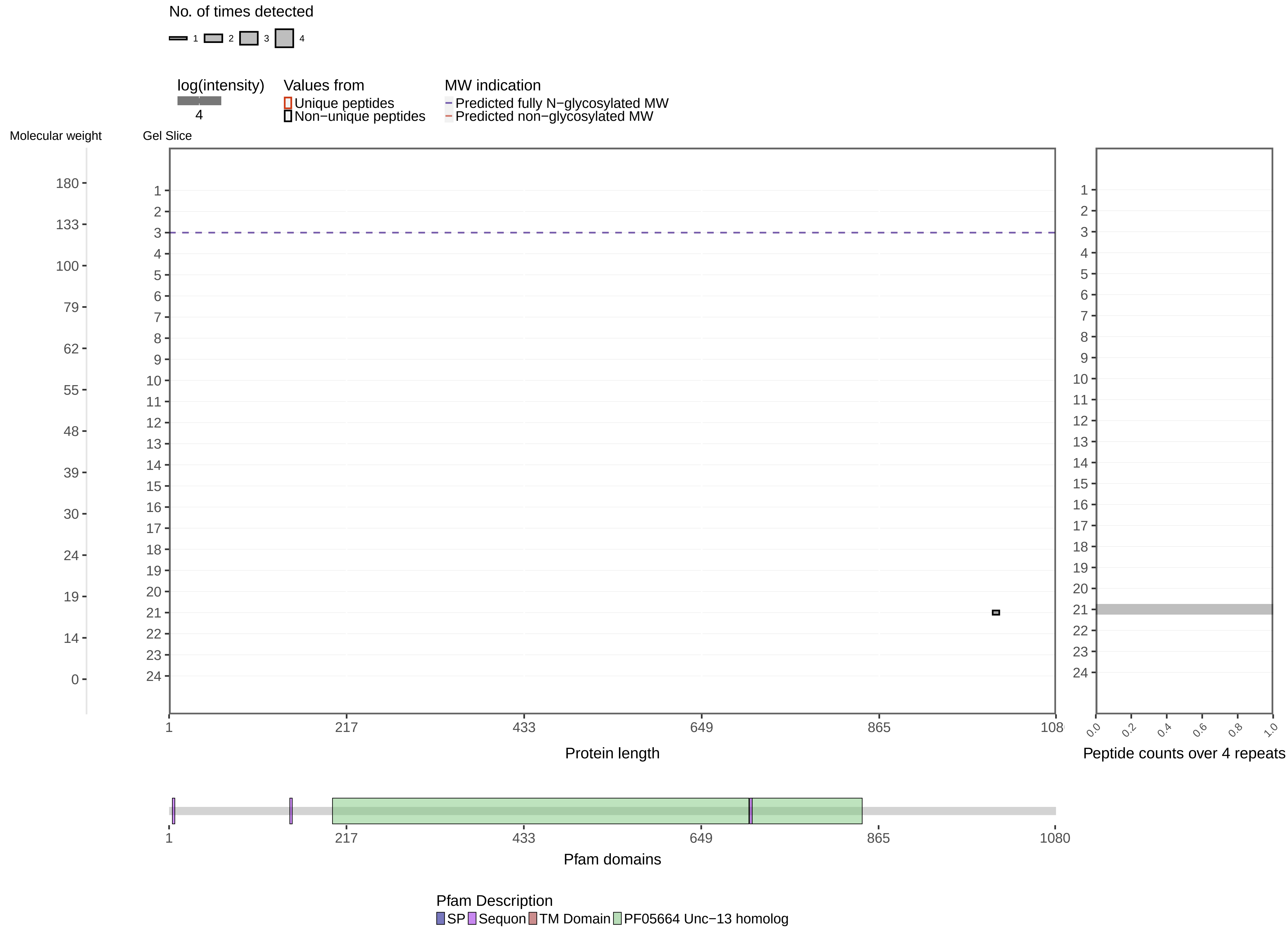


Protein sequence:

.PISKWADKQLGDYHLNYAEGSAMMENTVAVAMLVRRLLLEEPEMAMQYAAISDTEQIEFYVTSSIKNAFTRIIQDVEATAHATNEHPLALLAEHTKELLHRDNTIYMPILSQRHRNAAAVSASILHKLYGIKLRPFLENVEHLTEDTIAVFPAADSLEQYIMQVILSTCADGTSDGYCRKLNVSKRWTPVSPQQRHGSSIVEVYRIVEETVDQFFALKVPMRPGELSSLFRGIDNAFQVYAKTILDKIANKEDVVPPVPILTRYSREHGIKAFVKKELKDTRIPDVVKSVEIDVVATSTLC

NbL11g17110.1 protein_MAINTENANCE_OF_PSII_UNDER_HIGH_LIGHT_1 [MW = 24.53 kDa]

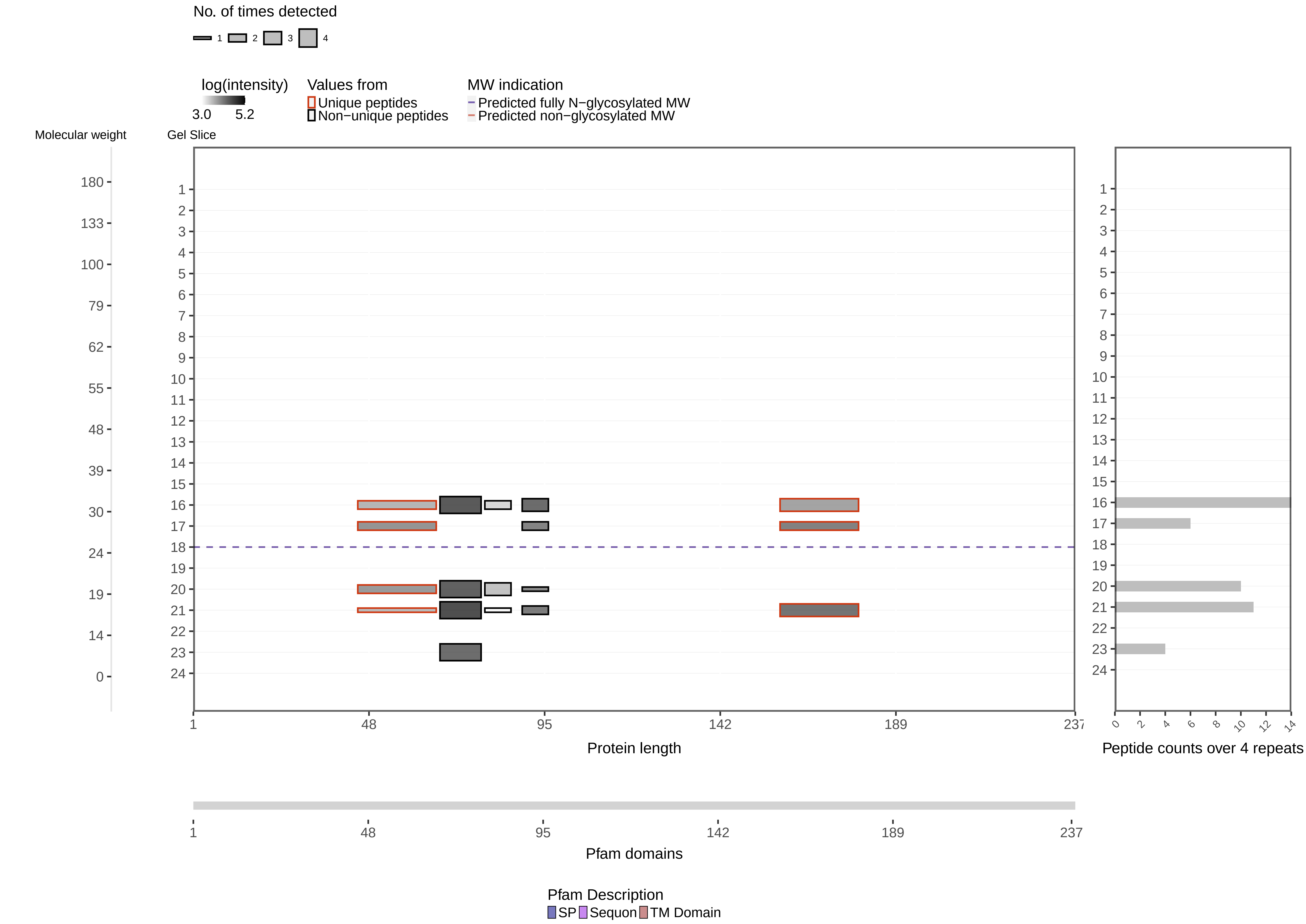


Protein sequence:

NIYSVEPAVYVAESAISSGSAGSTSDGAENTAAIAAGIALISIAAASSILLQVGKNSPPPVQTIEYSGPSLSYYINKFKPAEILQASVAAETEAPTAPEVVQASVAPETEAPNAPEVESSAPQVEVQSEAPQDT*

NbL11g17070.1 dnaJ_homolog_subfamily_C_member_2 [MW = 74.08 kDa]

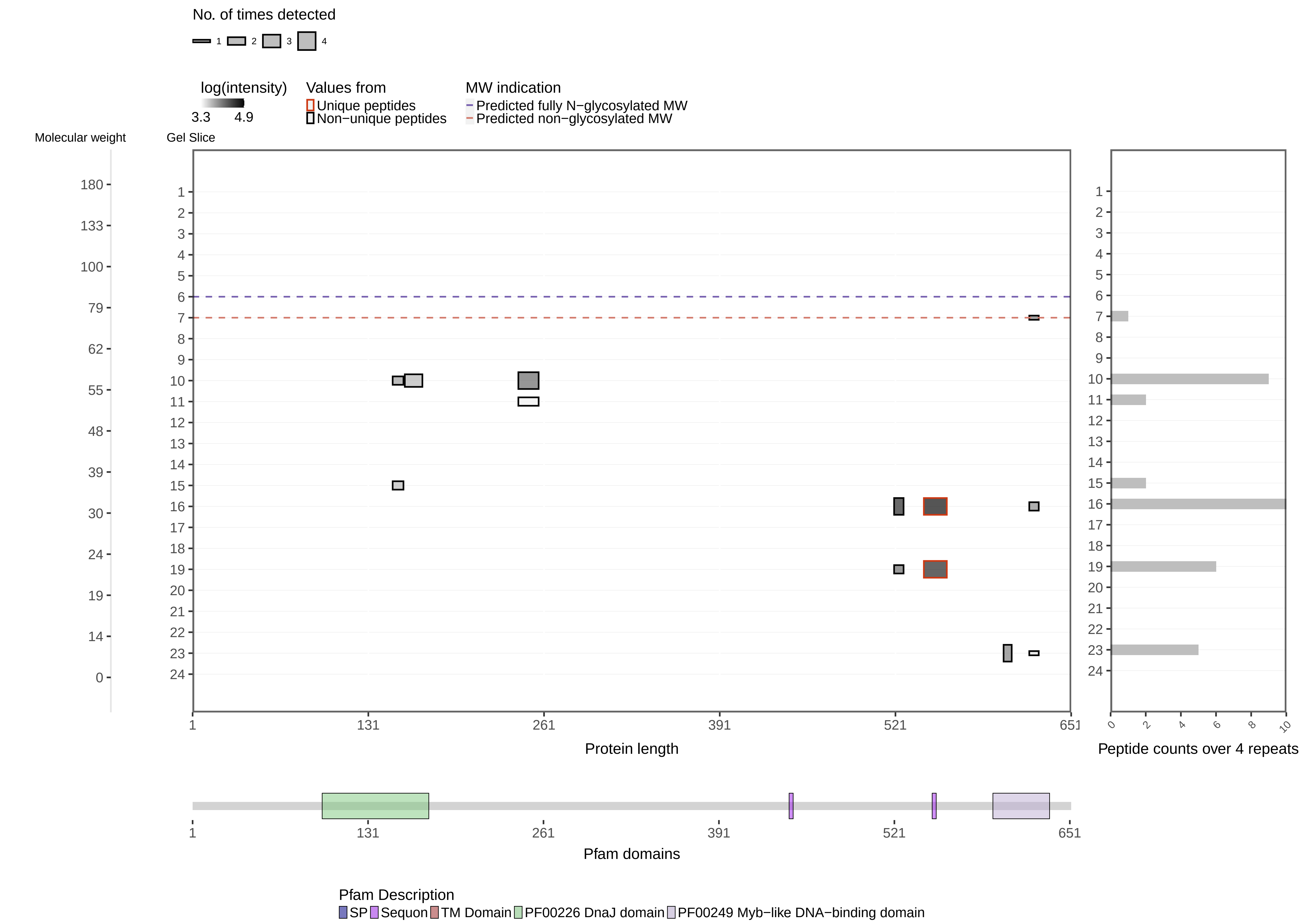


Protein sequence:

### NbL11g17020.1 SUN_domain−containing_protein_3−like [MW = 50.55 kDa]

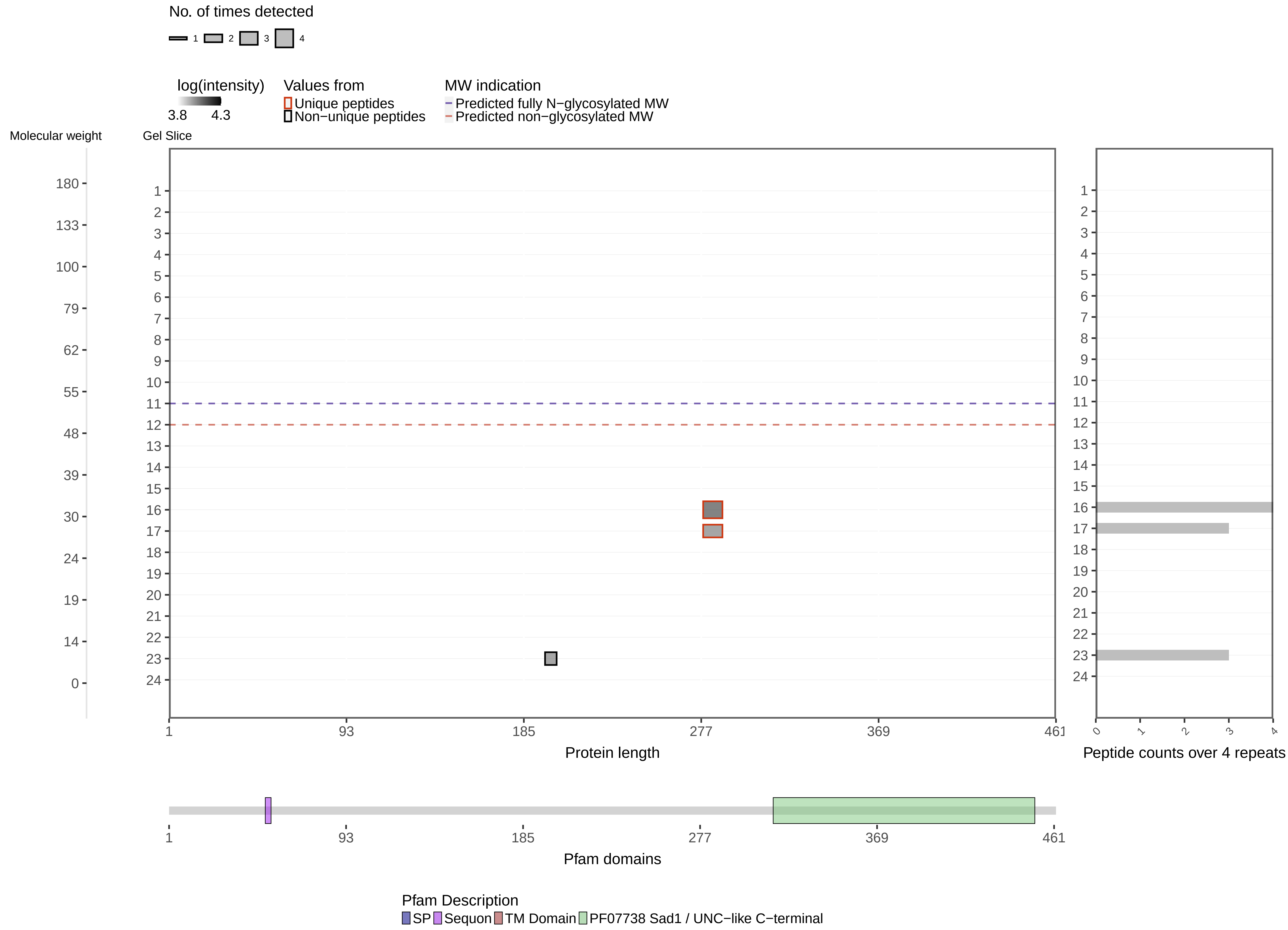


Protein sequence:

SEFREKDWVSKDELTHLFEEYRKSKGDTEADDKSLDEFRAYAREVVEKEIEKHAADGLGRVDYALASGGGRIVKHSEPFITKSGGGDVWSLLNRNAVSNDAQKILTPSFGEPGQCFPLKGDRGFVQIRLRTSIIPEAVTLEHVAKSVAYDRSSAPKNCRISGWLHDKKETDLAADSEKMFLLTEFIYDLDKSNAQTFNVLESAGSSVIDTIRFDFTSNHGMPHTCIYRLRVHGREPNSLVMQS*

NbL11g16990.1 peter_Pan−like_protein [MW = 39.71 kDa]

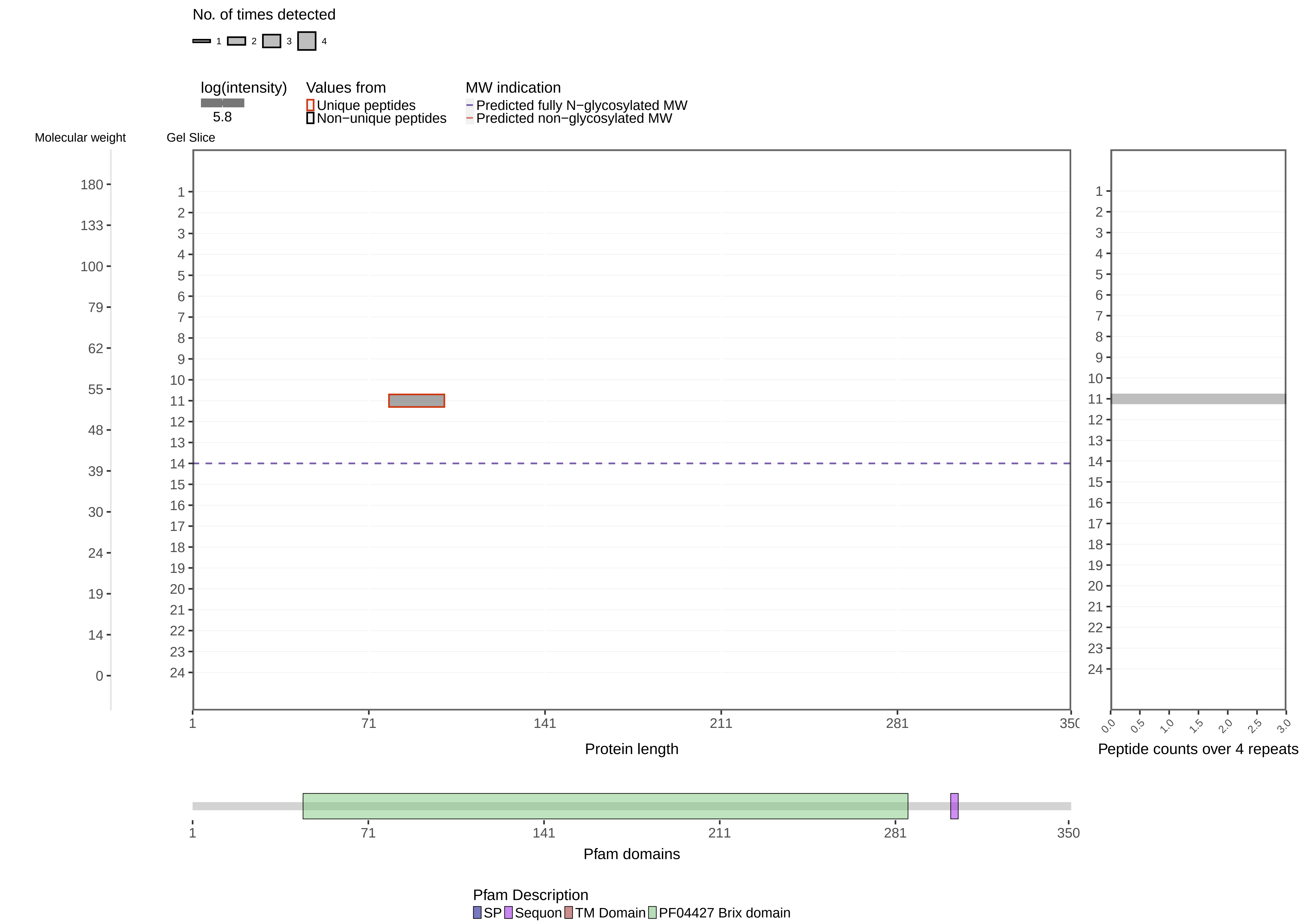


Protein sequence:

MFQNIFPAIDINTVKLPSCQRIVLLNYNKETKLIDLRHYSIRLQPVGVSRRIRKFVQNHQVPDLRSLQDVSDFVTKAGYGSESEPDDEAATVSLSSDIGRVNRASTKSAVKLQEIGPRMTLQLVKIEEGLCTGRVIFSEYGNTSAKKLGNEEEDEQESEEEMDTDGQENADEEEQEIEDEEEQEIEQD*

NbL11g16900.1 proline−−tRNA_ligase [MW = 61 kDa]

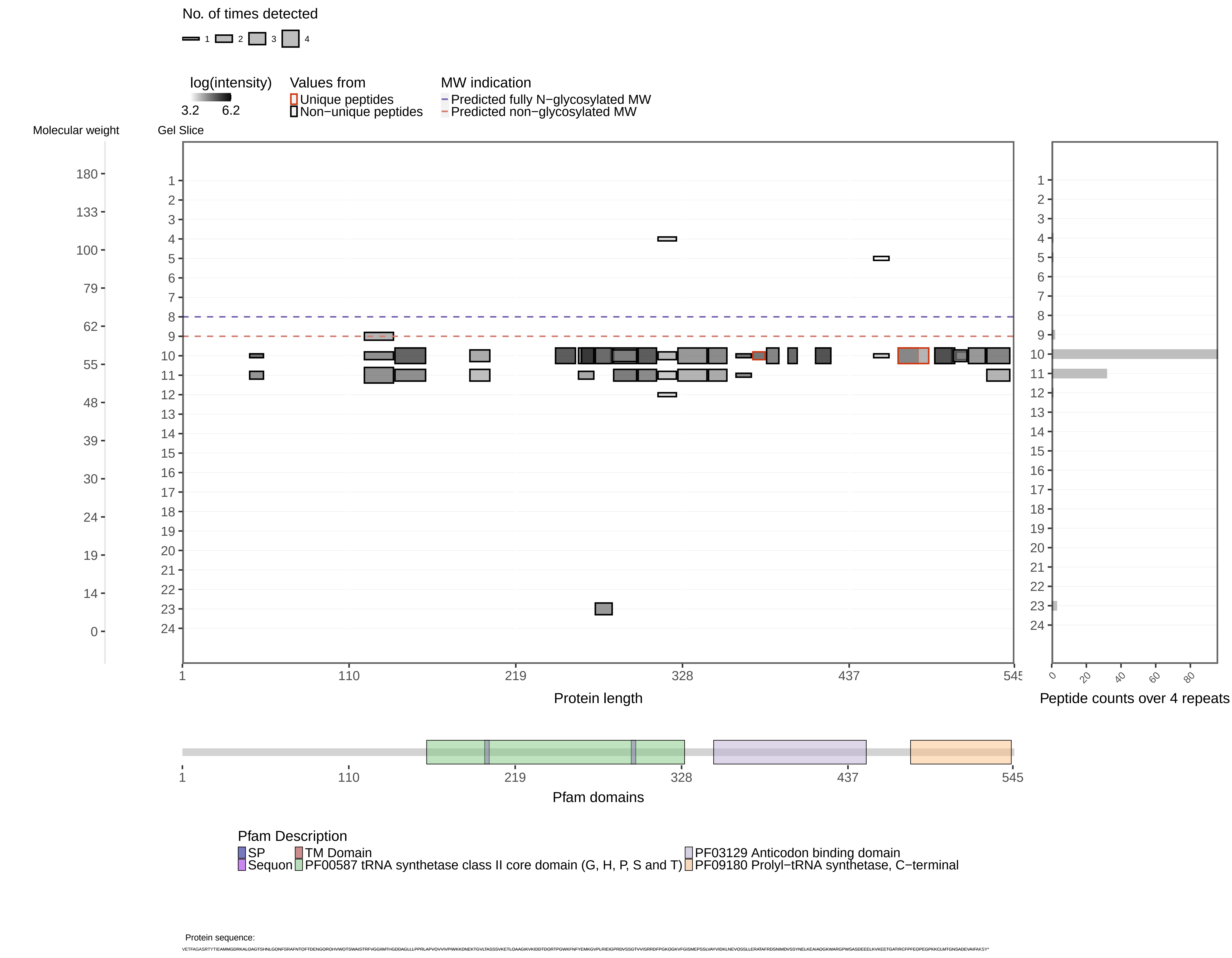

NbL11g16880.1 DNA_gyrase_subunit_A_chloroplastic/mitochondrial [MW = 104.82 kDa]

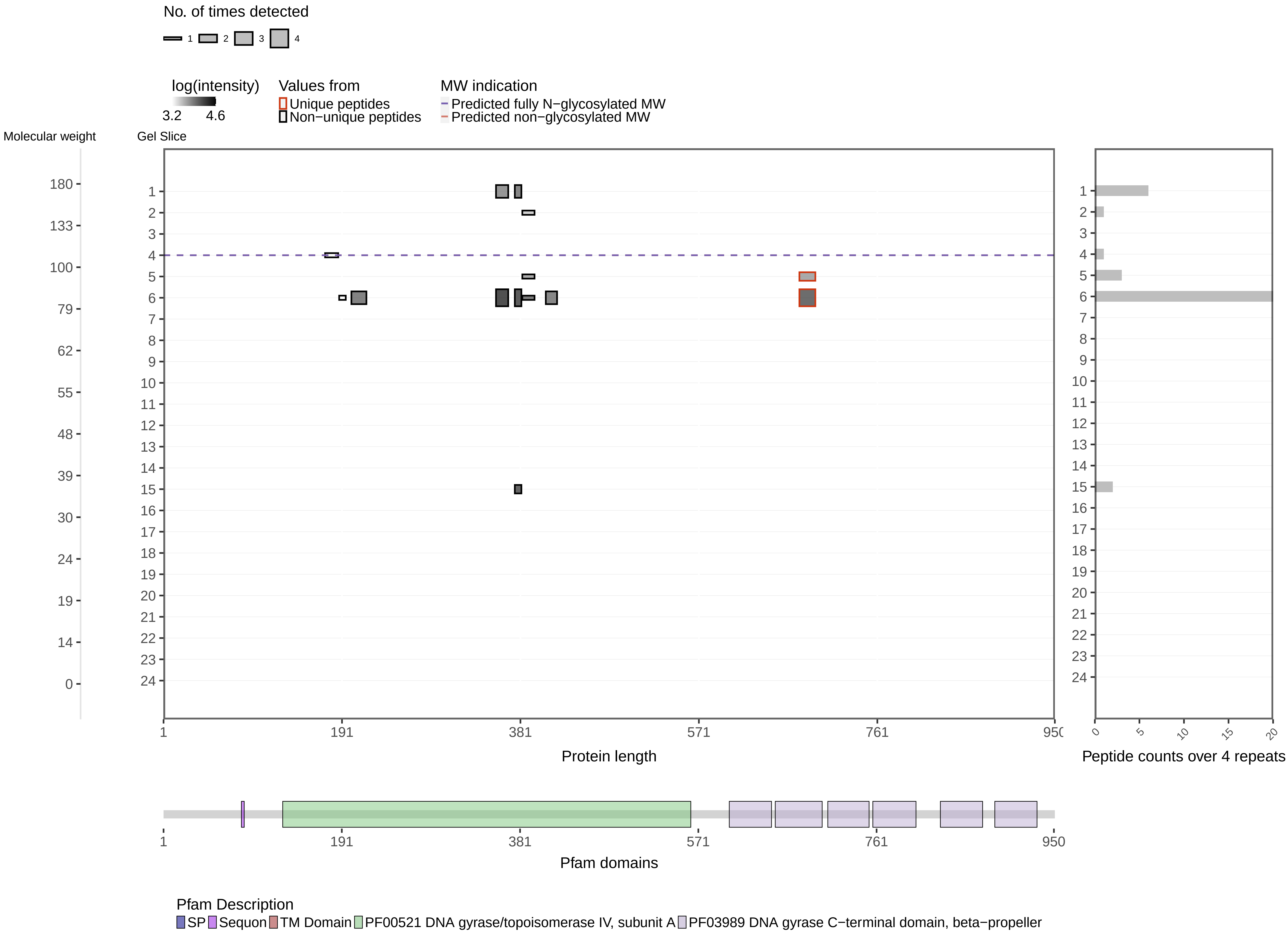


Protein sequence:

VERRAKFKLSQAQERNHIVEGIIVGLDNLDEVINTIRKASSNALAAASLRKEFELSEKQAEAILDISLRRLTALERNKFVEEGKSLRTQISKLEELLSSKKQILQLIEEEAIEIKNKFFTPRRSMLEDTDSGEVEDIDVIPNEEMLLAISEKGYVKRMKSDTFNLQNRGTIGKSVGKLRVNDAMSDFLVCRAHDKVLYFSDKGTVYSSPAYKIPECSRTAAGTPLVQILSLSDGERITSIIPVSEFAGDQYLVMLTVNGYIKKVSLNYFASIRCTGIIAIQLVPGDELKWVKCCSNNDFVAMA

### NbL11g16800.1 beta−glucosidase_BoGH3B−like [MW − SP = 69.53 kDa]

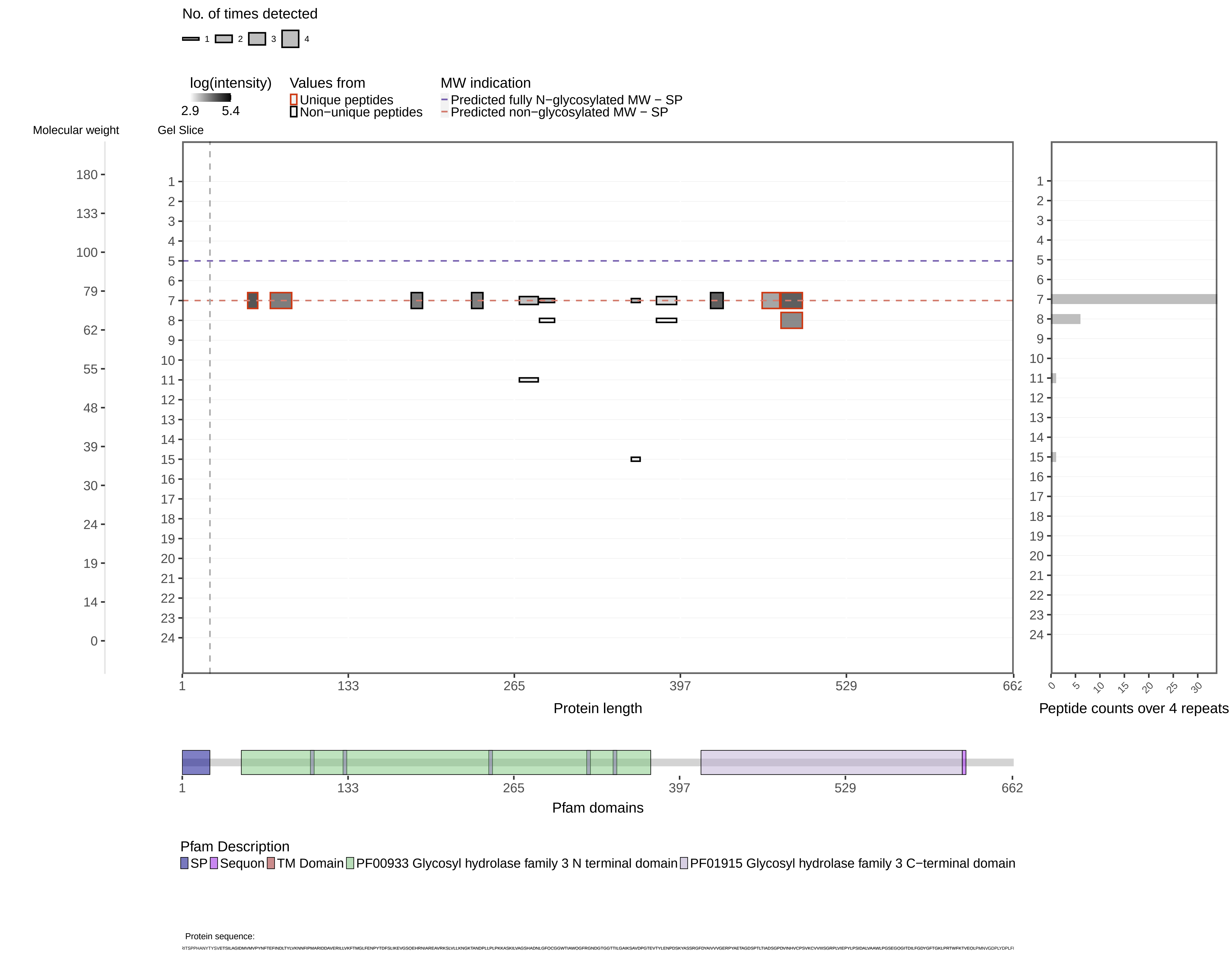

### NbL11g16470.1 PITH_domain−containing_protein_At3g04780−like [MW = 19.66 kDa]

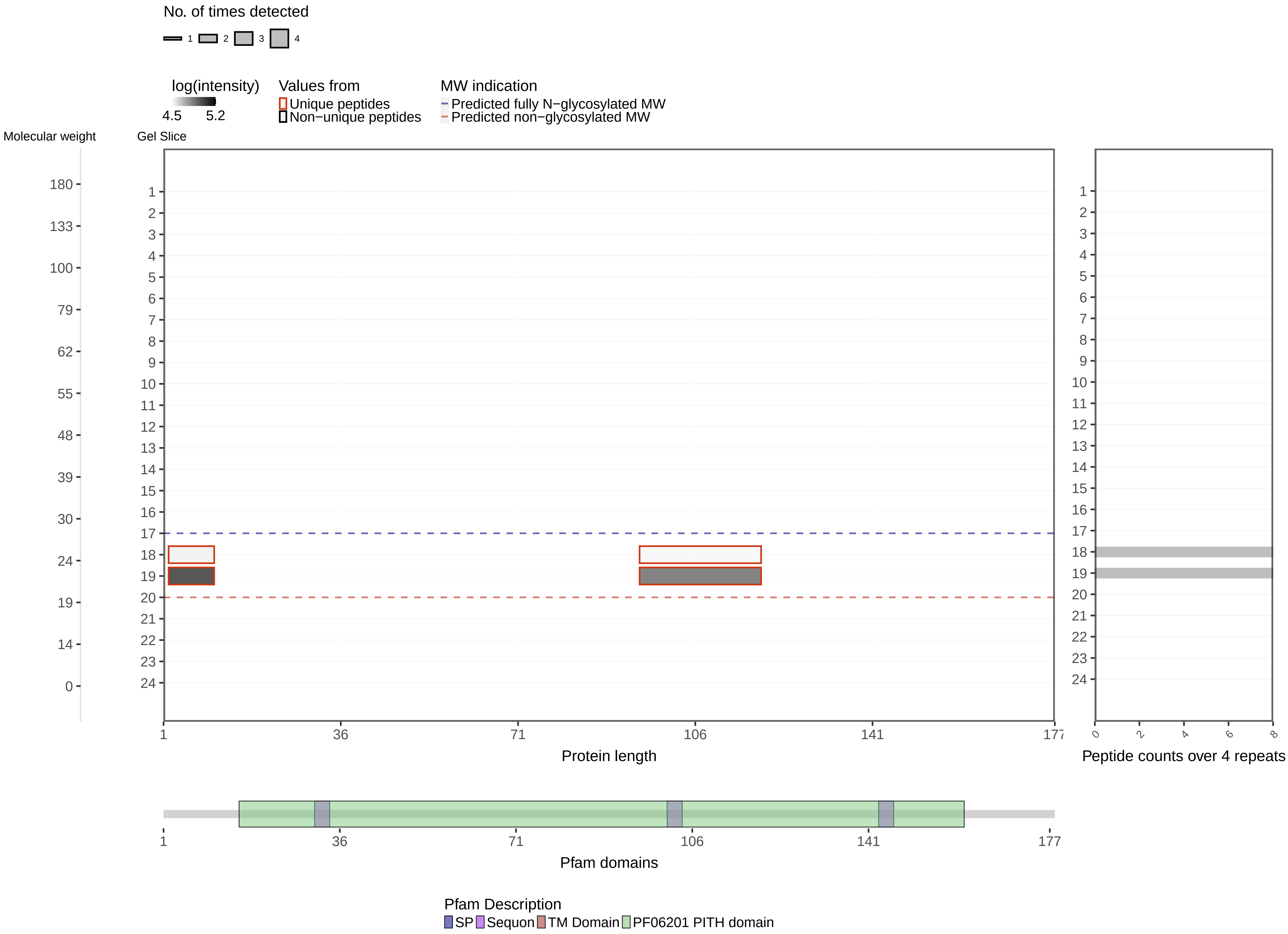


Protein sequence:

VIKGPEEEGPKTVKLFSNKDHMGFSNVSDYPPSDTAVLSEDNLKGKPVILKYVKFQNVRSVTIFIEDNQSDSELTRVQKIVLYGTTVETTDMKGLKKIEDH*

### NbL11g16460.1 probable_ribose−5−phosphate_isomerase_3_chloroplastic [MW = 30.42 kDa]

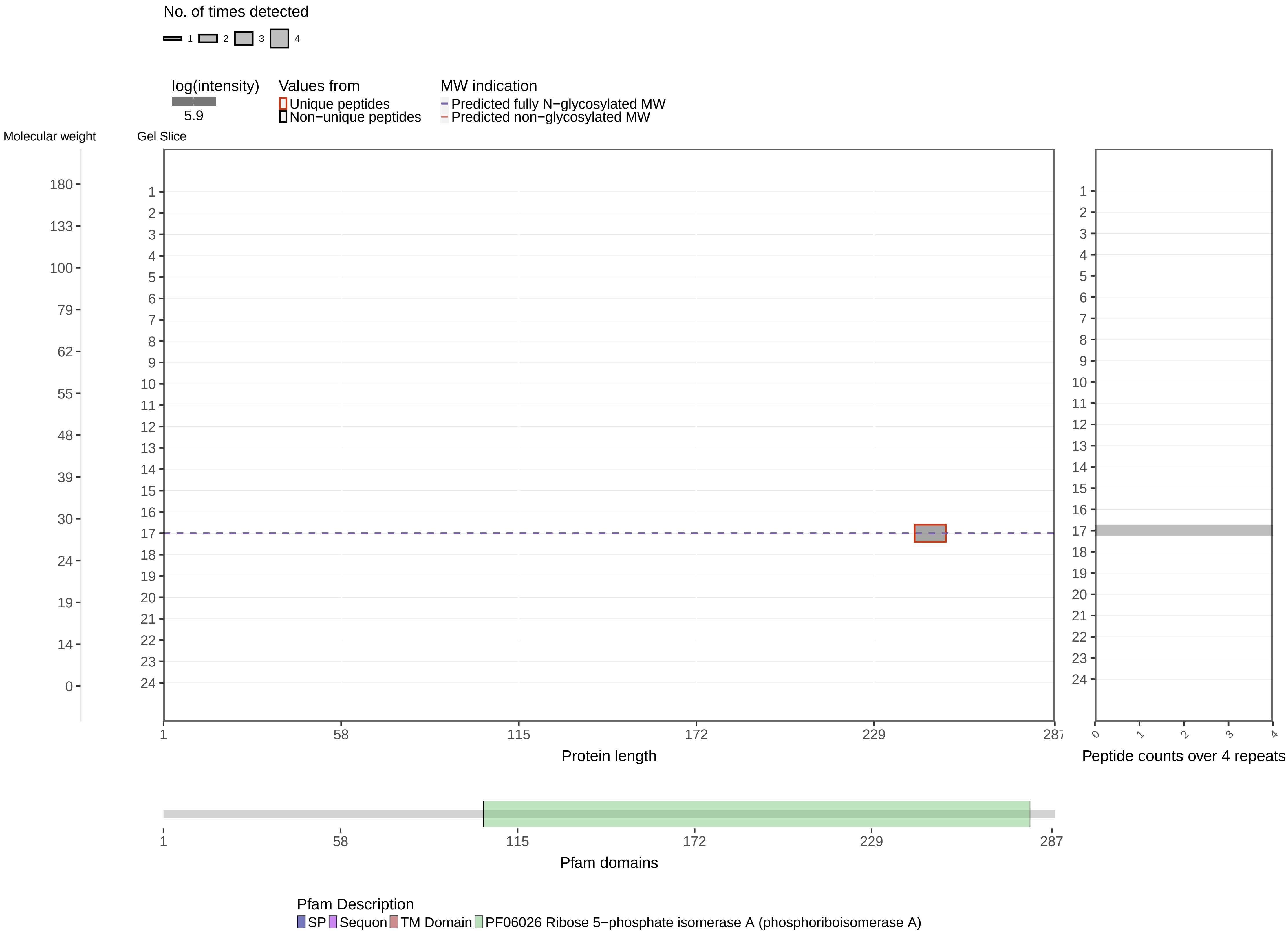


Protein sequence:

DLAIDGADEVDPNLDLVKGRGGALLREKMVEAASDKFVVVVDDSKLVSGLGGSGLAMPVEVVQFCWKYNLVRLQELFKAEGVDAKLRVDGNGKPYVTDNSNYIVDLYFKTPIRDSAVAGKEIAALEGVVEHGLFLDMTTAVIIAGKEGVSVQSK*

NbL11g16430.1 annexin_D4−like [MW = 40.12 kDa]

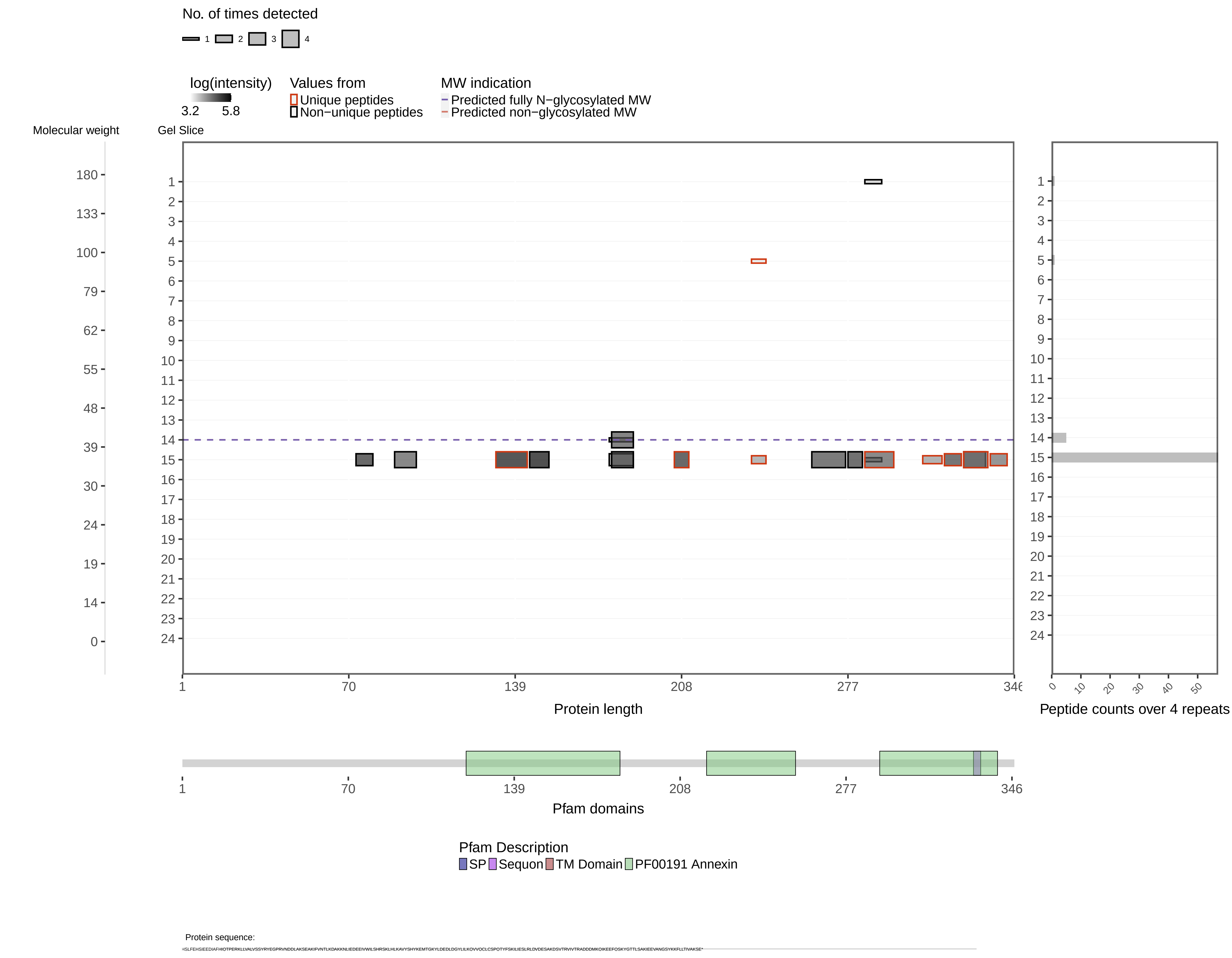

### NbL11g16310.1 phototropin−2−like [MW = 107.81 kDa]

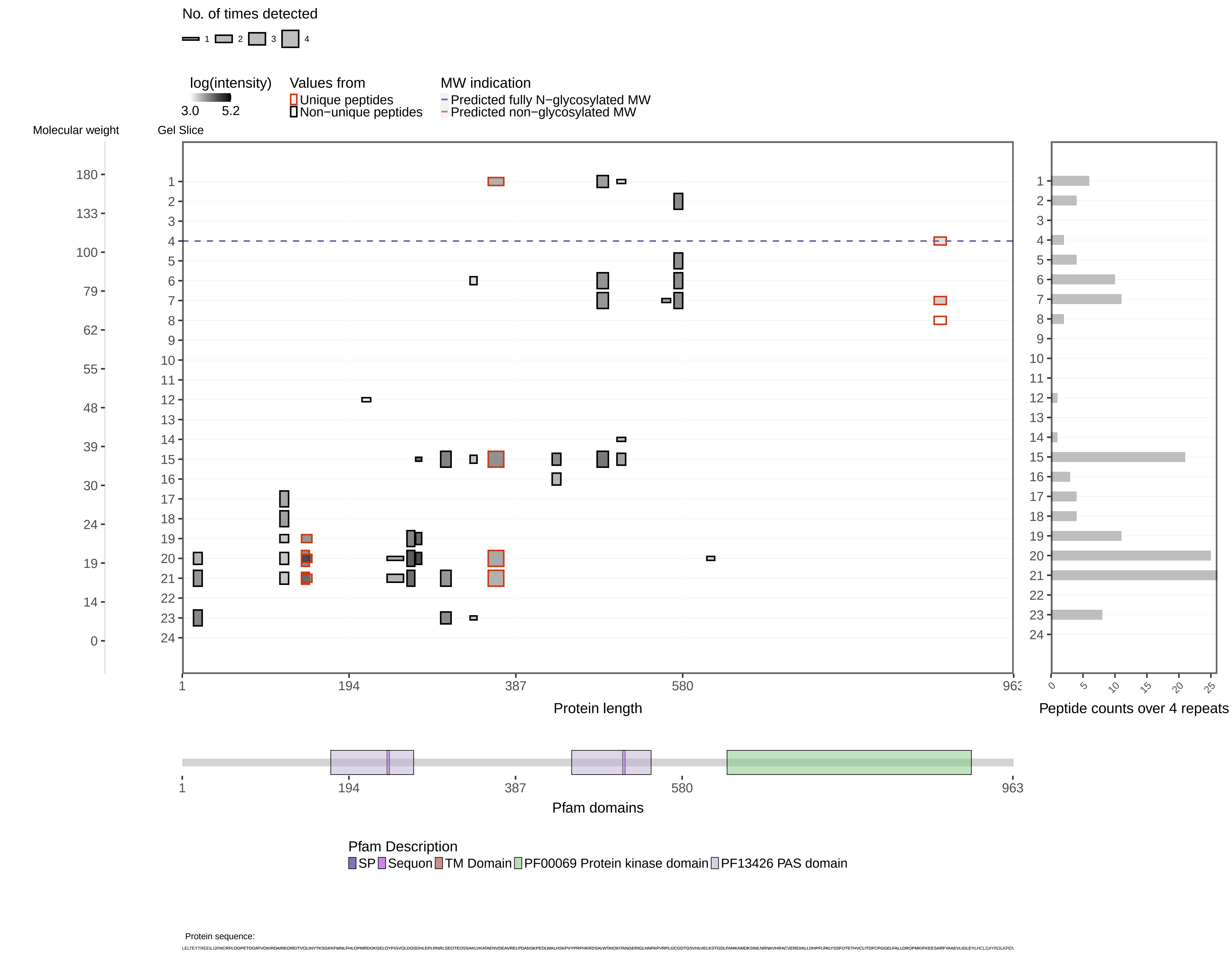

### NbL11g16300.1 DNA_damage−repair/toleration_protein_DRT102 [MW = 33.8 kDa]

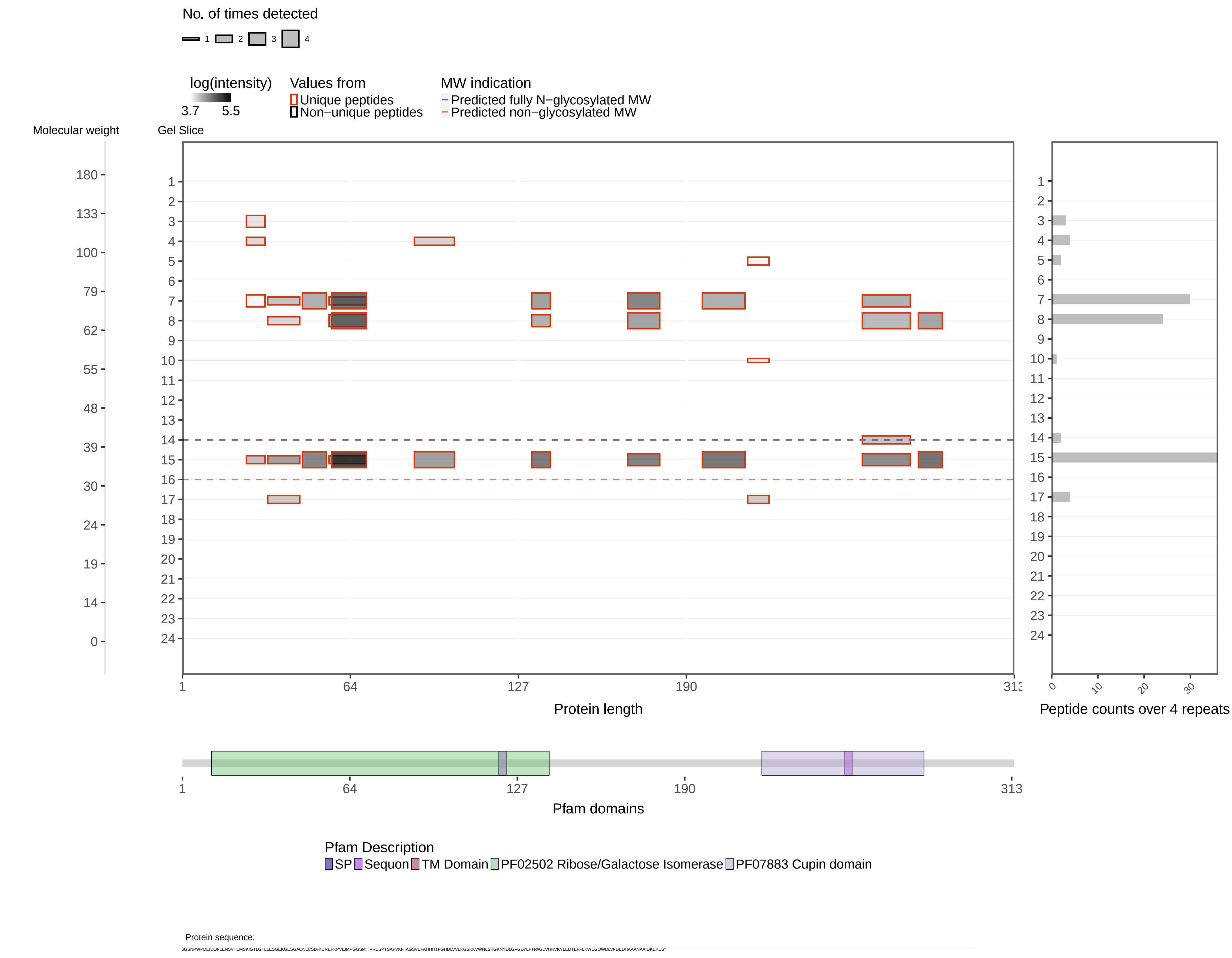

### NbL11g16210.1 rubredoxin−like [MW = 22.73 kDa]

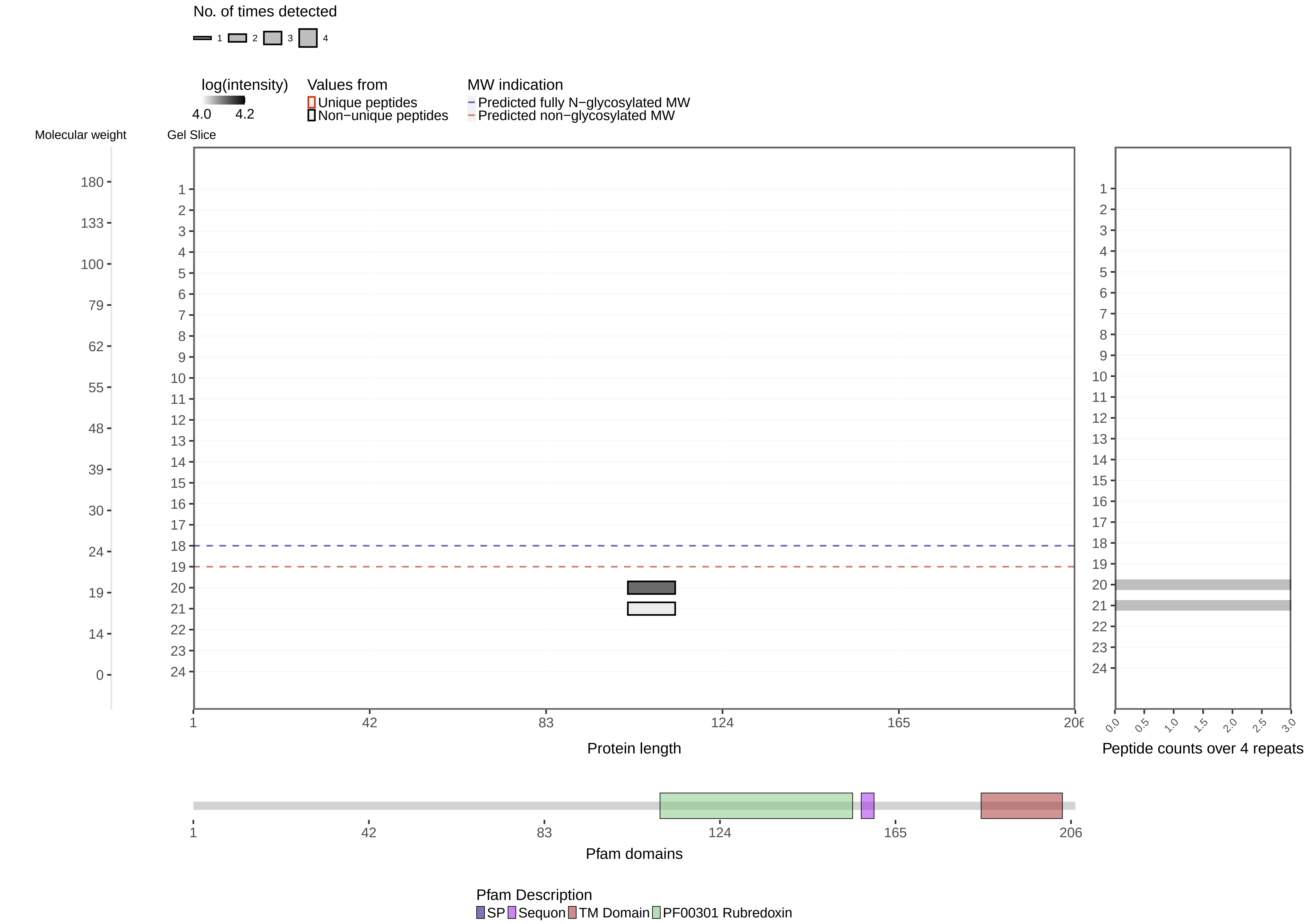


Protein sequence:

KNFDKRRLEEKFAVLNTGIYECRSCGFKYNEAAGDPSYPIPPGLPFDQLPSDWRCPTCGAAKSFFDNKSVEIAGFAQNQQYGFGGNTLTSGQKALLIYGGLLLGFVFFLSGYFLQ*

### NbL11g16160.1 translation_initiation_factor_IF−2−like [MW = 52.65 kDa]

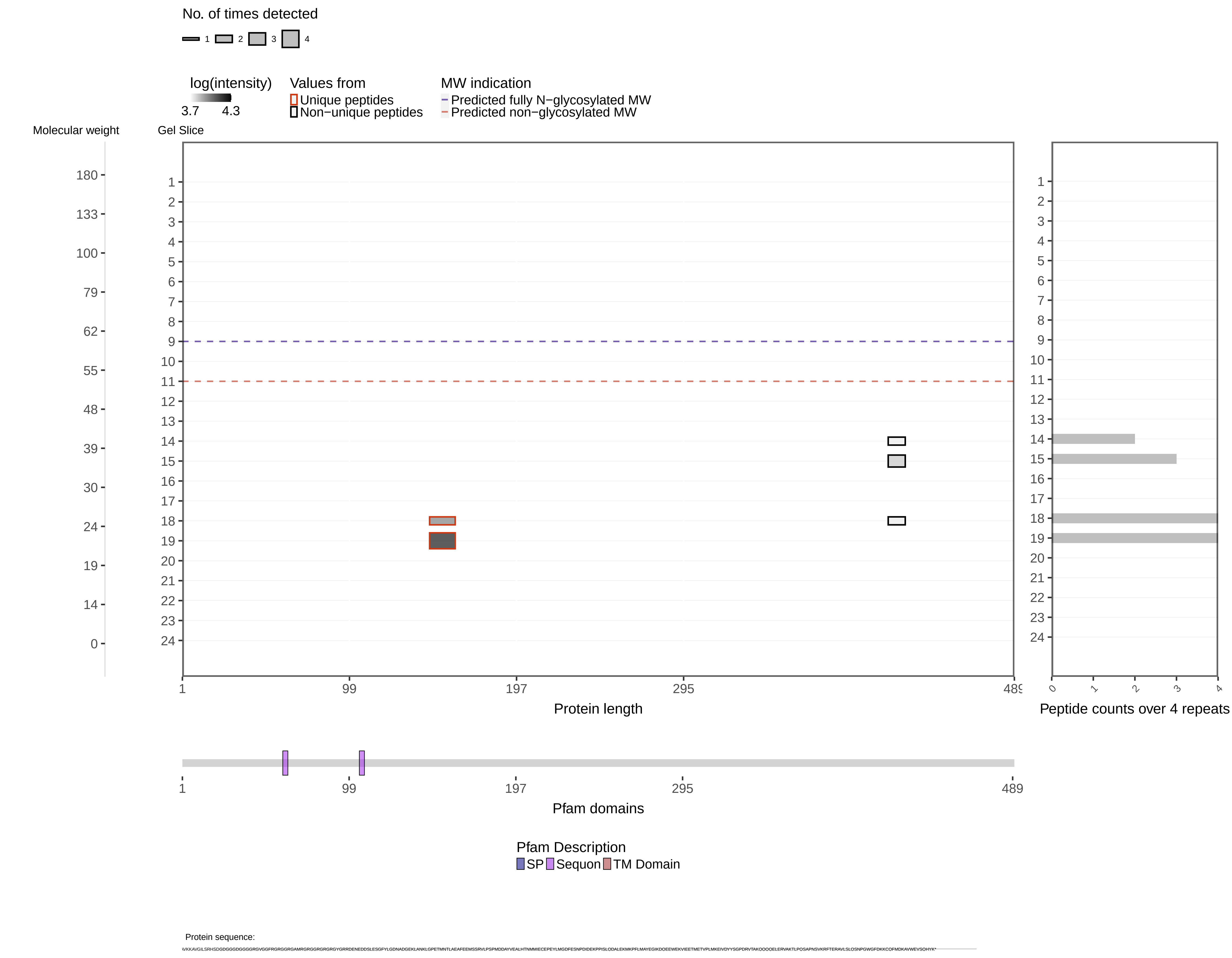

NbL11g16030.1 4−hydroxy−tetrahydrodipicolinate_reductase_1_chloroplastic−like [MW = 36.54 kDa]

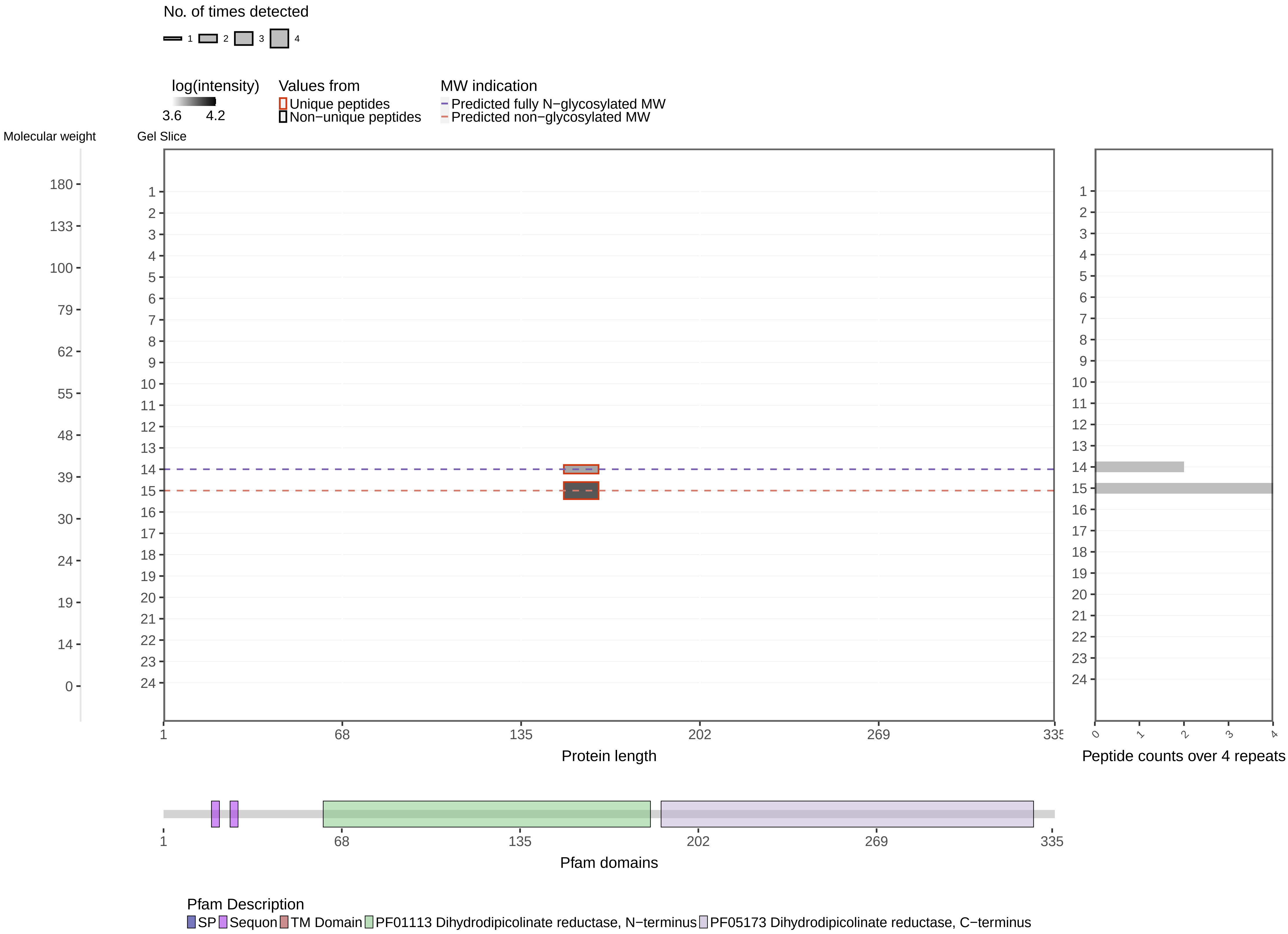


Protein sequence:

FVLGTTGGDRERLYKTVSDSKVYAVISPQMGKQVVAFLAAMEIMAEQFPGAFSGYTLQVMESHQASKLDTSGTAKAVISCFQKLGVSFDLDEVQLIRDPKQQVEMVGVPEEHLLGHAFHMYHLTSPDGTVSFEFQHNVCGRSIYAEGTVDAILFLAKKVKLKEDKRIYDMIDVLREGNMR*

NbL11g15870.1 tryptophan_synthase_alpha_chain−like [MW = 33.81 kDa]

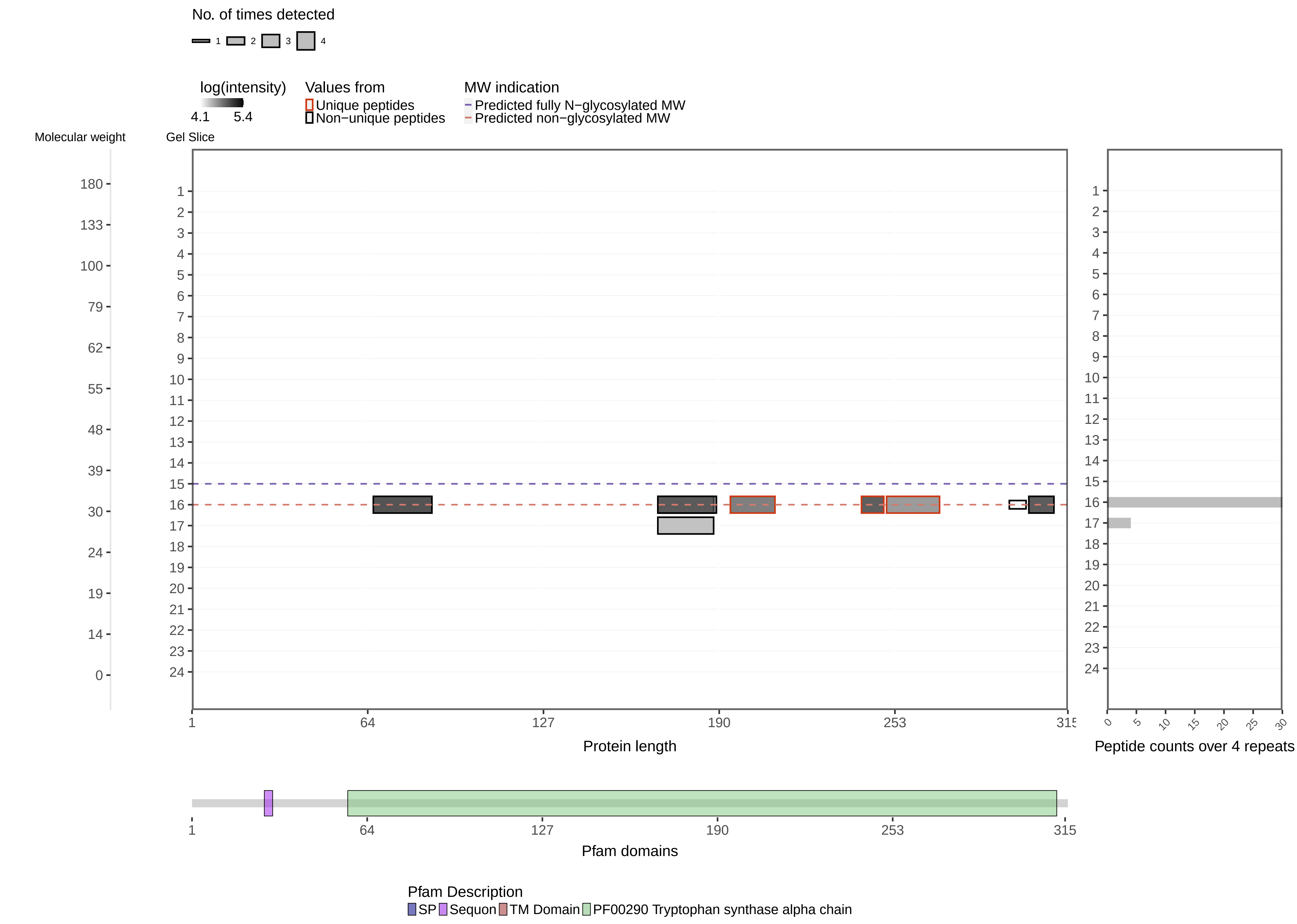


Protein sequence:

CPIALFTYYNPILKRGTEKFMATVRDAGVHGLVVPDVPLEETEILRKEAARHNIELVLLTTPTTPTVRMKAITEASEGFVYLVSSIGVTGARESVSSKVOSLLIDIKEATSKPVAVGFGISKPEHVKOVAGWGADGVIIGSAIVRILGEAKSPEEGLKELEVFATSLKSALS*

### NbL11g15800.1 glutathione_synthetase_chloroplastic [MW = 61.62 kDa]

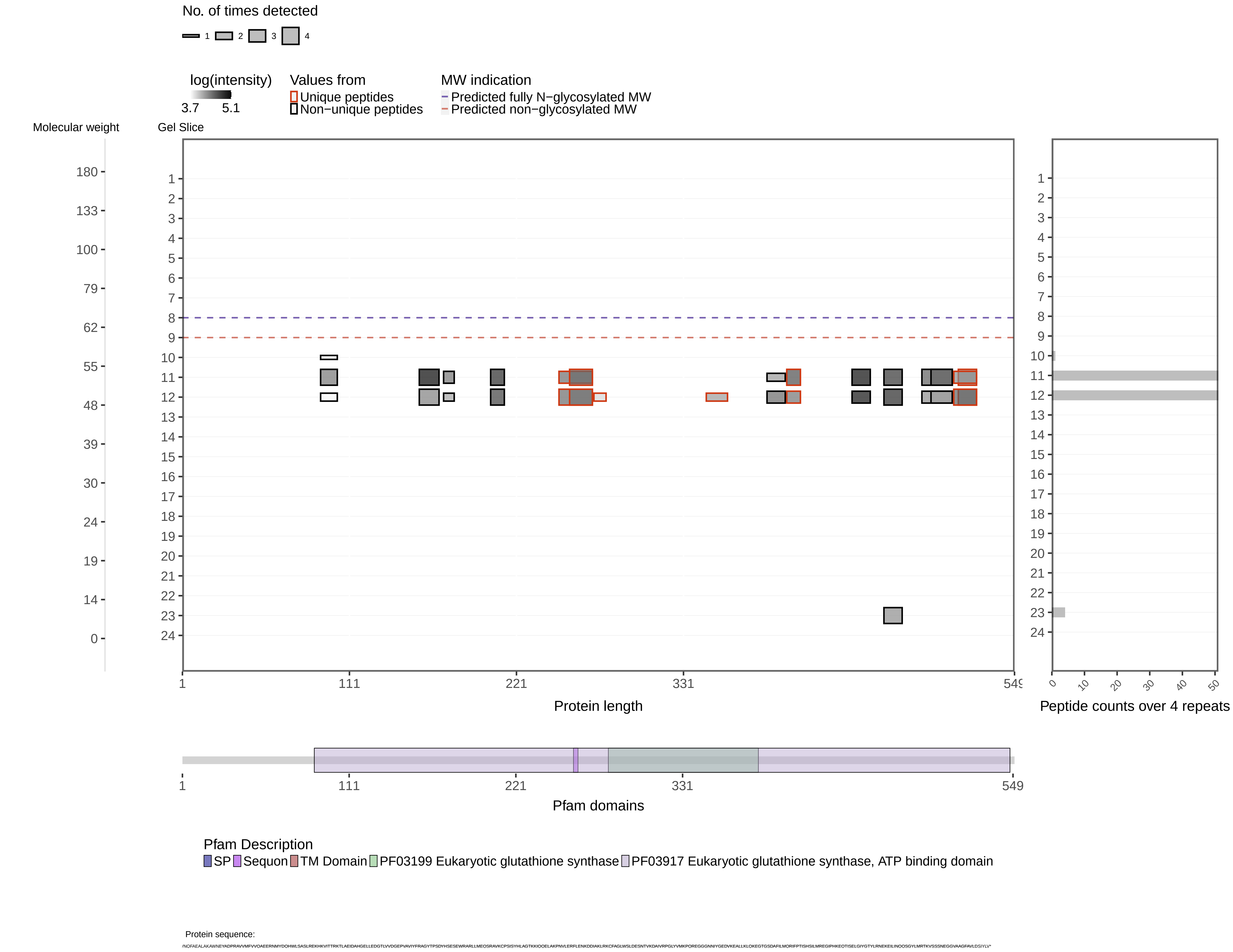

### NbL11g15780.1 UPF0603_protein_At1g54780_chloroplastic [MW = 31.99 kDa]

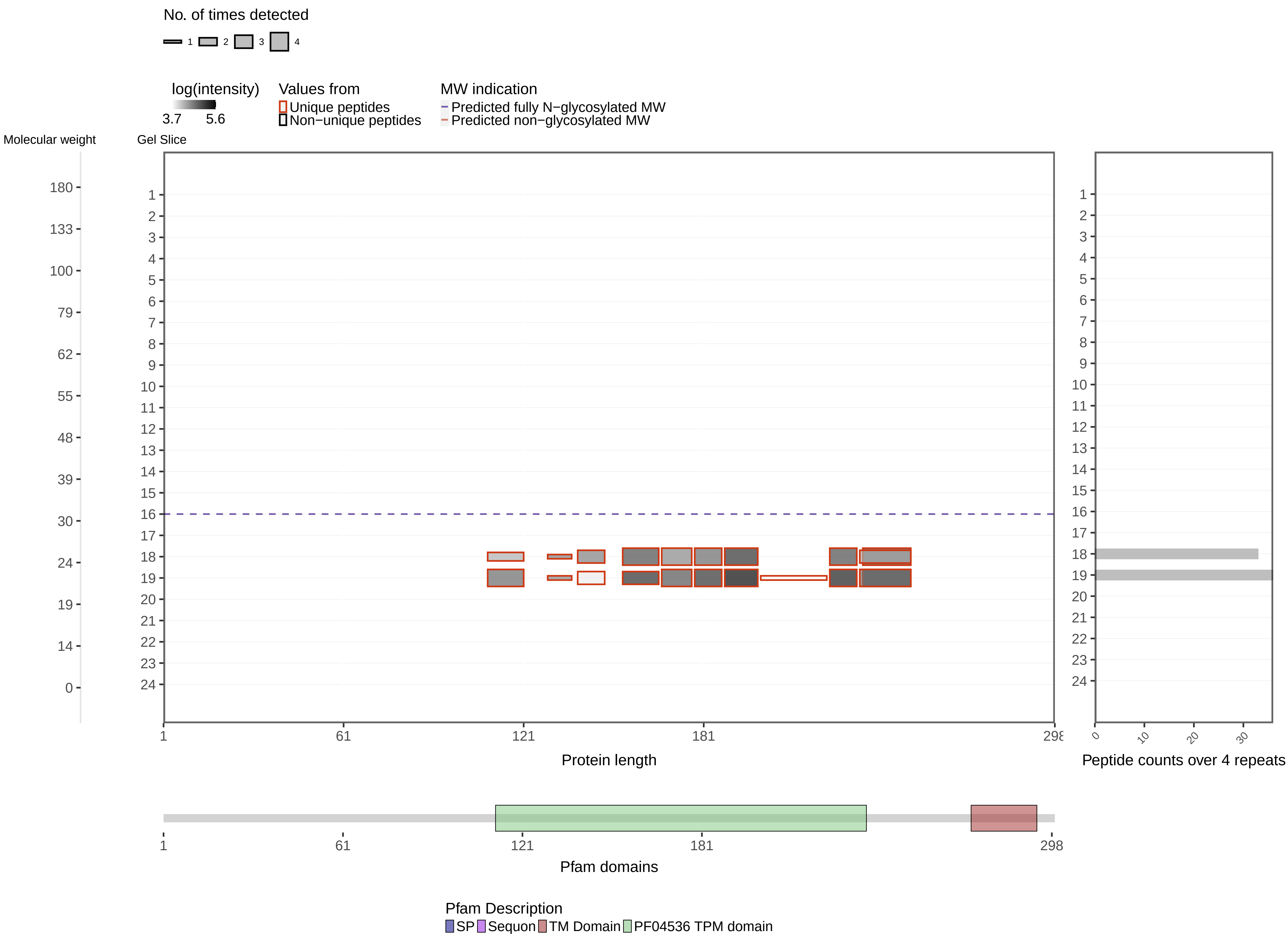


Protein sequence:

RKGFHINFITVRKLTSKADAFEYADTVLEKWYPSVEQGNDKGIVVLVTSQKEGAITGGPDFVKAVGDTVLDAAVSENLPVLATEEKYNEAVYSTAKRLVAAIDGLPDPGGPQLKDNKRESNFKSREETDEKRGQFTLVVGGLLVIAFVVPMAQYYAYVSKK*

NbL11g15760.1 branched−chain−amino−acid_aminotransferase−like_protein_1 [MW = 62.33 kDa]

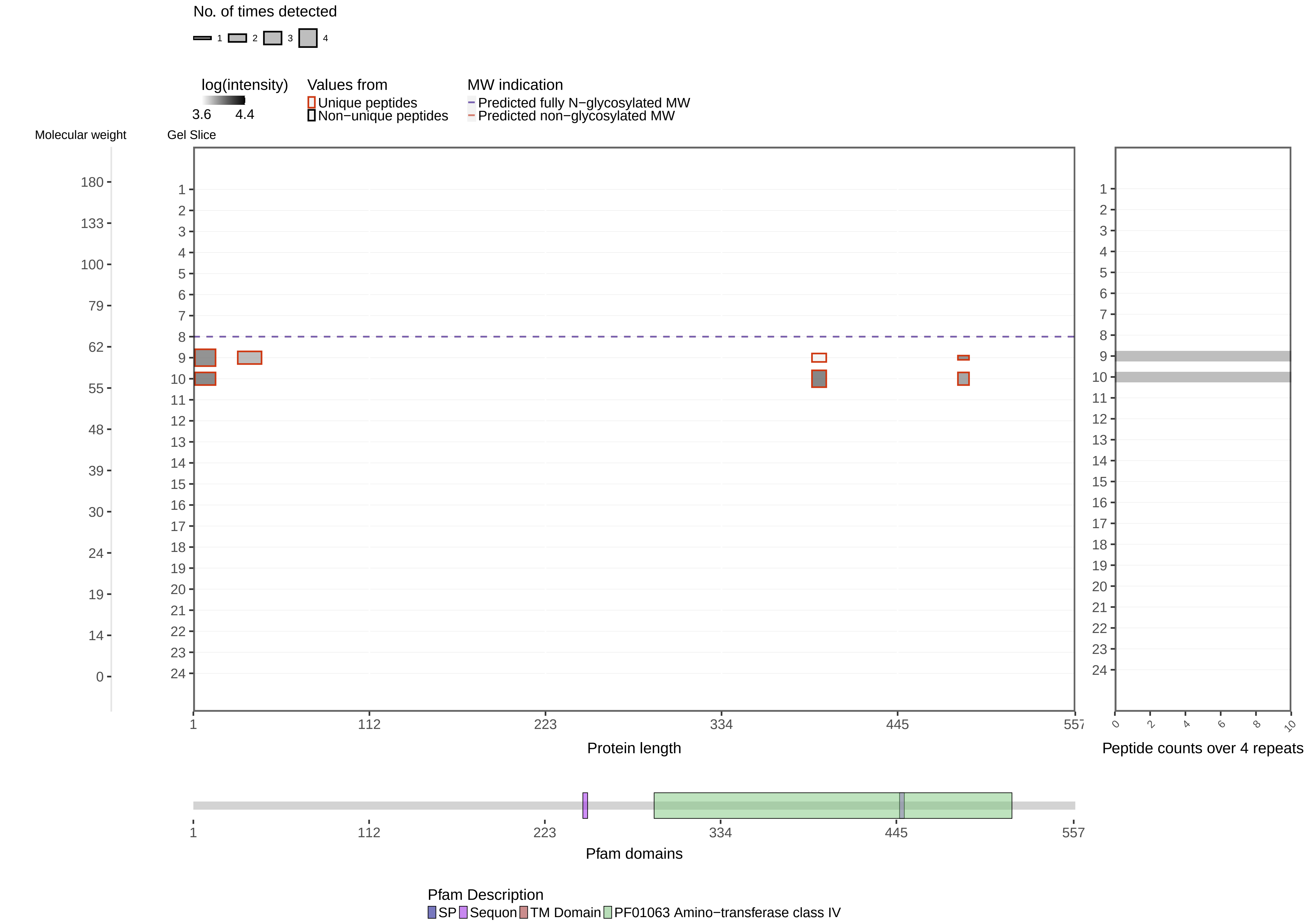


Protein sequence:

AWVGDEIVPRETAKVSVFDSVVQGGDAVWEGLRVYSGKLFKLEEHLDRLFDSAKALAFSNVPTREEVKEAIFKTLLLTNGMYDNAHIRLTLTRGKKVTSGMSPAFNQYGCTLIVLAEWKPPVYDNEKGIILVTATTRRNSPNNLDSKIHHNNLLNNILAKIEGNNAGADDAVMLDKDGYVSETNATNIFLVKKGRVVTPHADYCLPGITRATVMELVLKENLVLEERRISLSEFHTADEVWTTGTMGELSPVVKIDGRIIGDGRVGPVTQRLQNAYKKLAEDSGVSIPTYQKS*

NbL11g15610.1 mevalonate_kinase−like [MW = 40.83 kDa]

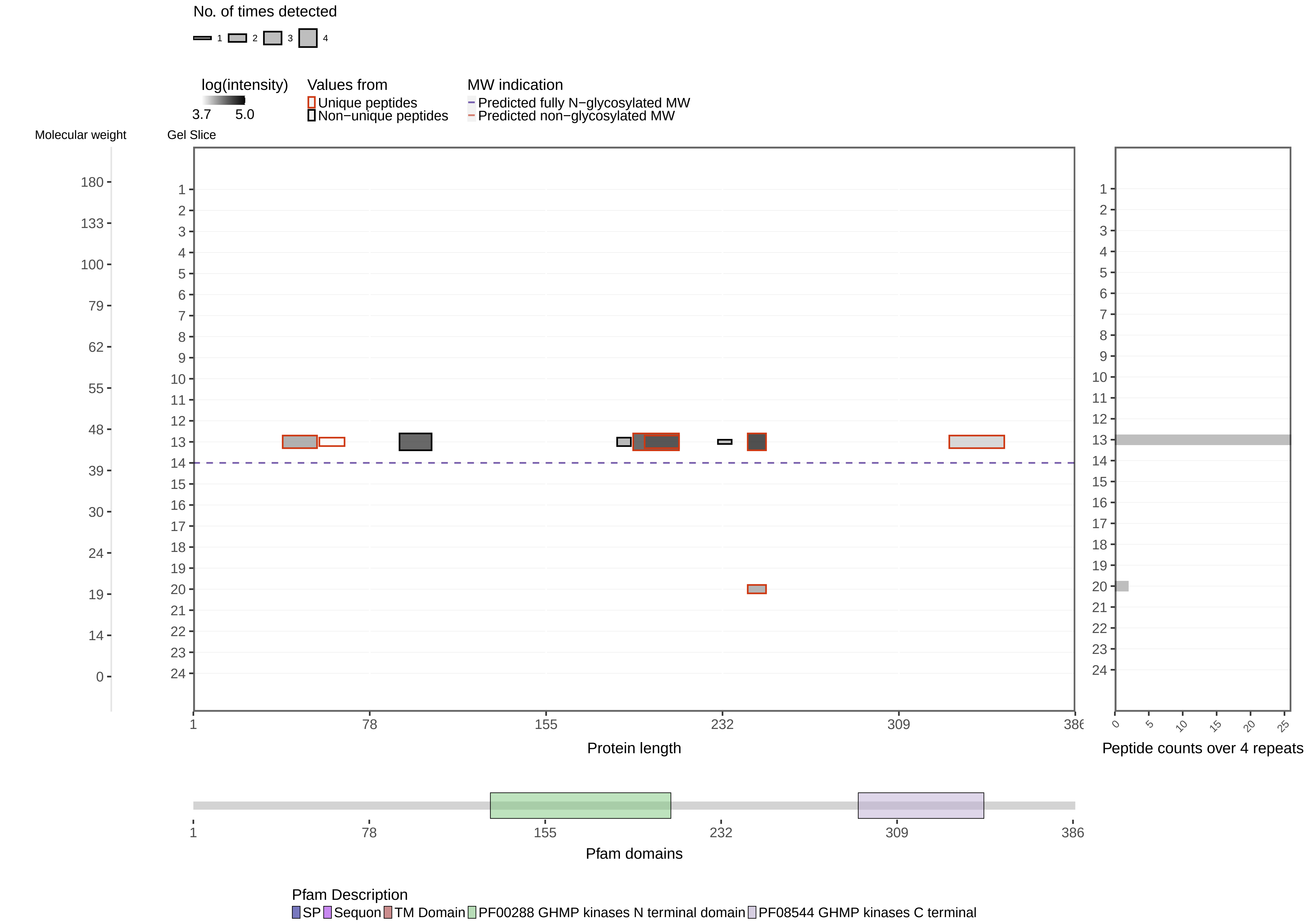


Protein sequence:

LELVNKWAFEGEKIIHGKPSGIDNTVSTYGNLIKFKSGDLTRLKTNMPLKMLITNTKVGRNTKALVASVSERTLRHPTAMASVFTAVDSISSEVAAIIQSPVPDDLAITEKEEKLEELMEMNQGLLQCMGVSHASIETVLRTTLKYKLSSKLTGAGGGGCVLTLLPTLLSGTVVDKVIAELETCGFQCLLAGIGGNGVEILFSGTS*

### NbL11g15580.1 serine−−tRNA_ligase [MW = 51.09 kDa]

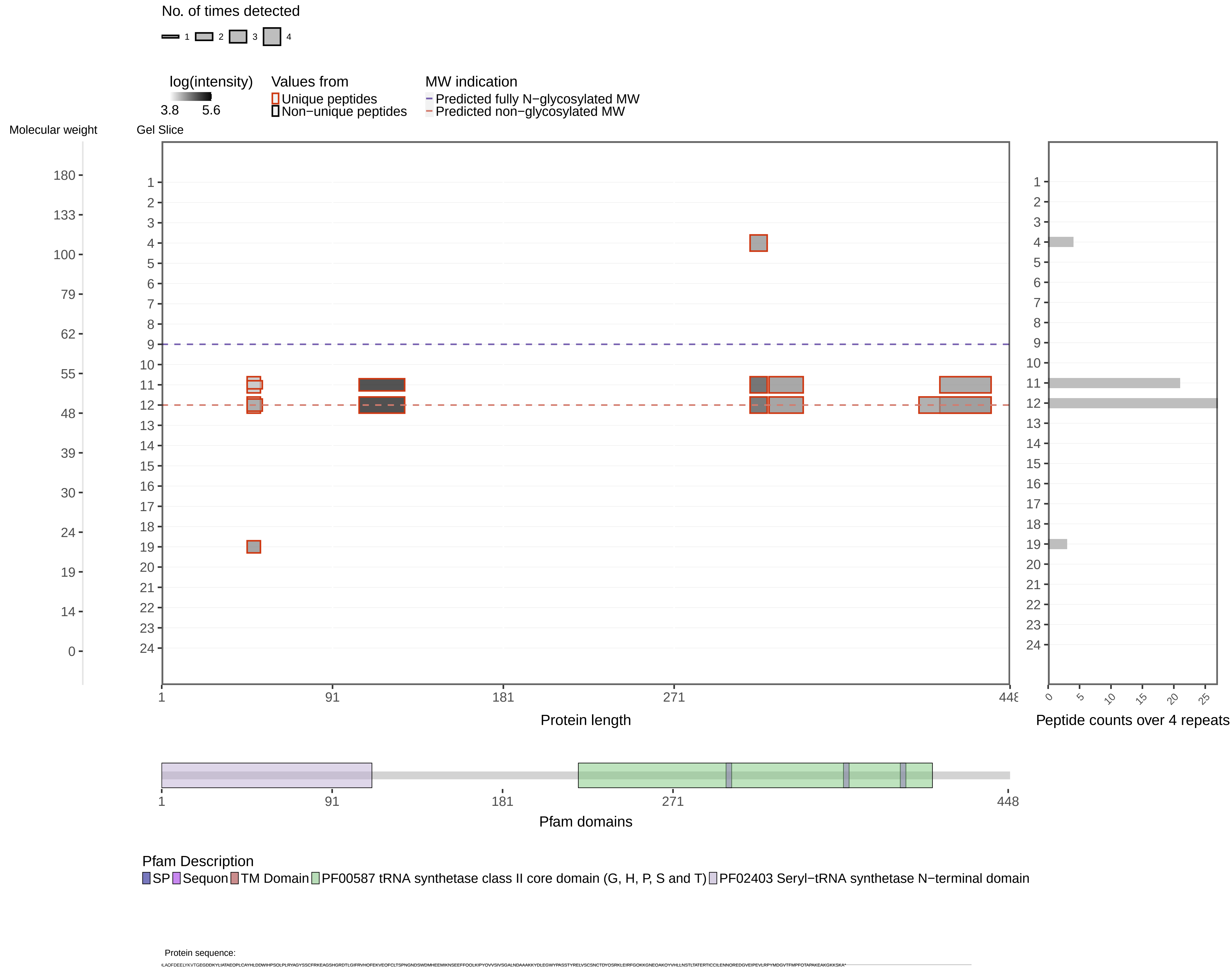

NbL11g15450.1 acetylajmalan_esterase−like [MW − SP = 39.6 kDa]

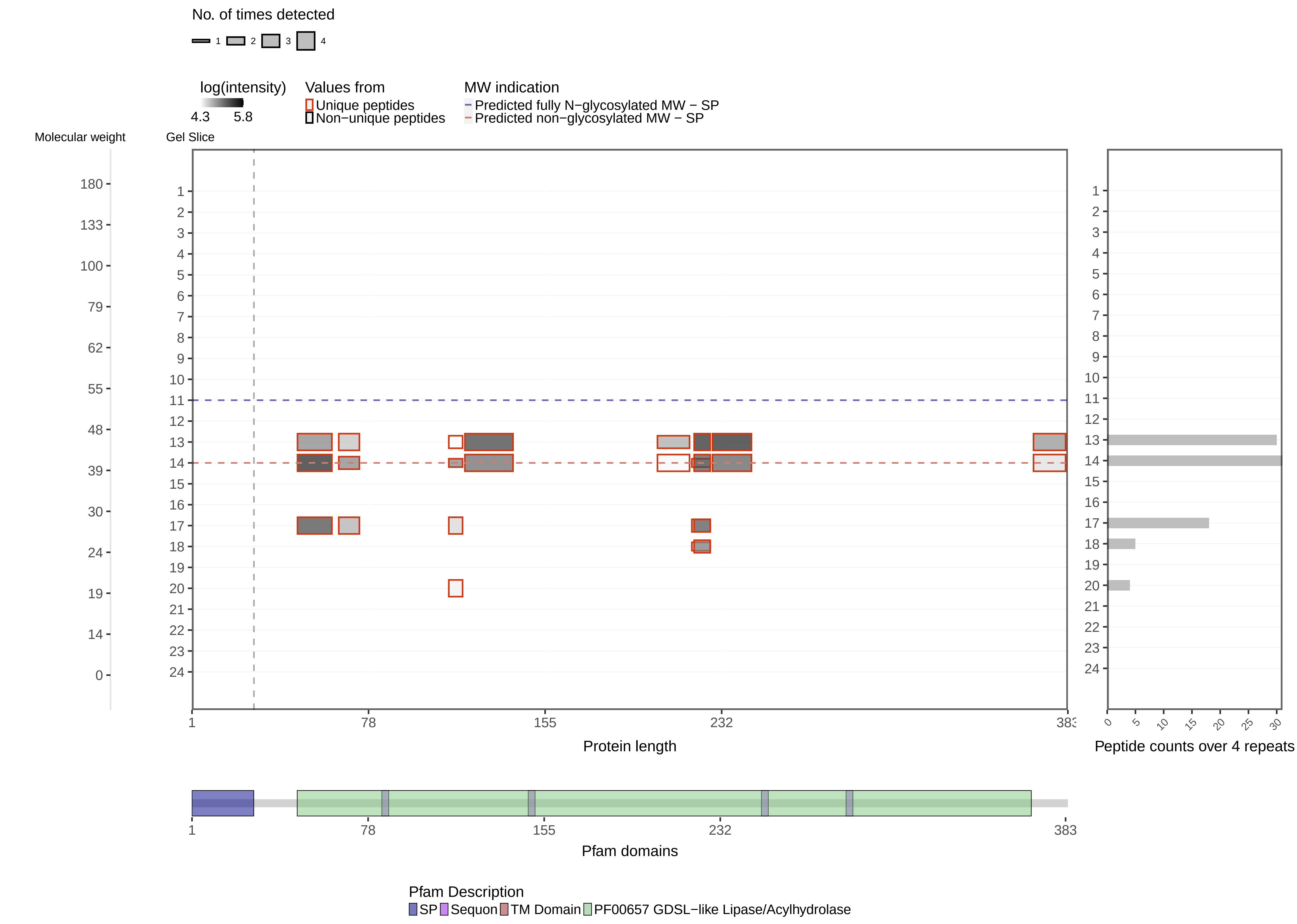


Protein sequence:

ALFLVGGIGGNEFNFALSQGKTLEESRTIVPDIVEAIIHGVRRVISFGATRIVVPGNFPGGCFPIILTRFMTDNSSAYDEHHCLKDFNSLIIFFNDHLQQAILDLKKEYPNITLIYGDYYNAYLWLLQNAVSLGFDKNSLHKACCGIGGDYNYNIHNQCGTIGVPVCTDPTTYISWDGAHLTQEAYSWLARWLIDNNILPQLNCQA*

### NbL11g15410.1 calcium−dependent_protein_kinase_5 [MW = 63.88 kDa]

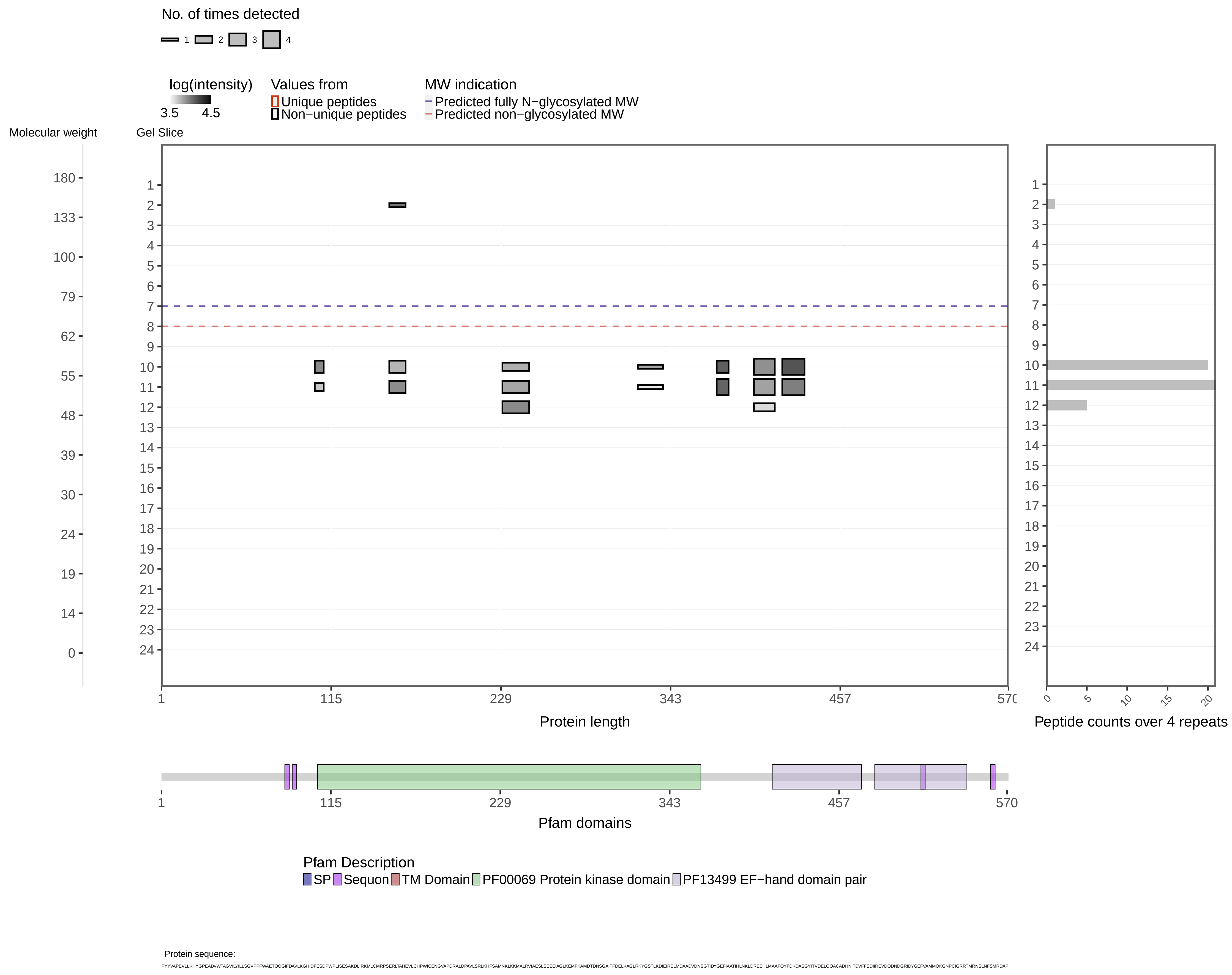

### NbL11g15220.1 lysM_domain−containing_GPI−anchored_protein_2 [MW − SP = 35.25 kDa]

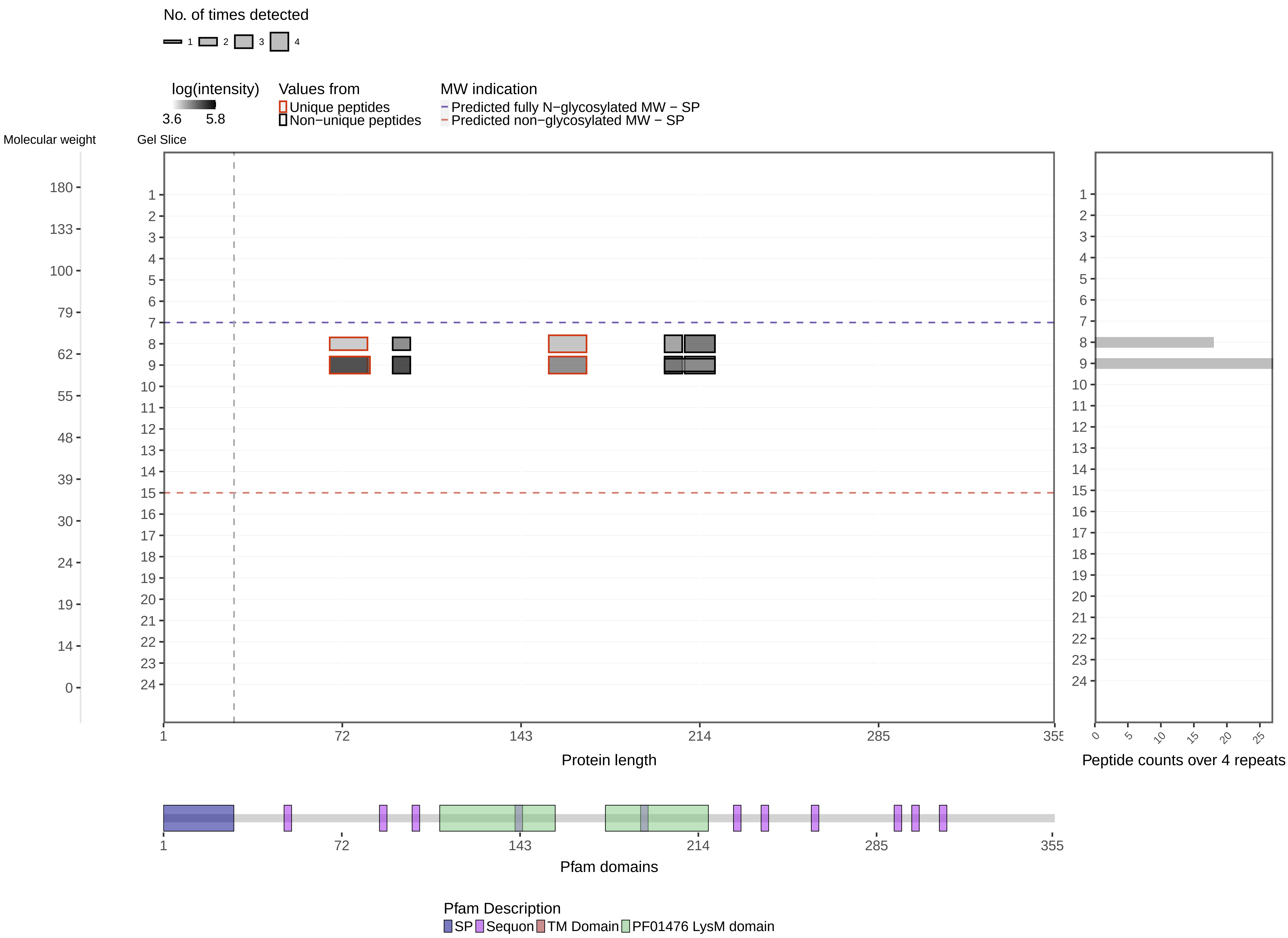


Protein sequence:

GEKVVHYGRLVAAGNTVDDIAOOYNVSODTLLKLNGLSDPKELIAGSVLDVPLKACOSMVSNTSLDYPLLVPNDTYVFTAANCVACKCDAAKNWTLOCOPSOIKSSLWKSCPSMOCOGLDSFYIGNVTSSDCNSTACAYAGYSNOTIFTTSTOLTCPASTASEMTGTWRWNIILVAIHLALLSASYMR*

### NbL11g15150.1 expansin−like_A2 [MW − SP = 26.47 kDa]

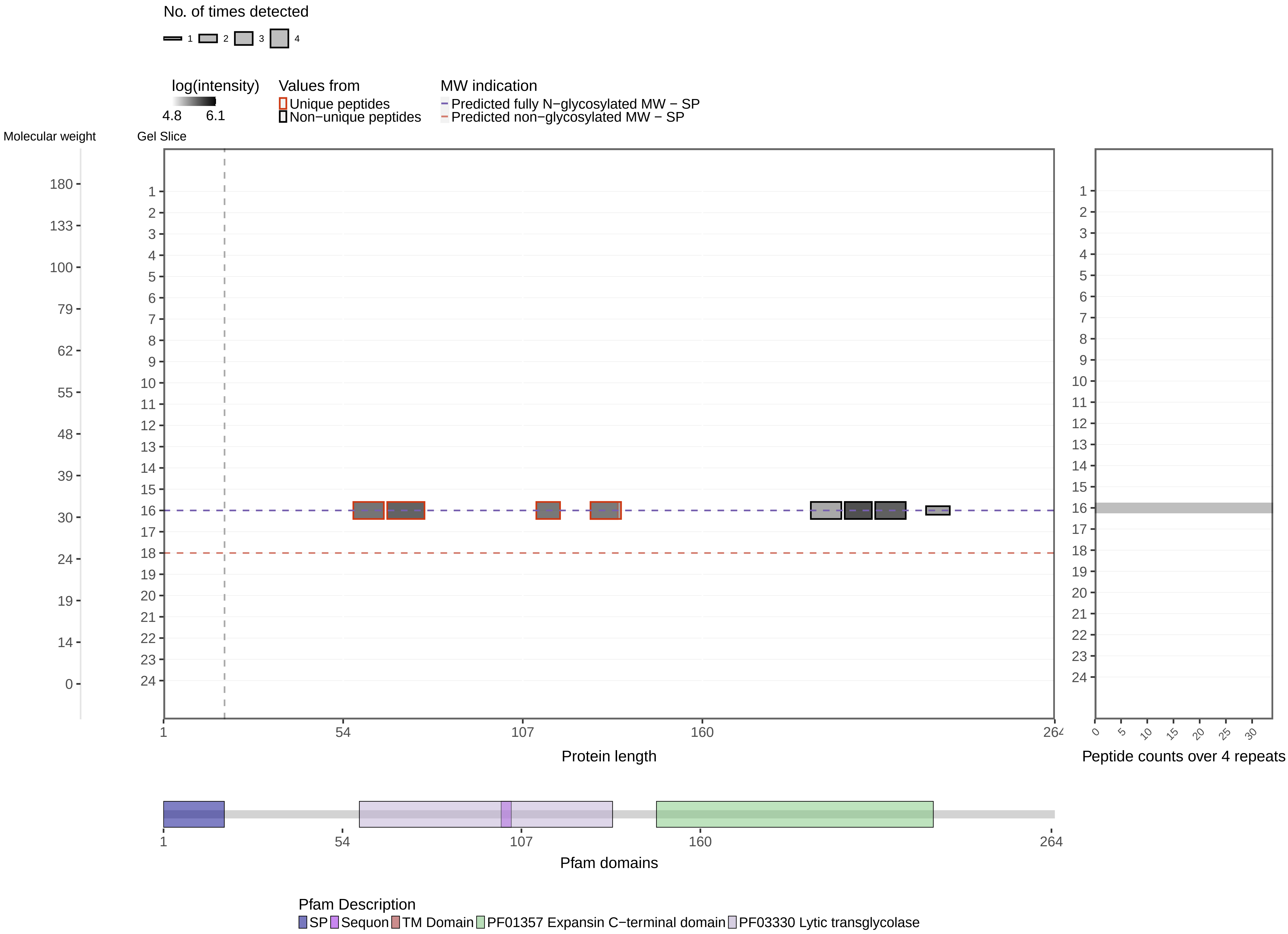


Protein sequence:

KAKDVLKLGIADVEYKRVPCDYKSKNLAIRVEESSQKPNYLAISFLYQGGQTEIVAVDVAQVGSSTWNFLSRNHGAIWDTSRVPTGALQFRFVVTAGYDGKWYWAKSVLPADWKNGVIYDTGLQITDIAQEACSPCDDGNWKLH*

### NbL11g14930.1 V−type_proton_ATPase_subunit_B2 [MW = 60.01 kDa]

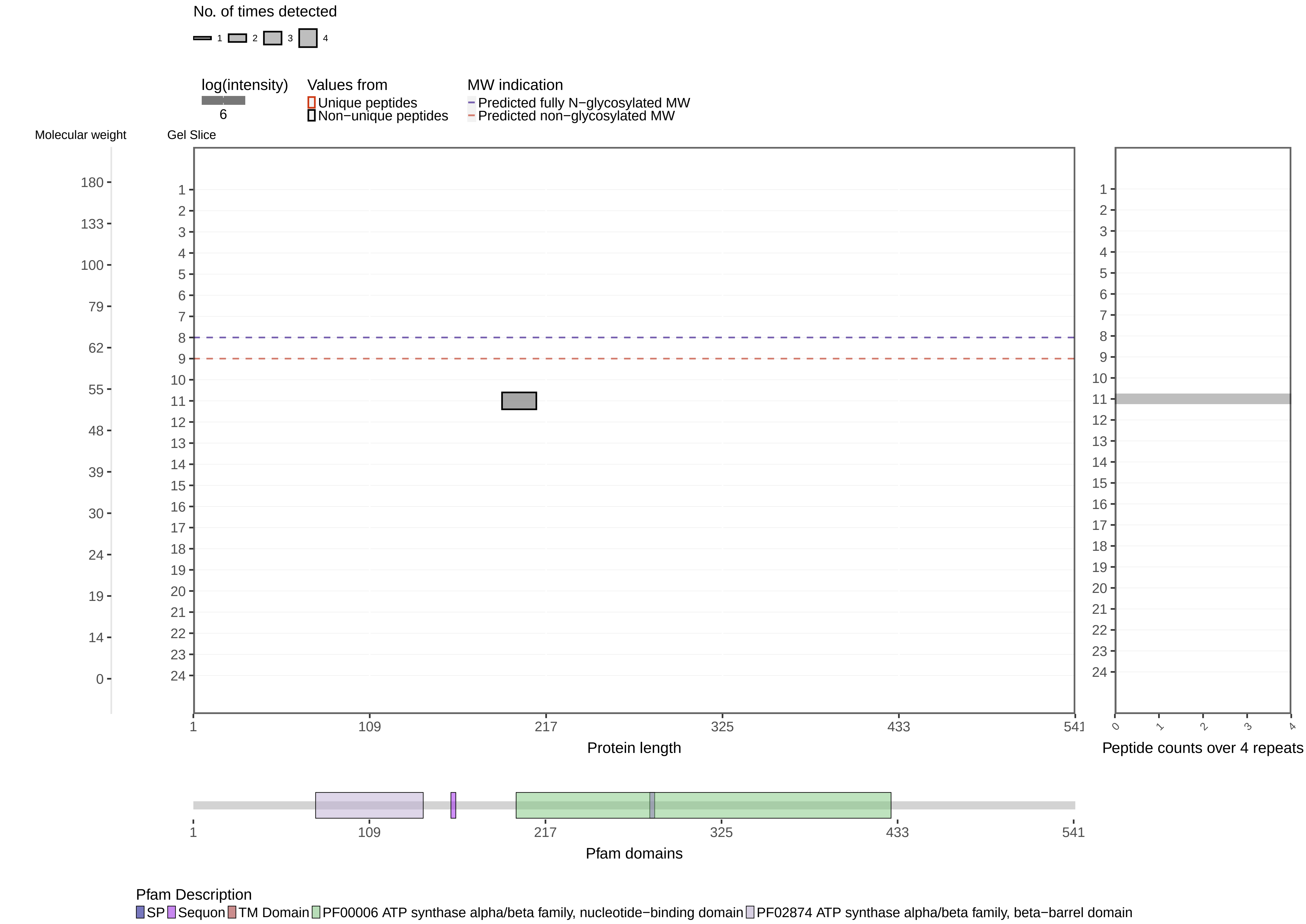


Protein sequence:

NFAIVFAAMGVNMETAQFFKRDFEENGSMERVTLFLNLANDPTIERIITPRIALTTAEYLAYECGKHVLVILTDMSSYADALREVSAAREEVPGRRGYPGYMYTDLATIYERAGRIEGRTGSITQIPILTMPNDDITHPTPDLTGYITEGQIYIDRQLHNRQIYPPINVLPSLSRLMKSAIGEGMTRRDHSDVSNQLYANYAIGKDVQAMKAVVGEEALSSEDLLYLEFLDKFERKFVAQGAYDTRNIFQSLDLAWTLLRIFPRELLHRIPAKTLDQYYSRDASN*

### NbL11g14870.1 probable_26S_proteasome_non−ATPase_regulatory_subunit_3 [MW = 55.48 kDa]

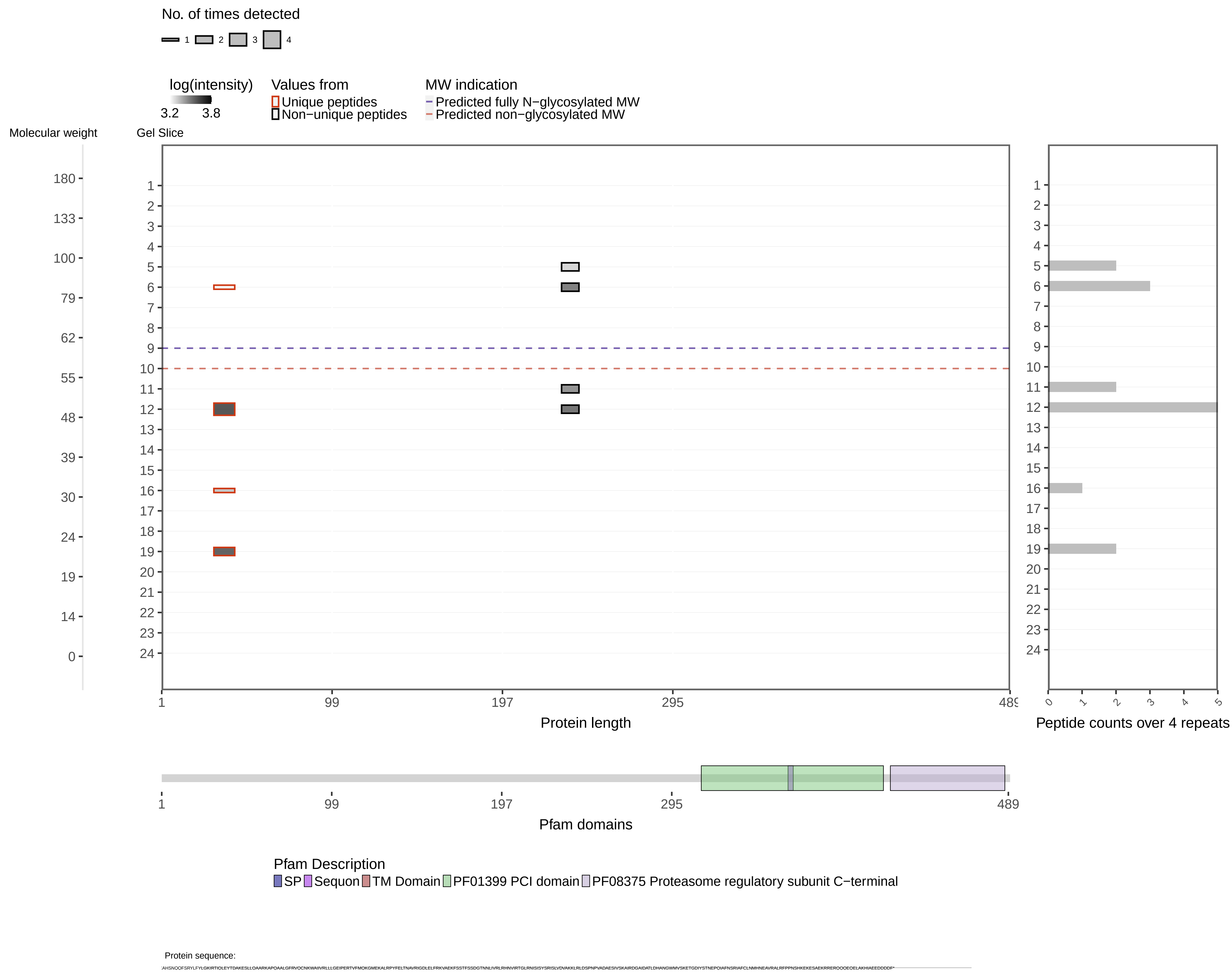

NbL11g14840.1 PIP2−7 [MW = 30.47 kDa]

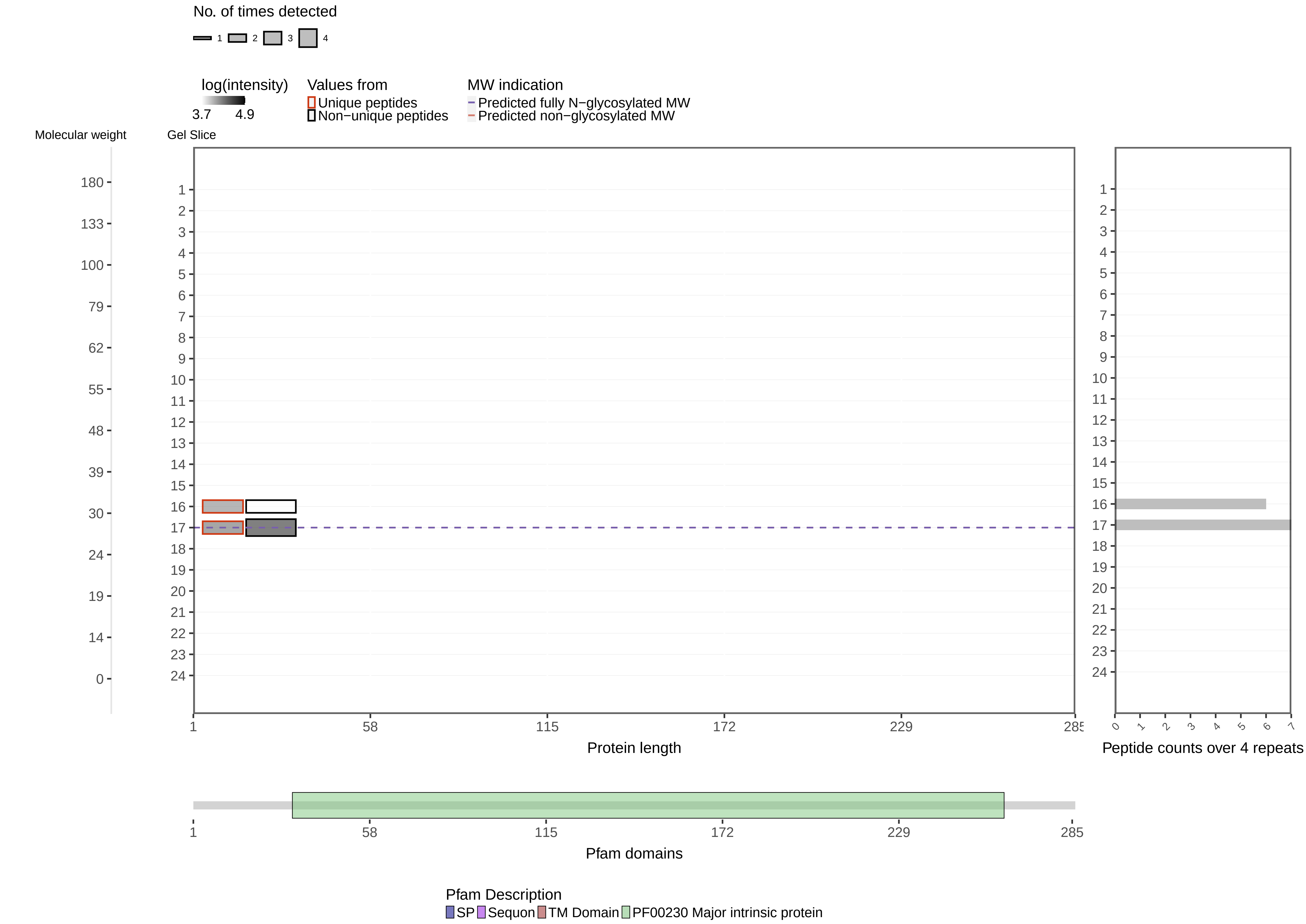


Protein sequence:

LGAICGVGLVKGFMKHYYNTLGGGANFVQPGYNKGTALGAEIIGTFVLVYTVFSATDPKRSARDSHVPVLAPLPIGFAVFMVHLATIPITGTGINPARSFGAAVIYNTDKVWDDQWIFWVGPFVGALVAAVYHQYILRGSAIKALGSFRSNPTN*

### NbL11g14820.1 glycerate_dehydrogenase [MW = 42.31 kDa]

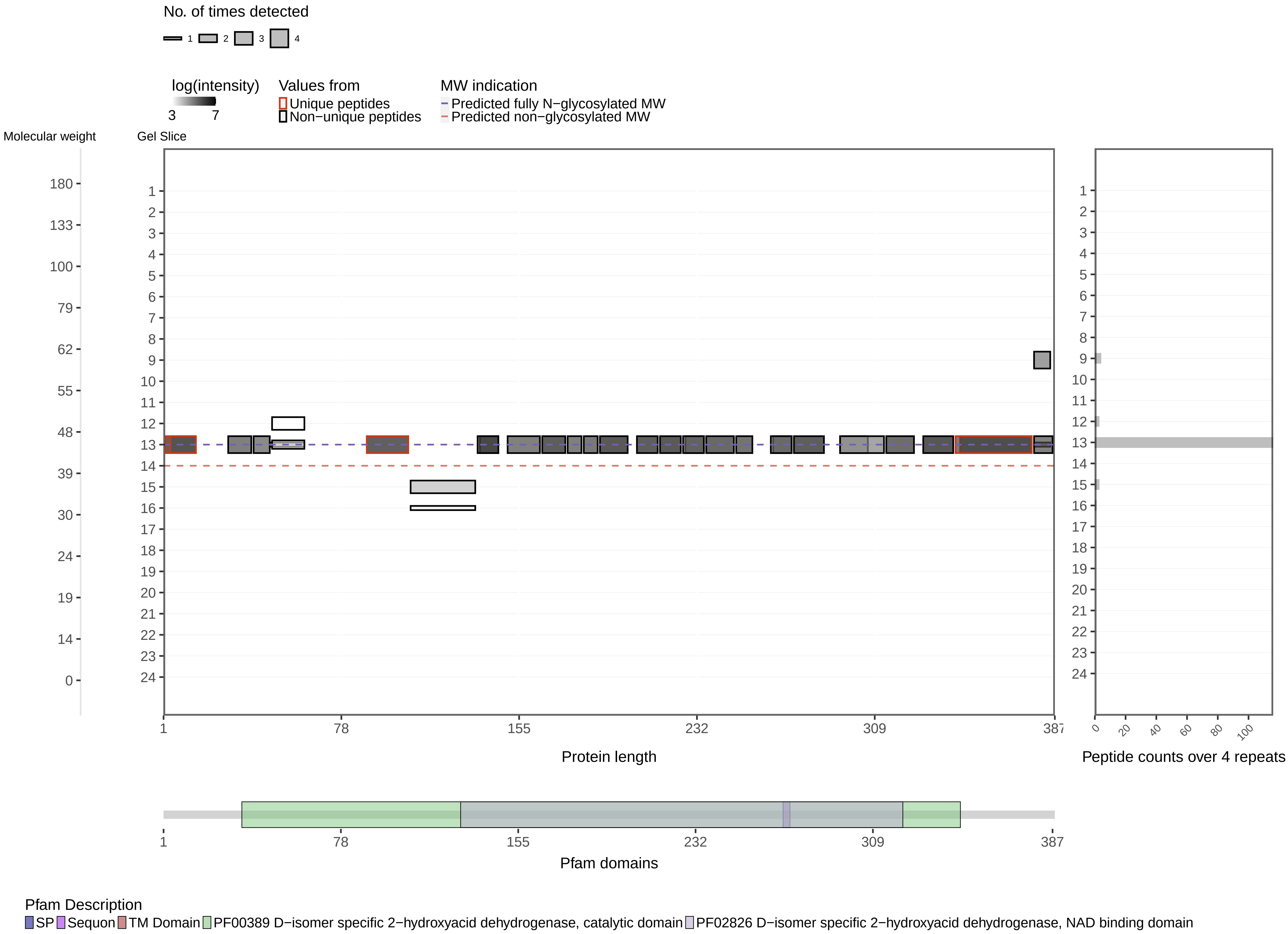


Protein sequence:

ARMMIEGFKMNLIYYDLYQATRLEKFVTAYGQFLKASGEQPVTWKRAASMEEVLREADVISLHPILDKTTYHLVNKERLAMMKKEAILVNCSRGPVIDEVALVEHLRENPMFRVGLDVFEDEPYMKPGLADMKNAVVVPHIASASKWTREGMATLAALNVLGKIKGYPVWGNPNSVDAFLNENSQPPAACPSIVNSKTLGLPVSKL*

NbL11g14690.1 L−ascorbate_peroxidase_3_peroxisomal [MW = 31.6 kDa]

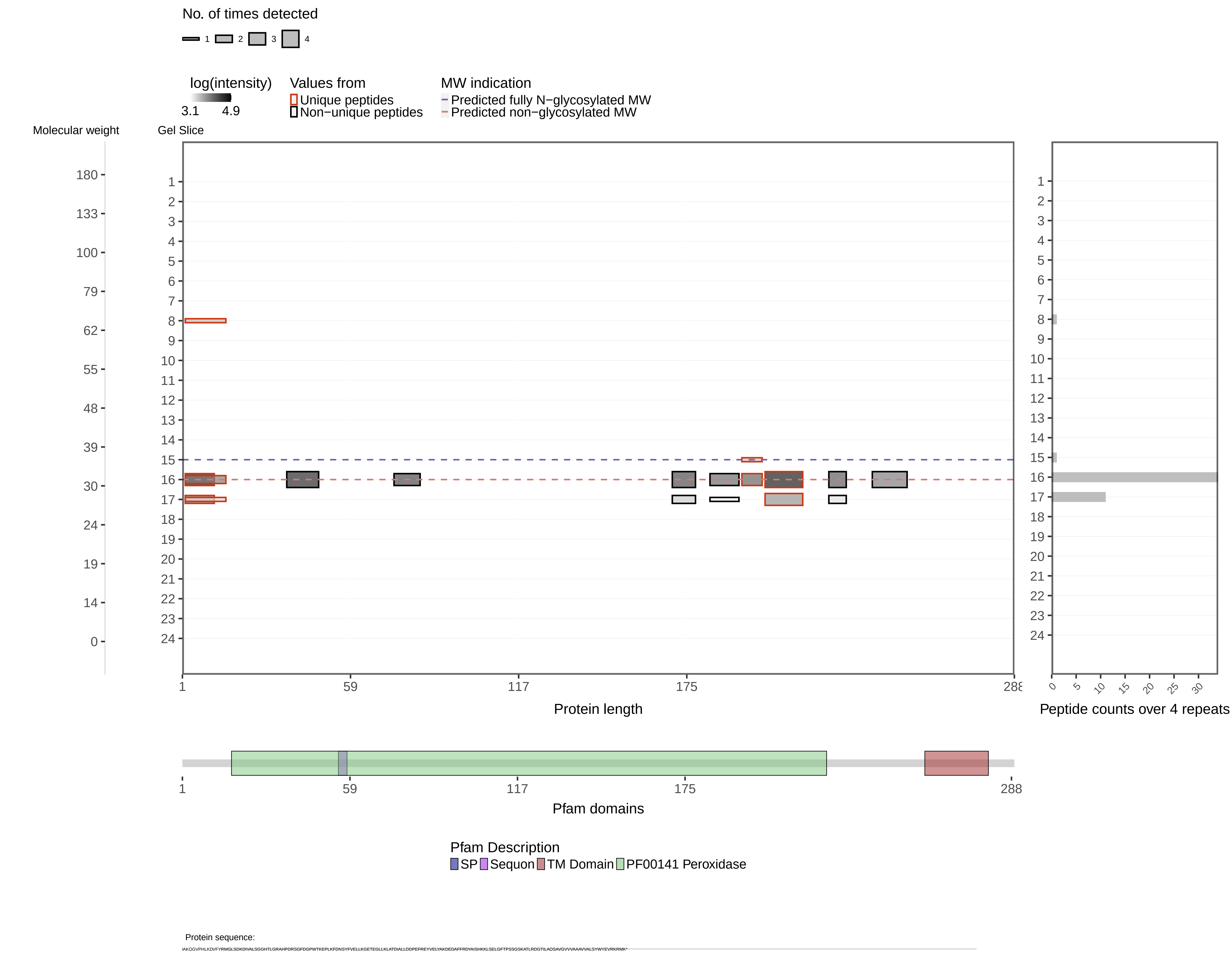

### NbL11g14670.1 proteasome_subunit_alpha_type−7 [MW = 27.03 kDa]

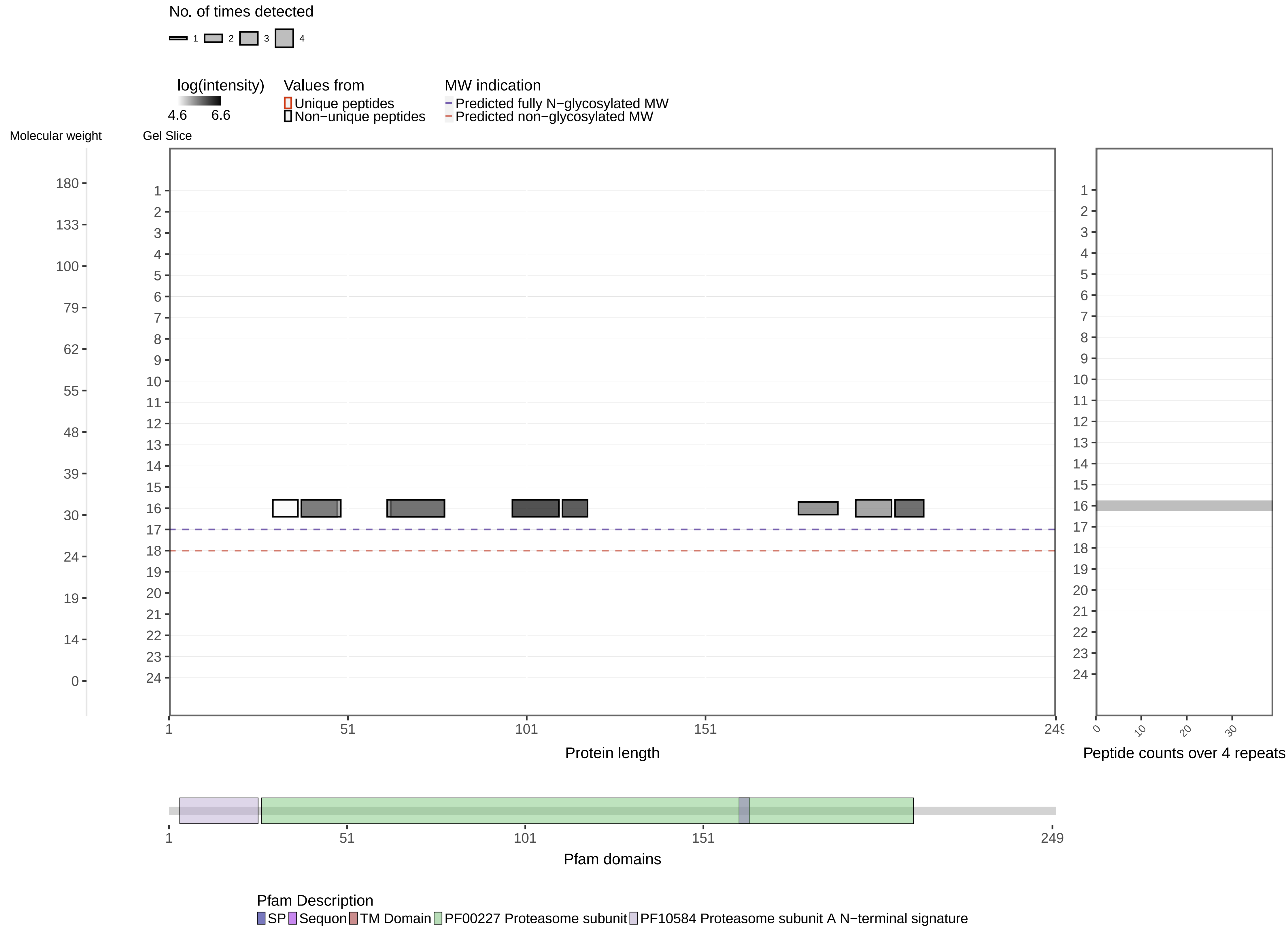


Protein sequence:

YIAGLQQKYTQSGGVRPFGLSTLIIGFDPYTGVPALYQTDPSGTFSAWKANATGRNSNSLREFLEKNYKETSGQETVKLAIRALLEVVESGGKNIEVAVMTKEHGLKQLEEAEIDAIVAEIEAEKAAAEAAKKAPKET*

### NbL11g14620.1 subtilisin−like_protease_SBT1.6 [MW − SP = 78.88 kDa]

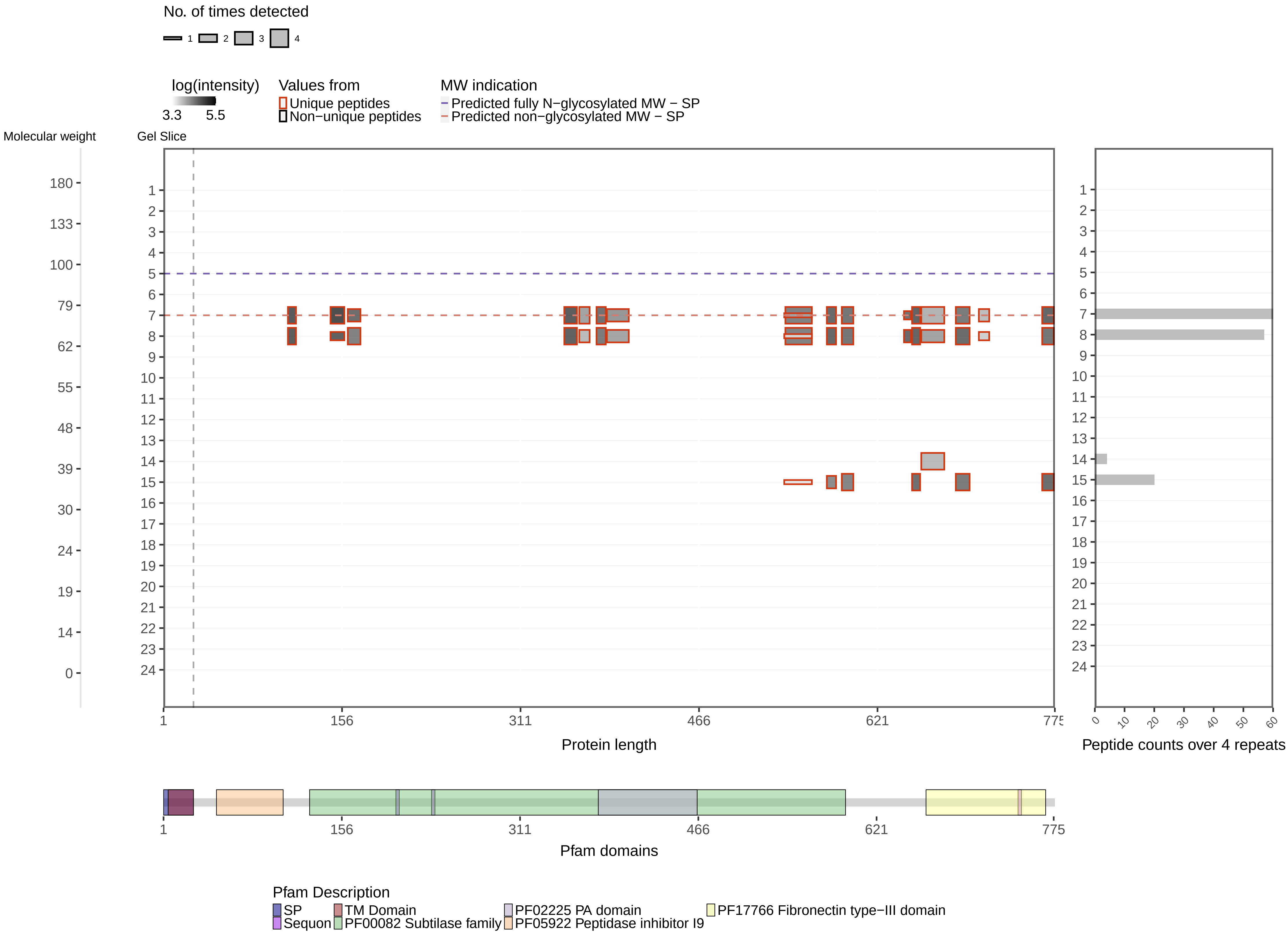


Protein sequence:

GKMYPIVYPGKSGVLSASLCMENSLDPHLVRGKIVICDRGSNPRVAKGMVVNKAGGVGMILANGVSNGEGLVGDAHLIPTCAVGANEGDAIKSYIASHPTASATINFHGTVIGVKPAPVVASFSGRGPNGLNPEILKPDLIAPGVNILAAWTDAVGPTGLDLDNRKAEFNILSGTSMACPHVSGAAALLKSAHSDWSPAAIRSAMMTTANRVDNRLQPMTDEATGKPATPYDYGAGHLNLDLALDPGLVYDLANEDYVSFLCAIEYDPKTIQVITKSPVDCPMKKPLPENLNYPS

NbL11g14550.1 protein_2_(GRP−2)_mRNA_X60007.1_N.sylvestris_mRNA_for_glycine_rich_protein_2 [MW = 19.45 kDa]

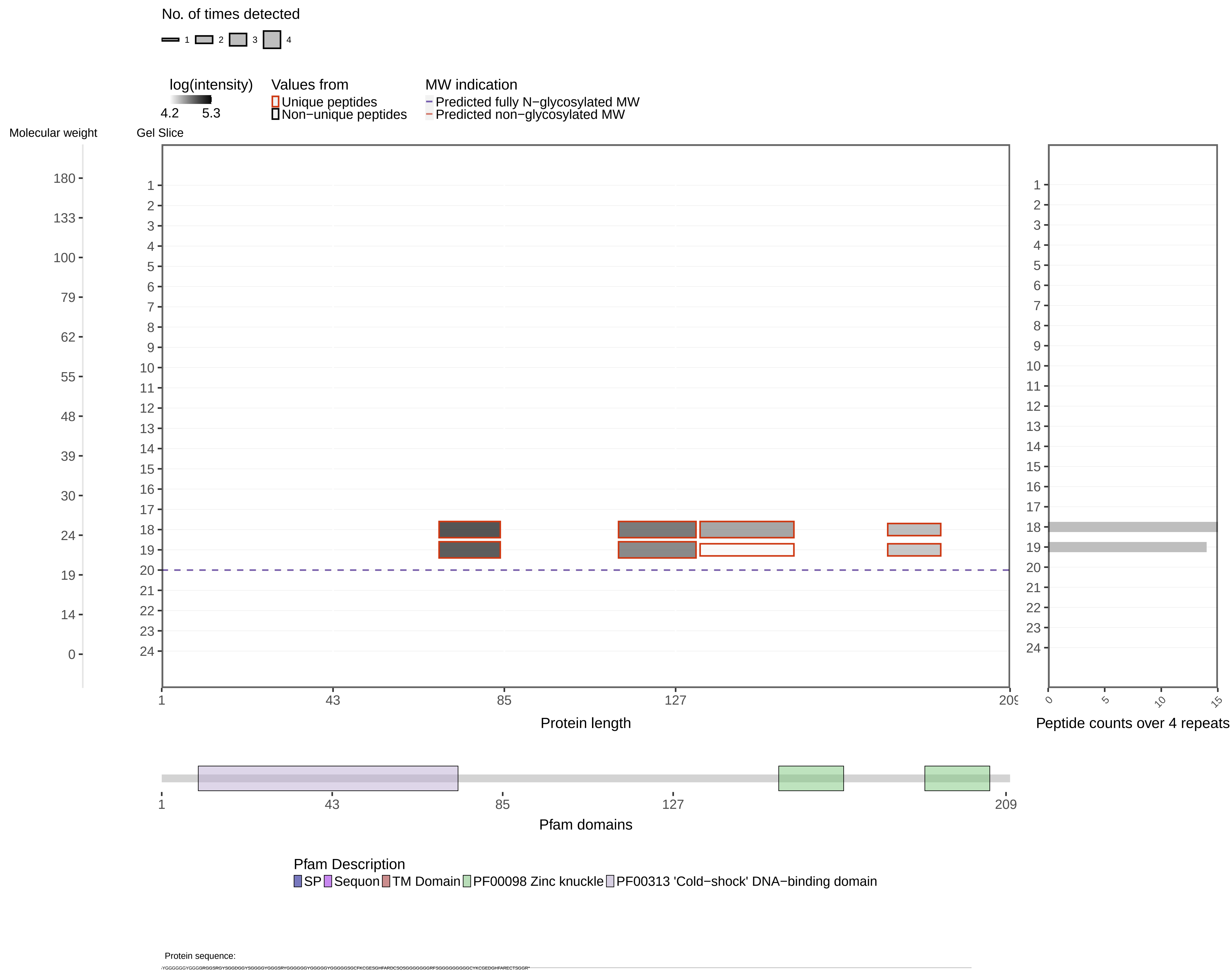

NbL11g14520.1 PI−PLC_X−box_domain−containing_protein_DDB_G0293730−like [MW = 36.84 kDa]

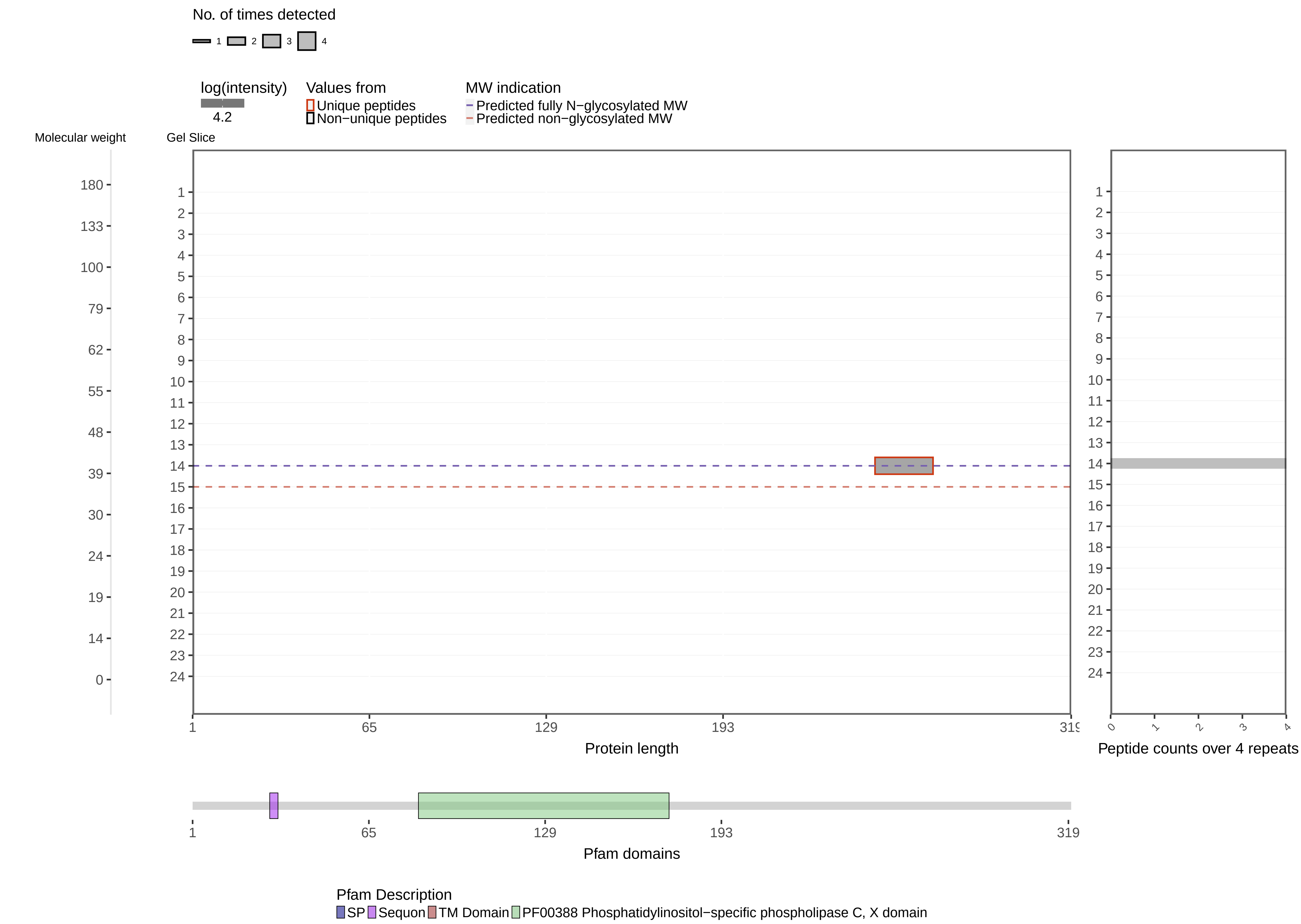


Protein sequence:

GHEDPPEFDKYLEDQLGEYLIHQDDSVFHKTVAELLPKRVICVWKPRKSPQPKHGSPLWSAGYLKDNWIDTDLPETKFESNMKHLSEQPPVTSRKYFYRVENTVTPQADNPVVCVKPVTNRIHPYARLFITESISRGYGDRLQIYSTDFIDEDFVDACVGLTNARIEDKV*

### NbL11g14480.1 probable_amidase_At4g34880 [MW = 53.92 kDa]

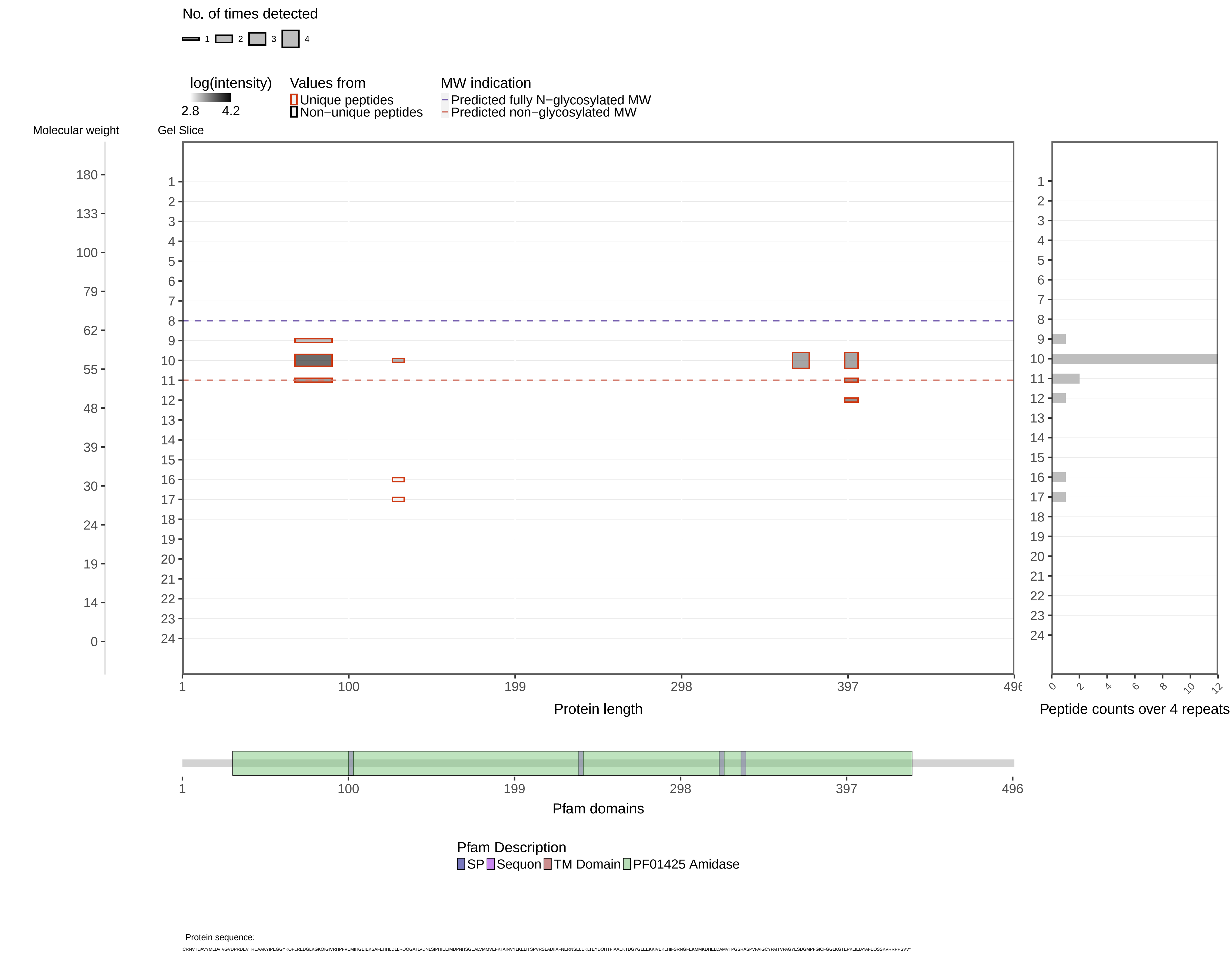

### NbL11g14460.1 eukaryotic_translation_initiation_factor_4B3 [MW = 44.75 kDa]

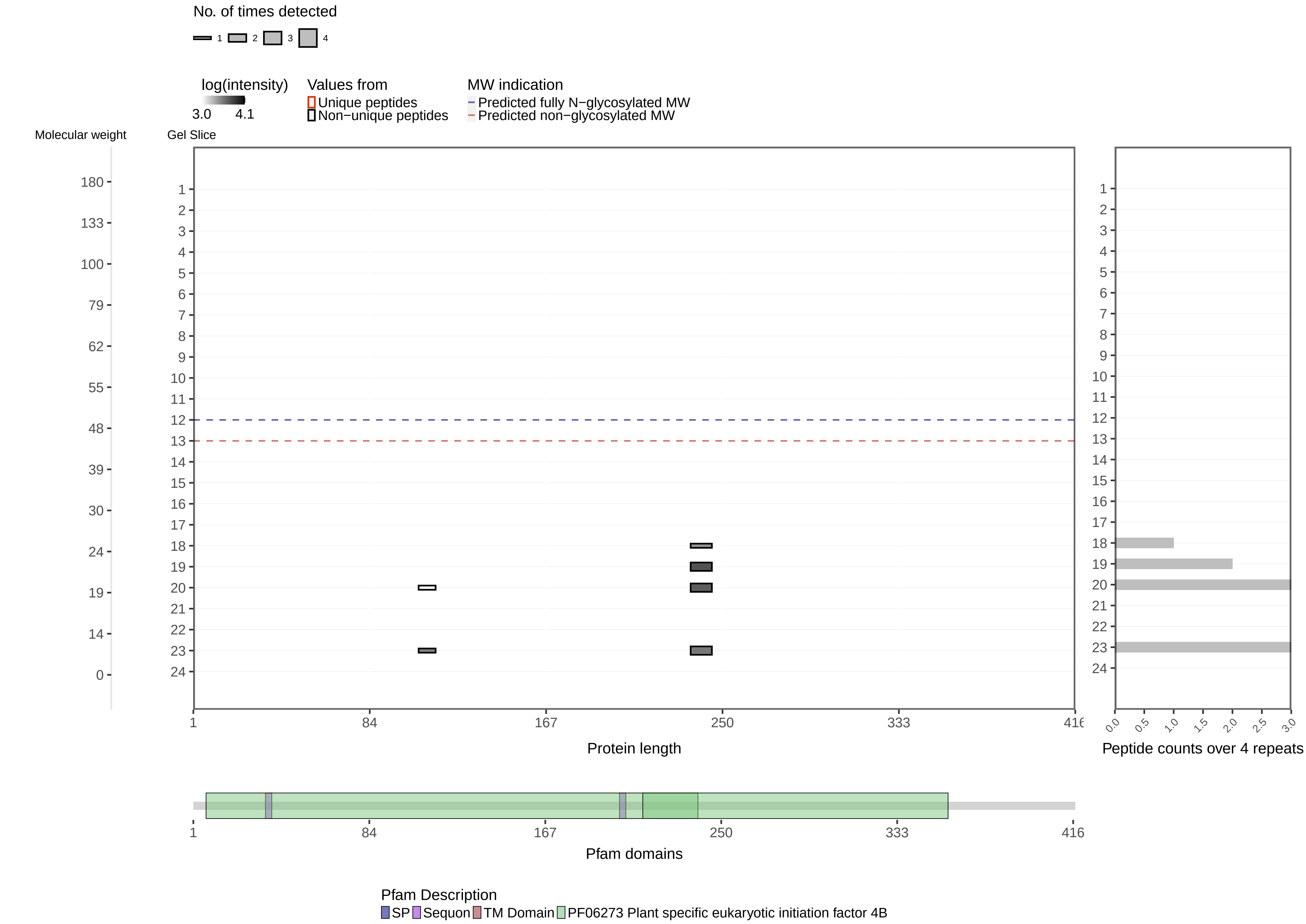


Protein sequence:

ONWGANKSFVPSSGRRFDRRGSFGSNGGDSDSDKWTKRSGGGGAFDSLRERRGGGYESYNSGVDSENWGRGKREETLGGGGGGRPKLNLQPRTLPITEVQQQNGGTNEATVVVKPKGSNNPFGAARPREEVLKEKGQDWKEIDQKLESLKVKEVVSESSDKKAWGNGKPSFMREESTEKSWRKPQLNEVRPLSAEVTVNGAVEETAQGAAKESGGEPKI*

NbL11g14440.1 peptidyl−prolyl_cis−trans_isomerase−like [MW = 18.04 kDa]

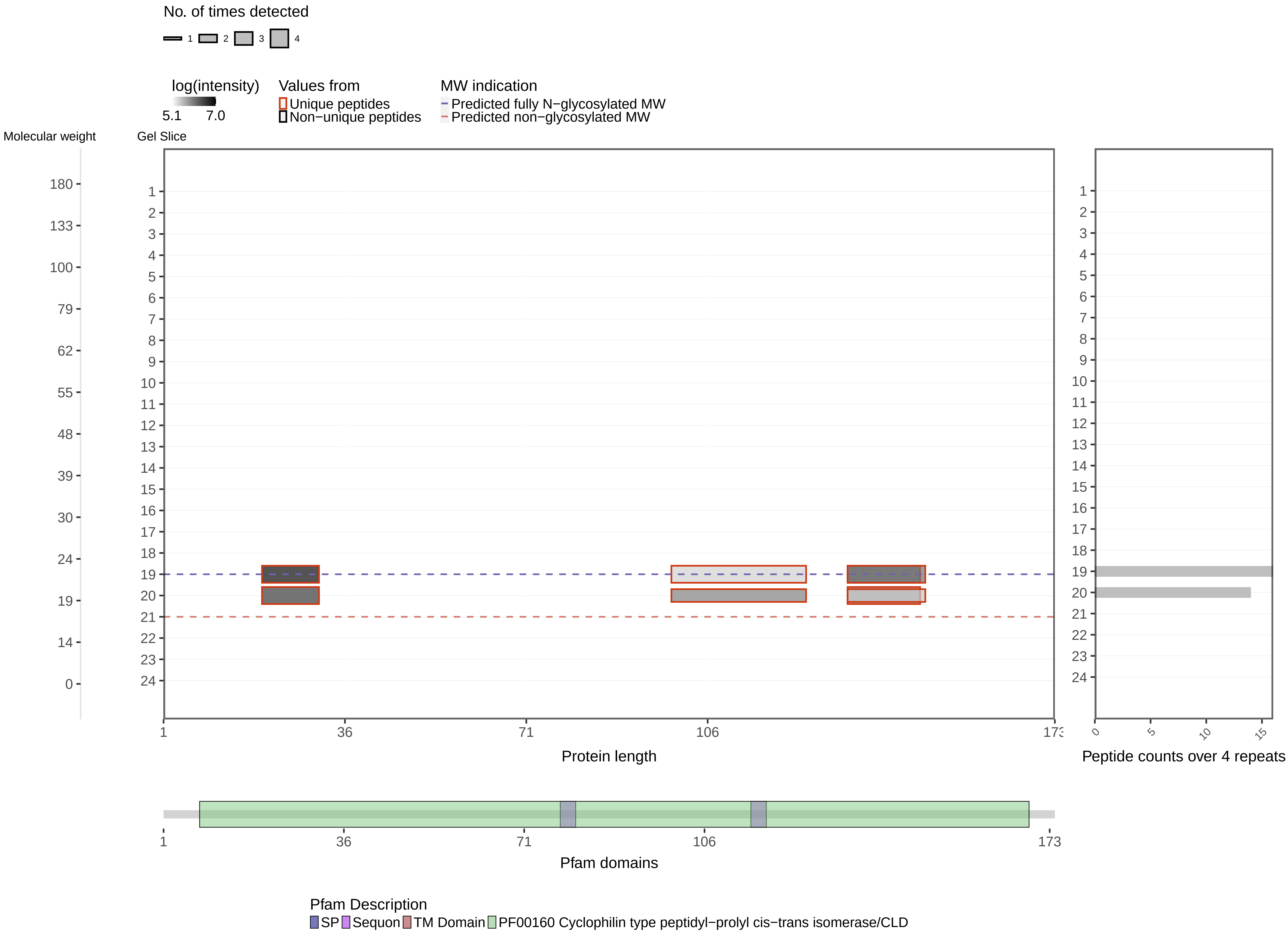


Protein sequence:

FTAGNGTGGESIYGAKFADENFKRKHTGPGVLSMANAGPGTNGSQFFICTAKTEWLDGKHVVFGQVVEGYDAIKKAEAVGSGSGRCSKPVVIADCGQLC*

NbL11g14390.1 triosephosphate_isomerase_chloroplastic−like [MW = 34.81 kDa]

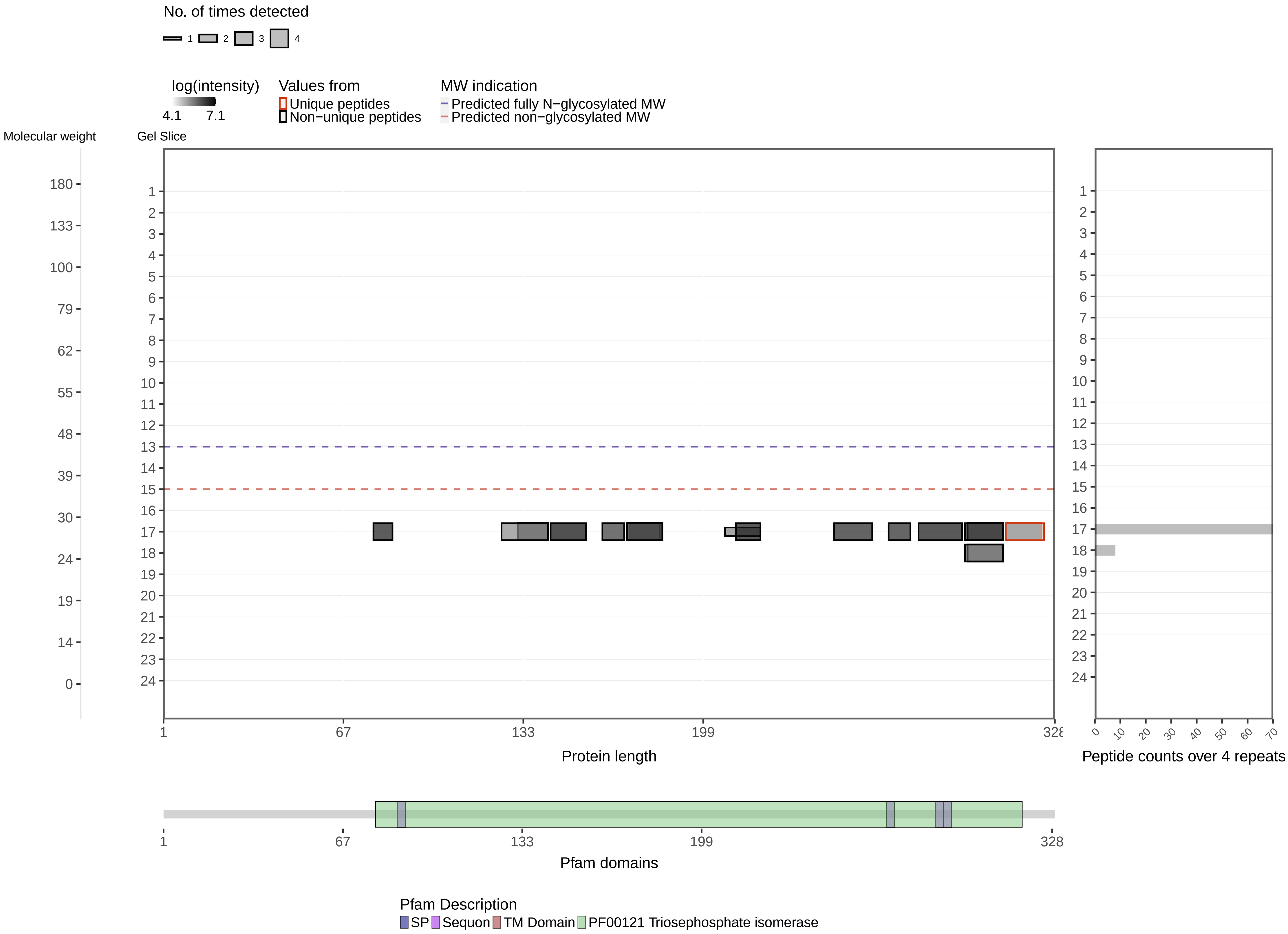


Protein sequence:

EISVEOLKDLGCKWVILGHSERRHVIGENDEFIGKKAAYALSOGVGVIACIGELLOEREAGKTFDVCFOOLKAFADALPSWDNVVIAYEPVWAIGTGKVASPEOAOEVHAAVRDWLTKNVSAEVASKTRIIYGGSVNGSNSSDLAKKEDIDGFLVGGASLKGPEFATIINSVTSKKVAA*

### NbL11g14370.1 5'−methylthioadenosine/S−adenosylhomocysteine_nucleosidase_2 [MW = 27.95 kDa]

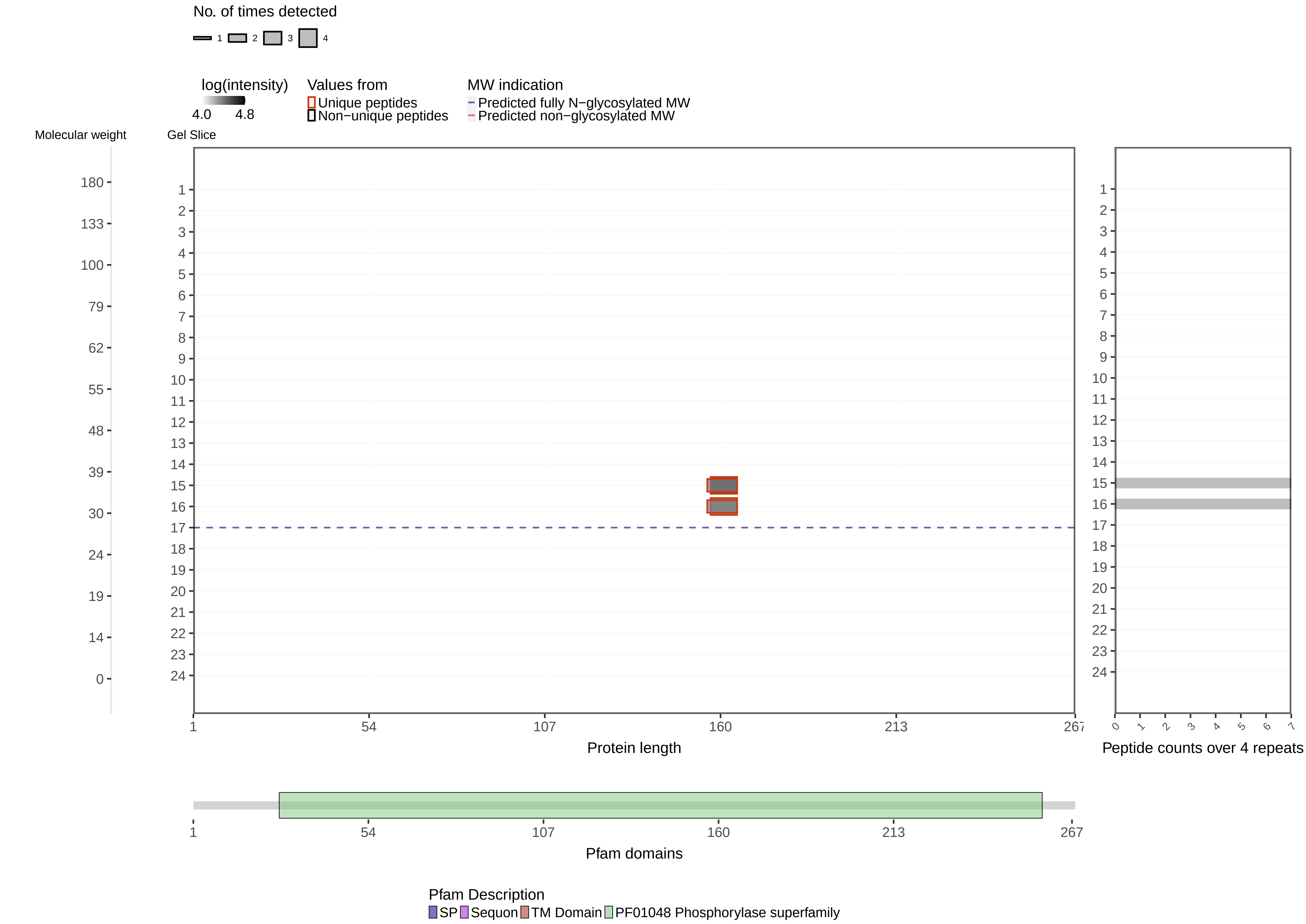


Protein sequence:

GAKGASIGDVFLASEVAFHDRRIPIPVFDIYGVGSRKAFATPYLLKELNLKIGKLSTGDSLDMSPMDEASIVANDATVKDMEGAAVAYVADLLKVPVIFLKAVTDIVDGDKPTAEEFLLNLAAVTAALDQTATLVVDCISGKCLSEL*

### NbL11g14320.1 uncharacterized_LOC104087214 [MW = 39.48 kDa]

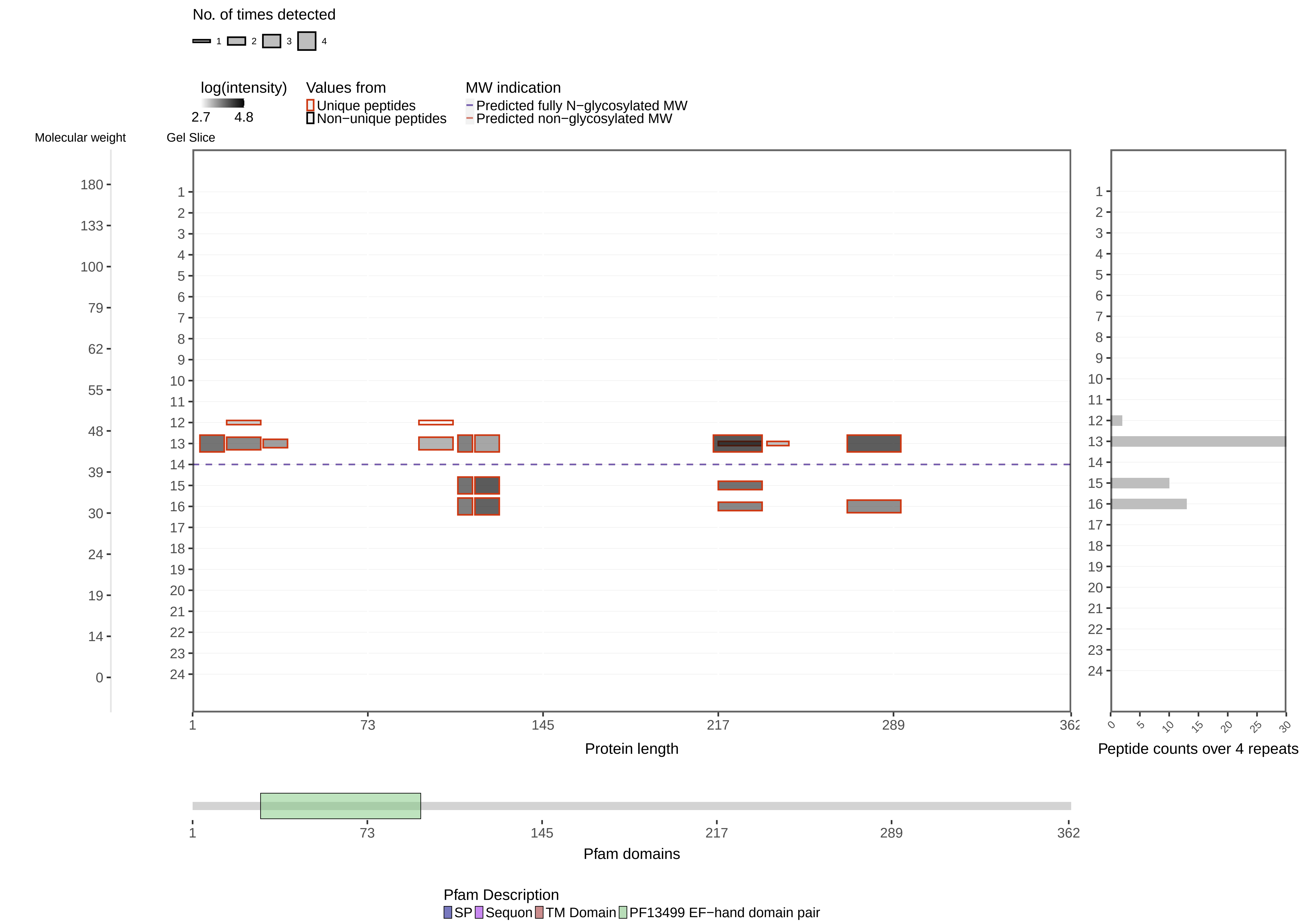


Protein sequence:

SDSWVMSNVVEPVVDSCIGASNEQPVSQETFLAEFKKVAESAAQRLKEQPVIVAHSENTFDGSGIRRLLSNKFELDKALDSAVKTVPKDRHGKMSKEYLRVALDLLAPSAGLPPIGAVDQMDKIIIEACKMLDADDGKMVKEEEFKKLLTEVLGSMMLQLEGNPVSVSTNSVVHEPLASASTLLQPPSNSEM*

NbL11g14290.1 26S_protease_regulatory_subunit_S10B_homolog_B [MW = 44.6 kDa]

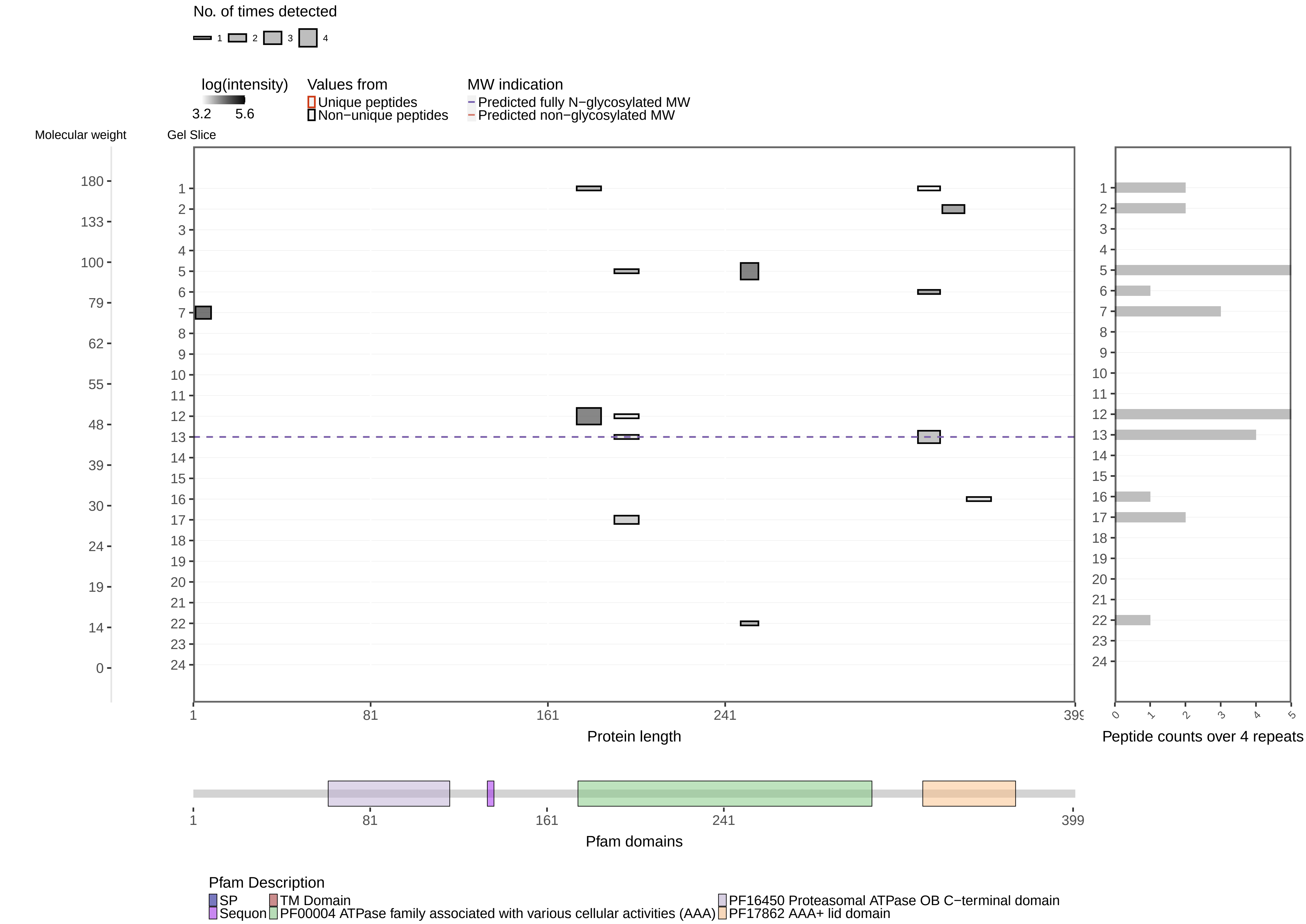


Protein sequence:

TLLARAIASNIDANFLKVVSSAIIDKYIGESARLIREMFNYARDHOPCIIFMDEIDAIGGRRFSEGTSADREIORTLMELLNOLDGFDOLGKVKMIMATNRPDVLDPALLRPGRLDRKIEIPLPNEOSRMEVLKIHAAGIAKHGEIDYEAVVKLAEGFNGADLRNVCTEAGMSAIRAERDYVIHEDFMKAVRKLNDAKKLESSAHYSADFGKE*

NbL11g14220.1 receptor_protein_kinase_TMK1−like [MW − SP = 98.86 kDa]

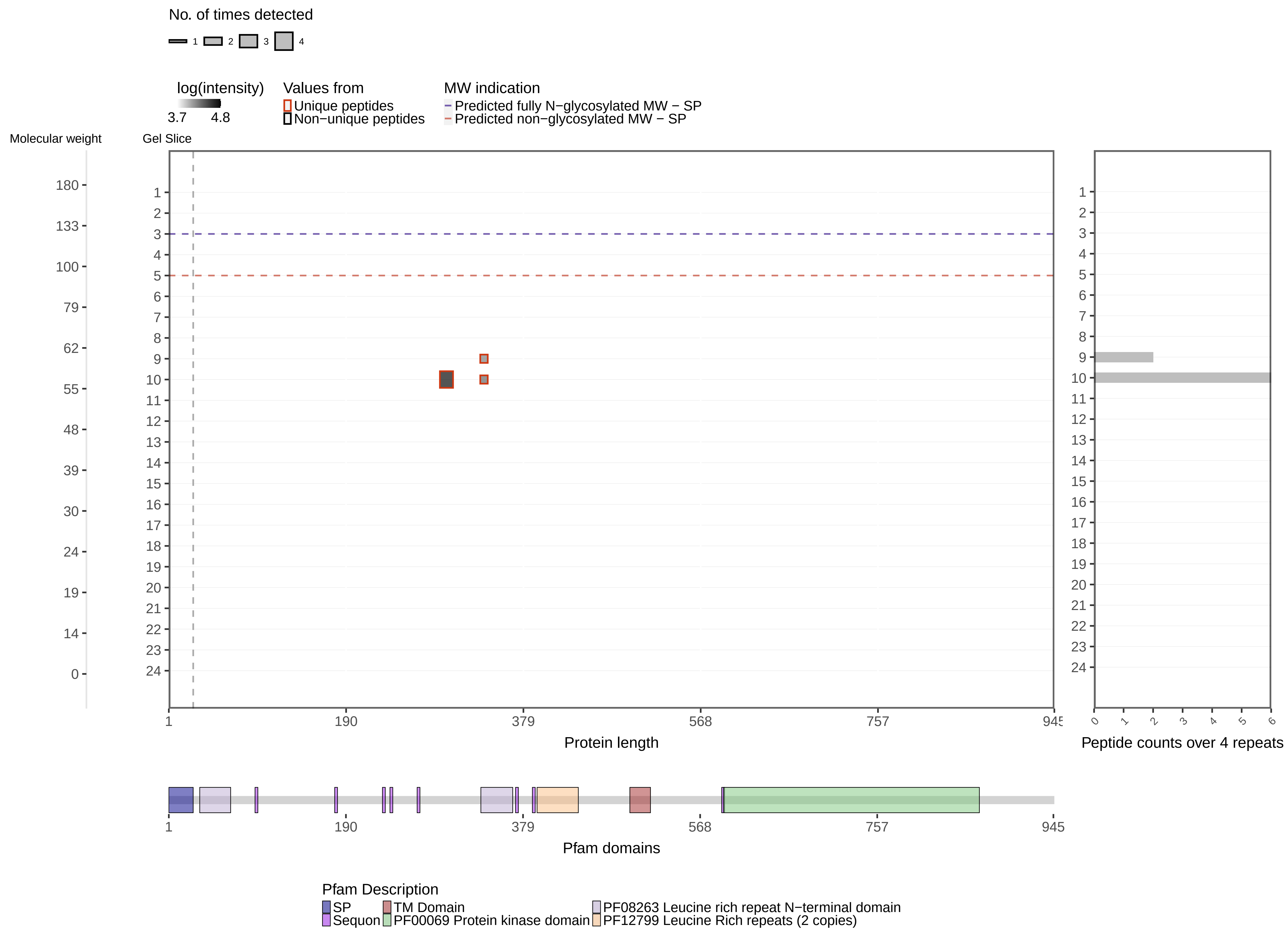


Protein sequence:

TPSGSTPGSSDGGGGGQTHANGGKKSSTGVVVGSVIGGVCAAAVLAGLIVFCLYRTKHKRSGRVQSPHTVVIHPHHSGSDQDAVKITIAGSSVNGGDSCGSSSAPGDLHIVEAGNMVISIQVLRDVTNNFSEVNILGRGGFGTVYKGELHDGTKMAVKRMESGVMSEKGLDEFKSEIAVLTKVRHRHLVTLLGYCLDGNERLLVYEYMPQGTLSRYLFNWKEEGLKPLEWTRRLTIALDVARGVEYLHGLAQQSFIHRDLKPSNILLGDDMRAKVADFGLVRLAPDGKAS[illegible]

NbL11g14190.1 purple_acid_phosphatase_2−like [MW − SP = 50.85 kDa]

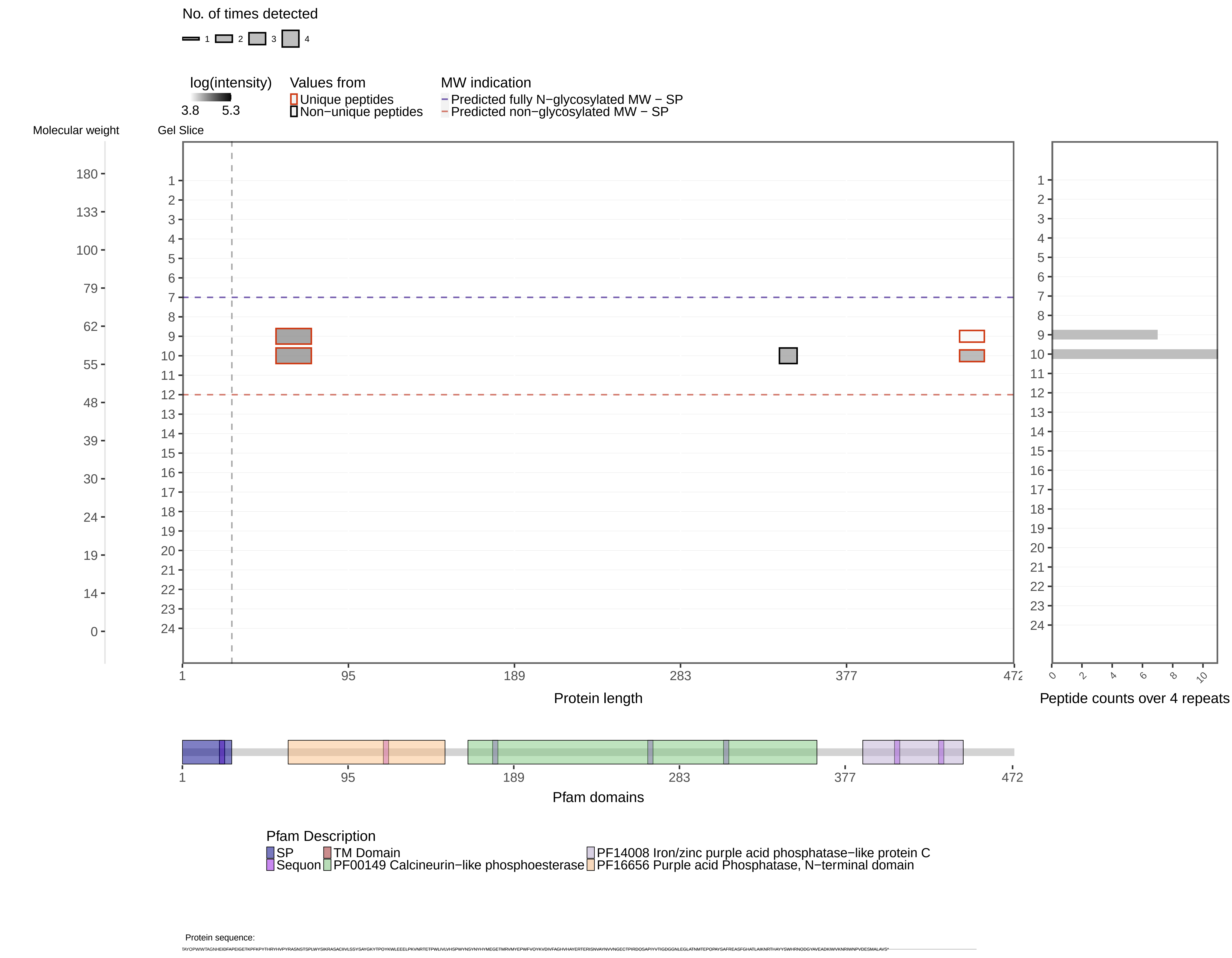

### NbL11g14110.1 cysteine_protease_RD19A−like [MW − SP = 38.04 kDa]

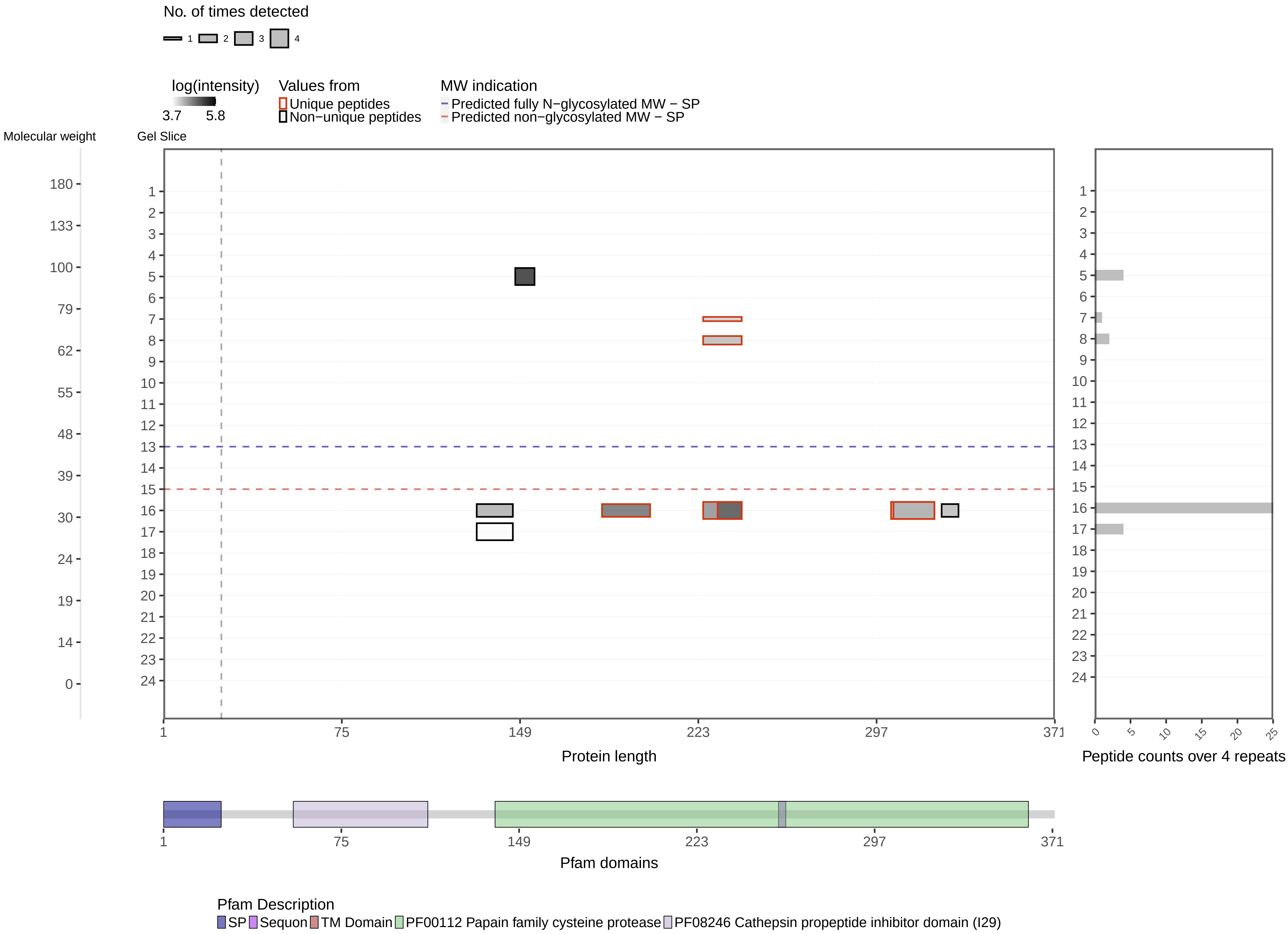


Protein sequence:

### NbL11g14100.1 V−type_proton_ATPase_subunit_a2−like [MW = 92.44 kDa]

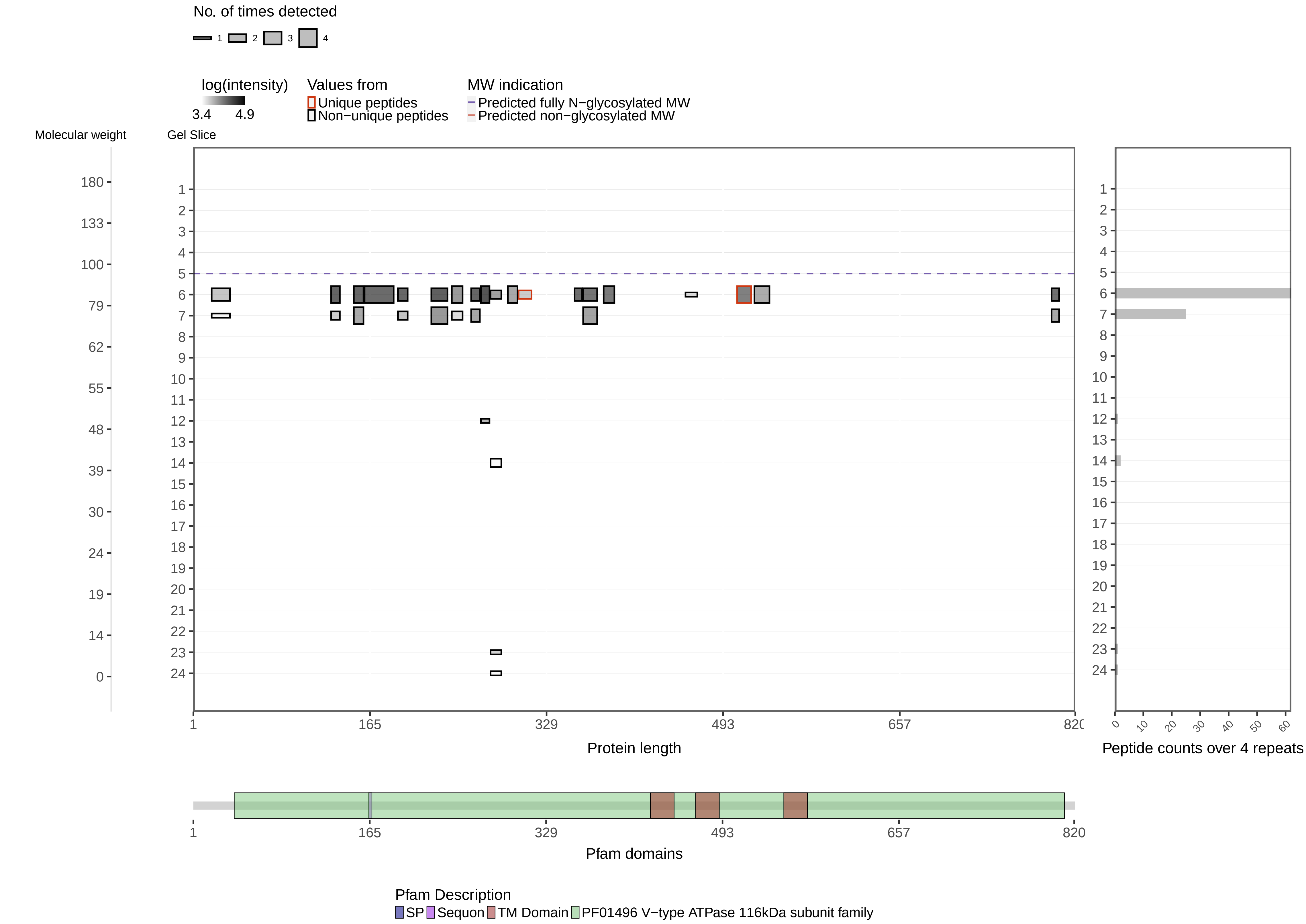


Protein sequence:

SSFQEIVDAYGVAKYQEANPGVYTIVTFPFLFAVMFGDWGHGICLLLATLFLLIREKKLSSQKLGDIMEMTFGGRYVIFMMSLFSIYTGLIYNEFFSVPFELFGRSAYGCRDPSCGDSTSAGLIKVRDTYPFGVDPAWHGTRSELPYLNSLKMKMSILLGVAQMNLGIFLSFFNALFFRSGINIWCQFVPQIIFLNALFGYLSILIIVKWCTGSQADLYHVMIYMFLSPTDELGENQLFAGQKTTQLVLLFSALVAVPWMLLPKPFLLKAQHERHQGQSYTALQEAEESLLVESSDDS

### NbL11g14060.1 uncharacterized_LOC104113315 [MW = 43.25 kDa]

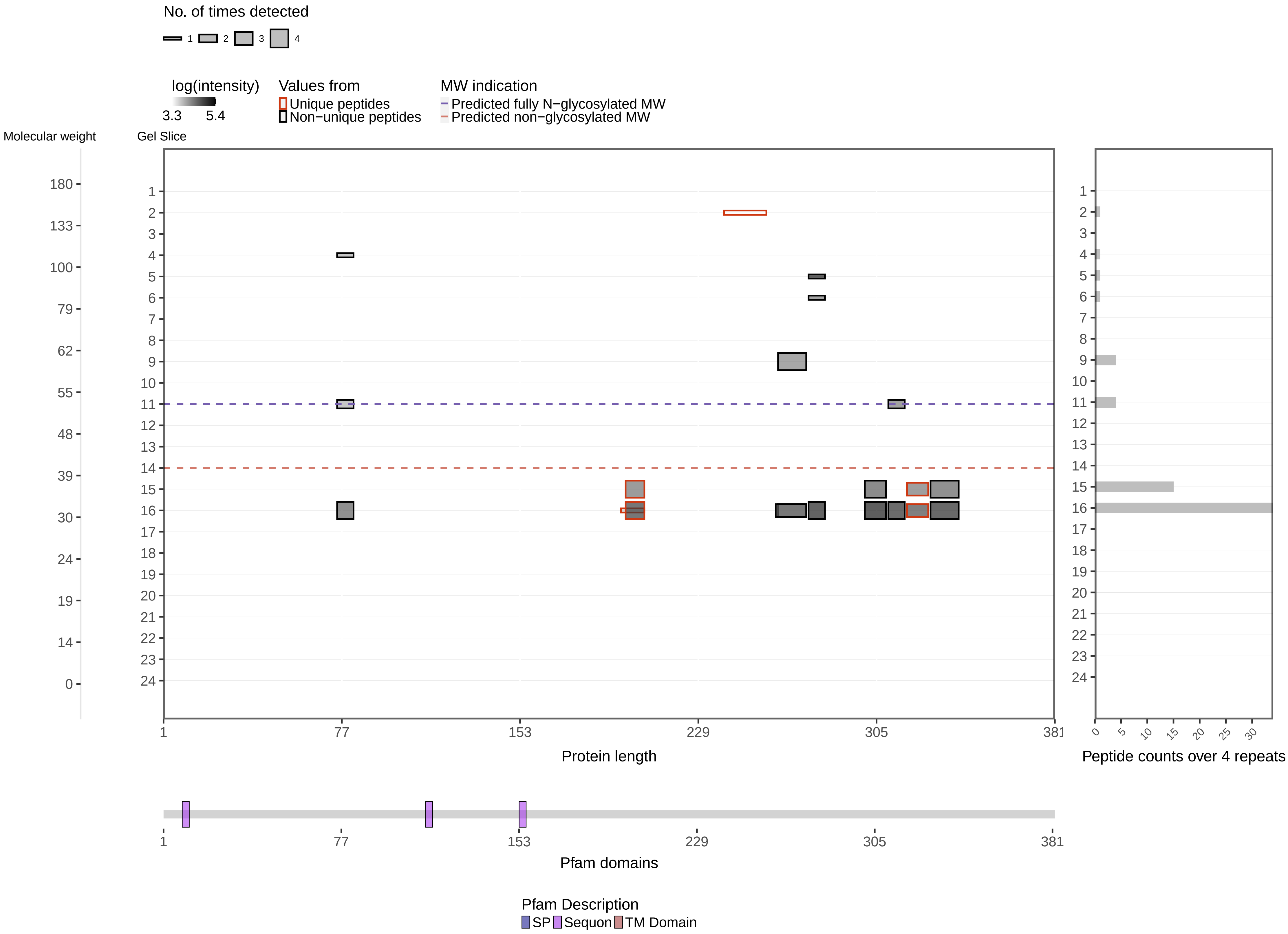


Protein sequence:

LALSVMNIVDRLVHKTKEKIDSSTGVLKAILEPVLDEVEEISWPVSDSEALSLMEKEINQREQDGQLDEGFLSEVNAQLRQAKKDGDKPGLEAMLQKVLQLYASRVLSKRSYAKKGNEVLKAEQFLESIIKAPEEEWNKLLINGMTVGKGDISPEELYAVIKKRMERTLIRTEGGSYQQRILIEYLKGIEARAGEIVKVLQG*

NbL11g13940.1 fructose−bisphosphate_aldolase_1_chloroplastic−like [MW = 42.81 kDa]

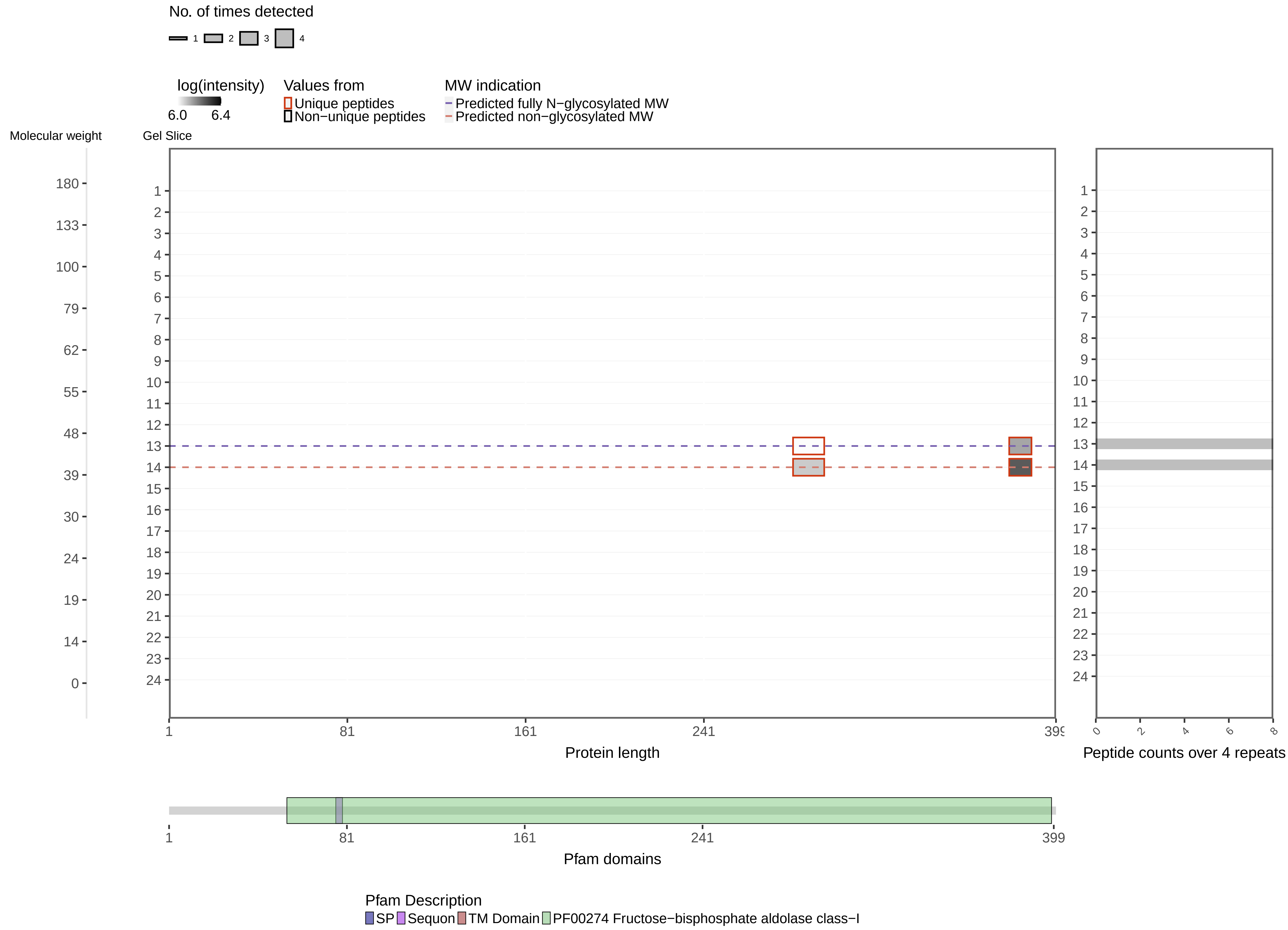


Protein sequence:

TVVSIPNGPSALAVKEAAWGLARYAAISQDSGLVPIVEPEILLDGEHGIDRTFEVAQKVWAEVFFYLAENNVMFEGILLKPSMVTPGAECKDRAIPQQVADYTLSLLKRRIPPAVPGIMFLSGGQSEVEATLNLNAMNQAPNPWHVSFSYARALQNTCLKTWGGQPENVKAAQDALLTRAKANSLAQLGKYTGEGESDEAKQGMFVKGYVY*

NbL11g13910.1 NADH_dehydrogenase_[ubiquinone]_1_beta_subcomplex_subunit_9−like [MW = 14.65 kDa]

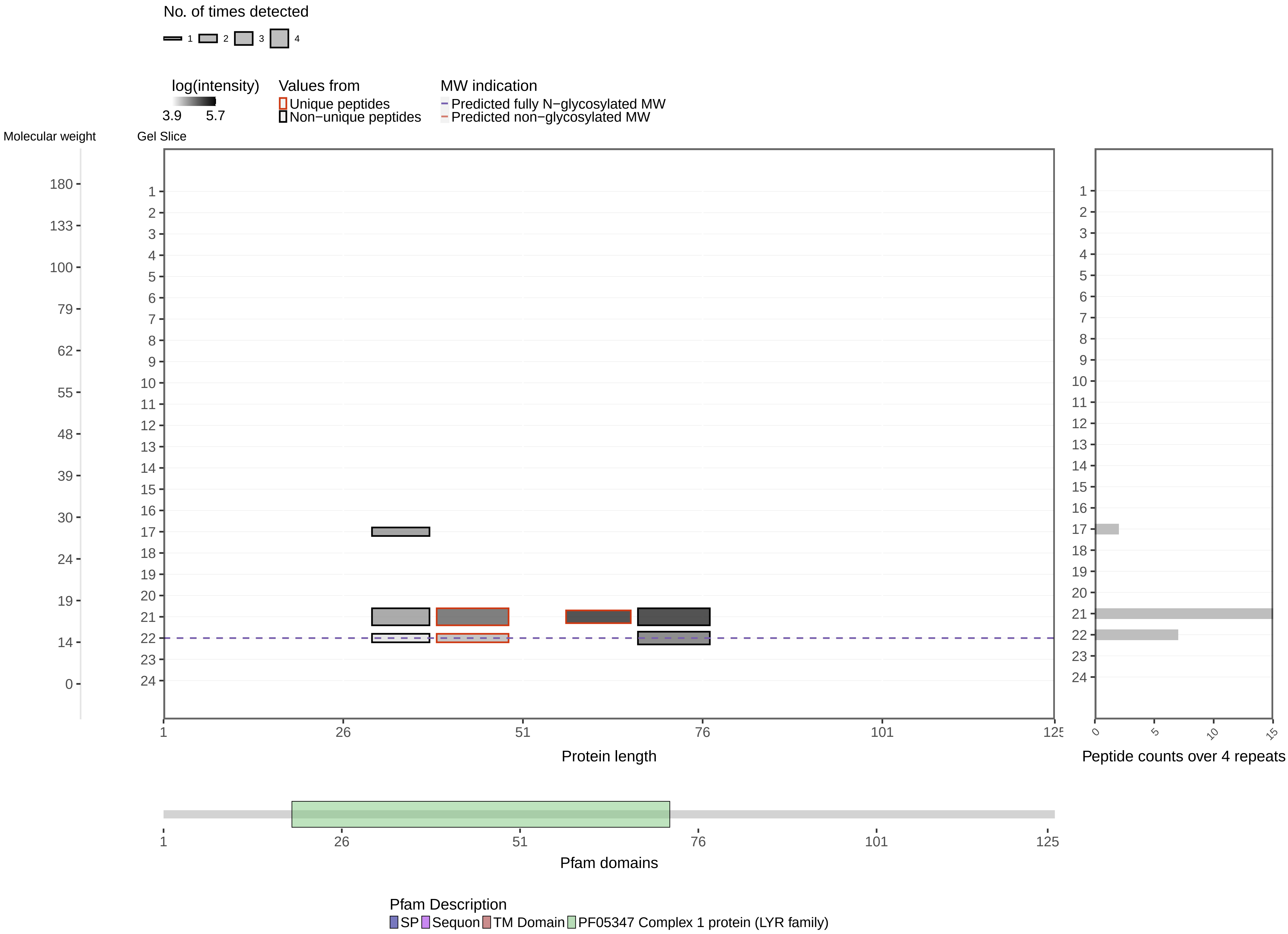


Protein sequence:

RERFEANRNVEDVETIDRLIADGEASYNKWRHPDPYIGMFLYEFHNIOLCLNPKOVESAIRILIVHVFFCEFHACS*

NbL11g13850.1 NADP−dependent_D−sorbitol−6−phosphate_dehydrogenase−like [MW = 34.76 kDa]

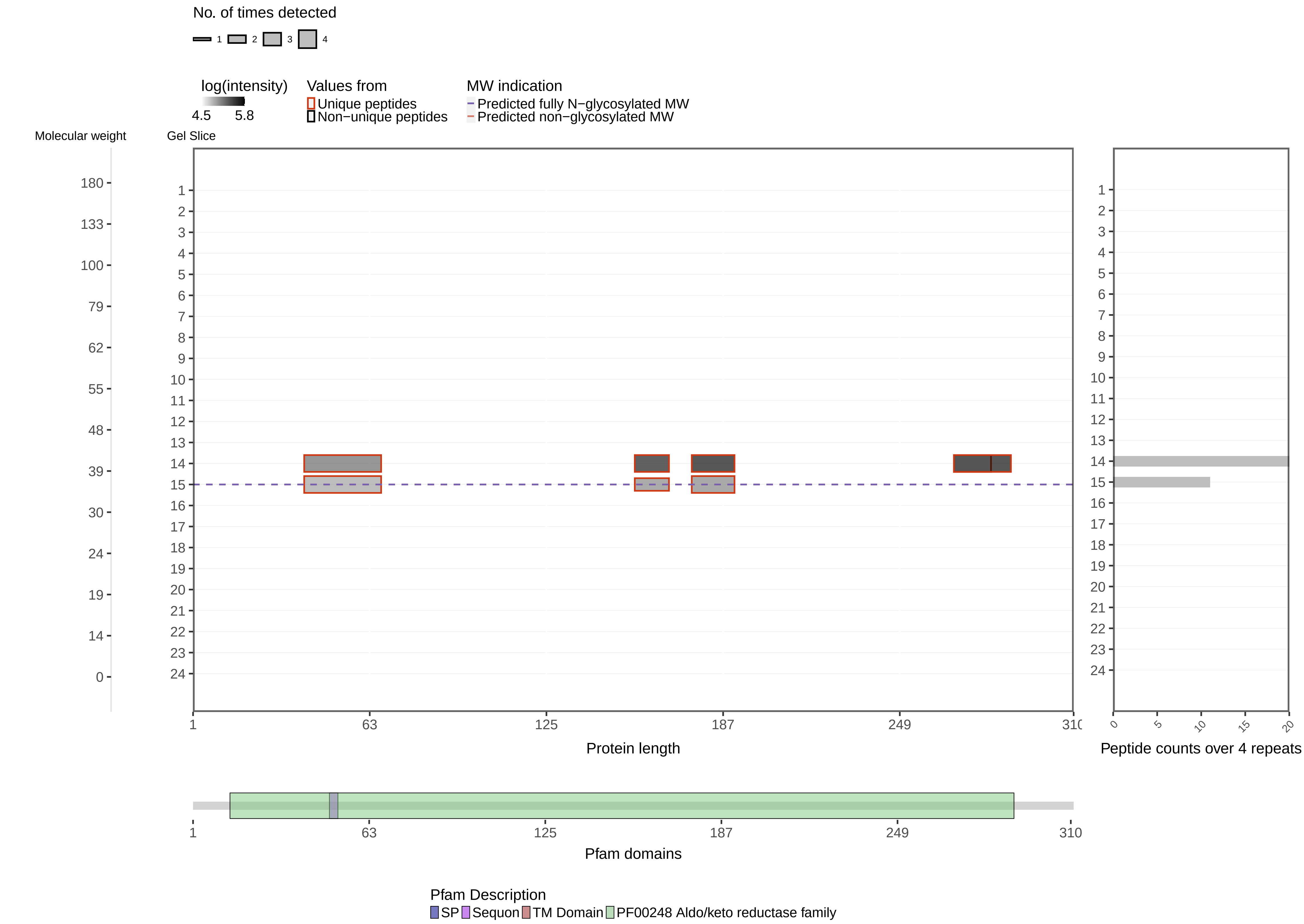


Protein sequence:

HAMENLVSLGLVRSIGISNYDIFLTRDCLAYSKVKPVVNOIETHPYFORESLVKFCOKHGICVTAHTPLGGAAANTEWFGTVSCLEDPALKGLAEKYKKTVAOVVLRWGIORNTVVIPKSSKLERLOENFNVLDFELTKEDMDLVKSLDRKYRTNOPAKFWGIDLYA*

### NbL11g13790.1 dihydrolipoyl_dehydrogenase_2_chloroplastic−like [MW = 64.18 kDa]

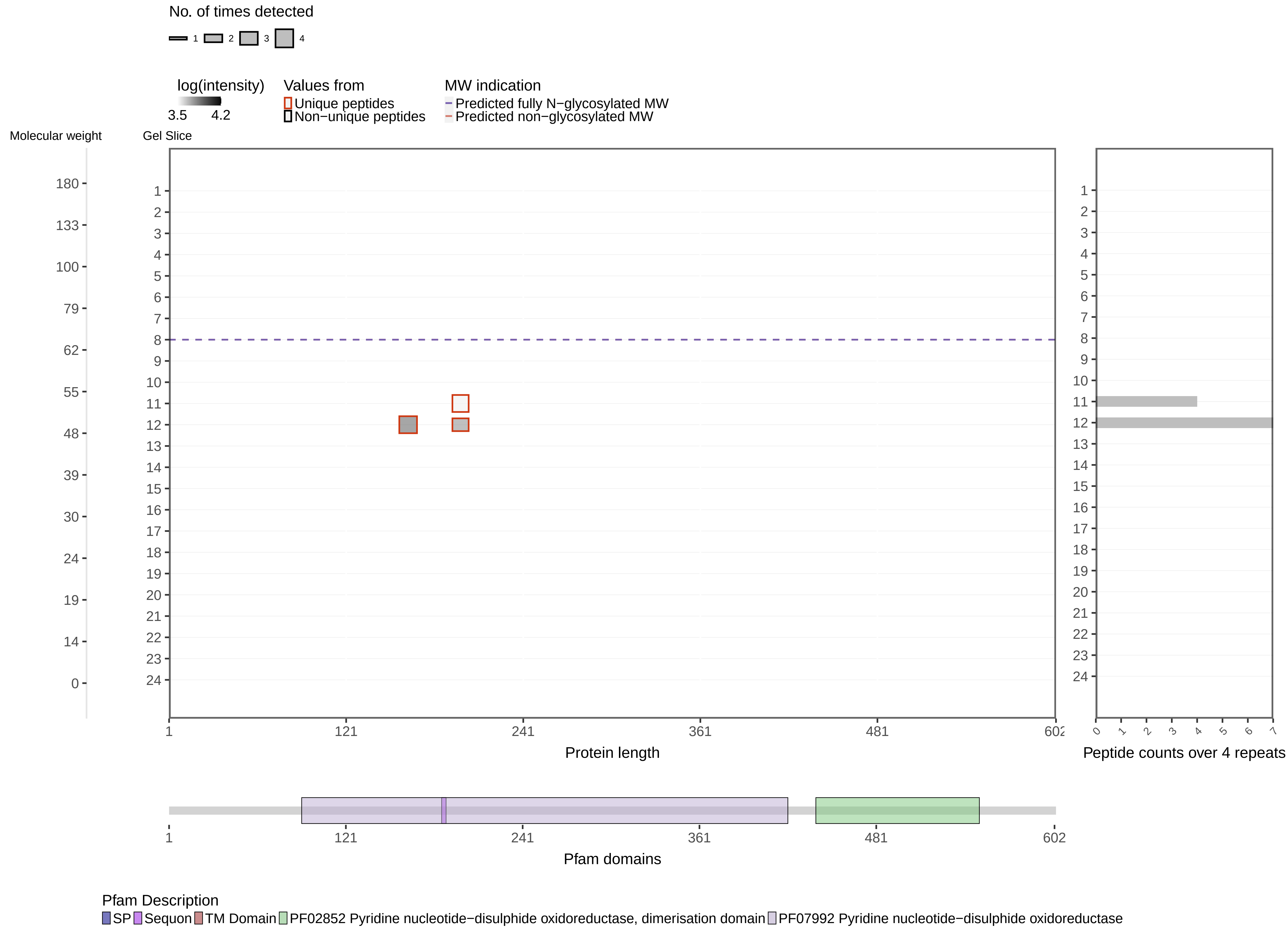


Protein sequence:

[illegible]

### NbL11g13750.1 calreticulin−like [MW − SP = 49.2 kDa]

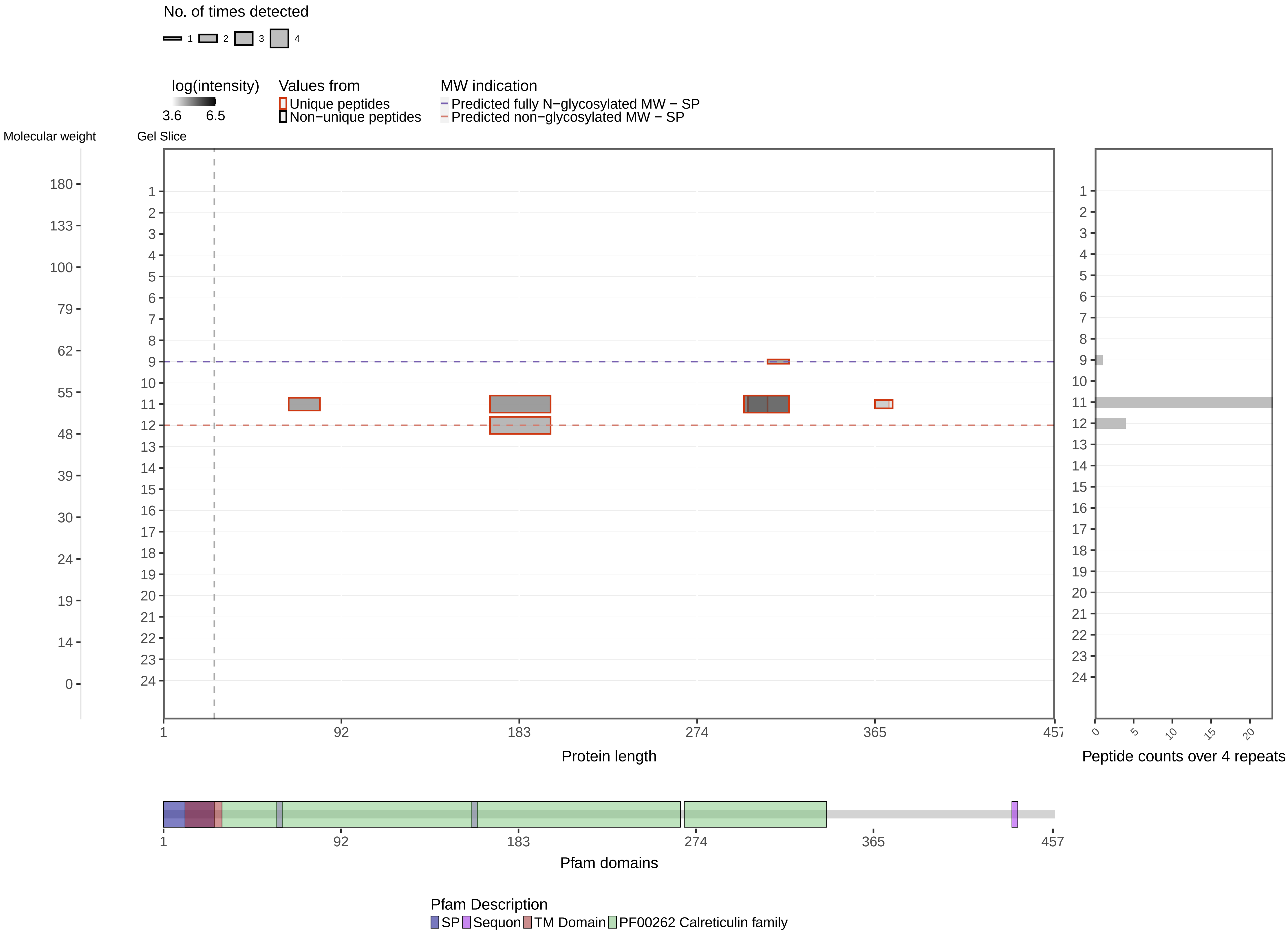


Protein sequence:
IKDPSAKKPEDWDEKEFIDDPEDKKPEGYDDIPEEITDPDAKKPEDWDDEEDGEWTAPTIPNPEYKGPWKPKKIKNPNYKGKWKAPLIDNPDFKDDPDLYVFPNLKYVGVELWQVKSGTLFDNIVICDDPEYAKAIAEETWGKQKDAEKASFDEEEKKREEEESKAAPADSDAEEDDDADDDSDDADENSESKDDAAHVRIVSCLSFKQRGSSRTFLVVHNLTAYILLNLFCRMNCKDQSF*

### NbL11g13720.1 54S_ribosomal_protein_L19_mitochondrial−like [MW = 16.8 kDa]

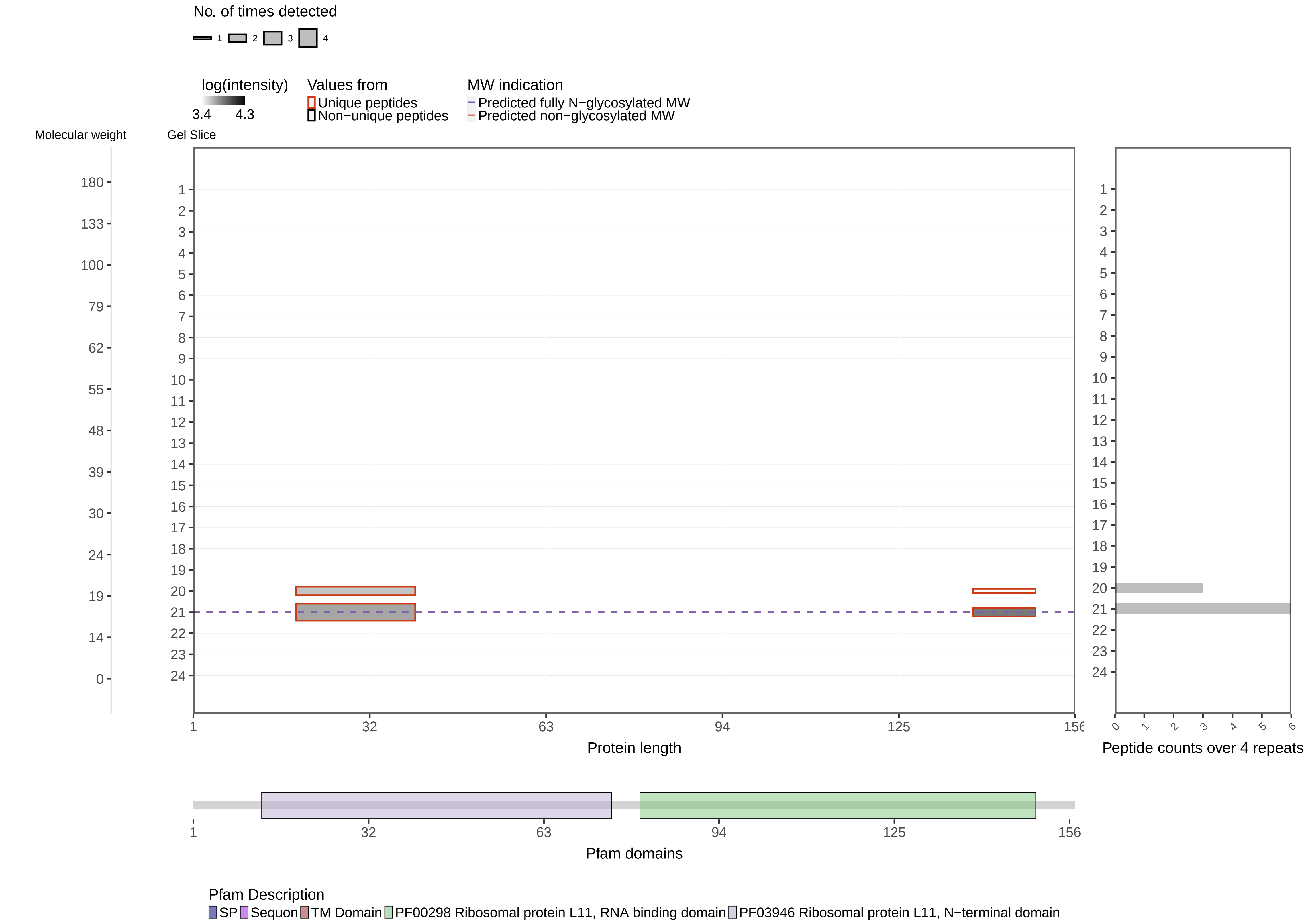


Protein sequence:

[illegible]VTIMAFKDGTFEFTVKSPSVTWYLKQAAGIELGSGRPGHVTASTLTLKHVYEIAKIKQSDPFCQYMPLESICKSIIGTANSMGIKVQKELD*

### NbL11g13630.1 ATP−dependent_Clp_protease_proteolytic_subunit_5_chloroplastic−like [MW = 32.26 kDa]

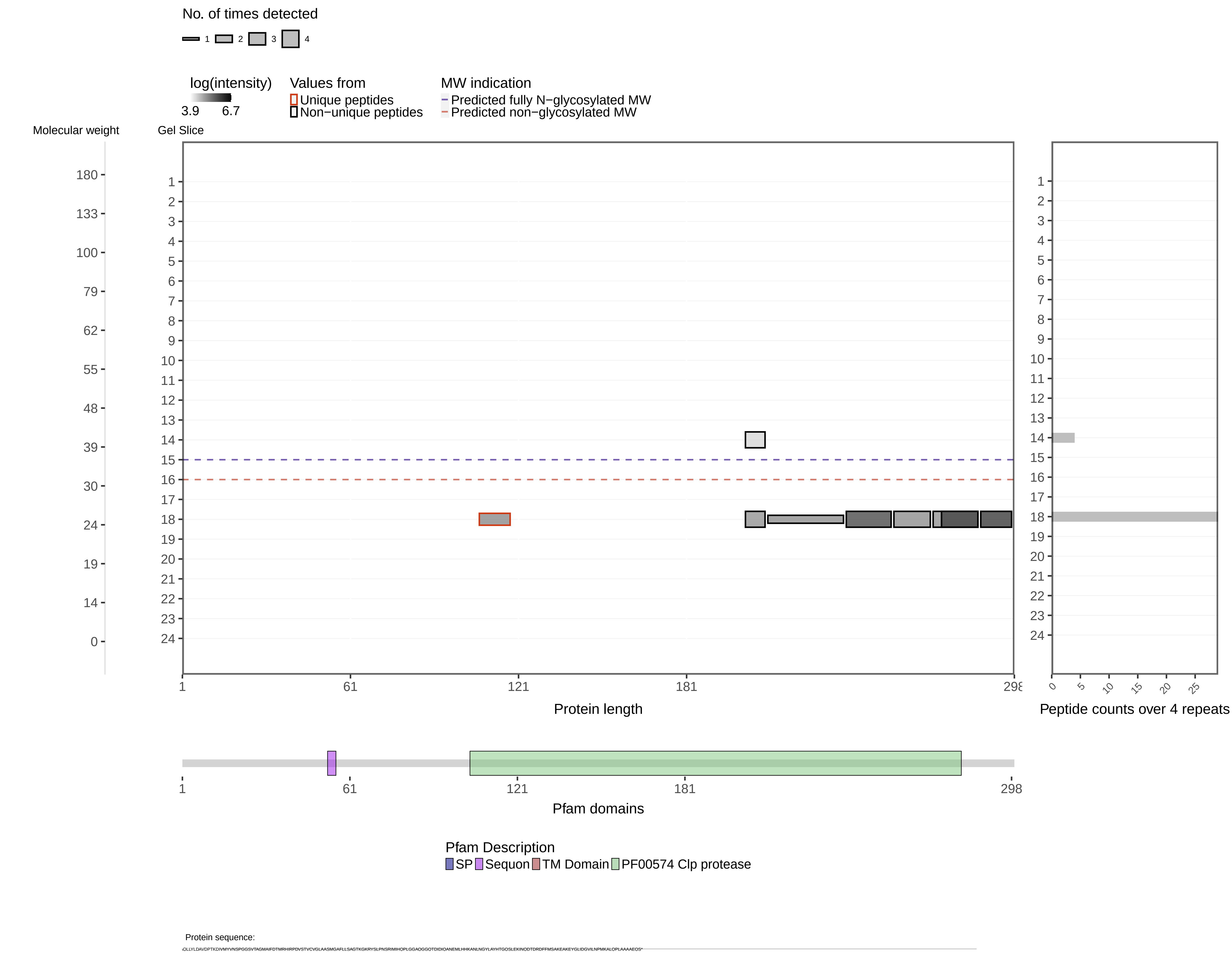

NbL11g13560.1 uncharacterized_LOC107804171 [MW − SP = 38.31 kDa]

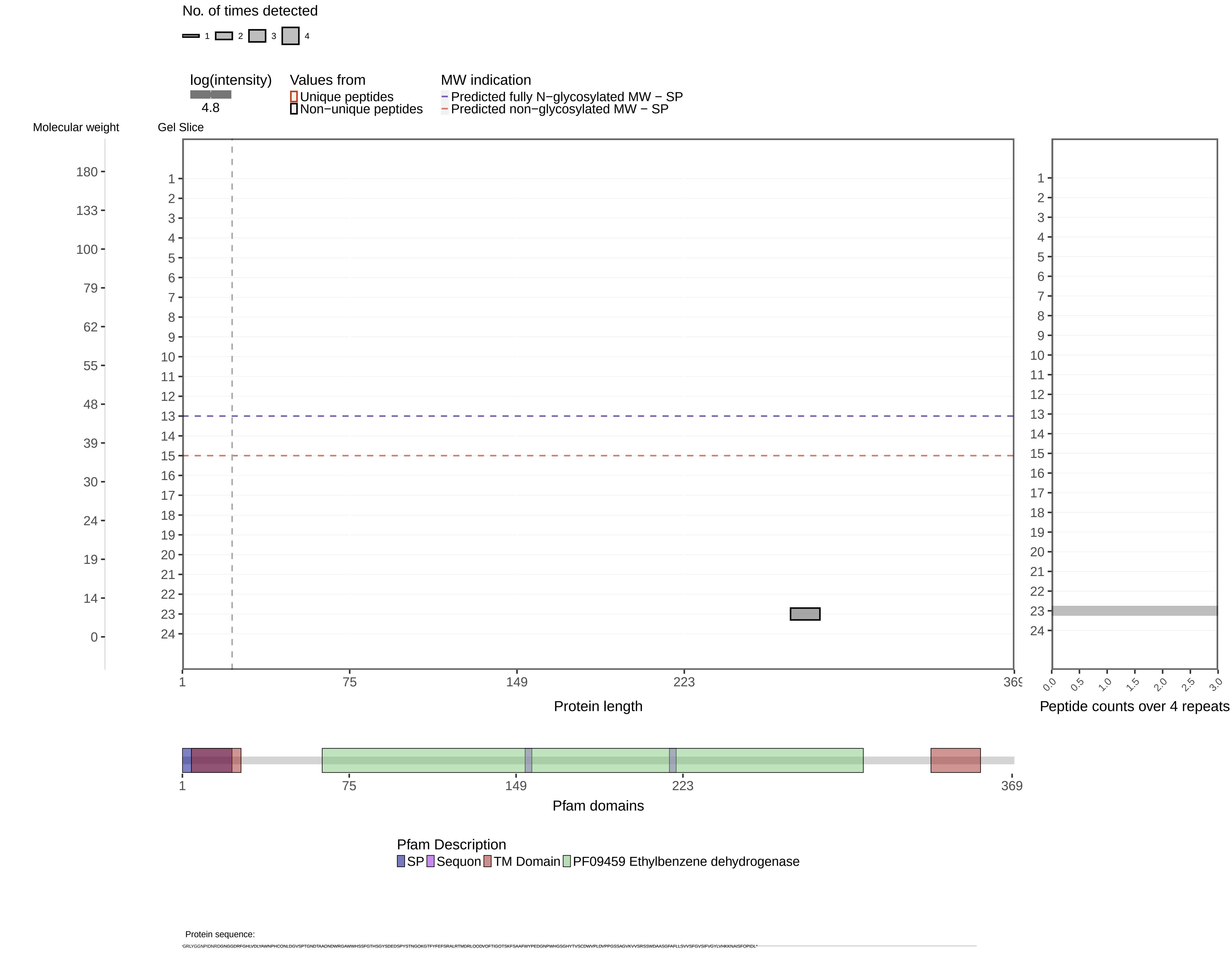

### NbL11g13530.1 NHL_repeat−containing_protein_2 [MW = 121.37 kDa]

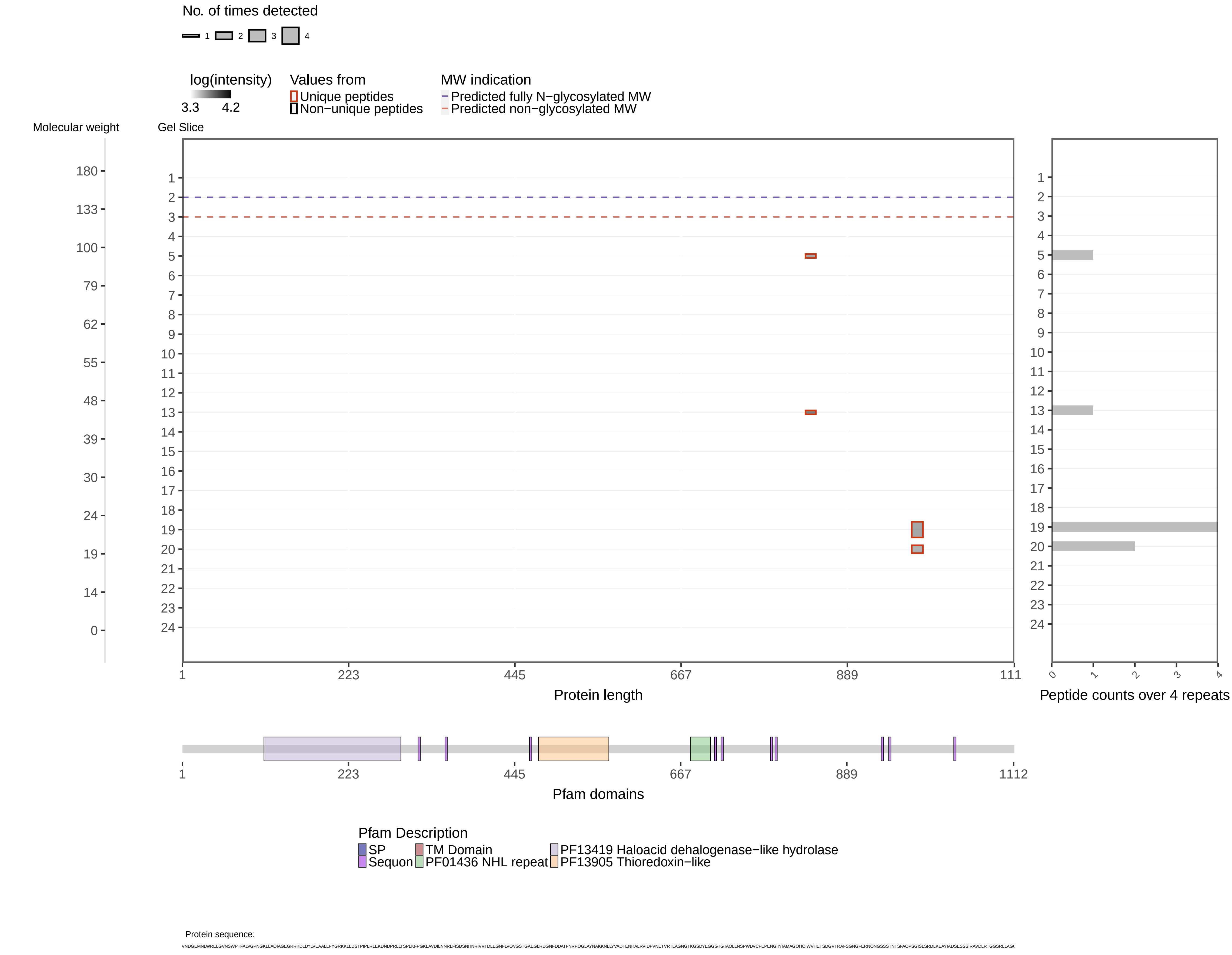

### NbL11g13450.1 probable_Xaa−Pro_aminopeptidase_3 [MW = 41.53 kDa]

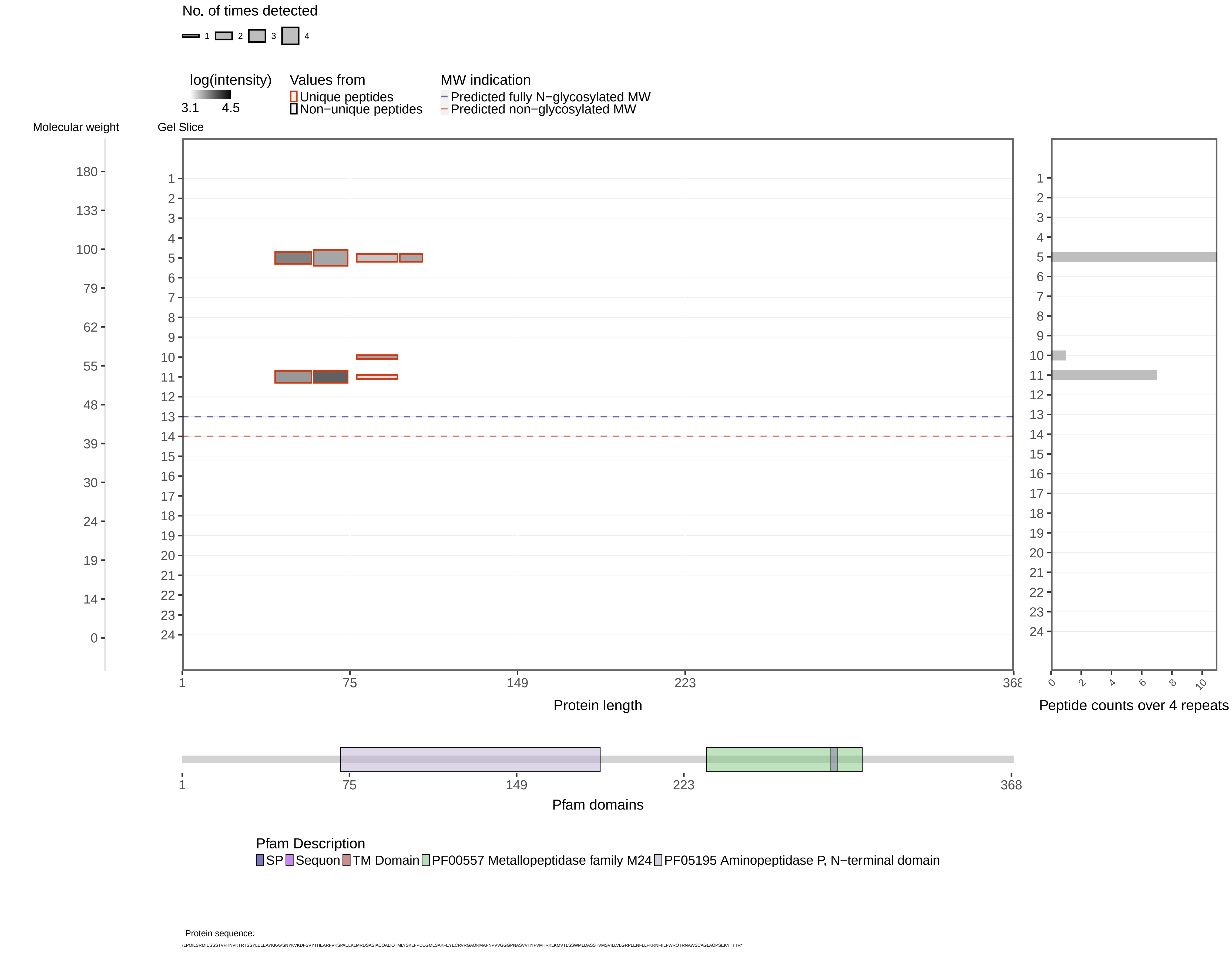

NbL11g13430.1 uncharacterized_LOC107799792 [MW = 23.01 kDa]

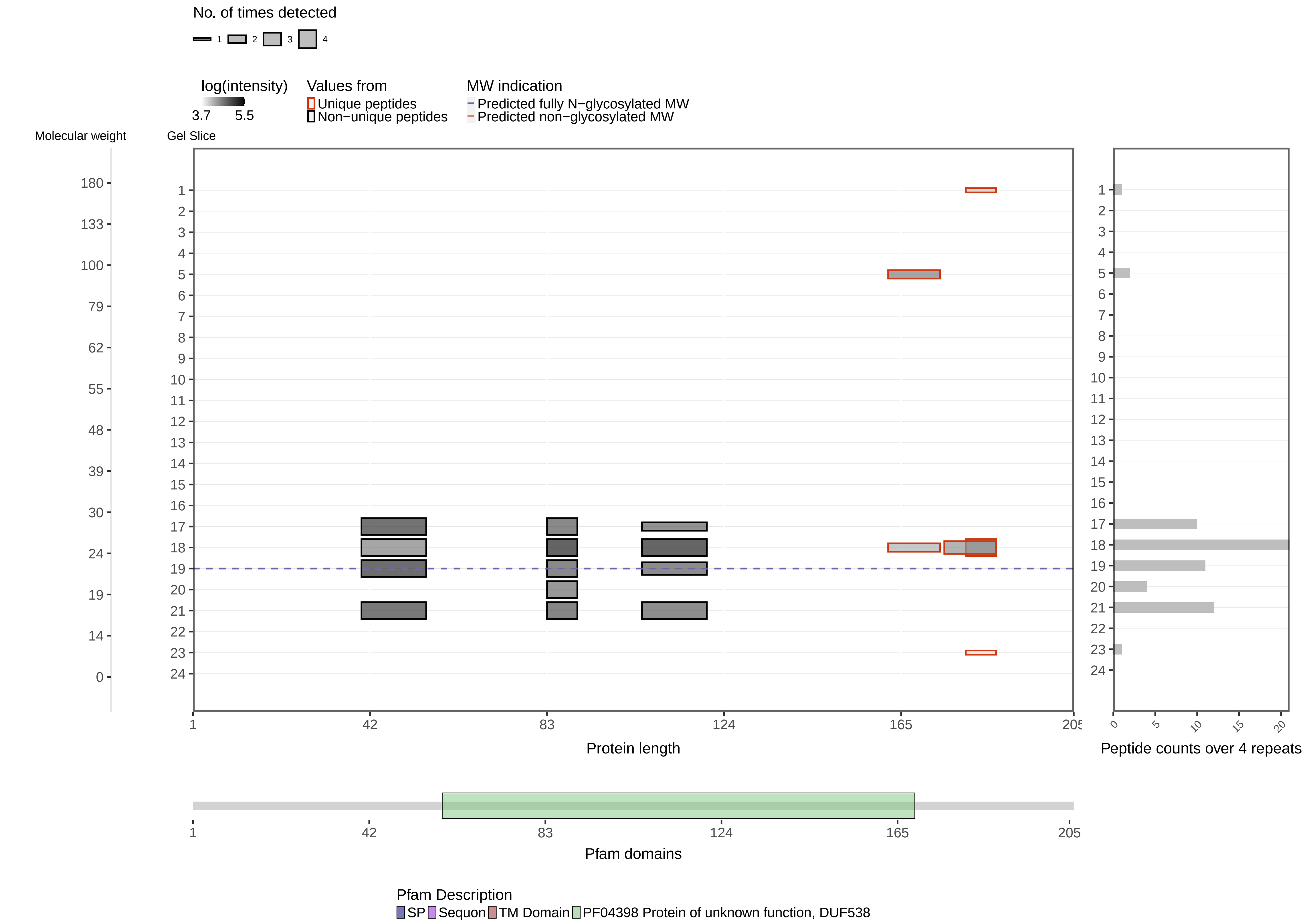


Protein sequence:

KSKKKTEHKFEKIGRQVQYGTEVTAIVEQNKIKKLTGVKAKELLMWLTLTEICVDDPPTGKIHFKATSGLSRTFPVSAFEIDVPKKEEVKDVNEVVNKENKEVIAAAAALEVKEV*

NbL11g13330.1 ABC_transporter_I_family_member_20−like [MW = 36.6 kDa]

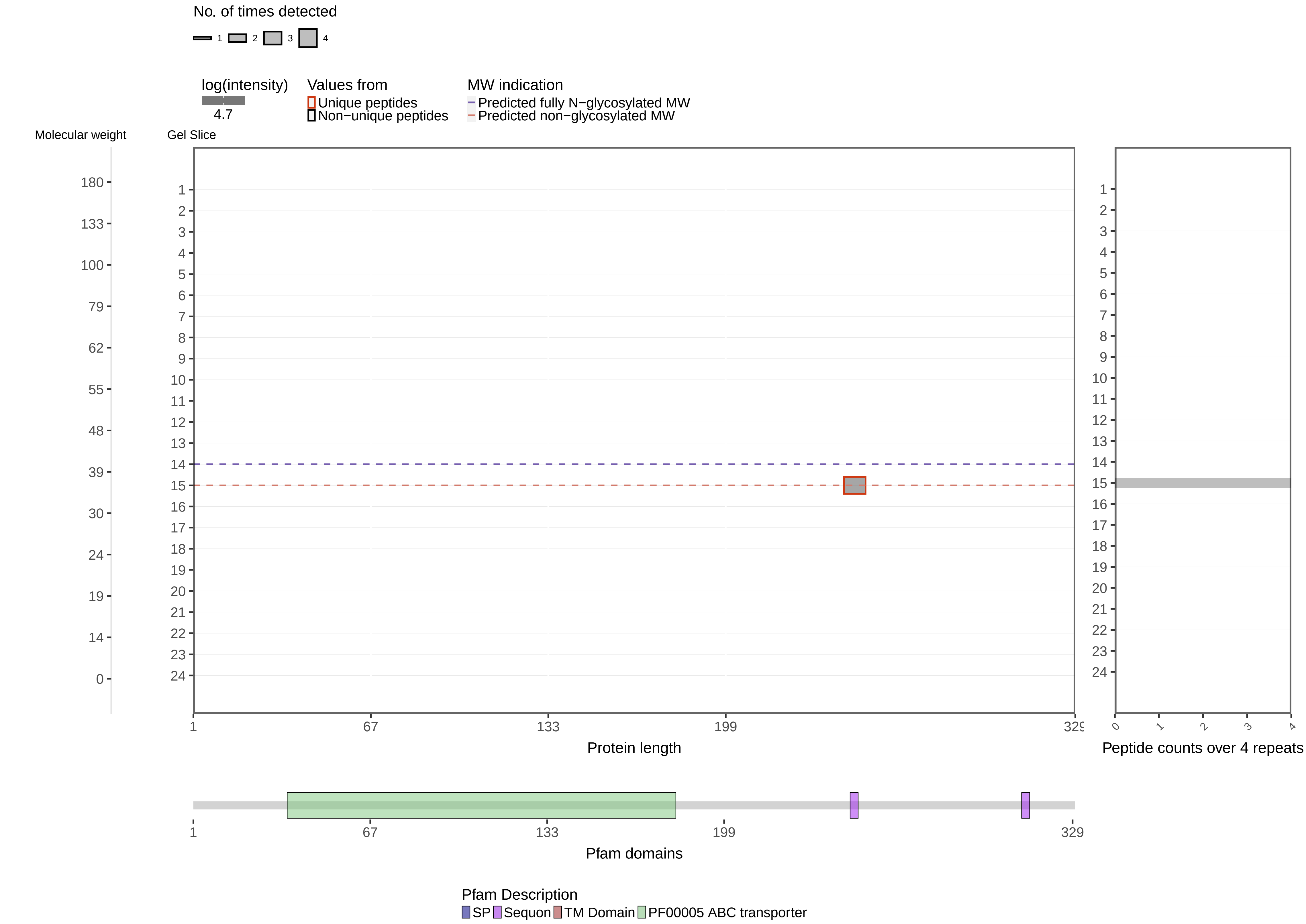


Protein sequence:

KASDGQRRRVQICMGLLKPFKVLLLDEITVDLDVLARADLLKFLSKECEERGATIIYATHIFDGLENWPSHMLYVAHGKLQLAMPMDKVKEISNLSLMRTVESWLRKERDEERKRRAERKAKGLPEYGKQVEGSRVVGDPARAAARPLNNGWAGGRLNSTIAGEENFVLSSNRVLR*

### NbL11g13150.1 ATP−citrate_synthase_alpha_chain_protein_2 [MW = 46.45 kDa]

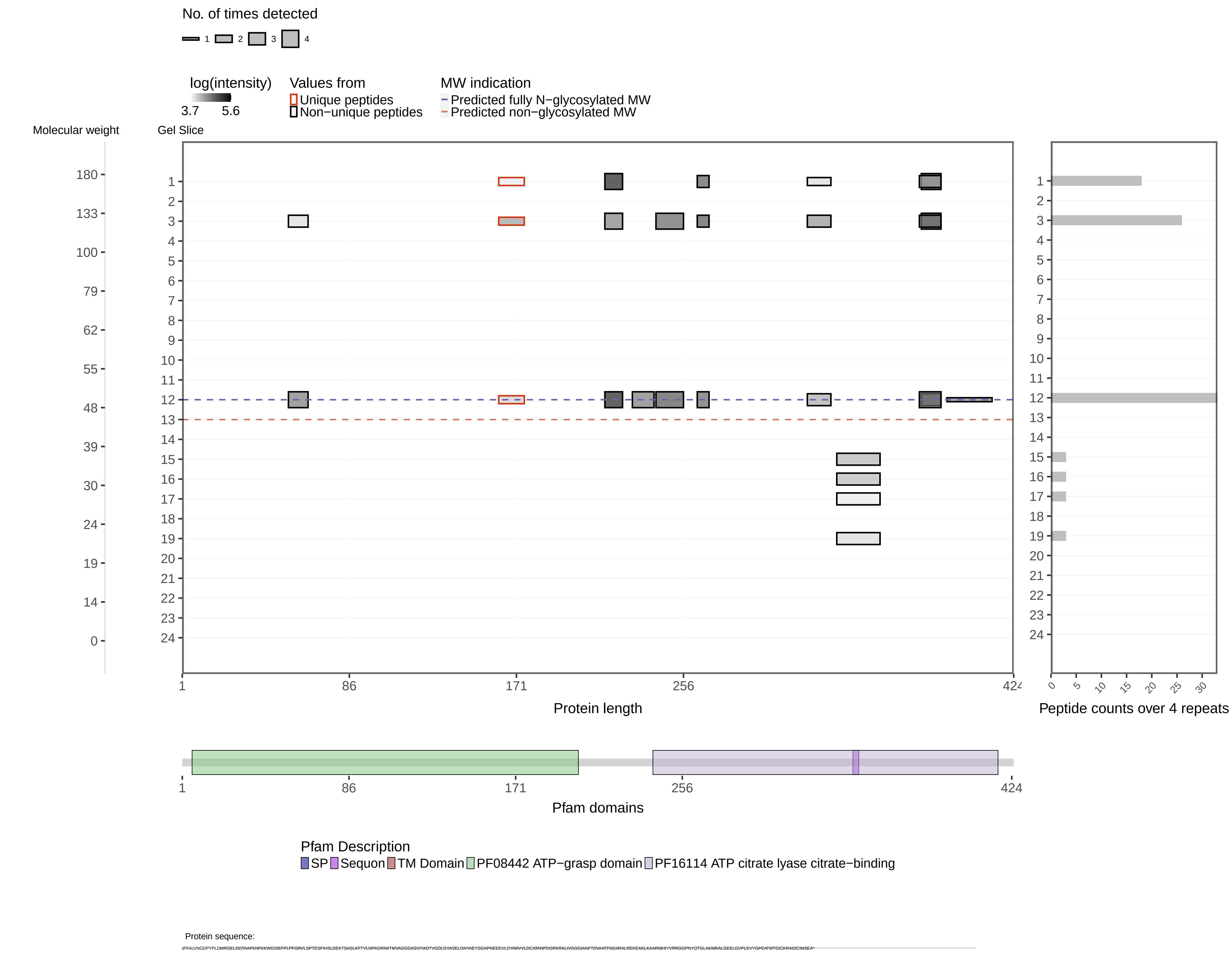

NbL11g13130.1 S−adenosylmethionine_synthase_1 [MW = 43.22 kDa]

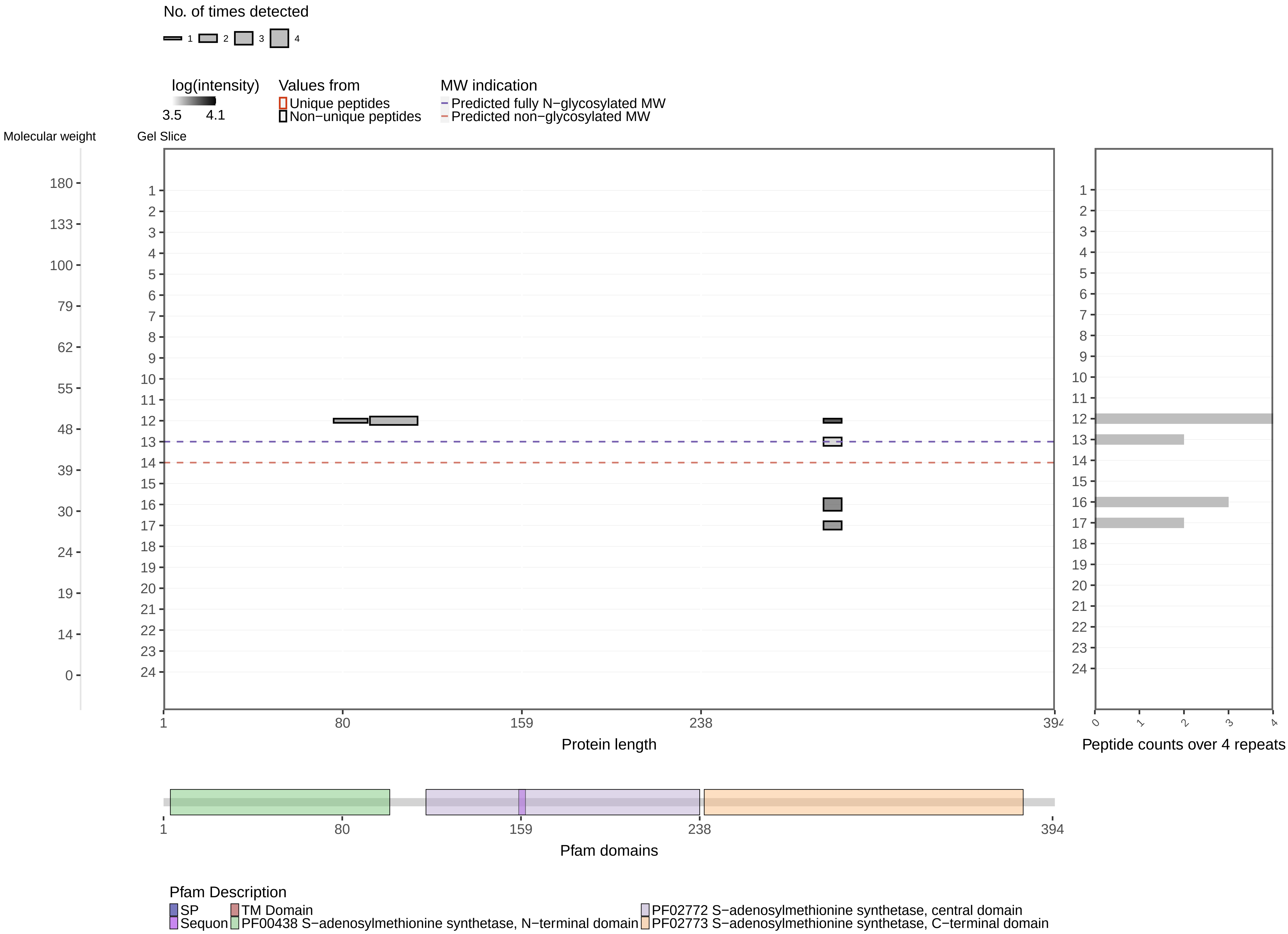


Protein sequence:

[illegible]MVPVRVHTVLISTQHDETVTNDEIARDLKEHVIKPVIPEKYLDEKTIFHLNPSGRFVIGGPHGDAGLTGRKIIIDTYGGWGAHGGGAFSGKDPTKVDRSGAYIVRQAAKSIVASGLARRCIVQVSYAIGVPEPLSVFVDTYGTGKIPDKEILKIVKENFDFRPGMISINLDLKRGGNGRFLKTAAYGHFGRDDSDFTWEVVKPLKREKPQD*

### NbL11g13000.1 cyprosin−like [MW − SP = 52.28 kDa]

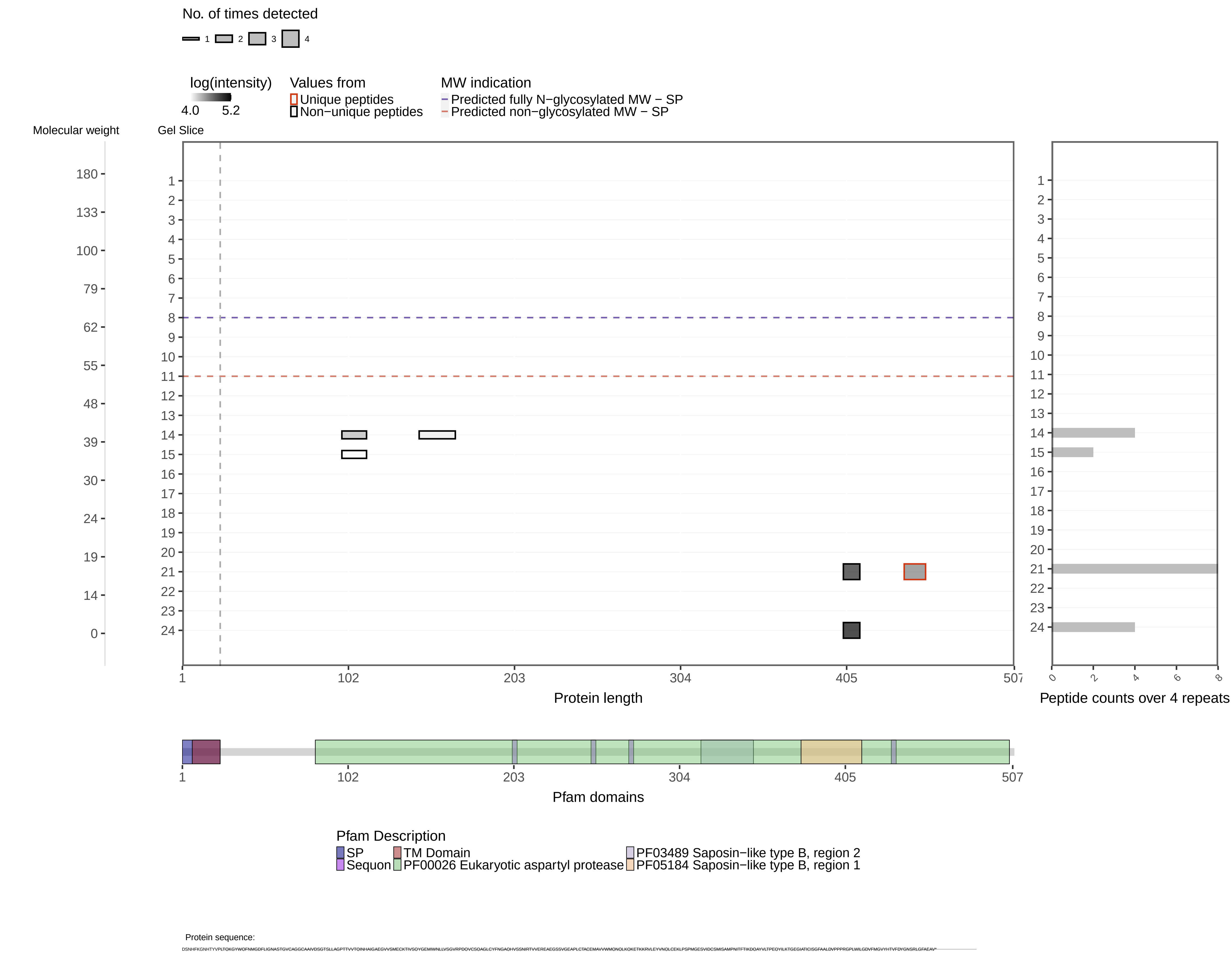

### NbL11g12940.1 FK506−binding_protein_5−like [MW = 82.85 kDa]

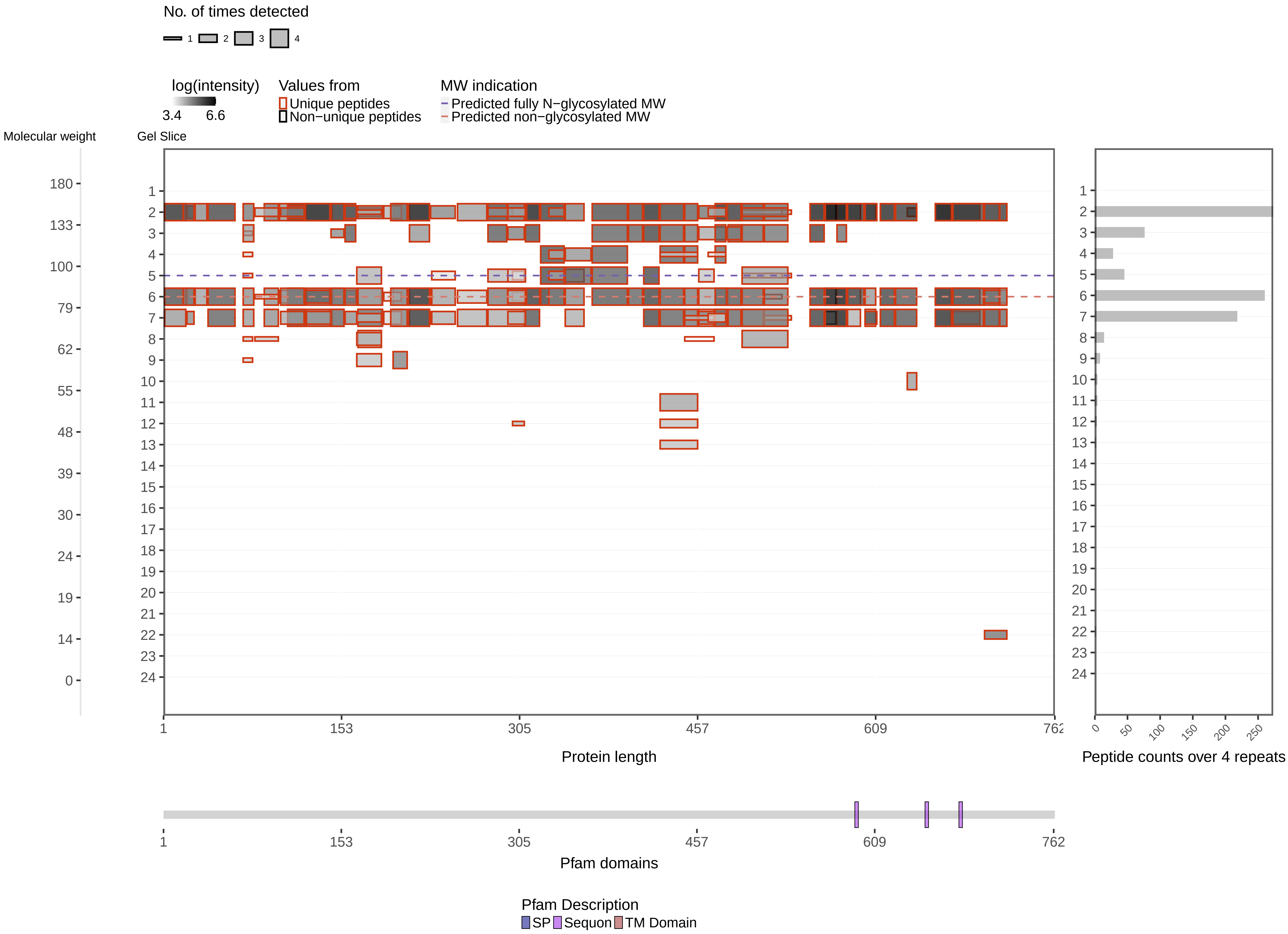


Protein sequence:

PVNEKPEEQPTTVDVVETPVGAVDNKNEQADLEASEPQKTVVTVSDVPVAEPKIEDISVSAPSIEEKPTEATEKFEQEVGEAVTGKEEAIPVDKVEEAPKGESLLAESSKKATSTTLGSLMQKTDEESPAEVAGDNAVVEEKEHESVPEATVAEAPAKEEVEKVERETKDVKTDDQDKGEKTVASEELAAPDAKVVETSEVAEKAPEVEISSRDMEPLAENESTKDEKIVTTNDQDEKREKSVATEELAAPEAKVVETSEVAEKVPELEIASRDIEPNAKNGSKKDEKIAASEAN

### NbL11g12930.1 probable_carbohydrate_esterase_At4g34215 [MW − SP = 27.28 kDa]

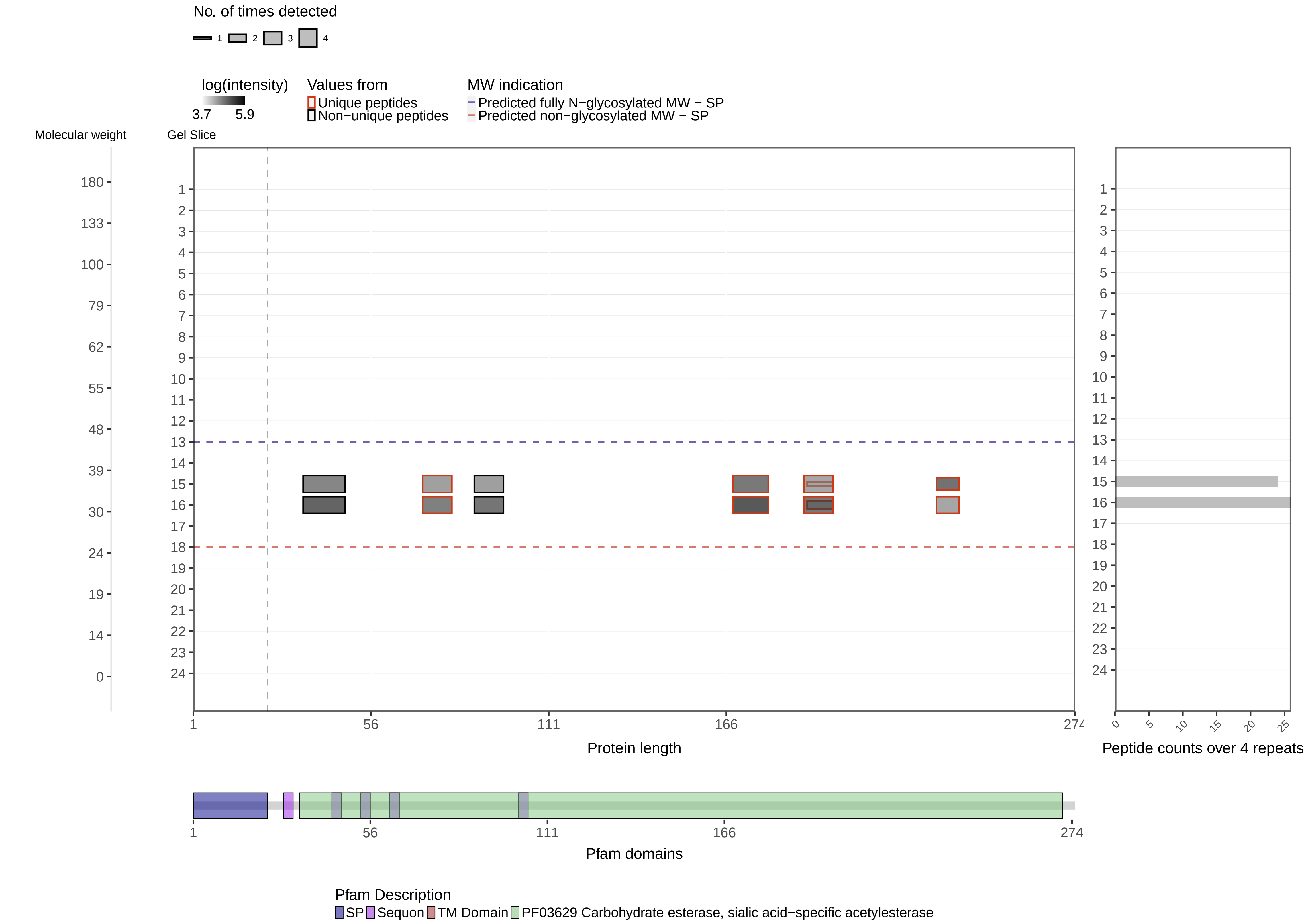


Protein sequence:

[illegible]VGVIGLVPCAVGGTSIIEWAPGGFLYKQMIRRTKVATRFGGRVRALLWYQGESDTKTLEDAKMYKRRLESFFNHVRHDLQLPTLPVIQVGLATALGPYMELVRKAQREINVGNVRYVDAKGLQVGPDYVHLTTAAQIQLGQMLADAFLRI*

### NbL11g12900.1 pectin_acetylesterase_12−like [MW − SP = 44.2 kDa]

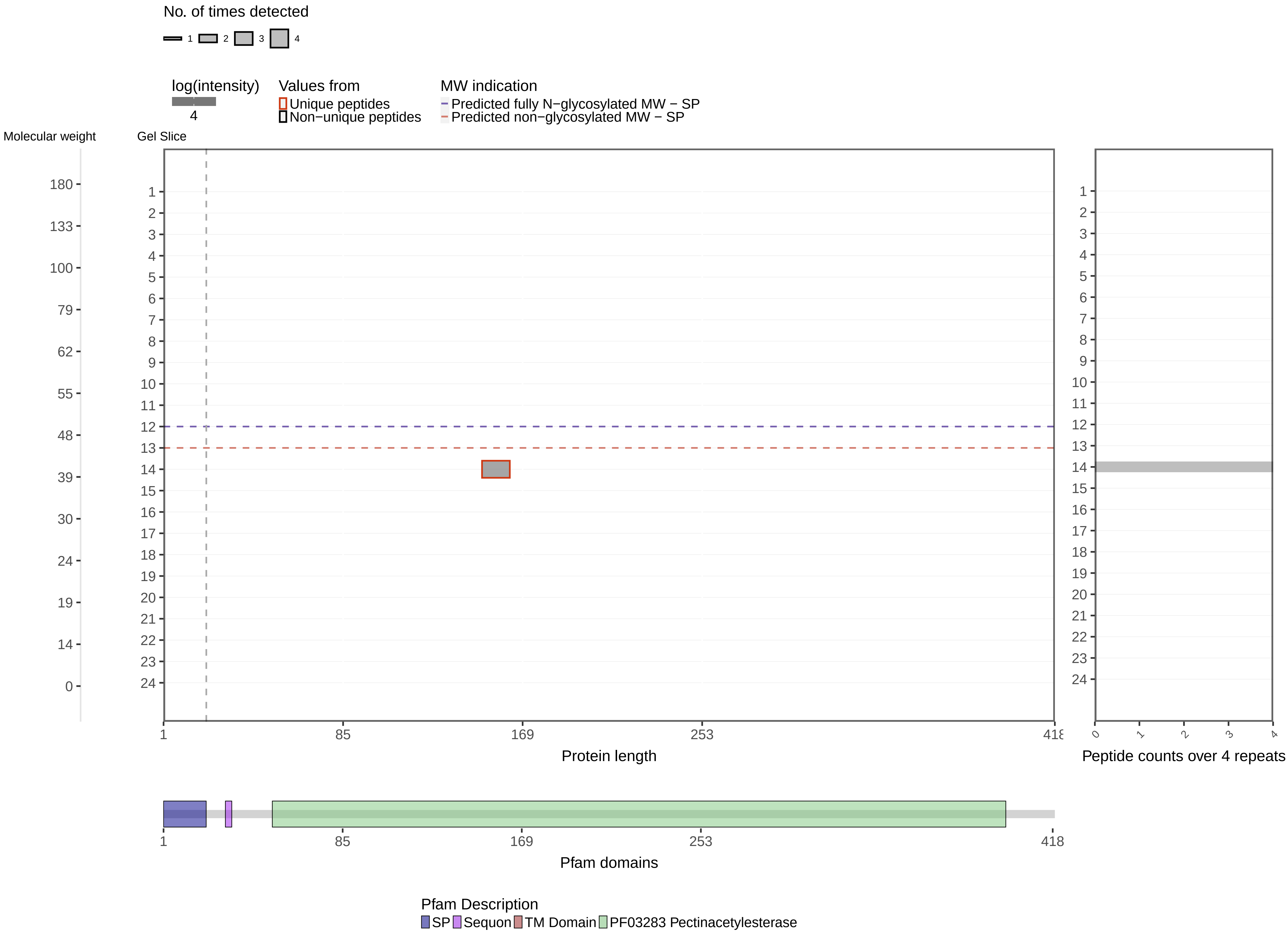


Protein sequence:

LSGCSAGGLASIMHCDDFRNLLPHSTKVKCLSDAGLFMDAVDVSGGRSLRNFFGGVVKLQGLHKTLPRTCTNHLDATSCFFPENLINNIKTPLFLLNAAYDSWQLQESLAPRGADPHGIWHDCTLNNERCSPSQIHFLQGFRIHMLNSIKRFAASRQNGLFINSCFAHCQSERQDTWFSDNSPMINNKPIALAVGDWYFDRSGMKAIDCAYPCDRTCHNLIFR*

NbL11g12890.1 flap_endonuclease_1 [MW = 45.89 kDa]

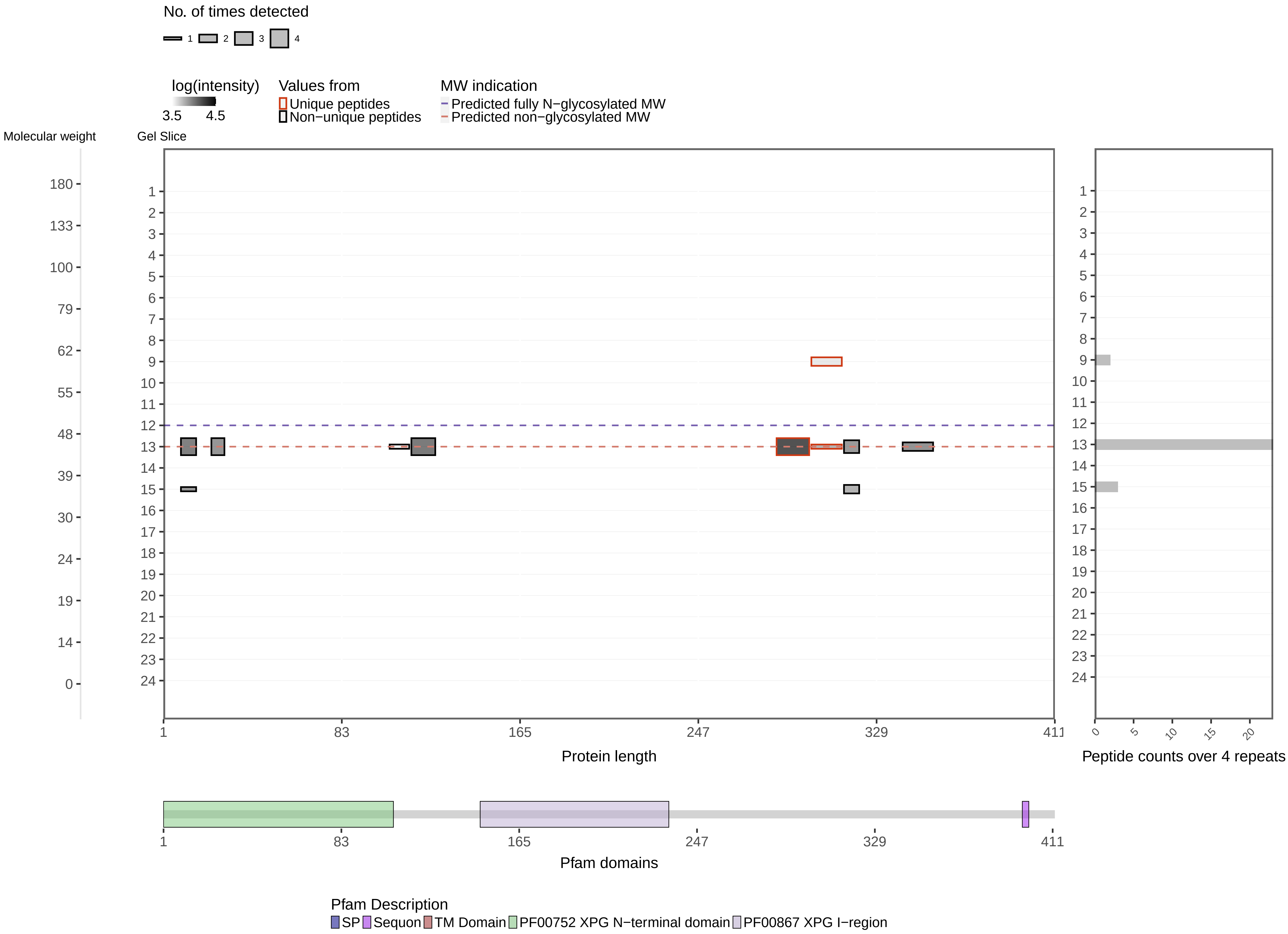


Protein sequence:

LRHLMDPSSKKIPVMEFDISQILEELELSMDQFIDLCILCGCDYCDSIRGIGPQTALKLVRQHGSMESILENINKERYQIPDDWPYEEARRLFKEPLVADDEQTELKWAAPDEEGLVNFLVKENGFNIDRVTKAVEKIKTAKTKSSQGRLESFFKPVSTSAPVKRKGTKCVLVSPNPVTKFKTLSLQSRPMVLGSQIFPSLDLGSNYSVPLSRHVCFGT*

NbL11g12850.1 nectarin−1−like [MW = 26.36 kDa]

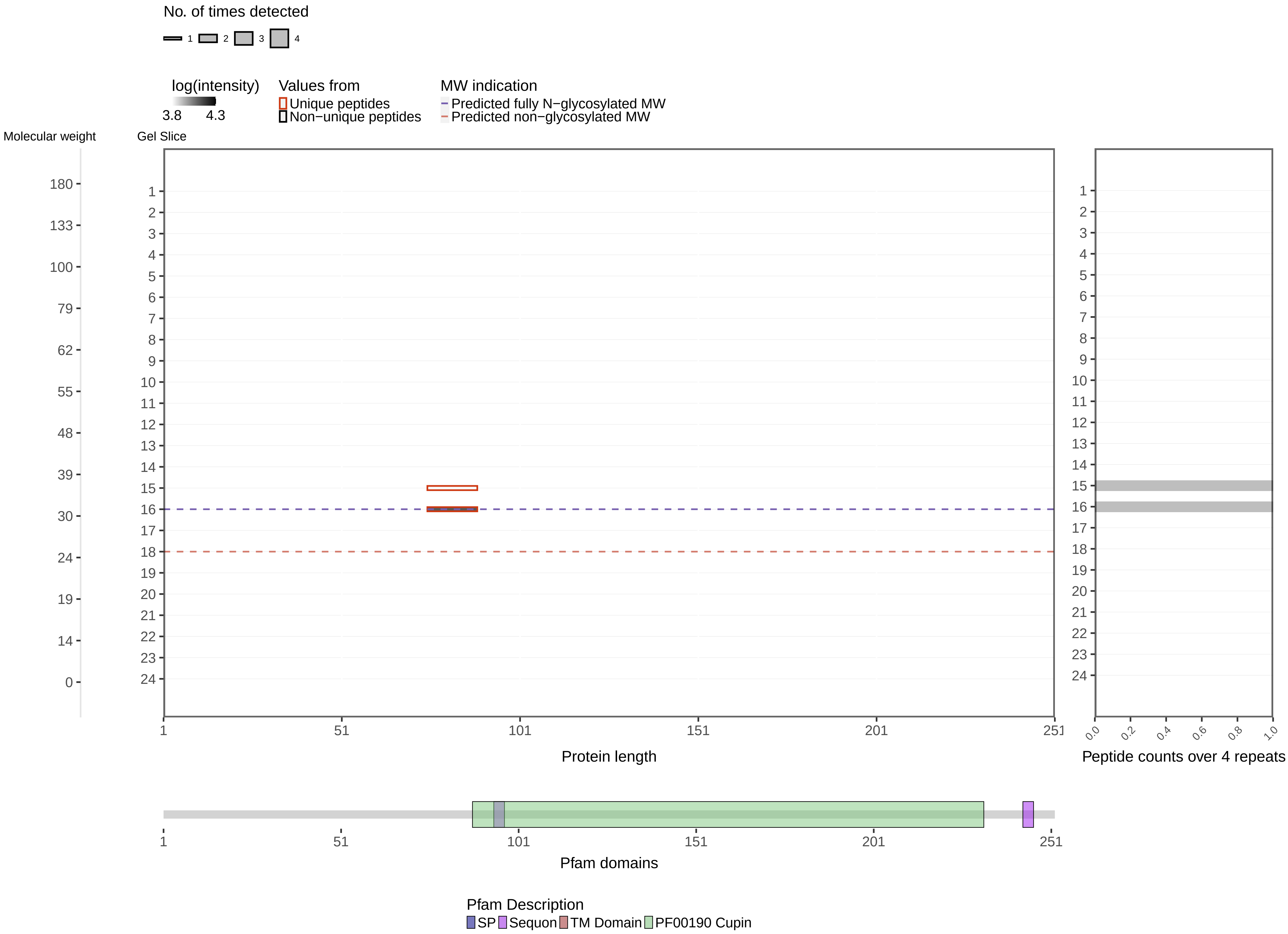


Protein sequence:

ITLGVSLARIDYAPGGINPPHVHPRATEMVYVLSGQLYVGFITTANVLVSKVIVQGEVFAFPRGLVHFQKNNGKVPAAVIAGFNSQLPGAQSMATTLFAASPSVPDDVLGQAFQINTMQVQQIKSRFAPGNRSSIFVC*

### NbL11g12840.1 nectarin−1−like [MW − SP = 20.65 kDa]

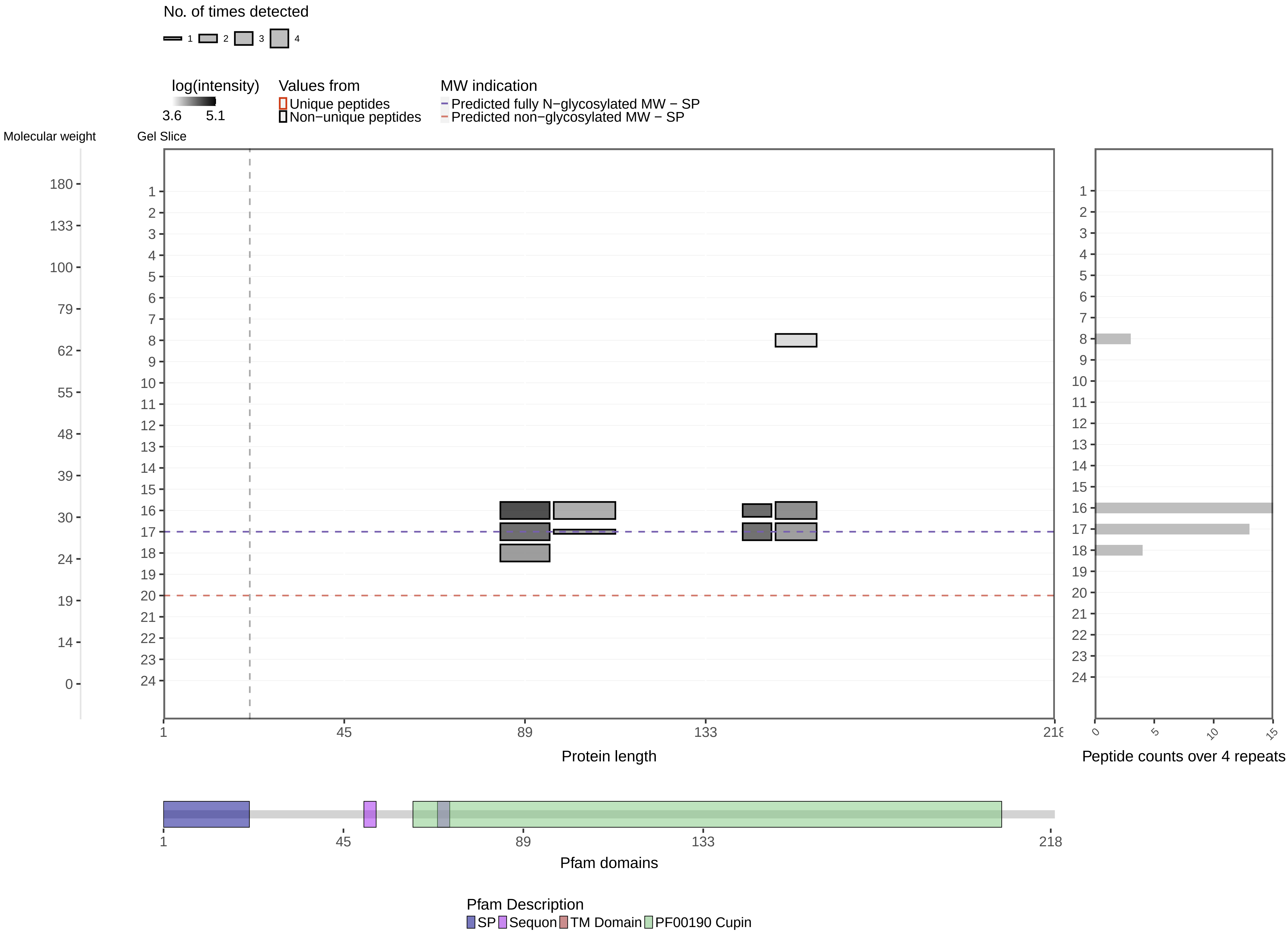


Protein sequence:

RIDYAPGGINPPHTHPRATEMIFVTEGELDVGFITTSNVLISKHIVKGEVFVFPKGLVHFQQNNGKVPAAVVAAFNSQLPGTQSIATTLFASTPTVPDHVLTKAFQIGTKQVEKIKSRLAPKK*

NbL11g12830.1 glutamate−−tRNA_ligase_cytoplasmic [MW = 86.8 kDa]

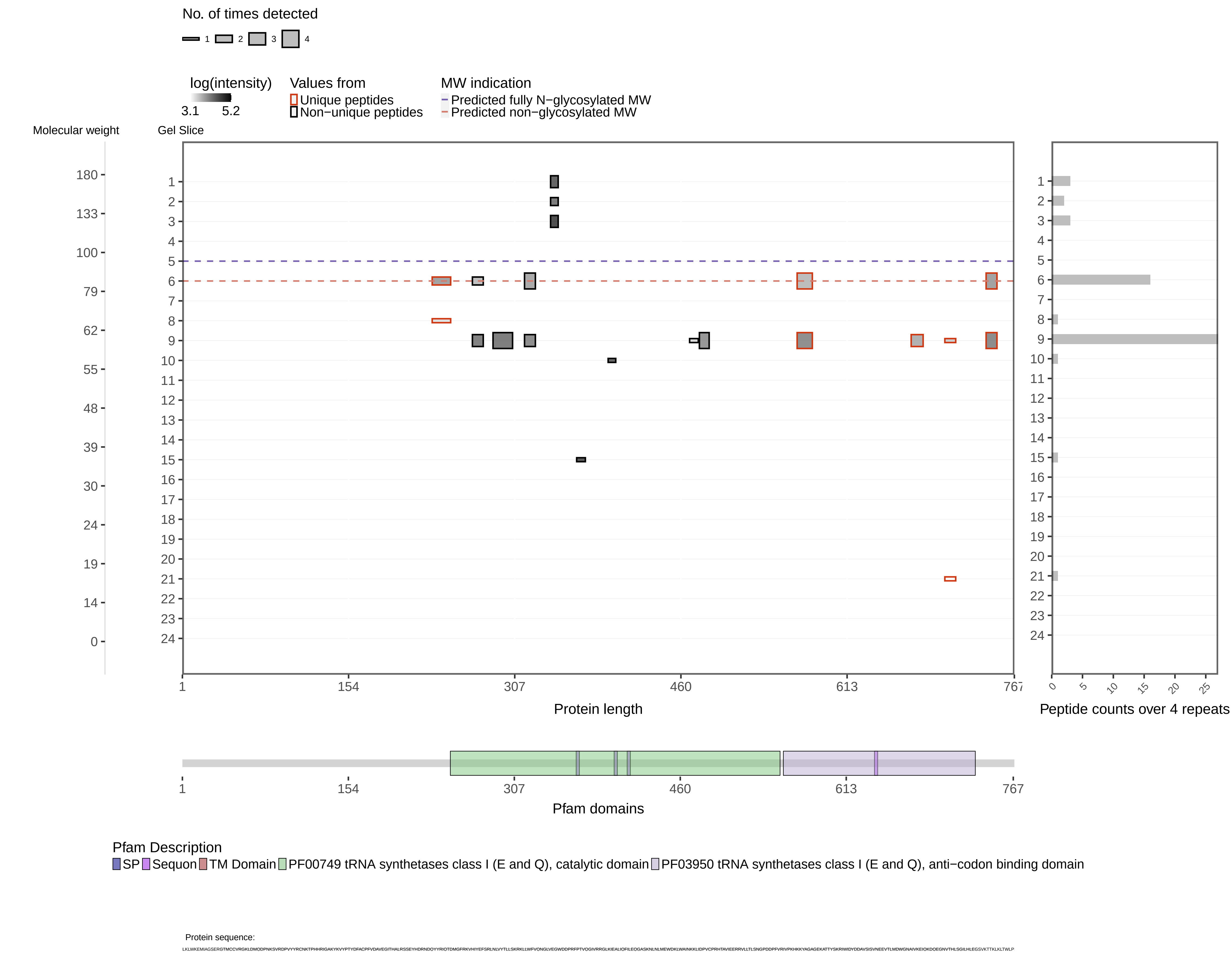

NbL11g12700.1 endoglucanase_25−like [MW = 68.69 kDa]

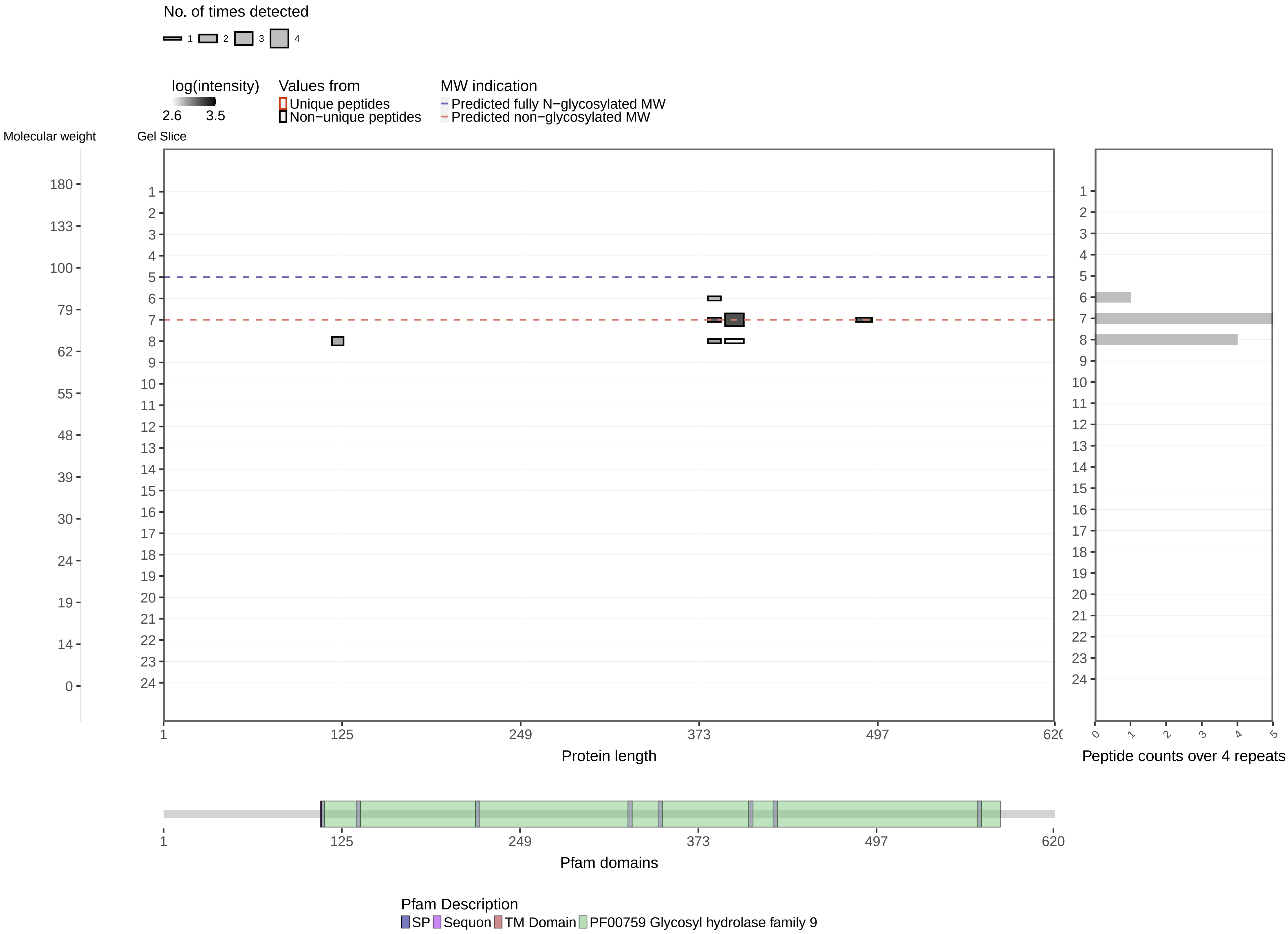


Protein sequence:

GAKTLFKFSRDQRGRYSVGNEAETFYNSTGYWDEFIWGAAWLYYATGNSSYLQLATTPGLAKHAGAFWGGPDYGVLSWDNKLTGAQVLLSRMRLFLSPGYPYEEILKTFHNQTSIIMCSYLPIFTSFNRTKGGLIQLNHGRPQPLQYVVNAAFLATLFSDYLEAADTPGWYCGPNFYSTDVLRRFAETQIDYILGKNPRKMSYVVGFGNHYPKRVHHRGASIPKNKVKHNCKGGWKWRDSSKANPNIIVGAMVAGPDKHDGFHDVRSNYNYTEPTLAGNAGLVAALVALS

NbL11g12640.1 probable_leucine−rich_repeat_receptor−like_protein_kinase_At5g49770 [MW − SP = 101.86 kDa]

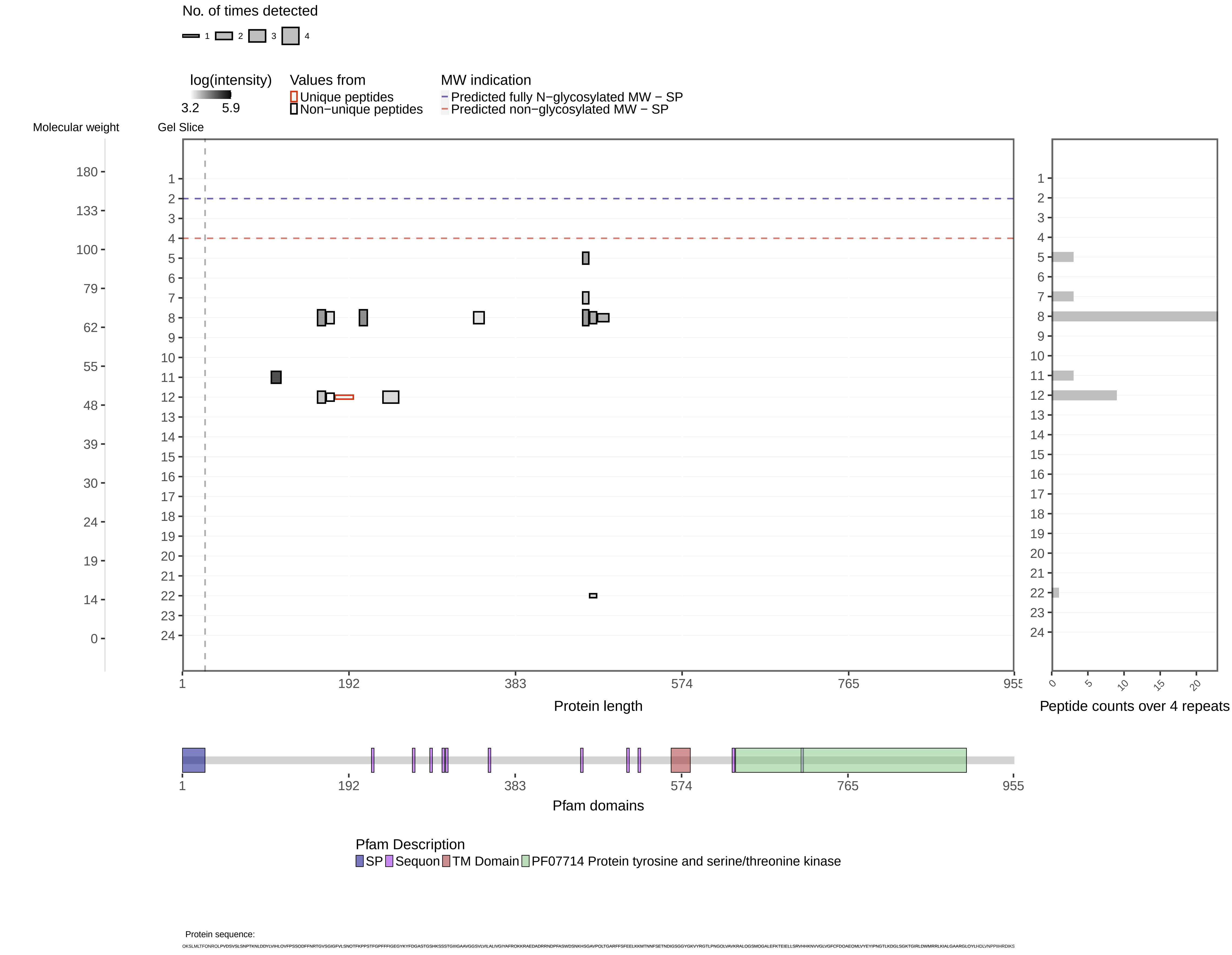

### NbL11g12600.1 cinnamoyl−CoA_reductase_1 [MW = 35.78 kDa]

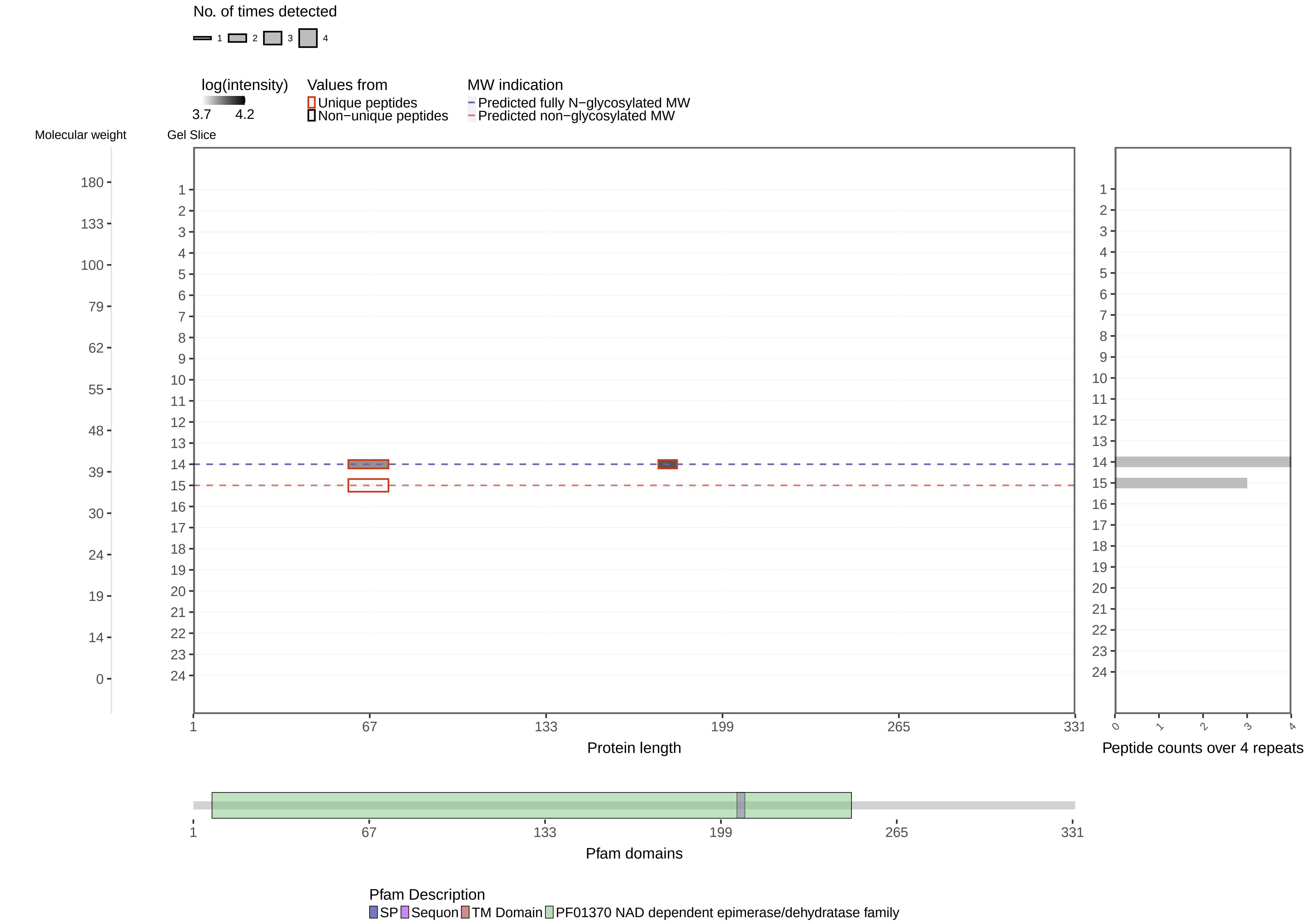


Protein sequence:

TDLDYCKSROKWYPVSKTLAEKTAWEFAEKNGLDVVAINPATCLGOLLOPGLNASCAVLOOLLOGSTDTOEYHWLGAVHVKDVAKAOILLFESPHASGRHLCTNGIYOFGDFAERVSKLYPEFPTHRFTGETOPGLVGCKDAAKKLIDLGLVFTSVEDAVRDTVESLKTKGFLKOOON*

NbL11g12420.1 cullin−associated_NEDD8−dissociated_protein_1 [MW = 134.57 kDa]

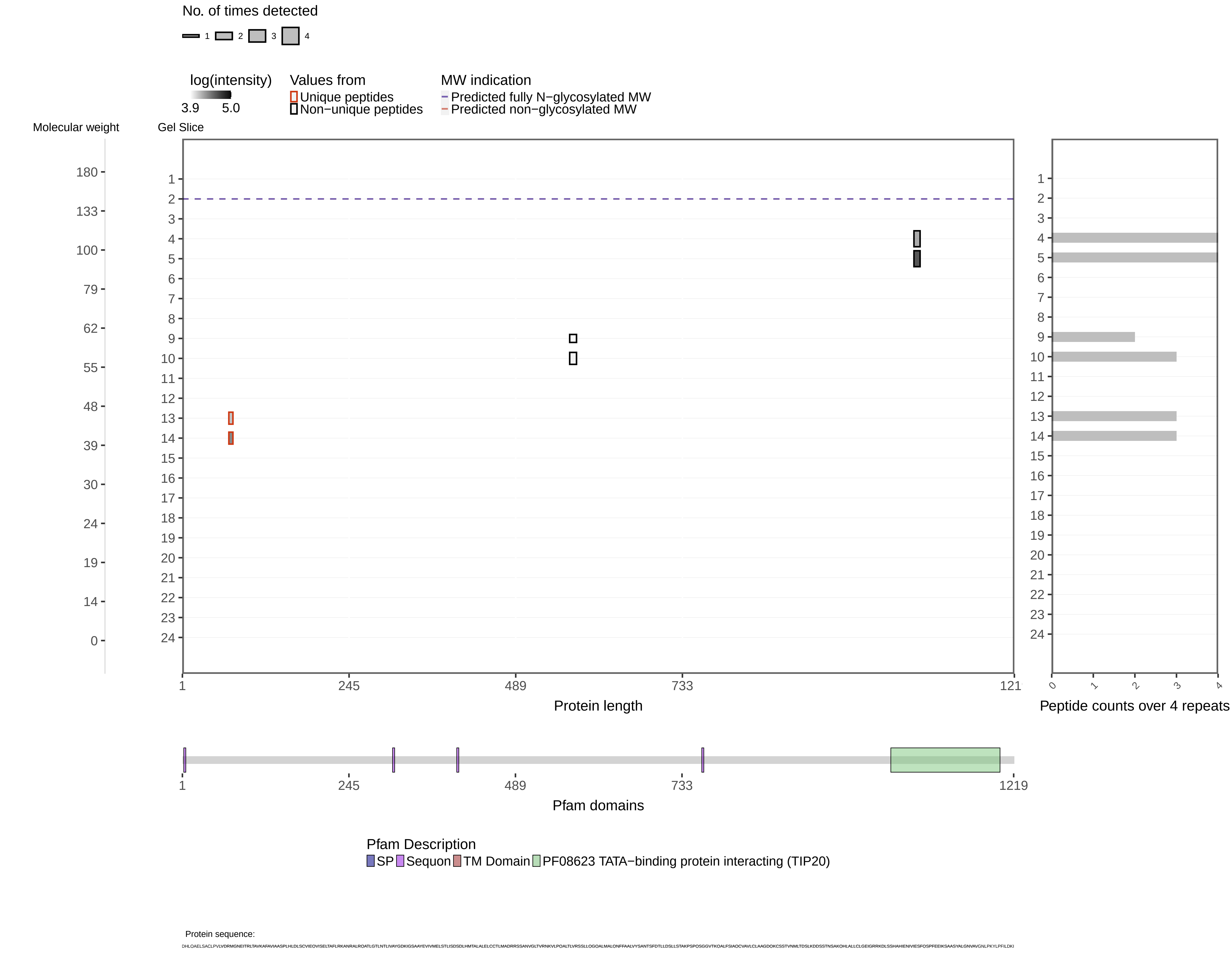

NbL11g12200.1 cytochrome_c [MW = 12.2 kDa]

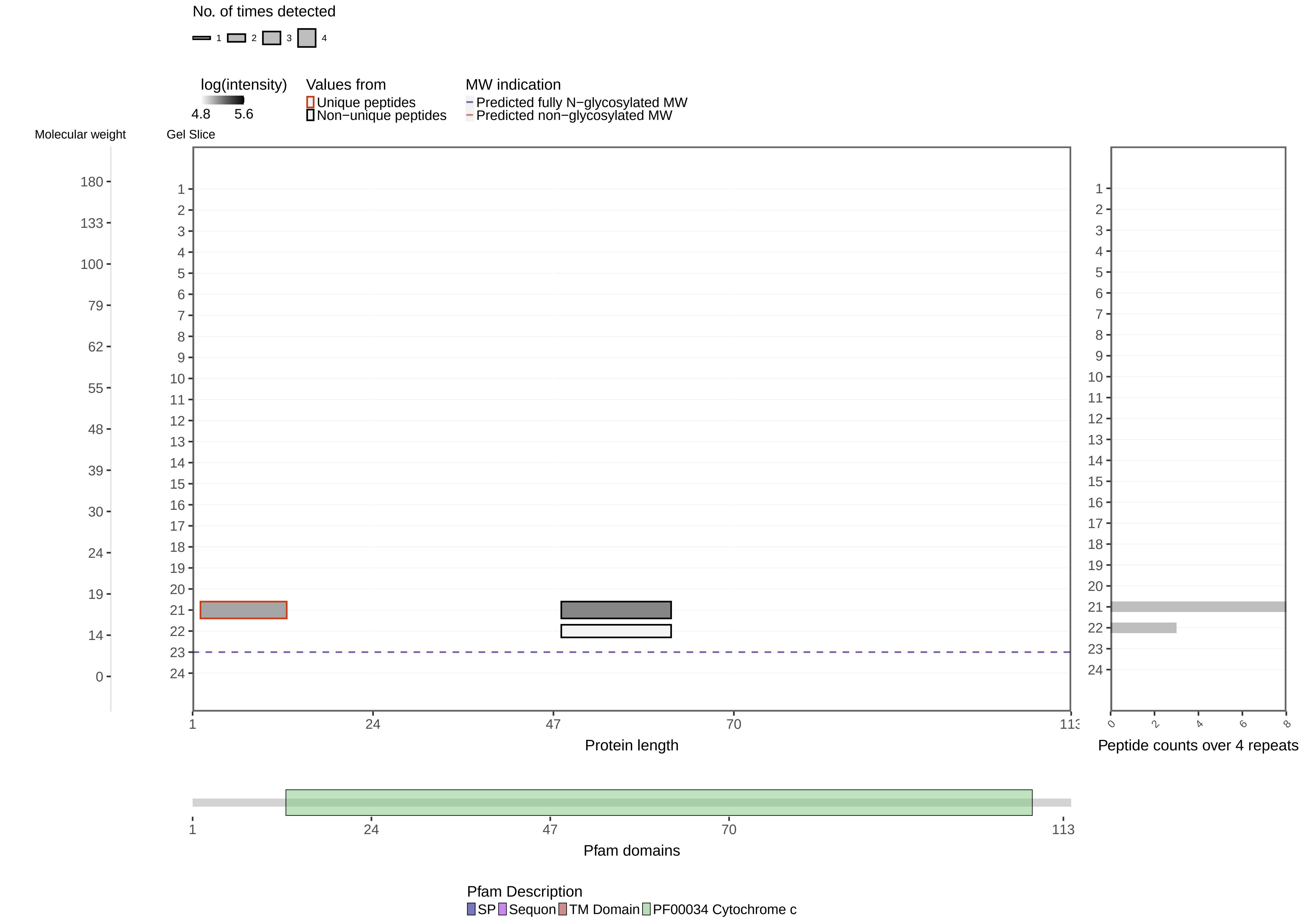


Protein sequence:

GLFGRQSGTTPGYSYSAANKNMAVIWGENTLYEYLLNPKKYIPGTKMVFPGLKKPQERADLIAYLKESTA*

NbL11g12120.1 ras−related_protein_RABD1 [MW = 22.43 kDa]

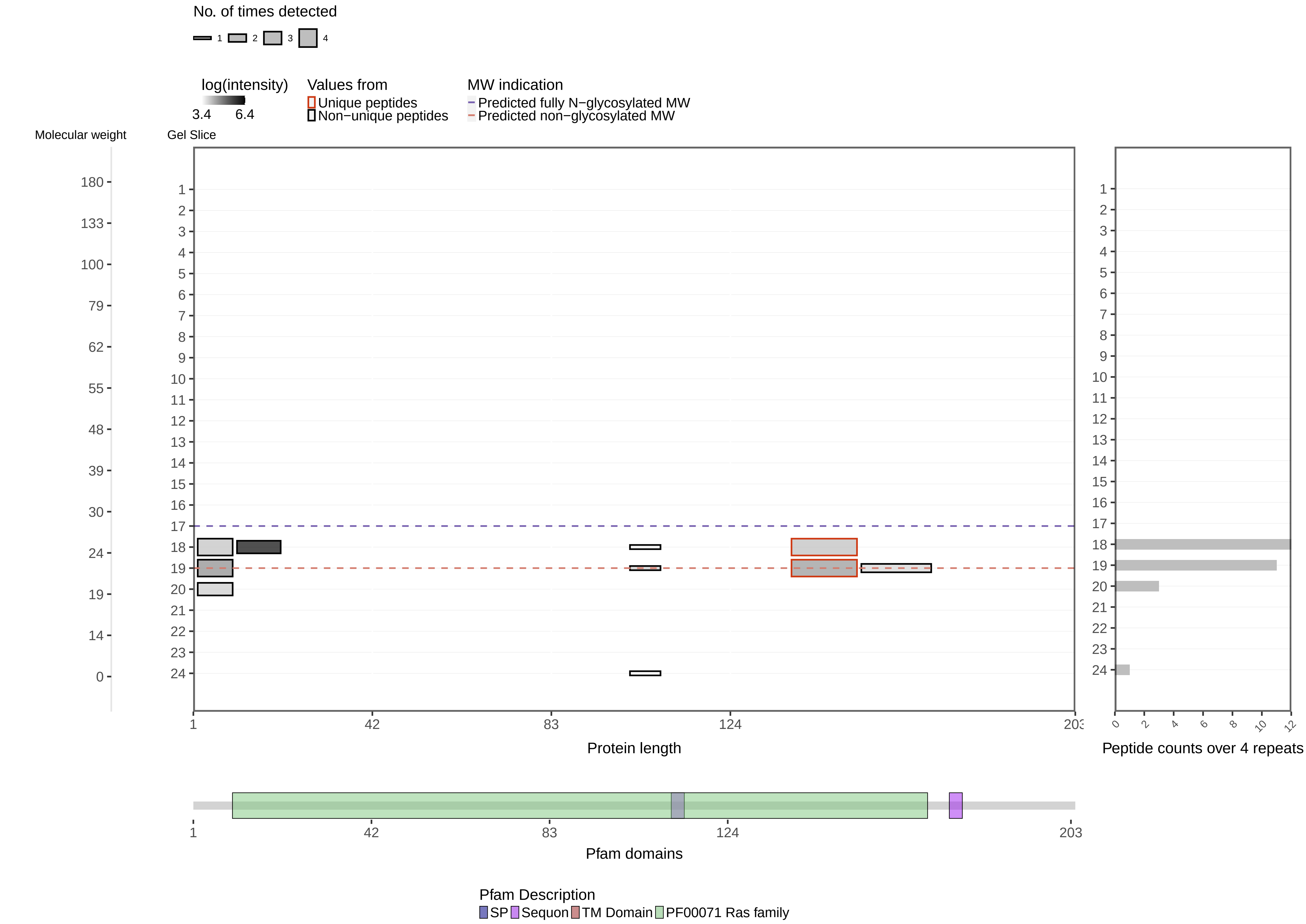


Protein sequence:

TERESFNNVKQWLNEIDRYANESVCKLLVGNKCDLVENKVVDTQTGKALADELGIPFLETSAKDSINVEQAFLTMAGEIKKKMGNQSAGAKKSGSTVQIKGQPIEQKSNCCG*

### NbL11g12110.1 acetolactate_synthase_1_chloroplastic [MW = 72.72 kDa]

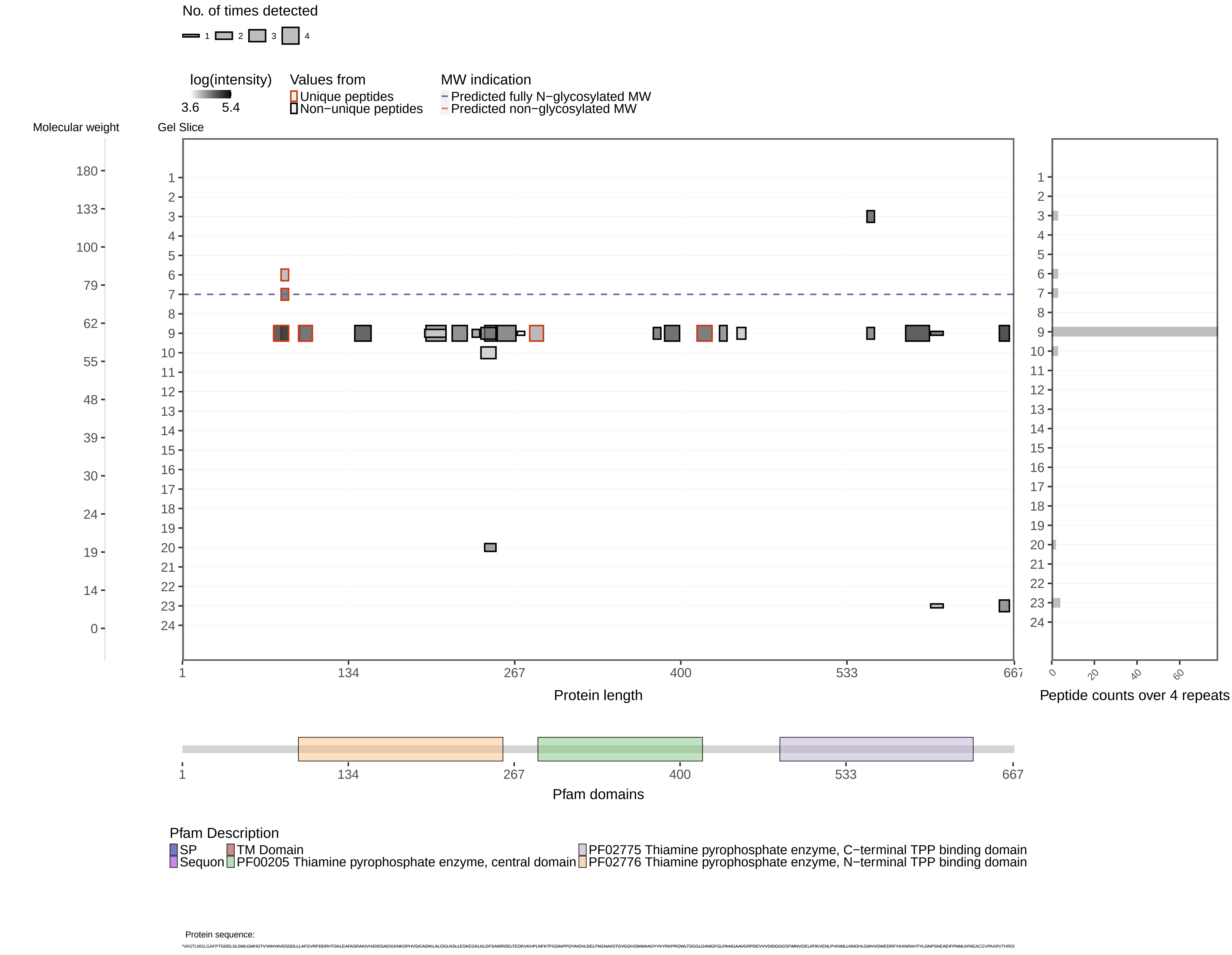

### NbL11g12090.1 ras−related_protein_RABA2a [MW = 23.92 kDa]

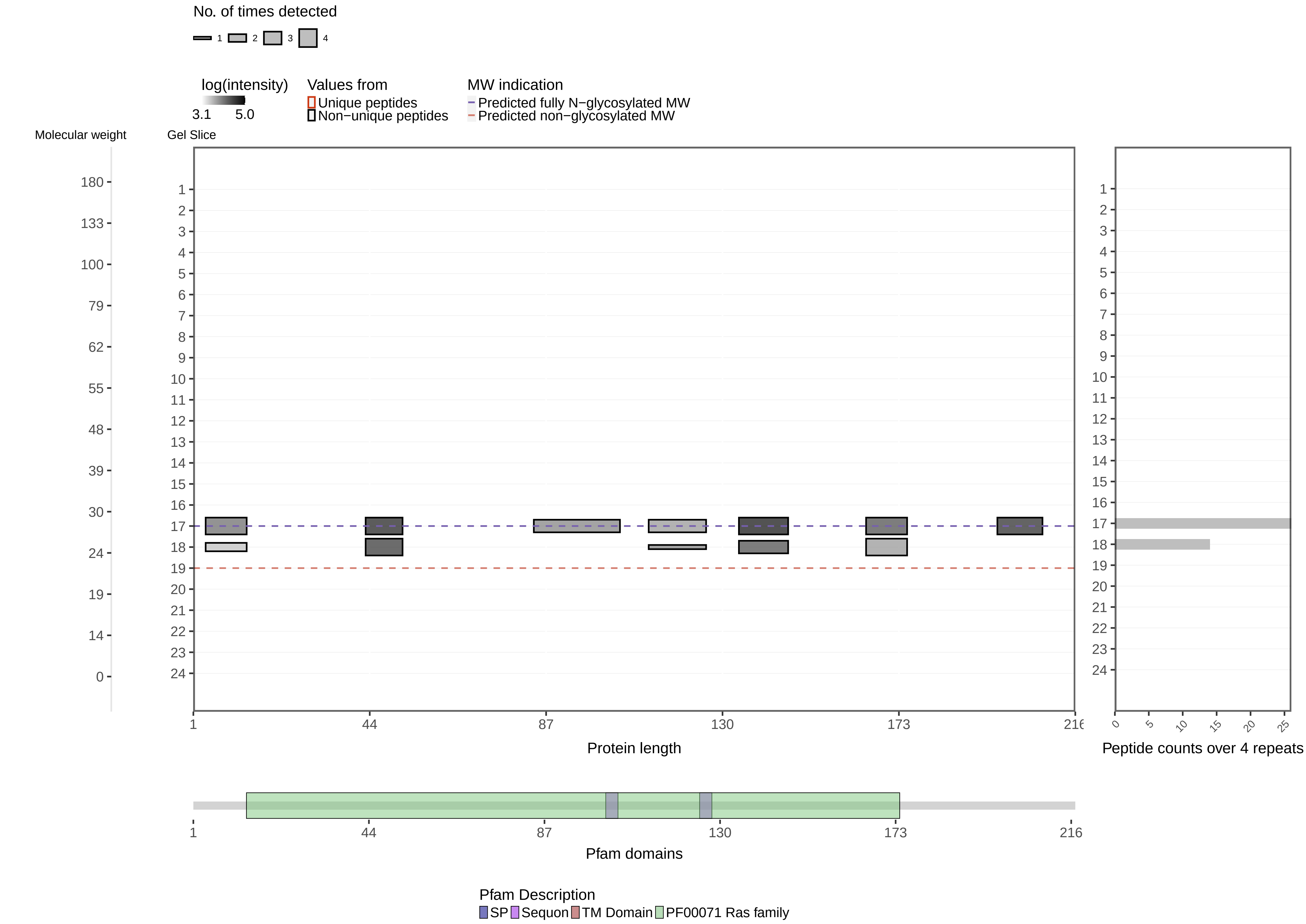


Protein sequence:

VTKPTTFENVSRWLKELRDHADSNIVIMLIGNKTDLKHLRAVDTEDAQSFAEKEGLSFIETSALEATNVEKAFQTILSEIYRIISKKPLSSEEPATANVKEGKTLVVGAEEPAPKKACCSSS*

### NbL11g11990.1 coatomer_subunit_delta−like [MW = 58.07 kDa]

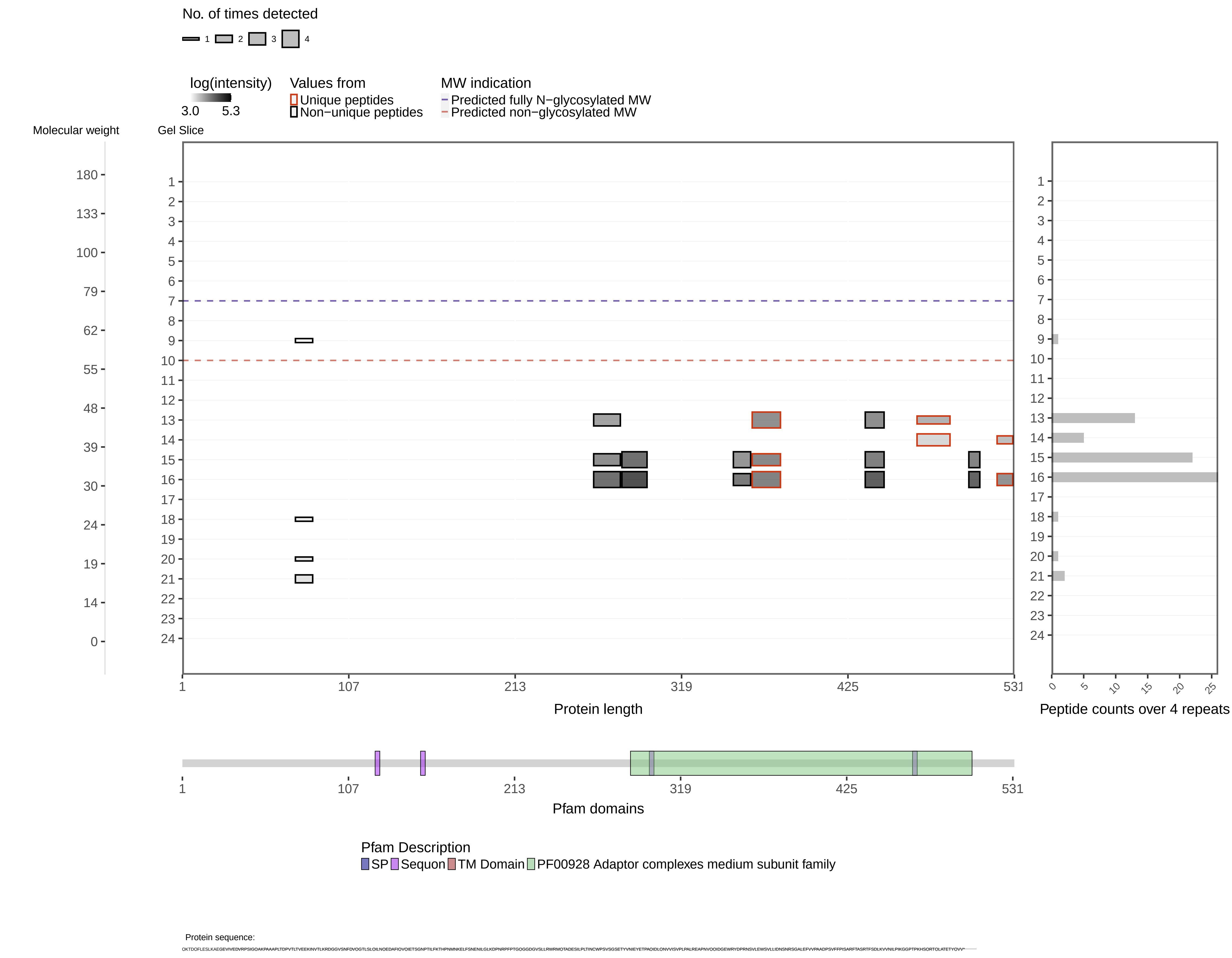

### NbL11g11950.1 60S_ribosomal_protein_L3−2 [MW = 44.48 kDa]

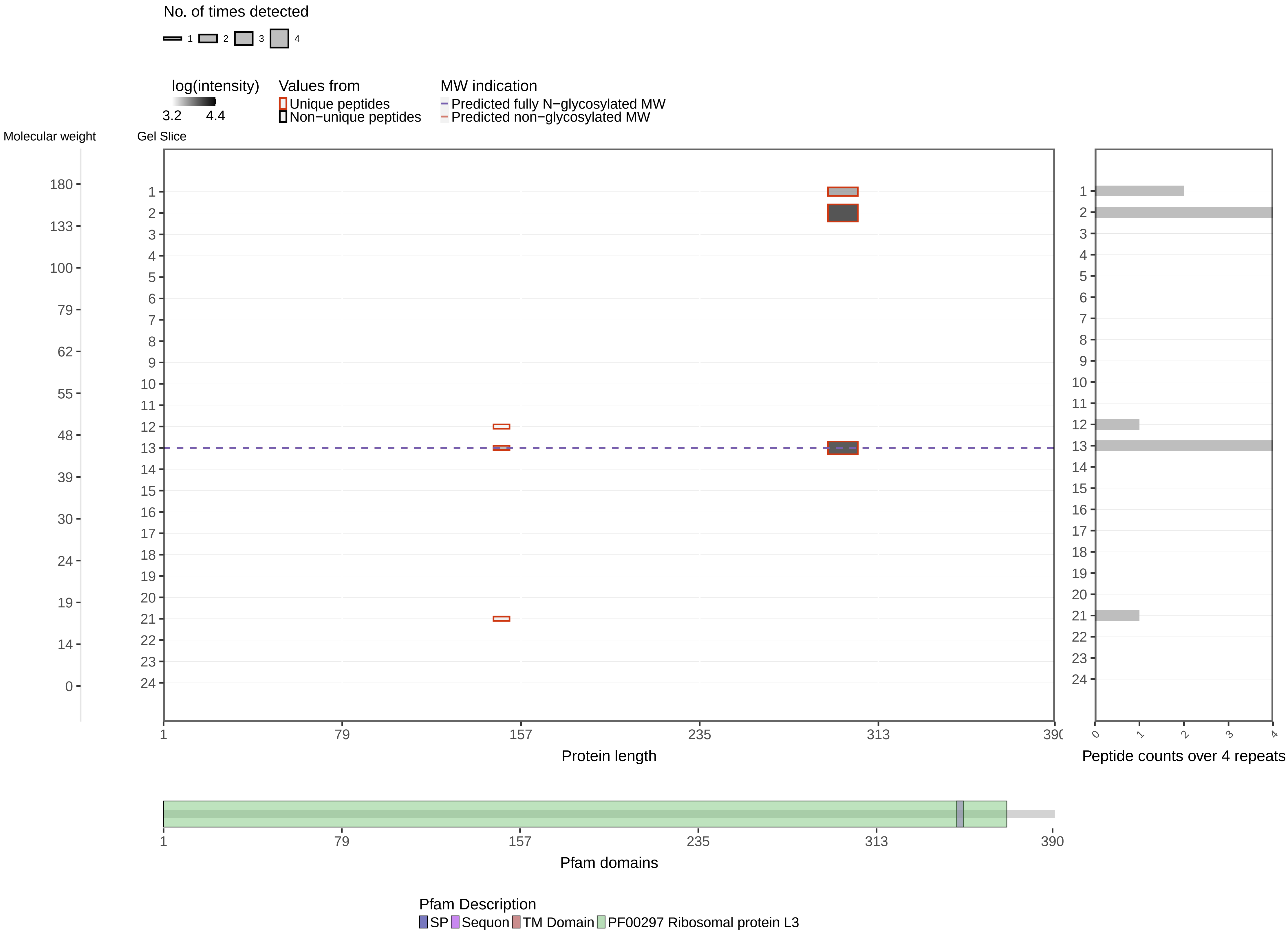


Protein sequence:

LMEIQVNGGDISQKVDYAYGFFEKQIPVDAIFQKDEMIDIIGVTKGKGFEGVVTRWGVTRLPRKTHRGLRKVACIGAWHPARVSYTVARAGQNGYHHRTELNKKVYRLGKAGQESHSAITEFDRTEKDITPMGGFPHYGIVKEDFLLIKGCCVGPKKRVVTLRQSLLNQTSRVALEEIKLKFIDTSSKFGHGRFQTTQEKAKFYGRLKA*

### NbL11g11690.1 40S_ribosomal_protein_S12−like [MW = 15.13 kDa]

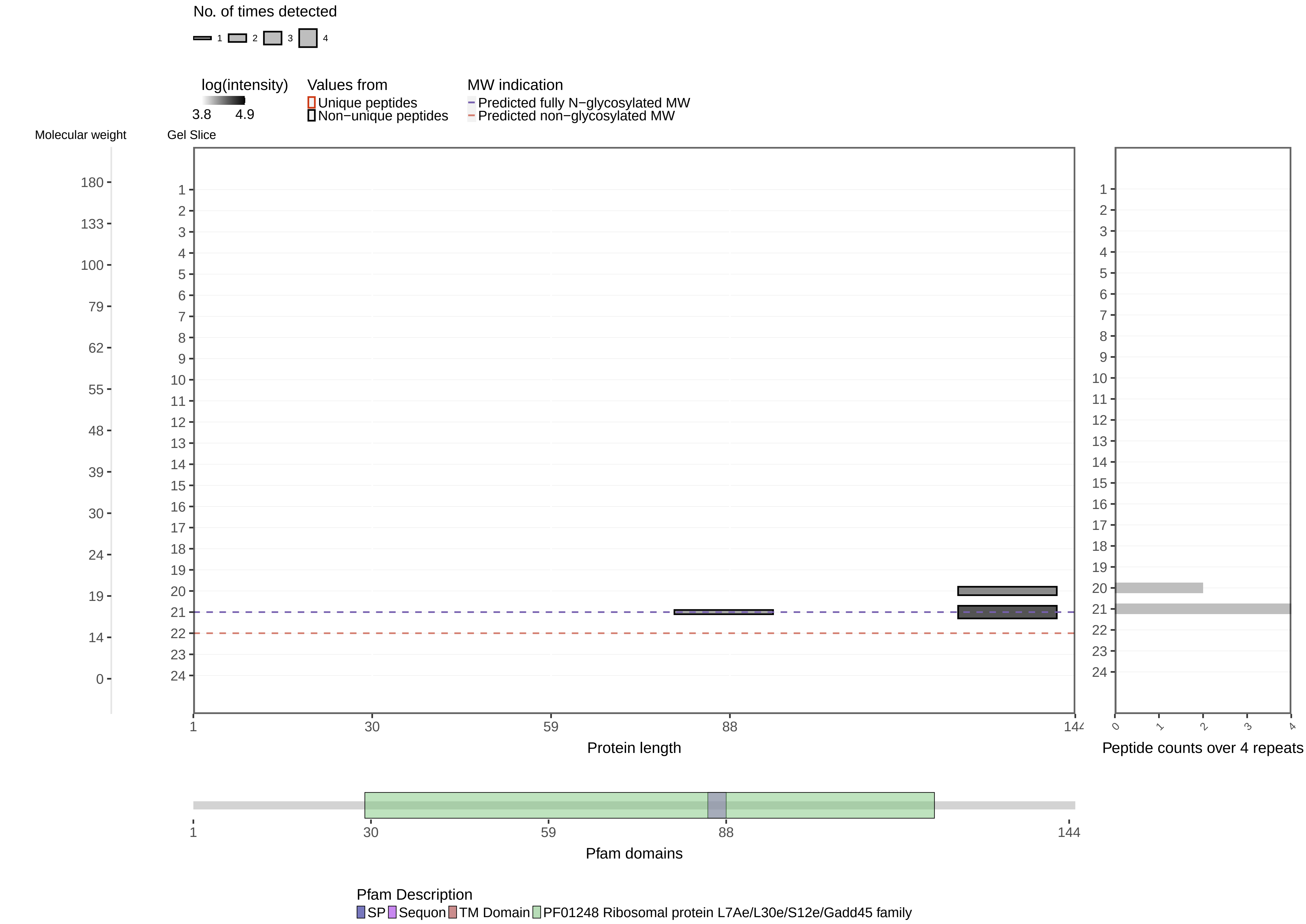


Protein sequence:

QLCVLAEDCDQPDYVKLVKALCADHNVSLITVPNAKTLGEWAGLCKIDSEGKARKVVGCGCVVVKDYGEETEGLHIVQEYVKSH*

NbL11g11640.1 ATP_phosphoribosyltransferase_2_chloroplastic−like [MW = 42.93 kDa]

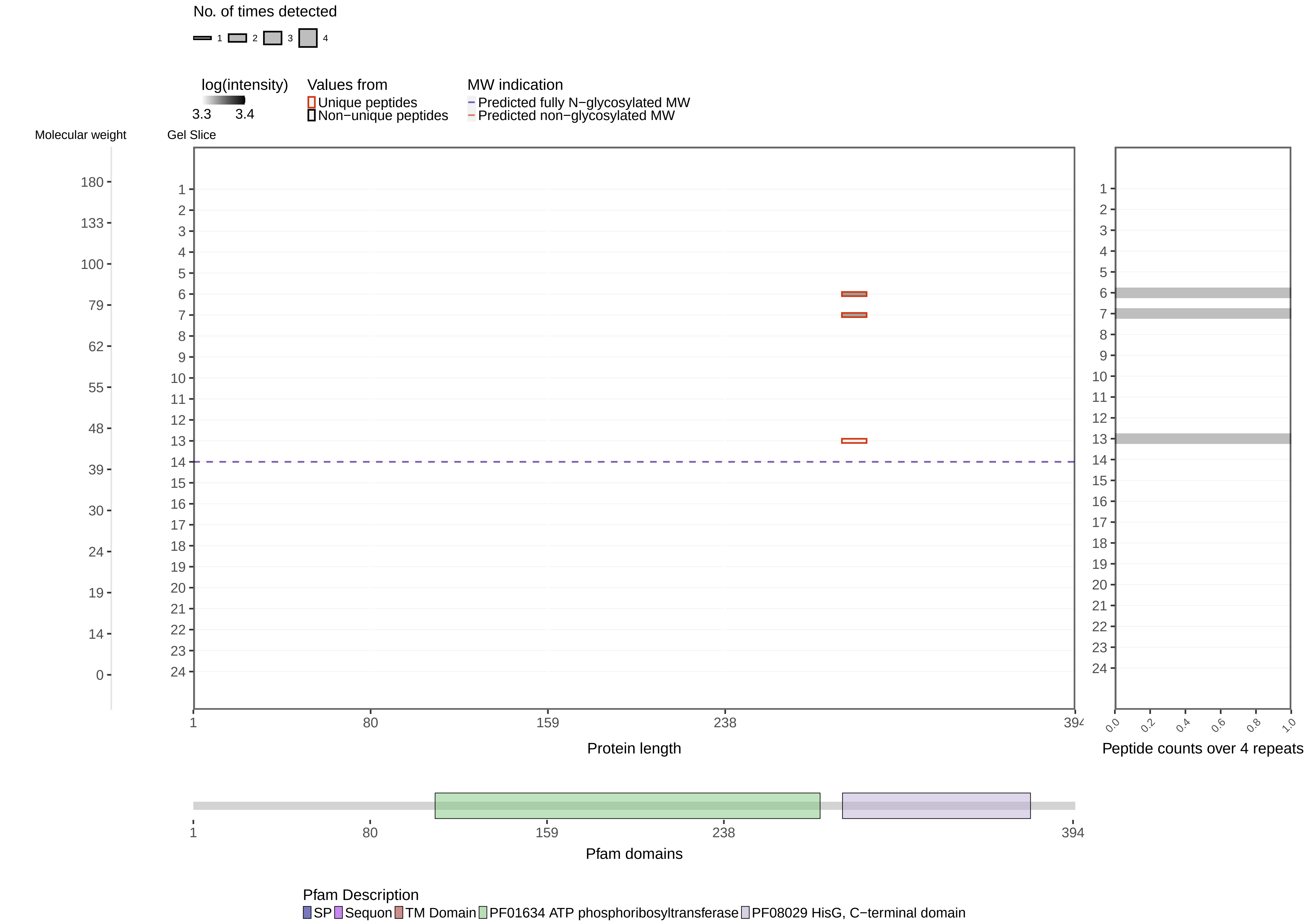


Protein sequence:

SERPLRVATGFTYLGPKFMKDNGLKHVTFSTADGALEAAPAMGIADAIVDLVSSGTTLRENNLKEIEGGVILESQAVLVAGRKSLMQRKGVLDITHEMLERLEAHLRAVGQFTVTANMRGSSAEEVAERILSQTSLSGLQGPTVSPVFCKRDGRVTADYFAIVICVPKKALYKSVQQLRAIGGSGVLISPLTYIFDEETPRWRQLLSMLGL*

### NbL11g11630.1 UDP−glucose:glycoprotein_glucosyltransferase [MW − SP = 183.1 kDa]

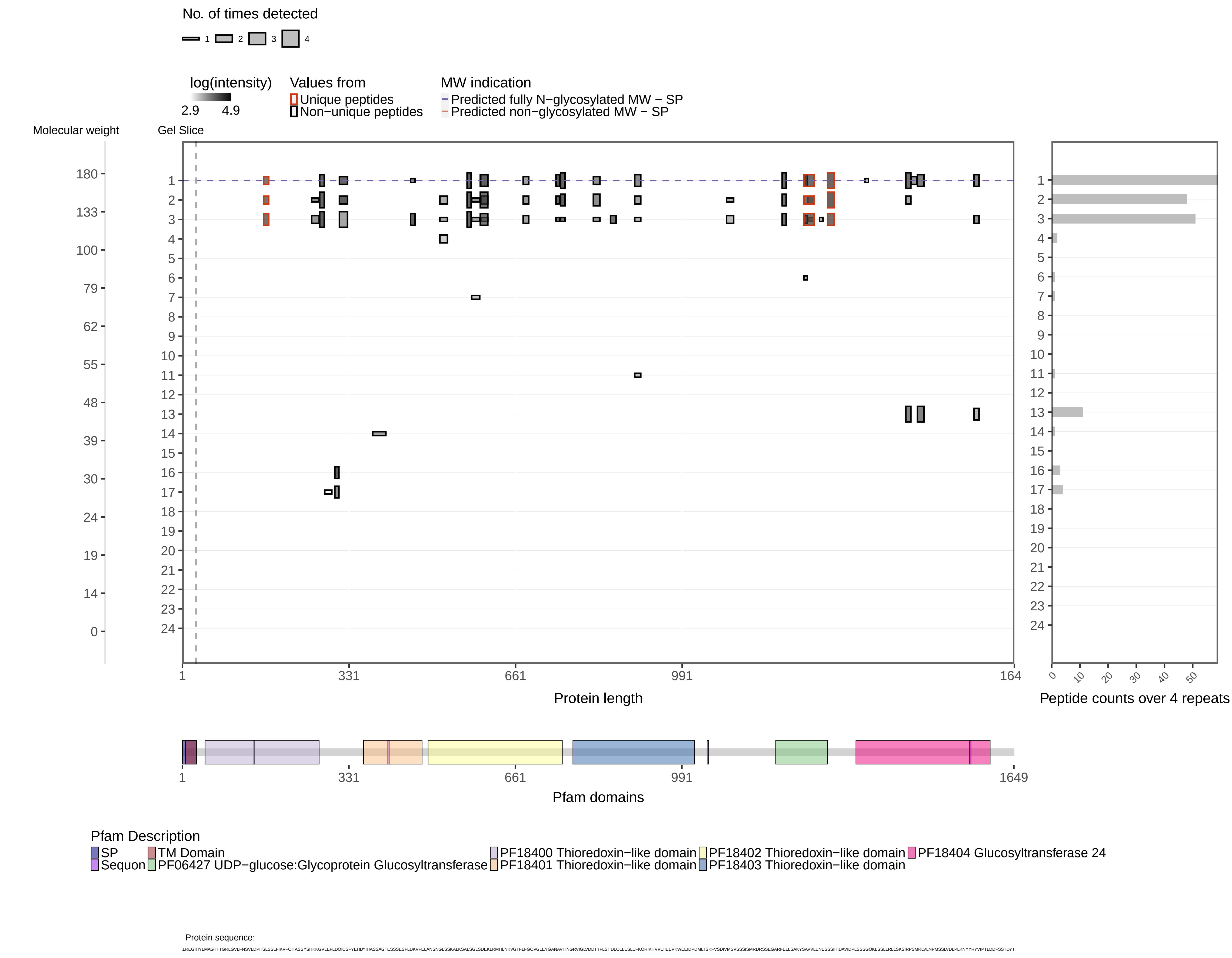

NbL11g11530.1 ankyrin_repeat_domain−containing_protein_2B−like [MW = 37.3 kDa]

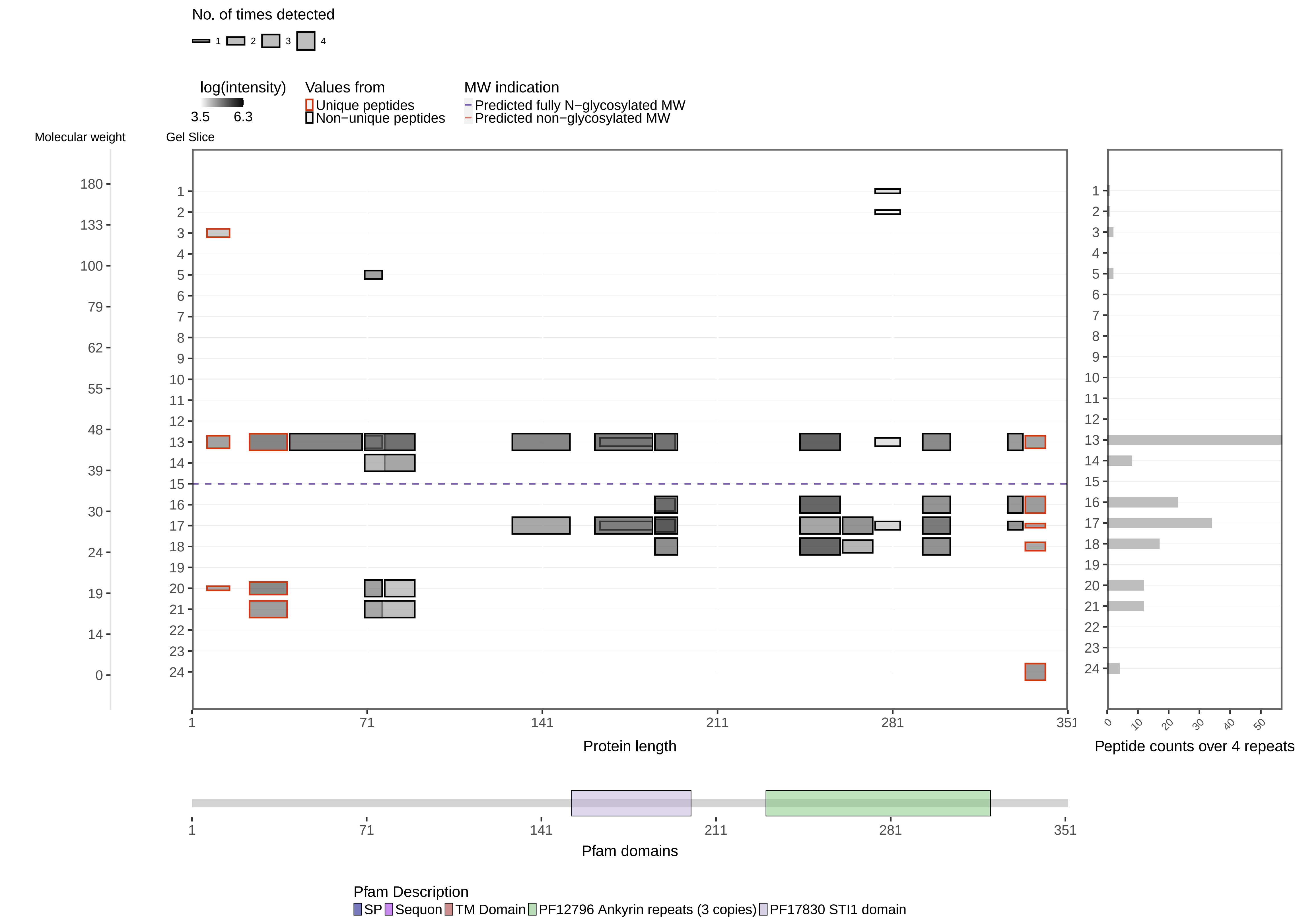


Protein sequence:

RIKEDPSLKPILEEIESGGPAAMMRYWNDOETLKKIGEAMGFAAGGEGATSSAIPGTDETEEANEDESVVHOCASVGDAEGLKAALTAGADKDEEDSEGRTALHFACGYGEVKCAOILLEAGAKVDALDKNKNTALHYAAGYGRKECVALLLENGAAVTLONLDGKTPIDVAKLNNOOEVLKLLEKDVFL*

NbL11g11420.1 60S_acidic_ribosomal_protein_P1−like [MW = 11.43 kDa]

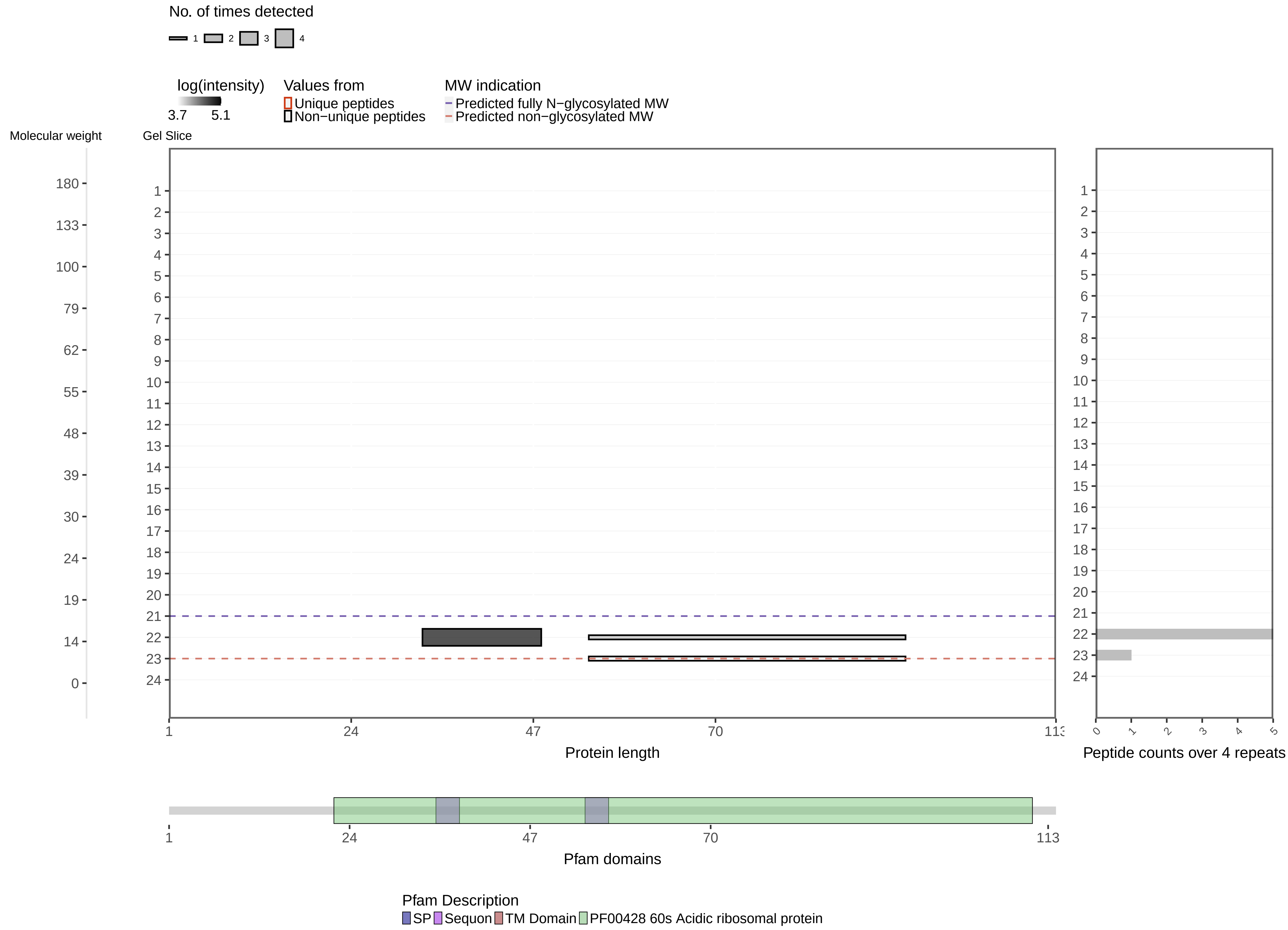


Protein sequence:

FAKLAEKRNLSDLIMNVGAGGGAAAVAVAAPTGGAAAGGAAAAPAAEEKKEEPKEESDDDMGFSLFD*

### NbL11g11400.1 basic_blue_protein [MW − SP = 10.41 kDa]

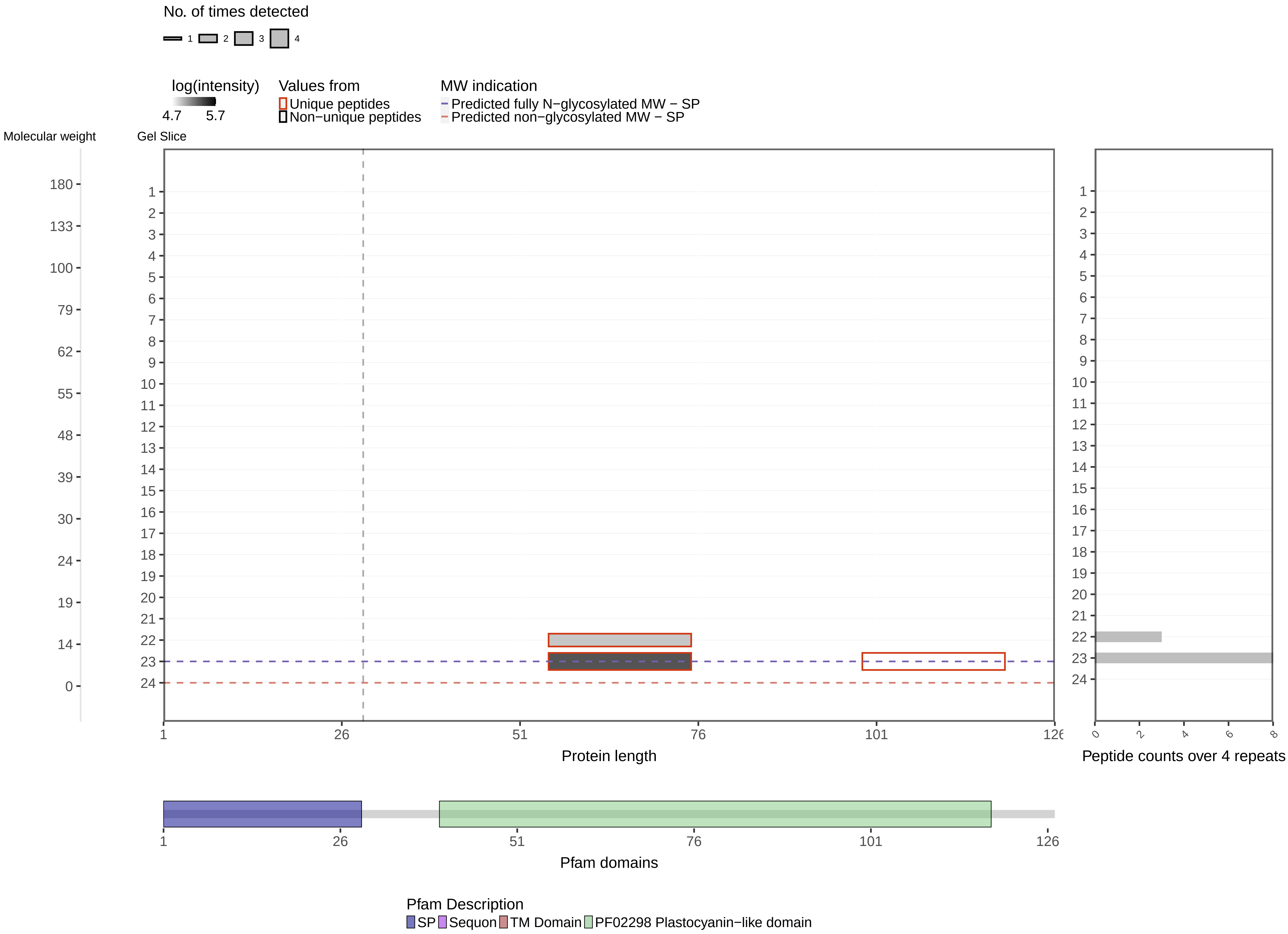


Protein sequence:

:GKRFRAGDILAFNYPSGAHNVVAVDKAGYSNCRTPRGAKVYTKGGDKIKLVNGQNYFICNFPGHCQSGMKISVFAS*

### NbL11g11370.1 partner_of_Y14_and_mago [MW = 24.98 kDa]

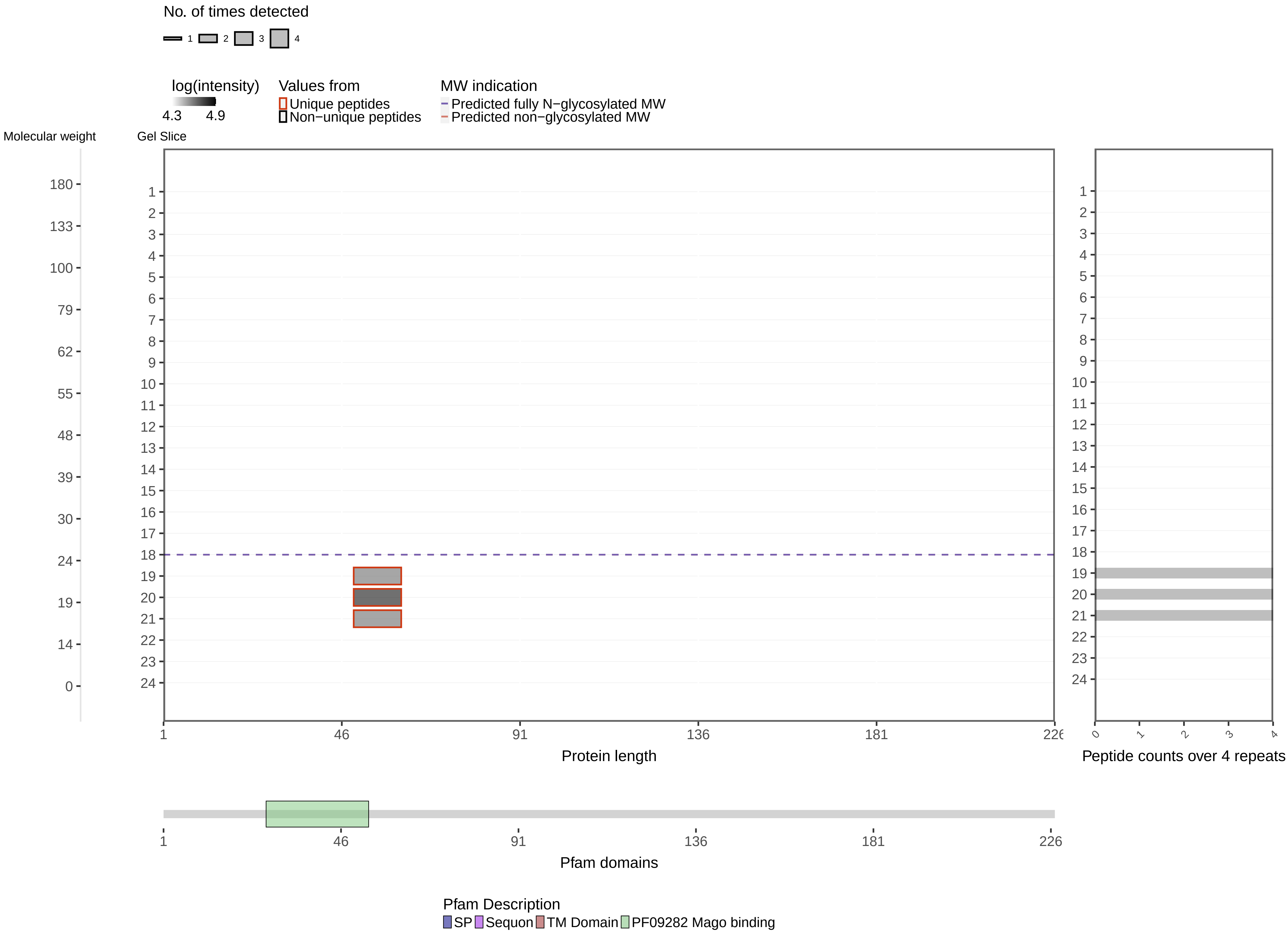


Protein sequence:

RKKEKRQQAAVEKGTNPESGEISSAENSVGVPEQVESVMSQINSLAISANPVVPLSNSIESSAVGDSVQDIDKRIRALKKKIRLTEAQQQKTEEKDMKPEQLEKVAKLESWRKELKVLEDKKAELEA*

### NbL11g11360.1 mRNA_turnover_protein_4_homolog [MW = 28.22 kDa]

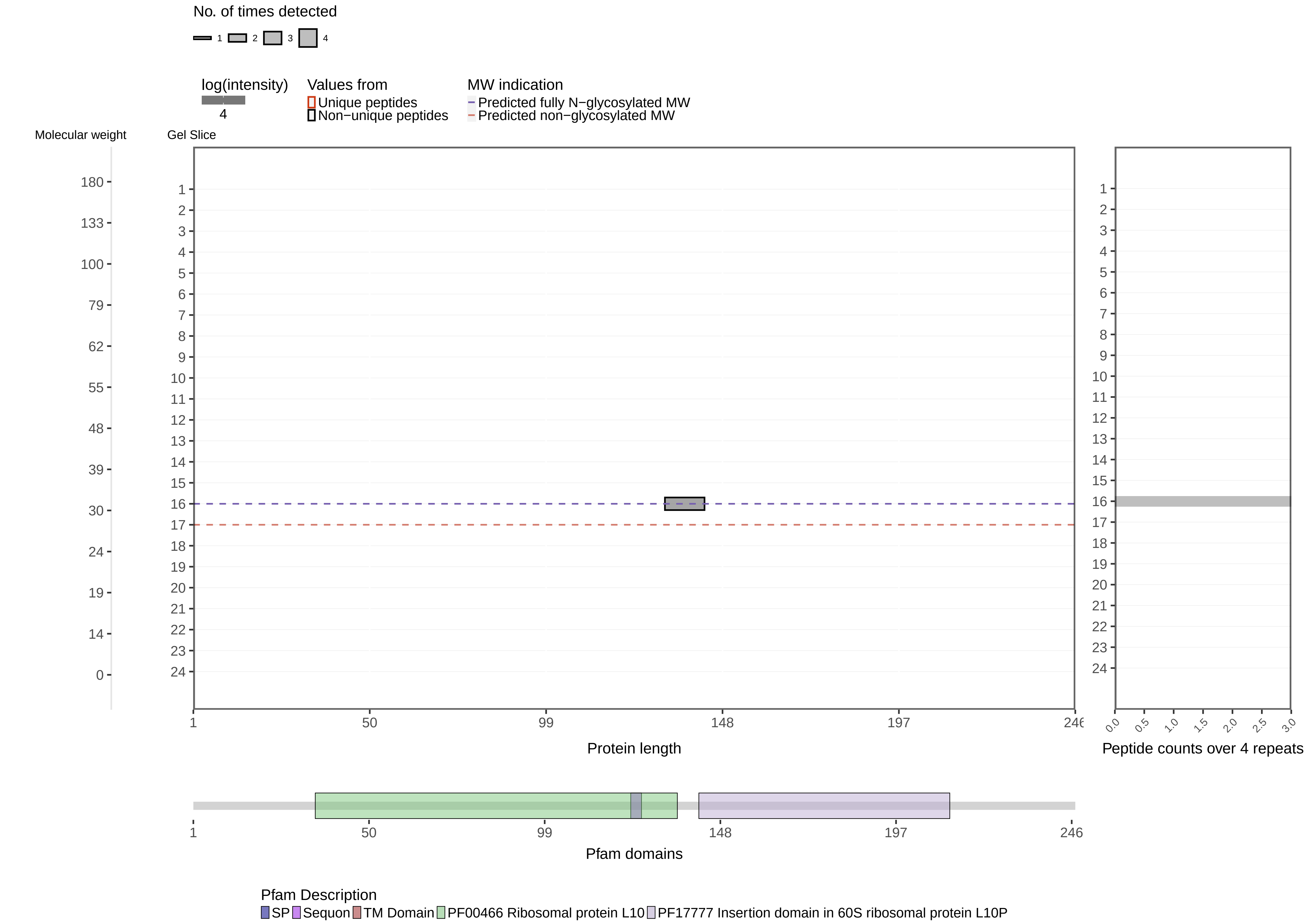


Protein sequence:

SKLVRGDSGICFTNLSKEEVQRLFNEFEGYDFARTGSAATEKVELQEGPLDQFTHEMEPFLRKQRMPVRLNKGVVELVSDFVVCEEGKALSPEAARILRLLGIKMATFRLNLICRWTPDEFEIYREGLEESDIESS*

### NbL11g11260.1 60S_ribosomal_protein_L3 [MW = 44.41 kDa]

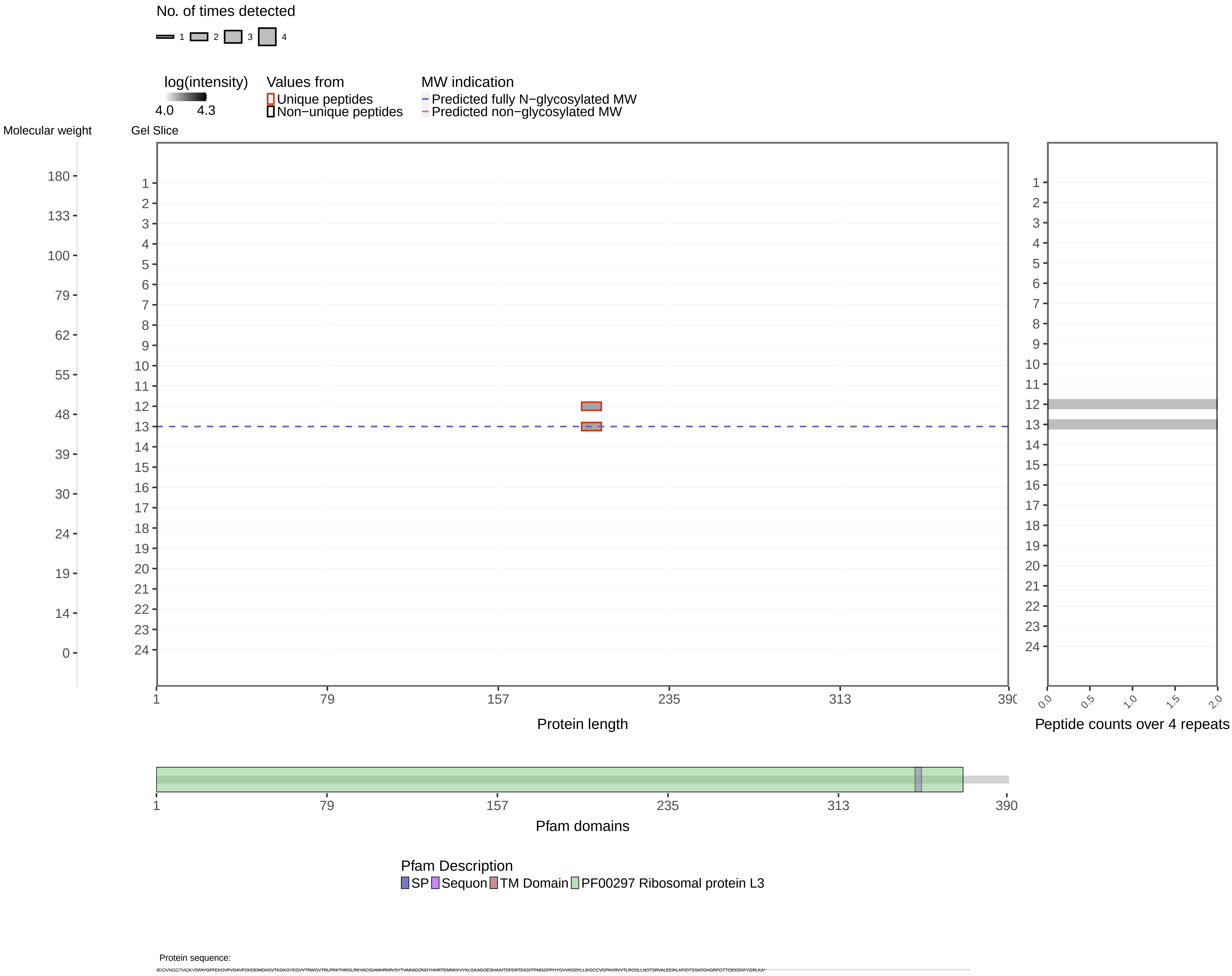

### NbL11g11250.1 COP9_signalosome_complex_subunit_8 [MW = 22.59 kDa]

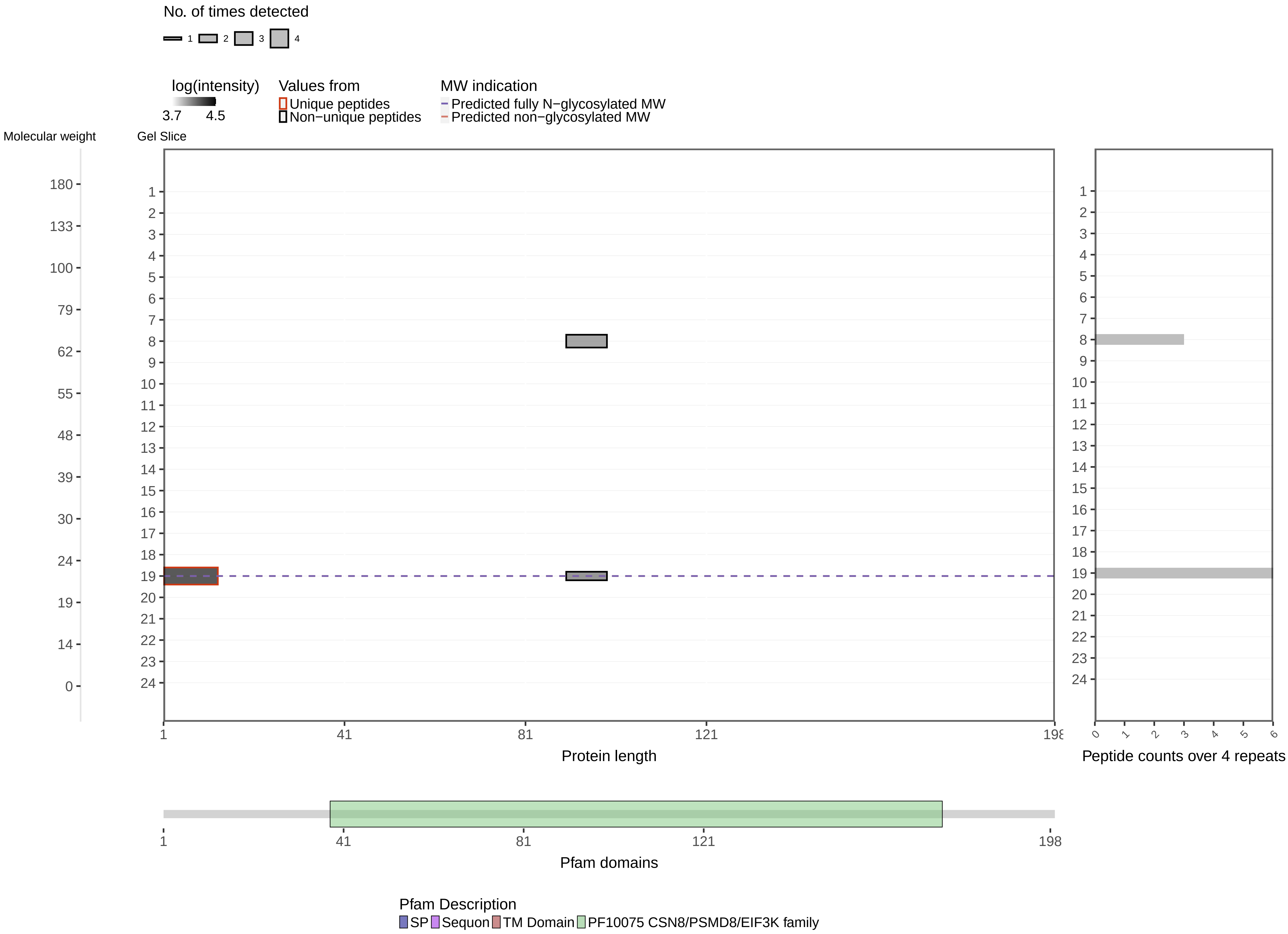


Protein sequence:

WTRDYVGVYEAIREFSWTPDVQPIVASFAELYRKRMFELLLSAYSTISTQDTAQFLGMNENDATNYVLQQGWVLDPASQMLTVKKQAVVTEQKLDPSKLQRLTEYVFHLEH*

### NbL11g11160.1 actin−101 [MW = 41.64 kDa]

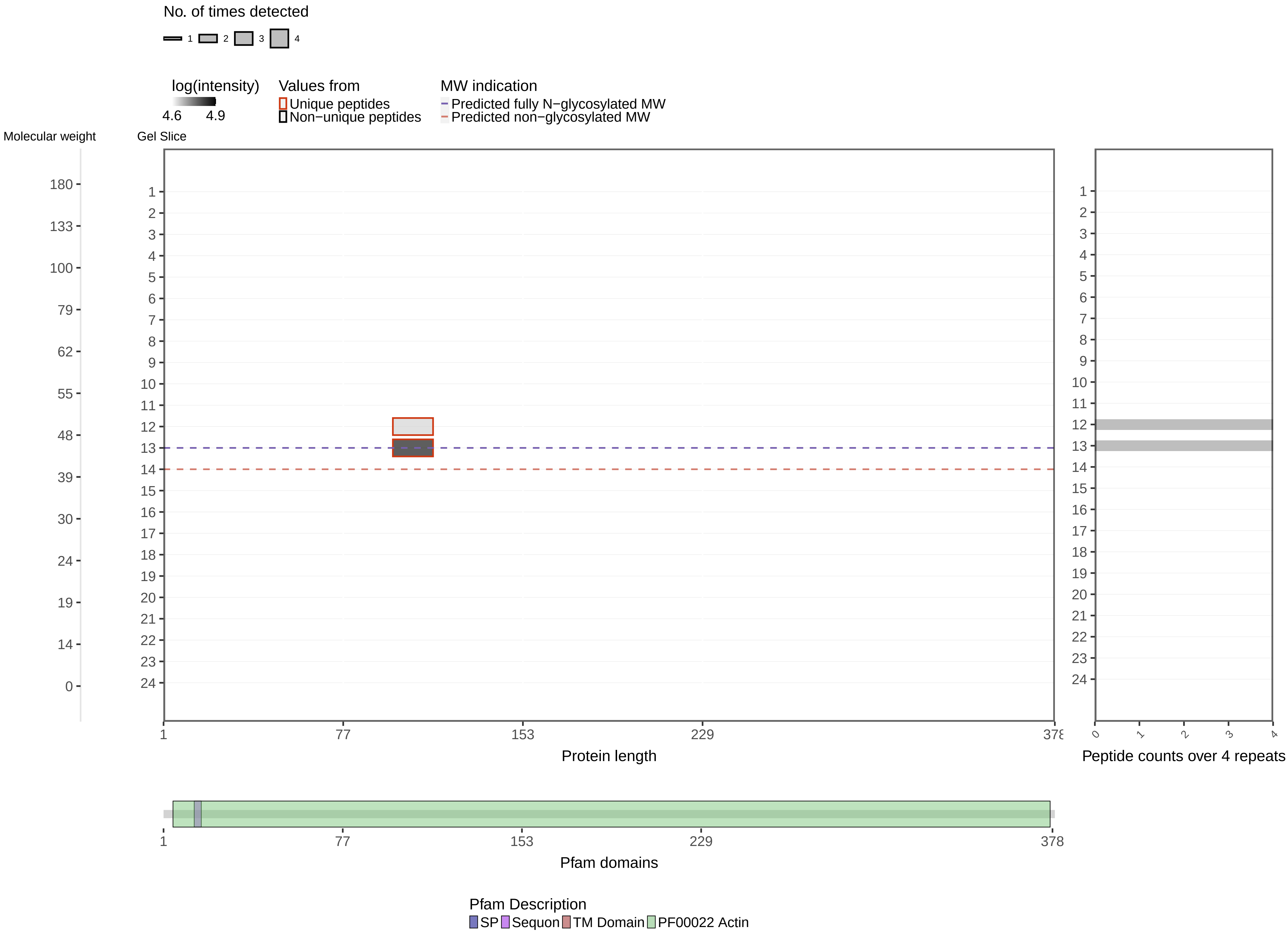


Protein sequence:

PHAILRLDLAGRDLTDCLMKILTERGYSFTTSAEREIVRDVKEKLAYVALDYEQELETAKSSSSVEKSYELPDGQVITIGSERFRCPEVLFQPSLVGMEAAGIHETTYNSIMKCDVDIRKDLYGNIVLSGGTTMFPGIADRMSKEITALAPSSMKIKVVAPPERKYSVWIGGSILASLSTFQQMWITKGEYDESGPSIVHRKCF*

NbL11g10980.1 beta−xylosidase/alpha−L−arabinofuranosidase_2−like [MW = 89.57 kDa]

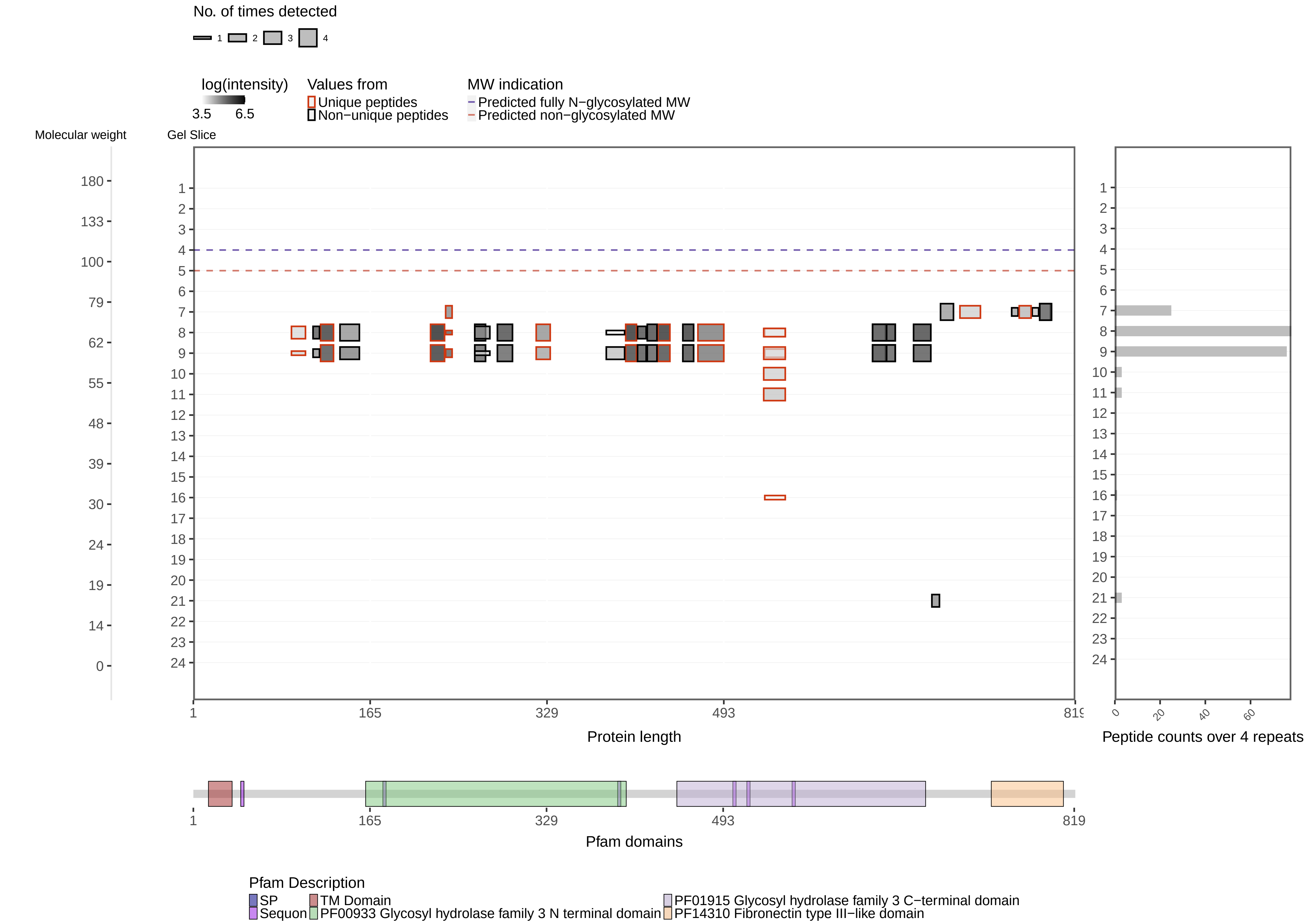


Protein sequence:

NbL11g10930.1 non−specific_lipid−transfer_protein−like_protein_At2g13820 [MW − SP = 15.77 kDa]

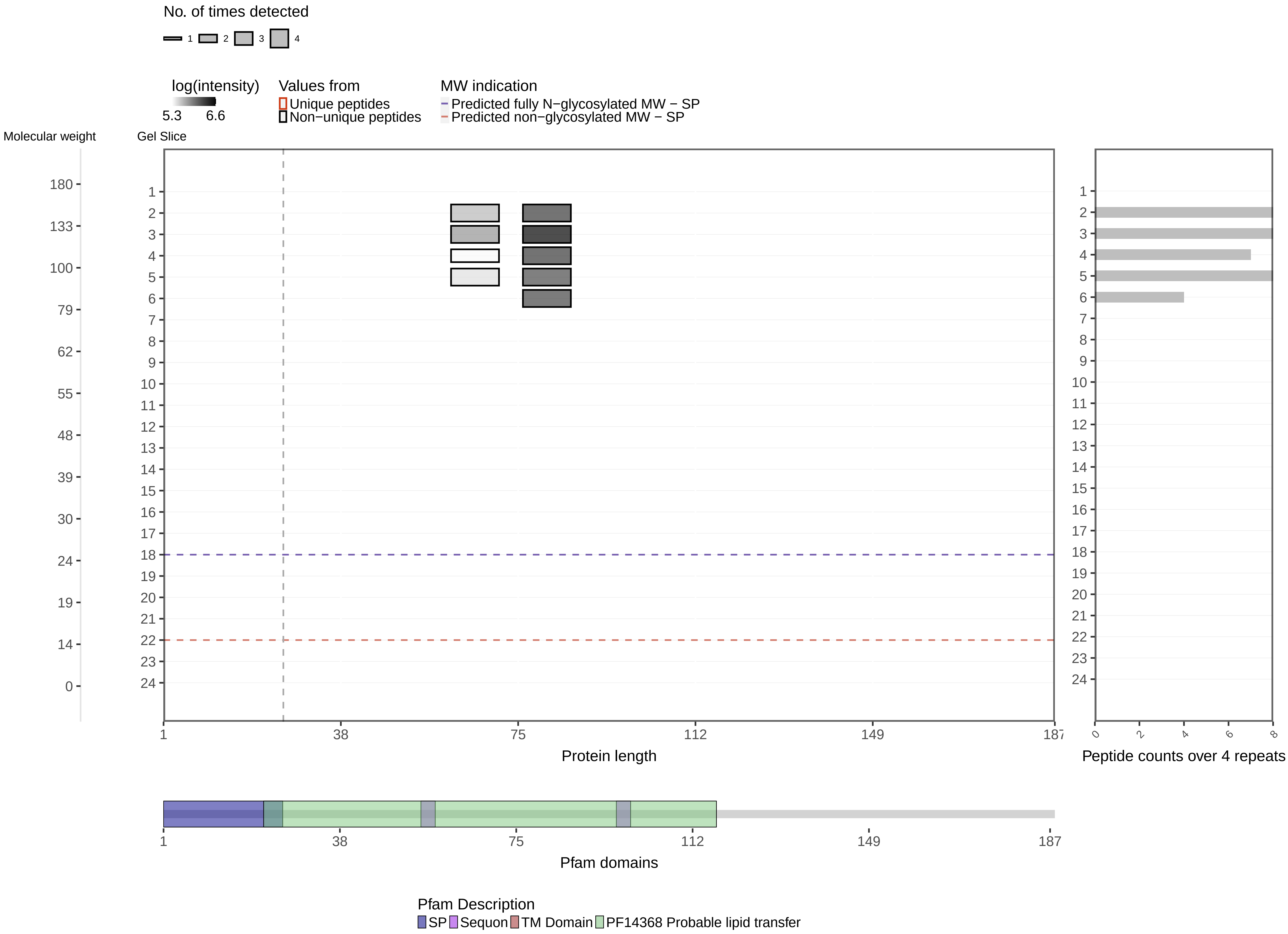


Protein sequence:

CLCEGFKNSAQLGVVLNITKAMALPAACHVSAPSVSSCGLSTDTGAAPALSPIAVSPTMAGAVAPTTVEGASEVAPAPAPGSSDSNTLGLSIGQLVLTAMAVAFCWF*

### NbL11g10920.1 probable_protein_phosphatase_2C_62 [MW = 86.78 kDa]

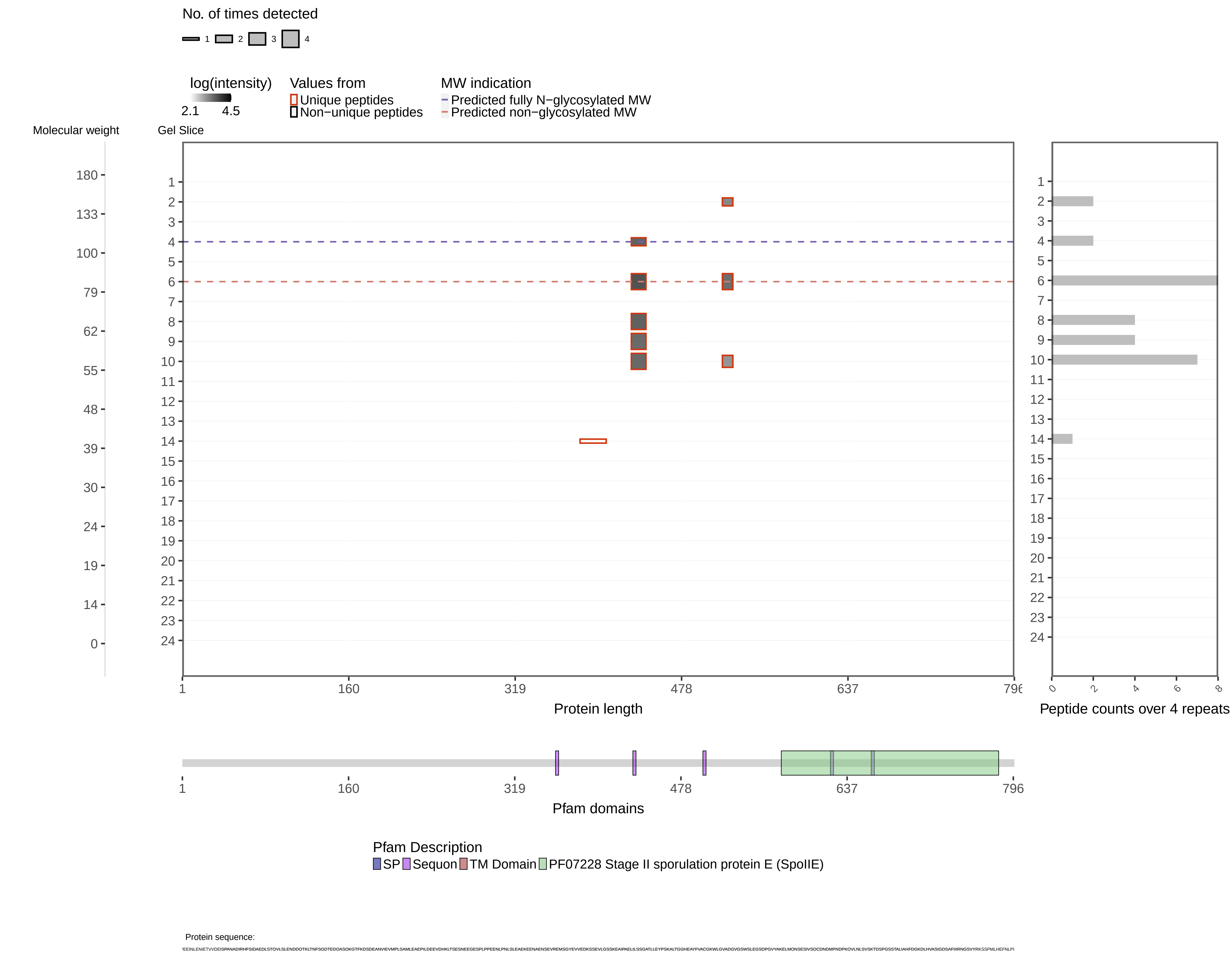

NbL11g10910.1 chlorophyll_a−b_binding_protein_CP24_10A_chloroplastic−like [MW = 27.34 kDa]

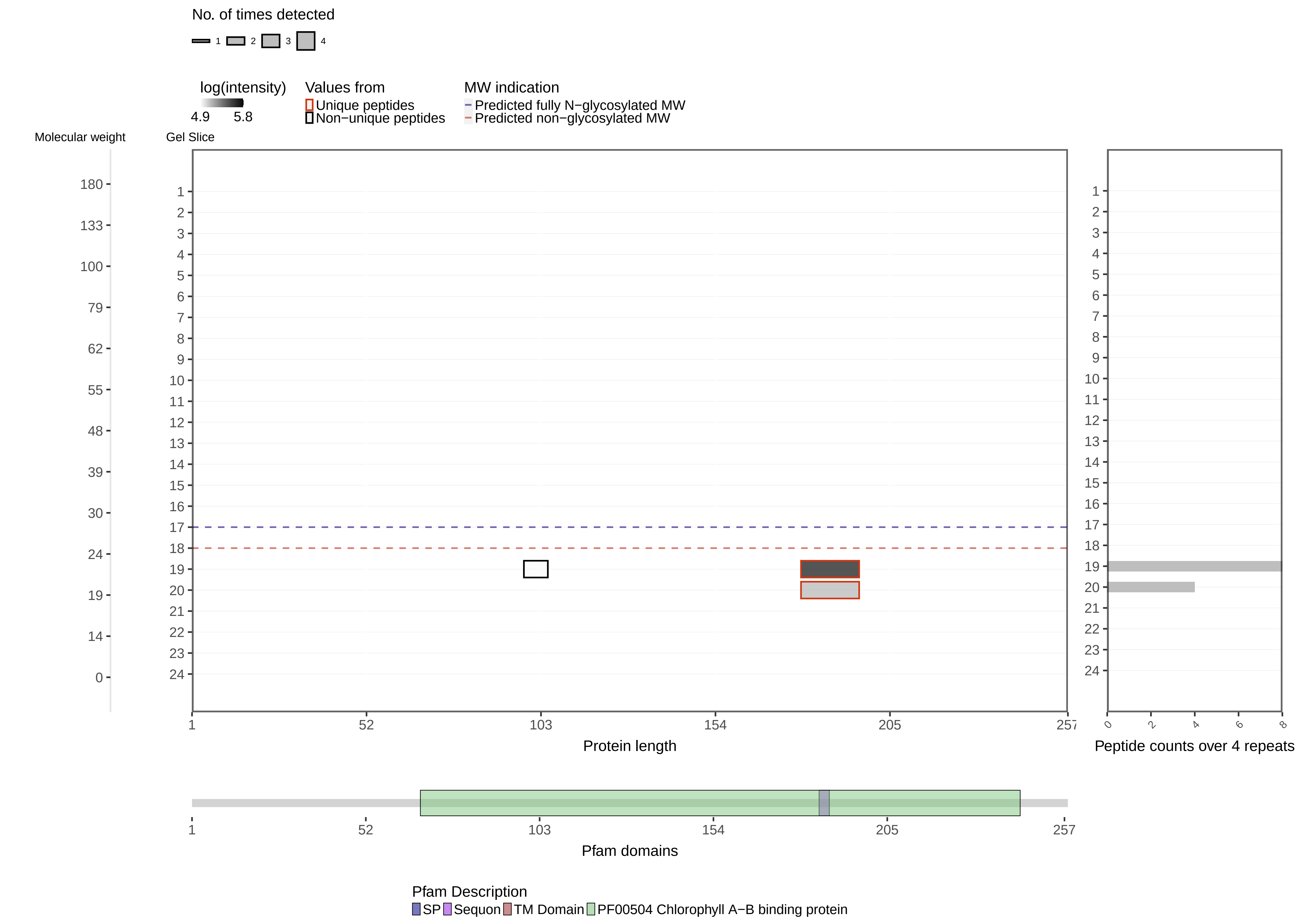


Protein sequence:

VGQAWSGIPWFEAGADPGAIAPFSFGSLLGTQLLLMGWVESKRWVDFFNPESQSVEWATPWSKTAENFANFTGEQGYPGGKFFDPLSLAGTLQDGVYIPDTEKLERLKVAEIKHARLAMLAMLIFYFEAGQGKTPLGALGL*

### NbL11g10880.1 peroxidase_N1 [MW = 38.62 kDa]

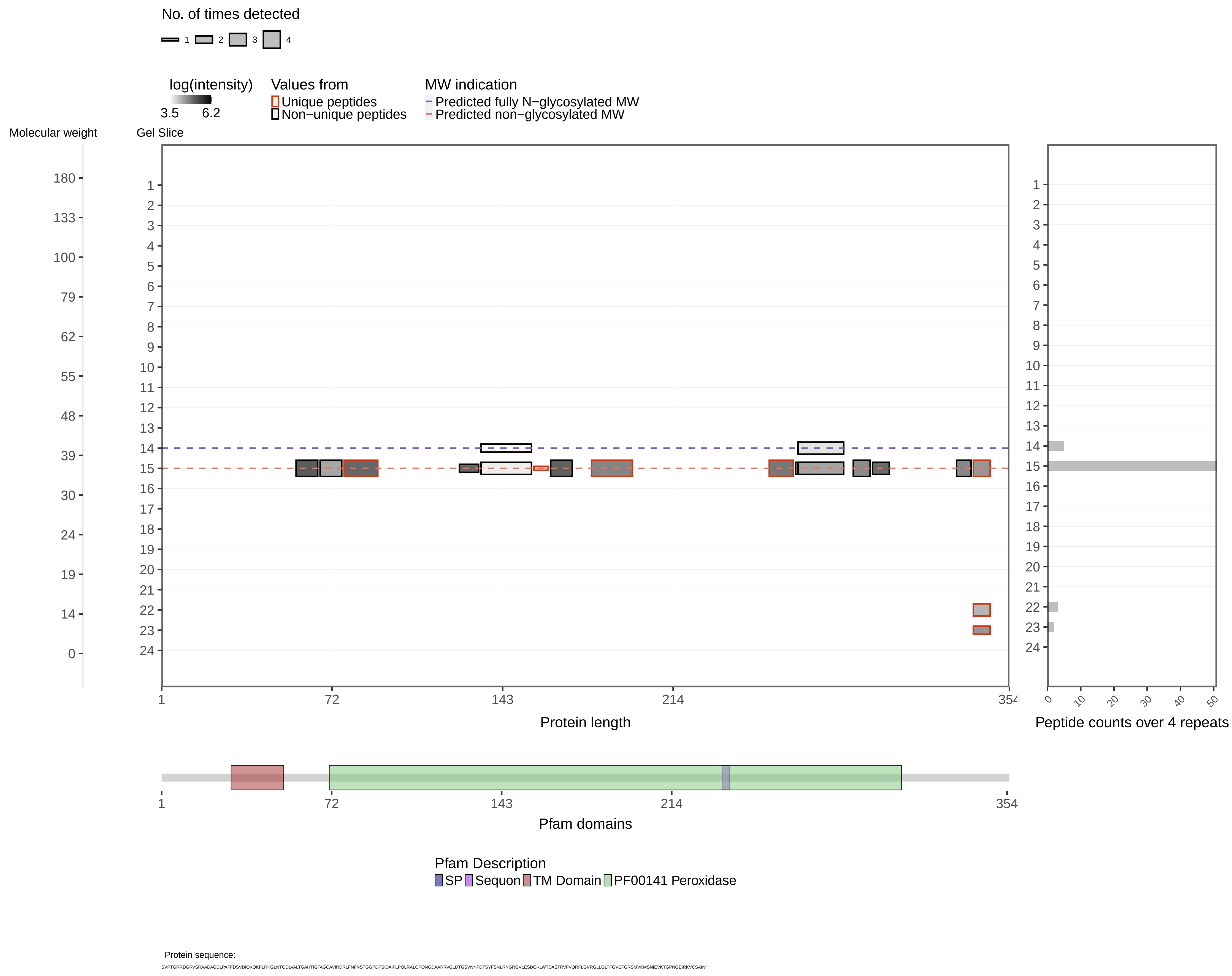

### NbL11g10690.1 nitronate_monooxygenase [MW = 36.12 kDa]

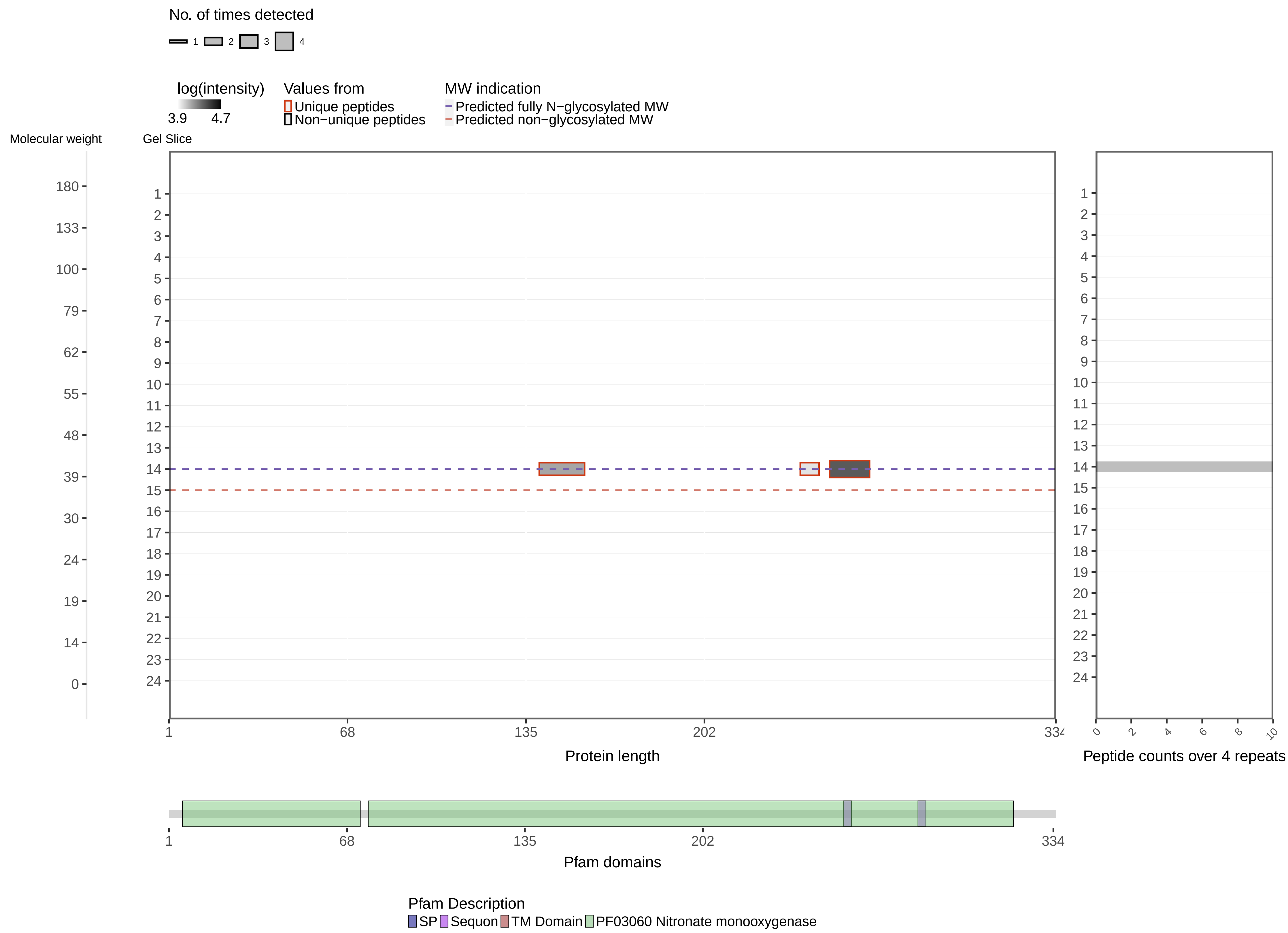


Protein sequence:

TLLPRVVDLVRGRDIAVIAAGGIVDARGYVAALALGAOGVSLGTRFLATEESYAHPTYKRKLIEFEOTEYTDLFGRARWPGAPHRVLATPFFKEWKALPSHENETNOPVIGRTIIHGKEREVRRFAGTVPNASTTGDIESMAMYAGOSIGLIKEILPAGEVIKOIVEGAORLIDOLFVPVN*

NbL11g10680.1 protein_EXORDIUM−like_2 [MW − SP = 29.09 kDa]

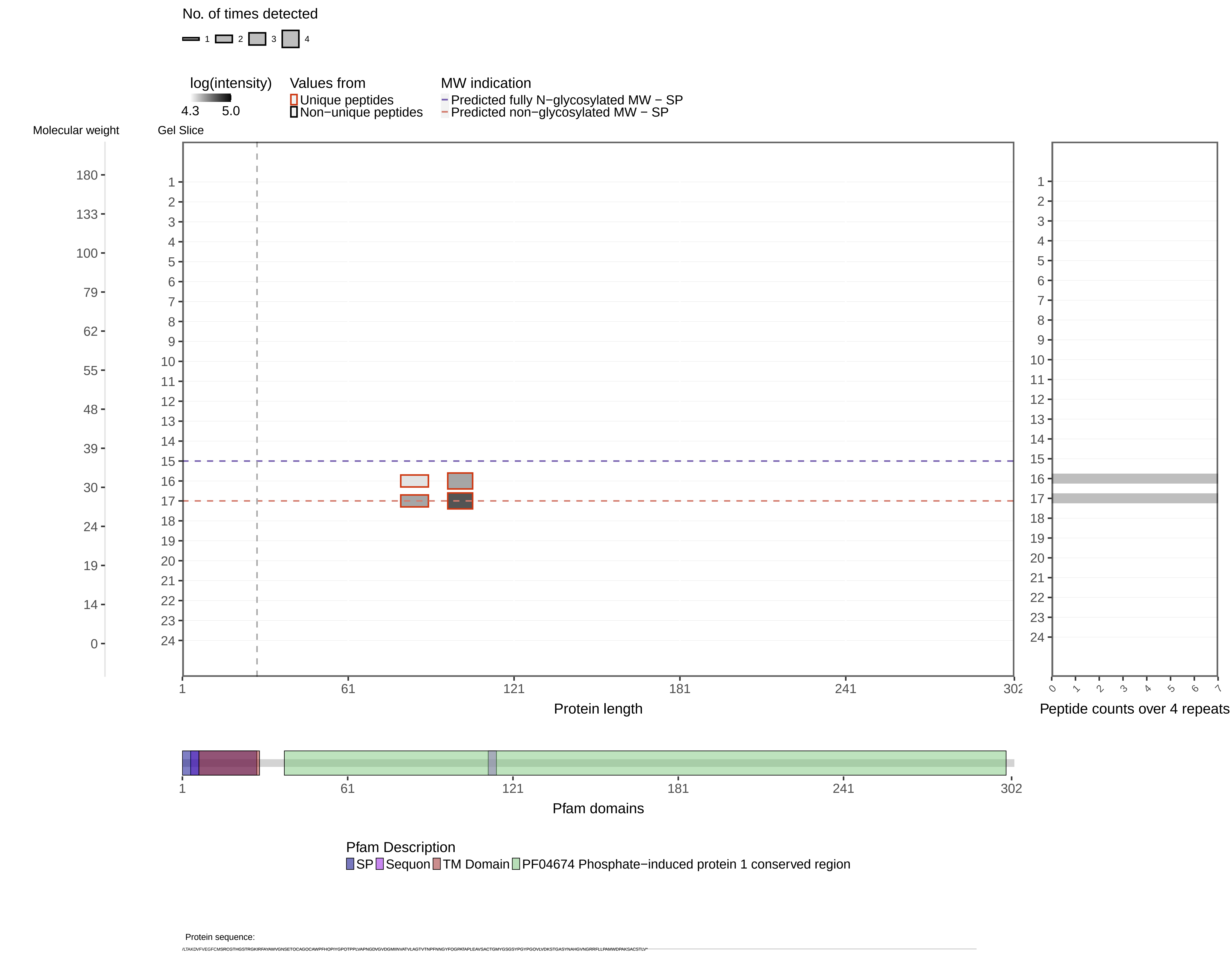

NbL11g10670.1 phospho−2−dehydro−3−deoxyheptonate_aldolase_2_chloroplastic−like [MW = 58.78 kDa]

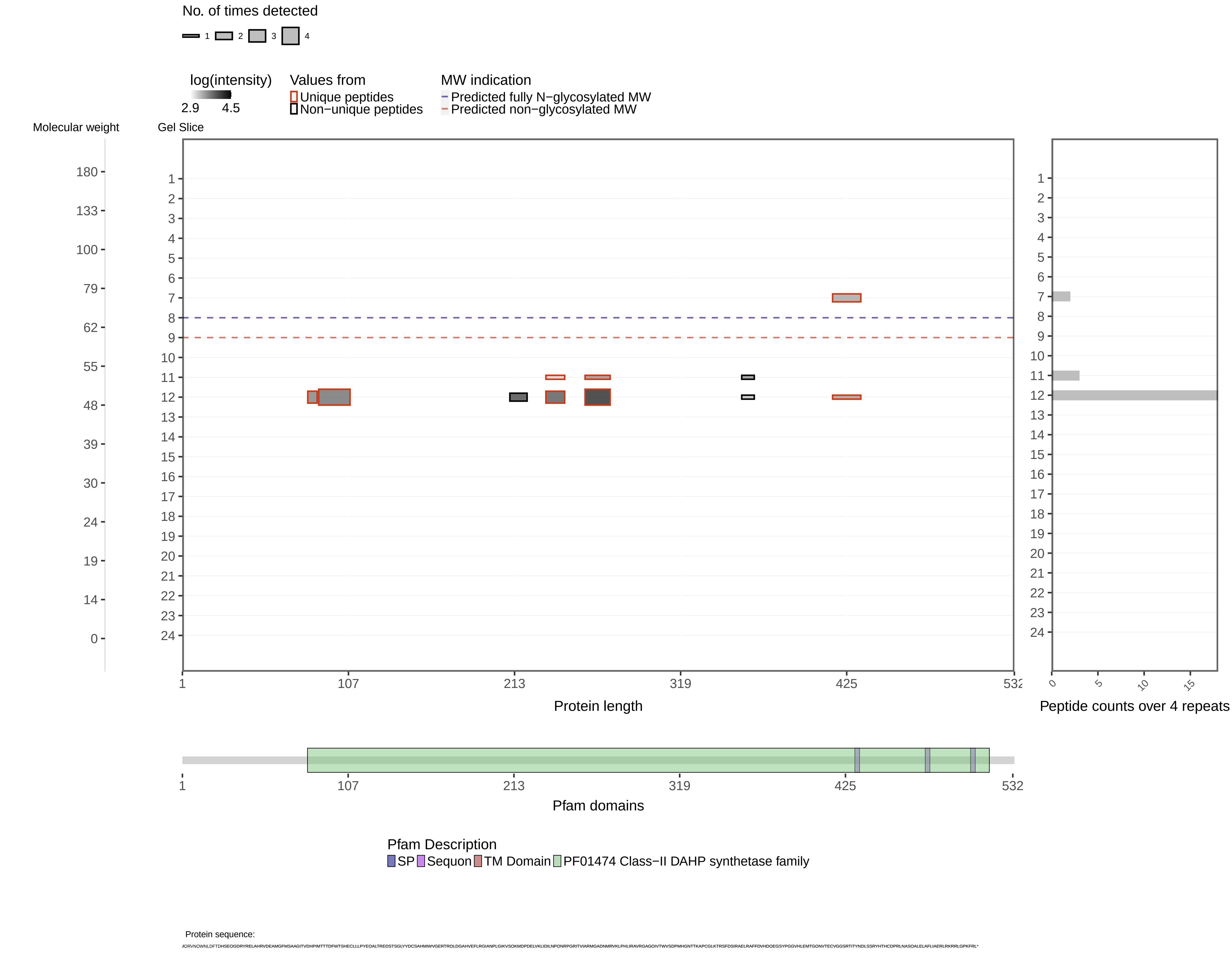

NbL11g10520.1 bifunctional_riboflavin_biosynthesis_protein_RIBA_1_chloroplastic−like [MW = 60.16 kDa]

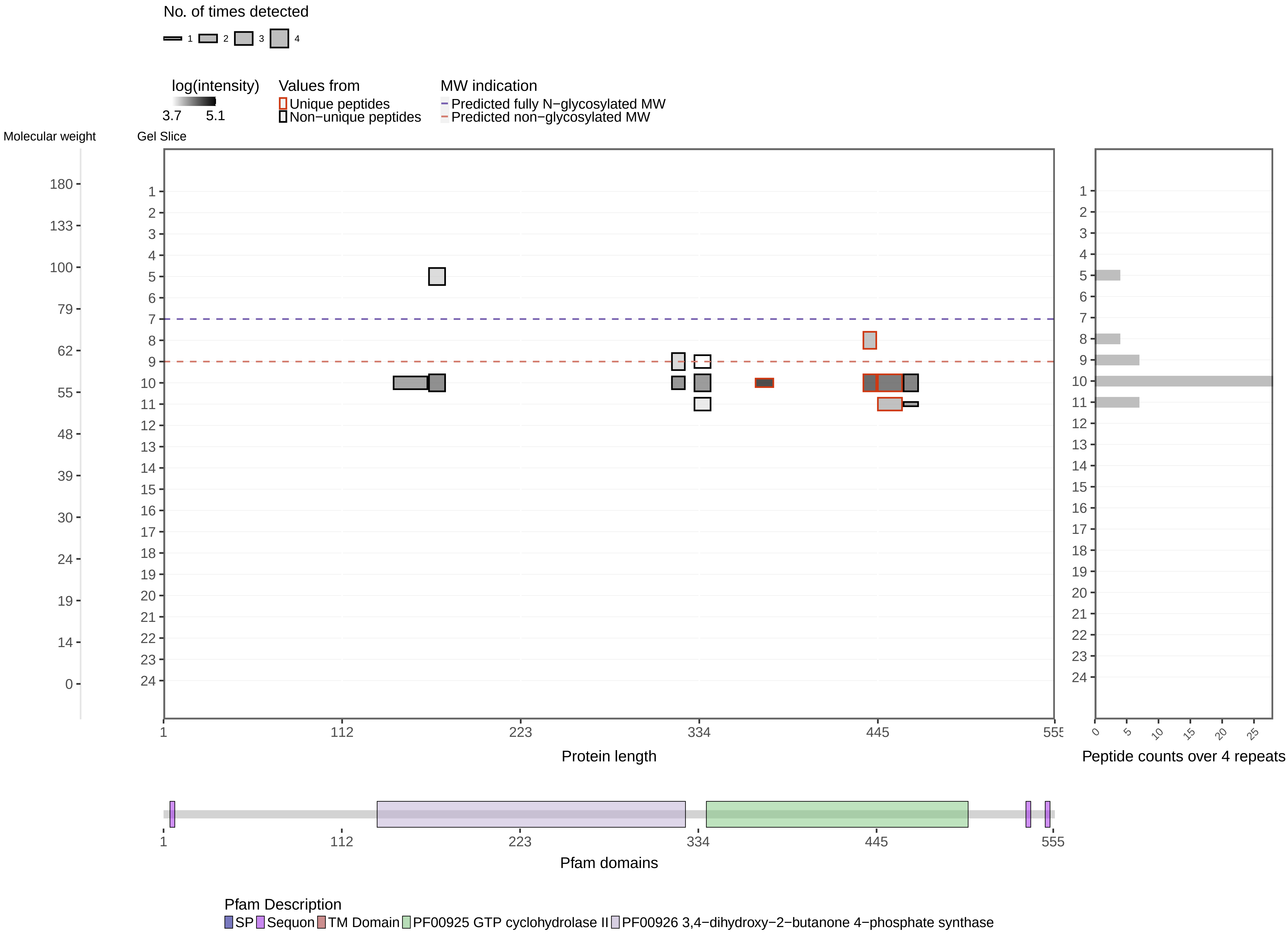


Protein sequence:

GVLKRAGHTEASVDLAMLAGLDPIGVICEIVDEDGSMARLPKLRQFAKEHNLKIISVADLIRYRRKRDQLVEHASAARLPTMWGPFTAHCFKSIIDGIEHIAMVKGDIGDGQDILVRVHSECLTGDIFGSARCDCGNQLALAMQQIEAAGKGVLVYLRGHEGRGIGLGHKLRAYNLQDAGRDTVEANEELGLPVDSREYGIGAQMLRDLGVRTMKLMTNNPAKYRGLKGYGLAISGRVPLLCPITKENKKYLETKRAKMGHVYGLDLIRPATSSNTSTTNGKPNVENISTI*

NbL11g10290.1 guanosine_nucleotide_diphosphate_dissociation_inhibitor_At5g09550 [MW = 49.66 kDa]

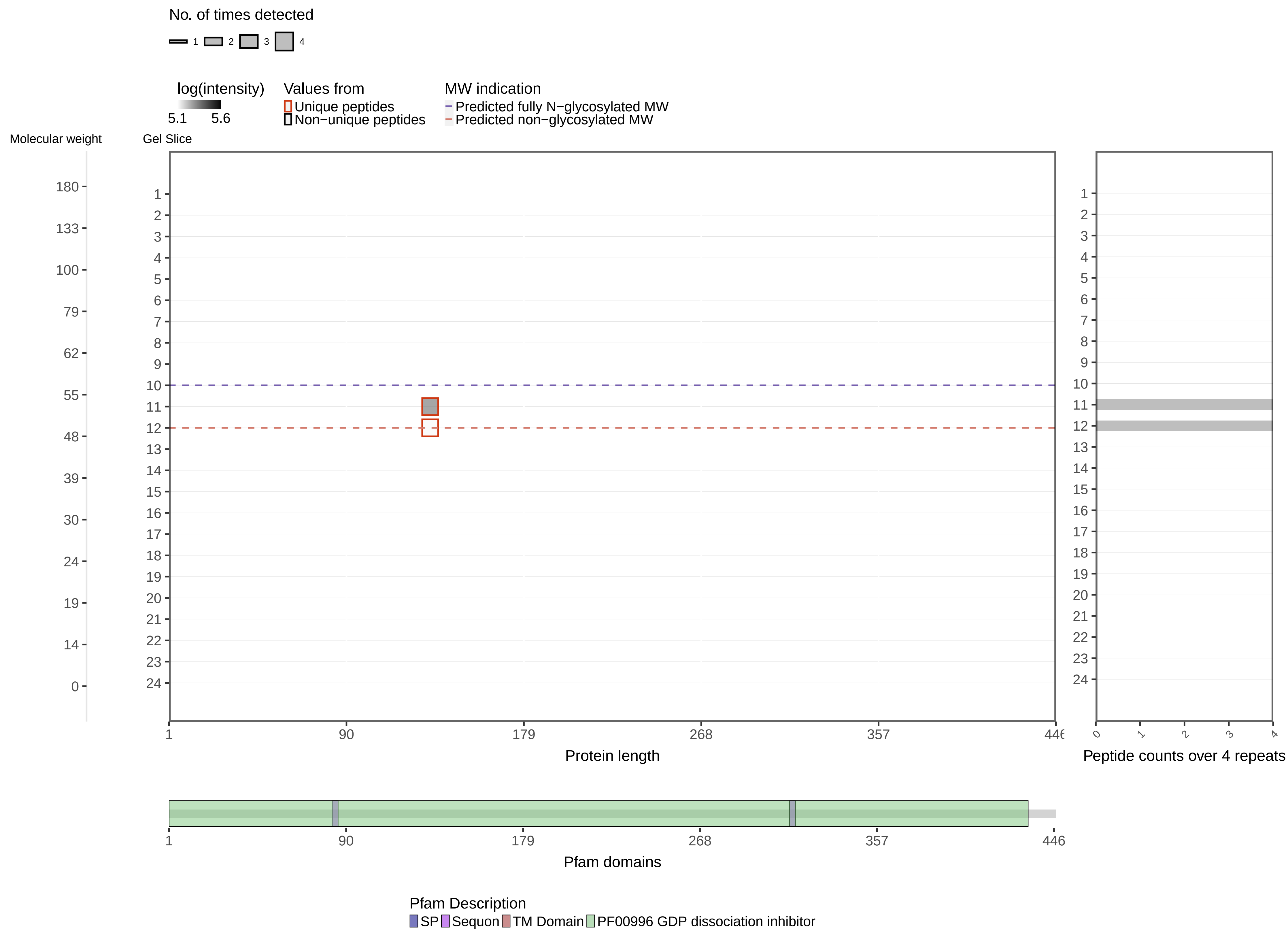


Protein sequence:

AESLARFQAGSPYIYPMYGLGELPQAFARLSAVYGGTYMLNKPQCKVEFDENGKVVGVTSEGETAKCKKVVCDPSYLPDKVQKVGKVARAICIMSHPIPDTNDSHSAQVILPQKQLGRKSDMYLFCCSYSHNVAPKGKYIAFVSTEAETDDPETELKPGVGILGPVDEIFYETYDRFAPSHDSAADNCFISKSYDSTTHFESTVMDVLDMYSKITGKVVDLSVDLSAASATAEE*

NbL11g10180.1 T−complex_protein_1_subunit_alpha [MW = 59.11 kDa]

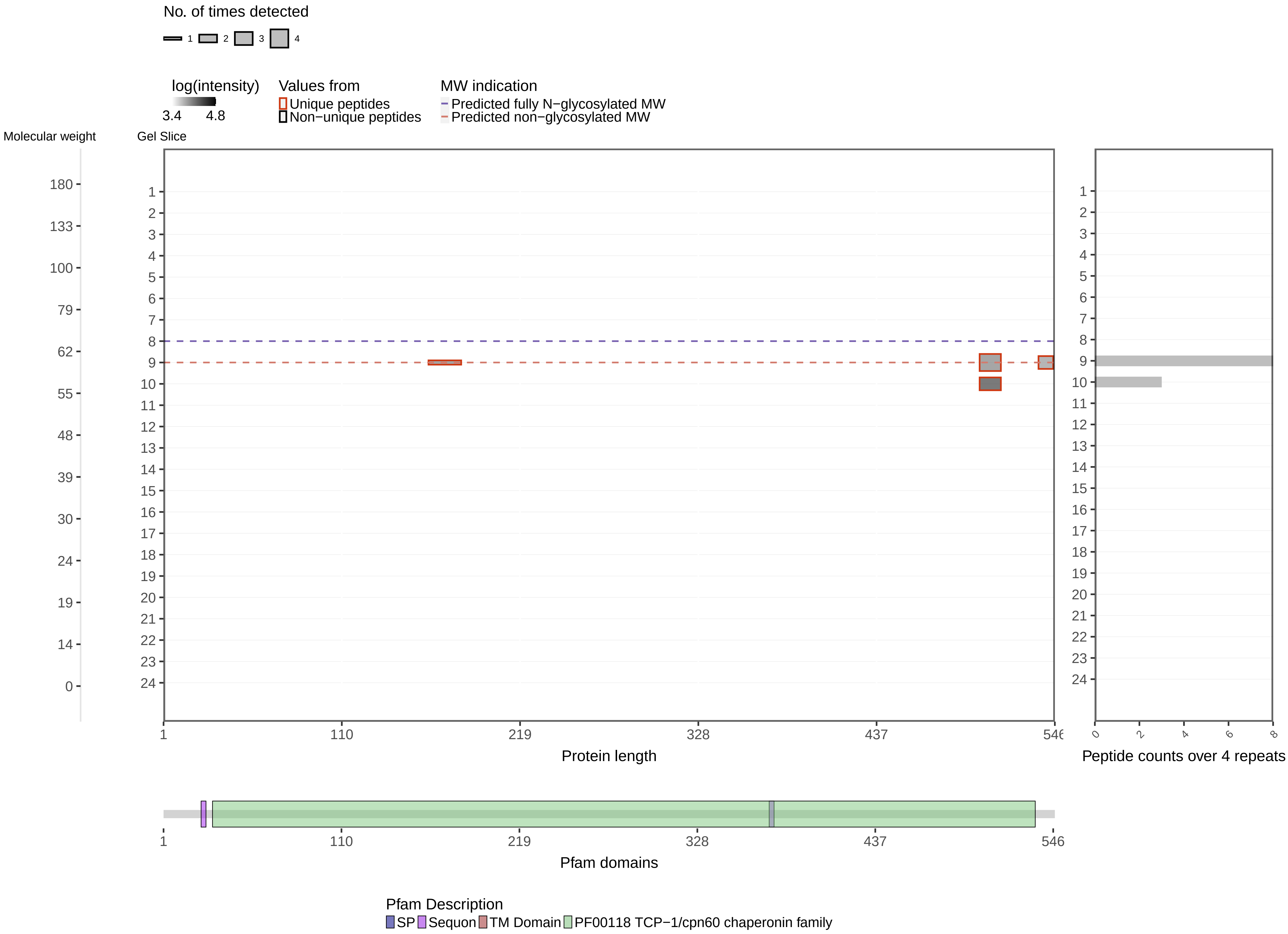


Protein sequence:

DPRELERIRQREADMTKERIQKLLSAGANVVLTSKGIDDMALKYFVEAGAIAVRRVRKEDLRHVAKATGATVVSTFADMEGEETFEPSLLGHAEEVVEERIADDDVIMVKGTKNSSSVSLILRGANDFMLDEMERALHDALCIVKRTLESTTVVAGGGAVEAALCVYLEYLATTLGSREQLAIAQFAESLLVIPKVLANNAAKDSSDLVSKLRSCHYLAQTKADKKHLSSMGLDLAKGTVRNNLGAGVIEPAMSKVKIIQFATEAAITILRIDDMIRLVKDESQNDD*

NbL11g10140.1 malate_dehydrogenase_cytoplasmic−like [MW = 39.49 kDa]

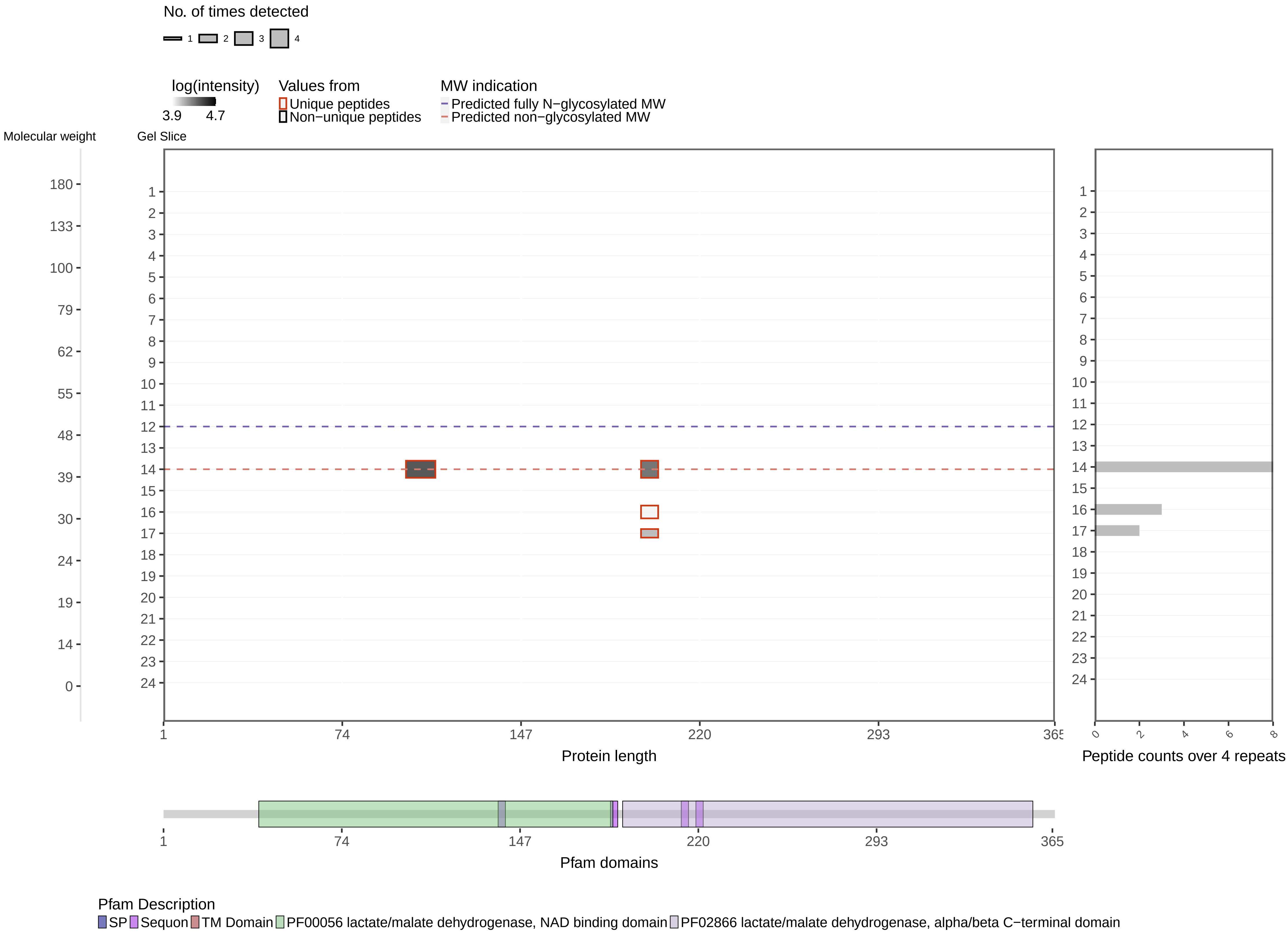


Protein sequence:

LILKEFAPSFPEKNITCLTRLDHNRALGQVSERLNVHVGDVKNVTIWGNHSSTQYPDVNHATVKTTSGEKSVRELVANDQWLNTEFITTVQQRGAAIIKARKLSSALSAASSACDHIRNWVLGTPKGTWVSMGVYSDGSYGIPPGLIYSFPVTCEKGEWSIVQGLKIDEFSRAKMDASAKELIDEKSLAYSCLN*

### NbL11g10110.1 elongation_factor_G−2_mitochondrial [MW = 84.29 kDa]

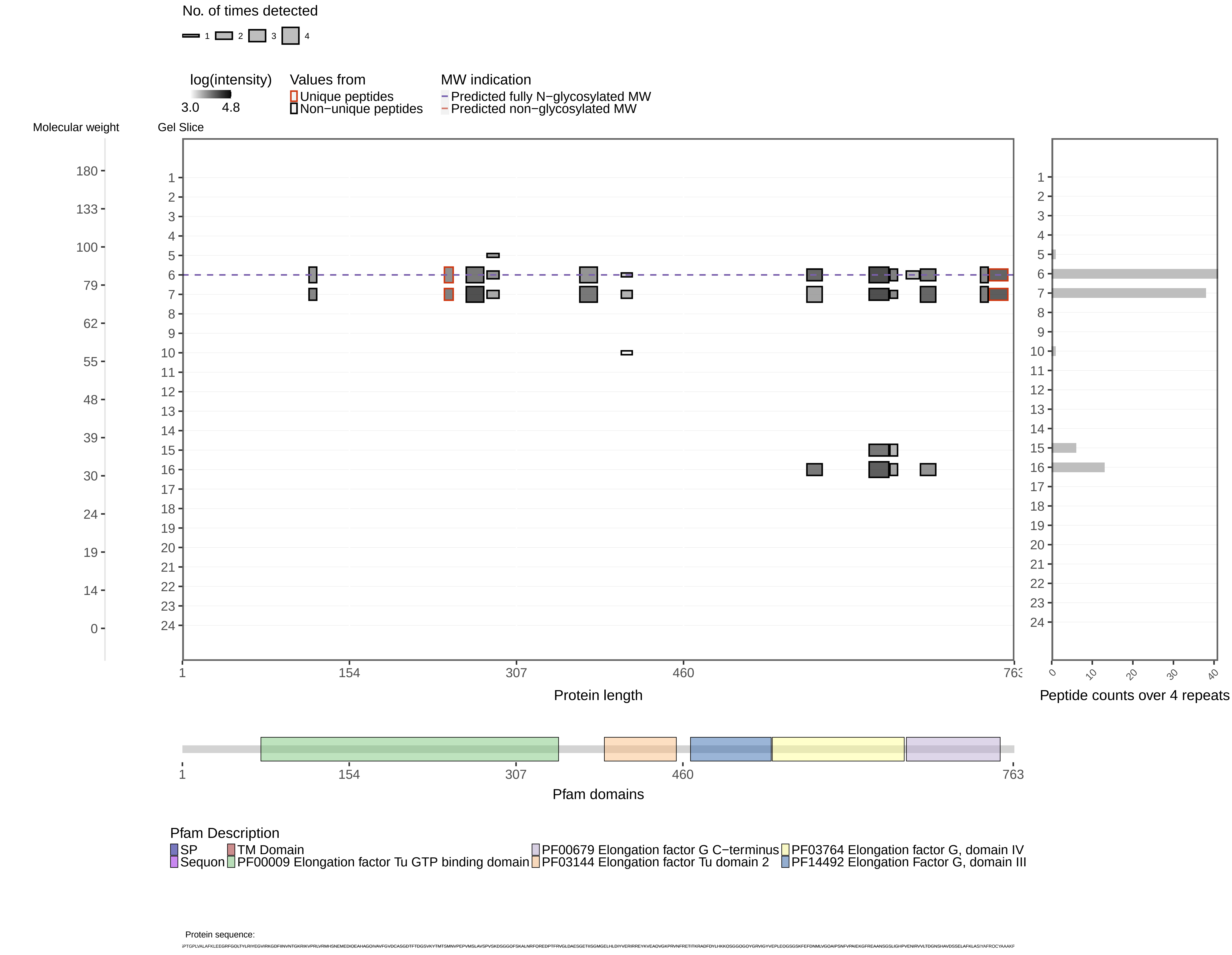

### NbL11g10000.1 phosphoribosylamine−−glycine_ligase−like [MW = 54.82 kDa]

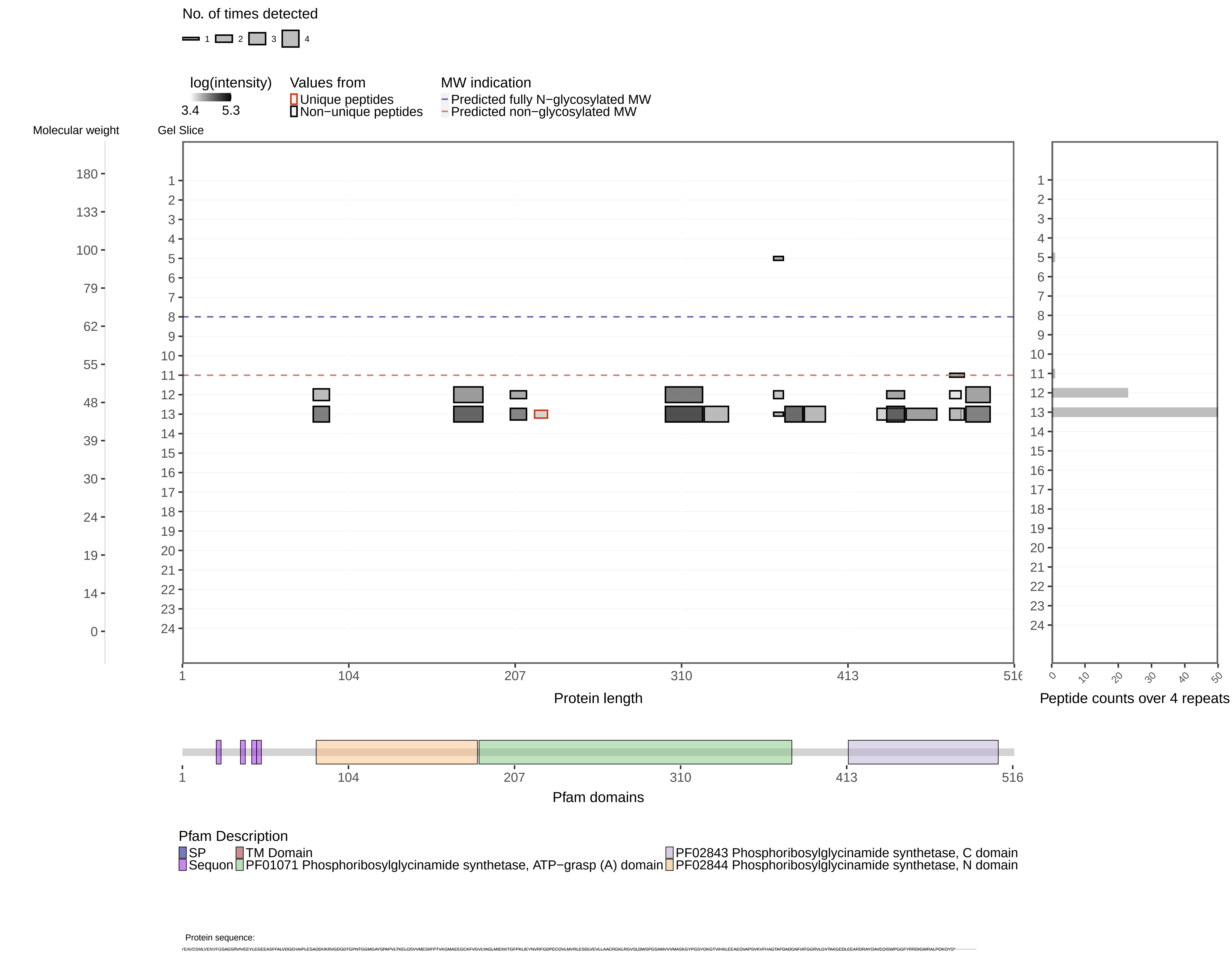

### NbL11g09790.1 peptidyl−prolyl_cis−trans_isomerase_CYP40 [MW = 40.32 kDa]

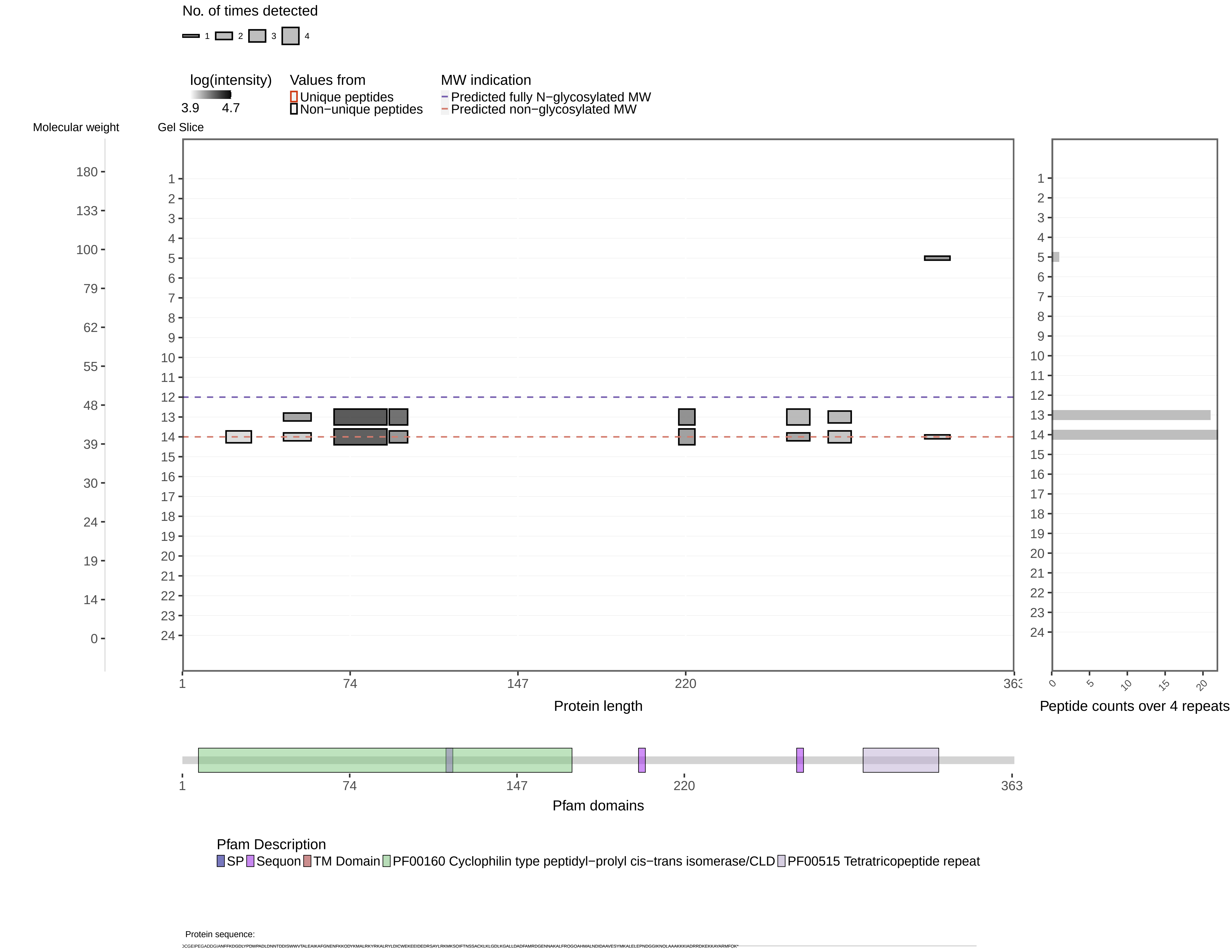

NbL11g09670.1 probable_N−acetyl−gamma−glutamyl−phosphate_reductase_chloroplastic [MW = 44.22 kDa]

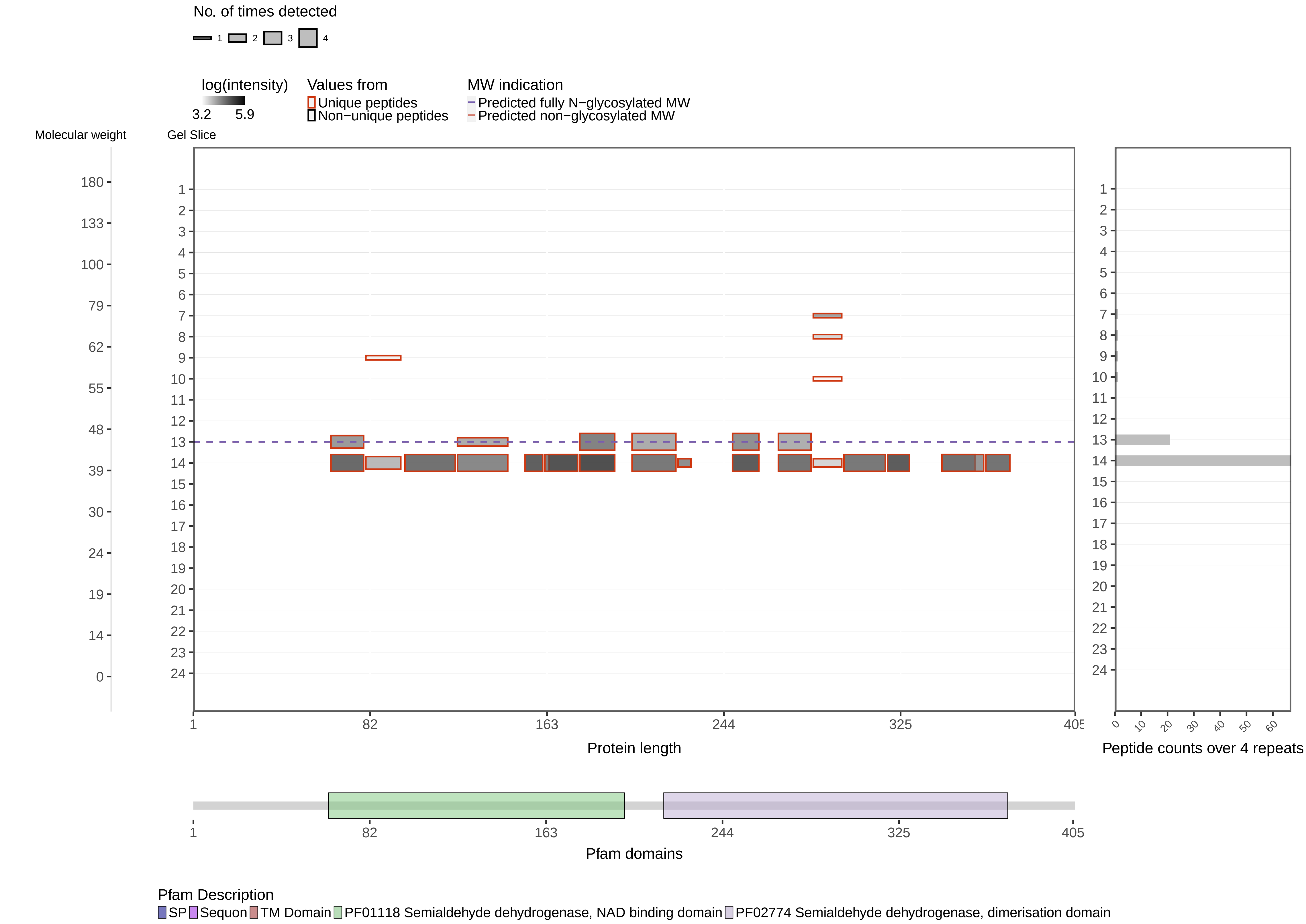


Protein sequence:

.TEISRREIQSARLVANPGCYPTSVQLPLIPLIKSNLIEVRNIIIDSKSGVSGAGRAAKEANLFTEIAEGIRSYGITRHRHVPEIEQGLSEAANSKVTVSFTPHLMPMSRGMQSTIYVEMAPGVTTQDLKDHLVNYYEHEKFVVLLKNKEVPYTRHVTGSNYCLMNVFPDRIPGRAIIVSVIDNLVKGASGQALQNLNLMMGLPENLGLGSLPLFP*

NbL11g09490.1 2−hydroxyisoflavanone_dehydratase−like [MW = 35.05 kDa]

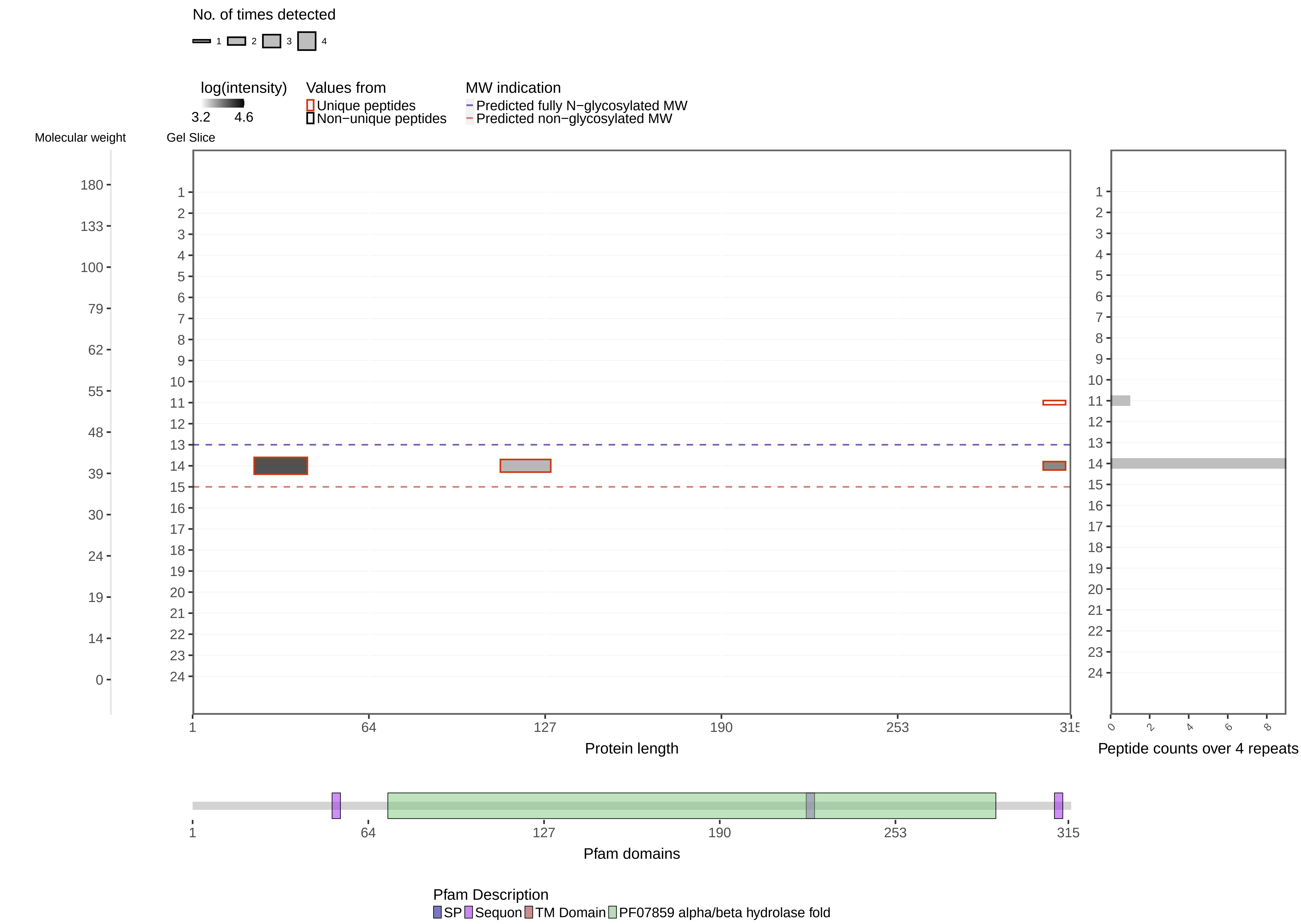


Protein sequence:

DHADLNRVYFGGDSAGANIAHNVAIRVGLEKLDGVKLDGIFLACPYFWGIDLIDGEAENLDCKNYIEKLWALAHPNSSGLDDPLMNPEKDPKLSSLGCDKVLVYVAAKDPLKYRGLYYKELLEKSGWOGTVEAVEIKDEEHVFHLFAPTSENAMVVLKKLVSFLNOSKA*

### NbL11g09420.1 ferredoxin−−nitrite_reductase_chloroplastic−like [MW = 67.84 kDa]

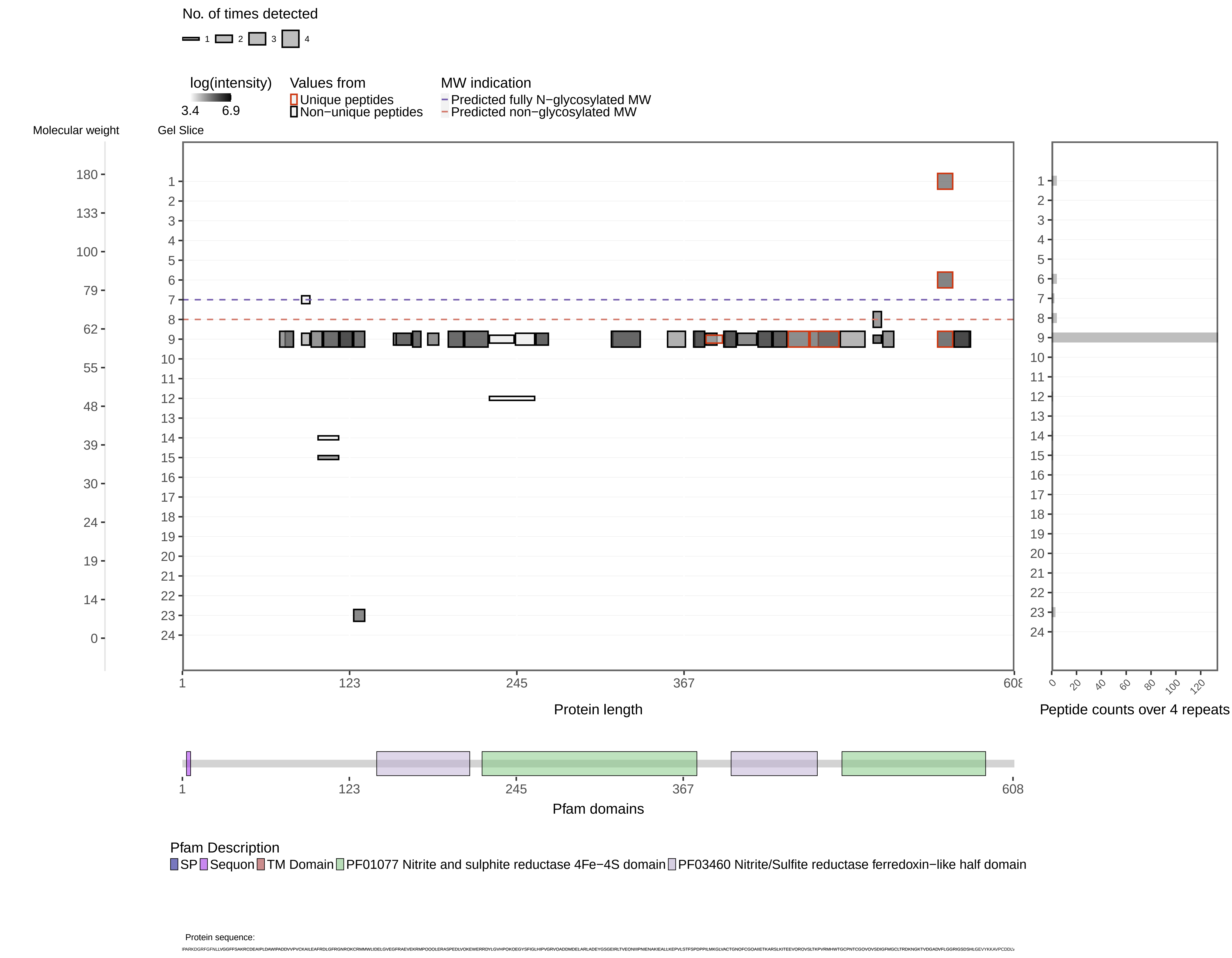

### NbL11g09310.1 probable_quinone_oxidoreductase [MW = 68.32 kDa]

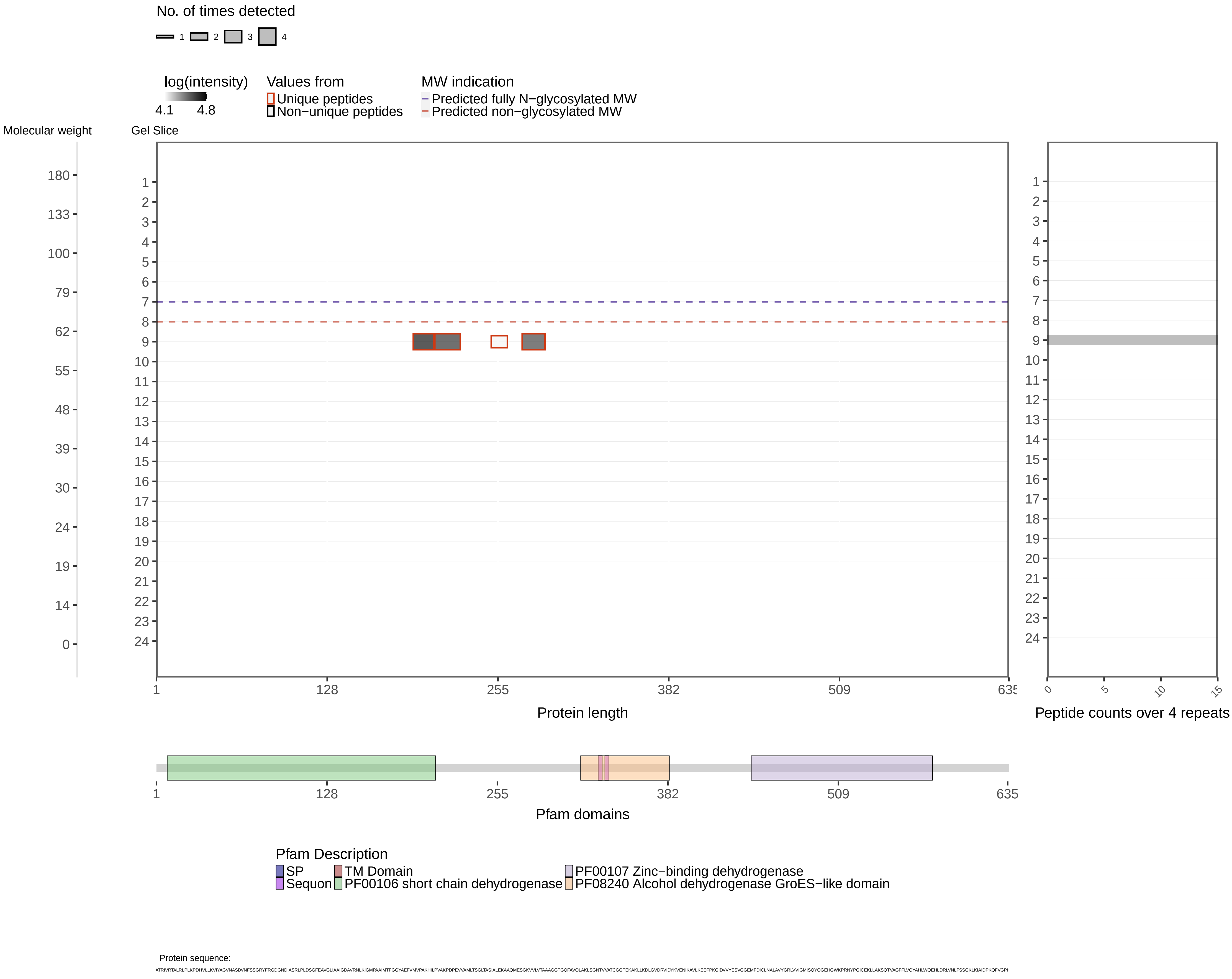

NbL11g09070.1 adenylate_kinase_chloroplastic [MW = 33.98 kDa]

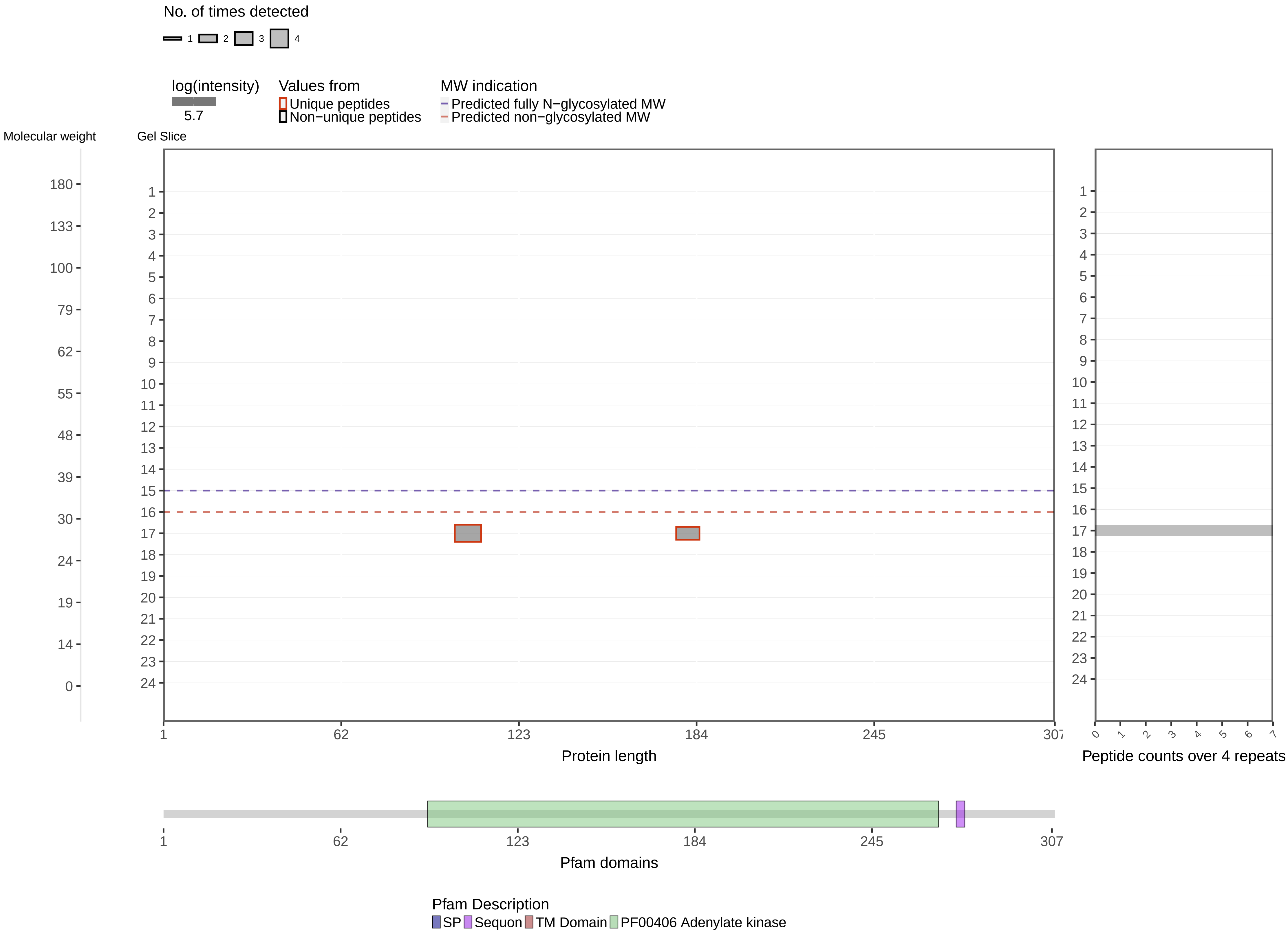


Protein sequence:
DKGKLVPDEIVVTMVKERLNGPDSREKGWLLDGYPRSSSQAIALKEFGFQPDIFILLEVPEEILVERVVGRRLDPVTGKIYHLKYSPPETEEIAARLTQRFDDTEEKVKLRLHTHHQNVEAVLSLYKDITLKVNGSASKQDVFAQIDDALTQLREQKQGKLGTVAA*

### NbL11g09050.1 protein_TIC_40_chloroplastic [MW = 49.23 kDa]

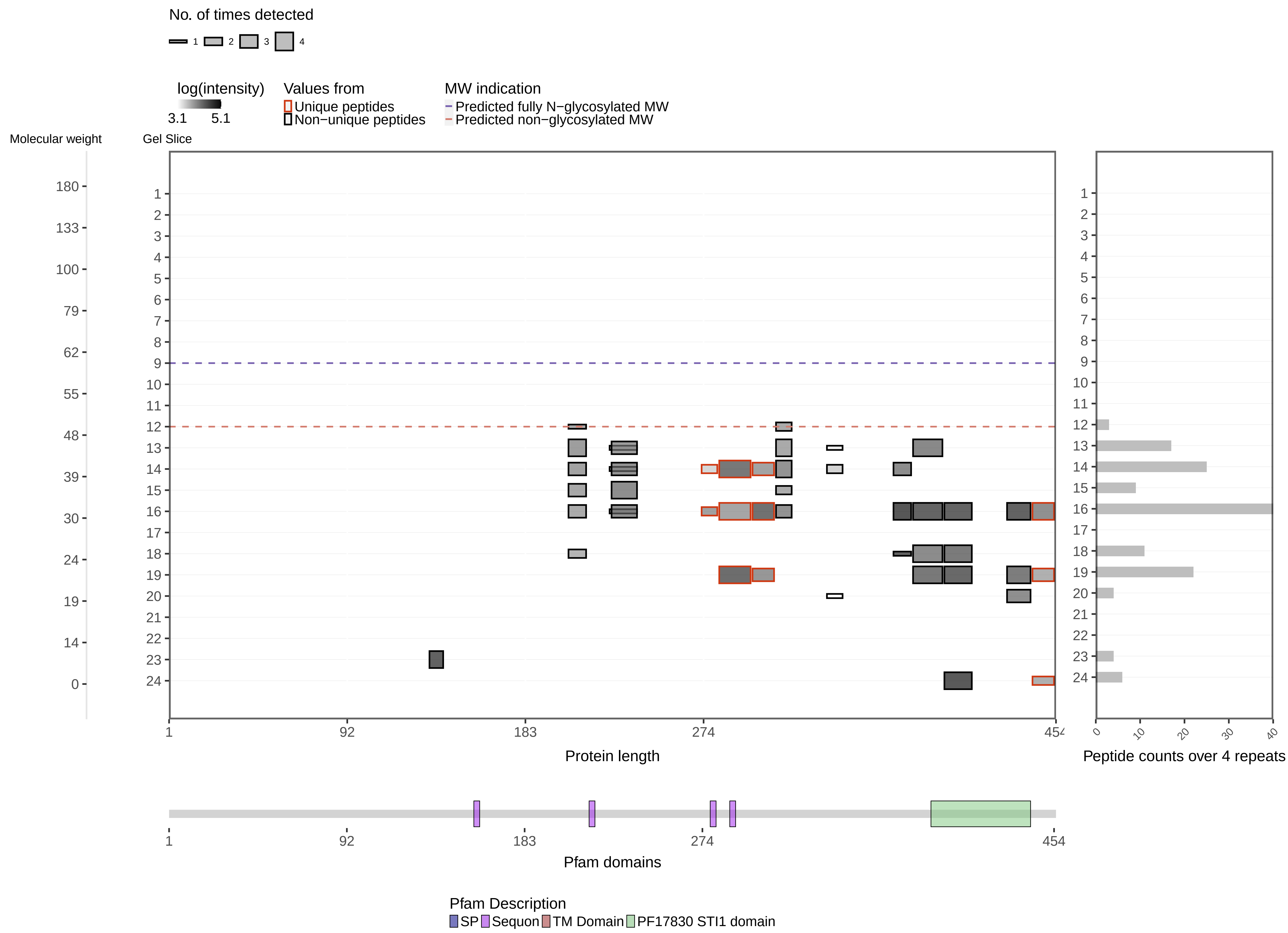


Protein sequence:

NDTEAERPPKKNAFVDISPDETFQKGAFENFKESTETASVTVDVTVDQVTQNGAASKSEFGANTSKSTGFGANTSDSTSSTEKSNPPMSVEAVEKMMEDPTVQKMVYPYLPEEMRNPTTFKWMLQNPQYRQQLQDMLNNMGGNPVWDNRMVDSLKNFDLSSPEIKQQFDQIGLTPEEVISKIMANPDVAMAFQNPRVQAAIMDCSQNPLSIAKYQNDKEVMDVFNKISELFPGVSGSL*

### NbL11g08900.1 putative_pentatricopeptide_repeat−containing_protein_At5g37570 [MW = 44.07 kDa]

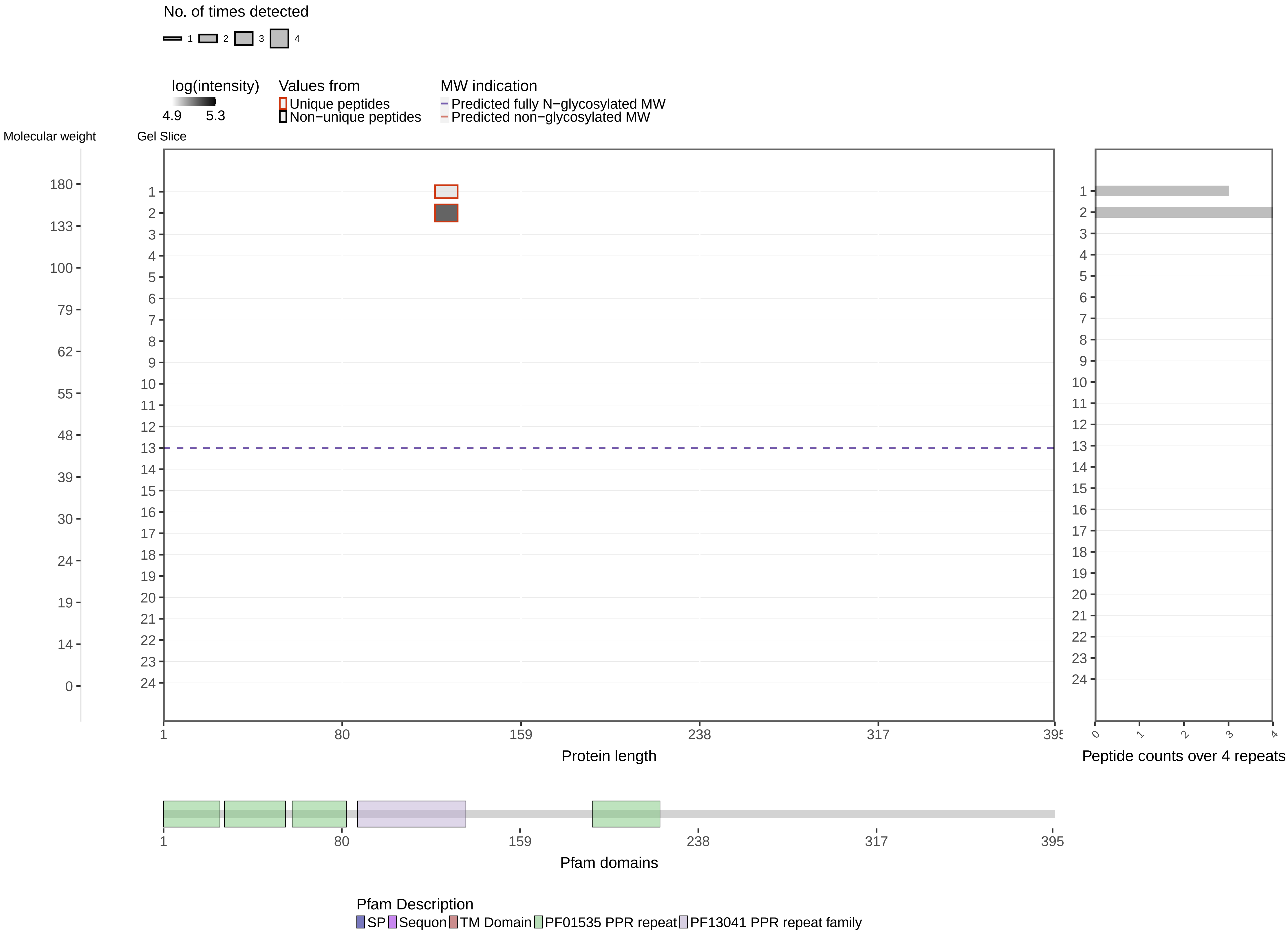


Protein sequence:

GTPKRDLVSYCSMIQGLSIHGCGSQAVVFFDRMLNEGLVPDDVSLKVIMTACSRAEEGCRIFNLVINKYHVKPSPDHYACVVDLLGRSGKLKDAYEFIRSMAIGPHAGAWGANLGACKLHCDIEVGEEVSHRLFELEPQNTGNYVLLSNIYAAANRWLDVALLRQQMSEKGLKKIPGCSWYNVMAICGHSLKLNKELQKEIKVDNSLLHDW*

### NbL11g08740.1 elongation_factor_1−gamma_2−like [MW = 15.29 kDa]

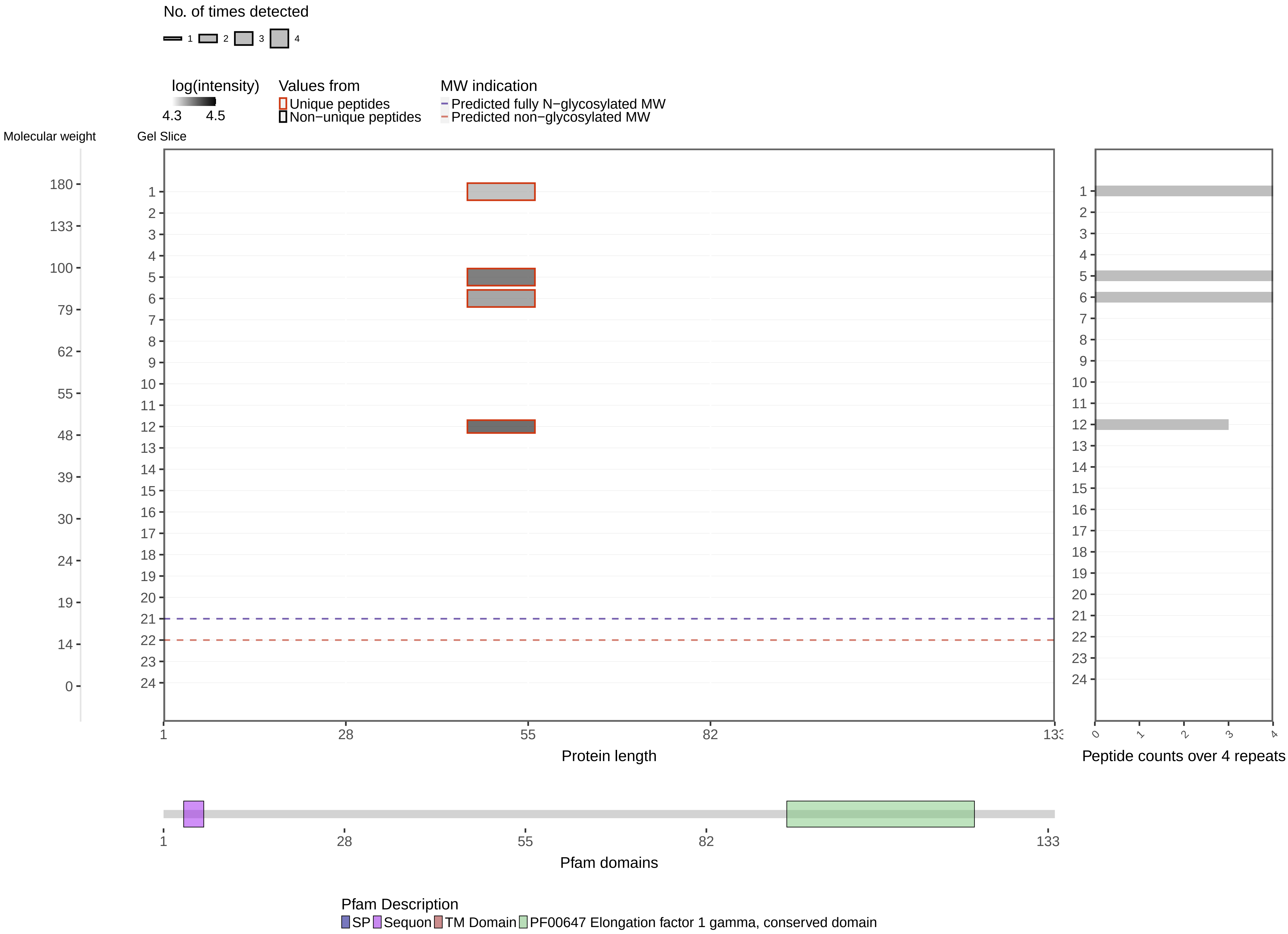


Protein sequence:

PPSKKPAPAKEPAKPKAKEEPKKEVKKEEPKFEEEEEAPKPKAKNPLDLLPPSKMILDEWKRLYSNTKTNFREVAIKGIF*

### NbL11g08730.1 elongation_factor_1−gamma_2−like [MW = 47.67 kDa]

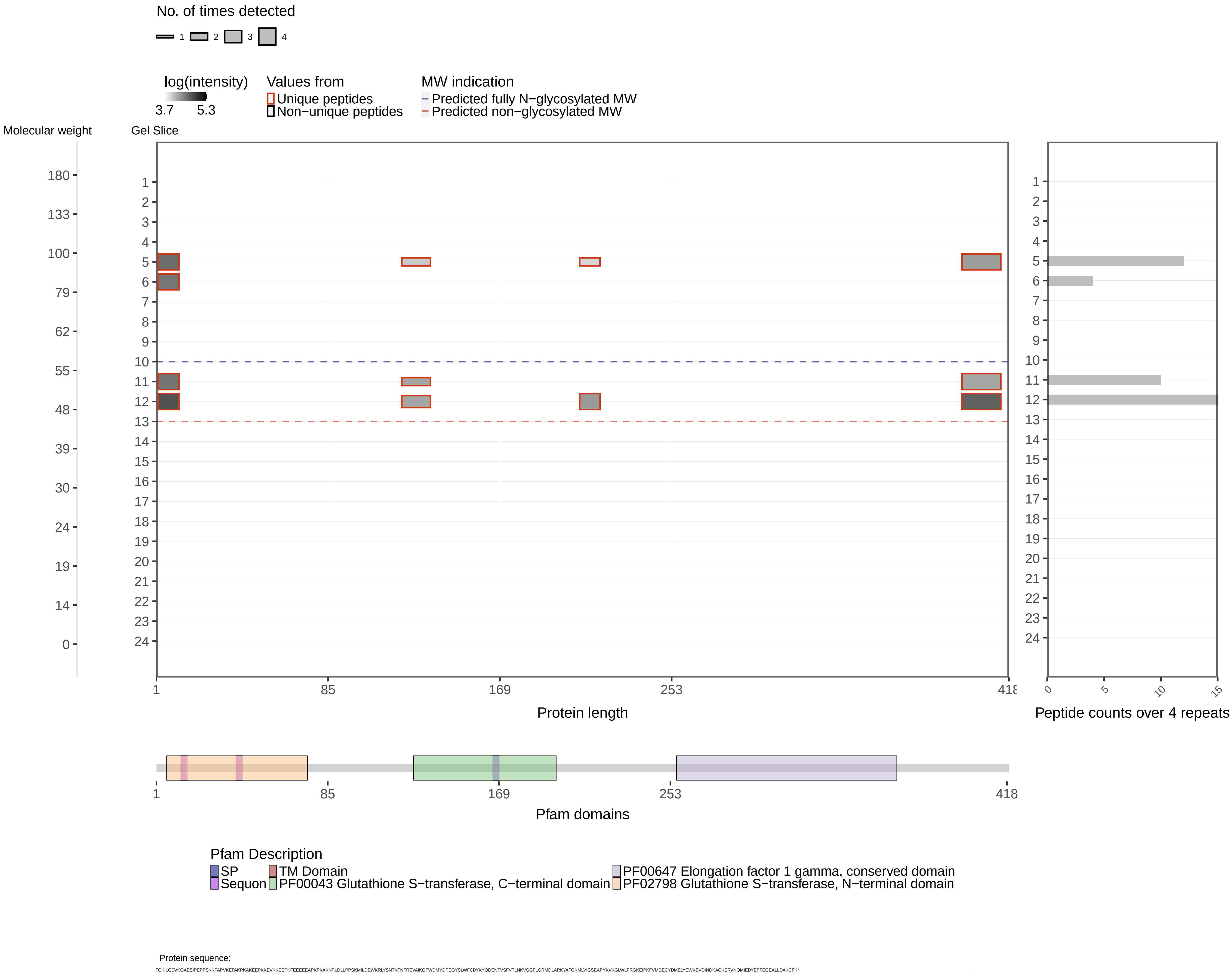

### NbL11g08700.1 patellin−3 [MW = 59.34 kDa]

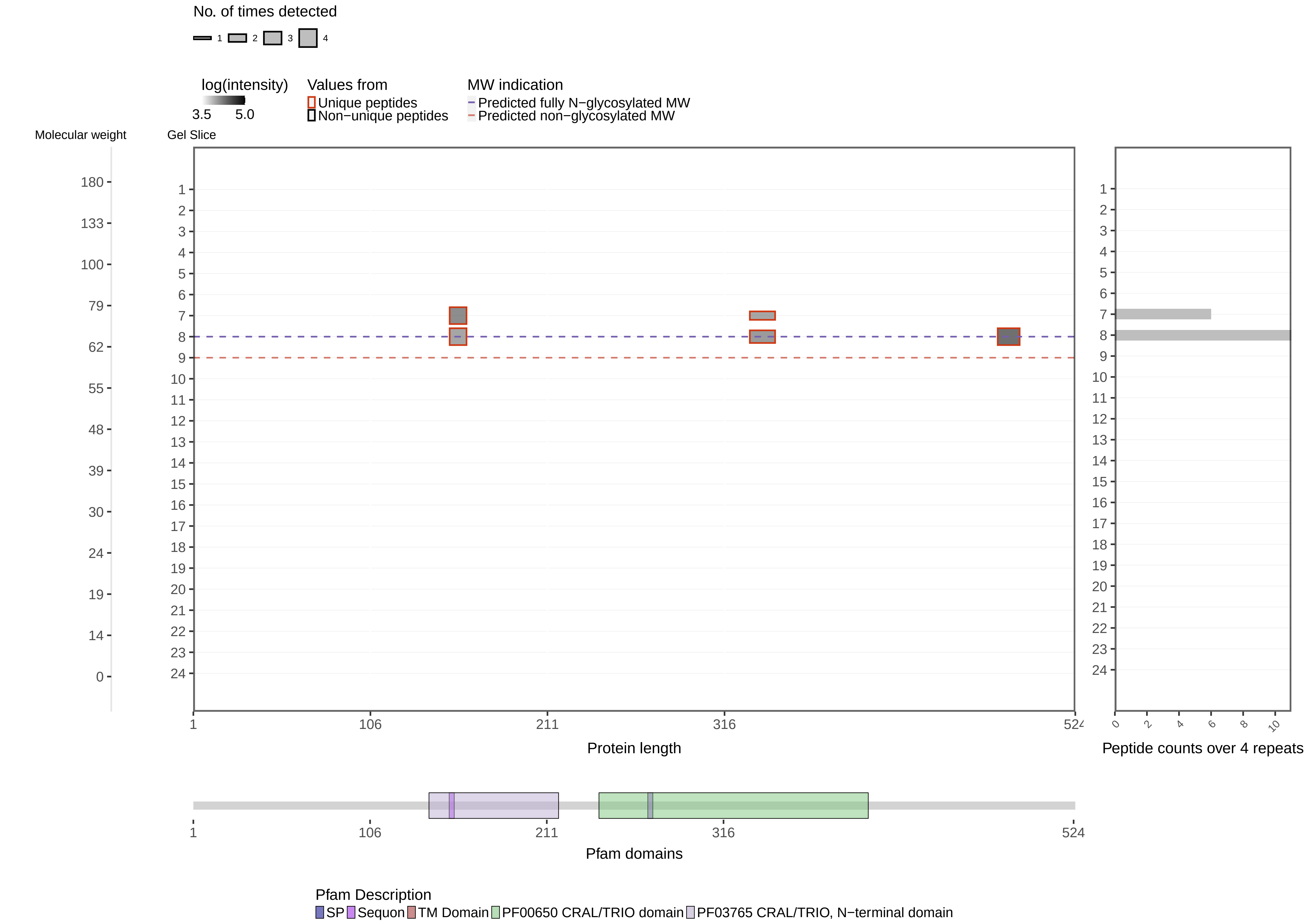


Protein sequence:

IDKEGHPVCYNVYGEFQNKELYNKTFGDEEKRNKFLRWRIQFLERSIRKLDFNPGGINTIFQVSDLKNSPGPGKRELRLATRQALHLLQDNYPEFVAKQVFINVPWWYLAFYTMISPFMTQRTKSKFVFAGPSKTAETLYKYVSPEQVPVQYGGLSVDYCECNPEFTVSDPVTEIIVKPATKQTVEIIVNEKCIIVWELRVLGWEVTYSAEYVPNTDSGYTVNIQKPRKMAPTDEPVVSSSFKIVELGKILLTIDNPTSKKKKLLYRYKDKPYST*

### NbL11g08690.1 uncharacterized_LOC104109700 [MW = 10.75 kDa]

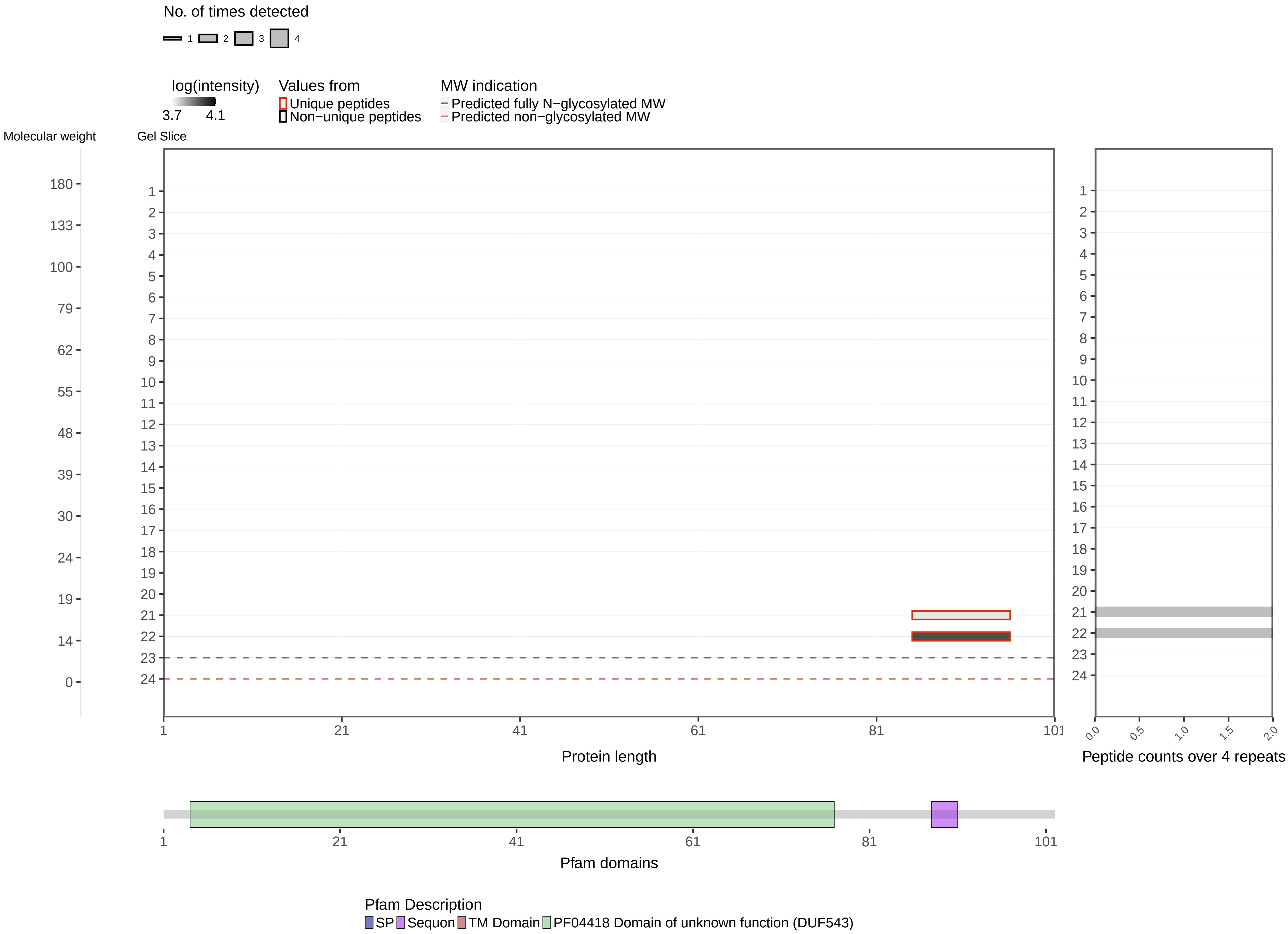


Protein sequence:

FSLIGGFAGLLLFRSPVTRWASTAFGAGTGLGSAYTECSQKLGGYPGKLTANVSEAPVTKDGEH*

### NbL11g08640.1 patellin−3−like [MW = 69.43 kDa]

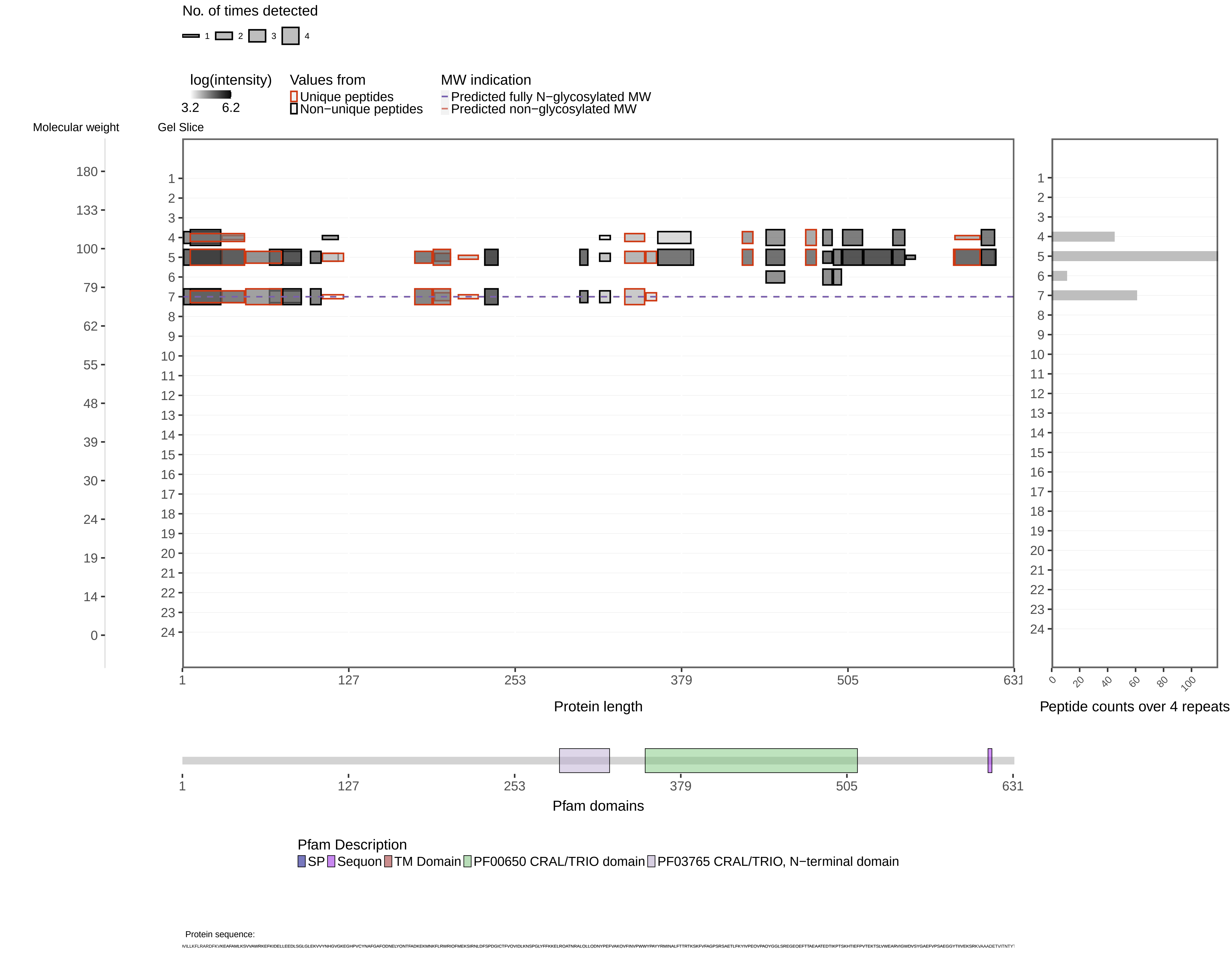

NbL11g08520.1 probable_polygalacturonase_At3g15720 [MW − SP = 49.76 kDa]

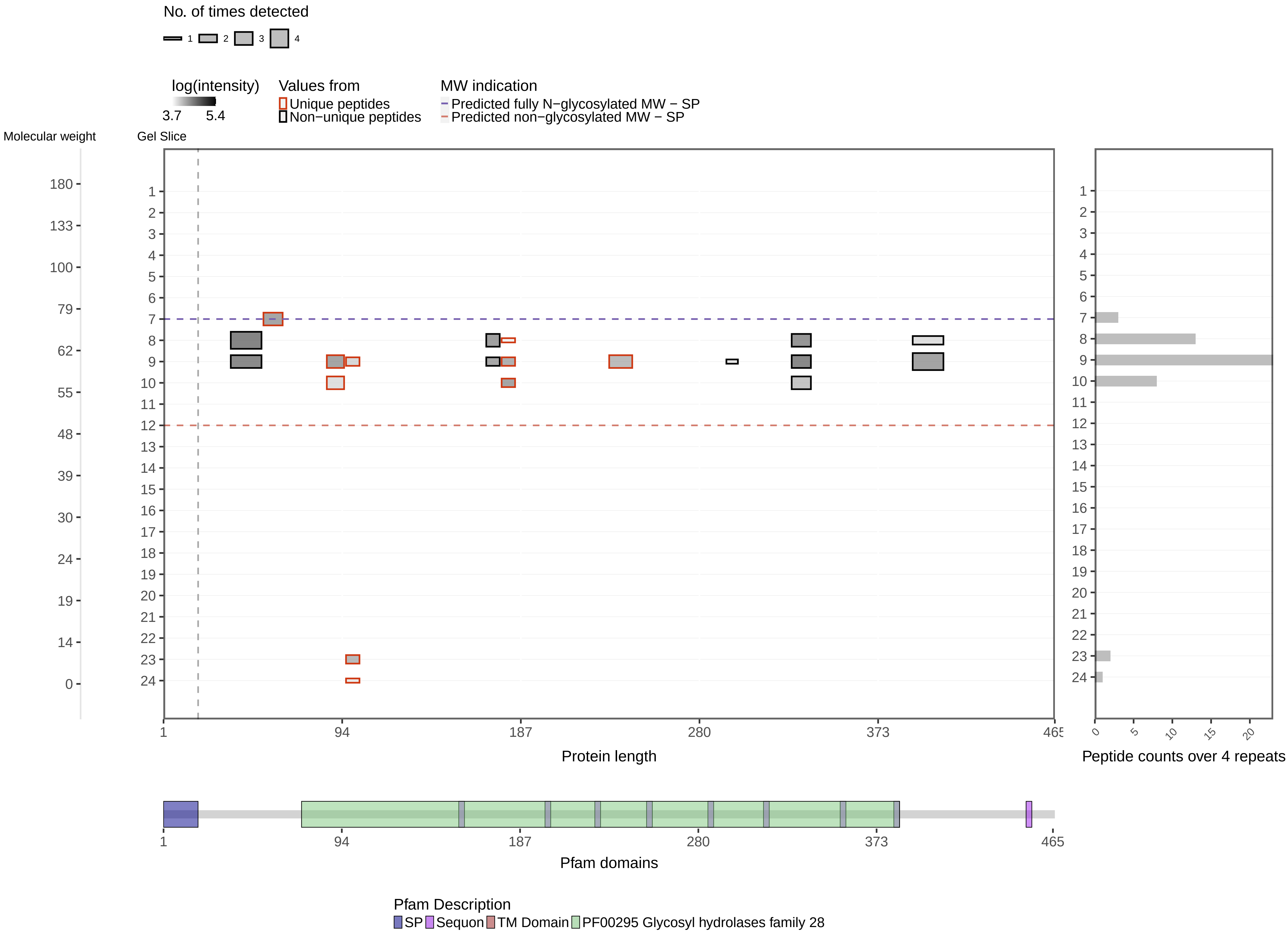


Protein sequence:

DIEDSNNTVITRCYINTGDDAICPKSSTGPVHNLTATNCWIRTKSSAIKLGSASFYTFNNFLFDNITIFESHRGLALQIRDGGDVSDITFSNINISTRYYHPSWWGRAEPIYVTTCPRDATSKAGSISNLQFVNITSTSENGIFLSGSGGGVLSNLKFINMNLTYKRWTKYPDGWVDYRPGCQELVKHQTAGFMMEHIDGLVFENVVMKWSNDESMRWNNPLDFRPSTVNNISMLNFYSRSYQQ*

NbL11g08390.1 60S_ribosomal_protein_L7a−1 [MW = 29.18 kDa]

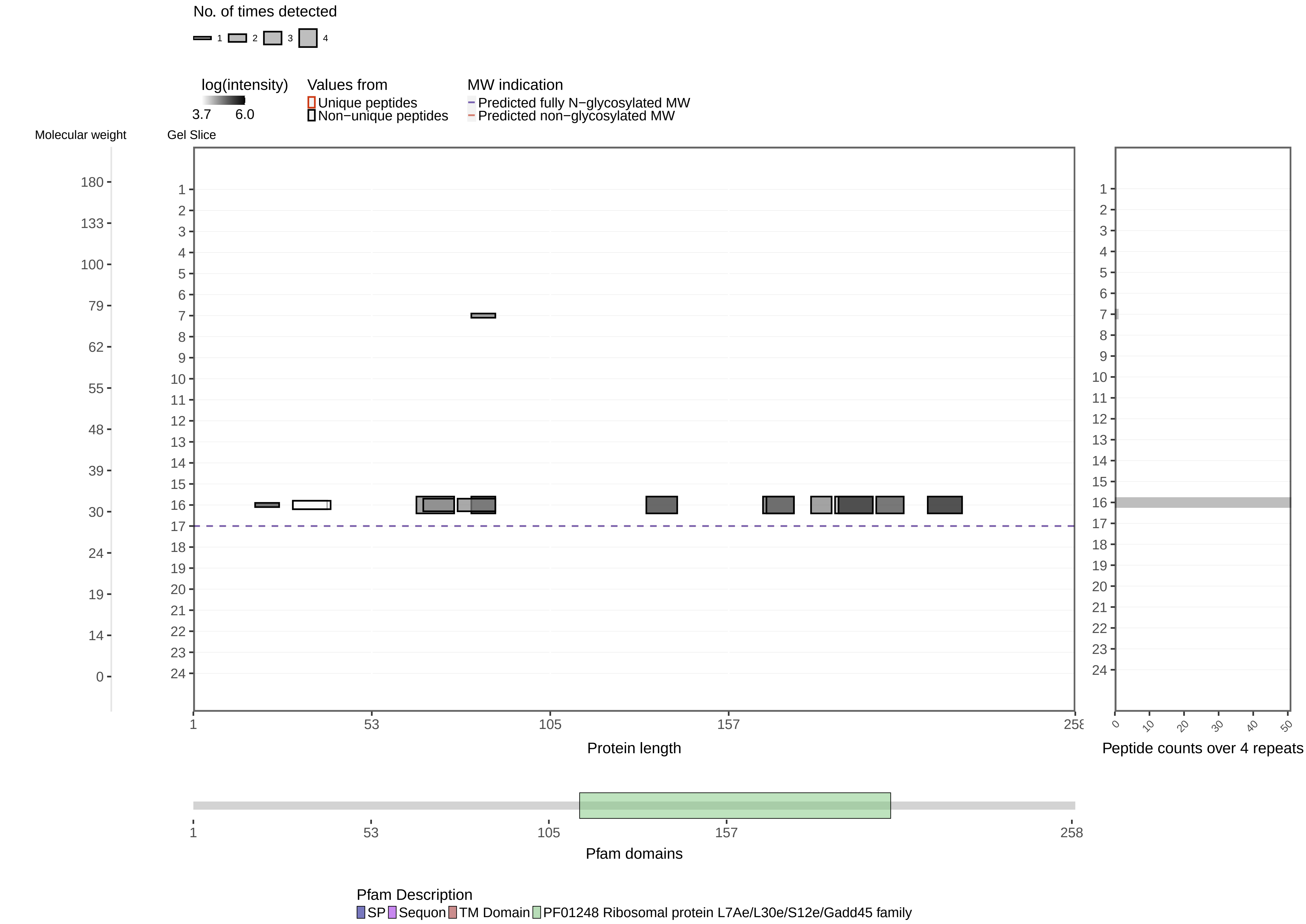


Protein sequence:

EGKTPETKKPIIVKYGLKHITYLIEQNKAQLVVIAHDVDPIELVVWLPALCRKMEIPYCIVKGKARLGSIVHKKTASALCLTTVKNEDKMEFSRILEAIKANFNDKYEENRKKWGGGVMGSKSQARTKAKERVLAKEAAQRMN*

### NbL11g08220.1 50S_ribosomal_protein_L13_chloroplastic [MW = 27.11 kDa]

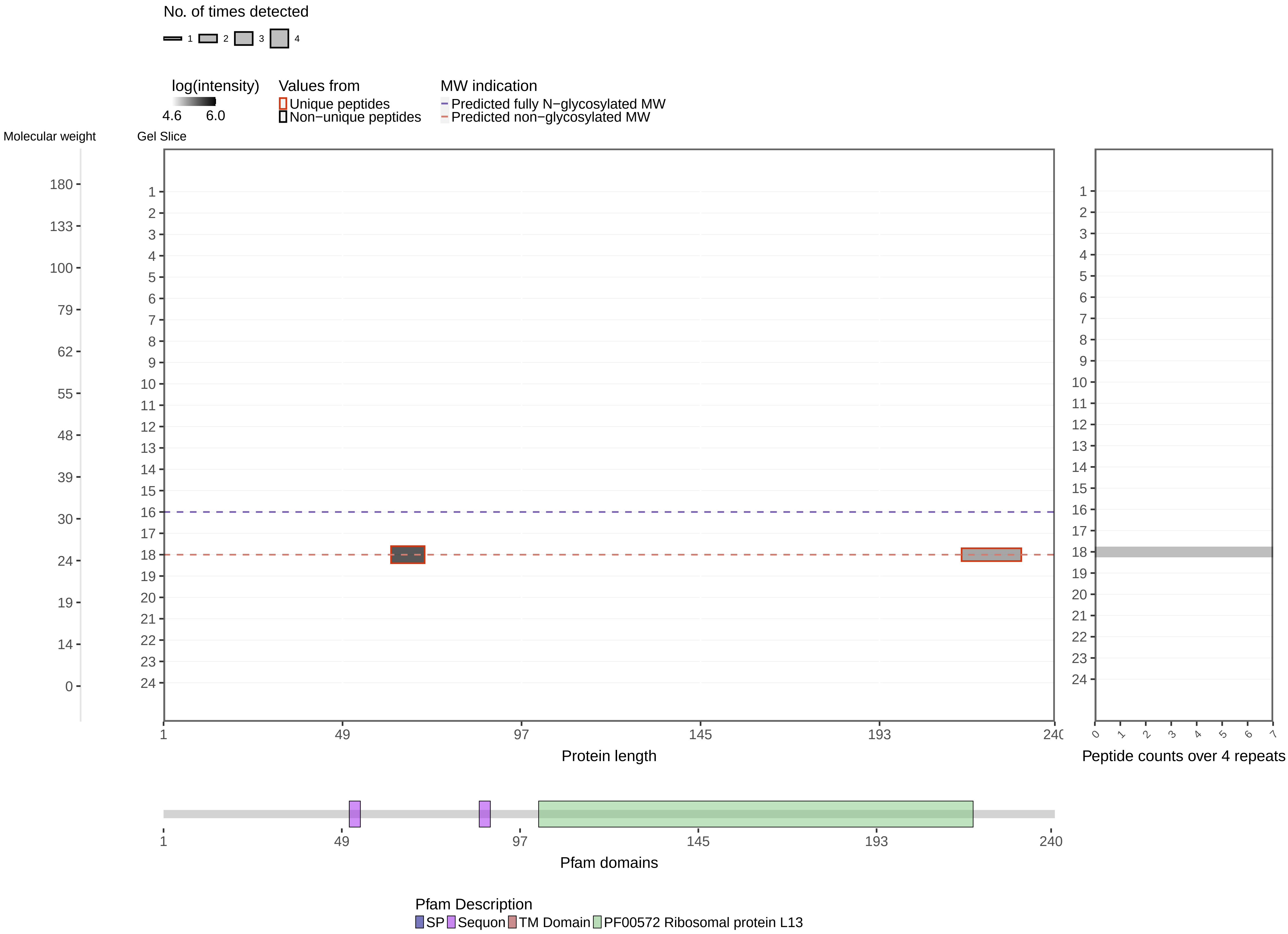


Protein sequence:

DASDLILGRMASTIAIHMRGKNLATYTPSVDMGAFVIVVNAEKVAVSGKKRTQKLYRRHSGRPGGMKVETFDQLQQRIPERIIEHAVRGMLPKGRLGRQLFTHLKVYKGPEHPHEAQKPIELPIRDKRIQKQR*

NbL11g08130.1 ubiquitin_carboxyl−terminal_hydrolase_14−like [MW = 89.37 kDa]

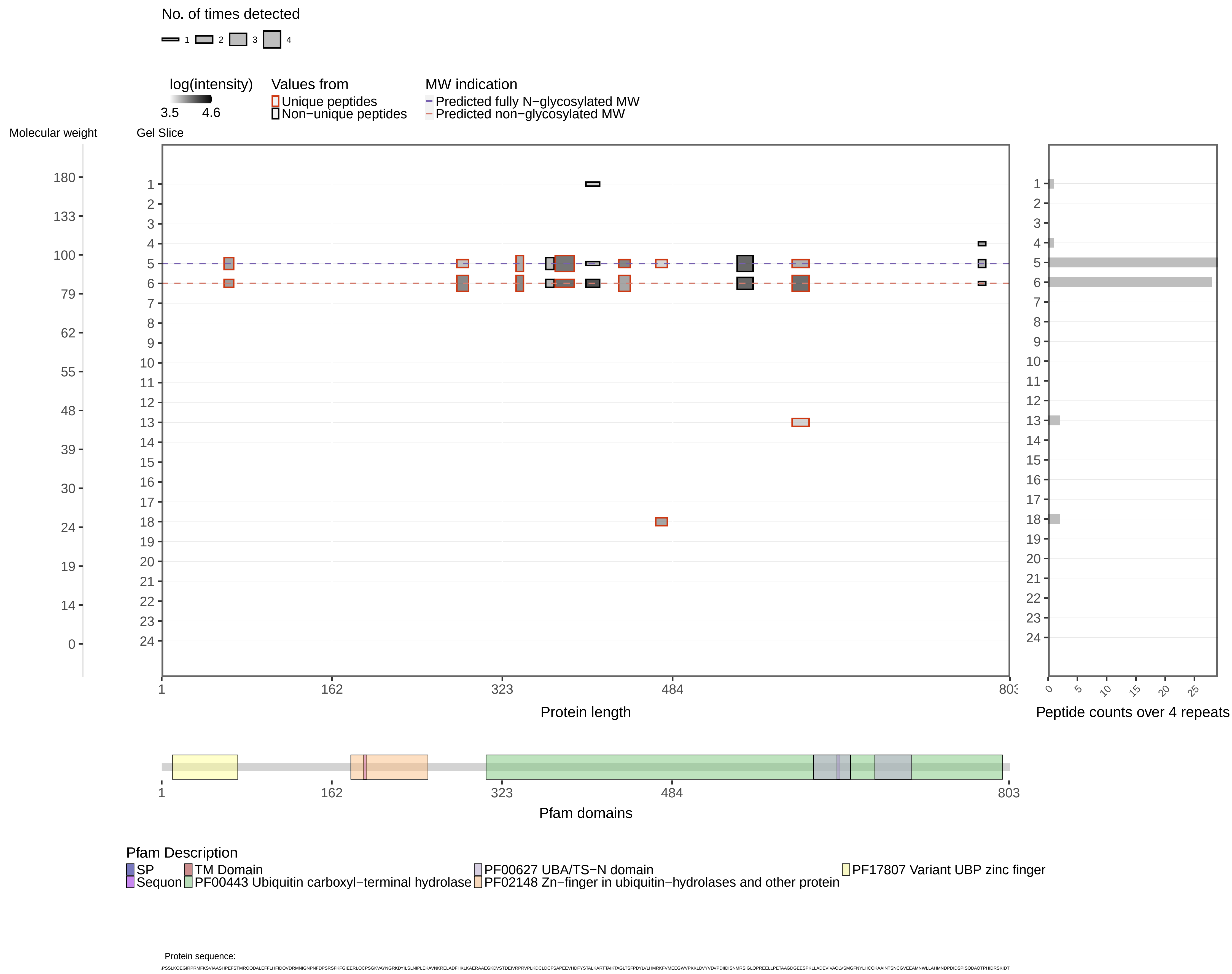

### NbL11g08100.1 60S_ribosomal_protein_L8−like [MW = 28.16 kDa]

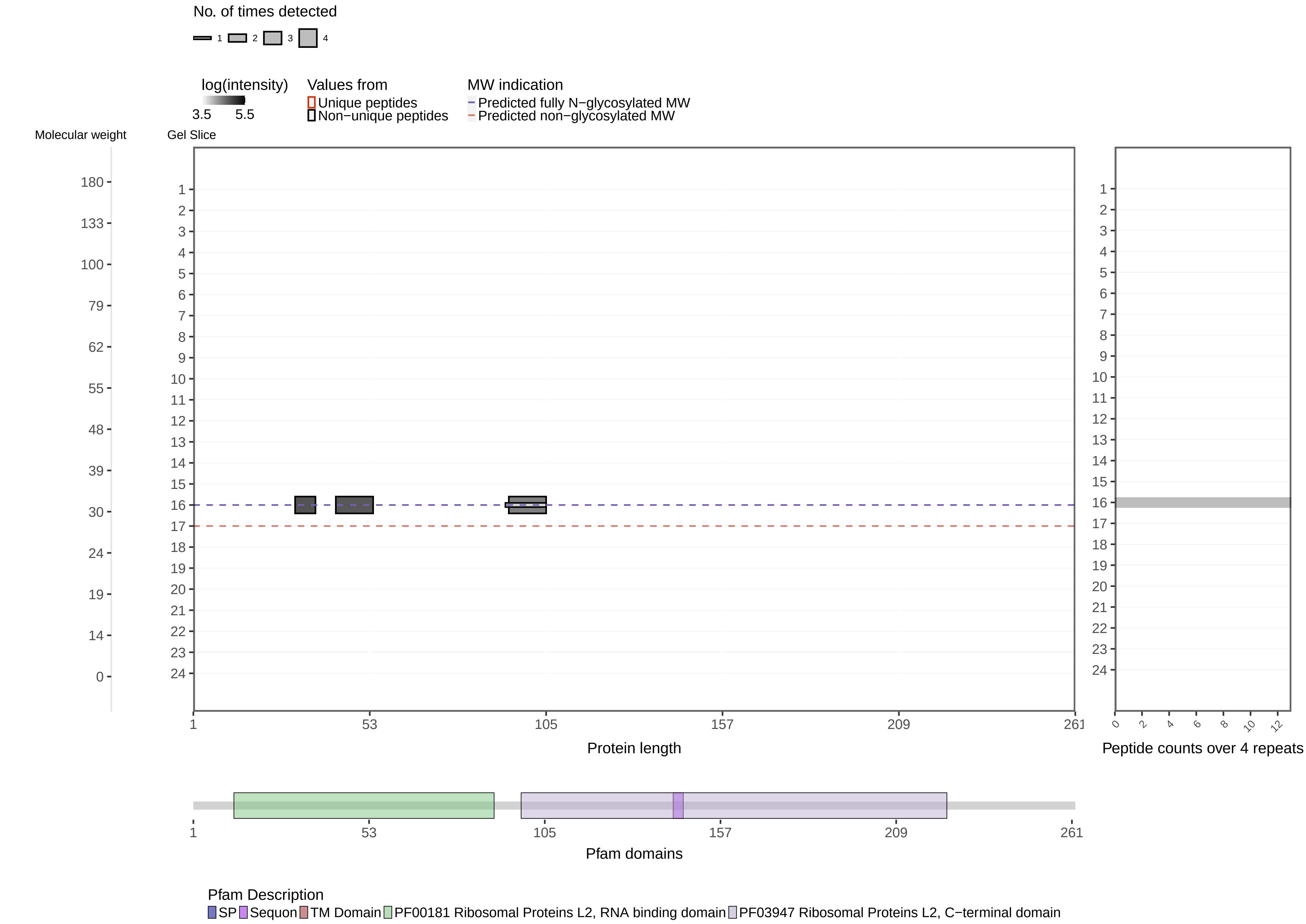


Protein sequence:

IKVGDRGVFARCSGDYAIVISHNPDNGTTRIKLPSGAKKIVPSGCRAMIGQVAGGGRTEKPMLKAGNAYHKYRVKRNCWPKVRGVAMNPVEHPHGGGNHQHIGHASTVRRDAPPGQKVGLIAARRTGRLRGQAAATAAKADKA*

### NbL11g08090.1 glycine−rich_RNA−binding_protein_3_mitochondrial−like [MW = 16.5 kDa]

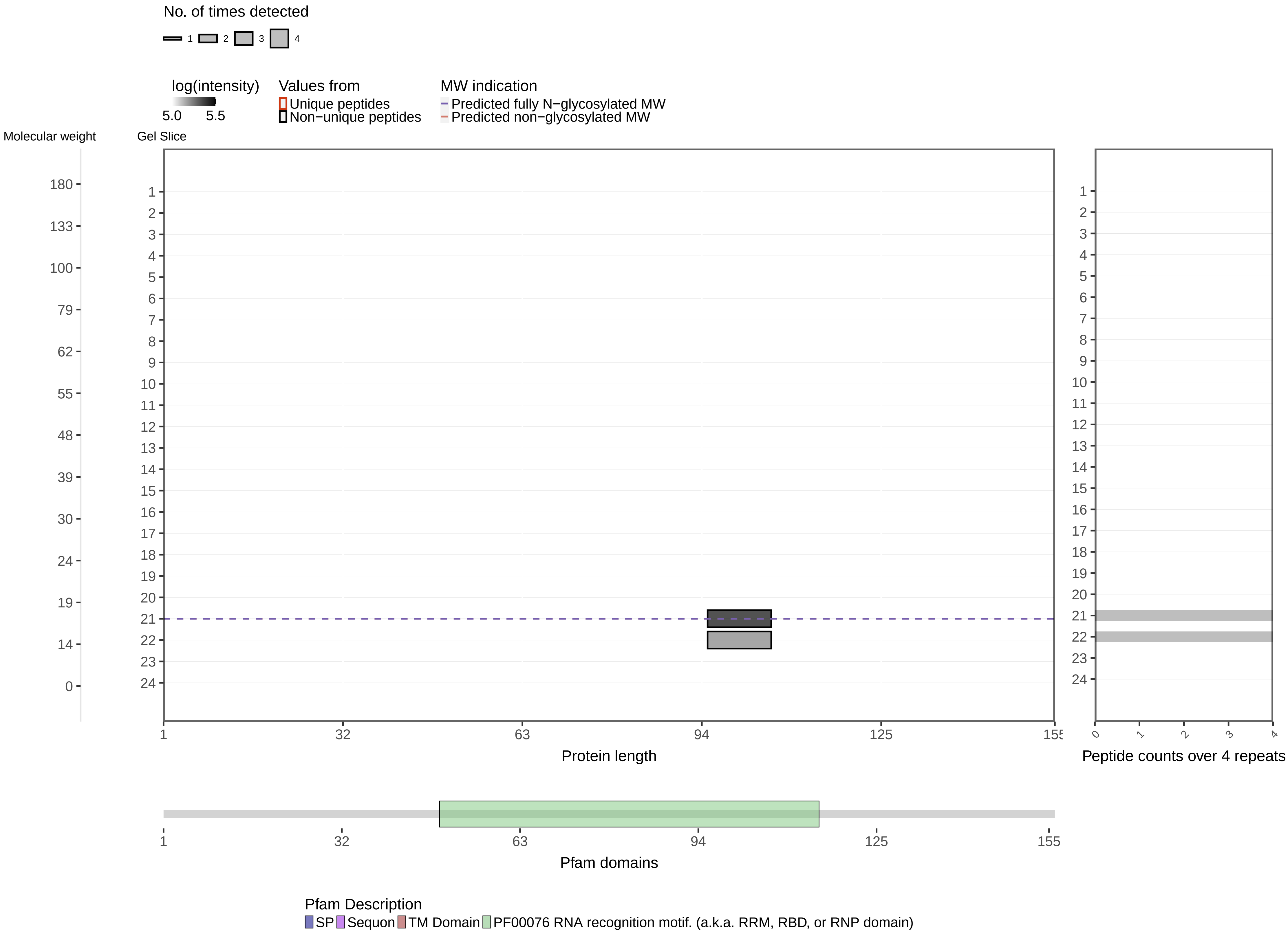


Protein sequence:

AFAKHGEVVNARVVTDRVSGYSKGFGFVRYATLEDAAAGIKGLDGQFLDGWVIFAEYARPRPPPPSLQNDNPAGGYGNYNGPSYGRNNM*

NbL11g07960.1 electron_transfer_flavoprotein_subunit_alpha_mitochondrial−like [MW = 37.18 kDa]

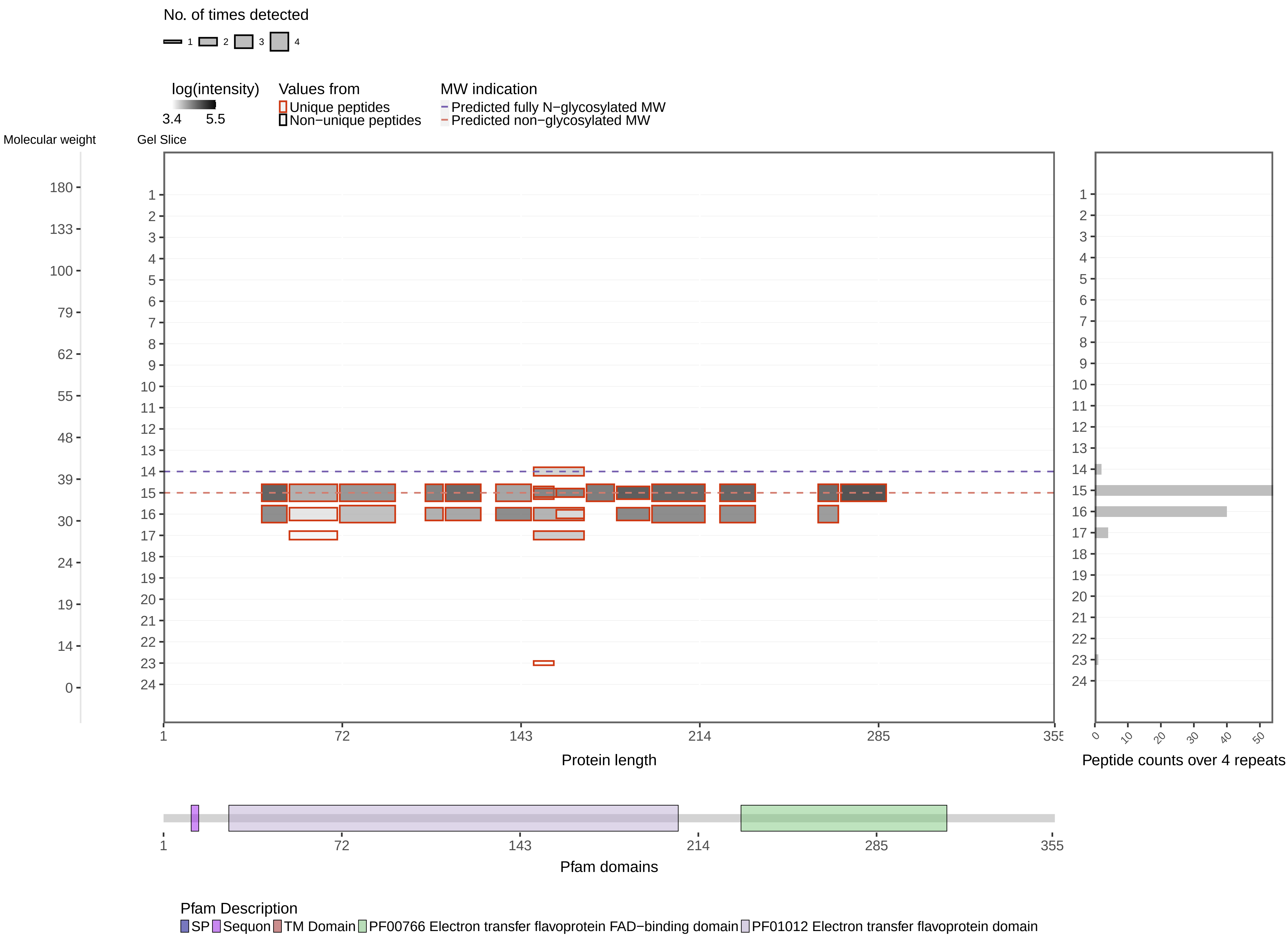


Protein sequence:

STVRYTGSSPCMLSIRATSFPIASLPADSKSDAASIDQVDLSTLDEDDSVGKSRYIKLSSQNVERPDLGNARIVVTGGRGVKSAENFKMIDKLAEKIGAAVGATRAAVDAGFVPNDLQVGQTGKIVAPELYMAFGVSGAIQHIAGMRDSKVIVAVNKDADAPIFQIADYGLVGDLFEVIPELLEKIPEKK*

### NbL11g07940.1 chloroplastic_lipocalin−like [MW = 37.26 kDa]

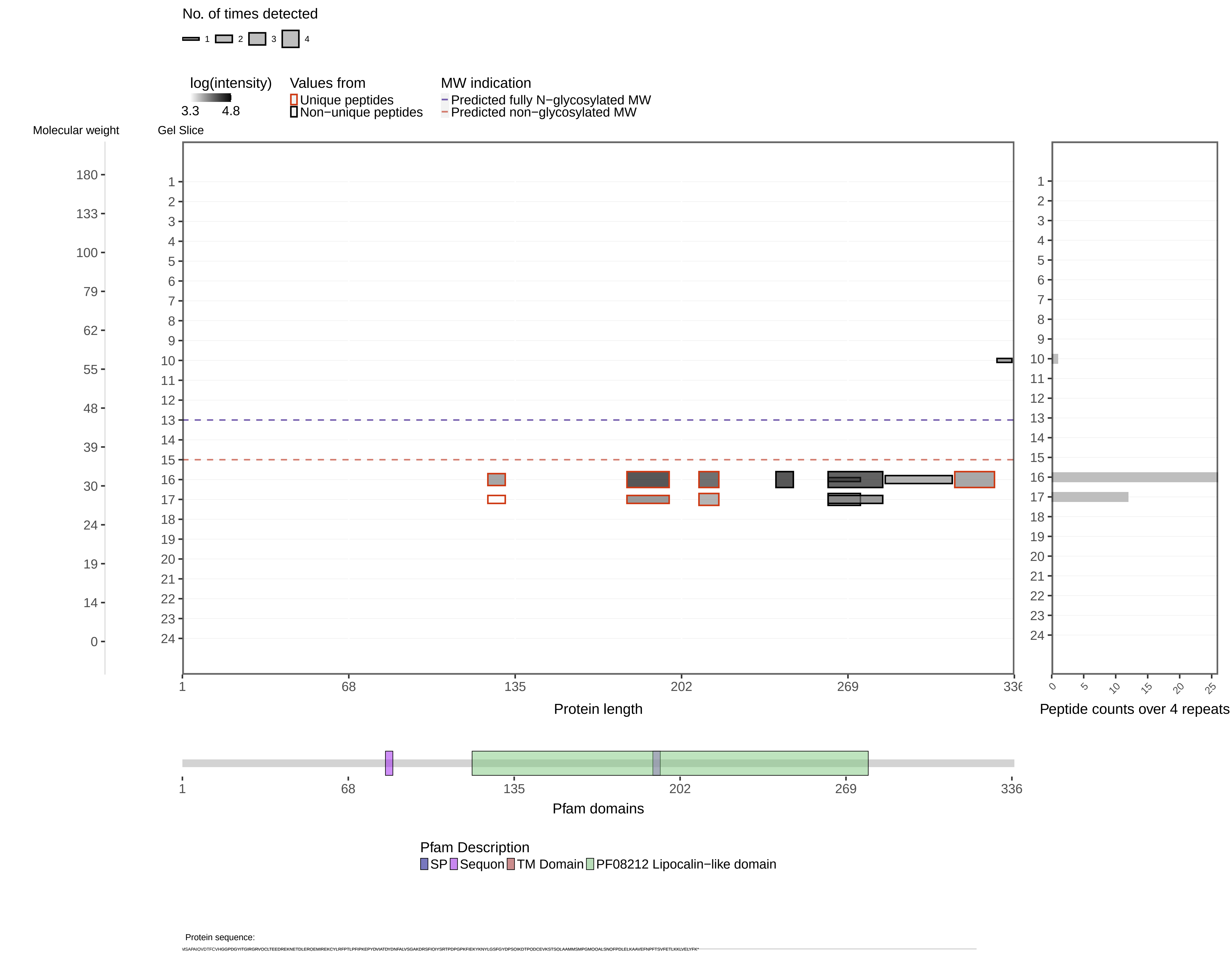

NbL11g07780.1
dihydrolipoyllysine−residue_succinyltransferase_component_of_2−oxoglutarate_dehydrogenase_complex_1_mitochondrial−like
[MW = 51.42 kDa]

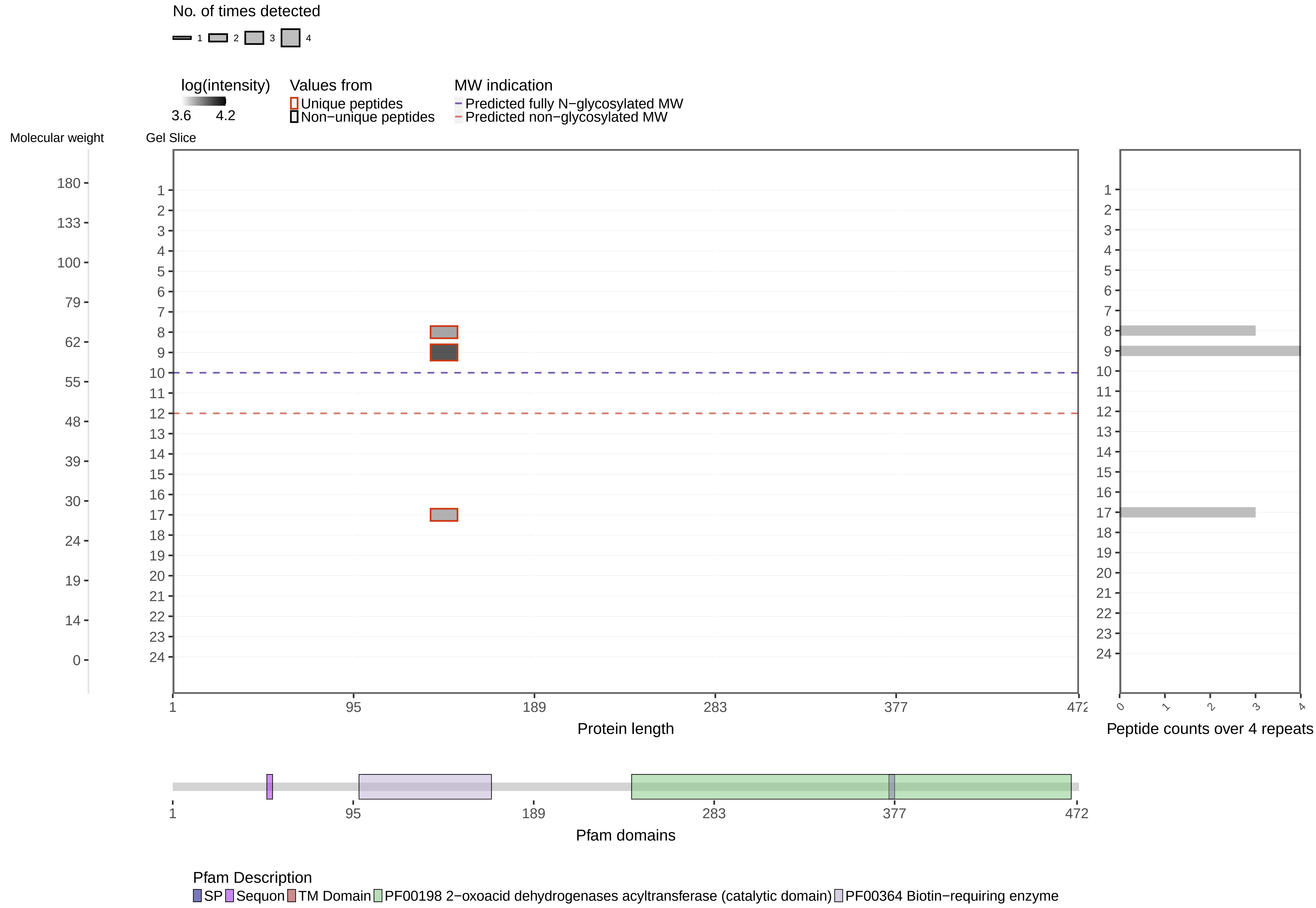


Protein sequence:

TSPPPPKRSATEPQLPPKERERRVPMTRLRKRVATRLKDSQNTFALLTTFNEVDMTNLMKLRSDYKDAFVEKHGVKLGLMSGFVKAAVSALQNQPIVNAVIDGDDIIYRDYVDISIAVGTPKGLVVPVLRNAEKMNFAEIEKTINGLAKKATNGTISIDEMAGGSFTISNGGVYGSLLSTPIINPPQSAILGMHSIVNRPMVVGGDVVSRPMMYIALTYDHRLIDGREAVYFLRRIKDVVEDPRRLLLDV*

NbL11g07710.1 uncharacterized_LOC109235633 [MW = 45.54 kDa]

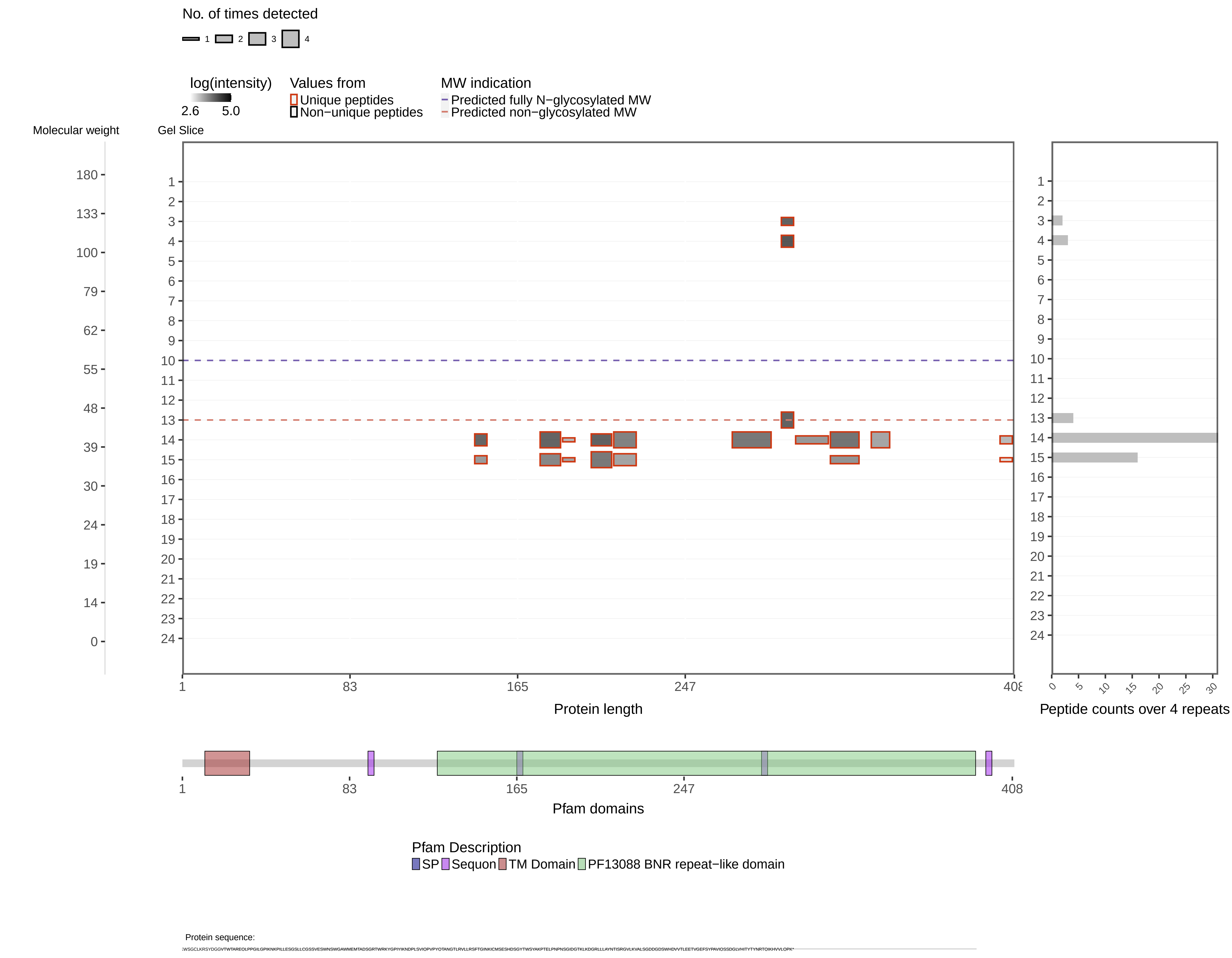

### NbL11g07590.1 glycerophosphodiester_phosphodiesterase_GDPDL3−like [MW − SP = 80.1 kDa]

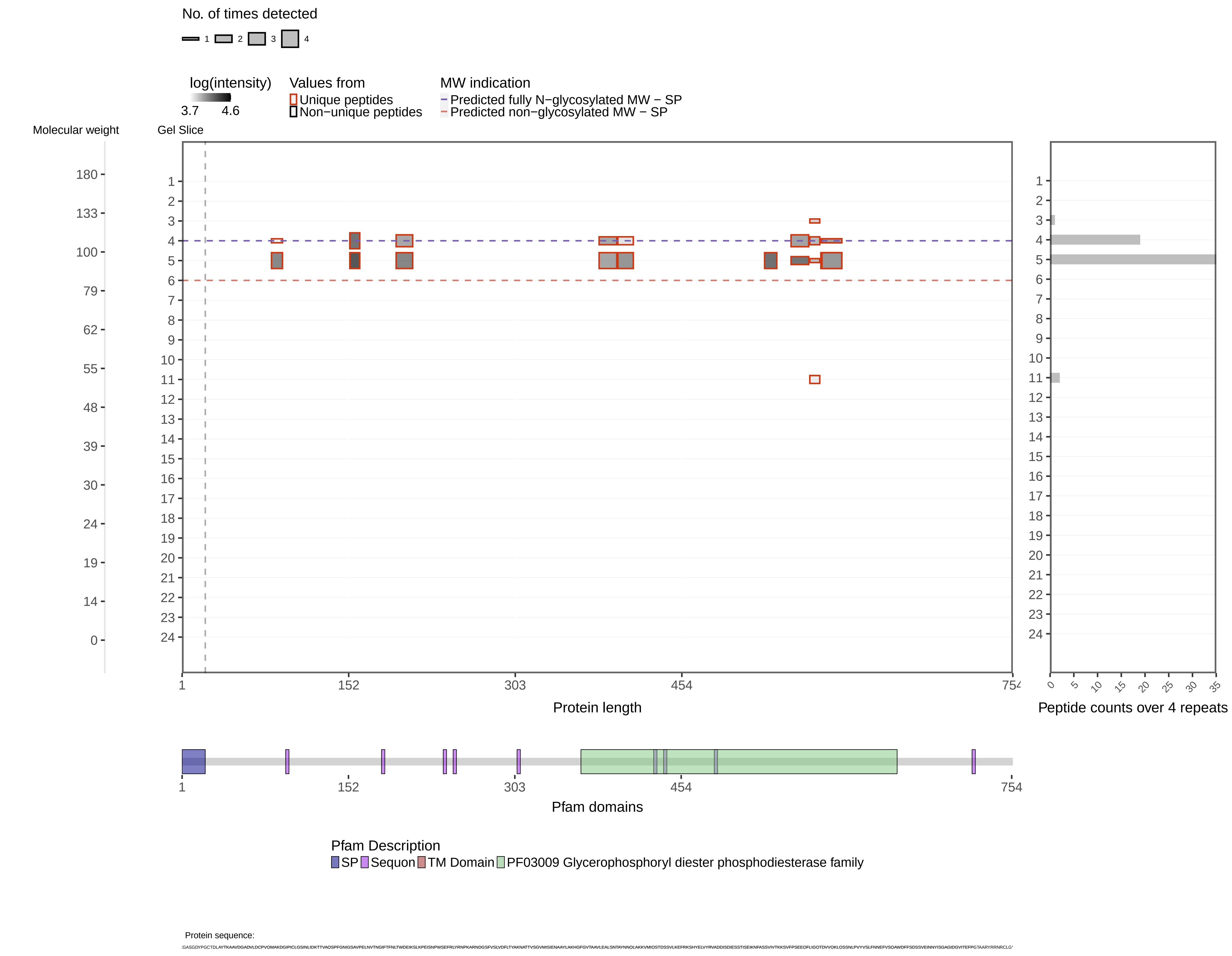

NbL11g07310.1 farnesyl_pyrophosphate_synthase_1−like [MW = 39.45 kDa]

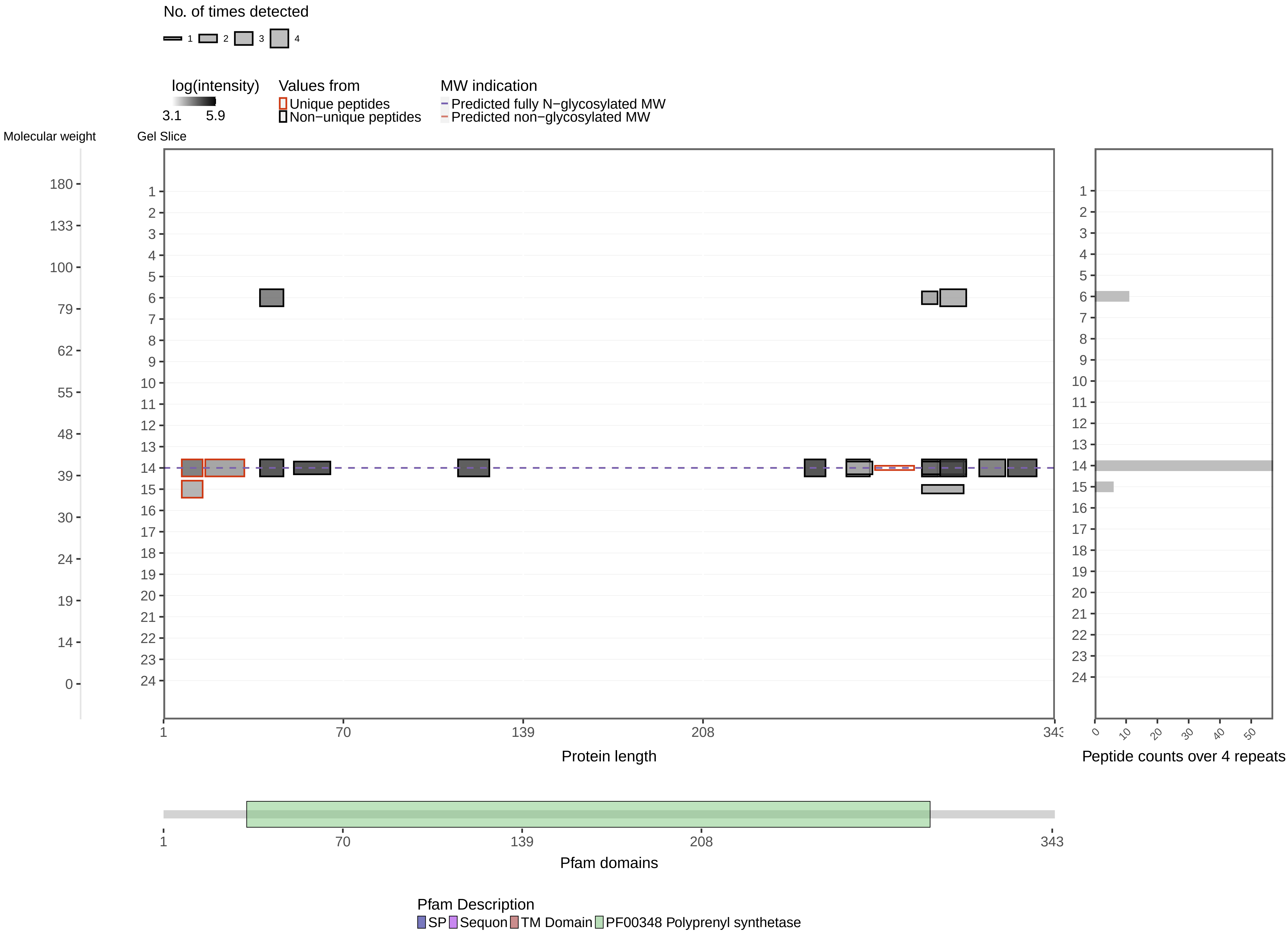


Protein sequence:

)TASGQMIDLITTLVGEKDLSKYSLPIHRRIVQYKTAYYSFYLPVACALLMAGDNLDKHVDVKNILVEMGIYFQVQDDYLDCFADPEVLGKIGTDIQDFKCSWLVVKALELCNEEQKKLLYENYGKDDSECIAKIKALYNDLKLEEVFLEYEKKSYEKLTSSIAAHPSKAVQAVLHSFLGKIYKRQK*

### NbL11g07290.1 heat_shock_cognate_protein_80−like [MW = 80.27 kDa]

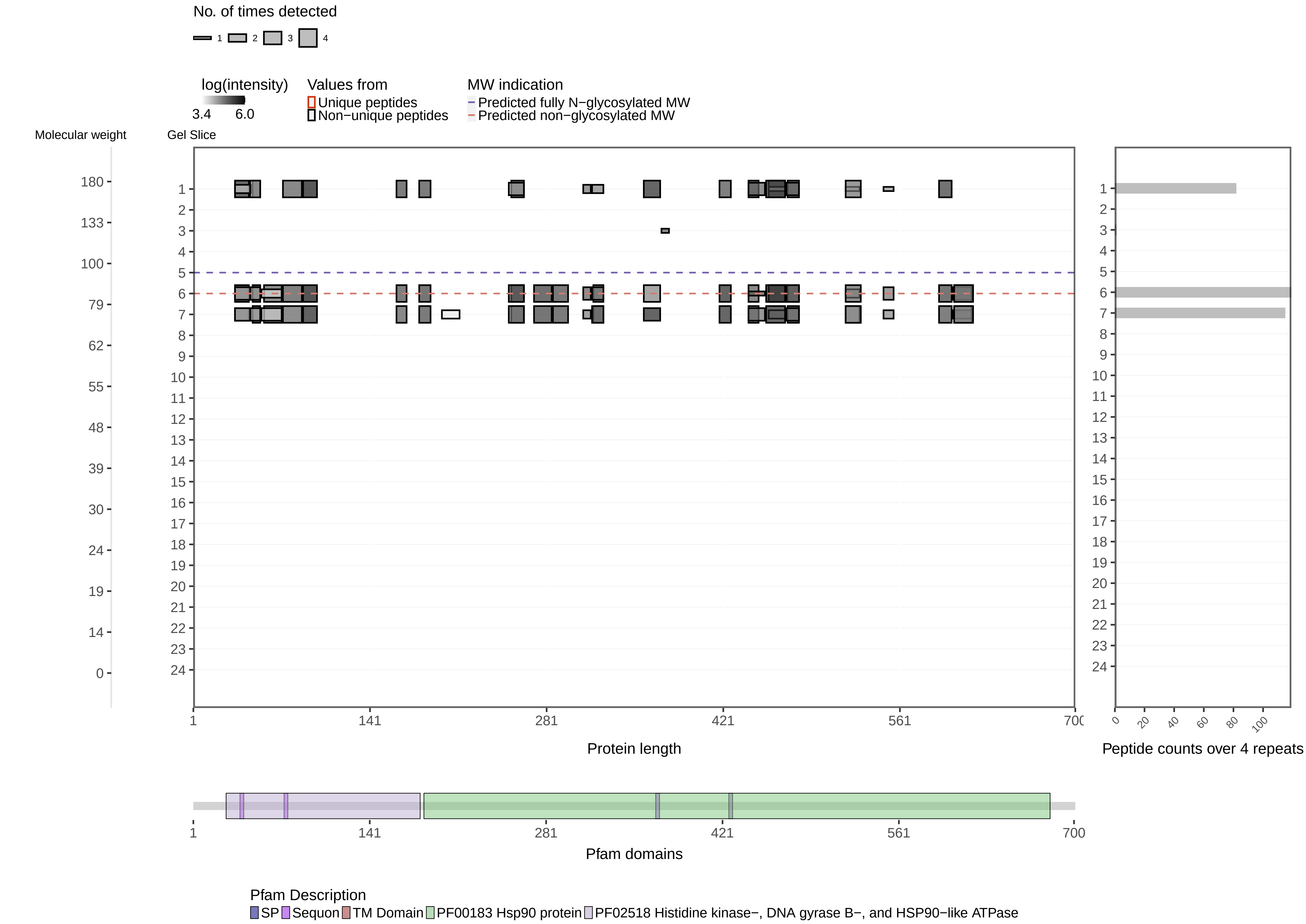


Protein sequence:

[illegible]

### NbL11g07240.1 GPI−anchored_protein_LORELEI−like [MW − SP = 16 kDa]

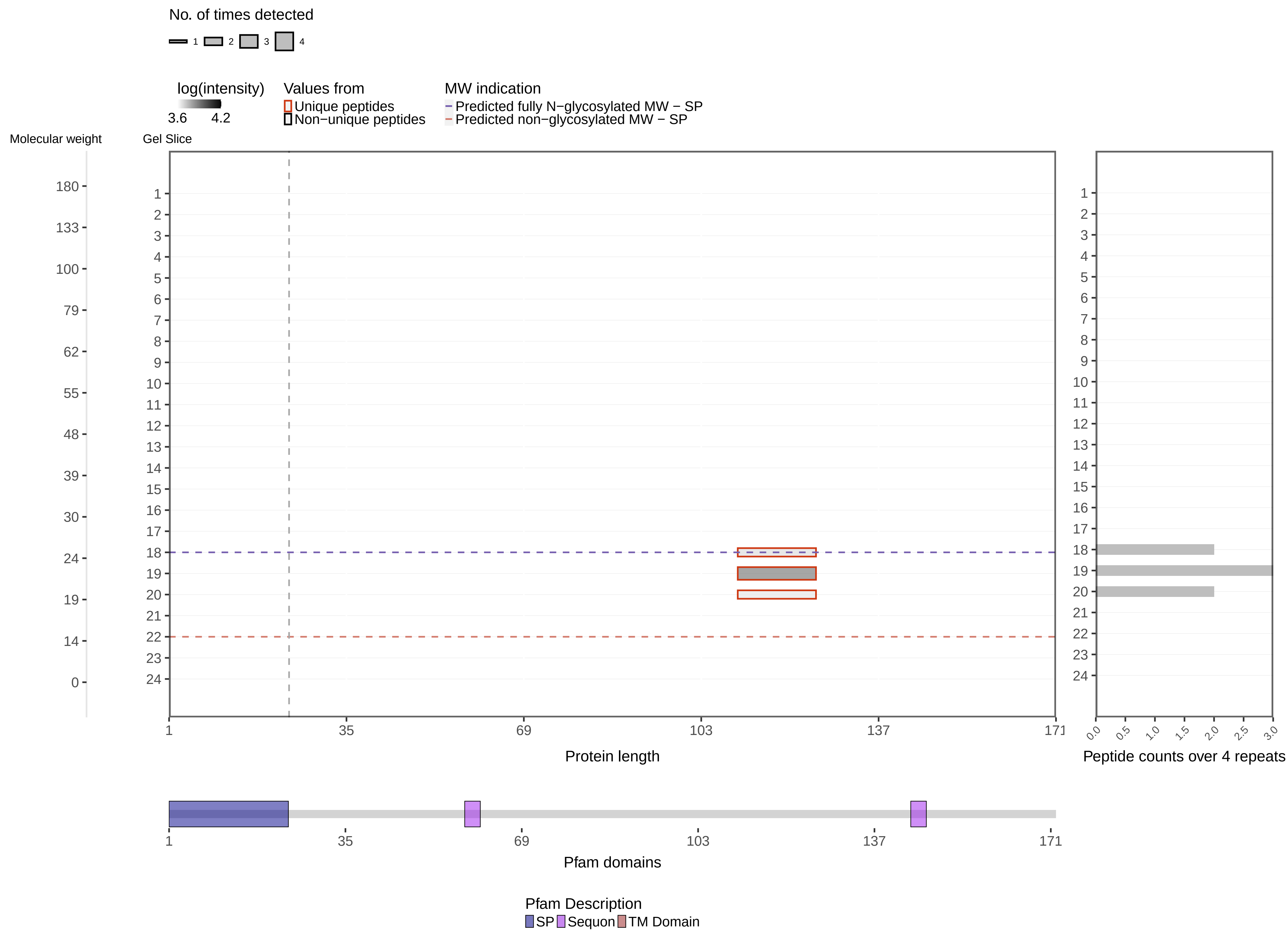


Protein sequence:

KQCCDAFVQFACPFADLLNDLSNDCSTTMFSYINLRGKYPPGLFGSMCTGTKEGLECPILAPSQSANDDANSSQNNCKLSPVQIMATAFMWMLFLVF*

### NbL11g07160.1 porphobilinogen_deaminase_chloroplastic−like [MW = 39.94 kDa]

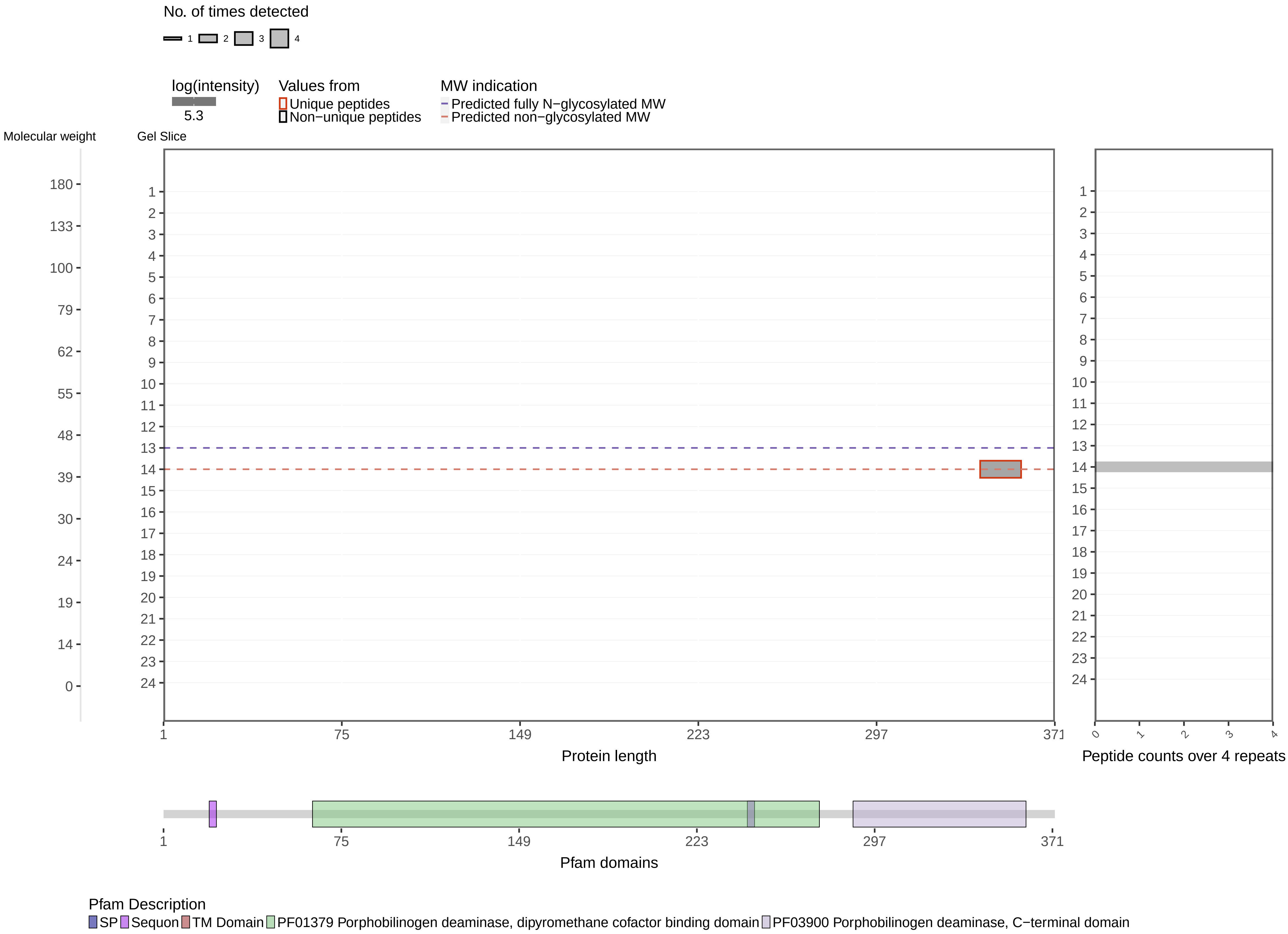


Protein sequence:

### NbL11g07140.1 thiosulfate/3−mercaptopyruvate_sulfurtransferase_1_mitochondrial−like [MW = 41.78 kDa]

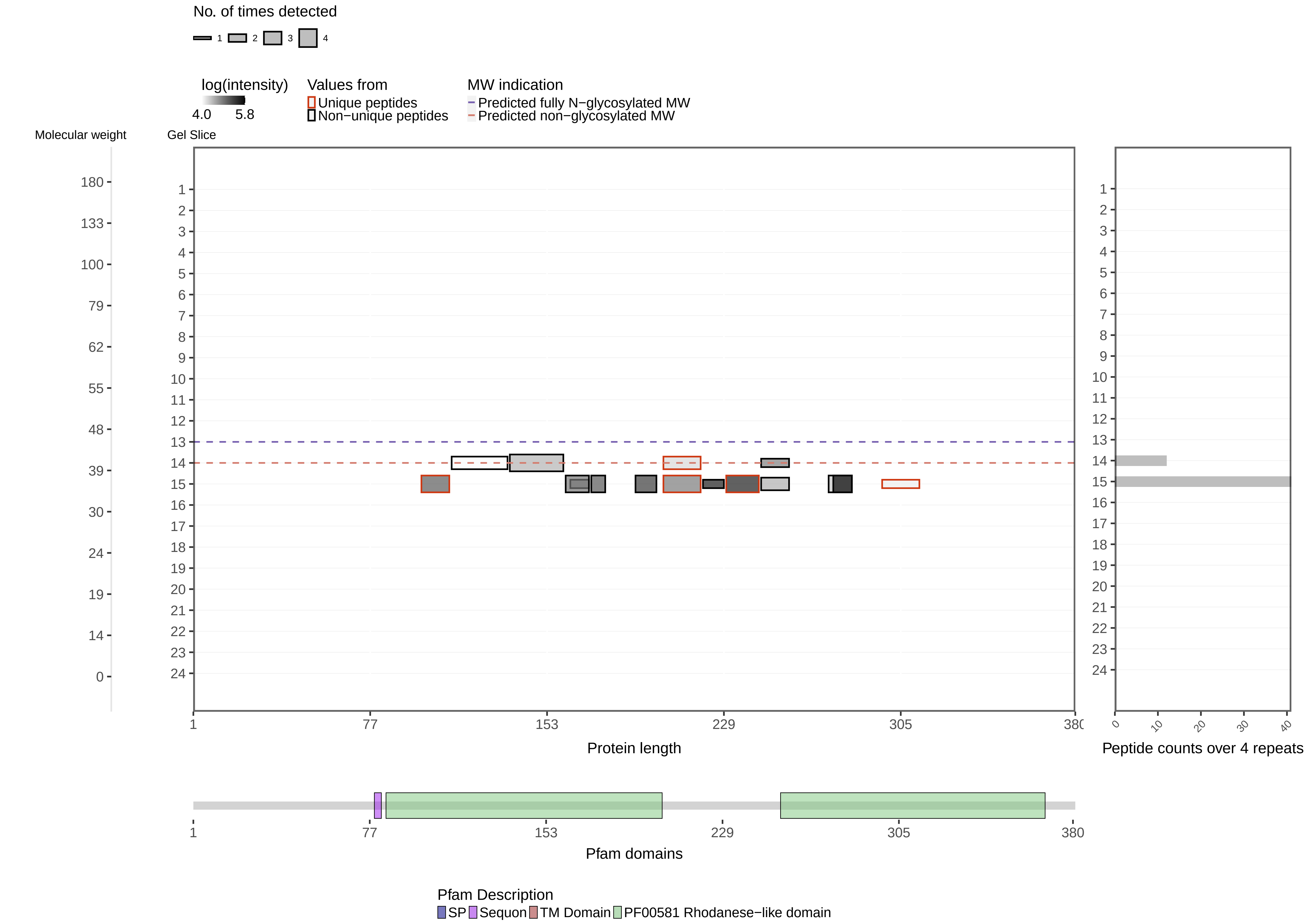


Protein sequence:

VWWMFRAFGHDKVWVLDGGLPRWRASEYDVESSASGDAILKASAASEAIEKVYQGQMVGPTTFETKFQPHLVWNLDQVKRNIEDKTYQHIDARSKARFDGVAPEPRKGIRSGHVPGSKCIPFPQMLDDSQTLLSREELKKKFDLEGISLDKPVGTSCGTGVTACILALGLHRLGKTDIPVYDGSWTEWGAHPDTPVSTSET*

### NbL11g07130.1 phosphoglycerate_kinase_cytosolic−like [MW = 42.49 kDa]

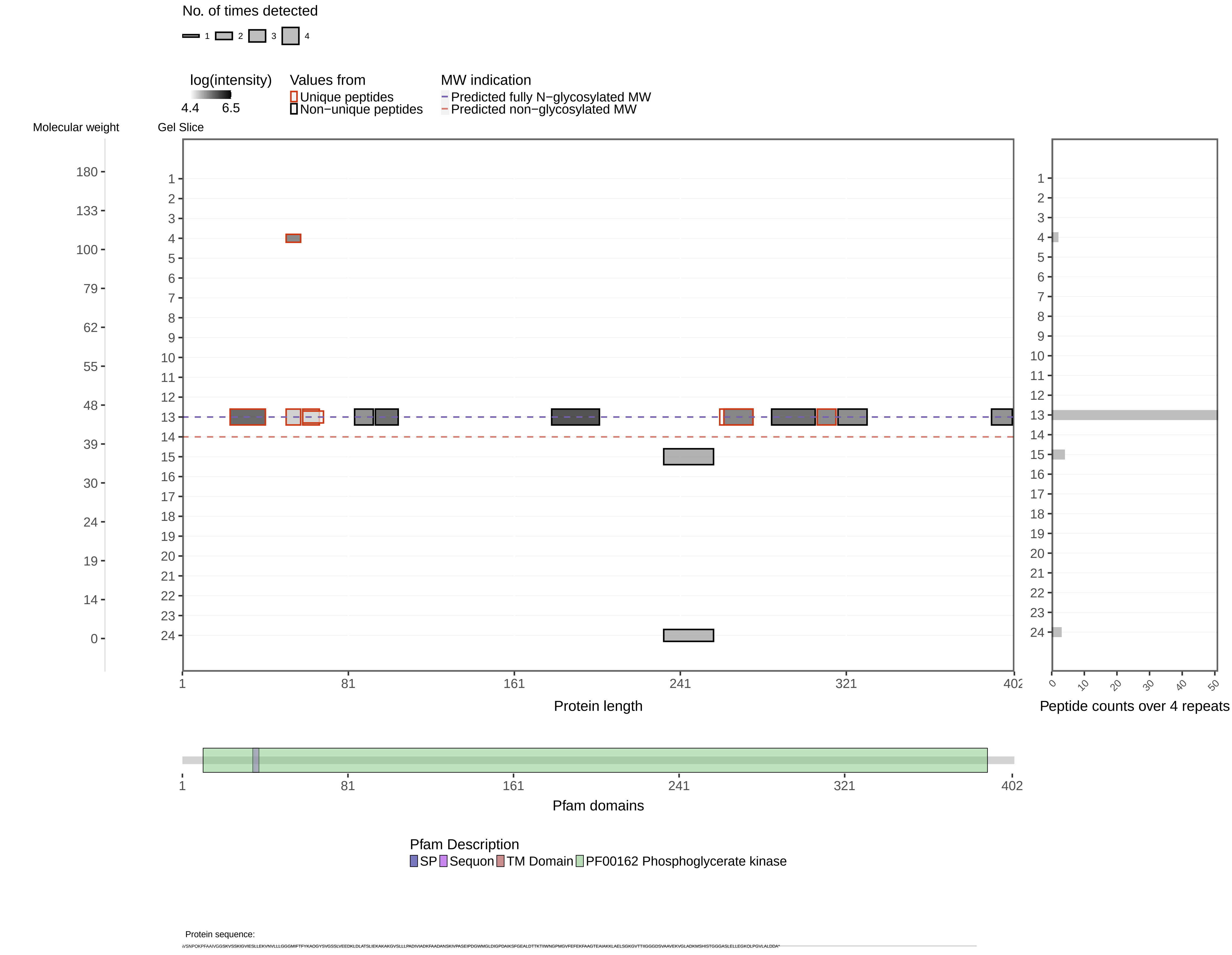

NbL11g06990.1 40S_ribosomal_protein_S8−like [MW = 25.03 kDa]

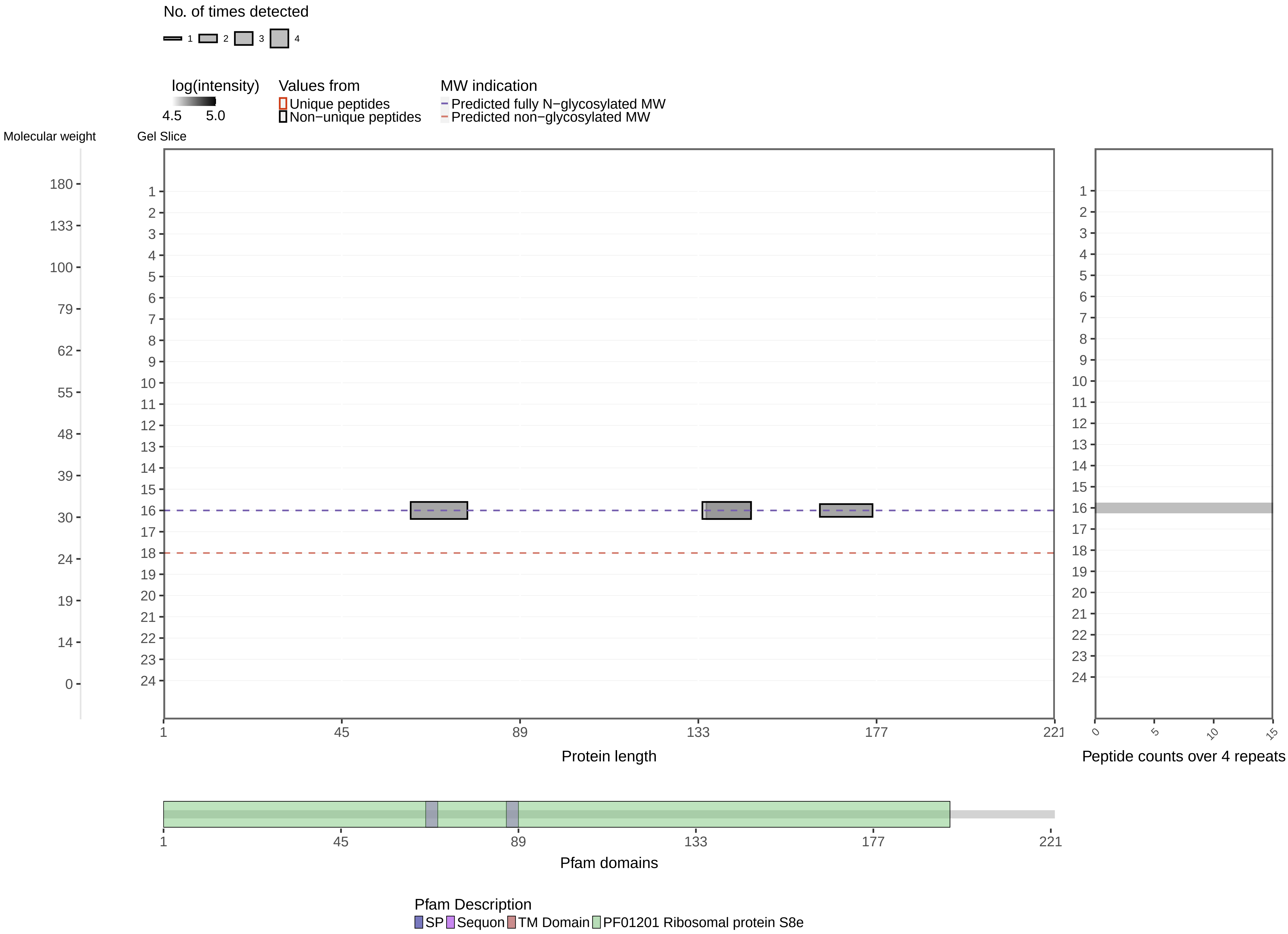


Protein sequence:
TLVKSAIVQVDAAPFKQWYLQHYGVDIGRKKKTAASKKEGEEAEAVPEEKKSNHVQRKLQKRQQDRKIDPHIEEQFASGRLLAAISSRPGQCGRADGYILEGKELEFYMKKLQRKKGKGASGAT*

NbL11g06860.1 guanosine_nucleotide_diphosphate_dissociation_inhibitor_2 [MW = 49.85 kDa]

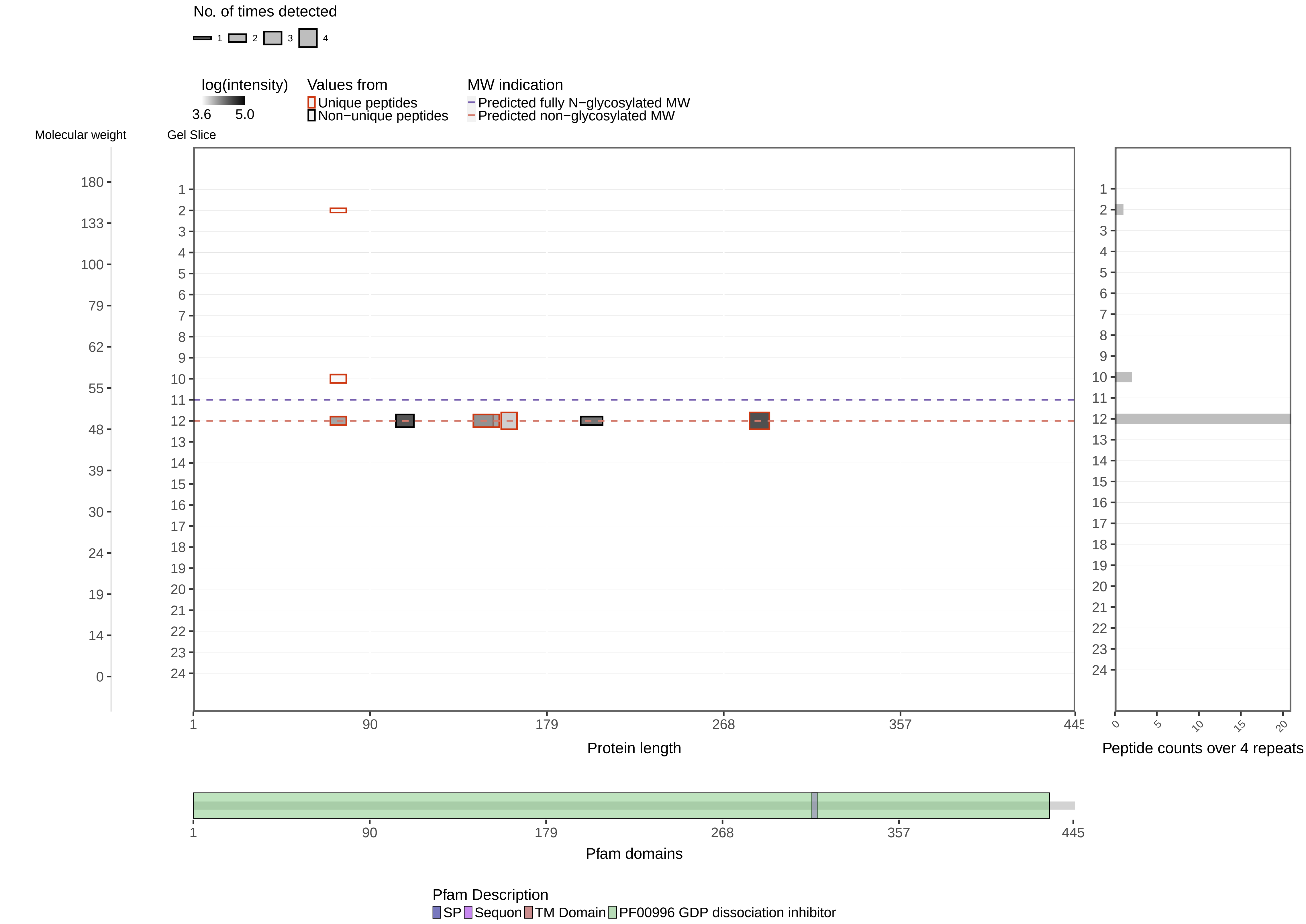


Protein sequence:

KLYAESLARFQGGSPYIYPLYGLGELPQAFARLSAVYGGTYMLNKPECKVEFDEEGKVCGVTSEGETAKCKKVVCDPSYLTNKVRKVGKVARAIAIMSHPIPNTNDSHSVQIILPQKQLGRKSDMYLFCCSYTHNVAPKGKFIAFVSTEAETDNPESELKPGIDLLGQVDEIFFEAYDRFEPVNEPSLDNCFISTSYDPTTHFESTVDDVLNMYTLITGKVLDLNVDLSAASAAEE*

### NbL11g06830.1 brefeldin_A−inhibited_guanine_nucleotide−exchange_protein_5 [MW = 196.42 kDa]

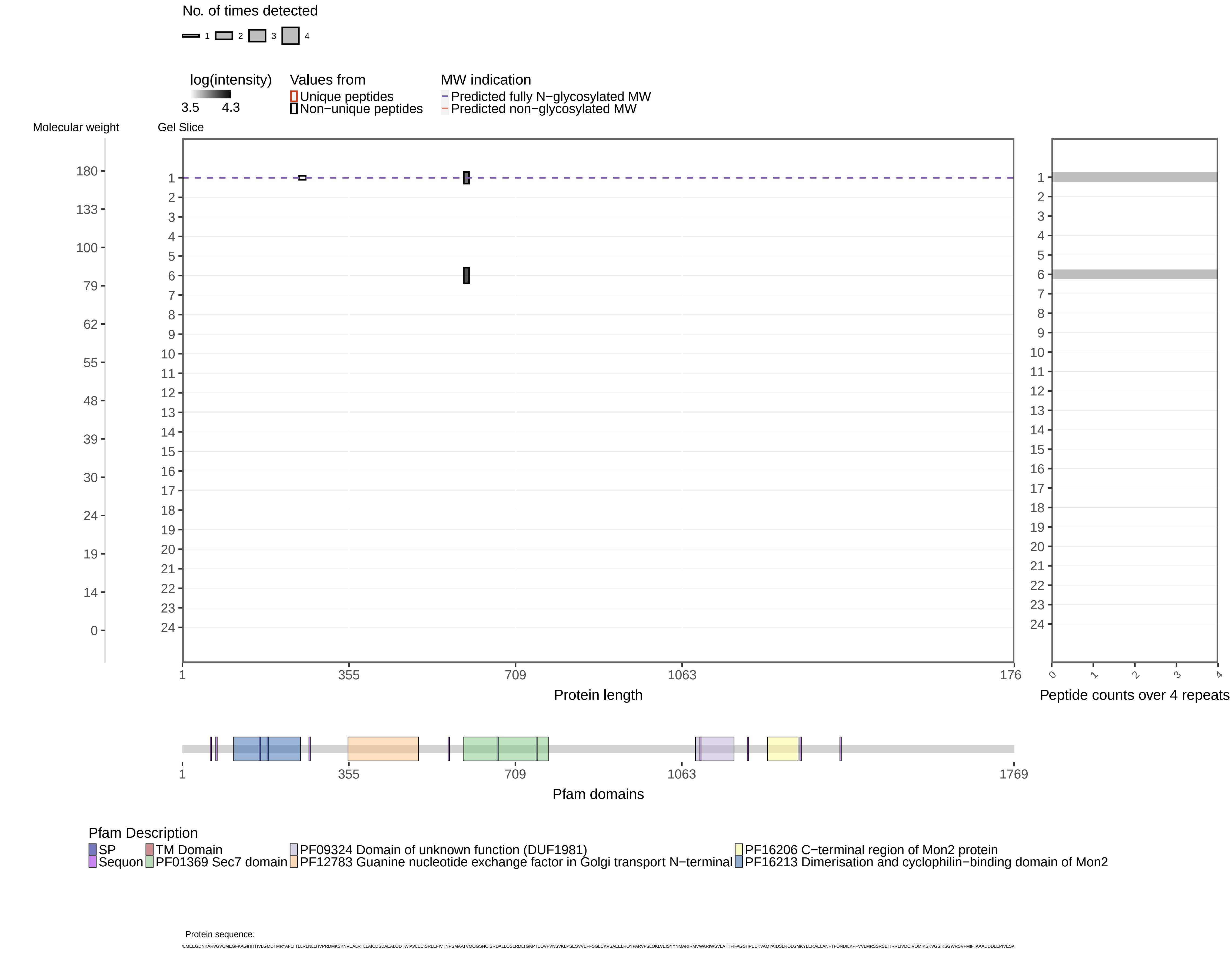

### NbL11g06800.1 DExH−box_ATP−dependent_RNA_helicase_DExH10−like [MW = 111.5 kDa]

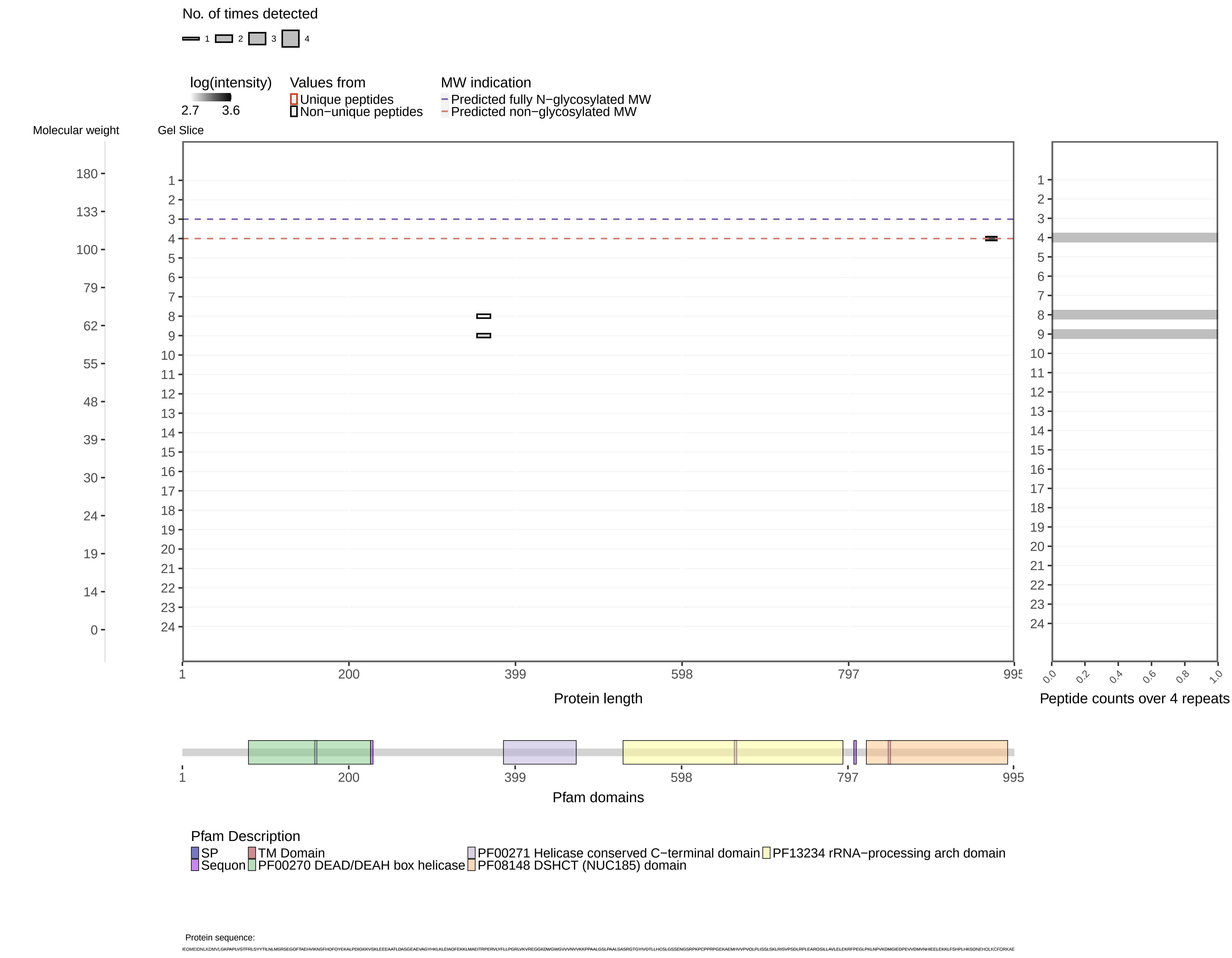

NbL11g06740.1 probable_uridine_nucleosidase_2 [MW = 34.29 kDa]

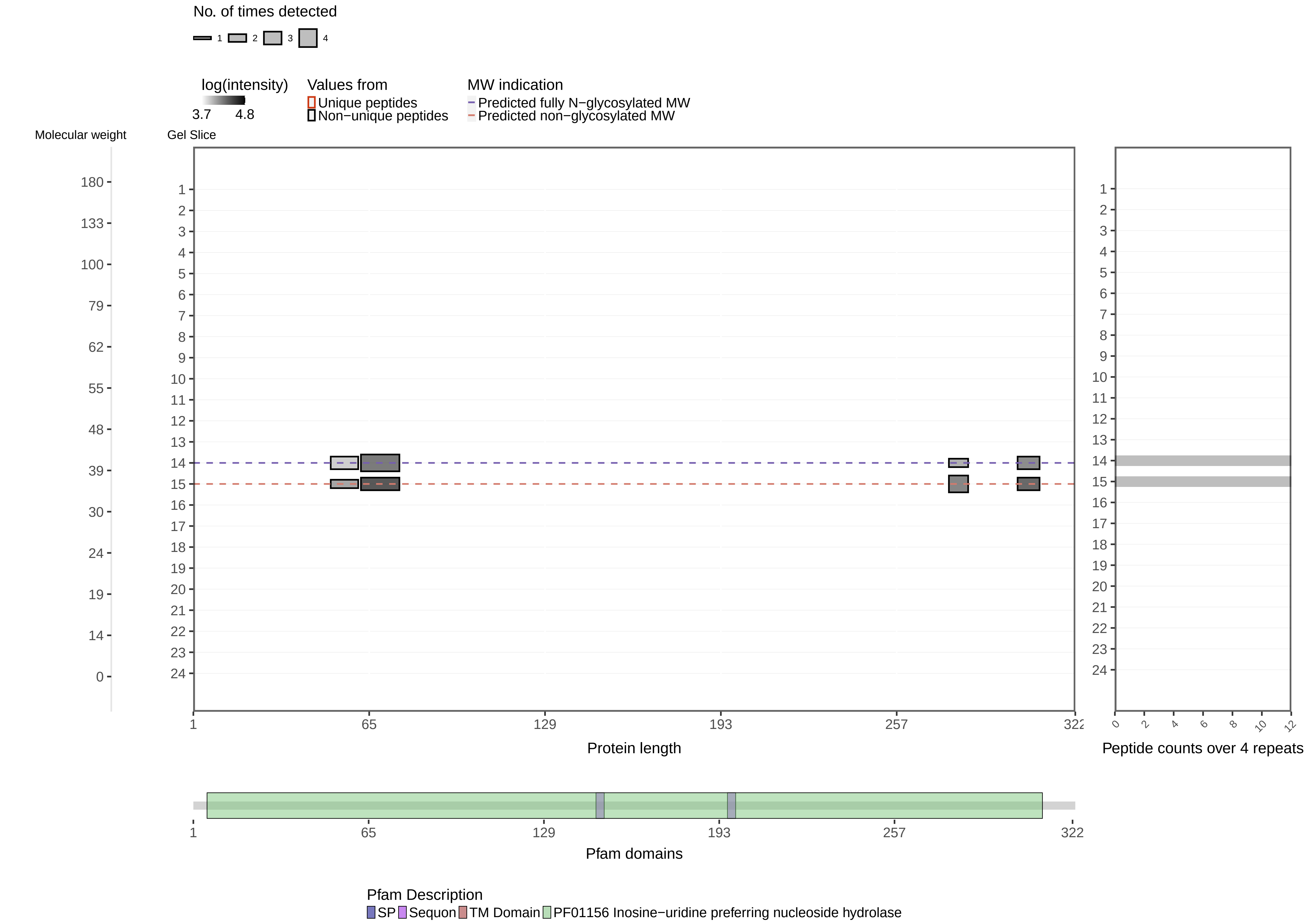


Protein sequence:
SQIVVLGGSFAVNGNVNPAAEANIFGDPDAADIVFTSGADVLAVGINVTHQVVLTDSDRNELAKSNGKFAKYLDKILGVYFGYHHDAYSTKGVYLHDPTALLAAVDPSLITYTEGAVRVQTIGITRGLTLFYNKQKRFGEVTEWCDKPSVKVAVTVDAPKVLKLVMERLINS*

NbL11g06640.1 valine−−tRNA_ligase_mitochondrial [MW = 109.55 kDa]

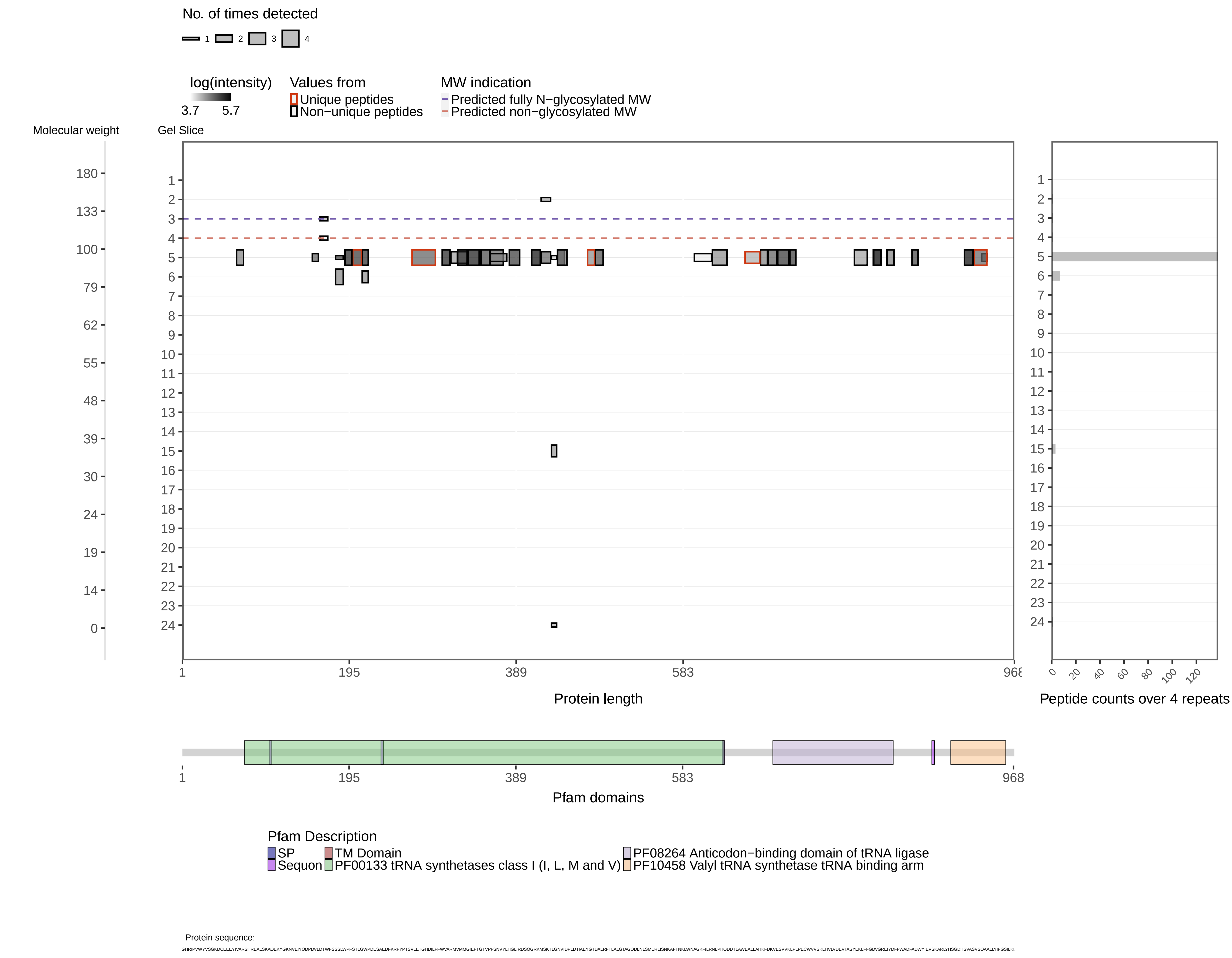

NbL11g06550.1 acetolactate_synthase_small_subunit_1_chloroplastic−like [MW = 53.34 kDa]

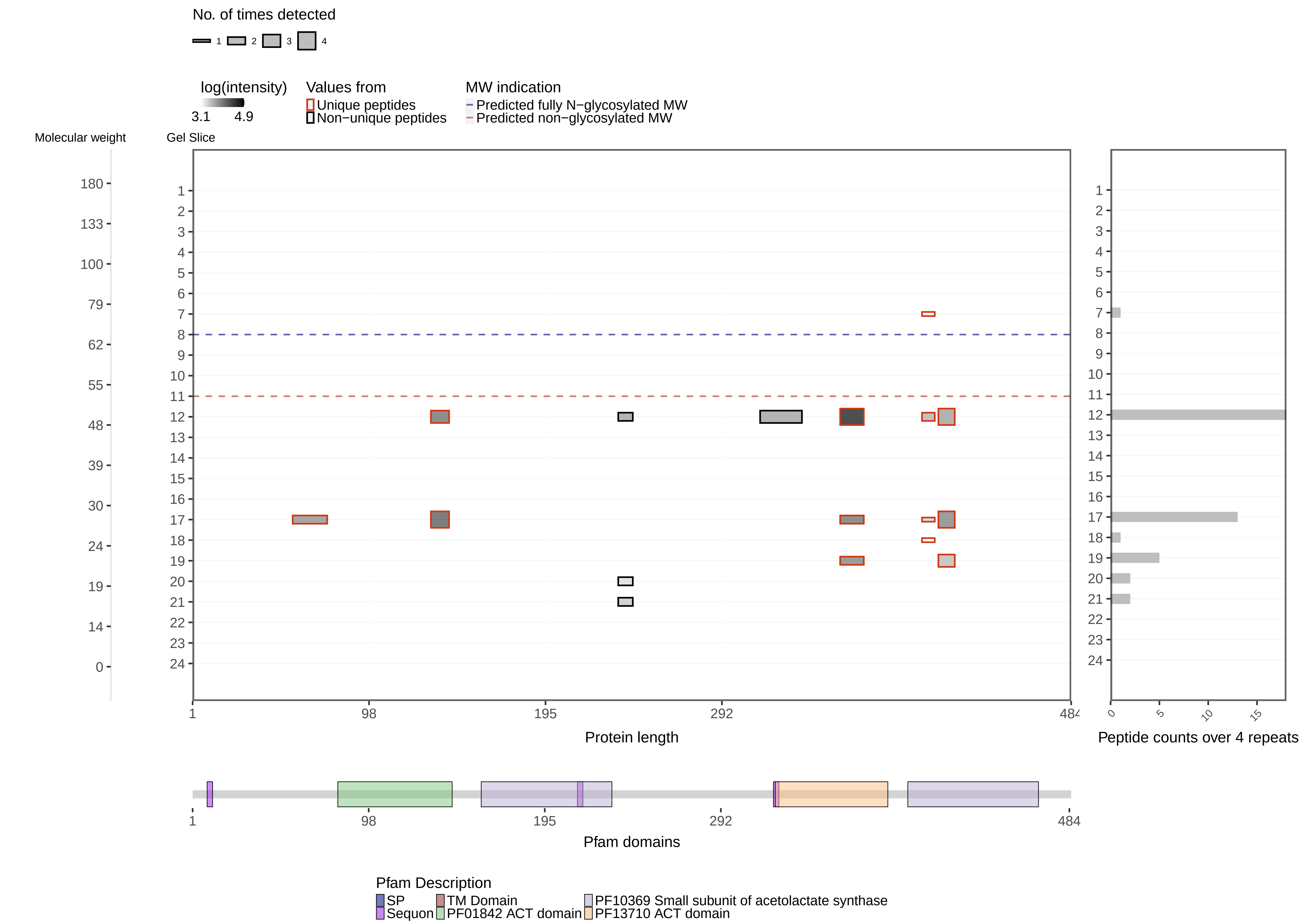


Protein sequence:

NbL11g06520.1 2−alkenal_reductase_(NADP(+)−dependent) [MW = 38.07 kDa]

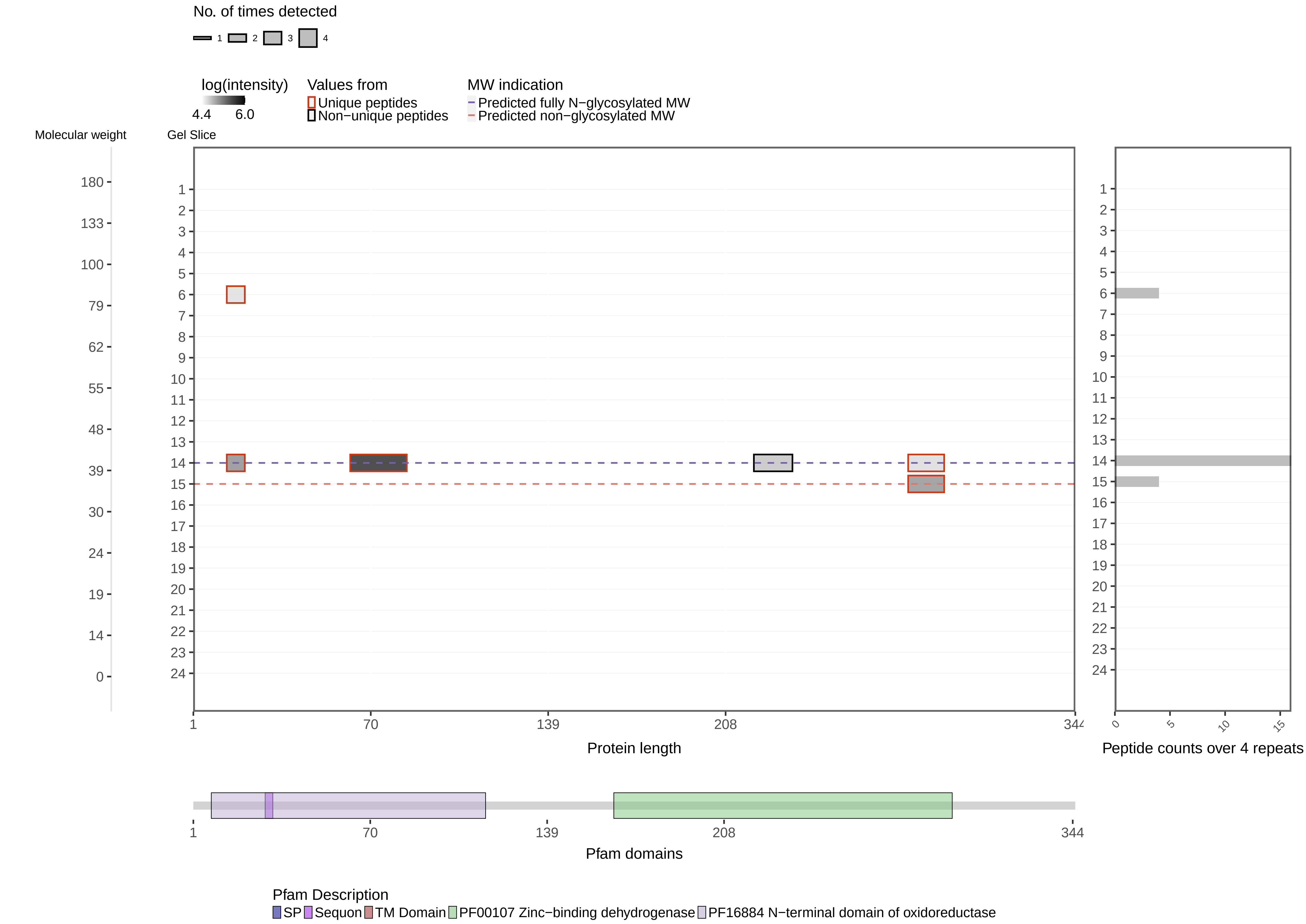


Protein sequence:

VSAASGAVGQLVGQFAKMLGCYVVGSAGSKEKVDLLKTKFGFDEAFNYKEEQDLNAALKRYFPDGIDIYFENVGGKMLDAVLLNMKLHGRIAVCGMISQYNLEQTEGVHNLFCLITKRIRMEGFLVFDYYHLYEKYLEMVIPQIKAGKVVYVEDVVDGLESAPSALVGLFSGRNIGKQVVIVSRE*

NbL11g06330.1 U5_small_nuclear_ribonucleoprotein_40_kDa_protein [MW = 41.32 kDa]

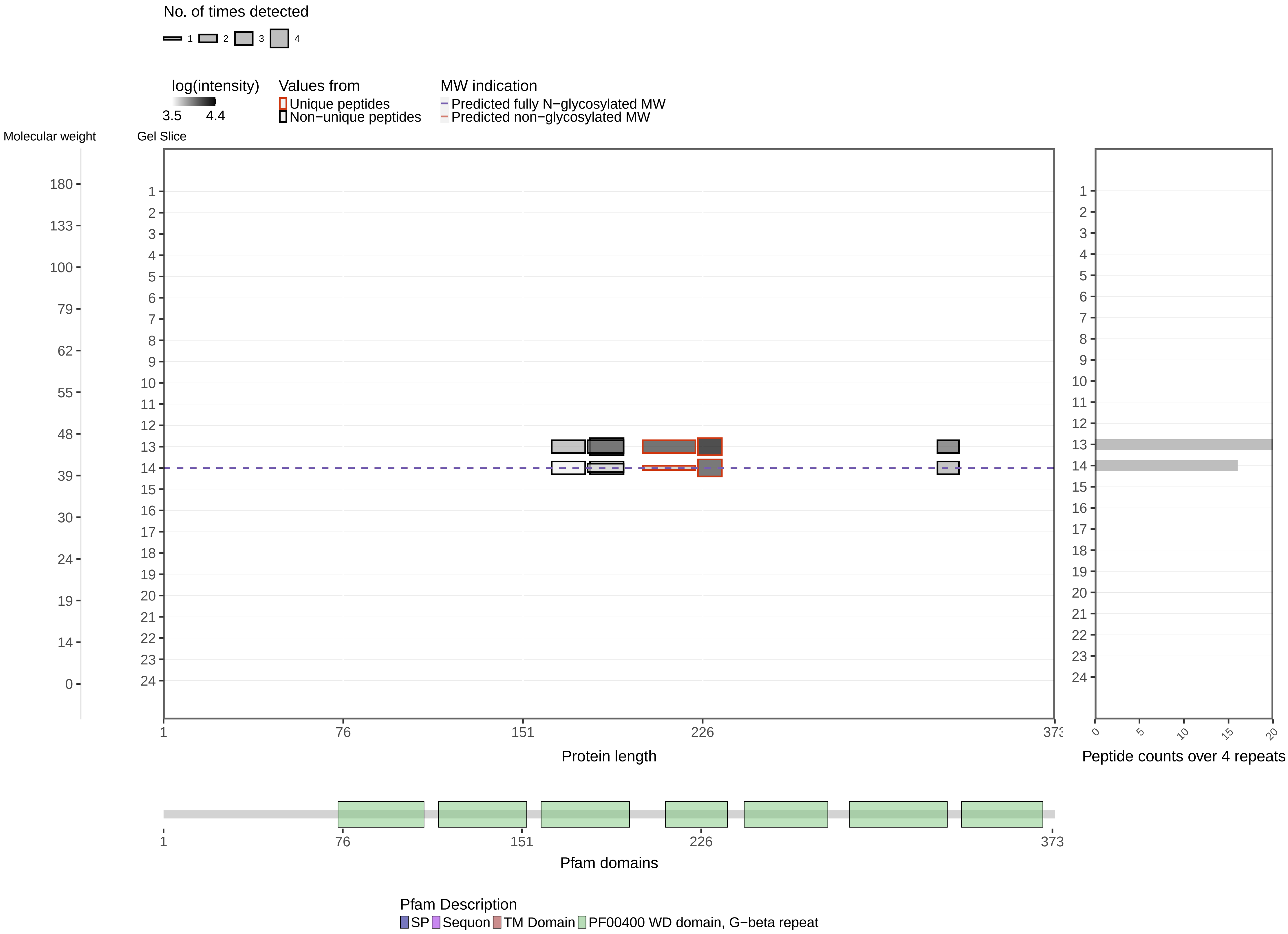


Protein sequence:

CPARRGPPLIVSGSDDGTAKLWDMRQRGAIQTFPDKFQITAVSFSDASDKIYSGGIDNDVKVWDLRRCEVTMTLQGHQDTITGMQLSPDGSYLLTNAMDCTLRIWDMRPYAPQNRCVKILEGHQHNFEKNLLKCSWSPDGSKVTAGSSDRMVYIWDTTSRRILYKLPGHTGSVNECVFHPNEPIVGSCSSDKQIYLGEI*

### NbL11g06190.1 VHS_domain−containing_protein_At3g16270 [MW = 76.05 kDa]

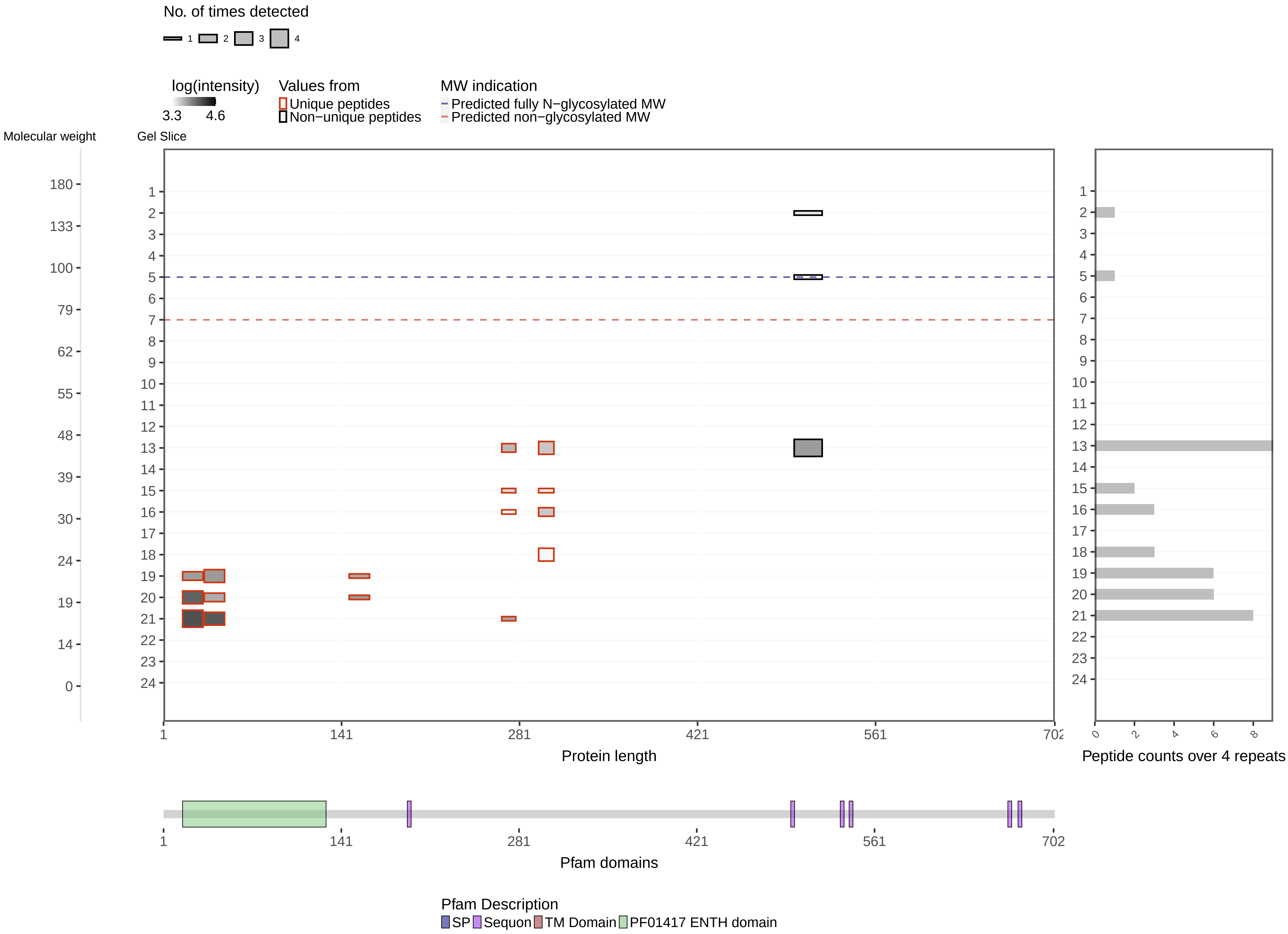


Protein sequence:

YFNENKDVVVKCSESPQASLREKANKVLSLLNDGQTADSVSHVDRSARASTPAVQMPDLIDTGNSDDLFGADDLANMQSGEGINIASTSAAPLVDDLFGDNLGGGVTSGQQKNDDDPFADVSFHTSNEKALEADLFSGMIFDKSDAAEVHSAVNRTGPELFDIFGPSVEVPQEPNNPRKEIHDLMNSFSLNGNDSSKKQNGSSRGTYQDMCQESTIDPHHQASNDALNSIFSSQAGGVNSNPMFPLGAMQYNLPPGFVLNPSFAPQALNYNAMGNMFAQQQLFATLS

### NbL11g05910.1 probable_pectinesterase/pectinesterase_inhibitor_34 [MW = 66.29 kDa]

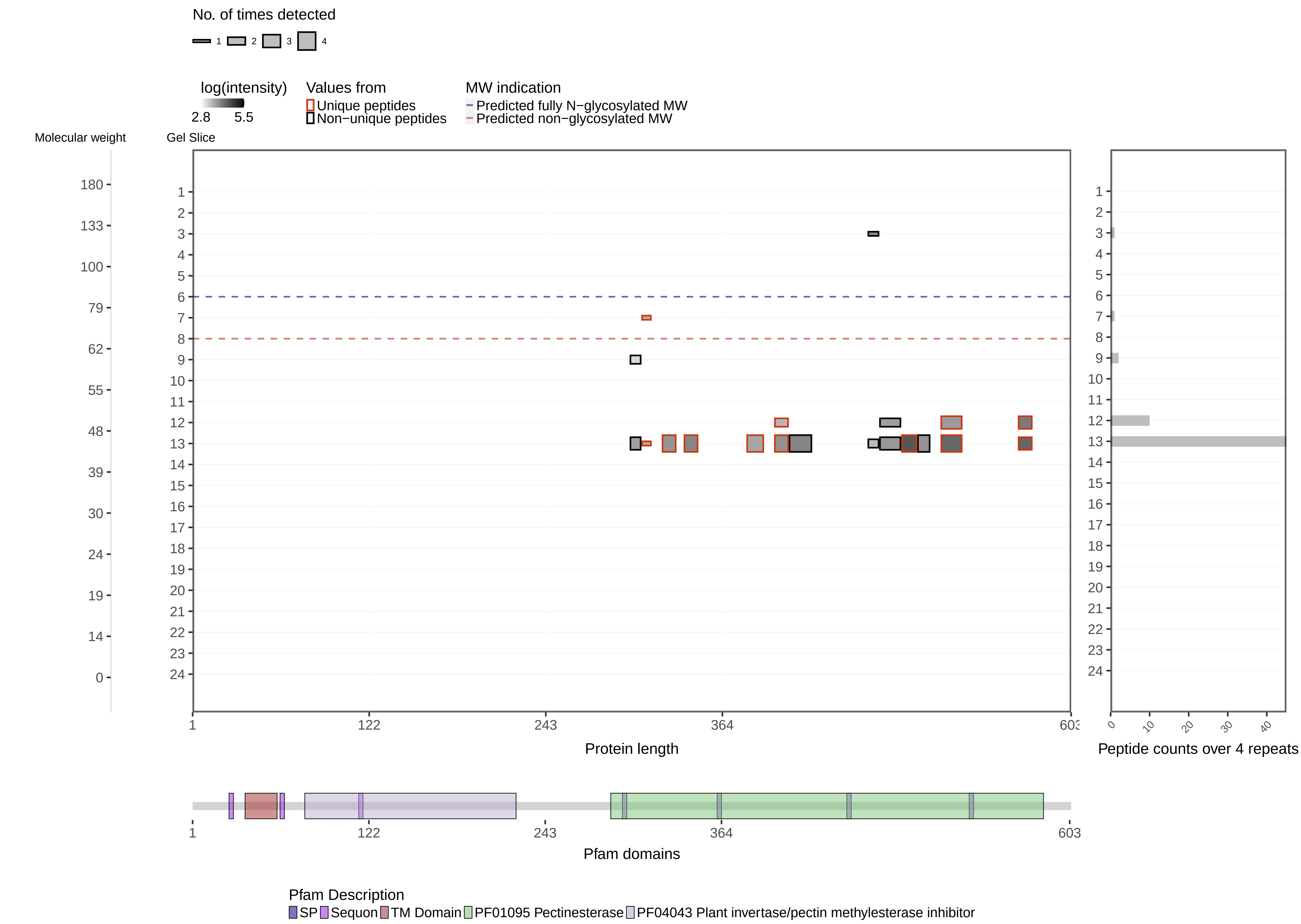


Protein sequence:

### NbL11g05840.1 binding_partner_of_ACD11_1−like [MW = 31.3 kDa]

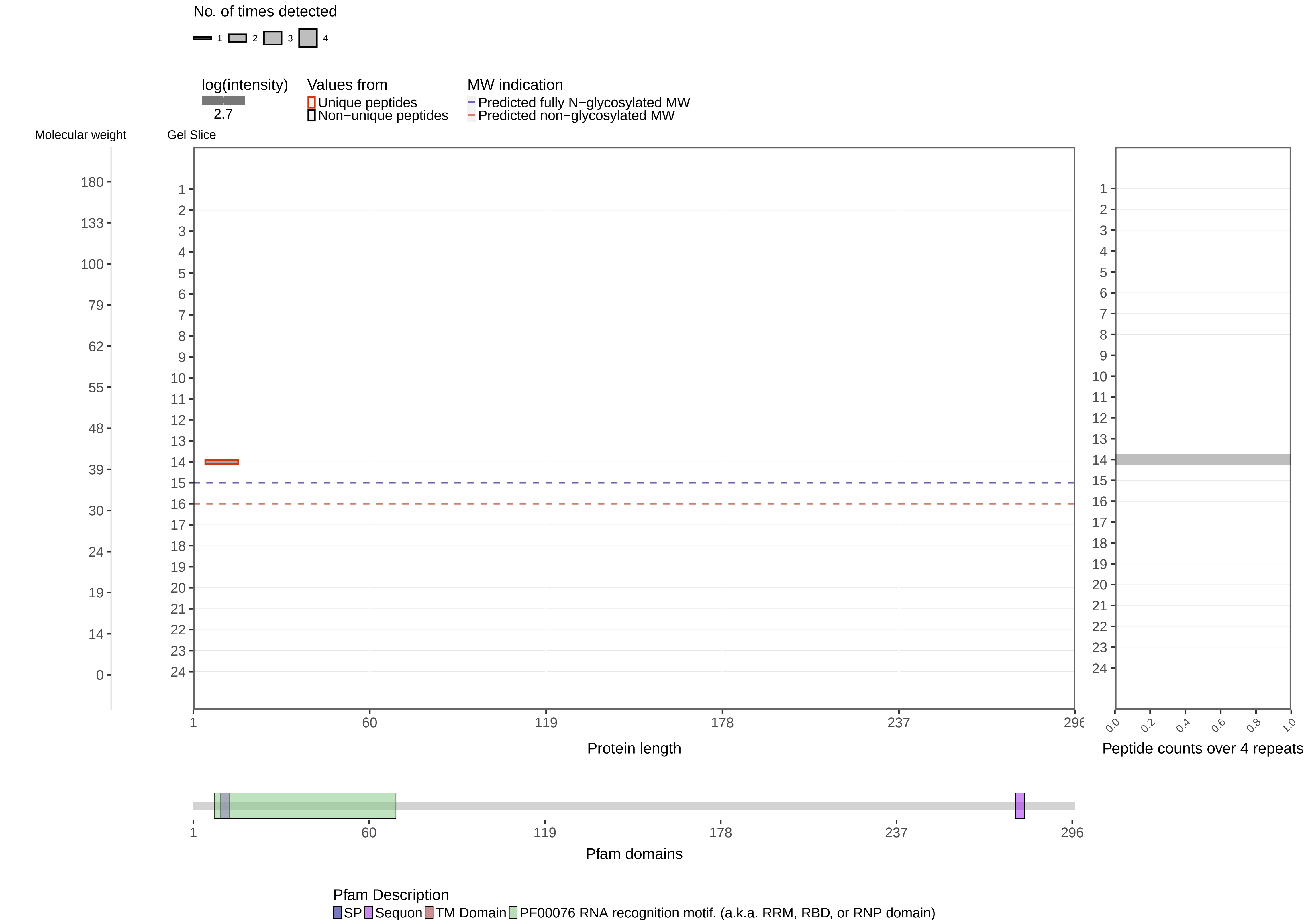


Protein sequence:

KHKITSTTSARVASLDQKYGLSEKITTGATIVNHKVKEMDQKFQVSEKSKSAFAAAEQTVSNAGSAIMKNRYALTGVTWAAGAFSRVTKAAGEVGQKTKEKLAEEEKGRNSAKGHVQEHAETQTGIKEPTIESQTGIKEPANDSLTQIQELTASSLPPKA*

### NbL11g05770.1 60S_ribosomal_protein_L13−1 [MW = 23.66 kDa]

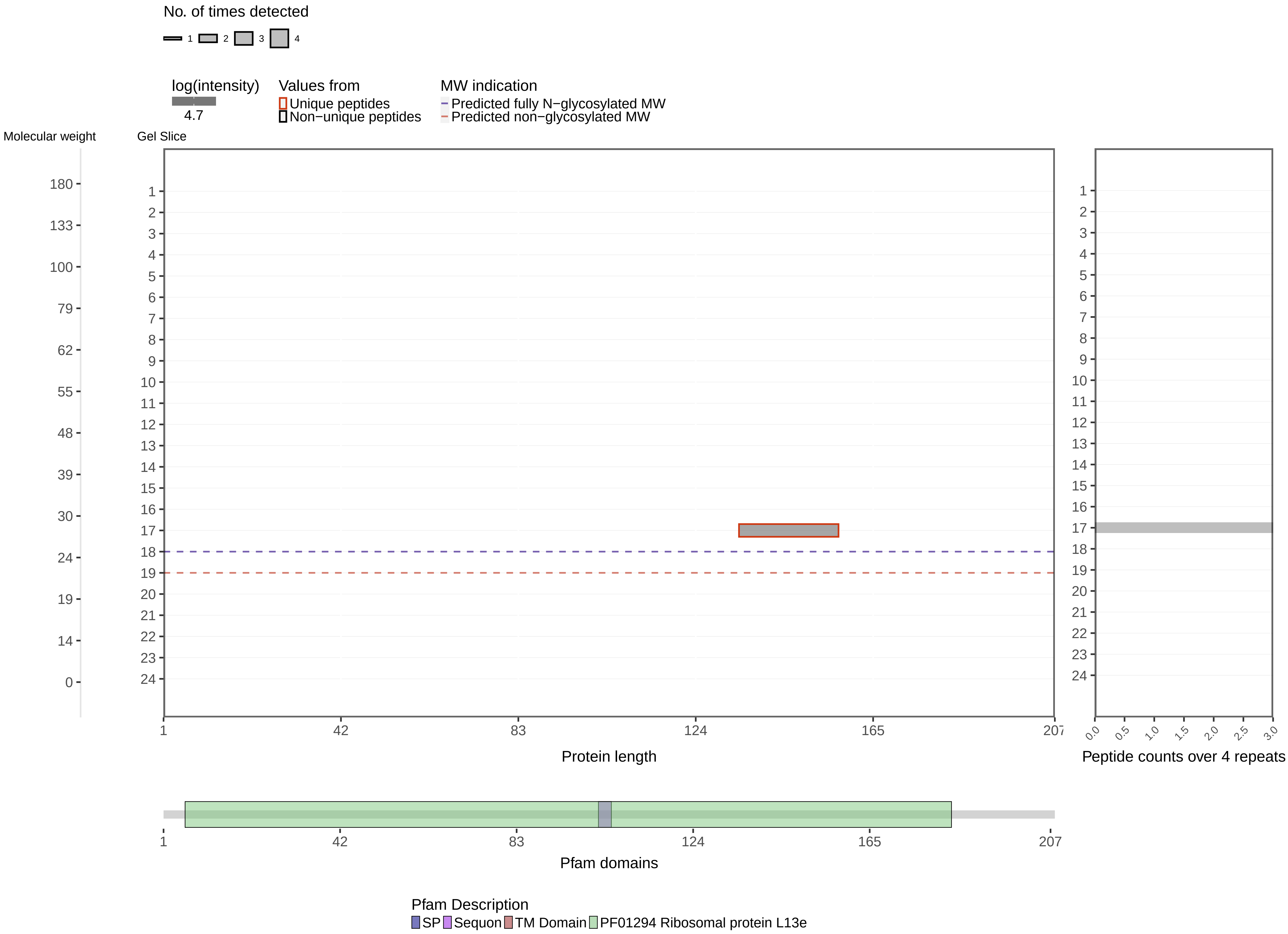


Protein sequence:

TIGIAVDHRRRNRSLEGLQTNVQRLKTYKAKLVVFPRRARKVKAGDSAPEELATATQVHGAYMPIAREKPSVDLVKVTEEMKSFNAYGKLRVERTNERHLGARMKRAAEAEKEEKK*

### NbL11g05610.1 ribosome−binding_factor_PSRP1_chloroplastic−like [MW = 33.83 kDa]

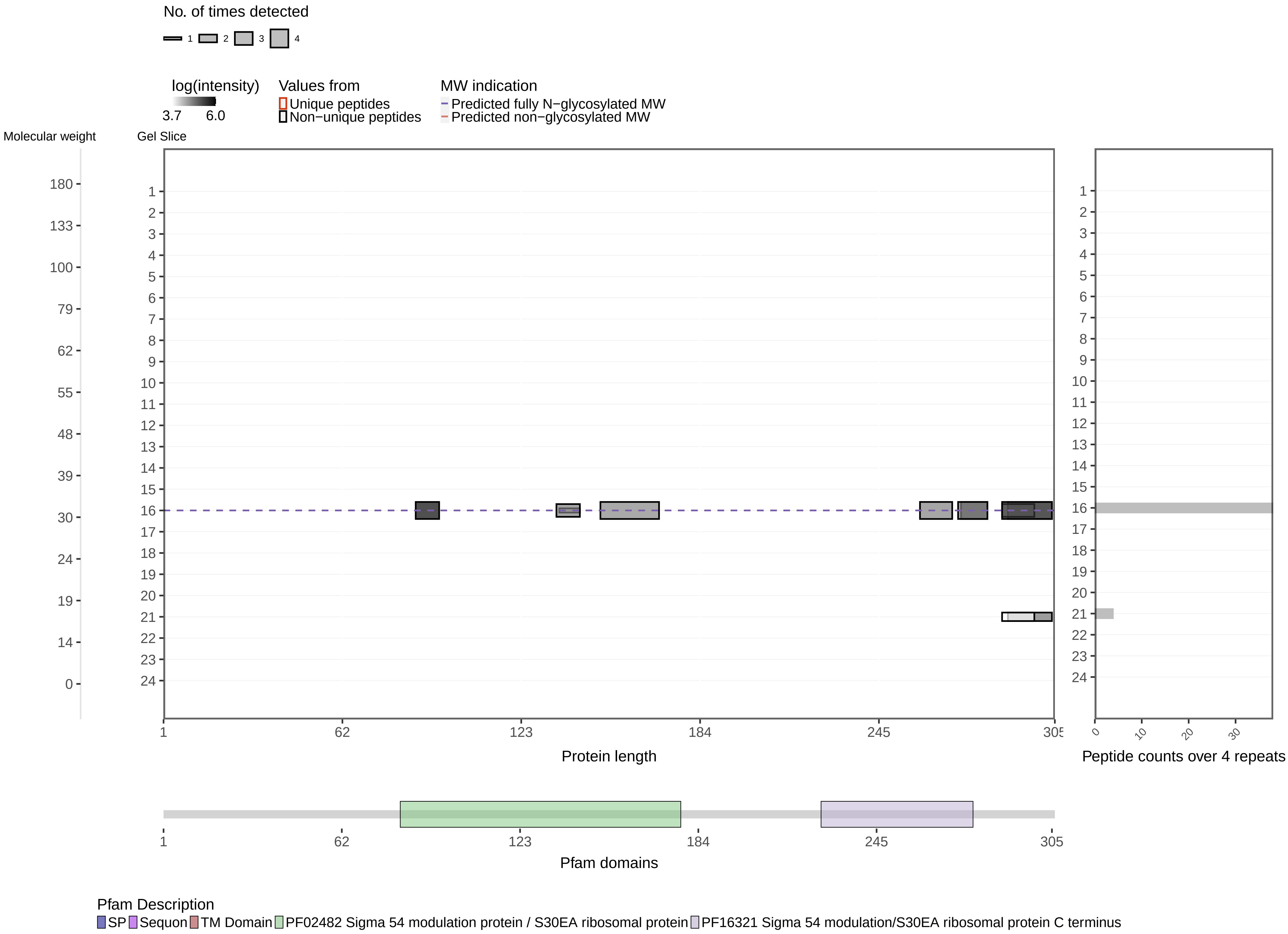


Protein sequence:

LFTKKHGVVRAEEDAESVYGSIDMVSSIIQRKLRKIKEKDSDHGRHMKGFDRLKVRDPELLAVQEDLGTLPQEEEVEDDESEGFVTEIVREKLFDMPPLTVTEAIEQMESVDHDFYGFRNEETGEINIVYRRKAGGYGLIIPKEDGKTKLEPVEVELEKETSMAE*

NbL11g05550.1 probable_6−phosphogluconolactonase_4_chloroplastic [MW = 28.58 kDa]

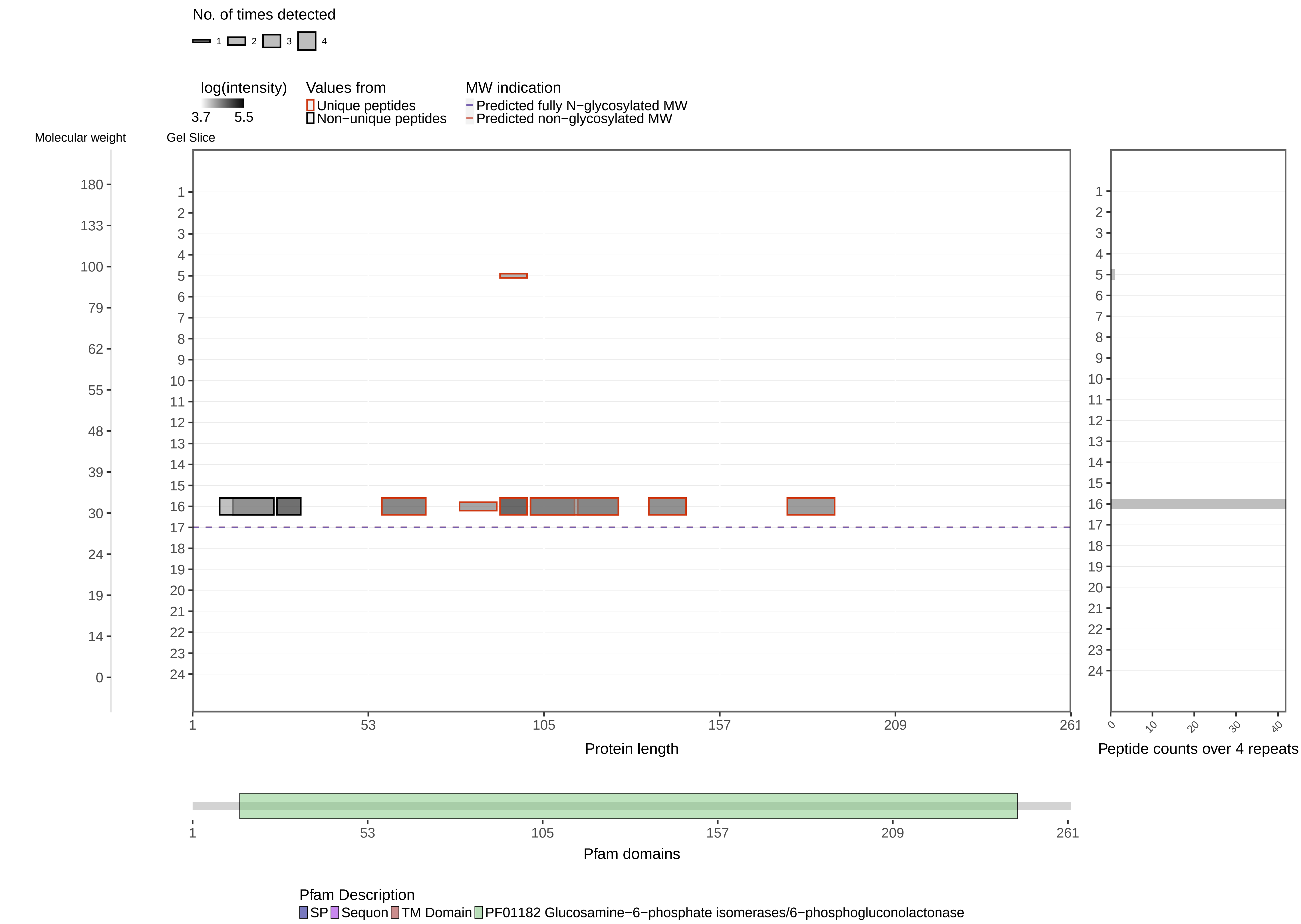


Protein sequence:

PEGAATDYEARVKQLVESKVLPLSAVSGFPKFDLMLLGMGPDGHVASLFPTRSQRHEKERLVTFITDSPKPPPPRITFTFPVINSASEIAMVITGAELANMVDVVLGSGLTADGIPPPATEVSAEEELTWFLDKDAASKLQTSE*

NbL11g05490.1 rhodanese−like_domain−containing_protein_14_chloroplastic [MW = 25.93 kDa]

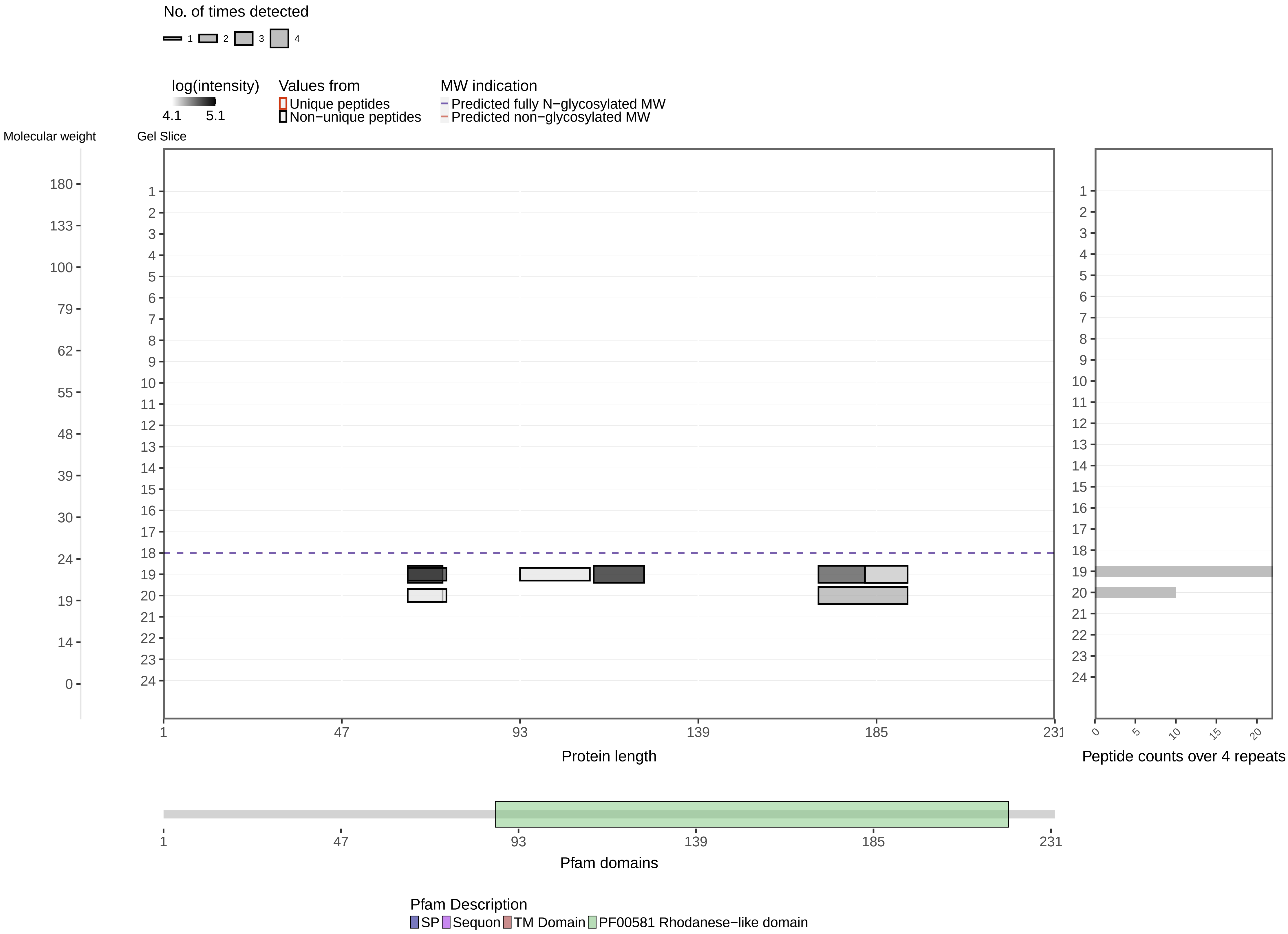


Protein sequence:

LDVRPEAEFKEAHPEGAINVQIYRLIKEWTAWDIARRVAFAFFGIFSGTEENPEFIQMVESKINKDAKIIVACSSGGTMKPTQNLPEGQQSRSLIAAYLLILNGYTNVFHLEGGLYNWFKQGLPAVSEE*

### NbL11g05440.1 luminal−binding_protein_4 [MW − SP = 70.41 kDa]

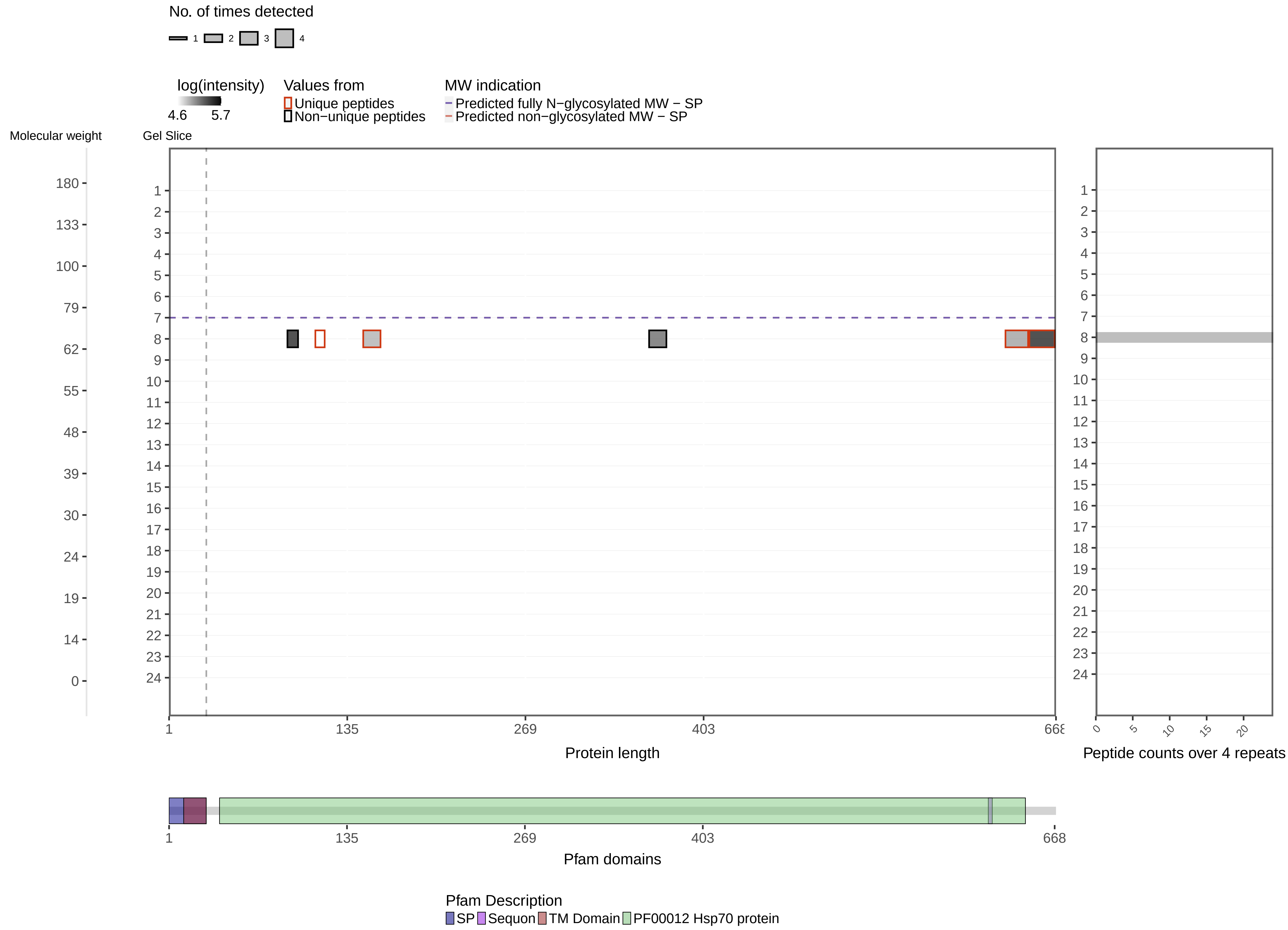


Protein sequence:

/DFSEPLTRARFEELNNDLFRKTMGPVKKAMDDAGLEKTQIDEIVLVGGSTRIPKVQQLLKDYFDGKEPNKGVNPDEAVAYGAAVQGGILSGEGGDETKDILLLDVAPLTLGIETVGGVMTKLIPRNTVIPTKKSQVFTTYQDQQTTVTIQVFEGERSLTKDCRLLGKFDLTGIAPAPRGTPQIEVTFEVDANGILNVKAEDKASGKSEKITITNDKGRLSQEEIERMVKEAEEFAEEDKKVKERIDARNSLETYVYNMRNQINDKDKLADKLESDEKEKIETATKEALEWLDDNQS/

### NbL11g05290.1 pyruvate_dehydrogenase_E1_component_subunit_beta−1_mitochondrial [MW = 39.53 kDa]

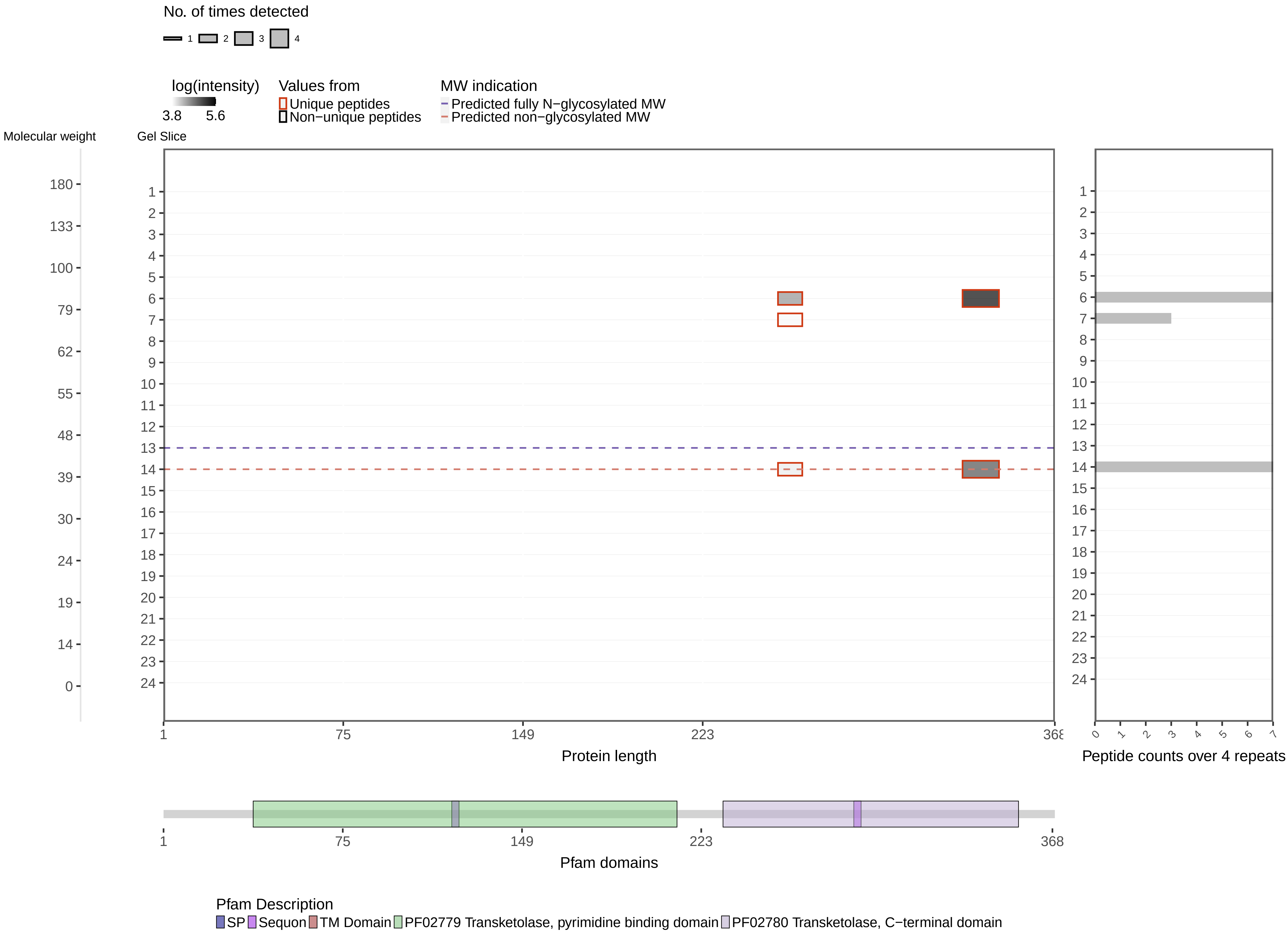


Protein sequence:

AWFGSCPGLKVLAPYSSEDARGLLKAAIRDPDPVVFLENELLYGESFPISAEALDSSFCLPIGKAKIEREGKDVTITAFSKMVRCALQAAEILAKEGISAEVINLRSIRPLDRPAINASVRKTNRLVTVEEGFPQHGVGAEICASVVEESFEYLDAPVERITGADVPMPYAANLERMAVPQVEDIVRAAKRACYRSST*

### NbL11g05280.1 ubiquitin−conjugating_enzyme_E2_27 [MW = 21.13 kDa]

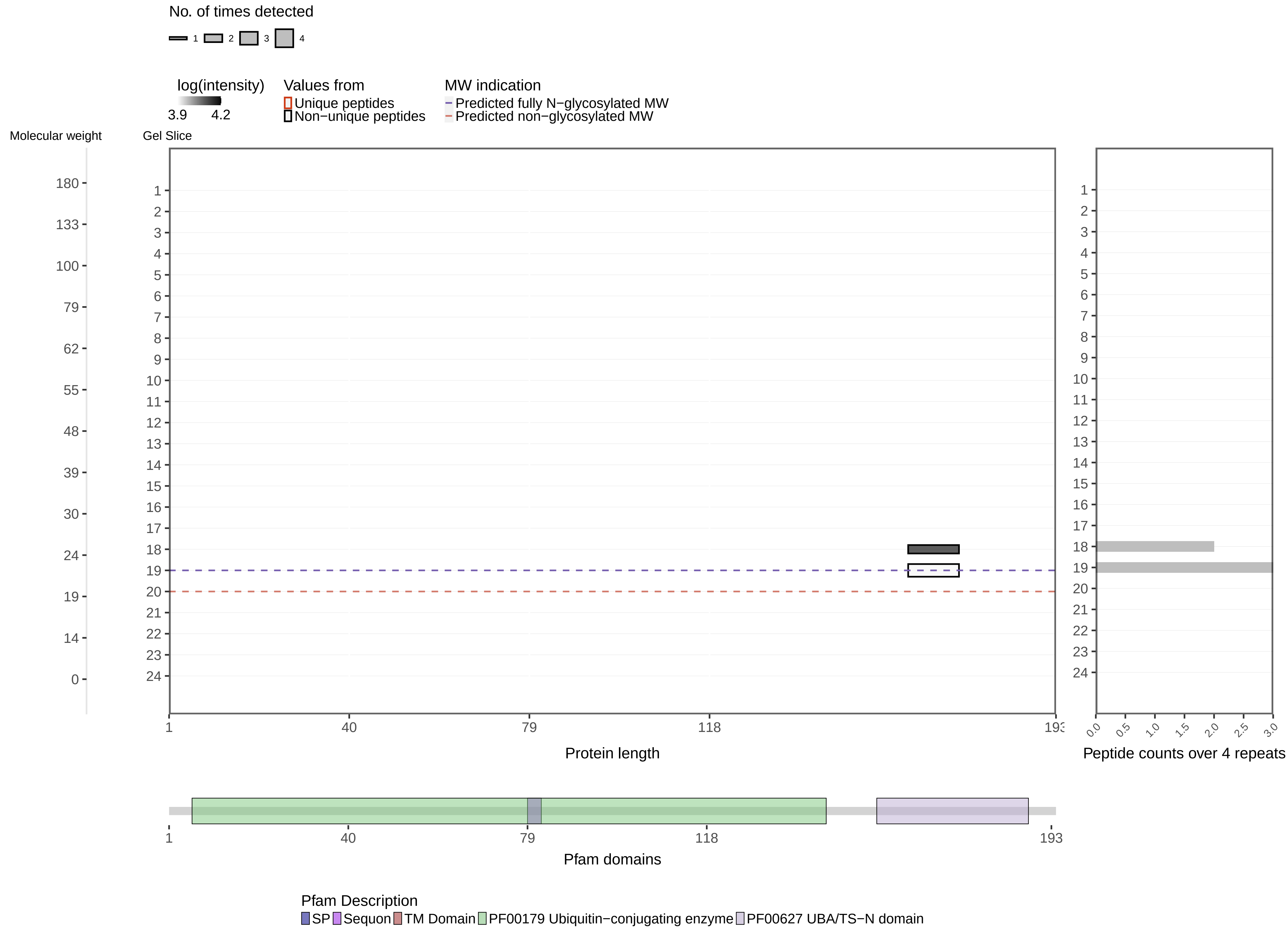


Protein sequence:

SGAICLDILKDQWSPALTLKTALLSIQALLFAPEPDDPQDAVVAQQYLRDHQTFVGTARYWTETFAKTLGTEEKIQKLVEMGFPEAQVRSTLEGTGWDENMALEKLCSG*

NbL11g05250.1 40S_ribosomal_protein_S3a [MW = 29.73 kDa]

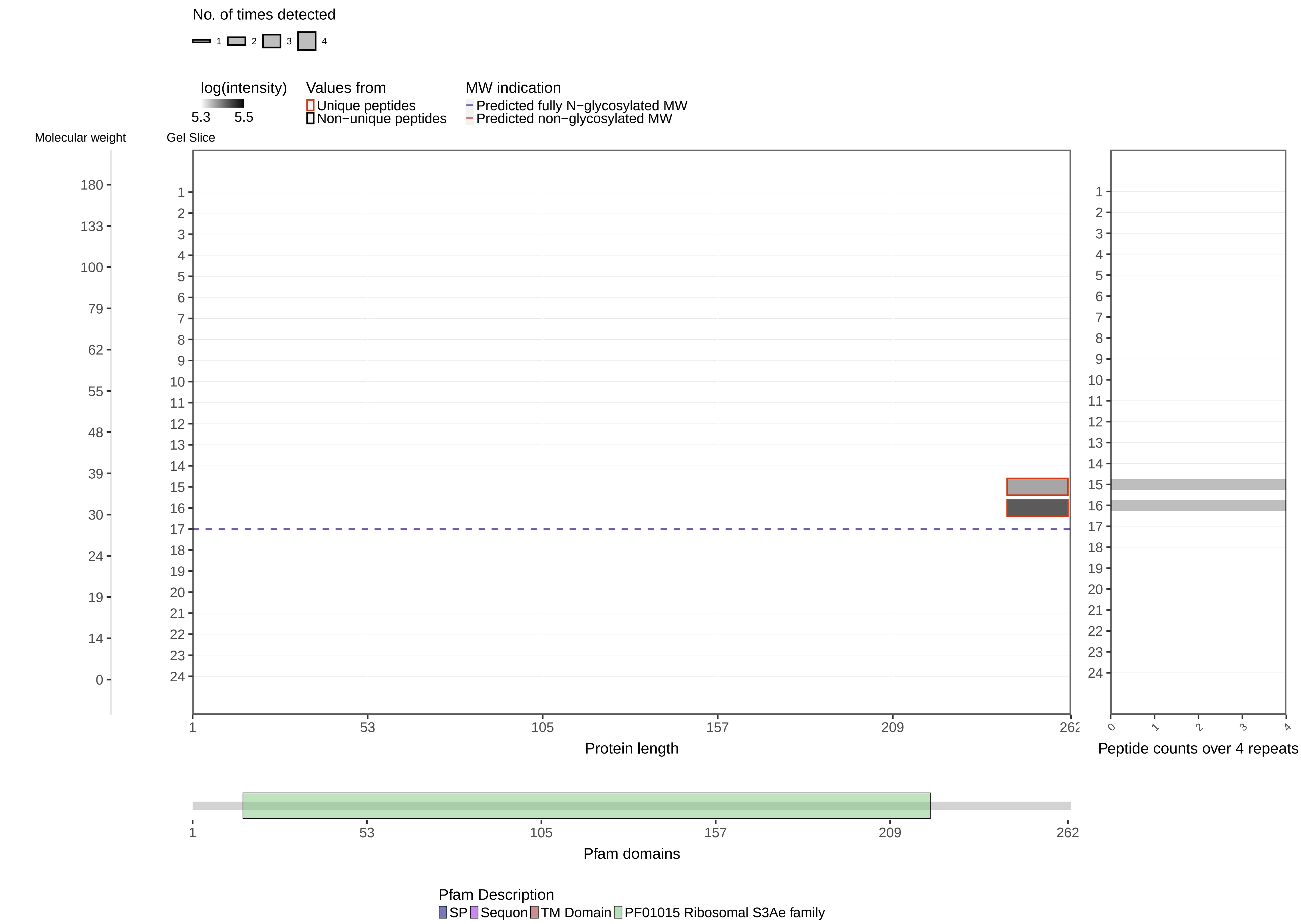


Protein sequence:

[illegible]WQTLIEAHVDVKTTDSYTLRMFCIAFTKKRPNQQKRTCYAQSSQIRQIRRKMVEIMRNQASSCDLKELVAKFIPESIGREIEKATSSIFPLQNVYIRKVKILKAPKFDIGKLMEVHGDYSEDVGVKLDRPADETVAEAEPEIPGA*

NbL11g05210.1 50S_ribosomal_protein_L29_chloroplastic−like [MW = 19.07 kDa]

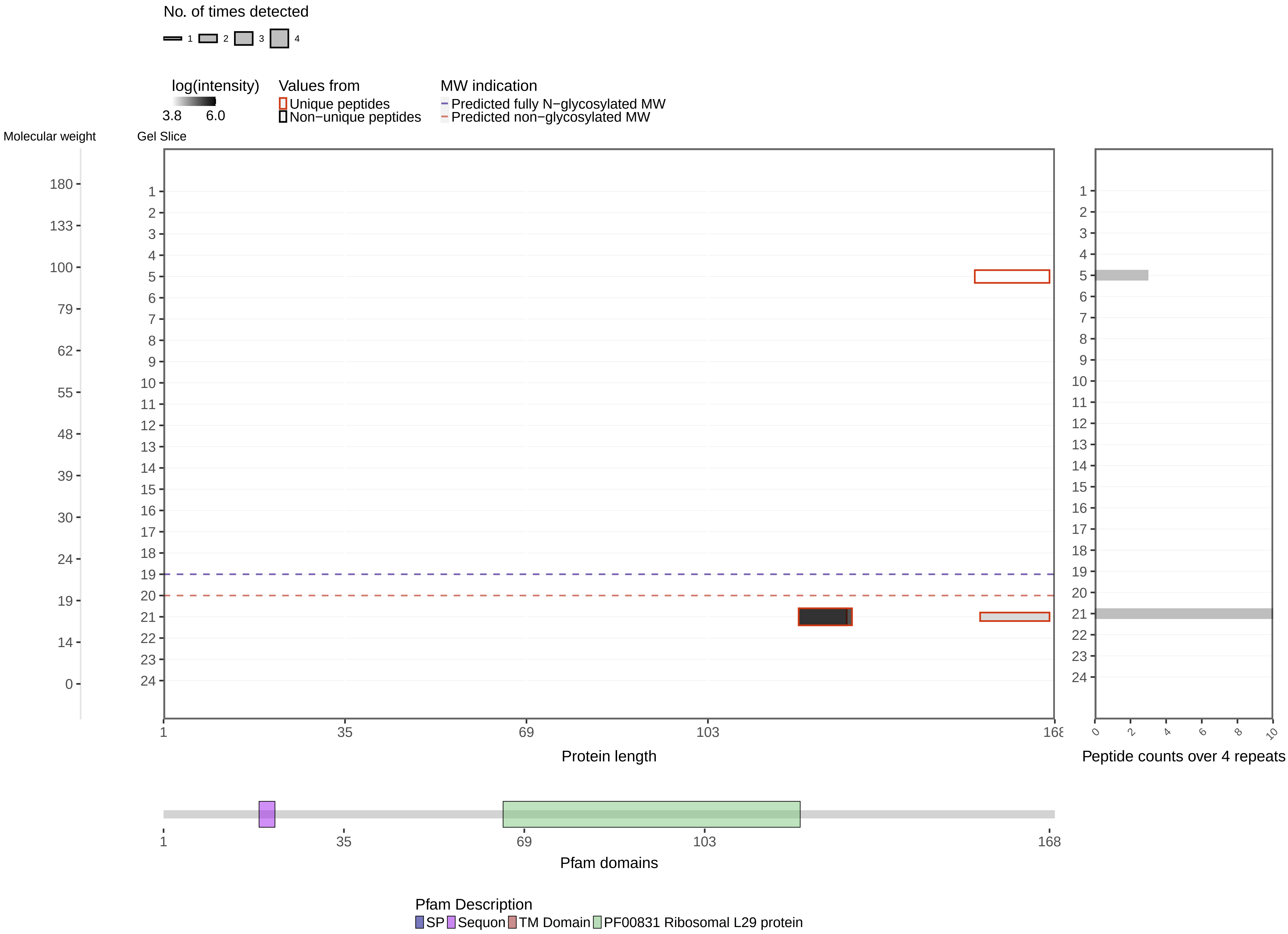


Protein sequence:

TEEIQEEIVDLKGELFMLRLQRSARNEFKSSEFLRMRKRIARMLTVKRERELEEGINKRISRKLDRKWKKSIVPRPPPSLKKLQEEEAAAEAKESA*

### NbL11g05150.1 putative_34−dihydroxy−2−butanone_kinase [MW = 61.85 kDa]

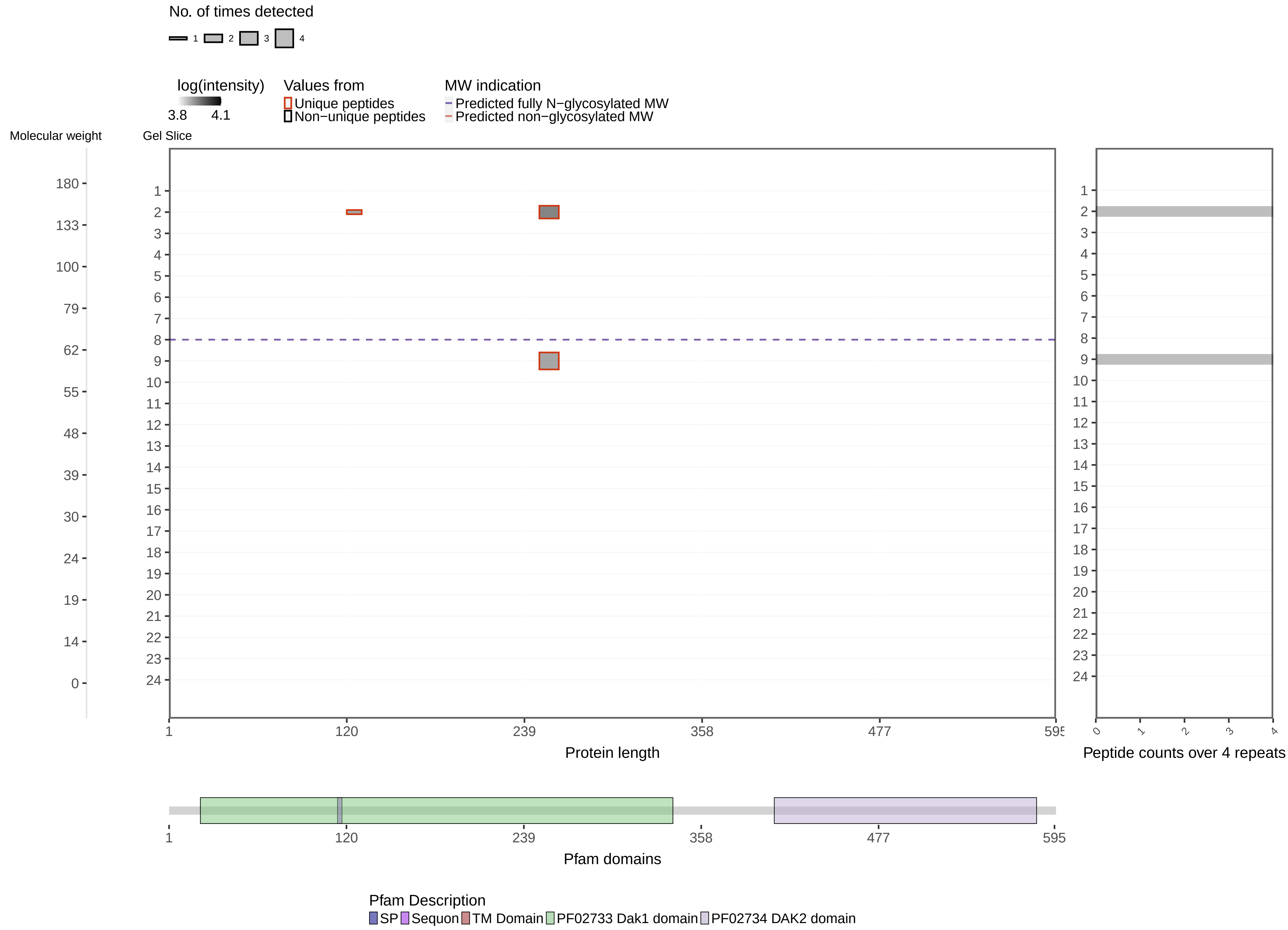


Protein sequence:

IAGKAVPGLQLEHGLAVDRVYTGSLMTSLDMAGFSISVMKADQAILDRLDAPTKAPNWPVGADGNRPPAKIPVPLPPPHSIQSDKTLSRPEKLSSLGHILEAAIEAAATELVNLRDNLNEWDSKVGDGDCGSTMFRGAVAILEDMKKYYPLNDPAETVNEIGSSIRRVMGGTSGILYDIFCKAAYAKLKADSELVITALHWADALDAAIAAVSKYGGARAGYRTLLDALIPALSALKERLSAGDNPVDAFVISAEAAVAGAESTKHMQAQAGRSTYIPGDVLASIPDPGAMAAAAW

### NbL11g05110.1 CASP−like_protein_4D2 [MW − SP = 16.77 kDa]

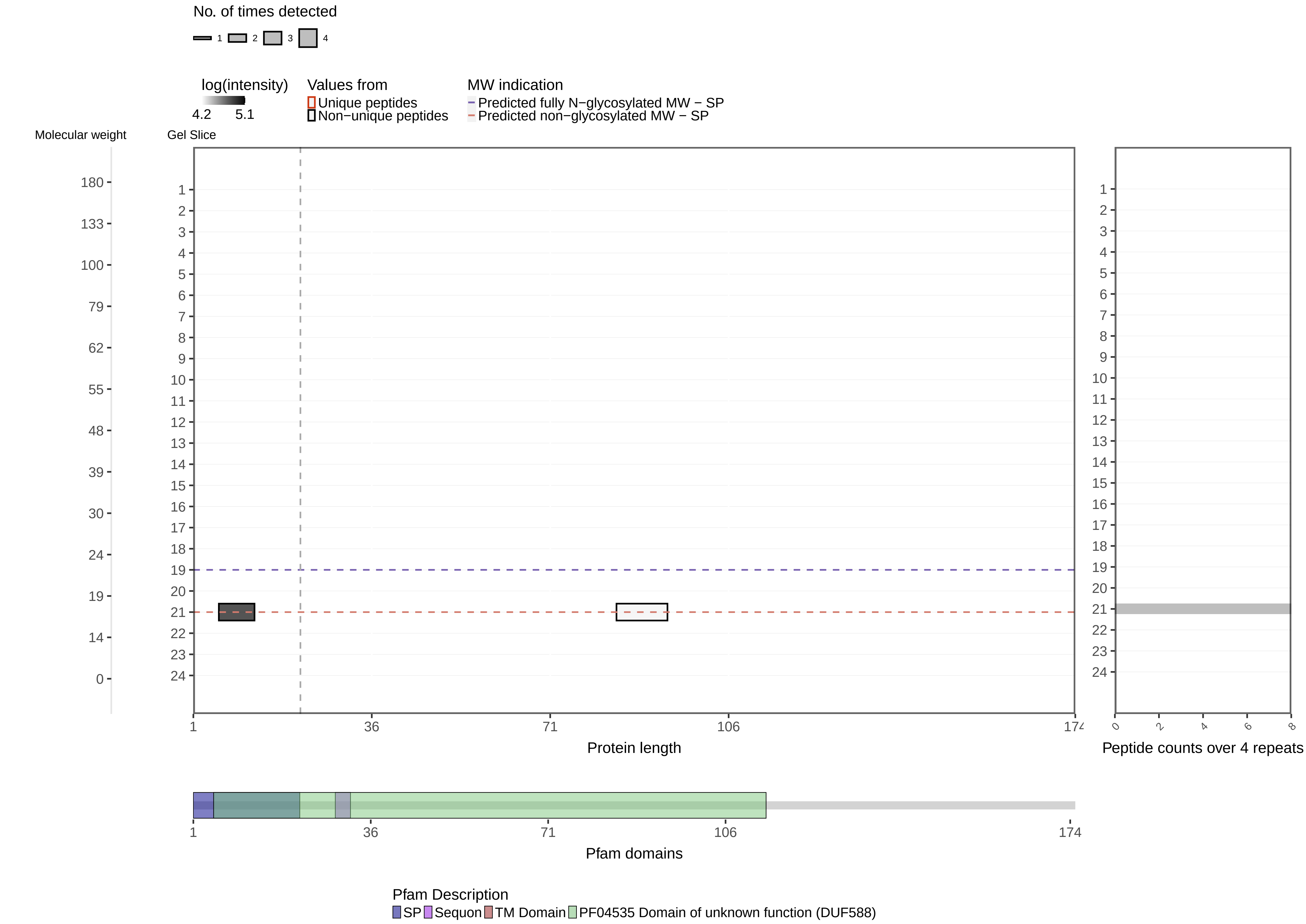


Protein sequence:

'AFTGEKVIKGRFLGMLDFYADKIITFFLASGLGVGFGVSSELKRYINGFVDTIETSGIDTFEELRTESKKFFDRGTLATTPLLAGFTTMAVLTIITSFHRK*

### NbL11g05000.1 miraculin−like [MW − SP = 21.27 kDa]

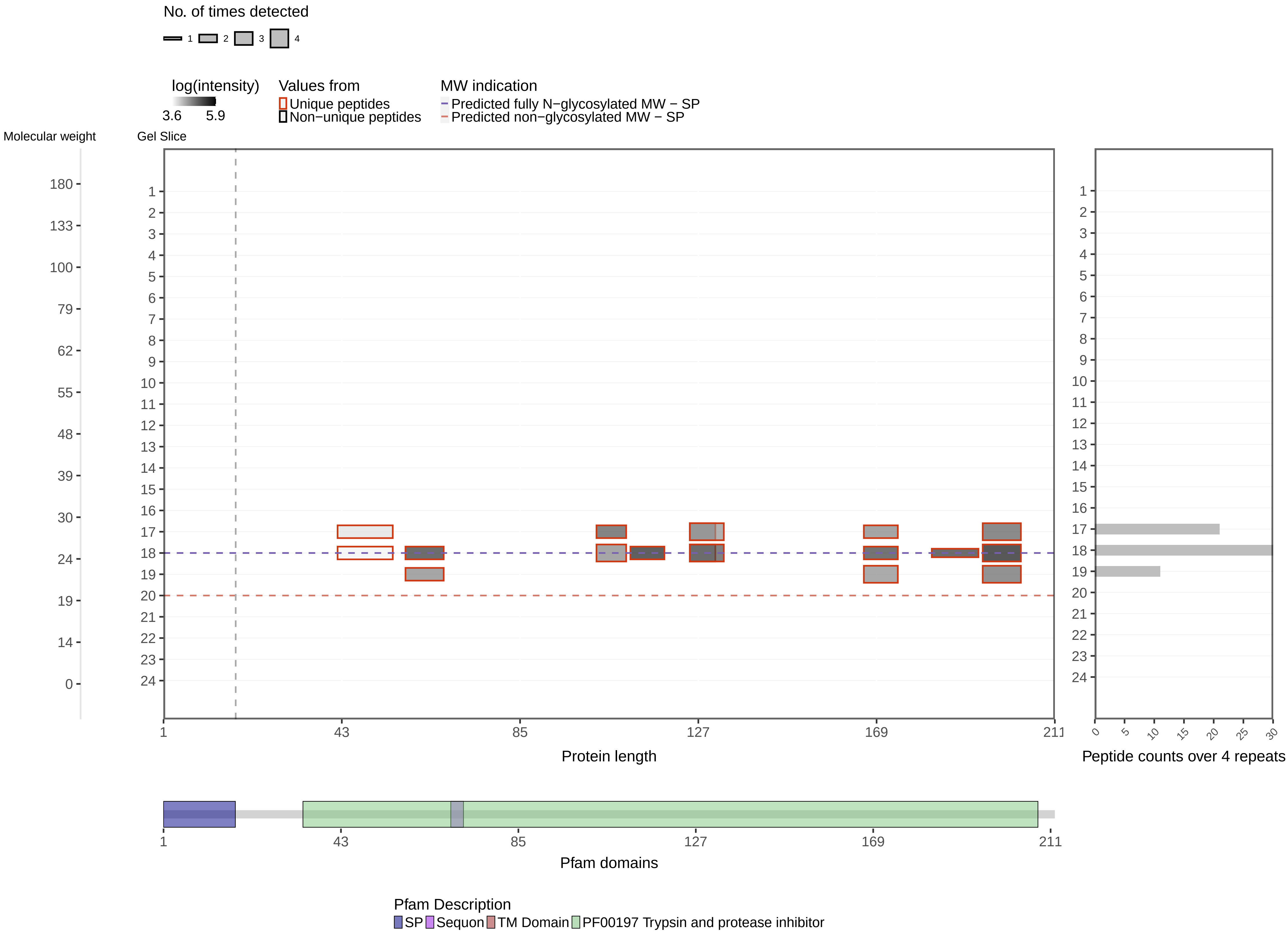


Protein sequence:

PFDAKKGVVRLSTDLNVRFFTPTICARETIWKLGTYDDKLKQYFIVTGGVEGNPGPQTLSSWFKIEKLGTDYKFVFCPSVCKICKVICKDVGIYSGNDGVRFMALSDTPFRVMFKKTF*

### NbL11g04780.1 glyceraldehyde−3−phosphate_dehydrogenase_cytosolic [MW = 36.65 kDa]

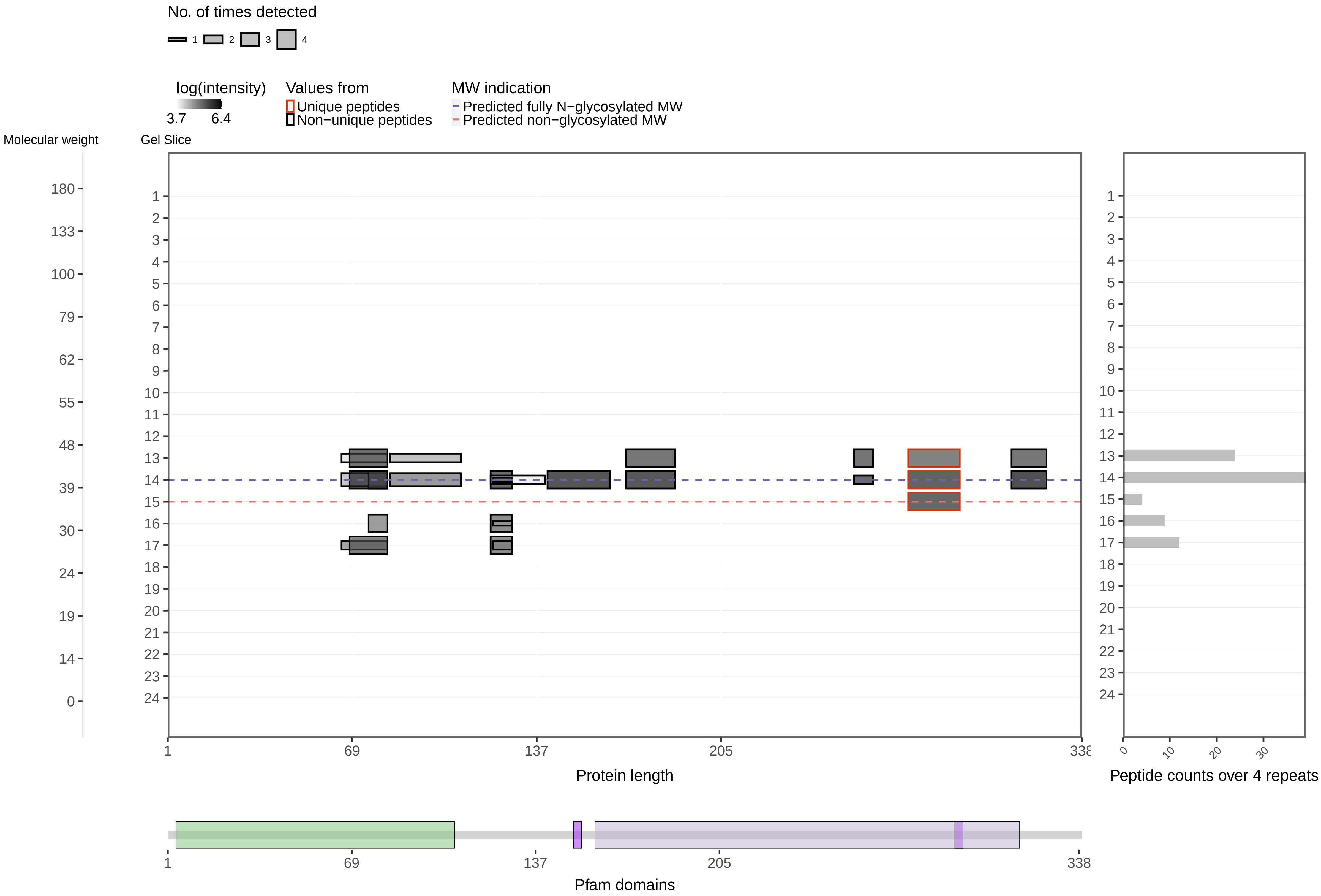


Pfam Description

SP | Sequon | TM Domain | PF00044 Glyceraldehyde 3−phosphate dehydrogenase, NAD binding domain | PF02800 Glyceraldehyde 3−phosphate dehydrogenase, C−terminal domain

Protein sequence:

TNCLAPLAKVIHDRFGIVEGLMTTVHSMTATQKTVDGPSAKDWRGGRAASFNIIPSSTGAAKAVGKVLPSLNGKLTGMAFRVPTVDVSVVDLTVRLEKEATYDEIKAAIKEESEGKLKGILGYTEDDVVSTDFIGDNRSSIFDAKAGIALSKNFVKLVSWYDNEMGYSTRVVDLIKHMASVQ*

### NbL11g04680.1 elongation_factor_TuB_chloroplastic [MW = 52.65 kDa]

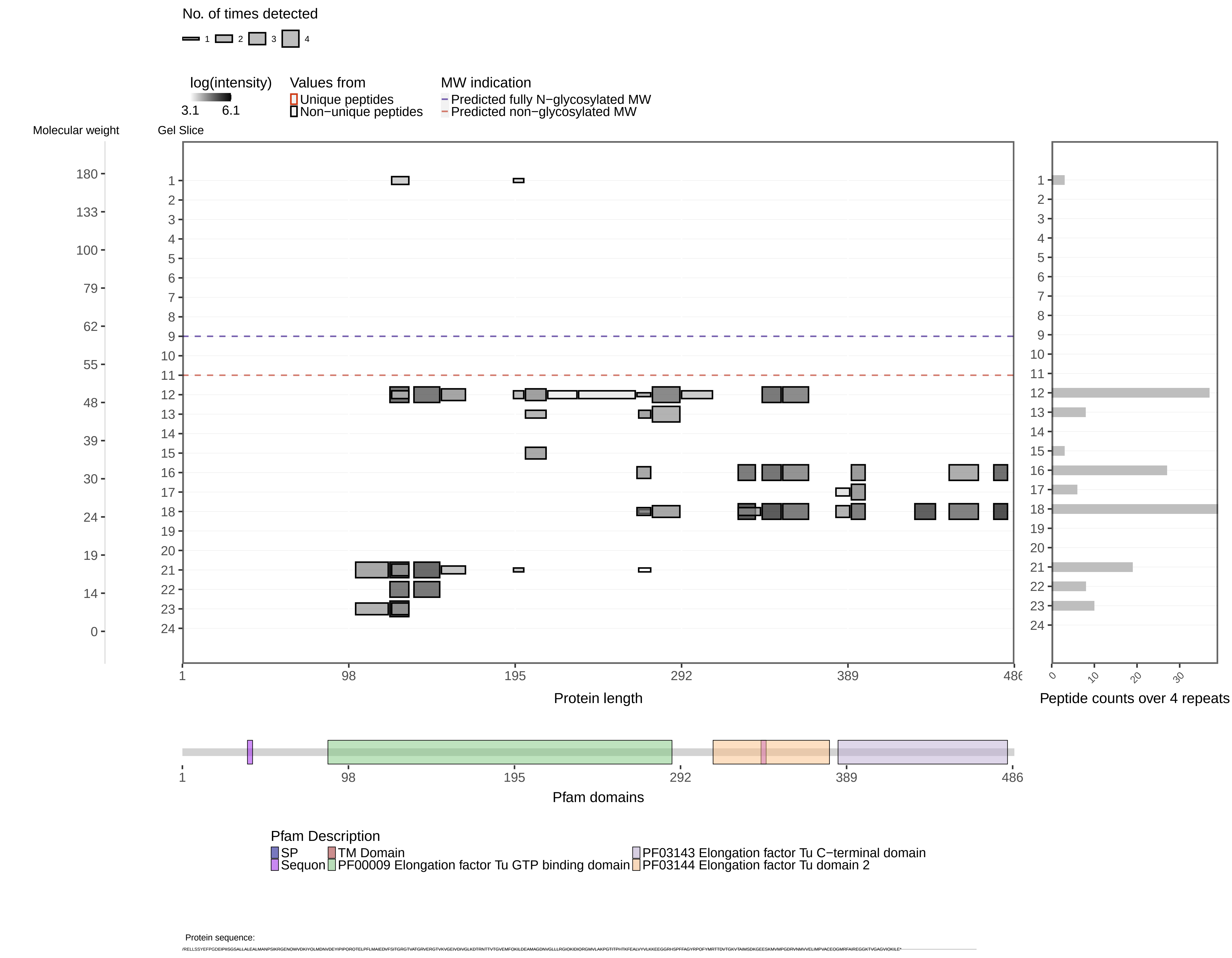

### NbL11g04530.1 anthranilate_phosphoribosyltransferase_chloroplastic−like [MW = 42.38 kDa]

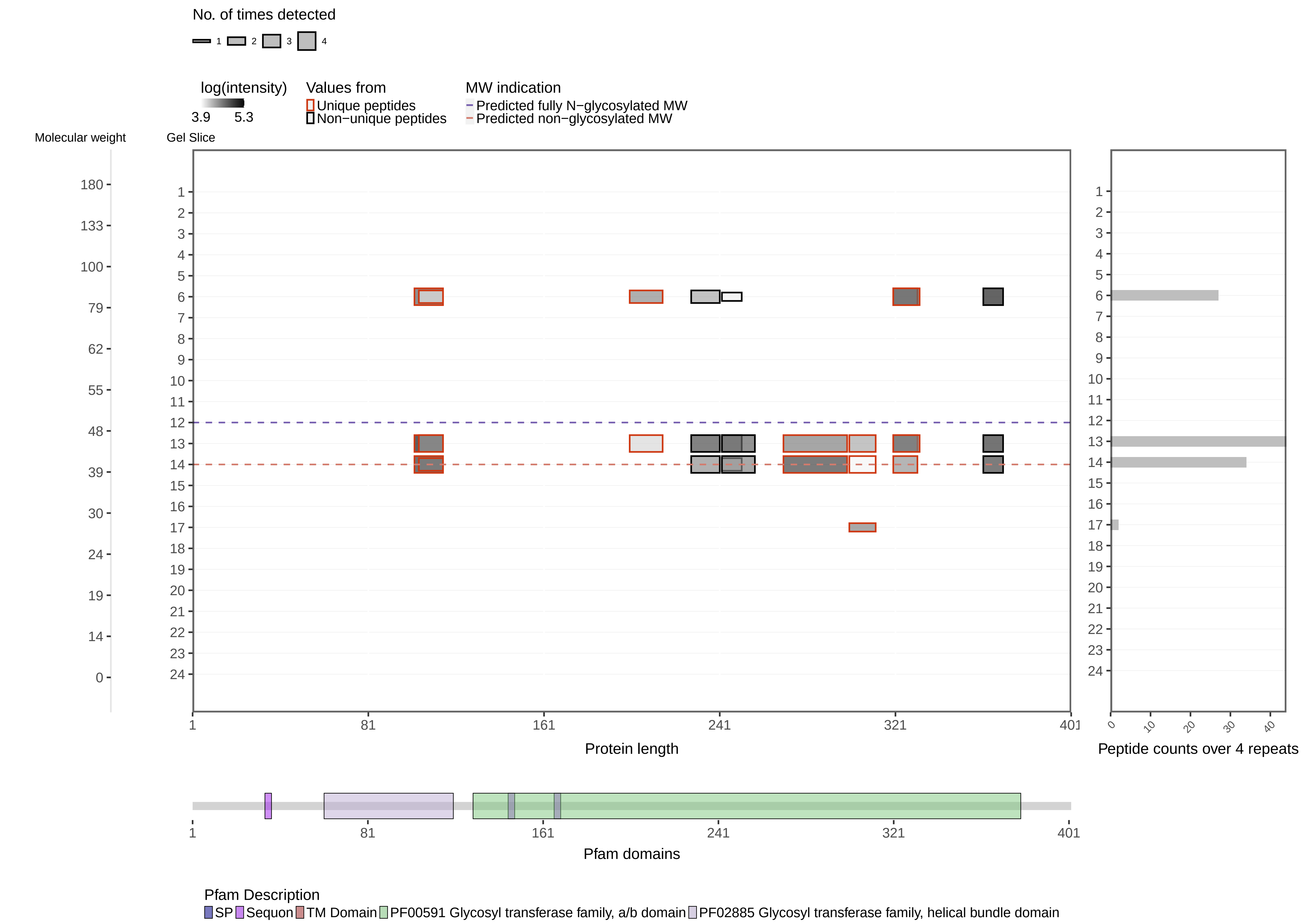


Protein sequence:

PEGVKECVTRAGIGFMMAPVYHPAMKIVRPIRKKLKVKTVFNILGPMLNPARVPFAVVGVYKEDLVHKMAKAVQRFGMKRALVVHSEGLDEMSPLGPGLVLDVTPDNIEKFSFDPLVDFGIPRCTLESLRGGGPDYNADMLKRVLSGEKGPIADAFALNAAAALLVSGLVKNLAEGVALARETLVSGKALKTLDLWVEVSNEVKAAVSQVA*

### NbL11g04490.1 GDSL_esterase/lipase_At5g62930 [MW = 27.35 kDa]

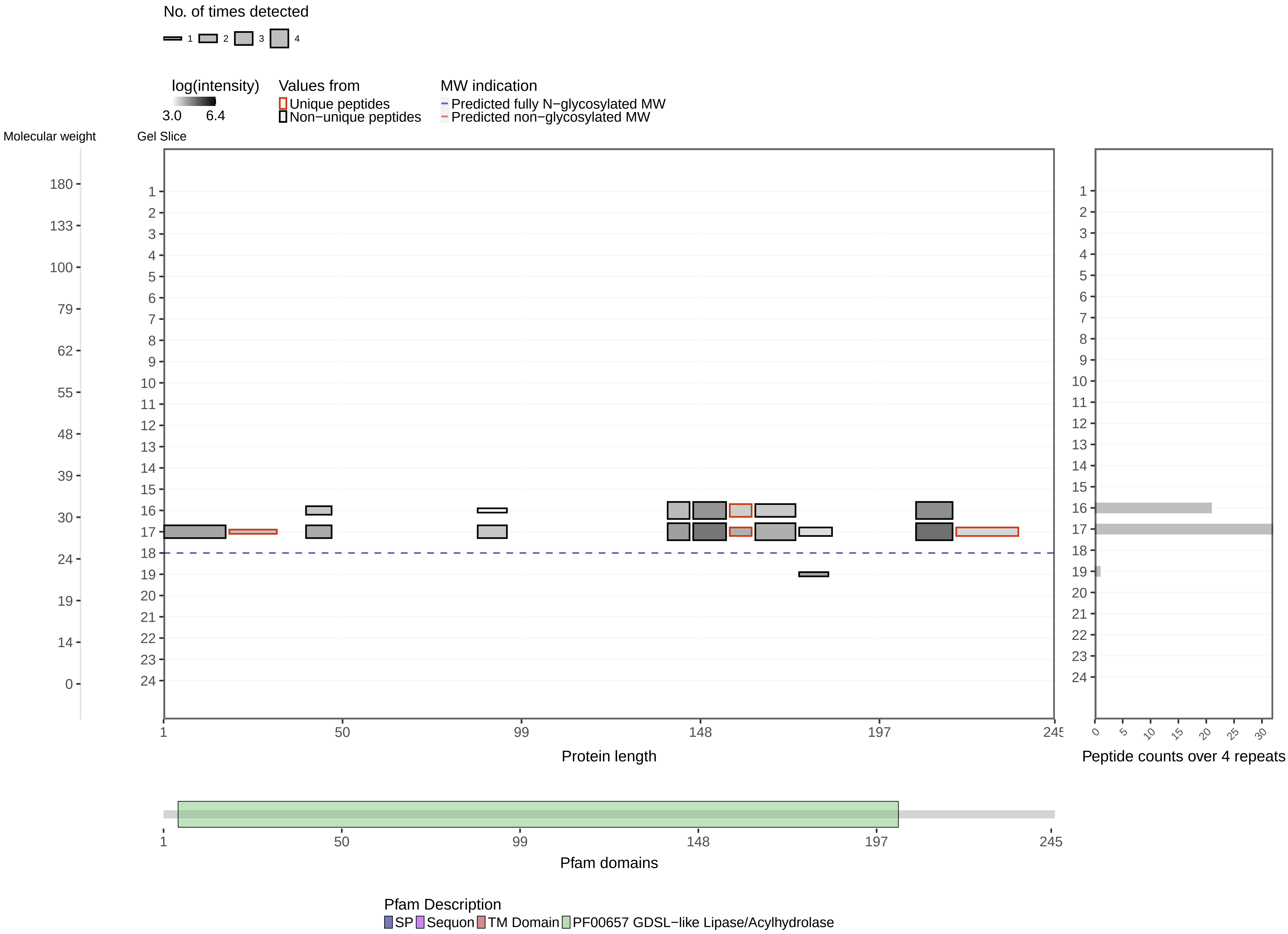


Protein sequence:

SILVVLITPPPVDEAGRFEQARSMYGDKAMELPERTNEVTGEYAKQCVQLARELGLPSINLWSKMQETEGWQKKFLSDGLHLTAEGNAIVHQEVVKAFNEAGFSAQKMPYDVPHHSEINGQNPEKTFQLQCPAV*

### NbL11g04430.1 peroxisomal_membrane_protein_PEX14−like [MW = 56.97 kDa]

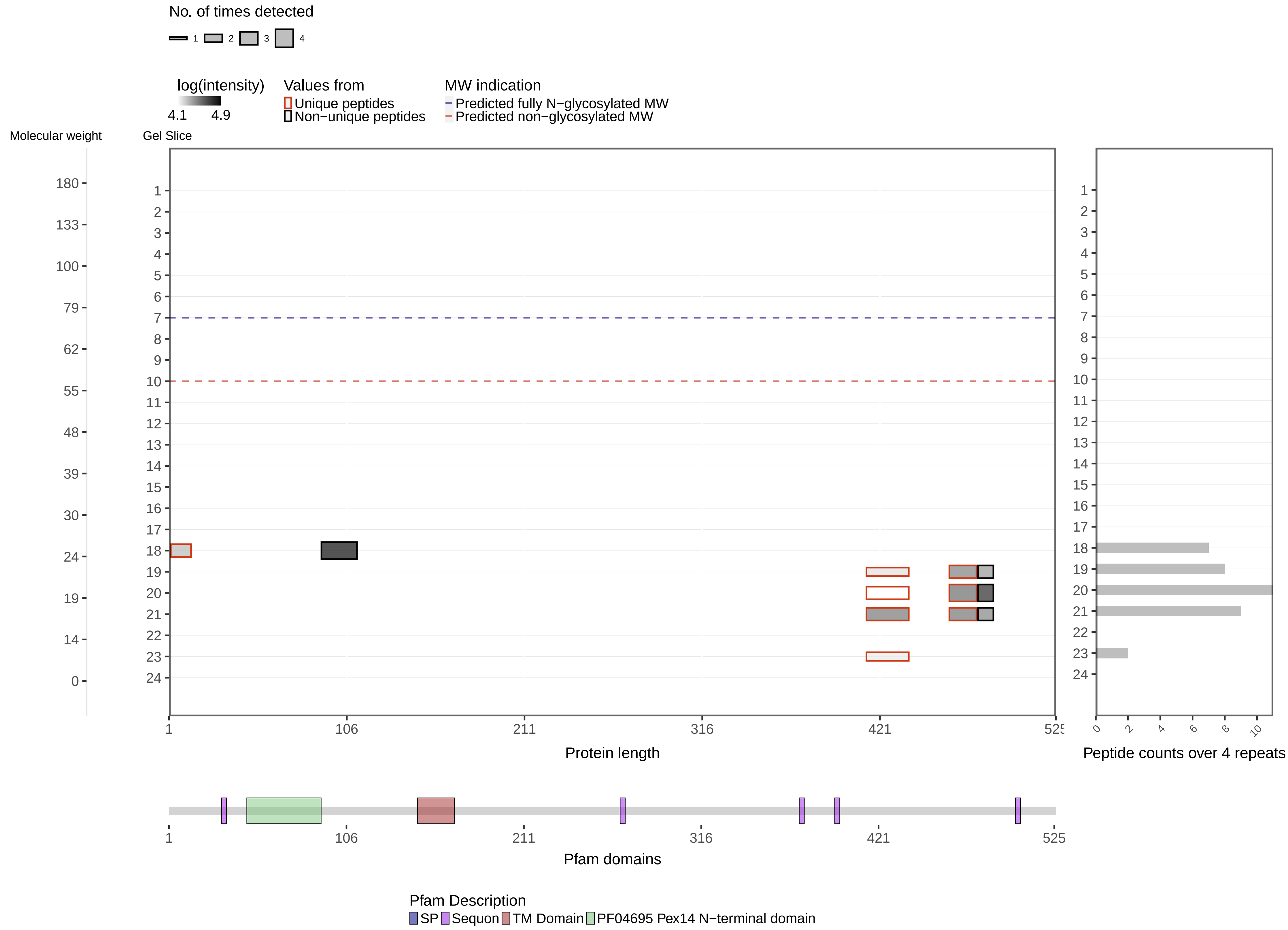


Protein sequence:

### NbL11g04370.1 E3_ubiquitin−protein_ligase_SIS3−like [MW = 42.28 kDa]

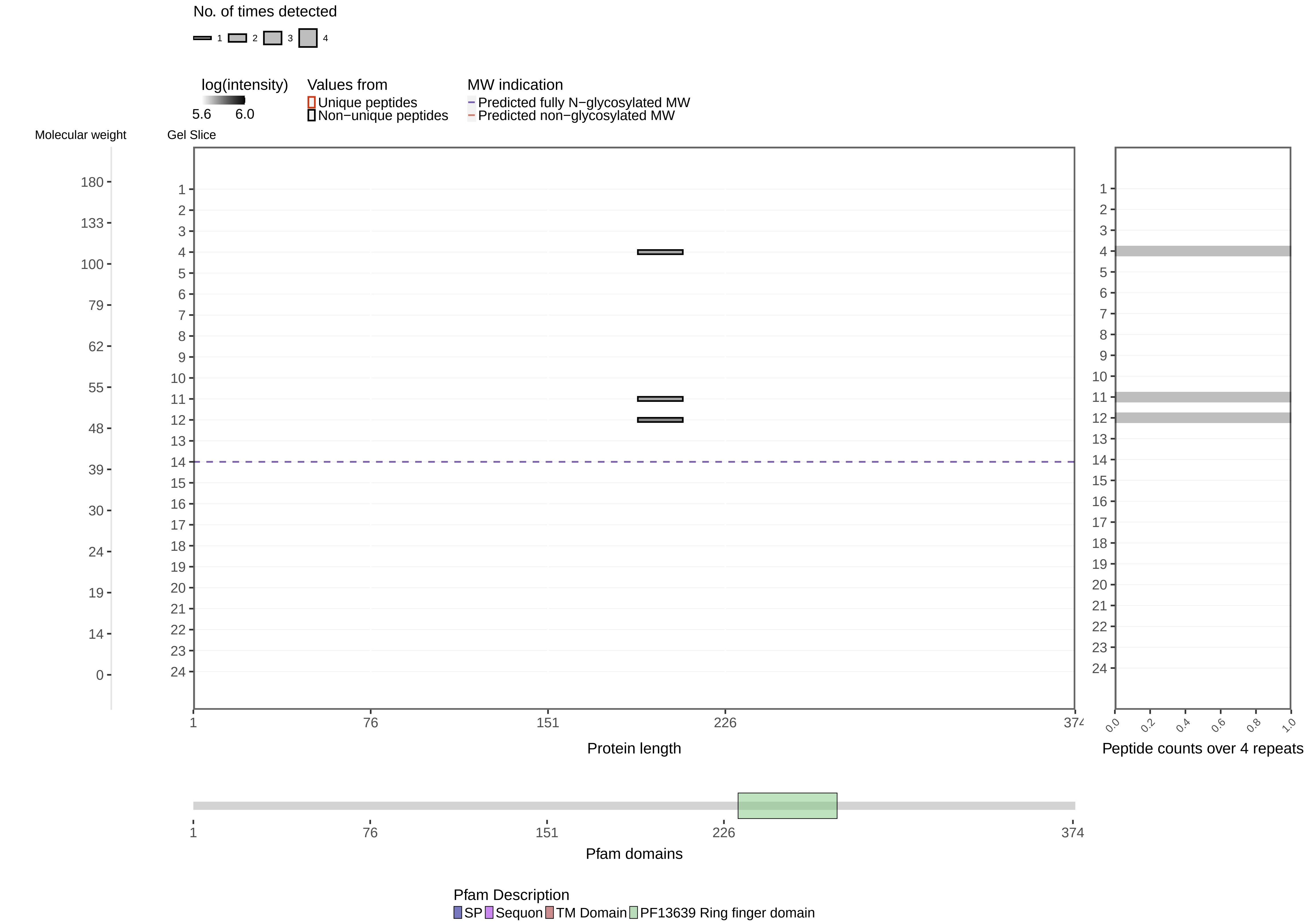


Protein sequence:

WTFAAAGOEMRGMGODATSYHPGVYLTOAOREAVEALIOELPKFMLKAVPTDCSECPICLEEFRVGNEVRGLPCAHNFHVECIDEWLRLNVKCPRCRSSVFPNLDLSALSNIRADIESSTADVVTEARYMRSOPSSOSYLLRLOGLLRPVRTODTGMGSEVDATSGTAESRNOALTPHHOGNMEPVOVVVEOSPRE*

NbL11g04350.1 probable_26S_proteasome_complex_subunit_sem1−2 [MW = 15.87 kDa]

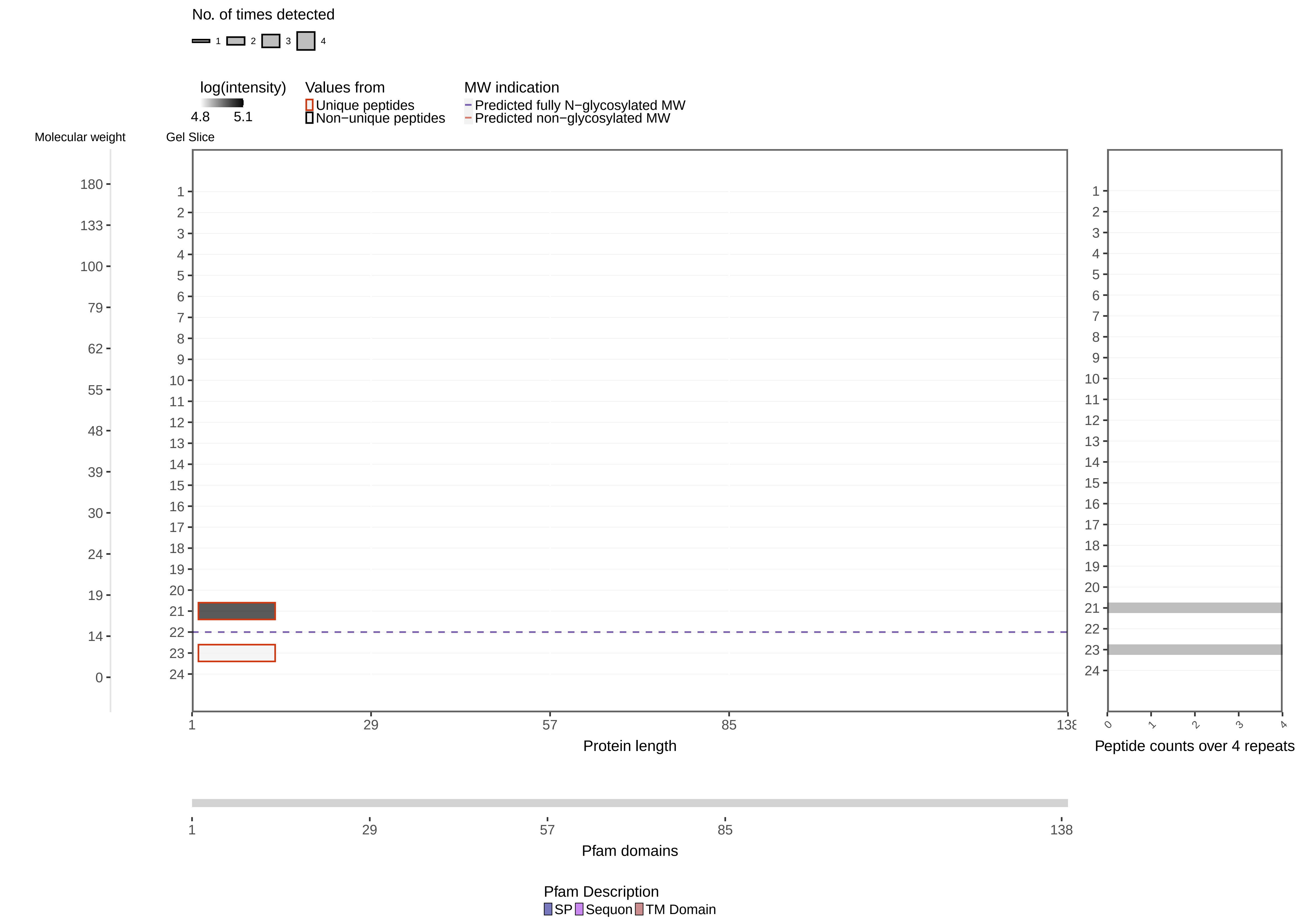


Protein sequence:

YWIIVVVKFENGRRKRKGKKLPSNGKMIGMMMMSMMTSLFSSKGNWKATRRRRNEPSLKGLFLFFKNELQSFCSVQMFPIM*

NbL11g04250.1 betaine_aldehyde_dehydrogenase_chloroplastic−like [MW = 55.4 kDa]

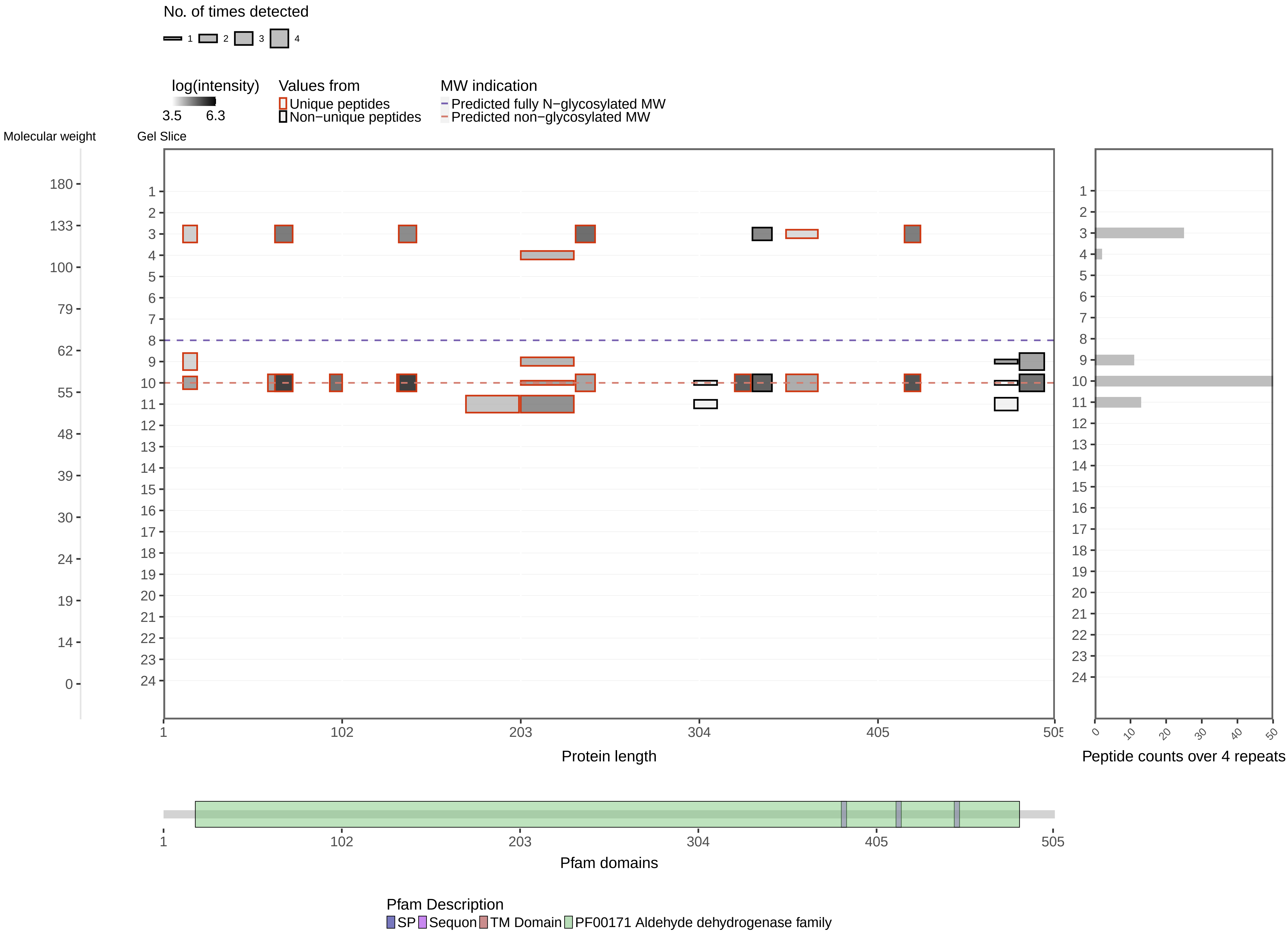


Protein sequence:

TGVKIMTAAAQLVKPVTLELGGKSPIVVFDDIDNLDLAVEWTLFGCFANAGQVCSATSRLILQESIASEFLERLLLWIKNIKLSDPLEEDCKLGPIVSSGQYEKVLKFISKAKDEGATILCGGERPQHMKKGYYVQPTIITDVNTSMEIWKEEVFGPVLCVKTFKTEEEAIELANDTKYGLASAVLSKDLERCERFTKAFQTGIIWLNCSQPTFTQLPWGGRKRSGFGRDLGEWGLENYLNIKQVTQYTSDEPWAFYKSPSKV*

### NbL11g04080.1 plasma_membrane_ATPase_2 [MW = 74 kDa]

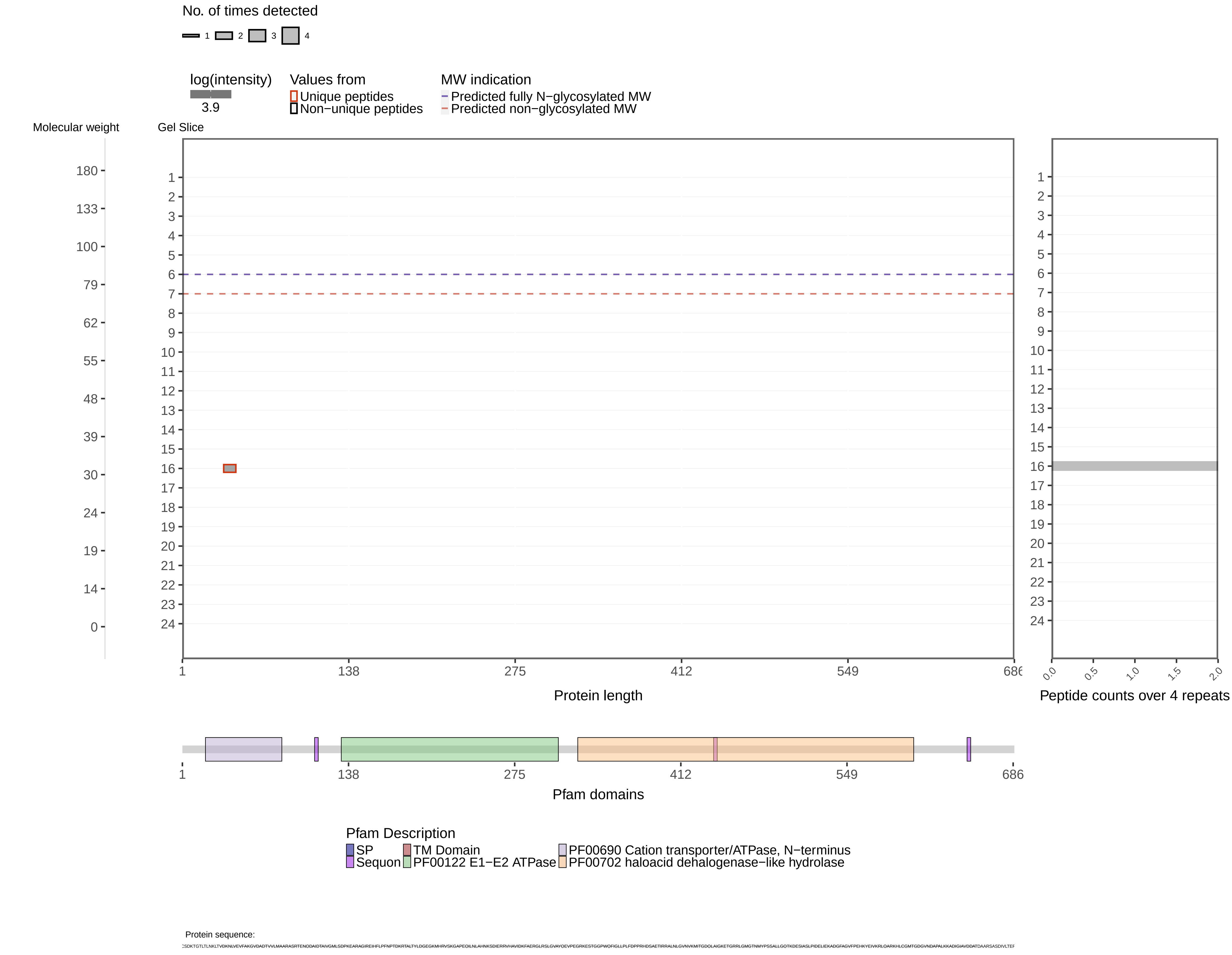

NbL11g04000.1 delta−1−pyrroline−5−carboxylate_dehydrogenase_12A1_mitochondrial−like [MW = 60.75 kDa]

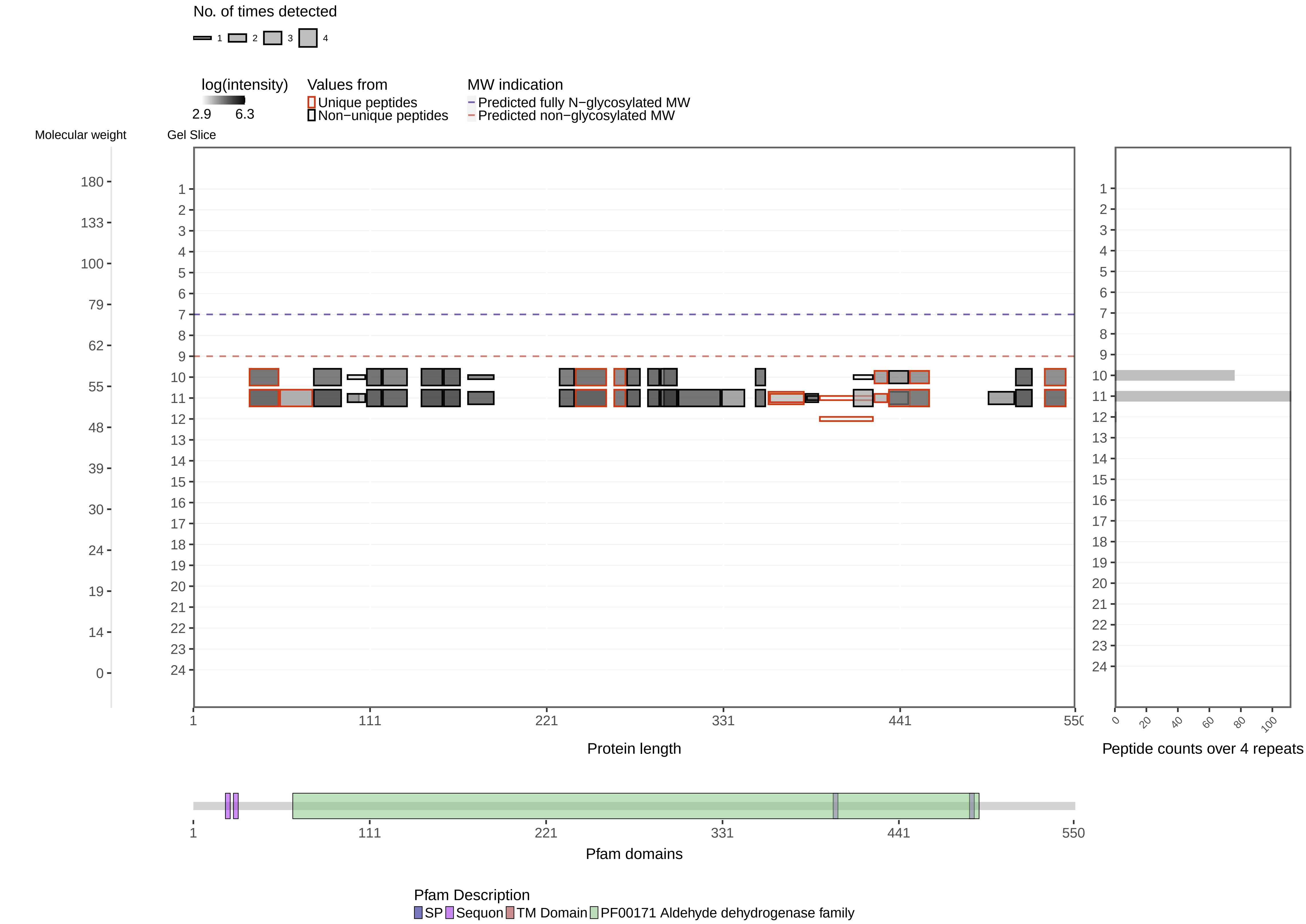


Protein sequence:

TMNKLLIEGKPRMTLFTGSSRVAEKLADDLSGRVKLEDAGFDWKILGPDVHEVDYVAWVCDQDAYACSGQKCSAQSILFMHENWGRSSLLDKMTELAARRKLDDLTIGPVLTVTTEAMLDHAKKLLQIPGSRLLFGGEPLQNHSIPAIYGAIKPTAIFIPLEEILKGQHYHLVTKEIFGPFQVVTEYKDNQLPLVLNALEKMHAHLTAAVVSNDILFLQEVIGNSVNGTTYAGLRARTTGAPQNHWFGPAGDPRGAGIGTPEAIKLVWSCHREVIYDVGPVPLGWKTPAST*

NbL11g03820.1 ran−binding_protein_1_homolog_b−like [MW = 28.76 kDa]

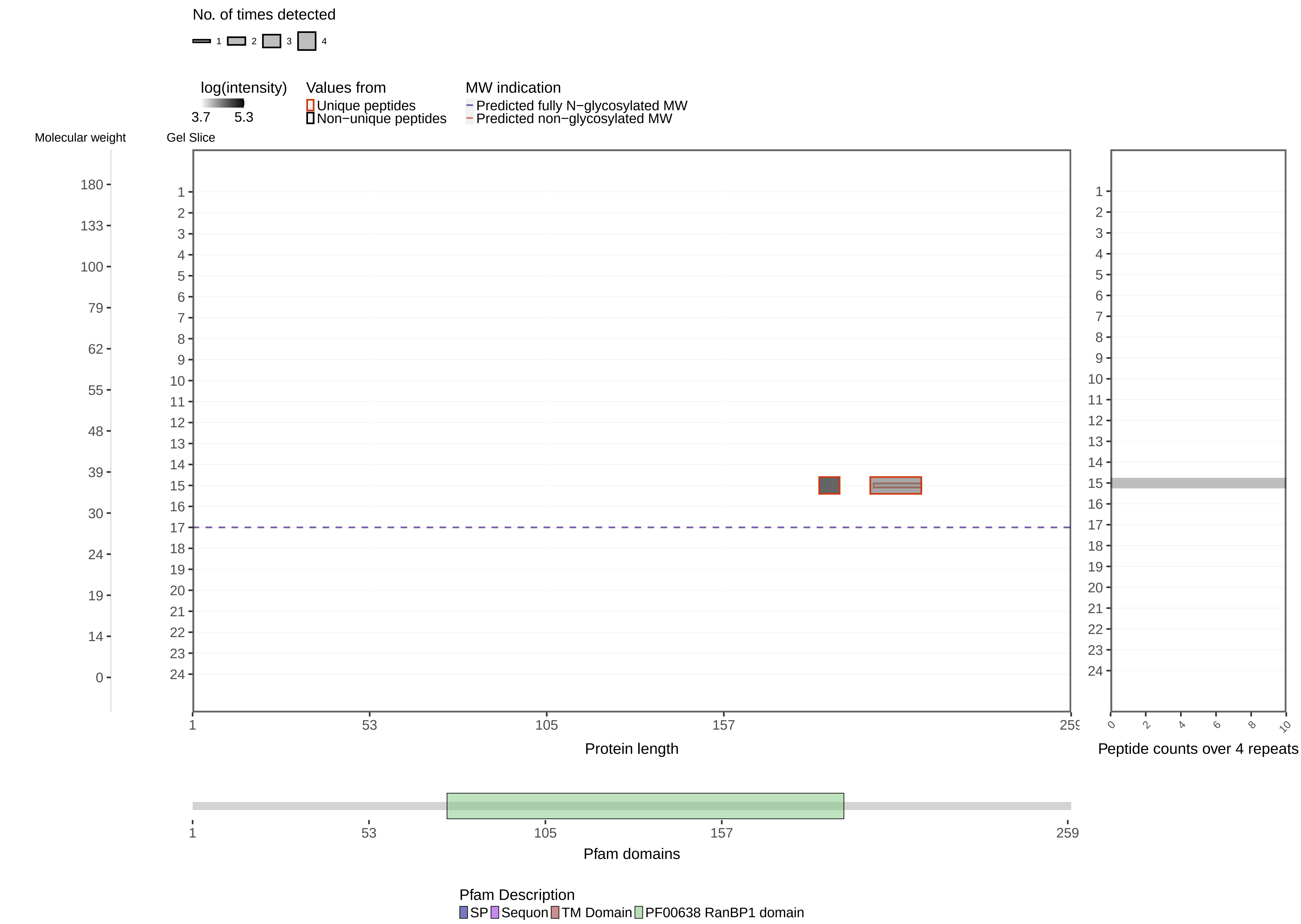


Protein sequence:

ETGKVRLVMROSKTLKICVNHLVIPTMTVOEHAGSEKSCVWHAADFADGELKDEFFCIRFASIENCKTFMETFREVAESOKKKKENEDASNAAGLLEKLSVEDKKSEEKSVEKAEEKTKEDDVKDAKPSDNAEKRDETASD*

NbL11g03790.1 ERBB−3_BINDING_PROTEIN_1 [MW = 42.75 kDa]

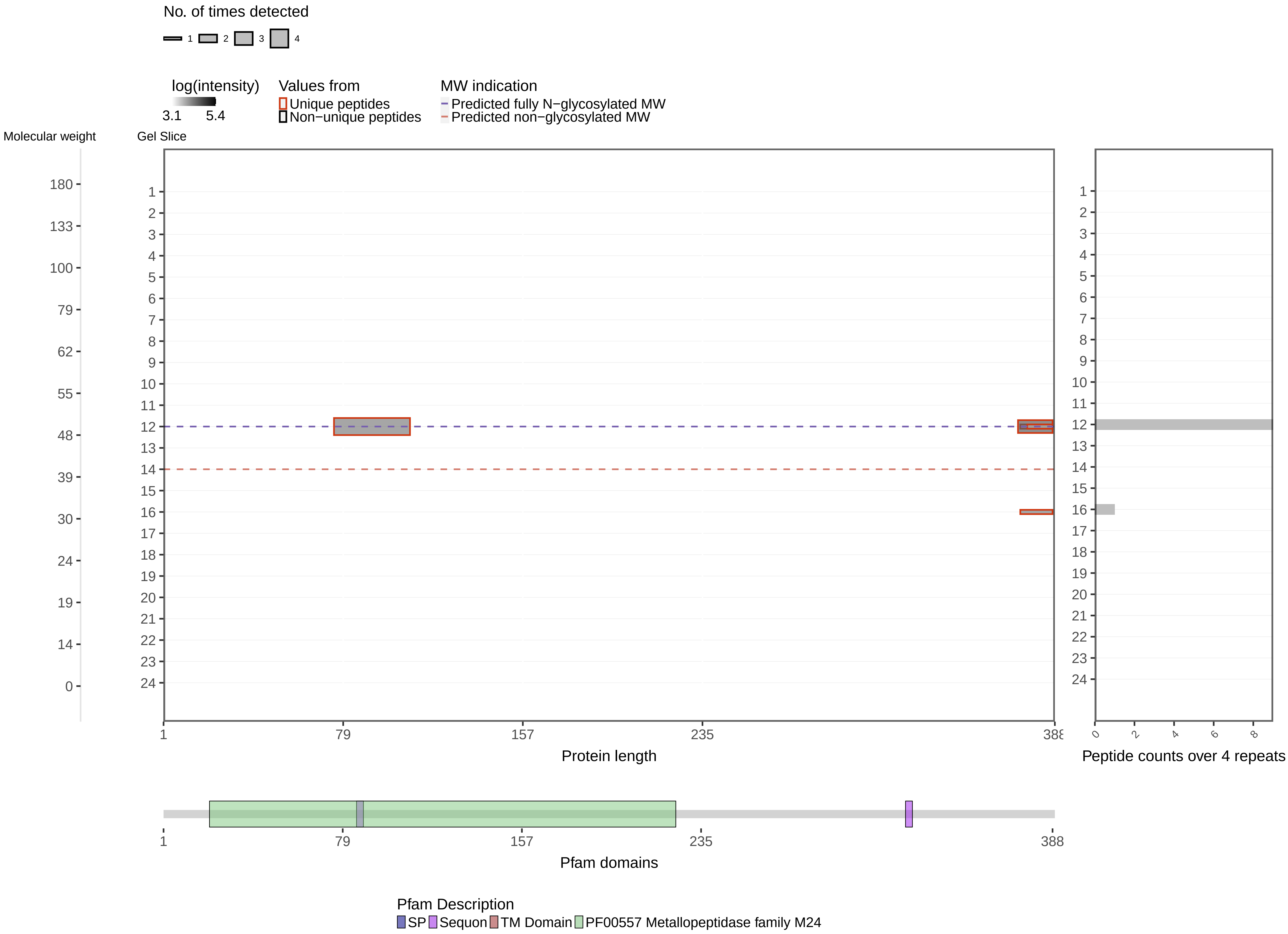


Protein sequence:

GVLSHQMKQFVIDGNKVVLSVTNPETRVDEAEFEENEVYSIDIVTSTGEGKPKLLDEKQTTIYKRAVDKSYNLKMKASRFIFSEISQKFPIMPFTARDLEEKRARLGLVECVNHELLQPYPVLHEKPGDLVAHIKFTVLLMPNGSDRVTSHALQELKTAKTIENEPEIKAWLALPVKTKKKGGGKKKKGKKGDKTEDSSQAEPTEG*

### NbL11g03670.1 probable_pyridoxal_5'−phosphate_synthase_subunit_PDX1 [MW = 33.12 kDa]

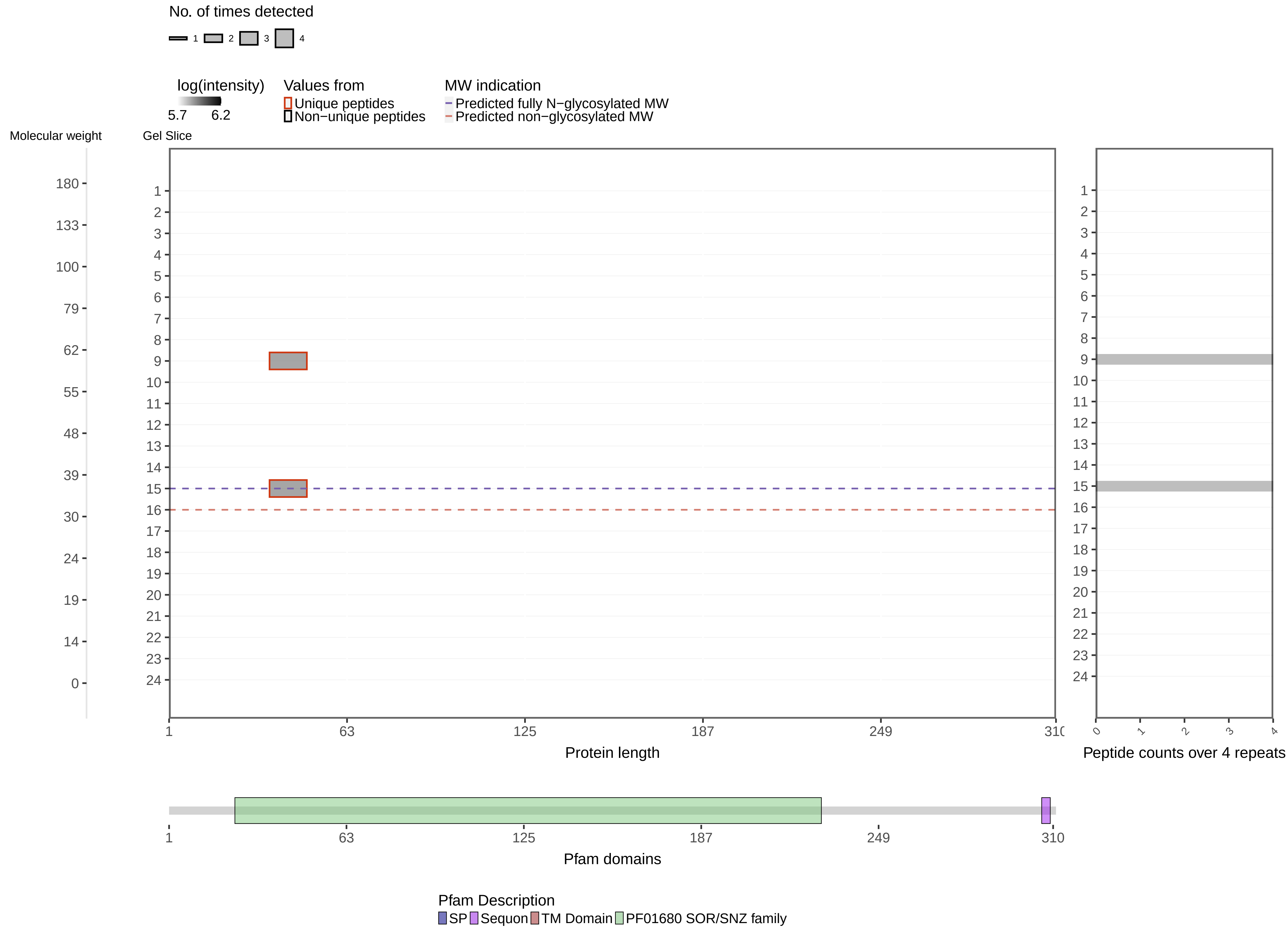


Protein sequence:

CGCRNLGEALRRIREGAAMIRTKGEAGTGNIIEAVRHVRSVMGDIRVLRNMDDDEVFTFAKKLQAPYDLVMQTKQLGRLPVVHFAAGGVATPADAALMMQLGCDGVFVGSGIFKSGDPAKRGRAIVQAVTHYSDPGLLAEISCGLGEAMVGINLDDKVERYANRSE*

### NbL11g03580.1 uncharacterized_LOC107793146 [MW = 12.17 kDa]

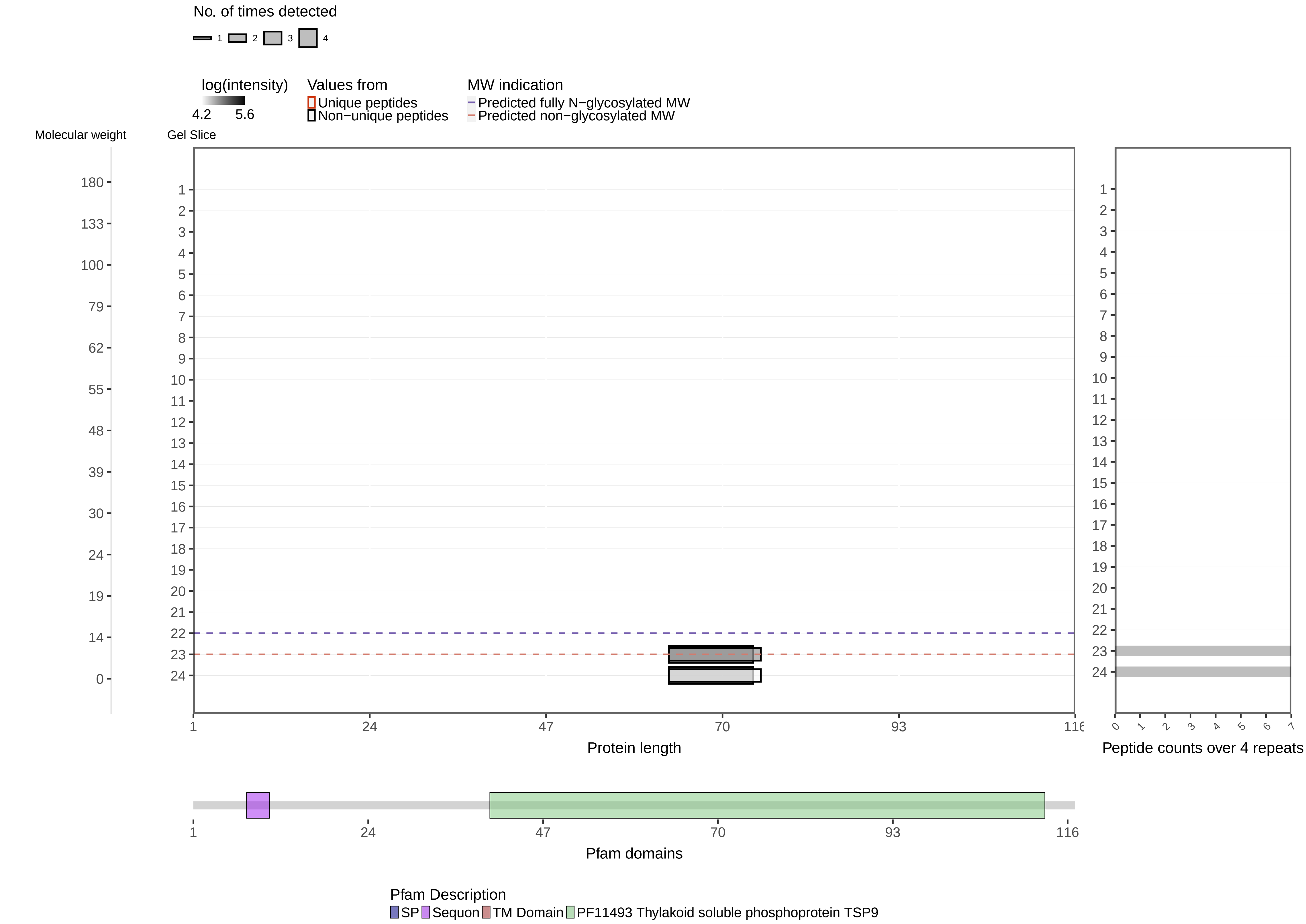


Protein sequence:

SSEEKSLLDFILGSLQKEDQLLEVDPILKKVEDKSVGTTSRGKNSVAVPPKKNSNGGGFGGFGGLFAKKD*

### NbL11g03560.1 beta−glucosidase_BoGH3B−like [MW = 65.76 kDa]

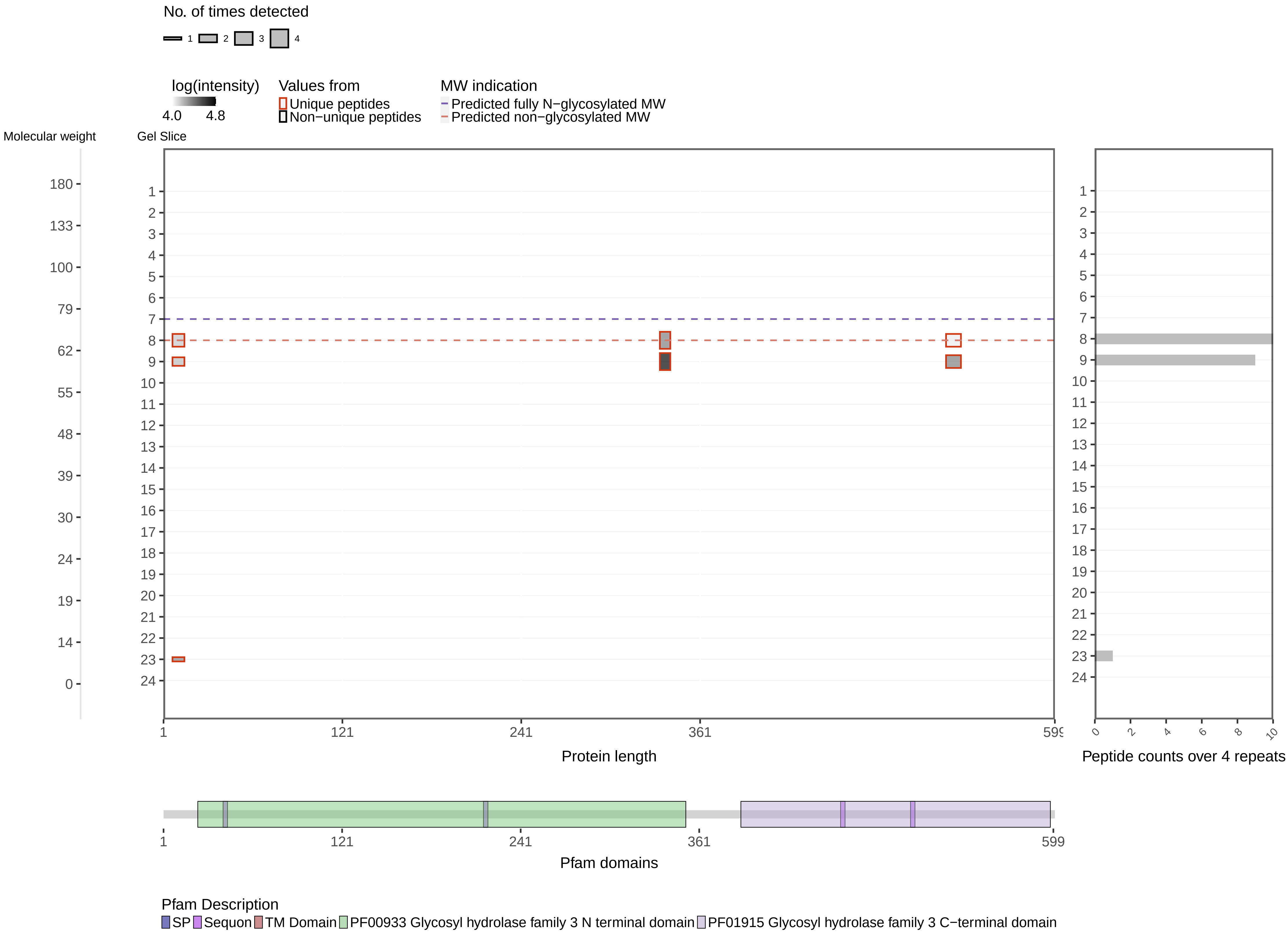


Protein sequence:

ITDWEALDRLTDPHGSDYROSIKSTINAGIDMVMVPFRYELFLEELLSLVKSGEISMTRIDDAVERILRVKFVAGLFEHPFTDRSLIELVGCEAHRELAREAVRKSLVLLKNGKDPKKPFLPLDRNAKKILVAGTHADDLGYOCGGWTATWTGESGRITVGTTLLEALRKLIGNGTEIVFEPNPTTETFANOEFSFAIVAIGEGPYCETGGDDPELKIPFNGTEIATFVADRVPTVTILISGRPMVLEPSLFEKVDAFVAAWLPGTEGTGITDVLFGDYPFOGKLPVTWFKTVDOLPMH[illegible]

### NbL11g03500.1 acyl−coenzyme_A_thioesterase_13−like [MW = 17.11 kDa]

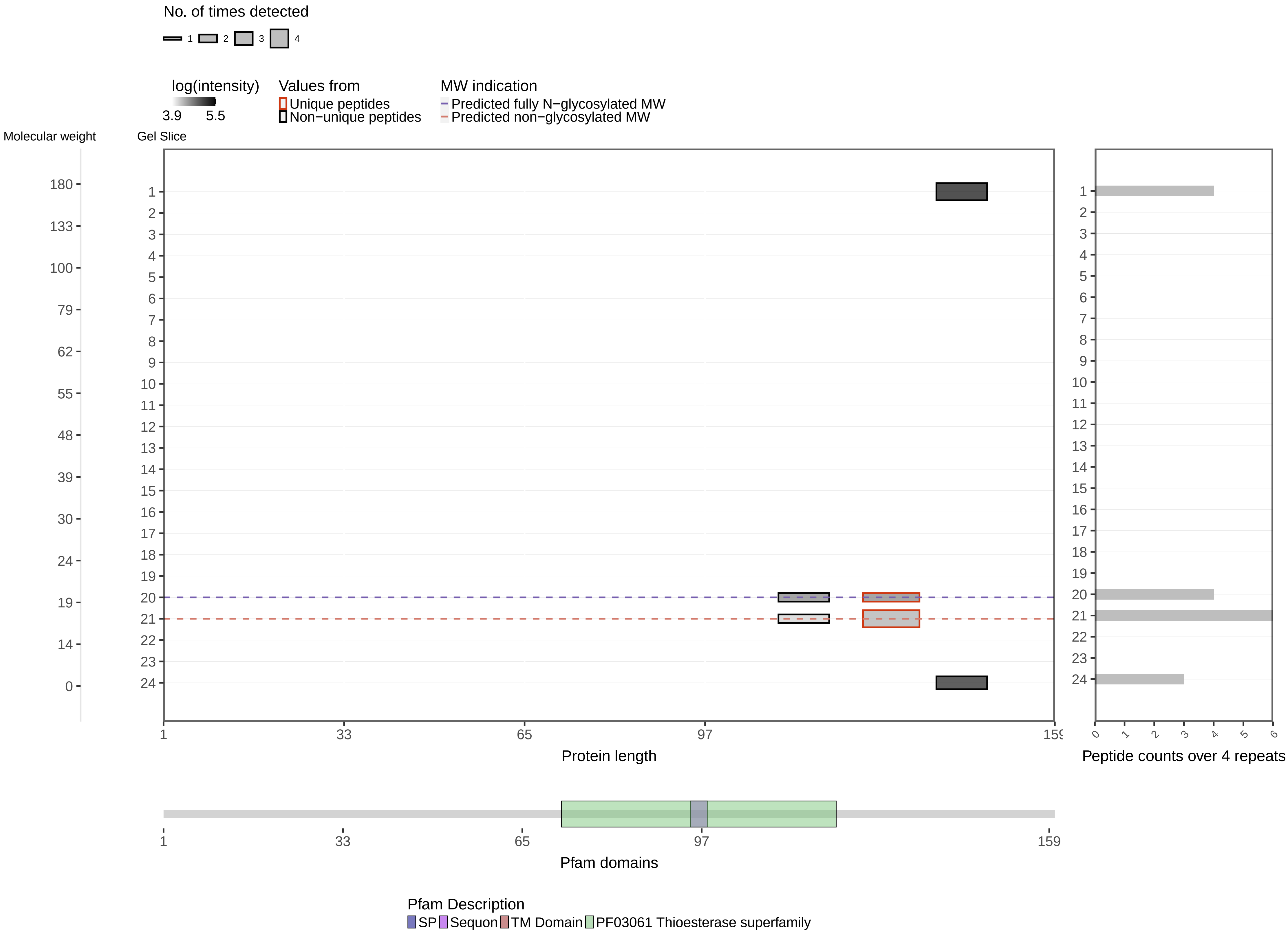


Protein sequence:

EGKLASGAIANLVDEVGGAVVFVEGLPMNVSVDMSISFLSSAKADDELEIIGRVLGQKGGYSGTNVLMKNKATGELIAEGRHSLFGKHASKM*

### NbL11g03460.1 proliferating_cell_nuclear_antigen [MW = 29.23 kDa]

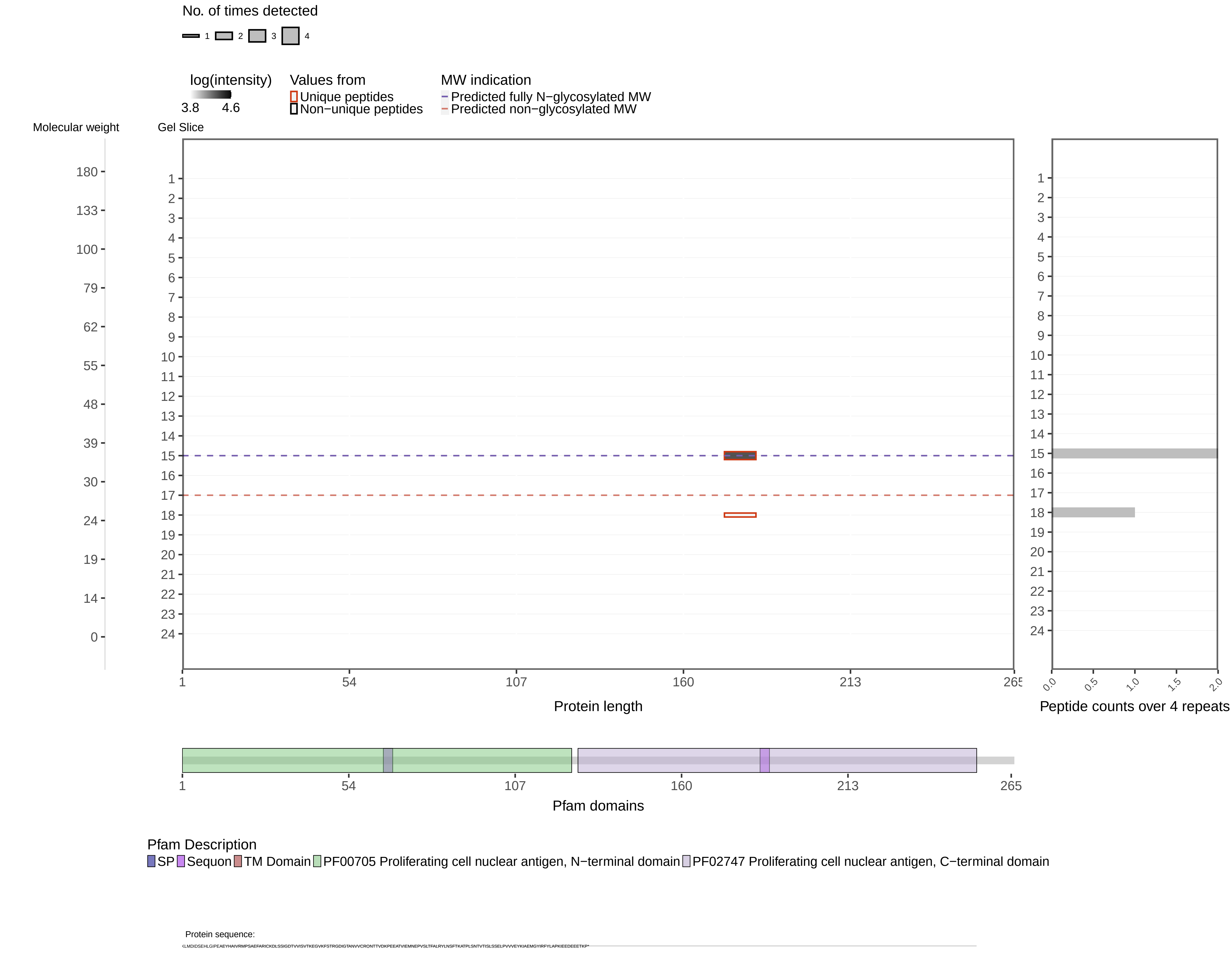

### NbL11g03450.1 cytosolic_enolase_3 [MW = 51.62 kDa]

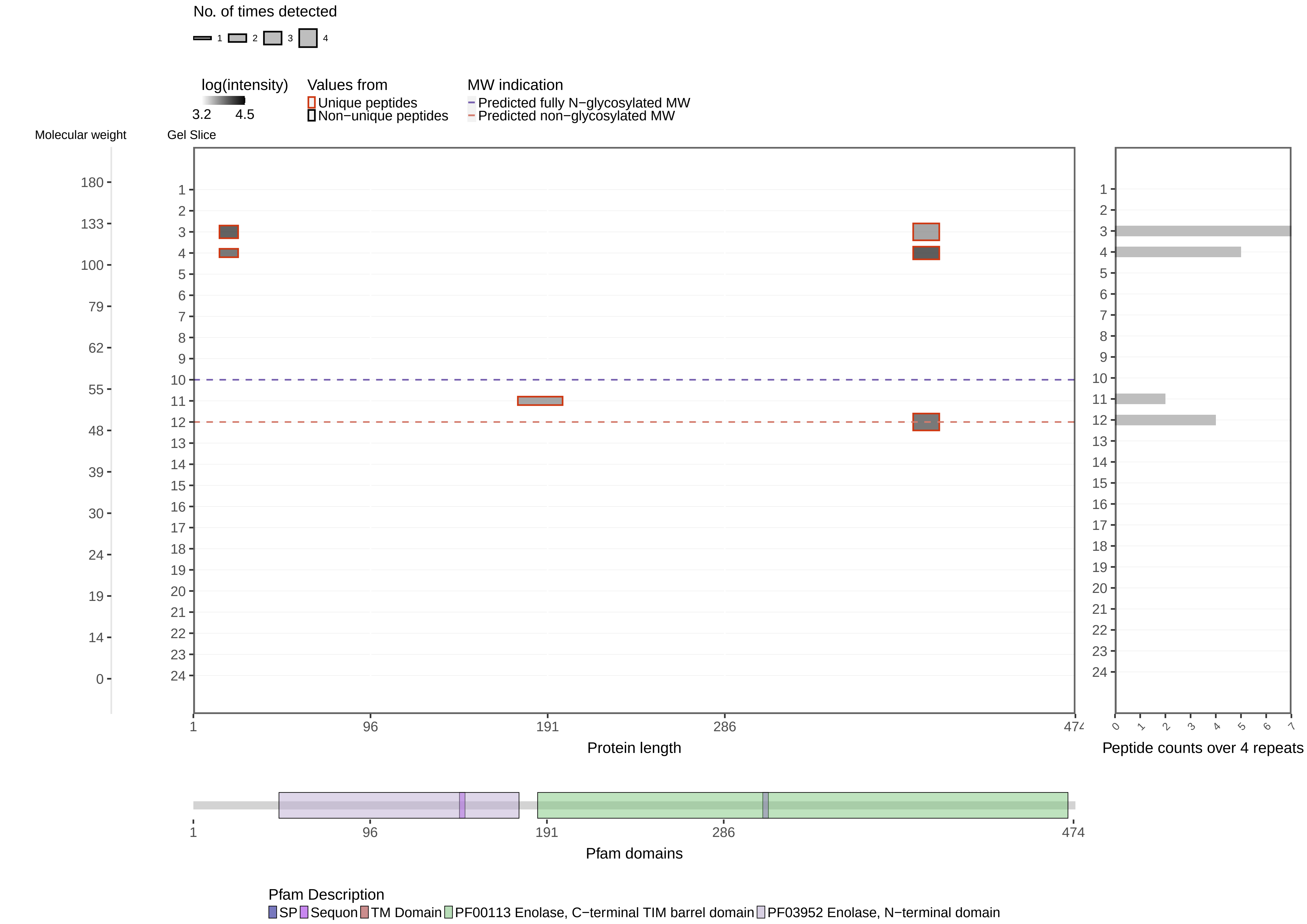


Protein sequence:

IGSETYHHLKVVITEKYGARGCNVGEDGGFAPDISSLRDGLDLVLEAIGRTGYKEKIKIAIGVAATEFCIGTKYDLDFKTPNKSGQNFKSGQDMIDMYKELCADYPVVSIEDPFDKEDWEHVKYFSSLGMCQVVGDDLLMSNPKRIERAIQENACNALLLKVNQLGTVTEAIEVVKMAKDAQWGVVISQRSGETEDSFIADLSVGLATGQIKAGAPCRGERLAKYNQLLRIEQELGDQAFYIGEKWRT*

### NbL11g03440.1 uncharacterized_LOC109226983 [MW = 17.6 kDa]

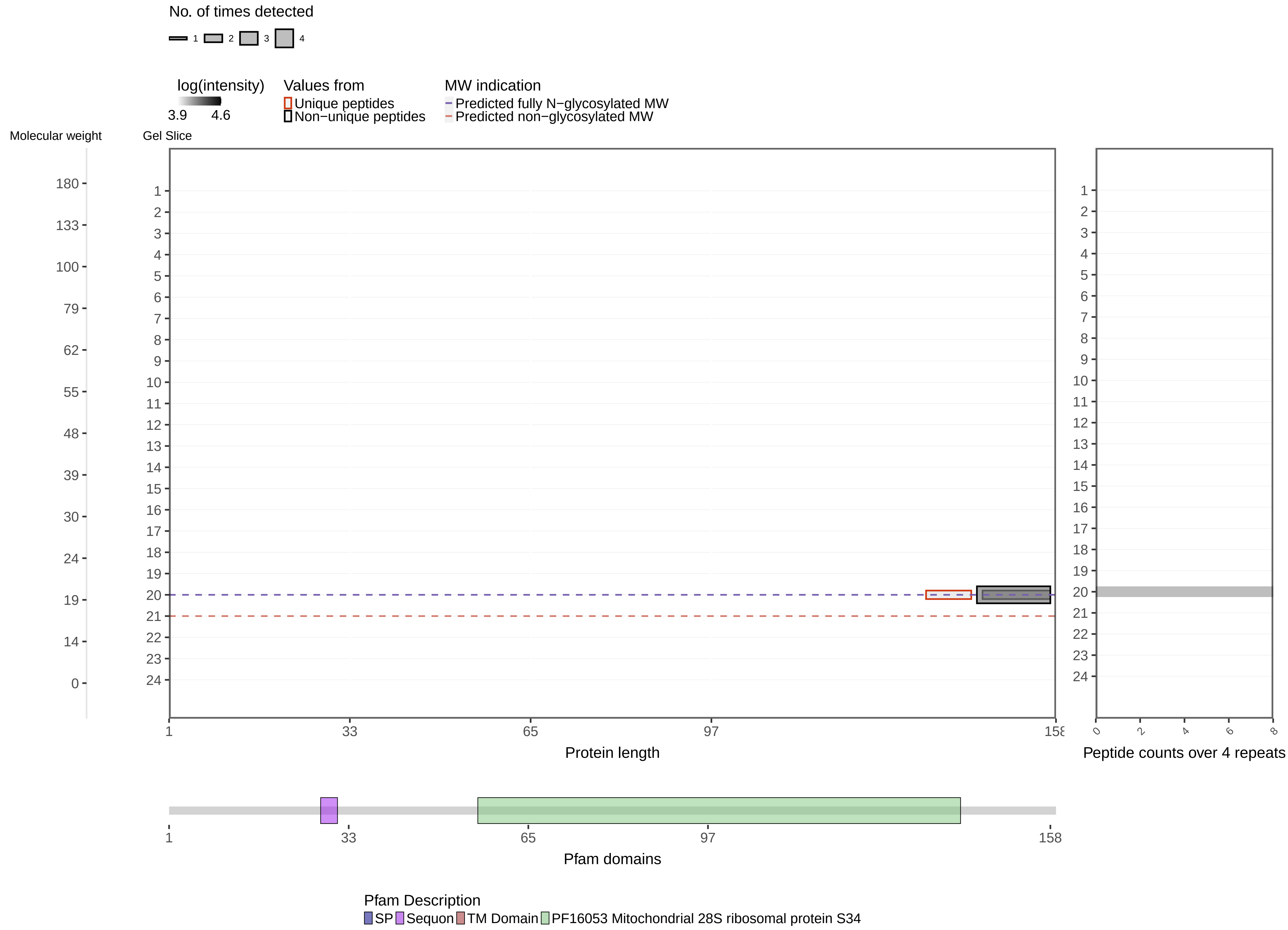


Protein sequence:

NWGIGYHMAKTHWTGISYEITKINLYKDGRHGKAWGLAYKDGLPIADAPKKMSGVHKRCWKYIANFKKAEGSLDQIKKAEESTPNSDVEAA*

NbL11g03380.1 phosphoserine_phosphatase_chloroplastic [MW = 31.94 kDa]

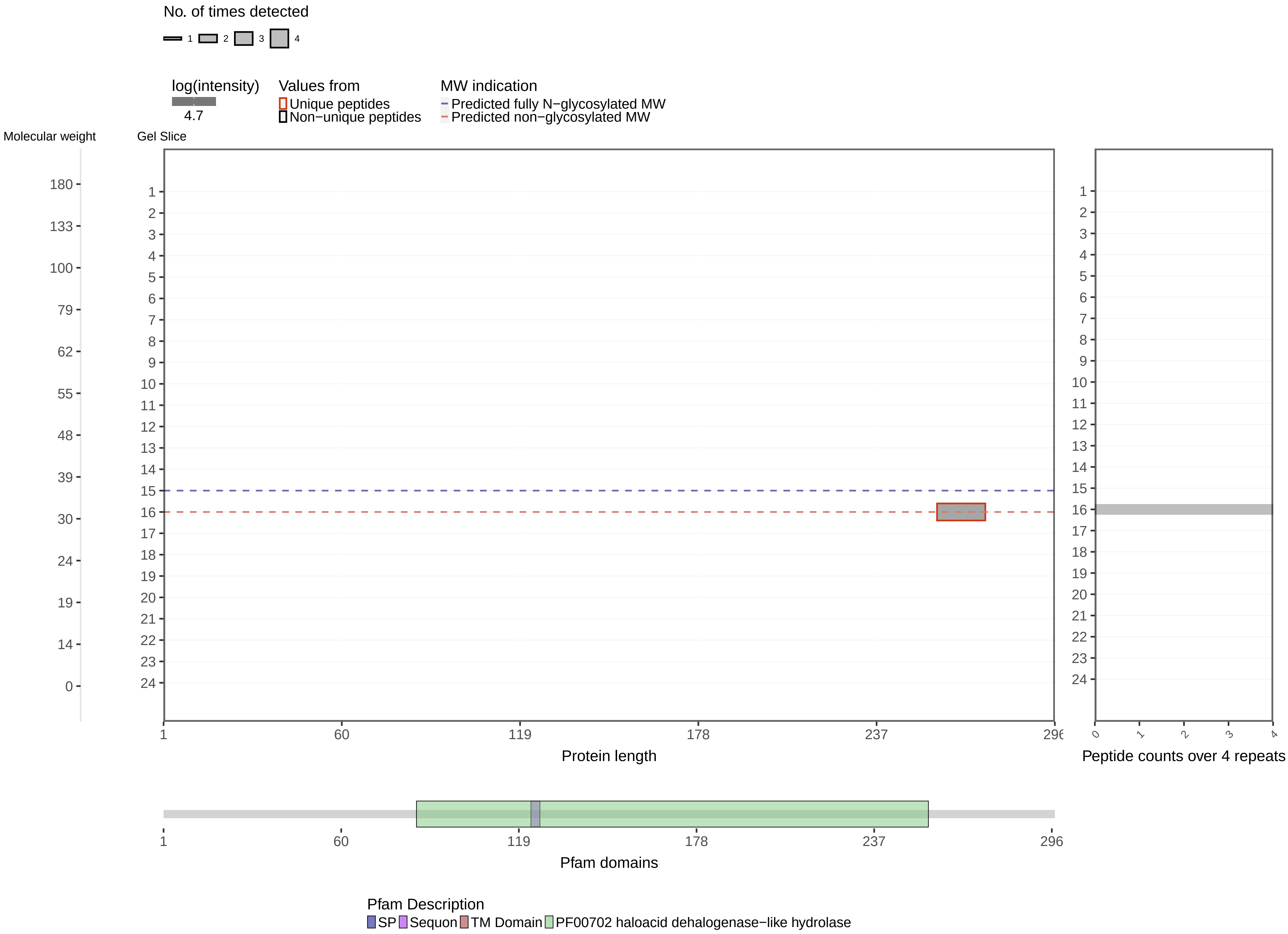


Protein sequence:

ARLSLINPSLSQLQDFLKRPPRLSPGIDLLVKKLKDKKKDVYLVSGGFRQMINPIASILGIPLENIFANQMLFGSDGECAGFDKNEPTSRSGGKPTAVQQIKKACGYKSVVMIGDGATDLEARMPGGADLFICYGGVQQREPVAAKADWLVFNFKDLINSLE*

### NbL11g03300.1 DNA−directed_RNA_polymerase_II_subunit_RPB7 [MW = 19.55 kDa]

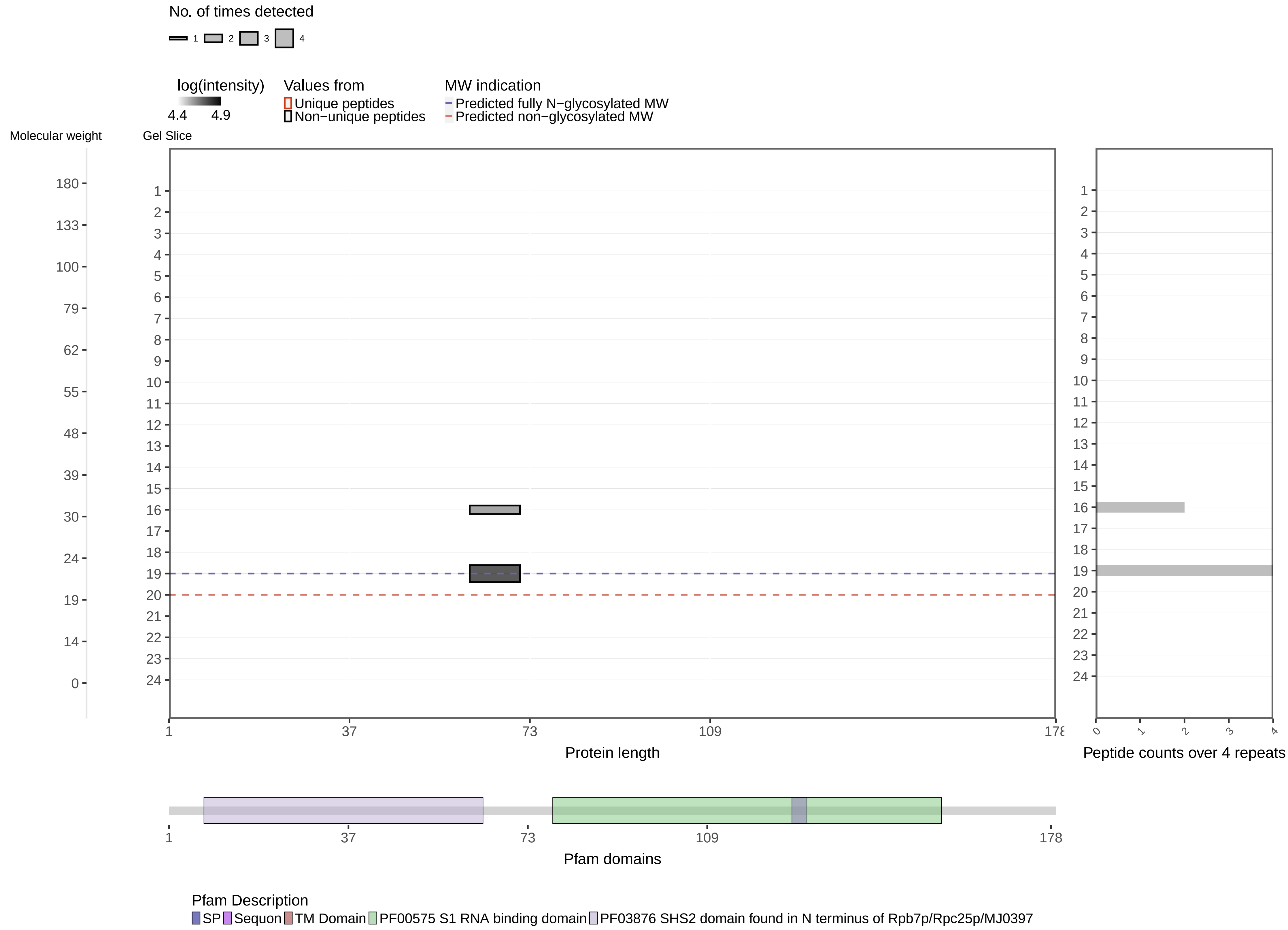


Protein sequence:

VVFRPFKGEILEAVVTVVNKVGFFAEAGPVQIFVSNHLIPDDMEFQSGDVPNYTTSDGSVKIQKESEVRLKIIGTRVDATEIFCIGTIKDDFLGVISDPGANA*

### NbL11g03240.1 carbamoyl−phosphate_synthase_large_chain_chloroplastic−like [MW = 131.8 kDa]

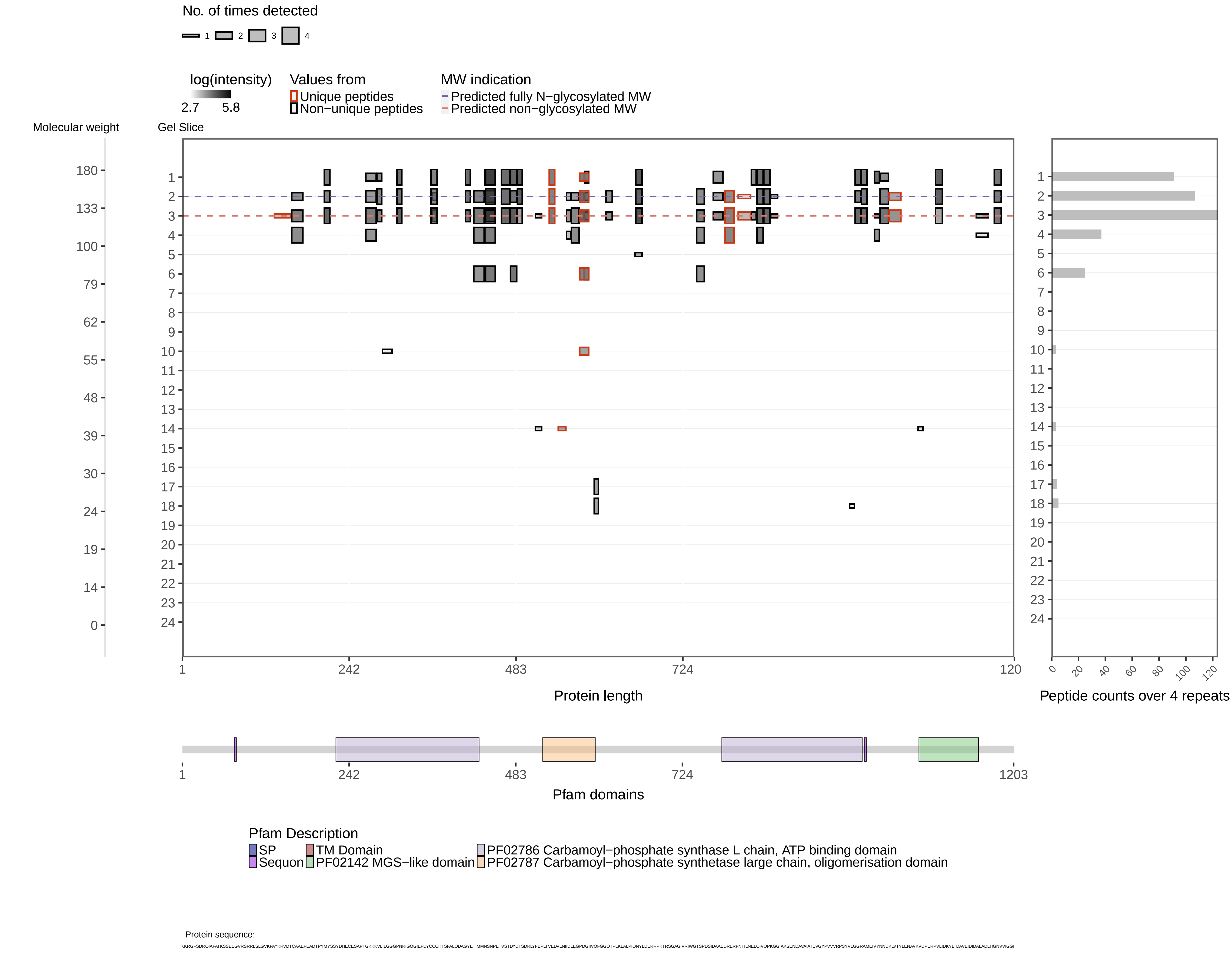

### NbL11g03170.1 L−ascorbate_peroxidase_1_cytosolic−like [MW = 27.94 kDa]

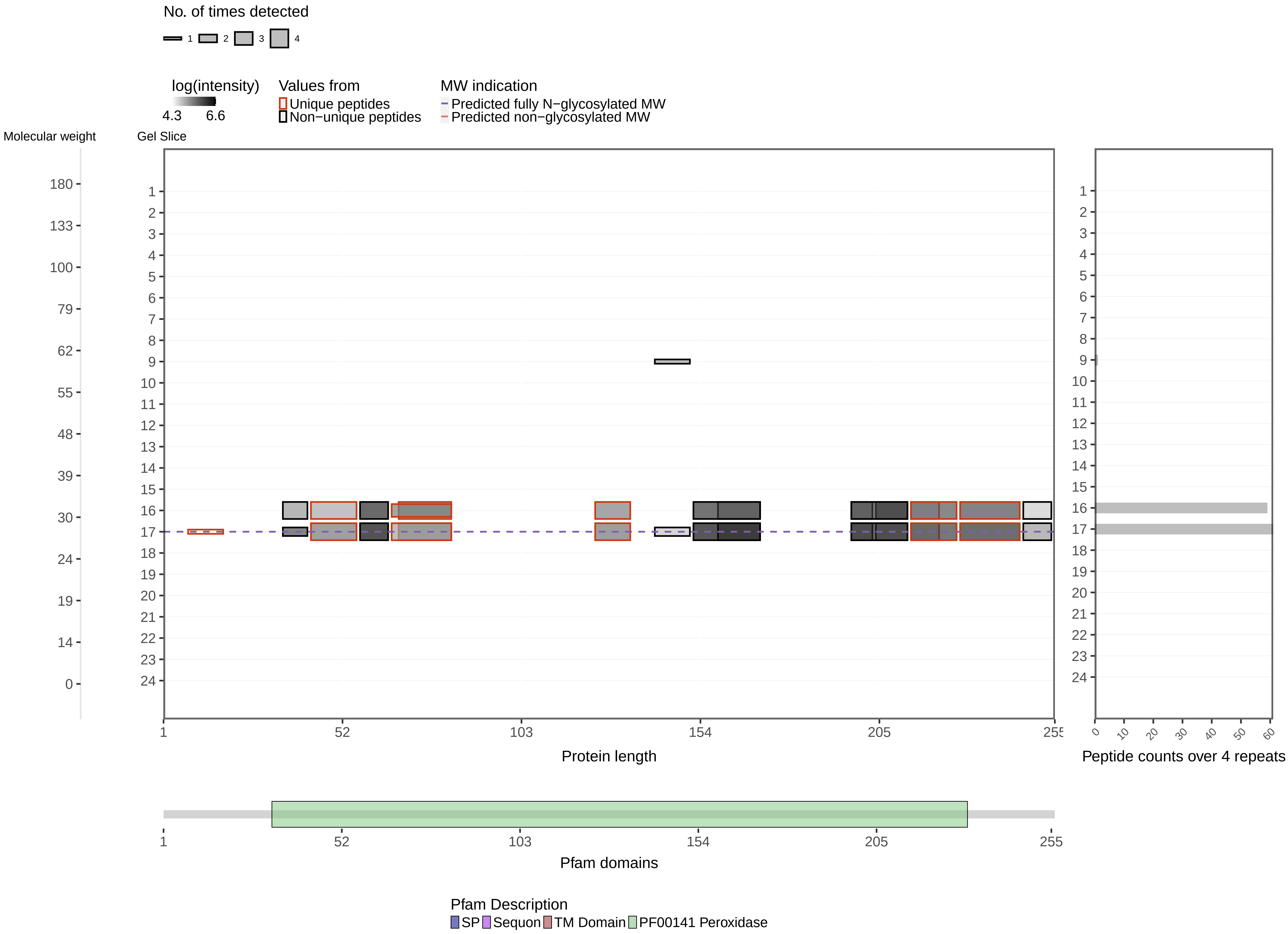


Protein sequence:

PDVPFHPGREDKTEPPLEGRLPDATKGSDHLRDVFVKQMGLSDKDIVALSGGHTLGRCHKERSGFEGPWTANPLIFDNSYFKELLSGEKEGLLQLPSDKALLSDPVFRPLVEKYAADEDAFFADYAEAHMKLSELGFAEA*

### NbL11g03120.1 uncharacterized_protein_YKR070W [MW = 45.34 kDa]

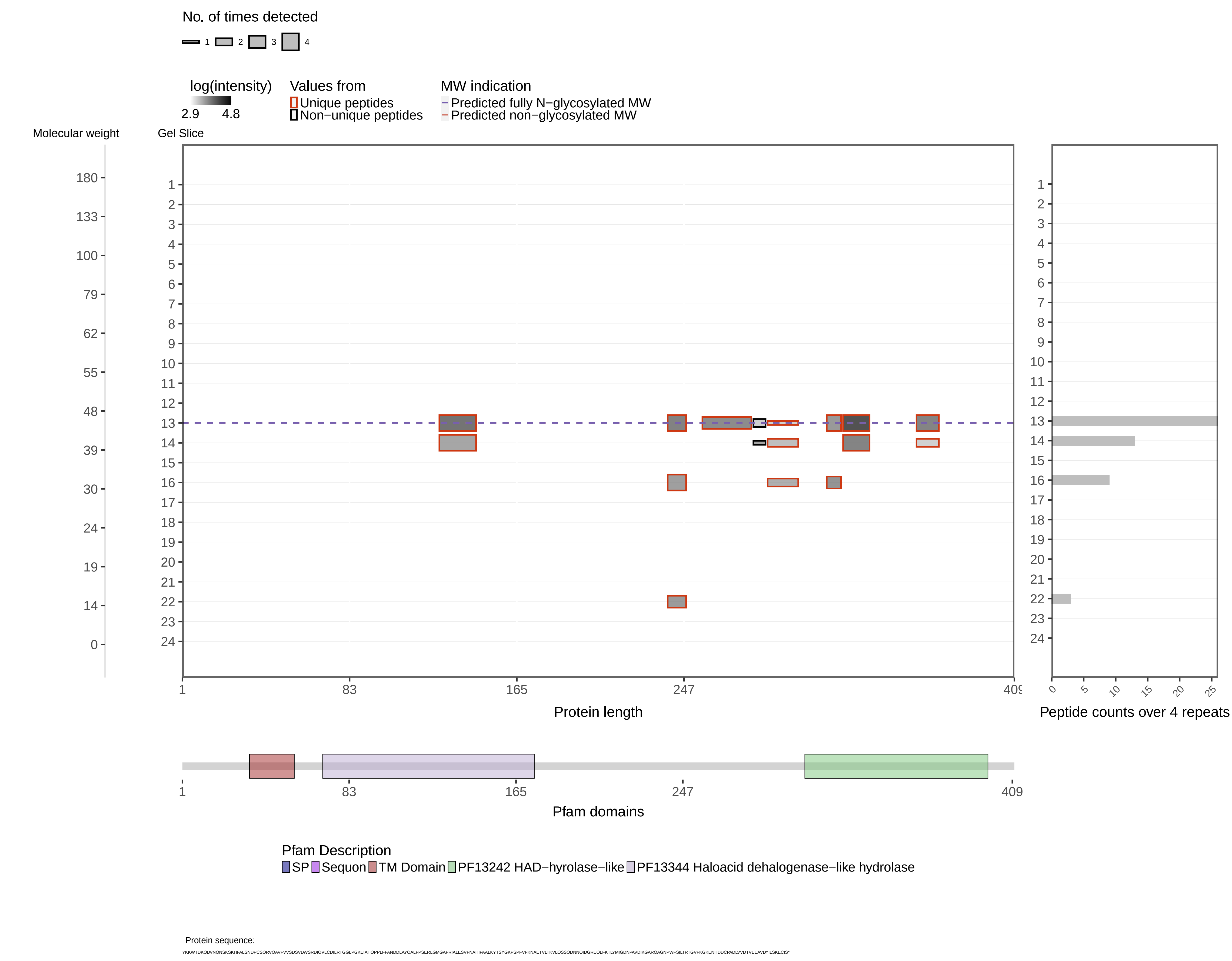

### NbL11g02980.1 aquaporin_TIP1−3 [MW = 25.89 kDa]

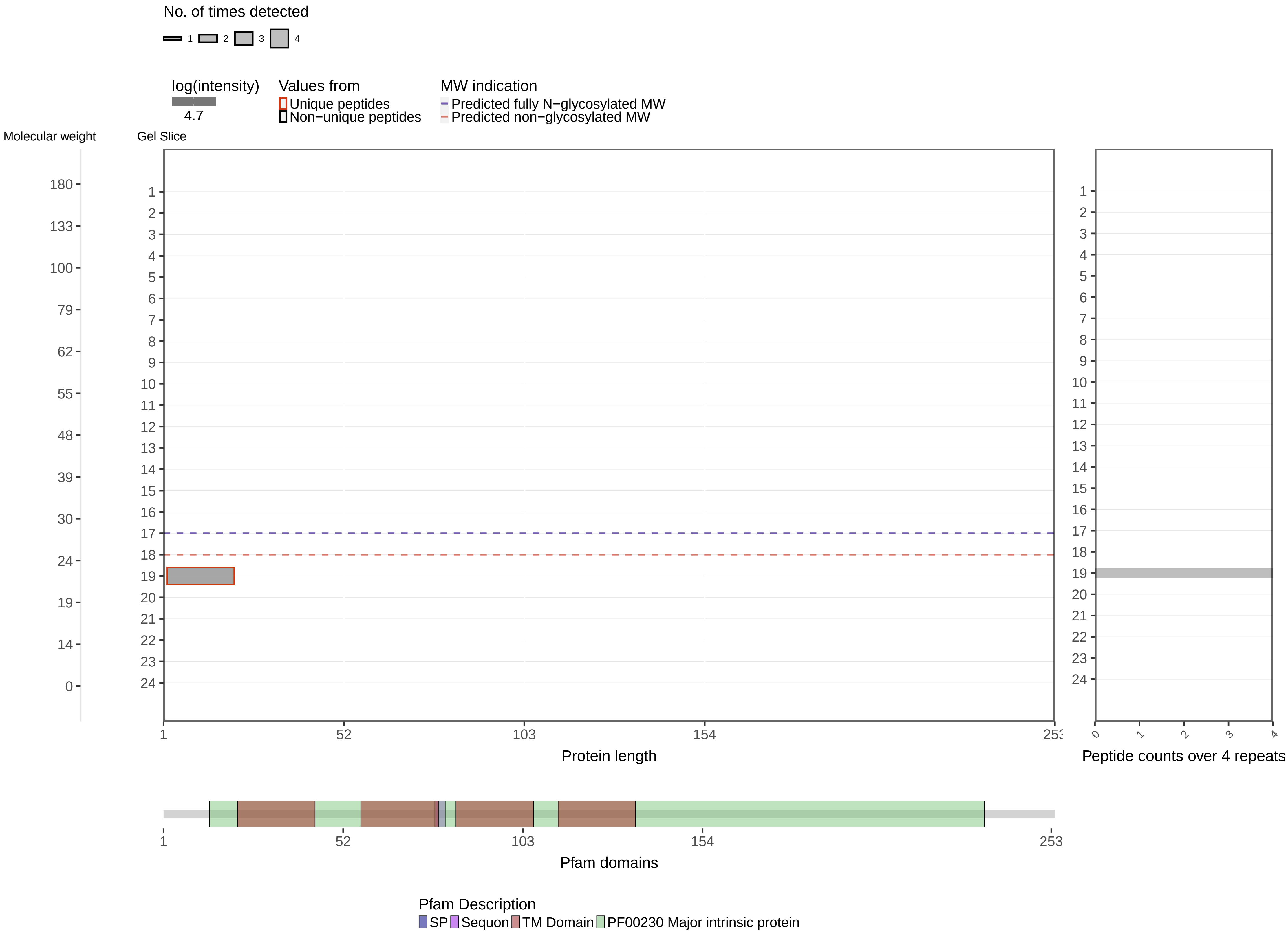


Protein sequence:

SVVACVLLKFATGGLETSAFALSTGVTPWNAVVFEIVMTFGLVYTVYATAVDPKRGNLGIIAPIAIGFIVGANILAGGAFDGASMNPAVSFGPAVVSWTWNCHWVYWLGPFVGAAIAALVYEIIFIGDNTHEOLPTADY*

NbL11g02900.1 uncharacterized_LOC104240093 [MW = 89.21 kDa]

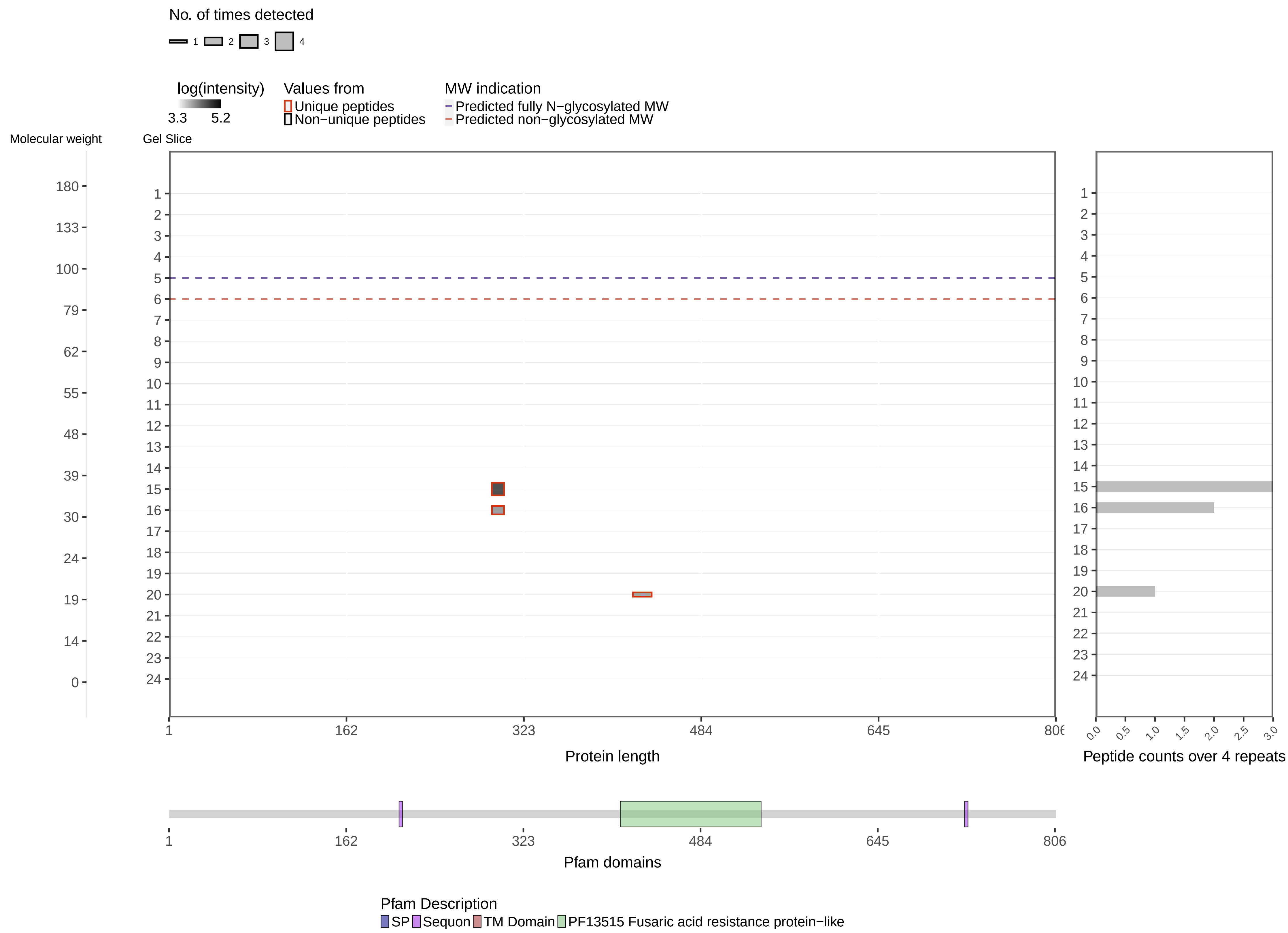


Protein sequence:
ITINNKRFMEAFKCSLSLGLAILFGSIYSKENGFWAGLPVAVSLAGAREASFKVANYKAQGTVLGSIYGILGCFVFEKYVQIRFLSLLPWFIVSSFLRQSKMYGQAGGISAIVGAVLILGRKGFGPPSEFAIARITETFIGLSCSIMVEILLQPTRASTLAKLQLSKSFEILHECIGSISFGSLGNLVETQNKLKFHVNKMGKFIGEAEAEPNFWFVPFNSACYRKLMGSLSKMVDCLYFGSQAILFLEQESGGIDNFVWKETLNKLDADFMLFKDFIGSFMKCFEEVSLVKSLAVLDK

NbL11g02880.1 uncharacterized_protein_Mb0911c [MW = 15.36 kDa]

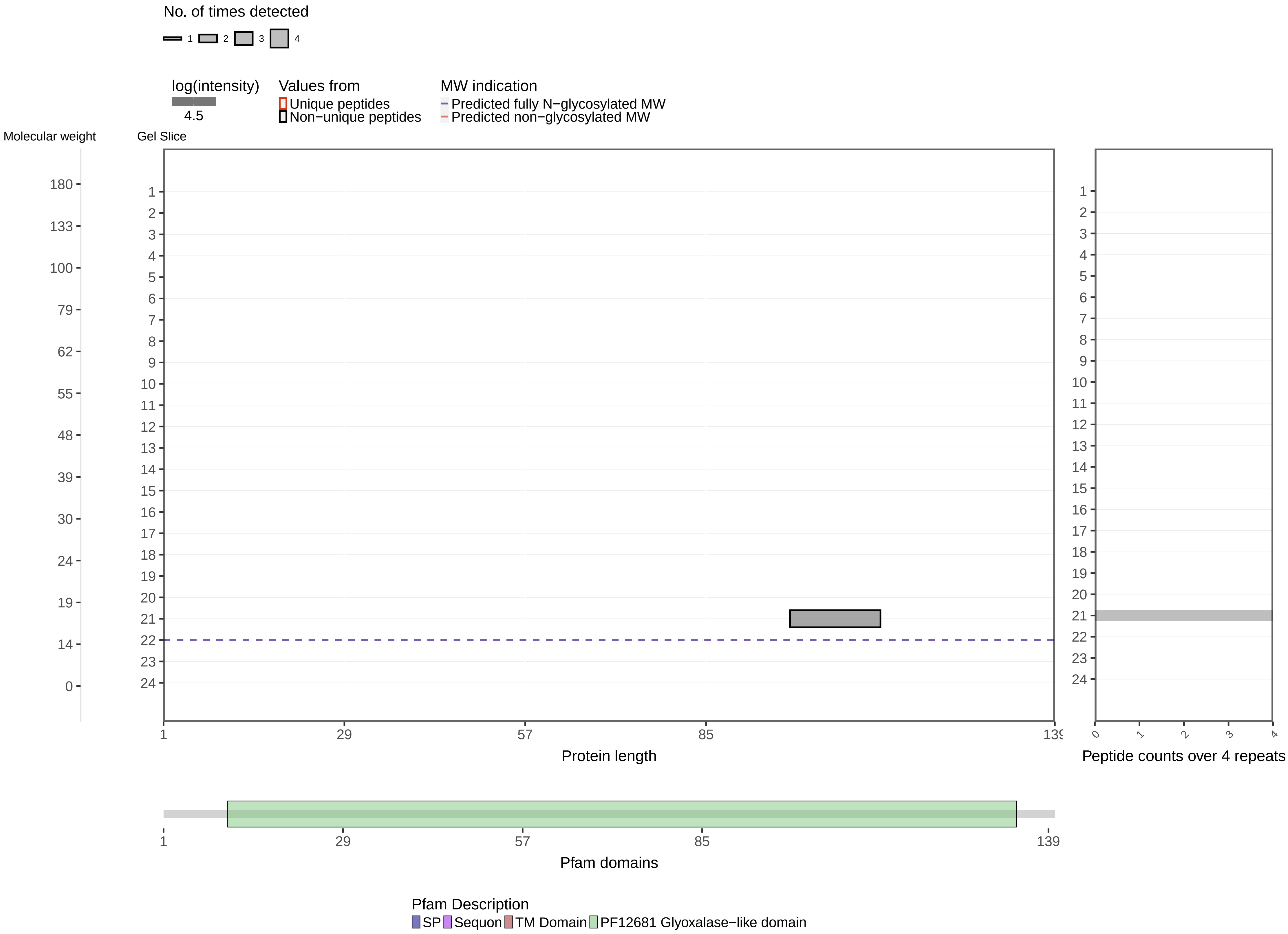


Protein sequence:

IHQHETDDLTGQVQTPHSRGDRQPVELCFDYVDIDAAYKRAVENGAVAVSQPEEKKWGQKVGYVRDLDGNIVRLGSHVDPK*

NbL11g02850.1 heat_shock_cognate_70_kDa_protein_1−like [MW = 61.37 kDa]

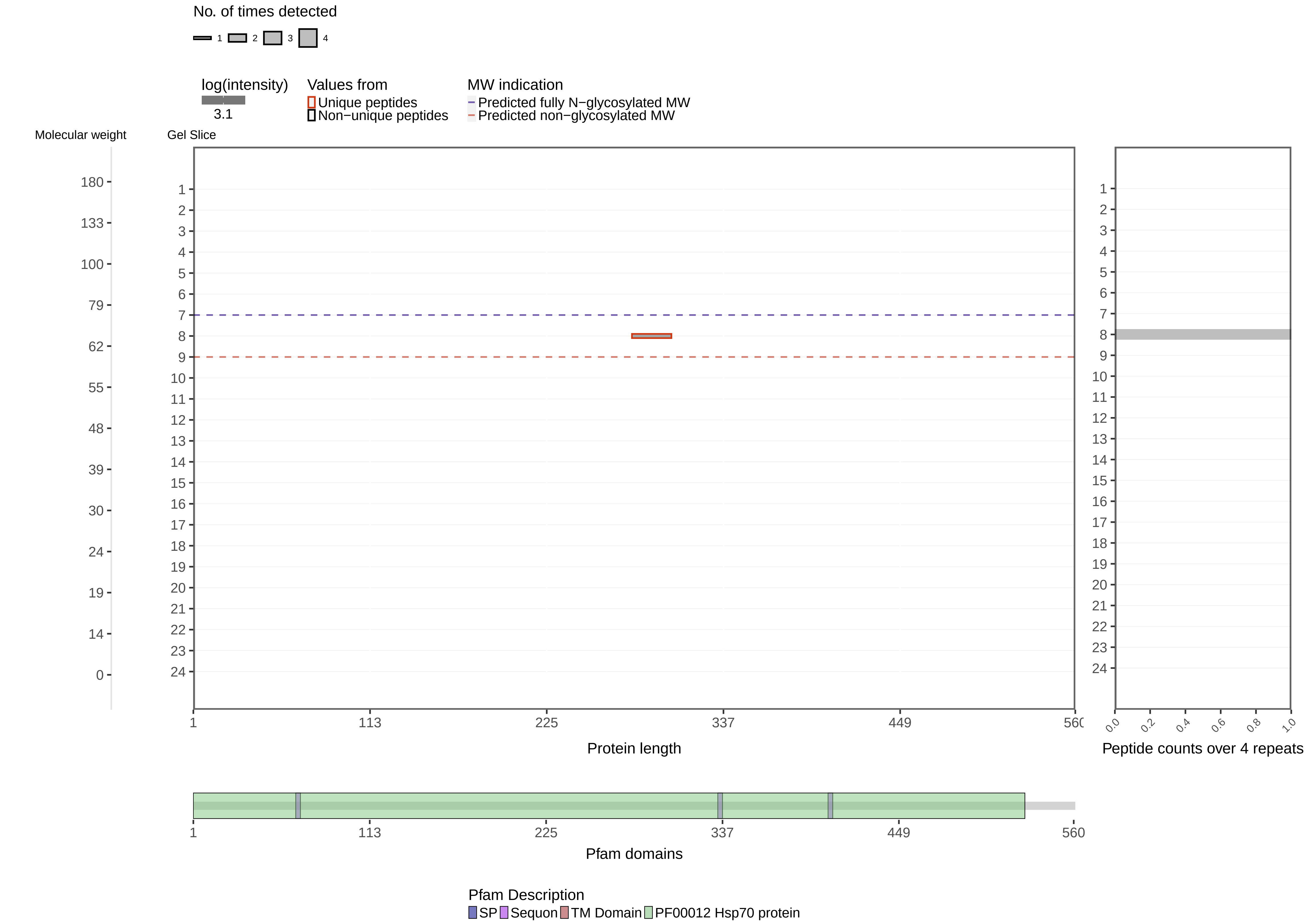


Protein sequence:

QDFFNGKELCKSINPDEAVAYGAAVQAAILSGEGDEKVQDLLLLDVTPLSLGLETAGGVMTVLIPRNTTIPTKKEQVFSTYSDNQPGVLIQVFEGERARTRDNNLLGKFELSGIPPAPRGVPQITVCFDIDANGILNVSAEDKTTGQKNKITITNDKGRLSKEEIEKMVQEAEKYKAEDEDHKKKVEAKNALENYAYNMRNTVKDEKIGSKLSTDDKKKIEDAIDQAISWLDSNQLAEADEFEDKMKELEGICNPIIAKMYQGAGGDAGAAMDDDAPPAGASSAGPKIEEVD*

### NbL11g02790.1 actin−like [MW = 41.73 kDa]

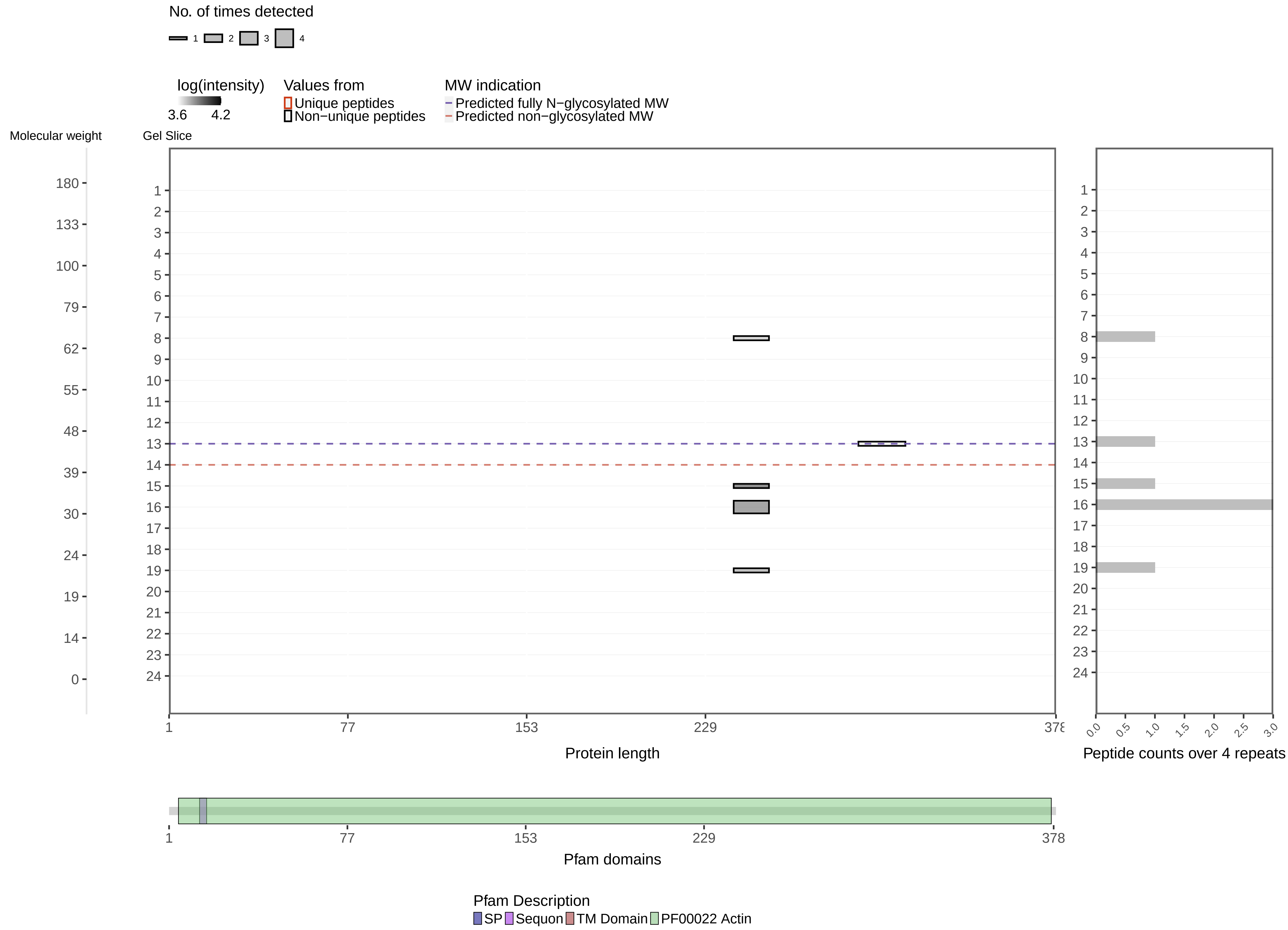


Protein sequence:

PHAILRLDLAGRDLTDHLMKILTERGYSFTTSAEREIVRDVKEKLSYIALDFEOELETSKTSSSVEKSYELPDGOVITIGSERFRCPEVLFOPSMIGMEAAGIHETTYNSIMKCDVDIRKDLYGNIVLSGGSTMFNGIADRMSKEITALAPSSMKIKVVAPPERKYSVWIGGSILASLSTFOOMWIAKAEYDESGPSIVHRKCF*

### NbL11g02430.1 nudix_hydrolase_19_chloroplastic [MW = 48.48 kDa]

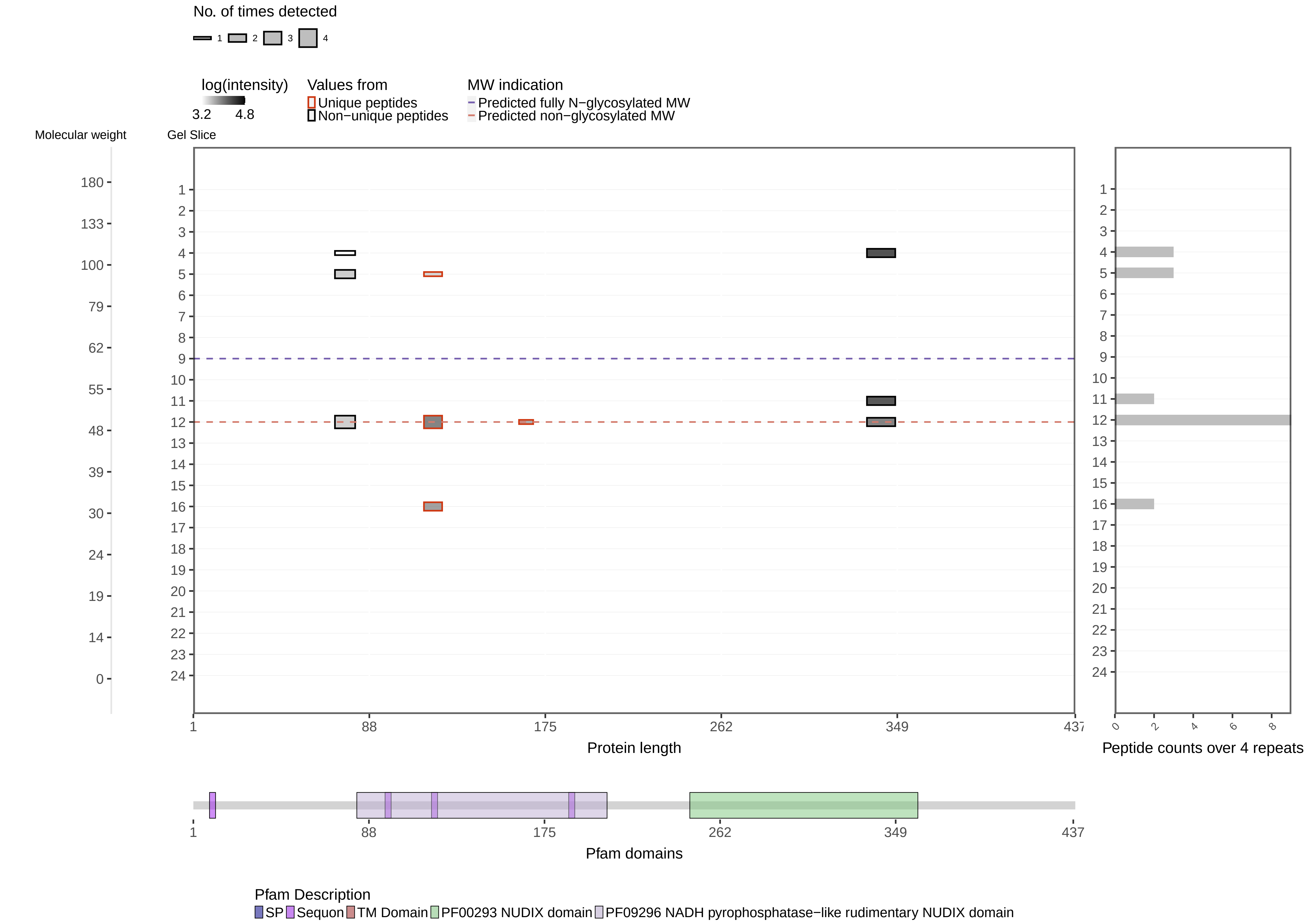


Protein sequence:

ITVSRFCGFCGGKNNLIDAGRRKQCSNELCKKKIYPRVDPVVIMLVIDKENDRVLLSRQSRFVPRMWSCLAGFMEPGESLEEAVRRETWEETGIEVGQVVYHSSQPWPVGPSSMPCQLMVGFFAYAKSLDINVDKVELEDAKWHGREDVKKALTFTEYKKAQRTAAGKVDQMCKGVERGQSLSSDFNVESGELAPMFIPGPFAIAHHLISSWVNGAEAQVKQLASSFSNL*

NbL11g02380.1 ruBisCO_large_subunit−binding_protein_subunit_alpha−like [MW = 62.37 kDa]

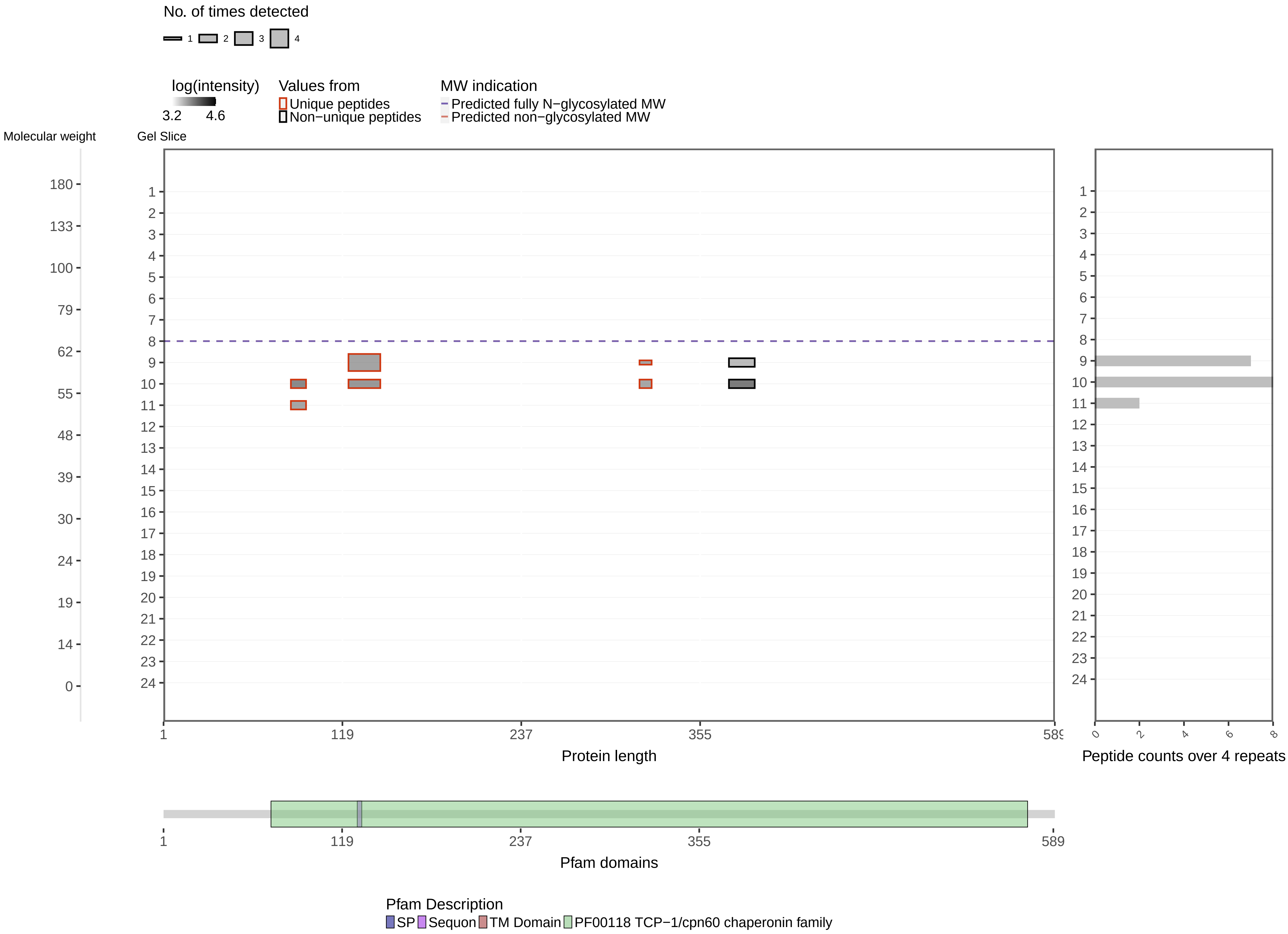


Protein sequence:

PLLEKSTELRAPLLIVAGDVTGEALATLVVNKLRGVLNVAAIRAPGFGQRRKALLQDIAIVTGAEYQATELGMLVENTPVEALGFARKVTISKDSTIILADDASKDEIQARISQLKKELDMTDSISDSEKLAERIAKLCGGVAVIKVGAATEAELEDRKLRIEDAKHATFAAIEEGIVPGGGAALVHLSTYVHAIKDNLEDPDEKLGADIMQKALVAPASLIAQNAGVEGEVVVEKIKASKWKMGYNAMTDKYENLVEAGVIDPAMVTRCALQNAASVAGVVLTTQAIVVDKHTPDVPAFA

### NbL11g02370.1 (R)−specific_enoyl−CoA_hydratase−like [MW = 18.36 kDa]

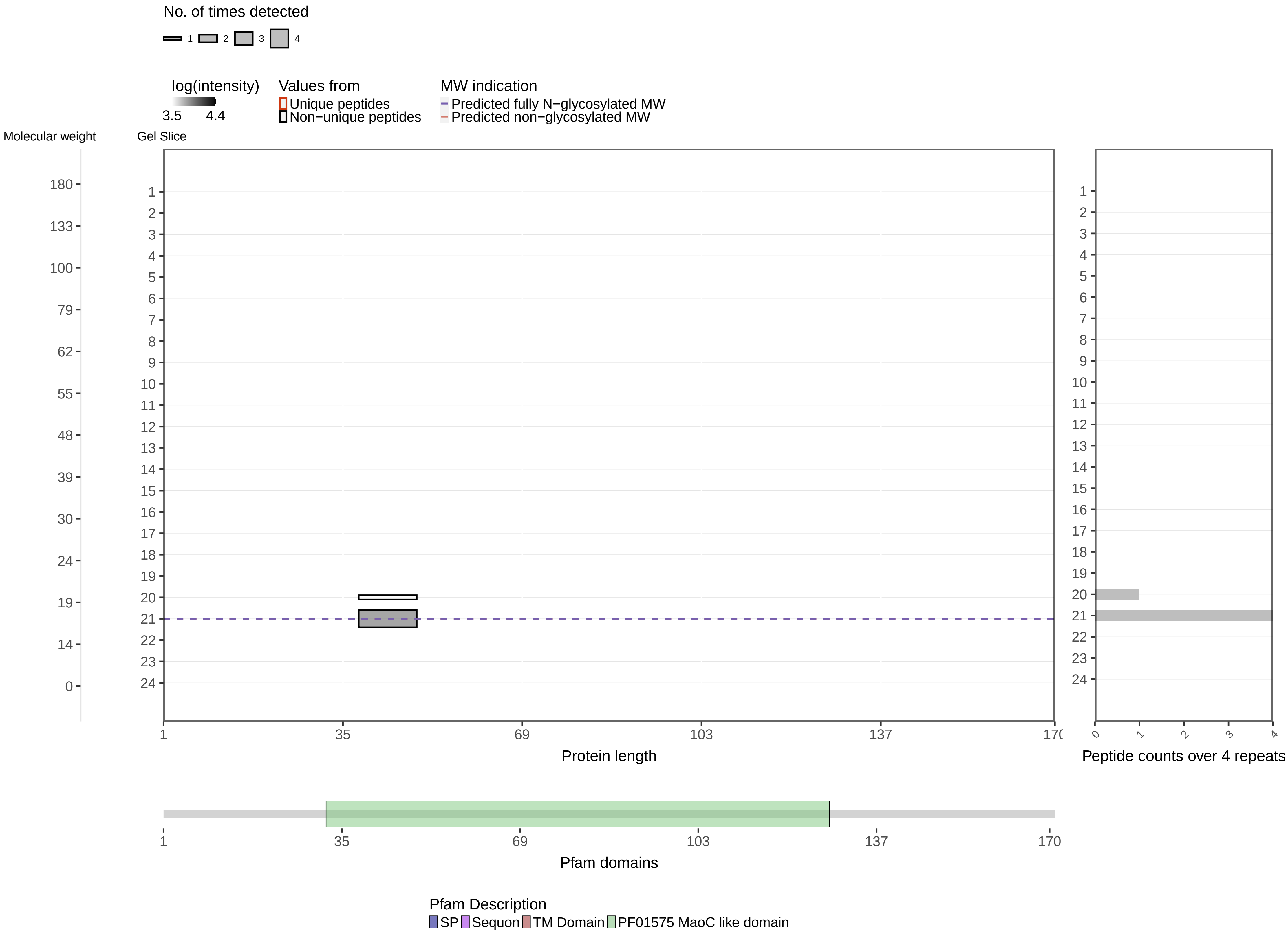


Protein sequence:

ORLVPGMLVASLFPRIIAAHFPGALYVSQTLHFKLPVYIGDEIIAEVQASSIRQMKNKYIAKLSTKCFKCDGPLVIDGEATAILPSLVMEPTHLASTSS*

NbL11g02310.1 cell_division_cycle_protein_48_homolog [MW = 89.37 kDa]

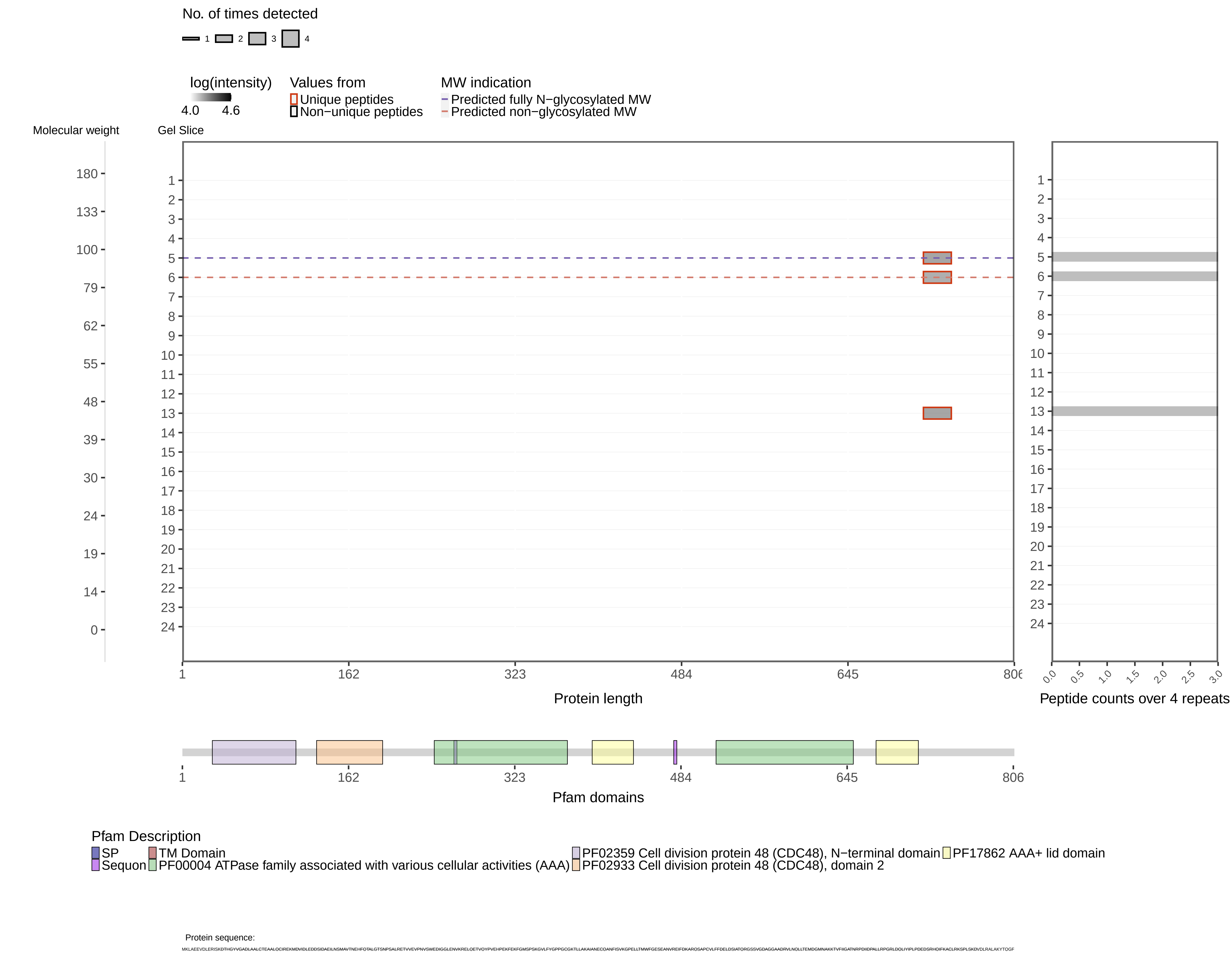

NbL11g02170.1 probable_aquaporin_TIP1−1 [MW = 25.78 kDa]

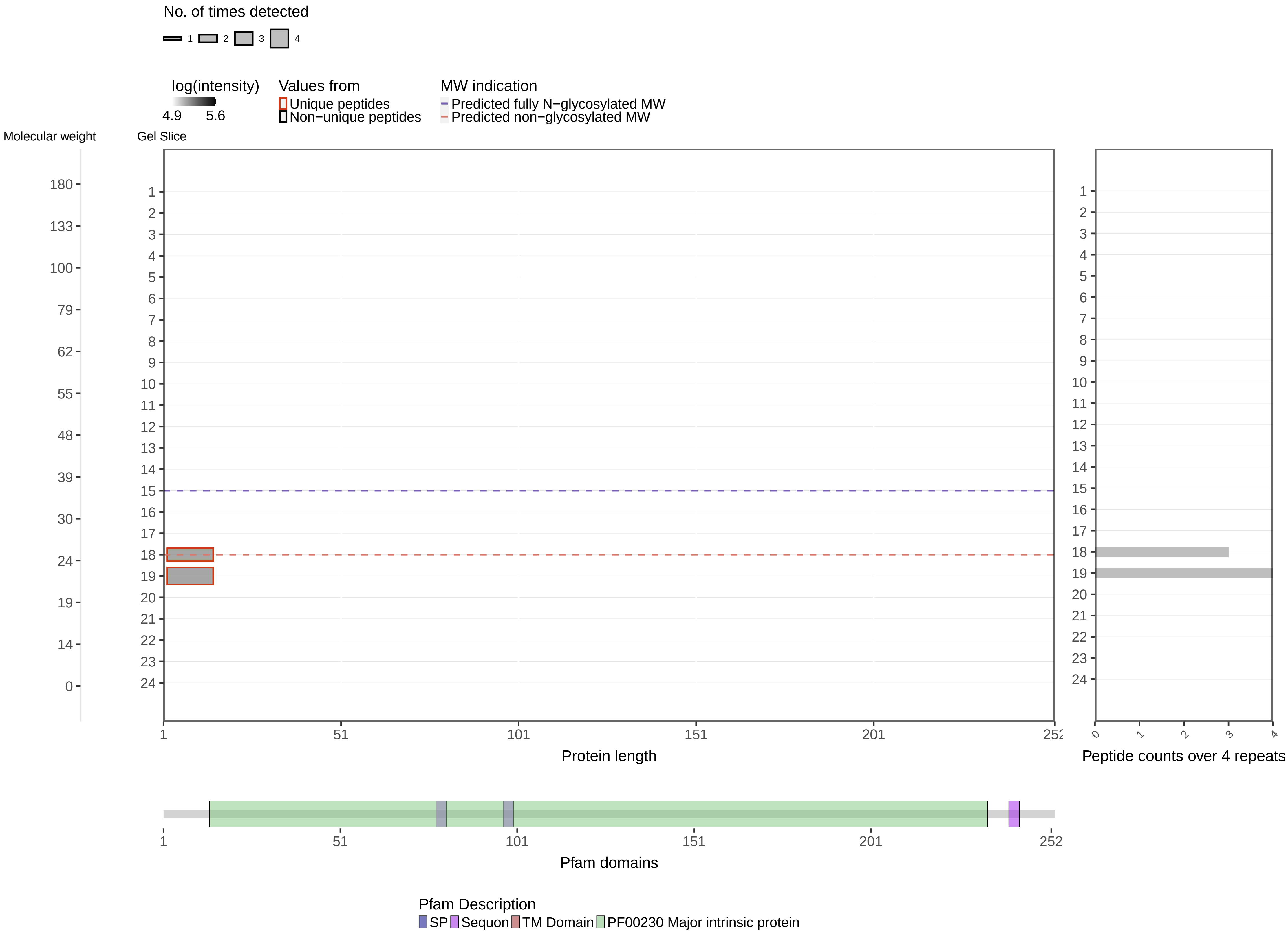


Protein sequence:

STVACFLLEFATGGMSTGAFALSAGVSVWNAFVFEIVMTFGLVYTVYATAVDPKKGDLGVIAPIAIGFIVGANILAGGAFTGASMNPAVSFGPALVSWTWTHQWVYWAGPLVGGGIAGVVYELIFINHSHEPLPSGDF*

### NbL11g01930.1 phospholipase_A−2−activating_protein [MW = 83.34 kDa]

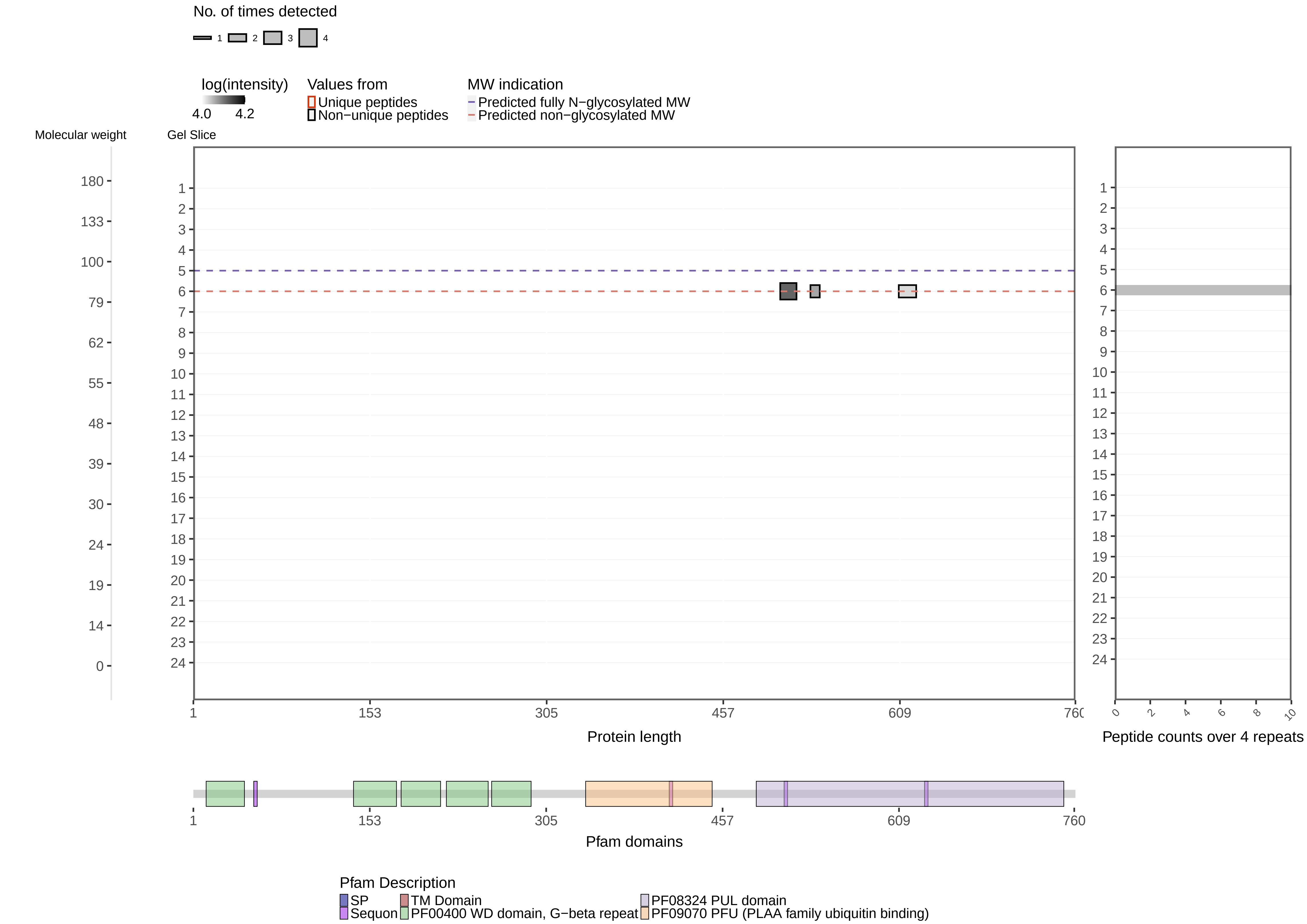


Protein sequence:

MEQKWDKIGEVVDGPDDGMTHPIHDGIQYDYVFDVDIGDGEPVRKLPYNRSDNPYDIADKWLLKENLPLSFREQIVEFILQNTGQRQFTPDPSFRDPYTGNSAYTPGQPSKPVATAAKPTFKHIPKKGMLVFDAALFDGILKKISEFNSTLLSGSEQKHMALTEAYMSRLASILNVLKDTSHYHSTKLSDDDVAMVLKLLKTWPLTMLFPVIDVLRMIMLHADGAAVLFKHVSNPNDVLMELIKKVTTSPPLPANLLTSVRALTNLFKNGSYHDWLLKHRGEVLDAFSTCYLSSNK

NbL11g01880.1 4−hydroxy−3−methylbut−2−enyl_diphosphate_reductase_chloroplastic [MW = 56.05 kDa]

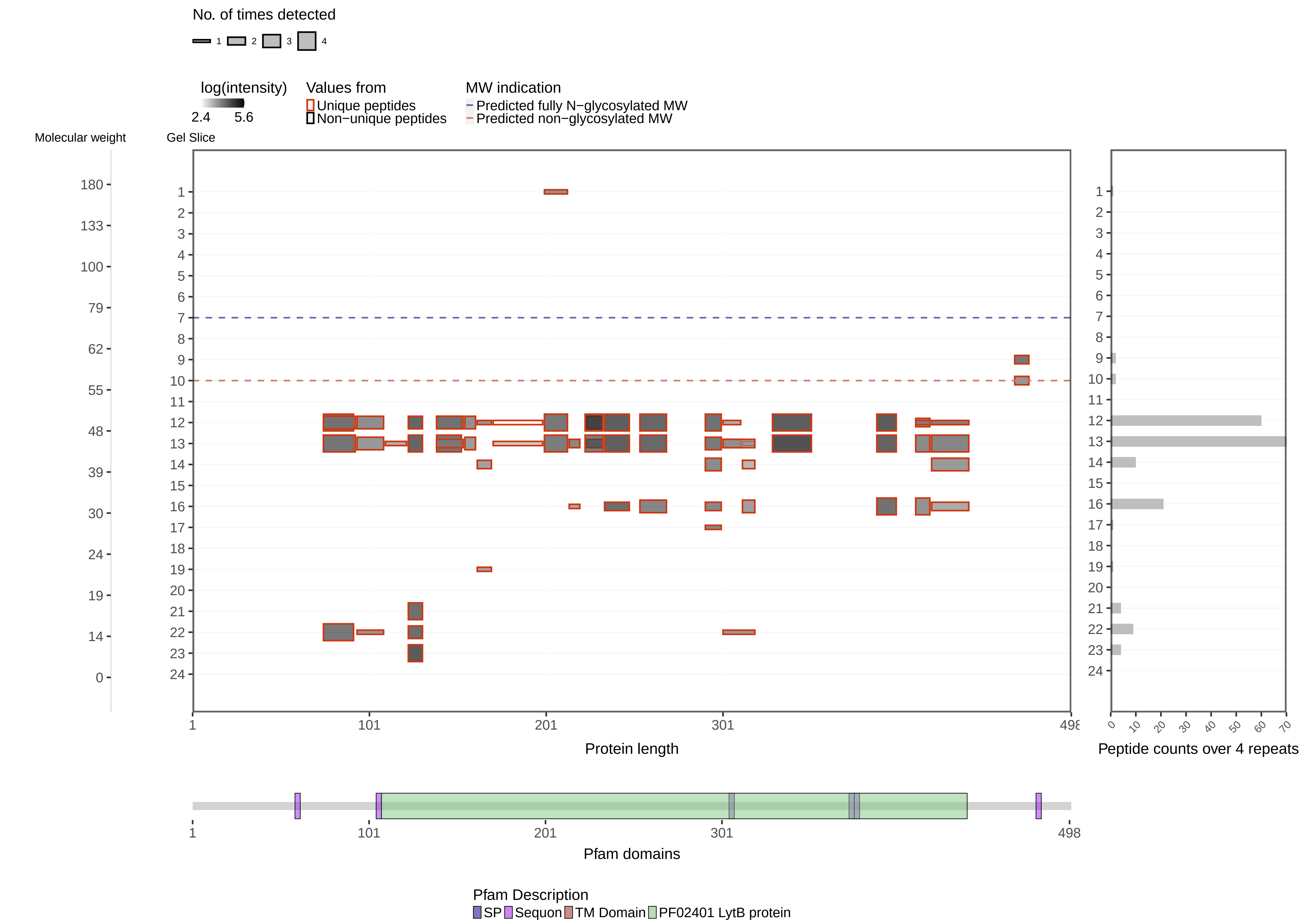


Protein sequence:

HEETVATASFAGKYIIVKNMAEATYVCDYILGGKLDGSSSTKEAFMQKFKNAVSKGFNPDVDLVKNGIANQTTMLKGETEEIGKLVERTMMQRYGVENINNHFISFNTICDATQERQDAMYKLVDQKLDLMLVVGGWNSSNTSHLQEIAEERGIPSYWIDSEQRIGPGNRISYKLLHGELVEKENFLPEGPITVGVTSGASTPDKNQMNGRLSYRKLCVLTIAGIIQQGKYIDIYNSIKVRCNNFSRHKSFCHDQETEPVYL*

NbL11g01680.1 coatomer_subunit_gamma [MW = 93.57 kDa]

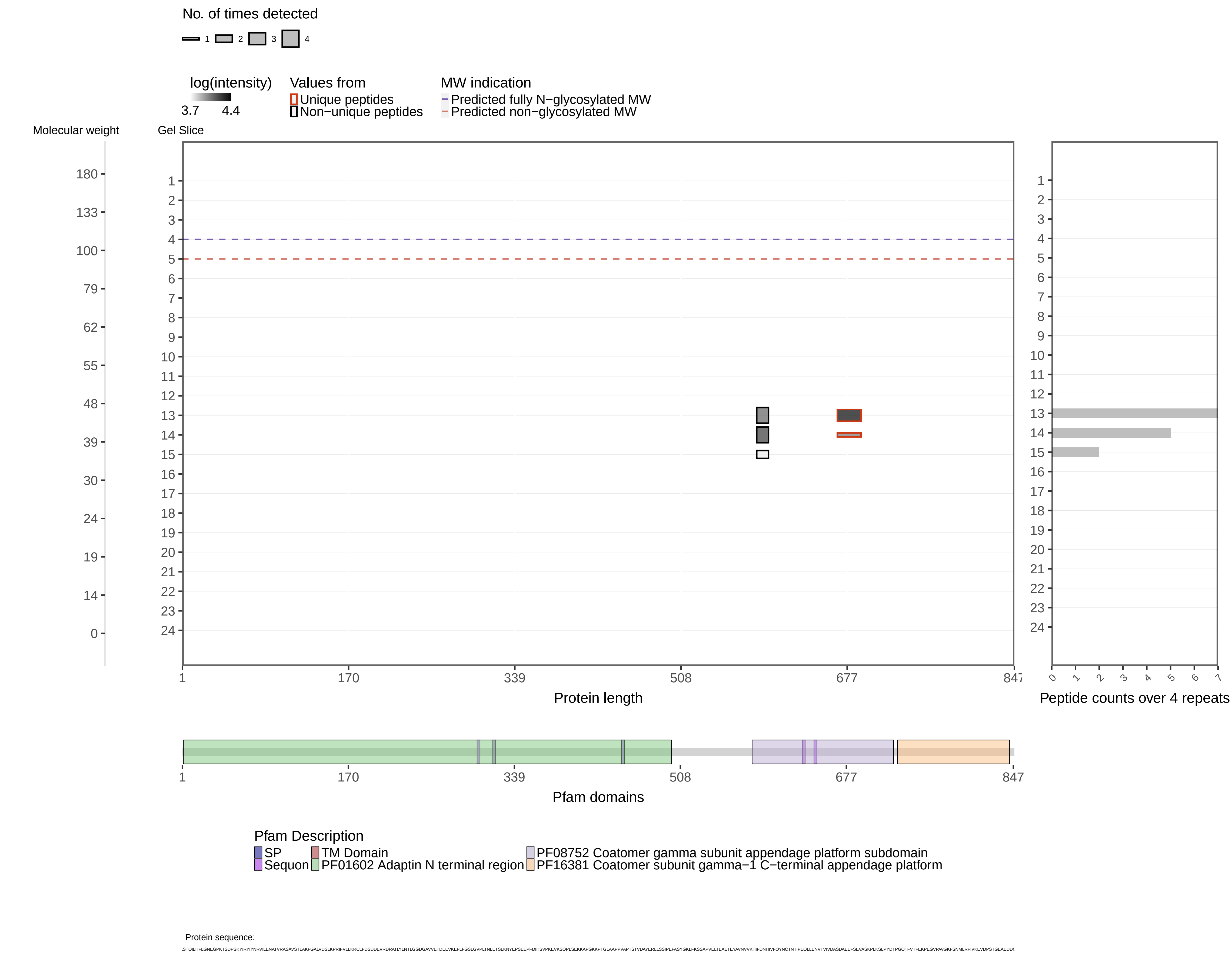

### NbL11g01550.1 glycine−rich_RNA−binding_protein_GRP1A−like [MW = 14.03 kDa]

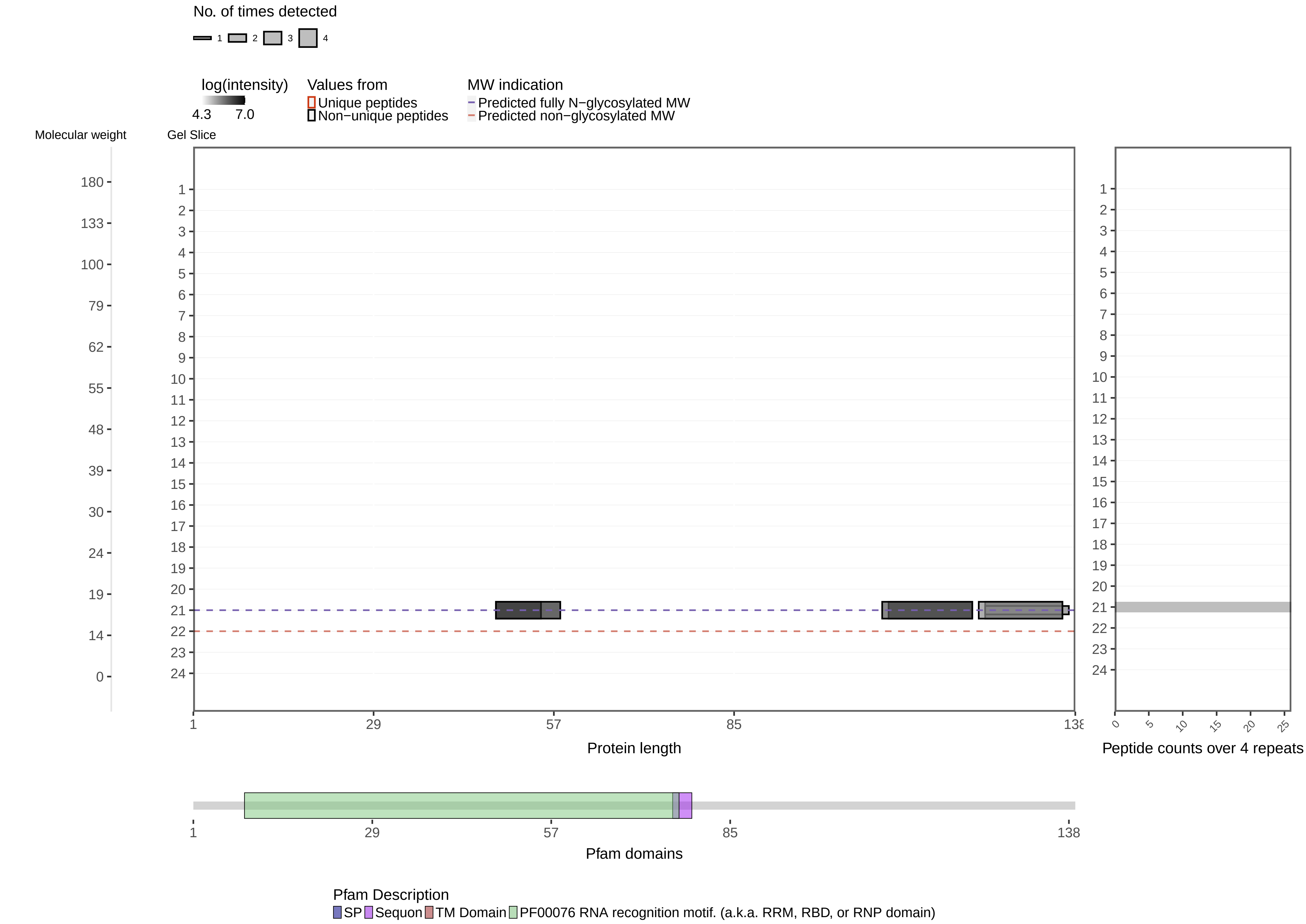


Protein sequence:

MRDAIEGMNGQDLDGRNITVNEAQSRGSGGGGGYRGGGGGGYGGGRREGGYGGGGGYGGGRREGGYGGGSEGNWRS*

### NbL11g01400.1 diaminopimelate_decarboxylase_1_chloroplastic−like [MW = 52.85 kDa]

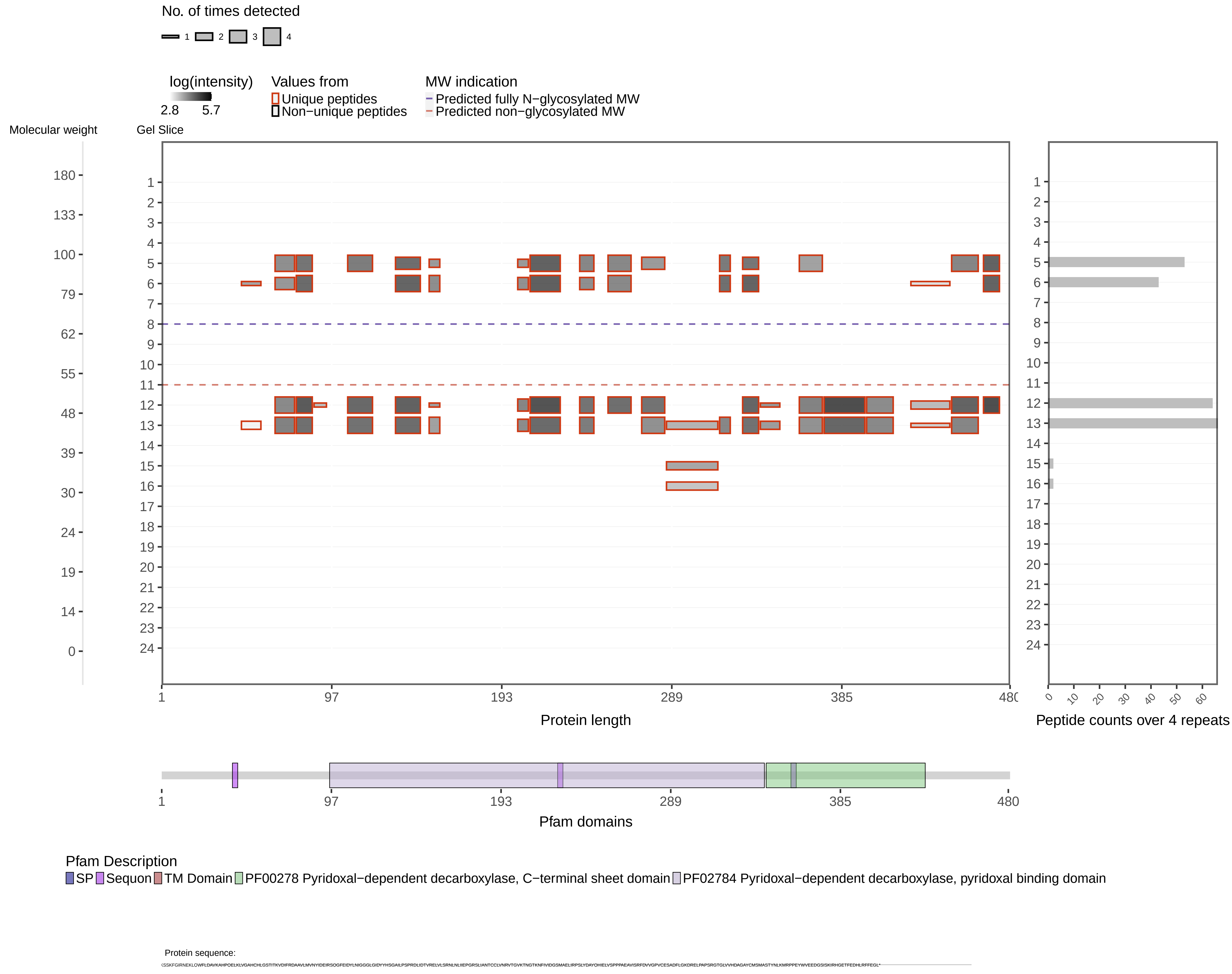

### NbL11g01340.1 chaperone_protein_dnaJ_10−like [MW = 18.28 kDa]

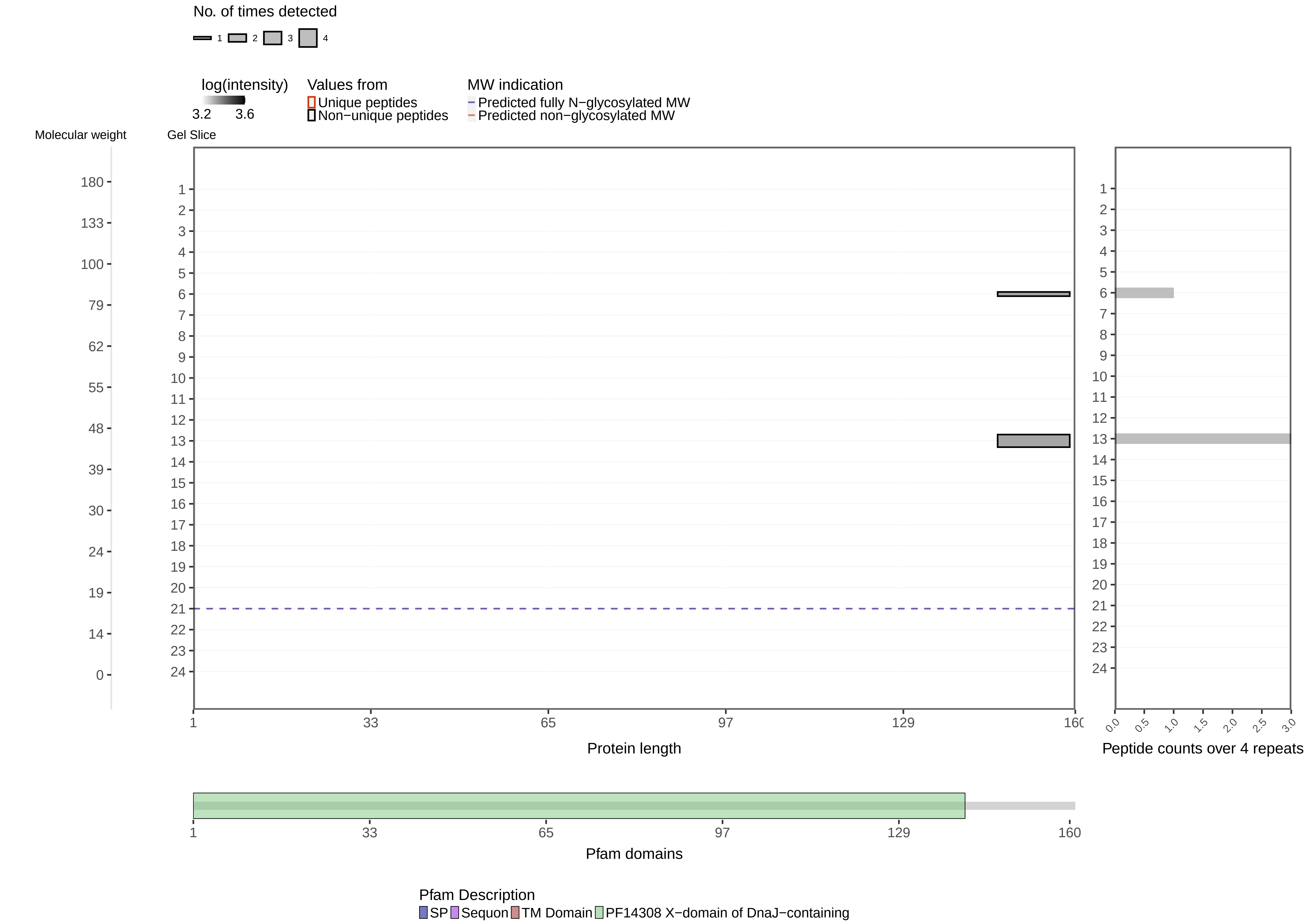


Protein sequence:

NKVEDAIKTMEEKKDIMIKSLWQINVVDIELTLSRVCQAVLKEPNVPRDVLRLRARAMKKLGTIFQGAKPMYRRESSLRSENTDMVDGGTSSK*

### NbL11g01070.1 amidophosphoribosyltransferase_chloroplastic−like [MW = 62.02 kDa]

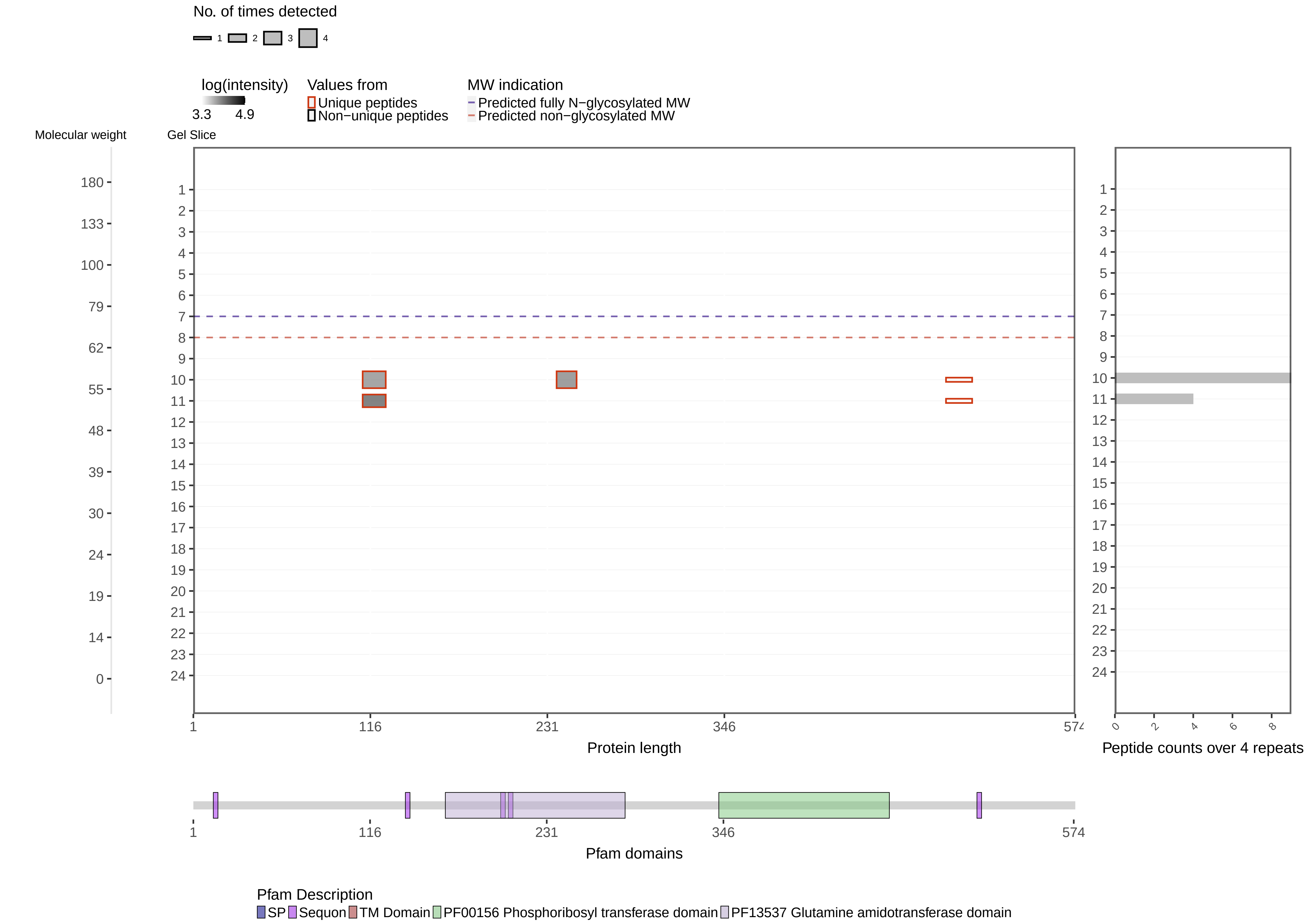


Protein sequence:

/VFASETCALDLIEATYEREVNPGEVVVVDKDGVQSICLMPHPERKSCIFEHIYFALPNSVVFGRSVYESRRAFGEILATEAPVECDVVIAVPDSGVVAALGYAAKAGVPFQQGLIRSHYVGRTFIEPSQKIRDFGVKLKLSPVRAVLEGKRVVVVDDSIVRGTTSSKIVRLLKEAGAKEVHMRIASPPIIASCYYGVDTPSSDELISNRMSVEEIKEFIGSDSLAFLPMDSLNKLLGNDSKSFCYACFSGNYPVEPTGKVKRIGDFMDDGLSGDMDSIDGGWLPGSSRVQKTILNE

### NbL11g01060.1 peptidyl−prolyl_cis−trans_isomerase_Pin1 [MW = 12.97 kDa]

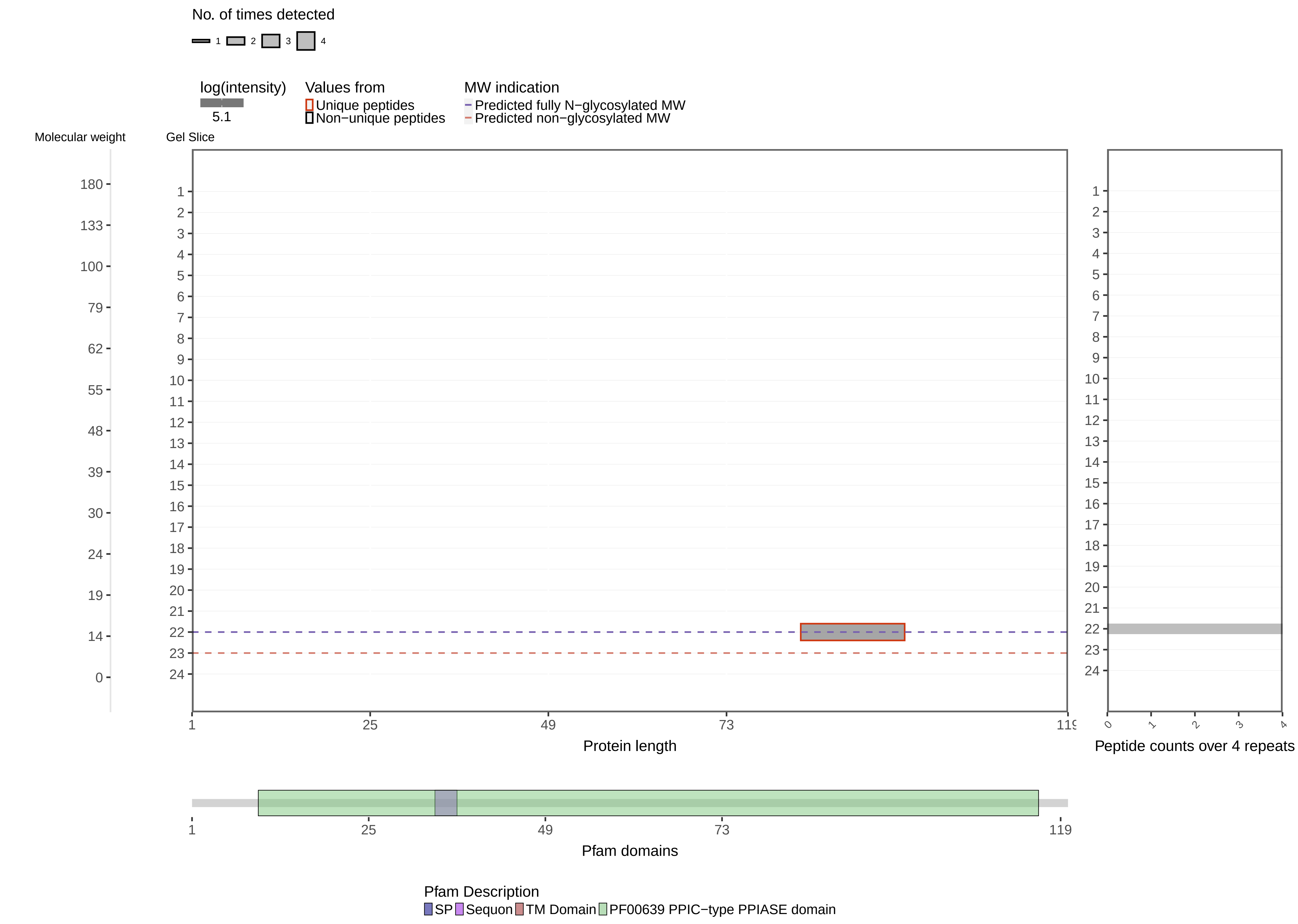


Protein sequence:

AFRDDIISGKSKFEDVASRFSDCSSAKRGGDLGPFCRGQMQKPFEDTTFALKVGEISDIIDTDSGVHIILRTG*

### NbL11g01030.1 probable_ribosome−binding_factor_A_chloroplastic [MW = 25.58 kDa]

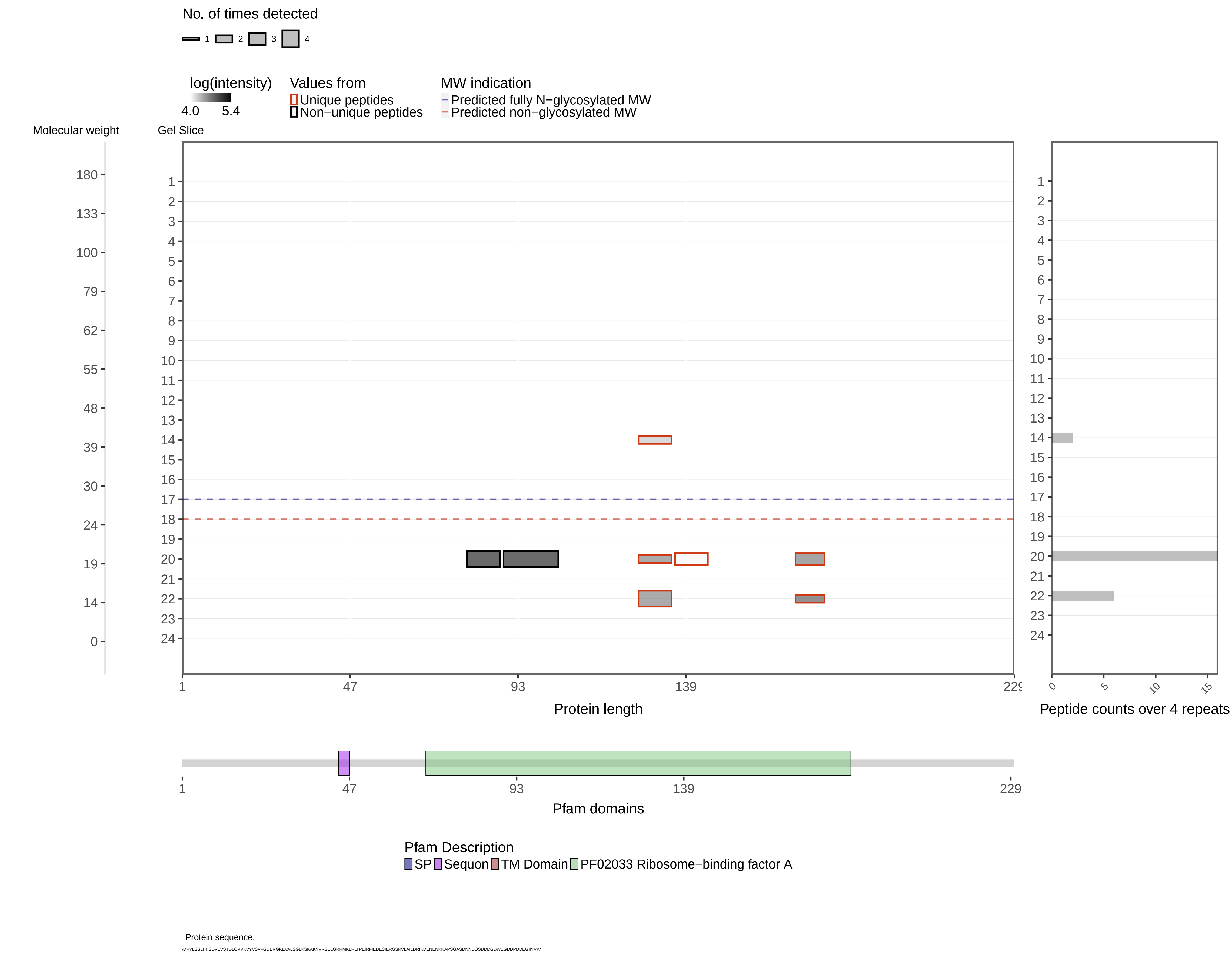

### NbL11g00880.1 single−stranded_DNA−binding_protein_mitochondrial [MW = 23.76 kDa]

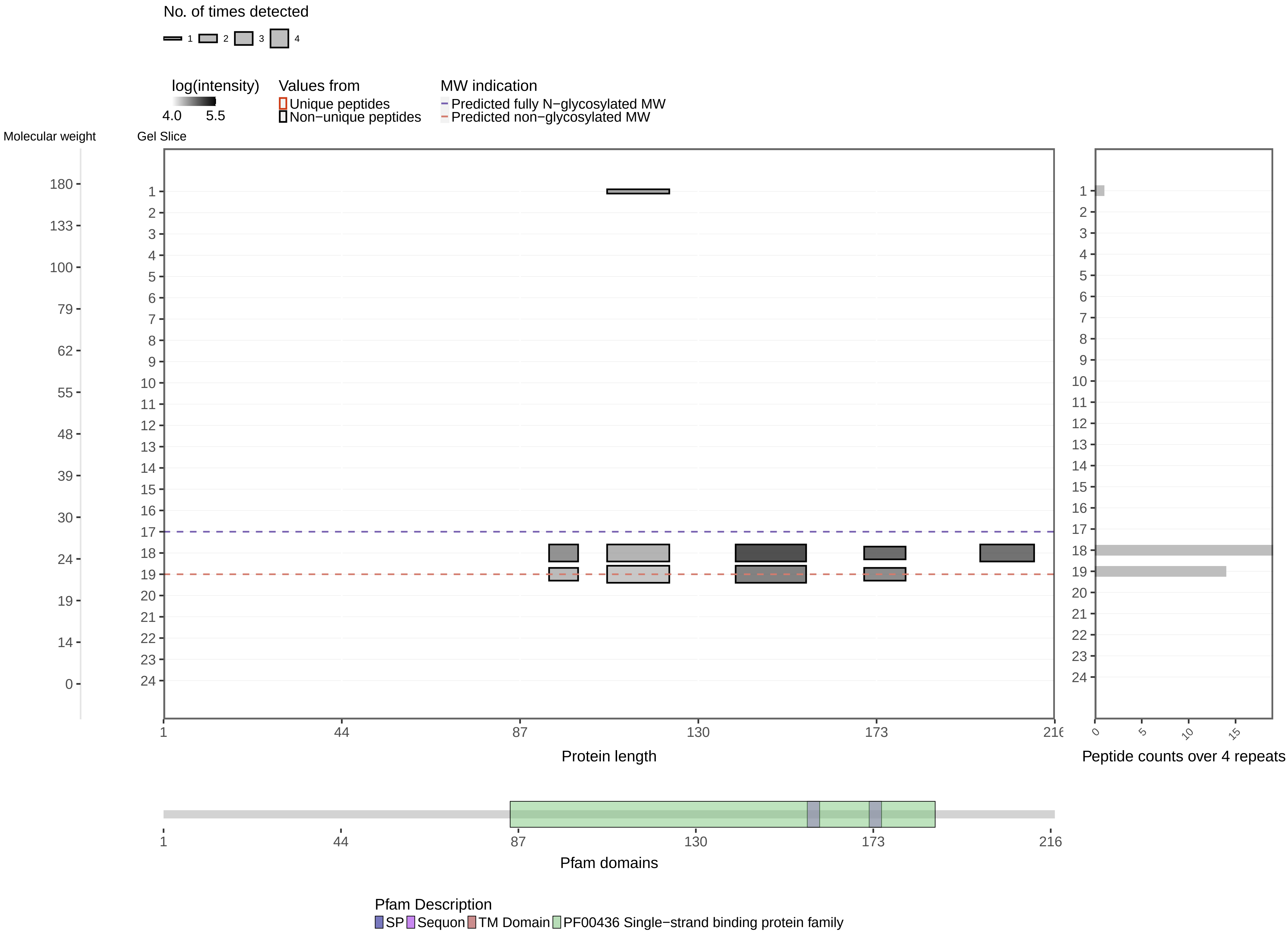


Protein sequence:

/GQSPVQKILRNGRTVTIFTVGTGGMFDQRITGDRNLPKPGQWHRVAVHNDMLGQYSVQQLTKNSSVYIEGDIETRVYNDSLSGEVKSIPEICVRRDGRIRLIKGGESVSSISIDELREGLL*

### NbL11g00760.1 deoxyuridine_5'−triphosphate_nucleotidohydrolase−like [MW = 18.17 kDa]

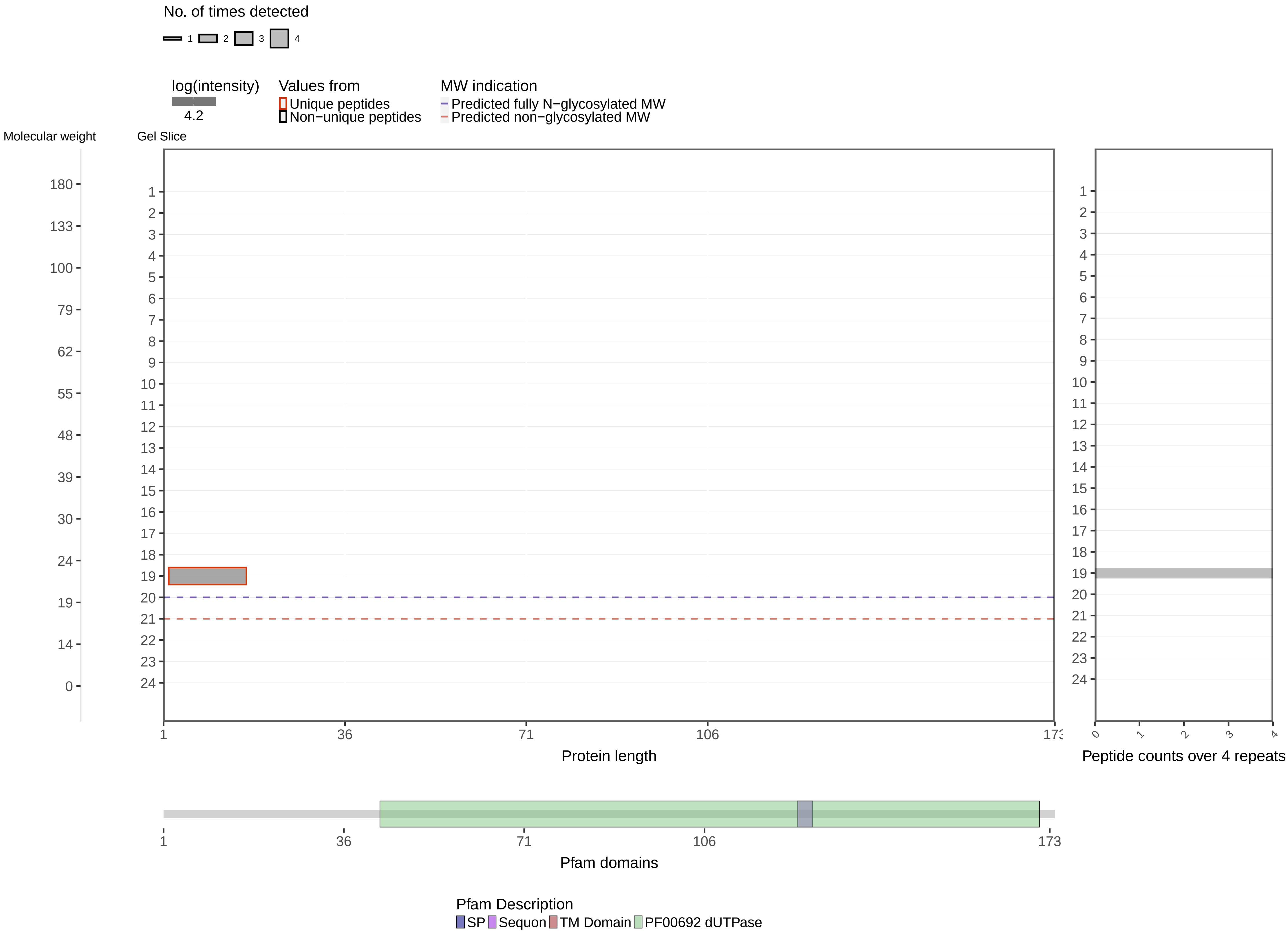


Protein sequence:

ALIPTDLSIAVPQGAYARIAPQSGLAWKHSIDVGAGVIDADYRGPVGVVLFNHSDVDFEVKAGDRIAQLIIQNVETPDVEEVDDLDSTVRGSGGFGSTGV*

NbL11g00660.1 bifunctional_epoxide_hydrolase_2−like [MW = 38.02 kDa]

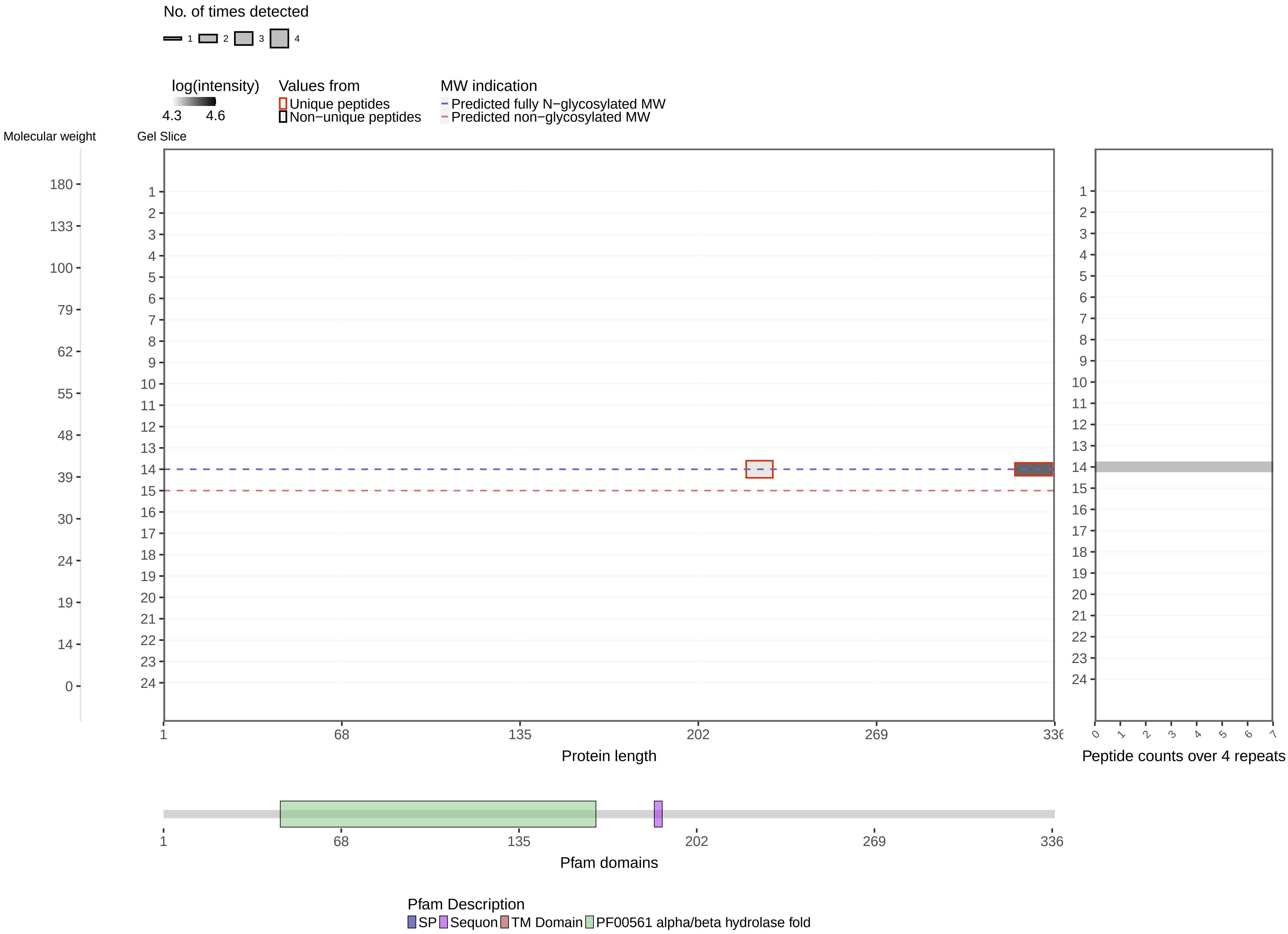


Protein sequence:

KPIPTMRAFFGDDYYMCRFQDPRMEDEIAKNGSEAVLKKILTDRKPGPPCLPKENPFGISPDSKLPSWLSQDDLKYYSTKFDQKGFTGGLNYYRALDLNWELTAAWTGAKAKVPVKFMVGELDLVYTTPGMKEYVHGGGFKKDVPMLDEDVVVMEGAAHFINQERAQETNSHIHNFINKF*

### NbL11g00650.1 60S_ribosomal_protein_L18−2−like [MW = 20.83 kDa]

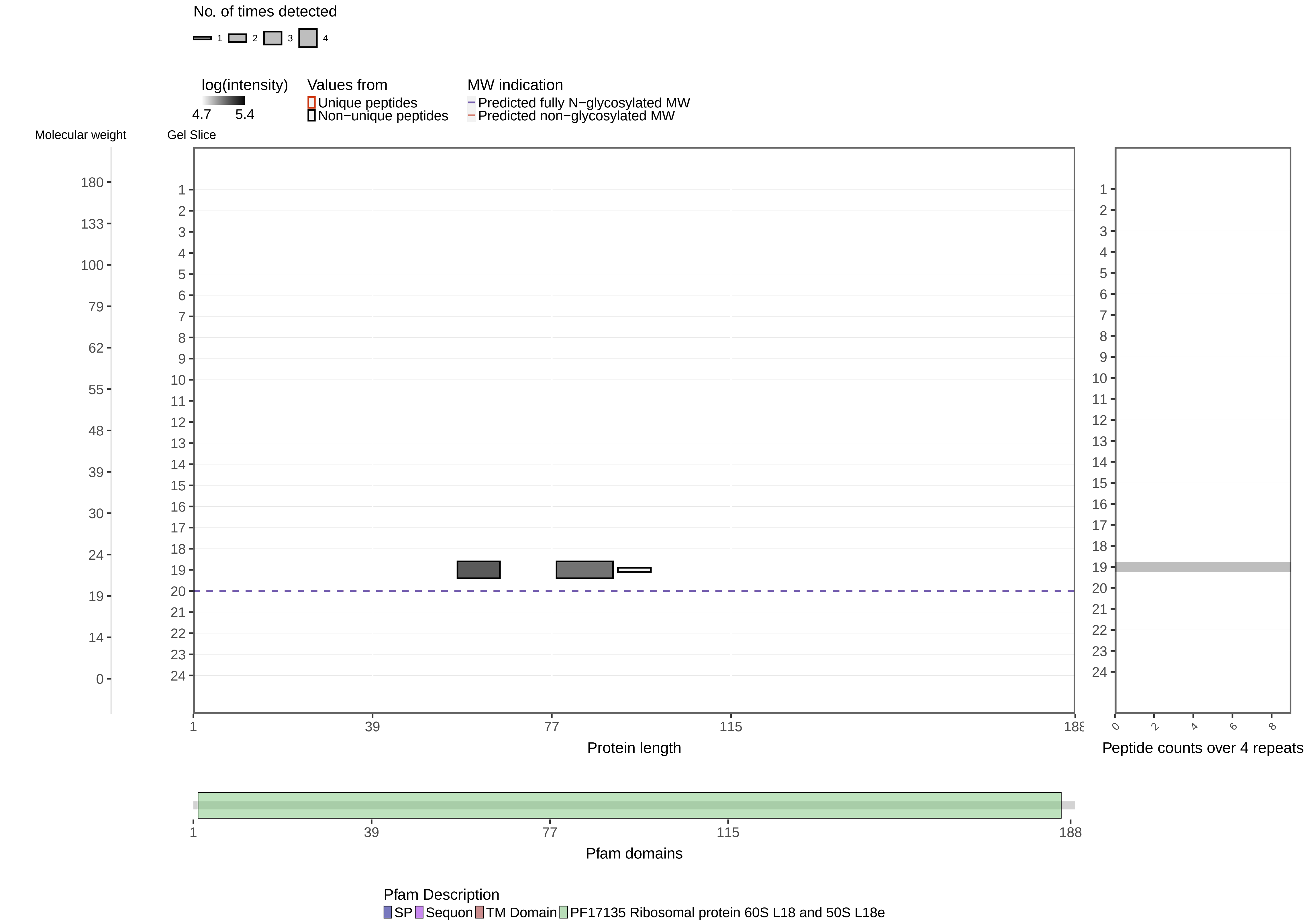


Protein sequence:

GTITDDVRAYEVPALKVCALRFTKTARARIEKAGGECLTFDQLALRAPLGQNTLLLRGPKNAREAVKHFGAAPGVPHSHTKPFVRSKGRKFERARGRRNSRGYKA*

NbL11g00590.1 auxin−repressed_12.5_kDa_protein−like [MW = 15.95 kDa]

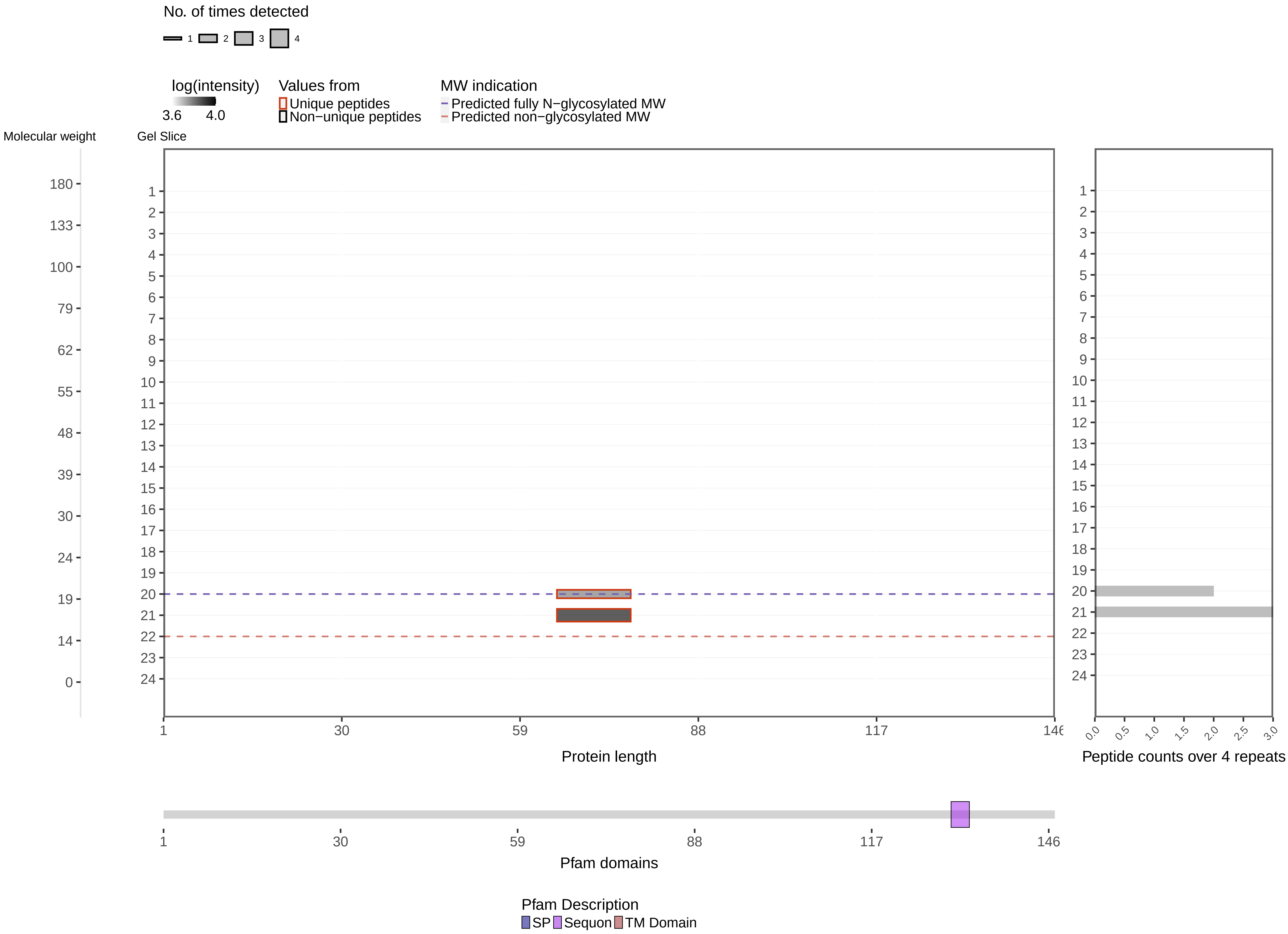


Protein sequence:
TPRSFNEETSSGDAVRVTRSIMIVKPPGSQNRDSPPVSPAGTTPPVSPFAGAGGREAYWSRRRSFAYENASGVGPRSPRPPYDL*

NbL11g00550.1 probable_Xaa−Pro_aminopeptidase_P [MW = 59.48 kDa]

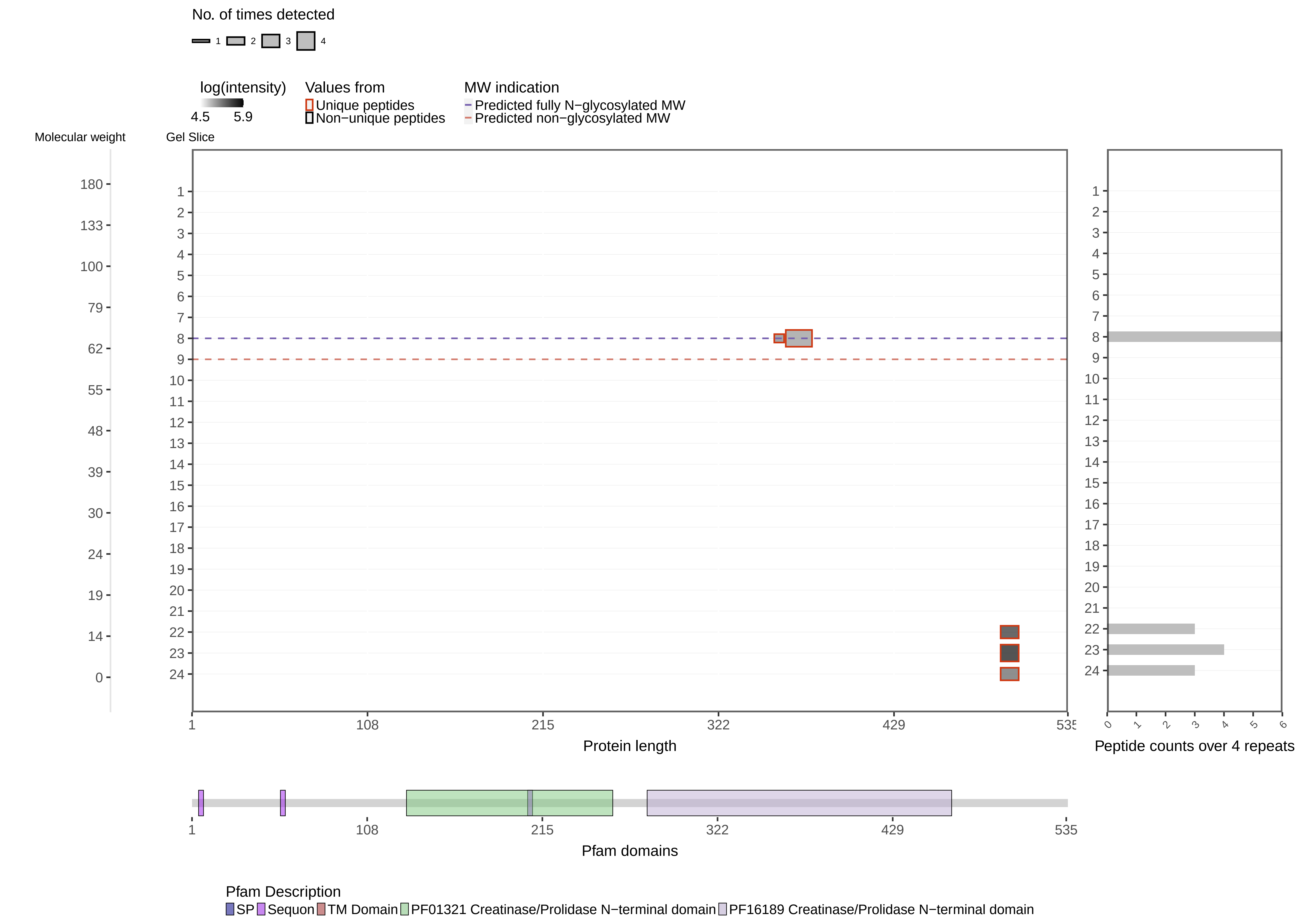


Protein sequence:

EDISKENHELVLLYDFNLVDEIWKESRPTPPKEPIRLHDLKYAGVDVSSKLSSLRSELKHAGASAIVISMLDEIAWLLNLRGNDVPRSPVMYAYLVVEIDGAKLFADDTKVTPEVMEHLTNAGIELRAYESILSEIESLAAKGANIWLDTSSVNAAILNAYESACYKYPWNSATKSRRKKTANLDSNGQSGGPCAVYRSSPVSLRKAVKNDAELEGMRNCHLRDAAALAQFWAWLEDEIQRNVALTEVEVADKLLEFRSNQDGFLDTSFDTISGILVSIIMI*

### NbL11g00400.1 ATP−dependent_Clp_protease_proteolytic_subunit−related_protein_3_chloroplastic [MW = 35.73 kDa]

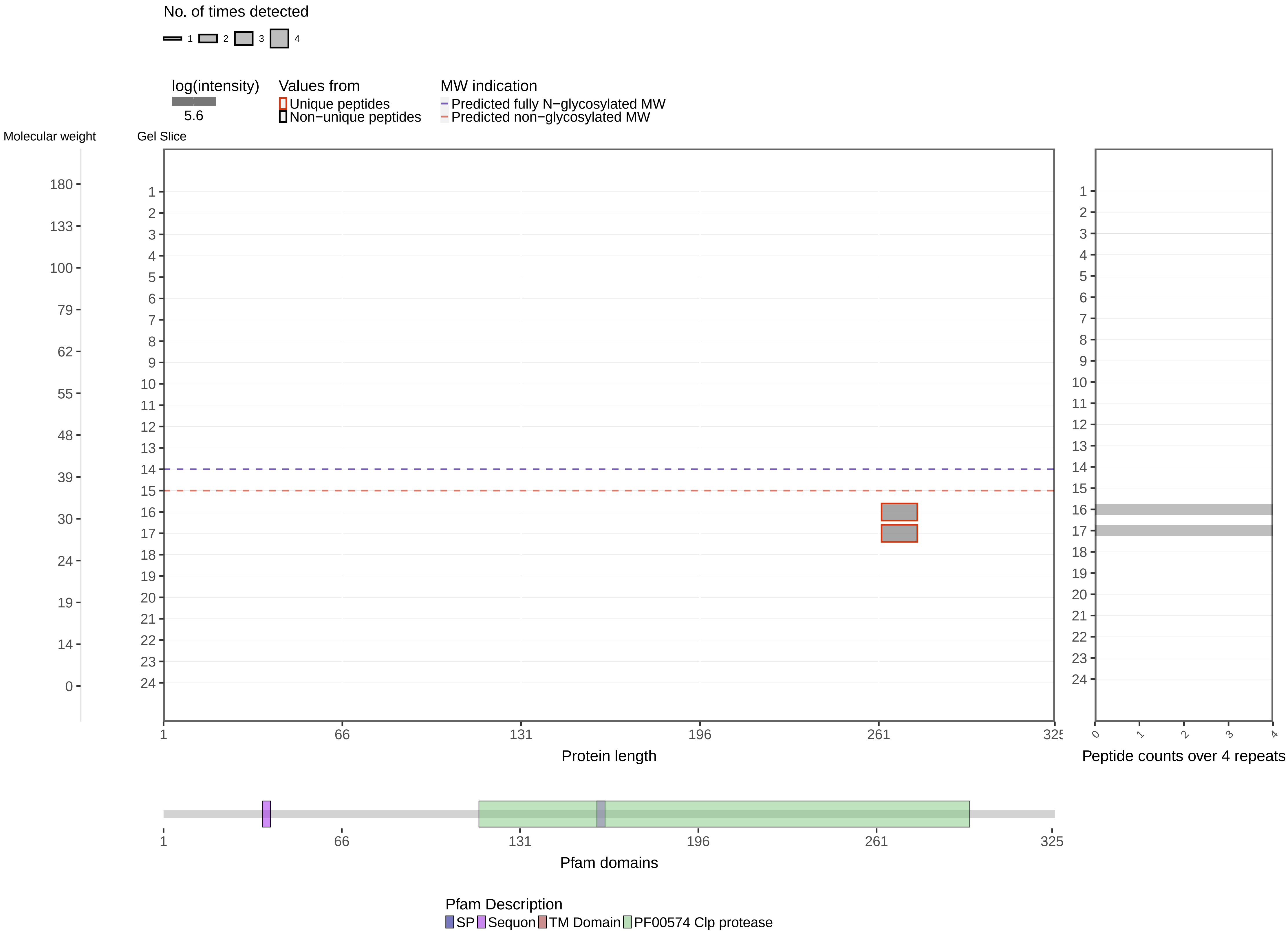


Protein sequence:

DPKEPIYLYINSTGTTRDDGETVGMETEGFAIYDAMMOLKNEIHTVAVGAAIGOACLLLAAGSKGKRFMMPHAKAMIOOPRVPSSGLMPASDVFIRAKEVIINRDTLVKLLAKHTENSEETVANVMRRPFYMDATRAREFGVIDKILWRGOEOIMADVSAPEOWDKNAGIKVADAI*

NbL11g00320.1 xyloglucan_endotransglucosylase/hydrolase_protein [MW − SP = 31.66 kDa]

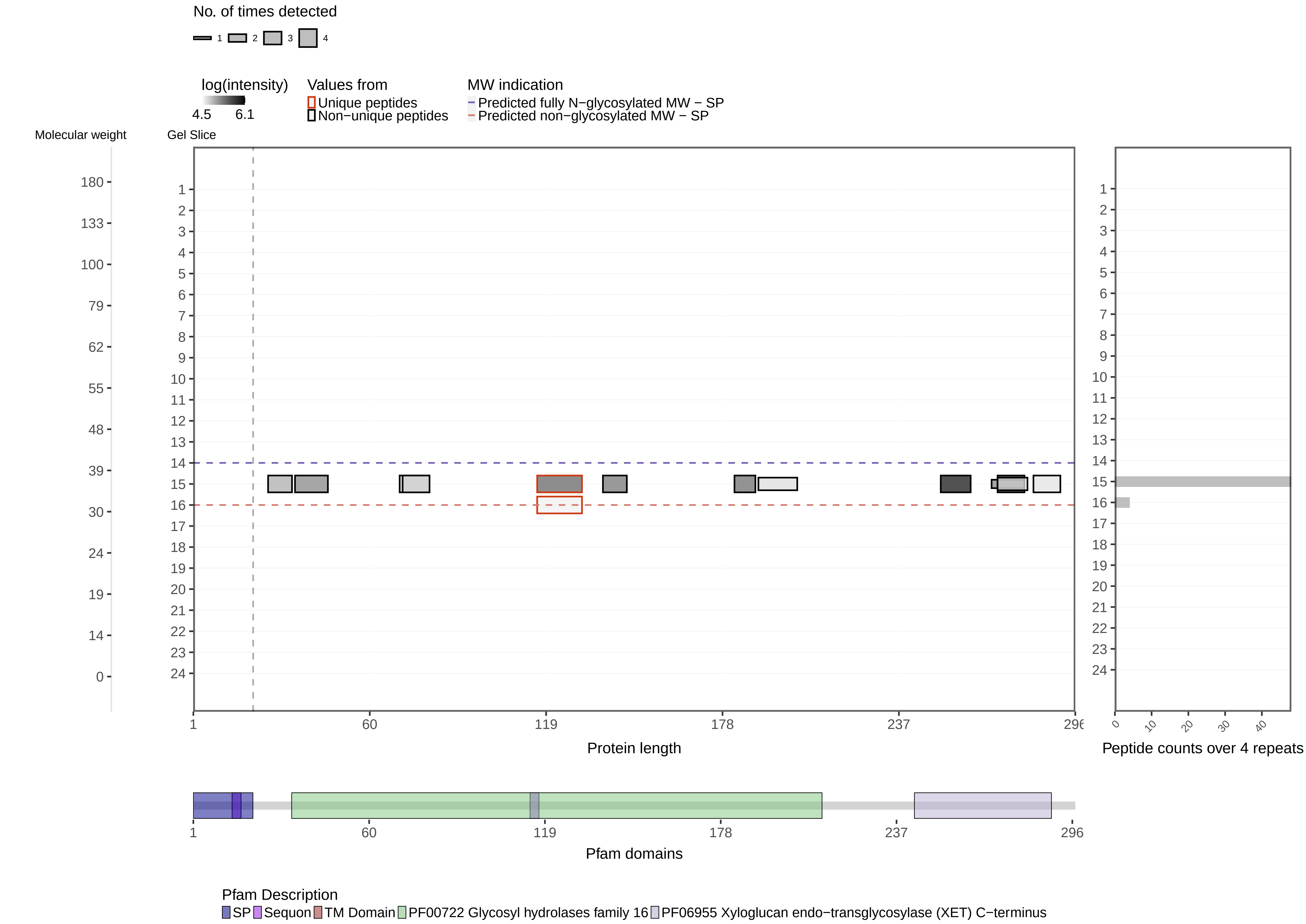


Protein sequence:

[illegible]RIYLWFDPTKGYHSYSVLWNTFQIVIFVDDVPIRAFKNSKDLGVKFPFNQPMKIYSSLWDADDWATRGGLEKTDWSNAPFSASYTSFHVDGCEAATPLEVQICNTKGMRWWDQKAFQDLDALQYRRLRWVRQKYTIYNYCTDRKRYPTLPPECTKDRDV*

### NbL11g00170.1 acyl−CoA−binding_domain−containing_protein_4 [MW = 73.99 kDa]

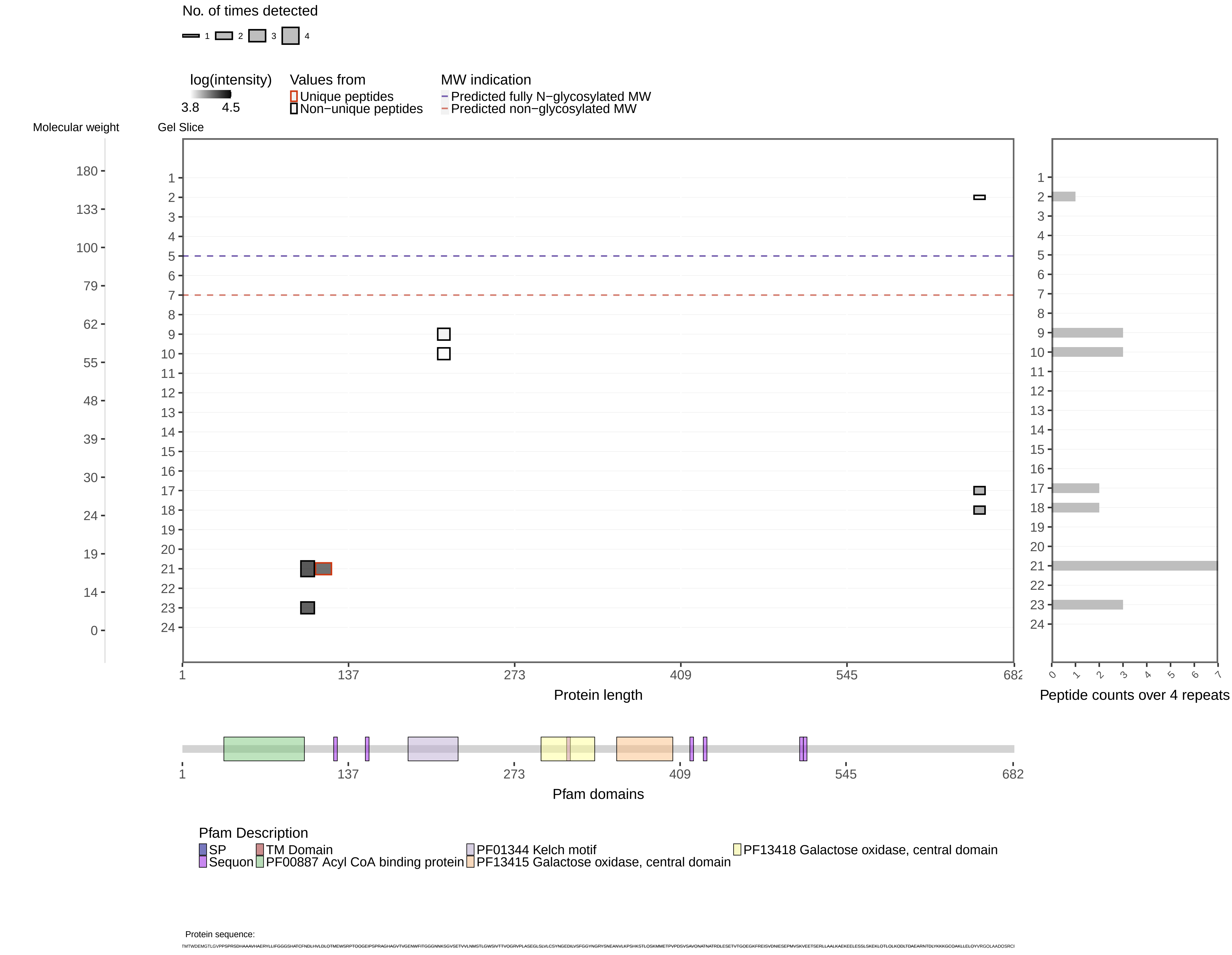

NbL11g00140.1 translationally−controlled_tumor_protein_homolog [MW = 18.91 kDa]

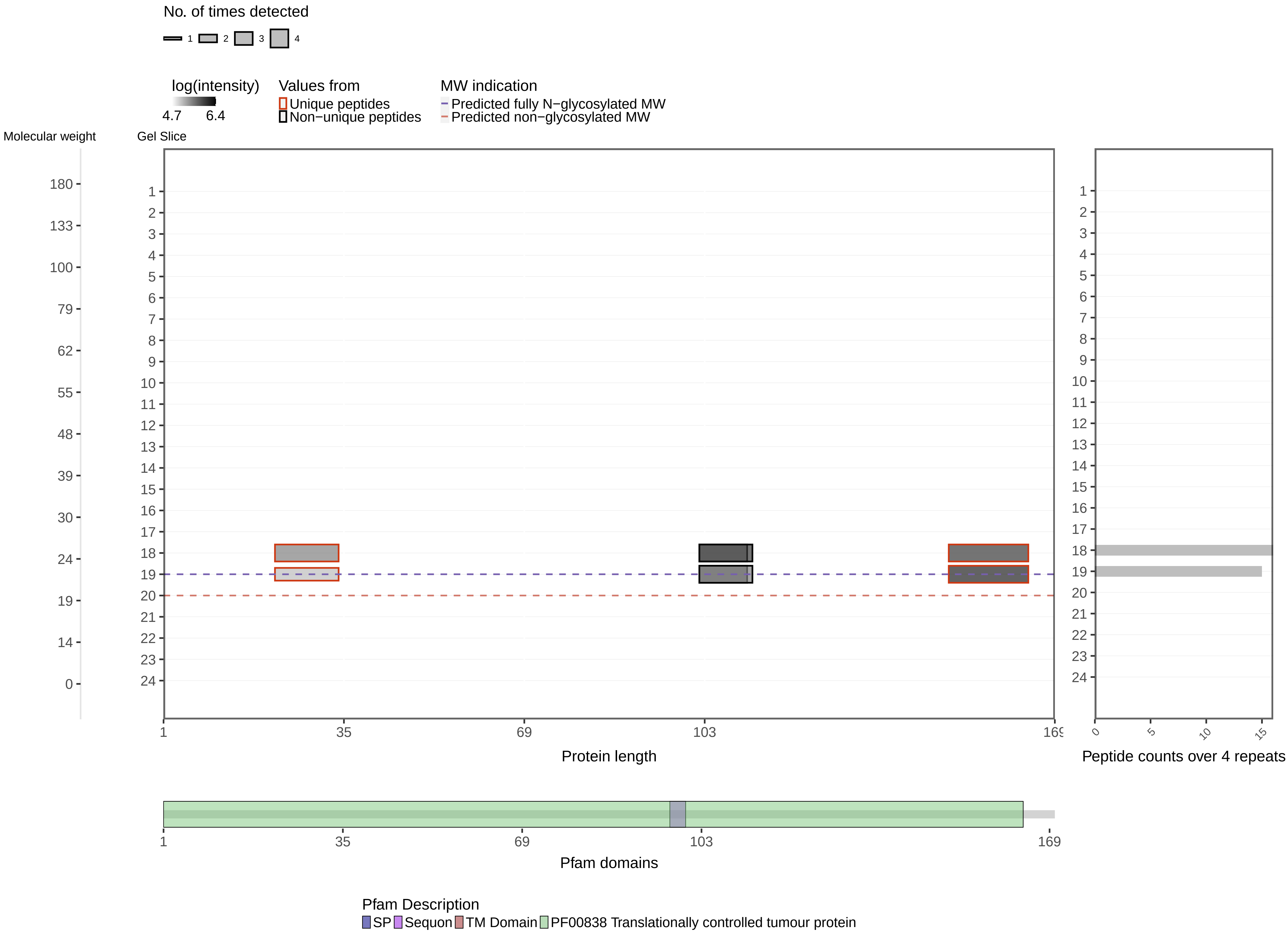


Protein sequence:

VDTFRLQEQPSFDKKQFVTYMKRYIKNLTPKLEGEAQEAFKKNIESATKFLMSKLKDLQFFVGESMHDDGALVFAYYKDGATDPTFLYLAPGLKEVKC*

### NbL11g00110.1 60S_ribosomal_protein_L22−3−like [MW = 14.06 kDa]

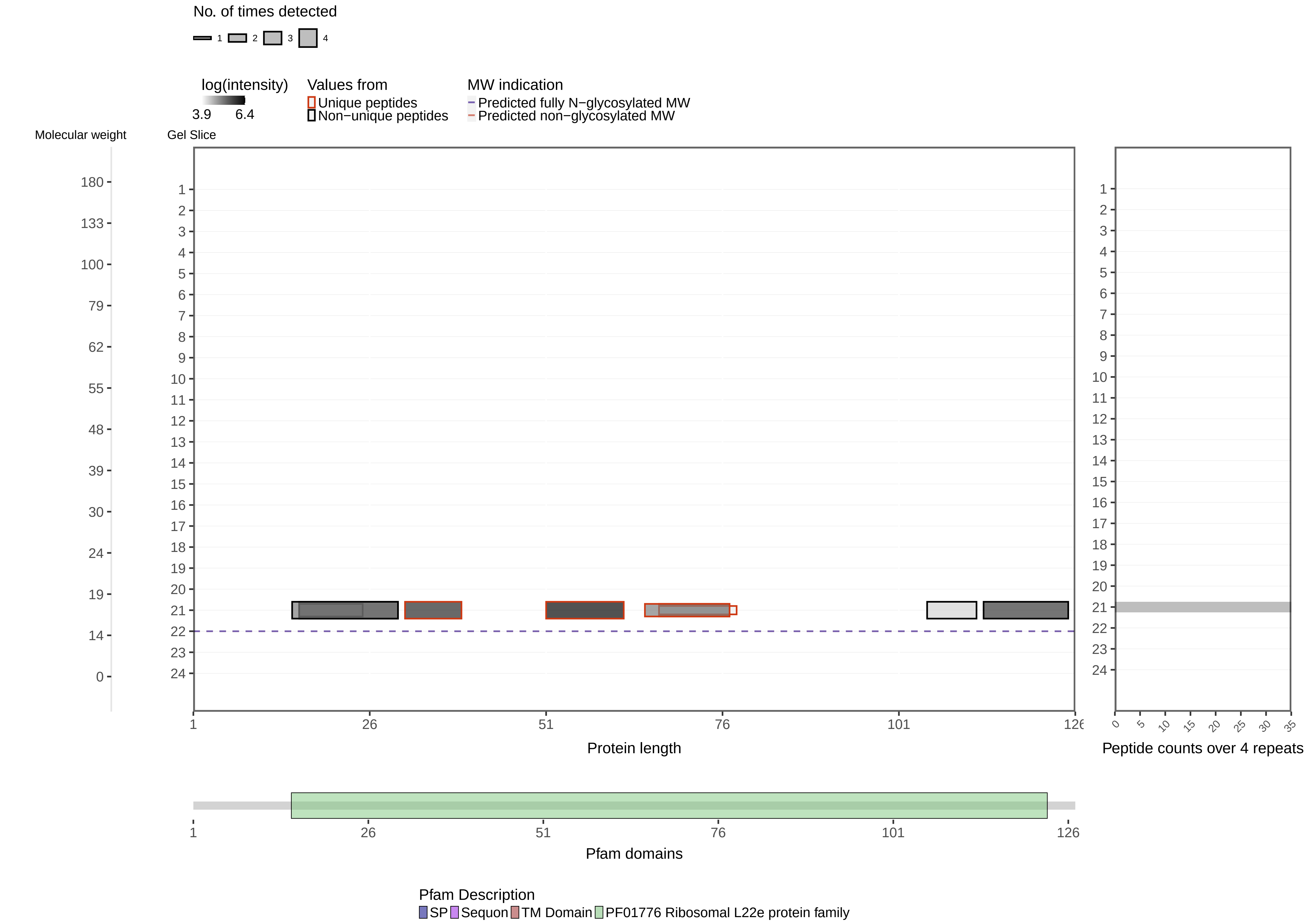


Protein sequence:

KAGALGDAVTVTRDKTKITVTSESTFSKRYLKYLTKKYLKKNNVRDWLRVISANKDRNVYELRYFNIAENEAEEED*

### NbL11g00080.1 acetylajmalan_esterase−like [MW − SP = 39.73 kDa]

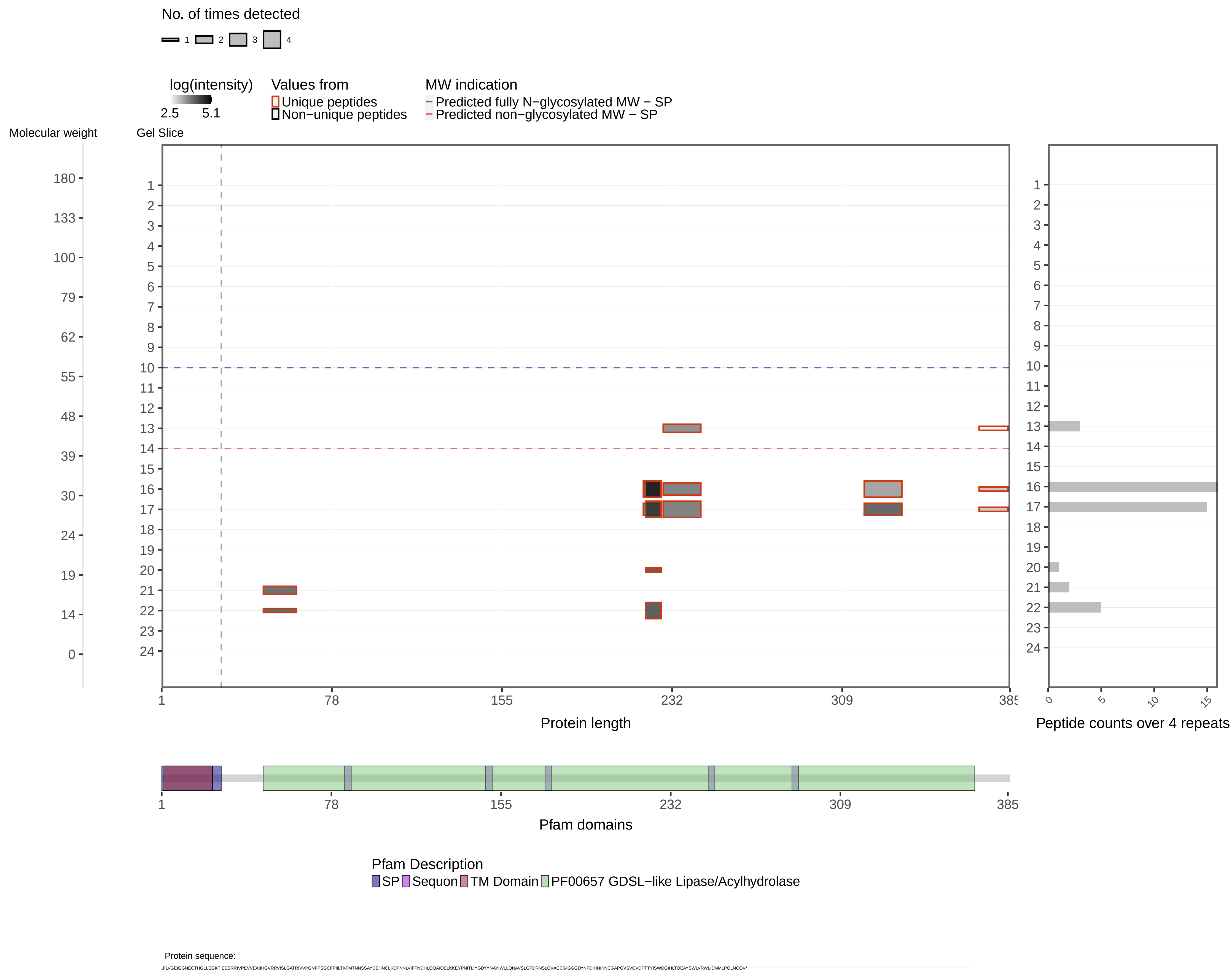

NbL11g00010.1 brefeldin_A−inhibited_guanine_nucleotide−exchange_protein_1 [MW = 98.41 kDa]

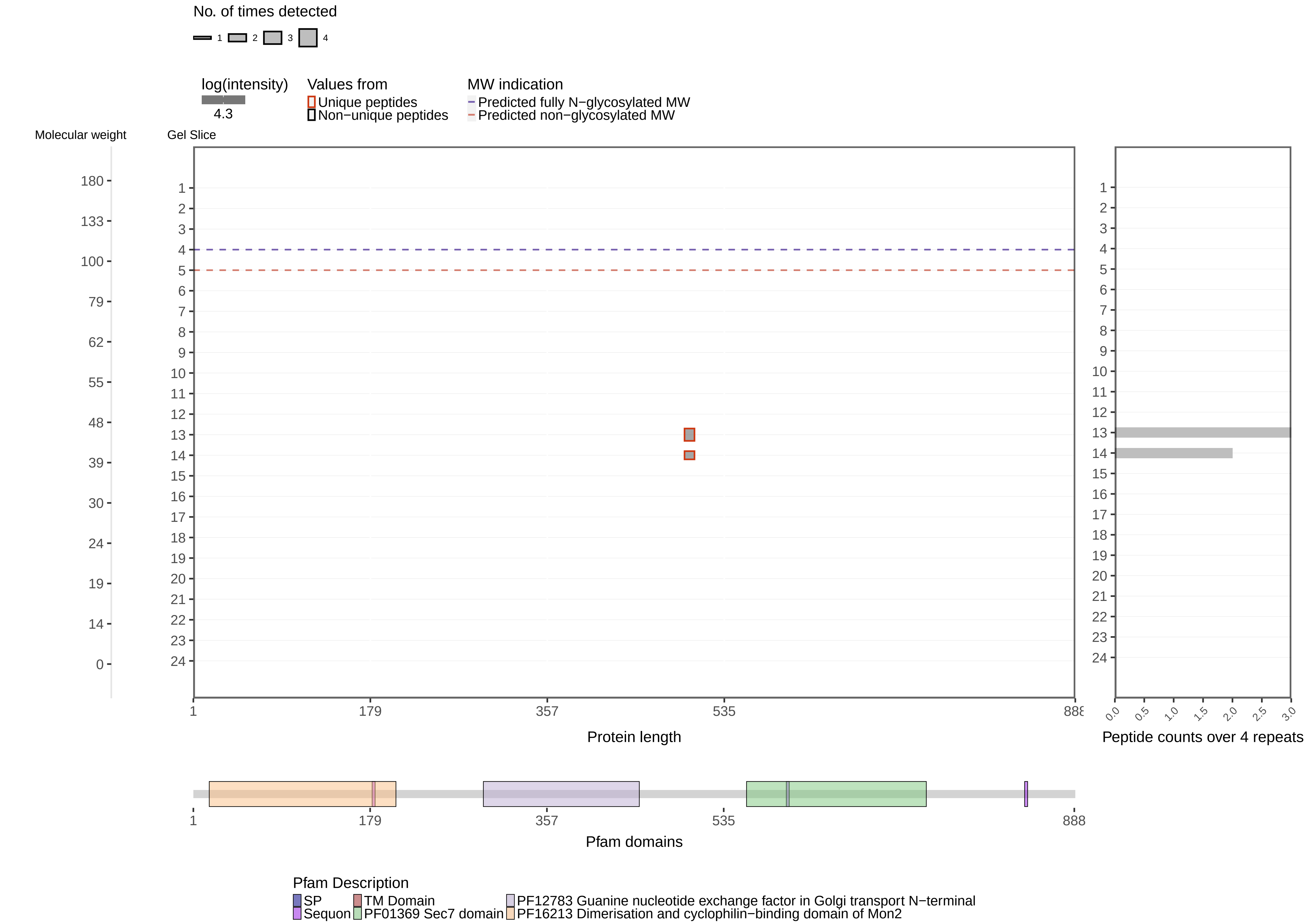


Protein sequence:

[illegible]

NbL10g24670.1 probable_inactive_purple_acid_phosphatase_27 [MW − SP = 66.7 kDa]

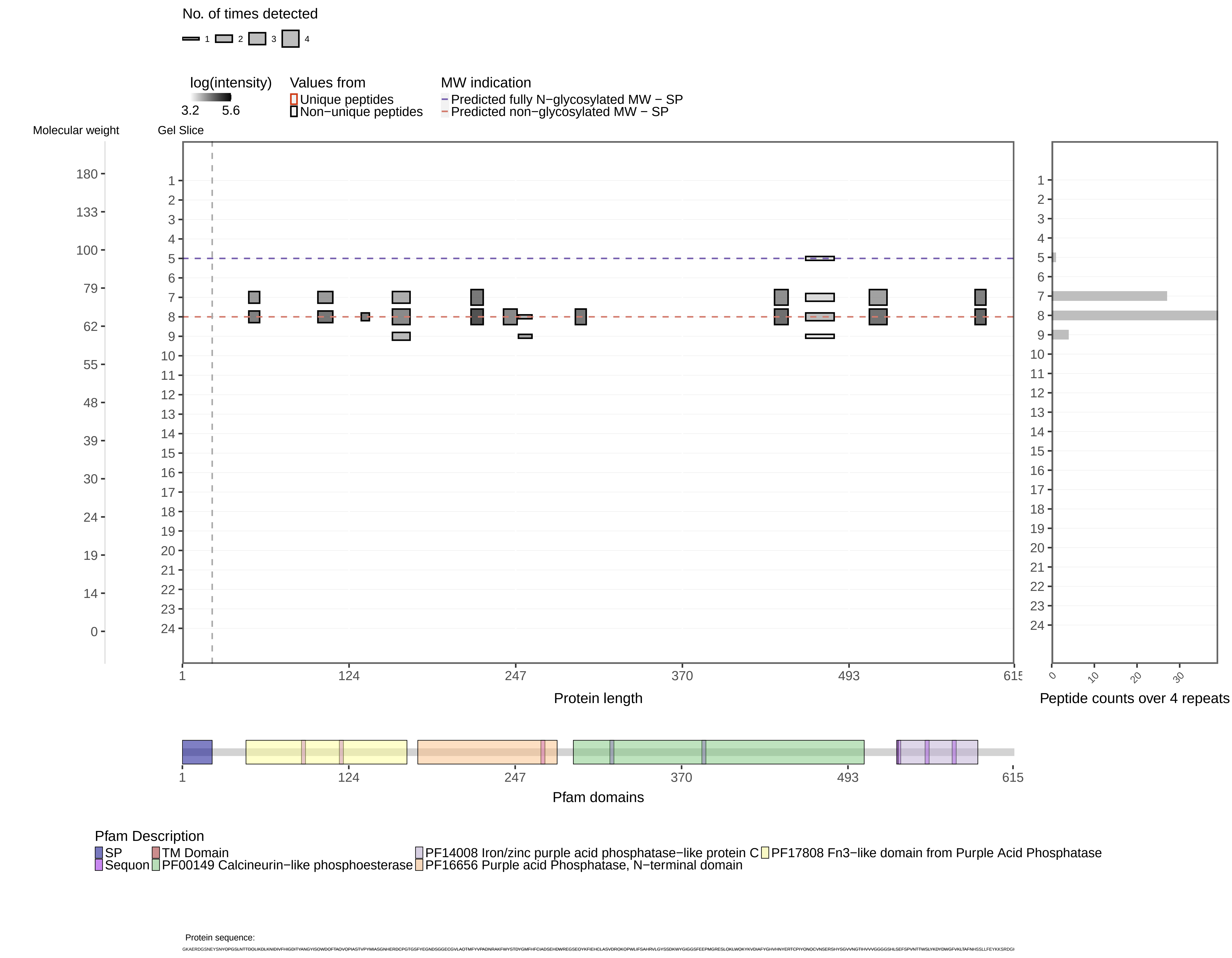

### NbL10g24660.1 snurportin−1−like [MW = 48.81 kDa]

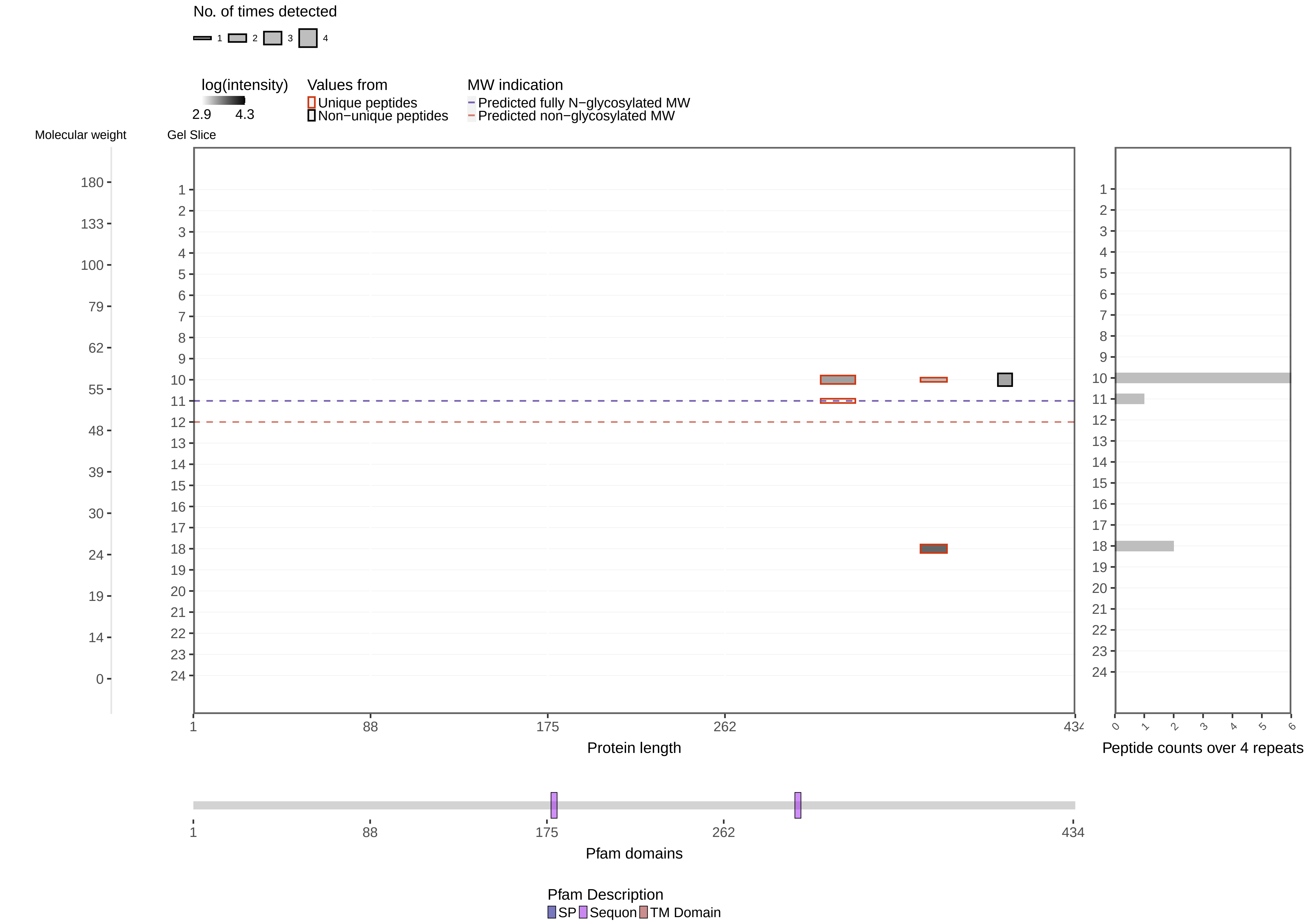


Protein sequence:

GVCWAGLSLYECTAEFRFFWLNSKLAETGACDAPSTCHRYKFSTLPVYNCDKEGLHTAYAGQVPYVKDGLLFYNKHAHYQTGNTPLTLVWKDENCSQYVIDTDNKGQVPSQQQVVLELLDDGRLATSDDPPVVFGCLLGDFIQKTELHRGDLIKFAIGEGGLVFGDSKLEKADLQYLGKSNRARAFADSYSKVLFQYAARHSPLRIEHLFASISSCVEDGRSTQDADMAG*

NbL10g24550.1 photosynthetic_NDH_subunit_of_subcomplex_B_3_chloroplastic−like [MW = 21.64 kDa]

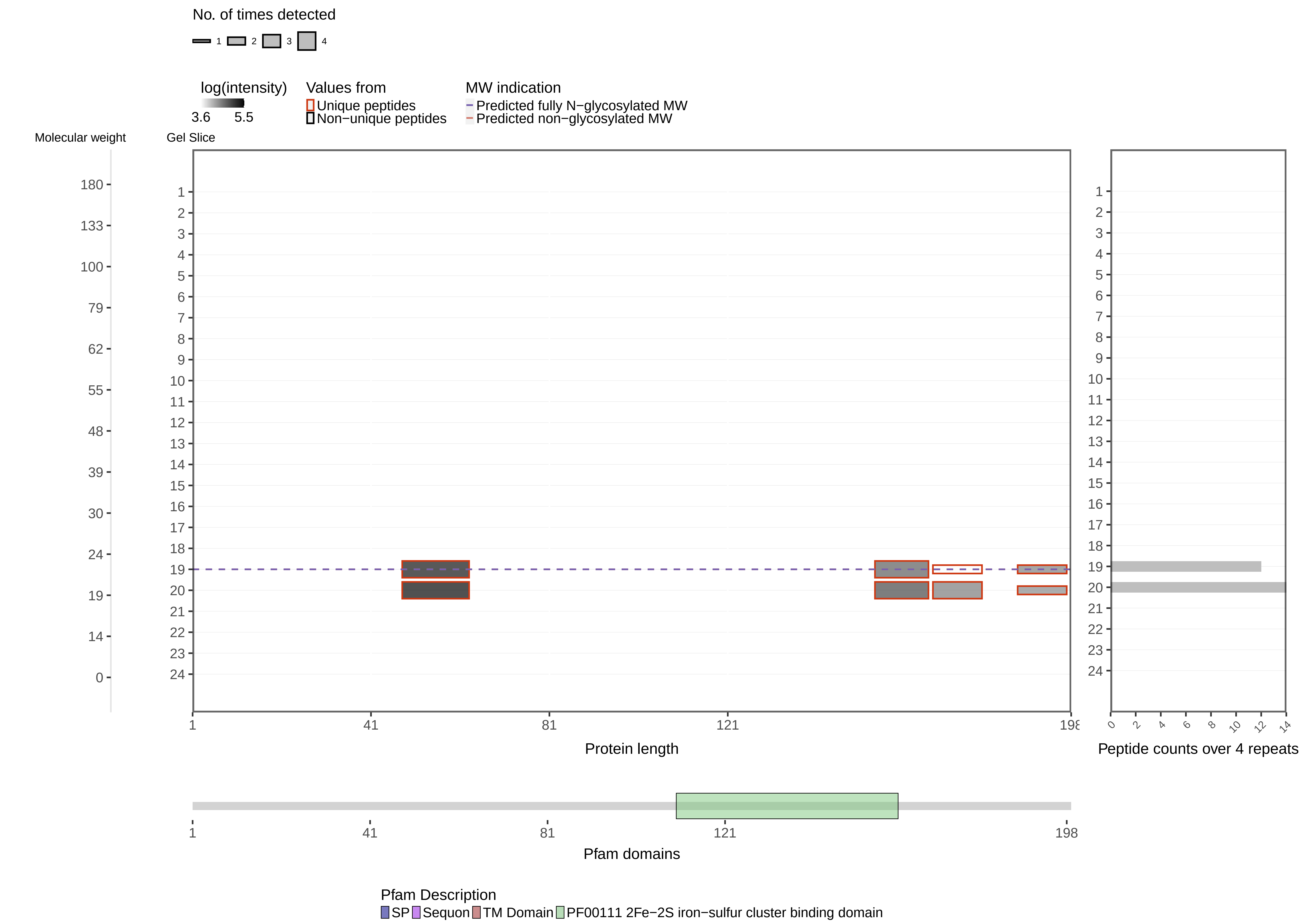


Protein sequence:

KLRDIMLDANVELYGPYARPLLNCGGGGTCGTCLVEVVEGKELLNPRTDKENNHEDLKRHPKNWRLACQAIVGKPDSRGMMVIQQLPEWKAHEWNYRGLPPIQEEEEKE*

### NbL10g24540.1 50S_ribosomal_protein_L11_chloroplastic−like [MW = 23.12 kDa]

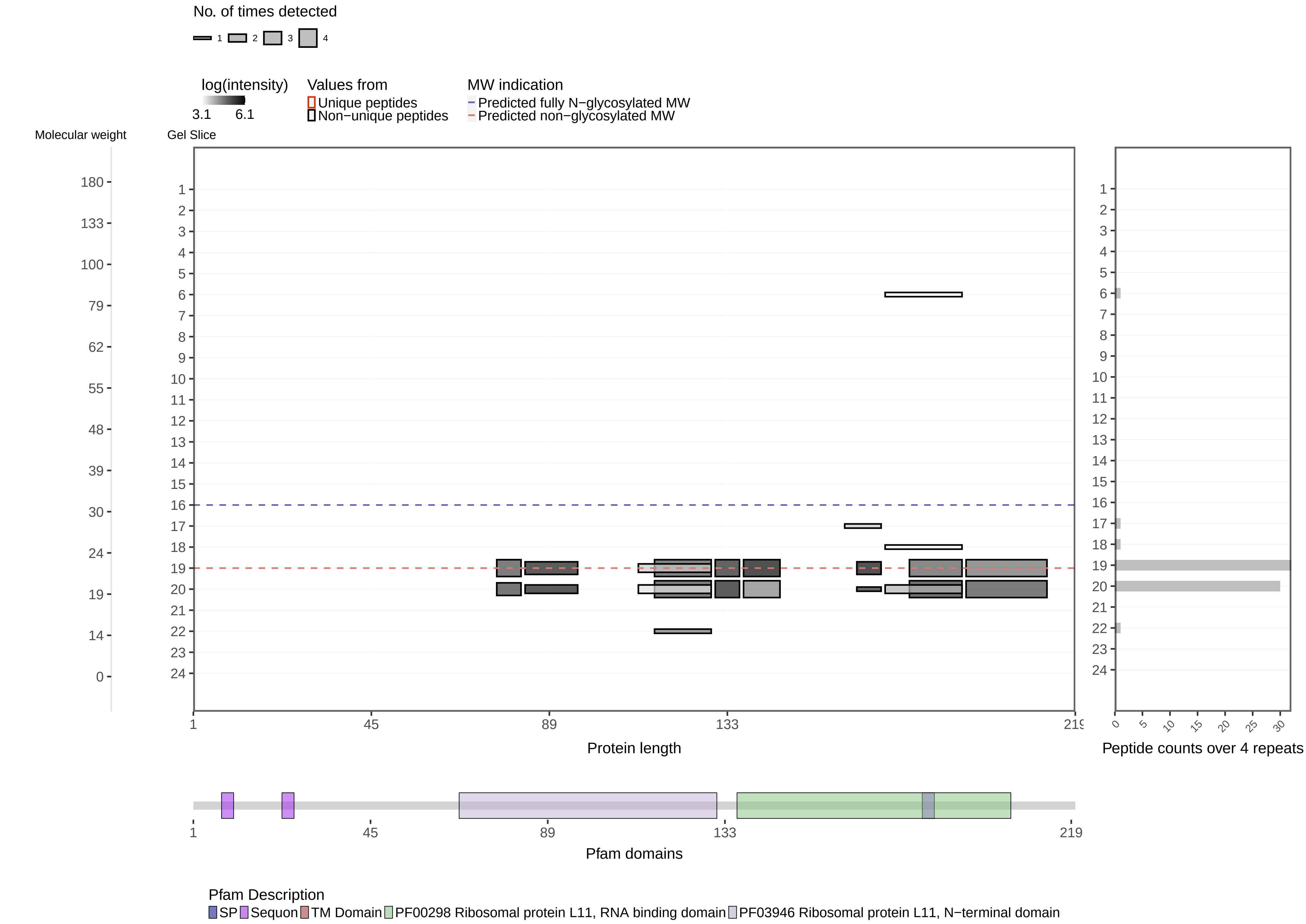


Protein sequence:

SKGVNIMAFCKDYNARTADKAGYVIPVEITVYDDRSFTFILKTPPASVLLLKAAGVEKGSKDPQRDKVGKITIDQLKTIAQEKLPDLNCTTIESAMRIIAGTAANMGIDIDPPVLEPKKKELIY*

### NbL10g24530.1 soluble_inorganic_pyrophosphatase_PPA1−like [MW = 24.07 kDa]

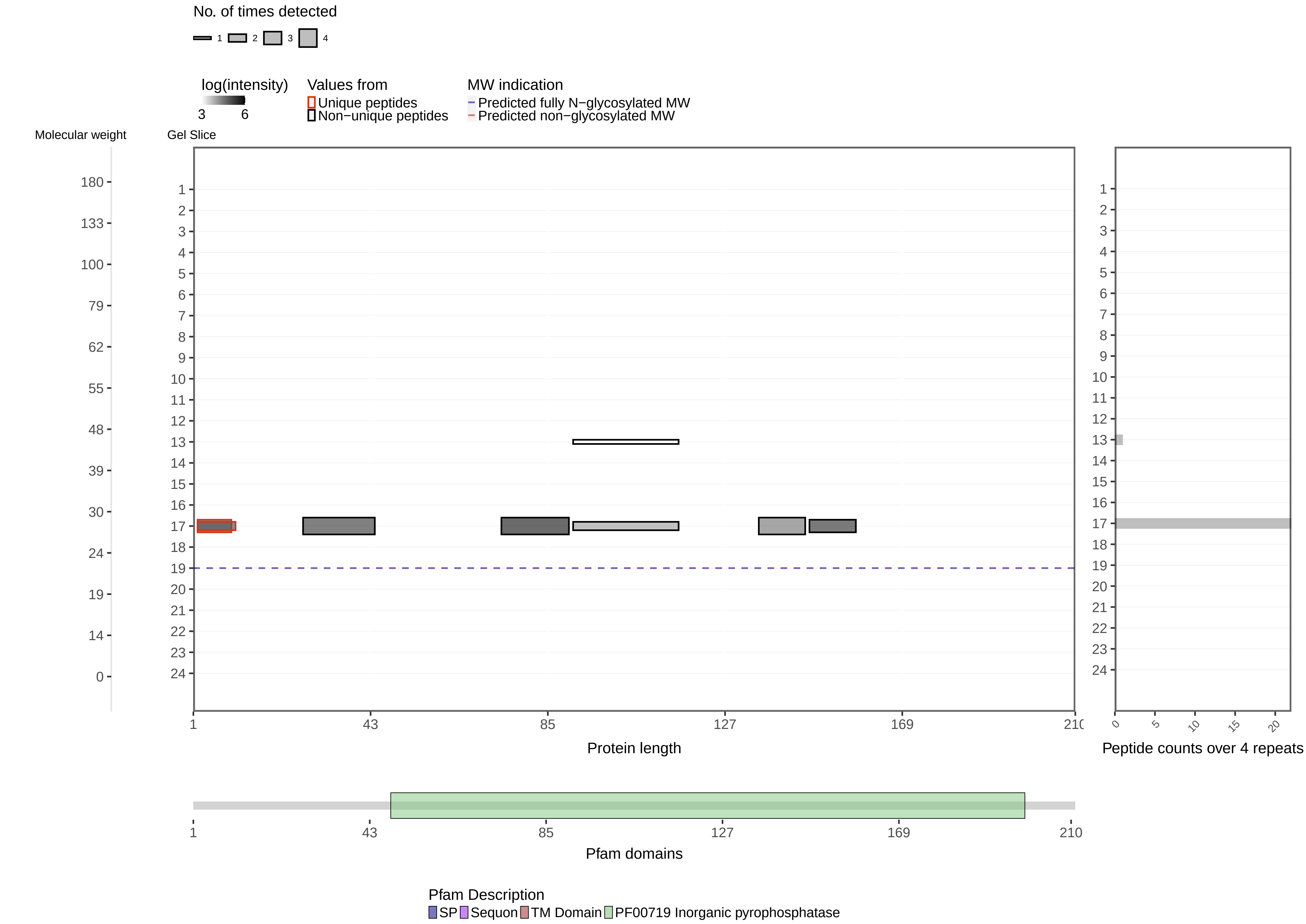


Protein sequence:

TLCEDNDPMDVLVLMQEPVLPGCFLRARAIGLMPMIDQGEKDDKIIAVCADDPEYRHYTDINQLAPHRLAEIRRFFEDYKKLENKEVAVNEFLPPNTAVDAIQYSMDLYAEYILETLRK*

### NbL10g24410.1 uncharacterized_LOC104214638 [MW = 61.16 kDa]

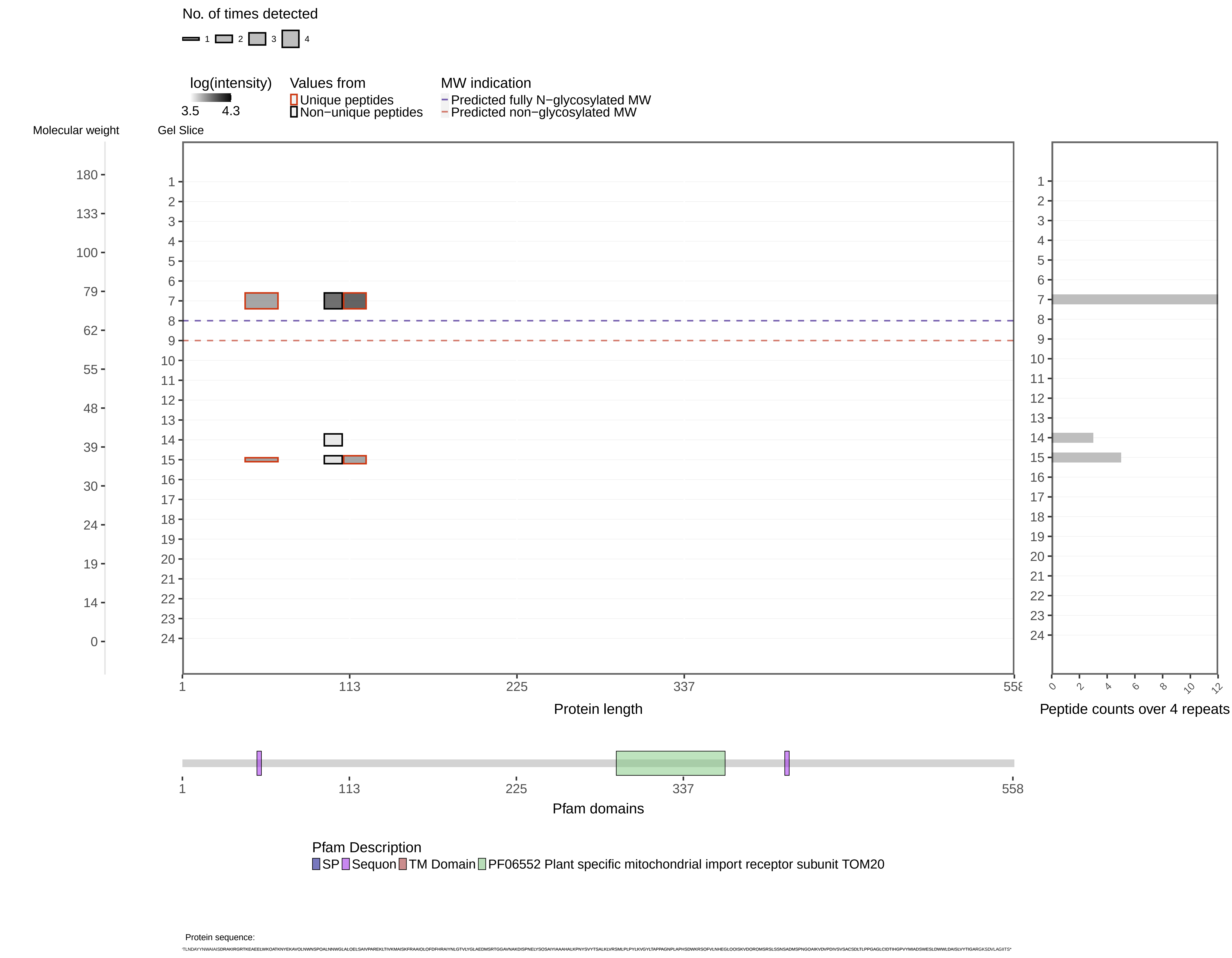

### NbL10g24170.1 probable_trehalase [MW = 65.95 kDa]

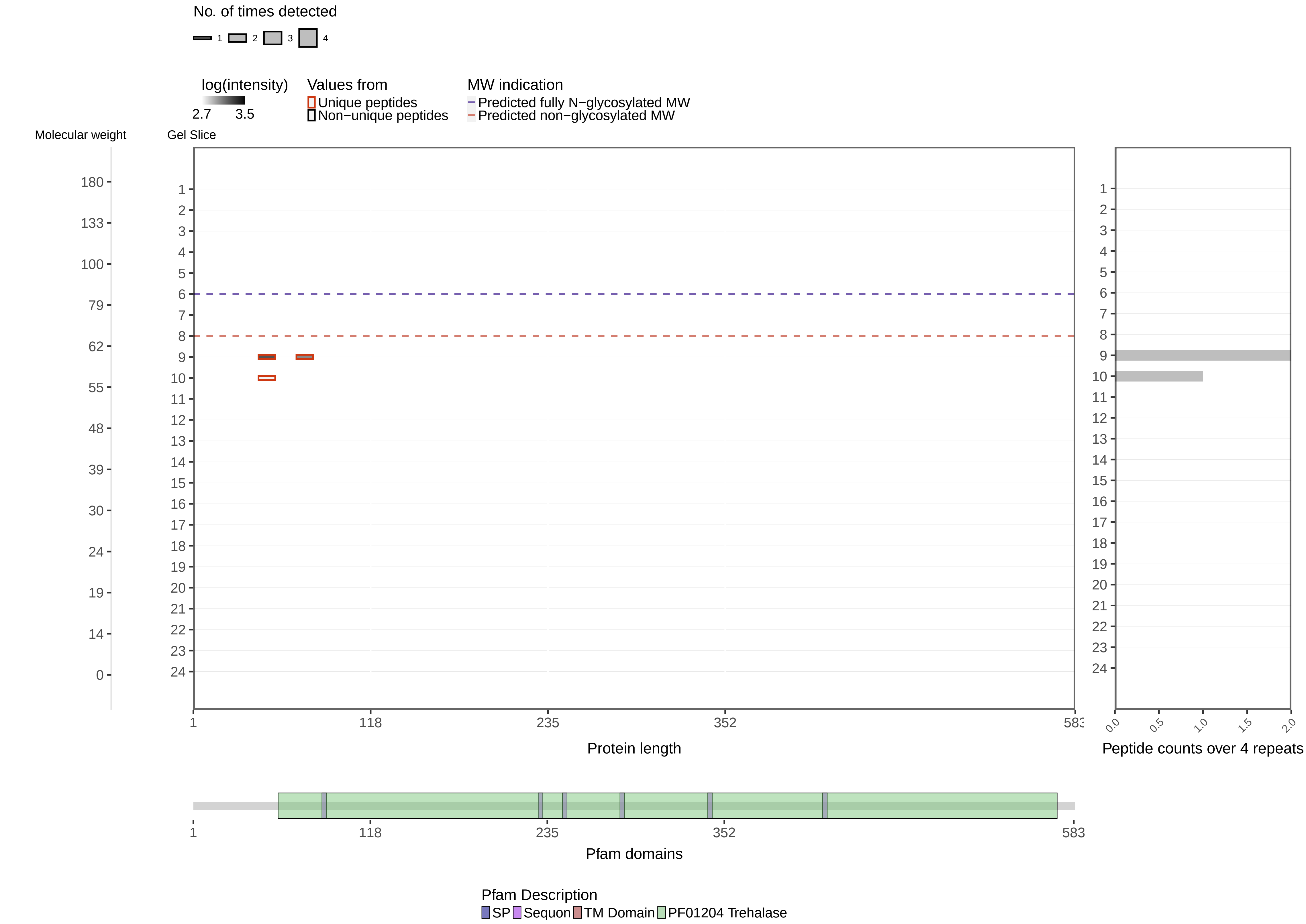


Protein sequence:

[illegible]

### NbL10g24160.1 ABC_transporter_F_family_member_3−like [MW = 80.39 kDa]

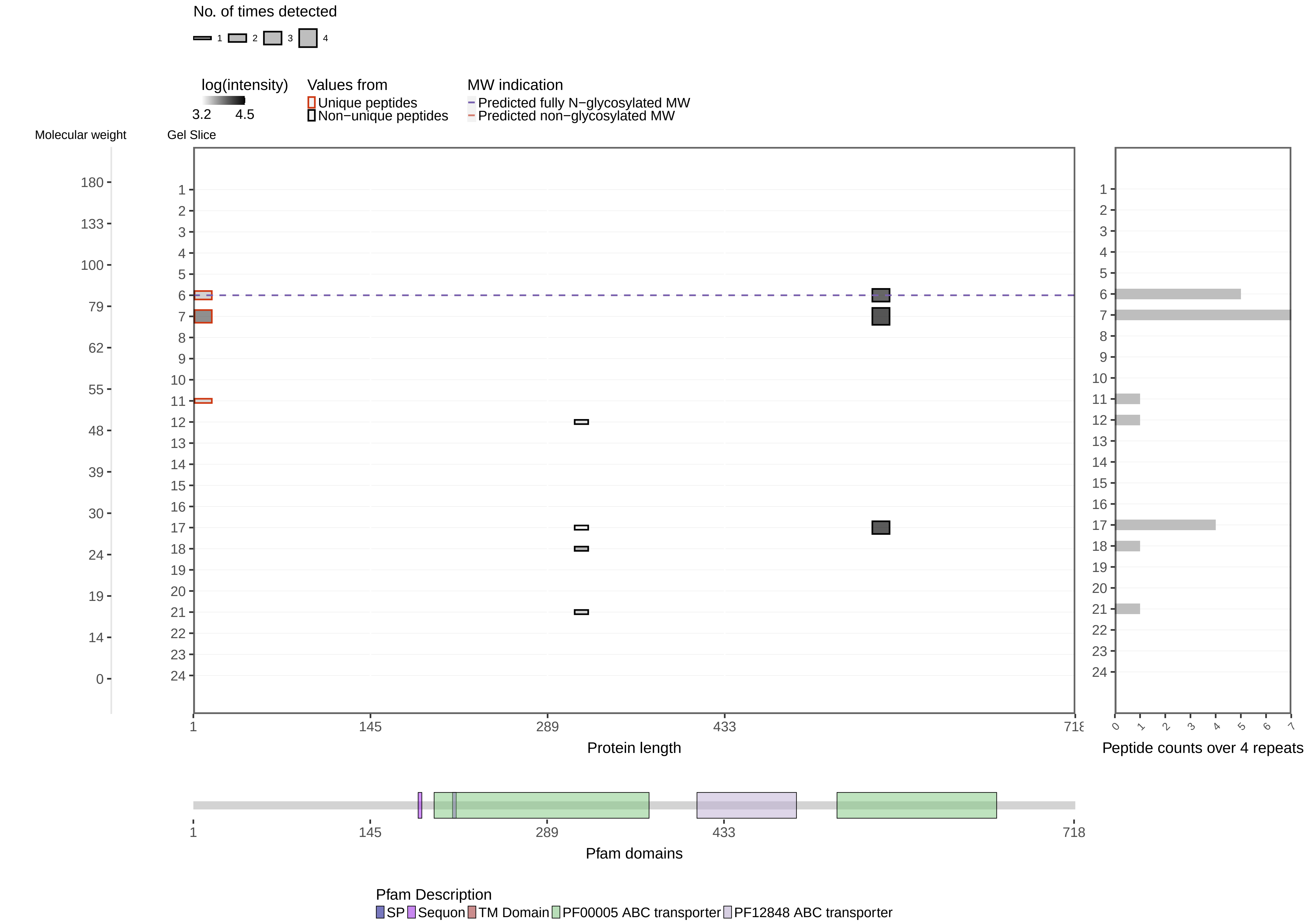


Protein sequence:

[illegible]

### NbL10g24150.1 glutamine−dependent_NAD(+)_synthetase [MW = 79.86 kDa]

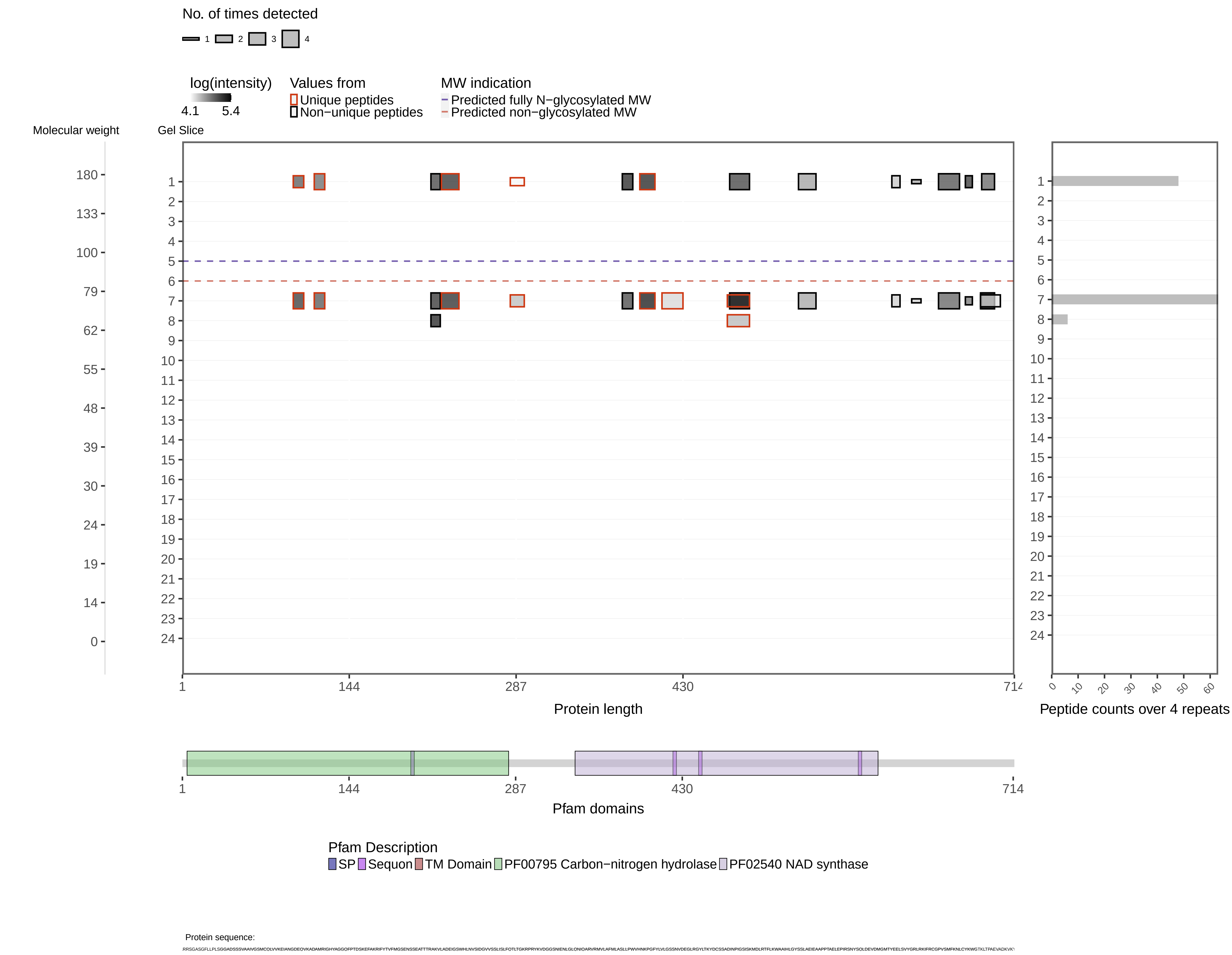

NbL10g24140.1 luminal−binding_protein [MW − SP = 70.64 kDa]

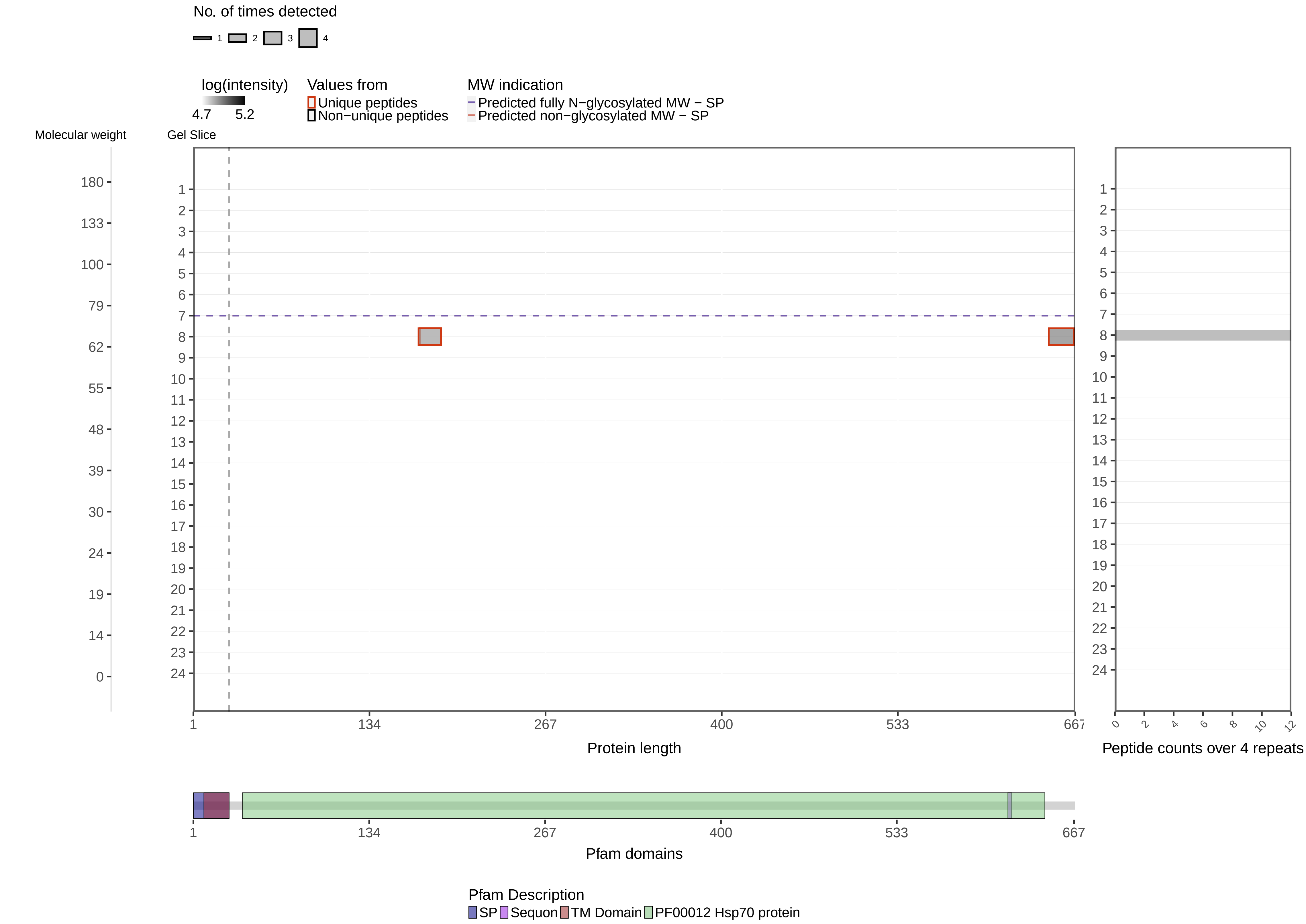


Protein sequence:

DFSEPLTRARFEELNNDLFRKTMGPVKKAMEDAGLEKNQIDEIVLVGGSTRIPKVQQLLKDYFDGKEPNKGVNPDEAVAYGAAVQGGILSGEGGDETKDILLLDVAPLTLGIETVGGVMTKLIPRNTVIPTKKSQVFTTYQDQQTTVSIQVFEGERSLTKDCRLLGKFDLTGIAPAPRGTPQIEVTFEVDANGILNVKAEDKGTGKSEKITITNDKGRLSQEEIERMVREAEEFAEEDKKVKERIDARNSLETYVYNMKNQINDKDKLADKLESDEKEKIETAMKEALEWLDDNQS

NbL10g24020.1 signal_recognition_particle_19_kDa_protein−like [MW = 18.44 kDa]

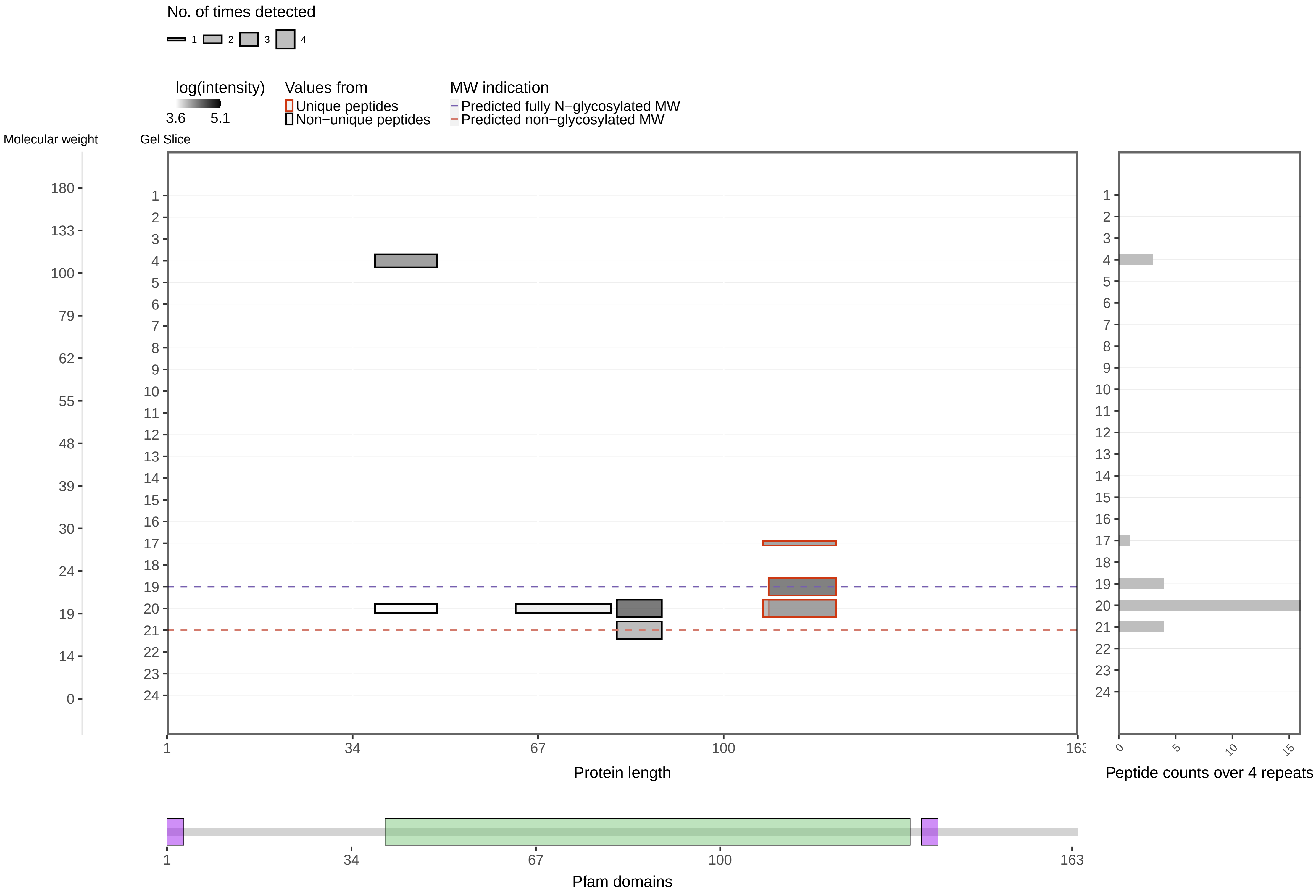


Protein sequence:

TEISDCCNYLKIPCAIELDKAYPRDFMQRGRVRVLLKKEDGTPYNPAISSRKQLMIHVAELVRRHPNRTKKQEPASSSAAGSSKSGKGGKKKR*

NbL10g23990.1 26S_proteasome_non−ATPase_regulatory_subunit_8_homolog_A−like [MW = 30.86 kDa]

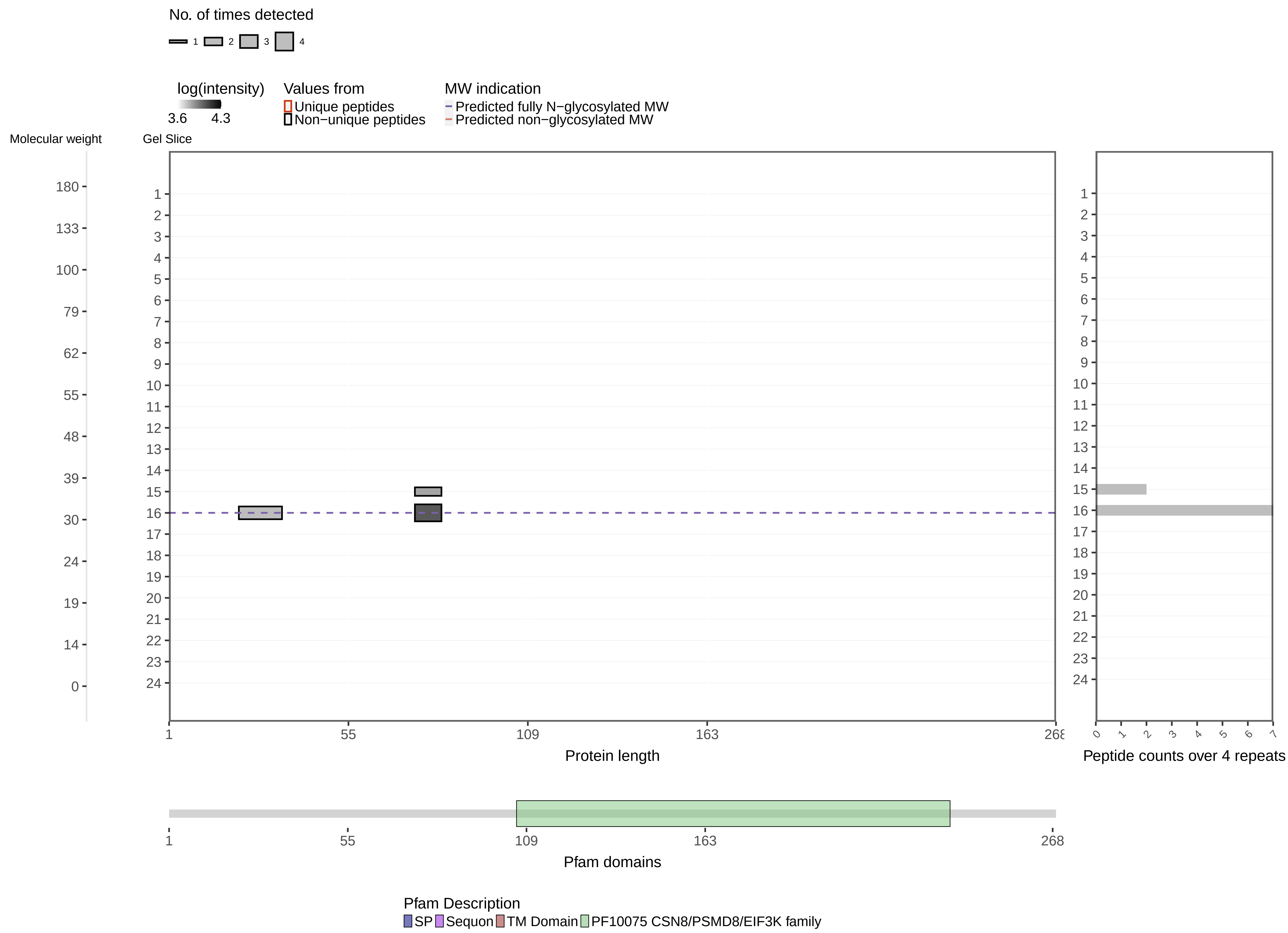


Protein sequence:

IRIAEFHTELELLSQSALDNPCIKHAVELEQSFMEGAYHRVLSARQTVPDATYAYFMDLLAKTVRDEIAGCSEKAYDRLSTSDARQMLLFSSDQELFEYVREEHPEWEIKDGFIVFQKVKDSVPCKEIPSLQLITQTLSYARELERIV*

### NbL10g23940.1 RNA_polymerase_II_transcriptional_coactivator_KELP [MW = 19.16 kDa]

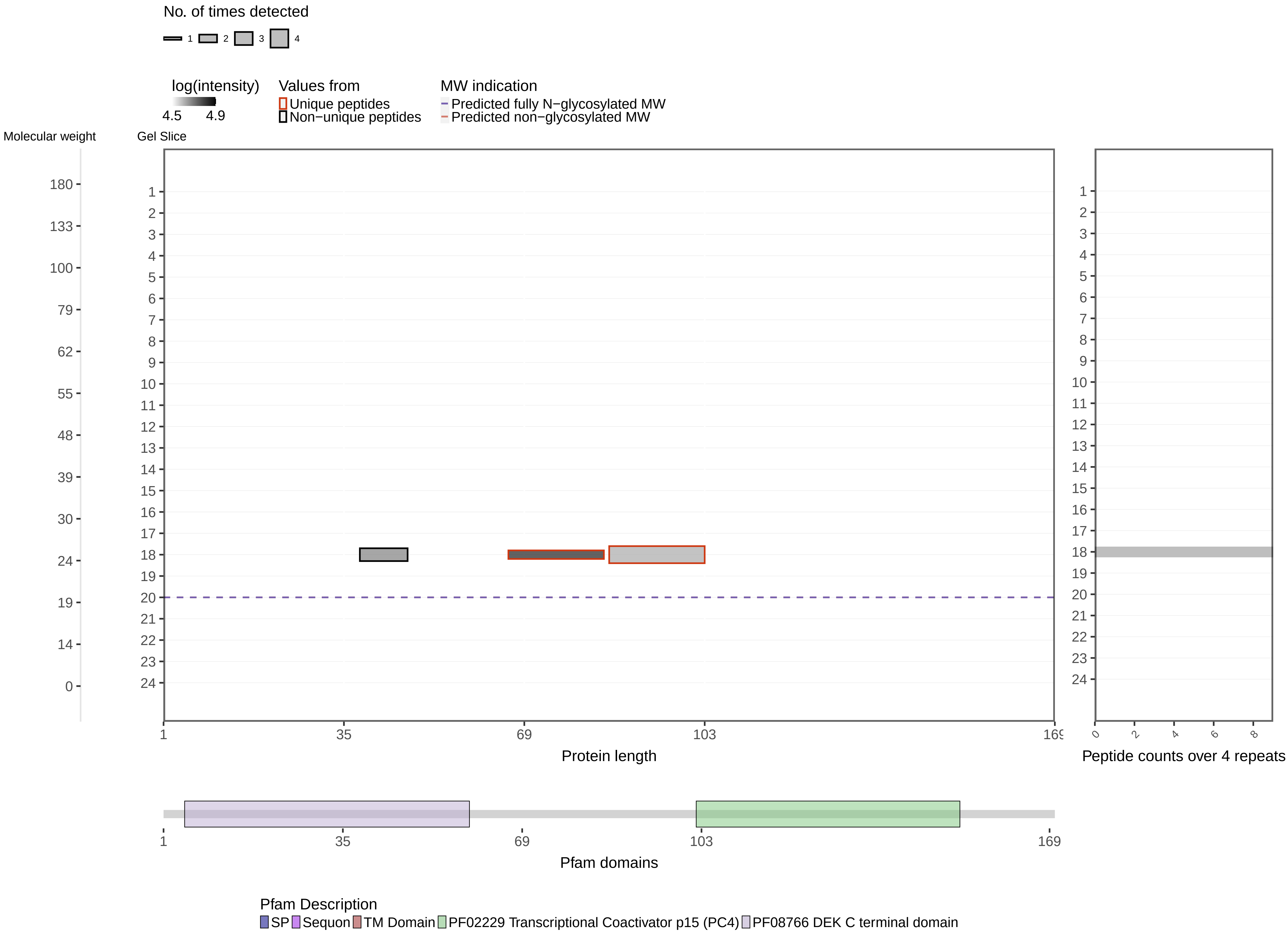


Protein sequence:

EEEEEEEEEDNKKKSSGAGDKEYDDDGDLIVCRLSHKRRVTITEFRGKTLVSIREYYNKDGKELPTAKGISLTAEQWATFNKNIPAVEKAIKKMESRA*

### NbL10g23840.1 UDP−glucose_4−epimerase_GEPI48−like [MW = 38.68 kDa]

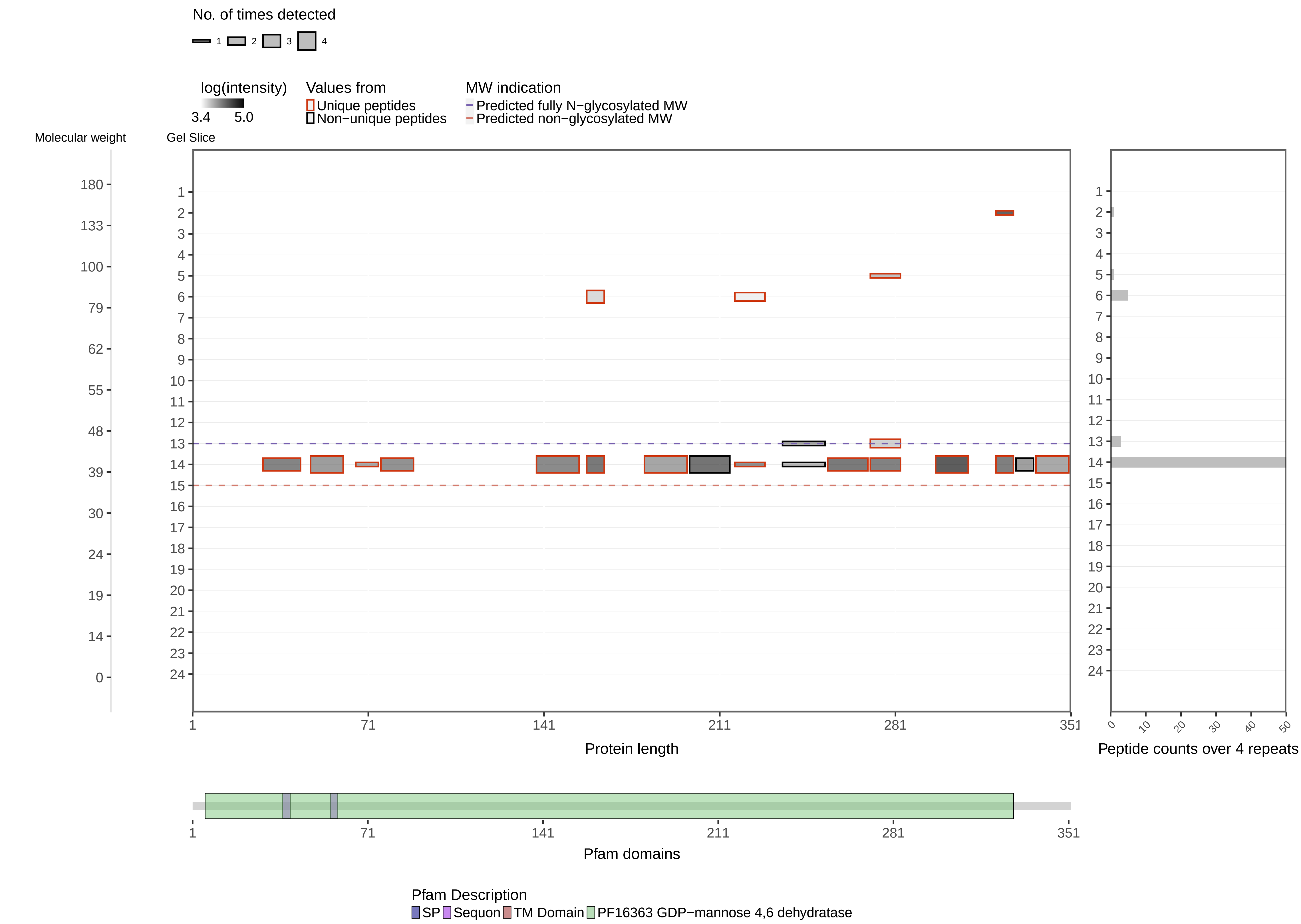


Protein sequence:

RDVRSSDSEWKIILLRYFNPVGAHSSGYIGEDPRGIPNNLMPFVQQVAVGRRLALTVYGTDYATKDGTGVRDYIHVVDLADGHIAALQKLSDPSIGCEVYNLGTGKGTSVLEMVAAFEKASGKKIPLVMSGRRPGDAEIVYAATQKAERELKWKAKYGIEEMCRDQWNWASKNPYGYEGIQESNCH*

NbL10g23810.1 NAD(P)H−quinone_oxidoreductase_subunit_S_chloroplastic−like [MW = 27.22 kDa]

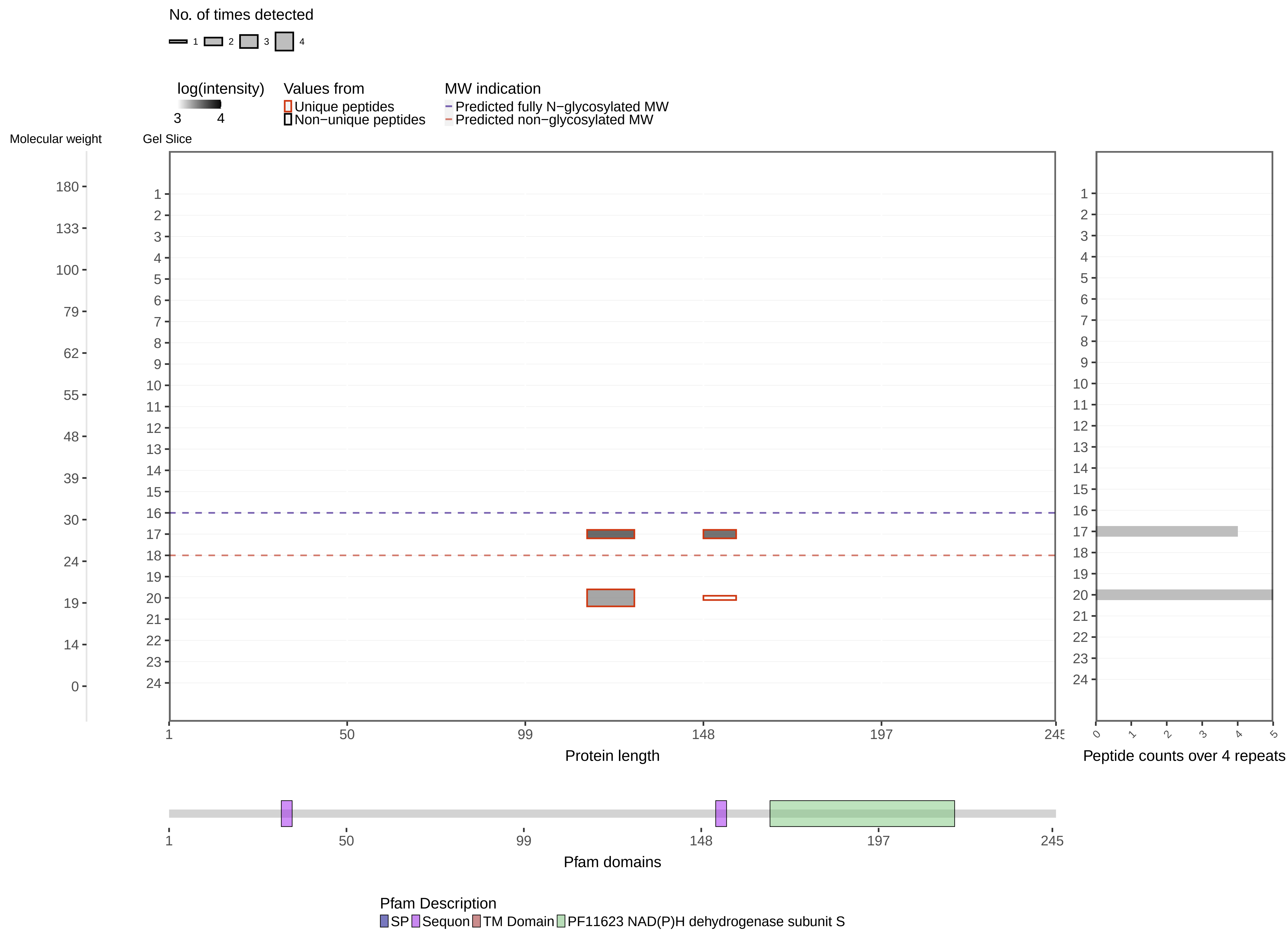


Protein sequence:

EDGFEKEMMGLVGGFPGGEKGLKKFIEKNPPPKKIESRIDGFNLSLVKKPKPPELPLLLPGMIAIVKNSNNPYYMYCGIVQRITDGKATVLFEGGNWDRLISFRLEELERREKGPPMVNPRSVILETMLEKSSES*

### NbL10g23720.1 long_chain_acyl−CoA_synthetase_4−like [MW = 73.41 kDa]

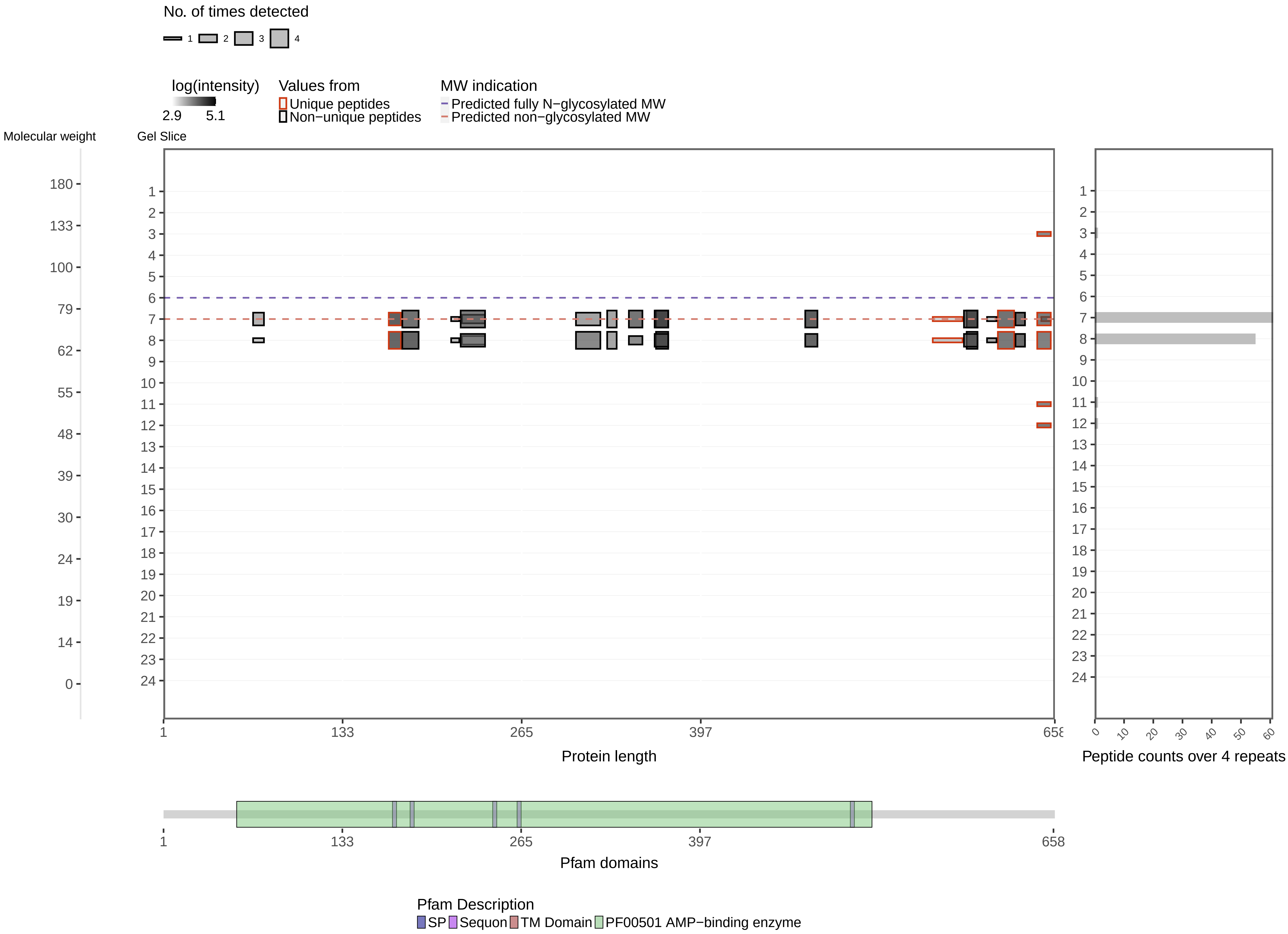


Protein sequence:

[illegible]

NbL10g23590.1 probable_polygalacturonase [MW − SP = 46.13 kDa]

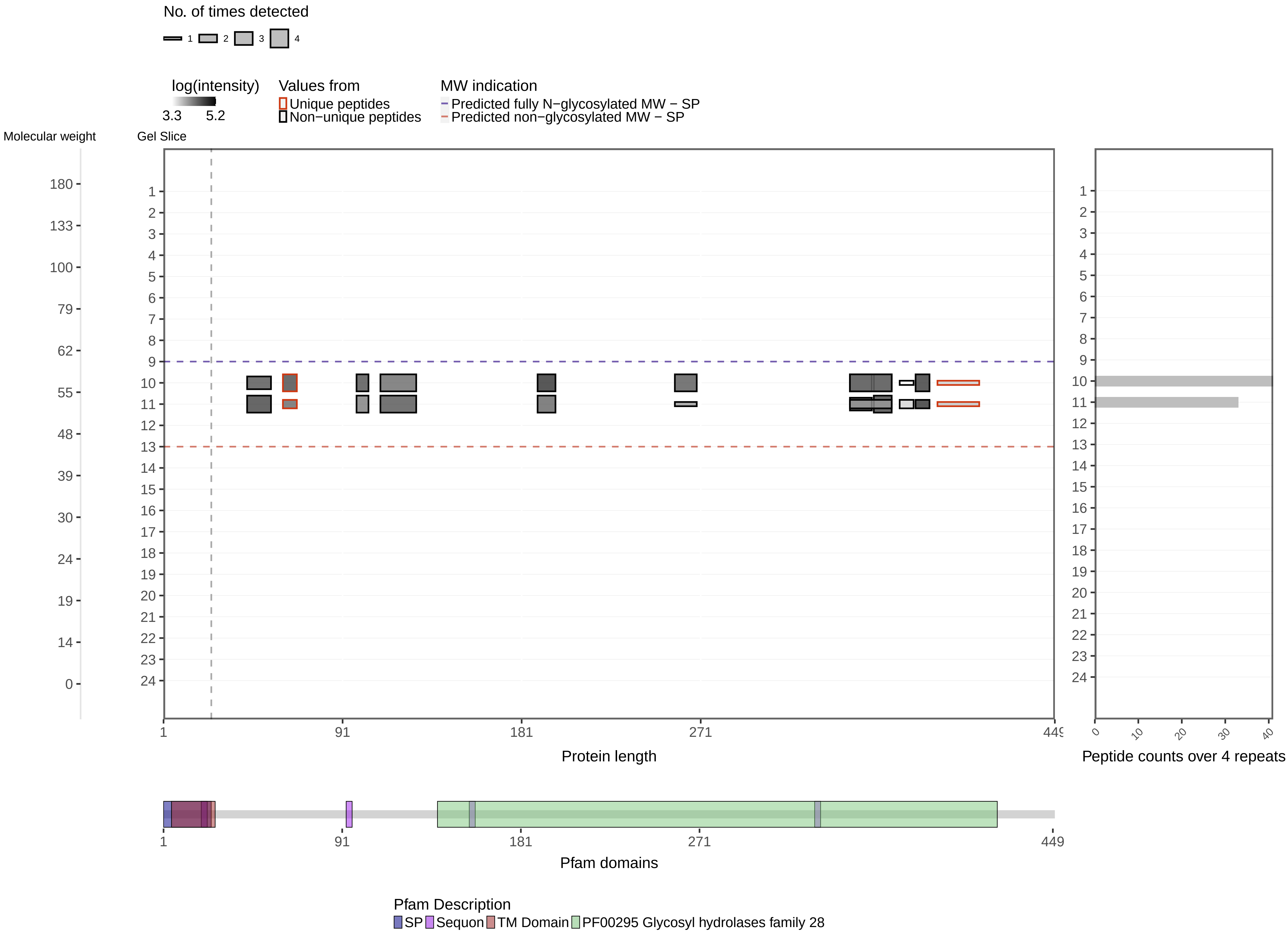


Protein sequence:

'CSNVVIRYVTILAPADSPNTDGIDPDSSSHVCIEDSYISVGDDLVAVKSGWDQYGIAYGRPSHGITIRRITGSSPFAGIAVGSETSGGVVDVLAEHINLFNMGVGIHIKTNIGRGGVIRNITVSNVYMENTRTGIKIAGDVGDHPDEKFNPNALPIVKGIKIKDVWGEKVLQAGLIRGLKNSPFSGICLSNINLNGNNPGPRKSPWKCSDVSGAAIQVSPWPCSELTSSQQTGECSTYF*

### NbL10g23460.1 myosin−11−like [MW = 158.7 kDa]

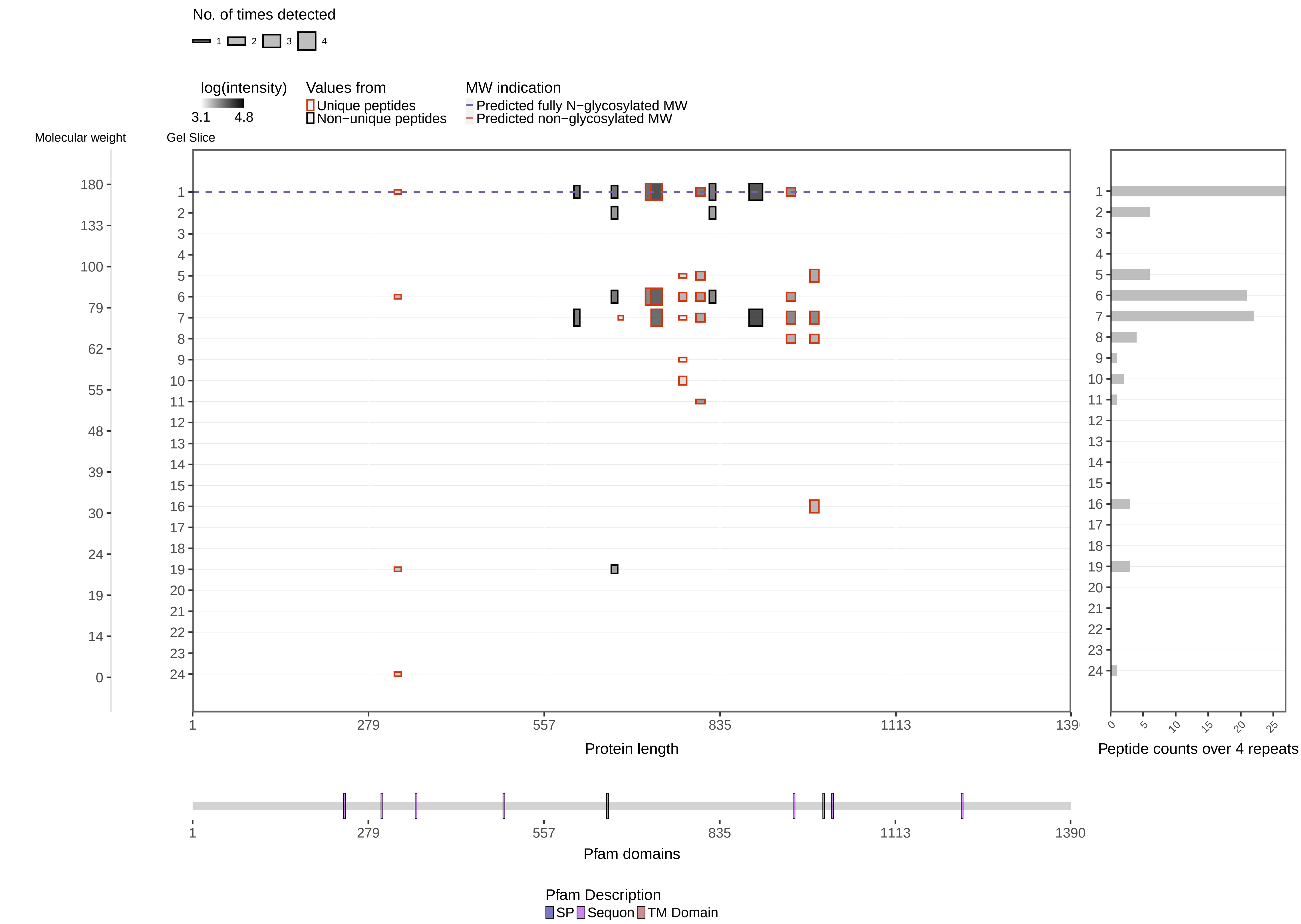


Protein sequence:

[illegible]

### NbL10g23400.1 putative_glucose−6−phosphate_1−epimerase [MW = 34.22 kDa]

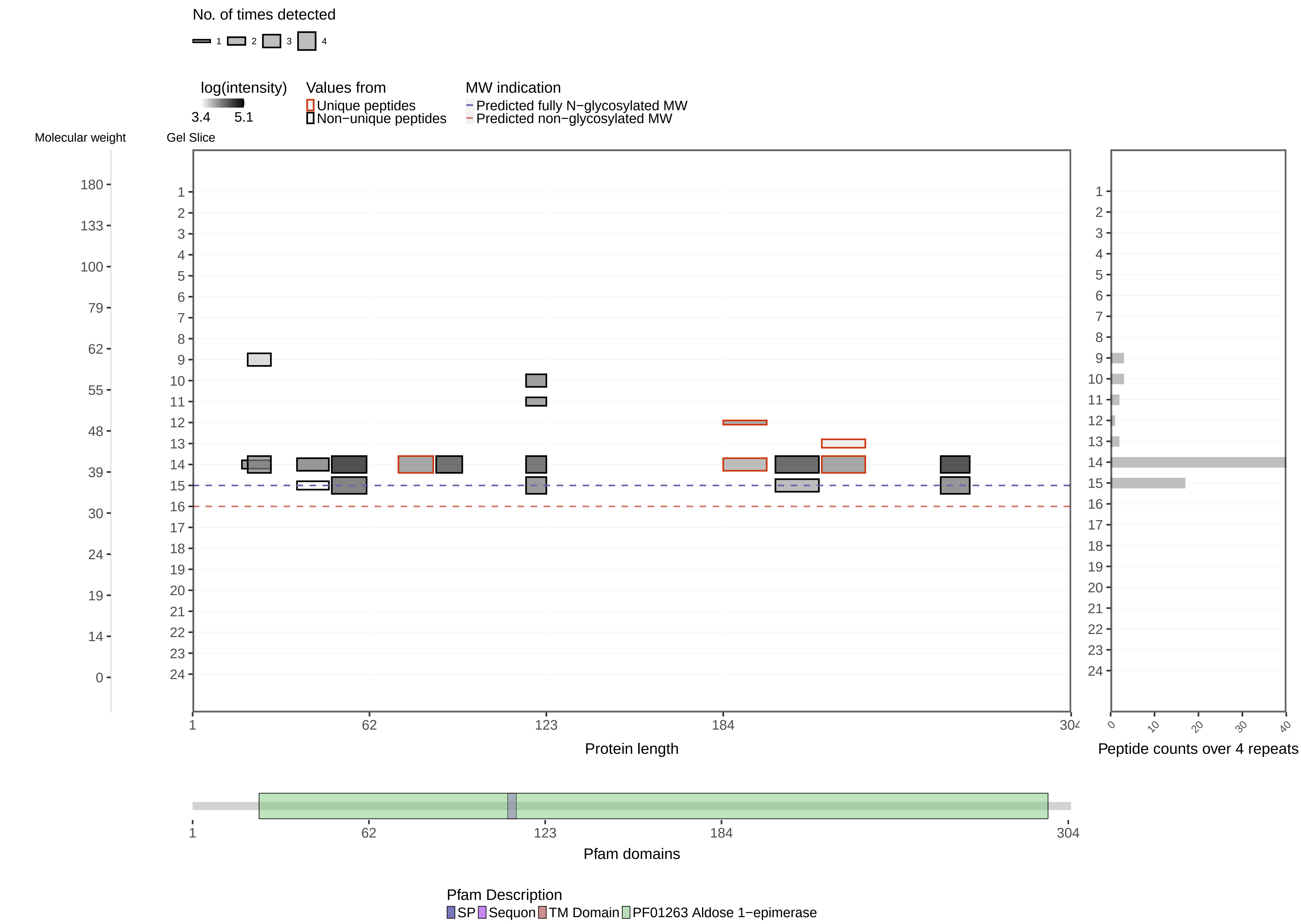


Protein sequence:

EFRLRVTLAFDGSLTLISRIRNVNCKPFSFSIAYHTYFALSDISEVRVEGLETLDYLDNLCNRERFTEQGDALTFETEVDRVYLSSSDVIAIFDHEKKRTFVIKREGLPDVVVWNPWEKKSKTIADFGDDEYRHMLCVDGAAIEKPITLKPGEEWTGRLELSVMPSS*

NbL10g23380.1 V−type_proton_ATPase_subunit_E−like [MW = 26.21 kDa]

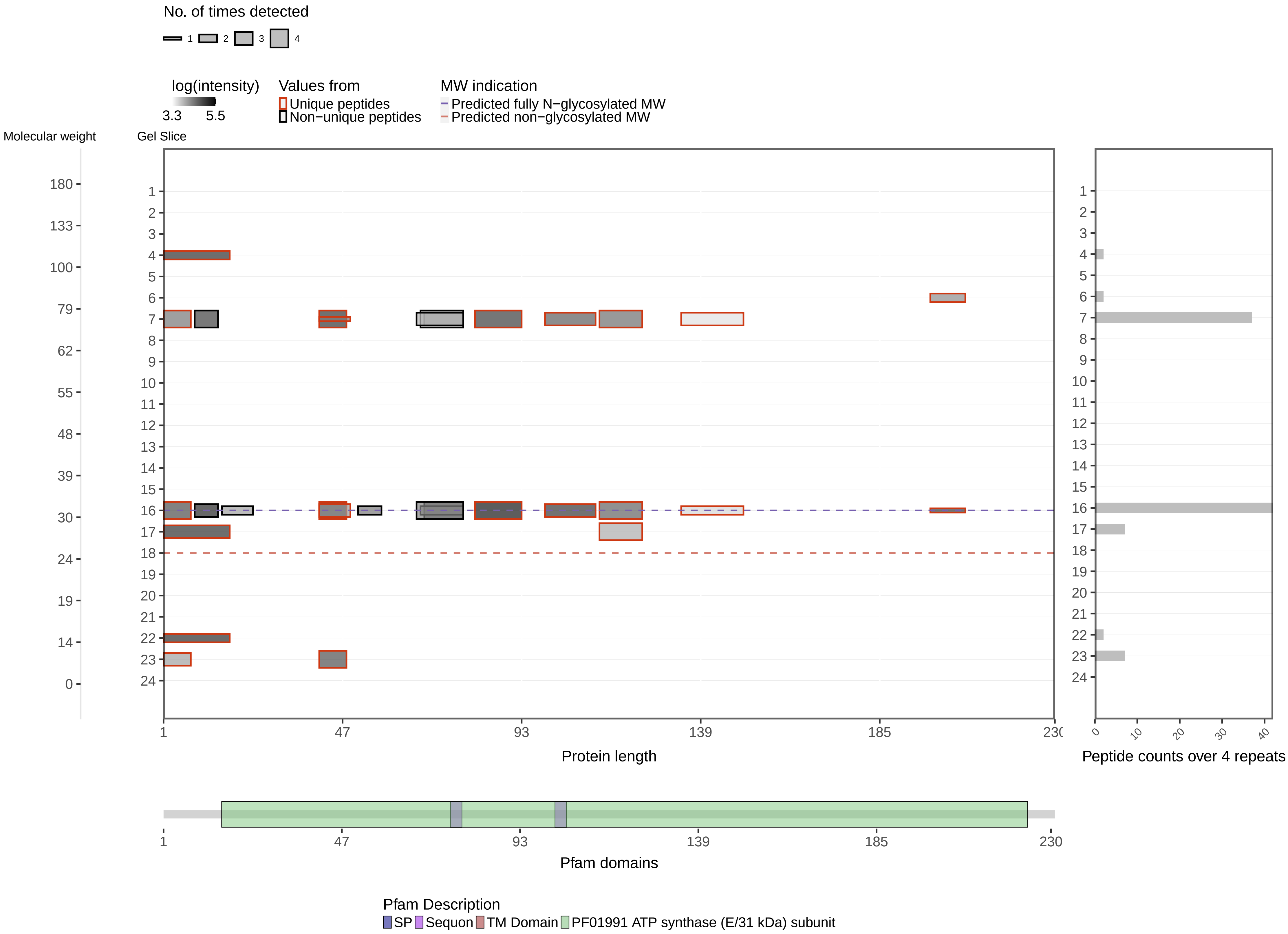


Protein sequence:

NVSQNHHIYEKLLQDLIVQSLLRLKEPCVLLRCREDDVSLVEGVLDAAKEEYAEKSQVHSPEVIIDQIYLPSAPSHHNAHGSSCYGGVVLASRDGKIVCENTVDARLEVVFRKKLPEIRKCLFGQIAA*

NbL10g23050.1 24−cyclodiphosphate_synthase_chloroplastic [MW = 24.3 kDa]

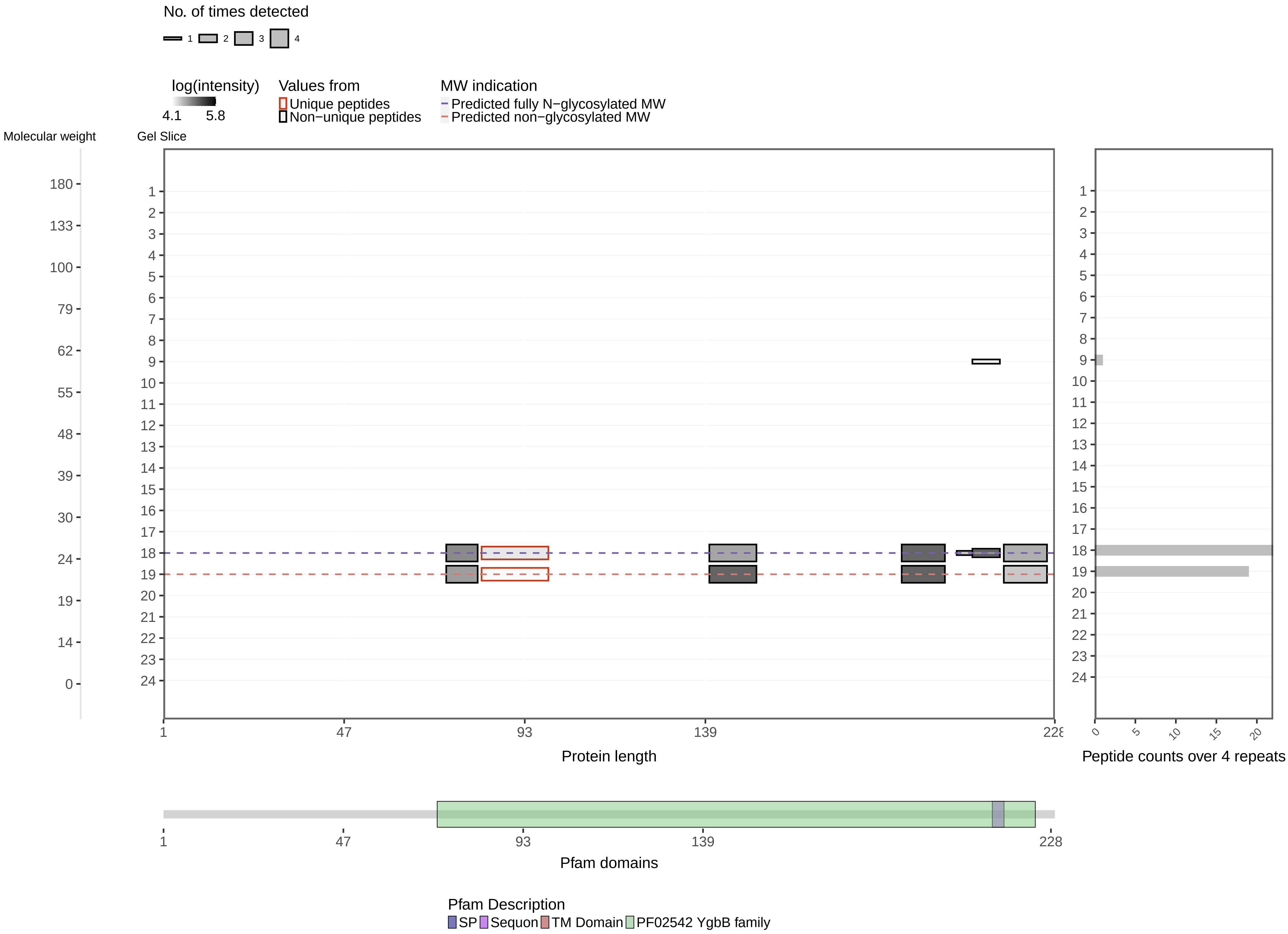


Protein sequence:

AHSDGDVLLHCVVDAILGALGLPDIGQIFPDTDPKWKGAPSSVFMEEAVRLMHEAGYELGNLDATLILQRPKVSPHKEAIRANLCKLLGADPSVVNLKAKTHEKVDSLGENRSIAAHTVVLLMKK*

NbL10g22980.1 monodehydroascorbate_reductase_chloroplastic [MW = 53.34 kDa]

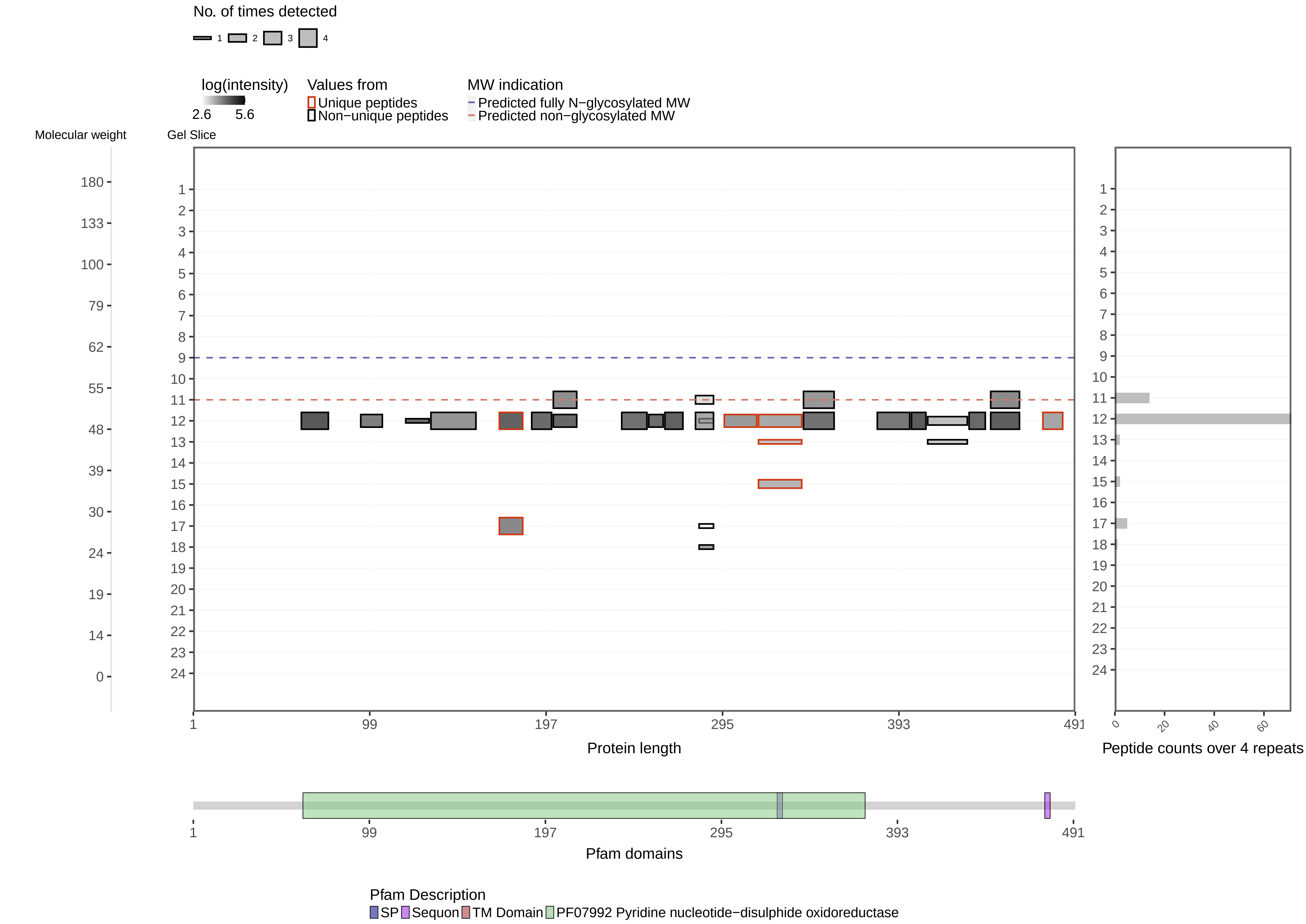


Protein sequence:

AAAVAWKLDTTIIFPEEHLLPRLFTPSLAQKYEQLYQDNGVKFVKGAKIKHLESGPDGRVVAVKLEDGSSIETDTVVIGIGAKPAVSPFDVVGLNSTVGGIAVDGQFRTSIPGIFAIGDVAAFPLKIYNRIARVEHVDHARKSAQHCIKSLLTAHTHTYDYLPYFYSRVFEYEGSPRKVWWQFFGDSVGETVEVGNFDPKVATFWIDSGKLKGVLLESGSPEEFQLLPKLARSQPSVDKSKLQNASSVEEALKIAQASL*

NbL10g22960.1 probable_polygalacturonase [MW − SP = 48.99 kDa]

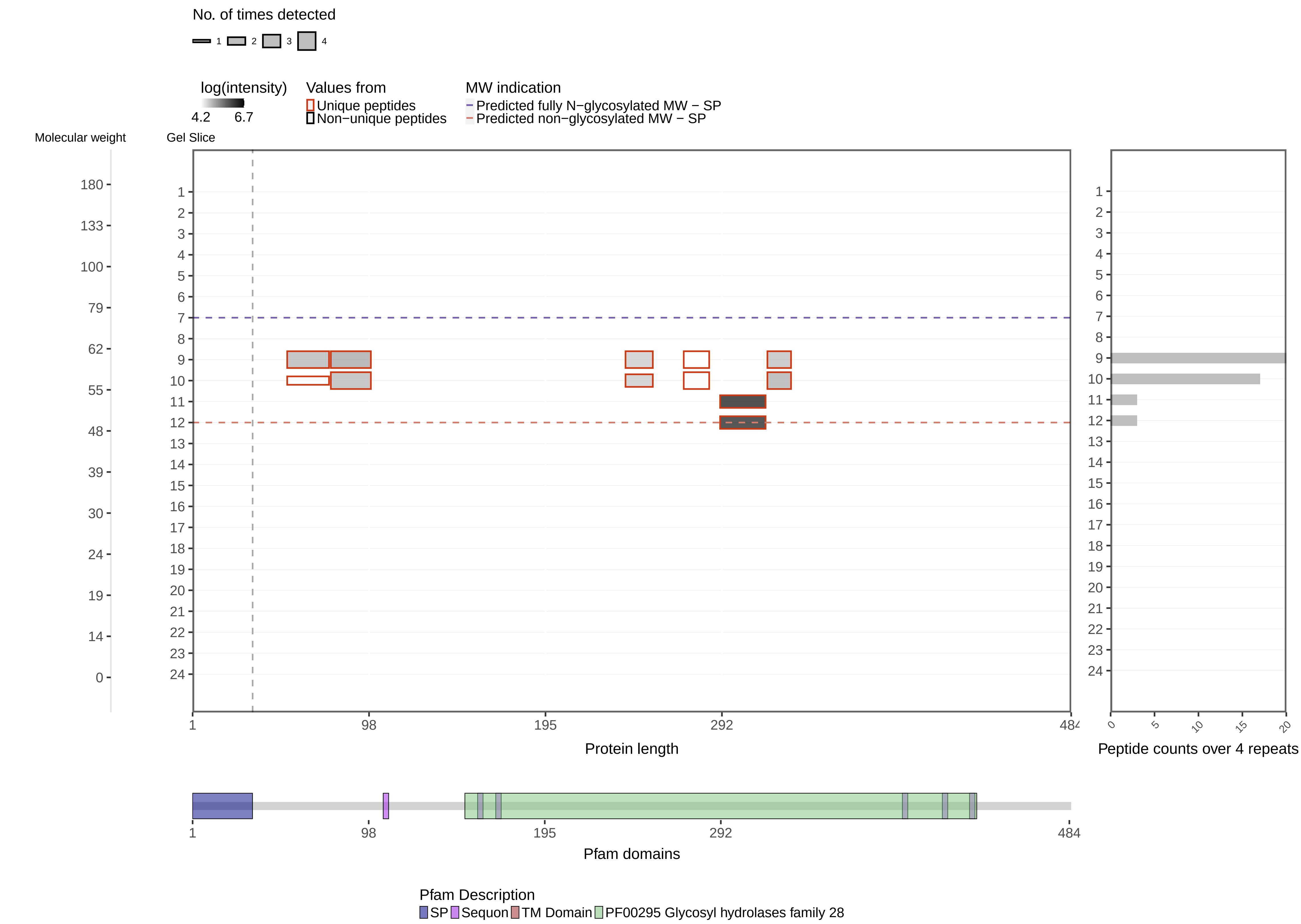


Protein sequence:

GITILAPVRSPNTDGINPDSCINTRIEDCYIVSGDDCIAVKSGWDEYGVAFGMPTKQLAIRRITCISPTSATIALGSEMSGGIQDVRAEDIVAINTESGVRIKTAVGRGGYVKDVYVKGVTMKTMKYVFWMTGDYGSHPDNNYDPNALPVIDNINYRDMVAENVTIAAKLAGISGDPFTGICISNVTIELAPKAKKLTWNCTDISGISSGVVPQPCELLPDQGTENAAPCNFPTENLPIDDMKVQTCSCRKKLY*

NbL10g22890.1 uncharacterized_LOC104246640 [MW = 39.62 kDa]

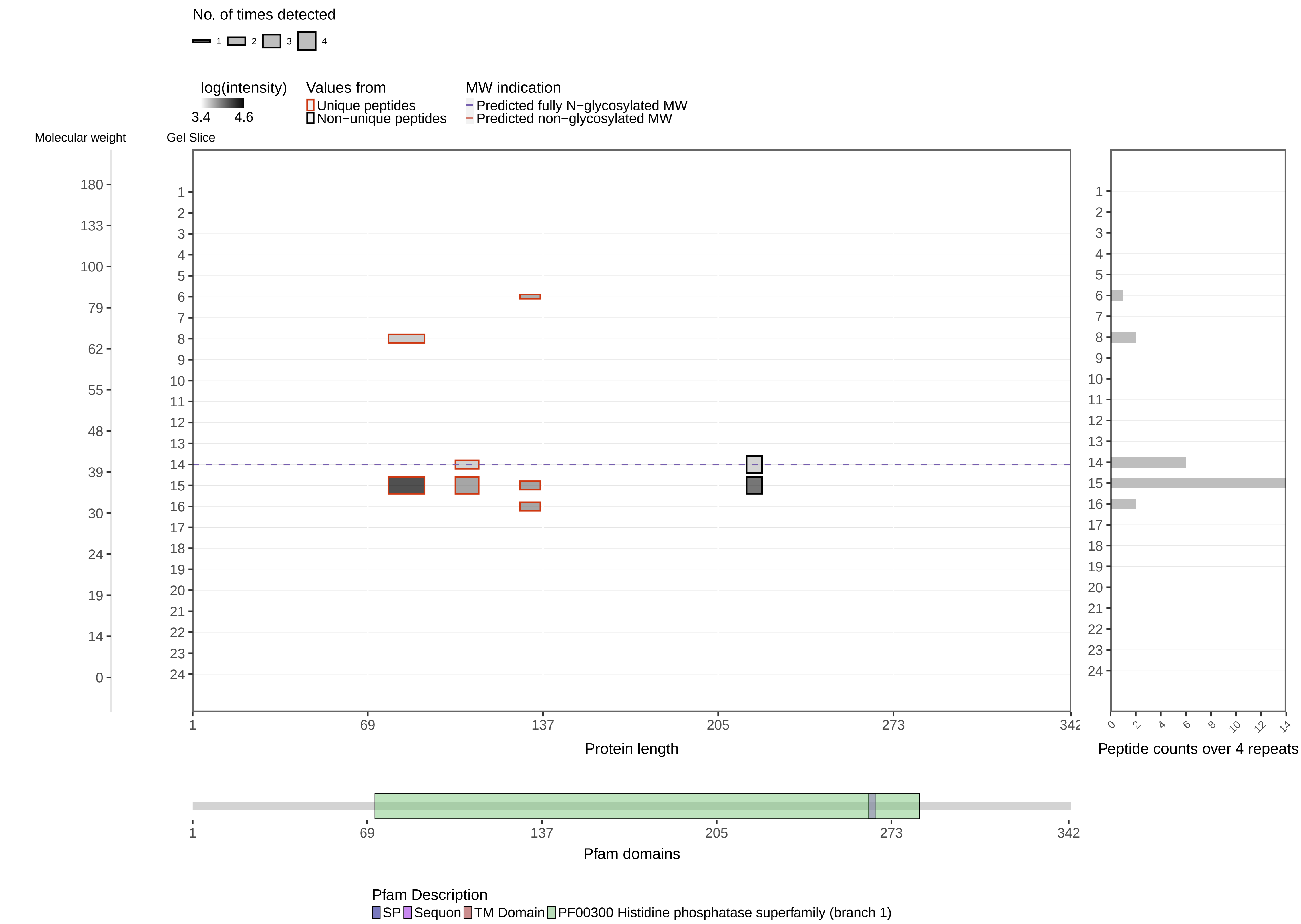


Protein sequence:

EPRLREQDFGNFODKEOMKIEKAVRARYGRFFYRFPNGESAADVYDRITGFRETLRSDIDIGRFOPPGOOSPNMNLVIVSHGLTLRVFLMRWYKWTVEOFEGLHNMSNGGMIVMERGYGGRYCLSVHHTKEELOKFGMTDEMLIDOEWOKIAKPGELNYDCLITGPSYFTHFDDDEEONFEL*

### NbL10g22730.1 vesicle−associated_protein_1−2−like [MW = 26.52 kDa]

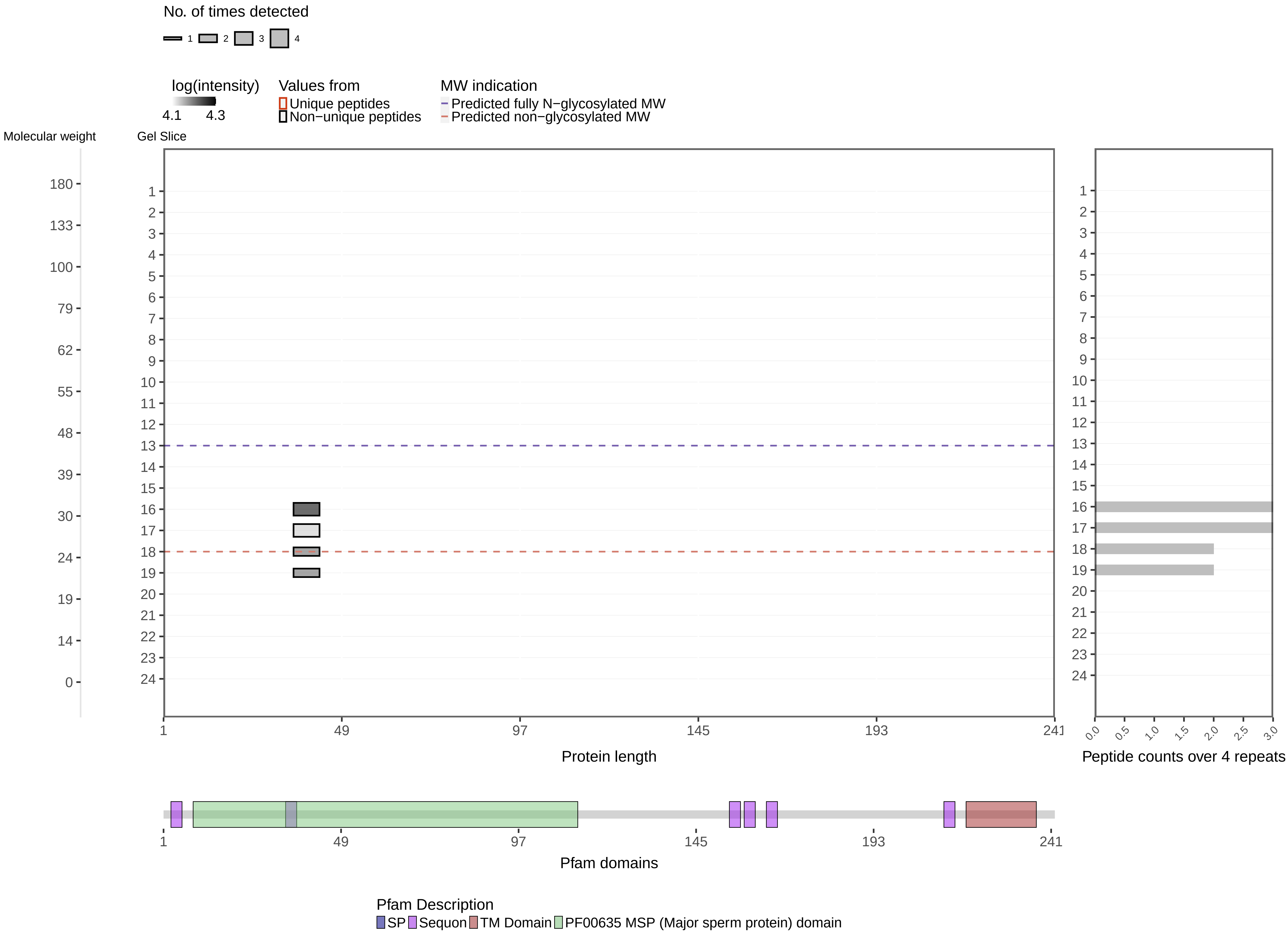


Protein sequence:

PEMFNKEEGNHVEDCKLRVVYVPPQQPPSPVQEGSEEGSSPRASVSENGTVNTSEFNNISRAYNEQQDSSSETKALISKLTEEKISVIQQNNKLQQELALLRRERNSSRGGIPTIYVLIIGLLGIILGYLLKKT*

NbL10g22420.1 uncharacterized_LOC107820933 [MW = 56.76 kDa]

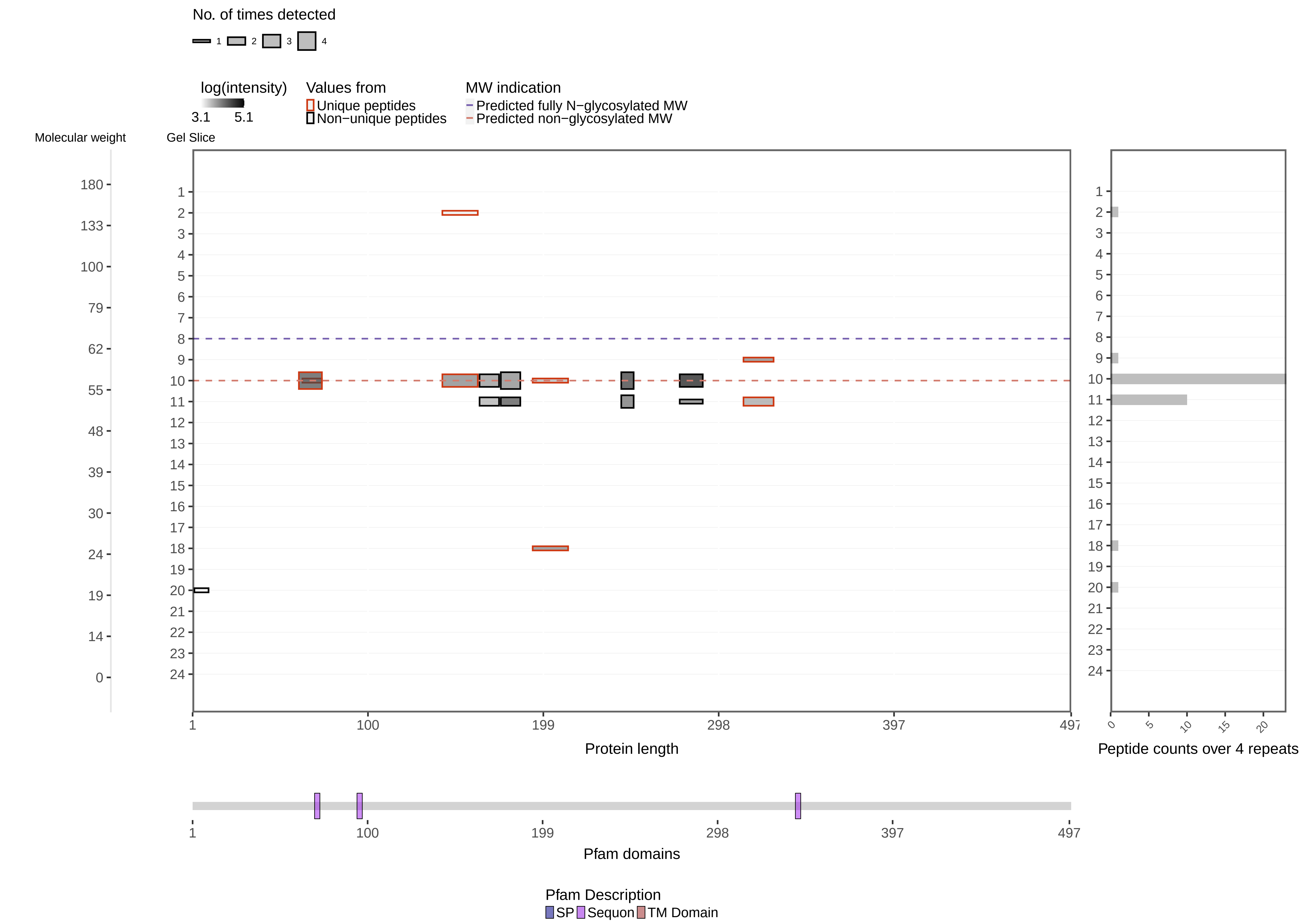


Protein sequence:

KSRHQKLPKFESFDFEKSDIMEEDIMGMEPGLVFFEDGSYSRGPVDIPVGELDESKYYISPTFKFEQCLVKGCHKRLRIVHTIEFSNGGSDIQILRVAVYEEQWISPANLSDESDTELDVKLFSQRRRLHPSELAGSWKVFEMSATPIYSEEGVTQETDDVPYVYLCTENLKKRSLPENPAYFGEEEMLDMQDVTVLWLPGGFTSYVDVNKDGILCIGVGWYSDEGVNLVMERDYEMDGKLKEVRWKSEMKRRWTNPPLIWLS*

NbL10g22330.1 acyl−CoA−binding_protein [MW = 13.74 kDa]

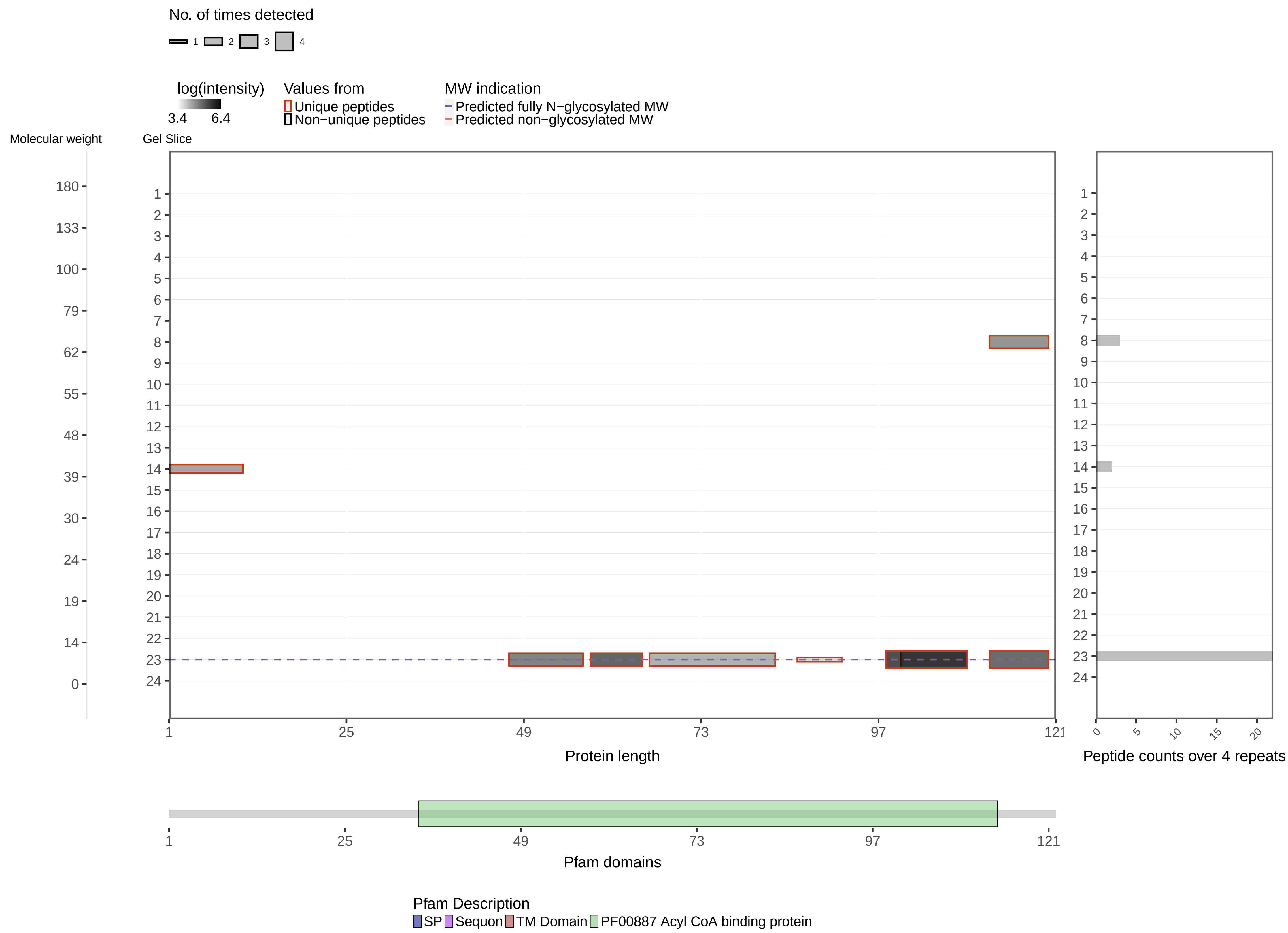


Protein sequence:

PESTSNENKLILYGLYKQATVGNVETCRPGIFNMRDRAKWDAWKAVEGKSKDEAMNDYITKVKQLKEEAASA*

### NbL10g22320.1 60S_ribosomal_protein_L13−2 [MW = 23.58 kDa]

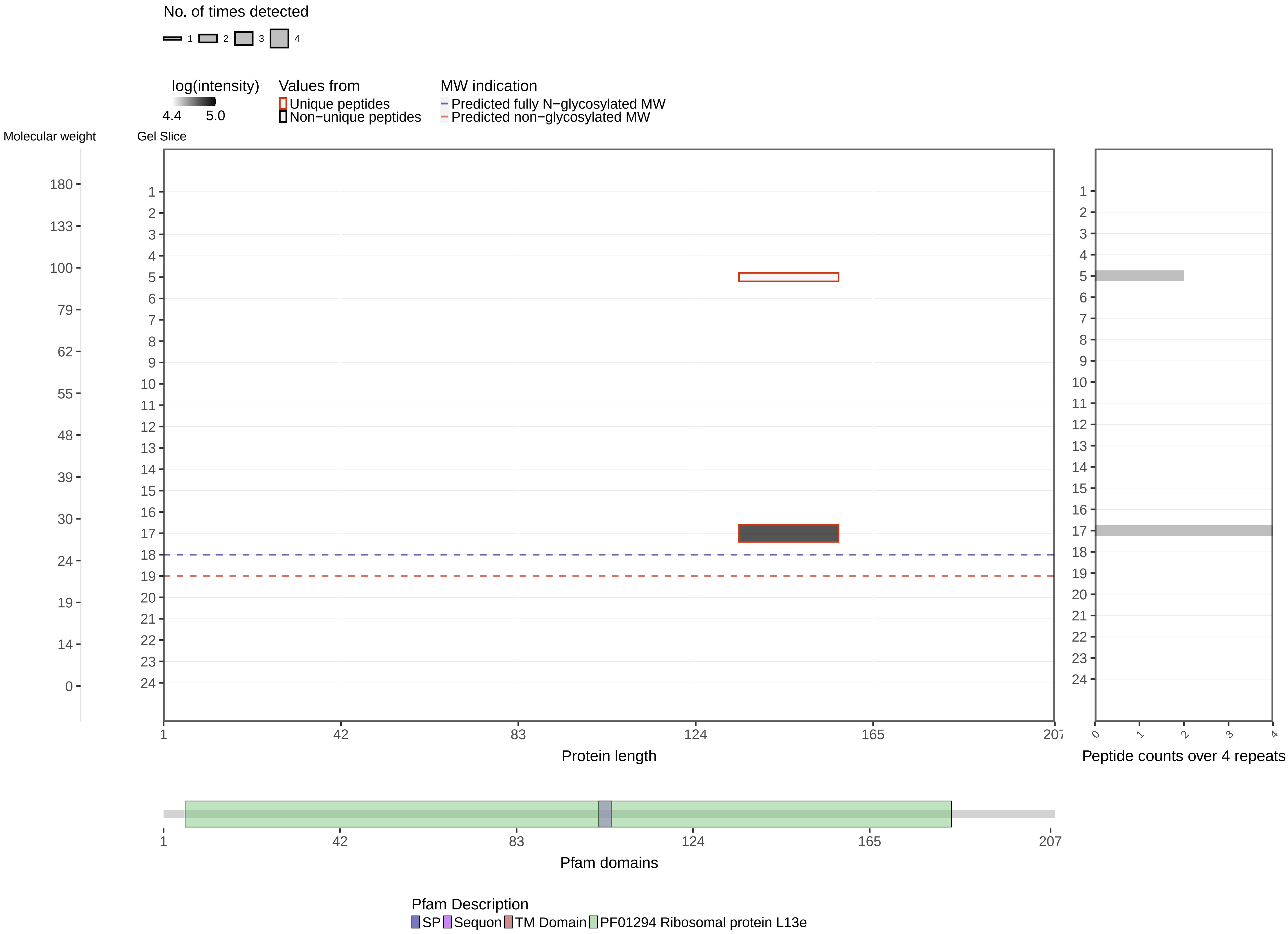


Protein sequence:

APTIGIAVDHRRRNRSLEGLQTNVQRLKTYKAKLVIFPRRAKKVKAGDSSVEELATATQVQGSYMPITREQPAVELVKVTDEMKSFNAYGKLRIERTNARHIGARLKRAAEAEKEEKK*

NbL10g22210.1 phosphate/phosphate_translocator_chloroplastic−like [MW = 43.91 kDa]

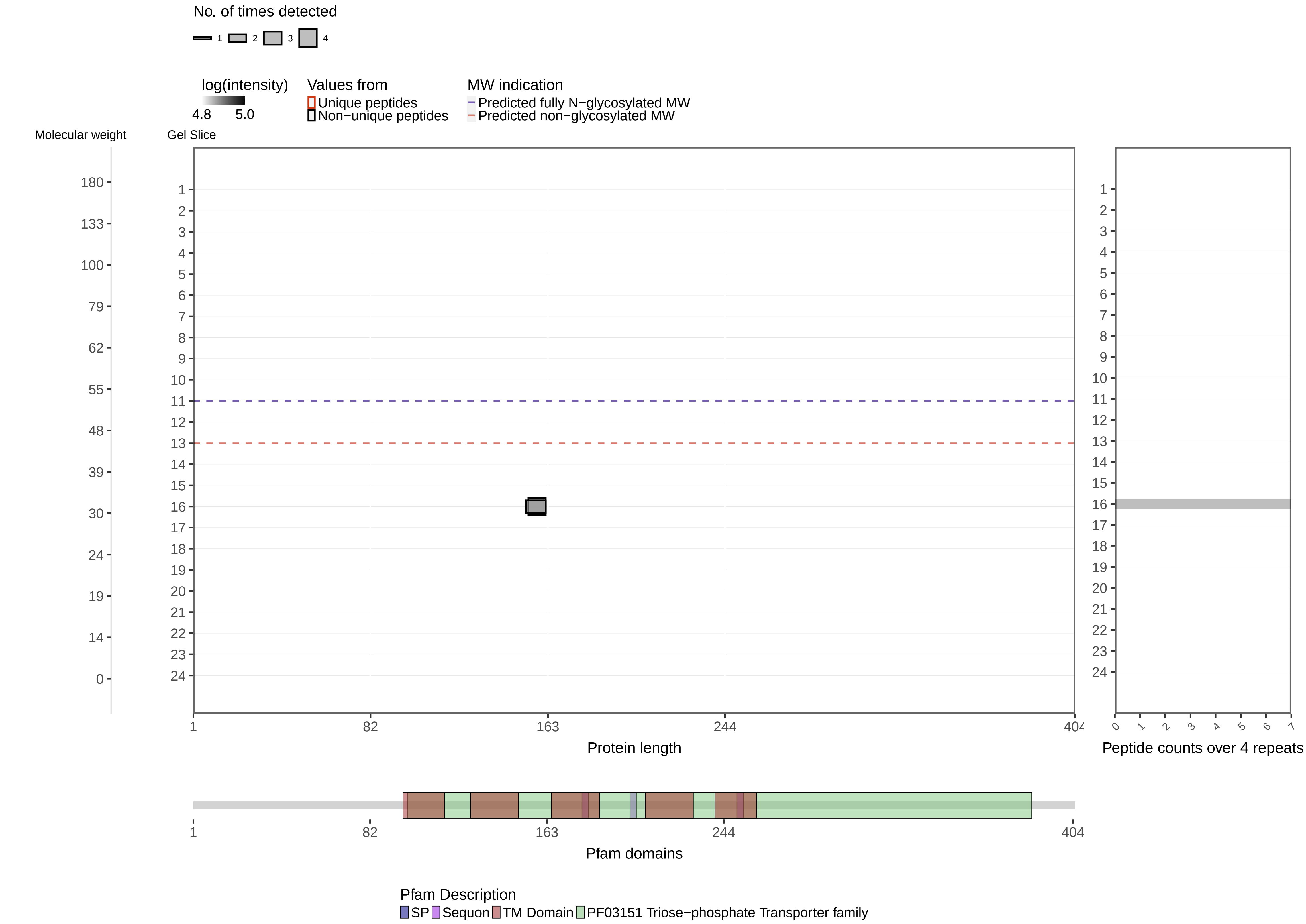


Protein sequence:

VSFTHTIKALEPFFNASASQFILGQQIPLALWLSLAPVVIGVSMASLTELSFNWLGFISAMISNVSFTYRSIYSKKAMTDMDSTNVYAYISIIALIFCIPPAIIIEGPQLLQHGFADAIAKVGLTKFVTDLFWVGMFYHLYNQVATNTLERVAPLTHAVGNVLKRVFVIGFSIIVFGNKISTQTGIGTCVAIAGVALYSFIKAKMEEEKRVSPTMQK*

### NbL10g22050.1 magnesium−chelatase_subunit_ChlI_chloroplastic [MW = 46.37 kDa]

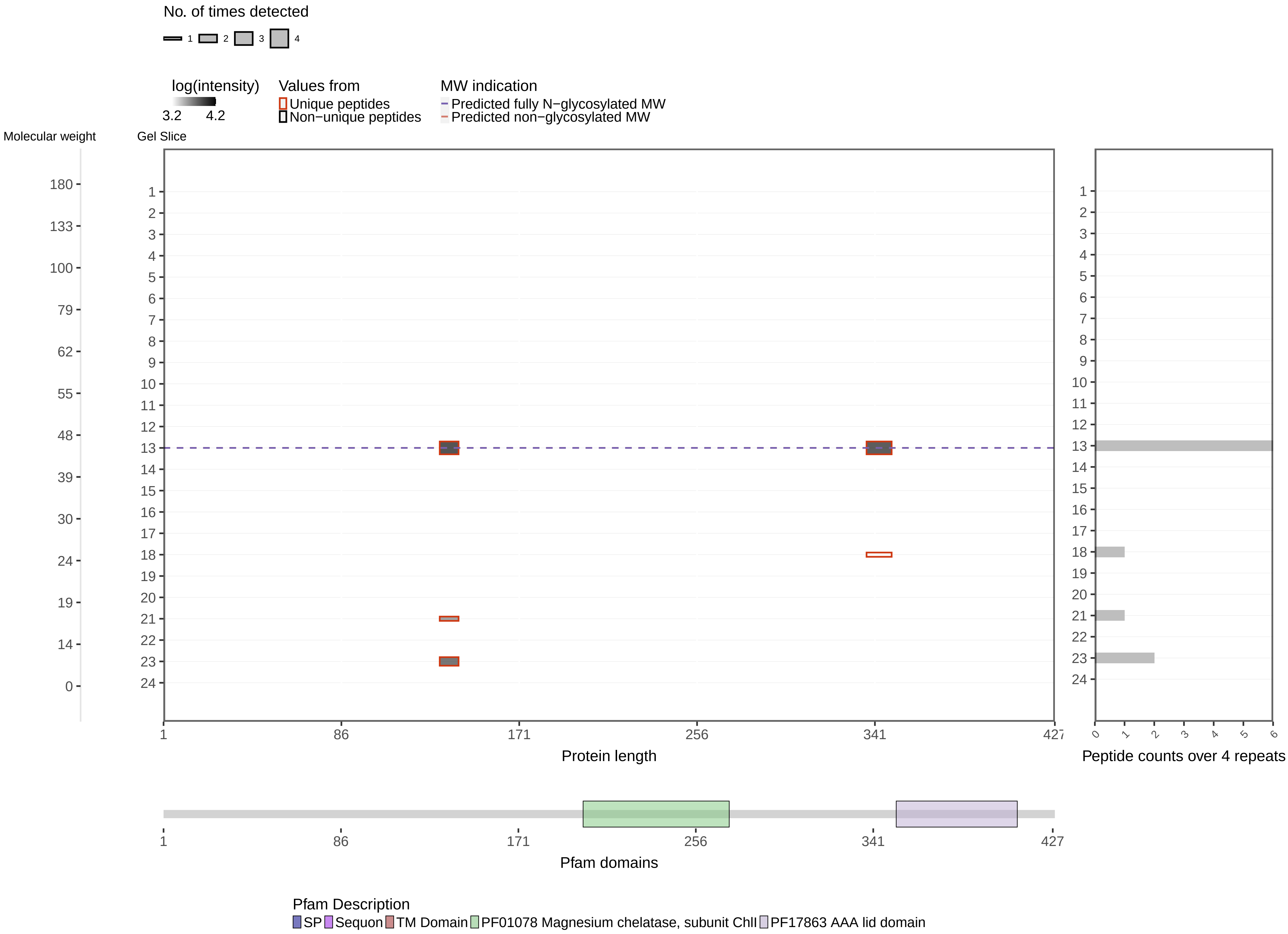


Protein sequence:

IEKALTEGVKAFEPGLLAKANRGILYVDEVNLLDDHLVDVLLDSAASGWNTVEREGISISHPARFILIGSGNPEEGELRPQLLDRFGMHAQVGTVRDAELRVKIVEERARFDKNPKEFRESYKAEQEKLQNQIDSARNALSAVTIDHDLRVKISKVCAELNVDGLRGDIVTNRAARALAALKGRDKVTPEDIATVIPNCLRHRLRKDPLESIDSGVLVVEKFYEVFA*

NbL10g22030.1 anthocyanin_5−aromatic_acyltransferase−like [MW = 51.04 kDa]

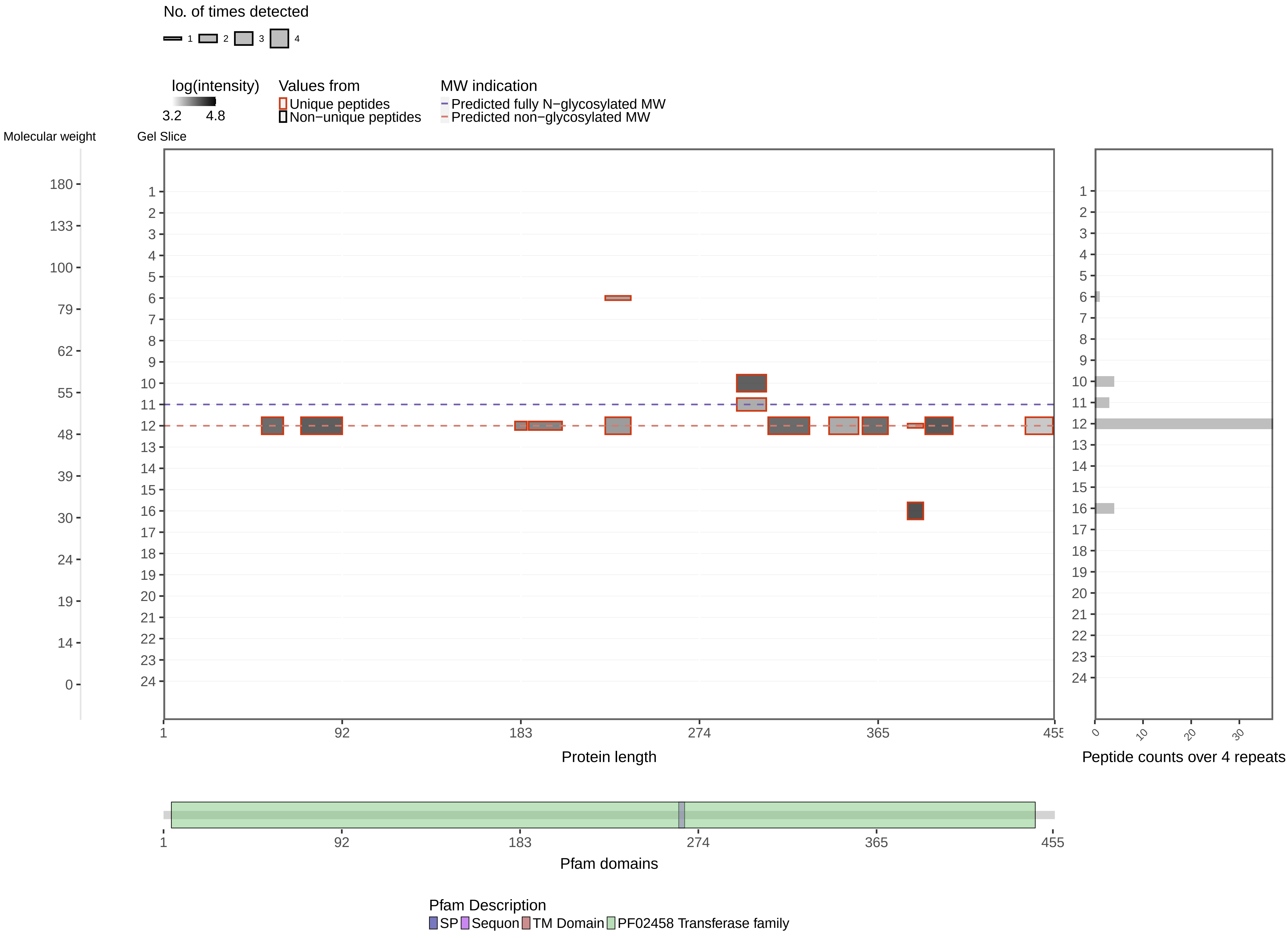


Protein sequence:

LIWDEMKKYKHMMTMSDIVTPPDKVRGTFIVSRDDILKFKNLILSRRPNLTHVTSFTVTCAYIWSCLIKSEAATGEKIDENGVEFFGFAADCRAQFNPPLPPSYFGNALVGYVARTRHVDLAGKEGFTIAAELIGEAIQKRMKDEEWILSGGWFKEYDKLDVKRSVSVAGSPKLDLYAADFGWGKPEKLEFVSIDSGDSISMSLSKYKDSDGDLEIGLSLSKTRMNAFAAMFTHGLSFL*

### NbL10g21990.1 diacylglycerol_kinase_5−like [MW = 39.6 kDa]

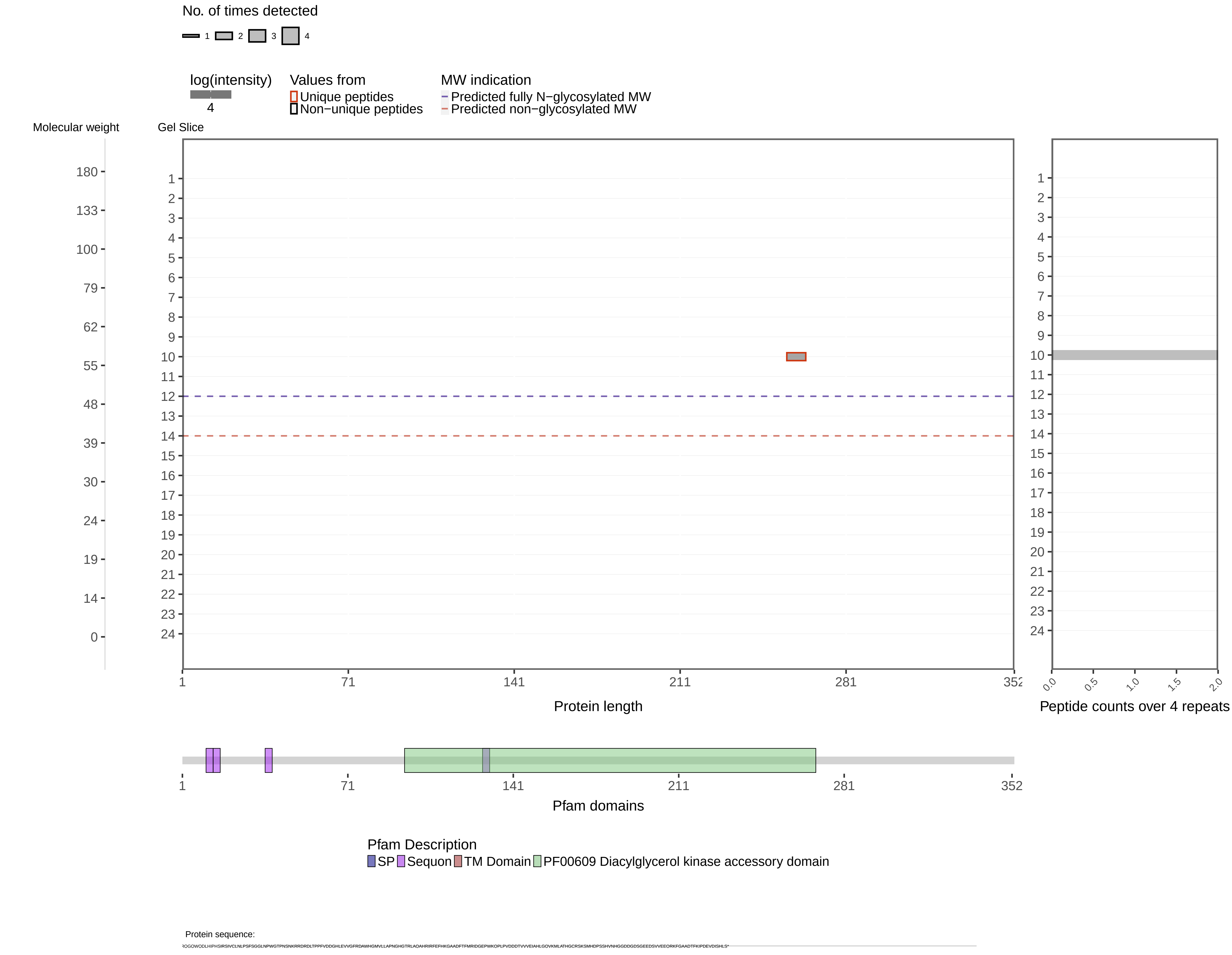

### NbL10g21950.1 deoxyribodipyrimidine_photo−lyase [MW = 44.08 kDa]

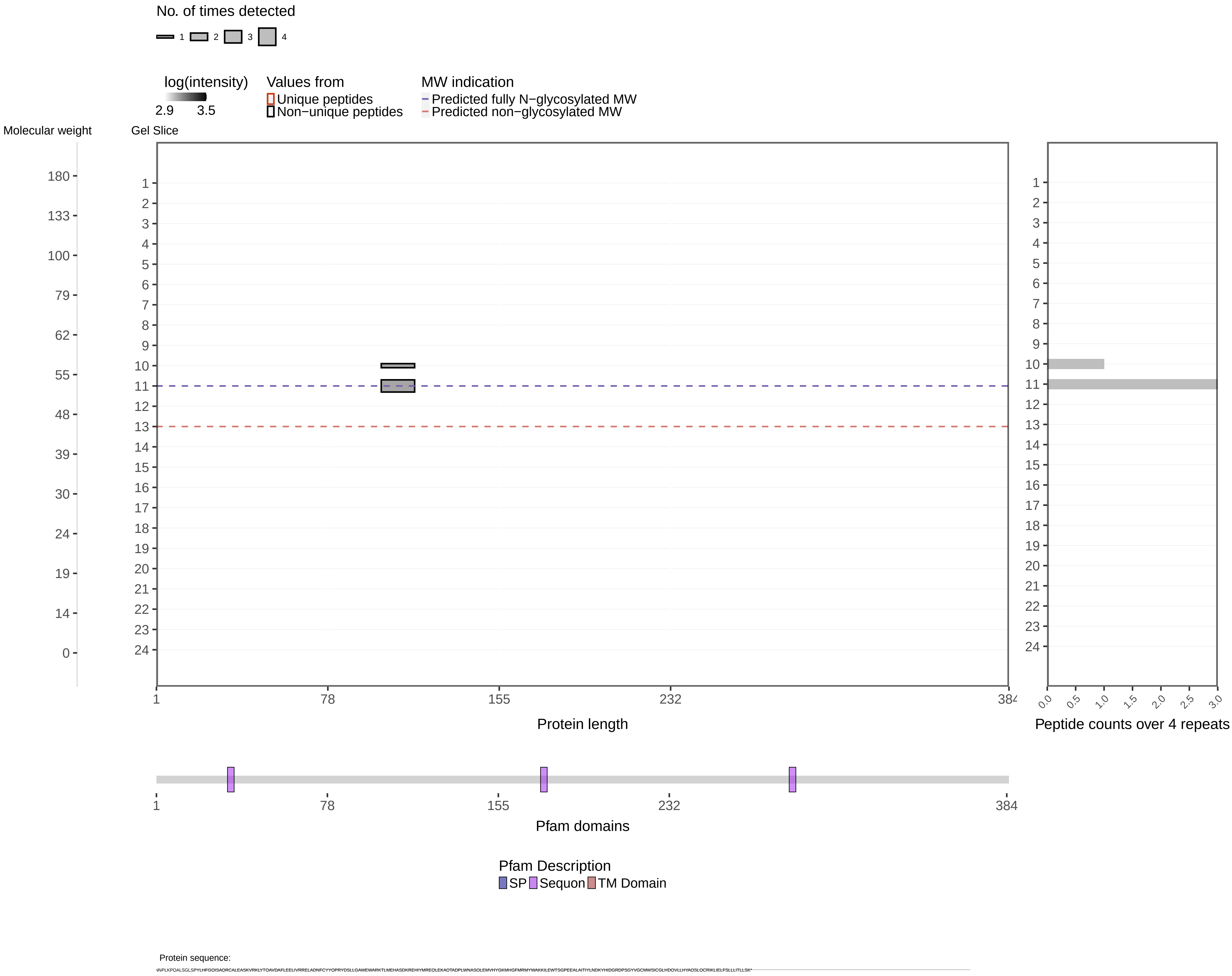

### NbL10g21940.1 uncharacterized_LOC109214416 [MW = 73.71 kDa]

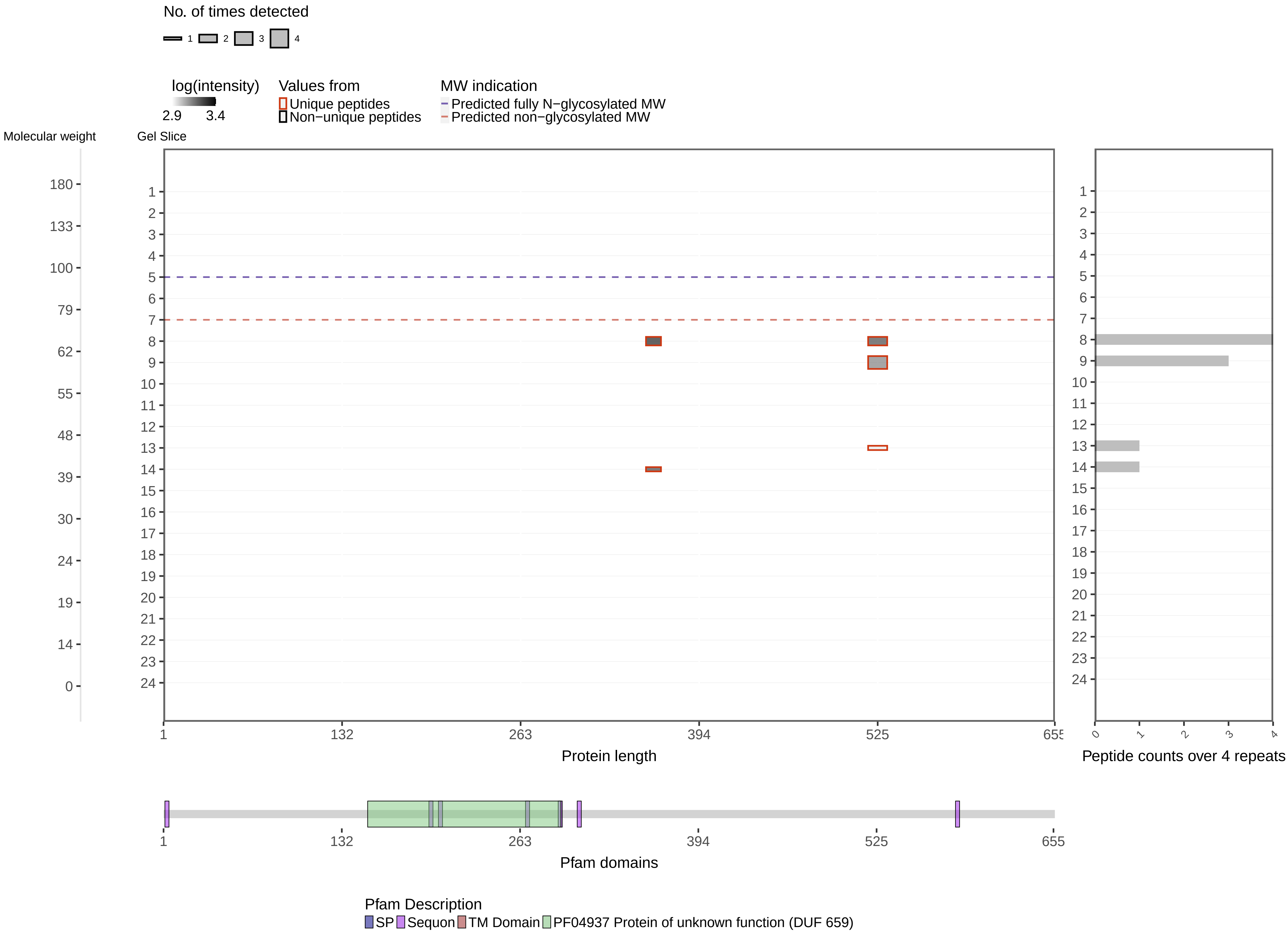


Protein sequence:

FHKYQLQEYGHASLLRVPLRSYERSDFEPVYTLVEDILSSARALQLVLLDESYKILSMEEQIARDIEEMMRSPHFWNELEAVHSLVKLIKTMAQDIQTEKSRVGQCLPLWEELRVKVKDWCLKFHIVEGPVEKVIERRFNKNYHPAWAAAFILDPLYLIRDNSGKYLPPFKCLTPEQEKDVDKLITRLVSREEAHIALMELMKWRTEGLDPVYAQAVQLKQRDASSGKMKIANPQSSRLVWETHLTEFKSLGKVAVRLIFLRASSCGYKCNWSFLRWANAHSHSRVGMDKAQKL

NbL10g21880.1 ATP−dependent_Clp_protease_proteolytic_subunit−related_protein_2_chloroplastic−like [MW = 35.66 kDa]

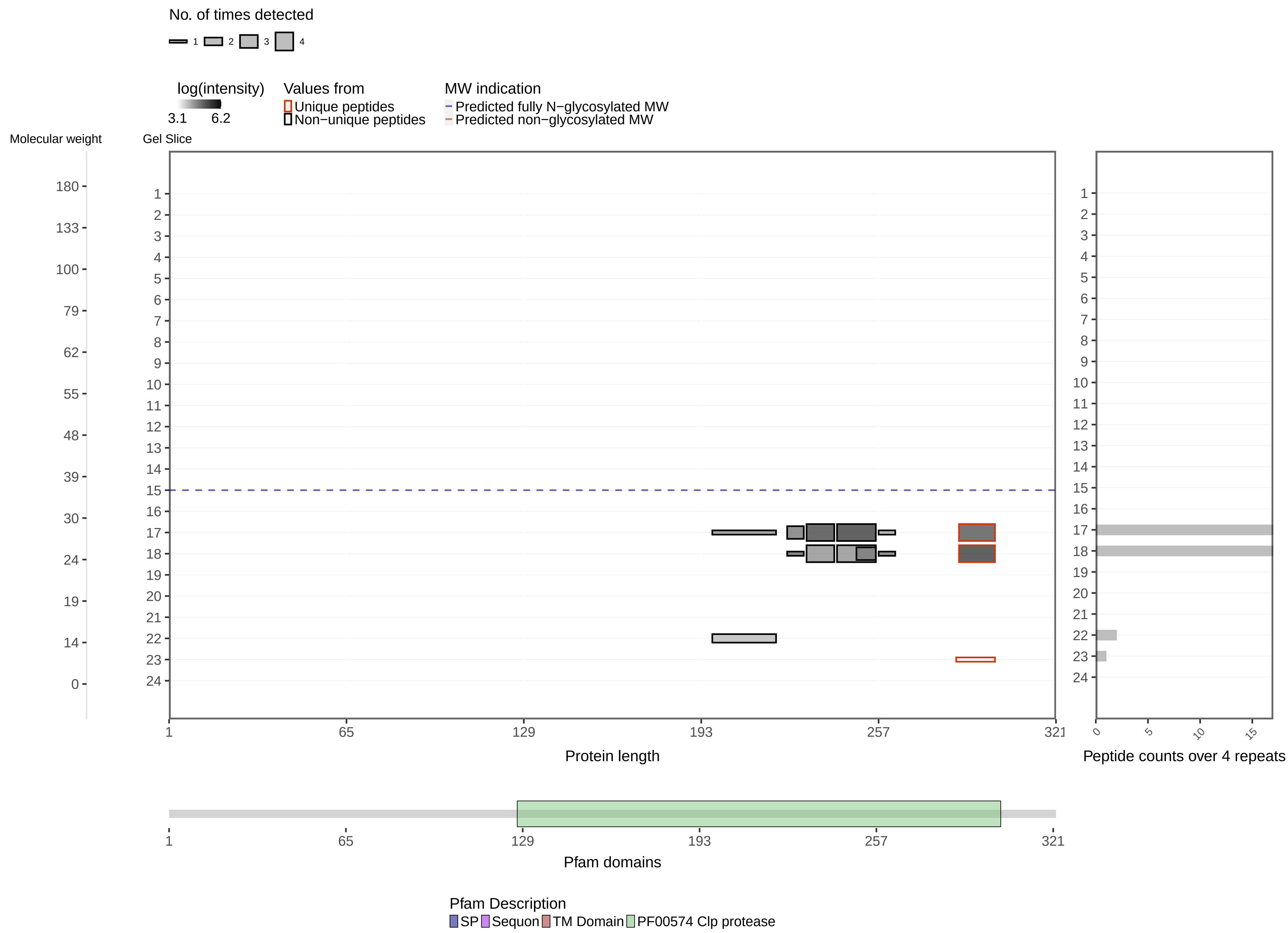


Protein sequence:

HIDEEFSNQILATMLYLDSIDDSKKLYLYINGPGGDLTPSMAIYDTMQSLKSAVGTHCVGYAYNLAGFLLAAGEKGNRFAMPLSRIALQSPAGAARGQADDIRNEADELLRIRDYLFKELAEKTGQPVEKVHKDLSRMKRYNAQEALEYGLIDRIVRPPRIKADAPRKDTTAGLG*

### NbL10g21850.1 ATP−dependent_Clp_protease_ATP−binding_subunit_CLPT2_chloroplastic−like [MW = 25.8 kDa]

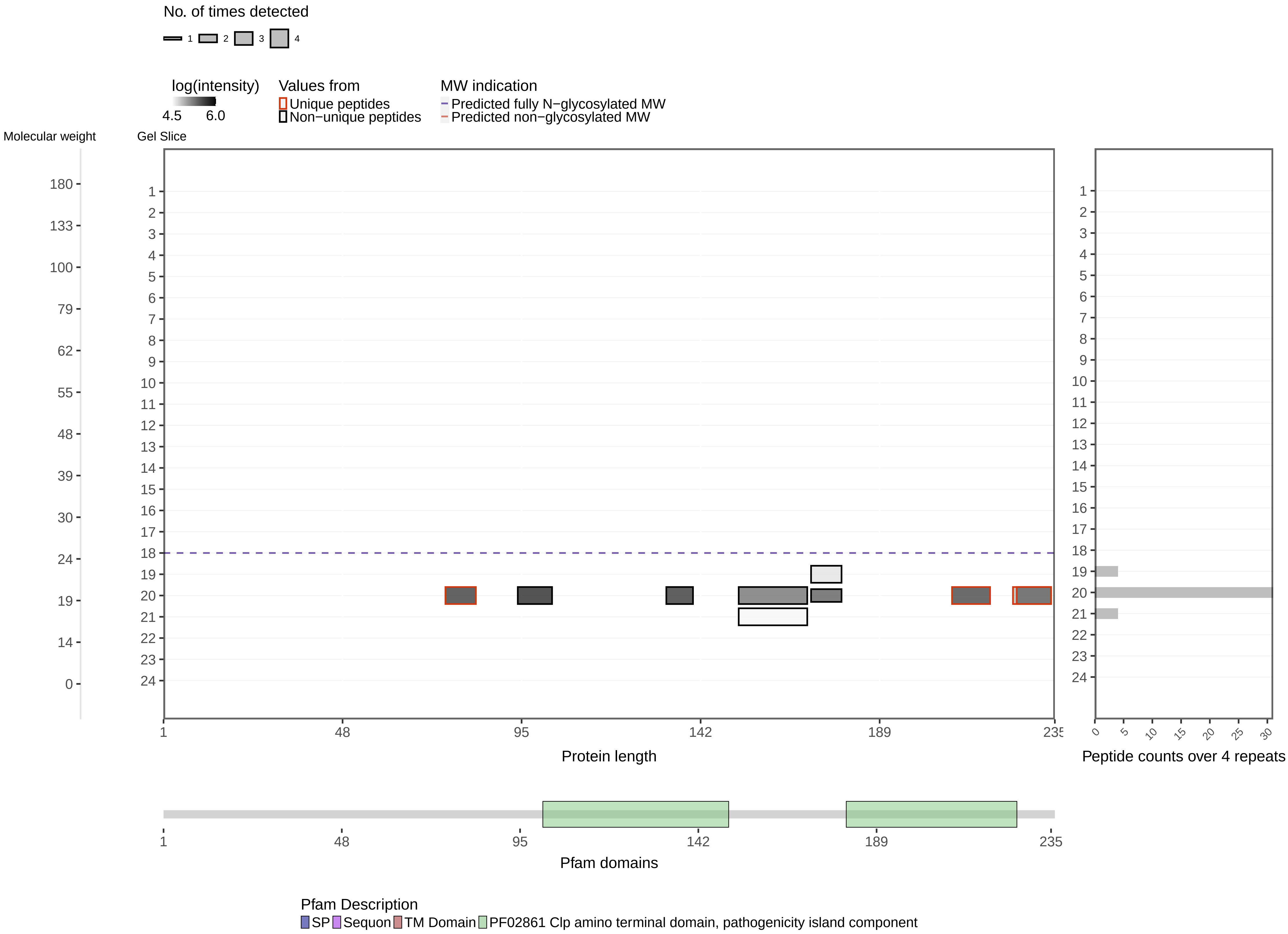


Protein sequence:

..KYPTTGTEALLMGILIEGTNFASKYLRANSITLFKVREETVKILGKADMWFFSPEHPPLTEDAQKALDWALDEKLKSSENGEITTTHLLLGVWSQVGSPGYKILSGLGFNDEKAQELKKVISEPGFVDD*

NbL10g21760.1 probable_LRR_receptor−like_serine/threonine−protein_kinase_At1g12460 [MW − SP = 92.56 kDa]

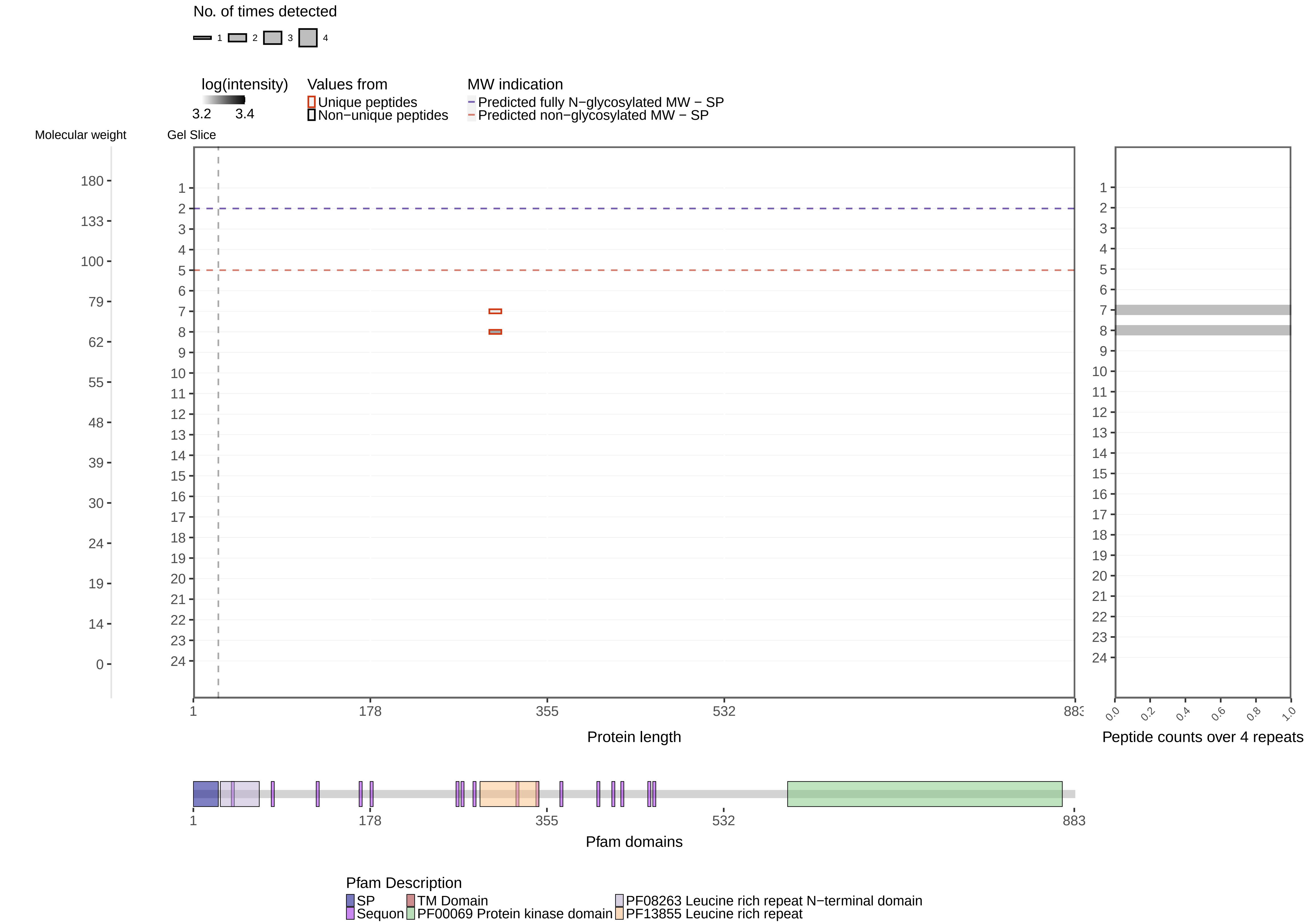


Protein sequence:

SNLHFLDLSENVLSGSIPVALENMHNLTHFNVSYNLLSGAIPSTESIQQFGPSAFFHNSGLCGAPLDVSCSPSGIAPAKGKPRLSASAIVAIVAAAVILTGVCVITIINMKARRRRRRADETIIVESTPLASTDSNVIIGKLVLFSKTLPSKYEDWEAGTKALLDKESLIGGGTIGSVYRTSFEGGVSLAVKKLETLGRIRNQDEFEHEIGRLGTLQHPNLVALQGYYWSSSMQLILSEFVPNGNLYDNLHGLGYPGTSIGAAGTPELNWPRRFQIAVGTARALAYLHHDCKPPILHLNV

NbL10g21690.1 enoyl−CoA_delta_isomerase_1_peroxisomal [MW = 26.84 kDa]

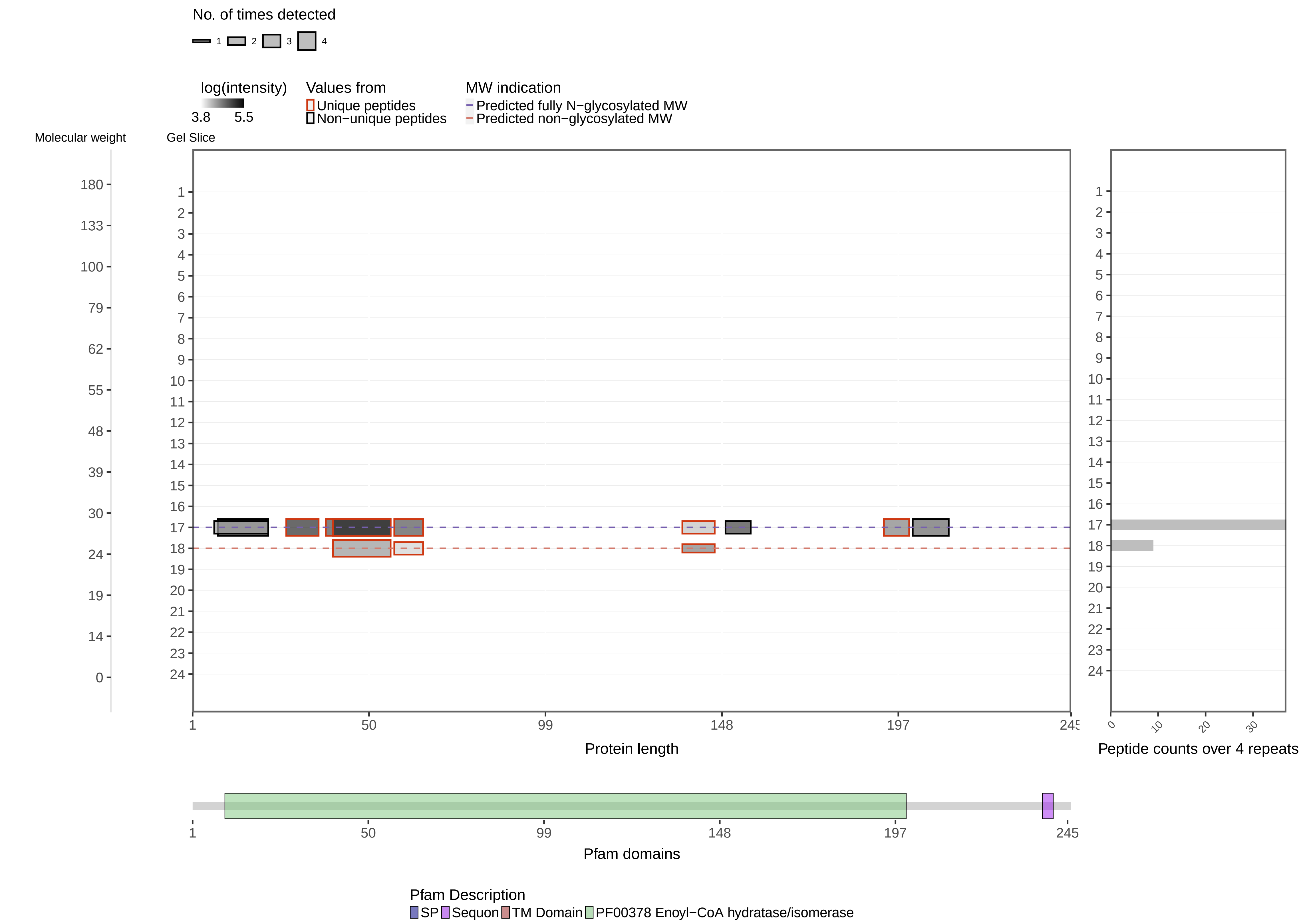


Protein sequence:

LSHDYILMRKDRGFLYMSELDIGLKVPIWFSALVKCKVGSPAVWREVVLKSAKMTAEMGVEMGIVDSAHSGAEETVEAAVRLAEELVSRNWDGNVYADCRKTLCVELLNSLGSDETVGDYGEEEKPNKTLSRL*

### NbL10g21670.1 copper_chaperone_for_superoxide_dismutase_chloroplastic/cytosolic [MW = 33.2 kDa]

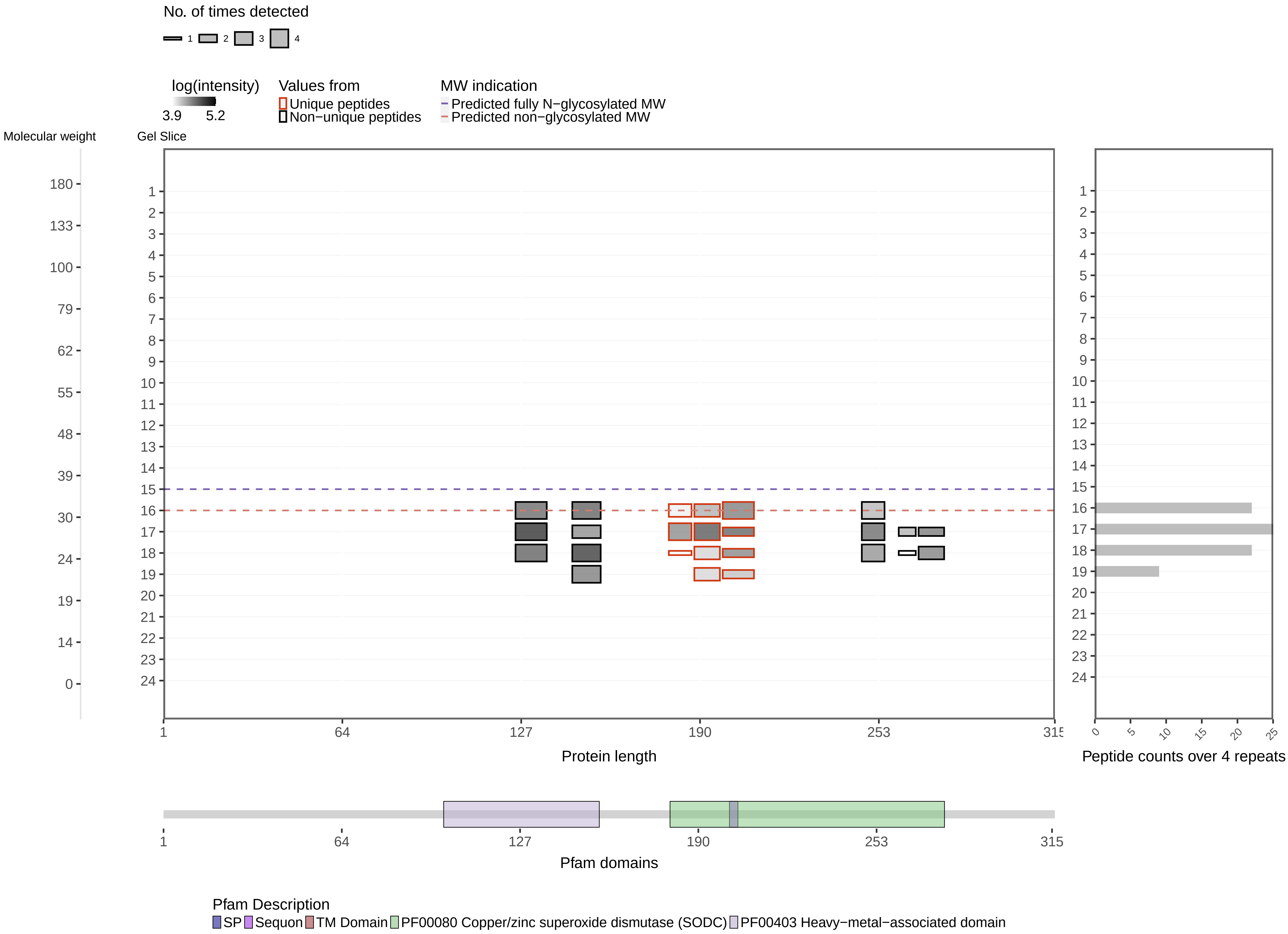


Protein sequence:
KILTEALEQTGRKARLIGQGVPADFLISAAVAEFKGPDIFGVVRLAQVNMELTRIEANFSGLSPGRHGWSINEFGDLTRGAASTGKLYSPPLGDLGTLEVDVKGEAFYSGAKEKLRVADLIGRAIAVYSTEDKSDPGLKAAVIARSAGVGENYKKLCTCDGTIIWEATNKL*

NbL10g21650.1 subtilisin−like_protease_SBT1.2 [MW − SP = 77.03 kDa]

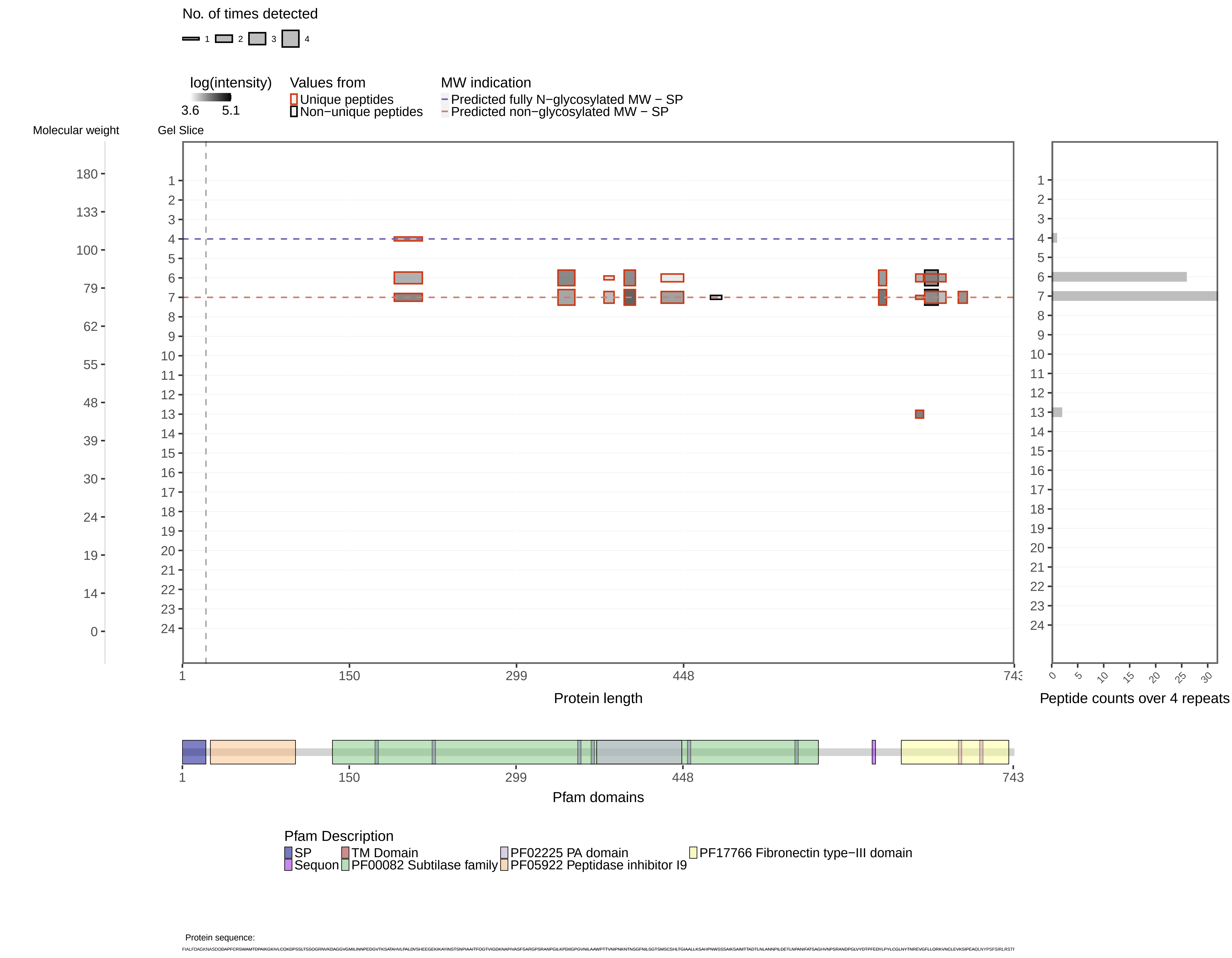

NbL10g21630.1 coiled−coil_domain−containing_protein_18−like [MW = 76.04 kDa]

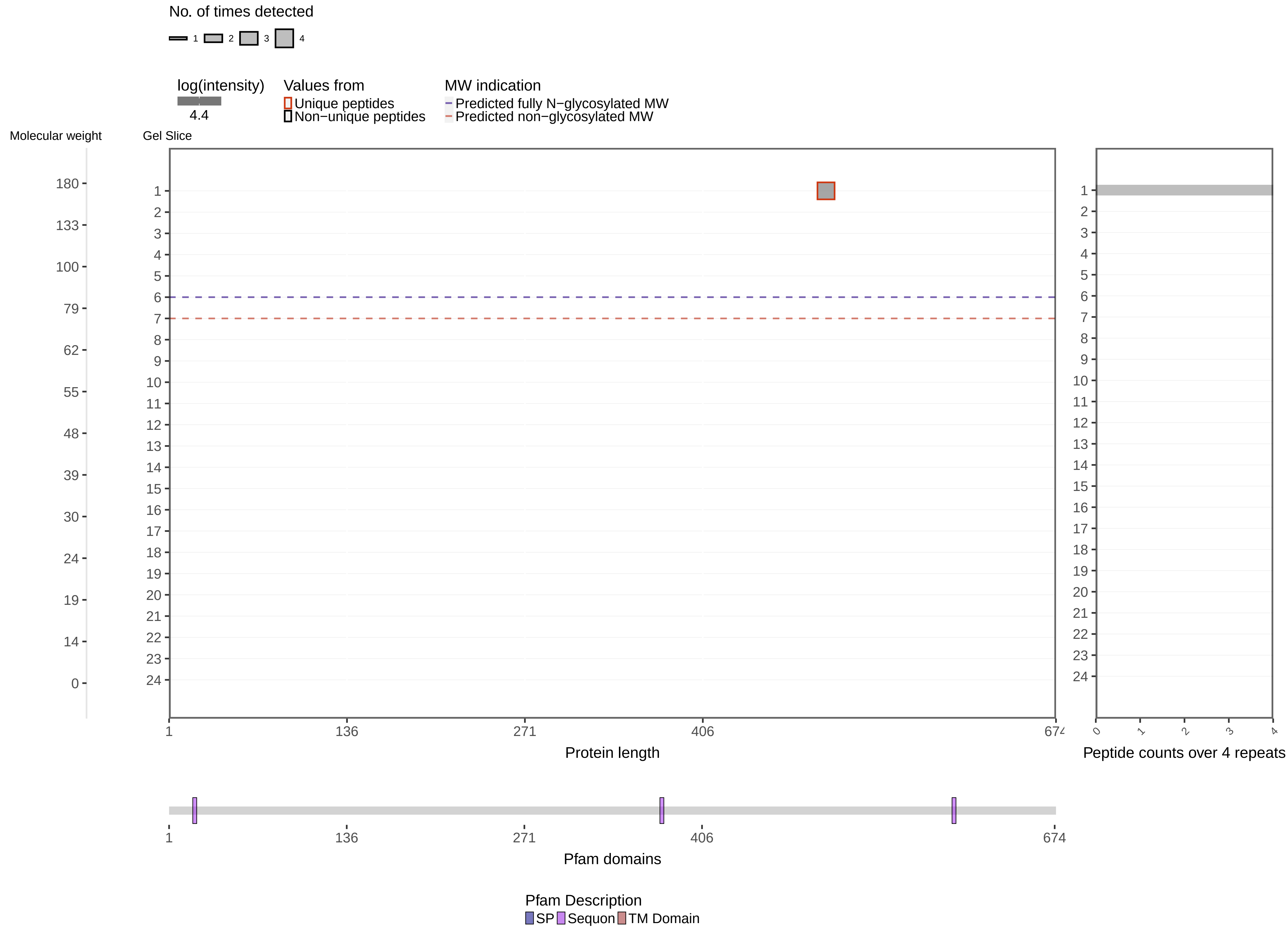


Protein sequence:

KISDHRIQKEEALKFRVTKTSEVGEIEKELATEVSALEKQRDEIEAQLKQVNISLAAANARLQNVREERDQFYEANDQIVAHLKTREDELTKSIGSCKVEANILGTWINFLEDTWNLQSSFTESKDKEVSVELEKHEDYFVNLVMQLLSAYEKELRPSIDRISKFVENLKTLGEGAAVGSGVSKSLSPRKSLEEEYLDYEAKIITTFSVVDNMREQFYSQQEKISRKDDPDMKKLFENIEKLRQEFESIERPNLEMEIPVQEGDDSSHETVDDNTSHPIKQAVETAIAGESKKNEPP

### NbL10g21580.1 glyoxylate/hydroxypyruvate_reductase_HPR3−like [MW = 37.76 kDa]

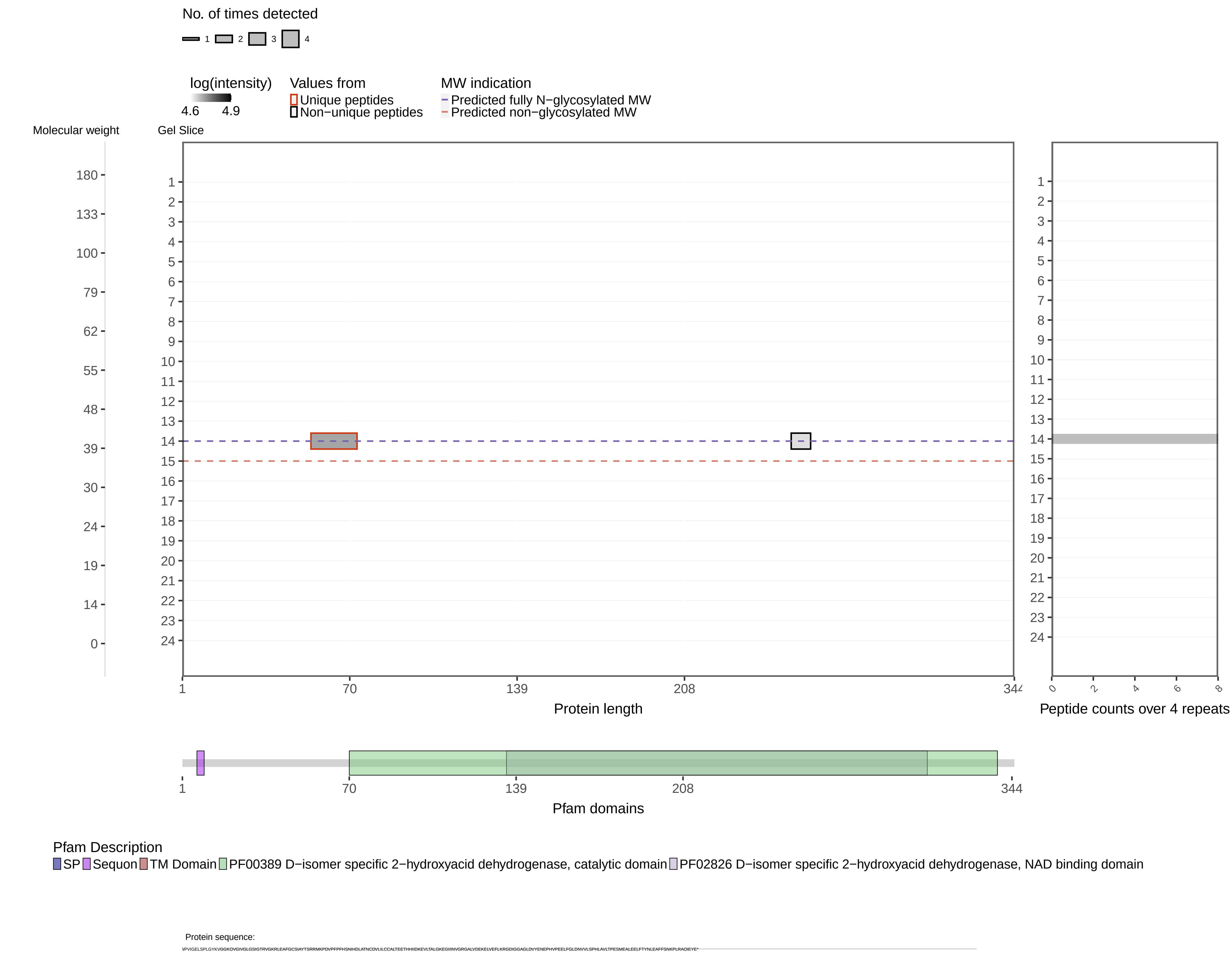

### NbL10g21560.1 PGR5−like_protein_1A_chloroplastic [MW = 36.29 kDa]

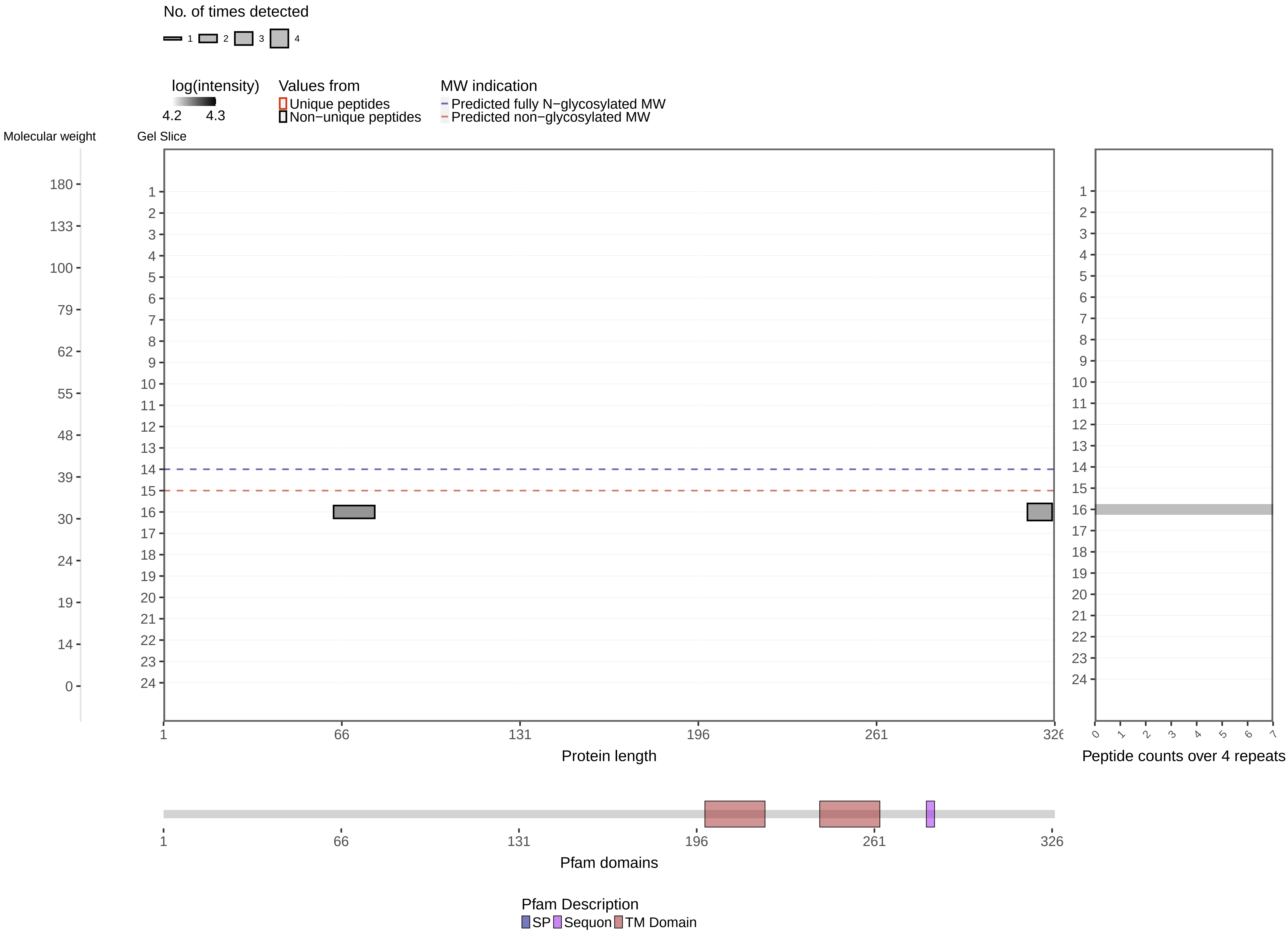


Protein sequence:

SMAYVSGNPIMTDKEFDKLKLKLKMDGSDIVVEGPRCSLRSRKVYSDLYVDYLRMFLLNVPAAVVALGLFFFLDDITGFEITYLLELPEPFSFIFTWFAALPFILWLSFTITNAIIKDFLILKGPCPNCGTENTSFFGTILSVSSGGNTNKVKCTNCGTDMVYDSGTRLITLPEGSSA*

### NbL10g21450.1 hydroxymethylglutaryl−CoA_synthase−like [MW = 51.25 kDa]

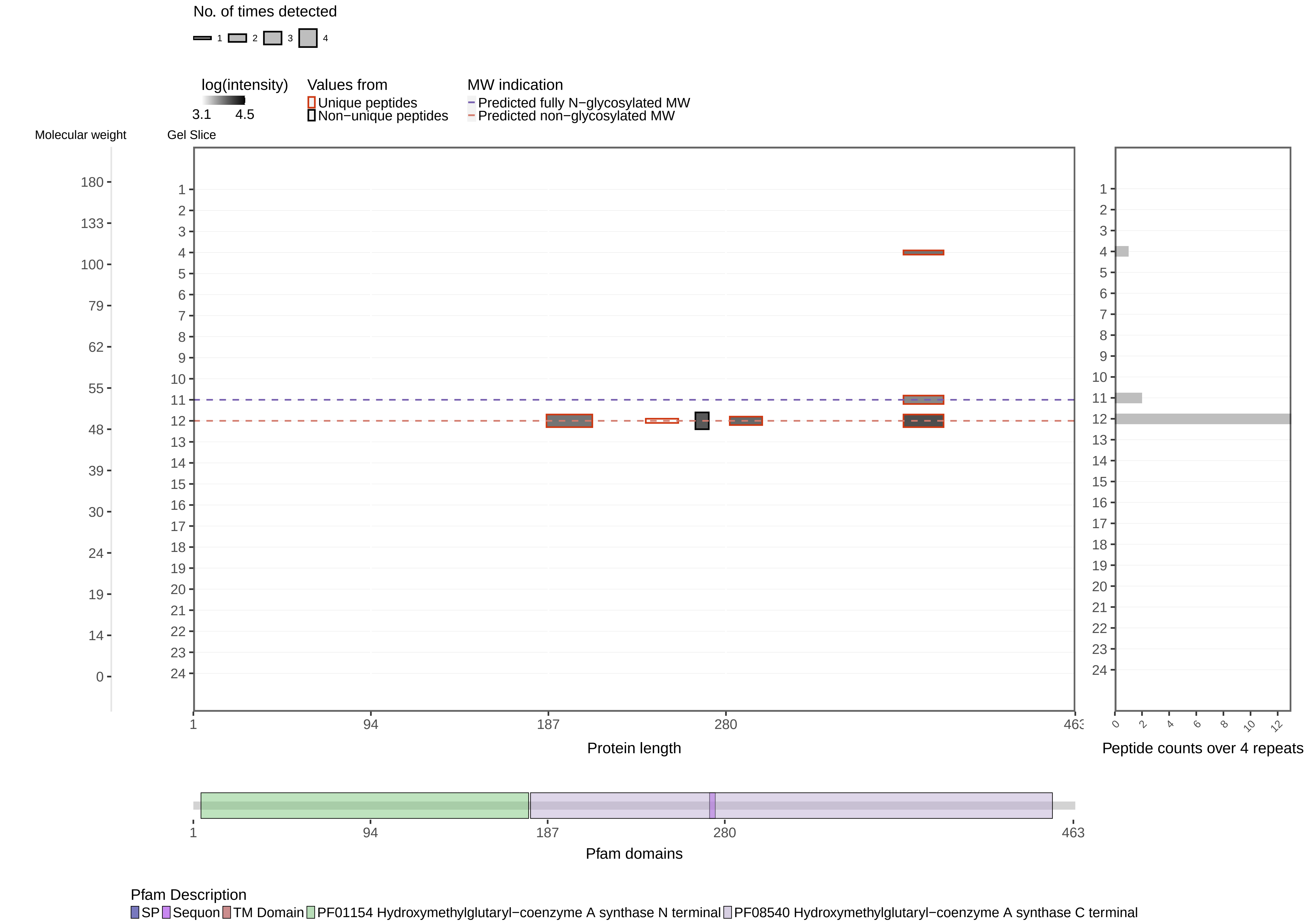


Protein sequence:

ALDSCYKSLCNKYEKLEGKQFSMADAAYFVFHSPYNKLVQKSFGRLLFNDFLRNASSIDESTKQILAPFESLTGDESYQSRDLEKASQQVAKPFYDEKVQPTTLIPKQVGNMYTASLYAAFASLIHNKHNTLAGQRVIMFSYGSGLTATMFSLKFNEGQHPFSLSNIASVMNVAEKLKSRHEFTPEKFVEIMKLMEHRYGAKDFVTSKDCSLLAPGTYYLTEVDSKYRRFYAQKAPEHGLINGH*

### NbL10g21430.1 protein_HOTHEAD [MW − SP = 60.11 kDa]

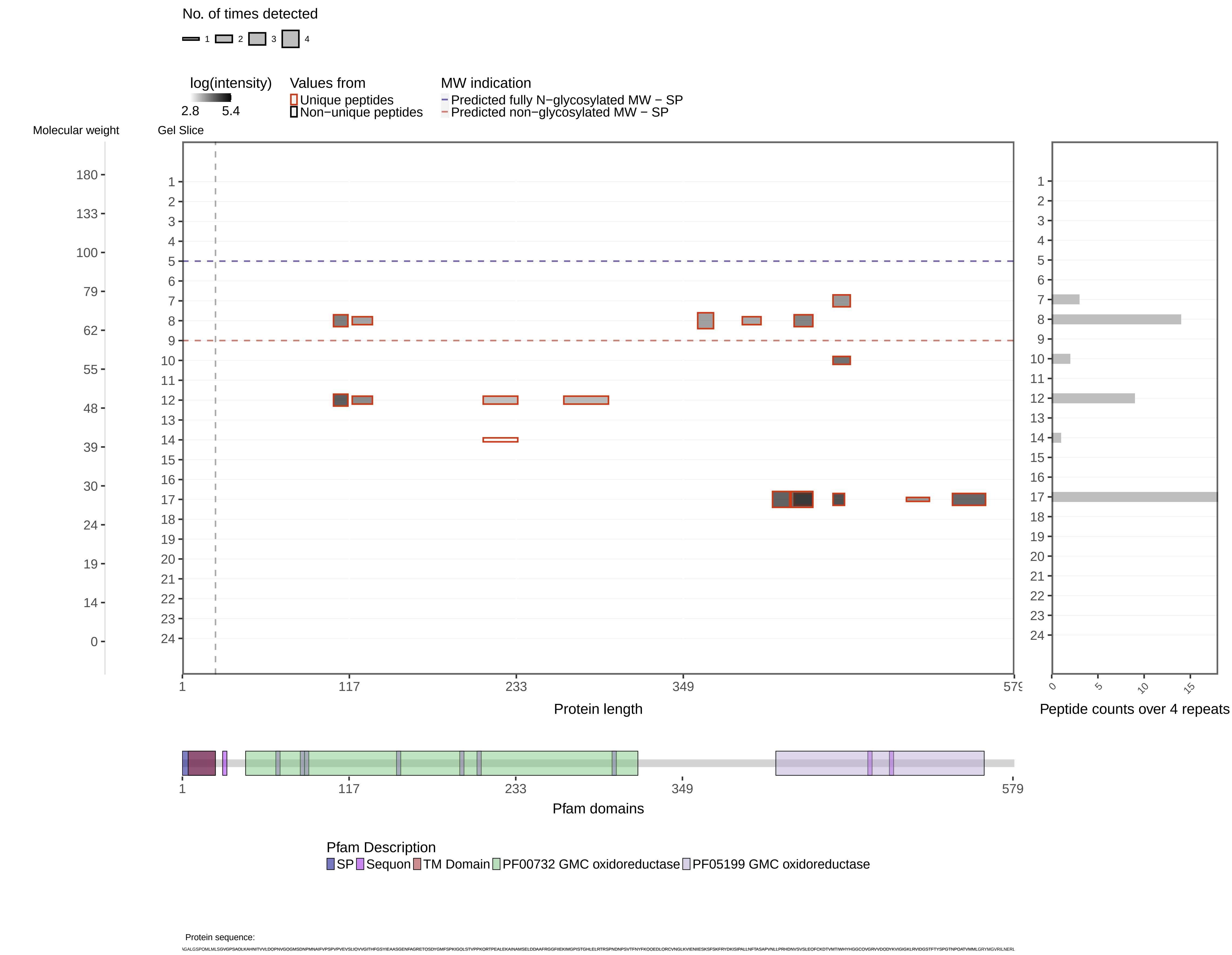

### NbL10g21380.1 NADH_dehydrogenase_[ubiquinone]_1_alpha_subcomplex_subunit_8−B−like [MW = 12 kDa]

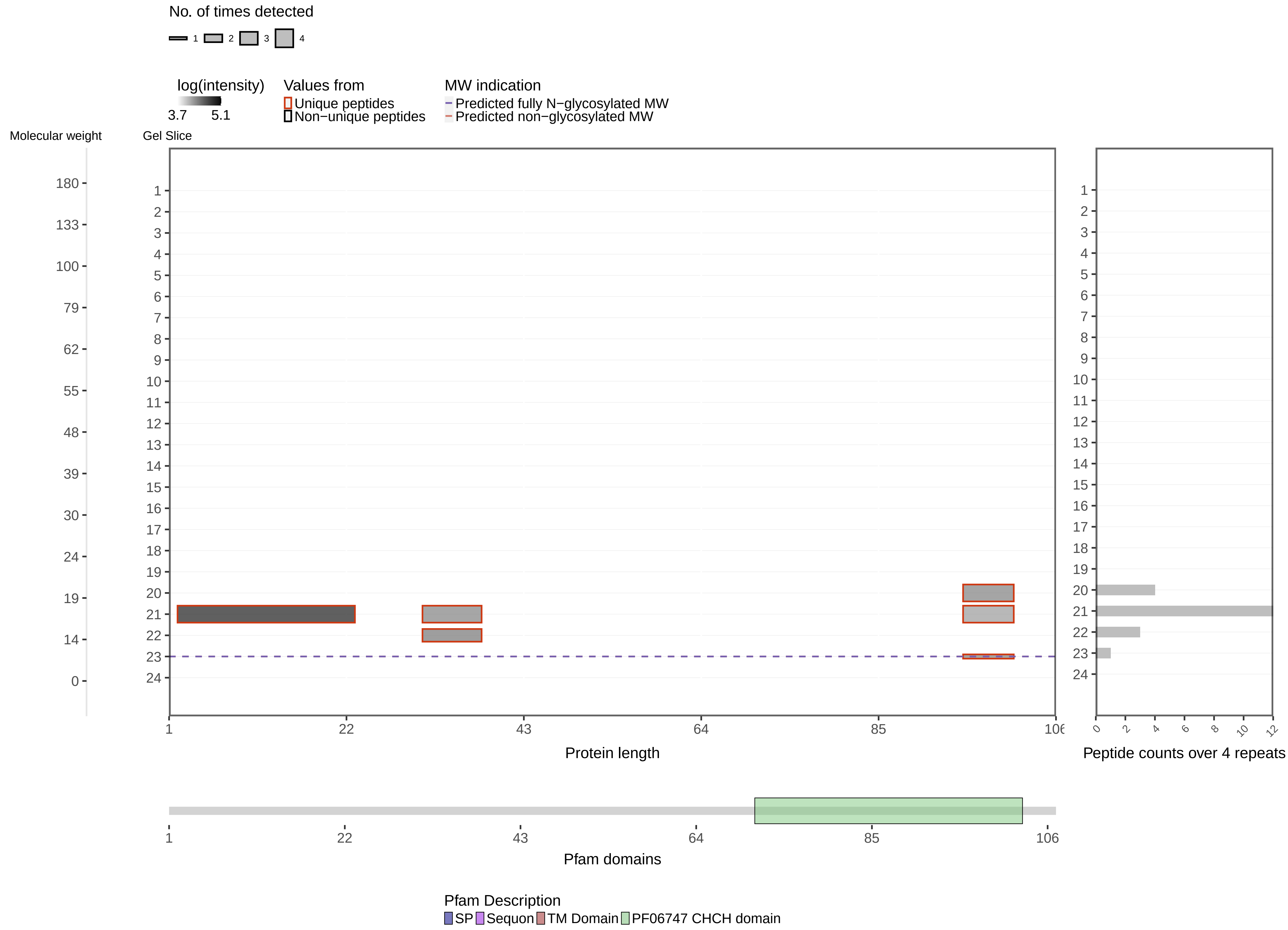


Protein sequence:

DDPNPEKCLHKGQQVTRCVLSLLKDLHQRCNKEMDAYAGCMYYNTNEFEMCRKEQQEFEKACPY*

### NbL10g21360.1 uncharacterized_LOC109212508 [MW = 39.74 kDa]

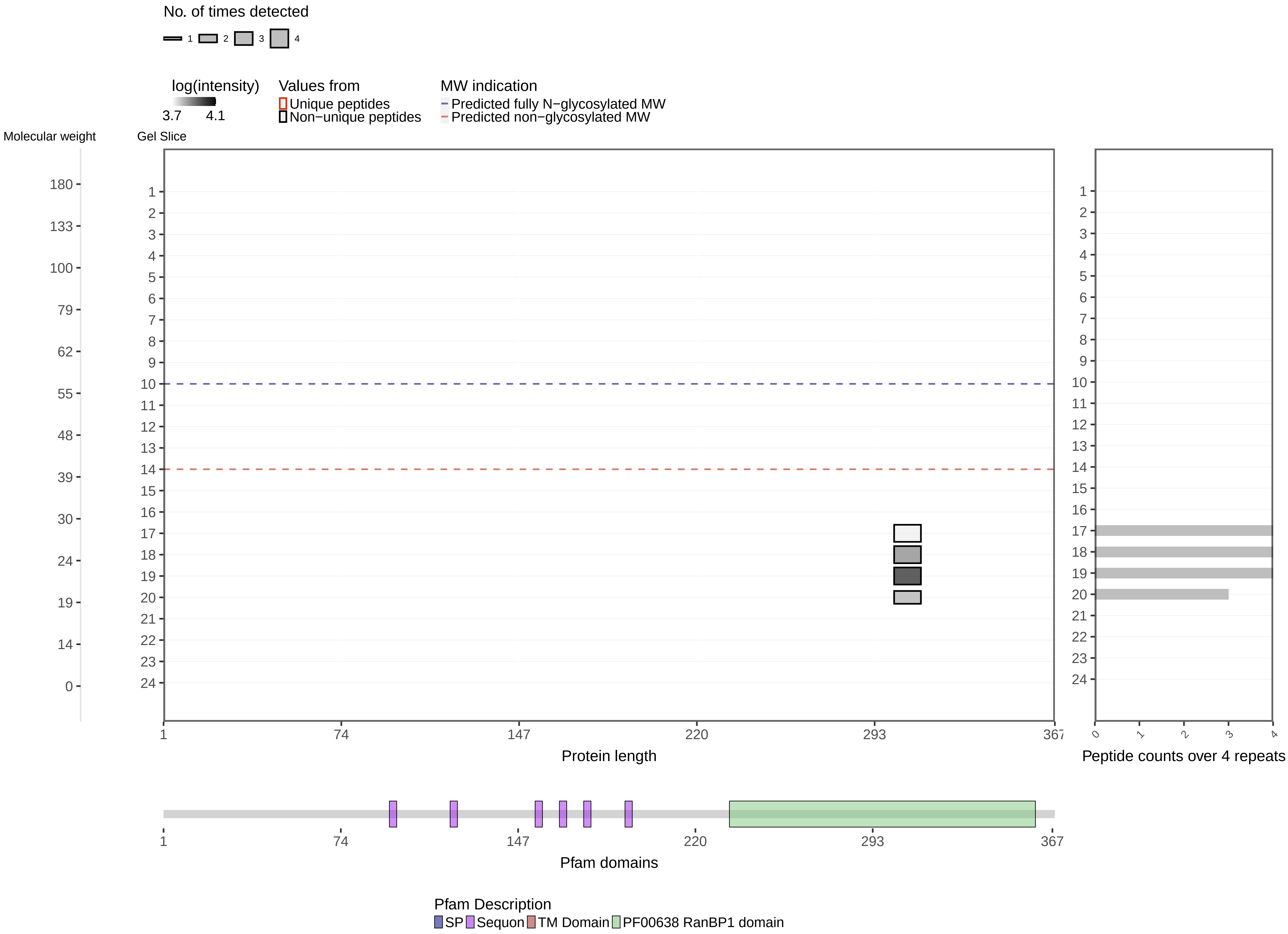


Protein sequence:

SFVGLANSSSSFGALWSSGGLFNNKSPFSFGLOSSVOTOSSVPVKSDAANDVODTEDEVEOPSSPSVKKTEEKGVTLVHEVKCKLYVKSTDPADKDAWKDRGTGOLSIKCKEGVSKGTRESKPTIVMRNDVGKILLNALLYPGIKTNMOKNSIVAIFHTSGDDDKDDAVVARTFLIRTKTGEDRDKLAAIIOEYAPAAA*

NbL10g21250.1 plant_UBX_domain−containing_protein_8−like [MW = 66.93 kDa]

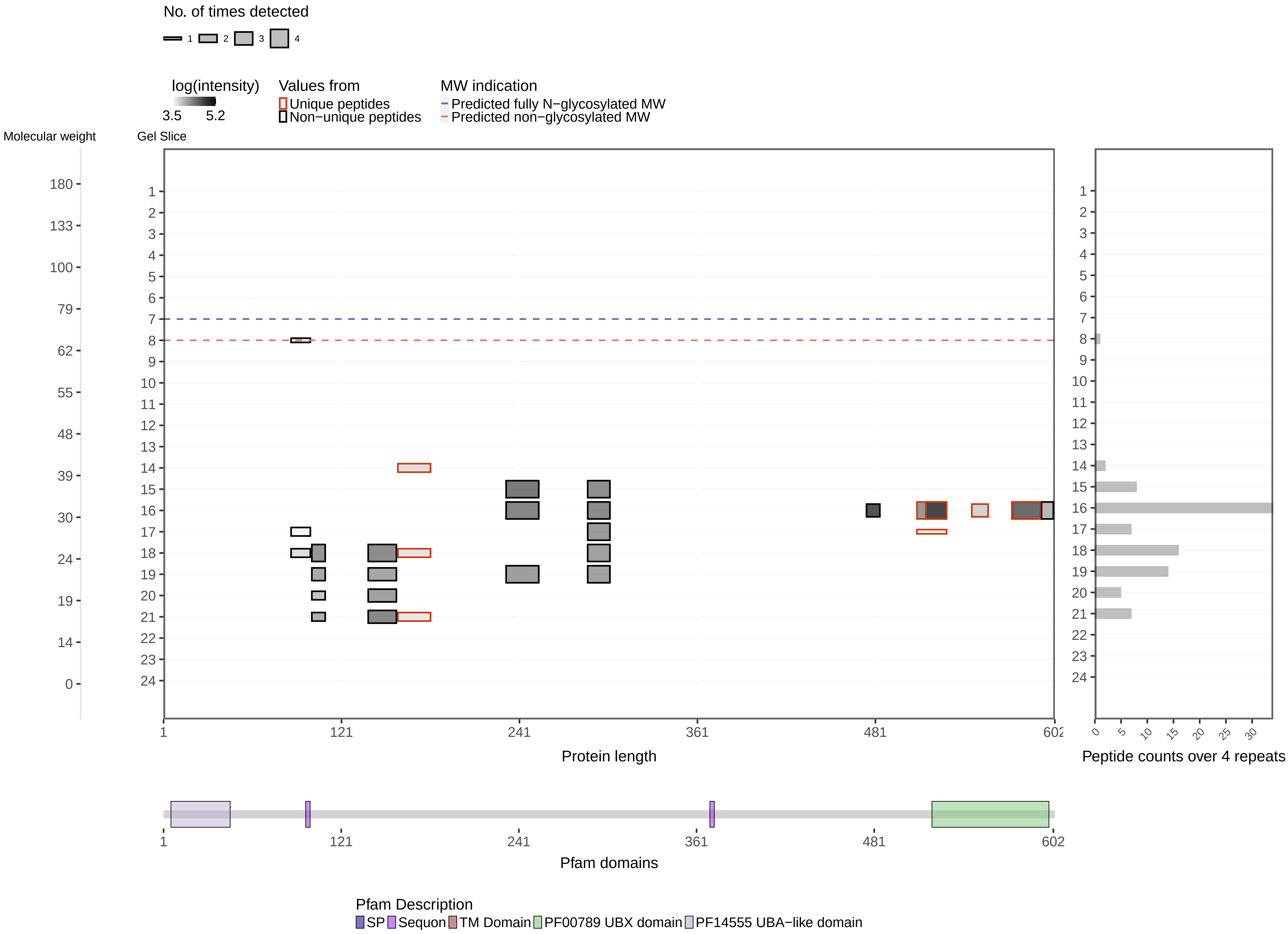


Protein sequence:

EGYKPAEEDERGKSITSNGRQFRSEVGSSFVPDEAEDLEEEPLVWRRRKPISSGSETAQDIEEHVVSPLSSPRQHDNLNHSPRNGADFPSDEWGGISSLEHDEAVMLEAALFGGIPEGSGYRLPNAPHQLMQNGVEGPMGPYQWRMPRPPSPSLVAQRLLREQQDDEYHAALQADREKELKAKEEAEAALEEKRQEEEESRRKAEEEKEMERQLAAKEASLPREPTTDNENAVNLVVRMPDGSRRGRRFLKSDRLQYLFDYIDVGRAVKPGTYRLVRPYPRRAFSI

NbL10g21180.1 protein_farnesyltransferase/geranylgeranyltransferase_type−1_subunit_alpha [MW = 40.27 kDa]

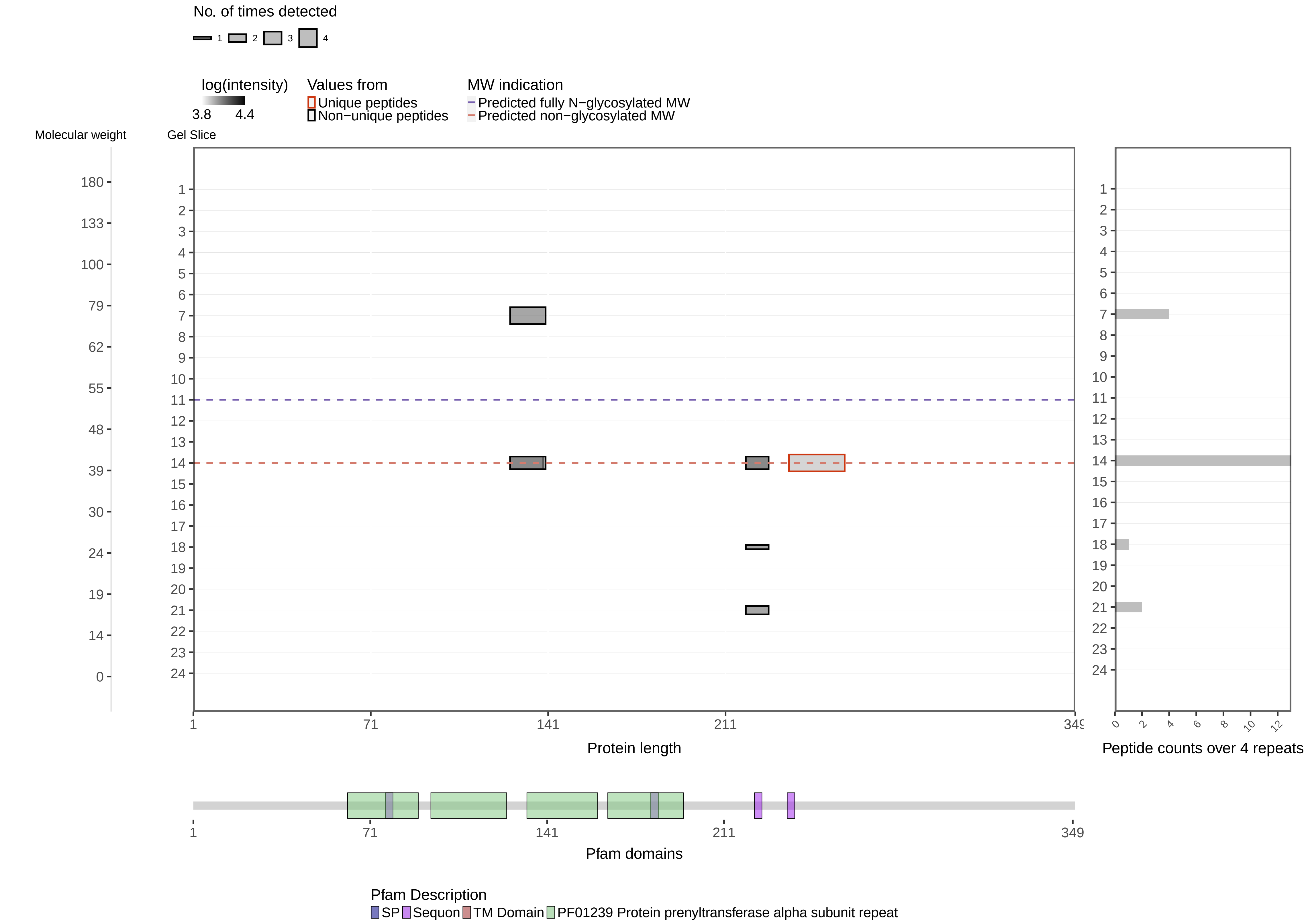


Protein sequence:

LGGWENELAYCQQLLEDDIYNNSAWNQRYFVVTRSPLLGGLVAMRELEVSYTVQALRASPENESPWRYLRGLYKNDTQSLGEDPQVASVLLDVLTSKNNHVHALSLLLDLLCHGFEPSEELKSAVDVLTPESCSMDLPLAKRICSILEHADPMRINYWNWRKSTILVQSAKCQNTDRLADLSMQDNL*

NbL10g21080.1 bifunctional_UDP−glucose_4−epimerase_and_UDP−xylose_4−epimerase_1 [MW = 48.11 kDa]

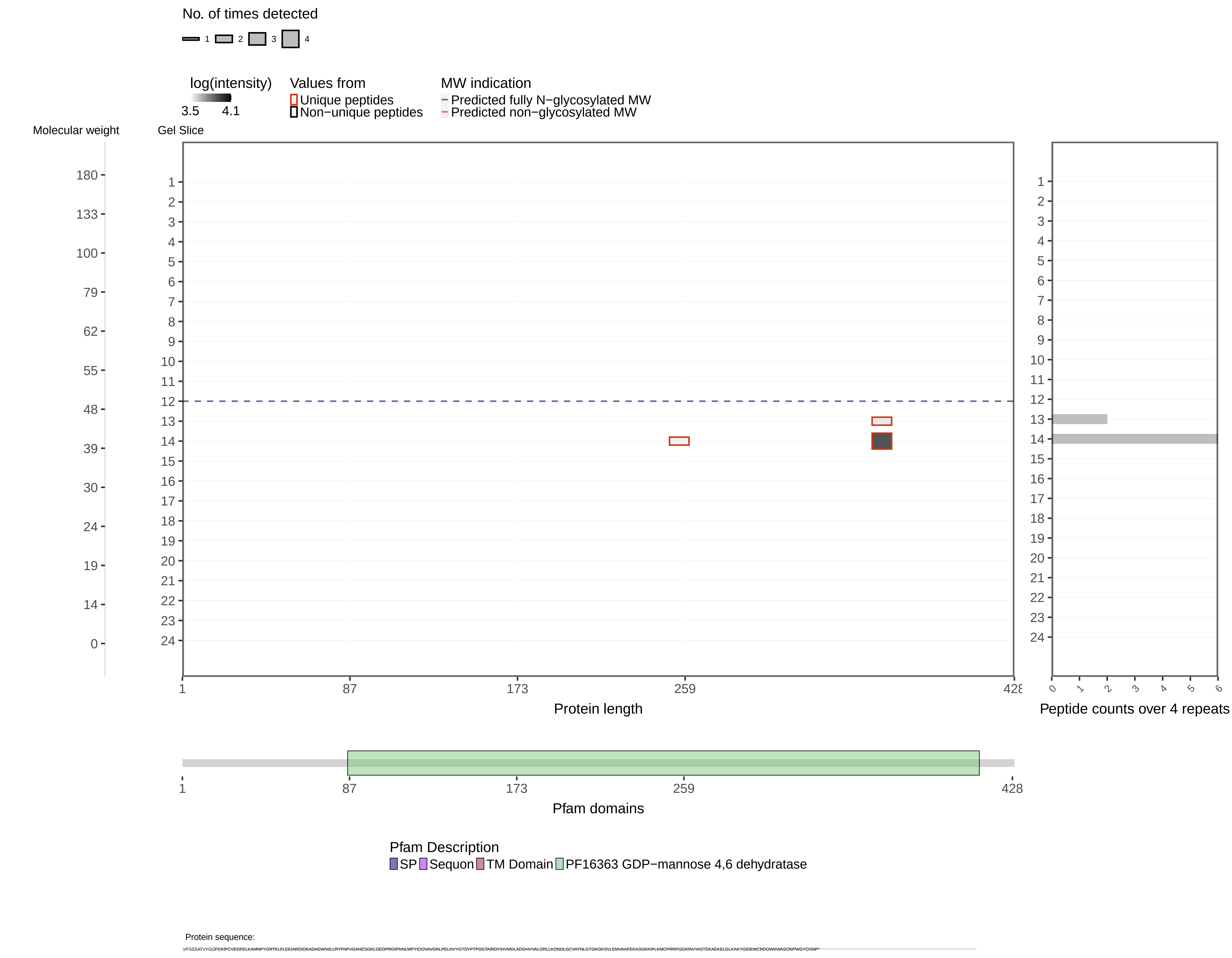

### NbL10g21060.1 riboflavin_biosynthesis_protein_PYRR_chloroplastic [MW = 66.11 kDa]

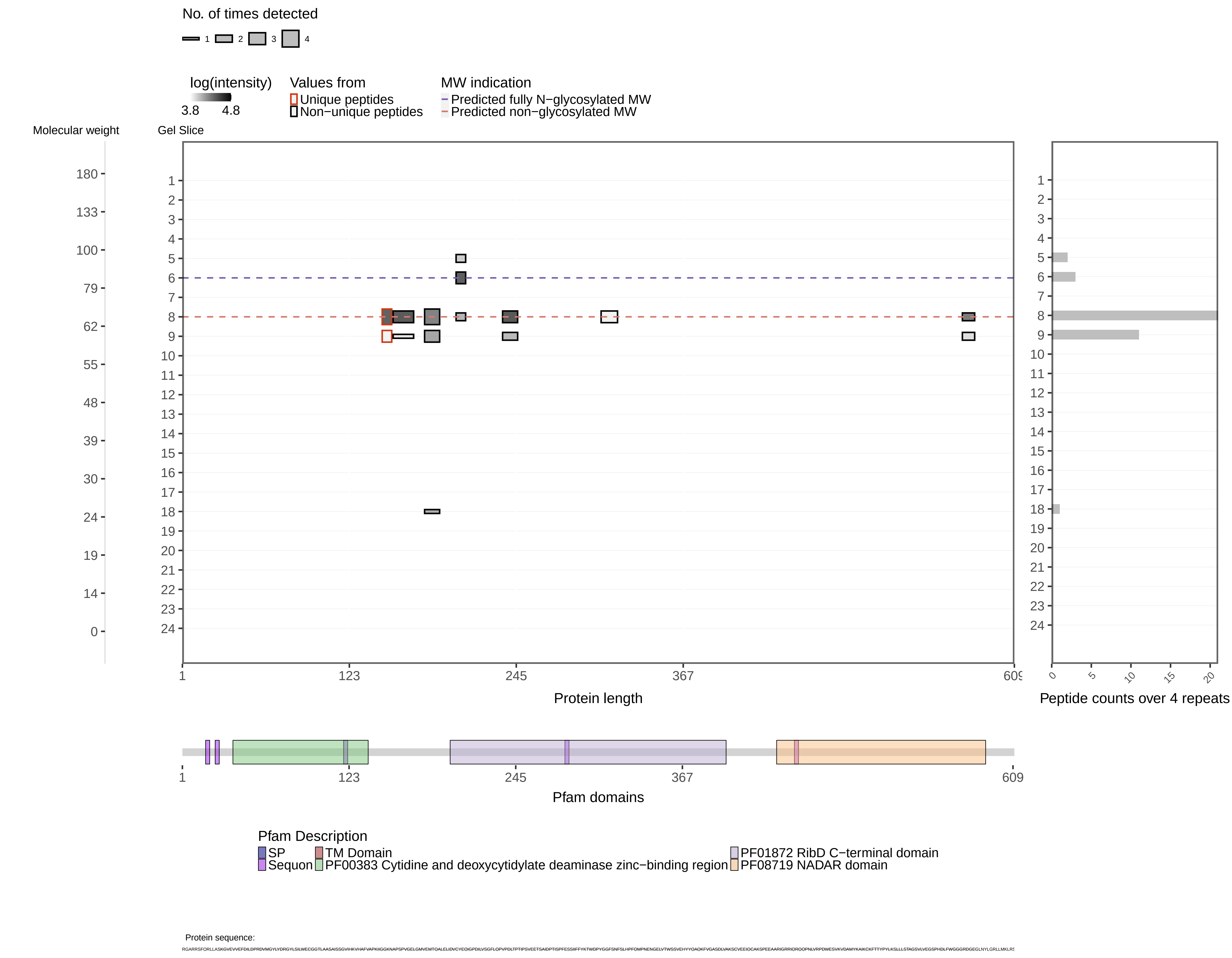

NbL10g21040.1 osmotin [MW − SP = 24.21 kDa]

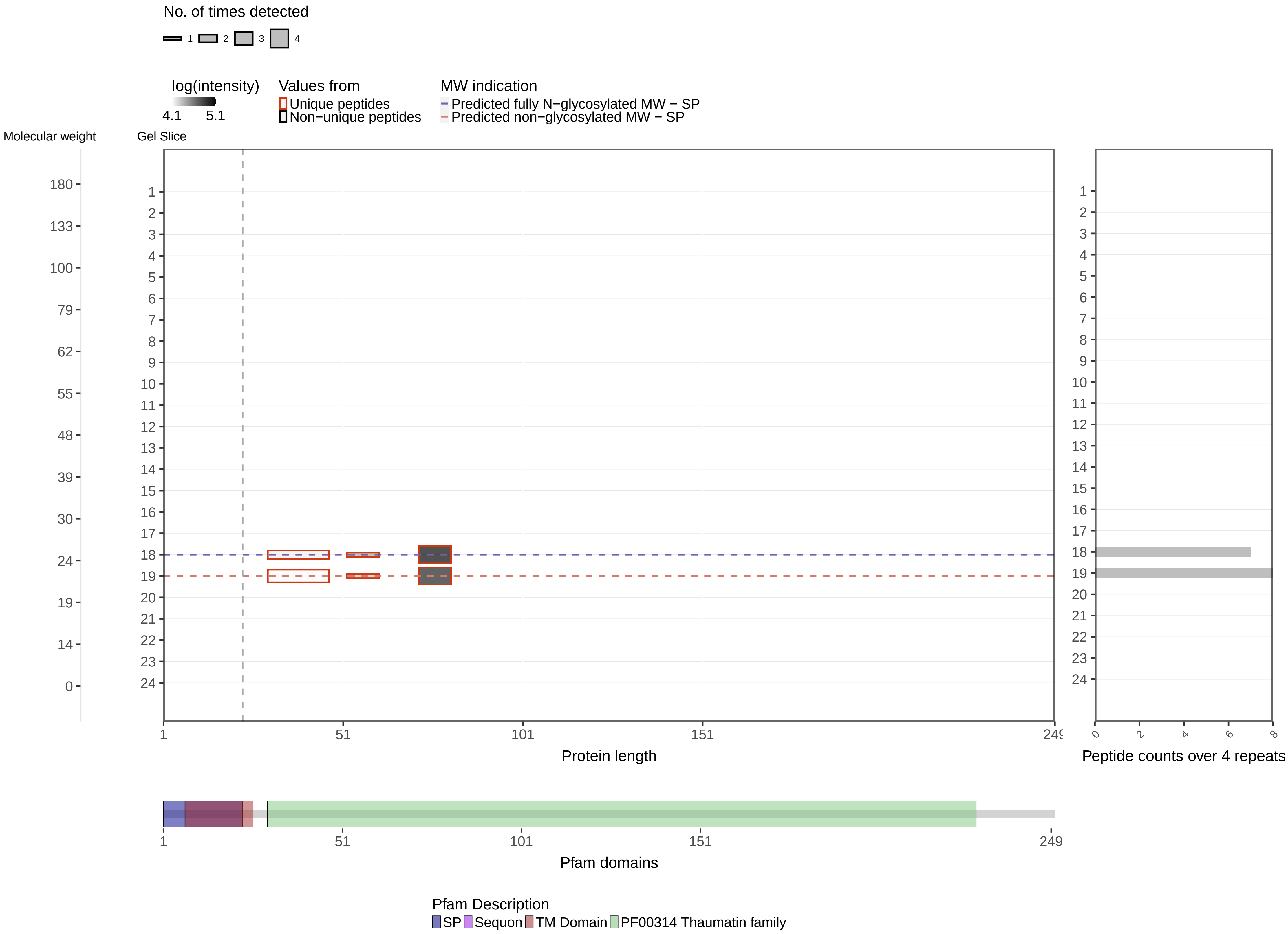


Protein sequence:
QFSGLDFWDISLVDGFNIPMTFAPTNPSGGKCHAIHCTANINGECPRELRVPGGCNNPCTTFGGQQYCCTQGPCGPTSFSKFFKQRCPDAYSYPQDDPTSTFTCPGGSTNYRVIFCPNGQGHPNFPLEMPGSDEVAK*

NbL10g21030.1 osmotin−like_protein [MW − SP = 25.21 kDa]

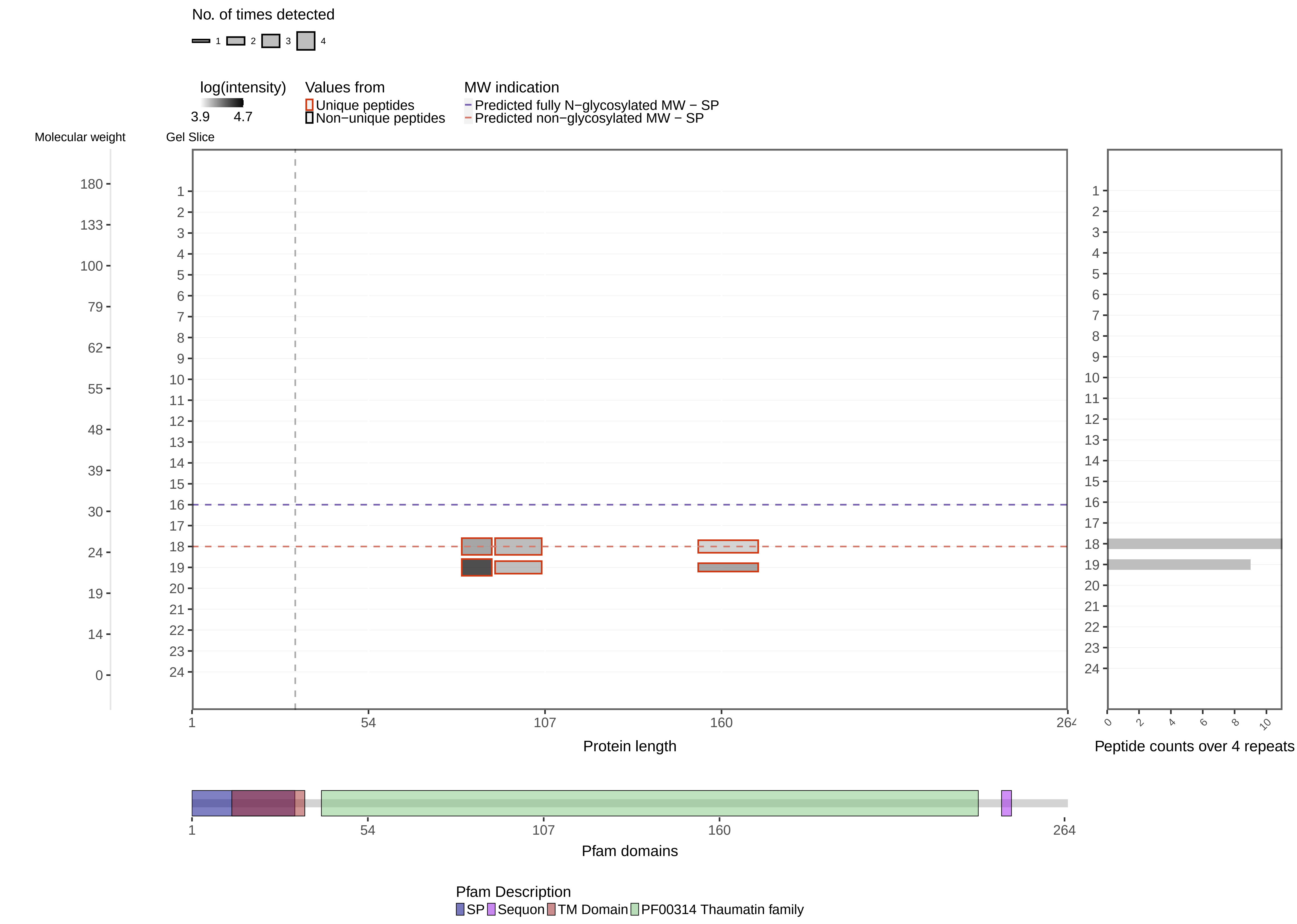


Protein sequence:
ALNQFSNLDFWDISVIDGFNIPMSFGPTKPGPGKCHGIQCTANINGECPGALRVPGGCNNPCTTFGGQQYCCTQGPCGPTELSRWFKQRCPDAYSYPQDDPTSTFTCTSWTTDYKVMFCPYGSAHNNNTTNFPLEMPSSTHEVAK*

### NbL10g20870.1 probable_phosphopantothenoylcysteine_decarboxylase [MW = 23.16 kDa]

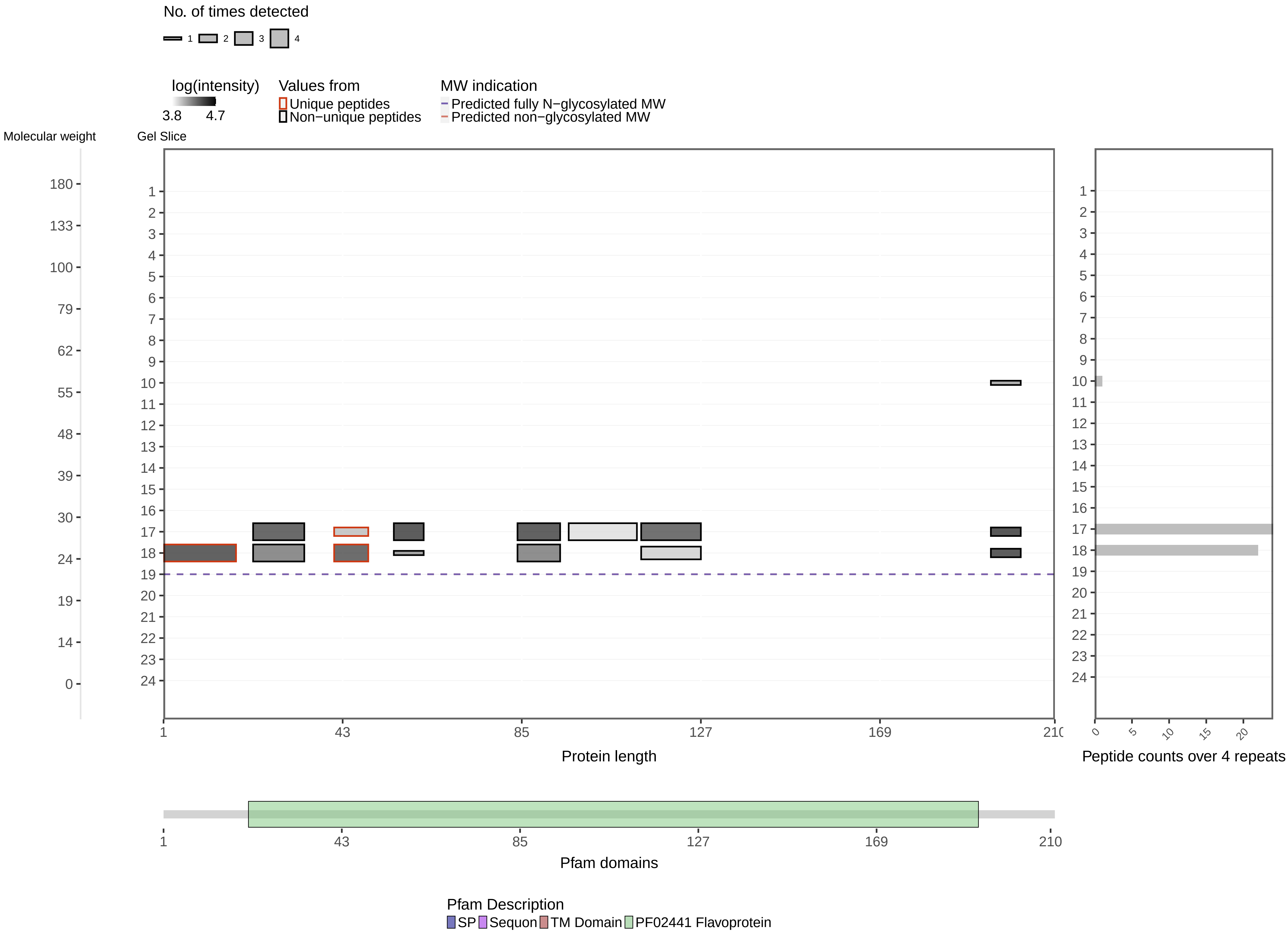


Protein sequence:

ELRRWADIMIIAPLSANTLGKIAGGLCDNLLTCIVRAWDYDKPLFVAPAMNTLMWNNPFTEKHLMVIDELGISLIPPVSKRLACGDYGSGAMAEPSLIYQAVRIYYDAQLRSGGSNVA*

NbL10g20860.1 glutathione_S−transferase_T1−like [MW = 28.62 kDa]

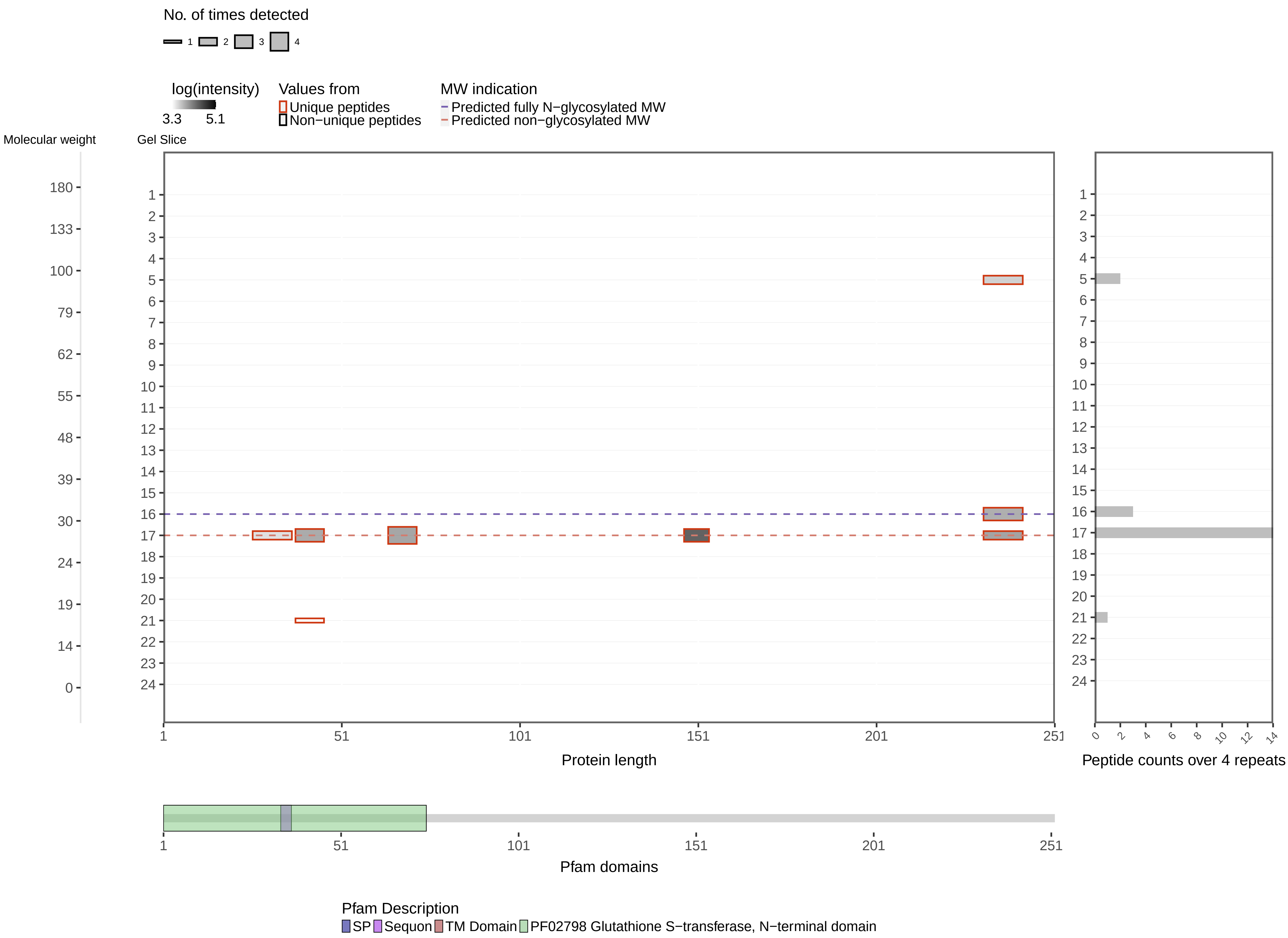


Protein sequence:

AVYVFSTVLAPAFGLPLNQQAAAEAEKVLLASLAKIESVWLQKKGRFLLGSGQPSIADLSLVCEIMQLEVLDEKDRKRIIGPYKRVLKWIDDTKNAMEPHFGEVHLILFKAKEKFQKQRDAVGSSIPQSSRKLDLHSKM*

NbL10g20720.1 glutamate−−cysteine_ligase_chloroplastic [MW = 58.93 kDa]

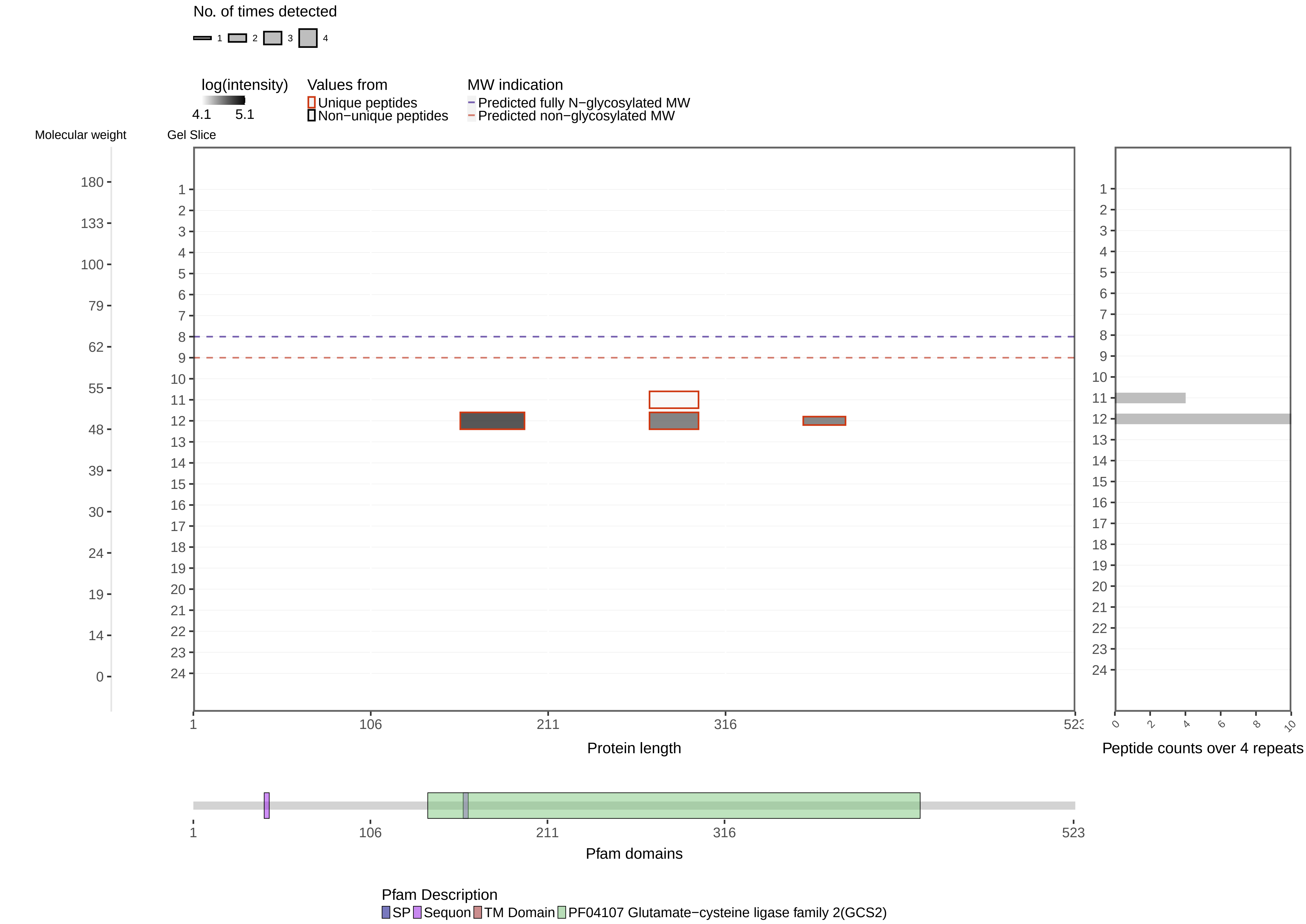


Protein sequence:

RTCTVQVNLDFSSEVDMIRKFRAGLALQPIATALFGNSPFTEGKPNGYLSMRSHIWTDTDNNRAGMLPFVFDDSFGFEQYVDYALDVPMYFVYRKKKYIDCAGMSFRDFMNGKLLPIPGDYPTLNDWENHLTTIFPEVRLKRYLEMRGADGGPWRRLCALPAFWVGILYDELSLQTVLDMTSDWTAEEREMLRNKVPTSGLKTPFRDGLLKHVAQDVVKLAKEGLERRGFKETGFLNEVTEVVRTGVTPAEKLLELYHGKWGRSVDPVFEELLY*

### NbL10g20560.1 aquaporin_PIP−type_pTOM75 [MW = 30.77 kDa]

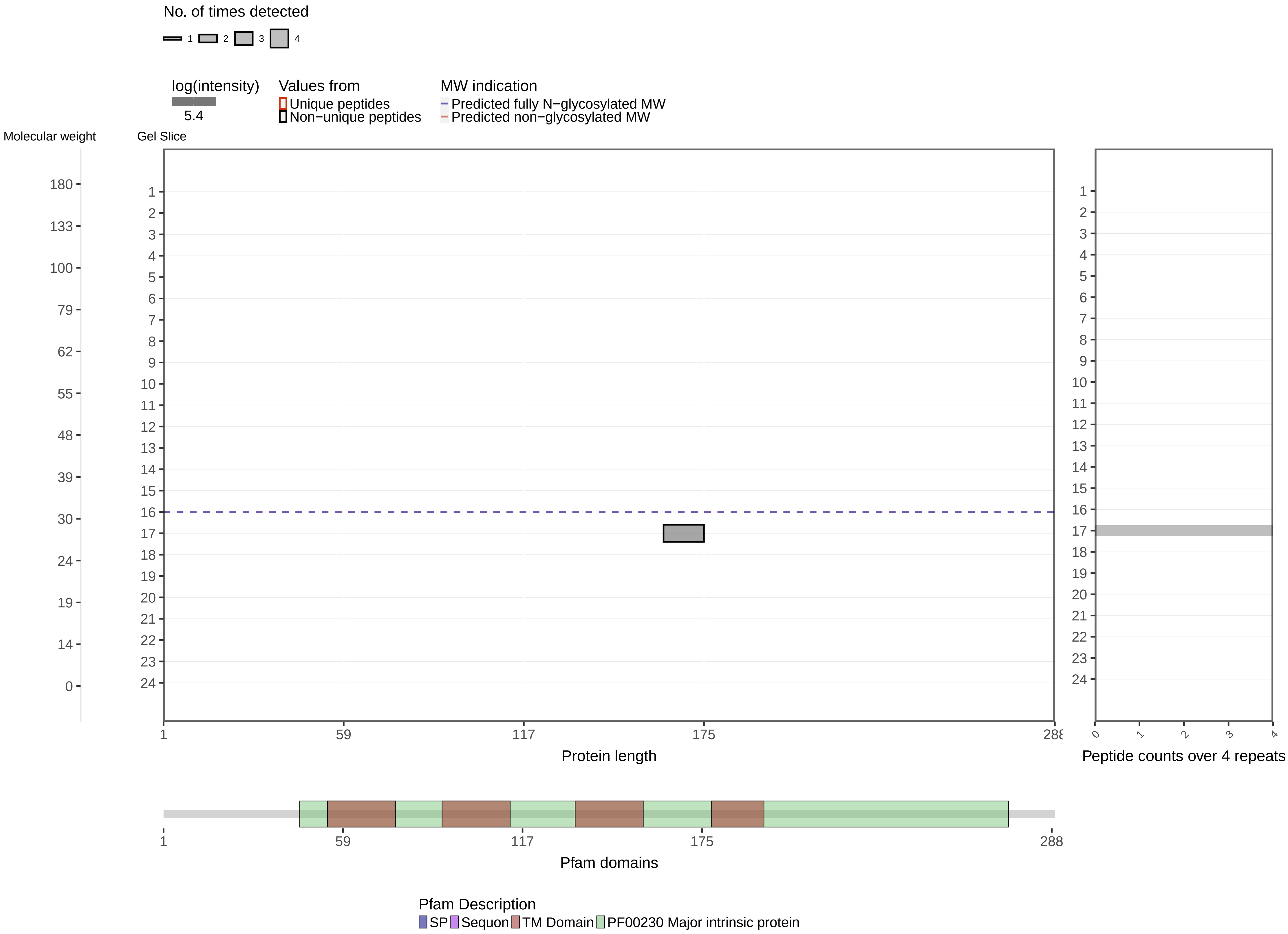


Protein sequence:

.TRAIFYMVMQCLGAICGAGVVKGFMVGPYQRLGGGANVVNPGYTKGDGLGAEIIGTFVLVYTVFSATDAKRNARDSHVPILAPLPIGFAVFLVHLATIPITGTGINPARSLGAAIIHNNDHAWDDHWIFWVGPFIGAALAAVYHQIIIRAIPFHKSS*

NbL10g20500.1 puromycin−sensitive_aminopeptidase−like [MW = 100.05 kDa]

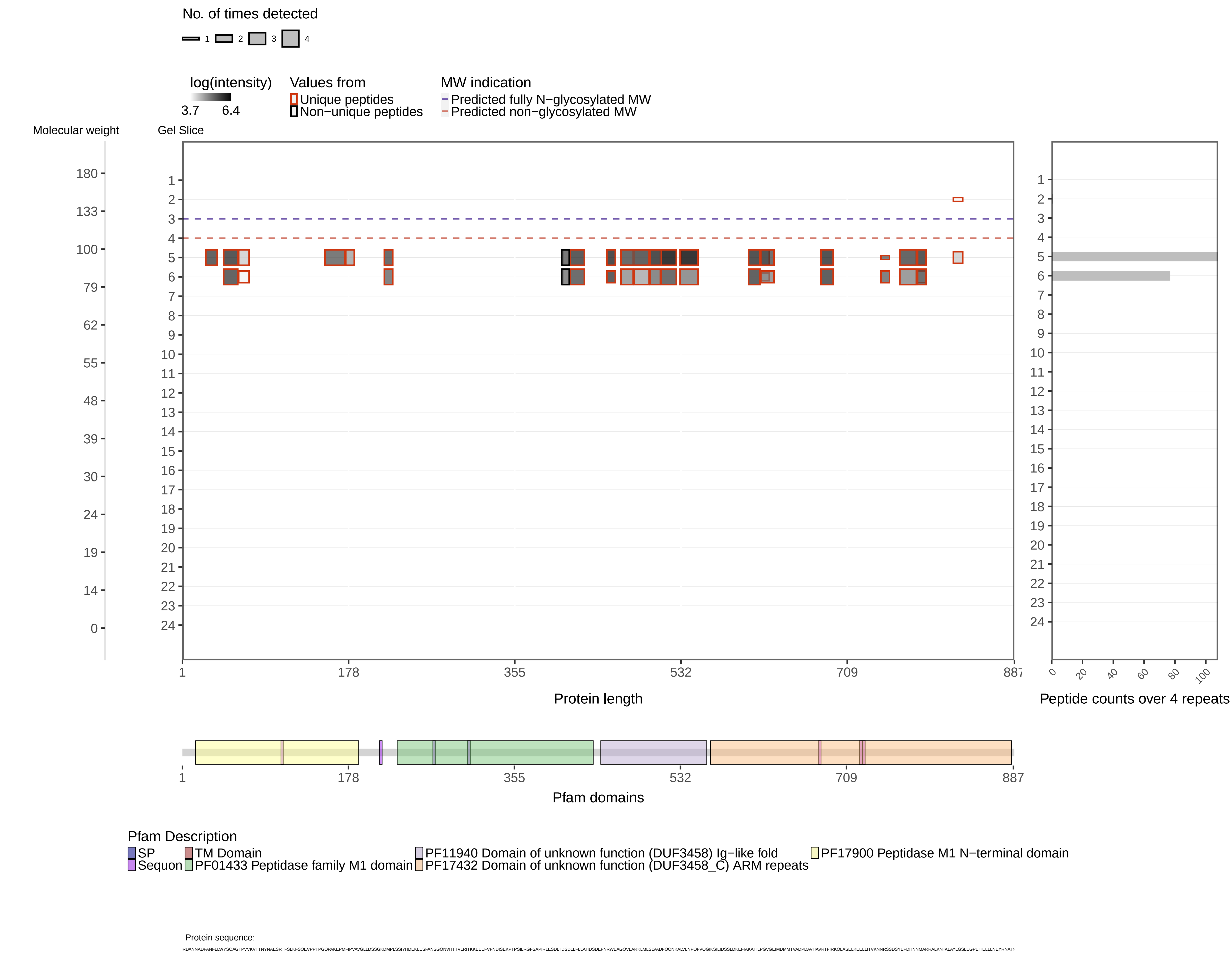

### NbL10g20150.1 uncharacterized_LOC109231095 [MW = 30.22 kDa]

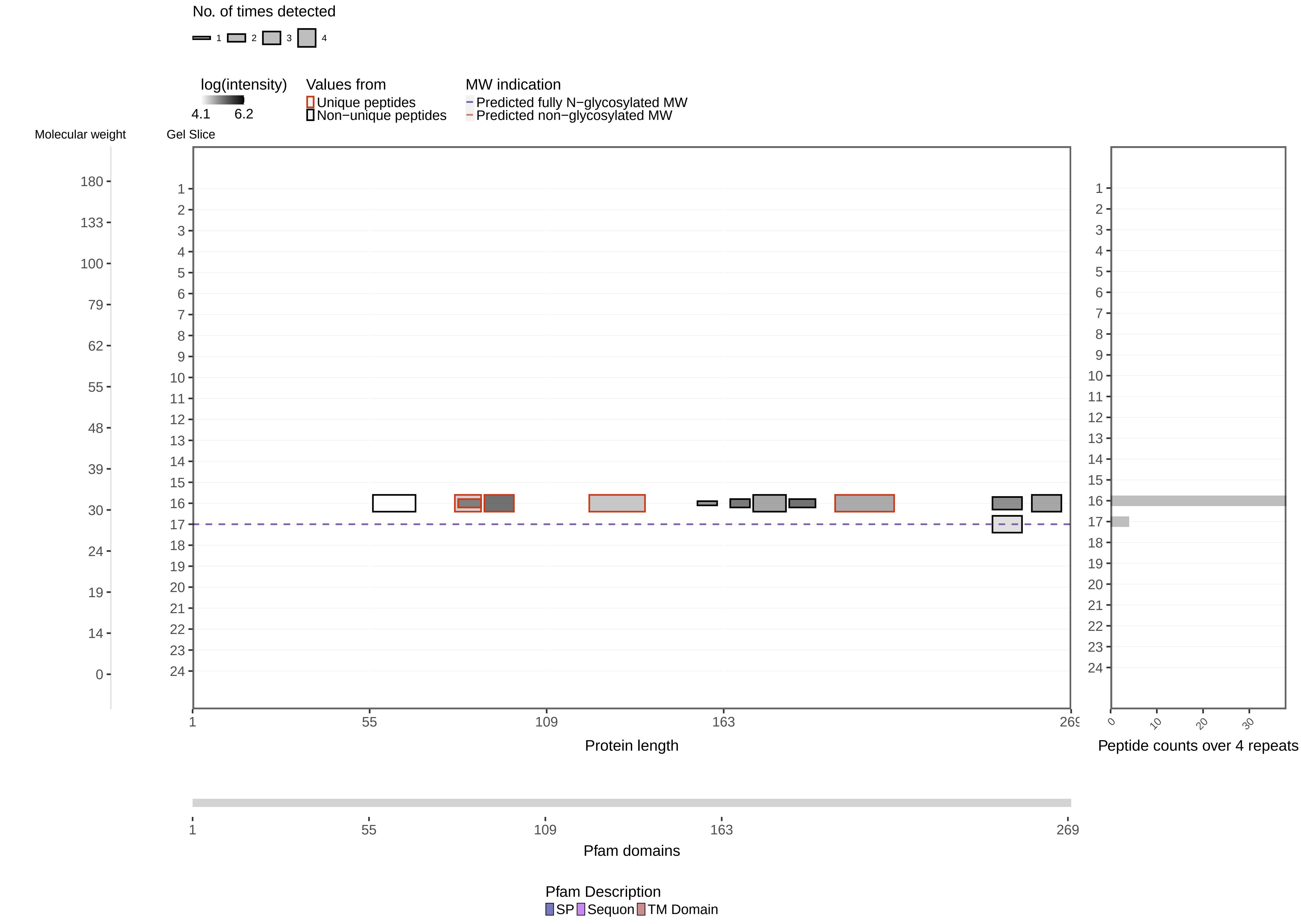


Protein sequence:

KVRDEWMDSDLATWIGANRFYPGVPDALKFASSKLYIVTTKQSRFADALLRELAGITIPAERIYGLGTGPKVKVLKLLQEMPEYQGLTLHFVEDRLATLKNVIKESELDGWNLYLGDWGYNTQKEREEAASIPRIRLLGLSDFSNKLK*

### NbL10g20130.1 bifunctional_L−3−cyanoalanine_synthase/cysteine_synthase_1_mitochondrial [MW = 40.28 kDa]

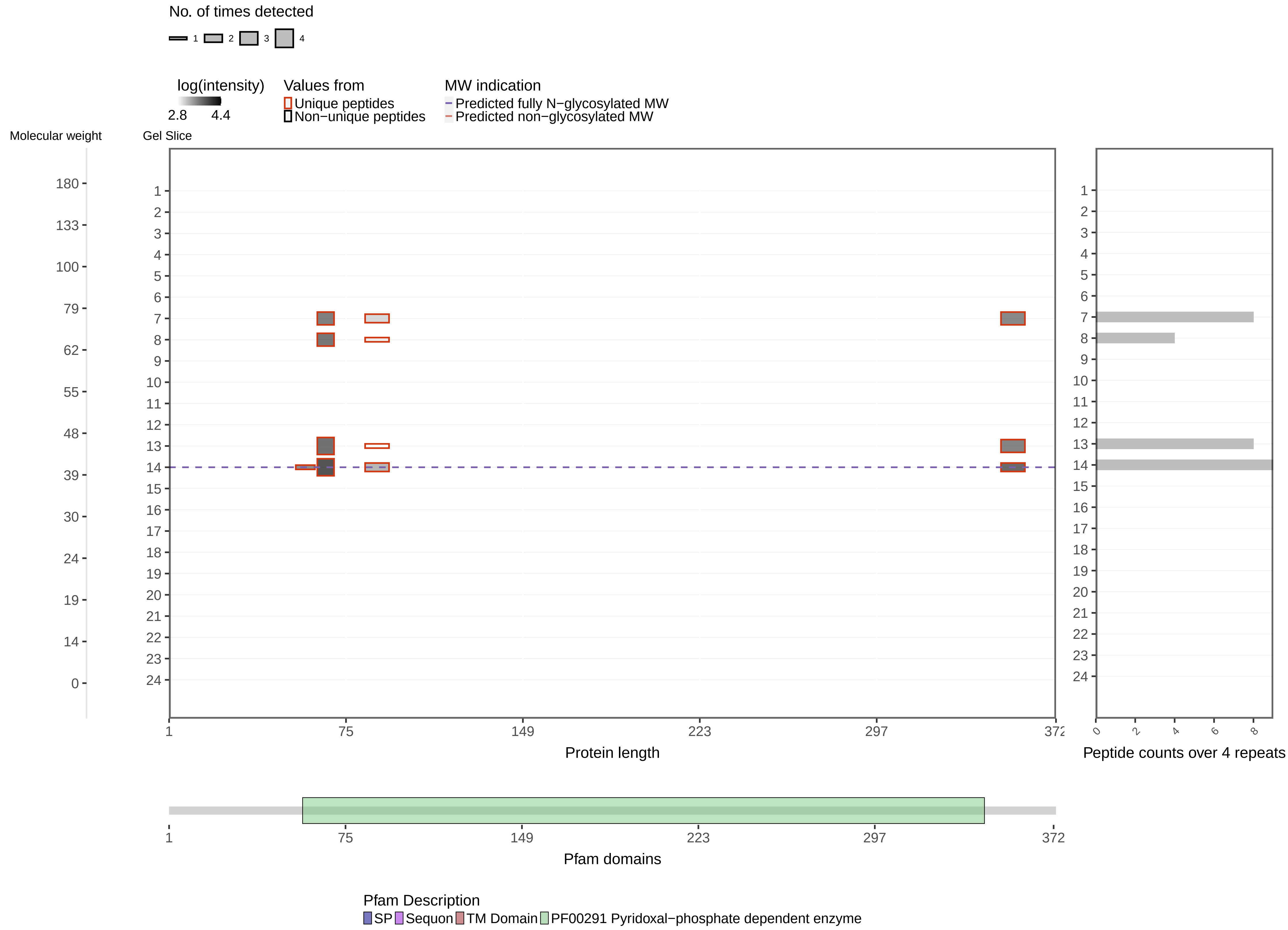


Protein sequence:

GTVKKAYDLLESTPNAFMLOOFSNPANTOVHFETTGPEIWEDTOGNVDIFVMGIGSGGTVSGVGOYLKSKNPNVKIYGIEPTESNVLNGGKPGPHHITGNGVGFKPDILDMDIMEEVLMVSSEEAVNMARELALKEGLMVGISSGANTVAALRLANKPENRGKLIVTIHPSFGERYLSSVLYOELRKEAENMOPVPVD*

NbL10g20020.1 uncharacterized_LOC107784094 [MW = 12.39 kDa]

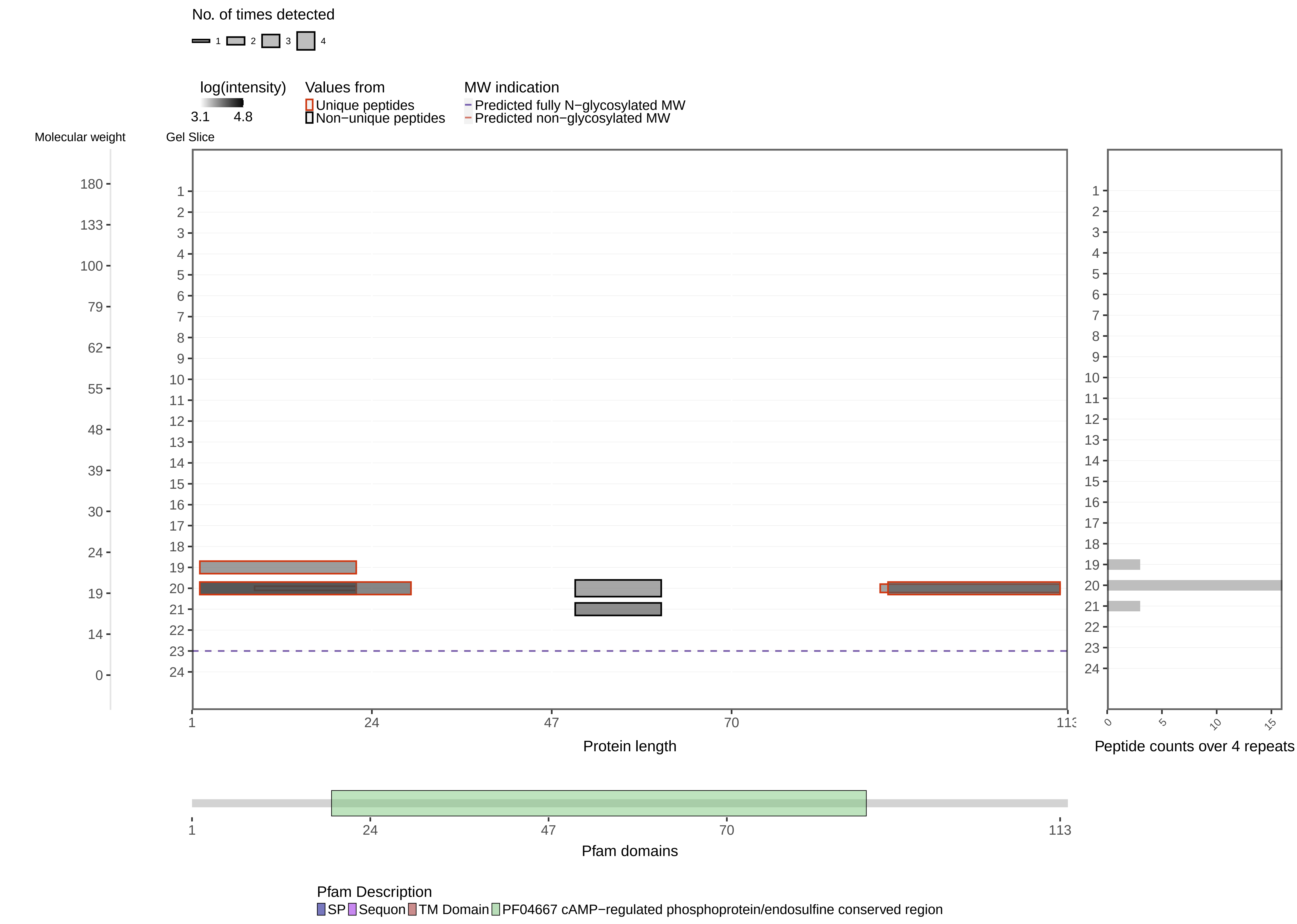


Protein sequence:
KDHERAYFDSADWALGKQGVAKPKGPLEALRPKLQPTNQQTRYRKSPYAPSEGEDGSNMPPEDATANE*

### NbL10g19910.1 ubiquitin−conjugating_enzyme_E2_28−like [MW = 15.44 kDa]

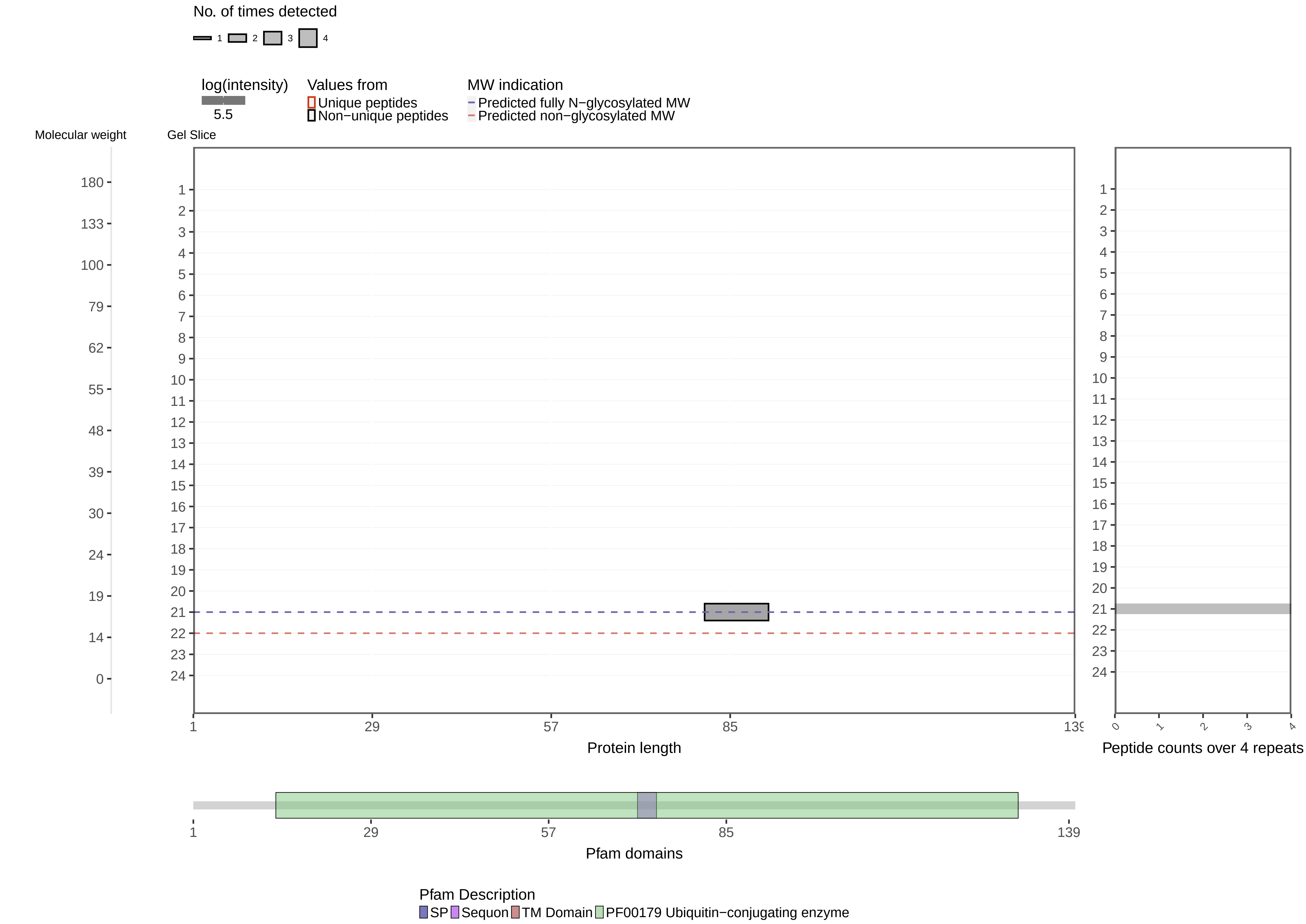


Protein sequence:

KVAFRTKVLHPNINSNGSICLDILKEQWSPALTISKVLLSICSLLTDPNPDDPLVPEIAHMYKTDRAKYETTARSWTQKYAMG*

### NbL10g19820.1 anthocyanidin_reductase−like [MW = 38.84 kDa]

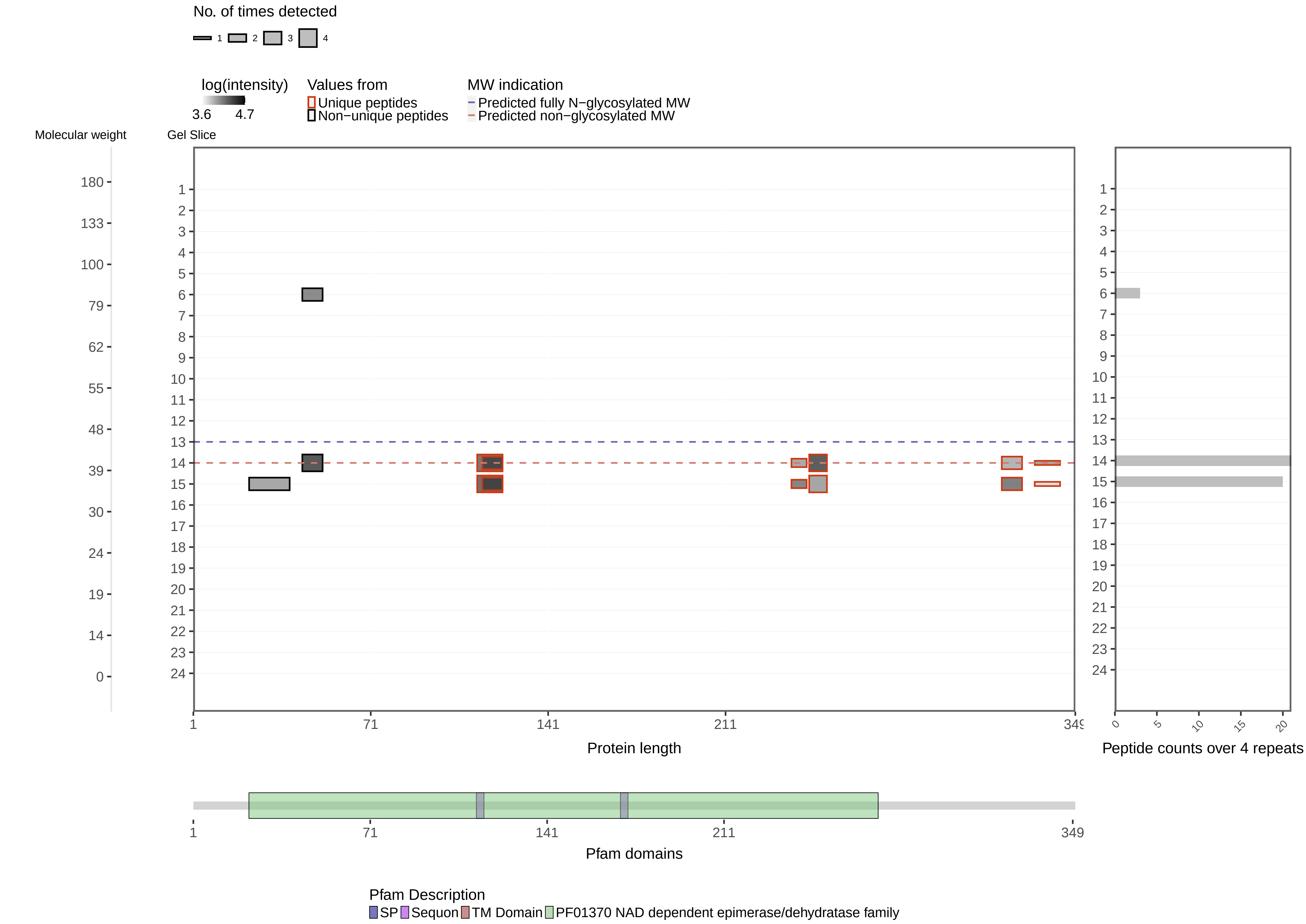


Protein sequence:

ETCWTPLNFSNPYANOGLKEYVESKMLAEKEMLNFGEEGLEVVTLCCGLVGGETYLPYTPTSTALFLSLFTODKSLYNSLKFLEELIGKVPIVHIEDVCEAHLLSMRAAVNGRFLCASSFISSAEIGCYYOONYPEFNLNOEYLNDPNREIKWGSKKLVDNGFVYKYGMKEILDDSVSCARKNGAF*

NbL10g19630.1 heterogeneous_nuclear_ribonucleoprotein_Q [MW = 52.27 kDa]

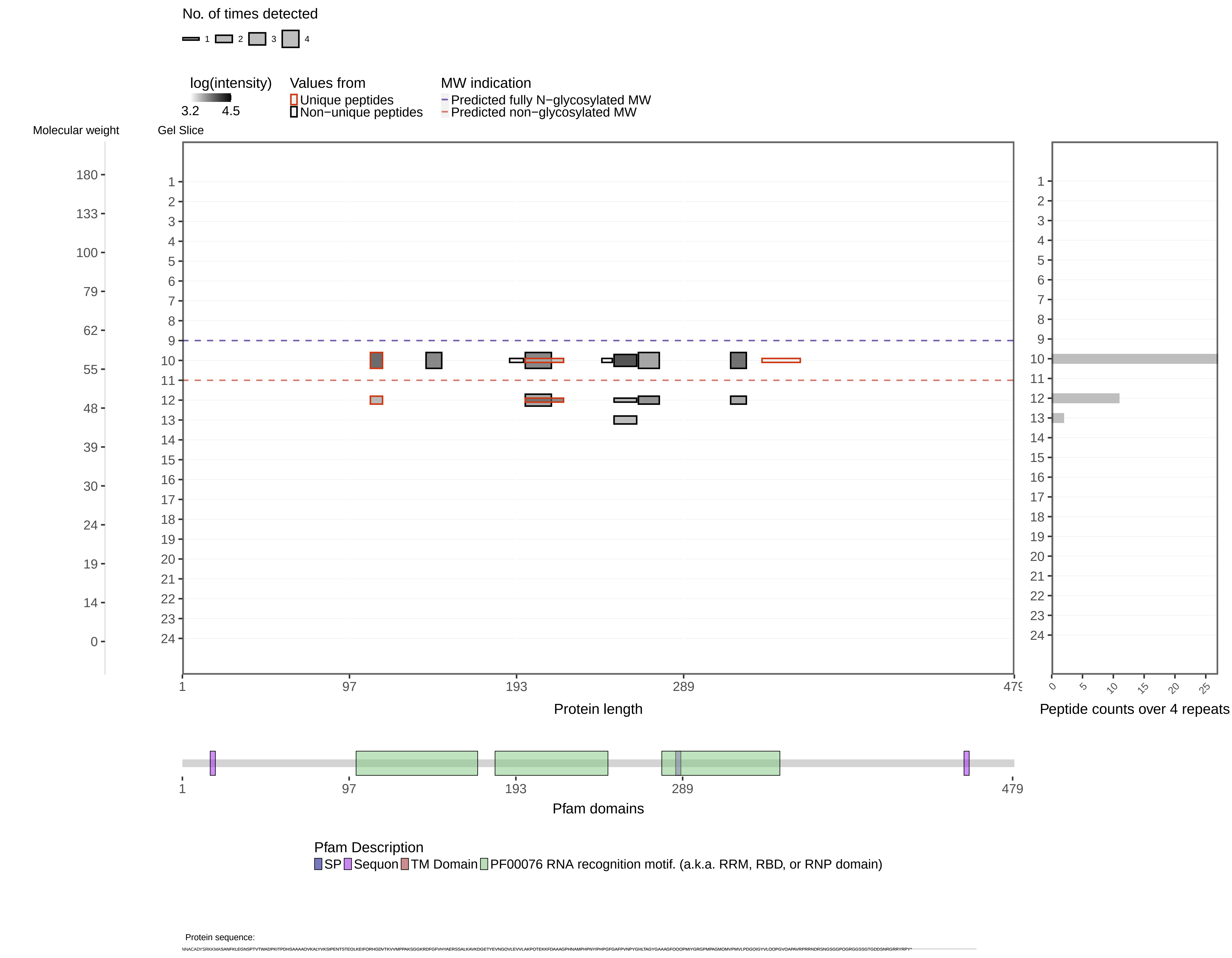

### NbL10g19560.1 protein_CURVATURE_THYLAKOID_1B_chloroplastic [MW = 19.22 kDa]

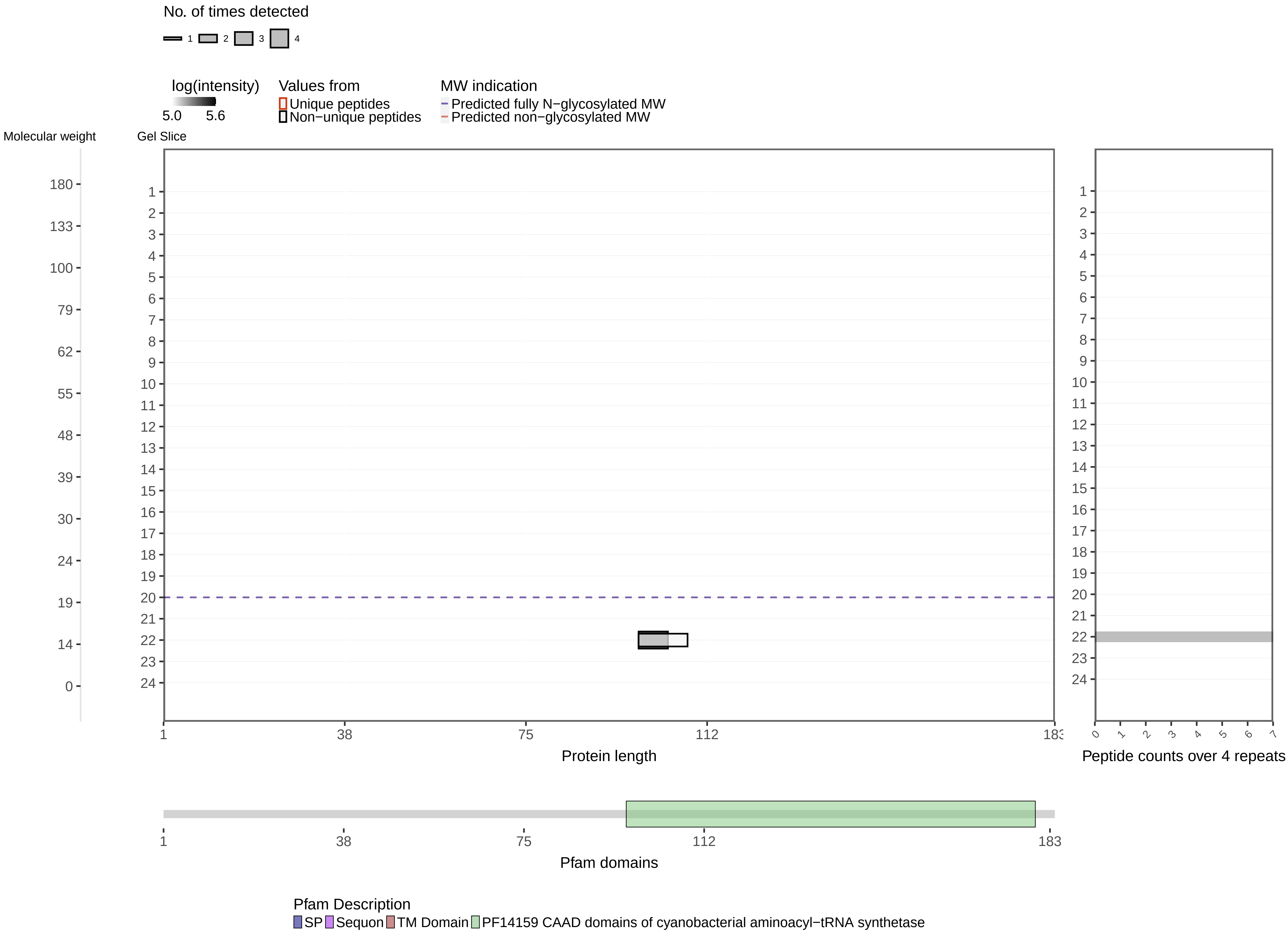


Protein sequence:

TTEASSADMATTELPSELLQKIQEAWDKLDDKYAVSSLGVAALLLLWSSTGVISAIDRLPLIPGVLELVGIGYTGWFAYKNLVFKPDREALISKIKDLYKEVIGSS*

### NbL10g19550.1 caffeic_acid_3−O−methyltransferase−like [MW = 40.19 kDa]

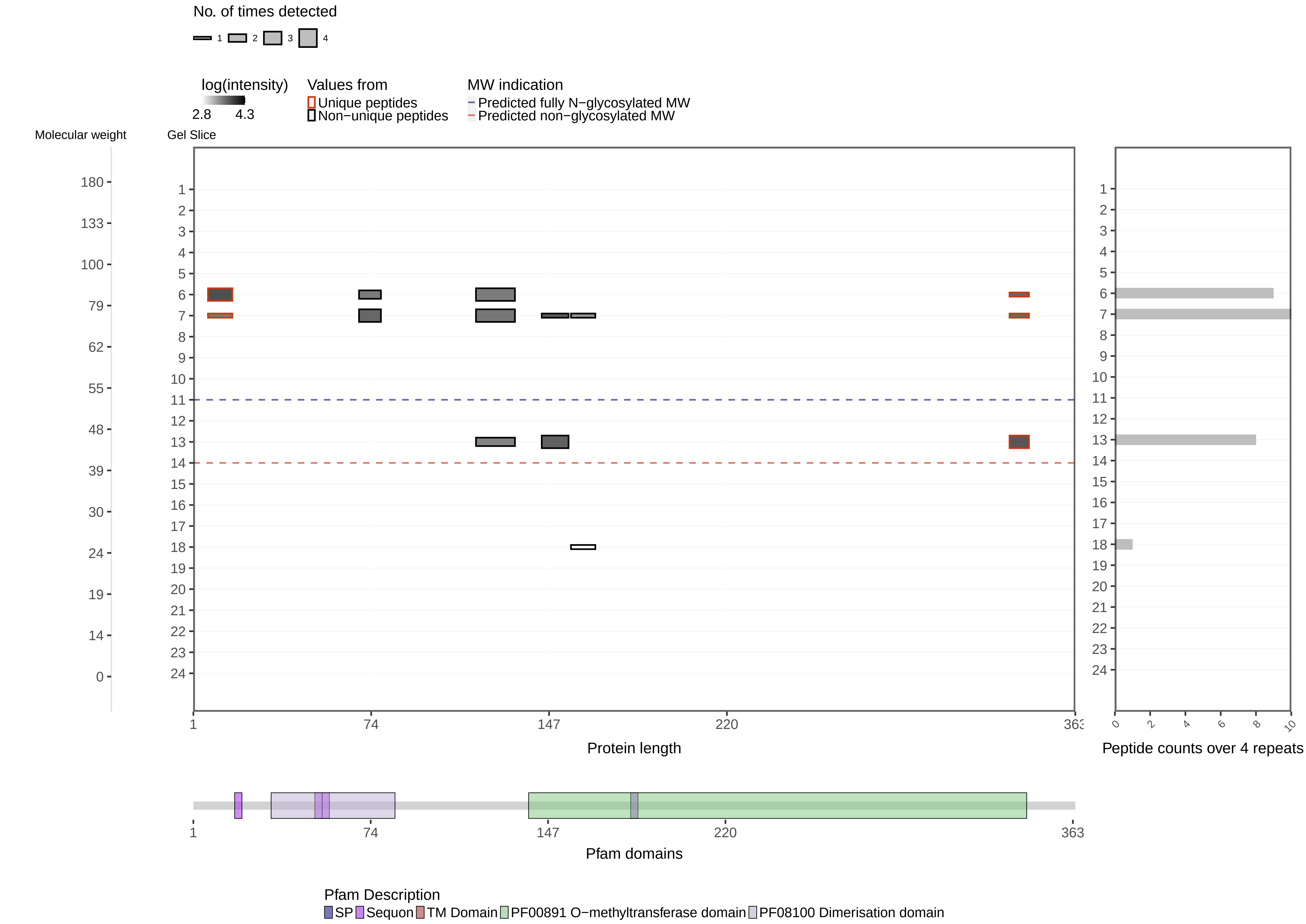


Protein sequence:

PKFNDVFNSAMLNHTTVAMKKILENYKGFDNLKTLVDVGGGLGVNLKMITAKYPTIKGTNFDLPHVVQHAPSYPGVEHVGGDMFESVPDGDAIFMKWILHDWSNSHNLKLLKNCYKALQENGKVIVVEAILPVKPDTDTAVIGVSQCDLIMMAQNPGGKERSEEEFRTLATEAGFKGVNLICCVCNFWVMEFFK*

NbL10g19490.1 probable_2−oxoglutarate/Fe(II)−dependent_dioxygenase [MW = 38.84 kDa]

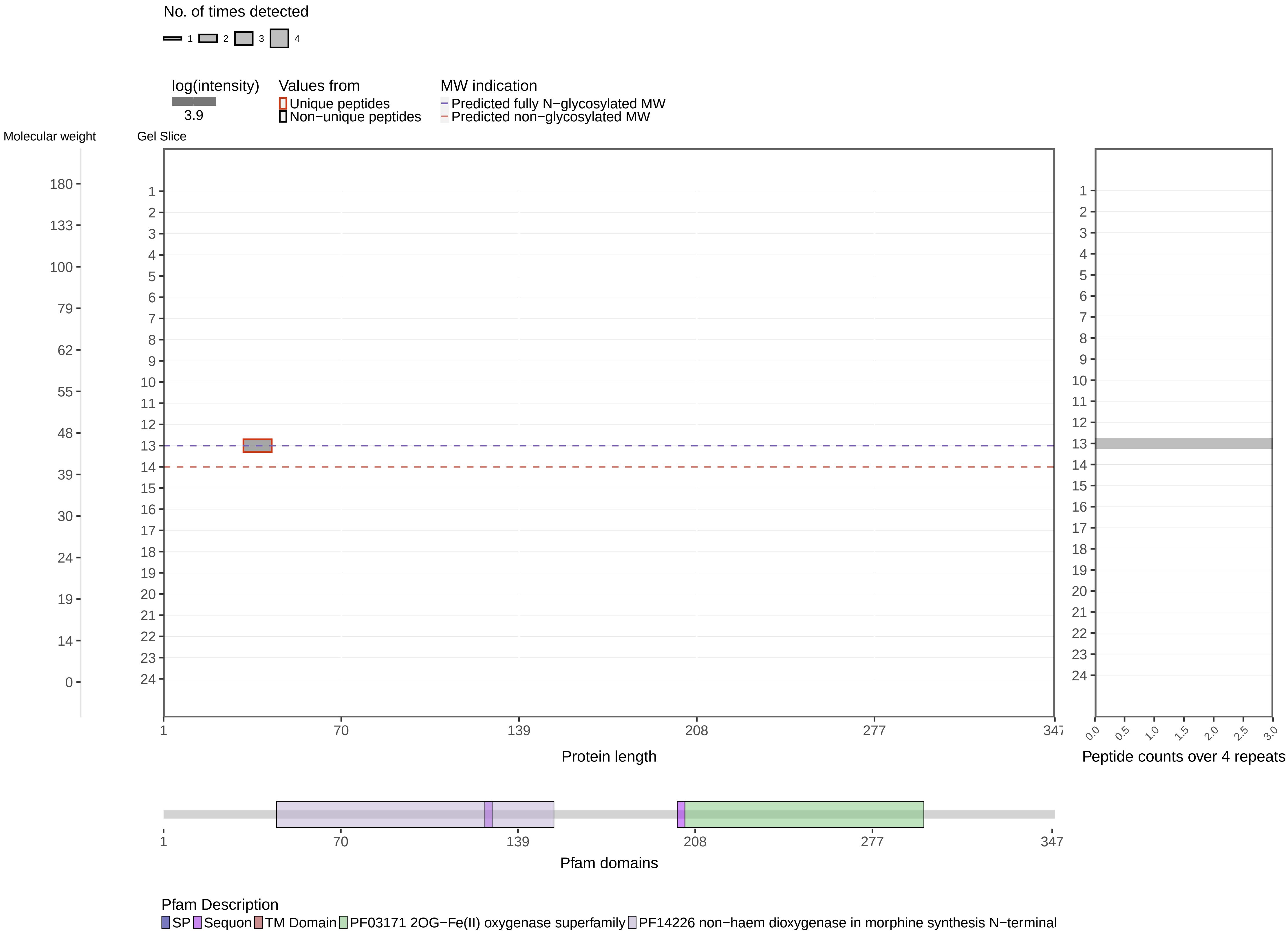


Protein sequence:

YRKVVTAYSDHMKALAQELLGLVSESLGLTCQSMENVVGEFYQNITISYYPPCPQPELTLGLQSHSDMGAITLLIQDDVGGLEVFNSGEWVTVNPMSDAICVILADQTEIISNGEYRSAQHRAVTNARKARLSVATFHDPAKTRQISPSPAFHPPKYRPVIYGDYVSSWYTKGPEGKRNLDALLNCQLN*

NbL10g19480.1 plasma_membrane−associated_cation−binding_protein_1−like [MW = 24.77 kDa]

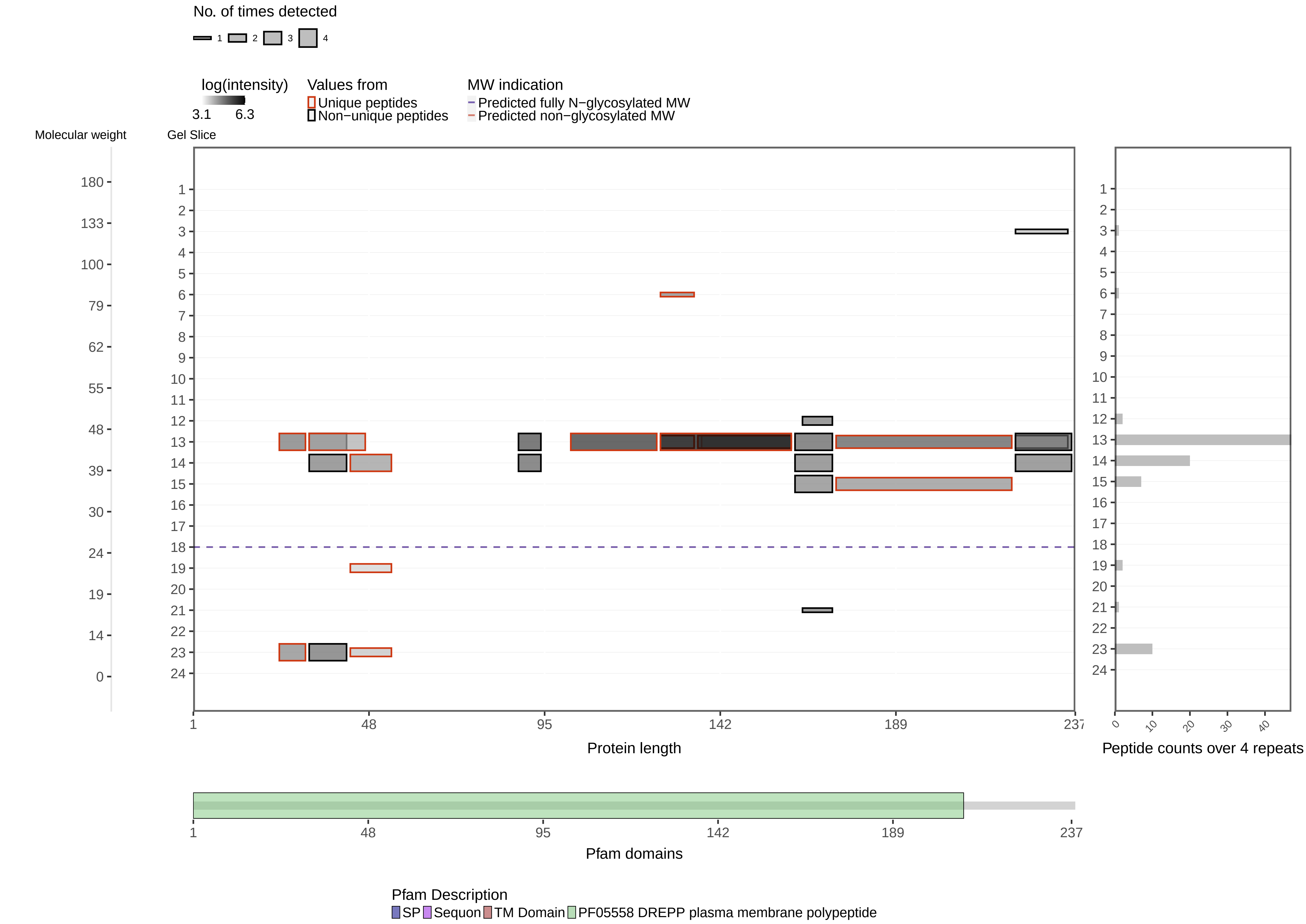


Protein sequence:

SEASSNFGPSYVSGPVLFVFEKVSTLIVTEDKKEEEAAIPVAAASEAEATTTAPVEVKEKEIVVEAEAKKEEEPAPAPAPAPAPAPAPAPAPAPAAAPAAEVAPAEVAPAADVAPAKVEEAPAAVAAPEPPKA*

NbL10g19370.1 polyadenylate−binding_protein_RBP45−like [MW = 52.28 kDa]

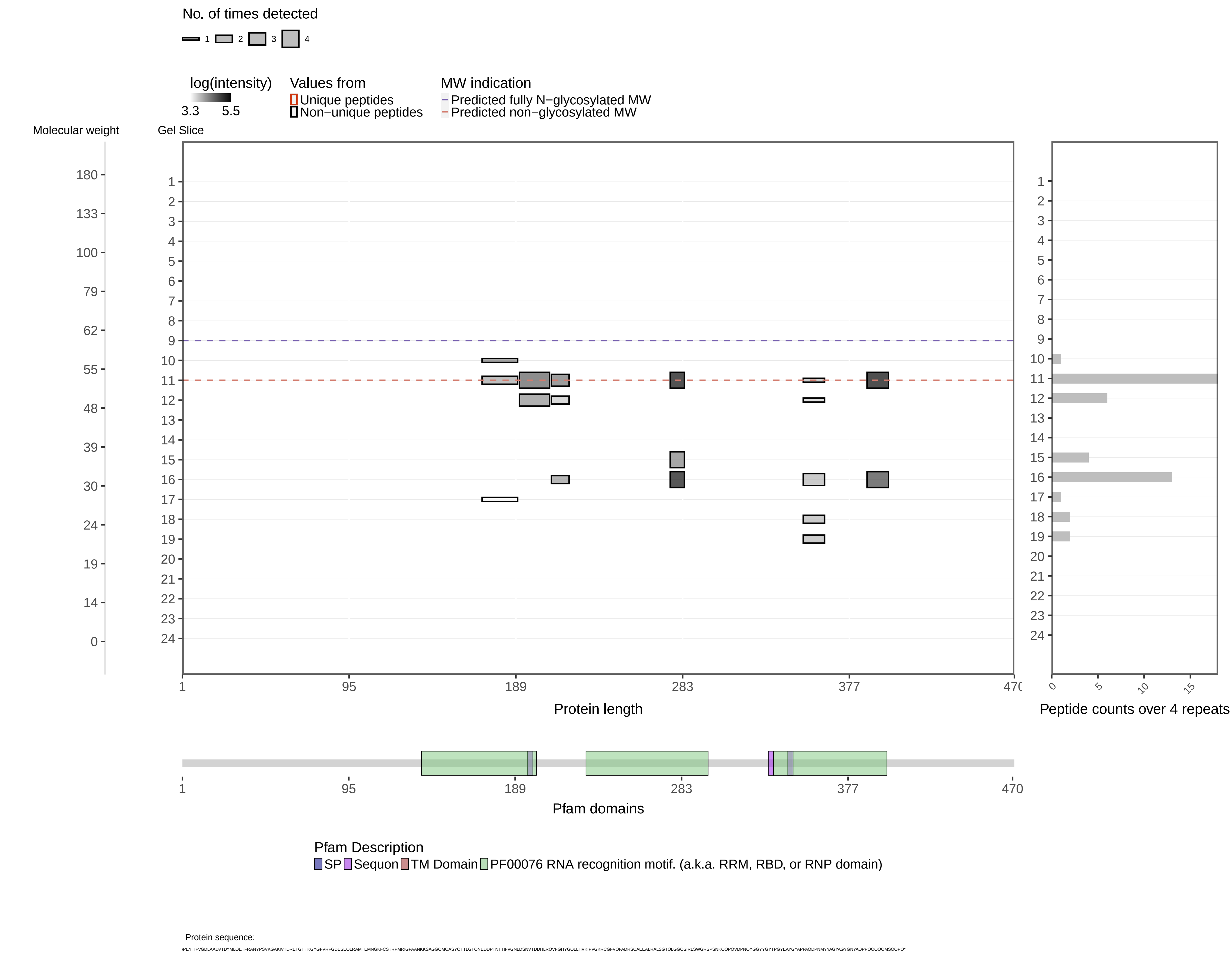

### NbL10g19320.1 adenosine_deaminase−like_protein [MW = 40.93 kDa]

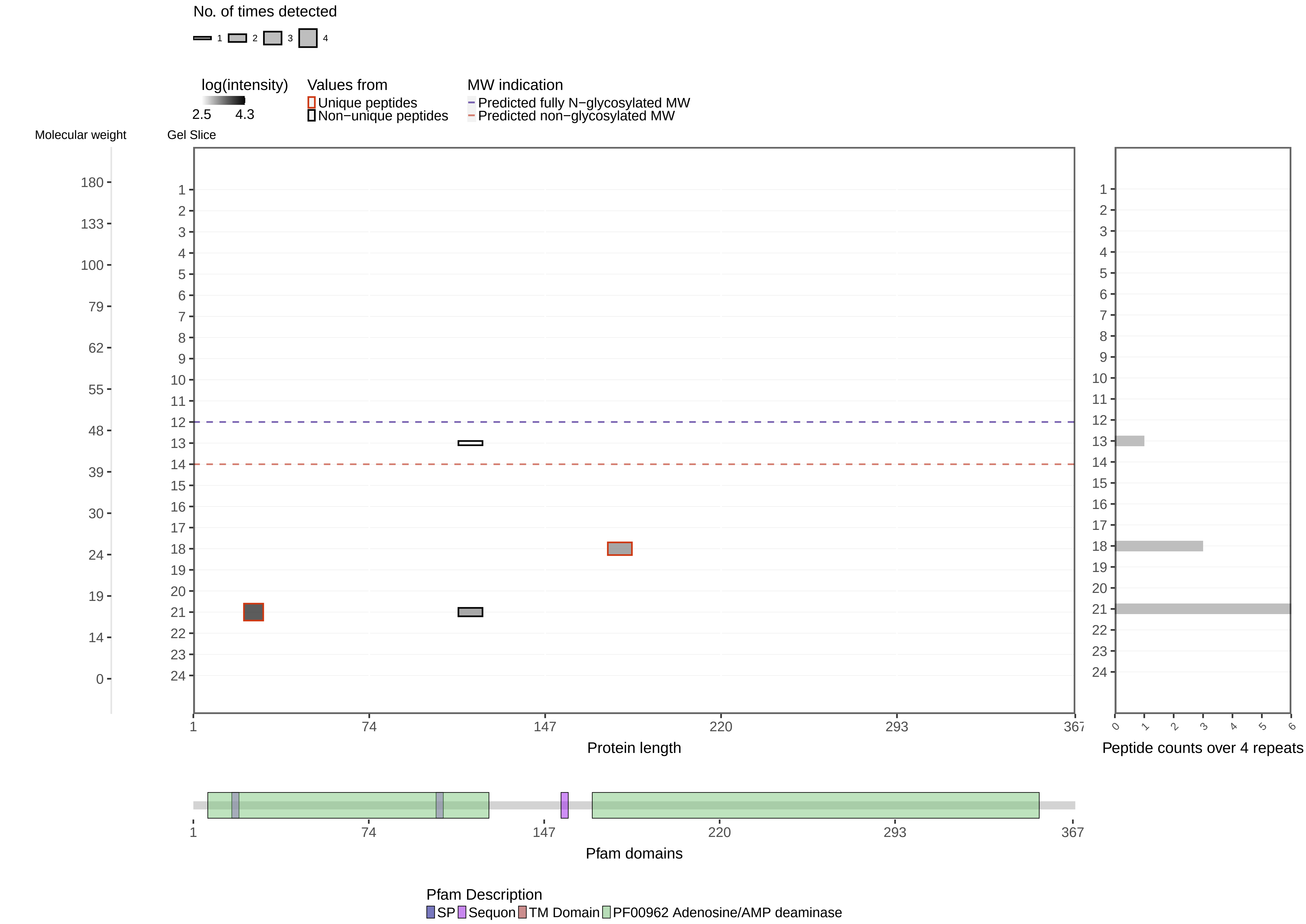


Protein sequence:

DRRESTEAAVETVKLALEMRHQGVVGIDLSGNPIIGEWLTFLPALEFAKEQGLLITLHCGEVPNPVEIHAMLDFLPARIGHACCFEEEEWAKLKSLKIPVEICLTSNIRTETISSLDIHHFADLYYSGHPIVLCTDDSGVFSTSVSGEYSLAASAFGIQKREMFQLARSAIKFIFAGDKVKLELEQLFDLAAKNLEF*

NbL10g19110.1 keratin_type_II_cytoskeletal_1−like [MW − SP = 9.35 kDa]

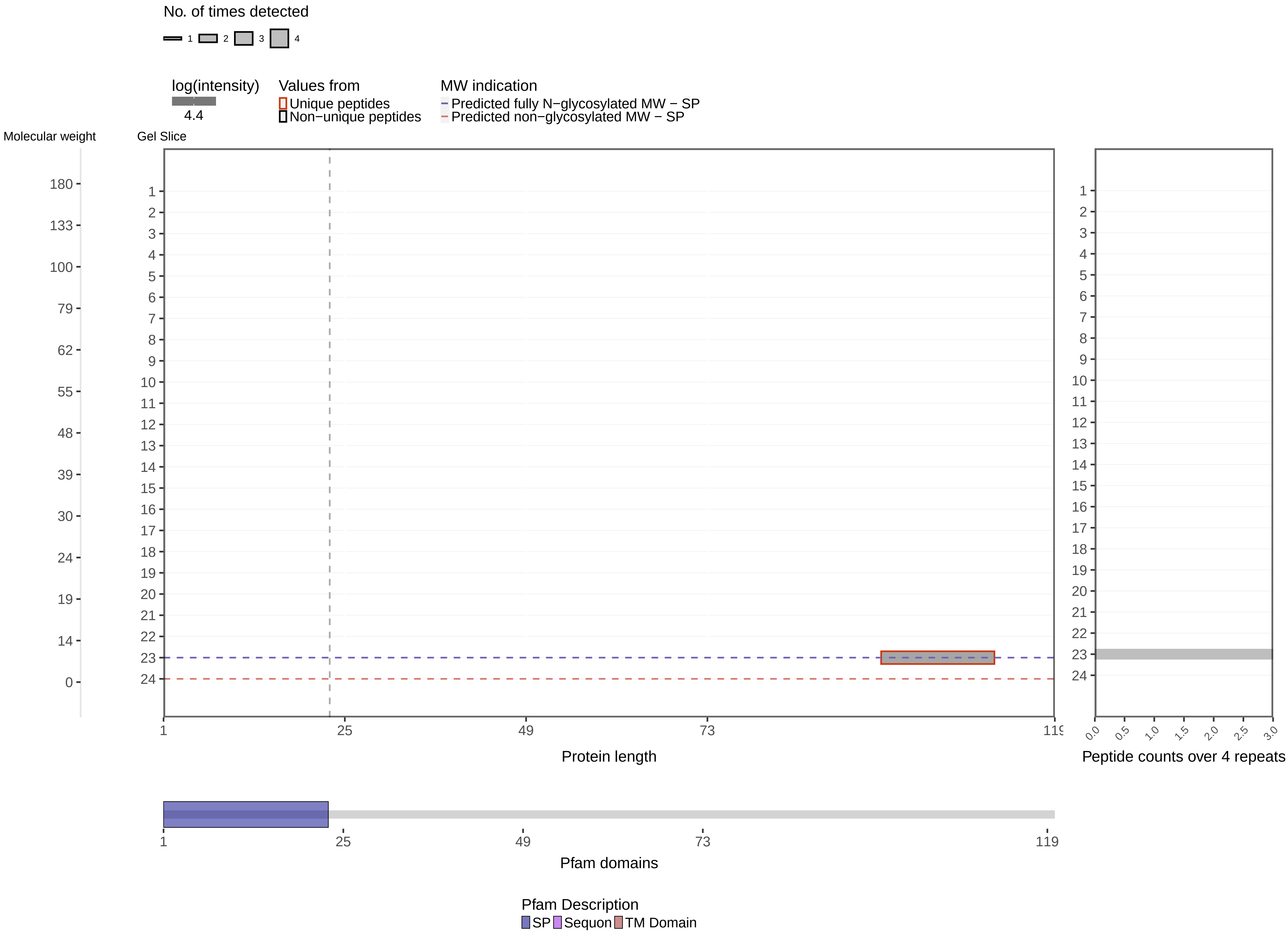


Protein sequence:
GFGGPKGGYAKGGIIRPTVVCKDKGPCLGKKLRCPAKCFKSFSRAGKGYGYGGGGGGCTMDCKKKCVAYC*

### NbL10g19080.1 granule−bound_starch_synthase_1_chloroplastic/amyloplastic−like [MW = 11.9 kDa]

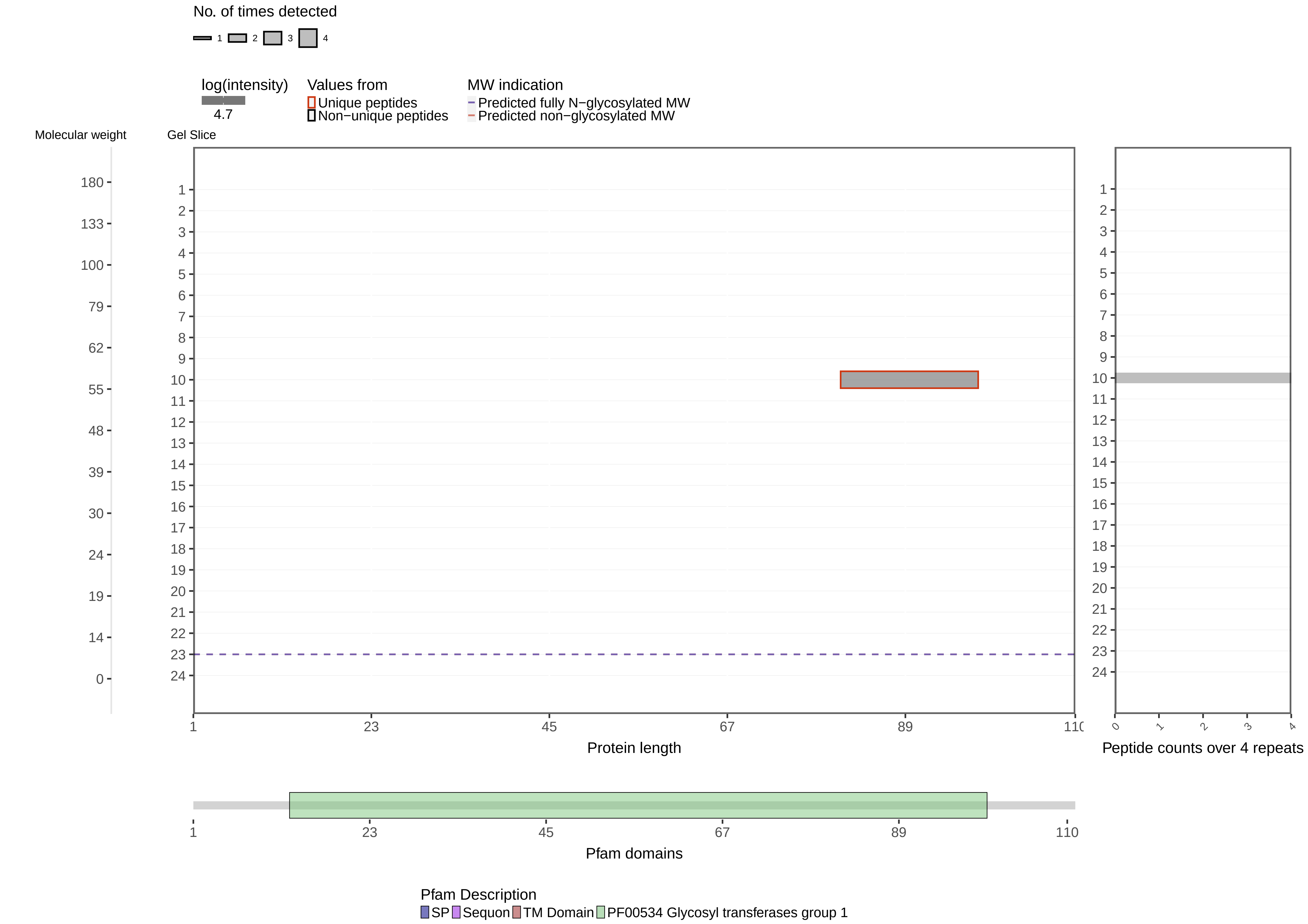


Protein sequence:

GVAKFNVPLAHMITAGADFMLVPSRFEPCGLIQLHAMRYGTVPICASTGGLVDTVKEGWIPYGSIQR*

### NbL10g18840.1 nudix_hydrolase_3 [MW = 88.9 kDa]

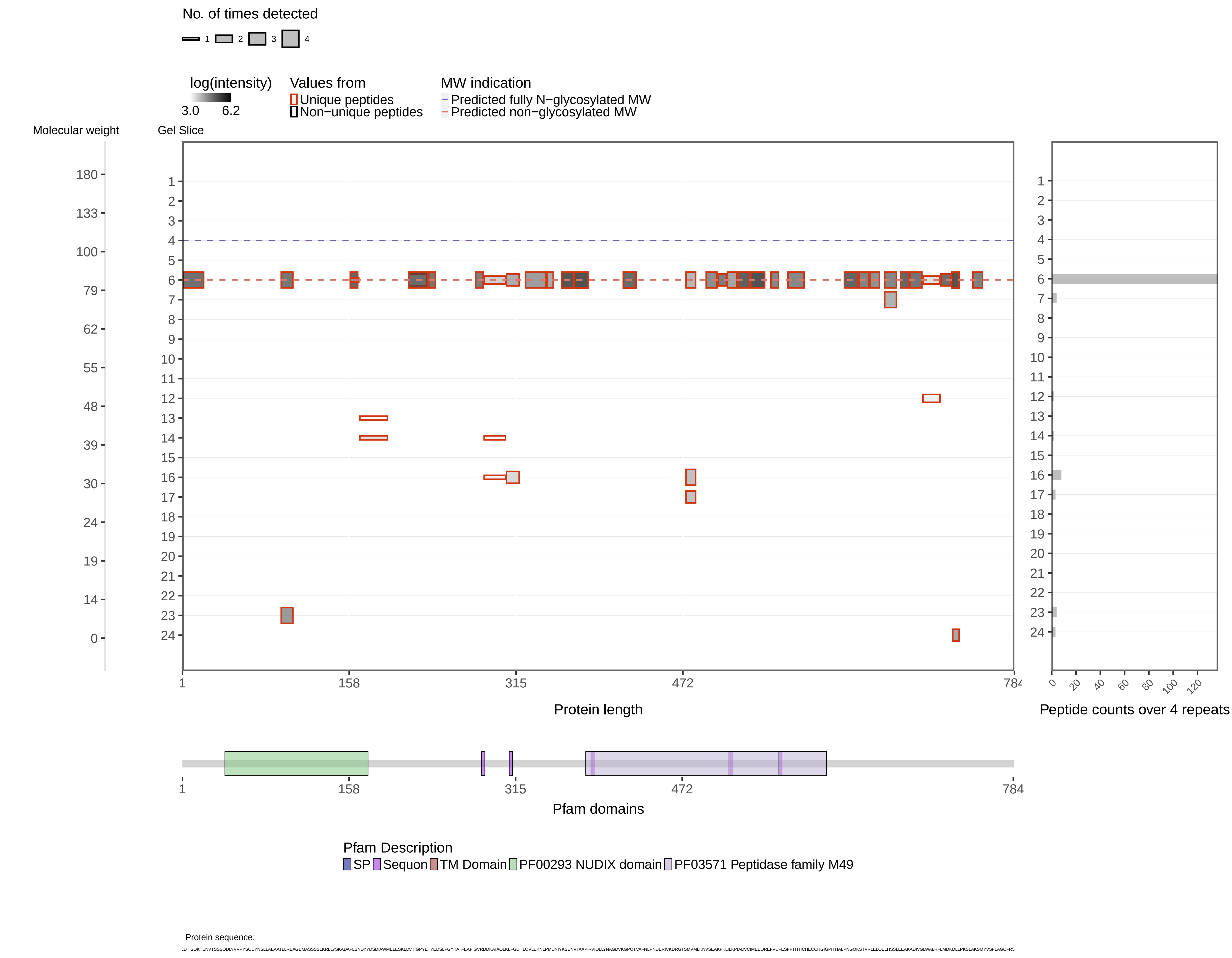

### NbL10g18810.1 ABC_transporter_E_family_member_2 [MW = 68.09 kDa]

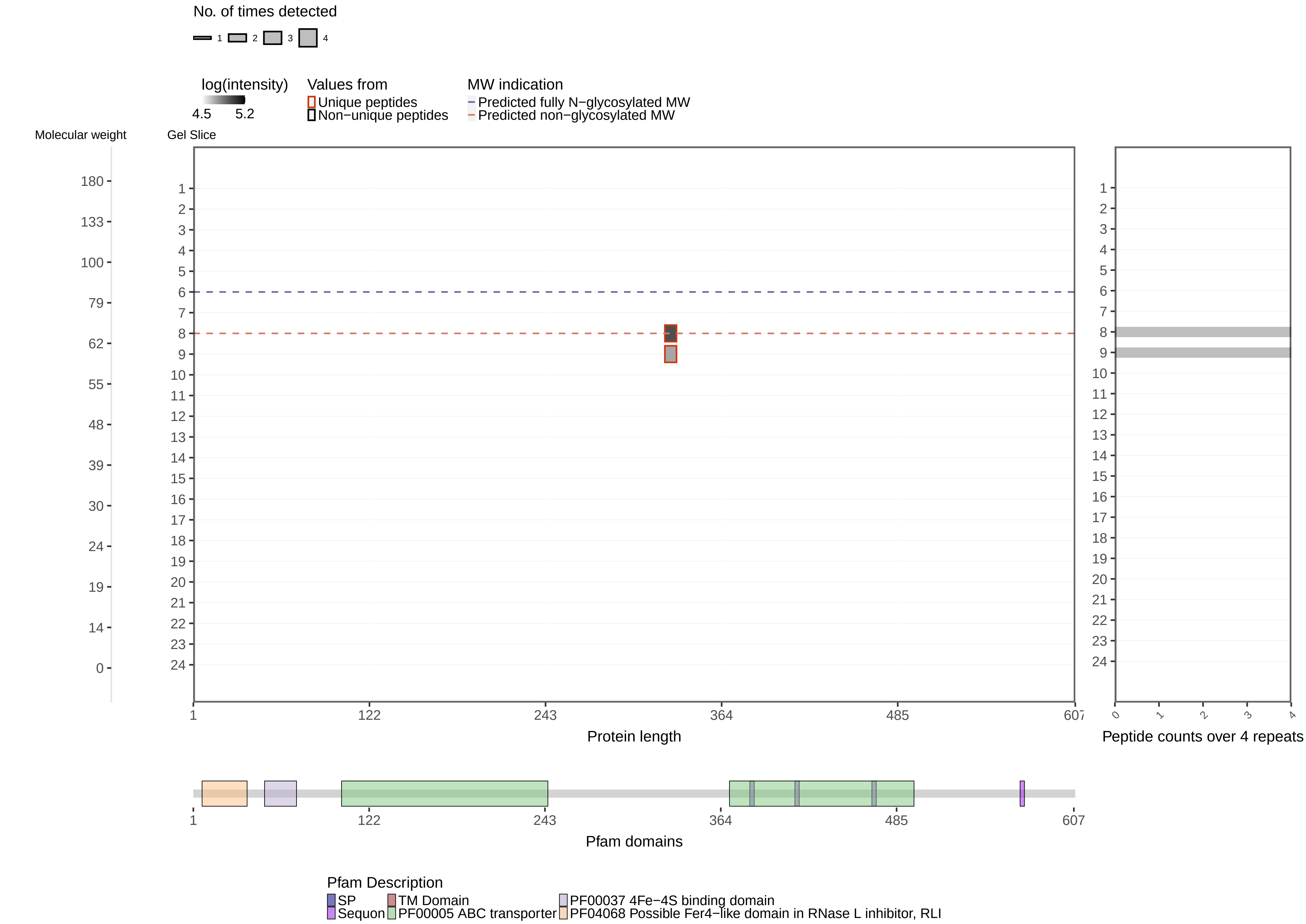


Protein sequence:

LYGKPGAYGVVTLPFSVREGINIFLAGFVPTENLRFRDESLTFKVAETPQEAAEEIETYARYRYPTMTKTQGNFKLKVAEGEFTDSQIIVMLGENGTGKTTFIRMLAGLLKPDVVEGSDTDIPEFNVSYKPQKISPKFPSTVRHLLHQKIRDSYMHPQFCSDVMKPLQIEQLMDQEVVNLSGGELQRVALALCLGKPADIYLIDEPSAYLDSEQRIVASKVIKRFILHAKKTAFIVEHDFIMATYLADRVIVYEGTPSIDCVANAPQSLLTGMNLFLSHLNITFRRDPTNFRPRINKLEST

NbL10g18560.1 NADP−dependent_malic_enzyme−like [MW = 65.39 kDa]

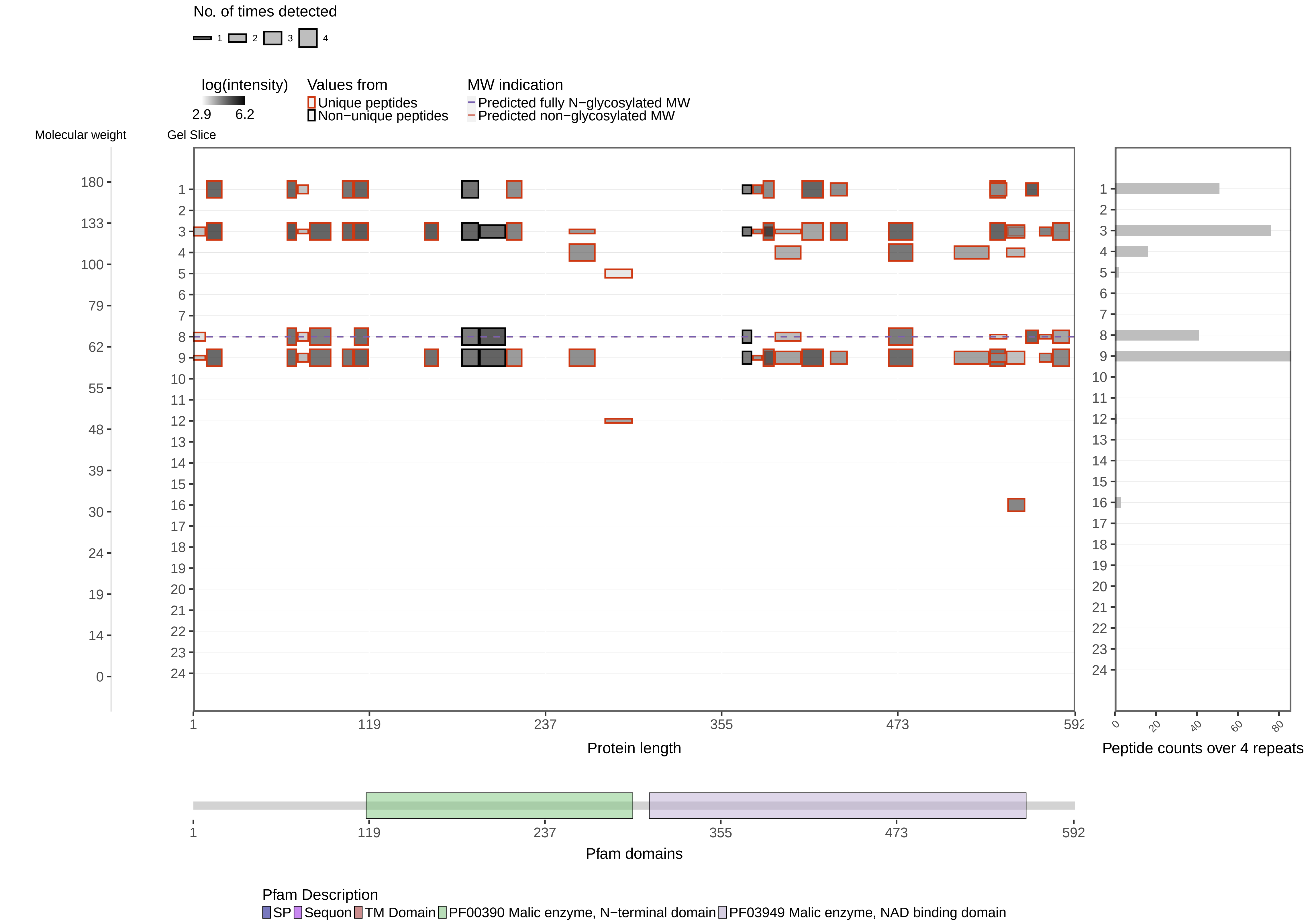


Protein sequence:

EDFANHNAFELLAKYGTTHLVFNDDIQGTASVVLAGLIASLKLLGGSLADHTFLFLGAGEAGTGIAELIALEISKKTKCPVEETRKKIWLVDSKGLIVSGRKETLQSFKKPWAHEHDPVNNLLDAVKSIKPTVLIGTSGVGRTFTKEVIEAMASTNERPLIMALSNPTSQAECTAEEAYTWSEGRAVFASGSPFPSFEYNGKLNIPGQANNCYIFPGFGFGLVMSGTIRVHDDMLLAASEALAAQVTEEHFAKGMIYPPFADIRKISAHIAASVATKAYELGVATRLPRPENLVKYAES[illegible]

NbL10g18390.1 polyphenol_oxidase_E_chloroplastic−like [MW = 67.64 kDa]

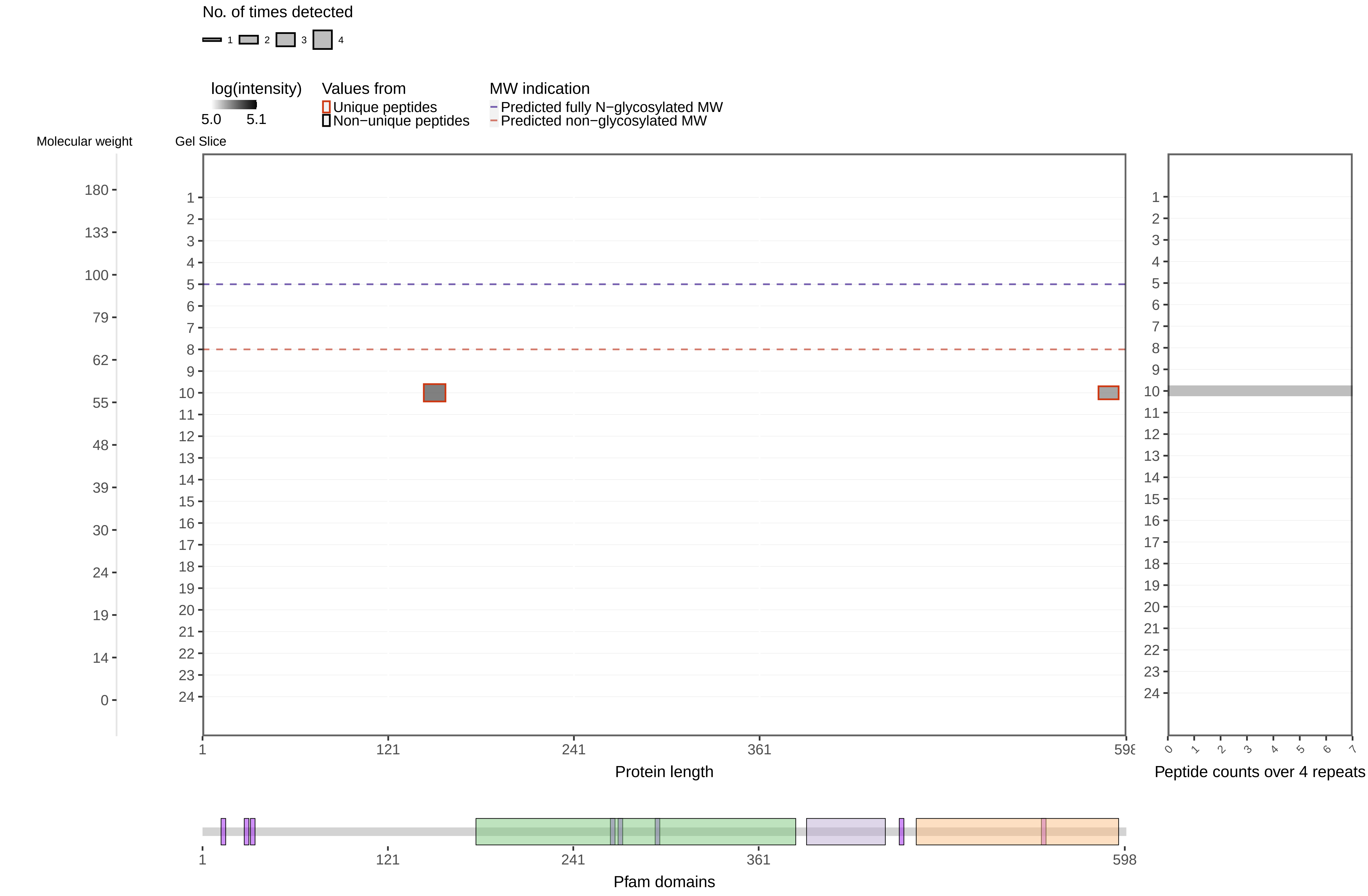


Protein sequence:

QLQQMTNNLTIMYRQMITNAPCPLLFFGQPYPLGTDPSPGMGTIENIPHTPVHIWVGSRPGENNVKHGEDMGNFYSAGLDPLFYSHHANVDRMWSEWKALGGKRRDLTHKDWLNSEFFFYDENRNPFRVKVRDCLHSKKMGFDYAPMPTPWRNFKPTRKSNAGKVNLSSVPPASKVFPLSKLDRAISFSIDRPSSSRTQQEKNEQEEMLTFNNIKYDDSKYIRFDVFLNVDKTVNADELDKAEYAGSYTSLPHVHGDNVSHVTSVTFQLAITELLEDIGLEDEDTIAVTV

NbL10g18300.1 tryptophan−−tRNA_ligase_cytoplasmic [MW = 45.92 kDa]

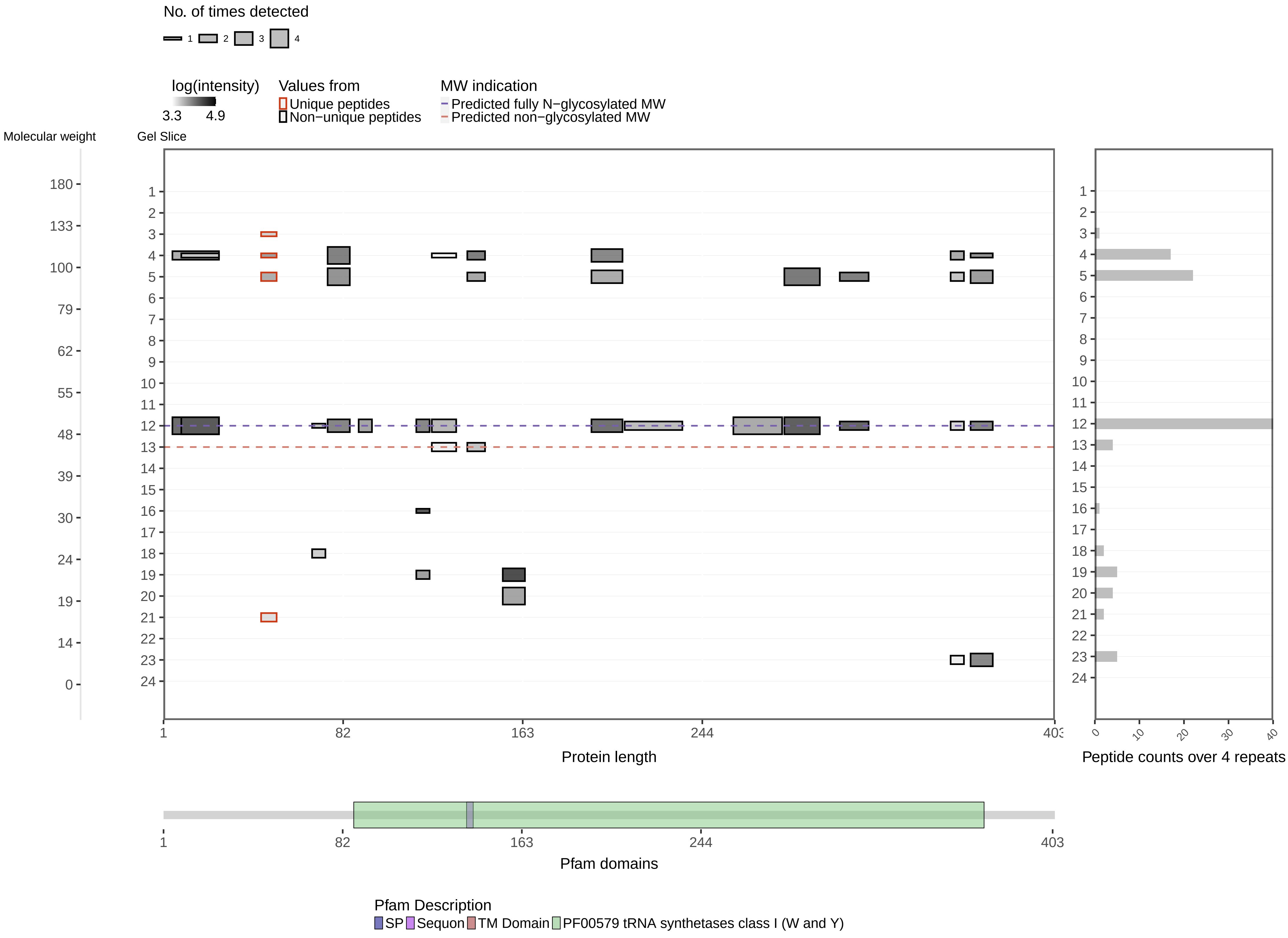


Protein sequence:

CVTYNKVVGIFGFTGEDHIGKVSFPPVQAVPSFPSSFPHLFSGNDNIRCLIPCAIDQDPYFRMTRDVAPRIGYHKPALIESSFFPALQGENGKMSASDPNSAIYVTDSAKEIKNKINRYAFSGGQDSIELHRKYGANLEVDIPFKYLGFFLDDDAELERIREEYGSGRMLTGEVKKRLTEVLTDLVERHRRARALVTDEMVDAFMAVRPLPNMFN*

NbL10g18290.1 glucan_endo−13−beta−glucosidase_5−like [MW − SP = 50.16 kDa]

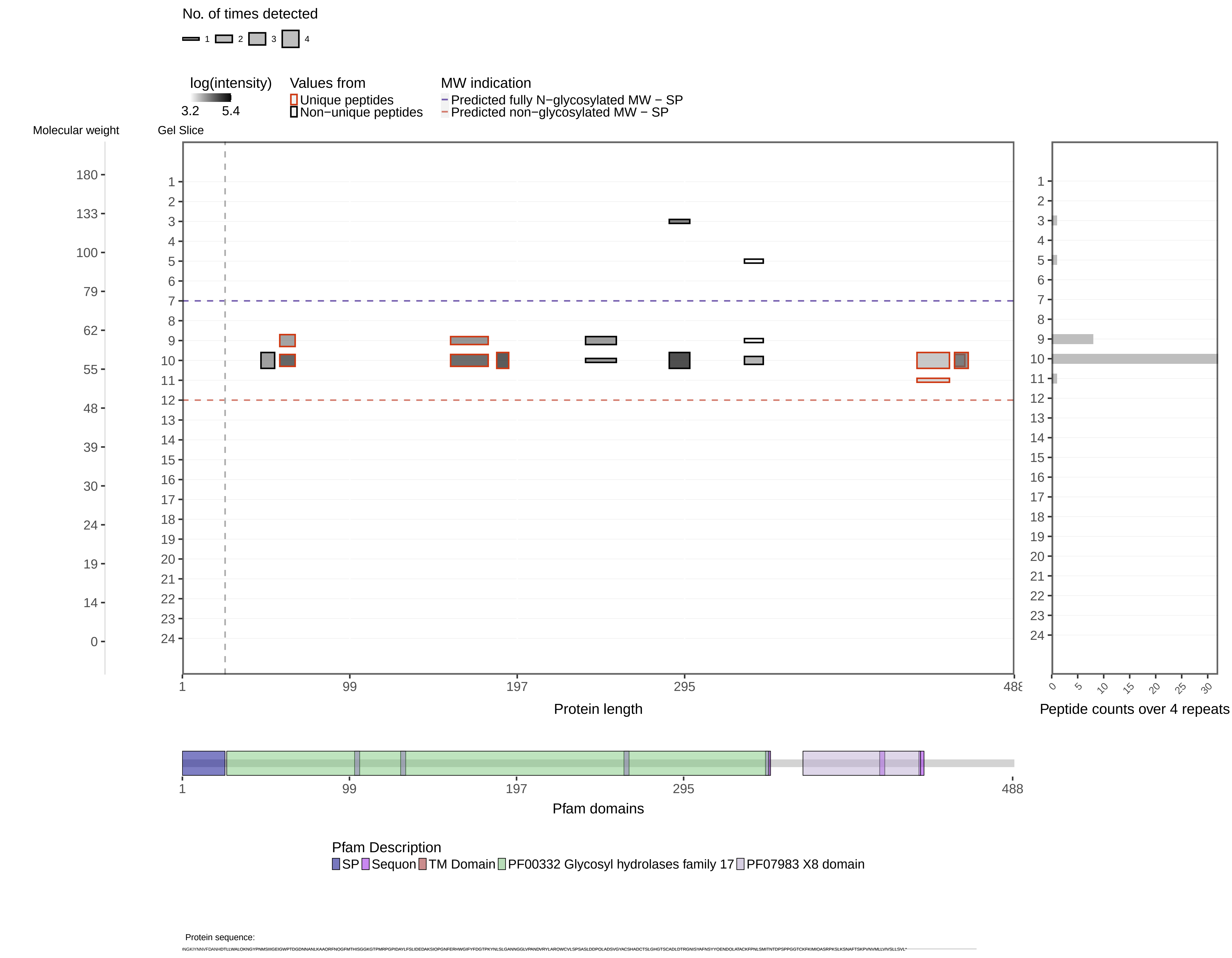

### NbL10g17910.1 phospholipase_D_Z−like [MW − SP = 54.64 kDa]

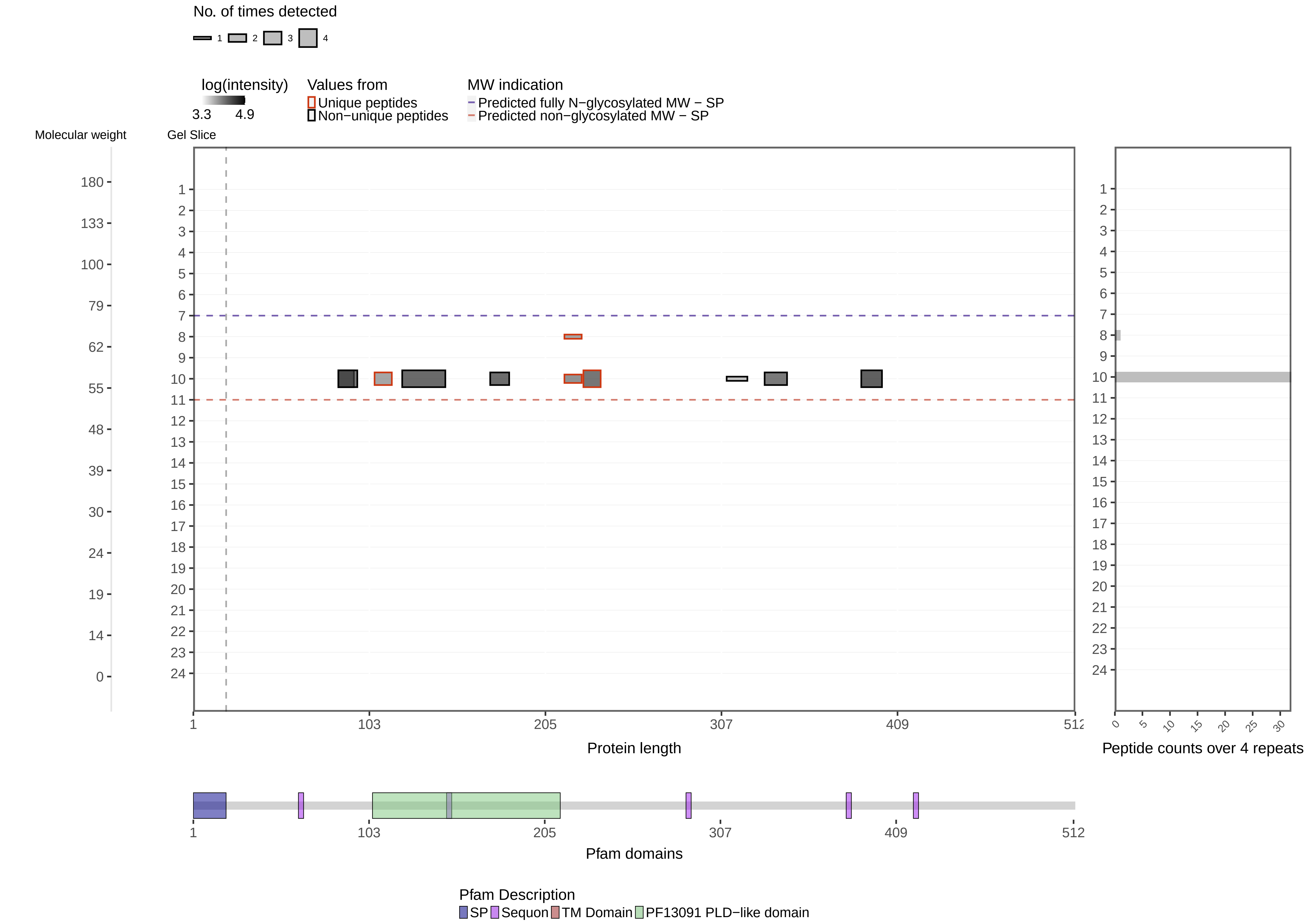


Protein sequence:

SHYVNSKDRCRSPLPKCMEVAHISGYPVLSDPFMFHIPIETPGVNDSFFHPQPSYLSFAPPELLFGKYYADEQAWVDTIKSVGNGGTVKINTMDWLGQSGYTKETVYYWASLSSAISEVVFSKSAKVQILVAKWAHFINNTDQYLKSLLYSNTLCSSSKYNHCSGKVEIKYYIVPGYNSTGPAALNGAATGNMYPGYTRVNHGKYAVSDTRAHISTSNLVWDYFYTTAGVSFGTYQPAIVLQLQEIFNADWNSPYAVPVEPLEEGHACSS*

NbL10g17900.1 tocopherol_cyclase_chloroplastic [MW = 58.96 kDa]

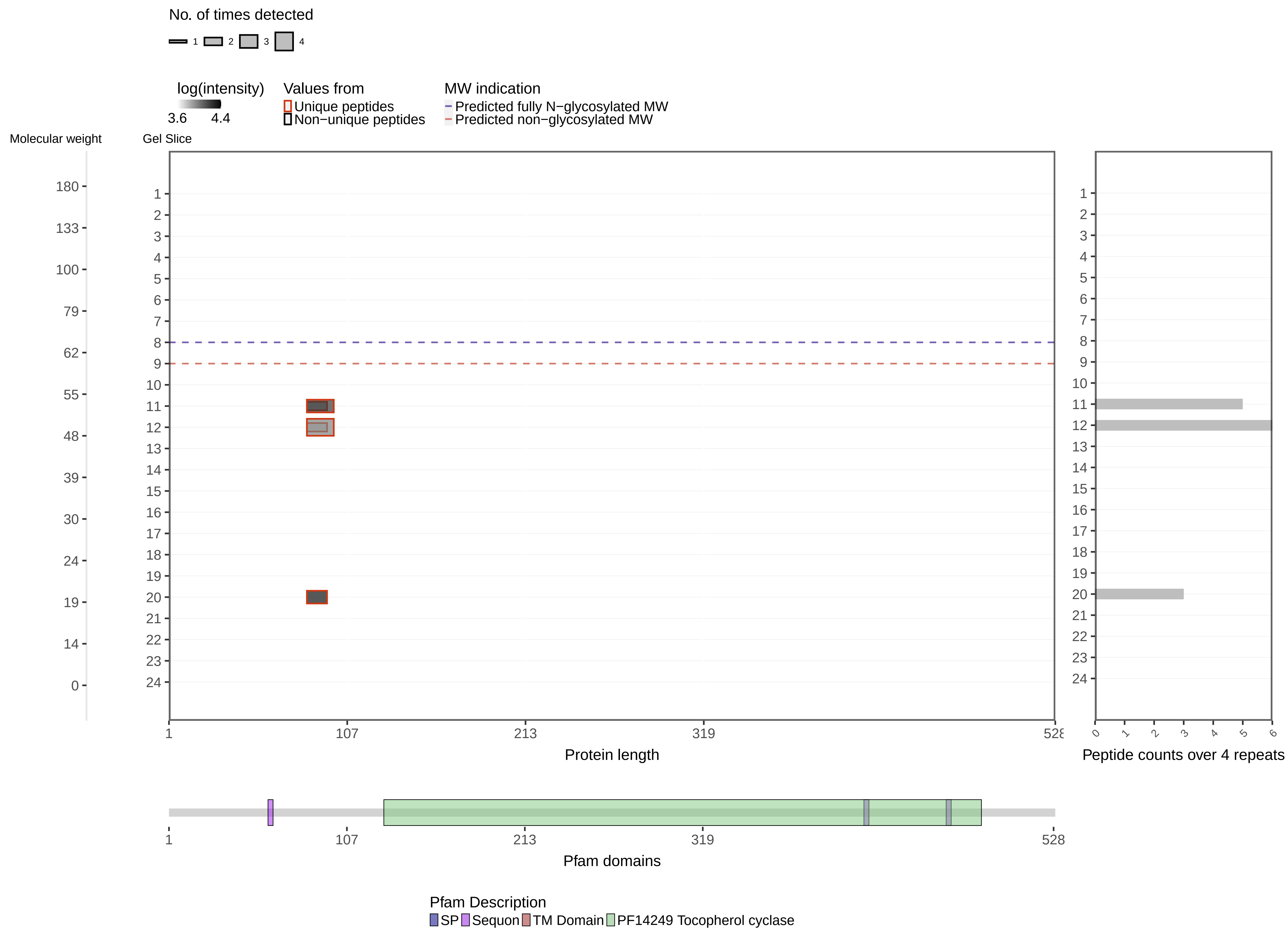


Protein sequence:

GRTDYTEIVETASWEYSMRPIYGWGDVNSKQKSTAGWPAAFPVFEPHWQICMAAGLSTGWMEWDGQRYEFQNAPSYSEKNWGGAFPRKWFWVQCSVFEGAIGDVALTAGGGLRQLPGLSQTFENAALIGIHYGGVFYEFVPWNACVSWEIAQWGKWHIFGENETHTVELEATAEDPGTTLRAPTDEMGLAPACRDTCFGDLRLRLWERKSNGSKGKVILDVTSNMAGLEVGGGPWFNTWKGKAQMPEIVSRAINVPVDLDAIFGLTPFLKPPGL*

NbL10g17850.1 peroxisomal_fatty_acid_beta−oxidation_multifunctional_protein_AIM1−like [MW = 82.3 kDa]

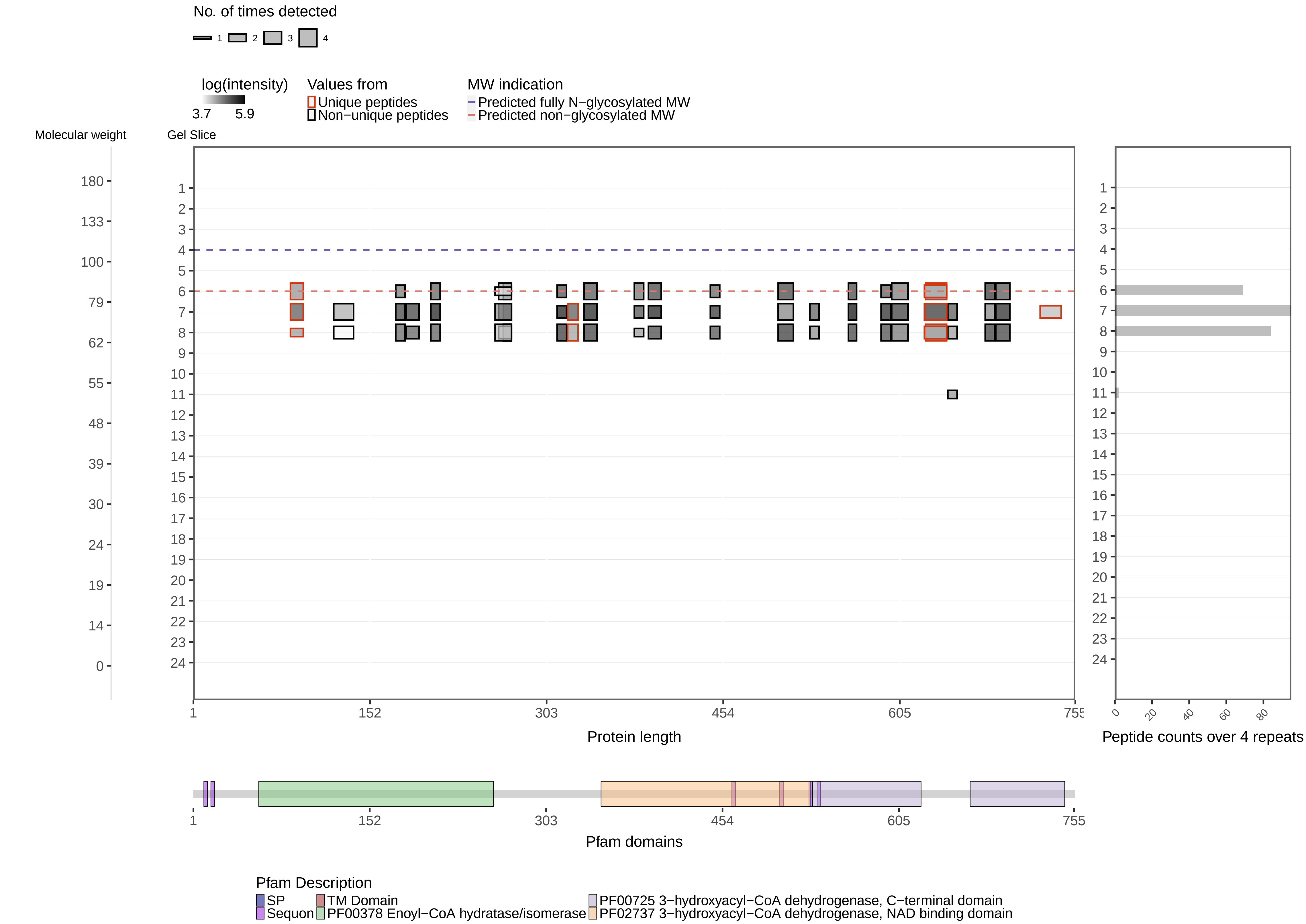


Protein sequence:

TALIMSNMNVVLKEINHEYLLKAVKSVEANLQGLVTRGQLNQDKMKRTLSLLKGTLDYEDLKDVDMVIEAVNEVVSLKQSIFEDIEKVCSSDCIFASNTSTISLDVIGERTSSQDRILGIHLFSPAHVMTLVEIVRTENTSPQVIVDVIRFTKTMKKVPIVVKNCTGFAVNRSFFPYMQGPDLLANLGVDIFRVDRVITEFGMRLGPFQLQDLSGYGIFLAGVKEFVAAFPDRAFQSPLVQLMVENGREGKKNGKGYYTYKKGSKPEADHSVIQVVEESMRFTNIAPGGKPISVTDEE

### NbL10g17840.1 peroxisomal_fatty_acid_beta−oxidation_multifunctional_protein−like [MW = 28.45 kDa]

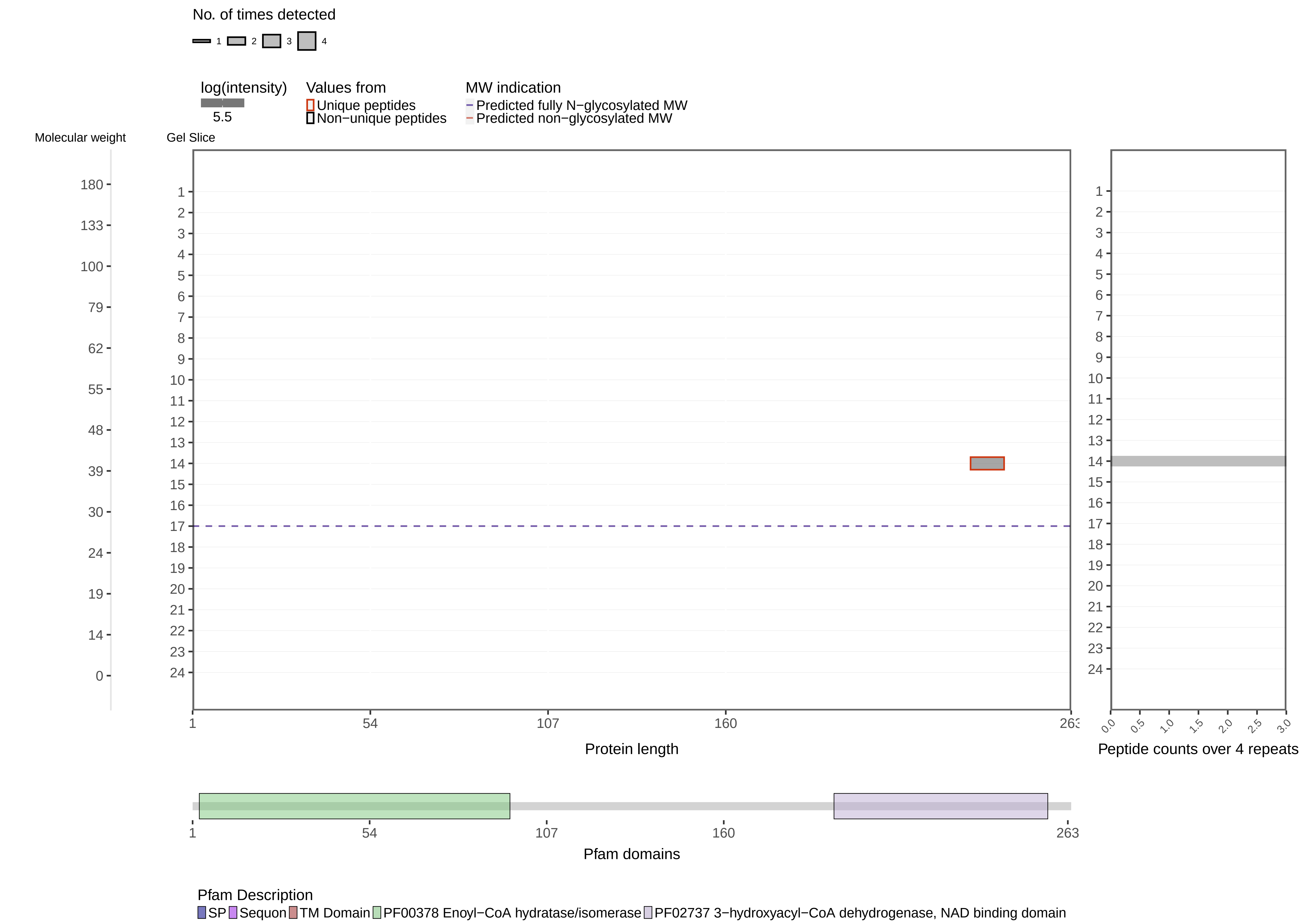


Protein sequence:

.KTSPSMPHYKACLDVIEVGIISGGYAGVLKEDKVLRELVLSKSSRALLHVYFAERATSKVPGITDCQLEHRKIEKVAVIGGGLMGSGIATALIISFVHVILKEIDYNCLQKSLKSVEVNLQSLVKGGHLVEEKMKKAFSFSKVL*

### NbL10g17810.1 eukaryotic_translation_initiation_factor_3_subunit_L−like [MW = 65.69 kDa]

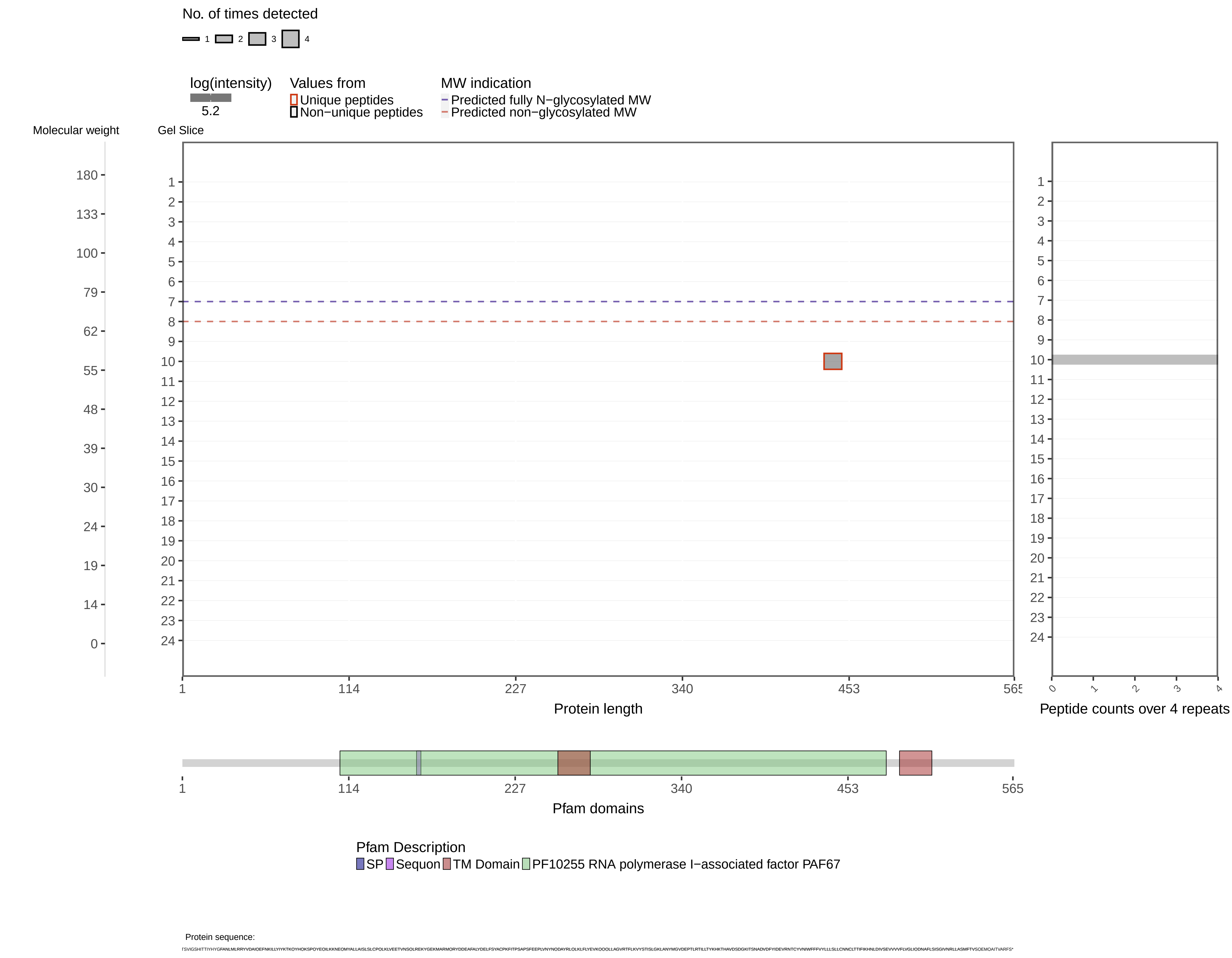

NbL10g17780.1 la_protein_1 [MW = 51.53 kDa]

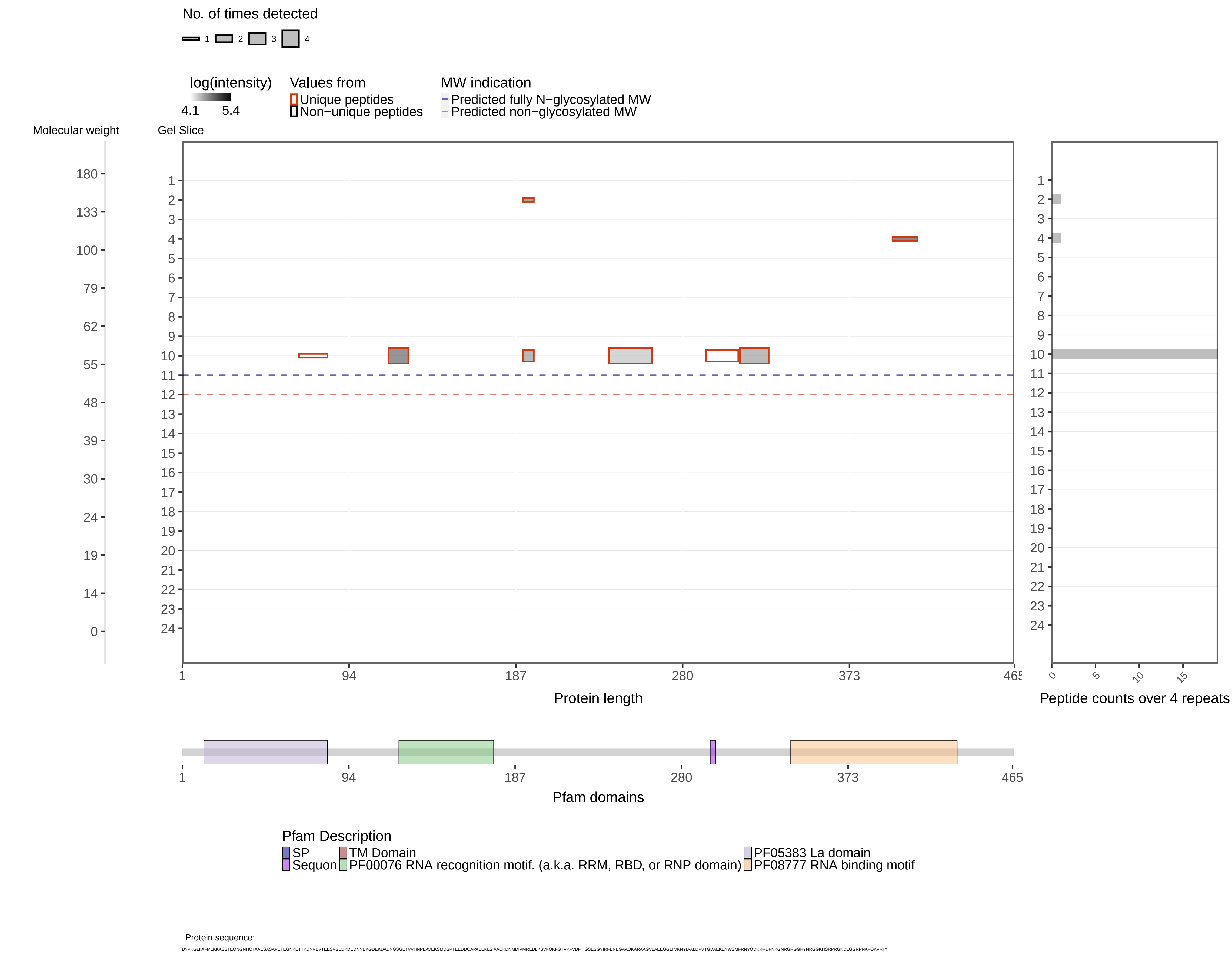

### NbL10g17770.1 coiled−coil_domain−containing_protein_25 [MW = 25.26 kDa]

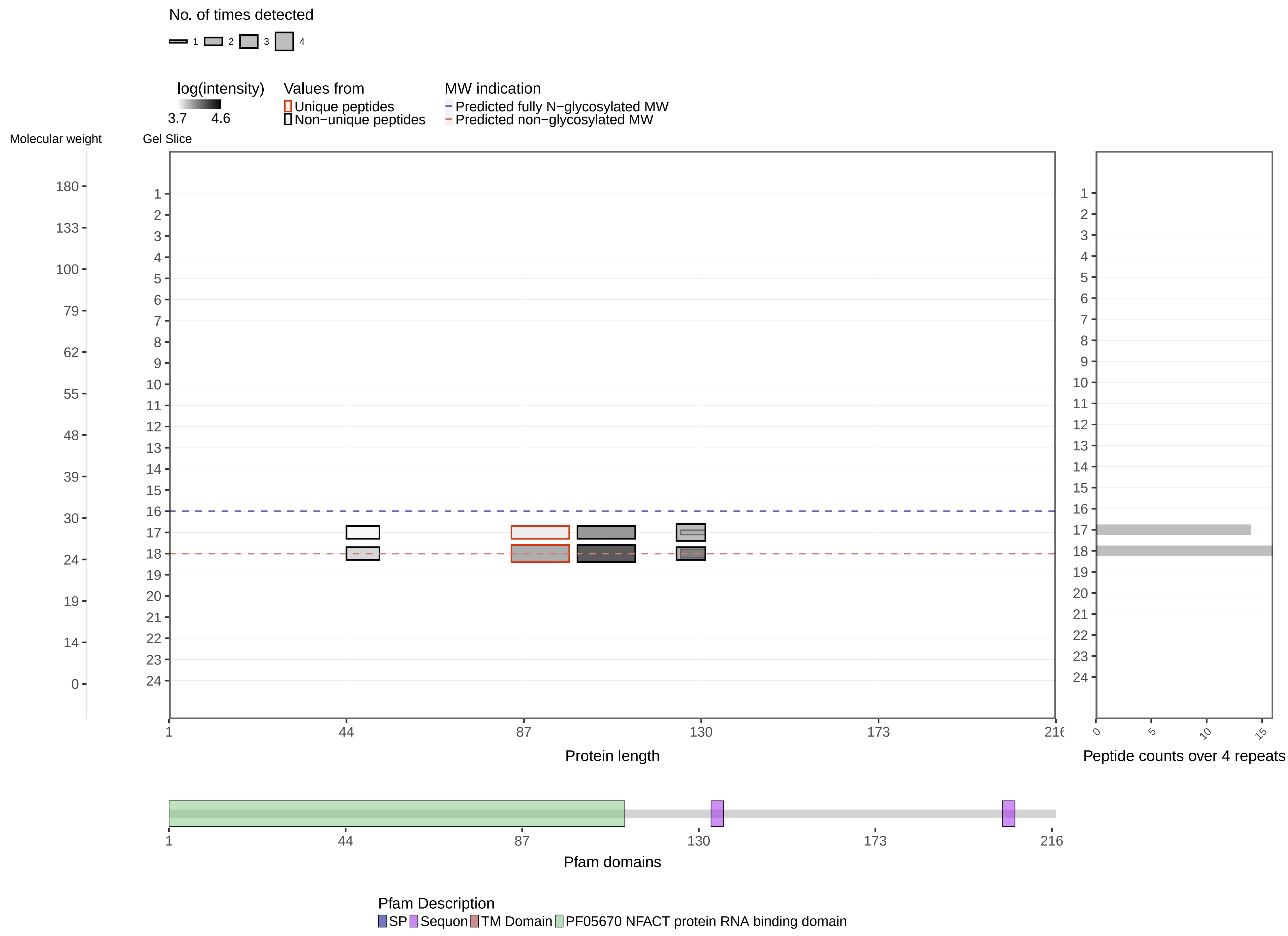


Protein sequence:

NLKKTASMDVGQVSFHNPKMVRTVKVEKRINEIVNRLNRTKVEKKPDLKAEREAVNAAERAERKQQLREKKRREEMDRLEKERQAEMRSYKNLMVADKMTSNKDIASTNKSLQELEEDFM*

NbL10g17660.1 uncharacterized_LOC109213855 [MW = 11.92 kDa]

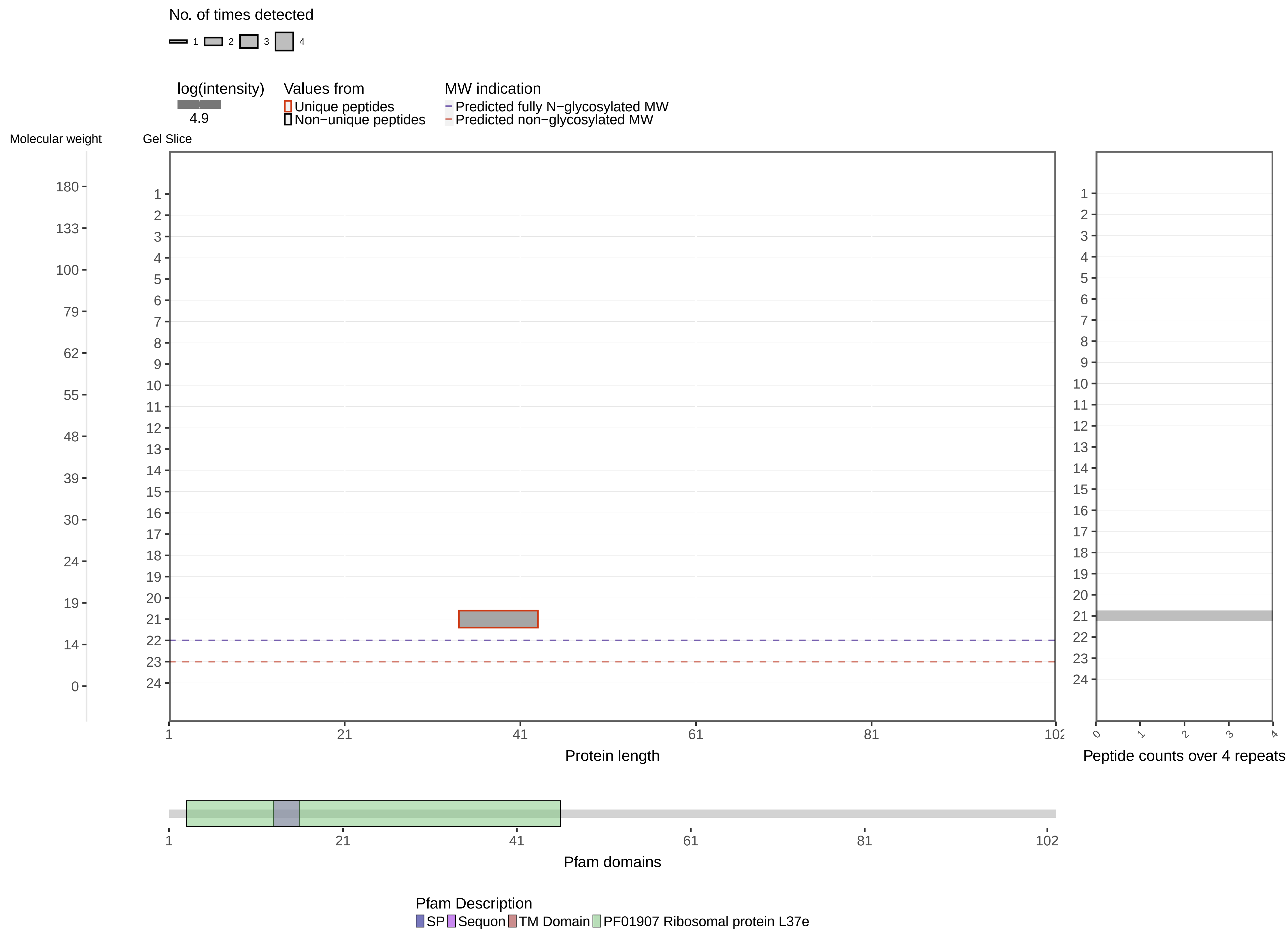


Protein sequence:

CAYPAARLRKCMCCFSLFGSPFFILIFACWNWYYREIFQCWKSMYYLVDYELVCFGLEKGLSLFK*

### NbL10g17630.1 BURP_domain_protein_RD22 [MW − SP = 37.03 kDa]

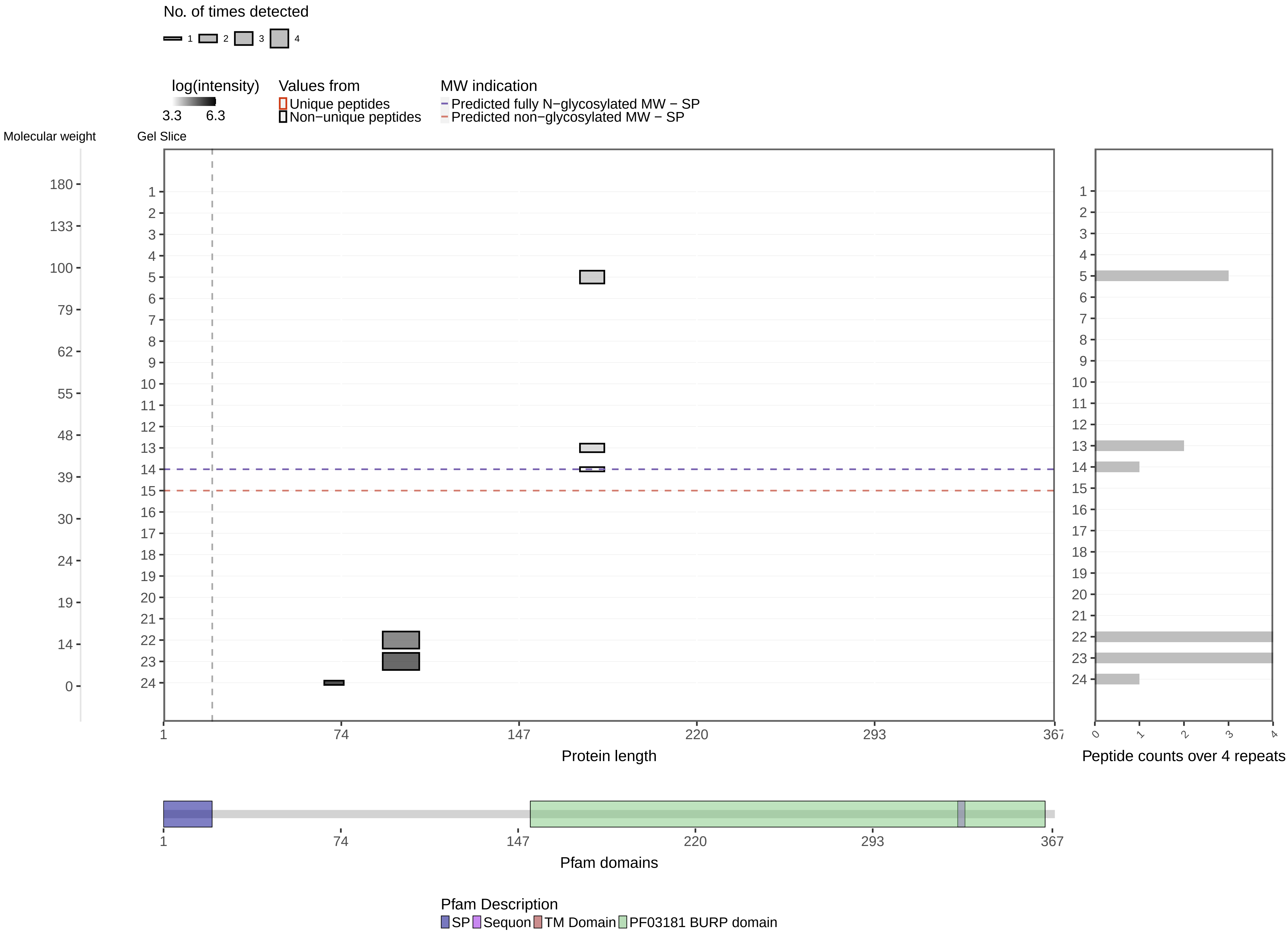


Protein sequence:

FVKTTNNAATFLPRQVANSIPFSSNEMPEILNQFSVKPDSEEAEILKQTVQECEEPGIKGEEKYCATSLESMVDFSTSKLGNKVQPVSTETEKETQMQKYTILGTKKMGNGKSDAVVCHKQNYAYAVFYCHKTETTQSYMVSLVGADGTKAKAVAVCHKNTSAWNPKHLAFKVLKVTPGSVPVCHFLPEDHIVWVPKN*

NbL10g17590.1 two−on−two_hemoglobin−3−like [MW = 15.3 kDa]

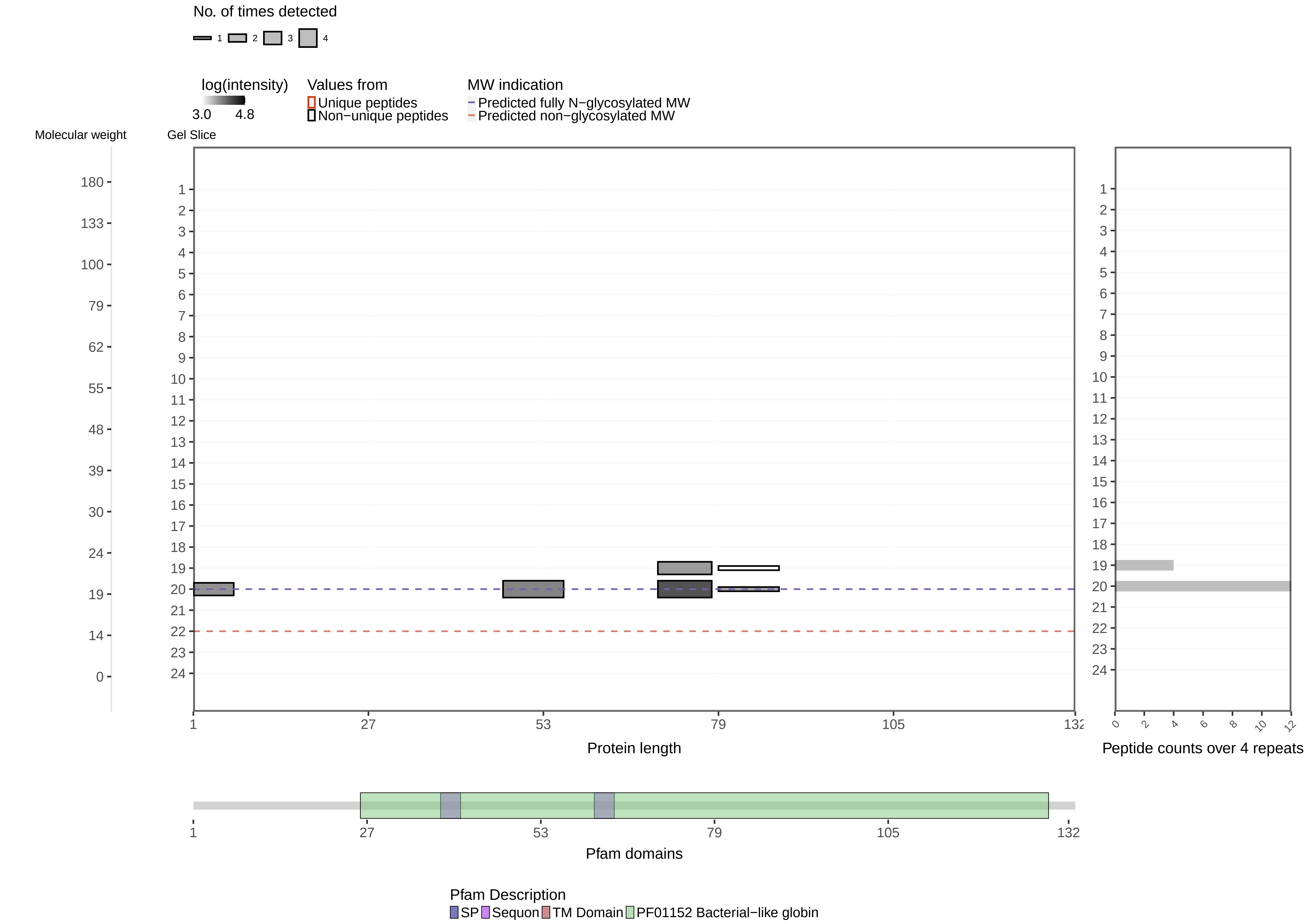


Protein sequence:

WFRSIFSNLSKEDAIRNQYEFFVQRMGGPPLYSQRKGHPALIGRHRPFPVTHKAAERWLQHMQLALDSVTVIDEDSQN*

### NbL10g17500.1 50S_ribosomal_protein_L17−like [MW = 18.46 kDa]

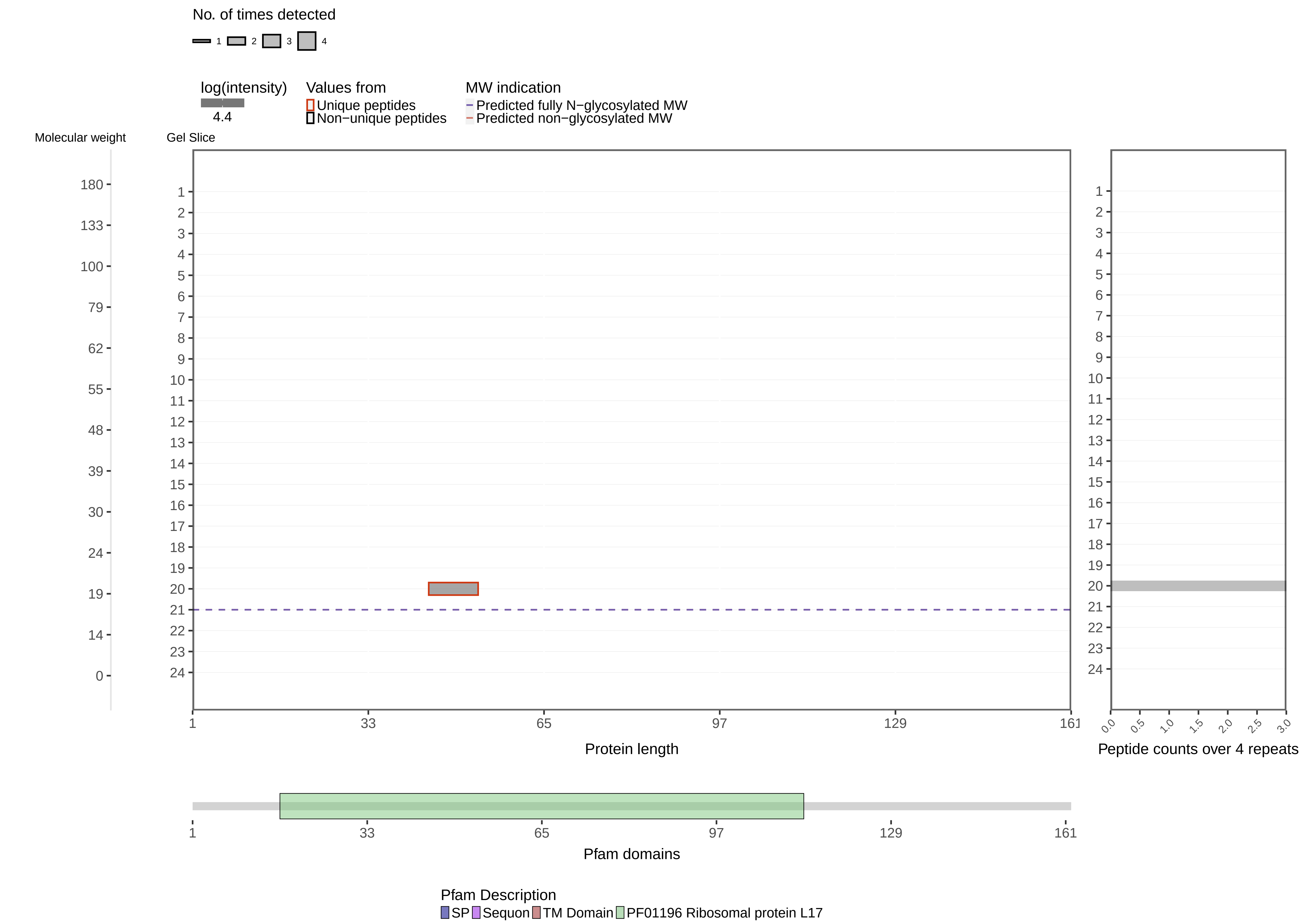


Protein sequence:

VRGDDVIHKLFTELAYRYKDRAGGYTRMLRTRIRVGDAAPMAYIEFIDRENELRQSKPPNPQPPQRPAMDPWTRSRLSRQFAPPKEEKSSDSDI*

NbL10g17410.1 psbP_domain−containing_protein_5_chloroplastic [MW = 33.49 kDa]

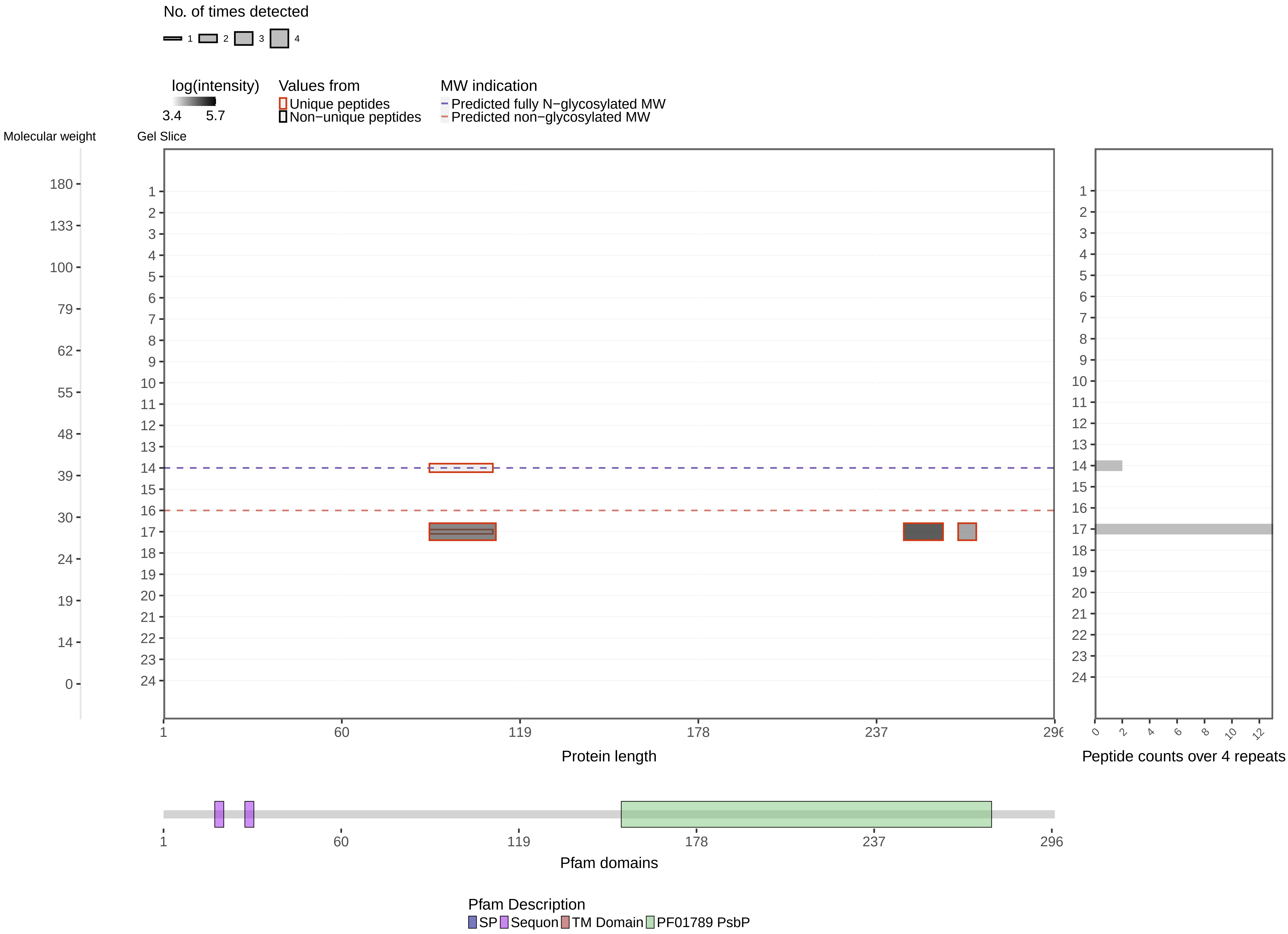


Protein sequence:

RLRIVSERVDIIDNLIISVSIGPPNSQFLKSEEKSTWTAKDVADSVLSDKSALRVTSTQRLAESSLLDAHTSEIDGEPYWYYEYLVRKSPTKTAQETNLFRHYAASTAERDGYLYTFSASTLGKQWDRMGPFLQKSVASFRLLPPTESYVPPYKDPWRFW*

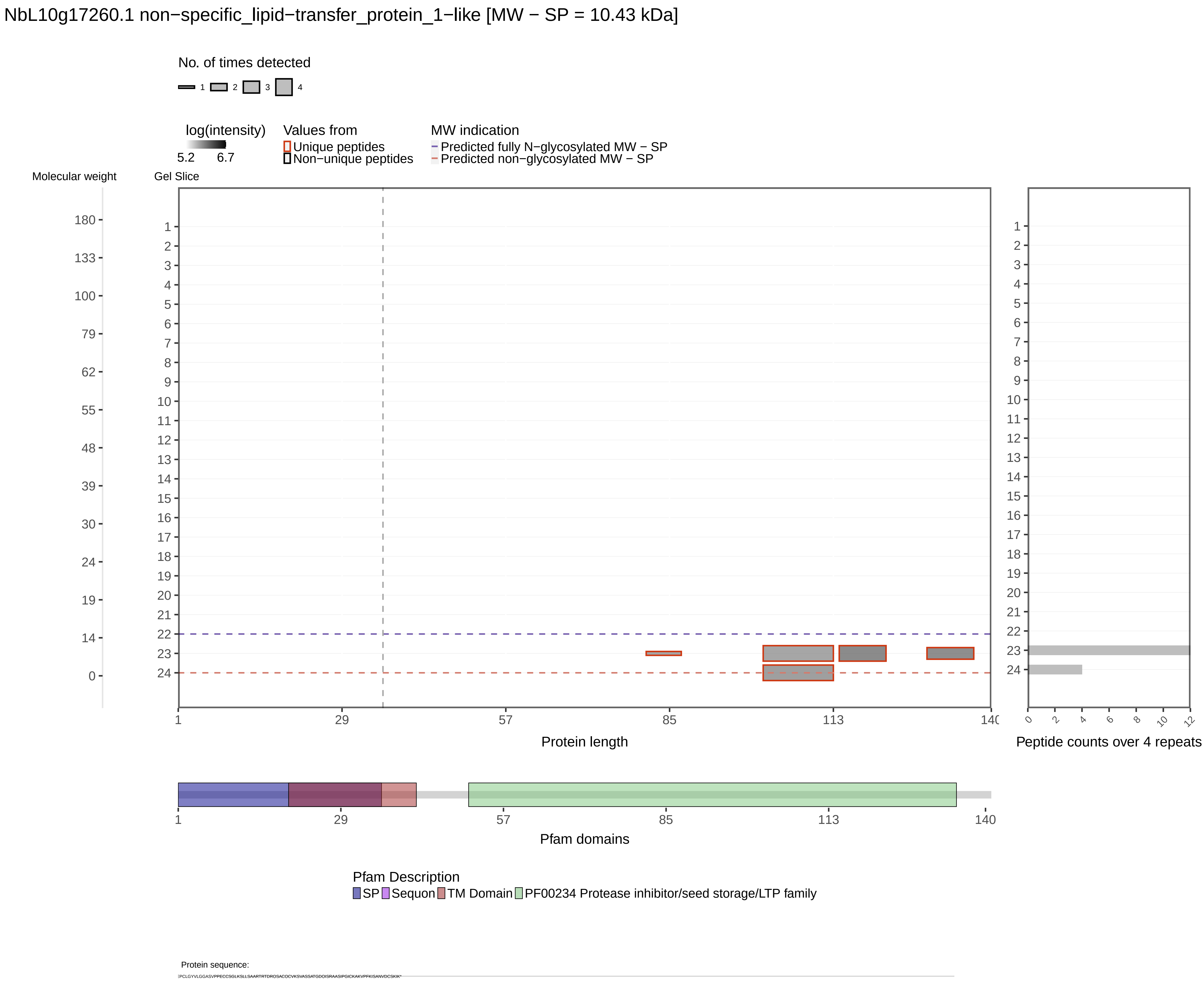

NbL10g17260.1 non−specific_lipid−transfer_protein_1−like [MW − SP = 10.43 kDa]
No. of times detected
1 2 3 4
log(intensity)
5.2 6.7
Values from
Unique peptides
Non−unique peptides
MW indication
Predicted fully N−glycosylated MW − SP
Predicted non−glycosylated MW − SP
Molecular weight
Gel Slice
Protein length
Peptide counts over 4 repeats
Pfam domains
Pfam Description
SP
Sequon
TM Domain
PF00234 Protease inhibitor/seed storage/LTP family
Protein sequence:

### NbL10g17250.1 non−specific_lipid−transfer_protein_1−like [MW − SP = 9.71 kDa]

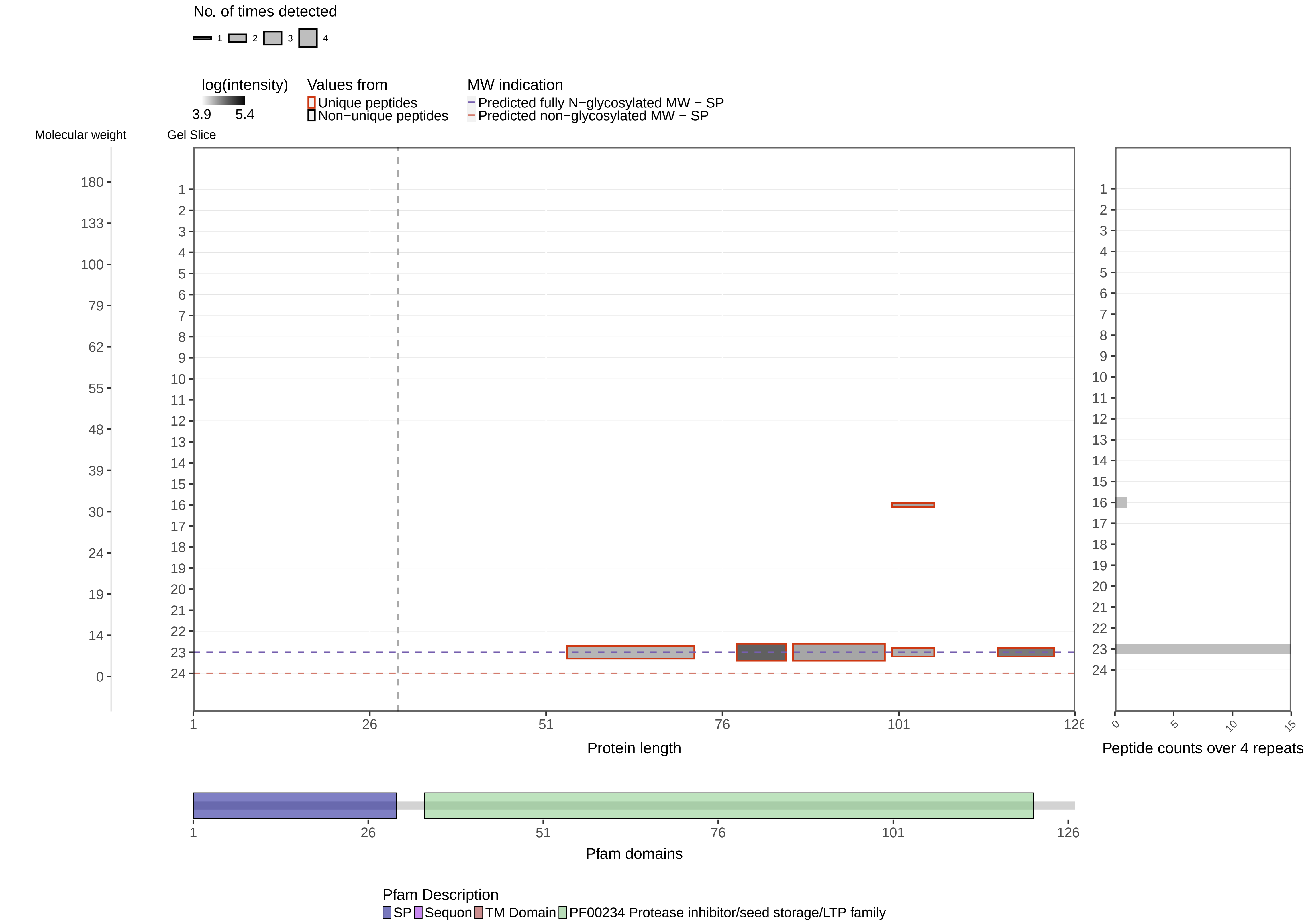


Protein sequence:

SSKQAAPSGCCSGIQSLYTTARTTVDRQSVCSCLKAGAASLGNSIDANKAAILLNKCGVTVPYKISPNIDCSKVQ*

NbL10g17230.1 6−hydroxynicotinate_3−monooxygenase [MW = 48.15 kDa]

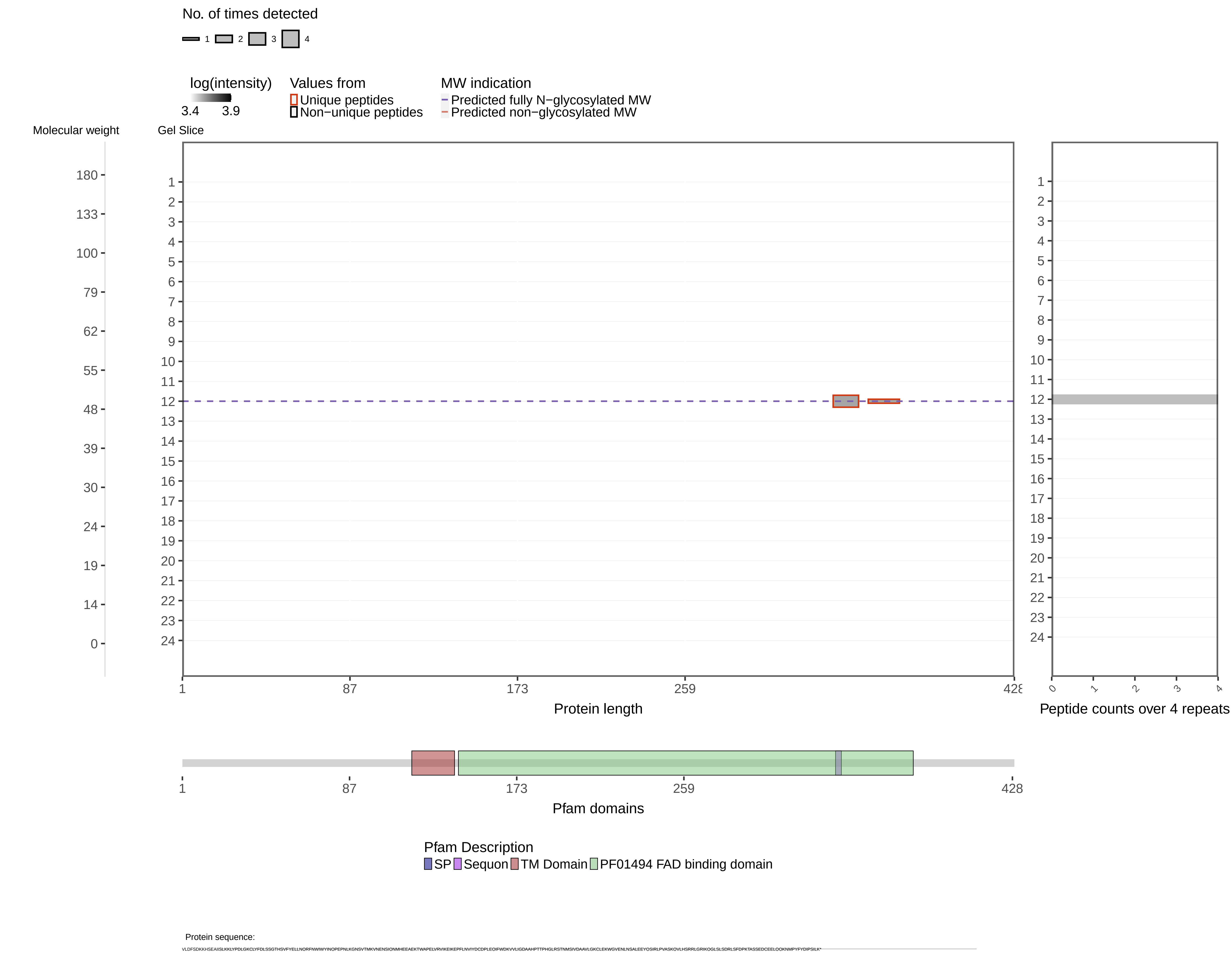

### NbL10g17200.1 EG45−like_domain_containing_protein [MW − SP = 12.03 kDa]

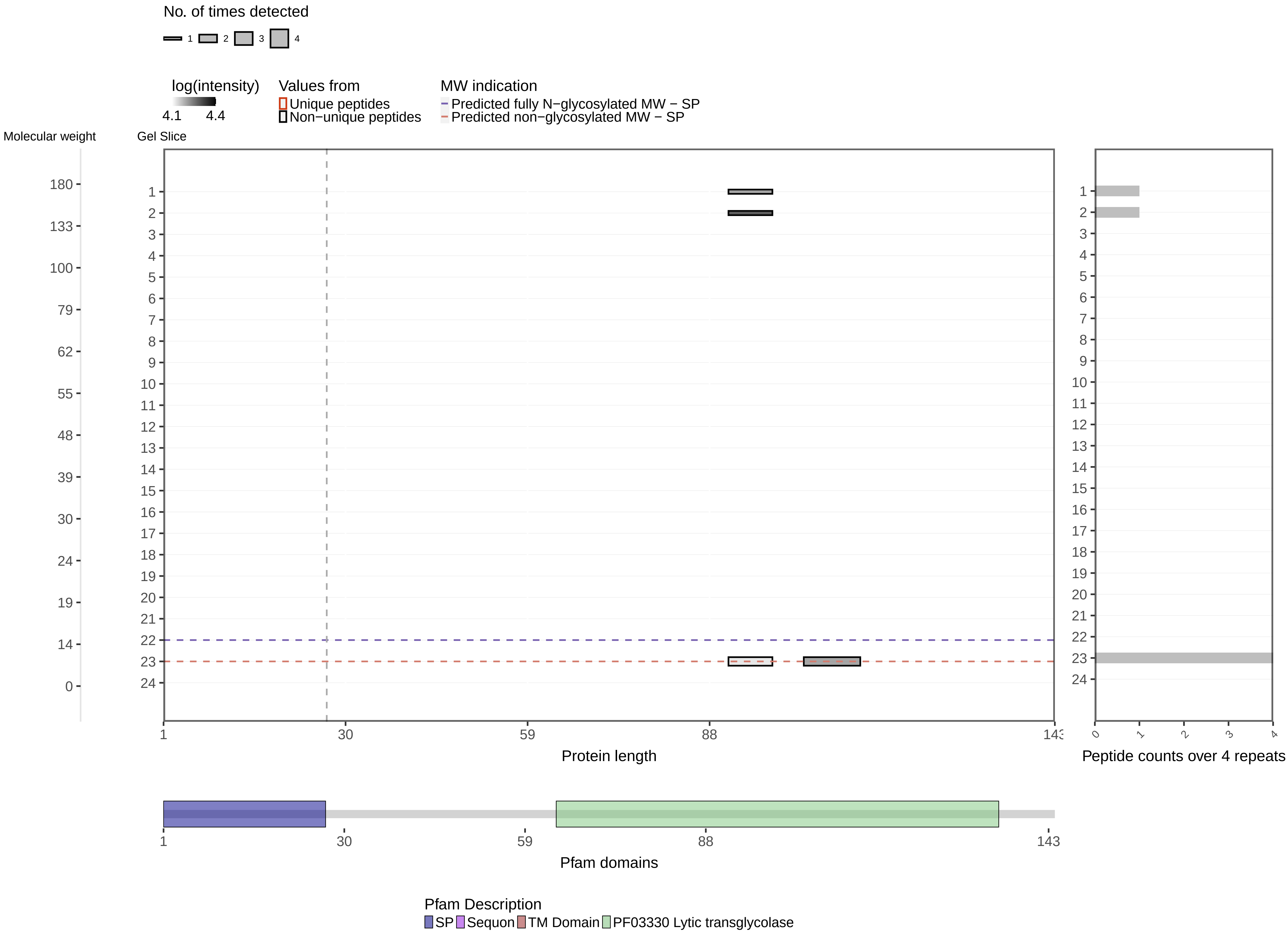


Protein sequence:

FAAAGEGIWDDGAACGRQYLISCISSILPRACKSGETIQIKIVDRAQTVASNPTRQGTTIVLSNAAFAAIADSNALSLNIDFQQV*

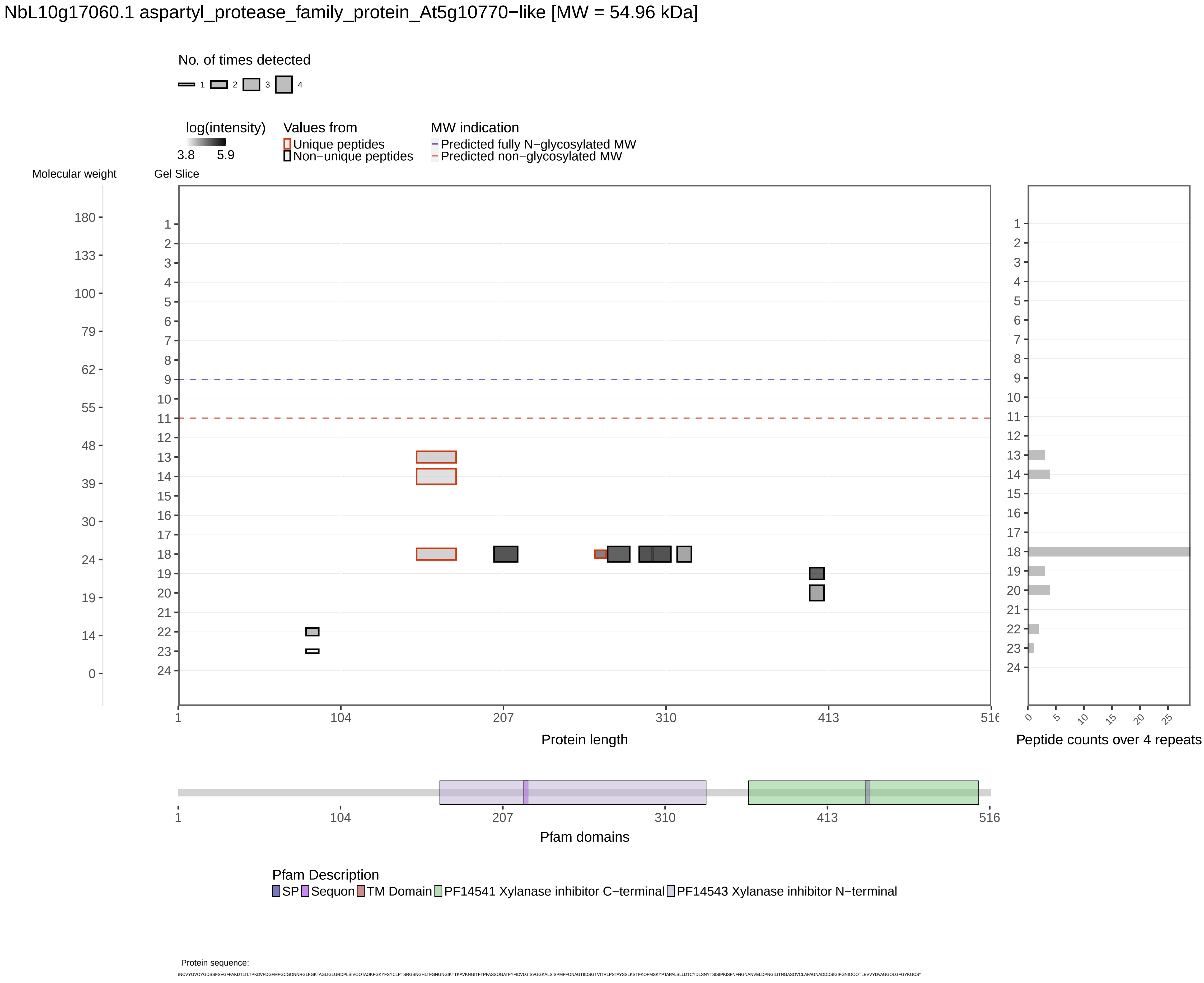
NbL10g17060.1 aspartyl_protease_family_protein_At5g10770−like [MW = 54.96 kDa]
No. of times detected
1 2 3 4
log(intensity)
3.8 5.9
Values from
Unique peptides
Non−unique peptides
MW indication
Predicted fully N−glycosylated MW
Predicted non−glycosylated MW
Molecular weight
Gel Slice
180 133 100 79 62 55 48 39 30 24 19 14 0
1 2 3 4 5 6 7 8 9 10 11 12 13 14 15 16 17 18 19 20 21 22 23 24
1 104 207 310 413 516
Protein length
0 5 10 15 20 25
Peptide counts over 4 repeats
Pfam domains
Pfam Description
SP
Sequon
TM Domain
PF14541 Xylanase inhibitor C−terminal
PF14543 Xylanase inhibitor N−terminal
Protein sequence:

### NbL10g16970.1 ATP_synthase_subunit_alpha_chloroplastic [MW = 32.94 kDa]

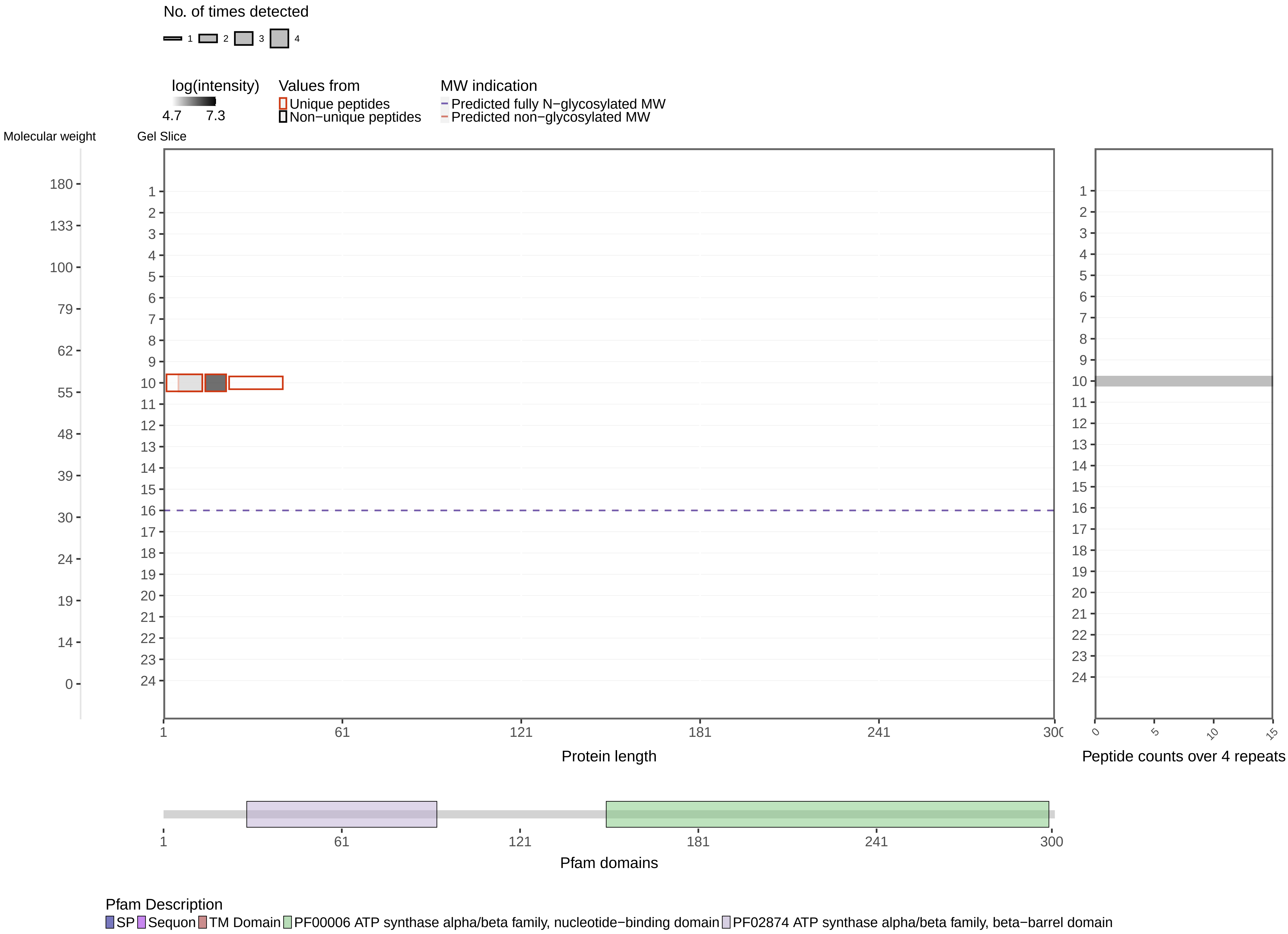


Protein sequence:

SRRSIYEPLQTGLIAIDSMIPIGRGQRELIIGDRQTDKTAVPTDTILNQQSQNVIYFYVAIGQKASSVAQVVTTLQERGAMEYTVVVAETADSPATLQYLAPYTGAALTEYFMYRERHTLIIYDDPSKQAQAYRQMSLLLRRPPDREAYPGDVFYLHSSLL*

### NbL10g16940.1 uncharacterized_LOC109237937 [MW − SP = 38.26 kDa]

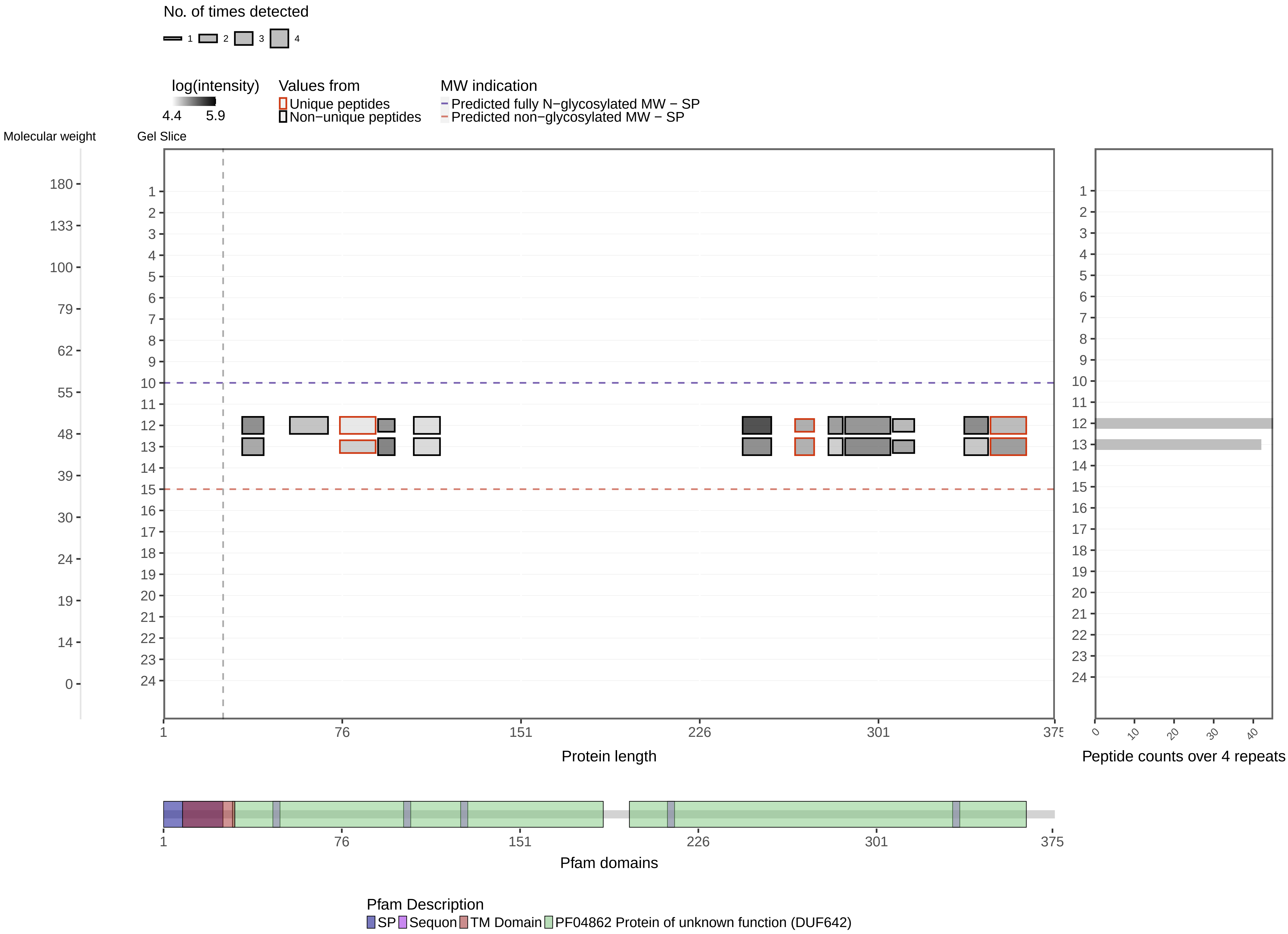


Protein sequence:

ACGPLIDSIAIRSLYPPRATNNNLLKNGDFEEGPYIFPNTSWGVLVPPNIEDDHSPLPAWMVESLKAVKYLDSDHFSVPQGKRAVELVAGKESAVAQVARTTIGKMYELSFKVGDASNSCEGSMIVEAFAGRDTLKVPYESKGKGGYKRAVLRFKAAANRTRIMFLSTYYHTRSDDFVSLCGPVVDDVKLLSVRNPRRFL*

NbL10g16820.1 early_nodulin−like_protein_1 [MW − SP = 18.51 kDa]

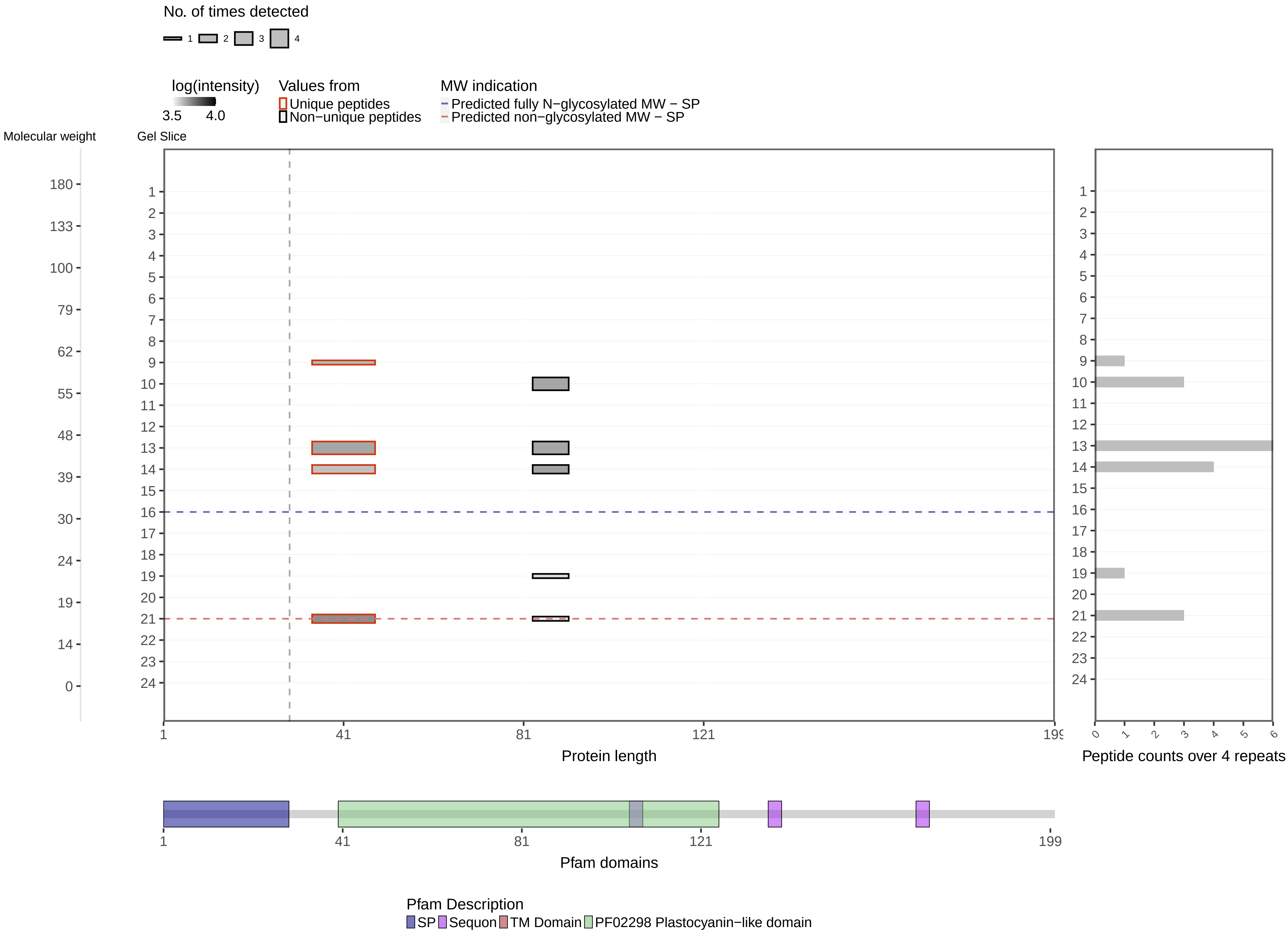


Protein sequence:

NSCNLKNPILYMNNGNSLFNITKPGEFYFTSGAEGHCEKSQKLHISIPGGNGTTYEEDSPAFAPSADSPAYPTVFGSIPVQSANQSSSGKLPISIFAAIGSLFFTLLIRSTIF*

### NbL10g16800.1 exosome_complex_exonuclease_RRP46_homolog [MW = 26.47 kDa]

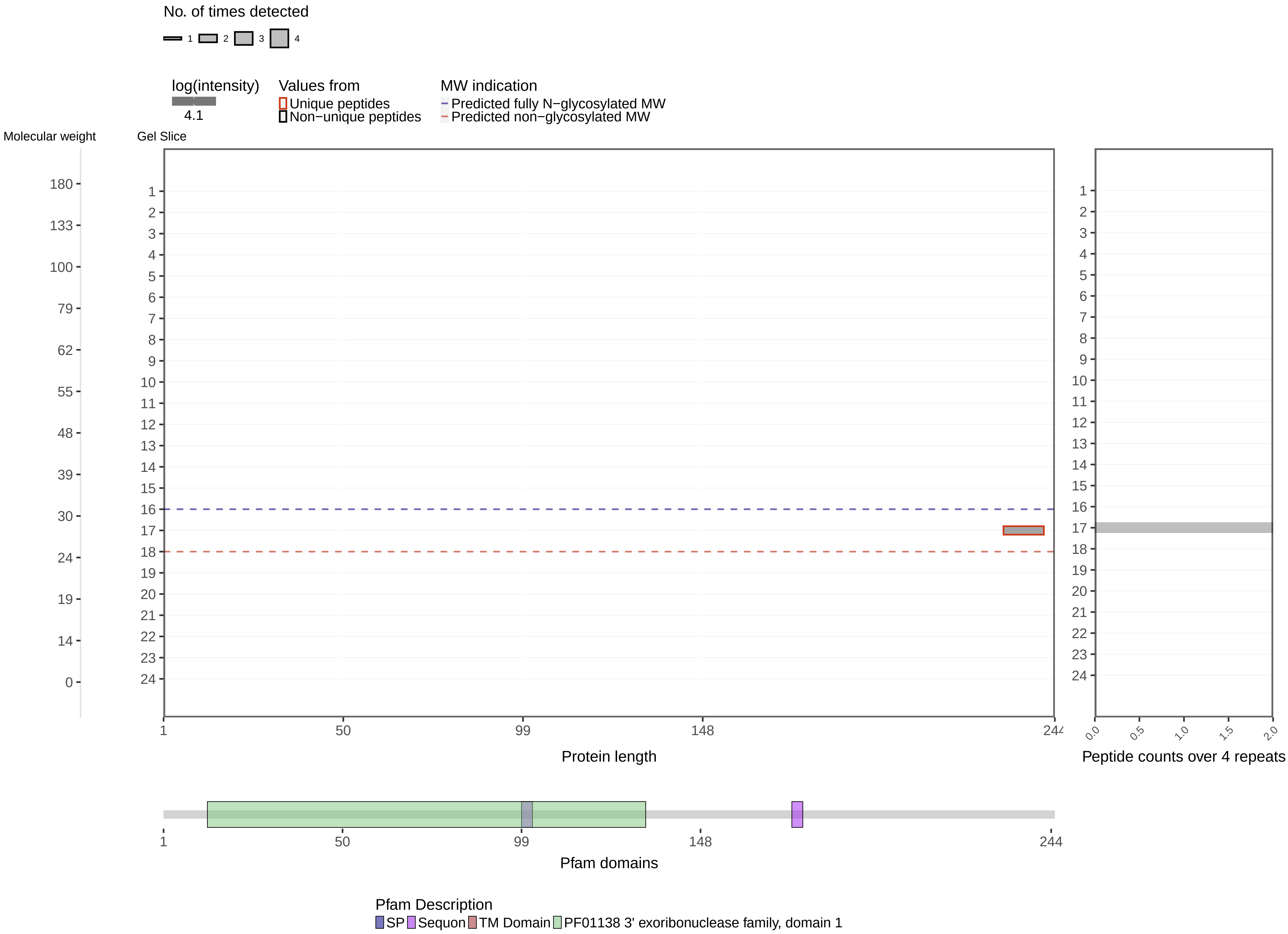


Protein sequence:

/VNDDGALLPCAINAVCAALVDAGIPLKHLAVAICCSLAESGHILLDPSKLEEQNMKAFVYLVFPNSTLSVLPEESLKVRGEATEHGLITSFTHGVMSVDDYIRCLERGRAATGKLSDFLRRSFQSQVQSDSSKLG*

NbL10g16490.1 NADP−dependent_malic_enzyme−like [MW = 69.3 kDa]

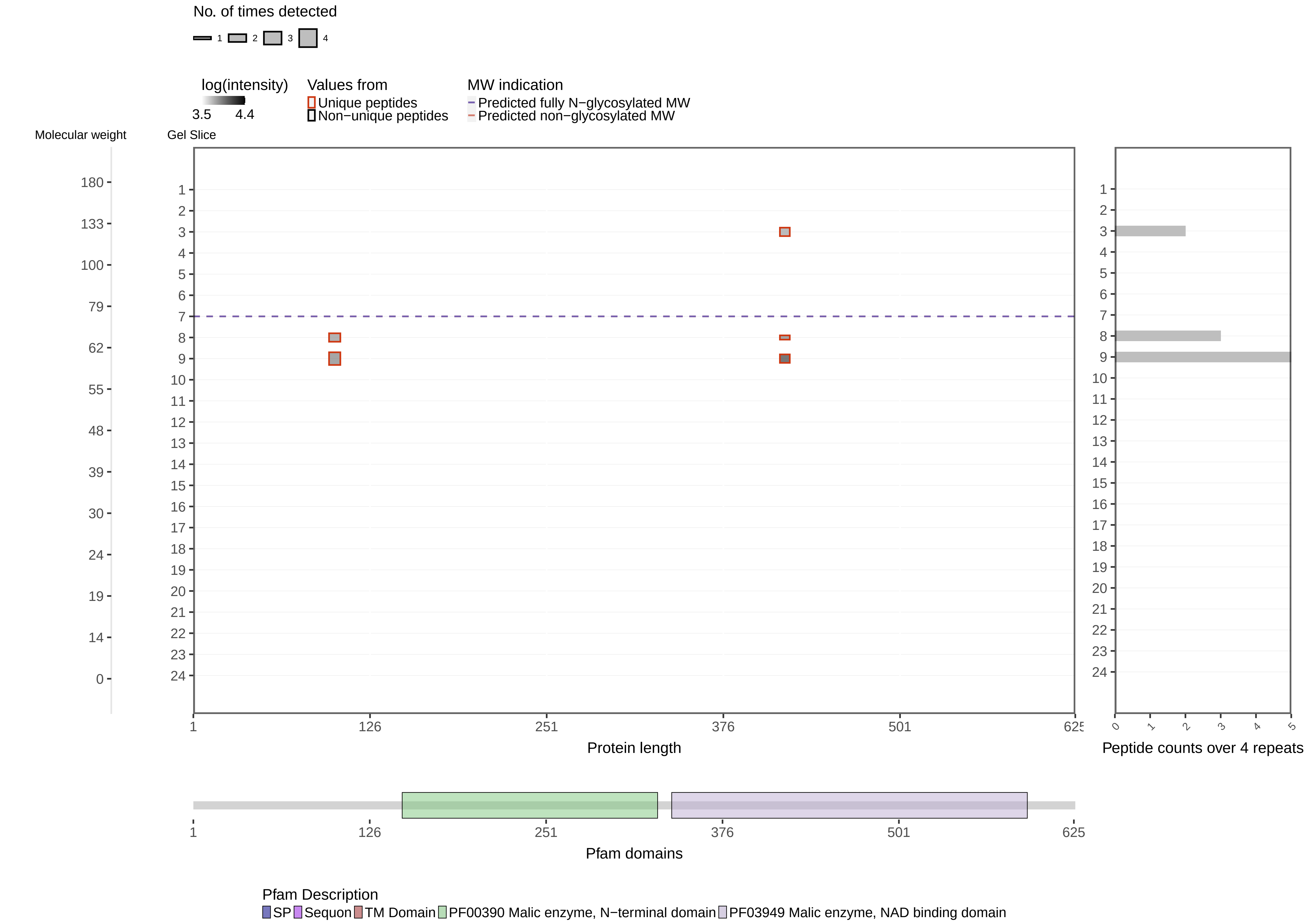


Protein sequence:

EFMSAVKQNYGEKVLIQFEDFANHNAFDLLAKYRTTHLVFNDDIQGTASVVLAGLIASLKLLGGTLADHTFLFLGAGEAGTGIAELIALEITKRTSVPLEEARKKIWLVDSKGLVVSSRLESLQPFKKPWAHGHEPVKELIDAVKAIKPTVLIGTSGVGKTFTKEVIETMASLNPKPLIMALSNPTSQAECTAEEAYTWSKGHAIFASGSPFDPFEYEGRTFVSGQANNAYIFPGFGLGIIMSGTIRVHDDMLLAASEALAAQVTEEDYLKGRIYPPFTNIRRISAHIAAEVAAKAYELGL

### NbL10g16470.1 cell_division_control_protein_2_homolog_A [MW = 33.83 kDa]

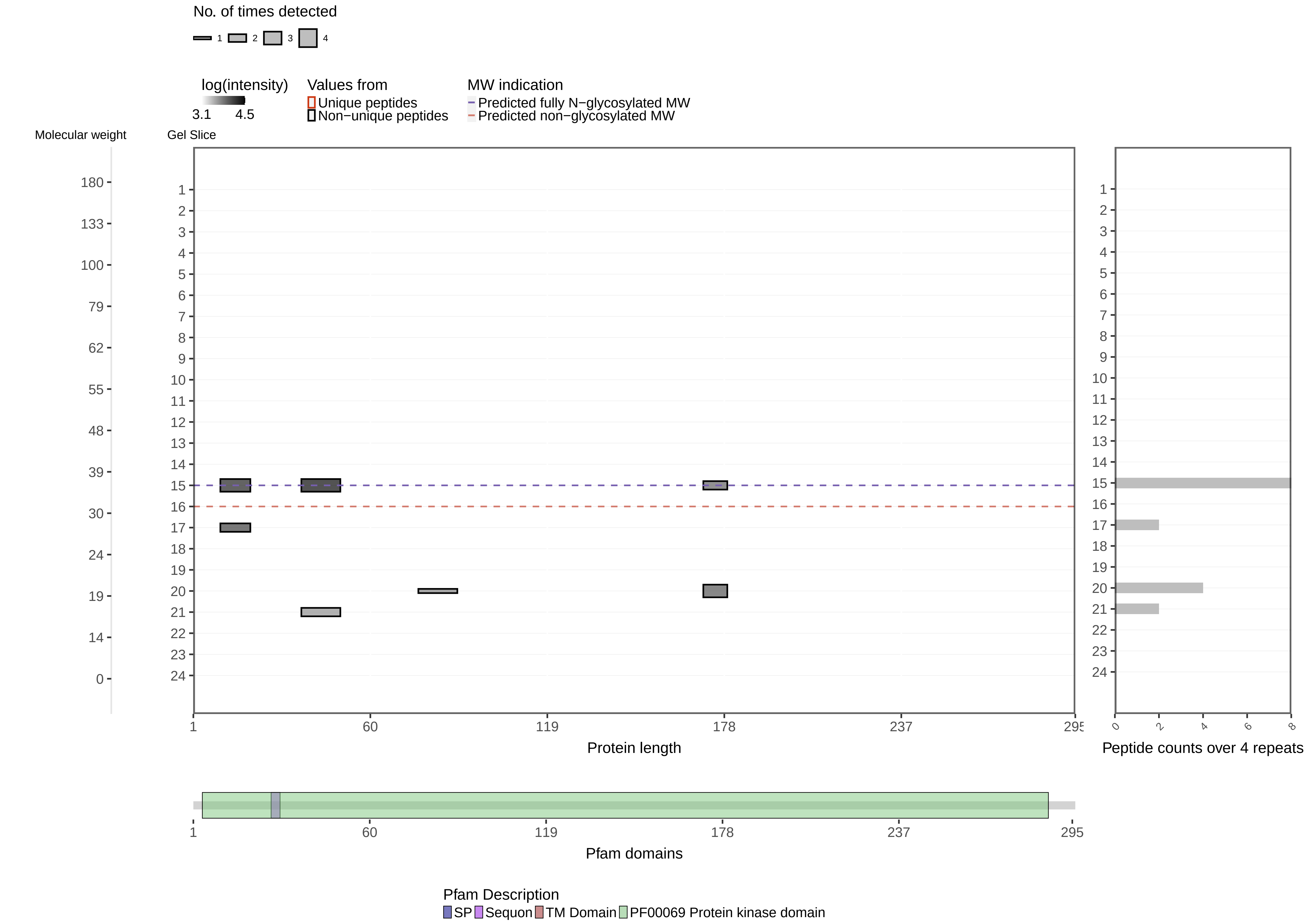


Protein sequence:

DRRTNALKLADFGLARAFGIPVRTFTHEVVTLWYRAPEILLGSRHYSTPVDVWSVGCIFAEMVTQRPLFPGDSEIDELFKIFRVMGTPNEDTWPGVTTLPDFKSAFPKWPSKDLAAIVPNLDGAGLDLLDKMLRLDPSKRITARNALEHEYFKGIGYVP*

### NbL10g16320.1 uncharacterized_LOC109241480 [MW − SP = 151.69 kDa]

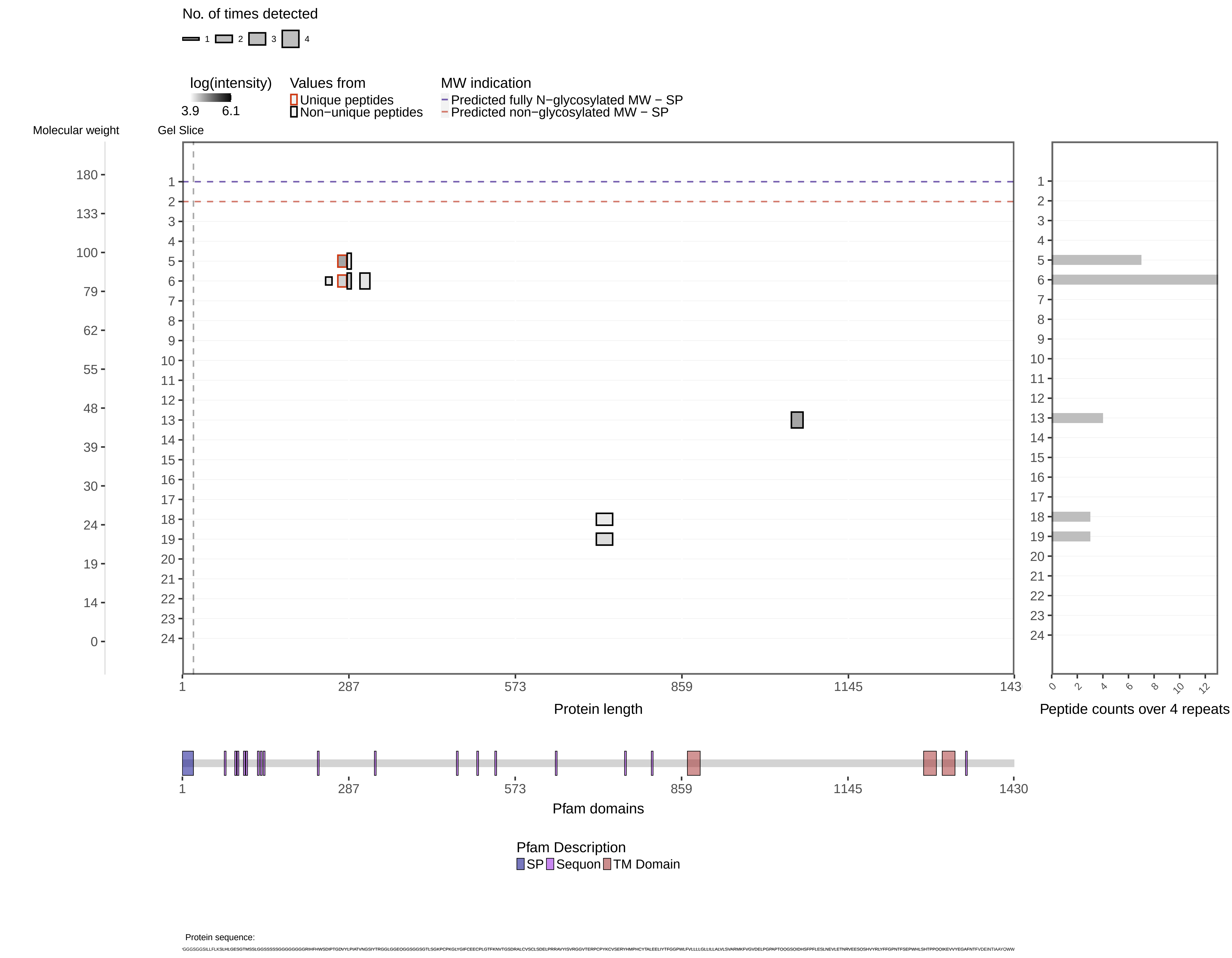

NbL10g16150.1 vacuolar−processing_enzyme−like [MW − SP = 50.88 kDa]

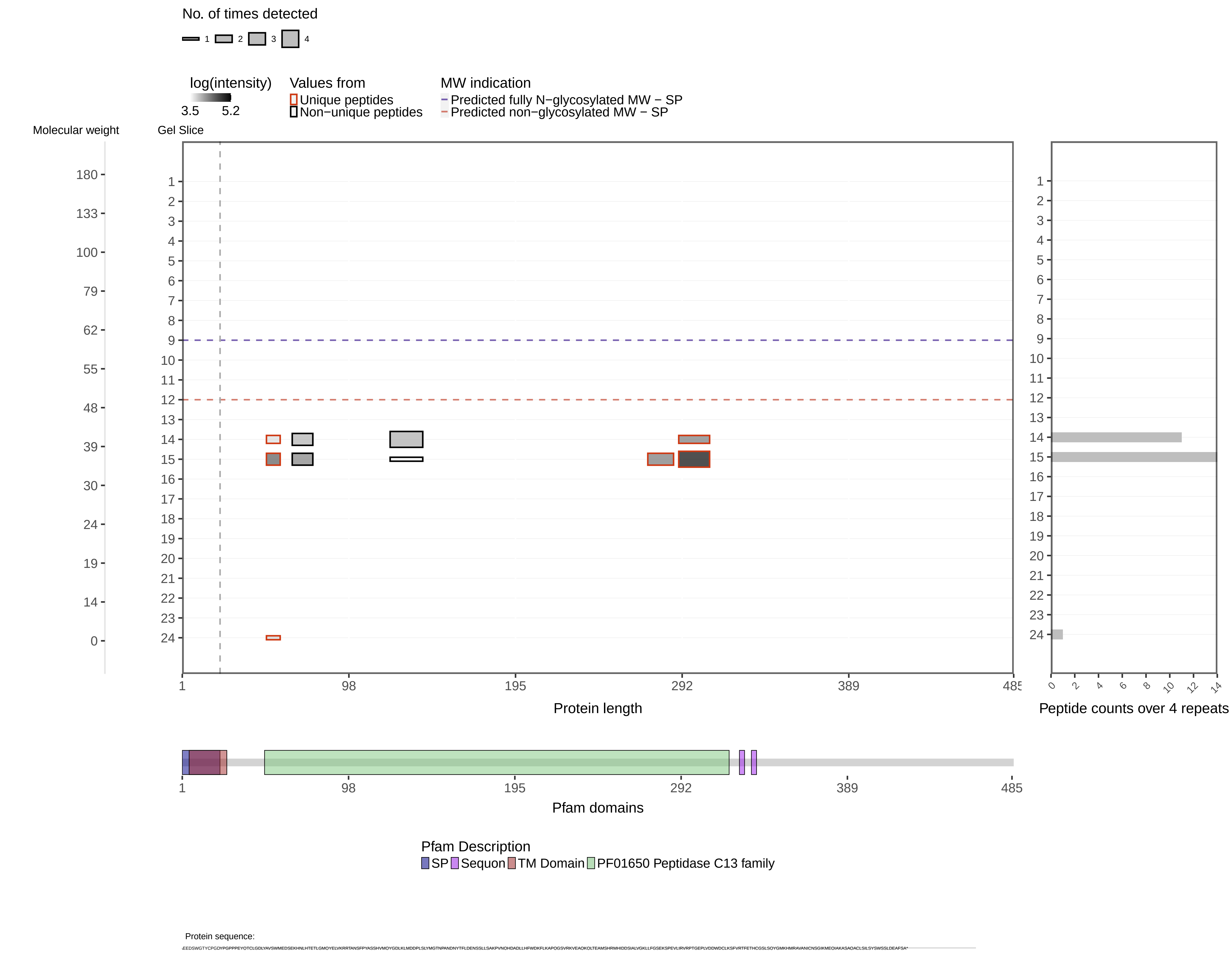

### NbL10g16050.1 serine_hydroxymethyltransferase_3_chloroplastic−like [MW = 56.08 kDa]

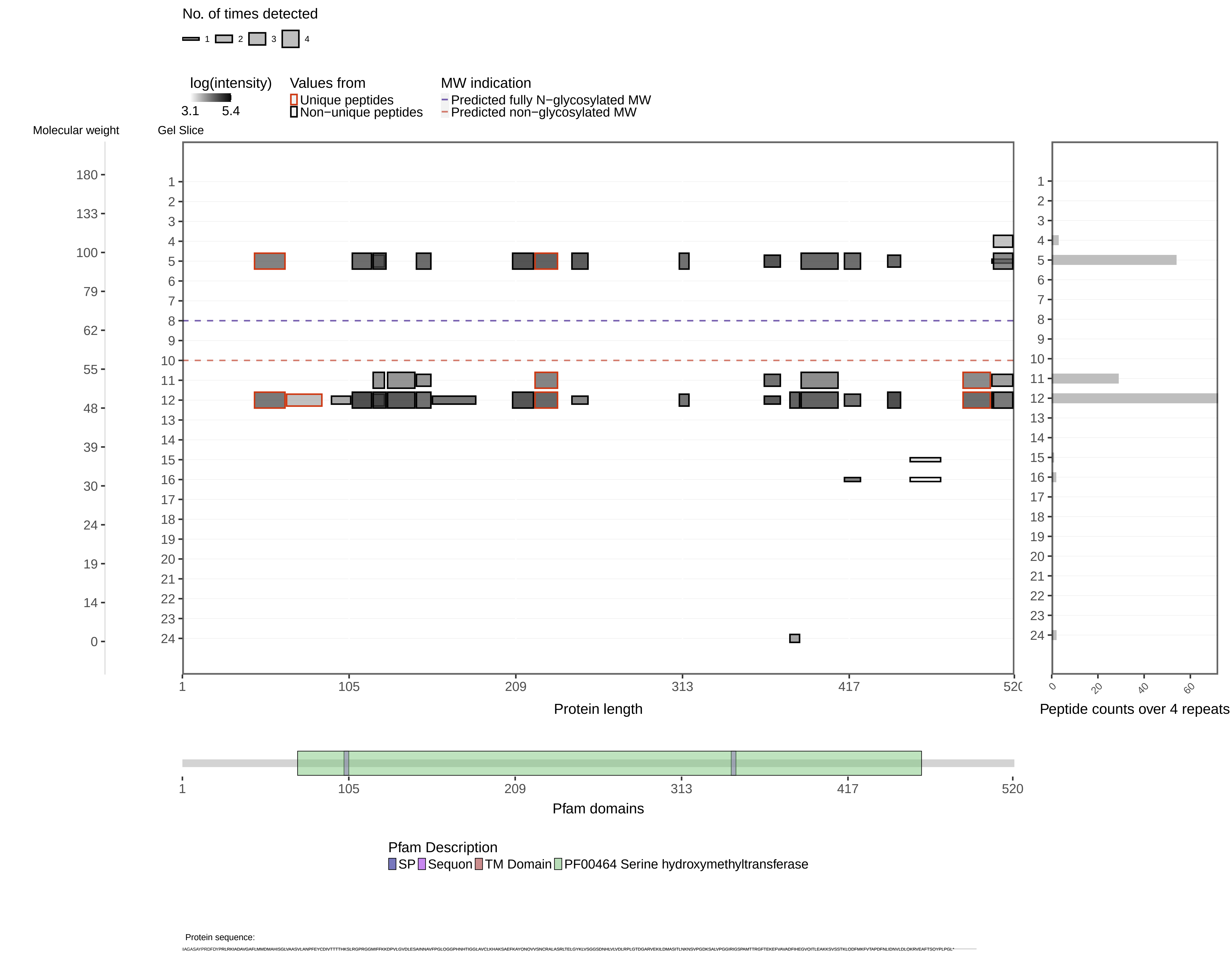

### NbL10g15970.1 abscisic_acid_receptor_PYL2−like [MW = 20.99 kDa]

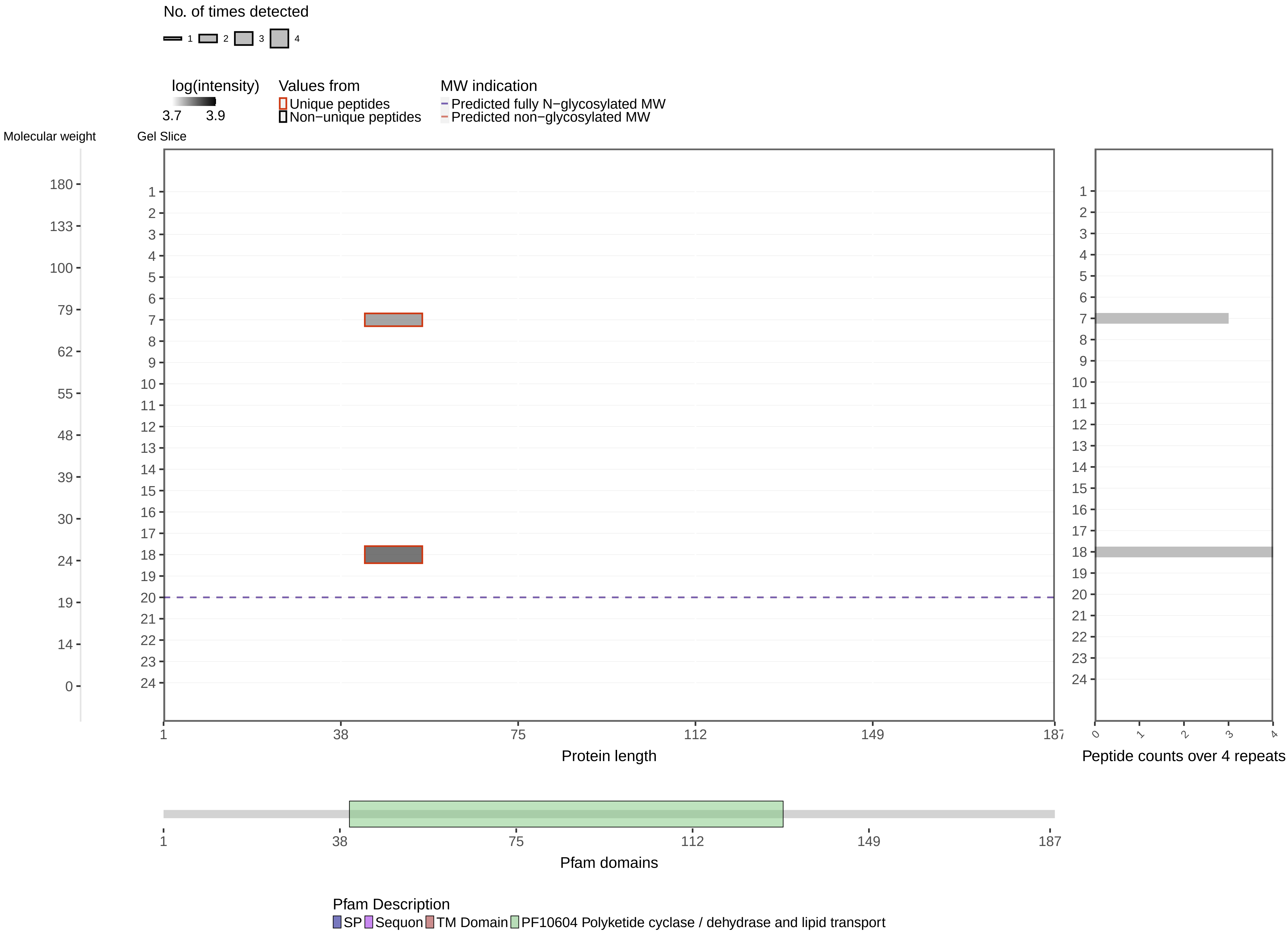


Protein sequence:

SMREVSVVSGIPALTSTERLEILDDEKHILSFKVVGGEHRLNNYRSVTSVNEFKKNGKPYTIVLESYIVDIPEGNTGEDTKMFTDTVVKLNLQKLGVVAMAALHGHE*

NbL10g15890.1 glycine_dehydrogenase_(decarboxylating)_mitochondrial [MW = 113.89 kDa]

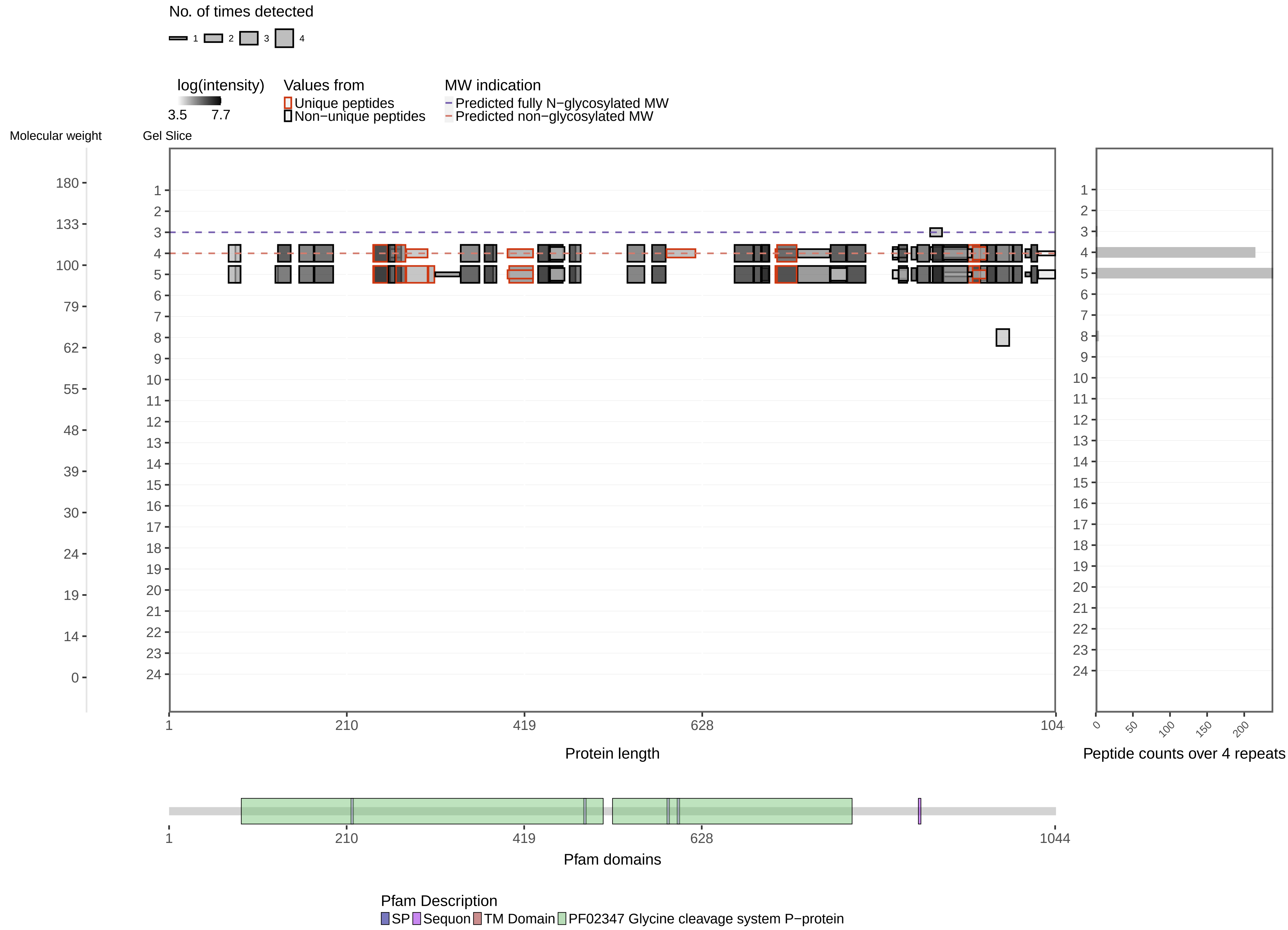


Protein sequence:

### NbL10g15840.1 UDP−sulfoquinovose_synthase_chloroplastic [MW = 53.35 kDa]

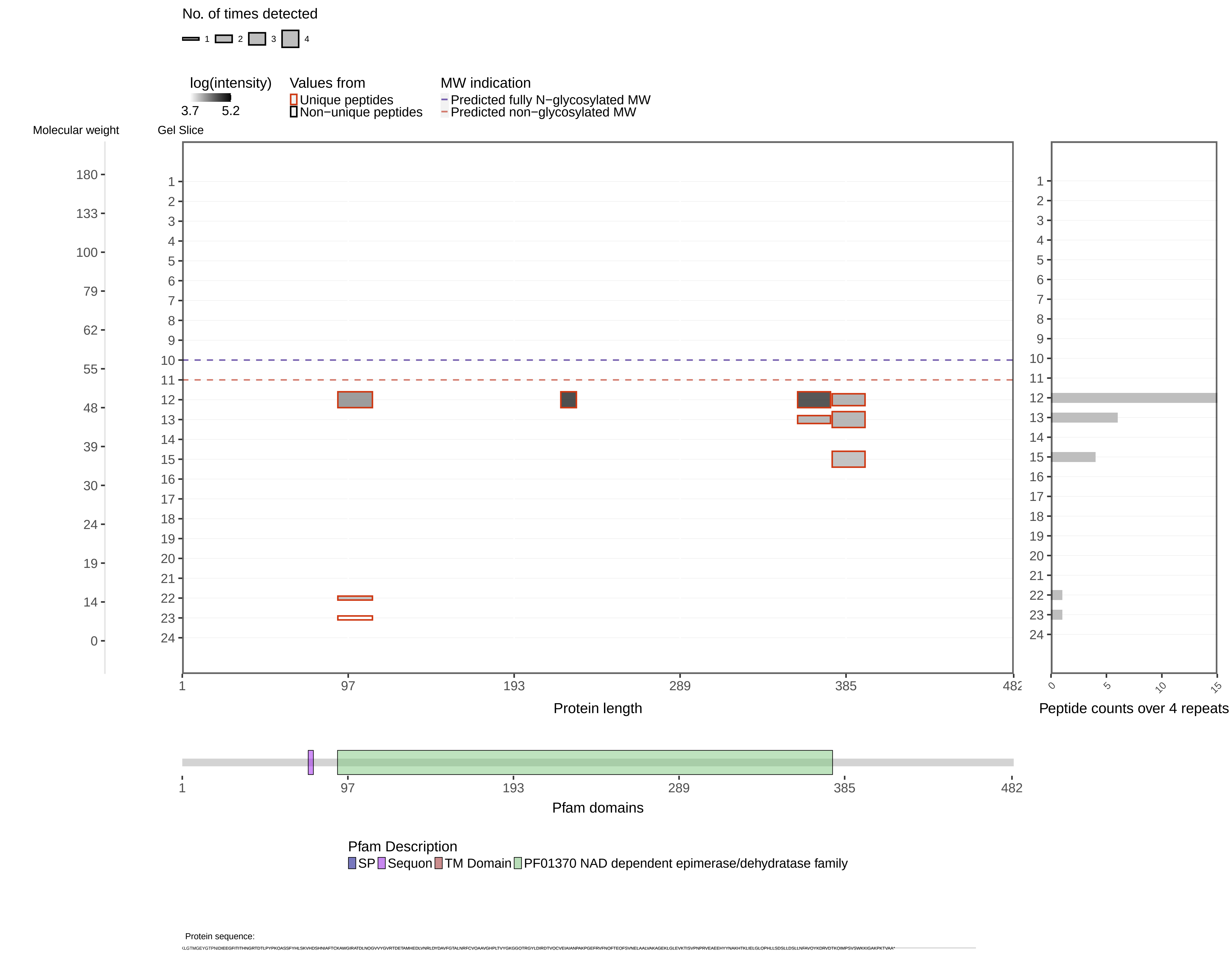

NbL10g15720.1 elongation_factor_2 [MW = 93.97 kDa]

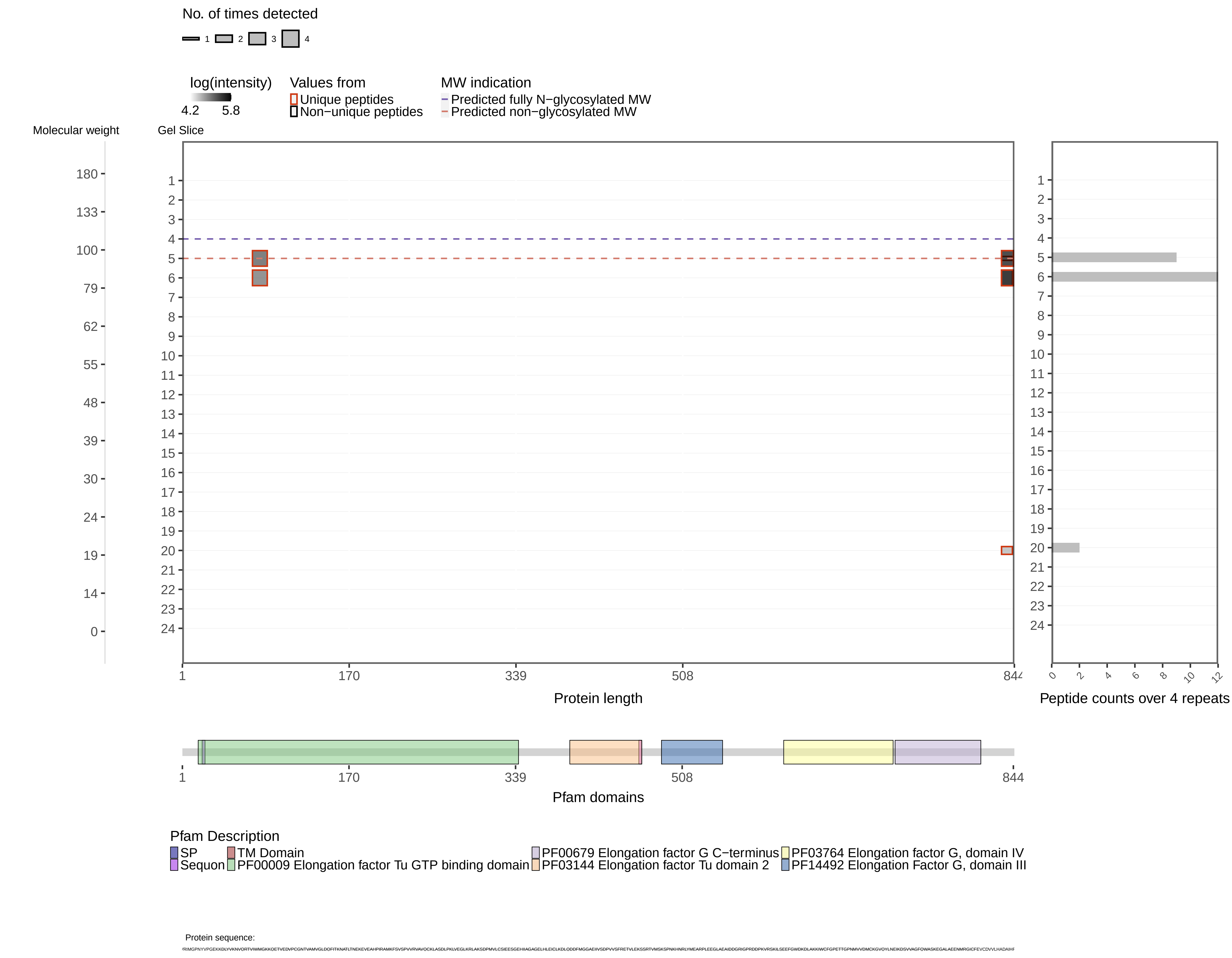

NbL10g15690.1 cytokinin_riboside_5'−monophosphate_phosphoribohydrolase_LOG8−like [MW = 23.36 kDa]

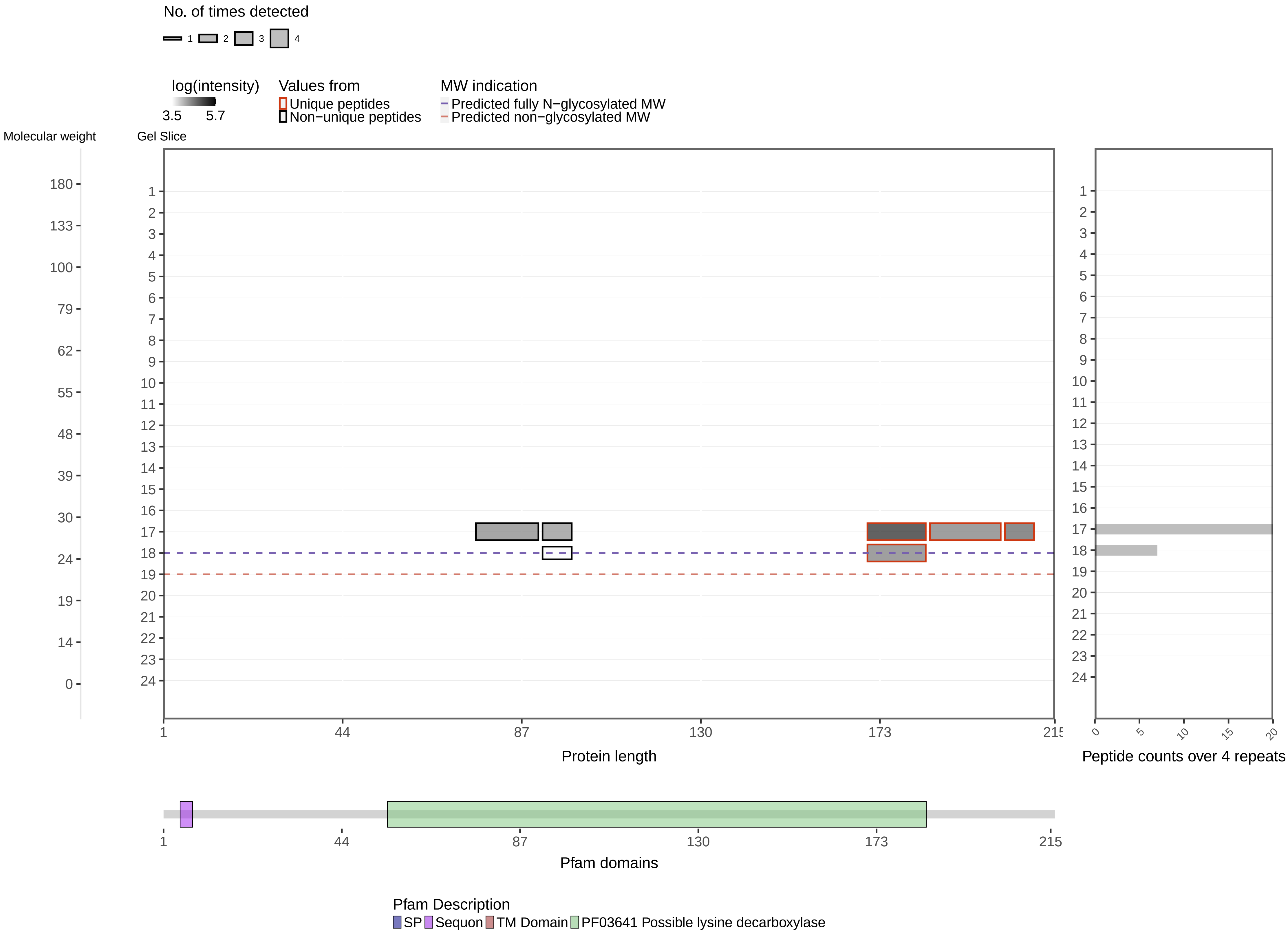


Protein sequence:

MHERKAEMASQADAFIALPGGYGTMEETLEMITWAQLGIHKKPVGLLNVEGYYNSLLTLFDNGVEEGFIKPGARDIVLAAPTAEELLSKMEQYTPSHDHVAPHDSWKMEELAYPKEQSP*

### NbL10g15540.1 ran−binding_protein_1_homolog_c−like [MW = 18.74 kDa]

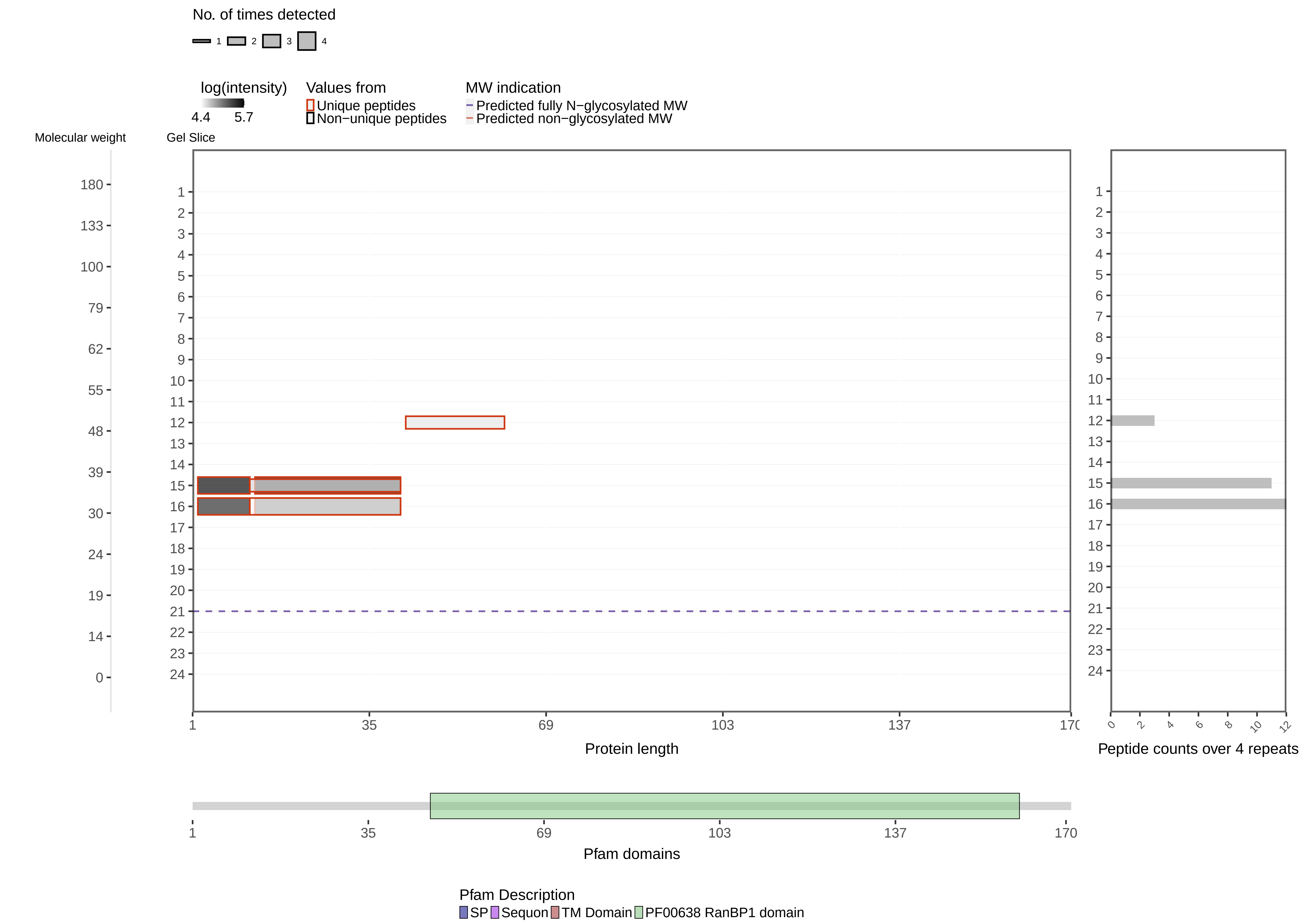


Protein sequence:

NAWKERGAGTVKLLKHKETGKVRLVMRQSKTLKICANHLVLPTMSIQEHAGNEKSCVWHAADFADGELKEETFCIRFASIESEYSVYASFGFCCSVFS*

### NbL10g15520.1 SNARE−interacting_protein_KEULE [MW = 75.56 kDa]

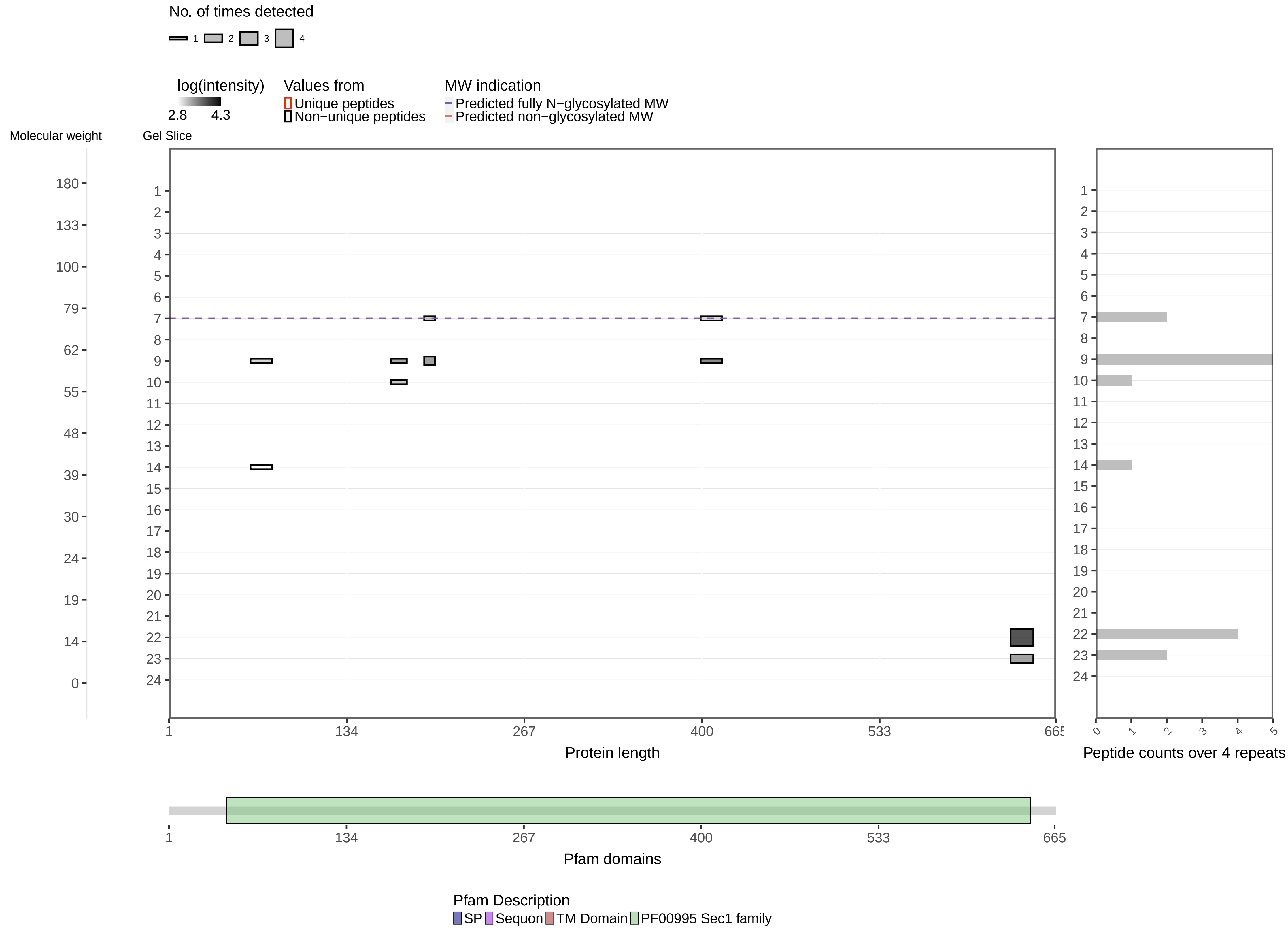


Protein sequence:

AHIADASERLHEKMTNFVSKNKAAQMHQSSRGGGQLSTRDLQKMVQALPQYSEQIEKLSLHVDIAGKLNRIIRESSLKEIGQLEQNLVFGDAGTKDLINFLKMDVTRENKLRLLMIYAAVHPEKLESDKLSKLMELARLPQDDMNAVYNMRLLEGSSDNKKSSLGPFSLKFDVHKKKHAARKDRTDEAATWQLSRFYPMIEELVEKLSKGELPKNDYPCMNDPNPTFHGTSQSASVQANQVPTPHSMRSRRTATWARPRNSDDGYSSDSILRHASSDFKKMGQRIFVFIVC

### NbL10g15500.1 thioredoxin_domain−containing_protein_9_homolog [MW = 24.26 kDa]

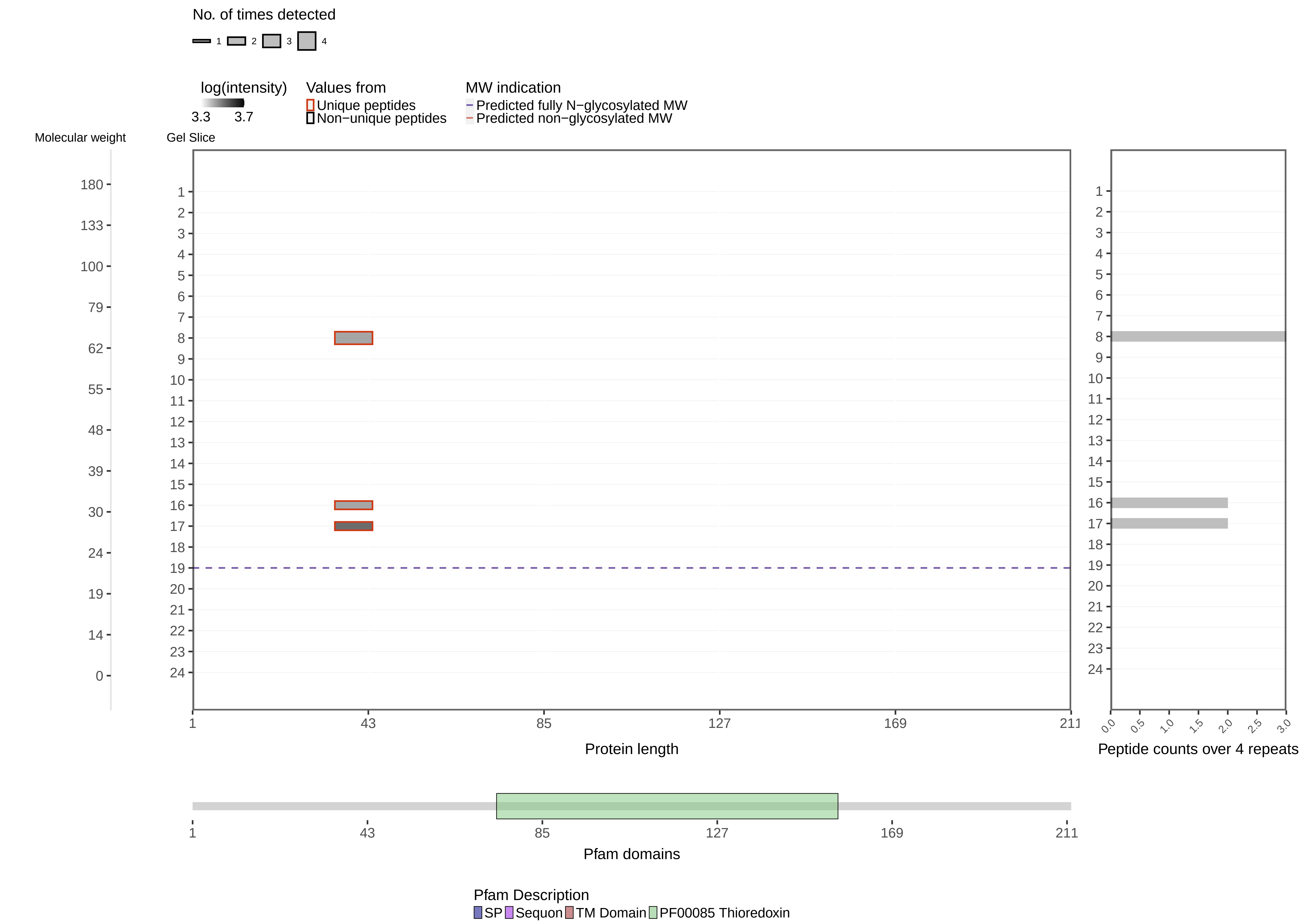


Protein sequence:

IFYRDNWPCKVMDKHLSILAKQHIETRFVKIHAEKSPYLAEKLRIVVLPTLALVKHAKVEDYVVGFDELGHTDEFSTEELEERLAKAEVIIFEGESSKLKSKAPSKSVRQSANPDSSDSE*

### NbL10g15460.1 putative_transferase_CAF17_homolog_mitochondrial [MW = 43.82 kDa]

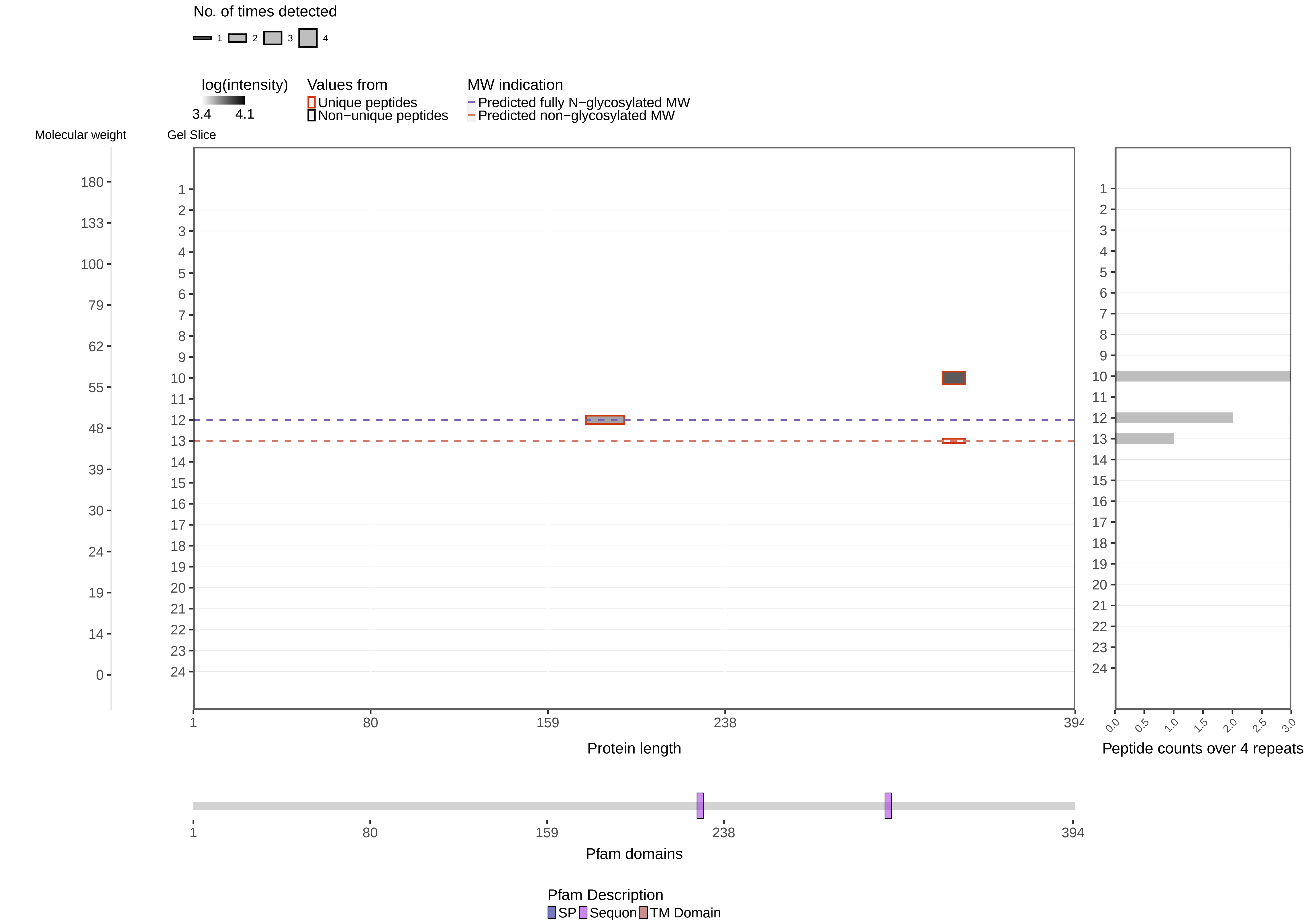


Protein sequence:

.GWGGGFDRSGQSSAQGNNRGWLWYRDPRLDCLGFRGIFPSNITPPLVEADKETNEENYLLWRLEKGVAEGSAEIPKGEAVPLEYNLAGLNGISFDKGCYVGQELIARTHHRGVIRKRLLPLKFVNDSGKEVEQKVAPRSEVVDASSGKKAGTVTTALGSRGLGLLRLDEAFKGSLTIRDLDDVKVQAIKPEWWRAEWLLDHQGQTAAA*

NbL10g15350.1 FAM10_family_protein_At4g22670−like [MW = 46.29 kDa]

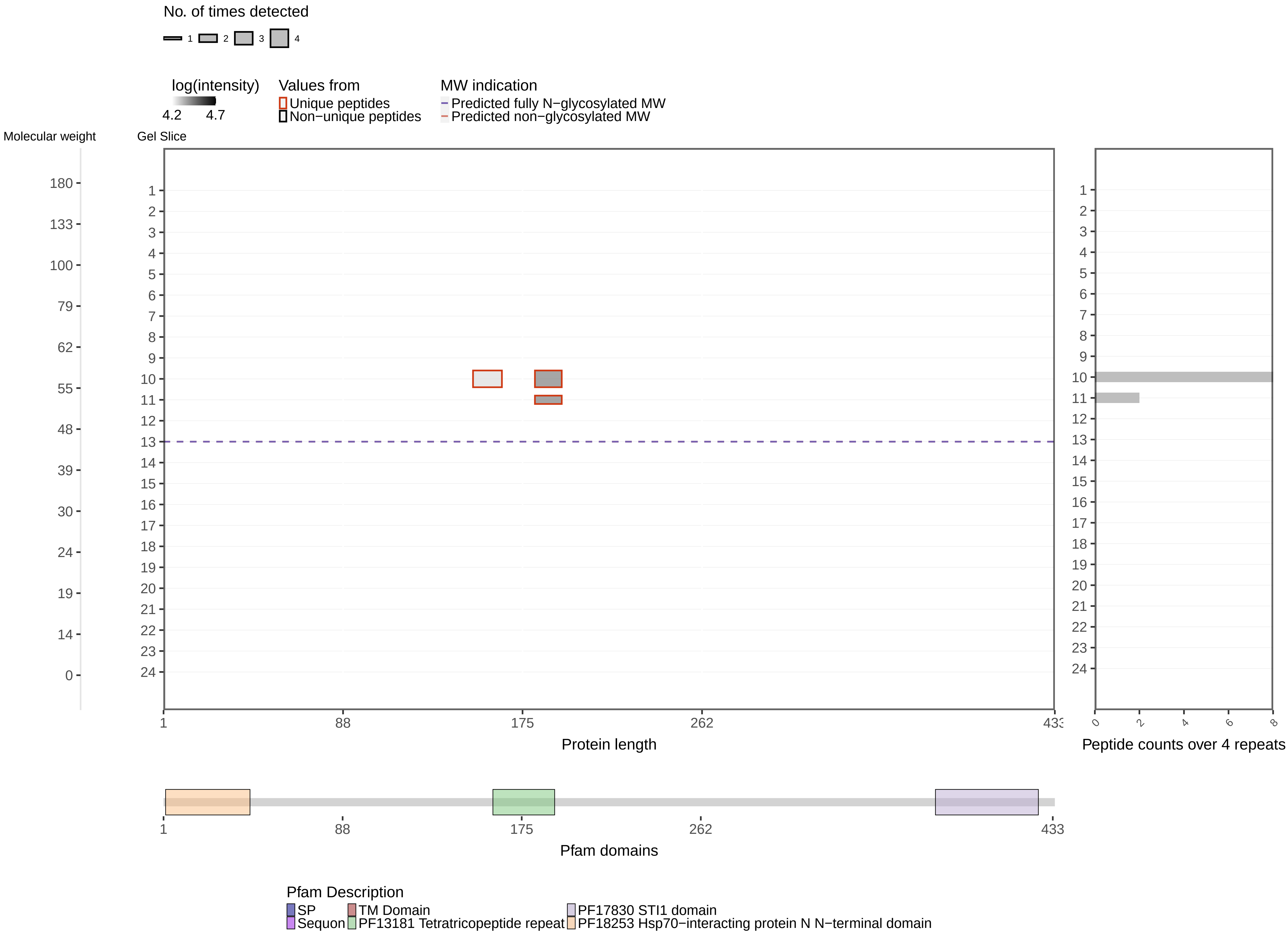


Protein sequence:

/EEAAKDLHLASKLDYDEEISAVLKKVEPNAHRIEEHHRKYDRLRKEREDRKNERERQRRKAEAQAAYEKAKKQEESSASRRAGAMPGGFPGGMPGDLPGGMPGGFPGGMPGGFPGGMPGGFPGGMPGGSPGAMPGGSPGAMPGGFPGAMPGGFPGAMPGGMPGNMDYRKILNDPELMAAFKDPDVMAALQDVMKNPANLAKHQANPKVAPIIAKMMSKFAGSN*

### NbL10g15330.1 uncharacterized_LOC107808286 [MW = 28.56 kDa]

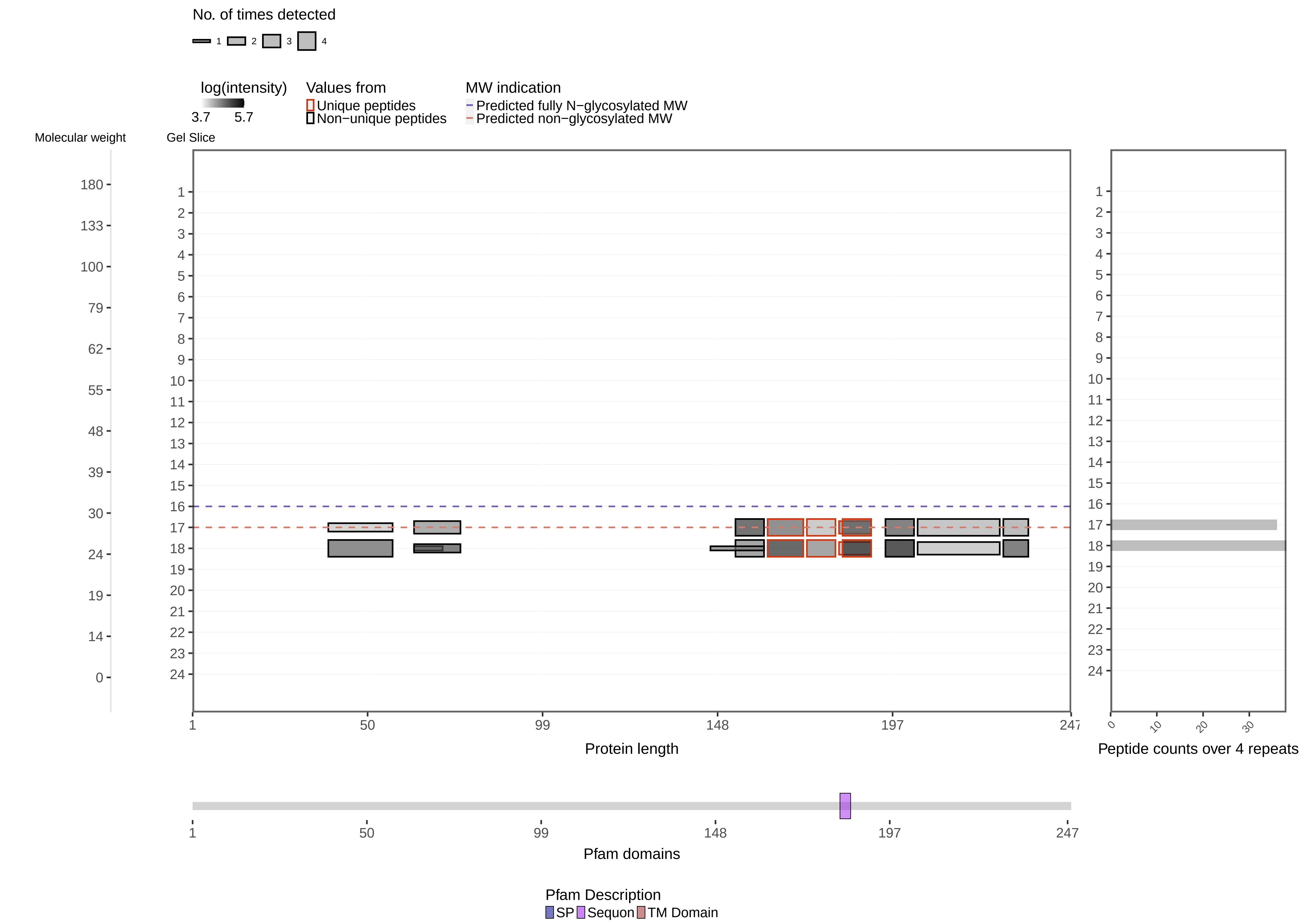


Protein sequence:
EAQEIEFEIARLRPRLNKEFFDHLQVEVGKLRFSMTKTQEVEDRLIELEALQKALHEATEAYDKMQADILATKKNLTDILTSKDVKATLLDLVERNELNRPLLTLLDENIATAHSVNQKQAAEYMEKLRGLVLKYLTV*

### NbL10g15310.1 elongation_factor_G−2_chloroplastic [MW = 86.07 kDa]

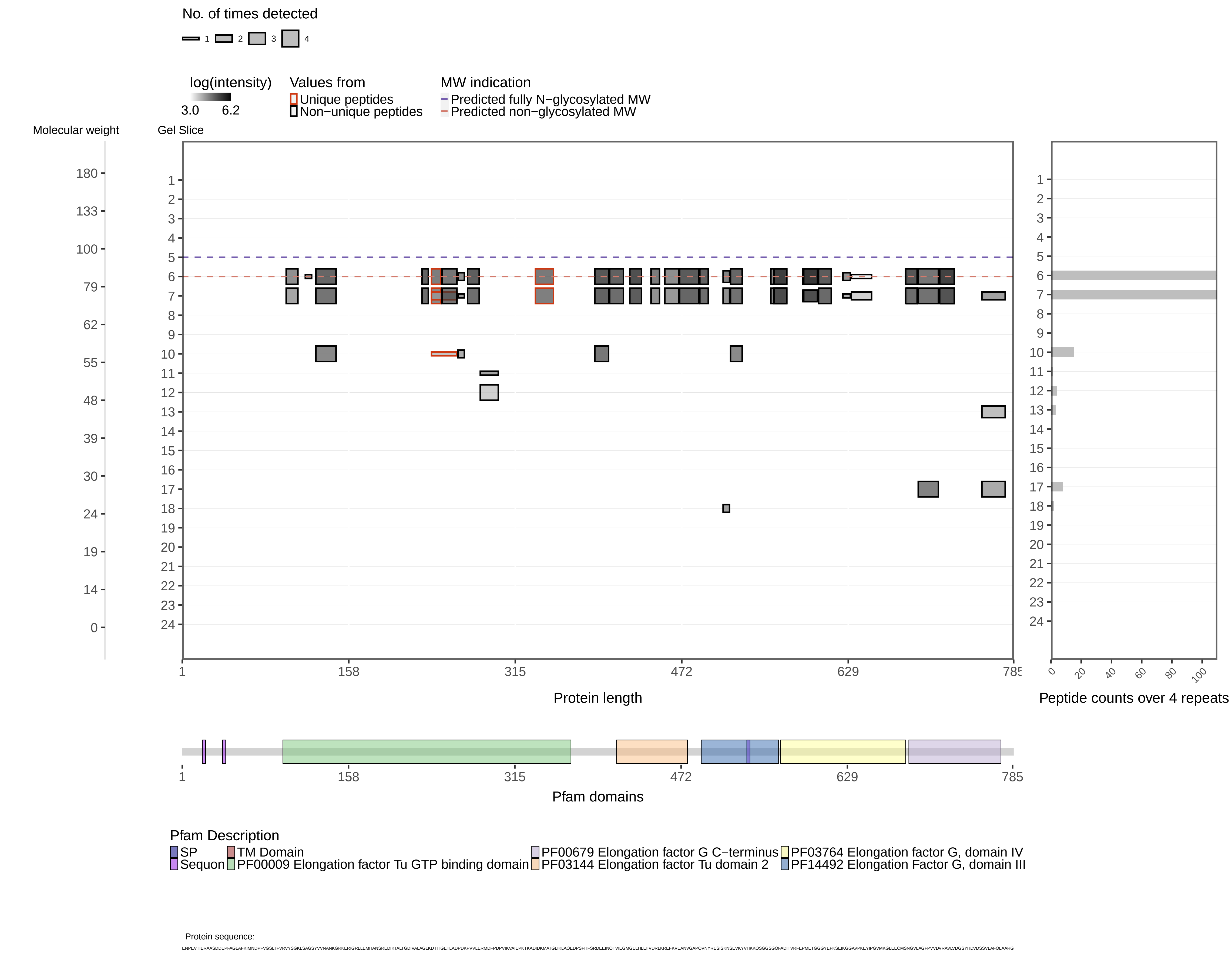

NbL10g15300.1 hsp70−Hsp90_organizing_protein_2 [MW = 65.01 kDa]

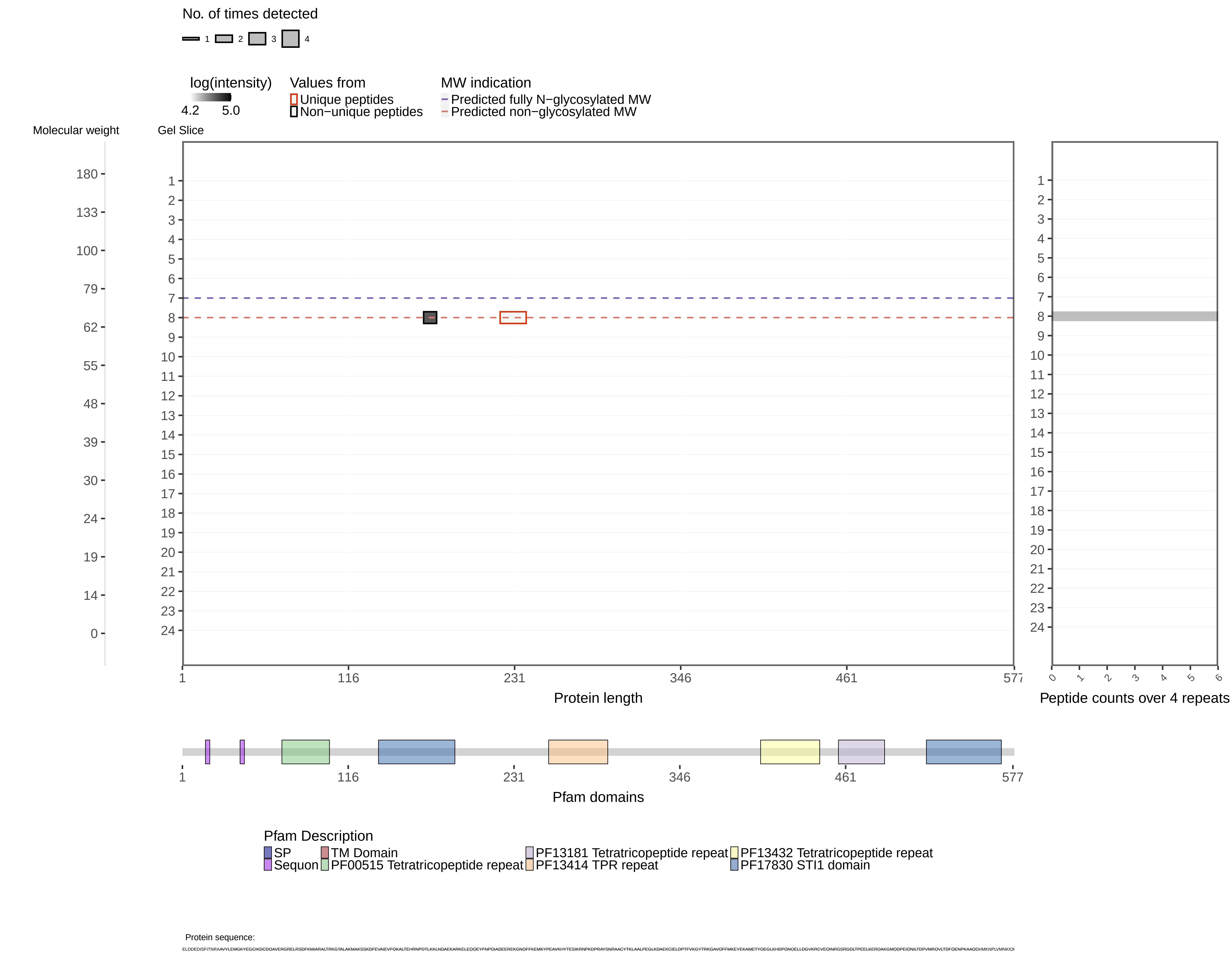

### NbL10g15230.1 monocopper_oxidase−like_protein_SKU5 [MW − SP = 63.88 kDa]

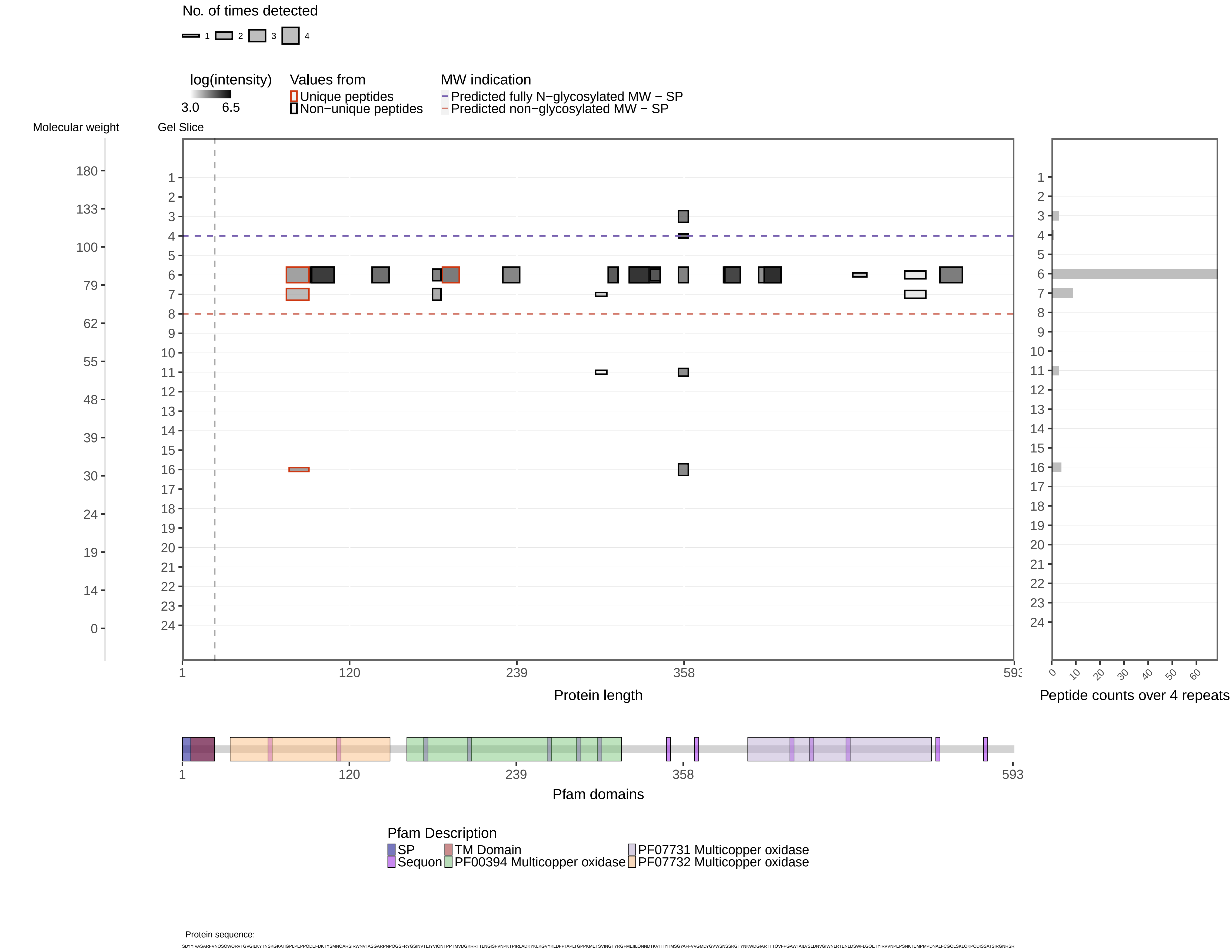

NbL10g15170.1 probable_xyloglucan_galactosyltransferase_GT19 [MW − SP = 48.29 kDa]

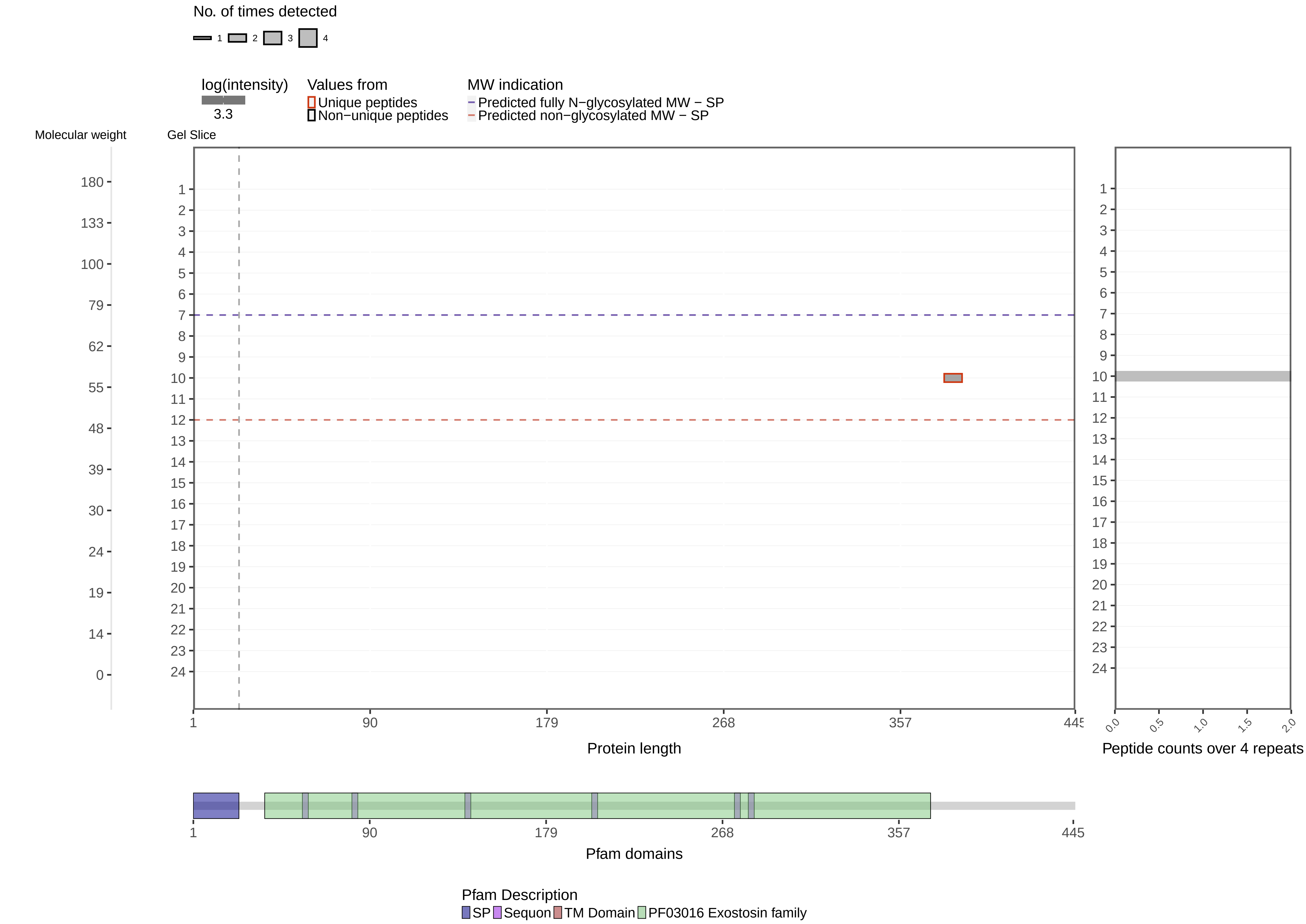


Protein sequence:

RAYPWQEQAIPYPTSFHPPNLAFFESWVNRVRRSRRNTLMLFAGGGGISANPYIRRSIRLECENVTSMSANGTGYQKLCEFVDCSNGICEHDPIRFMKPMLQASFCLQPPGDIPSRRSTFDGILAGCIPVFFEDLSAKKQYGWHLPAEKYEEFSVTIPKEDVVFKGLKVFDVLTSIPRAQVRRMREKVLEMMPRVMYRKHGSSLGLRSKKDAFDIAIEGTLERIKSRLQEVAAQ*

NbL10g15150.1 adenine_phosphoribosyltransferase_3 [MW = 20.07 kDa]

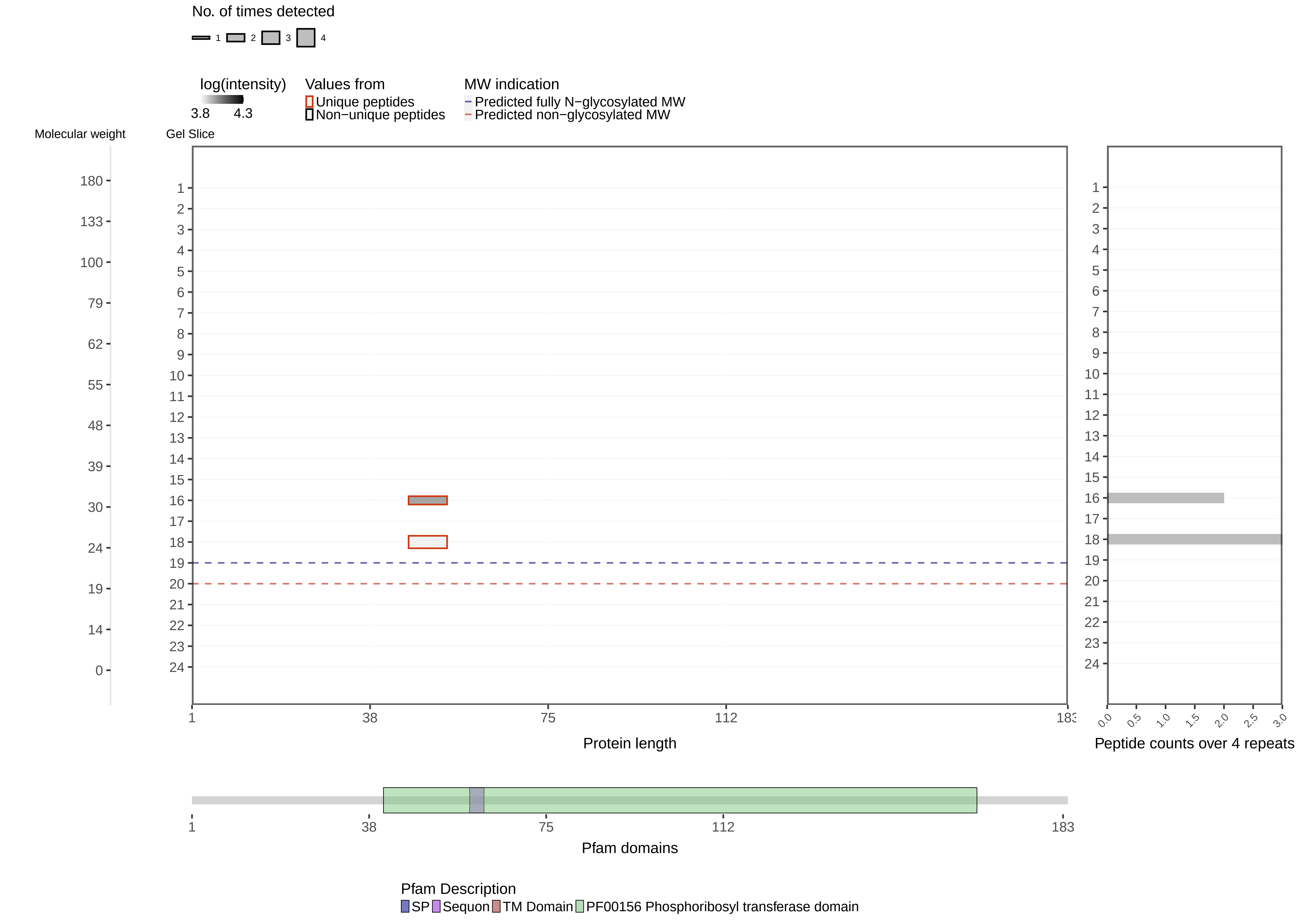


Protein sequence:

AIGAKFVPLRKPNKLPGKVFKQEYDLEYGSDCLEMHIGAVEAGERALVVDDLIATGGTLRAAMNLLERAGAEVVECACVIEIPDLKGKEQLNGKPLYVLVESQ*

NbL10g15080.1 kDa_chaperonin_chloroplastic−like [MW = 26.41 kDa]

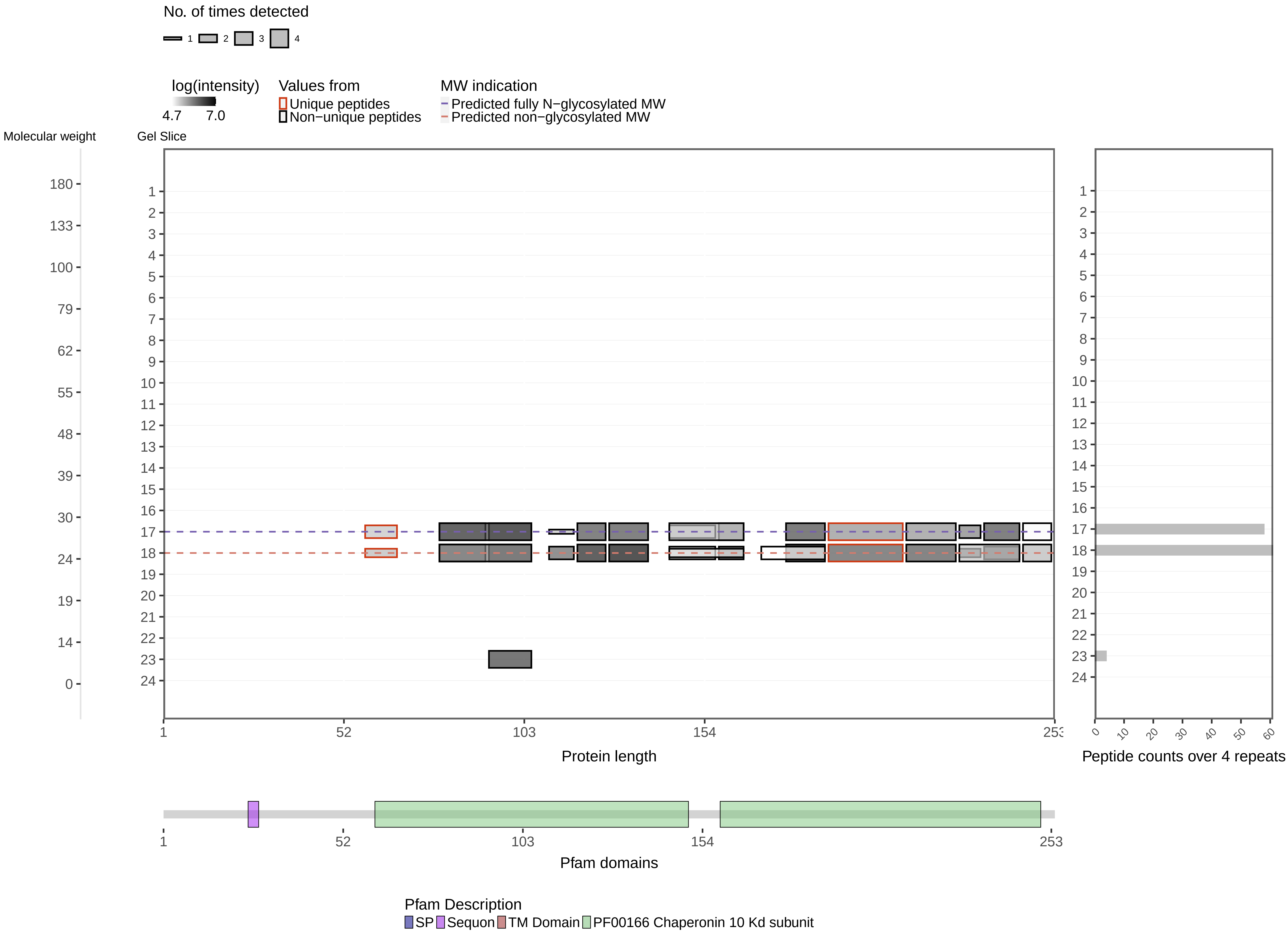


Protein sequence:

DISVKTGAQVVYSKYAGTEVEFDGSKHLILKEDDIVGVLETDDIKDLQPLNDRVLIKVAEAEEKTAGGLLLTEAAKEKPSIGTIIAVGPGPLDEDGNRKPLSVSPGNTVLYSKYAGSEFKGADGSDYITLRASDVMAVLS*

NbL10g14910.1 peroxisome_biogenesis_protein_19−1−like [MW = 27.92 kDa]

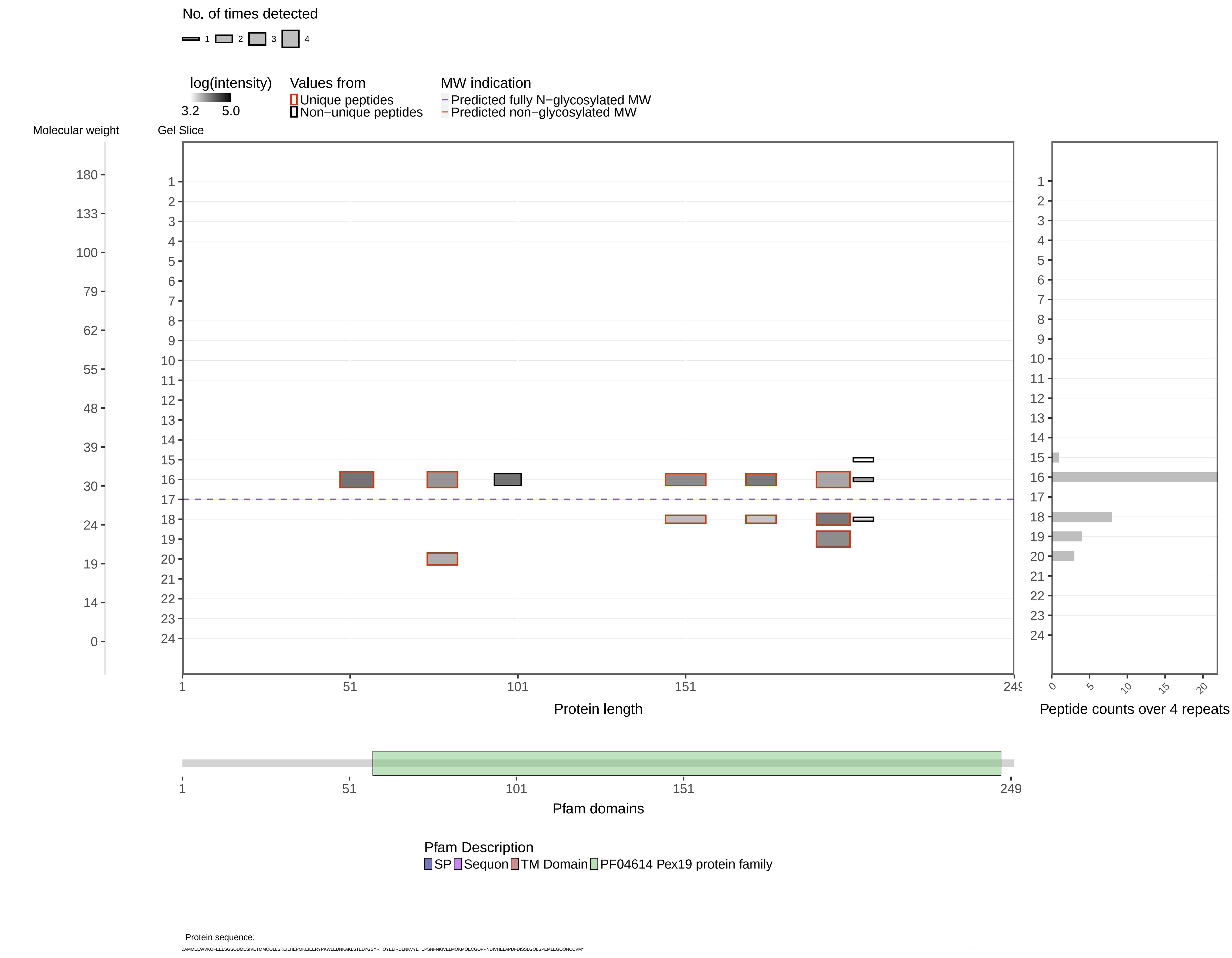

### NbL10g14800.1 uncharacterized_LOC107802887 [MW = 231 kDa]

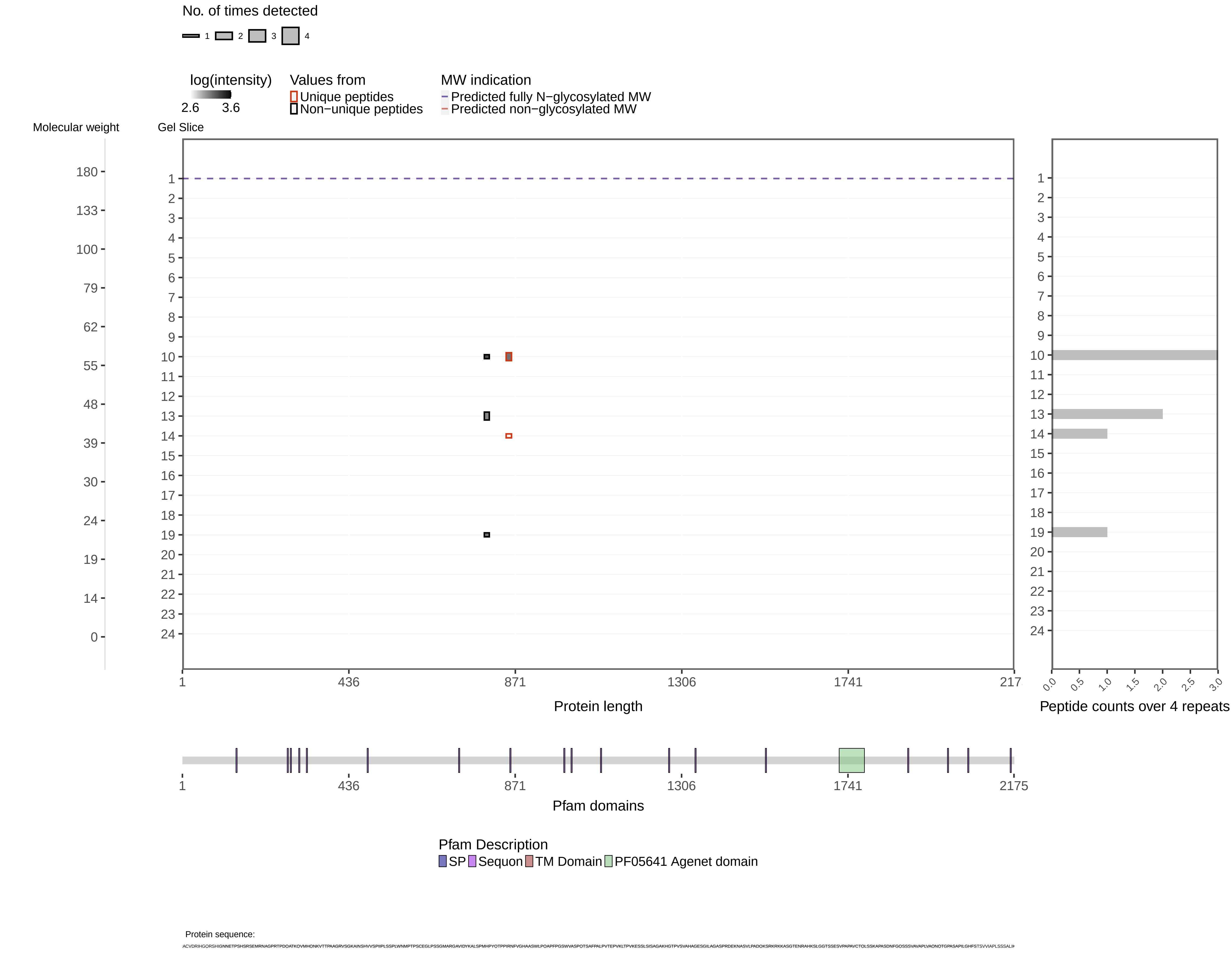

NbL10g14700.1 peroxisomal_acyl−coenzyme_A_oxidase_1 [MW = 74.33 kDa]

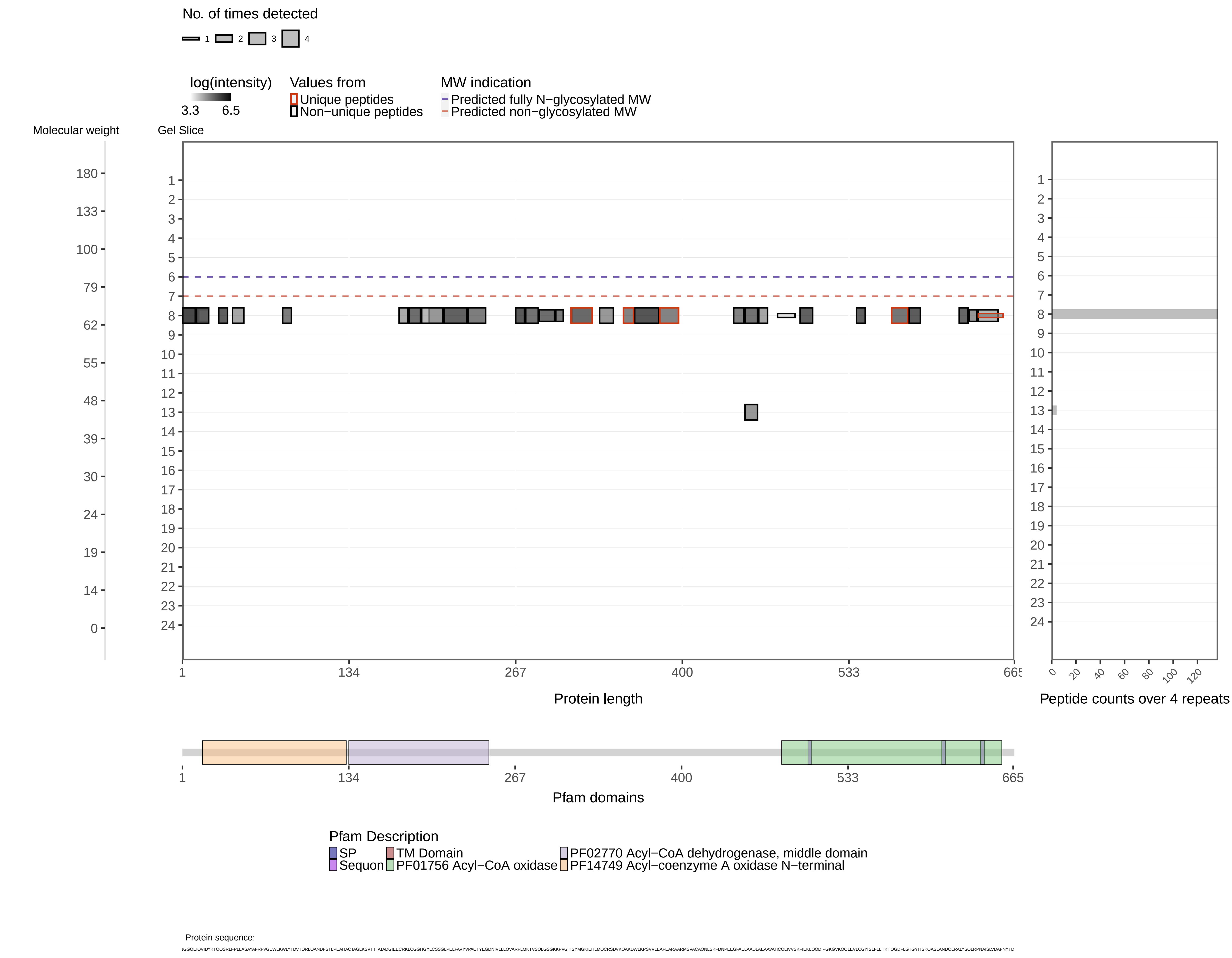

### NbL10g14380.1 plasminogen_activator_inhibitor_1_RNA−binding_protein−like [MW = 39.27 kDa]

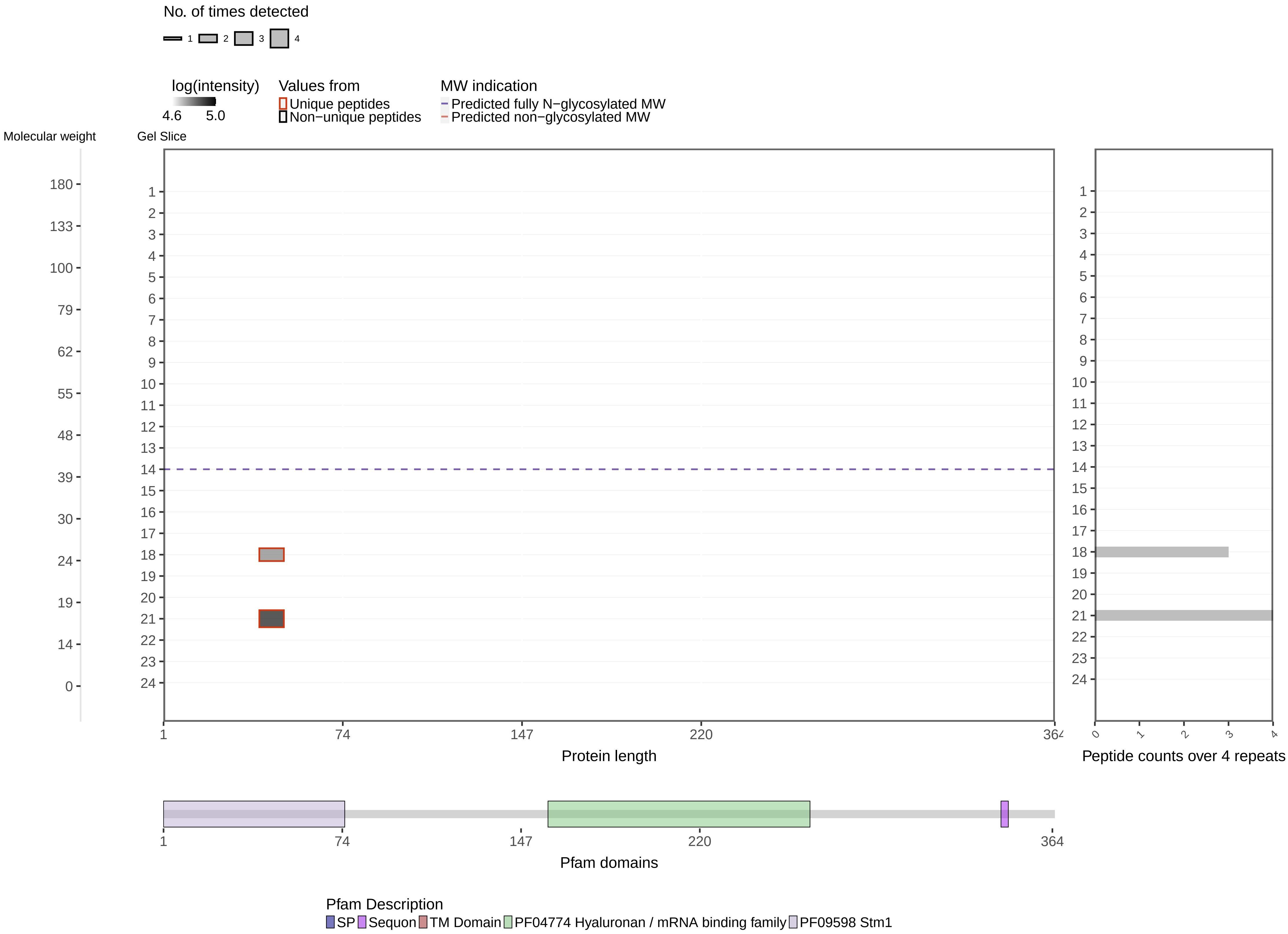


Protein sequence:

GTGRGNDFKREGAGRANWGTPTDEIAPETEEPVTEGEKIVDPEKPAGEEDAGDAKKDSSAAEPEEKEPEEKEMTLEEYEKLLEEKRKALLAOKPEERKVNLDKEFESMOLLSNKKNEDDIFIKLGPEKDKRKEAVEKAKKTOSINEFLKPAEGENYYRGRGRGGRGRGGYGGNGGNNYSVEAPKIEDVGOFPSLGAK*

### NbL10g14310.1 50S_ribosomal_protein_L19_chloroplastic−like [MW = 24.25 kDa]

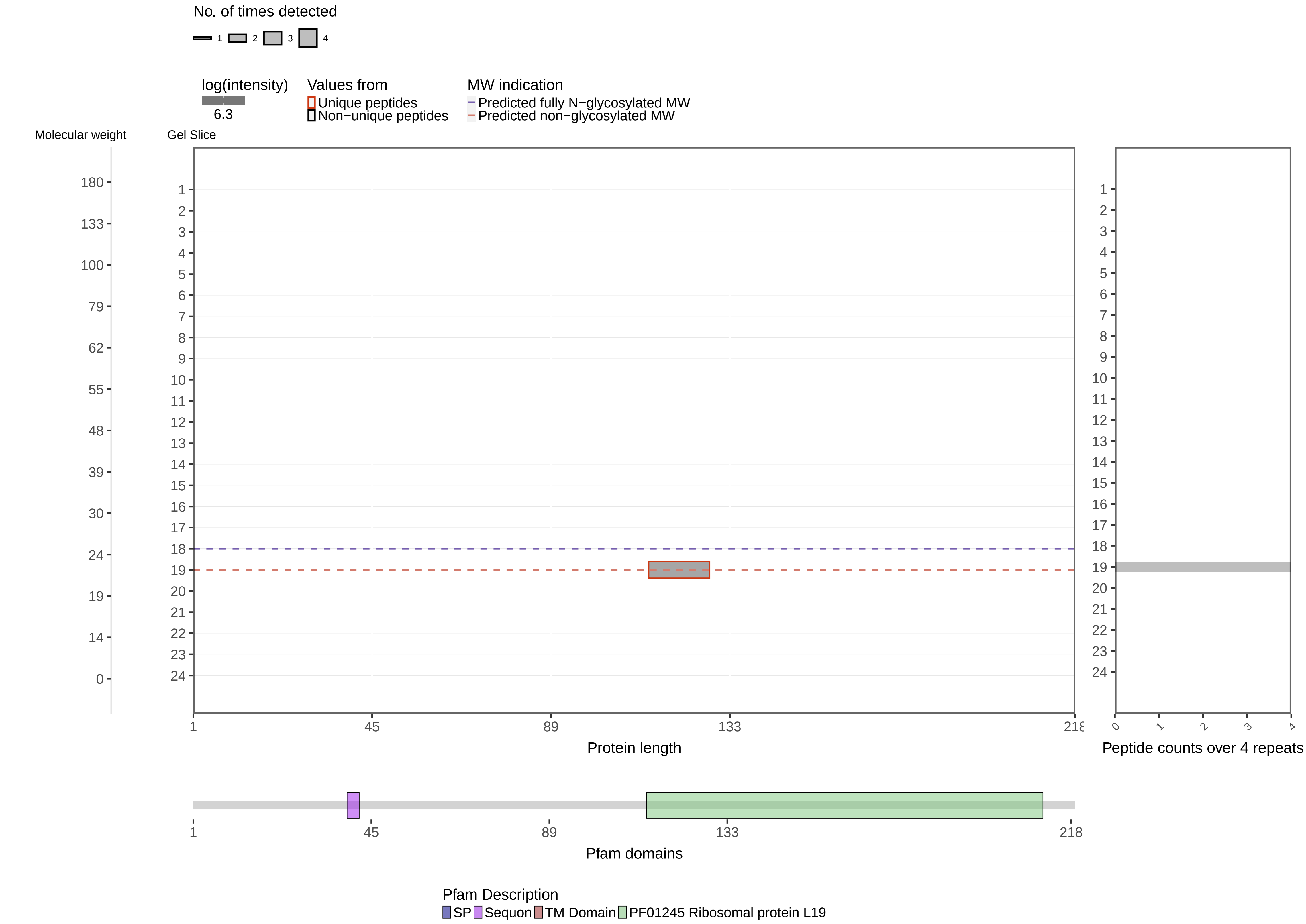


Protein sequence:

EERPPFKPRAKLGDIMGILNKQAIEASDAIRPIPDIRTGDVVEIKLEVPENRRRLSIYKGIVISKQNAGIHTTIRIRRIIAGIGVEIVFPVYSPNIKELKVVKHRKVRRARLYYLRDKLPRLSTFK*

### NbL10g14060.1 serine/threonine−protein_kinase_SAPK3−like [MW = 38.25 kDa]

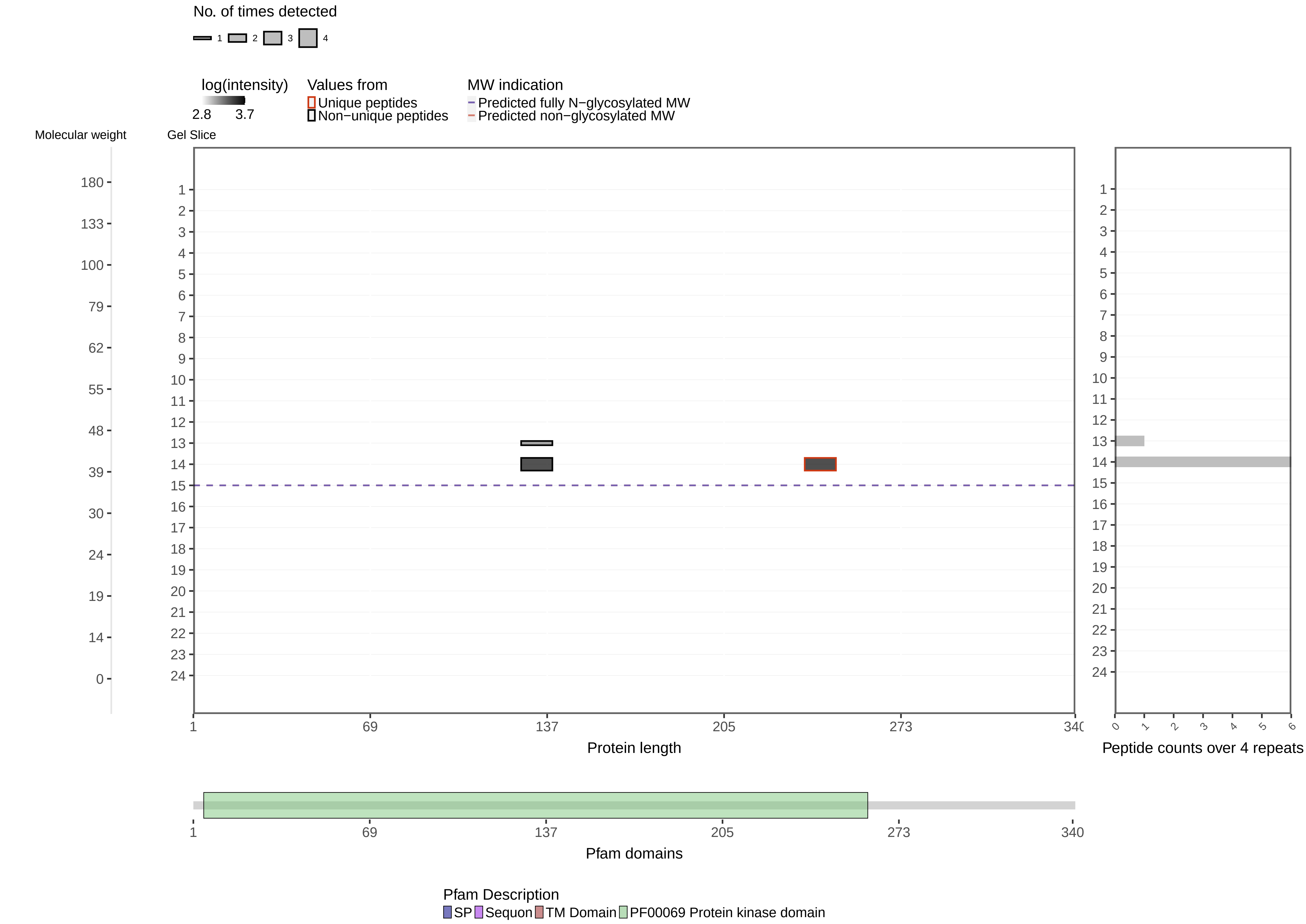


Protein sequence:

PKSTVGTPAYIAPEVLSRKEYDGKIADVWSCGVTLYVMLVGAYPFEDPEDPKNFRKTIGRIMSAQYSIPDYVRISADCKNLLSQIFVANPSKRITIPEIKKHPWFLKNLPKDLMDGKRSKFEEASEQQQQSVEEIMRIIQEAKIPGEVSKPEGQATAGTTEQDDTEDDLESEIDSSNDFAVYV*

### NbL10g13930.1 poly(U)−specific_endoribonuclease−B [MW = 49.45 kDa]

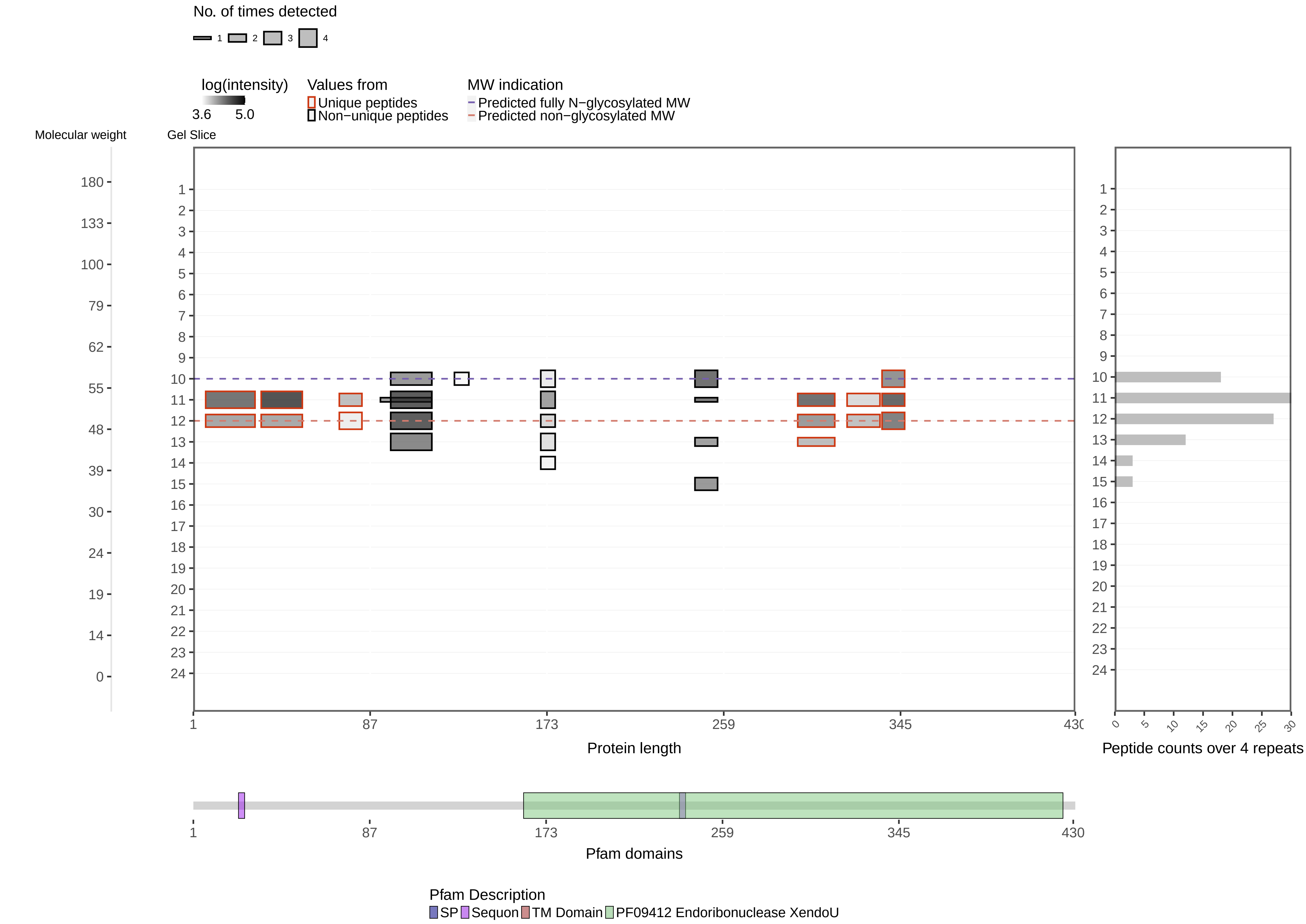


Protein sequence:

NbL10g13830.1 UMP−CMP_kinase_3−like [MW = 22.83 kDa]

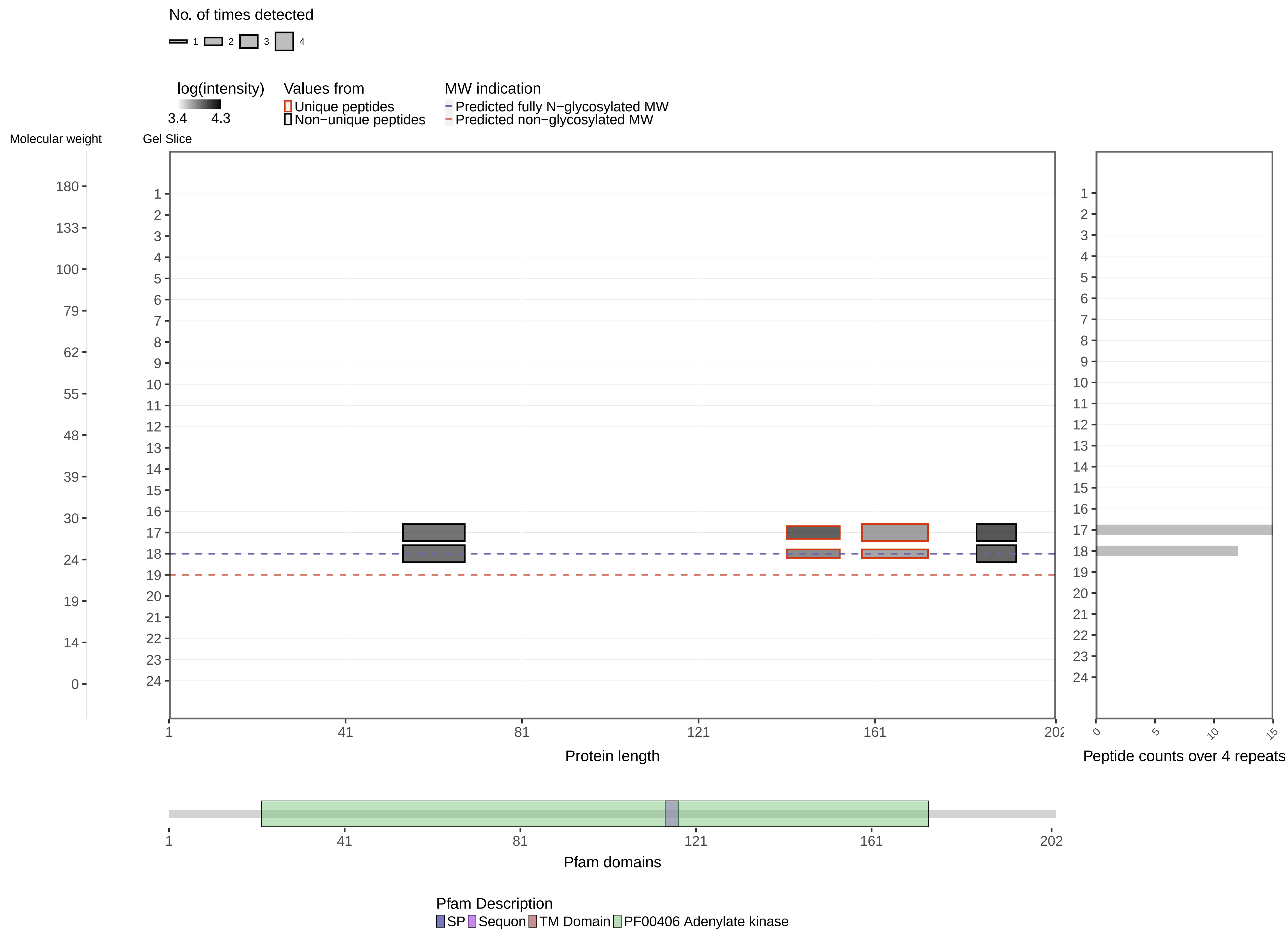


Protein sequence:

AMQGMDSDKFLIDGFPRNEENVKTFENLTKMEPEFVLYLDCPQDEMERRLLSRNEGREDDNTETIRKRFKVFLESTLPAVEYYESKGKVKKVDAGKSVDEVFESIKSILSRKRG*

NbL10g13800.1 haloacid_dehalogenase−like_hydrolase_domain−containing_protein_Sgpp [MW = 27.75 kDa]

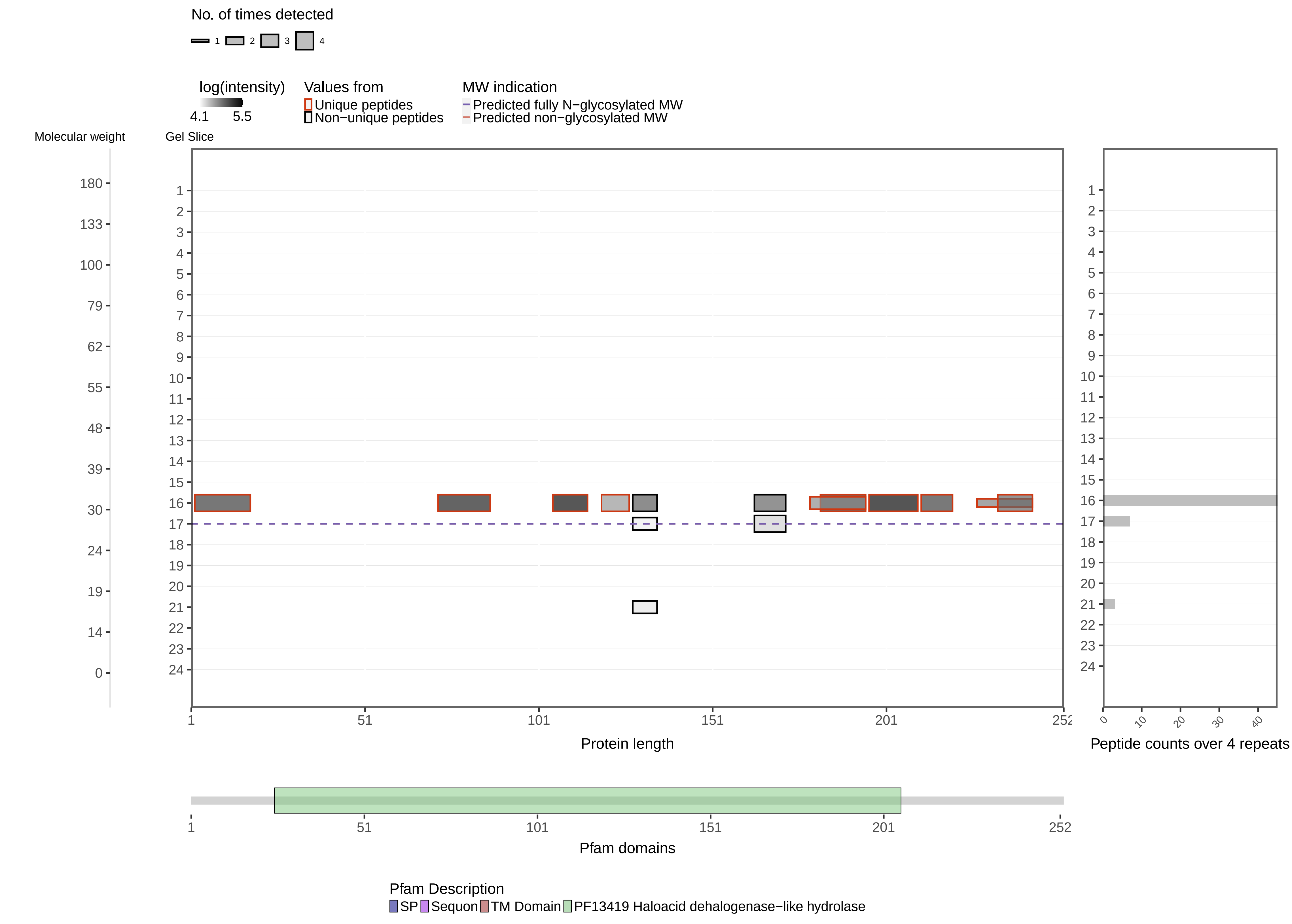


Protein sequence:

GLYKVKKWIEDHGLKRAAVTNAPRLNGELIIEILGLKDFFDVVIIGSECERAKPFPDPYLKALELLKVSKDHTFIFEDSVSGIKAGVAAGMSVVGLATRNPPHLLMEAKPAFLIKDYEDPKLWADLEEIDKKSGAGKATV*

### NbL10g13710.1 phosphoacetylglucosamine_mutase [MW = 65.93 kDa]

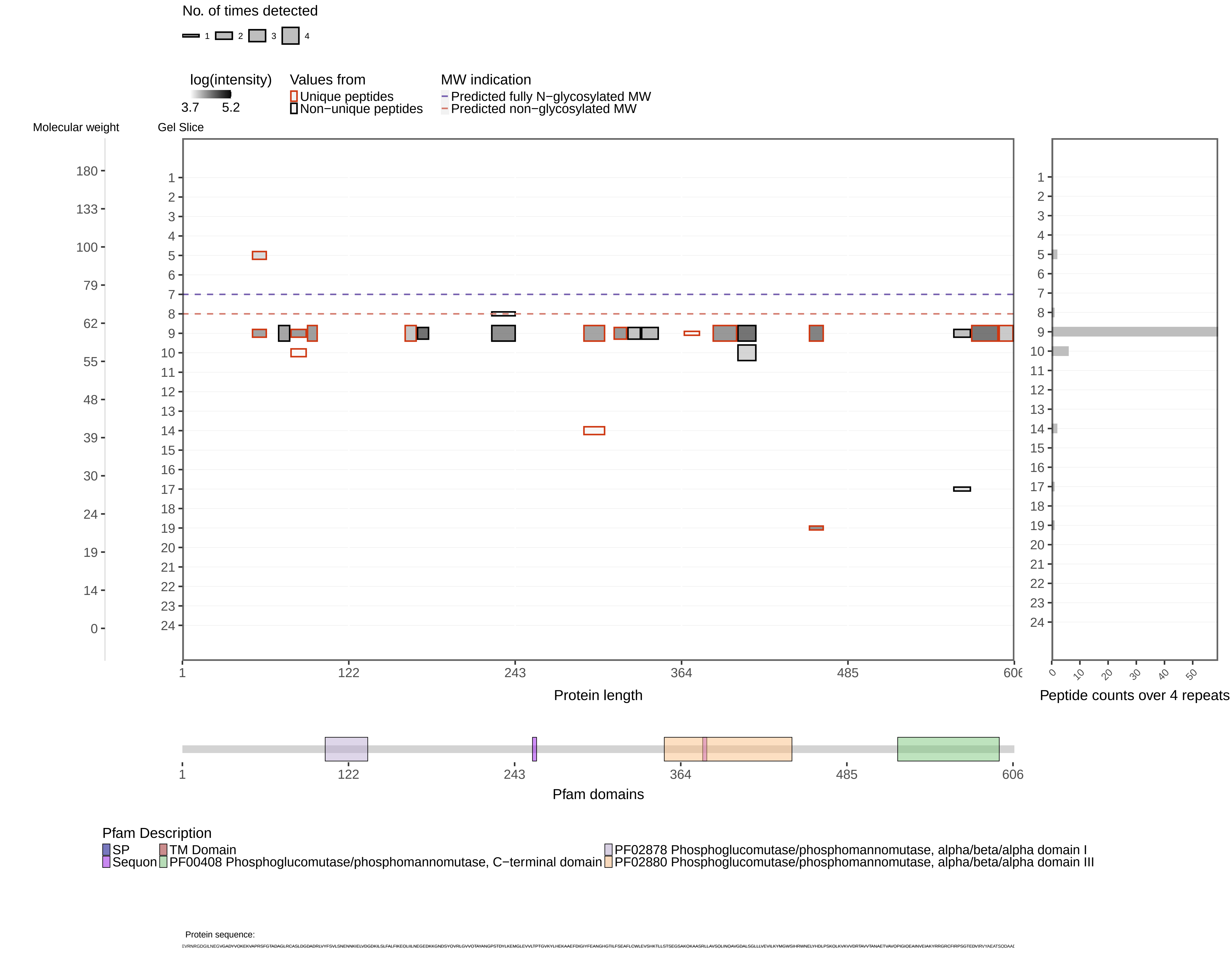

NbL10g13680.1 probable_methyltransferase_PMT14 [MW = 55.45 kDa]

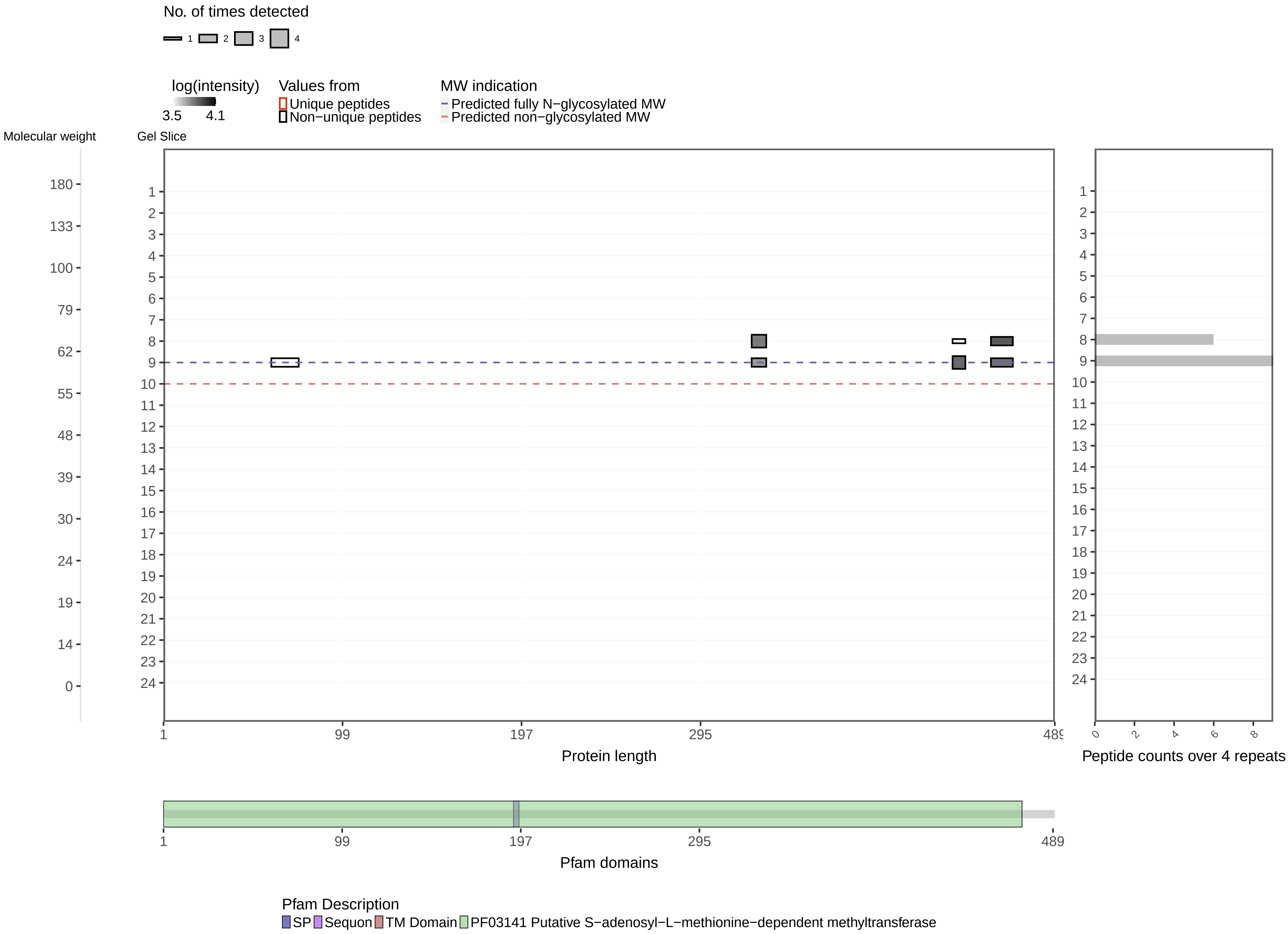


Protein sequence:

NGKSCSRRKSTNICQTKDTDNVWYKKMDACITPYPDVQSSDEVAGGELKKFPARLFAVPPRVANEMVPGVTIESYQEDNKLWKKHVTSYKRIVSLLGTTRYHNIMDMNAGLGGFAAALDSPKLWVMNVVPTIAENTLGVVYERGLIGIYHDWCEGFSTYPRTYDLLHANRLFTLYQDKCEFEDILLEMDRVLRPEGSVILRDGVEVLNKVRKIAAGLRWETKLVDHEDGPLVPEKIFVAVKQYHVEGDDNQNTQNDE*

### NbL10g13550.1 probable_xyloglucan_endotransglucosylase/hydrolase_protein_8 [MW − SP = 32.06 kDa]

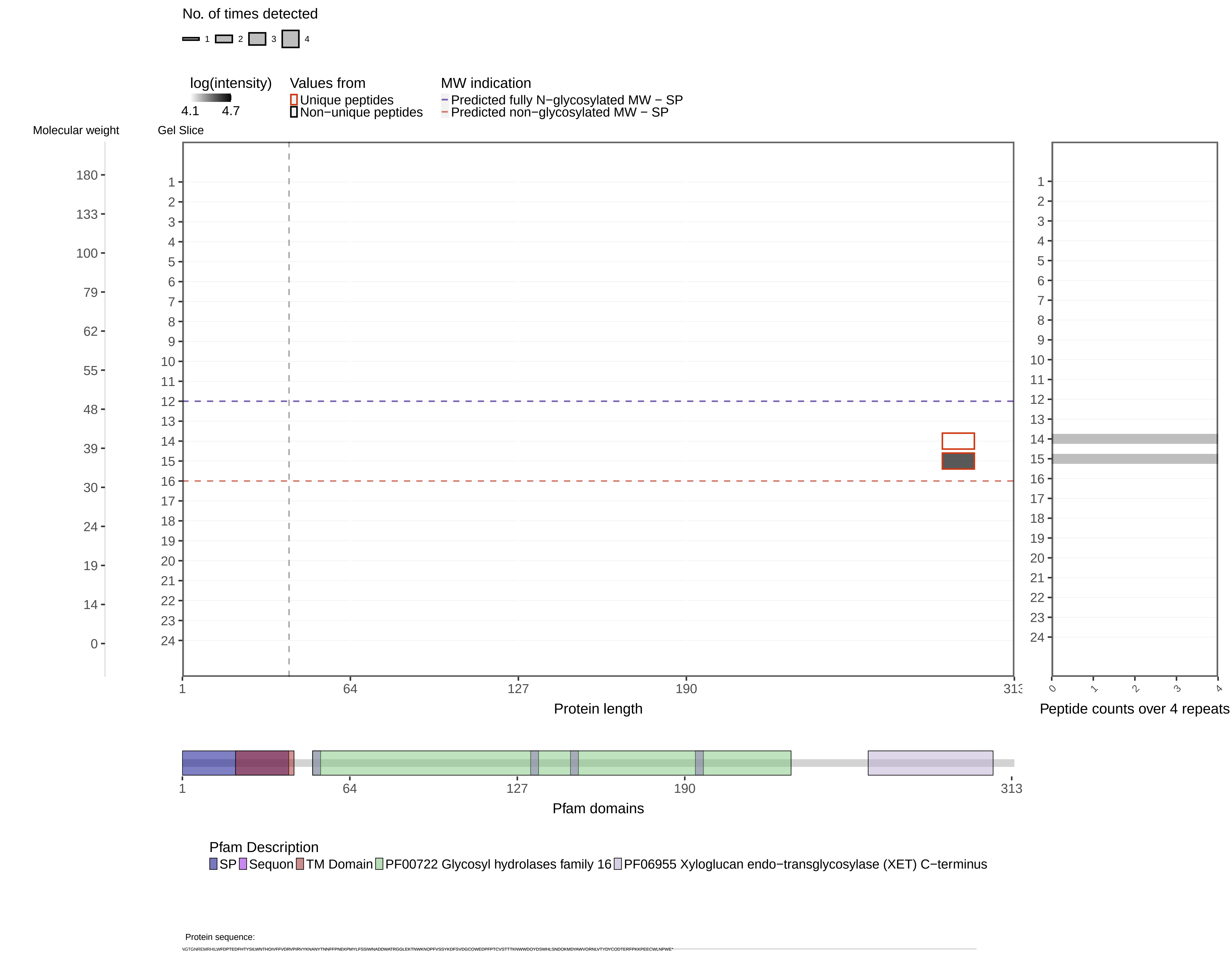

NbL10g13460.1 citrate_synthase_glyoxysomal−like [MW = 55.94 kDa]

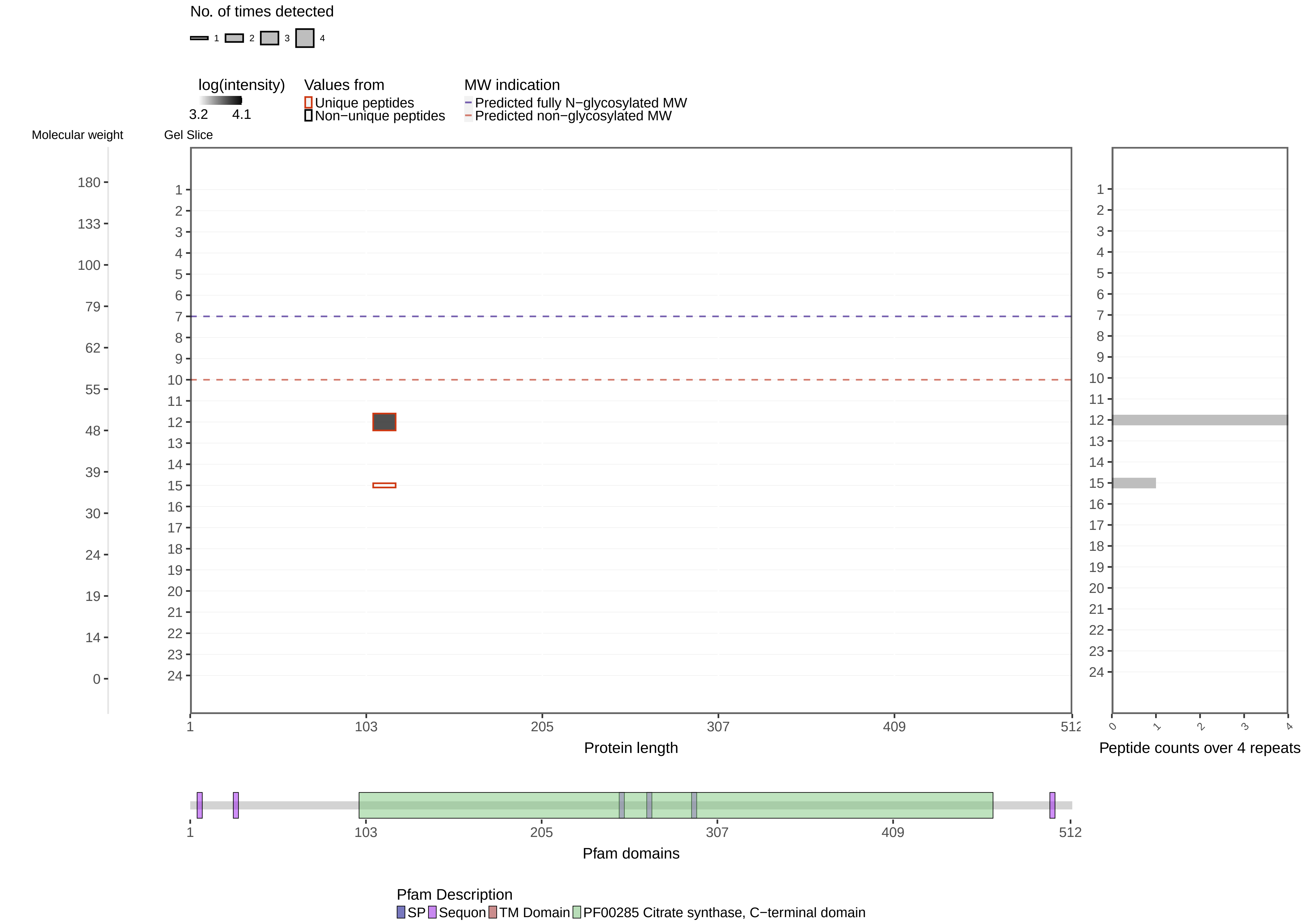


Protein sequence:

[illegible]VLPSNNLSYAENFLYMLDSLGNRSYKPNPRLARVLDILFILHAEHEMNCSTAAARHLASSGVDVYTAIAGAVGALYGPLHGGANEAVLKMLSEIGSVENIPEFIEGVKNRKRKMSGFGHRVYKNYDPRAKVIKKLADEVFSIVGRDPLIEVAIALEKAALSDEYFVKRKLYPNVDFYSGLIYRAMGFPTEFFPVLFAVPRMAGYLSHWRESLDDPDTKIMRPAQVYTGVWMRHYMPLKERSPYTETDKLGQVSVSNATKRRLAGSGA*

### NbL10g13430.1 ras−related_protein_RABD2a−like [MW = 22.54 kDa]

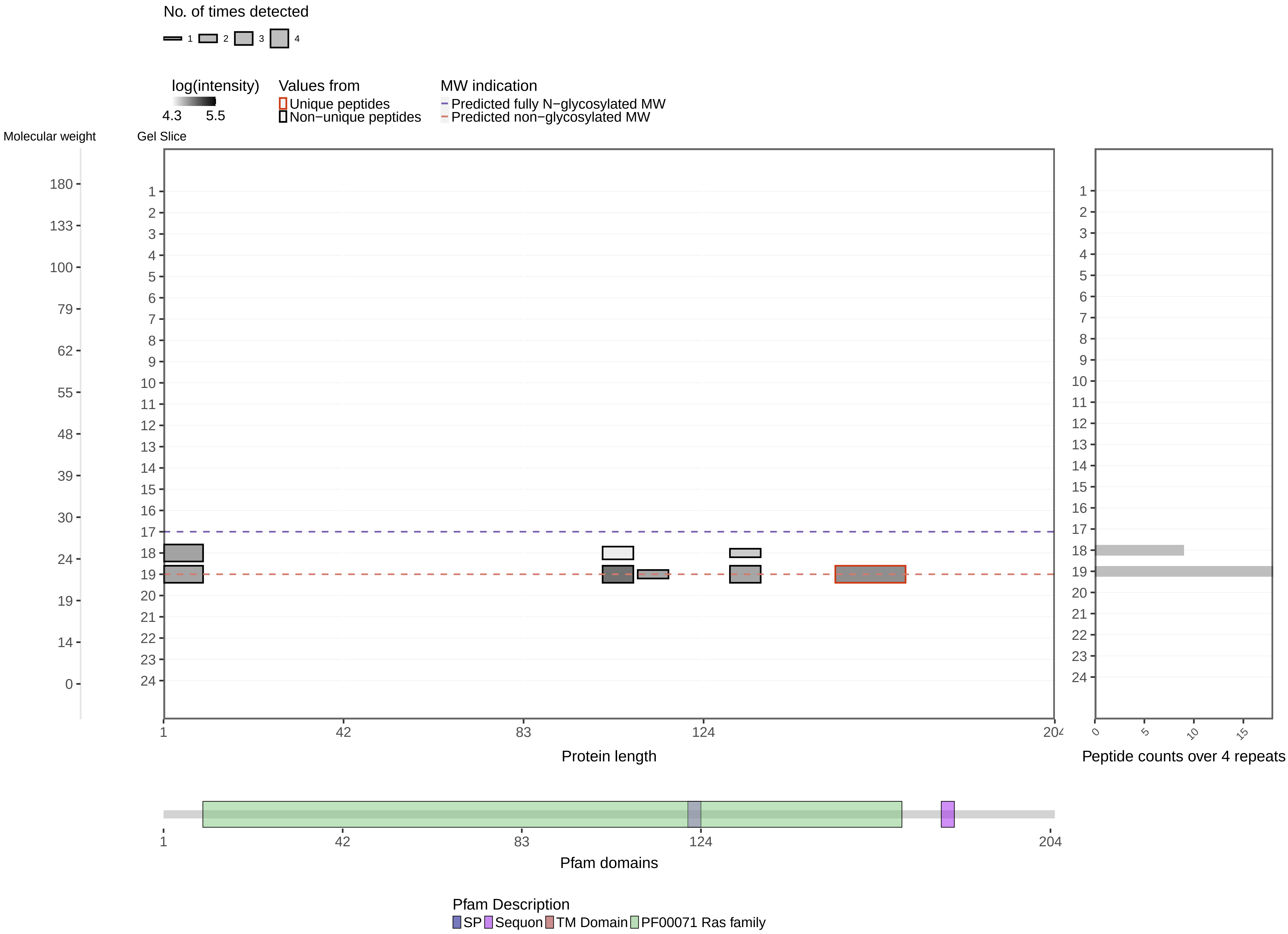


Protein sequence:

TDQESFNNVKQWLSEIDRYASDNVNKILVGNKSDLTANRVVSYETAKAFADEIGIPFLETSAKDATNVEQAFMAMTSAIKNRMASQPTNNSAKPPTVNIRGQPVTQSGGCCSS*

### NbL10g13420.1 nucleoside_diphosphate_kinase_1−like [MW = 16.16 kDa]

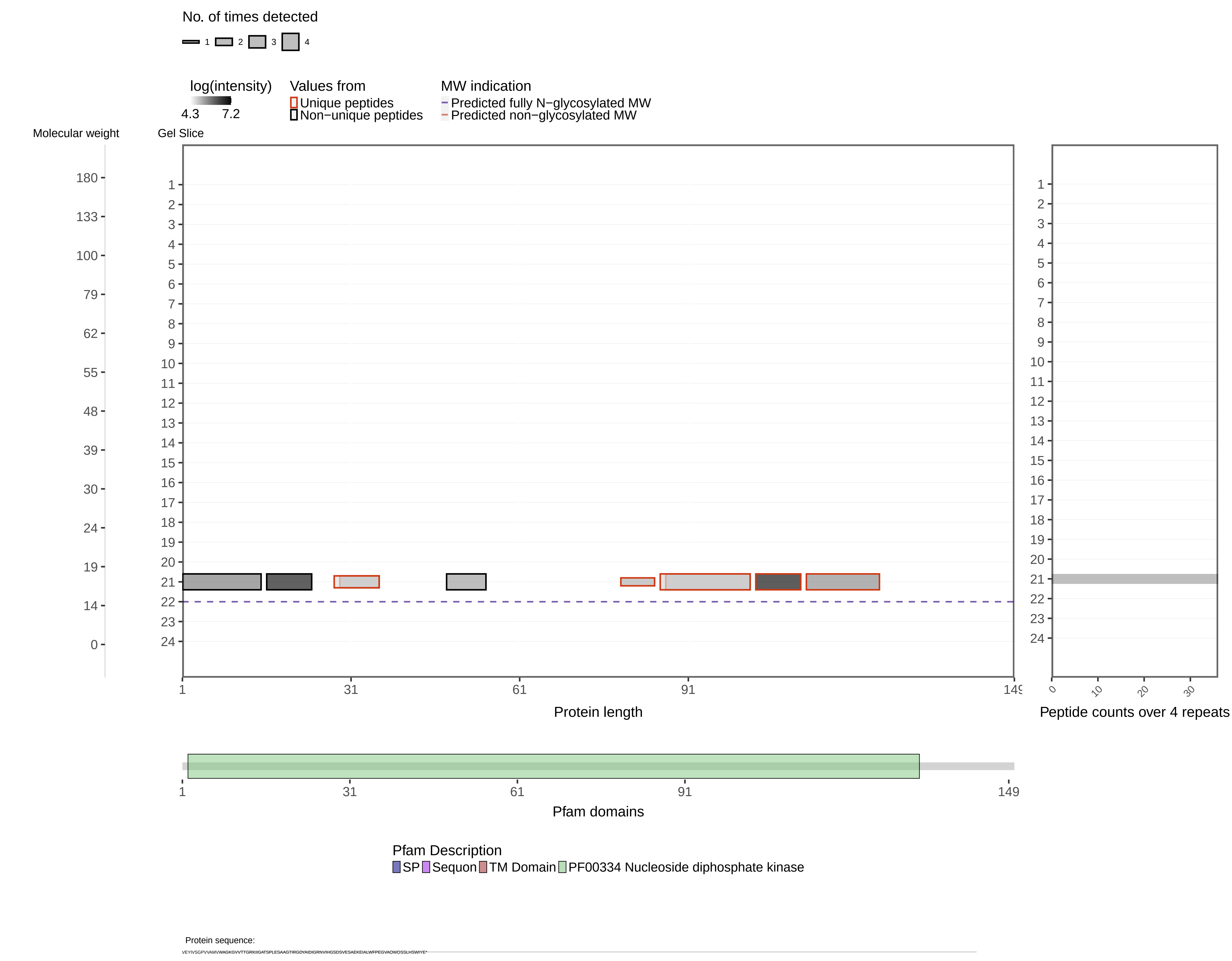

### NbL10g13330.1 ubiquitin−like_protein_5 [MW = 8.5 kDa]

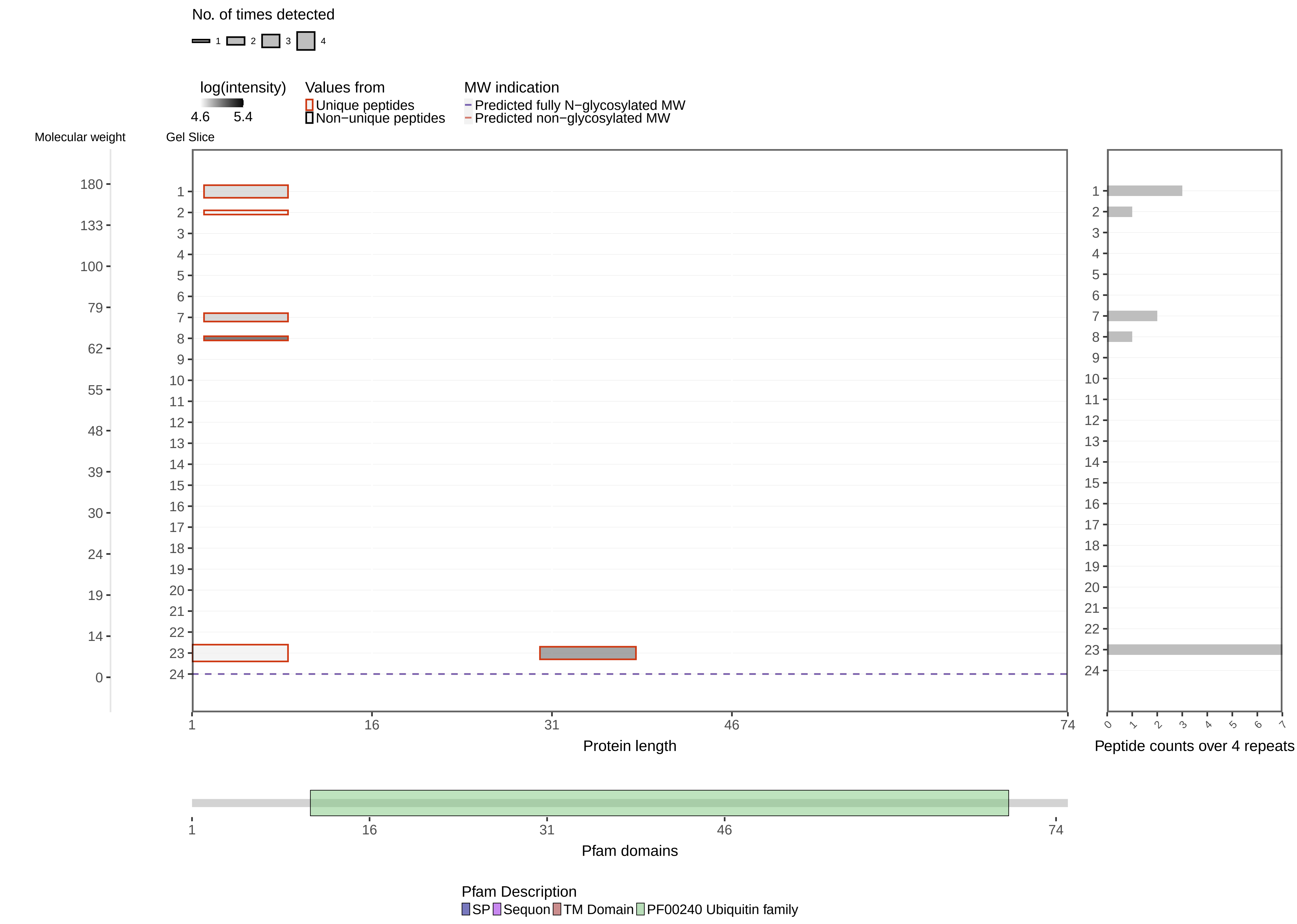


Protein sequence:

TIGDLKKLVAAQTGTRADKIRIQKWYNVYKDHIALKDYEIHDGMGLELYYN*

### NbL10g12800.1 protein_transport_protein_SEC23−like [MW = 85.22 kDa]

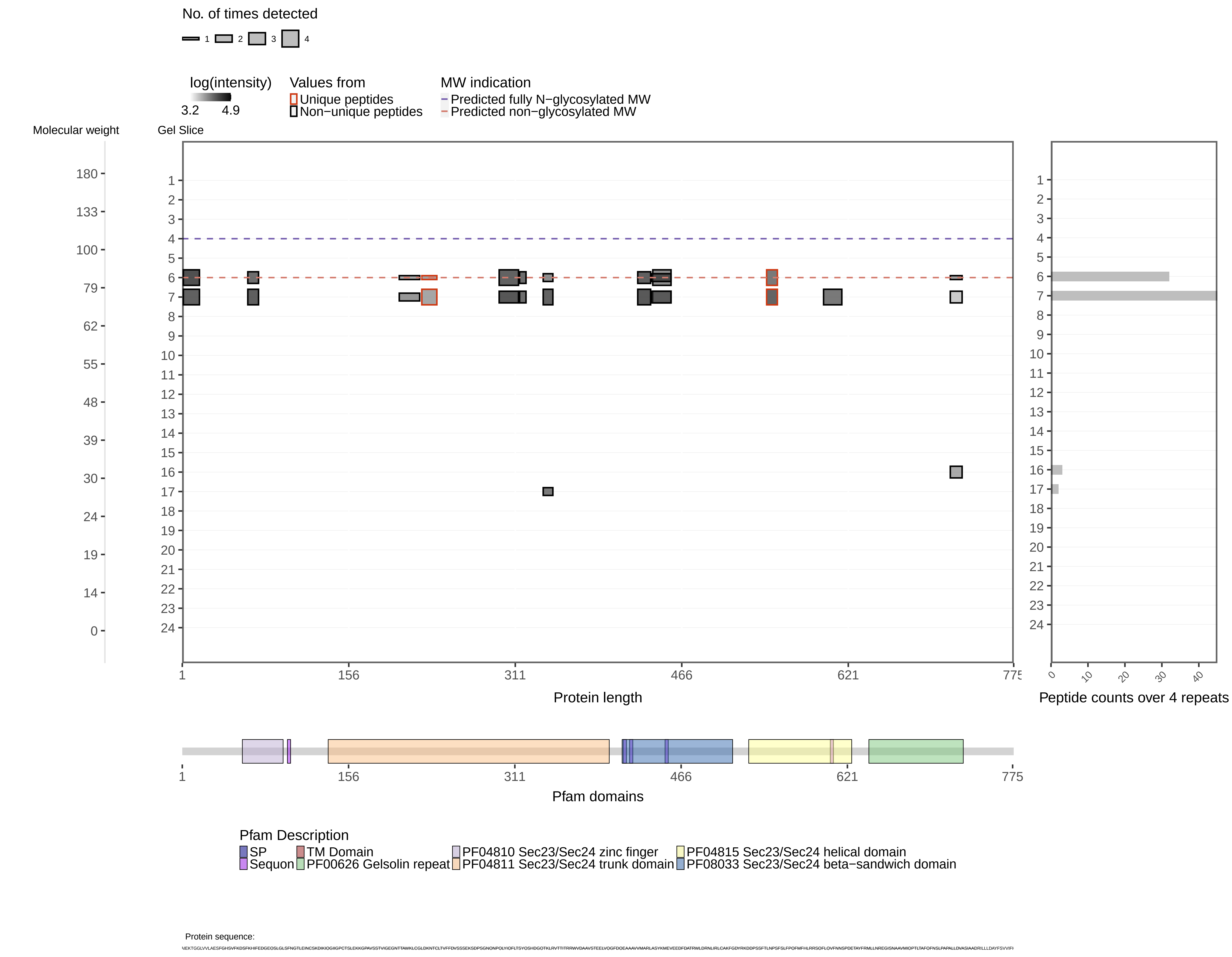

### NbL10g12780.1 adenosylhomocysteinase−like [MW = 53.14 kDa]

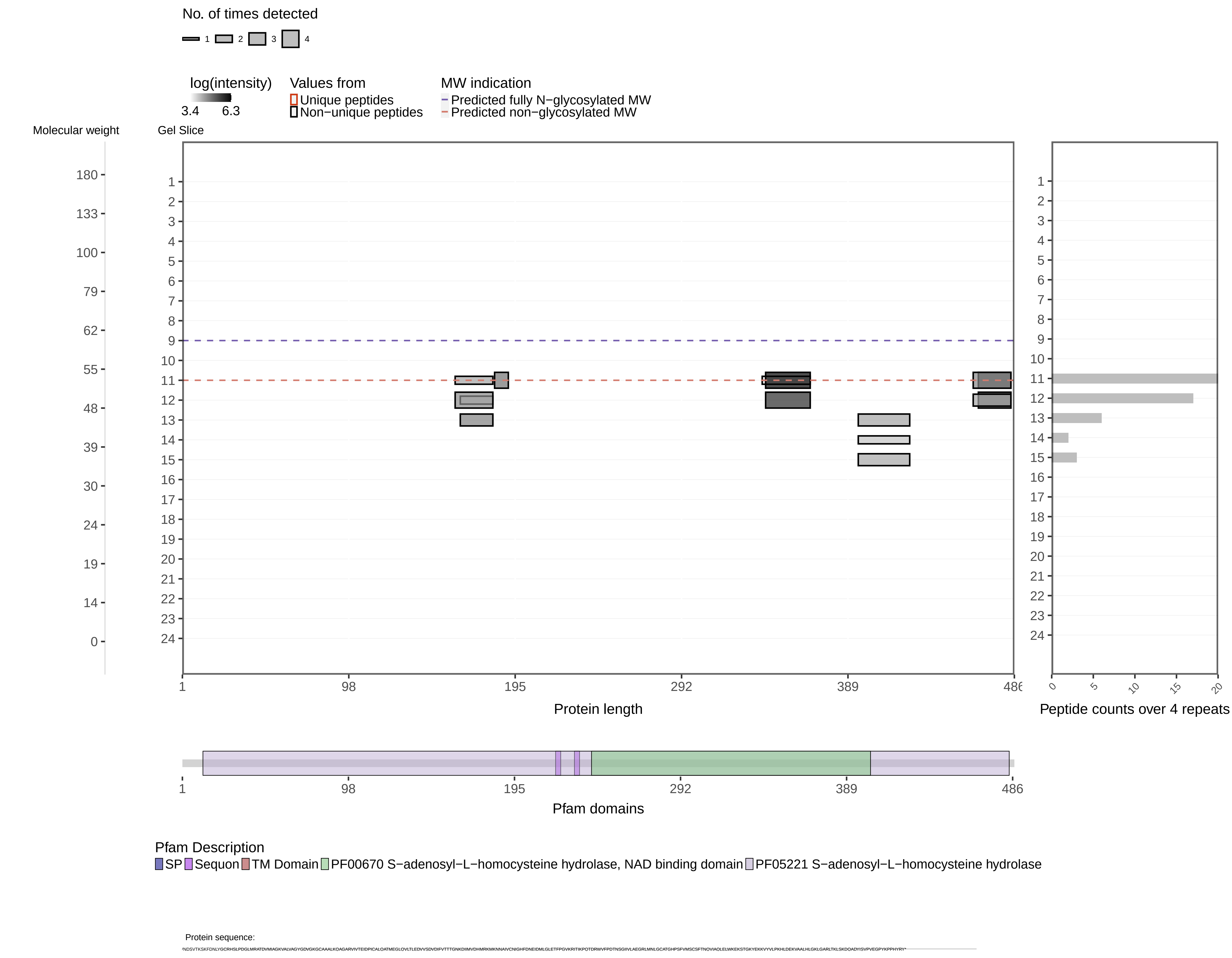

### NbL10g12770.1 serine_hydroxymethyltransferase_4−like [MW = 54.45 kDa]

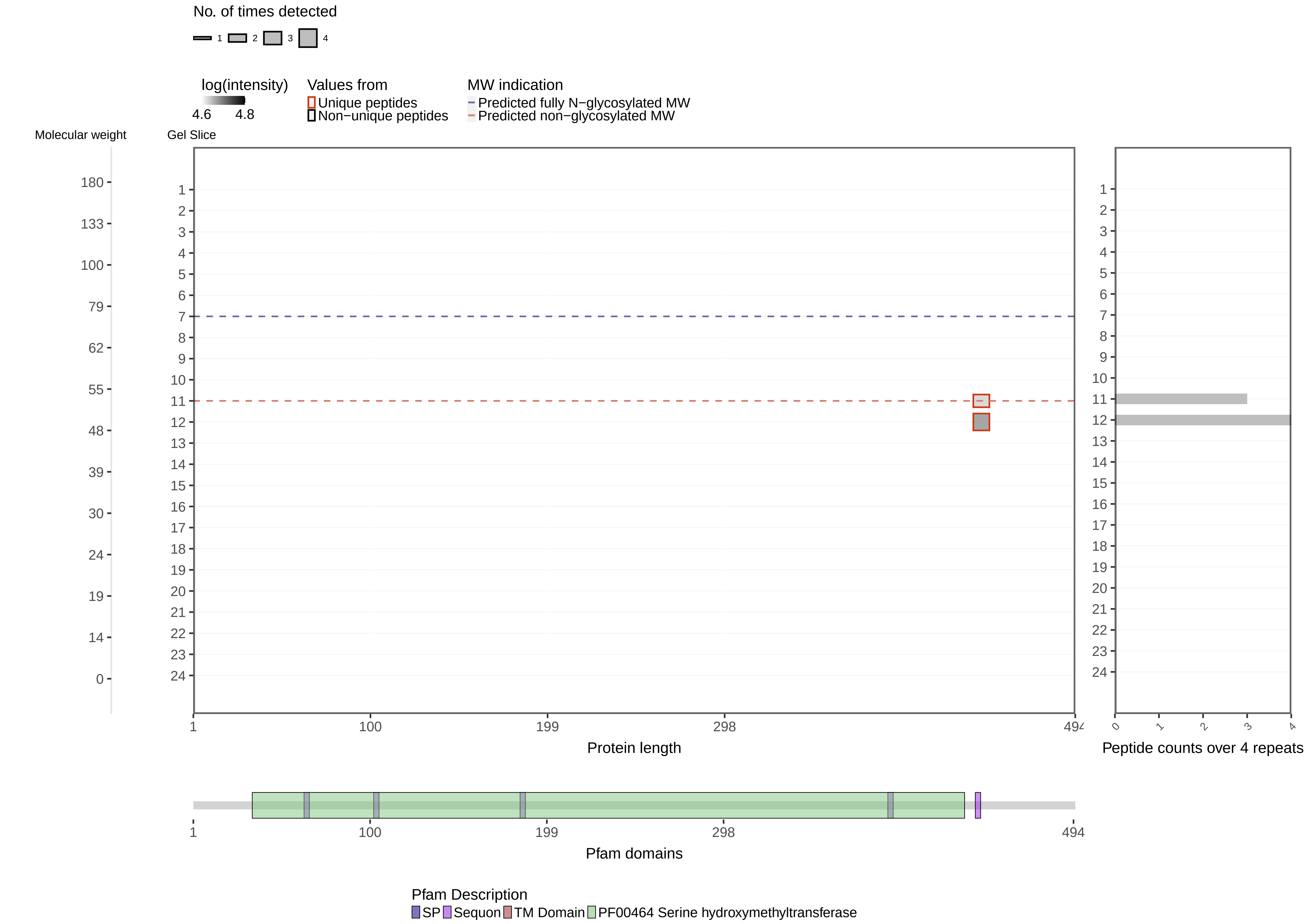


Protein sequence:

DMAHISGLVAAQEAADPFEYCDLVTTTTHKSLRGPRAGMIFYRKGPKPPKKGQPEDAVYDFEDKINFAVFPSLQGGPHNHQIGALAVALKQAATPGFKAYAKQVKANAVALGNYLMSKGYKLVTGGTENHLVLWDLRPLGLTGNKVEKLCDLANITVNKNAVFGDSSALAPGGVRIGTPAMTSRGLVEKDFEQIAEFLHRAVNITLNIQKEYGKLLKDFNKGLVNNKEIEELKADVEKFSSSFDMPGFKLSEMKYKD*

### NbL10g12670.1 60S_ribosomal_protein_L13a−4−like [MW = 23.52 kDa]

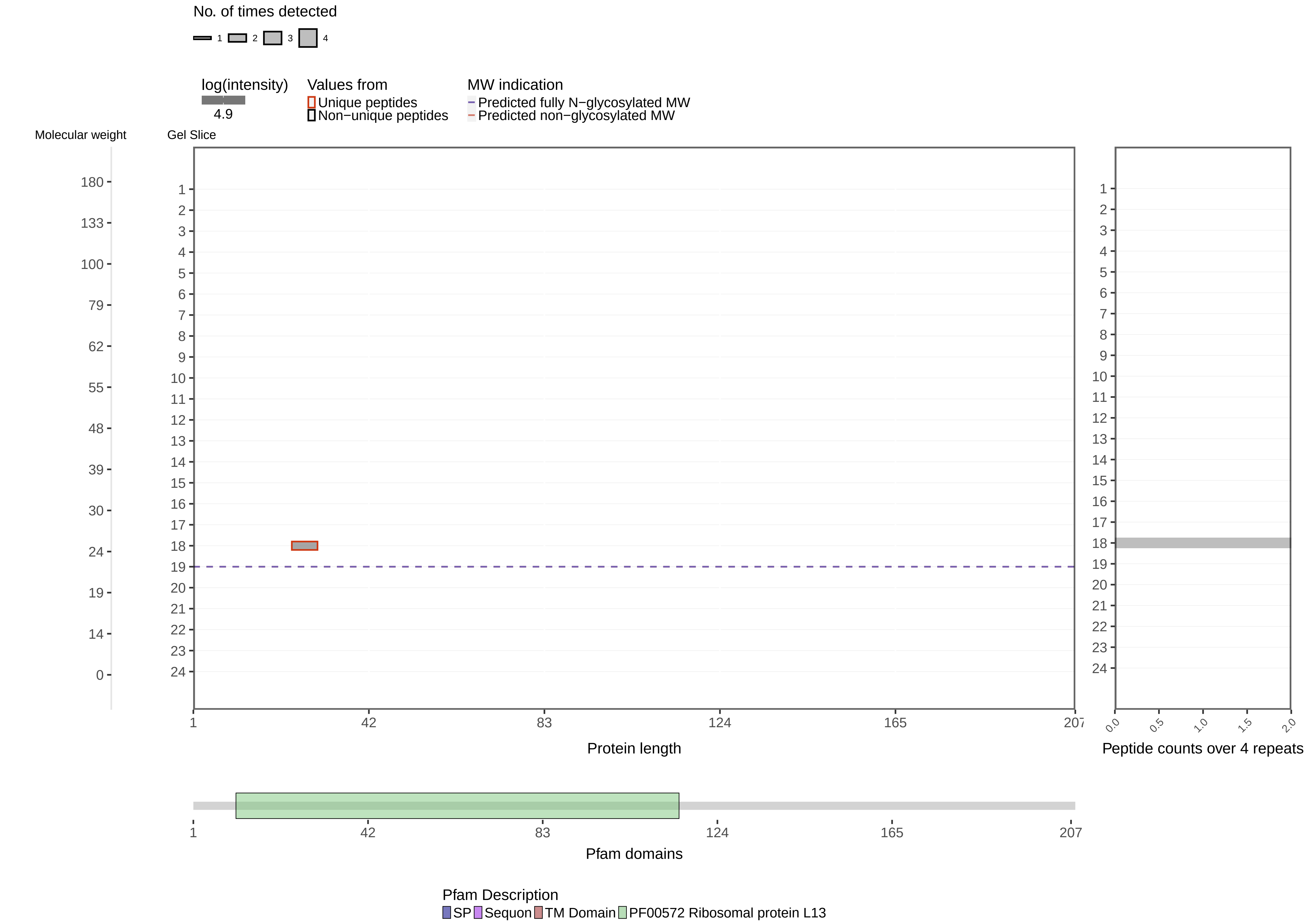


Protein sequence:

:GMIPHKTKRGAAALARLKVYEGVPPPYDKVKRMVIPDALKVLRLQAGHKYCLLGKLSSEVGWNHYDTIKELENKRKERAQVAYERRKQLAKLRVKAEKAAEEKLGPQLAVIEPIKY*

NbL10g12620.1 probable_methionine−−tRNA_ligase [MW = 90.66 kDa]

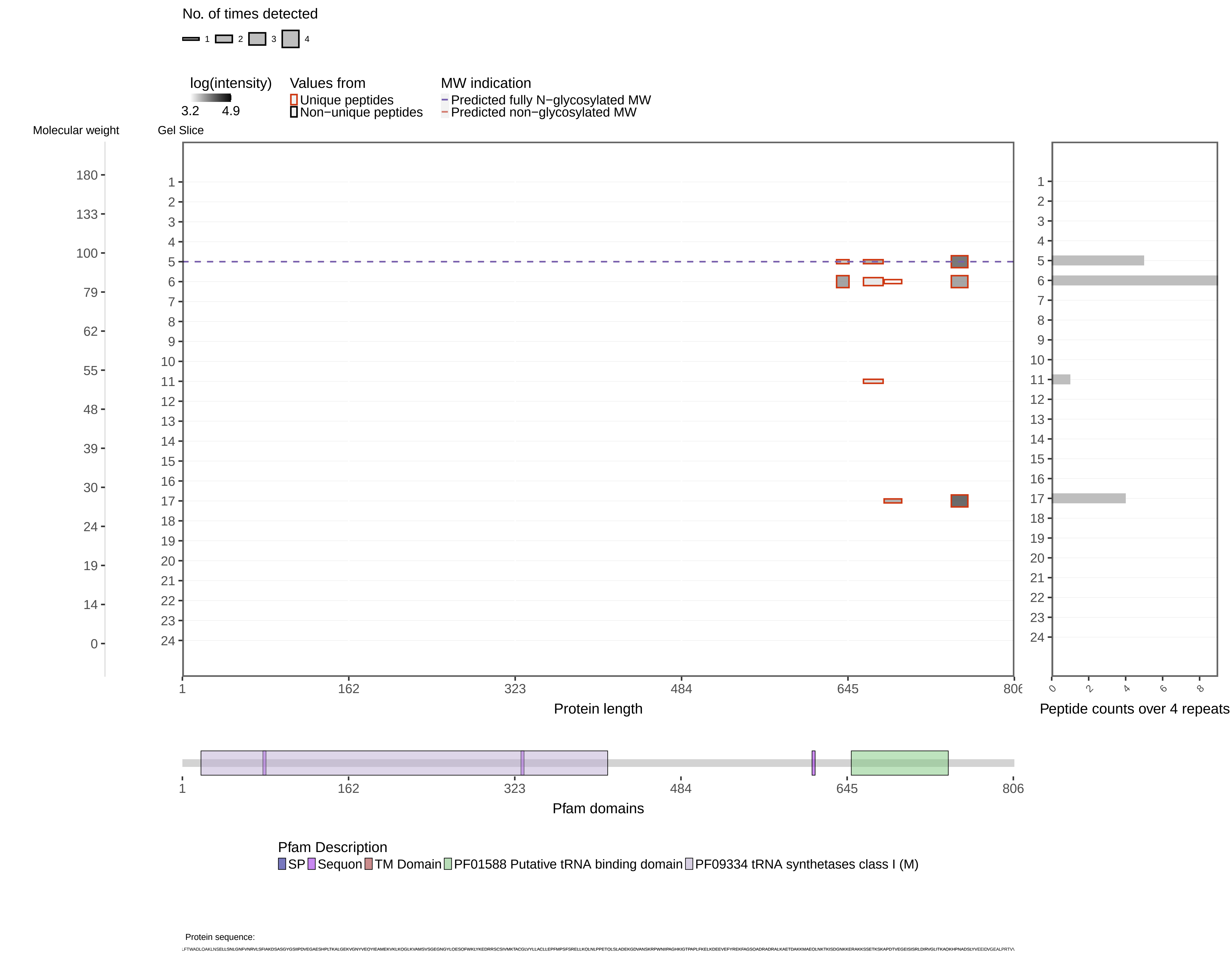

### NbL10g12550.1 dihydroxy−acid_dehydratase_chloroplastic [MW = 66.28 kDa]

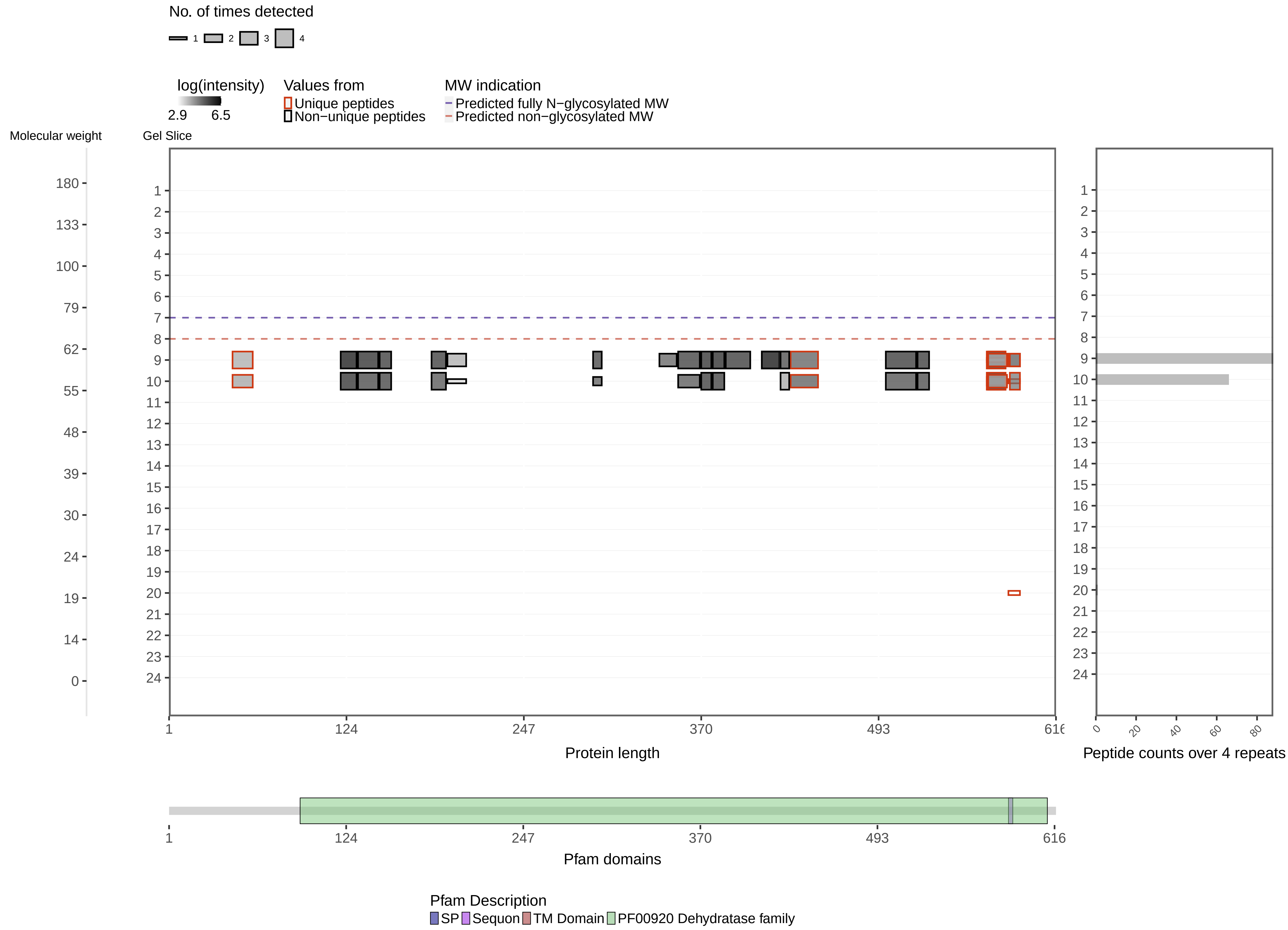


Protein sequence:

GKYLLELLKMDLKPRDIITKKSLRNAMVMVMALGGSTNAVLHLIAISRSVGLELTLDDFOKVSDAVPFLADLKPSGKYVMEDVHKIGGTPAVIRHLLELGYLDGDCMTVTGKTLAENAKLFPSLAEGOOIIRPLSNPIKETGHIOILYGNLAPEGSVAKITGKEGMYFSGPALVFEGEEAMIAAISEDPLRFKGKVVVIRGEGPKGGPGMPEMLTPTSAIMGAGLGKDCALLTDGRFSGGSHGYVVGHICPEAOEGGPIGLVODGDIITIDIOKRKMDVOLSDEEFEORRKNWTPPI

NbL10g12480.1 chaperonin_CPN60−2_mitochondrial−like [MW = 61.05 kDa]

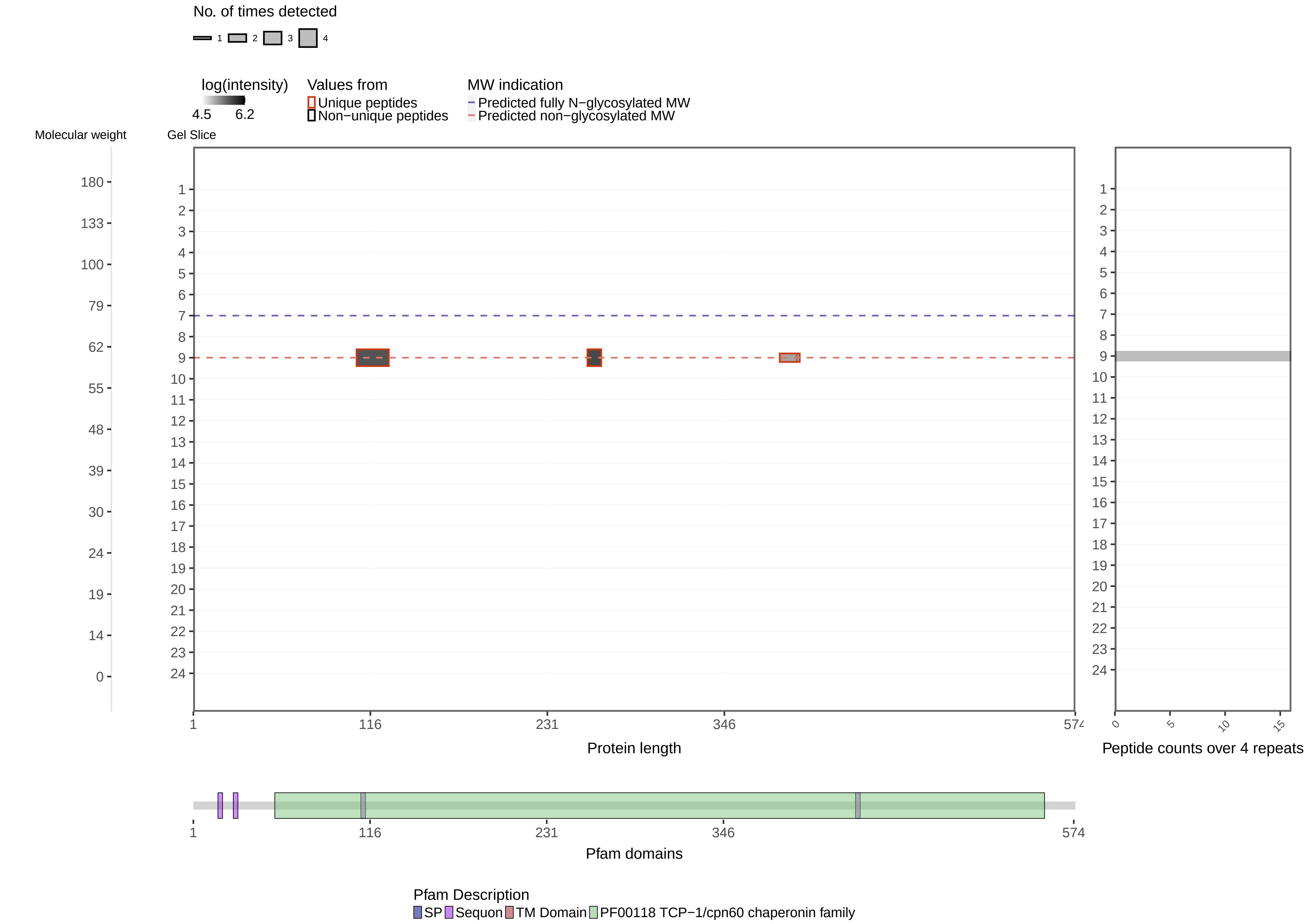


Protein sequence:

NbL10g12340.1 stress−response_A/B_barrel_domain−containing_protein_HS1 [MW = 12.15 kDa]

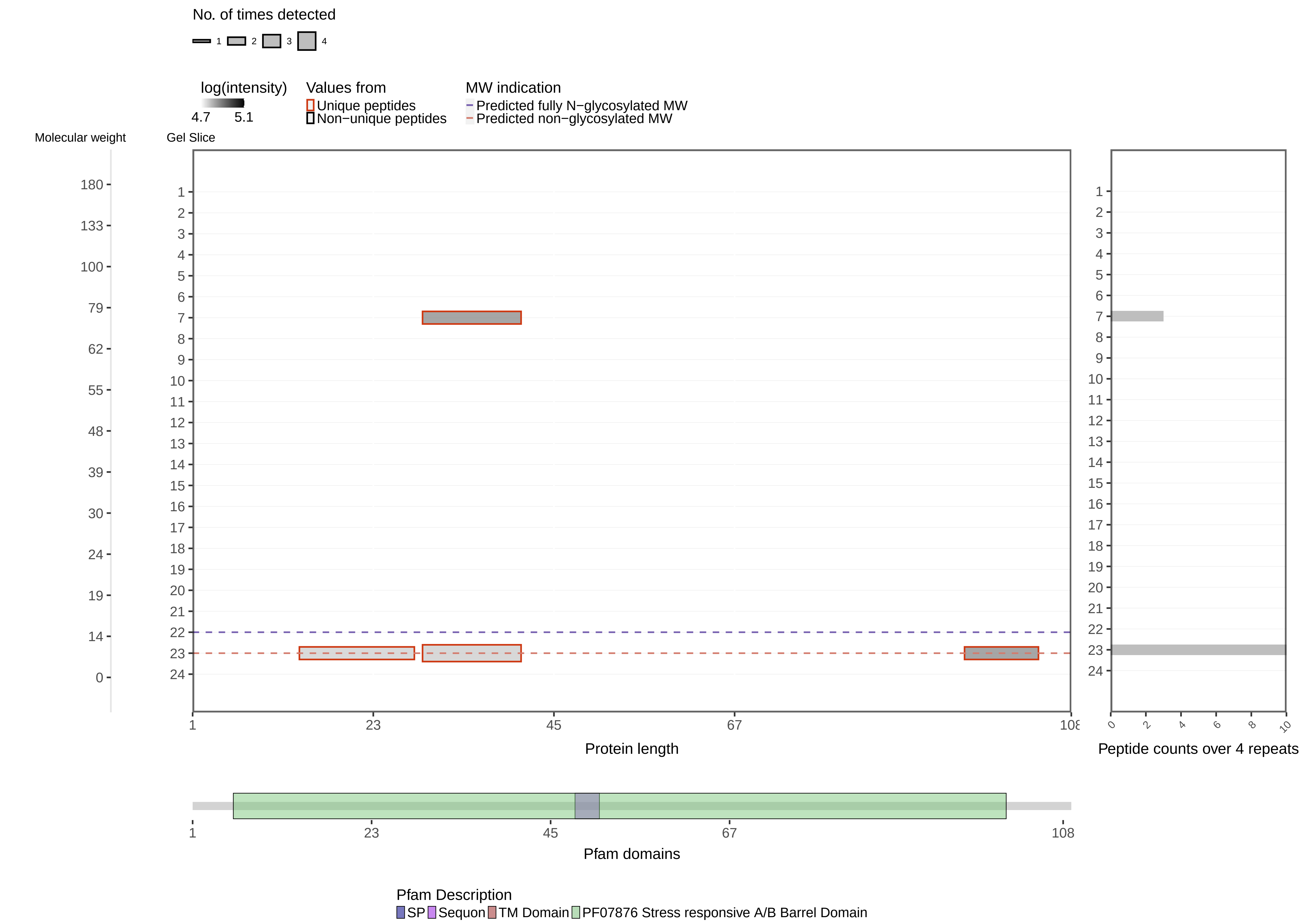


Protein sequence:

AFHWGENVSIENFHQGFTHVFESTFDSTEGIAEYIDHPAHVEYANTLLPQLEKVLVIDYKPEKVGP*

NbL10g12330.1 dihydrolipoyl_dehydrogenase_mitochondrial [MW = 53.4 kDa]

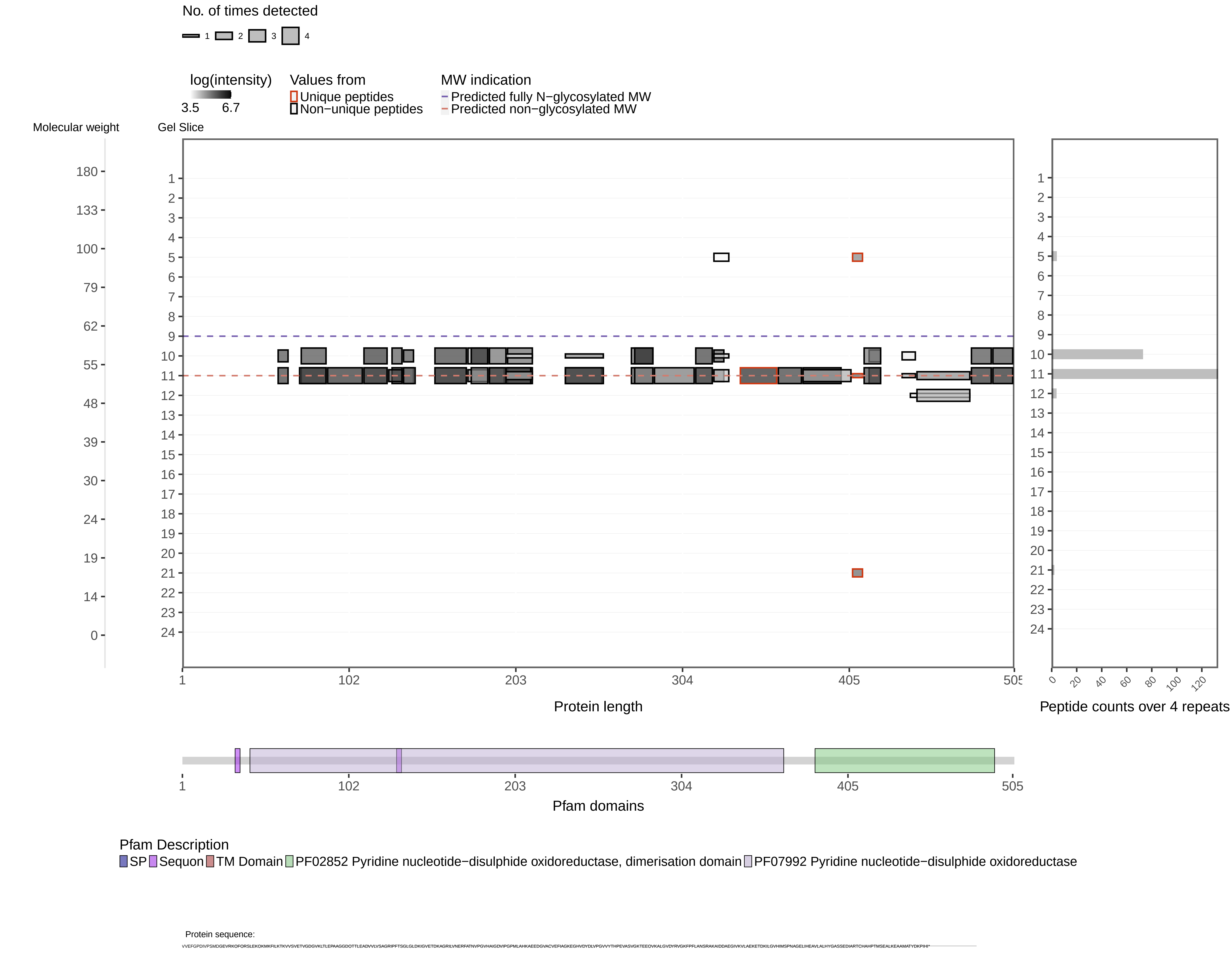

### NbL10g12200.1 dihydrolipoyl_dehydrogenase_2_chloroplastic−like [MW = 61.6 kDa]

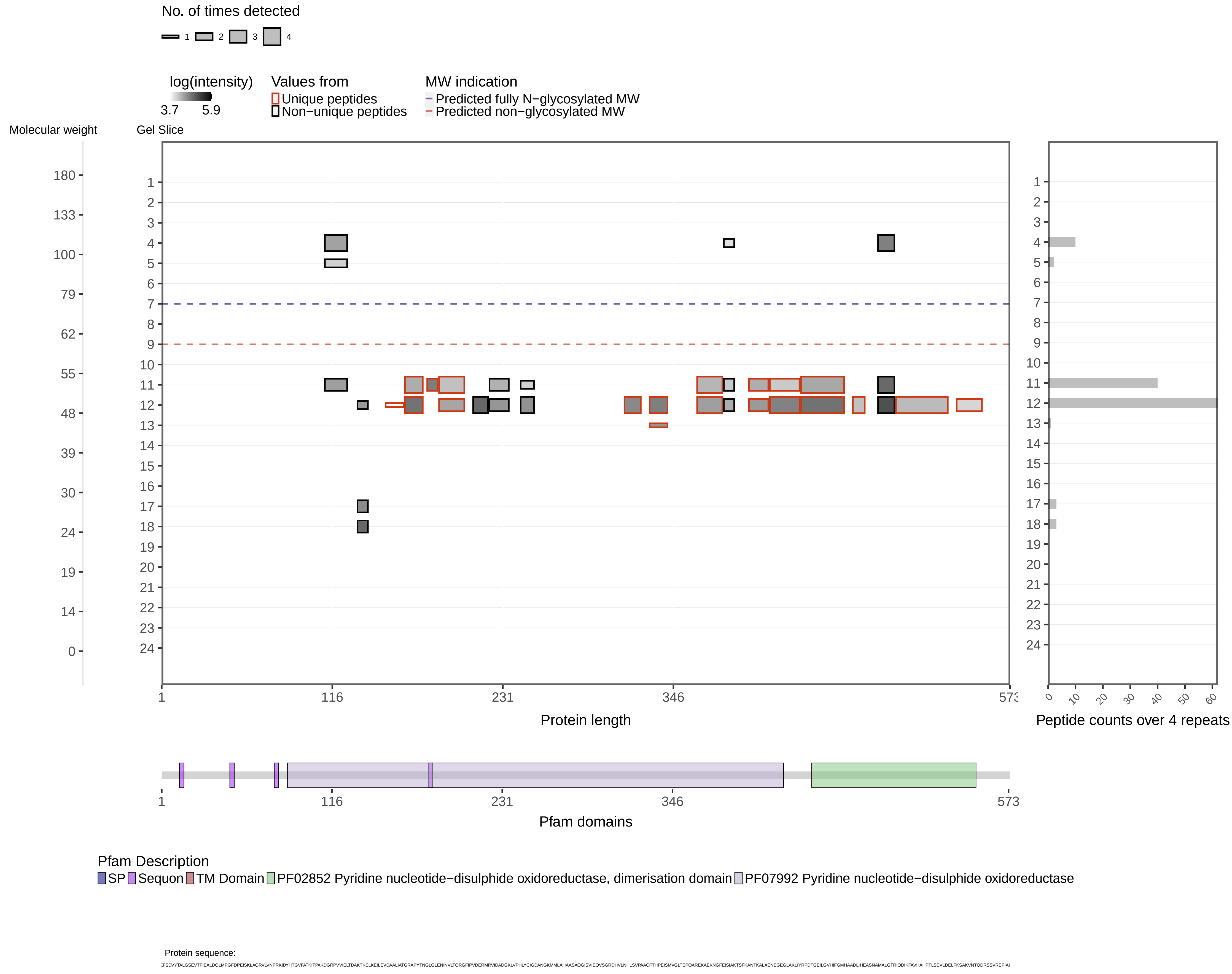

NbL10g12180.1 5−methyltetrahydropteroyltriglutamate−−homocysteine_methyltransferase−like [MW = 92.03 kDa]

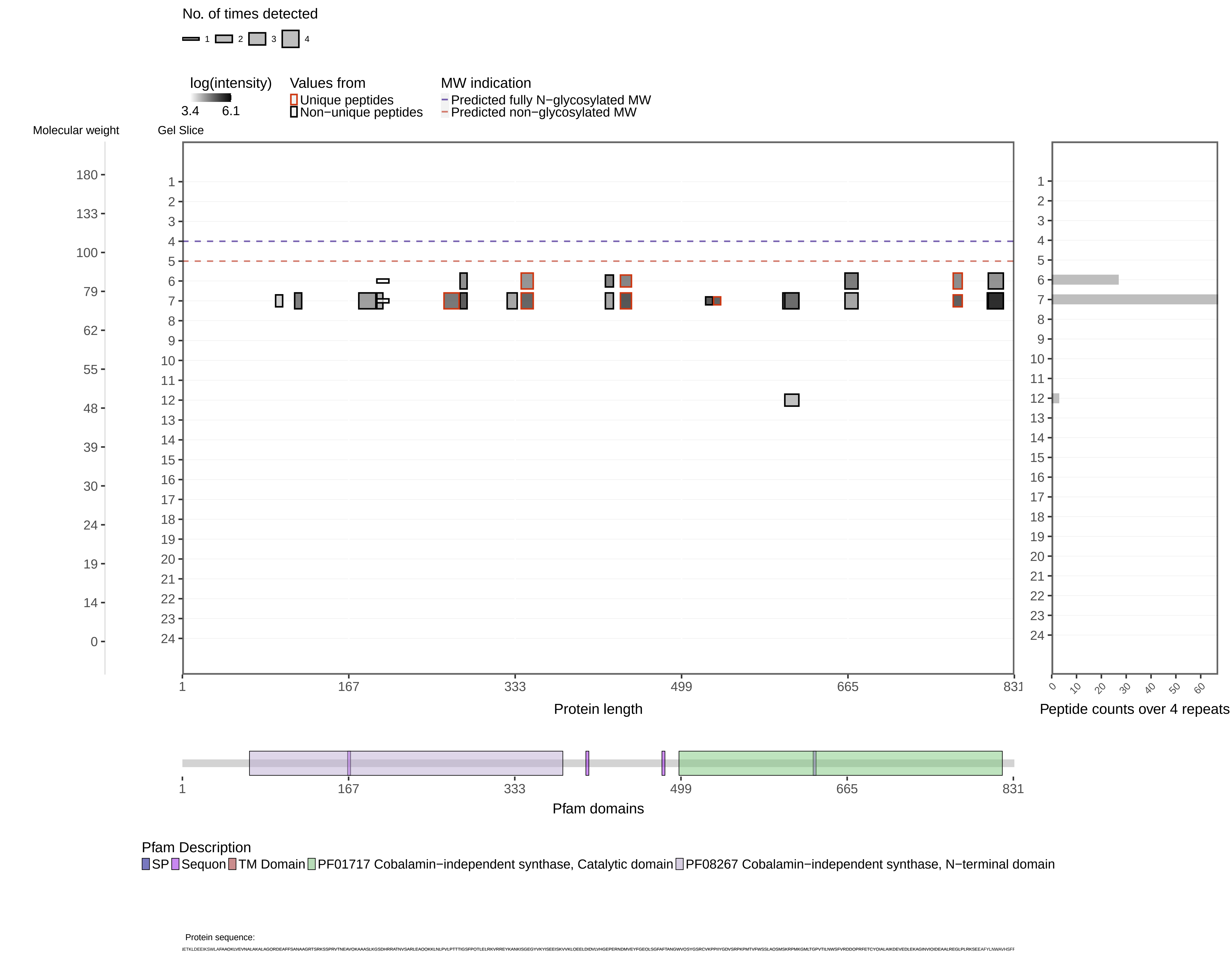

### NbL10g12050.1 folylpolyglutamate_synthase−like [MW = 68.15 kDa]

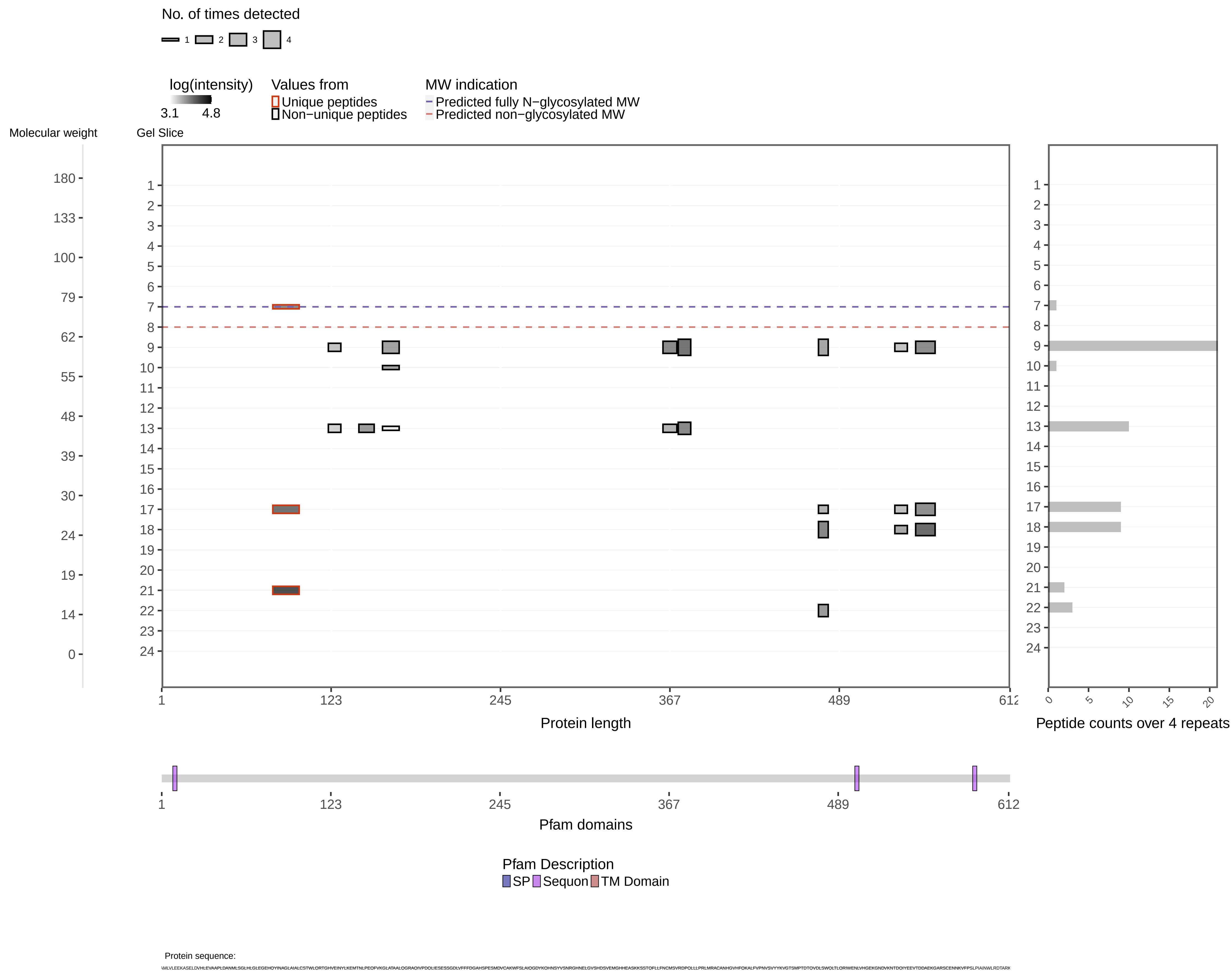

### NbL10g11880.1 pleckstrin_homology_domain−containing_protein_1−like [MW = 16.66 kDa]

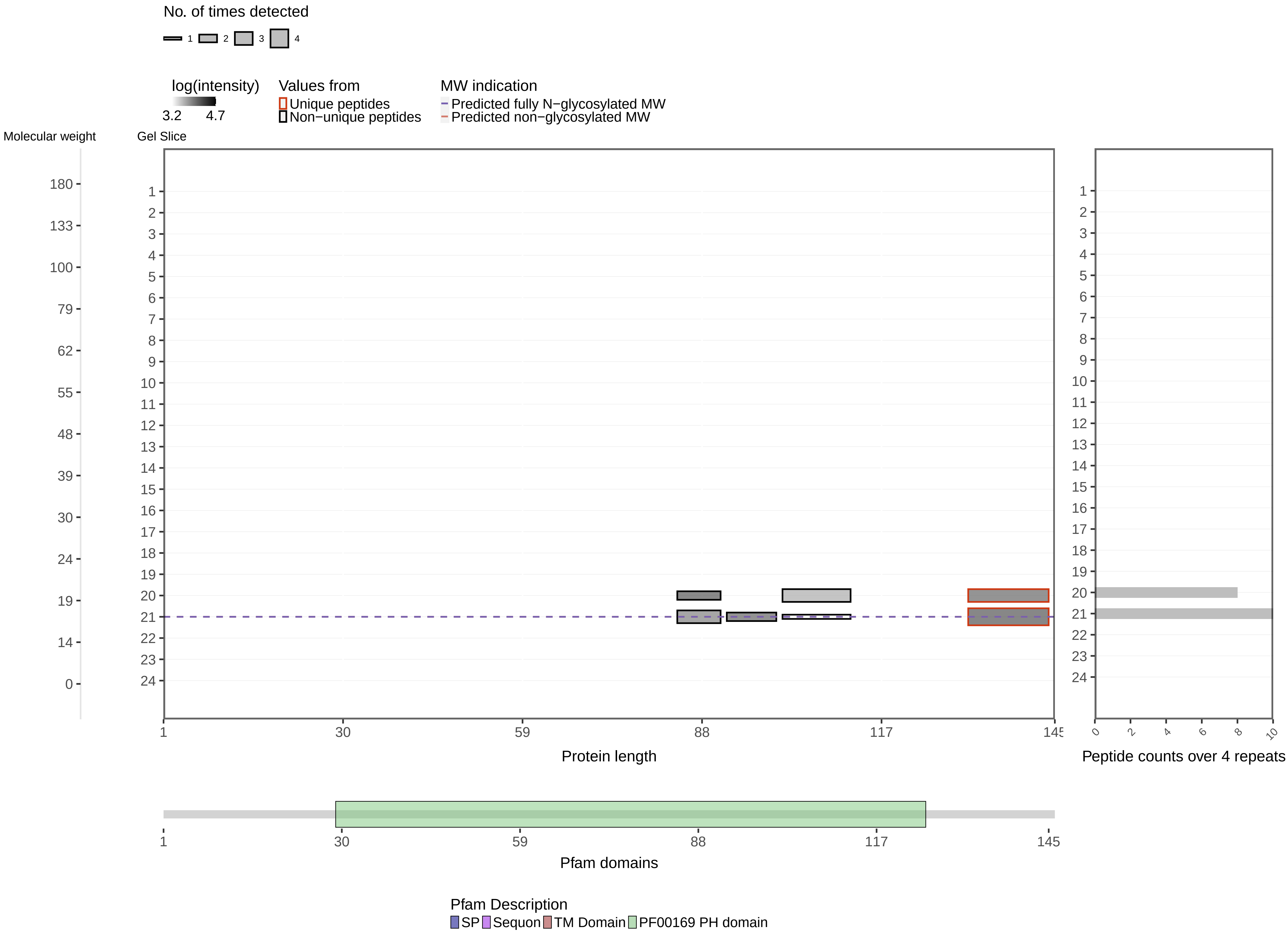


Protein sequence:

WFKEGSITGGSKPRGVIPVADCLTVKGAEDVLNKQFAFELSTRADTMYFIADSEKEKEDWINSIGRCIVQHSRSVTHDEILDYDSQK*

NbL10g11810.1 coronatine−insensitive_protein_1−like [MW = 68.81 kDa]

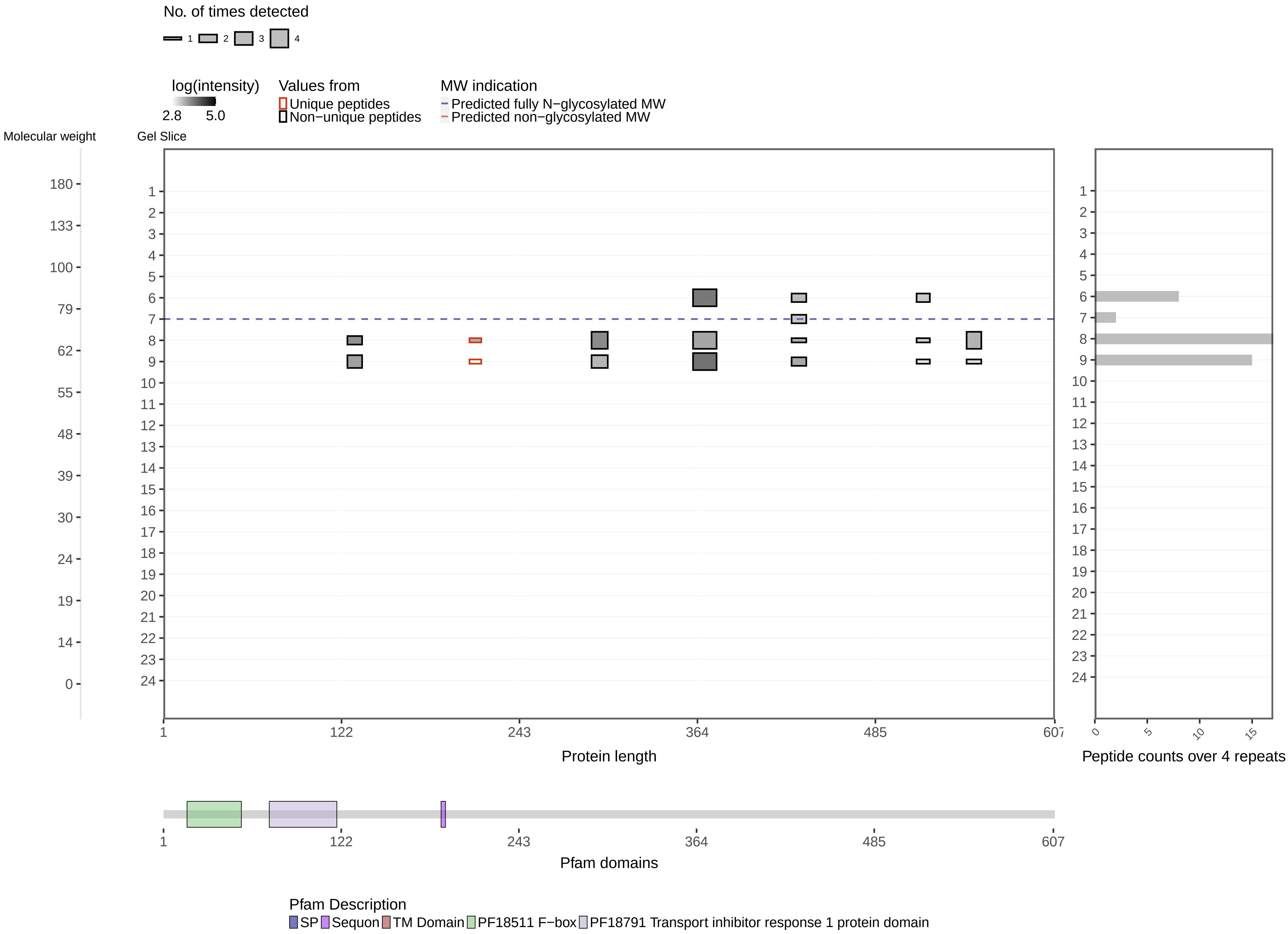


Protein sequence:

GKYEMPILFPIASRLTKLDLLYALLDTAAHCFLLQRCPNLEILETRNVVGDRGLEVLGQYCKRLKHLRIERGADDQEMEDEQGAVTHRGLTDLAKGCLELEYMAVYVSDITNEAFENIGTYLKNLCDFRLVLLDREERITDLPLDNGVRALLRGCYKLRRFALYVRPGGLTDVGLSYVGRYSPNVRWMLLGYVGESDEGLLEFSKGCPNLQKLEVRGCCFSERALALAAMQLKSLRYLWVQGYRASSAGRDLLAMARPFWNIELIPARRVVATEGNNGETVVAEHPAHILAYYSL

### NbL10g11790.1 sedoheptulose−17−bisphosphatase_chloroplastic−like [MW = 45.65 kDa]

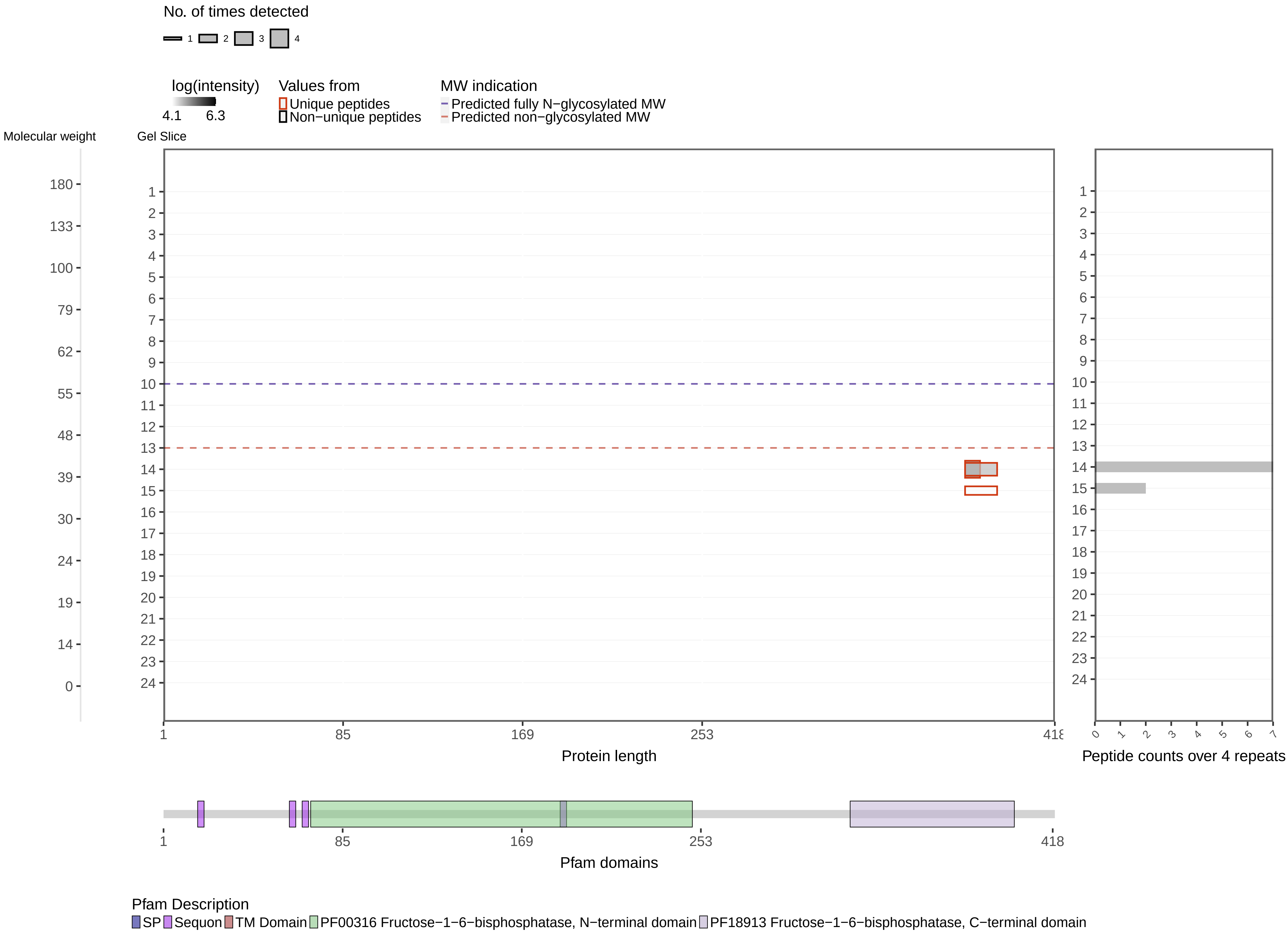


Protein sequence:

VWPGDKLTRITGRDQVAAAMGIYGPRTTYVLALKDVPGTHEFLLLDEGKWQHVKDTTEIGEGKMFSPGNLRATFDNPDYAKLIDYYVKEKYTLRYTGGMVPDVNQVFYLCTYSLKCHFEFRNSFLNFMQIIVKEKGIFTNVISPTTKAKLRLLFEVAPLGFLIEKAGGYSSDGKQSVLDRVVANLDDRTQVAYGSKNEIIRFEETLYGSSRLKSAEAVGAAA*

### NbL10g11700.1 clathrin_heavy_chain_1−like [MW = 198.65 kDa]

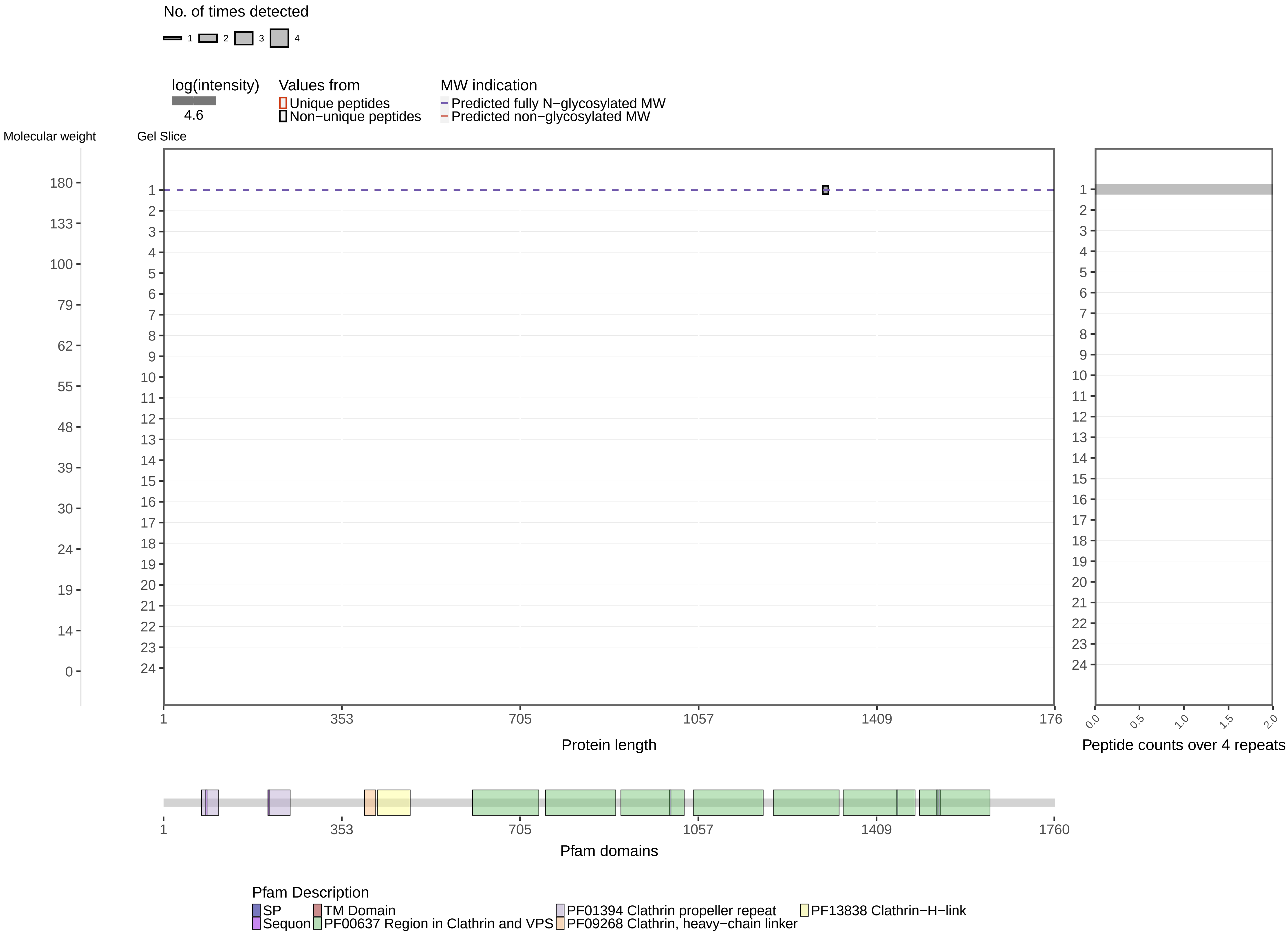


Protein sequence:

YVQKVNPGNAPLVVGQLLDDECPEDFIKGLILSVRSLLPVDPLVAECEKRNRLRLLTQFLEHLVSEGSQDVHVHNALGKIIIDSNNNPEHFLTTNPYYDSRVVGKYCEKRDPTLAVVAYRRGQCDEELINVTNKNSLFKLQARYVVERMDGDLWEKVLTPENEFRRQLIDQVVSTALPESKSPEQVSAAVKAFMTADLPHELIELLEKIVLQNSAFSGNFNLQNLLILTAIKADPSRVMDYINRLDNFDGPAVGEVAVEAQLYEEAFAIFKKFNLYVQAVNVLLDNIRDINRAVEFAFR

NbL10g11680.1 probable_methyltransferase_PMT23 [MW = 68.84 kDa]

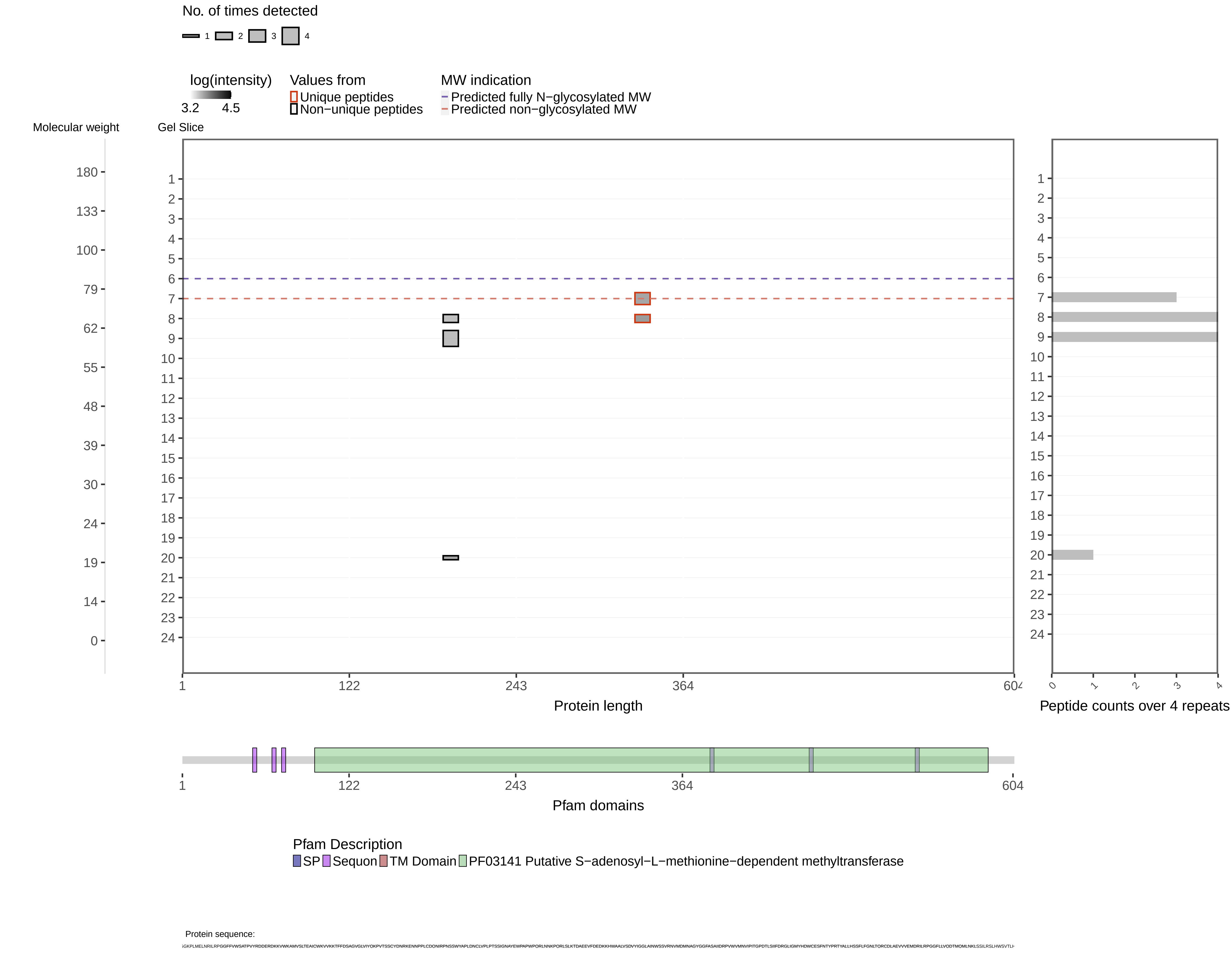

NbL10g11470.1 peroxidase_52−like [MW − SP = 31.95 kDa]

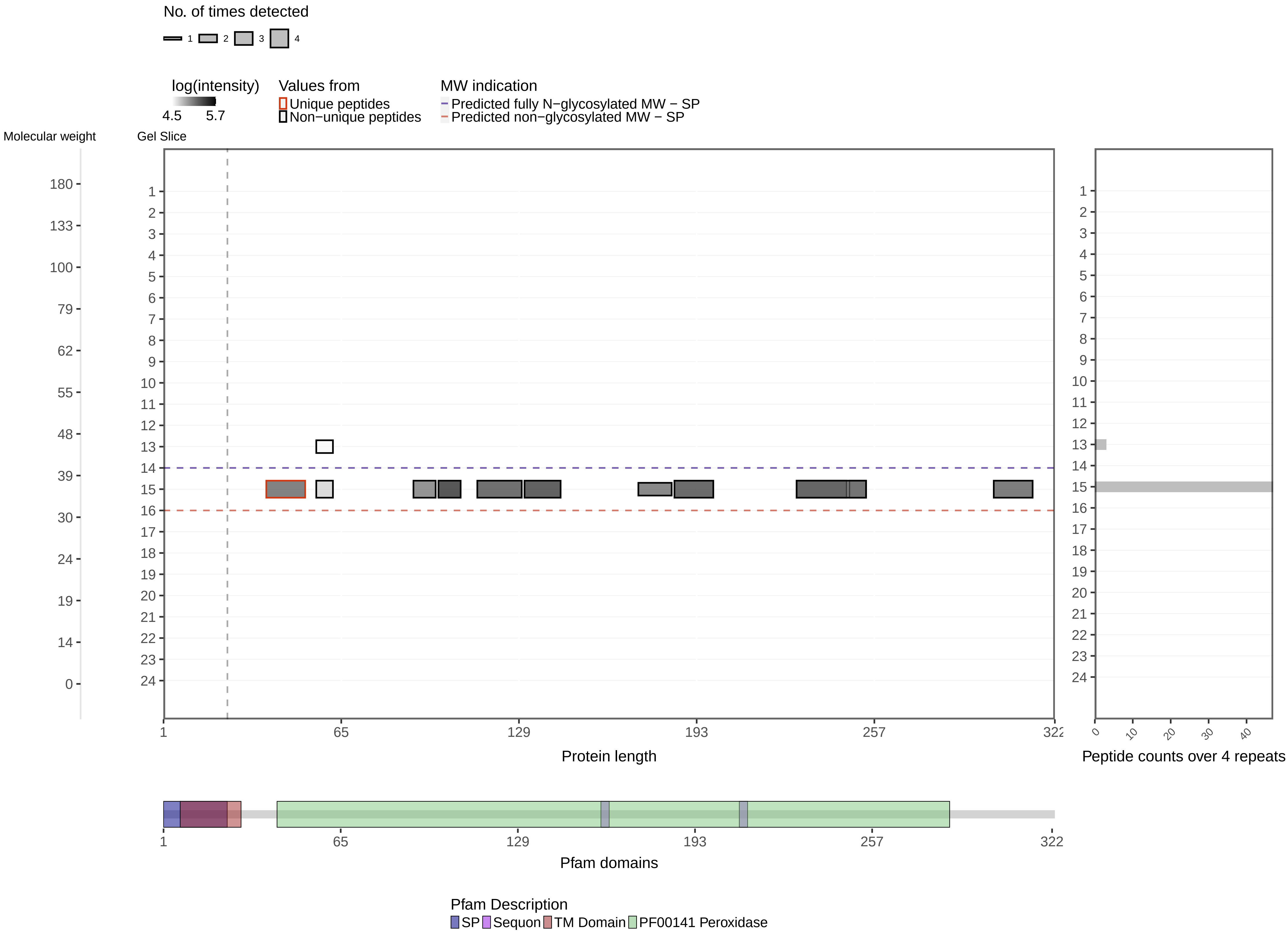


Protein sequence:

ARTASOSAANSSIPPASSNLNRLISSFSAVGLSTKDMVALSGAHTIGOARCTSFRARIYNETNNLDASFARTROSNCPRSSGSGDNNLAPLDLOTPNKFDNNYFKNLVNKKGLLHSDOOLFNGGSADSIVMSYSNNPSSFSSDFVTAMIKMGDIRPLTGSNGEIRKNCRRIN*

### NbL10g11440.1 superoxide_dismutase_[Mn]_mitochondrial−like [MW = 25.51 kDa]

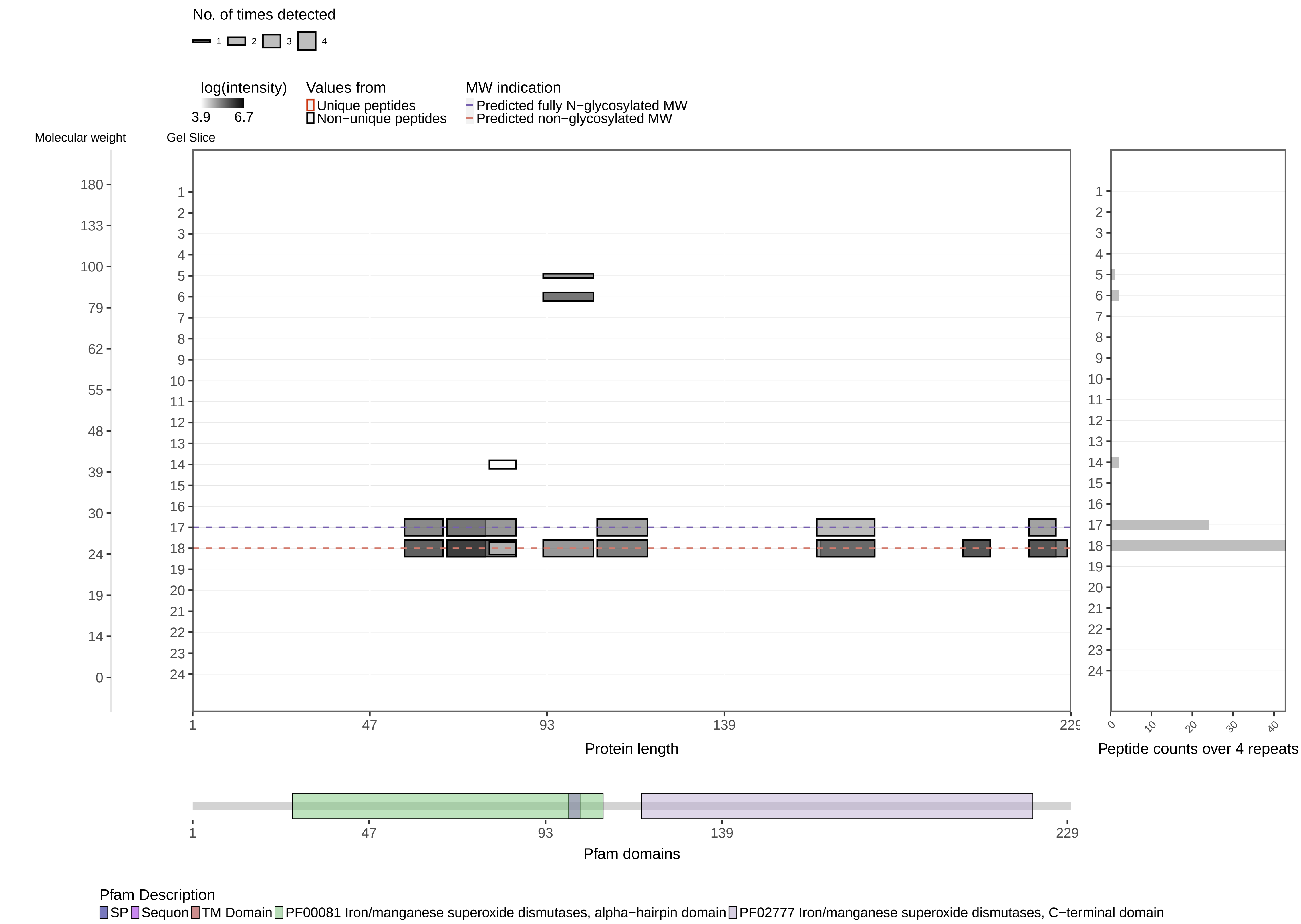


Protein sequence:

FWKNLAPIQEGGGEPPKGSLGSAIDTNFGSLEALIQKMNAEGAALQGSGWAWLALDKELKRLAIVTTSNQDPLVTKGPNLVPLFGIDVWEHAYYLQYKNVRPDYLKNIWKVMNWKYAAEVYEKECP*

### NbL10g11330.1 ATP_synthase_subunit_delta'_mitochondrial [MW = 21.51 kDa]

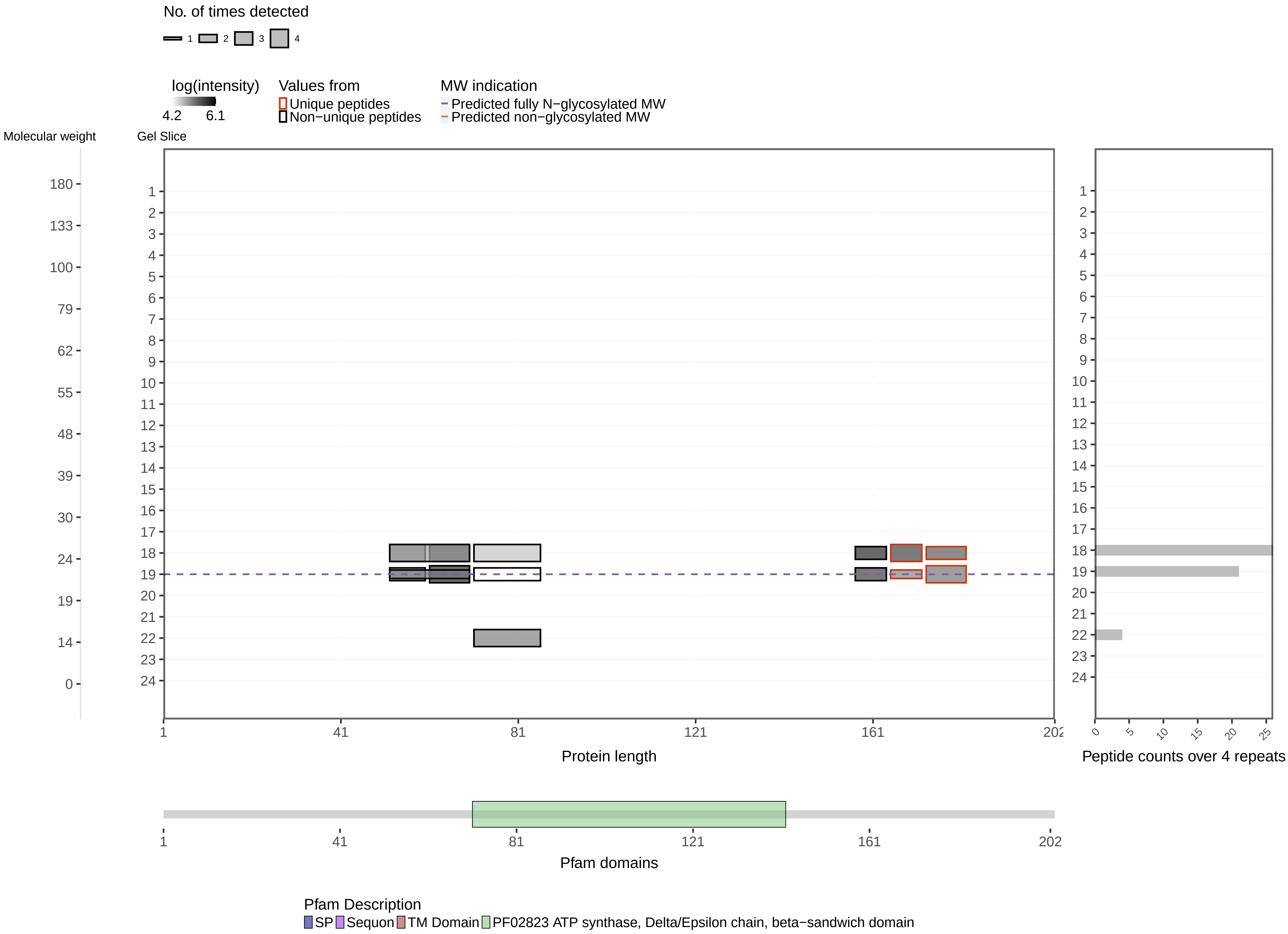


Protein sequence:

EVDMVIIPATTGQMGVLPGHVATIAELKPGVLSVHEGNDVTKYFVSGGFAFVHANSFADIIAVEAVPLDRIDPNLVQKGLAEFTQKLGTASTDVEKAEAQIGVDVHSALNAALTG*

### NbL10g10970.1 inositol−3−phosphate_synthase−like [MW = 63.13 kDa]

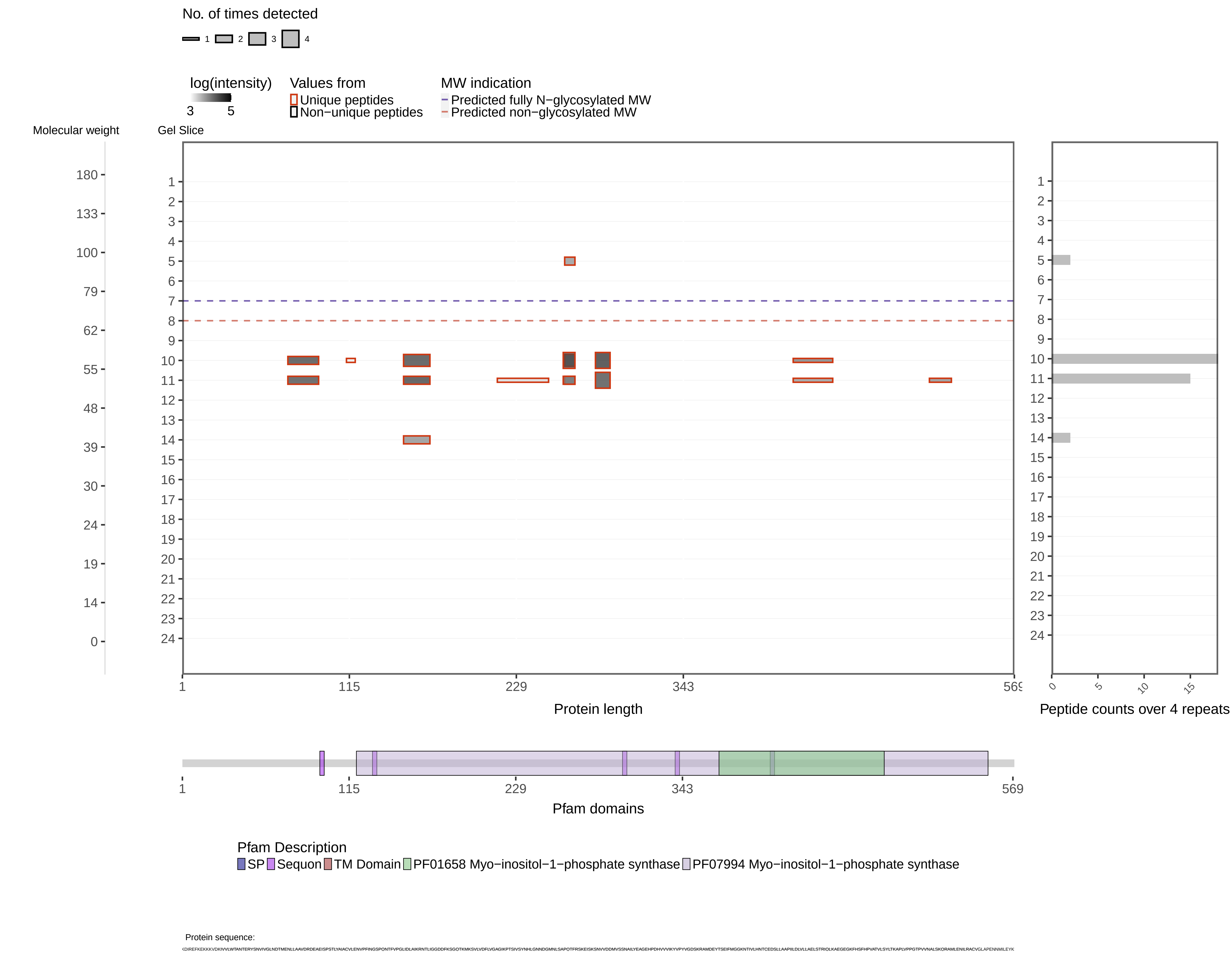

NbL10g10830.1 polyadenylate−binding_protein_2−like [MW = 24.81 kDa]

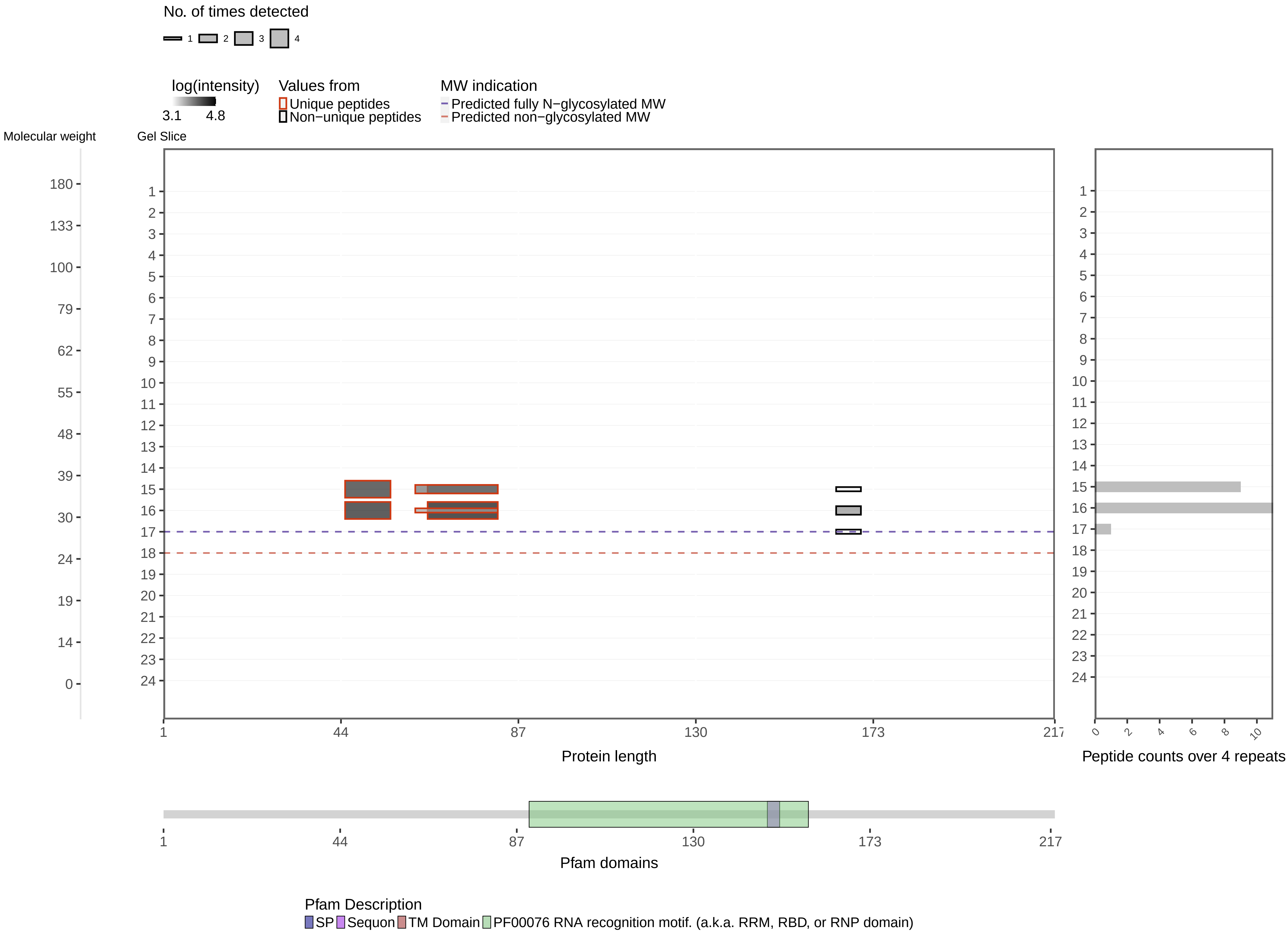


Protein sequence:

VDYACTPEEVQQHFQACGTVNRVTILTDKFGQPKGFAYVEFVEQEAIQVALQLNESELHGRQLKVMPKRTNVPGMKQFRGRRFNPYMAYGFRRPYTPPYFYSPYGFGKVPRFRRPTRFMPYY*

### NbL10g10770.1 histone_H3.3 [MW = 15.41 kDa]

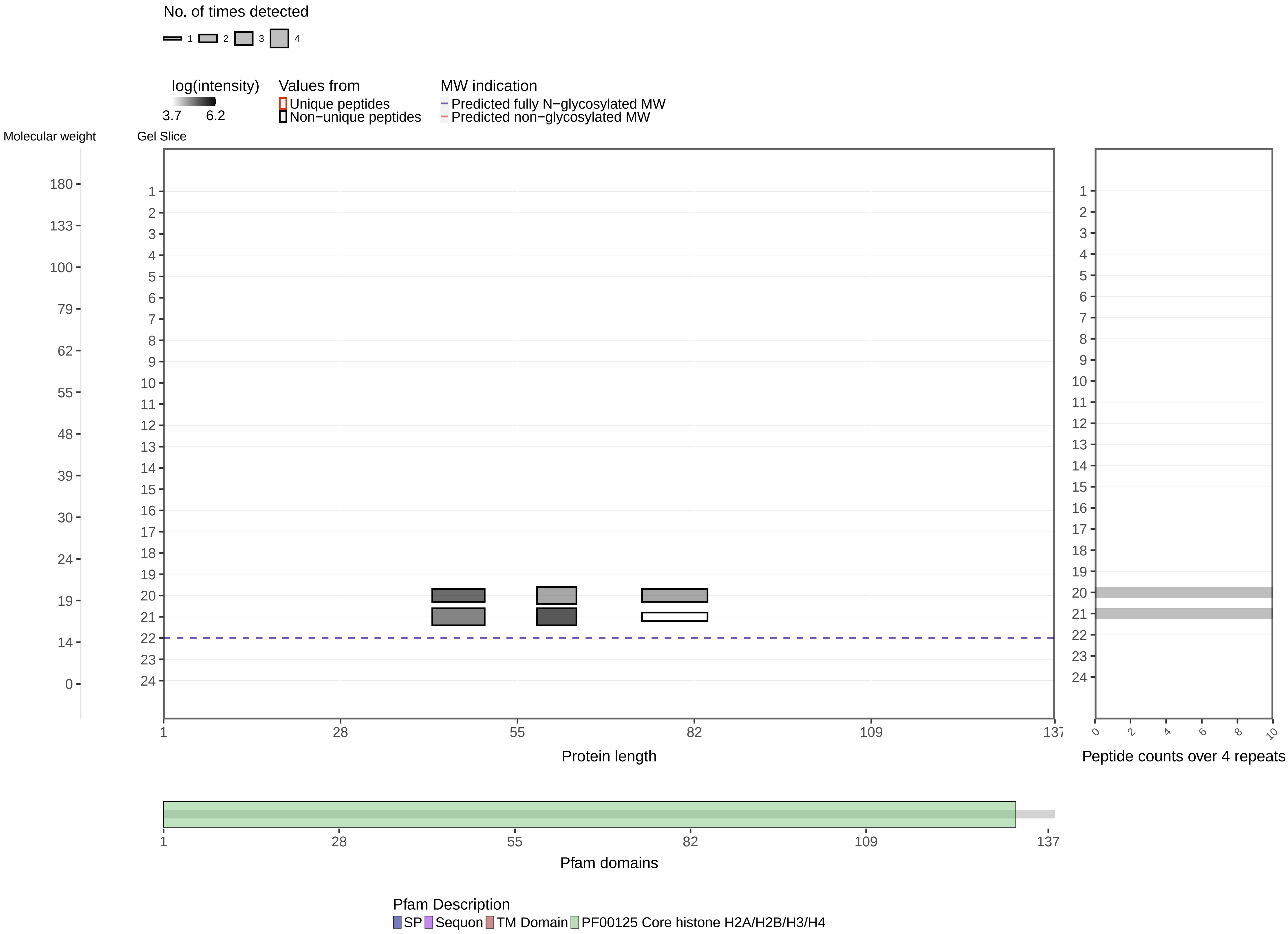


Protein sequence:

KYQKSTELLIRKLPFQRLVREIAQDFKTDLRFQSHAVLALQEAAEAYLVGLFEDTNLCAIHAKRVTIMPKDIQLARRIRGERA*

NbL10g10720.1 dihydrofolate_reductase−like [MW = 25.61 kDa]

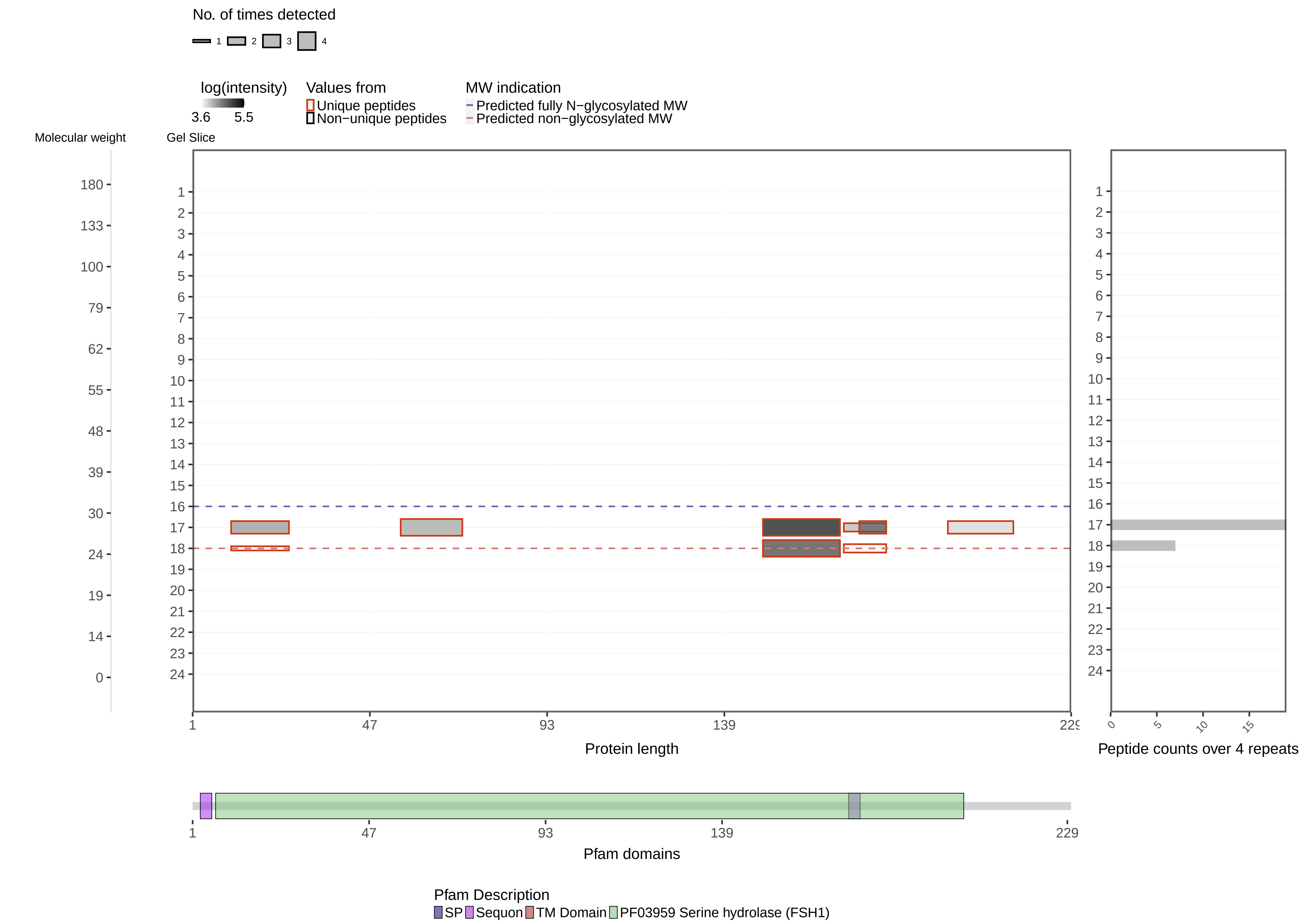


Protein sequence:

VLGFSQATIVTATMPGMQWERVALTKVPNIKFVIIISGLMFGAPGFGCPKLAANAYSSPIETPSLHFIGEKEANKTSEEELVKCFVNPVVIHHPEGHKVPNLDAESVETVITFINKVKKIKMALQGNSKM*

### NbL10g10650.1 O−fucosyltransferase_31−like [MW = 23.55 kDa]

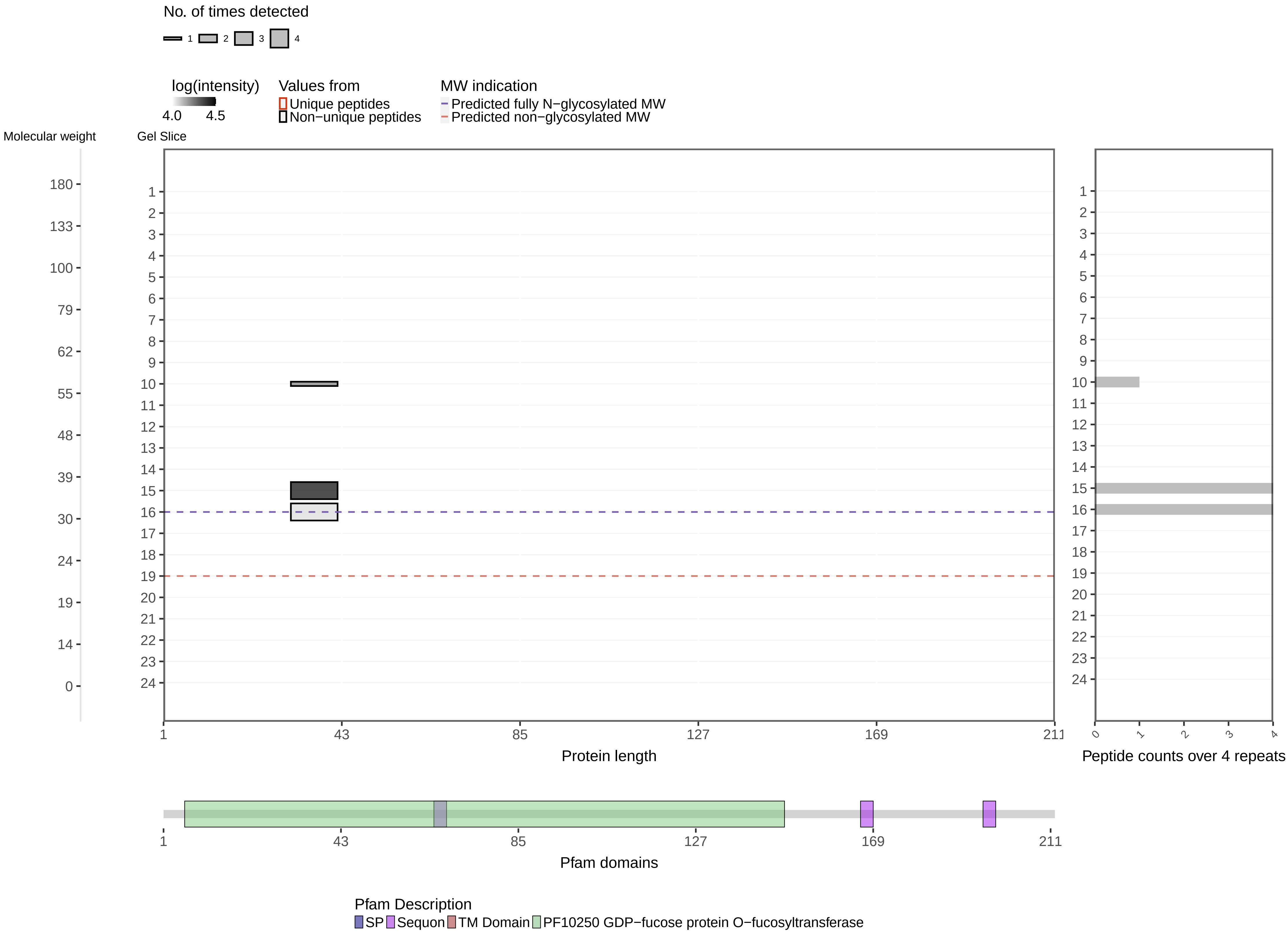


Protein sequence:

MEDKKSLASSDERALIKEKASLLAAVDYYVSMHSDIFISASPGNMHNAMVGHRTYYNMKTIRPNMALLGQLFLNKTLSWPEFKEAVVEGHKNRQGQIRLRKSNQSLYTYPAPNCMCQD*

NbL10g10570.1 HMG−Y−related_protein_A−like [MW = 17.75 kDa]

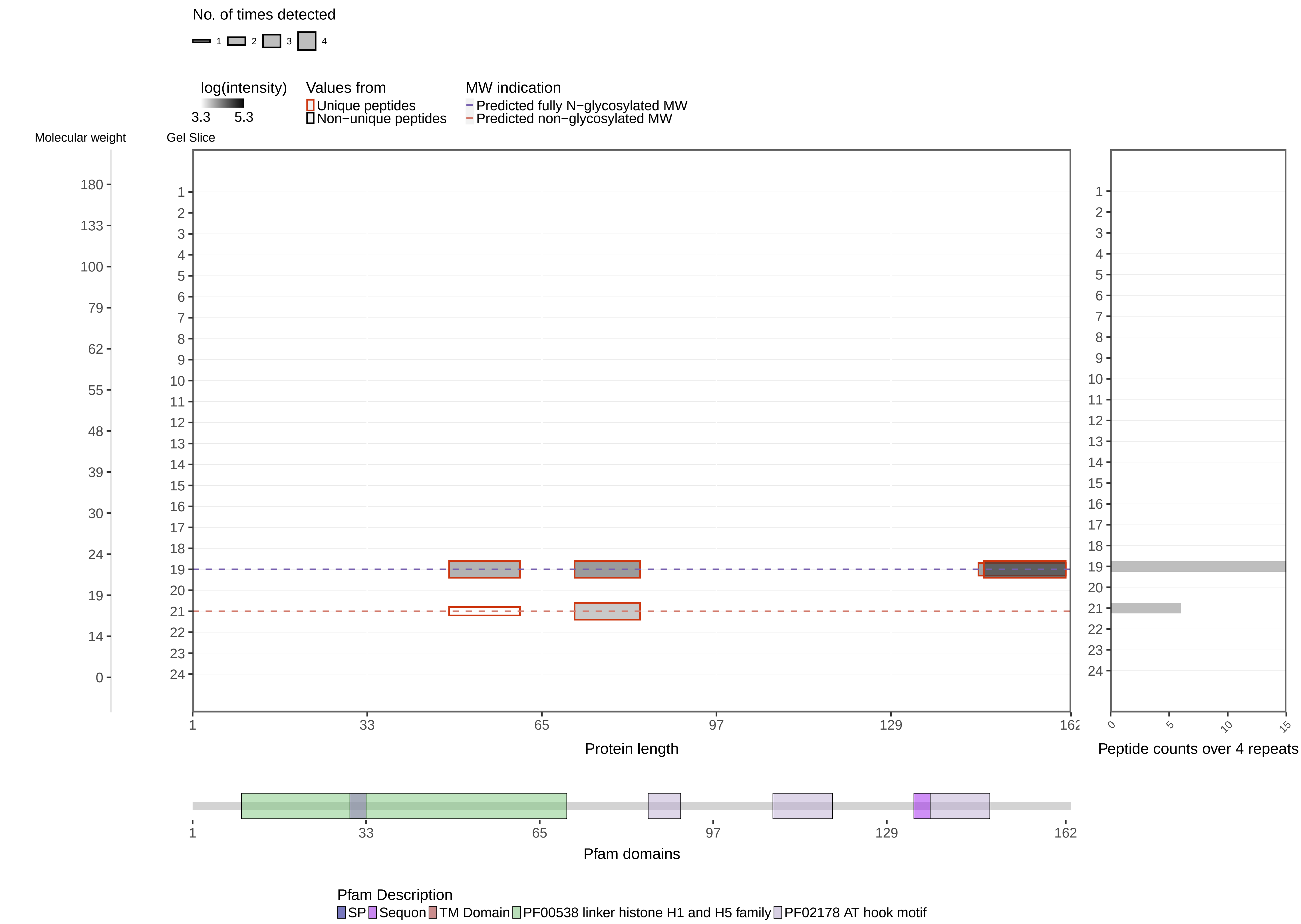


Protein sequence:

NNYMKPGTDVPQKWGRGRPPKPKVSLPPGAESGEIASPRPRGRPRKDPNAPPTSKKPKPTPAPNSSGKPRGRPRKVRPQPAQDDDDDVEES*

### NbL10g10340.1 major_latex_allergen_Hev_b_5−like [MW = 16.27 kDa]

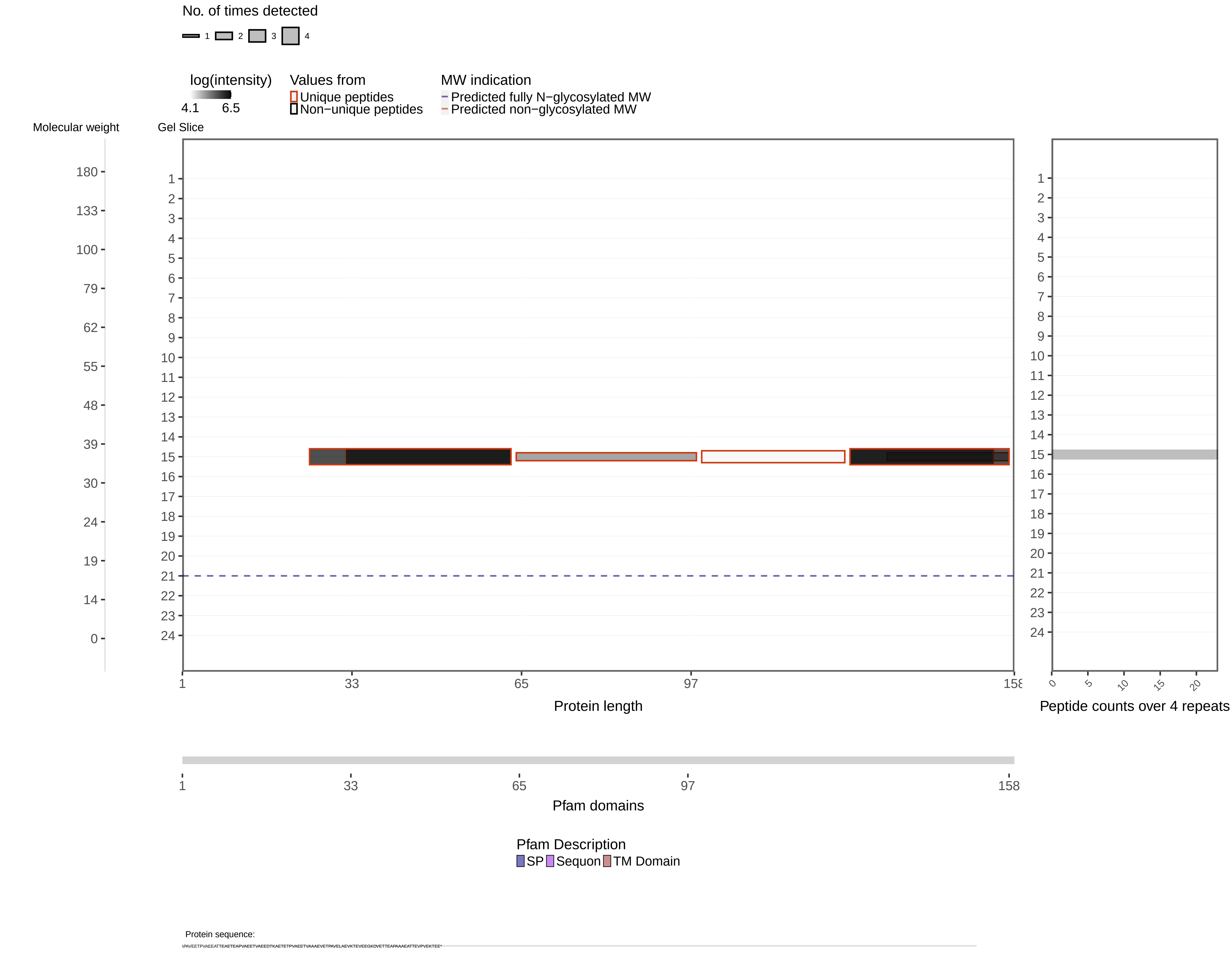

NbL10g10300.1 ras−related_protein_RABA5a−like [MW = 24.99 kDa]

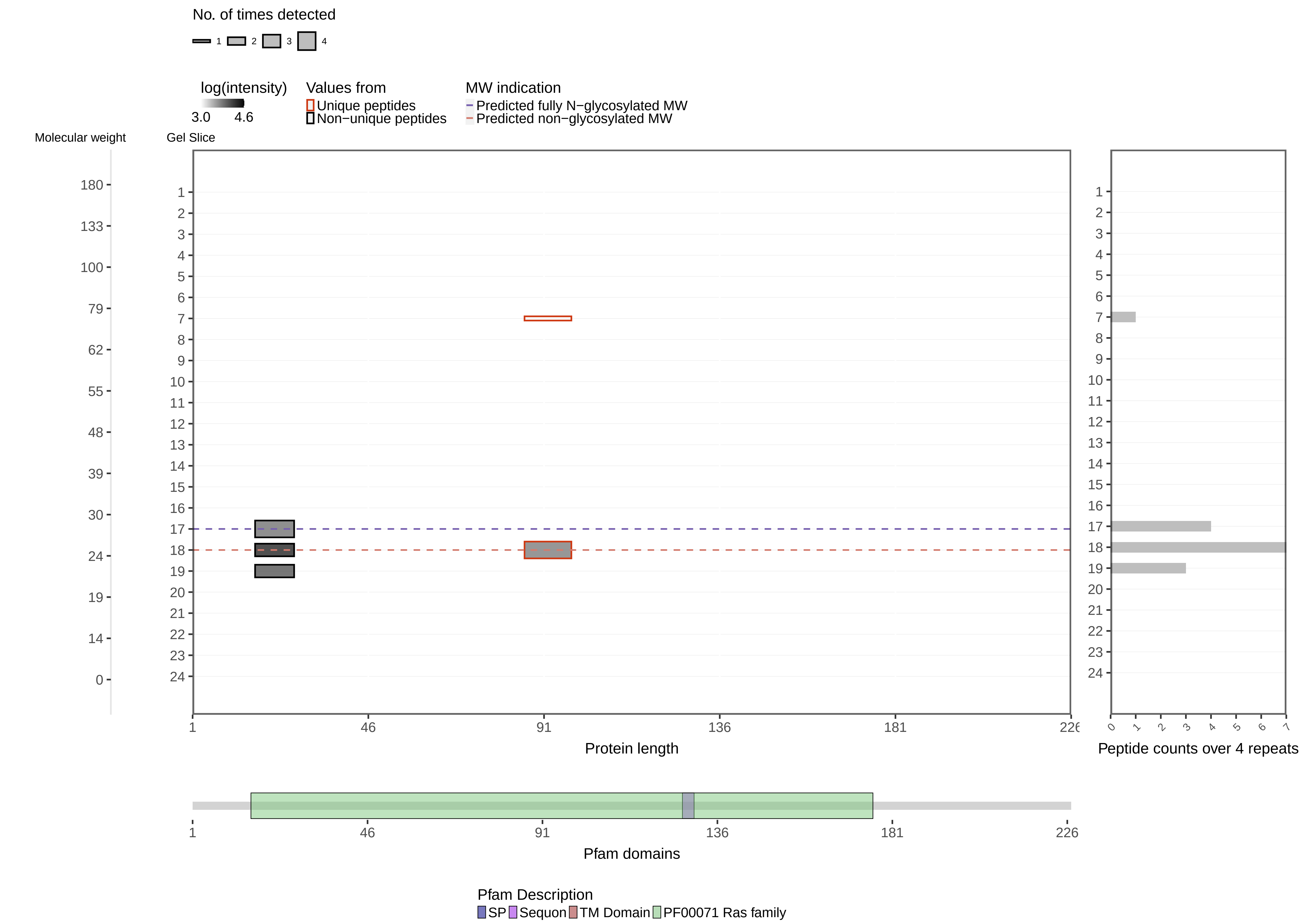


Protein sequence:

TFDSIGRWLNELQTHSDMNVVTILVGNKSDLKDAREVTTAEGKMLAEAKGLFFIETSALDSSNVASAFQTVVREIYNILSRKVIQSQELQKKDSGRLANGKTVVLQADGNQETDAETKKGGCCSS*

NbL10g10230.1 nudix_hydrolase_2−like [MW = 39.29 kDa]

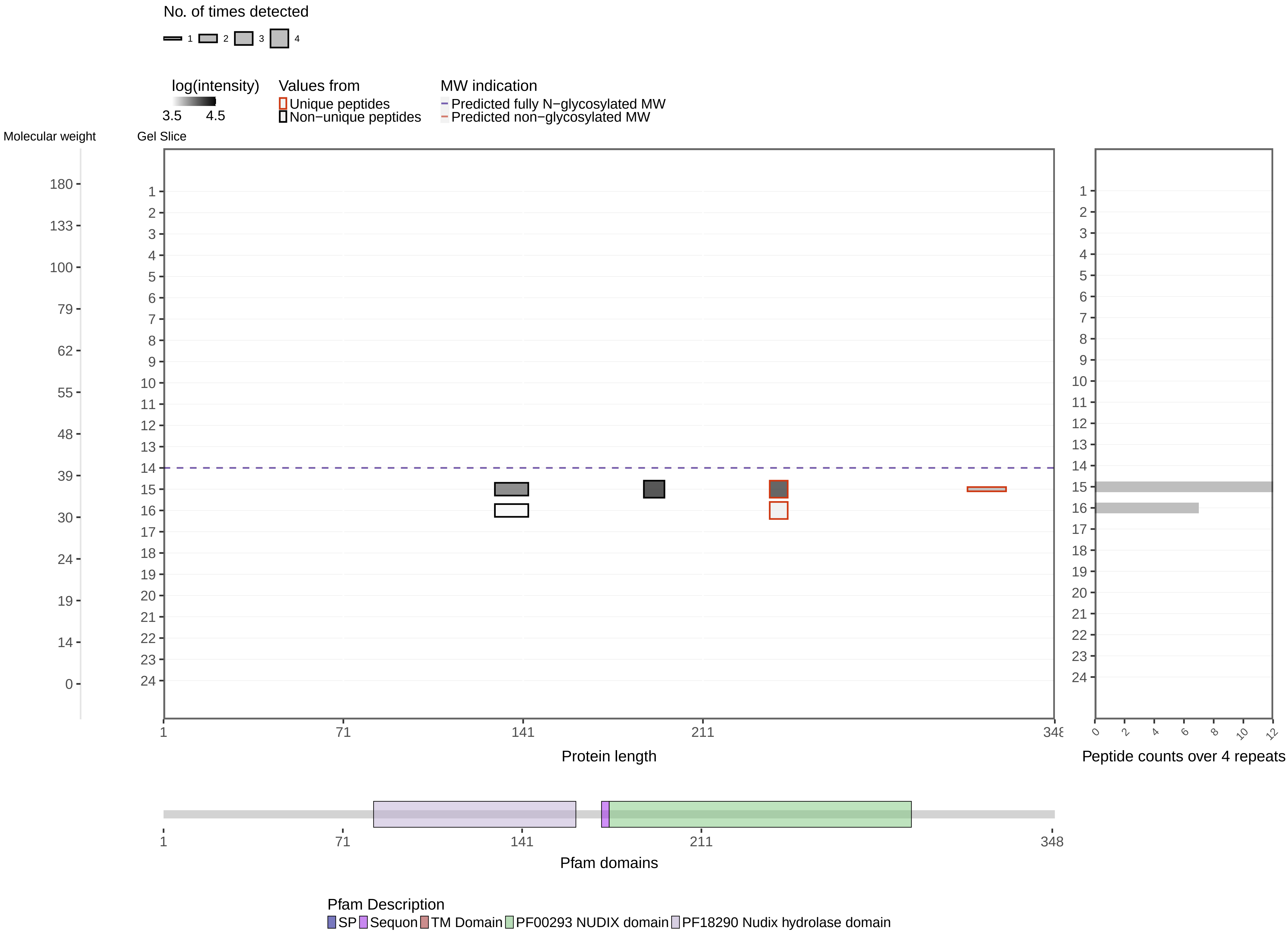


Protein sequence:

.PETANTIPANATHRVGIGAFVINERNEVLVVQEKNGRFRGTGVWKFPTGVVDEGEDICDAAVREVKEETGVNAKFVEILAFRQSHKSFFDKSDLFFVCMLQPLSFDIQKQDAEIEAAQWMPFEQYAAQPFVHGHELLRYISDICSAKMEGRYTGFSSVPTMTSFSQKNTYLYMNGNVRTTGSNDP*

### NbL10g10000.1 actin−97 [MW = 41.69 kDa]

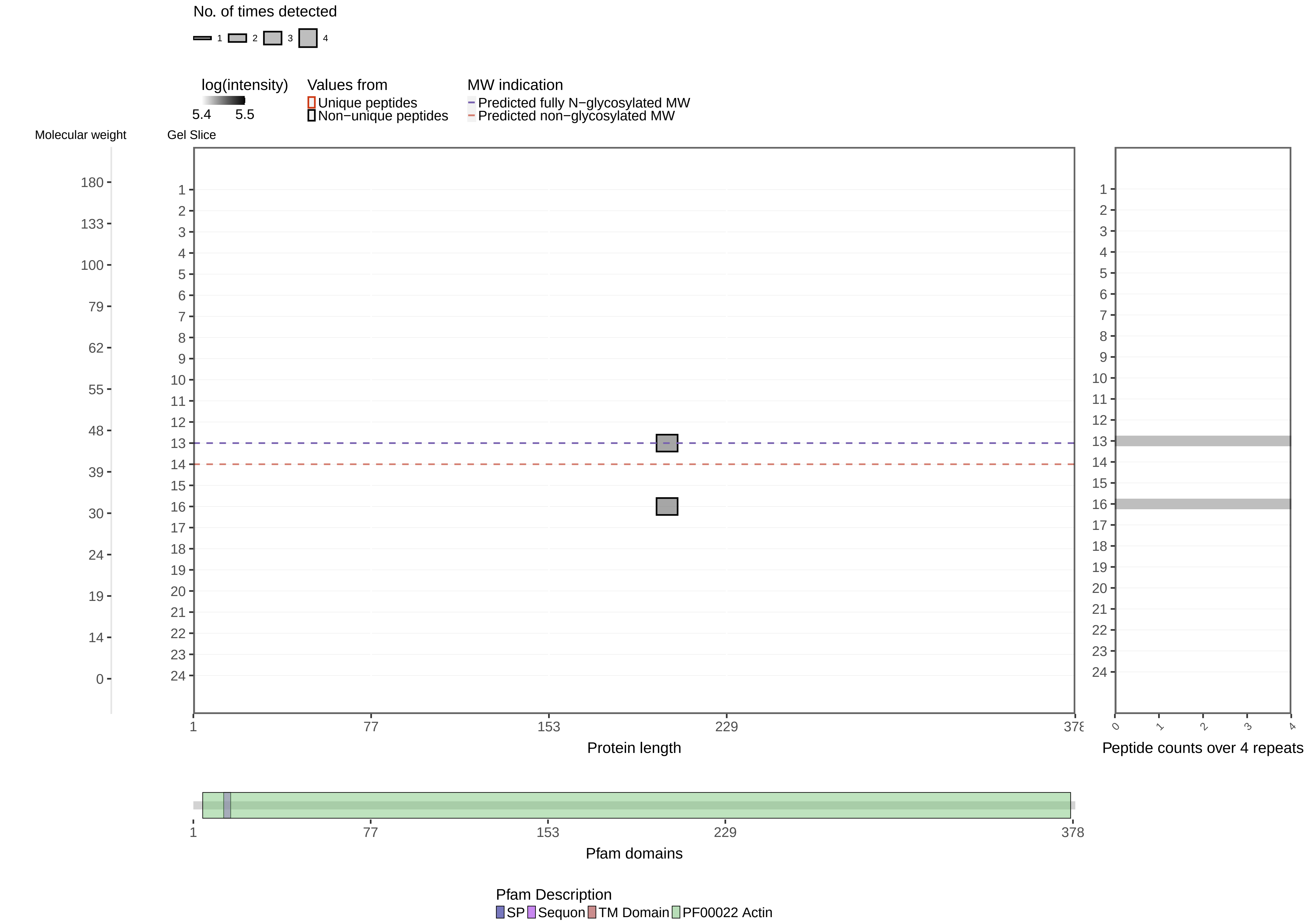


Protein sequence:

.PHAILRLDLAGRDLTDSLMKILTERGYSFTTTAEREIVRDVKEKLAYIALDYEQELETSKTSSSVEKSYELPDGQVITIGAERFRCPEVLFQPSMIGMEAAGIHETTYNSIMKCDVDIRKDLYGNIVLSGGSTMFPGIADRMSKEITALAPSSMKIKVVAPPERKYSVWIGGSILASLSTFQQMWISKAEYDESGPSIVHRKCF*

NbL10g09900.1 UDP−glucuronic_acid_decarboxylase_6−like [MW = 38.73 kDa]

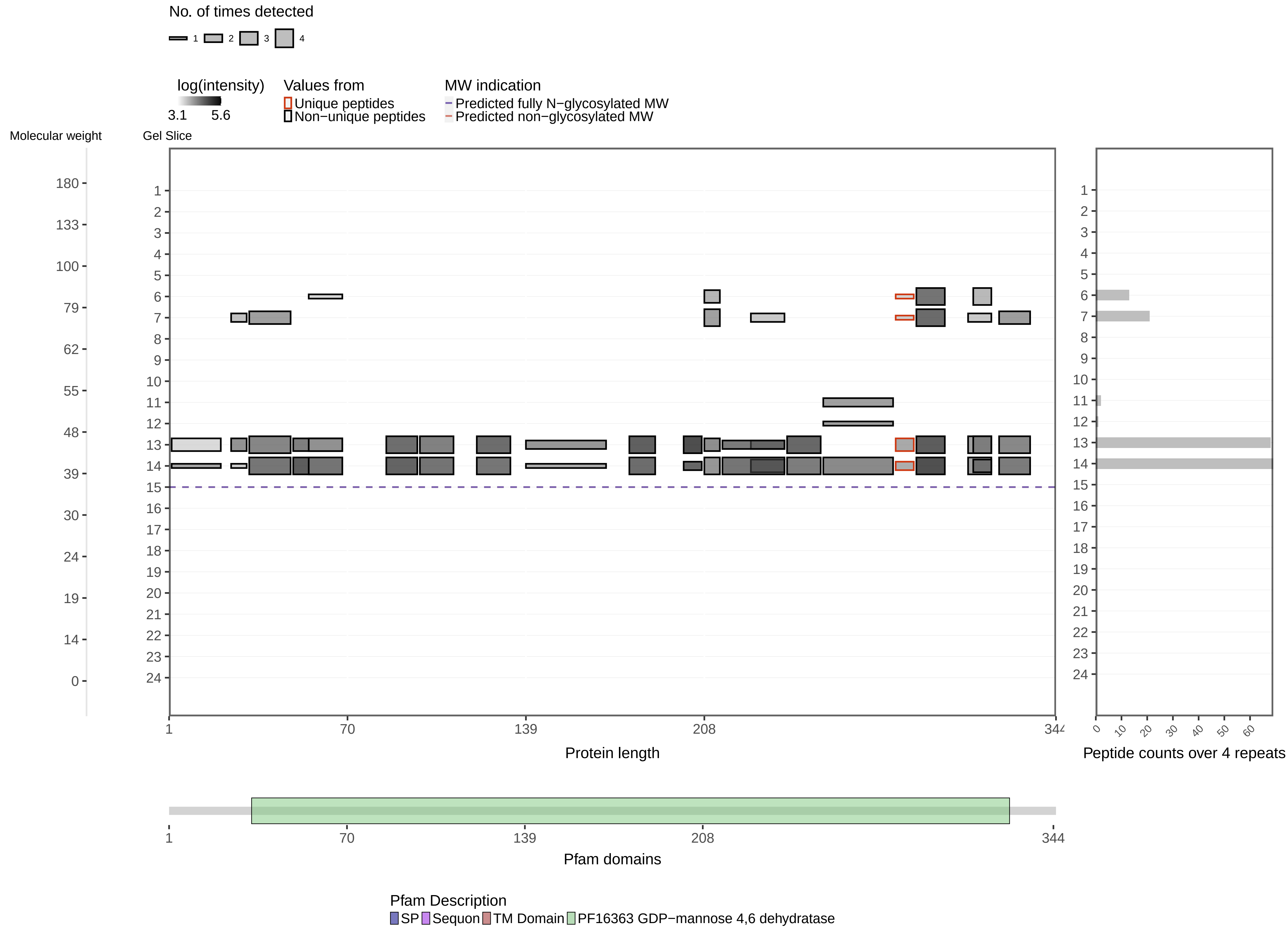


Protein sequence:

ESYWGNVNPIGVRSCYDEGKRVAETLMFDYHRQHGIEIRIARIFNTYGPRMNIDDGRVVSNFIAQAIRDEALTVQLPGTQTRSFCYVSDMVDGLIRLMEGDNTGPINIGNPGEFTMIELAENVKELINPEVKIITVENTPDDPRQRKPDITKAKELLGWEPKIKLRDGIPLMEEDFRGRLGISRKK*

### NbL10g09890.1 methionine−−tRNA_ligase_mitochondrial [MW = 68.46 kDa]

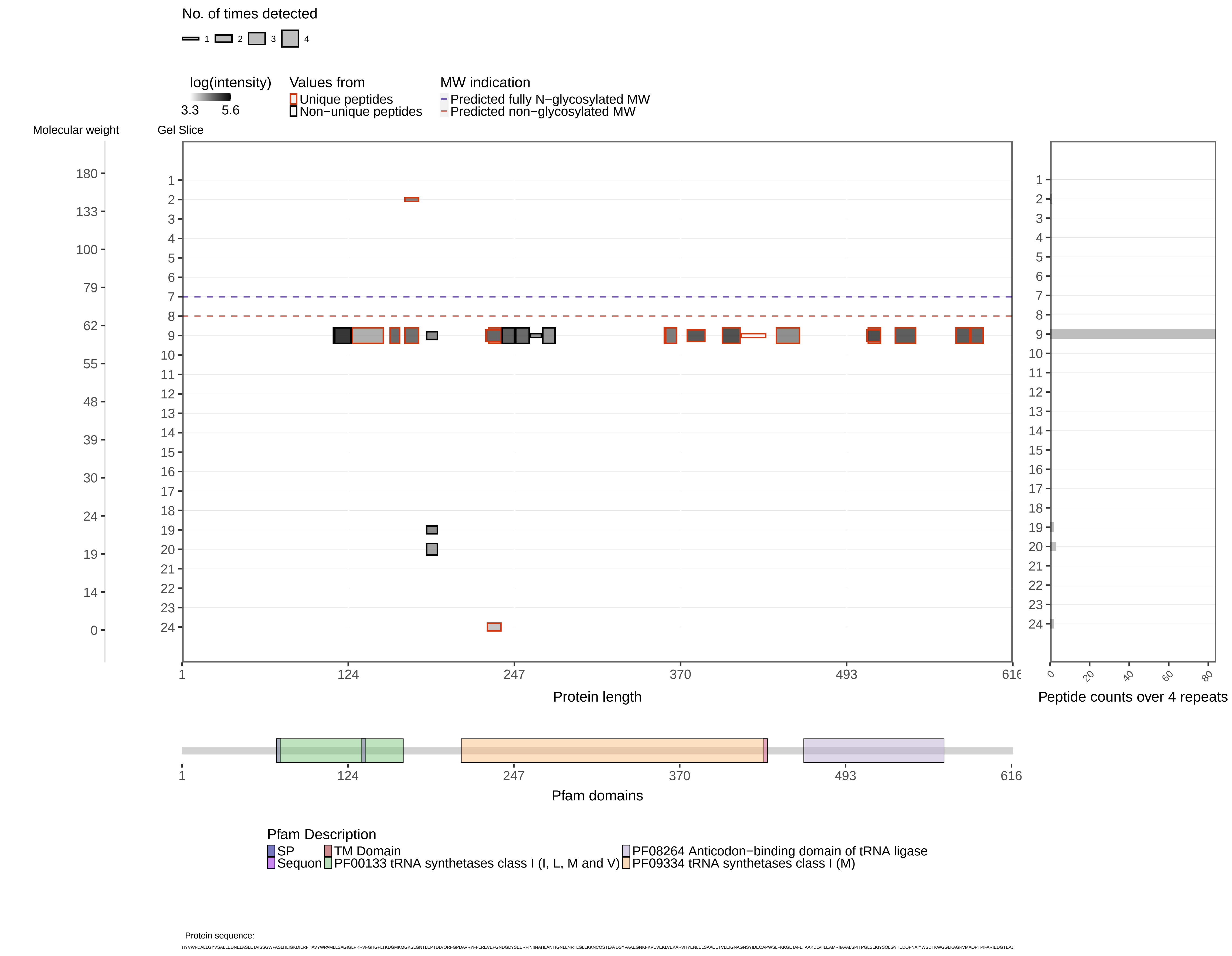

### NbL10g09850.1 2−oxoglutarate_dehydrogenase_mitochondrial−like [MW = 115.66 kDa]

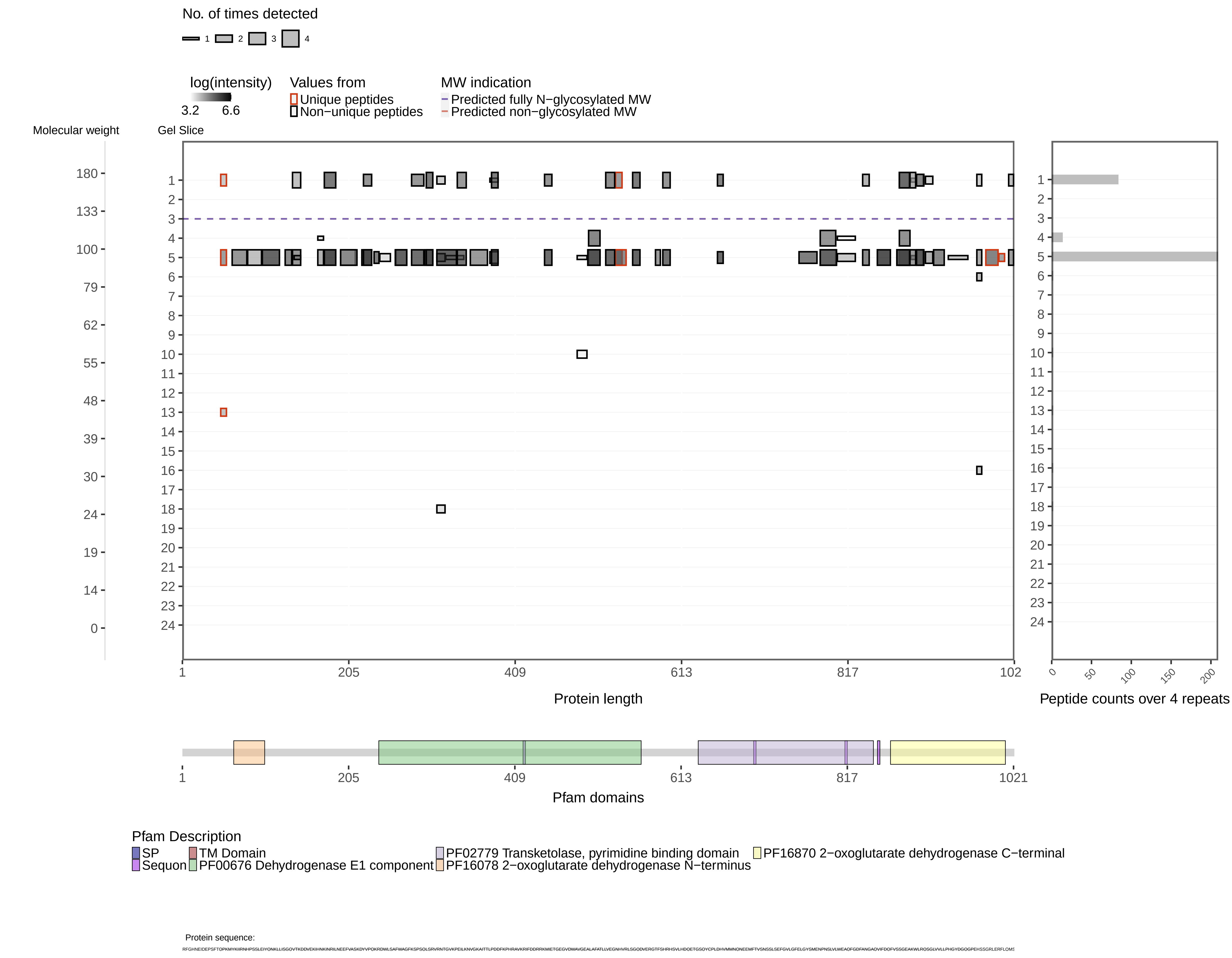

### NbL10g09770.1 beta−hexosaminidase_1−like [MW − SP = 58.98 kDa]

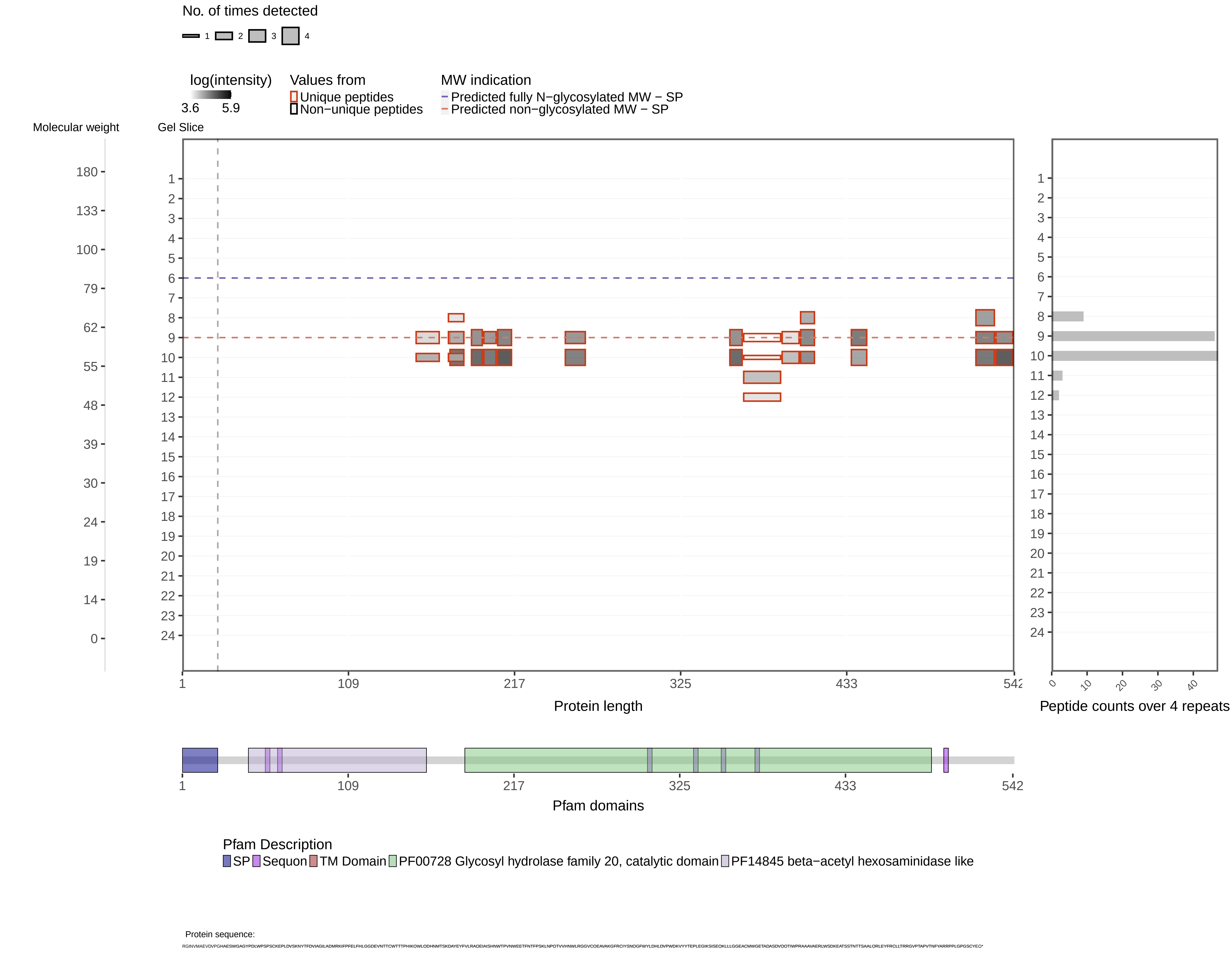

### NbL10g09750.1 glutathione_S−transferase_DHAR2−like [MW = 23.62 kDa]

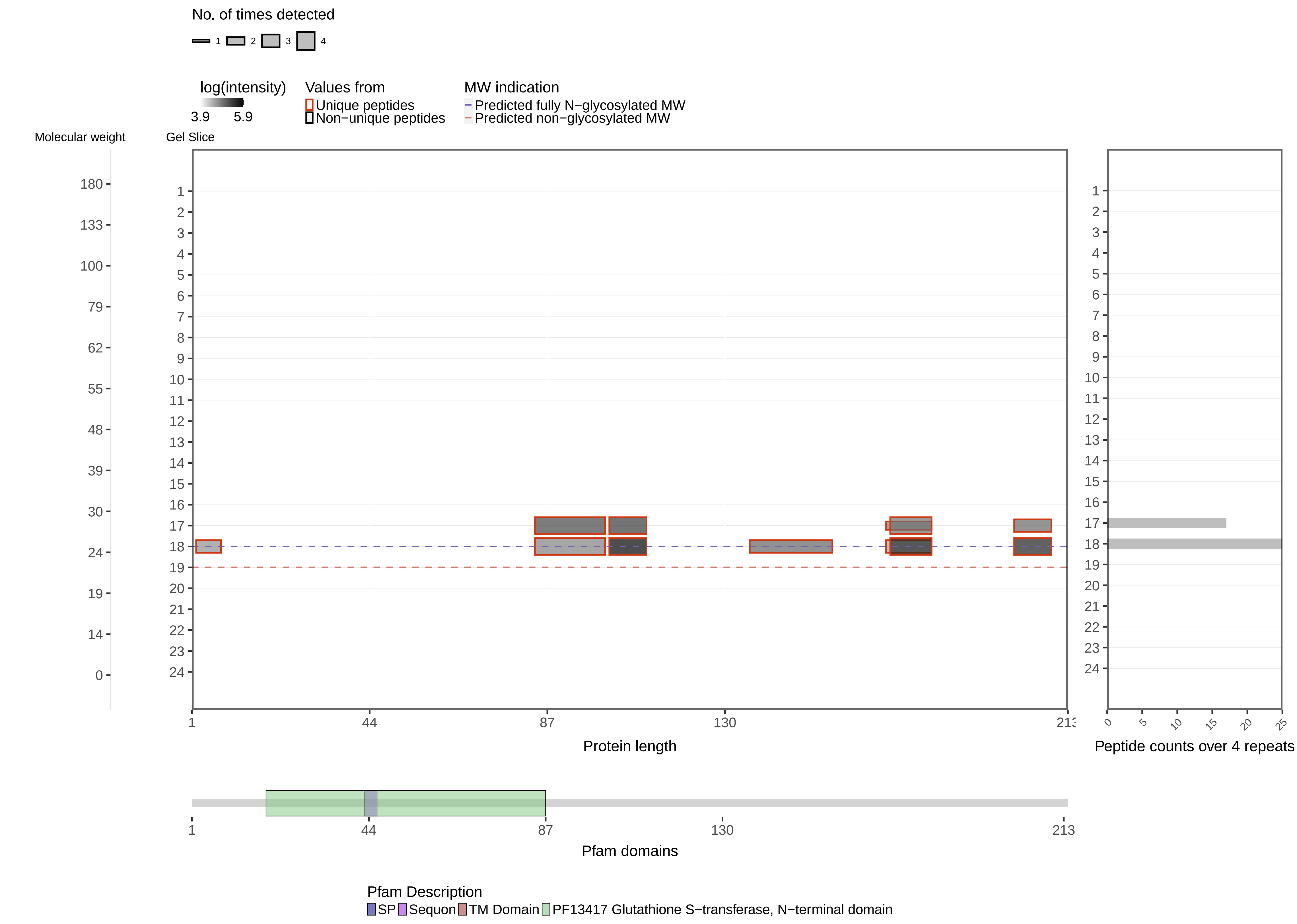


Protein sequence:

EFASVGSKIFPTFVTFLKSKDASDGTEQALLDELKALEEHLKAHGPYVNGANICSVDLSLAPKLYHLEVALGHFKKWSVPESLSHVRNYMKLLFERESFQKTKAAKEYIIAGWAPKVNP*

### NbL10g09530.1 multiple_organellar_RNA_editing_factor_2_chloroplastic−like [MW = 30.32 kDa]

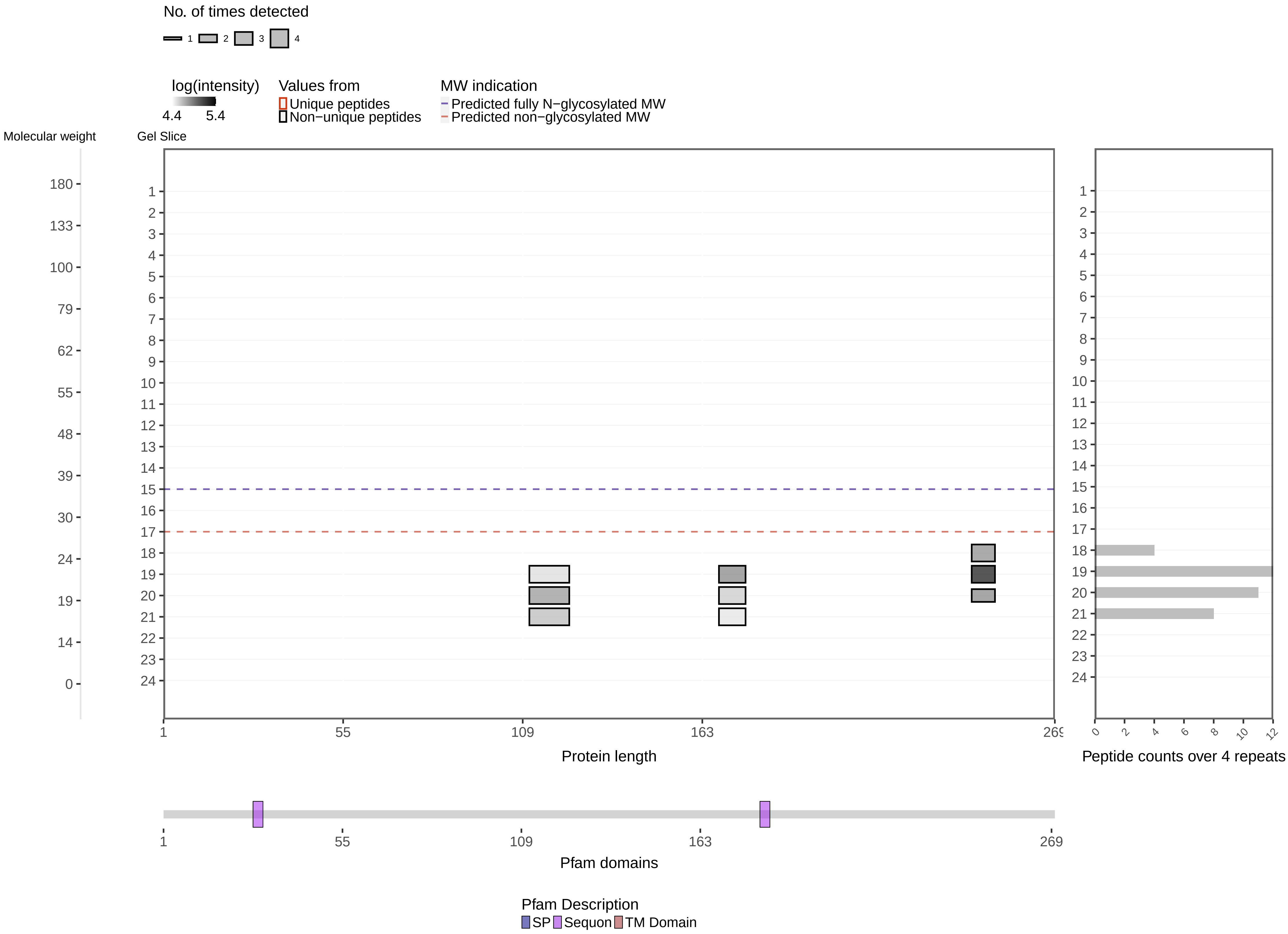


Protein sequence:

TEMAPLFPGCDYEHWLIVMDKPGGEGANKQQMIDCYIQTLSKVLGSEEEAKKKIYNVSCERYFGFGCEVDEETANKMEGLPGVLFVLPDSYVDPEYKDYGAELFVNGEIVQRSPERQRRVEPVPQRNDRIRPNRYNRRRDNMR*

### NbL10g09490.1 cysteine_desulfurase_1_chloroplastic−like [MW = 52.83 kDa]

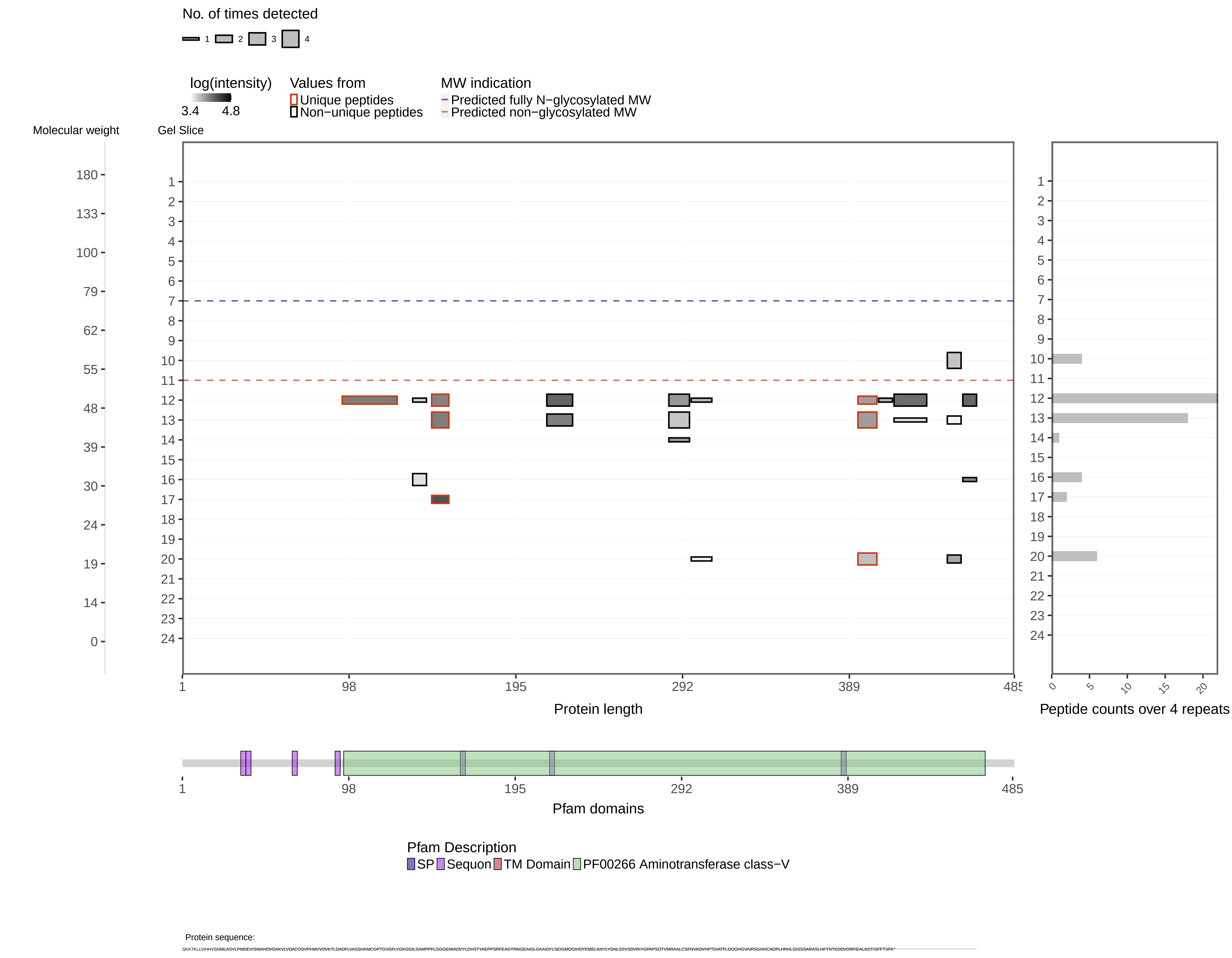

### NbL10g09400.1 ubiquitin_carboxyl−terminal_hydrolase_12−like [MW = 130.62 kDa]

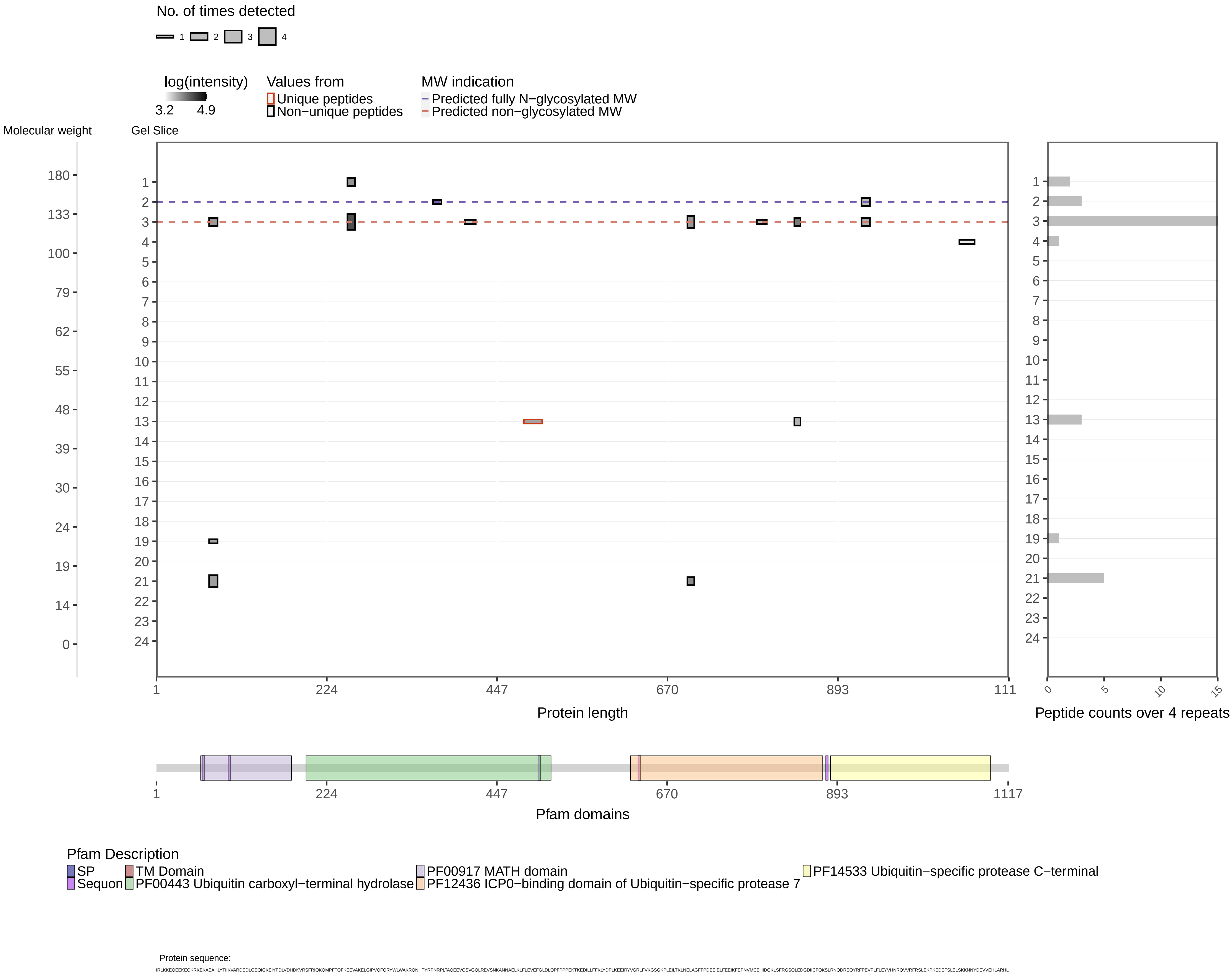

### NbL10g09300.1 40S_ribosomal_protein_S17 [MW = 16.22 kDa]

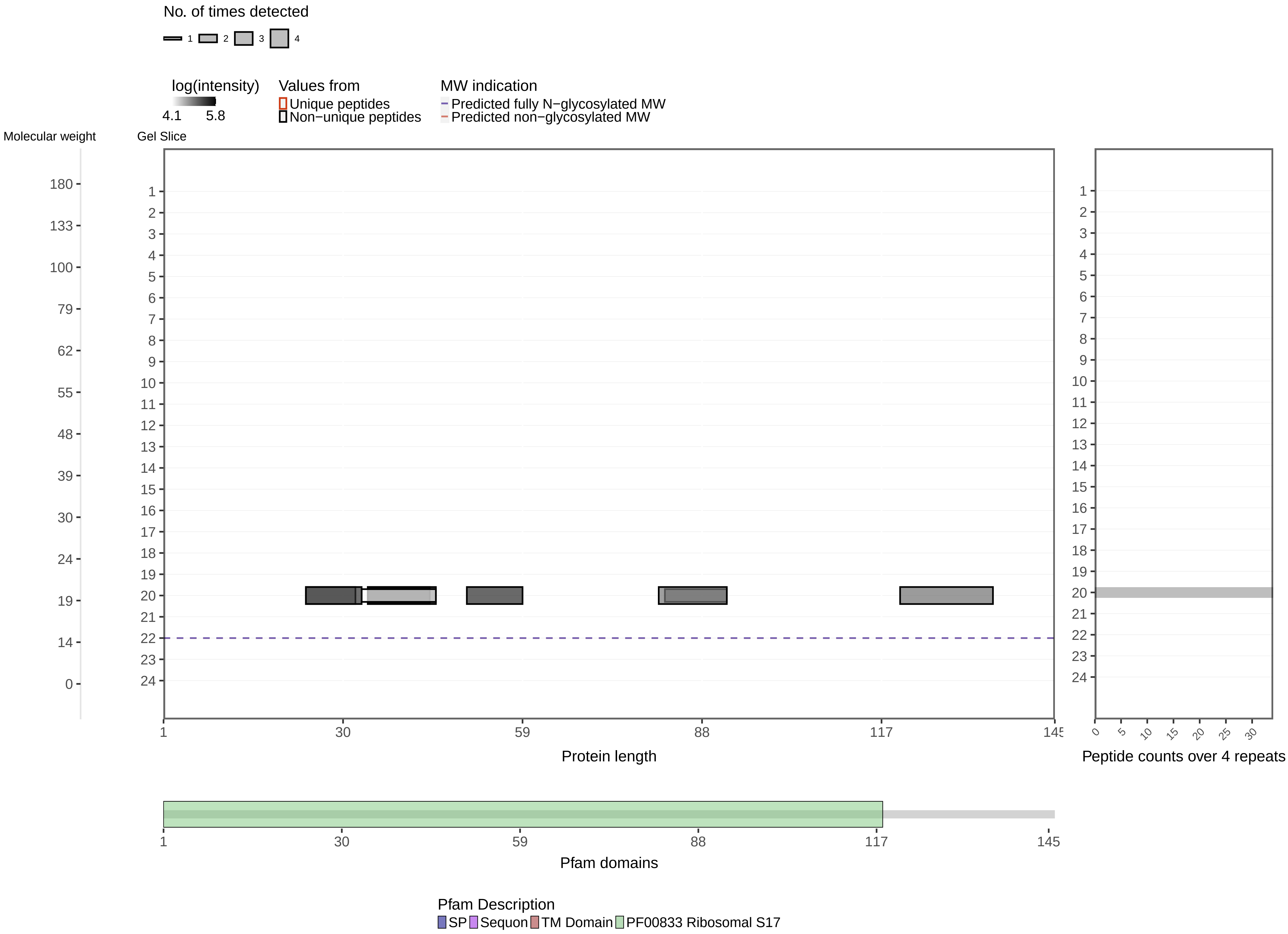


Protein sequence:

IQKGPVRGISLKLQEEERERRMDFVPDESAIKTDLIEVDKETLDMLSALGMSDLPGVVKQAAEPQAVAALPTYGRGAGGFGRKY*

NbL10g09250.1 copper_transport_protein_ATX1−like [MW = 13.63 kDa]

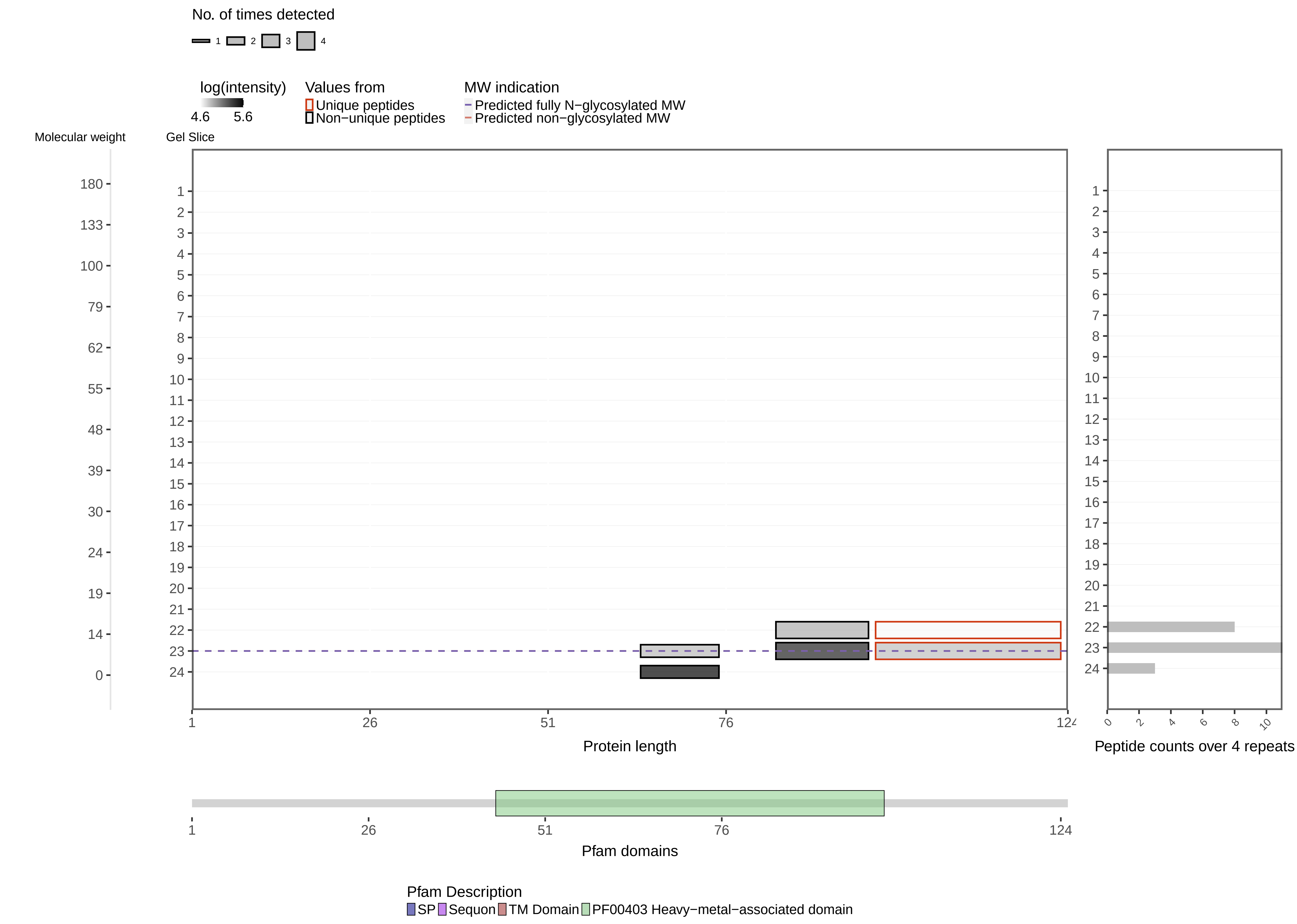


Protein sequence:

GCVGAVKRVLGKMEGVENFDIDLKEQKVTVKGNVQPEAVLQTVSKTGKPTSFWEAGEPAQTEEKPAEAVGTS*

### NbL10g09110.1 protein_BTR1−like [MW = 35.7 kDa]

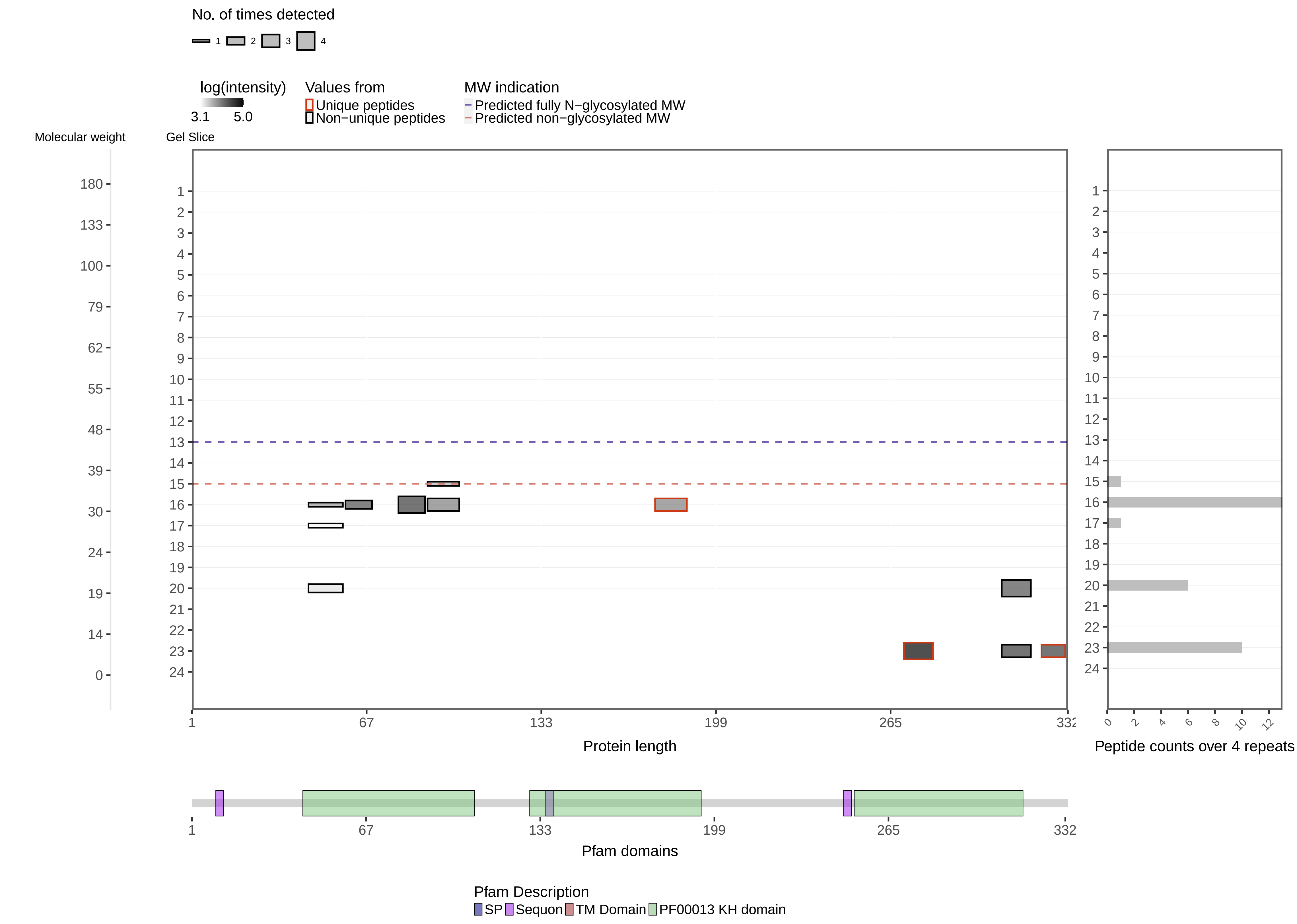


Protein sequence:
EDSHAGIKISSODDNFPGPHDRLVIVTGTLGEOMRAIELILHKLAEDSHYMONMNAPFPYAGYVGMNYGPRNGVGGRFPNNRYONNVOSNSEDRNNSVTIGVADEHIGLVLGRNGRSIMEISOOSGARIKISDRGDFISGTSDRKVTITGSORAISMAESMISOKVARVSESDVFVKG*

NbL10g08920.1 probable_methylenetetrahydrofolate_reductase [MW = 67.29 kDa]

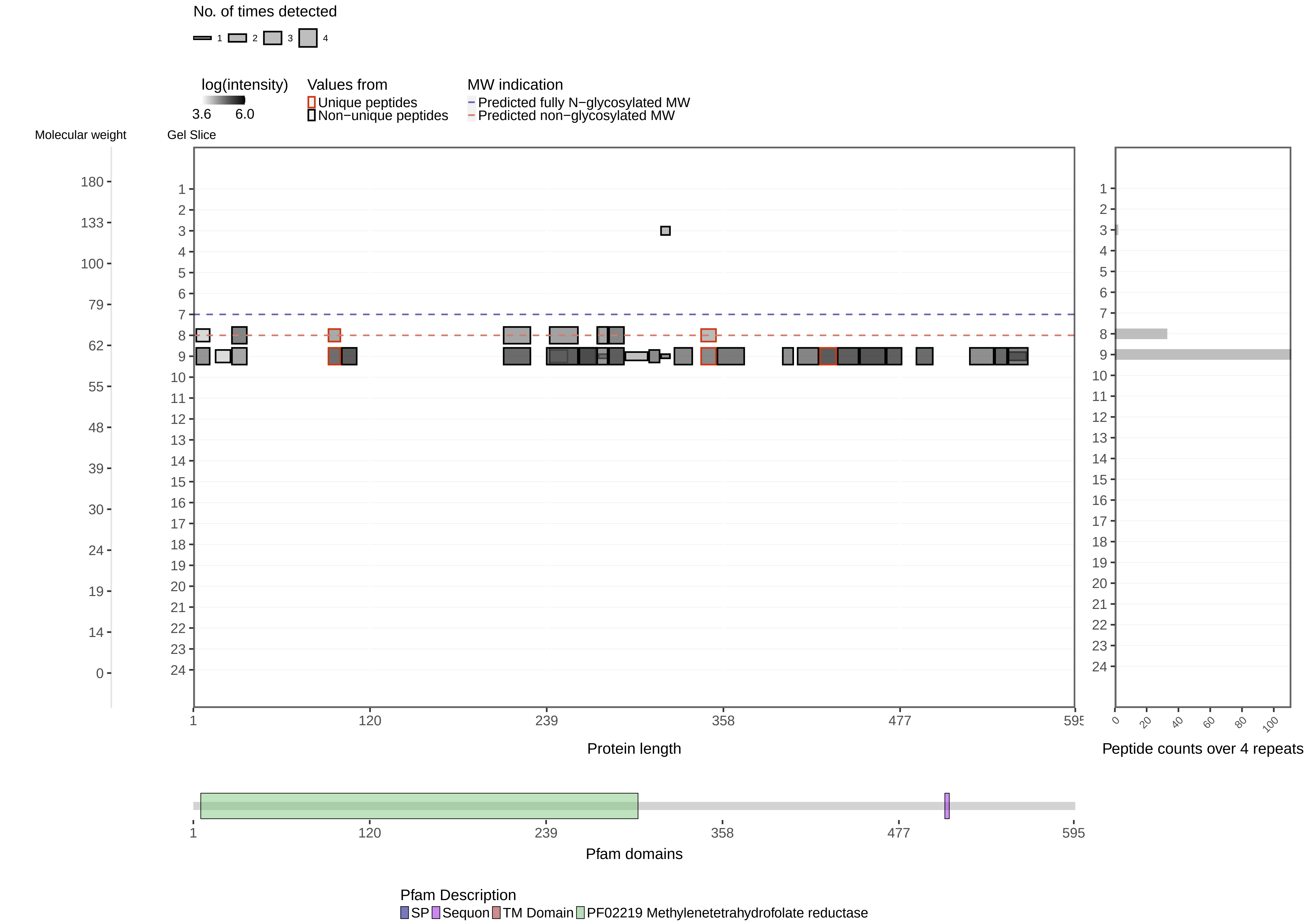


Protein sequence:

[illegible]MEKSALAILMNLGLIEESKISRPLPWRRPTNVFRVKEDVRPIFWANRPKSYISRTIDWDEYPHGRWGNAQNPSYGALSDYQFMRARSRDKKLQEEWAFALNSVEDIYERFMNYCLGKLRSCPWSELDGLQPETKIIDEHLSNINTKGFLTINSQPAVNGAKSDAPSIGWGGPGGYVYQKAYLEFFCSHEKLNALVEKCKAFPFLTYMAVNKEGNWISNANQTDINAVTWGVFPGKEIVQPTVVDPASFMVWKDEAFEIWSRGWAQLYQESDPSRKLLEQVQNSYFLVSLV

### NbL10g08790.1 uncharacterized_LOC109225451 [MW = 29.71 kDa]

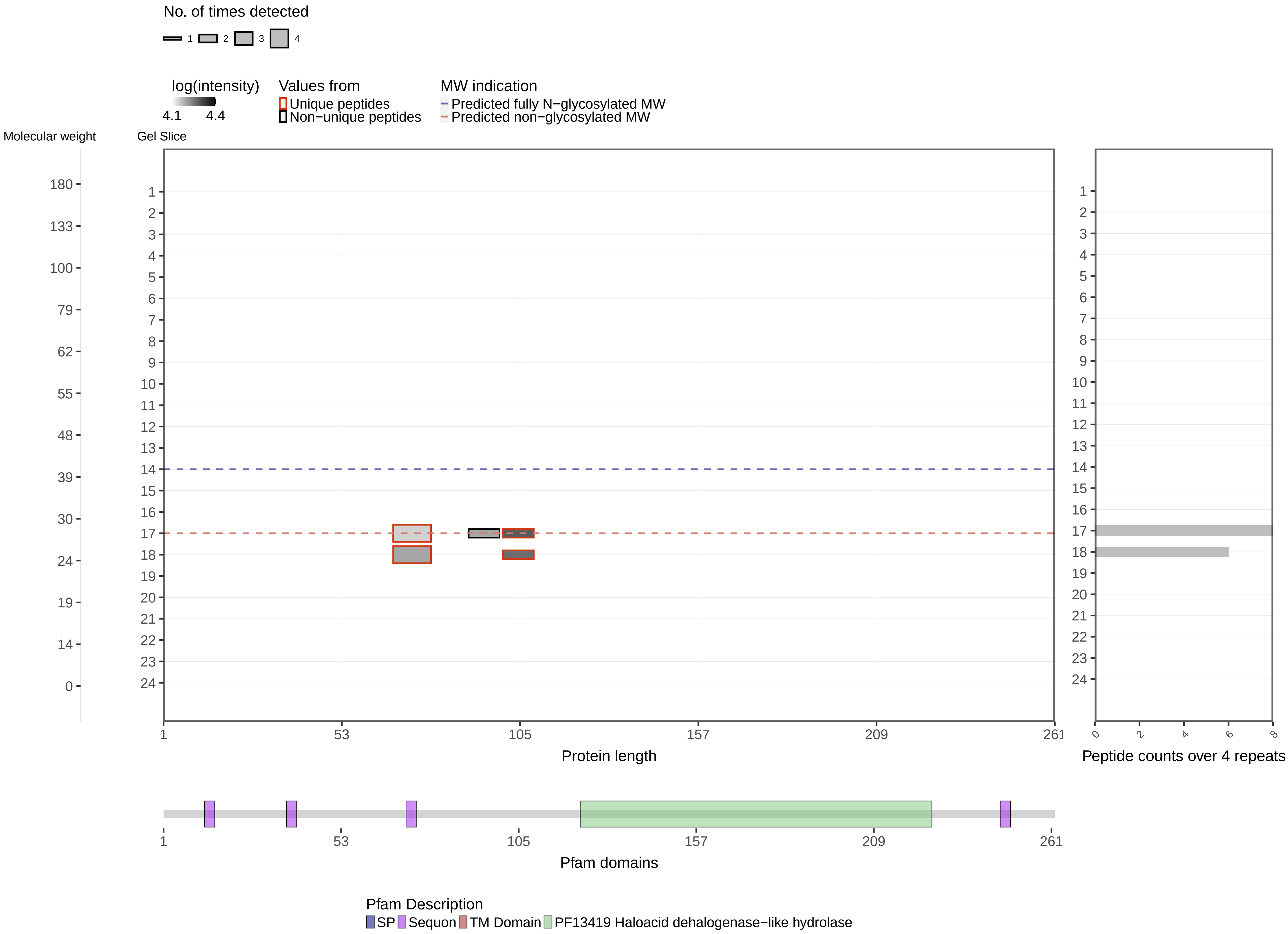


Protein sequence:

DLEGLKNCMRRAYSYIEGVEGLLNSLKENGYEIHAFTNYPIWYQMIEDKLKLSNYLSWTFCSCIFGKRKPDPDFYLEVVKHLNVEASSCIFVDDRMRNVEAAVEVGLNGLQFQNADLLRKDLSLLGVNISANENPDLAECSS*

NbL10g08470.1
1−(5−phosphoribosyl)−5−[(5−phosphoribosylamino)methylideneamino]_imidazole−4−carboxamide_isomerase_chloroplastic−like
[MW = 34.73 kDa]

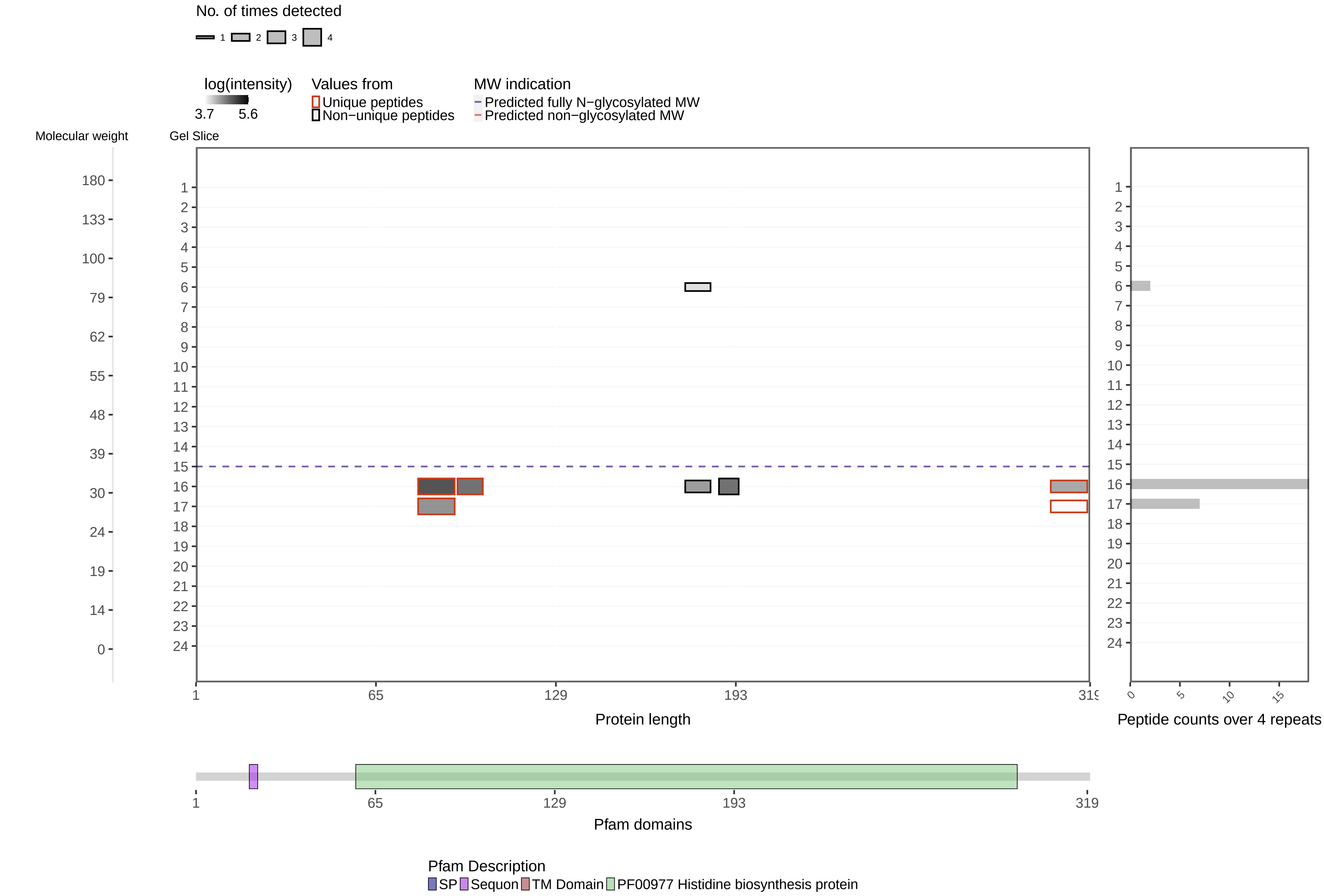


Protein sequence:
ALSYIDEGASHVIVTSYVFNNGQMDLERLKELASLVGRKRLVLDLSCRKKNMQEGEYVIVTDRWQKFTDVRLNEKVLNFLADYADEFLVHGVDVEGKKLGIDAELVALLGKYSPIPVTYAGGVTVMADLEKIKLAGMGRVDVTVGSALDIFGGNLAYKDVVAWHALQDPLAM*

### NbL10g08440.1 ammonia−lyase [MW = 78.73 kDa]

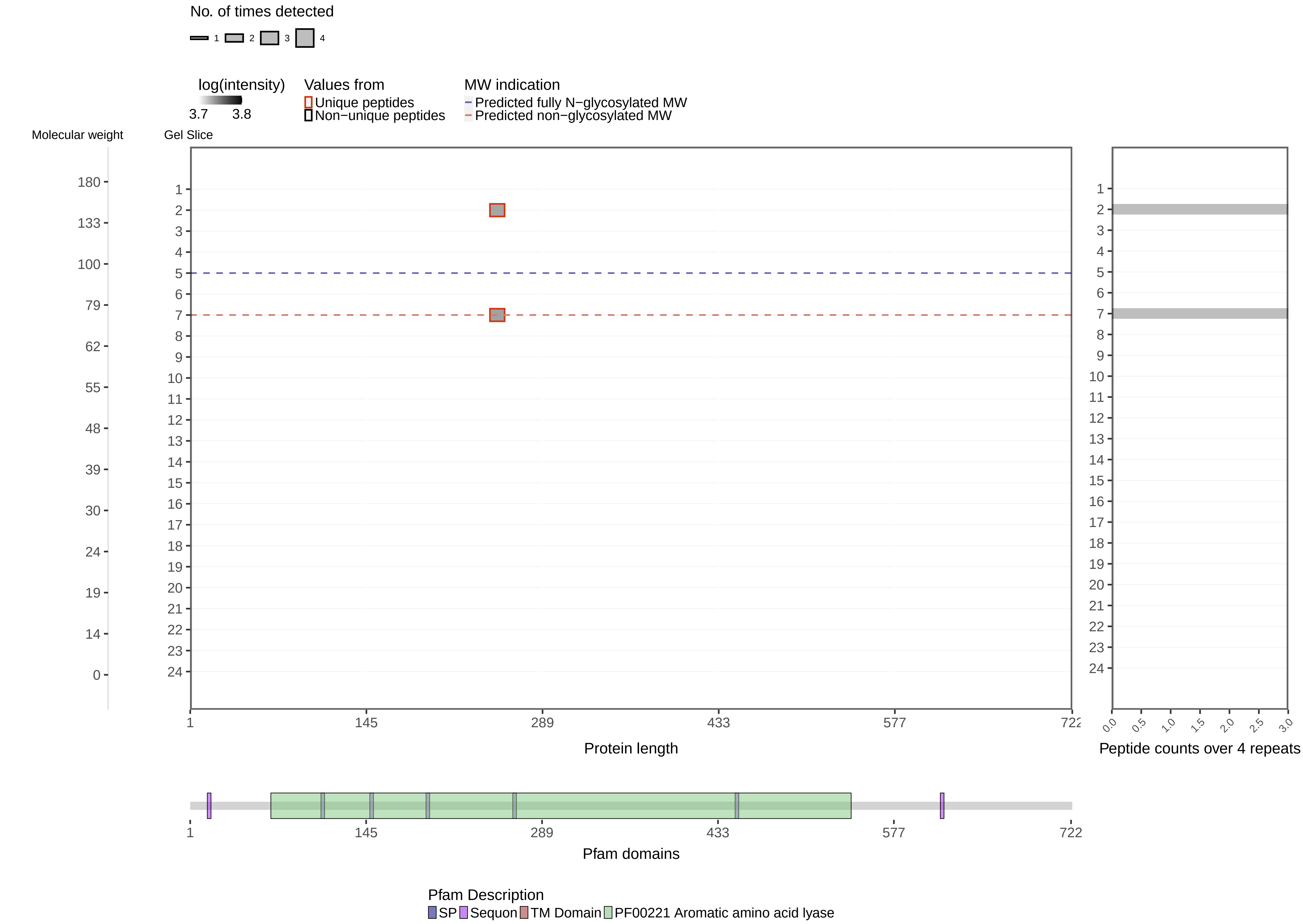


Protein sequence:

NbL10g08410.1 calreticulin−3−like [MW = 51.1 kDa]

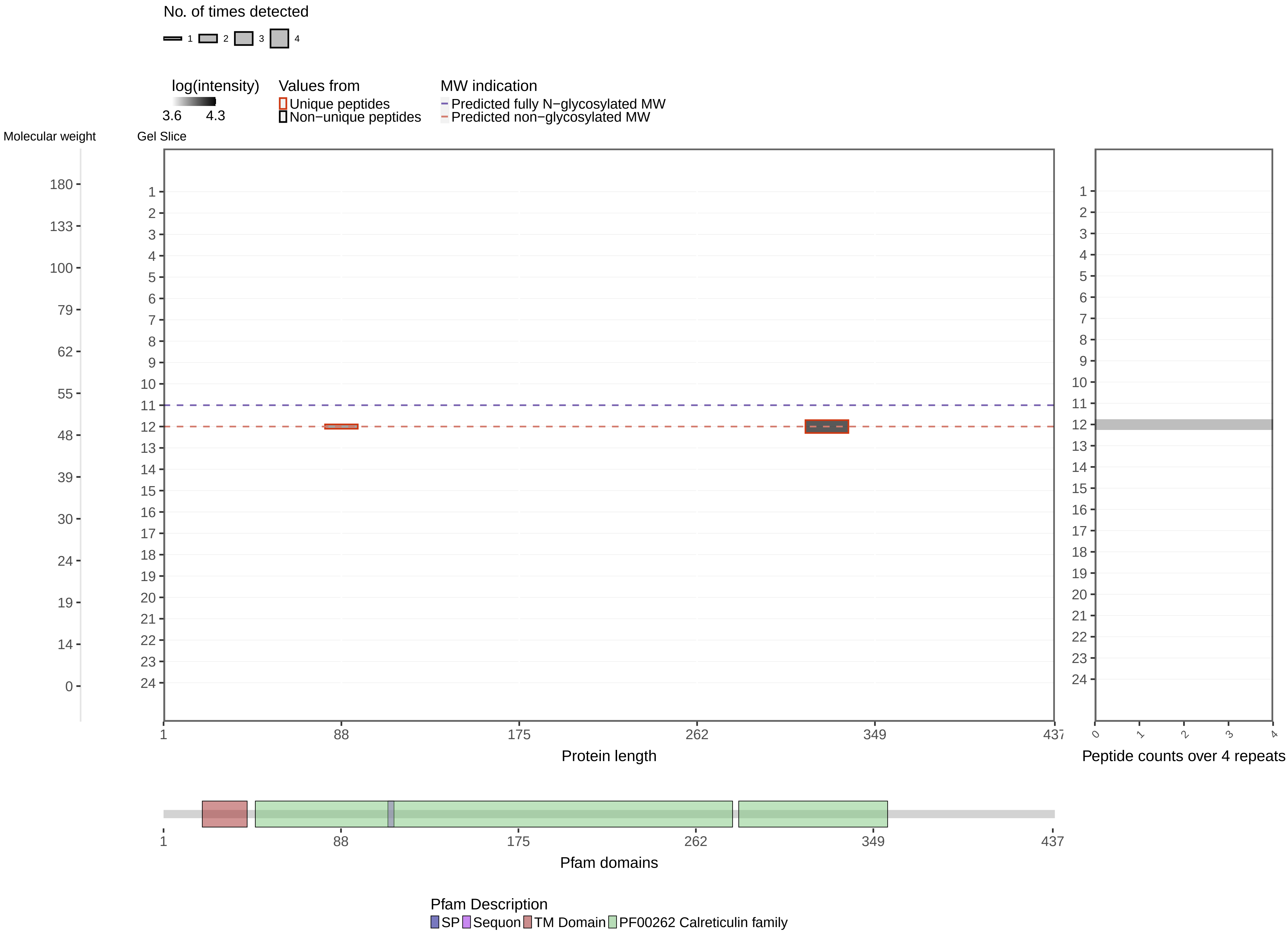


Protein sequence:

IWIDGRERDSGSMYTDWDILPPRKIKDVNAKKPADWDDSEYIEDPNKVKPEGYDSIPREIPDPKAKKPDHWDDEDDGIWRPPKIPNPAYKGPWKRKKIKNPNYKGKWRTPWIDNPEFEDDPDLYVLKPIKYVGIEVWQVKAGSVFDNILICDEPDYAKKVIEEVFANREAEKEAFEEAEKVRKAKEEEEARRAREEGERRRRERDRDRHRDRNKRRYRRDYDDDYHVSPE*

NbL10g08380.1 phenylalanine−−tRNA_ligase_chloroplastic/mitochondrial−like [MW = 48.59 kDa]

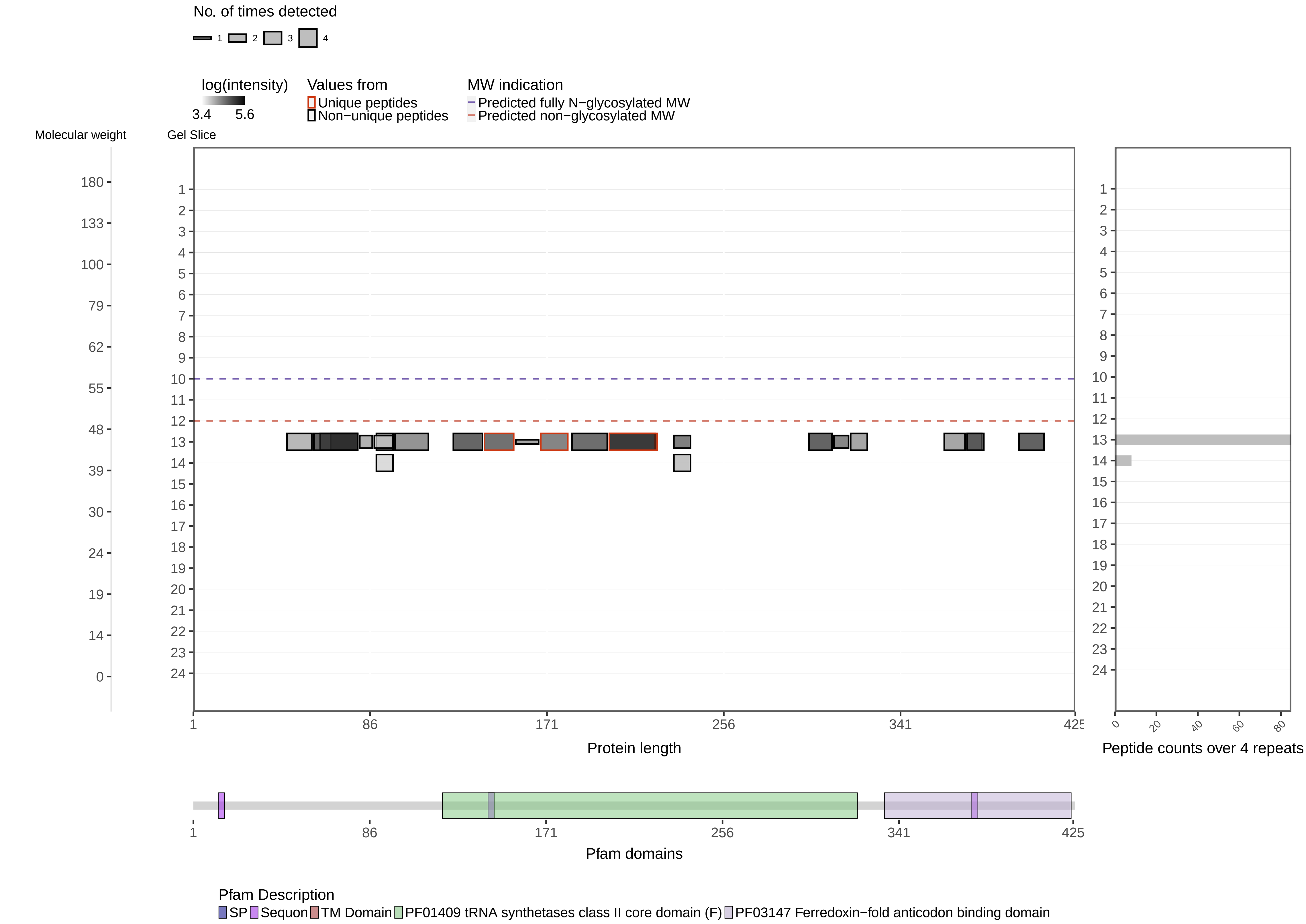


Protein sequence:

FAPNDWEGSGADATSHAAEDLKKCLEGLARHLFGGVEMRWVDTYFPFTNPSFELEIYFQNEWMEVLGCGVTEQEILKRSGRTDNVAWAFGLGLERLAMVLFDIPDIRLFWSTDERFTSQFSSGRLGLKFKPFSKYPPCYKDMSFWISDSFTENNLCEVVRGIAGDLAEEVKLIDNFTNKKGMTSHCYRIAYRSMERSLTDEEINDLQWKVREQVESKLKVVLR*

### NbL10g08370.1 23−dimethylmalate_lyase [MW = 54.41 kDa]

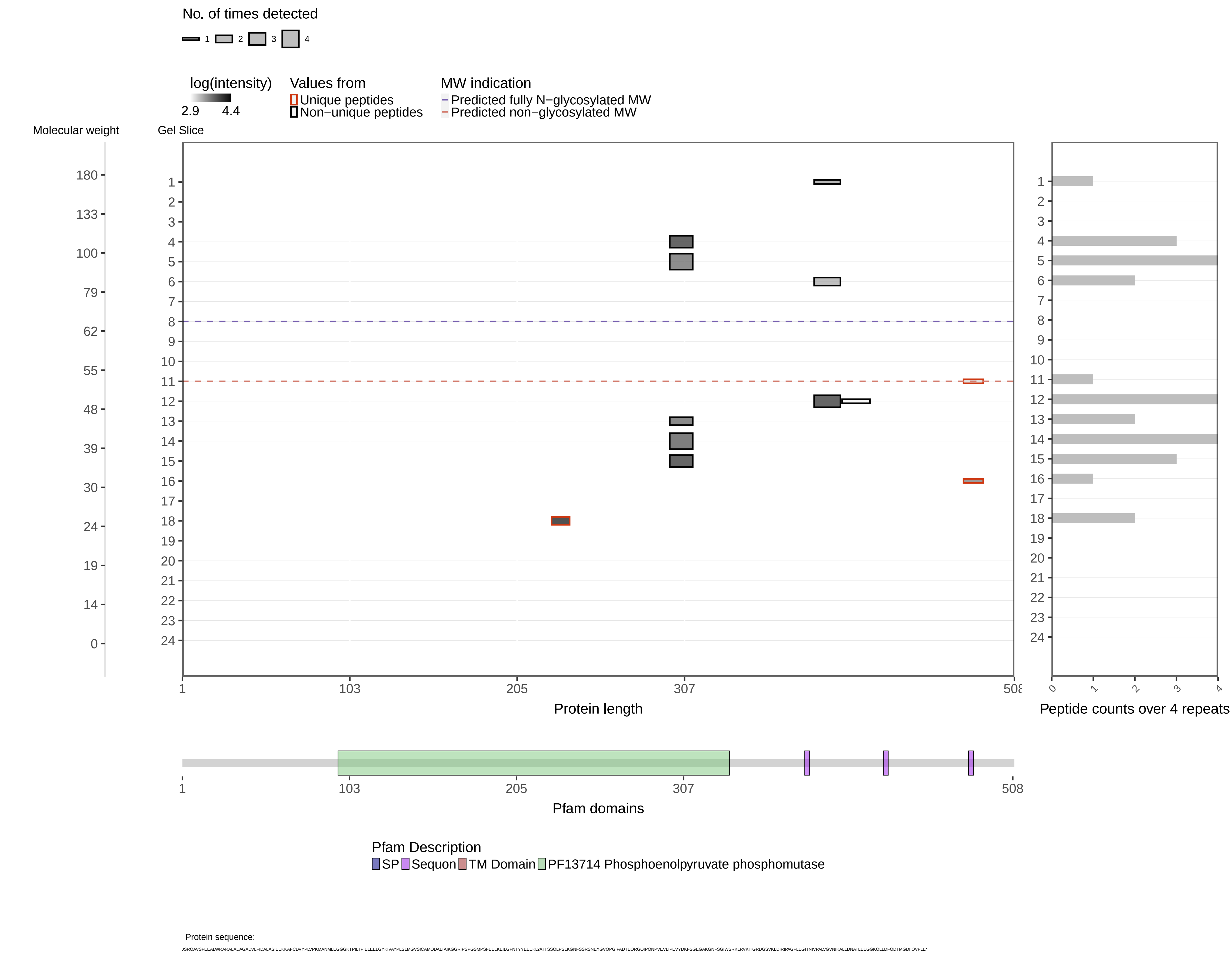

NbL10g08350.1 biotin_carboxyl_carrier_protein_of_acetyl−CoA_carboxylase_chloroplastic−like [MW = 34.36 kDa]

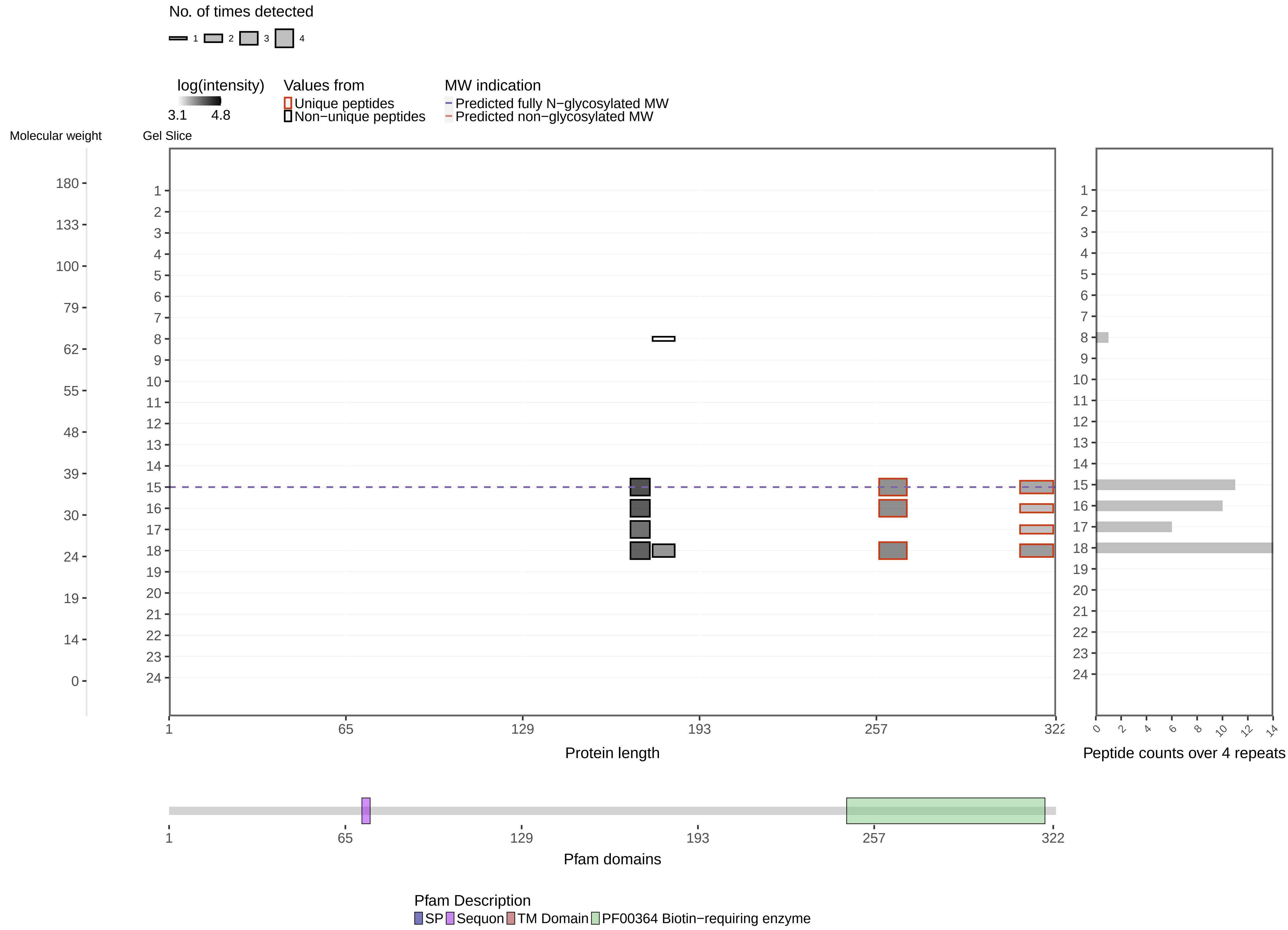


Protein sequence:

EESISNFISQVSSLVKLVDSRDVVELHLKQLDCEILIRKKEALPQPPPPAHTPLLQSYHVPSVQFSPPPPAPAPAPAAPPVQIPAPSPAAAKPSDSSVPPLKCPMAGTFYRSPGPGEPPIVKVGDKVQKGQVLCIIEAMKLMNEIEADQSGTIIDIFAEDGKSVSVDMPLFVIKP*

NbL10g08340.1 thioredoxin_F−type_chloroplastic−like [MW = 24.56 kDa]

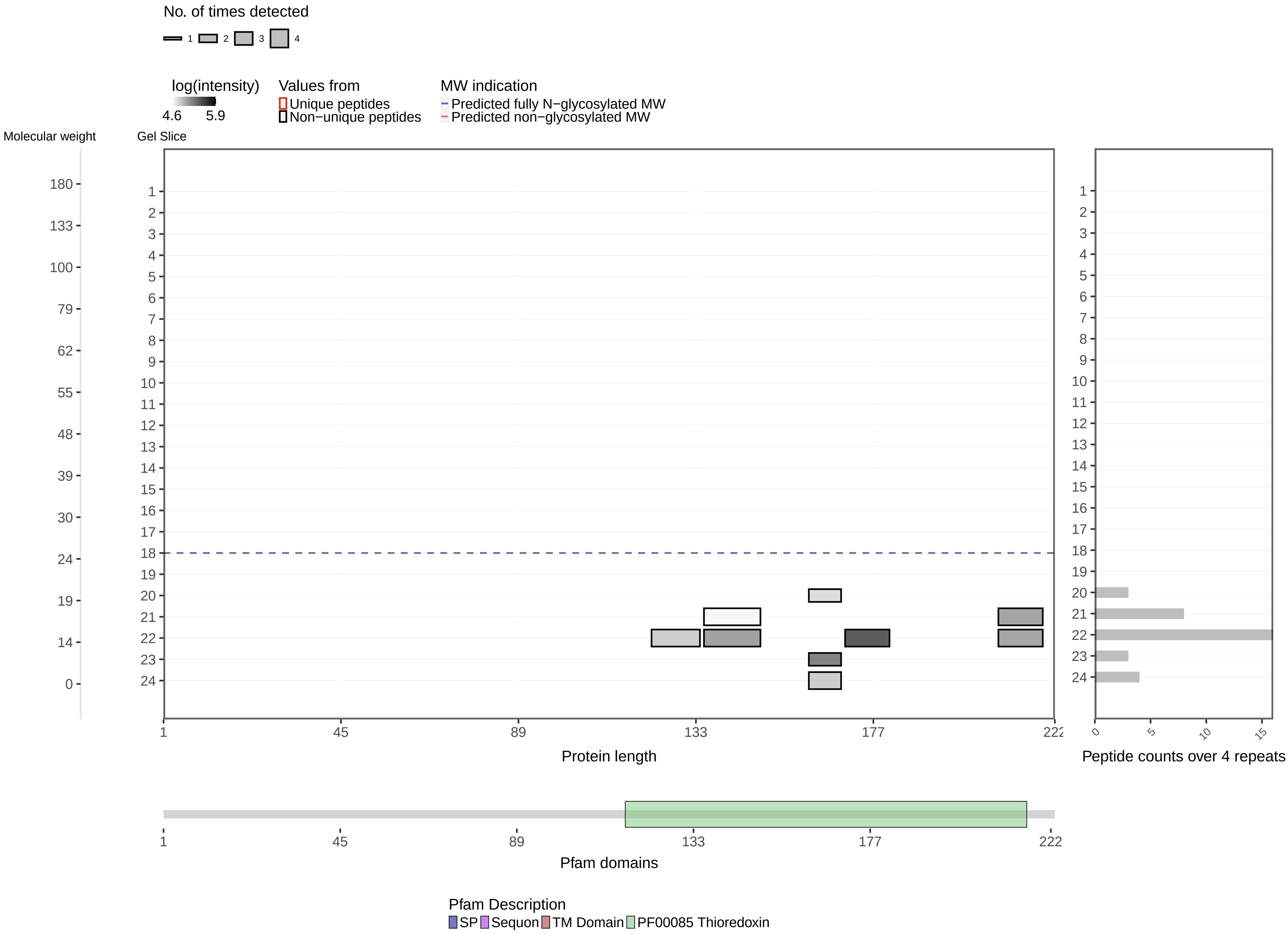


Protein sequence:

[illegible]VKCSSDATATTTVTLGQVTEVCKDTFWPIVEAAGDKTVVVDMYTQWCGPCKVIAPKFQELSKNYNDVVFLKLDCNQDNRPLAKELGIKVVPTFKILKNNKVVKEVTGAKLDNLIAAIEDVRSS*

### NbL10g08330.1 T−complex_protein_1_subunit_gamma−like [MW = 63.27 kDa]

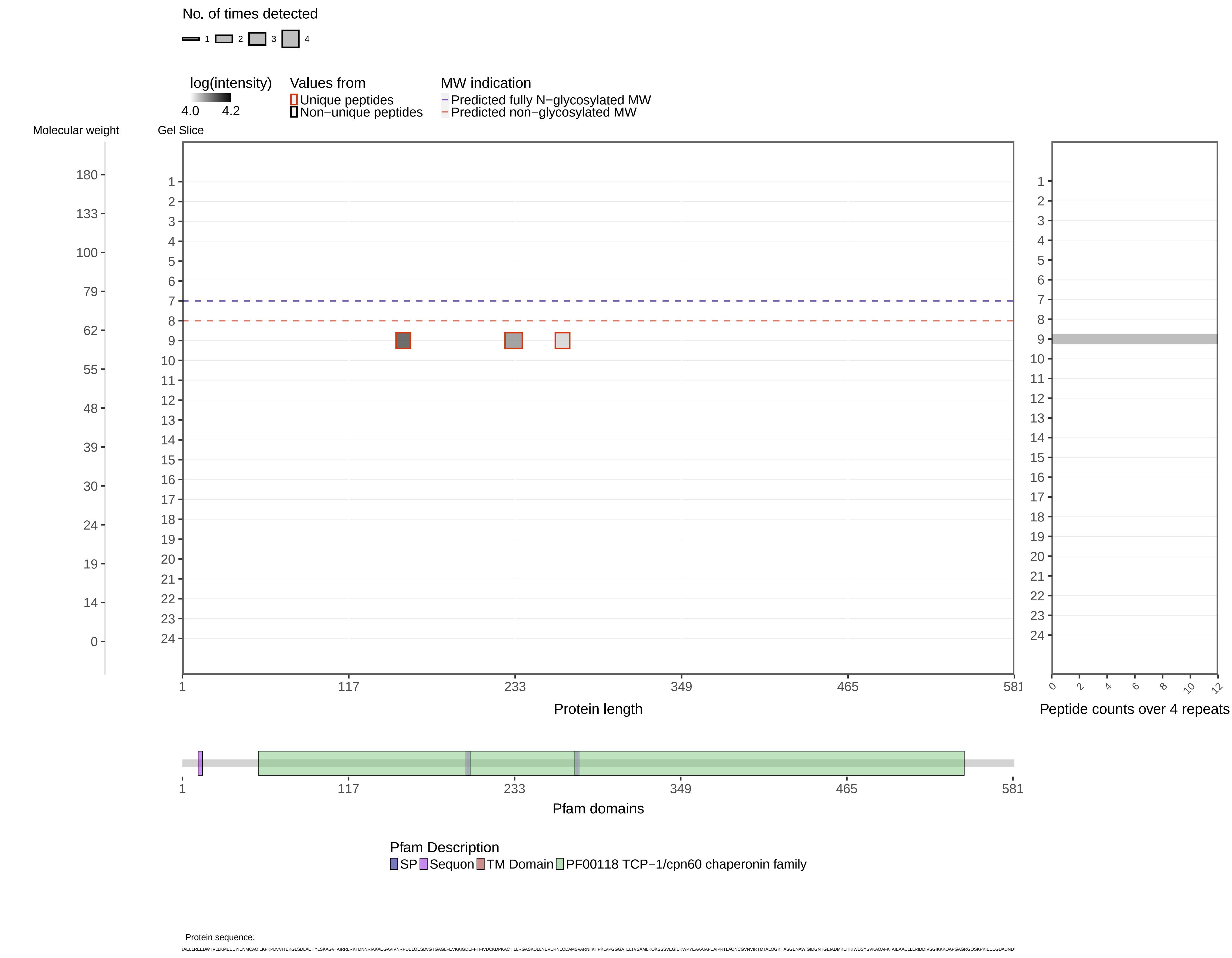

NbL10g08270.1 10_kDa_chaperonin_mitochondrial−like [MW = 10.46 kDa]

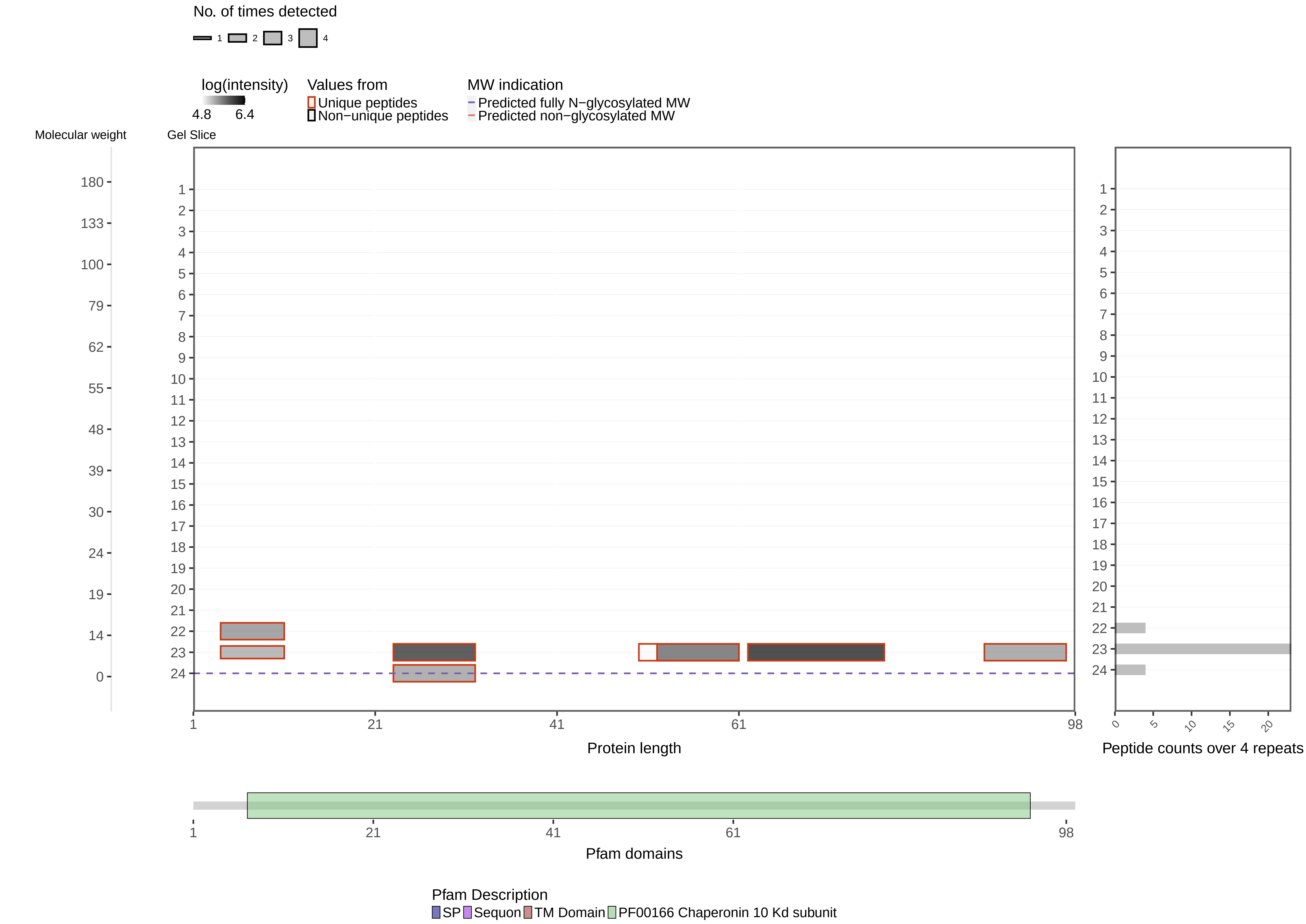


Protein sequence:

NSGKVVAVGPGLRDKSGNLIPTAVKEGDTVLLPEYGGTQVKLGEKEFHLYRDEDILGTLHD*

### NbL10g08140.1 SAL1_phosphatase−like [MW = 15.48 kDa]

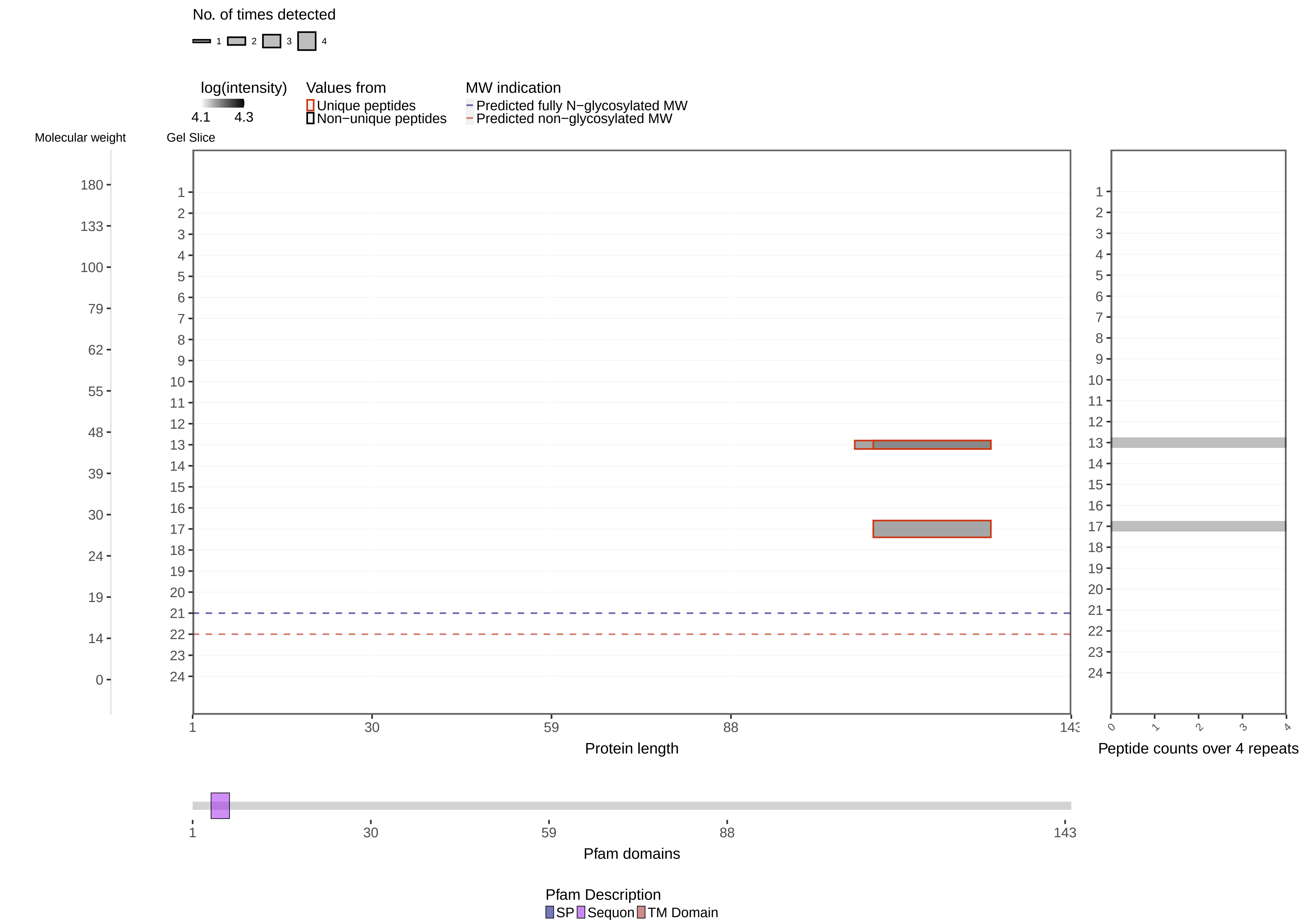


Protein sequence:

KASKTNLRTSSMSYDKQLAAAKKAASLAARLCQKVQKALLQADVQSKSDKSPVTVADYGSQAVVSIVLQKELDSASFSLVAEE

### NbL10g08030.1 acyl−coenzyme_A_thioesterase_13 [MW = 16.45 kDa]

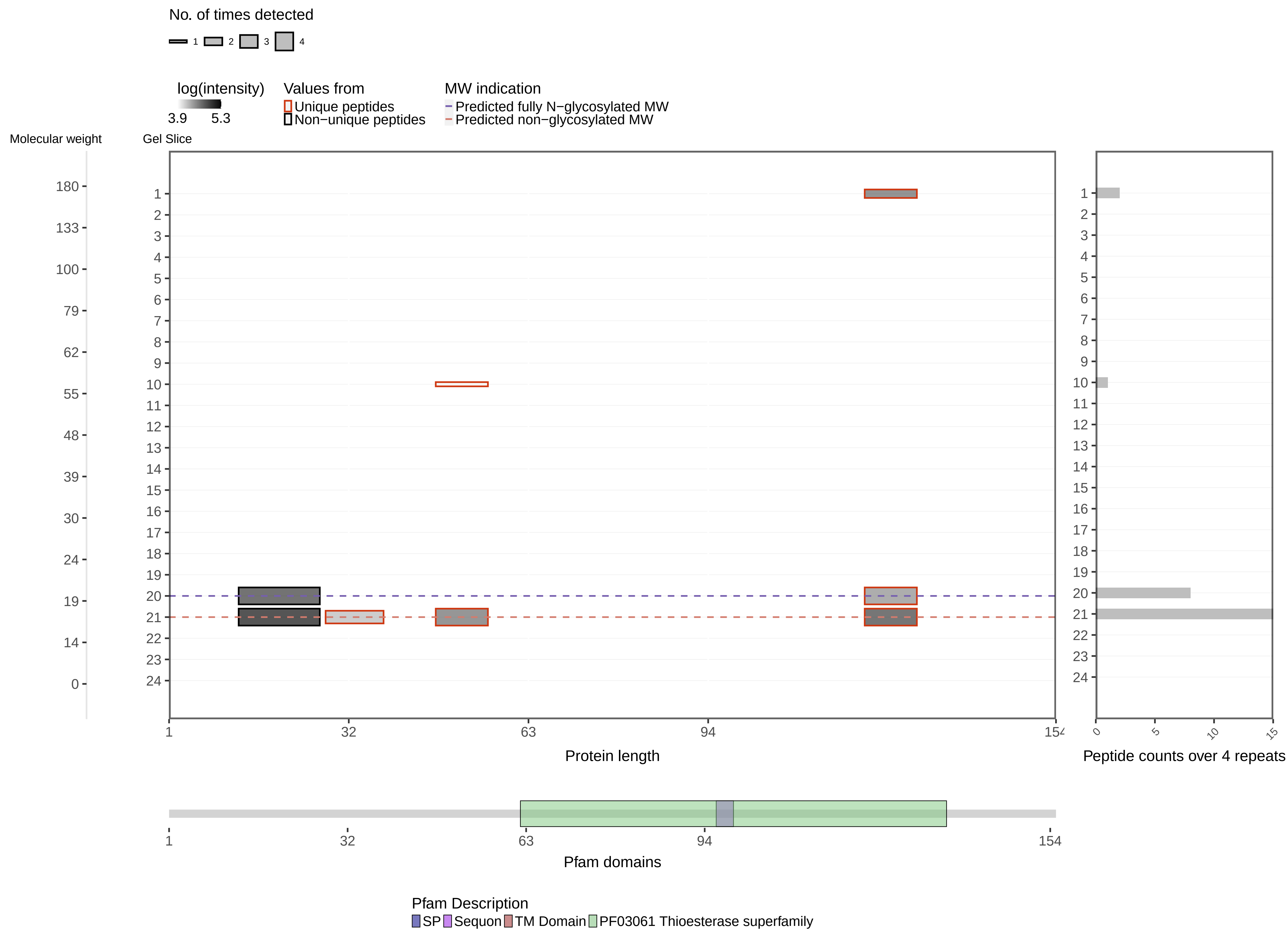


Protein sequence:

FLHGGATASLVDILGSAVIHTVGAAFTGVSVEINVSYLDAAFVGEEIEVEATVLRVGKTIGVVSVELRKKETGKIVARGRHTKYLAVNSKL*

NbL10g07990.1 4−hydroxy−tetrahydrodipicolinate_synthase_chloroplastic [MW = 39.91 kDa]

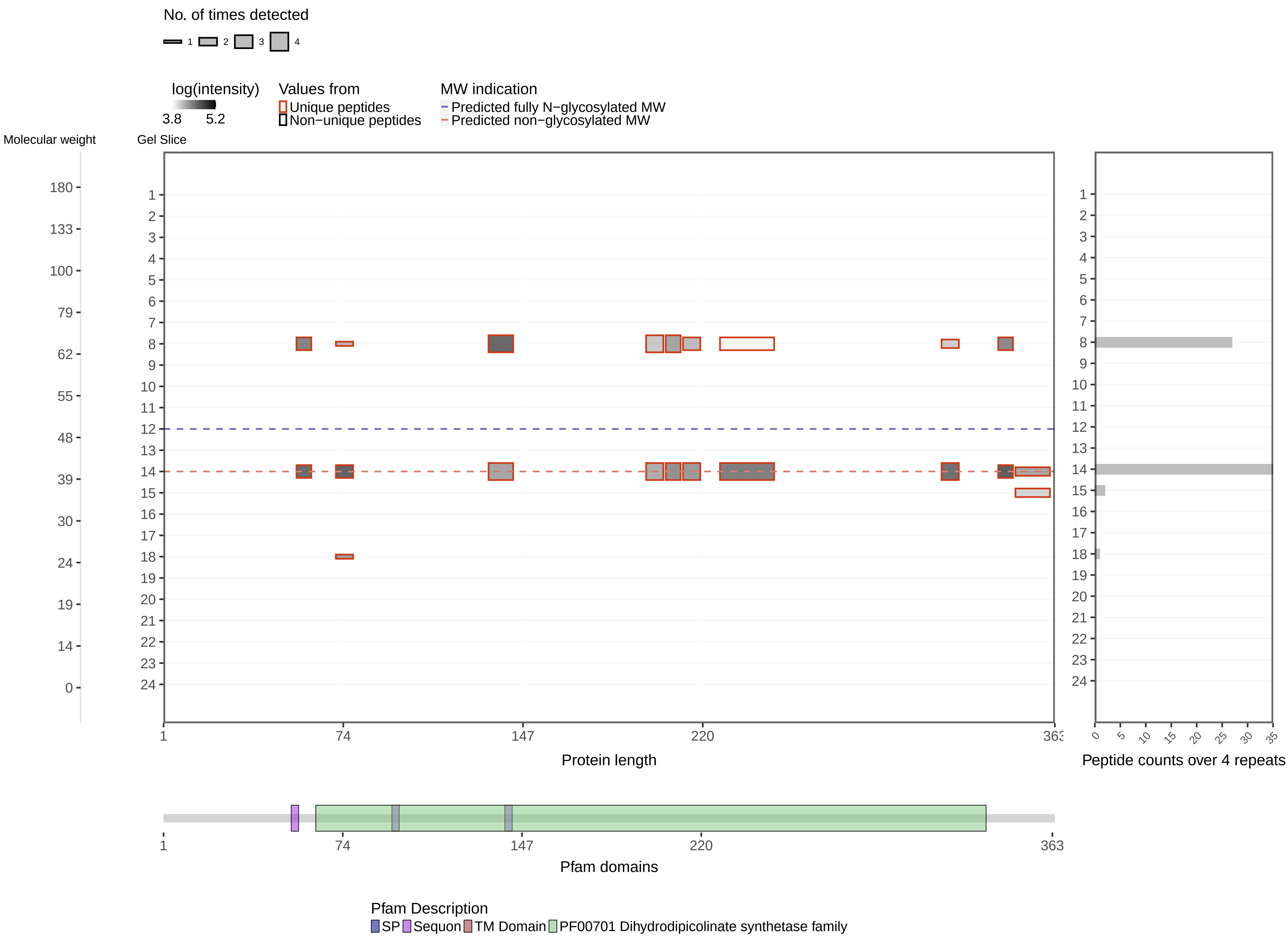


Protein sequence:

[illegible]TSLEGLISHFESVLPMGPTIIYNVPSRTGQDIPPRVIQTMAKNPNLAGVKECVGNDRVEQYTSNGIVVWSGNDDECHVSRWDYGATGVISVTSNLVPGLMRELMFGGKNLALNSKLMPLIEWLFHEPNPIALNTALAQLGVVRPVFRLPYVPLTKEKREEFVEIVKEIGRENFIGERDVQVLDDNDFILVARY*

### NbL10g07980.1 probable_methyltransferase_PMT26 [MW = 89.25 kDa]

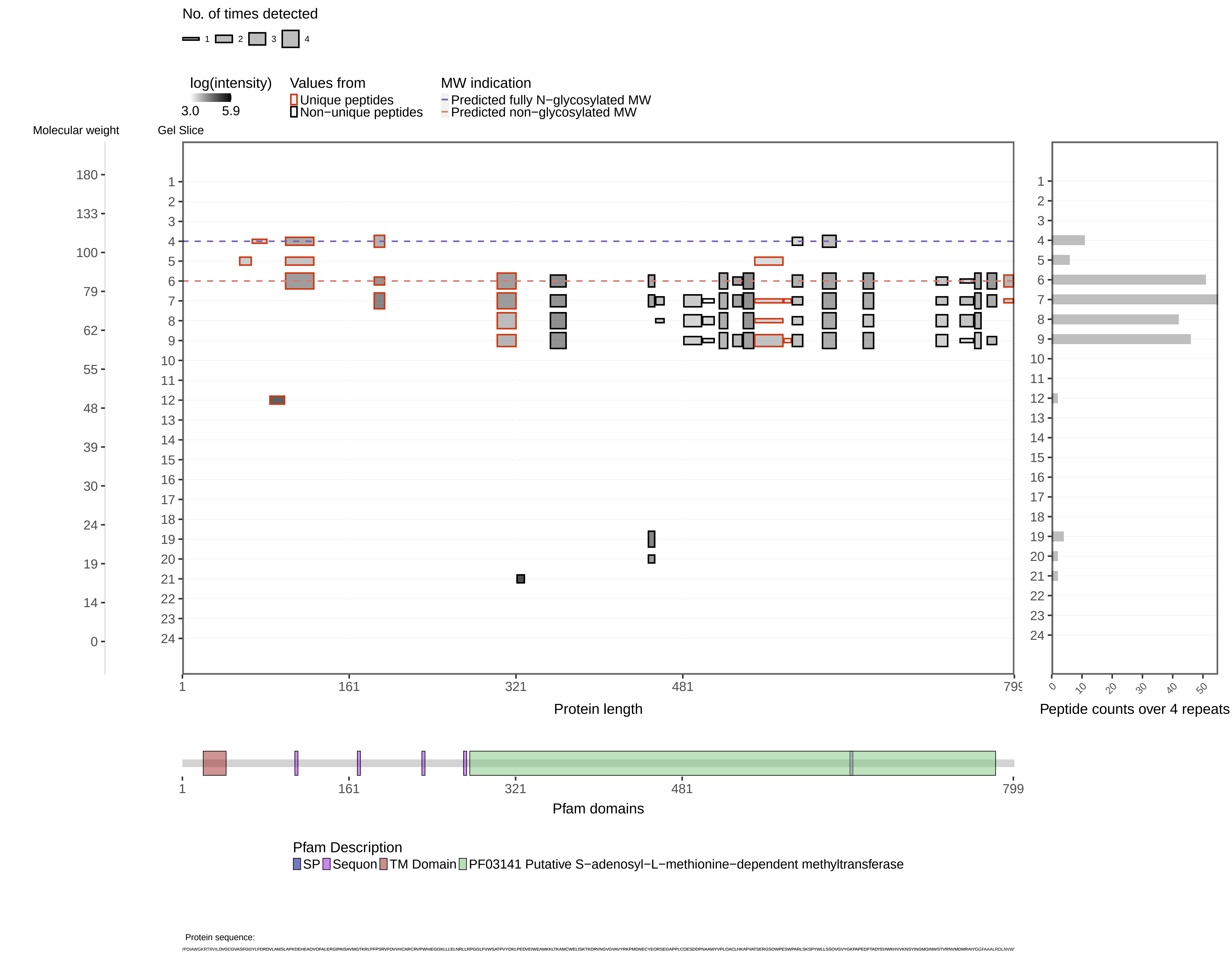

NbL10g07910.1 proteasome_subunit_beta_type−6−like [MW = 25.18 kDa]

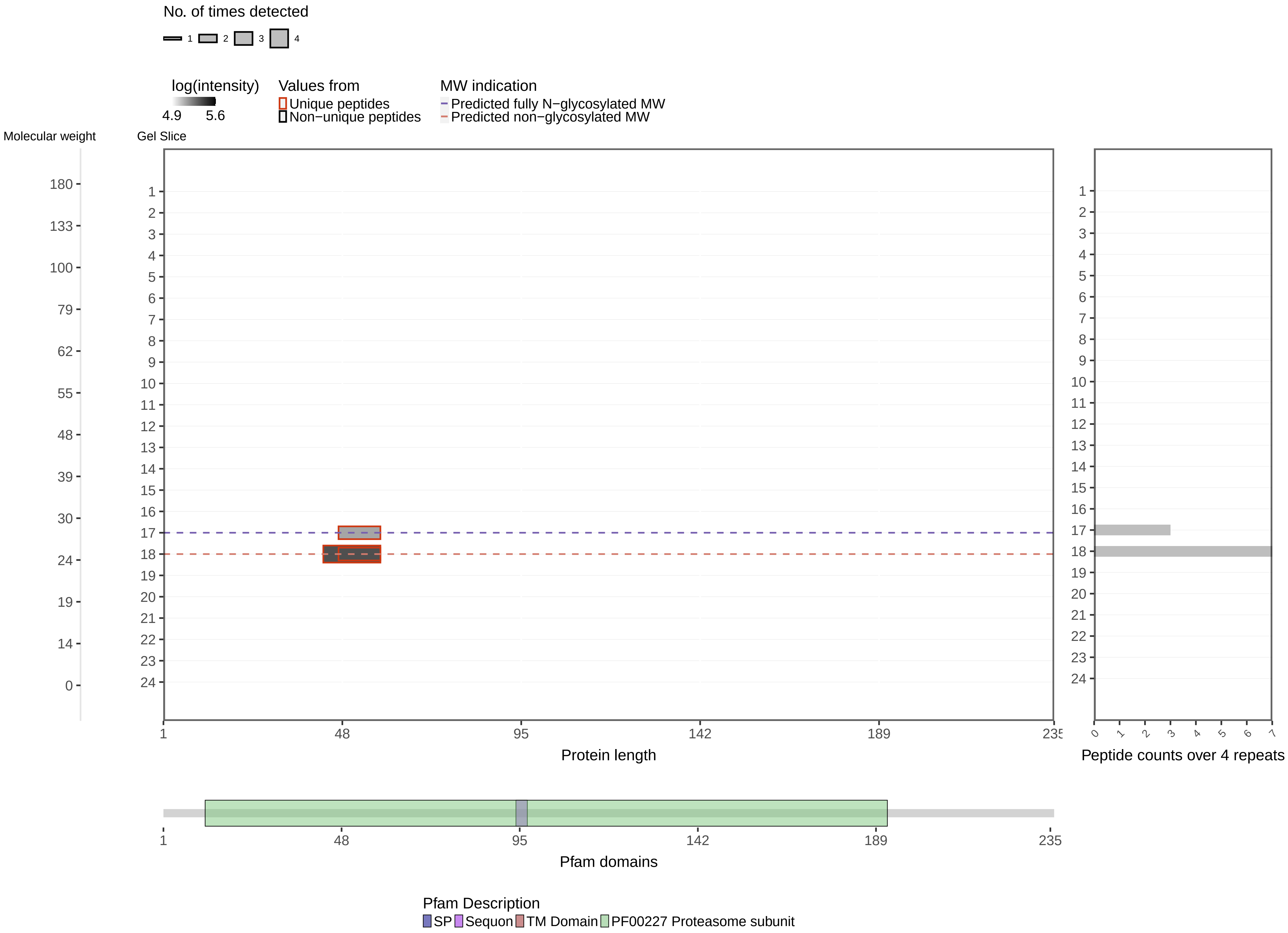


Protein sequence:

DRLQTGMIIGGWDKYEGGKIYGIPLGGTVLEQPFAIGGSGSSYLYGFFDQAWKEGMTQEEAEKLVVTAVSLAIARDGASGGVVRTVTINKDGATRKFYPGDSLQLWHEELEPVNSLLDVVSASSPDPMVS*

NbL10g07690.1 protein_usf−like [MW = 26.45 kDa]

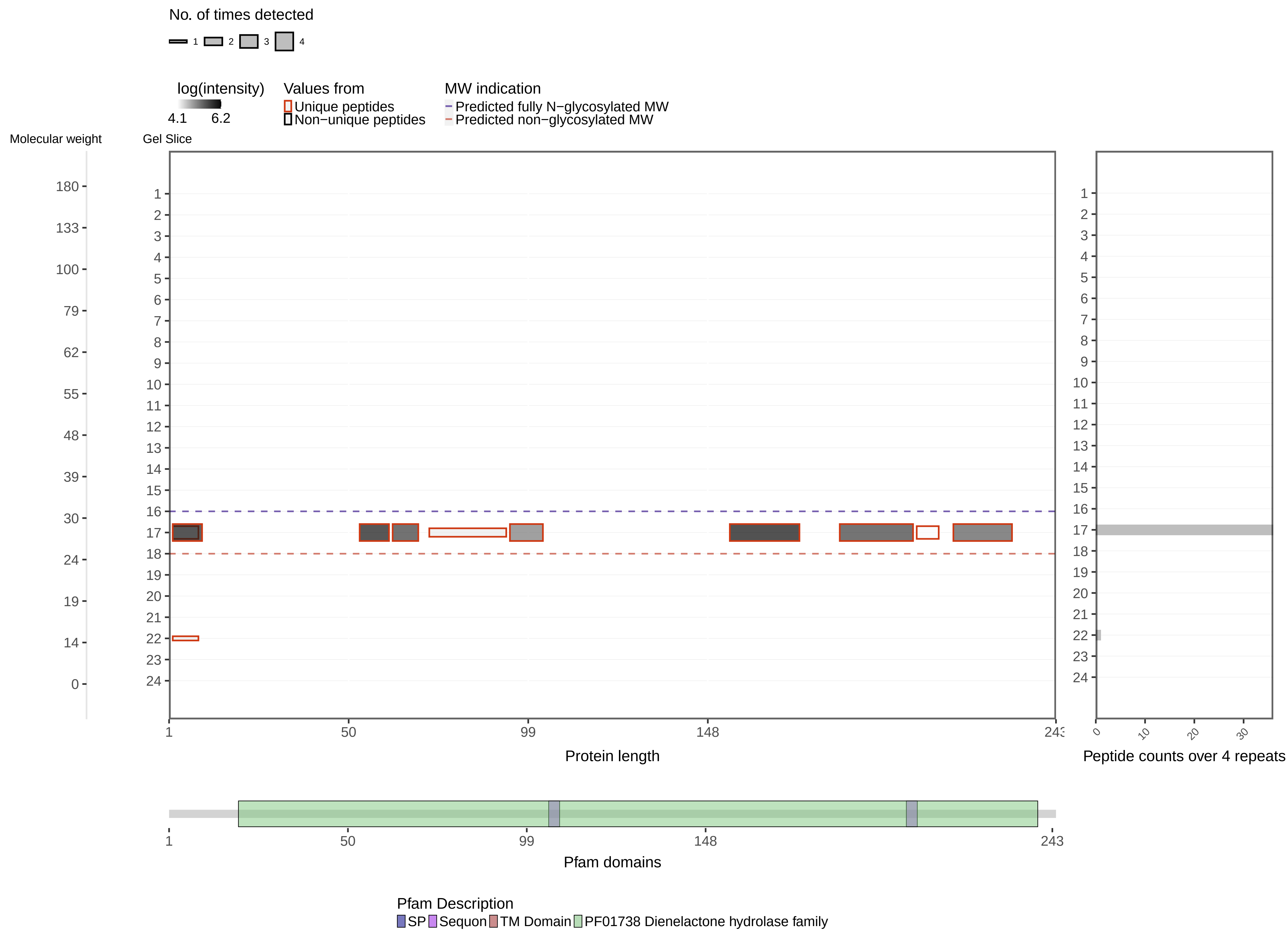


Protein sequence:

'GVTGYCMGGALAIASSVLVPGVDAVVAFYGVPSPELADPITAKAPVQAHFGEDDGFVGFSDVKTGKALEEKLKTSGVPHEVYFYPQTGHAFMNKSPEGVERRKKMGMPDVEDAAVEQAWTRFRSWMSRFLSA*

NbL10g07570.1 cytochrome_b561_and_DOMON_domain−containing_protein_At5g47530−like [MW − SP = 41.48 kDa]

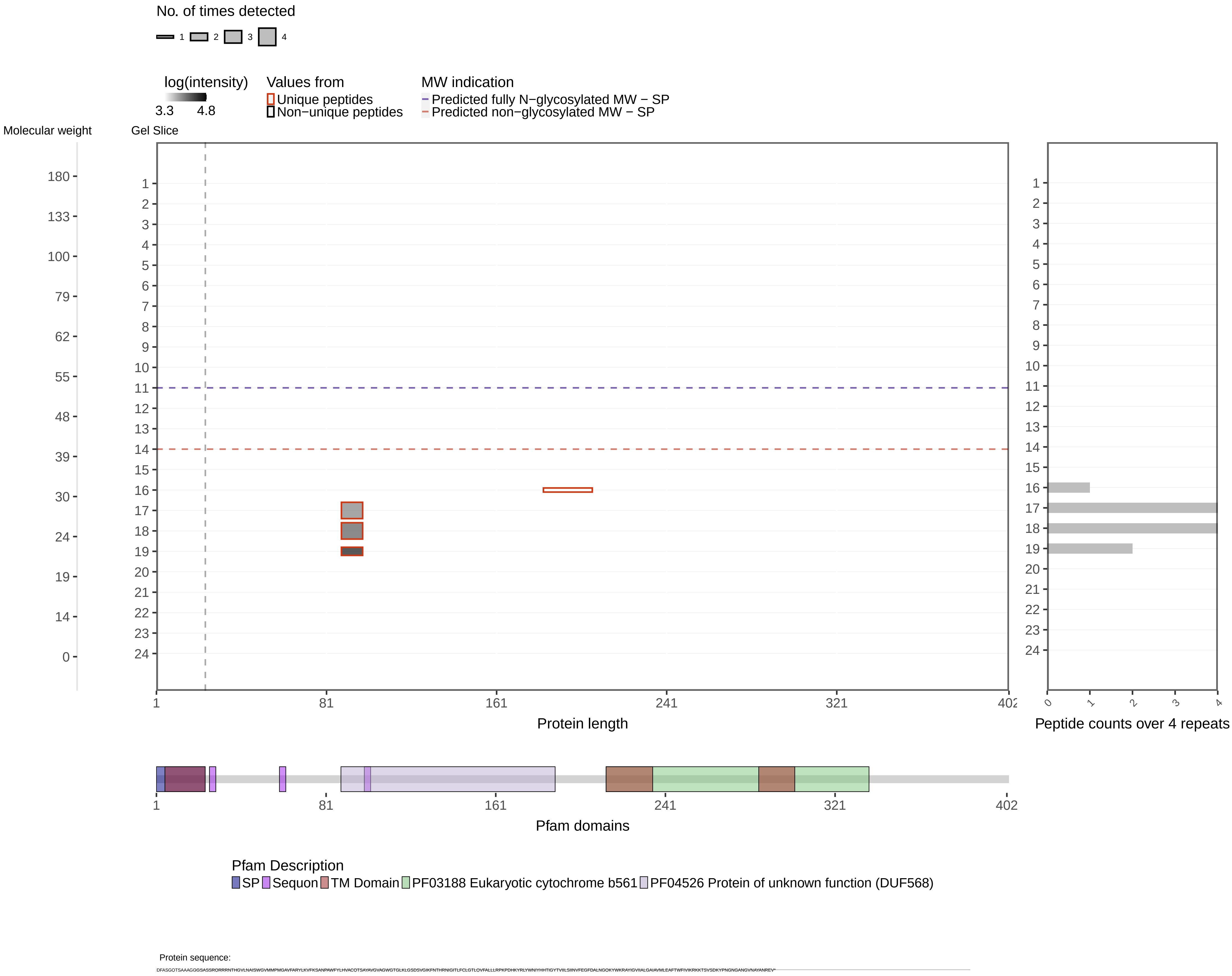

### NbL10g07550.1 thymidylate_kinase−like [MW = 28.98 kDa]

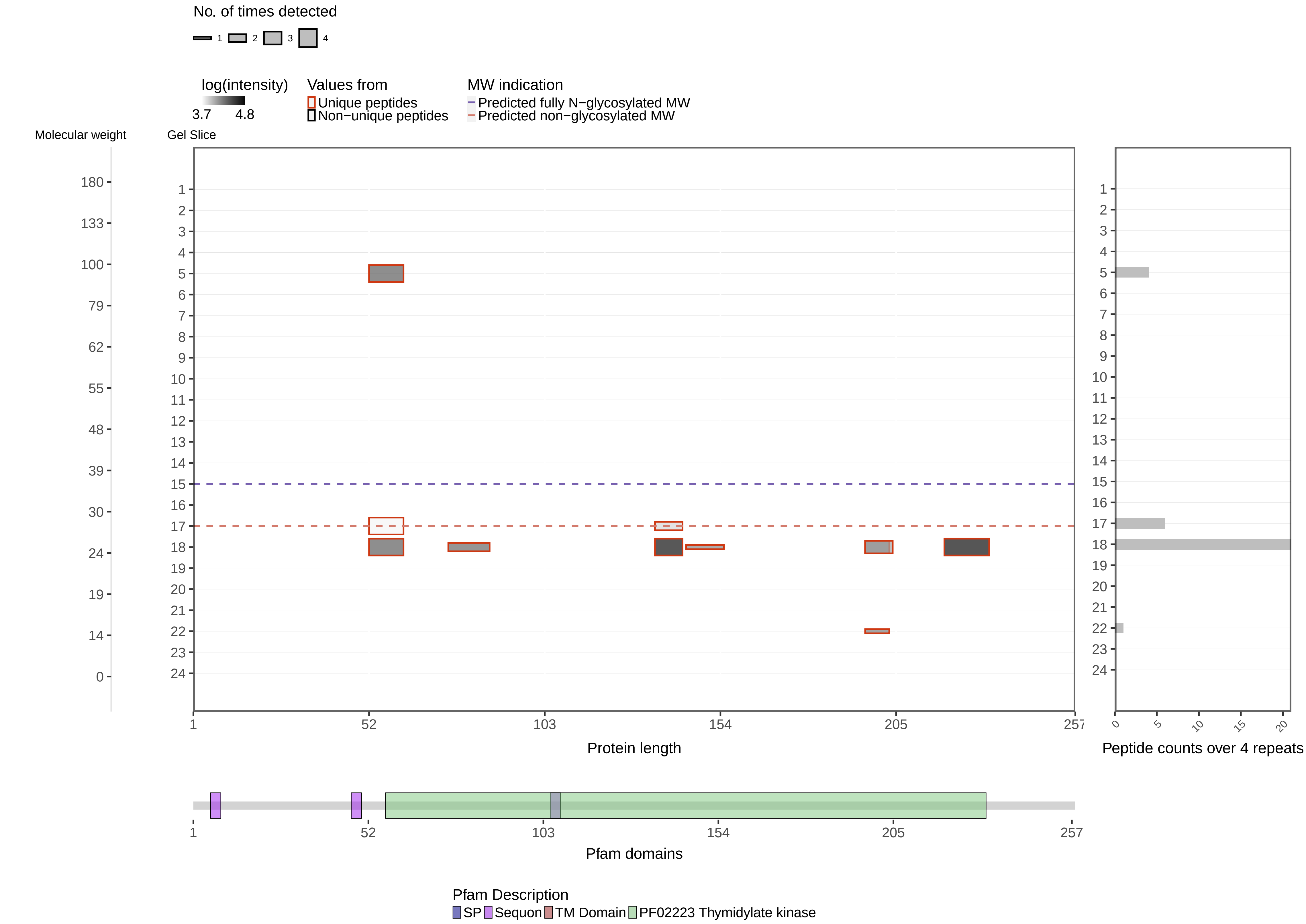


Protein sequence:

IHLLFSANRWEKRMLMEDKLRSGTTLIVDRYSYSGVAFSSAKGLDIEWCKAPEIGLLAPDLVIYLDISPEKAAERGGYGGERYEQLEFQKKVAKSYLTLHDSSWKVIDATLPVEDIEEKLREIVLDSMTICQKGKPLSQLWSS*

### NbL10g07470.1 uncharacterized_LOC109241615 [MW = 13.13 kDa]

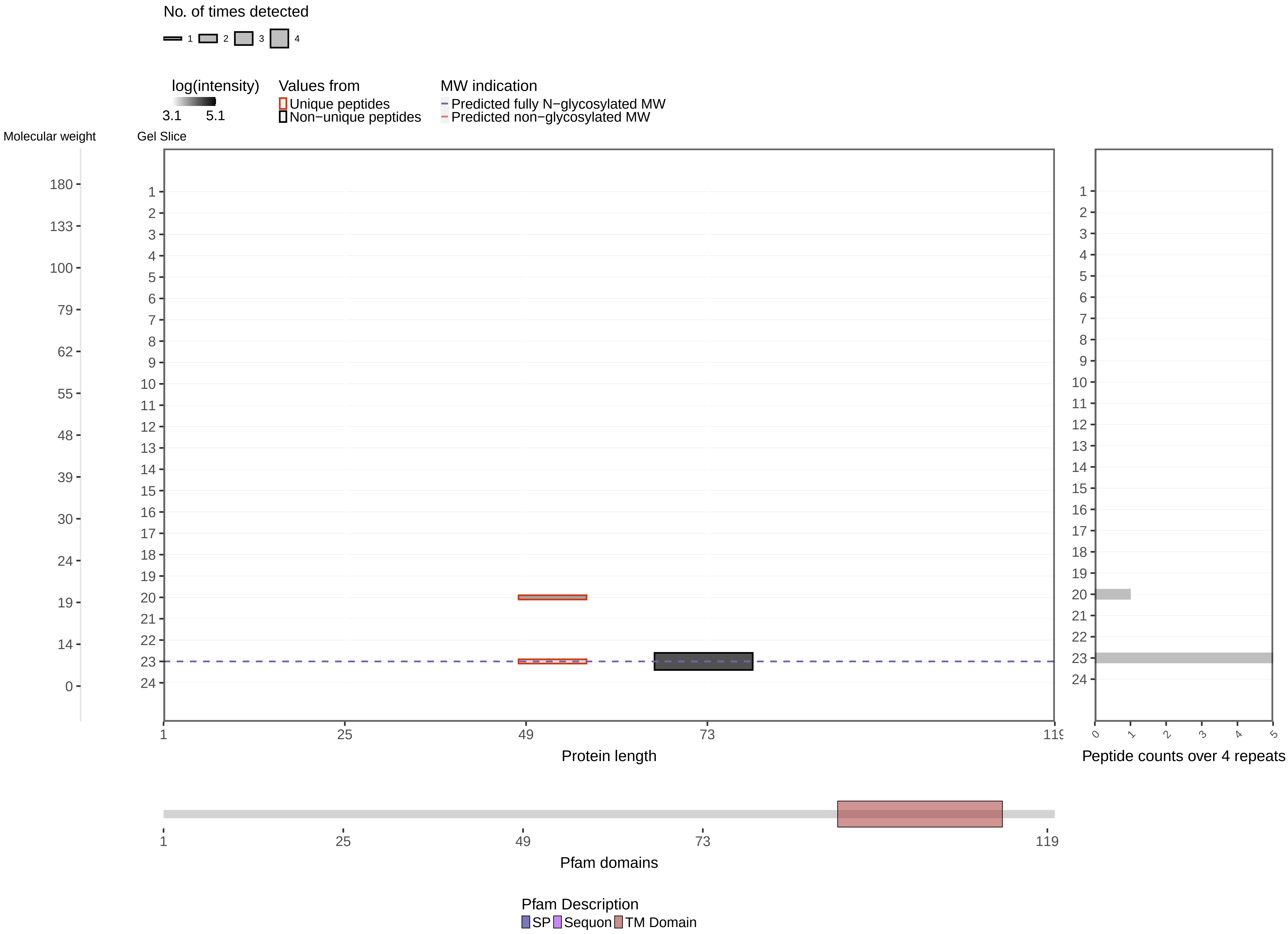


Protein sequence:

AKLGGEEGDVKKDKKKFITKEQEPEQYWQTAGERAGENPMMTPLPYIIIFGMSTPFVILAVAFANGWIKVPIR*

### NbL10g07460.1 nitrile−specifier_protein_5 [MW = 35.43 kDa]

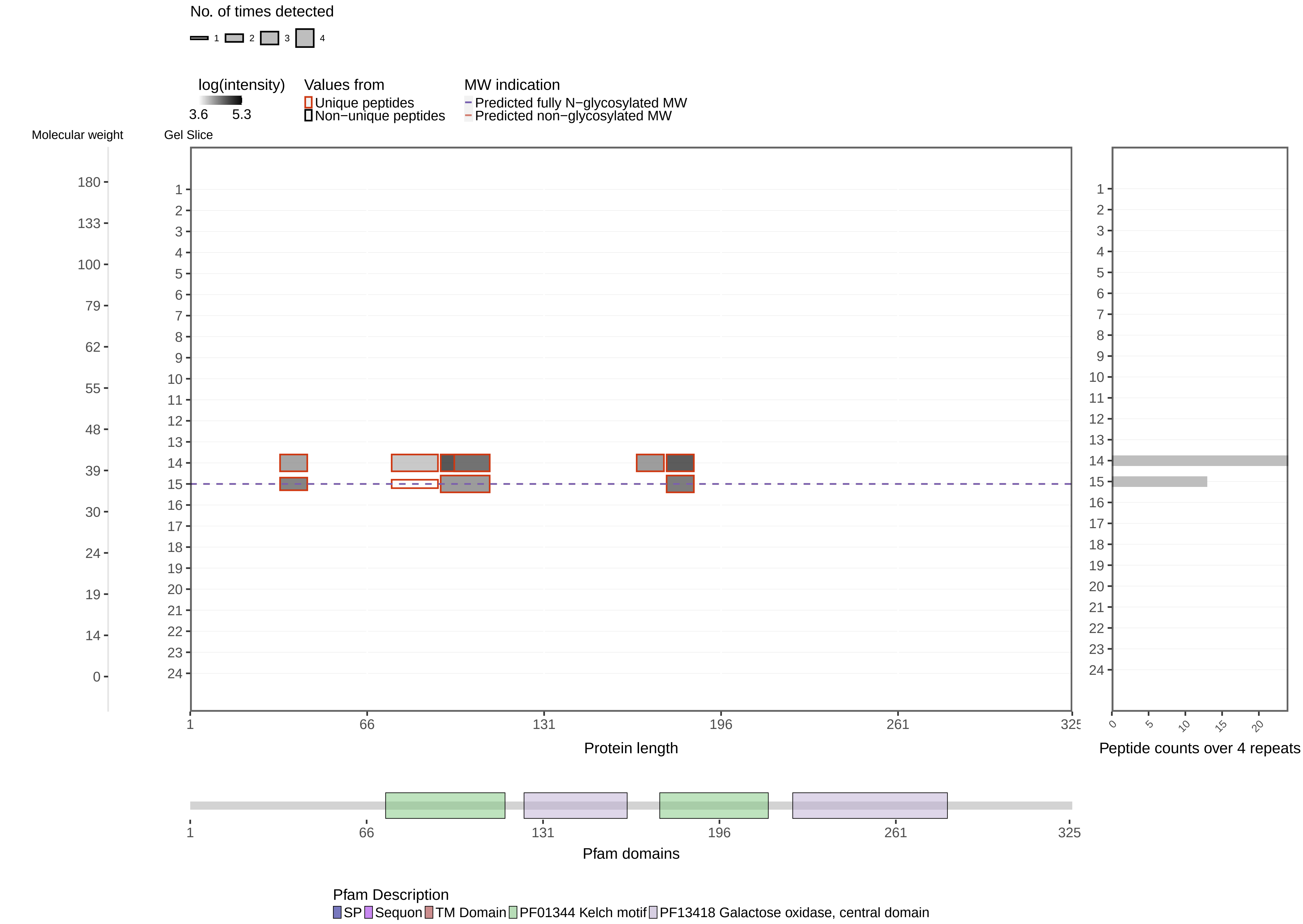


Protein sequence:

SDLWAYDAVDKKWIKFPVPGENCKPRGGPGLTVILGKIWVVYGFSGDELDDVHYFDAIEEKWVOVETNGEKPTARSVFSTVGIGKYIFIYGGEIDPSDLGHLGAGKFSGDVYALDTETLVWKOWIDSGDHPGPRGWCAFASGRRDGEEGLLVYGGNSPSNDRLGGIFFFTPLVET*

### NbL10g07330.1 nitrile−specifier_protein_5−like [MW = 38.52 kDa]

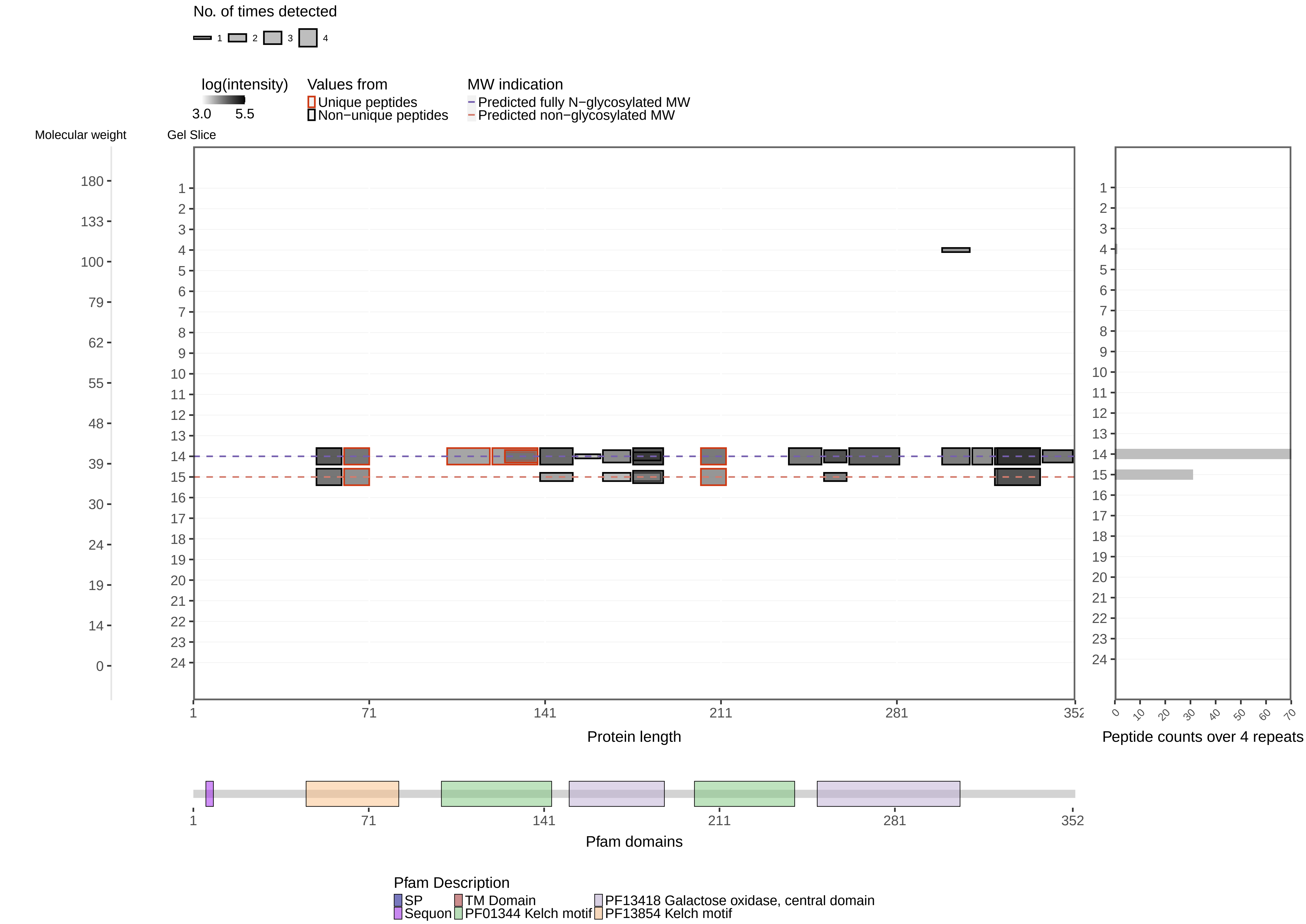


Protein sequence:

RIYVFGGCGNAGRLSDLWAYDAVDKKWIEFPVPGENCKPRGGPGLAVILGKIWVVYGFSGDELDDVHYFDAIEEKWVQVETNGEKPTARSVFSTVGIGKYIFIYGGEIDPSDLGHLGAGKFSGDVYALDTETLVWKQWIDSGDHPGPRGWCAFASGRRDGEEGLLVYGGNSPSNDRLGGIFFFTPLVET*

### NbL10g07320.1 4−hydroxyphenylpyruvate_dioxygenase [MW = 48.97 kDa]

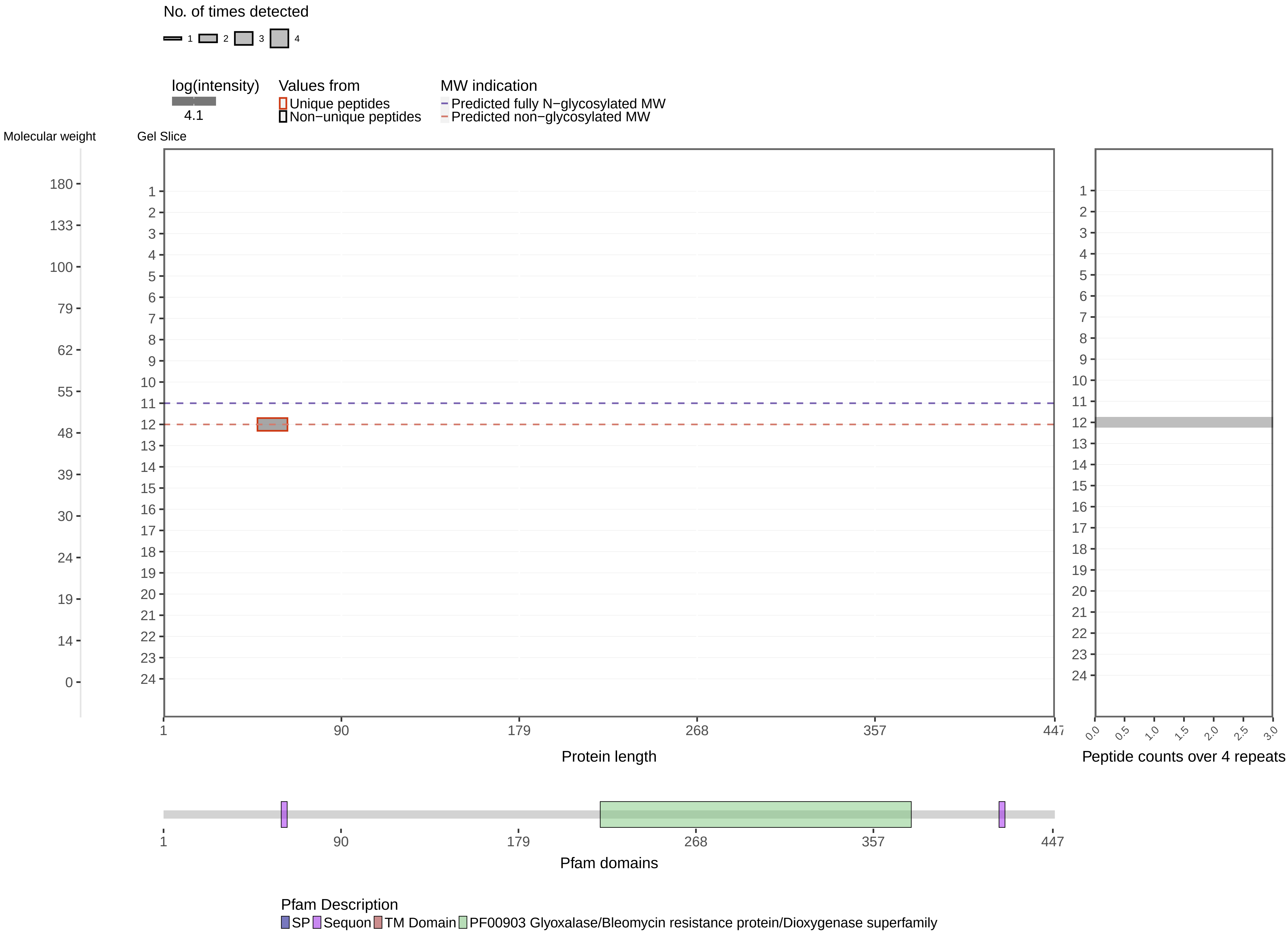


Protein sequence:

YKELDYGIRRLDHAVGNVPELGPAVDYIKRFTGFHEFAEFTSEDVGTAESGLNSMVLANNDETVLLPLNEPVYGTKRKSQIQTYLEHNEGAGVQHLALVTEDIFKTLKEMRKRSGVGGFEFMPSPPPTYYKNLKNRAGDVLTDEQIQACEDLGILVDRDDQGTLLQIFTKPVGDRPTIFIEIIQRIGCMLKDEKGQVYQKGGCGGFGKGNFSELFRSIEEYEKTLEAKQANQVAAA*

### NbL10g07270.1 oxygen−evolving_enhancer_protein_2−2_chloroplastic [MW = 28.72 kDa]

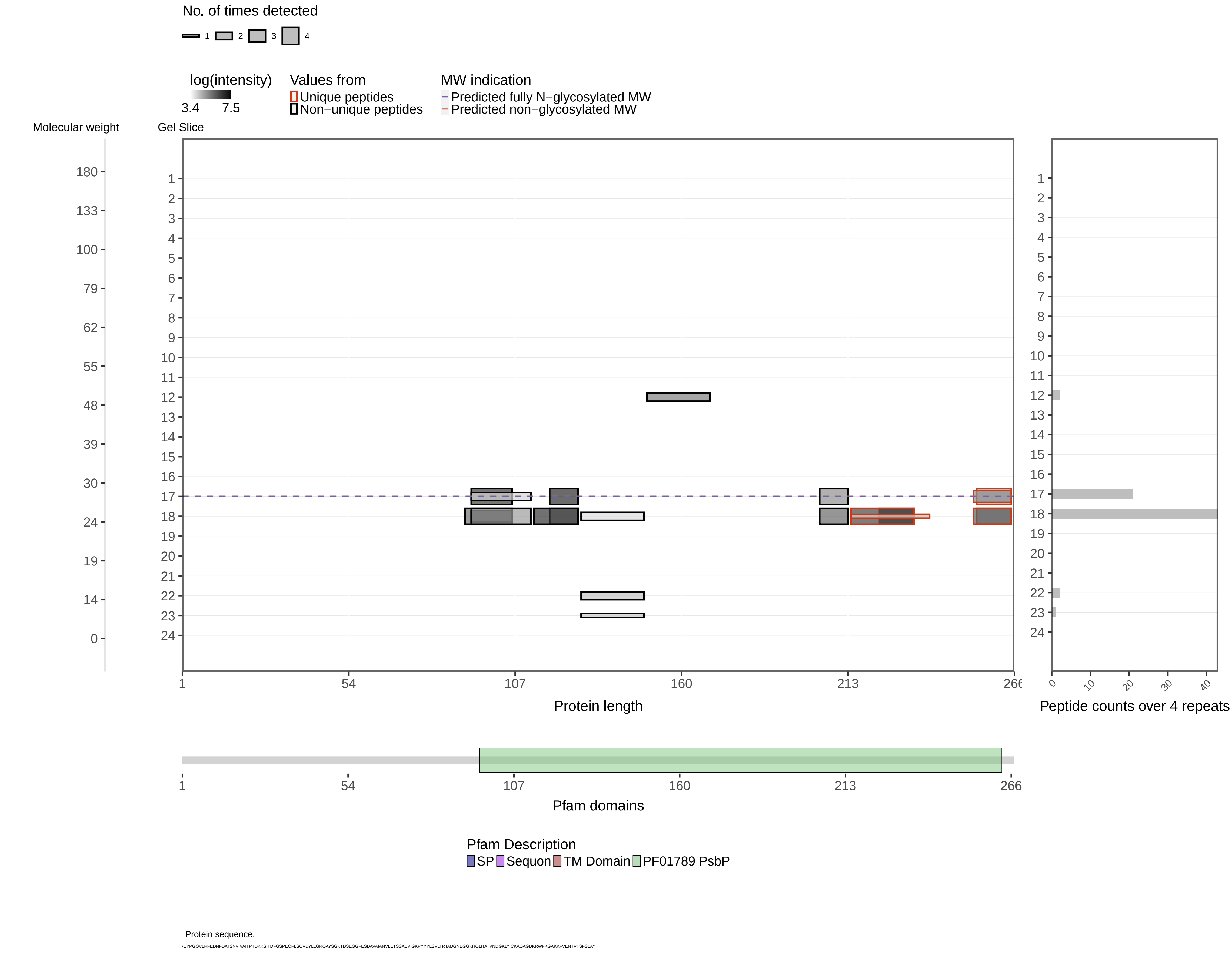

### NbL10g07210.1 switch−associated_protein_70 [MW = 61.47 kDa]

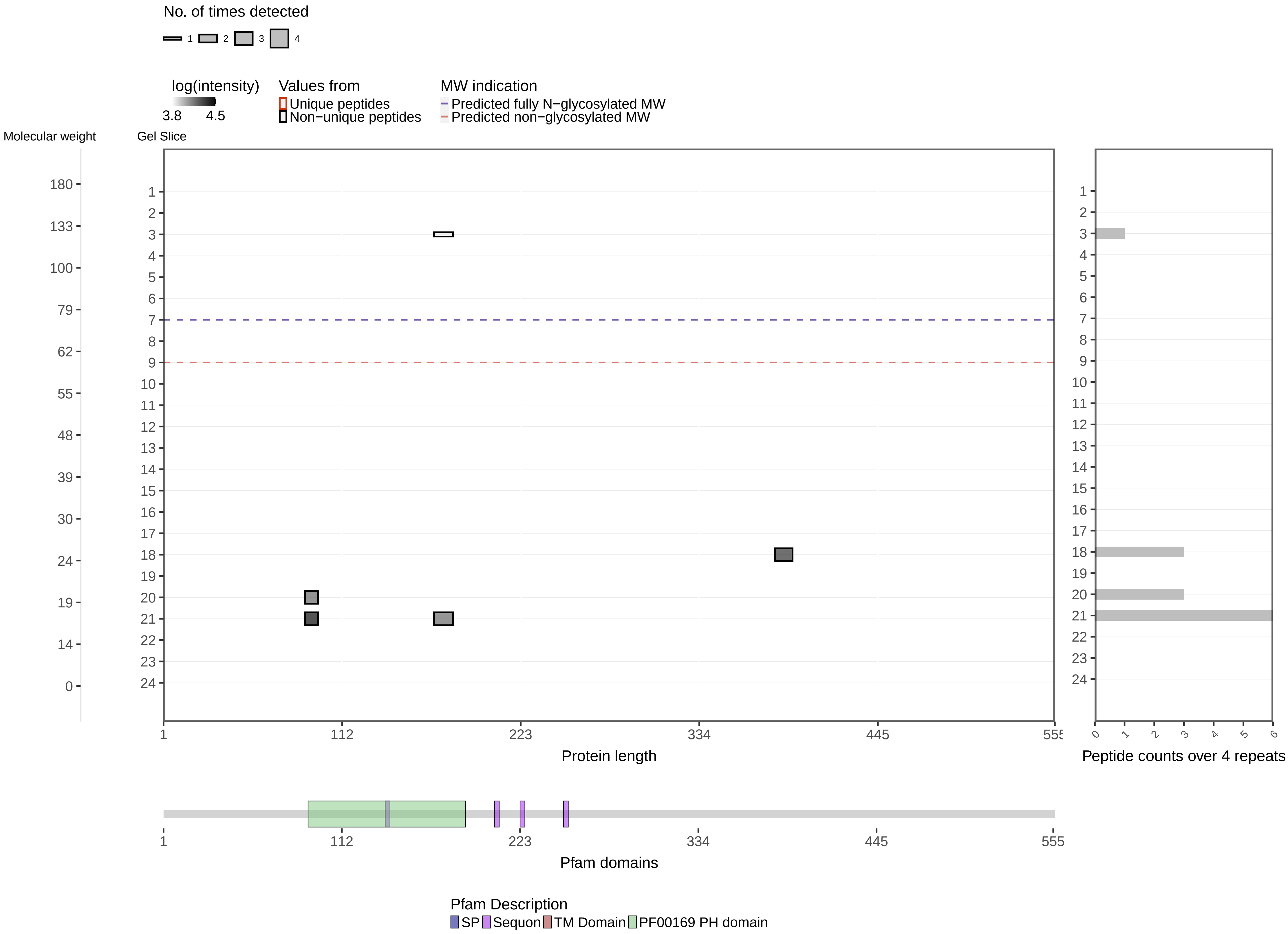


Protein sequence:

TLRVKDDELQNLAREVRARDSTIKELVDKLSETAEAAEAAASAAQTMNEQRKIACAEVERLKGDSEKQLESSKSKLREFEEKFMTLSKEMDQLIKQRDSAIQEAHLWRSELAKAREQAVILEGAAVRAEERVRVTEADAETRIKEAAQTEAASAKEKQELLAYVNMLQAQLTRQQVDMNQVFEKTESCSSSINNLPQTKHVDLSEENVDKACLSVSRAIDPENVVDLAVDQTNHQSSGHGEWSDIQPTESRIADVREVASETERSSLDIPVVISPSNTHQDHGHNSHLP*

NbL10g07170.1 cytochrome_P450_97B2_chloroplastic [MW = 64.65 kDa]

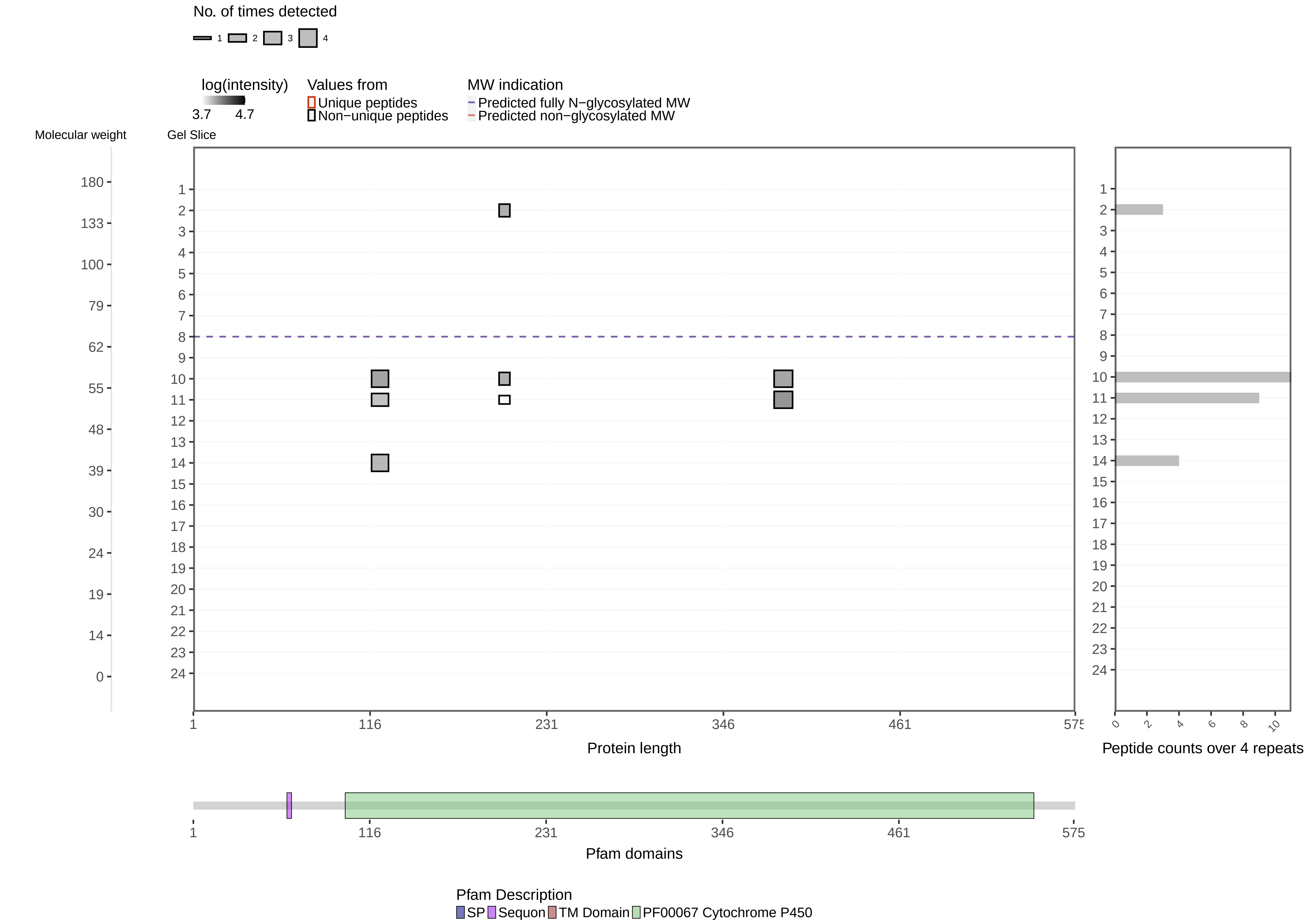


Protein sequence:

.VPRQRKFQNDLKVINDCLDGLIQNAKETREEADVEKLQQRDYLNLKDASLLRFLVDMRGVDVDDRQLRDDLMTMLIAGHETTAAVLTWAVFLLAQHPVKMKKAQSEIDAVLGQGRTTFESLKKLEYLRLIVVESLRLYPQPPLLIRRSLKSDKLPGGYNGDKDGYEIPAGTDVFLSVYNLHRSPYFWDKPDEFEPERFLVPKESSVEGWAGFDPSRSPGALYPNEIISDFAFLPFGGGPRKCVGDQFALMESTIALAMLLQKFDVELRGSRESVELVTGATIHTKTGLWCKLKK

NbL10g07150.1 probable_pectinesterase_8 [MW = 35.35 kDa]

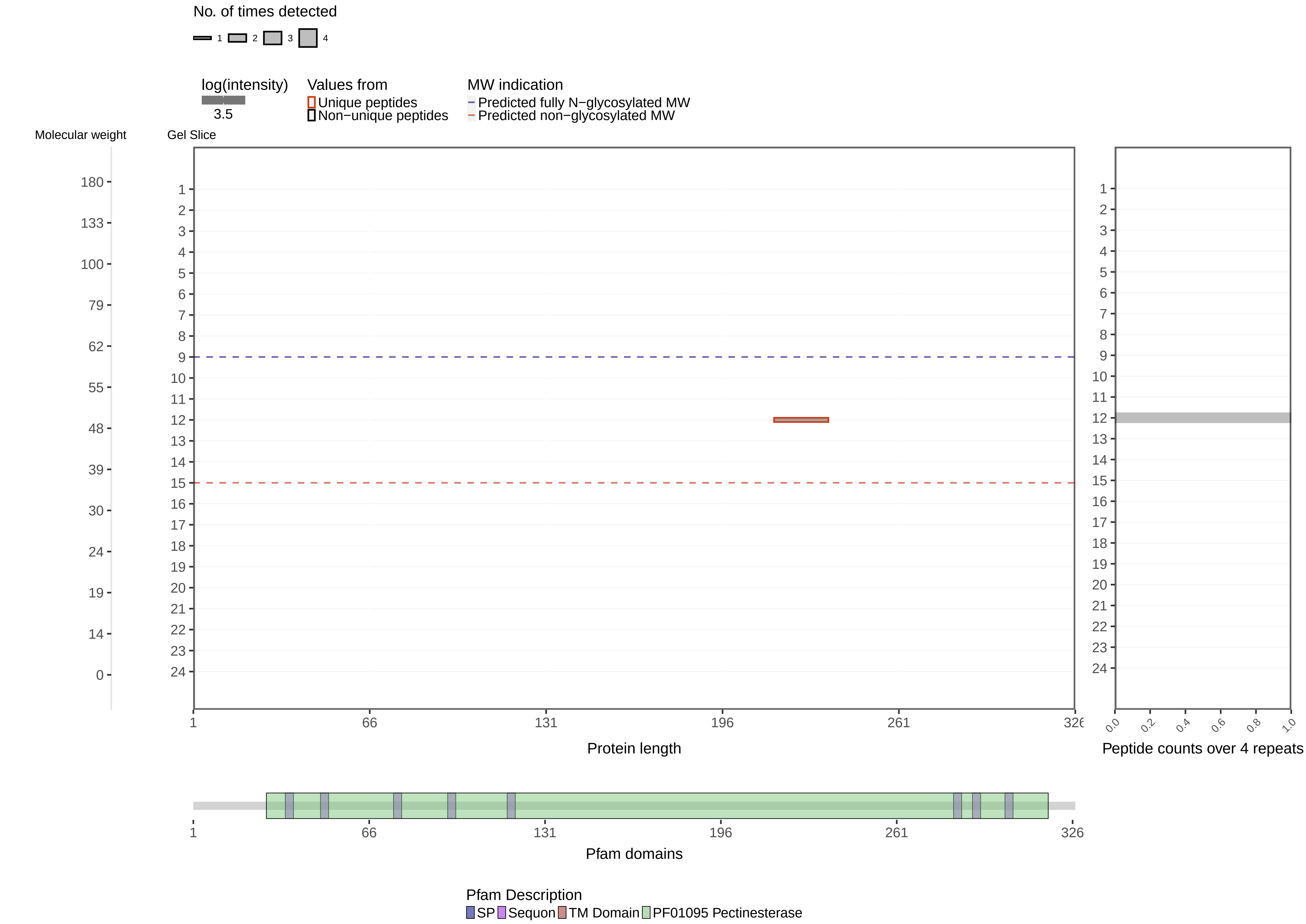


Protein sequence:

FGAQDTLHDDRGRHYFKECYIQGSIDFVFGNGKSLYENCELISMASPGWRSINGAVTAHGRASKEEDSGFAFVNCNVGGSGRIWLGRAWRPFSTVVFANSFMTNIISPDGWNDLNDPSRDQTIFYGEYNCTGAGSNMSQRAPYVQKLNDTQALPFLSSSFIDADQWLQSFSN*

NbL10g07080.1 glucan_endo−13−beta−glucosidase_6−like [MW − SP = 50.11 kDa]

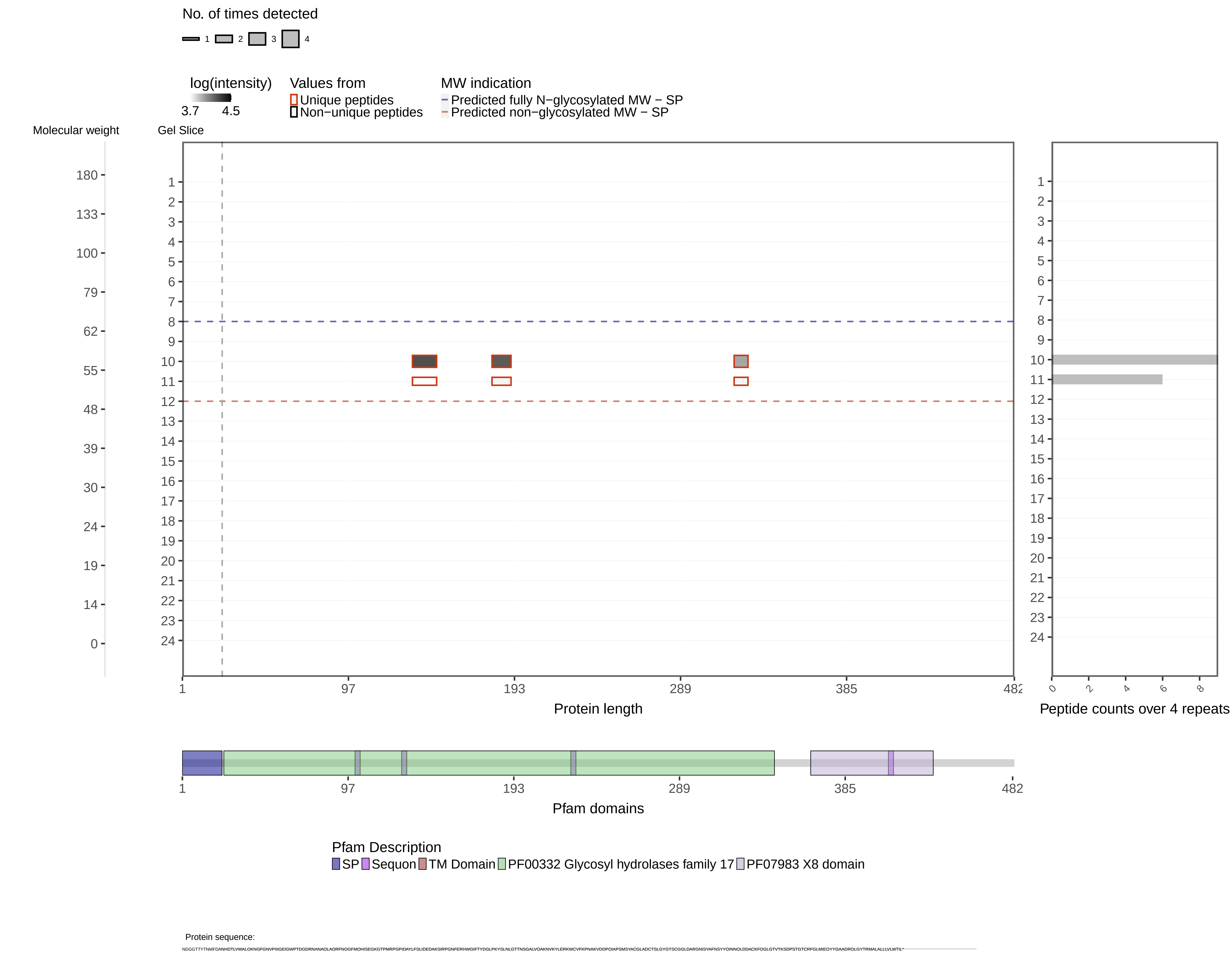

### NbL10g07050.1 uncharacterized_LOC104212881 [MW = 66.97 kDa]

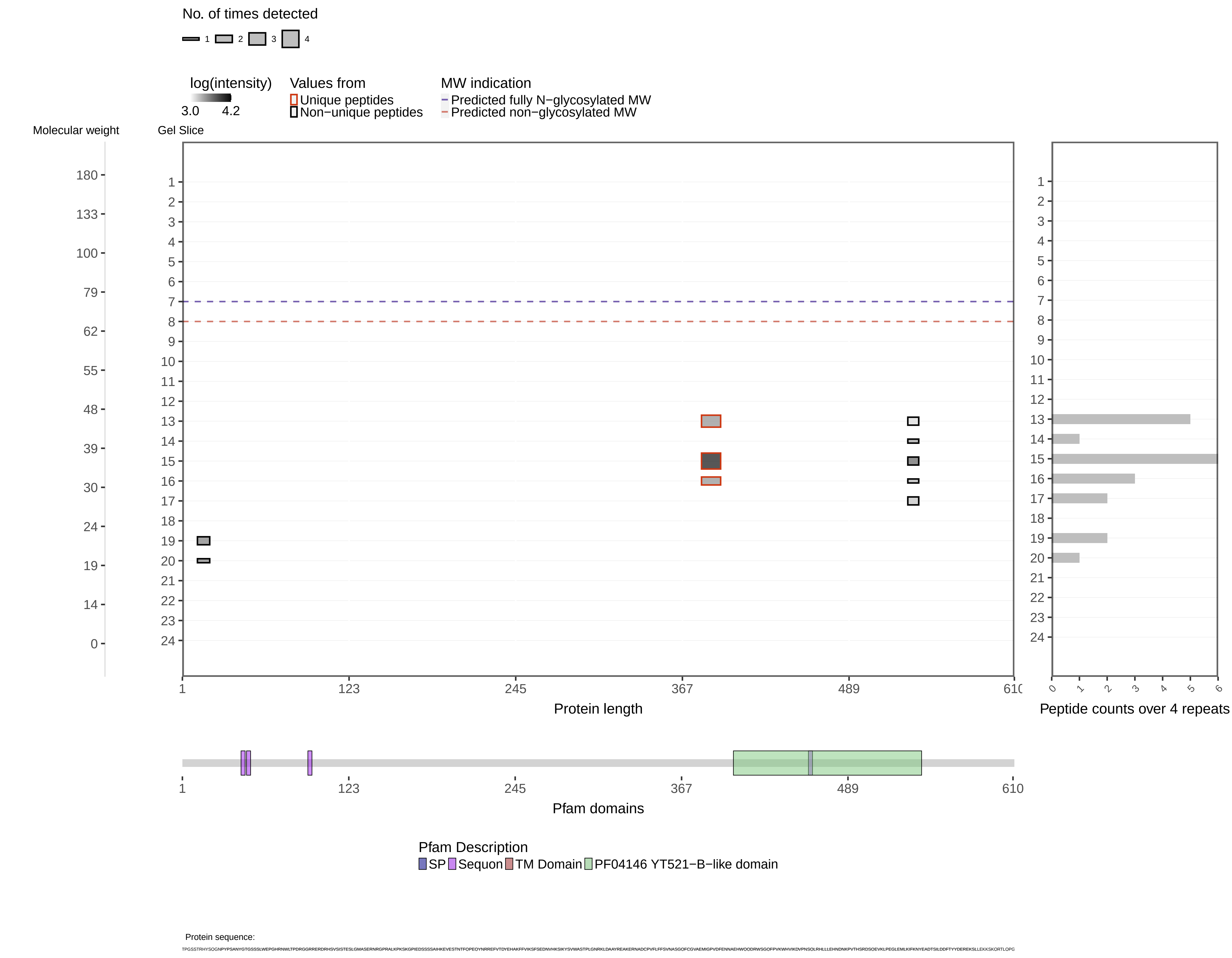

NbL10g06950.1 20_kDa_chaperonin_chloroplastic−like [MW = 26.22 kDa]

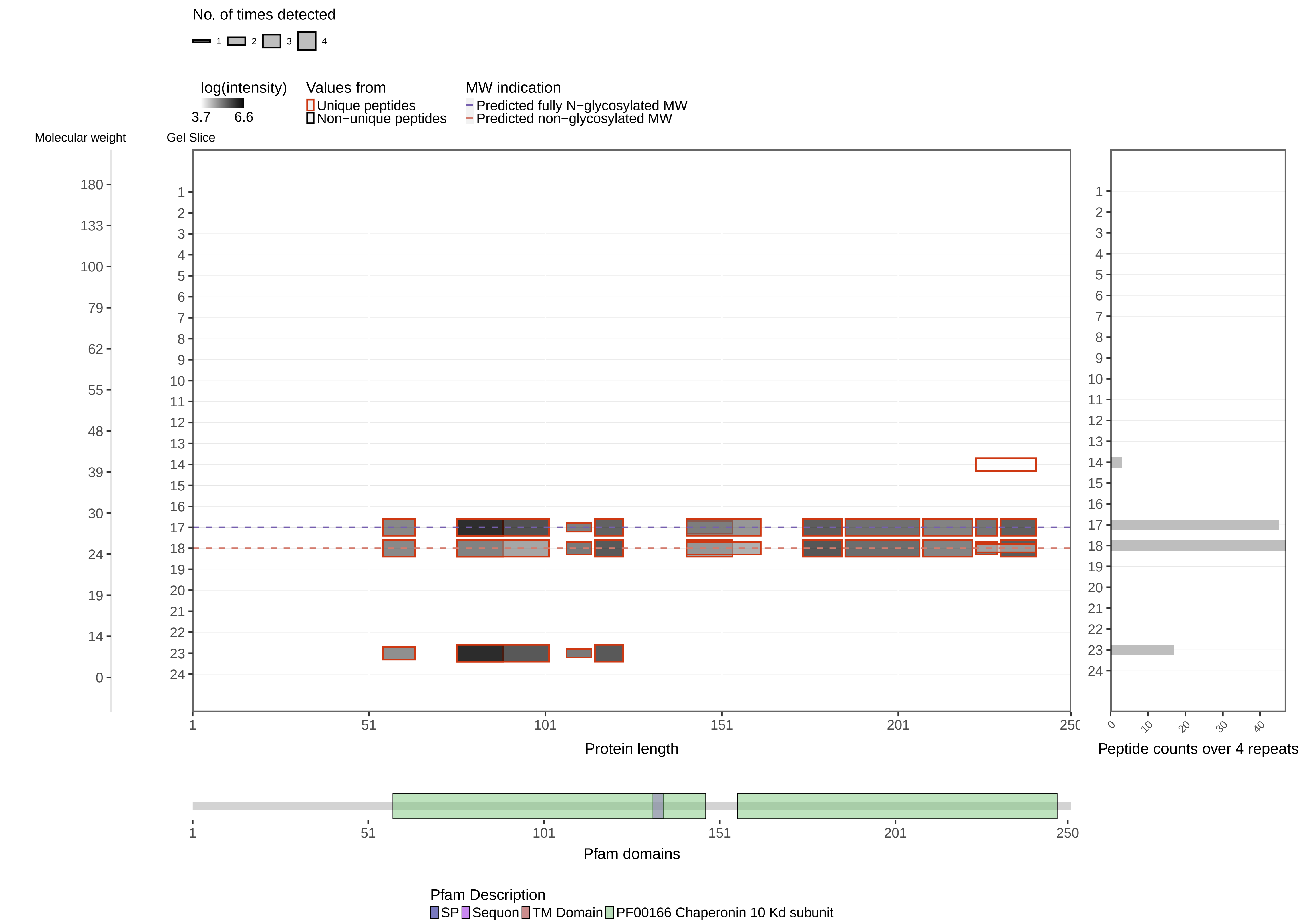


Protein sequence:

SVKTGAQVIYSKYAGTEVEFNGSNHLILKEDDIVGTLETDEIKDLQPLNDRVLIKVAEAEEKTAGGLLLTEASKEKPSIGTVIAVGPGPLDEEGNRKPLSVSTGNTVLYSKYAGNEFKGADGSEYIALRASDVMAVLS*

### NbL10g06900.1 pollen_receptor−like_kinase_3 [MW = 70.29 kDa]

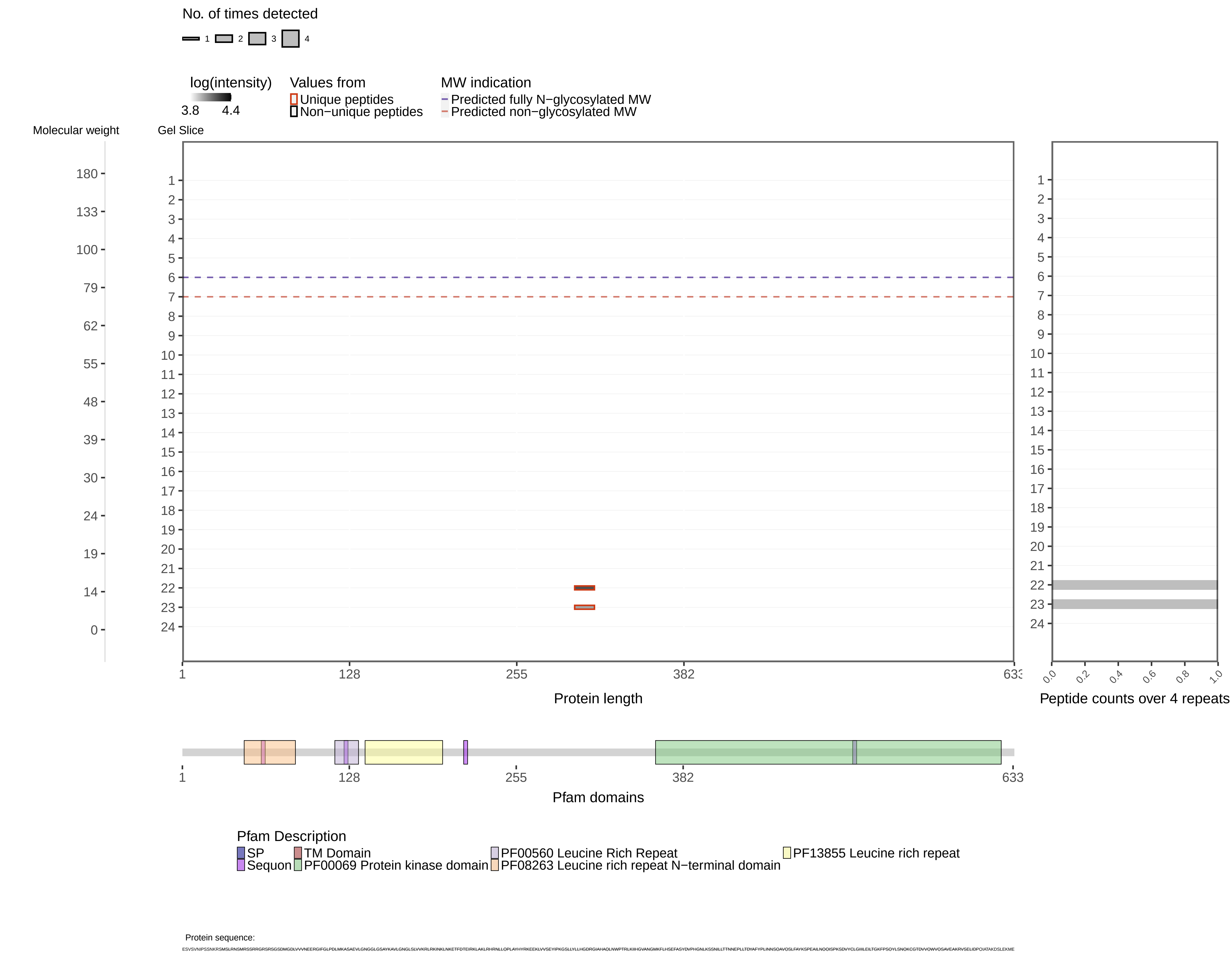

NbL10g06850.1 cysteine_proteinase_3−like [MW − SP = 37.62 kDa]

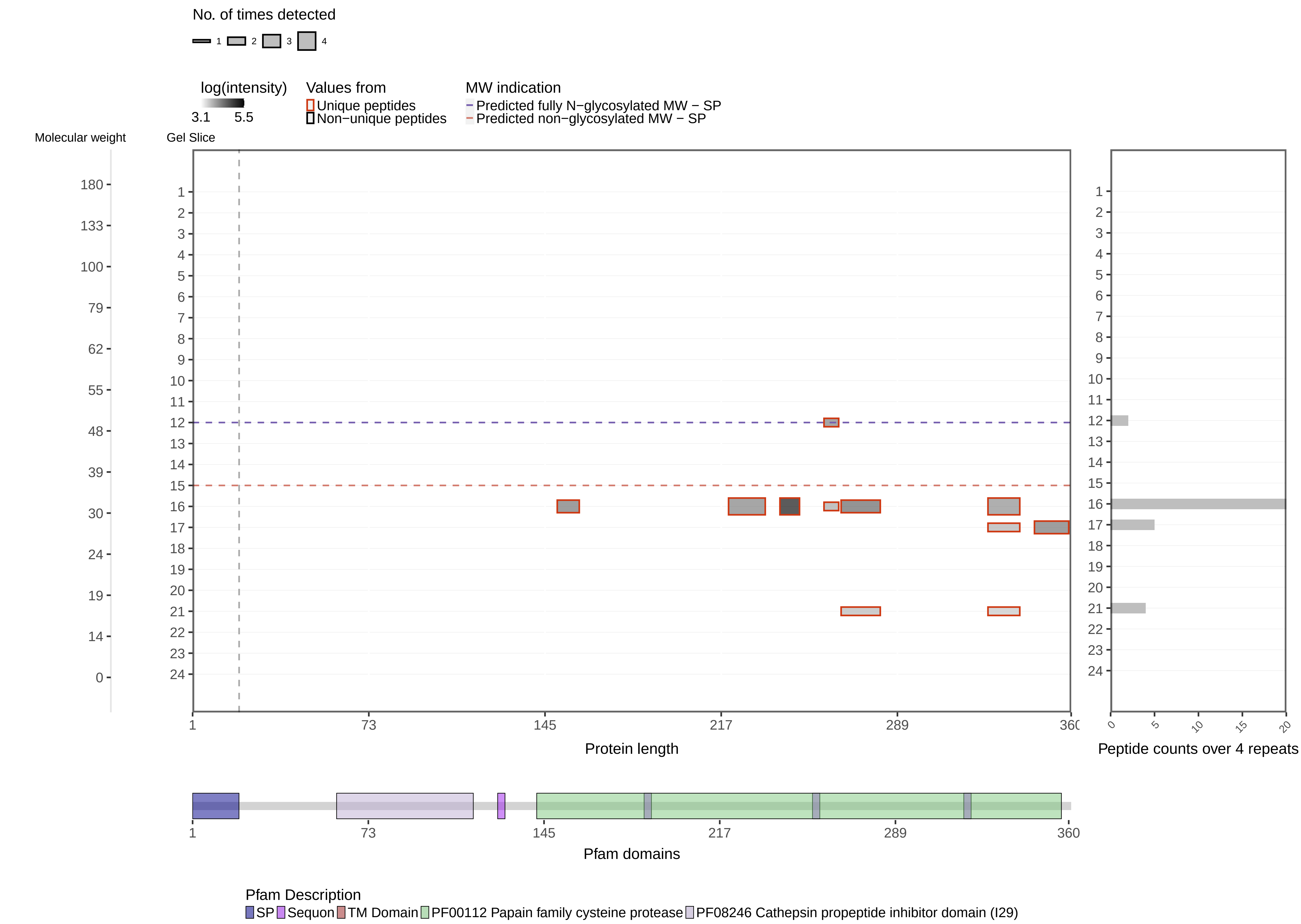


Protein sequence:

TFSTTGALEAAYAQAFGKNISLSEQQLLDCAGAFNNFGCHGGLPSQAFEYIKYSGGLDTEEEYPYAGKAGVCKFSSENIAVKVVGSVNITKDAEDELKYAVAFIRPVSVAYQVVKGFKQYKGGIYSSTVCGNTPQDVNHAVLAVGYGVENGTPYWLIKNSWGAEWGDNGYFKMEMGKNMCGIATCASYPIVA*

NbL10g06840.1 uncharacterized_LOC109225622 [MW − SP = 16.17 kDa]

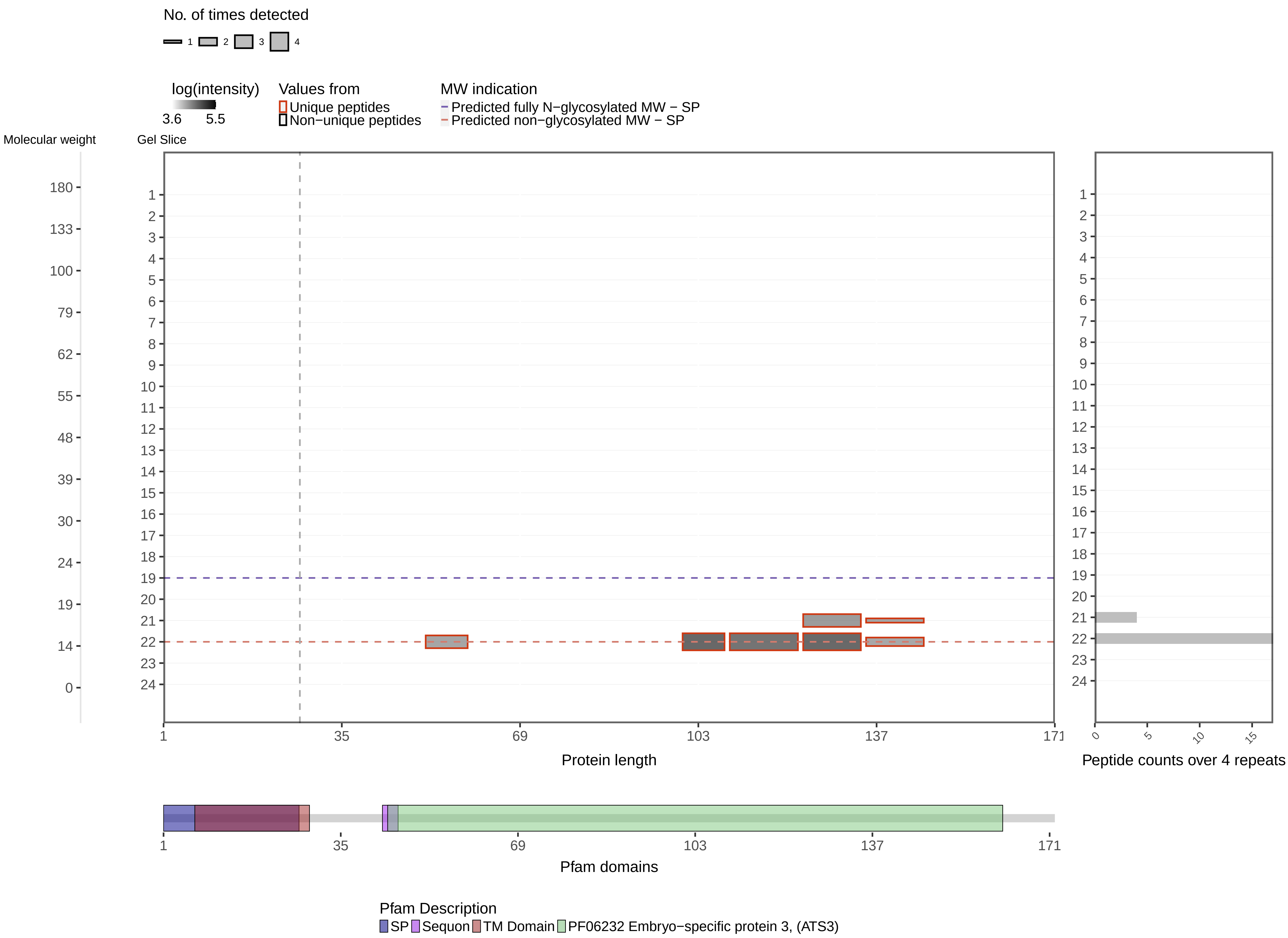


Protein sequence:

IAFGDAYGFEVYAPRLDNPSSRIFERCSTDTFQIRGPCTYEICYLNLLRVGSDGWKPESVKVYGPNRPVITFKYNKFLPNGVWFGFNHCRKVFGSVI*

### NbL10g06740.1 casein_kinase_1−like_protein_2 [MW = 53.02 kDa]

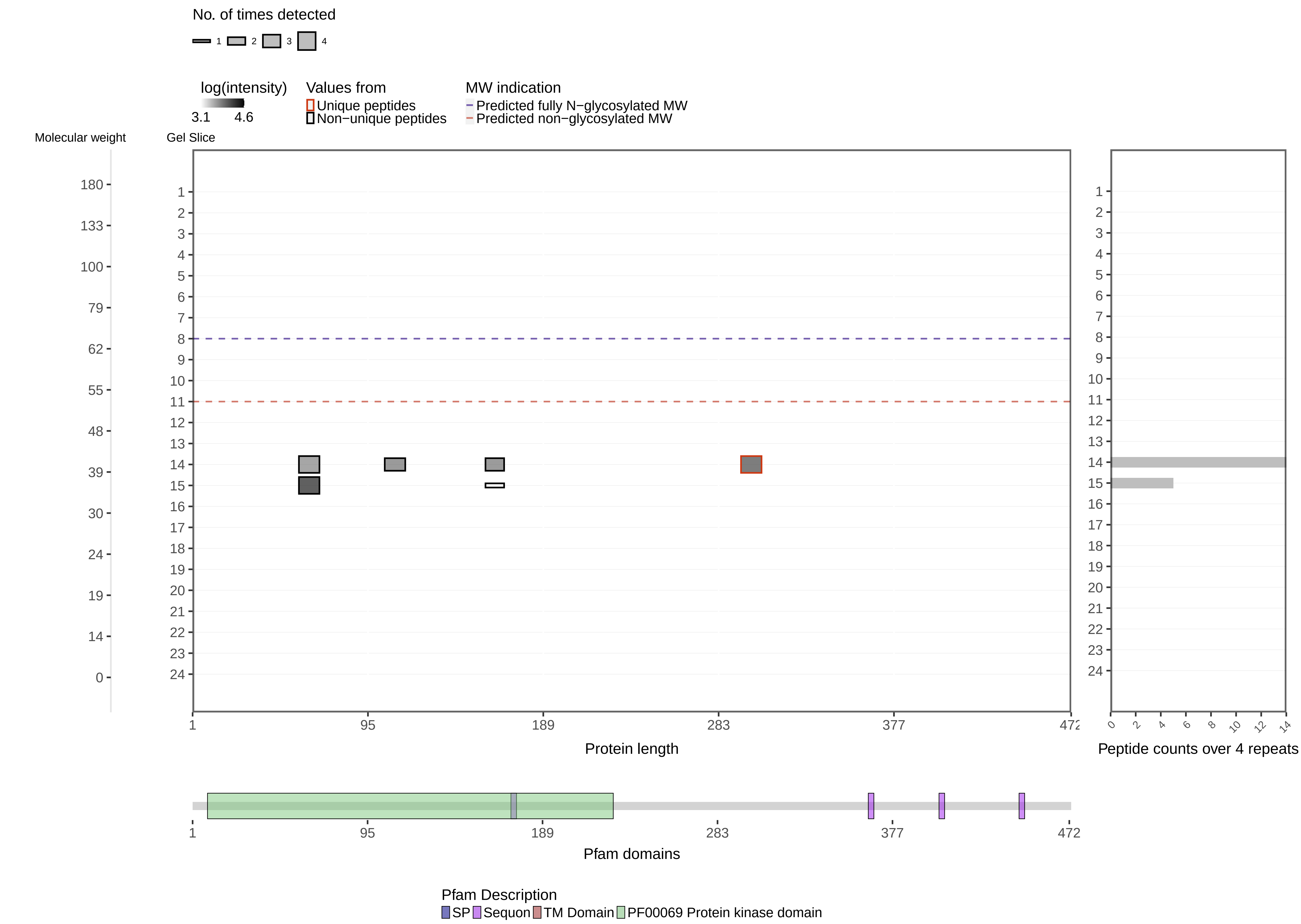


Protein sequence:

KYEKISEKKVSTSIEALCRGYPTEFASYFHYCRSLRFEDKPDYAYLKRIFRDLFIREGFQFDYVFDWTILKYQQSQIAAPTSRPLGAGAGTSSGMPPAIPNAERQSGEEEGSQPADPSRRRNSGPLINAGSFSKQKSPLRNDSTSKDAMLSSSTFLGRSSGSLRRGLVSGSRETFTMGNDSDPTRSRTPEASPATMHKISSGQRSSPLVGSSDAKHASSGRNTSGIKNYETTLKGIERLHFDEEDRSH*

### NbL10g06610.1 probable_polyamine_oxidase_5 [MW = 58.1 kDa]

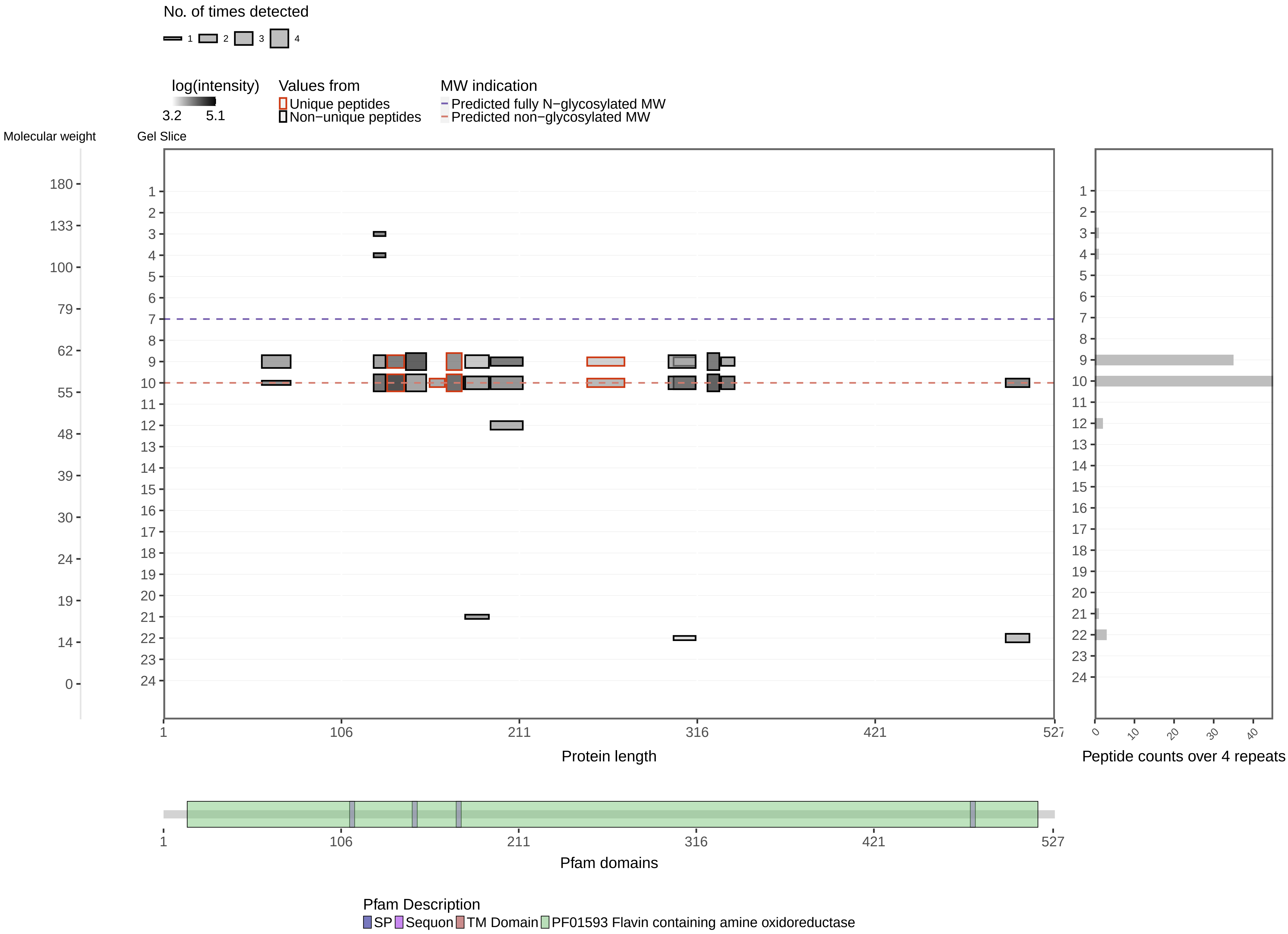


Protein sequence:

WQPDDCGPLDVENGDGTKPVKIHFYDGSIMYADHVIVTVSLGVLKREIRDDSGMFSPPLPSFKTQAISKLGFGVVNKLFLQLSPTKDEDGMKFPYLQMVFHQSDSRLRHPKIPWWMRRTASLCPIYGSSNVLLSWFAGAEALELESLGNEEIIDGFSTMISDFLANSKHVNKLSNGHAESSFKFDKVLKSQWGTDPLFLGSYSYVAVGSSGDDLDTMAEPLPTRSNYSCISPPLQILFAGEATHRTHYSTTHGAYFSGLREANRLLQHYHCIDI*

### NbL10g06280.1 rRNA_methyltransferase_1_mitochondrial [MW = 57.66 kDa]

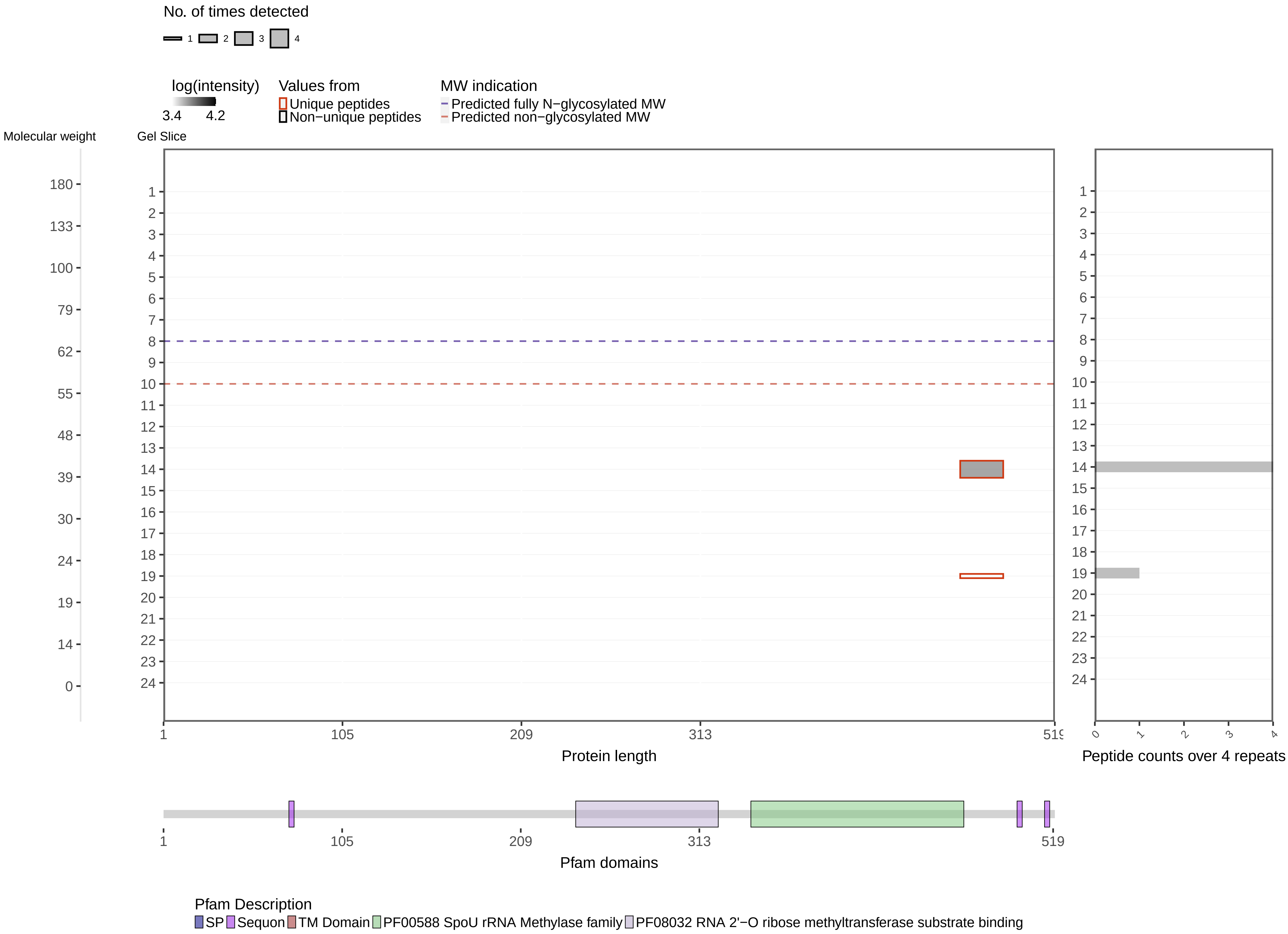


Protein sequence:

GPVLAALSAGRREFYSLYVQEGLDLSGNNRKKKDKKGFERVLRMAEKIGLSKKEVSKHDLNMFADNRPHQGLVLDASPLEMVGIKELEPVSTEEESTPLWVALDEVTDPQNLGAIIRSAYFFGAAGIVLCAKNSARLSGVVSKASAGSLELVELRSCKNMMQFLTSSAENGWRVIGGSVSSRAVPLHEIVPGAPTILVLGSEGTGLRPLVERSCTQLVKIPGNIPVDIIVGEDEDDHSSSGQNFRSFMAVESLNVSVAAGVLLHHLIDSNCSKN*

### NbL10g06190.1 probable_esterase_KAI2 [MW = 30.07 kDa]

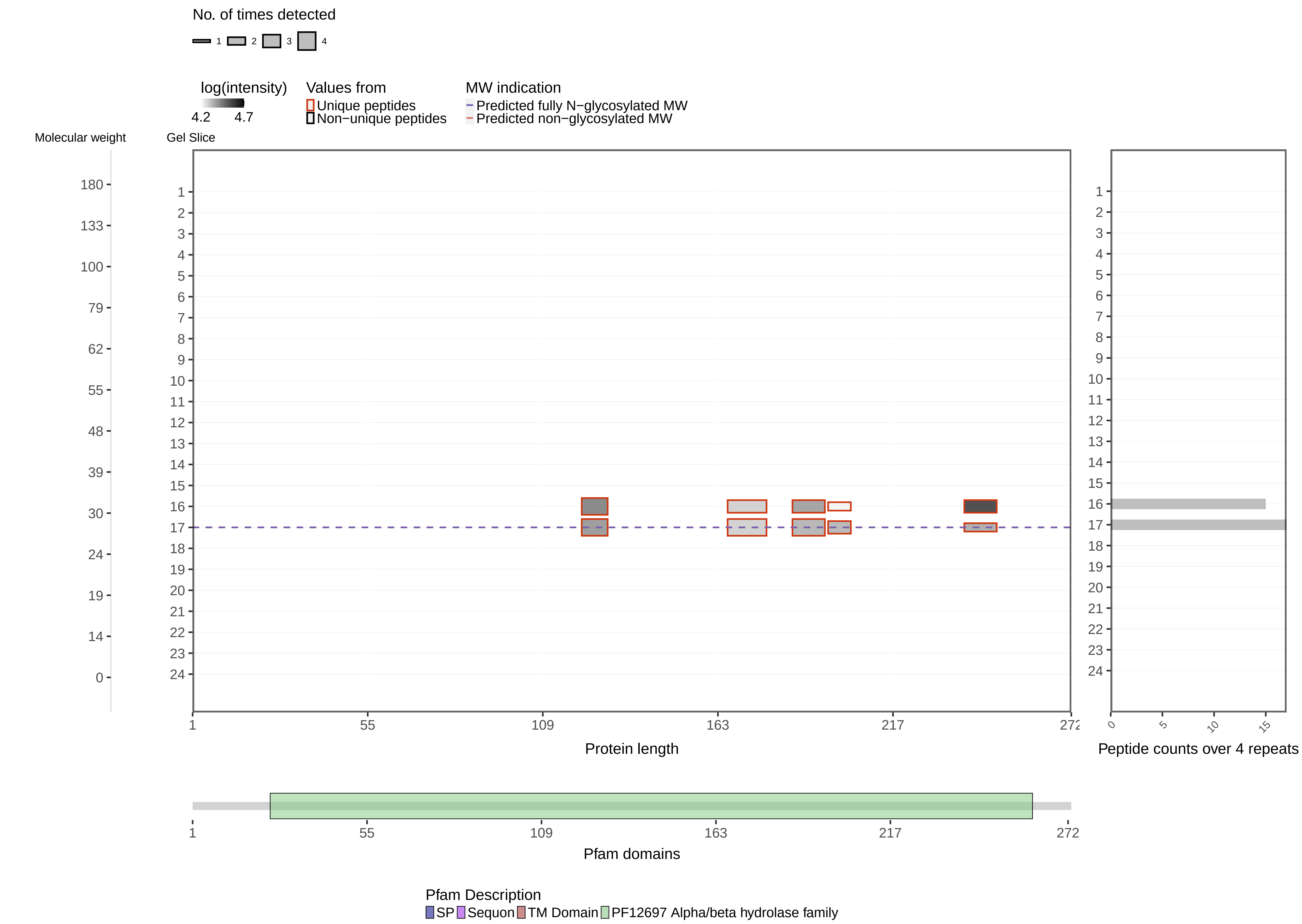


Protein sequence:

LILVASSPRFINLEDYVGGFEMTDIEQMFQNIGENYGIWSSGFARIAVDPSDPPSVEKFEKSLKRMGVEVALPLAKTVFLSDYRGILEKVVTPCTIIQCKIDFAVPNSVAIFMHTNIKGASTVEIINTRGHFPQLTAHEEFLNVIHRVLTS*

### NbL10g06040.1 thylakoid_lumenal_15_kDa_protein_1_chloroplastic−like [MW = 23.32 kDa]

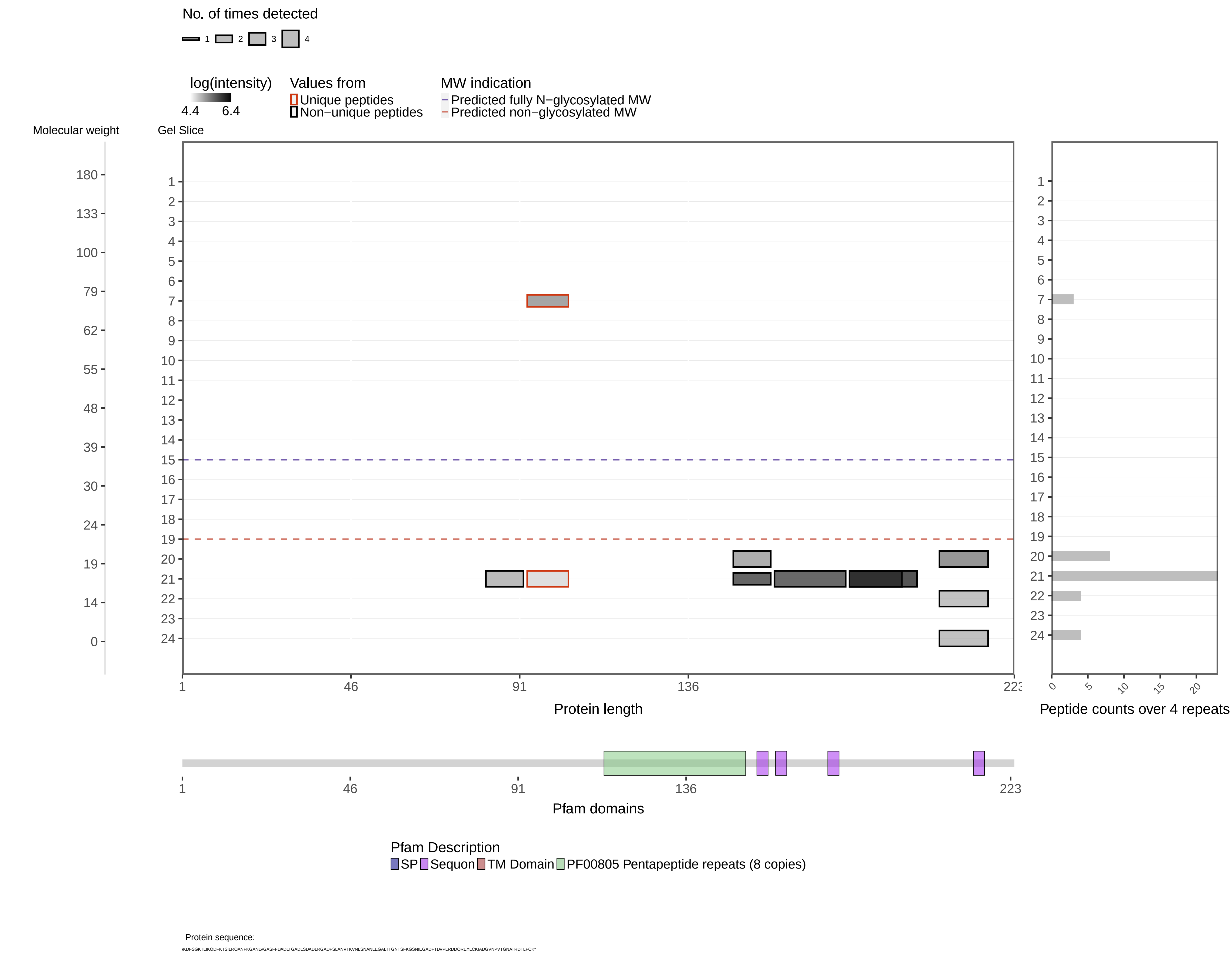

### NbL10g05890.1 tropinone_reductase_homolog_At5g06060−like [MW = 31.64 kDa]

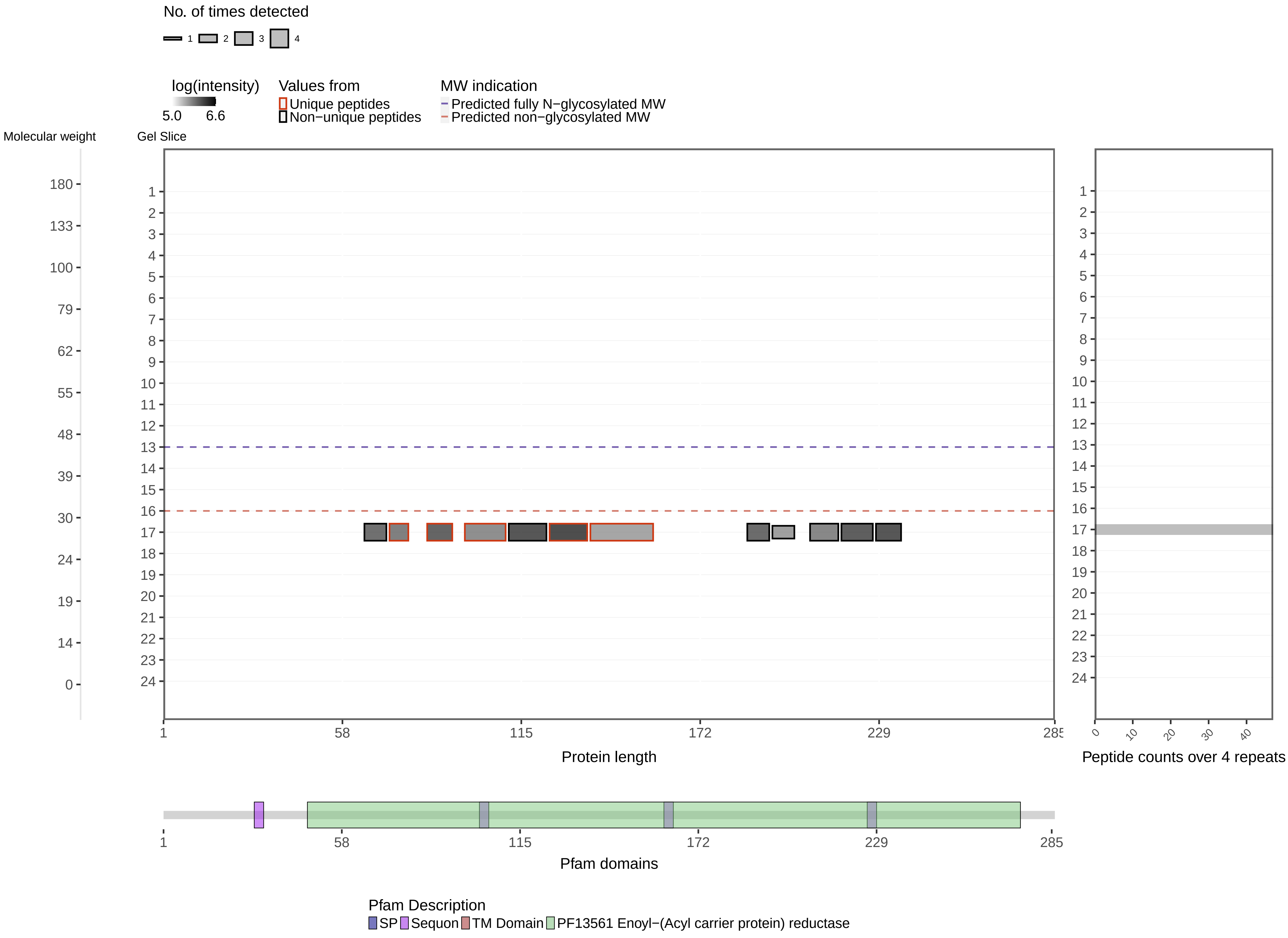


Protein sequence:

DYTAEEYARLLSTNLESAYHLCQLAHPLLKASGNGSVVFISSVAGLVHLSSGSVYGATKGAMNQLTRNLACEWAKDSIRVNGVAPWYIRTSLVEHLLENKTFLDGVISRTPLRRPGESQEVSSLVAYLCLPGASYITGQVIAVDGGFTVNGFEMTSY*

### NbL10g05880.1 uncharacterized_LOC107798942 [MW = 23.31 kDa]

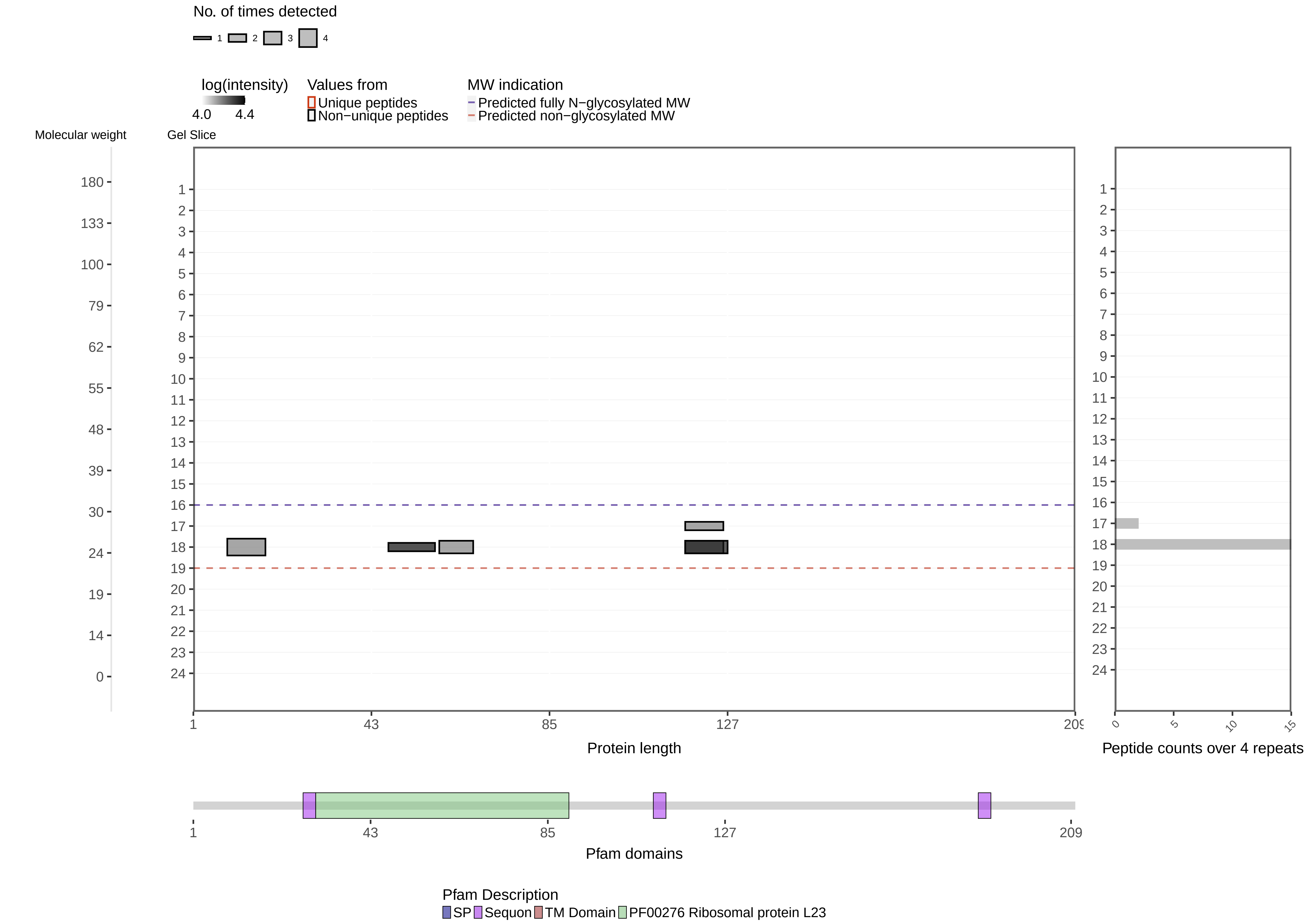


Protein sequence:

SRDLYPIRMIEEEKKNMSKKSKSSIVEDDEPKKMHWLDEKRDTRSSSERRFTRGRDREHGGERSSDVGAPAAKFPWSSMRSYAPRGSVGGFLNSSKVQGLLTVQVTVAGVGECC*

### NbL10g05780.1 protein_DOWNSTREAM_OF_FLC−like [MW = 28.24 kDa]

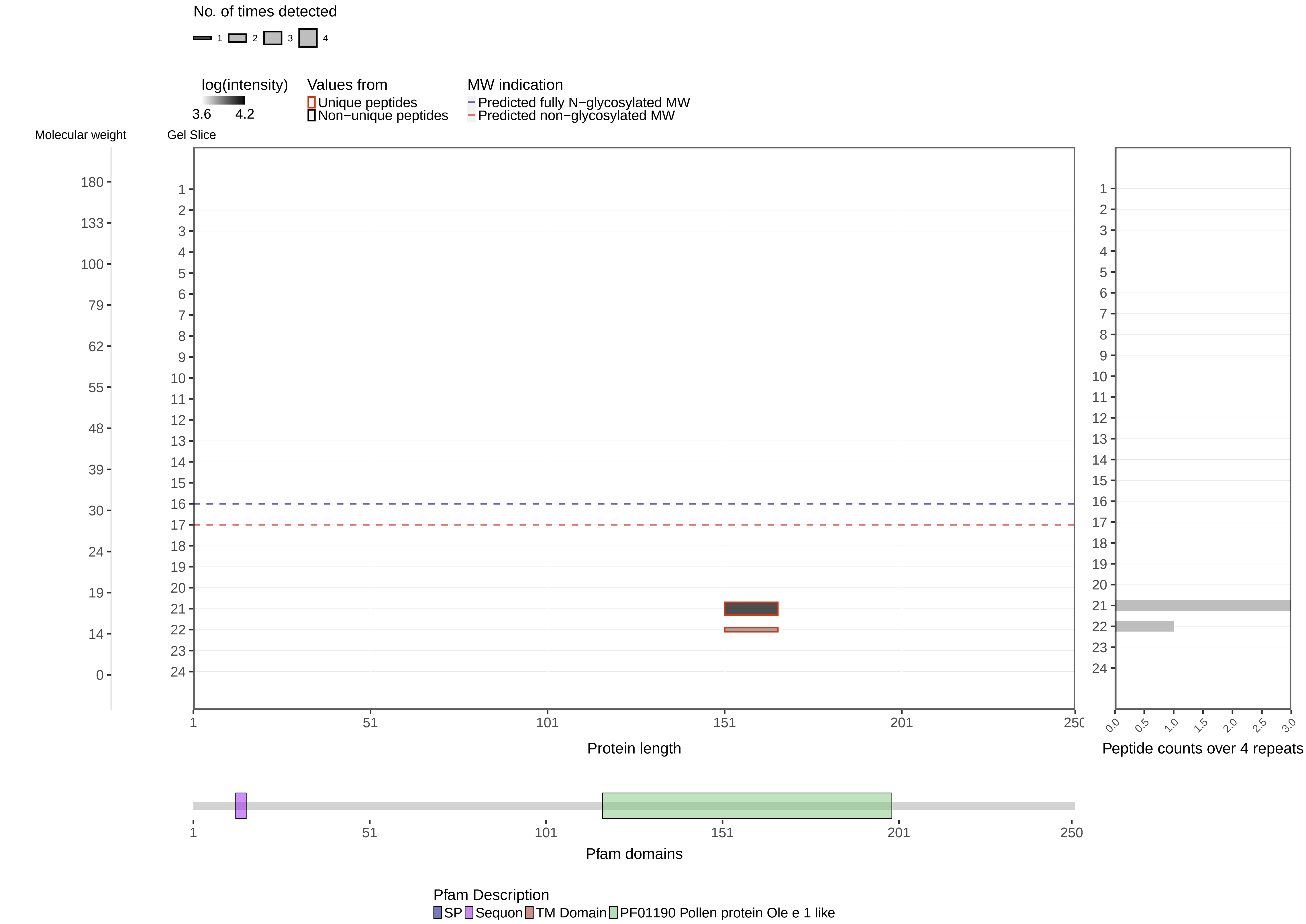


Protein sequence:

FLVKGKAYCDTCRCGFETPATKYLAGAKVRIECRHRVTEAPTYTVEGVTNSKGEYNILVNSDRGDDVCDVILVKSPDRTCAKPNAGRDRSRVILTRNNGMISNTRFANAMGFLRDEPLSSCTQILQQYQLTDDQV*

NbL10g05540.1 peptide−N4−(N−acetyl−beta−glucosaminyl)asparagine_amidase_A−like [MW − SP = 63.94 kDa]

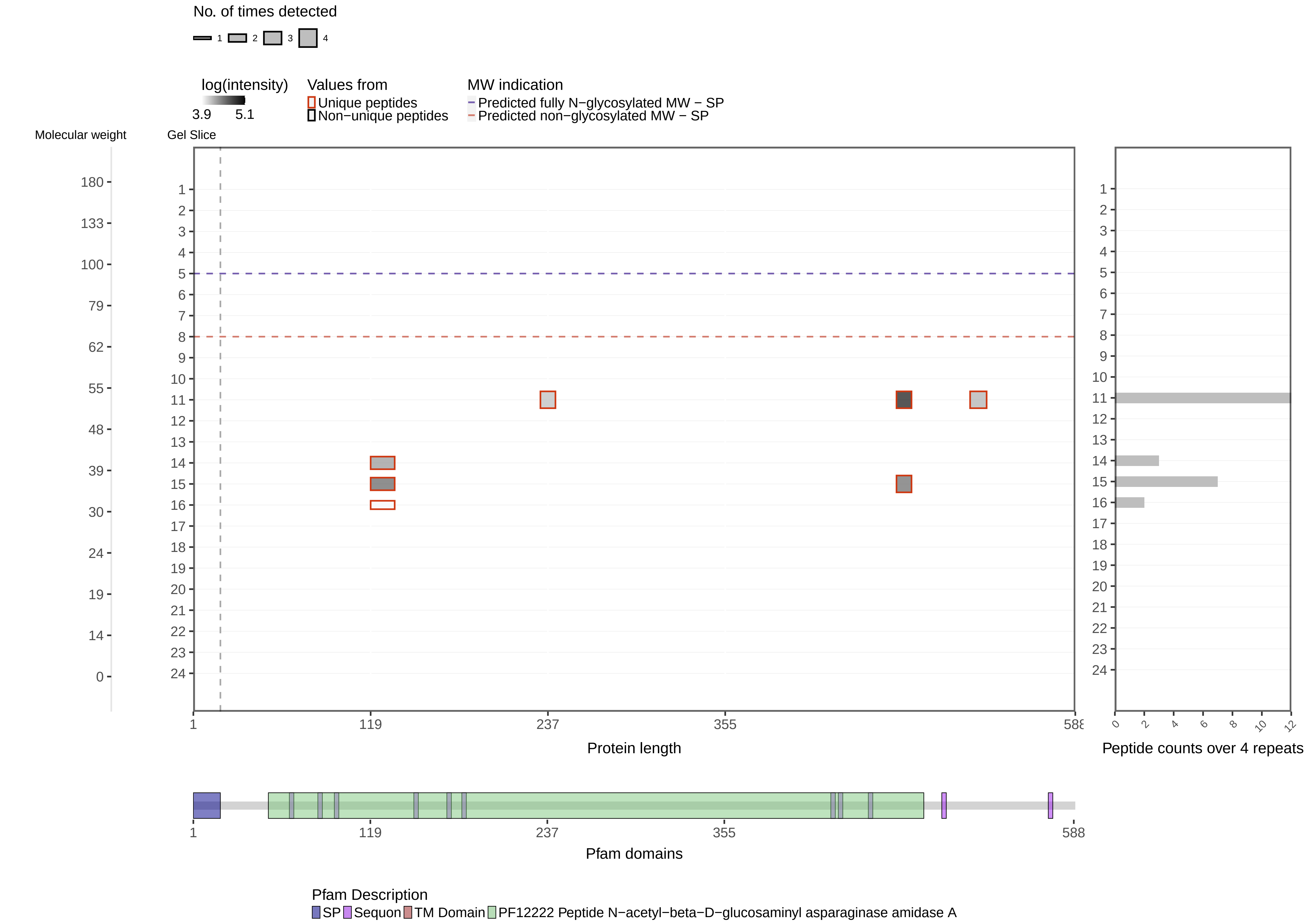


Protein sequence:

### NbL10g05530.1 eukaryotic_translation_initiation_factor_2_subunit_alpha_homolog [MW = 38.55 kDa]

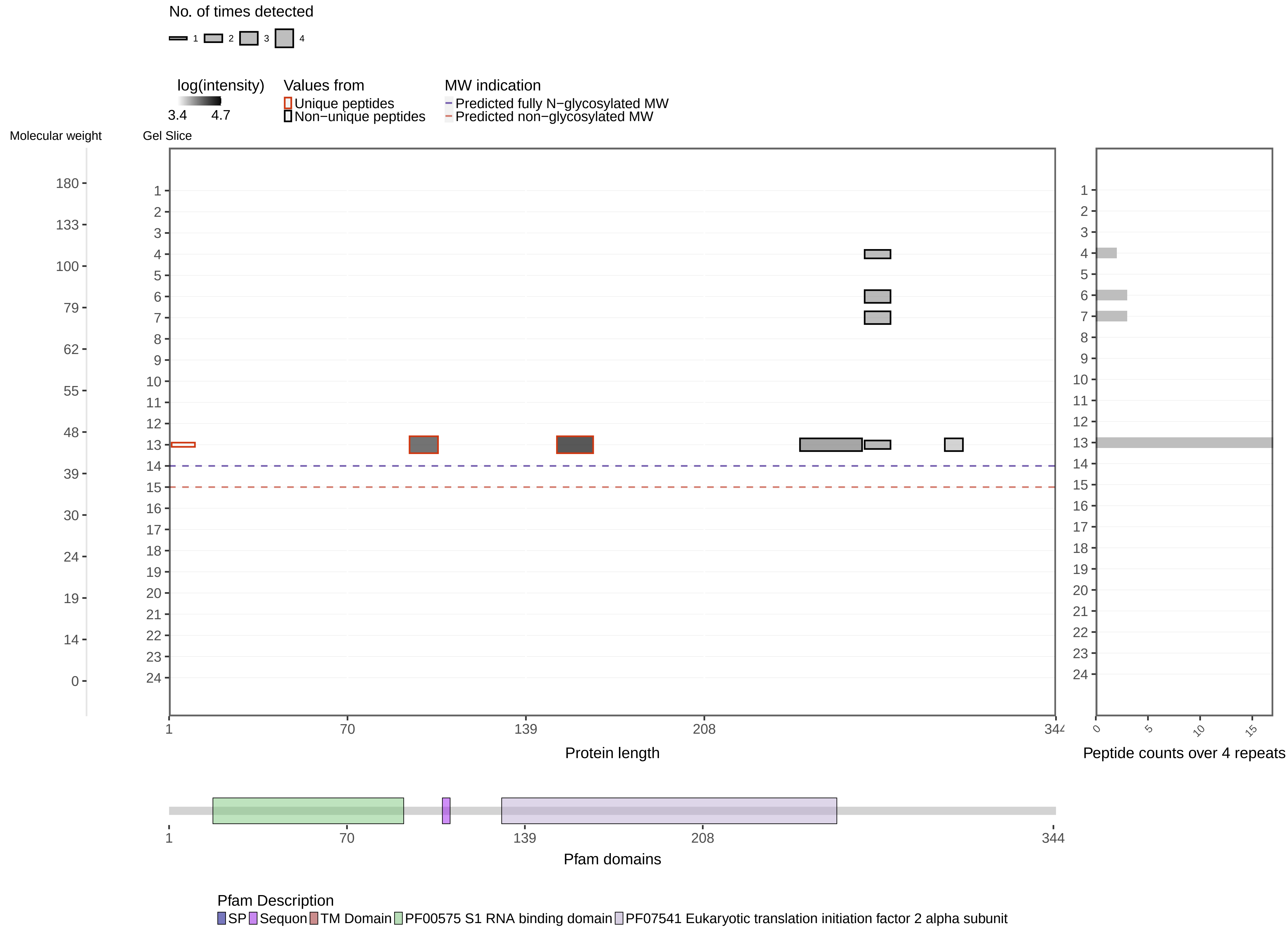


Protein sequence:

ILNSLTREVKENGPDGKEVTKVAPALSEEVKDALVKNIRRRMTPQPLKIRADIEMKCFQFDGVLHIKEAMRKAEAAGNDDCPVKIKLVAPPAYVLNTQTLDKEQGIAILTKAIAACTEEIERHKGKLAVKEAPRAVSEREDKLLAEQMAKLGRENEEISGDEDSEEEEDTGMGEIDVDSGAGIRE*

### NbL10g05400.1 protein_HOTHEAD−like [MW − SP = 60.23 kDa]

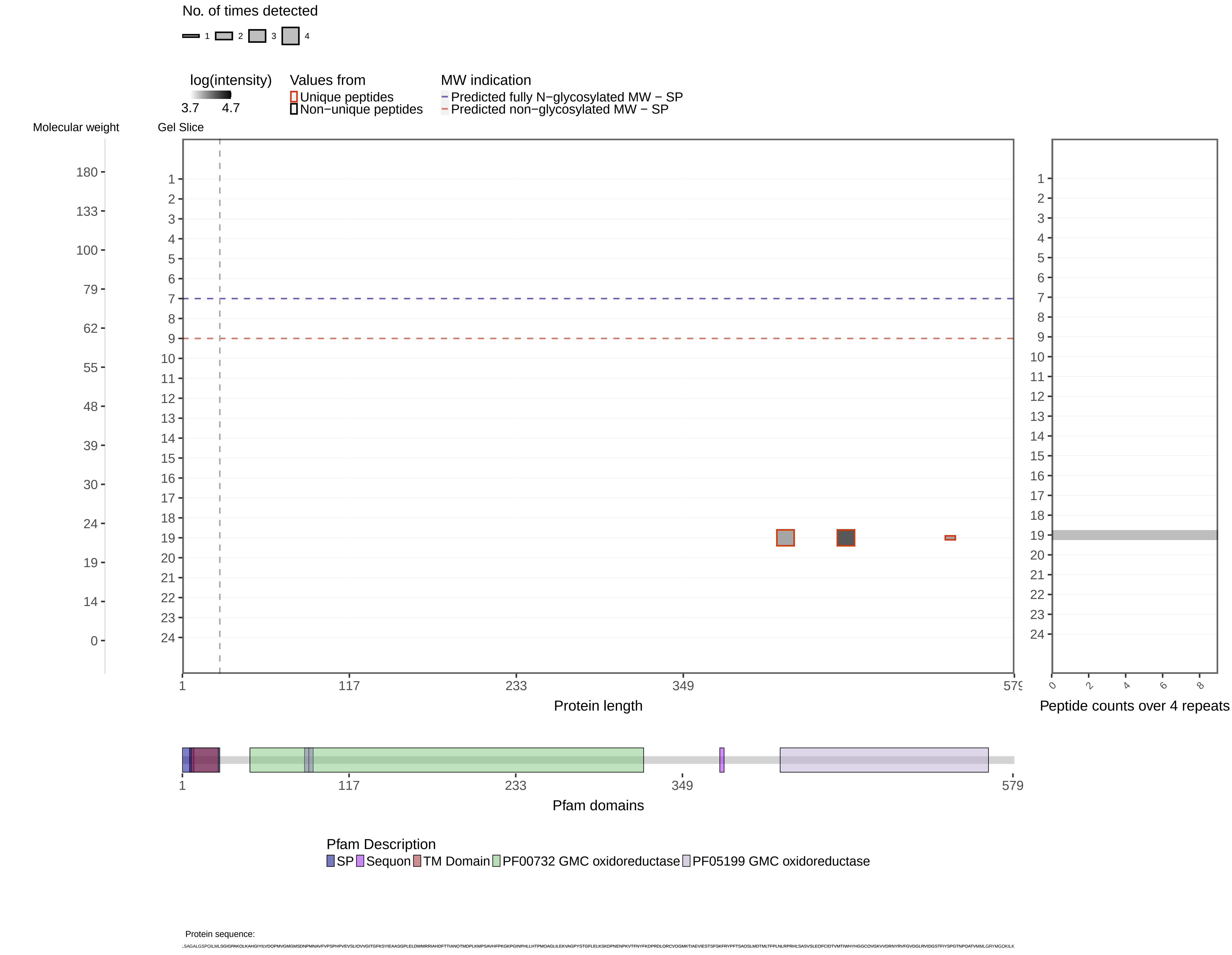

NbL10g05350.1 probable_calcium−binding_protein_CML13 [MW = 16.54 kDa]

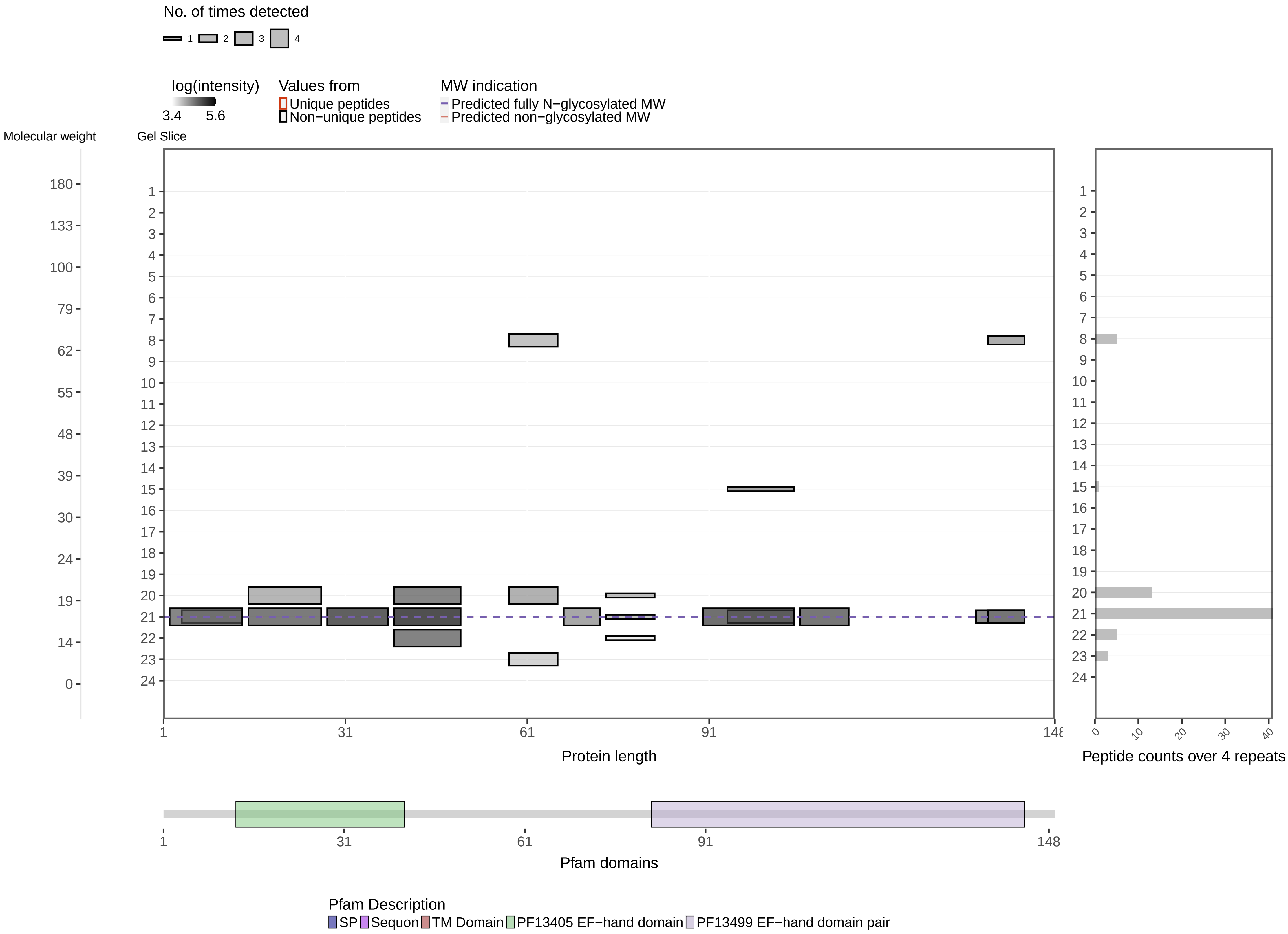


Protein sequence:

PFDFNRFLELMSKHLKPEPFDRQLRDAFKVLDKDGTGFVVVSDLKHILTSIGEKLEAAEFDEWIREVDVGSDGKIRYEDFIARMCAK*

### NbL10g05250.1 quinone_oxidoreductase−like [MW = 41.47 kDa]

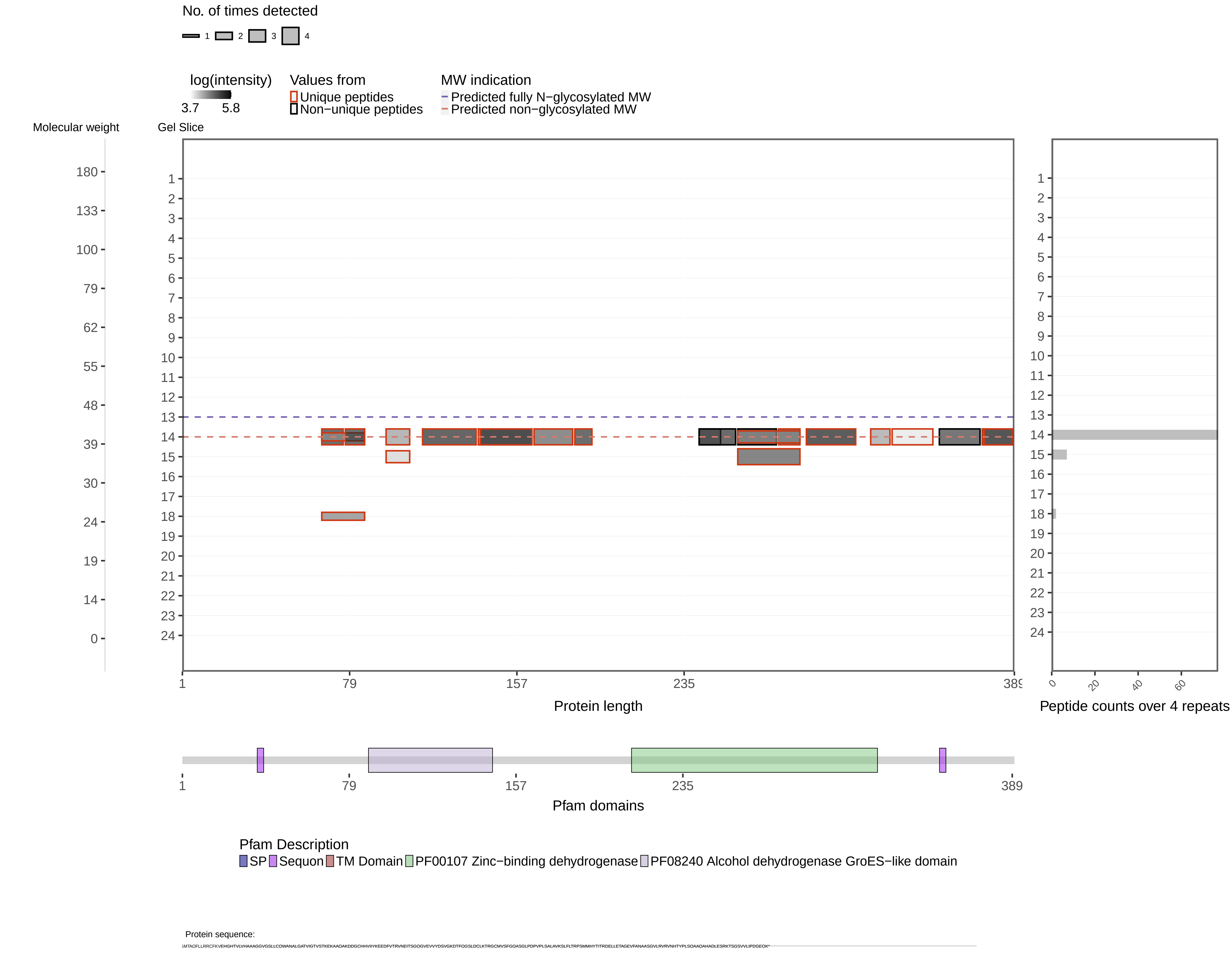

### NbL10g05190.1 protein_mago_nashi_homolog [MW = 20.74 kDa]

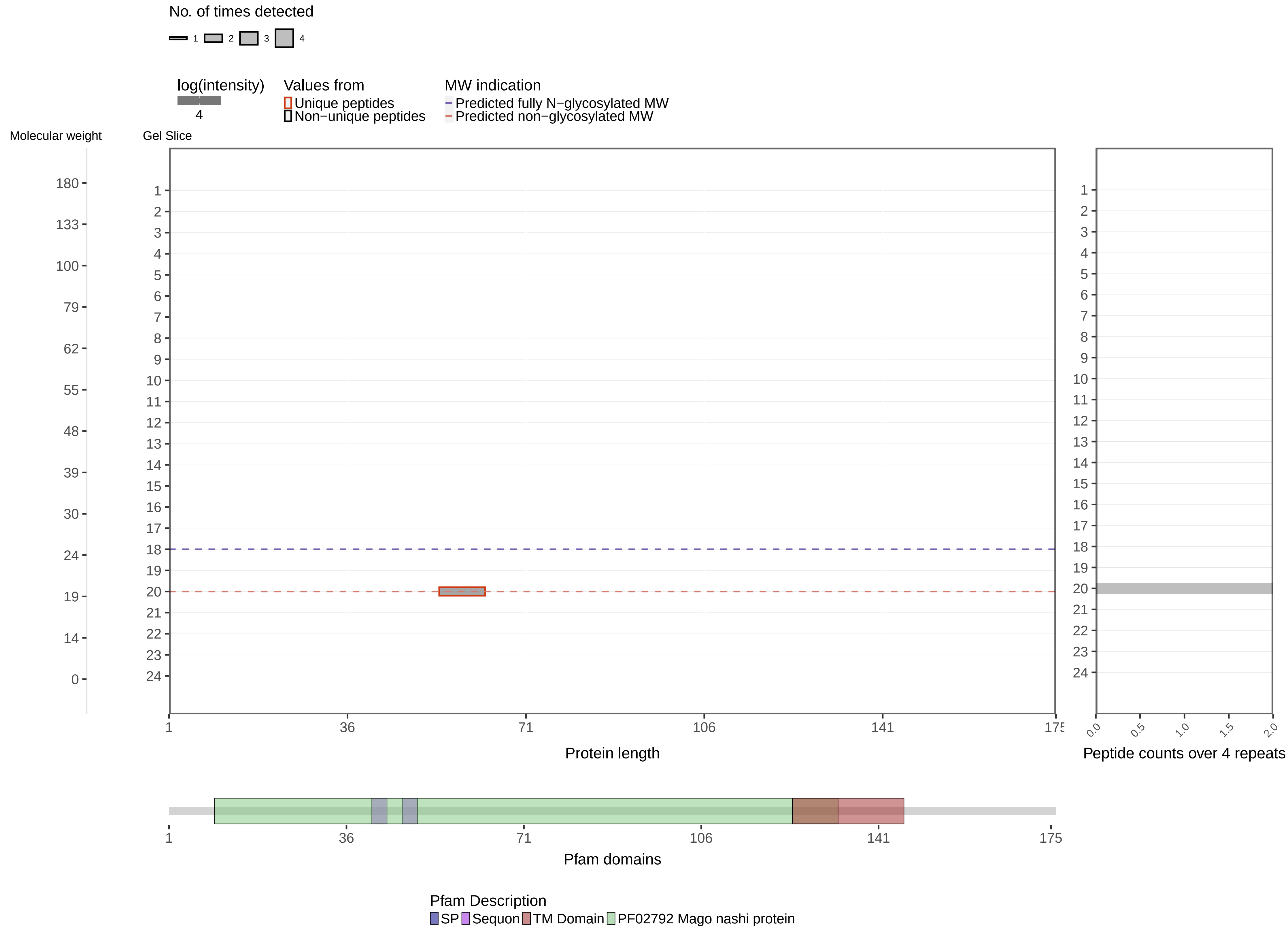


Protein sequence:

IMKEDDNNWPEPDRVGRQELEIVMGNEHISFTTSKIGSLMDVQTSKDPEGLRIFYYLVQVFIASFICPYFMFWMKFFKLIRCIVILHQVVAKIMWSNLYTW*

### NbL10g05170.1 glutamate_dehydrogenase_A [MW = 44.87 kDa]

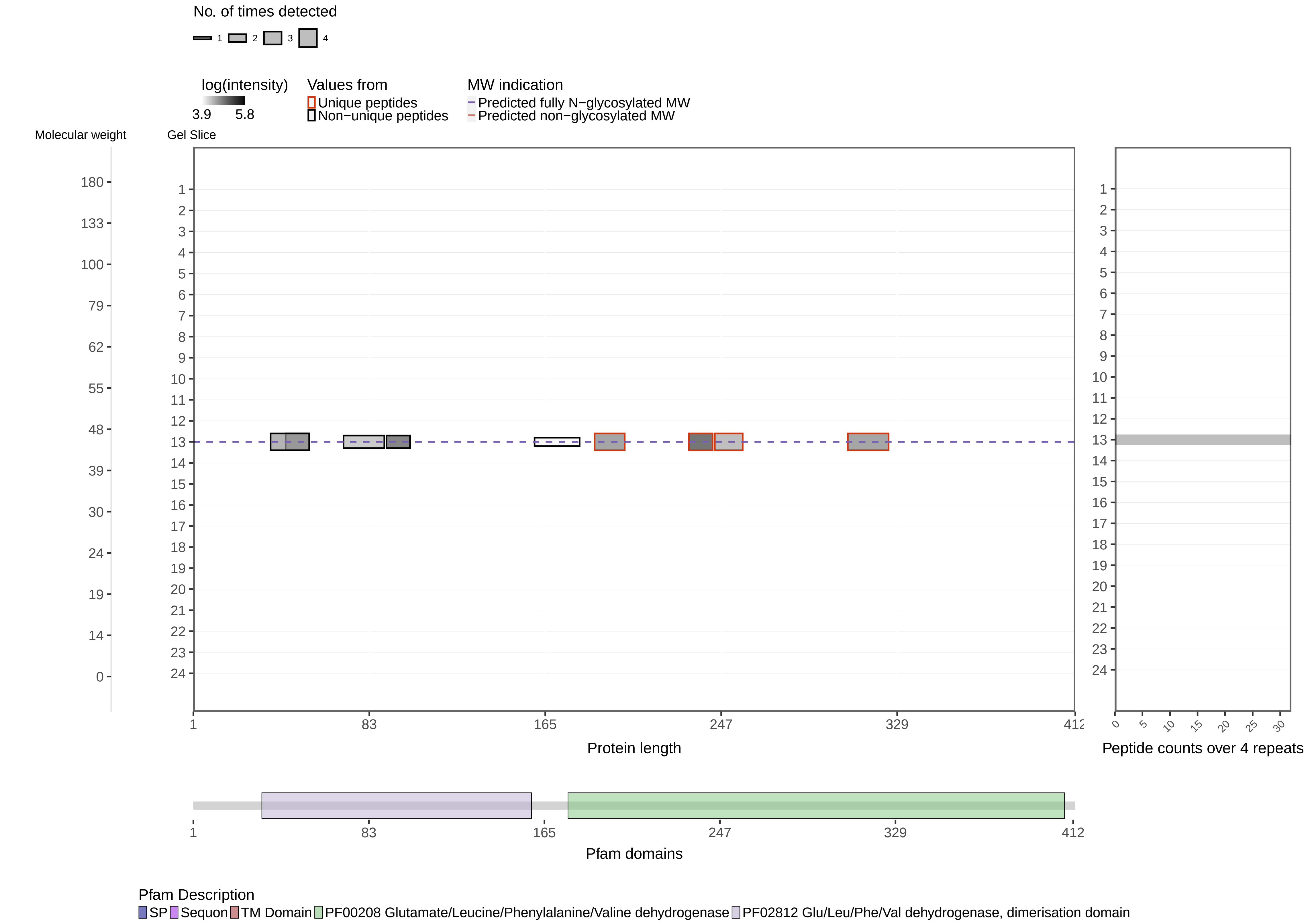


Protein sequence:

EALLAEYGKNIKDLTFAIQGFGNVGAWAAKLIHERGGKVIAVSDITGAVKNPDGLDIPALLNHKEATGKLIDFSGGDVMNSDEVLTHECDVLIPCALGGVLNRENSDNVRAKFIIEAANHPTDPEADEILSKKGIVILPDIYANAGGVTVSYFEWVQNIQGFMWDEEKVNRELKKYMTKAFHKLKNMCQSHDCNLRMGAFTLGVNRVARATTLRGWEA*

NbL10g05160.1 superoxide_dismutase_[Fe]_chloroplastic [MW = 28.04 kDa]

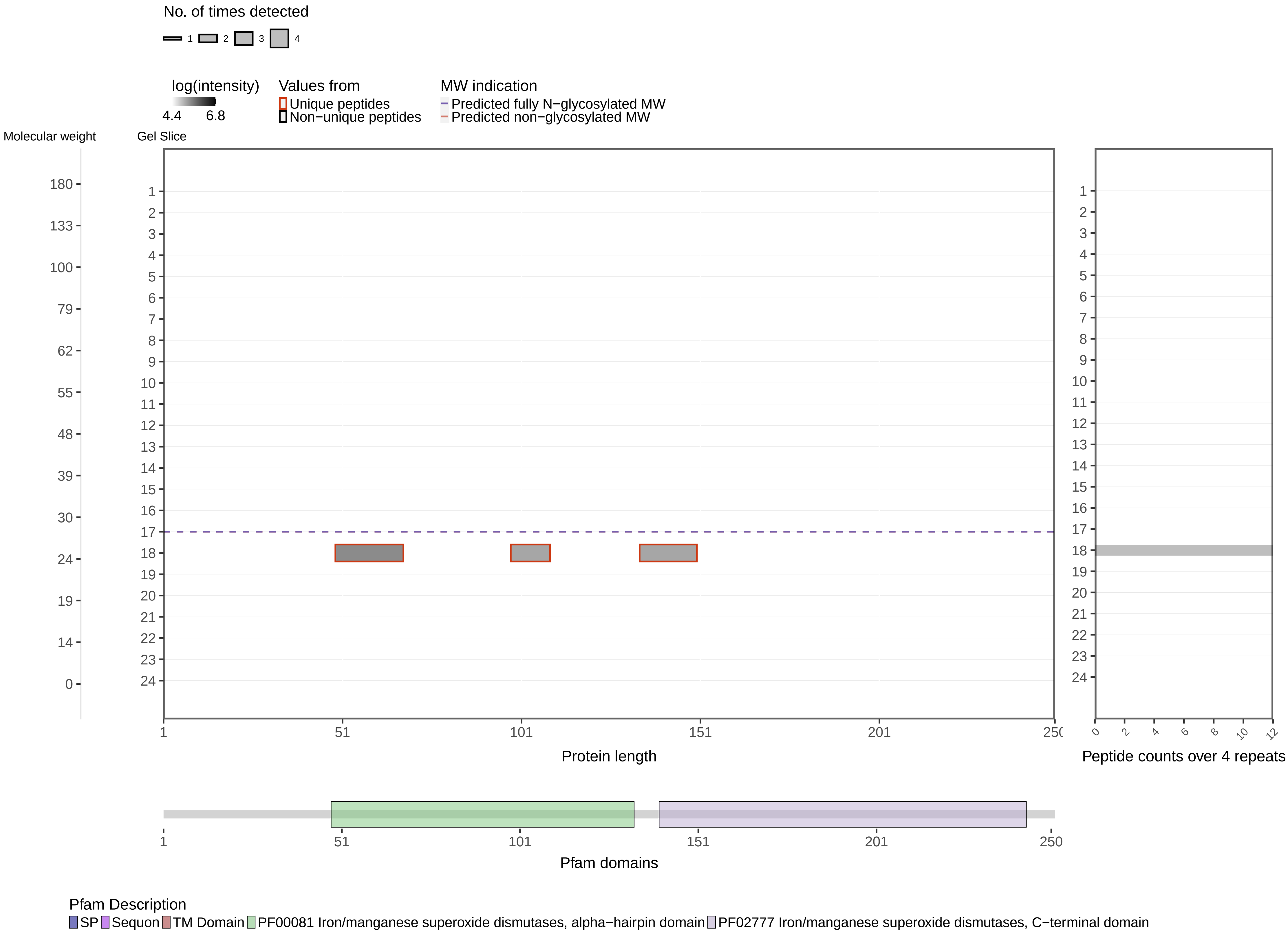


Protein sequence:

PLPAFNNAAQAWNHQFFWESMKPNGGGEPSGELLELINRDFGSYDAFVKEFKAAAATQFGSGWAWLAYKPEEKKLALVKTPNAENPLVLGYTPLLTIDVWEHAYYLDFQNRRPDYISIFMEKLVSWEAVSSRLKAATA*

NbL10g05150.1 ATP_synthase_subunit_delta'_mitochondrial−like [MW = 21.42 kDa]

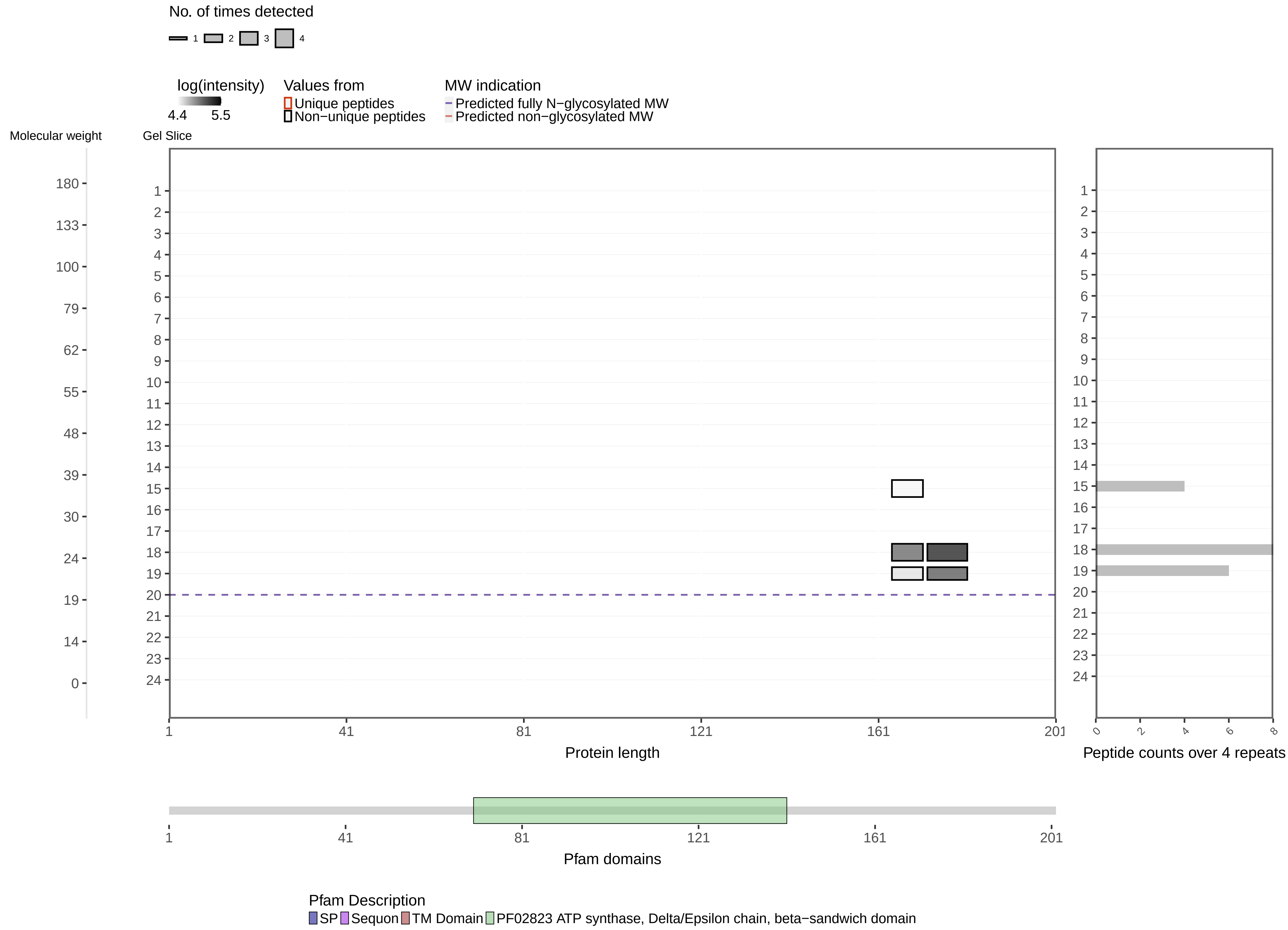


Protein sequence:

:VDMVIIPATTGQMGVLPGHVATIAELKPGVLSVHEGNDVSKYFVSGGFAFVHANSFADIIAIEAIPLDRVDPNLVQKGLTEFTQKLSTASTDVEKAEAQIGVDVHSALNAALTG*

NbL10g05100.1 hydroxyacylglutathione_hydrolase_cytoplasmic [MW = 28.86 kDa]

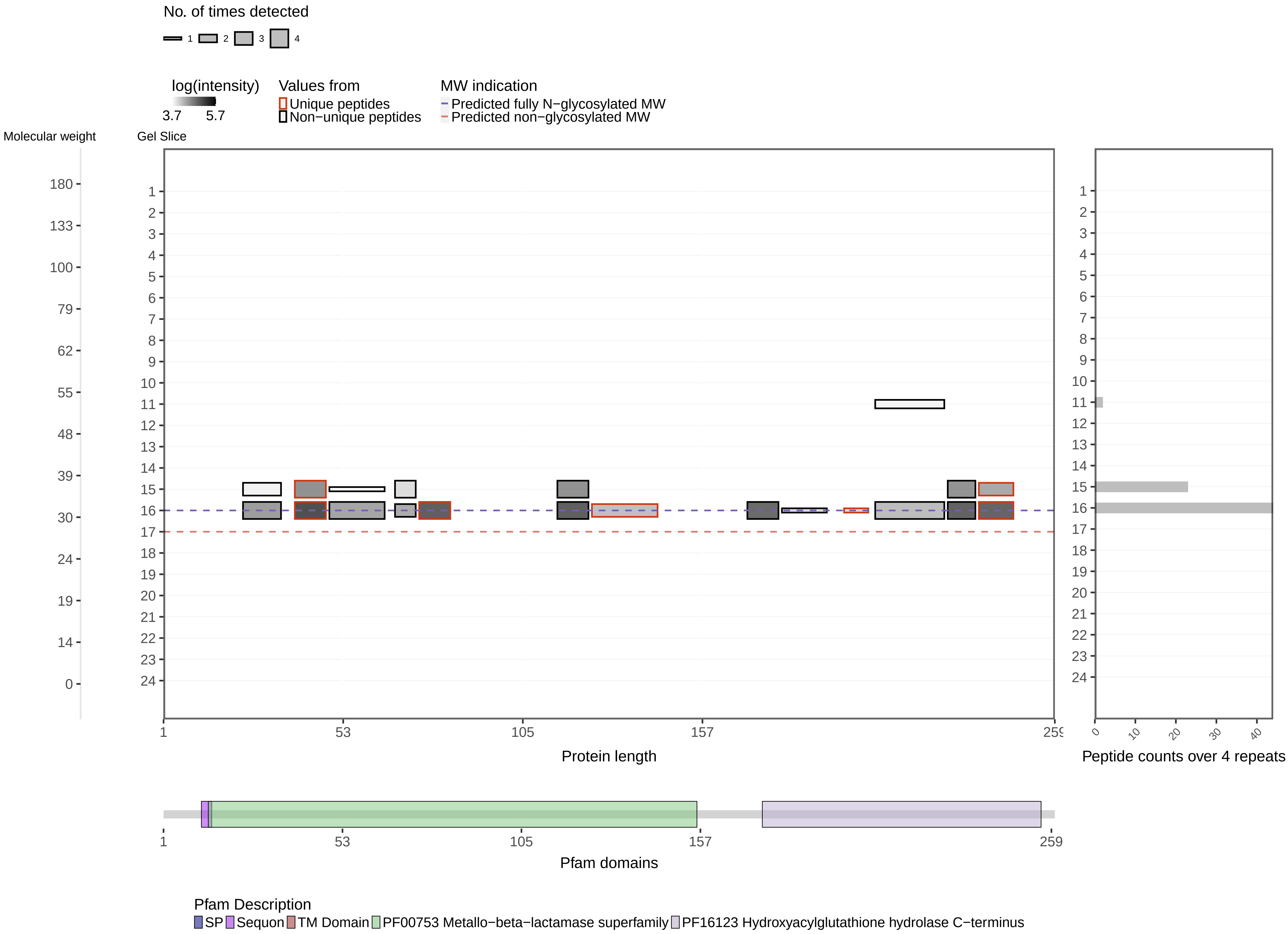


Protein sequence:

SYYVTGKEGEDPAVFTGDTLFIAGCGKFFEGTAEQMYQSLCVTLGSLPKPTRVYCGHEYTVKNLQFALTVEPDNAKIAQKLAWAEHQRRVGLPTIPSTIEEEMETNPFMRADLPEVQEKVGFTSPVEALREIRQRKDNWKS*

### NbL10g05090.1 ABC_transporter_I_family_member_6_chloroplastic−like [MW = 34.95 kDa]

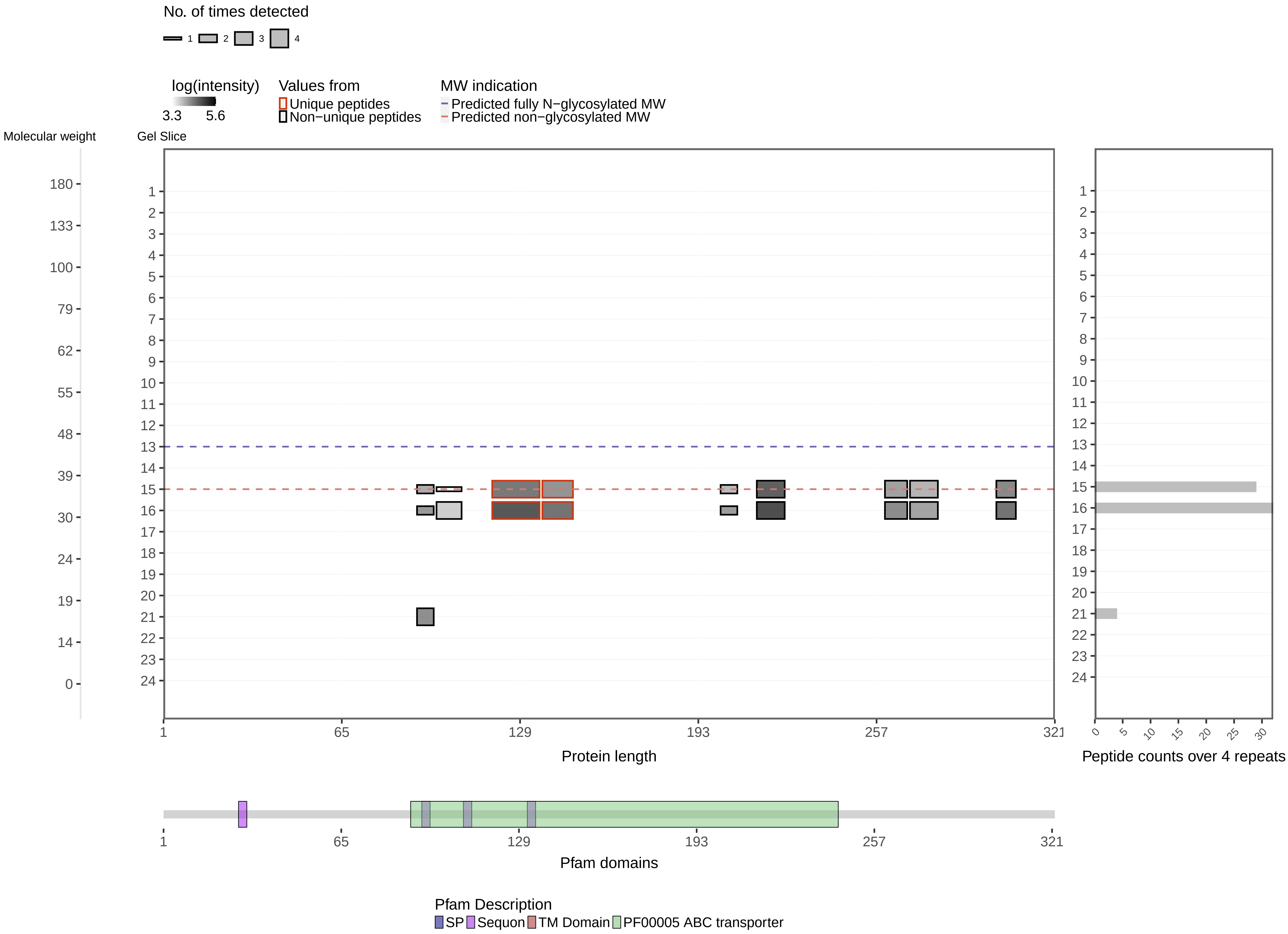


Protein sequence:

PEERSLAGLFMSFQSPVAIPGVSNIDFLNMAYNAQRRKLGLPELGPIEFYGYIAPKLELVNMKIDFLNRNVNEGFSGGERKRNEILQLAVLGADLAILDEIDSGLDVDALRDVAKAVNGLLSPKNSVLMITHYLRLLEFIKPTYIHIMENGRIVKTGDISIAKVLEKEGYKAISGA*

NbL10g05080.1 methylesterase_17−like [MW = 31.05 kDa]

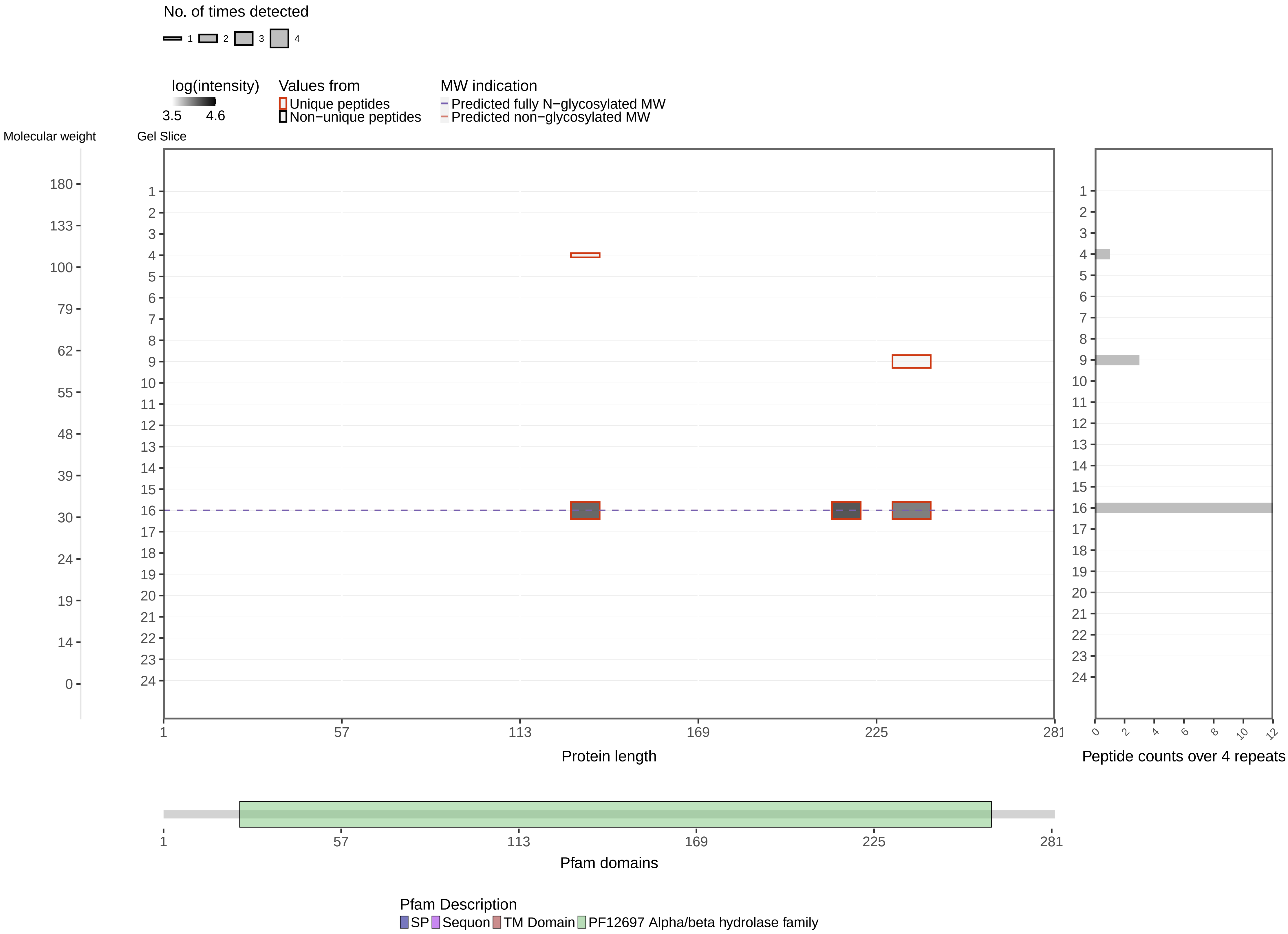


Protein sequence:

RTGFVTEQDVKDGIPDLSEFGEPIDVYDMGFGLGPEQPPSSAVIKRSLQRKIIYQTSPLEDSTLAAMLLRPGPIQALKSAQFKEGEGAEEVPRIYIRTAYDRVVKPEQQDAMITRWPPQNVYTLESDHSPFFSAPFLLFGLIINAASSLGCTD*

### NbL10g05010.1 GDSL_esterase/lipase_EXL3−like [MW − SP = 37.3 kDa]

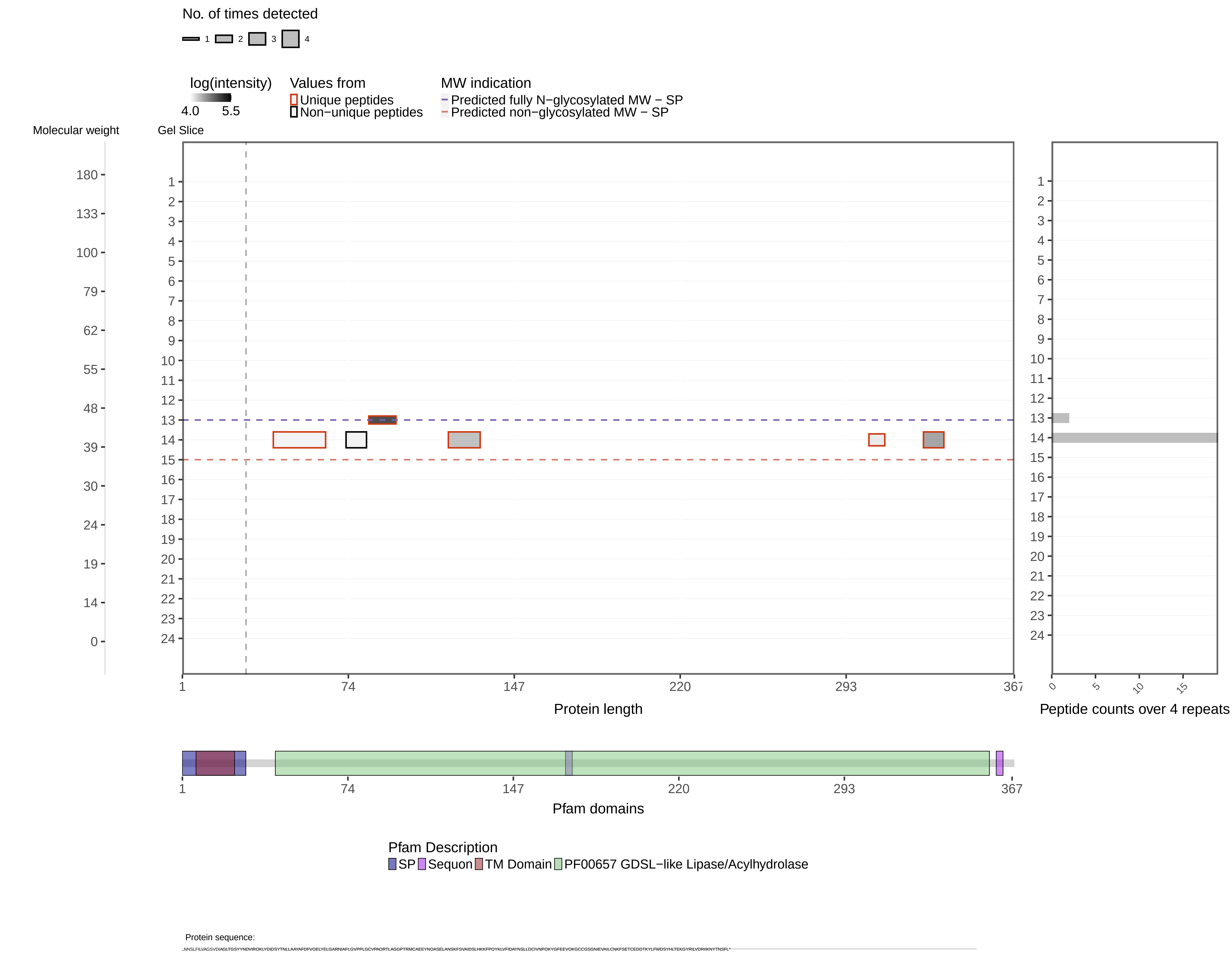

### NbL10g04930.1 membrane_steroid−binding_protein_2−like [MW = 24.84 kDa]

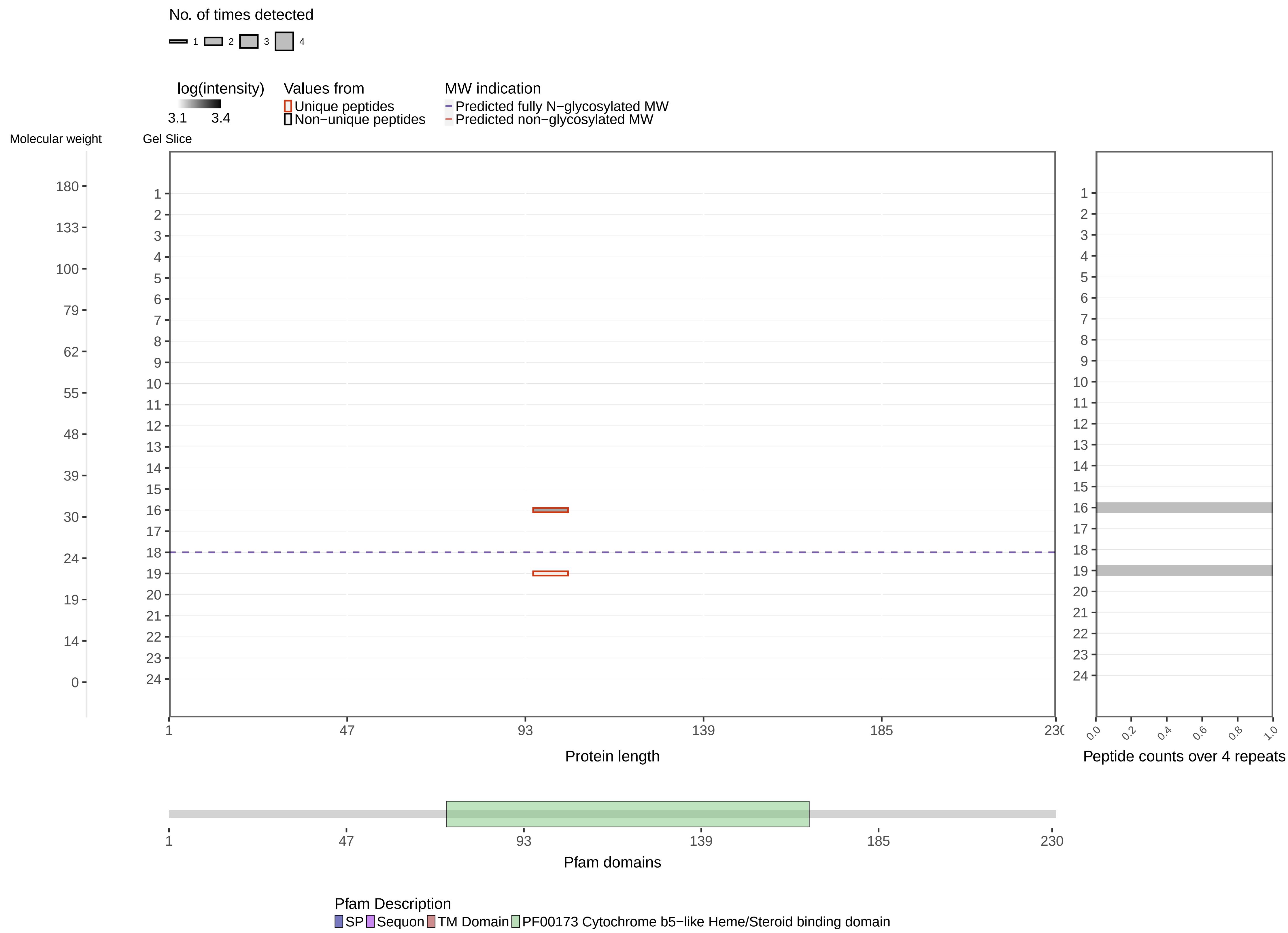


Protein sequence:

RMFYGPGGPYALFAGKDASRALAKMSFEEKDLTGDISGLGPFELDALQDWEYKFMSKYVKVGTIKQAVPVTDGAANGESVEATDREAKPAEDGASASSKPLEDDVPSASVAAETVEKSDVDVDKKD*

### NbL10g04920.1 tubulin_beta−1_chain [MW = 50.32 kDa]

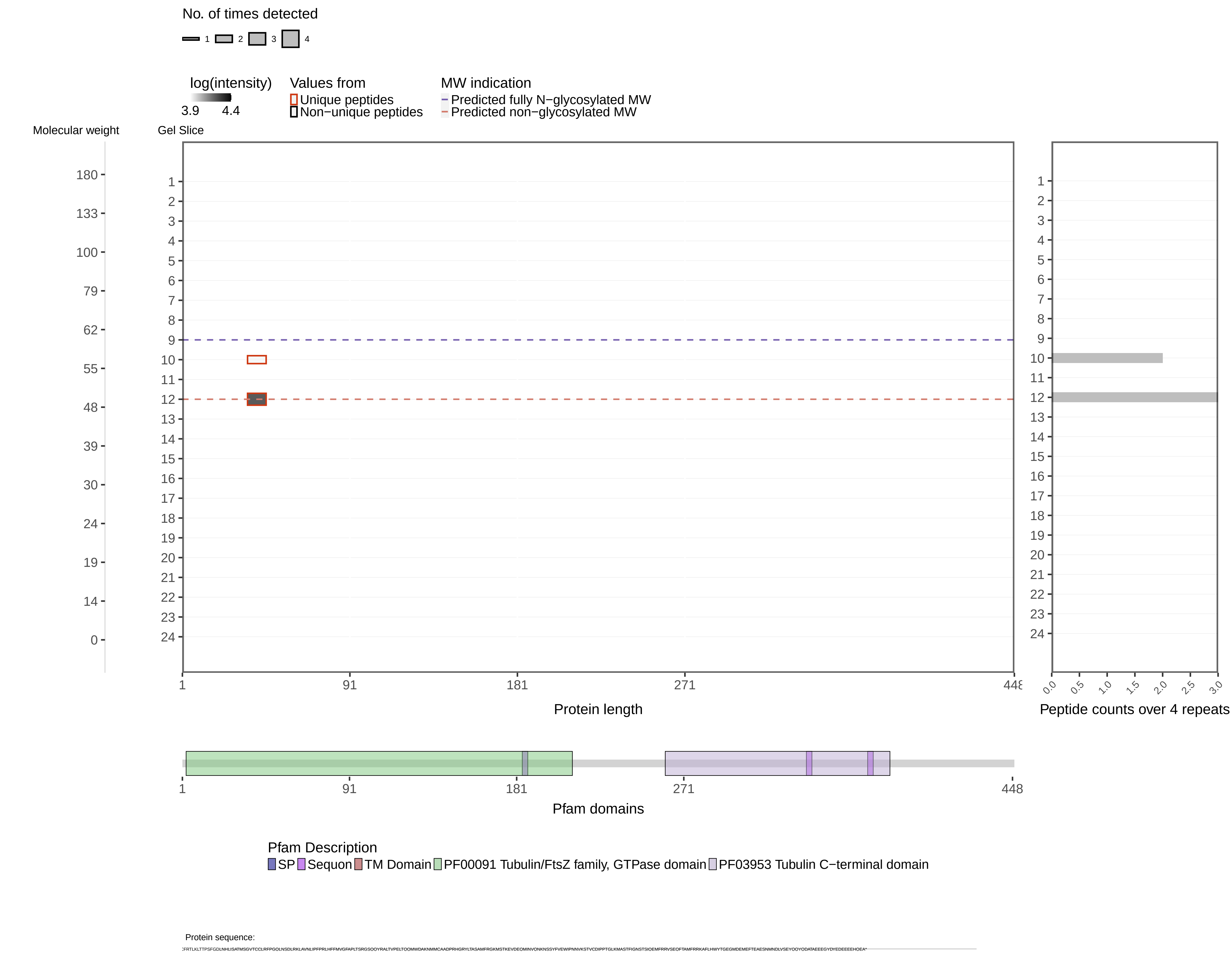

NbL10g04810.1 probable_glucan_13−alpha−glucosidase [MW − SP = 103.04 kDa]

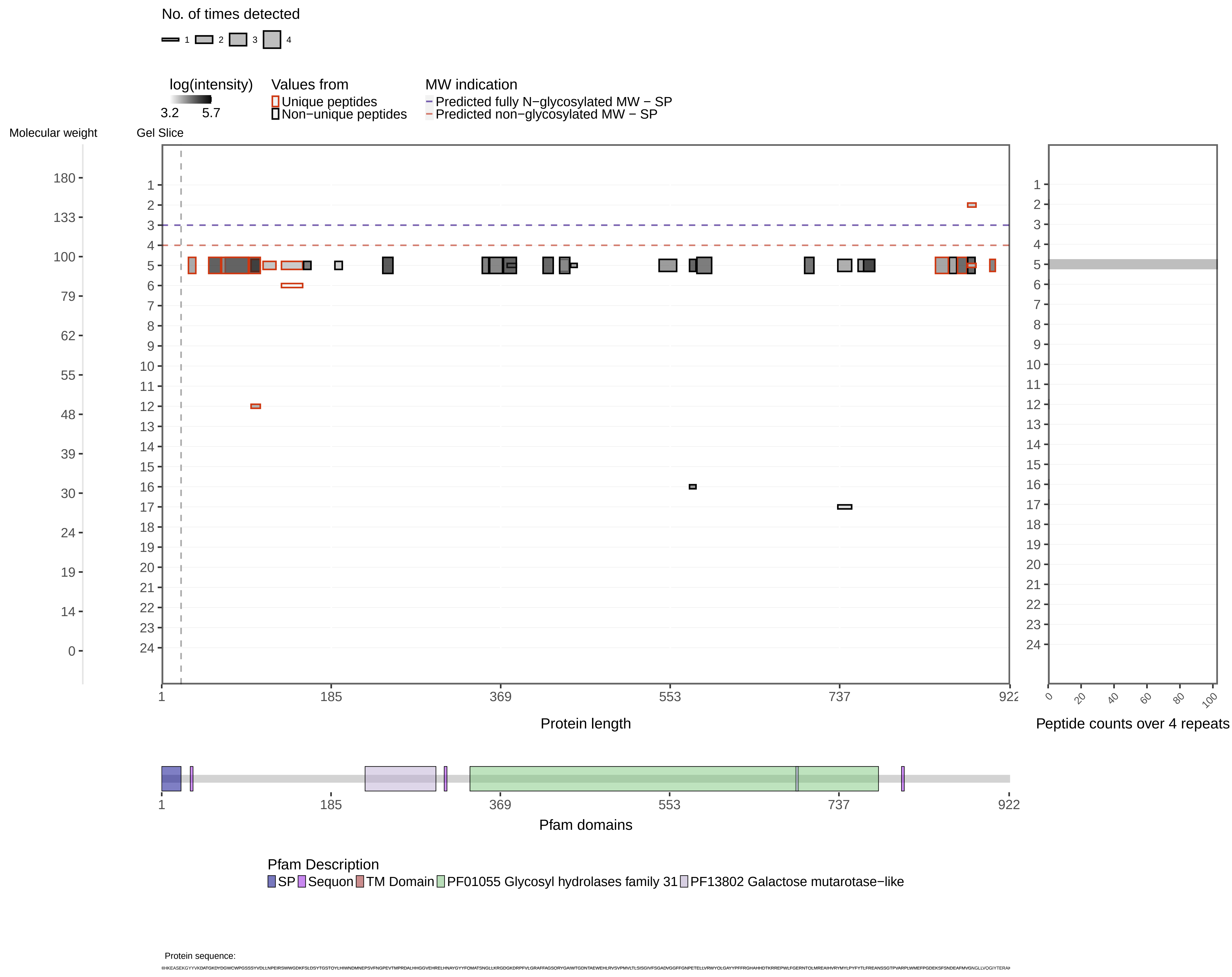

NbL10g04760.1 aspartate−−tRNA_ligase_2_cytoplasmic−like [MW = 60.57 kDa]

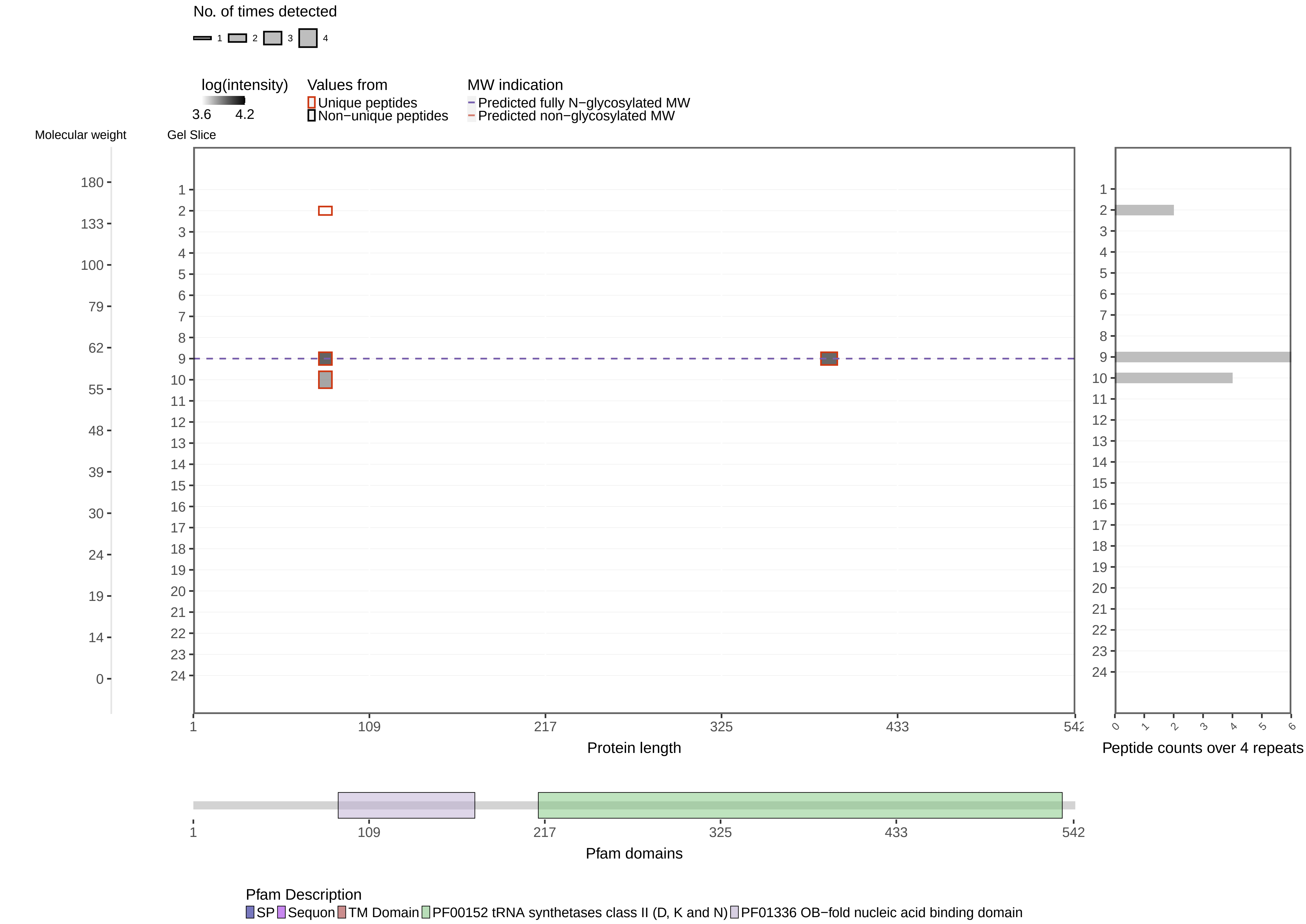


Protein sequence:

PKLIGGSSEGGSAVFRLDYKGQPACLAQSPQLHKQMAICGDFGRVFEIGPVFRAEDSFTHRHLCEFTGLDVEMEIKEHYSEVMDIVDRLFVTIFDSLNDKCKKELEAIGRQYPFEPLKYFRGDKKTLRLTFEEGVQILKEAGIEVDPFGDLNTESERKLGQLVAKKYGTDFYILHKYPLAVRPFYTMPCYDNPAYSNSFDVFIRGEEIISGAQRVHVPEFLAKRAEECGIDVKTISTYIDSFRYGAPPHGGFGVGLERVVMLFCALNNIRKTSLFPRDPQRIAP*

### NbL10g04740.1 acyl_carrier_protein_2_mitochondrial−like [MW = 18.69 kDa]

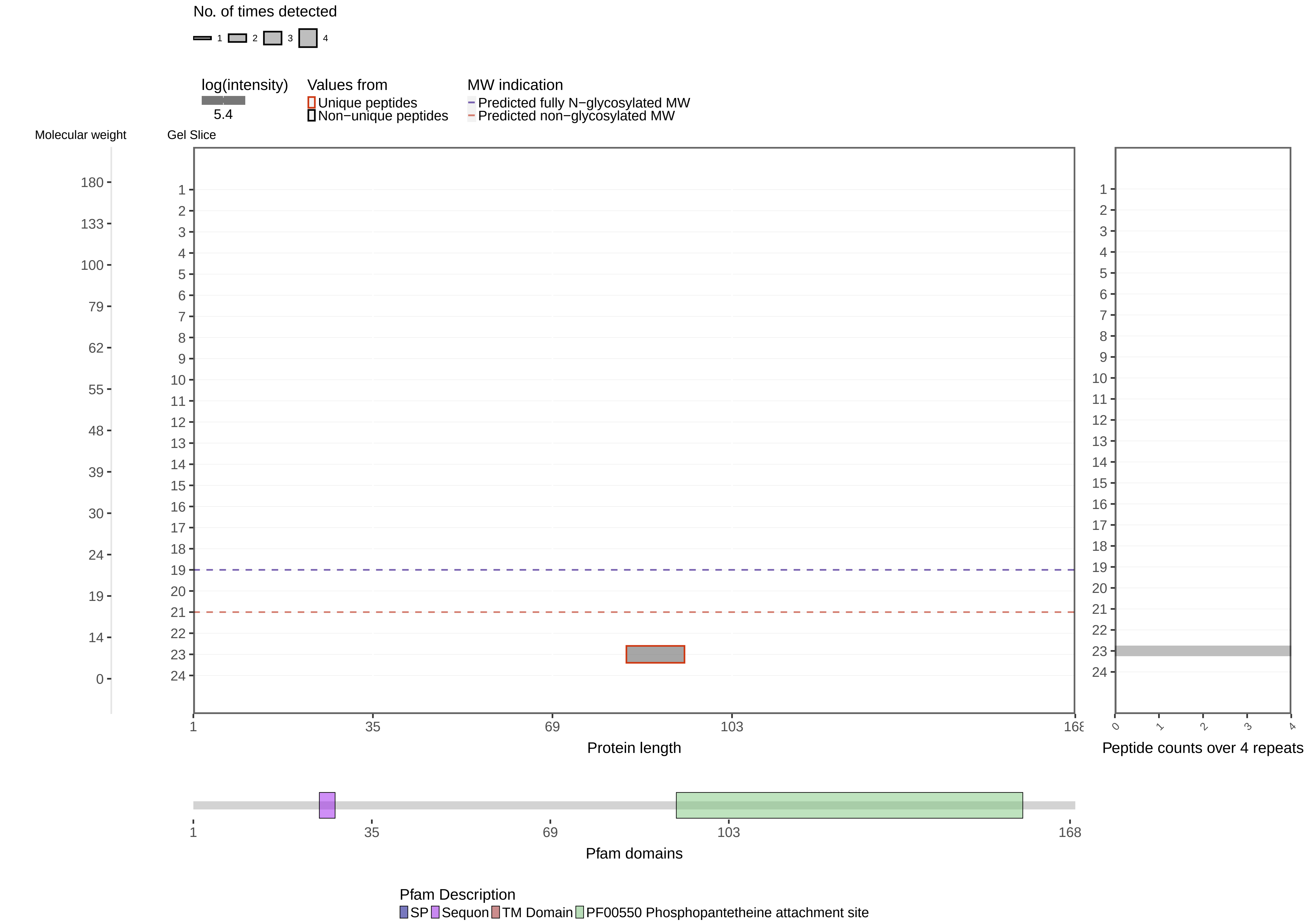


Protein sequence:

'FRRHFSEEVRGSFLDKSEVTDRVINCVKNFQKVDPAKVTPNAHFQNDLGLDSLDTVEVVMALEEEFGFEIPDNEADKINSINLAVDFIASHPQAK*

### NbL10g04730.1 calreticulin−3−like [MW − SP = 43.01 kDa]

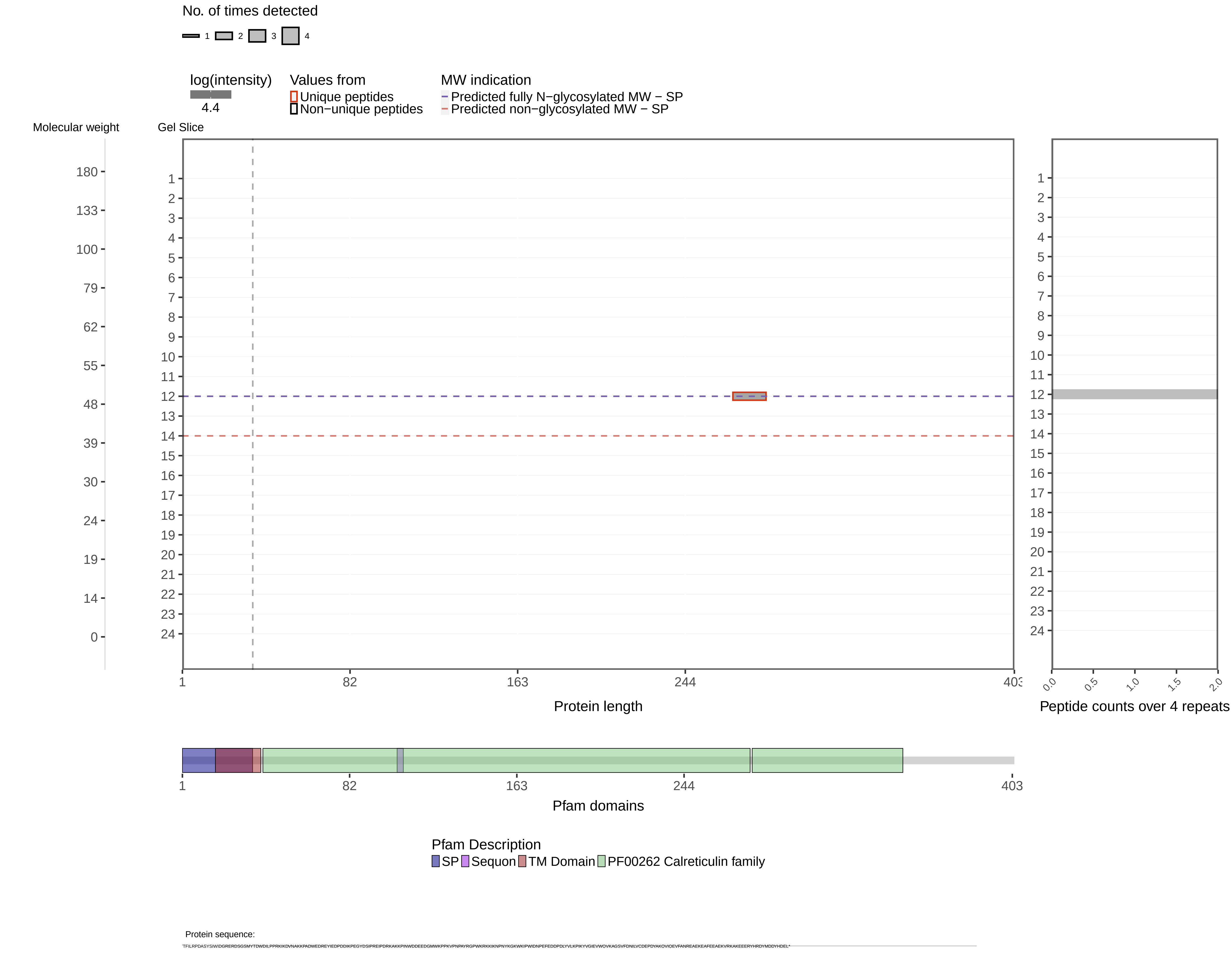

### NbL10g04670.1 phenylalanine_ammonia−lyase−like [MW = 78.25 kDa]

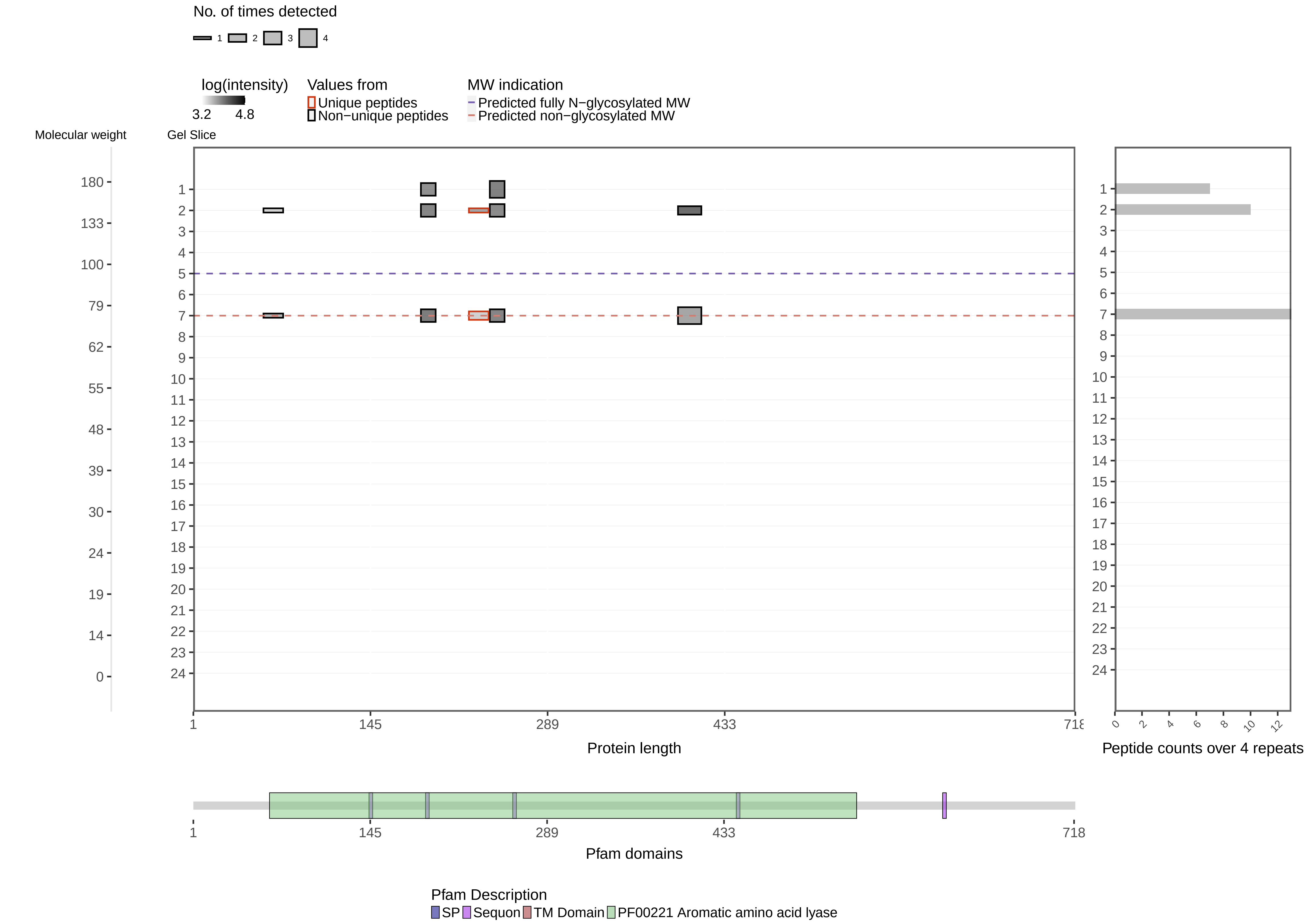


Protein sequence:

KPKQDRYALRTSPQWLGPQIEVIRTATKMIEREINSVNDNPLIDVSRNKALHGGNFQGTPIGVSMDNTRLALASIGKLMFAQFSELVNDYYNNGLPSNLTGGRNPSLDYGFKGAEIAMASYCSELQFSANPVTNHVQSAEQHNQDVNSLGLISSRKTAEAVDILKLMSSTYLVALCQAIDLRHLEENLKASVKNAVSLVAKKVLTIGQNGELHPSRFCEKDLLKVVDREYAFAYADDPCSCNYPLIQKLRQVMVDHALLNGEGEKNYSTSIFEKIGAFEEELKAILPKEVESARC

NbL10g04490.1 acetylornithine_aminotransferase_mitochondrial [MW = 54.01 kDa]

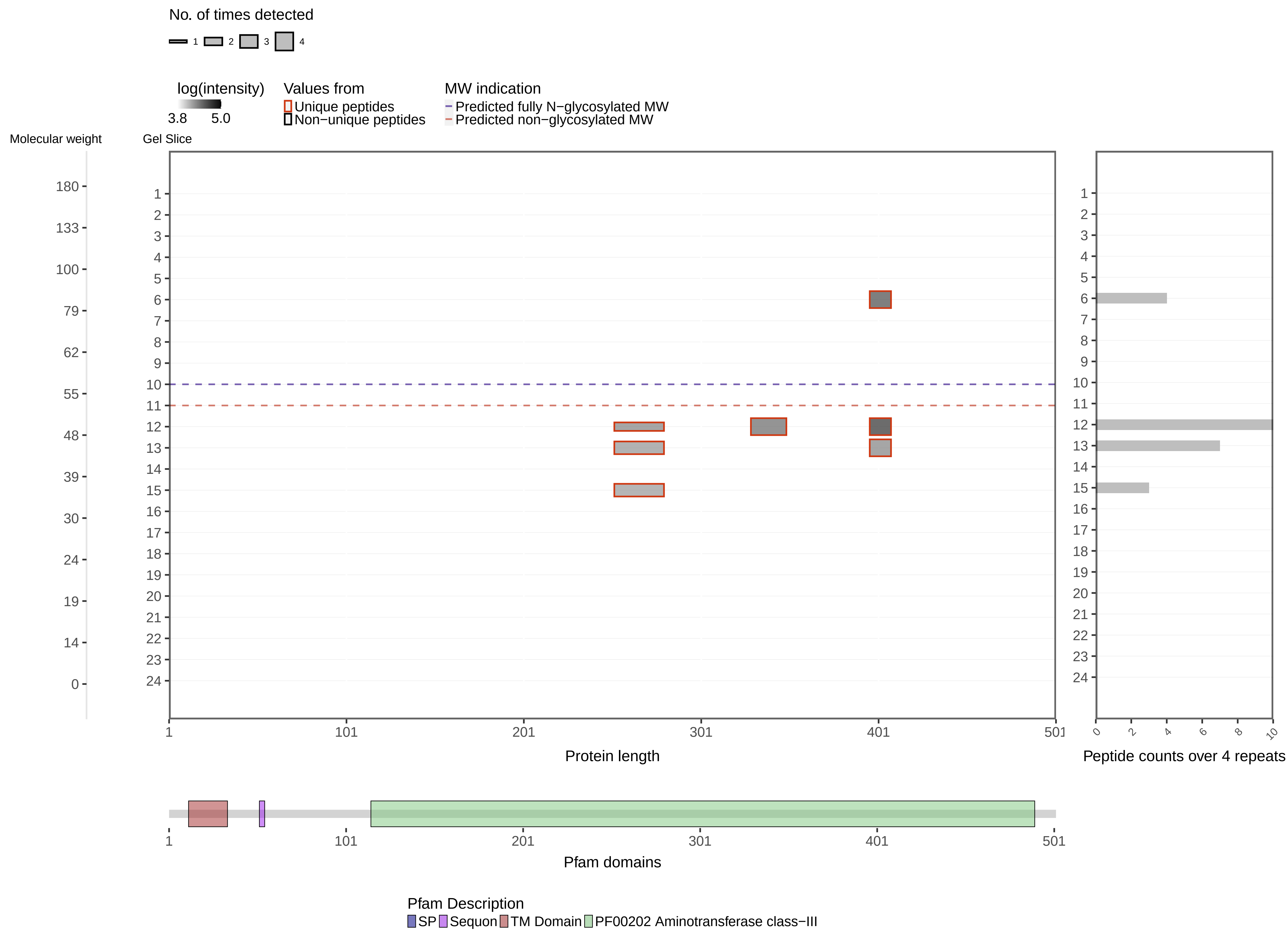


Protein sequence:

IGRTMGAVALTSKEHYRSPFEPVMPGVTFLEYGDIQAATELIQSGKIAAVFVEPIQGEGGIYSATKEFLQALRTSCDSVGSLLVFDEVQCGLGRTGHLWAHDAYGIYPDIMTLAKPLAGGLPIGAVLVTERVAAAINYGDHGSTFAGGPLVCNAAVAVLNKIAEPSFLASVTMKGQYFRELLVKKLGGNSHVKEVRGVGLIIGIDLDVPASPLVDACQQSGLLVLTAGKGNVVRLVPPLTITEQELDHAAEILFNCLPVLDKTNQN*

### NbL10g04410.1 uncharacterized_LOC109220890 [MW = 34.7 kDa]

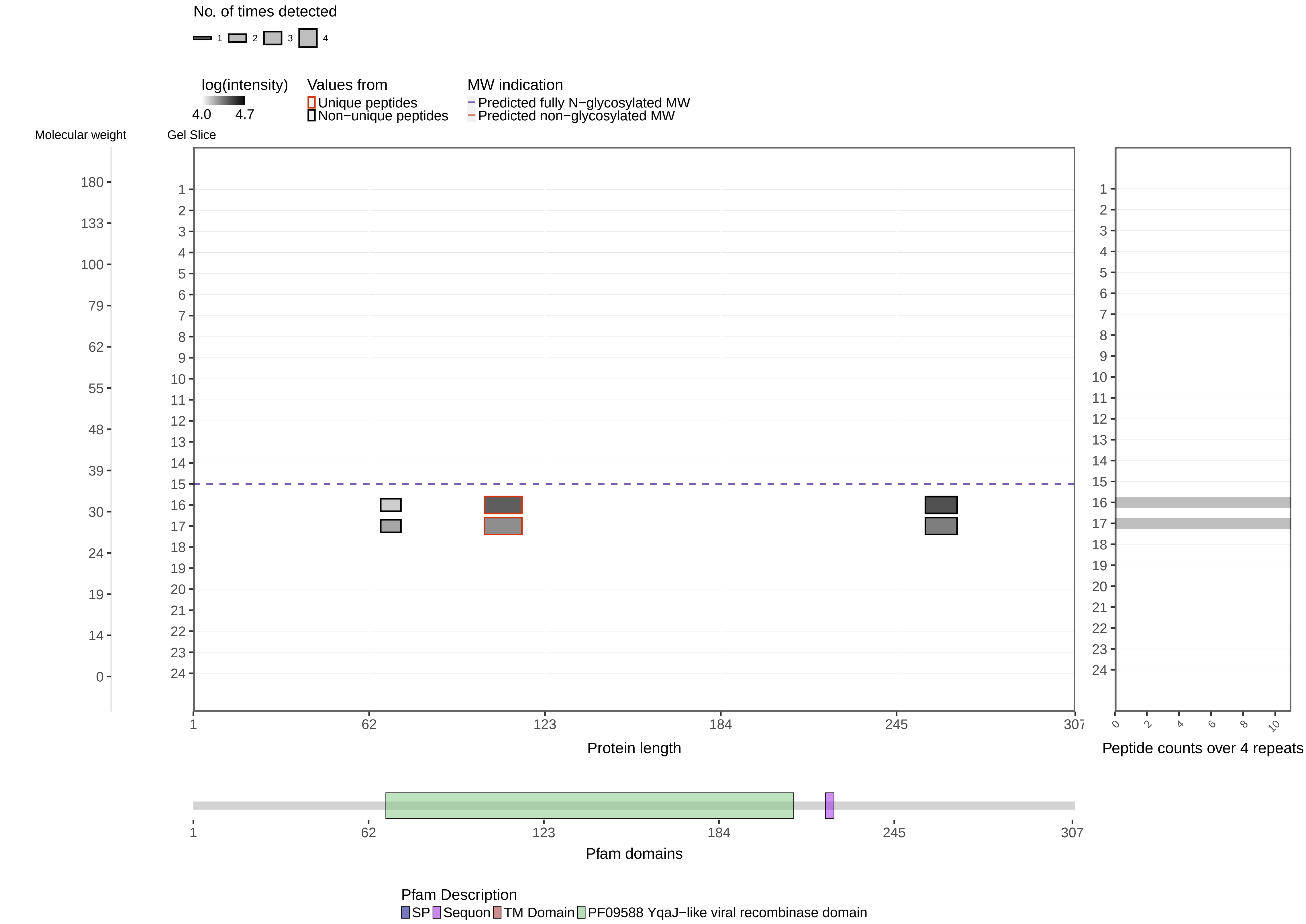


Protein sequence:

GFAVHSDERLGWVGASPDGLLGCLPGGGILEVKCPYNKGKPEKGLPWNTMPFYYMPOVOGOMEIMDRDWVDVYCWTPNGSTIFRVYRERSYWELMHEILWEFWWYNVVPAREALSLGNEEDAKSYKPTPTHKKTGLVISRSLKLAGEAKMLCRDIAGHVEFFR*

### NbL10g04270.1 chorismate_synthase_1_chloroplastic [MW = 47.68 kDa]

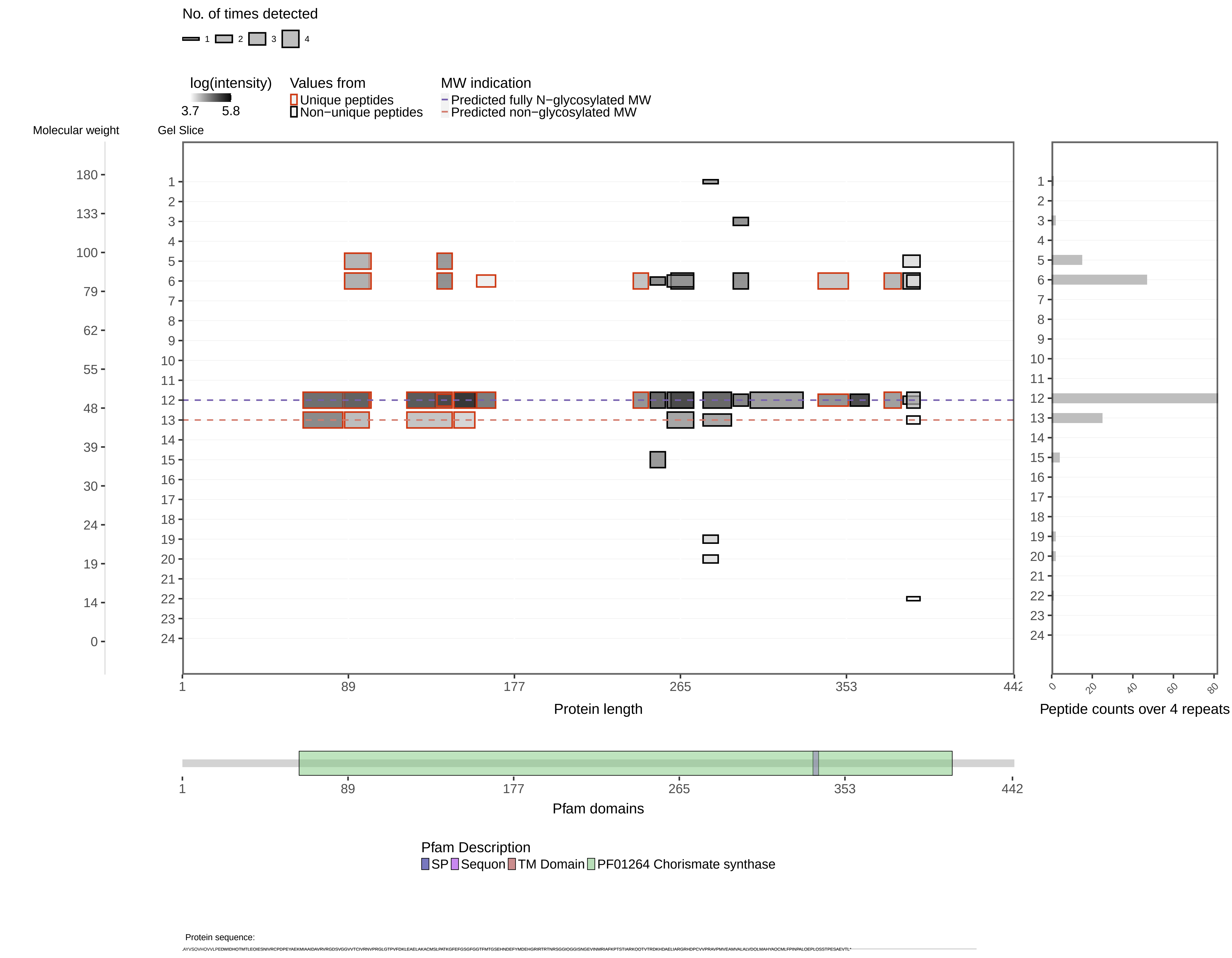

NbL10g04190.1 3−ketoacyl−CoA_thiolase_2_peroxisomal [MW = 48.95 kDa]

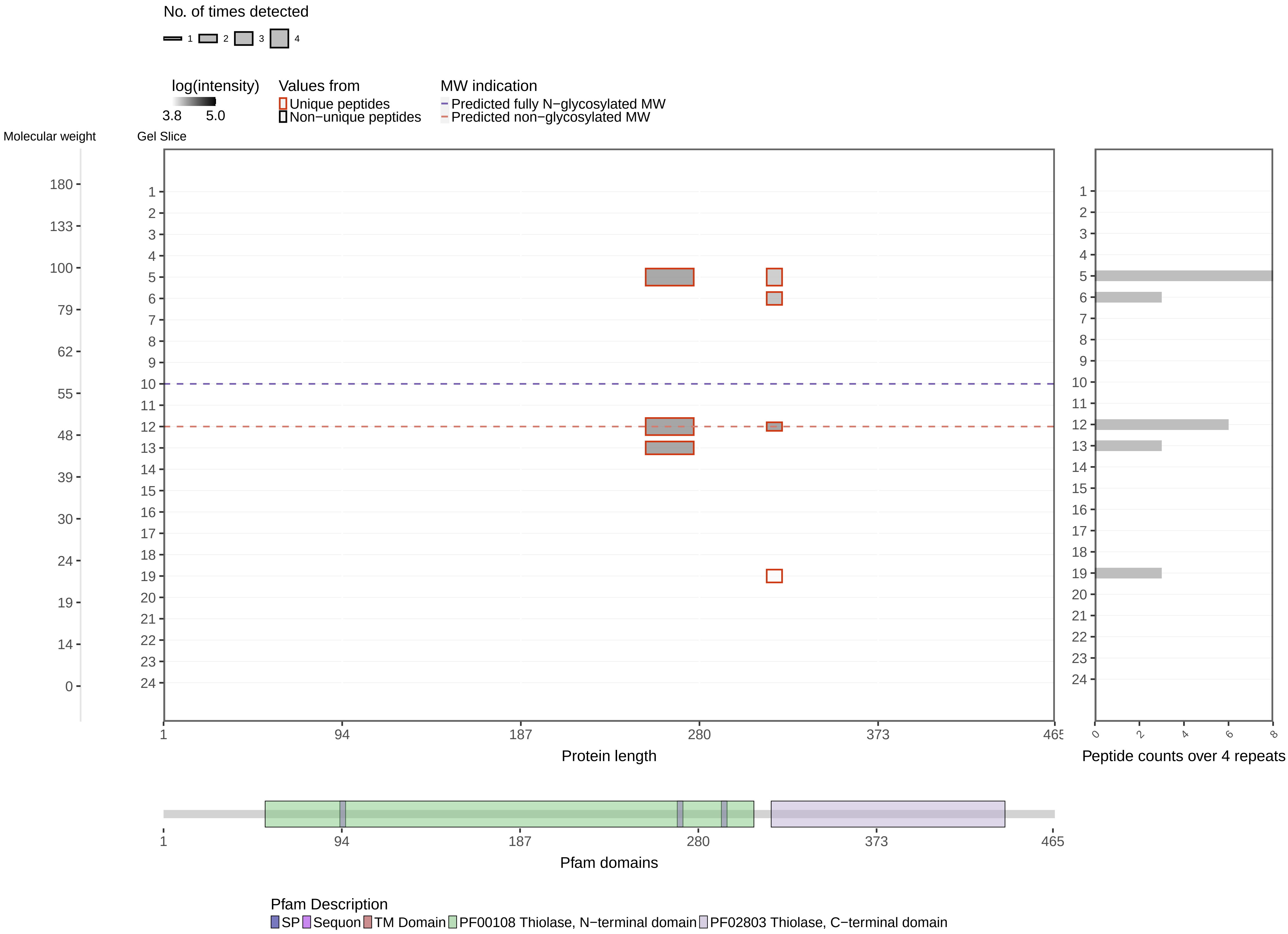


Protein sequence:

QAAVDSHRKAAAASASGKFKDEIIPVPTKIVDPKTRDETPVTISVDDGIRPNASVSDLAKLKPVFKKSGTTTAGNSSQVTDGAGAVLLMKRSLAMQKGLPVLGVFRTFAAVGVDPAIMGIGPAVAIPAAVKSAGLELEDIDLFEINEAFASQYVVYCRKKLELDPEKINVNGGAMALGHPLGATGARCVATLLHEMKRRGKDCRFGVVSMCIGTGMGAAAVFERGDSCDELCNARKIGSHNLLSKDVL*

### NbL10g04140.1 probable_pectate_lyase_8 [MW = 53.83 kDa]

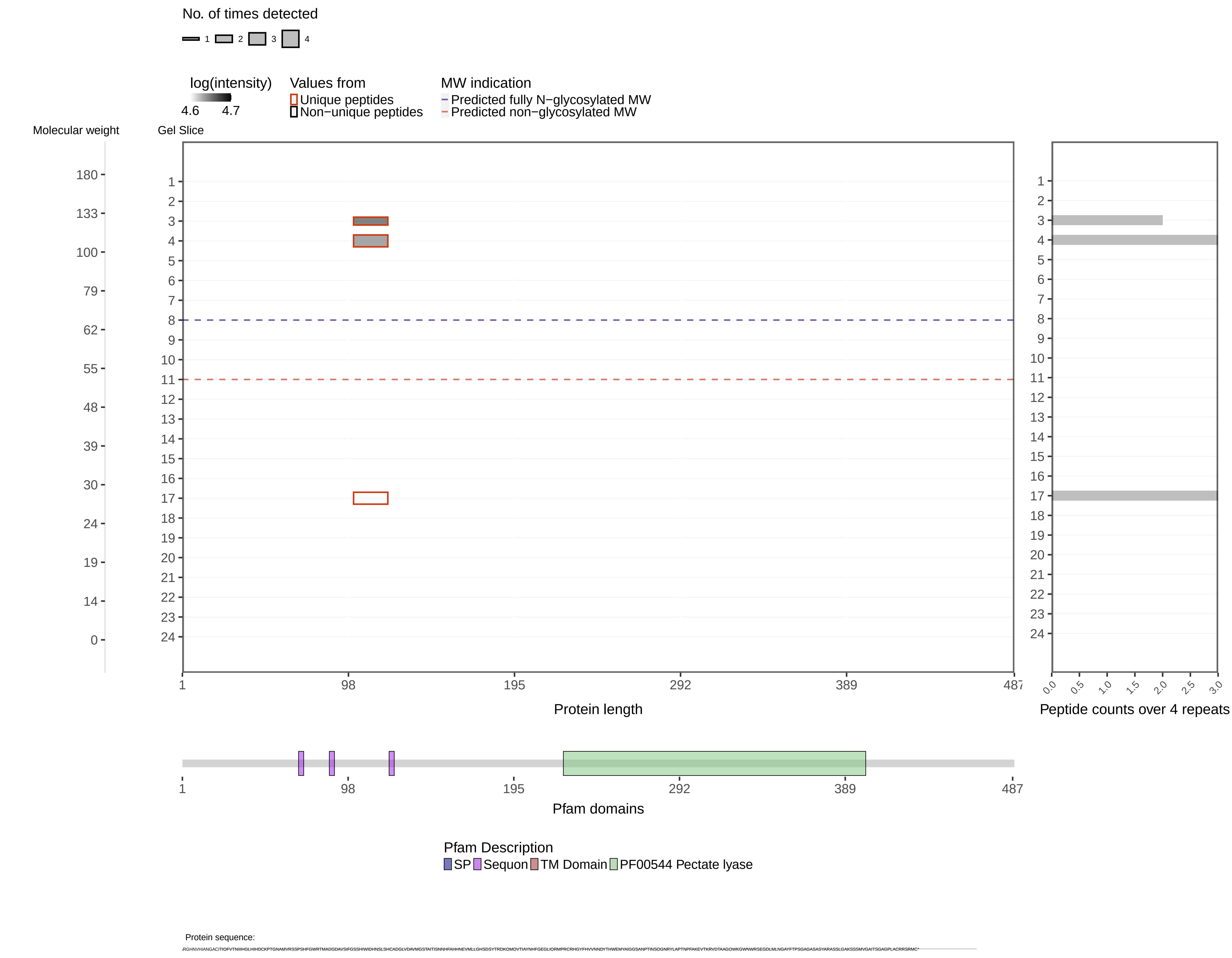

NbL10g03900.1 20_kDa_chaperonin_chloroplastic−like [MW = 26.37 kDa]

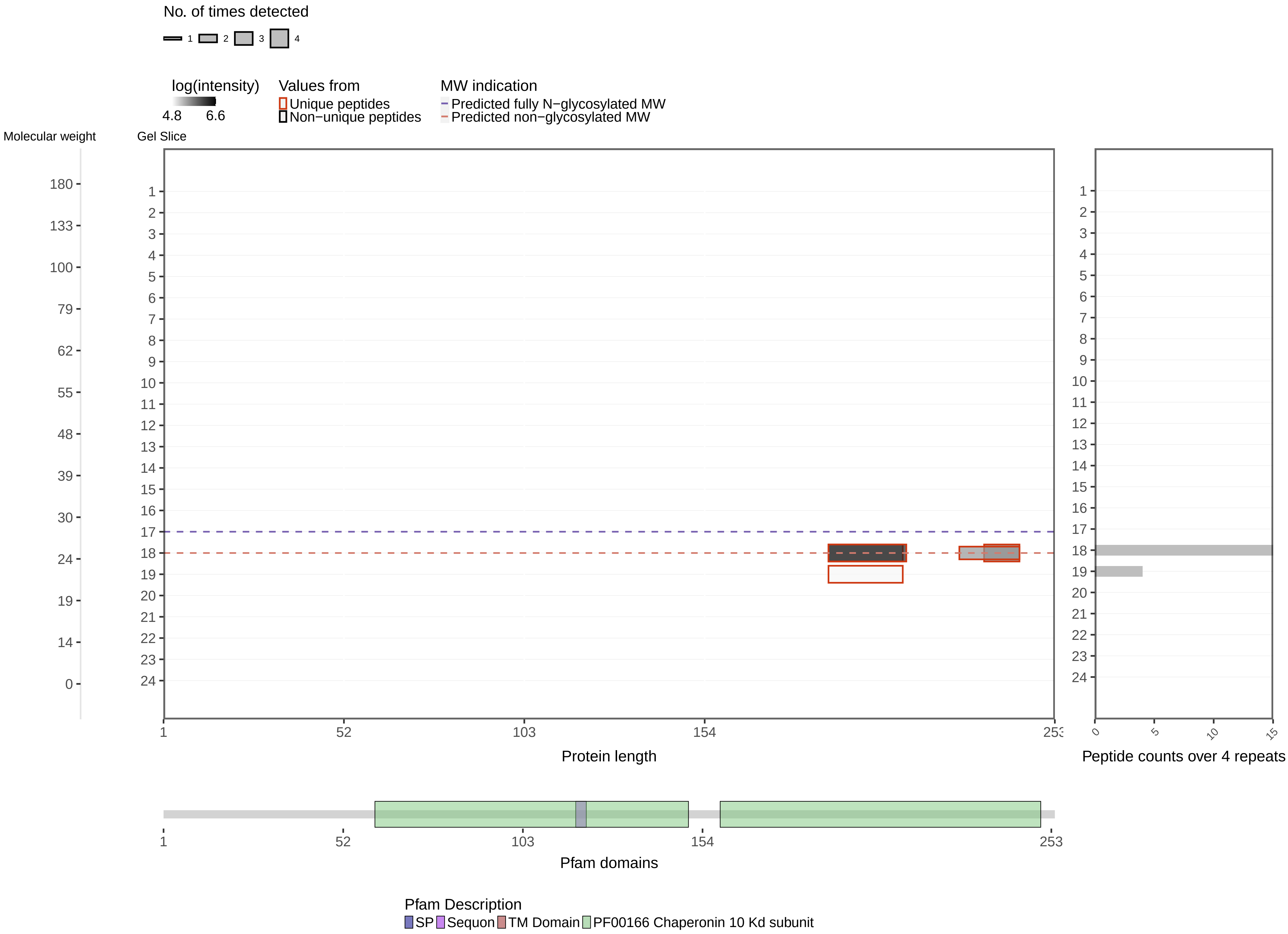


Protein sequence:

EISVKNGTQVLYSKYAGTEVLFDGSTHLILKEDDIVGILETDDIKDLQPLNDRILIKVAEVEEKTAGGLFLSEAAKEKPSIGSVIAAGPGPLDDEGNRKSLSVSPGNTVLYSKYAGNDFKGADGADYVTLRASDVIAVLS*

### NbL10g03850.1 sucrose_synthase [MW = 92.47 kDa]

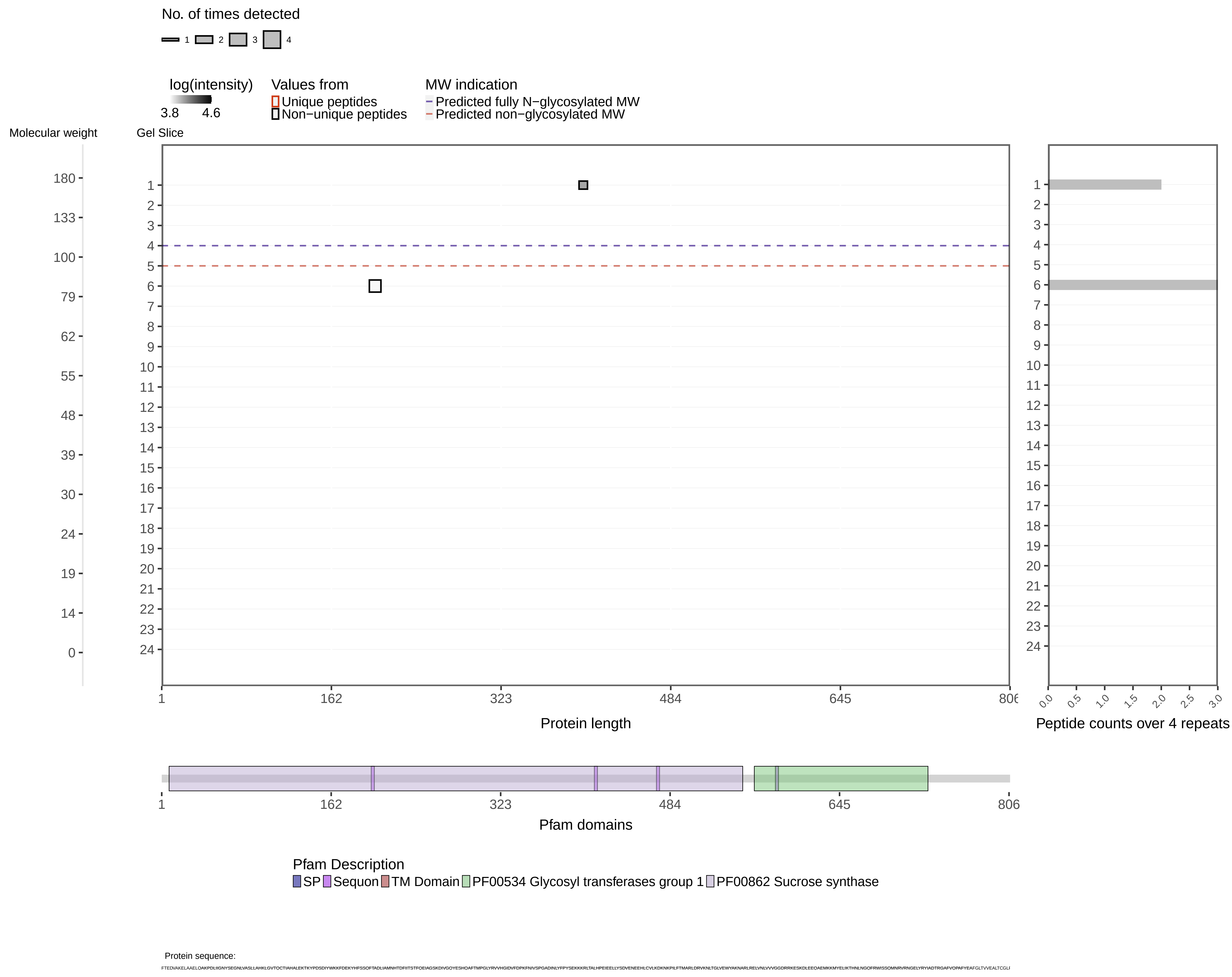

### NbL10g03670.1 yrdC_domain−containing_protein_mitochondrial [MW = 30.4 kDa]

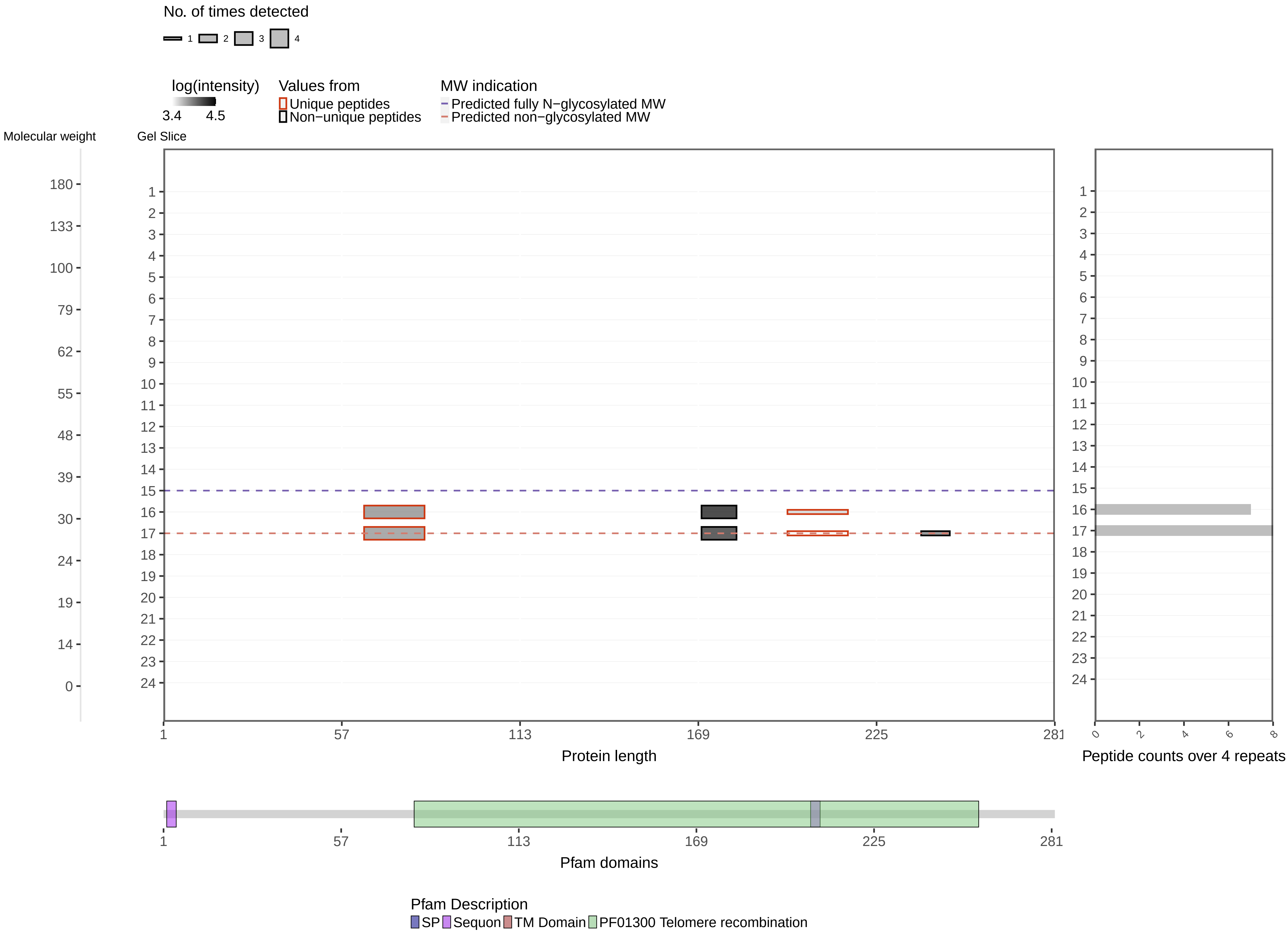


Protein sequence:

GDVHDIQRYAVTDHLPHGLLDCLLPGPVTVVLRRGESSILEKSLNPGLESIGVRVPDYNFIRVIARGSRSALALTSANLSGQPSSINIKDFENLWEHCAYVYDGGVLQDGRAGSTVVDLTKLGKYKILRPGSAKEETVAILERHSLLEDGTGG*

### NbL10g03470.1 fasciclin−like_arabinogalactan_protein_2 [MW − SP = 40.47 kDa]

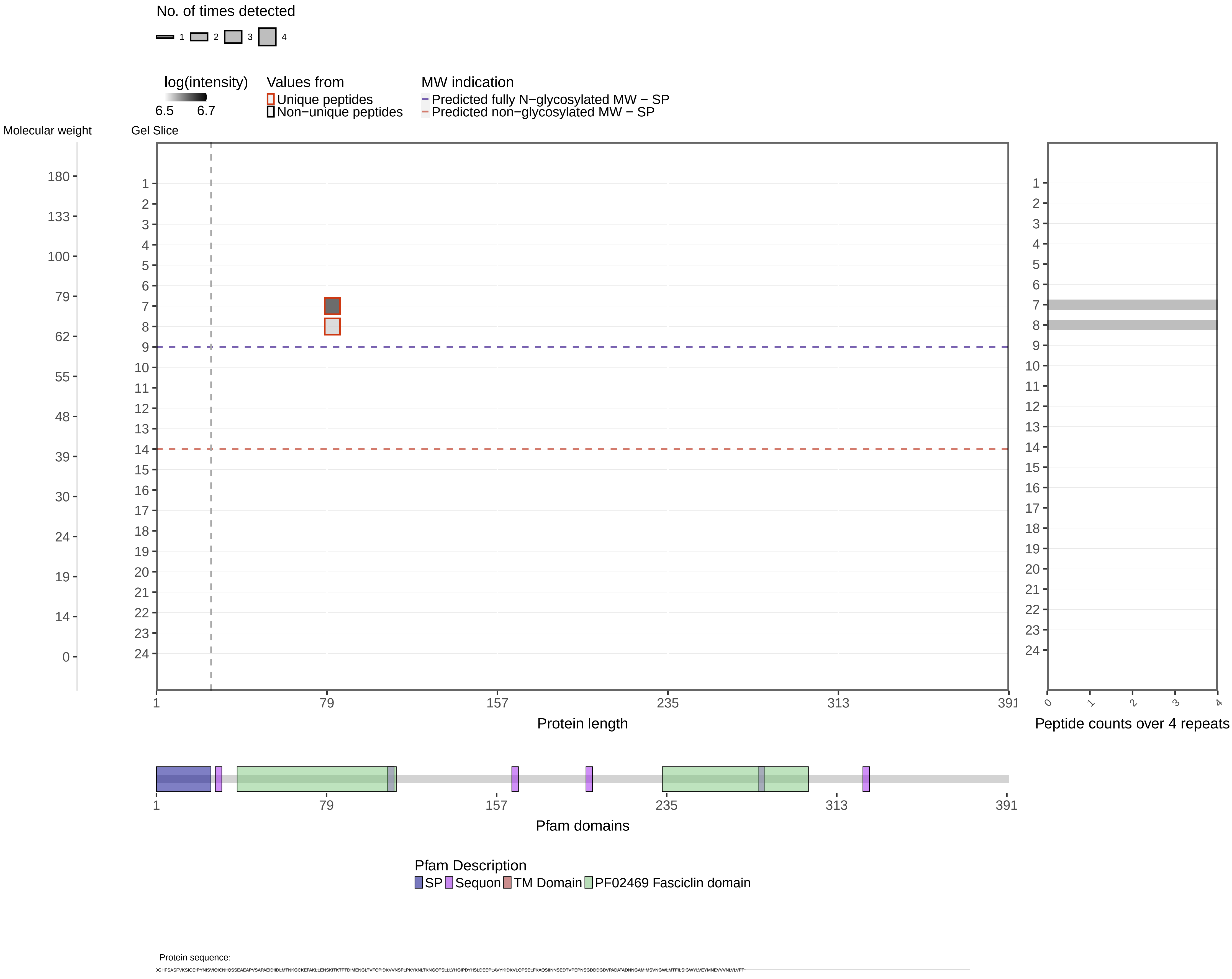

NbL10g03250.1 NEDD8−activating_enzyme_E1_regulatory_subunit_AXR1−like [MW = 58.13 kDa]

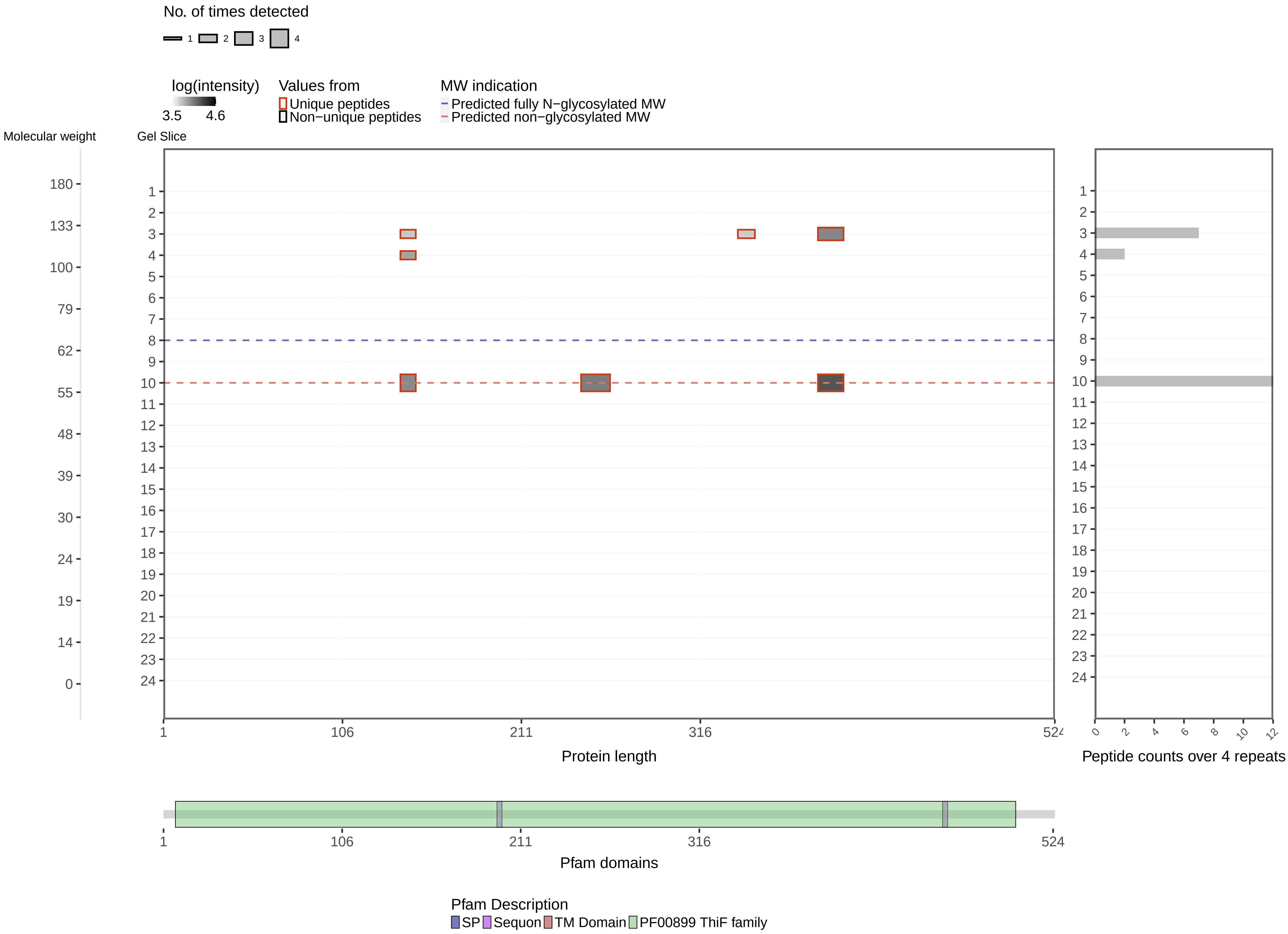


Protein sequence:

SDEENYKEAVEASYKLFSPQGTGPNLQKIIDDSCTEVDSSSSDFWVMVAALKEFIANEGGEETPLEGSIPDMTSSTELYINLQKTYQAKAEADFVIMEQKVRNLLKKIGRDPASIAKAHIKSCCKNARKLAVCRYRLIEDEFNSPVQPELQKYLIDEDYGTAVGLYILLRAADRFAANYNKFPGQFDGEMDEDISRLKTTAVGLLNDLGCNGSALSEDLINEMCRYGASELHAVAAFVGGVASQEVIKLLTRQFIPMSGTFIFNGIDHKSQLLLL*

### NbL10g03240.1 aconitate_hydratase_cytoplasmic−like [MW = 108.25 kDa]

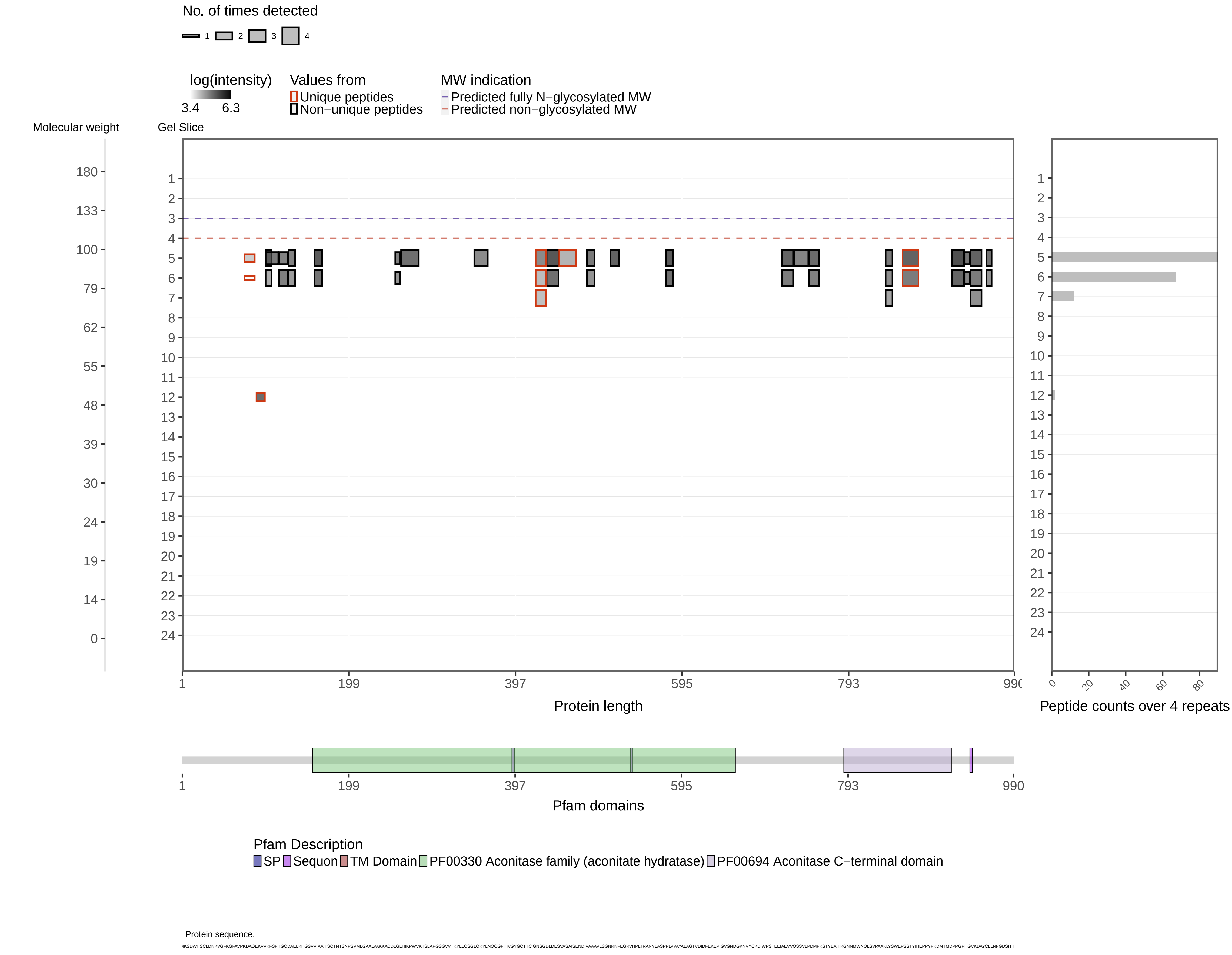

### NbL10g03070.1 cytochrome_b6−f_complex_iron−sulfur_subunit_2_chloroplastic [MW − SP = 22.53 kDa]

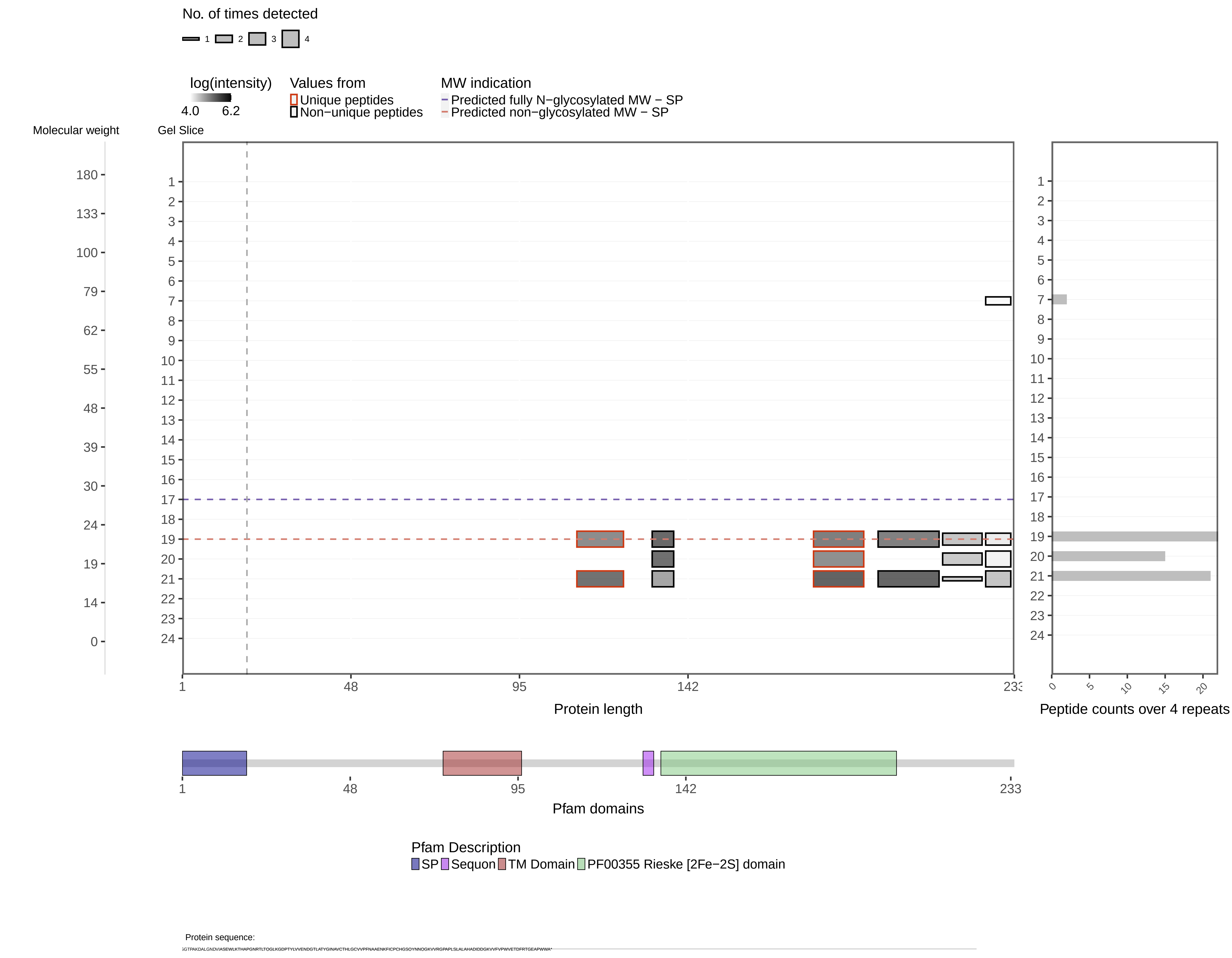

### NbL10g02760.1 ketol−acid_reductoisomerase_chloroplastic−like [MW = 63.7 kDa]

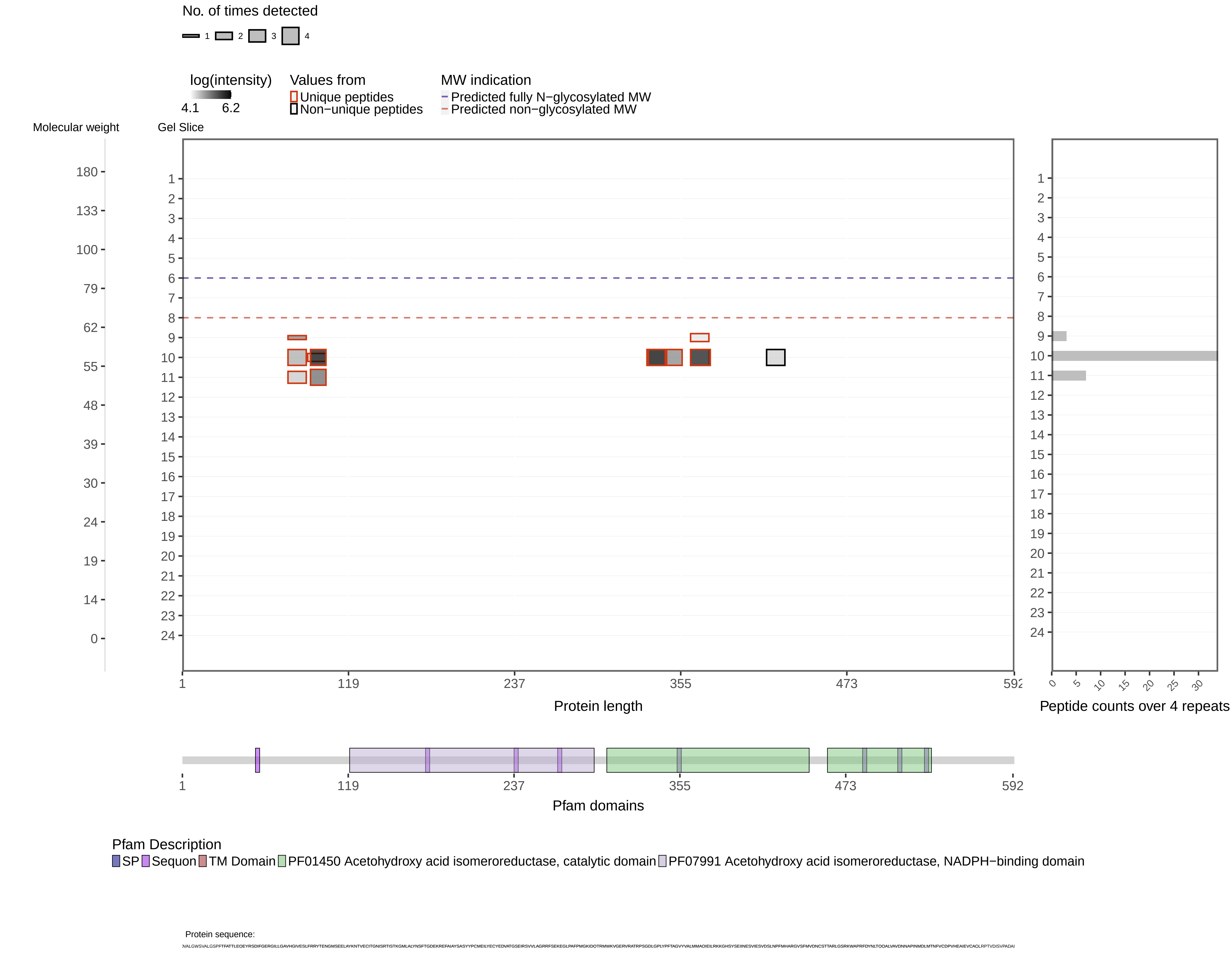

### NbL10g02650.1 ras−related_protein_RABA1f−like [MW = 24.1 kDa]

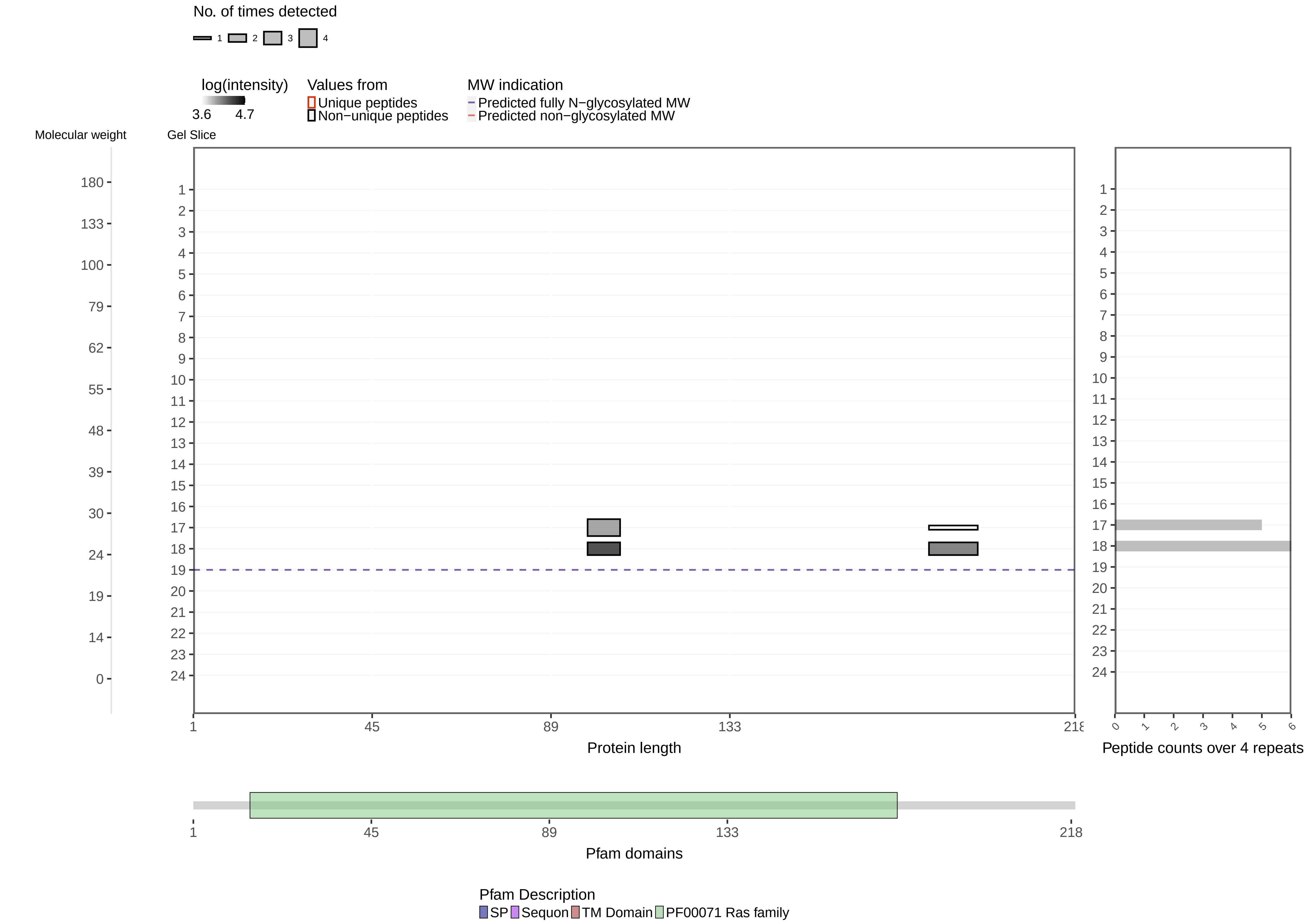


Protein sequence:

RHVTFENVERWLKELRDHTDSSIVIMLVGNKADLRHLRAVSTEDAKAFAEKESTFFMETSALESMNVDNAFTEVLTQIHRVVSRKALEVGDDPAALPKGQTINVGGKDDVSEVKKAGCCSS*

NbL10g02440.1 vacuolar_protein_sorting−associated_protein_35A−like [MW = 89.47 kDa]

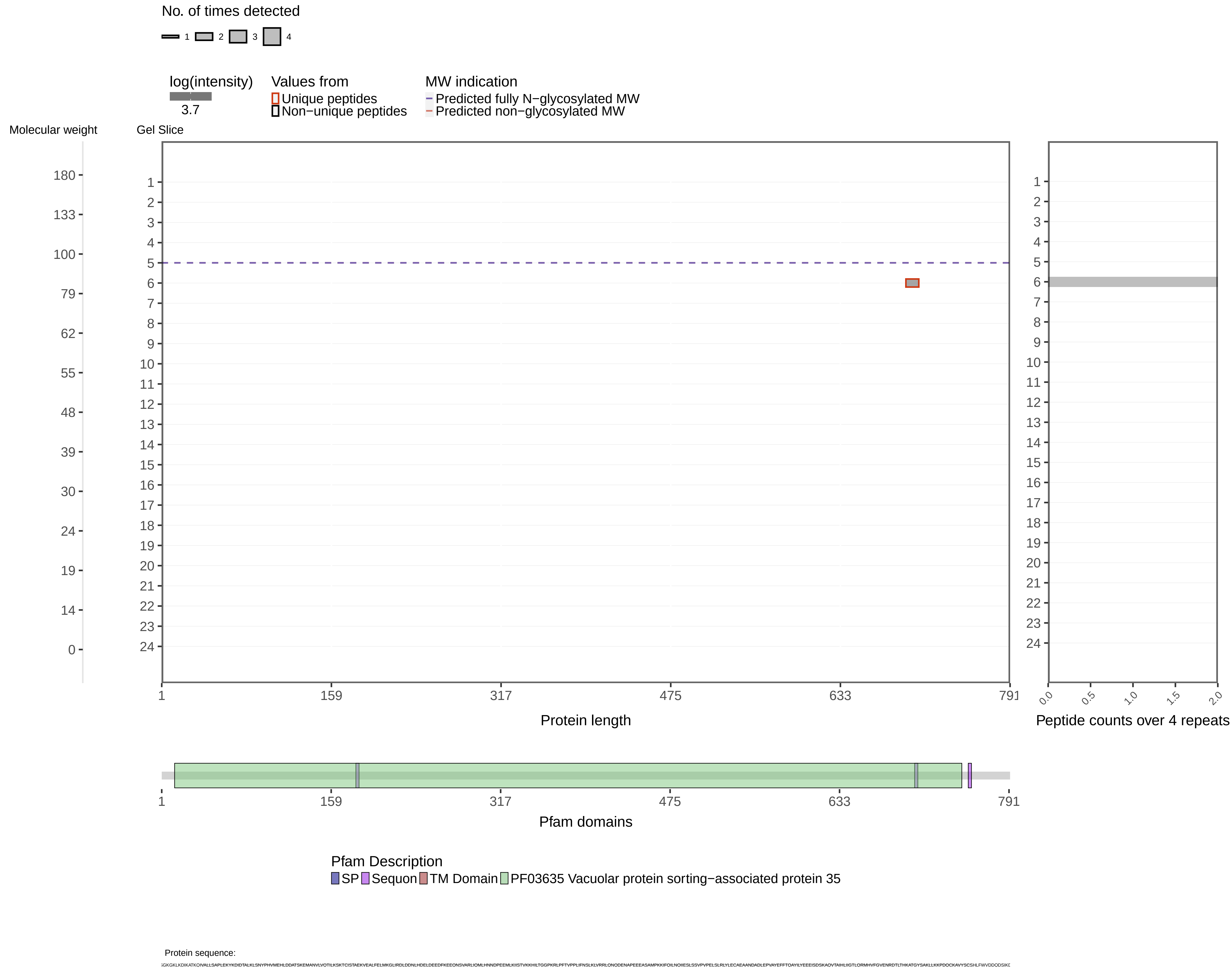

### NbL10g02420.1 proteasome_subunit_alpha_type−6 [MW = 28.42 kDa]

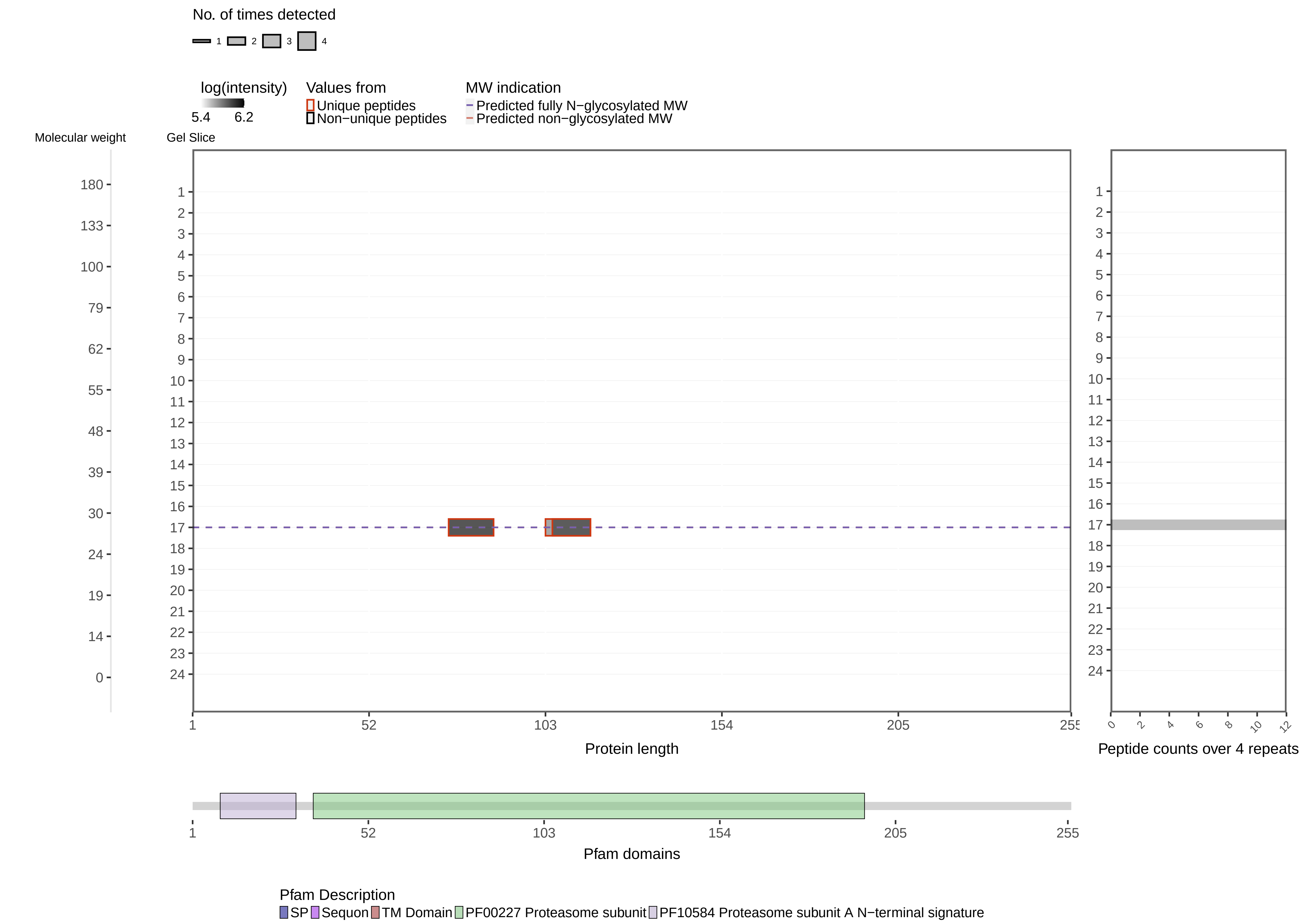


Protein sequence:

LSRWIADKSQVYTQHAYMRPLGVVAMILGIDEEKGPQLFKCDPAGHFFGHKATSAGSKEQEAINFLEKKMKNDPAFSYEETVQVKGKGVFRRGLRLPFSLYPIFSSFTFLSLQVSATAGIPVSRLYWYCIEYLCKILWALN*

NbL10g02260.1 probable_inactive_receptor_kinase_At2g26730 [MW − SP = 68.71 kDa]

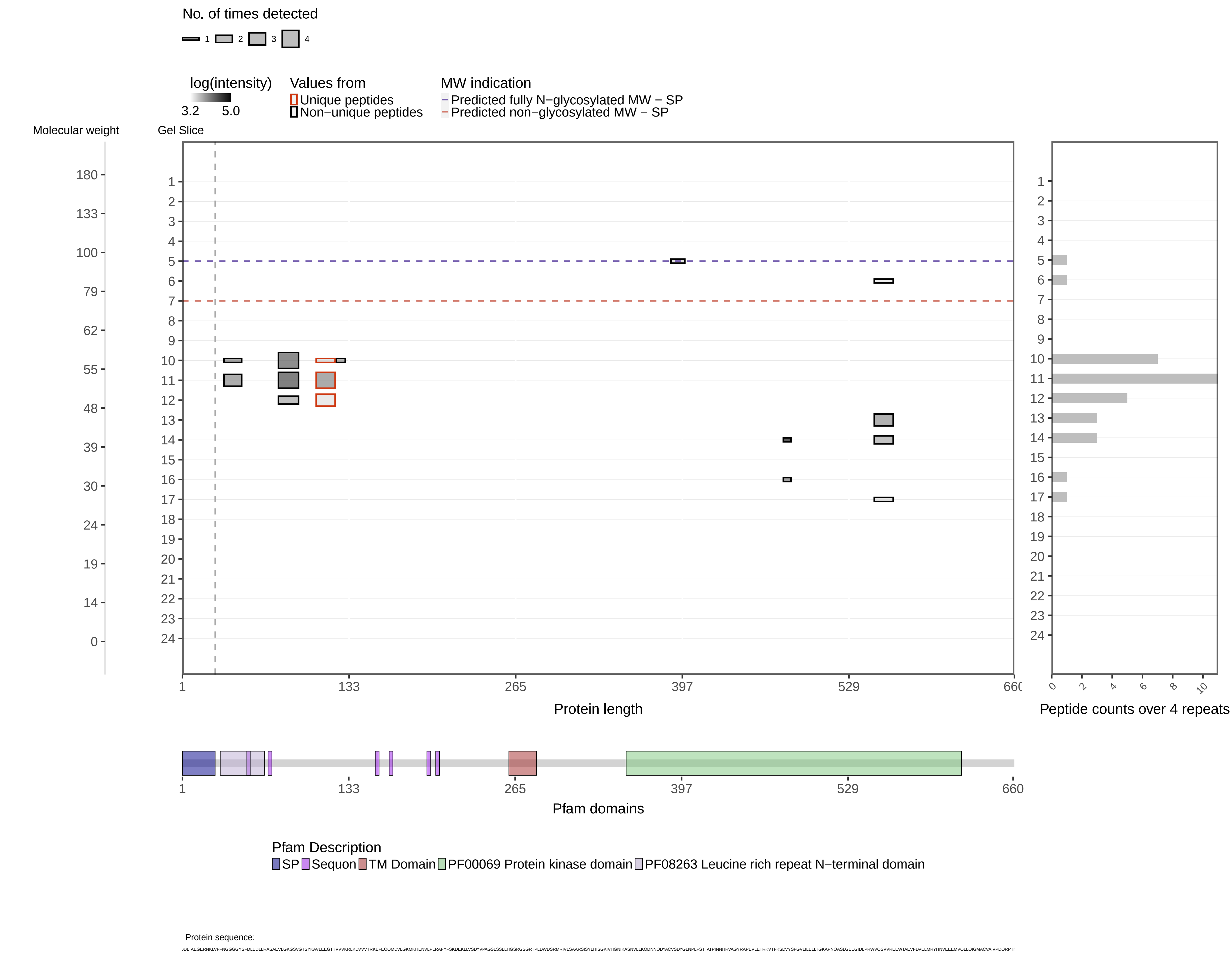

### NbL10g02160.1 carbamoyl−phosphate_synthase_small_chain_chloroplastic [MW = 47.22 kDa]

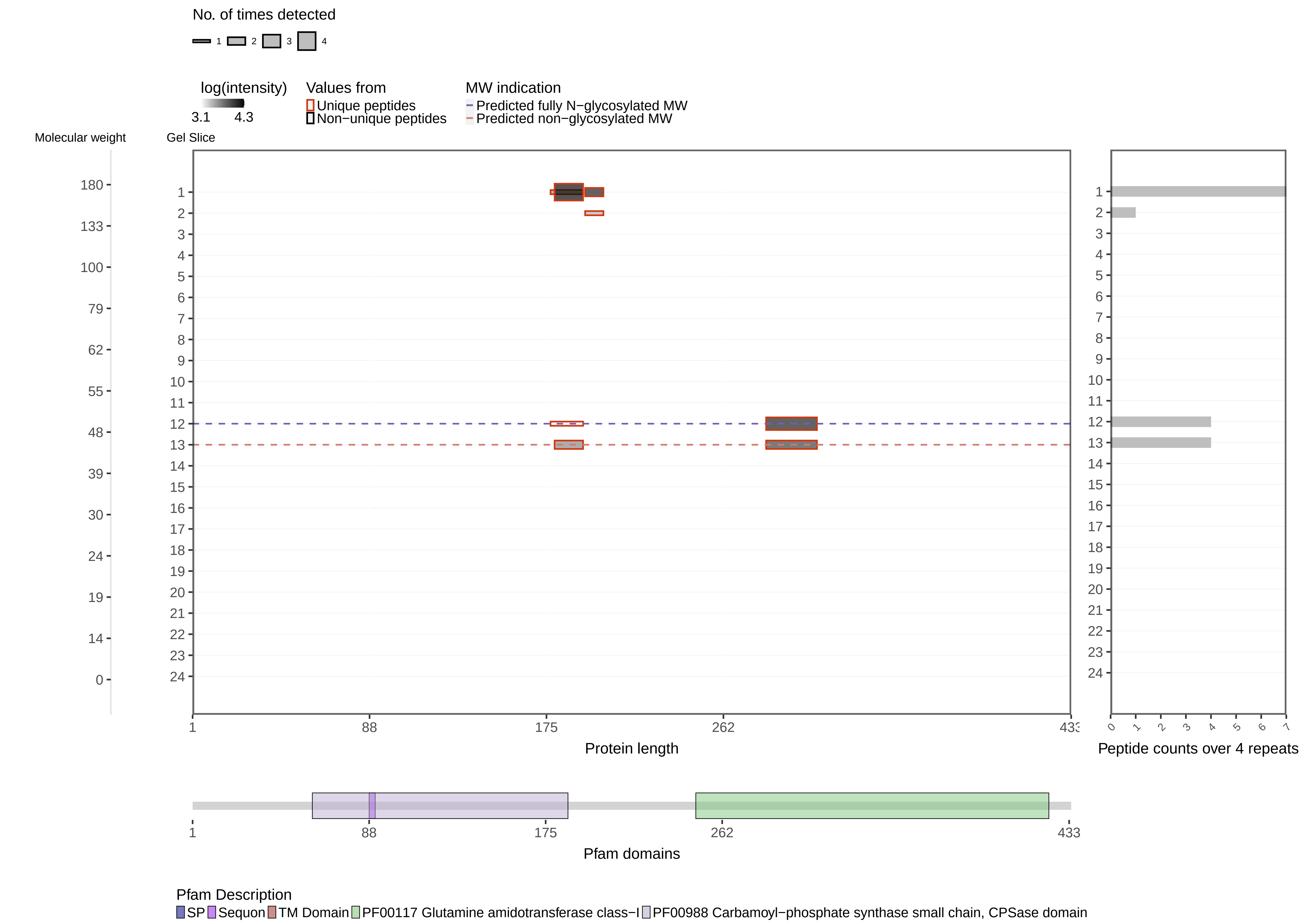


Protein sequence:

DIVGVDLITGVSCKSPYEWVDSTASDWDFNDNGRNKETFNVVAYDFGIKHNILRRIASYGCKITVVPSTWPASETLKMKPDGVLFSNGPGDPSAVPYAVEAVKELIGKIPVFGICMGHQLLGQALGGKTFKMKFGHHGGNHPVRNLRNGCVEISAQNHNYAVDPESLPEGVEVTHVNLNDGSCAGLASSKMKLMSLQYHPEASPGPHDSDPVFGEFIQLMKQEKVKA*

### NbL10g02120.1 probable_protein_arginine_N−methyltransferase_1.2 [MW = 45.1 kDa]

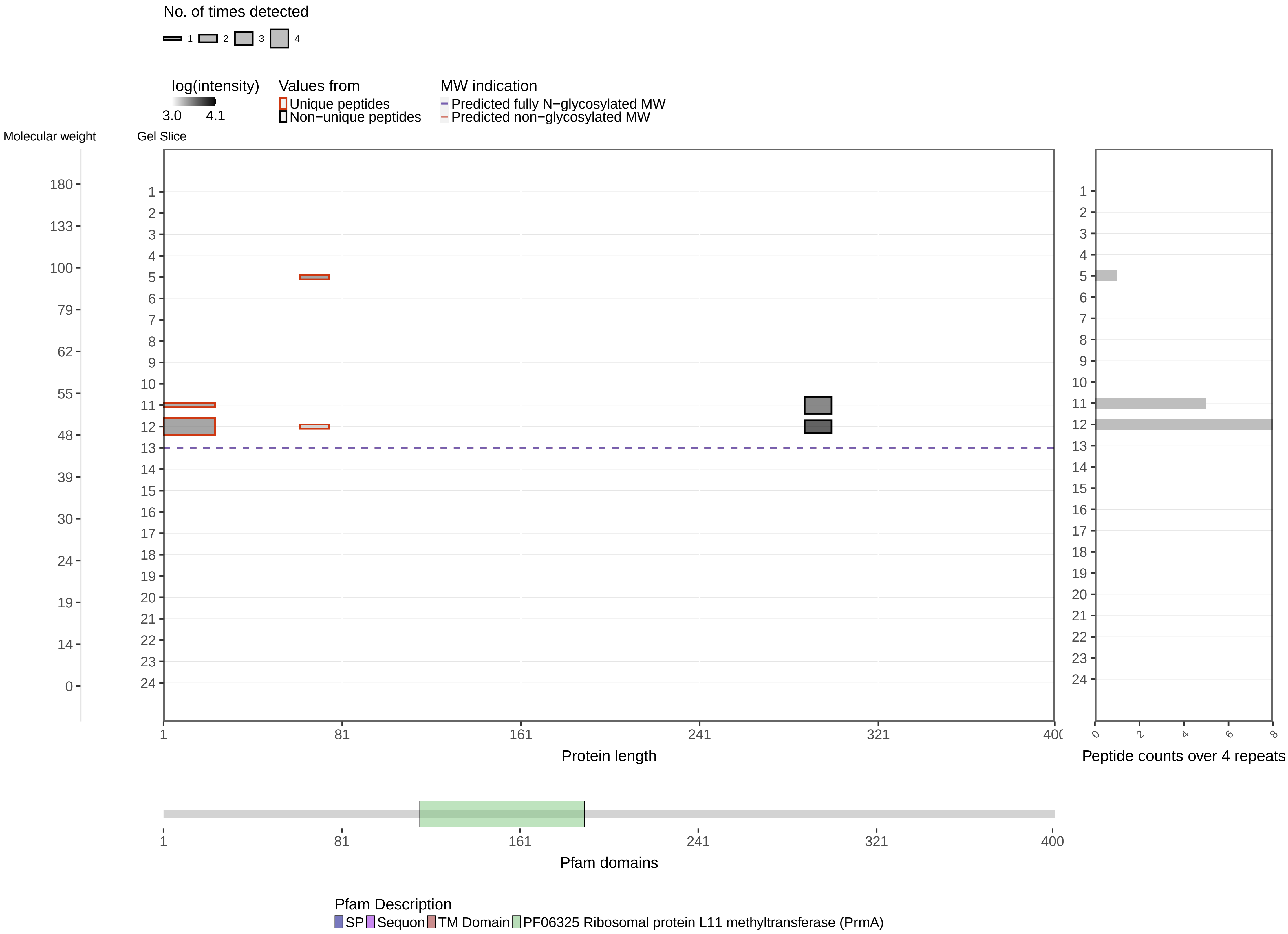


Protein sequence:

IISEWMGYFLLYENMLDTVLYARDKWLVKDGLVLPDKASLYLTAIEDADYKTDKIEFWNEVYGFDMSCIRKQAMMEPLVDTVDQNQIVTNCQLLKTMDISKMTSGDASFTAPFKLIAERDDYIHALVAYFDVSFTKCHKLMGFSTGPKSRGTHWKQTVLYLEDVITICQGEAVVGSITVAPNKKNPRDVDIMLKYSVNGRHCRVSRTQYHRMR*

### NbL10g01940.1 gamma−glutamyltranspeptidase_3 [MW = 69.4 kDa]

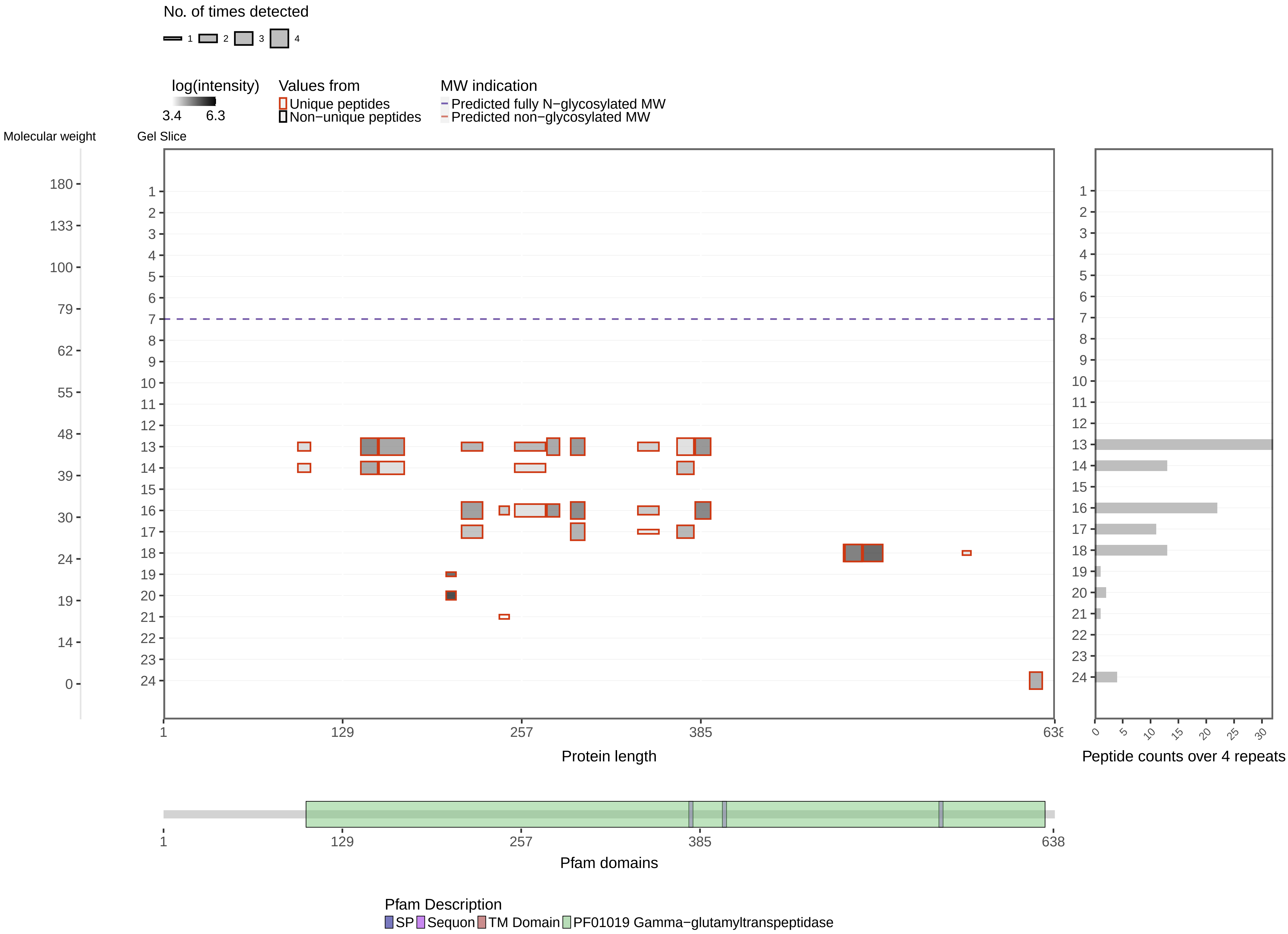


Protein sequence:

YKVETPEAVTFNAMGYTIVGMLPPSSGTLGISLILKILESYNTAEGYLGLHRLIEAMKHMFAVRMDLGDPDFVNISKTVSDMLSPSFAKEIQRKIFDNTTFPSEYYMPRWSQLRDHGTSHFCIVDSDRNAVSVTTTVNYAFGAGVLSPSTGIVINDEMGDFSTPNDISPDELPPSPANFIQPKKRPLSSMAPIIVLKDNQLAGVIGGSGGMKIIPAVVQVFINHFILGMDPLAAVQSPRVYHELIPNVVLYENWTCIDGDHIELSDEKKHFLEERGHQLEAHNGGAICQLVVQNLPNS

### NbL10g01890.1 arginine−−tRNA_ligase_chloroplastic/mitochondrial−like [MW = 73.4 kDa]

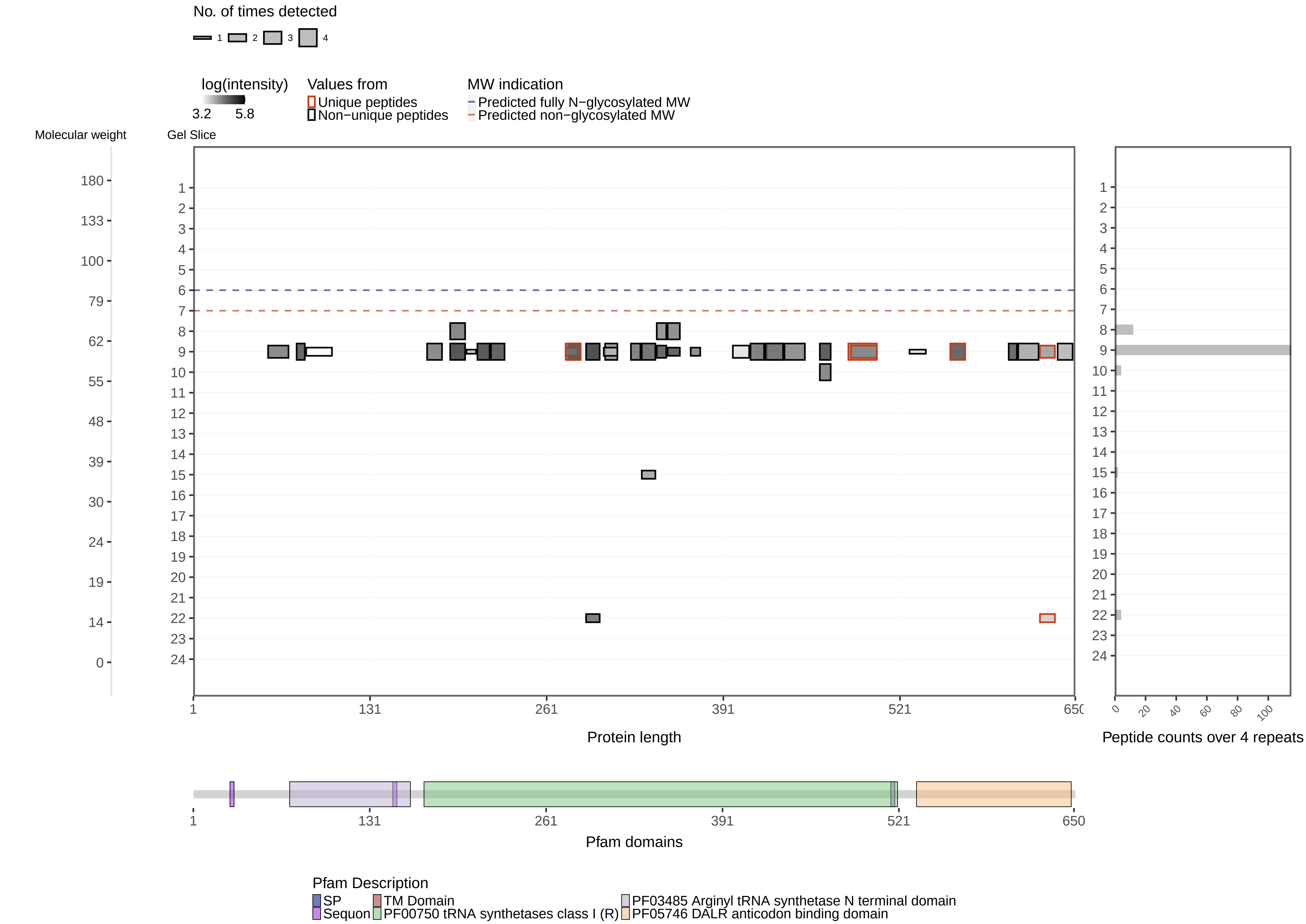


Protein sequence:

SRKEFQKVYERLGVHLEEKGESFYNPYIPKALELLNEKGLIEESEGARVIFIEGKKIPLIVVKRDGGFNYASTDLTALWYRLNEEKADWMIYVTDVGQREHFEMFFSAAKRAGWLPSNENDYPKASHVGFGLVLGEDGKRFRTRSTEVVKLVDLLDEAKSRCKLALEERGKAKEWTADELDQTAEAVGYGAVKYADLKNNRLTNYTFNFDLMLSDKGNTAVYLLYAHARICSIIRKSGRDIEELKKSGAVDLAHQDERTLGLHLLQFPEIVEEACTNLLPNLLCEYLYKLSEDFTK

NbL10g01690.1 probable_lactoylglutathione_lyase_chloroplastic [MW = 40.16 kDa]

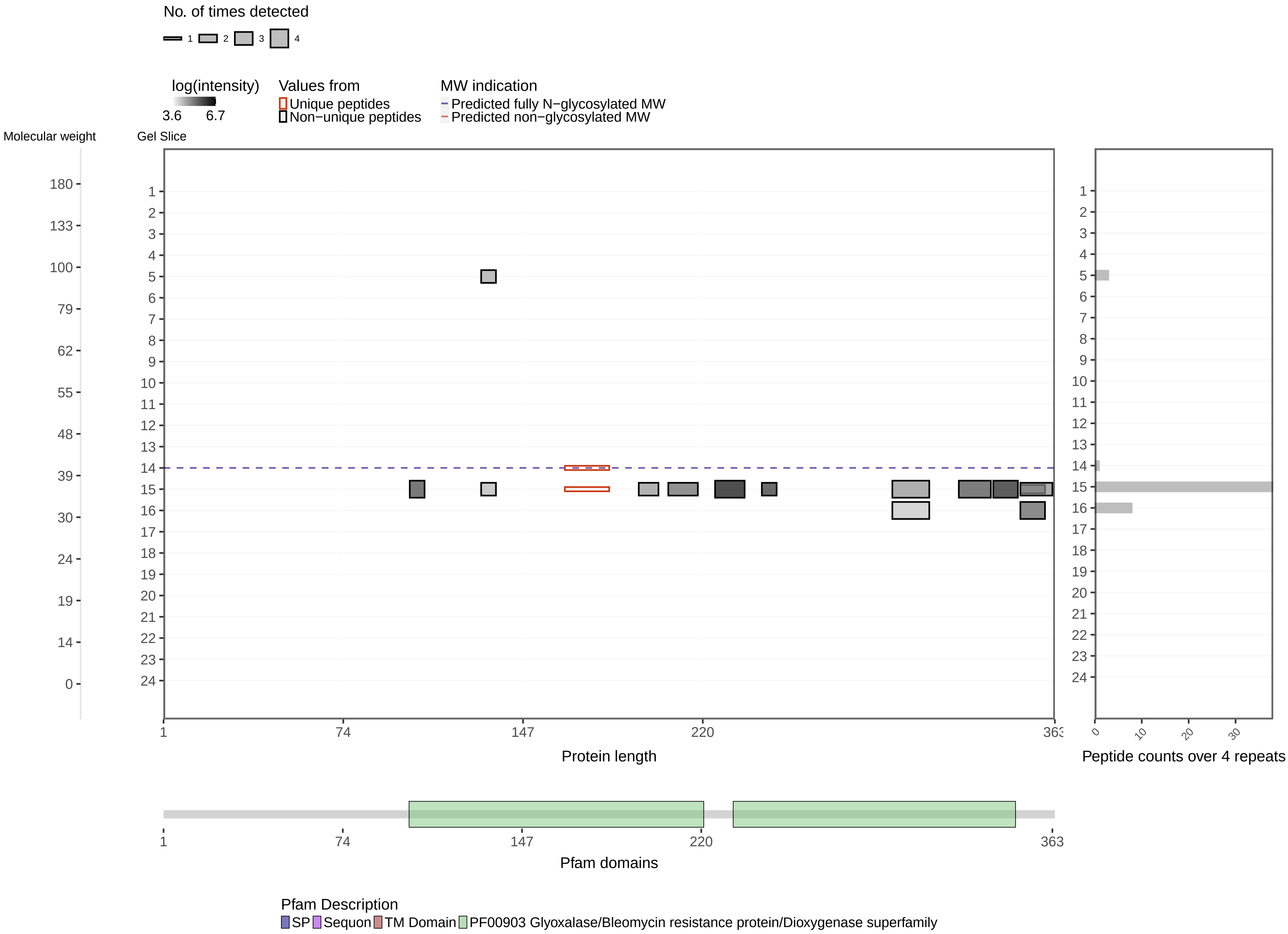


Protein sequence:

SGFGHFGIAVEDVAKTVELVKAKGGKVSREPGPVKGGKTVIAFIEDPDGYKFELLERGPTPEPLCQVMLRVGDLERAINFYEKAYGMKLLRTRDNPEYKYTIAMLGYGPEDSSAVMELTYNYGVTEYDKGNAYAQIAIGTDDVYKTAEAIRLCGGKITREPGPLPGISTKIVACLDPDGWKSVFVDNVDFLKELE*

NbL10g01430.1 aldehyde_dehydrogenase_family_2_member_C4 [MW = 54.67 kDa]

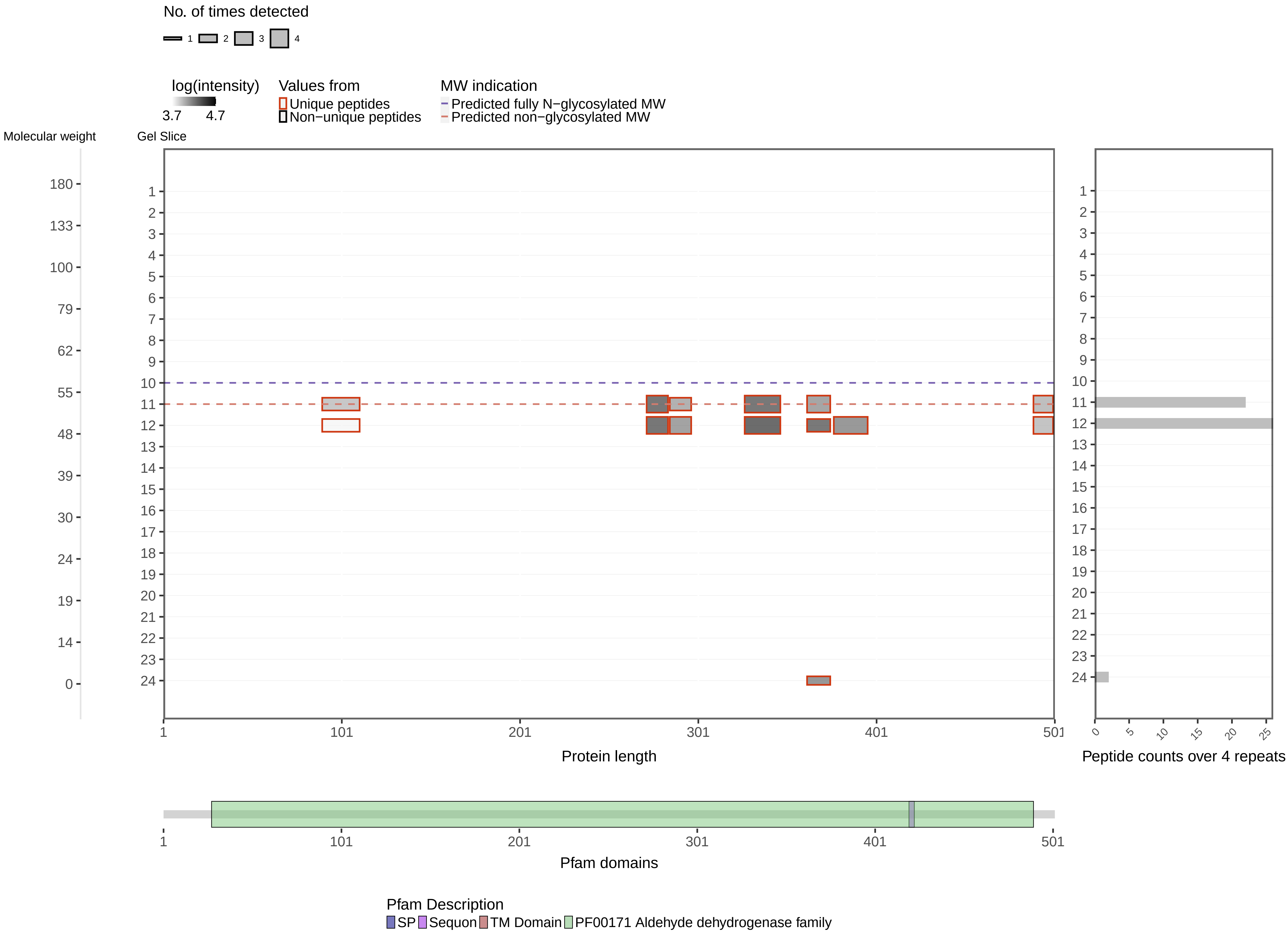


Protein sequence:

DKISFTGSTEVGRLVMQAAALSNLKPVSLELGGKSPFIVFDDVDVEKVAPLALIGILYNKGEVCVAGSRLFIQEGIYDKFVKKLVEITKTWVVGDTFDPNSHQGPQVDKKQFERVLSYIEHGKREGATVLTGGNALDRKGYFIEPTIFTDVEDHMIIAKEEIFGPVLSVMKFKTVEEVIKRANCTQYGLAAGVMTNNLNIANTVSRSIRAGVIWINCYFAFDPDCPYGGYKSSGFERDLGMEGLHKYLQVKSVATPIYNSPWL*

### NbL10g01040.1 probable_ADP−ribosylation_factor_GTPase−activating_protein_AGD14 [MW = 65.84 kDa]

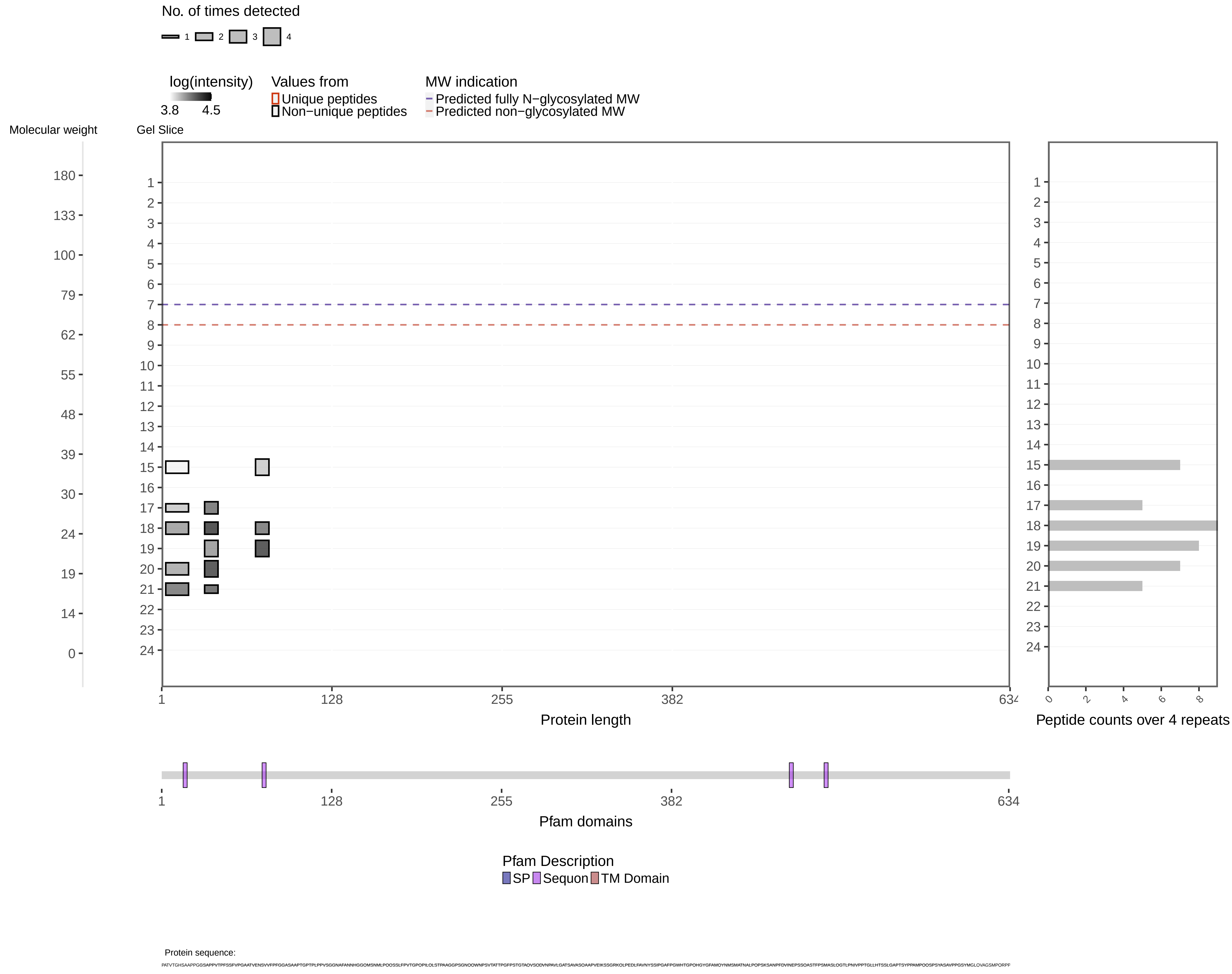

NbL10g00910.1 protein_BUNDLE_SHEATH_DEFECTIVE_2_chloroplastic−like [MW = 14.02 kDa]

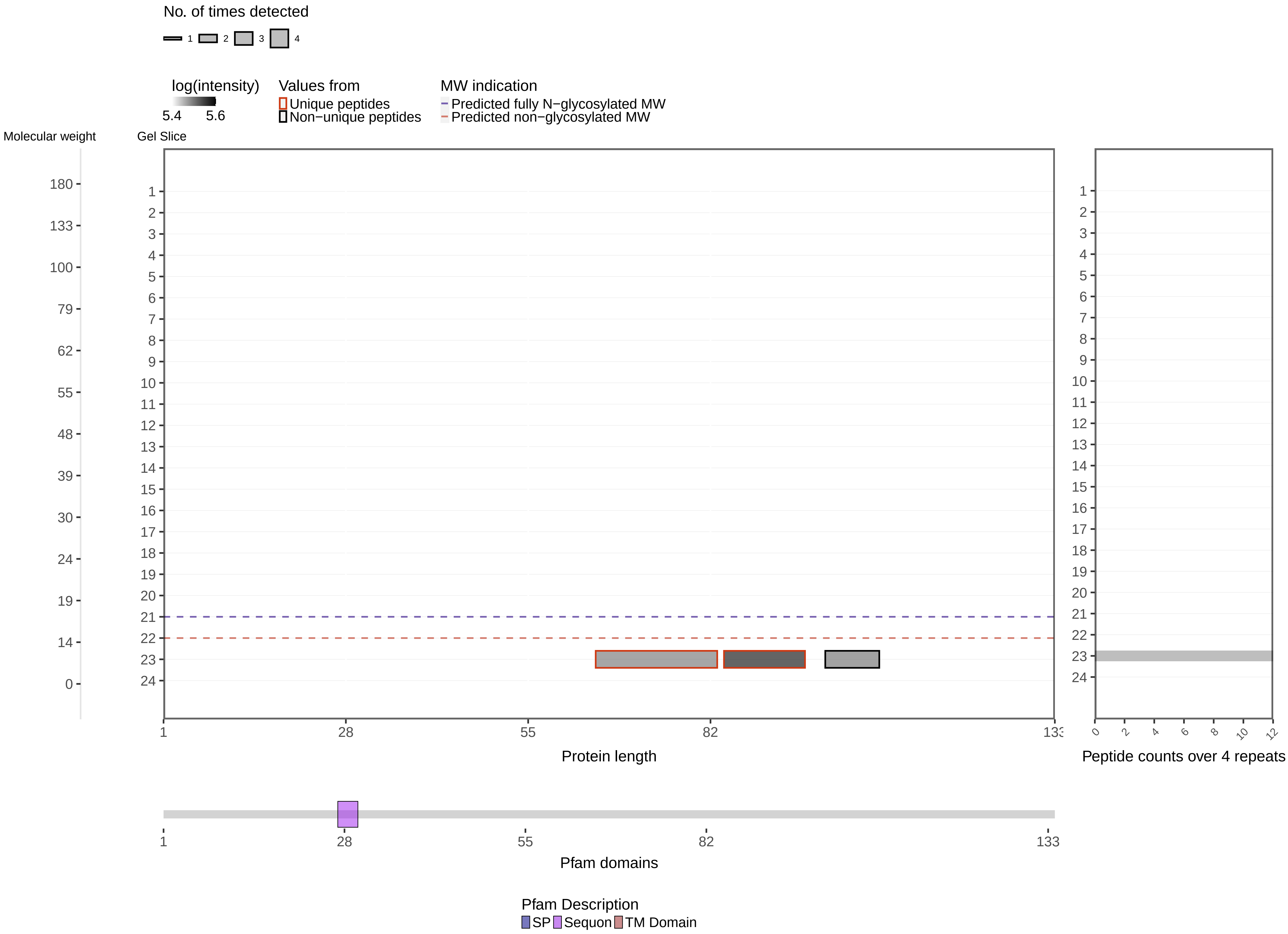


Protein sequence:

DSDKDTKVRSIVCQNCDGNGAVSCSQCKGTGVNSVDHFNGRFKAGGLCWLCRGKKDMLCGDCNGAGFLGGFMSTFDE*

NbL10g00880.1 oxygenase_1_chloroplastic−like_(LOC107792958)_mRNA_HQ690822.1_Nicotiana_tabacum_heme_oxygenase_1 [MW = 32.08 kDa]

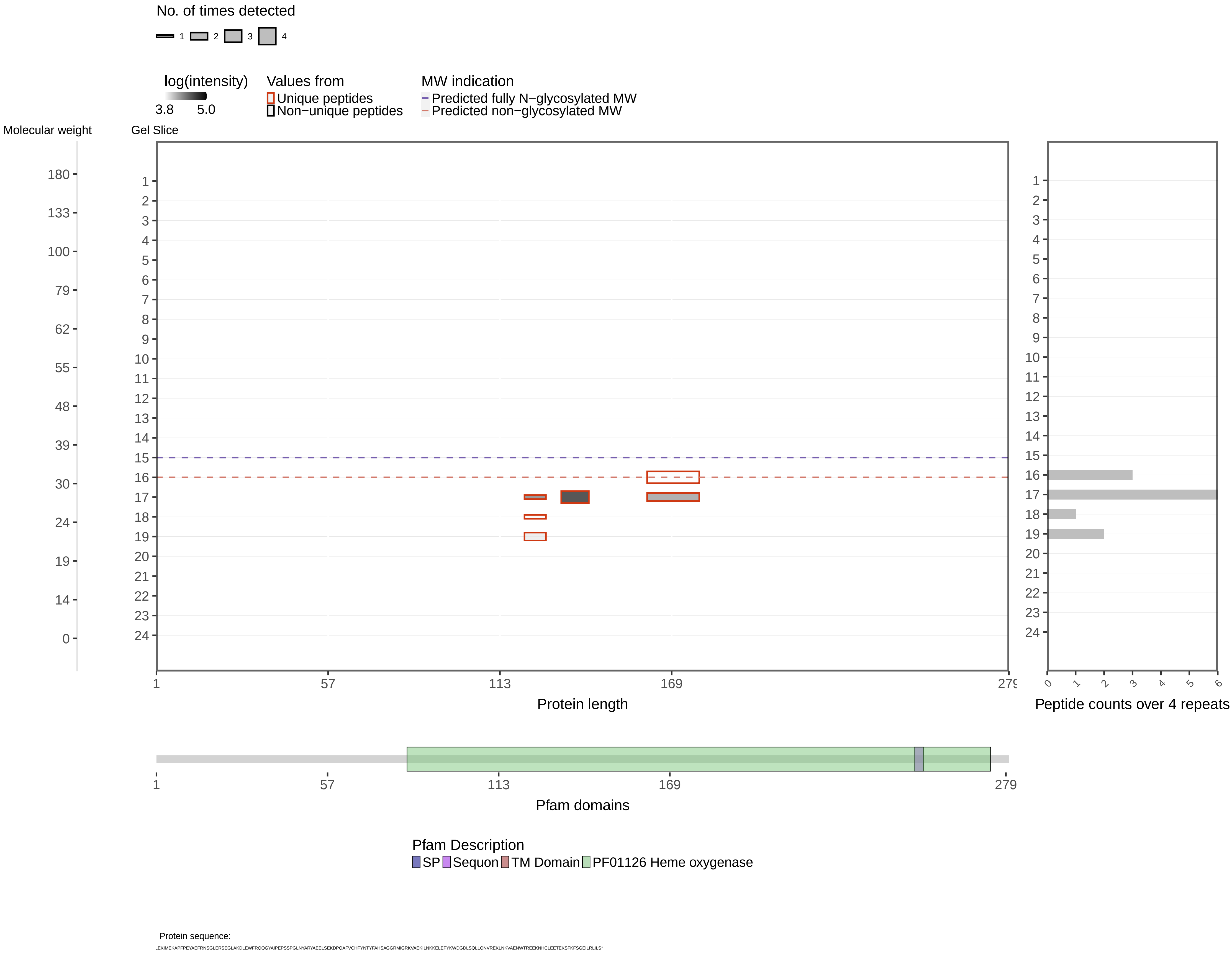

### NbL10g00790.1 repetitive_proline−rich_cell_wall_protein_1−like [MW − SP = 25.88 kDa]

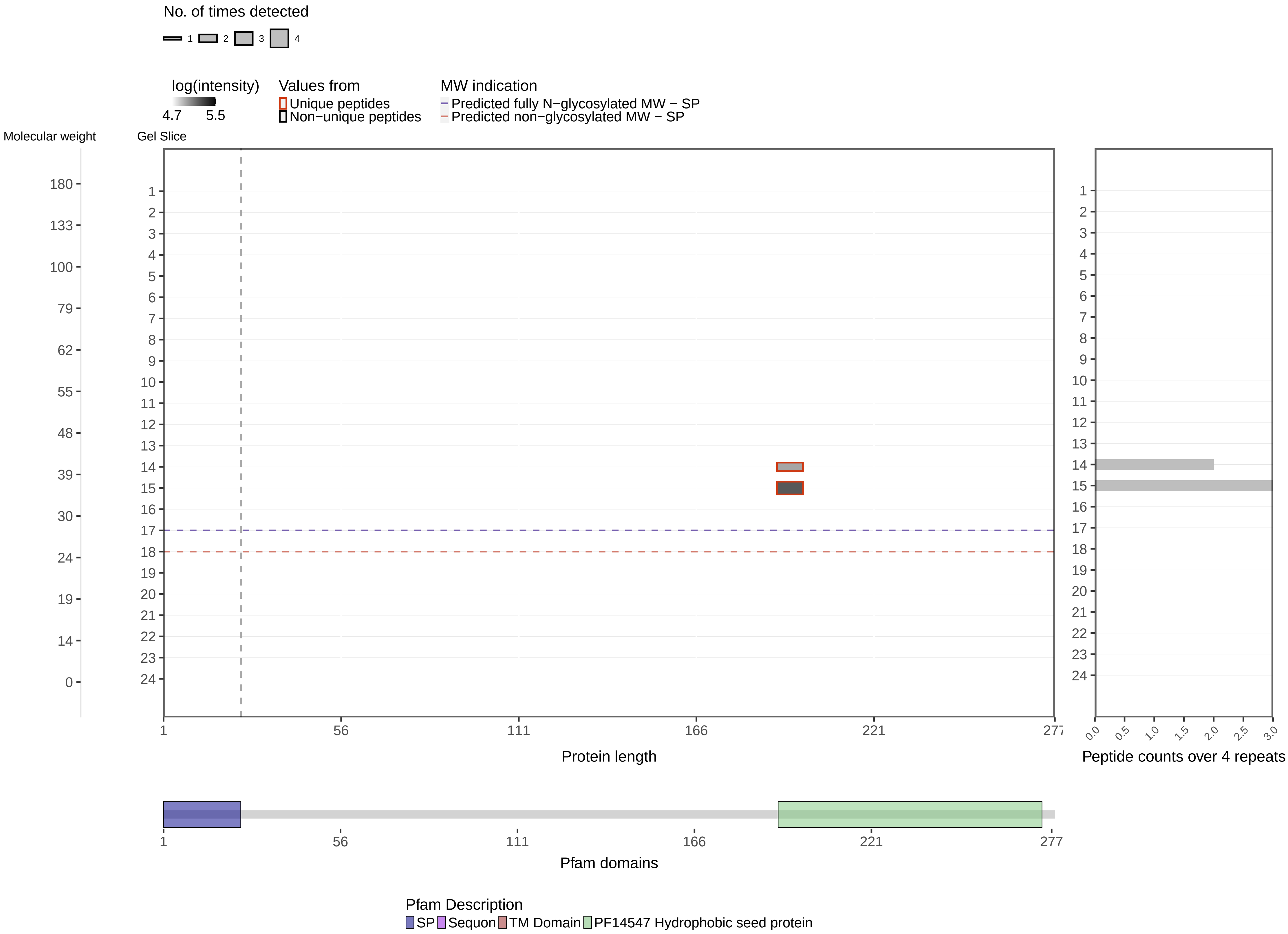


Protein sequence:

PPVIKPPIGIPPVTVPPVLKPPIGLPPIVKPPINLPPIGIPPVTIPPVITPSPKGKPCSPPPTTKATCPIDTLKLGACVDLLGGLVHVGIGDPVVNECCPILHGLVELEAAVCLCTTLKVKLLNLKIFVPLALQLLVTCGKTPPPGYTCSL*

### NbL10g00780.1 zinc_finger_BED_domain−containing_protein_DAYSLEEPER−like [MW = 89.66 kDa]

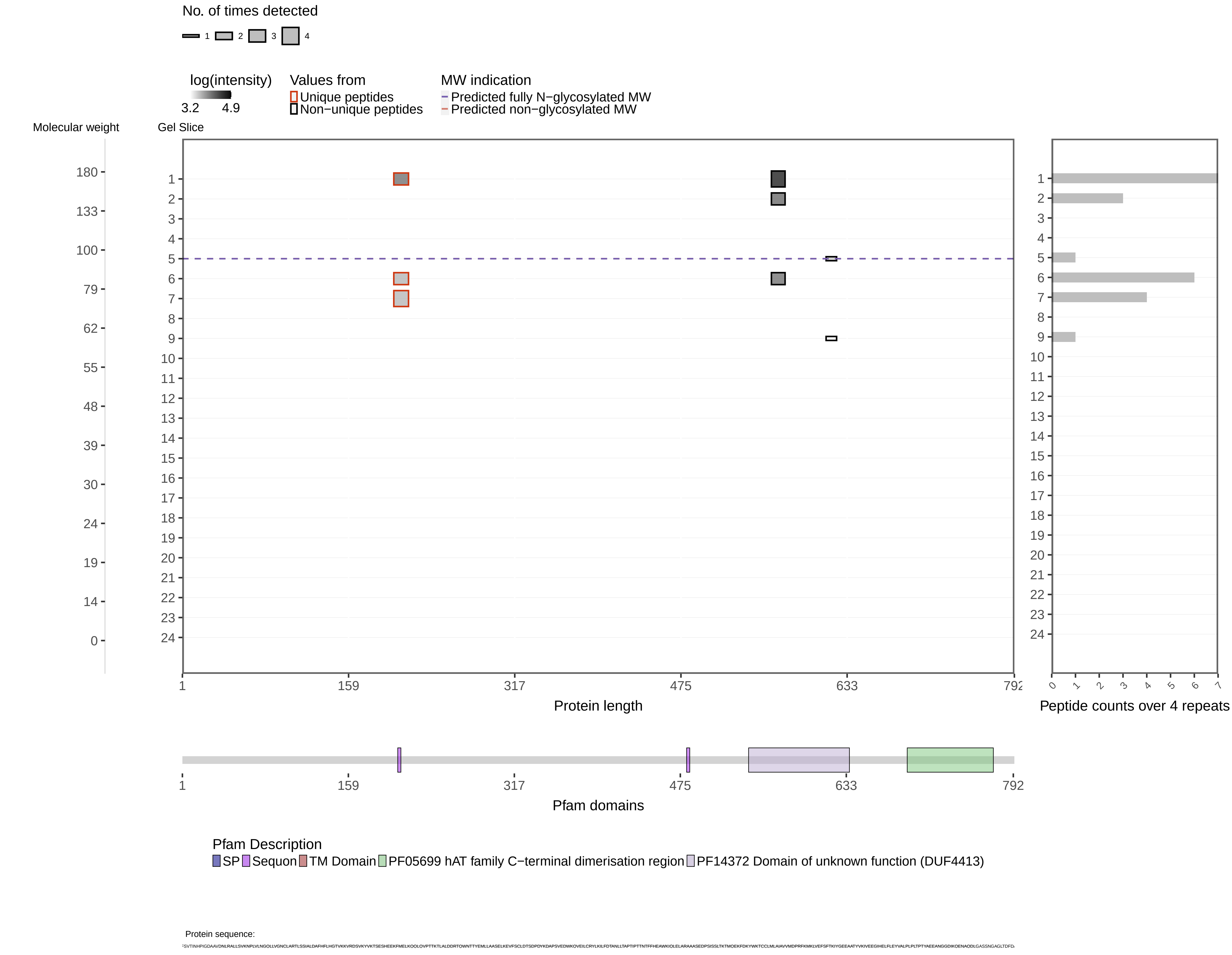

### NbL10g00730.1 methyl−CpG−binding_domain−containing_protein_11−like [MW = 35.9 kDa]

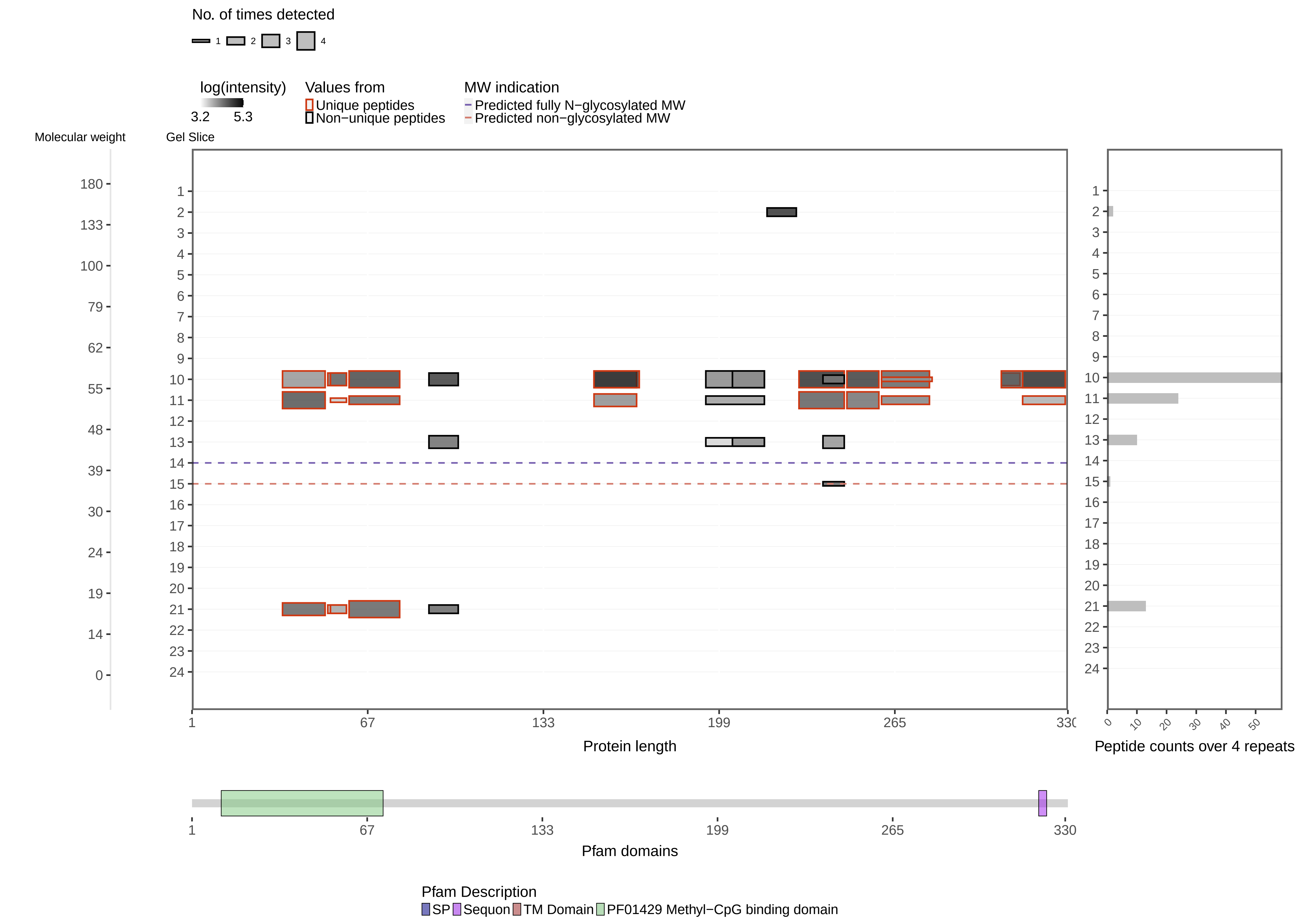


Protein sequence:

DENQDETQDGESGVEKKEEAEATEKDVSKKDEIKSSEIDGKKDESQNADGKVEDAPSEETQVEKDVEMADNVGHSKDVEEAAGDKVADDLEATKTNEEEKDVQVEEKVENMPTETVTLDSIAAEEKKHQAEGEQKDEQKTSAATTGTEDQHKLMNSEISKVEGEVTENGSNTNEAKP*

### NbL10g00550.1 shikimate_O−hydroxycinnamoyltransferase−like [MW = 48.95 kDa]

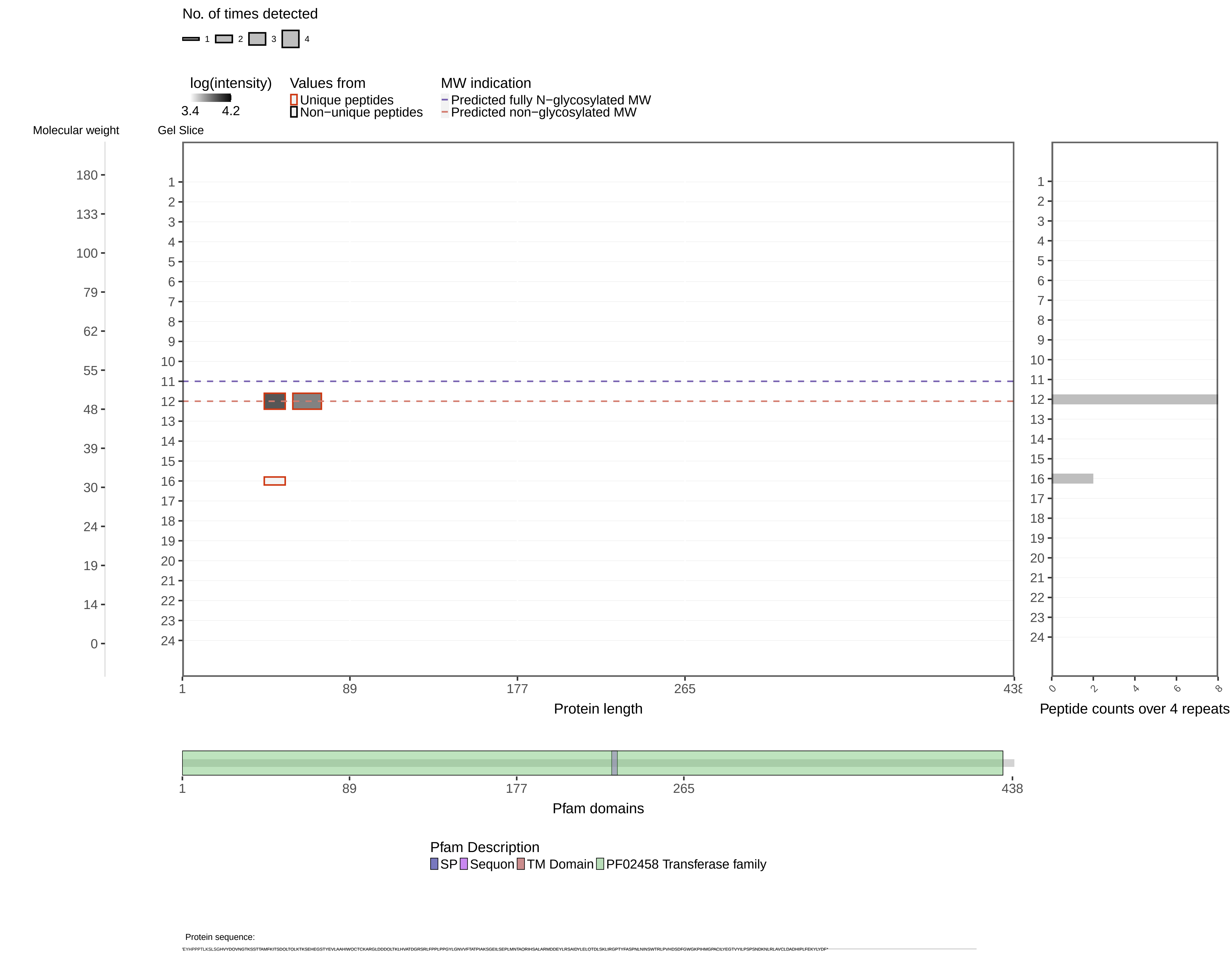

NbL10g00540.1 nucleoid−associated_protein_At4g30620_chloroplastic−like [MW = 20.01 kDa]

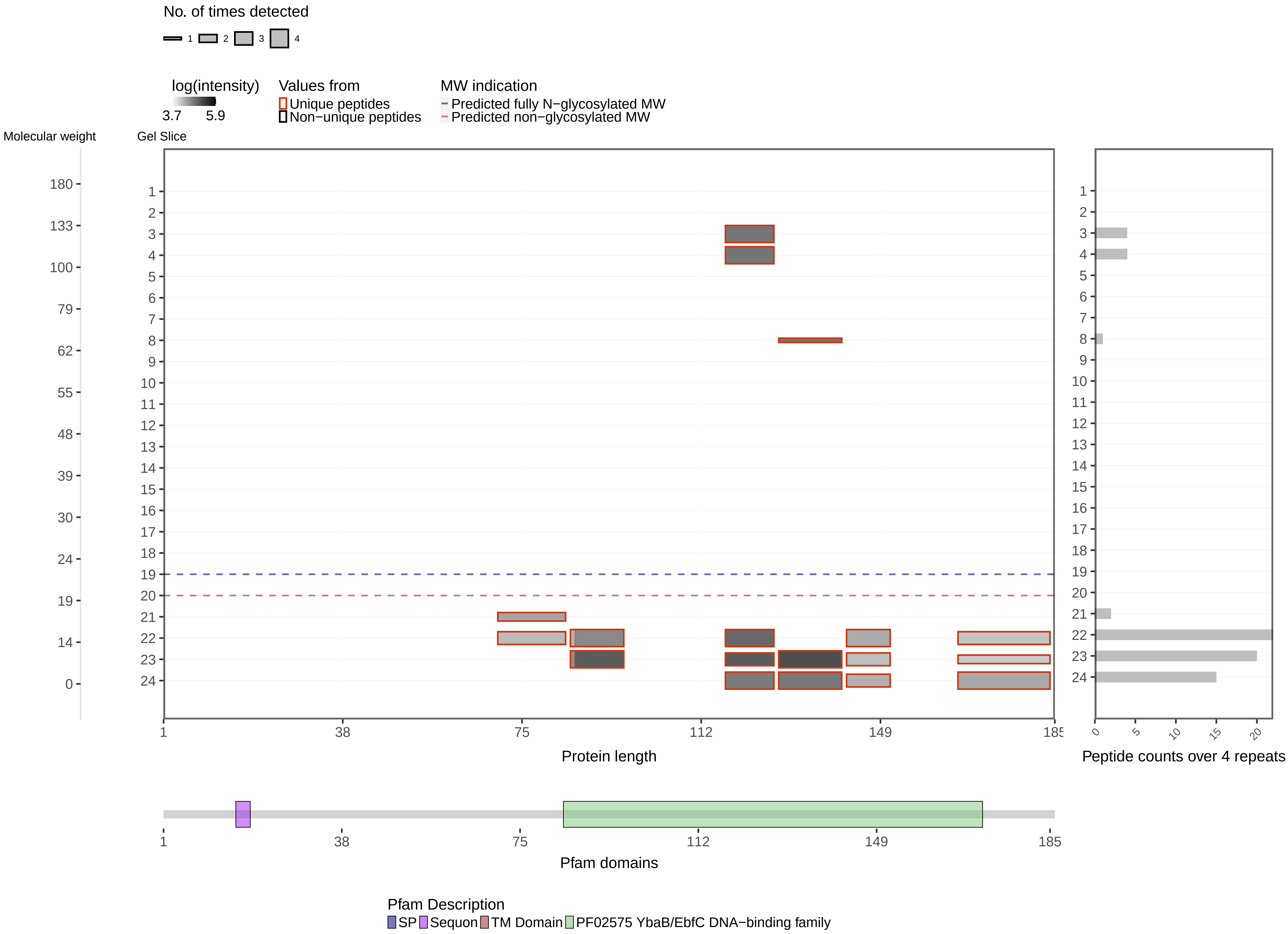


Protein sequence:

NLYETVKKAQNVVQVEAVRVQKELALAEFDGYCEGELIKVTLSGNQQPIRTEITEAAMELGPEKLSLLITEAYKDAHQKSVLAMKERMSDLAQSLGMPAGLGEGFKQ*

### NbL10g00510.1 leucine_aminopeptidase_2_chloroplastic [MW = 60.69 kDa]

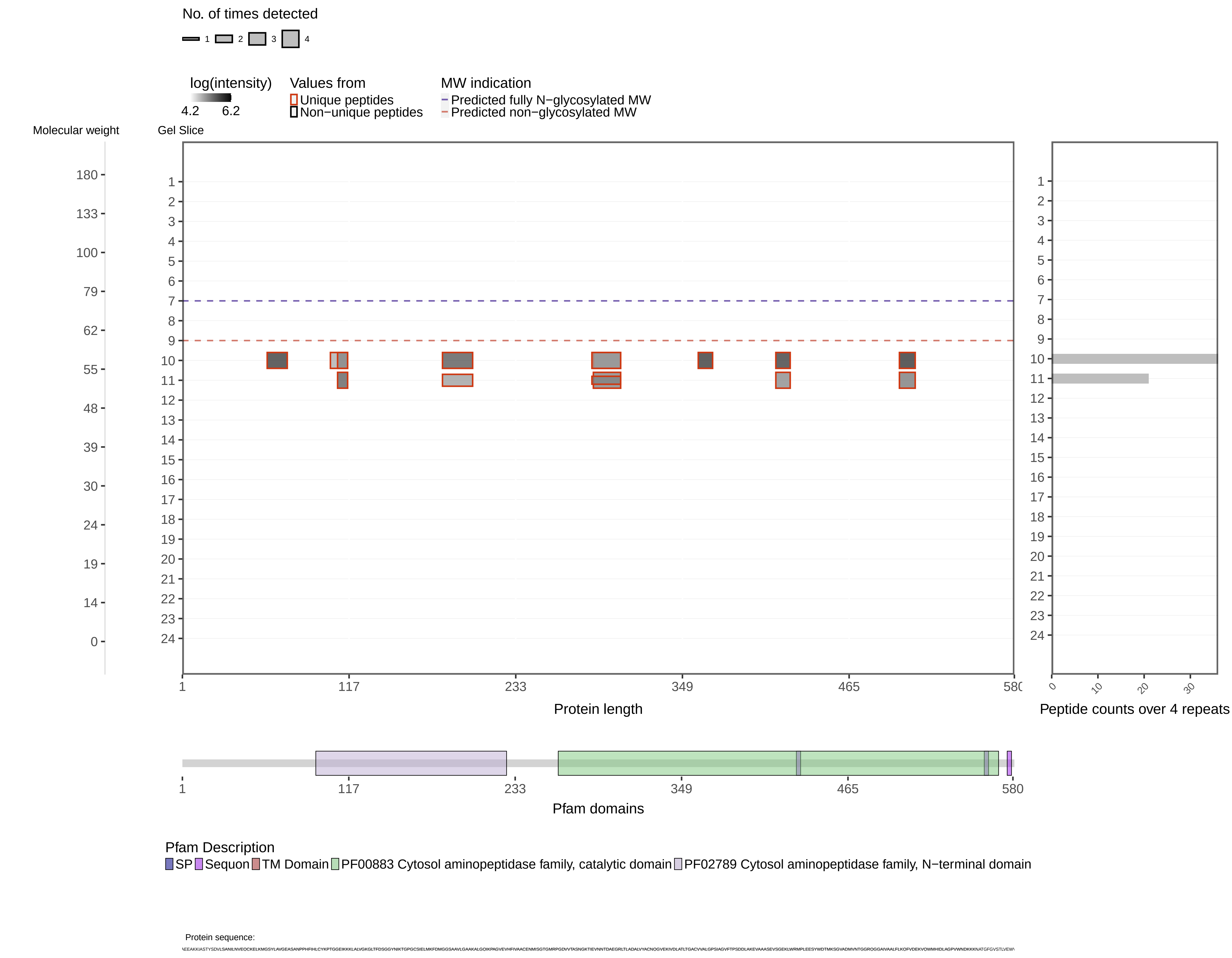

NbL10g00500.1 probable_GMP_synthase_[glutamine−hydrolyzing] [MW = 41.05 kDa]

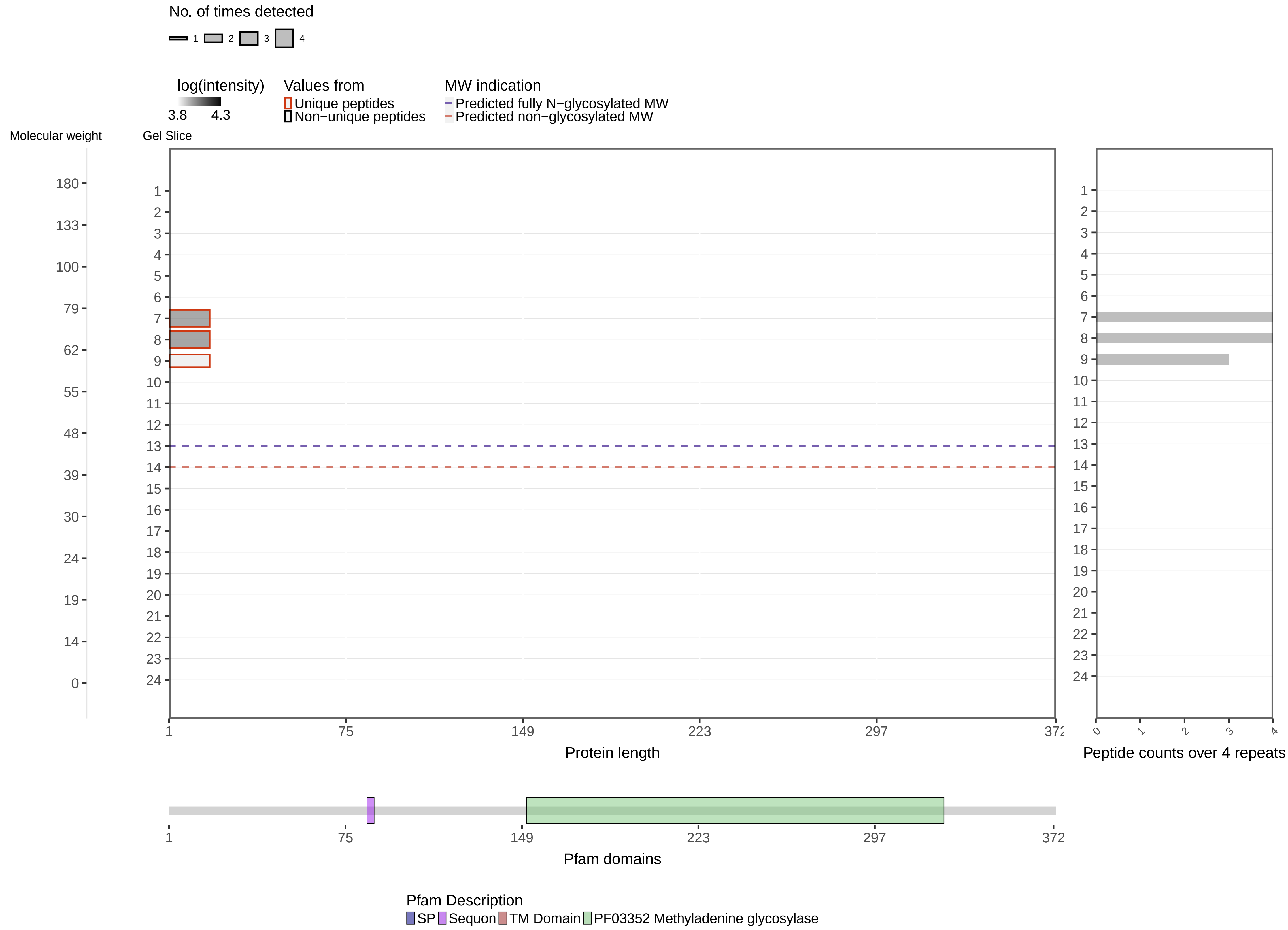


Protein sequence:

PVHDDKKLFELLVLCGALAELSWPSILSKRNIFREVFADFDPIVVAKLNEKKILAPGSTACSLLSELKLRAIIENAROMSKVIDEFGSFDKYIWSFVNNKPIVSGFRYPROVPVKTAKADLISKDLIRRGFRGVGPTVVYSFMOVAGITNDHLISCFRFHDCVESAEAKEKDSNKDETEATKVDEAIEAEICRSIDDLSFSSE*

### NbL10g00490.1 eukaryotic_translation_initiation_factor_5A−1/2 [MW = 17.51 kDa]

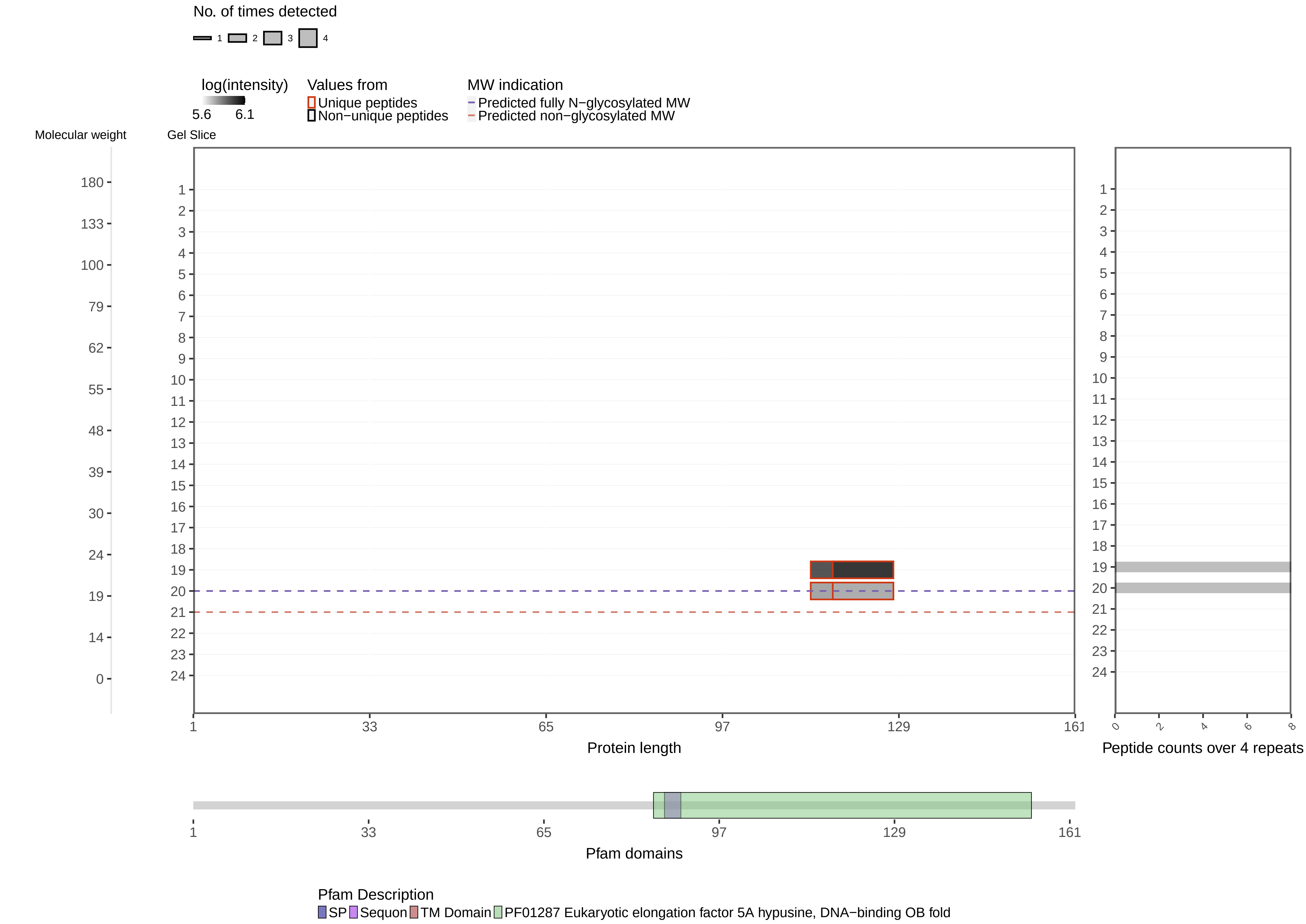


Protein sequence:

IGKKLEDIVPSSHNCDVPHVNRTDYQLIDISEDGFVSLLTENGNTKDDLRLPTDENLLNQVKGGFEEGKDLVLSVMSAMGEEQIAAVKDIGTKN*

NbL10g00420.1 uncharacterized_LOC109225199 [MW = 11.96 kDa]

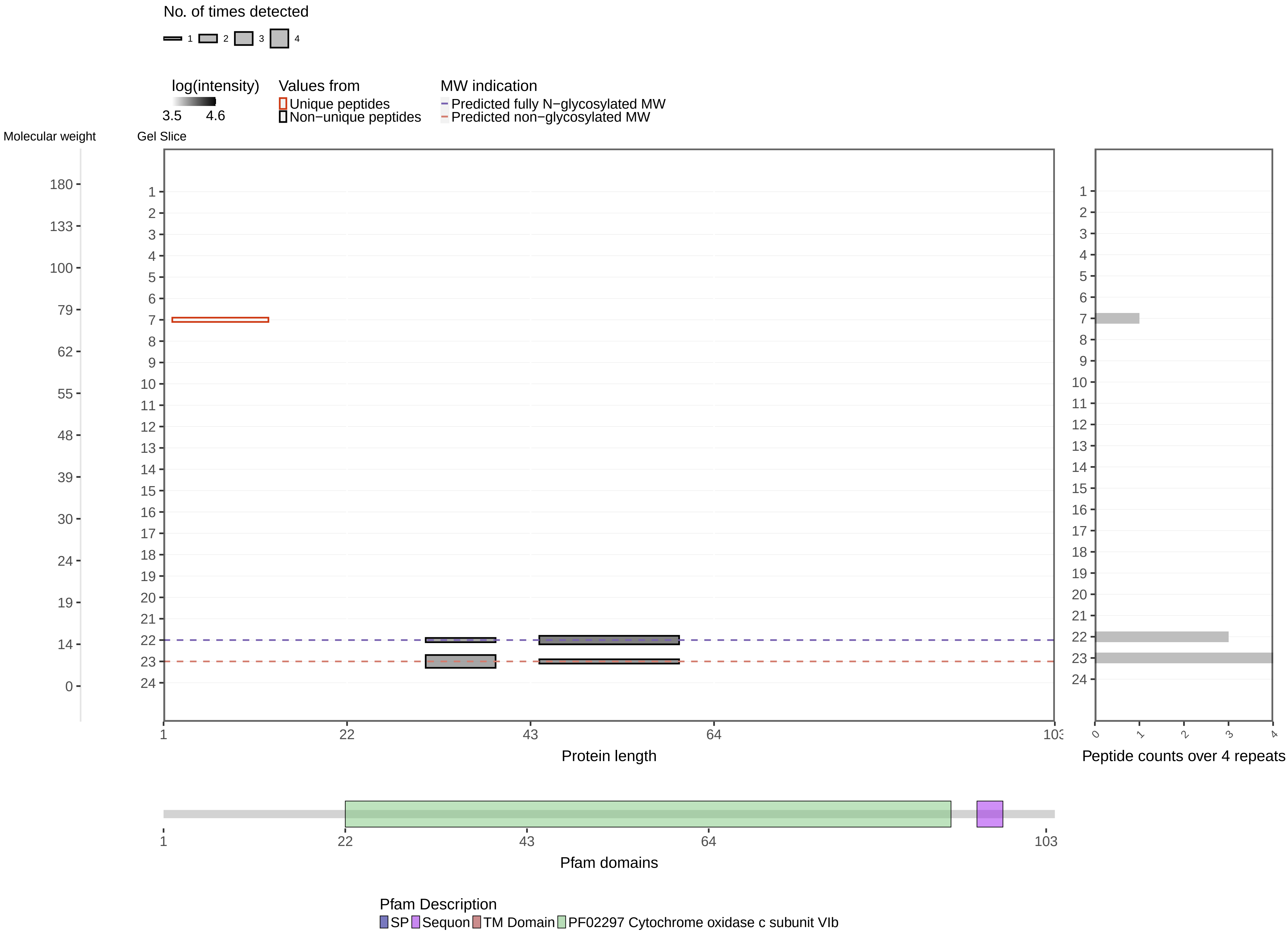


Protein sequence:

ESSKKPTEIATVGLLYPVECKKTREEYVKQCRPTWVKHFDRQYCAKMRVQRLLDDNDSRRECL*

### NbL09g23590.1 uncharacterized_LOC107760951 [MW = 17.44 kDa]

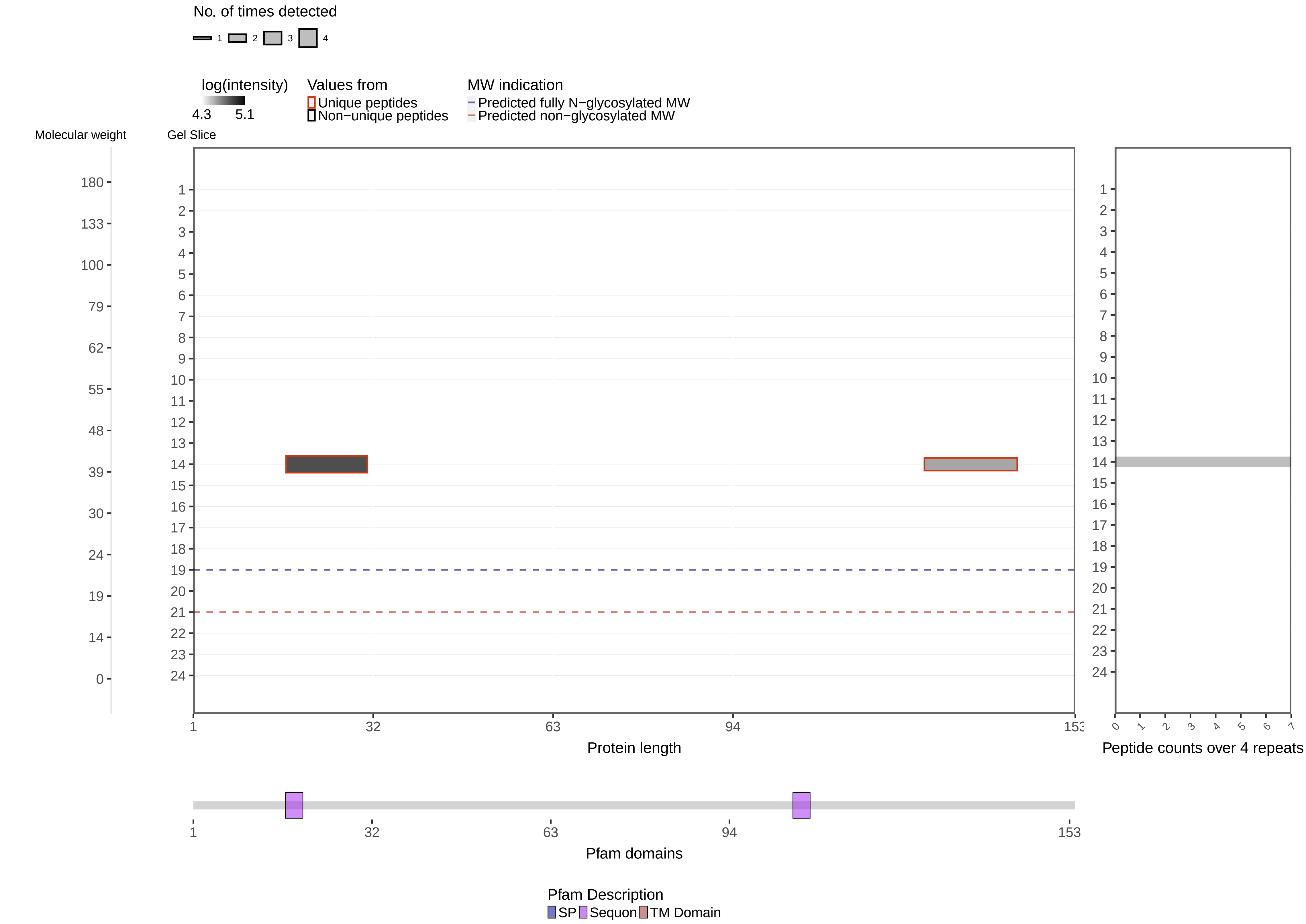


Protein sequence:

PTLAQDDDAFSFMYKSIRLLPTVGCEADAATRHSIDERSIVDDNVSAFAYQSSGCYVAAWSNENKANPYISWELEHCLIDPREKESRVRIIQ

### NbL09g23480.1 endochitinase_A [MW − SP = 32.73 kDa]

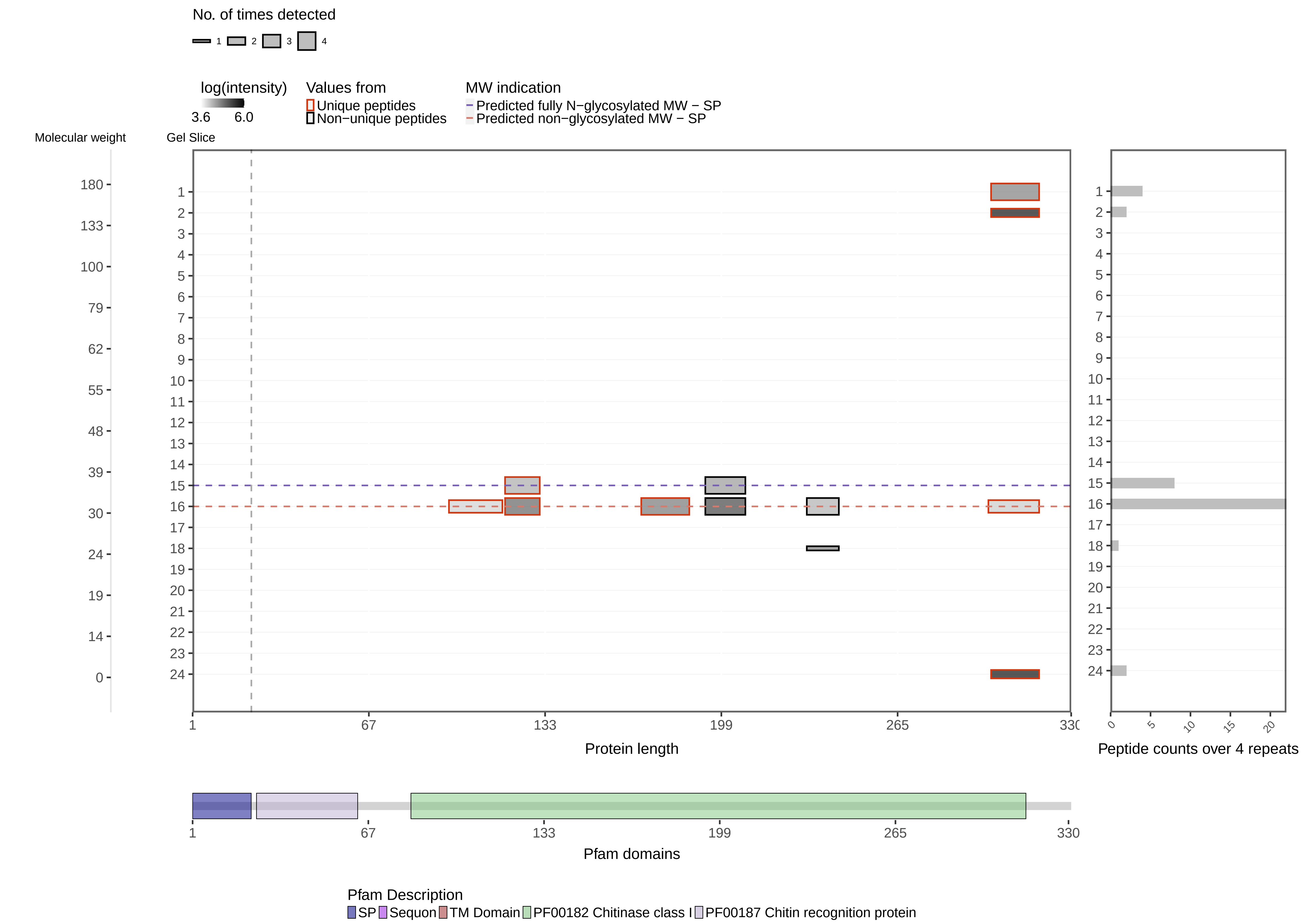


Protein sequence:

TAPDGPYAWGYCWLRERGSPGDYCTPSGQWPCAPGRKYFGRGPIQISHNYNYGPCGRAIGVDLLNNPDLVATDPVISFKSALWFWMTPQSPKPSCHDVIIGRWQPSSADRAANRLPGFGVITNIINGGLECGHGTDSRVQDRVGFYRRYCSILGVSPGDNLDCGNQRSFGNGLLVDTM*

### NbL09g23370.1 V−type_proton_ATPase_subunit_B2 [MW = 61.05 kDa]

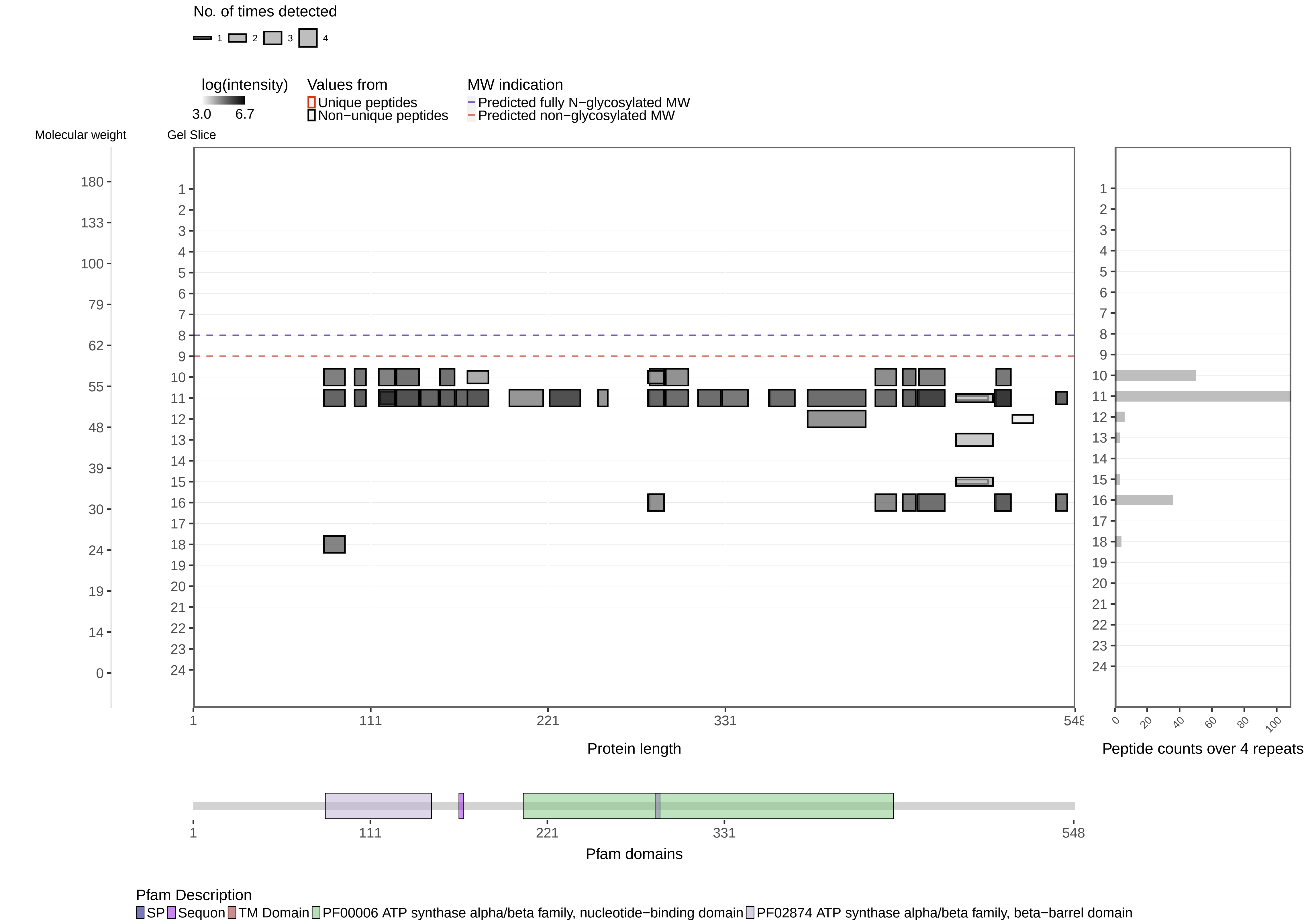


Protein sequence:

### NbL09g23340.1 aquaporin_PIP2−7−like [MW = 30.4 kDa]

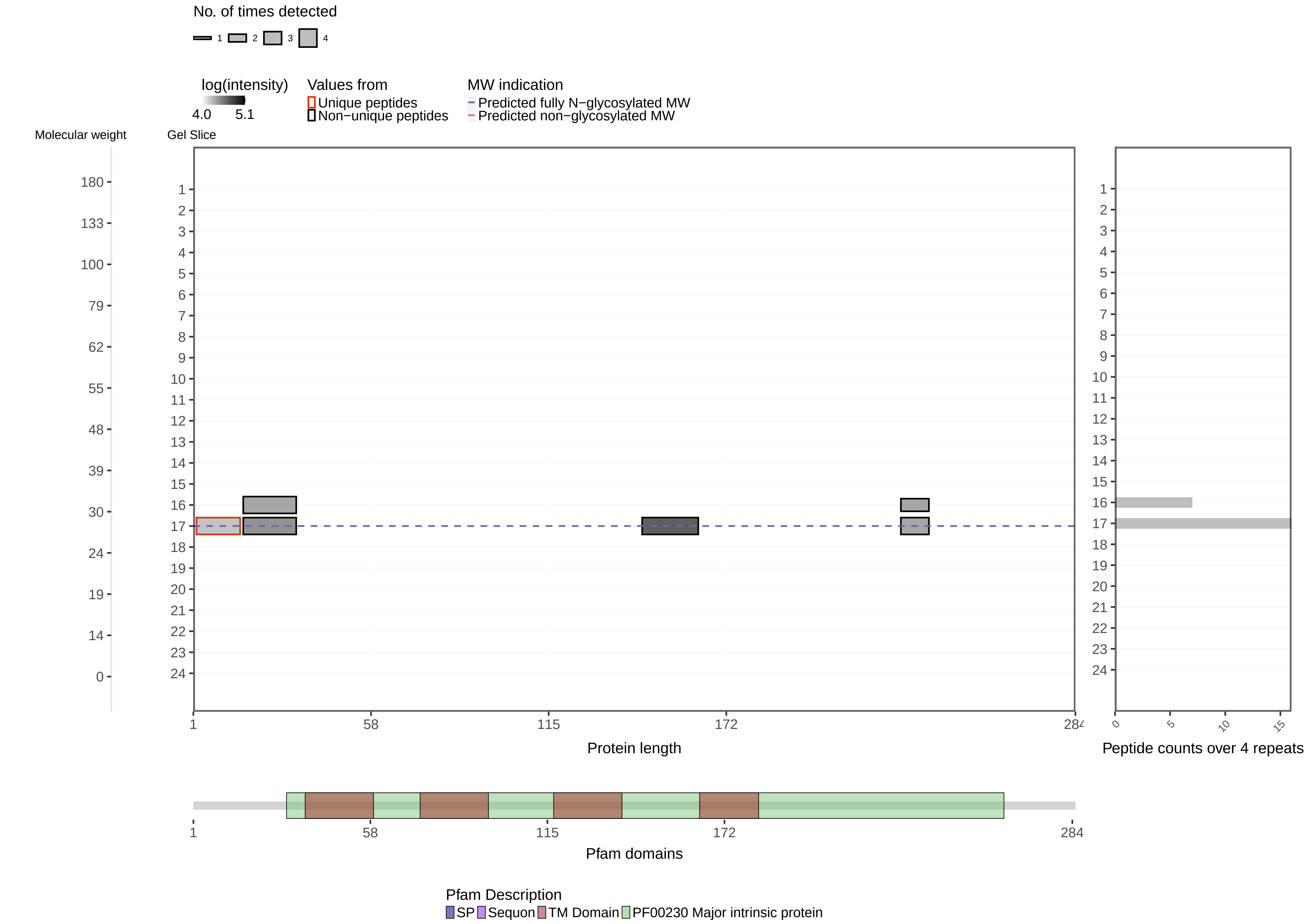


Protein sequence:

.GAICGVGFVKAFMKHYYNLEGGGANFVQPGYNKGTALGAEIIGTFVLVYTVFSATDPKRSARDSHVPVLAPLPIGFAVFMVHLATIPITGTGINPARSFGAAVIYNKEKIWDDQWIFWVGPFVGALIAAIYHQFILRAGAVKALGSFRSNPTN*

NbL09g23300.1 glycine−rich_protein_2−like [MW = 21.74 kDa]

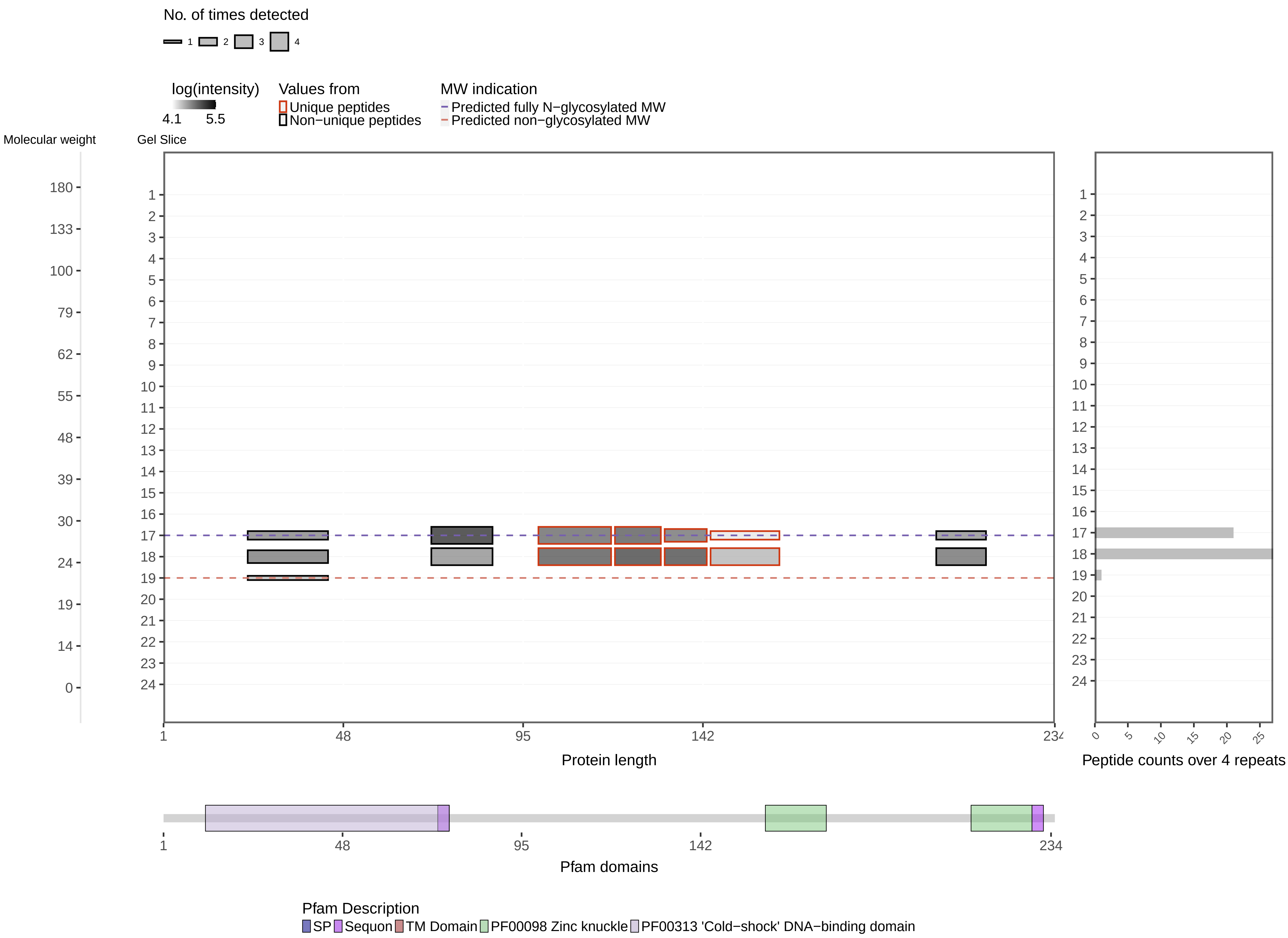


Protein sequence:
GGYGGGGGGRYGGGDGGYGGGGRYGGDGGYGGGGRYGGSGGYGGGGGGGSGCYKCGEEGHFARECTQGGGGGGYGSGGGGYGSGGGGYGGGGGRYGGSGGGGGGGCFKCGEEGHFARECPNSSGR*

### NbL09g23170.1 peptidyl−prolyl_cis−trans_isomerase−like [MW = 18.05 kDa]

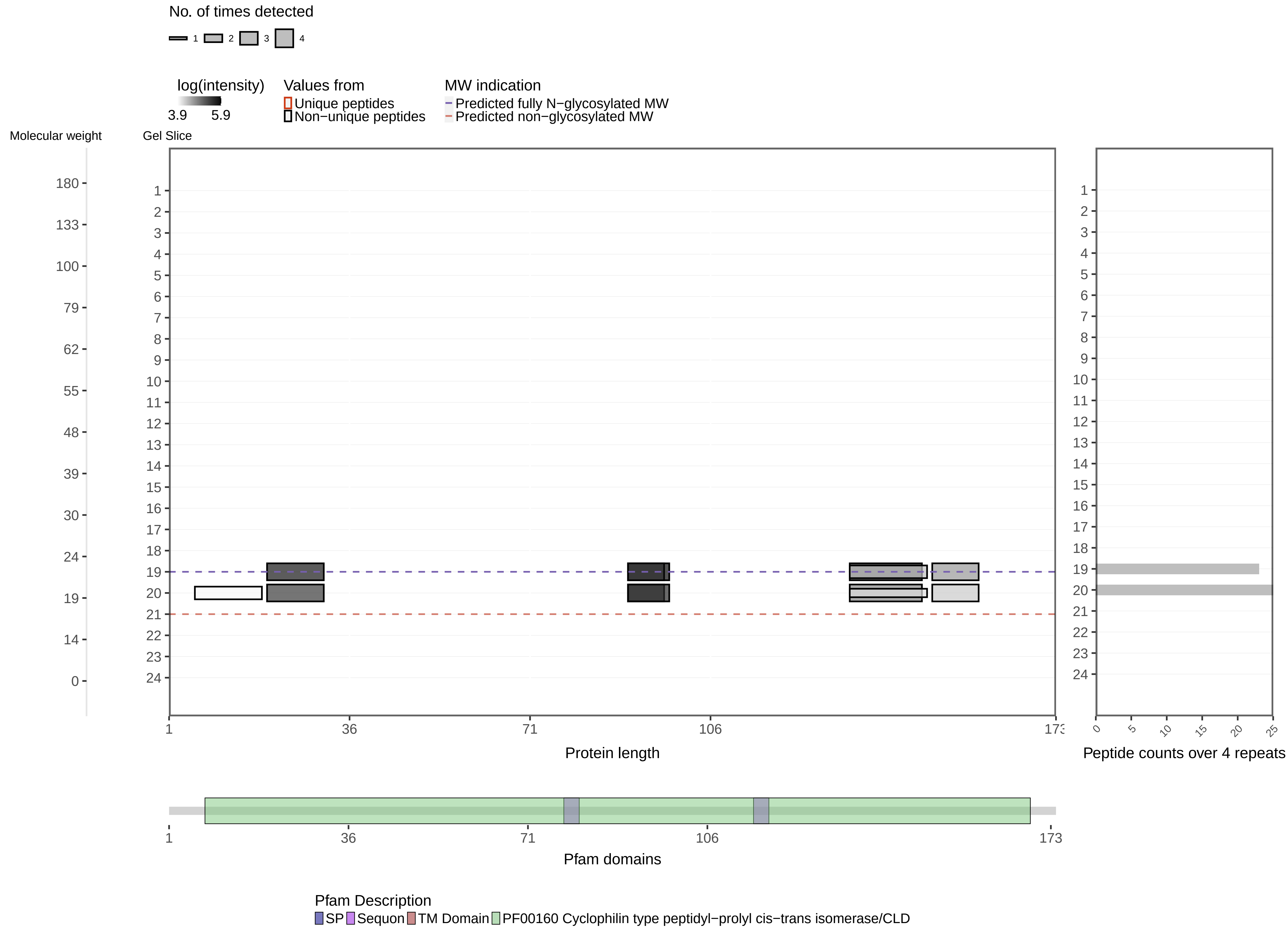


Protein sequence:

NbL09g23080.1 NADH_dehydrogenase_[ubiquinone]_1_beta_subcomplex_subunit_9−like [MW = 13.77 kDa]

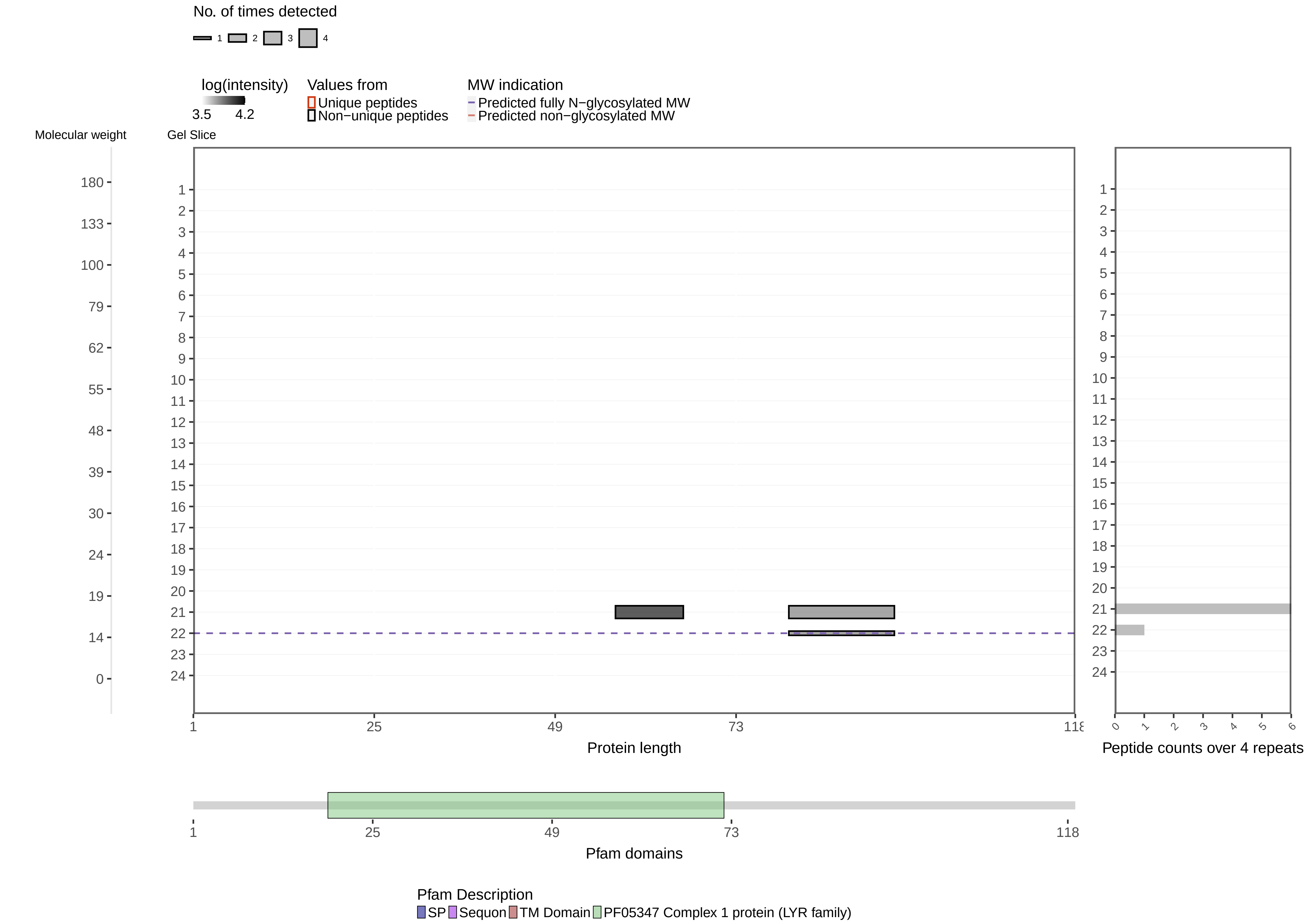


Protein sequence:

DALRERFEANKHVEDVETIDRLIADGEASYNKWRHPDPYIVPWAPGGSKFNRNPVPPEGLYFSRFMHQLLVE*

### NbL09g22930.1 isoflavone_reductase_homolog [MW = 33.89 kDa]

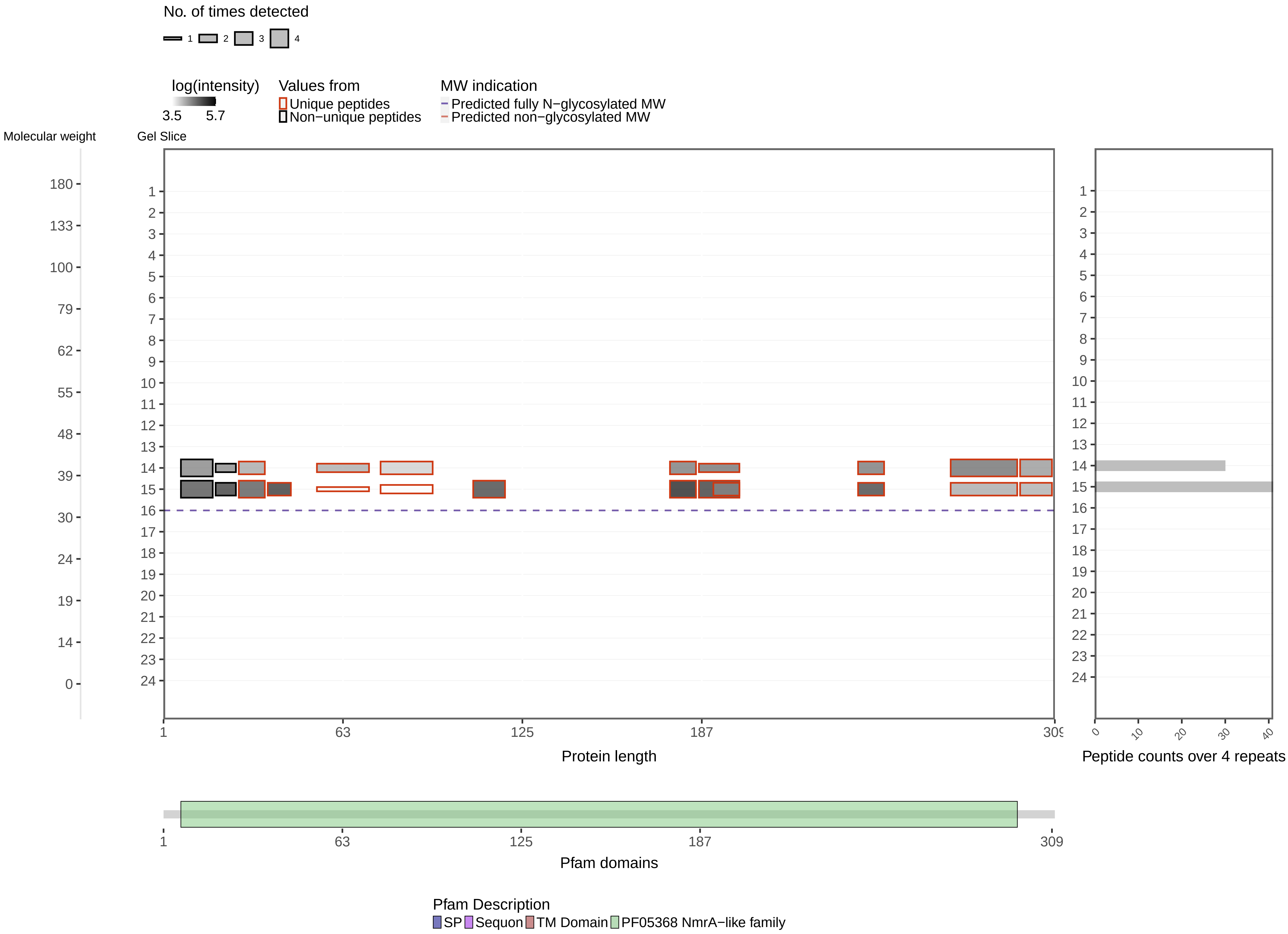


Protein sequence:

:AEGIPFTYVANFFFAGYFLPNLAQPGASGPPTDKVVILGDGNTKAVFNKEEDIAIYTIKAVDDPKTLNKILYIRPPHNIVTLNDVISLWEKKTGKNLERVYVPEEQVLKNIQEASFPLNVVLSISHTAFVKGDHTNFAIEPSFGVEASEAYPDVKYTPVDEILNQYV*

### NbL09g22900.1 glycine−rich_RNA−binding_protein−like [MW = 14.88 kDa]

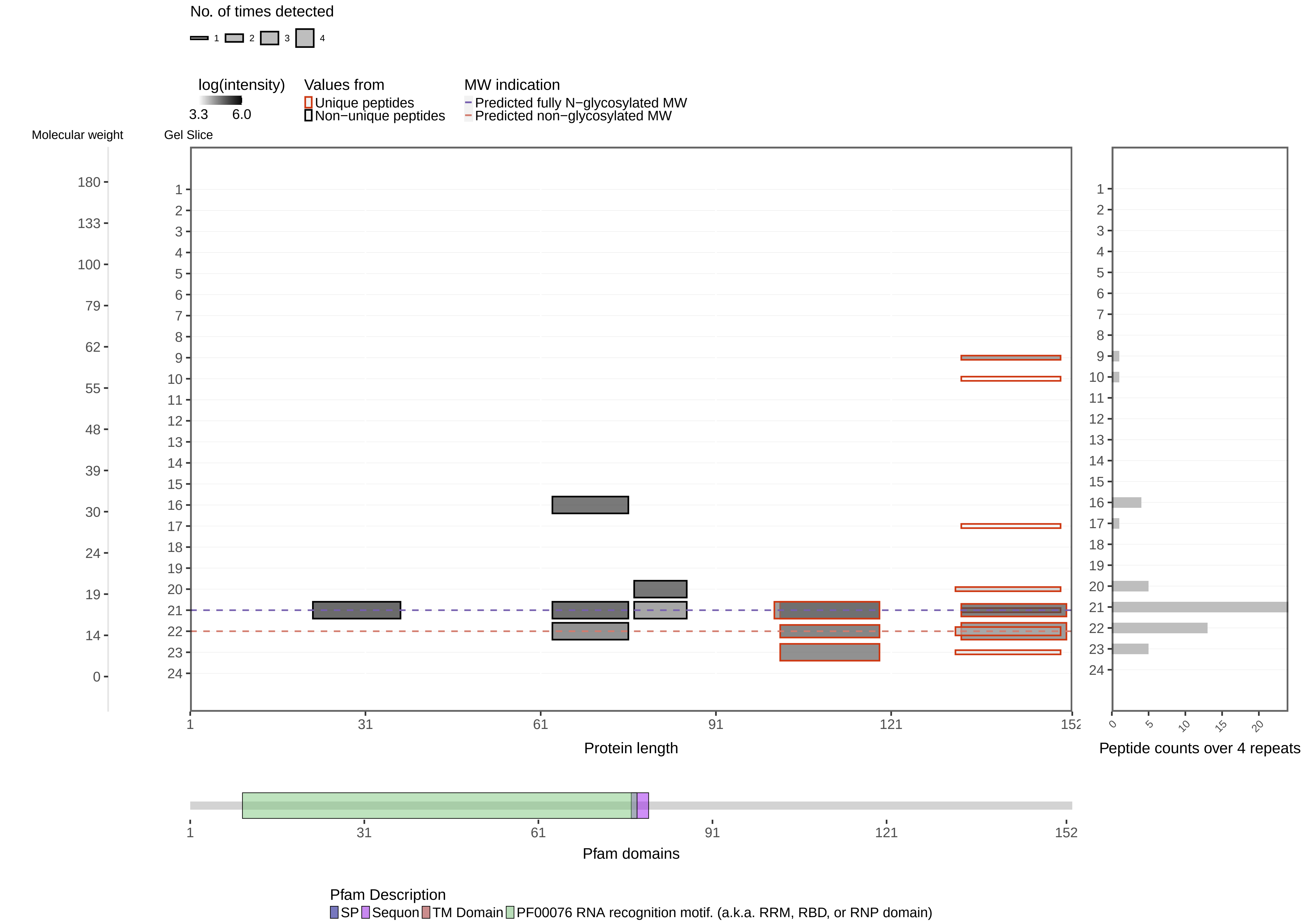


Protein sequence:

MNGQELDGRNITVNEAQARGSGGGGGGGYGGGRREGGGGGYGGGGGGYGGSRRDGGGGGYGGGRREGGGGGYGGGGGYGGGGRY*

NbL09g22880.1 glycine−rich_RNA−binding_protein−like [MW = 17.32 kDa]

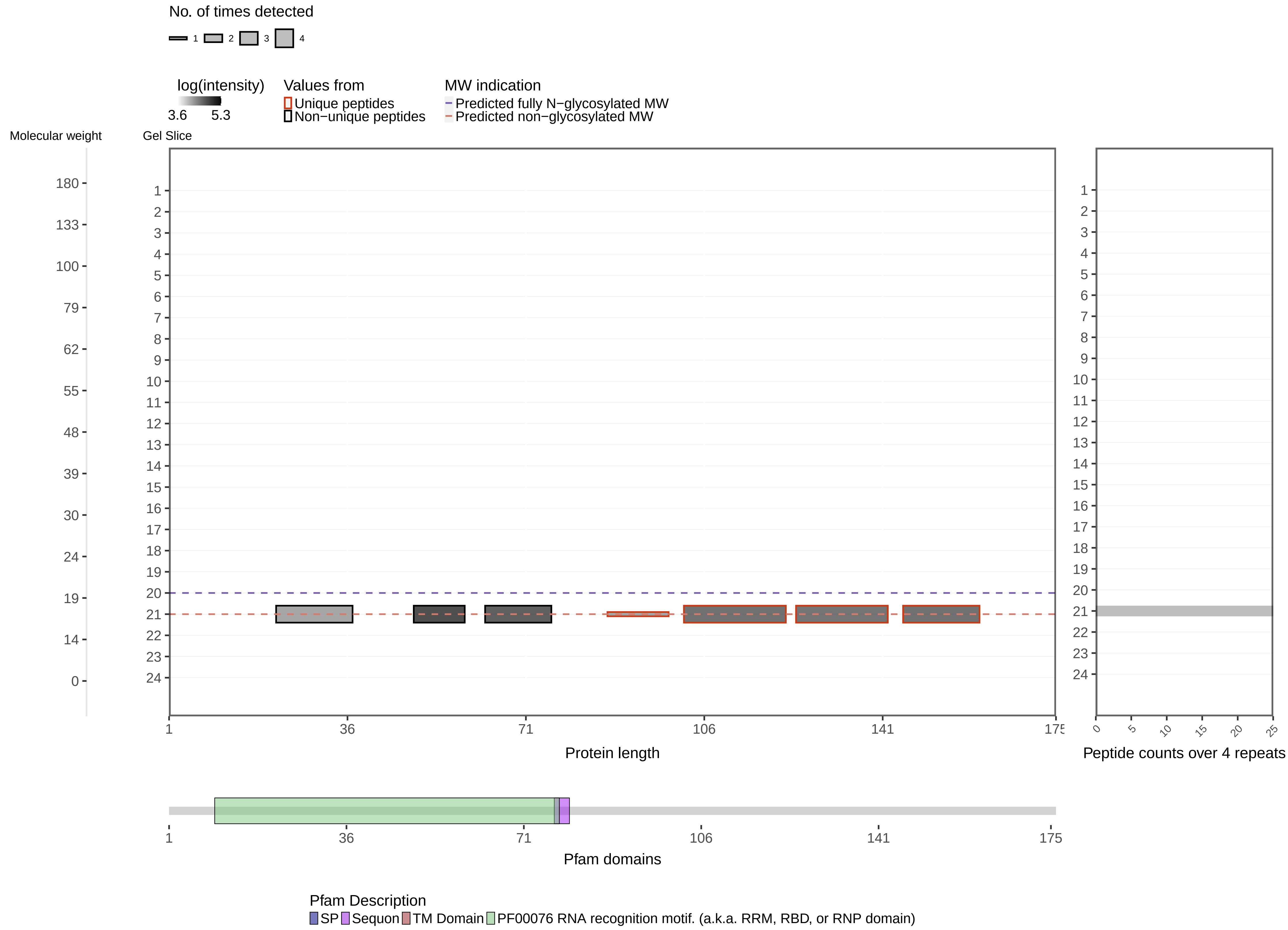


Protein sequence:

VNEAQSRGSGGGGGGFGGGRRREGGYSGGGGYGGGSGGYGGGRREGGYSGGGGGYGGGYGGGRNRGYGGGYGGGGGDGGSRYSRGGGASEGSWRN*

NbL09g22800.1 protein_TIC_62_chloroplastic [MW = 61.87 kDa]

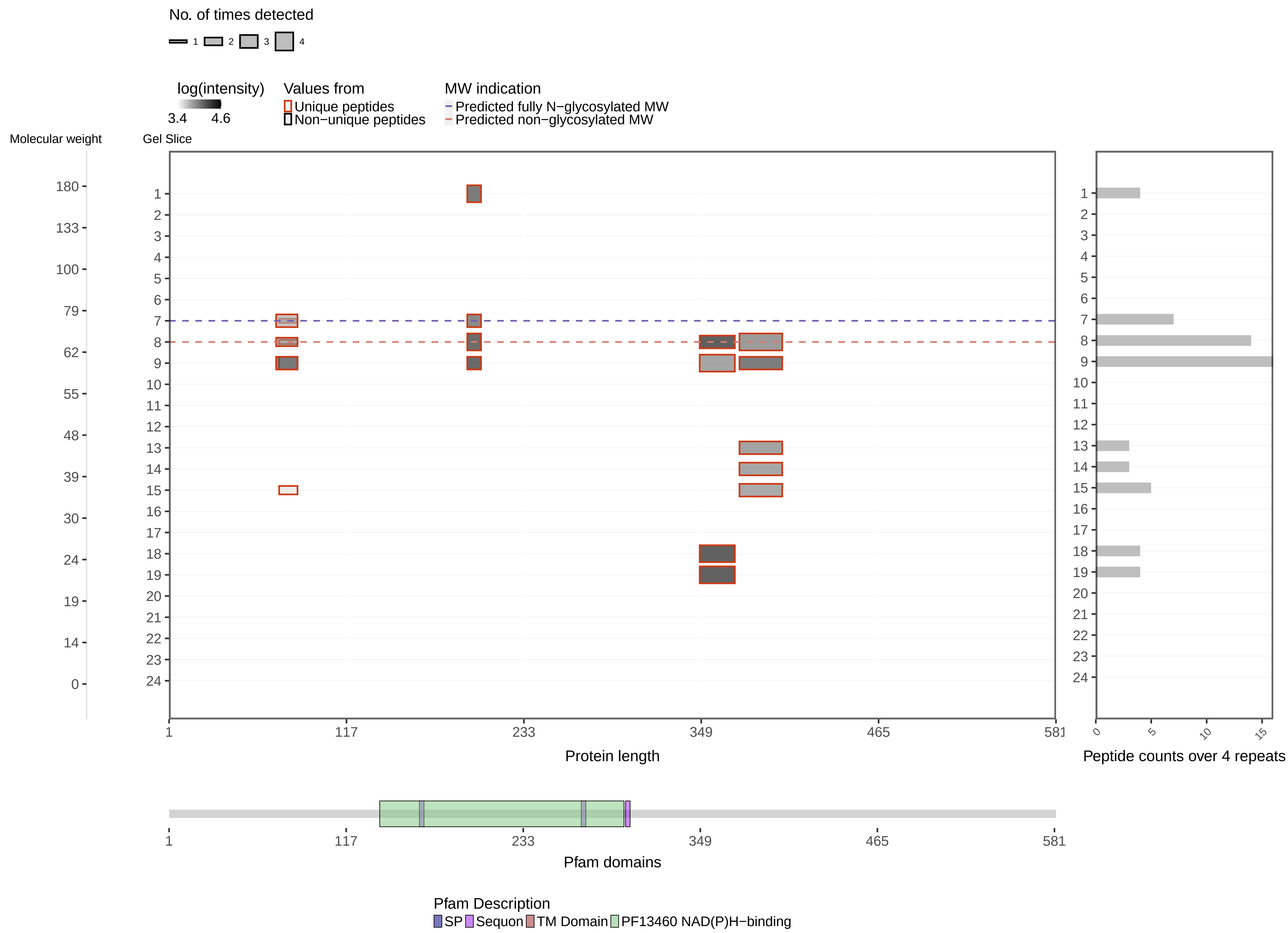


Protein sequence:

TLFGGQVSNLQVAELLAVMAKNLSLSYCKVVEVIAETTAPLTPMGDLLAKIPPQRVEVSPAKEAGDLQKLAASSSITSEIPSTSIEKEPAETKLKAAAPLSPYAAYEDLKPPTSPTPNPPGGSRVSASGAEDGKTTMFASVVVDGTSDKAELLTTRPLSPYTAYEDLKPPASPSPTPSGPKASISAETISQLAGGNDISPSITTISAEKDPSHTVAFCHSPFPFYEDLKPPTSPTPTPSGPQASTTVEVTSPLIGGNDVAIGSAADSTKEDTSKSLASCRSPYPMYEDLKPPTSPTPS

NbL09g22750.1 ferredoxin−−nitrite_reductase_chloroplastic−like [MW = 65.61 kDa]

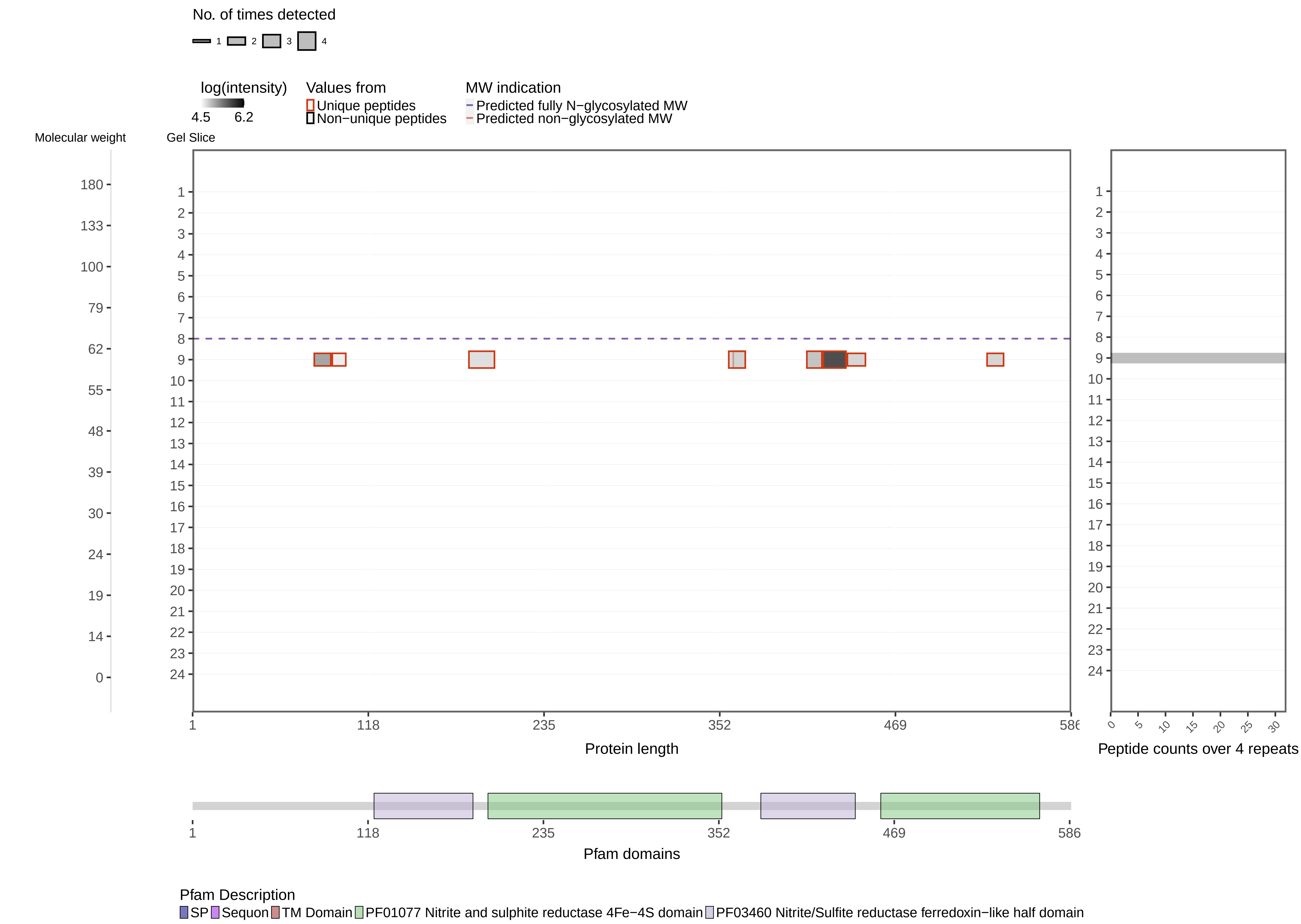


Protein sequence:

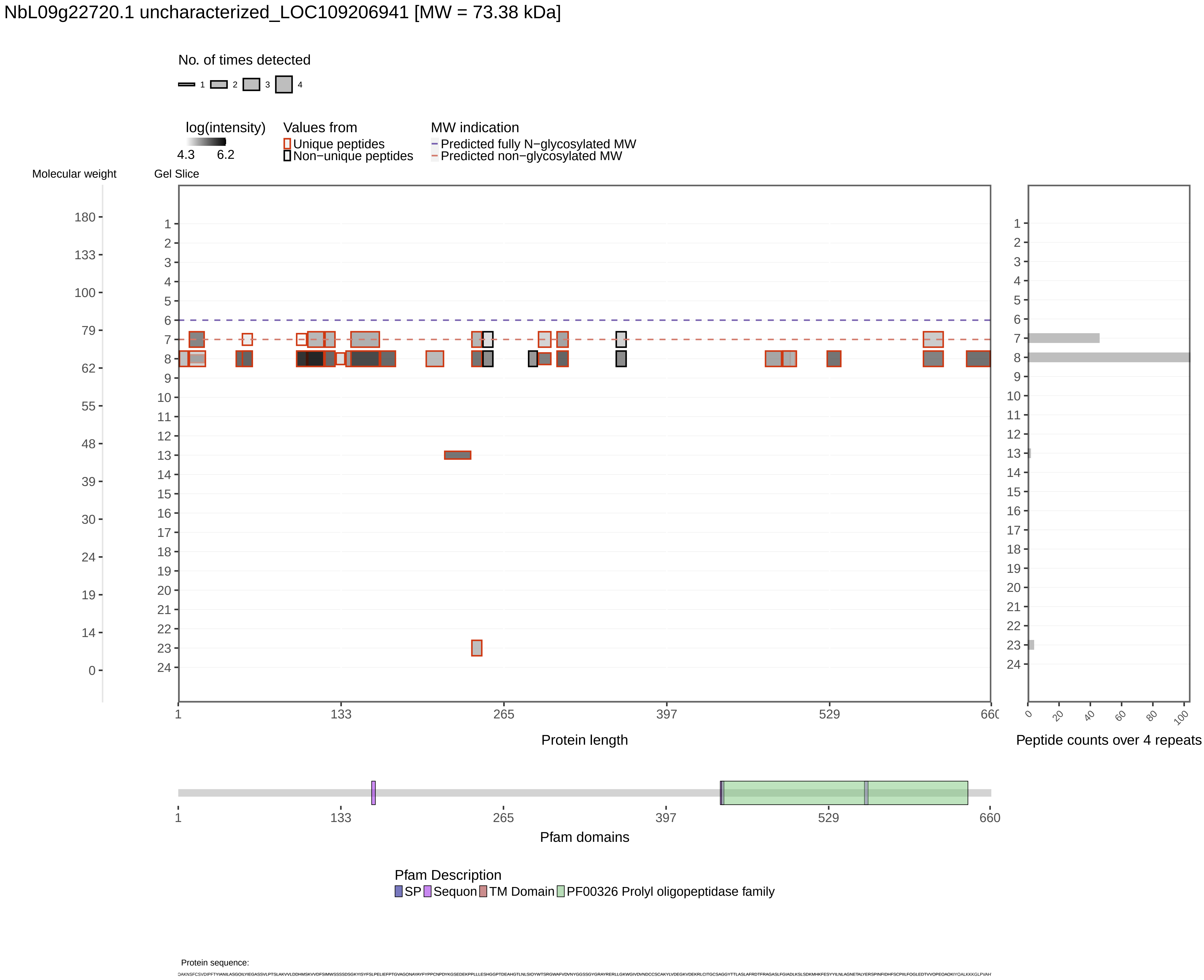

NbL09g22720.1 uncharacterized_LOC109206941 [MW = 73.38 kDa]
No. of times detected
1 2 3 4
log(intensity)
4.3 6.2
Values from
Unique peptides
Non−unique peptides
MW indication
Predicted fully N−glycosylated MW
Predicted non−glycosylated MW
Molecular weight
Gel Slice
Protein length
Peptide counts over 4 repeats
Pfam domains
Pfam Description
SP
Sequon
TM Domain
PF00326 Prolyl oligopeptidase family
Protein sequence:

### NbL09g22520.1 D−3−phosphoglycerate_dehydrogenase_1_chloroplastic−like [MW = 63.39 kDa]

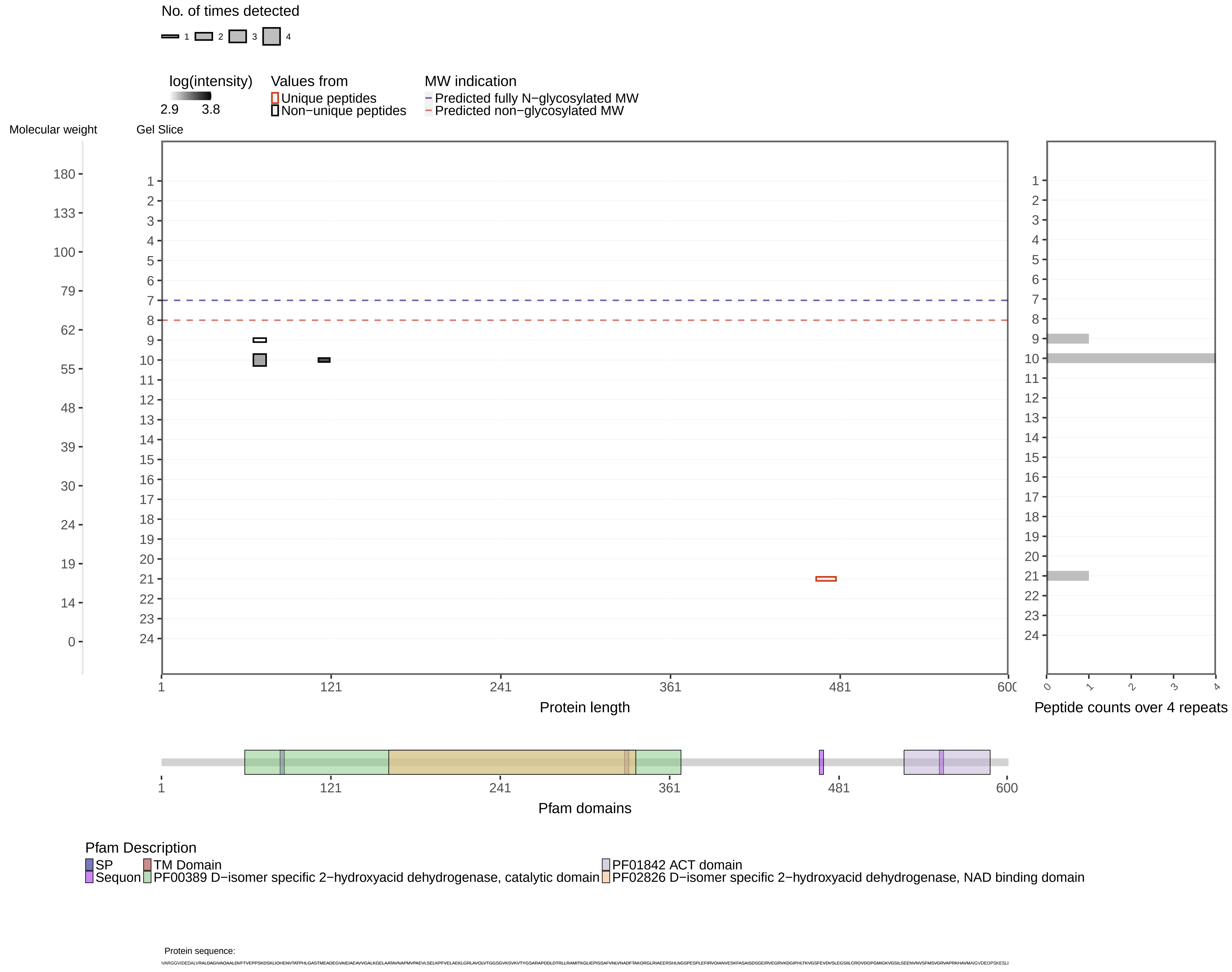

### NbL09g22370.1 heat_shock_70_kDa_protein_mitochondrial [MW = 73.02 kDa]

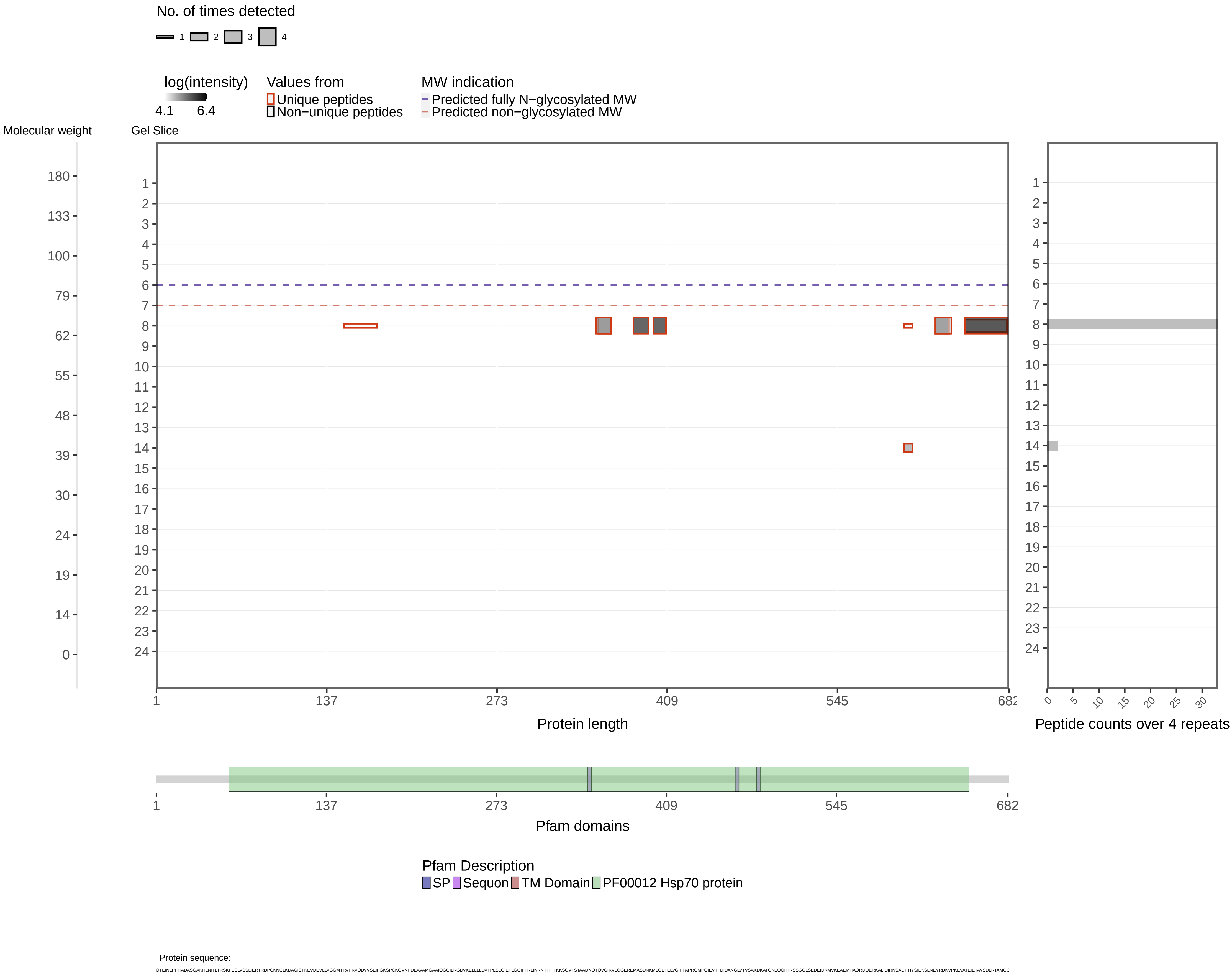

### NbL09g22320.1 elongation_factor_1−delta_1−like [MW = 28.98 kDa]

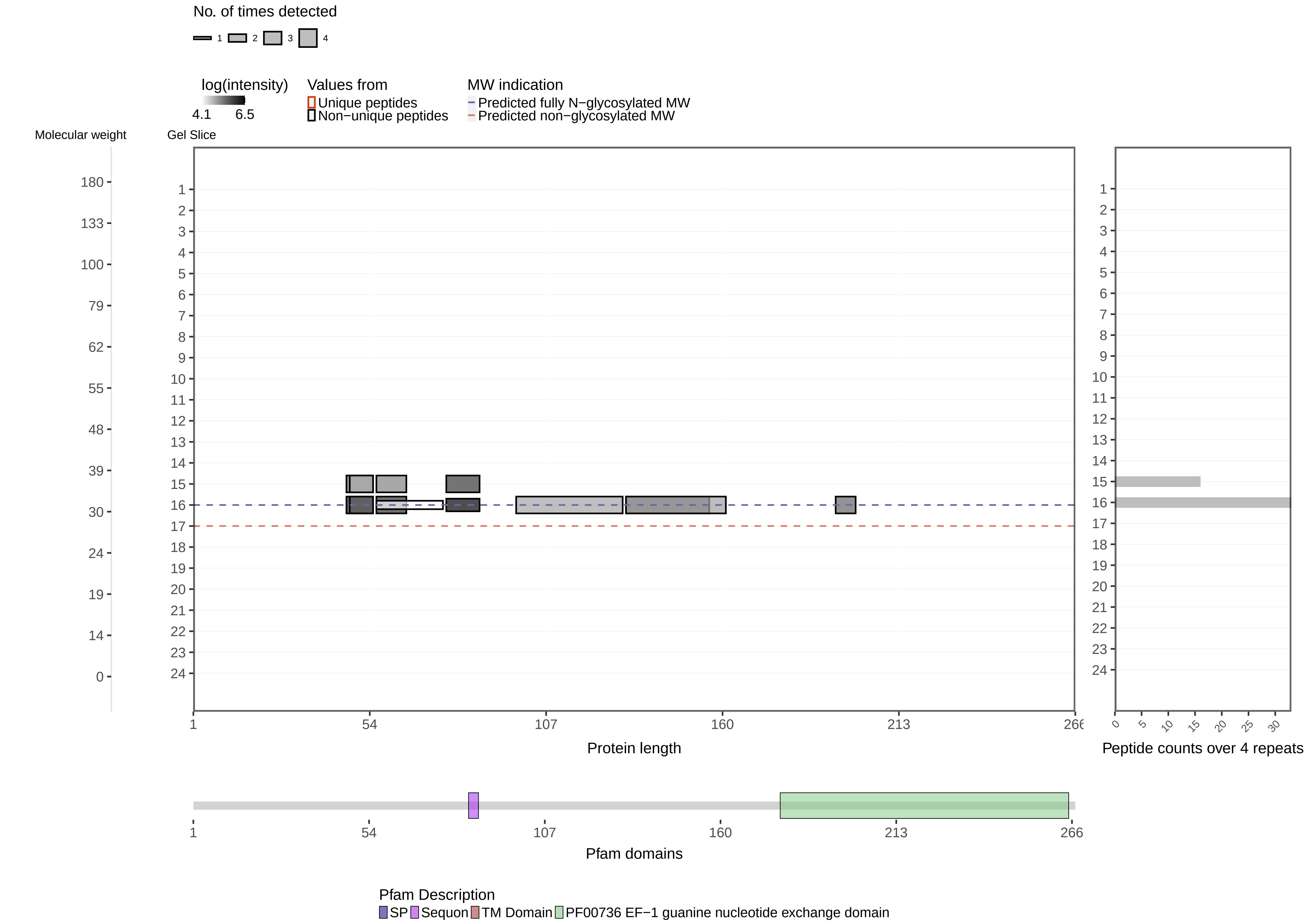


Protein sequence:

VATPPAADTKASAAEEEEEDDDDVDLFGEETEEEKKAAEERAAAVKASGKKKESGKSSVLLDVKPWDDETDMKKLEEAVRSVQMEGLTYGASKLVPVGYGIKKLQIMLTIVDDLVSVDDLVENYFMAEPINEYVQSCDIVAFNKI*

NbL09g22150.1 L−galactono−14−lactone_dehydrogenase_mitochondrial [MW = 66.94 kDa]

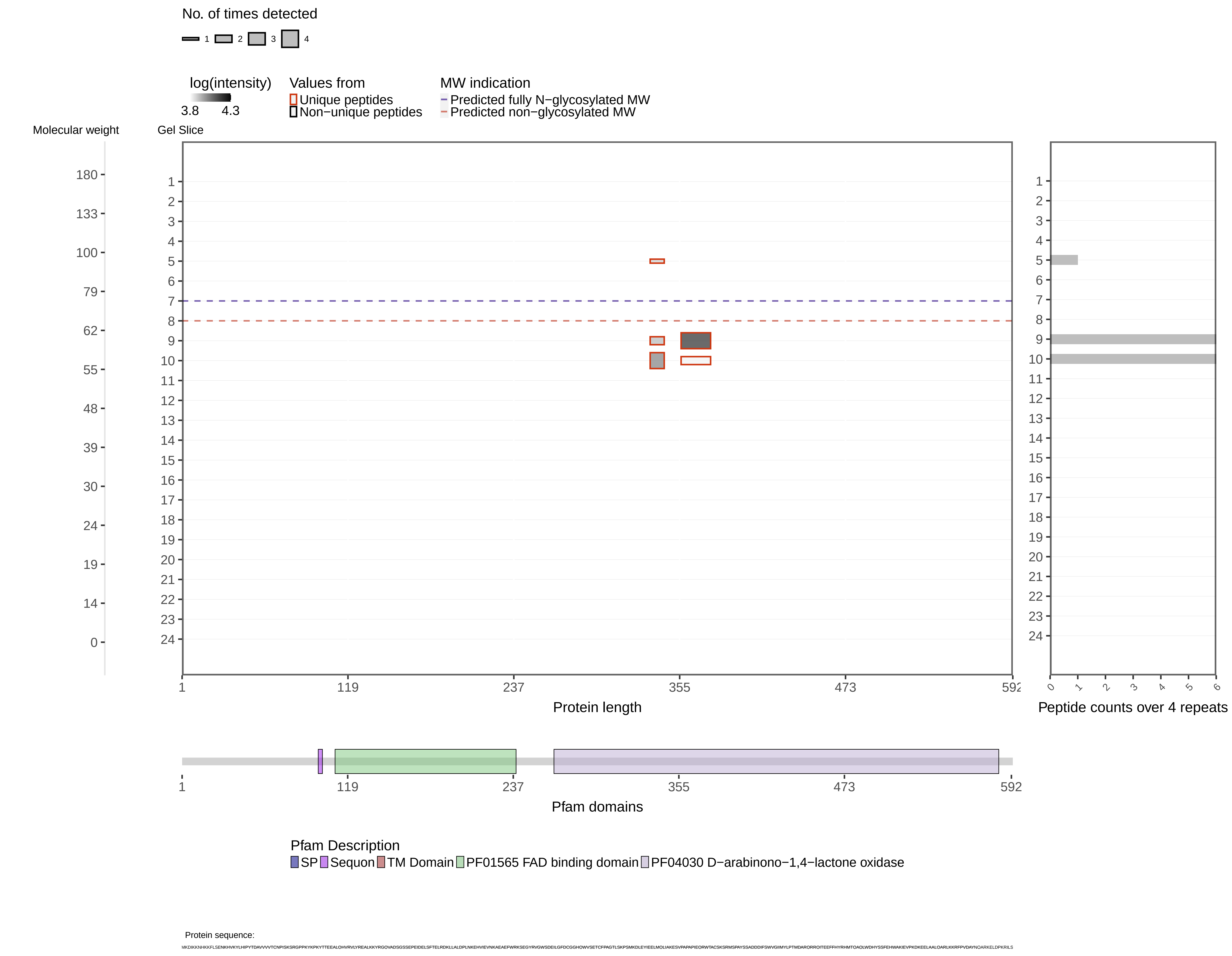

NbL09g21940.1 cystathionine_beta−lyase_chloroplastic [MW = 51.41 kDa]

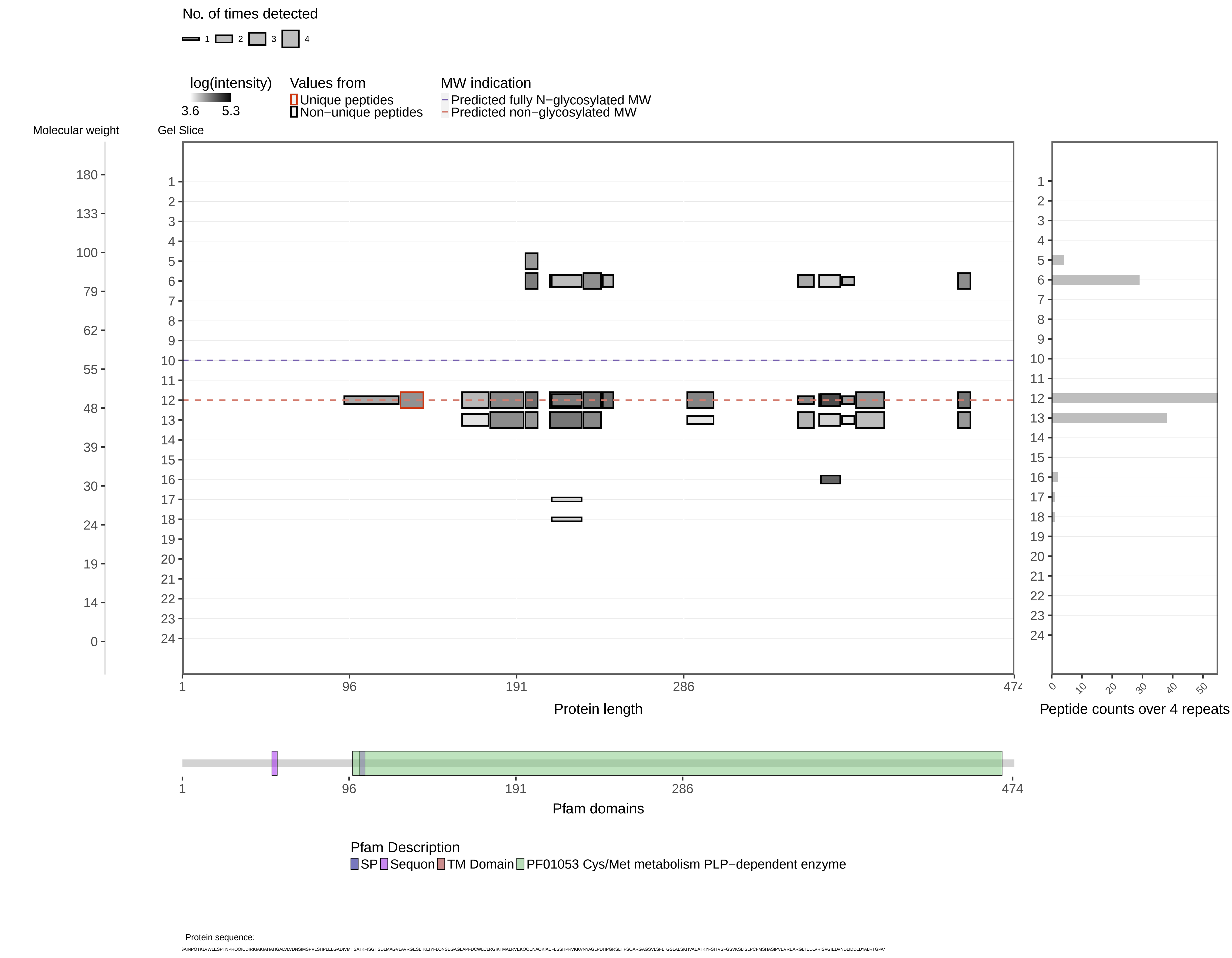

NbL09g21920.1 SNW/SKI−interacting_protein−like [MW = 69.59 kDa]

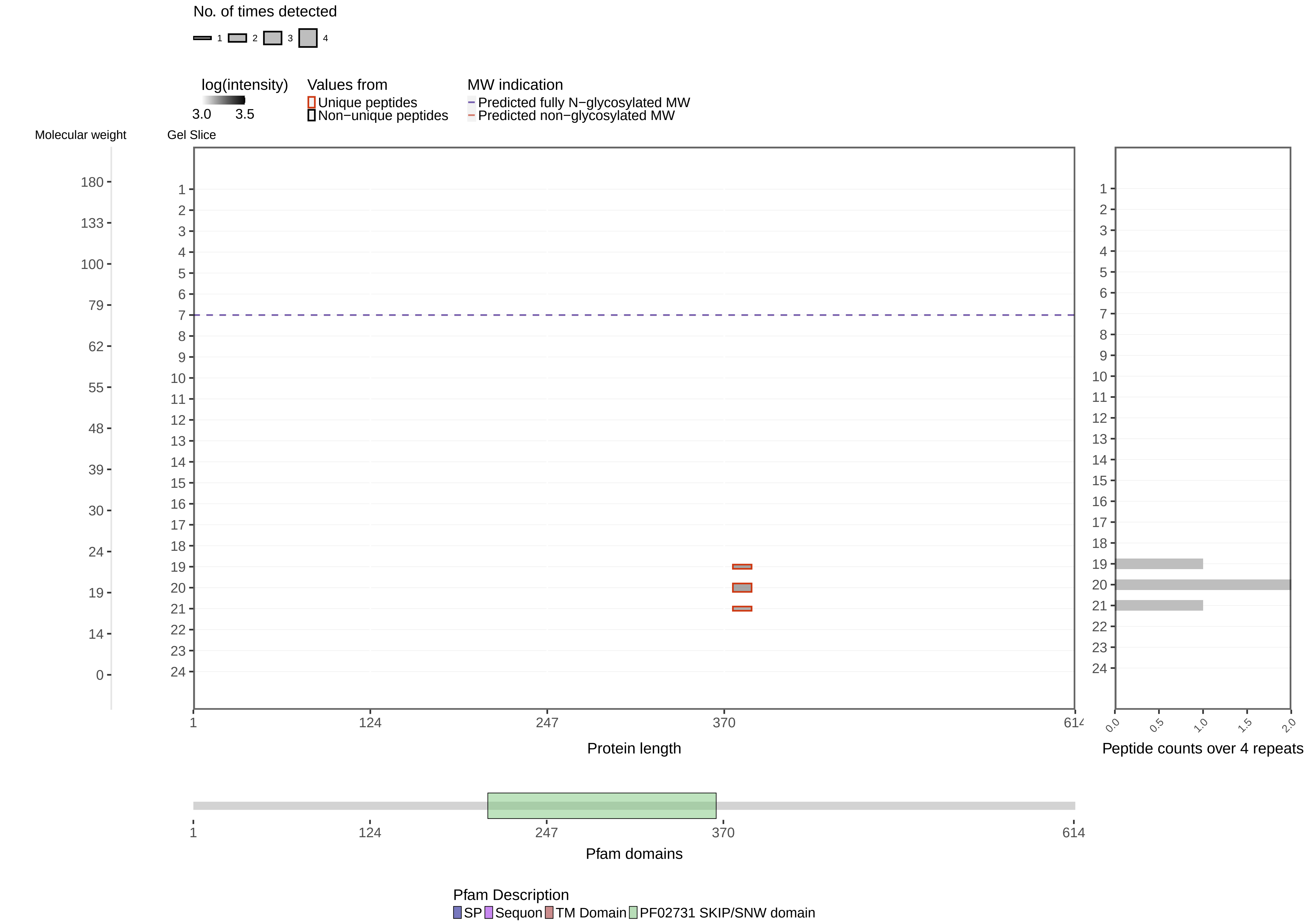


Protein sequence:

LAADGRGLQEVQINDNFAKLSEALYVAEQKAREAVAMRSKVQKEMMMKEKEKKEMELRELARKARSERTGVAPPAAERGTMNDDDMSVDYERARDFPKESREDREERSQREKMREERRRERERERRLEAKDAAMGKKSKITRDRDRDVSEKVALGMASTGTSRGEVMYDQRLFNQEKGMDSGFATDDAYNVYDKGLFTAQPTLSTLYRPKKDADSEMYGGADEQLEKIMKTERFKPDKAFAGTSERAAPRDGPVQFEKEVEEADPFGLDQFLTEVKKGKKAMAS

NbL09g21630.1 thioredoxin_reductase_NTRC−like [MW = 59.43 kDa]

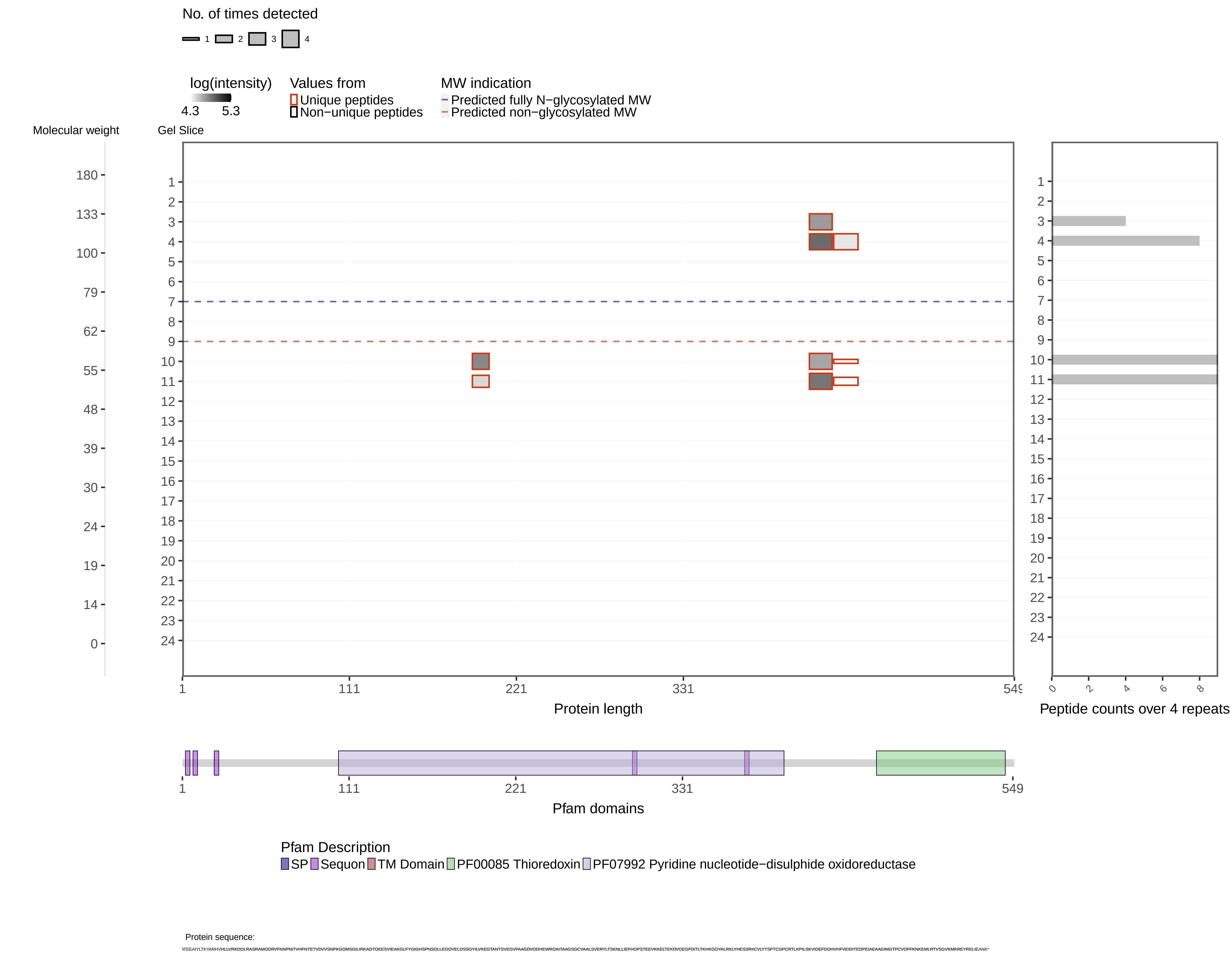

### NbL09g21500.1 adenylosuccinate_synthetase_chloroplastic [MW = 55.1 kDa]

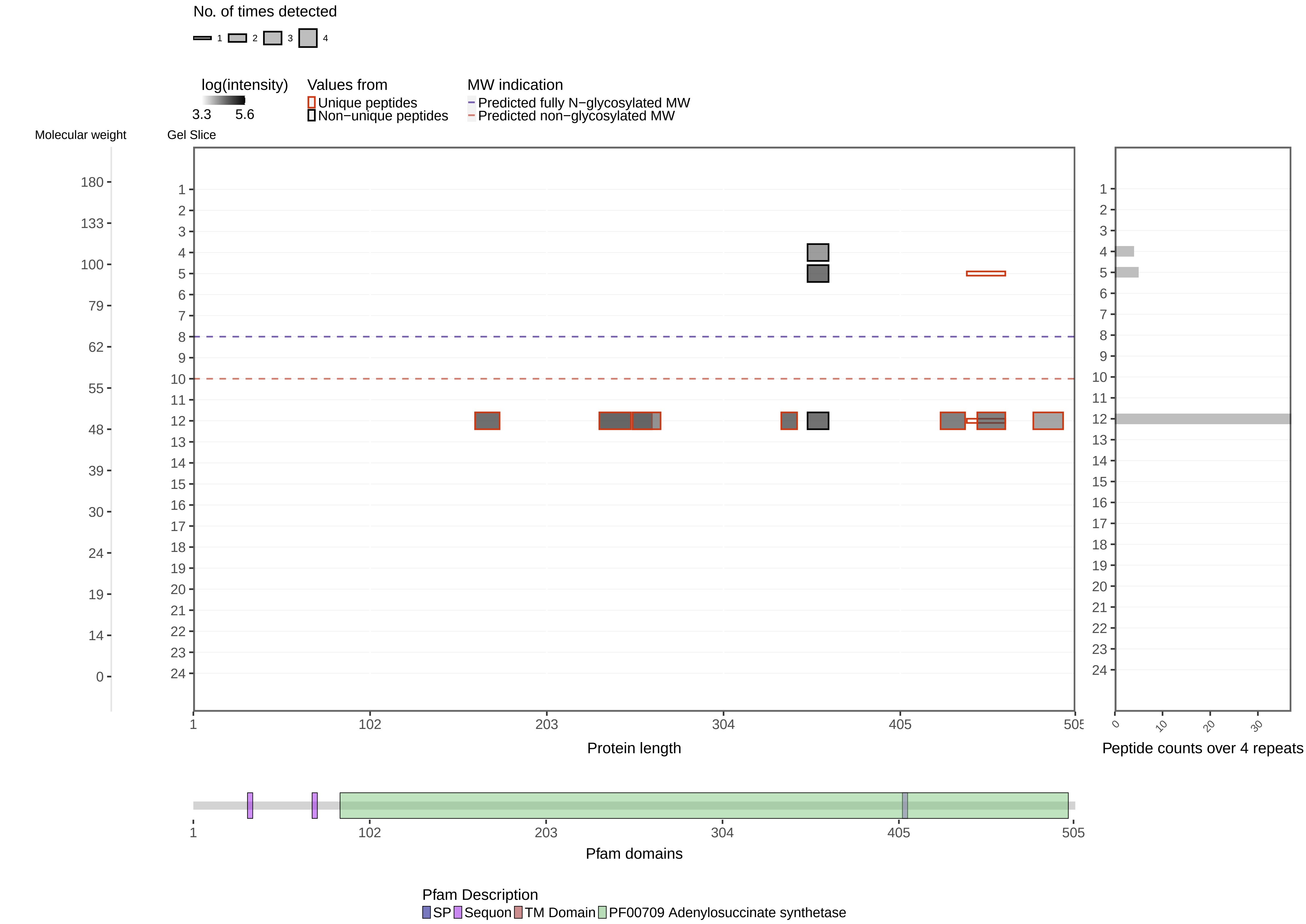


Protein sequence:

QLDLLLSDAASRFQGFNYGPDMLREEVERYKKFAERLEPFITDTVHFMNDAISQKKKILVEGGQATMLDIDFGTYPFVTSSSPSAGGICTGLGIAPRVVGDLVGVVKAYTTRVGSGPFPTEIMGKGGDLLRFAGQEFGTTTGRPRRCGWLDIVALRFCCQINGFASLNLTKLDVLSDLSEIQLGVTYRHPDGSTLNSFPSDLRLLEQIKVEYEVMPGWQSDISSVRKYSDLPKAAREYVERIEELVGVPVHYIGVGPGRDALIYK*

### NbL09g21360.1 zinc_finger_CCCH_domain−containing_protein_14−like [MW = 30.66 kDa]

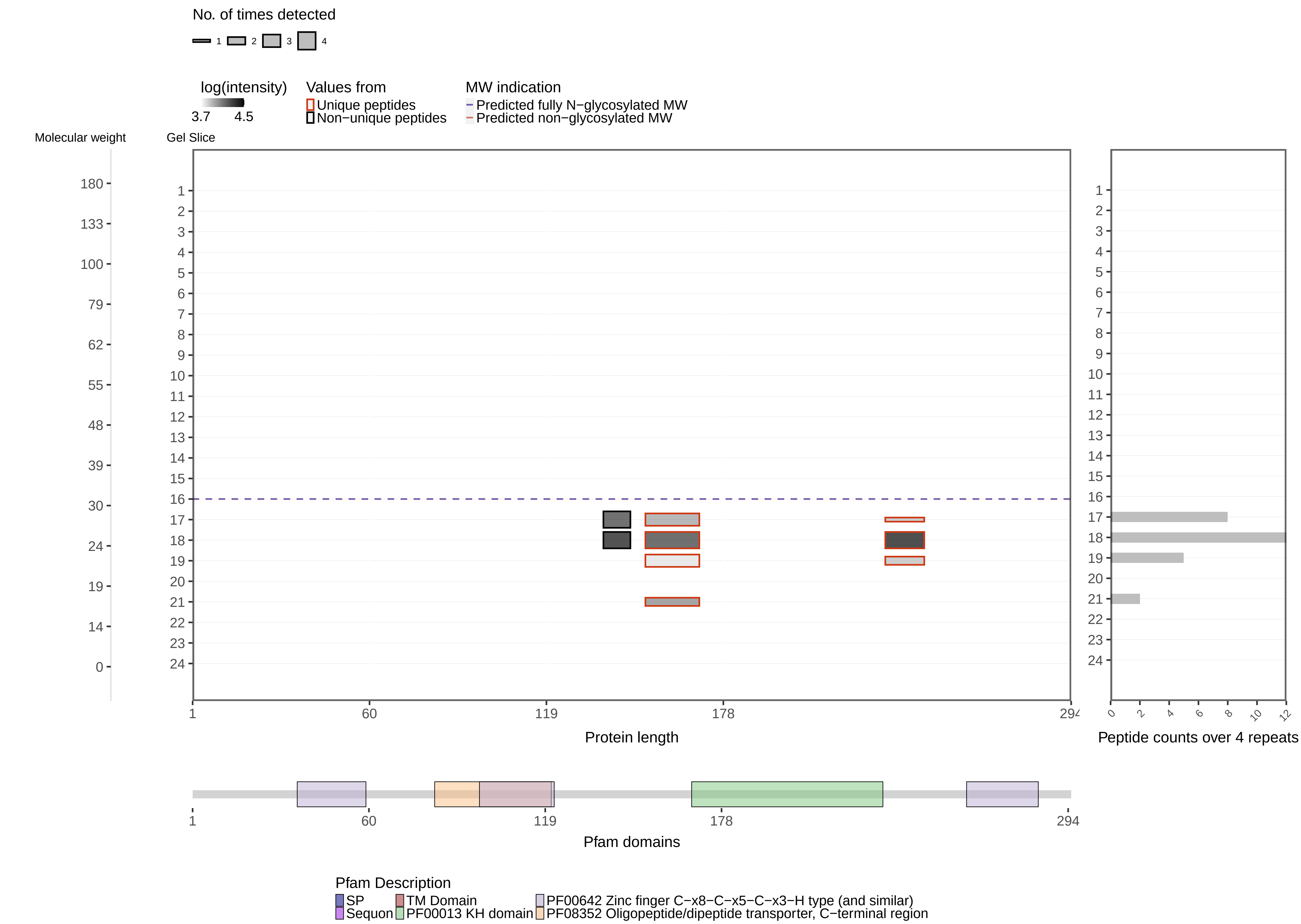


Protein sequence:

SQEDPRAMGVGPVPGRFGGRIEPAVSGPVASFGTSATAKISVDASLAGAIIGKGGVHSKKICRQTGAKLAIRDHETDKNLRNIELEGTFEQISQASAMVRELINSLGSVGGPGRTPAIPGGAAPPGSNFKTKLCENFAKGSCTFGDRCHFAHGAAELRKTGV*

NbL09g21340.1 probable_plastid−lipid−associated_protein_13_chloroplastic [MW = 35.9 kDa]

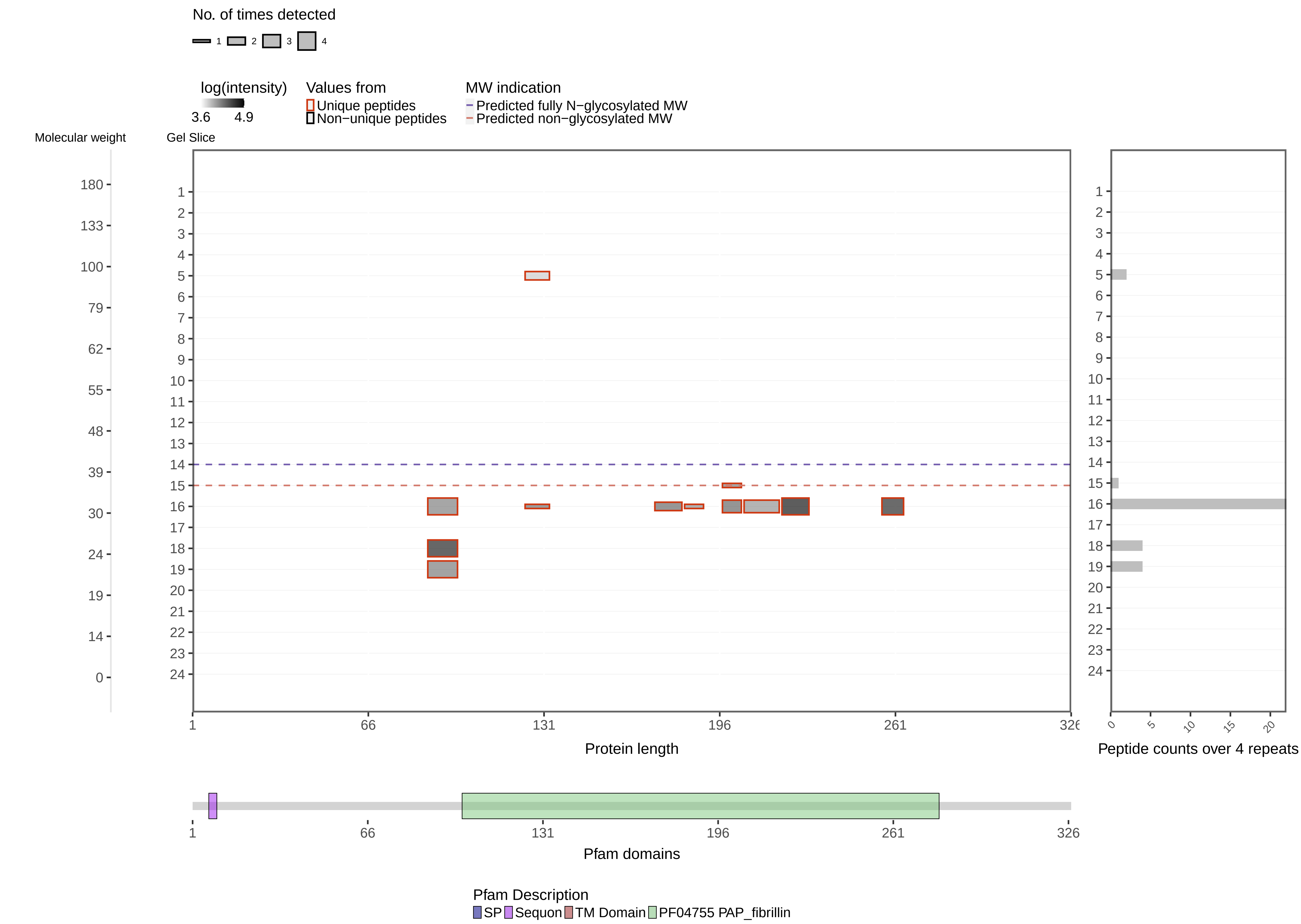


Protein sequence:

LATRLLFGRIPPTLASLSKLDLLIRNVYGTAIAHVKLLNSIESKFVLSSKFSVEGPLRMKEEYIEGIFETPKVNEDTMPEQLRGALGQAVSTLQQLPVPIKDTVSSGLKIPLGEAFQRLIMISYLDEEILVSPLQLVLQAALLIQDGYQTRNIPISGHECSNTFINAKTKPQPLVLLNY*

### NbL09g21330.1 actin−like [MW = 41.7 kDa]

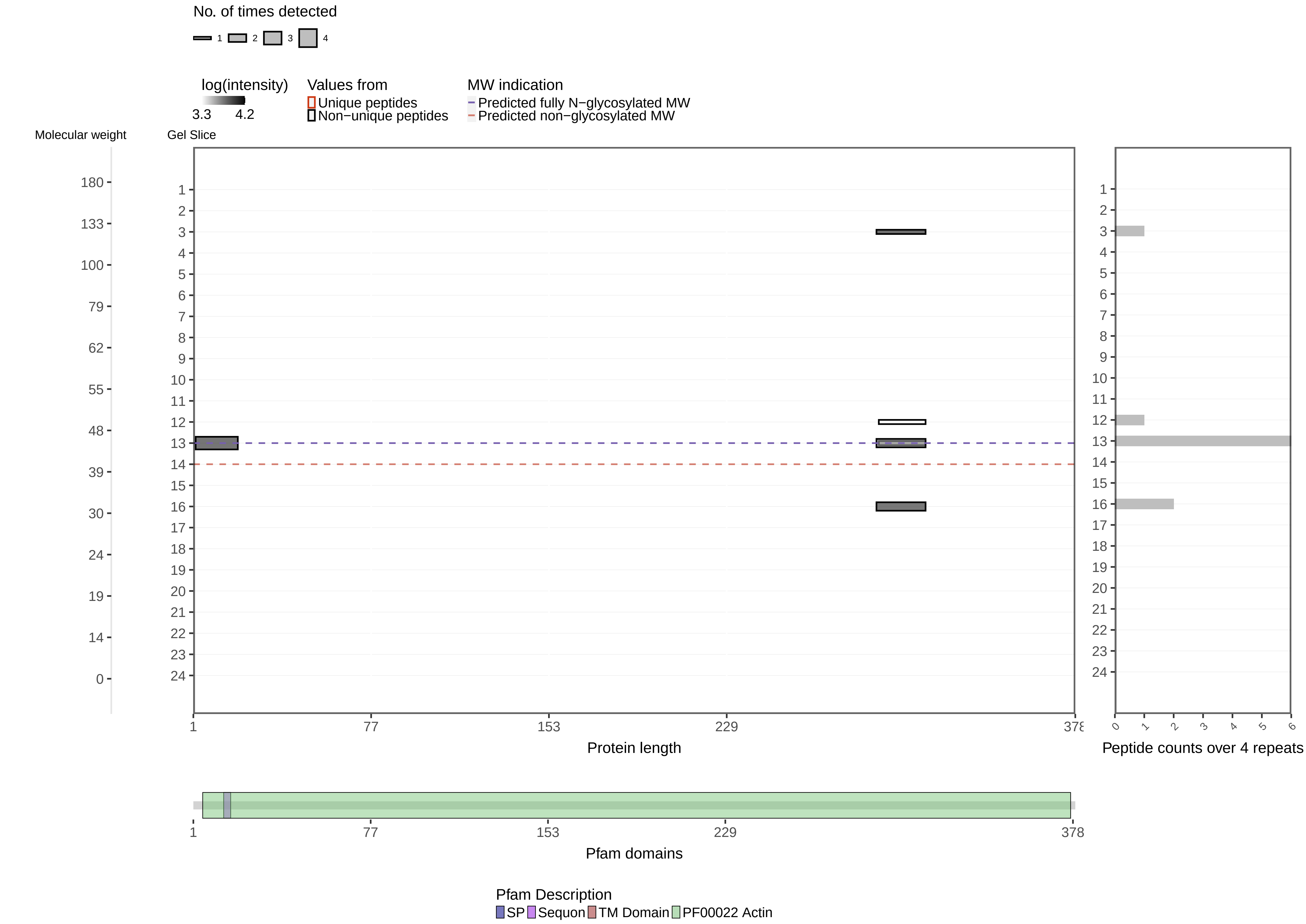


Protein sequence:

.PHAILRLDLAGRDLTDHLMKILTERGYSFTTTAEREIVRDVKEKLAYIALDYEQELETAKTSSSVEKSYELPDGQVITIGAERFRCPEVLFQPSMIGMEAAGIHETTYNSIMKCDVDIRKDLYGNIVLSGGTTMFPGIADRMSKEITALAPSSMKIKVVAPPERKYSVWIGGSILASLSTFQQMWIAKAEYDESGPSIVHRKCF*

### NbL09g21180.1 translation_initiation_factor_NCBP−like [MW = 24.47 kDa]

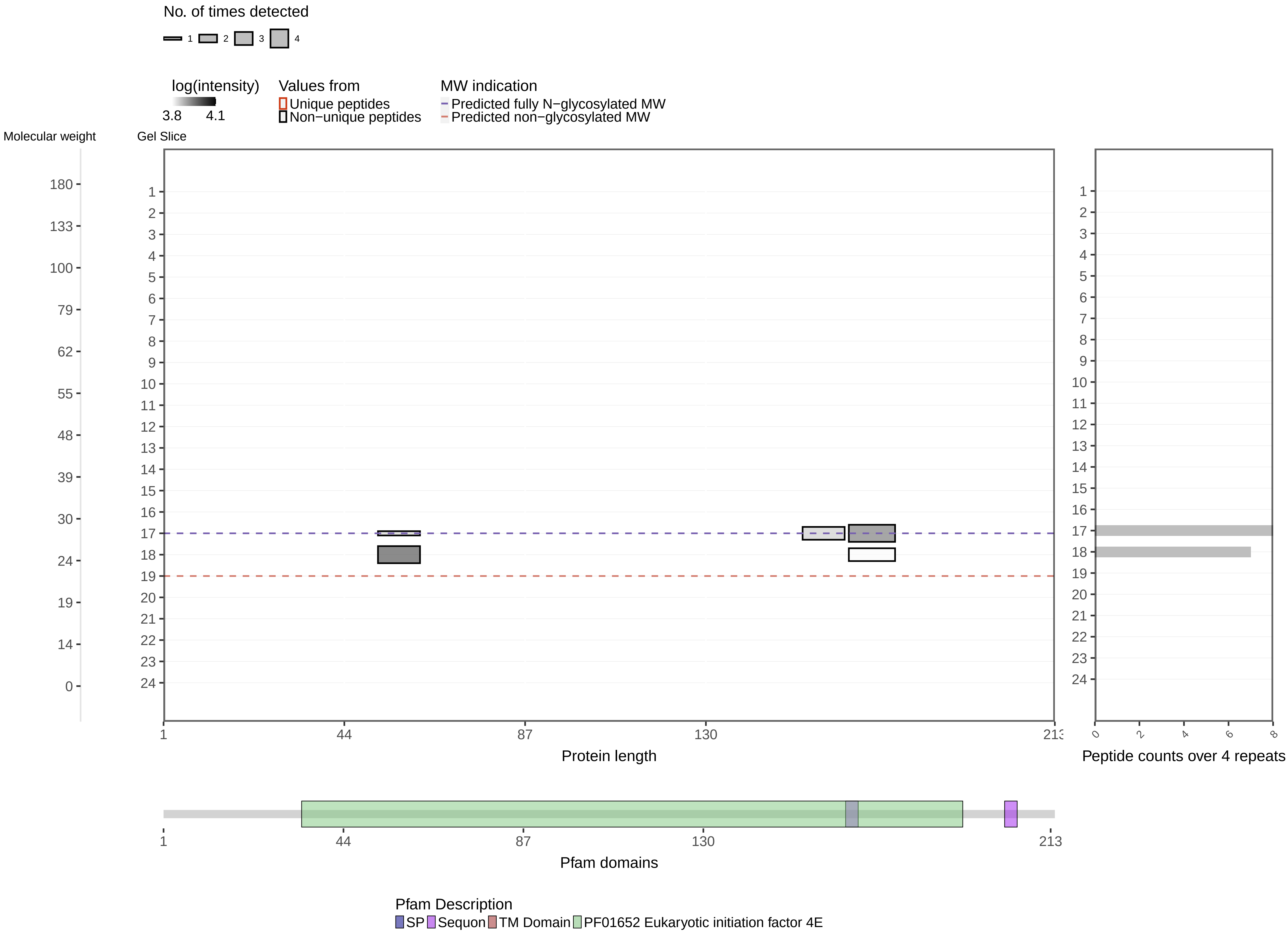


Protein sequence:

FKEGIRPLWEDAANCNGGKWIIRFKKAVSGRFWEDLVLALVGDQLDYGDNICGAVLSIRFNEDILSVWNRNASDQQAVMALRDAIKRHLKLPNGYVMEYKAHDASLRDNSSYRNTWIRG*

### NbL09g21110.1 asparagine−−tRNA_ligase_cytoplasmic_1−like [MW = 64.92 kDa]

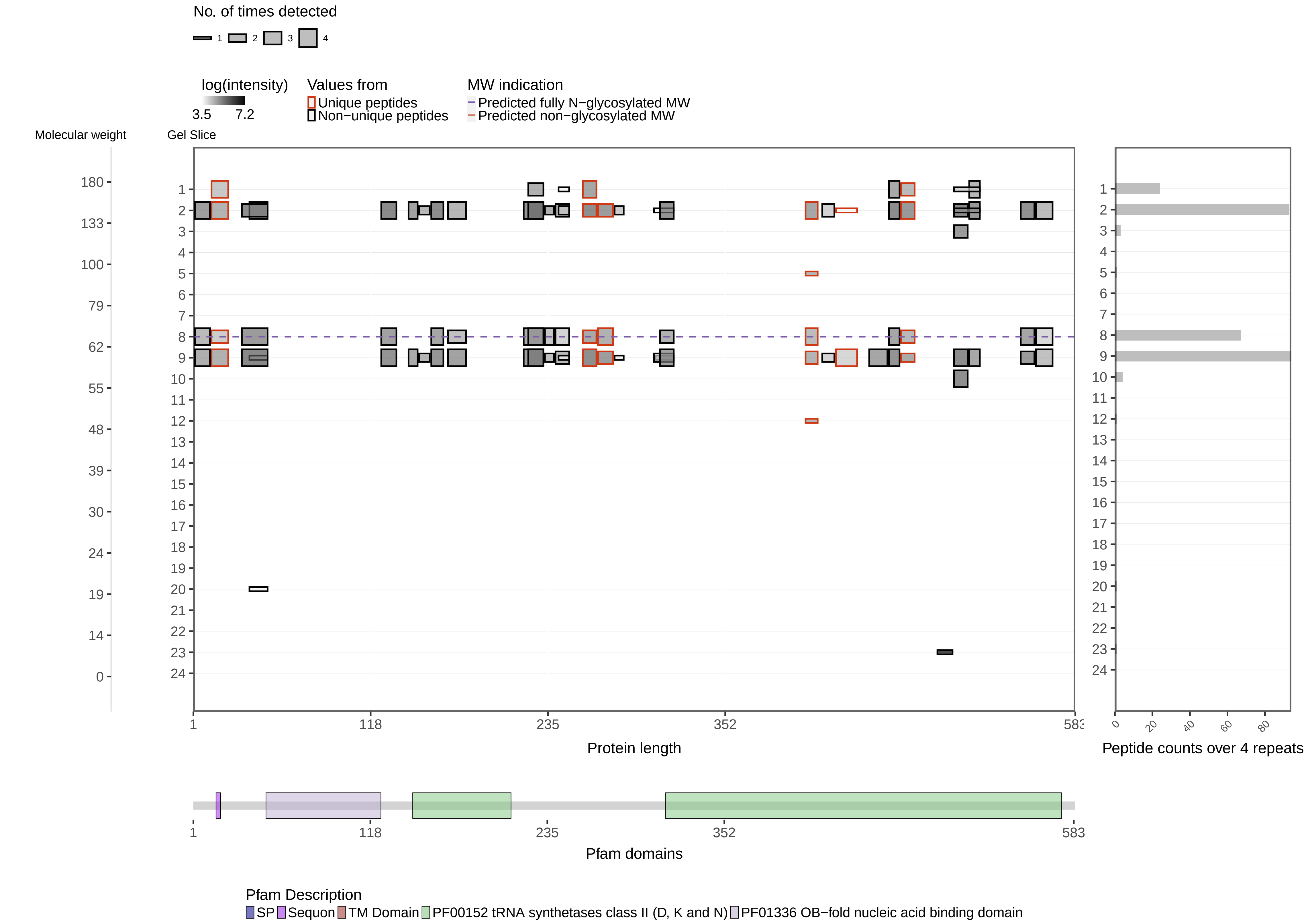


Protein sequence:

### NbL09g21080.1 protein_SMG7−like [MW = 108.07 kDa]

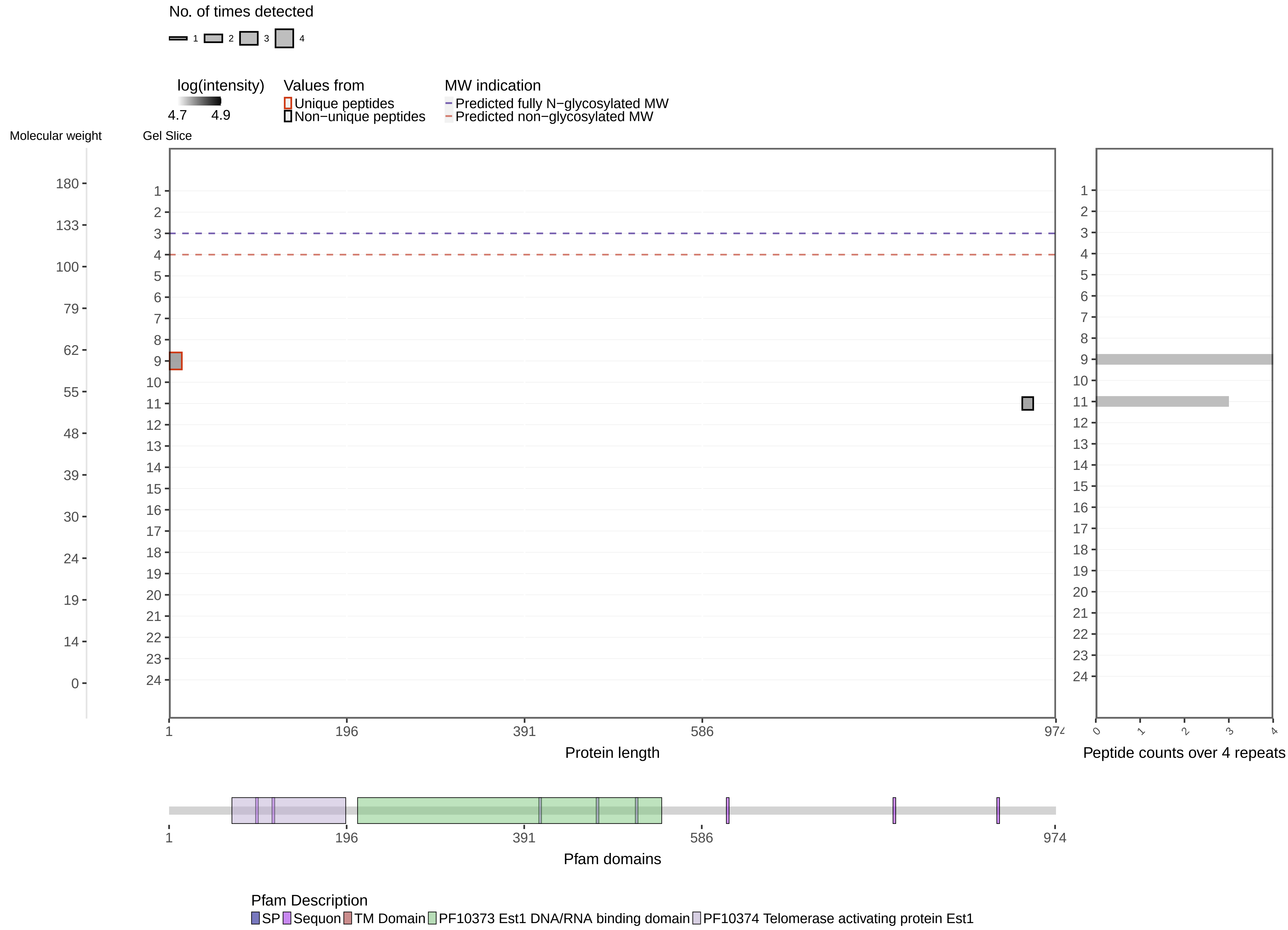


Protein sequence:

ARSFFWKNCITFFNKLLATGSKFVDEDEDETCFFNMSRYDESESGNRLALPEDFELRGFVPLLPAOLILDFSRKHSFGGDSGNKEKKARLORMIAAGKALANVVRVGEEGIYFDTRGKKFVIGLEPOASDDYOLNGSREVAKLSGIELENPDAGLMNVGDLOPKOOLYVECEEEDEVIVFKPSVMEKVNGISSNTMTLAVPVSVISASSVPSGVSMASINICSEMGPFSSALDGLSLONAWSANVRORTSIAHTNTOYVOPIOTSASMWSVEODAVMNGLVGGLNLIGNRLT

### NbL09g20930.1 3−oxoacyl−[acyl−carrier−protein]_reductase_FabG−like [MW = 31.22 kDa]

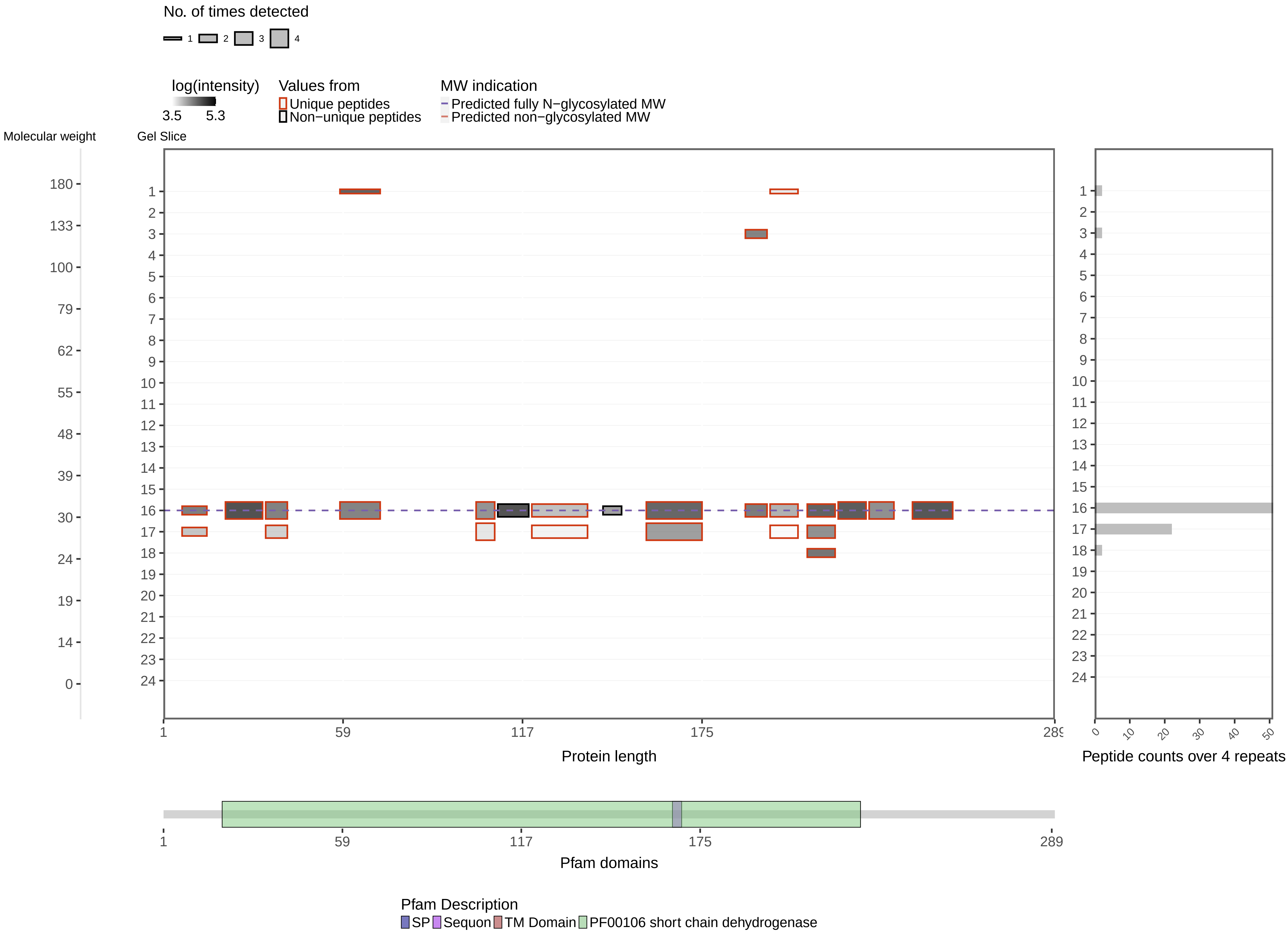


Protein sequence:

EEWDNIYRTNLKGAWLVSKYVCIRMRDANQGGSVINISSIAGLNRGQVPGSLAYSSSKTALNTVMKMMAIELGPYKIRVNSISPGLFRSEITEGLMQKDWLPNIALRTIPLRTHGTSDPALTSVVRYLIHDSSEYVSGNIFIVEAGATLPGVPIFSSL*

### NbL09g20870.1 tubulin_beta−1_chain [MW = 50.19 kDa]

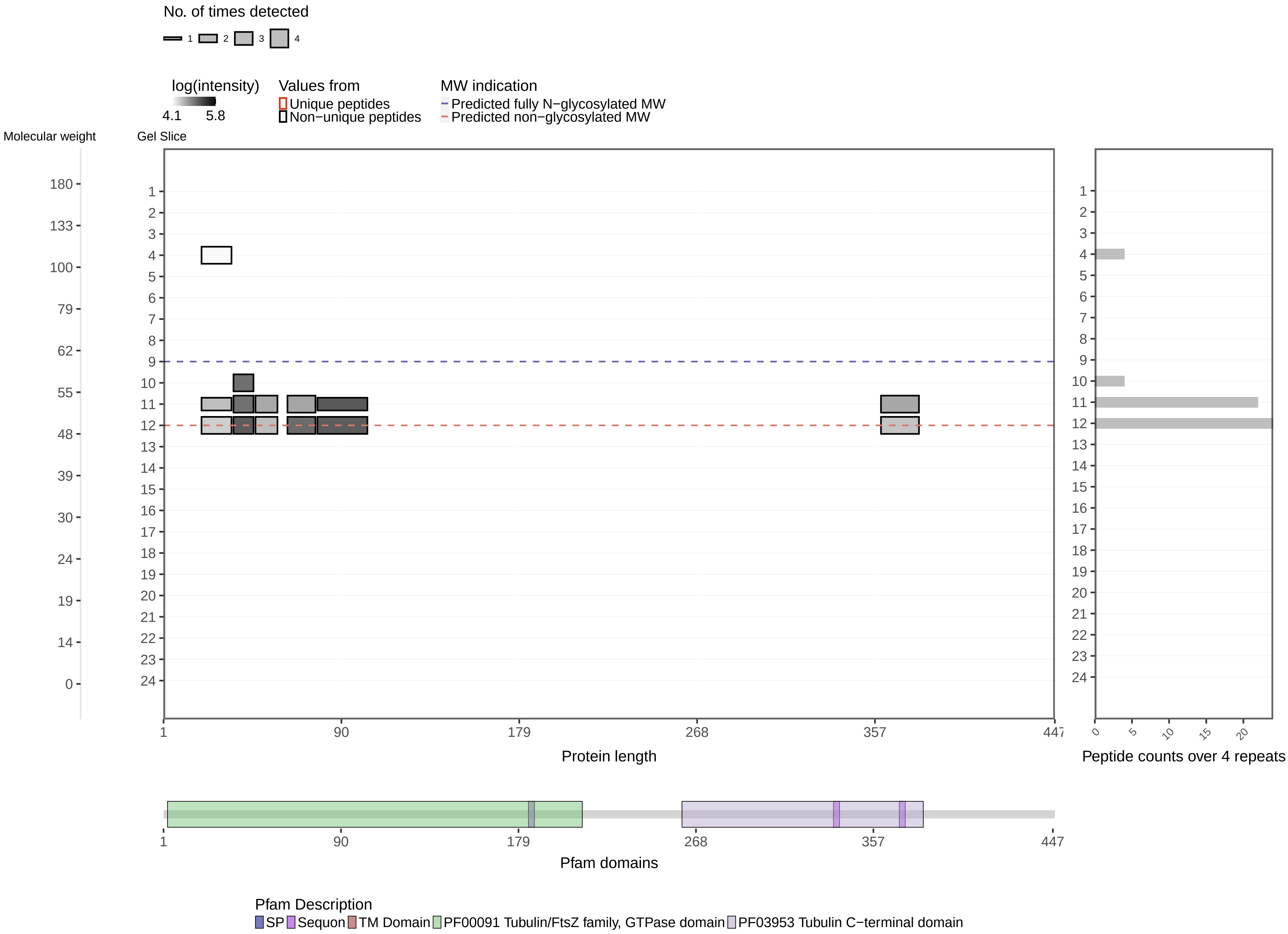


Protein sequence:

CFRTLKLTTPSFGDLNHLISATMSGVTCCLRFPGQLNSDLRKLAVNLIPFPRLHFFMVGFAPLTSRGSQQYRALTVPELTQQMWDAKNMMCAADPRHGRYLTASAMFRGKMSTKEVDEQMINVQNKNSSYFVEWIPNNVKSSVCDIPPRGLSMASTFIGNSTSIQEMFRRVSEQFTAMFRRKAFLHWYTGEGMDEMEFTEAESNMNDLVSEYQQYQDATADDEGEYEEEEYDHEGN*

### NbL09g20840.1 bifunctional_protein_FolD_2−like [MW = 32.2 kDa]

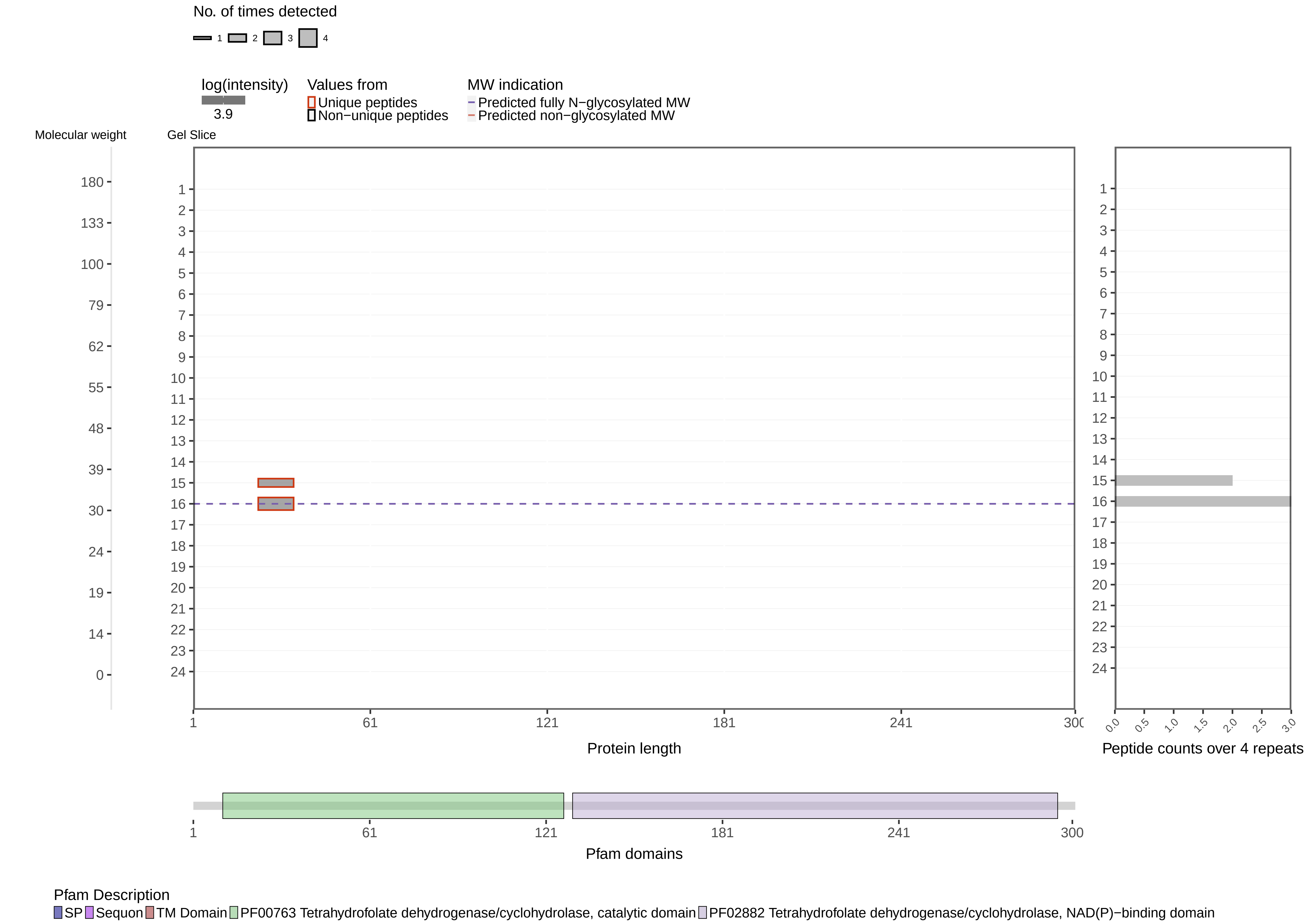


Protein sequence:

AMKGRQPLFLPCTPKGCIELLHRSGISIKGKNAVVVGRSNIVGLPVSLLLLKEDATVTIVHSRTKEPEKIVREADIIIAAAGQAMMIKGSWIKPGAAVIDVGTNAVDDPTRKSGYRLVGDVNFQEACKVAGWITPVPGGVGPMTVAMLLKNTLDGAKRVIEK*

### NbL09g20780.1 transcription_elongation_factor_SPT6 [MW = 186.02 kDa]

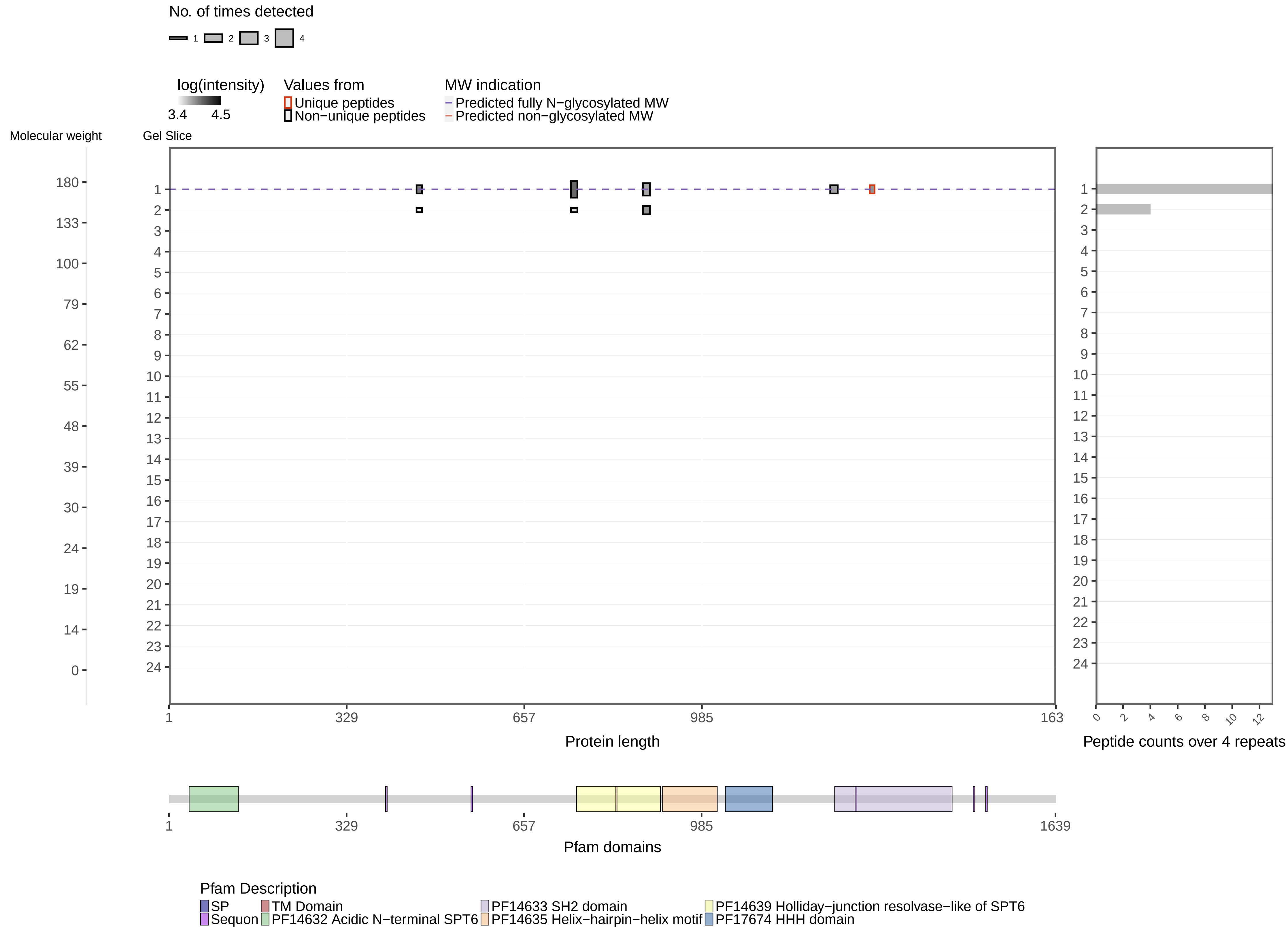


Protein sequence:

LLKFMMDHQPHVVVLGAVNLSCTRLKEDIYEIIFKMVEDNPRDVGHEMDNLNIIYGDETLPHLYENSRISVDQLPGQSGIVRRAVALGRYLQNPLAMVATLCGPGREILSWKLSTLDIFLTPDEKYGMVEQVMVDVTNQVGVDLNLAISHEWLFAPLQFISGLGPRKAASLQRSLVRQQTIFTRKDILTEHRLGKKVFVNAVGFLRVRRSGYTANSNTYIDLLDDTRIHPESYGLAQELAKDIYLNDIGEENNDDDELLEMAIEHVREKPHLARLLKVPNYAKAKDRQDKEETLND

NbL09g20770.1 nascent_polypeptide−associated_complex_subunit_alpha−like_protein [MW = 21.92 kDa]

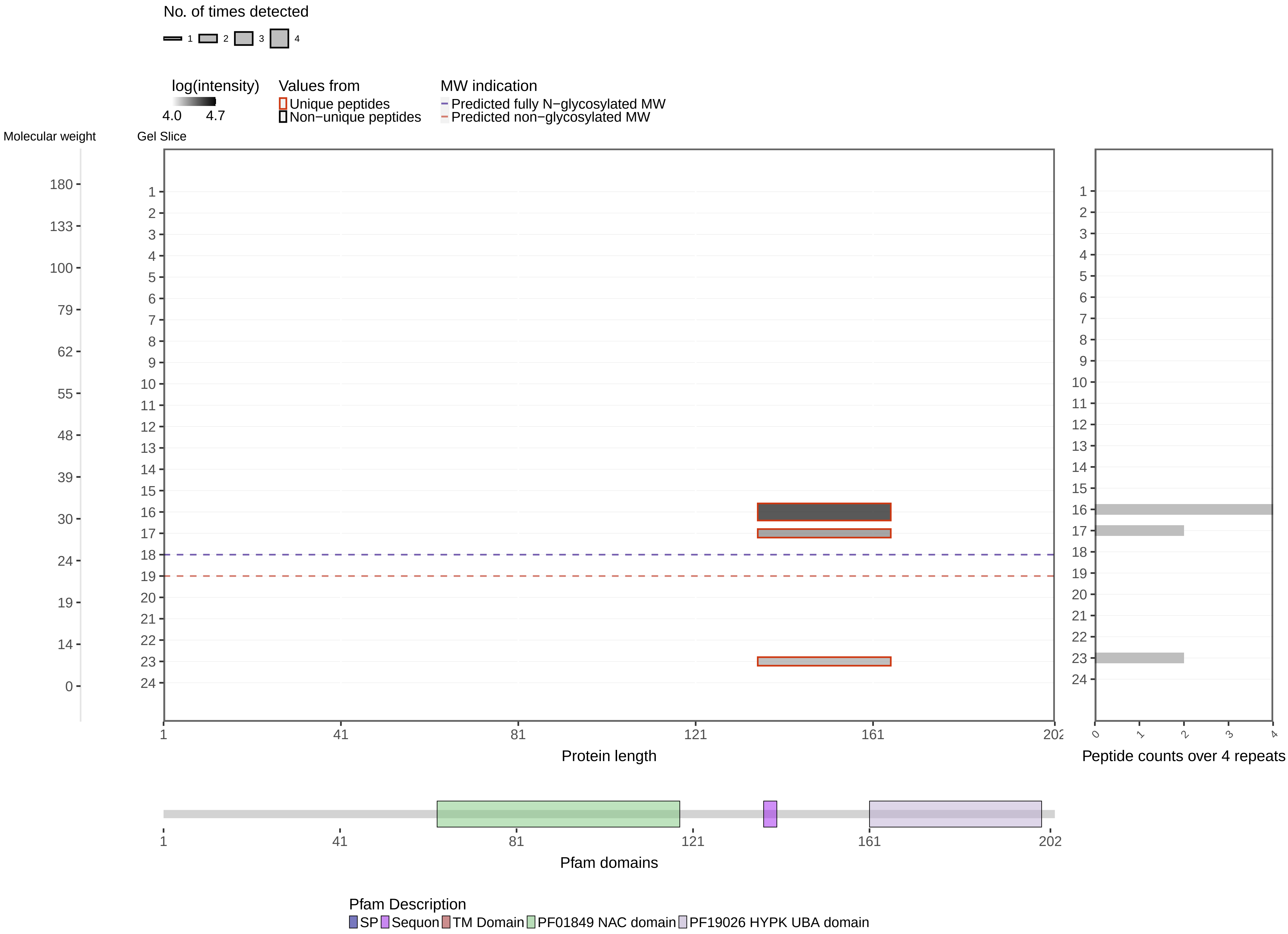


Protein sequence:
/KKSKNILFVISKPDVFKSPNSDTYVIFGEAKIEDLSSQLQTQAAEQFKAPNLSNVISKPEPSTVAQDDEDVDDTGVEPKDIELVMTQAGVSRAKAVKALKAADGDIVSAIMELTN*

NbL09g20720.1 alpha−L−arabinofuranosidase_1−like [MW − SP = 70.83 kDa]

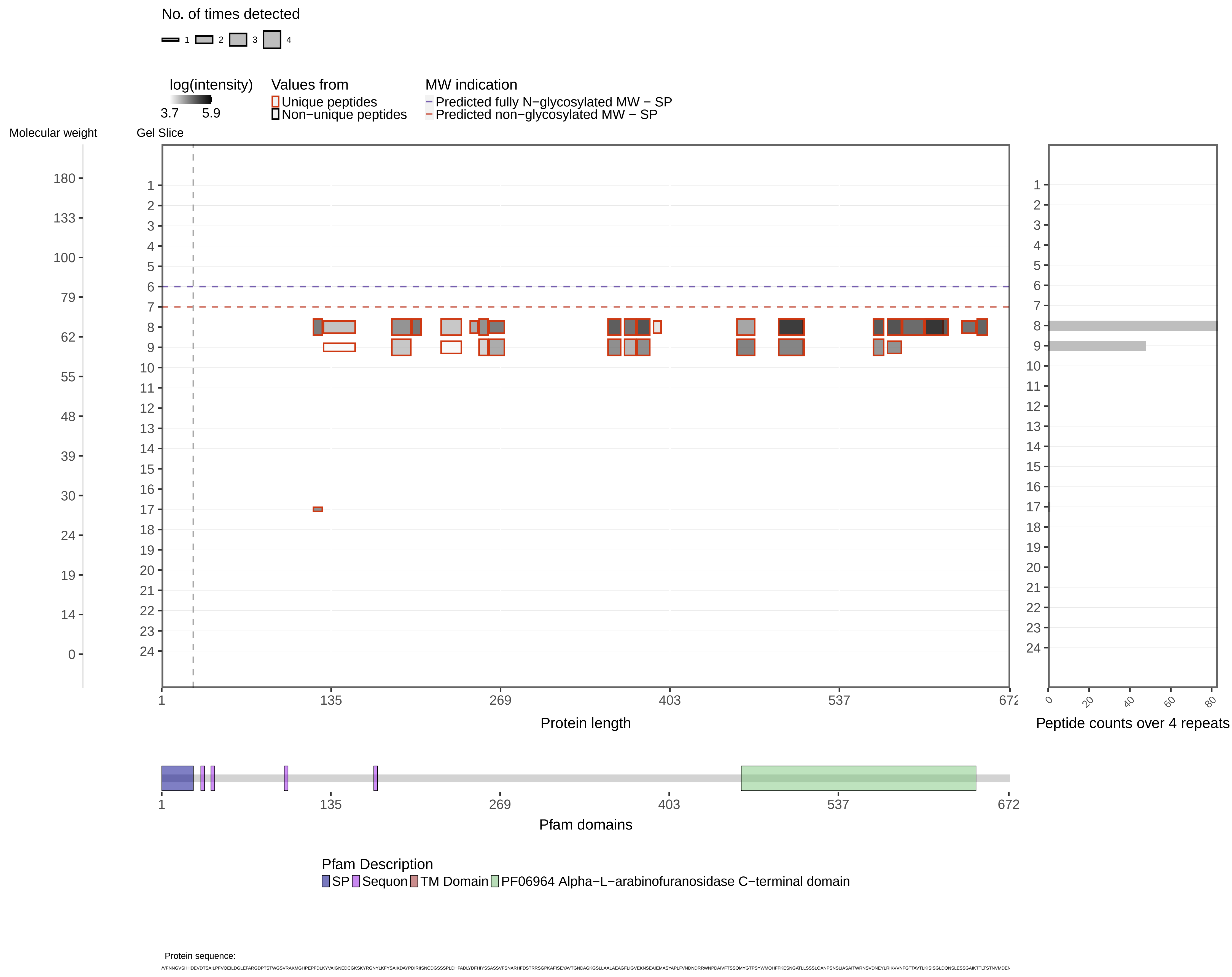

NbL09g20710.1 proteasome_subunit_alpha_type−3−like [MW = 27.2 kDa]

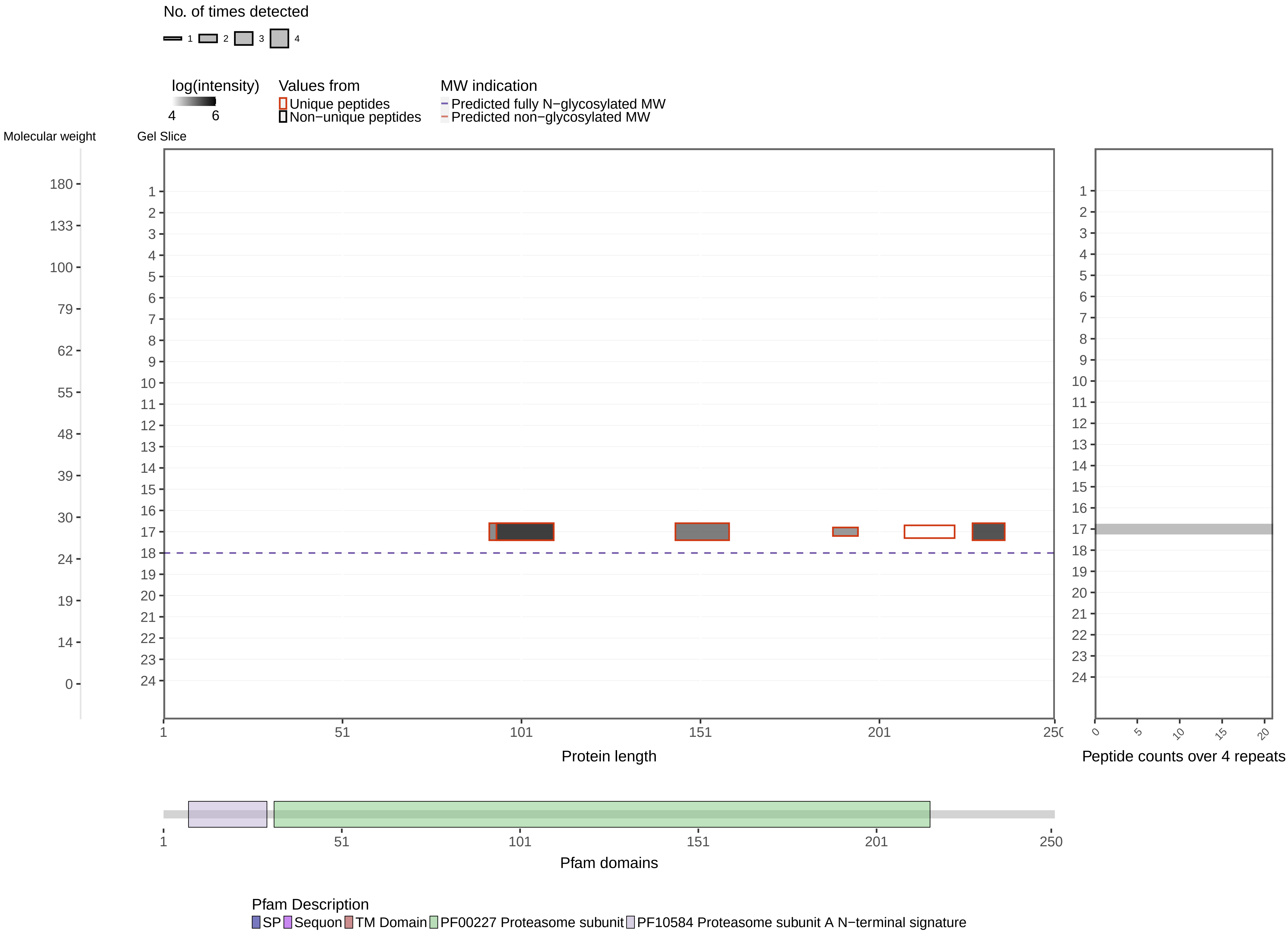


Protein sequence:

AERVASYVHLCTLYWWLRPFGCGVILGGYDRDGPQLYMVEPSGVSYRYFGAAIGKGRQAAKTEIEKLKLSEMTCRQGVIEIAKIIYGVHDEAKDKAFELEMSWVCDESNRQHQKVPADLLEEAKAAAKAALEEMDAD*

NbL09g20660.1 calmodulin−7 [MW = 16.83 kDa]

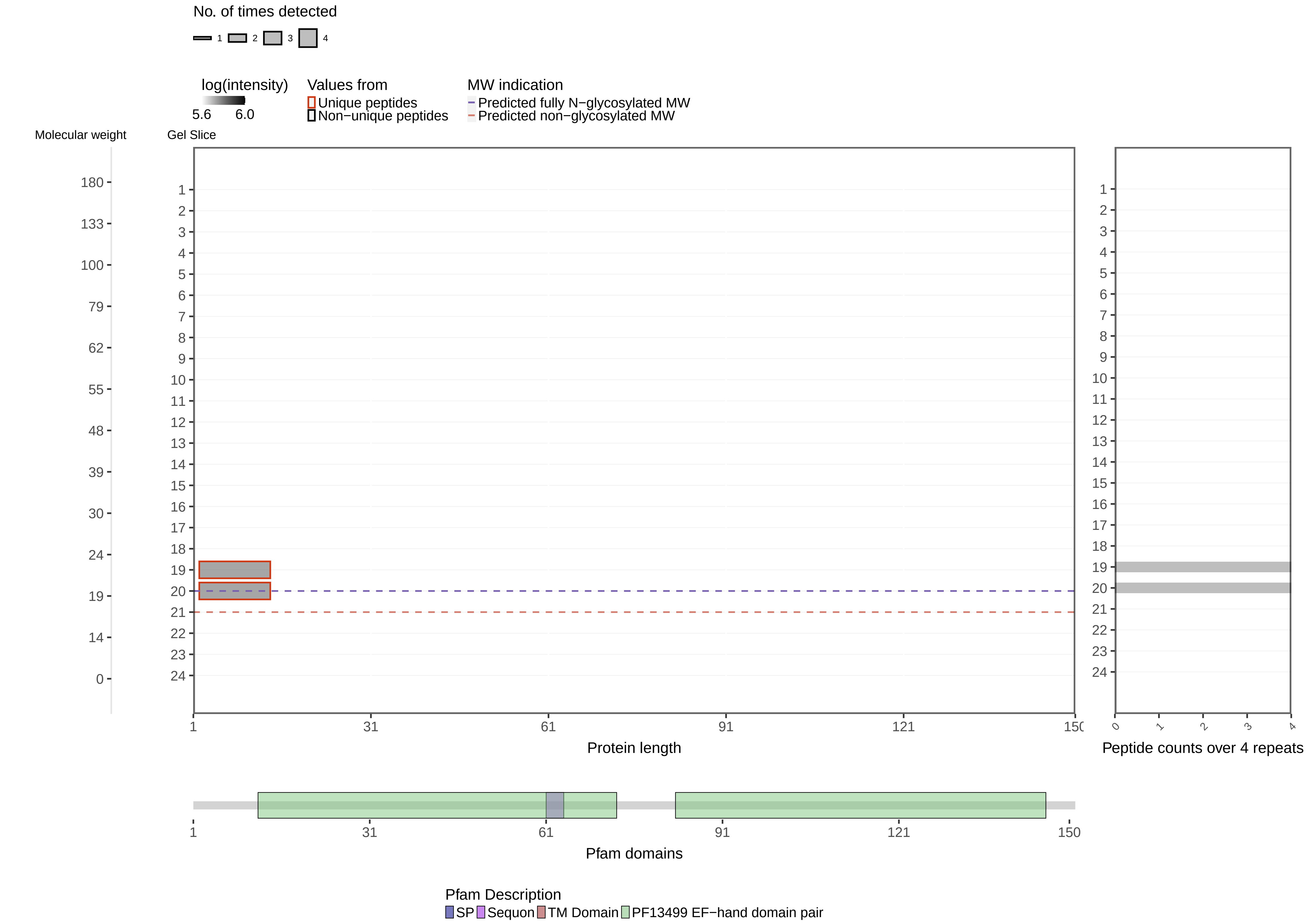


Protein sequence:

GTIDFPEFLNLMARKMKDTDSEEELKEAFRVFDKDQNGFISAAELRHVMTNLGEKLTDEEVDEMIREADVDGDGQINYEEFVKVMMAK*

### NbL09g20650.1 glycine−rich_RNA−binding_protein_2_mitochondrial−like [MW = 35.79 kDa]

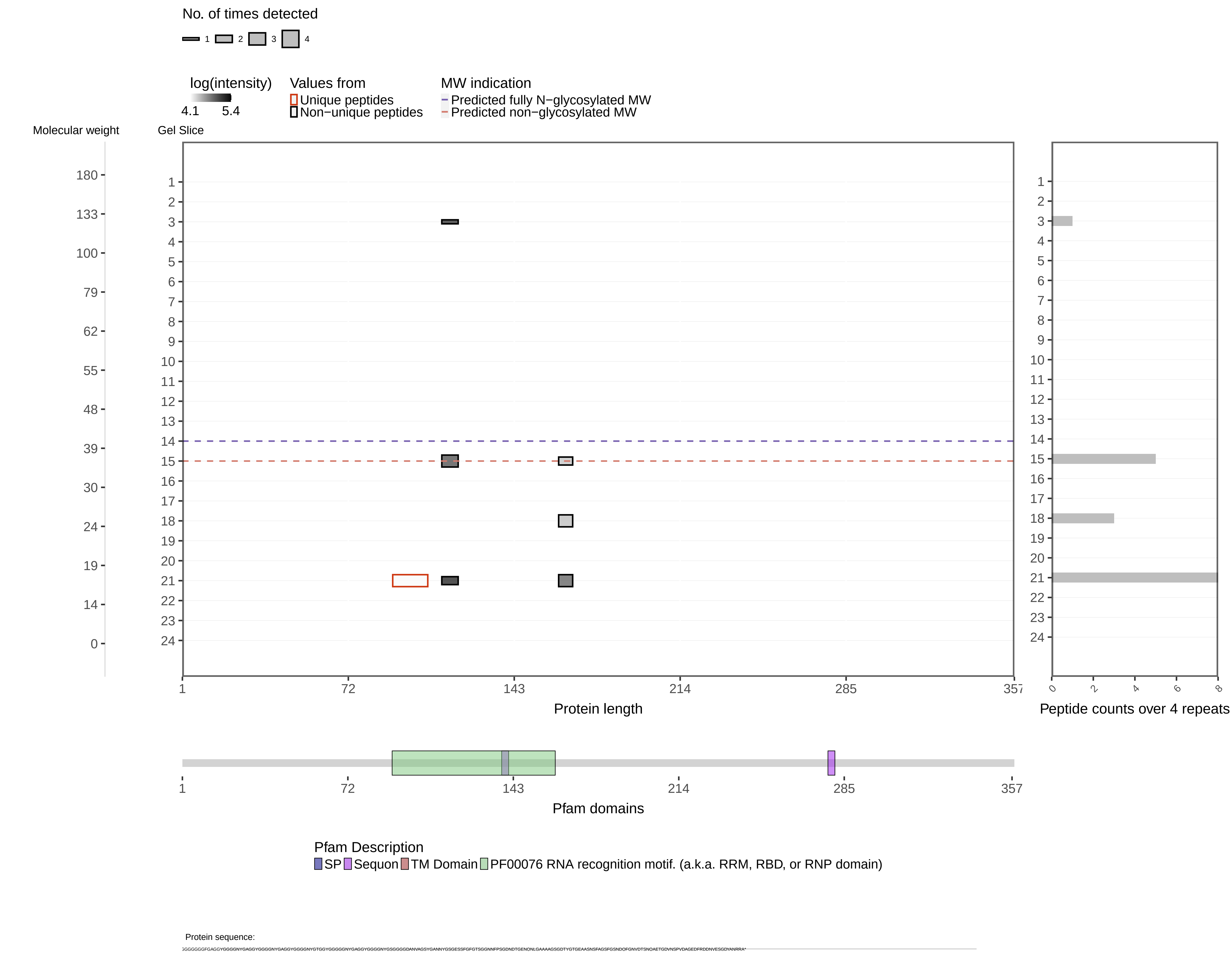

### NbL09g20620.1 protein_GrpE−like [MW = 42.71 kDa]

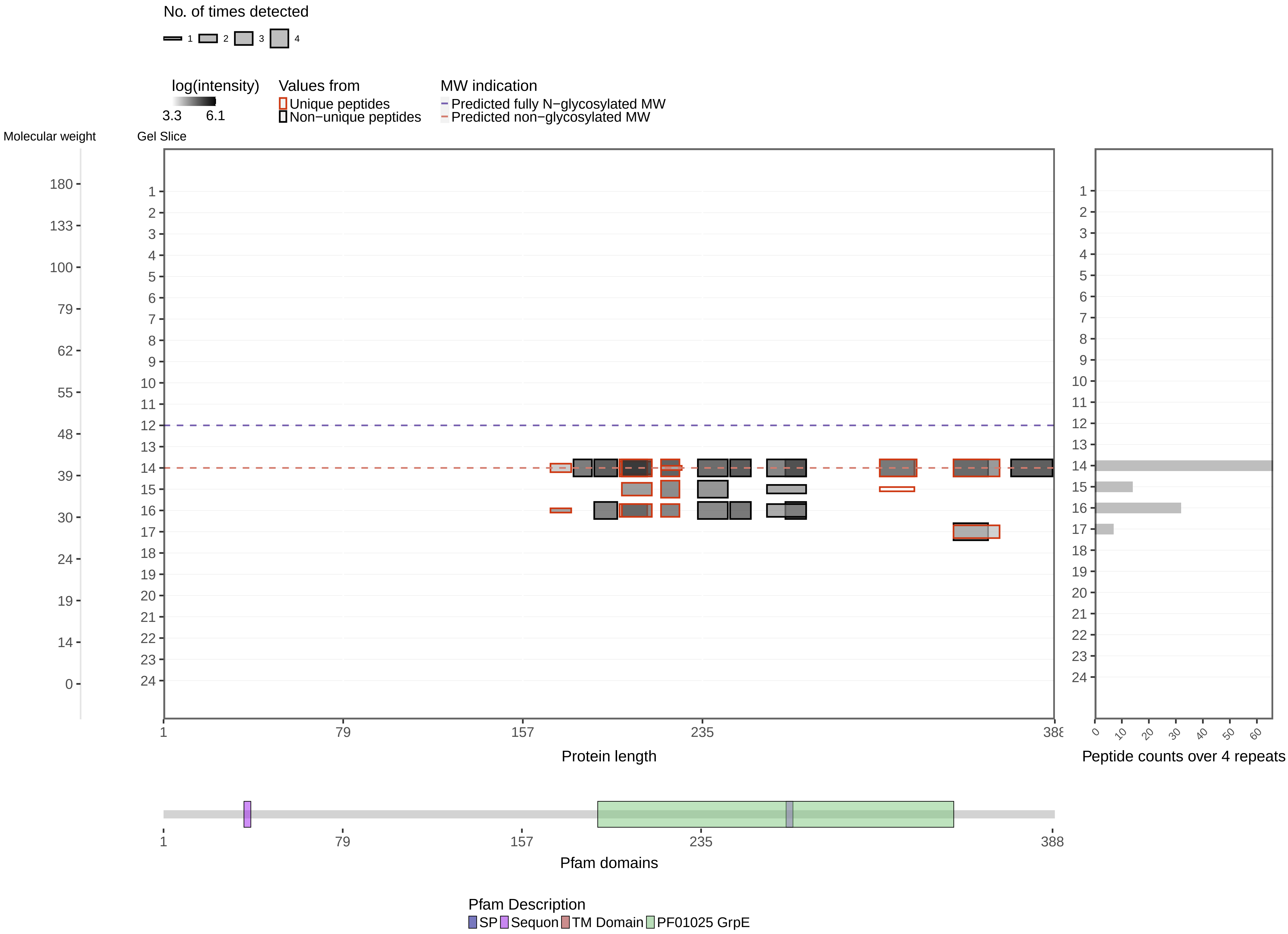


Protein sequence:

IEISLKSIENEKIELQRRVASLTEELSNERDRVLRVSADFDNFRKRTERERLSLVTNAQGEVVEKLLSVLDNFERAKMQIKVATEGEEKINNSYQSISKQFGEILGSLGVETVETVGKPFDPLLHEAIMREDSEEFEEGVVLEEYRKGFKLGDRLLRPSMVKVSAGPGPAKPETAEPKEEQNKVQEKSEEGTAETAGDEGTGEGGN*

NbL09g20510.1 14_kDa_zinc−binding_protein [MW = 17.62 kDa]

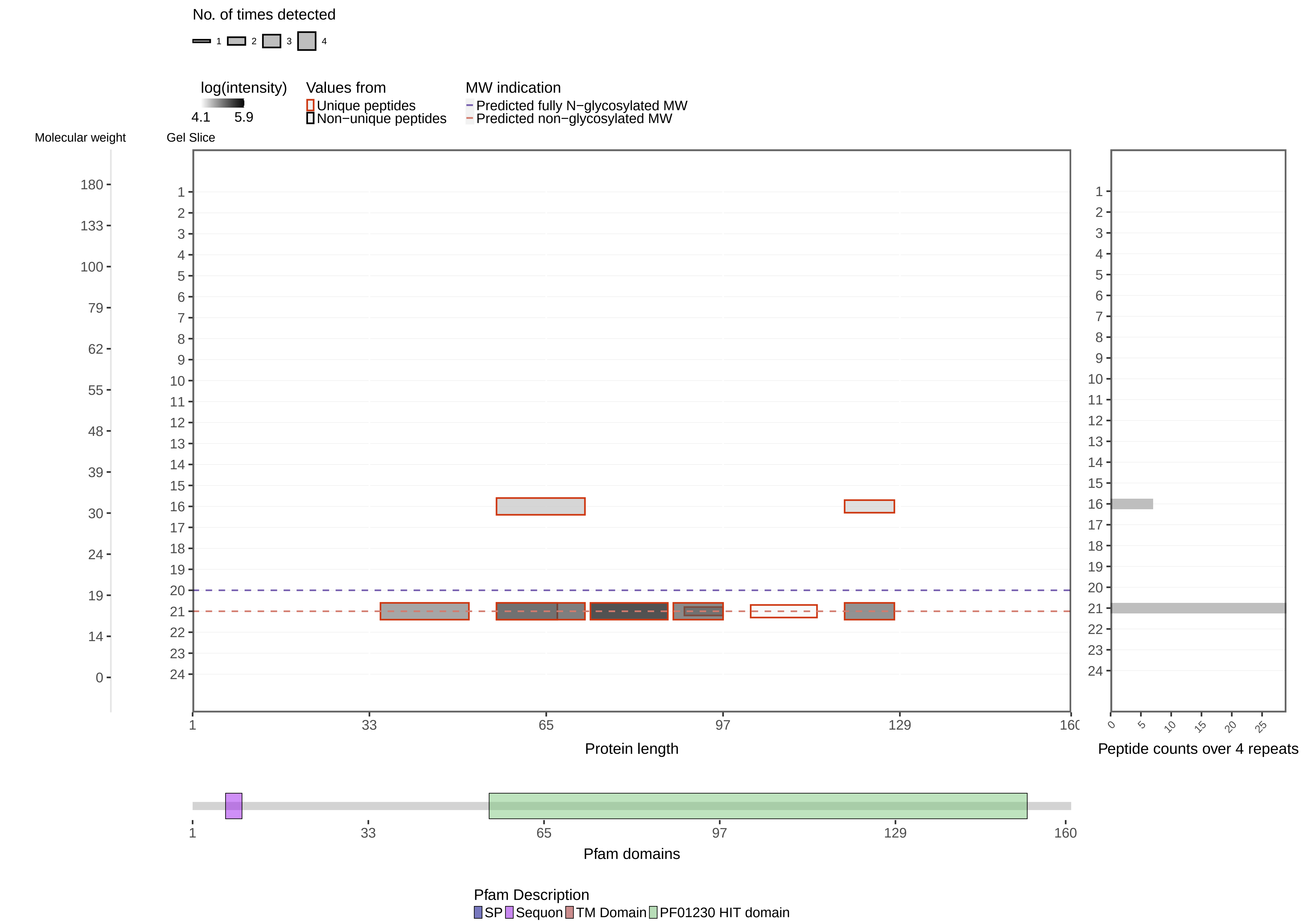


Protein sequence:

VLAFRDINPQAPVHILLIPKVRDGLTGLSKAEEKHCEILGQLLYTAKLVAKQEGLLENGFRLVINDGPSGCQSVYHLHLHLLGGRQMNWPPG*

### NbL09g20450.1 secoisolariciresinol_dehydrogenase−like [MW = 29.45 kDa]

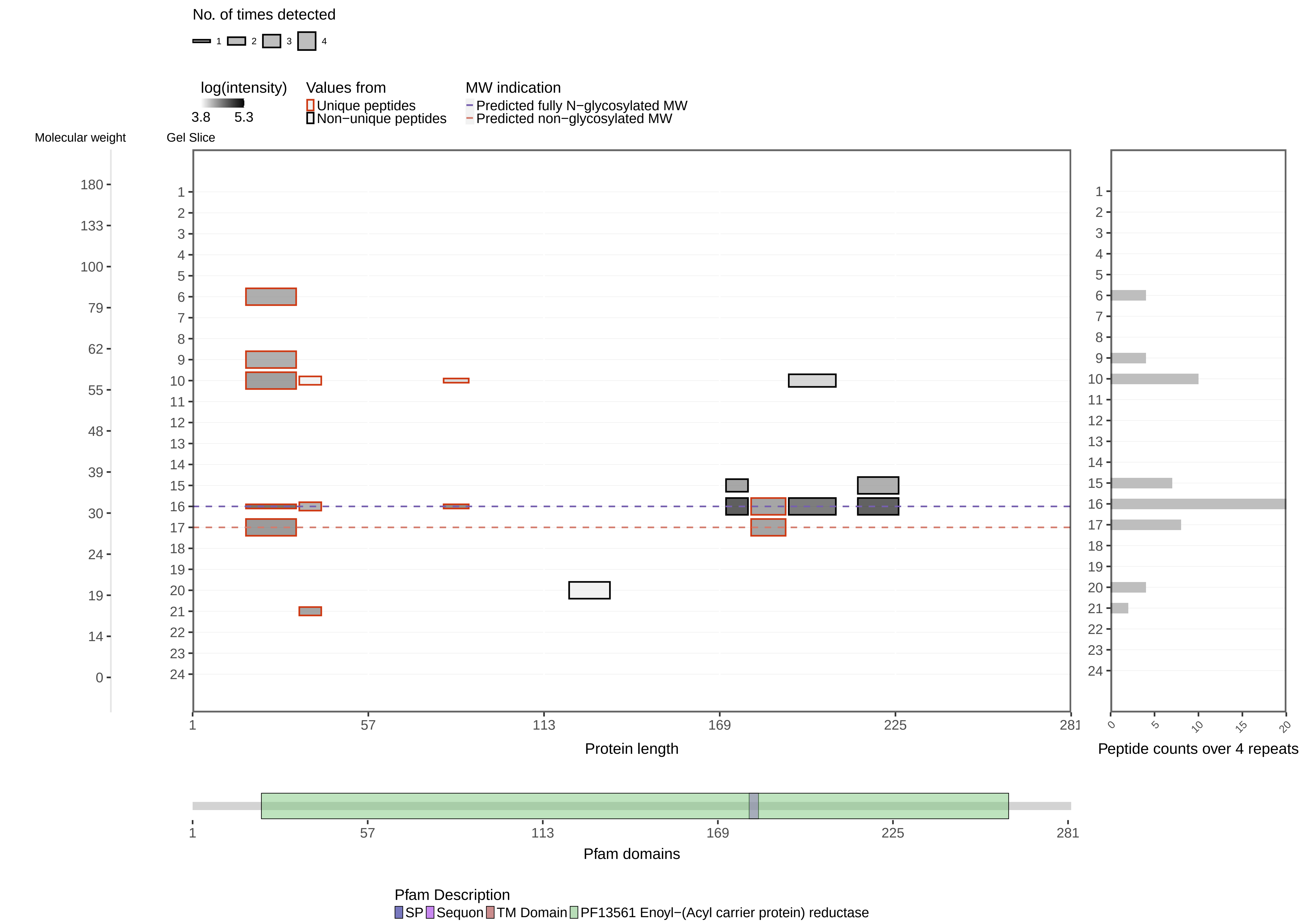


Protein sequence:

YGSFLGAKYAARVMIPAKKGVILFTSSAASVNAGEAPHSYTVSKHAVVGLTKNLSAELGQYGIRVNCVSPCAVATPLLVKAMGVDKSVVDGIICSSANLKGAVPTAEDVAEAALYLASDESKFVSGLNLVIDGGYSTTNQAYMKTIQSVLFSSV*

### NbL09g20210.1 sucrose−phosphatase_2−like [MW = 47.99 kDa]

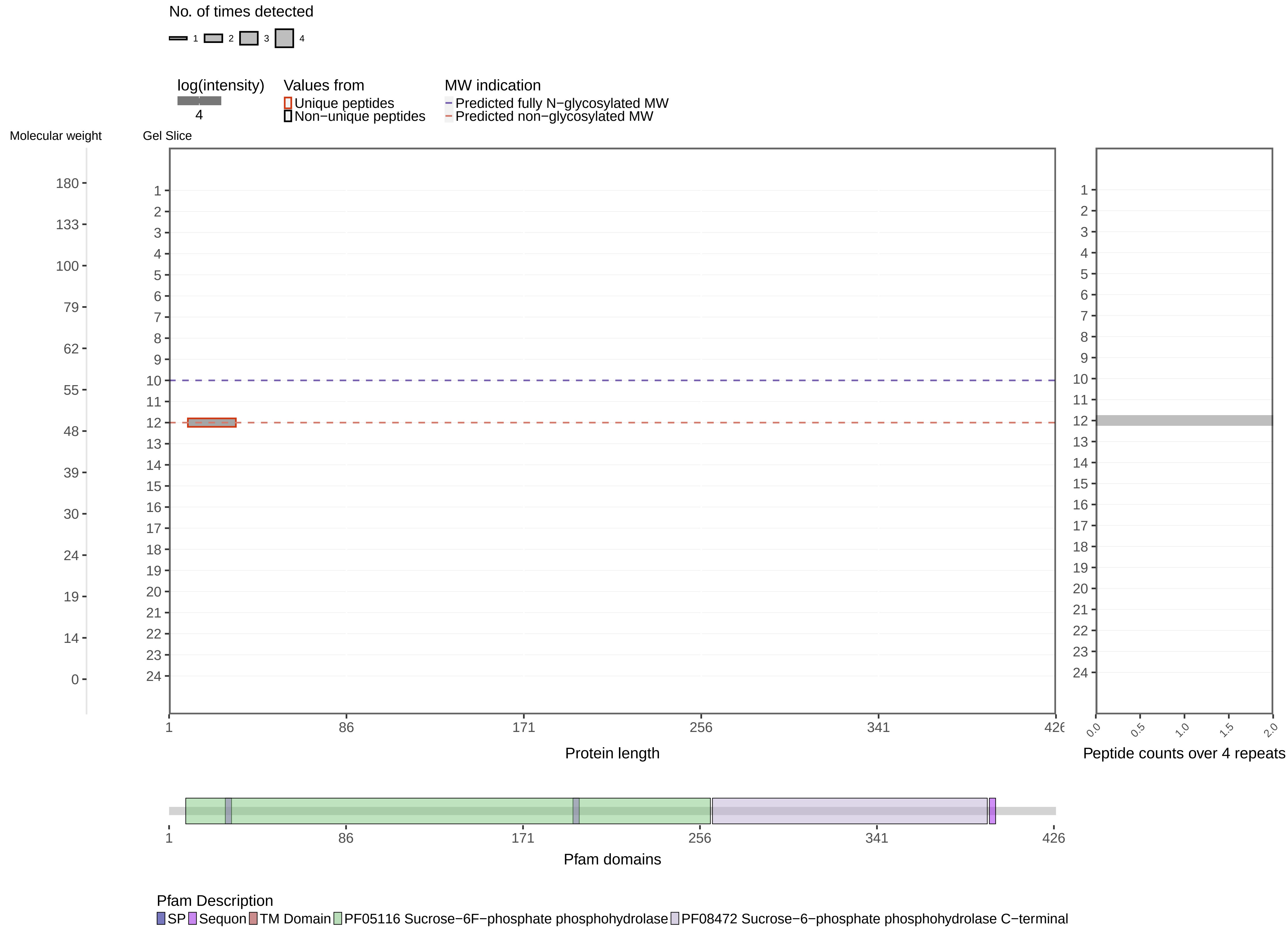


Protein sequence:

DSGNDAELFSIPDVYGVMVANAQEELLQWHAANAKNNPKVIHATERCAAGIIQAIGHFNLGPSTSPRDVMDLSDCKMENFVPAYEVVKFYLFFEKWRRGEIEHSEHYLSNLKAVCRPSGTFVHPFGVEKSLQECVTLFRTCHGDKQGKQFRIWVDQVLPVQVGSDSWLVSFKKWELSGEDRHCCITTVLLSSKNKTVADGLTWTHVHQTWLNGAAASDSTTWFF*

NbL09g20090.1 protein_WVD2−like_4 [MW = 48.52 kDa]

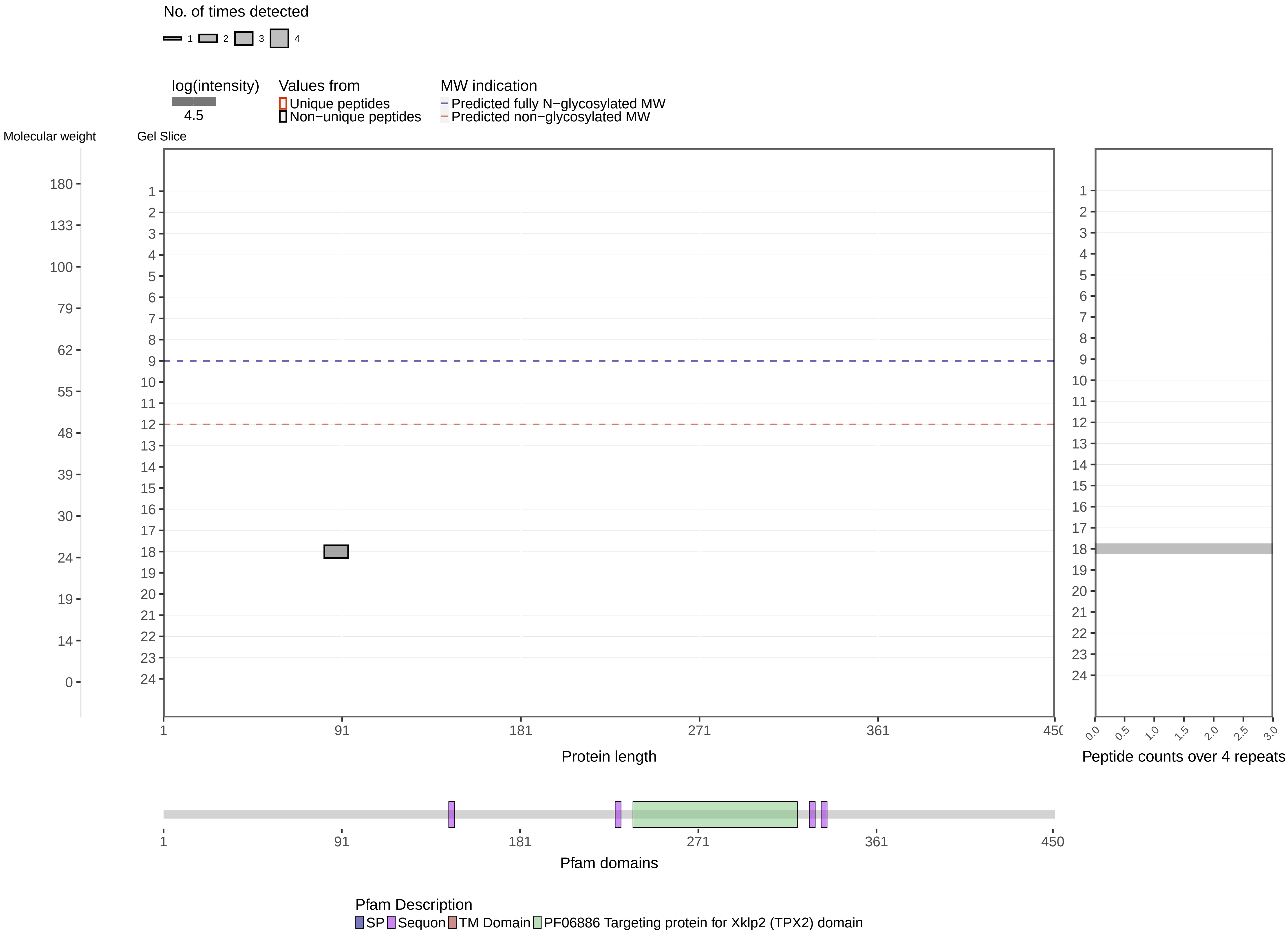


Protein sequence:

ATSPLKDDEDARSATSSNTTRTSTGGFSFRLEERAEKRKEFLAKMEERIQAREEEKNNLLAKSKENQEEEVKQFRKSLTFKATPMPNFYKEPPPKVELKKIPITRAISPKLGRNKNSTSATNGSESGGSSFSPKVIKDRSKSPRTSKDTIATKGKPVEIKKPVKNSHTISQSPKISTTKGKSAETKLKPAEAKGSDEKTEEMDIQPDSVPKPEMNSVEDNVPRPSNPVVVQAEVTVEV*

NbL09g20000.1 ubiquitin_carboxyl−terminal_hydrolase_12 [MW = 131.5 kDa]

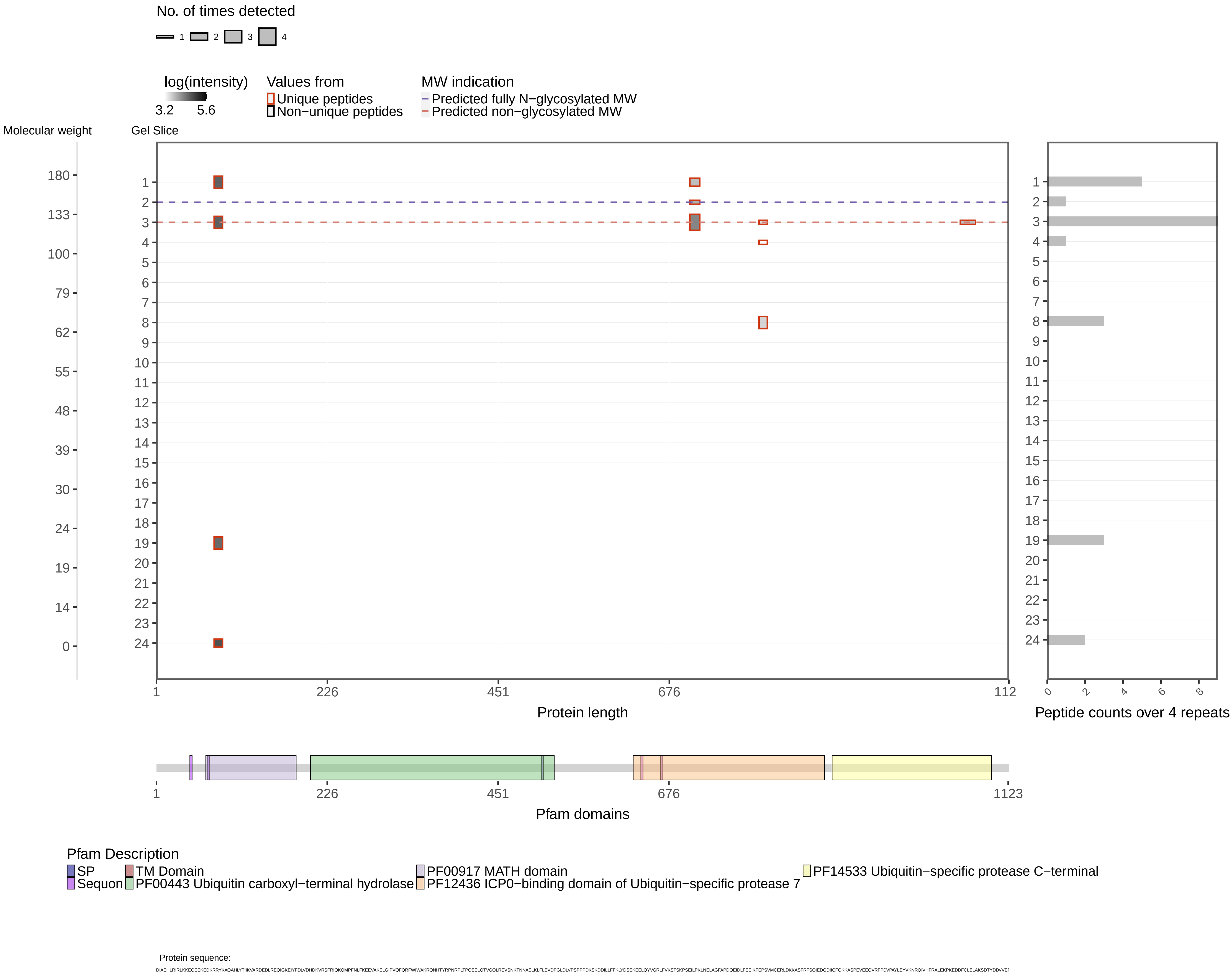

NbL09g19970.1 syntaxin−132−like [MW = 35.38 kDa]

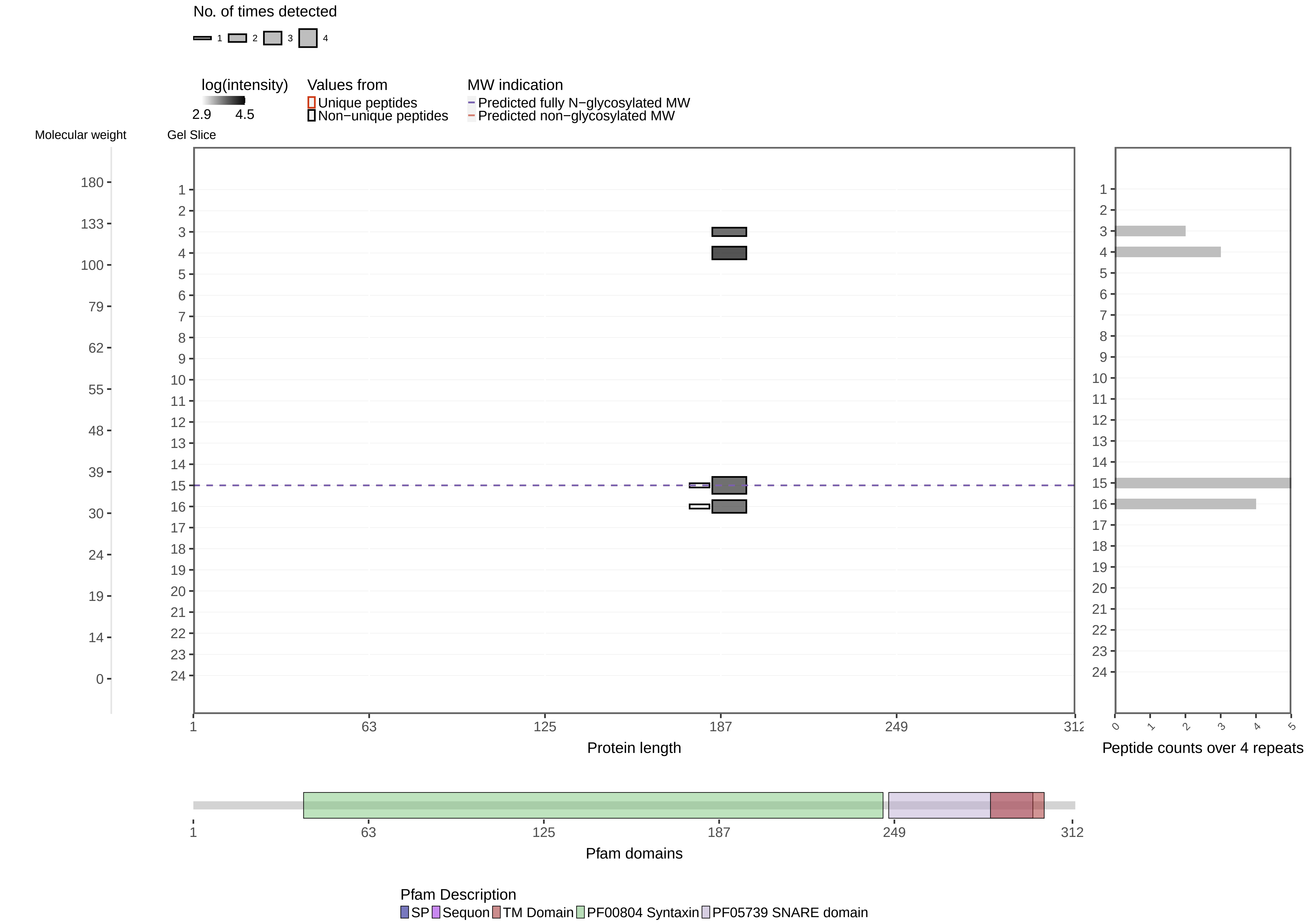


Protein sequence:

KFKDKMAEFQTLRENIQHEYREVVERRVFTVTGNRADEETIDRLIETGDSEQIFQKAIQQQGRGQIMGTLAEIQERHDAVRELERKLLELQQIFLDIAVLVDAQGDMLDNIESQVSTAVDHVQSGNTALQKAKKLQRNSRKWMCFAIIILLIIVAIIVVGVIKPWNSNKGA*

### NbL09g19950.1 tropinone_reductase_1−like [MW = 29.95 kDa]

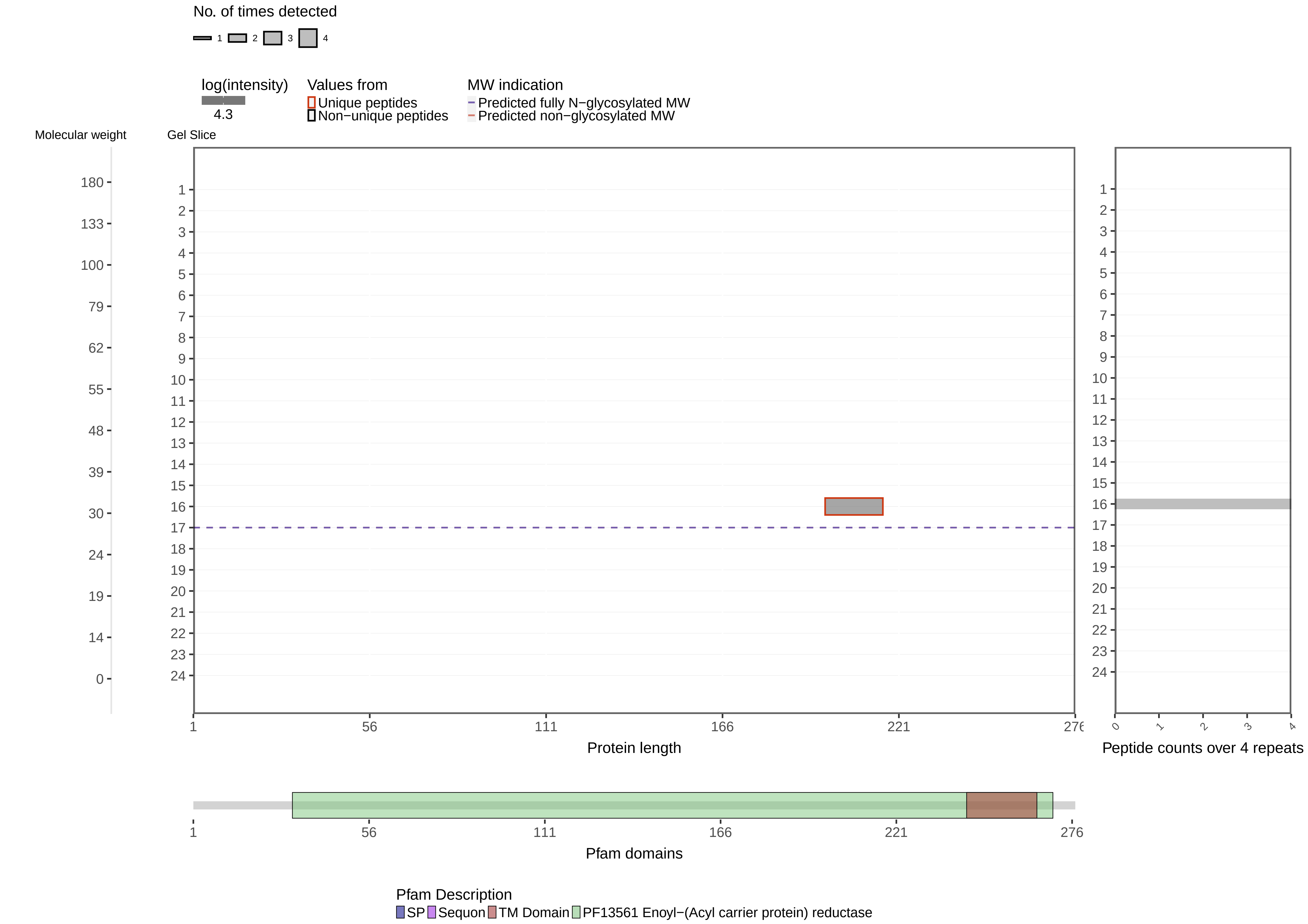


Protein sequence:

EDHDIVMGTNFEAGFHLSQLAYPLLKASENGNVIFLSSIAGFSALPSLSLYSASKGAINQMTKSLACEWAKDNIRVNSVAPGVIMTPLVETAIKKNPHQKKDIDHFIAKTPMRRTGKPEEISALMAFLCFPAASYITGQIIWVDGGFTANGGF*

NbL09g19720.1 protein_ABHD11−like [MW = 37.15 kDa]

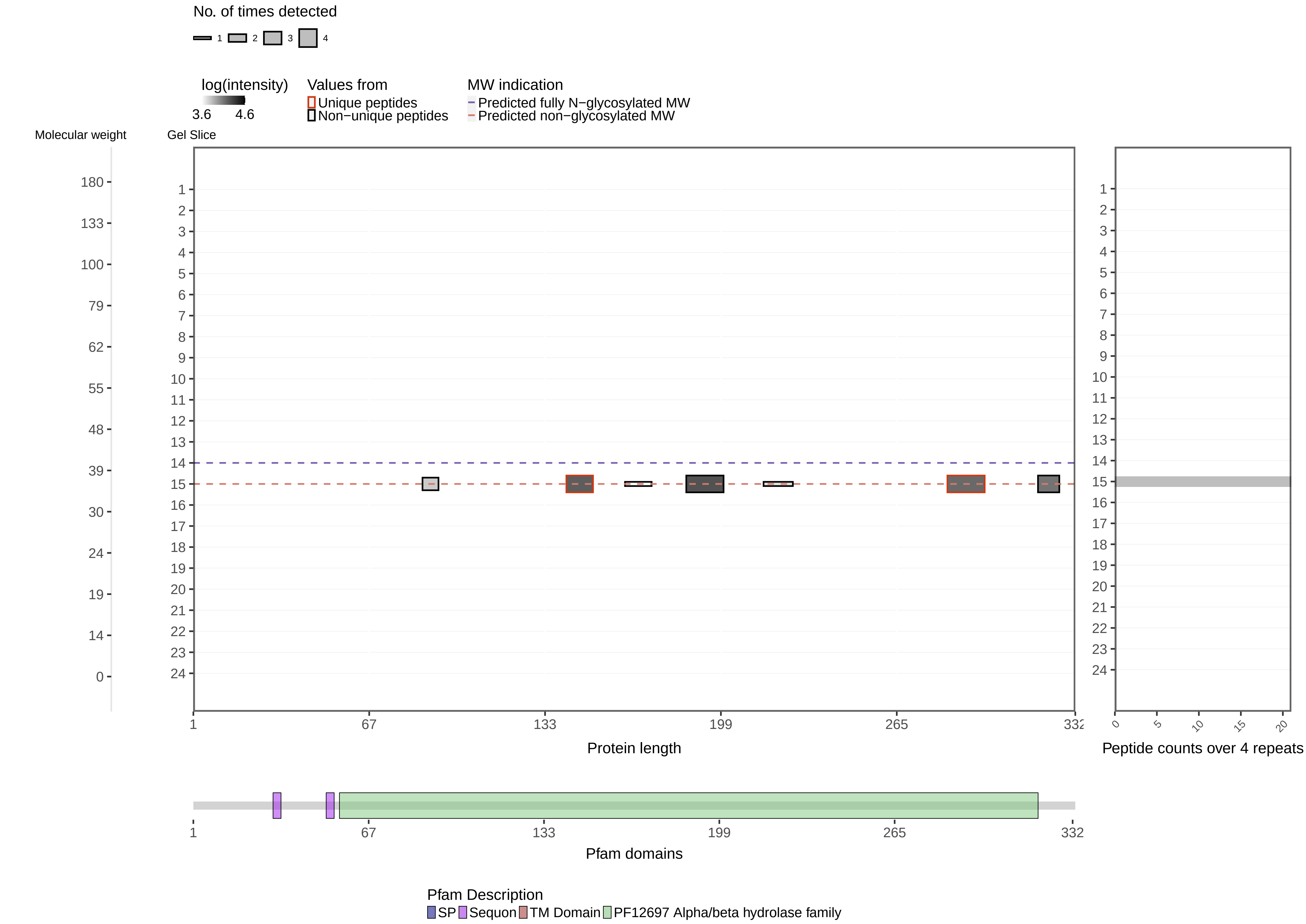


Protein sequence:

'YGOSARLPKOLWVLDSVPGKVDREDSSREVEKVMOTLOSLPSSIPSRKWLMDOMLKLGFSKALSEWLGSNLOKSGDOMKWTFNIEAAIEMFNSYREKDYCPLLEHPPKGTEIAIVRAEKSDRWDPVTVOKLESLASNGGGESEGKMSYHILPNSGHWVHVENPRGLLEIITPKLASLV*

### NbL09g19700.1 beta−fructofuranosidase_insoluble_isoenzyme_1−like [MW − SP = 63.54 kDa]

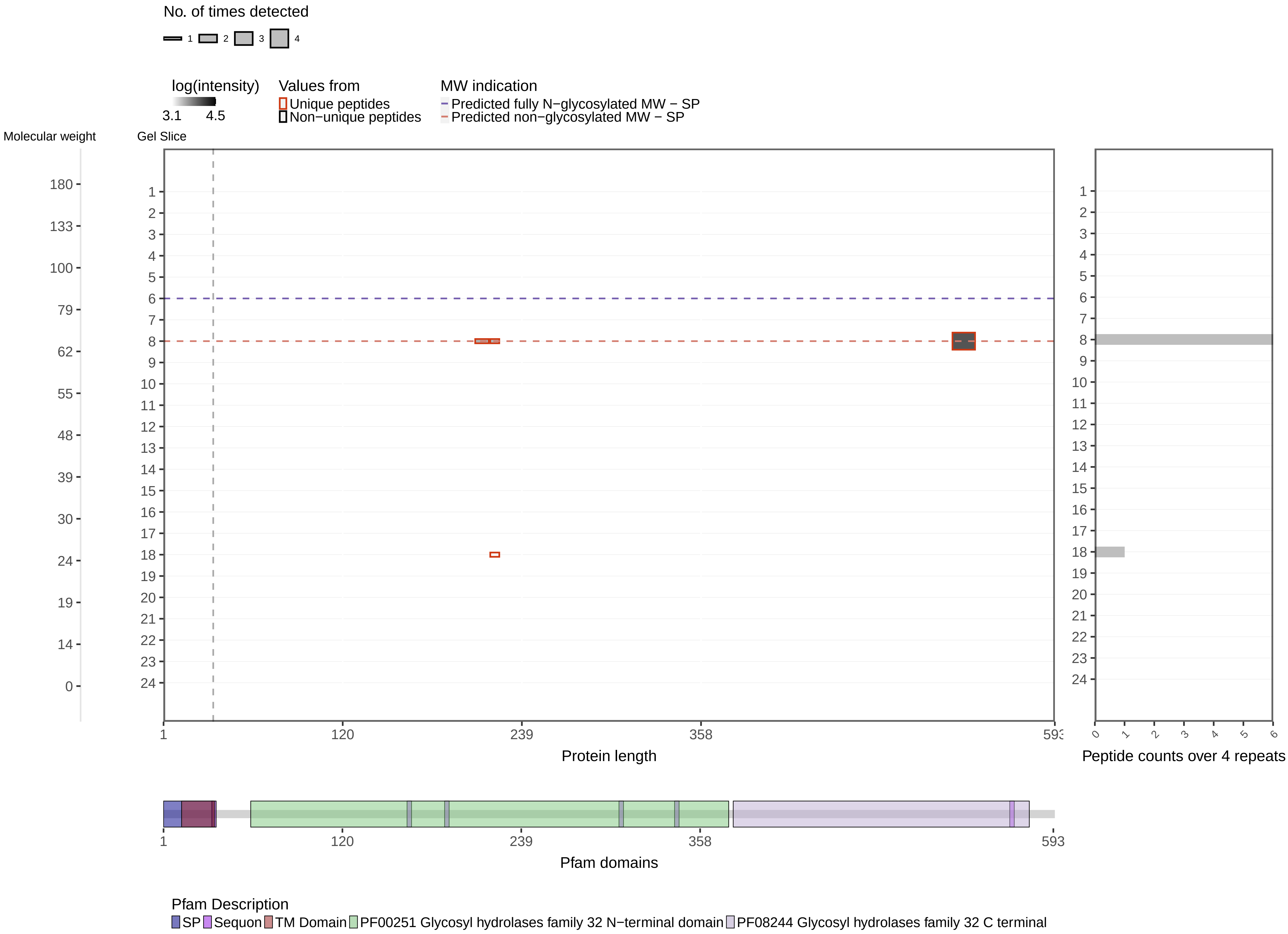


Protein sequence:

RFEYYTVGTYDIKKDRYIPDNTSIDGWKGLRLDYGNYYASKSFYDPCKKRRIIWGWANESDTVDDDVRKGWAGIQTIPRKLWLDPSGKQLVQWPVKELDTLREQKVQLKNHKLKKGENIEVKGITPAQADVEVTFAFSSLDKAEPFDLSWTDLYAQDVCAIKGSTVQGGIGPFGLLLLASEKLEEYTPVFFRVFKAQNKYKVLMCSDASRSSLENDKIMYKPSFAGYVDVDLAKNKKLSLRSLIDHSVVESFGAGGKTCITSRVYPTLAIYDNAHLFAFNNGTETIKIELKAWSM

### NbL09g19500.1 enolase [MW = 27.92 kDa]

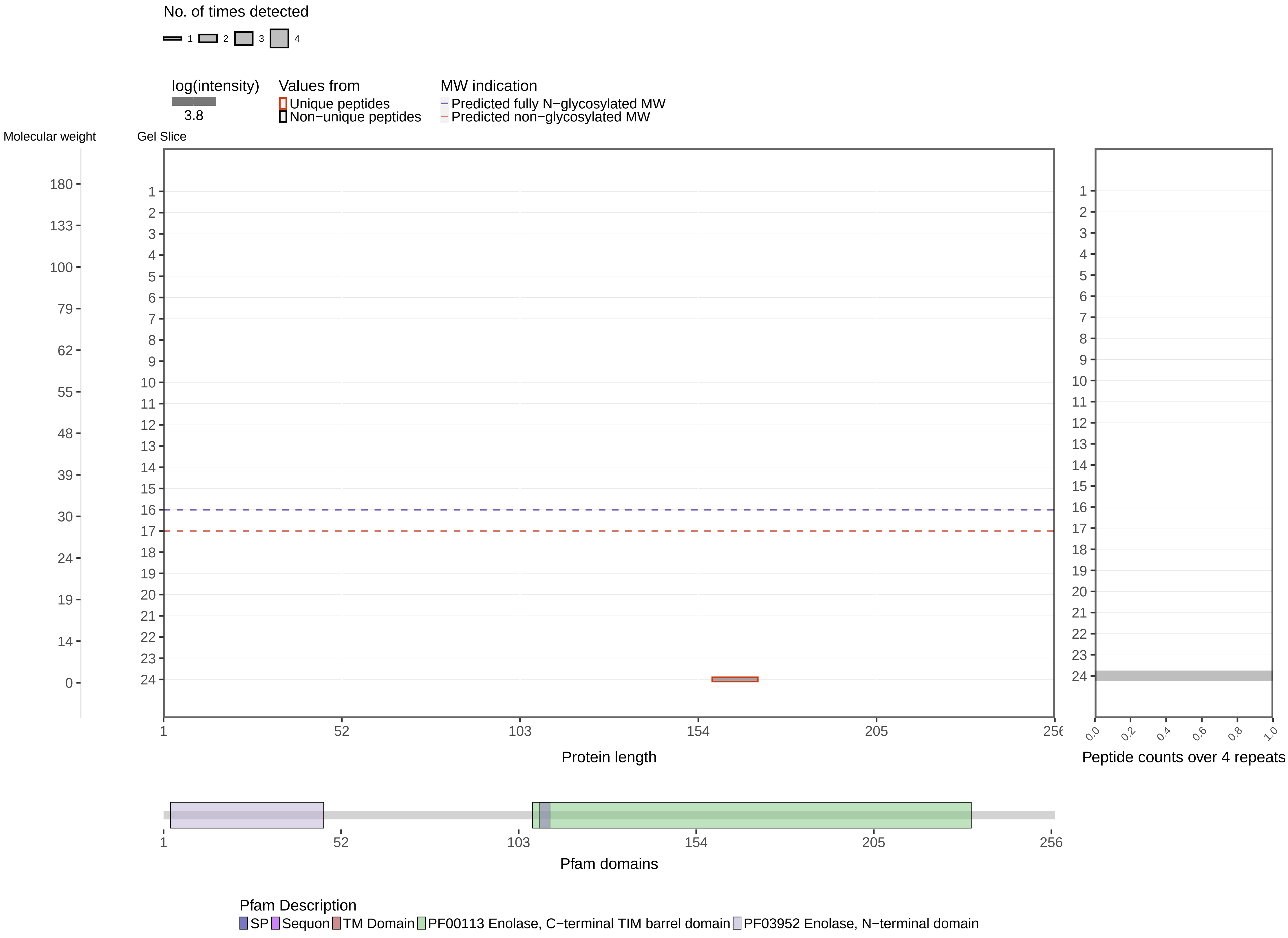


Protein sequence:

KISGDQLKDLYKSFVSEYPIVSIEDPFDQDDWETYAKLTAEIGEKVQIVGDDLLVTNPKRVAKAIEGMTCNALLLKVNQIGSVTESIEAVKMSKRAGWGLMTSHRSGETEDTFIADLAVVYQLDKSRLELPVGQSVSQSIARF*

NbL09g19490.1 peroxiredoxin−2E−2_chloroplastic−like [MW = 28.71 kDa]

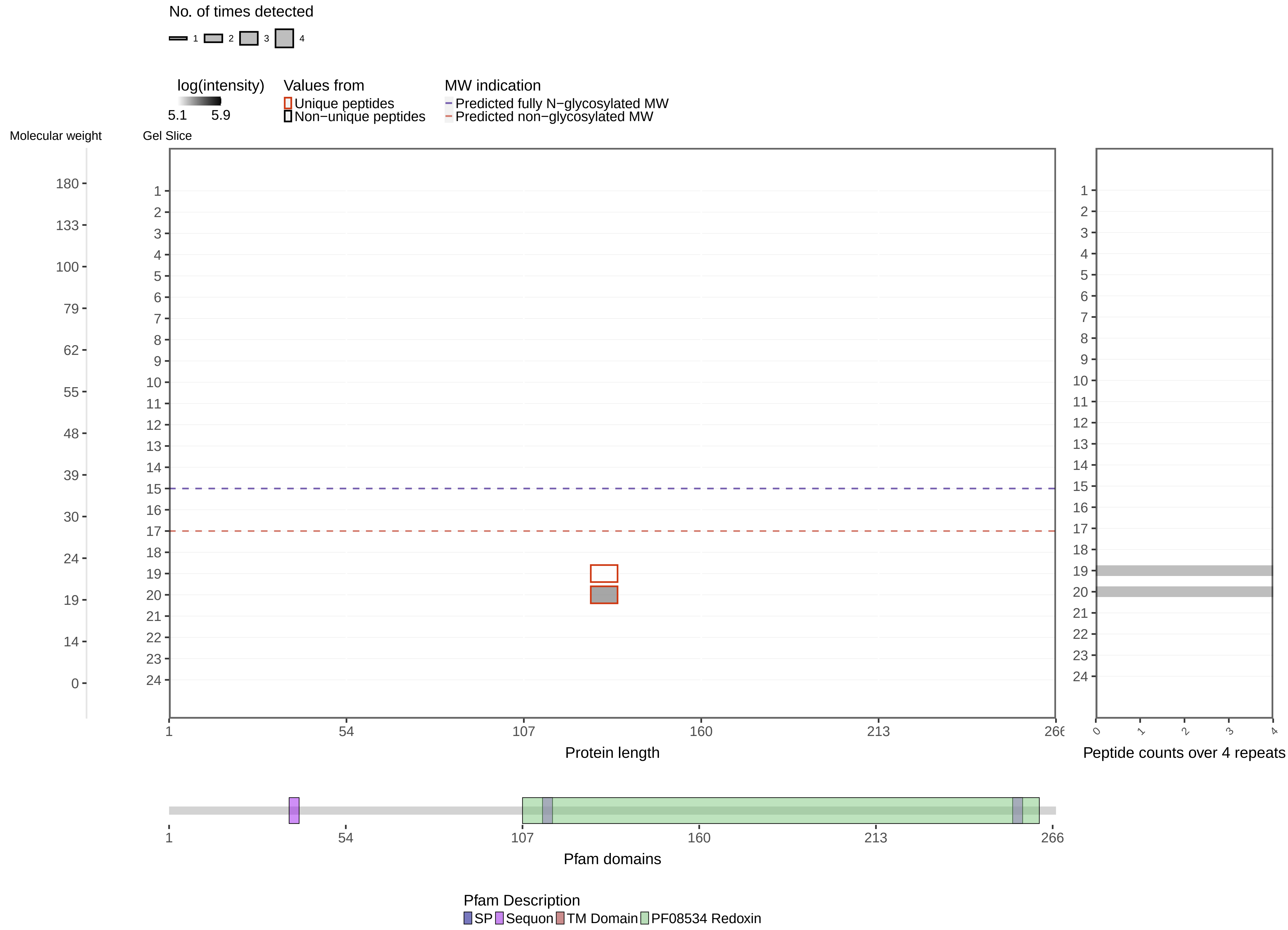


Protein sequence:
SDELKTLSISDLTKGKKVVLFAVPGAFTPTCSQKHLPGFVEKSKELKSKGVDTIACISVNDAFVMKFWKENLKINDEVLLLSDGNLEFTKAIGCELDLSDKPIGLGVRSRRYAMLVDDGVVKVLNLEEGGAFNVSSAEDILKTL*

### NbL09g19470.1 pyruvate_kinase_cytosolic_isozyme−like [MW = 63.09 kDa]

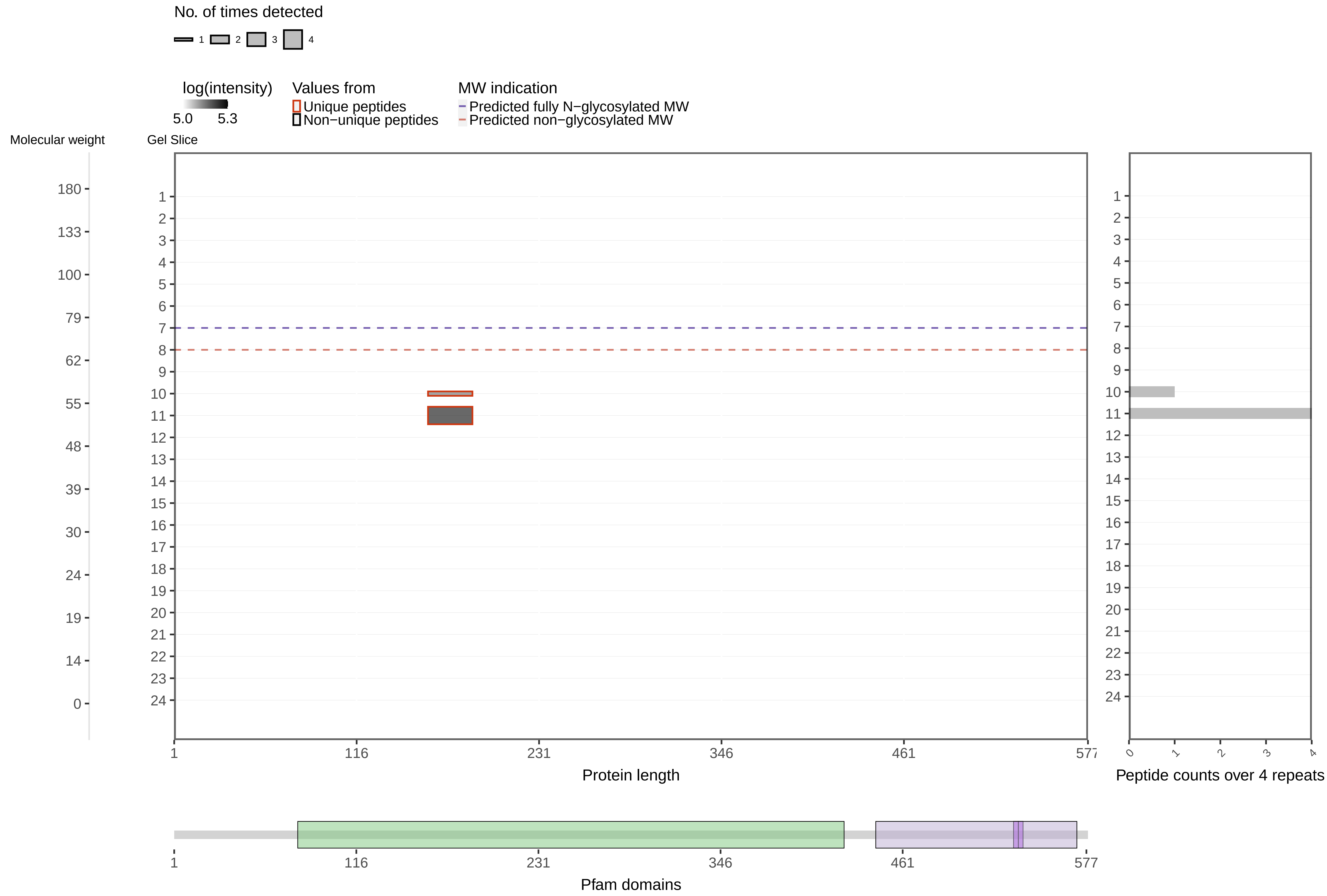


Protein sequence:

.SYTRHAEDVREAREFLSKLGDLSOTOIFAKIENVEGLTHFDEILKEADGIILSRGNLGIDLPPEKVFLFOKAAVHKCNMAGKPAVVTRVVDSMTDNLRPTRAEATDVANAVLDGTDAILLGAETLRGLYPVETISTVGRICFEAEKVFNODLYFKKTVKFVGEPMTHLESIASSAVRAAIKVKASVIICFTSSGRAARLIAKYRPTMPVLSVVIPRLKTNOLKWSFSGAFEAROSLIVRGLFPMLADPRHPAESTNASNESVLKVALDHGKASGVIKSHDRVVICOKVGDASVVKIIELI

### NbL09g19410.1 threonine_dehydratase_biosynthetic_chloroplastic−like [MW = 66.37 kDa]

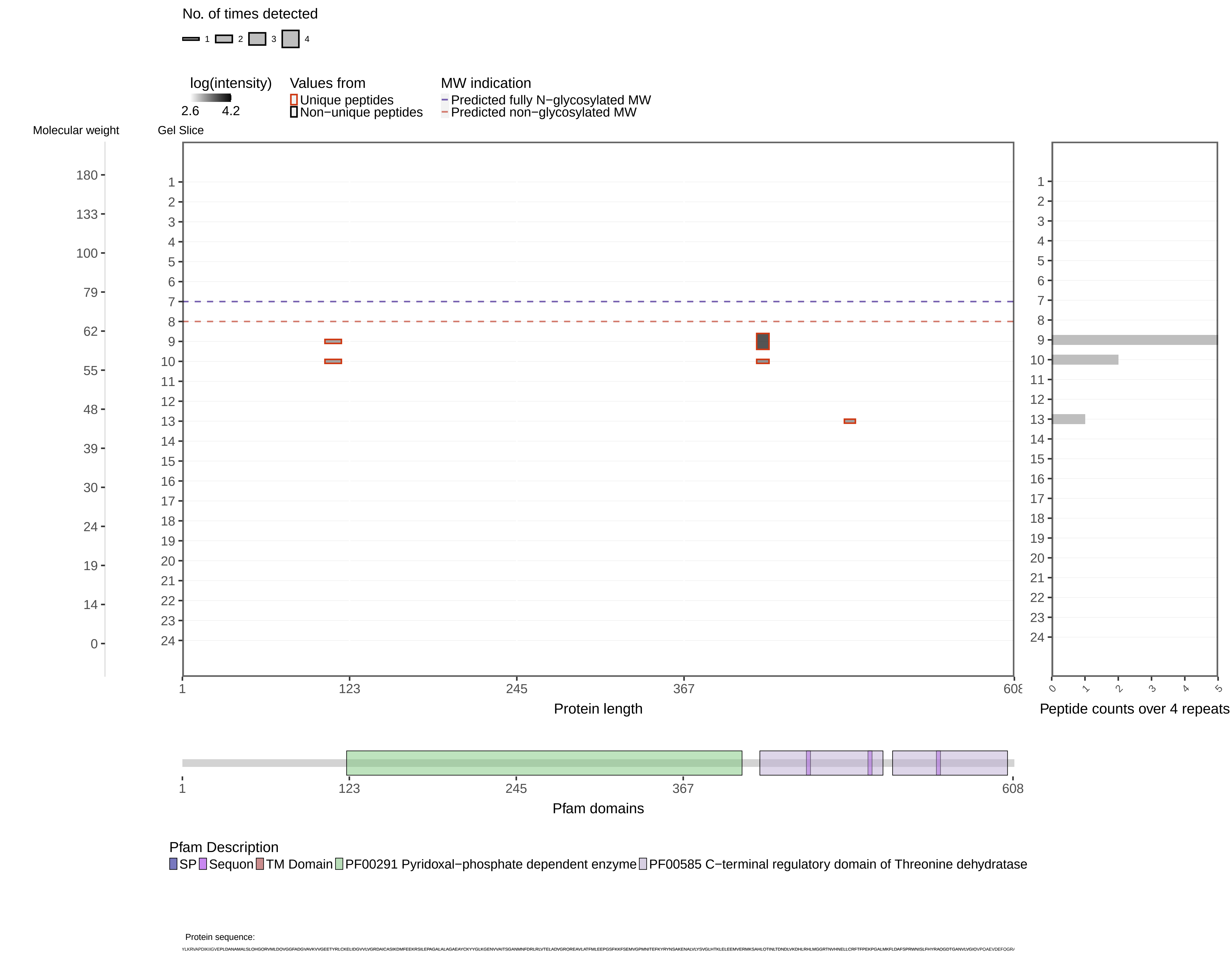

NbL09g19390.1 protein_fluG [MW = 93.43 kDa]

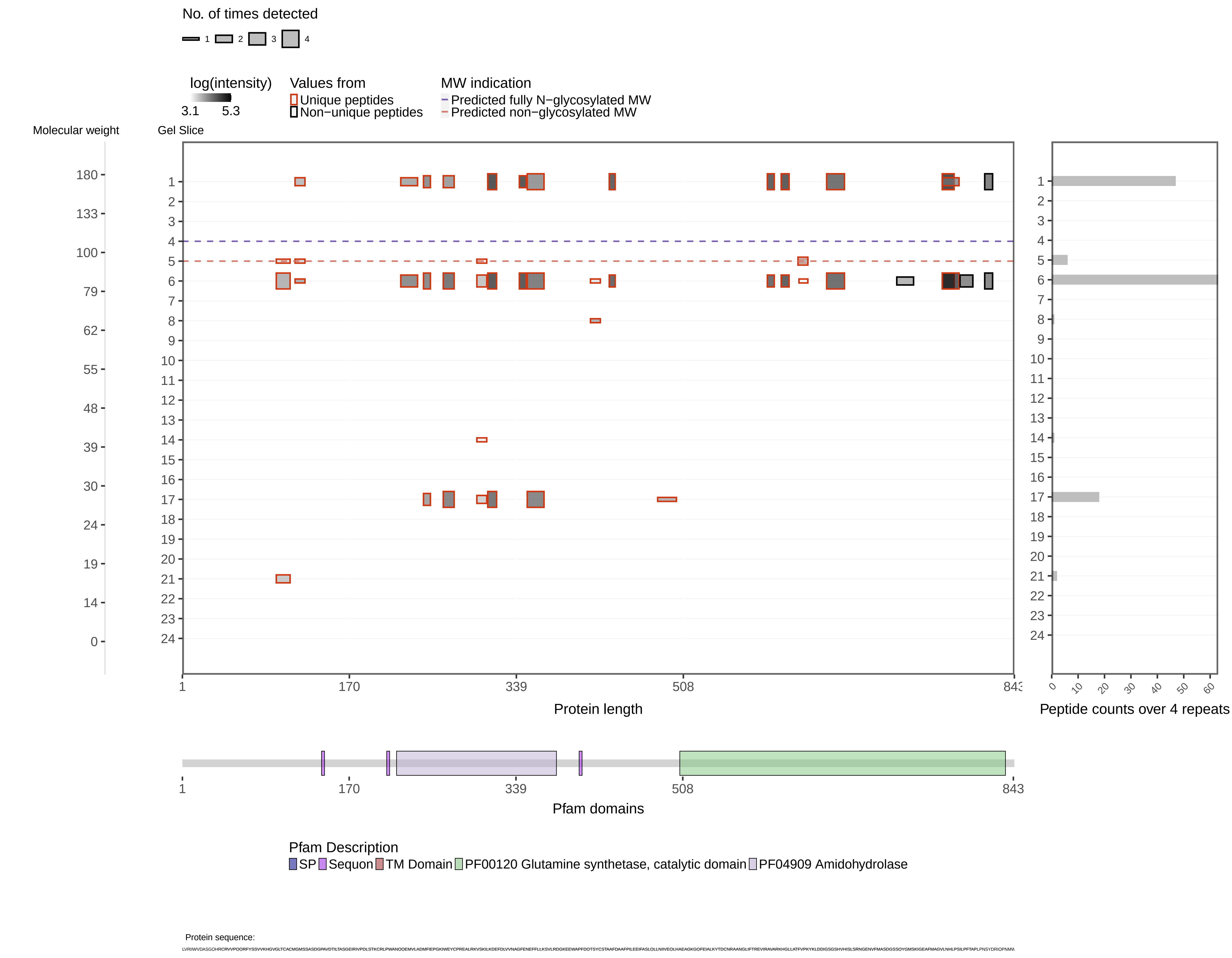

### NbL09g19180.1 synthase [MW = 34.22 kDa]

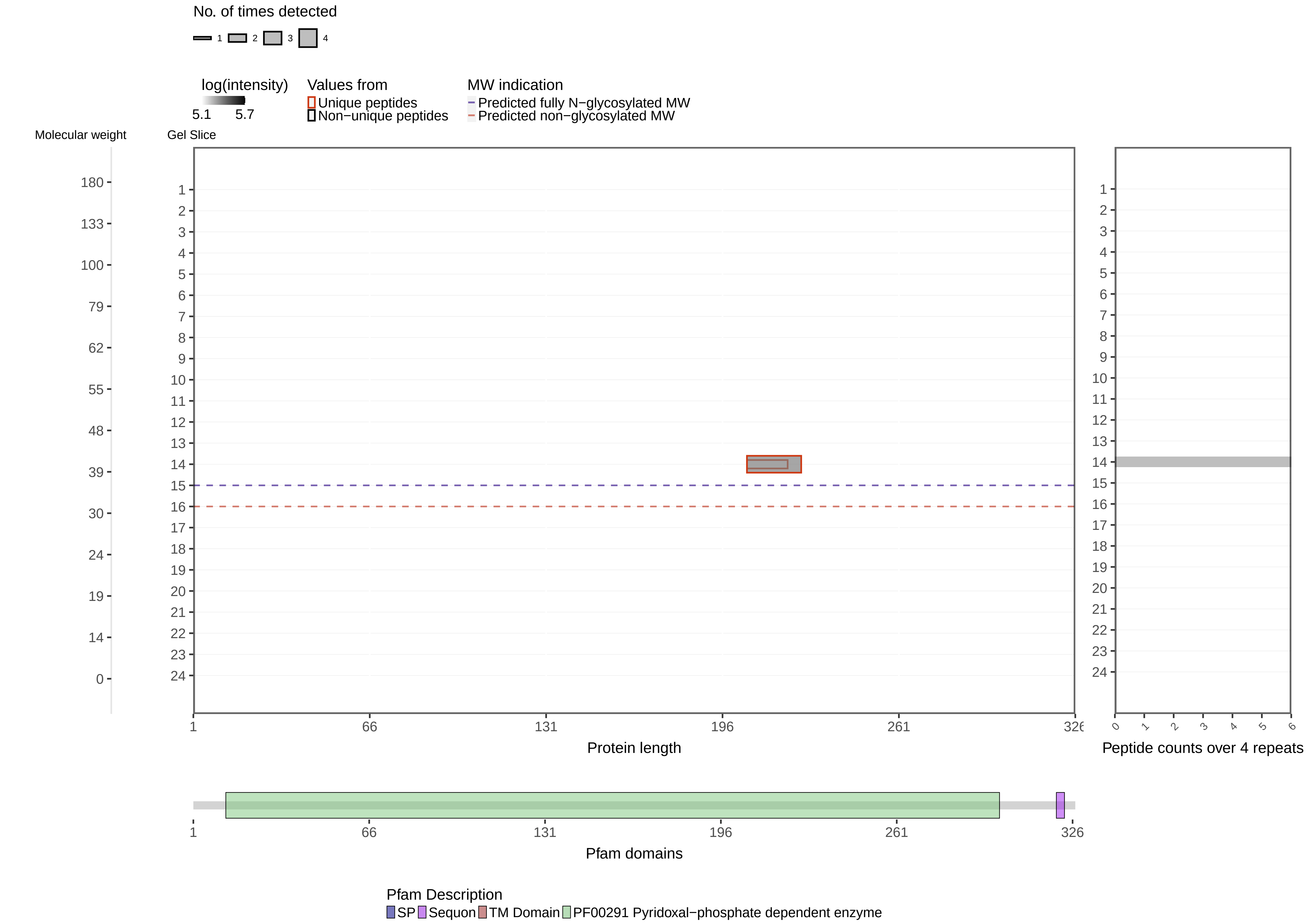


Protein sequence:

[illegible]FENPANPKVHYETTGPEIWKGSNGKVDALVSGIGTGGTITGSGKYLREQNPDIKLYGVEPVESAILSGGKPGPHKIQGIGAGFIPGVLEVGLIDEVIQVSSEEAIETAKLLALKEGLLVGISSGASAAAAIKLAKRPENAGKLIVVVFPSFGERYLSSVLFESVRREAENMTVEP*

NbL09g19060.1 proteasome_subunit_beta_type−2−A [MW = 22.6 kDa]

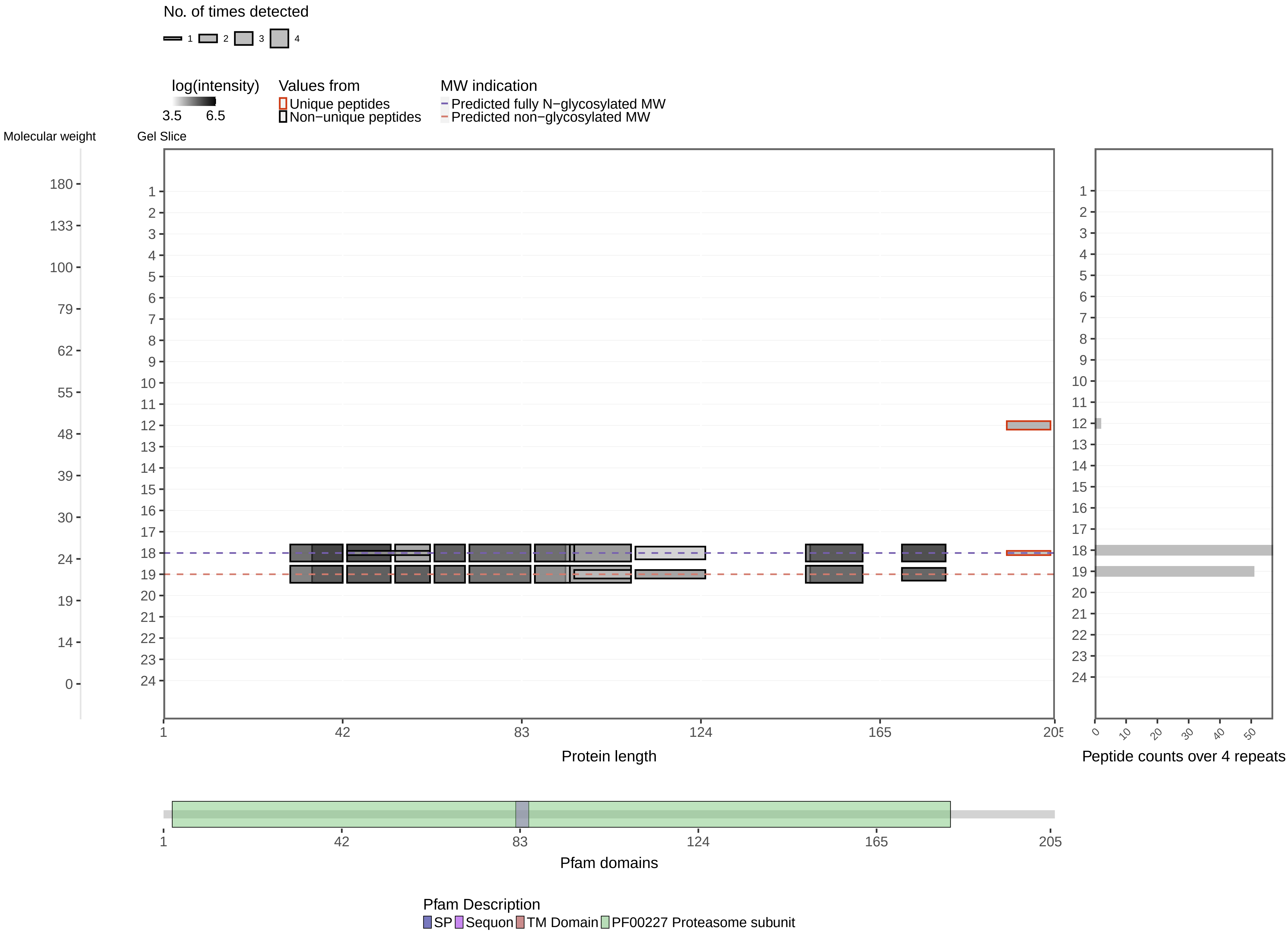


Protein sequence:

ATALRKNPYMVNIILAGYDKETGPSLYYVDYIATLHKVDKAAFGYGSYFSLAMMDRHYRKDMTVEEAVDLADKCIMEIRSRLVVAPPNFVIKIVDKDGAREYAWRQSVKDAPVSNS*

NbL09g18860.1 12−dihydroxy−3−keto−5−methylthiopentene_dioxygenase_2 [MW = 25.62 kDa]

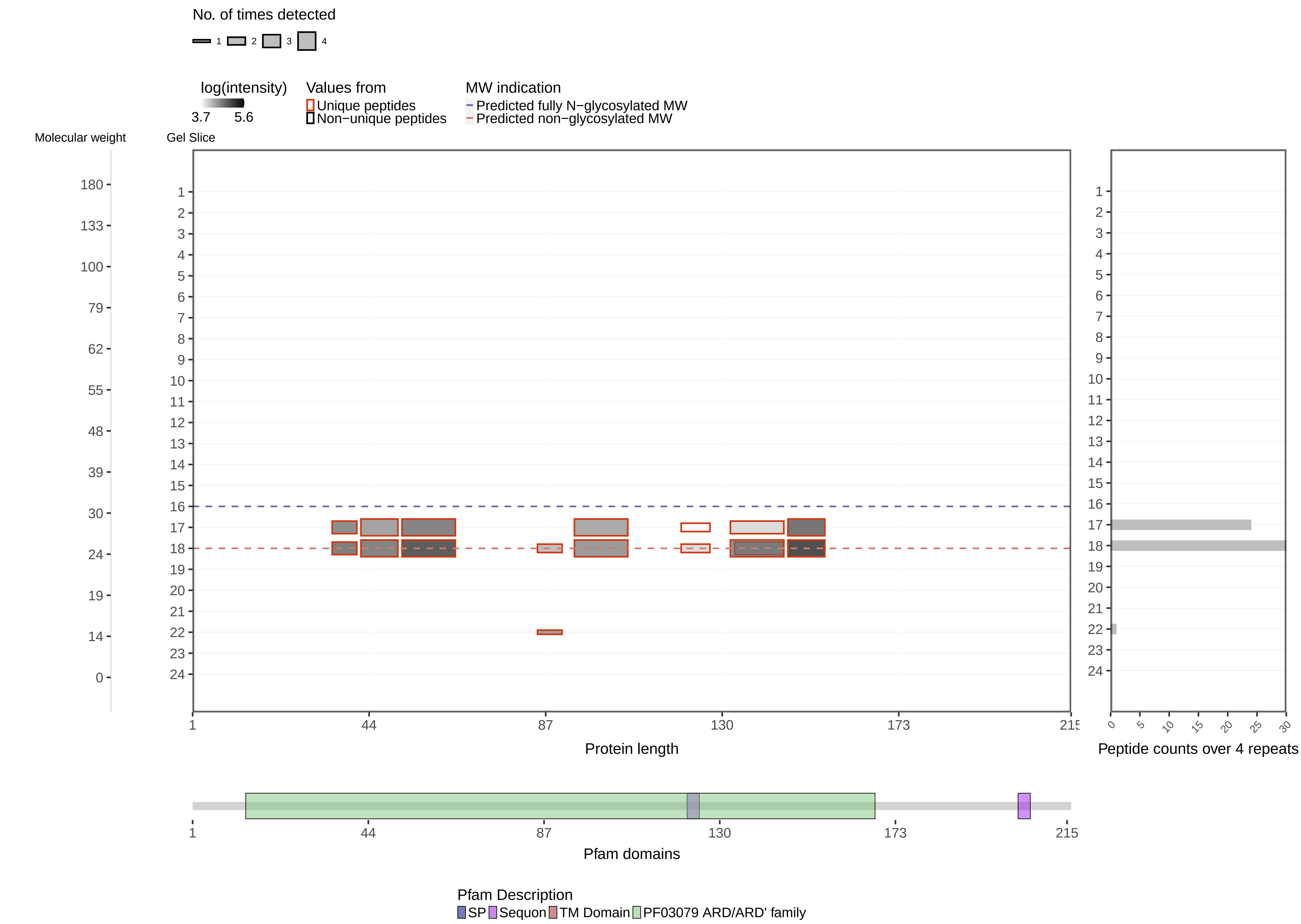


Protein sequence:

NFFEEHLHTDEEIRYCVAGSGYFDLRDWNDTWIRVWVKKGGMIVLPAGIYHRFTLDSDNYIKAMRLFVGDPIWTPYNRPHDHLPARSVSRHLLYSIFTLDNQKWSWVEANESFLIMQERIY*

### NbL09g18800.1 stem−specific_protein_TSJT1−like [MW = 27.29 kDa]

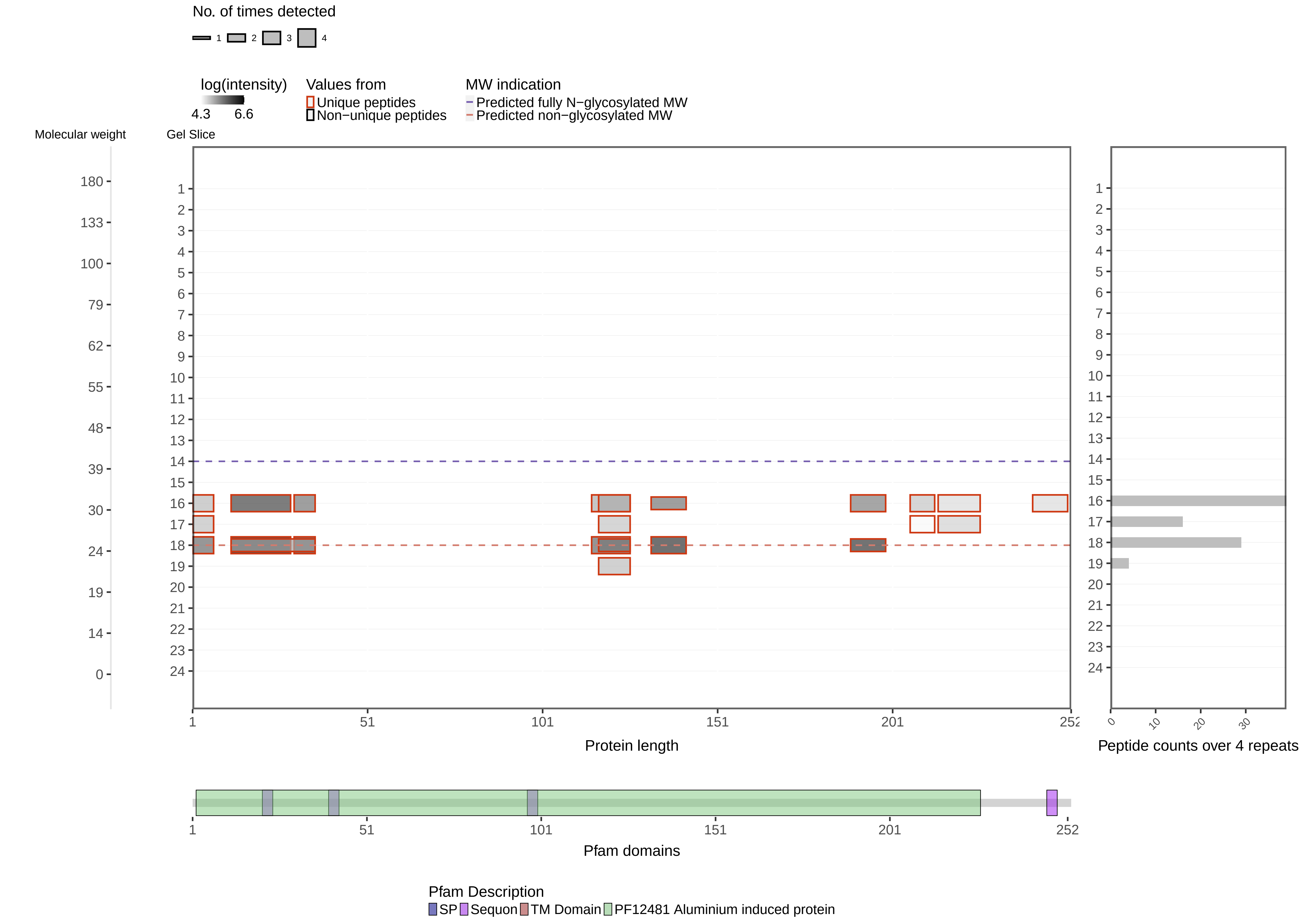


Protein sequence:

RGPYPPDQVVRDIHGKFAFVLYDSSSKTSFLASDVDGSVPFFWGTDSEGHLVLSDDADIAKQGCGKSFAPFPKGCFFTSSGGLRSYEHPRNELKPVPRVDSSGEVCGSNFKVDAESKKGGTGMPRVDSAANWSQHY*

### NbL09g18660.1 peroxiredoxin−2F_mitochondrial−like [MW = 20.97 kDa]

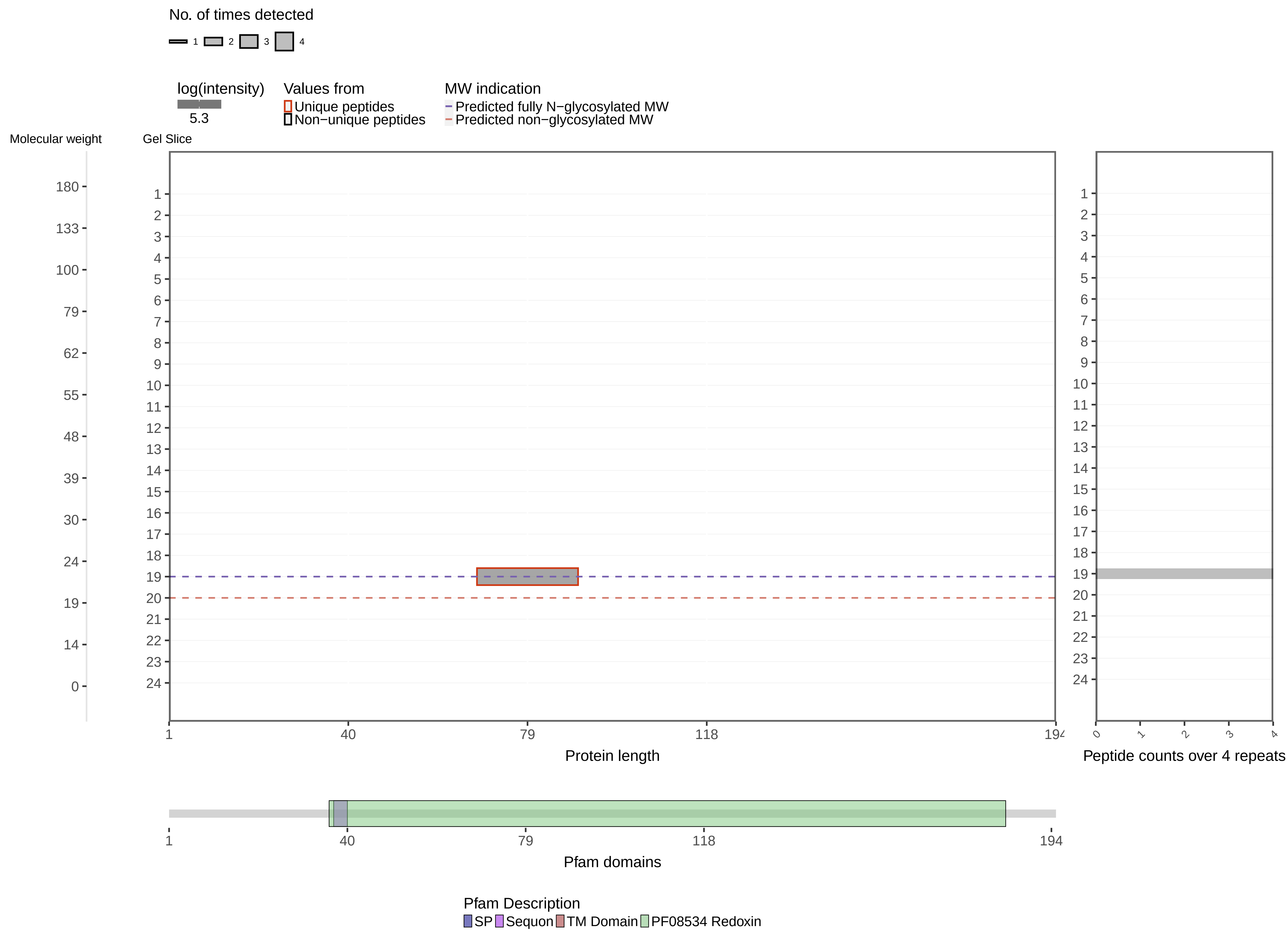


Protein sequence:

QHVPSYKNNIDKFKAKGIDAVICVAVNDPYVMNGWAEKLQAKEAIEFYGDFDGSFHKSLDLTIDLSAALLGPRSHRWSAYVDDGNVKVLNVEEAPSDFKVSGGDVILGQI*

### NbL09g18640.1 glucose−1−phosphate_adenylyltransferase_large_subunit_3_chloroplastic/amyloplastic−like [MW = 57.32 kDa]

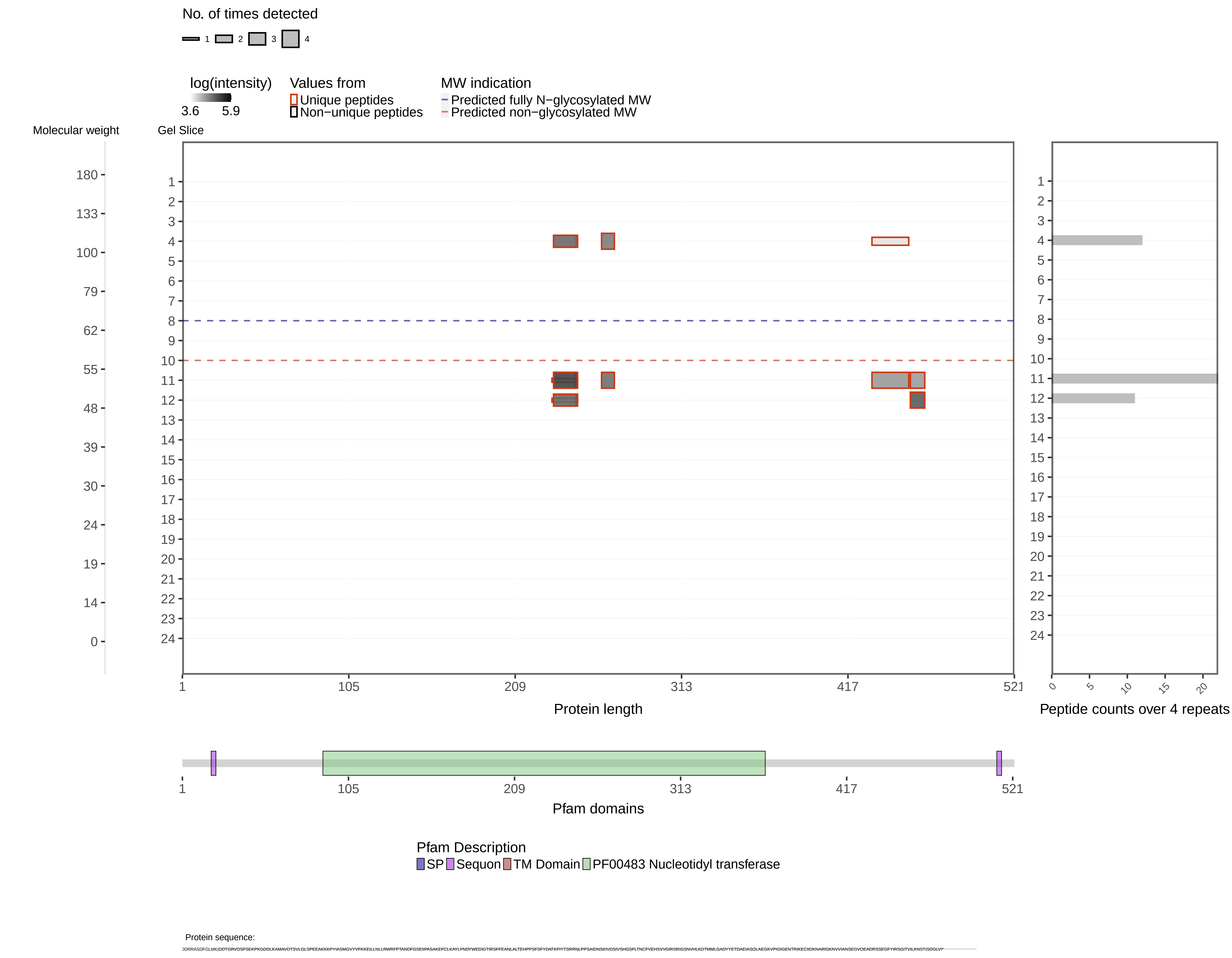

NbL09g18630.1 uncharacterized_GPI−anchored_protein_At3g06035−like [MW − SP = 18.34 kDa]

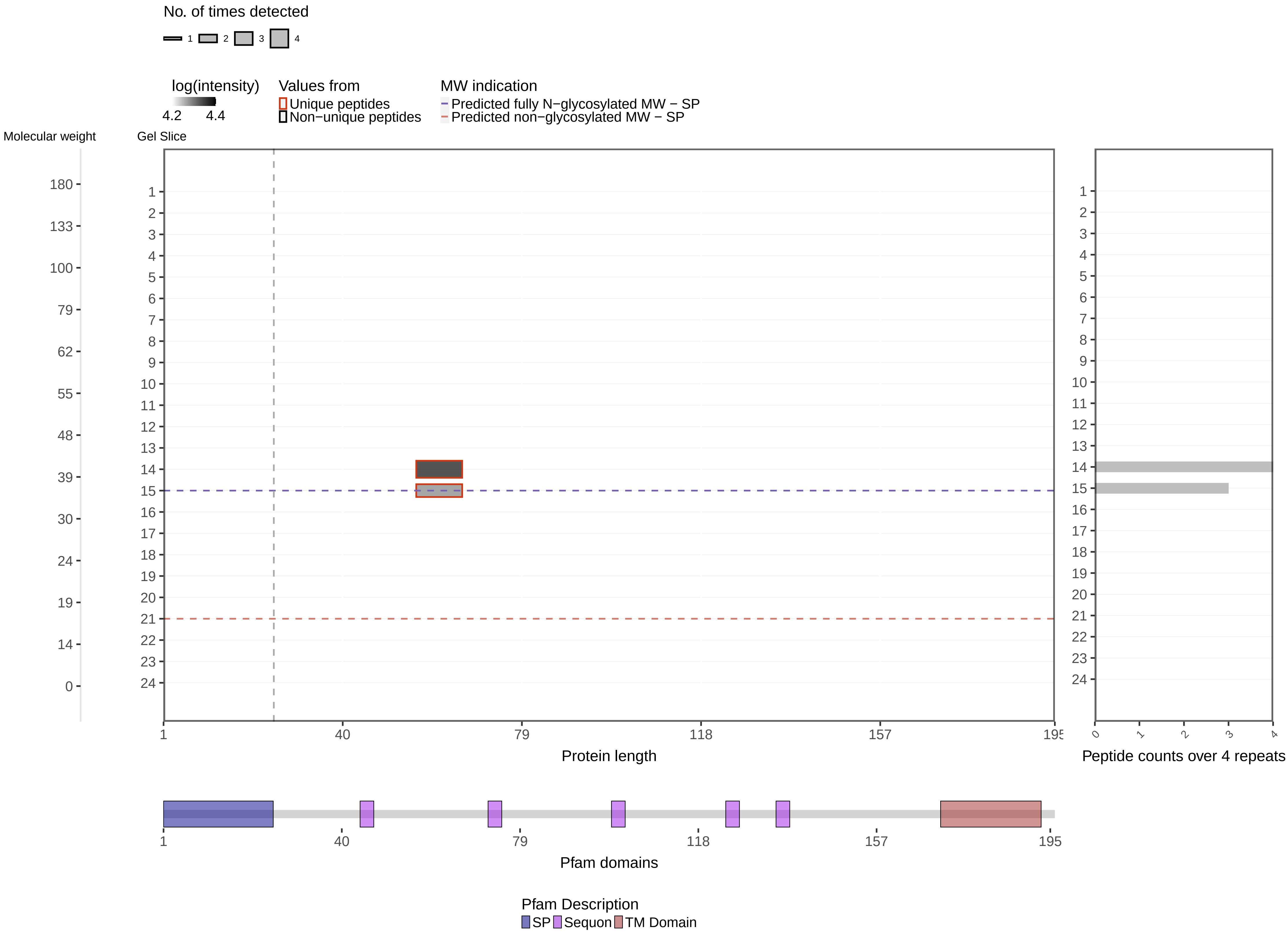


Protein sequence:

PQFSDYPKLLSKCDLNATTTRDGMIMPACVPNLVSDLVLSNYTKSQYSNYLNDTKFSGVGIGSDDNWVVVVVLTTNNPEGSFTPDTSSANLIFGLGLISHAVFILVAFVVLL*

### NbL09g18530.1 RAN_GTPase−activating_protein_2−like [MW = 67.48 kDa]

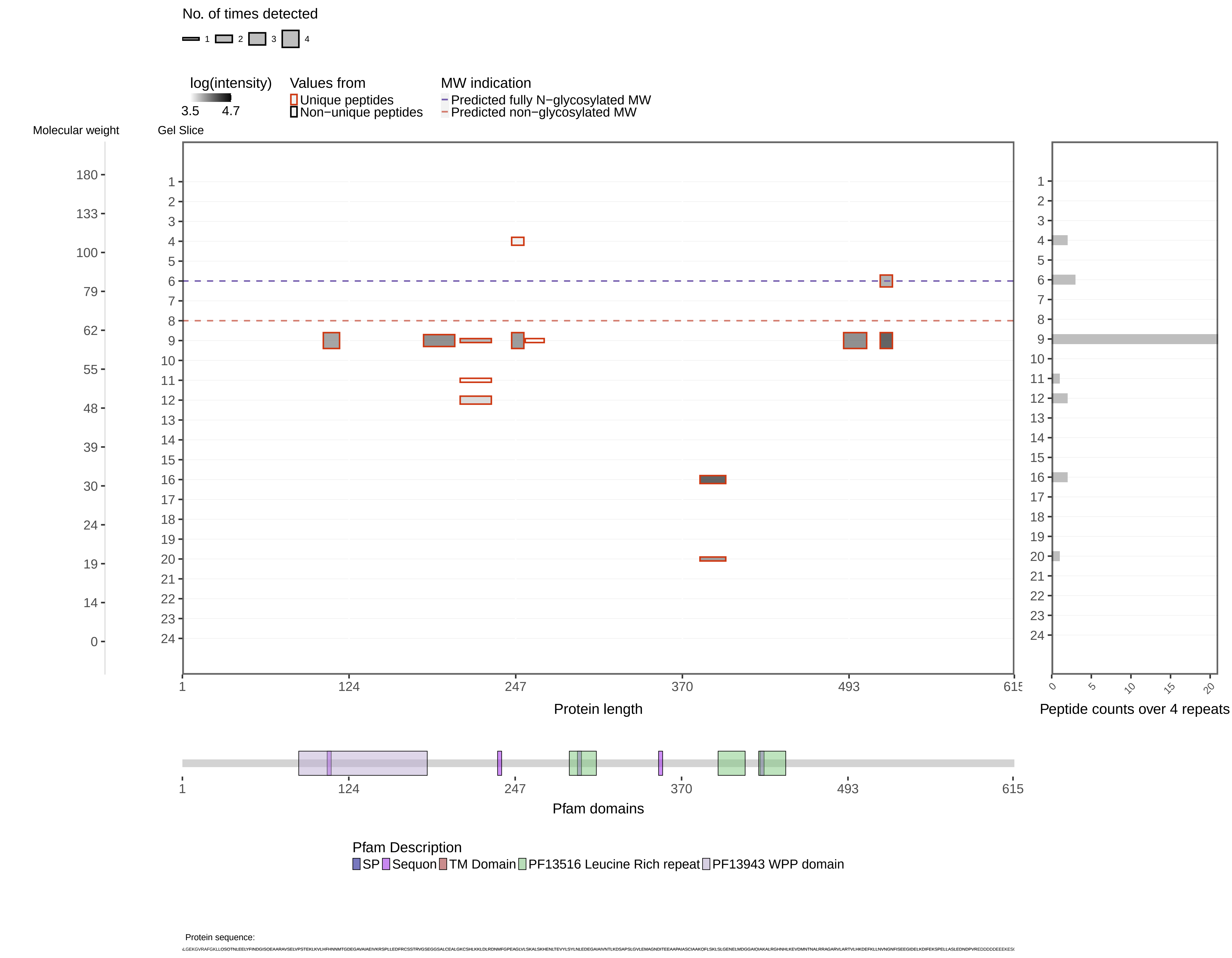

### NbL09g18400.1 heat_shock_70_kDa_protein_8 [MW = 62.45 kDa]

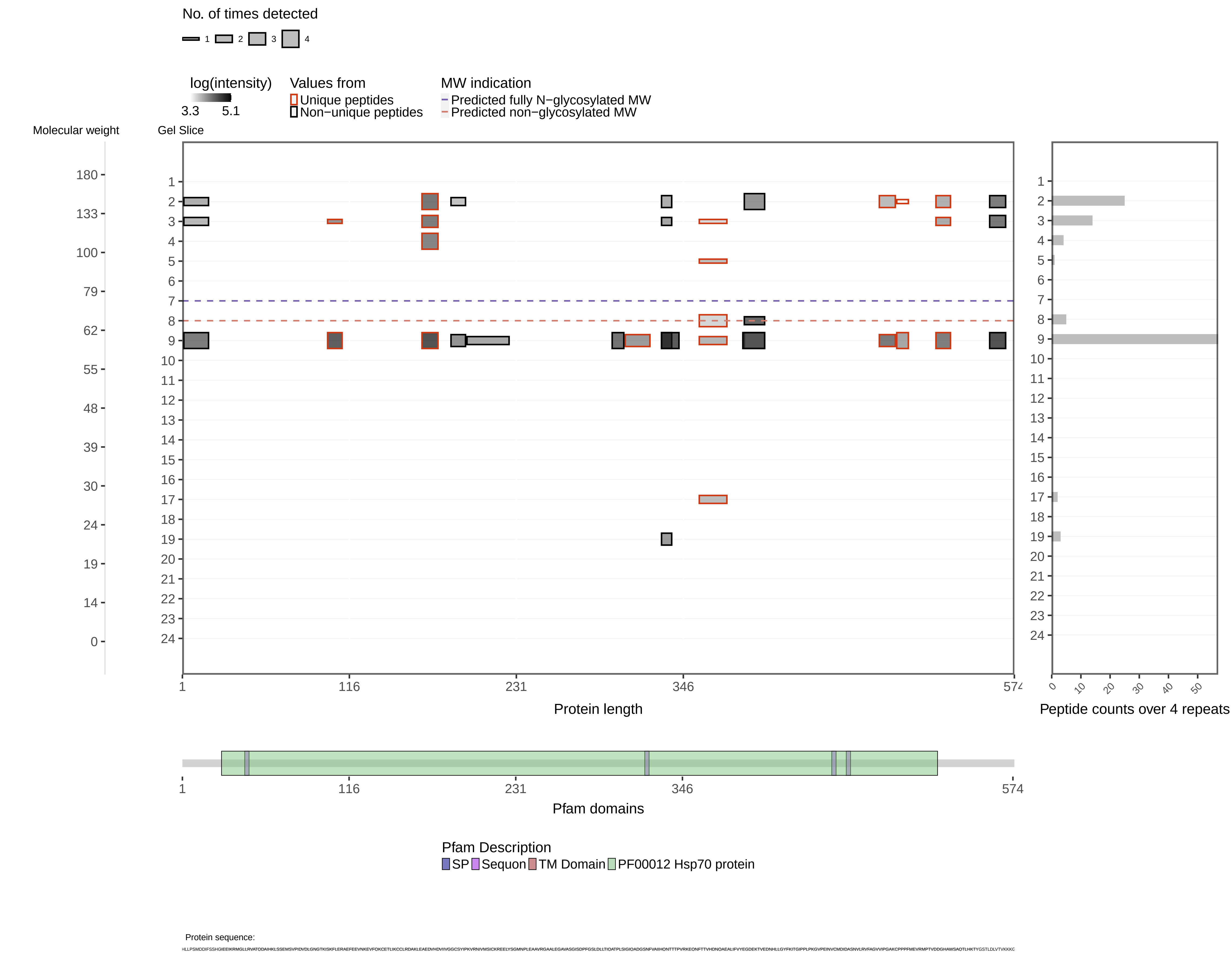

NbL09g18360.1 probable_serine/threonine−protein_kinase_WNK9 [MW = 83.02 kDa]

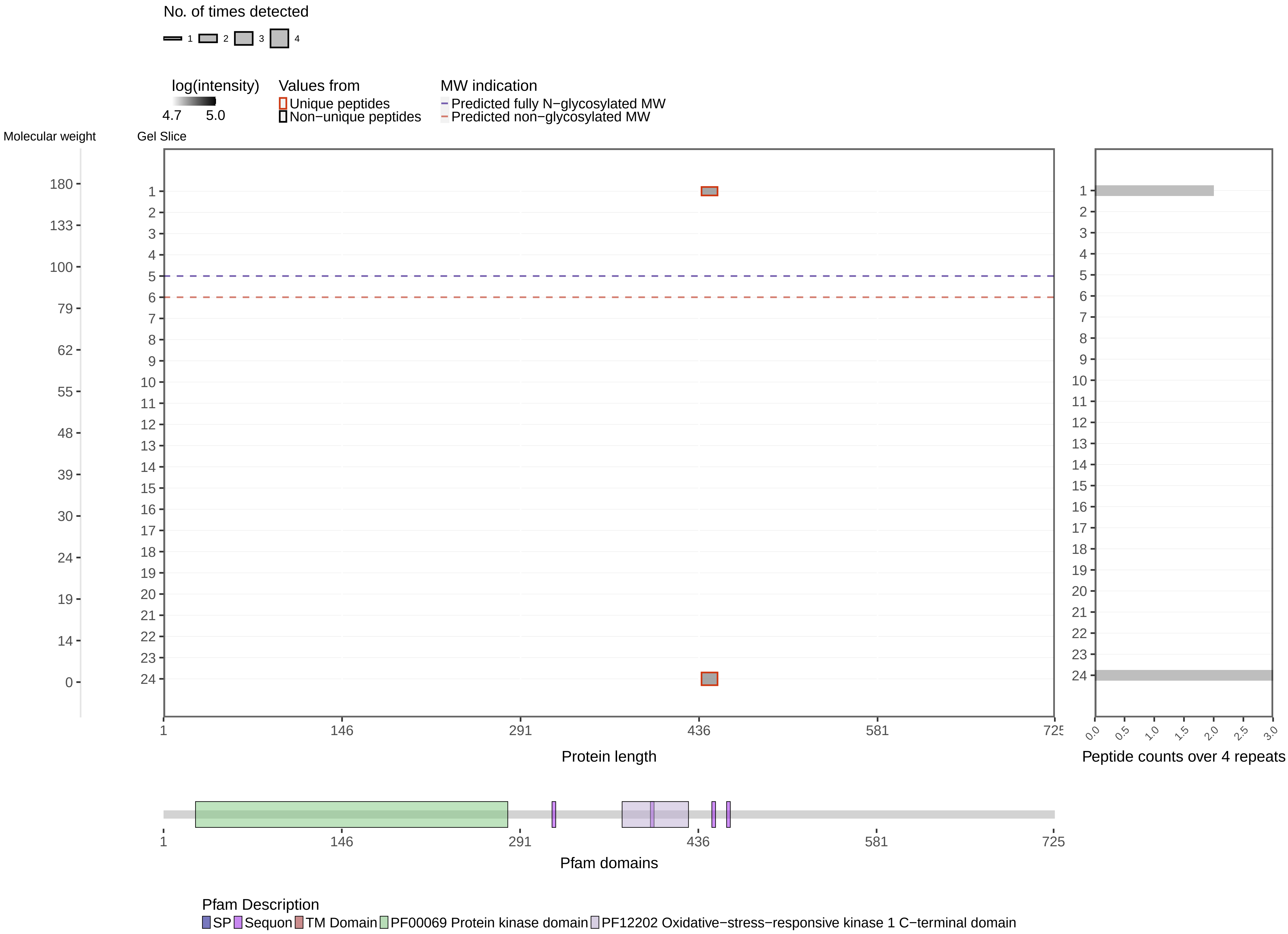


Protein sequence:

HEDDDMENVDISIKGRRREDDGIFLRLRIADKEGRVRNIYFPFDVENDTALTVATEMVAELDITDQDVTKIADMIDGEIASLVPVWKKGFGMEENLNHTNRGCHNCASNGSLVNYLLPDGPRNLQVLHCSKHGCGAMHGRFEEIMYHLDRSEQCITDAPEGIWESDGVHYTDIWAQHDKTKSRARGSVDYHSDVKDKLVDESIDGGEEKLTSVDKDDKSHVRLQDSPSSKSSGARTPVEDYENEIRRELRWLKAKYQMQLRGLKDQRLGVDSKYSRQSSSNKIPRKDKVS

### NbL09g18340.1 imidazoleglycerol−phosphate_dehydratase [MW = 30.22 kDa]

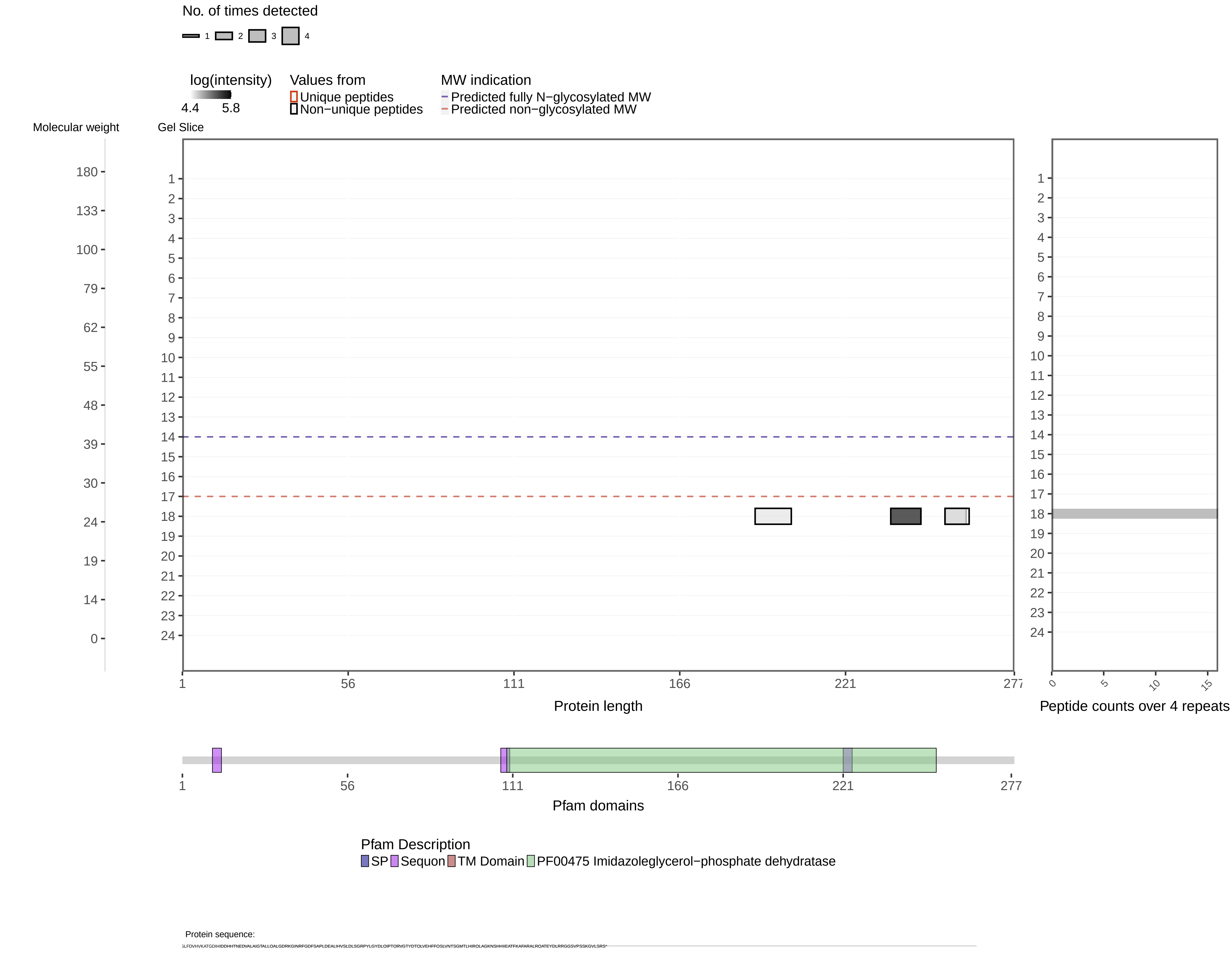

NbL09g18330.1 photosystem_II_D1_precursor_processing_protein_PSB27−H2_chloroplastic [MW = 22.64 kDa]

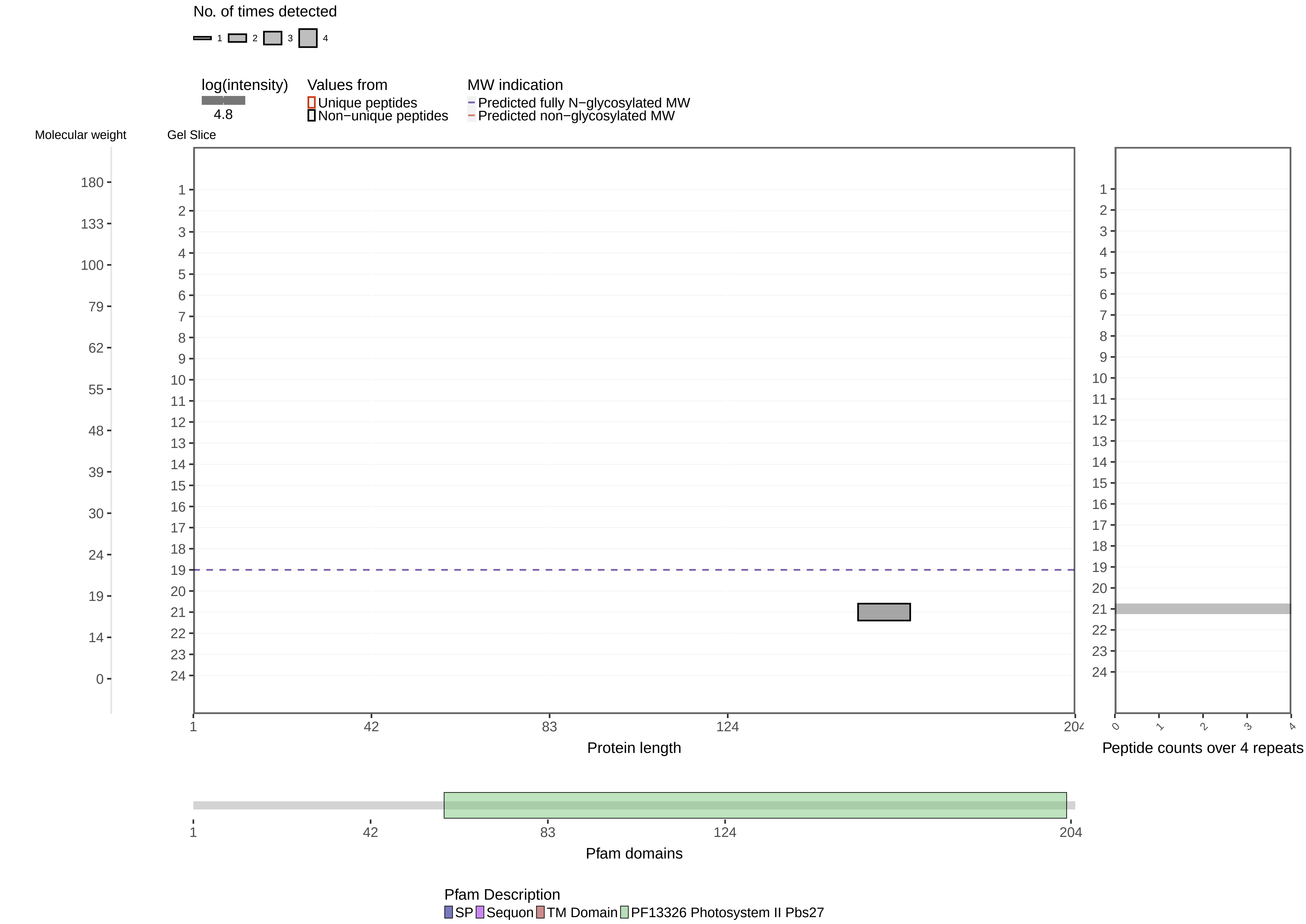


Protein sequence:

DPNEKTKSGKVLPKDYLKSAREVVKTLRESVKEDPNDMSKFRRTADAAKESIREYLGGWRGQKSVVNEESYVMLEKAIRSLASFYSKAGPSAPLPEEIKLEILDDLNKAEEAL*

NbL09g18200.1 protein_TPLATE−like [MW = 129.42 kDa]

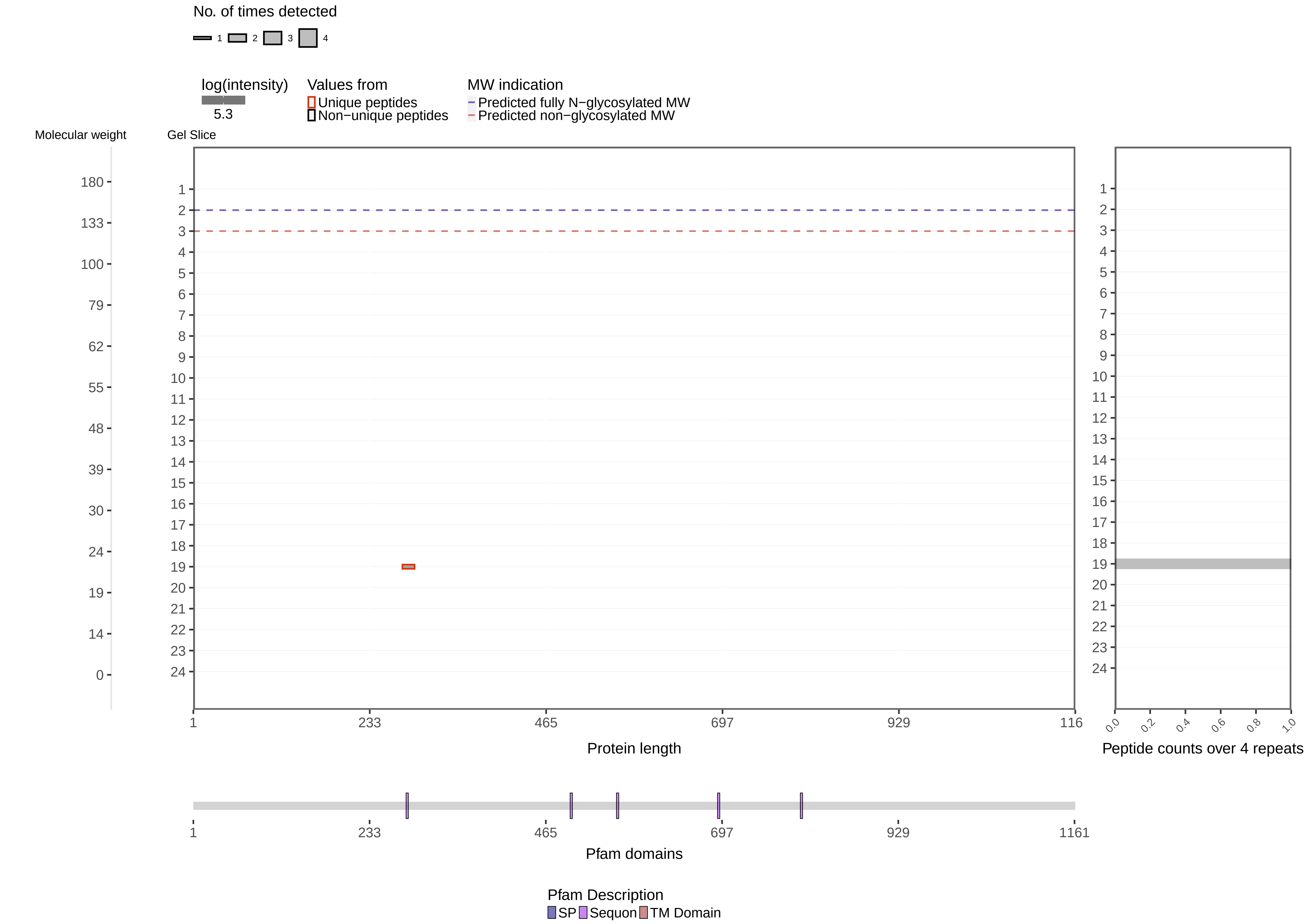


Protein sequence:

RDLREMNTPRVCTRLIWAISEHIDLEGLDPLLADDPEDPLNIIISNIHKVLFNIDSSASTTNRLQDVQAVLLCAQRLGSRNARAGQLLIKELEDFRSNAFADSVNKHQCRLILQRIKYVSNHSESKWAAVGEARGDYPFSHHKLTVQFYEAAAAQDRKLEGLVHKAILELWRPDPSELALLLAKRVDSTLLKVPPSAYTLTGSSDPCYVEAYHLTDPSDGRITLHLKVLNLTEIELNRVDLRVGLSGGLYFMDGSPQAVRQLRNLNSQEPVLSSVTVGVAHFERCDLWVQVLYYPFY

### NbL09g18190.1 uncharacterized_protein_At4g15545−like [MW = 35.9 kDa]

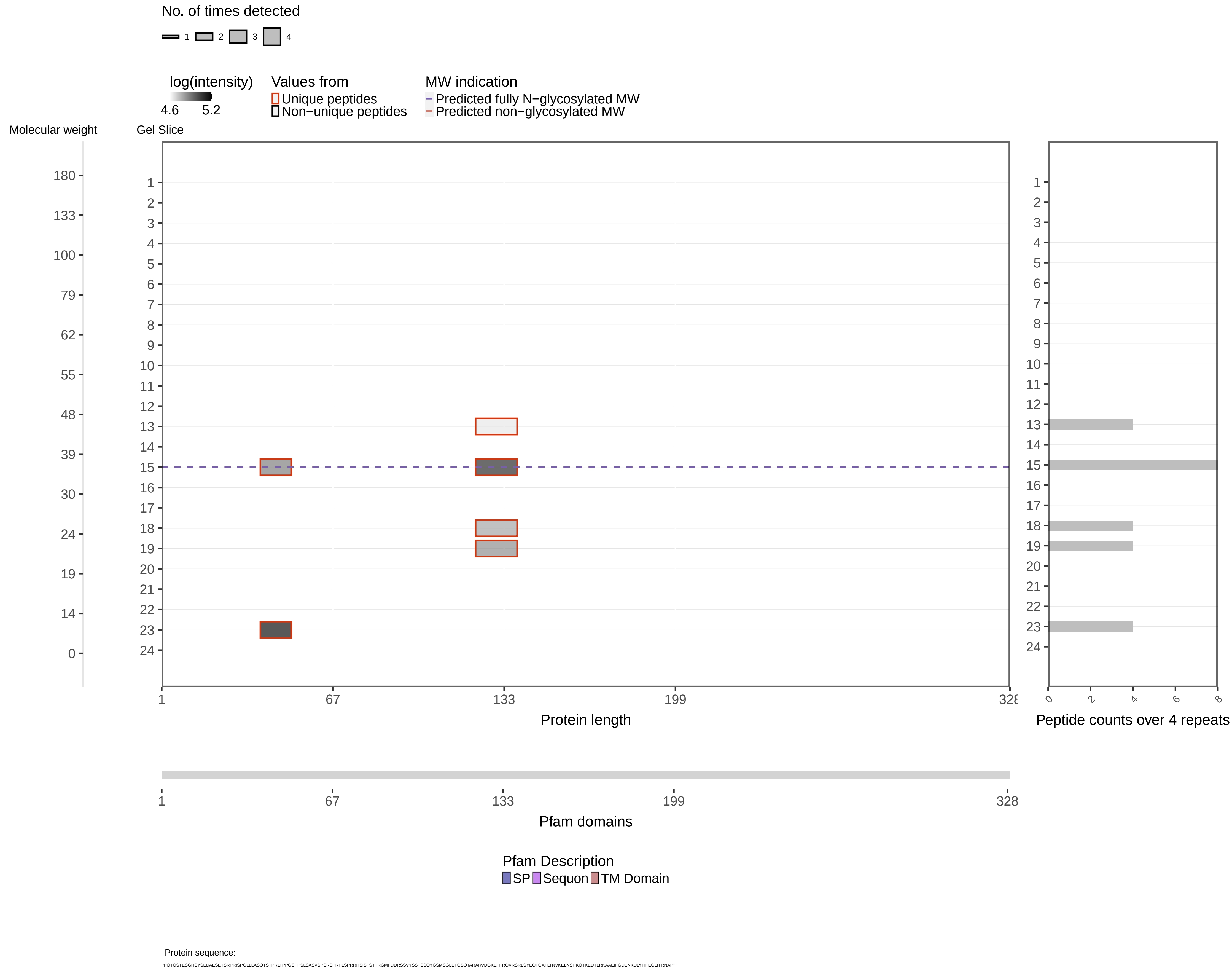

### NbL09g18170.1 probable_prolyl_4−hydroxylase_4 [MW = 35.62 kDa]

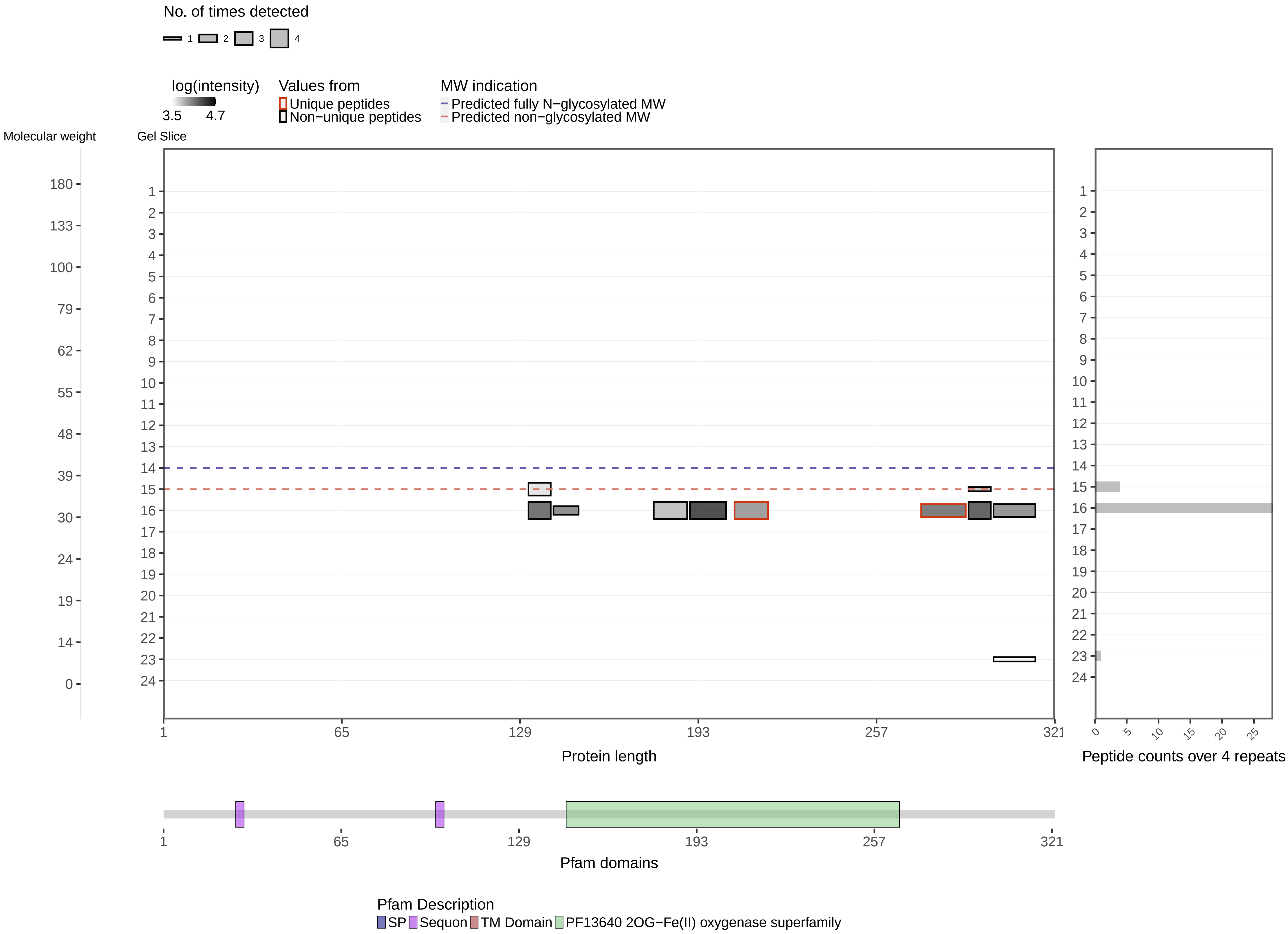


Protein sequence:

EEGQKYEPHYDYFSDKVNIVHGGHRLATVLMYLSDVEKGGETVFPEAEESTRRRSMAADDSLSECARKGIAVKPRKGDALLFFSLHPNAVPDPMSLHGGCPVIEGEKWSATKWIHVDSFDKIVGSGESCADQNENCERWAALGECTKNPEYMVGSSDLPGSCRKSCKAC*

### NbL09g18090.1 ABC_transporter_C_family_member_14−like [MW = 169.55 kDa]

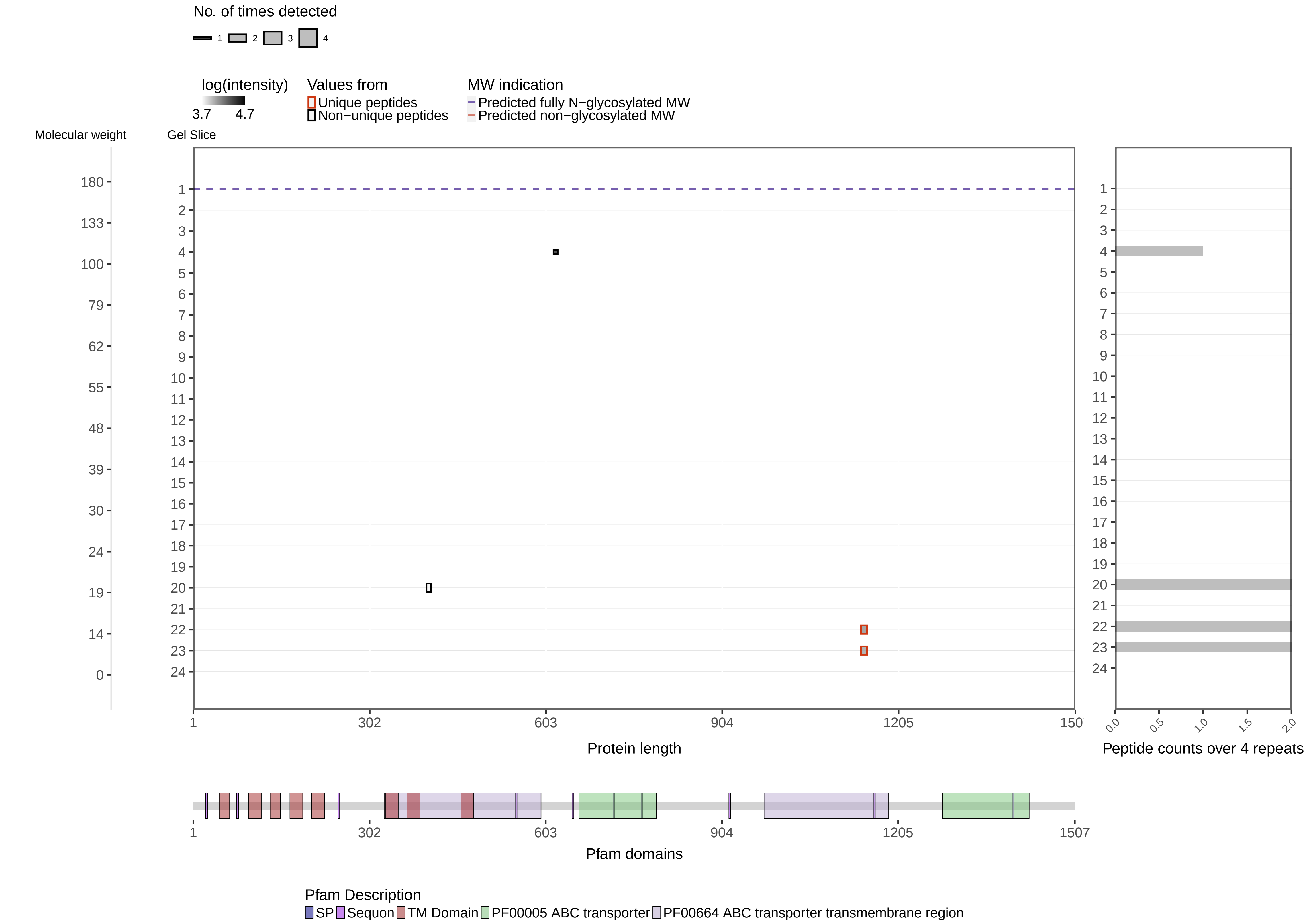


Protein sequence:

### NbL09g17990.1 isoleucine−−tRNA_ligase_chloroplastic/mitochondrial [MW = 123.94 kDa]

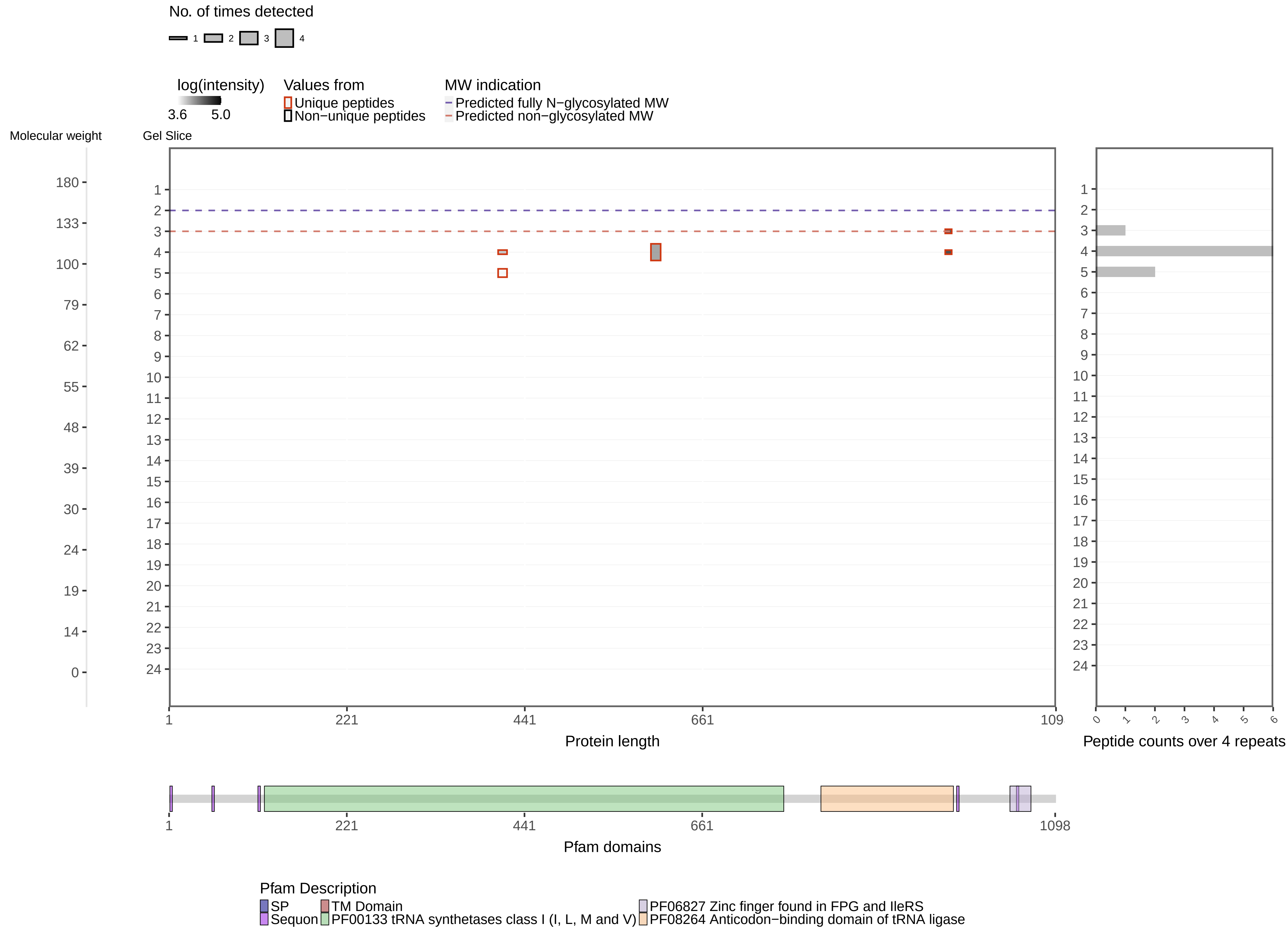


Protein sequence:

WFASVEGFRDAAMDAINQVTWTPTQAVNRISAMTSGRSEWCISRQRTWGVPIPVFYHVESKEPLMNEETIDHIKSIISQKGSDAWWYMSVEELLPEKYCDKASNYEKGTDTMDVWFDSGSSWAAVLDKRESLNYPADLYLEGTDQHRGWFQSSLLTSIATKGMAPYRGVITHGFVLDERGLKMSKSLGNVVDPRMVIEGGKNQKESPPYSADVLRLWVSSVDYTGDVLIGPQVLRQMSDIYRKLRGTLRFLLANLHDWKADCAVPYSDLPMIDQYALFQLANVVNNIRE

NbL09g17950.1 probable_1−deoxy−D−xylulose−5−phosphate_synthase_chloroplastic [MW = 77.4 kDa]

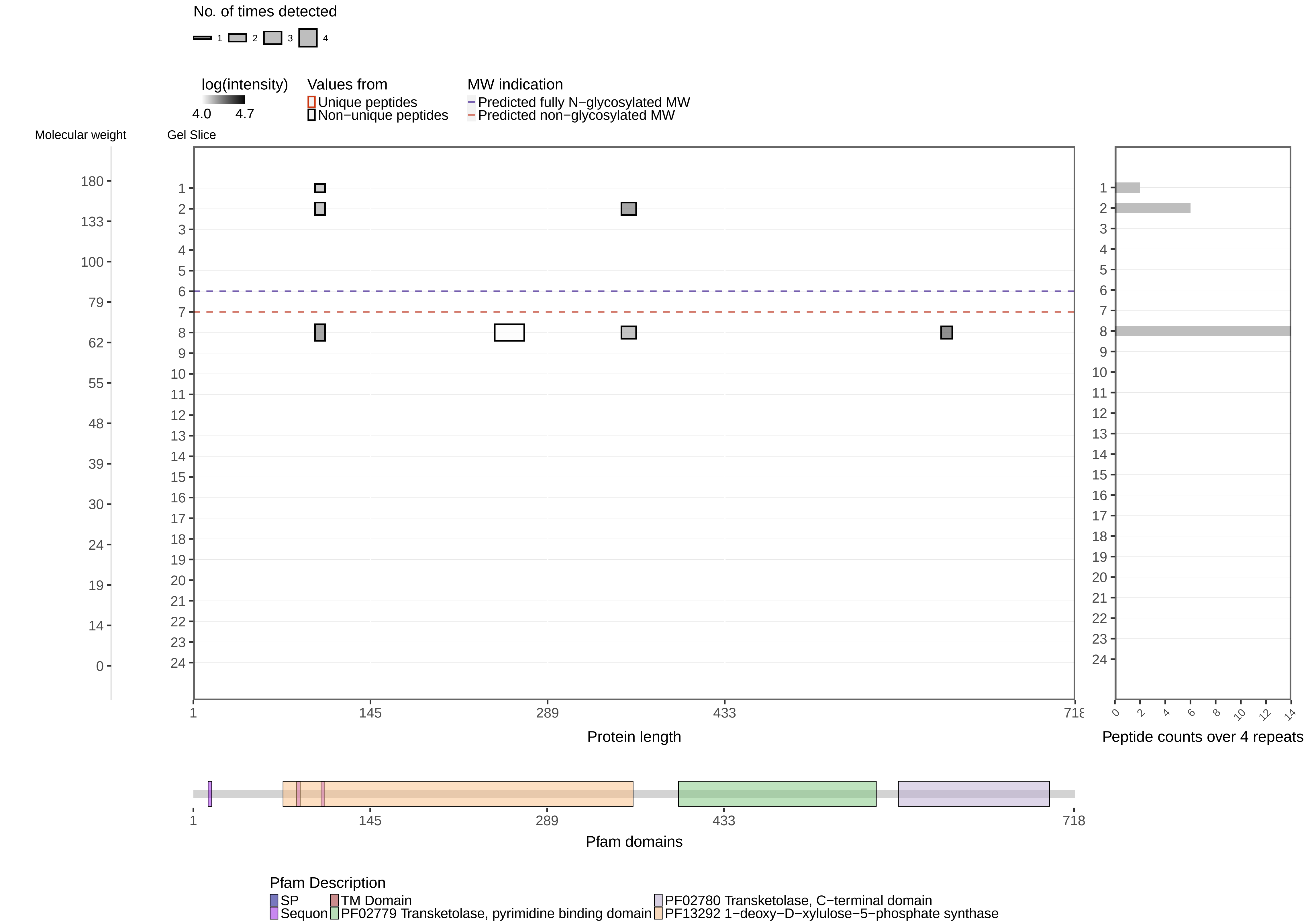


Protein sequence:

STKTTGPVLIHVITEKGRGYPYAERAADKYHGVAKFDPATGKQFKVSAKTQSYTTYFAEALIAEAEADKDIVAIHAAMGGGTGMNLFHRRFPNRCFDVGIAEQHAVTFAAGLACEGLKPFCAIYSSFLQRAYDQAMHDVDMQKLPVRFAMDRAGLVGADGPTHCGAFDVTFMACLPNMVVMAPSDEAELFHMVATAAAIDDRPSCFRYPRGNGIGVELPVENKGIPLEVGKGRILVEGERVALLGYGSAVQNCLTAAAVLETRGLQVTVADARFCKPLDGALIRSLAKSHEVL

NbL09g17860.1 bifunctional_3−dehydroquinate_dehydratase/shikimate_dehydrogenase_chloroplastic−like [MW = 57.53 kDa]

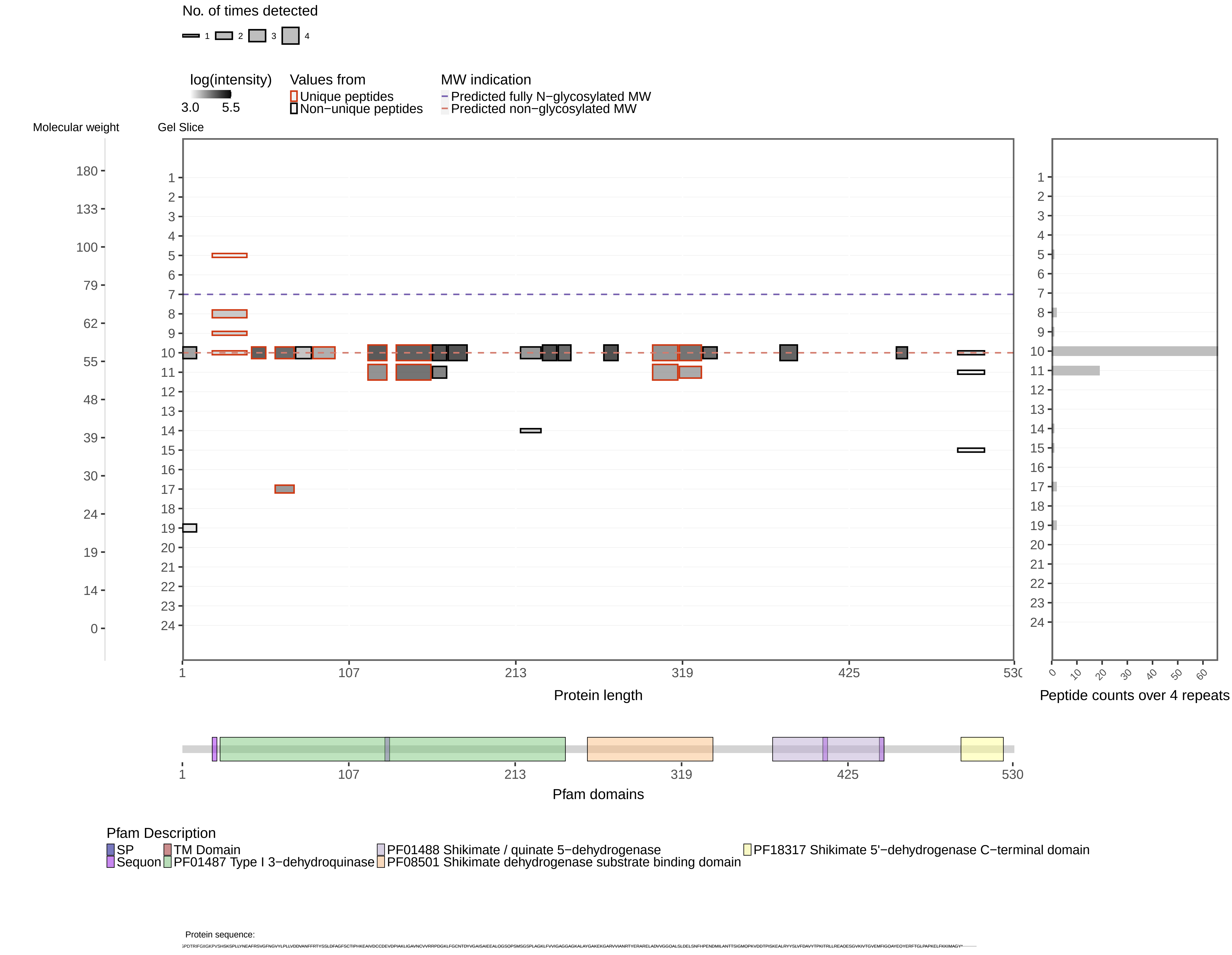

### NbL09g17850.1 superoxide_dismutase_[Cu−Zn] [MW = 15.27 kDa]

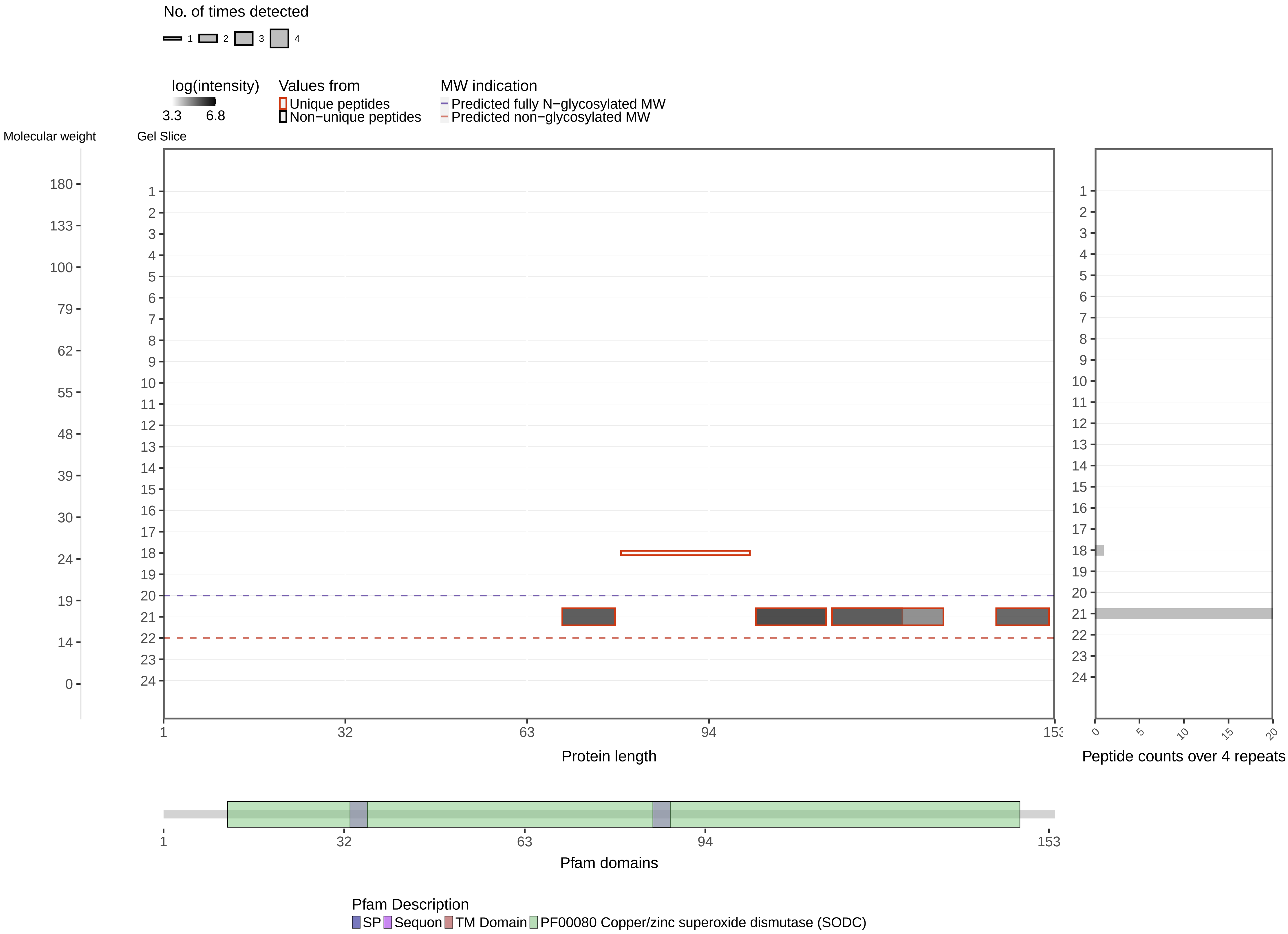


Protein sequence:

YNPAGKEHGAPEDEVRHAGDLGNITVGEDGTASFTITDKQIPLAGPQSIIGRAVVVHADPDDLGKGGHELSKTTGNAGGRVACGIIGLQG*

### NbL09g17840.1 acyl_carrier_protein_1_chloroplastic−like [MW = 13.99 kDa]

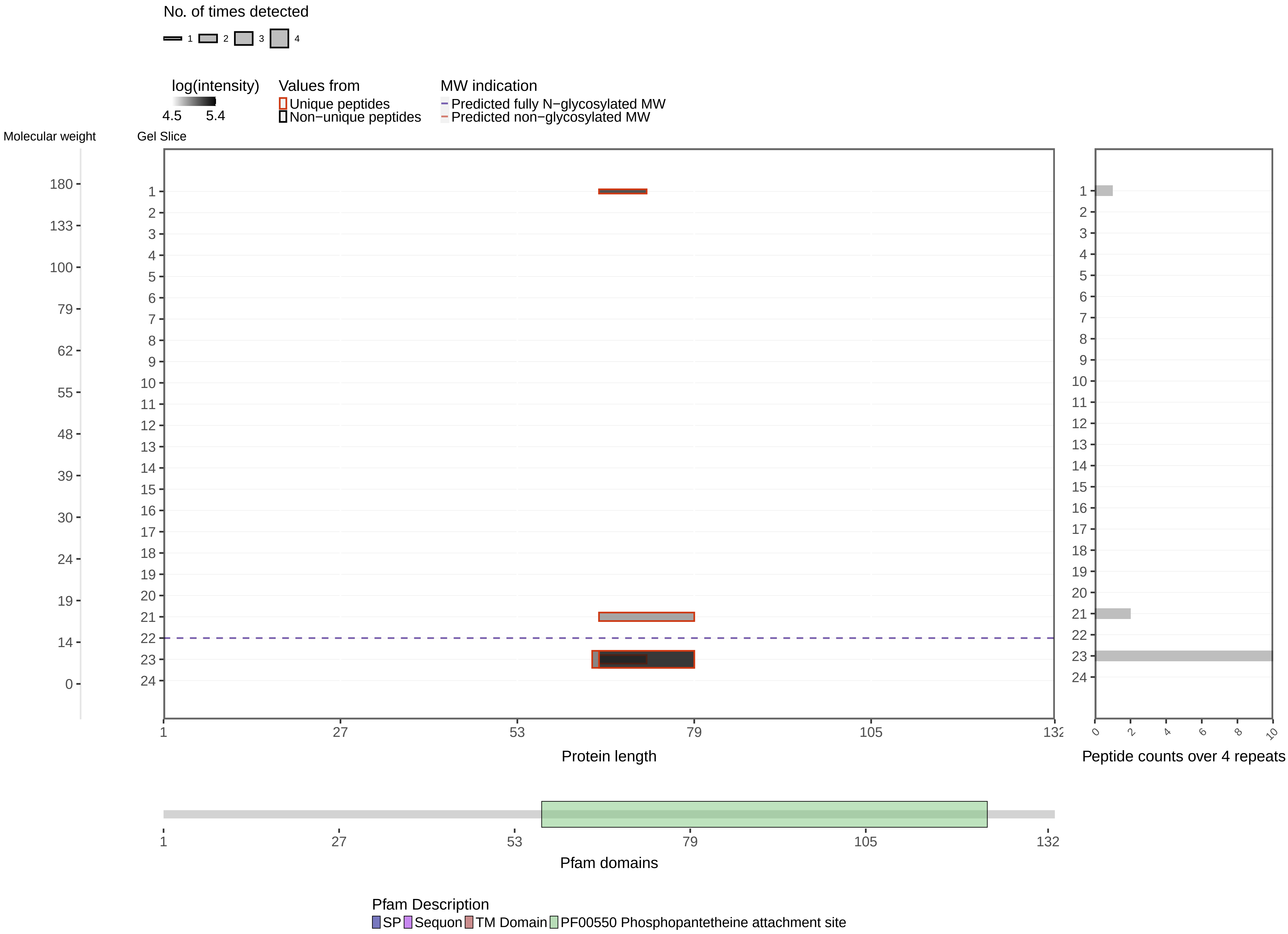


Protein sequence:

ETVNKVCEIVKKQLALTDDKVVNGESKFTALGADSLDTVEIVMGLEEAFGITVEEENAQTIATVQDAADLIEDLISKKA*

NbL09g17760.1 uncharacterized_LOC109208031 [MW = 26.81 kDa]

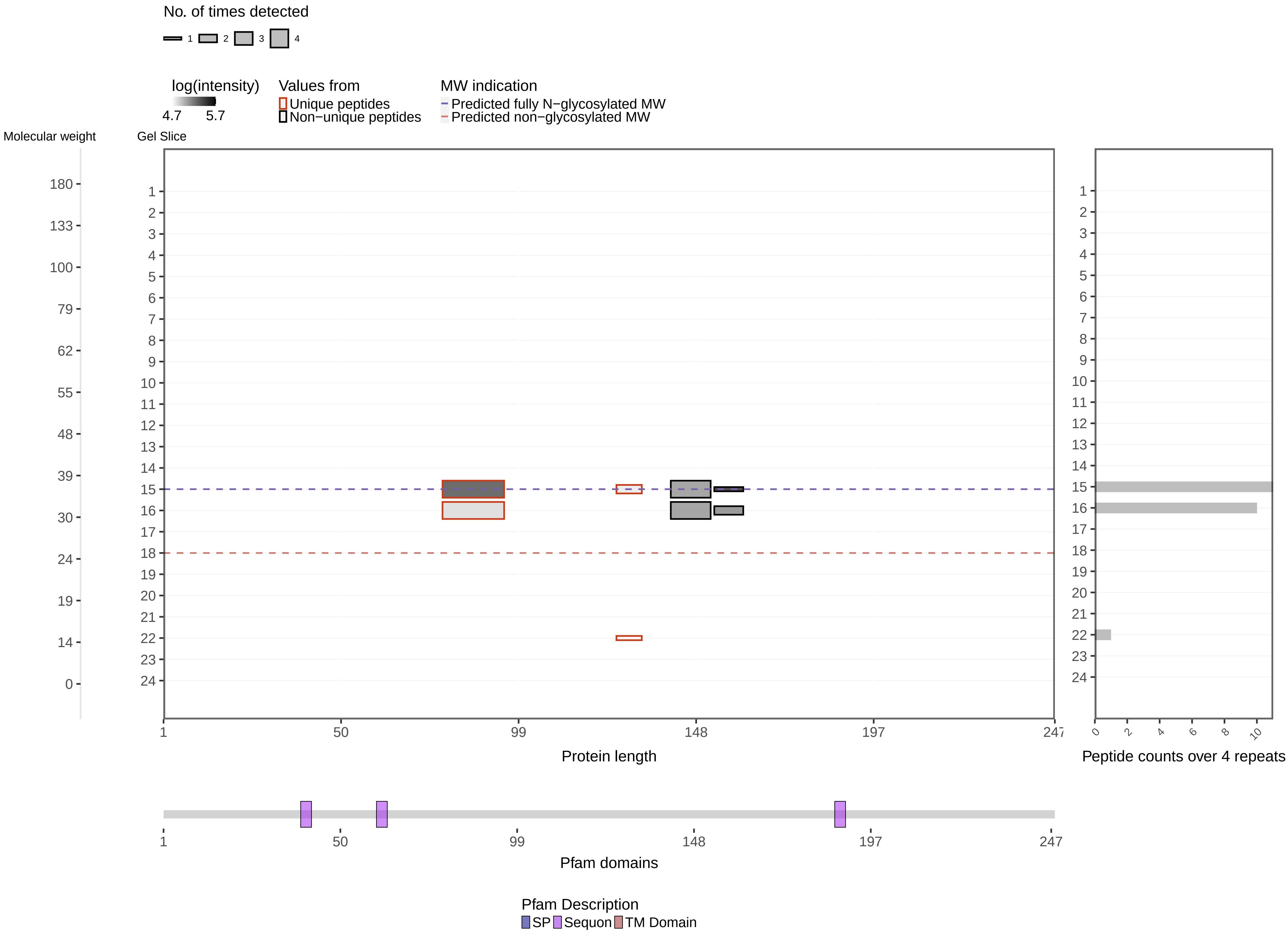


Protein sequence:

IFGOTMTTDHHDELIKMGISPSSRETKFTSKFPYVIFCAPPSRTEDYPGDIREAALKWSGEGSFLFTSSSAPYDCSDNGSVDEDGPIVPIGRSPRTDILLKAEEVVLEFGGCVIRLAGLYISLDMDFKELGFFLCSL*

### NbL09g17320.1 uncharacterized_LOC109205690 [MW = 17.24 kDa]

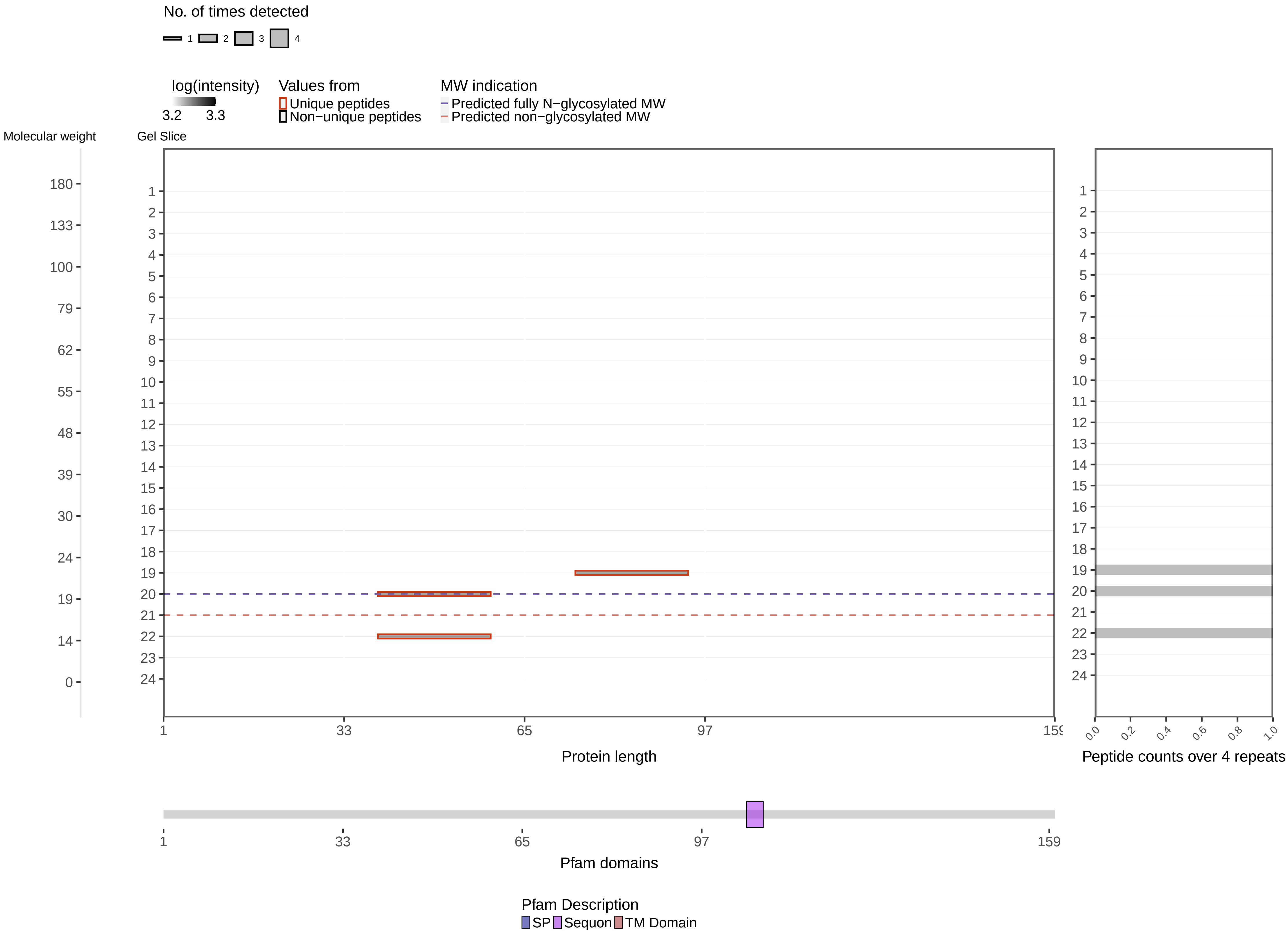


Protein sequence:

VGASMRGEMGEKPGLEGVSPGENEALKTYVNSNFQSINNSIMLGGSYCTNDPGVHLDISDYVDQETPTPKGHGKKKERGSSKHYHHSQHSE*

### NbL09g17090.1 acylpyruvase_FAHD1_mitochondrial [MW = 26.79 kDa]

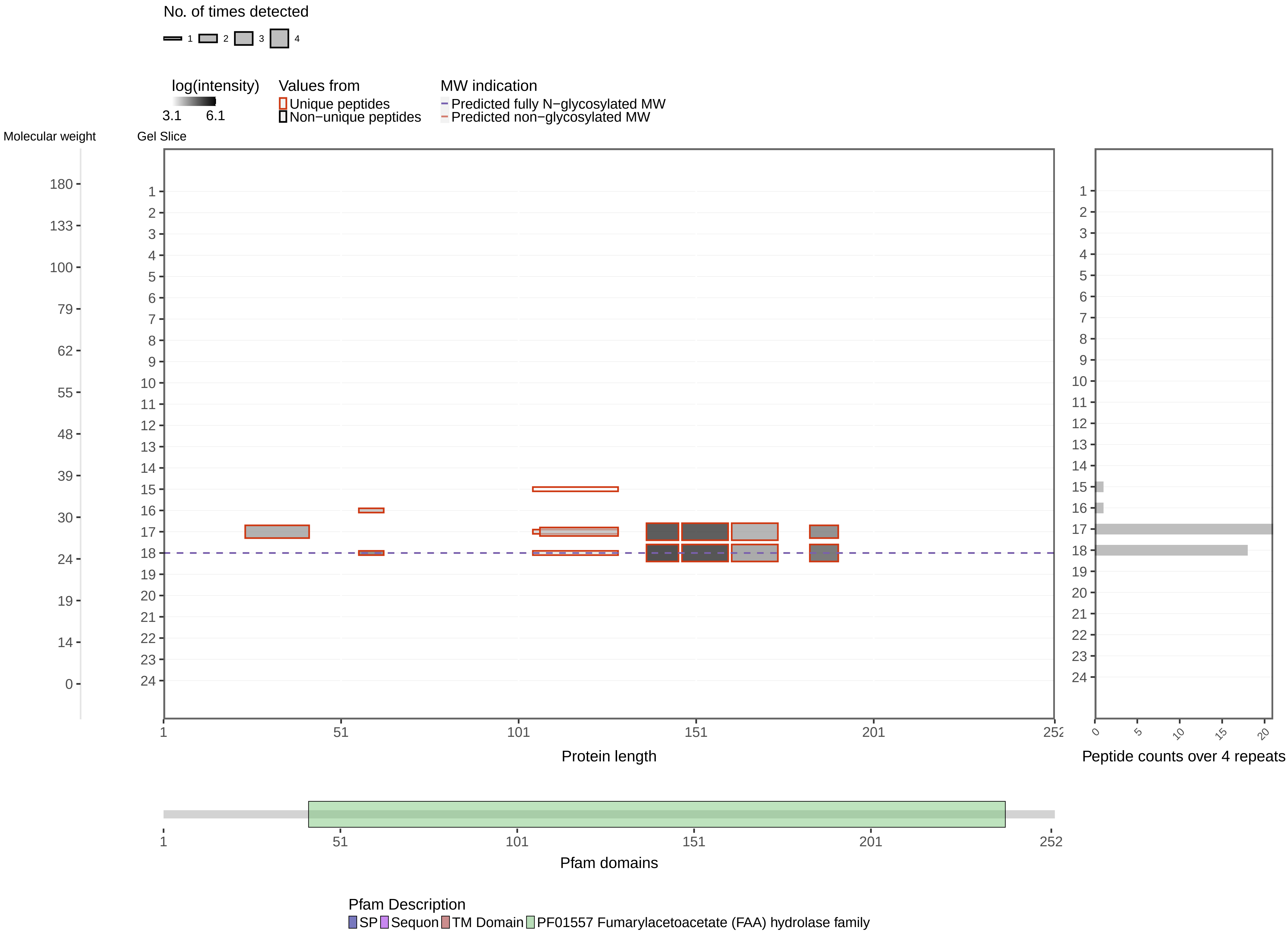


Protein sequence:

AMDYVGGYALALDMTAREIQATARSAGLPWTVAKGQDTFTPISSVLPKSMVPDPHDIELWLKVNGEIRQKGSTSDMIFKIPFLISHISSIMTLFQGDVILTGTPKGVGPVKVGQEIDAGITGLLDVHFDVGRRPGAKQQ*

NbL09g17020.1 probable_acyl−CoA_dehydrogenase_IBR3 [MW = 91.27 kDa]

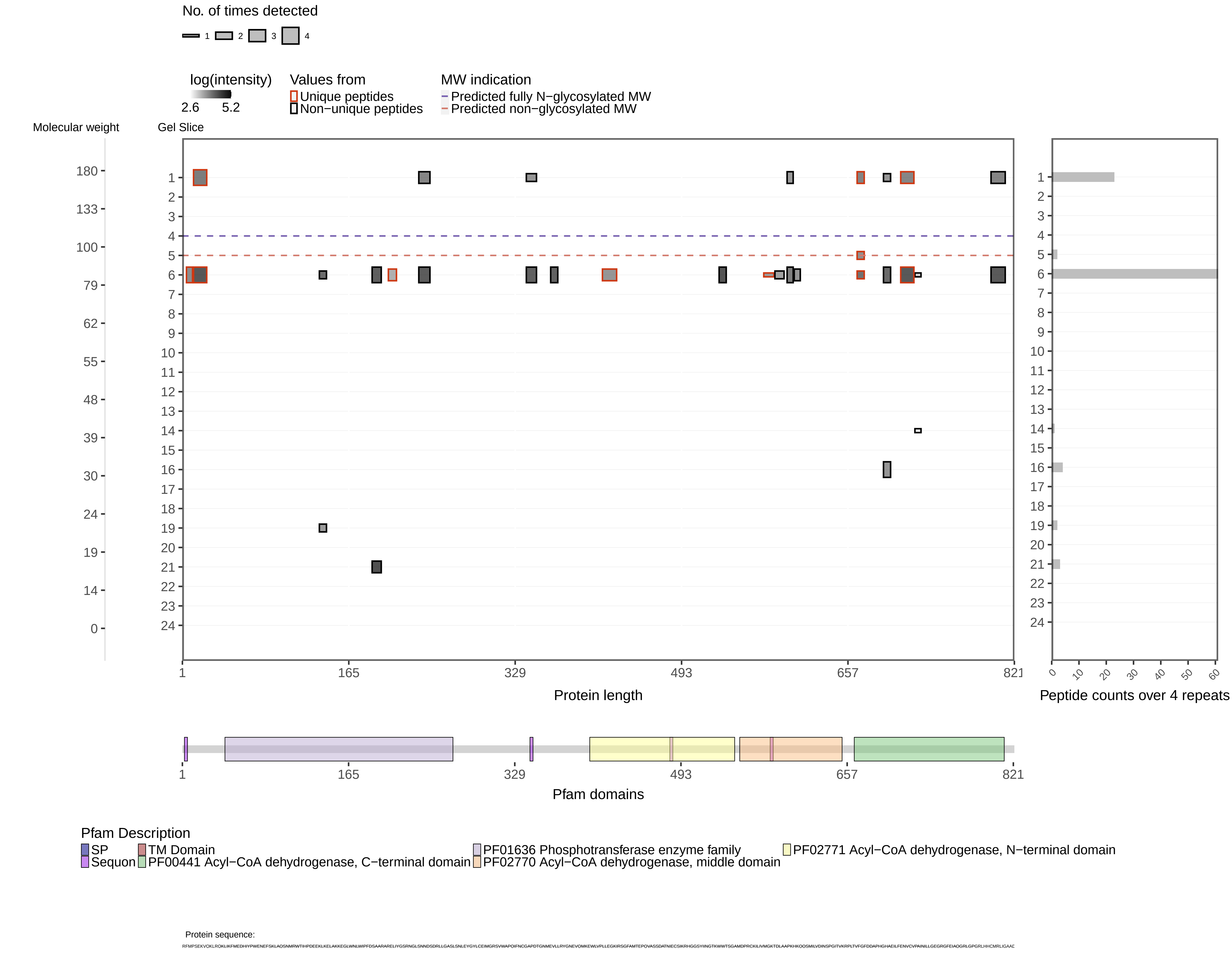

### NbL09g16800.1 probable_UDP−3−O−acylglucosamine_N−acyltransferase_2_mitochondrial [MW = 33.8 kDa]

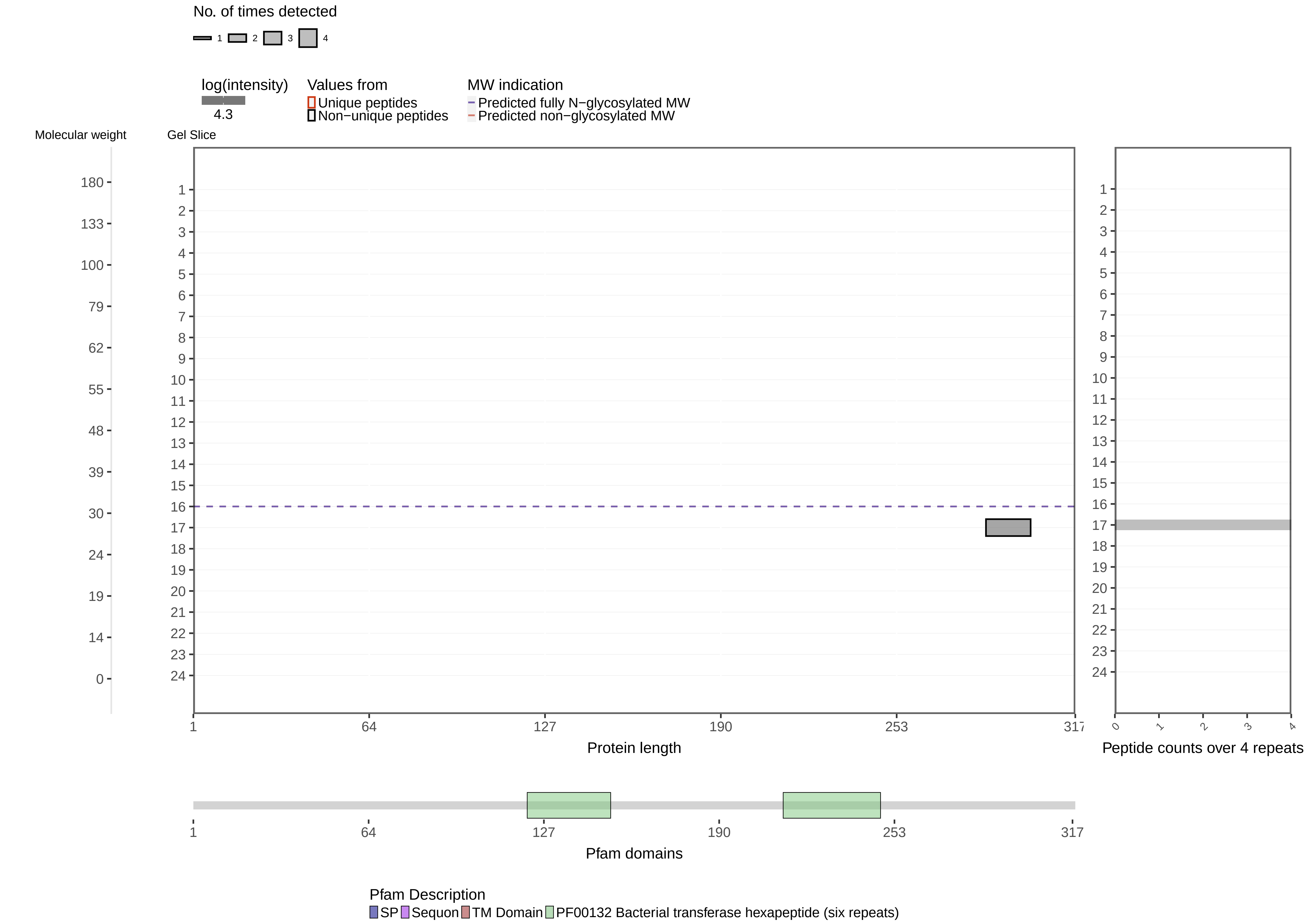


Protein sequence:

SYNVALANCMVGDFCVIHSGVCIGQDGFGFFVDEQGNMVKKPQTLNVRIENHVEIGANTCIDRGSWRNTVVGEHTKIDNLVQIGHNVVIGRSCIICGQVGVAGSVTIGDYVTFGGRVGIRDHVNIASKVRLAANSCVTKDISEPGDYGGFPAIPIRDWRRQIAMHRQSLKS*

### NbL09g16730.1 UPF0051_protein_ABCI8_chloroplastic−like [MW = 61.26 kDa]

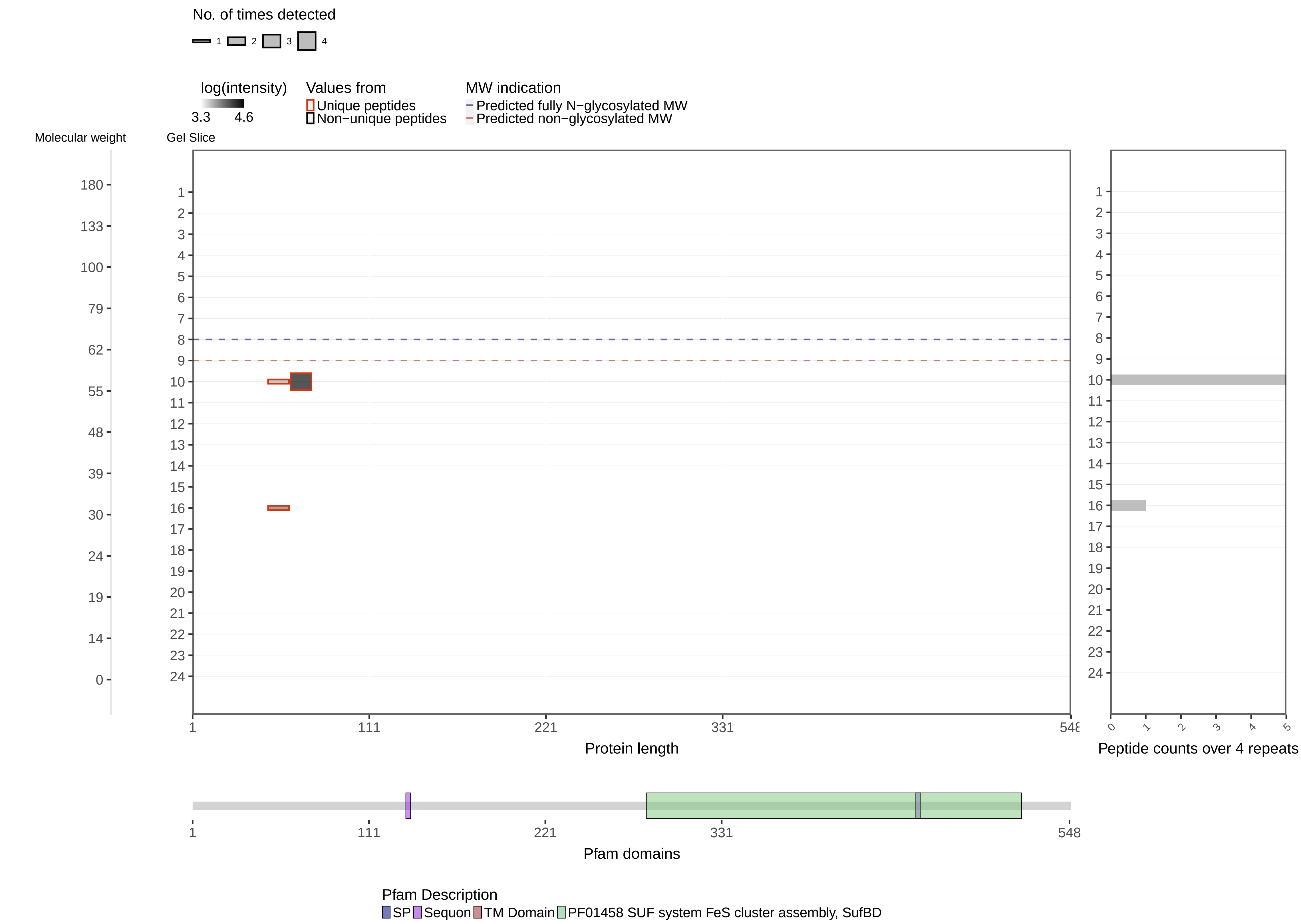


Protein sequence:

KNTRCPMQISTYFRINAMETGQFERTVIIAEEGSFVEYLEGCTAPAYDTNQLHAAVVELYCDEGAEIKYSTVQNWYAGDEEGKGGIYNFVTKRGLCAGARSKISWTQVETGSAITWKYPSVVLEGDESVGEFYSVALTNNYQQADTGTKMIHKGKNTRSRIISKGISAGHSRNCYRGLVQVLSNAENAKNSSQCDSMLIGDTAAANTYPYIQVSKNPTARIEHEATTSKIGEDQLFYFQQRGIDYEKAMAAMISGFCRDVFNELPDEFGAEVNQLMSLKLEGSVG*

### NbL09g16550.1 ubiquitin_domain−containing_protein_DSK2b−like [MW = 58.79 kDa]

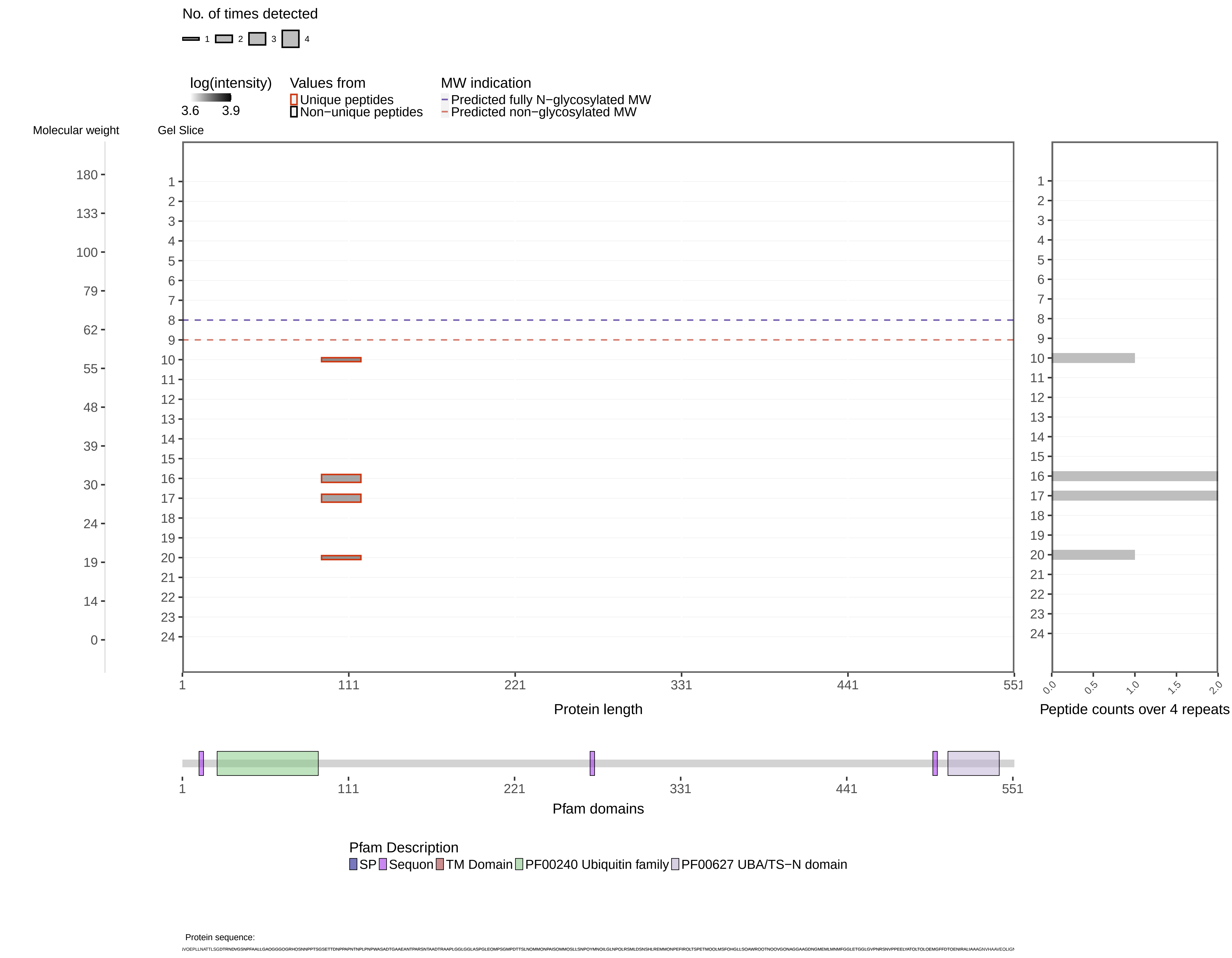

NbL09g16530.1 heterogeneous_nuclear_ribonucleoprotein_H2 [MW = 30.17 kDa]

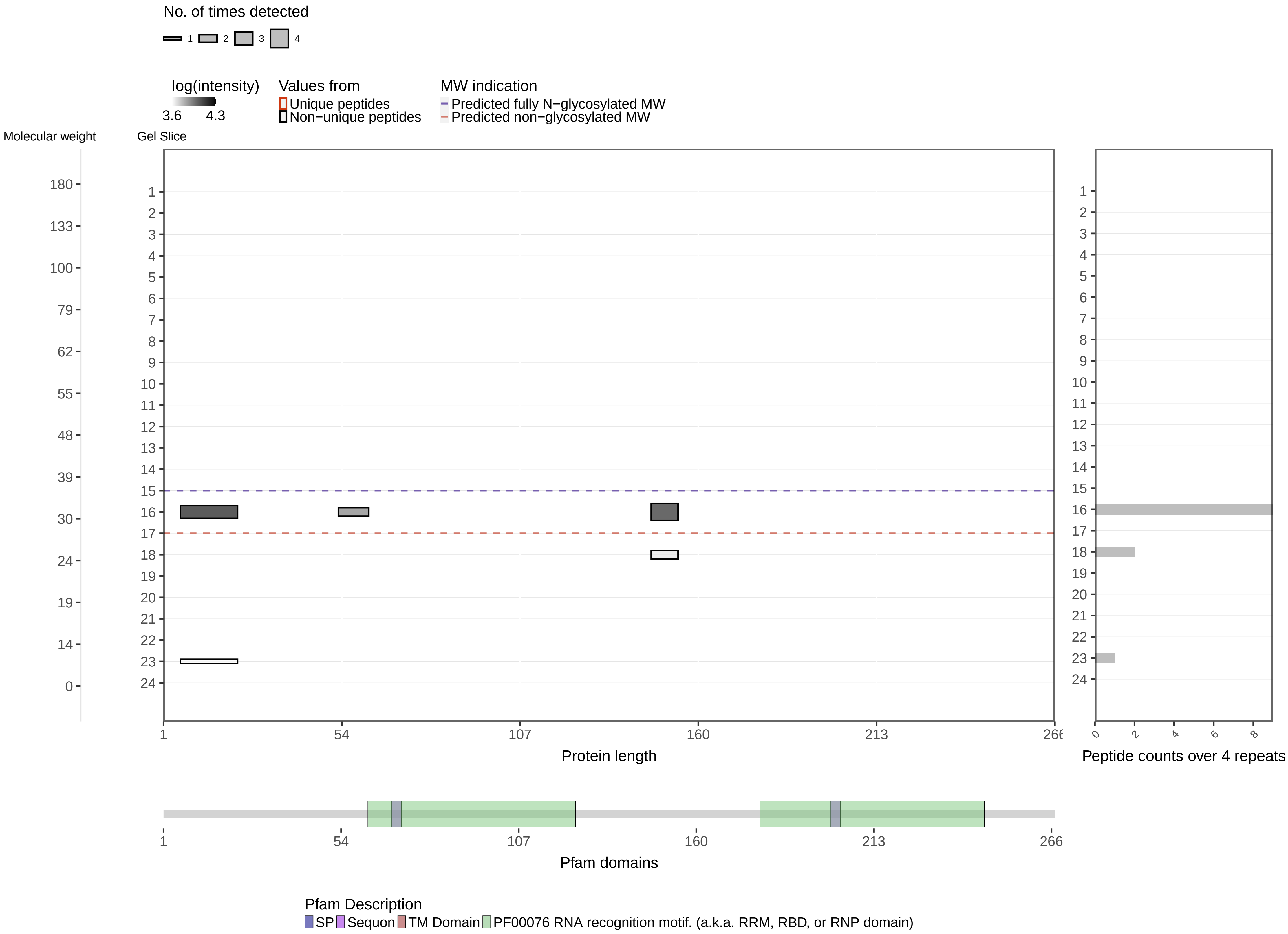


Protein sequence:

NMGRRYVEVFSCKKEEYYEAIAADVKEGGYDYEYRTSPPPSRPKRSMNKDQMEYTEILKLRGLPYSVRRTDIARFFGEEFNVSADKVHIAYRPDGKATGEAYVEFASSEEAKKAMCKDNMKIGSRYIELFPSHPDEARRAESRSRQ*

### NbL09g16340.1 fasciclin−like_arabinogalactan_protein_4 [MW − SP = 42.1 kDa]

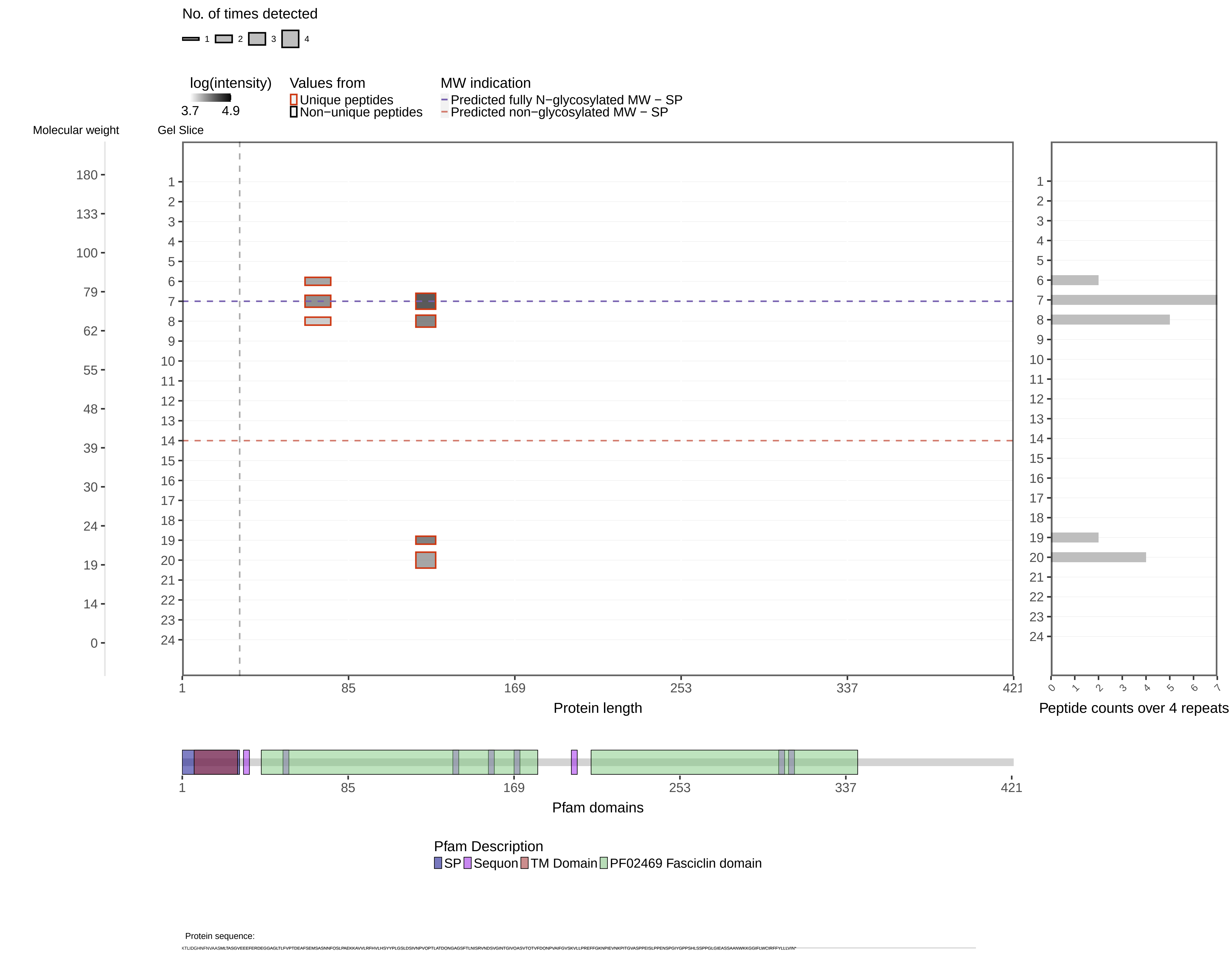

NbL09g16260.1 heat_shock_cognate_70_kDa_protein_1−like [MW = 47.96 kDa]

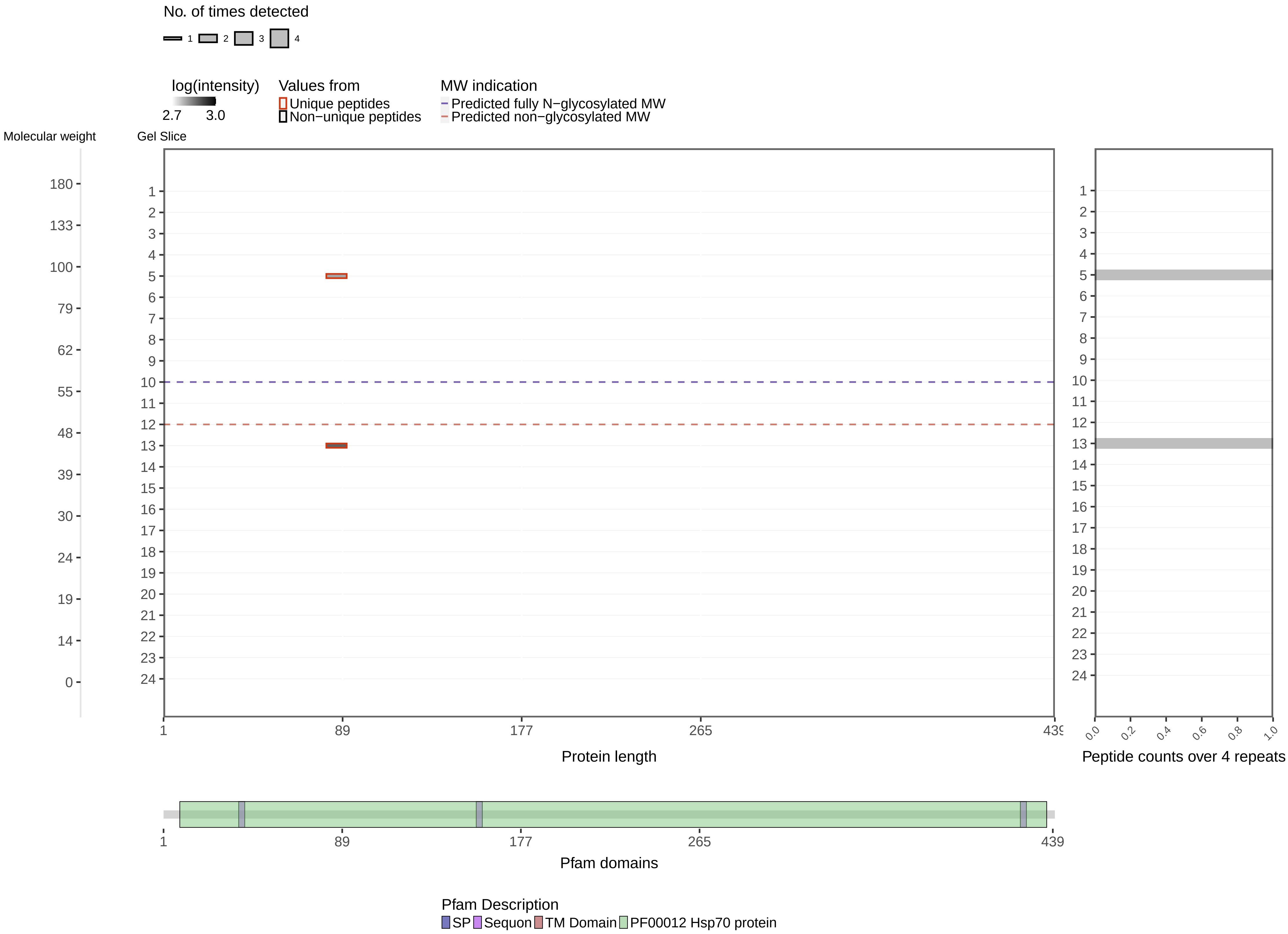


Protein sequence:

LGGGTFDVSLLTIEEGIFEVKATAGDTHLGGEDFDNRMVNHFVQEFKRKHKKDISGNPRALRRLRTSCERAKRTLSSTAQTTIEIDSLYEGVDFYSTITRARFEELNMDLFRKCMEPVEKCLRDAKMDKSTVHDVVLVGGSTRIPKVQQLLQDFFNGKELCKSINPDEAVAYGAAVQAAILSGEGNEKVQDLLLLDVTPLSLGLETAGGVMTVLIPRNTTIPTKKEQVFSHIL*

NbL09g16200.1 NADH_dehydrogenase_[ubiquinone]_1_alpha_subcomplex_subunit_2−like [MW = 10.97 kDa]

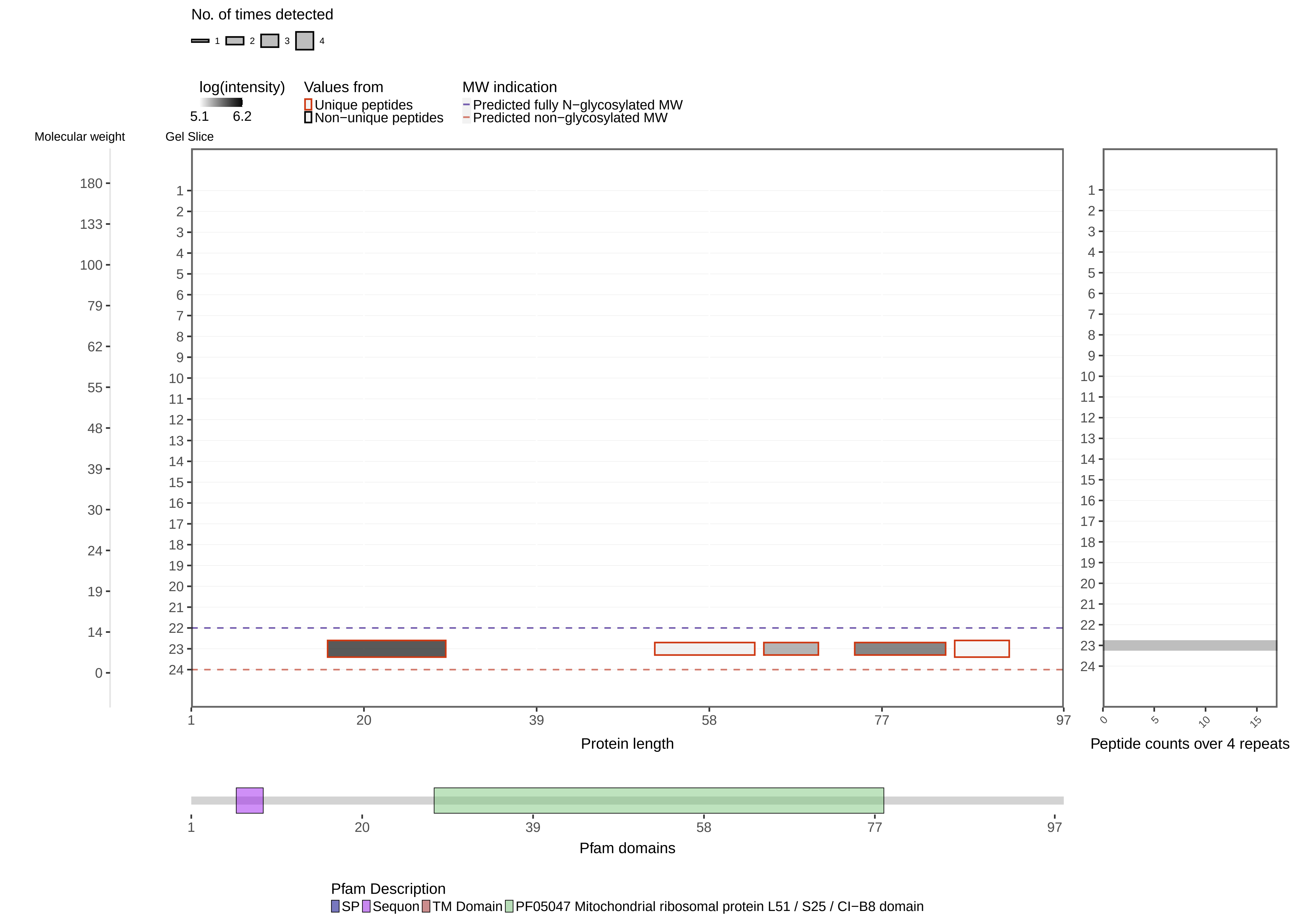


Protein sequence:

NYRDLKTHNPKLPILIREATSIEPQLWARYDLGVERGIRLEGLTEEQISKALEDLVKVGASR*

### NbL09g16180.1 probable_histone_H2B.1 [MW = 16.22 kDa]

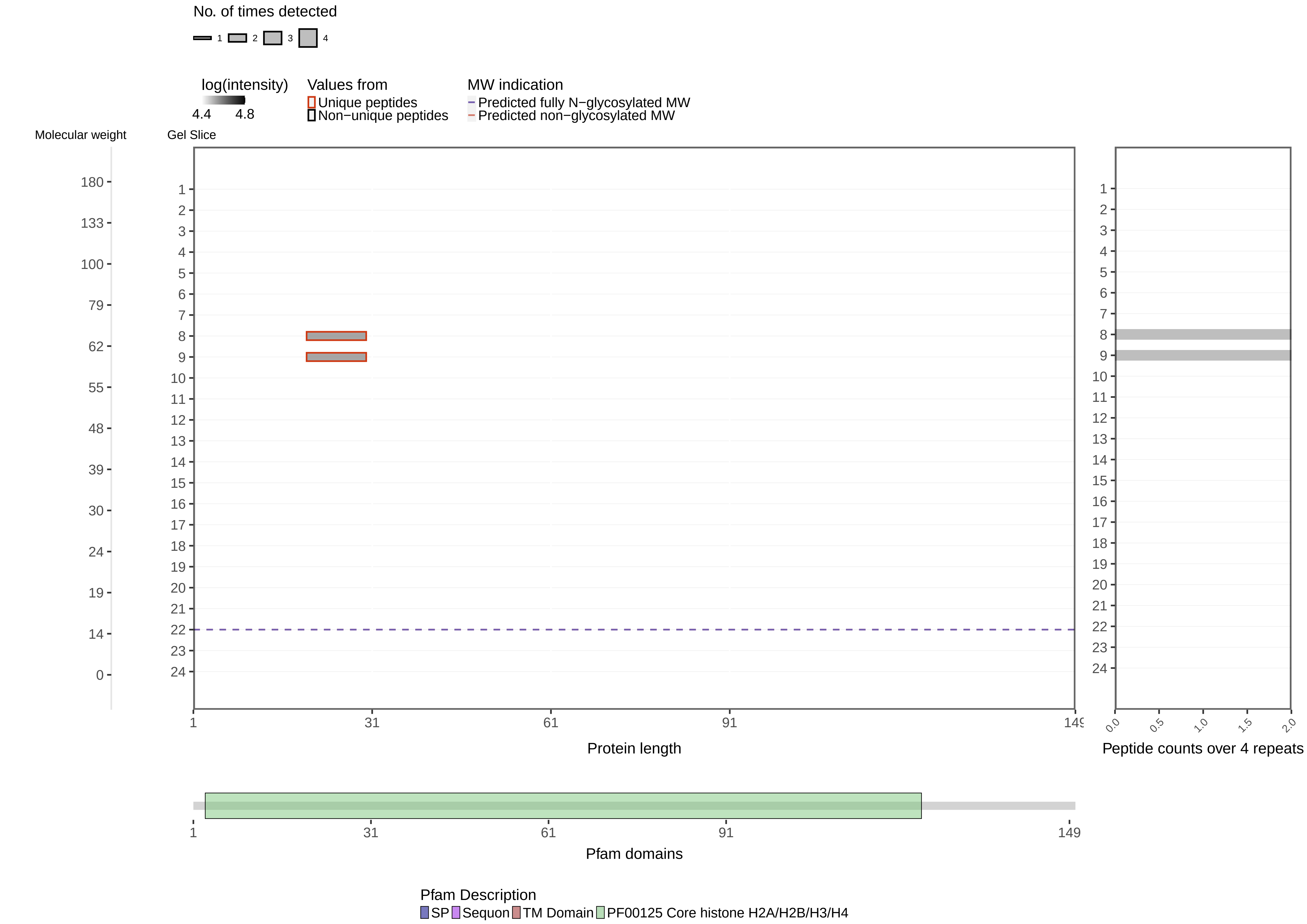


Protein sequence:

ETYKIYIFKVLKQVHPDIGISSKAMGIMNSFINDIFEKLAQESSRLARYNKKPTITSREIQTAVRLVLPGELAKHAVSEGTKAVTKFTSS*

### NbL09g16100.1 aquaporin_TIP1−3−like [MW = 25.9 kDa]

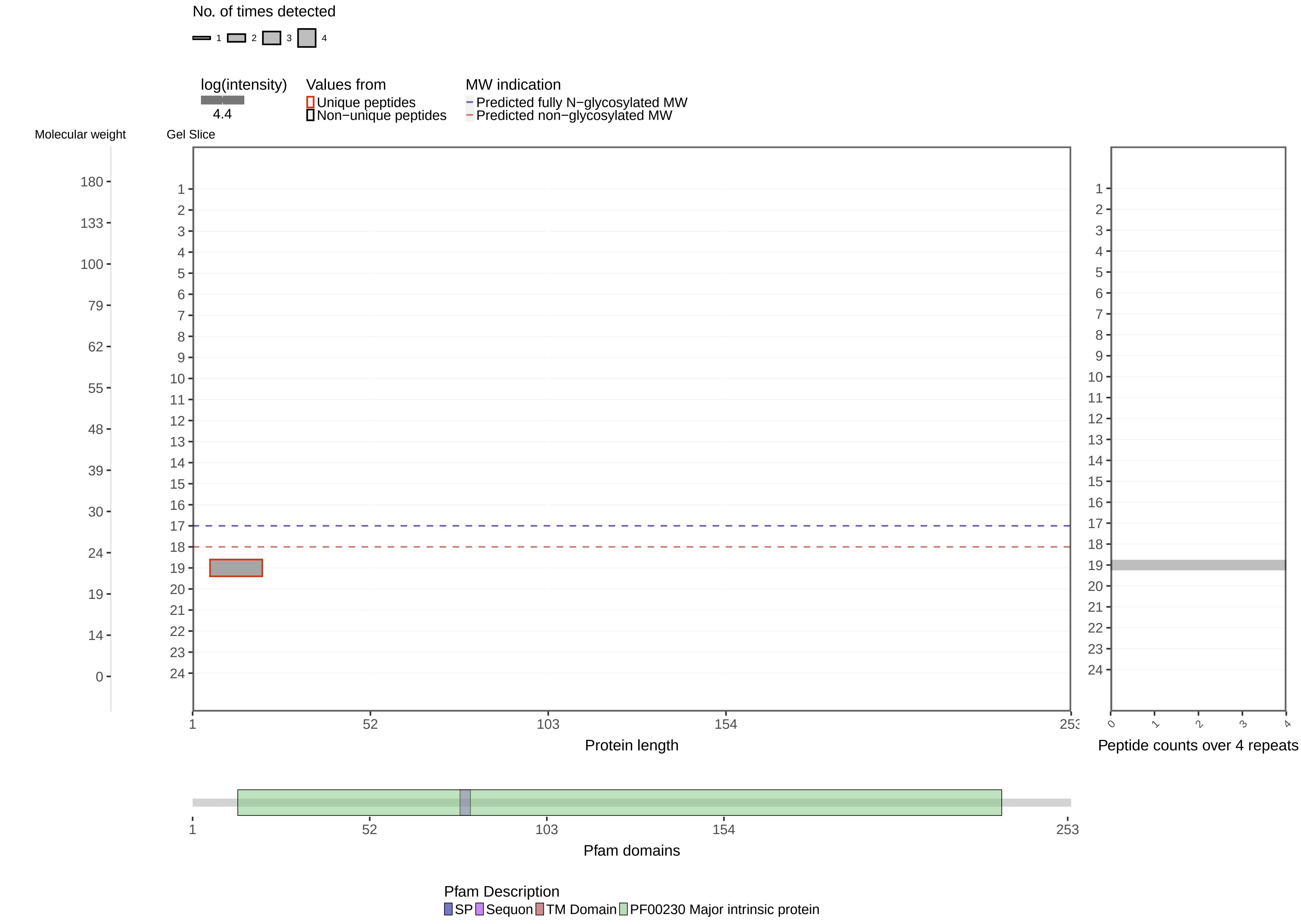


Protein sequence:

SVVACVLLKFATGGLETSAFALSTGVTPWNAVVFEIVMTFGLVYTVYATAVDPKRGNLGIIAPIAIGFIVGANILAGGAFDGASMNPAVSFGPAVVSWTWNCHWVYWLGPFVGAAIAALVYEIIFIGDNTHEQLPTADY*

### NbL09g15940.1 carbamoyl−phosphate_synthase_large_chain_chloroplastic−like [MW = 132.09 kDa]

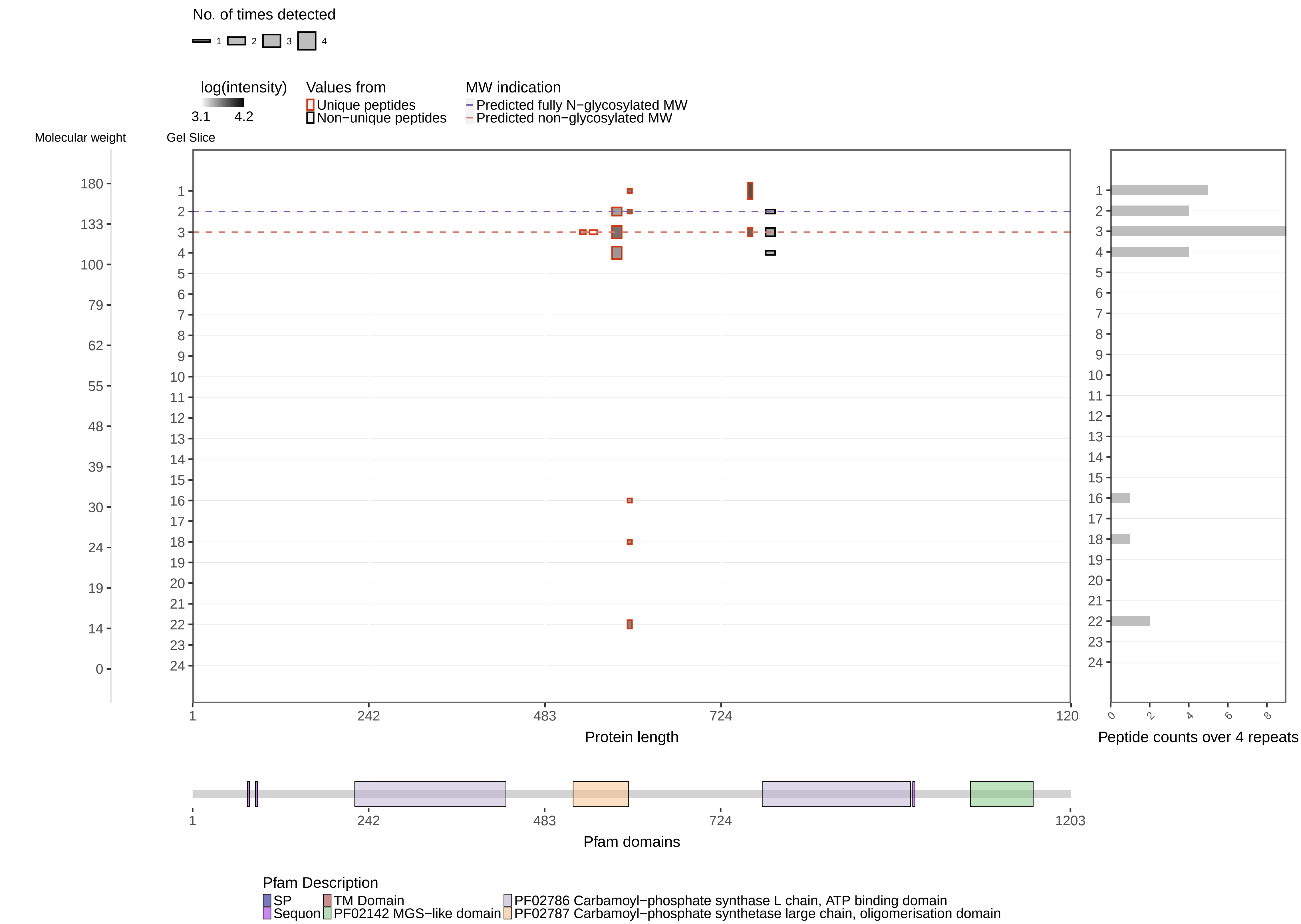


Protein sequence:

[illegible]

### NbL09g15770.1 cell_division_cycle_protein_48_homolog [MW = 89.38 kDa]

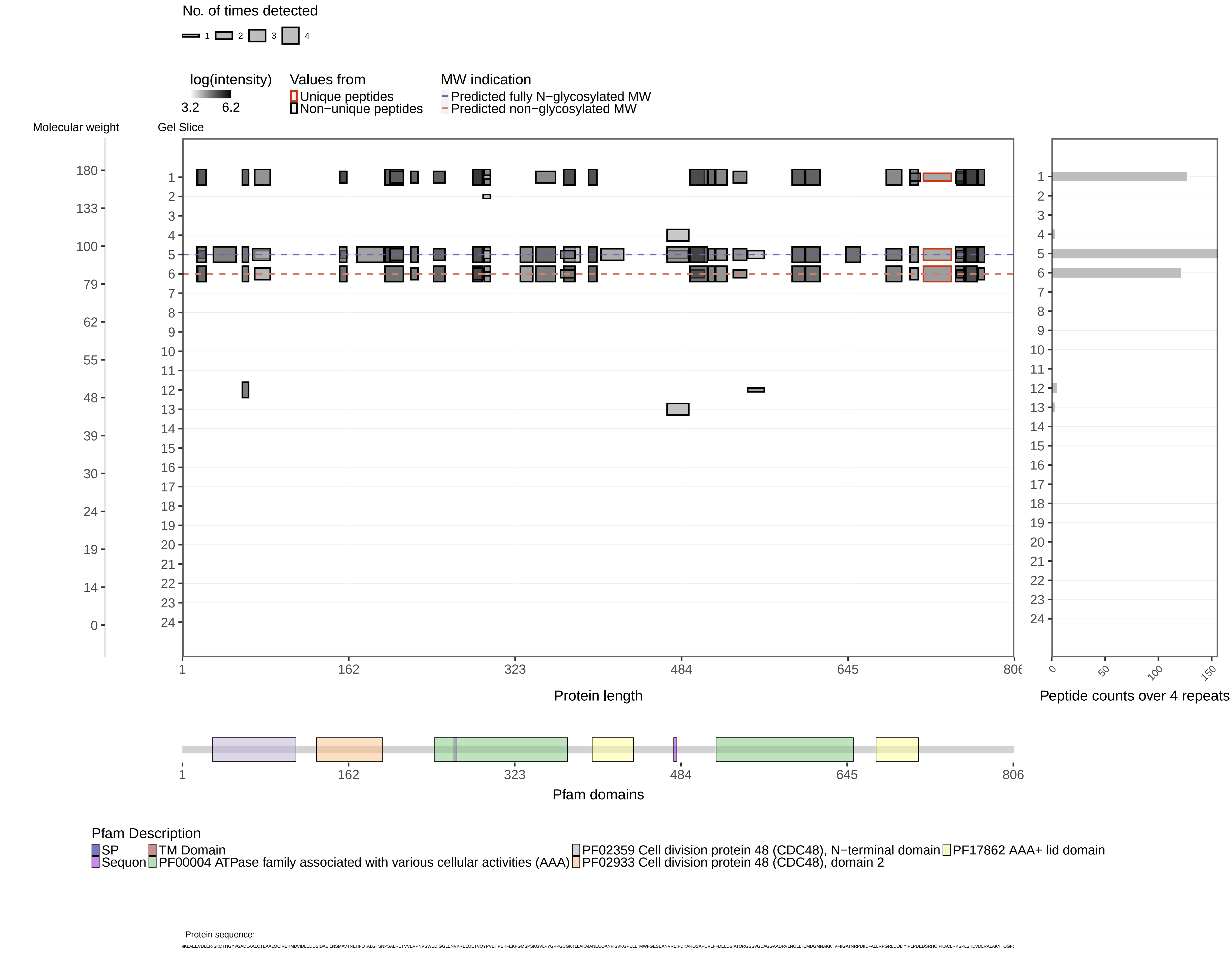

### NbL09g15740.1 ABC_transporter_F_family_member_1−like [MW = 67.55 kDa]

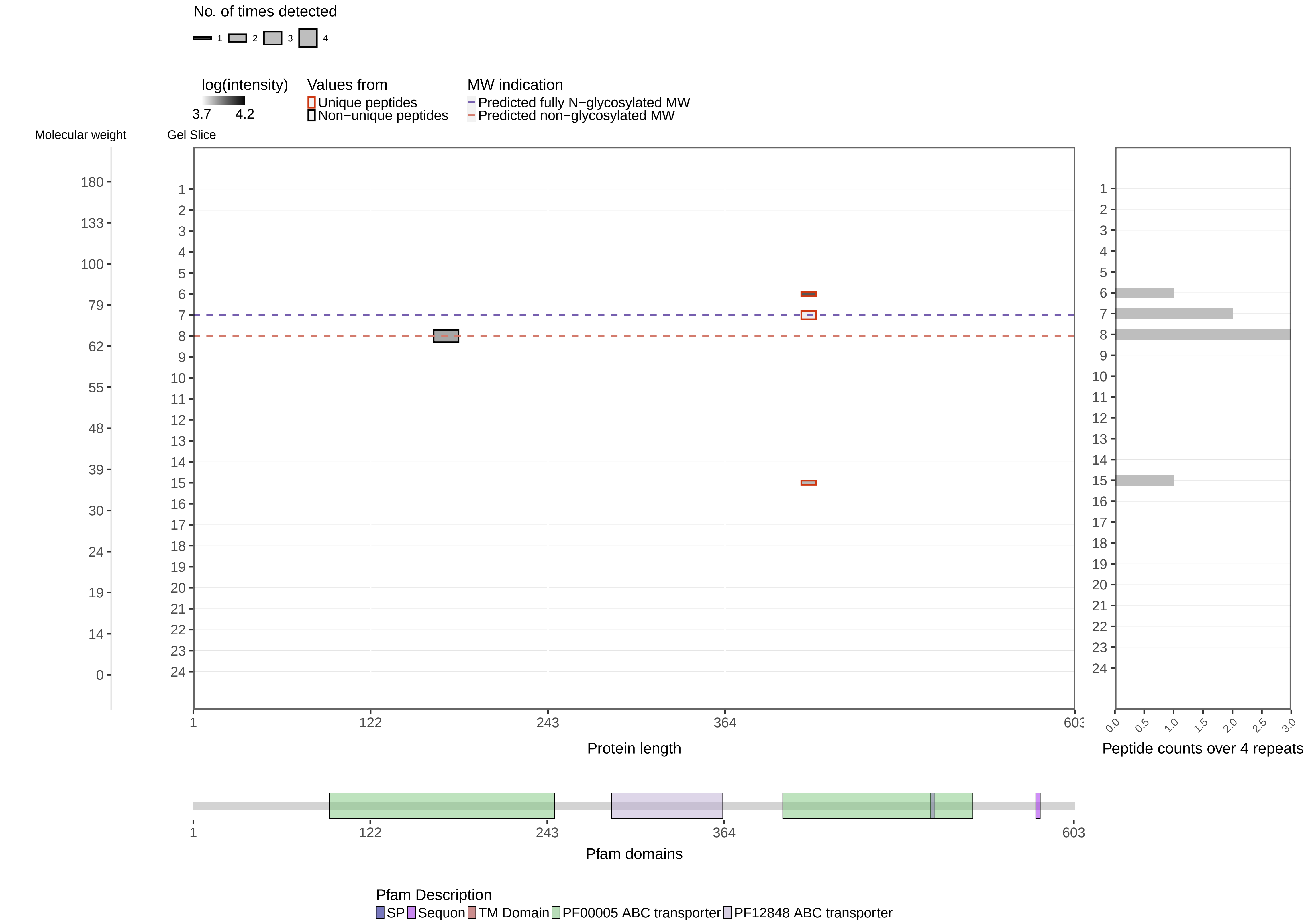


Protein sequence:

KKLKLYTGNYDQYVQTREELEENQMKQFRWEQEQIASMKEYIARFGHGSAKLARQAQSKEKTLAKMERGGLTEKVVKDKVLVFRFPDVGKLPPPVLQFVEVTFGYTPDNLIYKMLDFGVDLDSRVALVGPNGAGKSTLLKLMTGDLVPLDGMVRRHNHLRIAQFHQHLAEKLDMEMSALQYMIKEYPGNEEEKMRAAIGRFGLTGKAQVMPMKNLSDGQRSRVIFGWLAFRQPHILLLDEPTNHLDIETIDSLAEALNEWDGGMVLVSHDFRLINQVAHEIWVCENQTV[illegible]

### NbL09g15690.1 probable_aquaporin_TIP1−1 [MW = 25.74 kDa]

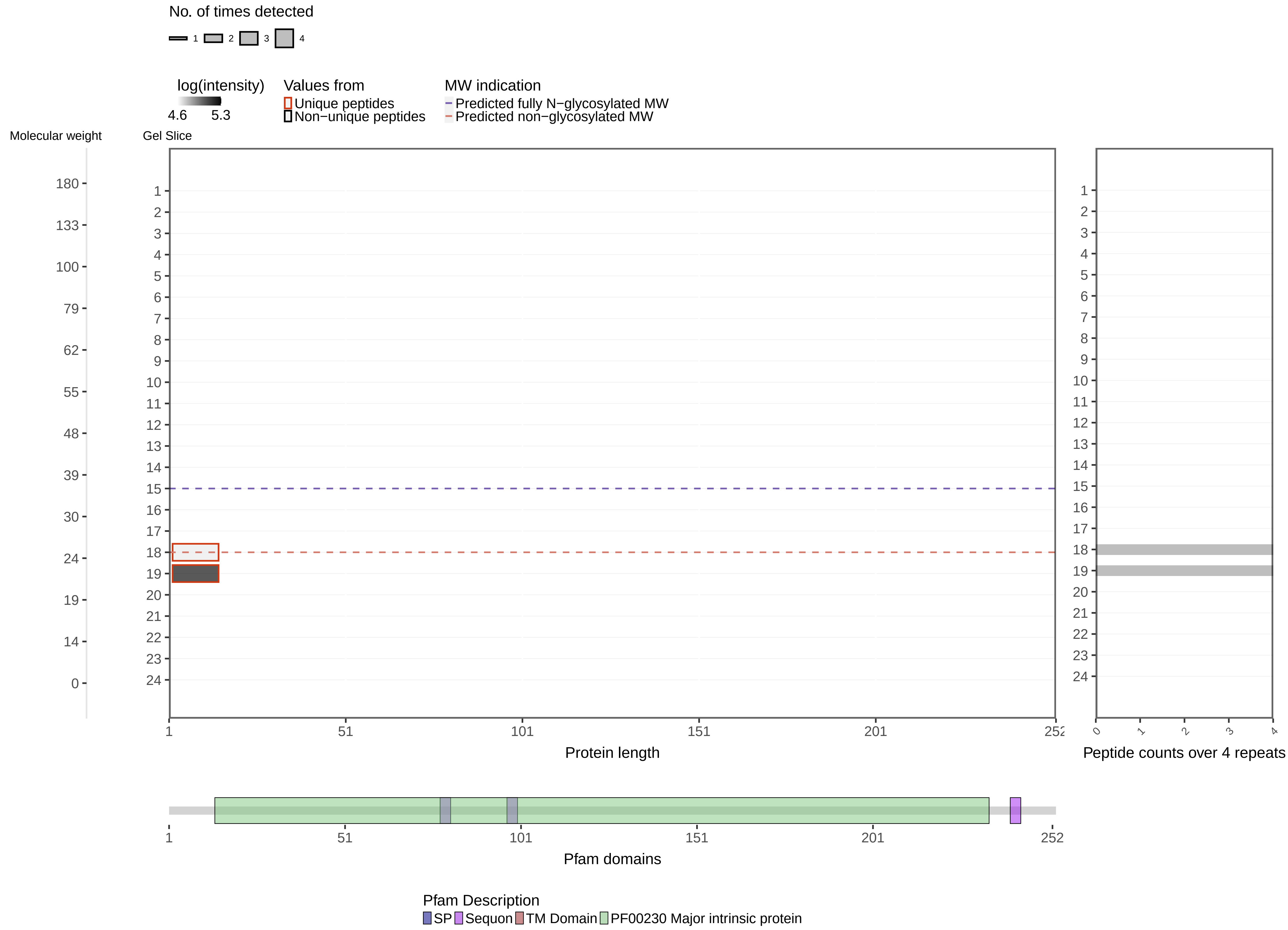


Protein sequence:

TVACFLLEFATGGMSTGAFAVSAGVSVWNAFVFEIVMTFGLVYTVYATAVDPKKGDLGVIAPIAIGFIVGANILAGGAFTGASMNPAVSFGPALVSWTWTNQWVYWAGPLVGGGIAGVVYELFFINHSHEPLPSGDF*

NbL09g15610.1 UDP−D−apiose/UDP−D−xylose_synthase_2−like [MW = 43.26 kDa]

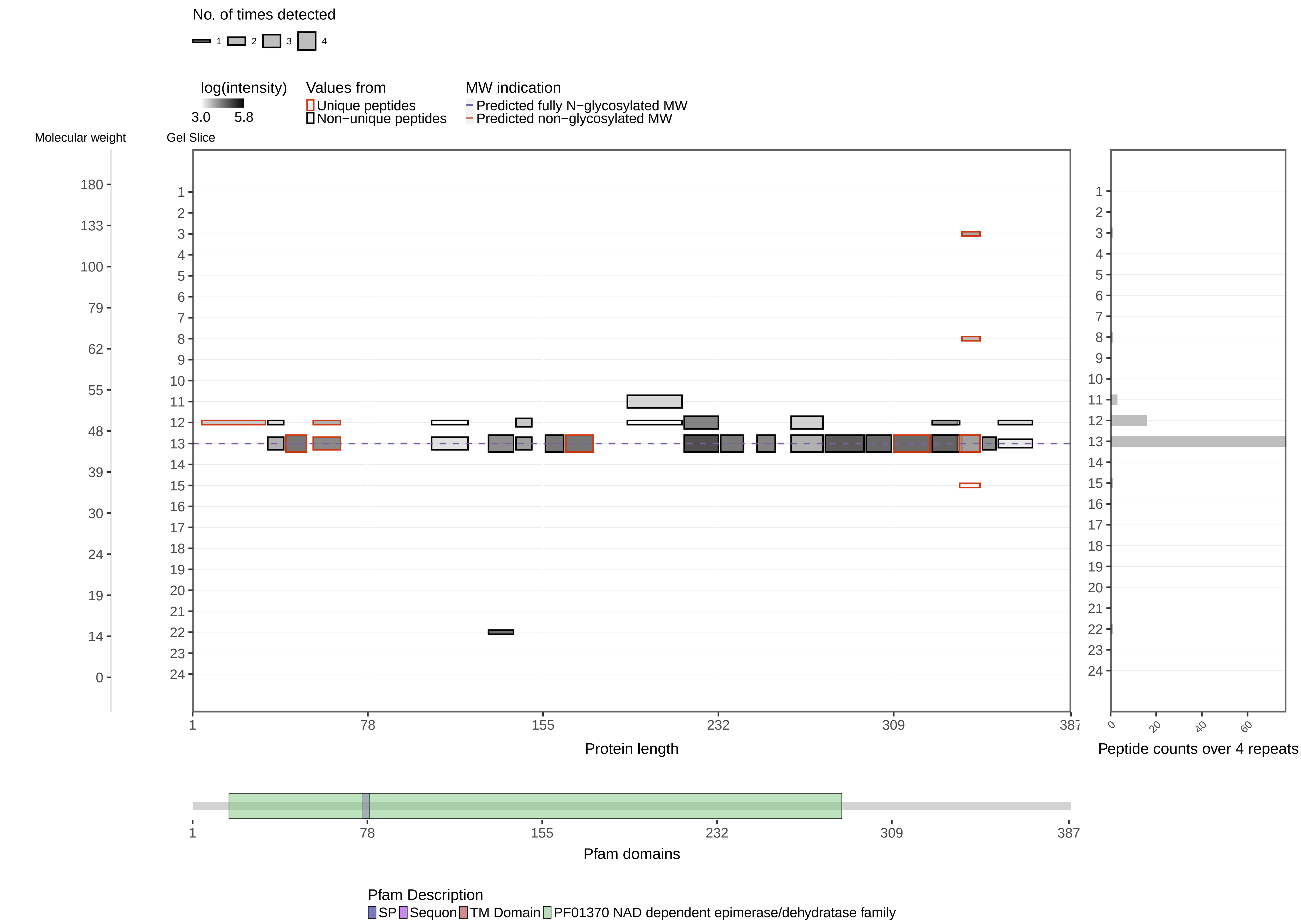


Protein sequence:

ACAKQLIERLIYAEGAENGLEFTIVRPFNWIGPRMDFIPGIDGPSEGVPRVLACFSNNLLRREPLKLVDGGESQRTFIYIKDAIEAVLLMIENPARANGHIFNVGNPNNEVTVRQLAEMMTQVYSKVSGESSLETPTVDVSSKEFYGEGYDDSDKRIPDMTIINKQLGWNPKTSLWDLLESTLTYQHRTYAEVIKQAMSKTTAN*

### NbL09g15540.1 arogenate_dehydratase/prephenate_dehydratase_6_chloroplastic−like [MW = 47.27 kDa]

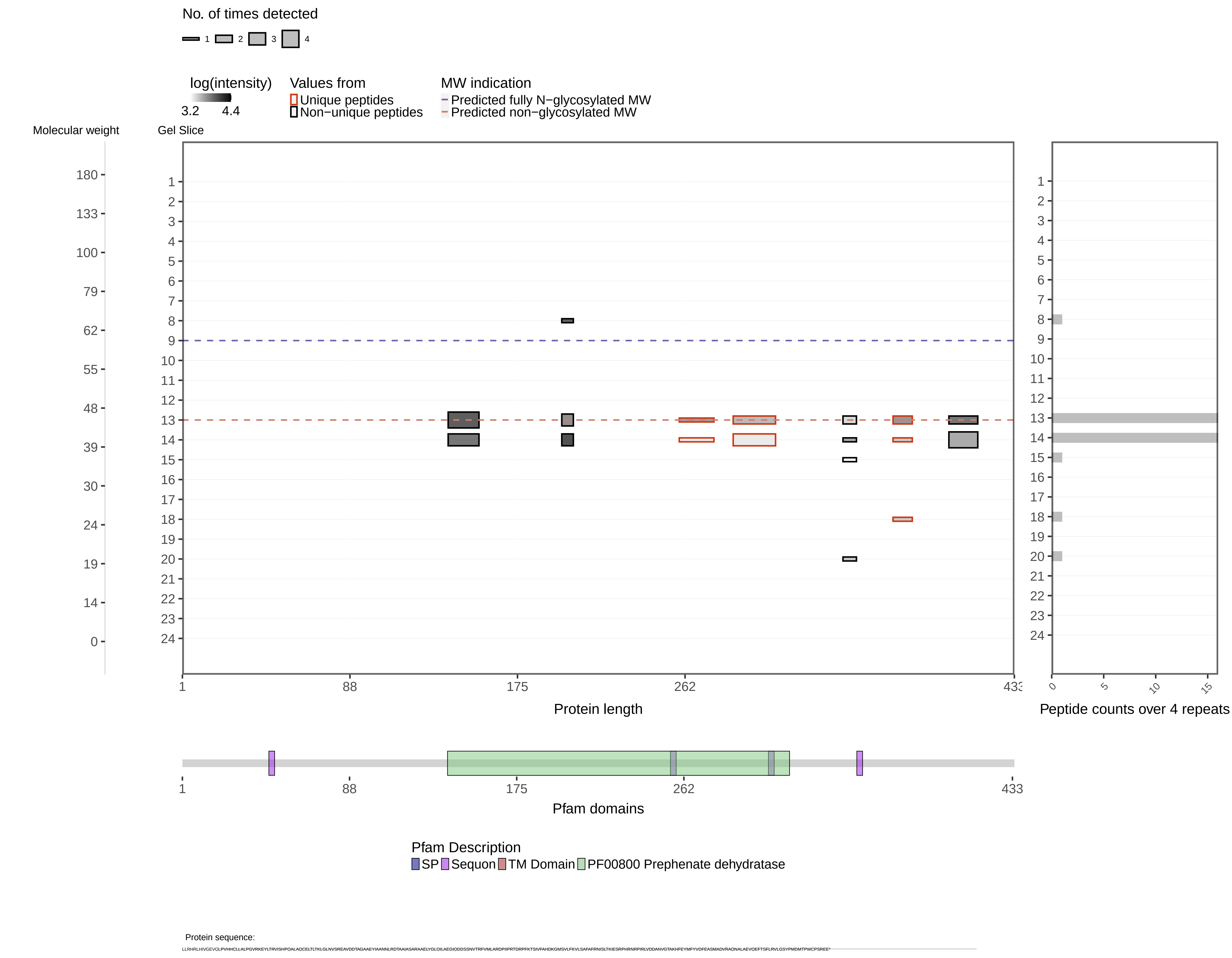

### NbL09g15530.1 probable_2−carboxy−D−arabinitol−1−phosphatase [MW = 55.89 kDa]

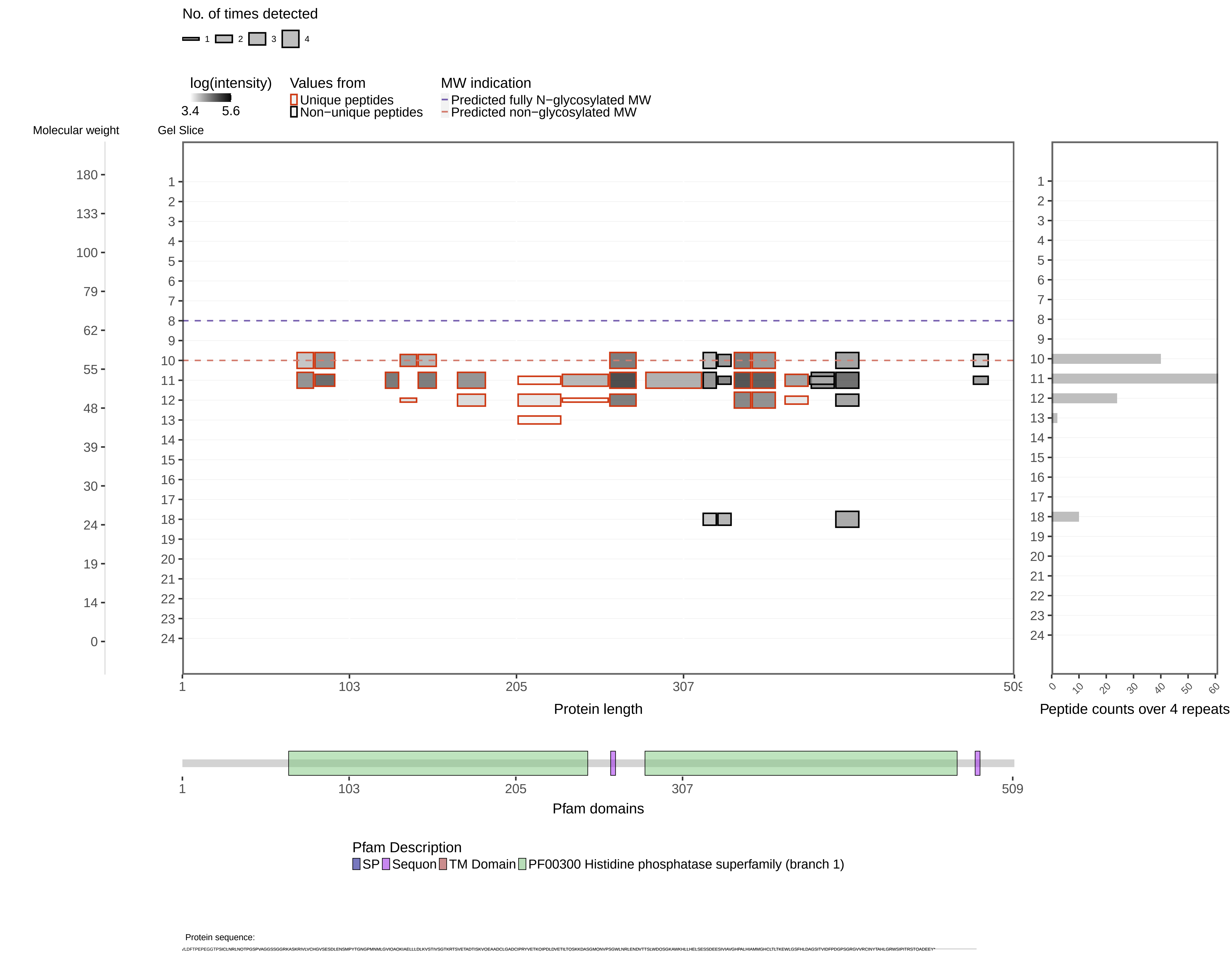

NbL09g15470.1 60S_acidic_ribosomal_protein_P2A−like [MW = 15.01 kDa]

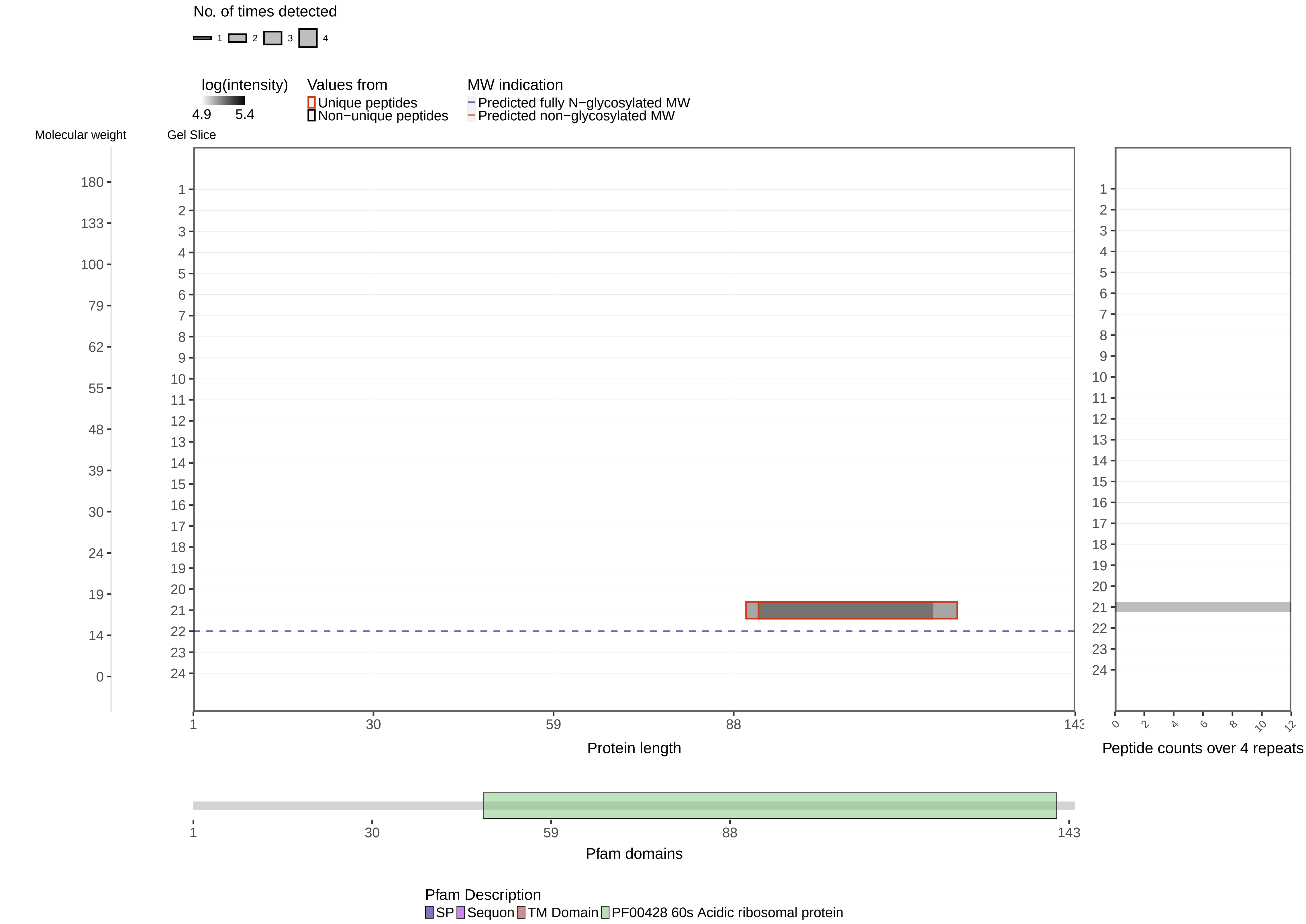


Protein sequence:

VGAEADDDRIELLLSQVEGKDITELIAAGREKLASVPAGGGGVAVAAPTSGAAAAPAAEEKKEEKKVEEKEESDDDMGFSLFD*

### NbL09g15210.1 protein_CDC73_homolog [MW = 45.37 kDa]

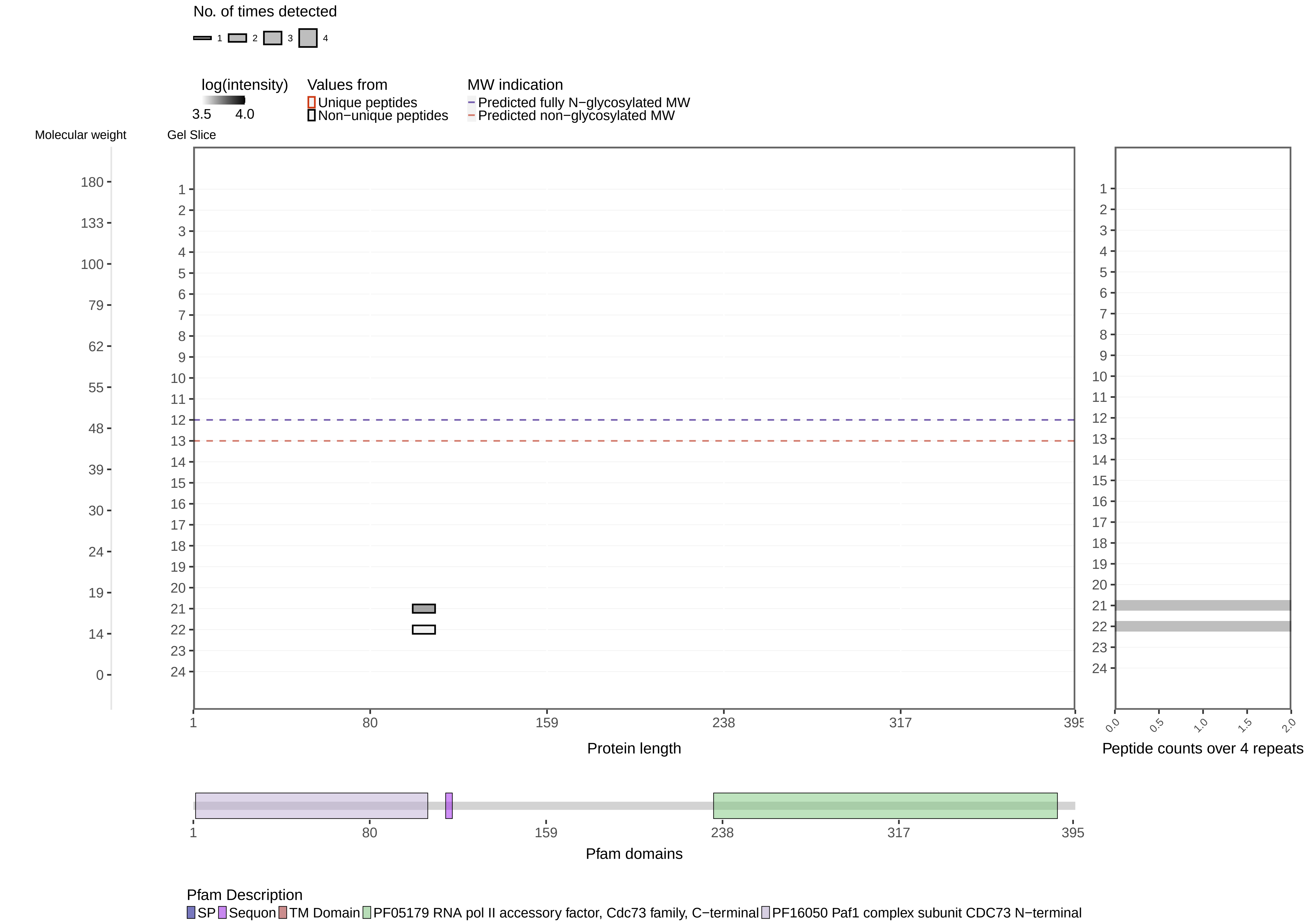


Protein sequence:

ESLQRKDGLVAKNRIERGGFGGGDELGYDGGPKAKMHLKGSKIGEGVPIILVPSAFSTLITIYNVKEFLEDGVFIPTDVKLKQMKGMKPDCITVQKKFSRDRVVTAYEVRDKPSALKAEDWDRVVAVFVLGKEWQFKDWPFKNHVEIFNKIVGFFLRFEDDSVESAKTAKQWNVKIISISKNKRHQDRAAALEVWEKLEEFMRSRSH*

### NbL09g15110.1 uncharacterized_LOC109240313 [MW = 65.51 kDa]

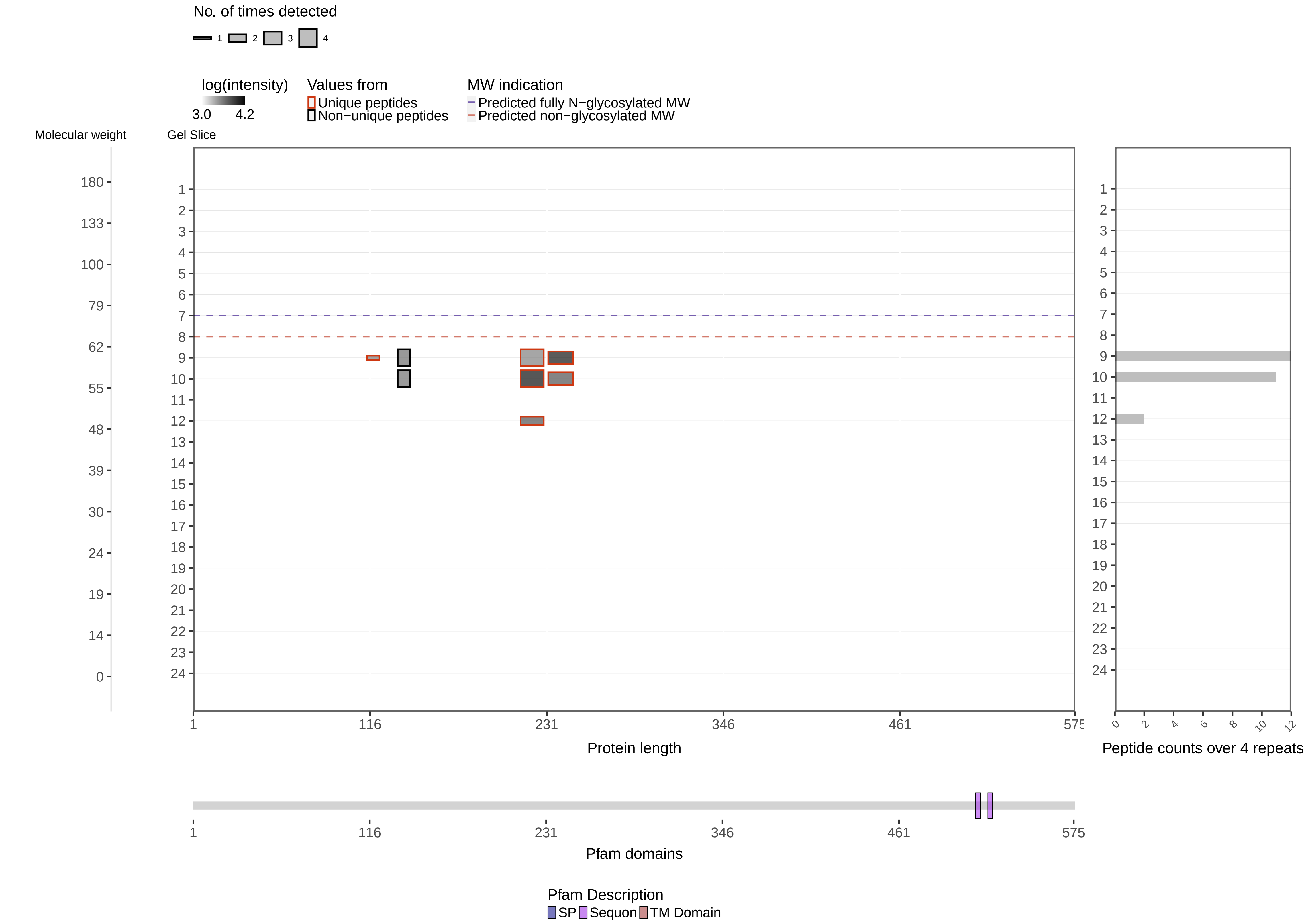


Protein sequence:

FSTNAAEAAYALSQENEASSQGINPDGSRWWKETGTEKRPDGVVCRWTLTRGVSADKTVEWEDKYWEAADEFGHKELGSEKCGRDAAGNVWHEFWKESMWQSGGLVHMEKTADKWGRNKKGEEWQEKWWEHYGAGGQAEKWAHKWCSIDPNTPLEAGHAHVWHERWGEKYDGKGGSTKYTDKWSERFEGDGWAKWGDKWDEHFDLNAHGVKQGETWWAGKSGERWNRTWGEGHNGSGWVHKYGKSSSGEHWDTHINEETWYERFPHYGFYH[illegible]

### NbL09g14990.1 60S_ribosomal_protein_L19−3−like [MW = 24.56 kDa]

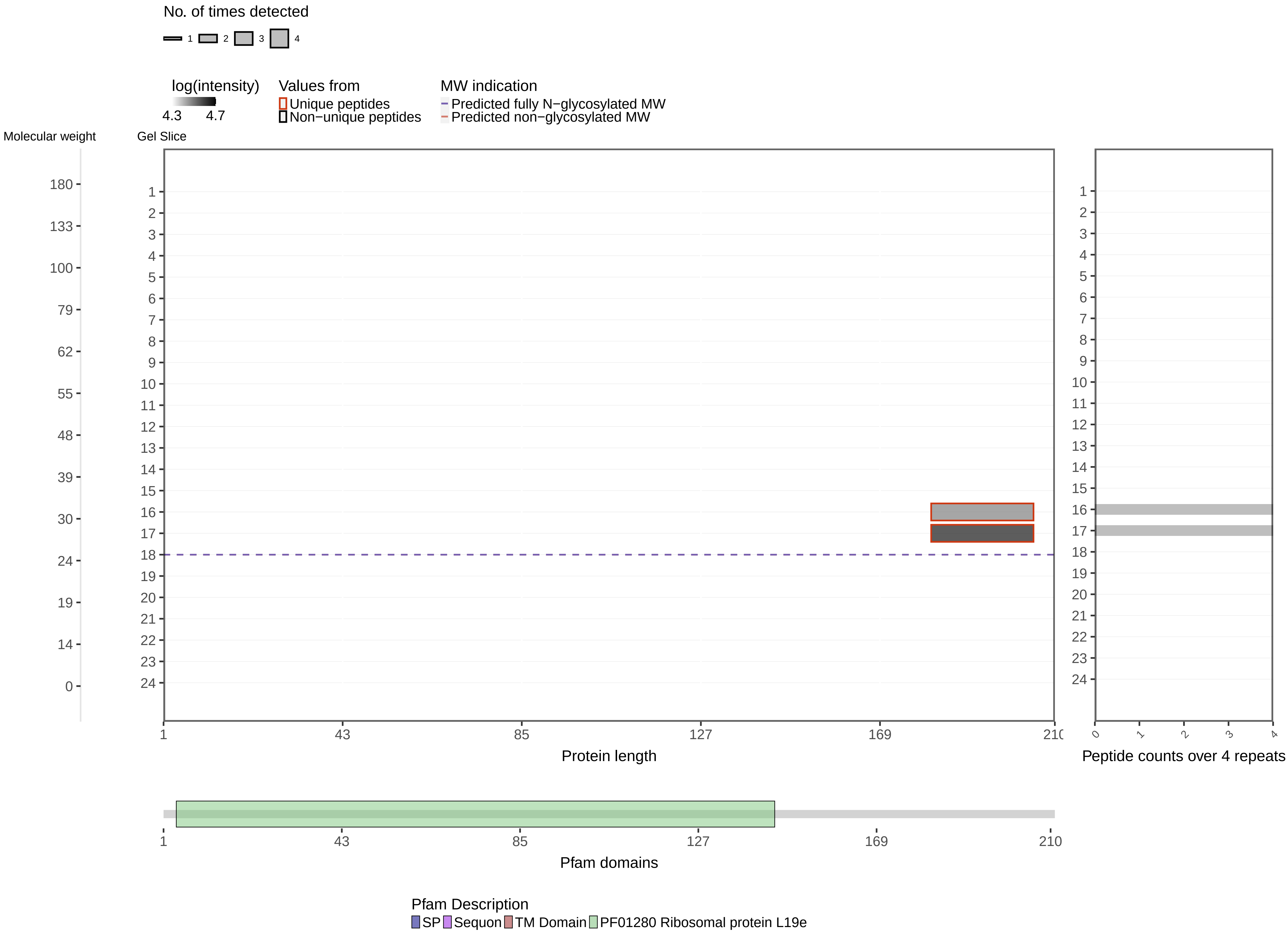


Protein sequence:

/LWMRRMRVLRRLLRKYRESKKIDRHMYHDMYMKVKGNVFKNKRVLMENIHKTKAEKAREKTLSDQFEARRAKNKASRERKFARREERLAQGPGERPVQPAAATEAPQAQGSKKSKK*

NbL09g14980.1 uncharacterized_protein_At5g01610−like [MW − SP = 15.98 kDa]

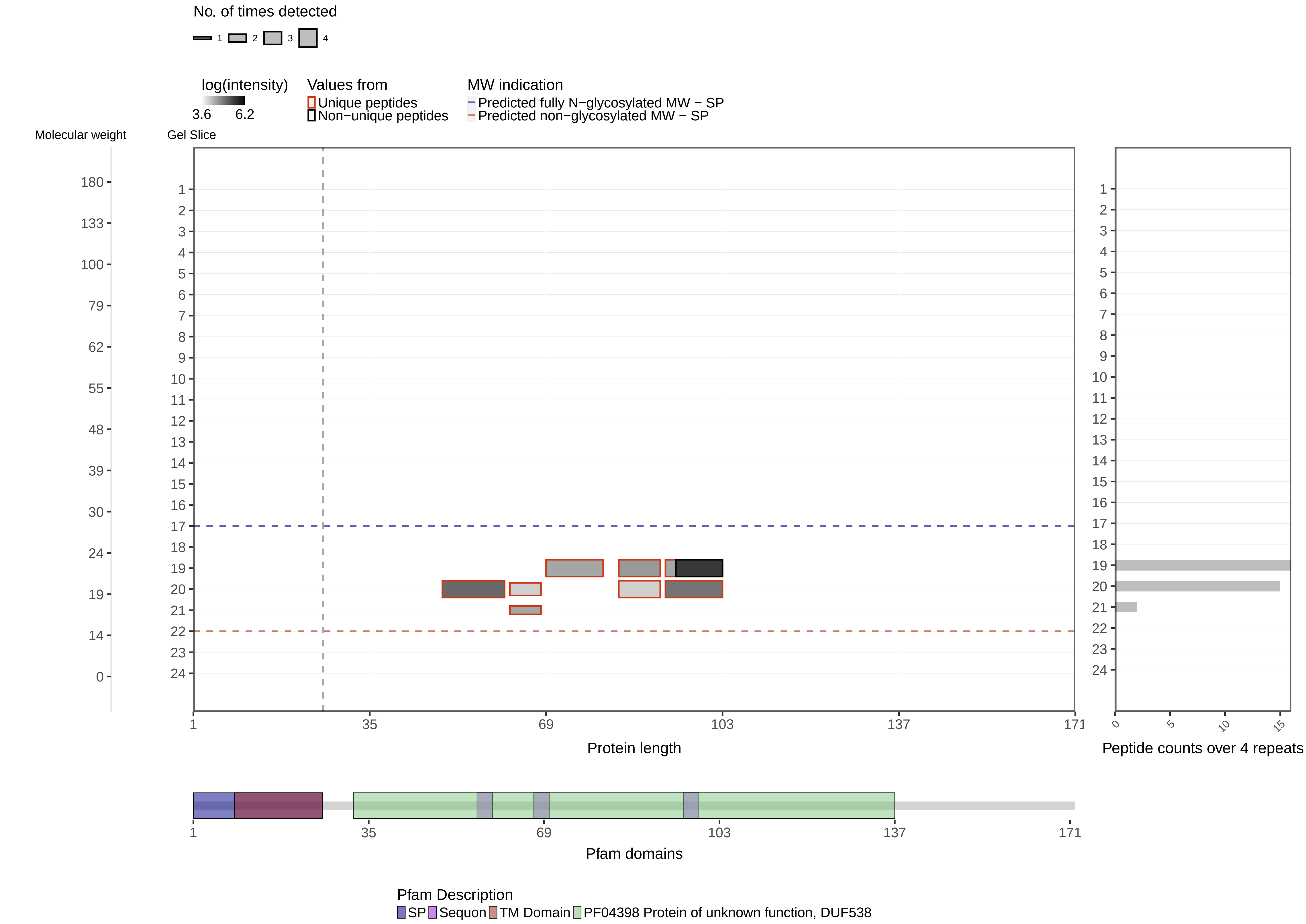


Protein sequence:

EIESYSLRYKSKITGTLSKDKLSNLSGVQVKVLFFWVNIGEVARVDDELDFTVGIVSADFPIDNFYESPQCGCGFDCVNSGGKSTGEIRGNIEMKEHK*

NbL09g14800.1 short−chain_dehydrogenase_reductase_3c−like [MW = 31 kDa]

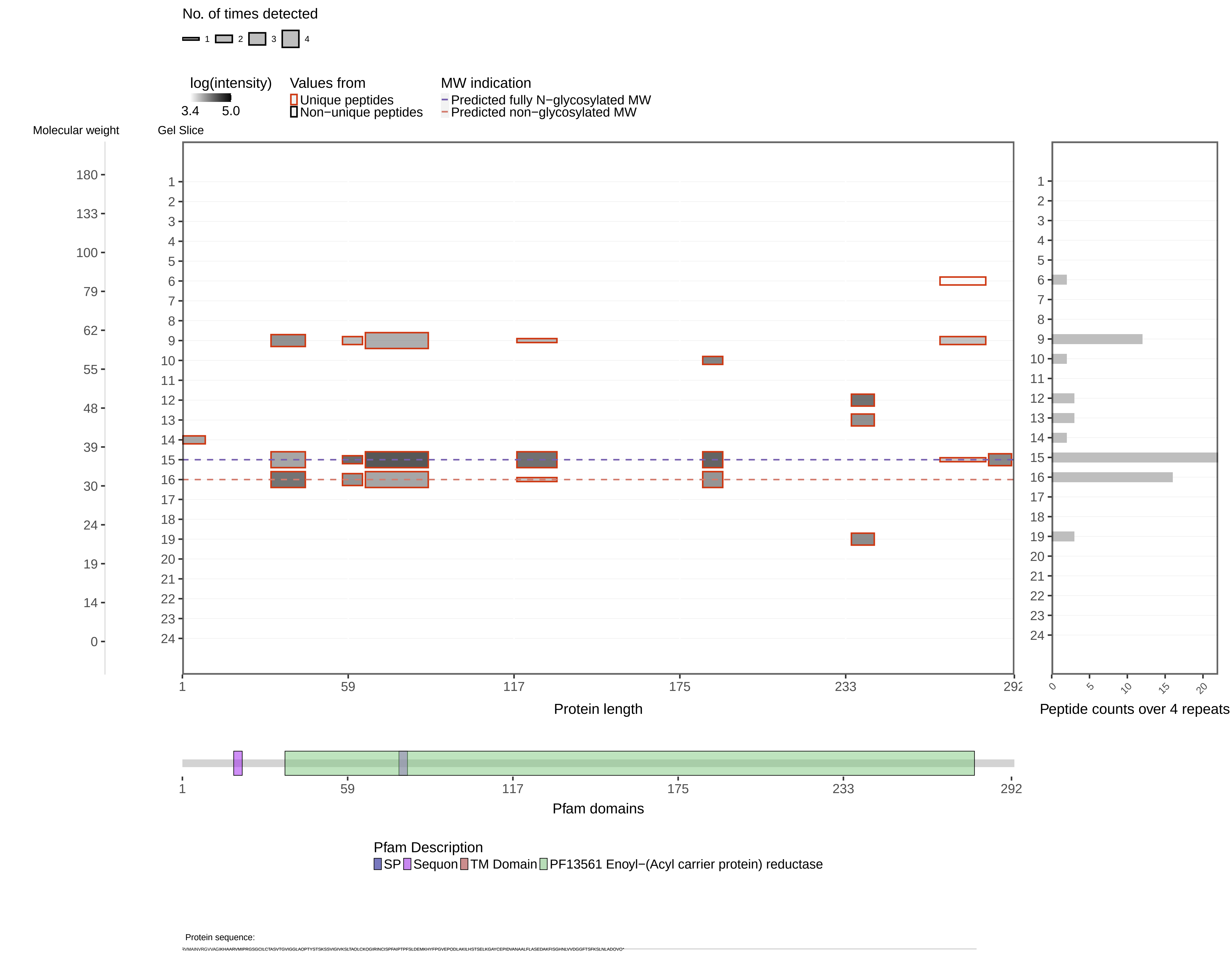

NbL09g14740.1 NPL4−like_protein_1 [MW = 46.36 kDa]

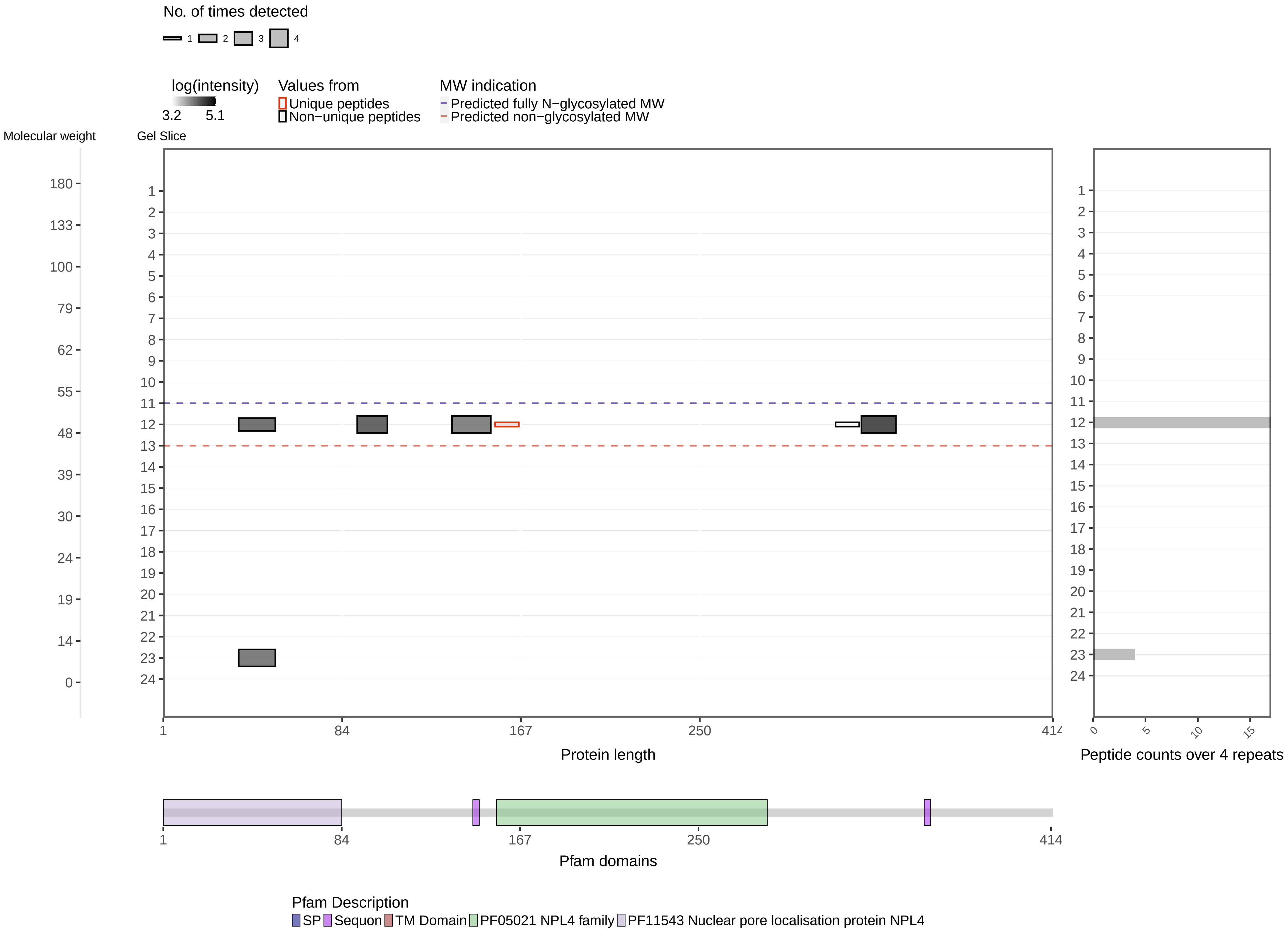


Protein sequence:

EKVVDALAIGLGMRRVGFIFTOTISODKKDYTMSTAEVLOAAELHAEGDLKEWVTAIVKLEVNEDGAADVHFEAFOMSDMCVRLFKEGWFETEVNKDEMIEPKLSKMKKDVVLGVKDTREVDNDFFLVVVKIADHOGPLSSSFPIENRNIPVSMRALKDHFNRTKSLSFVKRISDFHLLLLLAKFLDINADVPALAECVHTOTAVPEGYOLLIESMASAS*

### NbL09g14580.1 small_nuclear_ribonucleoprotein_Sm_D1−like [MW = 12.61 kDa]

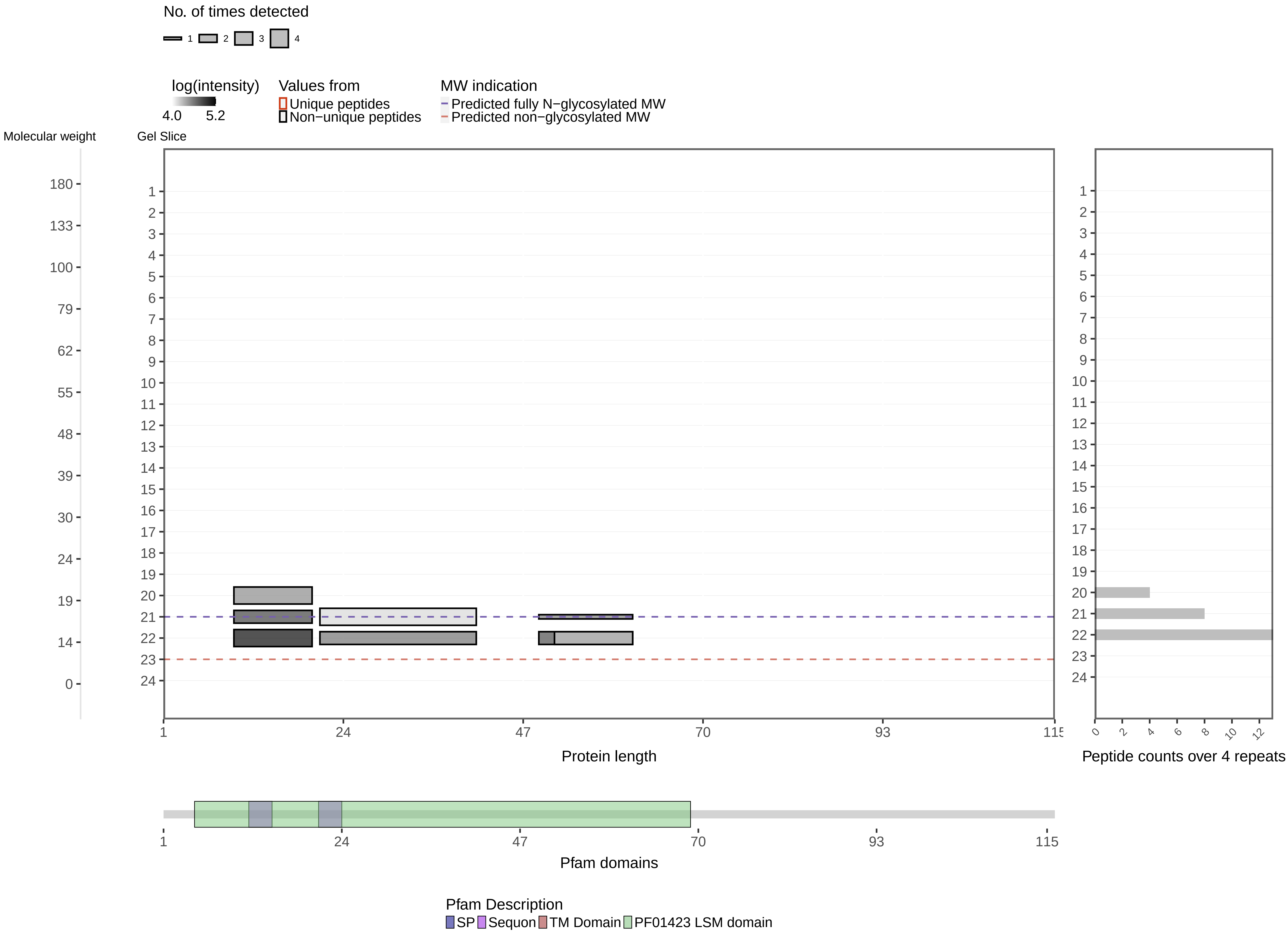


Protein sequence:

LTLKGKNPVTLDHLSVRGNNIRYYILPDSLNLETLLVEETPRVKPKKPTAGKPLGRGRGRGRGRGRGRGR*

NbL09g14530.1 fatty−acid−binding_protein_1 [MW = 32.3 kDa]

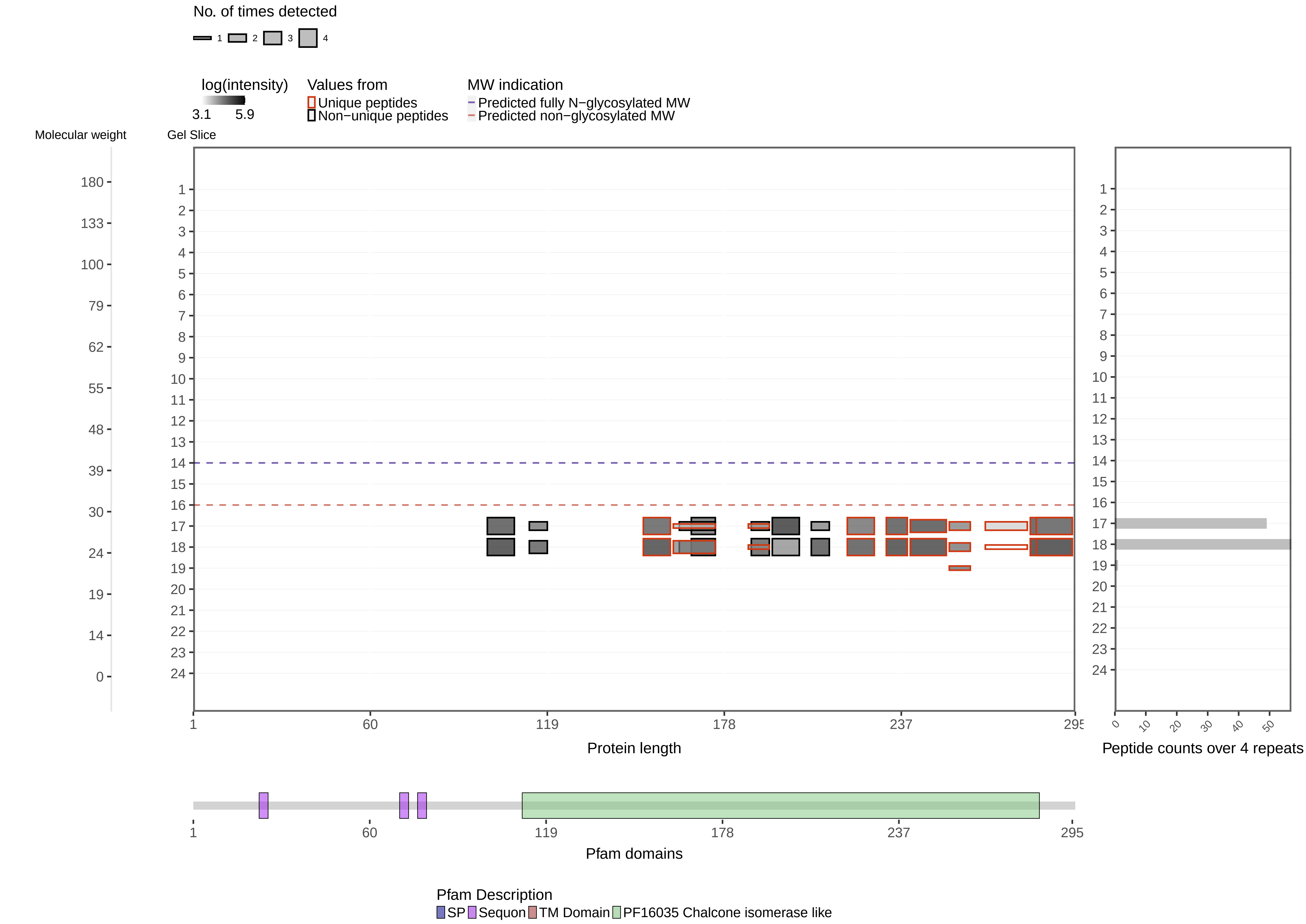


Protein sequence:

FGVYADDGEVKKCLAQKHVGPSDSELKSKEELRDHLMENDIRMTVRLQIVYRRLSIGSVRSAFEESVGSRLLKFGGYDNKELLHRFTSQFKDDIKIPRGSIIELSRDHGYILHTTIDGKEVGSIQSKLLCRSILDLYIGEESFDQKAKEDIESNLASLLHK*

### NbL09g14420.1 non−specific_lipid_transfer_protein_GPI−anchored_2−like [MW − SP = 13.62 kDa]

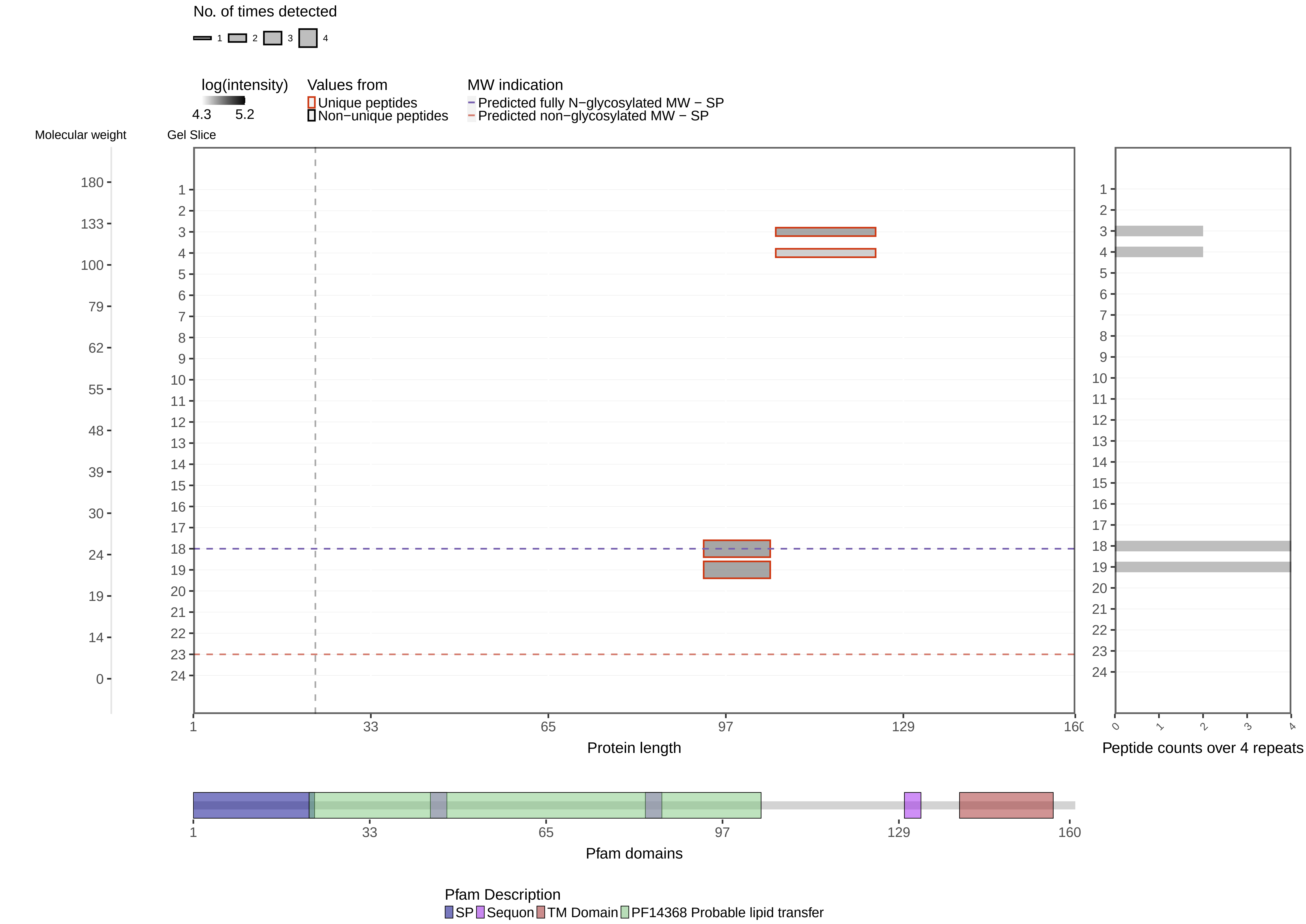


Protein sequence:

CQVVNGGGSNLGINVNQTQAMALPKACNVQTPSVSLCKGTTPSGSSGSSSNPSGGSKGEPSGNSSGNSVKMPYSLLFILVAIASLATVFTPI*

### NbL09g14140.1 glycolipid_transfer_protein_1 [MW = 24.68 kDa]

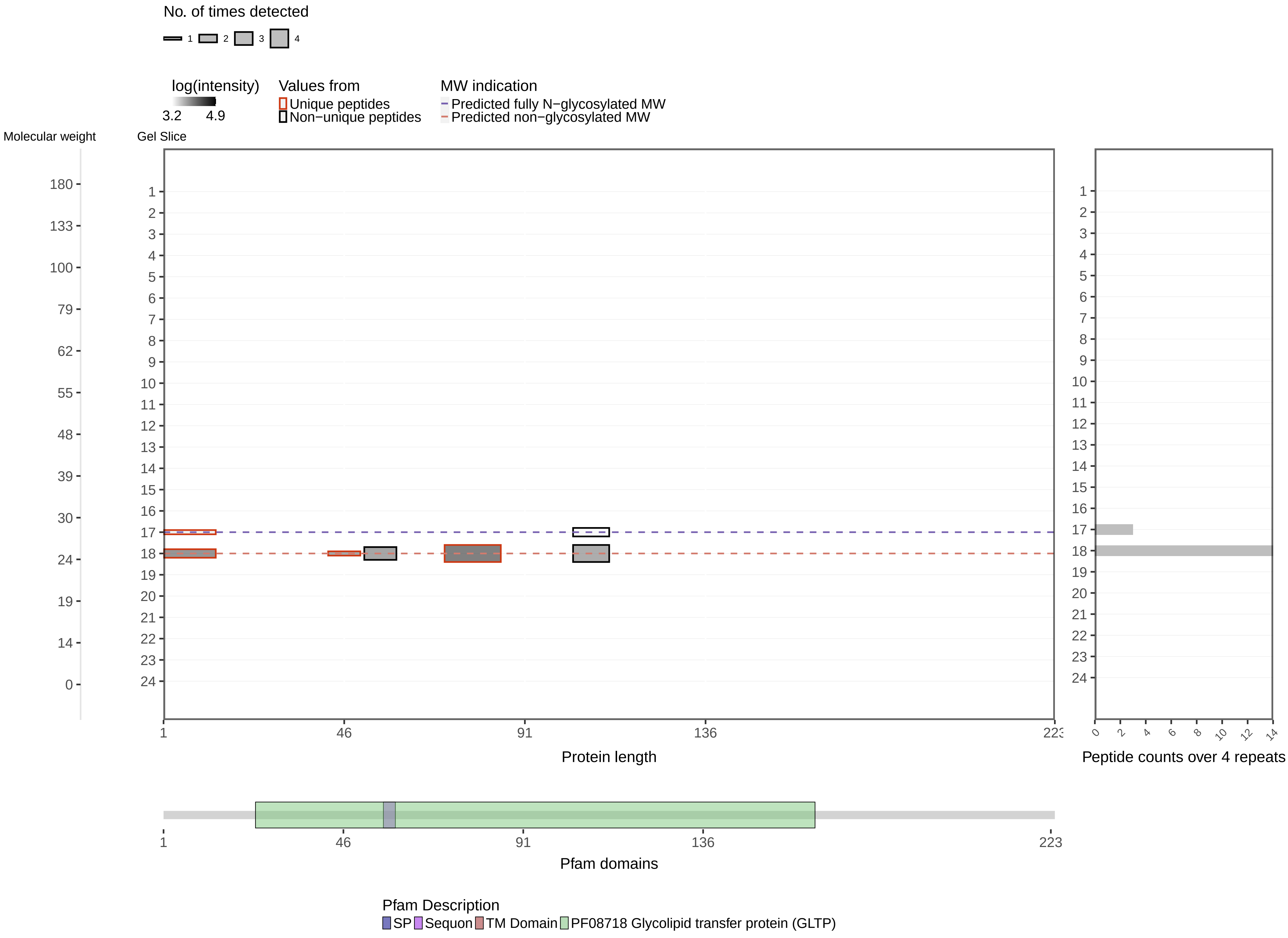


Protein sequence:

.TRAMDFLVELFRNLLEHGDWAMSQACSDSYSKTLKKFHGWLASSSFSVAMKLAPDRKKFMDVICGTGDIDSDIGQFCTTFSPLLEENHKFLVKCFLGGYSSDRCLDFFLGLHYCSCSFPAPT*

NbL09g14120.1 succinate−−CoA_ligase_[ADP−forming]_subunit_beta_mitochondrial [MW = 45.27 kDa]

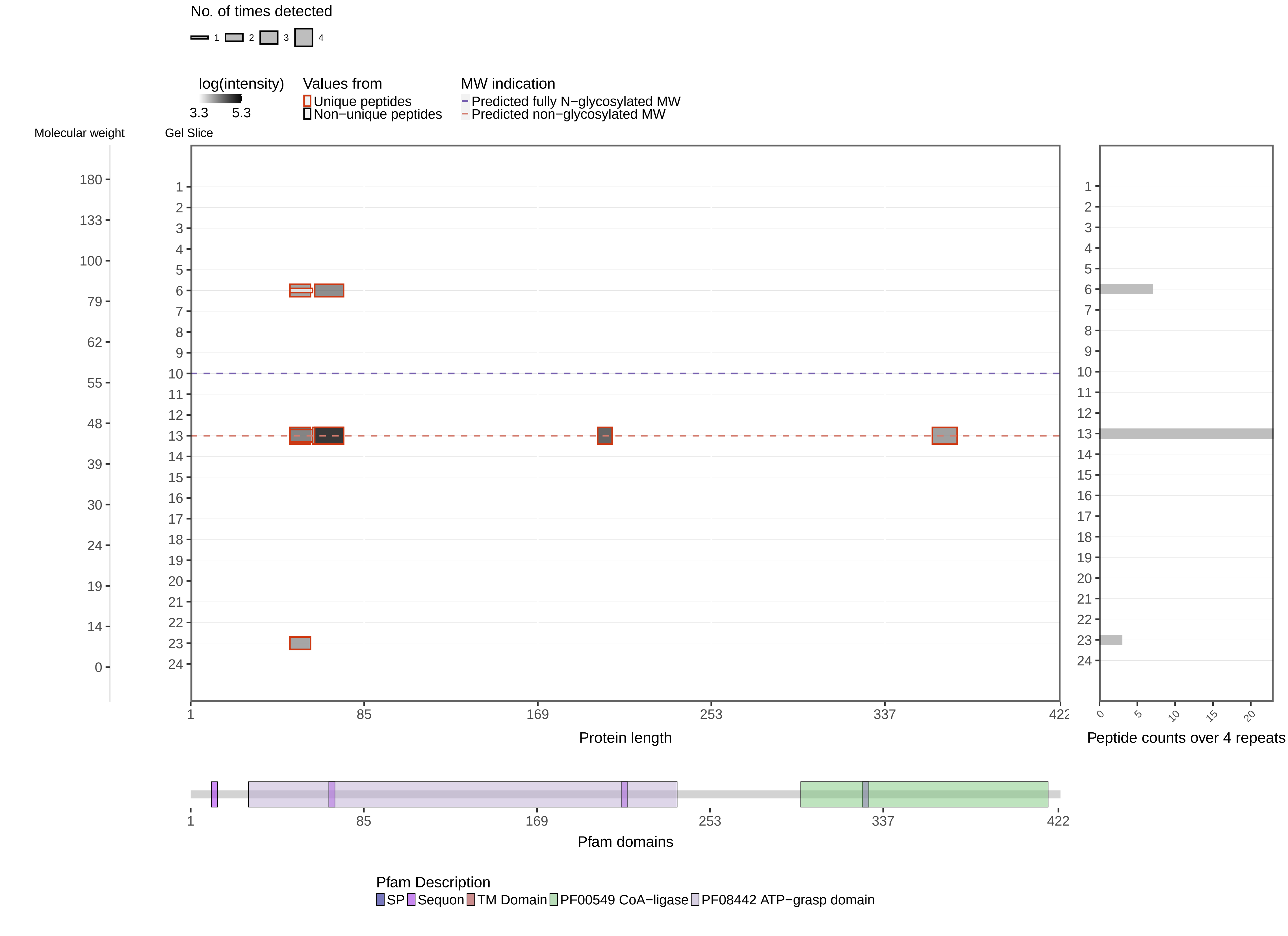


Protein sequence:

KVVDGFAPKVADRNDSIEQVKKLYQLFCESDCTMLEINPLAETSDNKLVAADAKLNFDDNAAYRQKEIFSLRDSSQEDPREVAAAKADLNYIGLDGEIGCMVNGAGLAMATMDIIKLHGGTPANFLDVGGNATEGQVVEAFKILTADEKVKAILVNIFGGIMKCDVIASGIVNAAKQVQLKVPVVVRLEGTNVEQGKRILKESGMTLITAEDLDDAAEKAVKALA*

NbL09g14000.1 monothiol_glutaredoxin−S10−like [MW = 28.11 kDa]

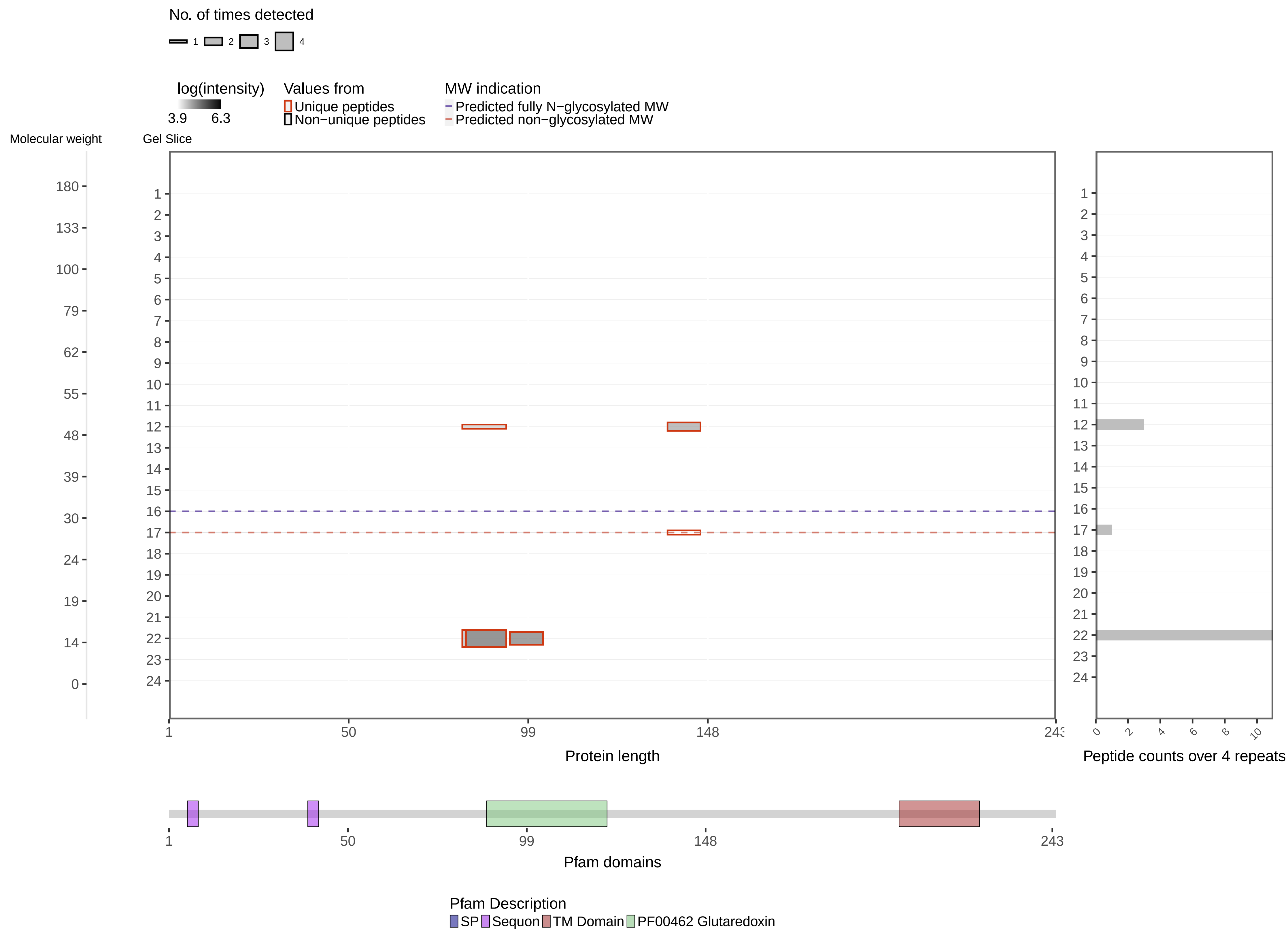


Protein sequence:

KLGVDPLVIELDEMVDMWFQVPKDHSCRKCWNGLLDNIRFPMYSLEPNTLVVVQILLSCIGKENLNLCCRKLMLEKQRAKPSWVSMLKRSYLMLSLYALFFQISLTYSYLIMLFLYKYILLLIIKLTVEAEIMLM*

### NbL09g13890.1 probable_pectate_lyase_8 [MW − SP = 46.28 kDa]

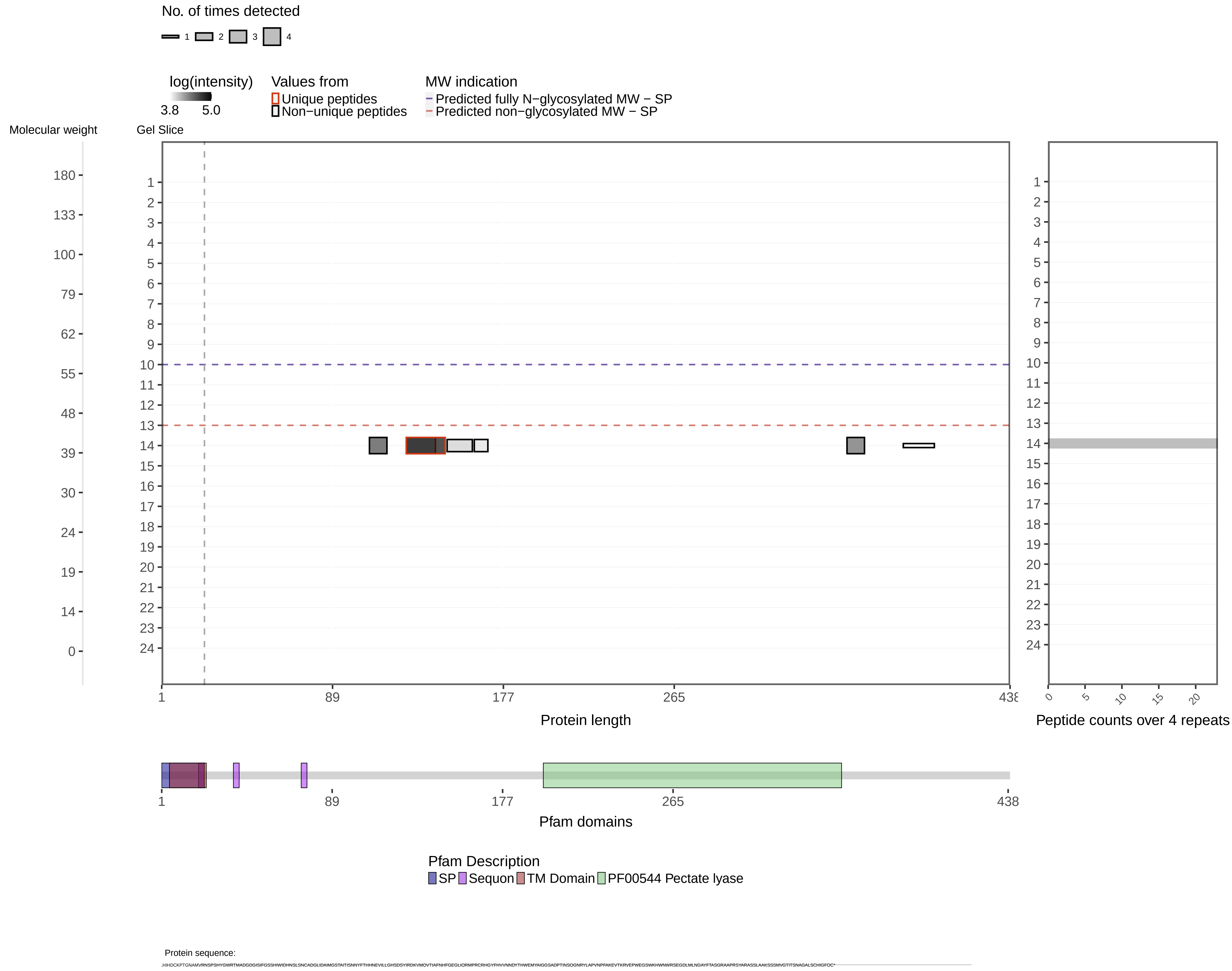

### NbL09g13540.1 peptidyl−prolyl_cis−trans_isomerase_FKBP62−like [MW = 63.37 kDa]

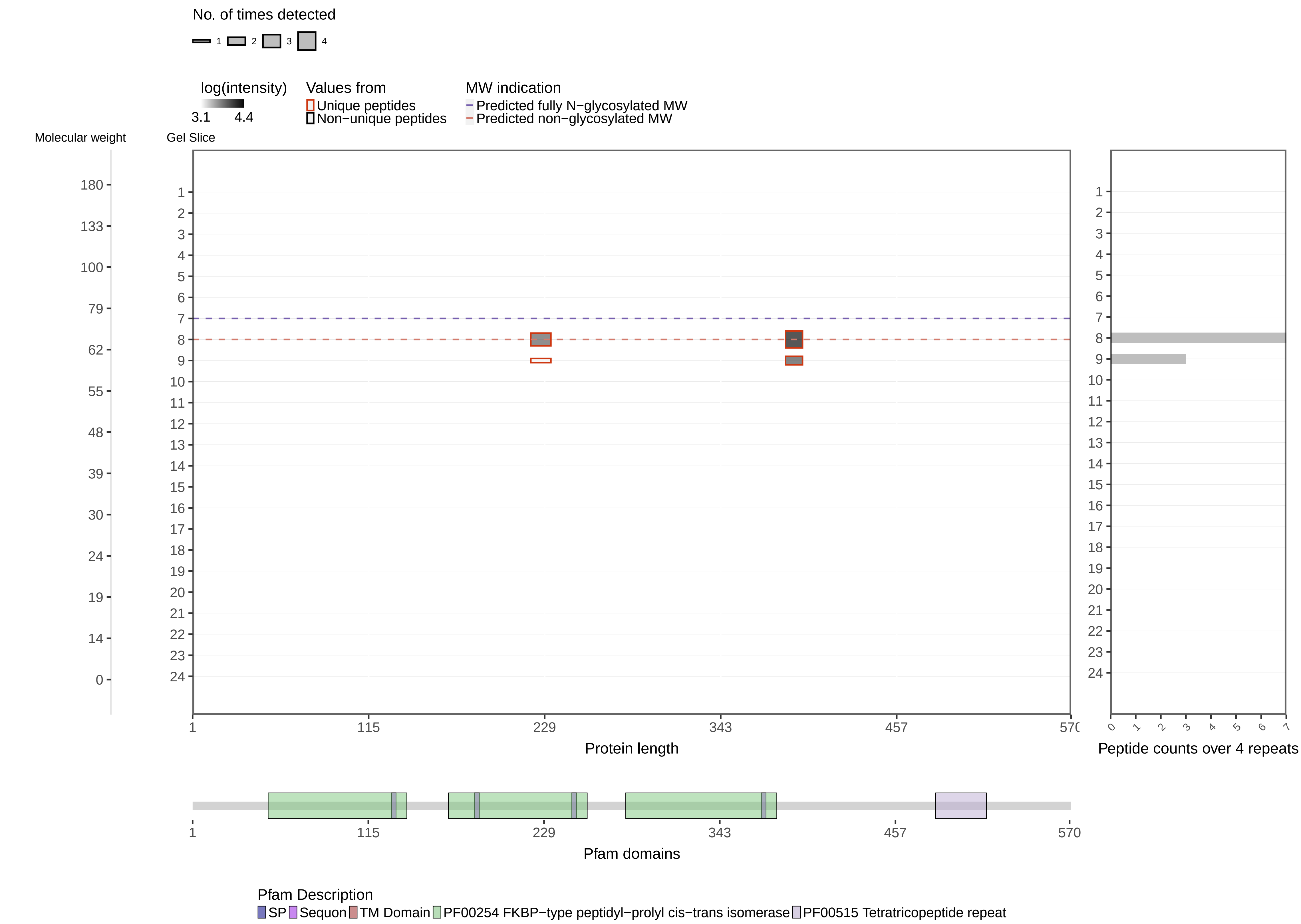


Protein sequence:

/VKKILKEGEGYEKPNEGAVVKLKLIGKLQDGTVFIKKGHDGENEDELFEFKTDEEQVIDGLDRAVMTMKKGEVALLTIAPEYAFGSSESKQDLAVIPPNSTVHYEVELVSFVKDKESWDMNTQGKIEAAGKKKEEGNALFKAGKYTRASKRYEKAAKFIEYDTSFSEEEKKQSKALKISCNLNNAACKLKLKDYKQAEKMCTKVLELESTNVKALYRRAQAYMNMADLDLAEFDIKKALEIDPDNRDVKLEYKALKGKVKDFNKKDAKFYGNMFAKLNKLESESAPKEPEPMS

NbL09g13440.1 serine_carboxypeptidase−like_20 [MW − SP = 52.86 kDa]

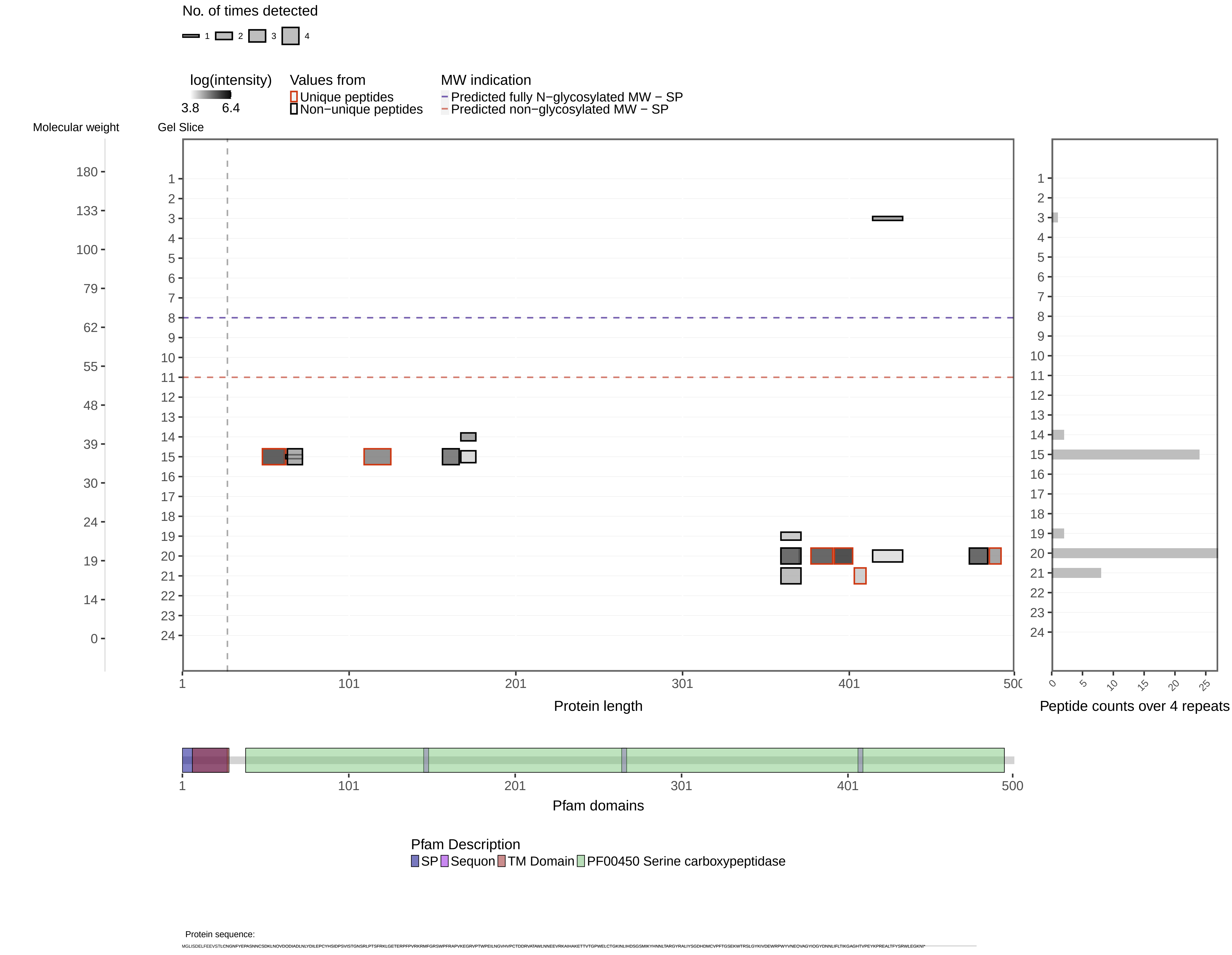

### NbL09g13430.1 dicamba_O−demethylase_ferredoxin_component−like [MW = 18.01 kDa]

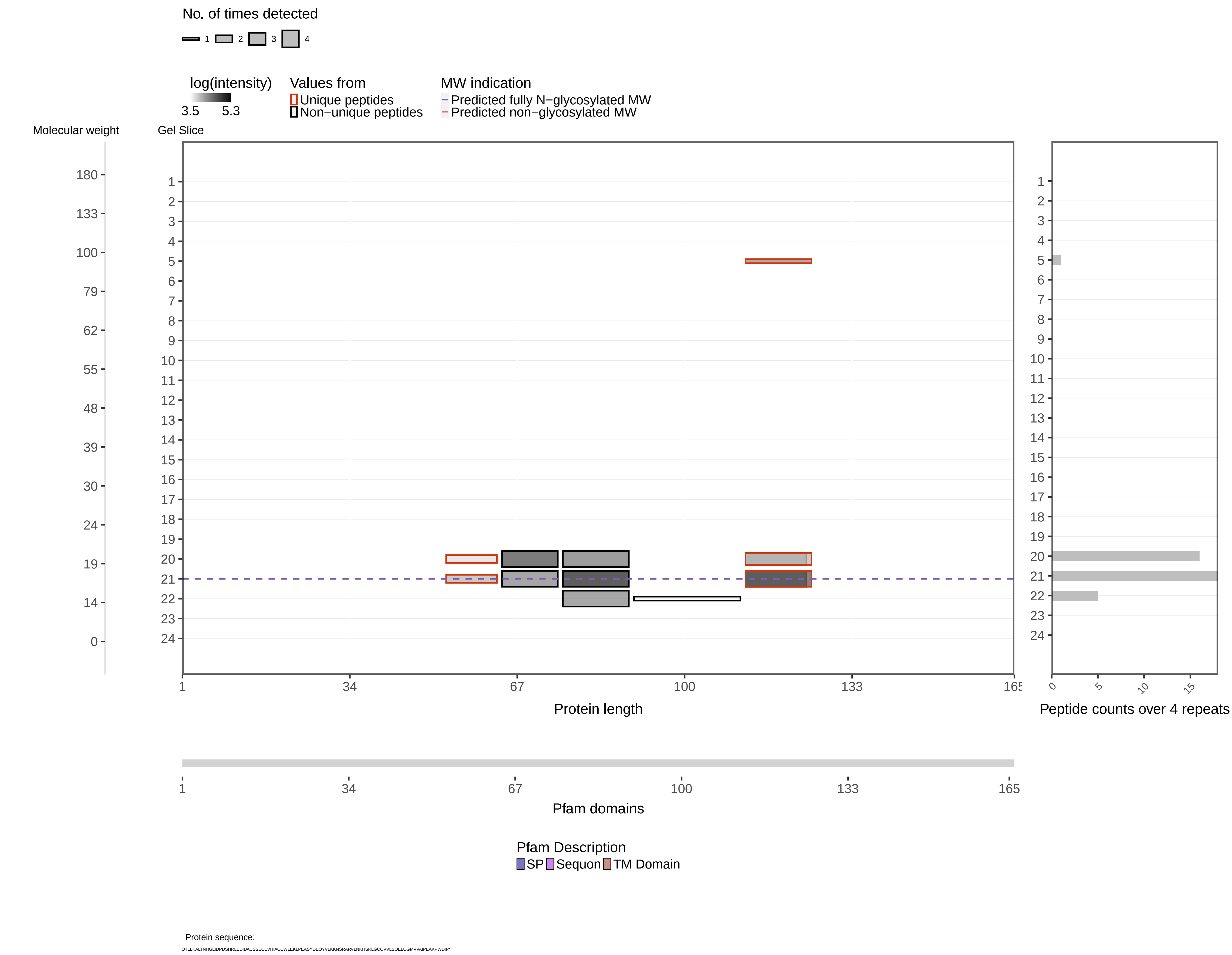

### NbL09g13400.1 photosystem_I_reaction_center_subunit_XI_chloroplastic−like [MW = 23.26 kDa]

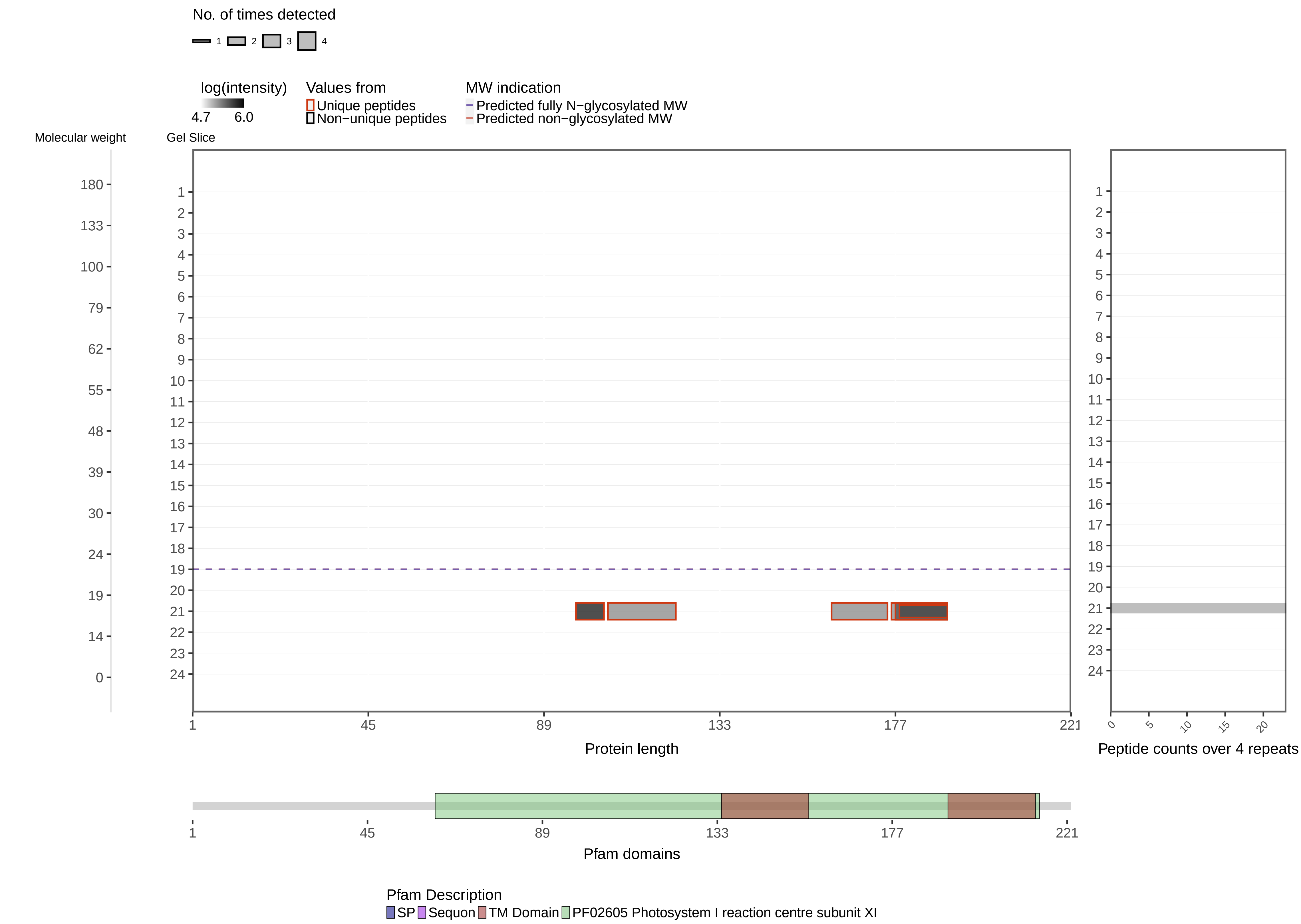


Protein sequence:

AVNPLLRGVEVGLAHGFLLVGPFVKAGPLRNTEYAGGAGSLAAAGLVVILSMCLTIYGISSFKEGDASTAPALTLTGRKKVPDQLQTAEGWSKFTGGFFFGGISGVTWAYFLLYVLDLPYYVK*

### NbL09g13380.1 probably_inactive_leucine−rich_repeat_receptor−like_protein_kinase_At5g48380 [MW = 68.4 kDa]

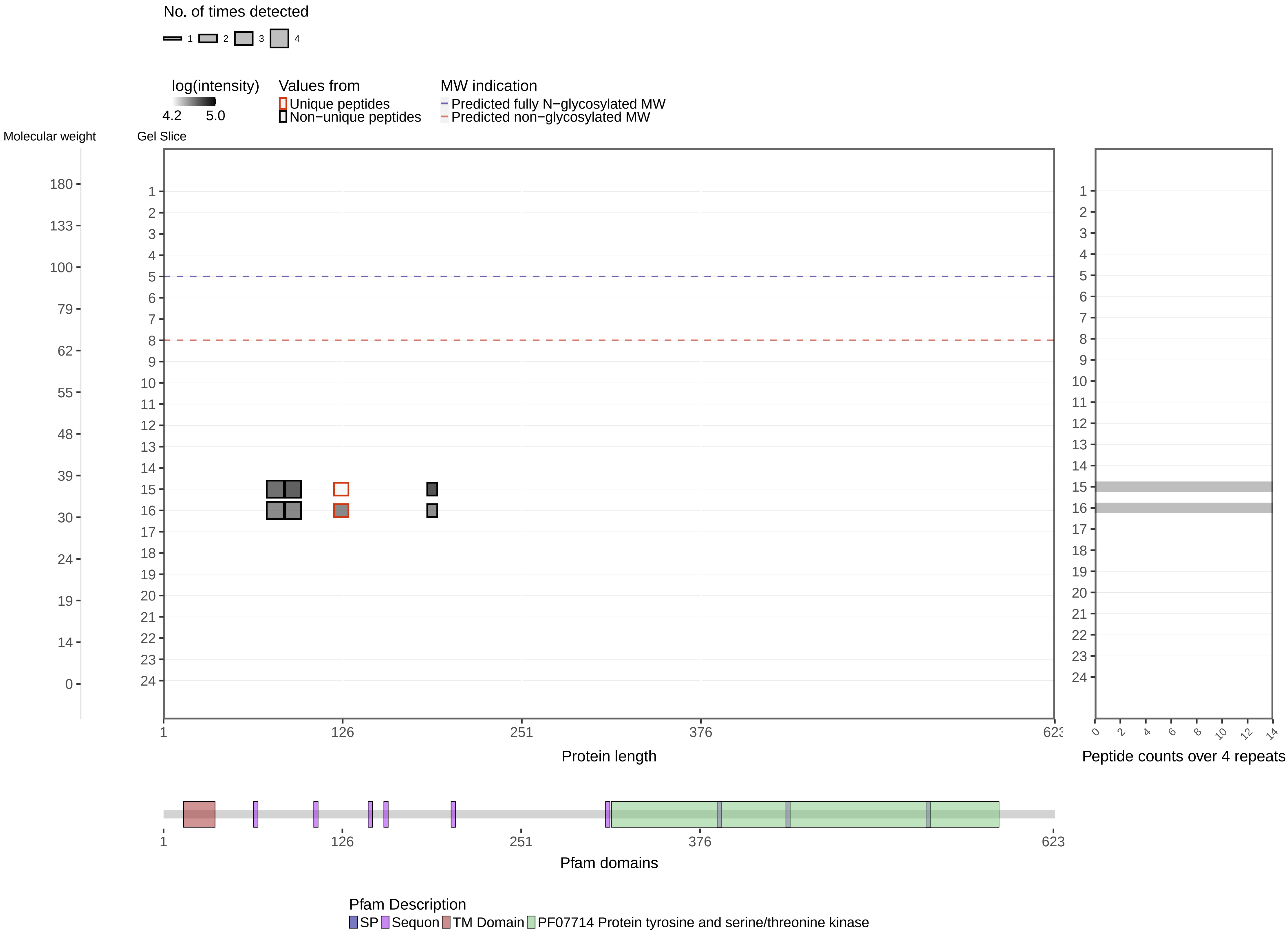


Protein sequence:

SDLMKATNNFSKSNIIGSASTGTFYKAVLDDGTSLMVKRLQNTQHSEKEFVSEMDTLGKVKHRNLVPLLGFCMAKKERLLVYKDMLNGTLHDKLHSVSEGEKTLEWPMRLKIGLGAAKGFAWLHHNCNPRIIHRNISSKCILLDVEFEPKISDFGLARLMNPIDTHLSTFVNGEFGDFGYVAPEYARTLMATPKGDVYSFGVVLLELVTGERPTSVTKAPETFKGNLVEWISNLSGESKLHDAIDHSLSGKGYDSEIFQVLKVACRCVLSAPKERPTMFEVYQFLRAIGERYHFT

### NbL09g13220.1 elongation_factor_1−delta_1−like [MW = 25.19 kDa]

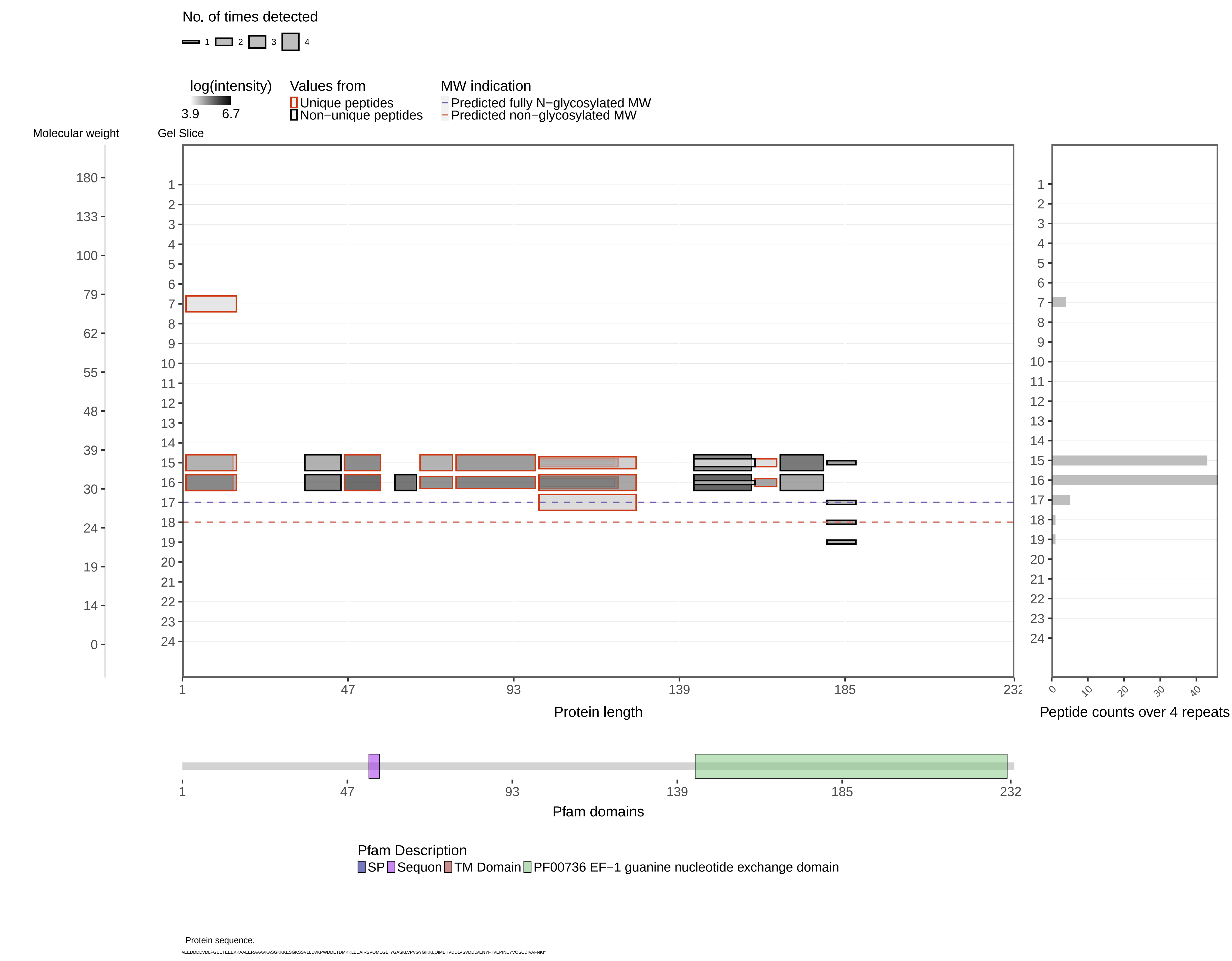

NbL09g12940.1 U2_small_nuclear_ribonucleoprotein_B" [MW = 26.14 kDa]

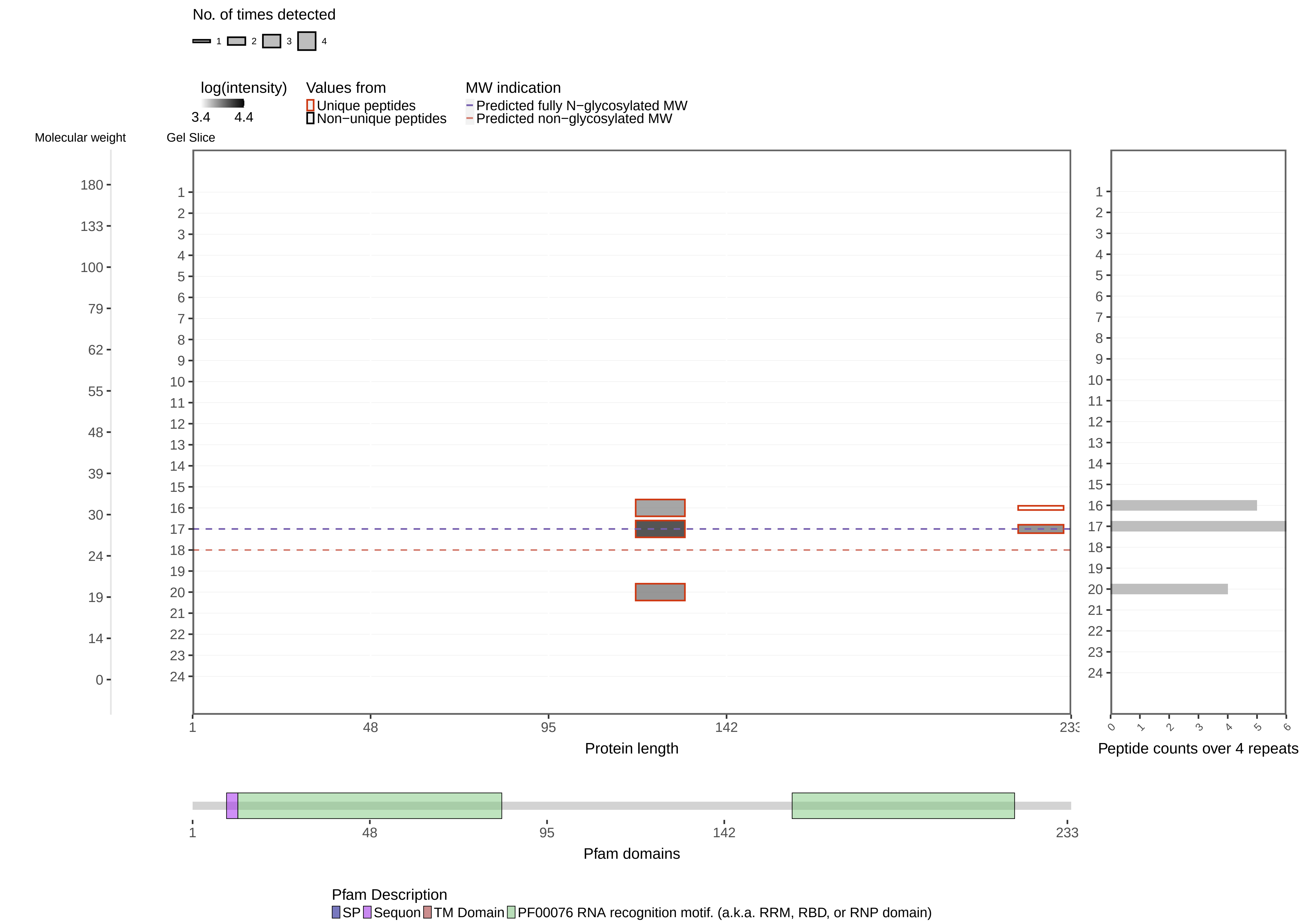


Protein sequence:

KKKQEEKVEKRKRTEETPQTGAANGPRADSNGGPAAAPRQGRPSAQETVAEPNNILFIQNLPHETTSMMLEVLFKQYPGFREVRMIEAKPGIAFVEFDDDVQSSVAMQALQGFKITPQNPMAITYAKK*

### NbL09g12890.1 UDP−glycosyltransferase_87A1−like [MW = 51.6 kDa]

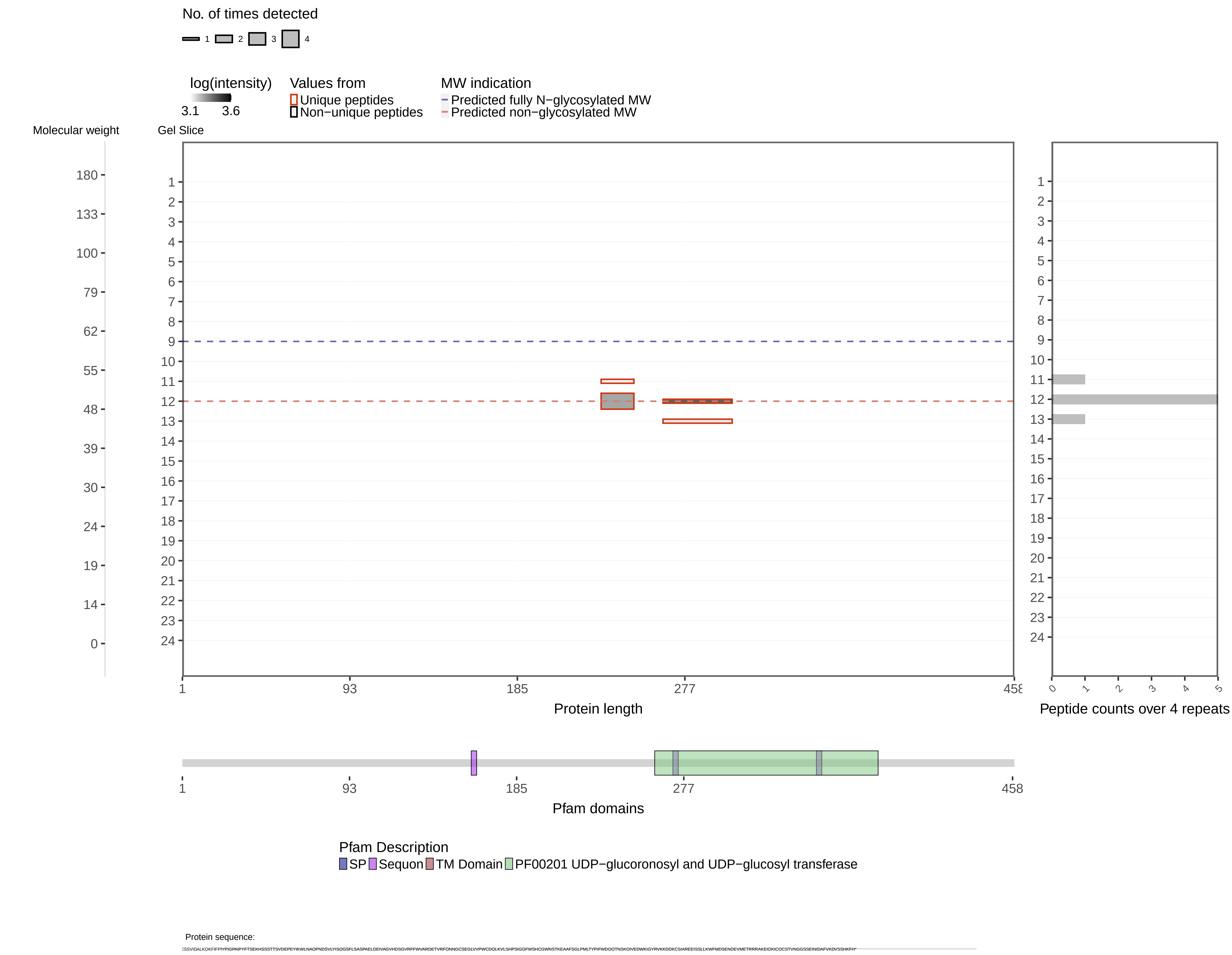

NbL09g12830.1 transcription_initiation_factor_TFIID_subunit_15b [MW = 43.64 kDa]

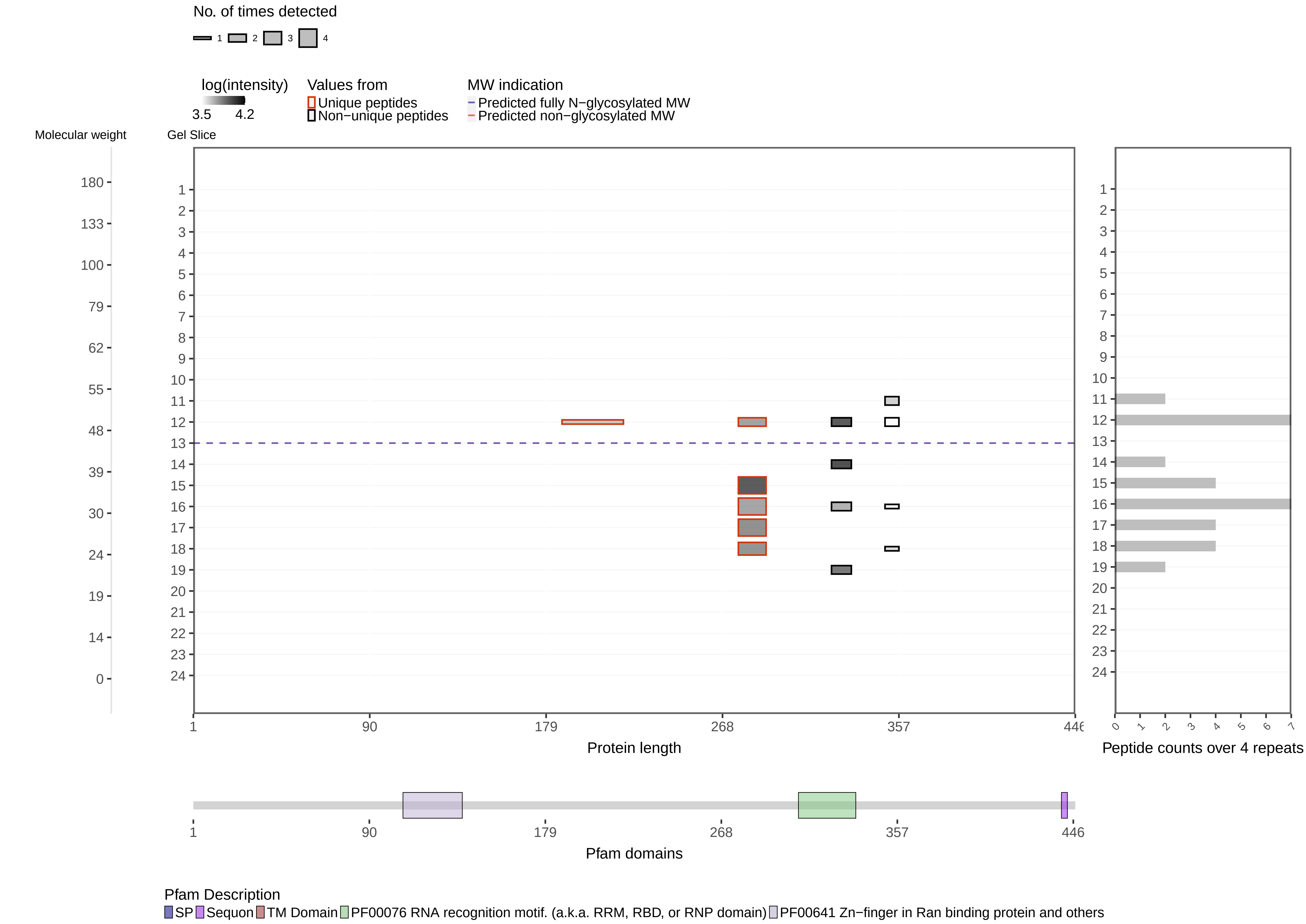


Protein sequence:

EADSNYPPPSNTYGRNPDAAPPPTSYVGGAASYPPSYGAPGGYGGDKTANARGGGRDGPPGGYDGGYGCGQRNAGGYDSSPAEAPEKVKQCDENCGDFCDNARIYISNLPPDVTIEELRELFGSIGQVARIKQKRGYKDQWPWSIKLYTDEQGKNKGDAVLSYEDPSAAHSAGGFFNNHDFRGYKISVAMAEKSAPKPAPAFGSGRGGRGGYGGGGRRDNYRDSGGSGPDRQSHGGNRSRPY*

NbL09g12780.1 protein_DCL_chloroplastic−like [MW = 26.54 kDa]

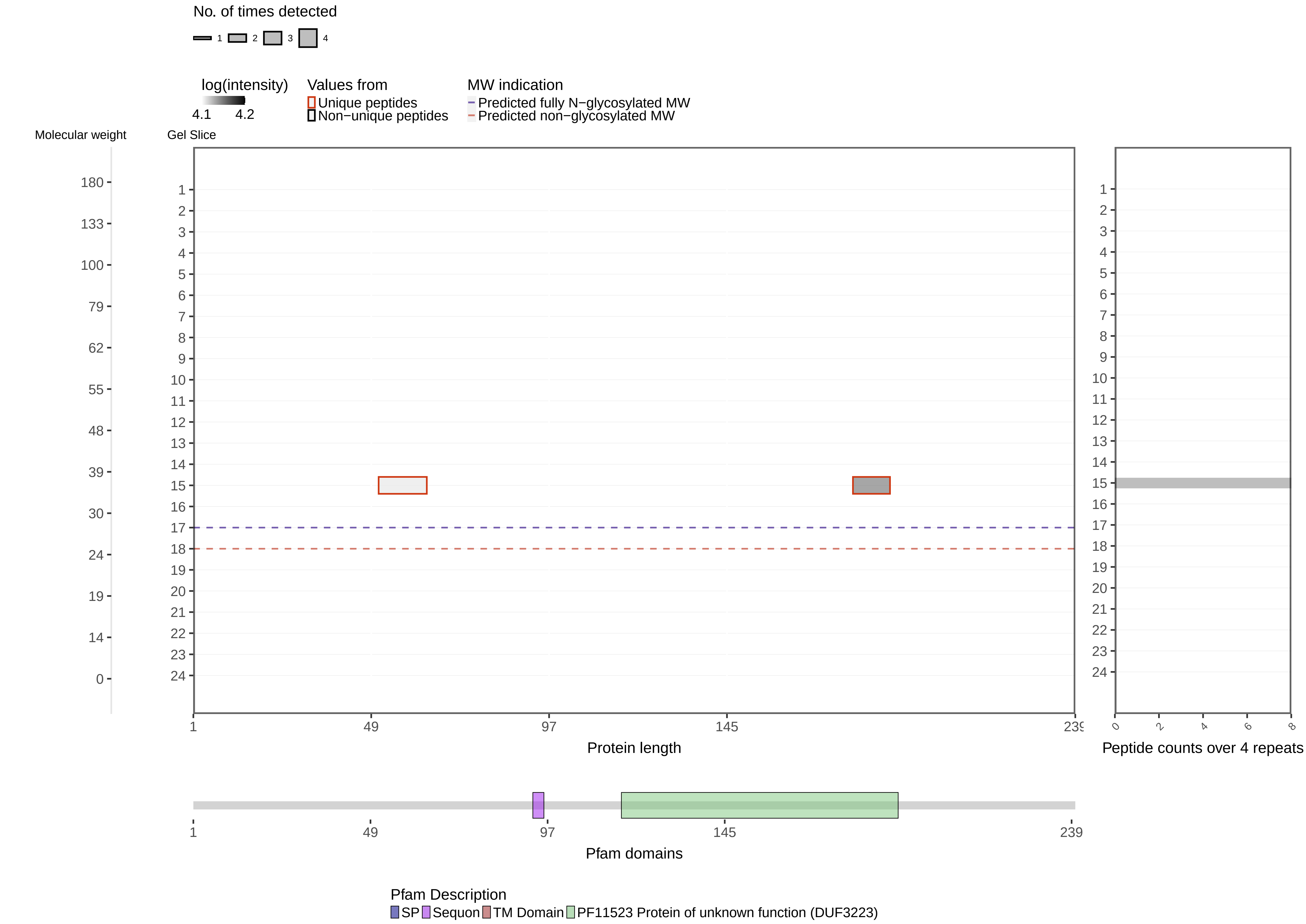


Protein sequence:

SSVEMFDYFYKLLHSWSPNLNLNKNLQYEHLVLLDLLKKGHLEPDRKIGTGIRAFQIRFHPQFKSRCFFVIREDDSVDDFSFRKCVDHILPLPENMQVKHEANKALAHSGKGRGRKGGGYGRDSGNGGKSRY*

### NbL09g12760.1 chlorophyll_a−b_binding_protein_4_chloroplastic−like [MW = 27.61 kDa]

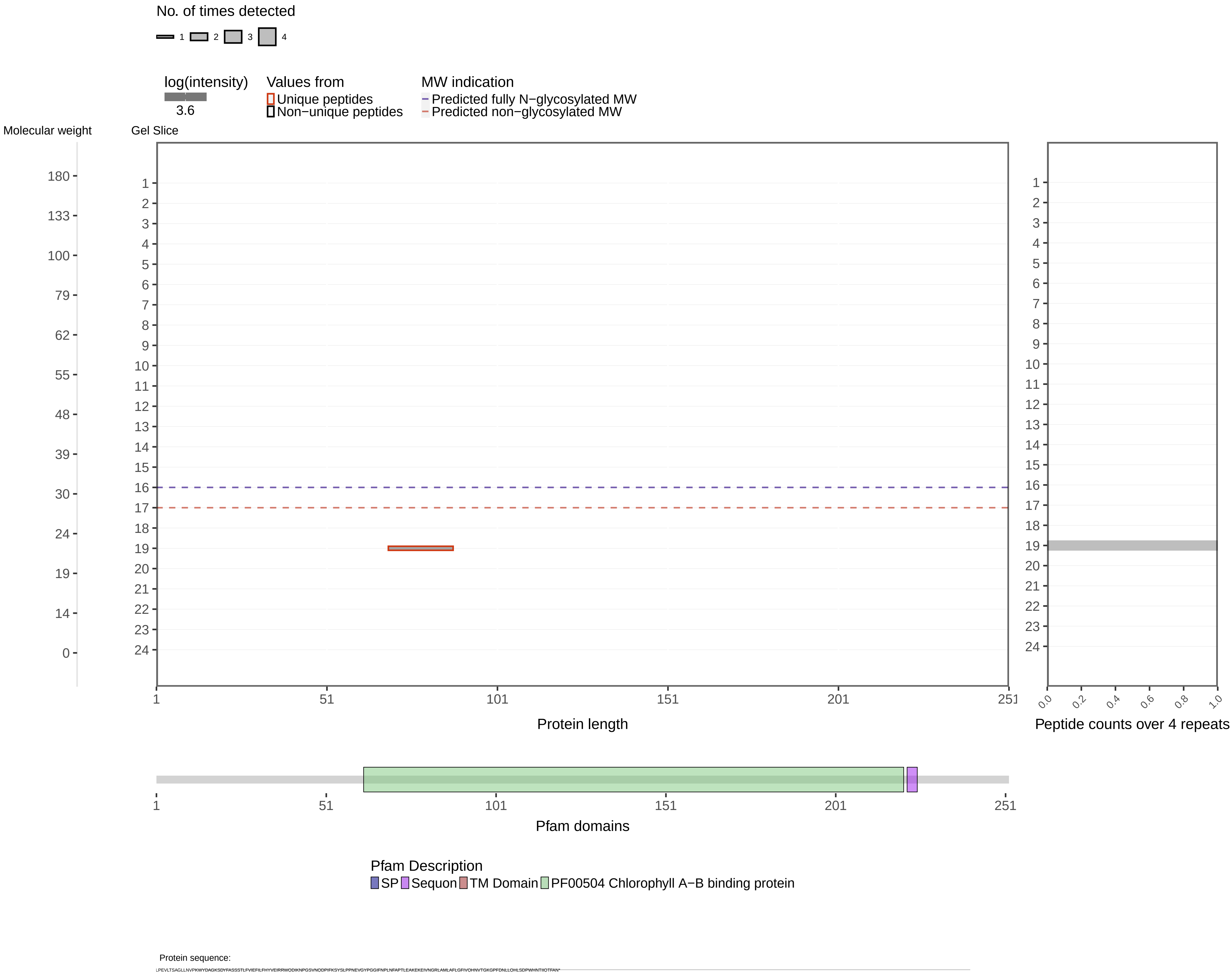

### NbL09g12590.1 biotin_carboxyl_carrier_protein_of_acetyl−CoA_carboxylase_1_chloroplastic−like [MW = 27.53 kDa]

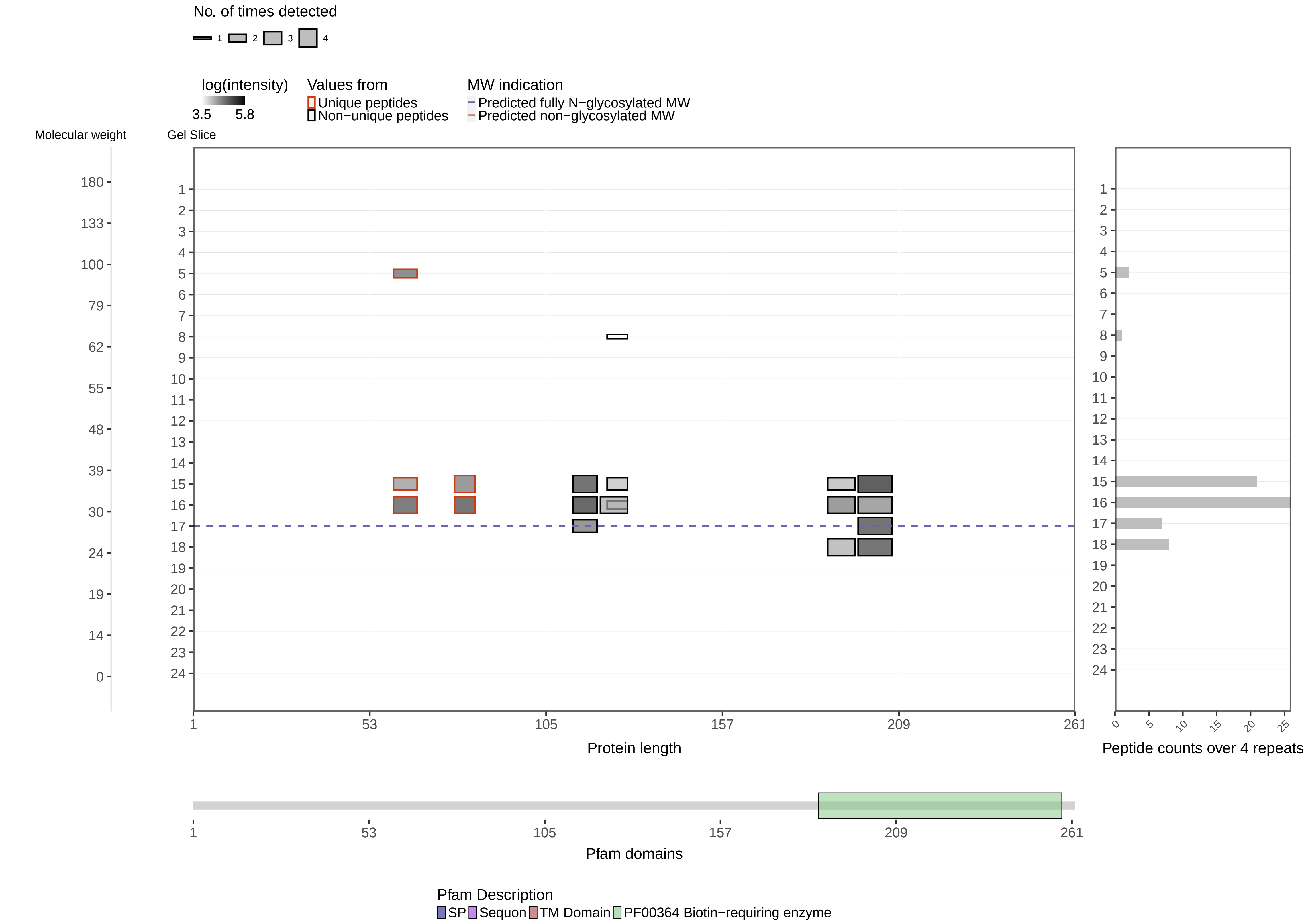


Protein sequence:

QLKQKDCEILIRKKAALPQPPVAMAPPSVQQGFYQPQLPPASAPAPAAPAAPALPPPAPSKPKSSHPPMKCPMAGTFYRSPAPGAPPFVKVGDKVQKGQVVCIIEAMKLMNEIEADSSGTIVDIIADDGKPVSVDTPLFVIEP*

### NbL09g12580.1 uncharacterized_LOC104224938 [MW = 237.3 kDa]

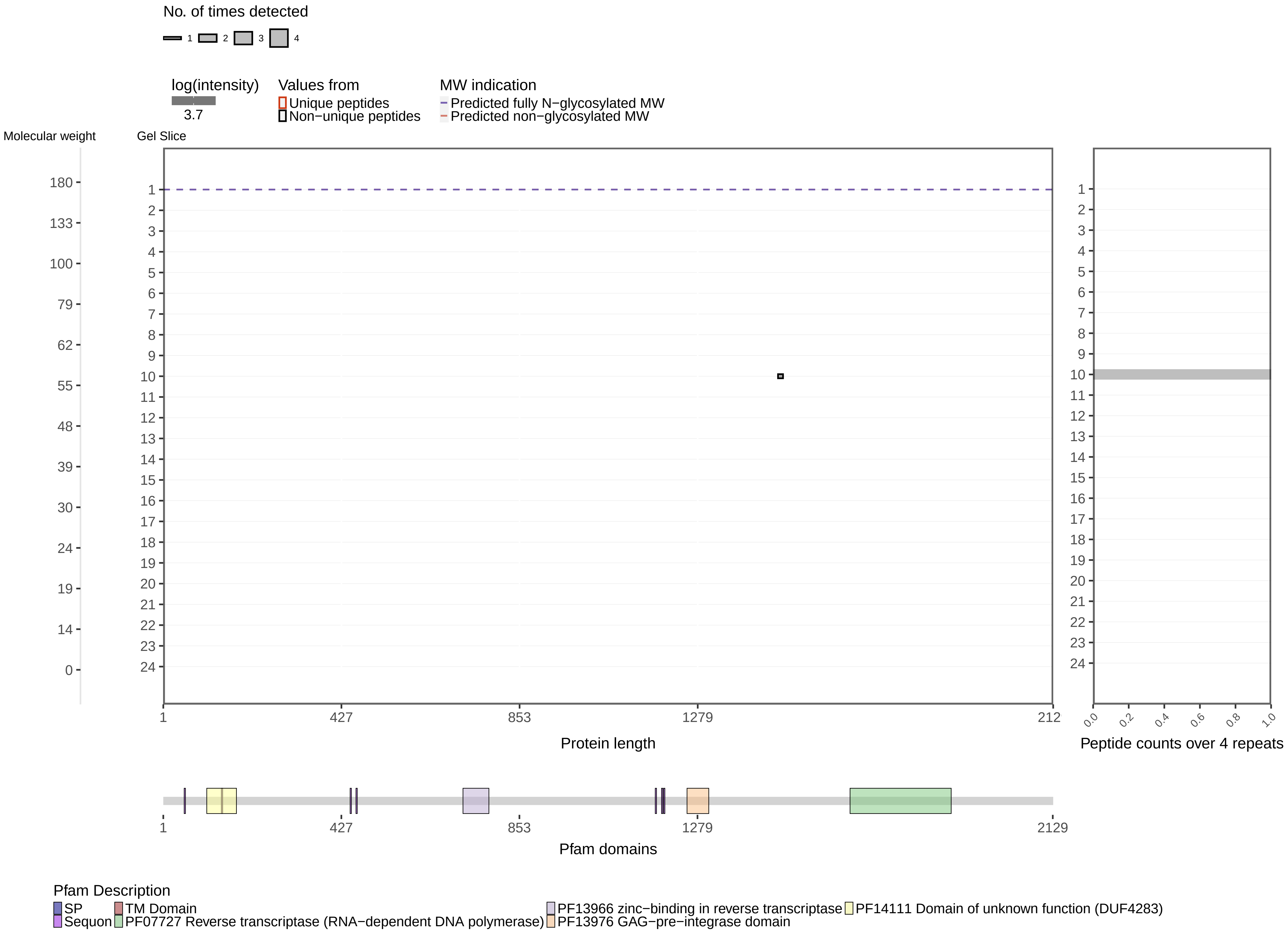


Protein sequence:

SQSVDNRASHTMEYRRGGGRFGRSRPKCSYCYKLGHTRDVCHSLHGRPTKNAYVAQTEPTGNQGFSVSEREYNEFLKYRASKQTSPQVATVAQPDTSVAGNSFACVSQSSPLEQWVVDSAASDHISGNKSLLSDITYSQFLPNVTLANGSQTKAKGVGQATPLPSVTLDSVLYVPGCPFNLASDRSTGQTIGTGLESDGIYYLNSLTSSKACLAADPPDLLHSRLGHPSLSKLQKMVPRLSSLSTLECESCQLGKHTRASFPRSIESRAESVFSLVHSDIWGPSRQFMS

NbL09g12510.1 glycerophosphodiester_phosphodiesterase_GDPD6 [MW − SP = 43.59 kDa]

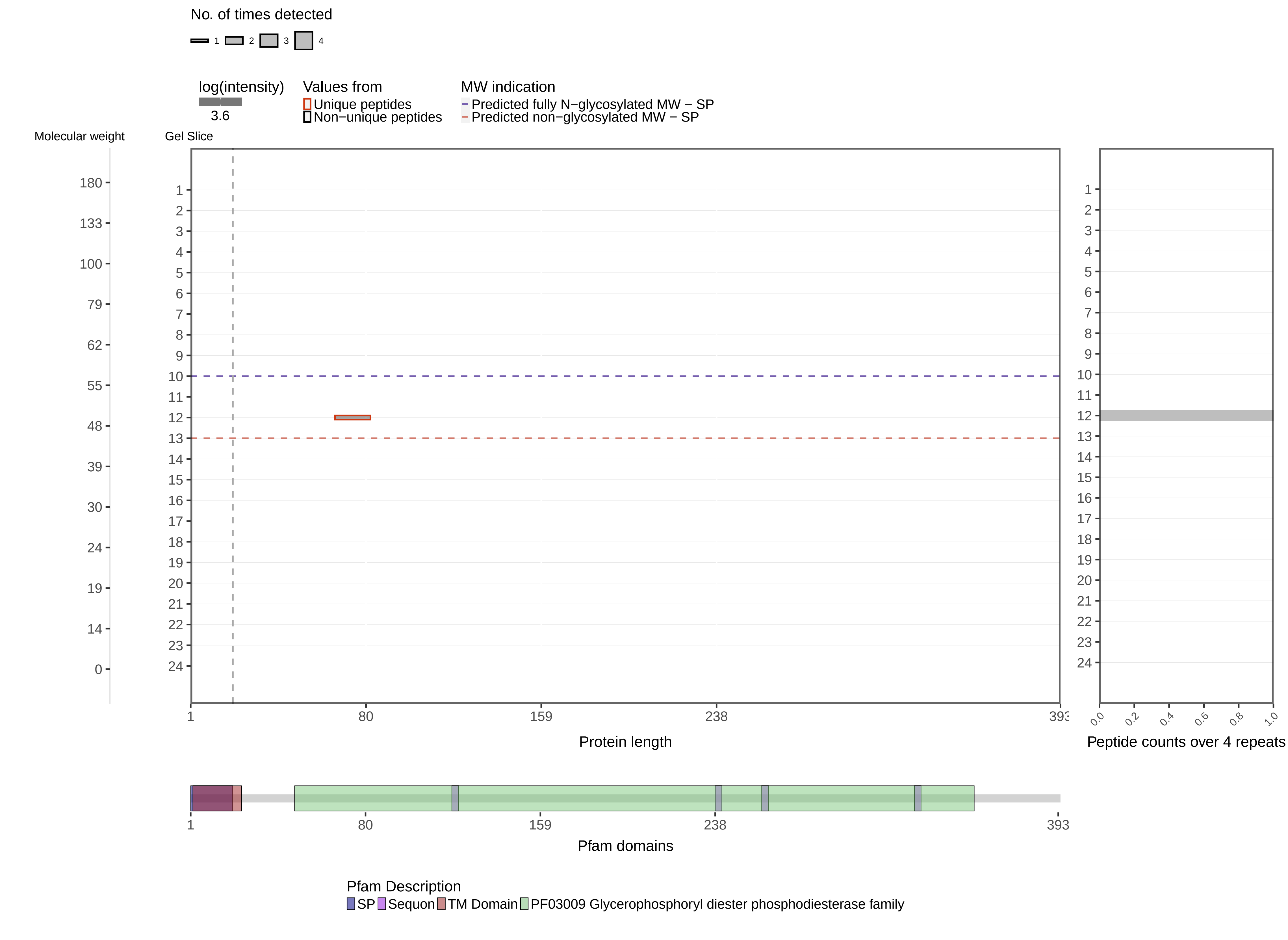


Protein sequence:
OVKWPGGKRFEDKFVETLKKYGYRGSYMSKOWLKOPAFIOSFAPTSLIYISNOTDLPKIFLIDDVDMPTODTNOSYWDITSDSYFDFIKEYVVGIGPWKDNIIPVSSNYLOTPSDLVARAHAHNLOVHPYTFRNENKFLHFNFSEDPYNEYDYWINKIGVDGLFTDFTGSLHHYOEWTNPFASGEKEATRLLHKIELMLSKFKYFN*

### NbL09g12470.1 uncharacterized_mitochondrial_protein_AtMg00810−like [MW = 13.71 kDa]

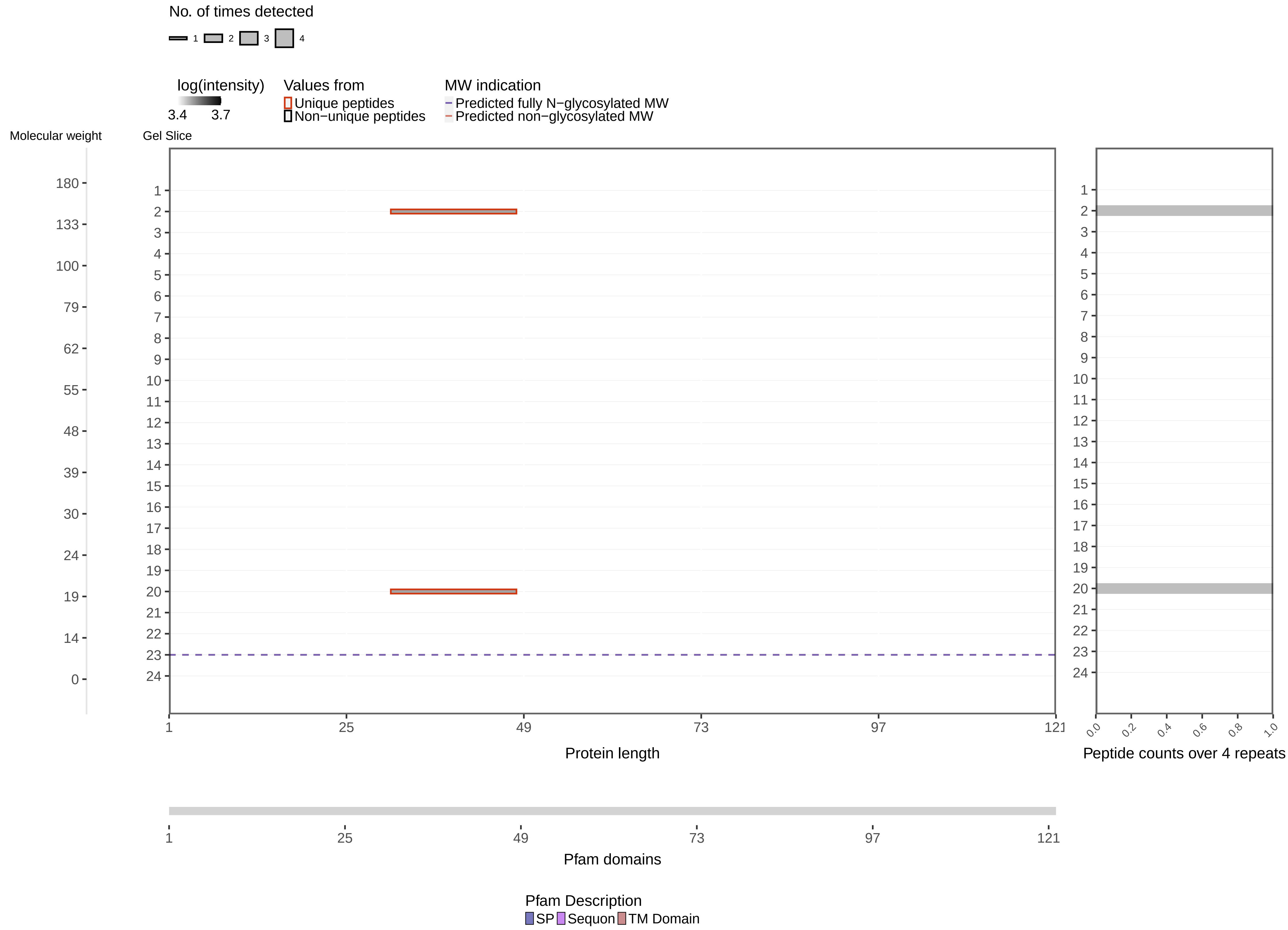


Protein sequence:

TKLSKFDKGEKVDPTFFISLVGSLRYLTCIKPDILFVVGVVSRSMEAPTSTQLKVTGRILHYLKGTIDFRLFYS*

NbL09g12250.1 serine/threonine−protein_phosphatase_2A_65_kDa_regulatory_subunit_A_beta_isoform−like [MW = 64.79 kDa]

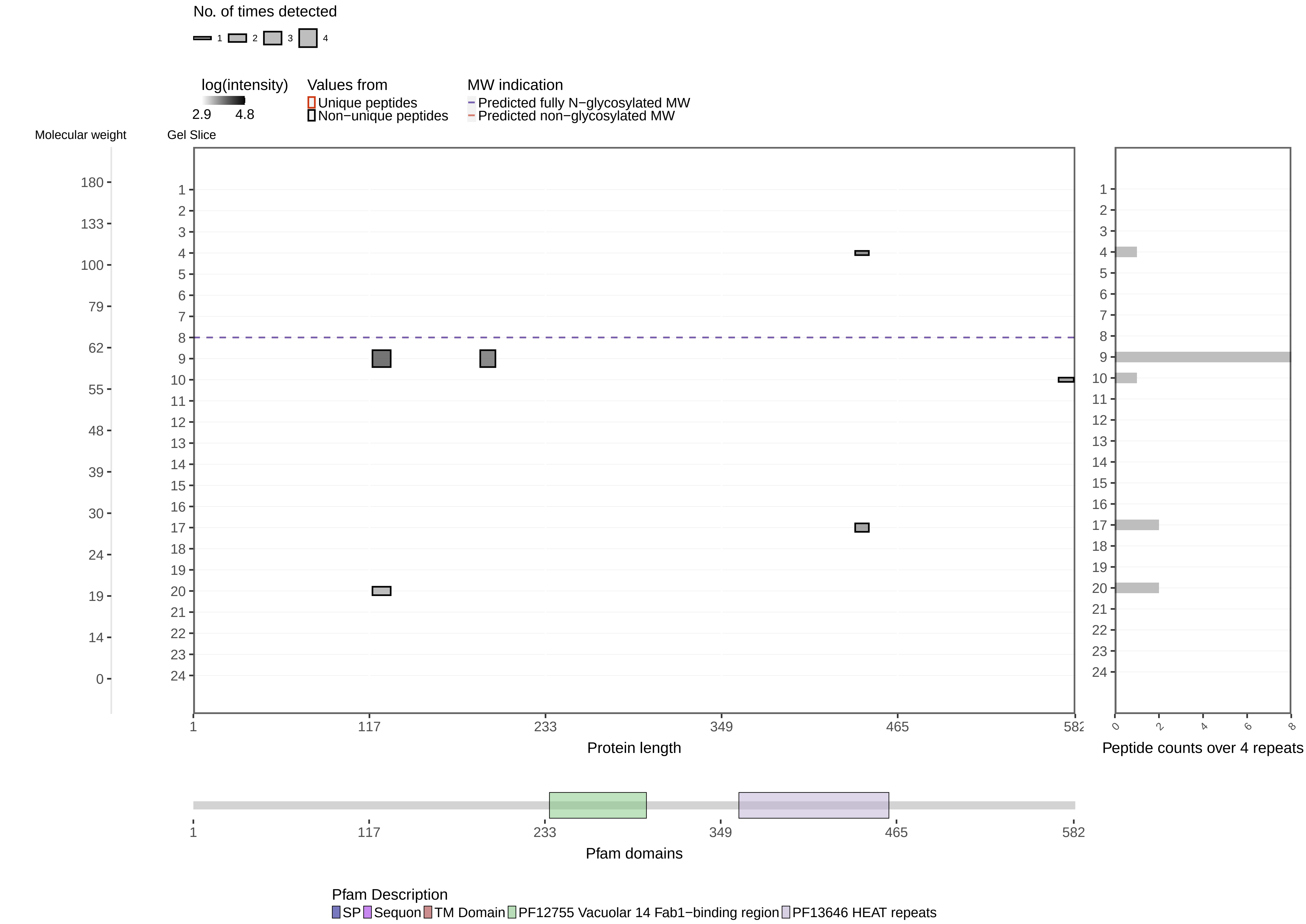


Protein sequence:

[illegible]

### NbL09g12160.1 40S_ribosomal_protein_S7−like [MW = 22.07 kDa]

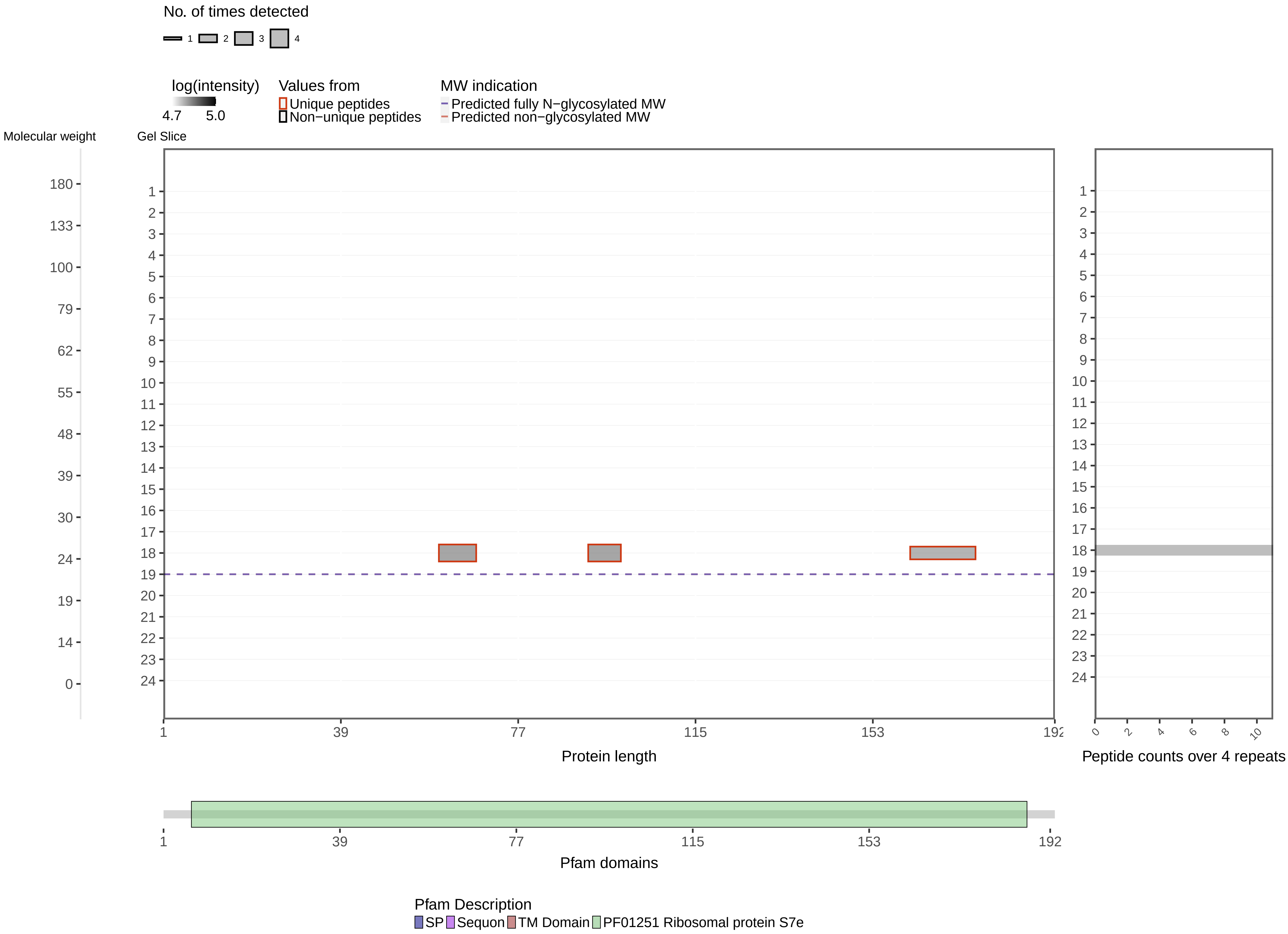


Protein sequence:

'RELEKKFSGKDIIFIATRRIARPPKRGSAAQRPRSRTLTSVHDAILEDLVVPAEIVGKRTRYRIDGSKIMKVFLDPKERTNTEYKLETFSAVYRKLSGKDVVFEYPITEA*

NbL09g12100.1 eukaryotic_peptide_chain_release_factor_GTP−binding_subunit_ERF3A−like [MW = 56.97 kDa]

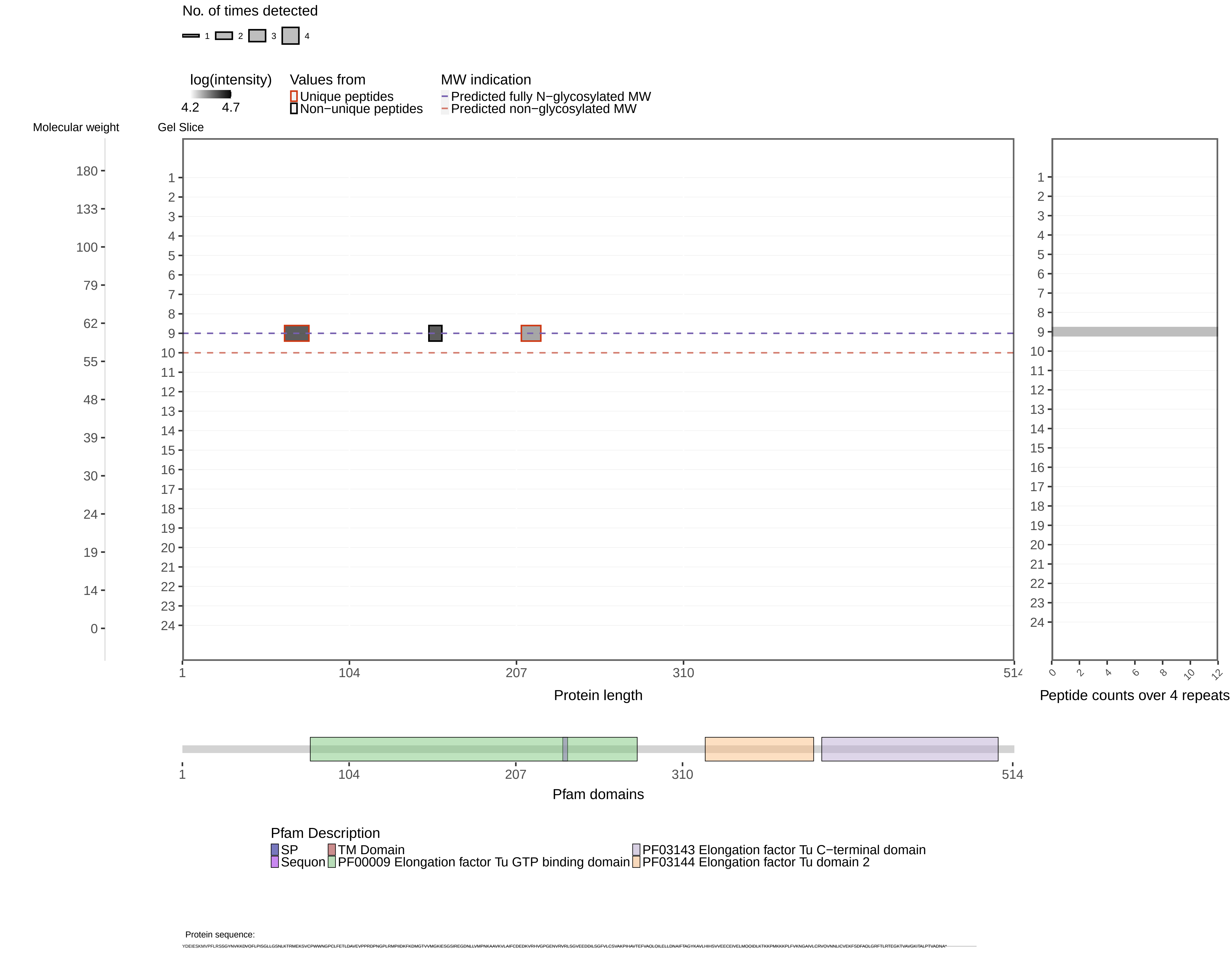

NbL09g11980.1 serine_carboxypeptidase−like_45 [MW = 39.28 kDa]

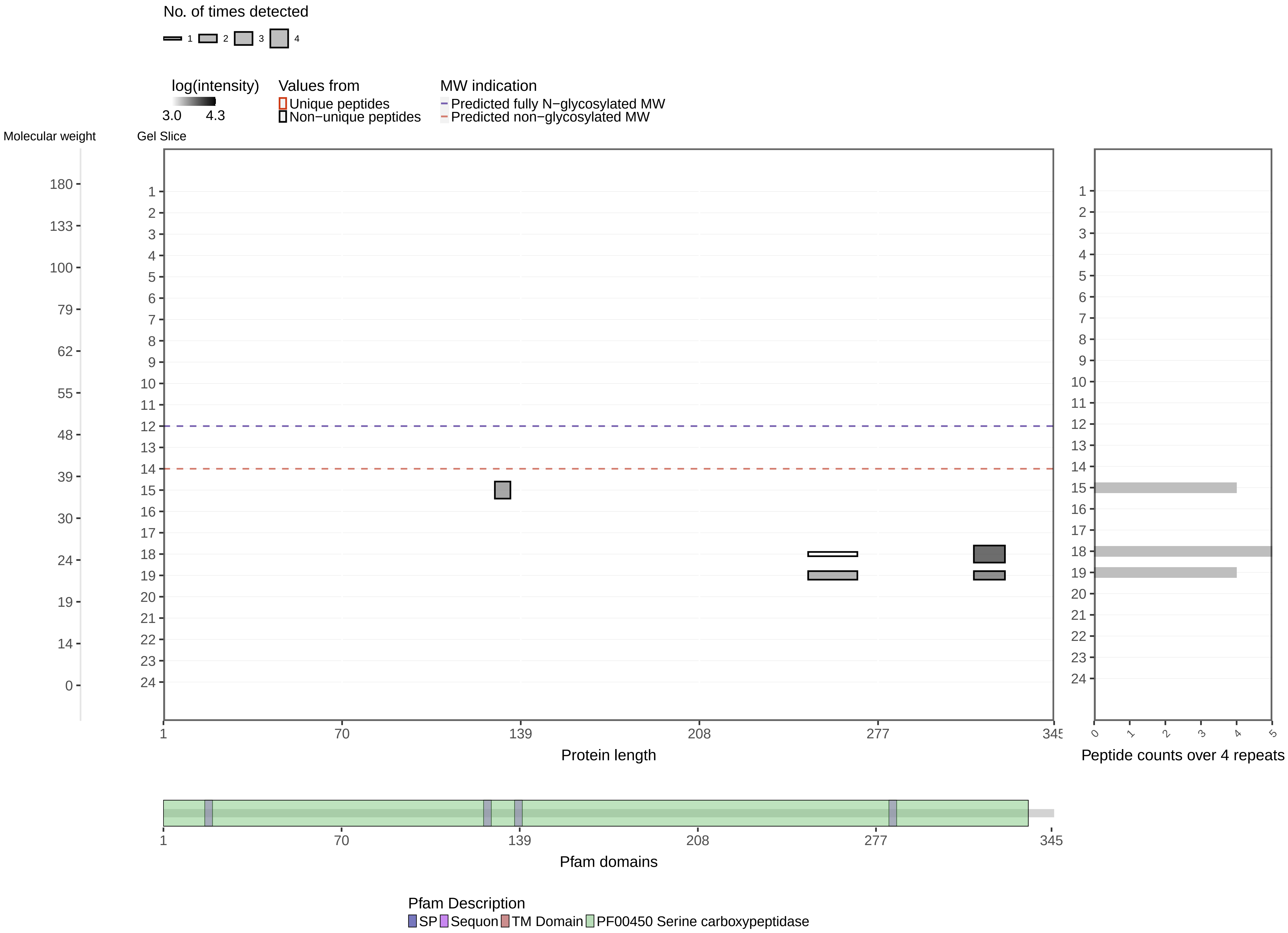


Protein sequence:

DKYDVTLDVCLSSVLSQSKIISPQENADAEKIDVCIDDKTVNYLNRQDVRRALHANLVGVRRWDVCSNILDYQLLDIEIPTISIVGLLVKERIPVLIYSGDQDSVVPLTGSRTVVHQLAKRMRLNTTVPYRVWFAGQQVGGWTVVYDNILSFATVRGAAHEVPFSQPERSFVLFKSFLEGRALPEVF*

### NbL09g11910.1 probable_beta−14−xylosyltransferase_IRX10L [MW − SP = 44.9 kDa]

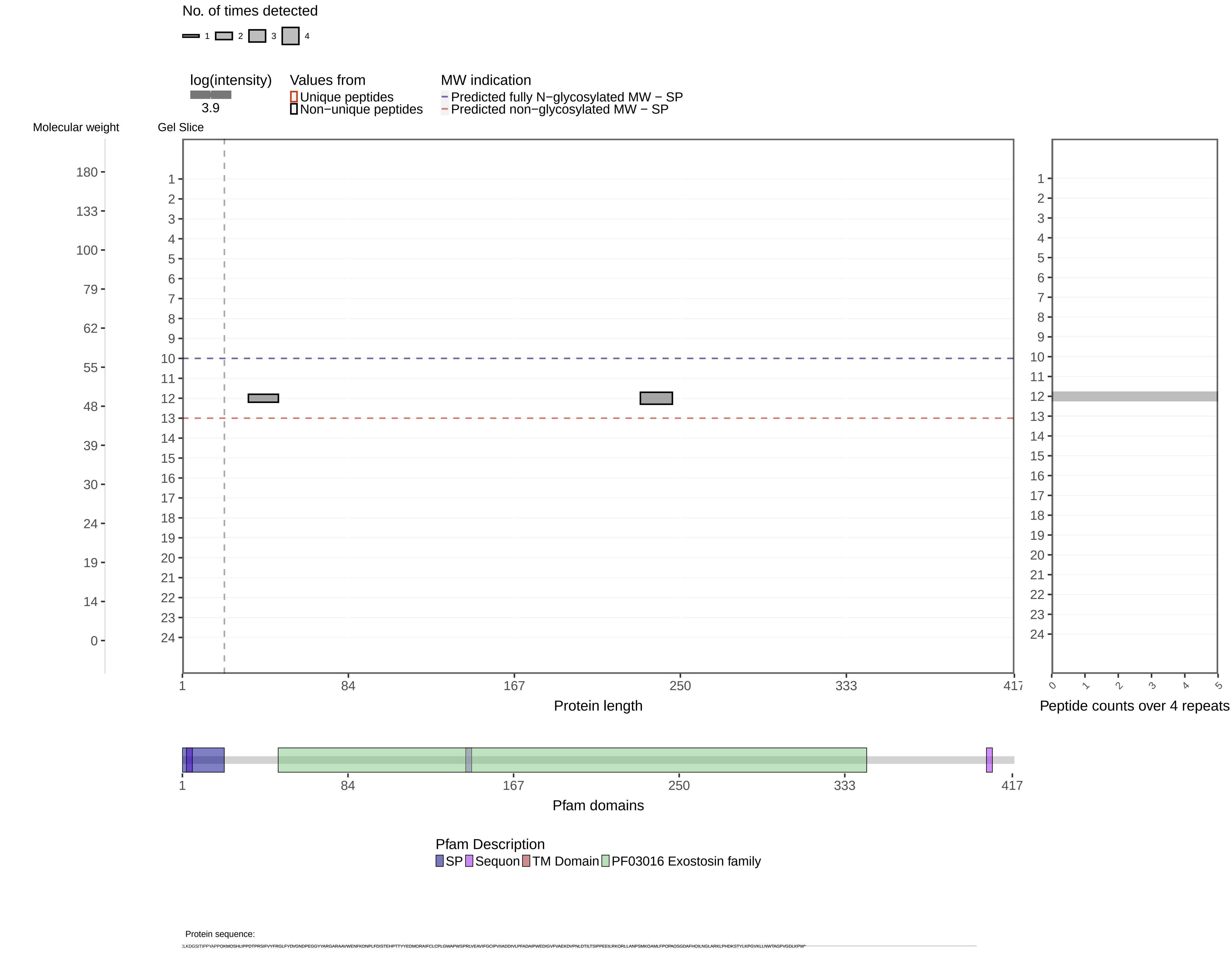

### NbL09g11890.1 uncharacterized_protein_At4g14100−like [MW − SP = 22.68 kDa]

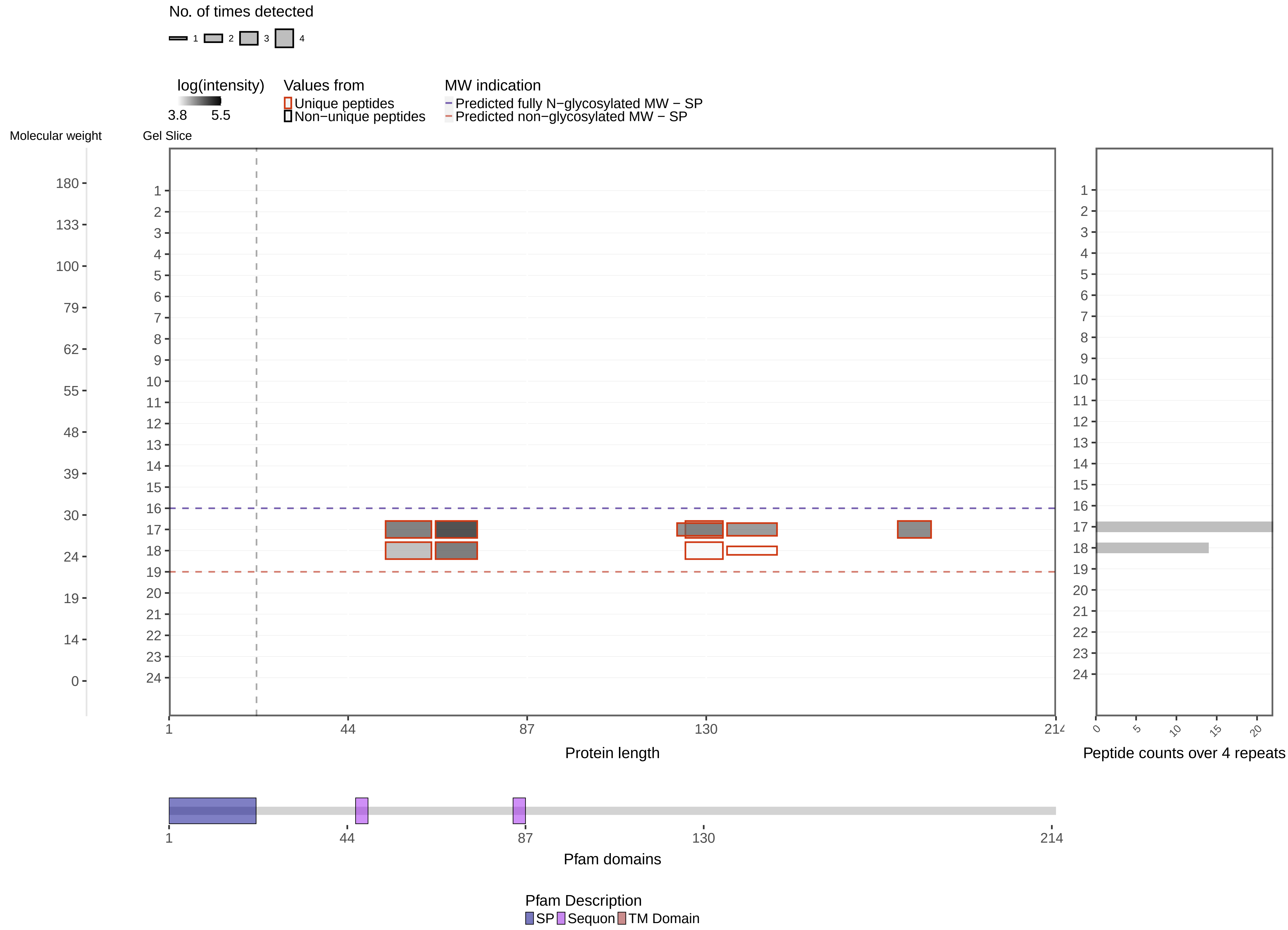


Protein sequence:

TQECTIRHFPVGILRPNWLEGSNYVGHKYKDGFLCNVWEKVGFLQYYEDAFTKIPVYWHFYDGLVEHIMTFEVGKVLEDSKWQAPAYCFKEVEKERISLLENLATTHFRDVVNGEVAEN*

### NbL09g11850.1 calnexin_homolog_1−like [MW = 65.68 kDa]

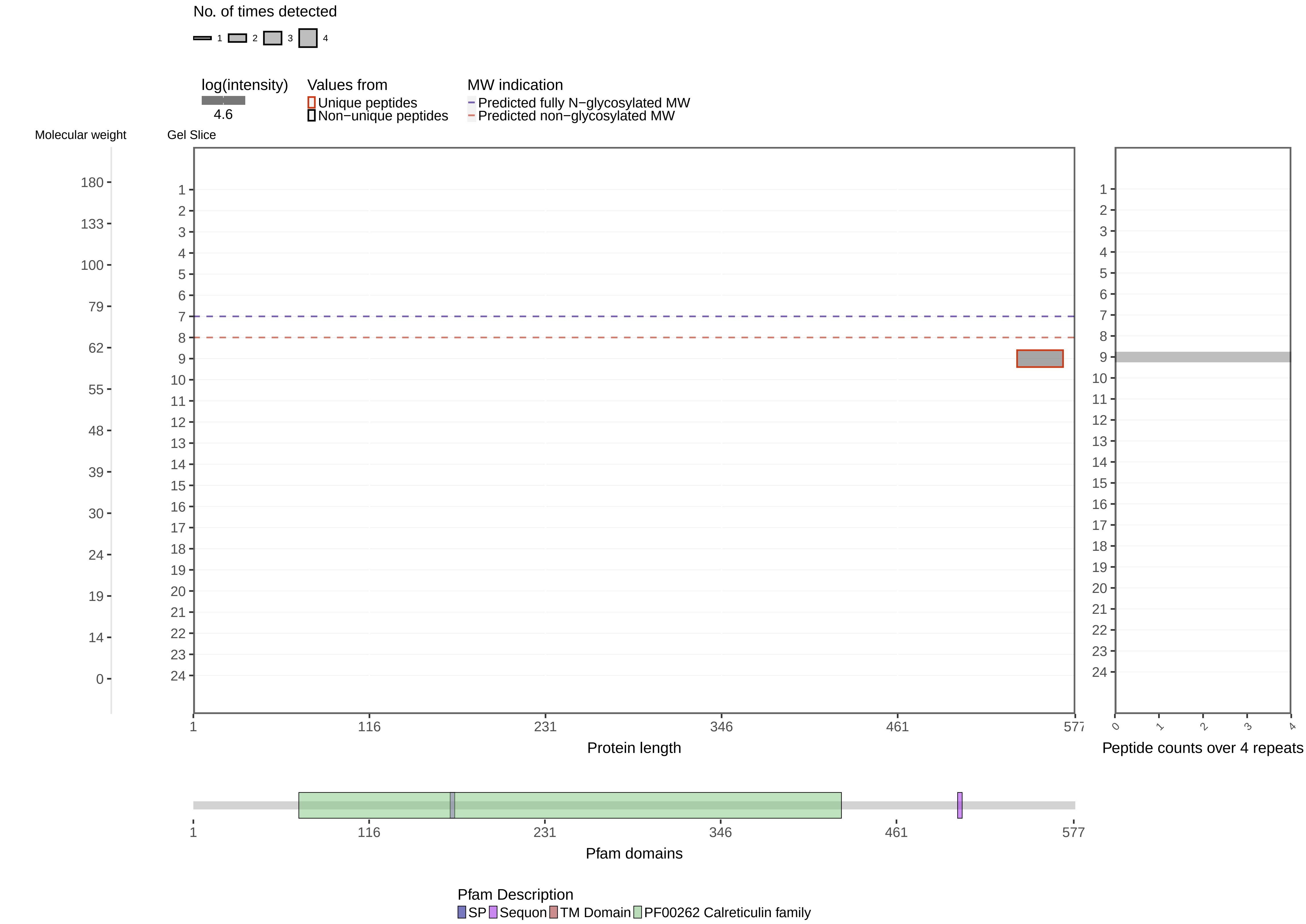


Protein sequence:

EKIPDPDAKKPDDWDEDAPMEIEDEEAEKPEGWLDDEPEEINDPEATKPEDWVDEEDGEWEAPNIDNPKCAEGPGCGEWRRPMKSNPAYKGKWHAPLIDNPNYKGIWKPRDIPNPNYFELNKPNFEPVAAIGIEIWTMQDGILFDNILIASDEKVAESYRKTAWKPKFDAEKEKQKAEEATISGGLKGFQKVVFDLLYKVADIPLLGEHKSKVLDLLEKAETQPNLTVGVVISIIVIIFTVLLKLIFGGKKQQPARATEEPKRSDVAEASTSNSQGATEEKEEQNEDAAAAPRRR

### NbL09g11830.1 calnexin_homolog_1−like [MW − SP = 57.94 kDa]

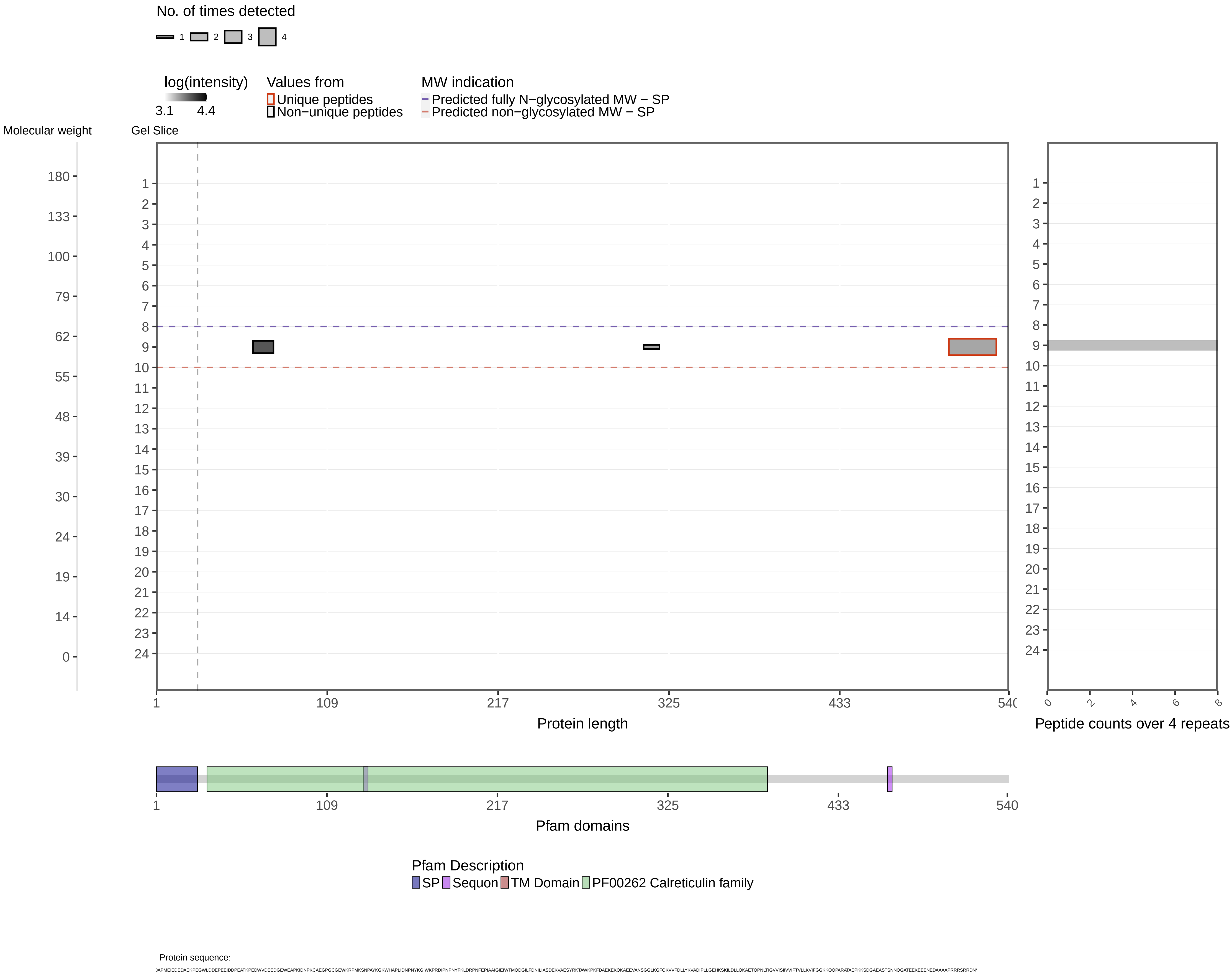

### NbL09g11290.1 beta−glucosidase_BoGH3B−like [MW − SP = 65.67 kDa]

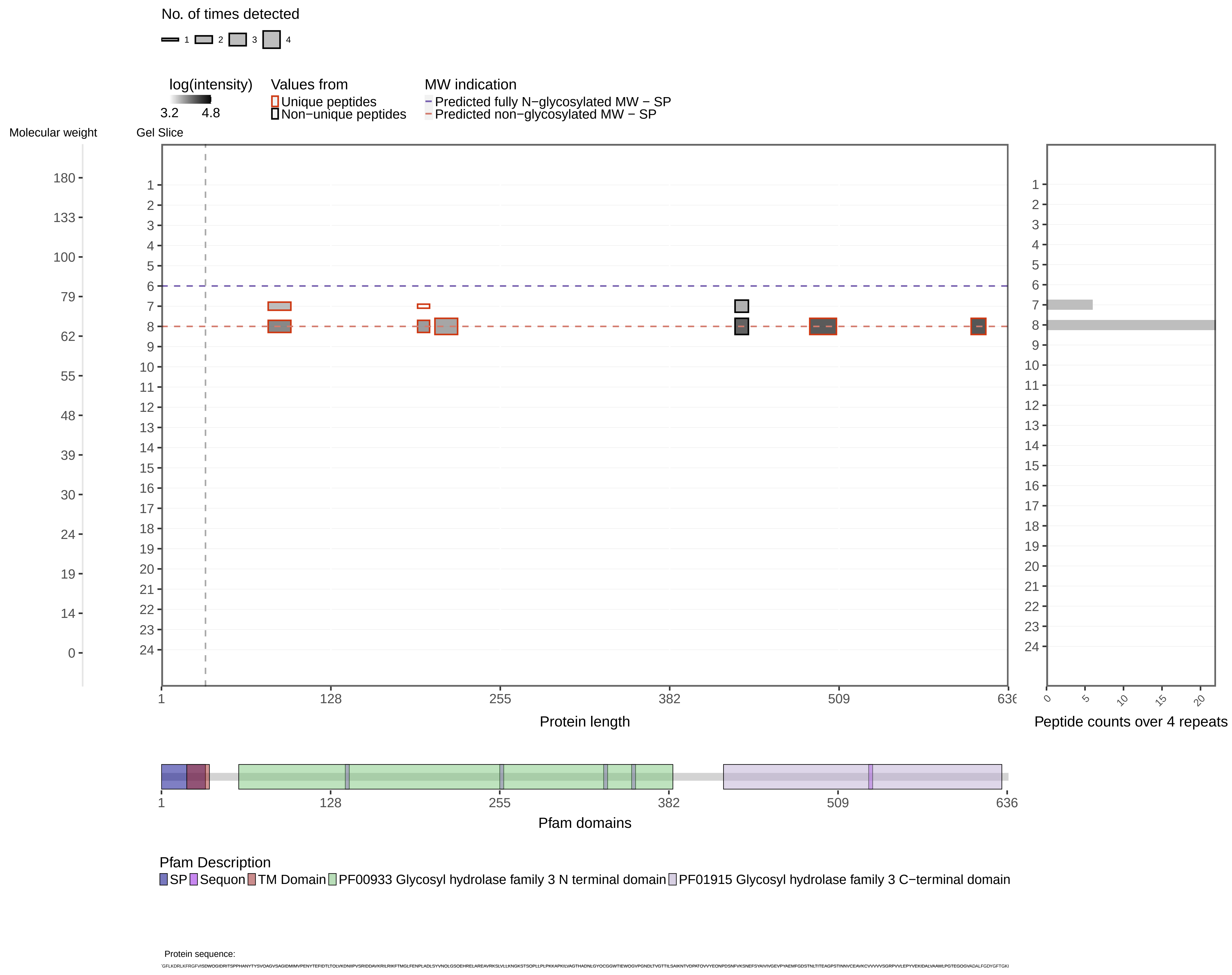

NbL09g11260.1 beta−glucosidase_BoGH3B−like [MW − SP = 65.7 kDa]

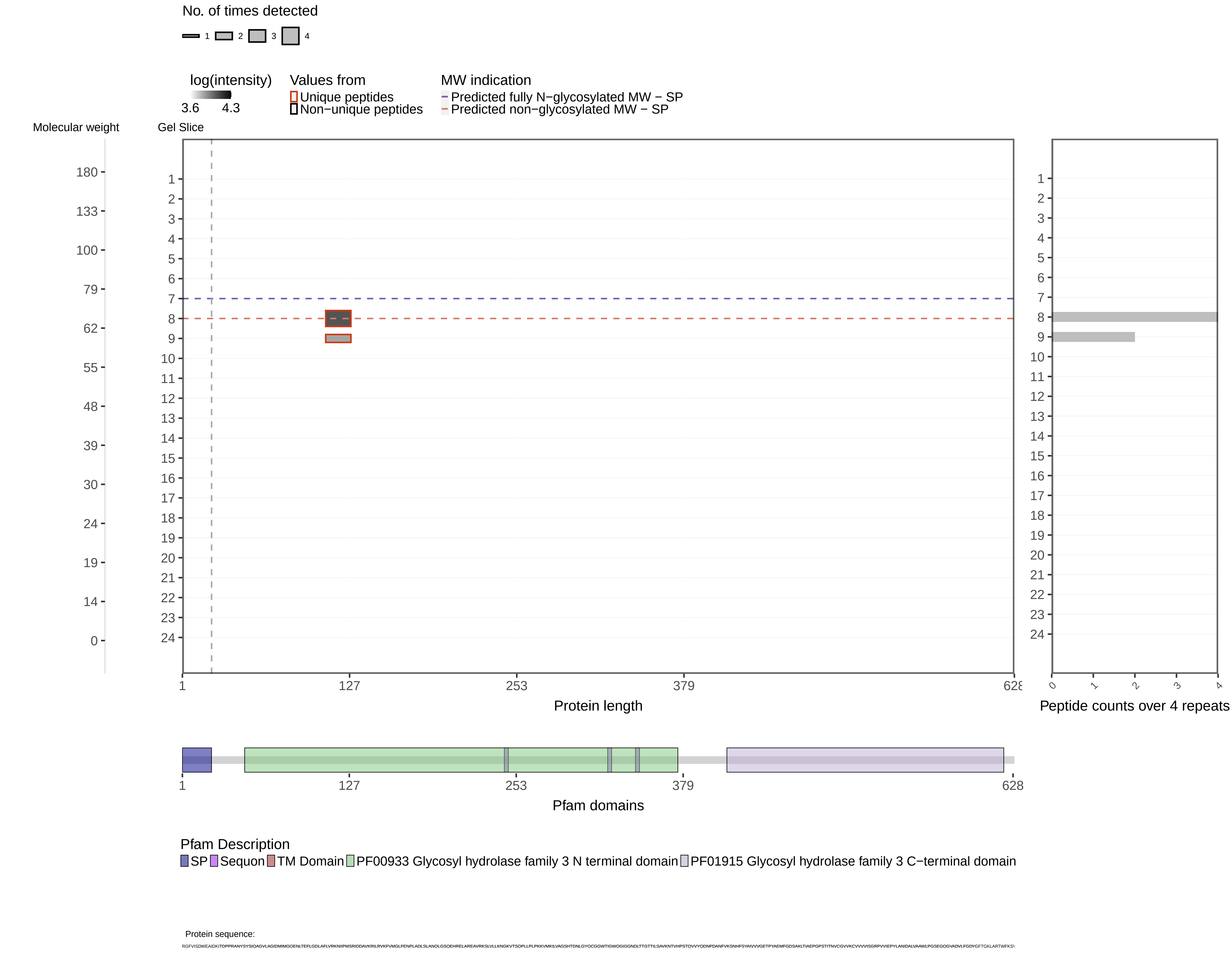

### NbL09g11200.1 60S_ribosomal_protein_L30 [MW = 12.38 kDa]

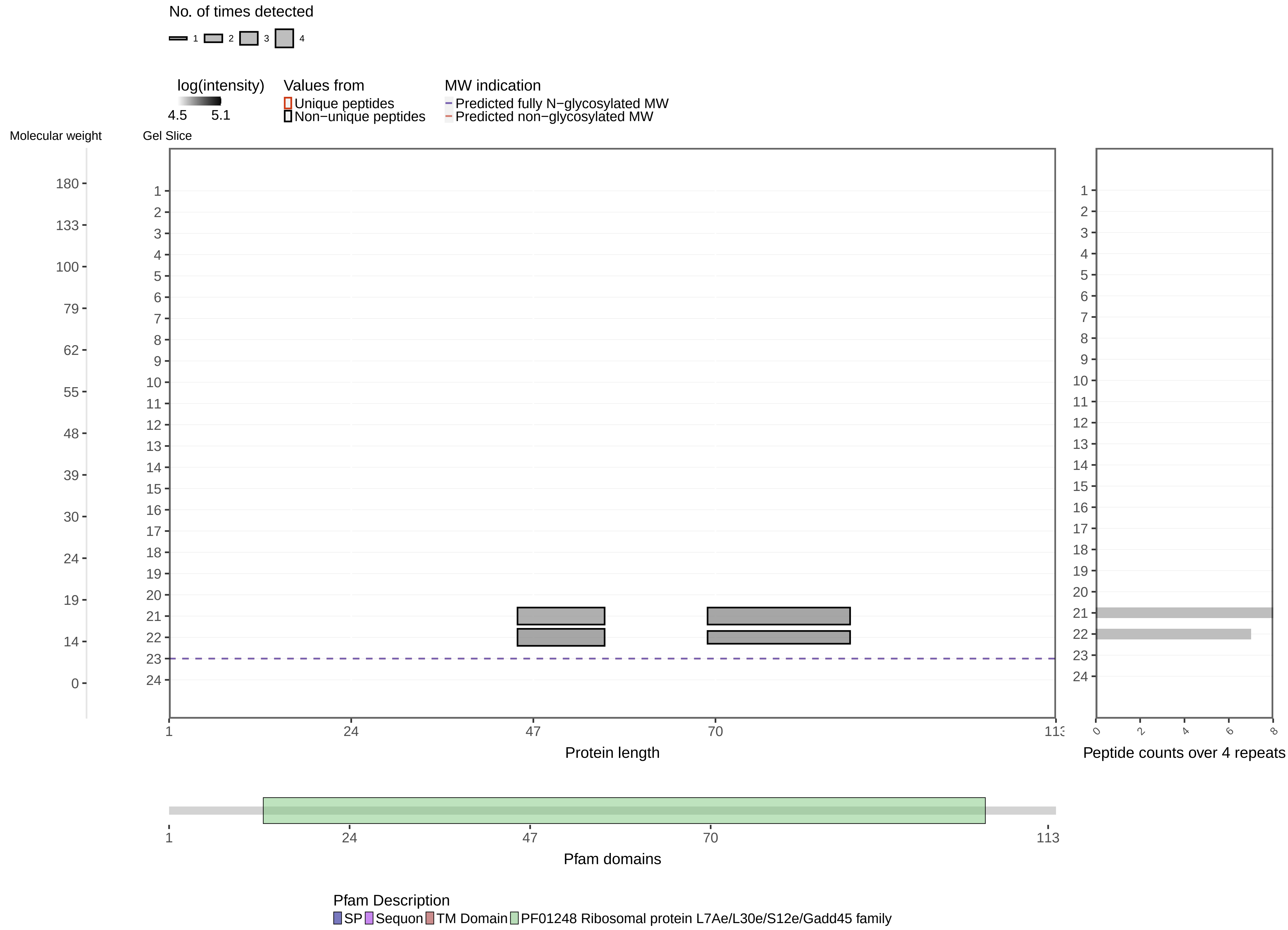


Protein sequence:

GKLVIIANNCPPLRKSEIEYYAMLAKIGVHHYNGNNVDLGTACGKYFRVCCLSIIDPGDSDIIKSMPGDO*

### NbL09g11180.1 NADH_dehydrogenase_[ubiquinone]_flavoprotein_1_mitochondrial−like [MW = 53.67 kDa]

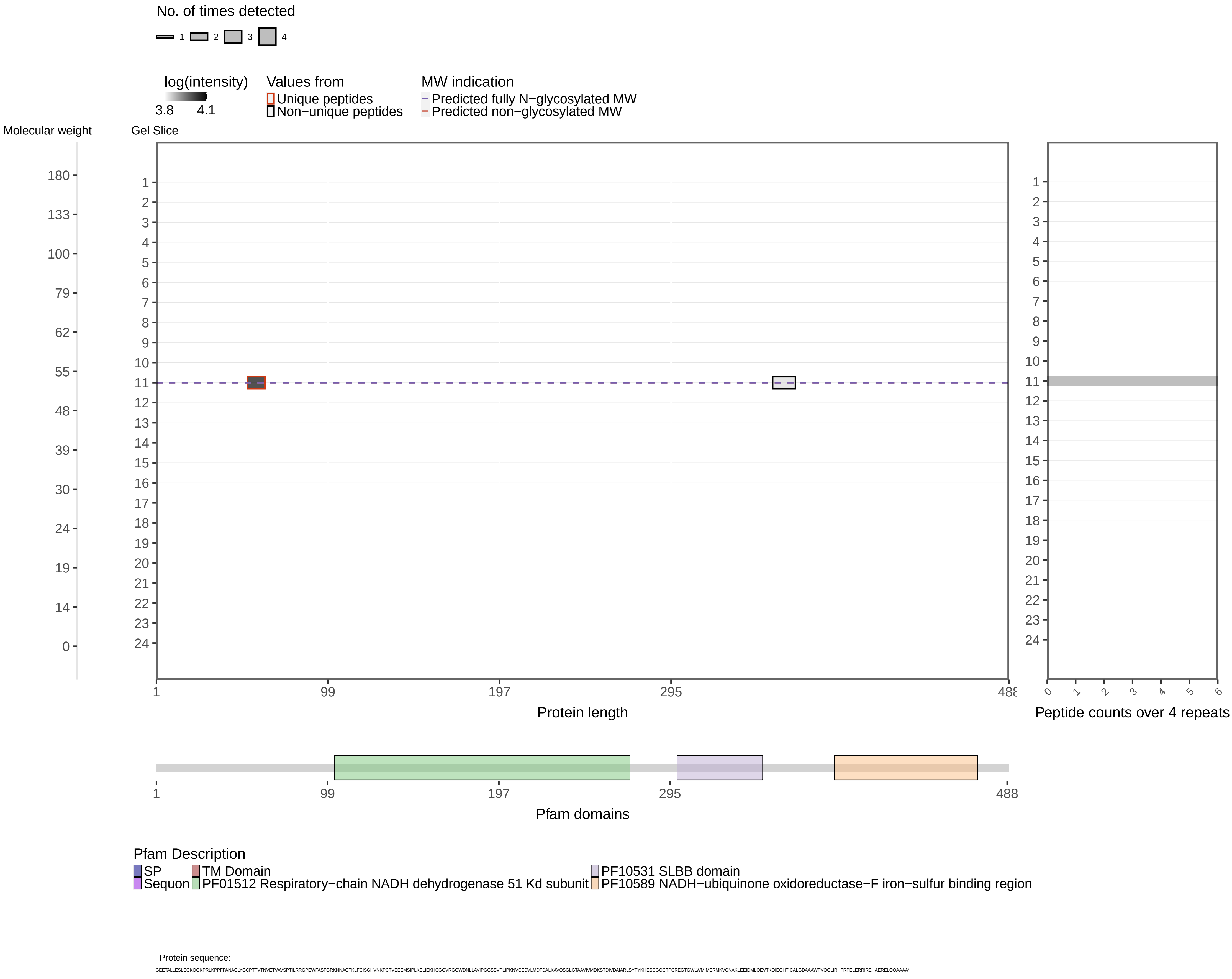

NbL09g11160.1 non−specific_lipid_transfer_protein_GPI−anchored_2−like [MW − SP = 14.64 kDa]

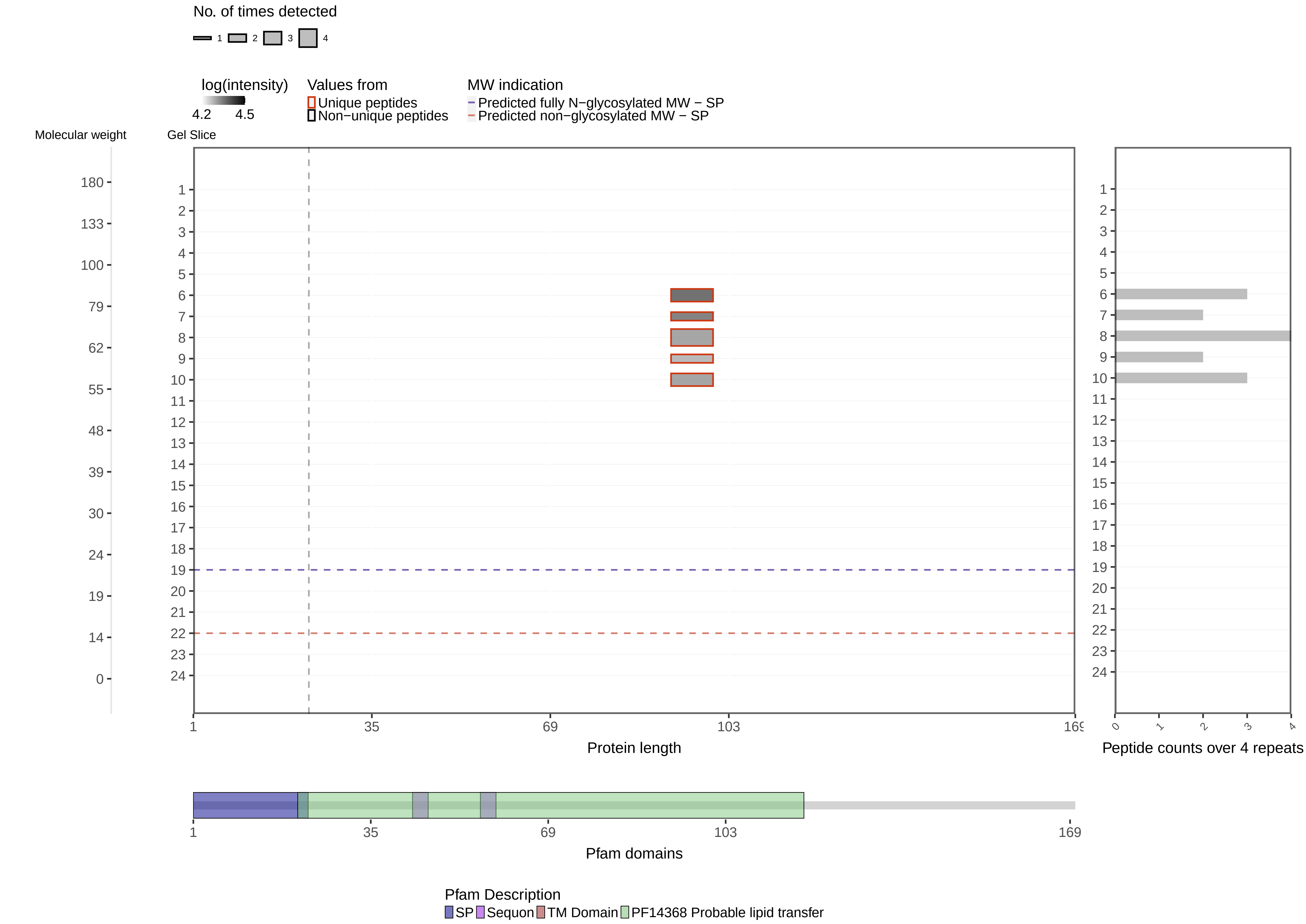


Protein sequence:

.DSNPICLCQMLGRASSGAKIGFNIDVDKALKLPSVCGLEIPPATTCSDLGVPIGAPIASDDGPAPLPGGFATSPTSDNKDNAASTMAGLMTVSSYE*

### NbL09g11050.1 protein_argonaute_4−like [MW = 101.85 kDa]

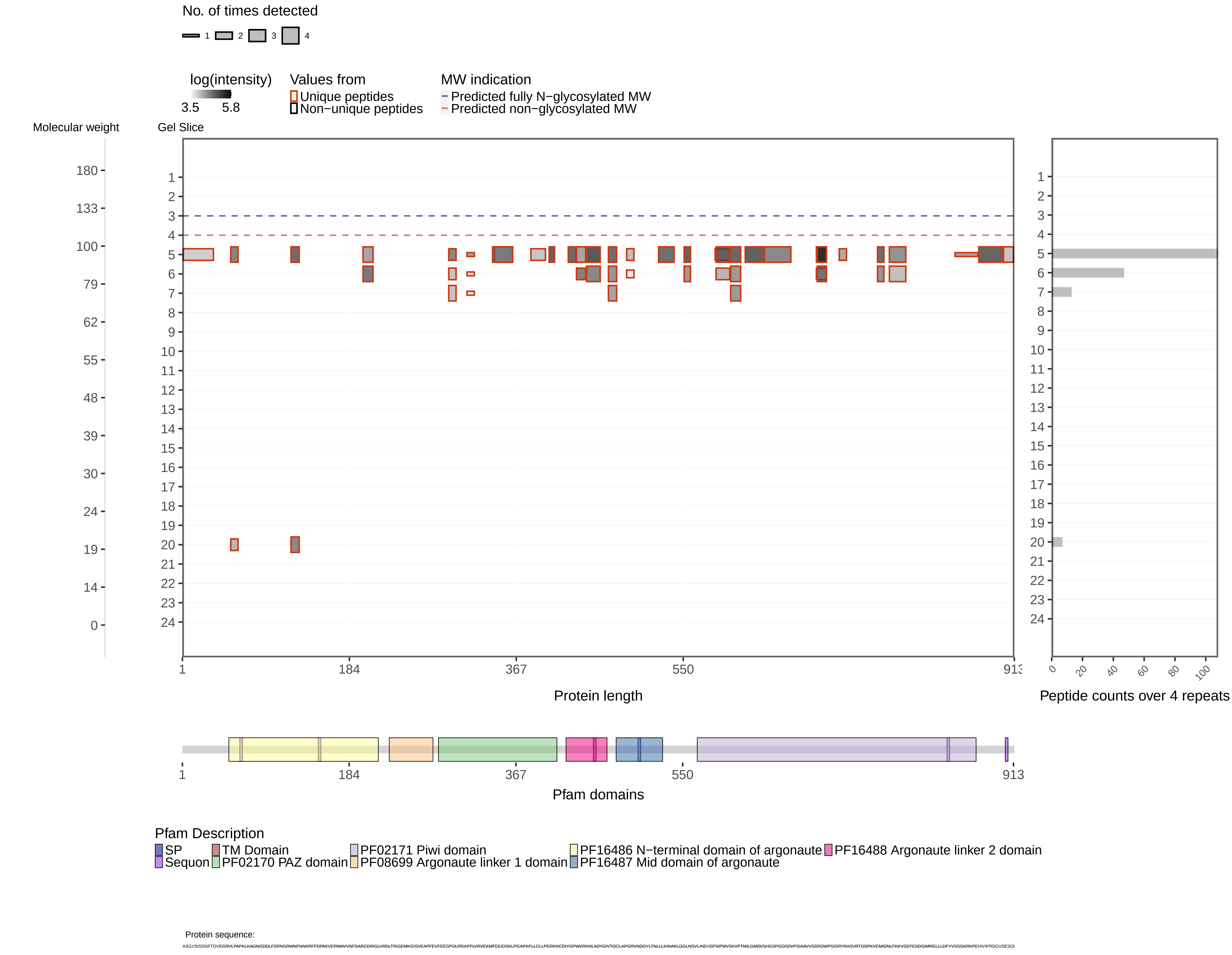

NbL09g11010.1 biotin_synthase [MW = 36.85 kDa]

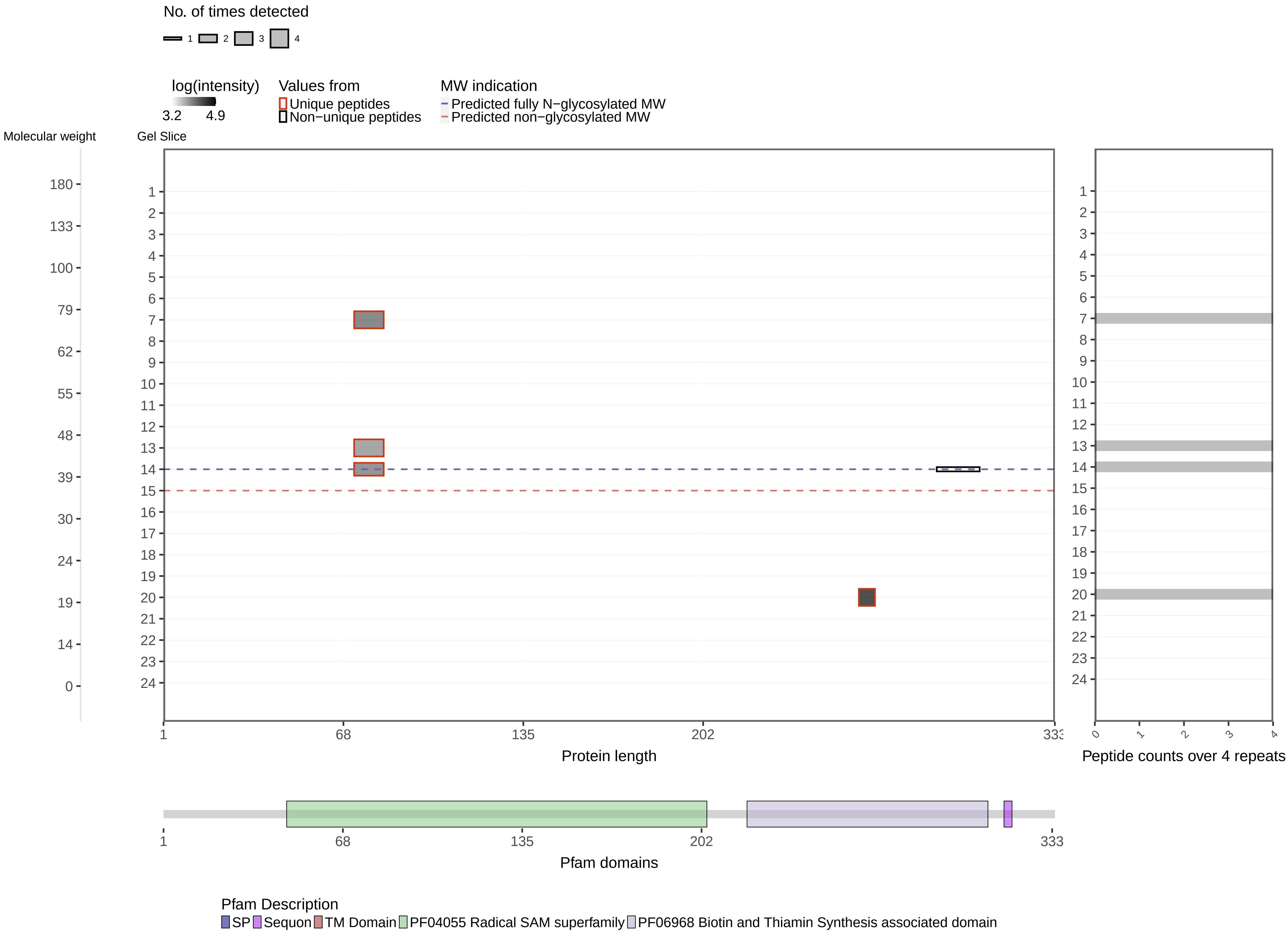


Protein sequence:

NHNLDTSREYYPNIITTRTYDERLETLKHVRKAGINVCSGGIIGLGEAEEDRIGLLHTLATLPTHPESVPINALIAVKGTPLEDQKPVEIWDMIRMISTARITMPSAMVRLSAGGVRFSIPEQALCFLAGANSIFTGDKLLTTPNNDYDADQYMFKLLGLVPKAPNFSADEKDSEDEKLEASI*

### NbL09g10960.1 nuclear_pore_complex_protein_NUP50B−like [MW = 47.6 kDa]

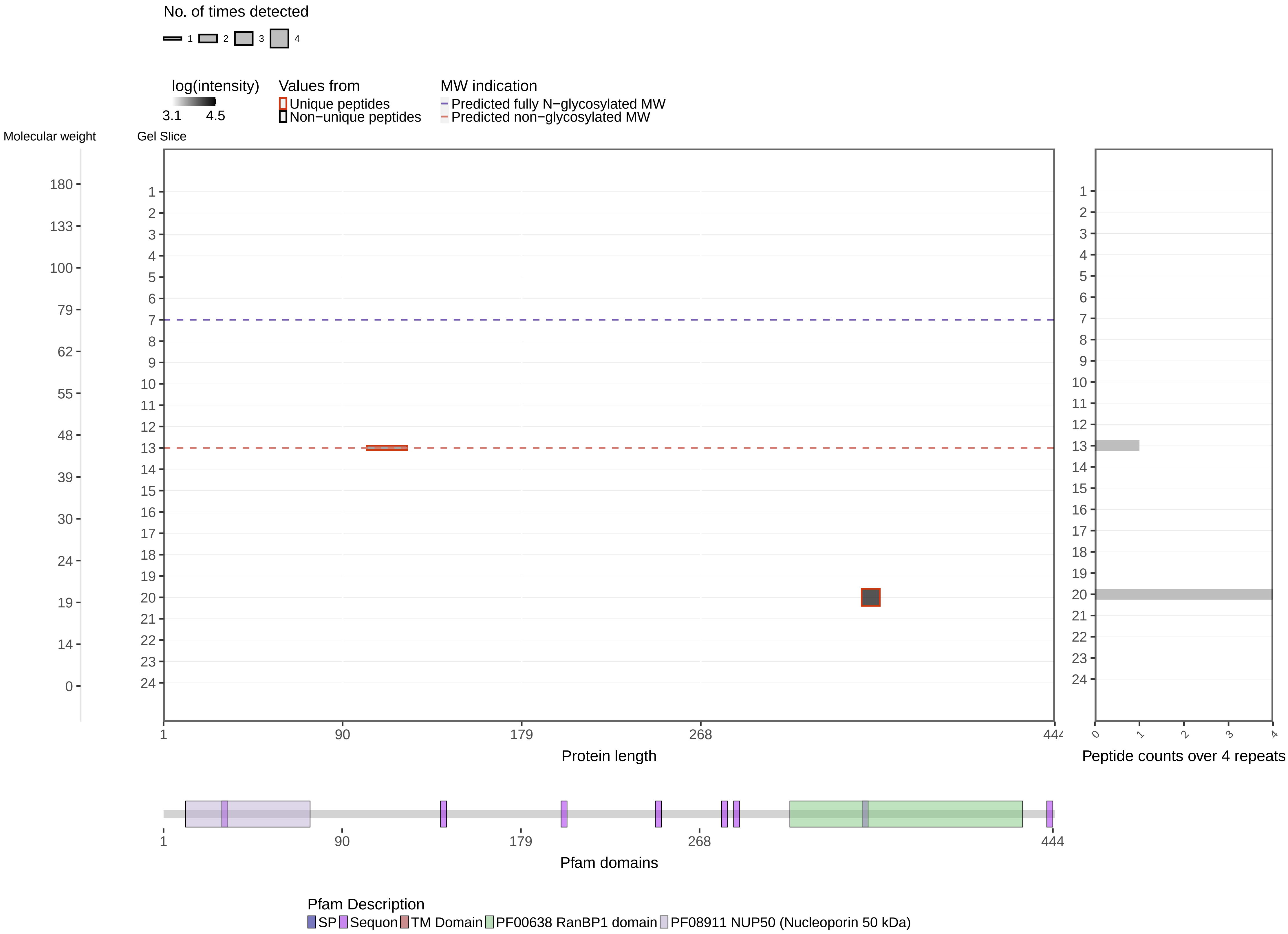


Protein sequence:

GNGENEKGAETASFSSFQQLSSSQNAFTGLAGTGFSNTTFSFGASSKEGSPLGSGSLFGVKSDQSPFGLNLSTNGNASLFGNSGSSRVNKGEGIEIPSMQEVPVETGEENEKAIFTADSALFEFLNGGWKERGKGELKVNISSTETGKARLVMRTRGNYRLILNANLYPEMKLTNMDKKGVTFACMNSAGDGKDGLSTIALKFKDASIIEEFRAVVMEHKDKMPGSLKTPQNSS*

### NbL09g10840.1 protein_CURVATURE_THYLAKOID_1C_chloroplastic−like [MW = 16.68 kDa]

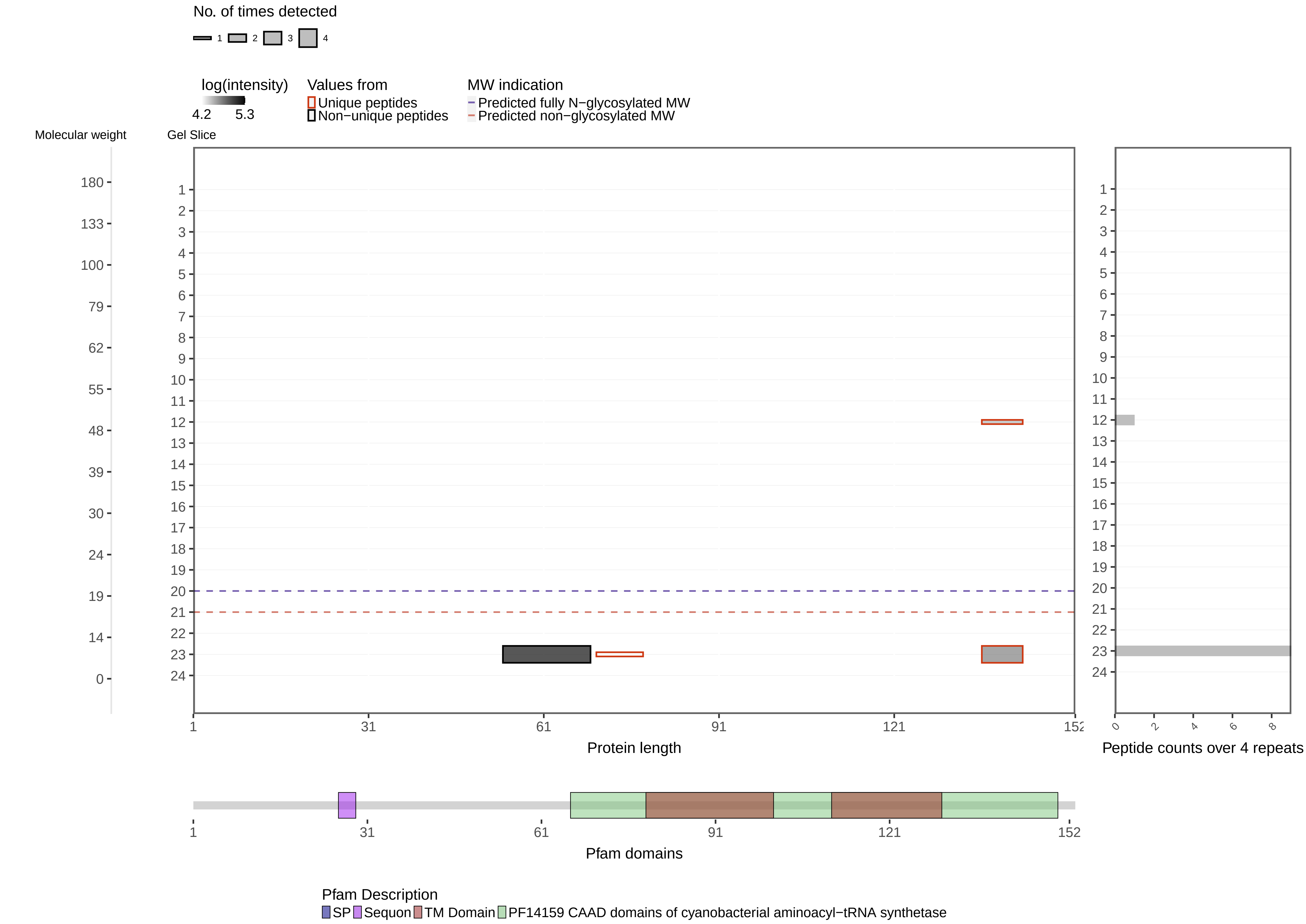


Protein sequence:

SSVVQSVKNIWDKPEDRVAVIGLGIAAIVGFWASSNLIAAIDRLPLIPSVFEFVGILFSSWFVYRYLLFKPDRKELFQAISKSISDILGQ*

### NbL09g10830.1 golgin_candidate_5 [MW = 112.09 kDa]

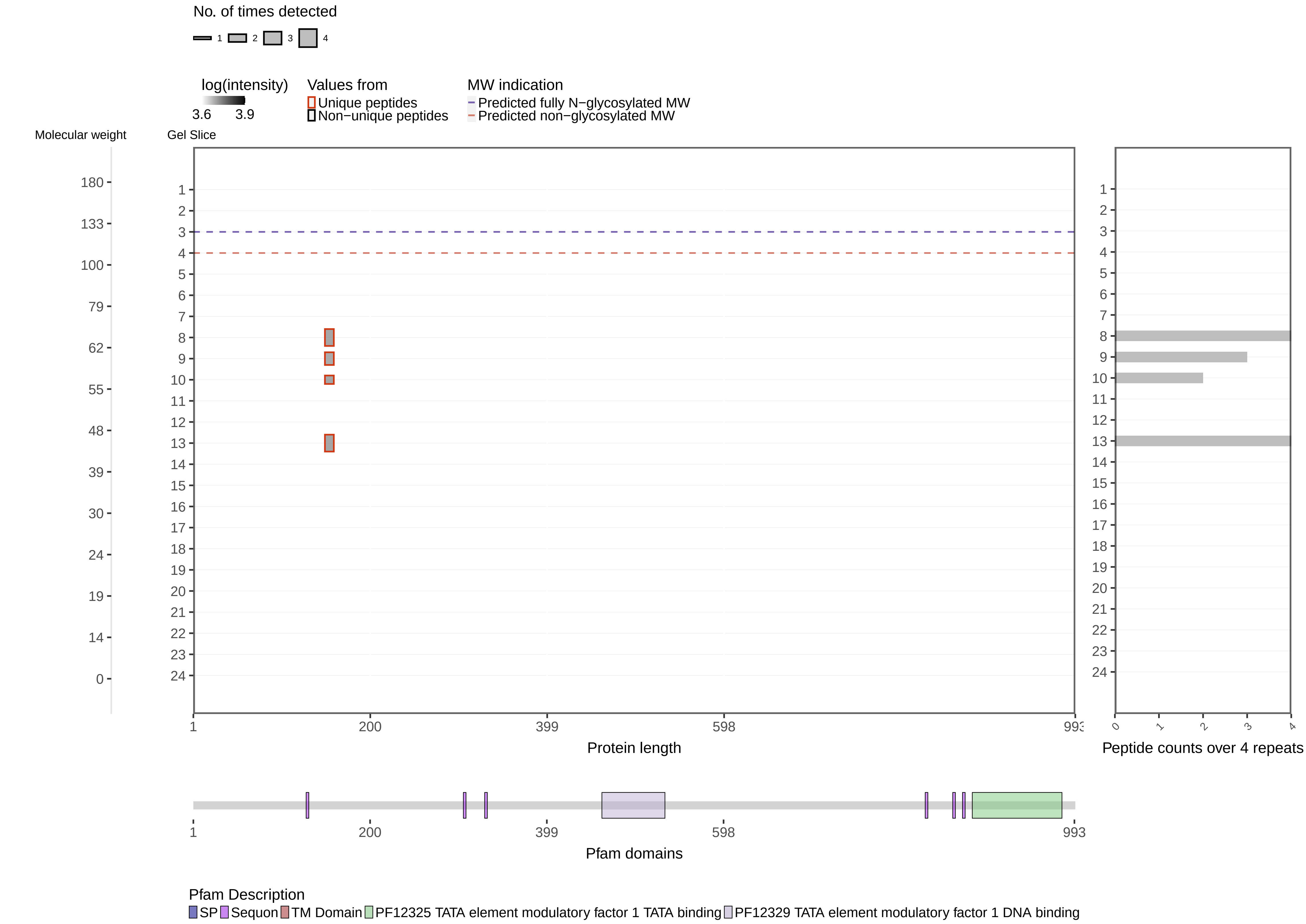


Protein sequence:

QLSKKQAAQEAQMRKLRAQIRELEEEKKGLLTKLEVEENKVESIKRDKAATEKLLHETVEKHQAELGTQKDYYTNALNAAREAEALAEARANNEARTQLEGRLREAEDREAMLVQALEELRQTLSRTEQQAVFREDMLRRDIEDLQKRYQASERRCEELITQVPESTRPLLRQIEAMQETNARKTEAWAAVERTLNSRLQEAEAKAATAEEKERSISERLSQTLSRINVLEAQISCLRAEQTQLTRSLDKERQRAGENRQEYLALKEEAETNEGRVNQLEEEIKEVRRKHK

### NbL09g10720.1 hexokinase−2 [MW = 54.34 kDa]

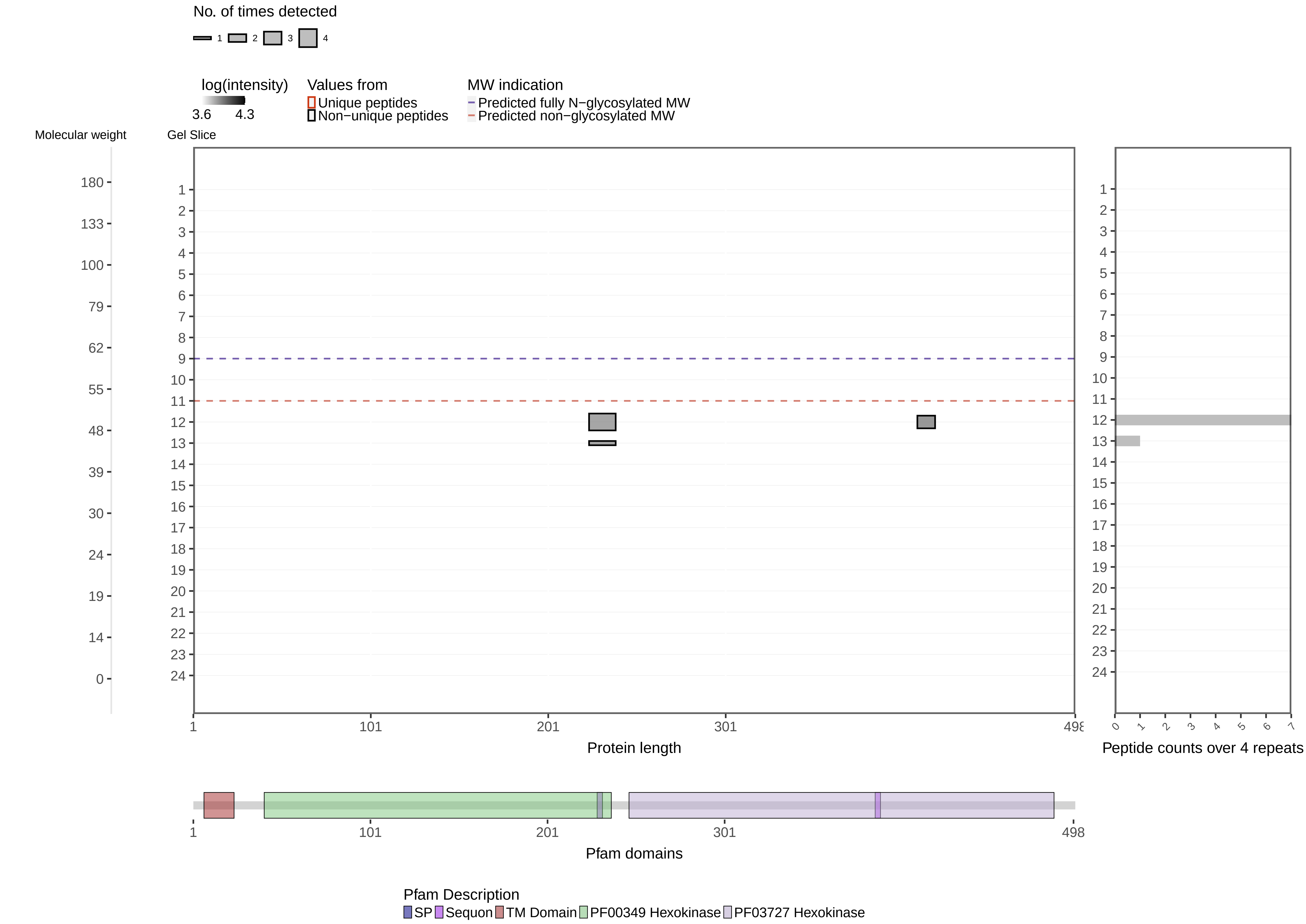


Protein sequence:

LAGGRFSNKDVSIAVILGTGTNAAYVERAQAIPKWHGPLPKSGEMVINMEWGNFRSSHLPLTEYDHAMDTDSLNPGEQIFEKICSGMYLGEILRRVLLRMAEEDAIFGDEVPPRLKNQFILRTPEMSAMHHDTSSDLRVVGDKLKDILEISNTSLKIRRLVVELCNIVATRGARLAAAGVLGIVKKMGRDTPRESGQEKIVVAMDGGLYEHYTEYRKCLENTLVELLGEEMATSIVFEHANDGSGIGAALLAASNSLYVEDKS*

### NbL09g10590.1 ESCRT−related_protein_CHMP1A−like [MW = 22.65 kDa]

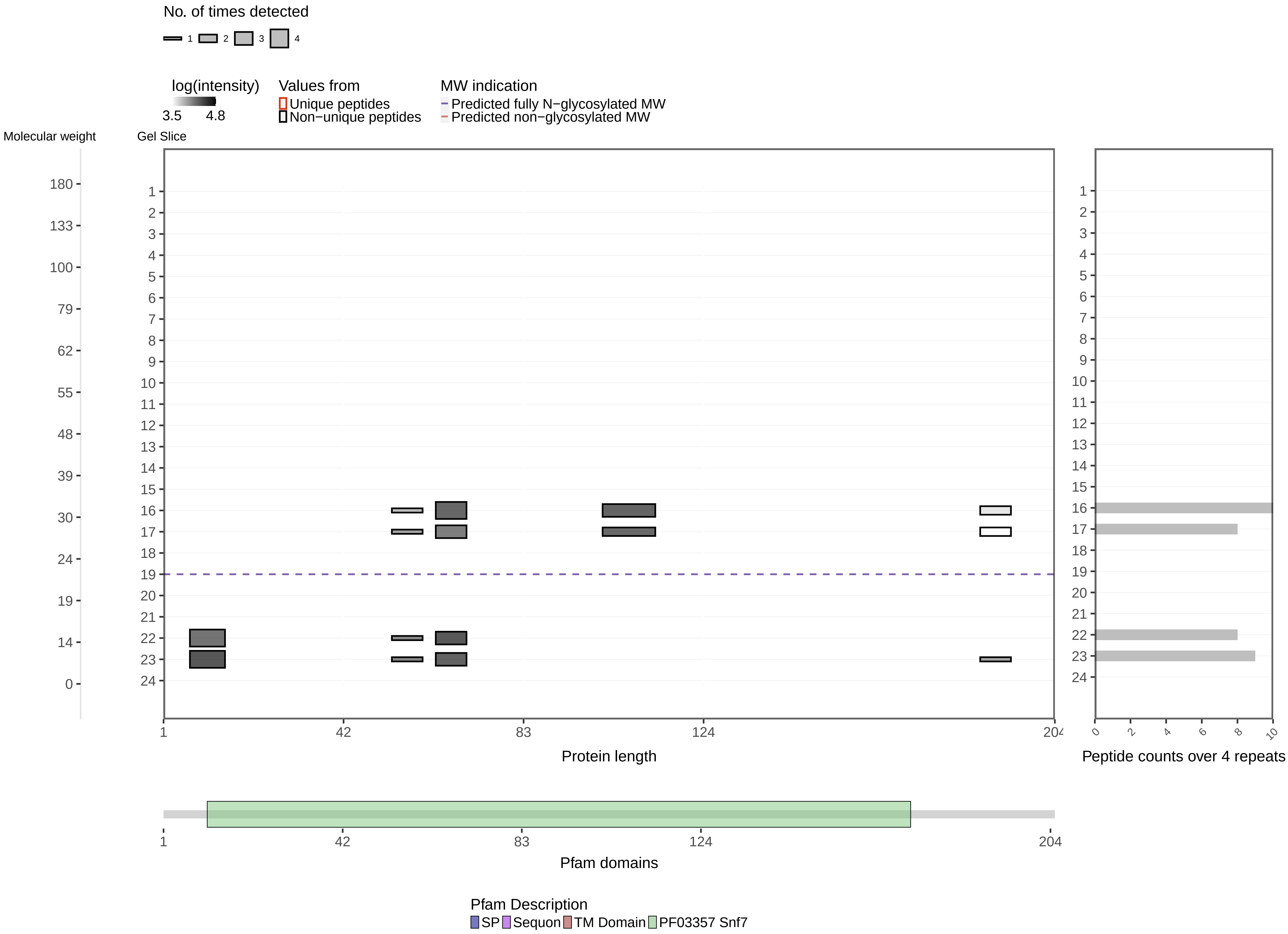


Protein sequence:

TISKSMSSIVKSLESSLNTGNLQKMSETMDSFERQFVNMEVQAEFMESSMAGSTSLSTPEDQVGSLMQQVADDYGLEVSVGLPQPAAHSISSKNEEKVNEDDLSRRLAELKARG*

### NbL09g10540.1 beta−galactosidase−like [MW = 93.89 kDa]

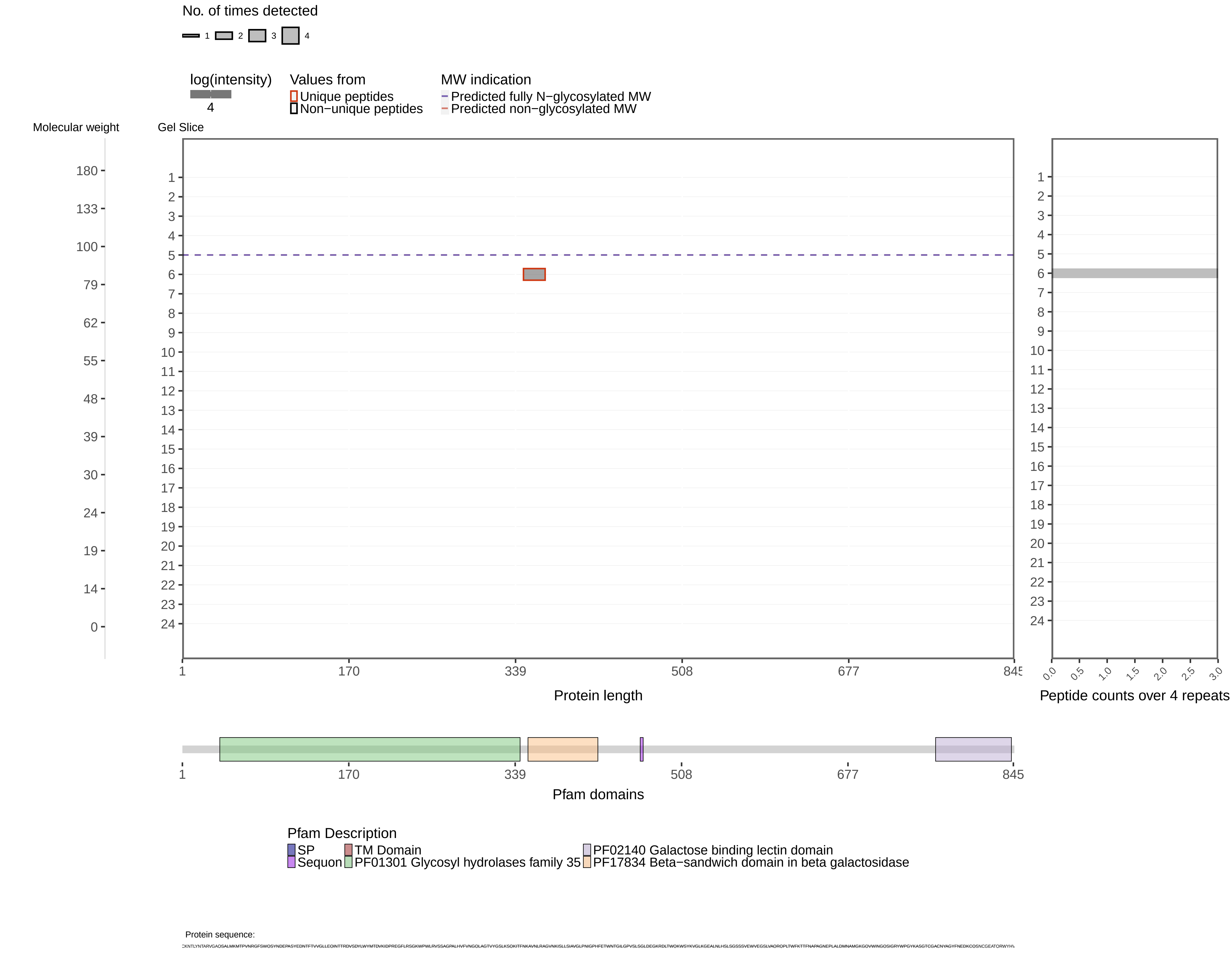

NbL09g10400.1 glycylpeptide_N−tetradecanoyltransferase_1−like [MW = 49.66 kDa]

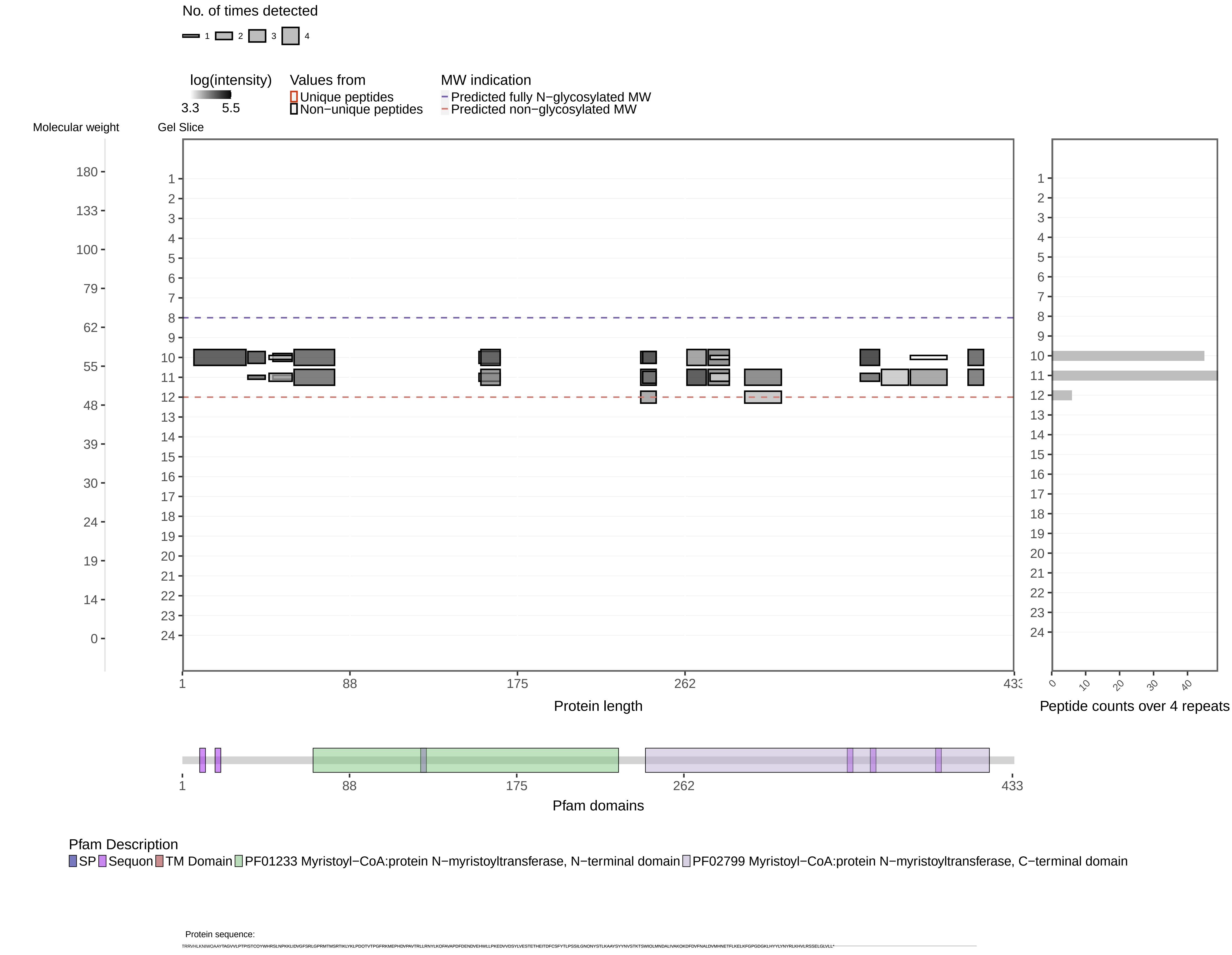

### NbL09g10390.1 glycylpeptide_N−tetradecanoyltransferase_1−like [MW = 49.76 kDa]

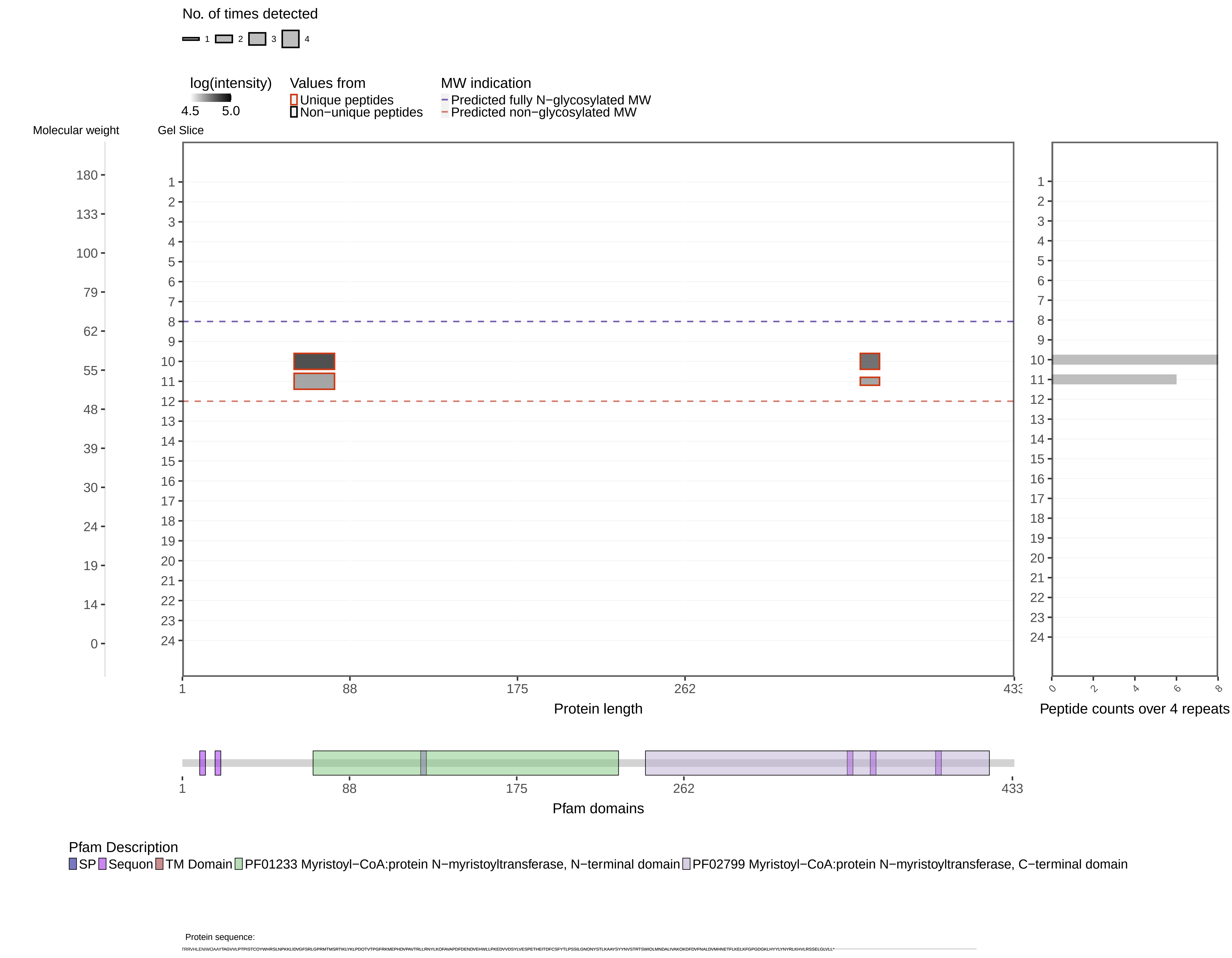

NbL09g10360.1 threonine_synthase_chloroplastic−like [MW = 60.4 kDa]

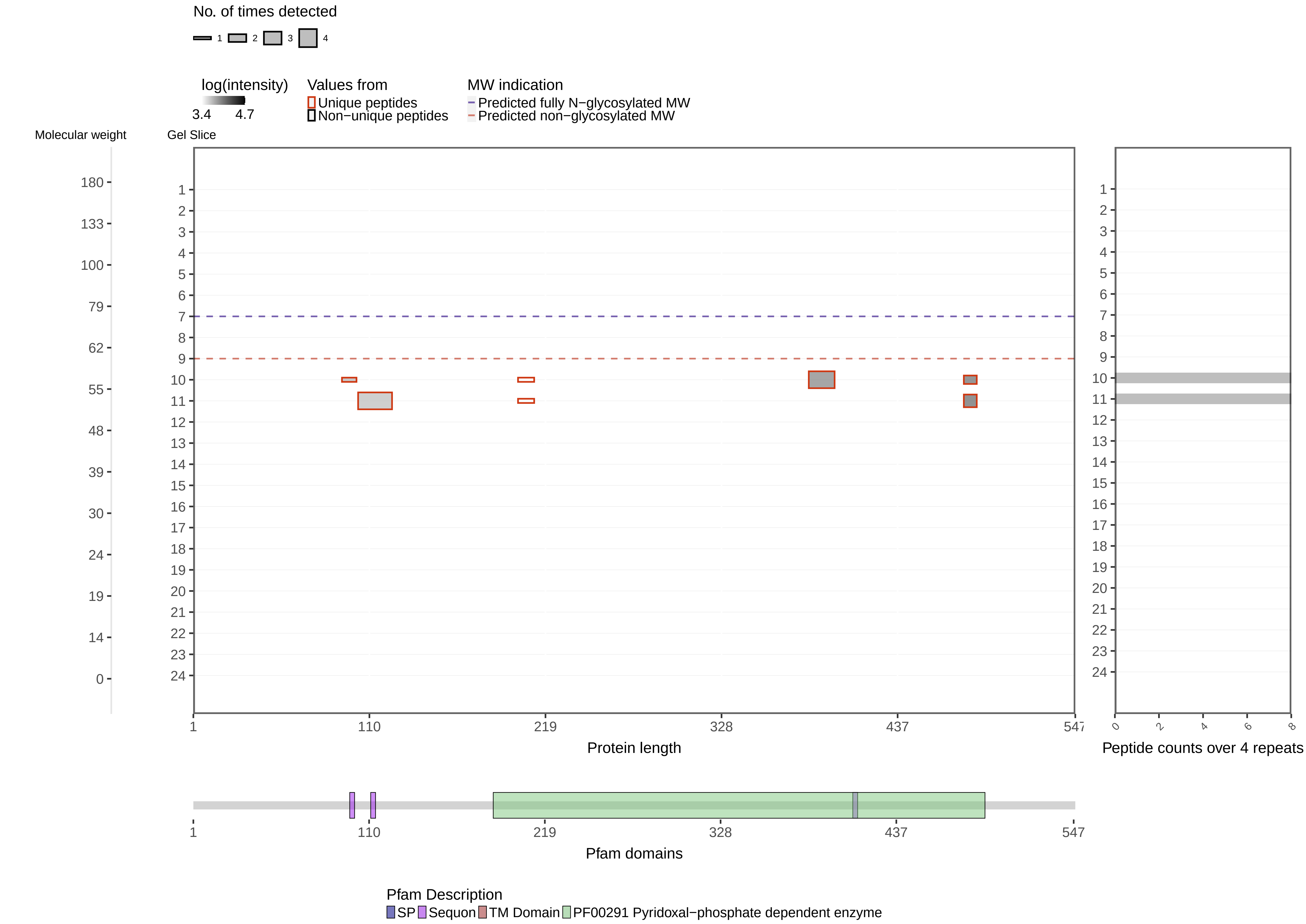


Protein sequence:

TSAALSAYCASAGIPSIVFLPANKISMAQLVQPIANGAFVLSLDTDFDGCMQLIREVTAELPIYLANSLNSLRLEGQKTAAIEILQQFDWEVPDWVIIPGGNLGNIYAFYKGFQMCKELGLVDRIPRLVCAQAANANPLYLHFKSGWKEYKSVKANTTFASAIQIGDPVSIDRAVYALKNSNGIVEEATEEELMDAMAQADSTGMFICPHTGVALTALSKLRKIGVIRPTDKTVVVSTAHGLKFTQFKVDYHSKEIKNMECRFANPPVQVKADFGSVMDVLKKYLLSKNSKF*

### NbL09g10160.1 alanine_aminotransferase_2−like [MW = 59.35 kDa]

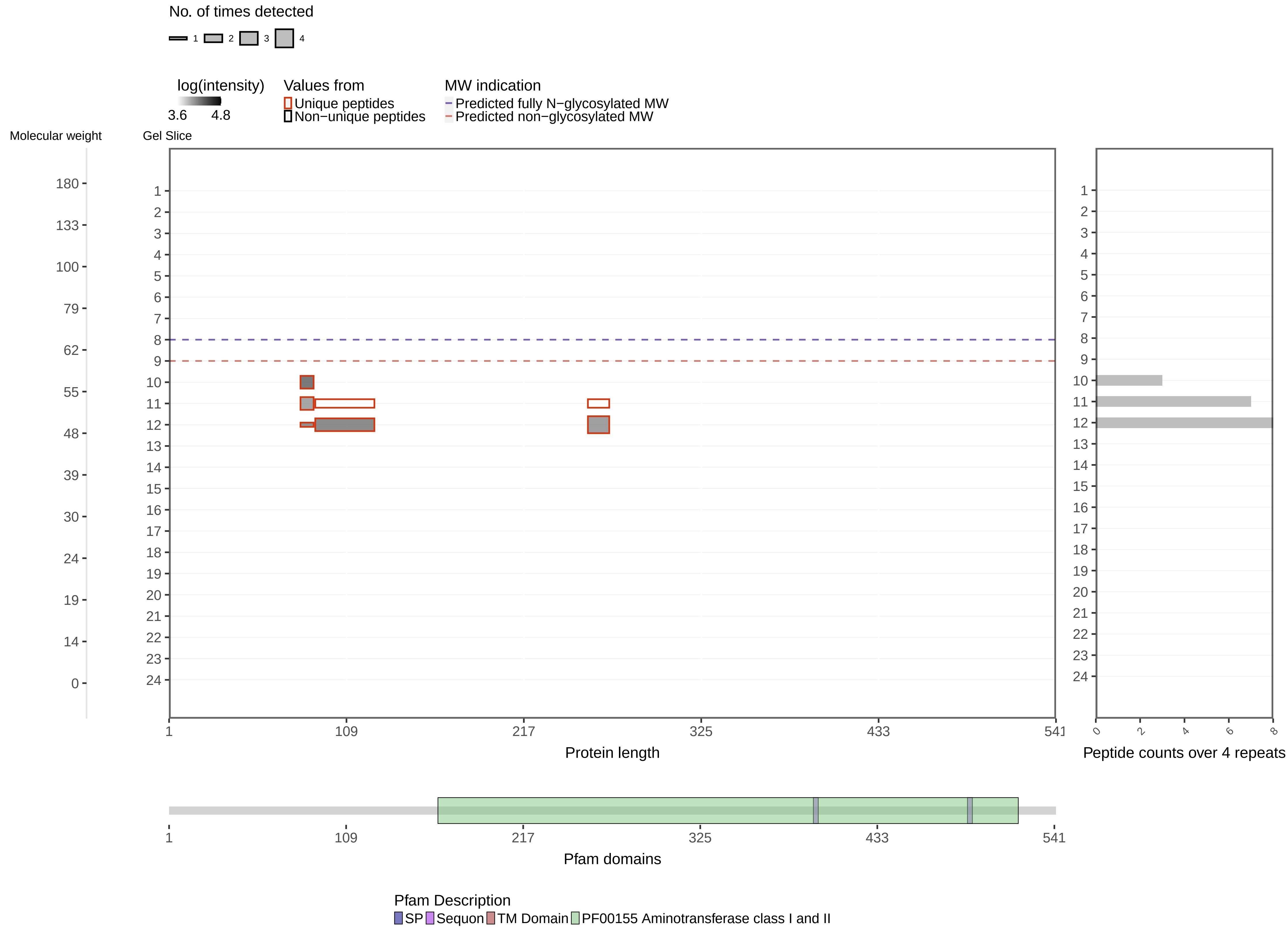


Protein sequence:

EISELEQQLKTAKSKGIDVRALVVINPGNPTGQVLGEANQREIVEFCKREGLVLLADEVYQENVYVPDKKFHSFKKVSRSMGYGEKDISLVSFQSVSKGFYGECGKRGGYMEVTGFSPEVREQIYKVASVNLCSNISGQILASLIMSPPKVGDESYESFSAEKEGILSSLARRAKTLEDALNSLEGVTCNRAEGAMYLFPRVNLPNKAVKAAEAAKTAPDAFYAKCLLNATGIVVVPGSGFRQVPGTWHFRCTILPQEEKIPAIVSRLTEFHEKFMNEFRG*

NbL09g10120.1 26S_protease_regulatory_subunit_7_homolog_A−like [MW = 47.77 kDa]

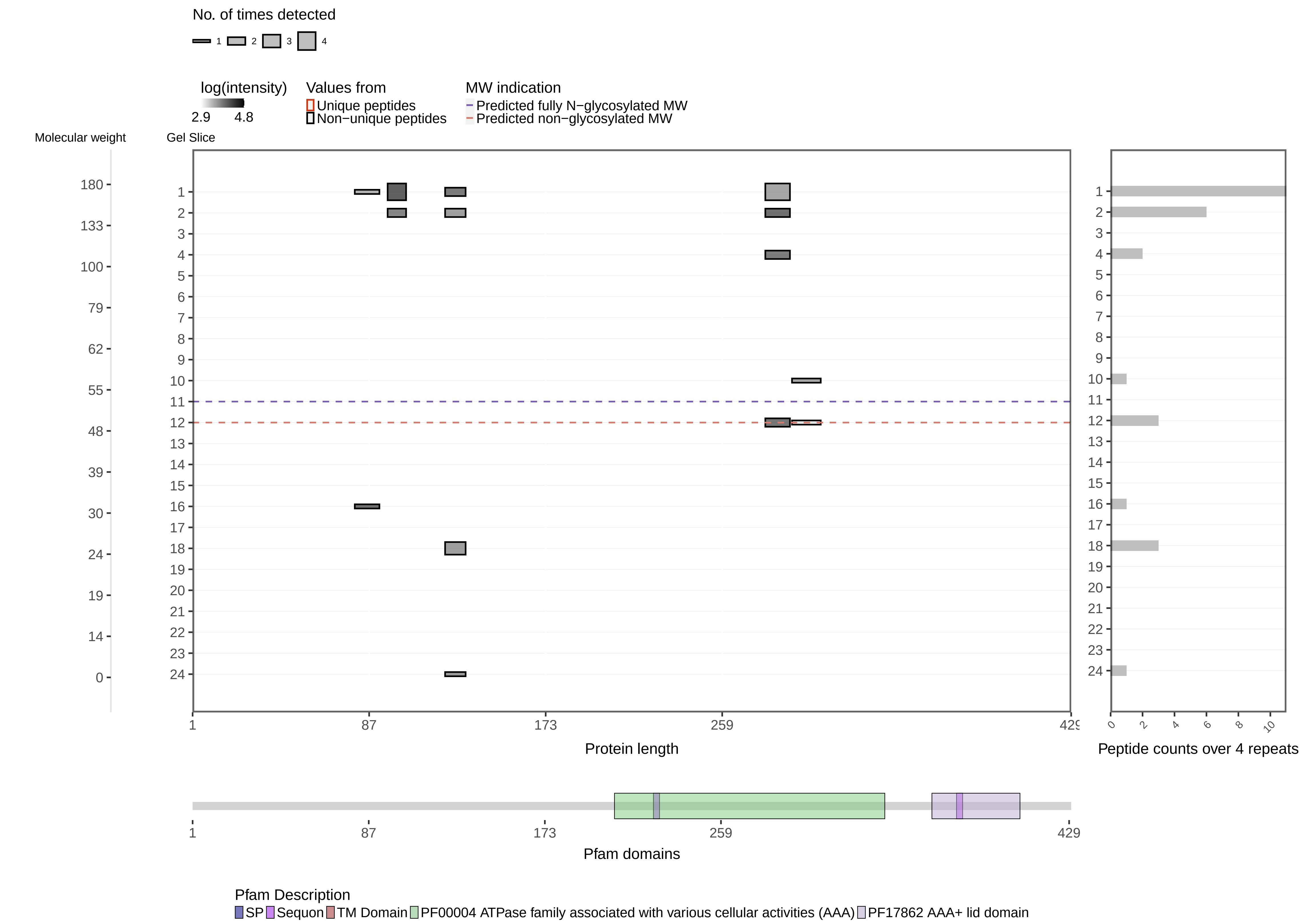


Protein sequence:

PPKGVLCYGPPGTGKTLLARAVANRTDACFIRVIGSELVQKYVGEGARMVRELFQMARSKKACIVFFDEVDAIGGARFDDGVGGDNEVQRTMLEIVNQLDGFDARGNIKVLMATNRPDTLDPALLRPGRLDRKVEFGLPDLESRTQIFKIHTRTMNCERDIRFELLARLCPNSTGADIRSVCTEAGMYAIRARRKTVTEKDFLDAVNKVIKGYQKFSATPKYMVYN*

NbL09g10030.1 mannosyl−oligosaccharide_12−alpha−mannosidase_MNS1−like [MW = 56 kDa]

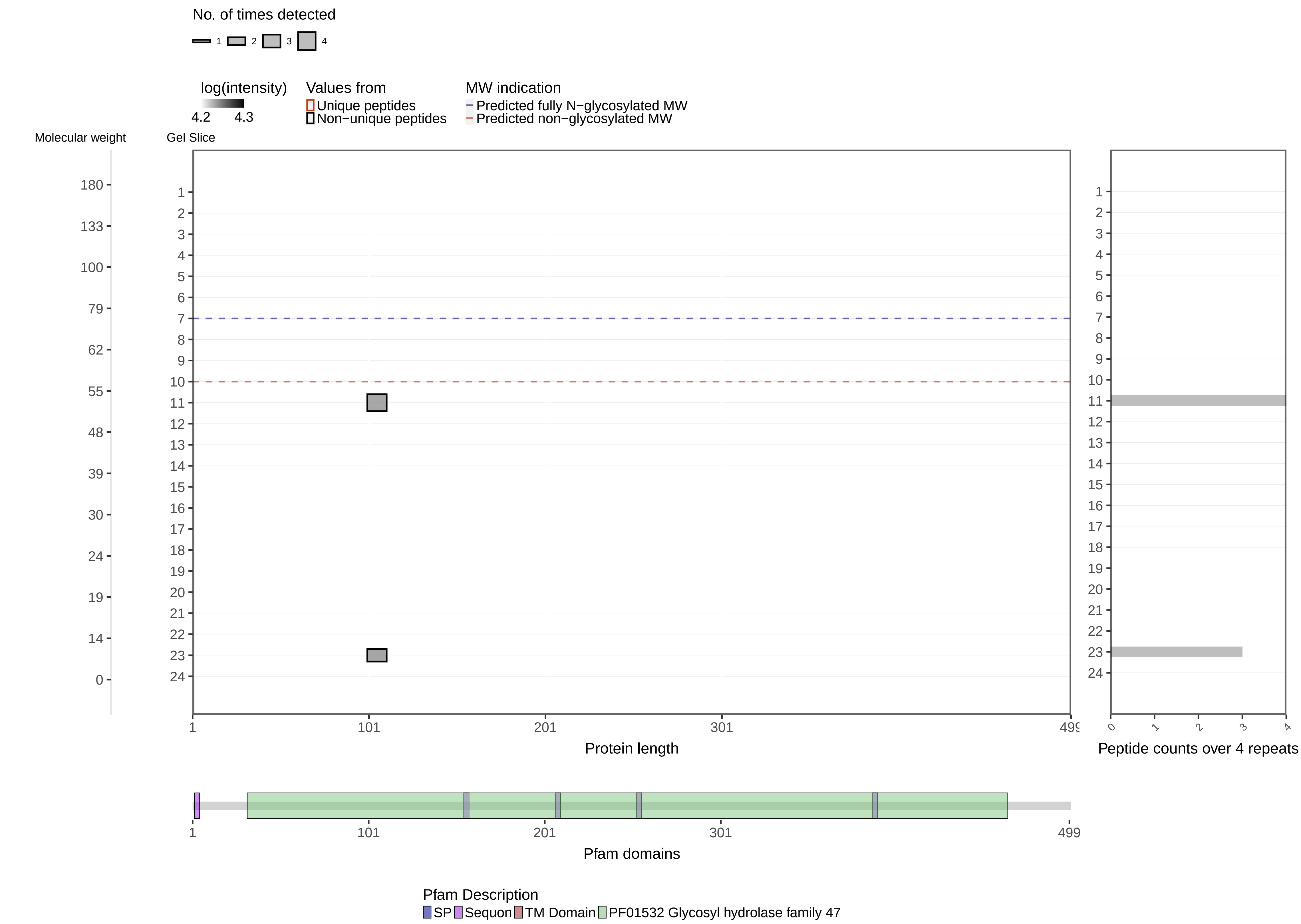


Protein sequence:

DSFYEYLLKAWIQGNQTAVVGHYRKMWETSMKGLLSLVRRTTPSSFAYIGEKIGSSLNDKMDELACFAPGMLALGSSGYGPEESQKFLSLAEELAWTCYNFYQSTPTKLAGENYFFNDDGQDMTVGTSWNILRPETVESLFYLWRLTGNKTYQEWGWNIFQAFEKNSRIESGYVGLKDVNTGVRDDMMQSFFLAETLKYLYLLFSPSSLIPLDEWVFNTEAHPIRIVSRNDPAVSSGRSVGQAKSYKQPRVRREGRFGNK*

### NbL09g10010.1 uncharacterized_LOC109227356 [MW = 44.98 kDa]

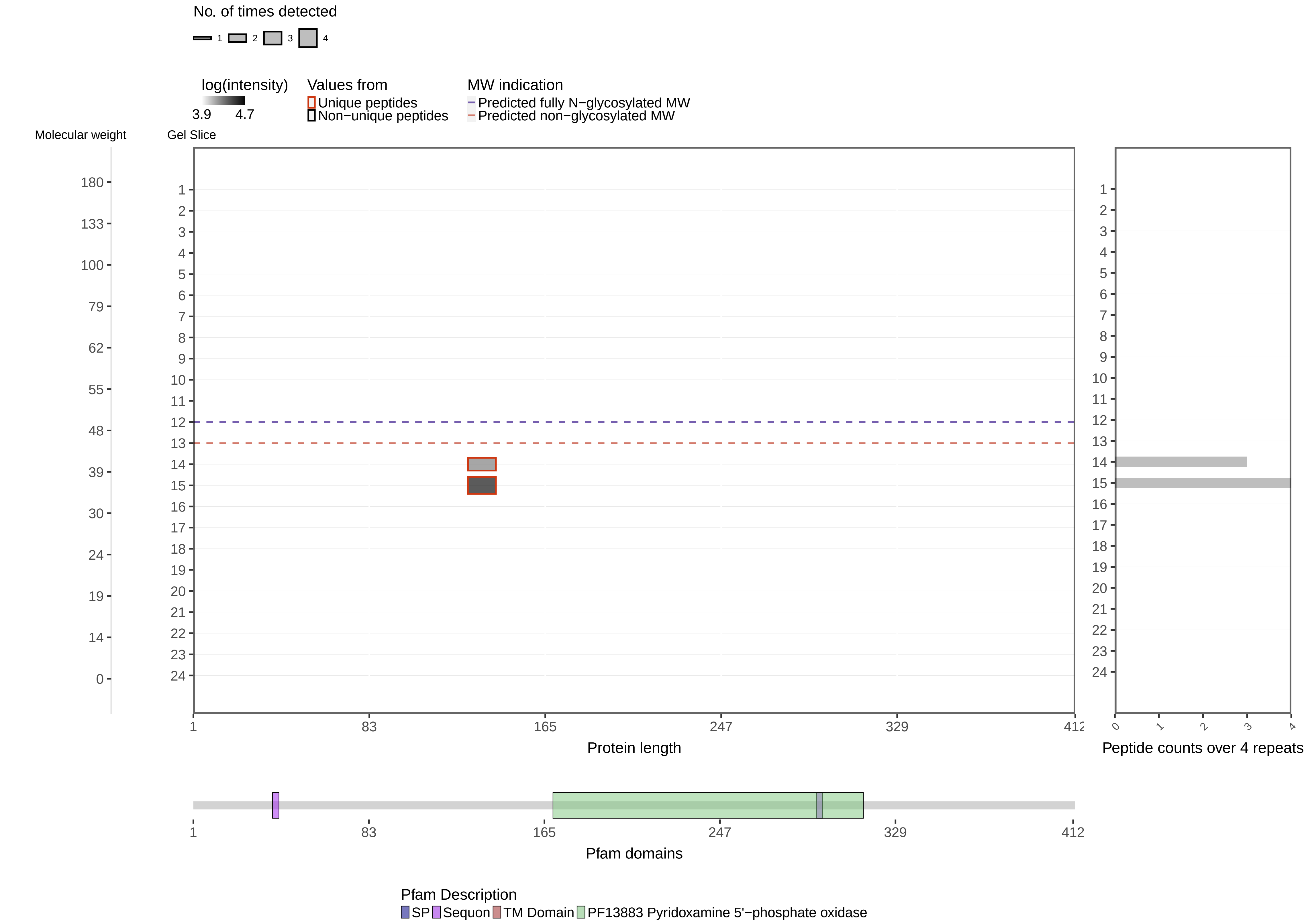


Protein sequence:

RREGYPFGSLVDFAPDSMGHPIFSFSPLAIHTRNLLADPRCTLVVOIPGWSGLSNARVTIFGDVYPLPEDOOAWAHKOYIAKHOOGPSOOWGNFFYFRMONISDIYFIGGFGTVAWVDVKEYETLOPDKIAVDGGEONLKELNAIFSKPLKDLLSOESEVDDAALISIDSKGTDVRVROGAOFNVORISFEVGHSVOTLEEAKAALWKLINGGOVYNLOK*

### NbL09g09970.1 V−type_proton_ATPase_catalytic_subunit_A [MW = 200.76 kDa]

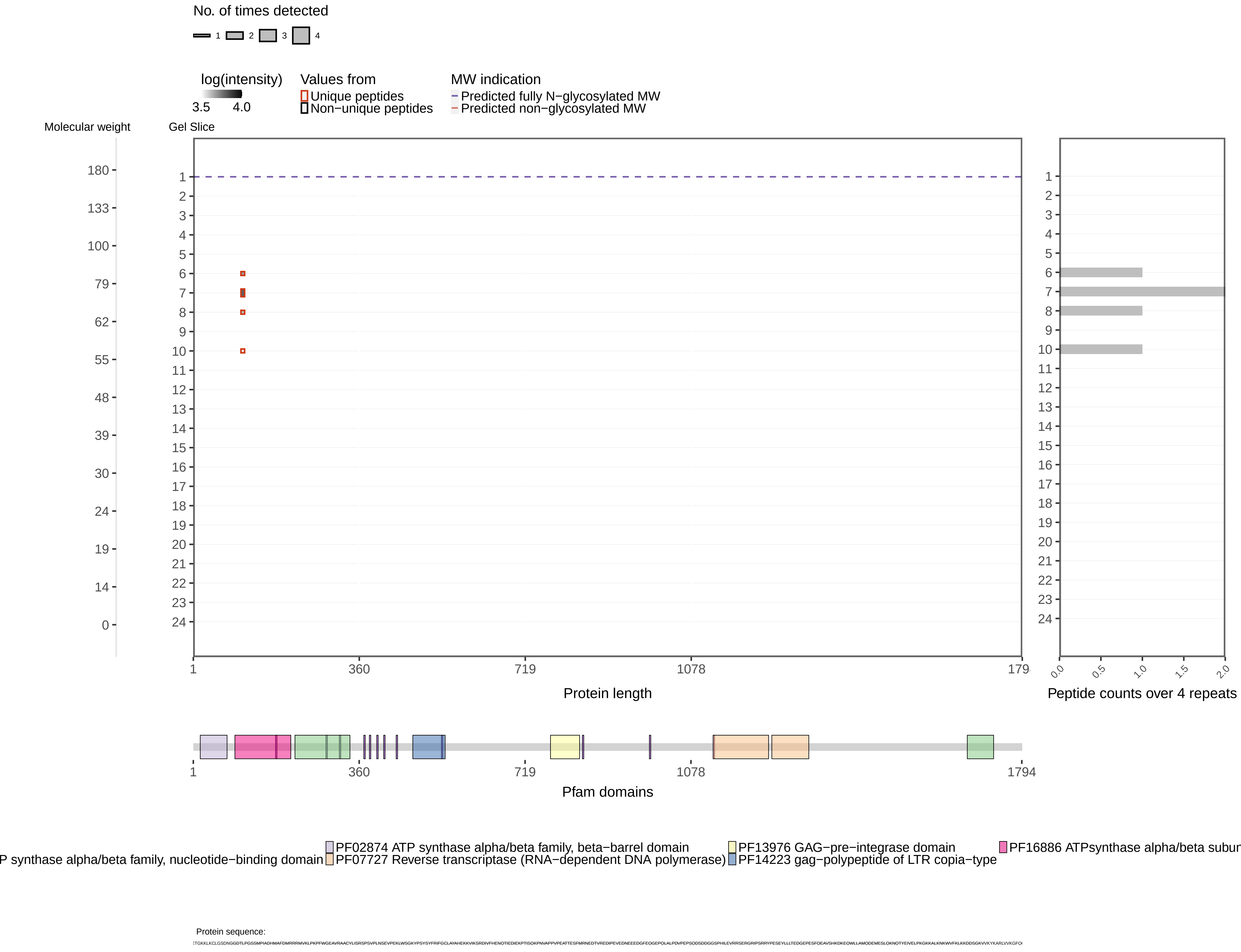

### NbL09g09910.1 chlorophyll_a−b_binding_protein_CP26_chloroplastic [MW = 30.41 kDa]

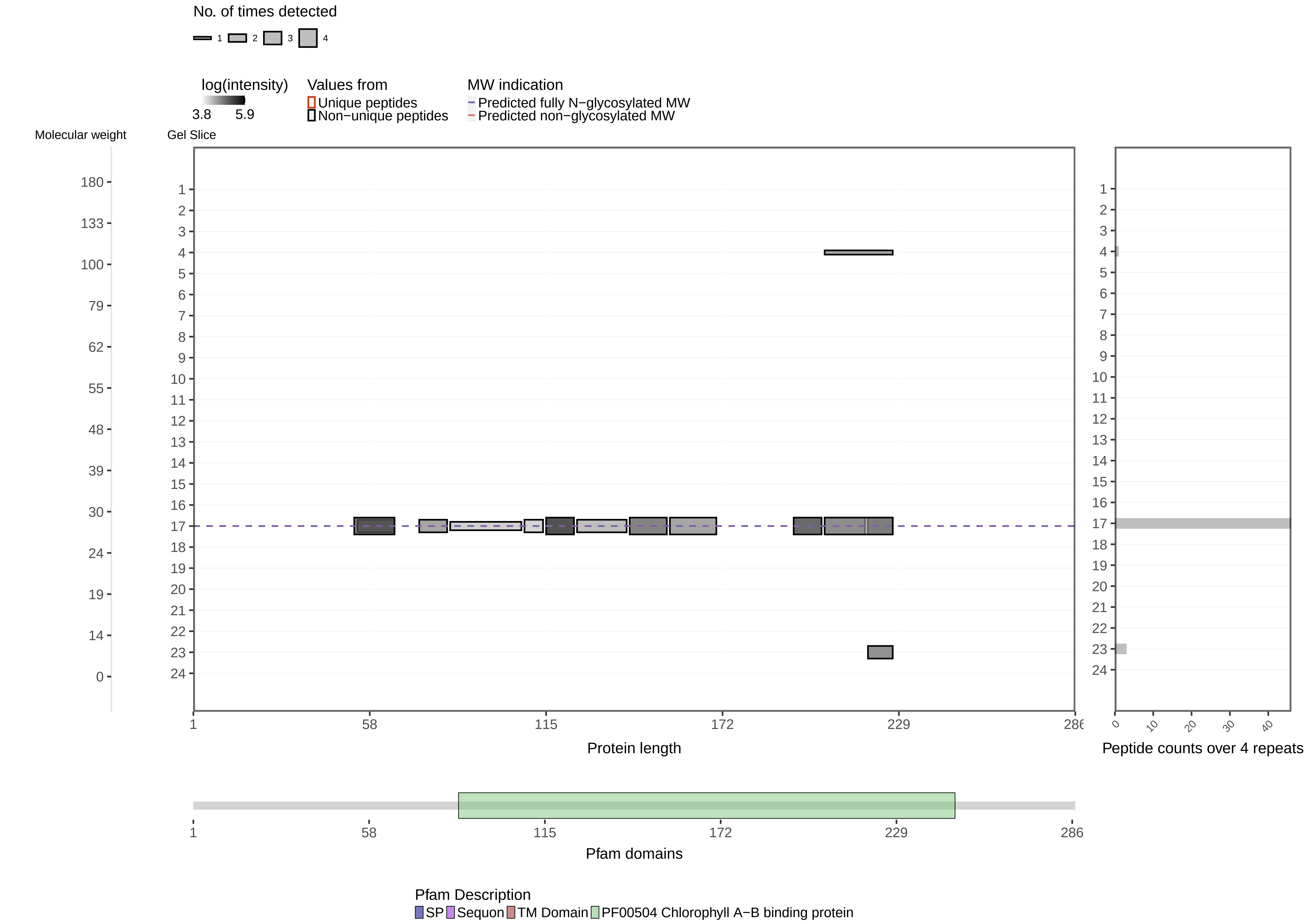


Protein sequence:

AAGFVIPEAFNKFGADCGPEAVWFKTGALLLDGNTLNYFGKNIPINLVVAVVAEVVLLGGAEYYRITNGLDLEDKLHPGGPFDPLGLAKDPDQAAILKVKEIKNGRLAMFAMLGFFIQAYVTGQGPVENLAAHLSDPFGNNLLTVIGGTAERVPTL*

### NbL09g09840.1 trans−resveratrol_di−O−methyltransferase−like [MW = 40.43 kDa]

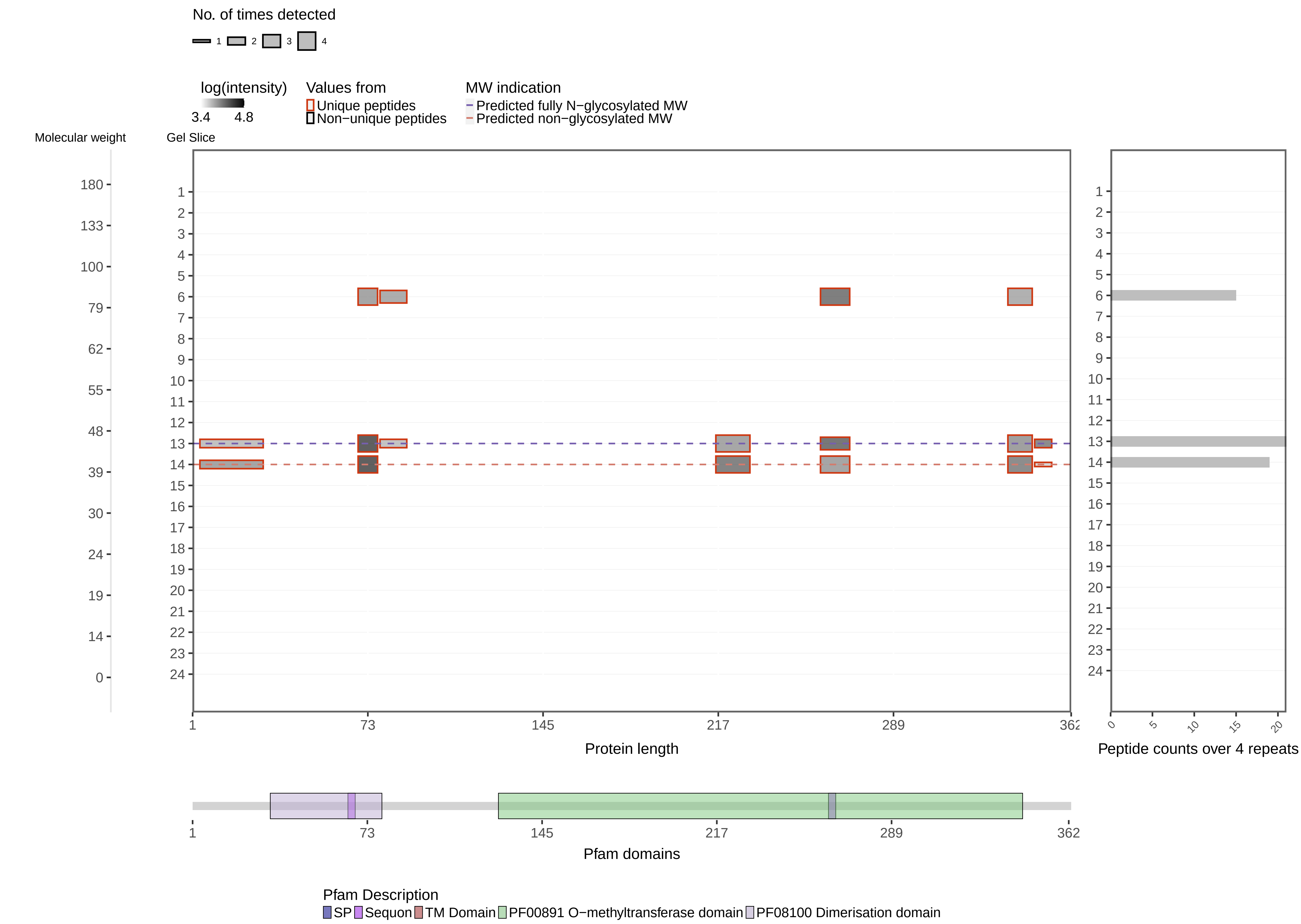


Protein sequence:

MASDSKLISHLLINSEQCKHVFEGLTSLVDVGGGTGTLAMAIANAFPRLNCIVLDLPHVIGDREGITTENVEFVAGNMFDNIPYANAVLLKYILHNWSDEDCVKLLKKCRESIPSKEKGGKVMVIDIVMEDYSSNNINEQLLQVQHLMHVVMKITYGAKERTKTEWEKLFLDAGFSEYKITANFGLRSLIEIYP*

NbL09g09570.1 patellin−3−like [MW = 63.9 kDa]

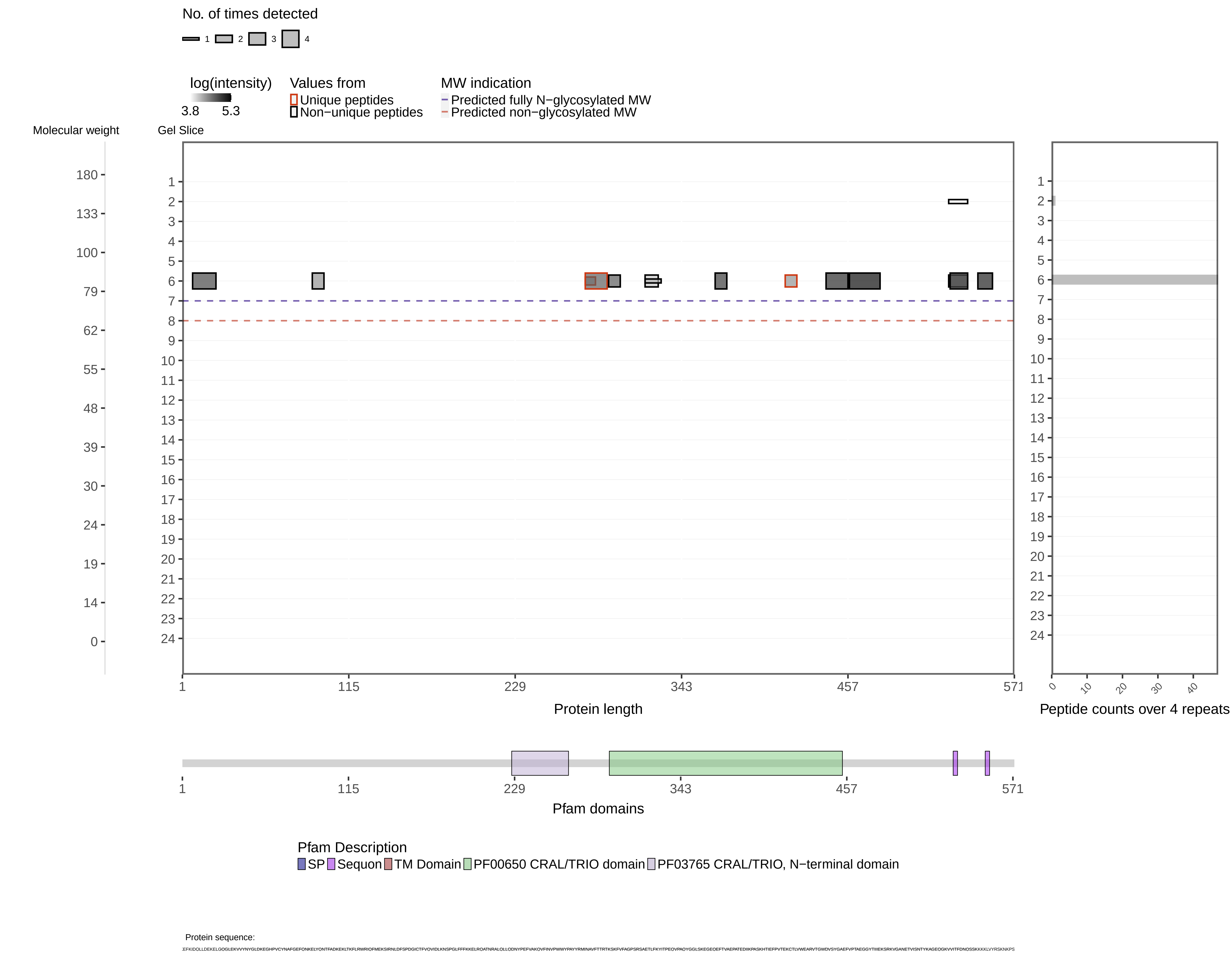

### NbL09g09440.1 uncharacterized_LOC104248794 [MW = 13.72 kDa]

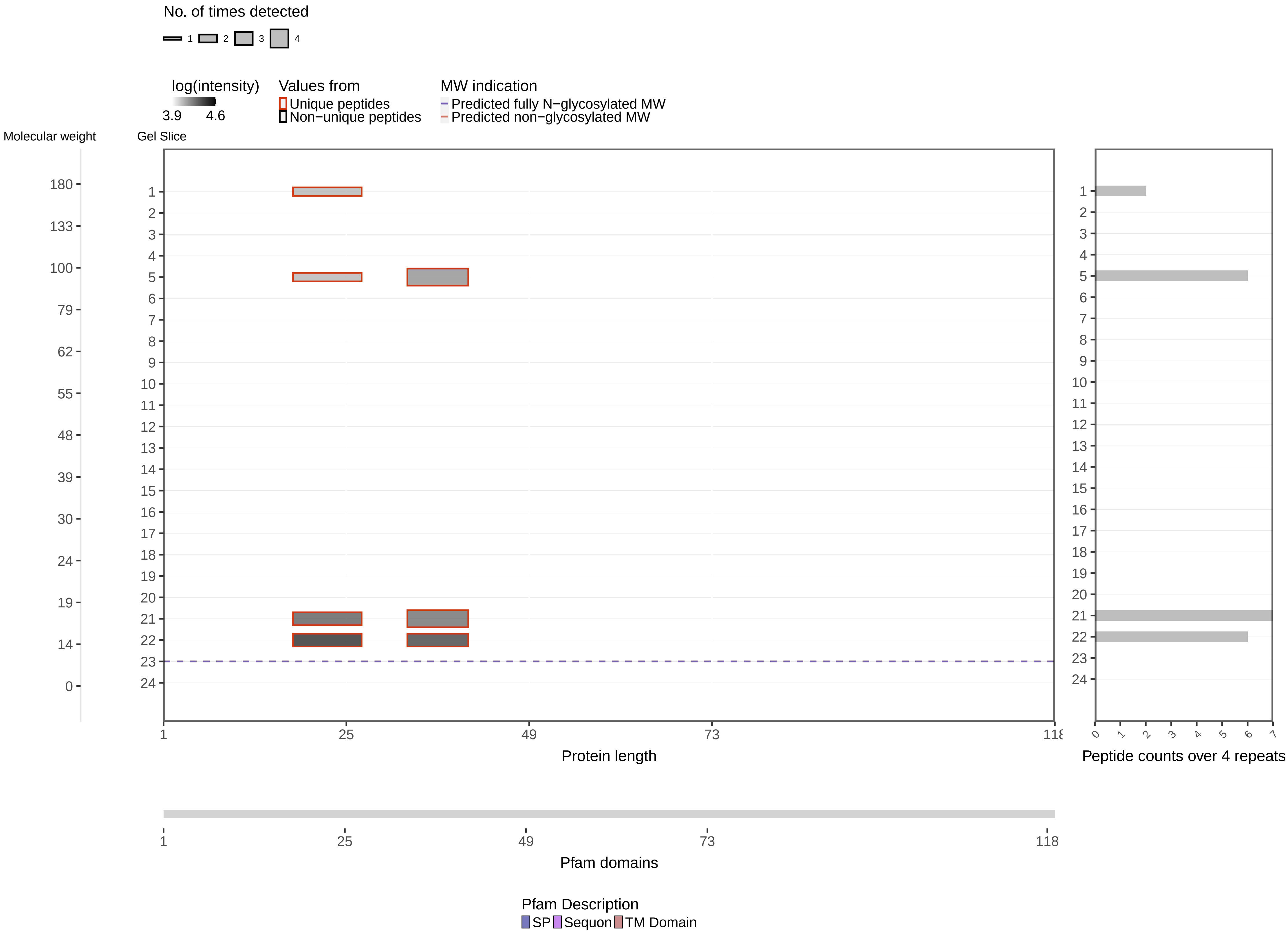


Protein sequence:

REAAWKAELSRREAEIARQEARLKMERENLEKEKSVLMGTASNQDNQDGALEITVSGEKYRCLRFAKAKK*

### NbL09g09310.1 50S_ribosomal_protein_L21_chloroplastic−like [MW = 23.46 kDa]

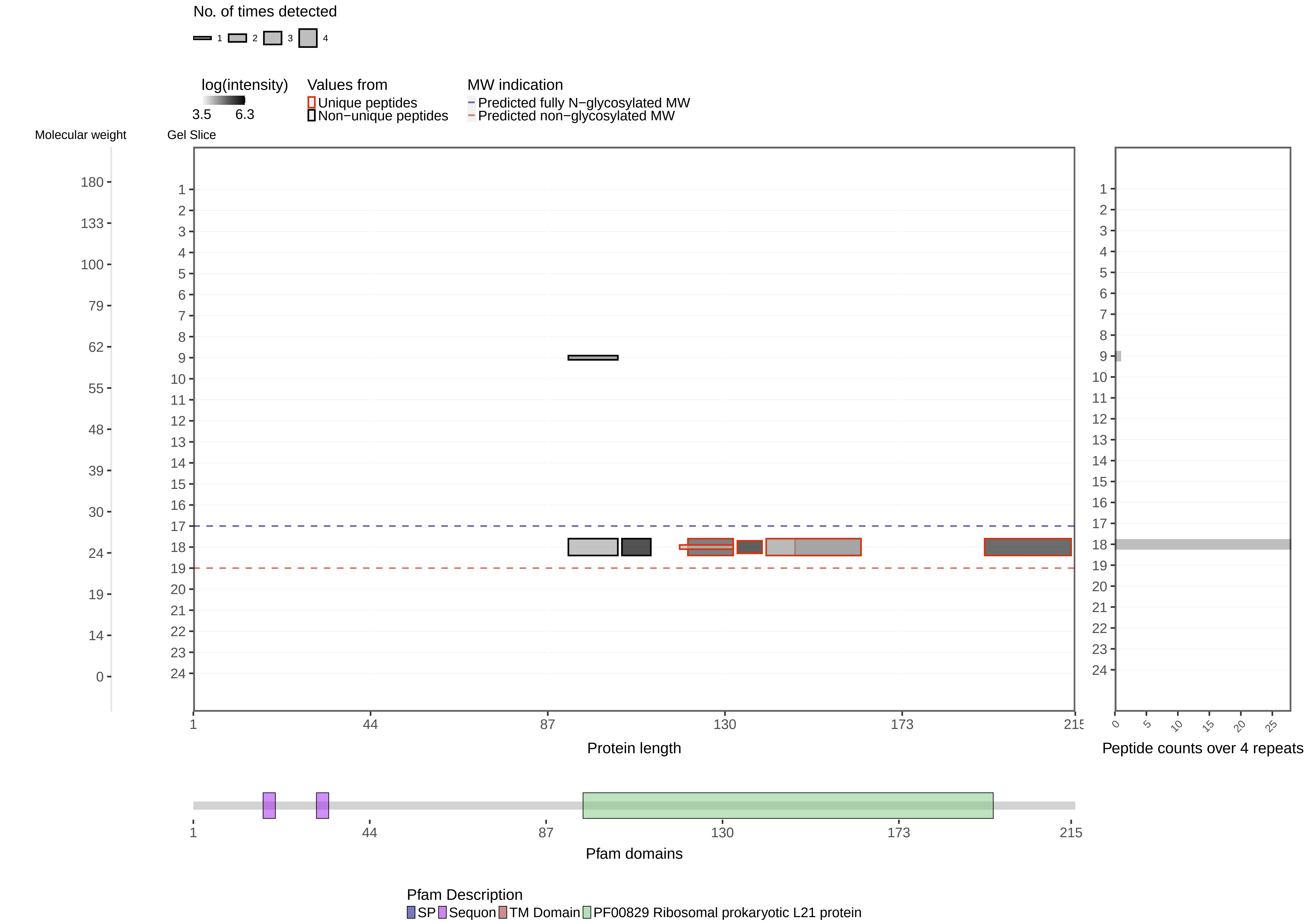


Protein sequence:

EVFAVVMIGSRQYIVFPGRYIYTQRLKGANVNDKIILNKVLLVGTKTSTWVGKPVVQNAAVHAVVEDQLKDKKVVVFKYKKKKNYRRNIGHRQPITRIRIMGITGYQDSPAVTLADLEAAK*

NbL09g09120.1 60S_ribosomal_protein_L18a−like [MW = 23.71 kDa]

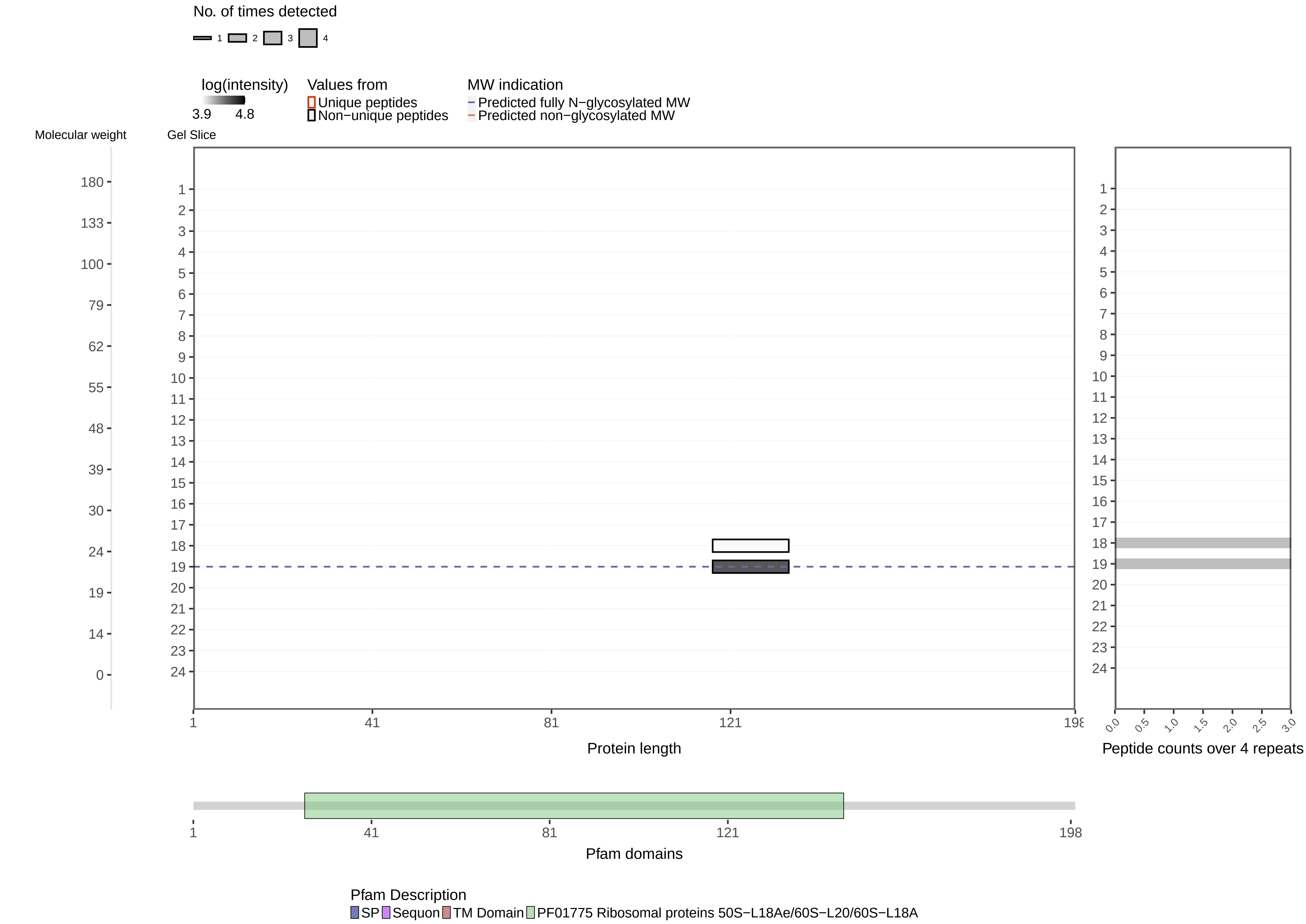


Protein sequence:

EIFEKNPTKIKNYGIWLRYQSRTGYHNMYKEFRDTTLNGAVEQMYTEMASRHRVRHHCIQIIKTATIPAKLCKRESTKQFHDSKIKFPLVFKKVRPPSRKLKTTYKATRPNLFM*

### NbL09g09040.1 T−complex_protein_1_subunit_eta [MW = 60.35 kDa]

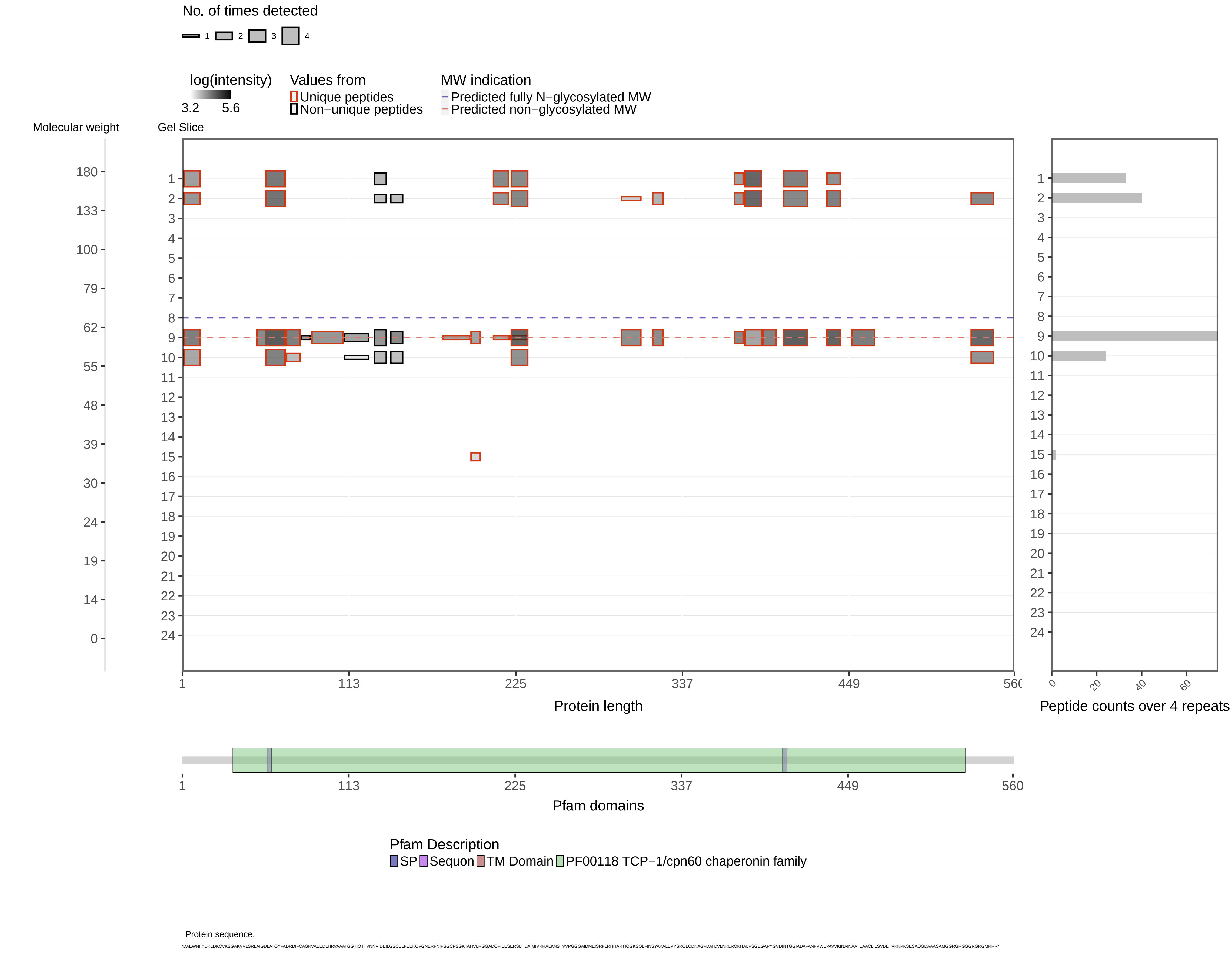

NbL09g08960.1 neurofilament_heavy_polypeptide [MW = 100.52 kDa]

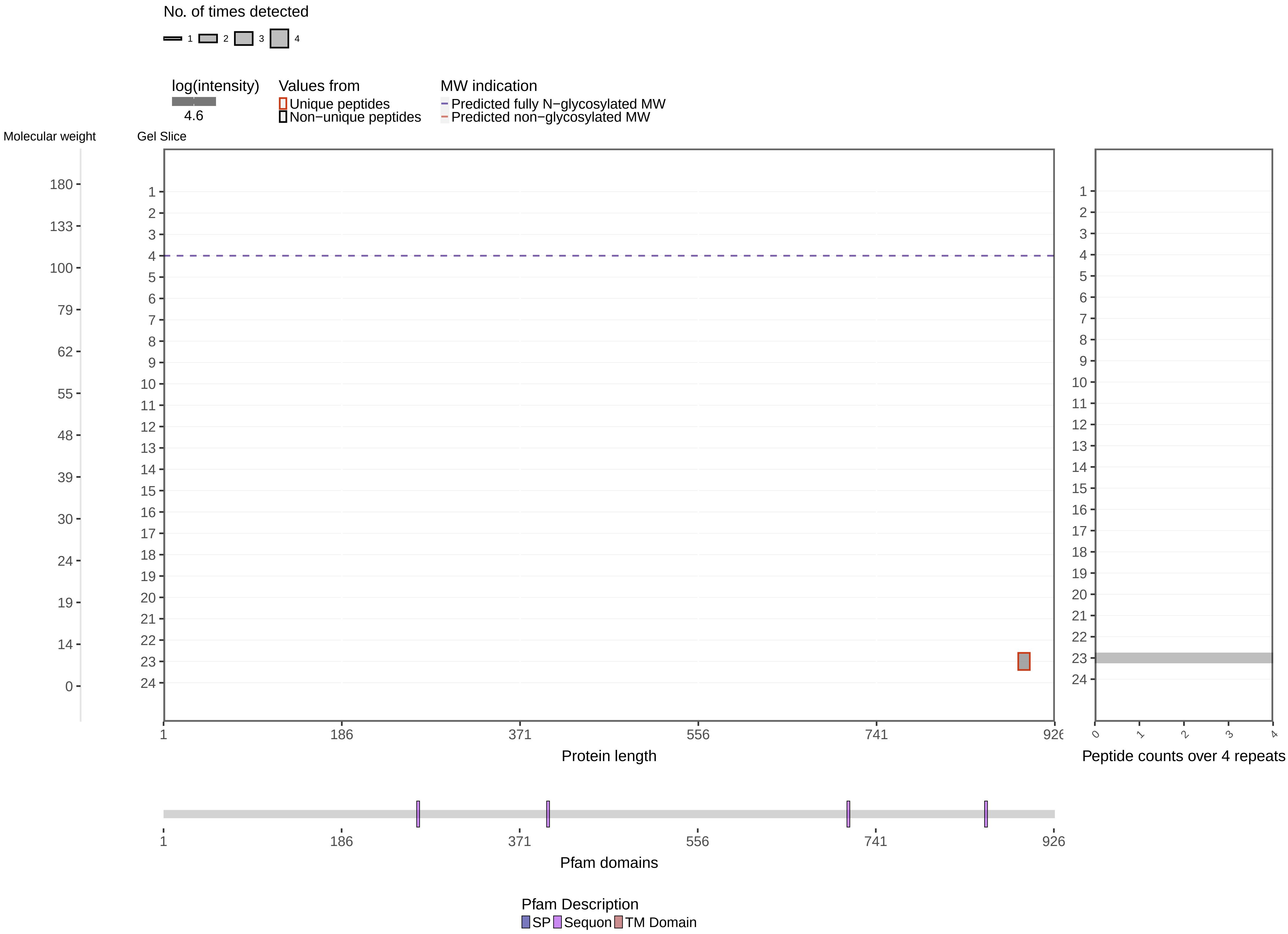


Protein sequence:

DSAPKESEGEAVNVAPLYTDLSLPDESAPKKGGRQKEDILNQEECVSEEEPEATSDLEVKQVRRPSKKAPAEPSHKENEGVSTSDVEAKKQKKSGKKIDTKNKNQVGPSVRNKEDSKKRGRGKASPETAPPQESPDDSAKHEFNEEDIPKTSAKKKPLSGKGRERVPSPVSLDNTAKHENNEEATPRTSAKRKQRSSGKDKVSGTVQHGESLVGKKIKVWWPLDKEFYEGVVEKFDPAKKKYRVAYTDGDVEIINLTEERWKLVEDDTMSEGEQIASADAASERQKKKK

NbL09g08800.1 14_kDa_proline−rich_protein_DC2.15−like [MW − SP = 10.71 kDa]

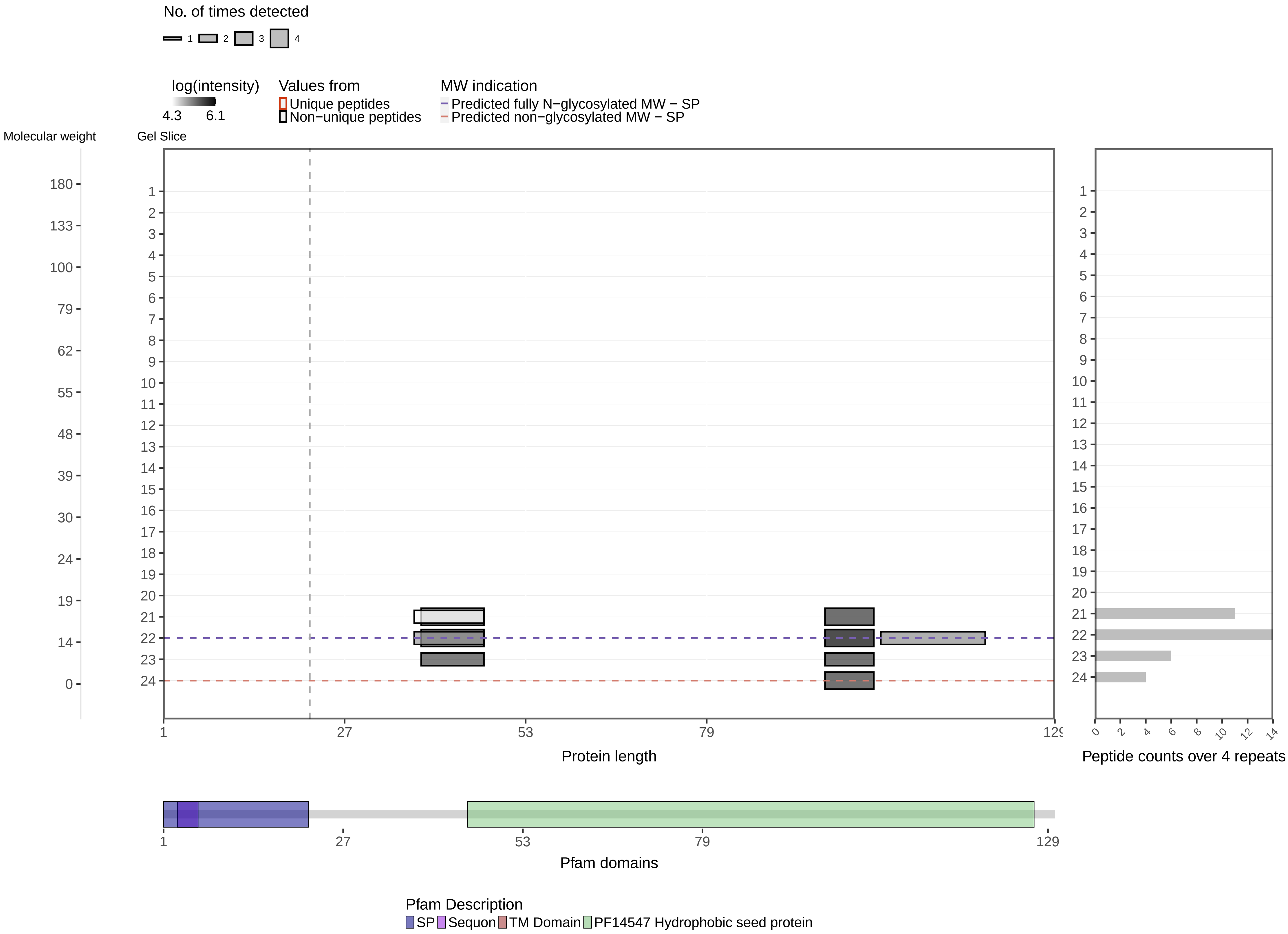


Protein sequence:

LGVCADVLGLVNVAVGSOVTSPCCSLLOGLADLEVAACLCTAIKANVLGIVKLDIPVALSALVSACAKTIPTGFKCN*

NbL09g08710.1 DEAD−box_ATP−dependent_RNA_helicase_56−like [MW = 48.42 kDa]

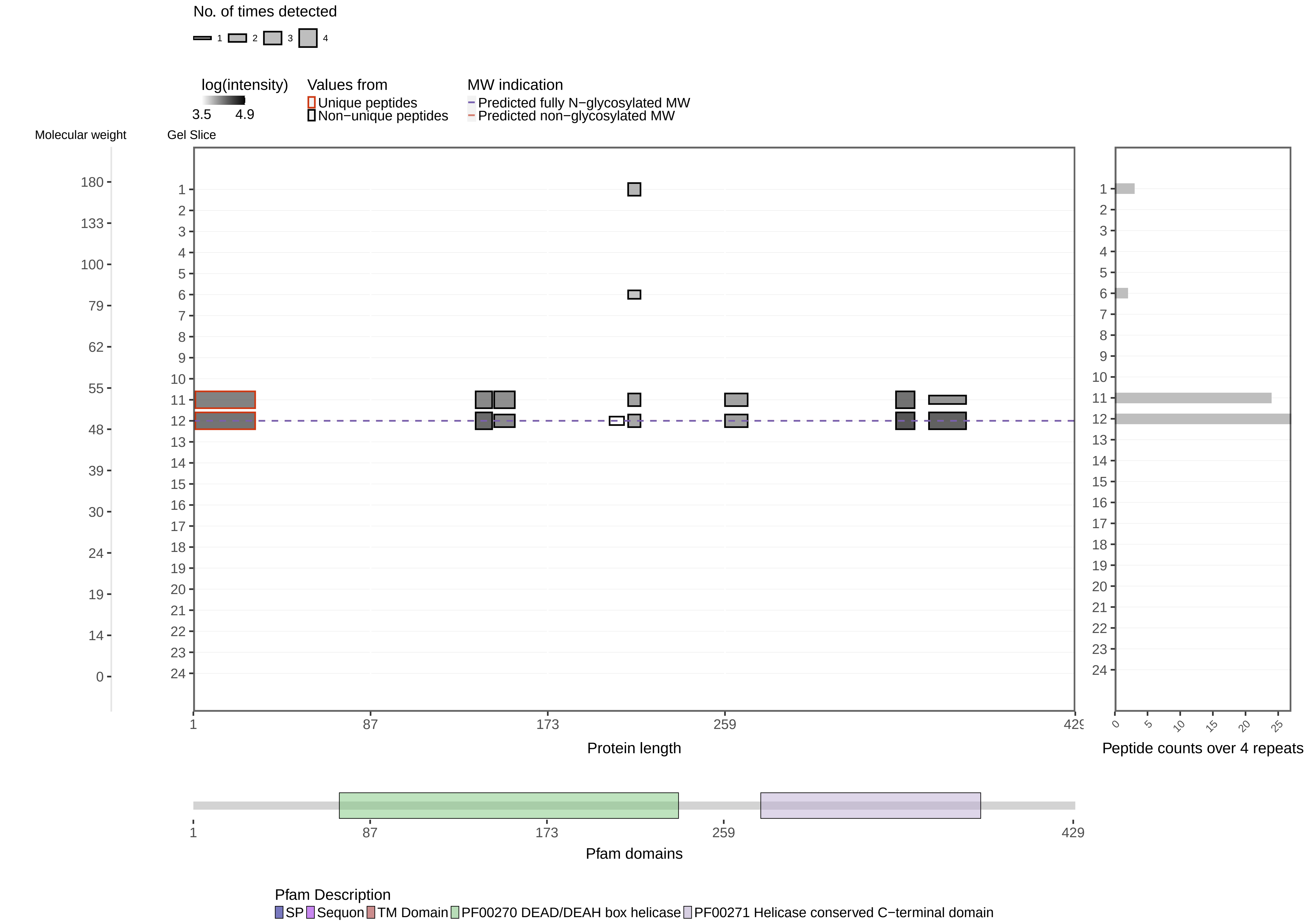


Protein sequence:

DKMLESLDMRRDVQEIFKMTPHDKQVMMFSATLSKEIRPVCKKFMQDPMEIYVDDEAKLTLHGLVQHYIKLSETEKNRKLNDLLDALDFNQVVIFVKSVSRAAQLDKLLVECNFPSICIHSGMTQEERLTRYKGFKEGHKRILVATDLVGRGIDIERVNIVINYDMPDSADTYLHRVGRAGRFGTKGLAITFVSSASDSDVLNQVQERFEVDIKELPEQIDTSTYMPS*

NbL09g08640.1 uncharacterized_protein_At2g34160−like [MW = 17.49 kDa]

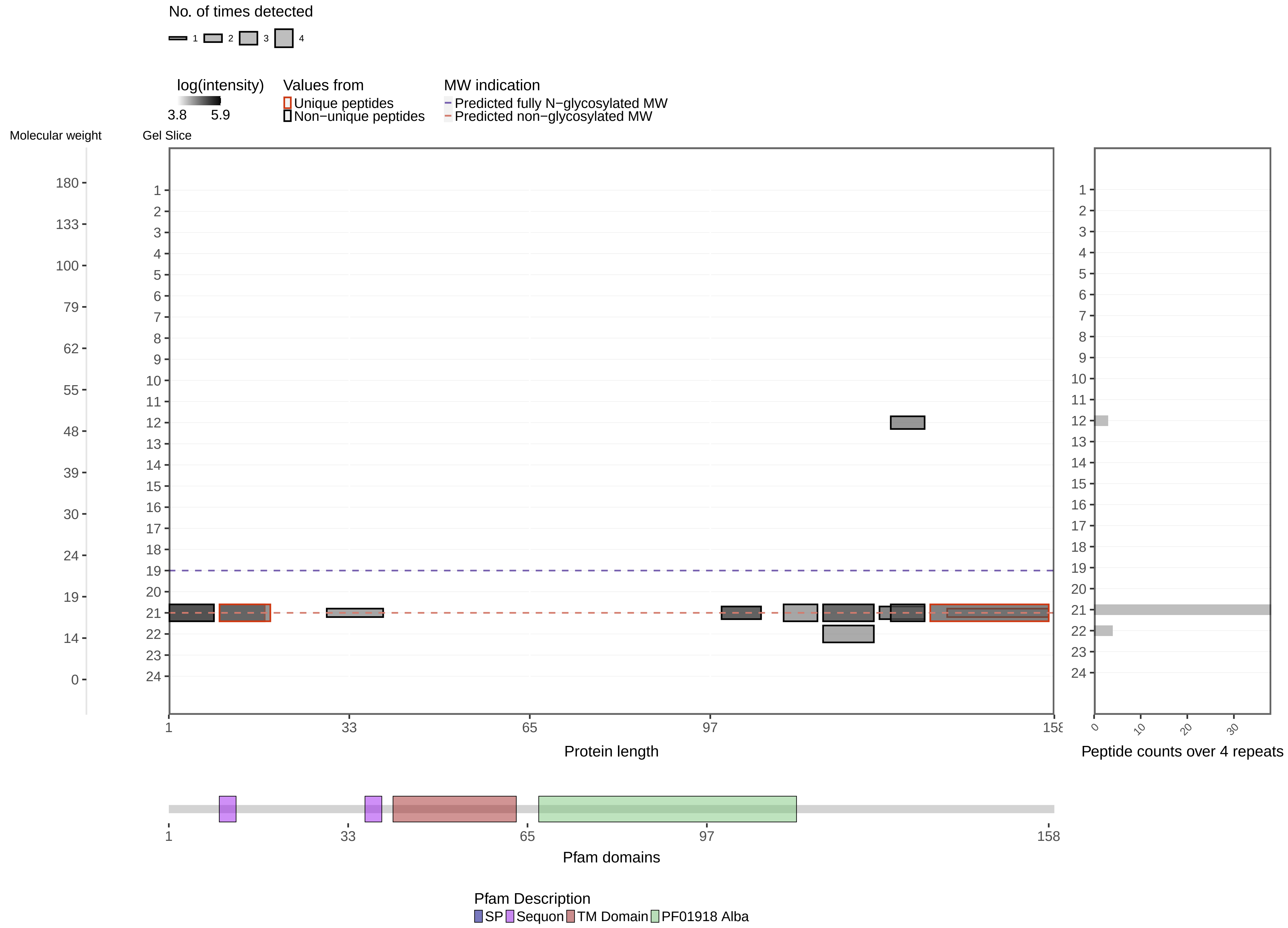


Protein sequence:

.IMQRYMQQYNEVELSALGMAISTVVTVAEILKNNGFAVEKKIRTLTIDMRDEPGARPIPKAKIEIVLGKTEKFDELMAAEAEQNGDSEEQT*

### NbL09g08500.1 protein_IWS1_homolog_1 [MW = 58.65 kDa]

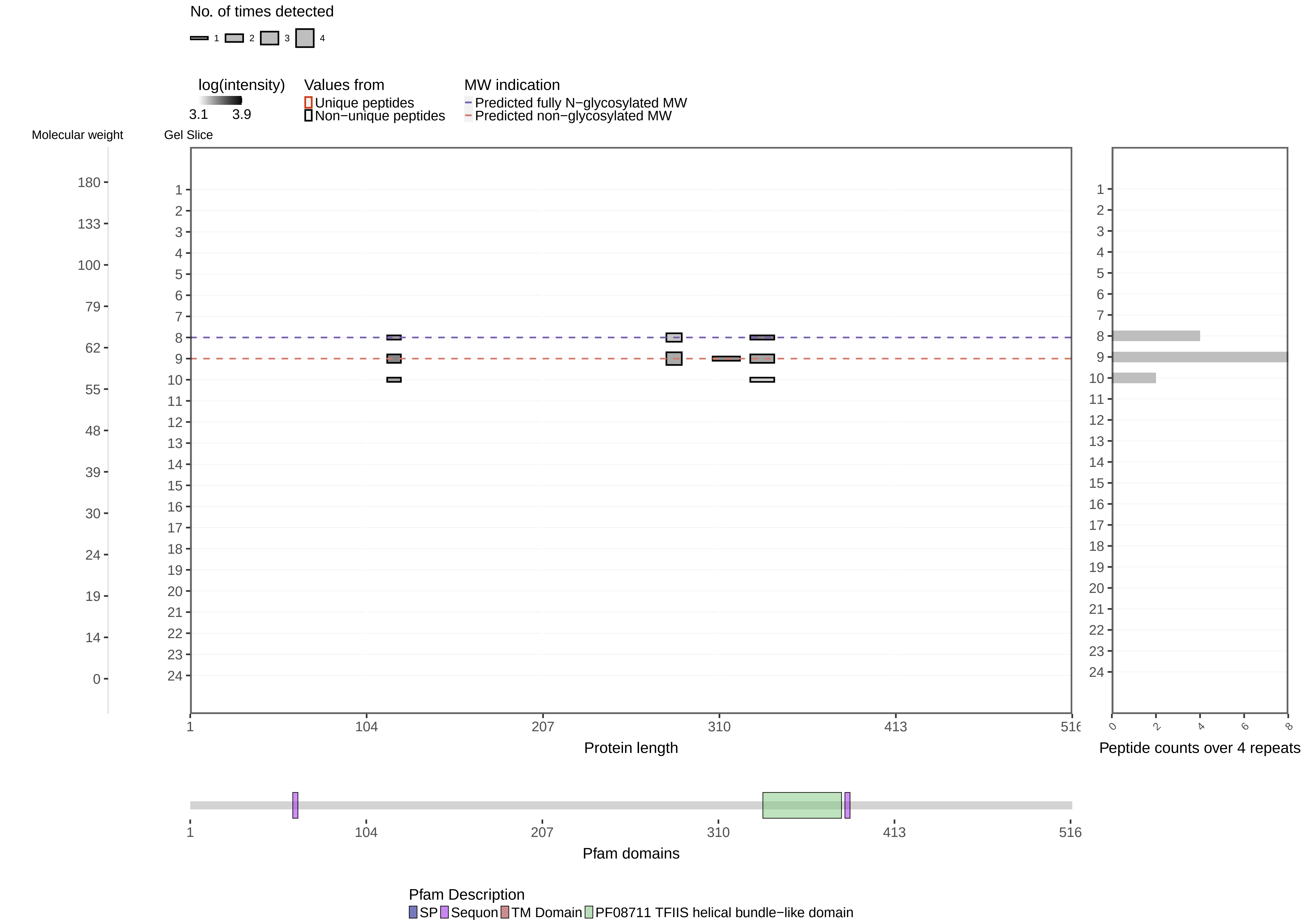


Protein sequence:

AEIALLVENVMAELEVVAEEDAELNRQSKPAINKLKKLPLLIDVLSKKQLQQEFLDHGVLTLLRTWLEPLPDGSLPNINIRAAILKILTDYPIDLEQYDRREQLKKSGLGKVIMFLSKSDEETTANRKLAKDLVDKWSRPIFNKSTRFEDMRNFEDERAAQFRRPSMKKPMNKASGMESRDDDLDLAGFSQGQKSGPSSSRQLTSRPEAMSMDFLVRPQSKIDPEEVRARAKAMVQDQRRMKMNKKLQQLKAPKKKKLQATKLSVEGRGMVKYL*

### NbL09g08360.1 uncharacterized_LOC109238939 [MW − SP = 6.01 kDa]

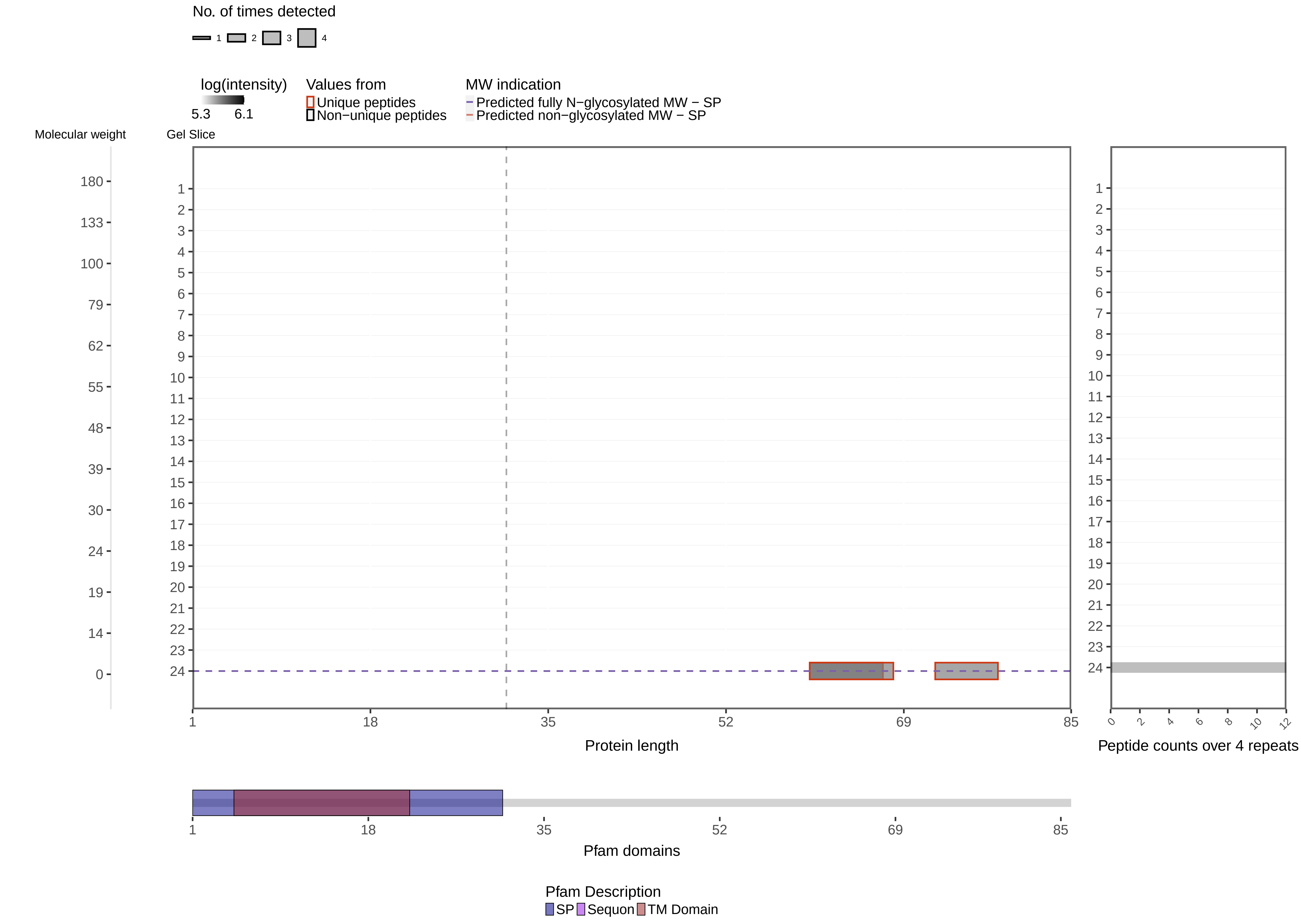


Protein sequence:

AARDVSRTIAELKSKLSDPTLMGGCGGYCRQASDCFDRRCPRCYYDVIKLTYGCR*

### NbL09g08340.1 abscisic_acid_receptor_PYR1−like [MW = 28.56 kDa]

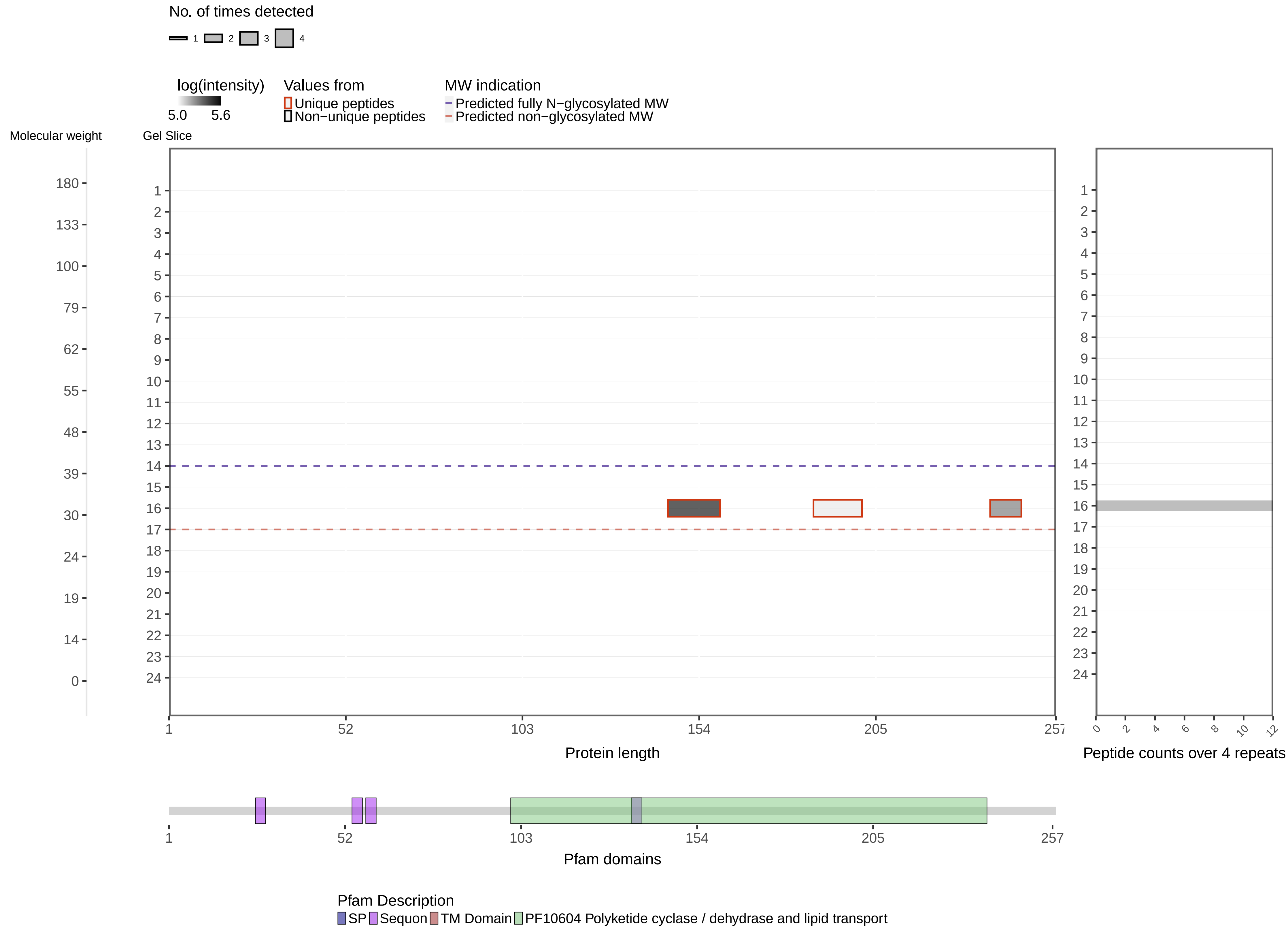


Protein sequence:

RFDKPQIYKHFIKSCSVGENFSIVVGATRYVNVISGLPADTSTERLDILDDEKYVTGFSIIGGEHRLRNYRSVTTVHGFYQQLDGKVSTVVLESYVVDVPEGNTEEDSRLFADTVVKLNLQKLGSVAEAIVRGGKGVVSEN*

### NbL09g08310.1 glycine_cleavage_system_H_protein_mitochondrial−like [MW = 17.43 kDa]

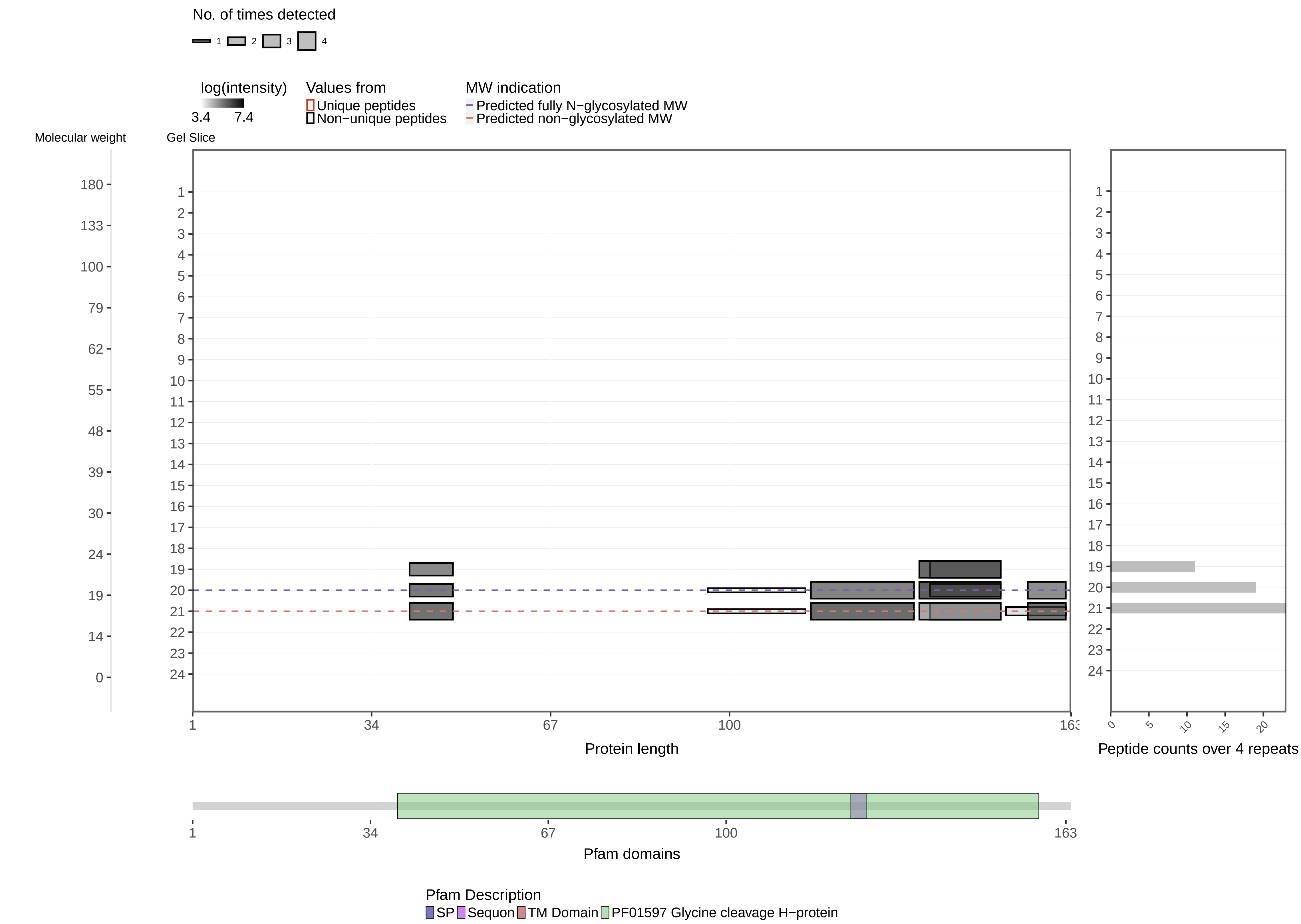


Protein sequence:

EVVFVDLPETGGSVSQGSSFGAVESVKATSDINCPISGEIVEVNTKLTETPGLVNSSPYEDGWMIKVKPSSPSELESLMGSKEYTKFCEEEDSH*

NbL09g07990.1 U3_small_nucleolar_RNA−associated_protein_25 [MW = 39.2 kDa]

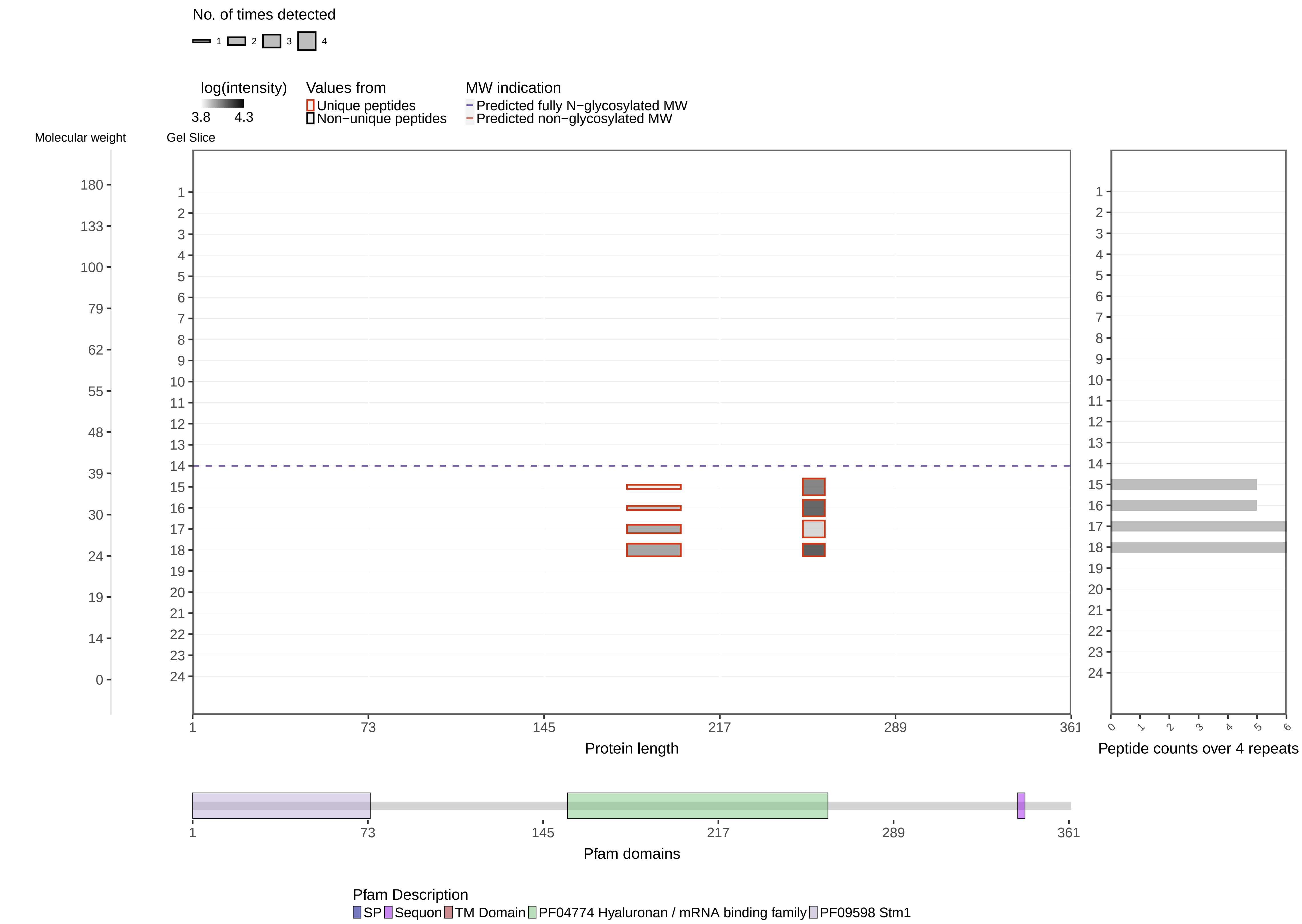


Protein sequence:

GRGNEFKREGAGRGNWGTPADEFAPEKEEPVNEGEKIVDTEKQAGQADAEDTSKDSPAAEPQEKEPEEKEMTLEEYEKLLEEKRKGLLALMAEERKVNLDKEFESMQLLSNKKSDDEVFIKLGSEKDKRKEAVEKAKKTKSINEFLLPADGESYYRQGGRGRGGRGRGRDGFVGNNSVSAPSIEDVRQFPSLGAK*

NbL09g07960.1 ubiquitin_carboxyl−terminal_hydrolase_3−like [MW = 25.84 kDa]

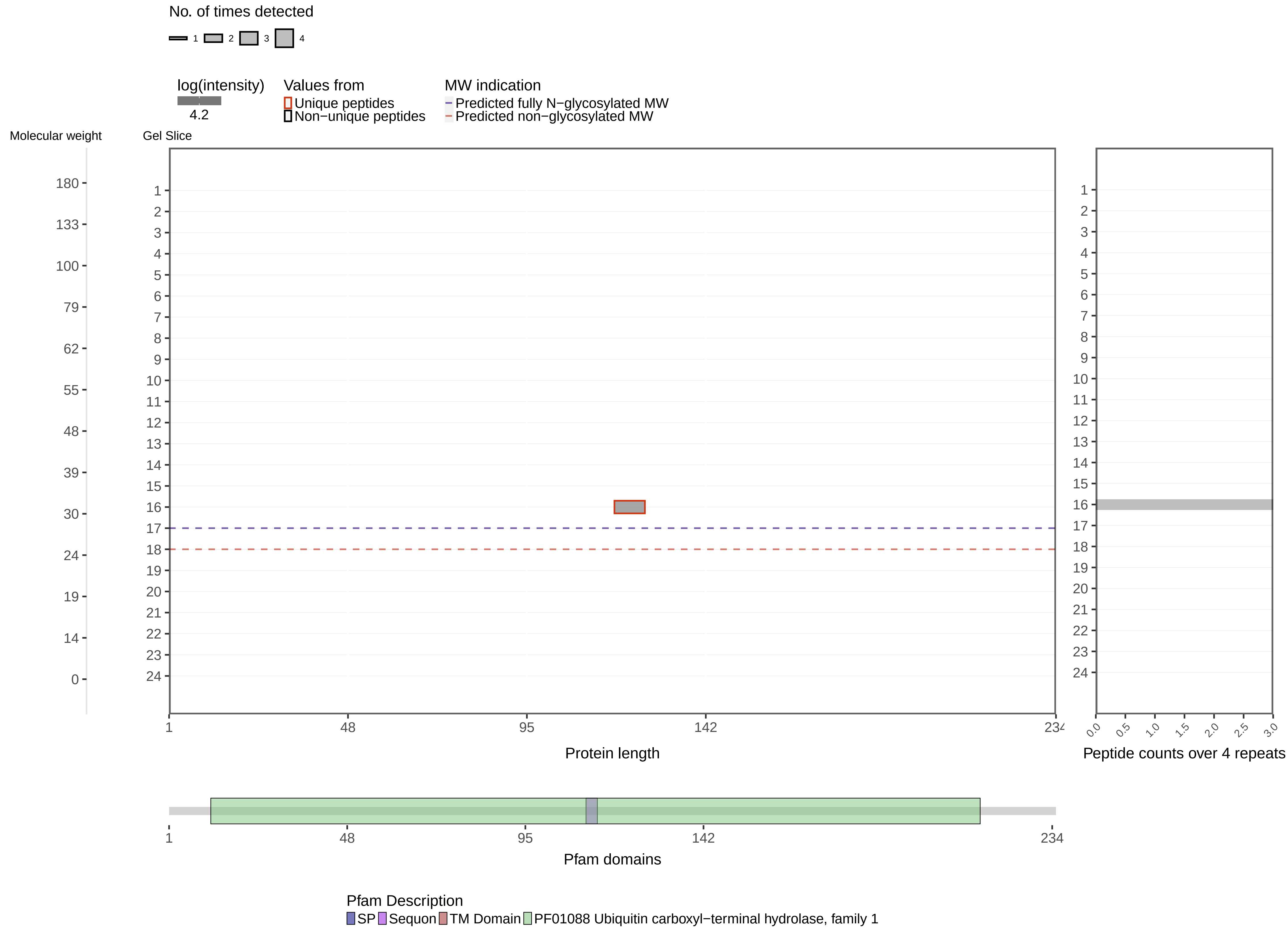


Protein sequence:

[illegible]TIGLLHAIGNITSQIKLTEESFLDRFFKSTASMDPMQRALFLENDREMEVAHSVAATAGETEASEDVNTHFICFTCADGQLYELDGRRSAPISHGASSPNNLLKDSAKVIQKMIEKNPDSINFNVIAISKRD*

NbL09g07880.1 formyltetrahydrofolate_deformylase_1_mitochondrial−like [MW = 36.4 kDa]

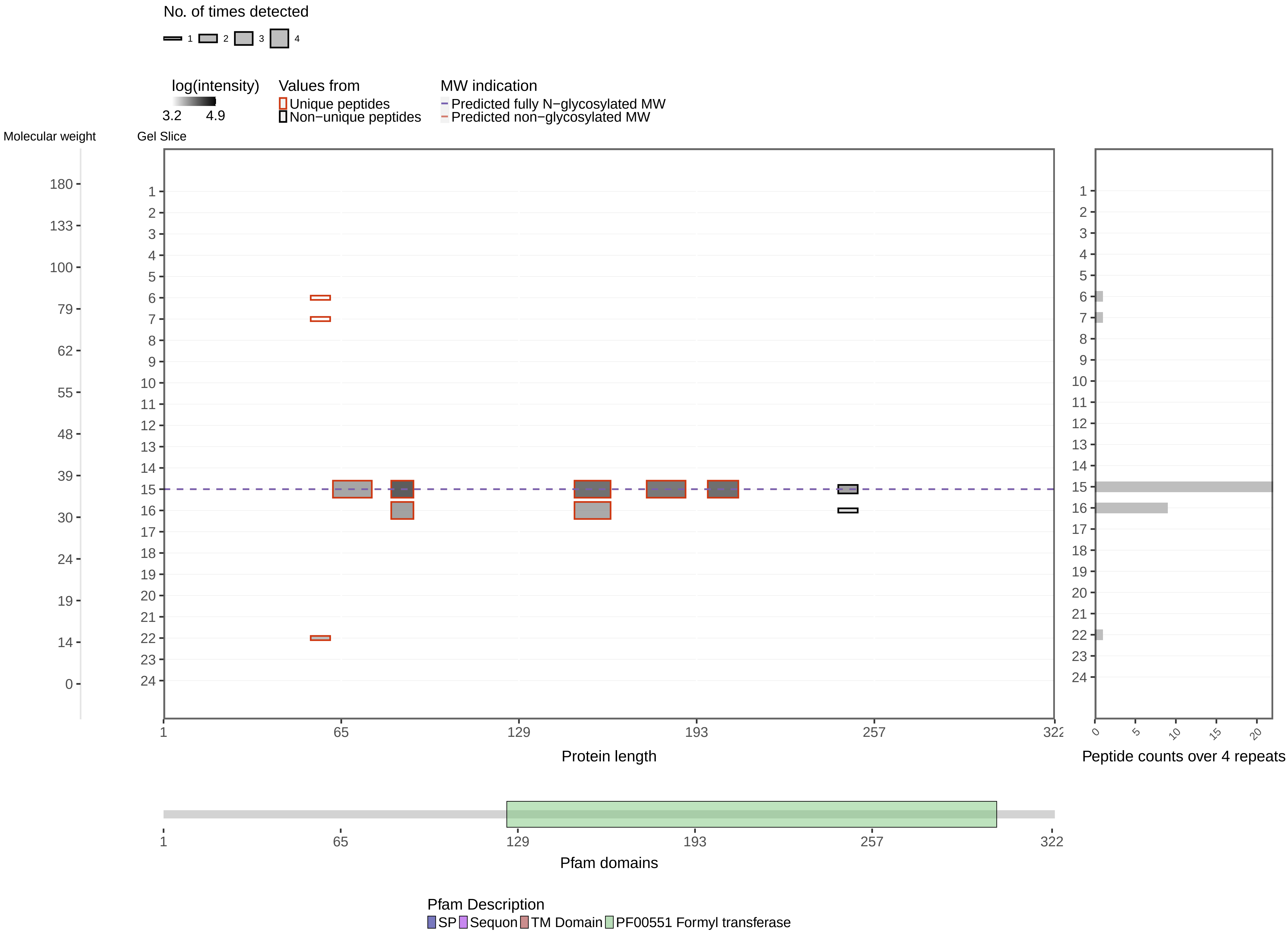


Protein sequence:

NGRLPVDITSVISNHDRGPDAHVIRLLERHGIPYYYLPTTMGTKREVEILKLVHDTDFLVLARYMQVFSAEFLESYGKDIINIHHGLLPSFKGGSPSKQAFEAGVKLIGATSHFVSEGLDAGPIIEQMVERVSHRDNLRSFIQKSEDLEKQCLAKAIKSYCELRVLPYYKDKTVVF*

### NbL09g07740.1 protein_disulfide−isomerase−like [MW − SP = 52.58 kDa]

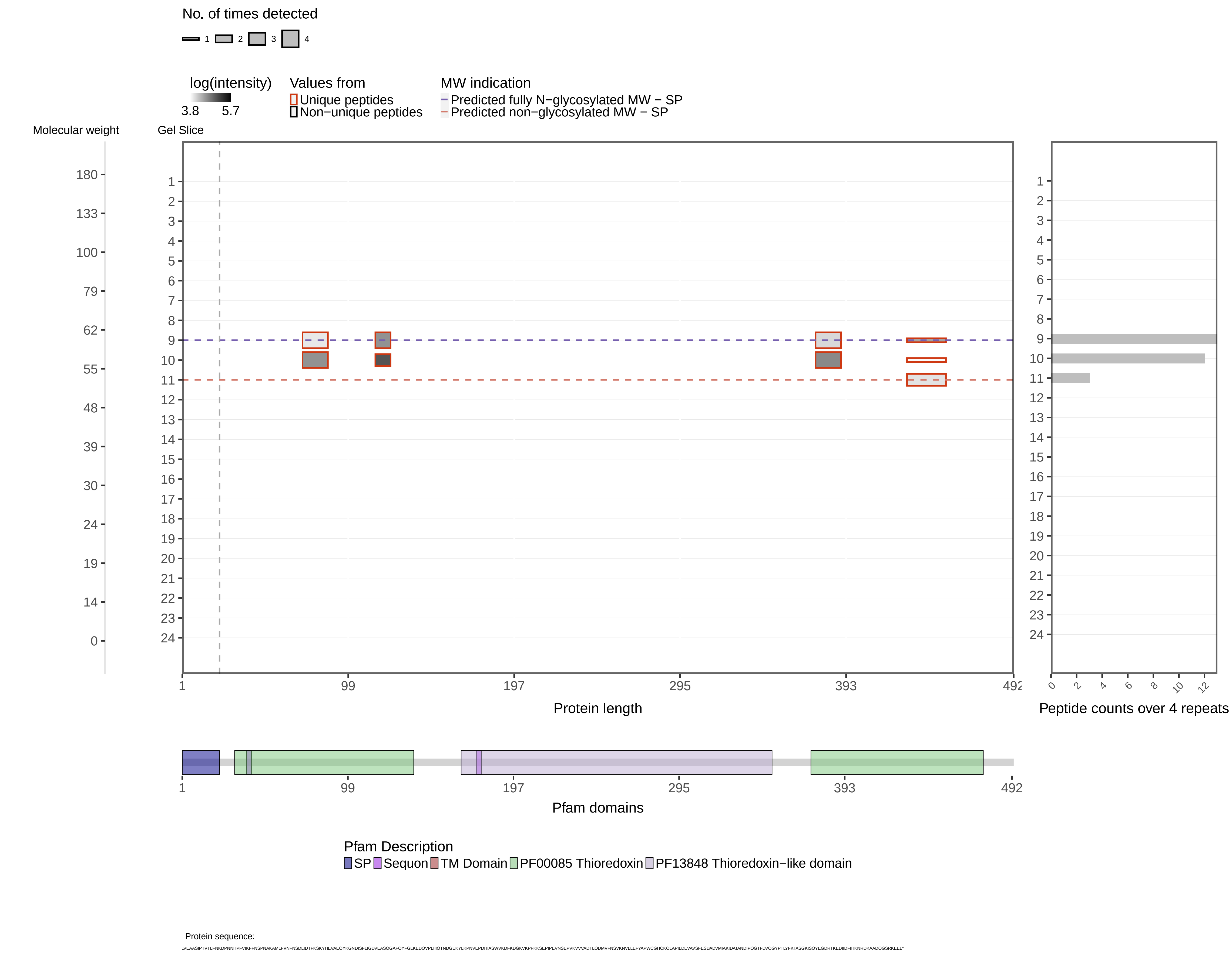

### NbL09g07690.1 photosystem_II_22_kDa_protein_chloroplastic [MW = 29.26 kDa]

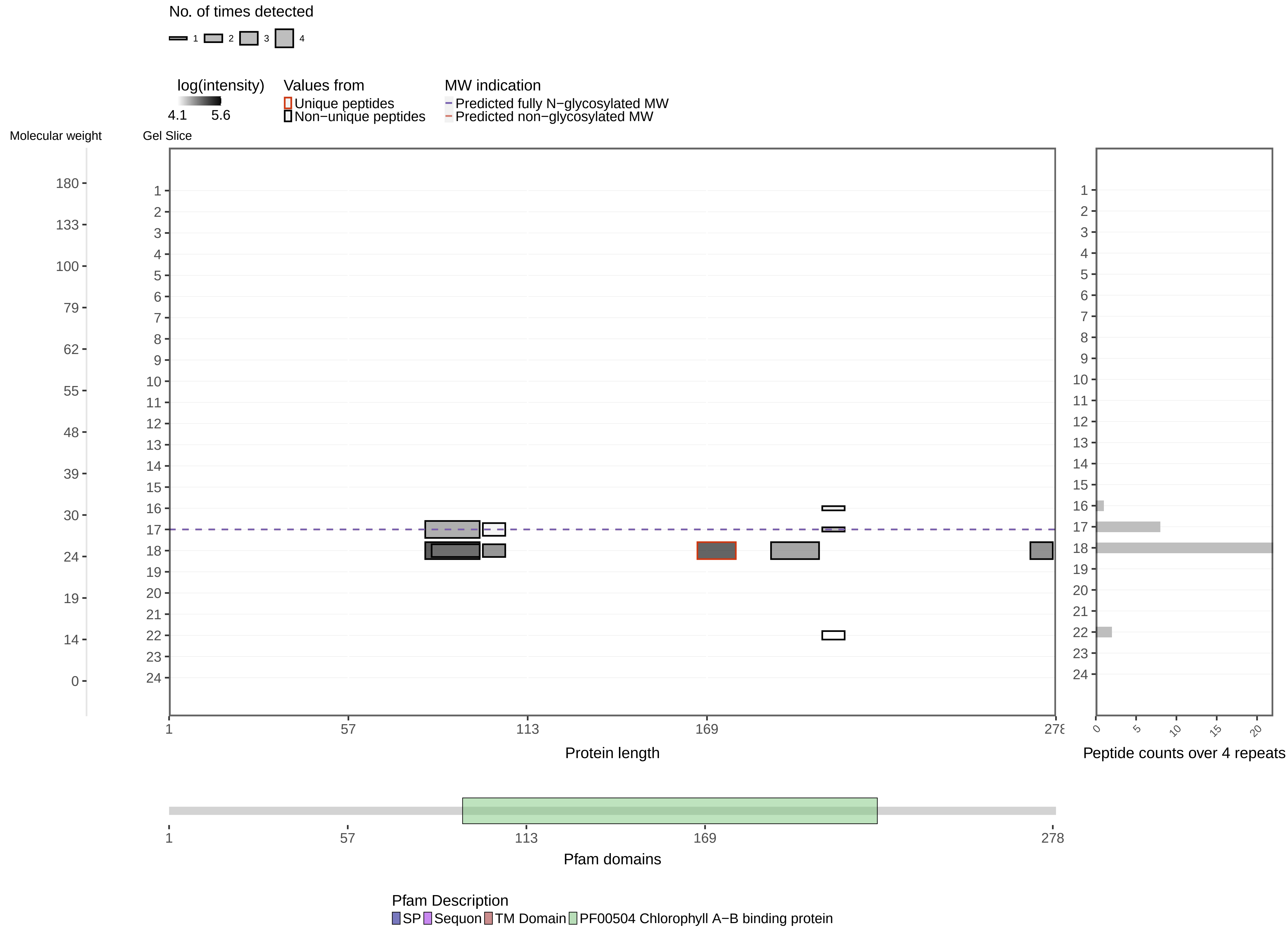


Protein sequence:

GKGILAQLNLETGIPIYEAEPLLLFFILFNLLGAIGALGDRGKFIDDPTPPTGLDKAVIPPGKGFKSALGLSEGGPLFGFTKANELFVGRLAQLGIAFSIIGEIITGKGALAQLNFETGVPINEIEPLLLFNIVFFFVAAINPGTGKFVTDEEEE*

### NbL09g07290.1 acetolactate_synthase_1_chloroplastic−like [MW = 71.78 kDa]

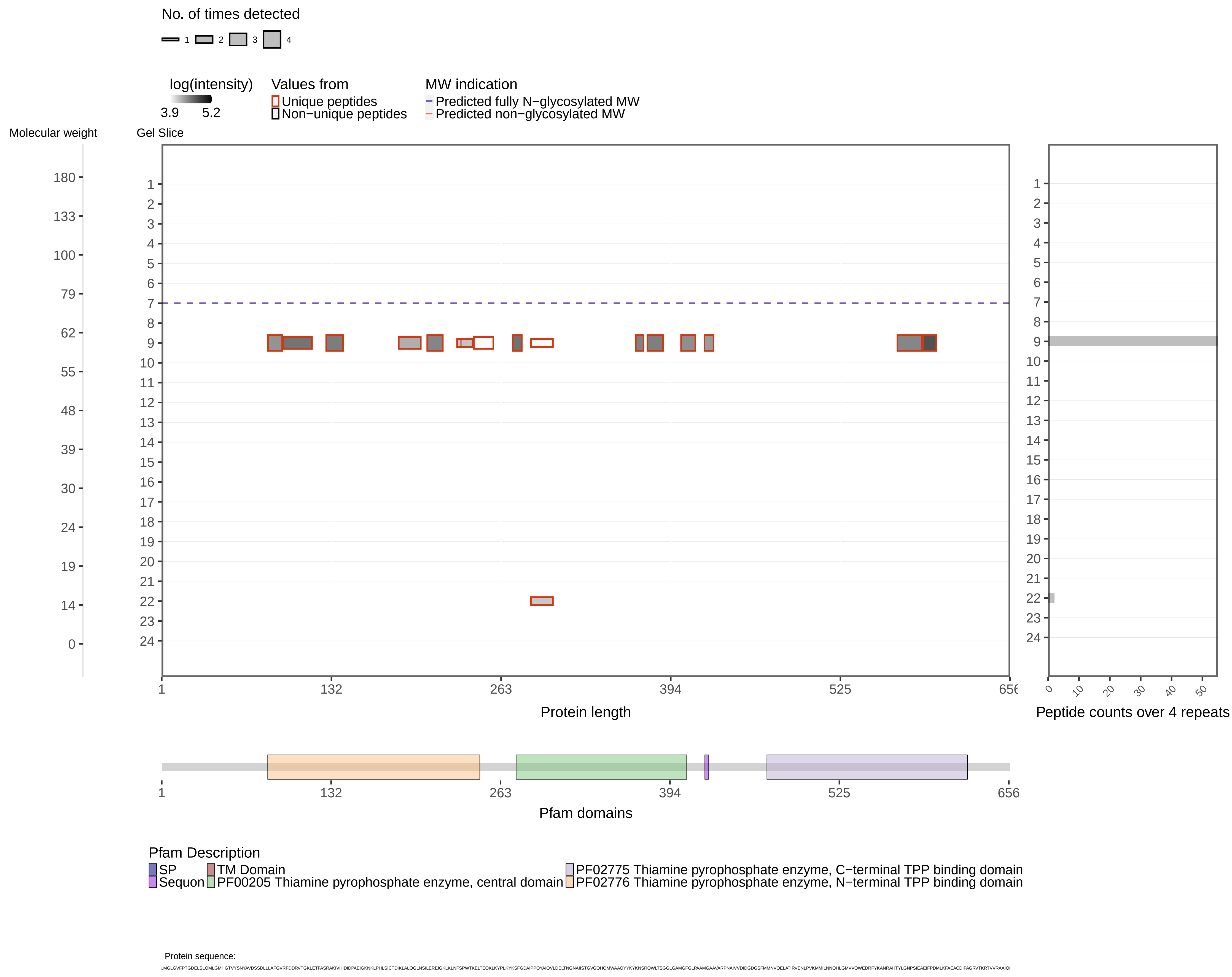

### NbL09g07260.1 40S_ribosomal_protein_S2−3−like [MW = 30.04 kDa]

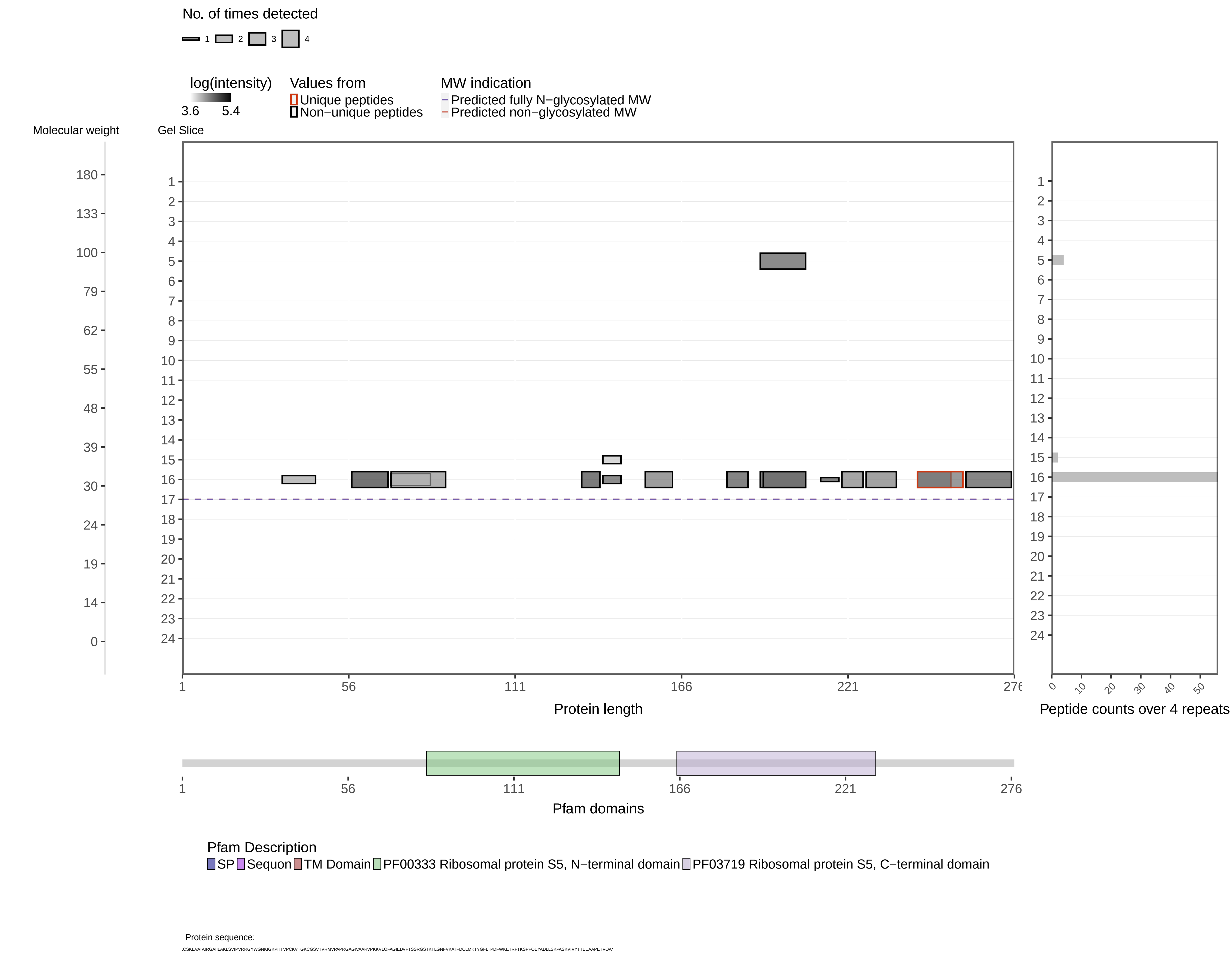

NbL09g06990.1 scarecrow−like_protein_14 [MW = 84.73 kDa]

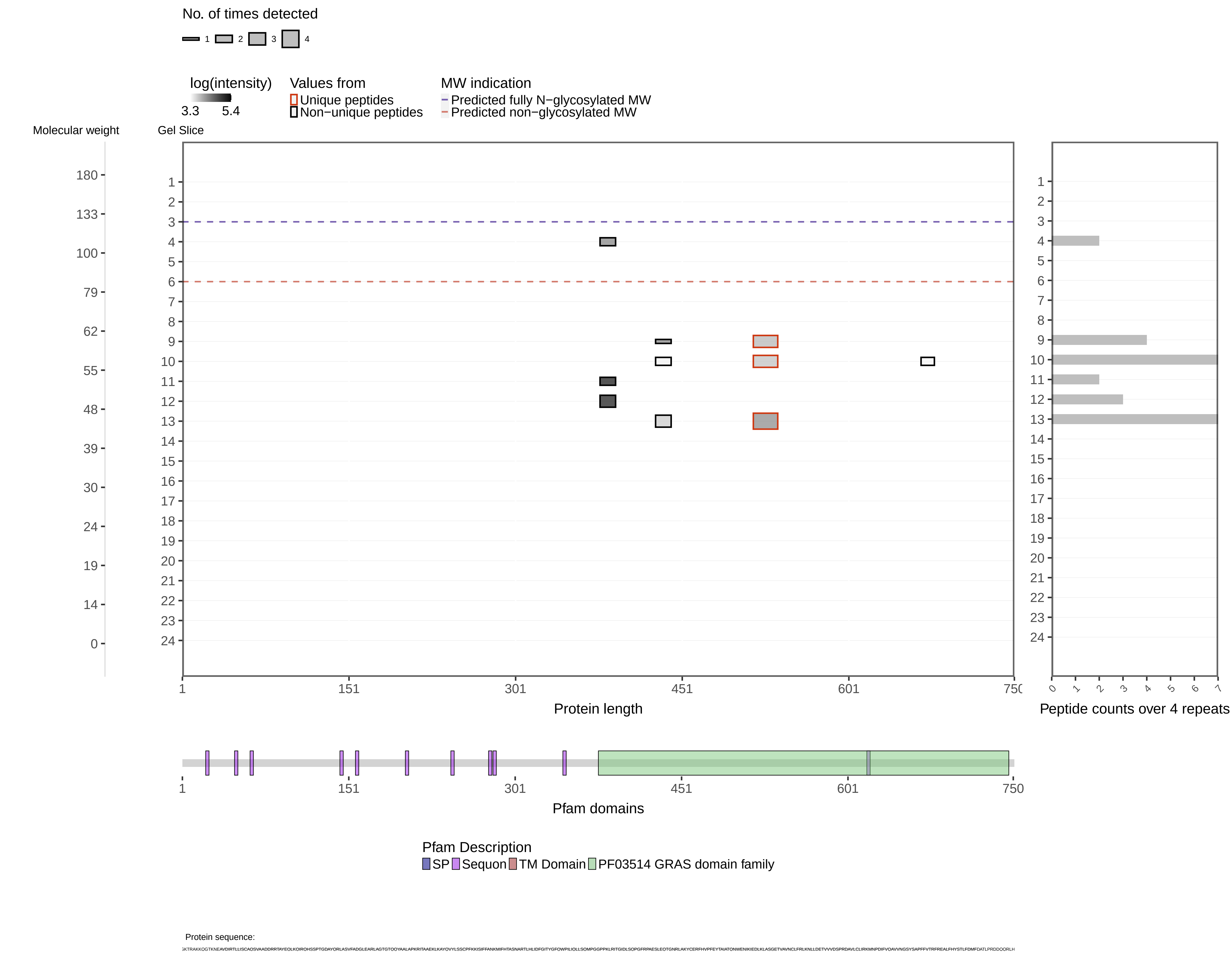

NbL09g06980.1 uncharacterized_protein_C24B11.05−like [MW = 33.12 kDa]

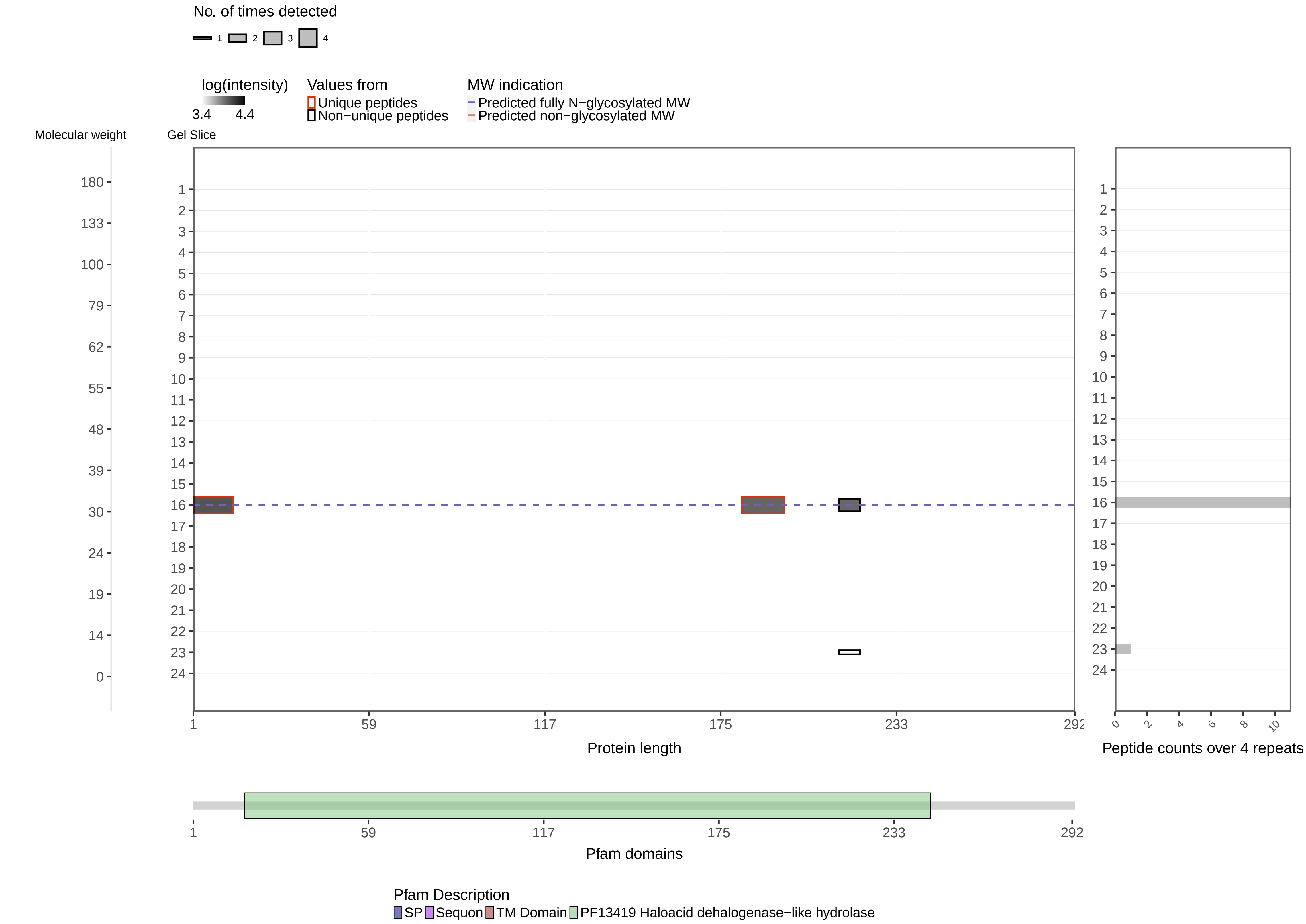


Protein sequence:

LRNLLHSLPFRKVIFSNANEAHVAKVLSRLGLEDCFDDIICFETVNPTRNSIVALDNINGESARTLVVCKPFEEAYERAFKIANINPQKTLFFDDSVRNLQTAKLMGLHTVWVGASHRTKGIDYALESIHNMKEALPELWEDVKPDVHISEKTAIEVRV*

NbL09g06800.1 phosphoserine_phosphatase_chloroplastic [MW = 35.31 kDa]

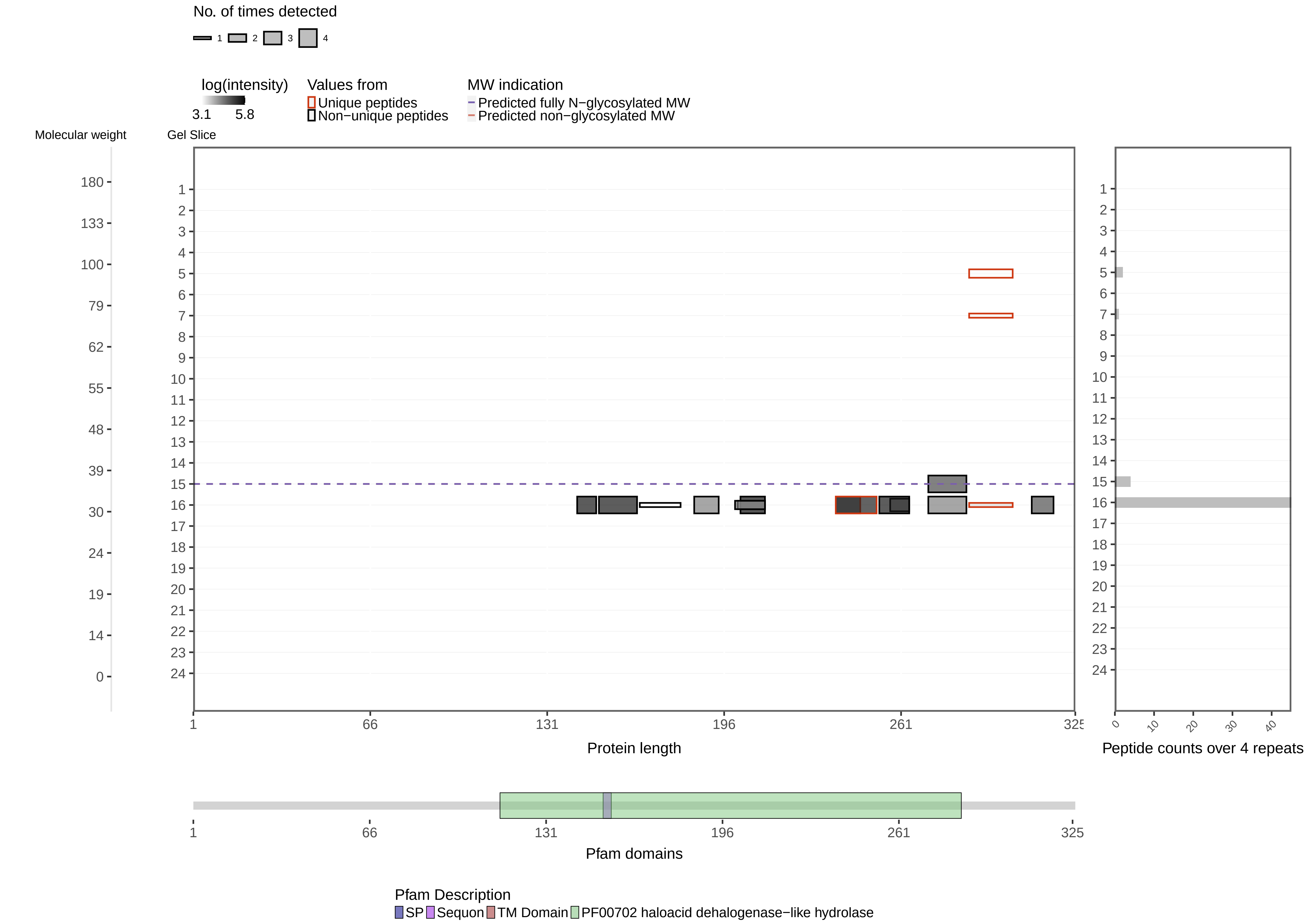


Protein sequence:

RAMNGSVPFEDALAARLSLINPSLSQLQDFLKRPPRLSPGIDLLVKKLKDKKKDVYLVSGGFRQMINPVASILGITLENIFANQMLFRSNGEFAGFDKNEPTSRSGGKPTAVQQIKKARGYKSVVMIGDGATDLEARLPGGADLFICYGGVQLREPVAAKADWLVFNFKDLINSLE*

### NbL09g06730.1 cytosolic_enolase_3 [MW = 51.71 kDa]

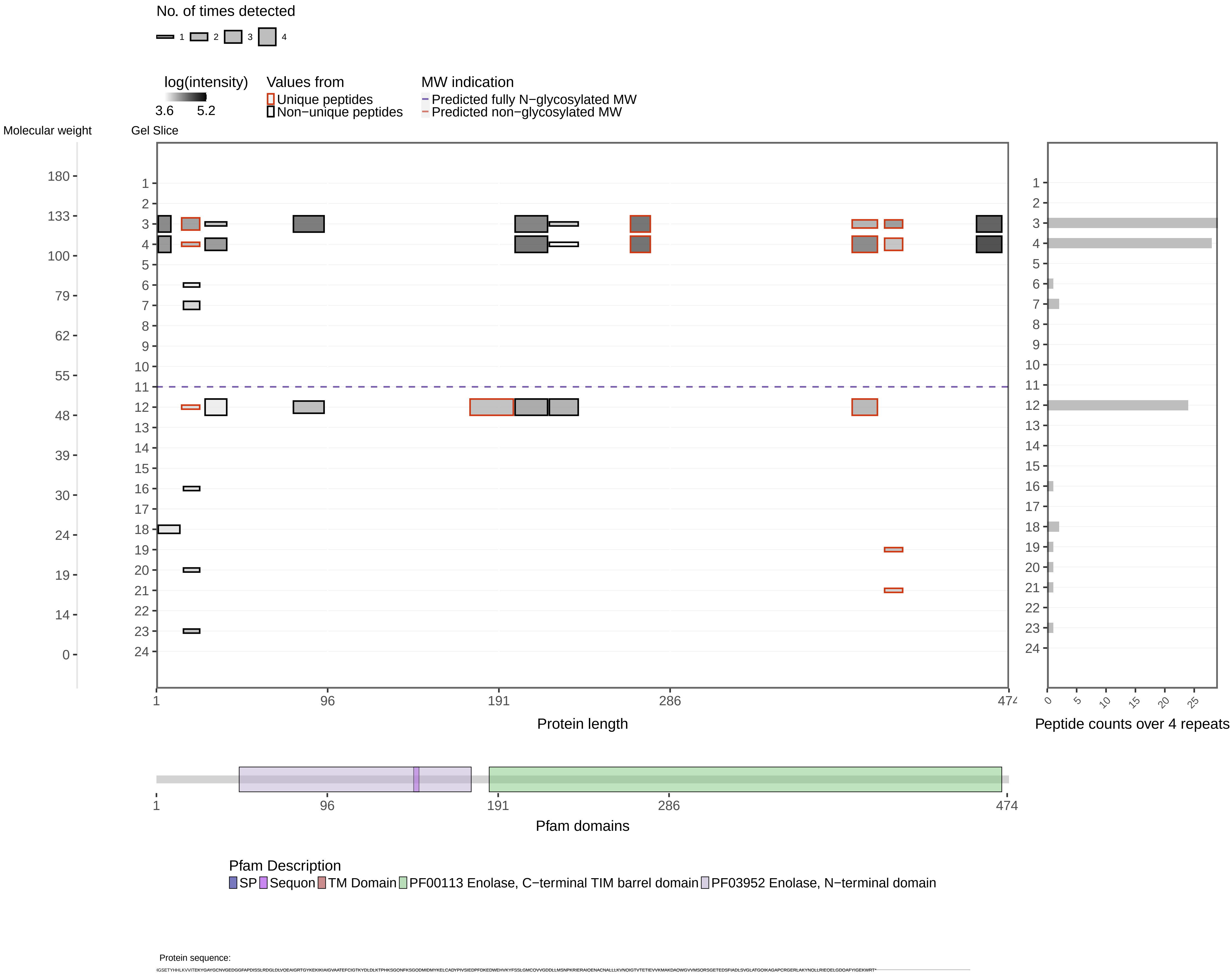

### NbL09g06720.1 proliferating_cell_nuclear_antigen [MW = 32.2 kDa]

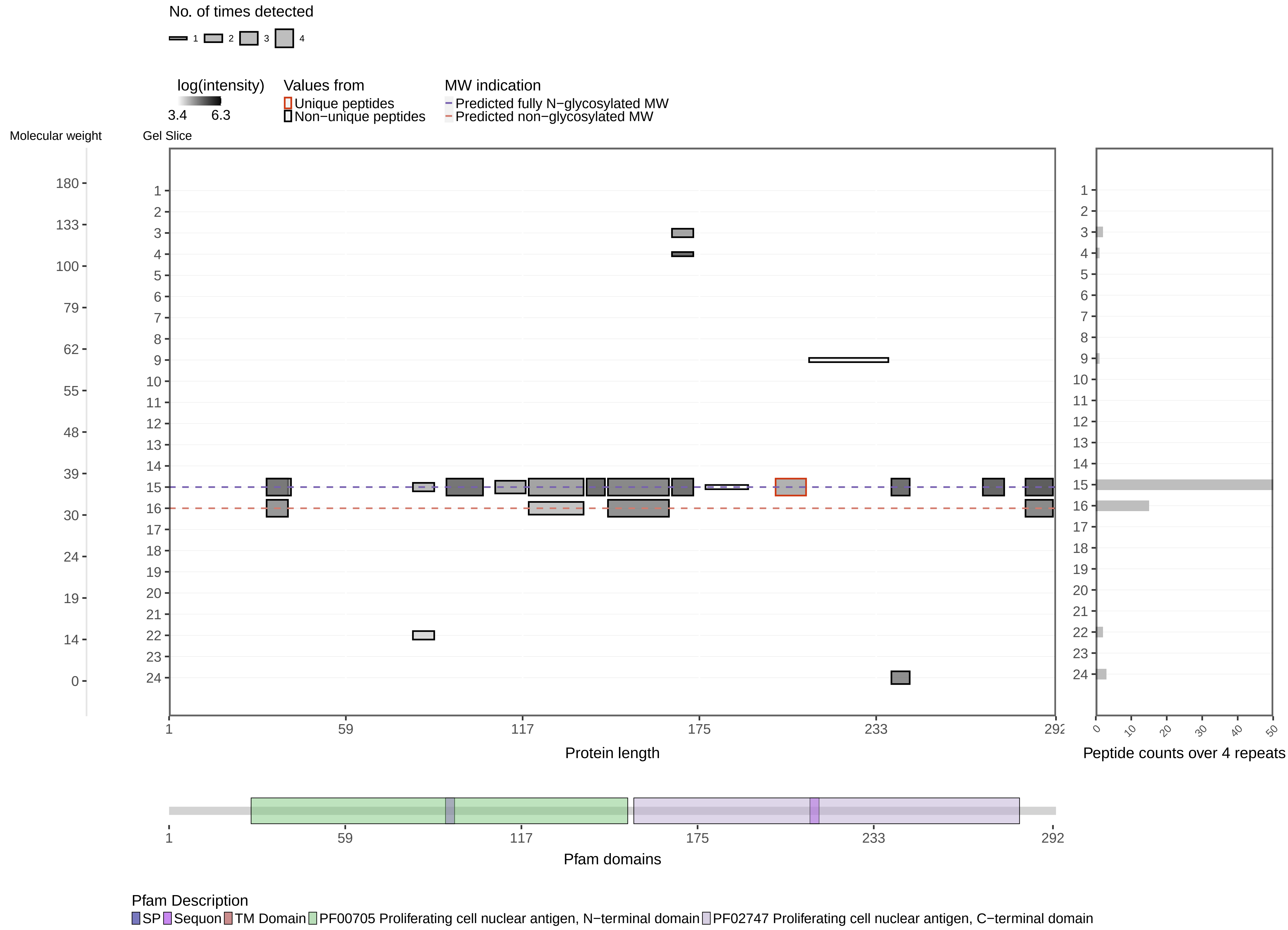


Protein sequence:

:ESPTQDKIADFEMKLMDIDSEHLGIPEAEYHAIVRMPSAEFARICKDLSSIGDTVVISVTKEGVKFSTRGDIGTANIVCRQNTTVDKPEEATVIEMNEPVSLTFALRYLNSFTKATPLSNTVTISLSSELPVVIEYKIAEMGYIRFYLAPKIEEDEEETKP*

### NbL09g06690.1 acyl−coenzyme_A_thioesterase_13−like [MW = 17.19 kDa]

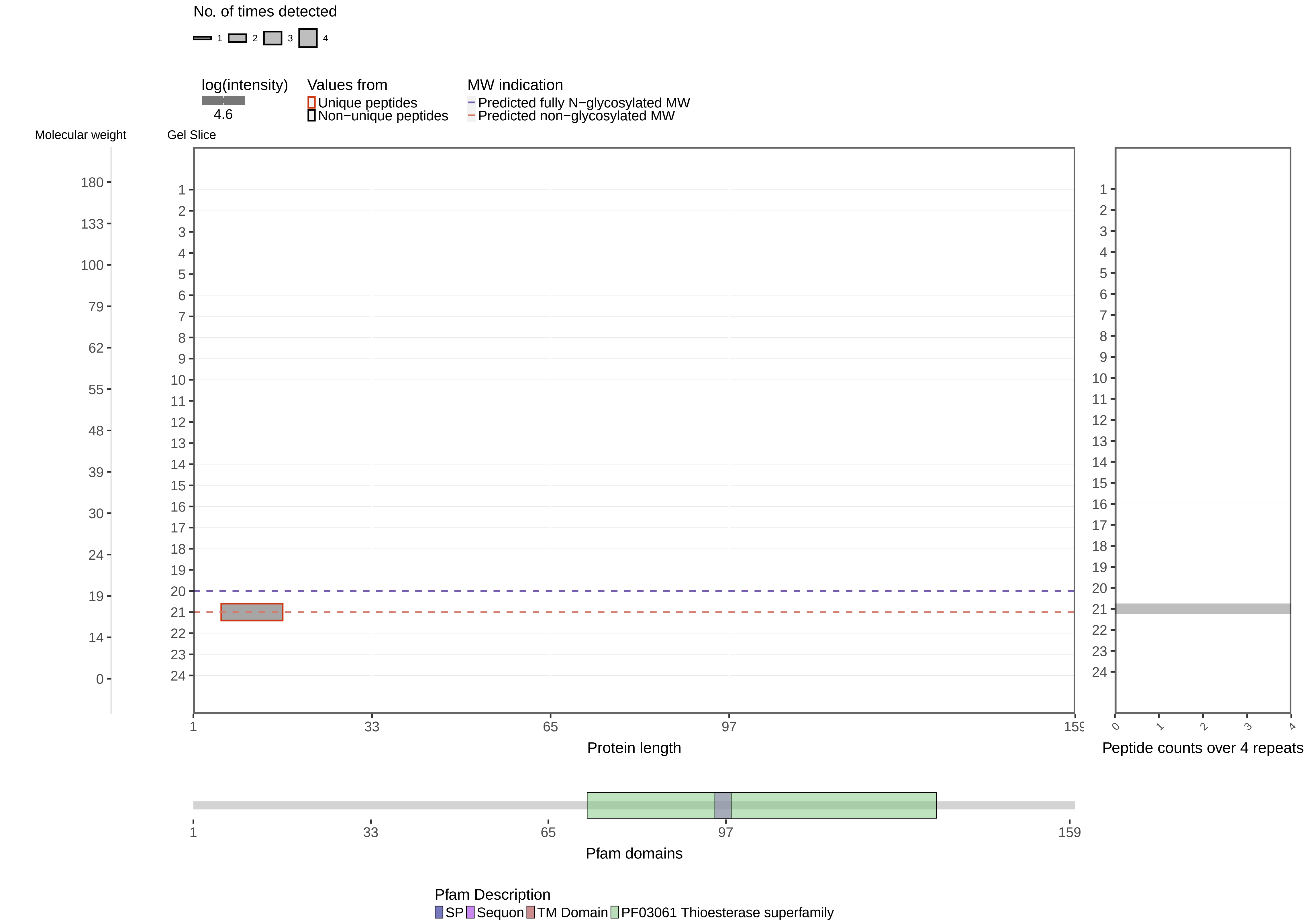


Protein sequence:

EGKLASGAIANLVDEVEGAVVFVEGLPMNVSVDMSISFLSSAKADDELEIIGRVLGQKRGYSGTSVLVKNKATGELIAEGRHSLFGKHASKM*

### NbL09g06630.1 beta−glucosidase_BoGH3B−like [MW = 65.71 kDa]

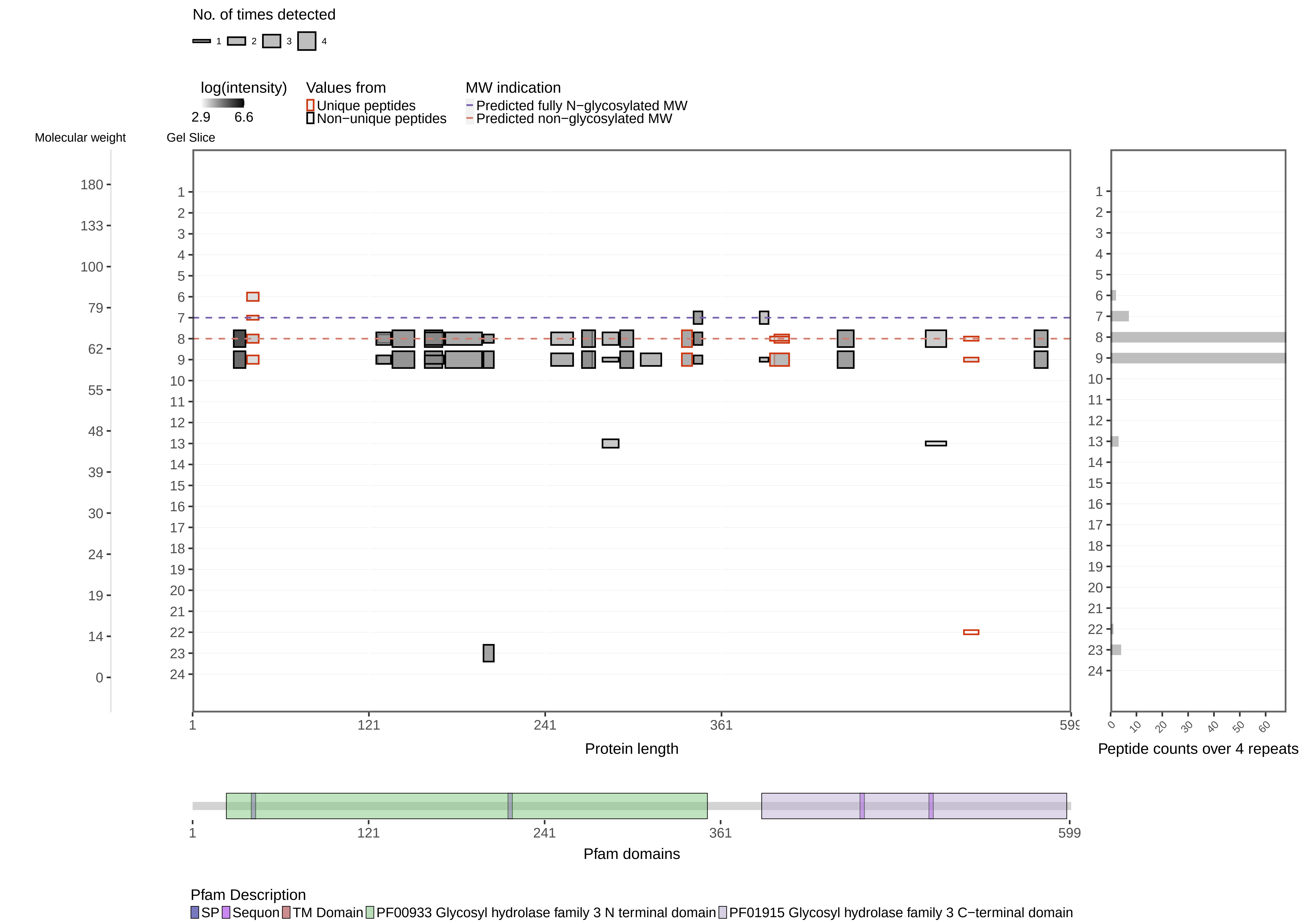


Protein sequence:

[illegible]

NbL09g06520.1 pleckstrin_homology_domain−containing_protein_1−like [MW = 16.64 kDa]

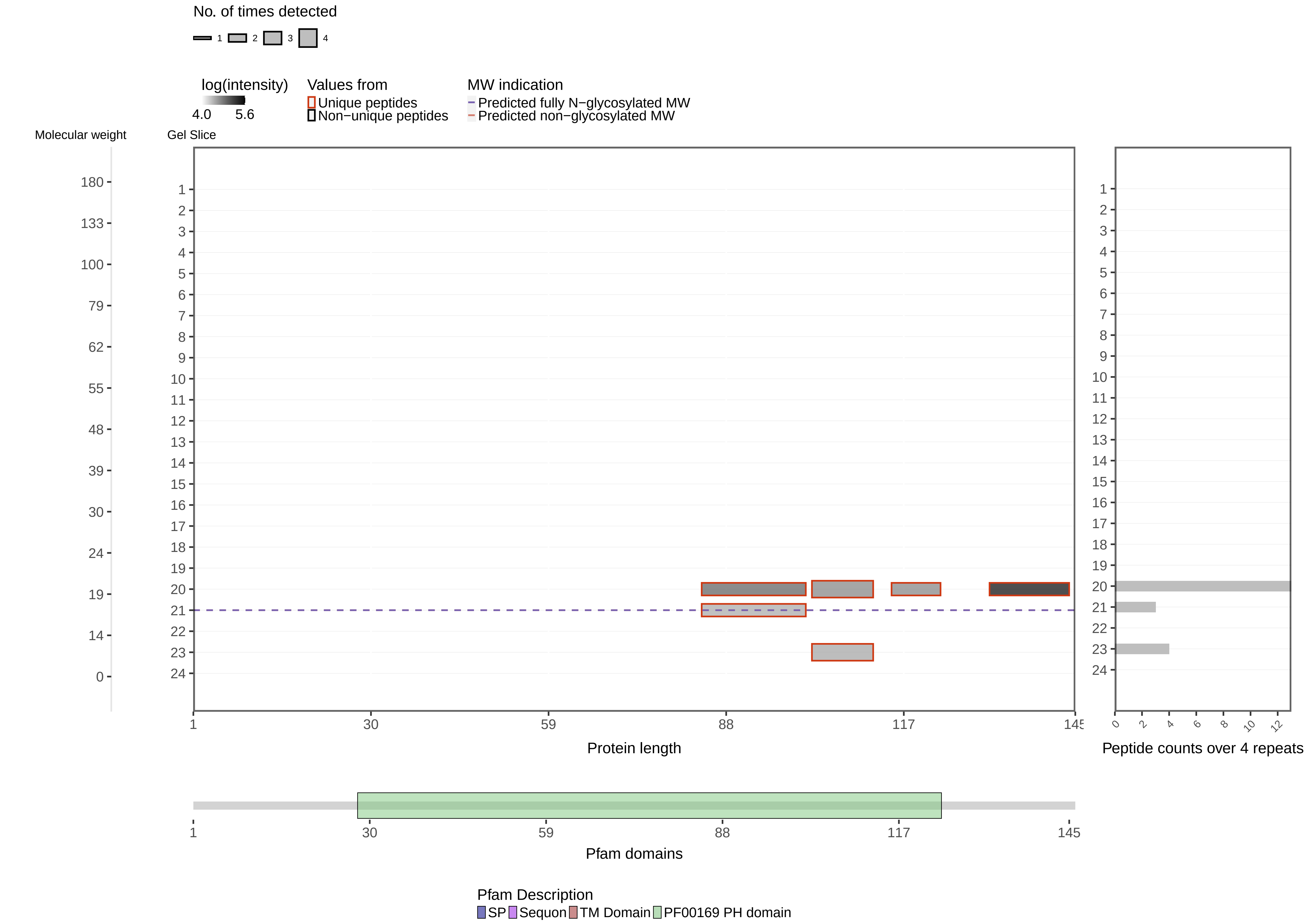


Protein sequence:

KDPSSVTRAAIPRGVVSVAECLTVKGAEDVINKPFAFELSTSRDTMYFIADSEKEKEEWINSIGRSIVQHSRSVTDSEVVDYDSRP*

NbL09g06430.1 peptidyl−prolyl_cis−trans_isomerase_CYP19−4−like [MW − SP = 19.54 kDa]

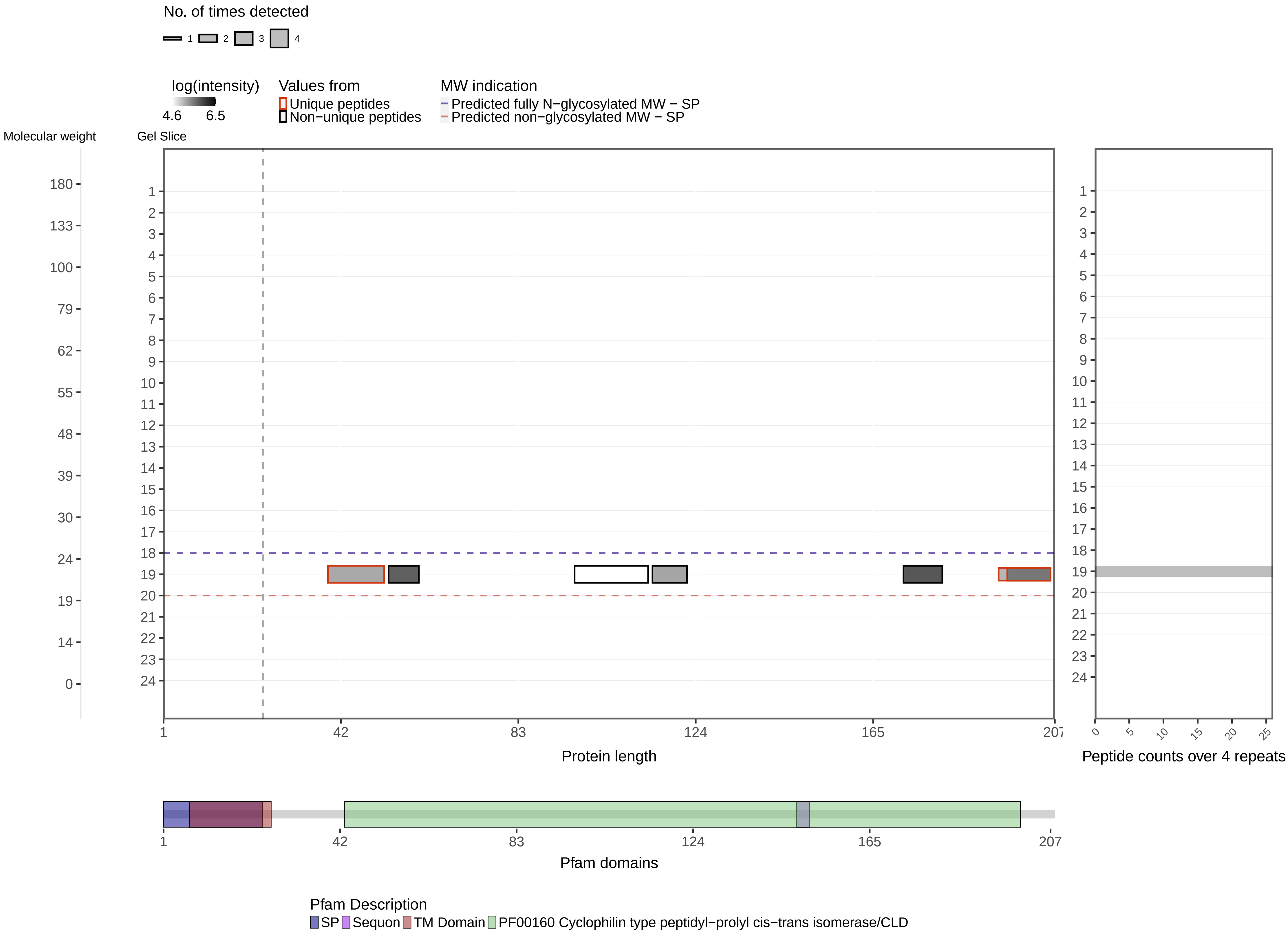


Protein sequence:

STFHRIIPSFMLQGGDFTLGDGRGGESIYGEKFADENFKIKHTGPGLLSMANAGPDTNGSQFFITTVTTSWLDGRHVVFGKVLSGMDVVYKIEAEGRQSGTPKSKVVIADSGELPL*

### NbL09g06410.1 probable_pyridoxal_5'−phosphate_synthase_subunit_PDX1 [MW = 33.09 kDa]

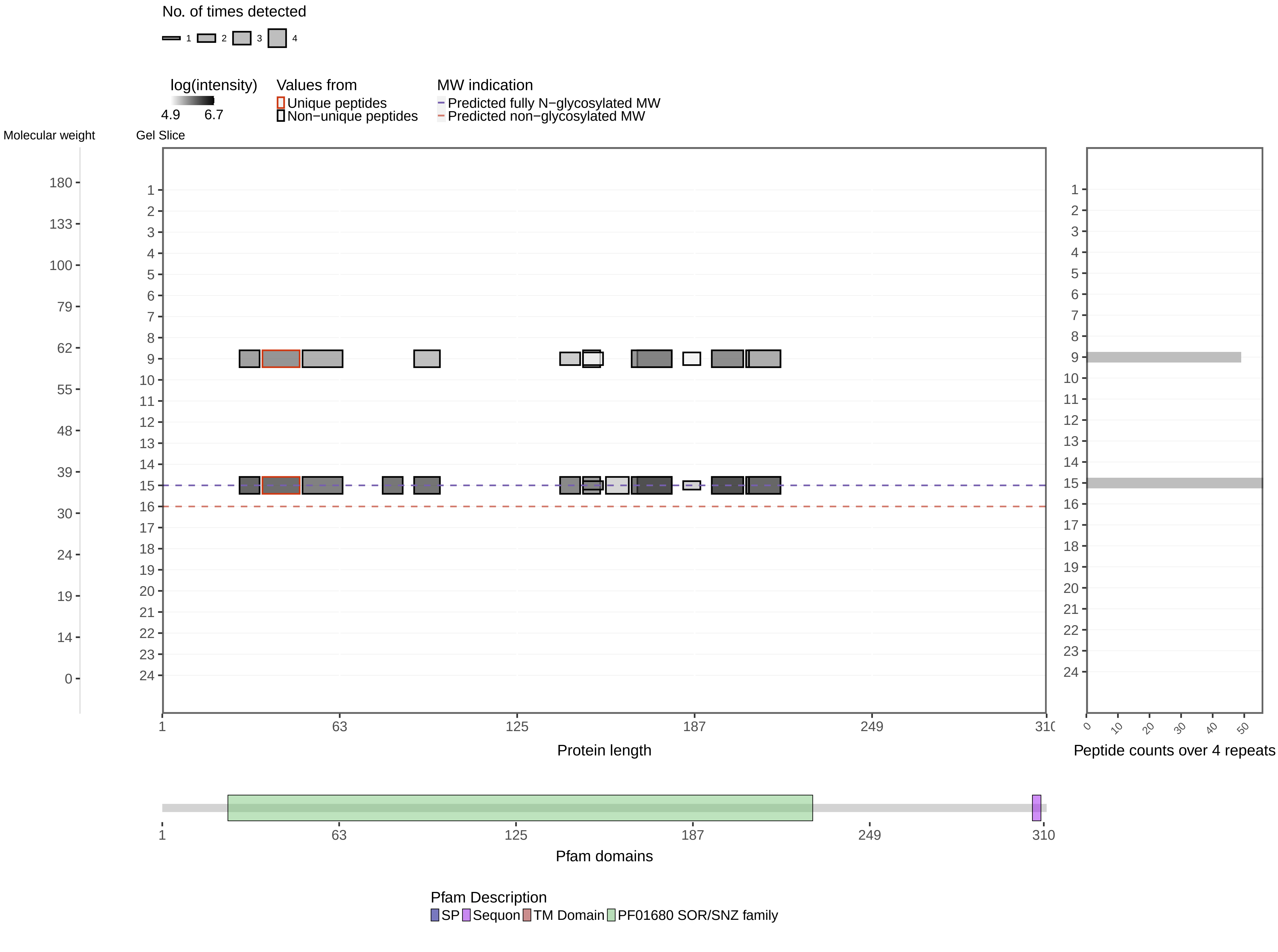


Protein sequence:

CGCRNLGEALRRIREGAAMIRTKGEAGTGNIIEAVRHVRSVMGDIRVLRNMDDDEVFTFAKKLQAPYDLVMQTKQLGRLPVVHFAAGGVATPADAALMMQLGCDGVFVGSGIFKSGDPAKRGRAIVQAVTHYSDPGLLAEISCGLGEAMVGINLDDKVERYANRSE*

NbL09g06350.1 ERBB−3_BINDING_PROTEIN_1 [MW = 42.78 kDa]

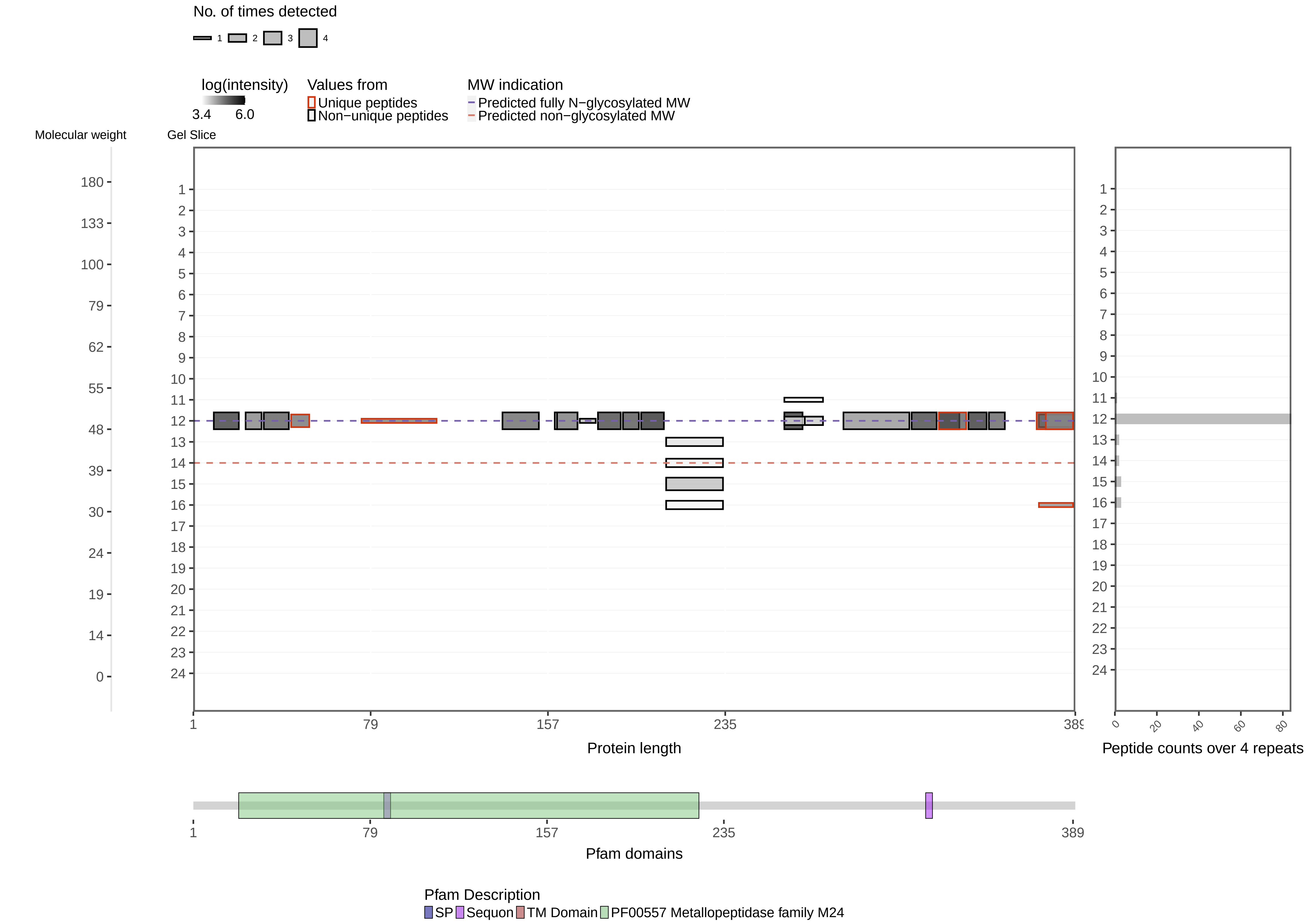


Protein sequence:

VLSHQMKQFVIDGNKVVLSVTNPETRVDEAEFEENEVYSIDIVTSTGEGKPKLLNEKQTTIYKRAVDKSYNLKMKASRFIFSEISQKFPIMPFTARDLEEKRARLGLVECVNHELLQPYPVLHEKPGDLVAHIKFTVLLMPNGSDRVTSHALQELKPAKTIENEPEIKAWLALPVKTKKKGGGKKKKGKKGDKTEDSSQAAEPTEG*

### NbL09g06340.1 U5_small_nuclear_ribonucleoprotein_200_kDa_helicase−like [MW = 247.4 kDa]

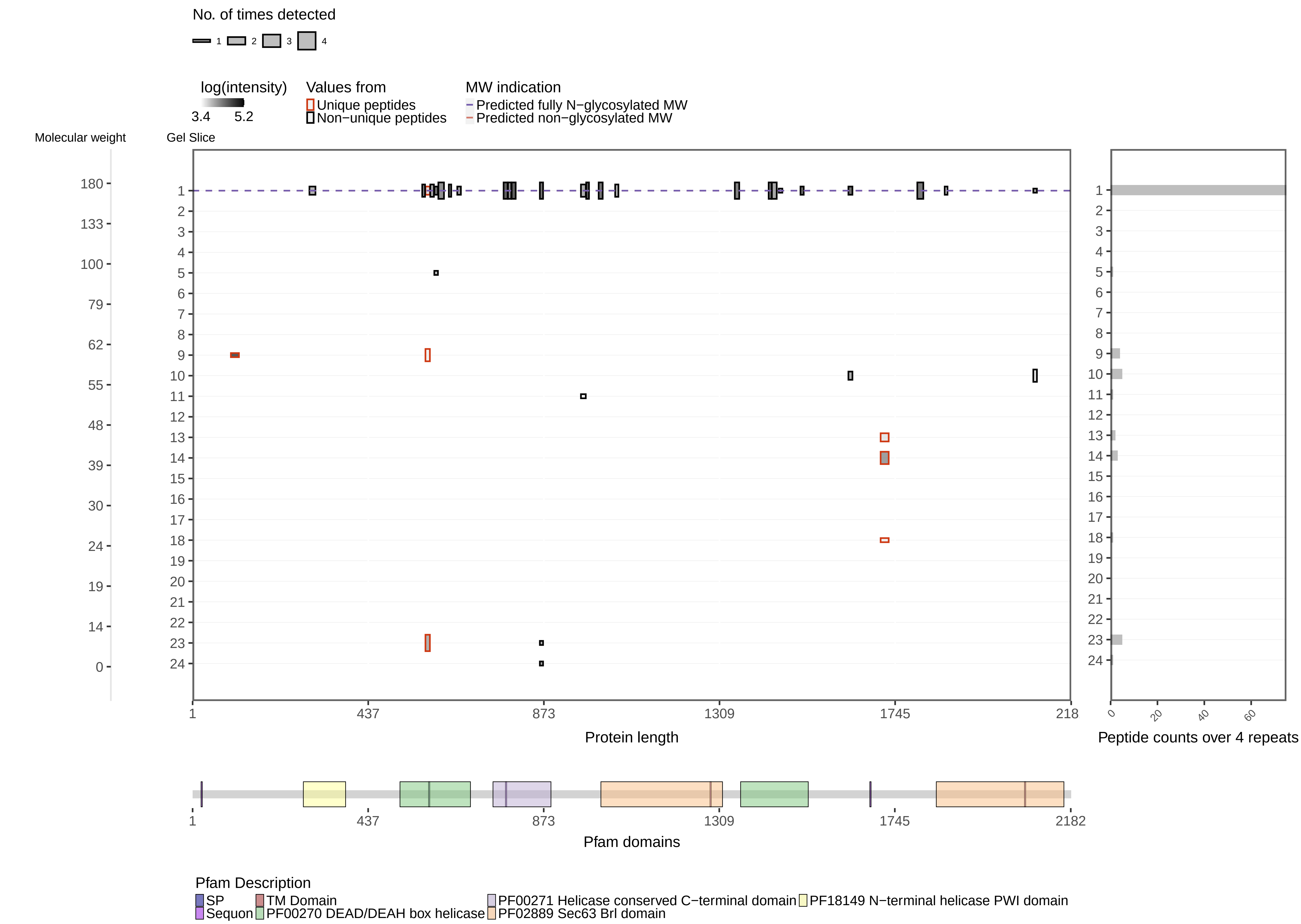


Protein sequence:

NbL09g06330.1 6−phosphofructo−2−kinase/fructose−26−bisphosphatase−like [MW = 83.16 kDa]

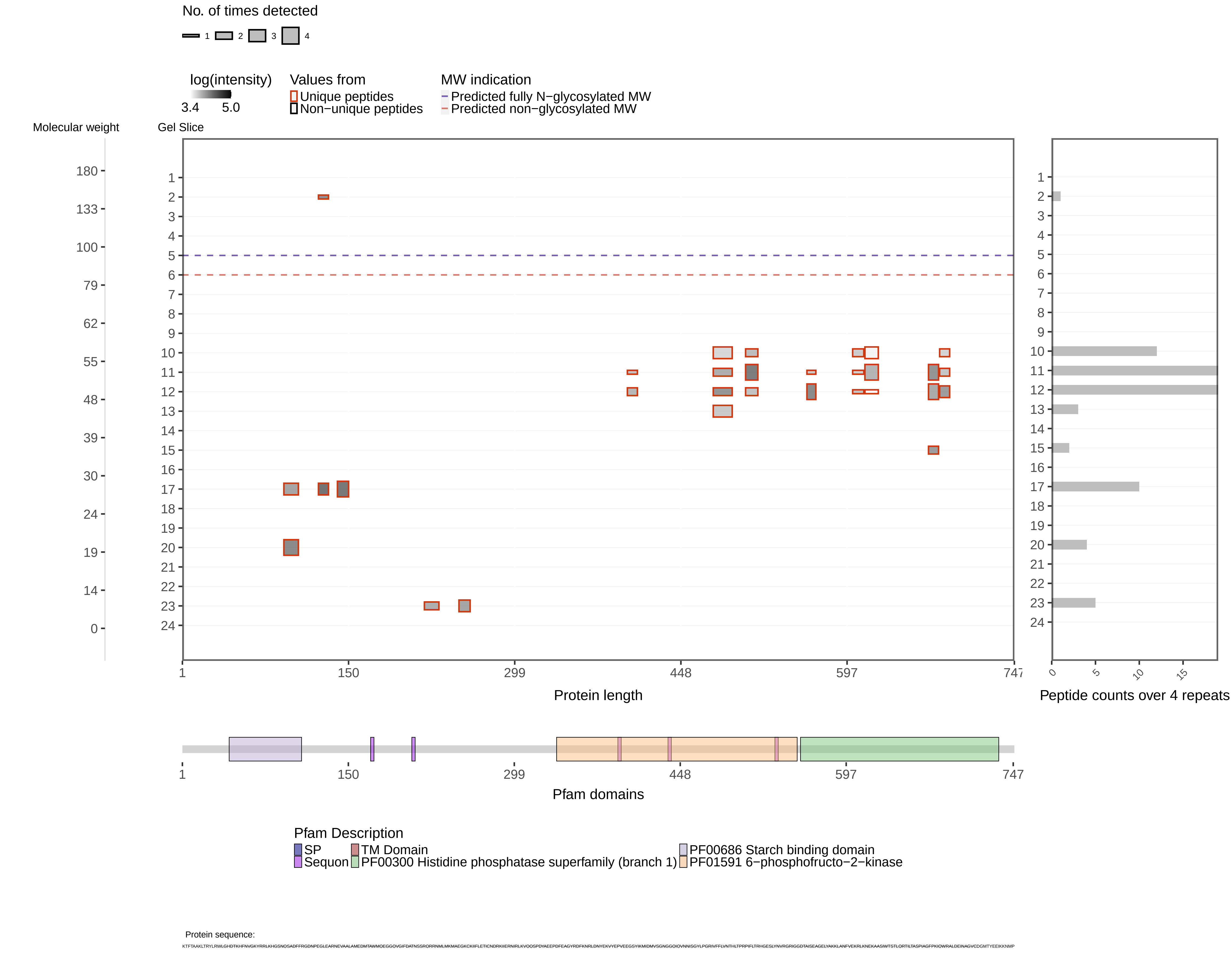

### NbL09g06280.1 NAP1−related_protein_2−like [MW = 28.94 kDa]

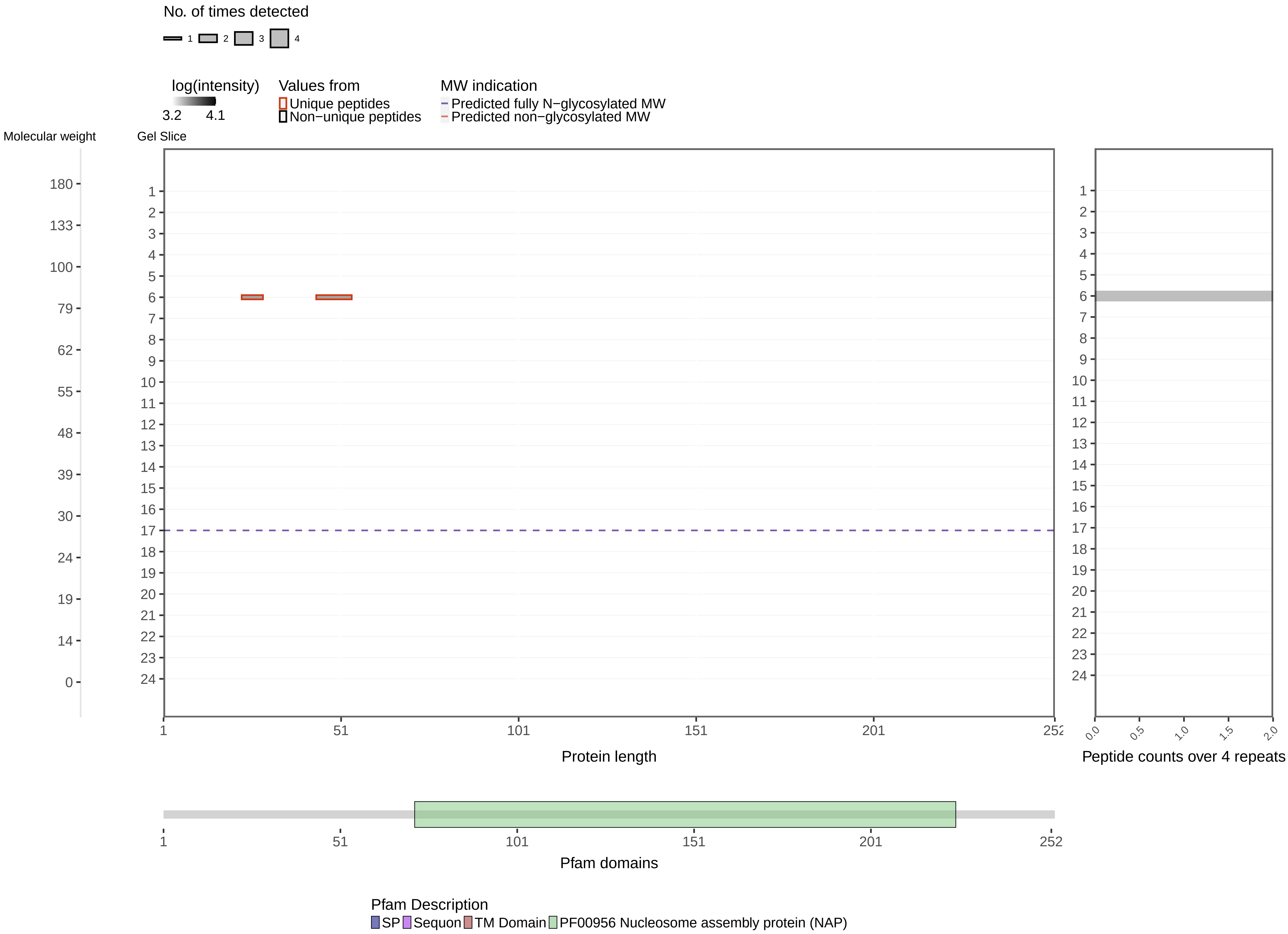


Protein sequence:

SGHSITFNFKPNPYFENSKLSKTYTFLEDGPTKITATTIKWKEGMAIPNGVADKNKGNKRSHAEESFFTWFSEVNQKGDVDDDENEILDIQDDEVAEIIKDDLWPNPLNYFDHDPDEEDIEGDEVQQHSCFIIAKLS*

NbL09g06210.1 delta−1−pyrroline−5−carboxylate_dehydrogenase_12A1_mitochondrial [MW = 60.84 kDa]

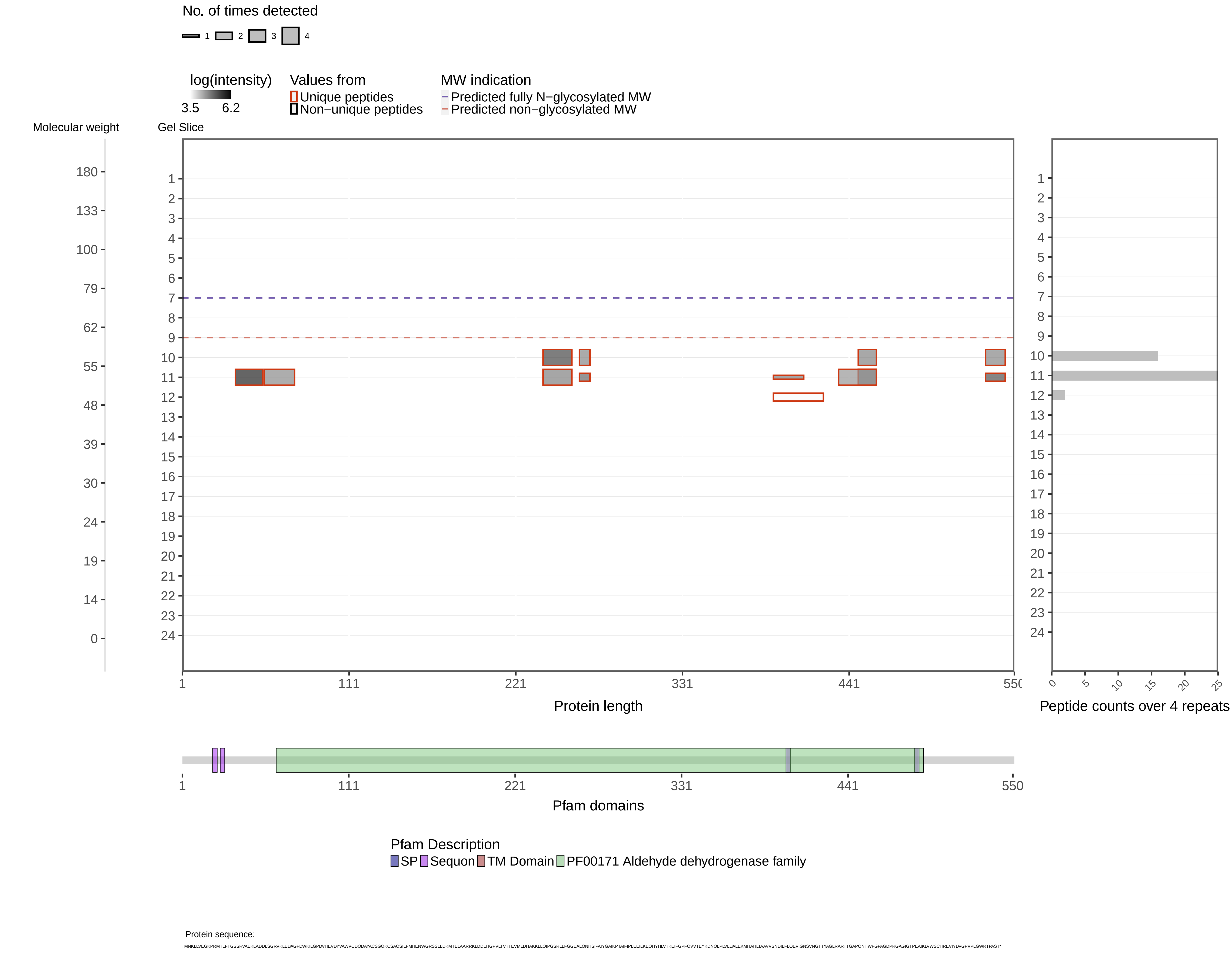

### NbL09g06120.1 plasma_membrane_ATPase_2 [MW = 70.41 kDa]

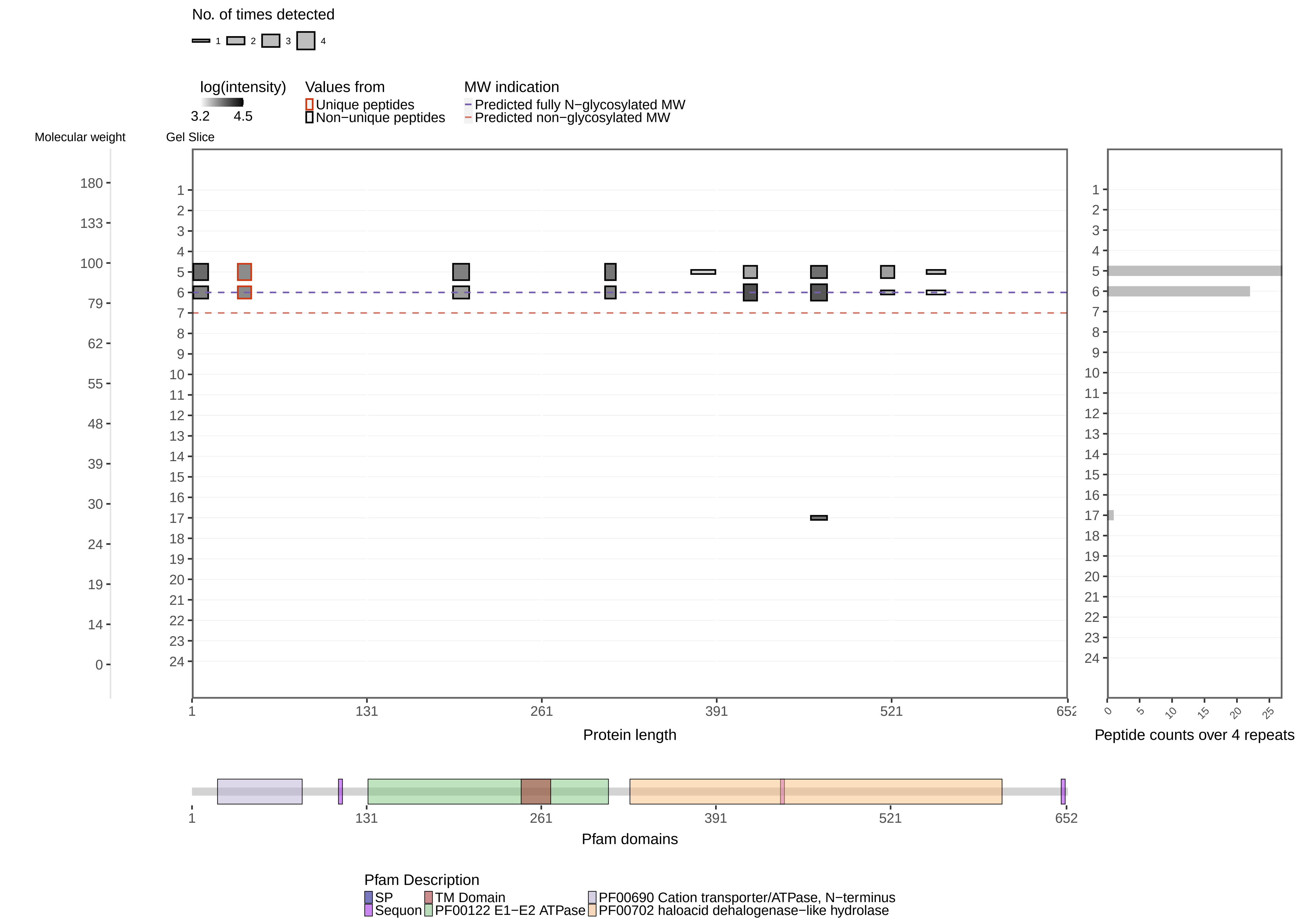


Protein sequence:

NbL09g05810.1 60S_ribosomal_protein_L13−1 [MW = 23.71 kDa]

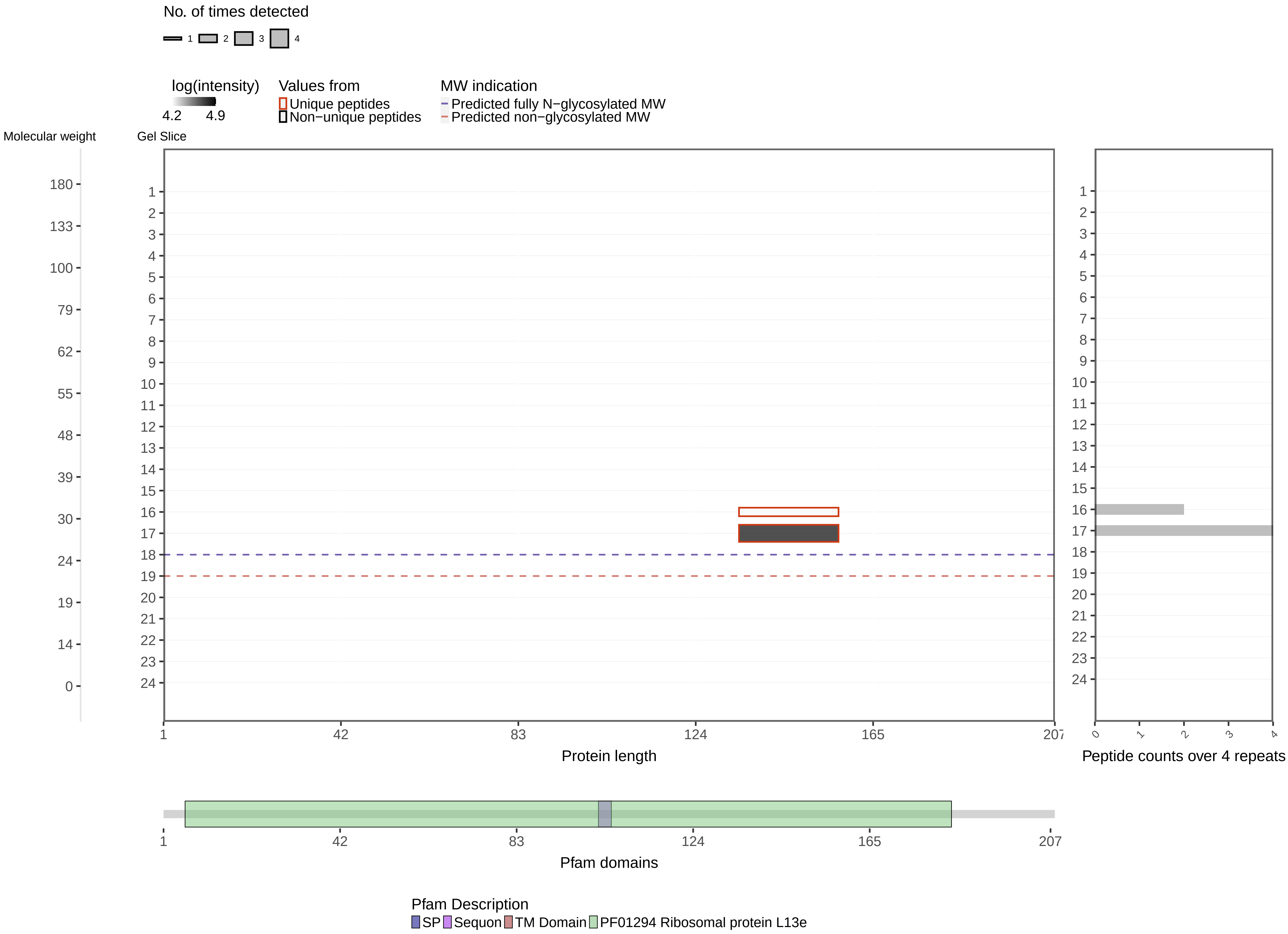


Protein sequence:

TIGIAVDHRRRNRSLEGLQTNVQRLKTYKAKLVVFPRRARKVKAGDSAPEELATATQVHGDYMPIAREKPSVDLVKVTEEMKSFNAYGKLRVERTNERHIGARMKRAAEAEKEEKK*

NbL09g05580.1 pyruvate_dehydrogenase_E1_component_subunit_beta−1_mitochondrial−like [MW = 39.42 kDa]

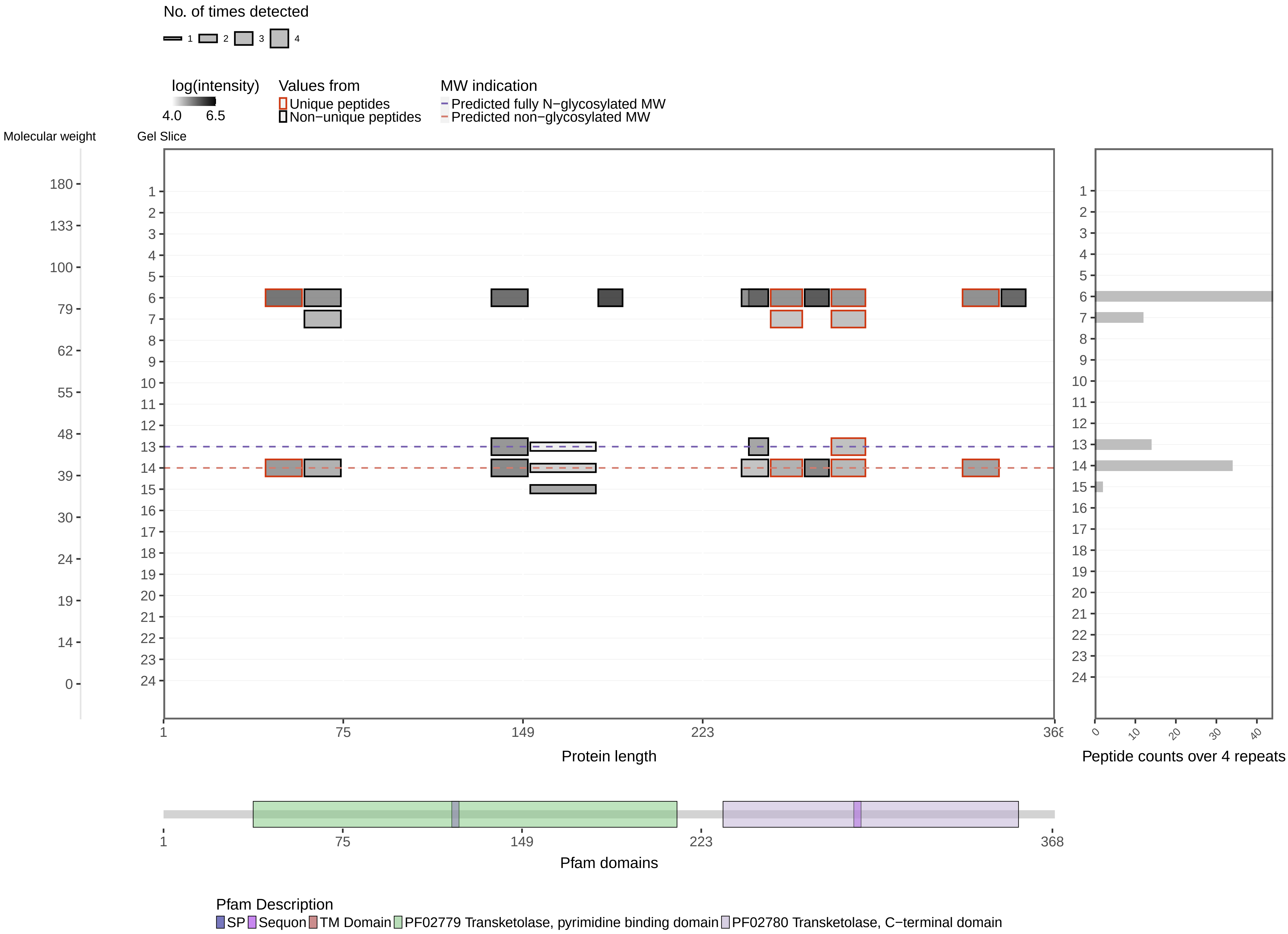


Protein sequence:

AWFGSCPGLKVLAPYSSEDARGLLKAAIRDPDPVVFLENELLYGESFPVSAEALDSSFCLPIGKAKIEREGKDVTITAFSKMVGYALQAAEILAKEGISAEVINLRSIRPLDRPAINTSVRKTNRLVTVEEGFPQHGVGAEICASVVEESFEYLDAPVERITGADVPMPYASNLERMAVPQVEDIVRAAKRACYRSST*

NbL09g05530.1 40S_ribosomal_protein_S3a [MW = 29.77 kDa]

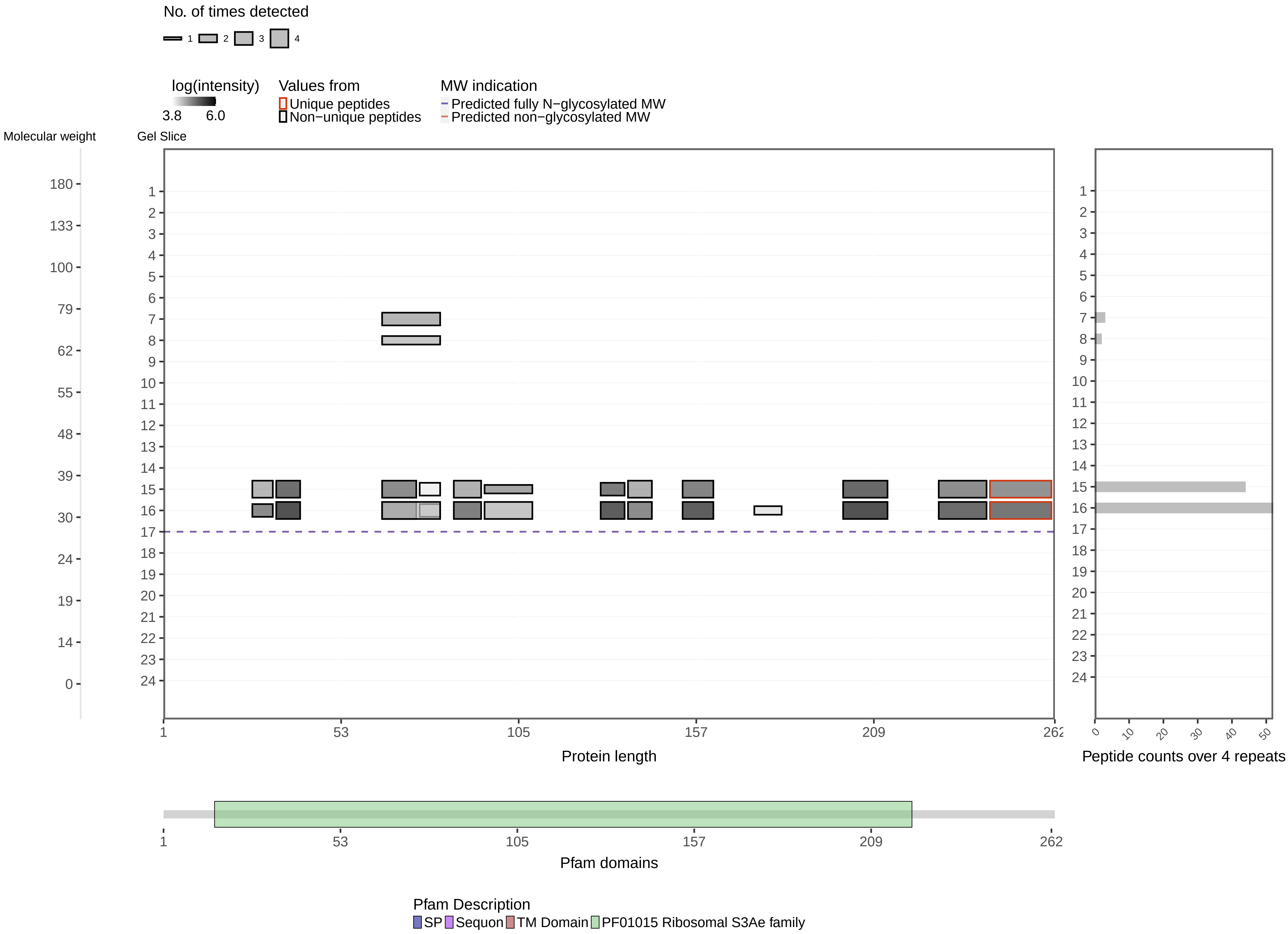


Protein sequence:

WOTLIEAHVDVKTTDSYTLRMFCIAFTKKRPNOOKRTCYAOSSOIROIRRKMVEIMRNOASSCDLKELVAKFIPESIGREIEKATSSIFPLONVYIRKVKILKAPKFDIGKLMEVHGDYSEDVGVKLDRPADENVAEAEPEIPGA*

NbL09g05510.1 50S_ribosomal_protein_L29_chloroplastic−like [MW = 19.06 kDa]

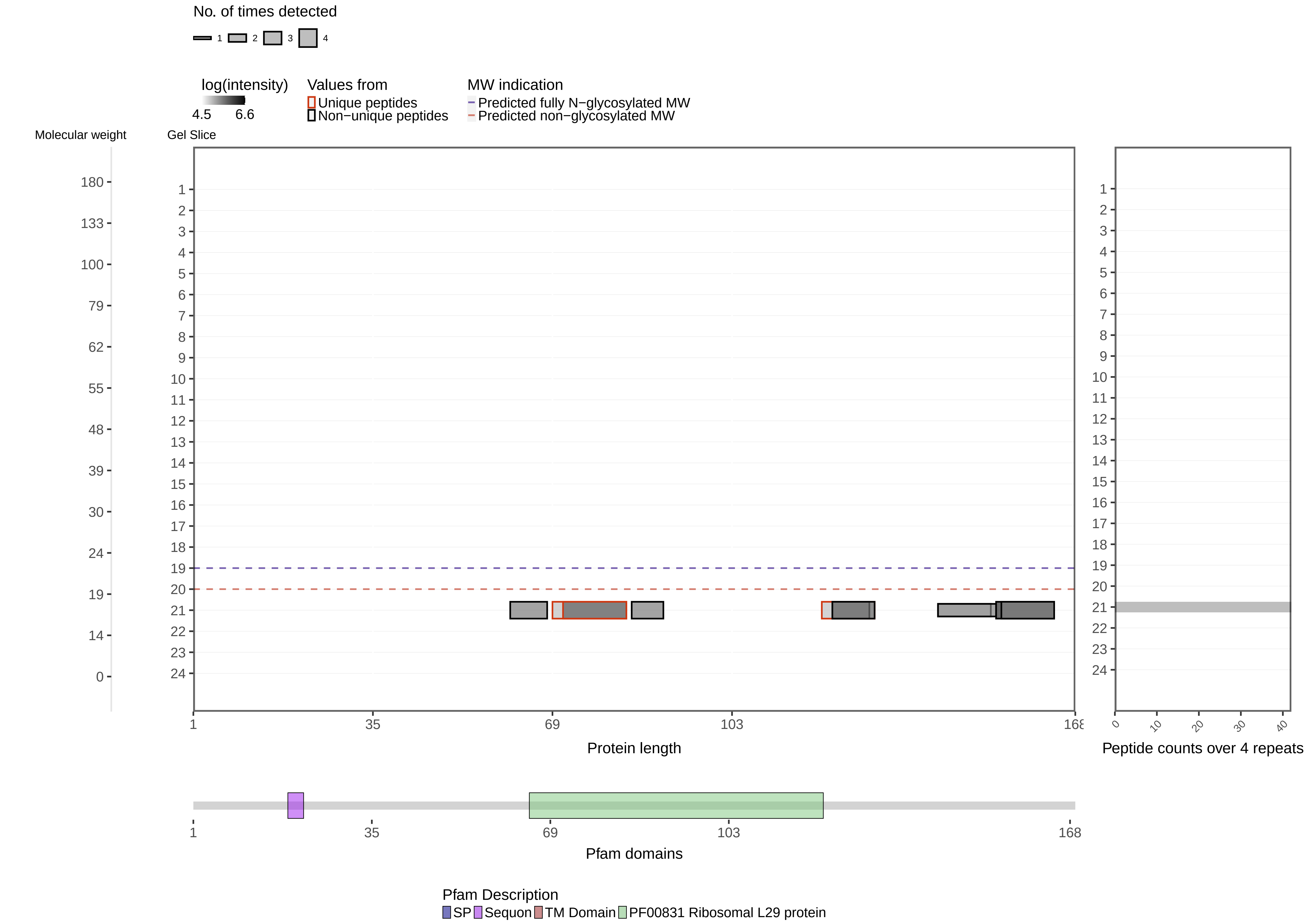


Protein sequence:

TEELQEEIVDLKGELFMLRLQRSARNEFKSSEFLRMRKRIARMLTVKRDRELEEGINKRISRKLDRKWKKSIVPRPPPSLKKLQEEEAAAEAKESS*

NbL09g05430.1 protein_argonaute_1−like [MW = 117.76 kDa]

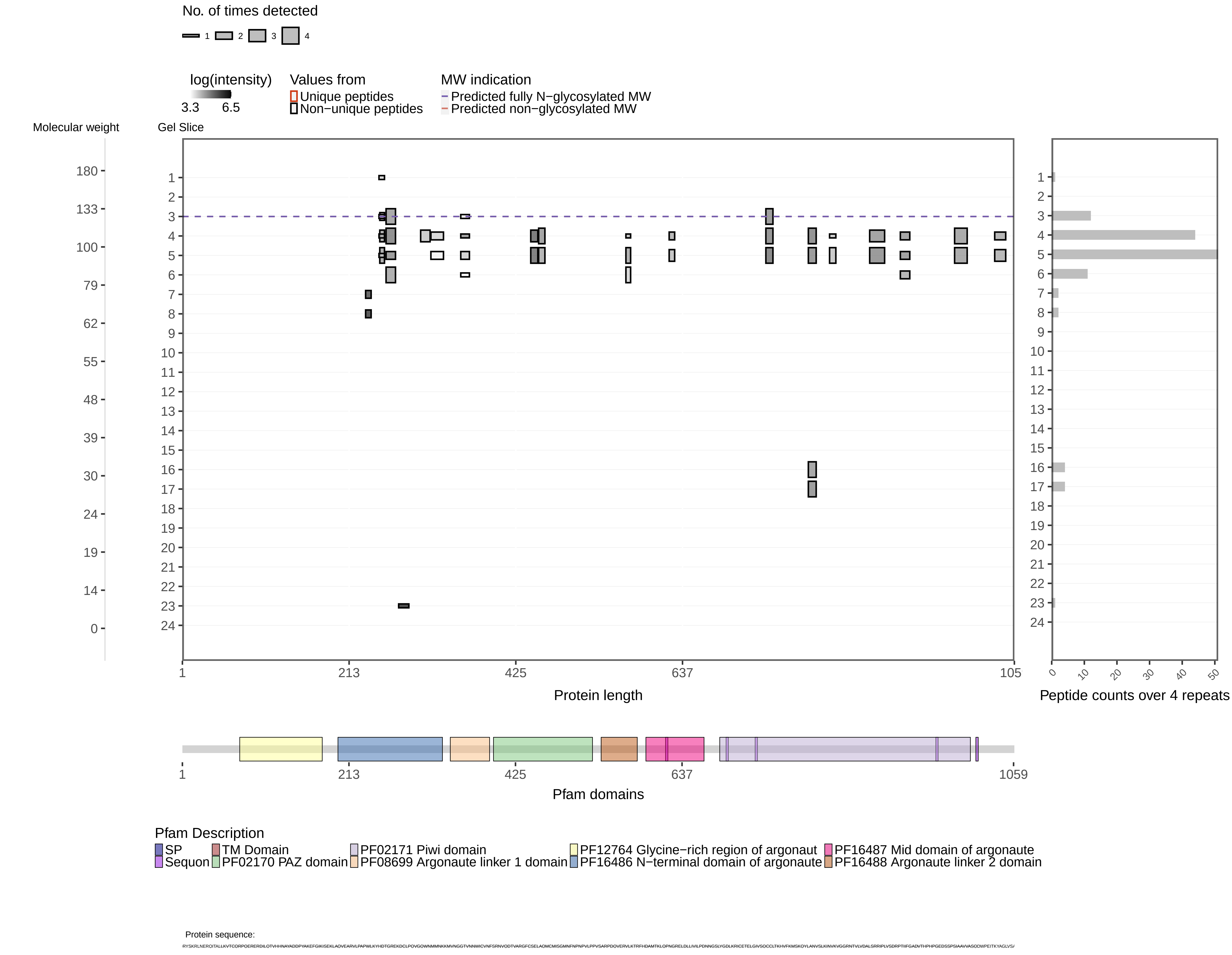

### NbL09g05410.1 pEARLI1−like_lipid_transfer_protein_1 [MW = 15.69 kDa]

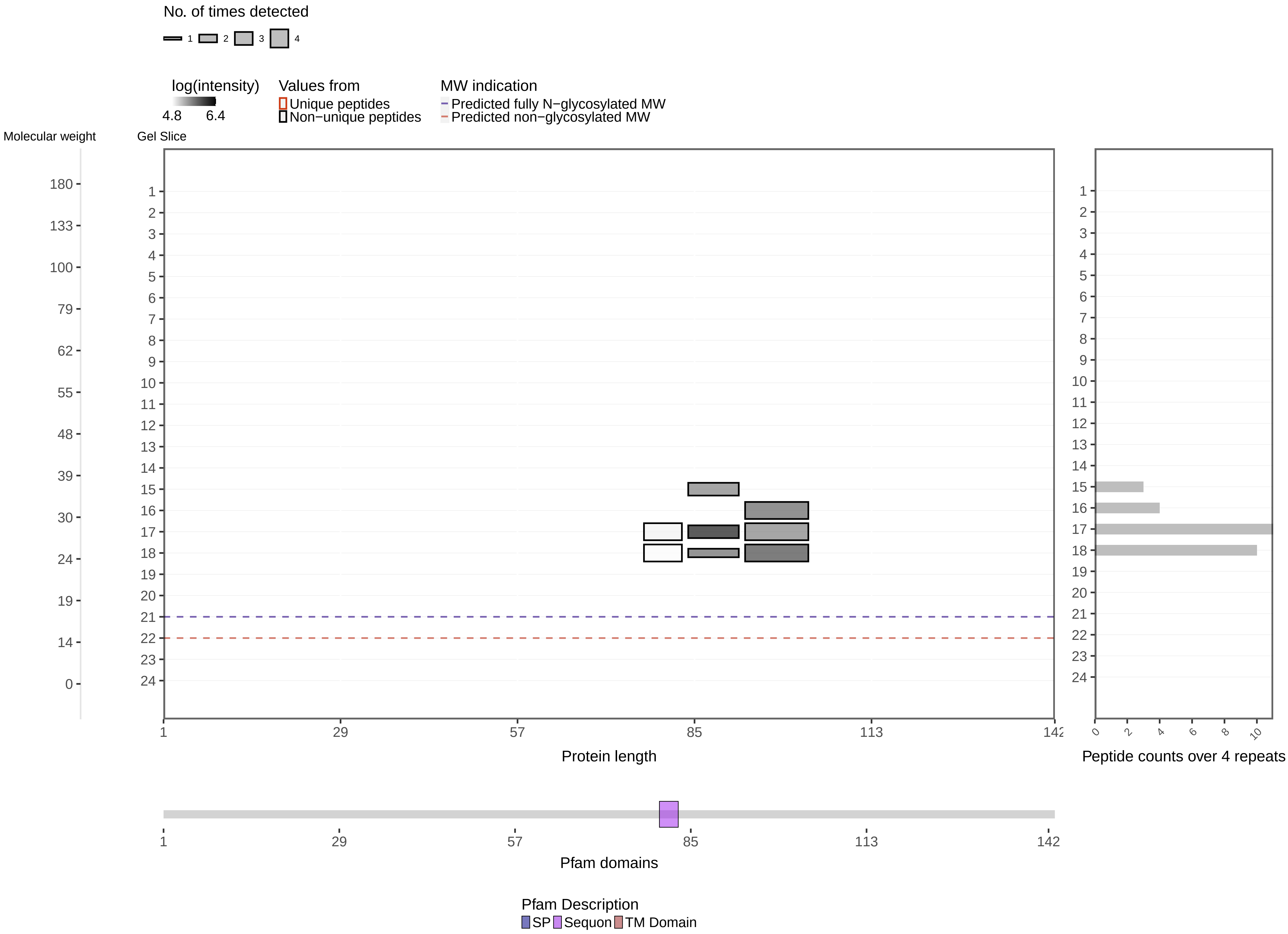


Protein sequence:

PPPPAPSSSCSASDEAKIYKCMFNETKIDPCCPTFKSILGTSCPCYKYAENLDNQVLITIEAYCDVYSPCKGLQAIKLSKEEEYK*

### NbL09g05350.1 phytochrome−associated_serine/threonine−protein_phosphatase_3−like [MW = 34.87 kDa]

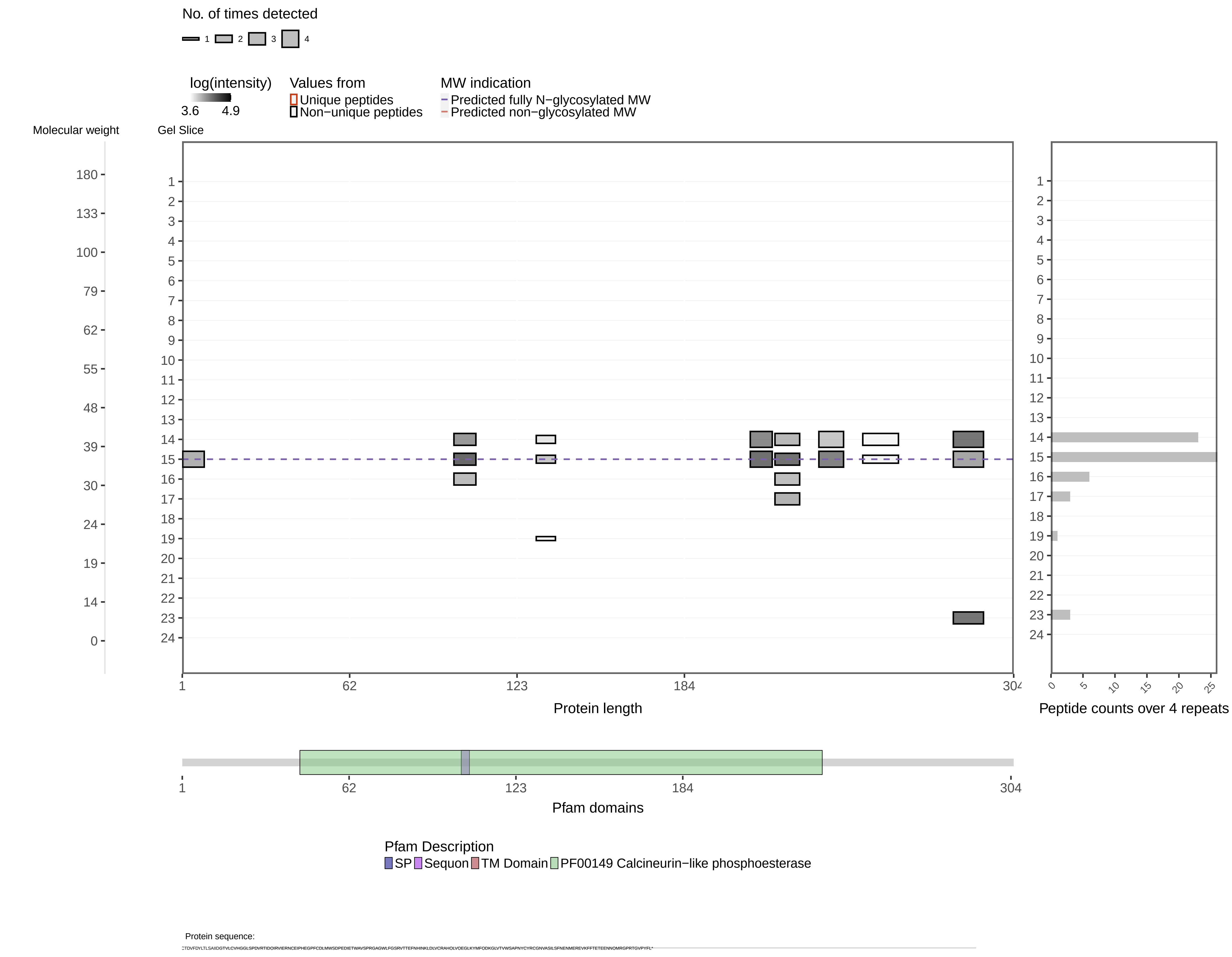

### NbL09g05330.1 40S_ribosomal_protein_SA−like [MW = 32.21 kDa]

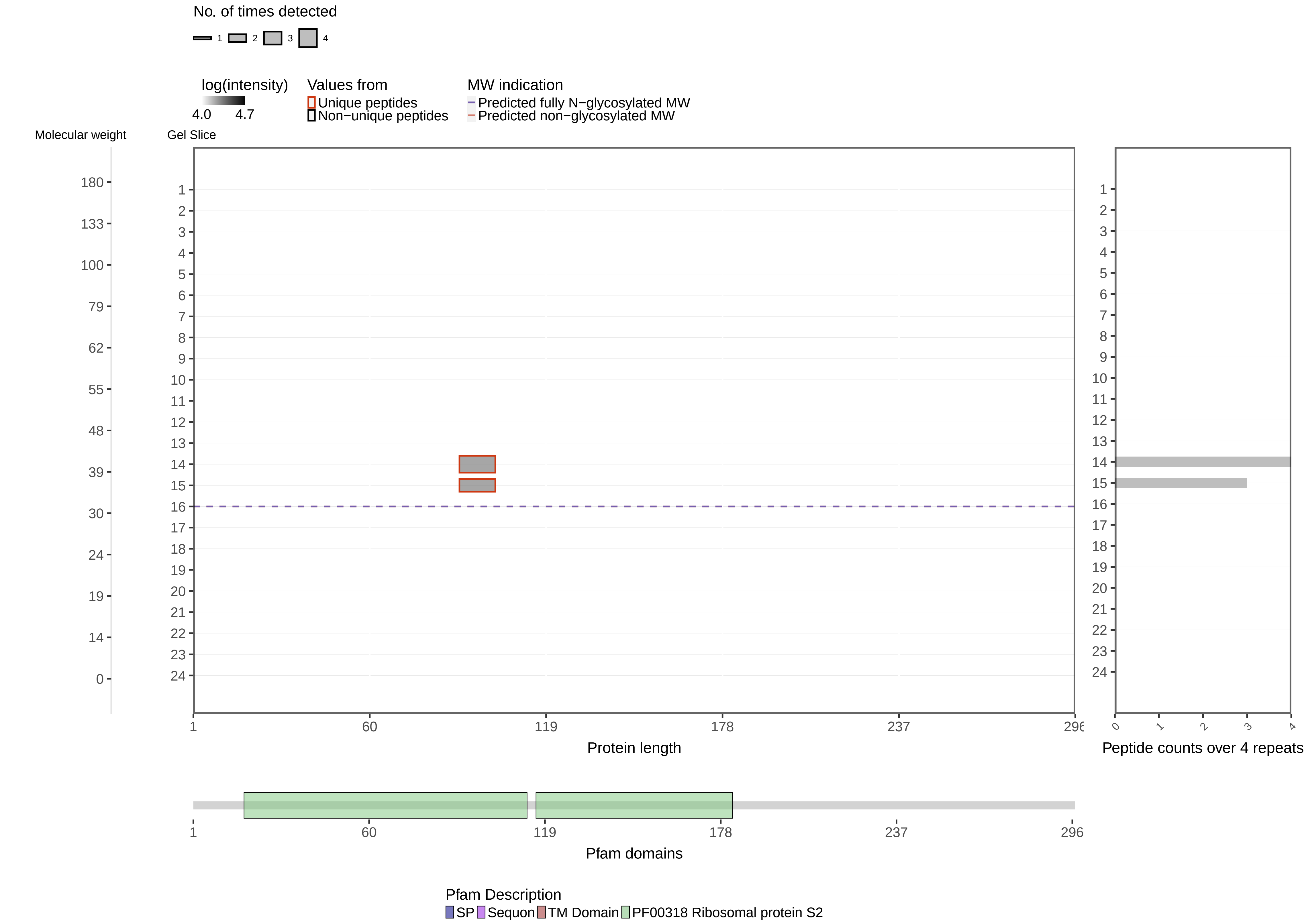


Protein sequence:

ALGNIPTIAFCDTDSPMRYVDIGIPANNKGKHSIGVLFWILARMVLQMRGSINQGHKWEVMVDLFFYREPEEAKEQEEEEAPAIDYADYSAGAALGGDWSSSQIPEAQWTGDAAPSGPVVASGWSGEGVAEGGWDTAAAPVPVPVPDAAPTAGATGWE*

### NbL09g05250.1 glyceraldehyde−3−phosphate_dehydrogenase_cytosolic [MW = 36.66 kDa]

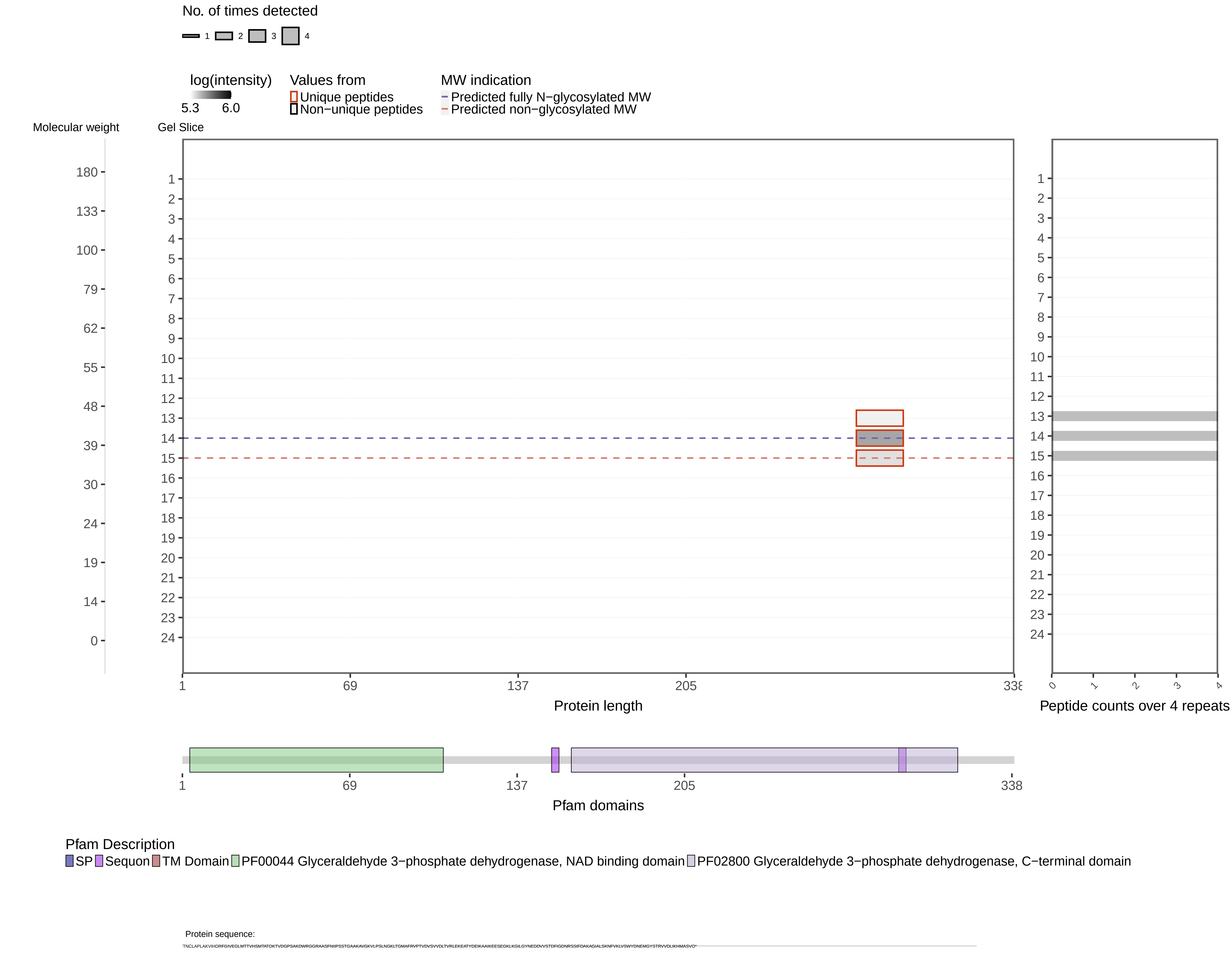

NbL09g05240.1 3−oxoacyl−[acyl−carrier−protein]_reductase_4−like [MW = 34.17 kDa]

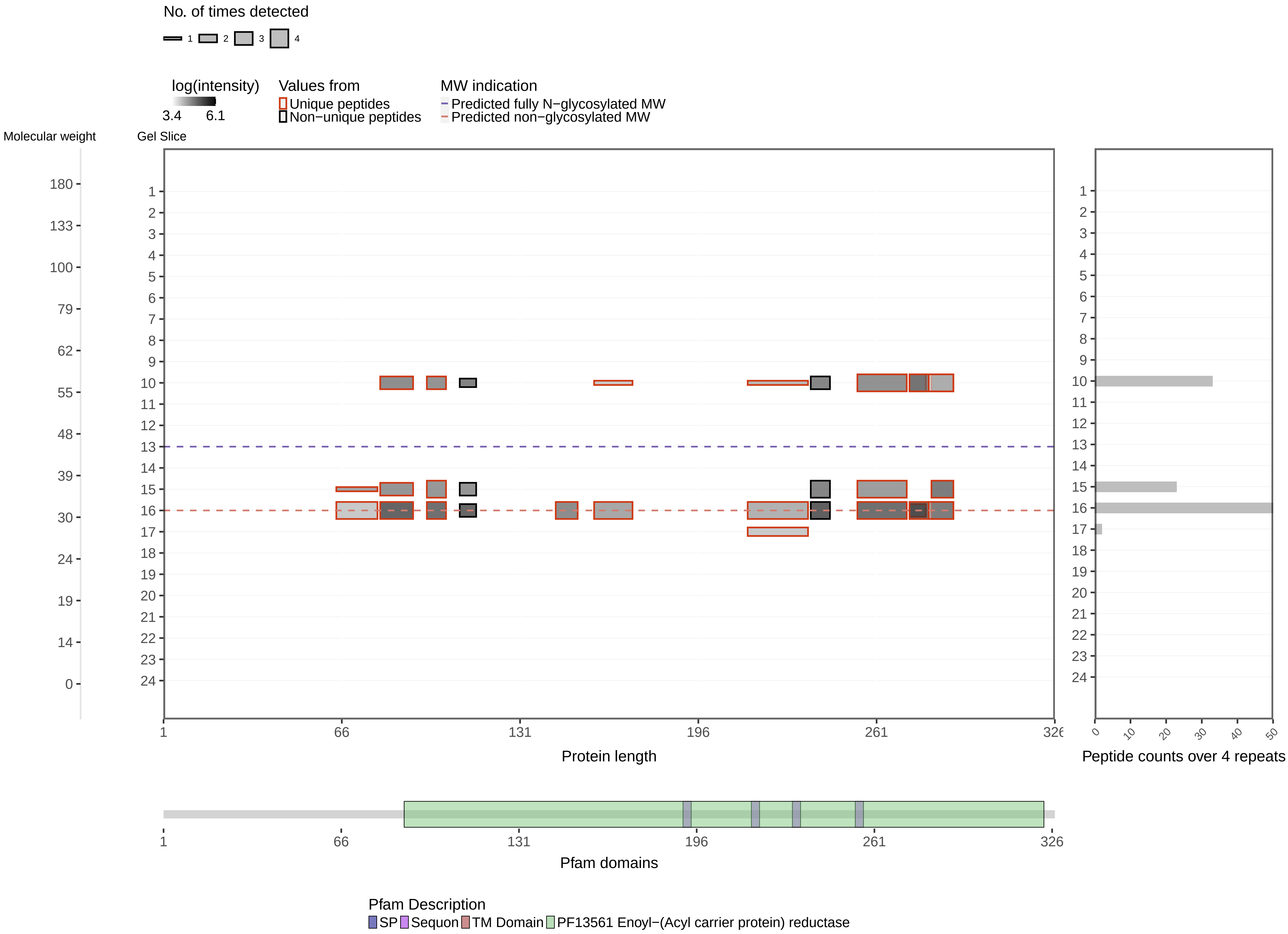


Protein sequence:

SMMKTAVDRWGTVDILINNAGITRDTLLMRMKKSQWQEVIELNLTGVFLCTQAAAKIMMKKKKGRIINISSVVGLVGNAGQANYSAAKAGVIGLTKSVAKEYASRNITVNAVAPGFIASDMTAKLSEDIEKKILQSVPLGRYGQPEEVAGLVEFLALNPAASYITGQVLTIDGGMVM*

### NbL09g05220.1 60S_ribosomal_protein_L17−2−like [MW = 20.09 kDa]

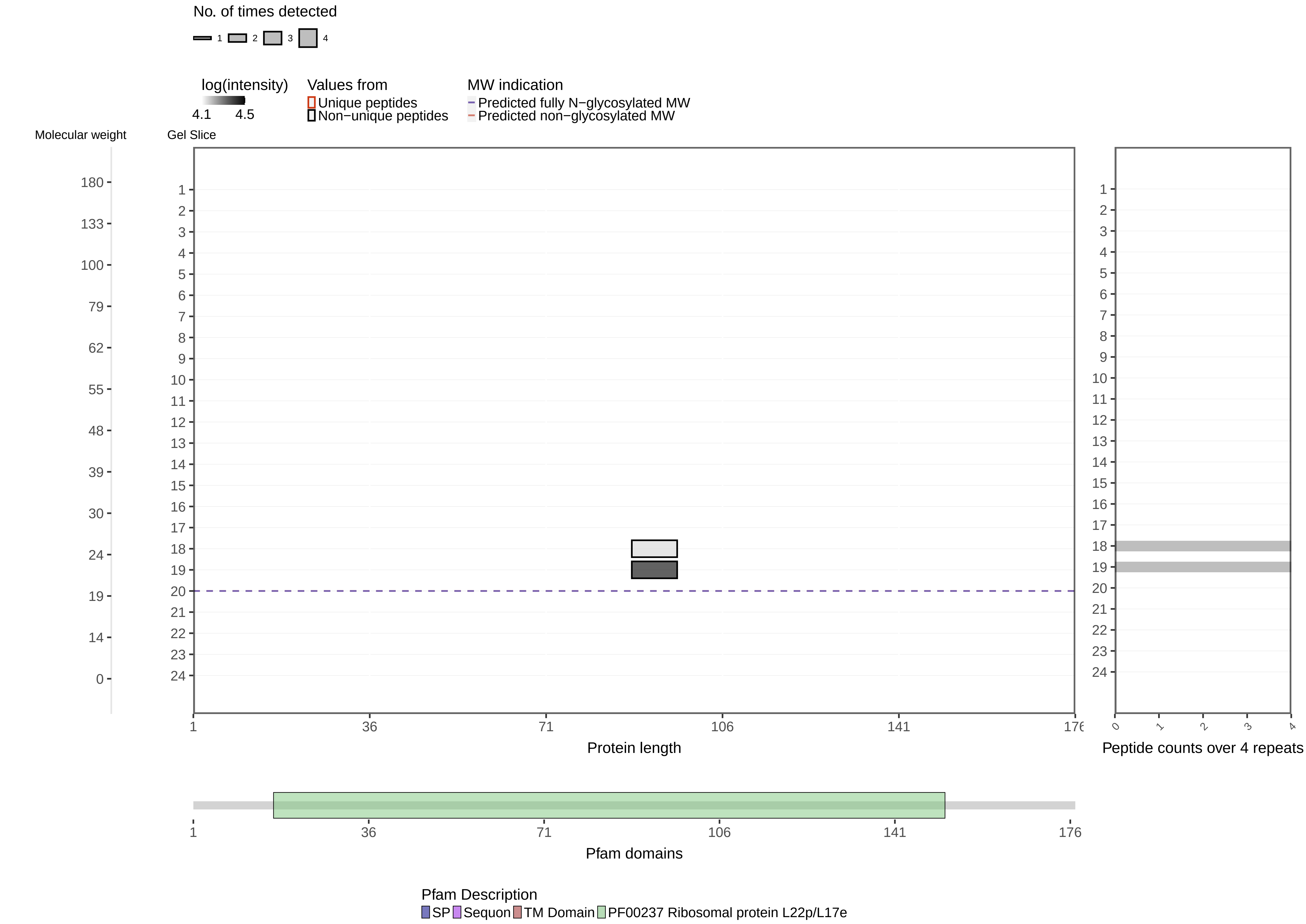


Protein sequence:

IRHSNGQGRWPVKSAGFILDLLKNAESNAEVKGLDVDSLYITHIQVNQAQKQRRRTYRAHGRINPYMSNPCHIELTLSEKEESVRKEPETQLAPRKTKASS*

### NbL09g05160.1 peroxisomal_membrane_protein_PEX14−like [MW = 57.29 kDa]

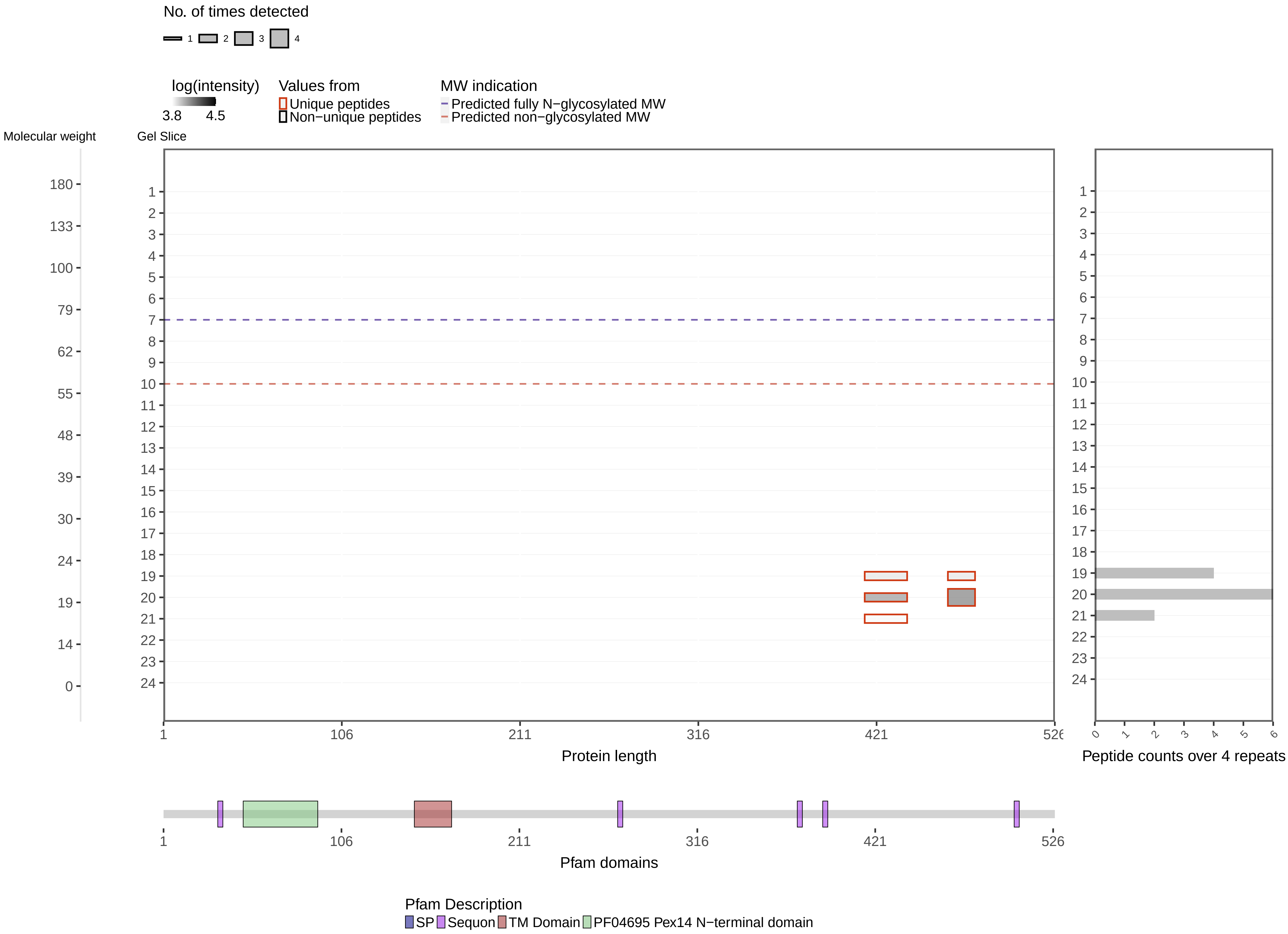


Protein sequence:

REMKSMTSAIEKLEGQSNTSGRIPITELDDRRISVTQSRQSYLNGKVDGDARSVRSLSLPASVEQSVAPHPKSYMEIMAMVQRGEKPSNIQDINDQPPNPNKPVSDPRVVPKPKPWEVGQIQSNSTNLLQFQGSGDGLNYSYQDSLTNGDSSAAWWQRRNARITEIESEGEQNFGSPAAPVQERPIQRSWVPPQPPPVAMAEAAAAIRQPRKPVLQNEQVTDDQLMARSSEITDELQRVTRFSESGDAPEANGSSSALHISETPIGEGDQILSS*

### NbL09g05110.1 GDSL_esterase/lipase_At5g62930 [MW = 27.43 kDa]

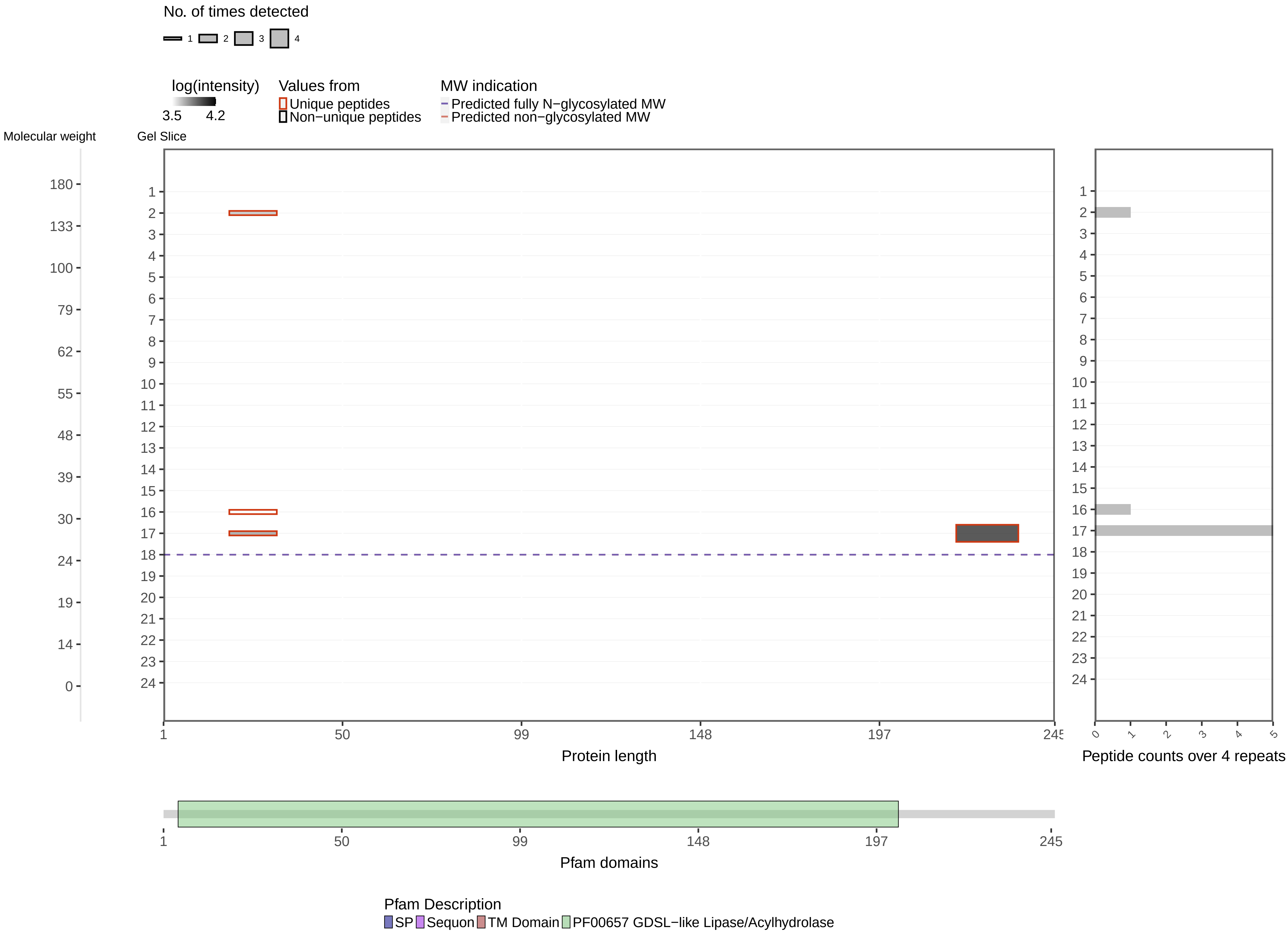


Protein sequence:

SILVVLITPPPVDEAGRFEOARSMYGDKAMELPERTNEVTGEYAKOCVELARELGLPSINLWSKMOETEGWOKKFLCDGLHLTAEGNAIVHOEVVKAFNEAGFSAOKMPYDVPHHSEIDGOHPEKAFOLOCPAV*

NbL09g05050.1 uncharacterized_LOC104247359 [MW = 39.88 kDa]

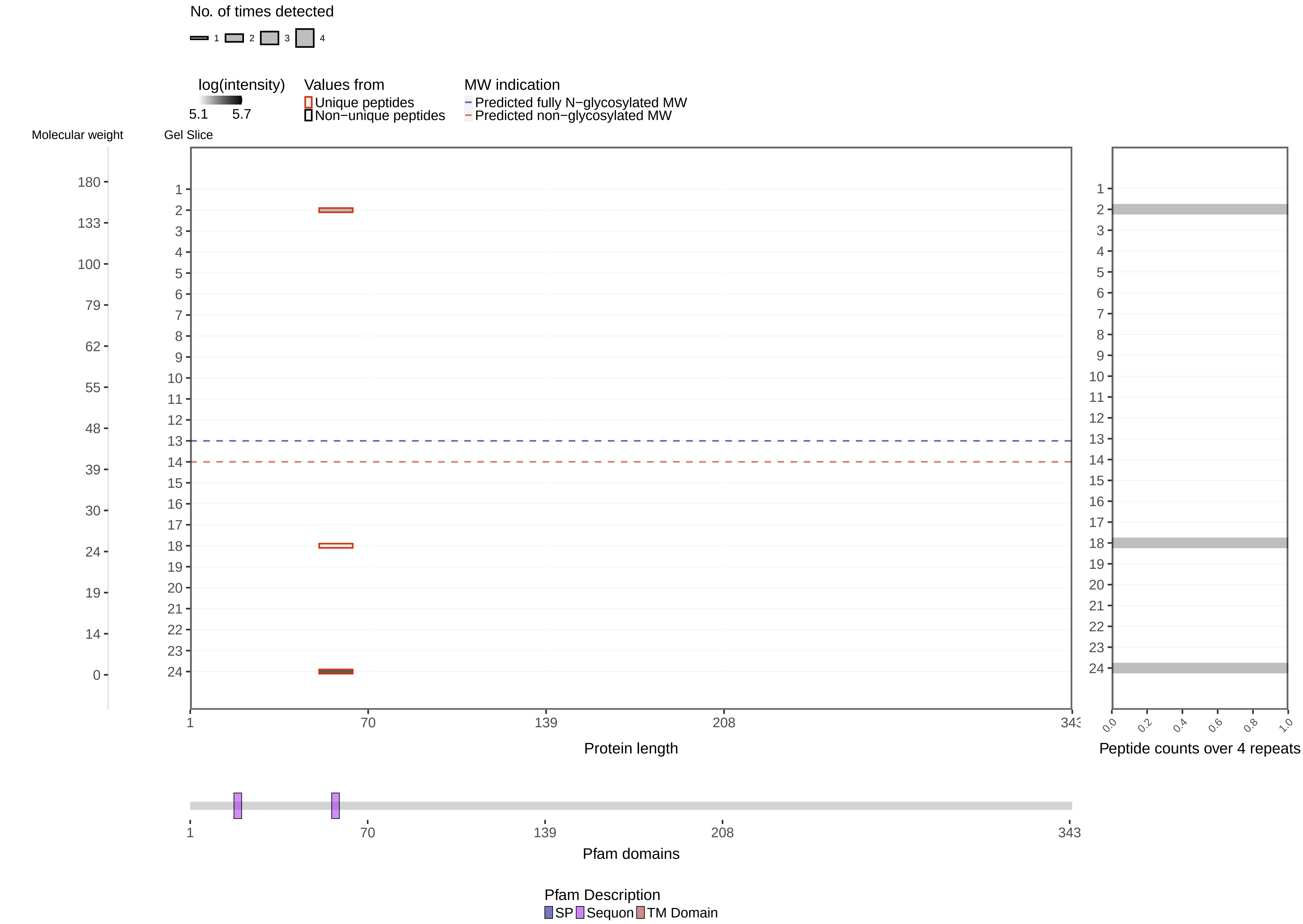


Protein sequence:

DAEYGSQGLAESSISSDAVKHIGEIAEQPYHRKTADFWGYVFQIIFQMNAGAVMLTDCVFWFILVPYLTIKGYKLNFWIINMHSVNVIFLLSDTALNCLRFPWFRIGYFFIWTTGYVIFQWAVHACISIWWPYPFLDLSSALAPLWYLLVAFMHIPCYGIFVLVMKLKHHLFSKWFPQSYQCAR*

### NbL09g04950.1 60S_ribosomal_protein_L27a−3−like [MW = 19.69 kDa]

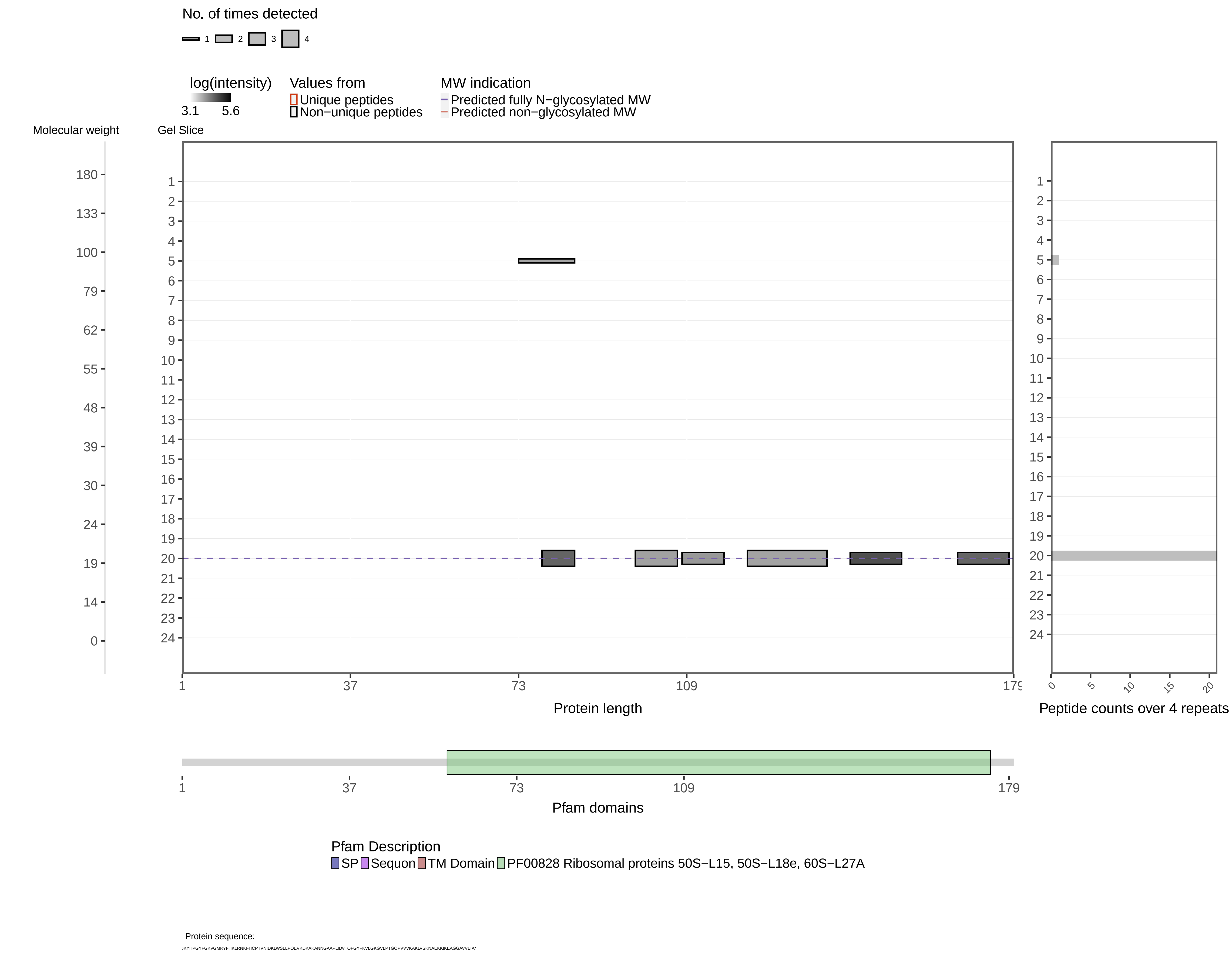

### NbL09g04760.1 IAA−amino_acid_hydrolase_ILR1−like_5 [MW − SP = 44.57 kDa]

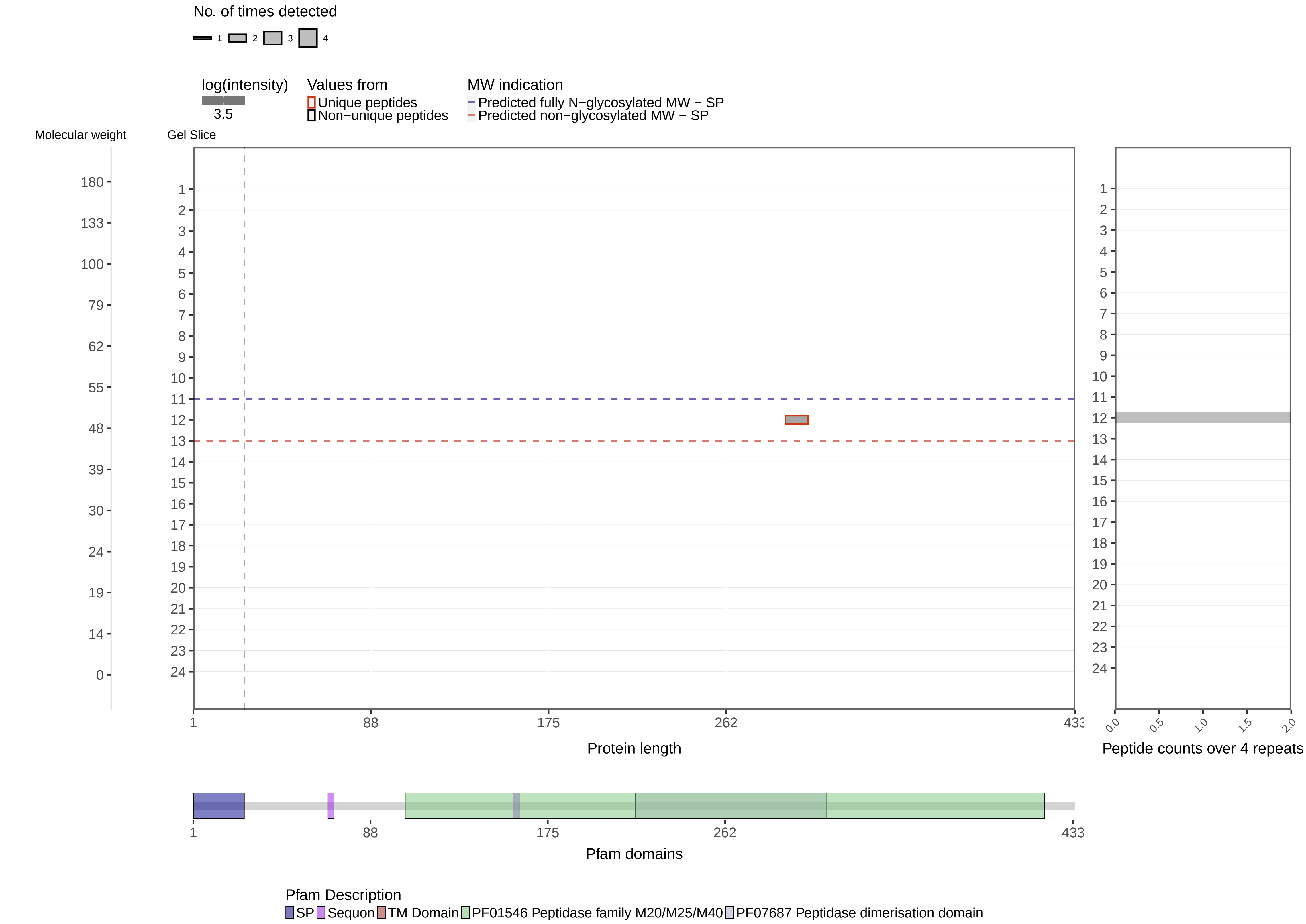


Protein sequence:

GSIGTSPGPILAAVSFFEAKIEGKGGHAAEPHATVDPILAASFAIVALQQLISREVDPLHSQVLSVTYVRGGSASNVIPPYVEFGGTLRSLTTEGLLQLQKRVKEVIEGQAAVHRCKAYIGMKEEDFPAYPACINDERLHQHVGRVGKLLLGSENIKEAEKVMAGEDFAFYQELTPGVMFQIGIRNEKVGSTHAPHSPHFFLDEDVLPIGVALHTAIAEMYLNDYQHPIAV*

### NbL09g04620.1 RNA−binding_protein_2−like [MW = 25.38 kDa]

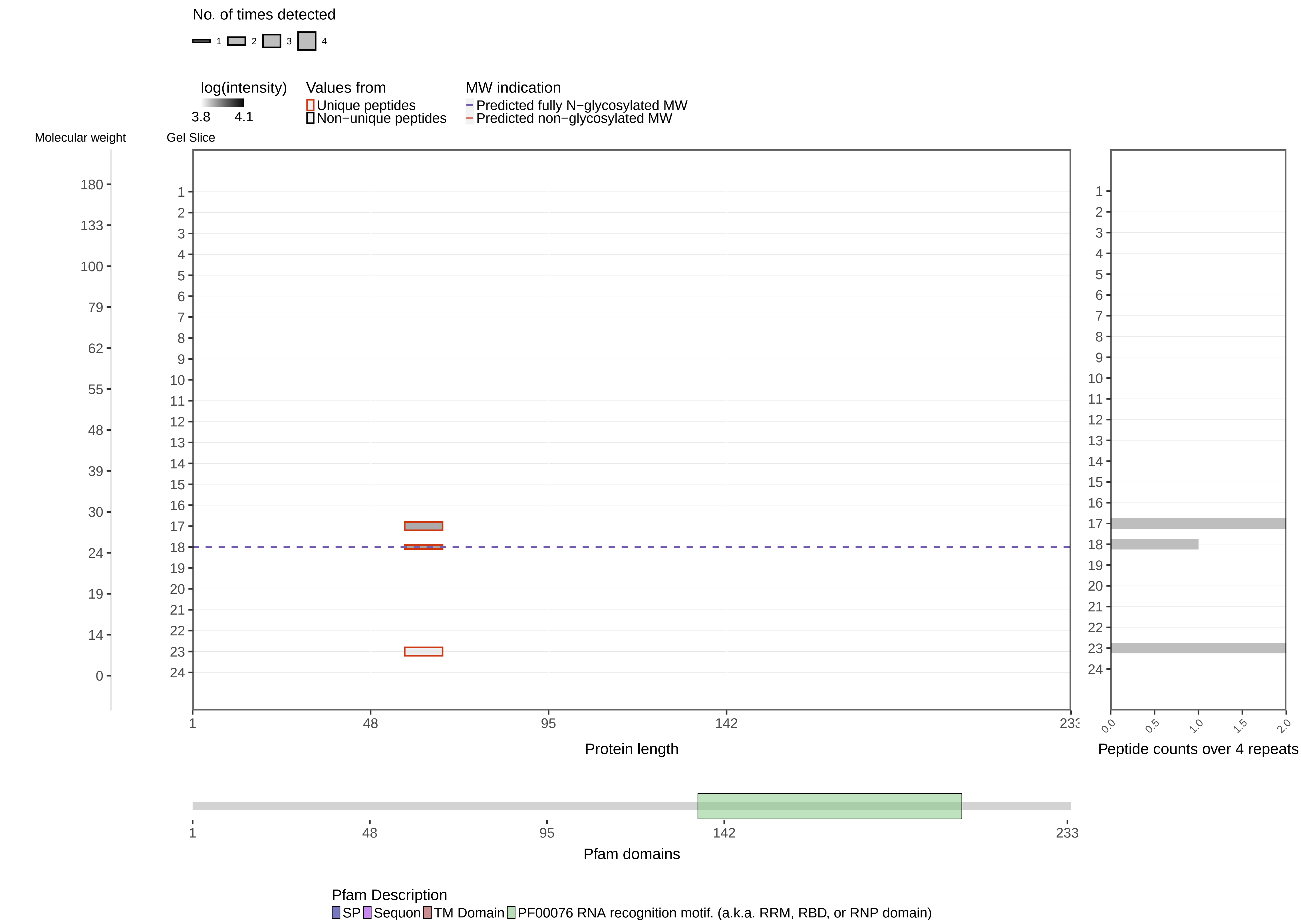


Protein sequence:

GPDPLALKSRNVGMVGGRPEISLPPDASSTLFVEGLPADCTRREVSHIFRPFVGYKEVRLVPKGSRHLGGDPLILCFVDFVTPLRAAAAMDALOGYKFDEHDRDSVSLRMOYARNPGARSGGGNRGKR*

### NbL09g04550.1 importin_subunit_beta−1−like [MW = 96.14 kDa]

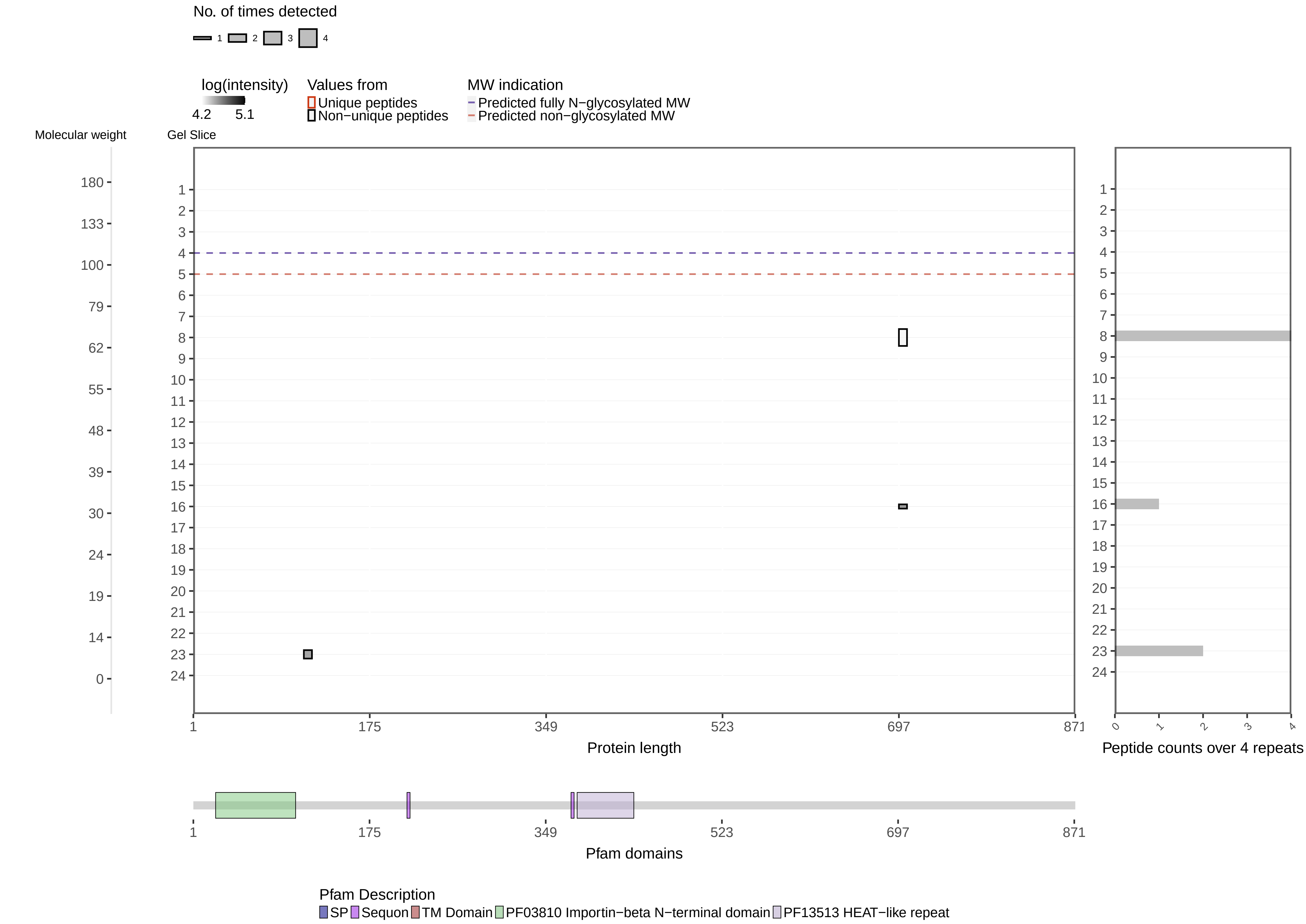


Protein sequence:

[illegible]

NbL09g04290.1 T−complex_protein_1_subunit_gamma−like [MW = 67.82 kDa]

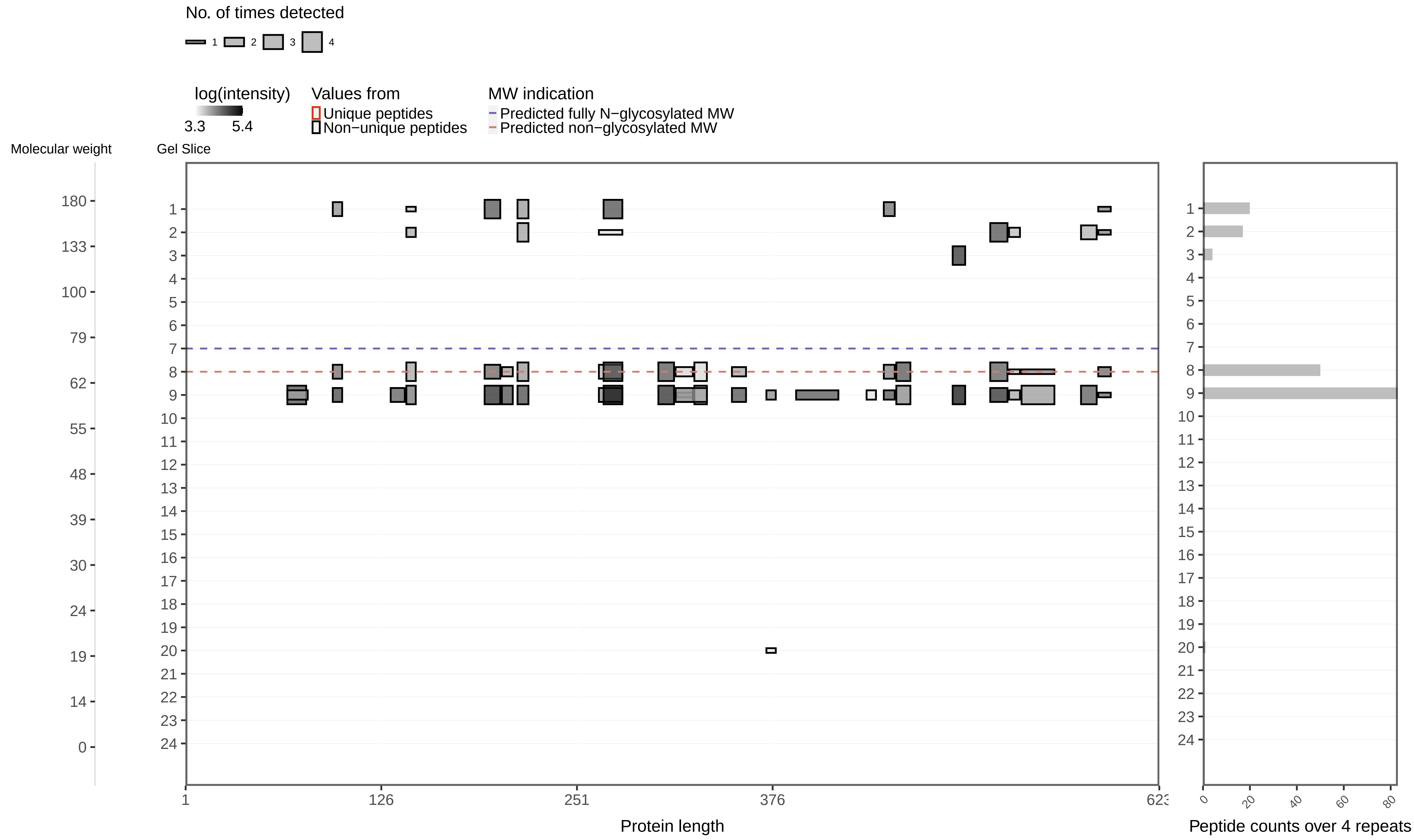


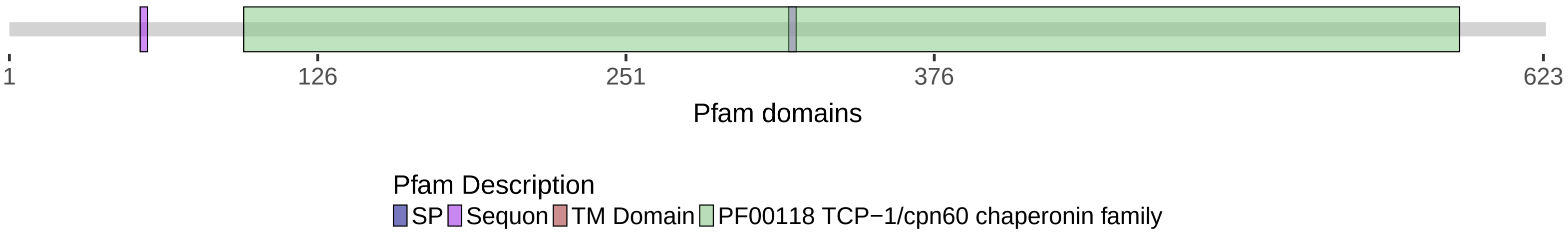


Protein sequence:

IPRIILLDSPLEYKKGENQTNAELLREEDWNVLLKMEEEYIENMCAQILKFKPDVVITEKGLSDLACHYLSKAGVSAIRRLRKTDNNRIAKACGAVIVNRPDELQESDVGTGAGLFEVKKIGDEFFTFIVDCKDPKACTVLLRGASKDLLNEVERNLQDAMSVARNIIKNPKLVPGGGATELTVSAMLKQKSSSIEGIEKWPYGAAAIAFEAIPRTLAQNCGVNVIRTMTALQGKHANGENAWIGIDGNTGEIADMKEQKIWDSYTVKAQAFKTAIEAACLLLRIDDIVSGIKKKQAPG

### NbL09g04030.1 probable_phosphoinositide_phosphatase_SAC9 [MW = 179.04 kDa]

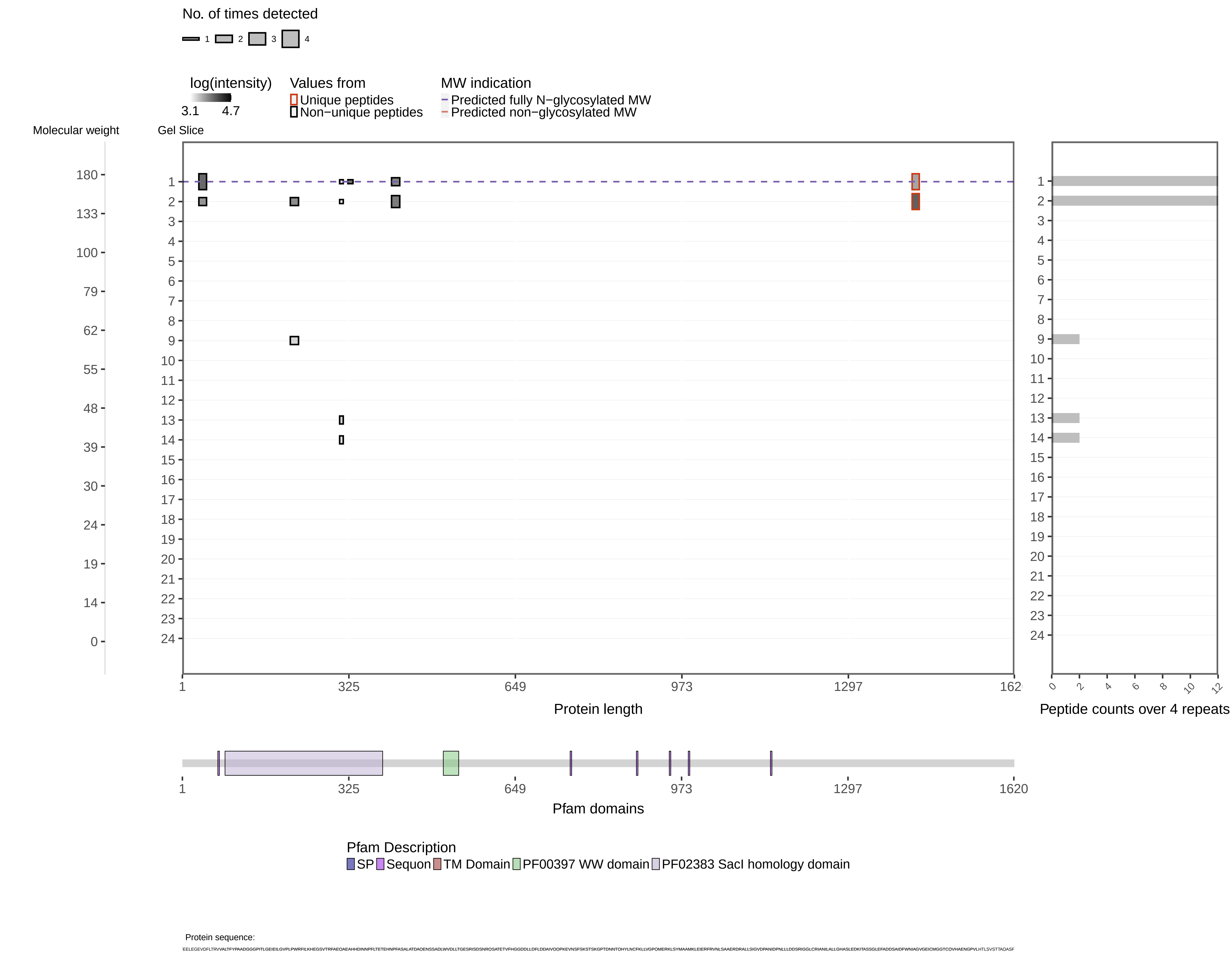

### NbL09g04010.1 peroxidase_47−like [MW − SP = 33.84 kDa]

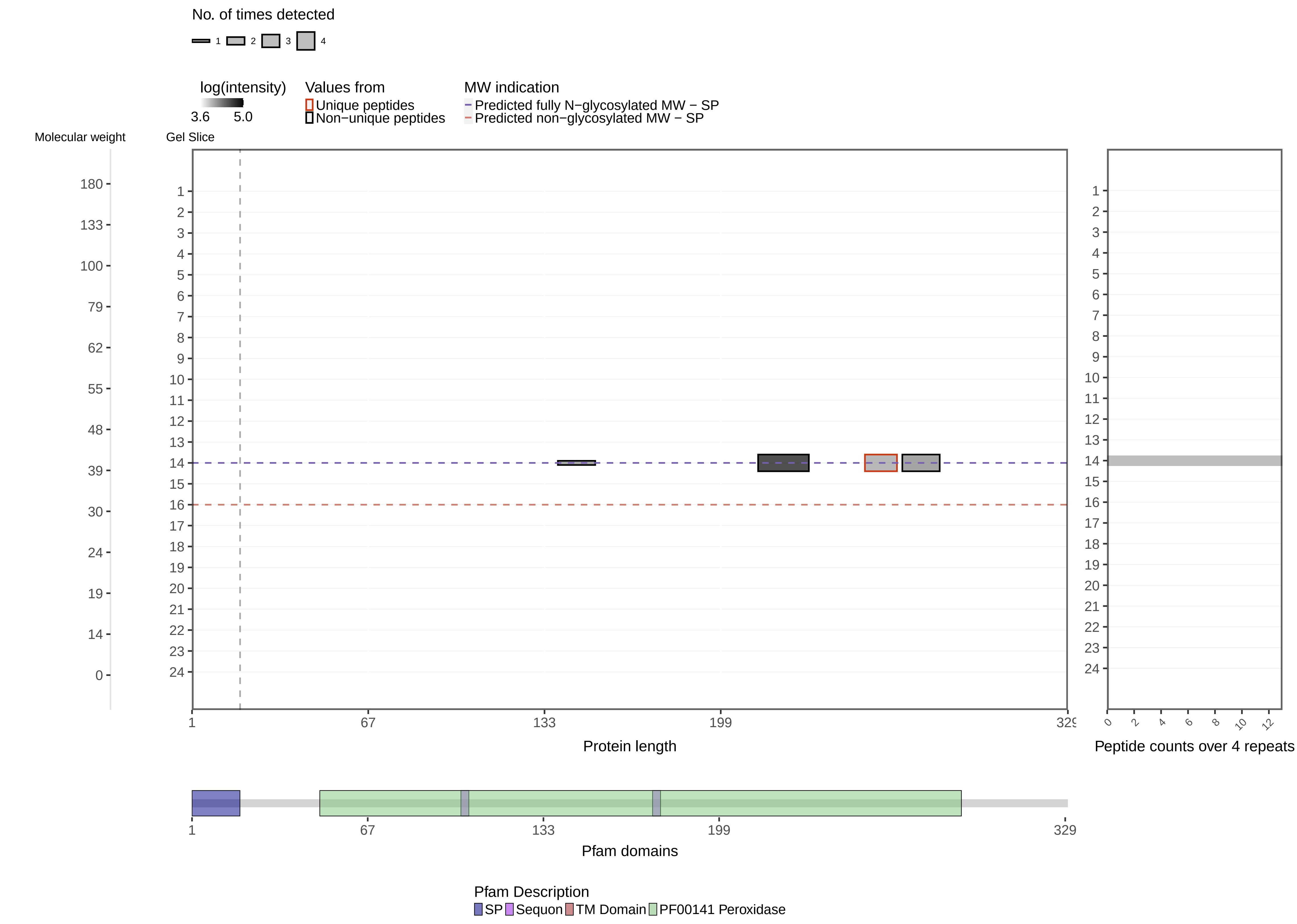


Protein sequence:

KGRKDGRRSRIEDTINLPSPALNSSELIRIFAQHGFSAQEMVALSGAHTLGVARCSSFKHRLSNFDSTHDVDPTLDAQFAKTLSKTCGGGDKSEQAFDTTKNDFDNNYFTALQRRSGVLFSDQTLYNSPRTRGIVNGYAFNQAMFFLDFQQAMVKMGLLDVKEGLDGEVRSNCRIIN*

NbL09g04000.1 probable_calcium−binding_protein_CML20 [MW = 21.22 kDa]

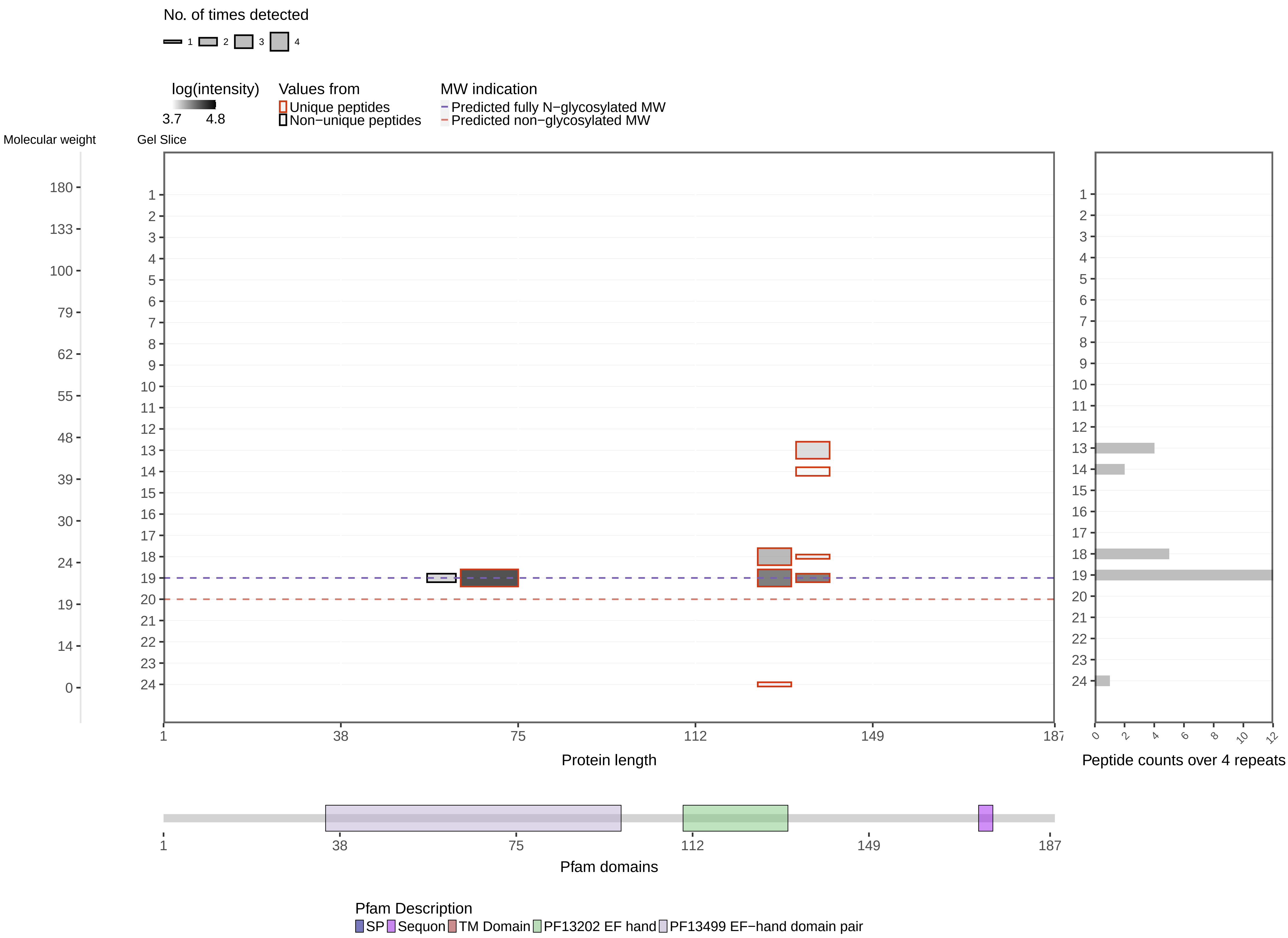


Protein sequence:

VDKDGSGAIDFDEFVHMMTAKFGERDTKEELKKAFDVIDQDKNGKISFADIQRIADELGERFTDREIQEMIEAADQDRRVCSLISTSLWMLLNLTITHSSAPLCSLP*

NbL09g03970.1 2−alkenal_reductase_(NADP(+)−dependent) [MW = 37.83 kDa]

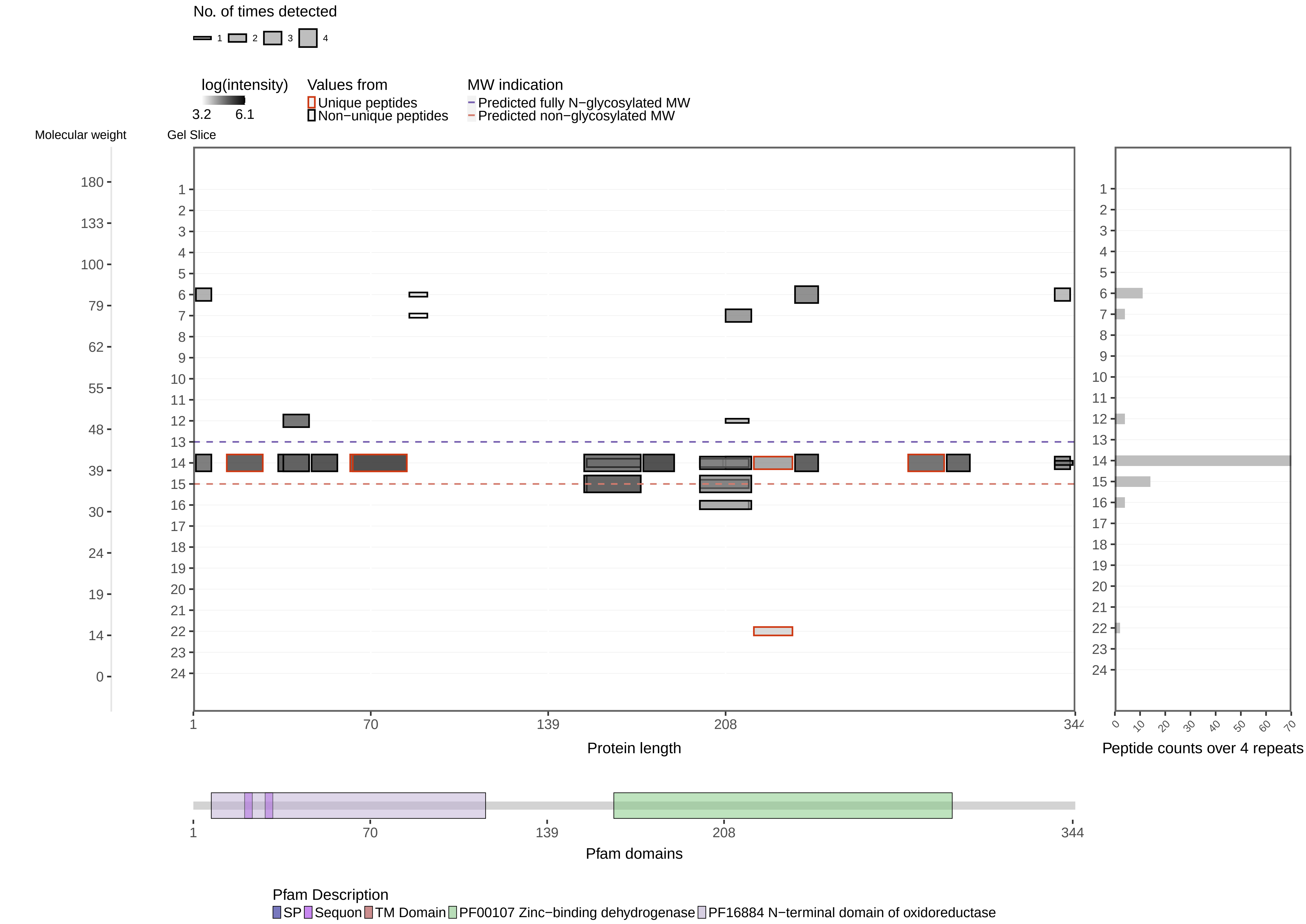


Protein sequence:

VSAASGAVGQLVGQFAKMLGCYVVGSAGSKEKVDLLKSKFGFDEAFNYKEEQDLNAALKRCFPDGIDVYFENVGGKMLDAVLLNMKLHGRIAACGMISQYNLEQNEGVHNLFCLITKRIRMEGLVVSDYYHLYPKYLEMVIPQIKAGKVVYVEDVADGLEIAPSALVGLFSGRNIGKQVVIVSRE*

### NbL09g03860.1 valine−−tRNA_ligase_chloroplastic/mitochondrial_2 [MW = 109.88 kDa]

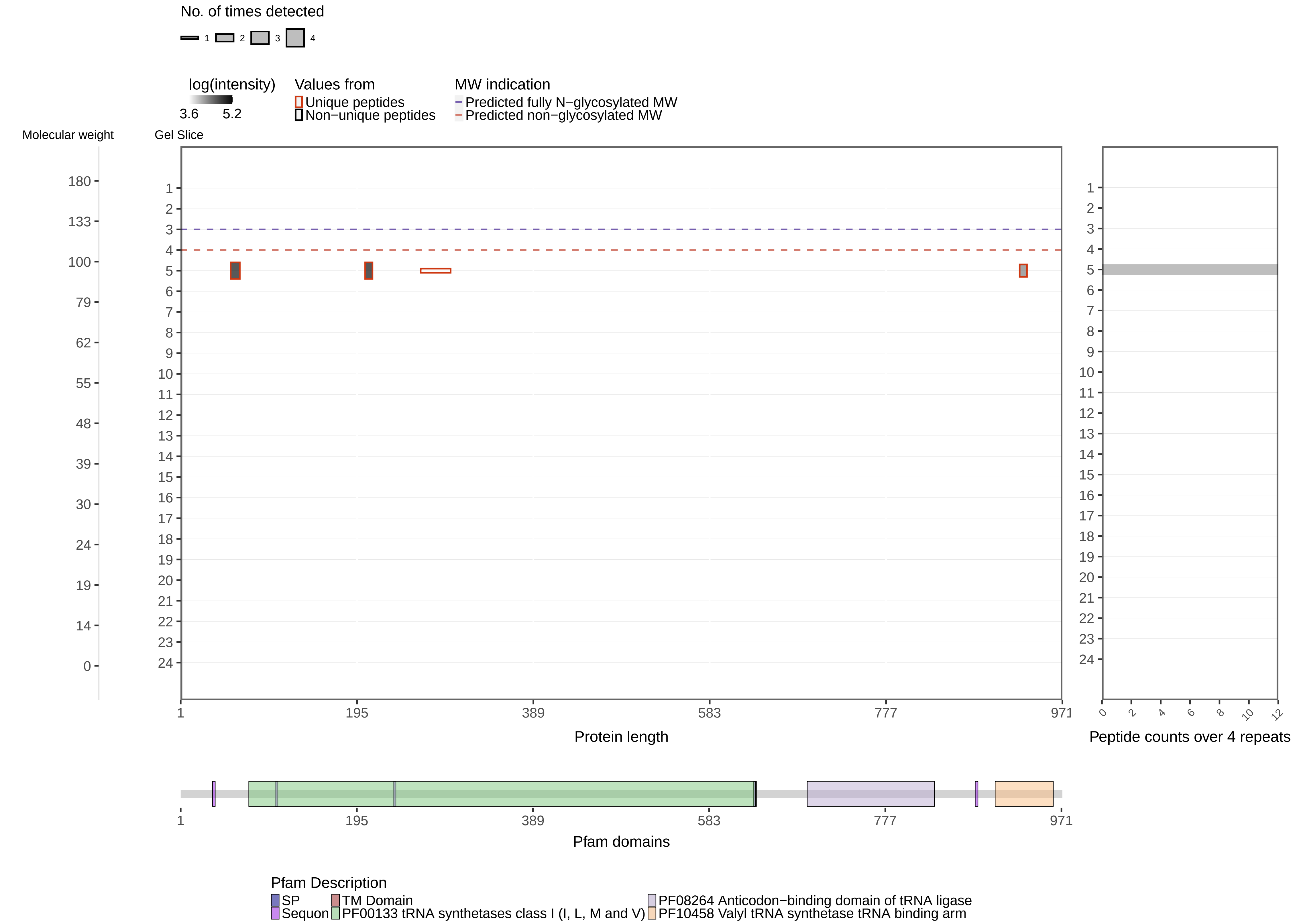


Protein sequence:

VGHRIPVWYASGKDCEEEYIVARSHREALSKAQEKYGKNVEIYQDPDVLDTWFSSSLWPFSTLGWPDESAEDFKRFYPTSVLETGHDILFFWVARMVMMGIEFTGTVPFSNVYLHGLIRDSQGRKMSKTLGNVIDPLDTIAEYGTDALRFTLALGTAGQDLNLSMERLTSNKAFTNKLWNAGKFILRNLPHLDDTVAWEALLAHKFDKVESVVKLPLPECWVVSKLHVLVDEVTASYEKLFFGDVGREIYDFFWADFADWYIEVSKARLYHSGDHSVSSVSQAALLYIFGTIL

NbL09g03640.1 ABC_transporter_D_family_member_2_chloroplastic [MW = 80.38 kDa]

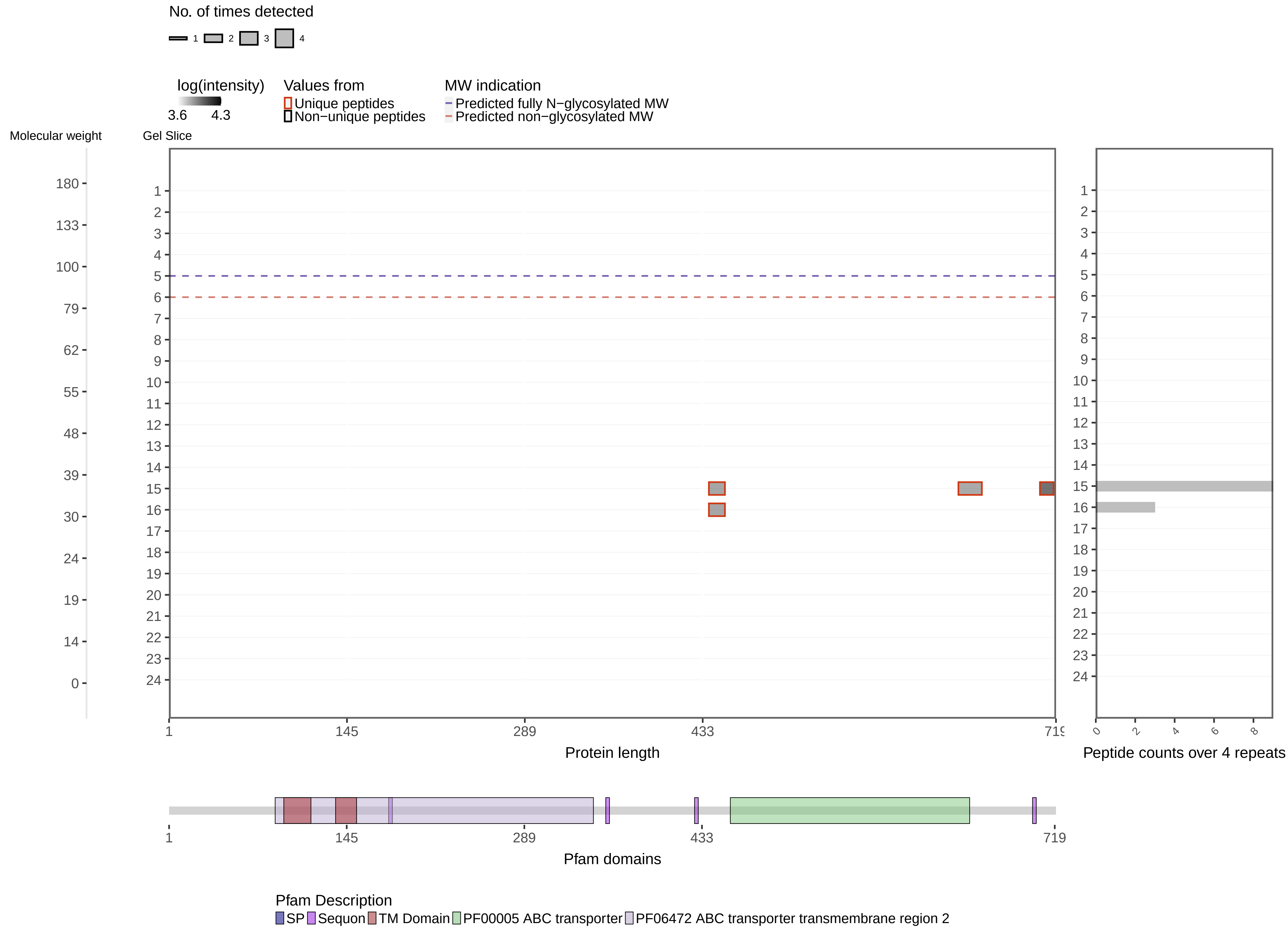


Protein sequence:

KIEFGVVNQSASAFNHILSDFSLIVYQFQAISAFSAVIDRLGEFDDVLDSSNRKKGDHDLKEEIQLNFCLISCSNGLQSNGSMPLSSRVKLLDVEQLTVQTPSKATLIKDLSFEIYEKDHLLITGPSGSGKTSLLRAVAGLWNFGSGAITIYVRPRGELDMPRSPDVISGQVTSTNEIIWDSTGRRNSEGVIFLPQKPYMVLGTLRQQLLYPTWTEDSDNVVDDAKQSGDVGCVNERPNQPTSDDLIRVLNDVSLRHLLSRFDGLDSTYEWSSVLSLGEQQRLAFARLLLAKPTLA

NbL09g03290.1 fasciclin−like_arabinogalactan_protein_1 [MW − SP = 26.26 kDa]

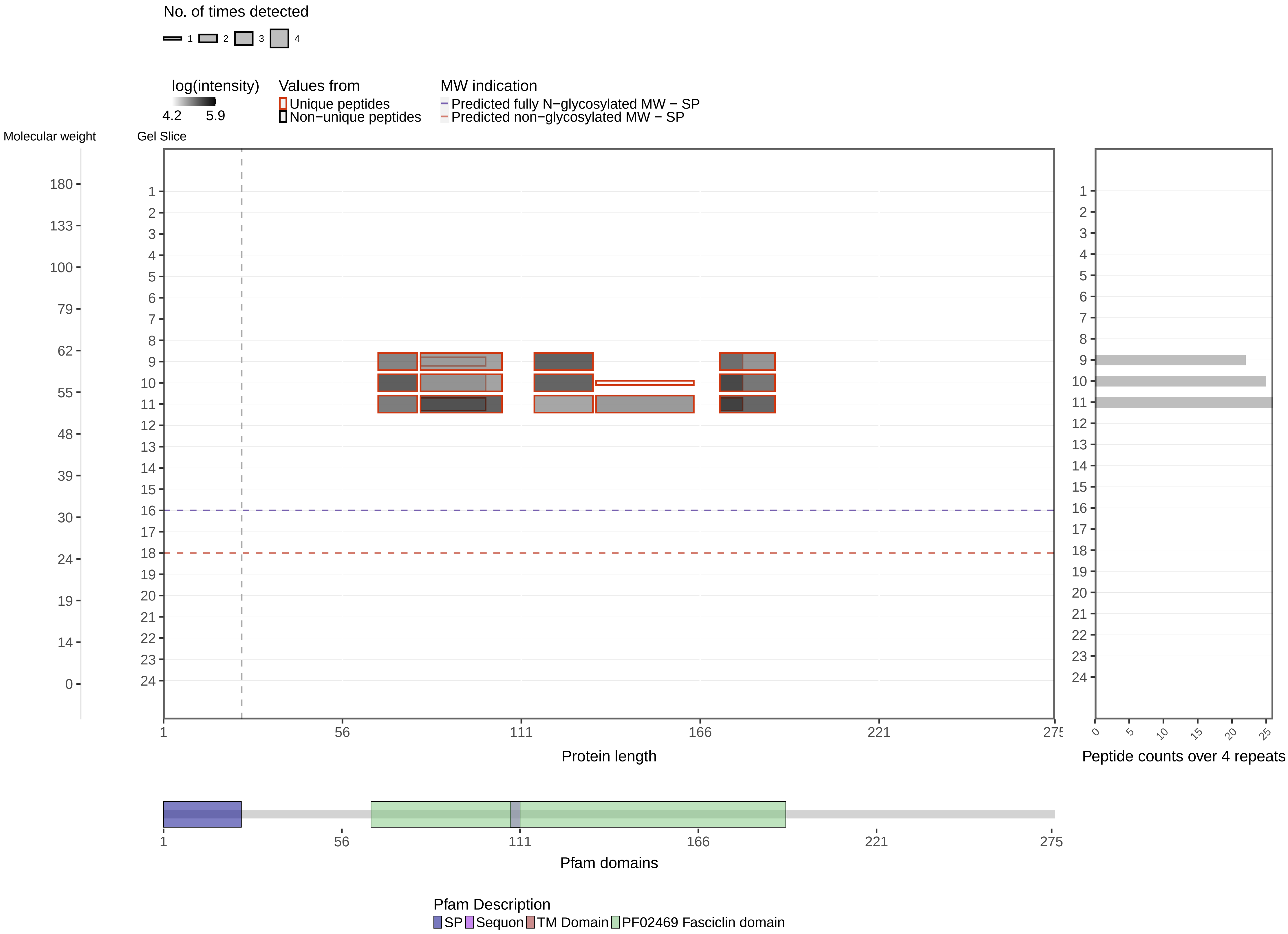


Protein sequence:

PVYQSLSSLRTSNGLMNTLATDGANNYDFVVQNDGQDVTLRTKIDTAKITGTVIDKEPLAIFSIDKVLEPKELFKGAPTPAPAPAPIAESPKPALKKHKSPPAPASPADSPSDGPTDDVADSKADKNGAFRYNAGALVVPVLCIWFGHLLL*

### NbL09g03050.1 porphobilinogen_deaminase_chloroplastic−like [MW = 40.08 kDa]

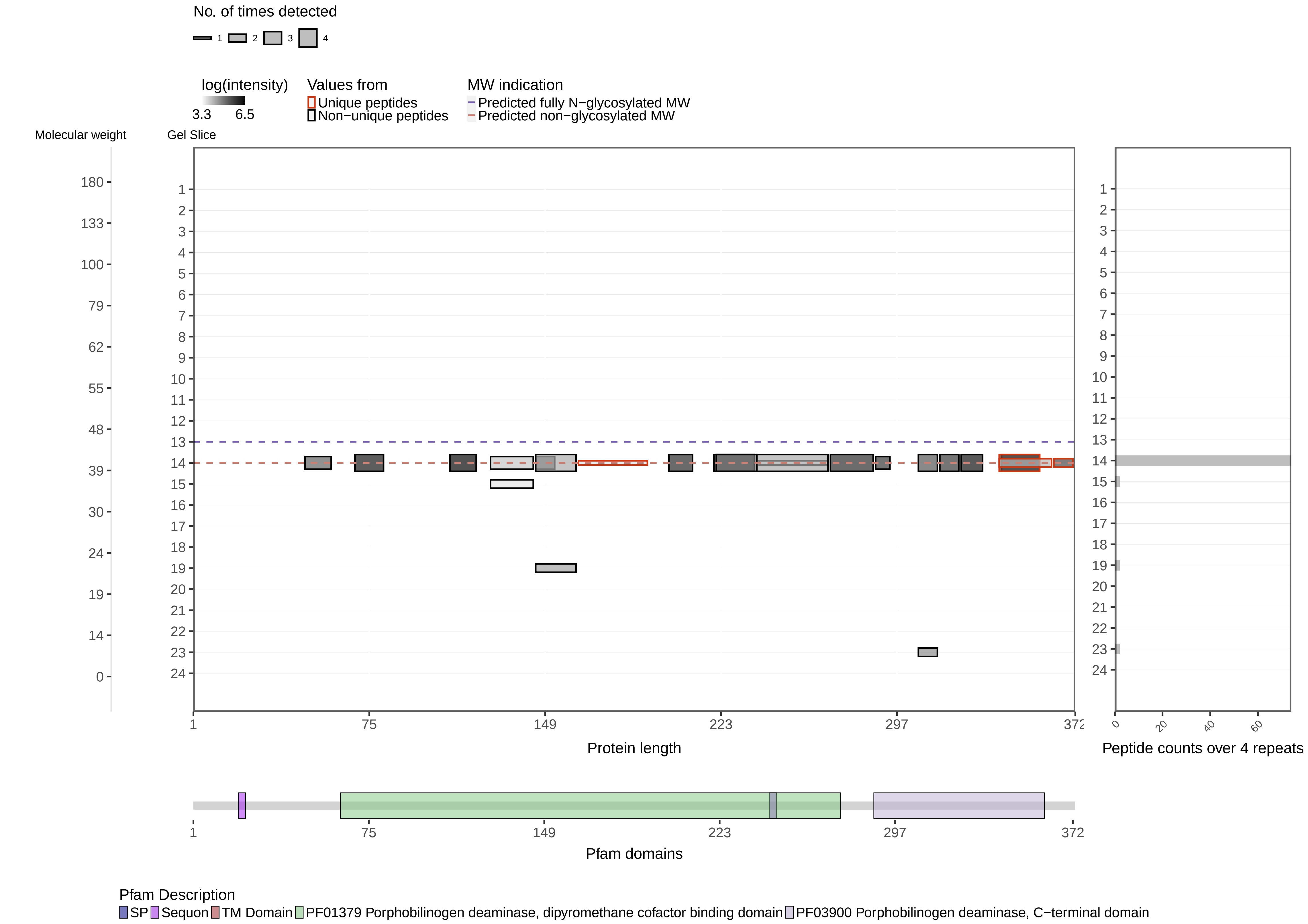


Protein sequence:

AQLPSGSTVGTASLRRKSQILHRYPSLNVLENFRGNVQTRLRKLNEGVVQATLLALAGLKRLDMTENVTSILPIEDMLPAVAQGAIGIACRSDDETMANYIALLNHEETRLAVACERAFLKTLDGSCRTPIAGYACRGEDGDCIFKGLVASPDGTRVLETSRKGPYSFEDMTVMGEDAGKELLSRAGPGFFGTK*

### NbL09g03010.1 phosphoglycerate_kinase_cytosolic−like [MW = 42.48 kDa]

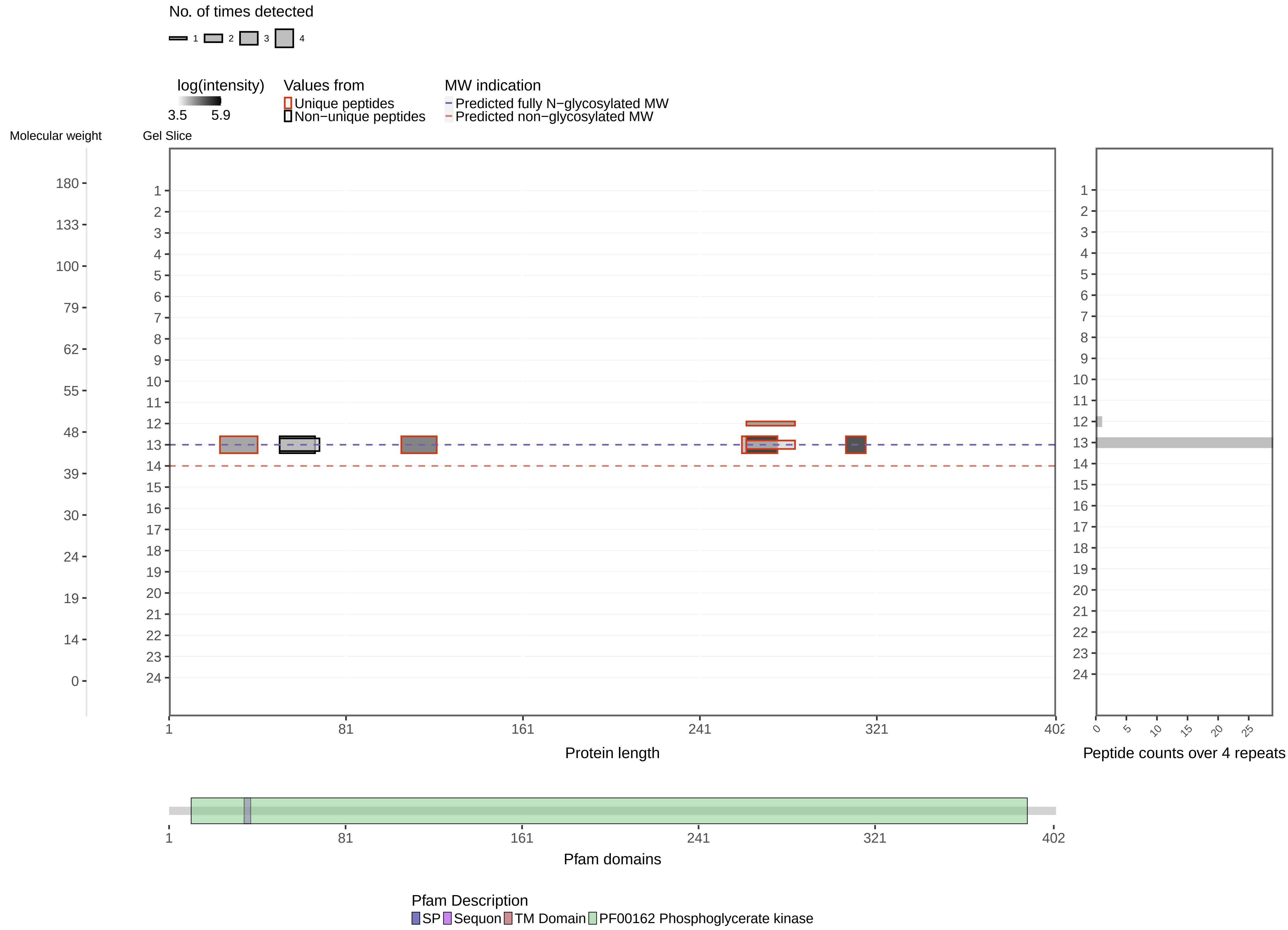


Protein sequence:

VSNPQKPFAAIVGGSKVSSKIGVIESLLEKVNVLLLGGGMIFTFYKAQGYSVGSSLVEEDKLDLATSLIEKAKAKGVSLLLPADVVIADKFAADANSKIVPASEIPDGWMGLDIGPDAIKSFGEALDATKTIIWNGPMGVFEFEKFAAGTEAIAKKLAELSGKGVTTIIGGGDSVAAVEKVGLADKMSHISTGGGASLELLEGKQLPGVLALDDA*

### NbL09g02980.1 glucan_endo−13−beta−glucosidase_12−like [MW − SP = 49.49 kDa]

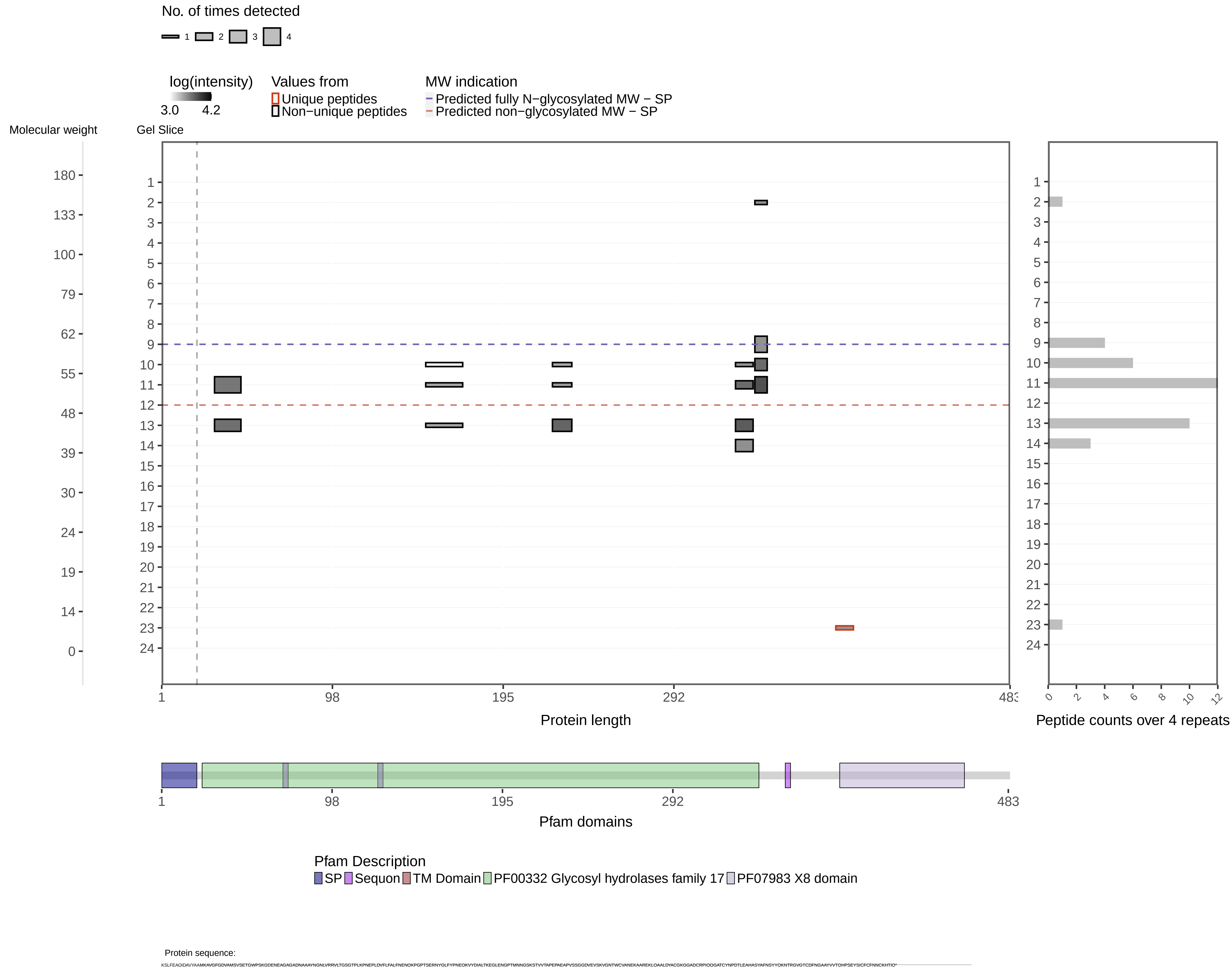

NbL09g02930.1
dihydrolipoyllysine−residue_succinyltransferase_component_of_2−oxoglutarate_dehydrogenase_complex_1_mitochondrial−like
[MW = 51.42 kDa]

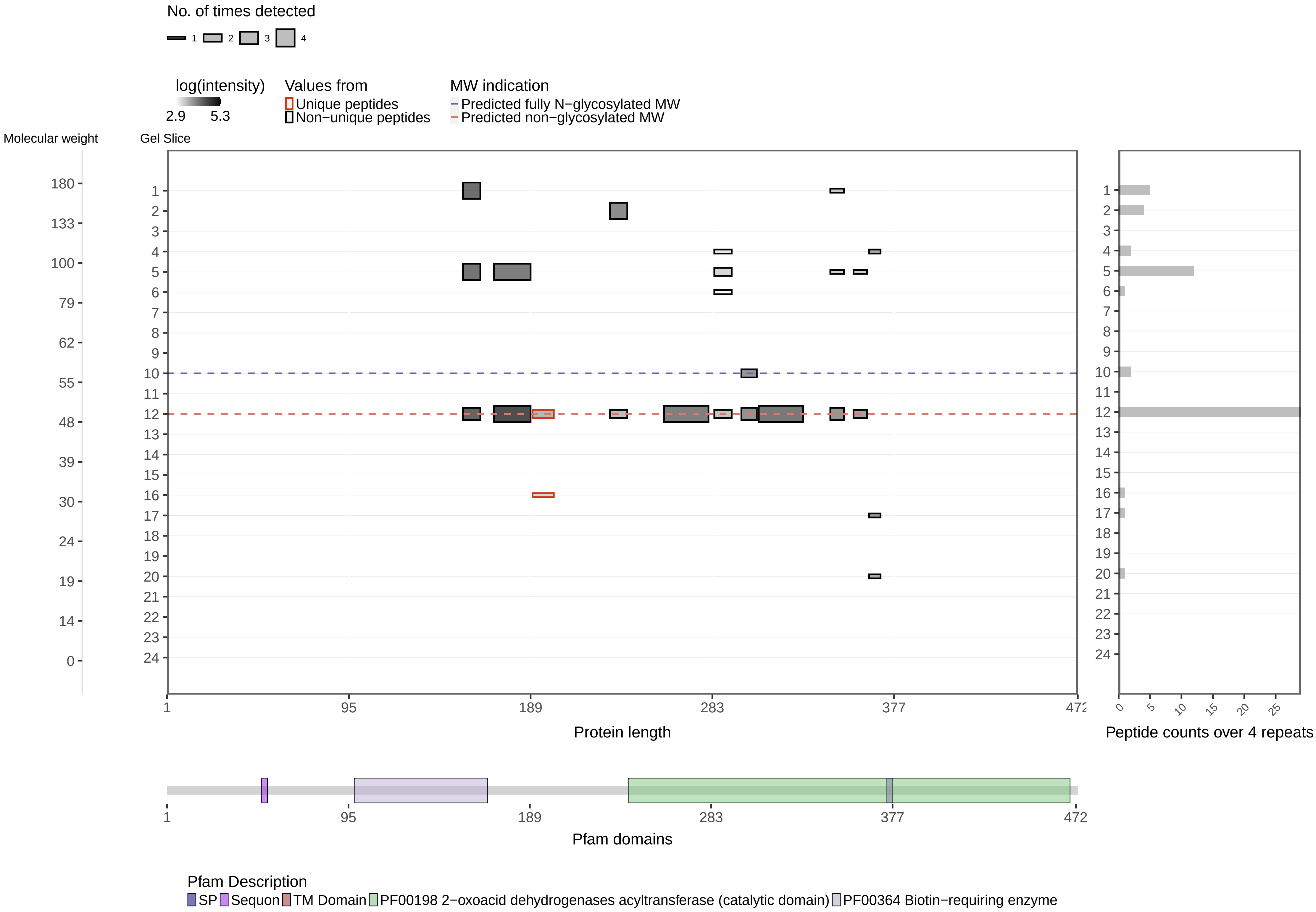


Protein sequence:
TSPPPPKRSATEPQLPPKERERRVPMTRLRKRVAIRLKDSQNTFALLTTFNEVDMTNLMKLRSDYKDAFVEKHGVKLGLMSGFVKAAVSALQNQPIVNAVIDGDDIIYRDYVDISIAVGTPKGLVVPVLRNAEKMNFAEIEKTINGLAKKATNGTISIDEMAGGSFTISNGGVYGSLLSTPIINPPQSAILGMHSIVNRPMVVGGNVVSRPMMYIALTYDHRLIDGREAVYFLRRIKDVVEDPRRLLLDV*

### NbL09g02790.1 ras−related_protein_RABB1c [MW = 26.66 kDa]

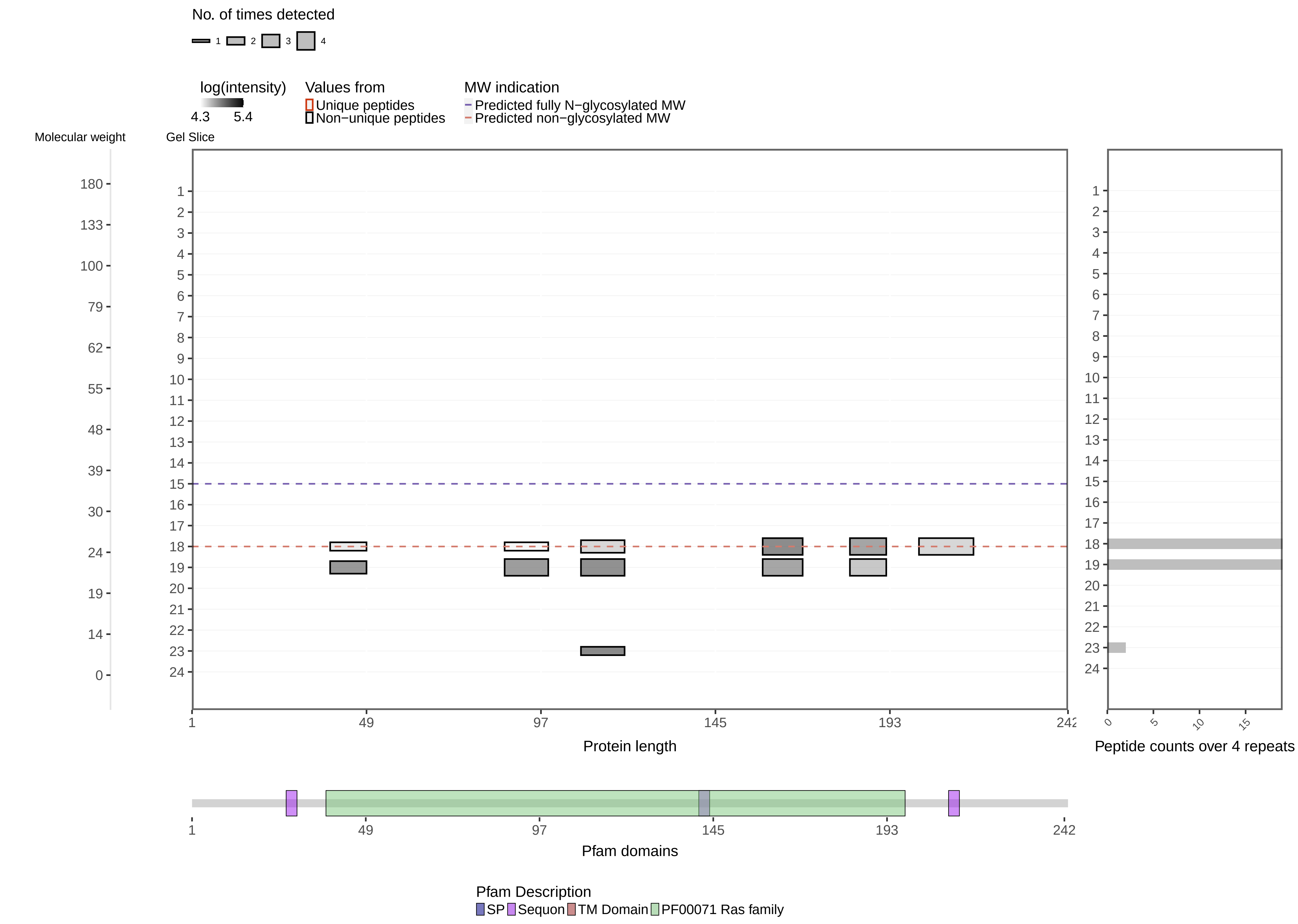


Protein sequence:

AGALLVYDITRRETFNHLASWLEDARQHANANMTIMLIGNKCDLAHRRAVSTEEGEQFAKENGLIFMEASAKTAQNVEEAFIRTASTIYKKIQDGVFDVSNESYGIKVGYGGIPGPSGGRDGVASQGGGCCS*

### NbL09g02750.1 elongation_factor_Ts_mitochondrial [MW = 43.29 kDa]

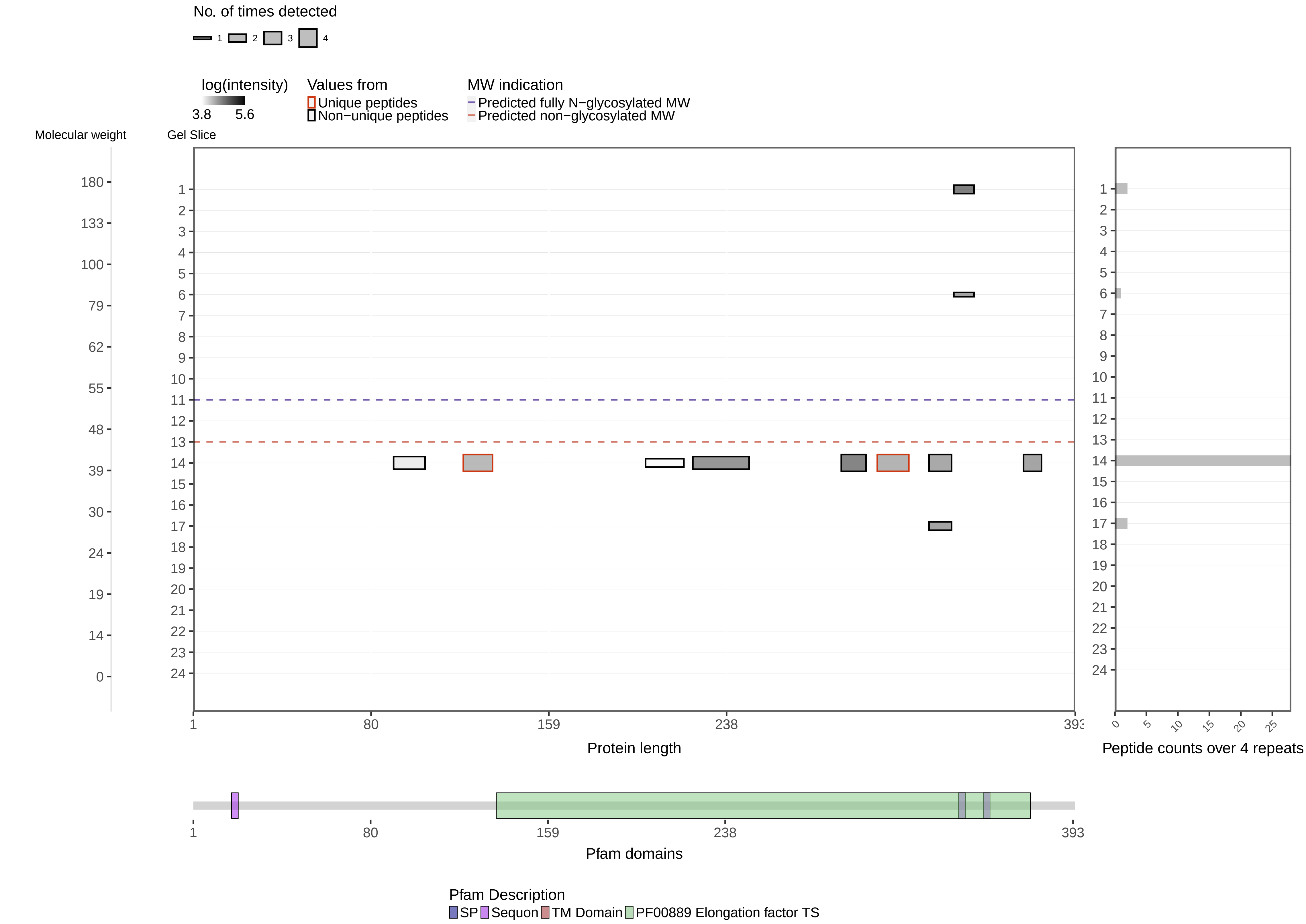


Protein sequence:

EHLNMNLDHPKLSGEKSVQNAITEVAAMMGENVKLRRGFAMSTPSIGVISTYLHTSPQPGVGRIAGILSLEVEDQNVPQDALQRVGSEIAMHVVAAKPLFLTKEDVSSDALSNEREILKSQAESLGRPQIAIEKMVEGRMRKYFEEVALMEQKFVVNDTINVKTLISNLSKEVGSPVKIGSFLRMEVGEGLQRLEASNENEPLANAA*

NbL09g02650.1 persulfide_dioxygenase_ETHE1_homolog_mitochondrial−like [MW = 33.04 kDa]

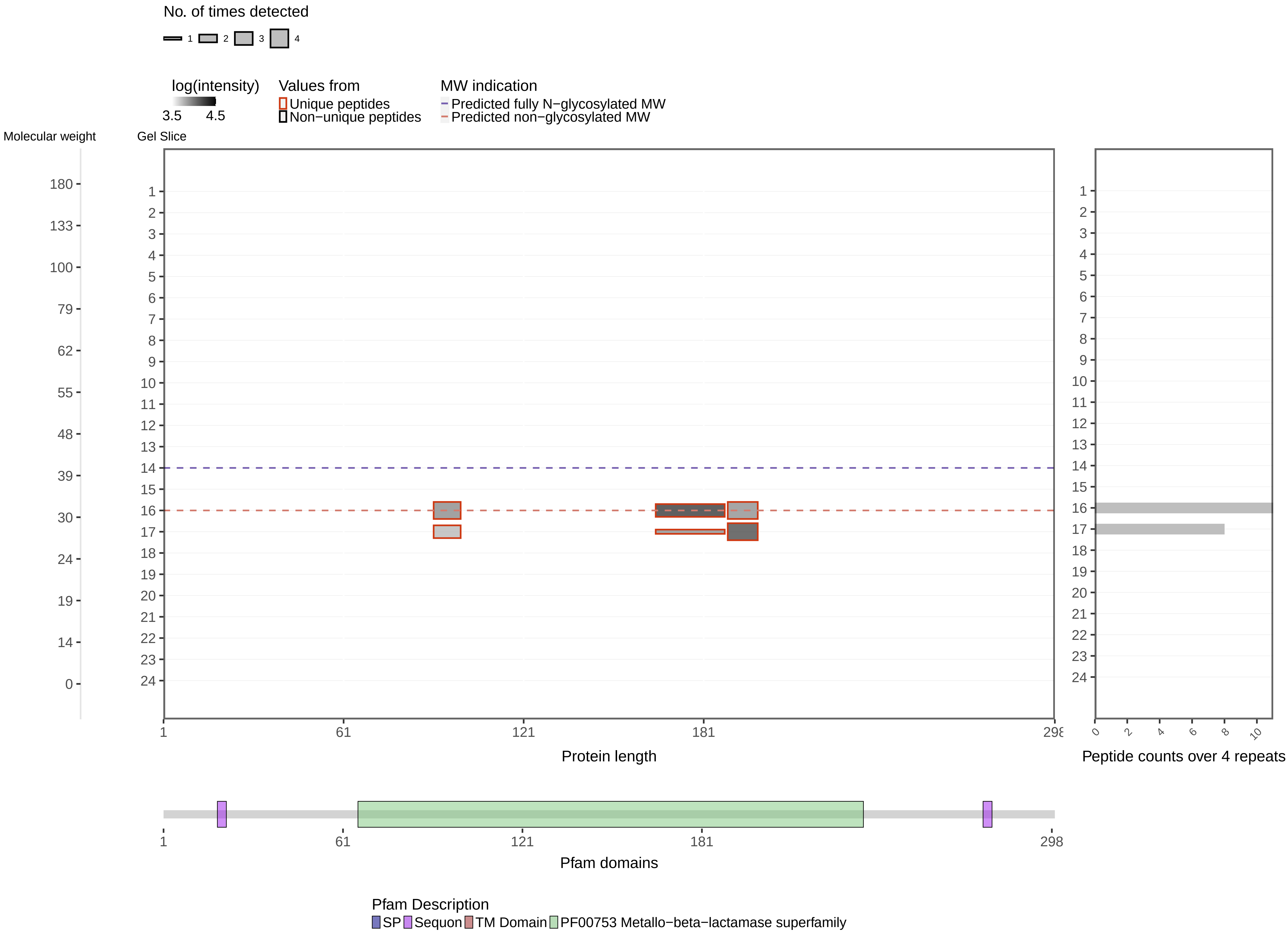


Protein sequence:
ASNAKADLFVEPGDKIYFGDIFLEVRATPGHTLGCVTYVTGDGPNQPQPRIAFTGDALLIRGCGRTDFQGGSSDKLYDSVHSQIFTLSKDTLVYPAHDYKGFTVSTVGEEMLYNPRLSKDKEQFLQETFNNIMQNLNLSYPKMMDVAVPSNMLCGLQET*

### NbL09g02630.1 a−b_binding_protein_13_chloroplastic [MW = 28.5 kDa]

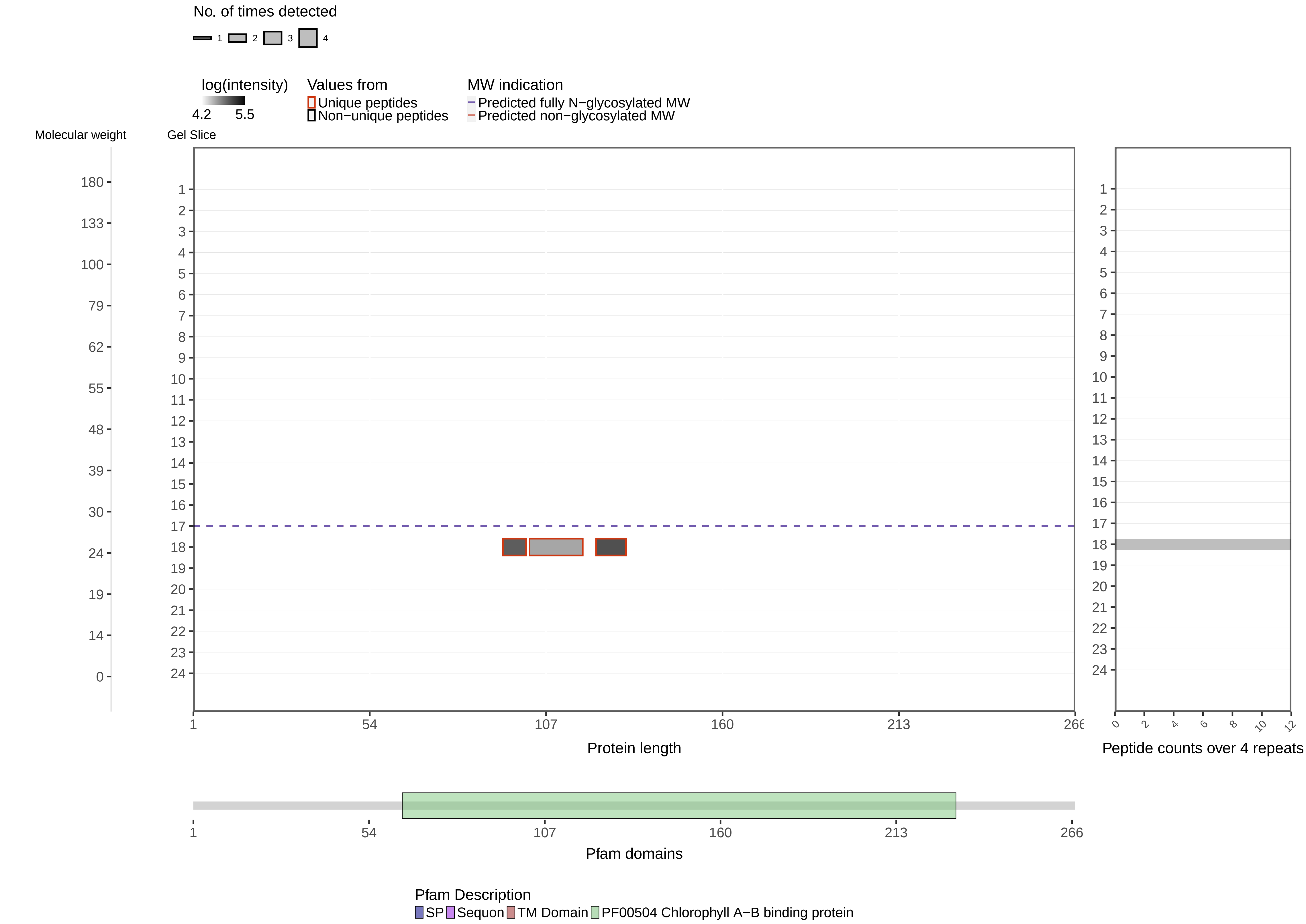


Protein sequence:

/VKVDFKEPVWFKAGAQIFSEGGLDYLGNPNLVHAQSILAVLGFQVVLMGLVEGFRINGLPGVGEGNNLYPGGQYFDPLGLADDPTTFAELKVKEIKNGRLAMFSMFGFFVQAIVTGKGPLENLLDHLDNPVANNAWVYATKFVPGA*

### NbL09g02580.1 peptidyl−prolyl_cis−trans_isomerase_CYP37_chloroplastic [MW = 53.15 kDa]

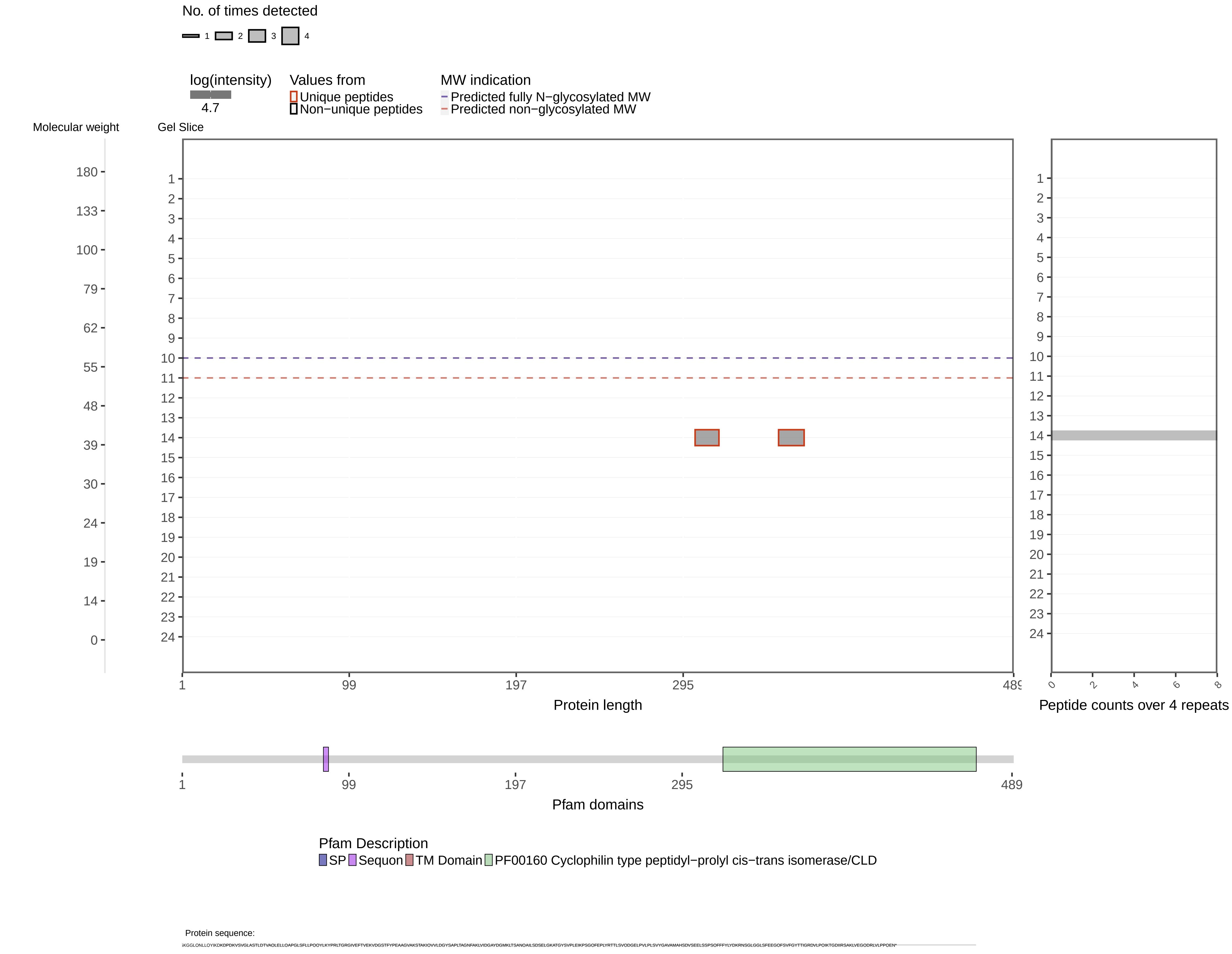

### NbL09g02350.1 early_nodulin−like_protein_2 [MW − SP = 34.11 kDa]

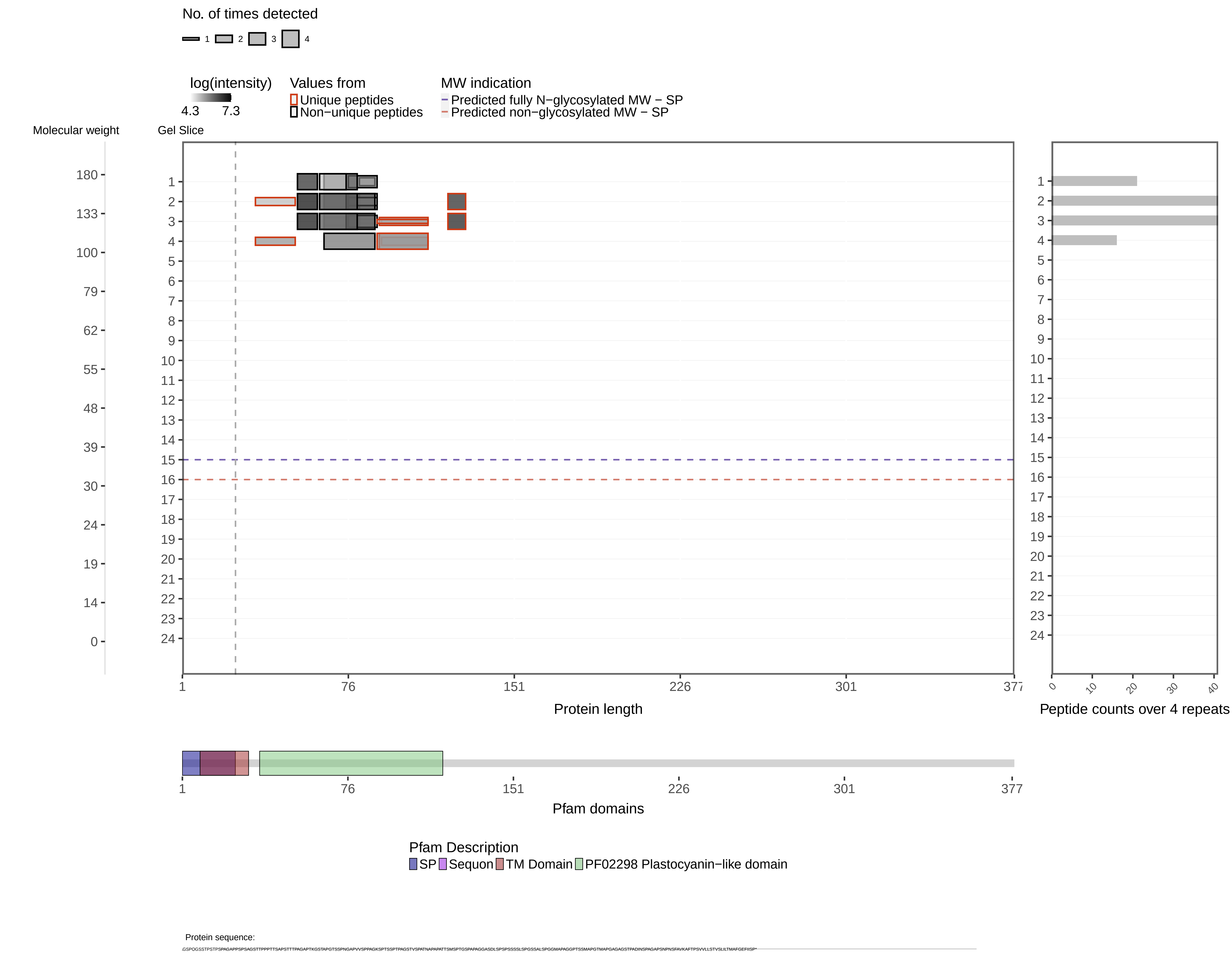

NbL09g02270.1 reticulon−4−interacting_protein_1_mitochondrial [MW = 39.65 kDa]

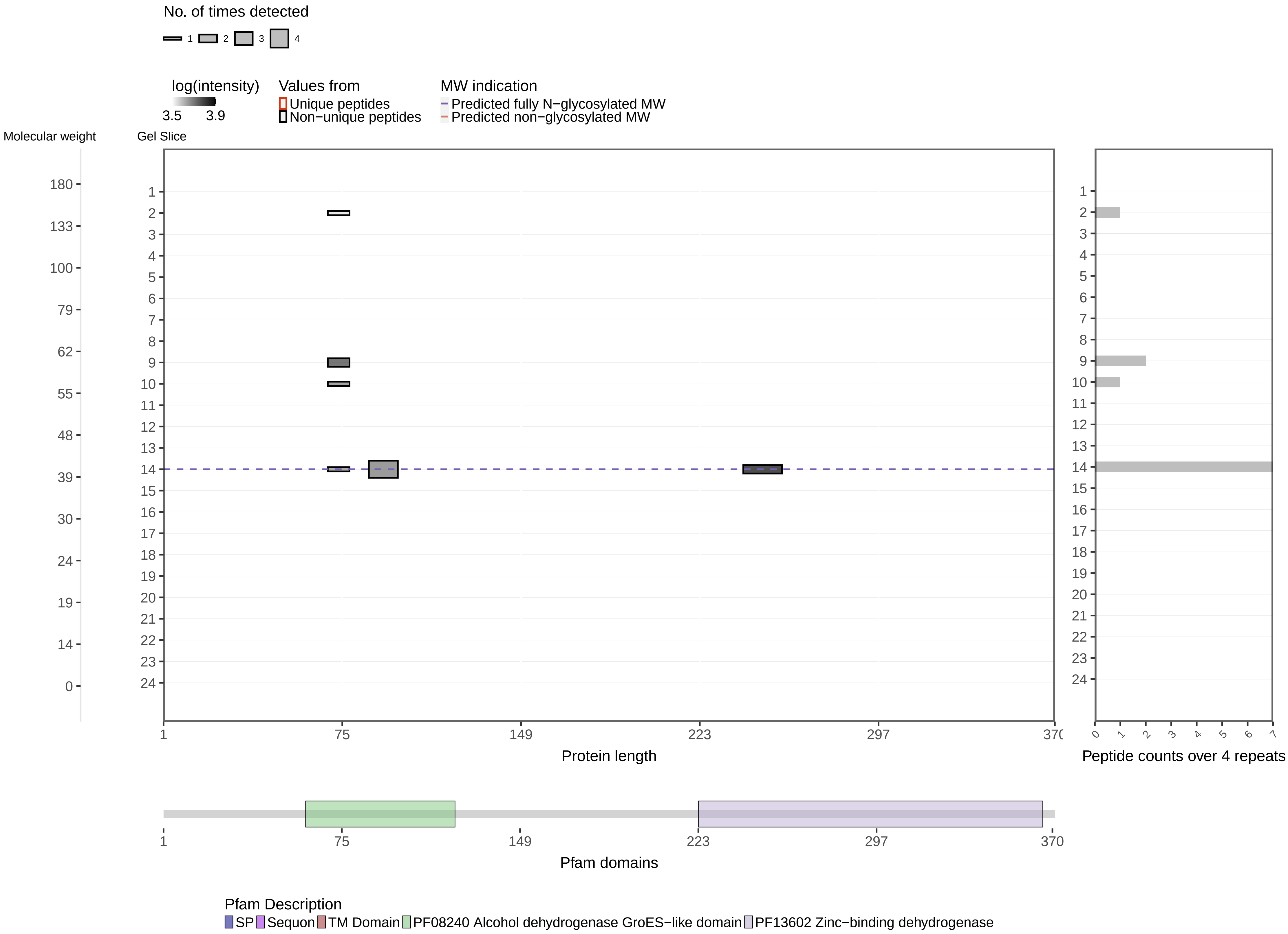


Protein sequence:

STARIREGHRVLVIGGGGAVGFSAIQLAVAAGCHVSSTCGESISISRILAAGAEQAVDYTTEDCEIALKGHFDAVLDTIGVPETERIGINLLKRGGHYMTLQGEAASLTDRYGLAFGLPLATAILLKKQIQYRYSHGIEYWWNYMRADAEGLHEIRRLSEAGKLRIPVDKTFPLSQAREAHEAKDRRQIPGKVVLEID*

### NbL09g02210.1 UBP1−associated_protein_2C−like [MW = 45.76 kDa]

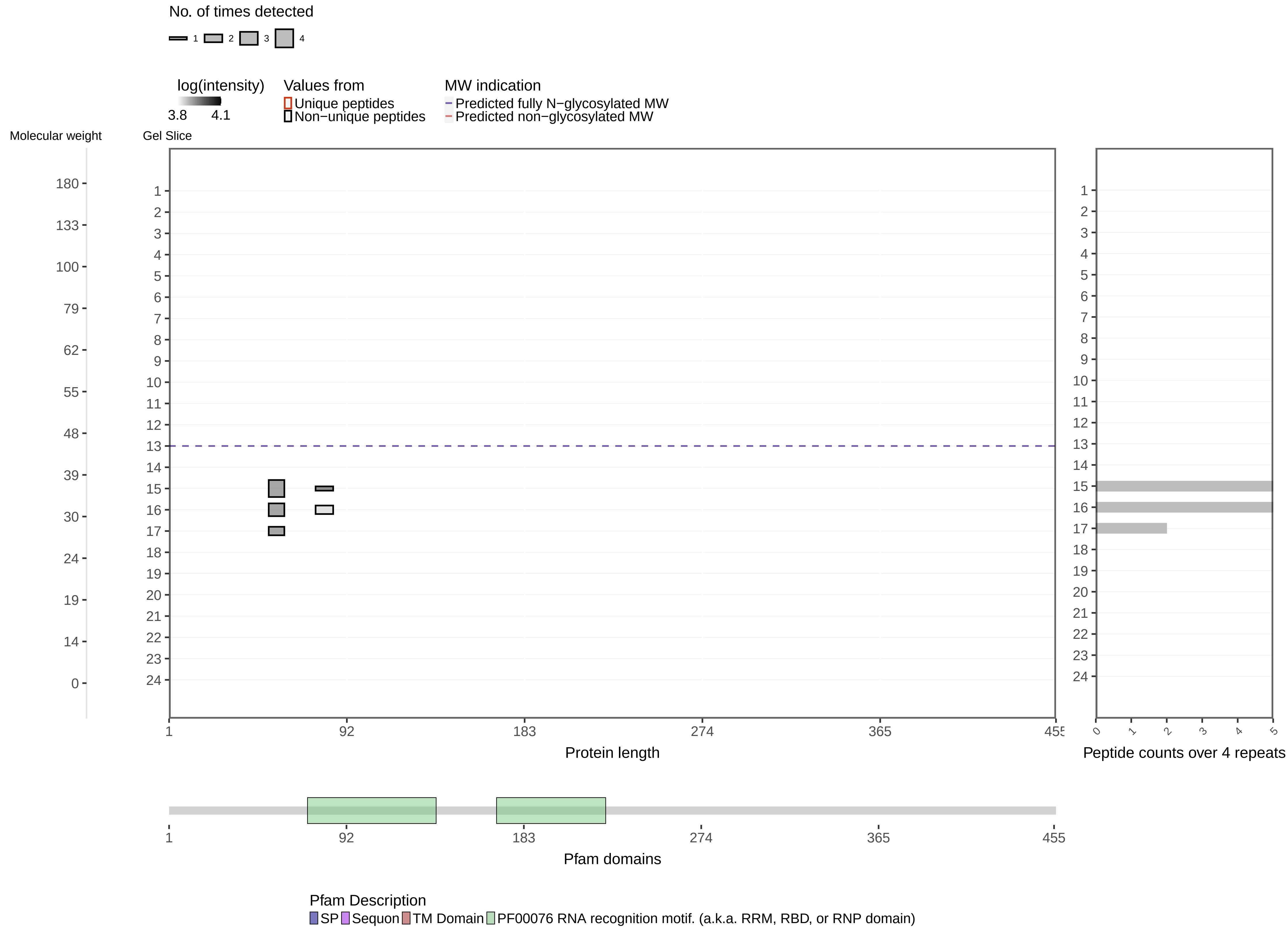


Protein sequence:

KASLVDPVKNIDGHQLNCKLAIDGKKGGKPGGGAQVQTDGHVYQVGPGQYGGPSGITGYGGFPGHSTFGGPGVHGLNSAYGSGNGIGVGGPGGGSSYGGSQPGGSVSGGAYGGPYGGGGGSHYGGPGSAGYGALASTATGGGGGYVSPGALGAGGYGSTGALGGGGLSGVGGALGGAGRGSSMYGLPPSSTGLASGDYPPQGPHYSQQNQVPGVSPAPRVAPGGMYQGMHSYY*

### NbL09g02180.1 malate_dehydrogenase_2_mitochondrial [MW = 35.63 kDa]

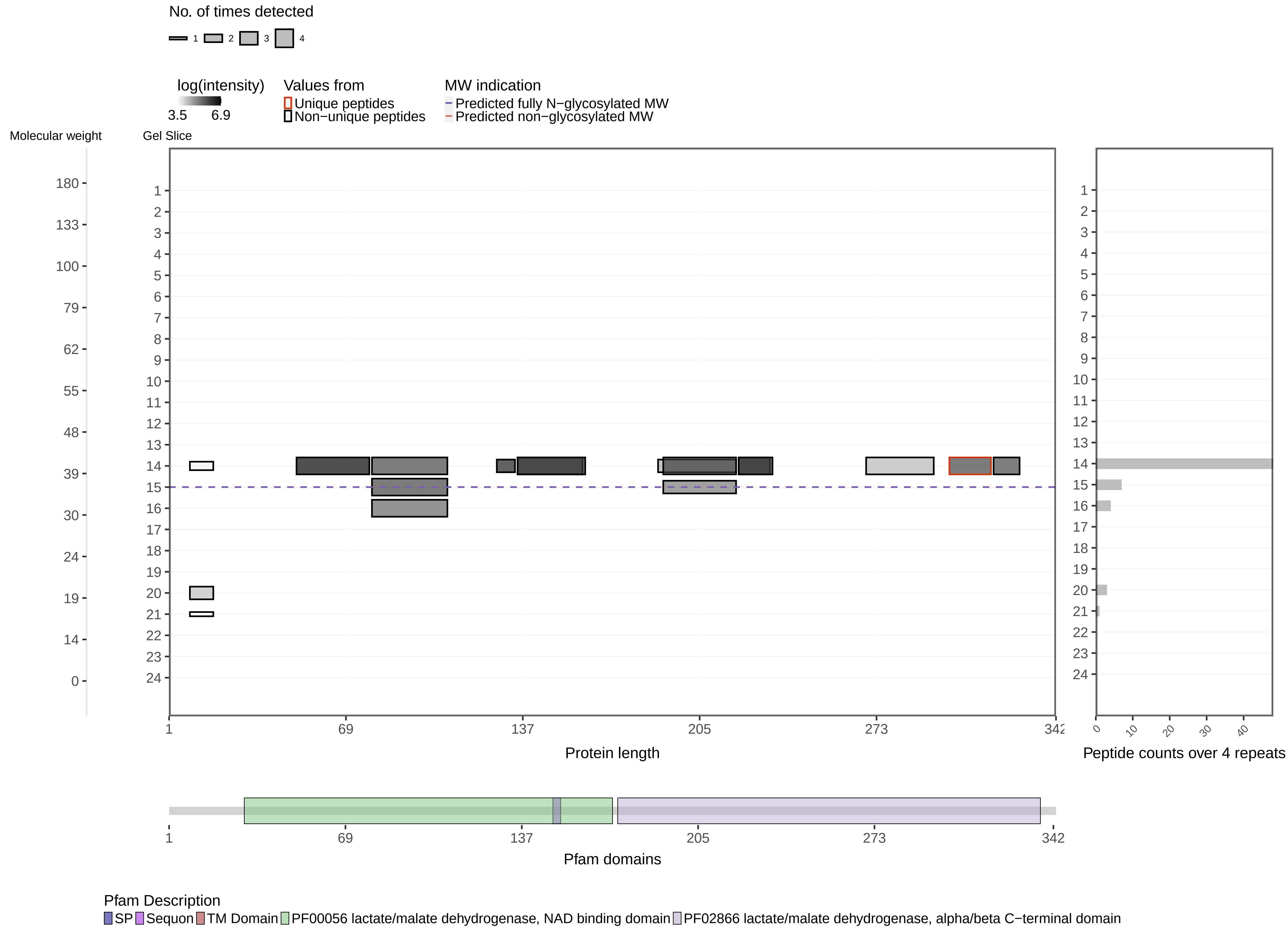


Protein sequence:

FKKAGTYDEKRLFGVTTLDVVRAKTFYAGKAKVNVADVIVPVVGGHAGITILPLFSQATPKANLGDEEIEALTKRTQDGGTEVVEAKAGKGSATLSMAYAGAIFADACLKGLNGVPDVVECSFVQSTVTDLPFFASKVRLGKNGVEEVLGLGPLSDFEKQGLEALKPELKSSIEKGIKFANDN*

NbL09g02100.1 homogentisate_12−dioxygenase−like [MW = 52.36 kDa]

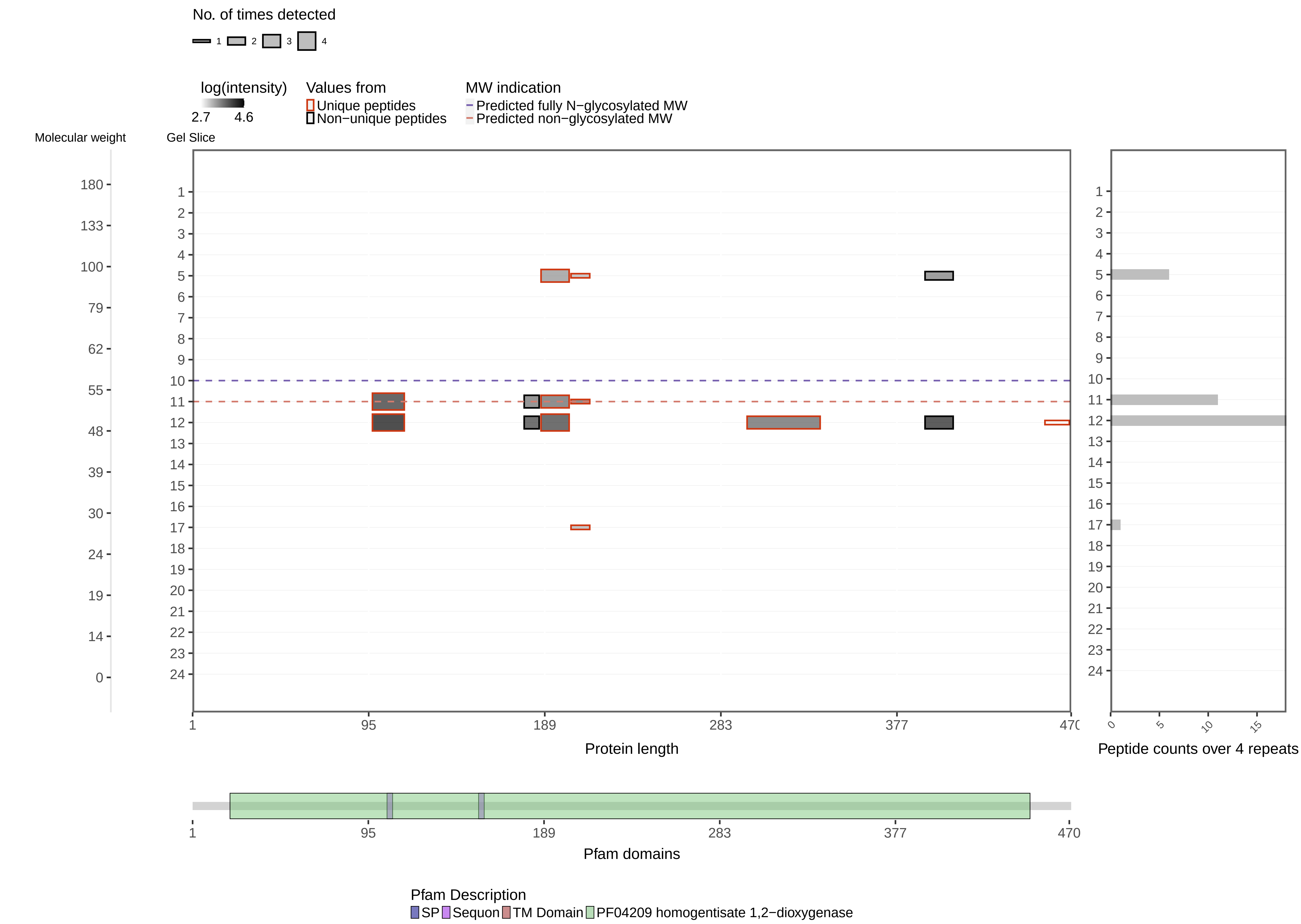


Protein sequence:

.QLPDLGPIGANGLAAPRDFLVPVAWYEDGSRPGYTIVQKYGGELFTAKQDFSPFNVVAWHGNYVPYKYDLSKFCPYNTVLIDHSDPSINTVLTAATDKPGVALLDFVIFPPRWLVAEHTFRPPYYHRNCMSEFMGLIYGGYEAKADGFHPGGASLHSCMTPHGPDTKTYEATIALGNEAGPHKIGDTMAFMFESCLIPRVCPWALESPFMDHDYYQCWIGLKSHFSGLSMKDDNRDFQNGKPVER*

### NbL09g02080.1 ras−related_protein_Rab11B [MW = 24.24 kDa]

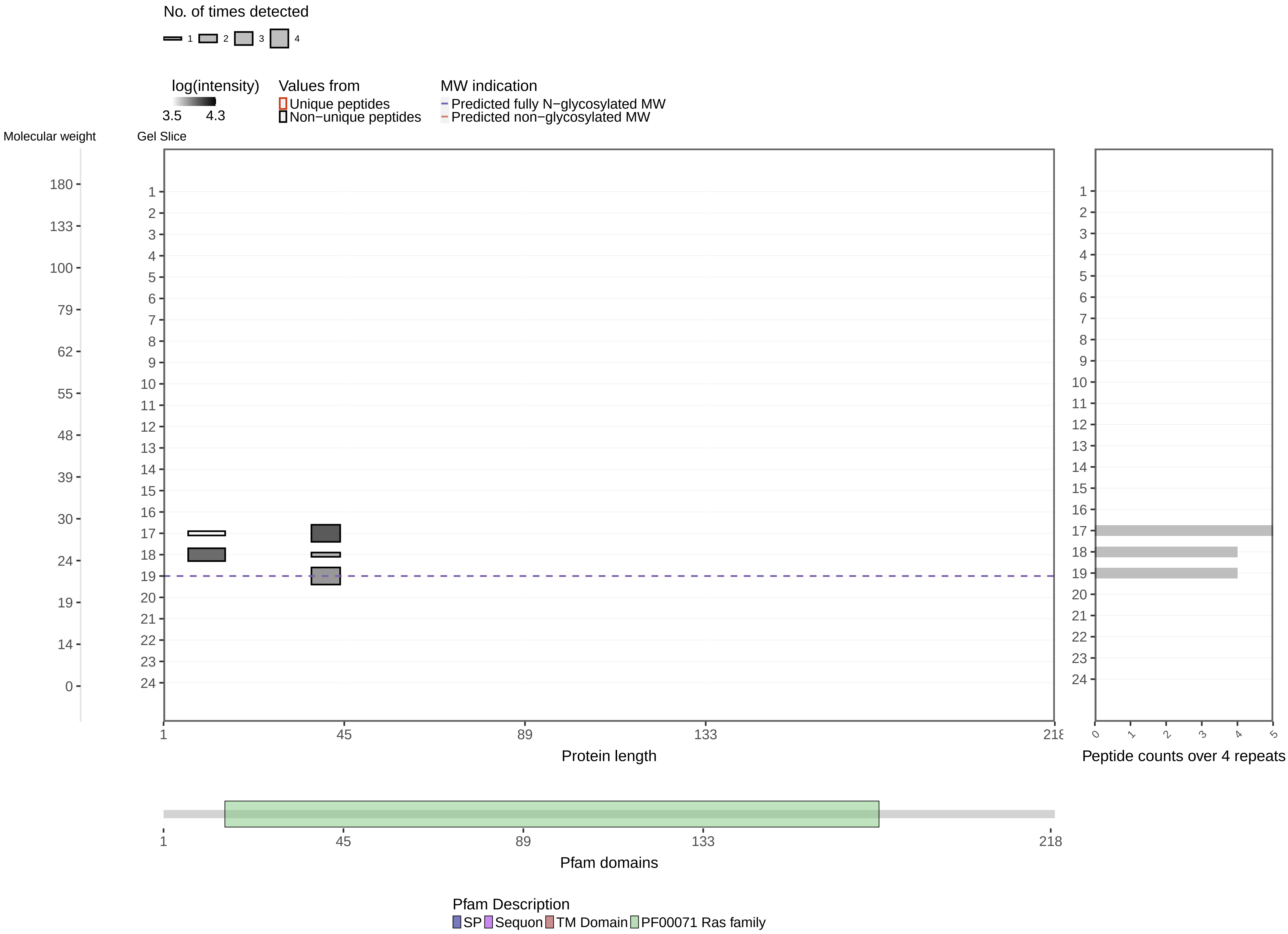


Protein sequence:

TRHVTFENVERWLKELRDHTDQNIVIMLVGNKADLRHLRAVSTEDAKAFAERESTFFMETSALESLNVENAFTEVLTEIYKVVCRKALEVGDDPAALPKGQTINVGKDDVSAVKKVGCCSV*

### NbL09g02050.1 multiple_organellar_RNA_editing_factor_8_chloroplastic/mitochondrial [MW = 47.2 kDa]

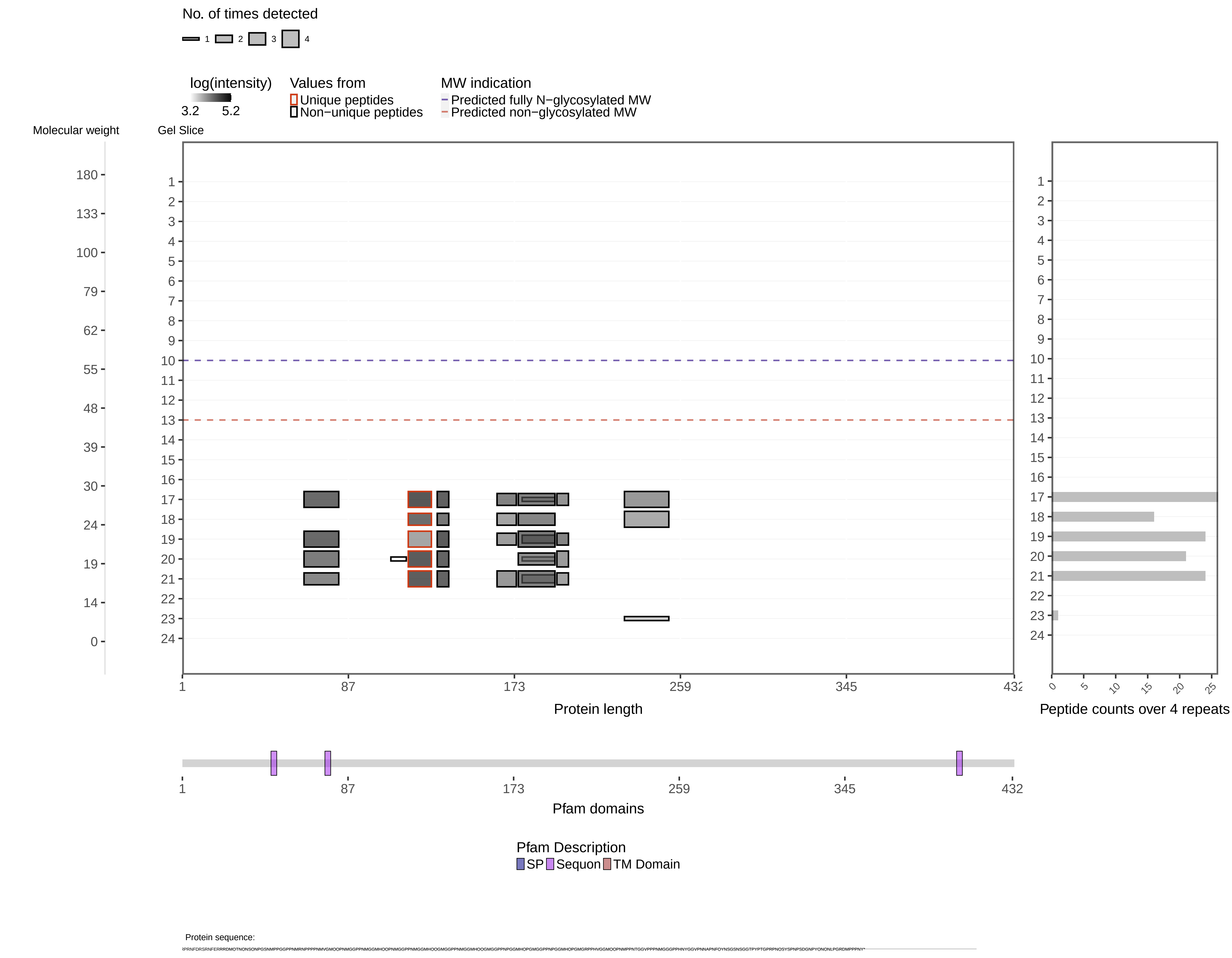

NbL09g02010.1 phosphoenolpyruvate_carboxylase [MW = 110.33 kDa]

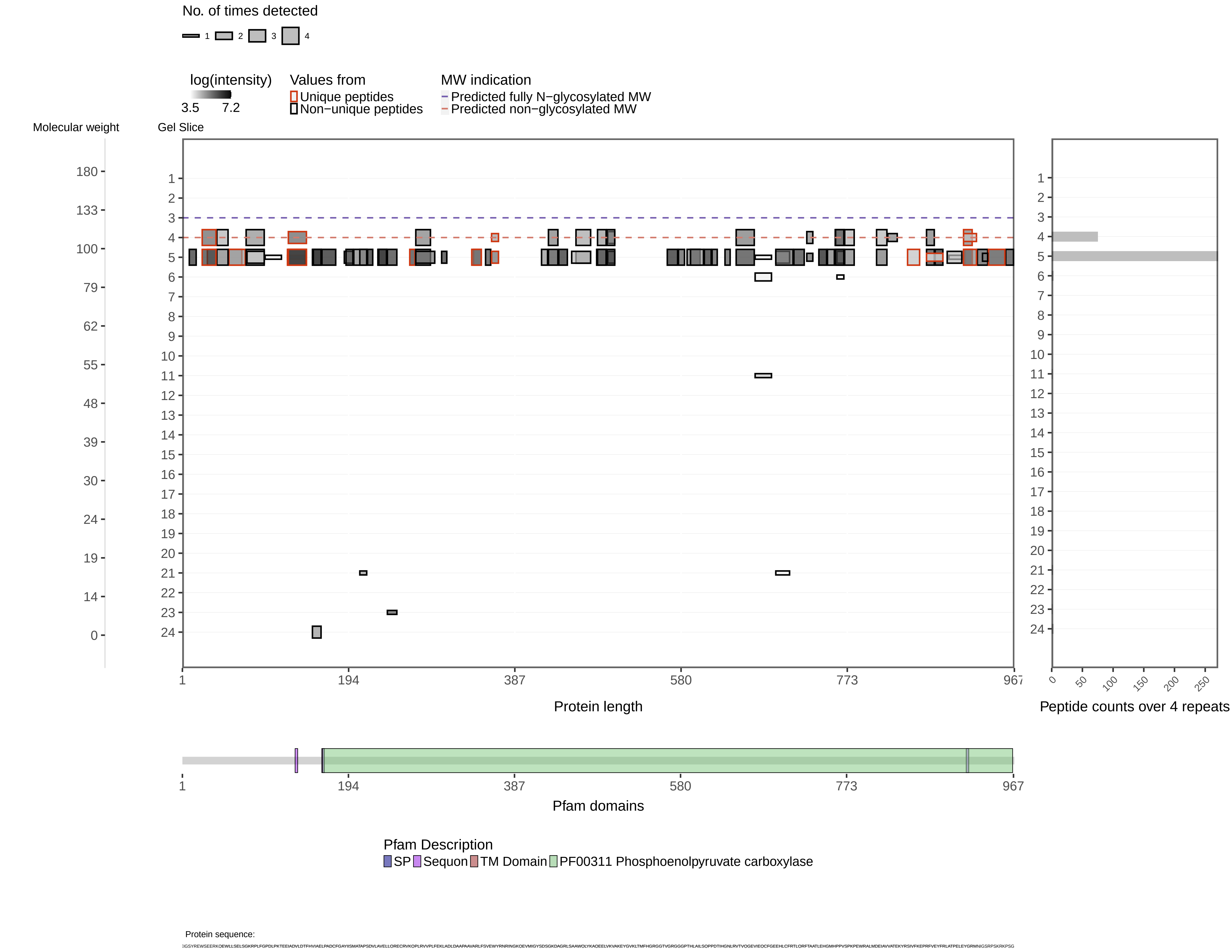

### NbL09g01990.1 multiprotein−bridging_factor_1b−like [MW = 15.27 kDa]

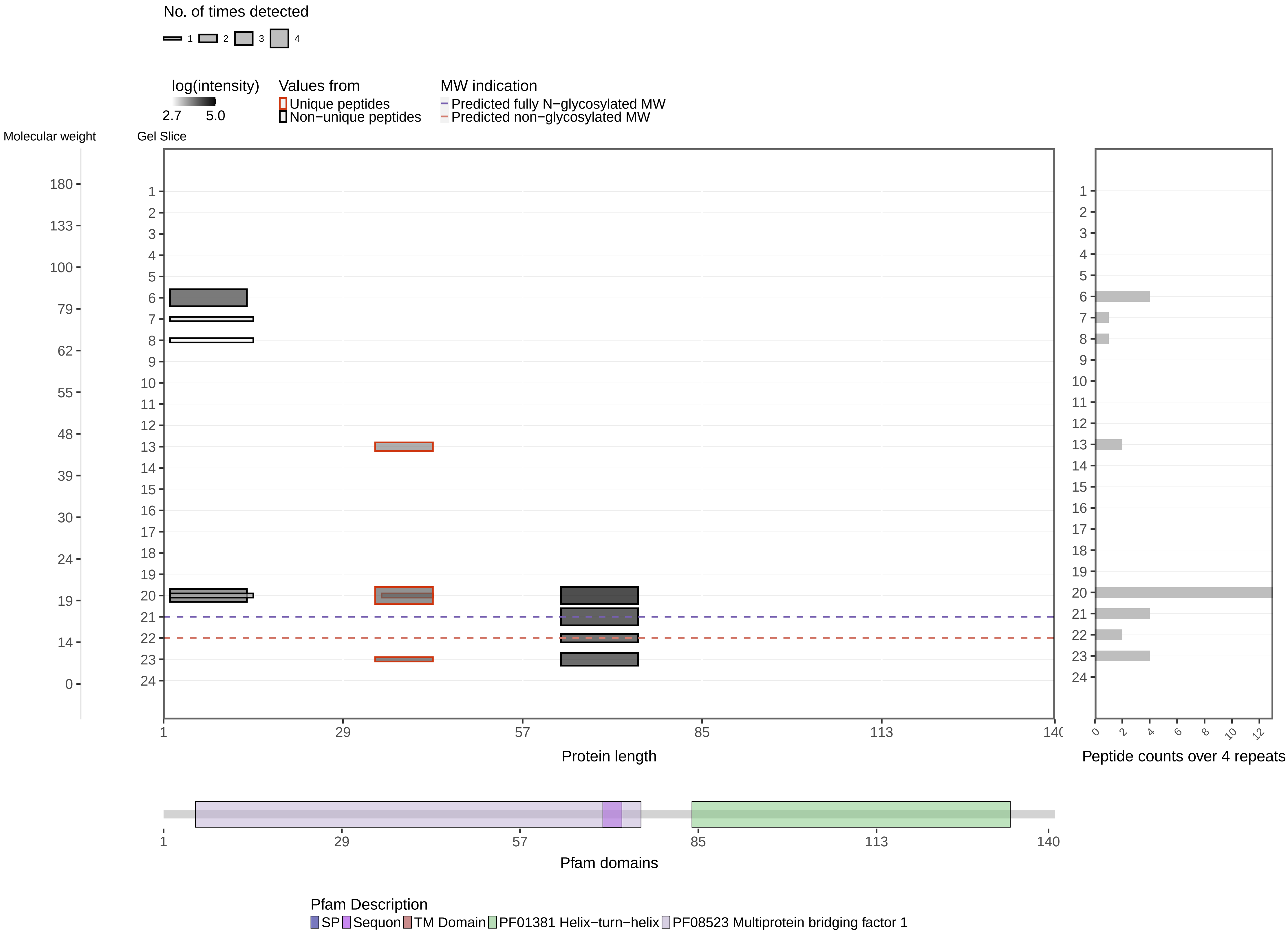


Protein sequence:

STTLNTRKLDEDTENLSHQKVPTELKKAIMQARQDKKLTQSQLAQLINEKPQIIQEYESGKAIPNQQIISKLERALGAKLRGKK*

### NbL09g01900.1 50S_ribosomal_protein_L13_chloroplastic [MW = 27.12 kDa]

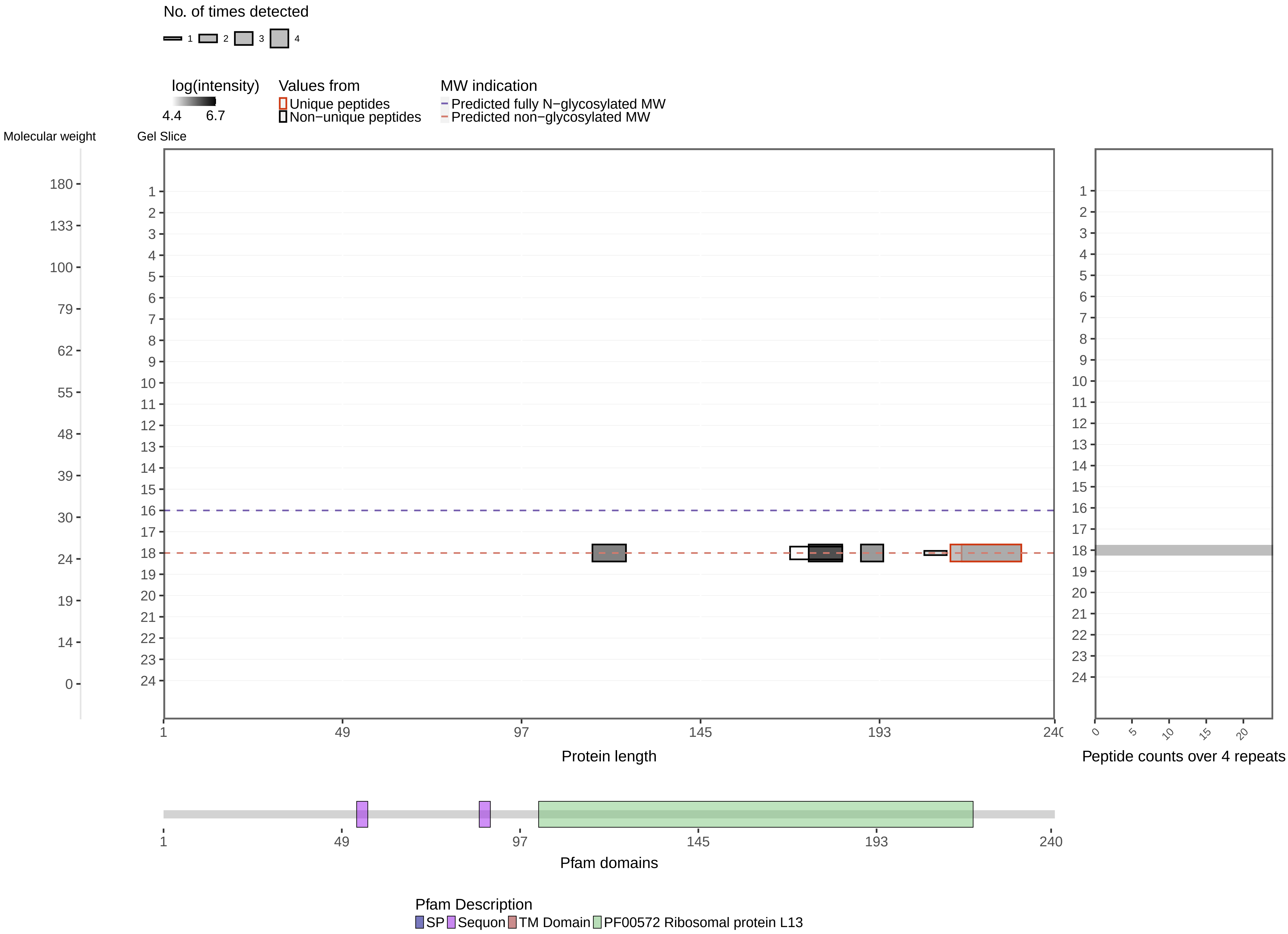


Protein sequence:

/DATDLILGRMASTIAIHMRGKNLATYTPSVDMGAFVIVVNAEKVAVSGKKRTQKLYRRHSGRPGGMKVETFDQLQQRIPERIIEHAVRGMLPKGRLGRQLFTHLKVYKGPDHPHEAQKPIELPIRDKRIQKQR*

NbL09g01710.1
putative_SWI/SNF−related_matrix−associated_actin−dependent_regulator_of_chromatin_subfamily_A_member_3−like_1 [MW = 99.39 kDa]

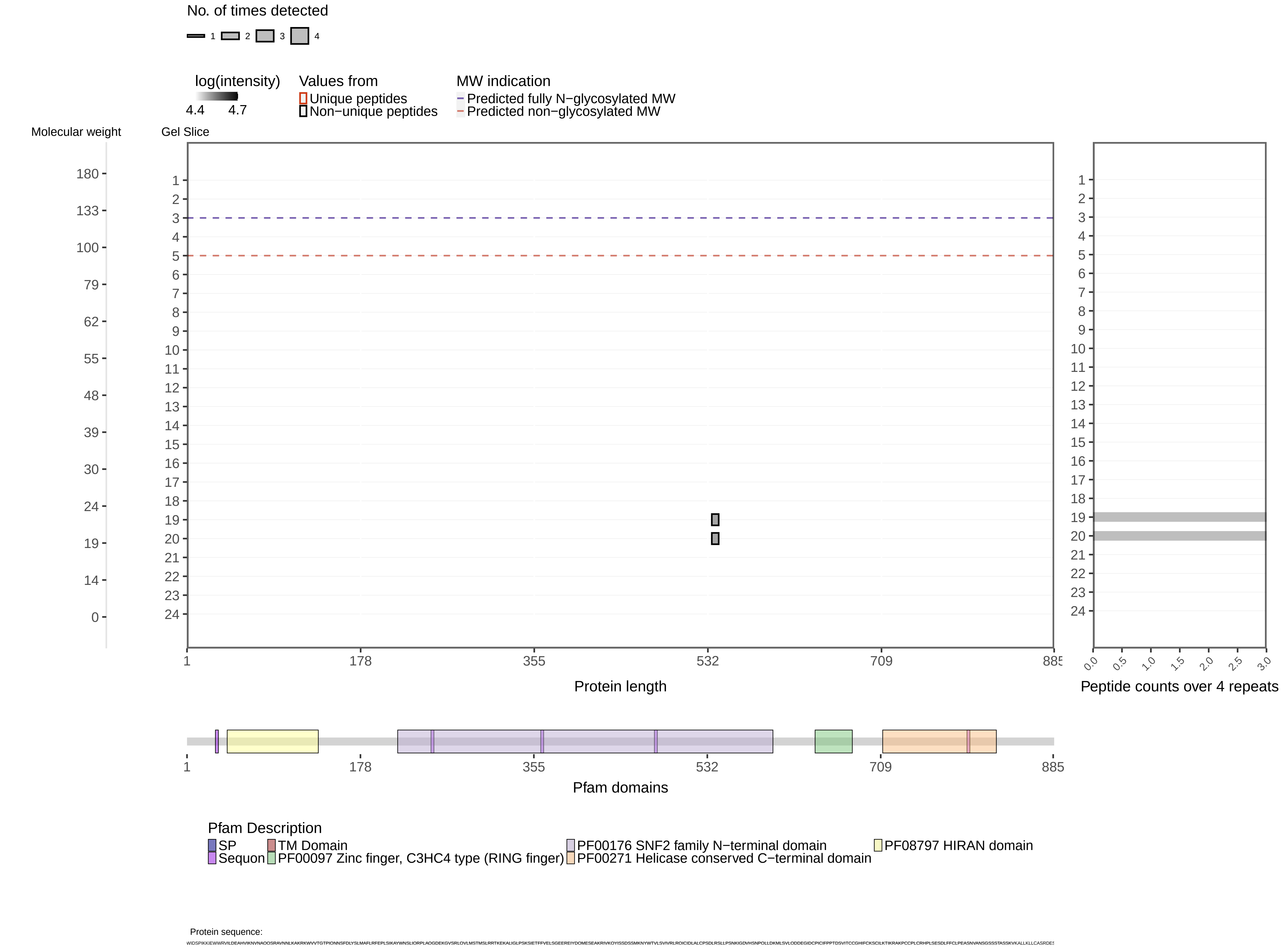

NbL09g01660.1 photosystem_I_reaction_center_subunit_II_chloroplastic [MW = 22.65 kDa]

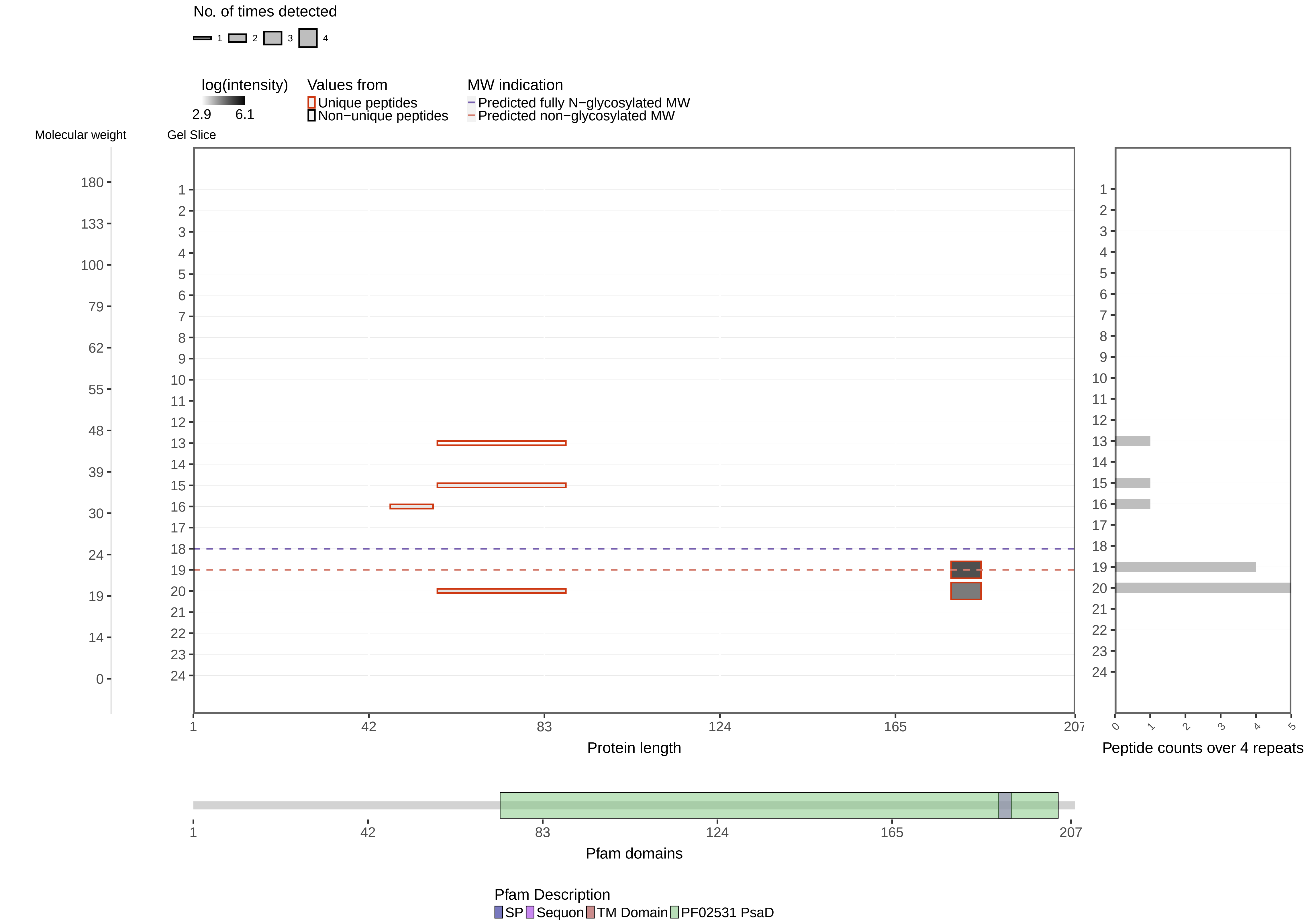


Protein sequence:

[illegible]VEEFYVITWESPKEQIFEMPTGGAAIMREGANLLKLARKEQCLALGTRLRSKYKINYRFYRVFPNGEVQYLHPKDGVYPEKVNAGRQGVGQNFRSIGKNKSPIEVKFTGKQVYDL*

NbL09g01630.1 protein_LOW_PSII_ACCUMULATION_3_chloroplastic [MW = 42.46 kDa]

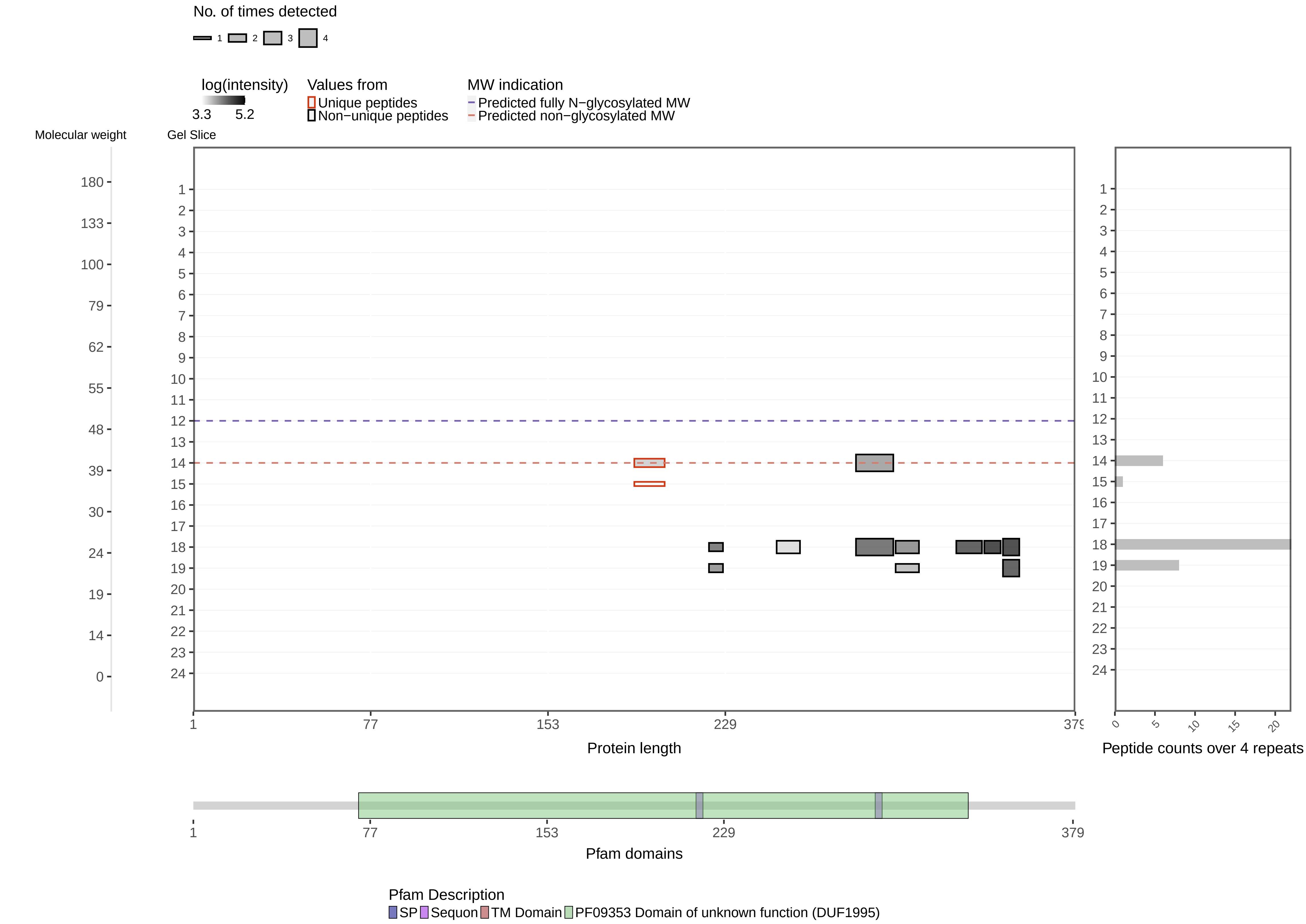


Protein sequence:

GAVTSFFRSVRNTLDFDFDDDNEGRWKSDKPPTLYIFINCSTRDLAYIEKYVEKIATSTPSLLFNLELDTLRADLGIFGFPPKELHYOFLSOFTPVFYIRTREYSKTVAVAPYIVNYSGALFROYPGPWOVMLKOTDGSFVCVAESATRFTLGETKEELLRVLGLOEEKGSSLEFLRRGYKTATWWEEDVDLEASSAWRS*

### NbL09g01500.1 triphosphate_tunnel_metalloenzyme_3−like [MW = 26.78 kDa]

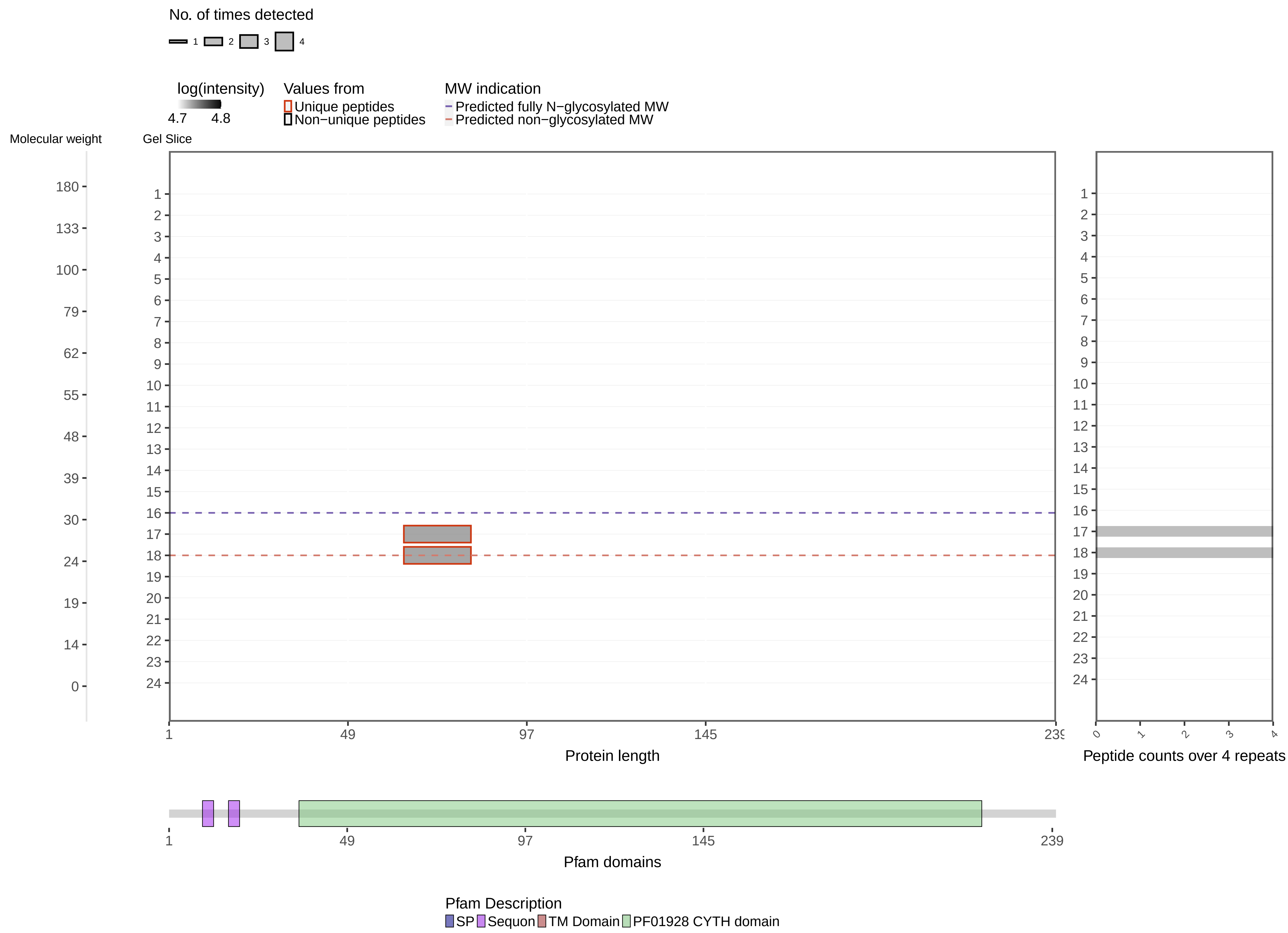


Protein sequence:

ISNGISRIEEDEEPIDPSIGRACVSDPWRLLLIDSSRVIQRVKDEYGFGEKQGLVCLGGFRNVRAVYEWKGLQLALDETHYEFGMNFEIECESSDPERAKDLLEELLKSHGIEYSYSTVSKFAIFRSGKLPQ*

NbL09g01470.1 clustered_mitochondria_protein_homolog [MW = 189.08 kDa]

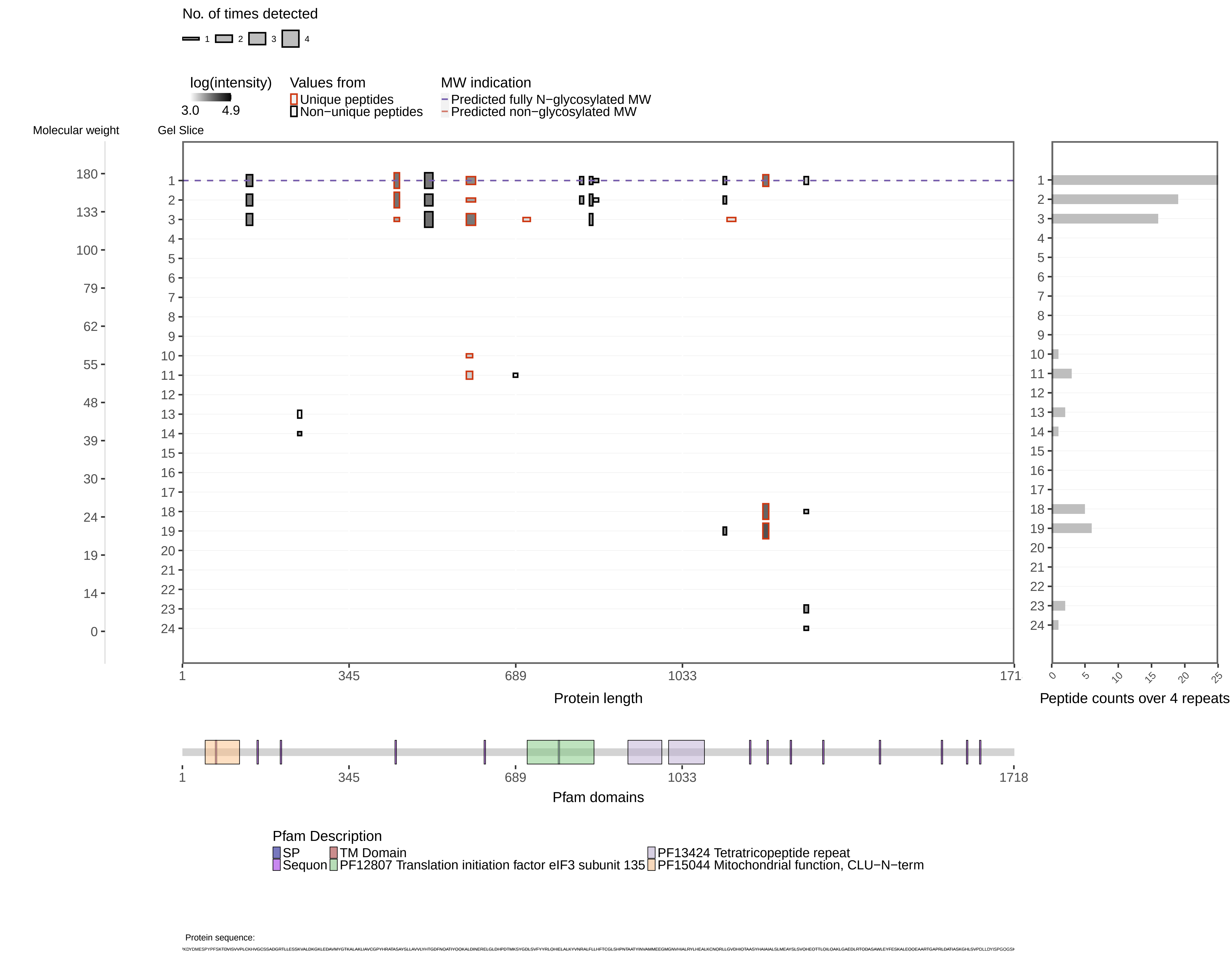

NbL09g01460.1 haloalkane_dehalogenase−like [MW = 35.75 kDa]

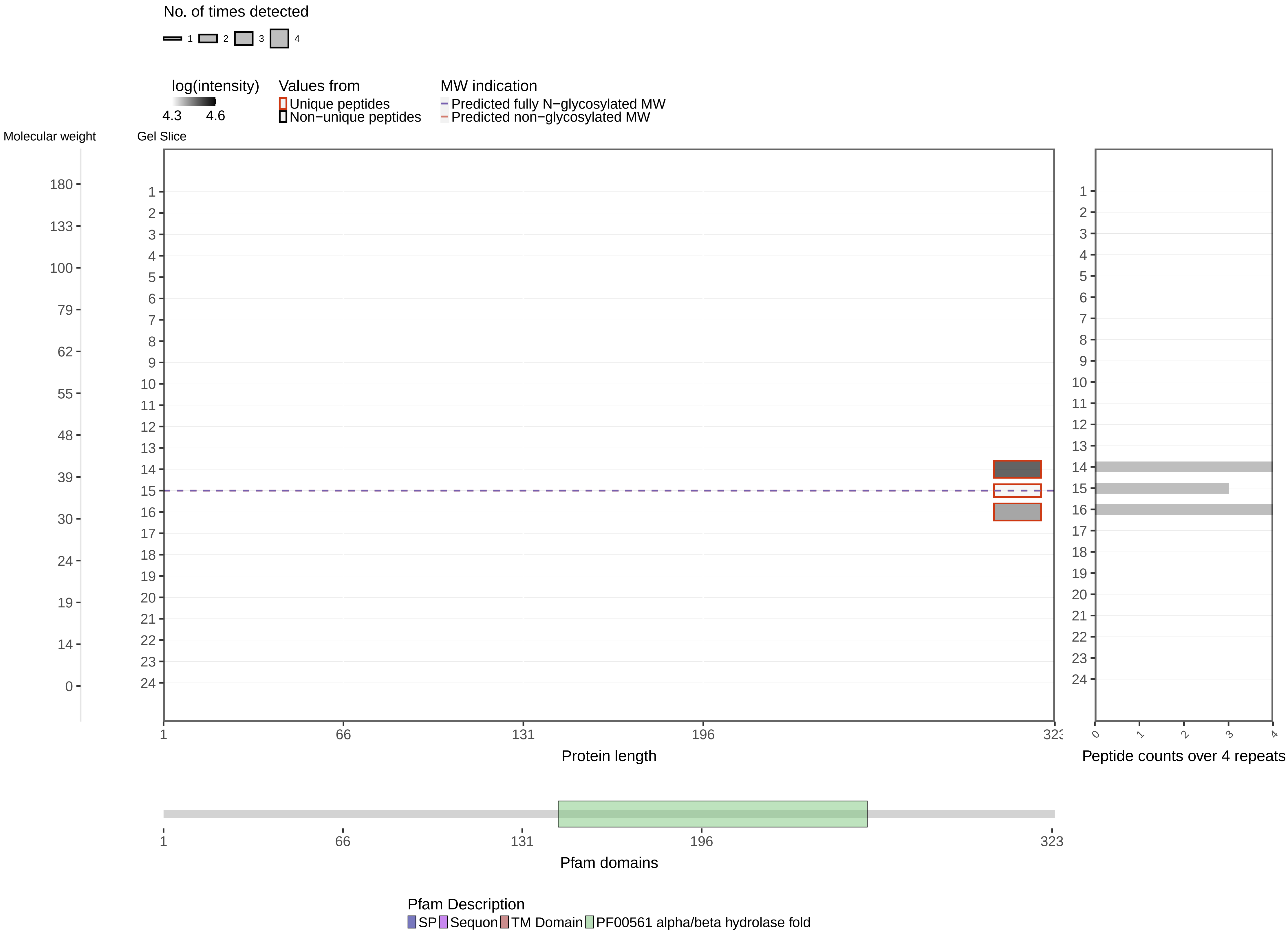


Protein sequence:

LHGAPTQSYSYRVVMSQMADTGFHCFAPGWIGFGFSDKPQPGYGFDYTEKEFHEEFEKLLEVLGVTSPFFLVVQGFLVGSYGLTWTLKNPSKISKLAILNTPLTVSSPIPGLFQQLRIPFFGEFTCQNAVMAERFIEAGSAYVLKLEKADVYRLPYLASSGPGFALLETARKLLQ*

NbL09g01410.1 methyl−CpG−binding_domain−containing_protein_11−like [MW = 37.56 kDa]

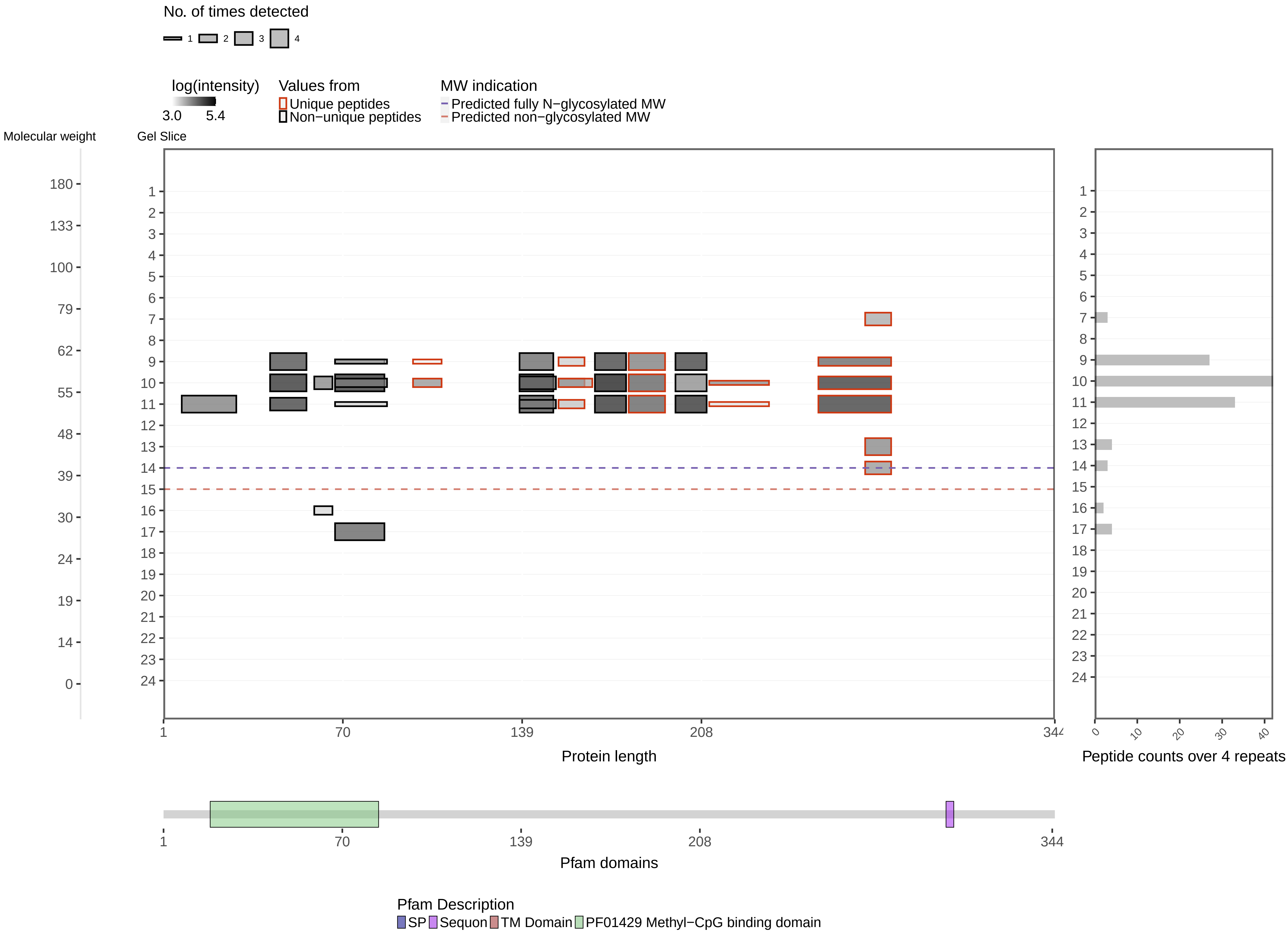


Protein sequence:

DDKGKKEVEAADEEAEGMKGAELHLEEKPOMDDKGKKEVEAANEEAEGMKGGELHPEEKPESEPEKVDSADDGKODKSENAKIEDKOVSGEKLESOSDKESPAADVNENKPEVVTAEGINKGVGGSSKDVEHAGNGESNNGENGSREEEKEMIGTDLVIGNNKINOPGPAHSOOHOSPAPISC*

### NbL09g01370.1 phospholipase_D_alpha_1 [MW = 92.1 kDa]

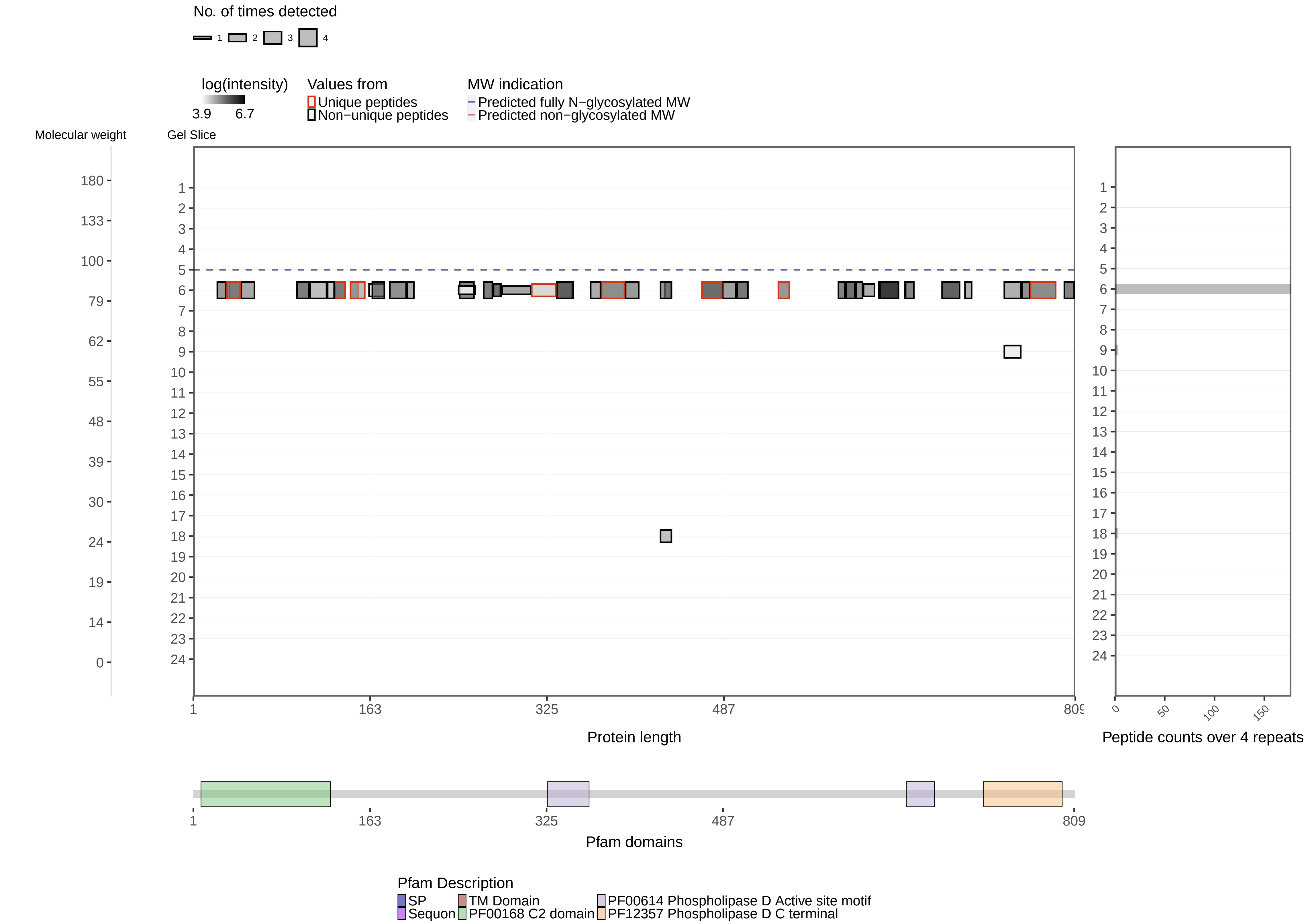


Protein sequence:

NbL09g01350.1 trafficking_protein_particle_complex_subunit_2−like [MW = 15.32 kDa]

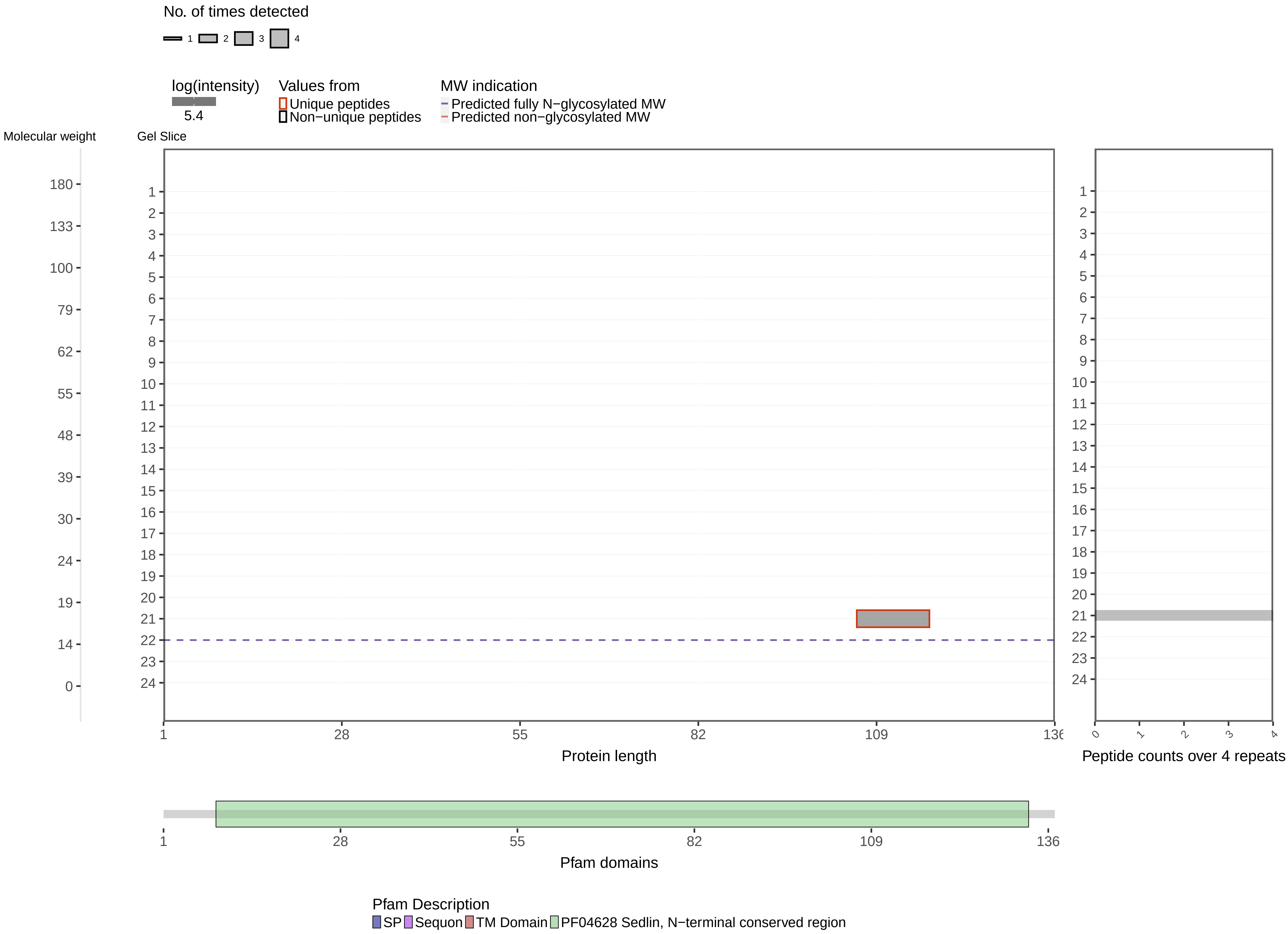


Protein sequence:

AMFLKSIDRFNDLVVSVYVTAGHTRLMLLHDSRNDDGIKSFFQEVHELYIKVLLNPLYLPGSRITSSHFDTKVRALARKYL*

NbL09g01330.1 uncharacterized_LOC107826806 [MW = 46.3 kDa]

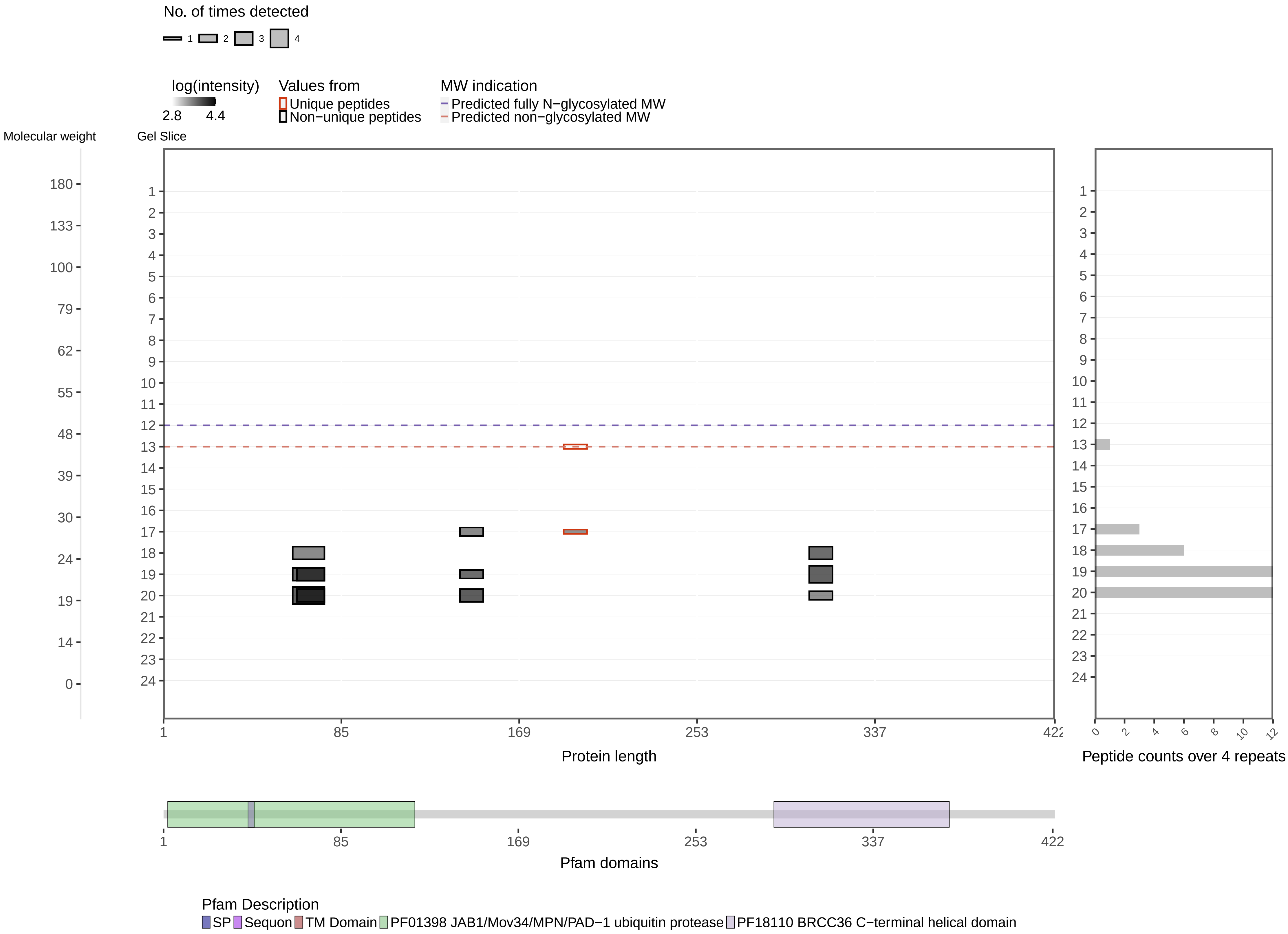


Protein sequence:

SRAAALASKGGARSTDLGNFFANVDNRVGENLQANSLSNAILDVDPMDMSDSMQEAMHRSNLEMSGAEYVRKEIPLLVLPSSSLLNLDAPLNSFTDLQRVLFEEERTAYNQAVVTNMREGKAHPLTFIHHTSTYQASMCKLMEYCLSPAISALQDHLRENEIRLKMLSEEAKVLETEITKGEPITSSPRRSSTSLGQRDLYSPTELSMKSVSGSGSRSRKAS*

### NbL09g01190.1 protein_CDI−like [MW = 29.42 kDa]

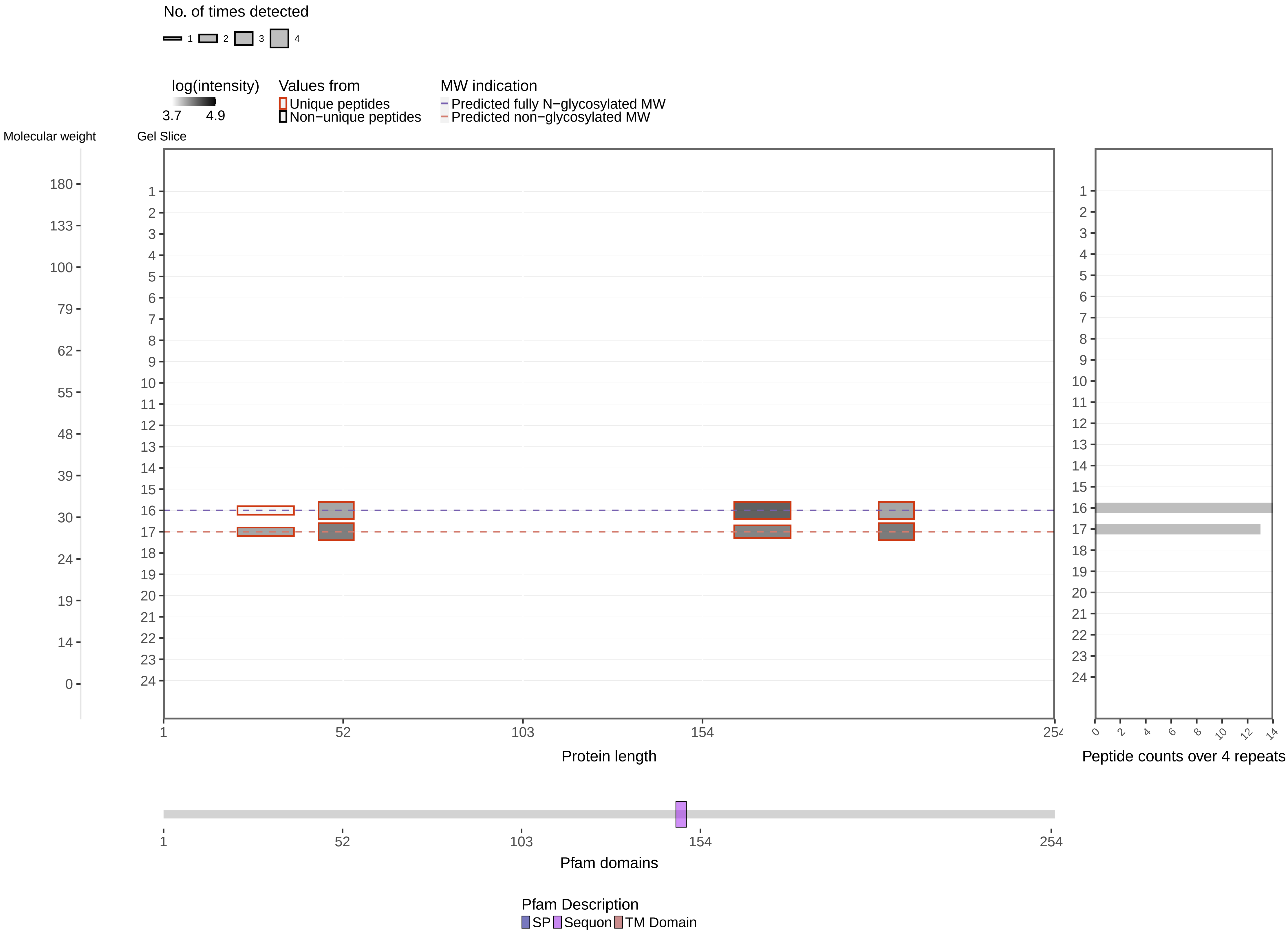


Protein sequence:

KYAVMCVHHDYAPKETTKMDGAVQTVYPRKNWSSMVLYNCGHPKNKVLTPEIVNTESGAFLHRFMWLEDNEIGEVPFVWNFLVGHNKVVEGDPNTFPKSIHYTLGGPWFEAWKDCQFGDLWIKELEEYKKSIDKKAD*

### NbL09g01110.1 chloroplast_stem−loop_binding_protein_of_41_kDa_b_chloroplastic [MW = 41.72 kDa]

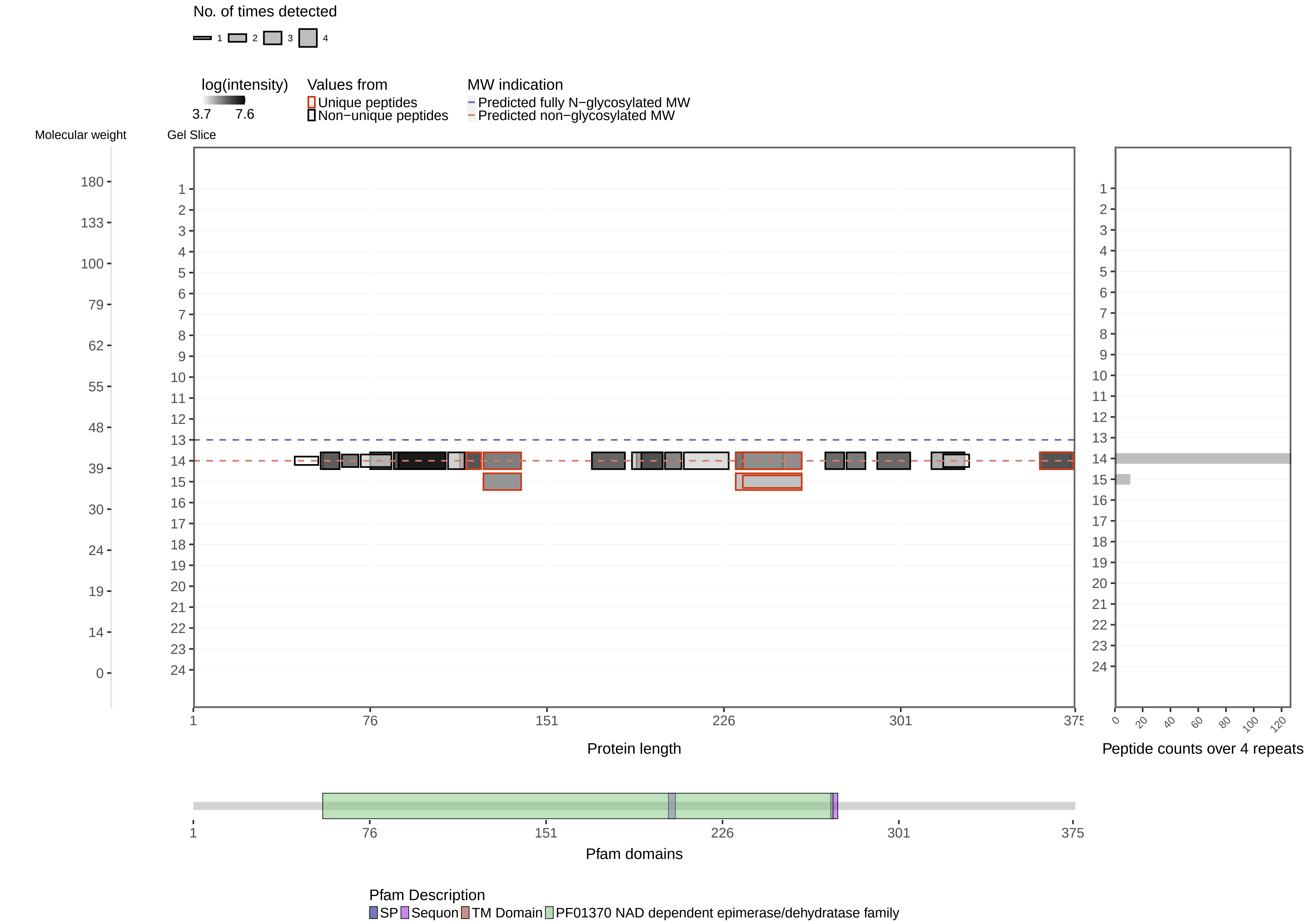


Protein sequence:

IFEADAVDPKSRHKGKLETESLLESRDVNWTSVRPVYIYGPLNYNPVEEWFFHRLKAGRPIPIPNSGLOITOLGHVKDLATAFIKVLGNEKASKOVFNISGEKYVTFDGLAKACAKAGGFPEPEIVHYNPKEFDFGKKKAFPFRDOHFFASVEKAKAVLGWKPEFDLVEGLTDSYNLDFGRGTFRKEADFSTDDLILGKN*

NbL09g01070.1 tetrapyrrole−binding_protein_chloroplastic [MW = 29.96 kDa]

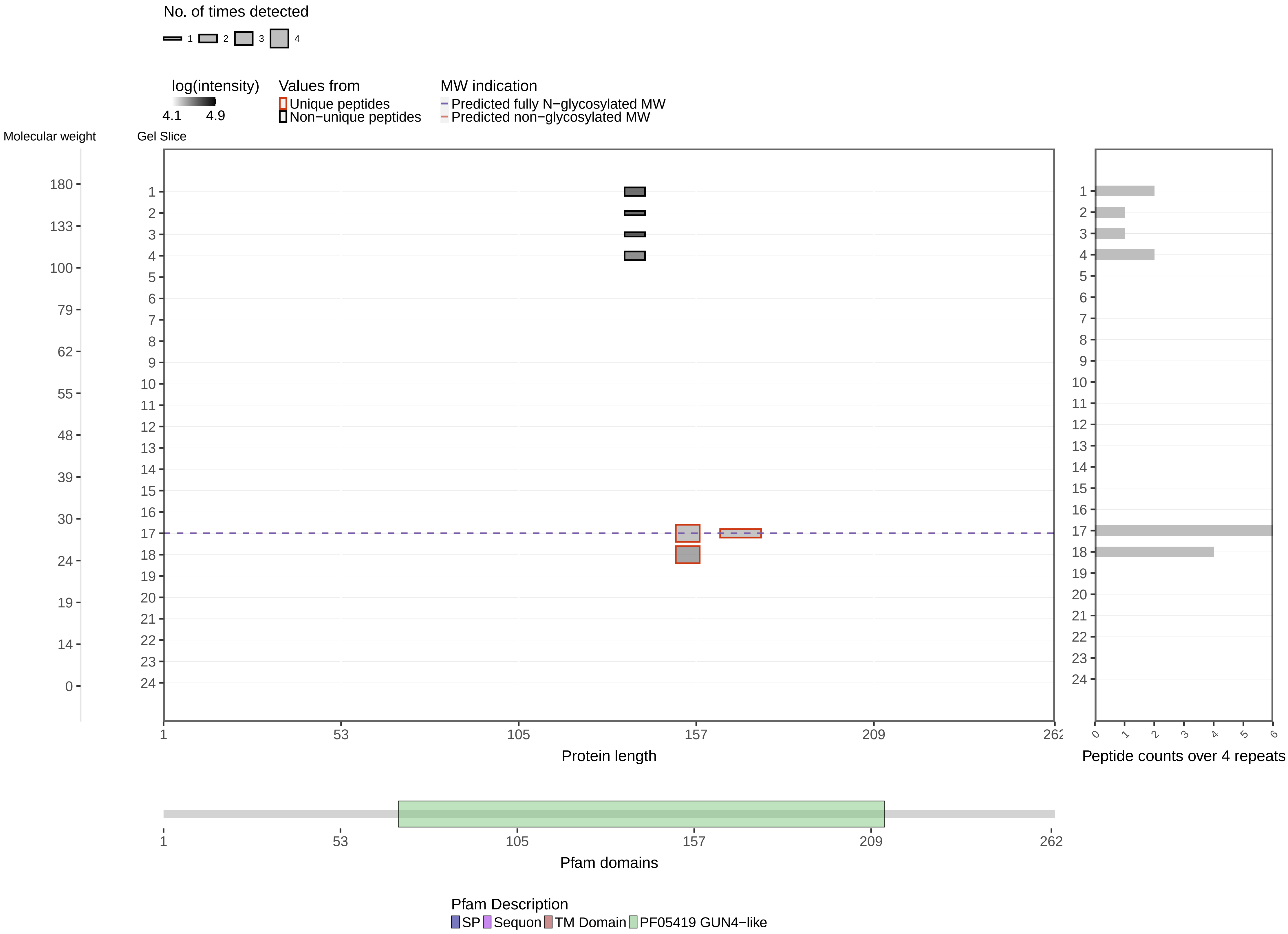


Protein sequence:
KEIDSLWKKYSDNKFGYSVQKKIWNKVNRDFTSFFIKVGWMKKLESSEVEQYNYRAFPNEFIWELNEETPEGHLPLTNALRGTQLLSSIFTHPAFVEQGEEEKDEEKQASFSGEDKSSVNKGGLLGGLRSKLFGKPDYSF*

### NbL09g01040.1 60S_ribosomal_protein_L9−1 [MW = 21.98 kDa]

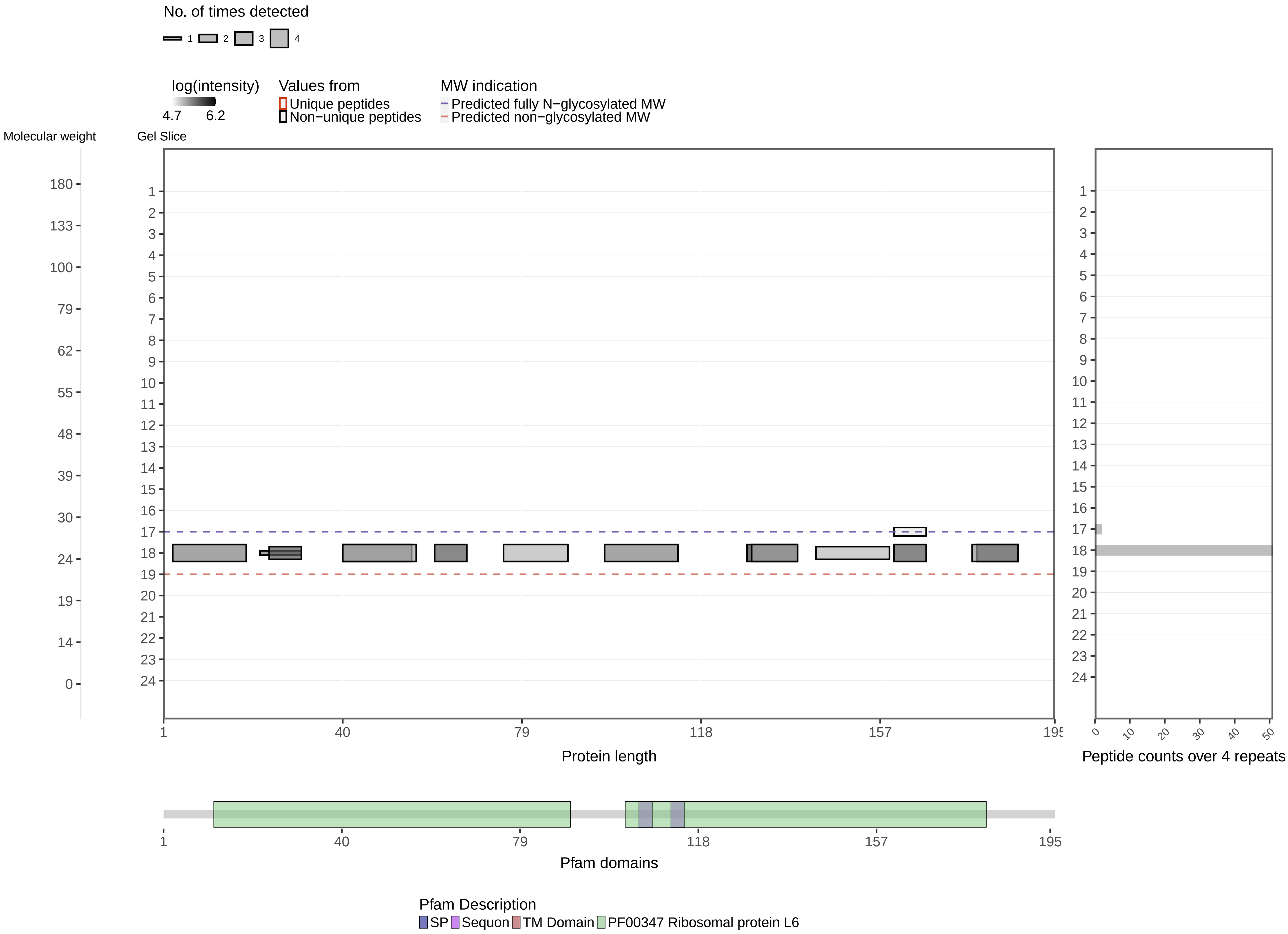


Protein sequence:

TGVTKGYRYKMRFVYAHFPINASITGGNKSIEIRNFLGEKRVRKVDMLDGVTVVRSEKVKDELVLDGNDIELVSRSAALINQKCHVKNKDIRKFLDGIYVSEKGRIAEEE*

NbL09g00870.1 nuclear−pore_anchor [MW = 229.22 kDa]

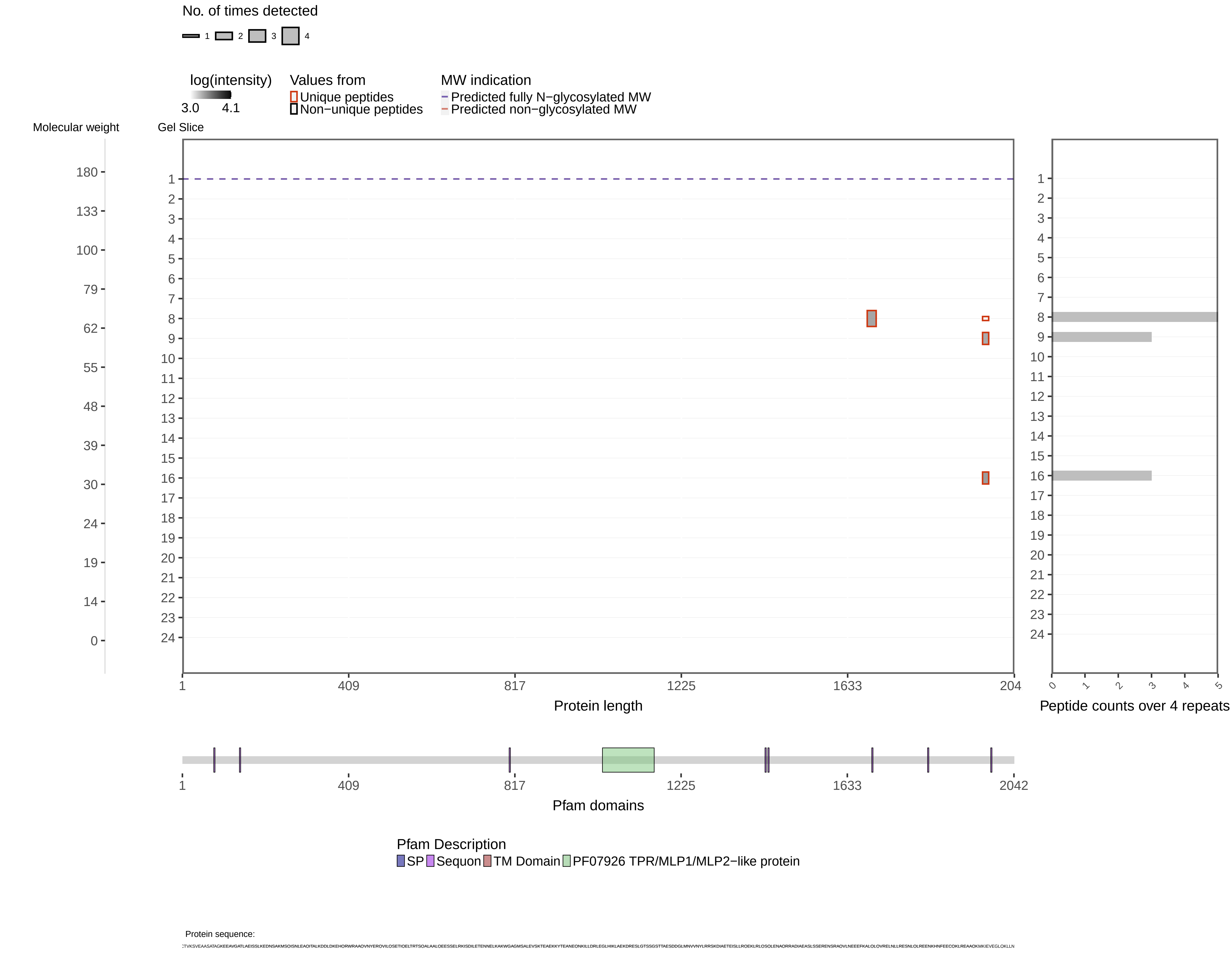

NbL09g00820.1 protein_FAM136A−like [MW = 17.02 kDa]

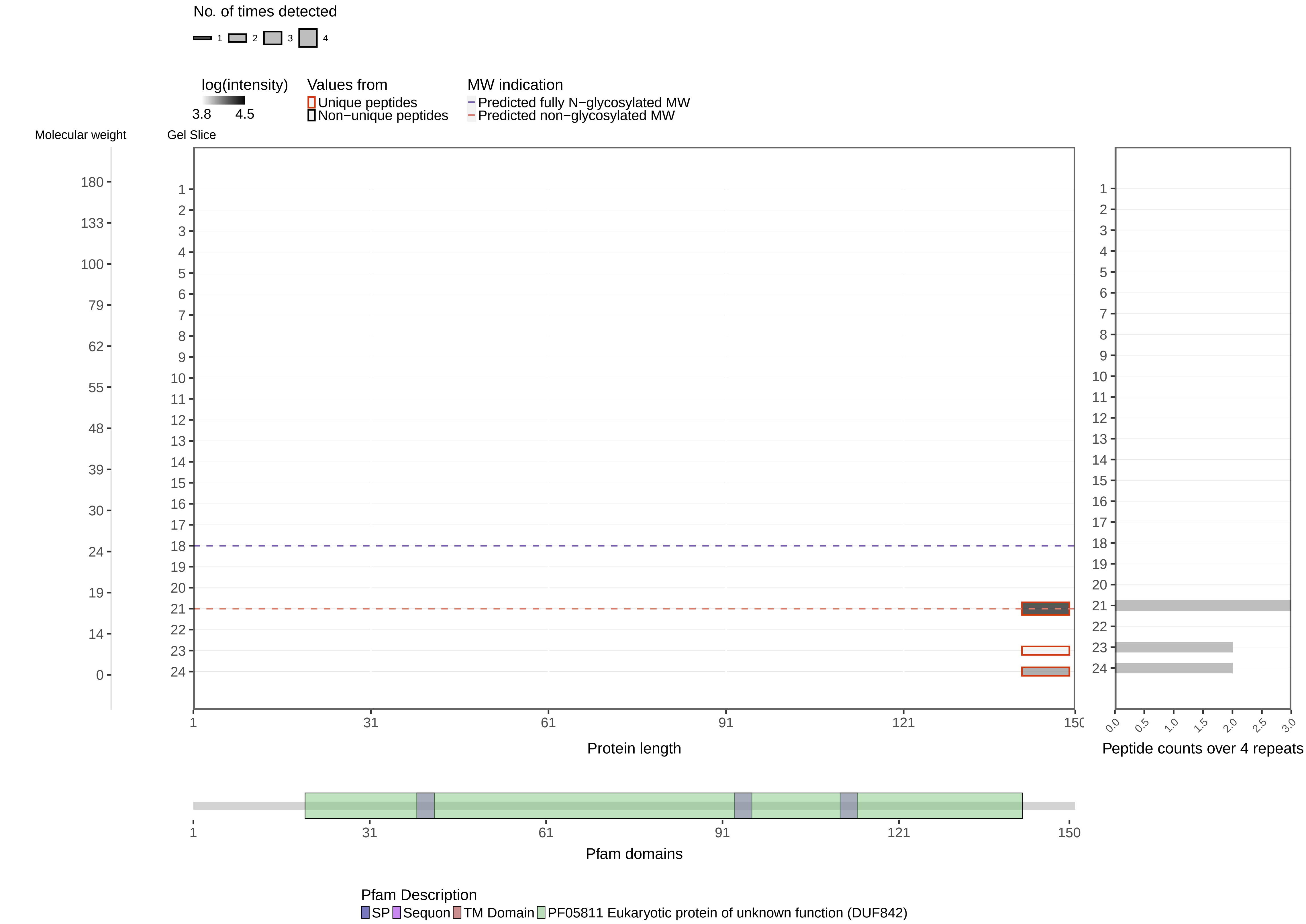


Protein sequence:

IGHCVEQCSVHVHNAQNLVQNEMAKFQERMNRSLMVCQDKYETAKLQKNRSDATKDLESCVDQSVHDSINTLPHLVARLKGSLGIAN*

### NbL09g00810.1 stearoyl−[acyl−carrier−protein]_9−desaturase_chloroplastic [MW = 47.78 kDa]

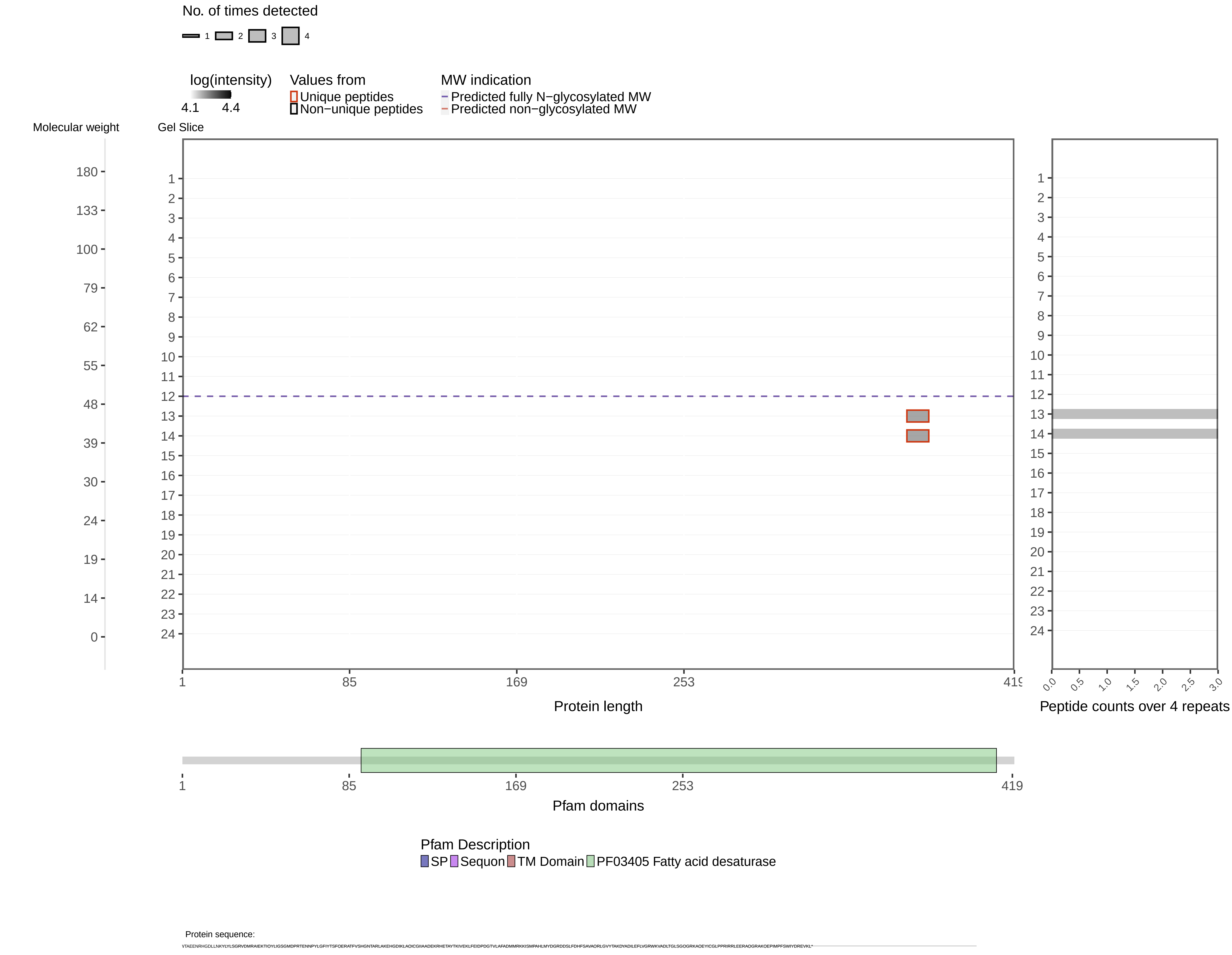

### NbL09g00740.1 fructokinase−2−like [MW = 39.05 kDa]

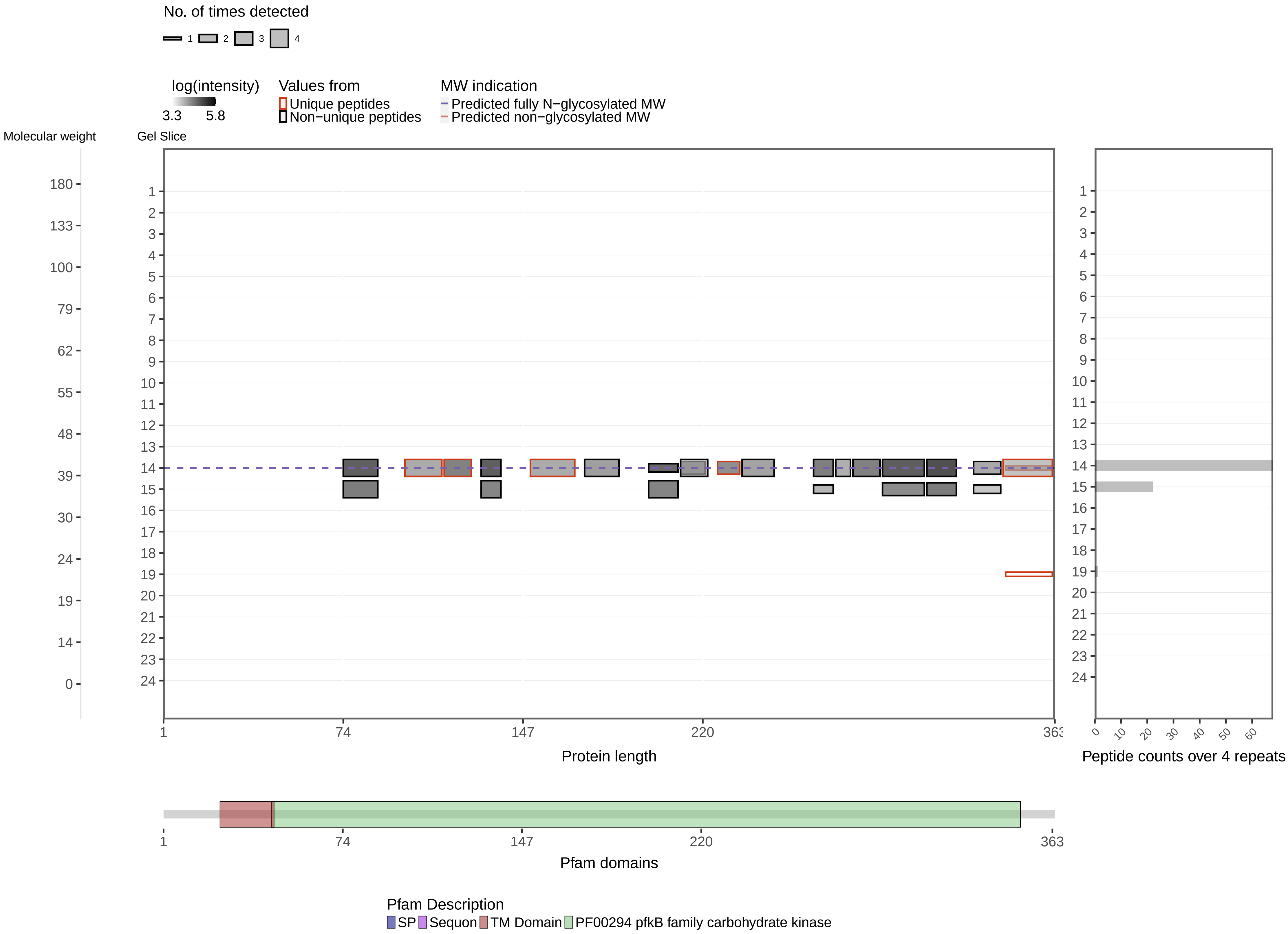


Protein sequence:

### NbL09g00610.1 probable_hydroxyacylglutathione_hydrolase_2_chloroplastic [MW = 36.4 kDa]

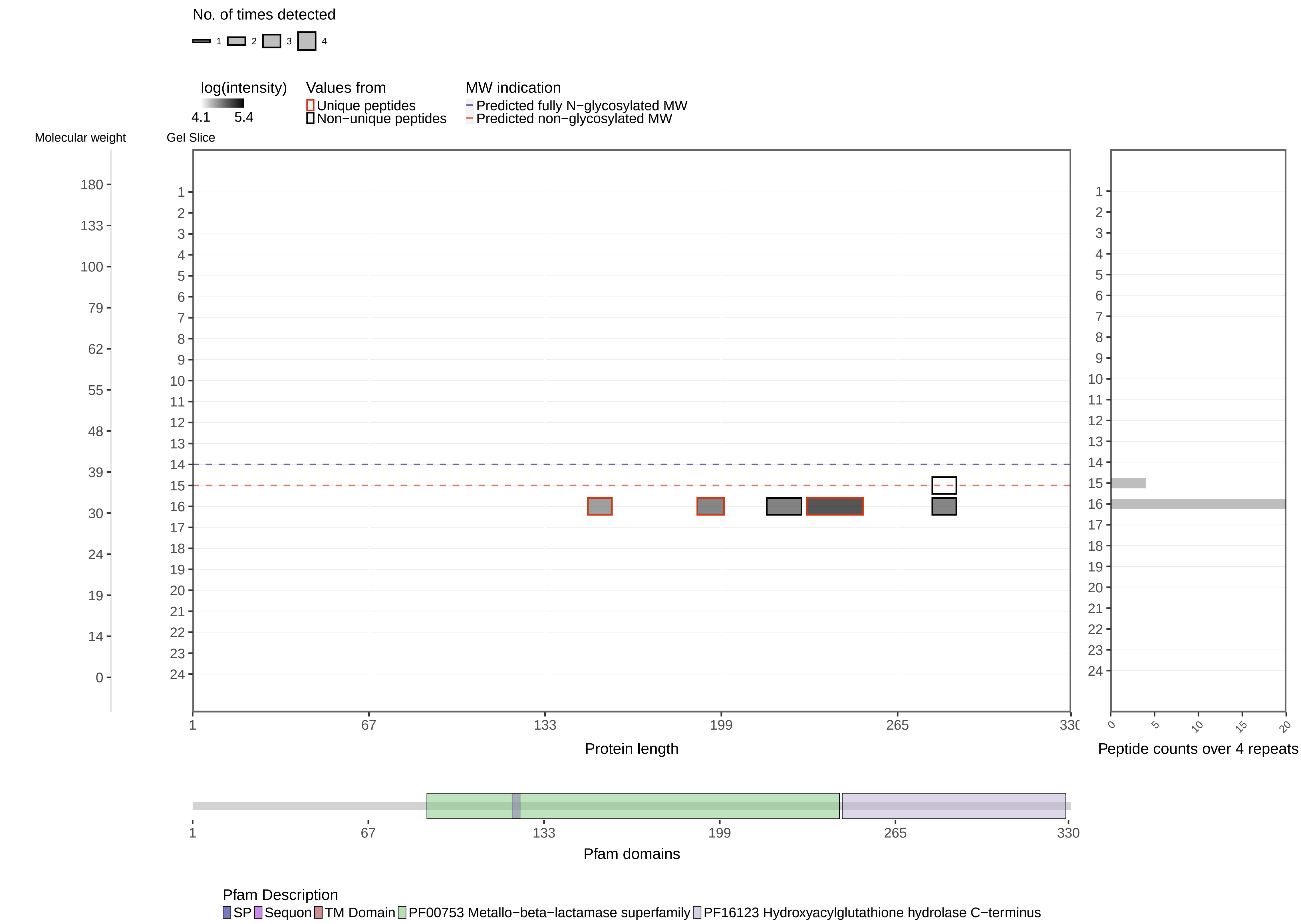


Protein sequence:

[illegible]SGVDSDRIPGIDIALNDGDQWMFAGHEVLVMETPGHTRGHISFYFPGSKAIFTGDTLFSLSCGKLFEGTPQQMLSSLRKITSLPDDTNVYCGHEYTLSNSKFALSVEPGNEELQSYSTQVVNLRKKGLPTIPTTLKAEKLCNPFLRTSSTEIRKLLNIPPTADDAEALGAIRRAKDNF*

### NbL09g00510.1 2−isopropylmalate_synthase_B [MW = 66 kDa]

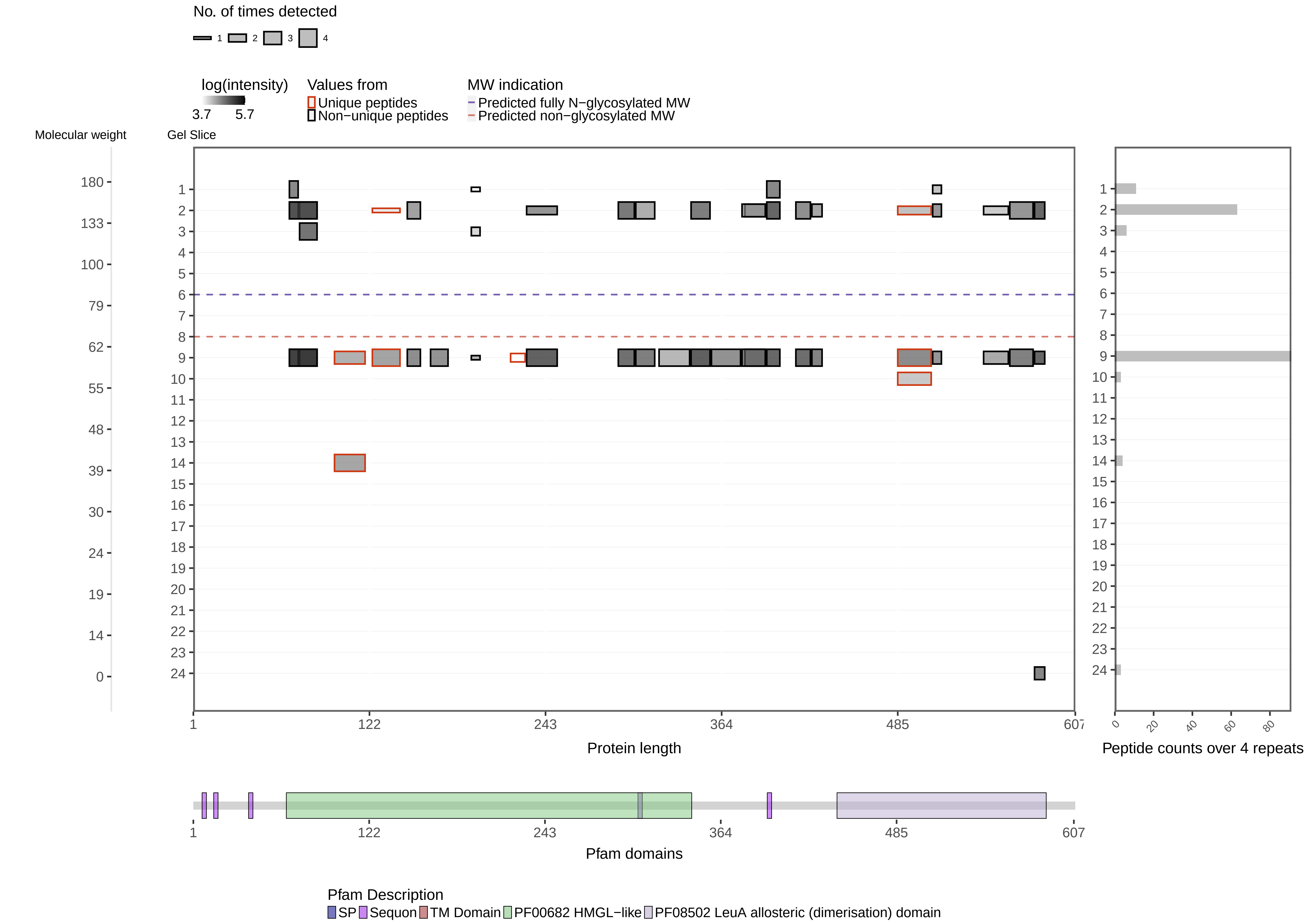


Protein sequence:

NbL09g00250.1 ATP_synthase_subunit_alpha_mitochondrial [MW = 49.53 kDa]

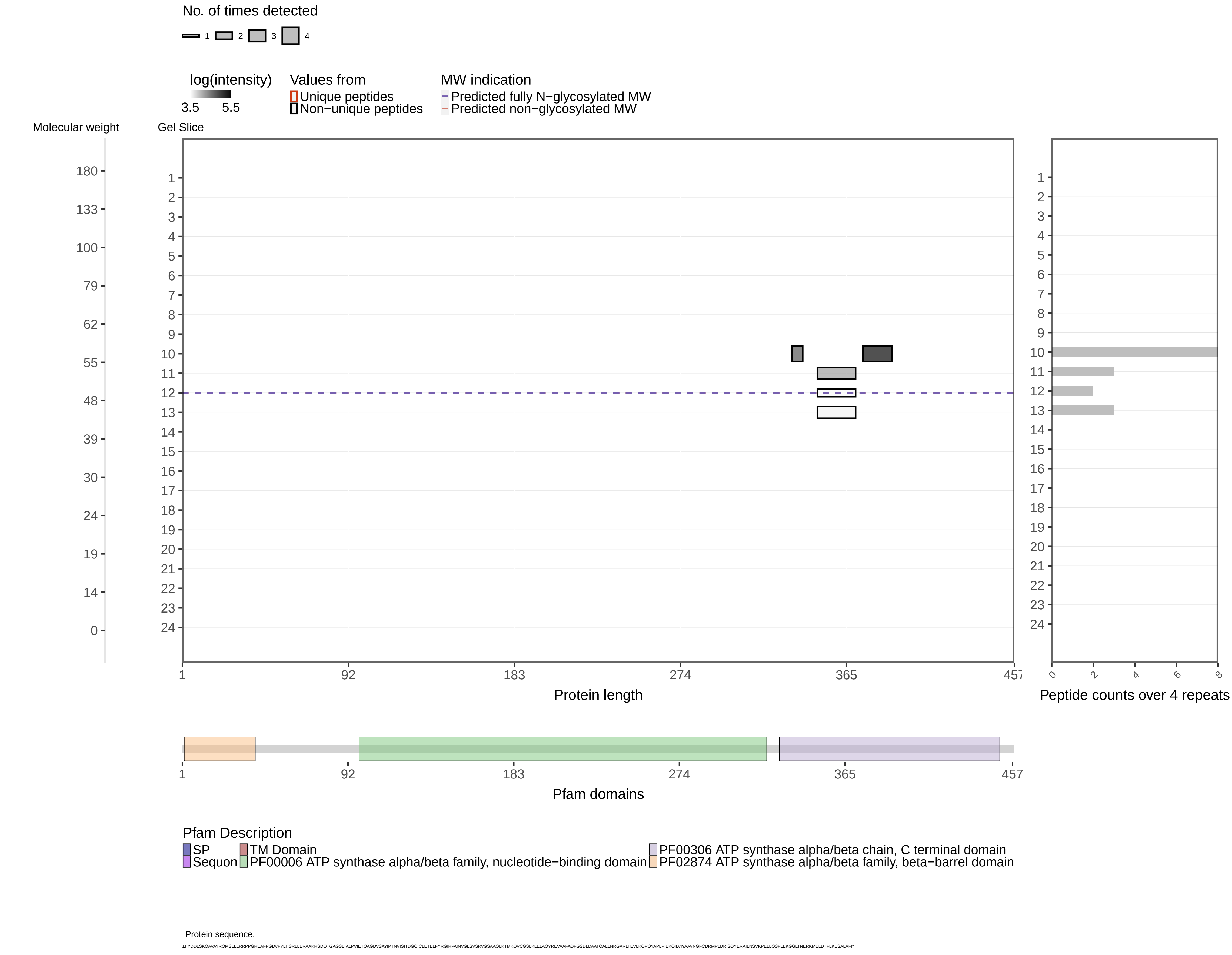

NbL09g00210.1 uncharacterized_LOC105138841 [MW = 49.25 kDa]

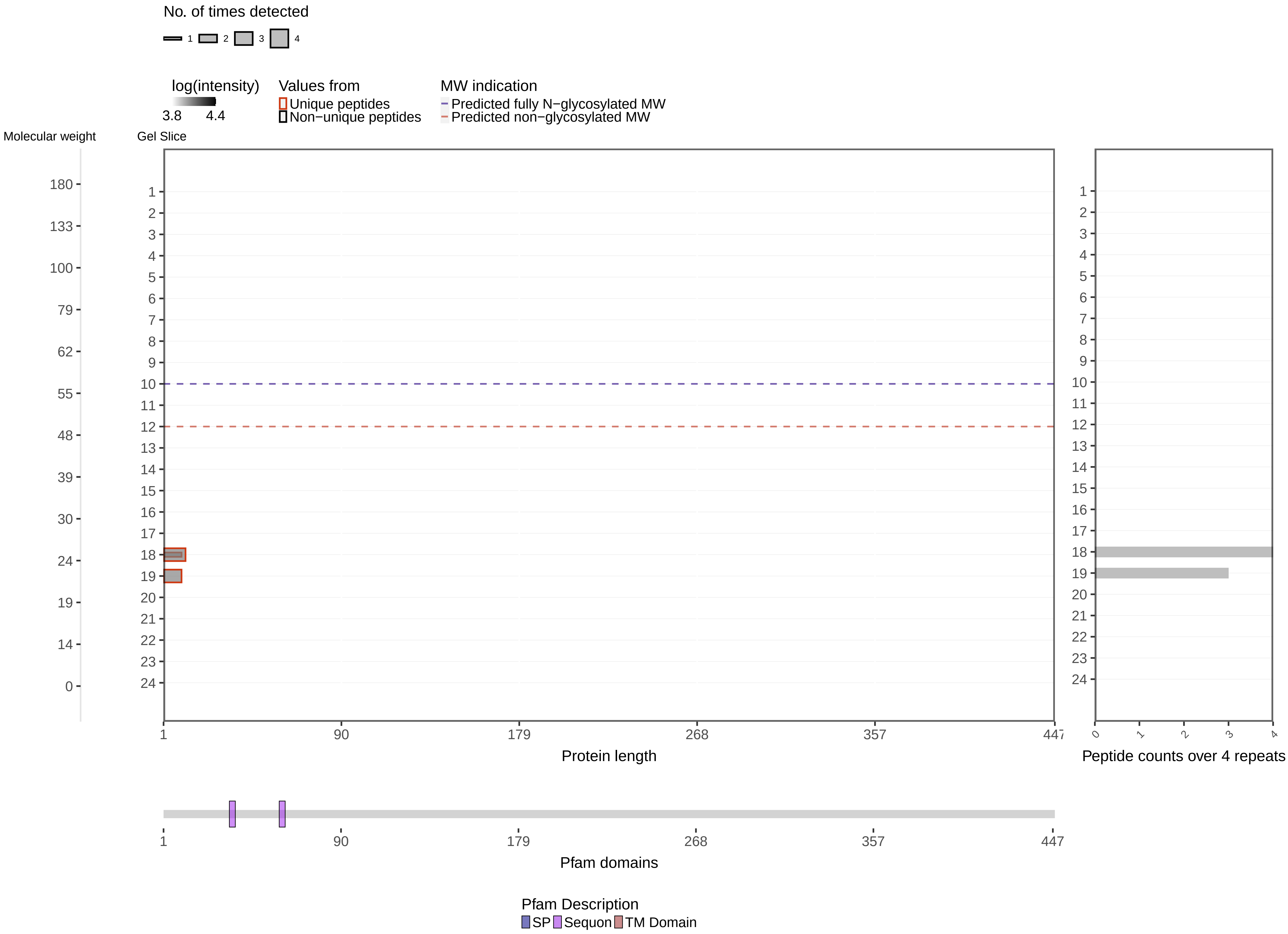


Protein sequence:

ELELAGCWRRHQSDLQEPGRSVFEPTGSGTSSGNSKEGLDTSCASGLTQLDRVSPNCFAVLRDPEQEEAKMPDLDTAEPEEISRDECLGNKAEEGVVKDRTLEESDLAQRSEDLEETVKYGSDEGDLFCDKQMTSNNDLRASKRGEPKKSSKSKRSSTRTMTNKVEDPLLESDKAALPSEKTKKGQTTPFFIDWLHFPLLPTLSSTNKIRVVTAYRHYFKGDCLHTGMVSPSTEL*

### NbL09g00040.1 40S_ribosomal_protein_S15 [MW = 17.17 kDa]

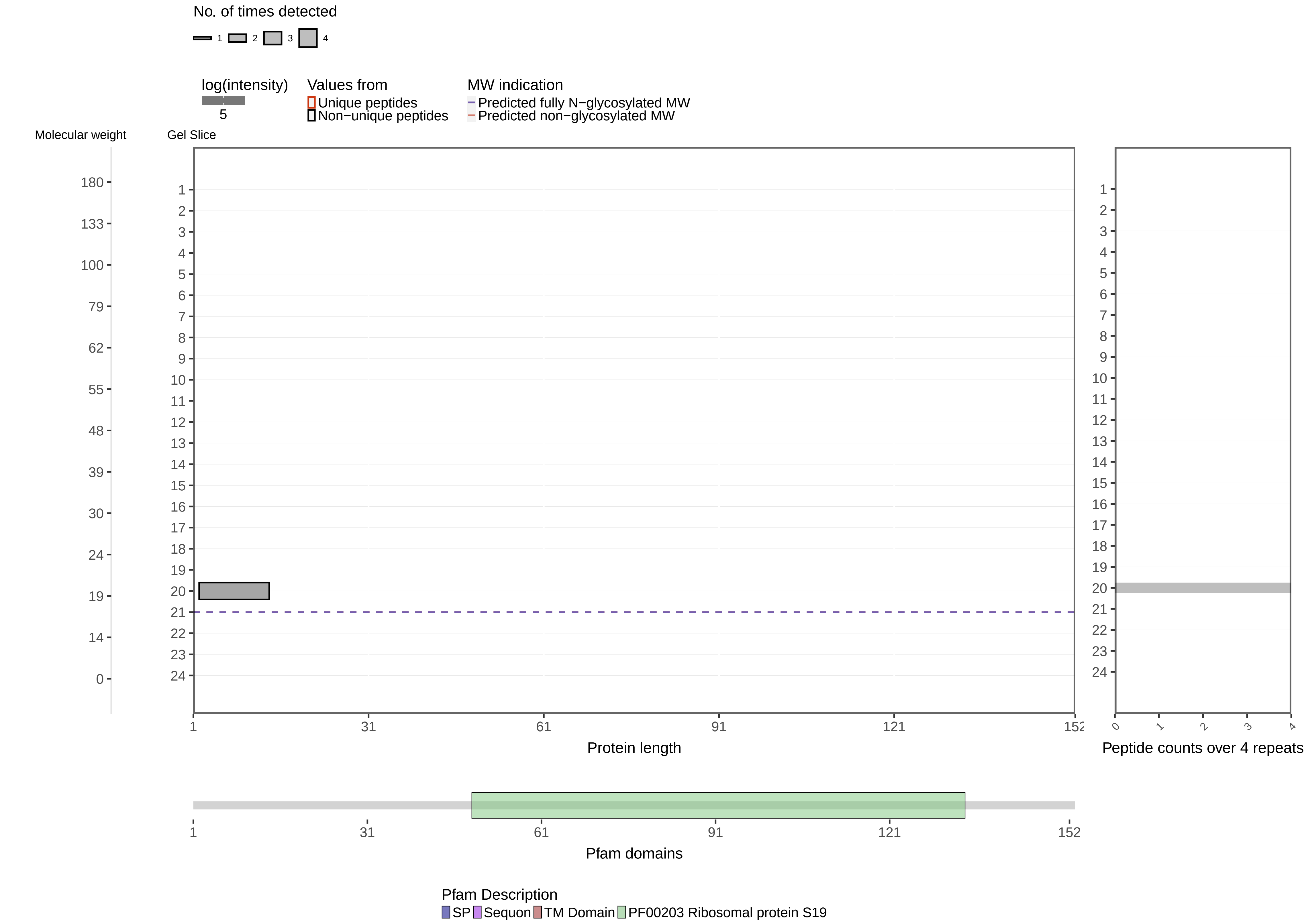


Protein sequence:

ALIKKLRKAKREAPPGEKPEPVKTHLRNMIIVPEMIGSVIGIYNGKTFNQIEVKPEMIGHYLAEFSISYKPVKHGRPGIGATHSSRFIPLK*

### NbL08g21680.1 phytanoyl−CoA_dioxygenase [MW = 32.27 kDa]

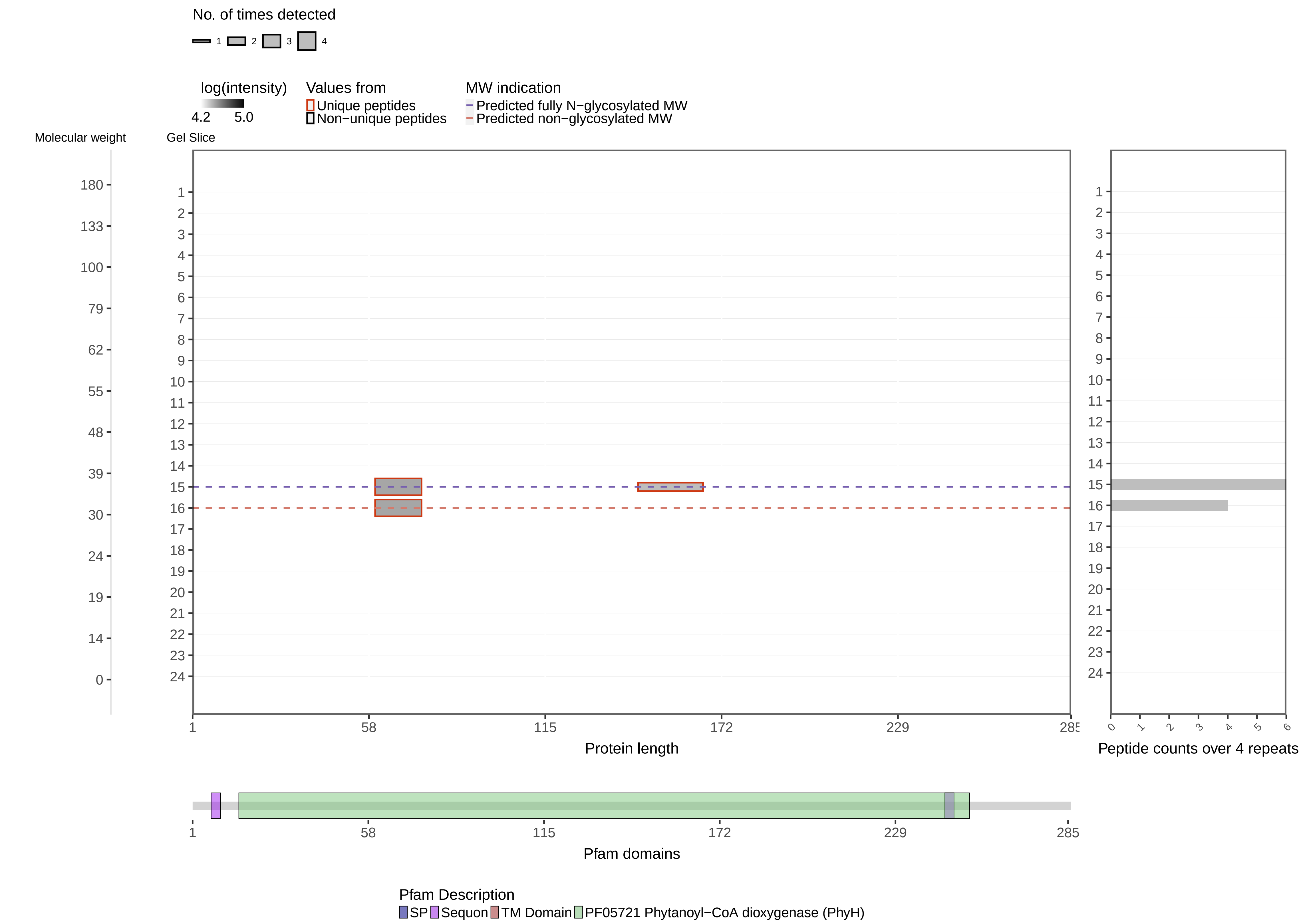


Protein sequence:

GYQRPAVIQSMYIFKQPGIGGEVVPHQDNSFLYTEPKTCTGLWLALEEATIVNGCLWAIPGSHKNGLVRRFLRDENGVHFDKPSPCYNQKDFVPLEVKAGSLVVIHGDLIHQSFENQSSKSRHAYSLHVVDTNGCKWAEDNWIRRNVDPEPLYSP*

NbL08g21650.1 putative_late_blight_resistance_protein_homolog_R1A−10 [MW = 102.56 kDa]

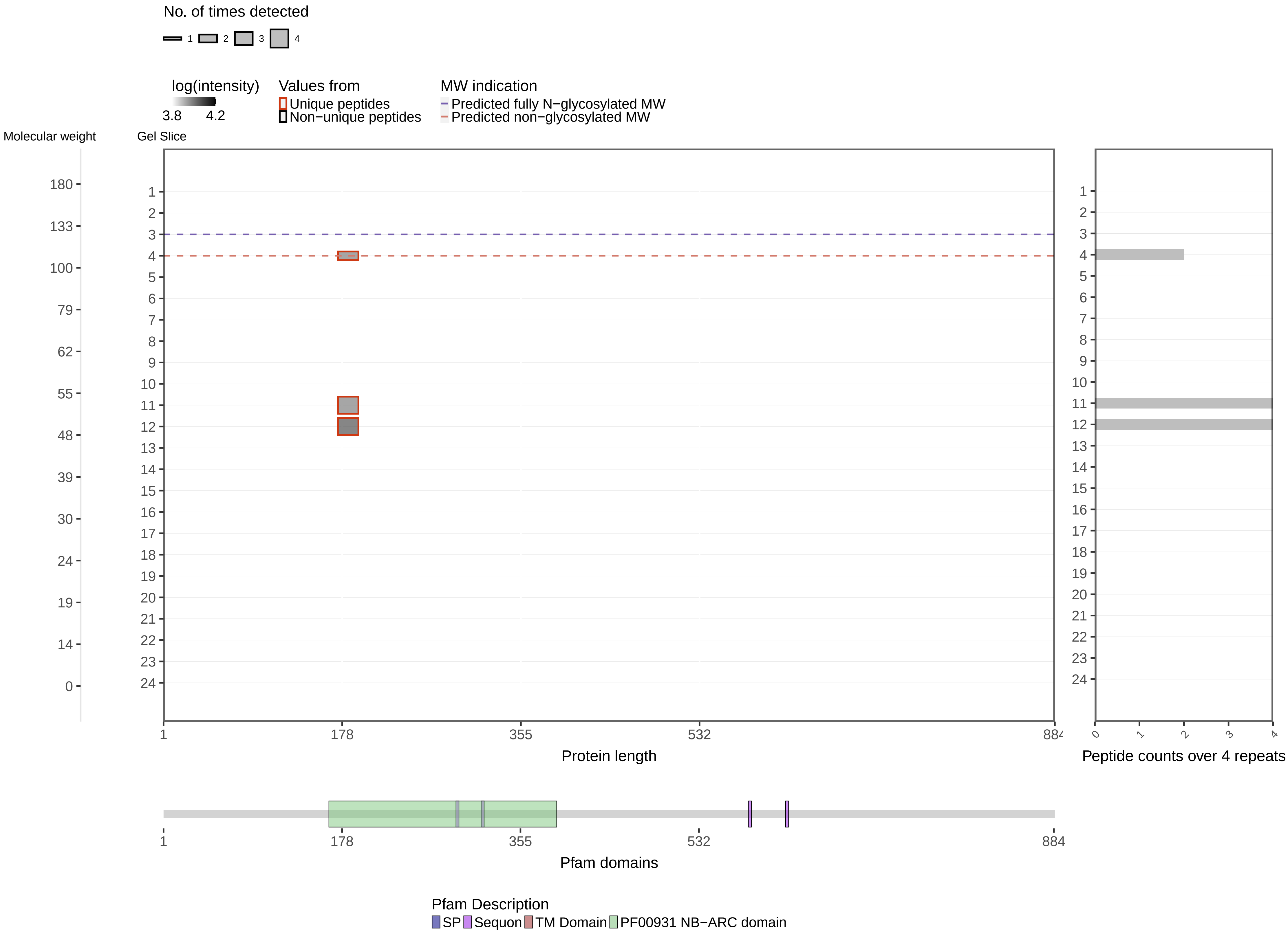


Protein sequence:

IGVSKLIKLWIAELFVRARSNKRLEMVAEEYLEELIDRSLILAGKRRANGRMKTCKIHDLLRQLCIREAQFENFVHVLSDIDHISSESIKYQRRAMLSLGNHWNHFYVPRHWRFITRTTNSLVFTGYSFQVSVMPQFVLQFKLLKVLEVSSVDYNFSWVIPKLVHLRYIAAKINIAPSLDKLWNLQTIILHNLTRRDFHLPLEIWTMSKIRHLDIDTGIHMRNPLEAENLCIIEQPLLLNNLQKLALPSSPFLAEILRRTPNLNKLQIVDMKHADWPAVLDSLILLSELETLAIKAERNIDRIV

NbL08g21570.1 nicotinate−nucleotide_pyrophosphorylase_[carboxylating]_chloroplastic [MW = 39.91 kDa]

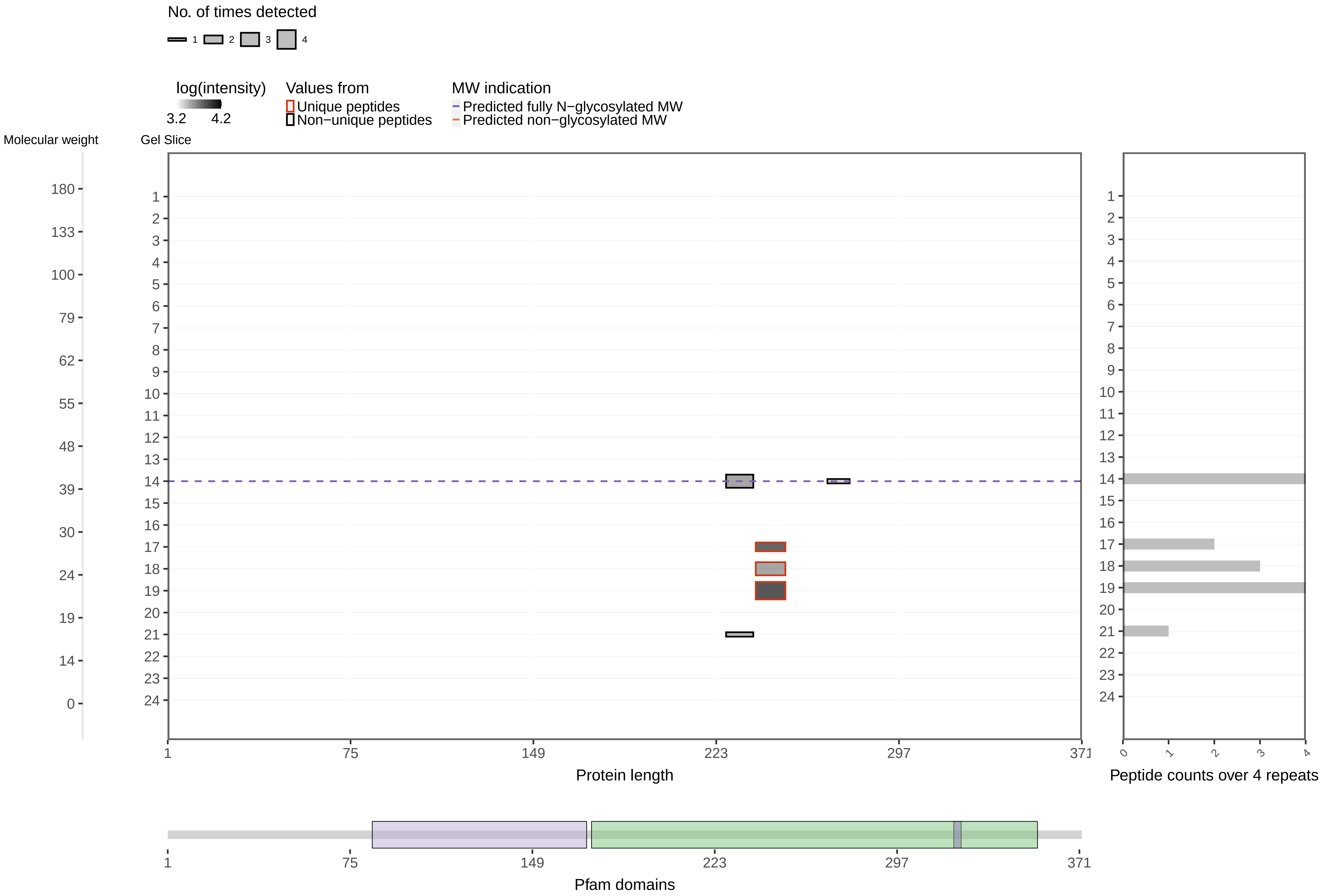


Pfam Description
SP Sequon TM Domain PF01729 Quinolinate phosphoribosyl transferase, C−terminal domain PF02749 Quinolinate phosphoribosyl transferase, N−terminal domain

Protein sequence:
ATLTKAMADAAHPATILETRKTAPGLRLVDKWAVLIGGGKNHRMGLFDMVMIKDNHISAAGGVSKALNSVDQYLEQNKLQMGVEVETRTIAEVREVLDYASQTKTSLTRIMLDNMVVPLSNGDIDVSMLKESVDLINGRFETEASGNVTLETVHKIGQTGVTYISSGALTHSVKALDISLKIDTELALEVGRRTKRA*

### NbL08g21480.1 multicopper_oxidase_LPR1−like [MW − SP = 62.95 kDa]

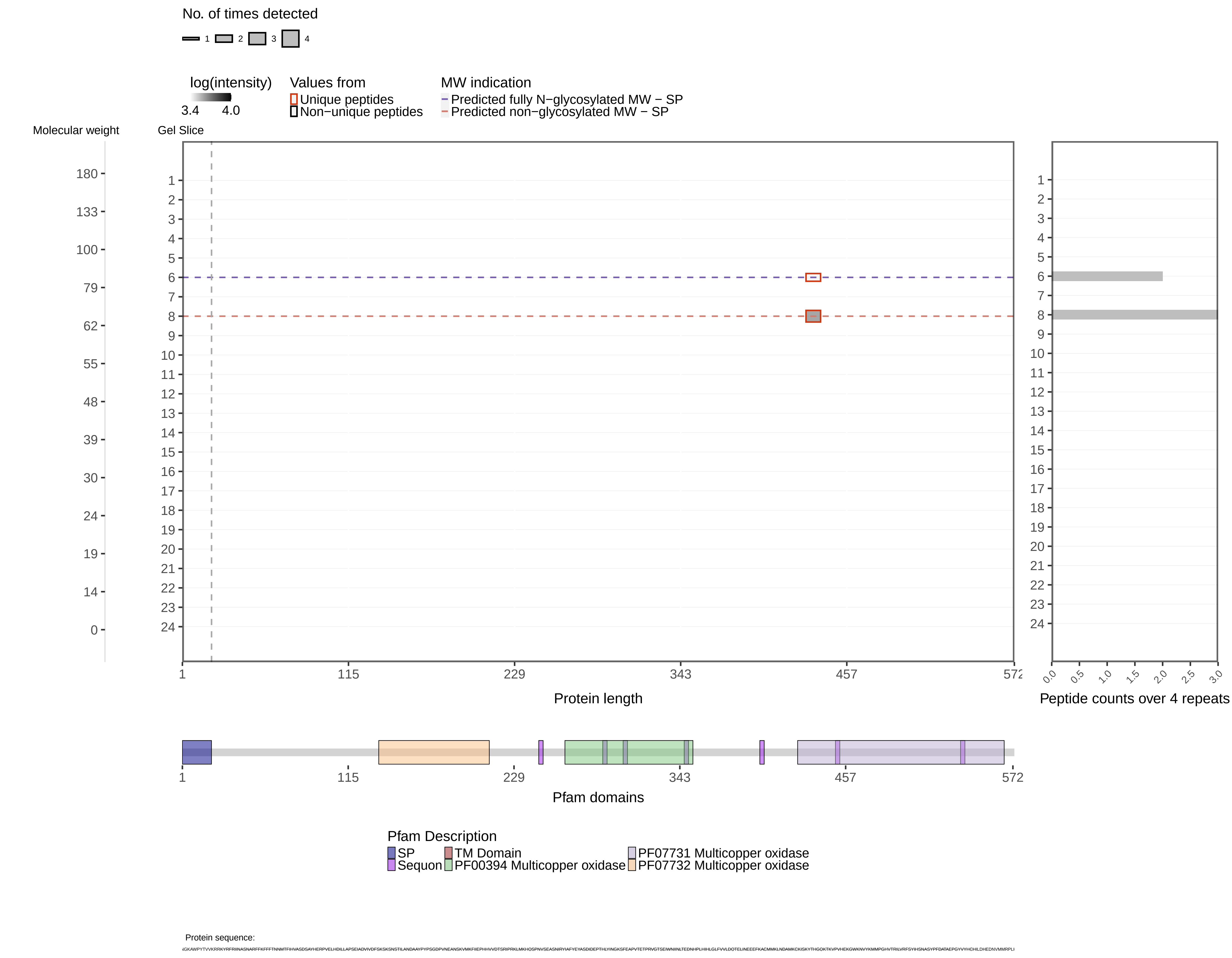

NbL08g21420.1 probable_ribose−5−phosphate_isomerase_2 [MW = 28.33 kDa]

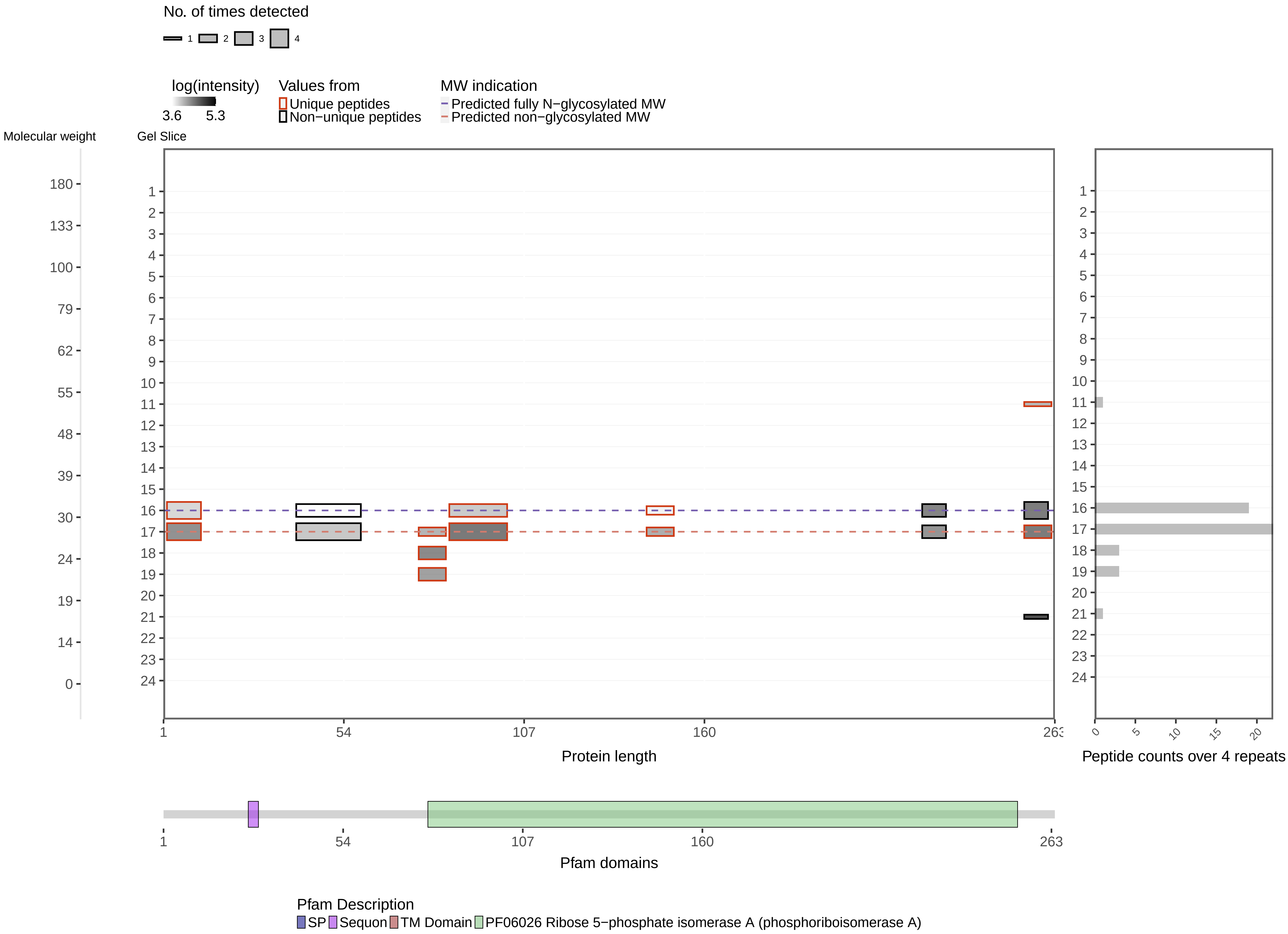


Protein sequence:

MNLVKGRGGSLLREKMVEAASKKFIVIIDESKLVNYLGGSGLAMPVEIVPFCWEFTLKRLEMLFIEAGCVGKLRTVGENGEVYVTDNGNYIIDLYFKKDMGDLKAASDAILRLAGVVEHGMFIDMATTVIVAGKLGVSVTNKE*

NbL08g21340.1 ATP−dependent_Clp_protease_ATP−binding_subunit_ClpA_homolog_CD4B_chloroplastic [MW = 102.26 kDa]

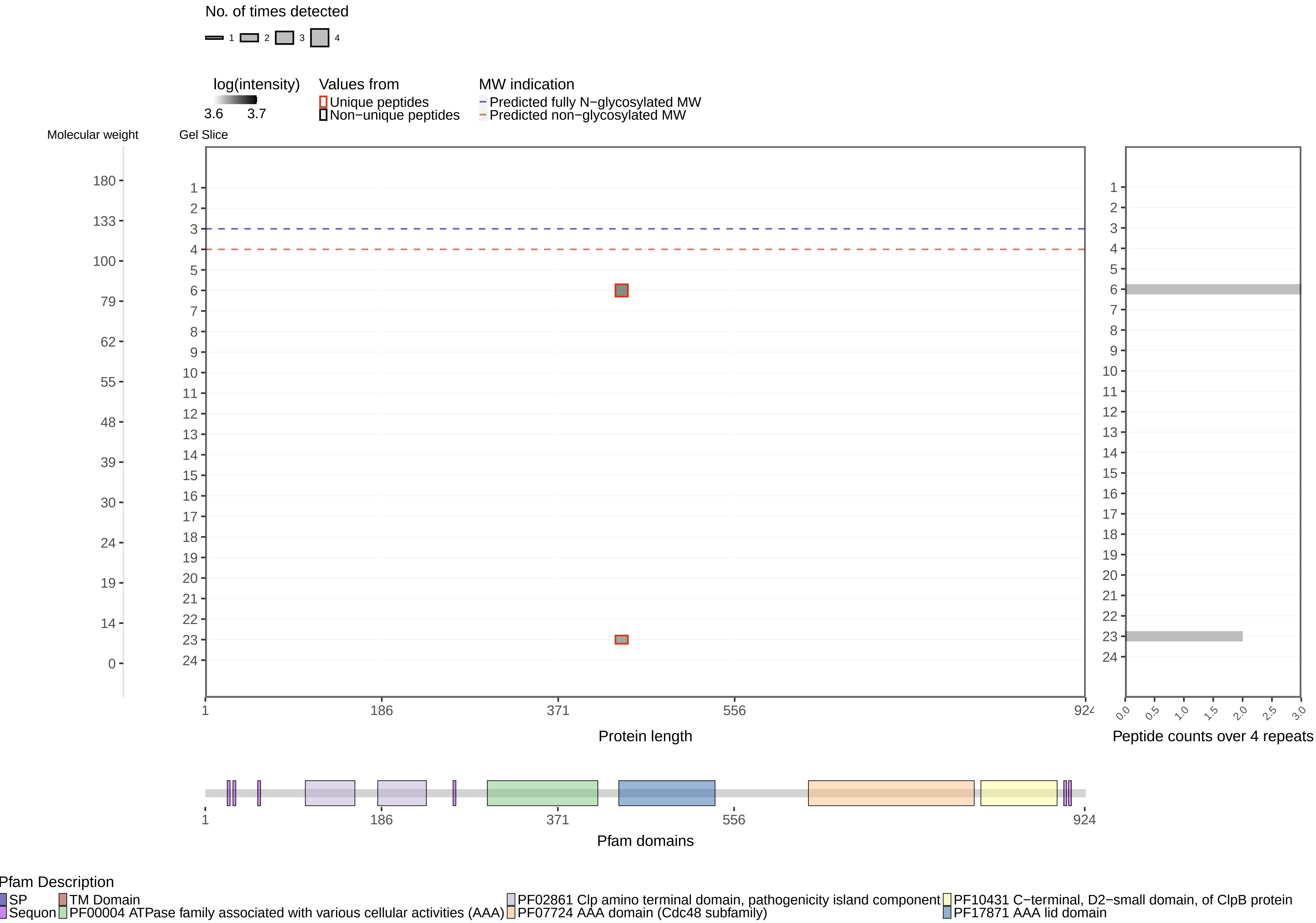


Protein sequence:

### NbL08g21220.1 uncharacterized_LOC107812854 [MW − SP = 48.81 kDa]

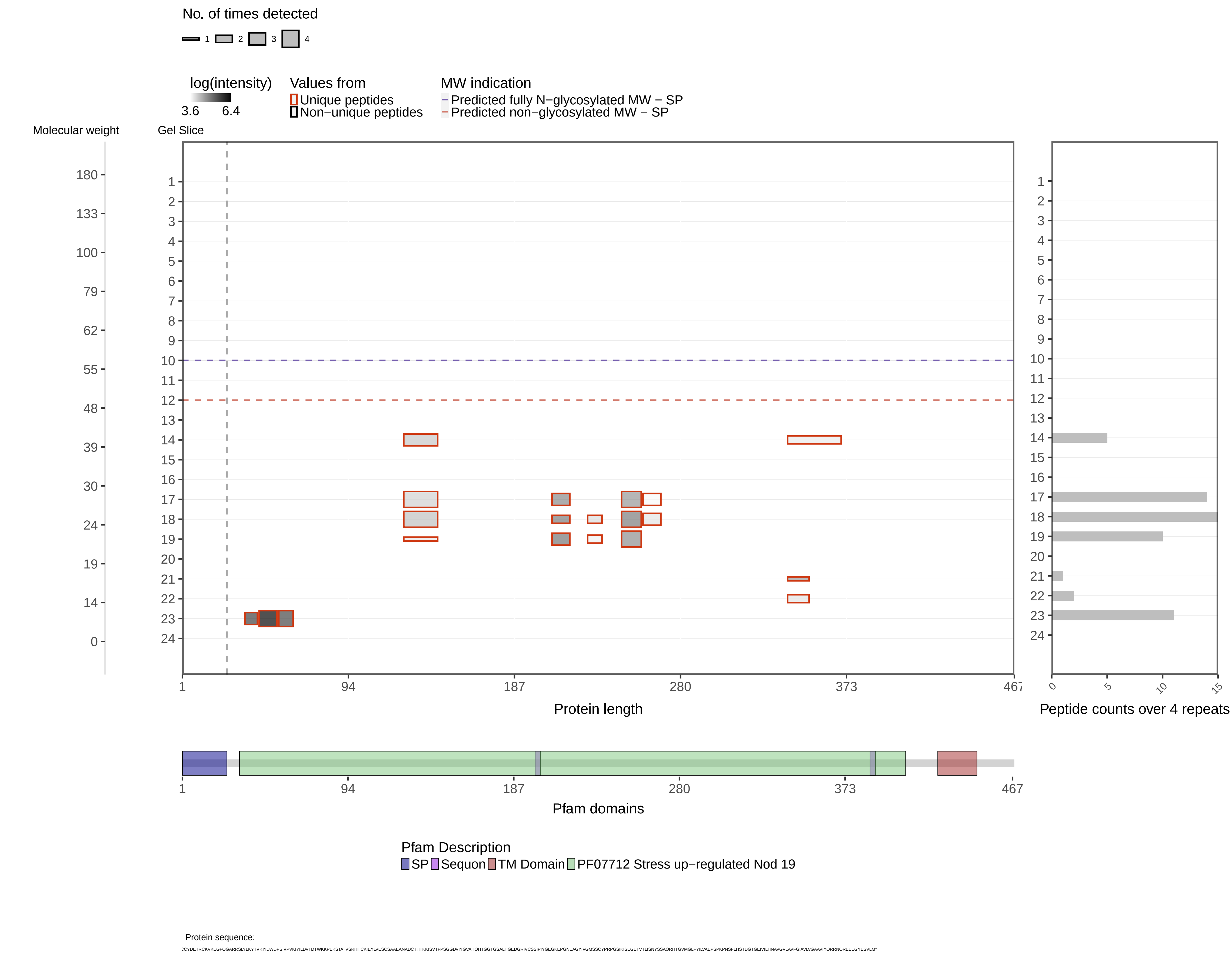

### NbL08g21110.1 shikimate_O−hydroxycinnamoyltransferase−like [MW = 48.09 kDa]

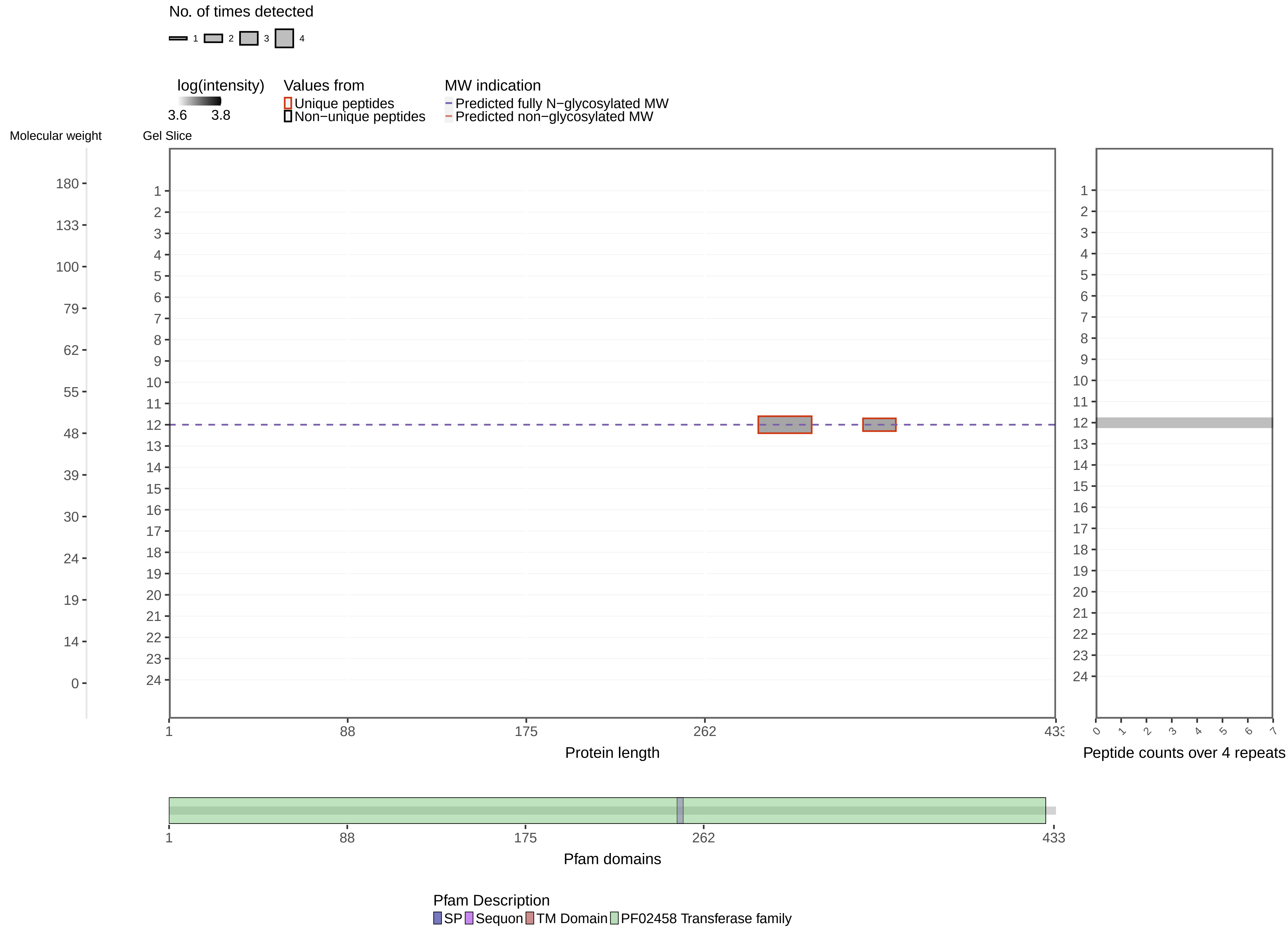


Protein sequence:

IEYQPPPTLKNIAKTEAVPETAVSIFKLTRDQIDALKAKSKEDGNTVNYSSYEMLAGHVWRSTCMARGLEEDQGSKLYIATDGRSRLRPTLPPGYFGNVIFTATPIAVAGDLKSKPIWYAASKIHDALARMDNDYLRSALDYLELQPDLTALVRGAHTFRCPNLGITSWVRLPIHDADFGWGRPIFMGPGGIAYEGLSFILPSPTNDGSLSVAISLQADHMKLFEKYLYDI*

### NbL08g21080.1 40S_ribosomal_protein_S12−like [MW = 14.95 kDa]

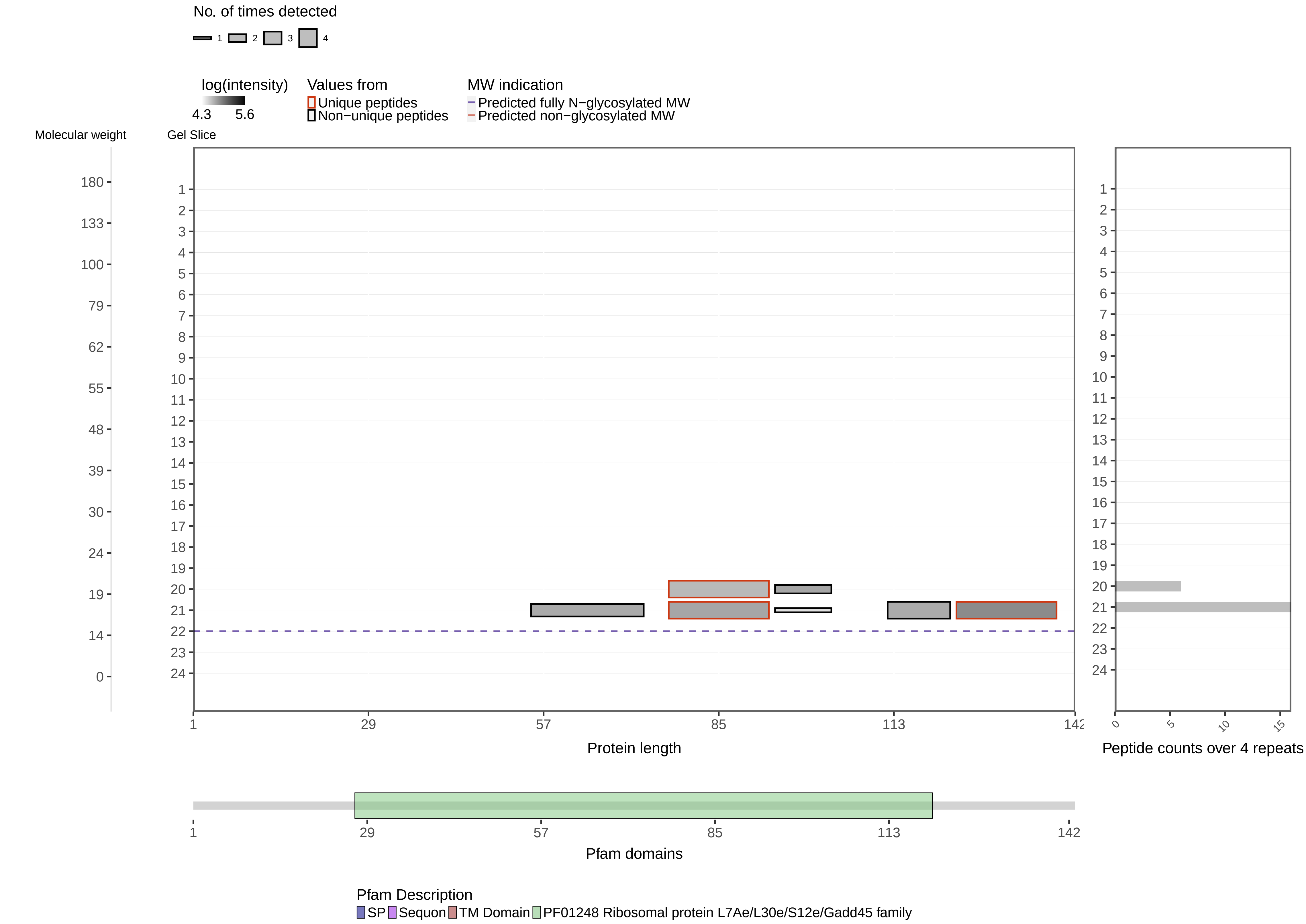


Protein sequence:

LCVLAEDCDQPDYVKLVKALCADHNVNLITVPSAKTLGEWAGLCKIDSEGKARKVVGCGCVVVKDFGEETEGLHIVQEYVKSH*

NbL08g20950.1 stress−response_A/B_barrel_domain−containing_protein_UP3 [MW = 34.43 kDa]

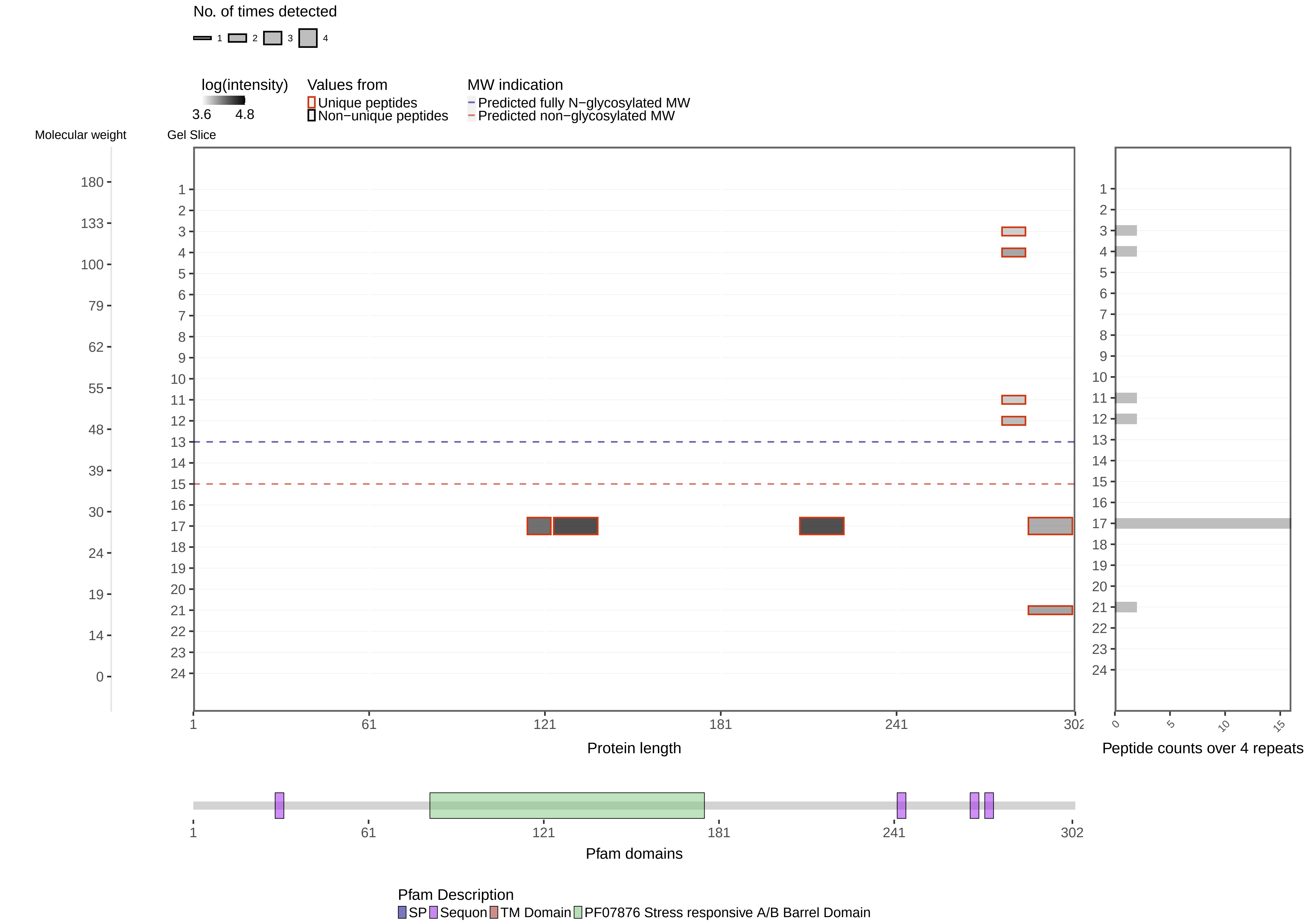


Protein sequence:

TYMRFQTKEDLSKFYENQFYVGVLRDHVLPYCHEIINADFESEVEDDILPIFRKGEEFNYGVELLLLIAFVKSSLDGPAEDALLVLSKLITEFPSLIVQATLGSIINISSVEYTHGVVIRFRSSEAFQMFINSSEYNNTWRSKFQPIVQKHVSVHFSVDPVGTELM*

### NbL08g20940.1 thaumatin−like_protein [MW − SP = 13.65 kDa]

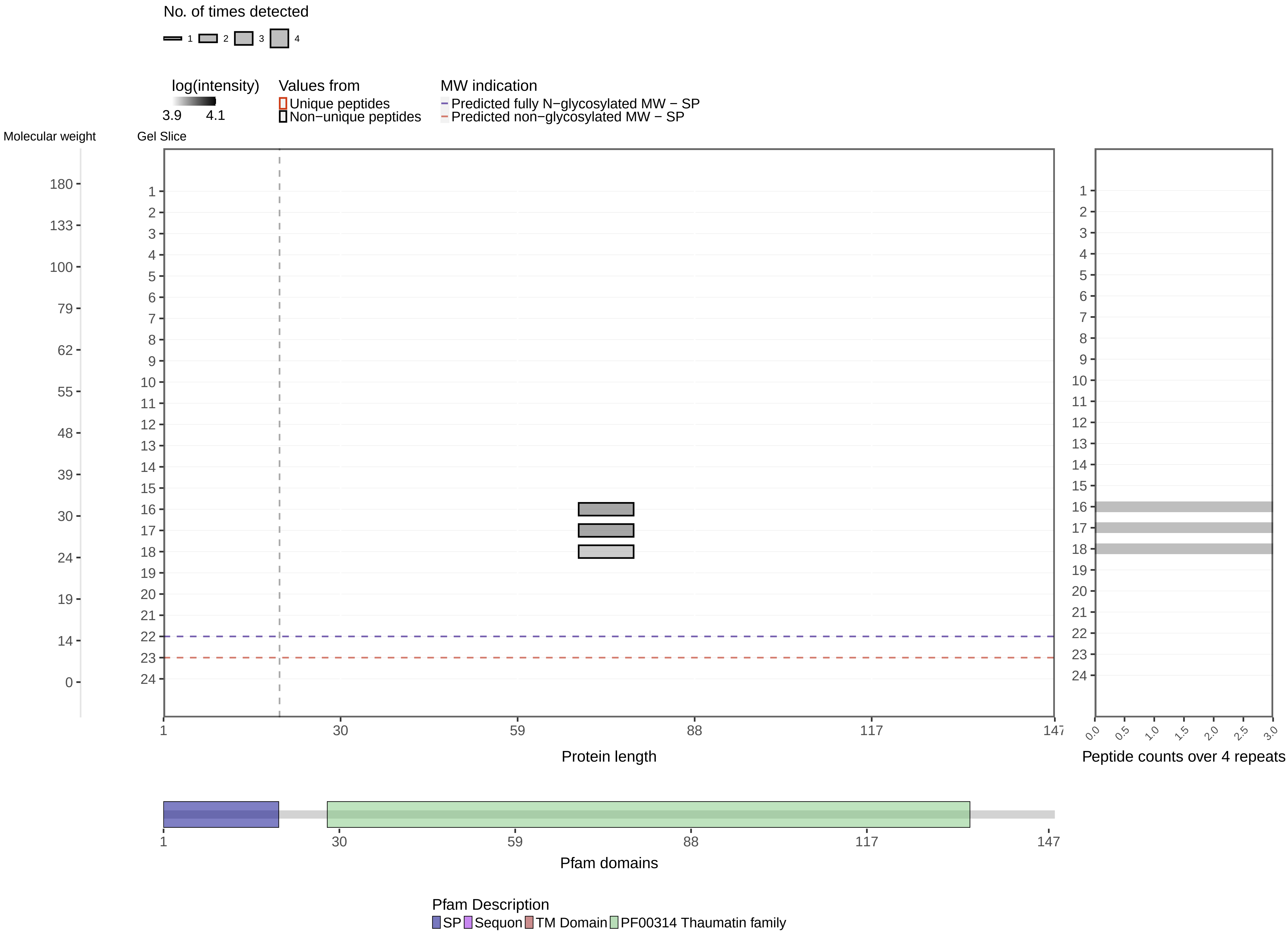


Protein sequence:

ITKRIWSRTNCNFDASGRGQCQTGDCNGVLECQAFGTPPNTLAEYSLKQFNNLDFFDISLVDGFNVPMEFSPTSGVLLRSAALPTS*

NbL08g20930.1 pathogenesis−related_protein_R_minor_form [MW − SP = 21.66 kDa]

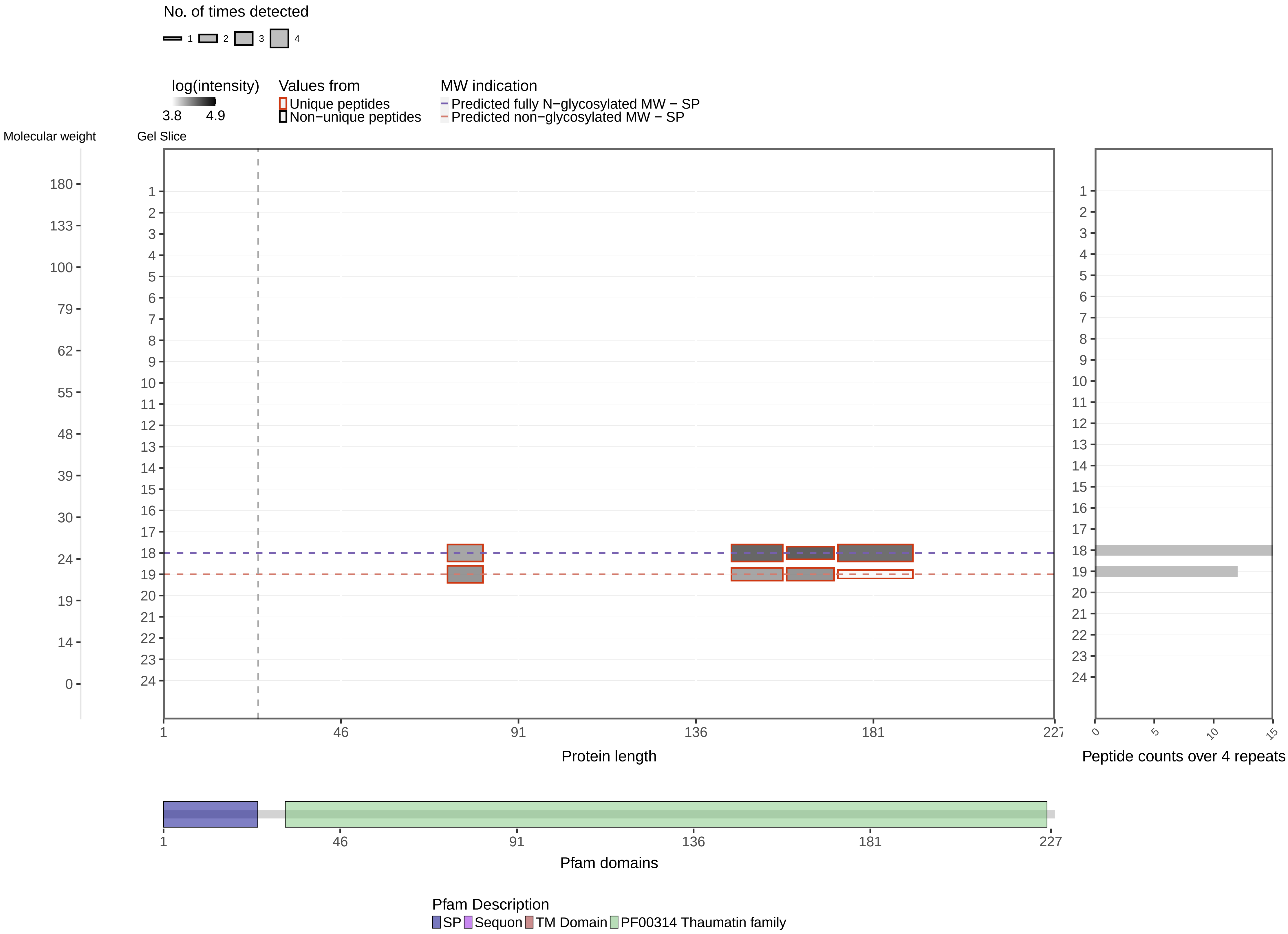


Protein sequence:

GKPPNTLAEFALNQPNQDFVDISLVDGFNIPMEFSPTNGGCRNLRCAAPINEQCPAQLKTQGGCNNPCTVIKTNEYCCTNGPGSCGPTDLSRFFKERCPDAYSYPQDDPTSLFTCPSGTNYRVVFCP*

### NbL08g20860.1 octanoyltransferase−like [MW = 26.43 kDa]

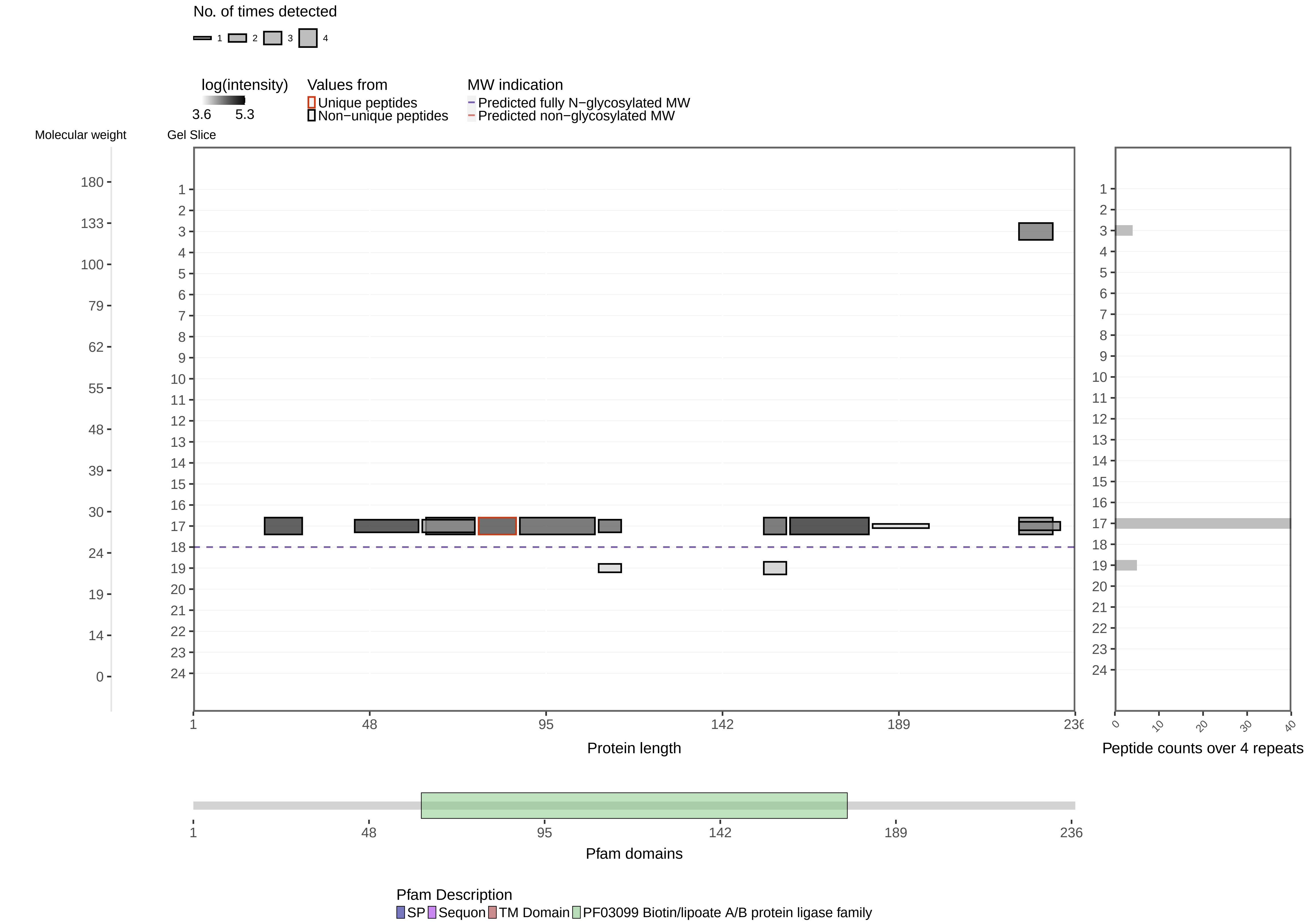


Protein sequence:

PIVSLRDIGLGARKYVEKLELTMIELASIYGVKAQVGKRCETGVWVEDRKIGAIGVRISSGITSHGLAFNMDPDLNYFKHIVPCGIADKDVTSLKKEADVELPAEEVIQEQLIACFVRIFGYNDVILKDKSVL*

NbL08g20690.1 ATP_synthase_delta_chain_chloroplastic−like [MW = 26.79 kDa]

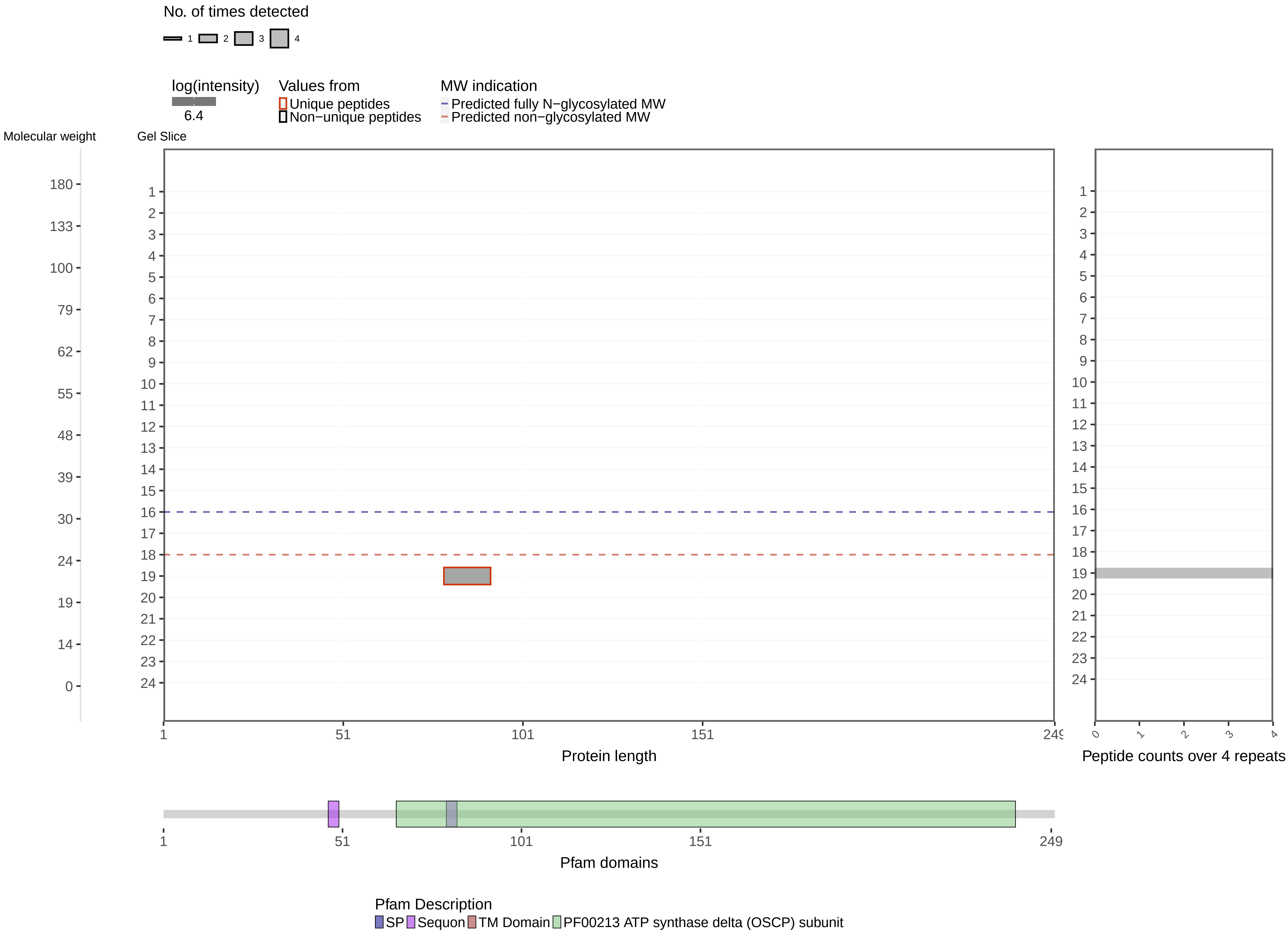


Protein sequence:

PIVGEEKKRELVDEIVSSSSIQPHVANFLNILVDMKRVELIKEIVKEFEKVYNTITDTELAVVTSVVKLESQHLAQIAKGVQRLTGSKNVRIKTAIDESLVAGFTIRYGNSGSKLIDMSVKKQLEDIAAQLEIGDIQLAV*

NbL08g20660.1 protein_tumorous_imaginal_discs_mitochondrial [MW = 32.85 kDa]

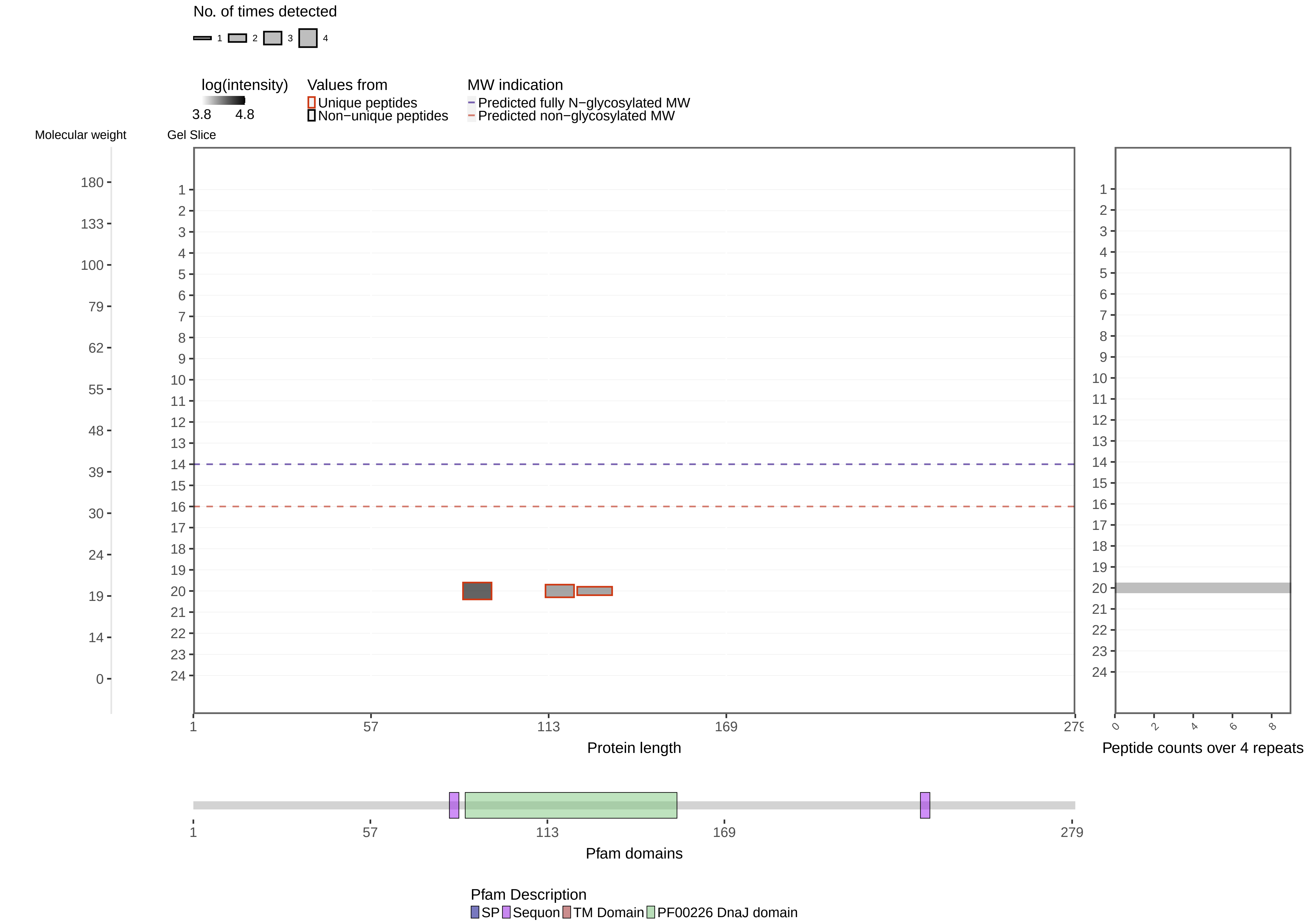


Protein sequence:

EEGETAEARFIKIQAAYELLIDEDERKKYDIANRVNPMKASEAWMEWLRKKRKAFDQRGDMAIAAWAEQQQLELNLKVRRLARSKMDPDEERRIIAKEKKASLENFNNTLKRHTLVLRKRDIMRRKAEEEKKKVISMLLAAEGLELEEEDDKNL*

NbL08g20630.1 probable_glucan_13−beta−glucosidase_A [MW = 36.26 kDa]

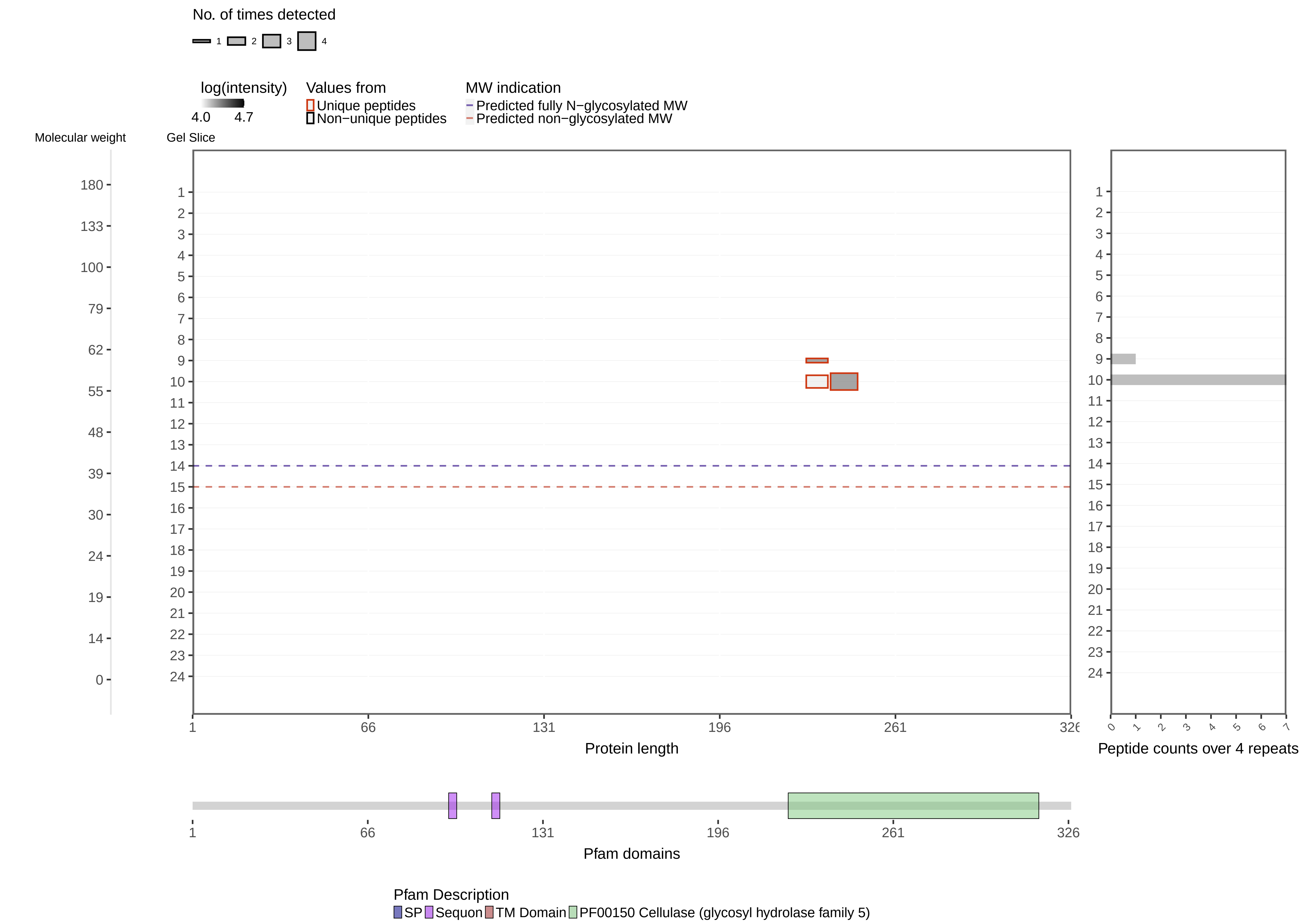


Protein sequence:

SDNPSRVRIKASNGFFLOVKTEELVTADNGGNGGWGDDDPSVFIMOTSGRLEGEFOITNGYGPIMAPOVMREHWKTFIVEEDFKFIASNGLNAVRIPVGWWIASDPTPPKPYVGGSLHALDNAFFWAKKYGLKVIIDLHAAPGSONPWEHSANRDGTIEWEOLMTTFNRLLOS*

NbL08g20540.1 7−deoxyloganetin_glucosyltransferase−like [MW = 54.34 kDa]

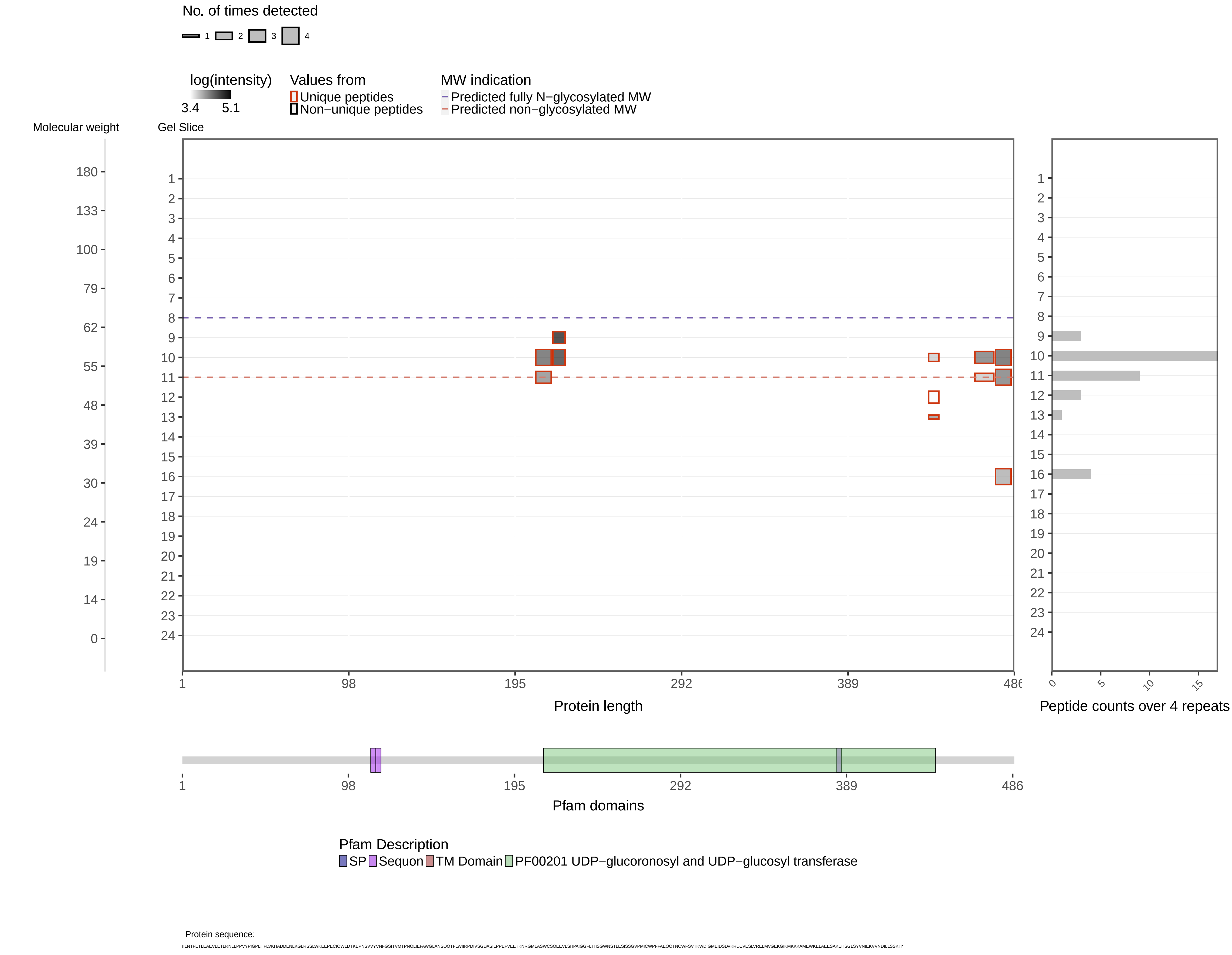

### NbL08g20530.1 14−3−3−like_protein_B [MW = 28.77 kDa]

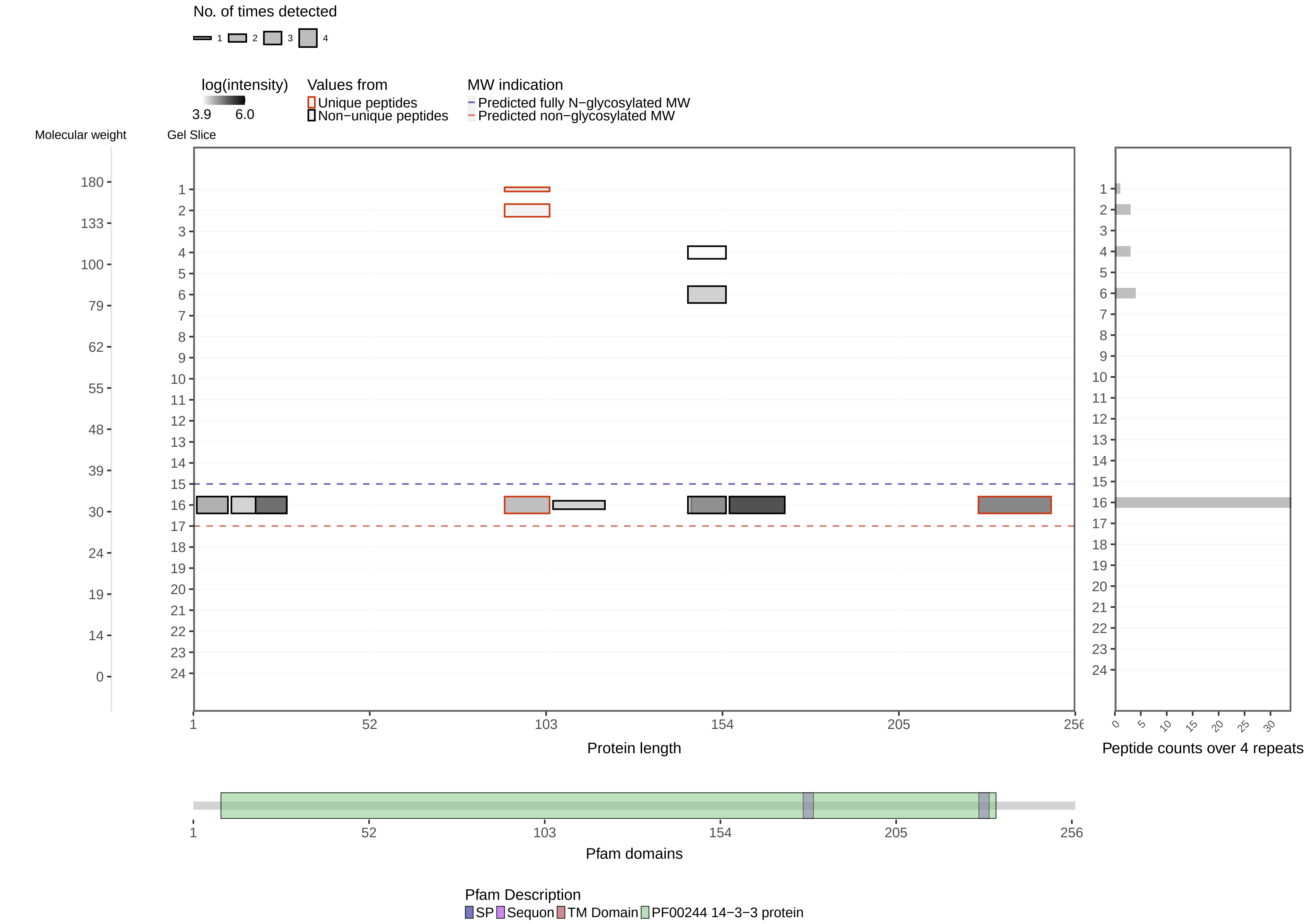


Protein sequence:

AGDSKVFYLKMKGDYHRYLAEFKTGAERKEAAESTLSAYKAAQDIANAELAPTHPIRLGLALNFSVFYYEILNSPDRACNLAKQAFDEAIAELDTLGEESYKDSTLIMQLLRDNLTLWTSDMQDDGADEIKETKADNEQQ*

### NbL08g20300.1 LIM_domain−containing_protein_WLIM1−like [MW = 21.43 kDa]

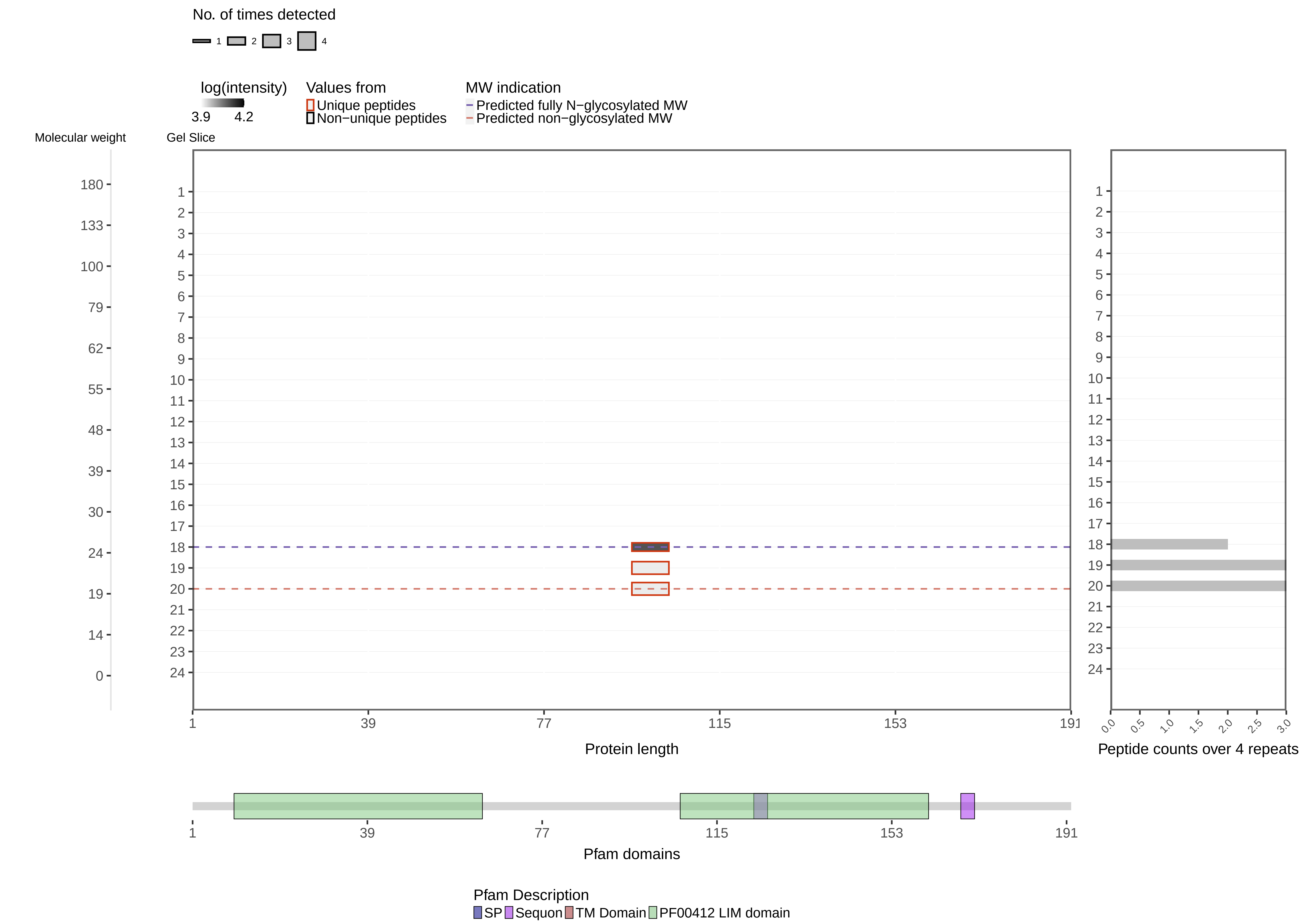


Protein sequence:

VKPEKDEKPHAAKVSSMFVGTREKCFGCKNTVYPTEKVSVNGTPYHKSCFKCSHGGCTISPSNYIAHEGRLYCKHHHIQLIKEKGNLSQLEGDHDKNTVRTAQVKAES*

NbL08g19980.1 low−temperature−induced_cysteine_proteinase−like [MW = 56.48 kDa]

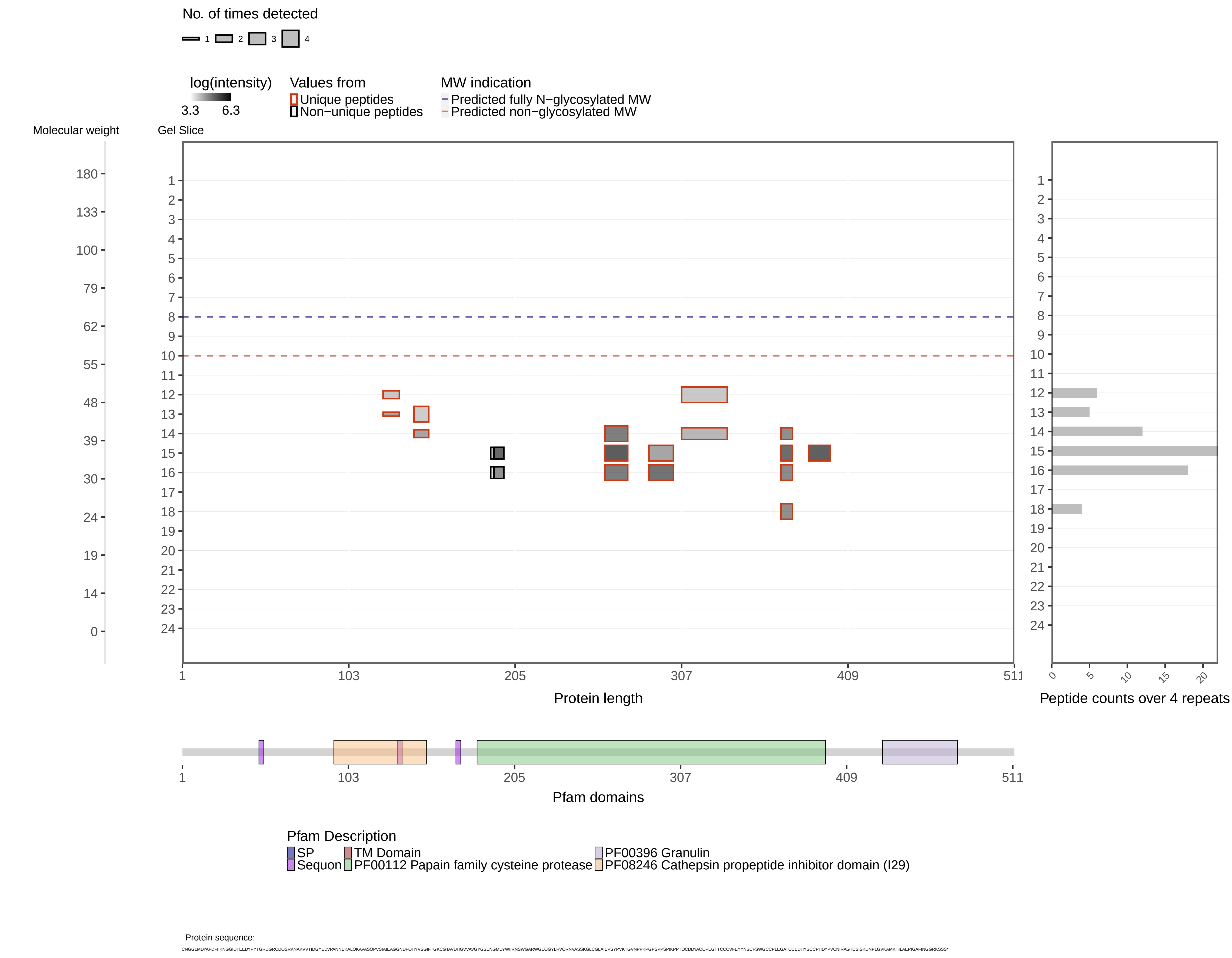

### NbL08g19960.1 polyadenylate−binding_protein_2−like [MW = 71.97 kDa]

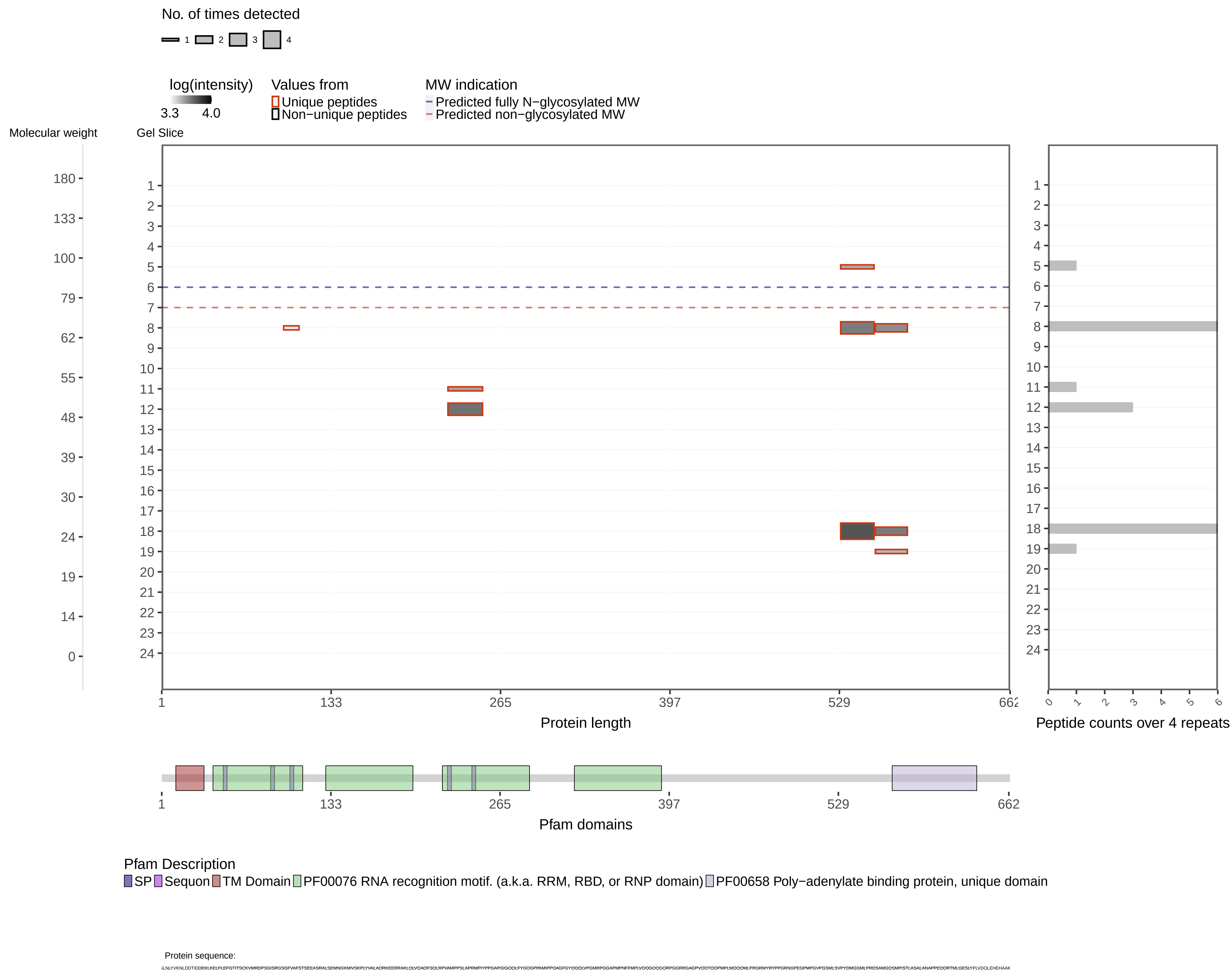

### NbL08g19950.1 50S_ribosomal_protein_L4 [MW = 34.3 kDa]

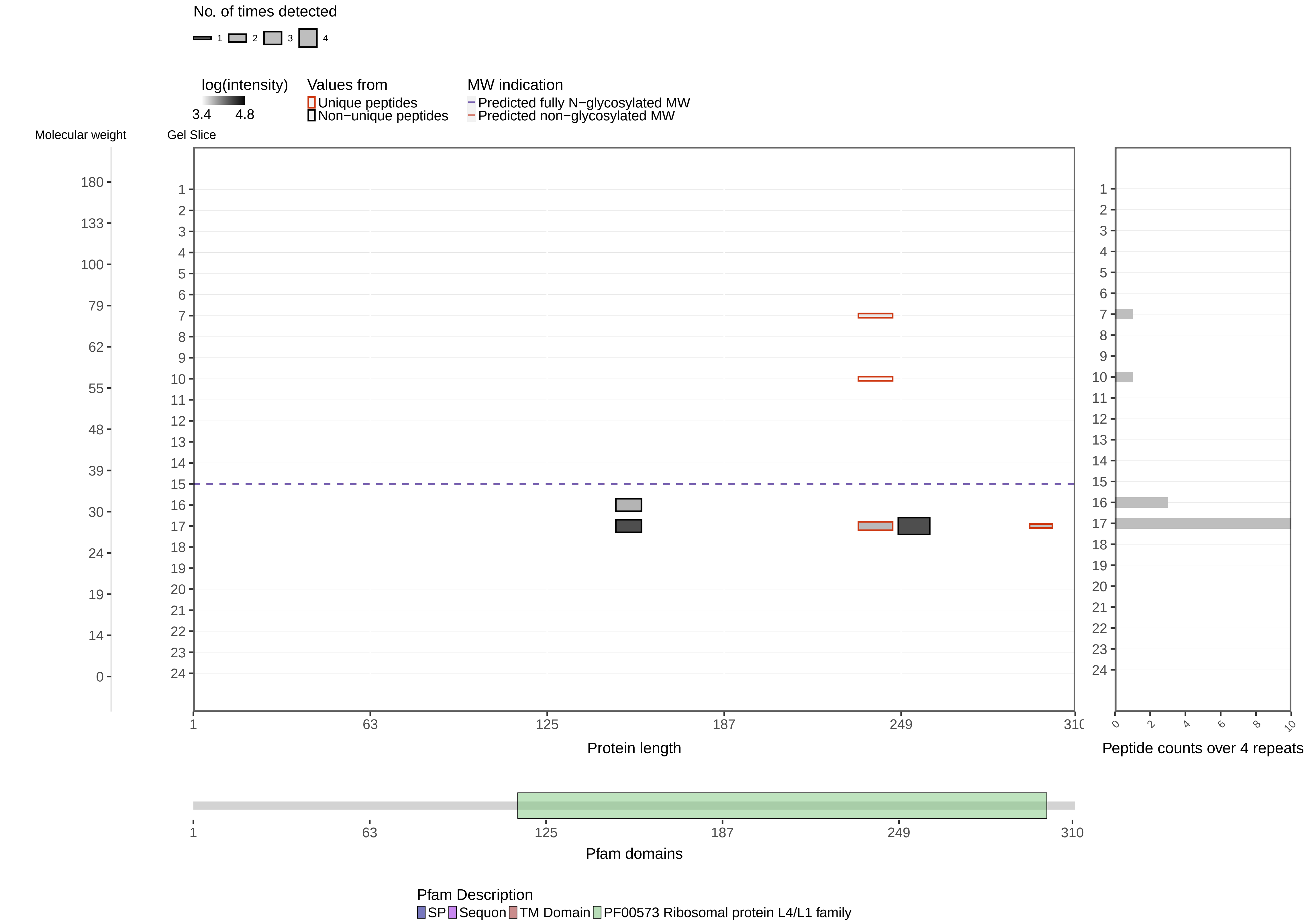


Protein sequence:

GTHSTKTISEVSGTGRKPWRQKGTGRARHGTLRGAQFRGGATMHGPKPRSHAIKLNKKVRRLGLKIALSARAAEGKLMVFEDLEISSHKTKNIVNYFNQLVNTKKLLLVDGGPISEKLKLATQNVHYVNVLPSIGLNVYSILLHDTLVMSRDAVNTIVERMHTPINR*

NbL08g19920.1 subtilisin−like_protease [MW − SP = 79.88 kDa]

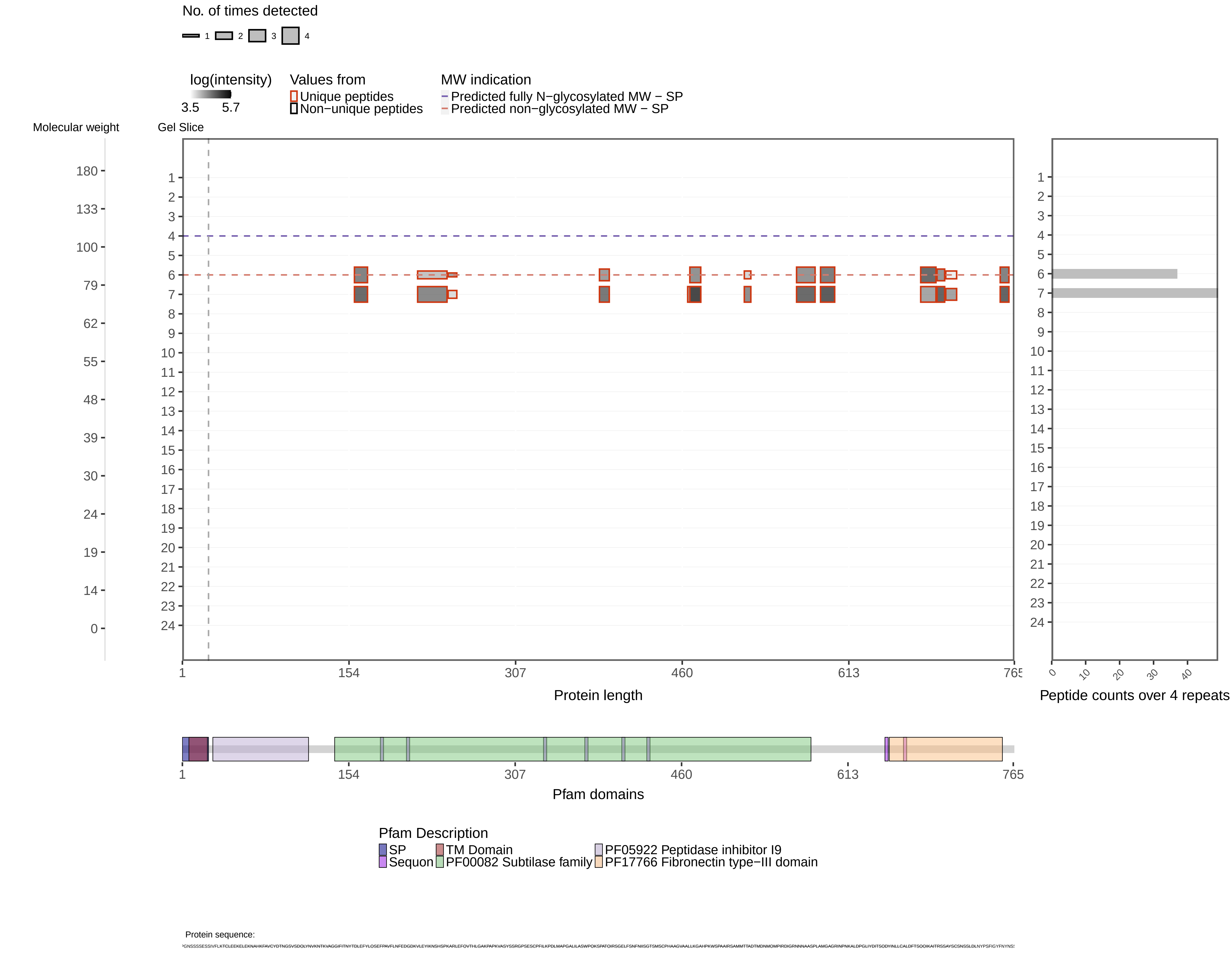

### NbL08g19570.1 ornithine_carbamoyltransferase_chloroplastic−like [MW = 41.83 kDa]

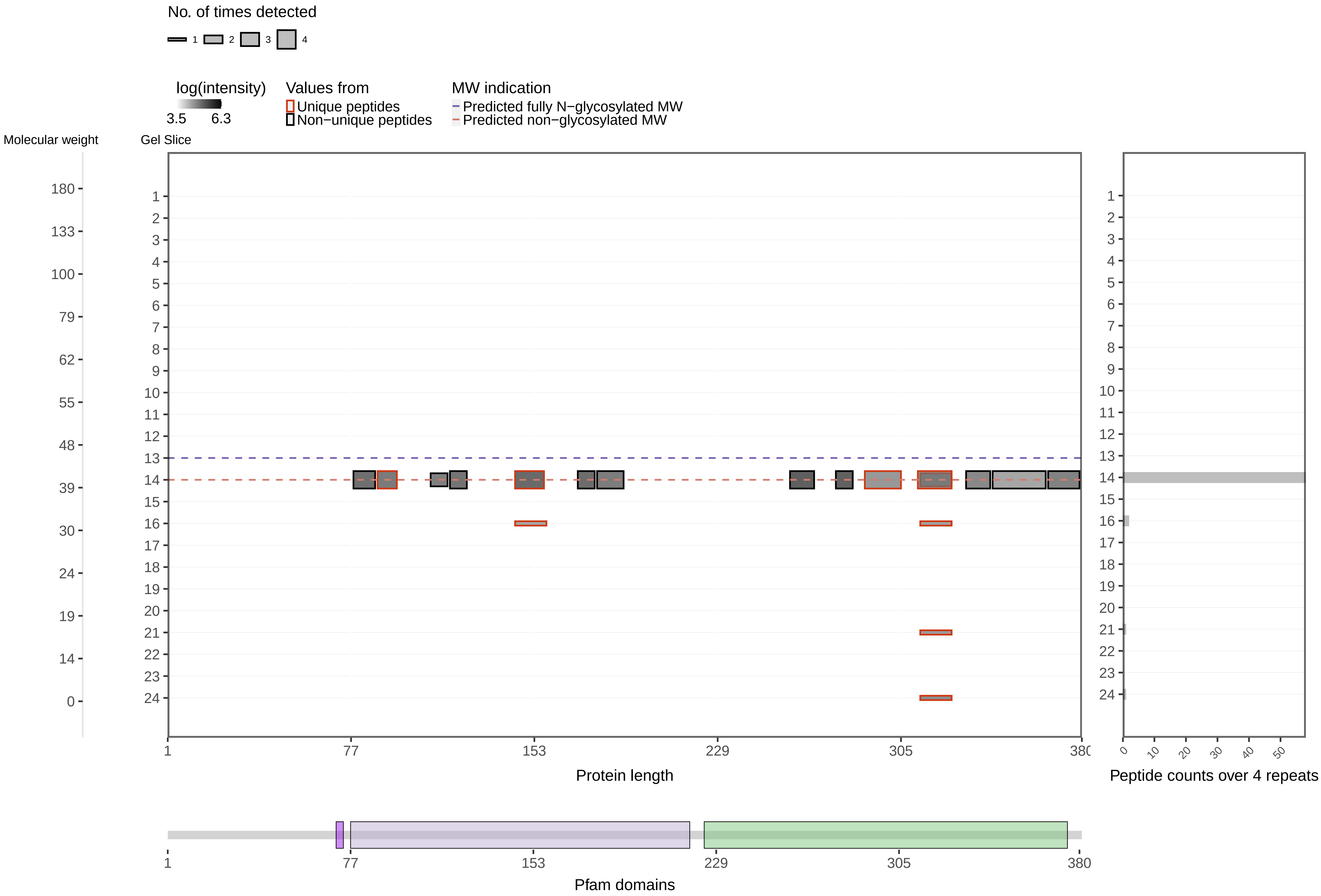


Pfam Description

SP Sequon TM Domain PF00185 Aspartate/ornithine carbamoyltransferase, Asp/Orn binding domain PF02729 Aspartate/ornithine carbamoyltransferase, carbamoyl−P binding domain

Protein sequence:

FAHQDILDLAKYATVPVINGLTDYNHPCQIMADALTMIEHVGQLEGTKVVYVGDGNNIVHSWLLLASVIPFHFVCACPKGFEPDQETVEKARQAGVSKIEITNDPKEAVRGADVVYADVWASMGQKEEAAHRRQVFQGFQVDEELMKLAGQKAYFMHCLPAERGVEVTDGVIEAPNSIVFPQAENRMHAQNAIMLHVLGL*

### NbL08g19380.1 uncharacterized_LOC104222813 [MW − SP = 22.33 kDa]

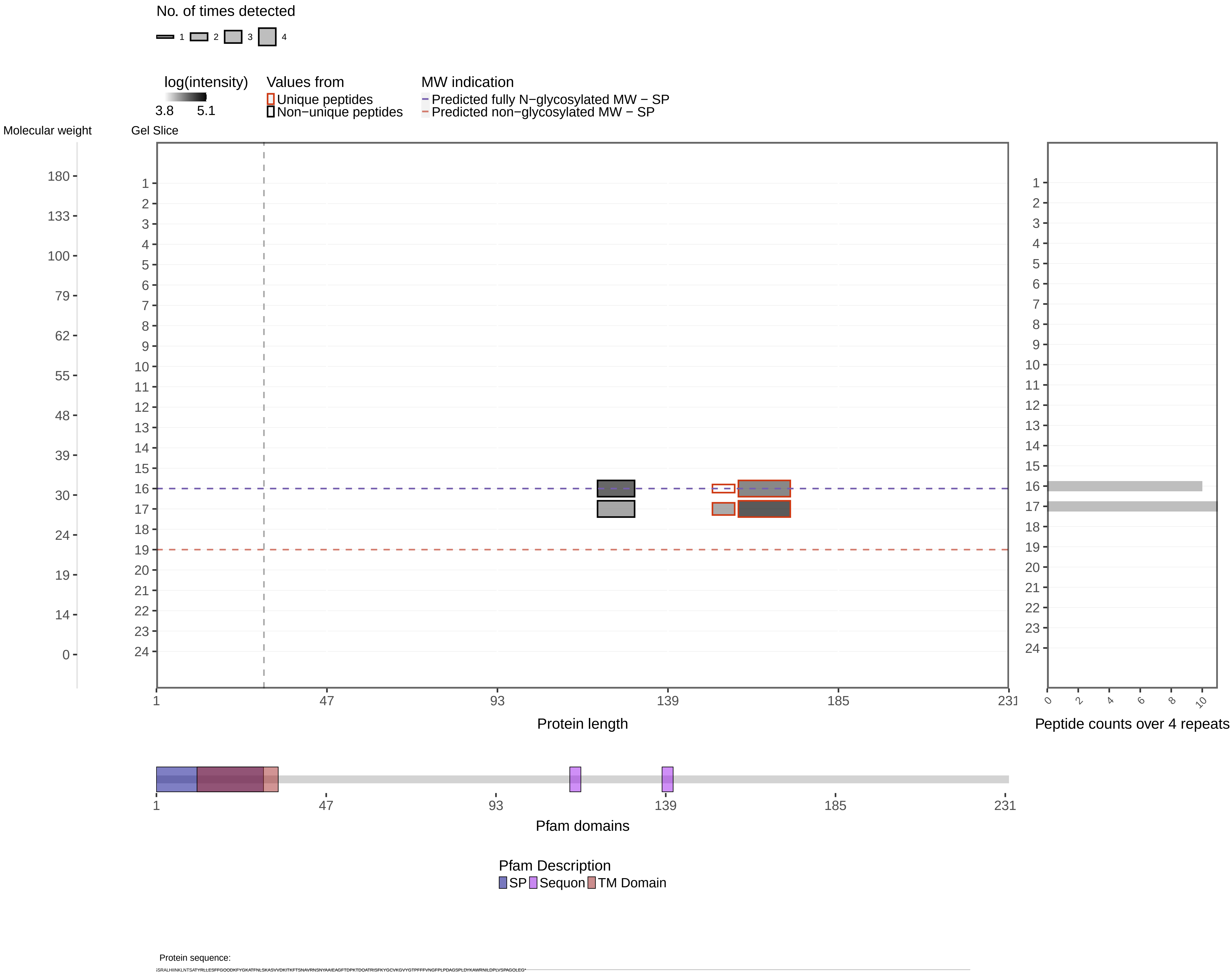

NbL08g19330.1 glutathione_S−transferase−like [MW = 25.47 kDa]

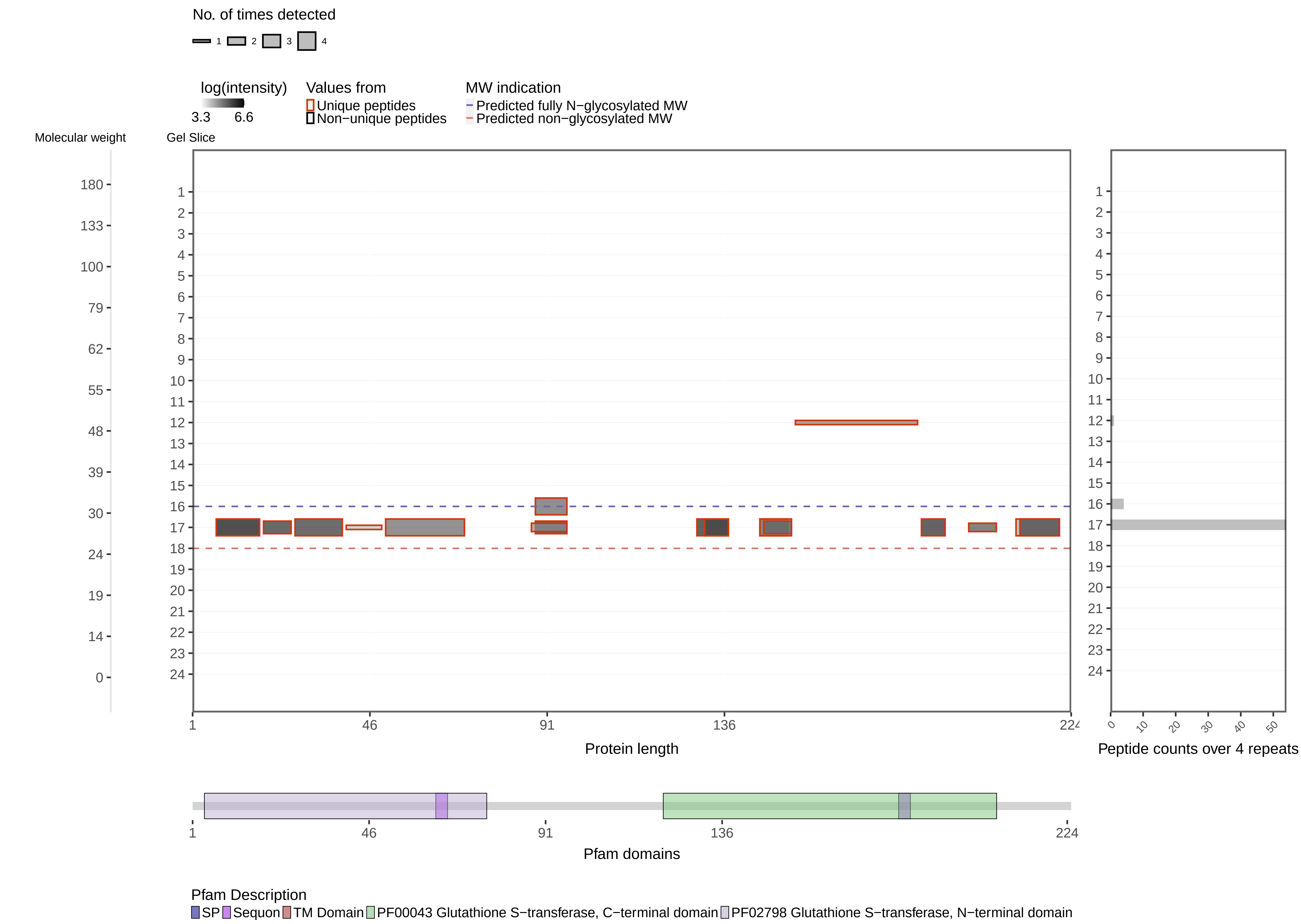


Protein sequence:

QWIEAEGQSFNPPSSILVFQLAFAPRMKIKQDENLIRQNEEKLKKVLDVYEKRLGESQYLAGDGFSLADLSHLPNIQYLVNGTDRAELFTSRKNVGRWWDEISSRESWKKVVEMQTSPPPKTC*

NbL08g19260.1 4−coumarate−−CoA_ligase−like_5 [MW = 60.65 kDa]

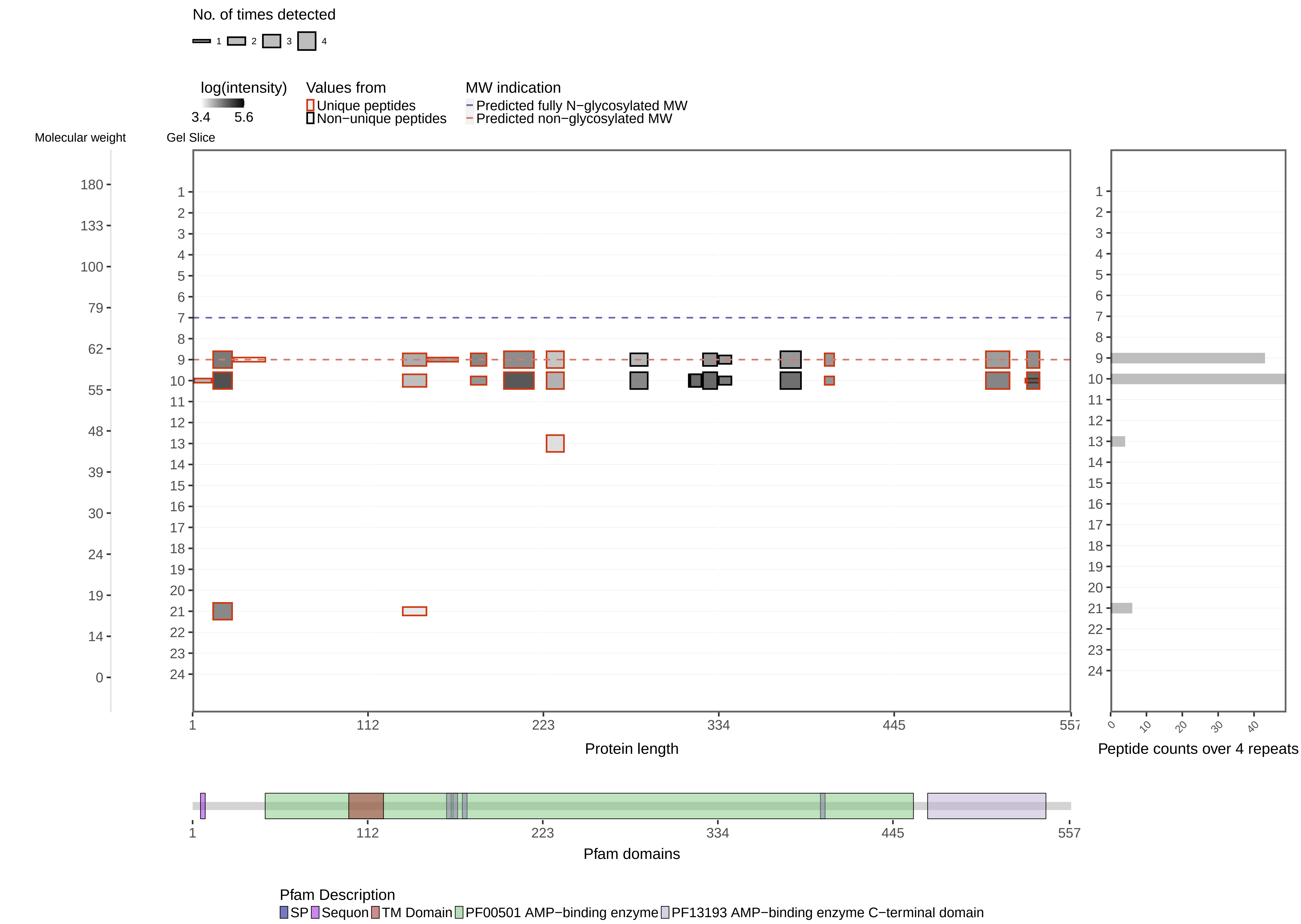


Protein sequence:

ASGSTVVVLSKFEMDEMLSSIHKYKATYLPLVPPILVALVNNAEMIKKKYDLSSLKKVLSGGAPLSKEVIEGFLEKYPSVTILQGYGLTESTAIGASTDSLEESRRYGTAGMLSPSMSAKIVDPESGKAFPVNKTGELWLRGPTIMKGYFSNEEATASTLDSEGWLRTGDLCYIDEDGFIFVVDRLKELIKYKGYQVPPAELEALLLTHPEISDAAVIPFPDKDVGQFPMAYVVRKSGSSLSESAVMDFIAKQVAPYKRIRRVAFVASIPKNPSGKILRKDLIKLATSKL*

NbL08g19150.1 glyceraldehyde−3−phosphate_dehydrogenase_B_chloroplastic−like [MW = 47.92 kDa]

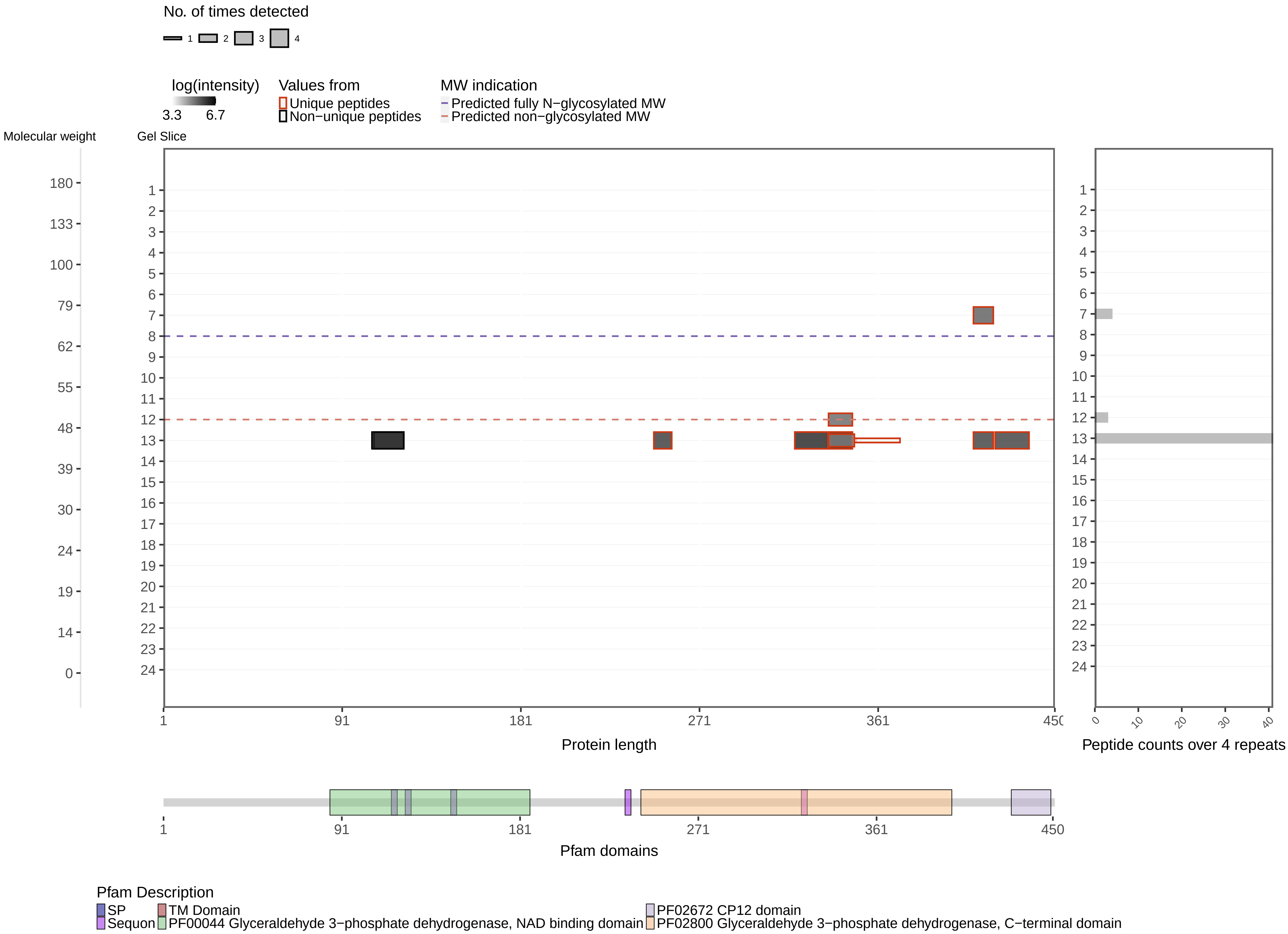


Protein sequence:

TYVVGVNEQDYSHEVANIISNASCTTNCLAPFVKVMDEELGIVKGTMTTTHSYTGDQRLLDASHRDLRRARAAALNIVPTSTGAAKAVSLVLPQLKGKLNGIALRVPTPNVSVVDLVVNVEKKGISAEDVNAAFRKAADGPLNGILAVCDVPLVSVDFRCSDVSSTIDSSLTMVMGDDMVKVVAWYDNEWGYSQRVVDLAHLVASKWPGSSVEGSGDALEDYCKTNPADKECKVYE*

NbL08g19050.1 pyrophosphate−−fructose_6−phosphate_1−phosphotransferase_subunit_alpha−like [MW = 67.17 kDa]

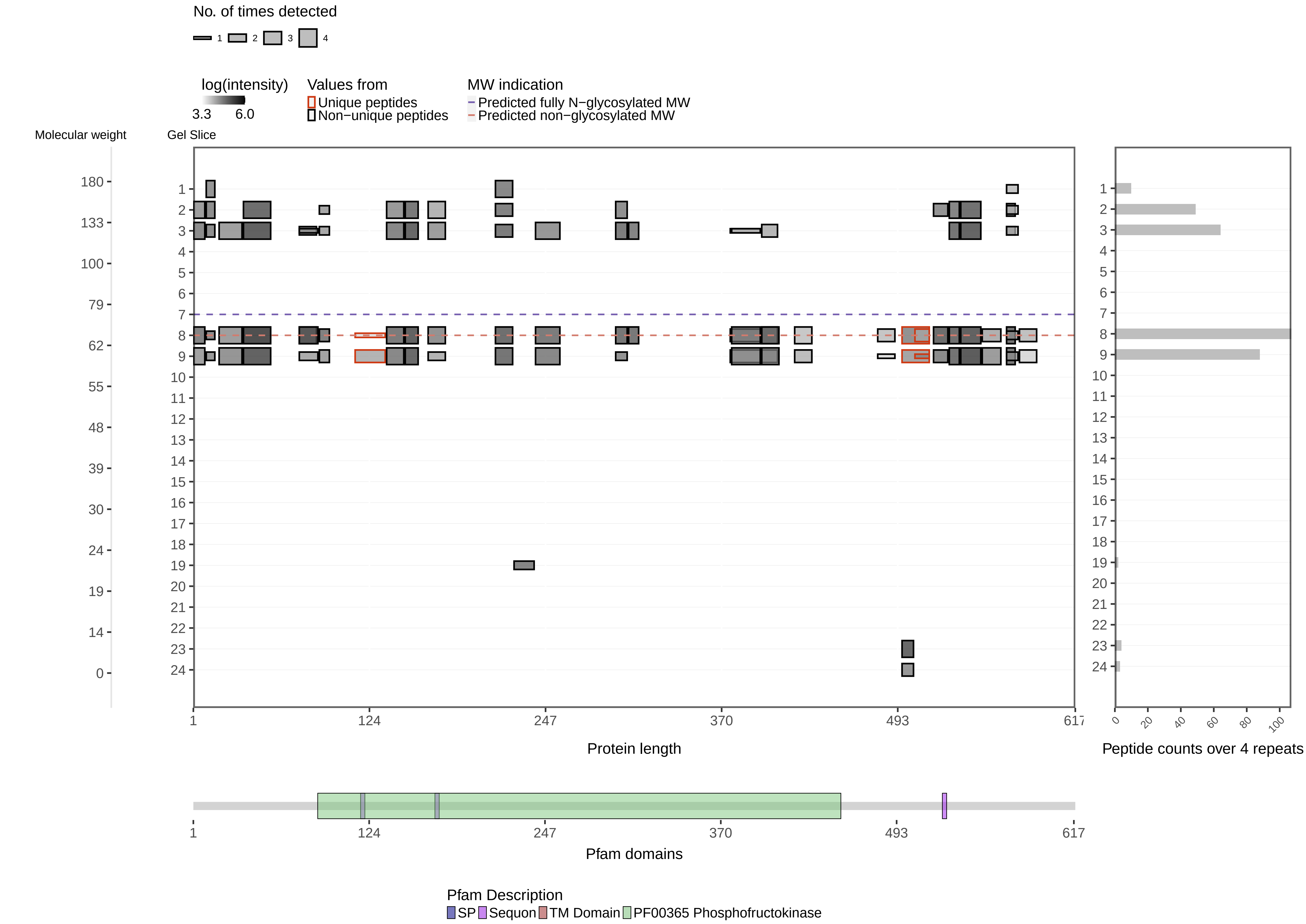


Protein sequence:

SKLTLFDITNKICDAVQARAEQDKNHGVILLPEGLIESIPEVYALLQEIHGLLRQGVSADKISSQLSPWASALFEFLPPFIKKQLLLYPESDDSAQLSQIETEKLIAHLVETEMNKRLKEGTYKGKKFNAICHFFGYQARGSLPSKFDCDYAYVLGHICYHILAAGLNGYMATVTNLKNPVNKWRCAAAPITAMMTVKRYGRGPGNAAIGKPAVHPATVDLKGKAYELLSQNATKFLLDDVYRNPGPLQFDGPGADAKAVTLCVEDQDYMGRIHKLQEYLDKVRTIVKPGCSQDVLKA

NbL08g19040.1 UPF0047_protein_YjbQ−like [MW = 20.84 kDa]

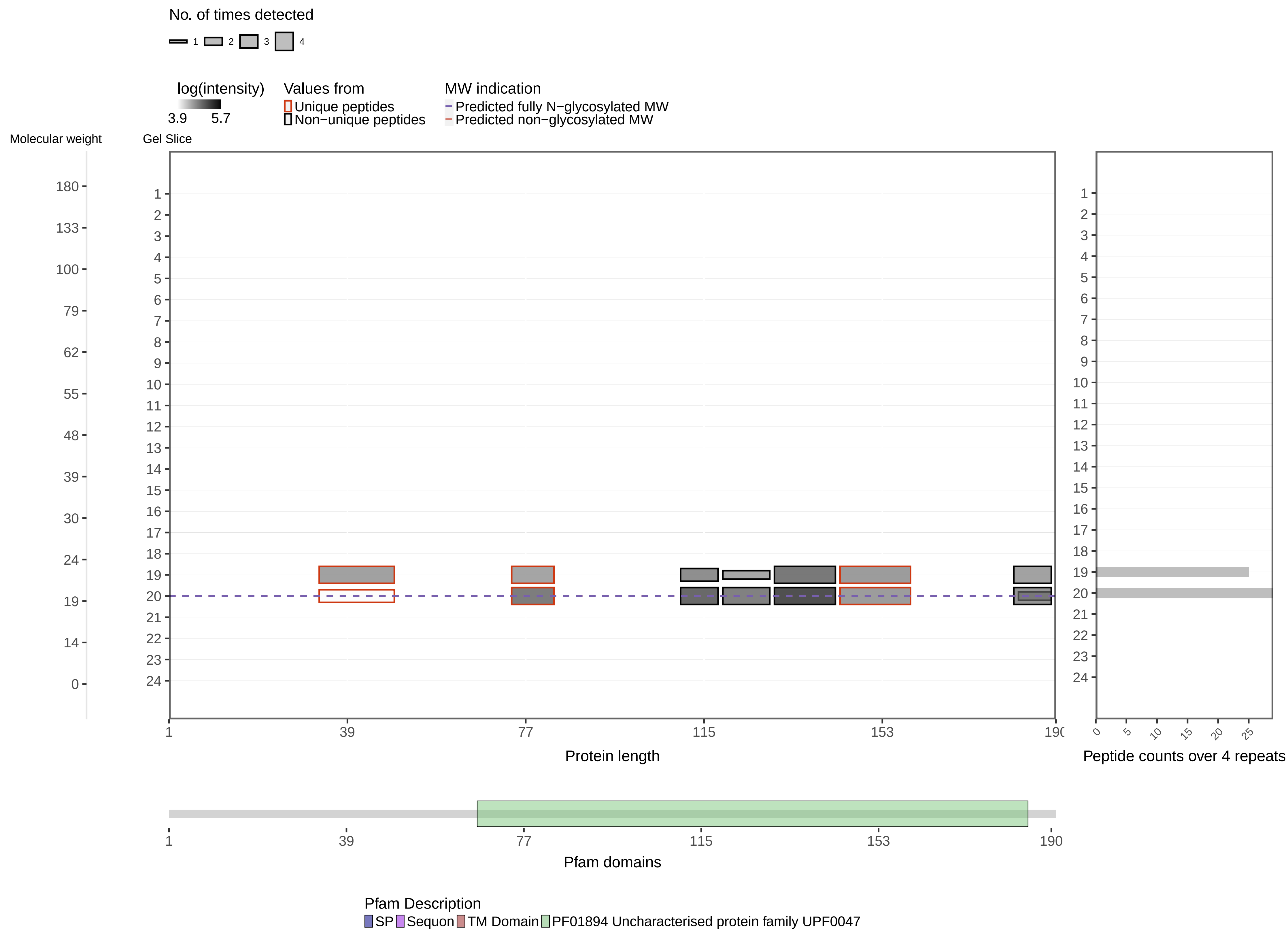


Protein sequence:

KCGLAHLFLQHTSASLTINENYDSDVRDDTETFLNRVVPEGISAPWKHTLEGPDDMPAHIKSSMFGCTLTIPITDGKLNMGTWQGIWLCEHRDAATPRRIVITLNGM*

### NbL08g18970.1 tryptophan−−tRNA_ligase_chloroplastic/mitochondrial [MW = 49.69 kDa]

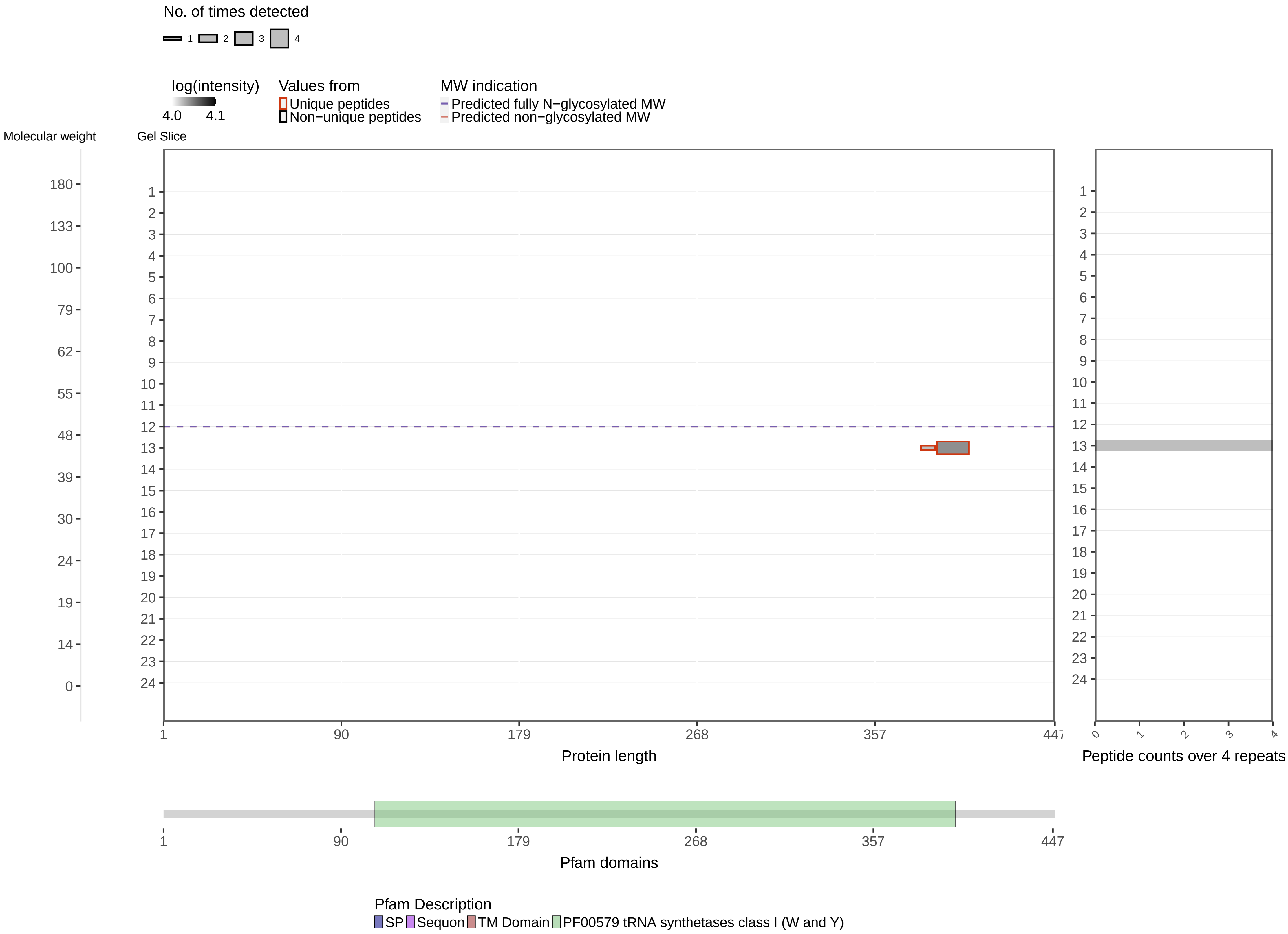


Protein sequence:

FKEKSRKAGDENVGVALLTYPVLMAADILLYQSDFVPVGEDQKQHLELTRELAERVNYLYGGRKWKKLGGRGGSIFKVPEPLIPPAGARIMSLTDGLSKMSKSAPSEQSRINLLDSKDVIANKIKRCKTDSFPGMEFDNPERPECNNLLSIYQLVTGKTKQEVAEECRHMNWGTFKLVLADALVDHLHPIQVRYEEIVSDASYLDGVLAEGARKAADVADVTVNNVFQAMGFLRR*

NbL08g18930.1 ATP−dependent_6−phosphofructokinase_6−like [MW = 57.05 kDa]

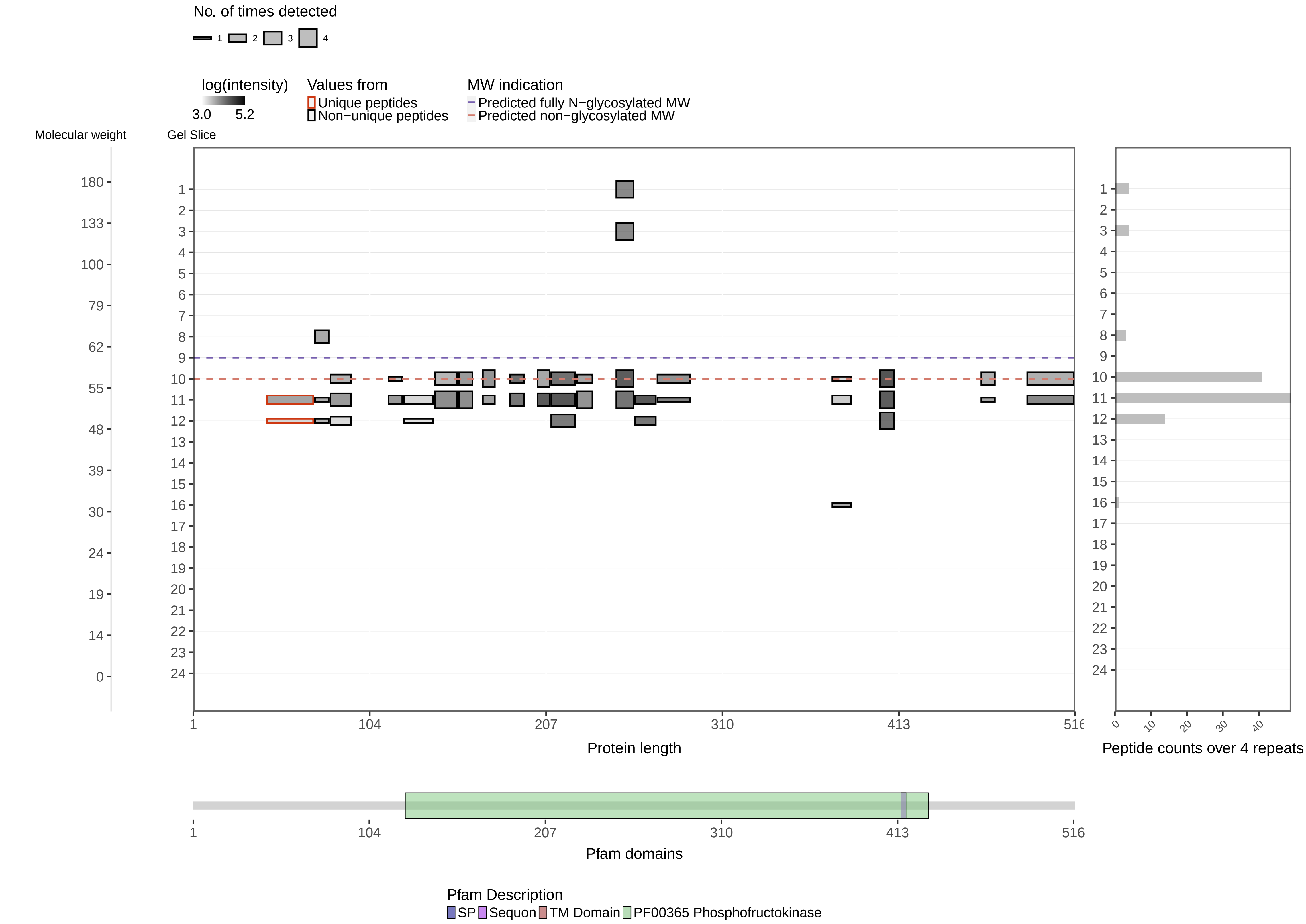


Protein sequence:

IPKTIDNDIPVIDKSFGFDTAVEEAQRAINAAHVEAESAENGIGVVKLMGRYSGFIAMYATLASRDVDCCLIPESPFYLEGEGGLYEYIEKRLKENGHMVIVIAEGAGQELLAAENSLAKNEQDASGNKLLQDVGLWISQKIRDHFAKNPKMAITLKYIDPTYMIRAVPSNASDNVYCTLLAQSCVHGAMAGYTGYTSGLVNGRQTYIPFNRITEKQNMVVITDRMWARLLSSTNQPSFLCPKDVEEVKNEEQPQTQLLDGDNNVHENSGH*

### NbL08g18900.1 vacuolar−processing_enzyme−like [MW − SP = 51.1 kDa]

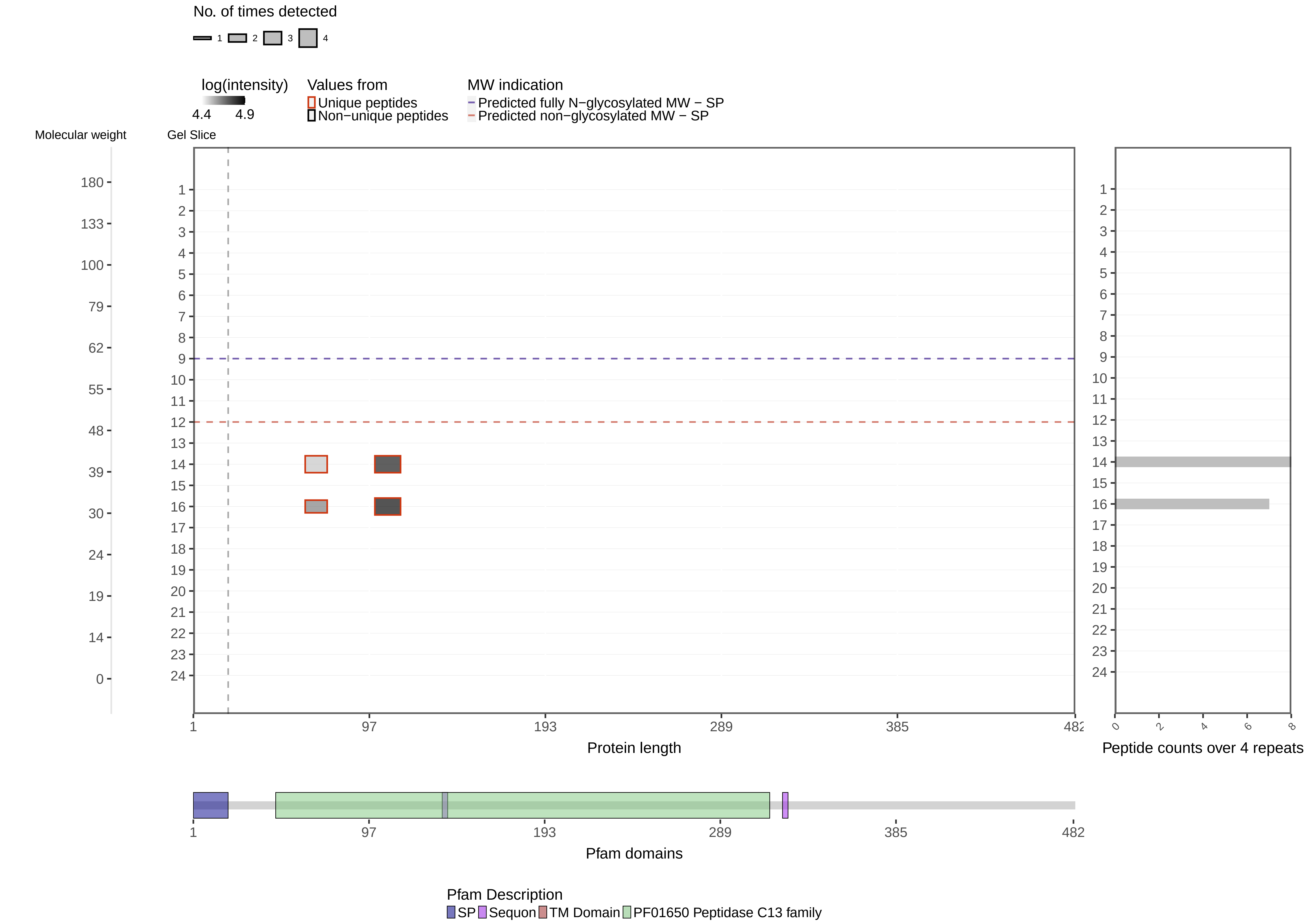


Protein sequence:

WGTYCPGDYPSLPPGYETCLGDLYAVSWMEDSEMHNLRTENLRQQYHLVKQRTANGNSAYGSHVLQFGDLQLSMDSLFMYMGTNPANDNYTYVDDNSLRASSKAVNQRDADLLHFWDKFRKAPEGSARKVEAQKQFTEAMSHRMHLDNSMALVGKLLFGIQKGPEALKHVRPVGQPLVDDWTCLKSFVRTFETHCGSLSQYGMKHMRSIANICNAGIKMEQMVEASAQACPSVPSNTWSSLHRGFSA*

### NbL08g18880.1 far_upstream_element−binding_protein_2 [MW = 67.55 kDa]

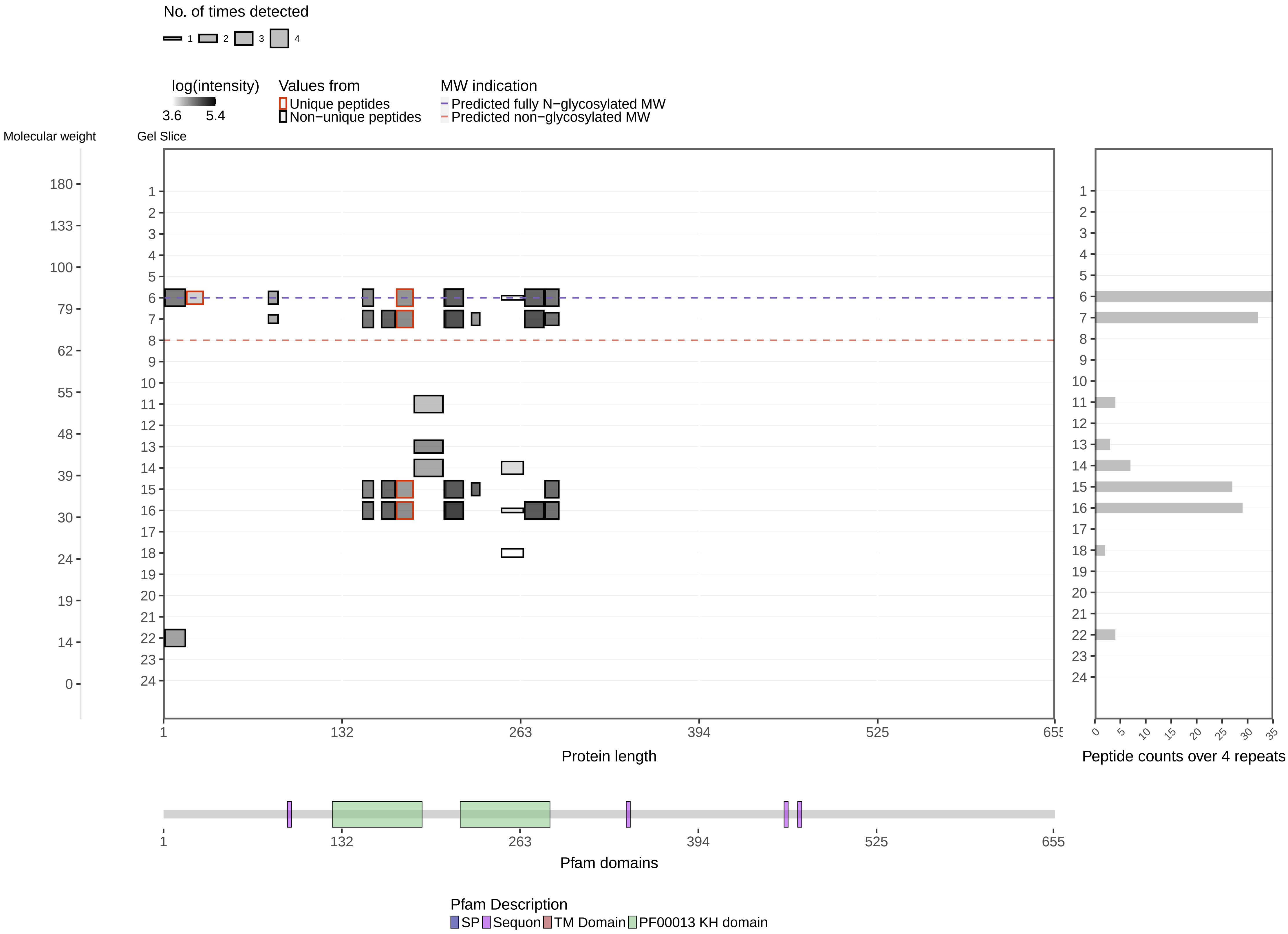


Protein sequence:

### NbL08g18800.1 60S_ribosomal_protein_L7−4 [MW = 32.93 kDa]

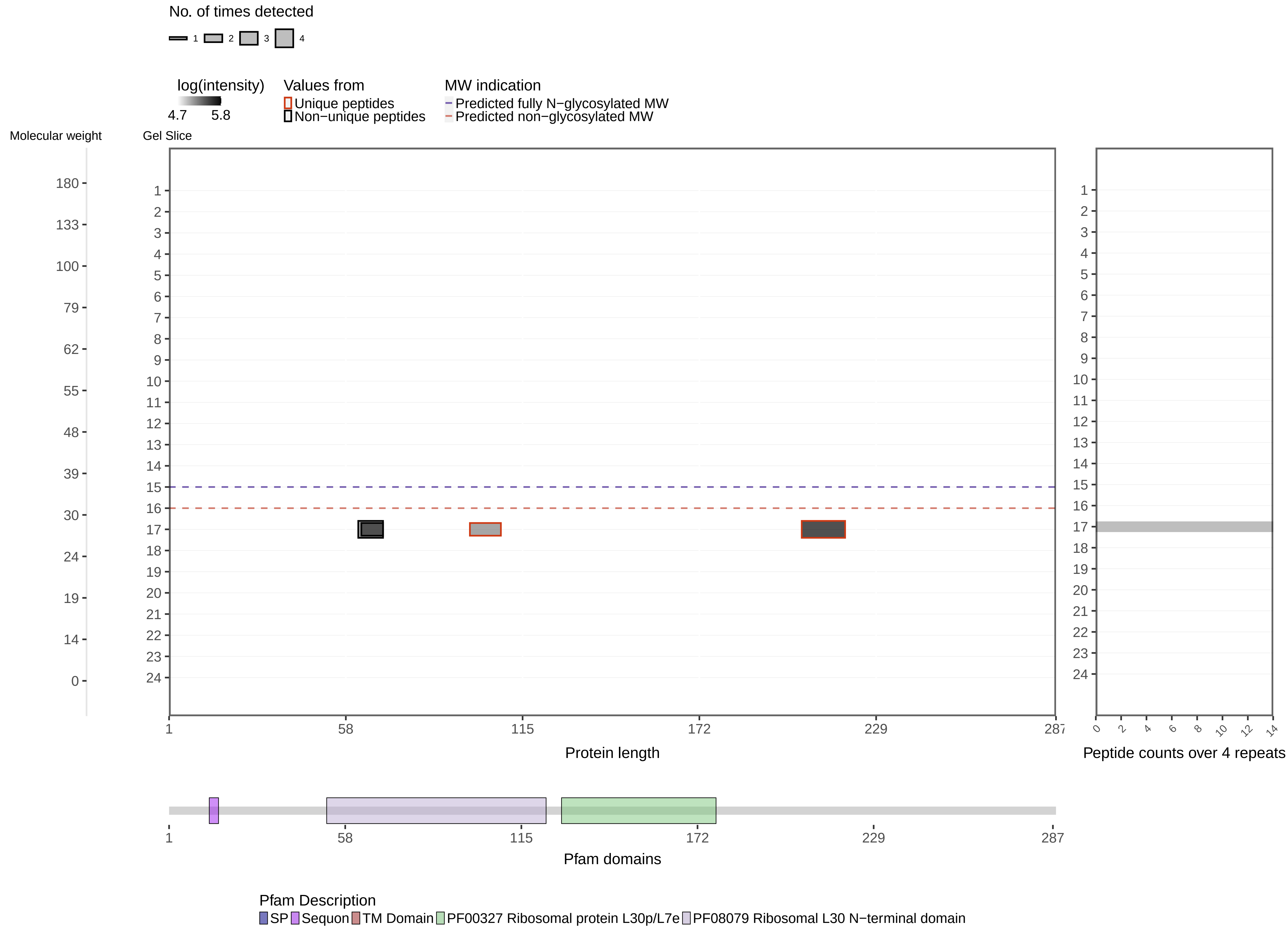


Protein sequence:

FIIRIRGINAMAPQTKKILQLLRLRQIFNGVFLKVNKATVNMLHRVEPYVTYGYPNLKSVRELIYKRGYGKLDKQRIALTDNSIIDQALGKHGIICAEDLVHEILTVGPHFKEANNFLWPFQLKAPLGGLKKKRNHYVEGGDAGNRENYINELIRRMN*

### NbL08g18750.1 tryptophan_synthase_beta_chain_2−like [MW = 54.1 kDa]

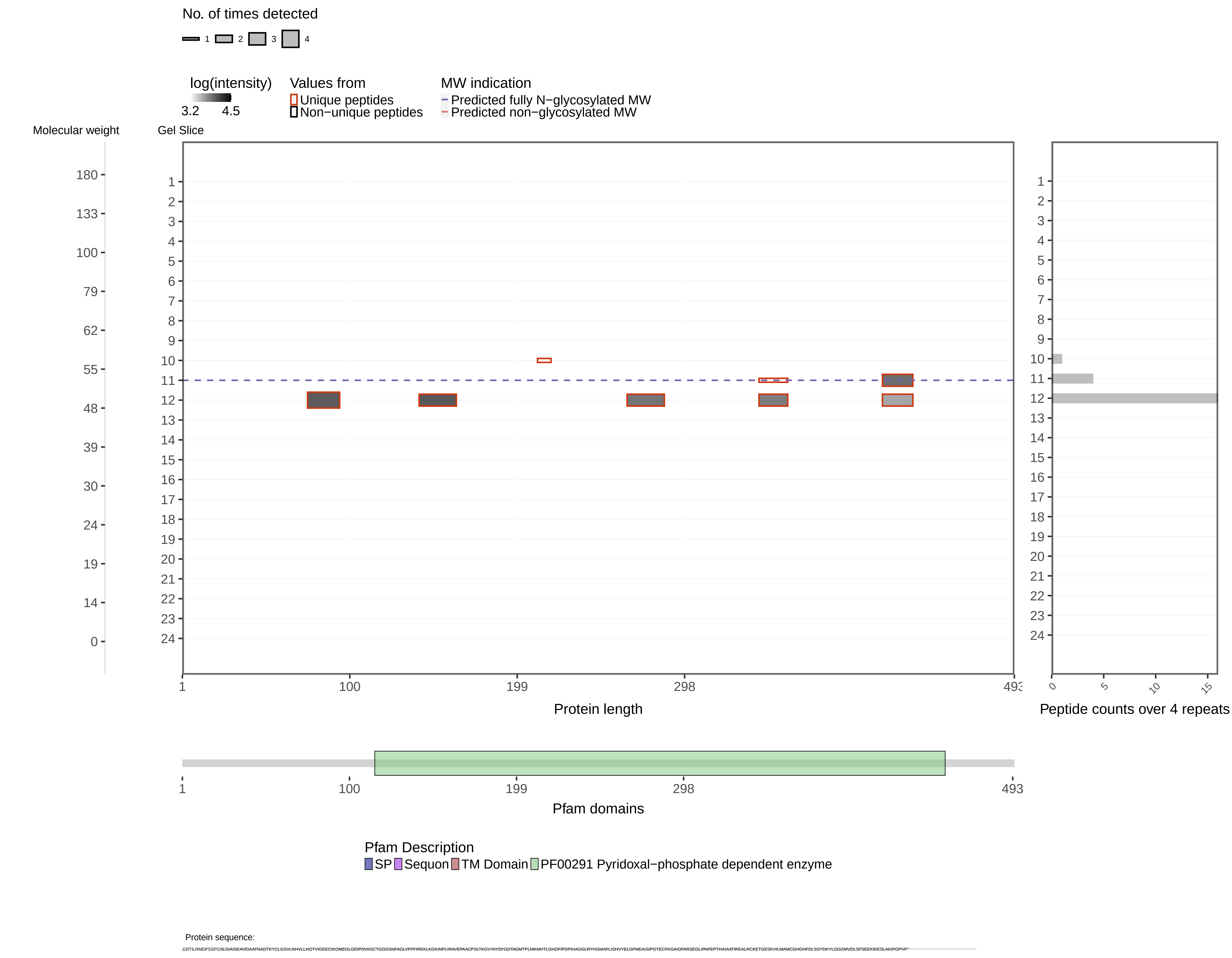

NbL08g18700.1 60S_ribosomal_protein_L13−1−like [MW = 23.73 kDa]

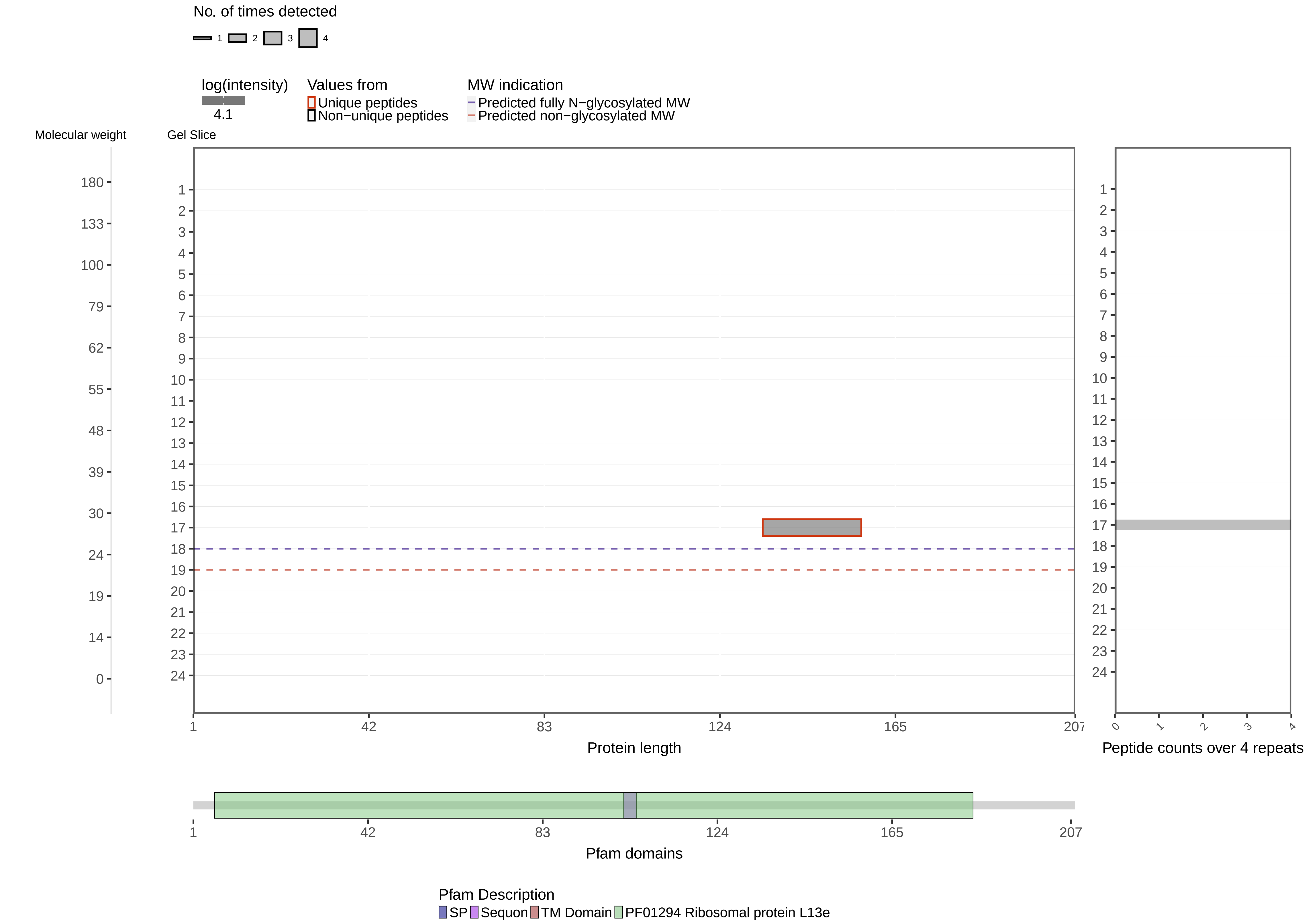


Protein sequence:

APTIGIAVDHRRRNRSLEGLQTNVQRLKTYKAKLVIFPRHAKIVKAGDSSPEELATATQVQGTYMPITREQSTIELVKVTDEMKSFNAYGKLRIERTNERHIGERLKRAAEAEKEEKK*

NbL08g18630.1 scopoletin_glucosyltransferase−like [MW = 39.54 kDa]

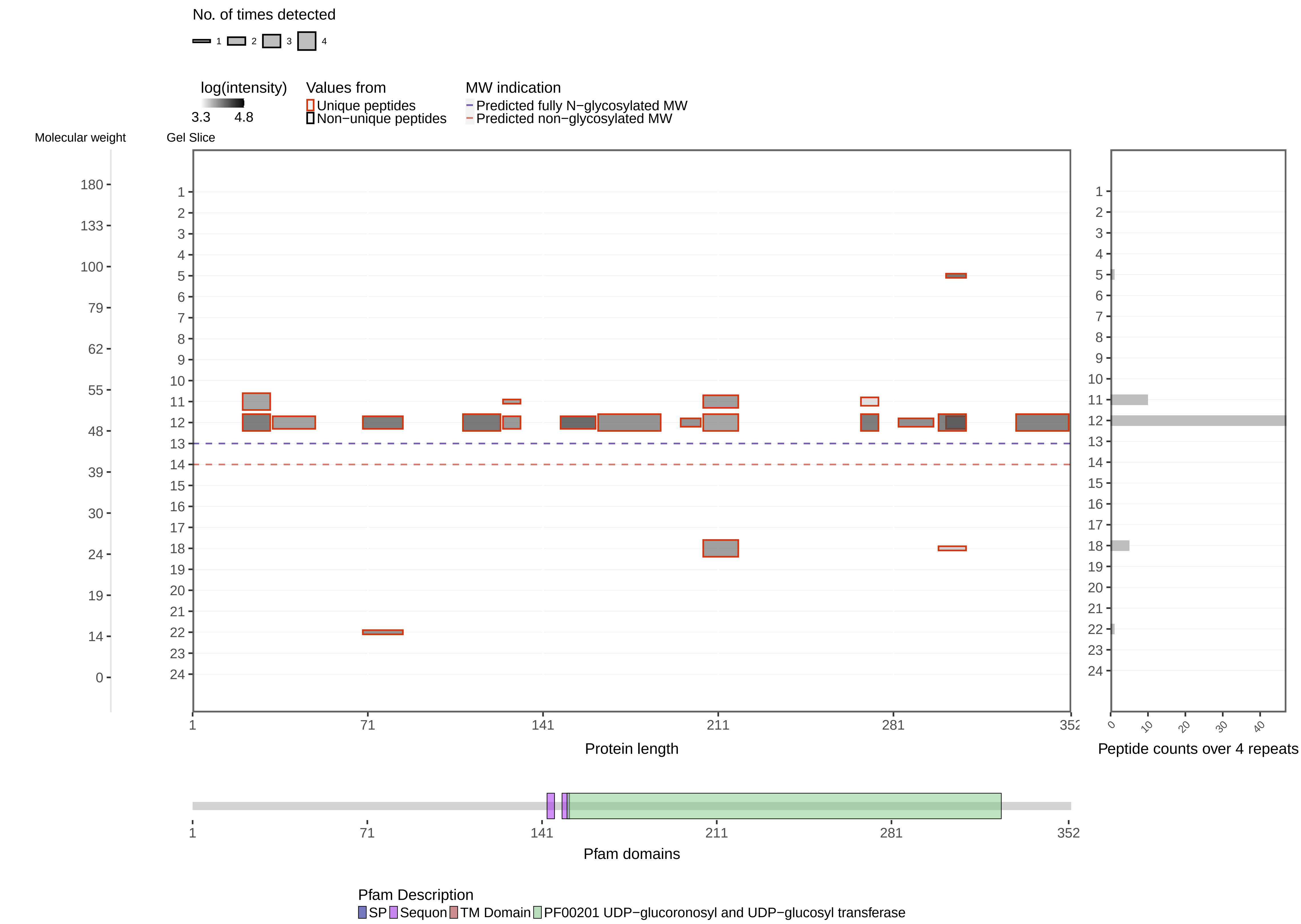


Protein sequence:

SKSPSSIIYIAFGSDVVPTLEEHEEIAKALEESSRQNFIWVISKNQDYYPNTLETQVGKRGLIINGWAPQLLILSHPSTGGFLTHCGWSSIMEAIGQGVPLLAWPIRGDQFYNAKLIVCYLKIGHMVSSIGENIGLKRVKKEEIIQGVQRLMEDKQVHNRVKDLSDKFRHGFPISSAASLNDFKDFIMQAK*

### NbL08g18120.1 peptidyl−prolyl_cis−trans_isomerase_NIMA−interacting_4 [MW = 15.13 kDa]

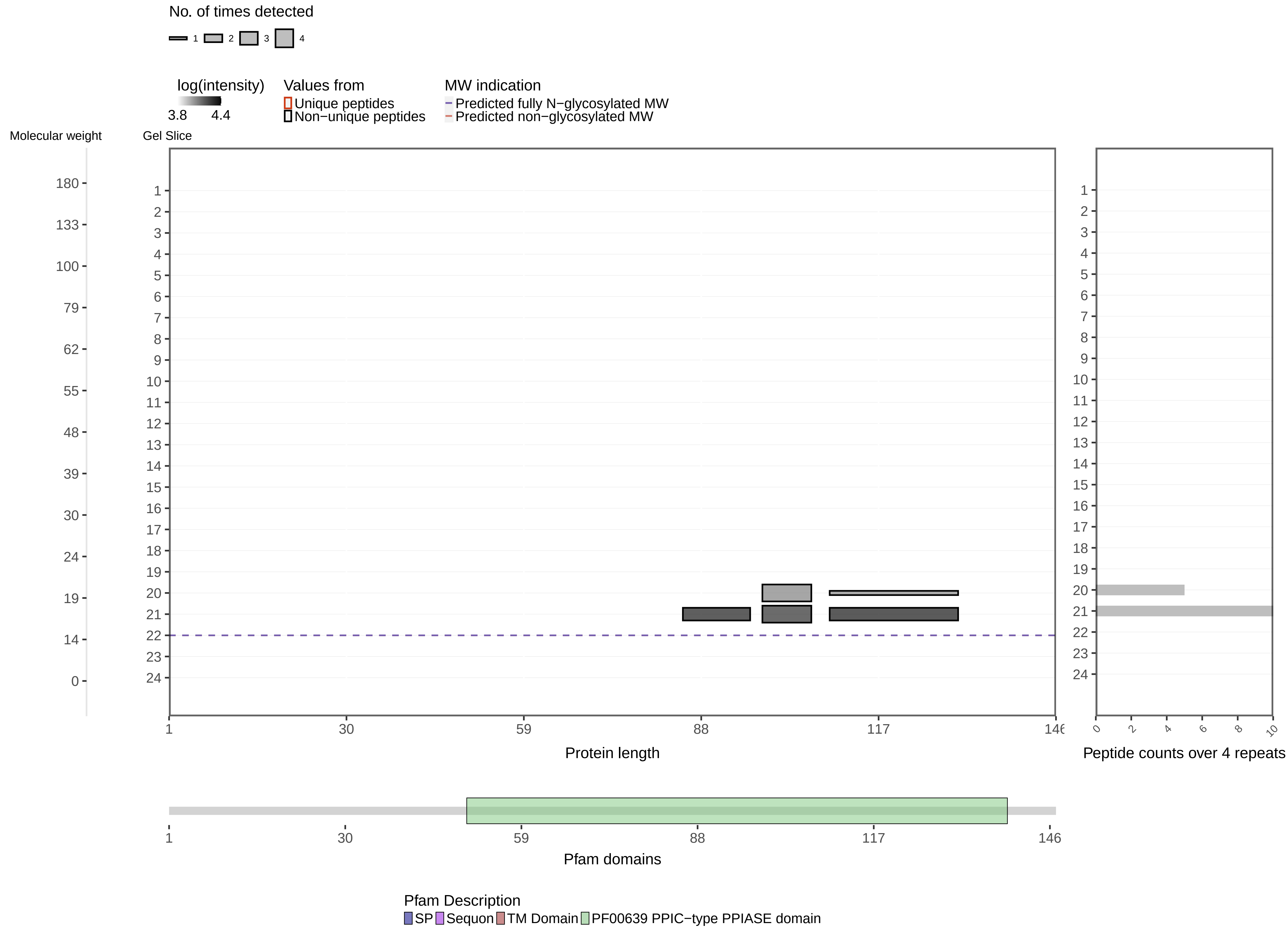


Protein sequence:

INEAYKKLQEGWLDIGDKVPPAEFAKVAAAYSECPSGKKGGDLGWFPRGKMAGPFQEVAFNTPVGVTSAPFKSTHGYHIILSEGRKN*

NbL08g17940.1 NAP1−related_protein_2−like [MW = 29.02 kDa]

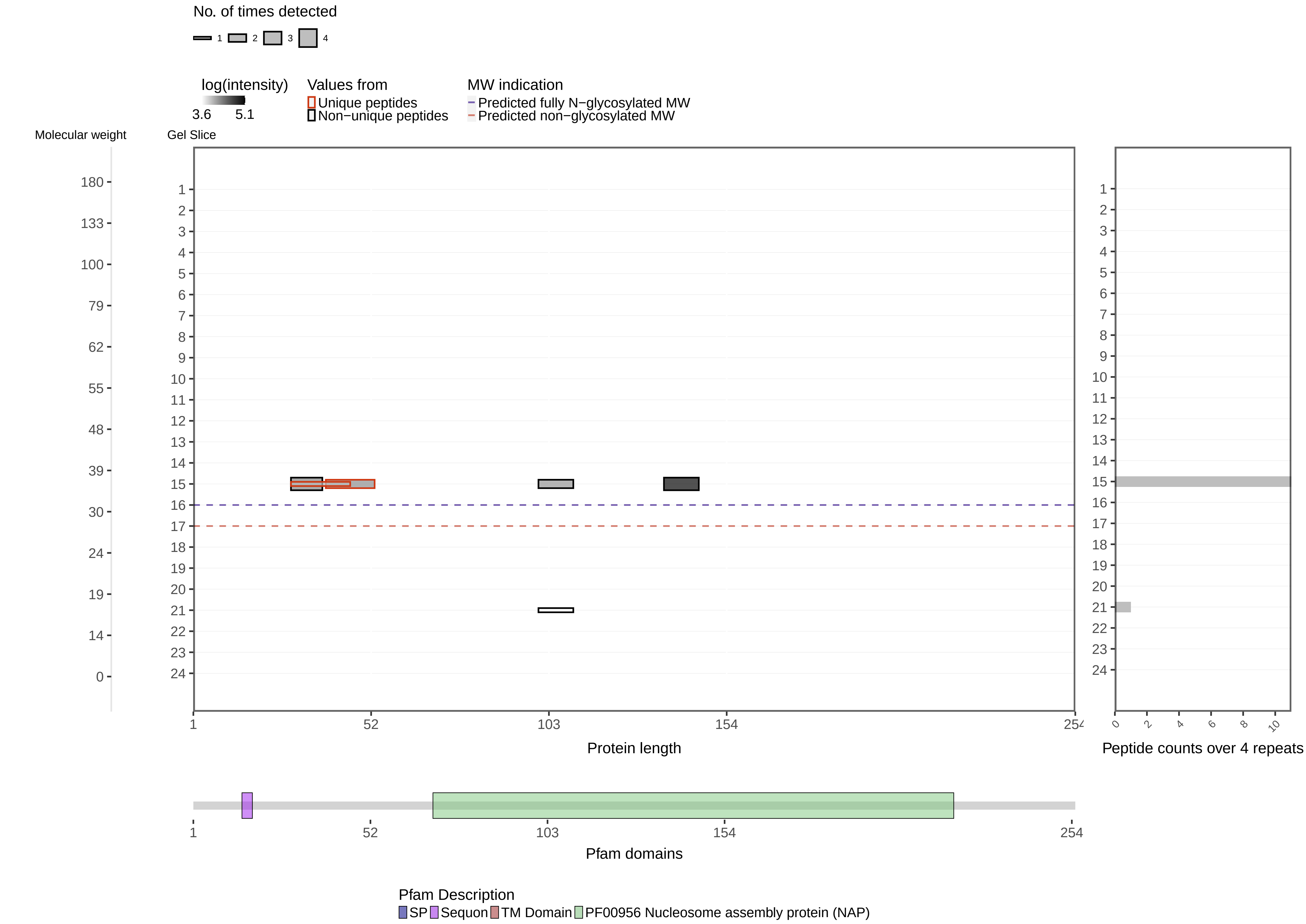


Protein sequence:

YSITFNFNANPYLENTKLAKTYTFLEDGPTKISATTIKWKEGTGIPNGVAHEKKGNKRSHAEESFFTWFSEVNOKDEDEALEIQDEVADIIKDDLWPNPLTYFNNEPDEEDFDGDEGKDSEGSEDEEEEEEDEDGDEE*

### NbL08g17820.1 plasma_membrane_ATPase_1 [MW = 105.24 kDa]

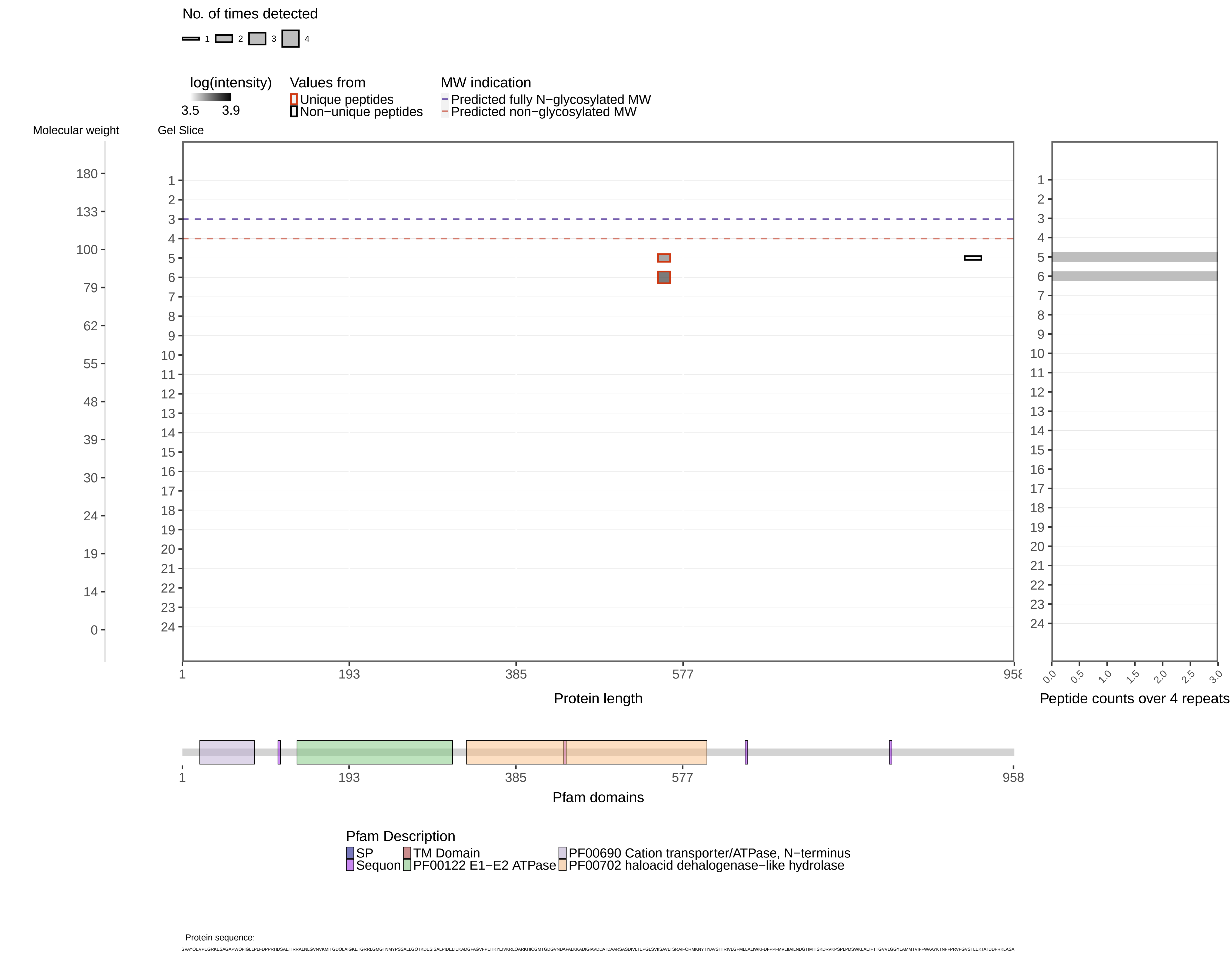

### NbL08g17690.1 protease_Do−like_9 [MW = 63.34 kDa]

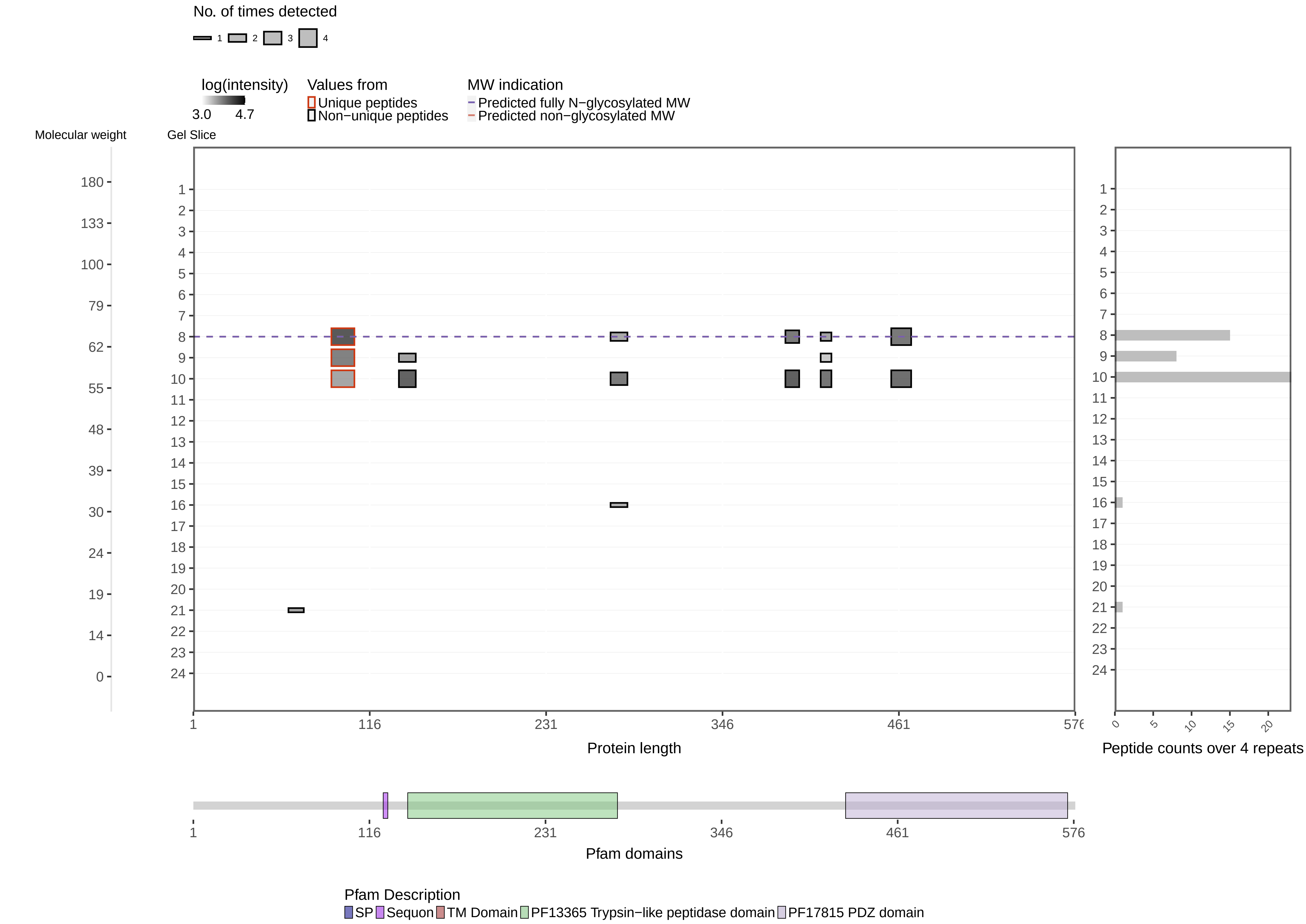


Protein sequence:

NCVGIAFQSLKHEDVENIGYVIPTPVIMHFIQDYEKNGAYTGFPILGIEWQKMENPDLRLSMGMKPDQKGVRIRRIDPTAPESMLLMPSDVLLSFDGVDIANDGTVPFRHGERIGFSYLVSQKYSGDSAAVKVLRRNSEILNFKIKLATQMRLVPAHNKGRPPSYYIIAGFVFTTVSVPYLRSEYGKDYEYEAPVKLLDKLLHEYRQSPDEQIVVVSQVLVADINIGYEEIVNTQVLAFNGKPVKNLKSLASMVENCNDEFLKLDLEYQQVVVLQTKTAKSATSDILTTHCIPSAMSE

### NbL08g17680.1 iron−sulfur_cluster_assembly_protein_1−like [MW = 18.19 kDa]

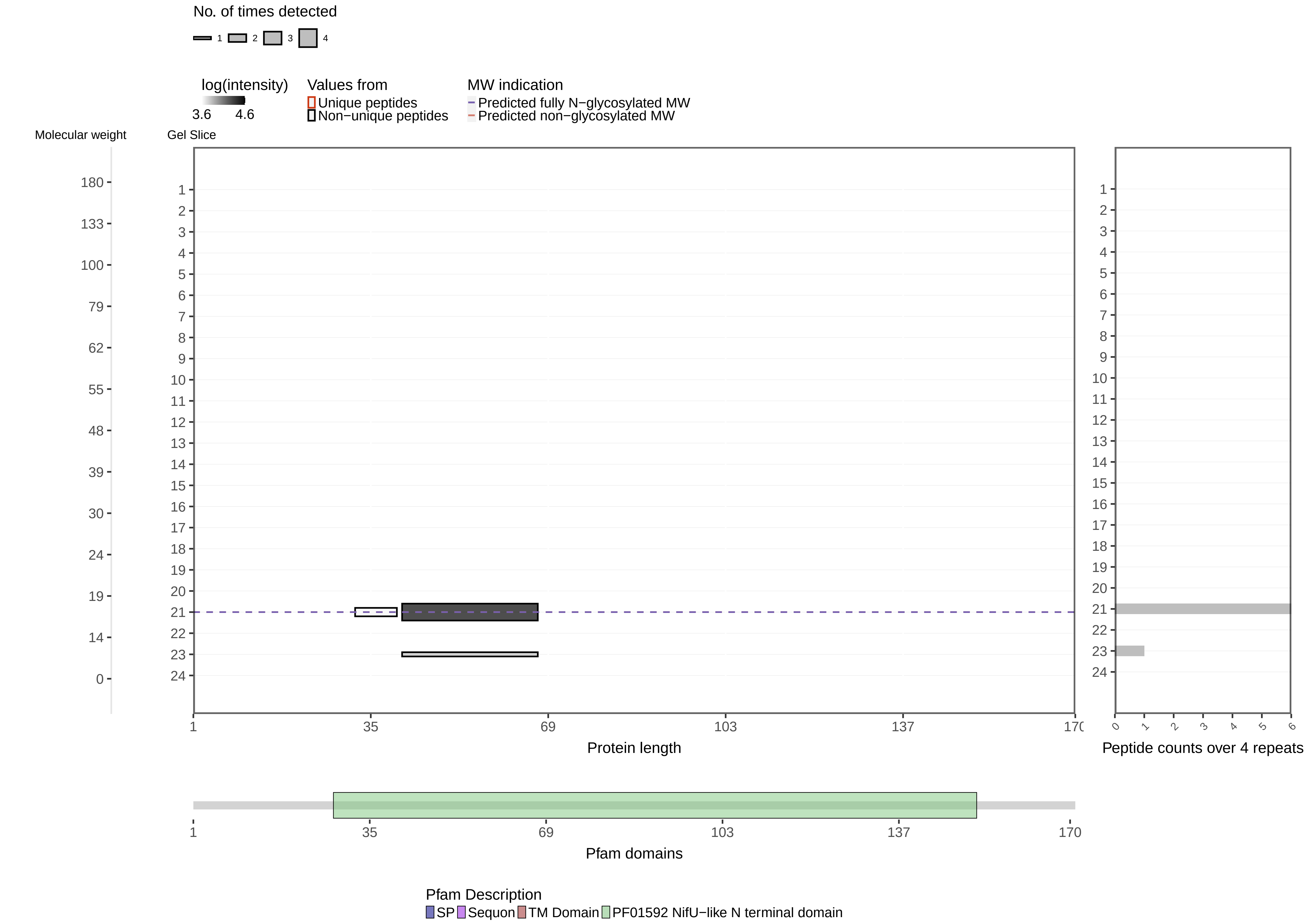


Protein sequence:

VDEETGKITDACFKTFGCGSAIASSSVATEWVKGKQMEEVLSIKNTEIAKHLSLPPVKLHCSMLAEDAIKAAVKDYSAKRAKLSDNAIDSQMGKAVDA*

NbL08g17510.1 31_kDa_ribonucleoprotein_chloroplastic [MW = 45.08 kDa]

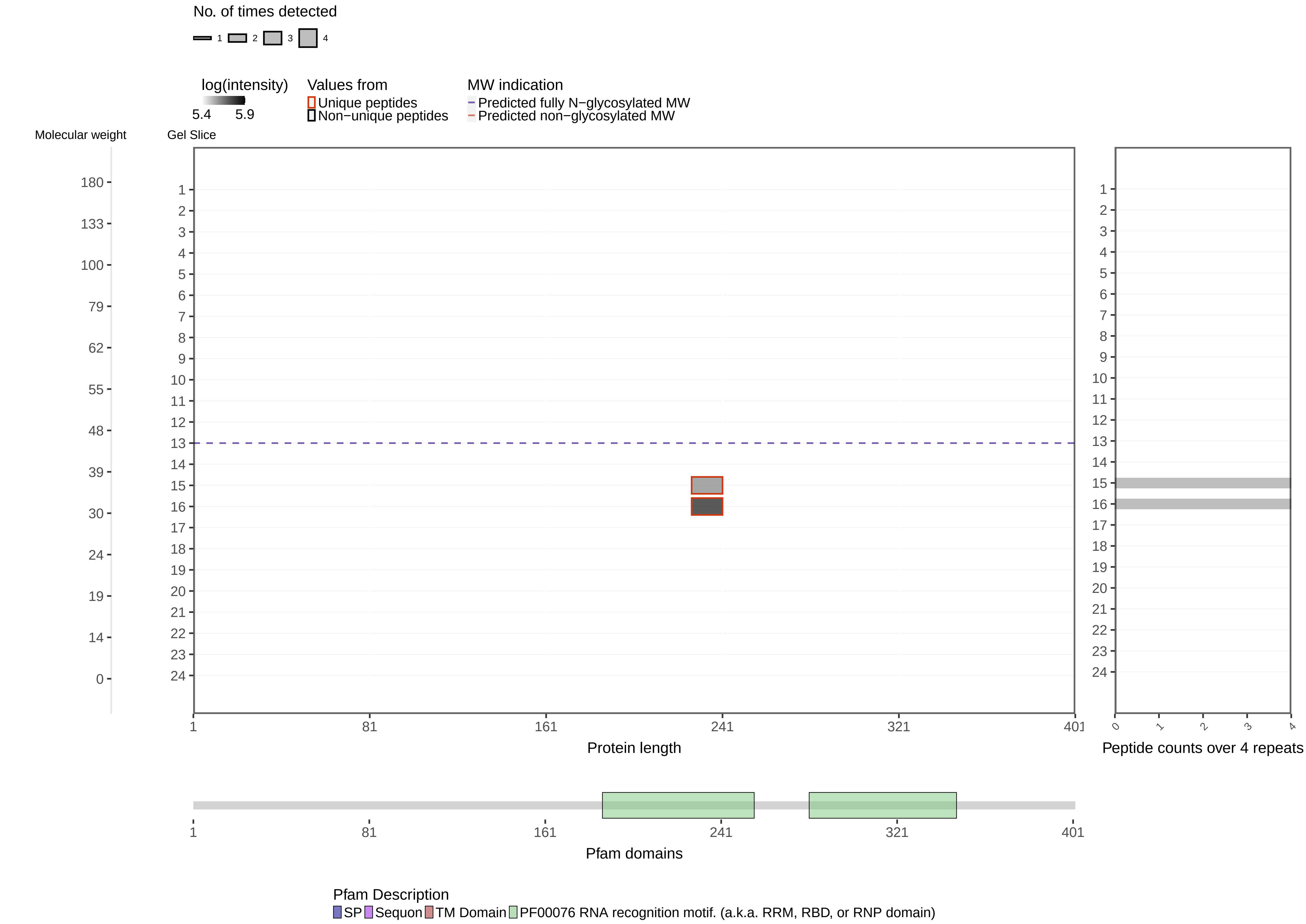


Protein sequence:

GNLPYDVDSEGLARLFEQAGVVEIAEVIYNRDTDQSRGFGFVTMSTVEEAQKAVEMYNRYDVNGRLLTVNKAARRGERPERPPQTFEQSYRIYVGNIPWGIDDARLEQLFSEHGKVVSARVVYDRETGRSRGFGFVTMASEAEMSDAIANLDGQVIHARRFLTHNQIHAHLLIHIVFEEFTRKGCVSILSIAFHIFHFLNFHFSFFGRSDF*

### NbL08g17390.1 glycine−rich_RNA−binding_protein_4_mitochondrial−like [MW = 18.32 kDa]

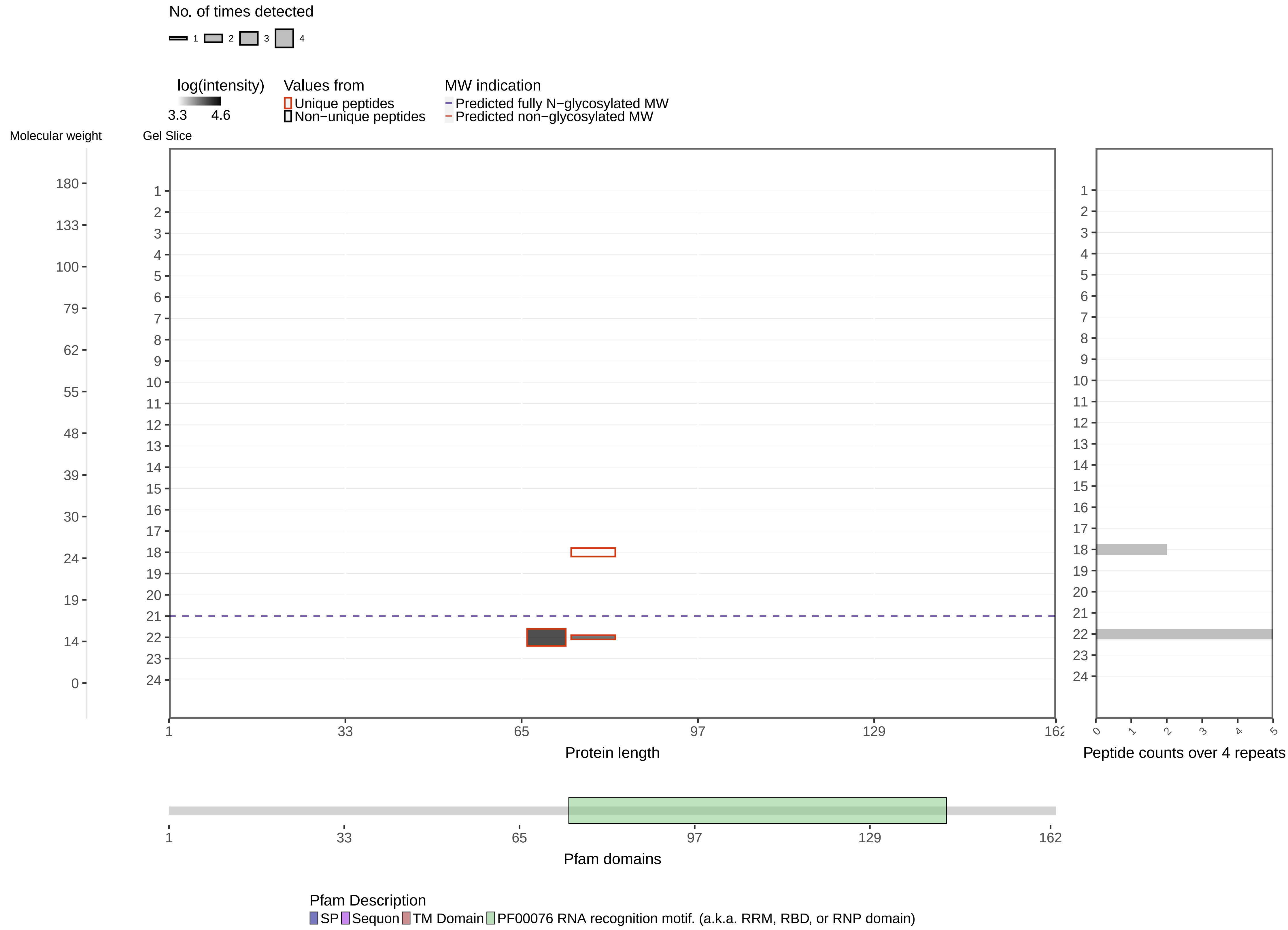


Protein sequence:

NPSTKLFVSGLSFRTTEESLKNAFNDFGELVEVNLVMDKIANRPRGFAFLRYATEEESQKAIEGMHGKFLDGRVIFVEVAKPRSELRRITARQ*

### NbL08g17080.1 GDSL_esterase/lipase_At5g03980−like [MW − SP = 38.77 kDa]

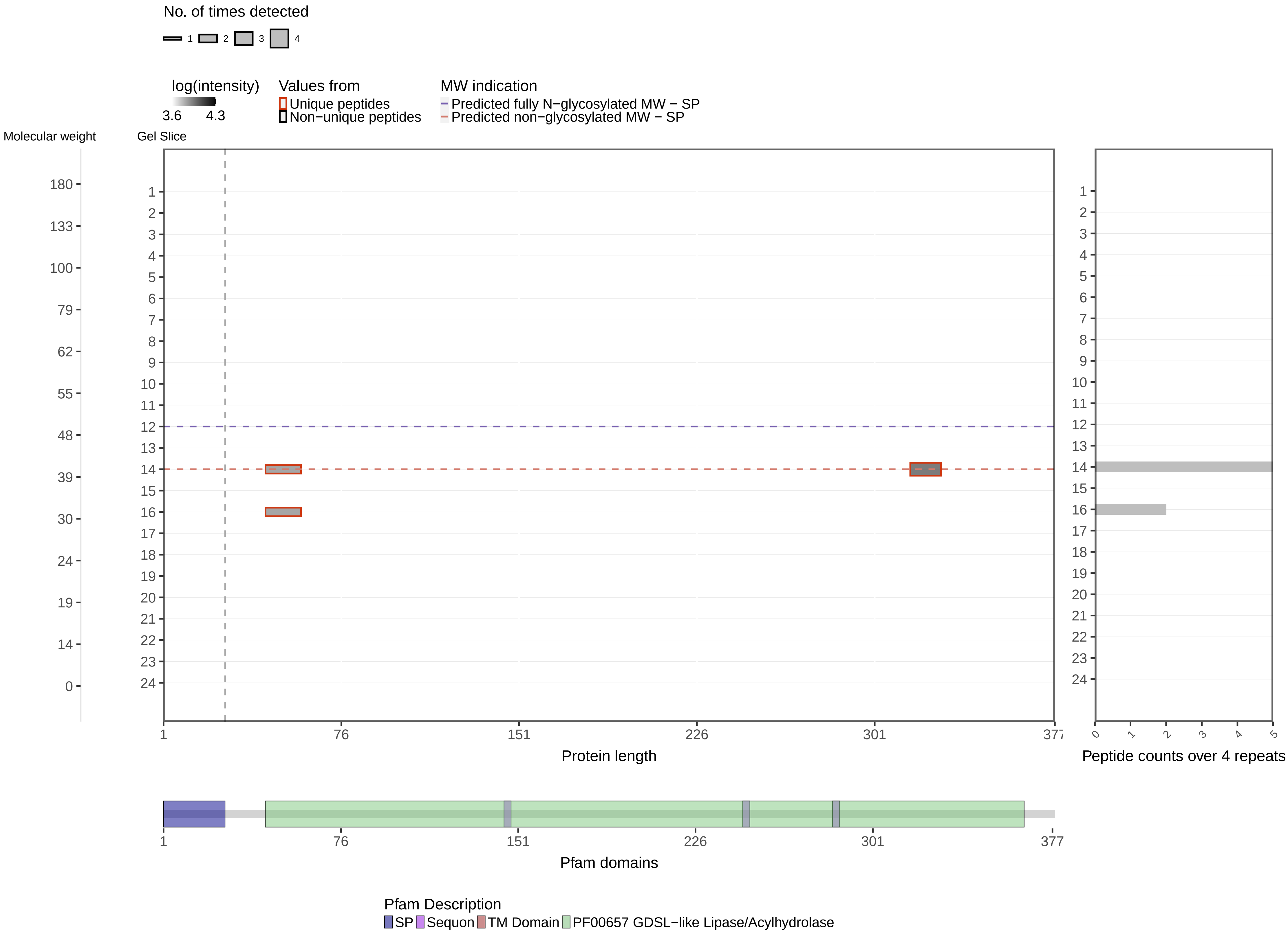


Protein sequence:

.FLVGEIGGNEFFYGLTOGKTMEELRNMVPEVVONIVEAVNTVTGFGAIRILIPGNFPIGCLPMTLTTFLTNNSMAYDENHCLKDLNNLAIFYNEHLOOAIEDLKKKNPNVTLIYGDYYNAYOWLLOHTVNOGFDONSLLKACCGEGGYYNYDAKROCGFPGVPVCADPGTYISWDGVHLTEKAYSLIATWLIDDILPOLKC*

### NbL08g17070.1 probable_U6_snRNA−associated_Sm−like_protein_LSm4 [MW = 15.76 kDa]

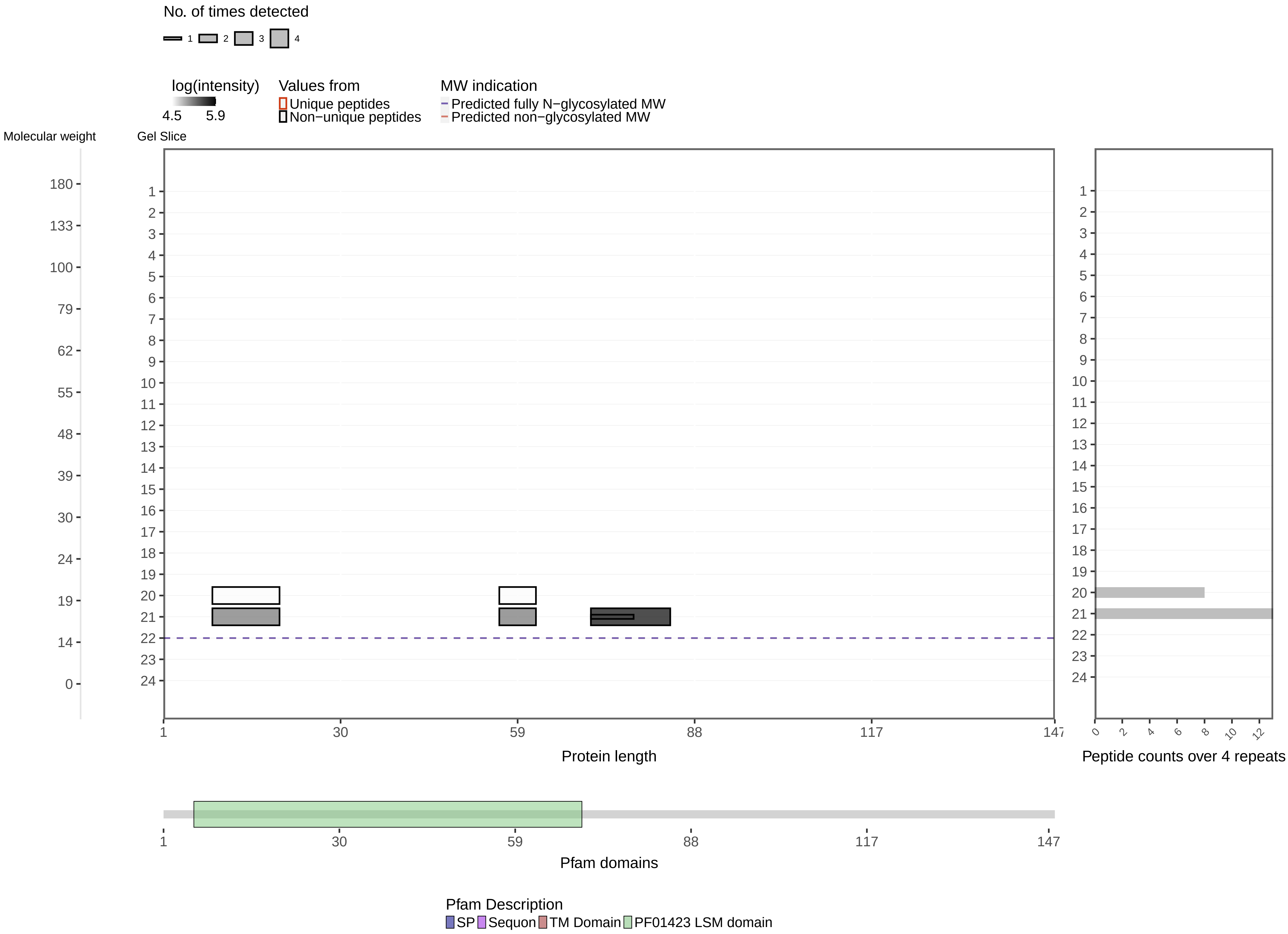


Protein sequence:

GNTIKYLRVPDEVIDKVQEEAKSRTDRKPPGVGRGRGRGGRDDSAAGRQAKGIGRGMDDGGAKGRGKGGPSAKSGGRGGGRGRG*

NbL08g17020.1 proteasome_subunit_beta_type−7−A [MW = 34.74 kDa]

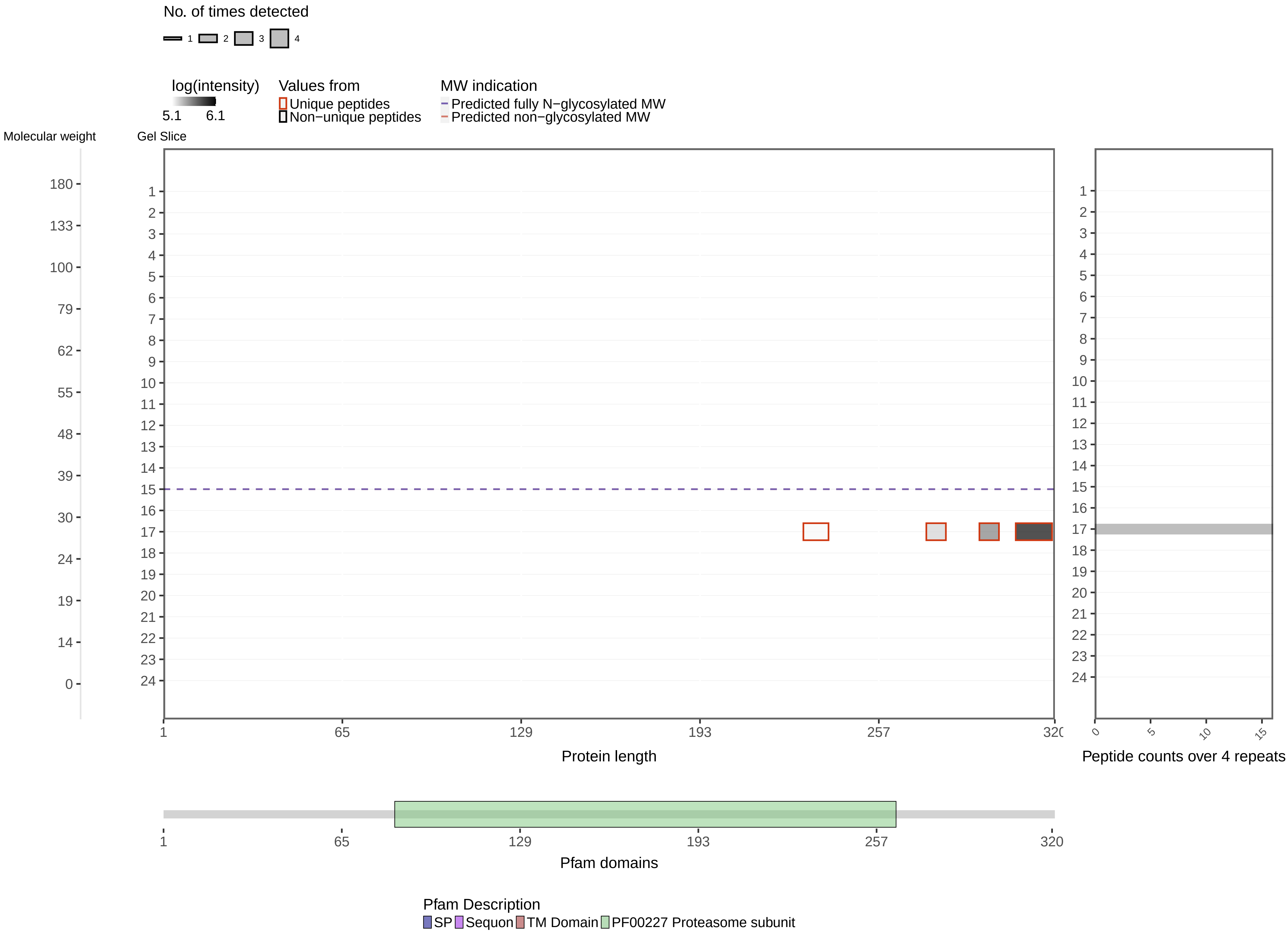


Protein sequence:

SQLKLHRFHTGRESRVVTALTLLKSHLFSYQGHVSAALVLGGVDVTGPHLHTIYPHGSTDTLPYATMGSGSLAAMAIFESKYREGMNRDEGIKLVAEAILSGVFNDLGSGSNVDICVITKGNTEYLRNHLSPNPRTYPQKGYSFPKKTEVLLTKITPLREIVQVIEGGDAMEE*

### NbL08g16870.1 peptidyl−prolyl_cis−trans_isomerase_CYP18−1 [MW = 26.82 kDa]

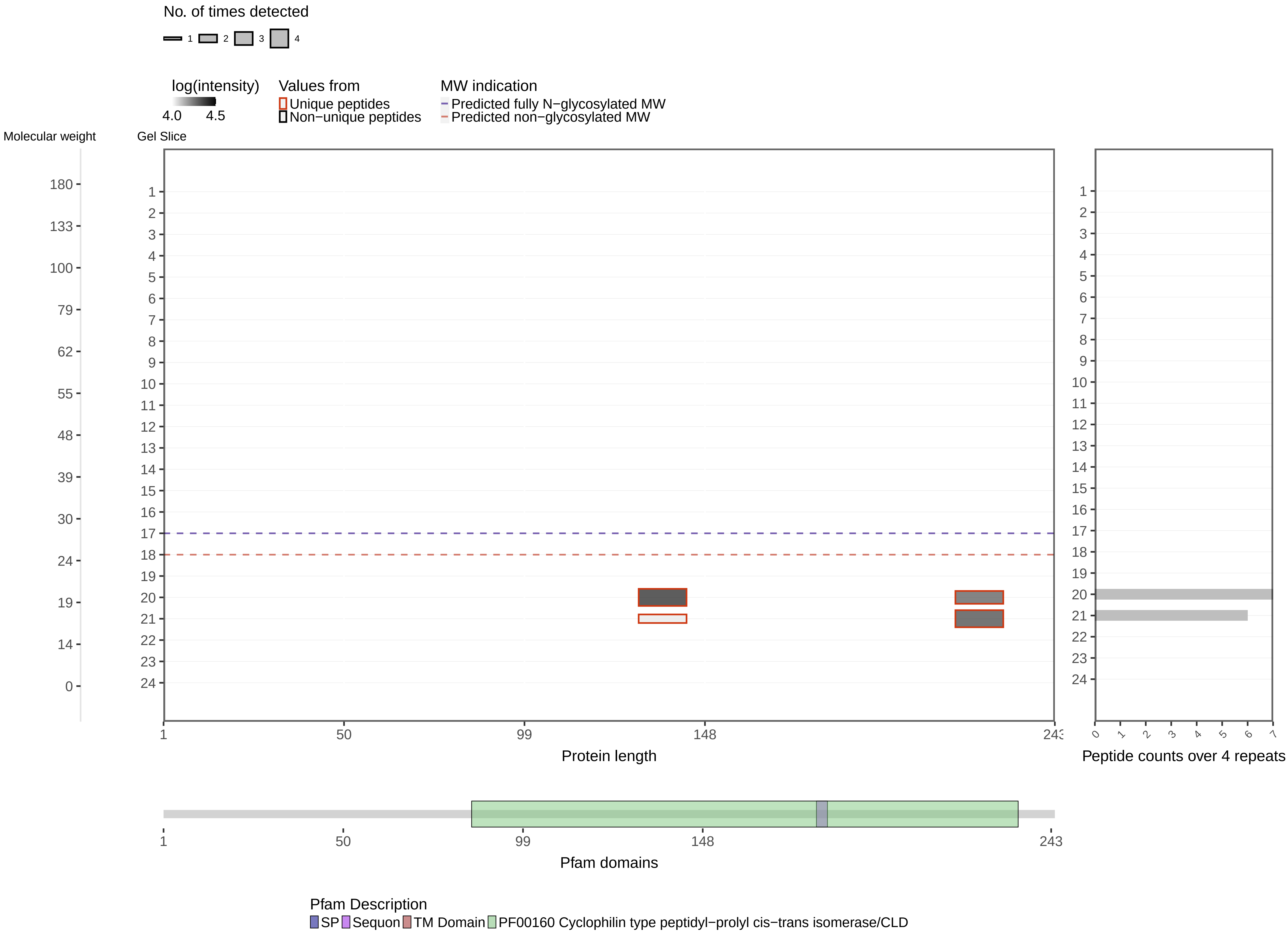


Protein sequence:

FLALCASGYYDGTIFHRNIKGFMIQGGDSTGTGRGGTSIWGKKFNDEIRESLKHNARGILSMANSGPNTNGSQFFMTYAKQPHLNGLYTIFGKVIHGFEVLDIMEKTPTGPGDKPLAEIRLNRVTIHANPLAG*

### NbL08g16540.1 D−3−phosphoglycerate_dehydrogenase_1_chloroplastic−like [MW = 63.48 kDa]

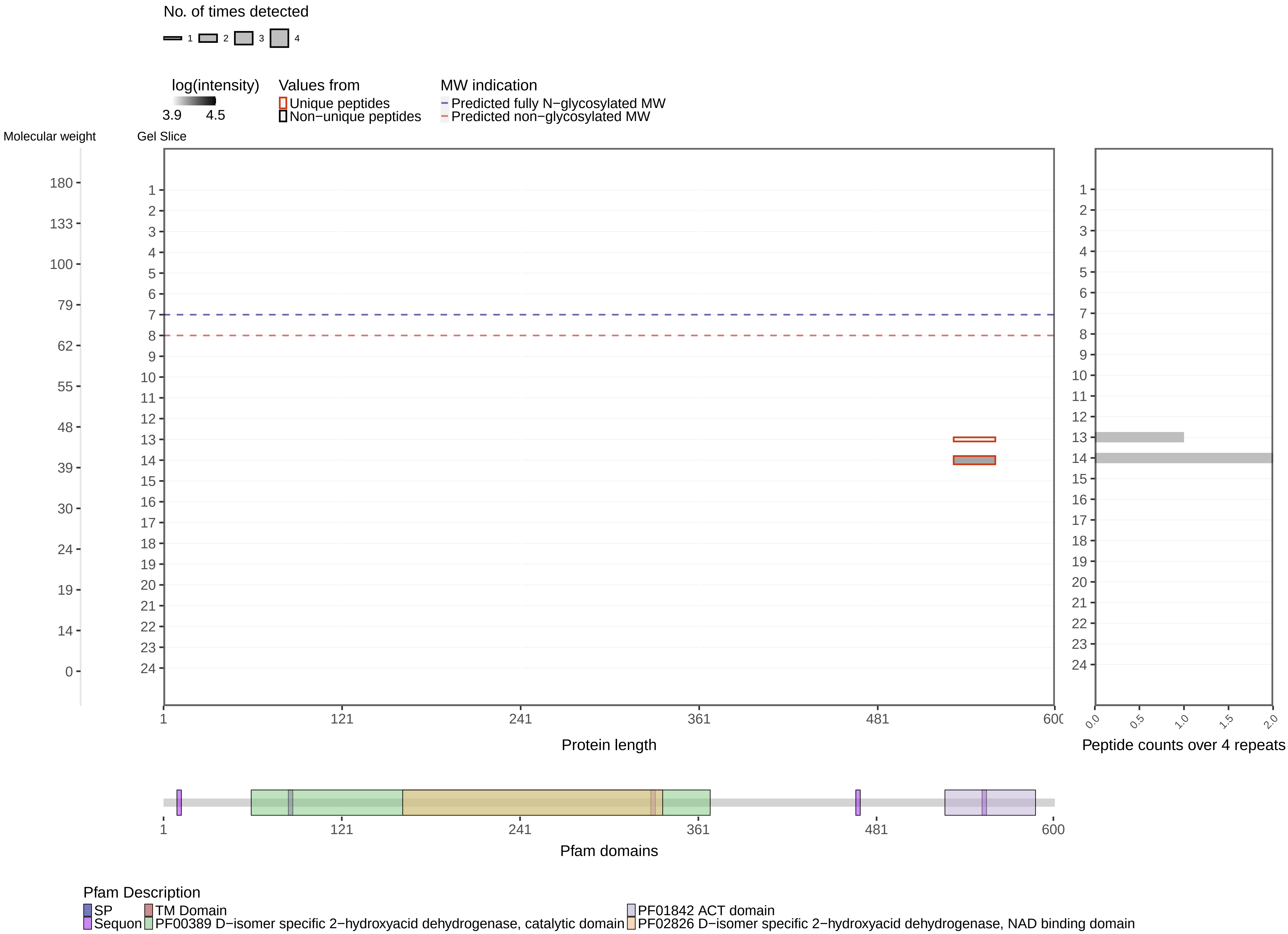


Protein sequence:

NbL08g16530.1 pathogenesis−related_protein_1A [MW − SP = 69.41 kDa]

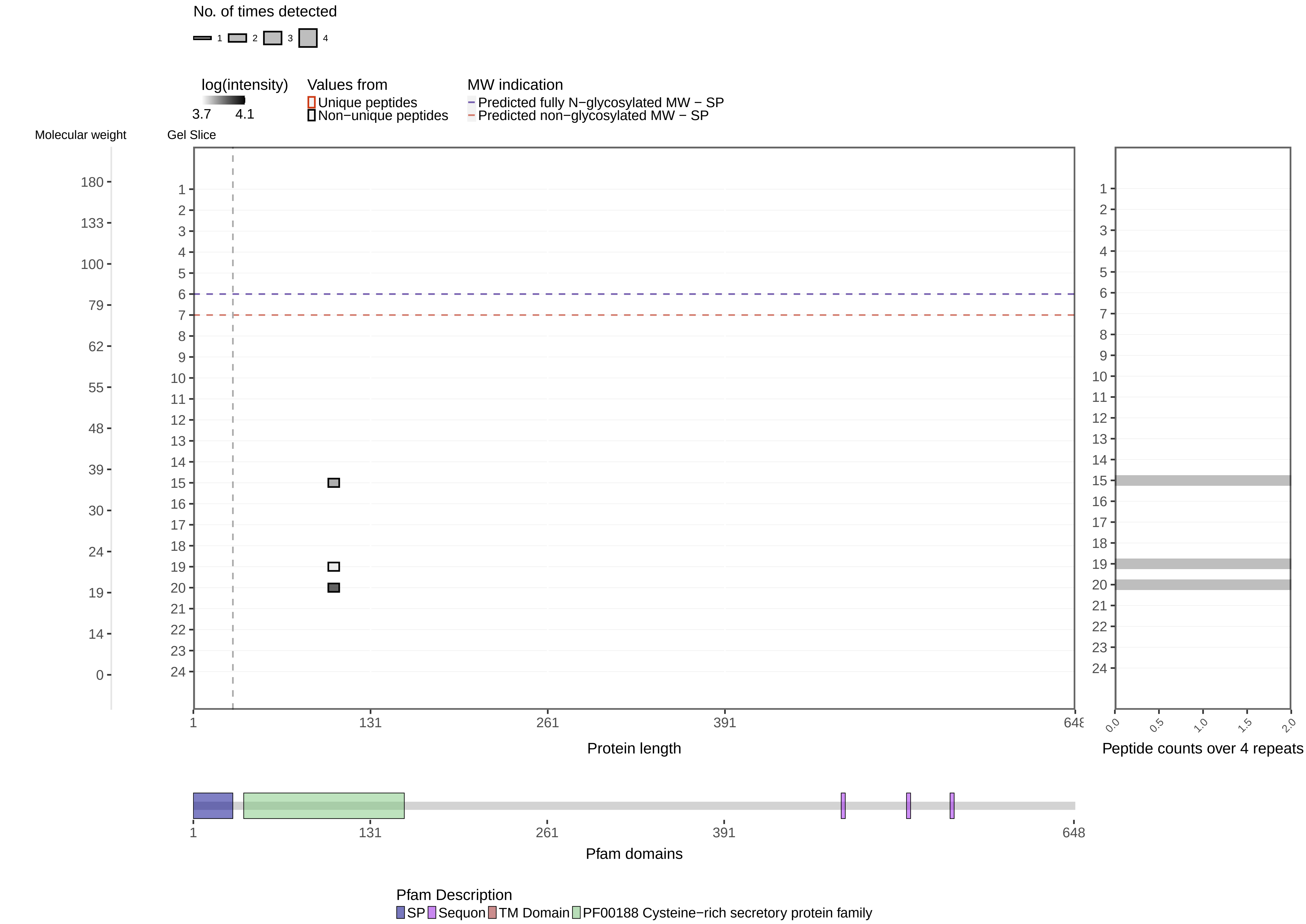


Protein sequence:

TREIDMGTRGYHOLLODSODVHTPIPSPOSSPRRSREGTPDGSHSNANAOFNNDEALONTNTLENPRSGLVNSGCGGISCESRDREPDHMTAFRTGVKGNDLTKODIESVLVKKFGEPLTKGAFTWYDHRSRNRESGSSSRFGNGRNRRESODDDRNLKARFGGYNFNVSTSELVVVSRRMGDKELWPKEIRSNPNRRNPNHWCEFHDDHRHKTANCSSNIILLRLLREMOAEDKLVPKEHTLSGFENASIVTMGEVILTTFTKCVIKDSKFOVVDMEIACNMILGRP

NbL08g16430.1 uncharacterized_LOC107772566 [MW − SP = 23.53 kDa]

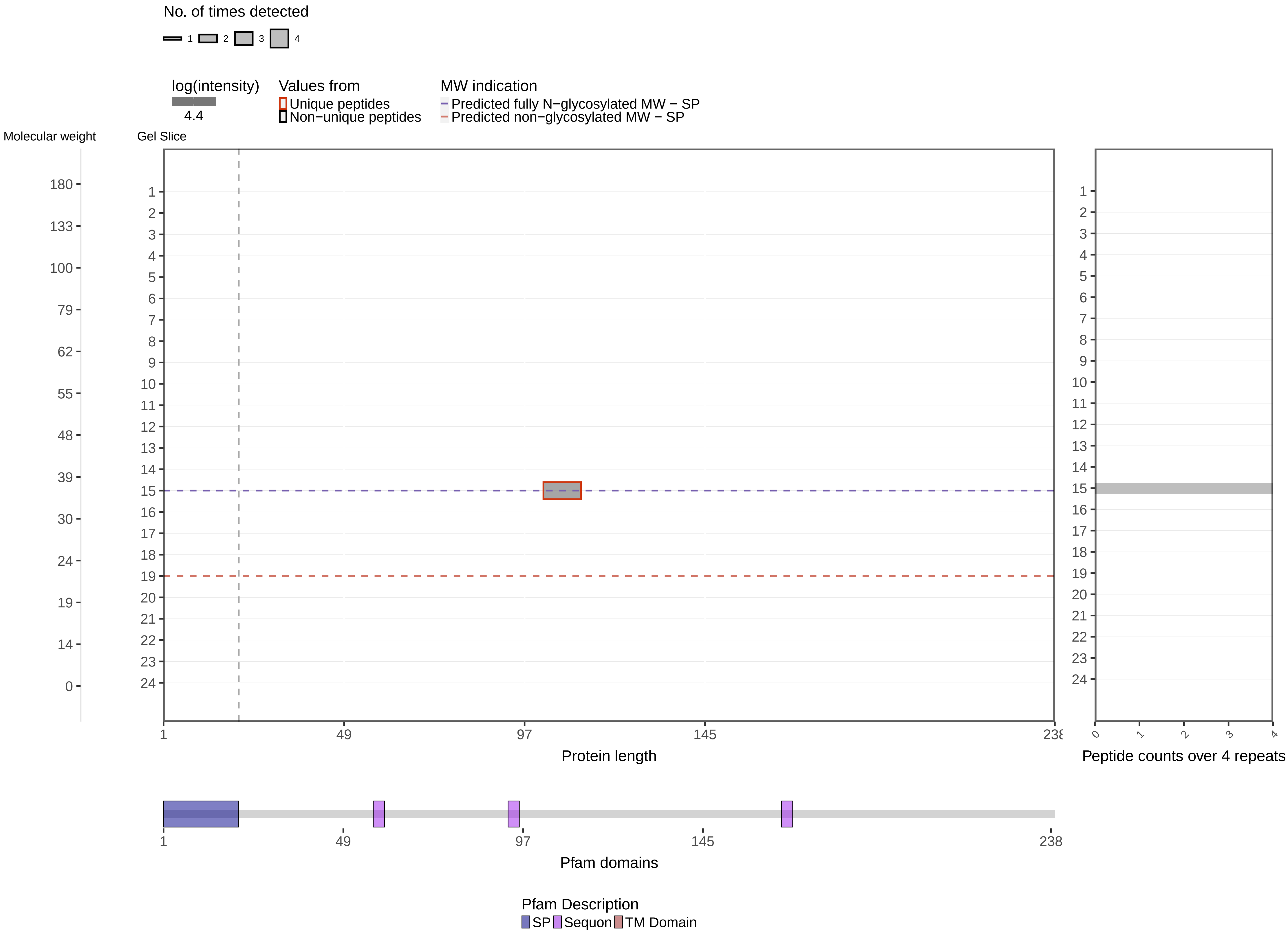


Protein sequence:
[illegible]VAFAGRKYAARSLPAFVADNTHTITSFTLVLEFQKGTLQNLFWKKFGCSSCTGGSFVCLNTTDCAVPNNKCNGNGGSVDCNMGIQLAFSGTDKNDDVLNSWYEVKNLRQYSLYGLFQDVSDSLTSPFKNLF*

NbL08g16390.1 isocitrate_dehydrogenase_[NAD]_regulatory_subunit_1_mitochondrial [MW = 40.48 kDa]

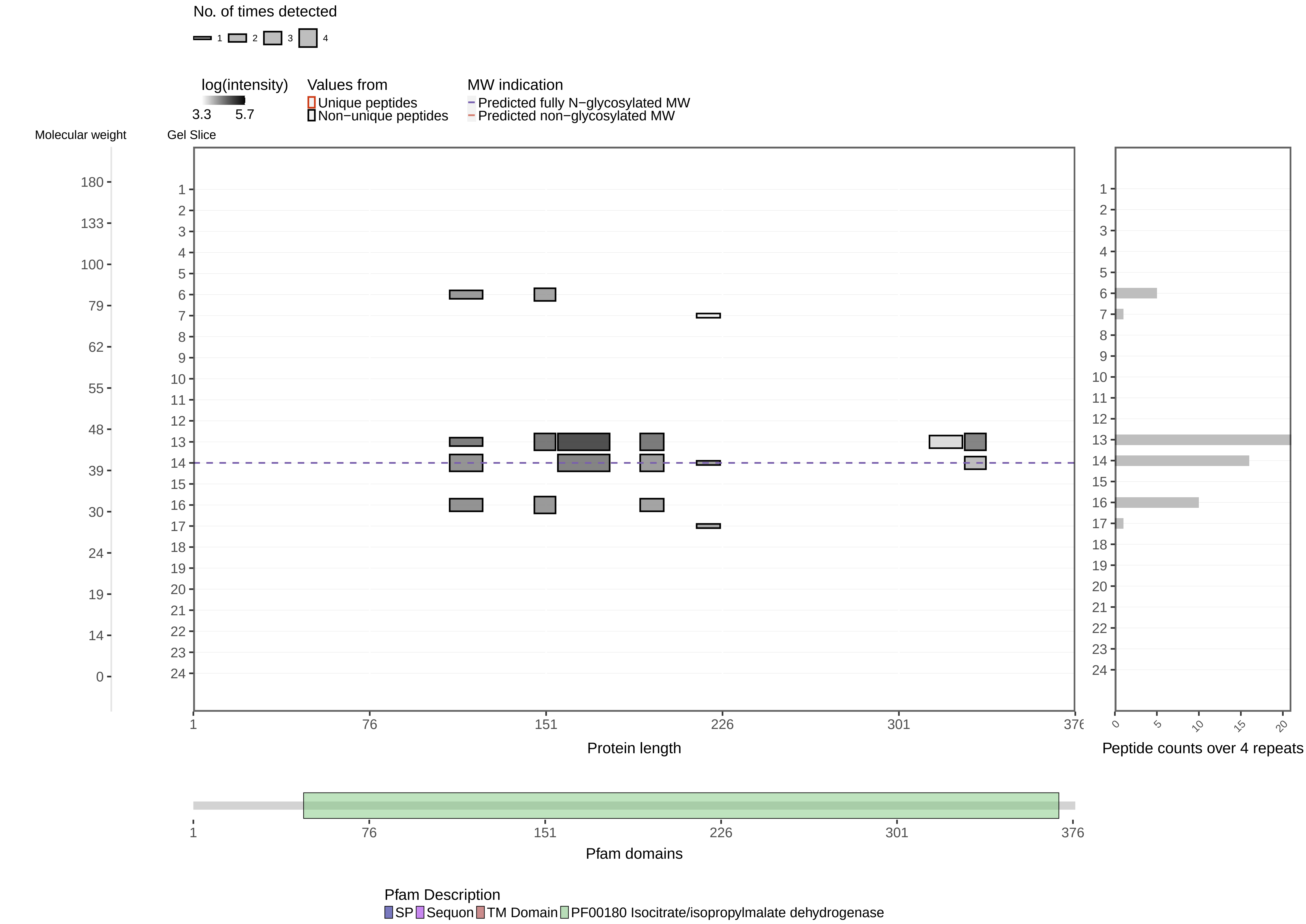


Protein sequence:

VESLKVMTKFCSERIAKYAFEYAYLNNRKVVTAVHKANIMKLADGLFLESCREVASKYPGIKYNEIIVDNCCMQLVSRPEQFDVMVTPNLYGNLVANTAAGIAGGTGVMPGGNVGADHAVFEQGASAGNVGNEKILEQKKANPVALFLSSAMMLRHLQFPSFADRLETSVKRVIAEGKCMTKDLGGGCTTQEITDAVIANLD*

### NbL08g16130.1 glycine−rich_protein_2−like [MW = 21.24 kDa]

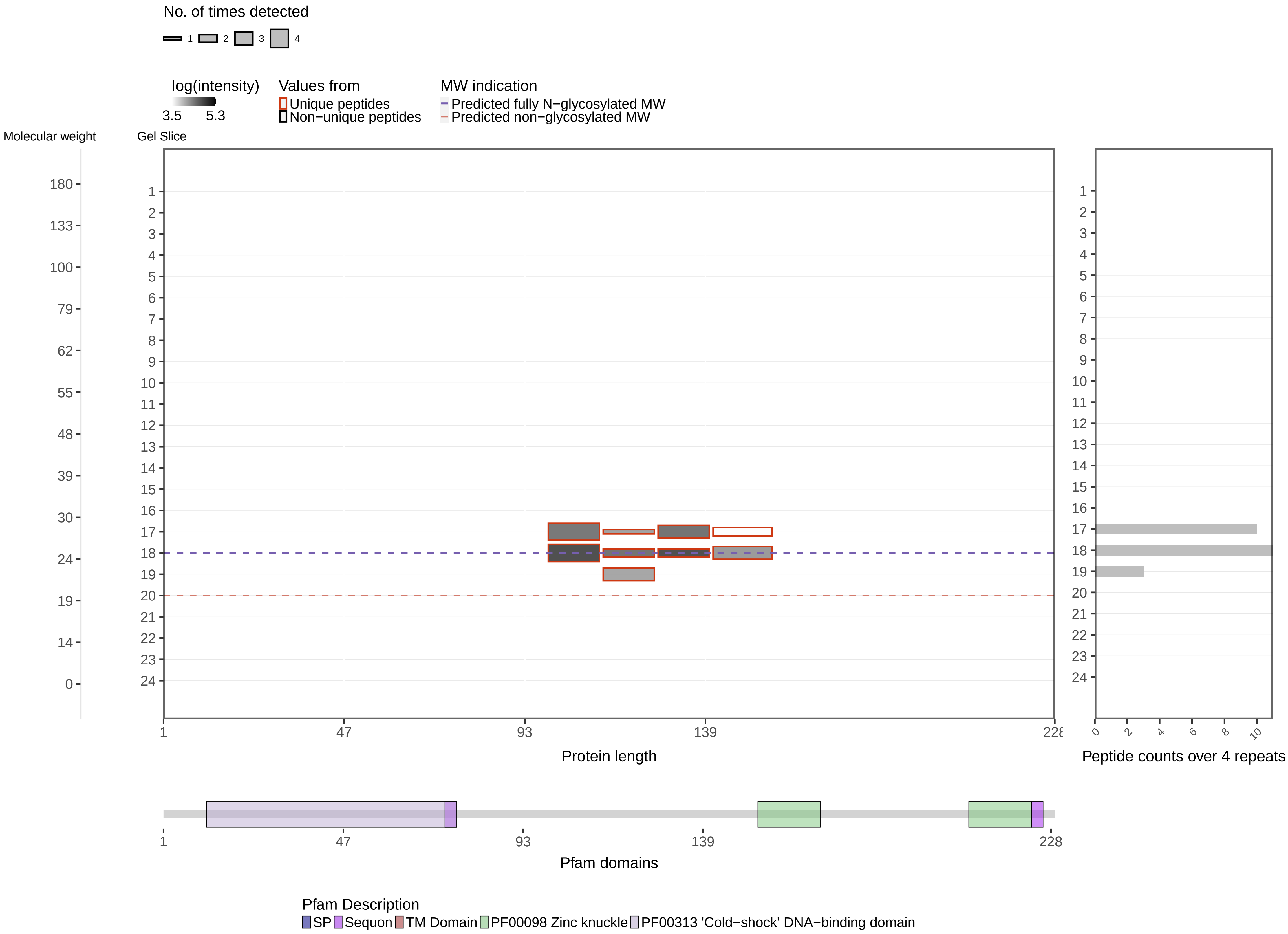


Protein sequence:

'GGGGGGRYGGGDGGYGGGGGRYGGDGGYGGGGGGRYGGGGYGGGGGSGCYKCGEEGHFARECSQGGGGGGYGSGGGGYGSGGGGYGGGGGRYGGSGGGGGGGCFKCGEEGHFARECPNSSGR*

### NbL08g16080.1 triosephosphate_isomerase_chloroplastic−like [MW = 34.16 kDa]

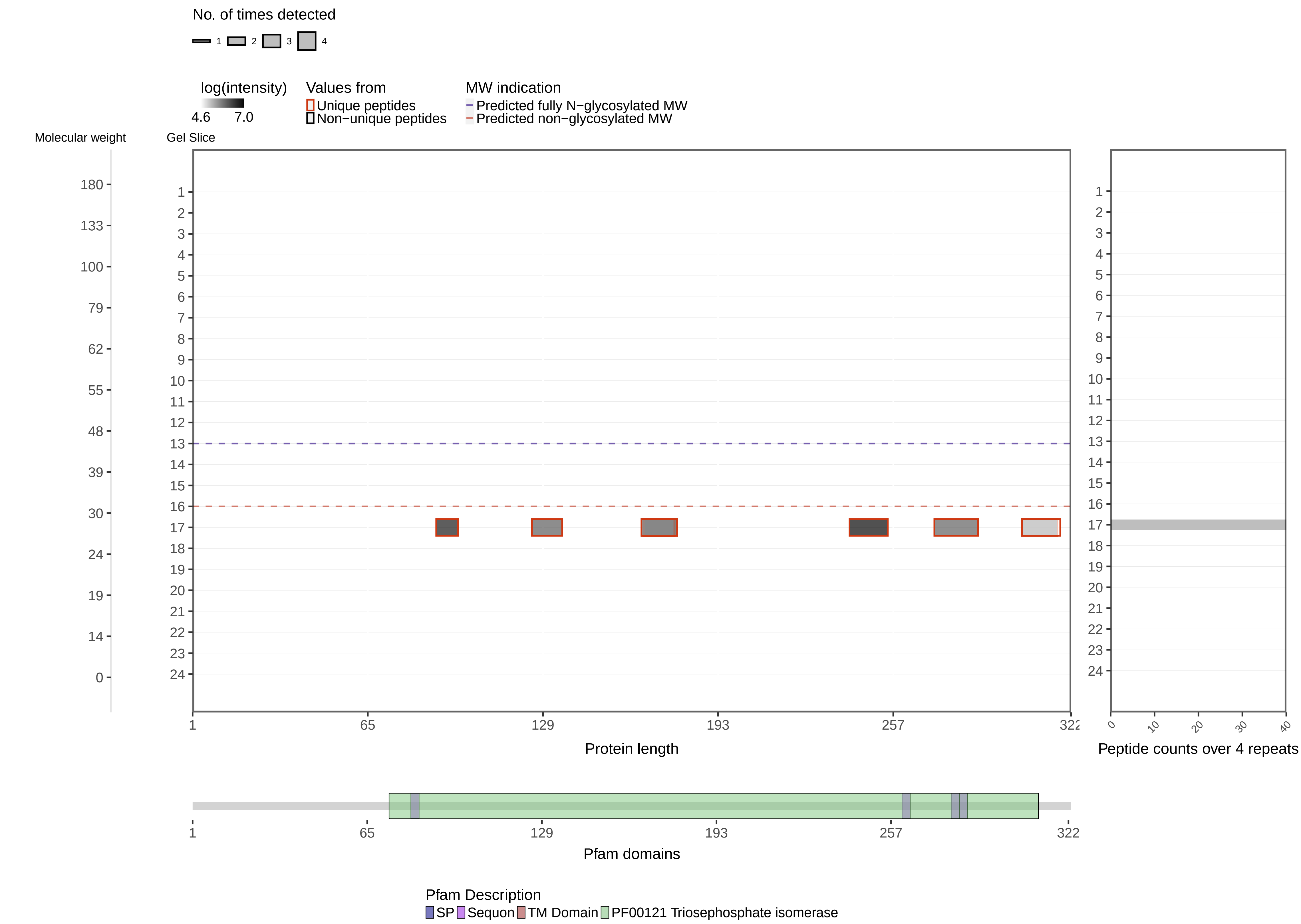


Protein sequence:

[illegible]QLKDLGCKWVILGHSERRHIIGENDEFIGKKAAYALSQNVGVIACIGELLEEREAGKTFDVCFQQLKAFADALPSWDNVVIAYEPVWAIGTGKVATPEQAQEVHAAVRDWLSKNVSAEVASKTRIIYGGSANGSNCSDLAKKEDIDGFLVGGASLKGPEFASIVNSVTSKKVAA*

NbL08g15890.1 60S_ribosomal_protein_L4−1−like [MW = 44.7 kDa]

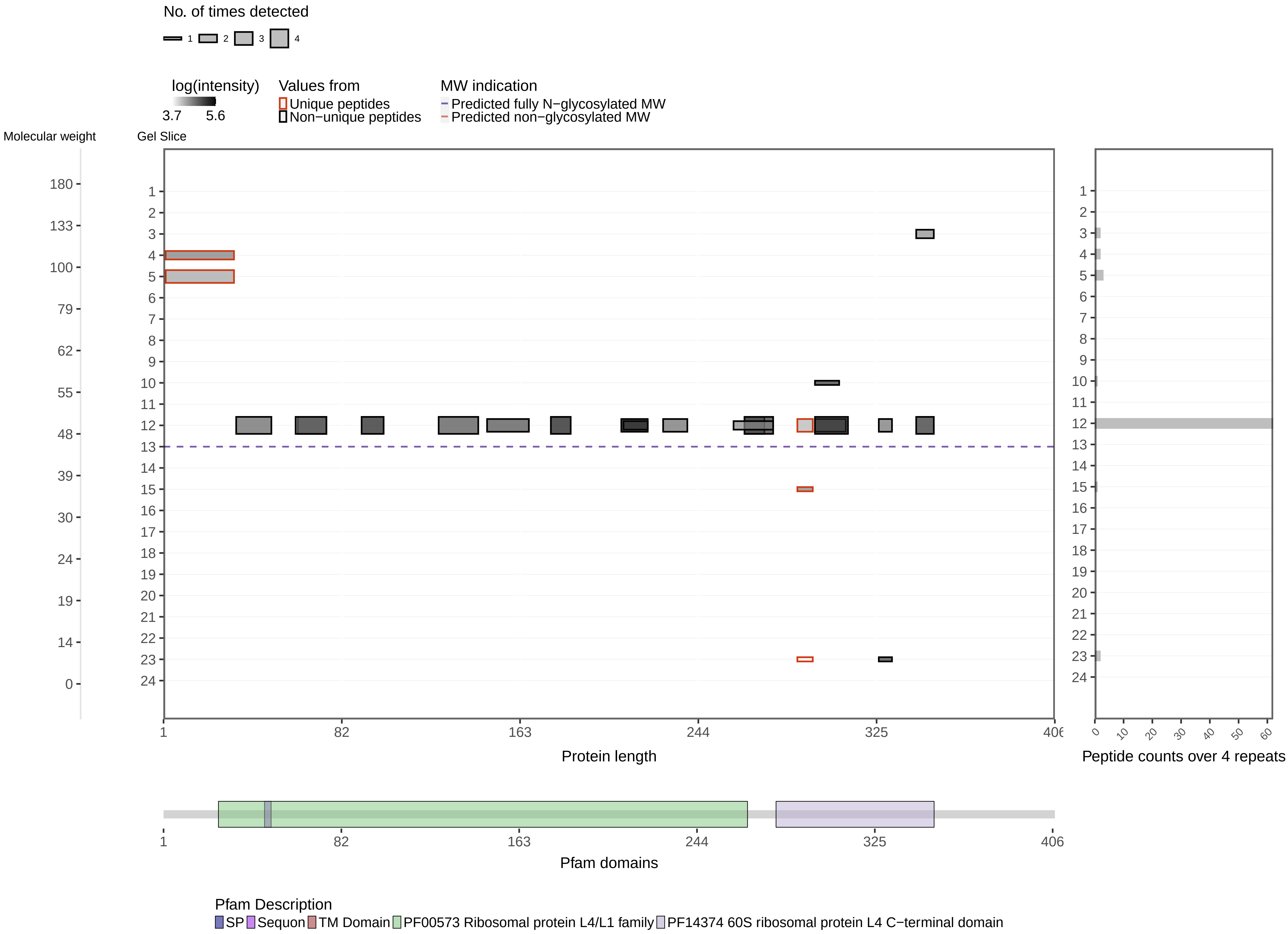


Protein sequence:

SHAIRPGKGKMRNRRYISRKGPLIVYGTEGAKLVKAFRNIPGVEICHVERLNLLKLAPGGHLGRFVVWTKSAYEKLDEIYGSFDKPSEKKKGYVLPRAKMANADIARIINSDEVQSVVRPIKKDVKRATLKKNPLKNLNVLLKLNPYAKTARRMSLLAEAQRVKAKKEKLDKKRHQVTKEEASAIRSASAAWYKTMISDSDYTEFDVFTKWLGVTQ*

### NbL08g15860.1 interactor_of_constitutive_active_ROPs_2_chloroplastic−like [MW = 52.84 kDa]

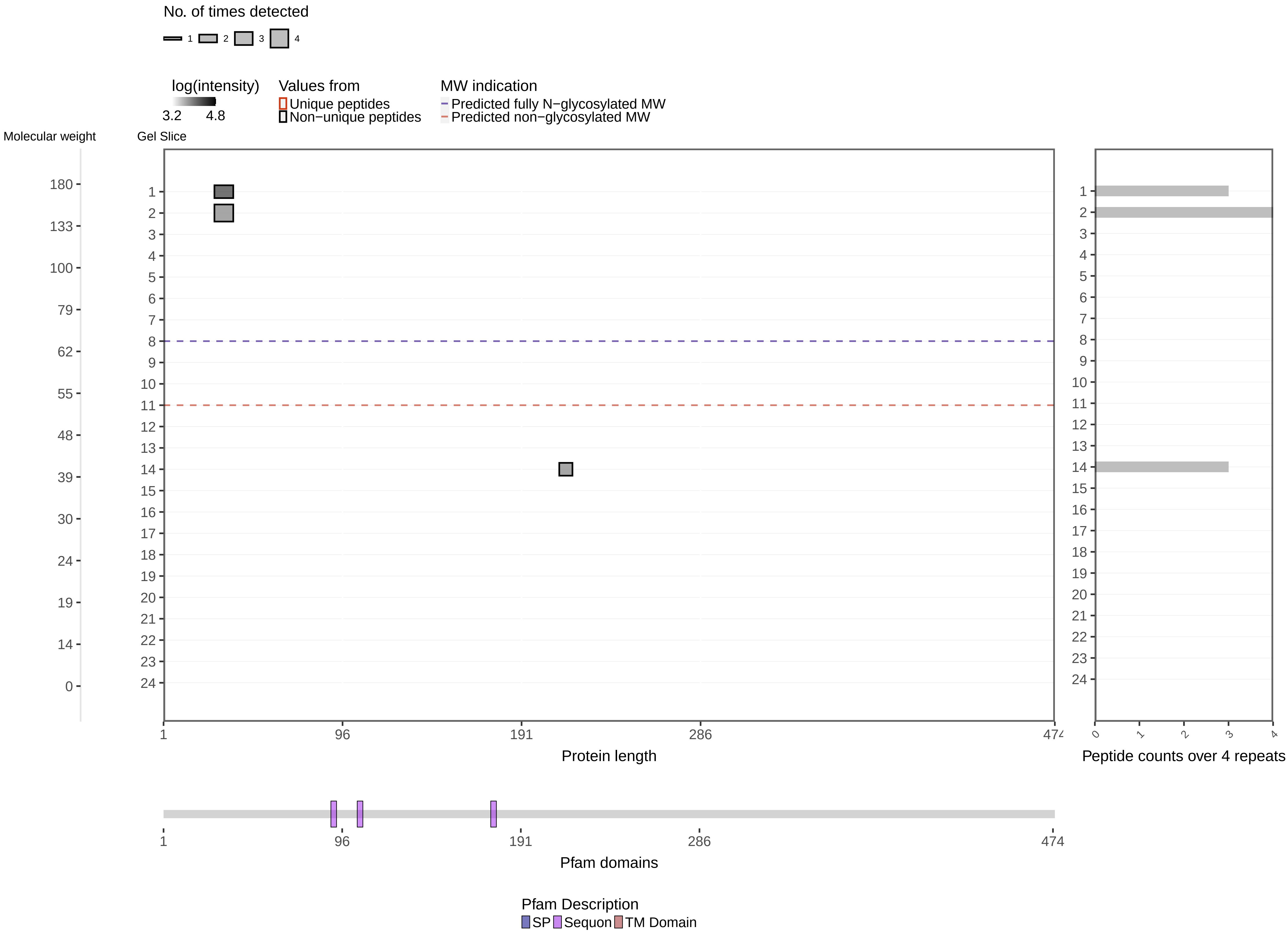


Protein sequence:

QSTLQITSAYEIVECTKSESIQRDTEWNSKLNAAKADVEELTGTLMNKEAELQKISDENKGLNLQVEEIQADDGETKLQAELKKSESILADLRASLFDKETEVQSTTEENEMLKSEMKKREMESTKVNDEVLALAEAAKTSEREALMKLGYLTEEADKRSRKAARVIEELDAAQAANSEMEAELRRLKVQSDQWRKAAEAAAAMLSTSNNGKYVERTGSLDYHTIGGKLRSPLSEDMDNDPRRRMAIC*

NbL08g15740.1 cytokinin_riboside_5'−monophosphate_phosphoribohydrolase_LOG3−like [MW = 24.01 kDa]

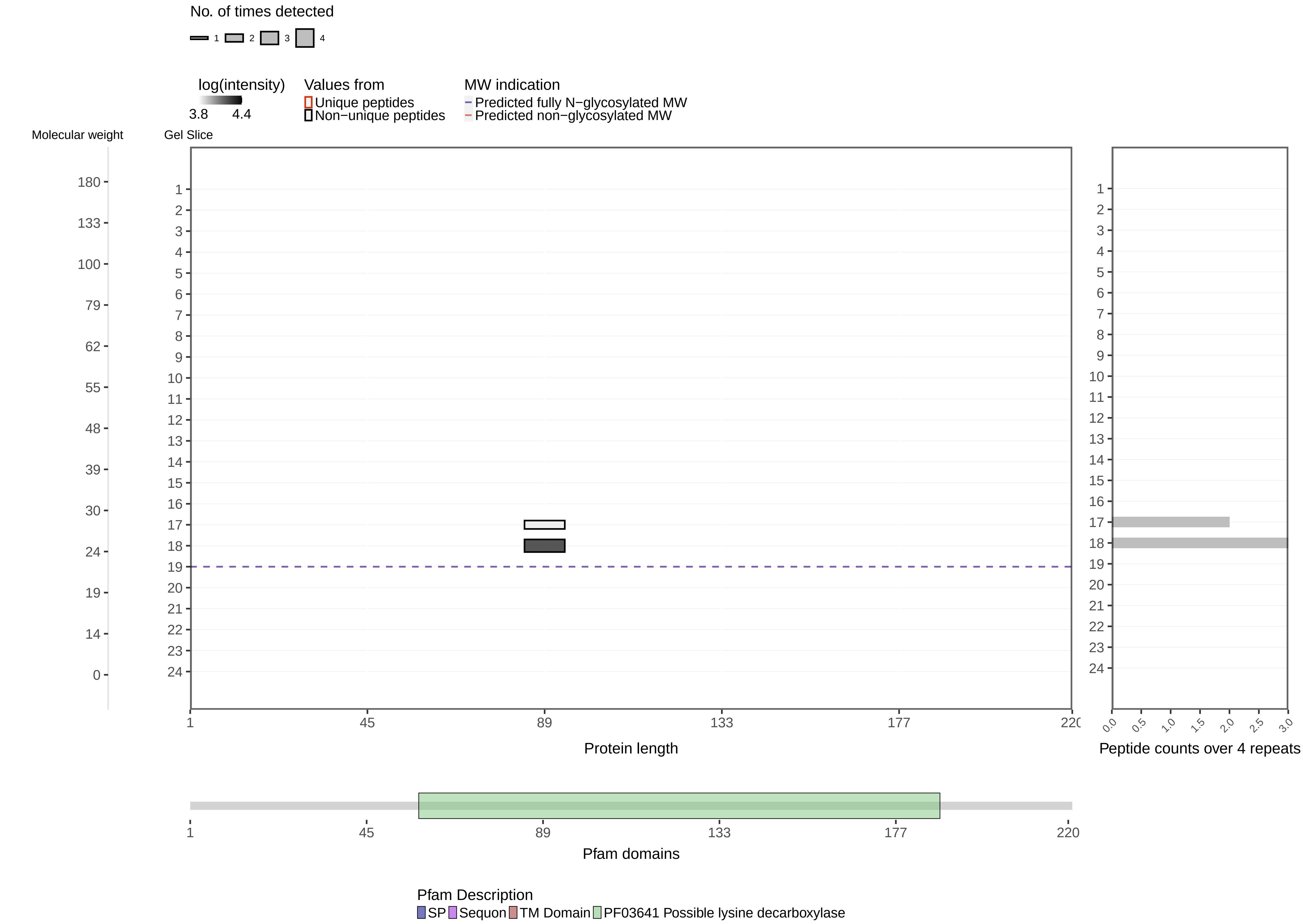


Protein sequence:

ADMHQRKAEMAKHSDAFIALPGGYGTLEELLEVITWAQLGIHDKPVGLLNVDGYYSSLLSFIDKAVEEGFICPNARHIIVSAPTPKELVNKLEEYVPCHERVASKLSWEMEQLGYPQAQEMAR*

NbL08g15640.1 thylakoid_lumenal_15.0_kDa_protein_2_chloroplastic−like [MW = 24.77 kDa]

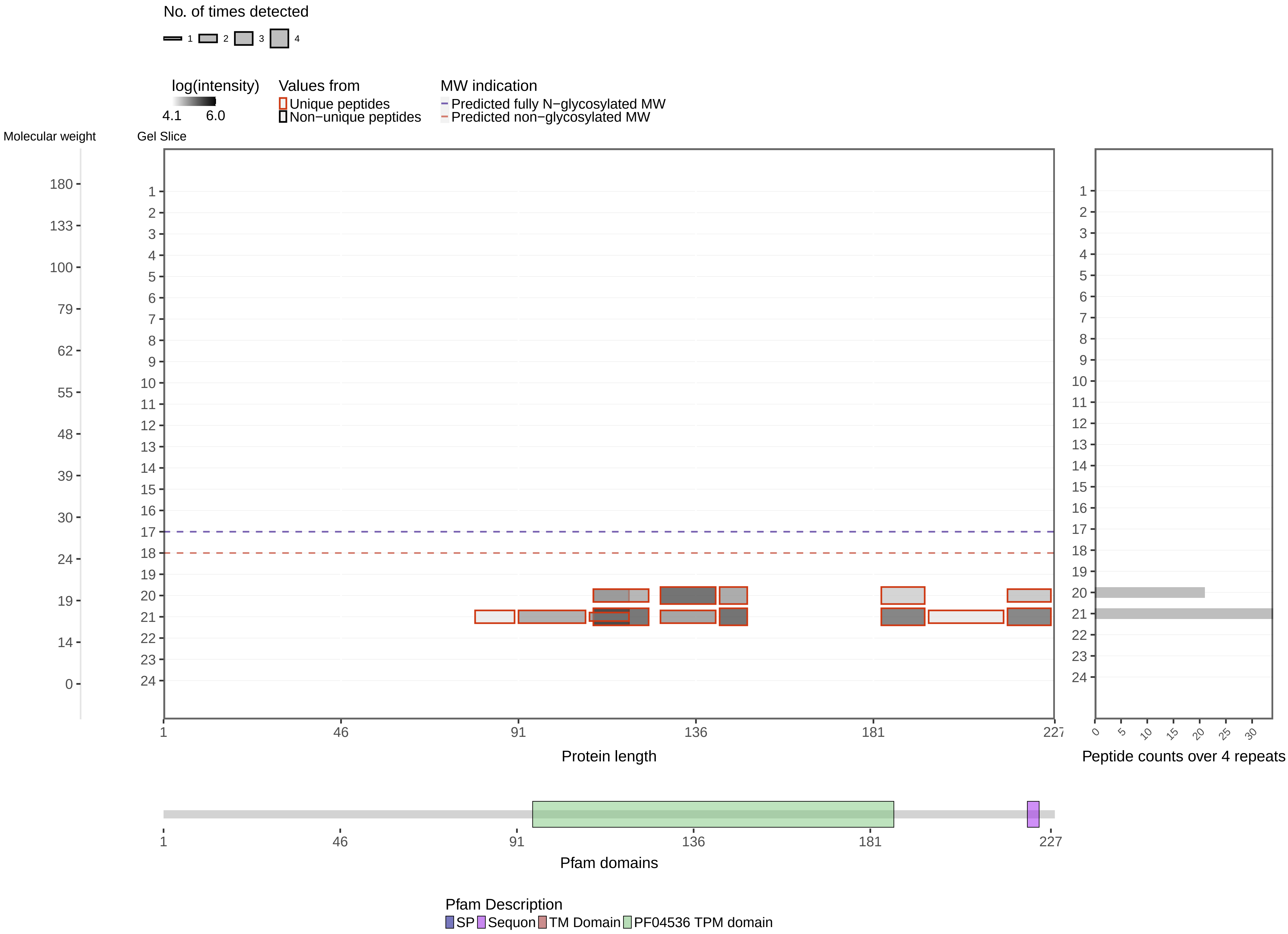


Protein sequence:

.SDGQEKRLAQEITDIEKDTGFKLRVLAQNYPDTPGLAIKDFWQVDDRTIVFVADPTFGNILNFNVGATVDLDIPRSFWSRLAGTYGNMFYWKEKGEDASIEAAVMAISTCLREPVGPNNCSEVK*

NbL08g15490.1 poly_[ADP−ribose]_polymerase_tankyrase−like [MW = 21.86 kDa]

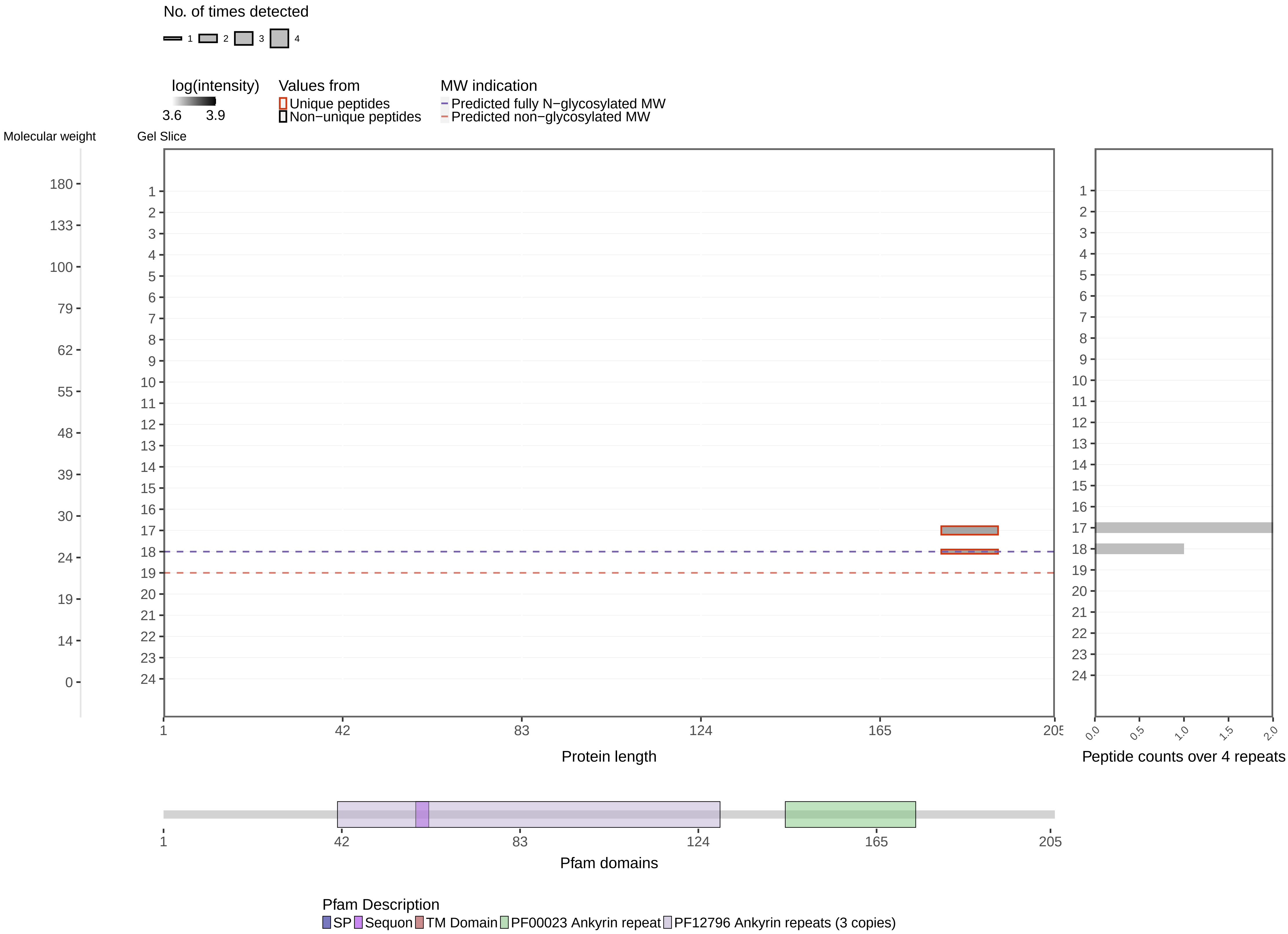


Protein sequence:

.LLERGASLEPKDEDGAIPLHDACAGGYTEIAQLLINSAPDPECVKRMLDSVDVEGDTPLHHAARGDHVTVVRLLLASGACPTRTNIYGKTPGELAEPDGEAQRILDETARAFPSH*

NbL08g15440.1 carotene_epsilon−monooxygenase_chloroplastic [MW = 60.82 kDa]

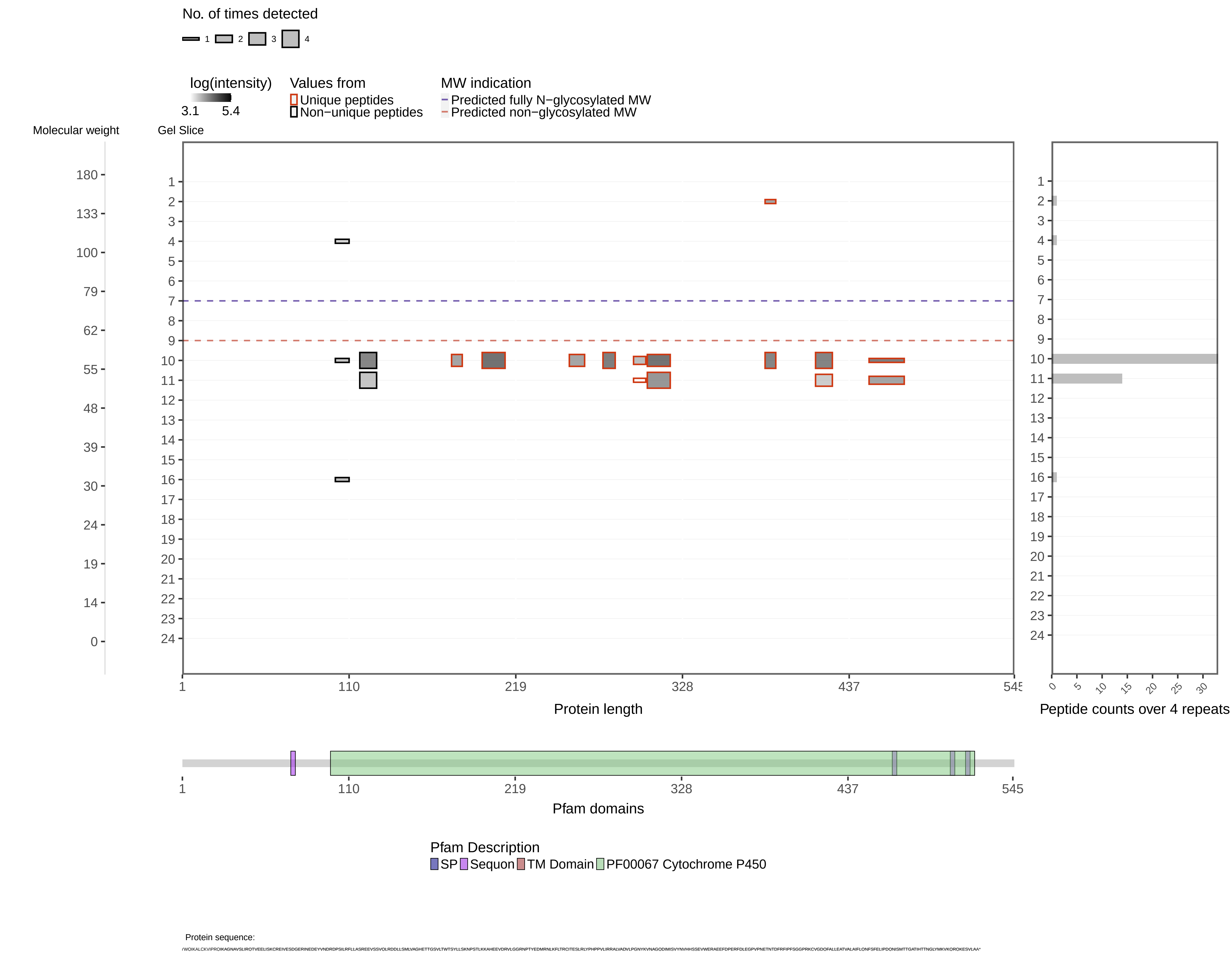

### NbL08g15400.1 threonine_dehydratase_biosynthetic_chloroplastic−like [MW = 66.07 kDa]

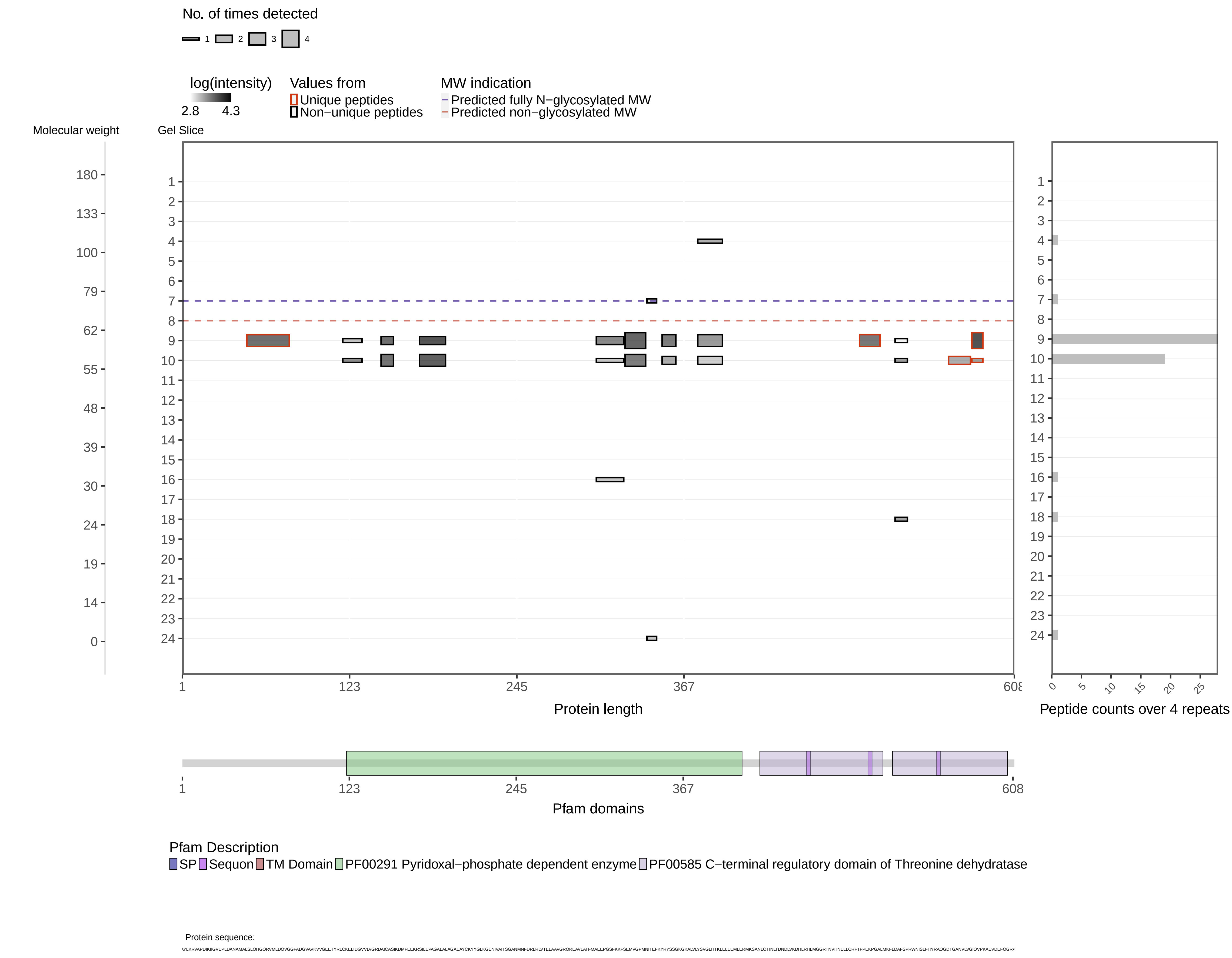

NbL08g15350.1 pyruvate_kinase_1_cytosolic [MW = 64.64 kDa]

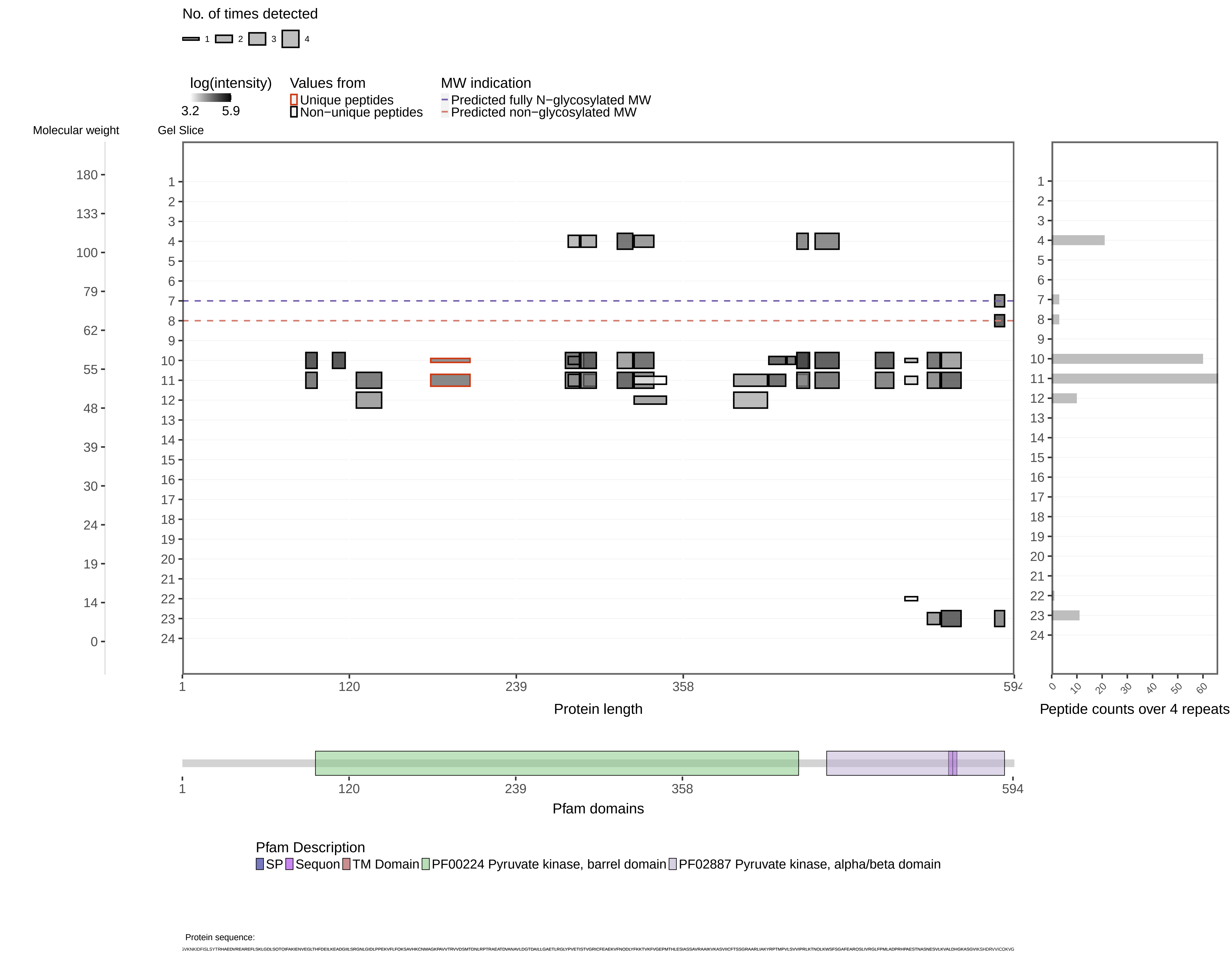

### NbL08g15300.1 peroxiredoxin−2E−2_chloroplastic−like [MW = 24.25 kDa]

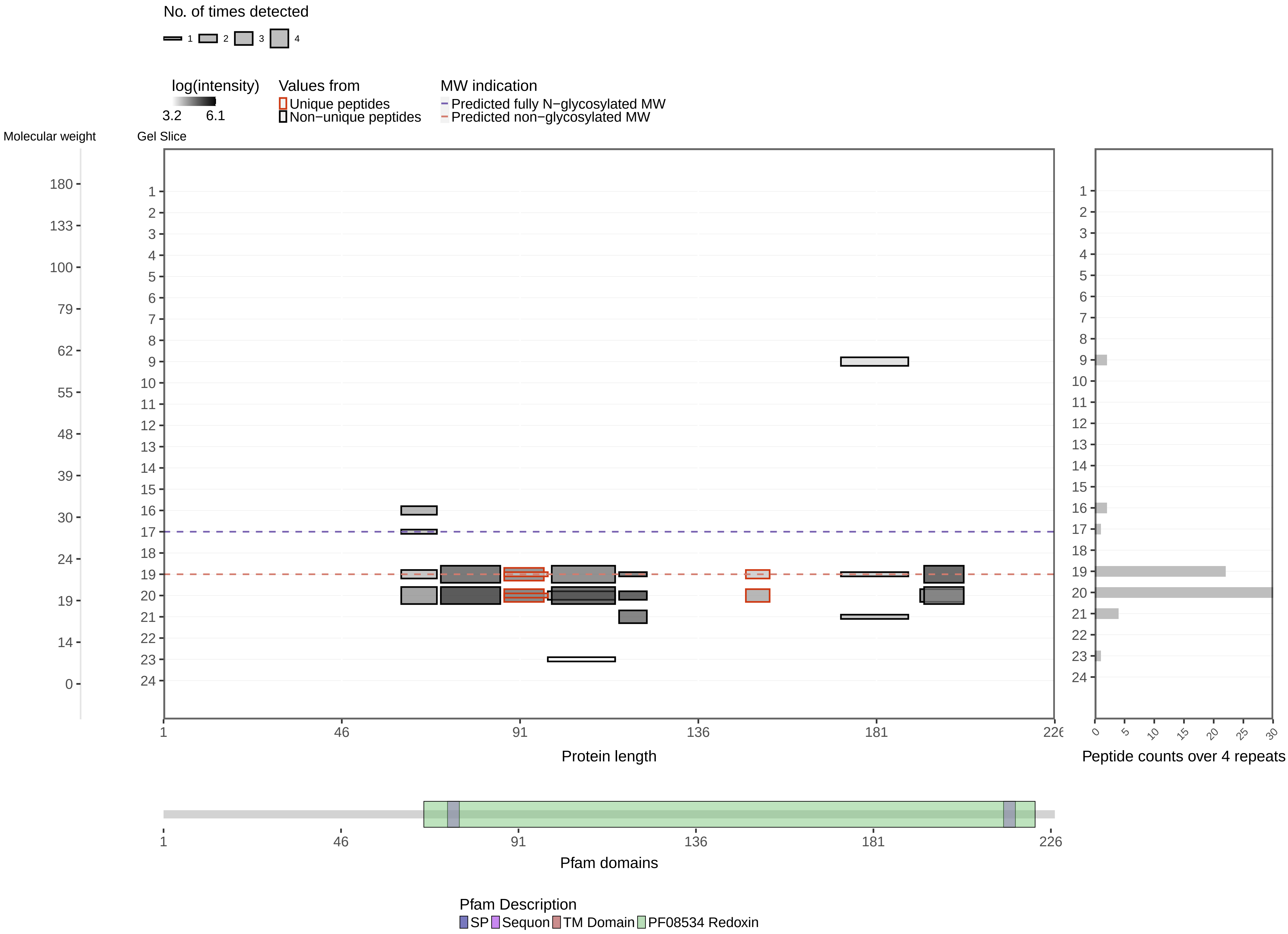


Protein sequence:

FAVPGAFTPTCSQKHLPGFVEKSKELKSKGVDTIACISVNDAFVMKSWKENLKINDEVLLLSDGNLEFTKAIGCELDLSDKPIGLGVRSRRYAMLVDDGVVKVLNLEEGGAFNVSSAEDILKTL*

NbL08g15270.1 calcium−dependent_protein_kinase_4 [MW = 61.78 kDa]

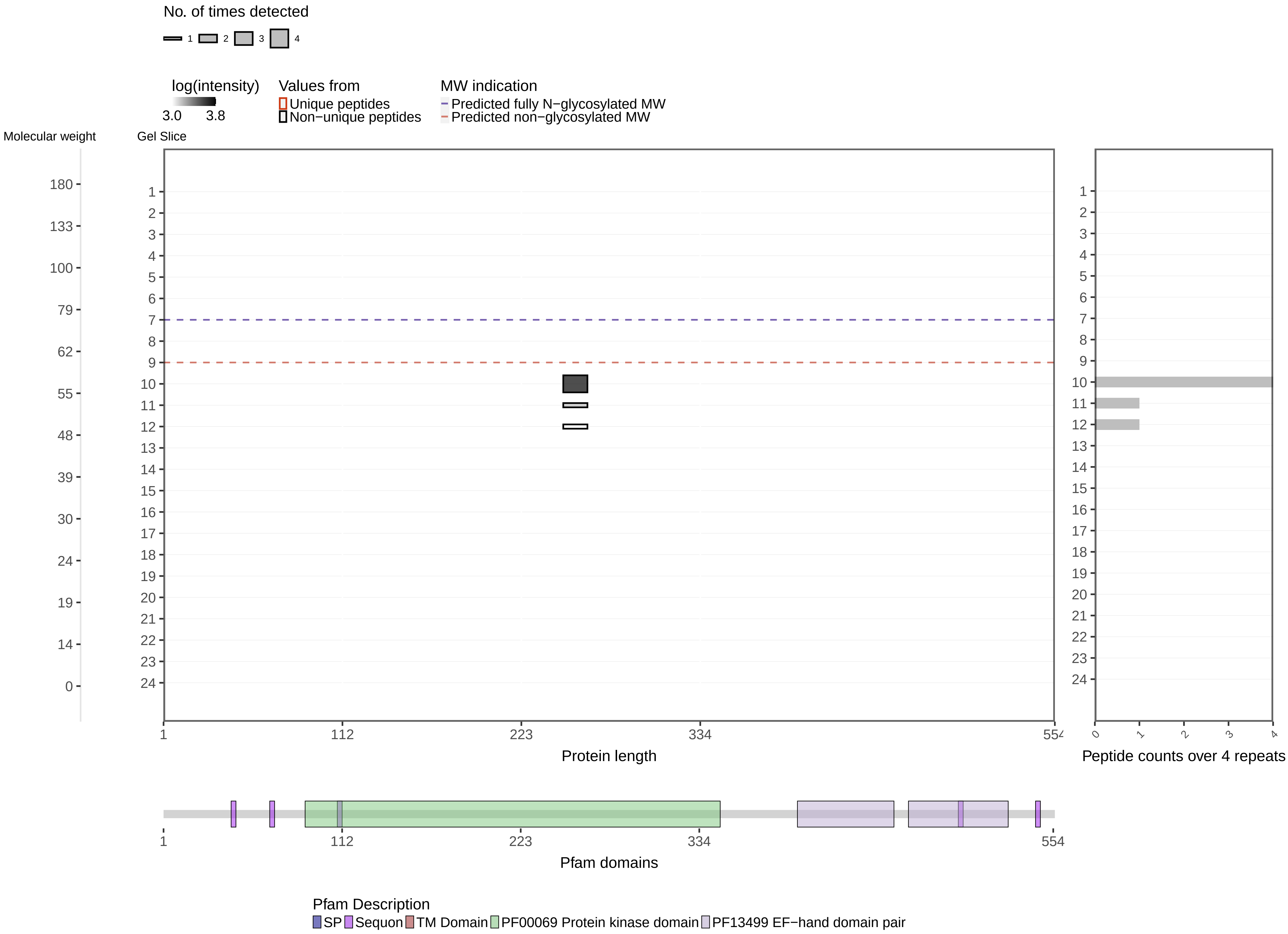


Protein sequence:

KHYGPEADVWTAGVILYILLSGVPPFWAETQQGIFDSVLKGHIDFDSDPWPLISESAKDLIRKMLCMQPSERLTAHEVLCHPWICENGVAPDRALDPAVLSRLKQFSAMNKLKKMALRVIAESLSEEEIAGLREMFKAMDTDNSGAITFDELKAGLRKYGSTLKDTEIRGLMNAADVDNSGTIDYGEFIAATVHLNKLEREEHLMAAFQYFDKDGSGYITVDEVQQACAEHNITDVYFEDIIREVDQDNDGRIDYGEFVAMMQKGNPCGVGRRTMRNSLNLSMRDAAGAH*

NbL08g15260.1 uncharacterized_LOC107760951 [MW = 34.16 kDa]

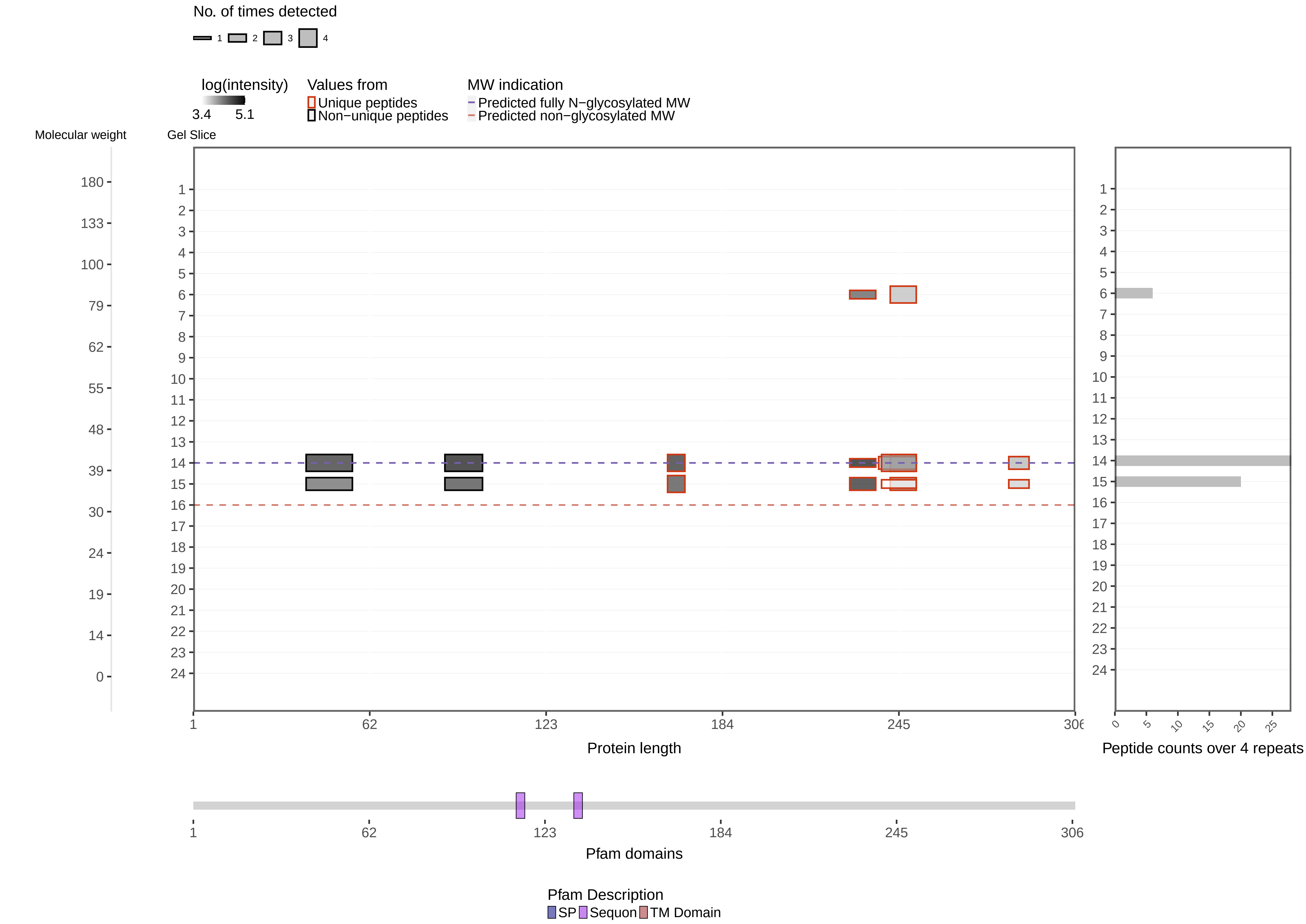


Protein sequence:

:HCLIDPRKKESRVRIIOVVRLEDSKLVLONVKVFCEHWYGPFRNGDOLGGCAIODSAFASTOALDPTOVIGVWEGKHAIFSFDNAPOKVOELVDGSTRKTVRDELDLVLLPROLWCCLKDVAGGETCCEVGWLFEKGRAMTSKCIFSDDGKLKEIAIACESAASAE*

NbL08g15250.1 uncharacterized_protein_At2g17340−like [MW = 44.41 kDa]

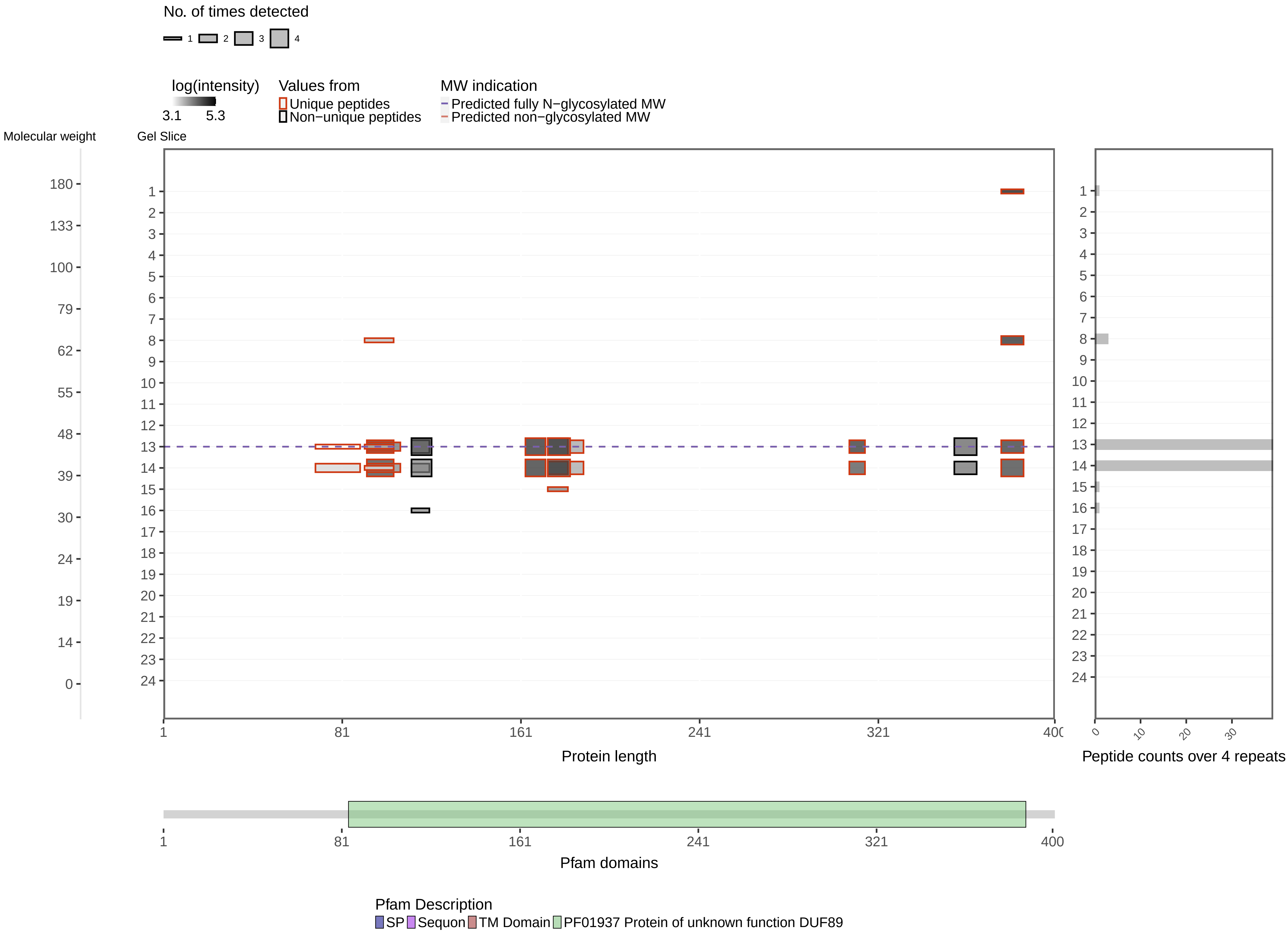


Protein sequence:

ENLVRGIFAGNIFDLGSAKLAELFSEDGMSFLASCQNLVPRPWVIDDVDTFITKWSRKTWKKAVIFVDNSGADIILGILPFARELLRQGTQVVLAANDLPSINDVTYTELVEILSKLKDEHGKLIGVDTFKLLVANSGNDLPVIDLTTVSQELAYLASDADLVILEGMGRGIETNLYAQFKCDSLKIGMVKHAEVAQFLGGRLYDCVFKFNEALS*

NbL08g15030.1 acetyltransferase_NSI [MW = 28.25 kDa]

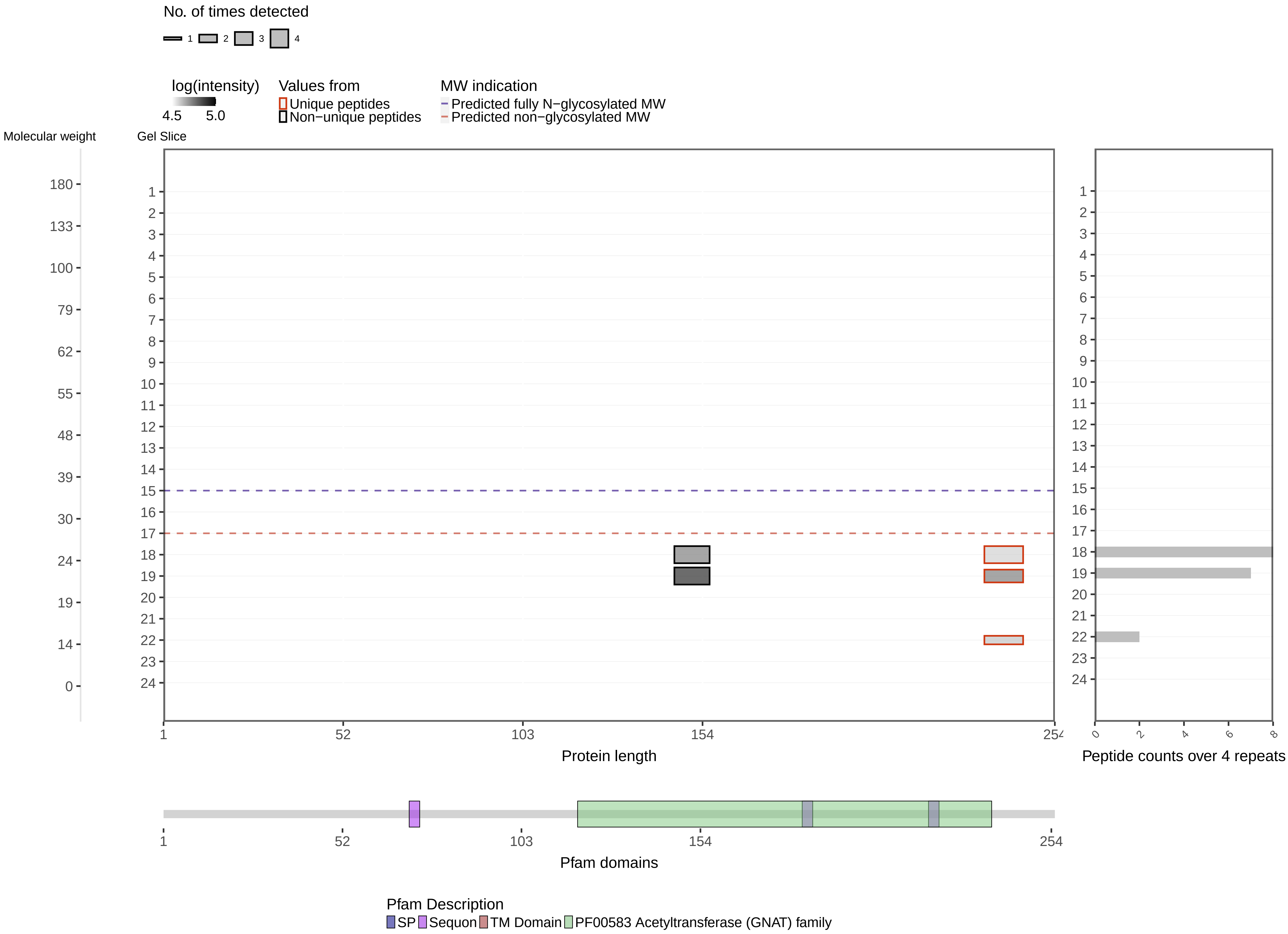


Protein sequence:

GGDVDVYDLQALCDKVGWPRRPLSKLAAALKNSYIVATLHSRKFSSGEEGNGEKKLIGMARATSDHAFNATIWDVLVDPSYQGQGLGKALIEKLIRTLLQRDIGNISLFADSQVVEFYKNLGFEPDPEGIKGMFWYPMY*

NbL08g15020.1 mannan_endo−14−beta−mannosidase_6 [MW − SP = 46.4 kDa]

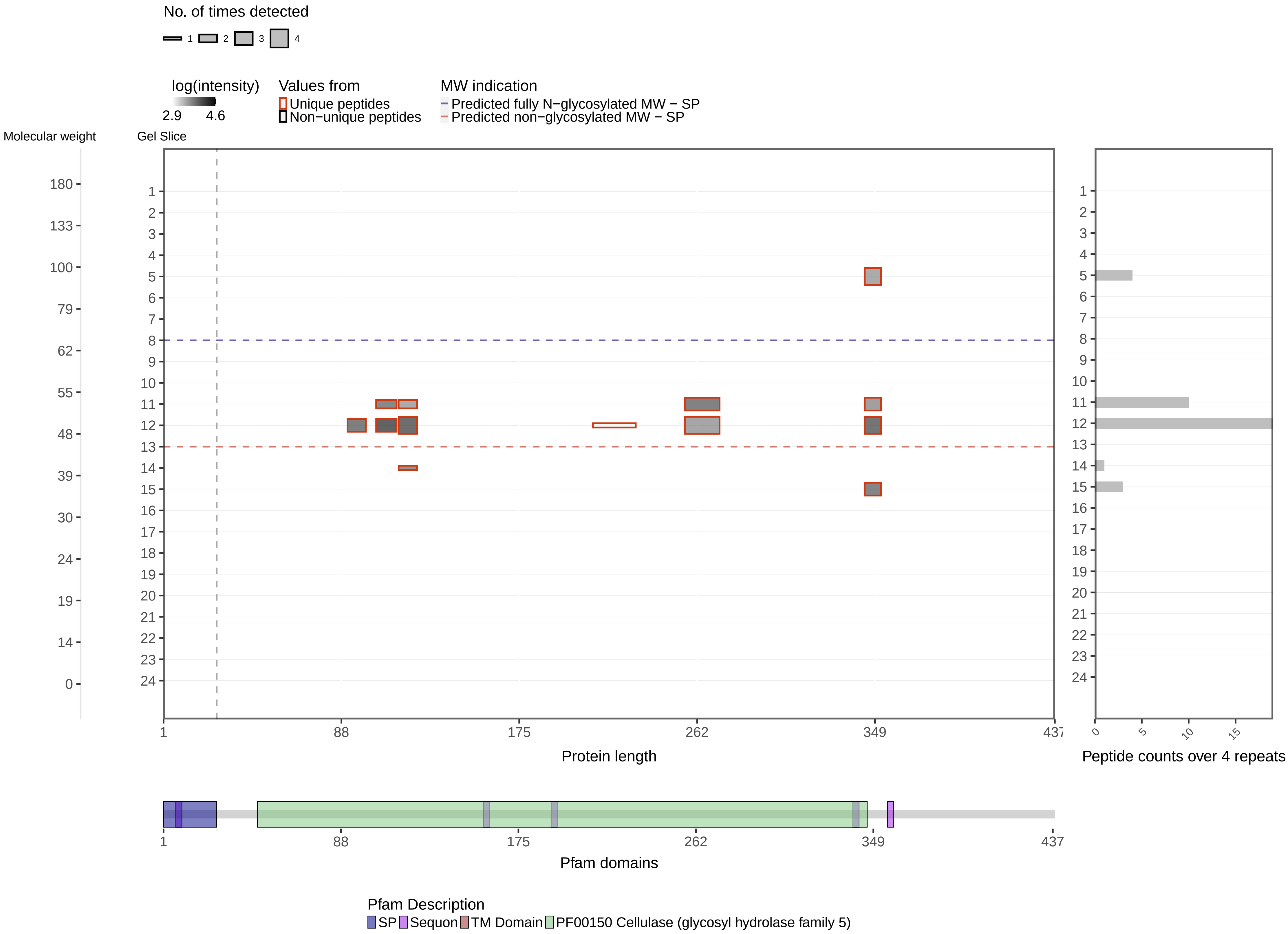


Protein sequence:

ELMNEPRCESDPSGDKLQAWIEEMAVYVKSIDPKHLVEIGLEGFYGLSTPNRAQVNPNSYAQQVGTDFIRNHQVLGVDFASVHIYPDSWISQSISEAHVEFTKSWMQSHIDDAEKYLNMPVVFAEFGVSTKDPGYNSTYRDTLIATVYKTILNSTRKGGAGGGSLLWQVFPEGTDYMDDGYAIVLSKSLSISNIVSLHSKKLNTFNSFCSWKCHWGCRKKHVLLDDFQSRDEL*

NbL08g14990.1 vacuolar−sorting_receptor_1−like [MW − SP = 67.2 kDa]

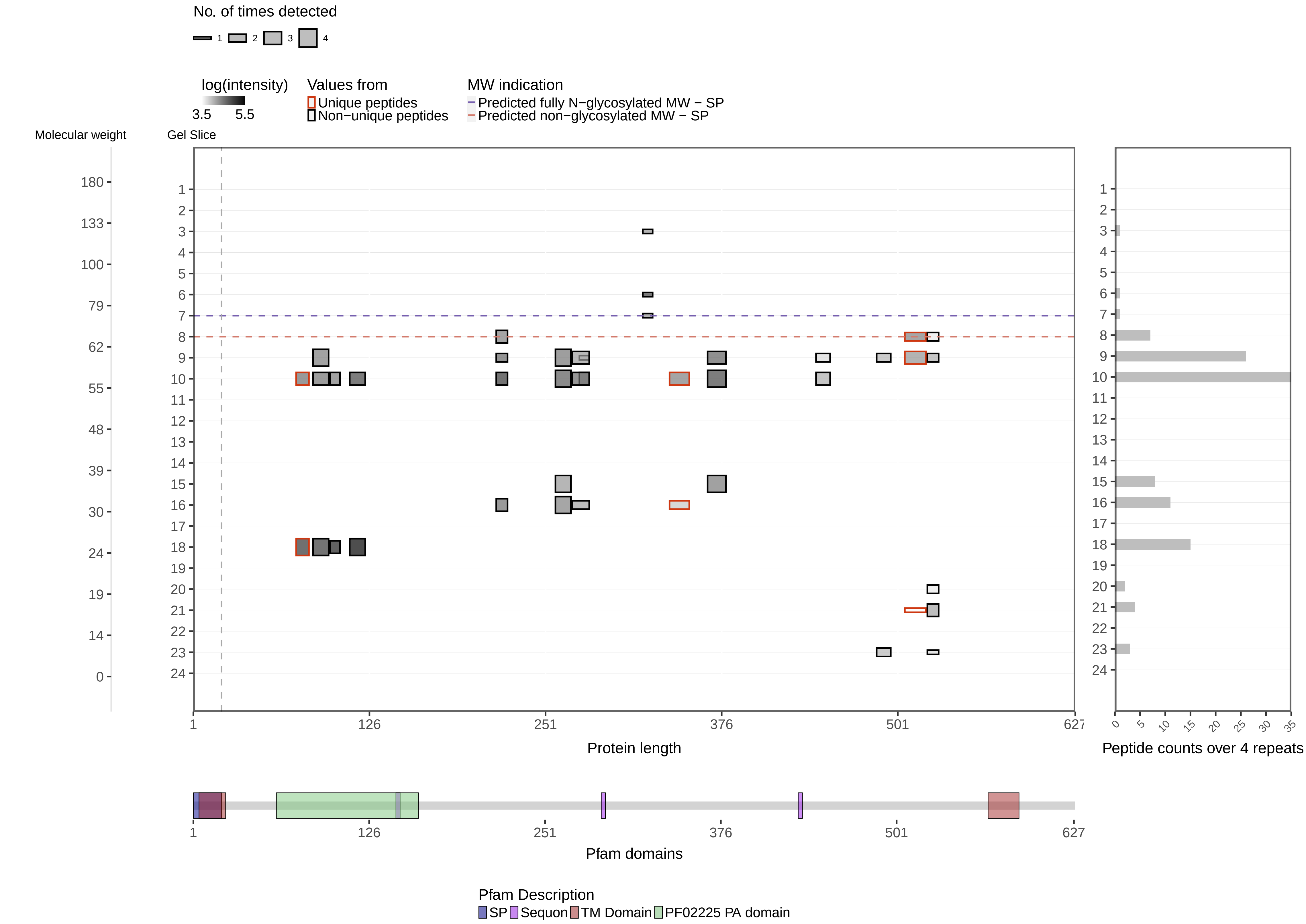


Protein sequence:

YVTDFAIRCPMKEKKYTKECADQVVKSLGFDVKQIDKCVGDPEADADNPVLKAEQEAQIGKDSRGDVTILPTLVINNRQYRGKLDKGAVLKAICSGFEETTEPAICLTDDIQTNECLESNGGCWQDKAANITACRDTFRGKVCECPMVQGVKFVGDGYTHCEASGALRCGINNGGCWKGTKDGRTFSACIDDHTKGCKCPPGFKGDGVNSCEDIDECNERLACQCPECKCKNTWGGYDCSCKGNLLYMHEHDTCINAGKDGKTEFSWGFVWVIILGLAAAGVGAYAVY

NbL08g14940.1 67−dimethyl−8−ribityllumazine_synthase_chloroplastic−like [MW = 24.65 kDa]

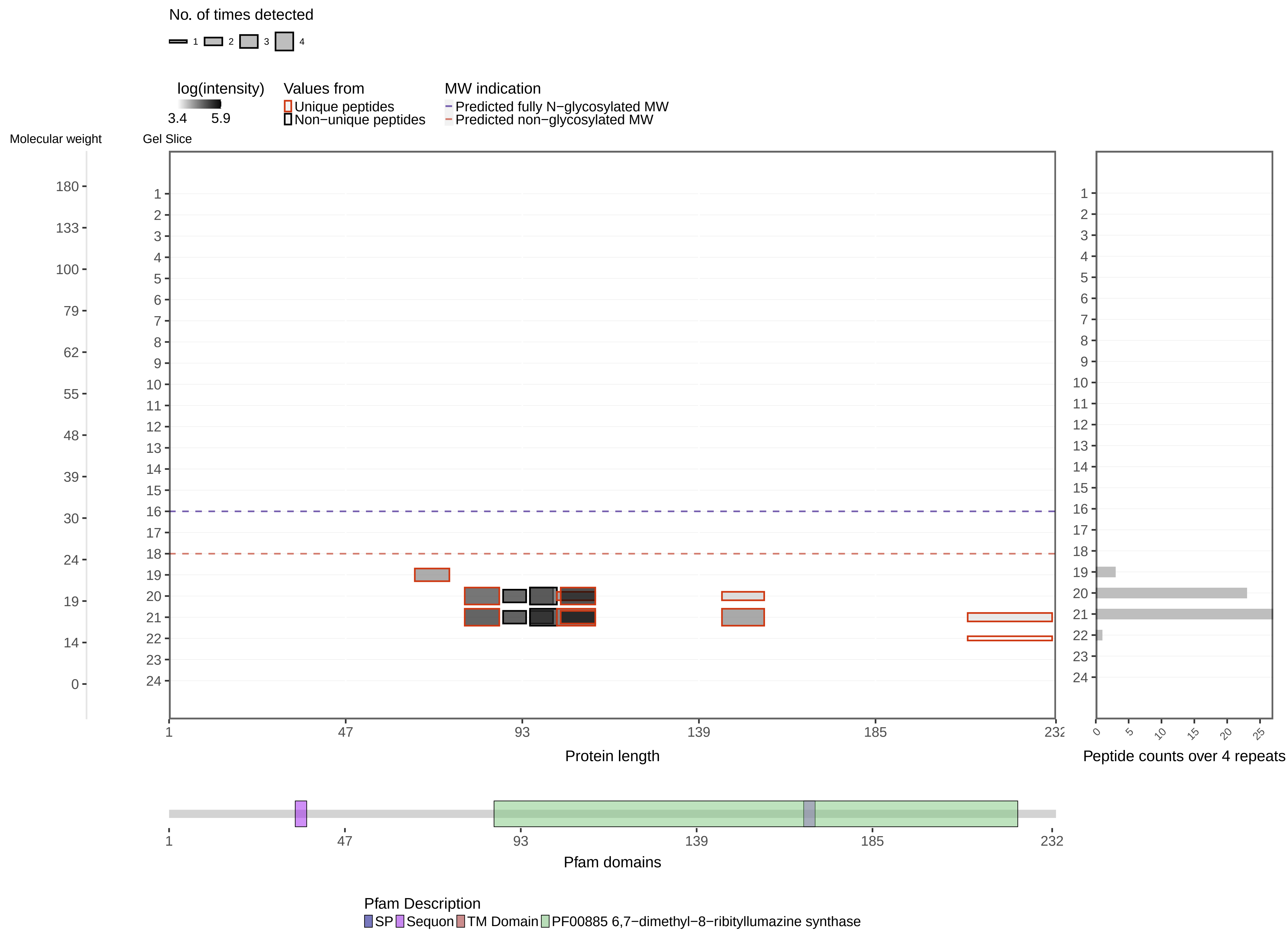


Protein sequence:
.LEGALDTFKSYSVREENIDVVWVPGSFEIGVVAQQLGKSQKYQSILCIGAVIKGDTSHYDAVVNATTSGVLSAGLNSGTPCIFGVLTCDTLEQAFNRAGGKSGNKGAEAALTAIEMASLFEHHLKPLE*

NbL08g14920.1 pyruvate_dehydrogenase_E1_component_subunit_beta−3_chloroplastic−like [MW = 44.49 kDa]

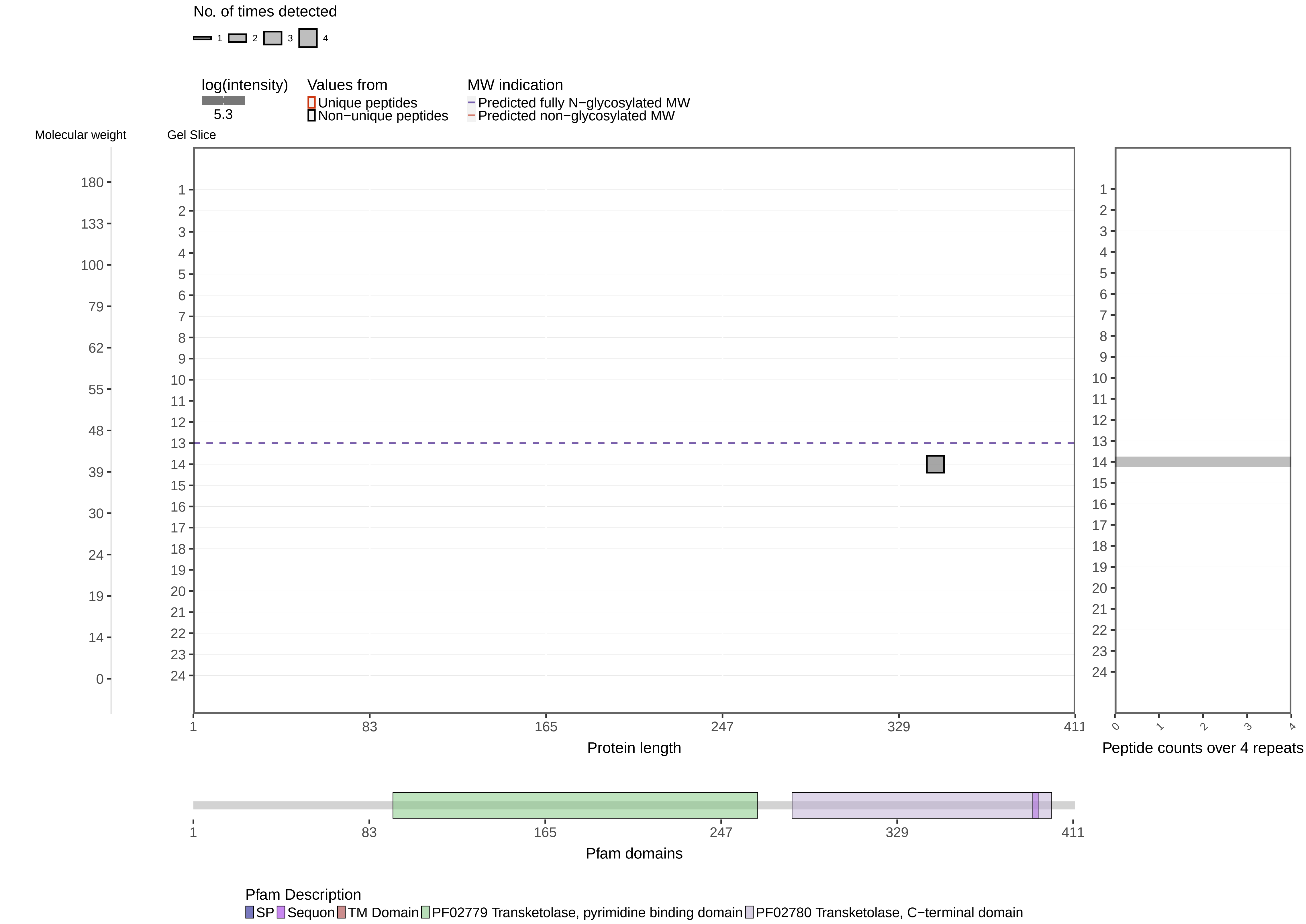


Protein sequence:

SGGQFKIPVVIRGPGGVGRQLGAEHSQRLESYFQSIPGIQMVACSTPYNAKGLMKAAIRSDNPVILFEHVLLYNLKERIPDEEYVLNLEEAEMVRPGEDVTILTYSRMRYHVMQAAKTLVDKGYDPEVIDIRSLKPFDLHTIGNSVKKTHRVVIVEECMRTGGIGASLTAAITENFHDYLDAPIVCLSSQDVPTPYAGTLENWTVVQPSQIVTAVEQLCR*

### NbL08g14850.1 uncharacterized_LOC107820649 [MW = 30.78 kDa]

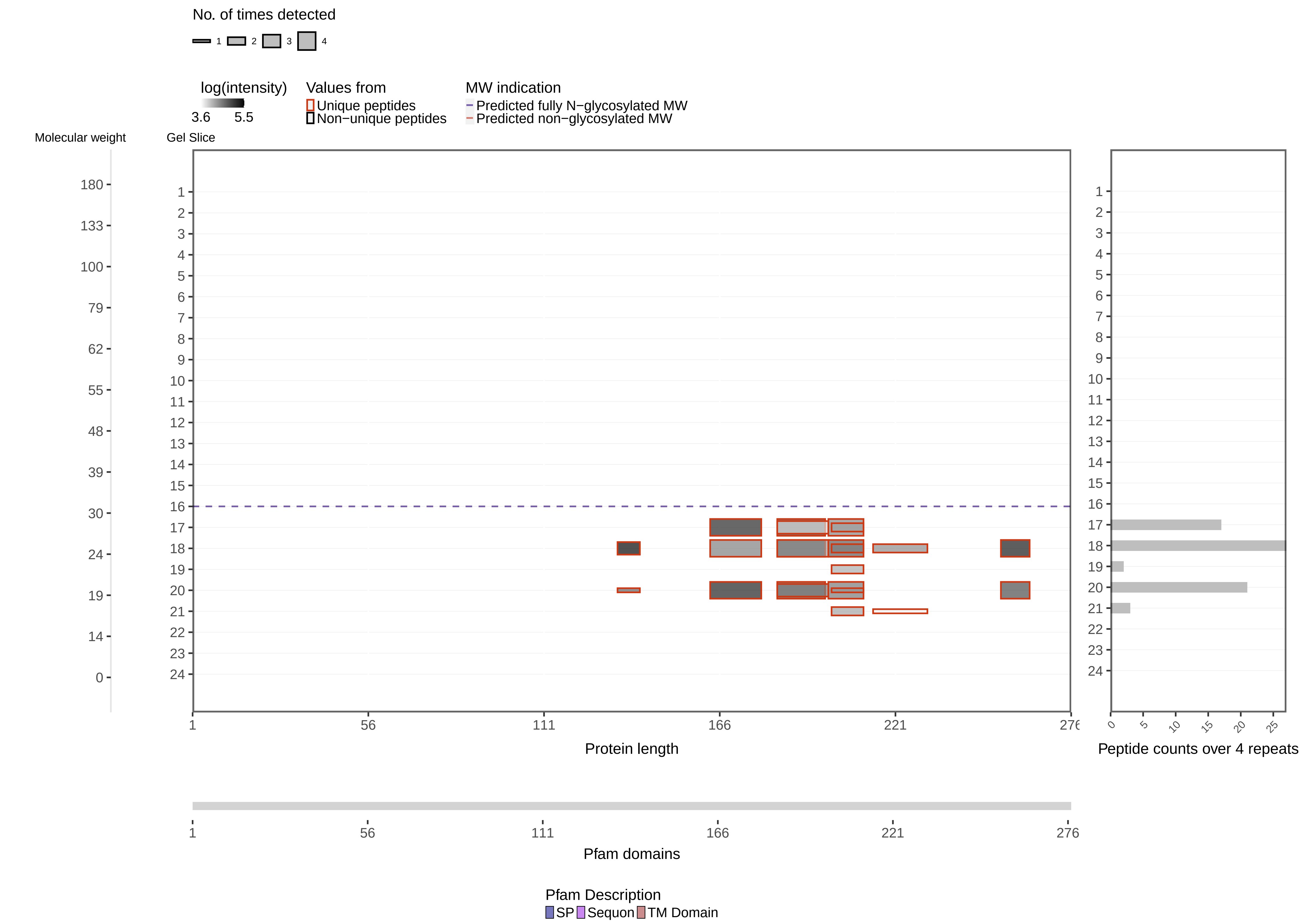


Protein sequence:

ELLAIQQQGPRNIGFFGTRNMGFMHQELIEILSYALVITKNHIFTSGAAGTNAAVIRGALRAEKPELLTVILPQSLKKQPPESQELLSKVKHVIEKPHNDHLSLLEASRLCNMDIIFQVQQVICFAFHDSRLLMETCQEAKNLRKIVTLFYLD*

### NbL08g14810.1 ATP_synthase_subunit_beta_mitochondrial−like [MW = 59.7 kDa]

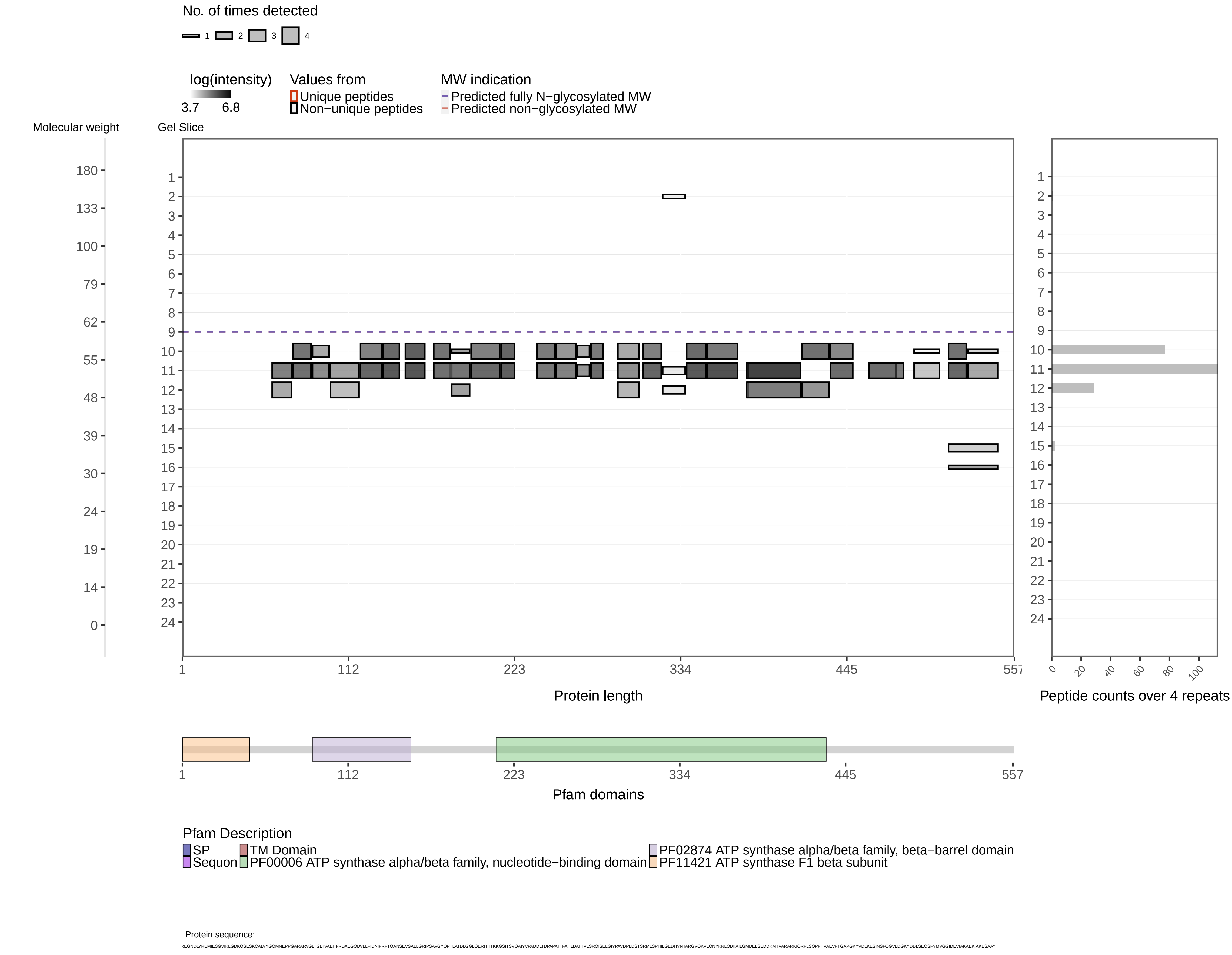

### NbL08g14800.1 fruit_protein_pKIWI502 [MW = 33.42 kDa]

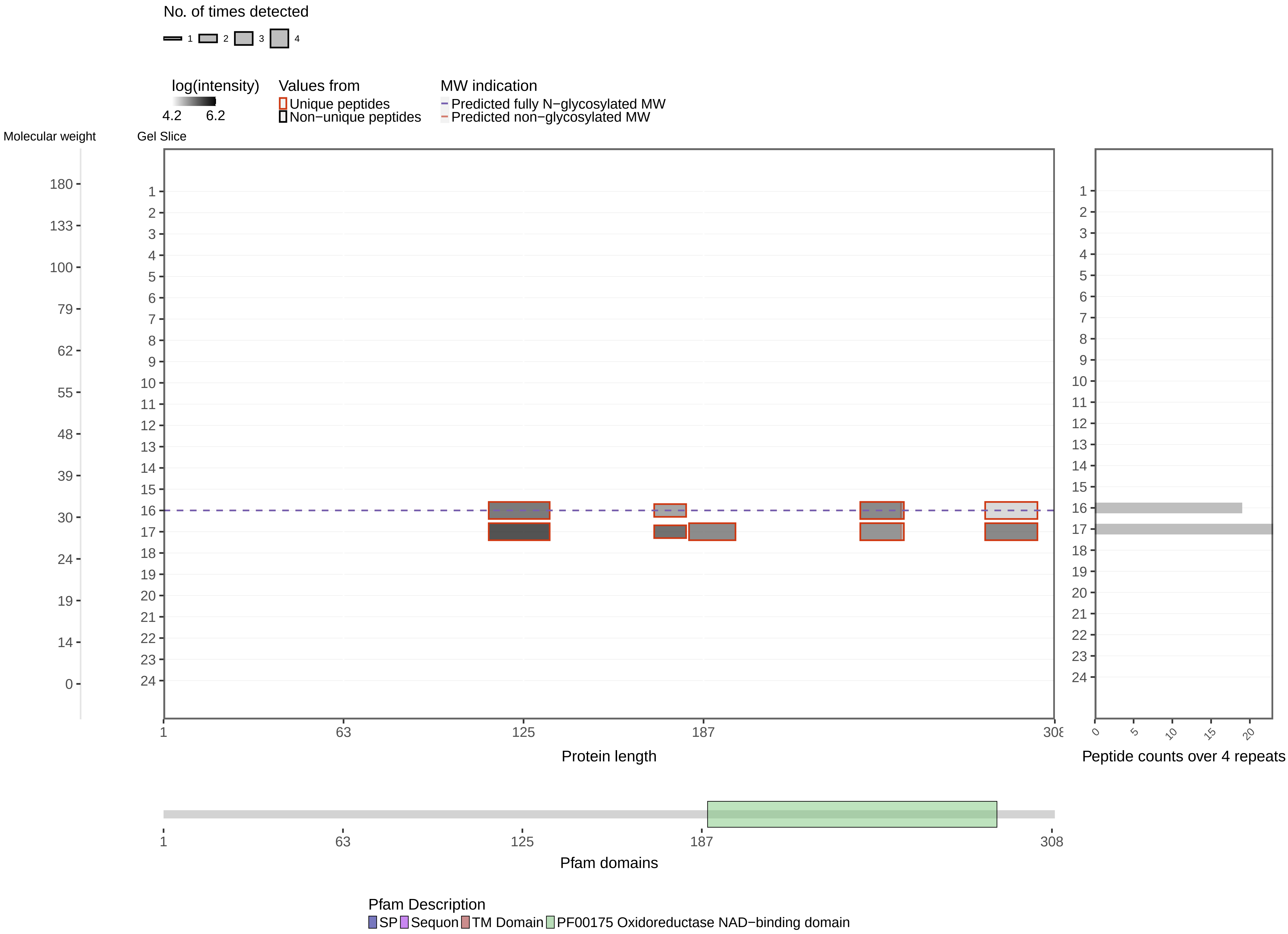


Protein sequence:

SISGSTAELLCGLQKGDVVELSQVMGKGFDLDQISPPEKYQTVLIFATGSGISPIRSLIEAGFSADKRSDVRLYYGARNLKRMAYQDRFENWASSGVKVVPVLSQPDDAWTGKRGFVQAAFARAKNIFSPQSTGAVLCGQKQMAEEVTALLVADGVSTEKILKNF*

### NbL08g14690.1 fructose−bisphosphate_aldolase_3_chloroplastic−like [MW = 42.91 kDa]

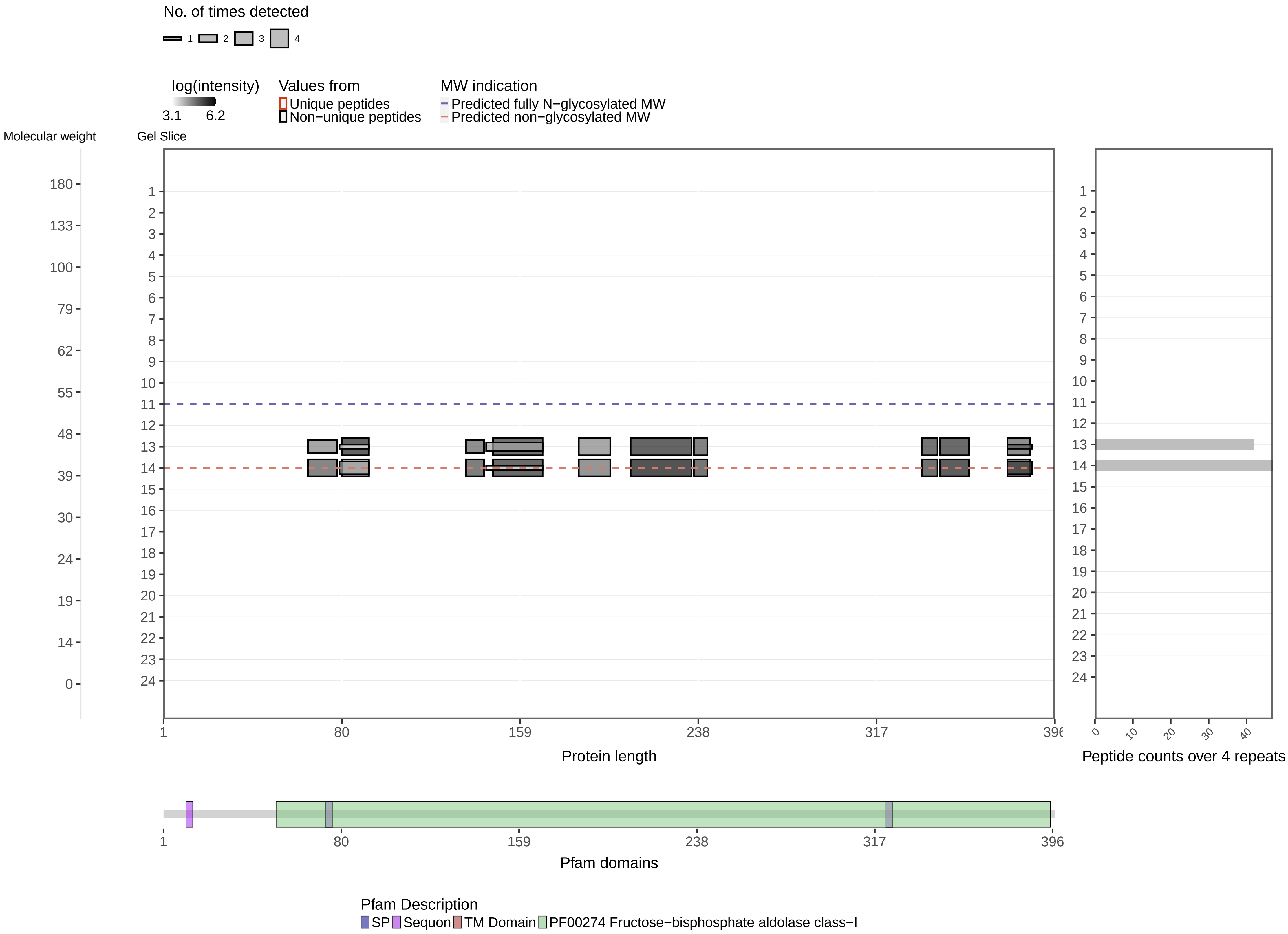


Protein sequence:

VVSIPCGPSALAVKEAAWGLARYAAISQDNGLVPIVEPEILLDGDHPIERTLEVAERVWAEVFYYLAENNVVFEGILLKPSMVTPAAEHKEKSSPDTIAKYTLKMLKRRVPPGVPGIMFLSGGQSEVEATLNLNAMNQSPNPWHVSFSYARALQNTVLKTWQGRPENVDAAQKALLIRAKANSLAQLGKYSAAGESEEAKKGMFVKGYTY*

NbL08g14320.1 uncharacterized_LOC107766948 [MW = 55.01 kDa]

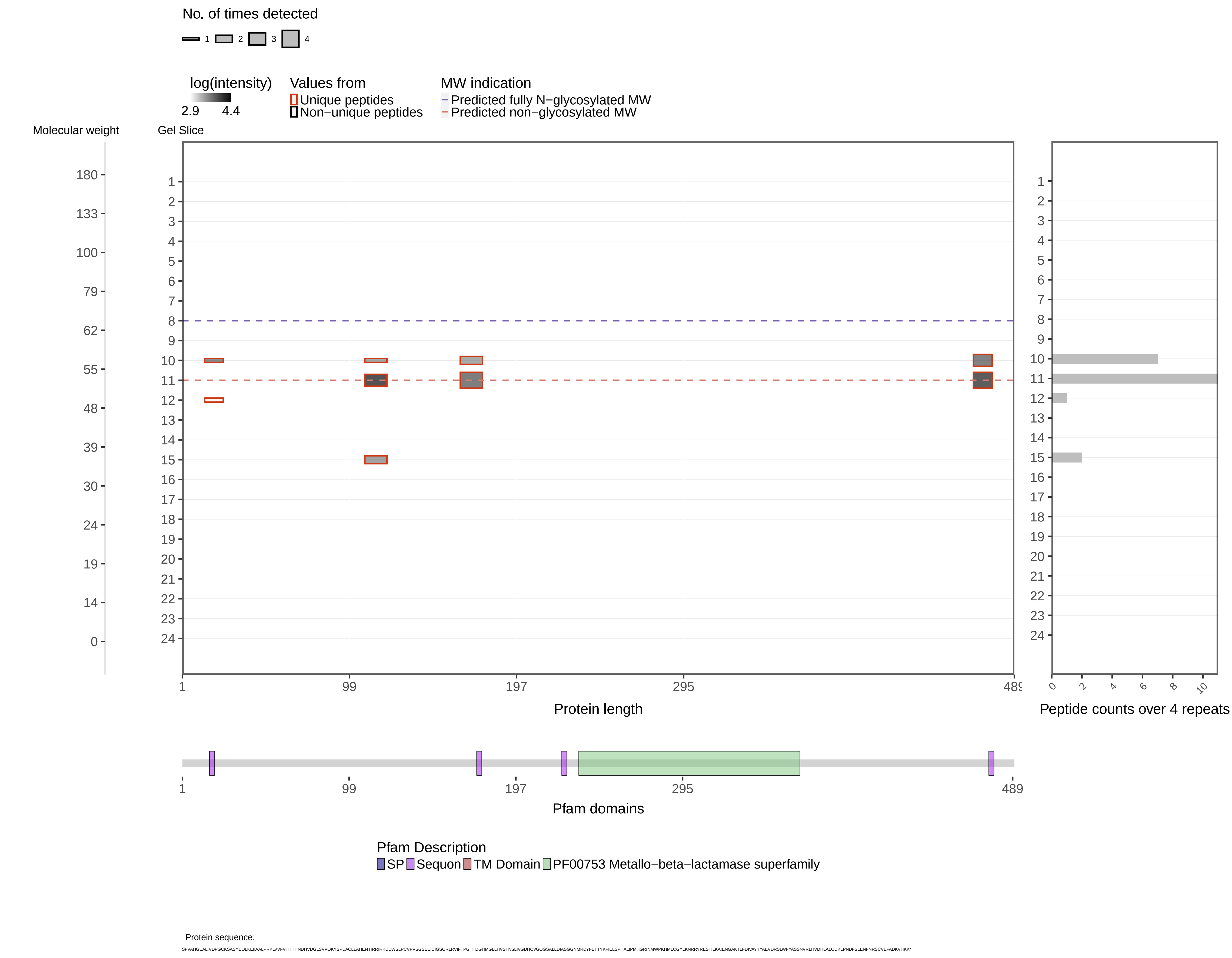

NbL08g14270.1 cysteine−−tRNA_ligase_cytoplasmic [MW = 66.62 kDa]

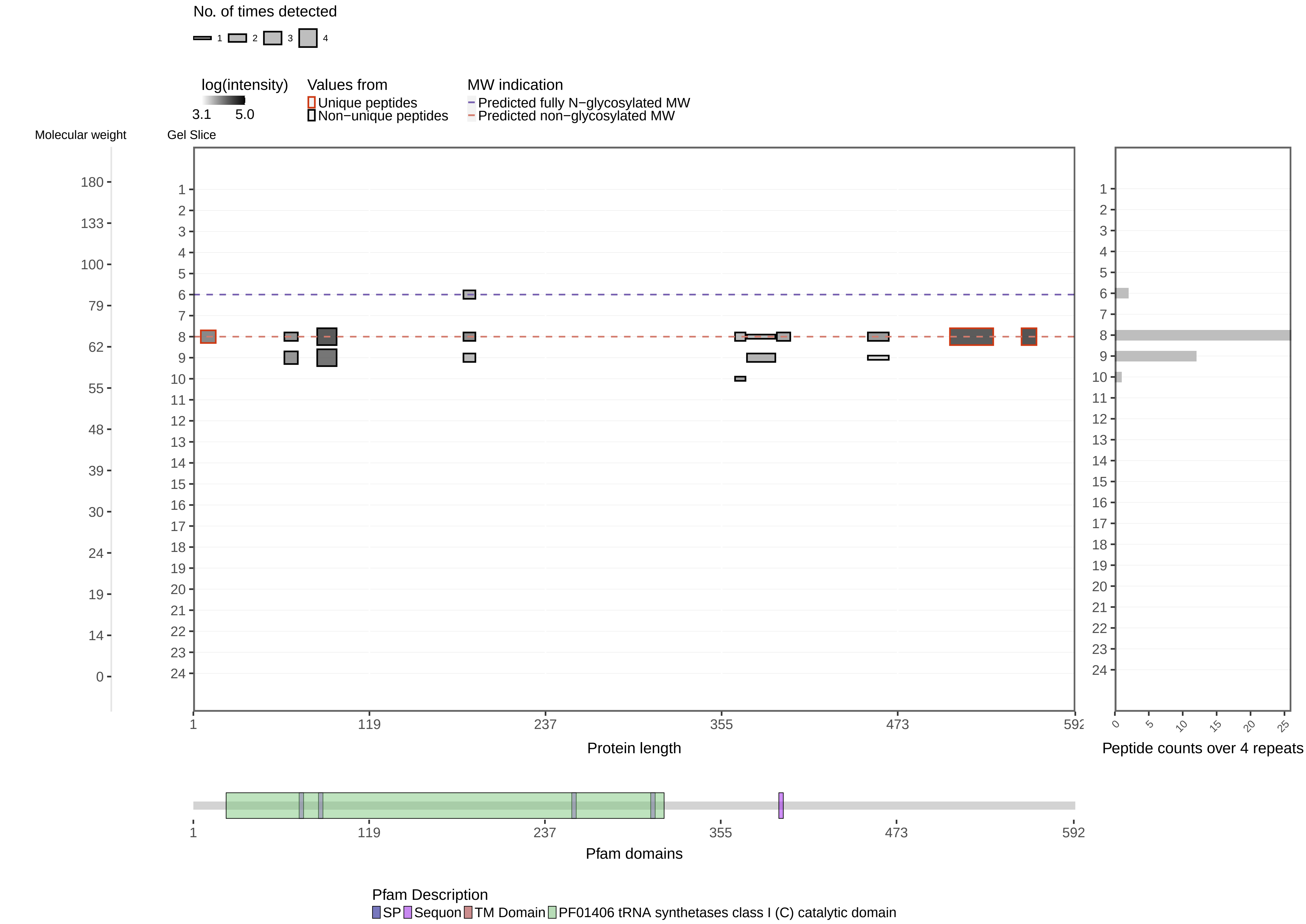


Protein sequence:

IREVTRLYHPIALRYFLLGMHYRSPVNYSISOIEIASESVFYLYOTLLDCEEALSVFOOGTETETEVRGVRVSSOVOECIKKLRNELETKLSDDLHTPTILNAALOEALRLMNSSLNMLKKKOOKKOOLSAVLSLAELLKEVKAVLDVLGLLSSSTCAEVLOOFKERALKRAELTEEDILHAIEERTLARKNKEFARSDOIRSDLVAKGIALMDIGAETVWRPCVRAOOEOPAAPAOOEOPAAPAOOEOSAAAMOKEKPVAPSOOKOSAGAPOKEOPVAAPSOPKHSAAAPOKE

### NbL08g14260.1 glutamine−−tRNA_ligase [MW = 95.97 kDa]

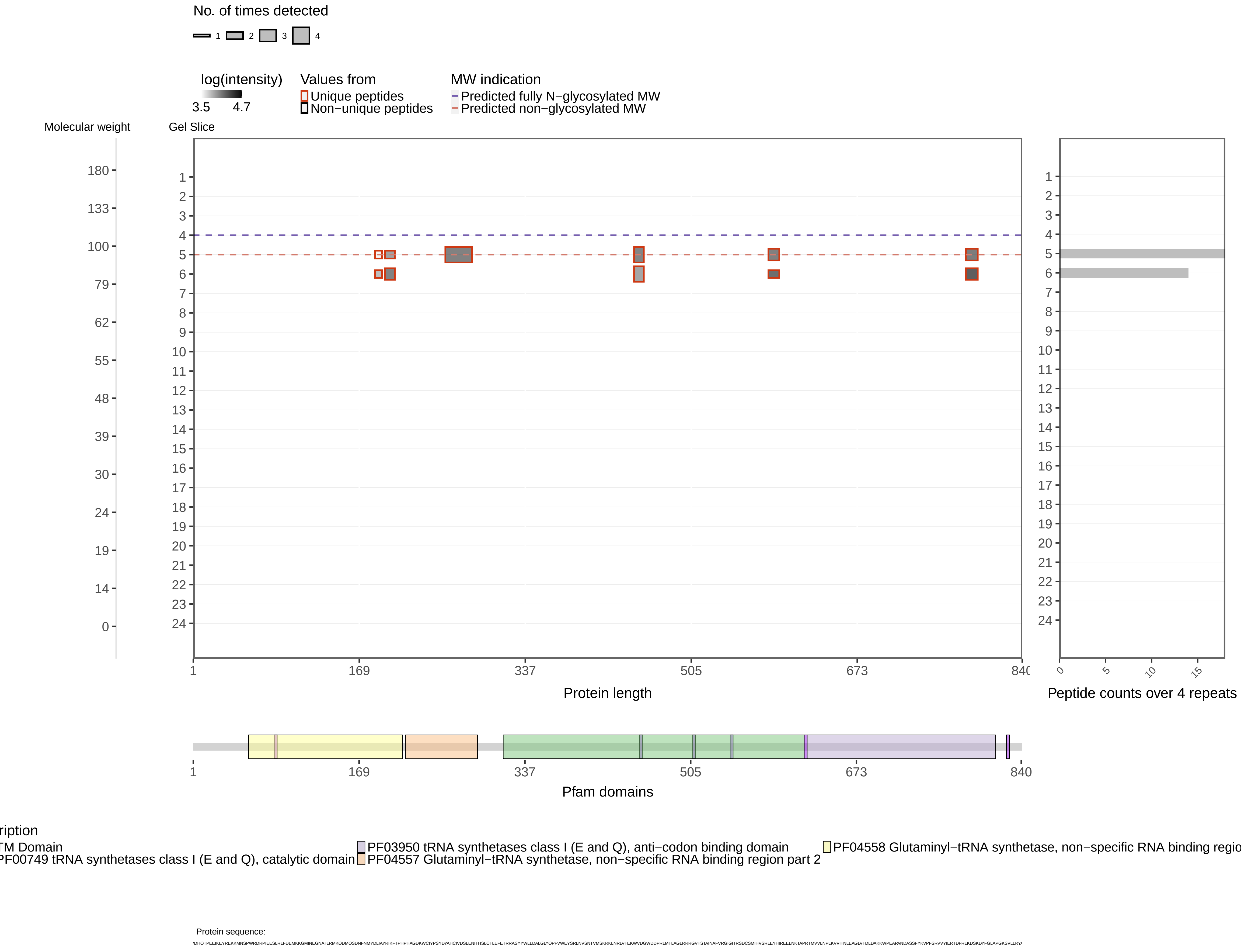

NbL08g14190.1 50S_ribosomal_protein_L15_chloroplastic−like [MW = 28.53 kDa]

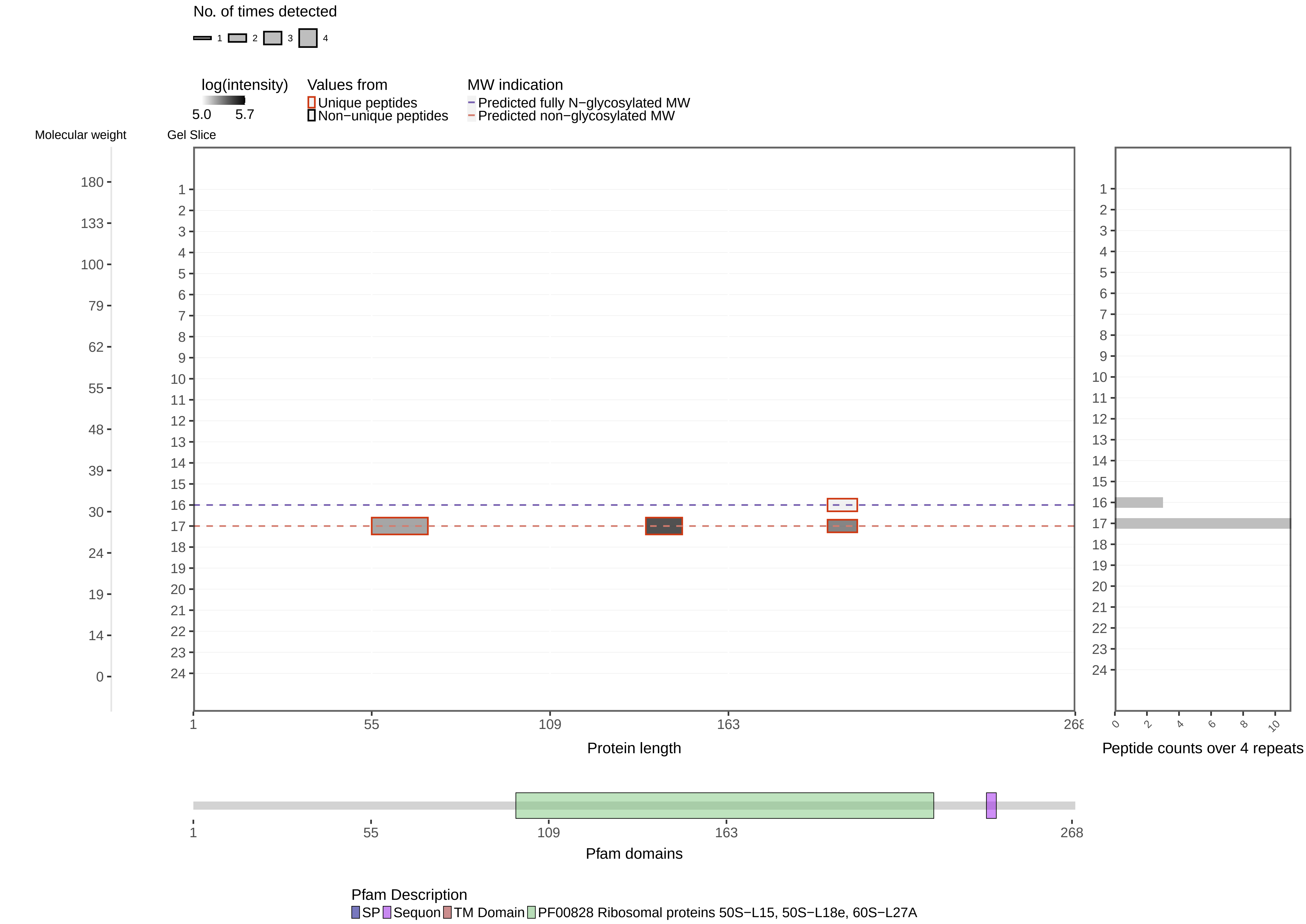


Protein sequence:

FEGGQMPLYRRLPKLRGIAGGMAAGLPKYVPVNLKDIEEAGFQEGEEVSLESLKQKGLINPSGRERRLPLKILGDGELSVKLNFKARAFSTSAKEKLEAAGCSLTVLPGRKKWVKPSVAKNLSRADEYFAKKRAAAASEAADPSSA*

### NbL08g14180.1 3−isopropylmalate_dehydratase_small_subunit_3−like [MW = 27.23 kDa]

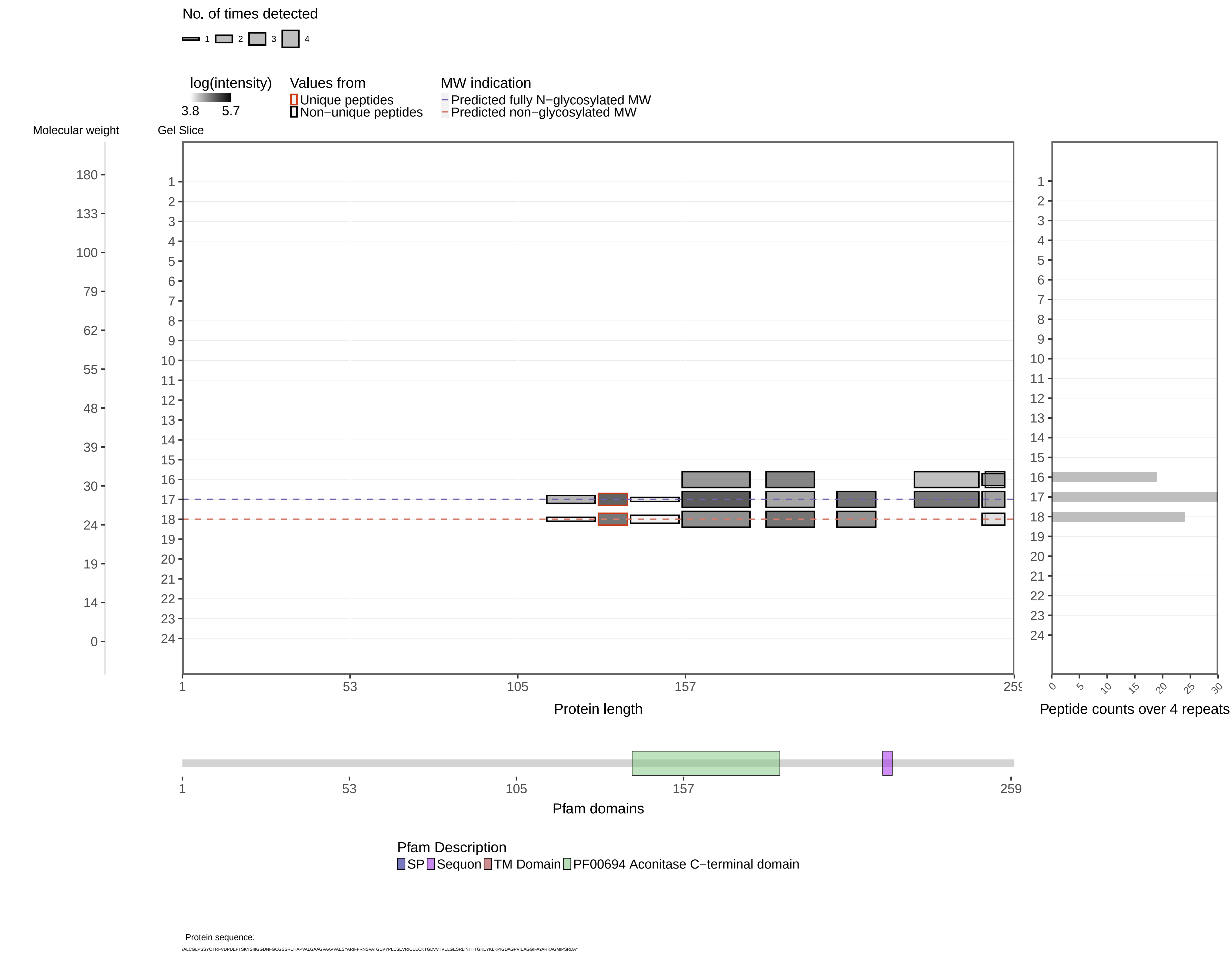

### NbL08g14010.1 serine/threonine−protein_phosphatase_2A_65_kDa_regulatory_subunit_A_beta_isoform [MW = 65.55 kDa]

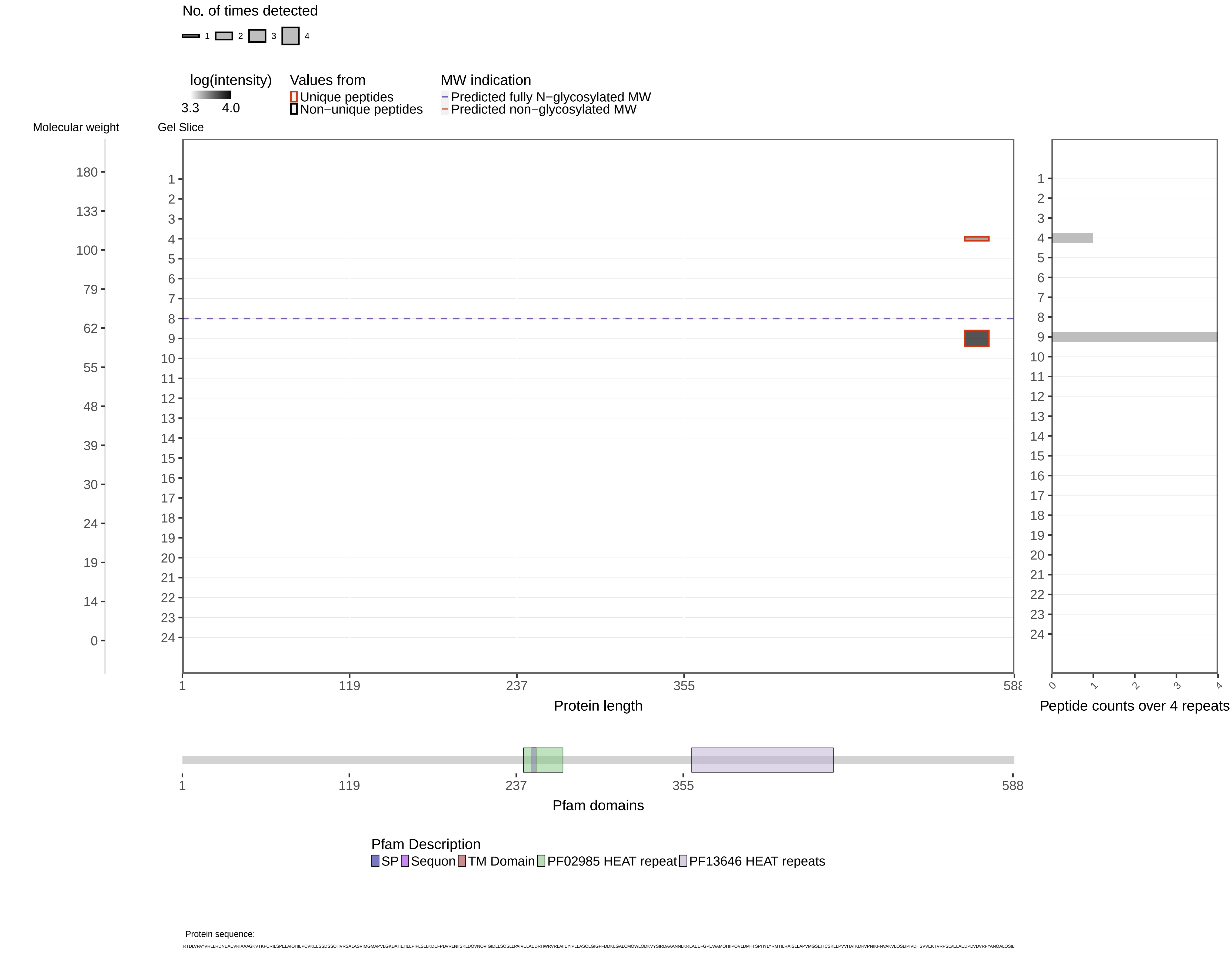

### NbL08g13930.1 fumarate_hydratase_1_mitochondrial−like [MW = 53.35 kDa]

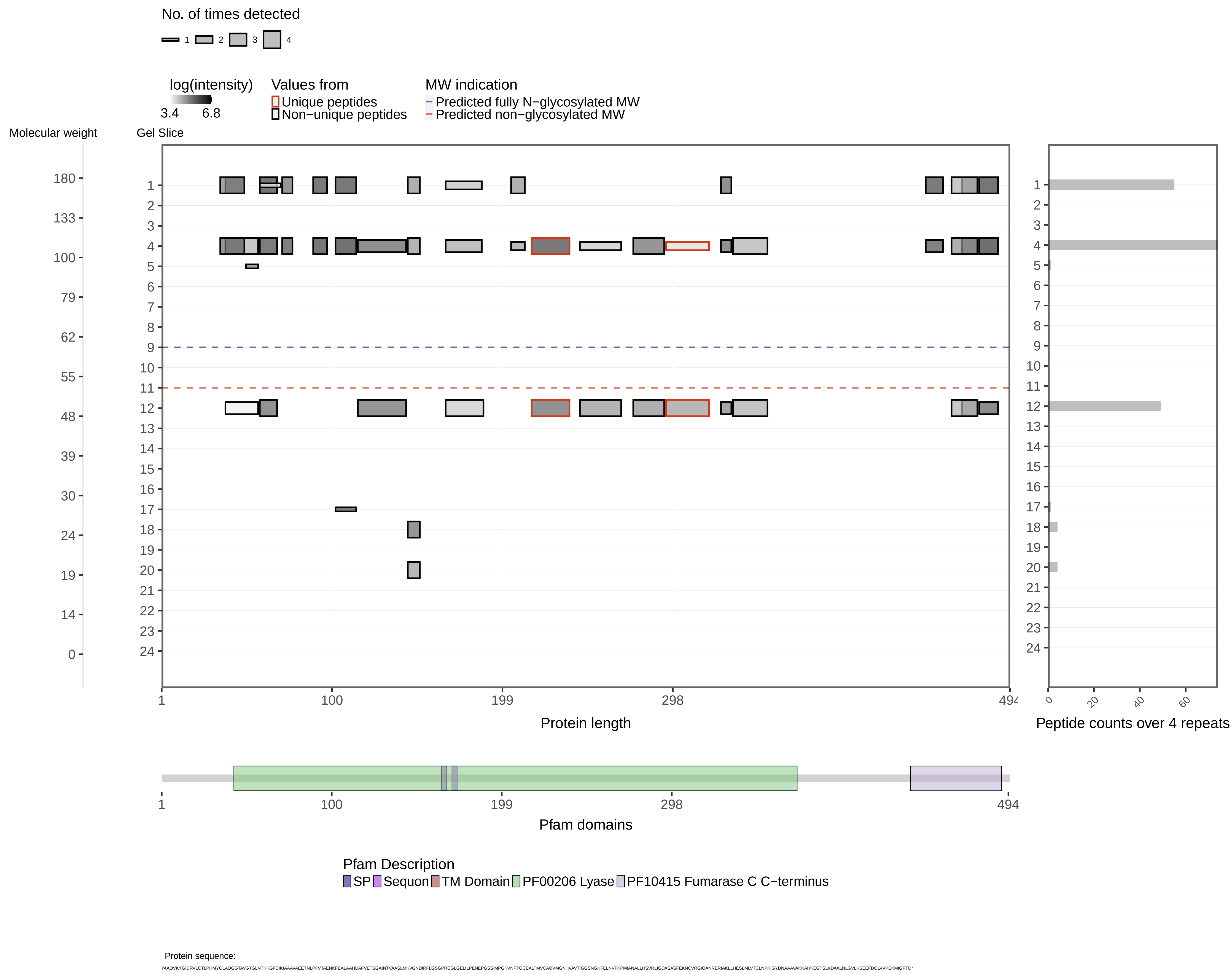

### NbL08g13660.1 protein_decapping_5 [MW = 62.31 kDa]

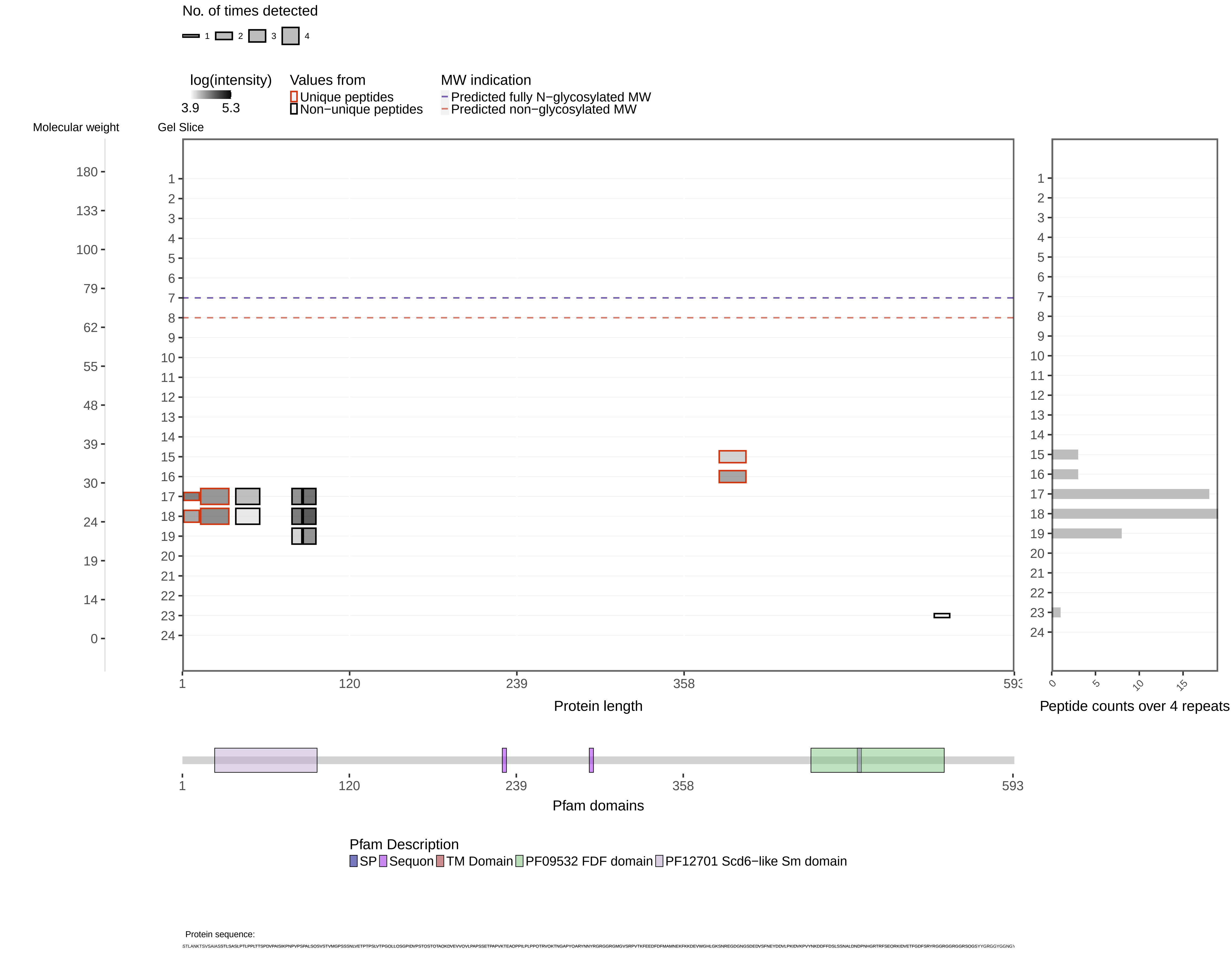

### NbL08g13560.1 HD_domain−containing_protein_2 [MW = 36.35 kDa]

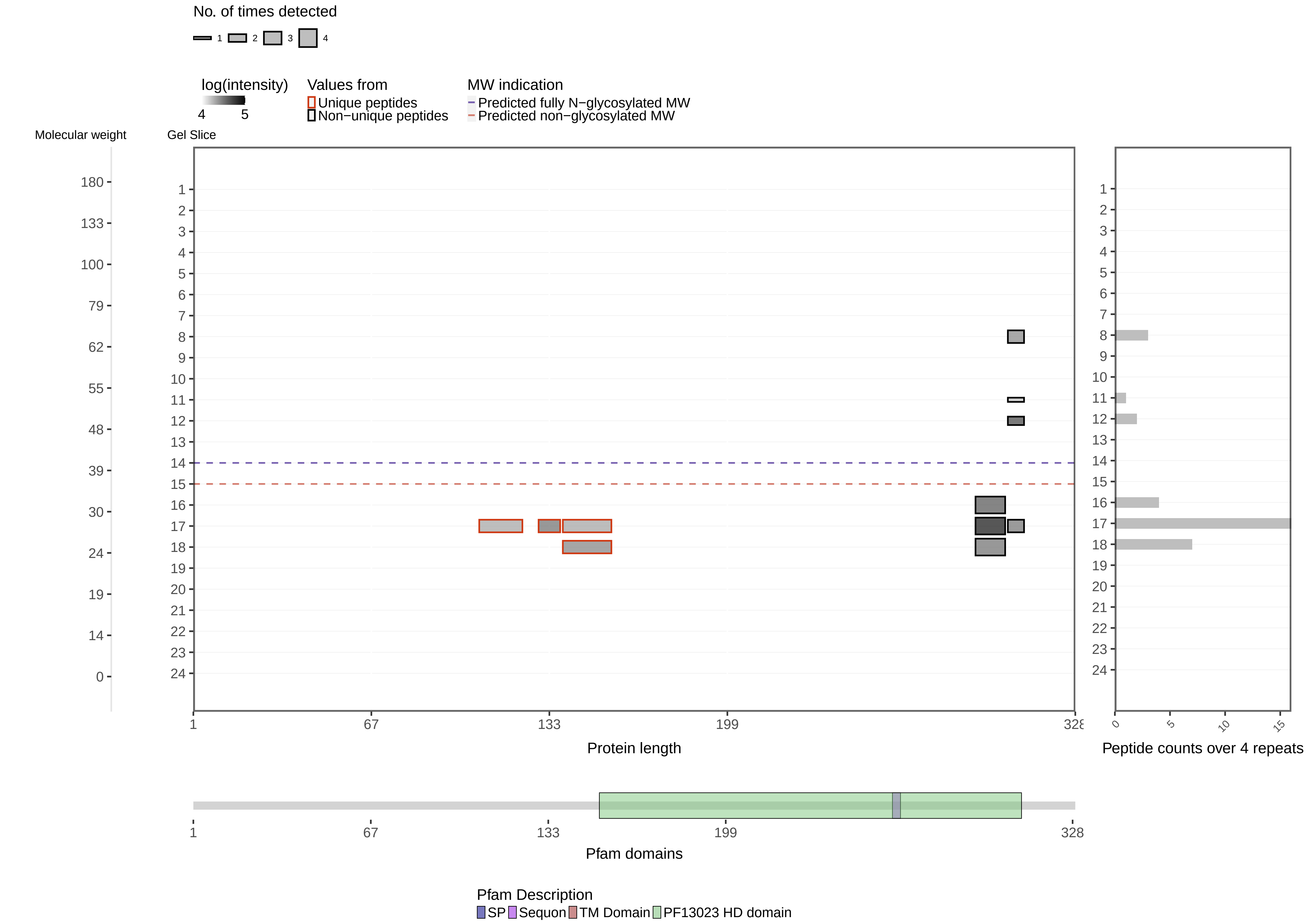


Protein sequence:

:HSLKTOKRKGWVNHGIKGPESIADHMYLMSVMALIASDLPGVNRERCIKMAIVHDIAEAIVGDITPSDGVPKAEKSRRELAALNEMCKILGGGMRAEEIKELWOEYENNASLEANLVKDFDKVEMILOALEYESEHGKVLDEFFLSTAGKFOTEIGKSWAAEIHSRRNSRSENRL*

### NbL08g13370.1 UDP−glycosyltransferase_89A2 [MW = 51.39 kDa]

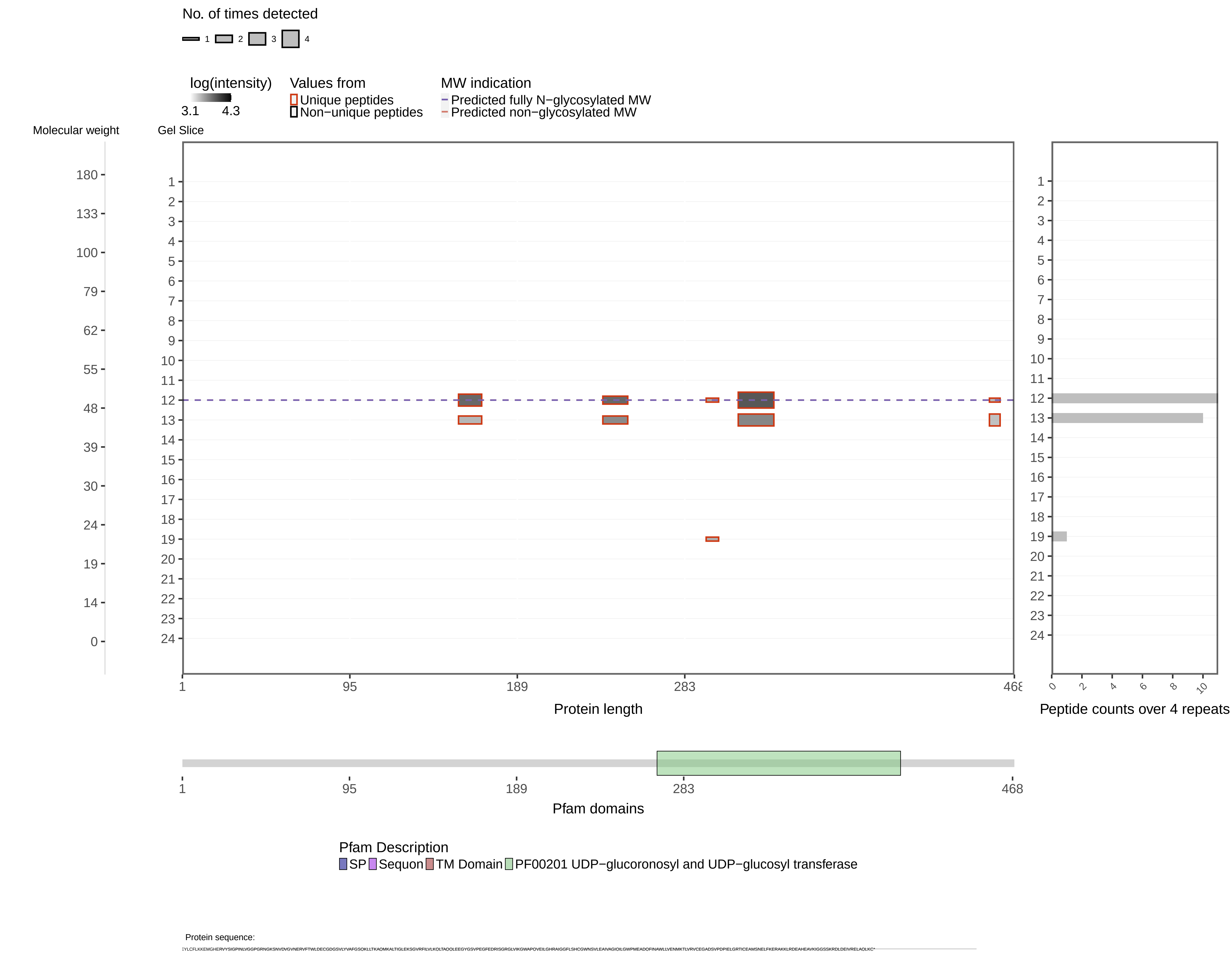

### NbL08g13350.1 heat_shock_protein_90−5_chloroplastic−like [MW = 91.01 kDa]

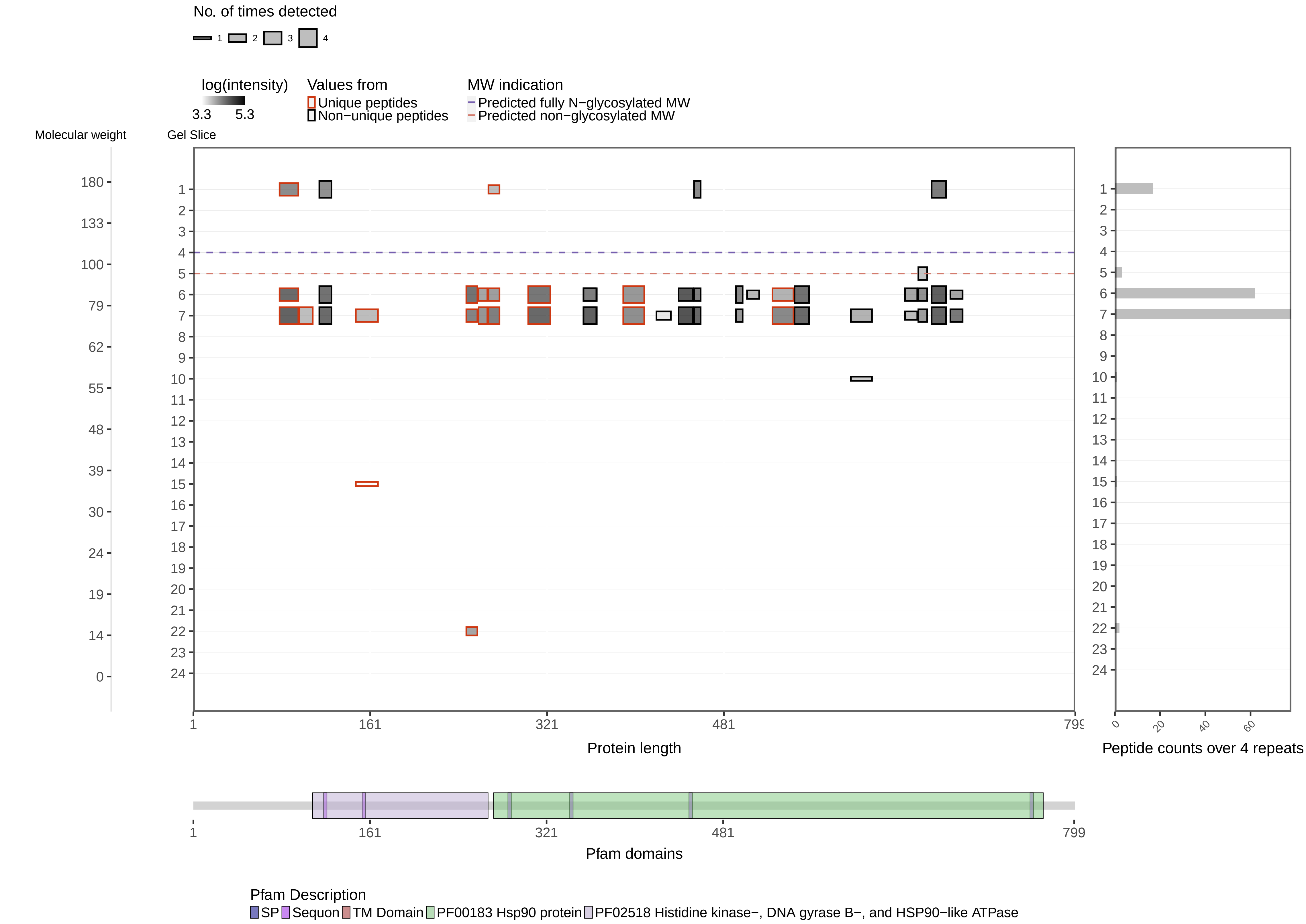


Protein sequence:

### NbL08g13240.1 chaperonin_60_subunit_beta_4_chloroplastic−like [MW = 65.73 kDa]

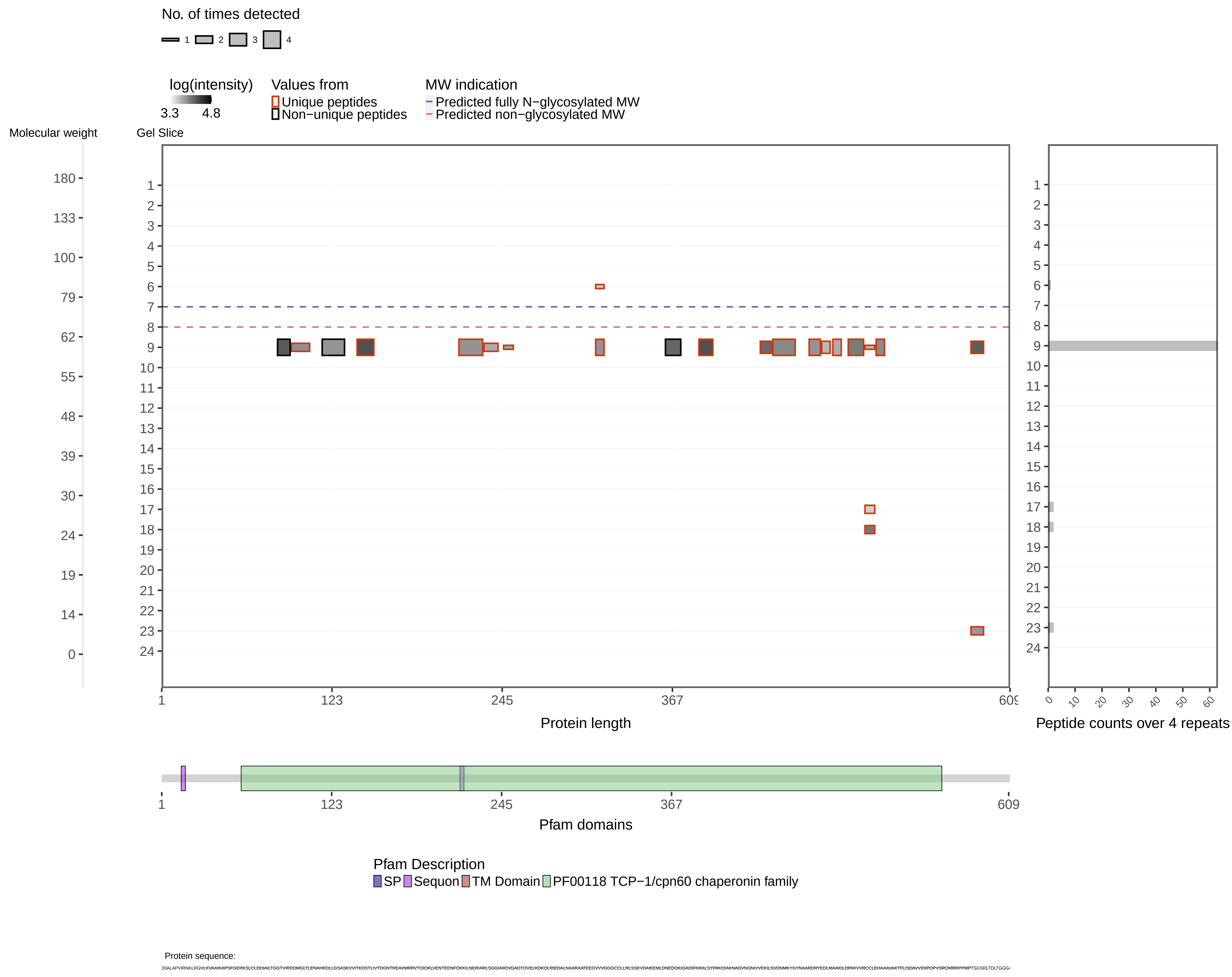

### NbL08g13220.1 serotonin_N−acetyltransferase_2_chloroplastic [MW = 21.69 kDa]

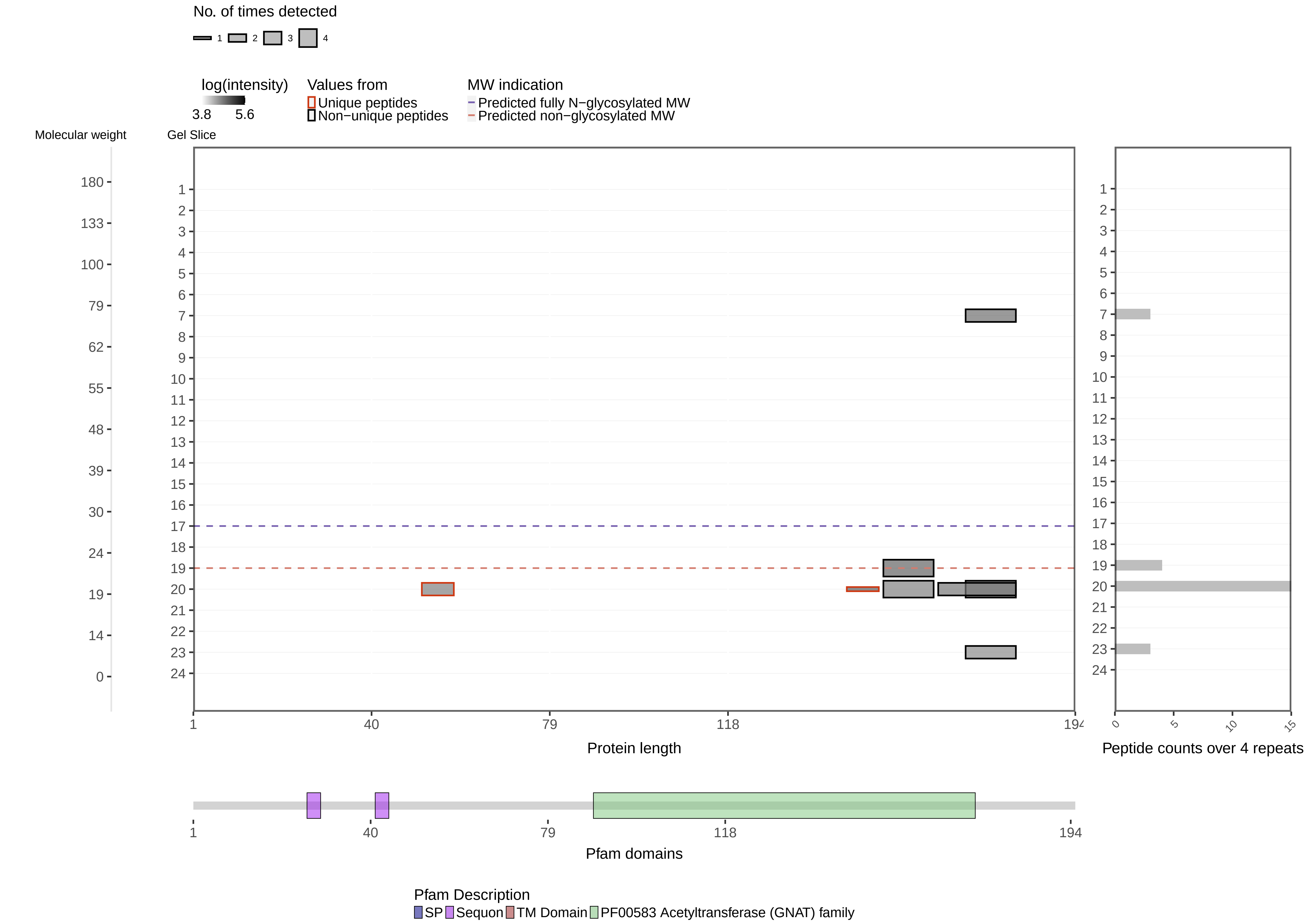


Protein sequence:

QLALENTDSLMWIEYEKTKRPVAFARATGDGVFNAIIWDVVVDPSFQGIGLGKAVMERLVTELLGKGINNIALYSEPRVLGFYRPLGFVADPDGIRGMVYSRKKKKNR*

NbL08g13210.1 6−phosphogluconate_dehydrogenase_decarboxylating_3 [MW = 54.04 kDa]

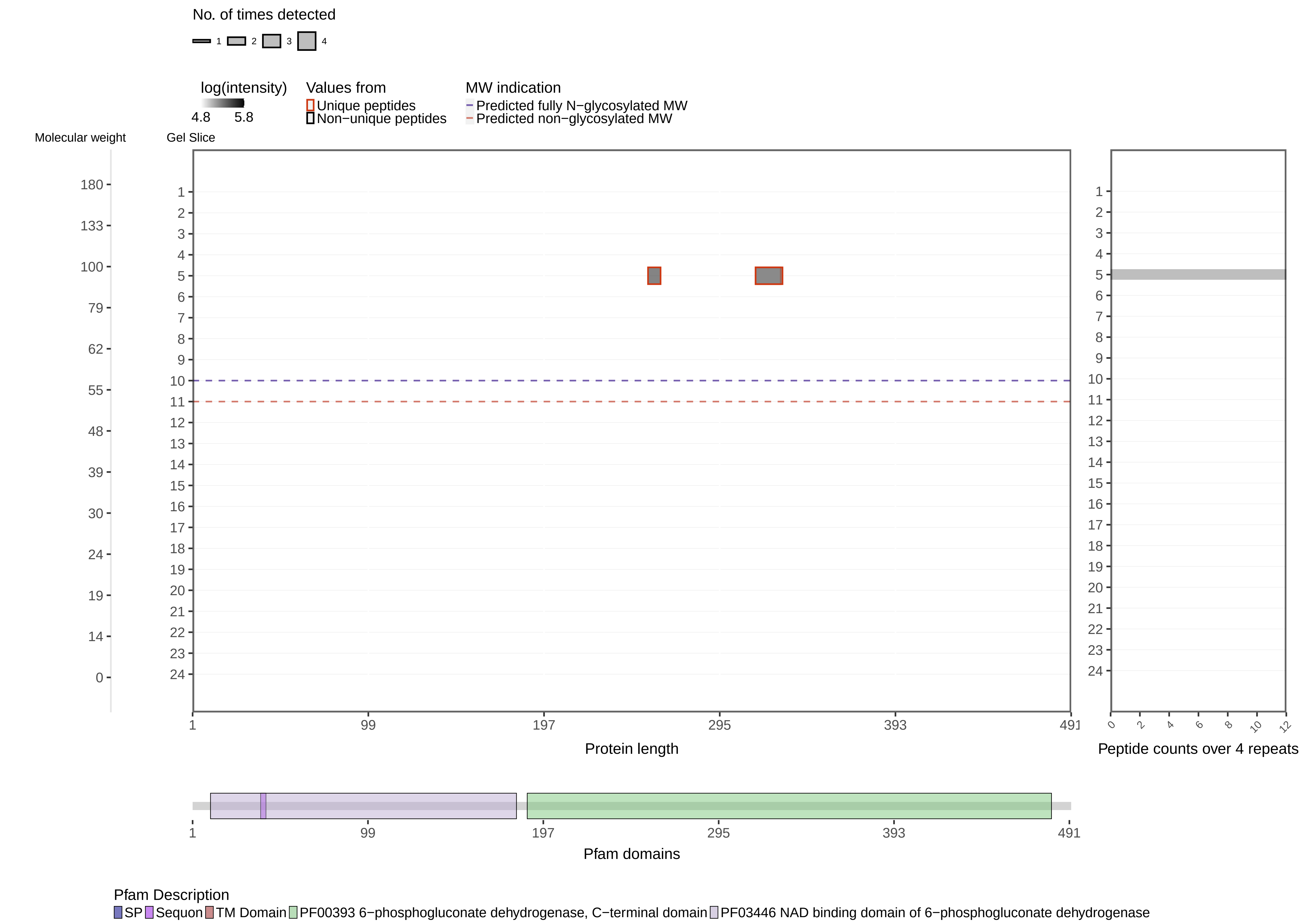


Protein sequence:

VNKGELLSFLIEITTDIFGIKDDKADGYLVDKVLDKTGMKGTGKWTVQQAAELSVAAPTIASSLDSRFLSGLKDERVQAAKVFKSSGVNDILVEQSVDKKQLIDDVRKALYASKICSYAQGMNLIRAKSVEKGWDLKLGELARIWKGGCIIRAIFLDRIKQAYDRNPDLANLLVDEEFAKEVVERQSAWRRVVCLAISSGISTPGMSSSLAYFDSYRRERLPANLVQAQRDYFGAHTYERTDMPGAFHTEWFKIAKQSKI*

NbL08g13170.1 uncharacterized_LOC104099690 [MW = 29.71 kDa]

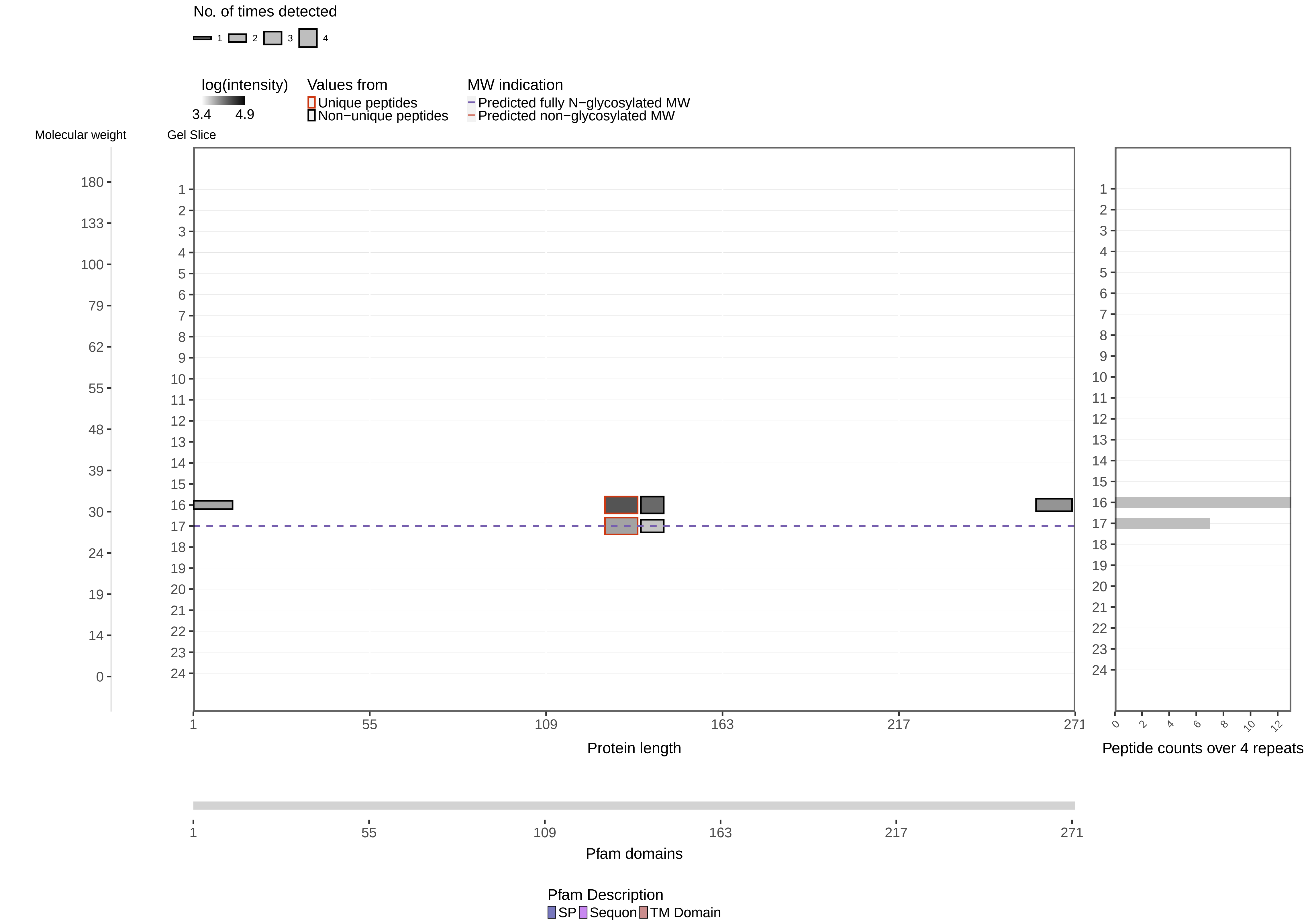


Protein sequence:

AVAKLLIGEQIIPEKVLILDSIRSNNFRGKLSPDEAFAFKLESSAERKAKADGHQDSPLVKCTDYLPSGSVVDGLAAALLSRCQLKKIRGTLCVSWPEAGISVISLVKSLLLKDVLSGVNHINVGDLEEEFLKLSRNKDFLLDSELYT*

NbL08g13160.1 DNA−directed_RNA_polymerases_II_IV_and_V_subunit_3−like [MW = 35.68 kDa]

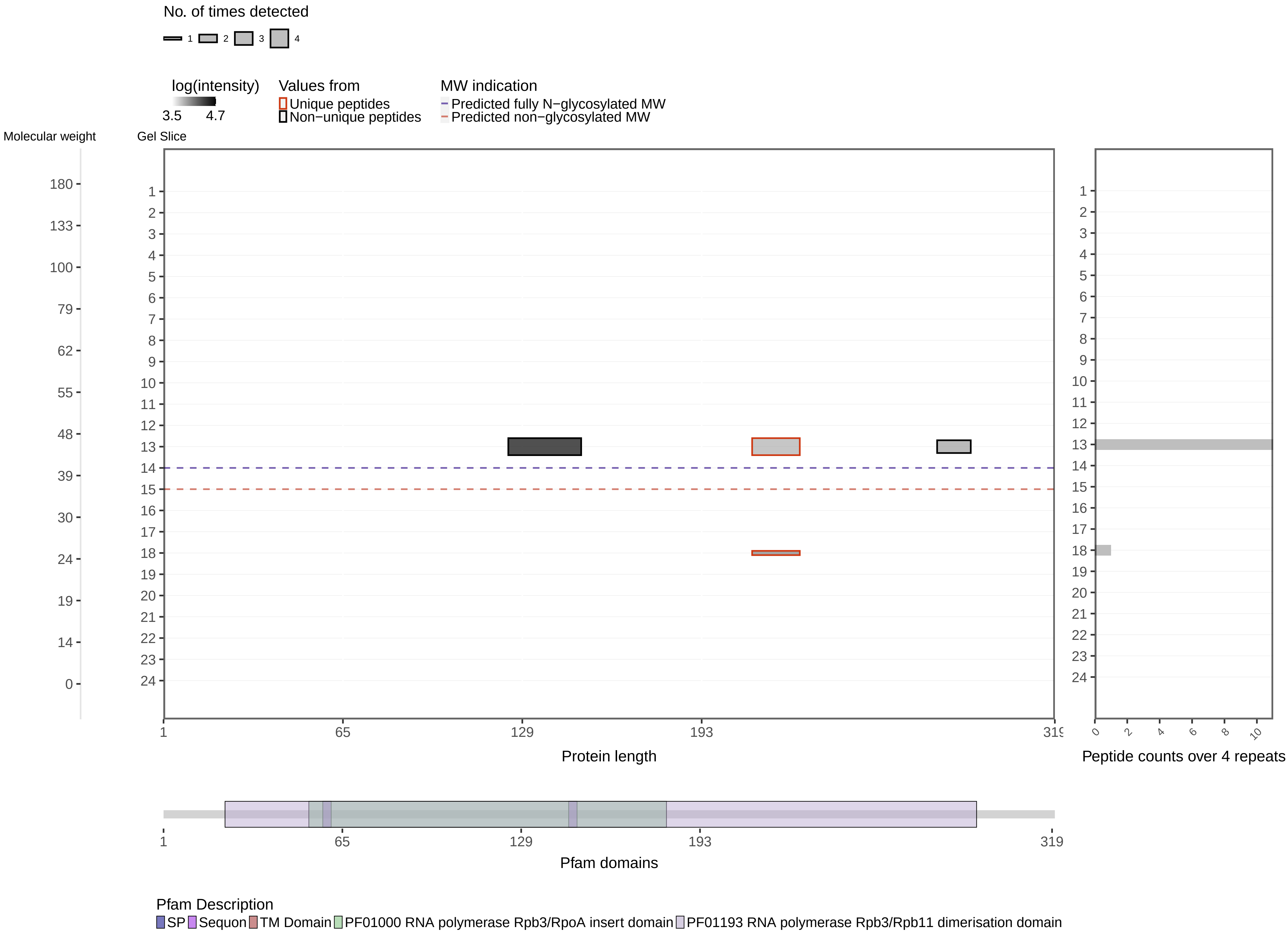


Protein sequence:

NSTNKGIIIVKLRRGQELRLRAIARKGIAKDHAKWSPAATVTFMYEPDIYINEDMMETLTLEEKRELVDSSPTEVFGIDPENKVIVVDPEAYTYDDEVLKKCEAMGKPGLIEIHAREDSFIFTVETTGAVKASQLVLNAIEVLKQKLDAVRLSDDTVEADDQFGELGAHMRGG*

### NbL08g13150.1 chalcone−−flavonone_isomerase [MW = 26.53 kDa]

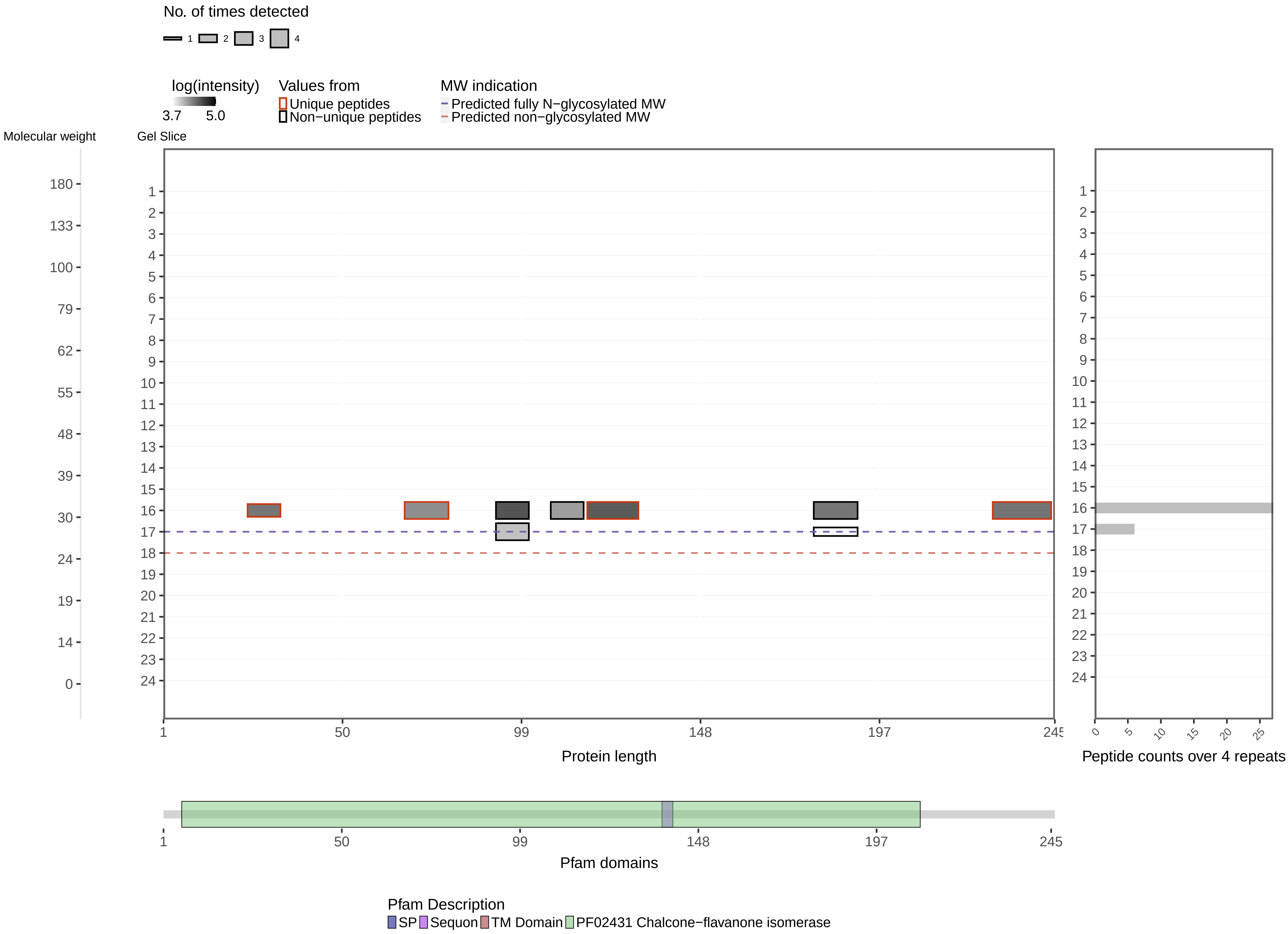


Protein sequence:
'AENCVAHWKAIGTYTNAESQAIEKFLNIFQNETFLPGASILFTQSPVGALTISFIKDDSITGTGNAVIENKQLSEAVLESIIGKHGVSPAAKCSIAERVSELFKKSYADASVCEKPGIEKSSDPVIEEKPTIPEIGV*

### NbL08g12840.1 probable_UDP−arabinopyranose_mutase_1 [MW = 41.13 kDa]

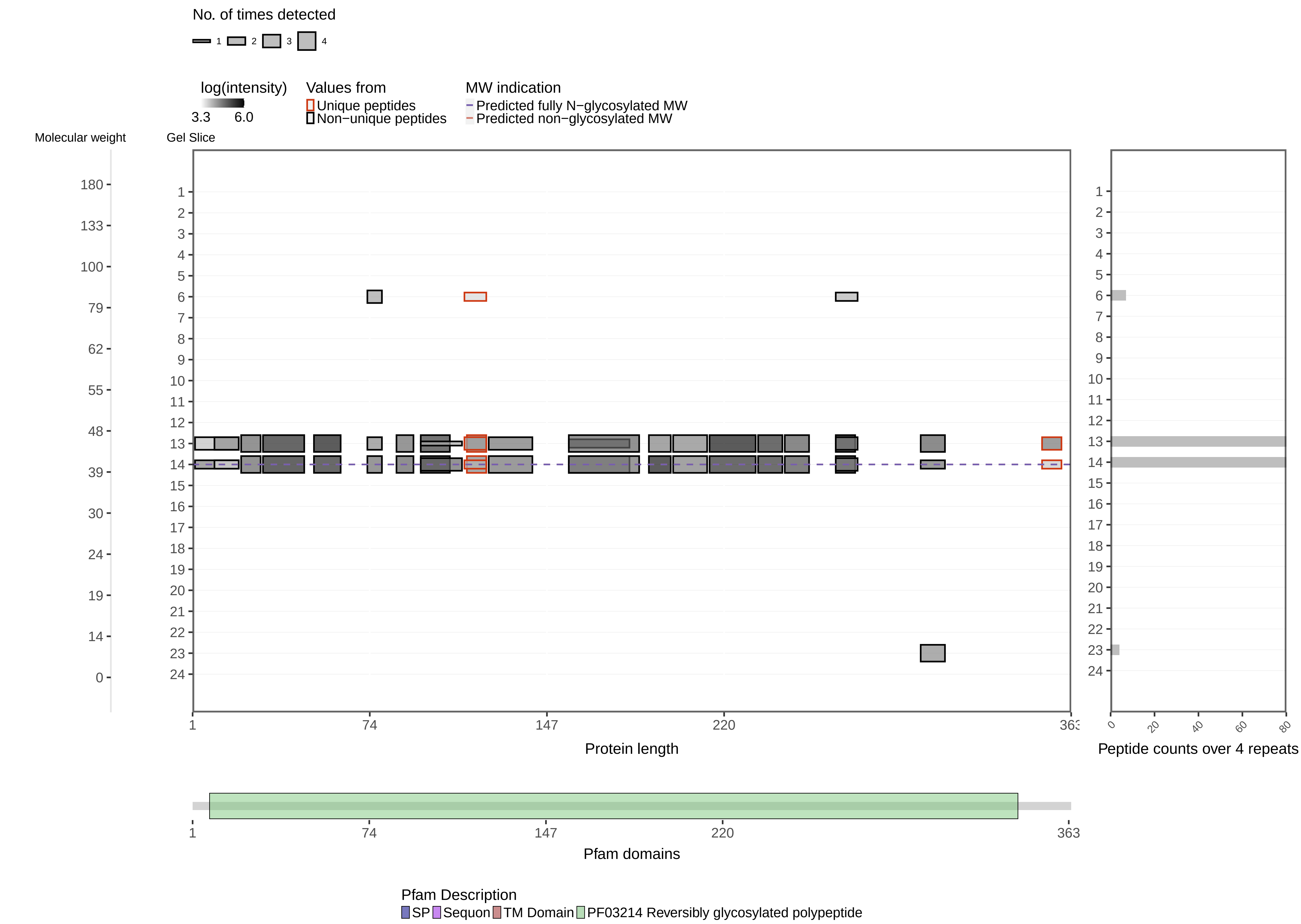


Protein sequence:

PDYDAPTQLVKPHERNTRYVDAVMTIPKGTLFPMCGMNLAFDRELIGPAMYFGLMGDGQPIGRYDDMWAGWCTKVICDHLGLGIKTGLPYIWHSKASNPFVNLKKEYNGIFWQEEIIPFFQAATLPKECKTVQQCYLELSKQVKDKLSKIDPYFTKLGDAMVTWIEAWDELNPTGESLAKLSISDGPAKAAK*

NbL08g12830.1 splicing_factor_YJU2 [MW = 37.43 kDa]

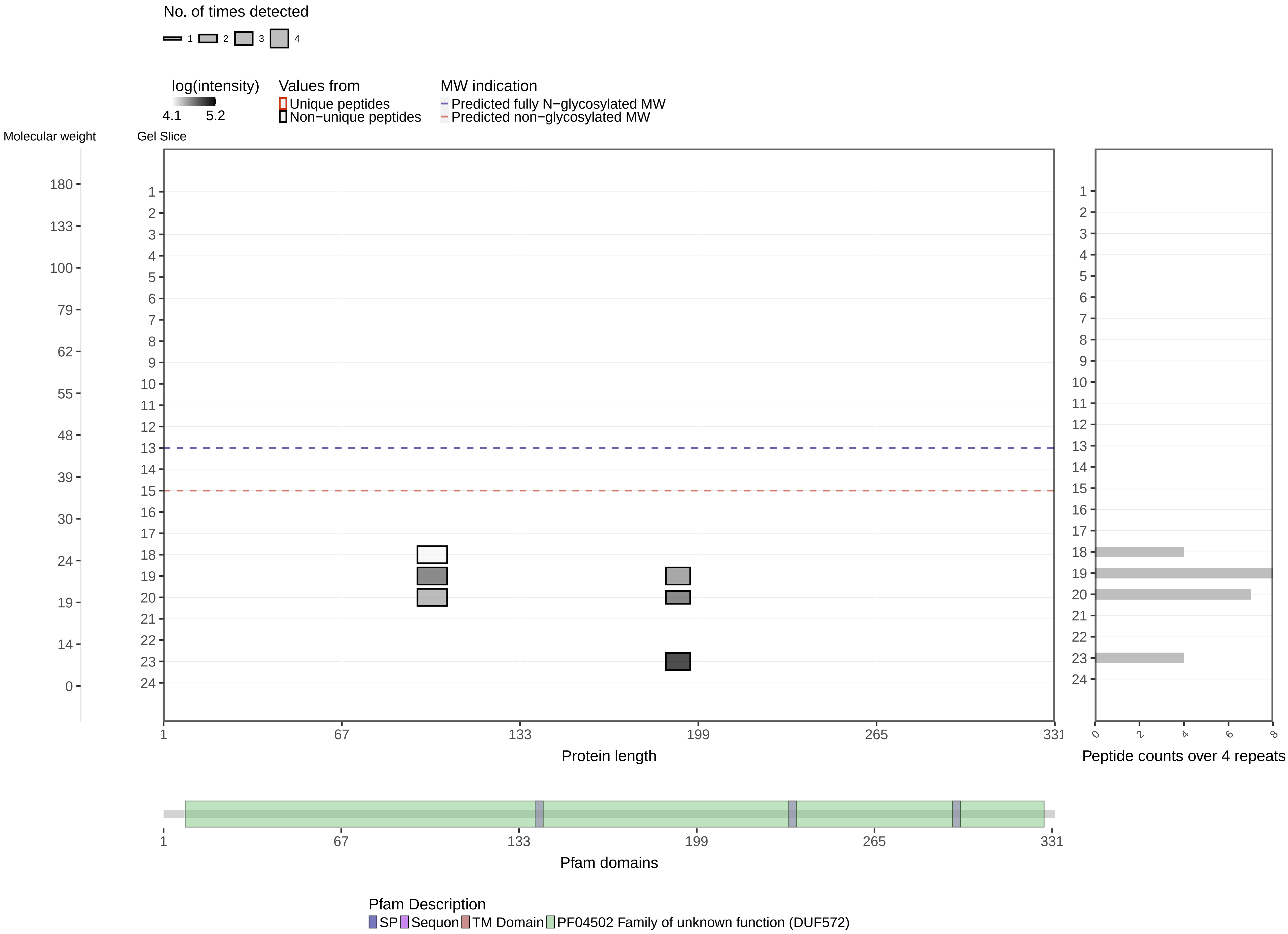


Protein sequence:

AALDEMKSMKSRHATVSVDAMLEALQRADEAKARKLEEKDEALIRSVFQGSRDIIKRIPDEELEEDDGDLAFFAPESSGSENNSLKRRKVSEESNEKPTNFLTKASVTDSSINKDKASSSDGKIIFKSSSVKISVVKKPSSDNNSKKTVEGNQQEAEKTGGTSNGLLSLCQQYESDEDD*

### NbL08g12790.1 fructokinase−like_2_chloroplastic [MW = 74.85 kDa]

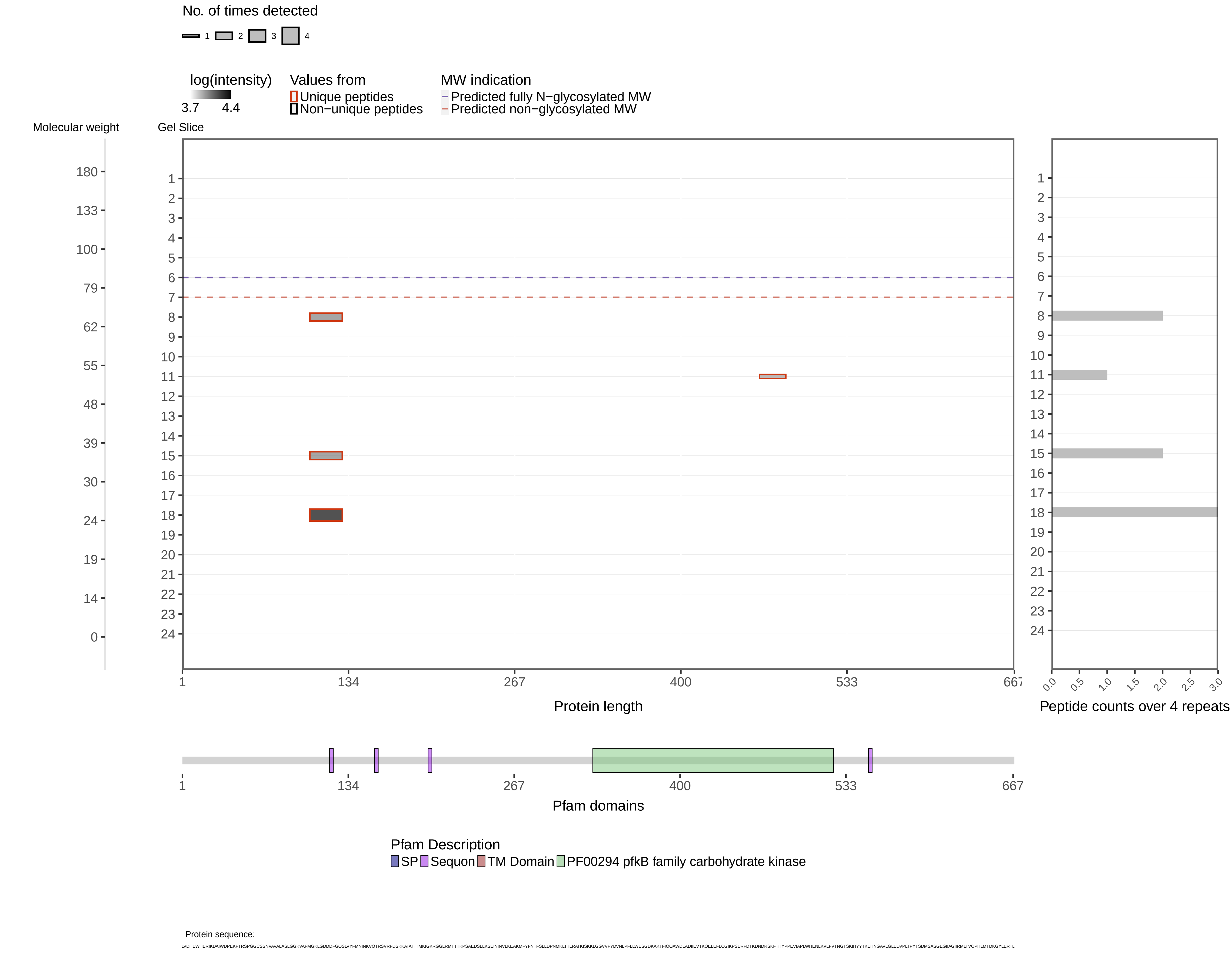

### NbL08g12630.1 probable_inactive_purple_acid_phosphatase_1 [MW − SP = 66.42 kDa]

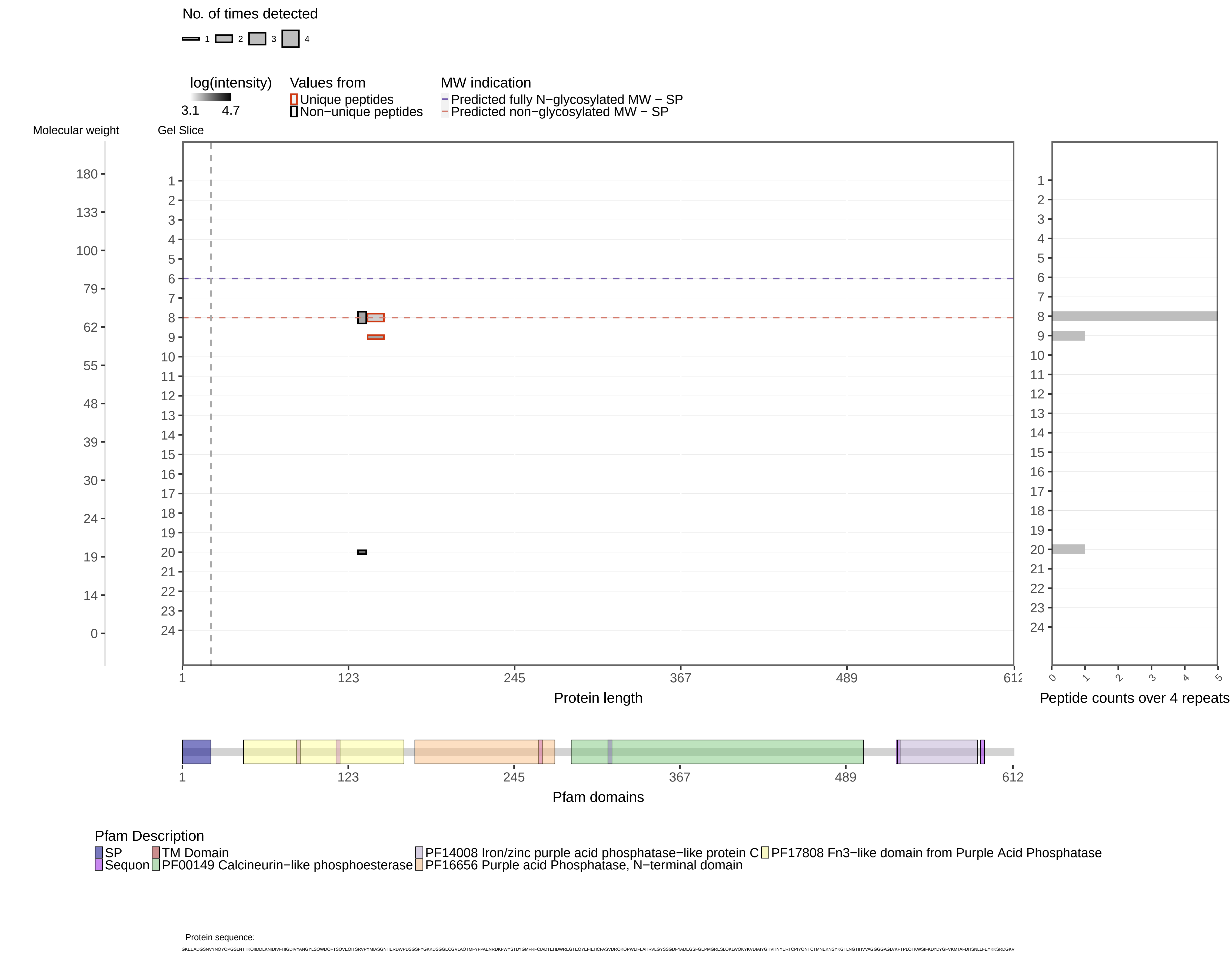

### NbL08g12620.1 argininosuccinate_synthase_chloroplastic [MW = 53.85 kDa]

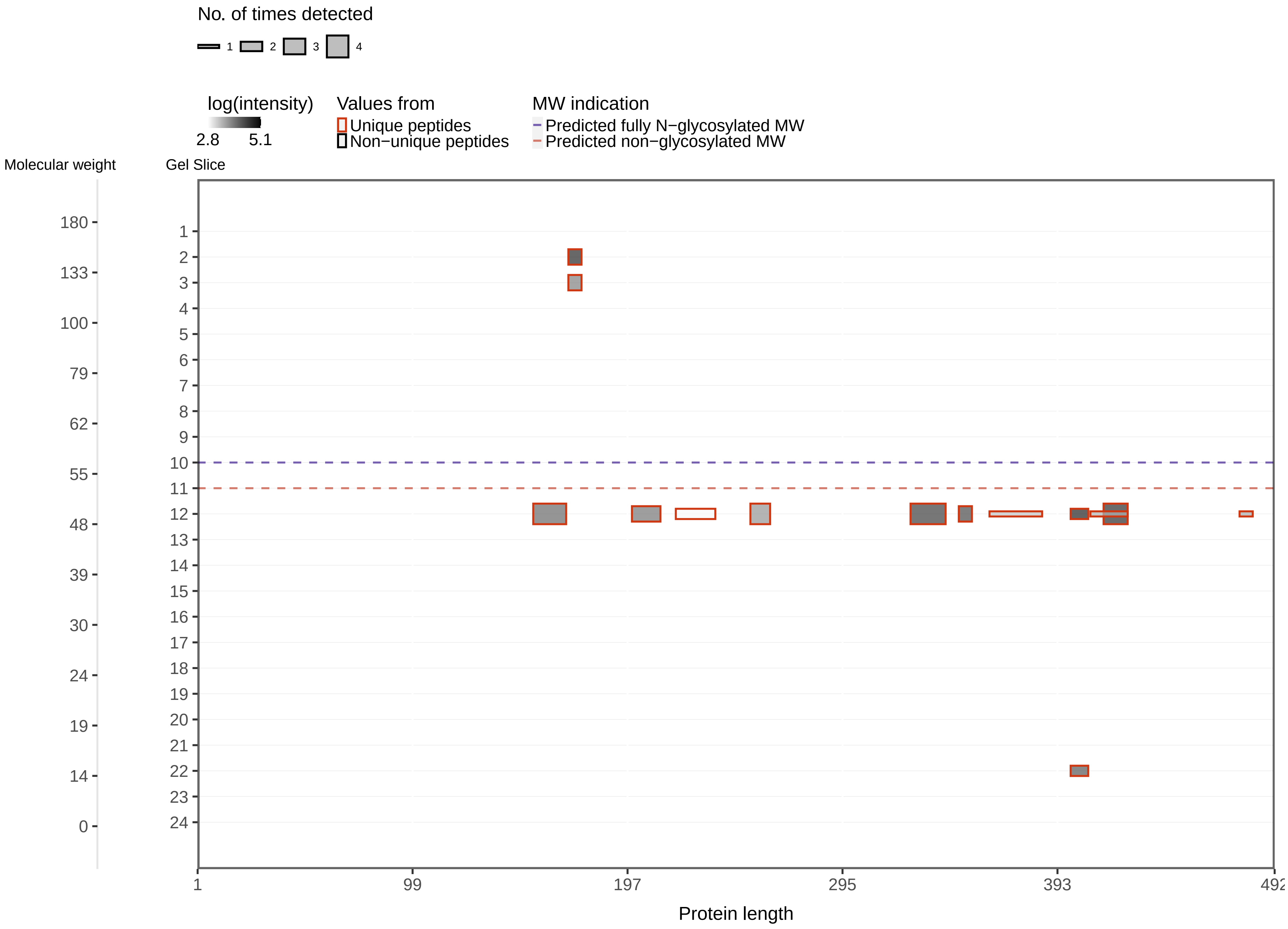


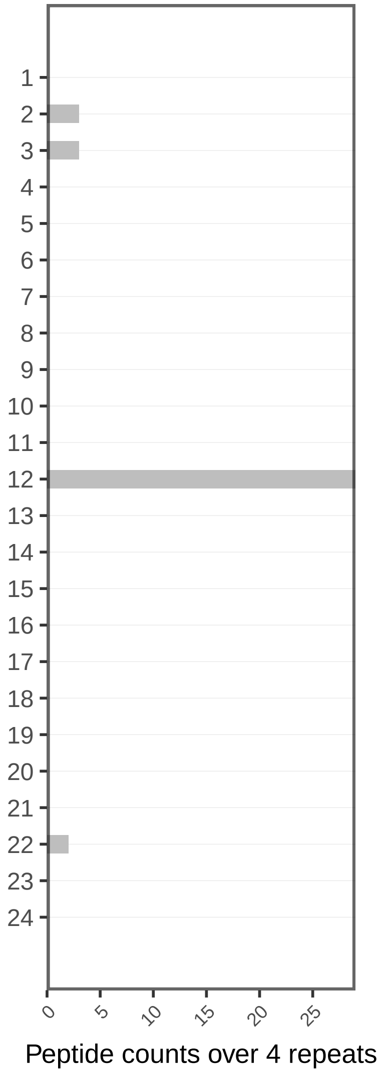


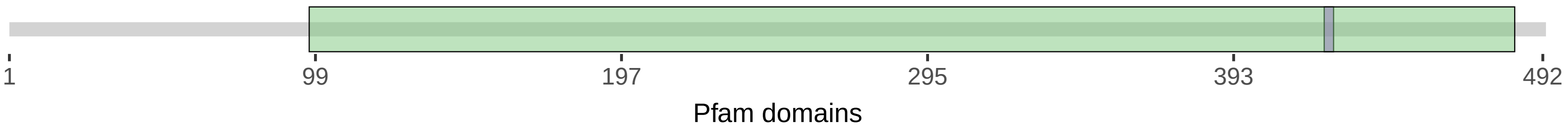


Pfam Description
SP Sequon TM Domain PF00764 Argininosuccinate synthase

Protein sequence:

VAPWREWEITGREDAIEYAKKHHVPVPVTKKSIYSRDRNLWHLSHEGDILEDPANEPMKDMYMMSIDPQDAPDQPEYVNIGIVAGVPVSVNGKELSPATLLSELNEIGGRHGIGRIDMVENRLVGMKSRGVYETPGGTILFTAVQELESLTLDRETIQVKDSLALKYAELVYAGRWFDPLRESMDAFMENITKTTTGSVTLKLYKGSVSVTGRQSPHSLYRQDISSFESGDIYNQADAAGFIRLYGLPMRVRAMLEKGL*

NbL08g12550.1 PLASMODESMATA_CALLOSE−BINDING_PROTEIN_3−like [MW − SP = 17.79 kDa]

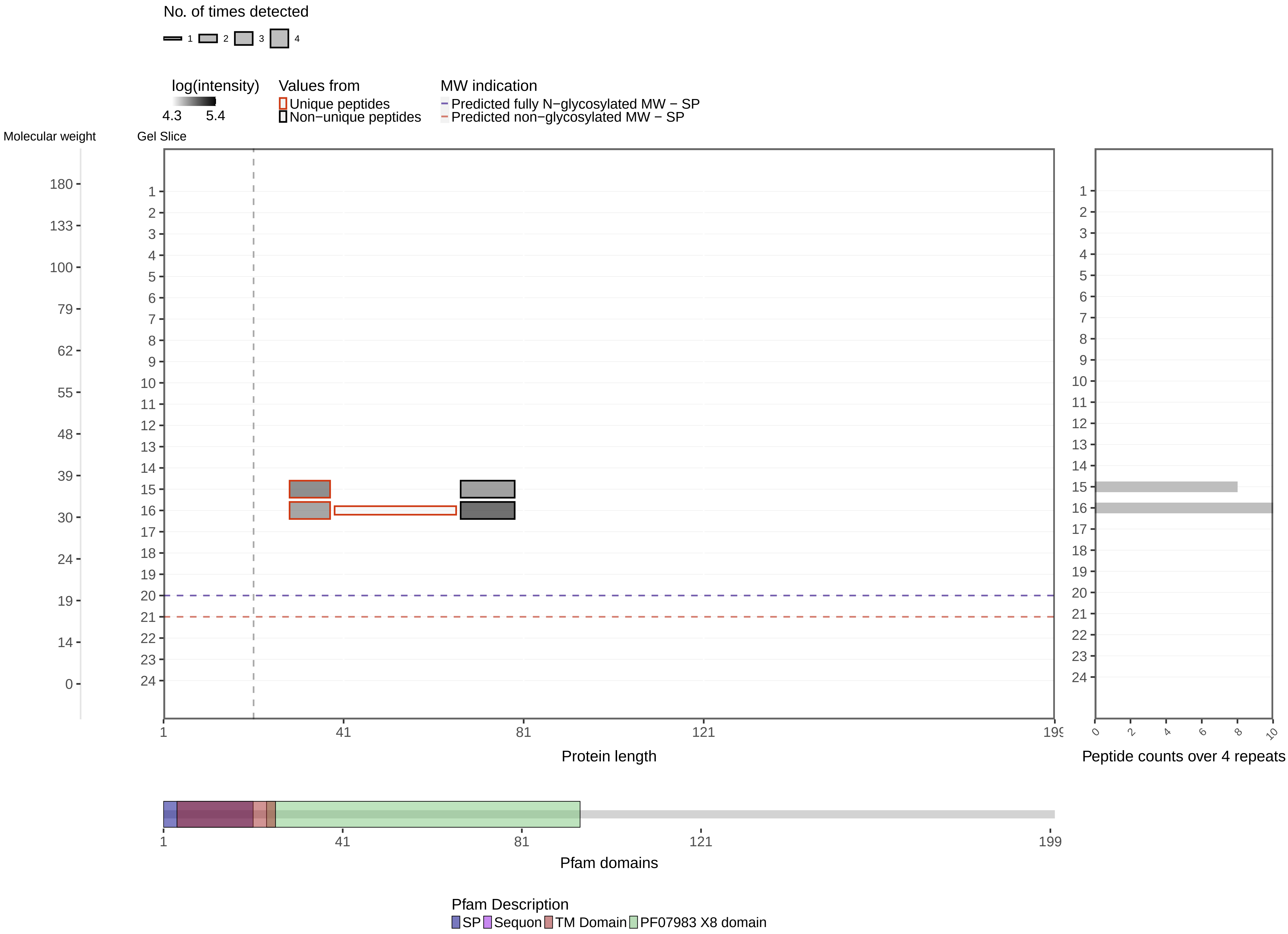


Protein sequence:

CDFSGTATLSSNPPSSSTSGCYQTTTGNTAGGTTTVPGTTTVPGTTTPGIGTGTGTGTGTGTTTGINNPTFGLGPTGSGGFNPDGSGTETLHQNTVFFTIAILFTSLVWLRL*

### NbL08g12520.1 14−3−3−like_protein_GF14_iota [MW = 29.94 kDa]

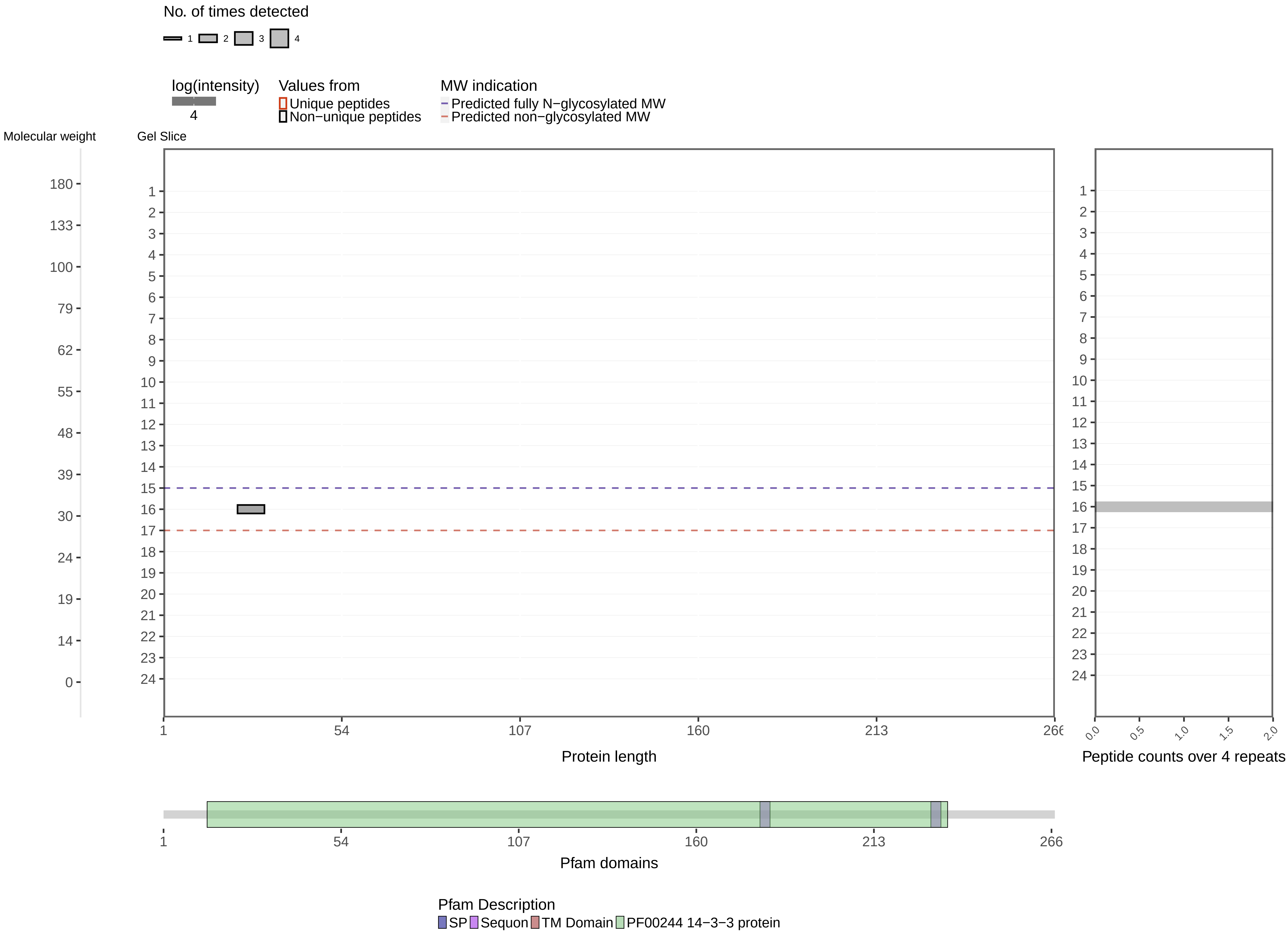


Protein sequence:

TVFYYKMKGDYYRYLAEFKTDSERKEAAEQSLKGYEAATATANTDLSSTHPIRLGLALNFSVFYYEIMNSPERACHLAKQAFDEAIAELDTLSEESYKDSTLIMQLLRDNLTLWTSDLPEDGGEENVKGDEPKGAEPKSADAKAC*

### NbL08g12460.1 probable_mitochondrial−processing_peptidase_subunit_beta_mitochondrial [MW = 59.43 kDa]

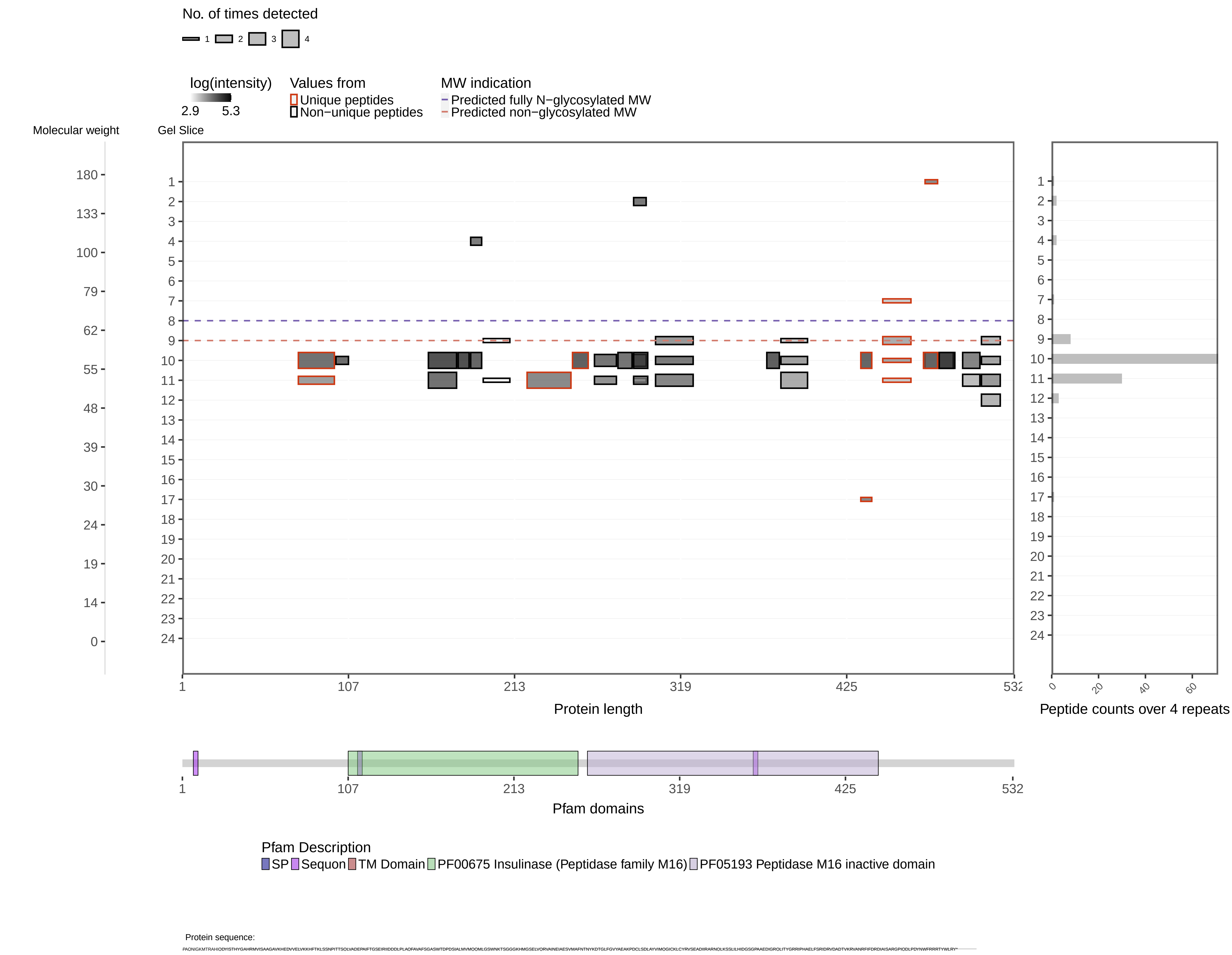

NbL08g12400.1 skin_secretory_protein_xP2−like [MW = 23.71 kDa]

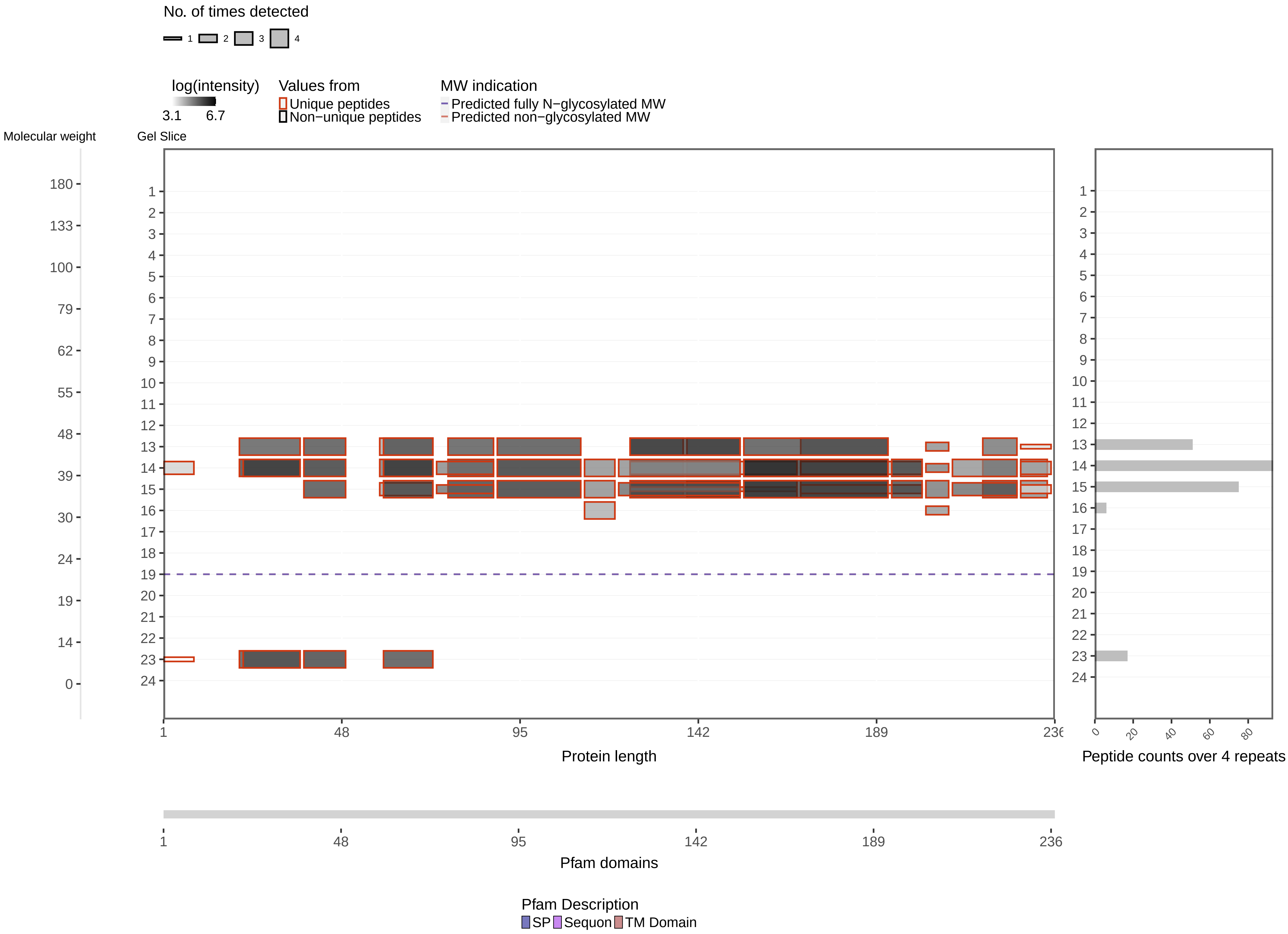


Protein sequence:

TTDAAAAKAPAATPDTKDTKAPTPAPAAADTDETKAPAPAPAAADTEETKASAPAPAAADTEETKAPATADTEEPPAPAPAPAPAPAPEEKGEGYGQYVKMAEGFLKSGDDESAKASEGGSDYLKMAGDFLGKK*

### NbL08g12390.1 26S_proteasome_non−ATPase_regulatory_subunit_10 [MW = 25.23 kDa]

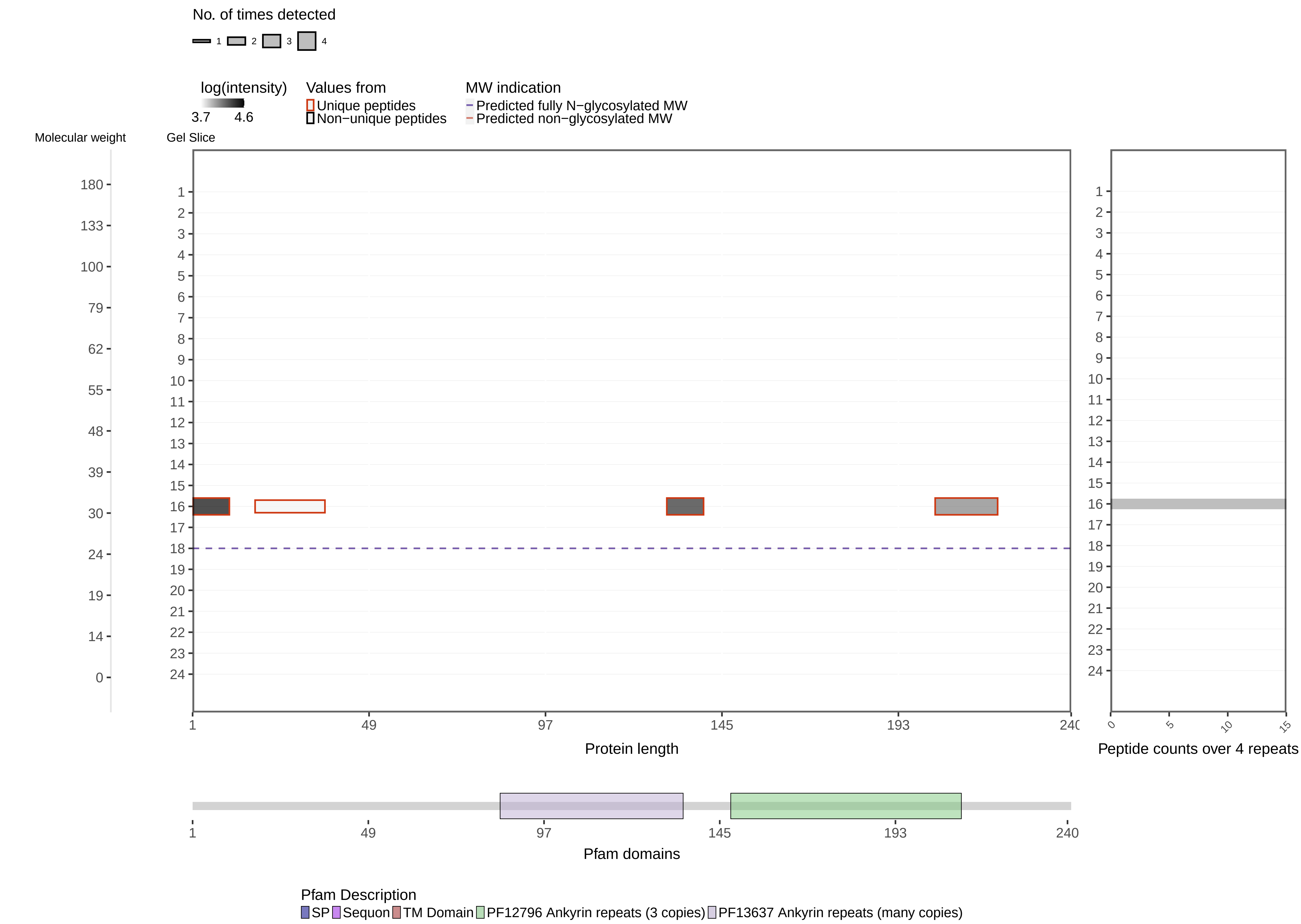


Protein sequence:

DANLKNDGGRTALHYAASKGRVKIAELLISHGAKINAKDKVGCTPLHRAASTGNSELCELLIEEGAEIDEVDKAGETPLMTAVVCGNKEVAMLLIRHGANVDAEDKEGYTVLGRASNDLRPALVDAAKTMLEG*

NbL08g12360.1 uncharacterized_LOC109212061 [MW = 37.96 kDa]

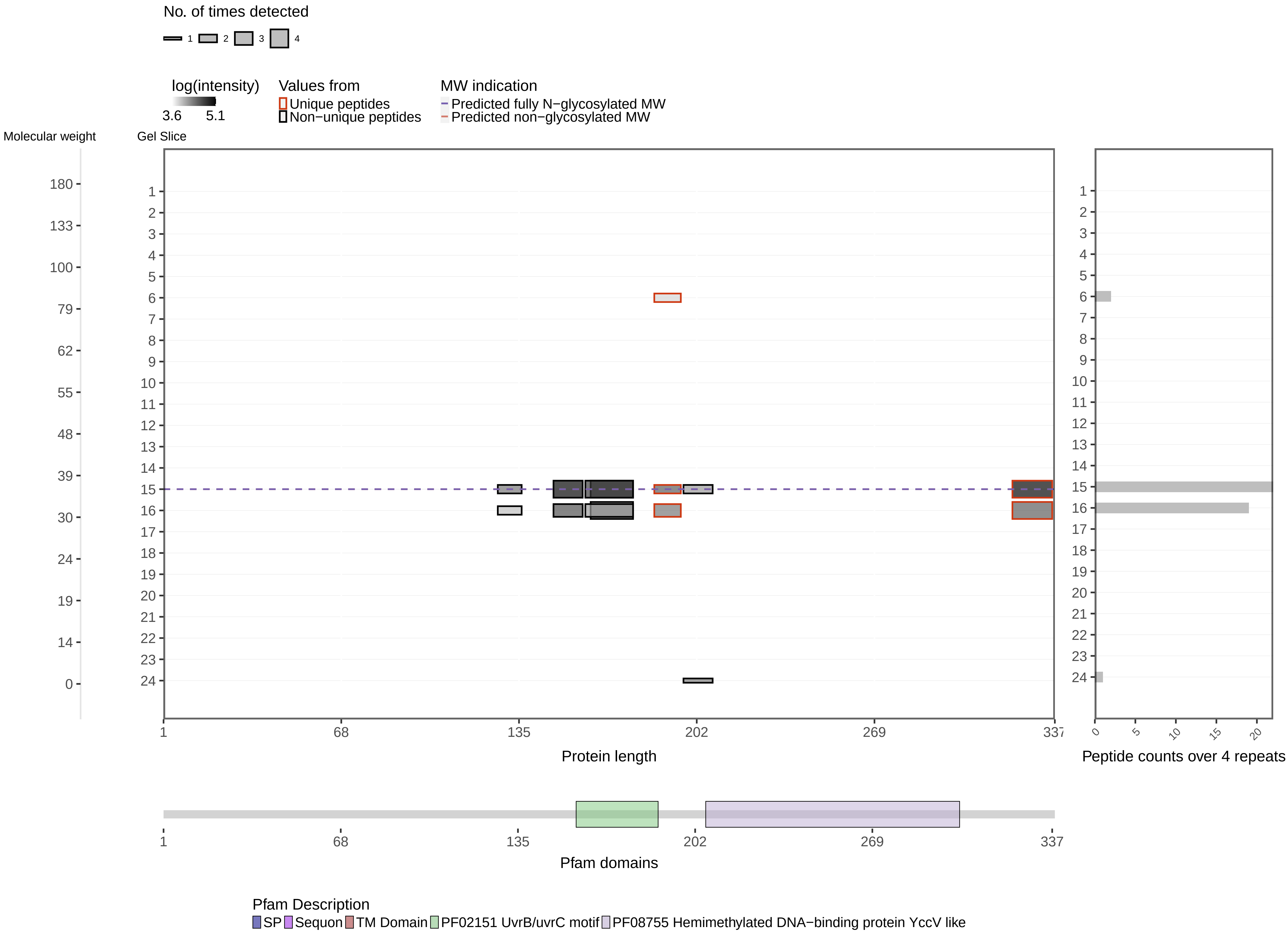


Protein sequence:

AISILRLRADLQNAVQSENYDLAAKLRDEISKLEAESLTASVRAQAYVNAQYAFRLGQKVRHKNFGYRGVICGMDPICCESSSWMETAQVDKLSRGPDQPFYQVLVDVHTDPNLLVAYVPEESLVAPSEPDKDRFDHPYTSFLFFGMDAAGDFIPIKQLREKYNRPRHEVPYDPEDEKSGEGS*

NbL08g12190.1 probable_L−cysteine_desulfhydrase_chloroplastic [MW = 49.09 kDa]

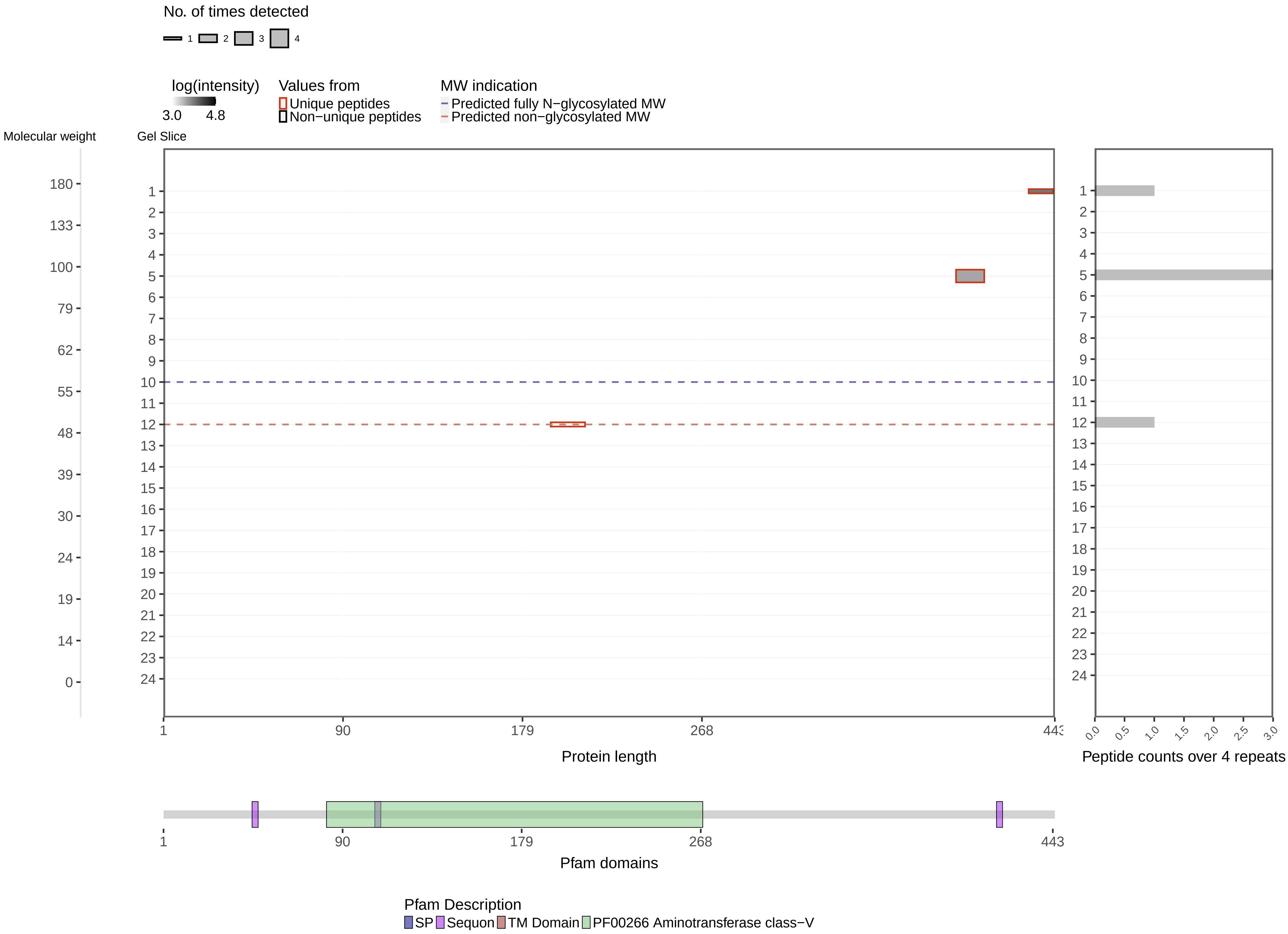


Protein sequence:

ELVKMCRDEGVDIIFVDGAHAIGNVQVDVGDIGADFYTSNLHKWFFSPPSAAFLYCKKSDKVLDLHHPVVSNEYGNGLPIESAWIGTRDYSAQLVIPEVLEFVNRFEGGIEGIRRRNHGKVVEMAEMLVKAWGTKLGTPPEMCSSMAMVGMPDCLGISSDSDALKLRTHLRVSFKVEVPIYYRAPLEGELNSITGYARISHQVYNTTEDYYRFRDAIIKLVNDGFTCAILSN*

NbL08g12150.1 uncharacterized_LOC109214568 [MW = 32.7 kDa]

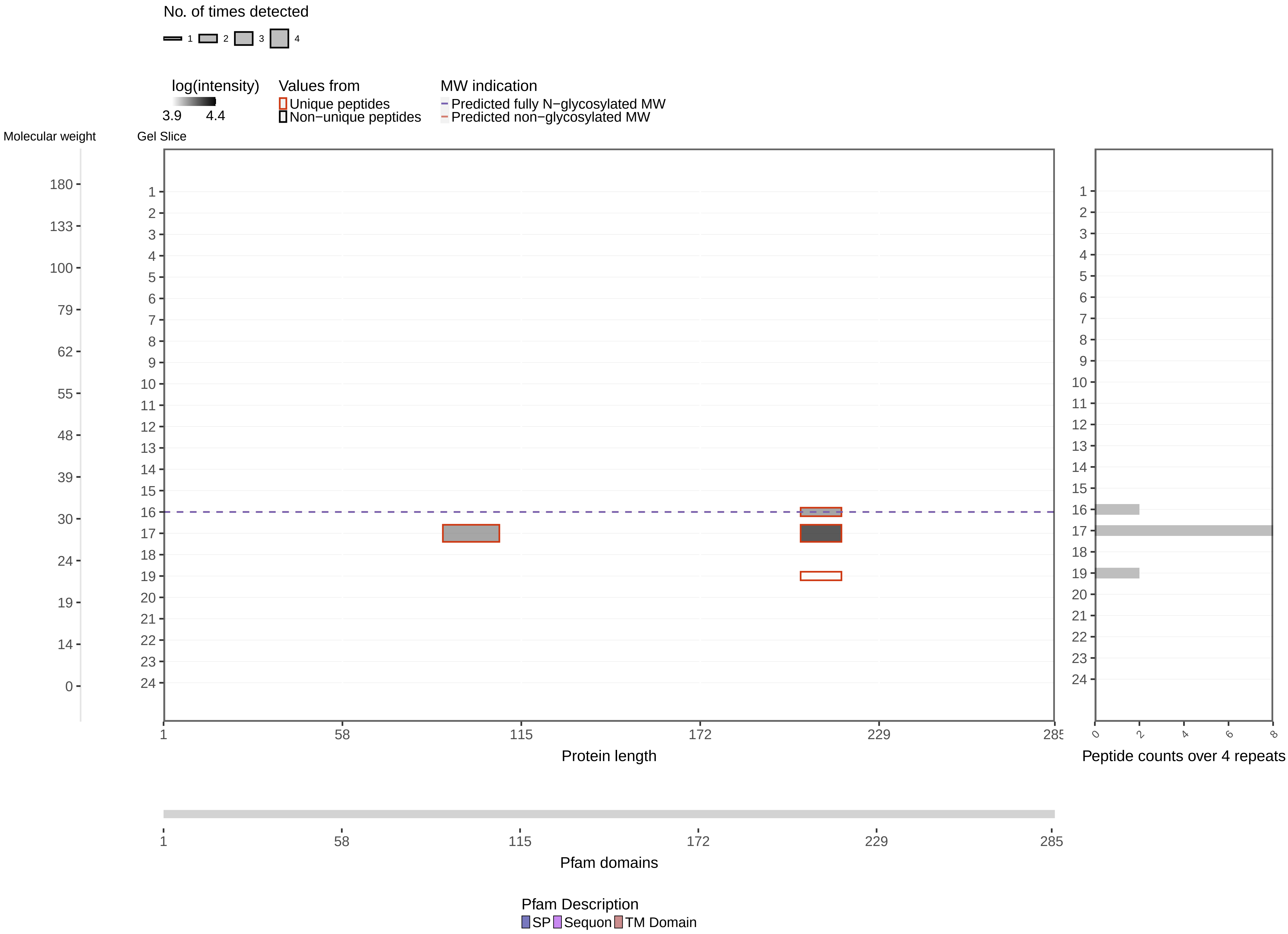


Protein sequence:

ANPWRETPKPVYVLTQRENQLCTMKTRKNQRFFEACSEVERELGMLFSKGGKWKNRAKQPETDTRFQMLVEDIREGVLVFEDENEAARYCDLLQGGGQGCEGVAEIEASSVFDLCRKMRALAVLFRRGRTPPLPESLKLNLRARKRSLEDQENLM*

### NbL08g12070.1 extracellular_ribonuclease_LE−like [MW − SP = 22.65 kDa]

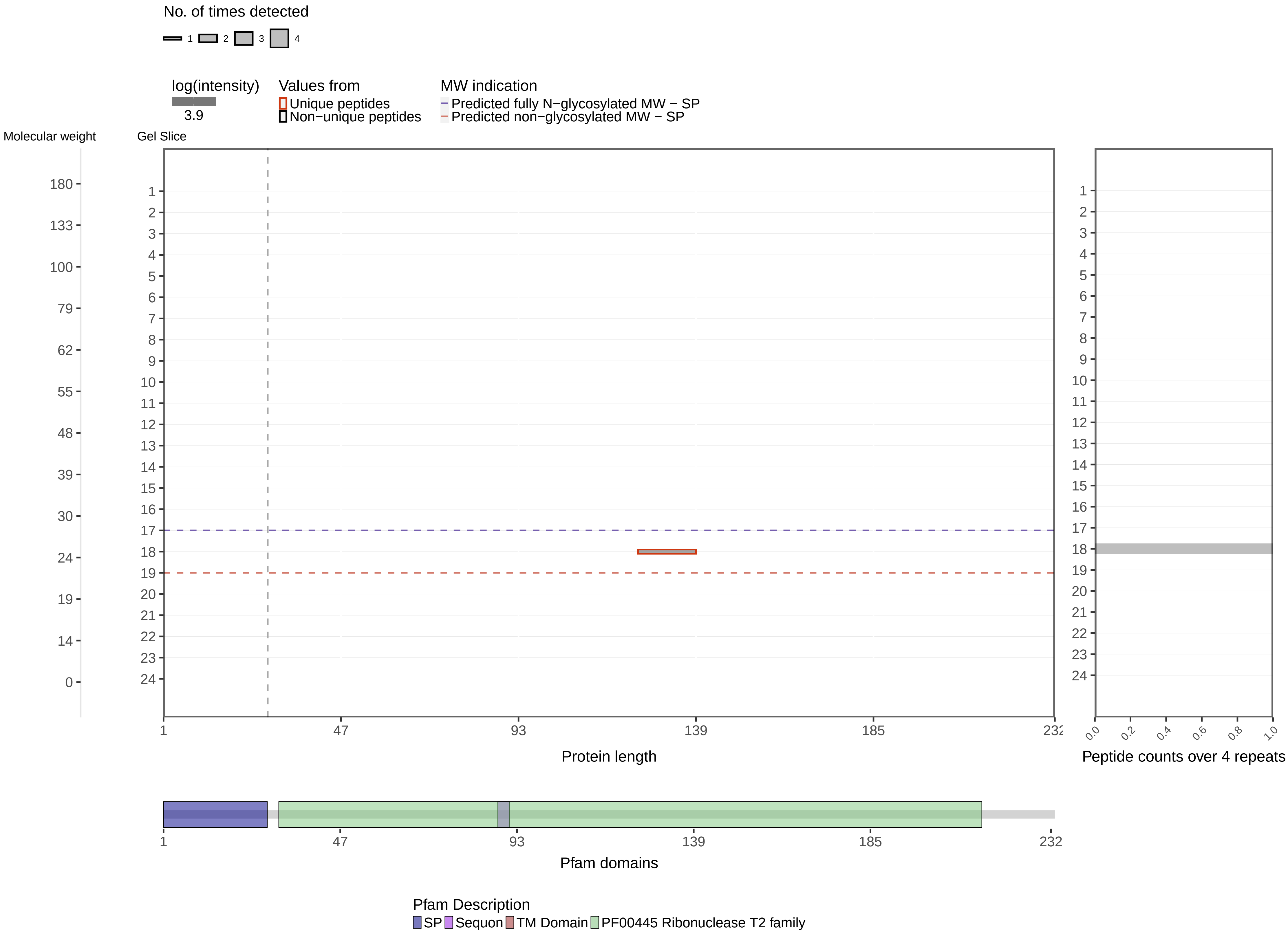


Protein sequence:

NWPTLACPSGTGSAFWSHEWEKHGTCSESIFDQHGYFKKALDLKNQINLLEILQGADINPDGGFYSLNNIKNAIRSAIGYTPGIECNIDESGNSQLYQVYICVDGSGSNLIECPVFPRGKCGSSIEFPTF*

NbL08g12060.1 intracellular_ribonuclease_LX−like [MW = 28.86 kDa]

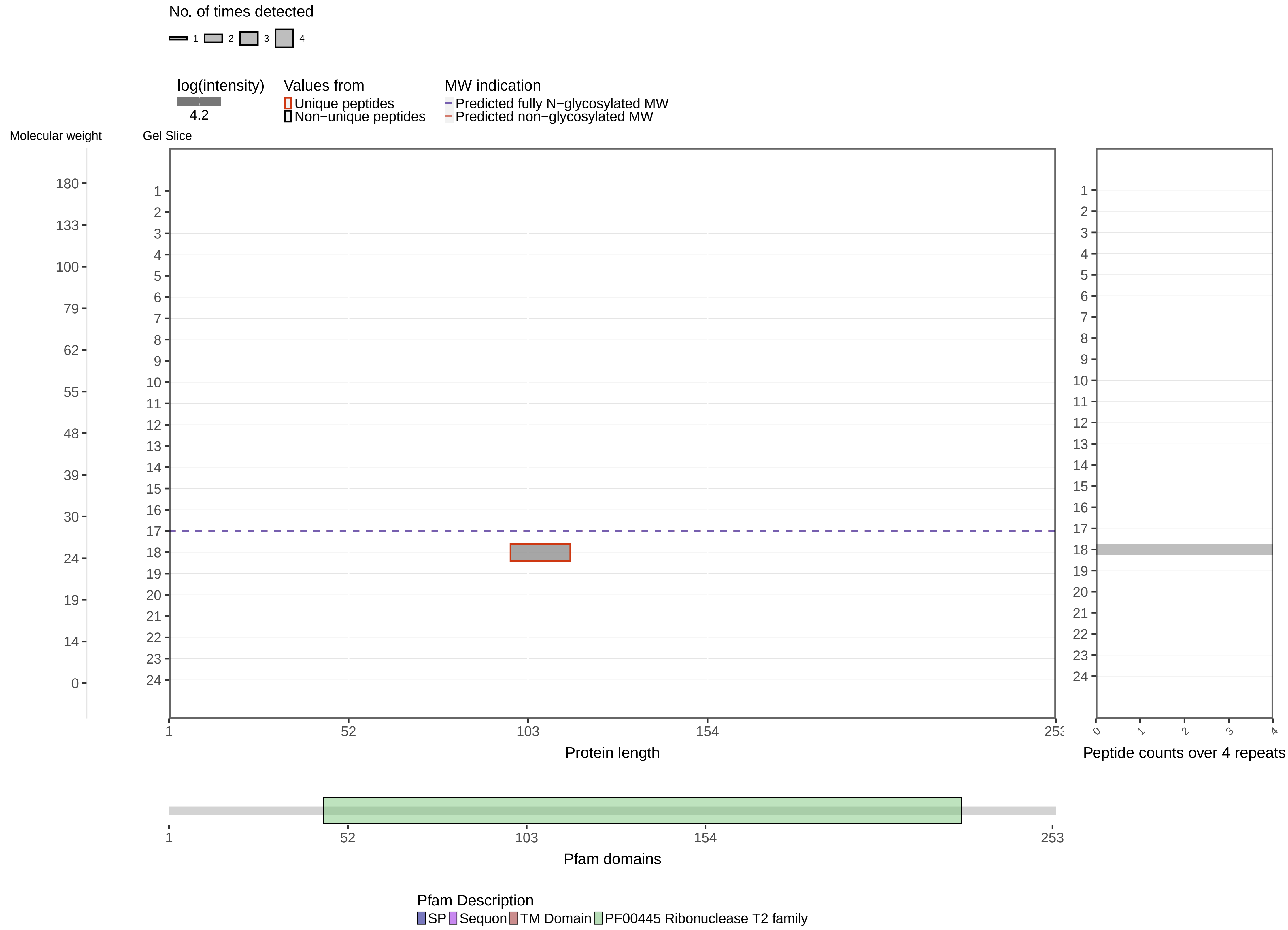


Protein sequence:

EKNWPSLACPSSDGVRFWSHEWLKHGTCSALGERAYFQAALDFRKKSNLLENLKNAGITPRNGEHYSLESIKKAIEEGVGHIPYIECNVDTQGNHQIYQVYLCVDKTASHFIDCPVFPHGRGCGAKIEFPPFSSDHDEF*

### NbL08g12030.1 methylecgonone_reductase−like [MW = 36.48 kDa]

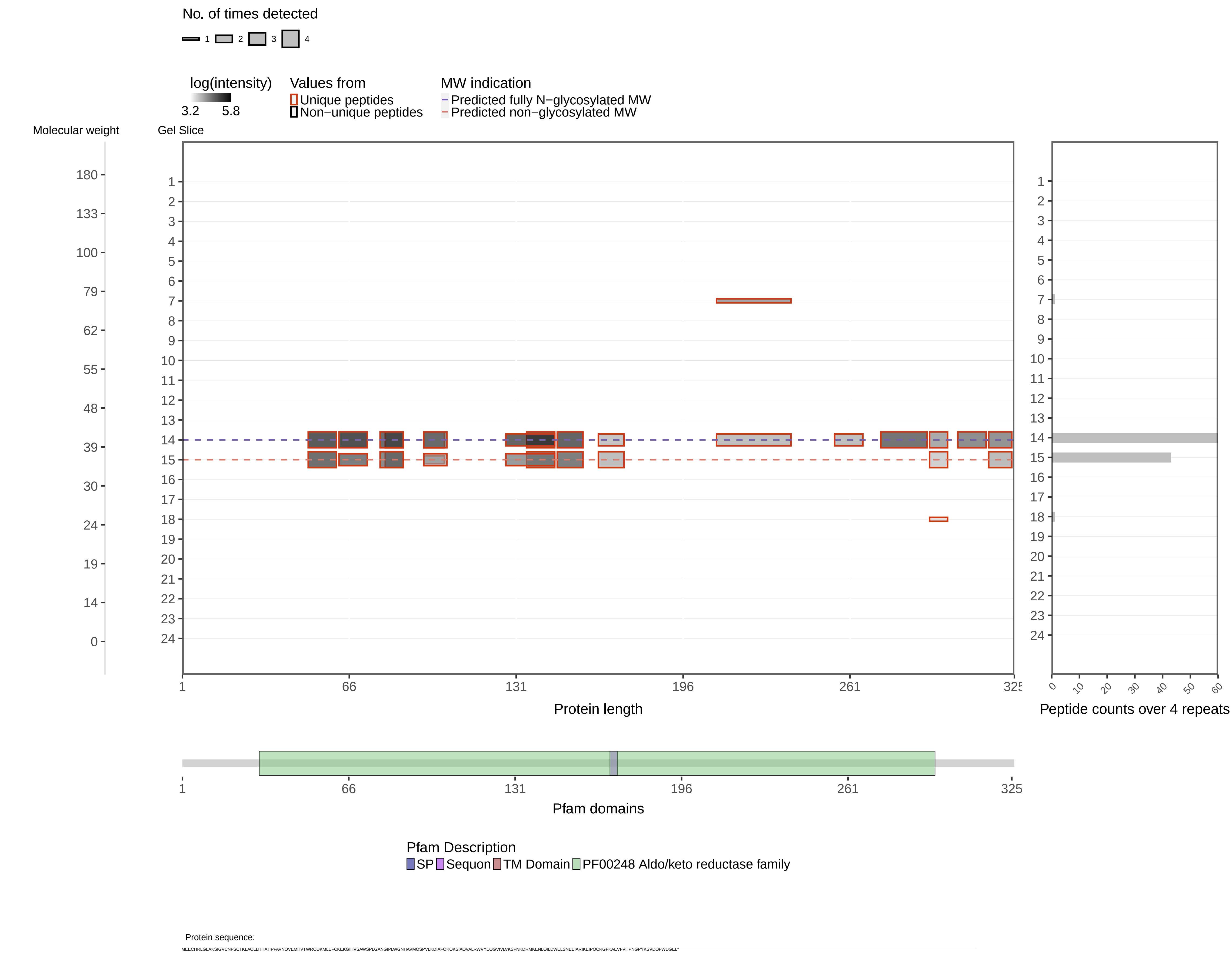

### NbL08g11930.1 KH_domain−containing_protein_At4g18375−like [MW = 57.42 kDa]

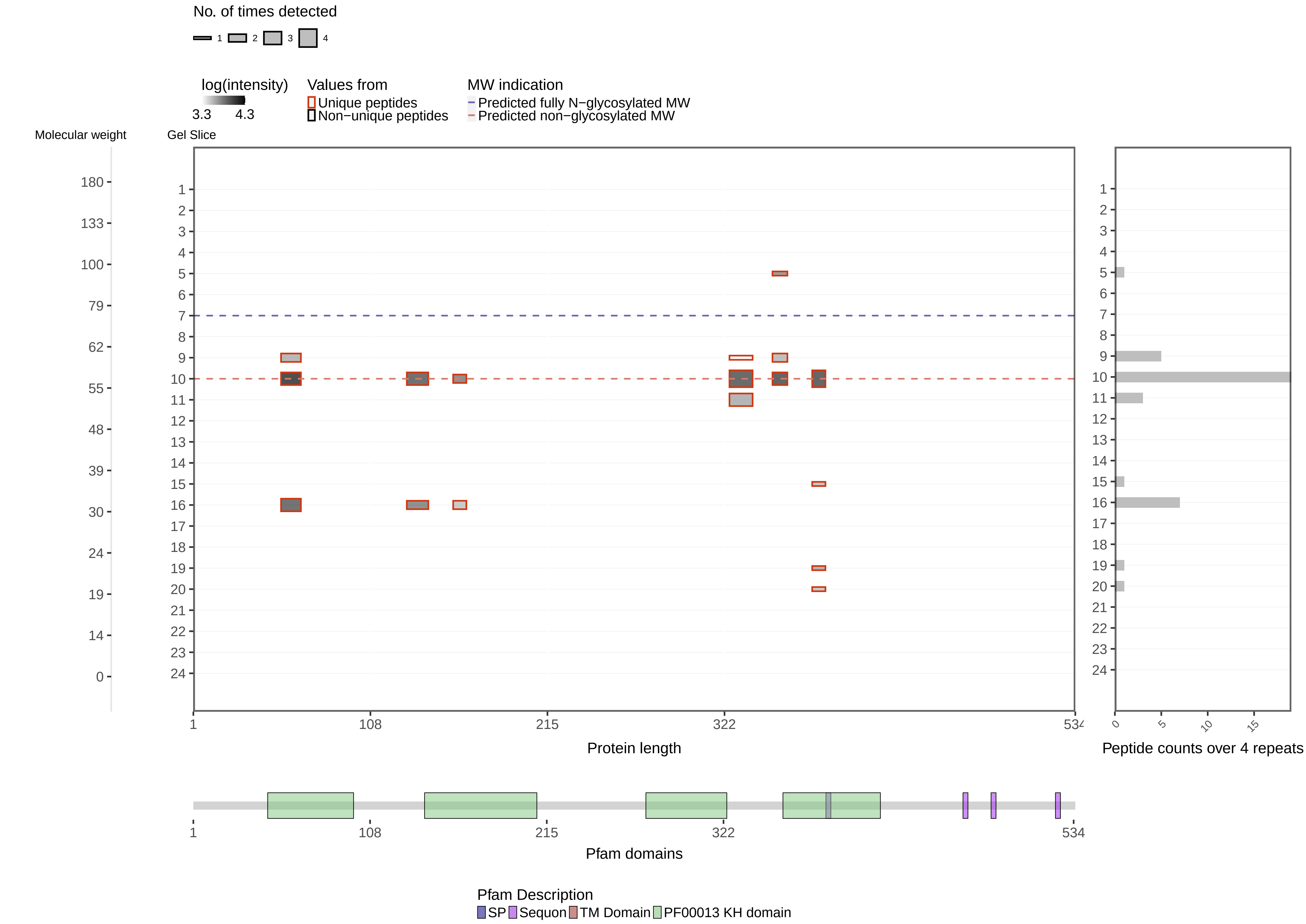


Protein sequence:

GSYKSDGRSRSASVKEFAVRLVCPTENIGAVIGKGGSIIKQLRQESGASIKVDSAAAEGDDCIIVVSAKEAYEDQSPTIDATMRLQPRSSEKTEKDSGDAILTTRLLVPSSRVGCLIGKGGSIVNEMRNTTRASIRILSKENLPKVASEDDEMVQITGDVNVAGNALLQVLMRLRANIFEMEGSFATFSPPVSYLPLSPSMSDGSRYANRDNKSRRHGYSSYSGGHDYNDSSQSDSYGGSQVGGGGGYAPYGVYSSGRSSSAGVSNHNSSAYGKSYRY*

NbL08g11820.1 uncharacterized_LOC107767450 [MW = 53.83 kDa]

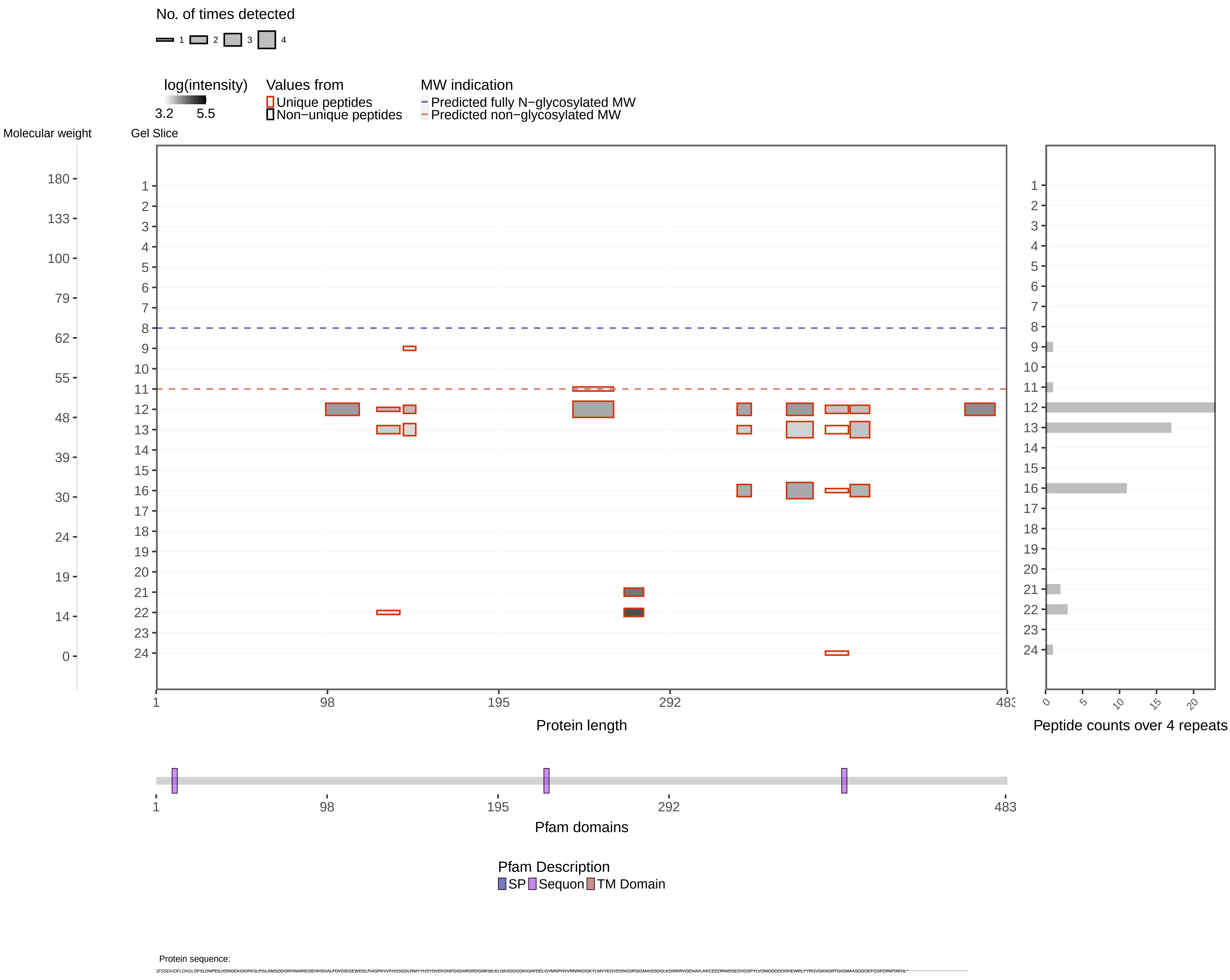

NbL08g11590.1 ribulose−phosphate_3−epimerase_cytoplasmic_isoform−like [MW = 25.7 kDa]

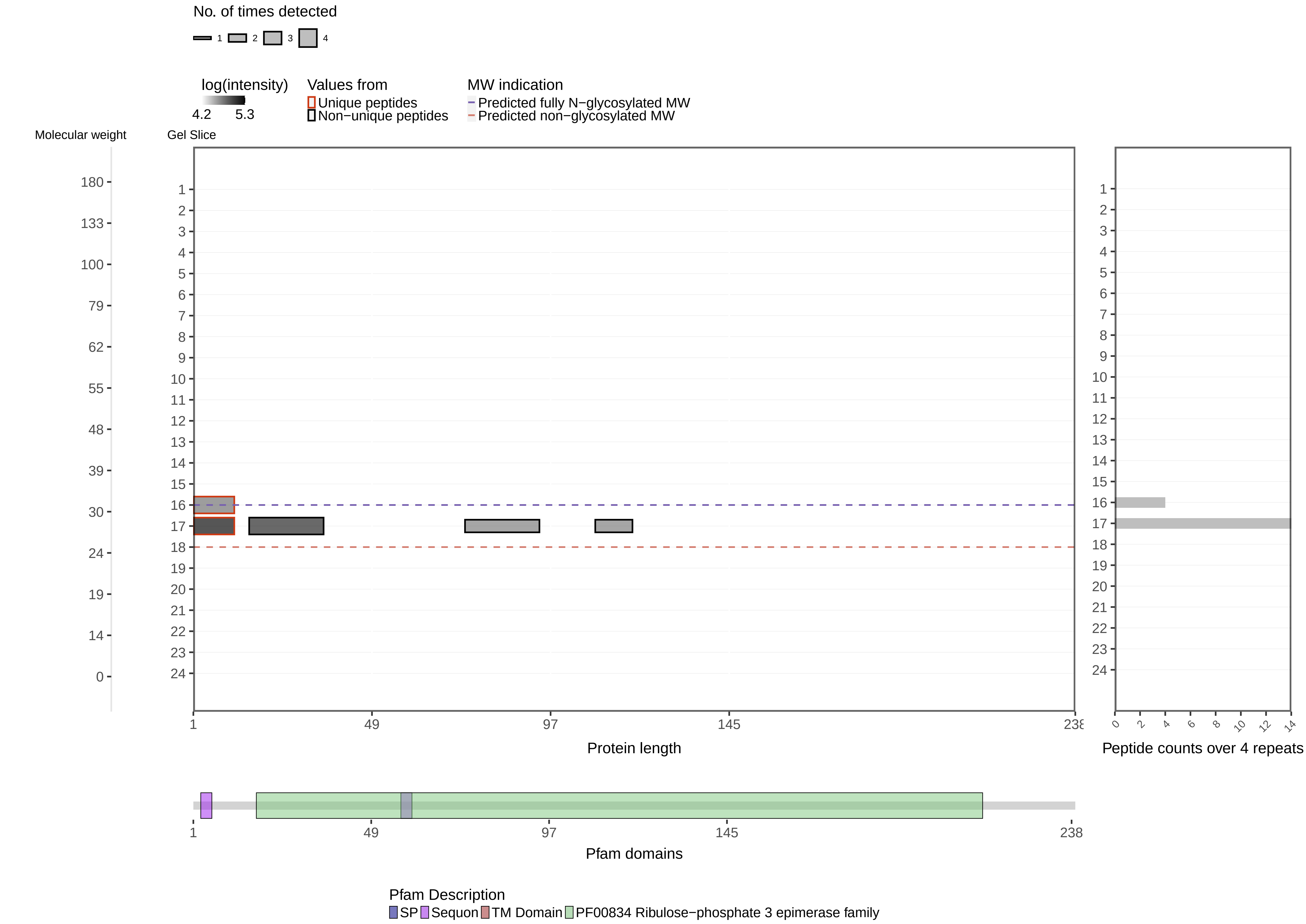


Protein sequence:

SRDNWQELVQQIKSKGMKPGVSLKPGTPIEEVYPQLDGENSVELVLVMTVEPGFGGQKFMPEMMDKVRTLRKKYPSLDIEVDGGLGPSTIEAASSAGANCVVAGSSVFGSPDPAQVISLMRNSVEEAQKIS*

### NbL08g11310.1 alpha−amylase_3_chloroplastic [MW = 101.8 kDa]

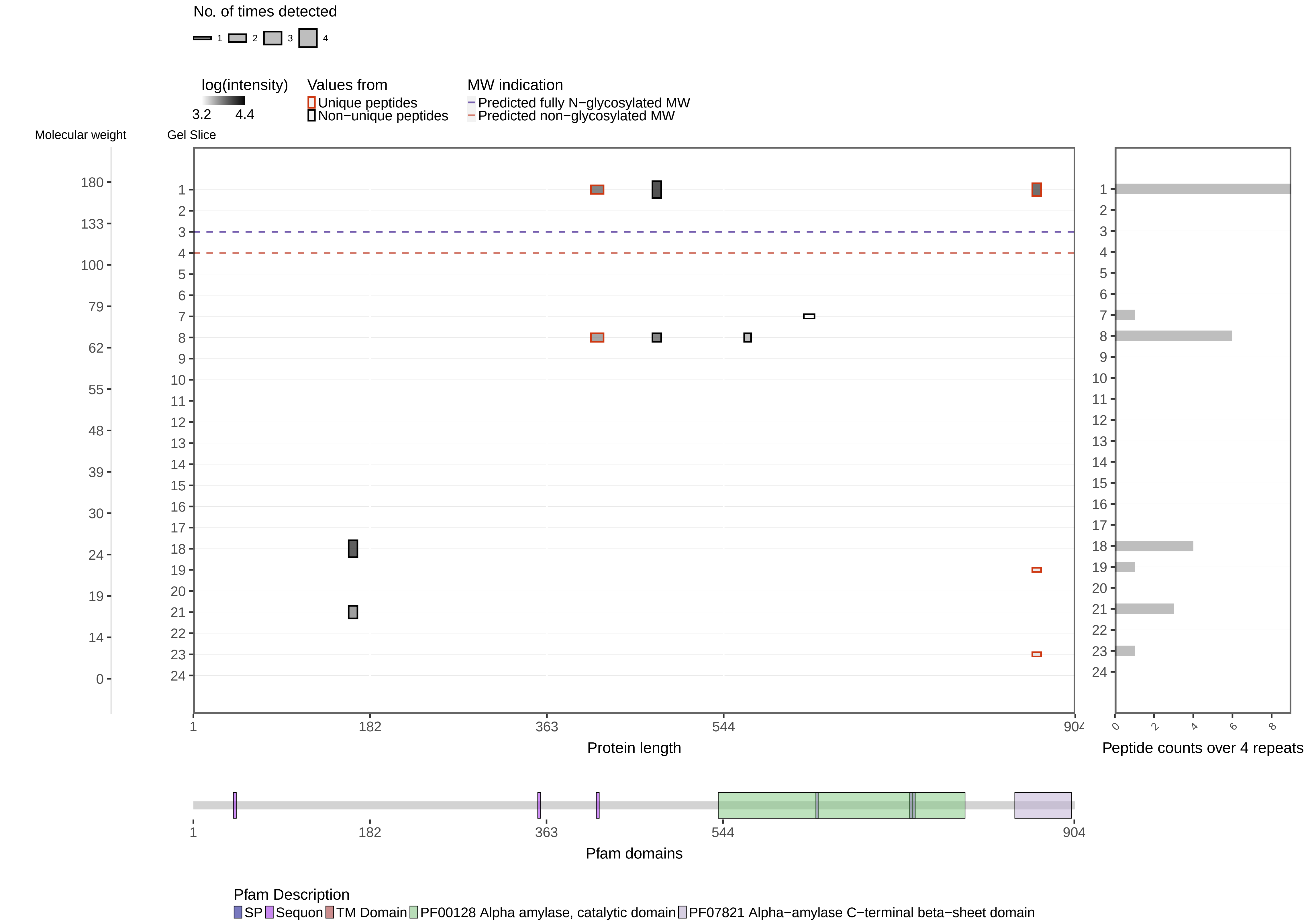


Protein sequence:

'TDISSEKSRKTKSKETQENILQEIEKLAAEAYSIFRNSIPTISEDVVSEAEVSQPDVKVSSGTGSGFEILCQGFNWESHKSGRWYVELHEKAAELSSLGFSVIWLPPPTDSVSPEGYMPRDLYNLNSRYGSFDELKVTVKKFHEVGIKVLGDVVLNHRCASERNQNGIWNIFGGRLNWDDRAVVADDPHFQGRGNKSSGDNFHAAPNIDHSQEFVRKDIKEWLRWLREEIGYDGWRLDFVRGFWGGYVKDYLDATEPYFAVGEYWDSLSYTYGEMDHNQDAHRQRIVD

NbL08g11290.1 uncharacterized_LOC109219735 [MW = 42.67 kDa]

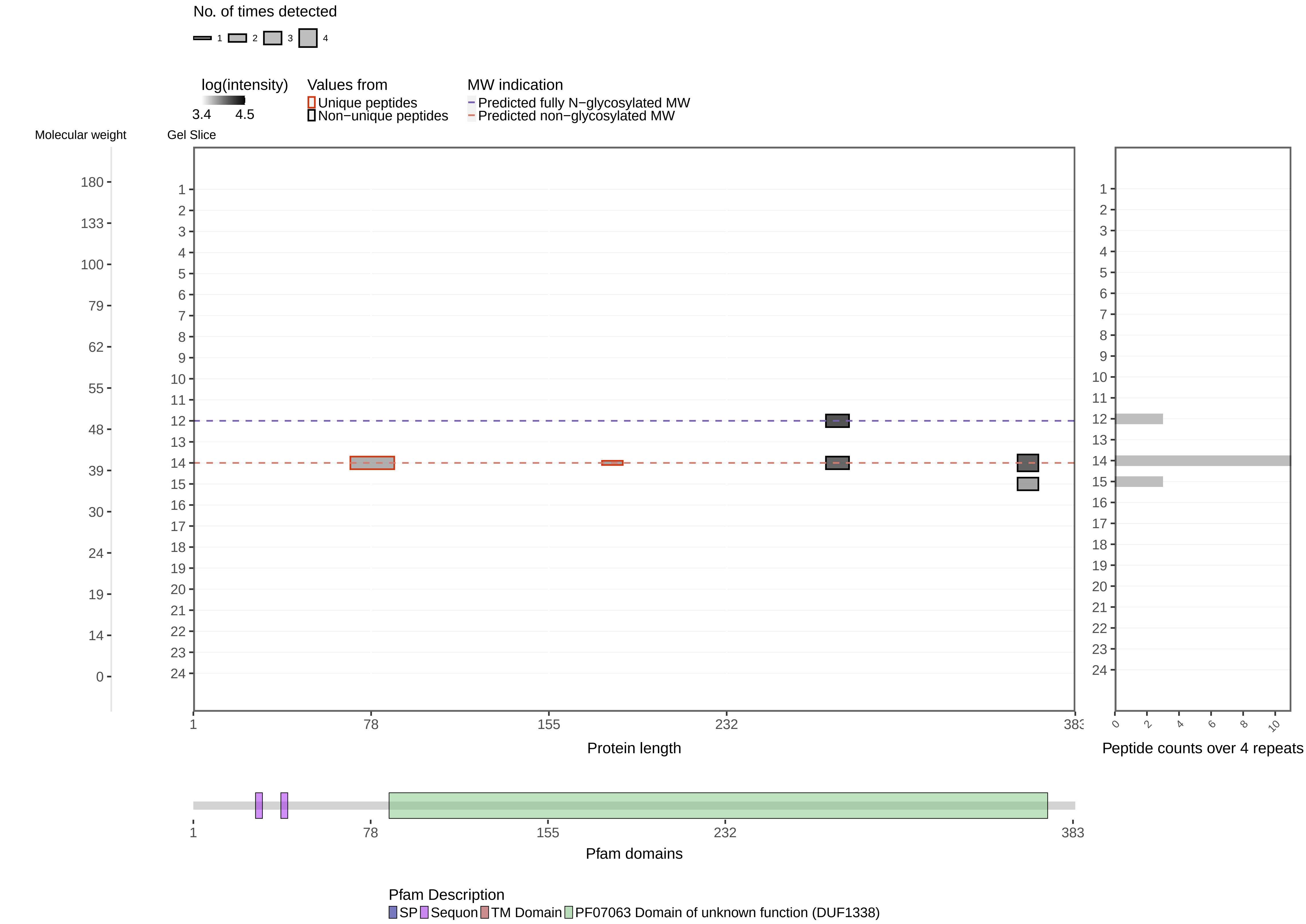


Protein sequence:

IRGSGVDGPLPRIFISELLVDQLSPEAQDIIKKYTDISRCGKEYAALASAFGVLTWERPSYSEFQQLARESEYAAWTLVNGYALNHVTISTHRLTSNLRSIGNLNQFIEKTGFNLNSEGGVLKVSPDGLLLQSSTVADSTSFQFSDGITEAVPCSYIEFAERLVHPQYKDLPAEEVEEFHRRDGFEVGNADKIFESTSKDQLTRRAA*

NbL08g10850.1 serine/threonine−protein_phosphatase_PP2A−2_catalytic_subunit−like [MW = 34.98 kDa]

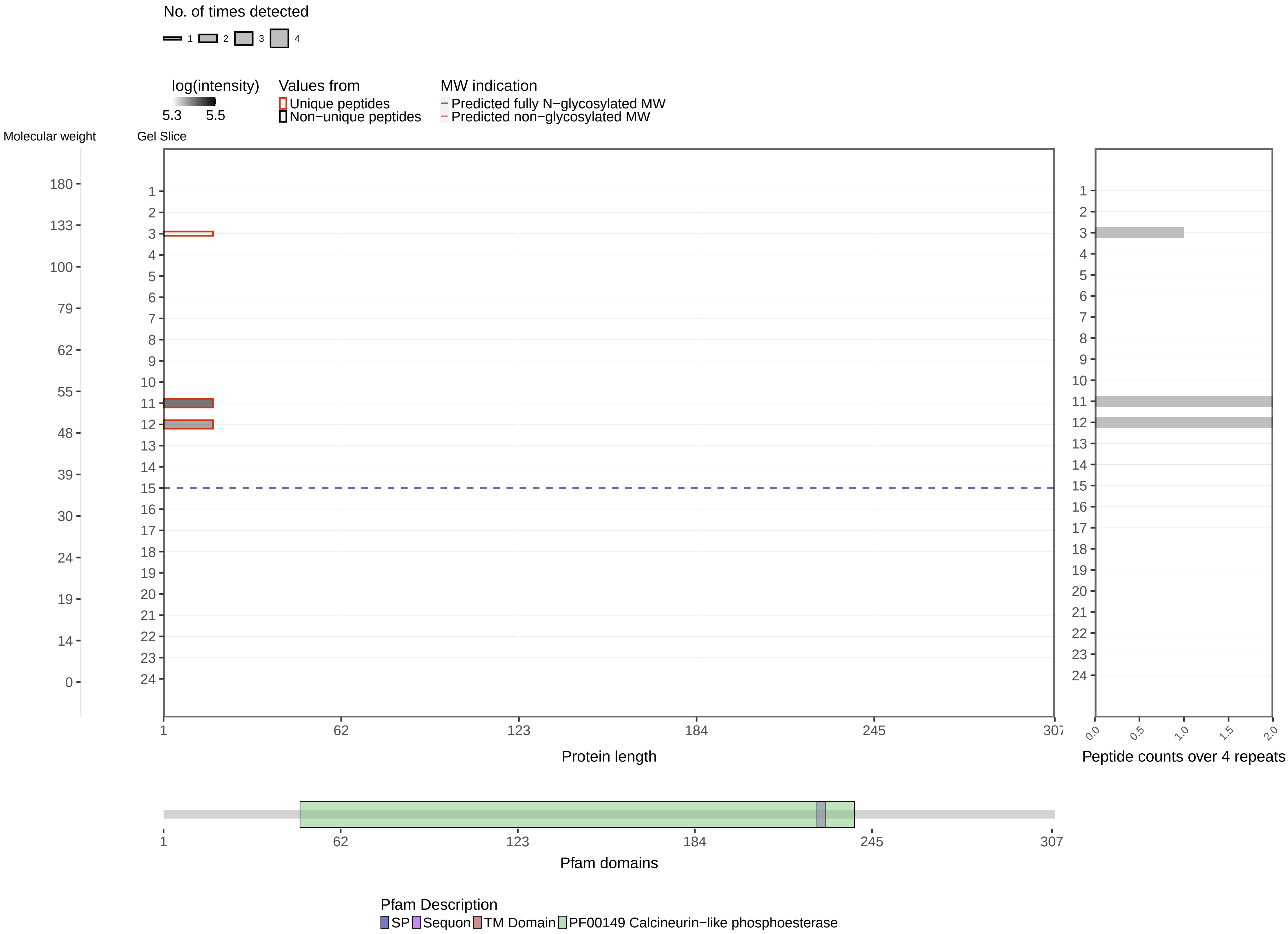


Protein sequence:

/KHFTDLFDYLPLTALIESOIFCLHGGLSPSLDTLDNIRALDRIOEVPHEGPMCDLLWSDPDDRCGWGISPRGAGYTFGODIASOFNHTNGLTLISRAHOLVMEGFNWCODKNVVTVFSAPNYCYRCGNMAAILEIGENMEONFLOFDPAPROIEPDTTRKTPDYFL*

### NbL08g10780.1 pyruvate_dehydrogenase_E1_component_subunit_alpha_mitochondrial−like [MW = 44 kDa]

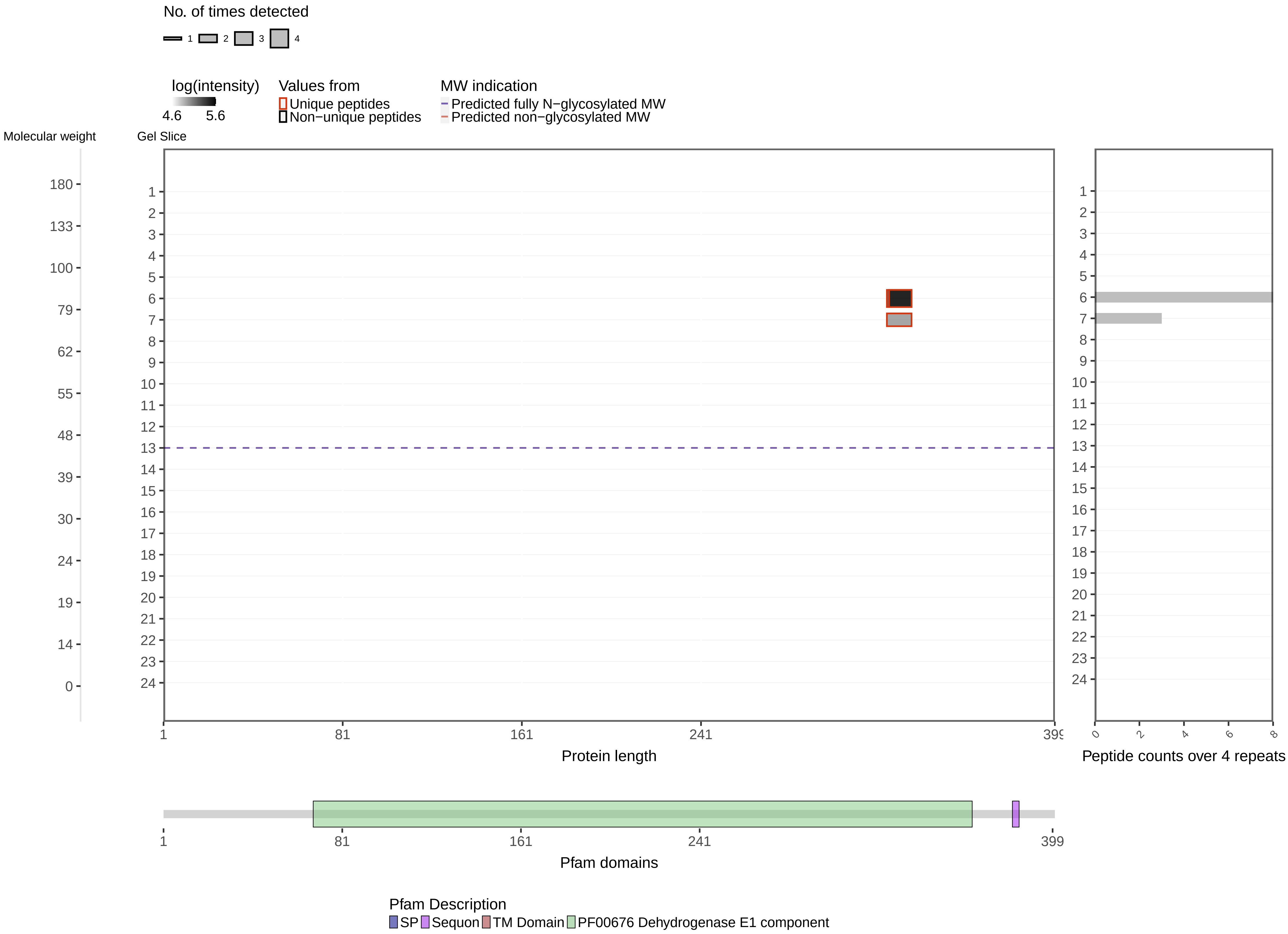


Protein sequence:

KEEHVSFAMYGDGAANQGQLFEALNMAALWDLPAILVCENNHYGMGTAEWRAAKSPAYYKRGDYVPGLKVDGMDALAVKQACKFAKEHALKNGPIILEMDTYRYHGHSMSDPGSTYRTRDEISGVRQERDPIERIRKLILAHDIATEKELKDIEKEKRKIVDEAIAKAKESPMPDPSELFTNVYVKGFGVEVCNCSNHQFIPSKLFTGCII*

### NbL08g10270.1 glyceraldehyde−3−phosphate_dehydrogenase_cytosolic [MW = 36.36 kDa]

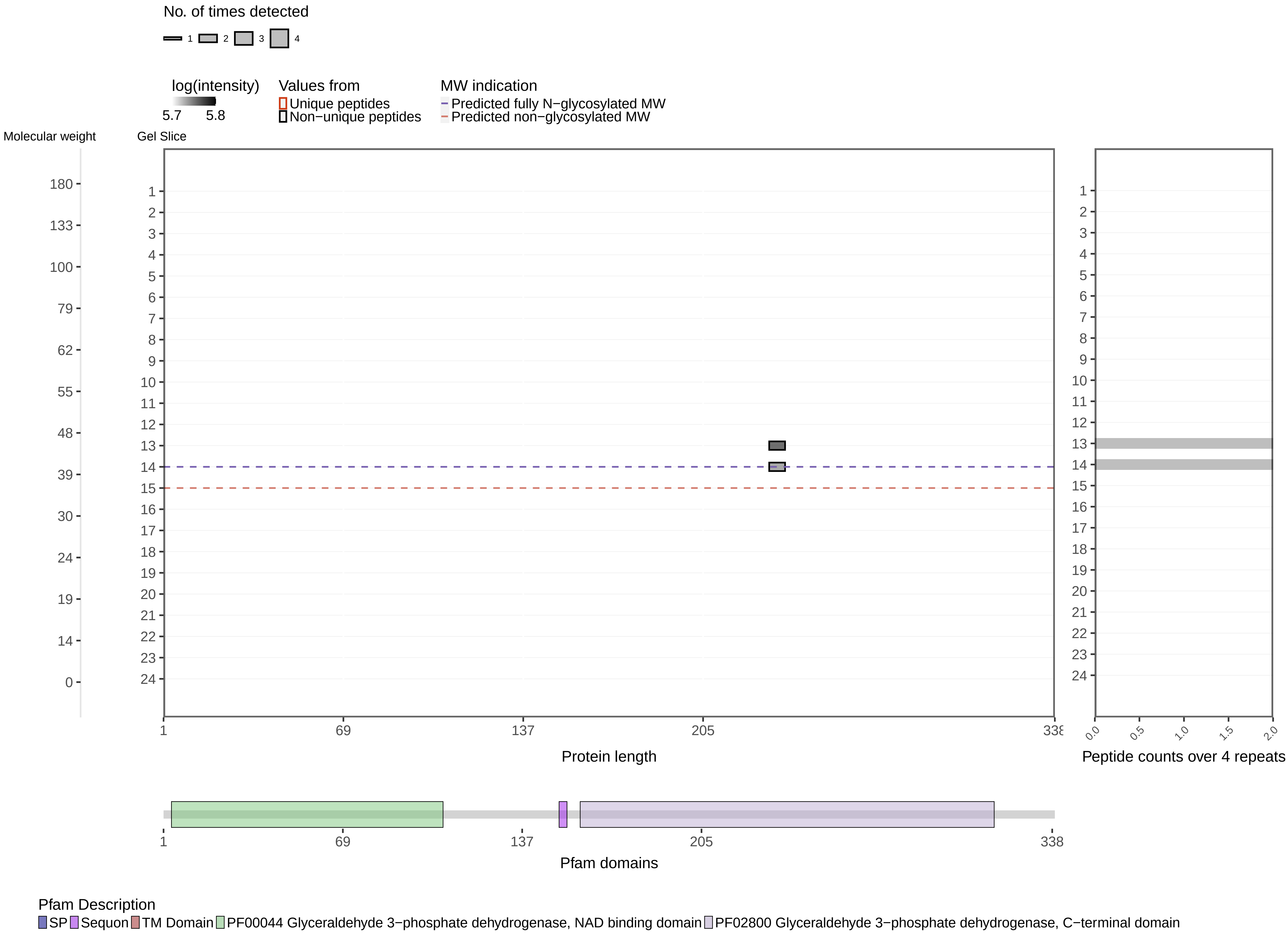


Protein sequence:

TTNCLAPLAKVINDKFGIVEGLMTTVHSITATQKTVDGPSMKDWRGGRAAAHNIIPSSTGAAKAVGKVLPALNGKLTGMSFRVPTVDVSVVDLTARLEKAATYDEIKAAIKAESEGKMKGILGYSEDDVVSSDFVGDCRSSIFDAKAGIALNGNFVKVVSWYDNEWGYSNRVIDLIRHMATVA*

### NbL08g10230.1 threonine−−tRNA_ligase_mitochondrial_1 [MW = 79.01 kDa]

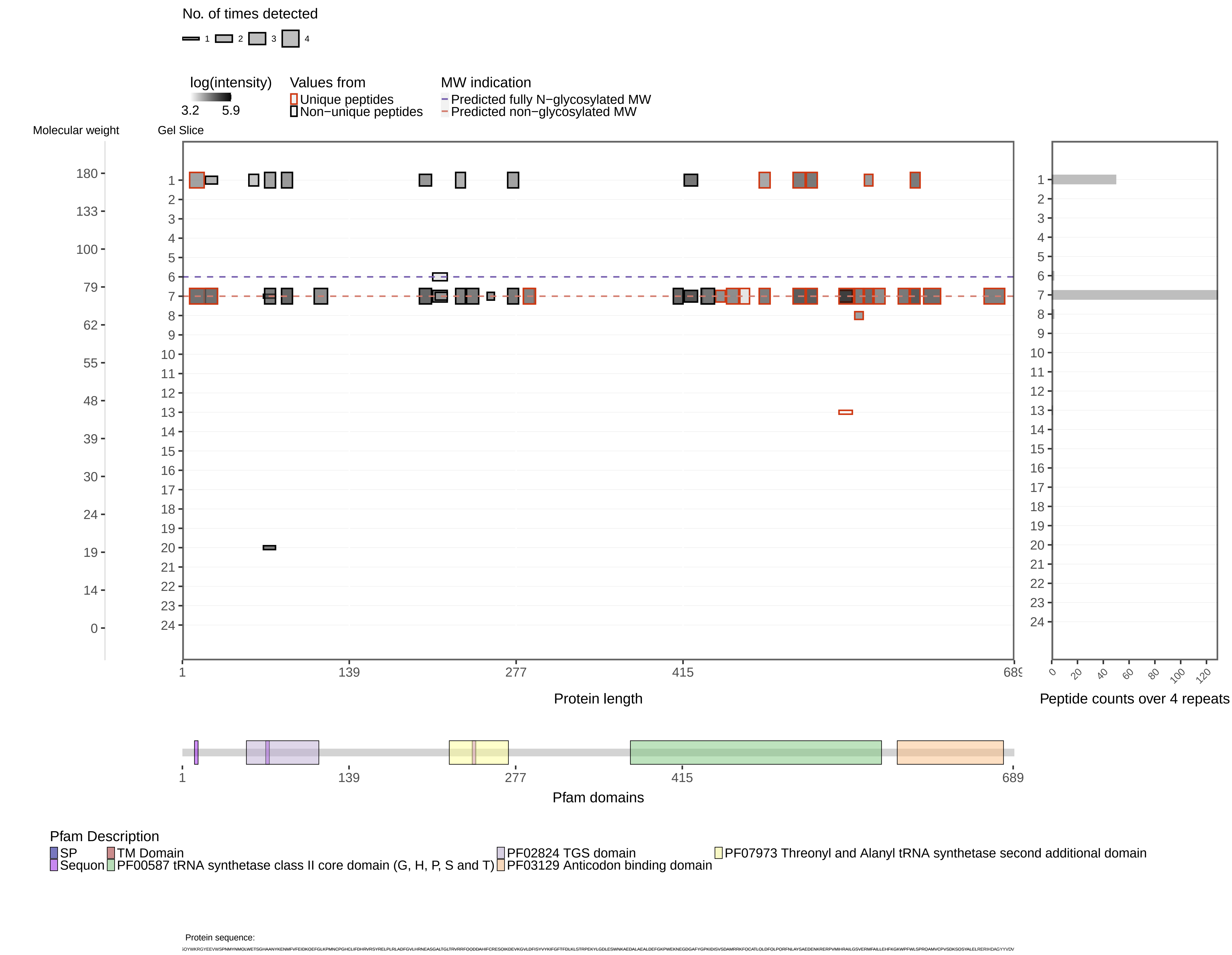

NbL08g10050.1 aldehyde_dehydrogenase_family_2_member_B7_mitochondrial−like [MW = 58.6 kDa]

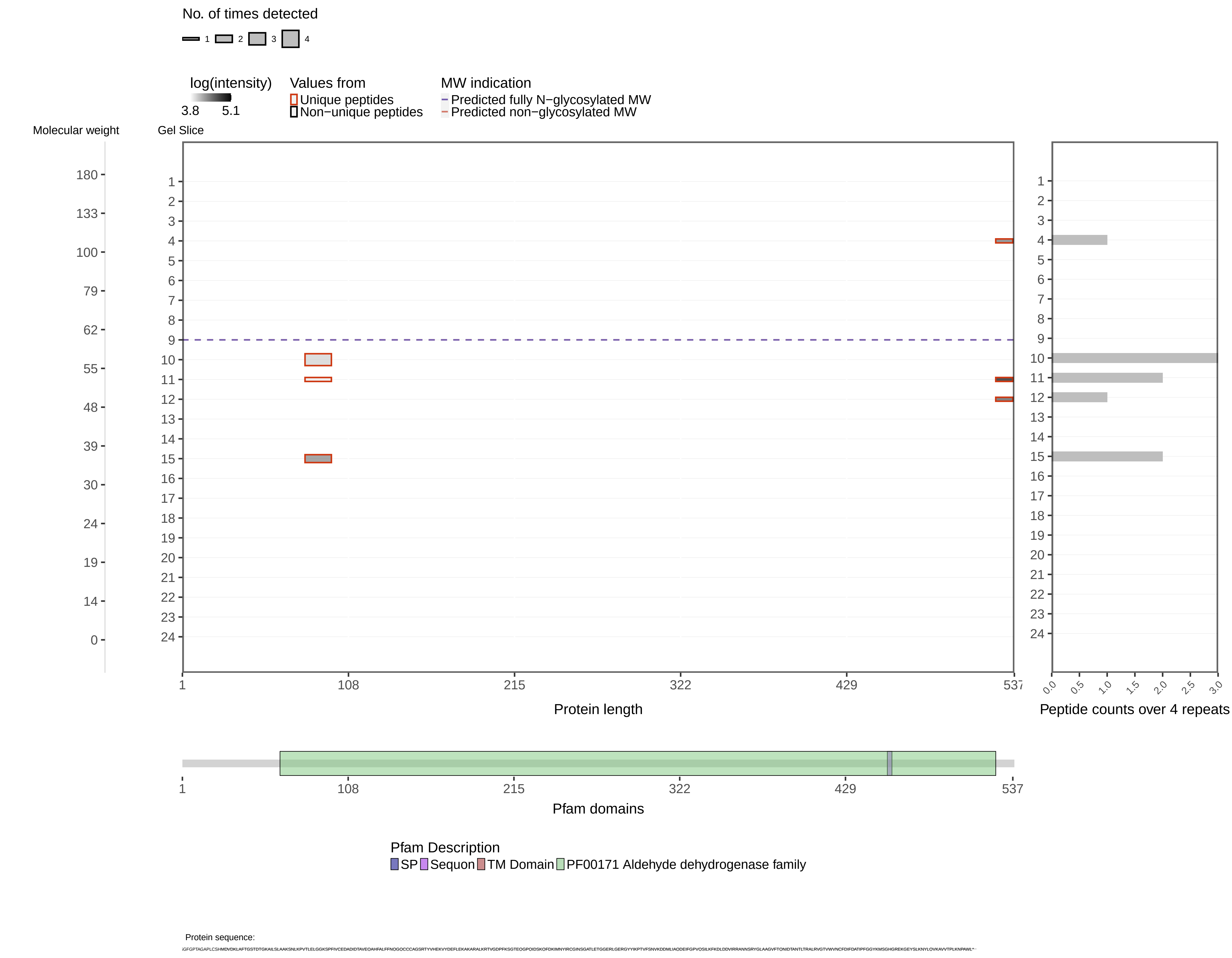

### NbL08g10040.1 40S_ribosomal_protein_S16 [MW = 16.7 kDa]

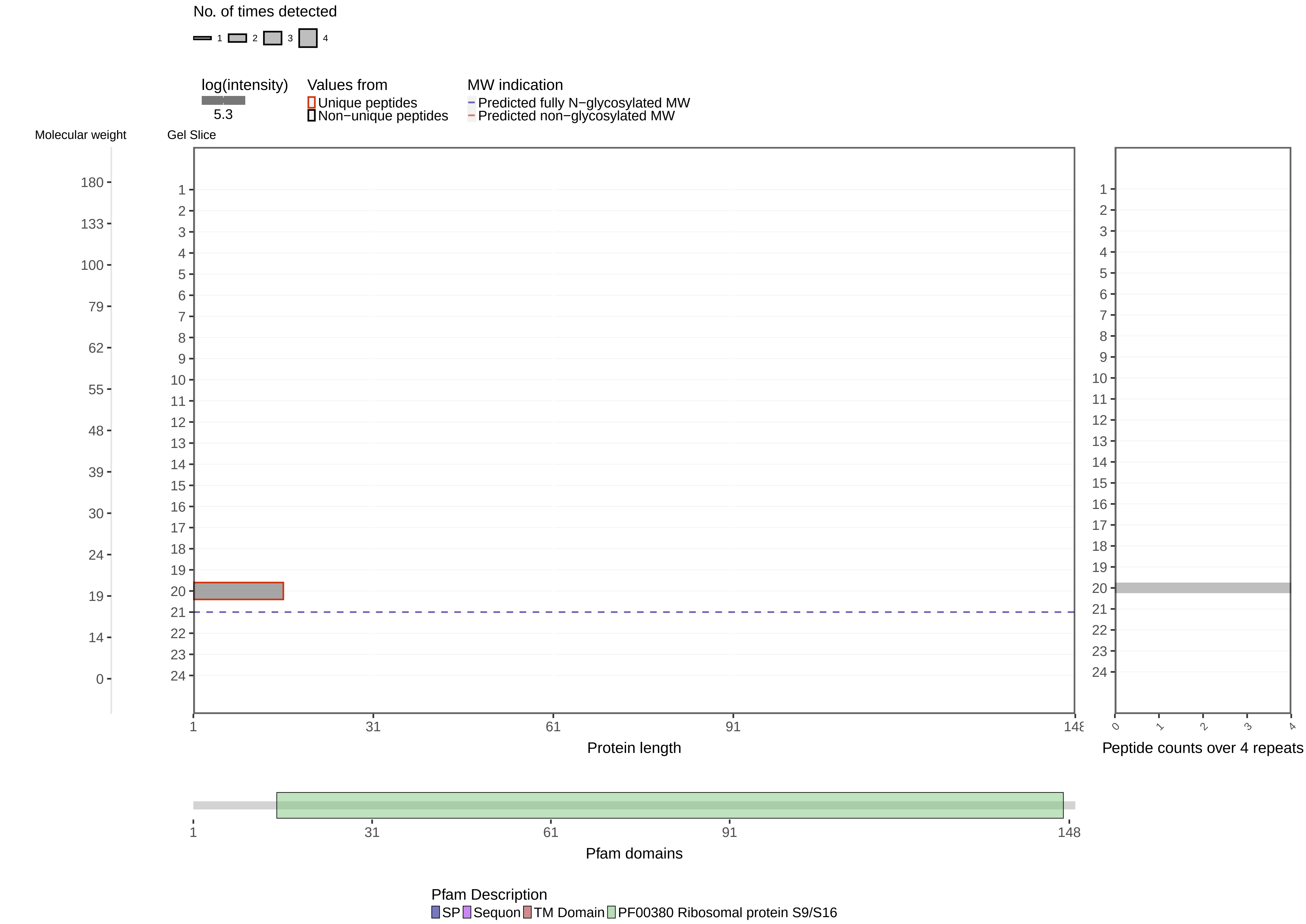


Protein sequence:

HRFAGVDMRIRVKGGGHTSQIYAIRQSIAKALVAFYQKYVDEQQKKEIKDILVRYDRTLLVADPRRCEPKKFGGRGARARFQKSYR*

### NbL08g09860.1 uncharacterized_protein_At4g28440−like [MW = 18.07 kDa]

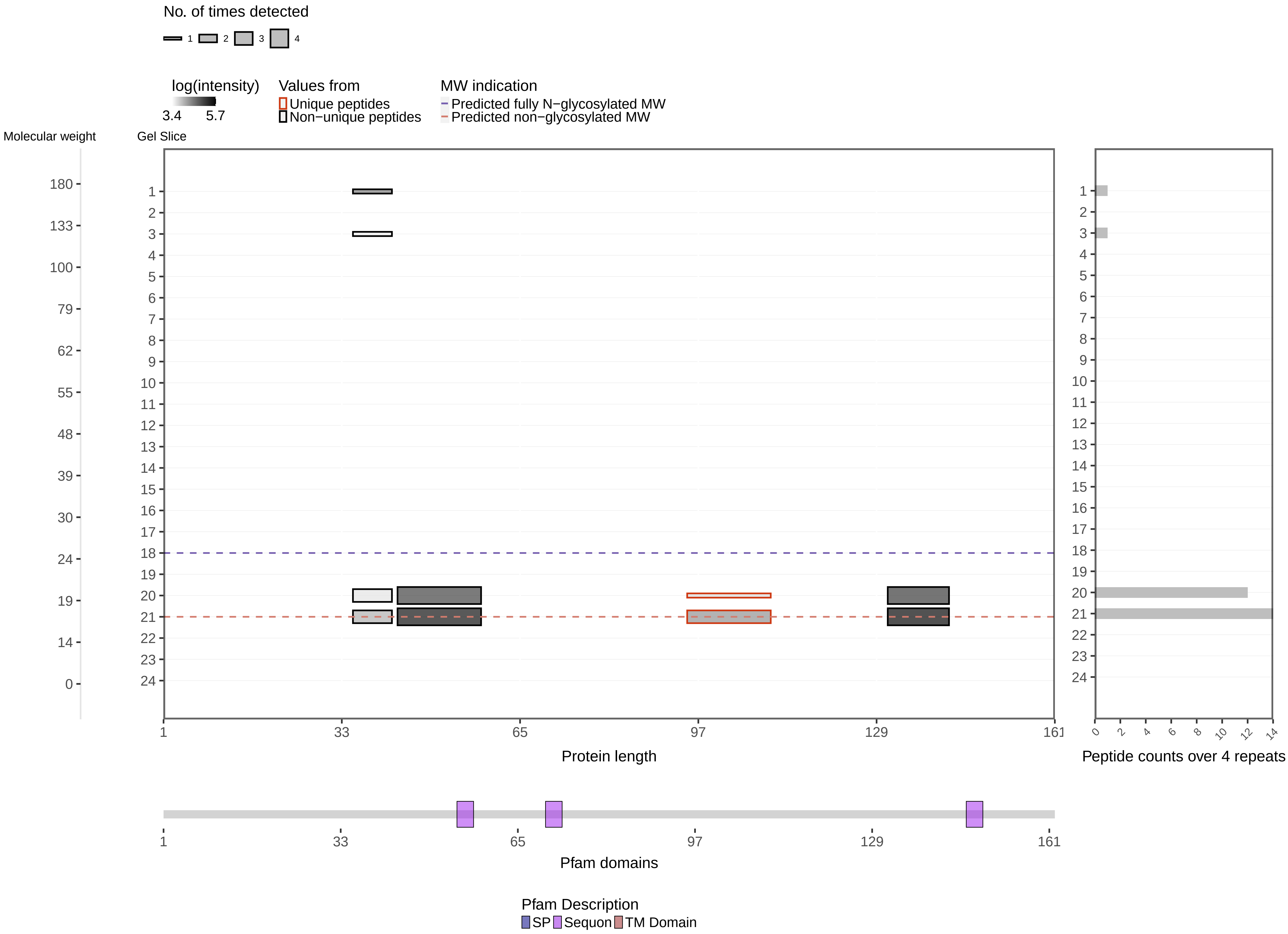


Protein sequence:

RGNQSRQMRLAECLVGDETGMIIFTARNDQVEMMTEGATVILRNAKIDMFKGSMRLAVDRWGRIEVTEPASFSVKEDNNLSLIEFEQVTVVEQ*

### NbL08g09850.1 carbonic_anhydrase_2−like [MW = 34.59 kDa]

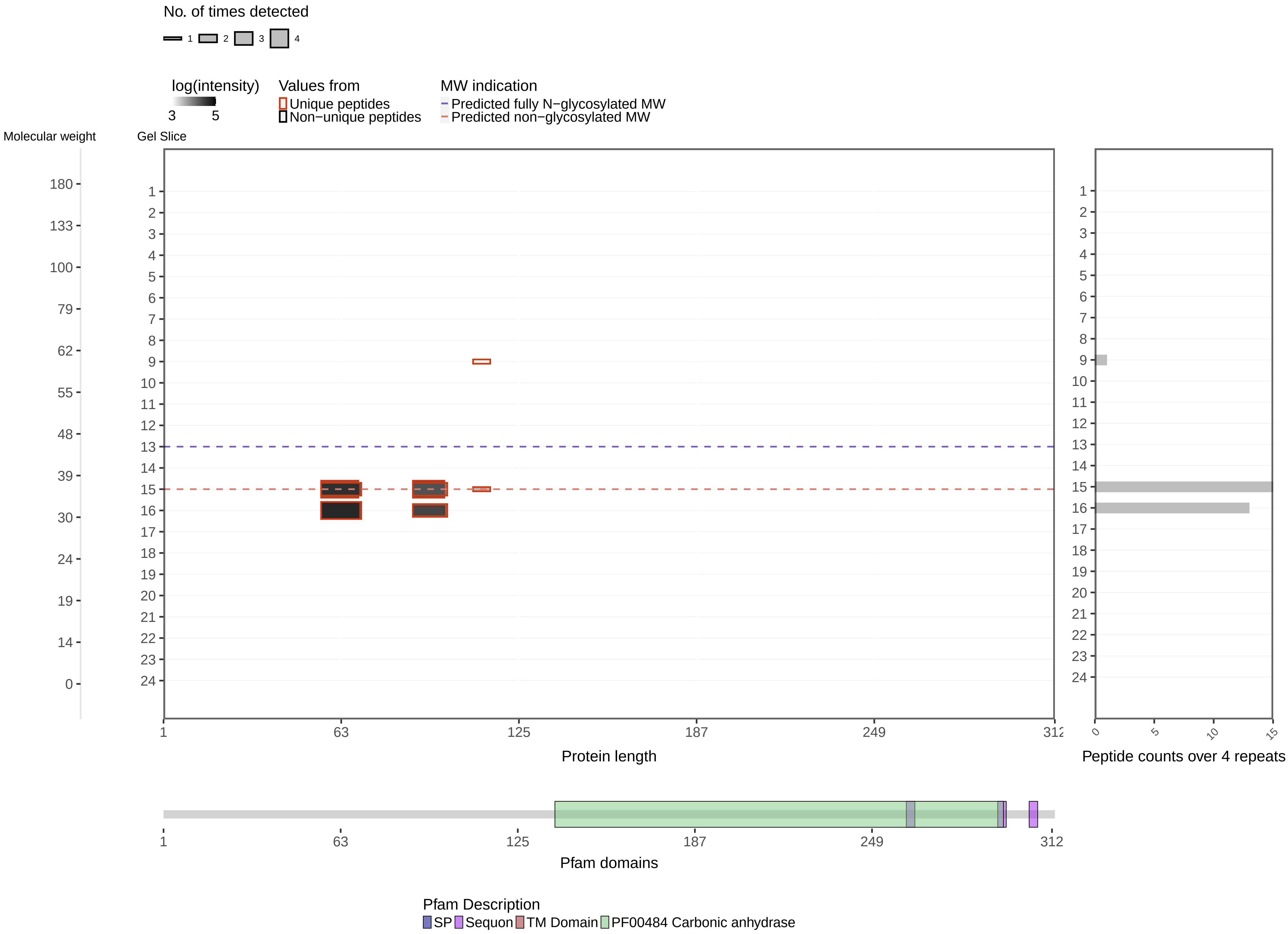


Protein sequence:

DSRVCPSHILNFQPGEAFVVRNIANMVPPFDQTKYSGVGAAVEYAVVHLKVENILVIGHSCCGGIKGLMSIPDDGSTNSDFIEEWVKICLPAKAKVKAECCHLDPTEQCTKLEKEAVNVSLGNLLTYPFVREAVVQKSLALKGGHYDFVNGSFELWNIDFNLTPSVSL*

### NbL08g09830.1 2−methylene−furan−3−one_reductase [MW = 41.2 kDa]

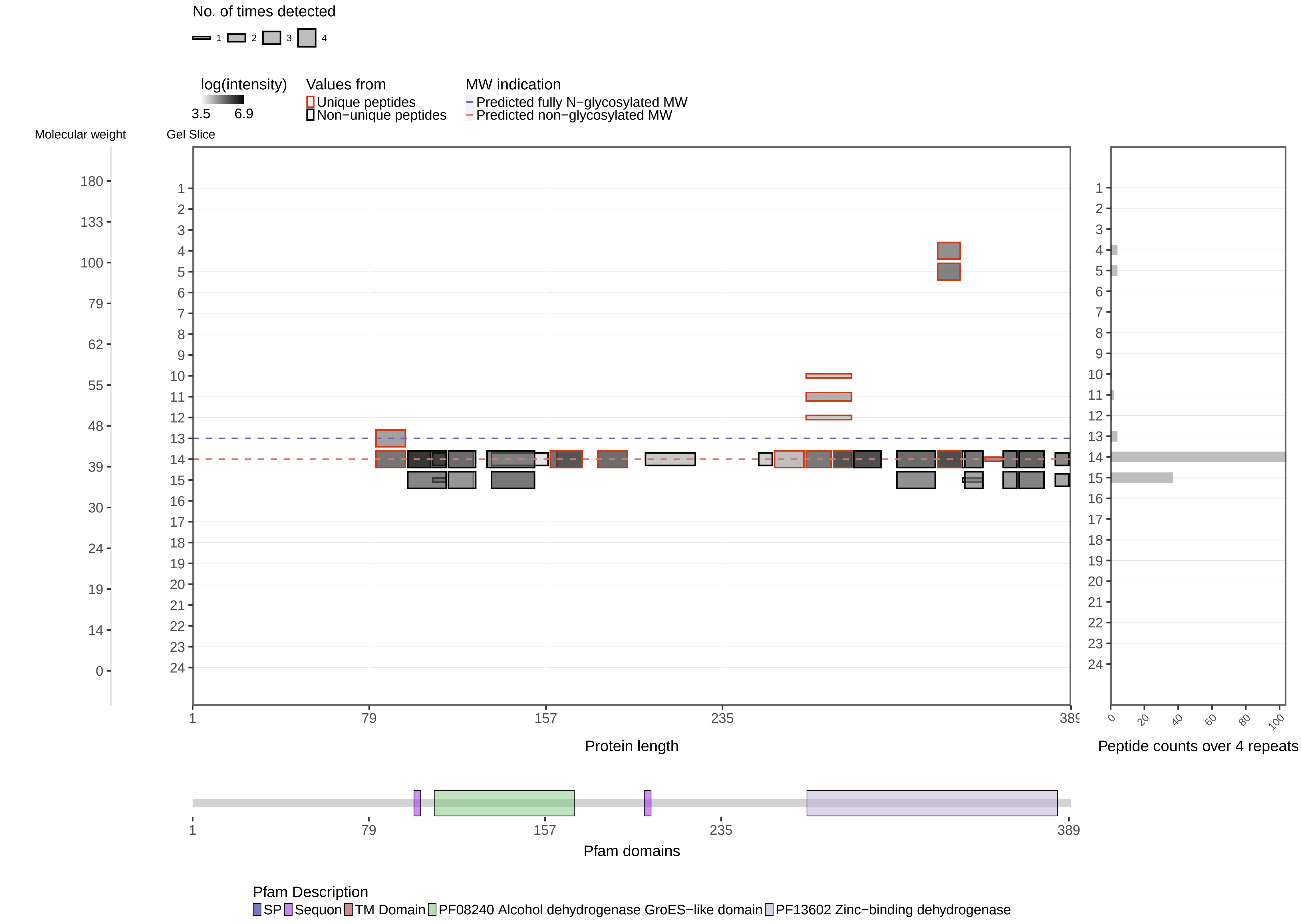


Protein sequence:

QFGSLAEYTAVEEKLVALKPKNLSFAEAASLPLAIETAYEGLEKAGFSAGKSILVLGGAGGVGSLVIQVAKHVFGASKVAATSSTGKLDLLKSLGADLAIDYTKENFEDLPEKFDVVYDTVGQGEKAVKAVKEGGSVVVLTGAVTPPGFRFVVTSNAEMLKKLKPYLESGKVTPVIDPKGPFPFDKVVEAFSYLETGRATGKVVIHPIP*

NbL08g09690.1 pyrrolidone−carboxylate_peptidase−like [MW = 23.64 kDa]

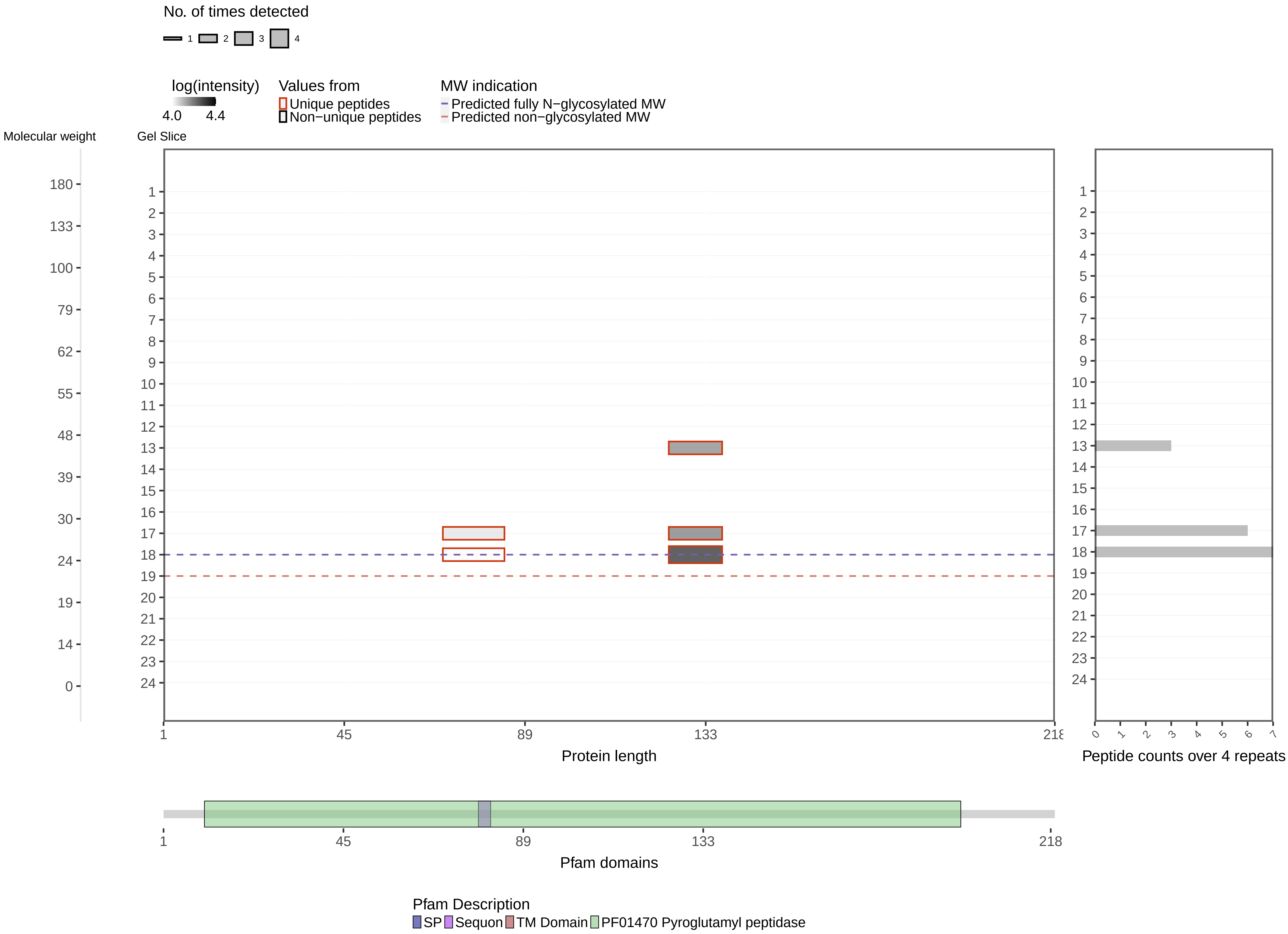


Protein sequence:

STKFAIERLAVNEATFRCADELGWQPQRLPIVPEDGGINQARKTSCSAEGILEFLKKKGFEVTMSDDAGRFVCNYVYYHSLRFAEKKGHKSIFVHVPTFNRVNQETQMEFVVALLEAIASTC*

NbL08g09670.1 14−dihydroxy−2−naphthoyl−CoA_synthase_peroxisomal−like [MW = 36.72 kDa]

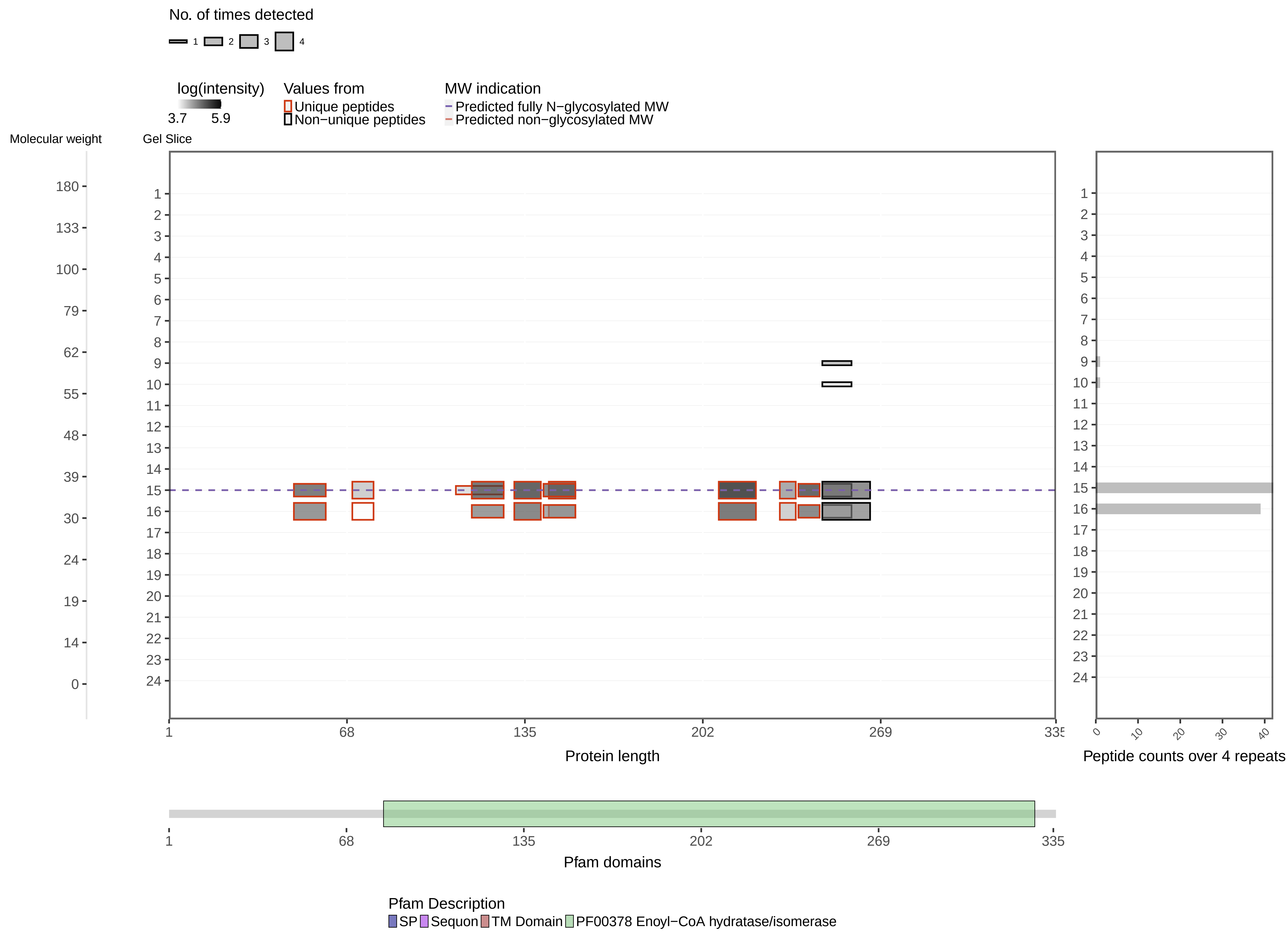


Protein sequence:

NVLDLQVQIRRLPKPVIAMVAGYAVGGGHILHMVCDLTIAADNAIFGQTGPKVGSFDAGYGSSIMSRLVGPKKAREMWFLTRFYTASEAEKMGLVNTVVPLDKLEQETIKWCREILRNSPTAIRVLKSALNAVDDGHAGLQGLGGDATLLFYGTEEGSEGKNAYNERRRPDFSKFPRLP*

NbL08g09640.1 ATP−citrate_synthase_alpha_chain_protein_2 [MW = 46.53 kDa]

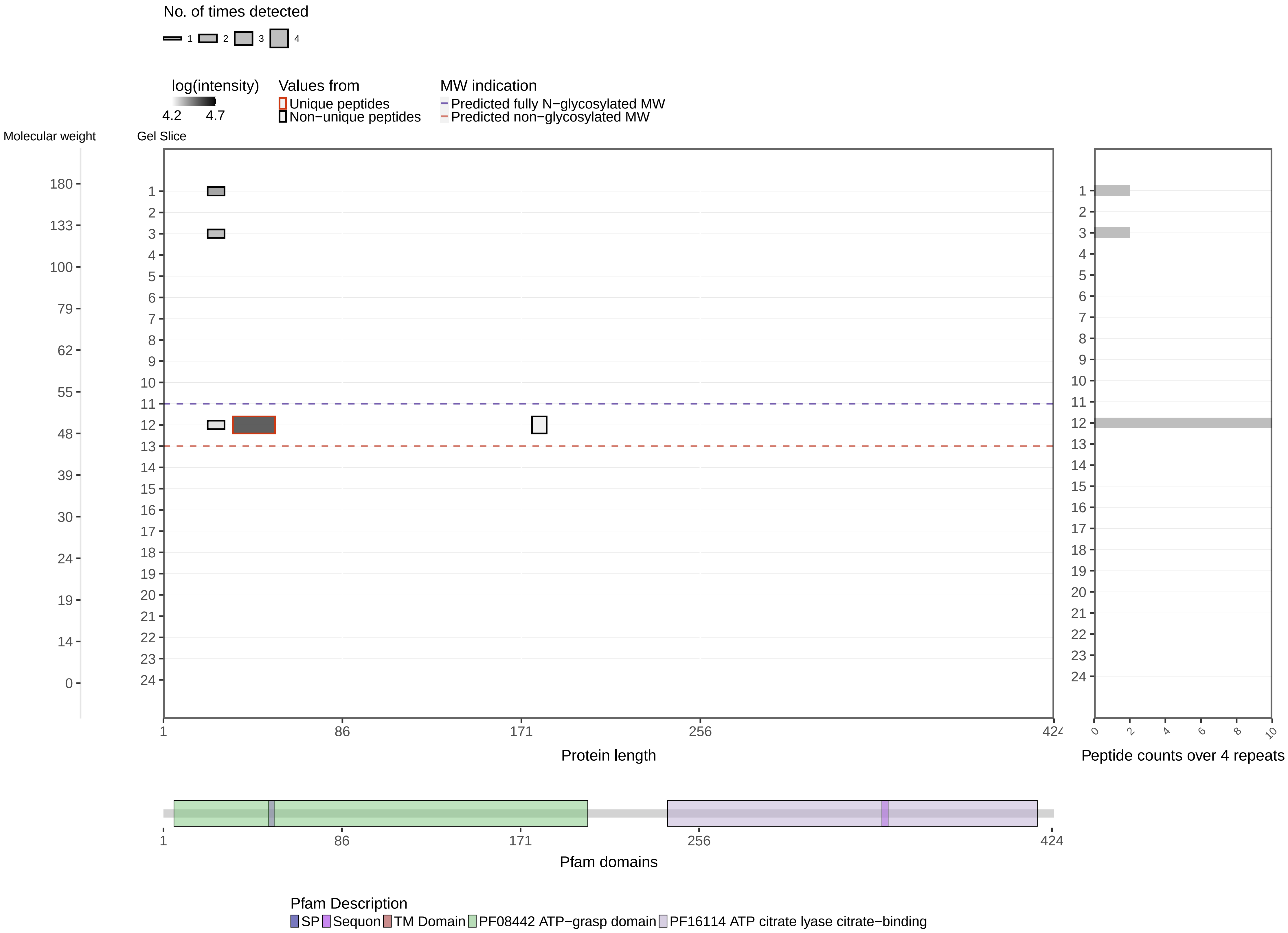


Protein sequence:

INPFALVEGKPYPLDMRGELDDTAAFKNFKKWGNIEFPLPFGRVMSATESFIHGLDEKTSASLKFTVLNPKGRIWTMVAGGGASVIYADTVGDLGYASELGNYAEYSGAPNEEEVLQYARVVIDCATANPDGRKRALVIGGGIANFTDVAATFSGIIRALKEKESKLKAARMQIFVRRGGPNYQKGLAKMRSLGEEIGIPIEVYGPEATMTGICKQAIECITPAA*

### NbL08g09520.1 alpha−glucan_water_dikinase_chloroplastic [MW = 166.69 kDa]

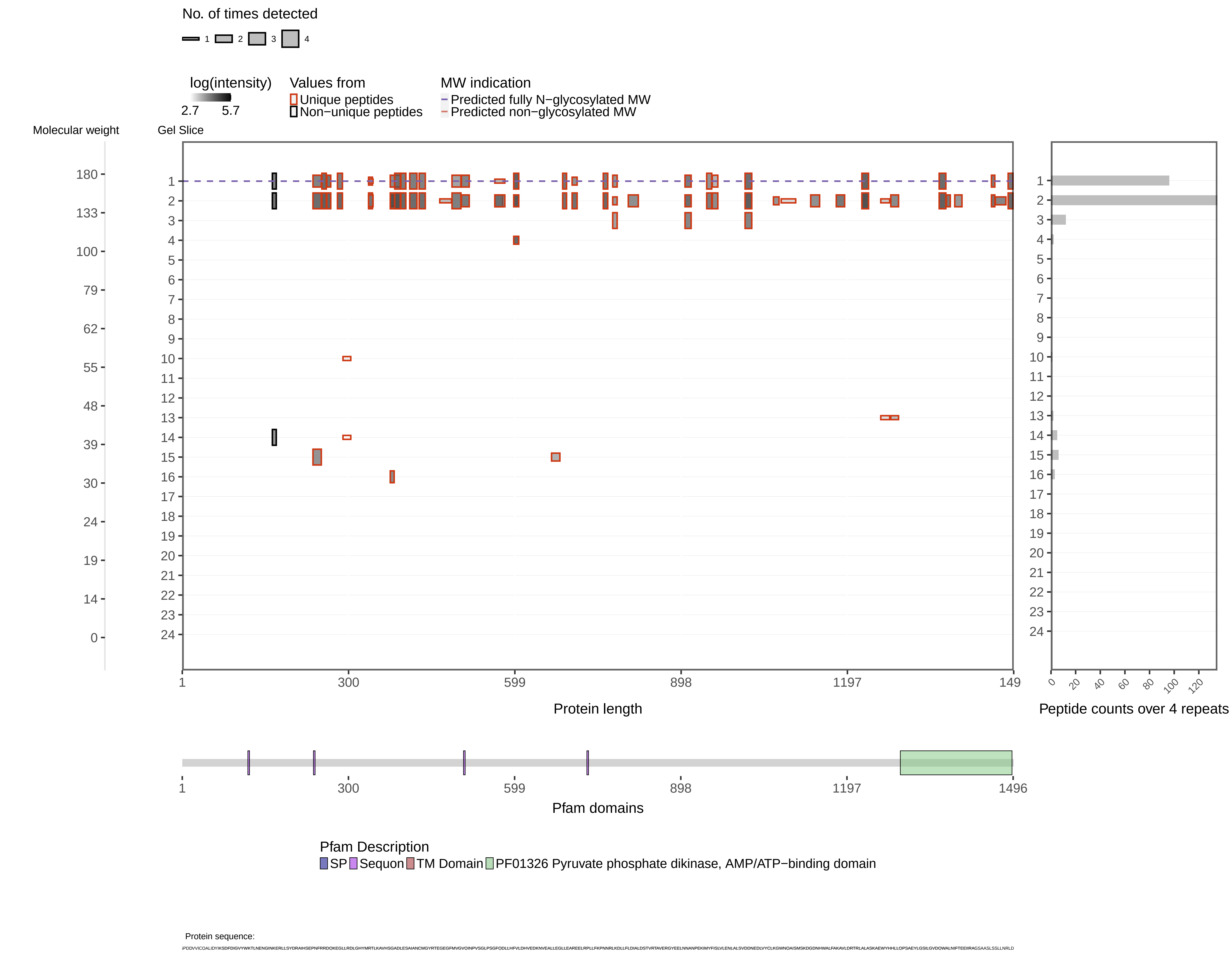

NbL08g09480.1 glutamate−−glyoxylate_aminotransferase_2 [MW = 53.5 kDa]

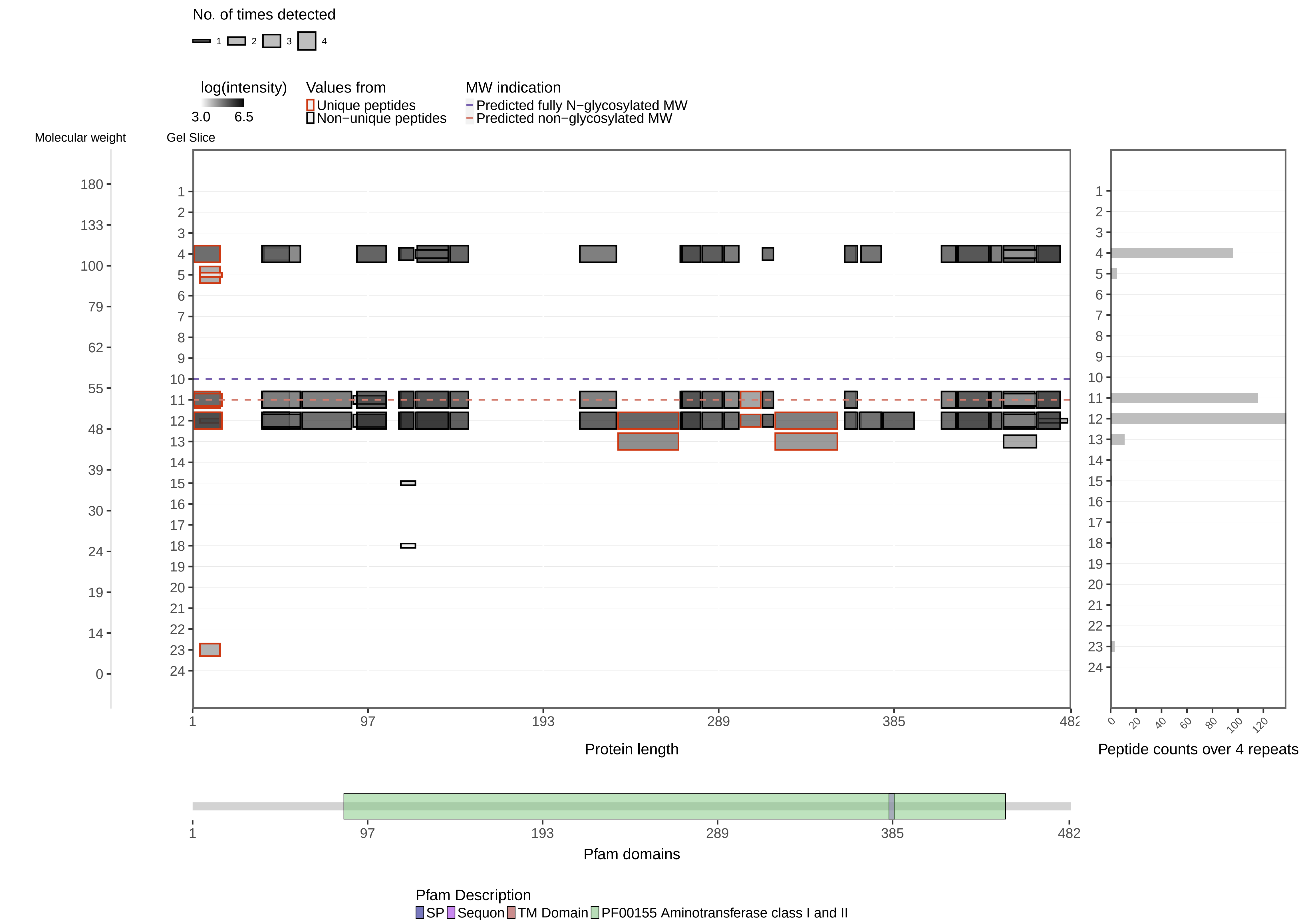


Protein sequence:

ANLKQIIQFCHQESLVLLGDEVYQQNIYQDERPFVSARKVLLDMGPPISKELQLVSFHTVSKGYWGECGQRGGYFEMTNIPPKSVEEIYKVASISLSPNVAGQIFMGLMVNPPKPGDISYDQYVRESKGILESLRKRAHMMTDGFNSCRNVVCNFTEGAMYSFPQIRLPPRAIEAAKKLGKVPDVFYCLRLLEATGISTVPGSGFGQKEGVFHLRTTILPAEEDMPAIMSSFKKFNDEFMEQYEDHRGYSRM*

### NbL08g09090.1 uncharacterized_acetyltransferase_At3g50280−like [MW = 49.43 kDa]

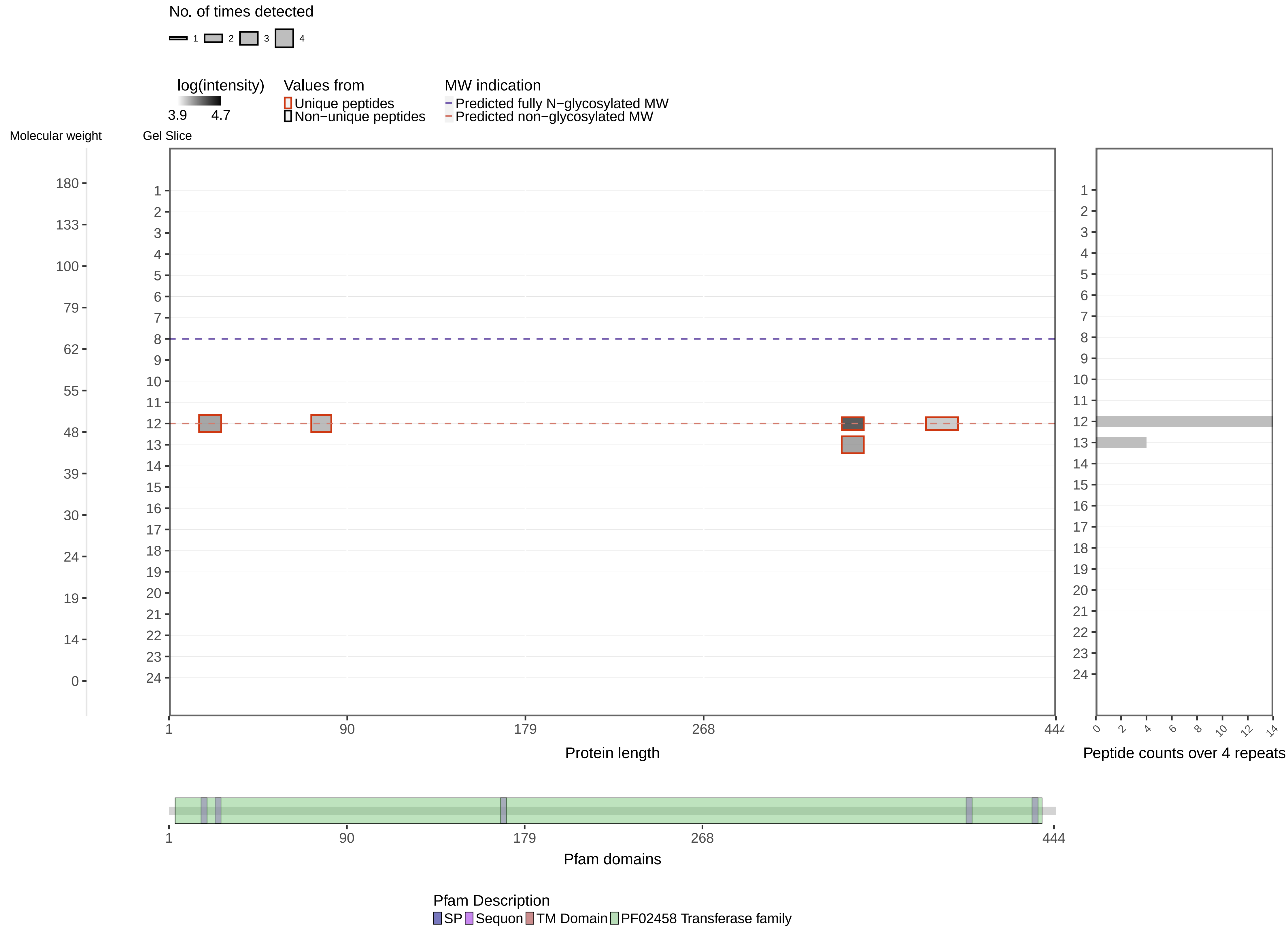


Protein sequence:

IHLPIKLDDEKLFEVFEIPHLKERIFHLSKESIAKLKEKANSEMGIKSISSLQAFLAHLWRSVTRCRKLDAKEEVVIGLIIGTRSRLNPPLPEGYFGNAIHAKRVKTTAGELLENGLGWAAMELNKVVNSQNSEEVVKIYKEWAENPVLFSKSTVFVGSKLAISSSPRYNIYGTDFGWGKPVAVRSGMANKSDGKITLFPGVEEGSVDIEVCSLPKTLQAMENDSEFMEAVTN*

### NbL08g09050.1 40S_ribosomal_protein_S6−like [MW = 28.36 kDa]

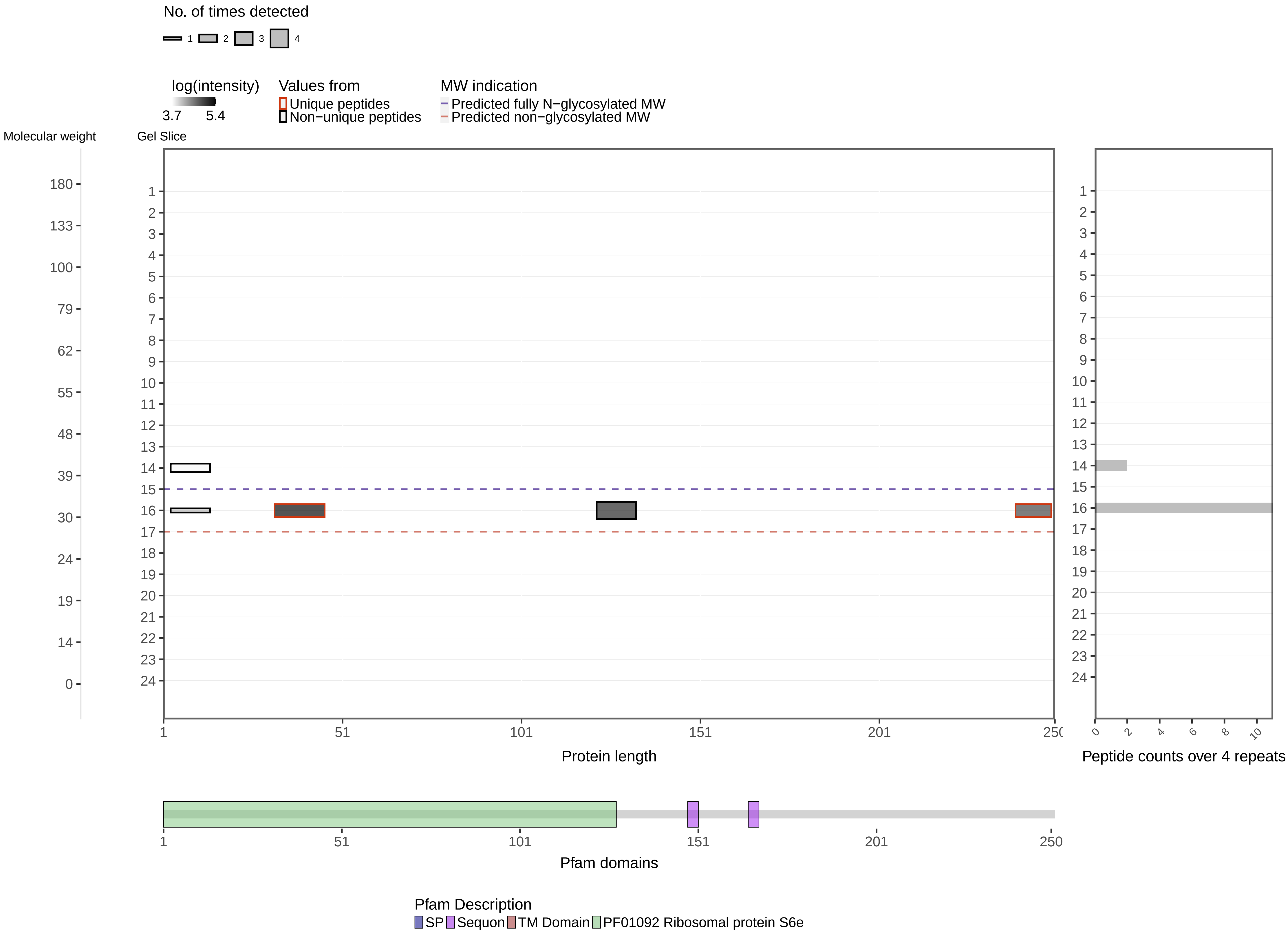


Protein sequence:

/LNLVIVKKGEKDLPGLTDTEKPRMRGPKRASKIRKLFNLSKEDDVRKYVNTYRRNFTTKSGKKASKAPKIQRLVTPLTLQRKRARIADKKKRIAKAKSEAADYQKLLATRLKEQRERRSESLAKKRSRLSAASKPSVAA*

### NbL08g08920.1 upstream_activation_factor_subunit_UAF30−like [MW = 13.22 kDa]

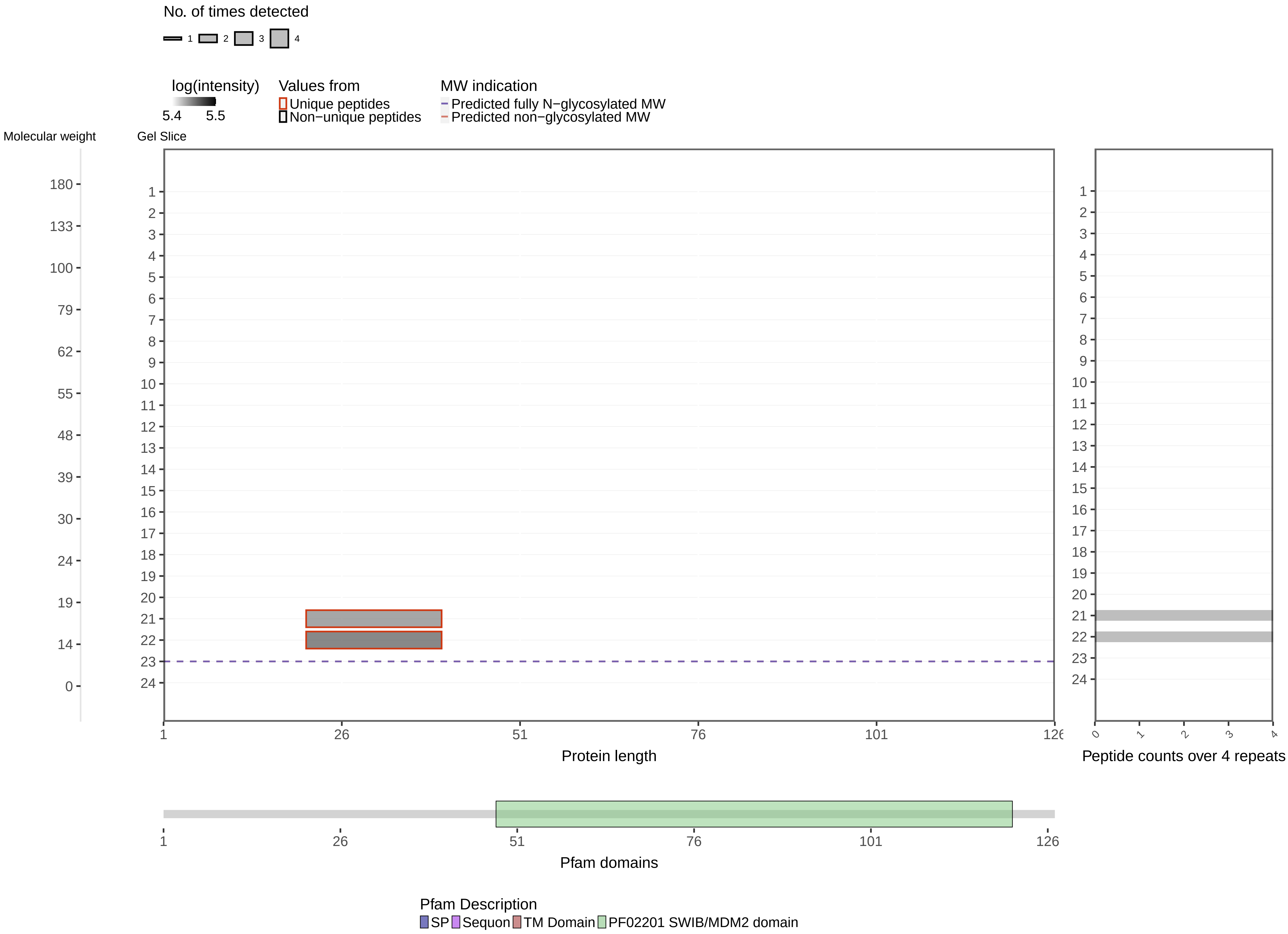


Protein sequence:

PLPVSPALGKFVGSSEISRTDAVKKVWDYIKTNNLQNPANKKEINCDDKLKTIFAGKDKVGFLEIAKLLSSHFQKAS*

### NbL08g08790.1 probable_methyltransferase_PMT20 [MW = 56.91 kDa]

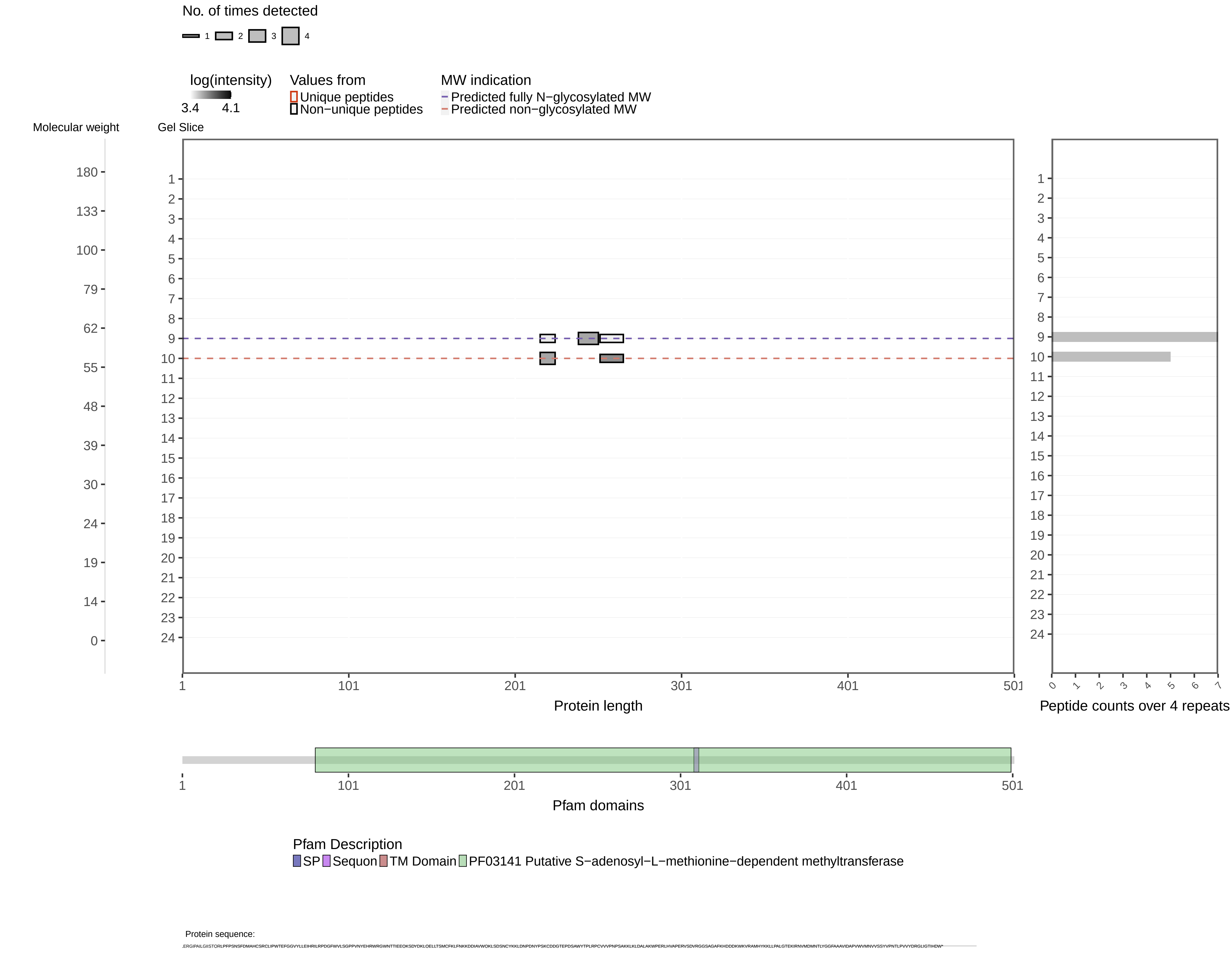

NbL08g08510.1 lipoyl_synthase_2_mitochondrial−like [MW = 42.51 kDa]

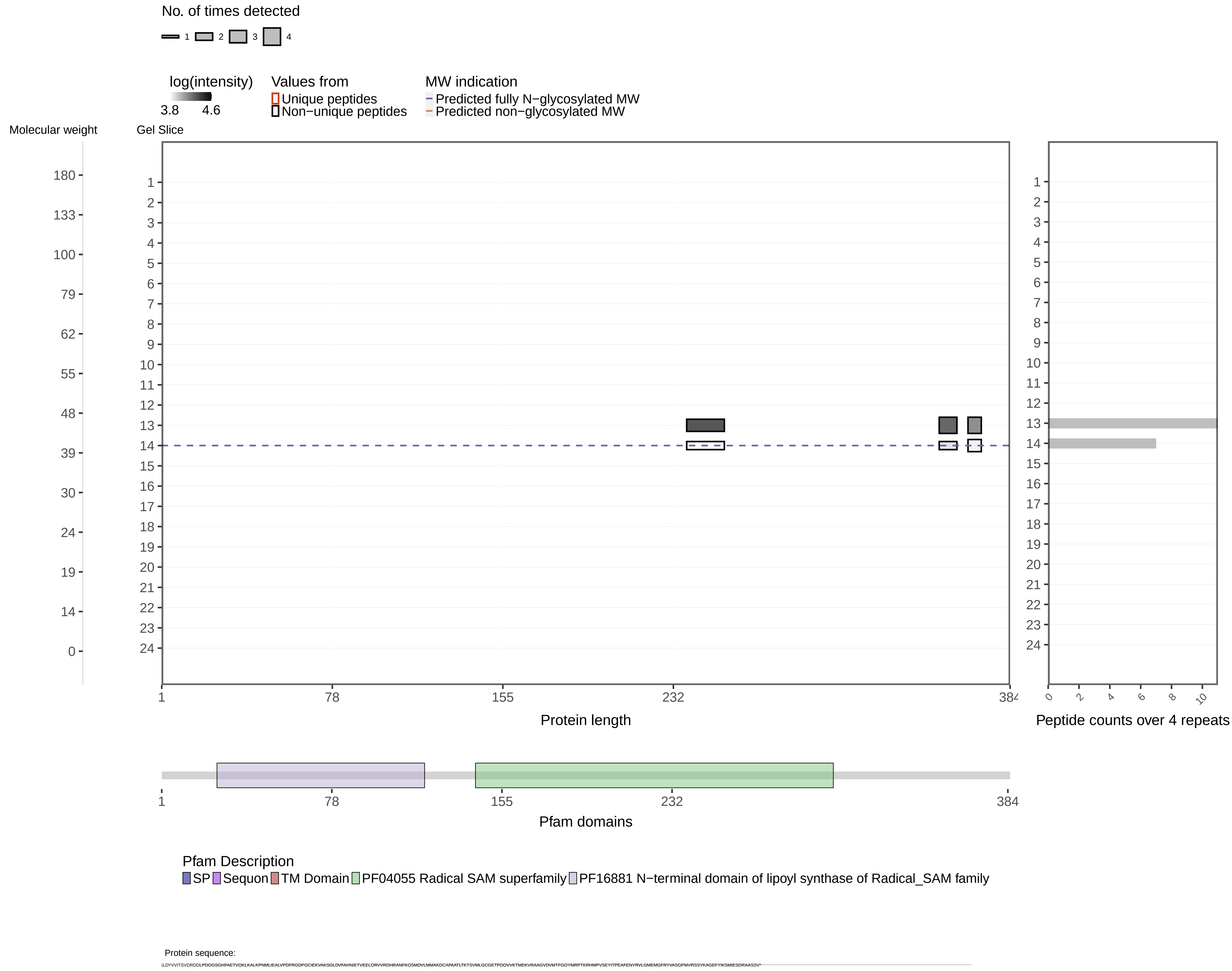

### NbL08g08460.1 5−aromatic_acyltransferase−like [MW = 50.22 kDa]

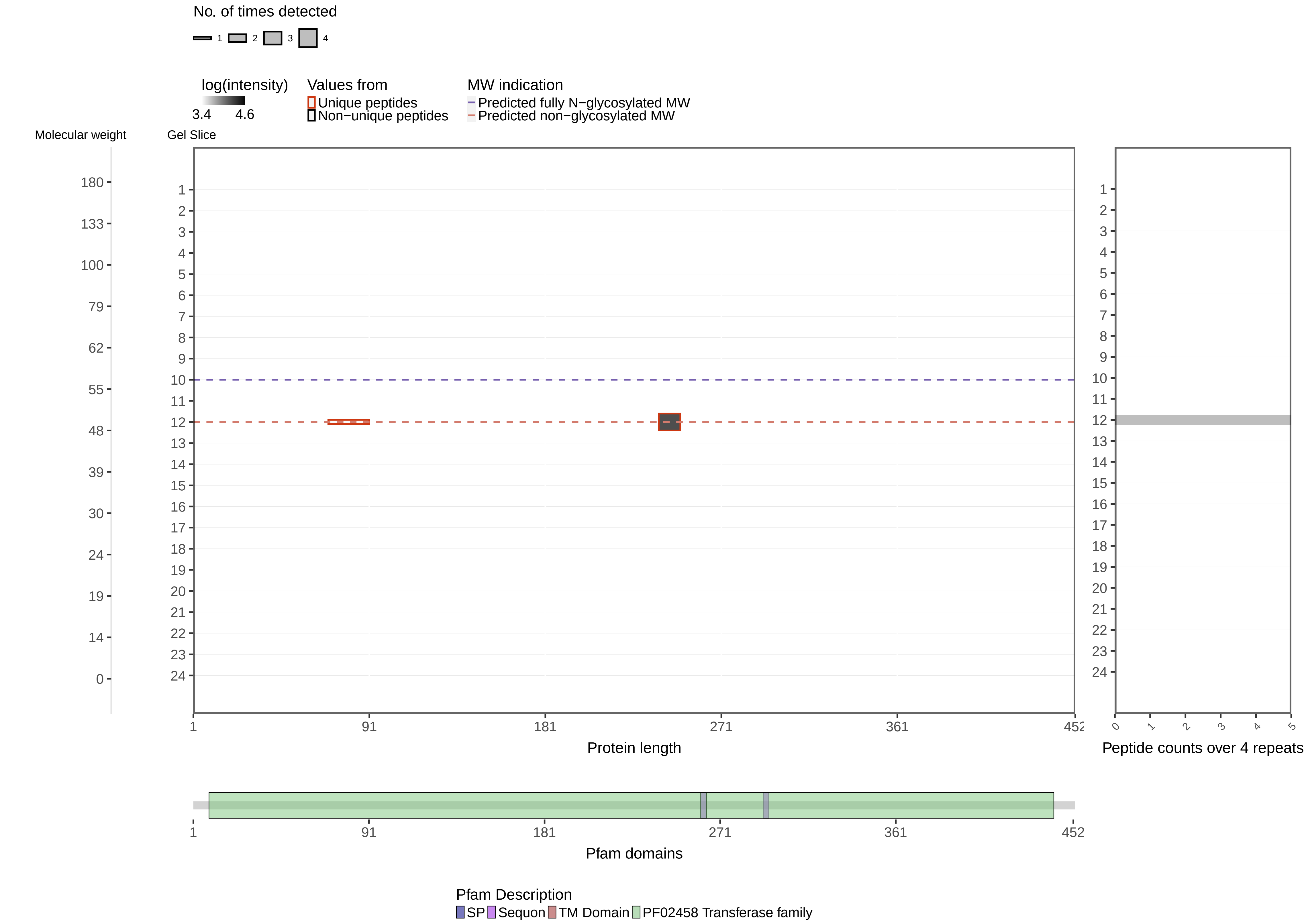


Protein sequence:

MSIWDEMQKFKLEMCDIVTPLDKVRGTFIITQDGITKLKNLMLSRRPNLSHVTSFTVTCAYVWTCLIKSEDSIREEIEENVTELFGCVADCRSRLNPPLPPSYFGNCLVPLVARAGHADLAGKEGFTIAVKVIGETIQKRMKDEDYILSGKWFKEFCSVDIKRTLSIAGSPKLDLYAADFCWGRPEKLEFVSIDNGDGVSMSLSKSKDSDGNLEIGLSLPKTRMNAFAALFTHGLSFL*

NbL08g08250.1 26S_proteasome_non−ATPase_regulatory_subunit_2_homolog_A−like [MW = 98.12 kDa]

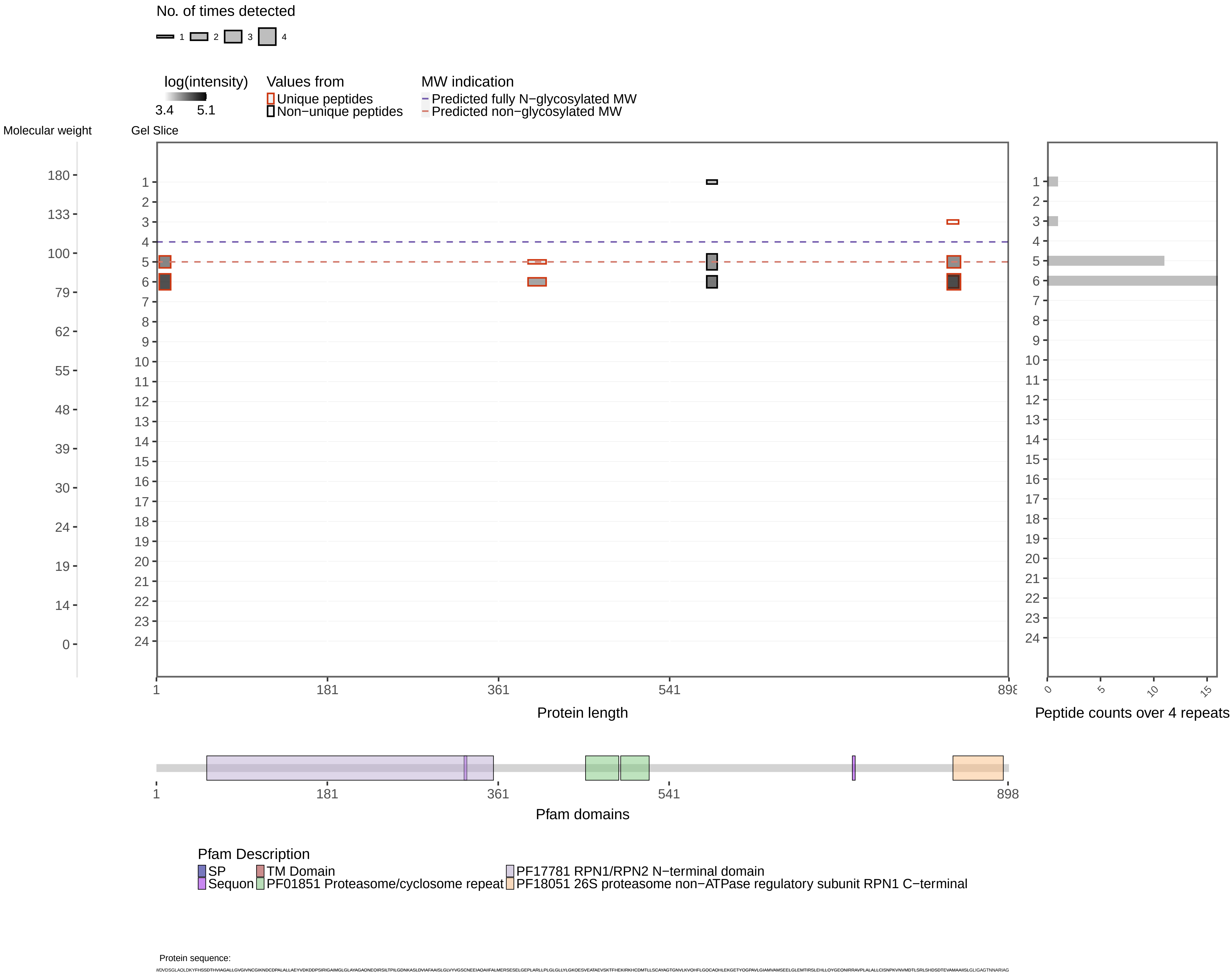

NbL08g08200.1 acyl−coenzyme_A_oxidase_3_peroxisomal−like [MW = 76.03 kDa]

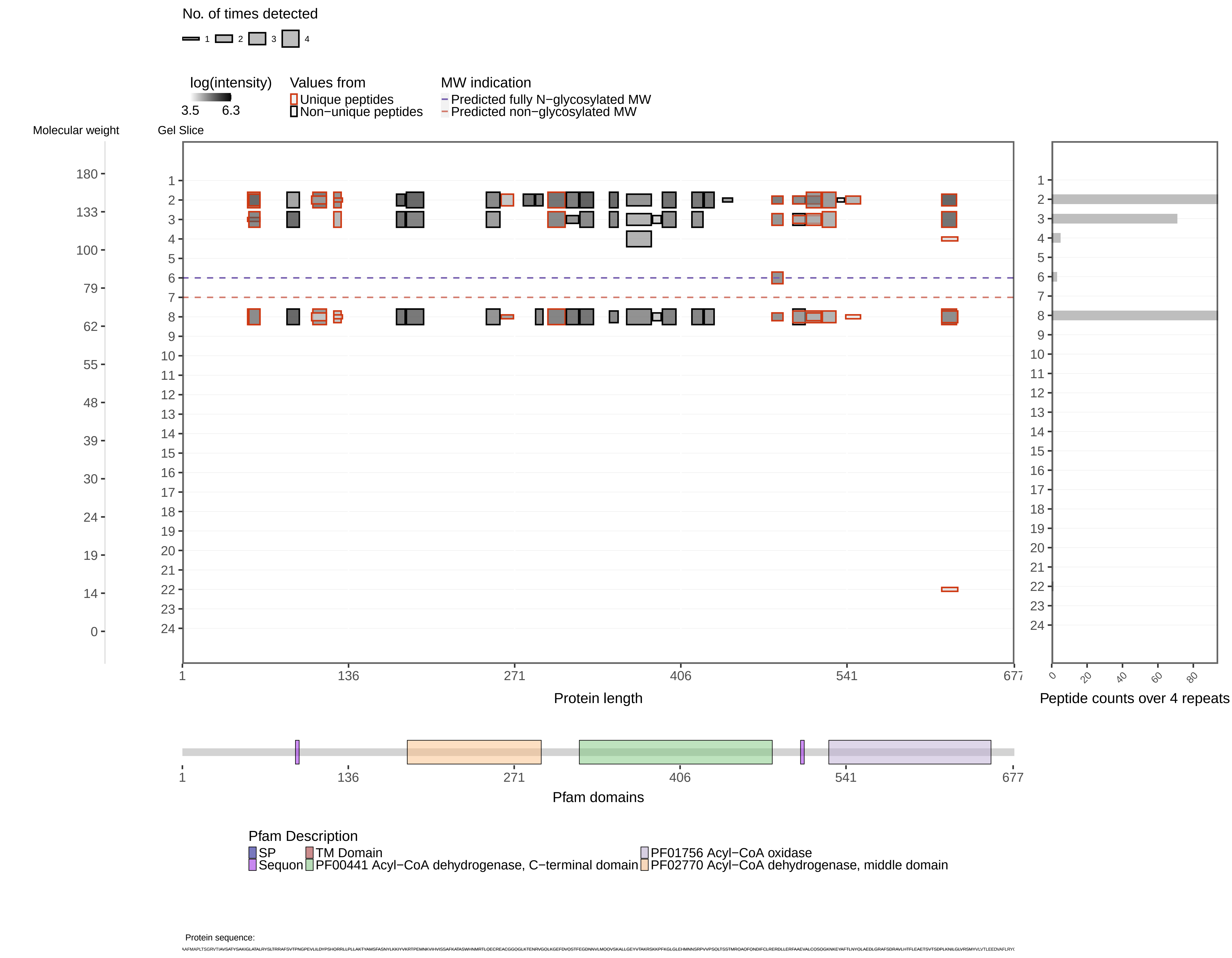

### NbL08g08030.1 ubiquitin−NEDD8−like_protein_RUB2 [MW = 17.09 kDa]

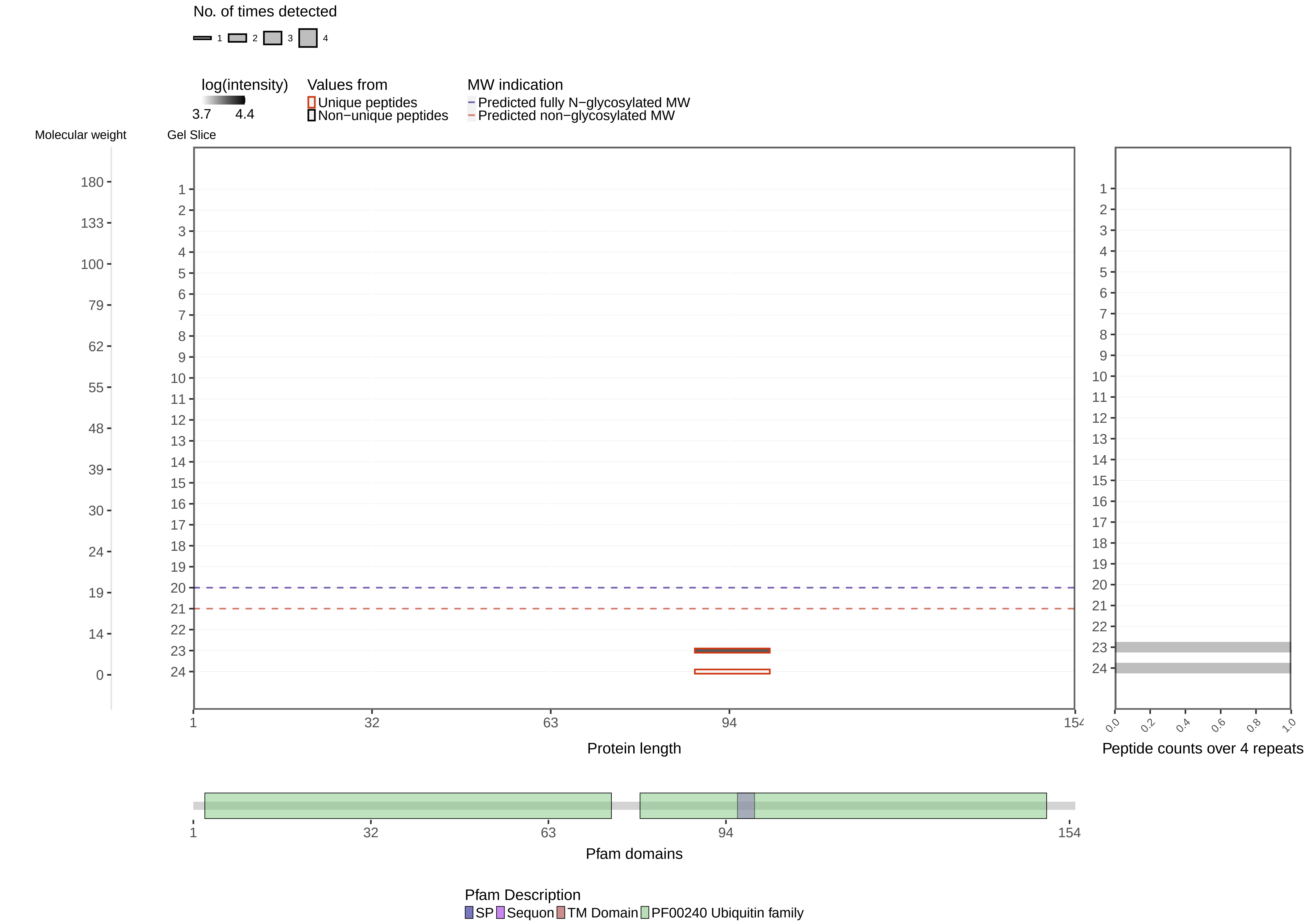


Protein sequence:

ESTLHLVLRLRGGTMIKVKTLTGKEIEIDIEPNDTIDRIKERVEEKEGIPPVQQRLIYAGKQLADDKTAKDYNIEGGSVLHLVLALRGGN*

NbL08g07930.1 putative_4−hydroxy−4−methyl−2−oxoglutarate_aldolase_3 [MW = 20.55 kDa]

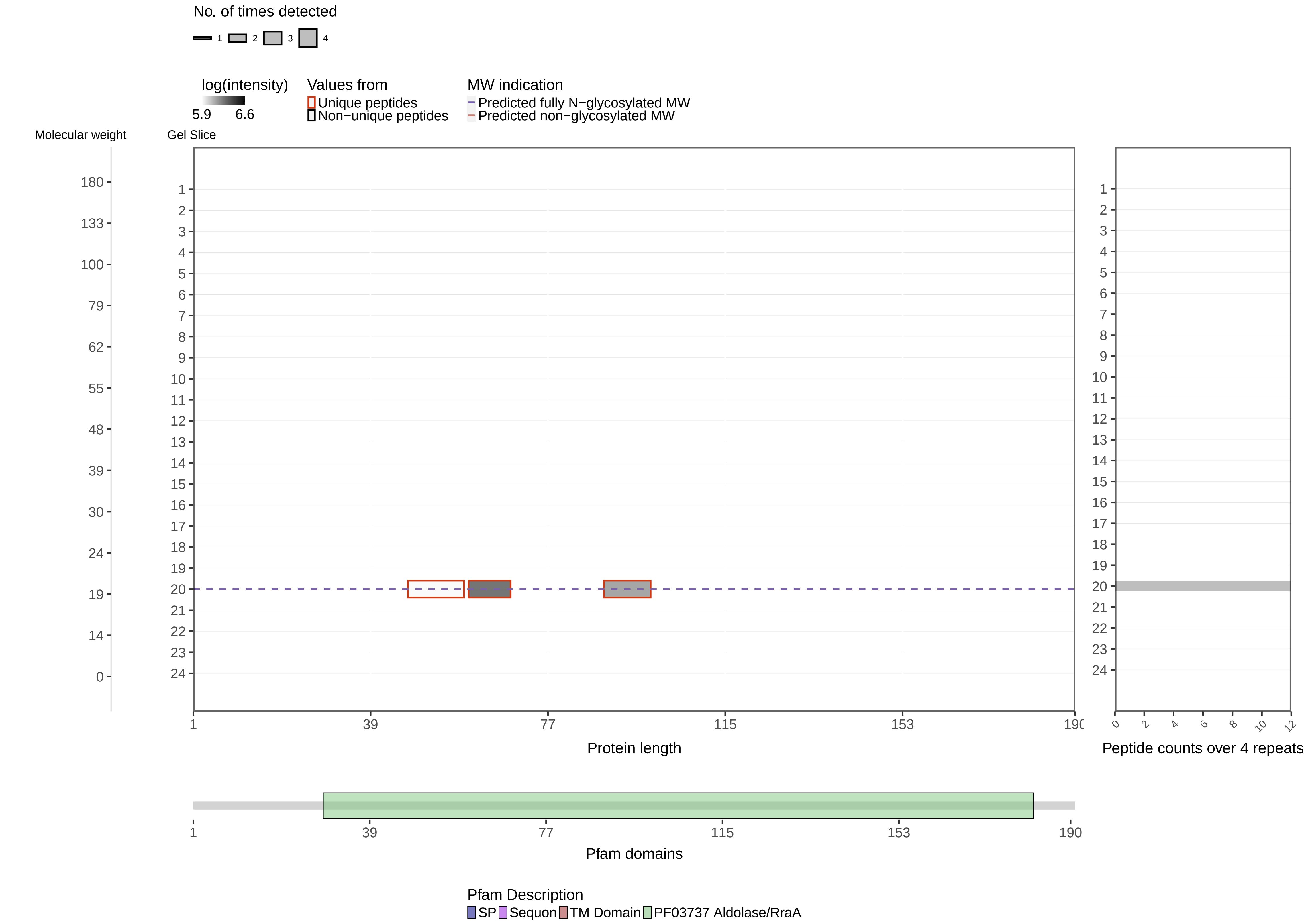


Protein sequence:

TRGEGRVLVIDGGGSMRFALVGGNLGQLAQNMGWAGIVVNGCIRDVDEINGCDIGVRALASHPQKSSKKGHGEKHVPVYIGGMLIREGEWLYADSDGILVSKTEVSV*

### NbL08g07880.1 photosystem_I_reaction_center_subunit_V_chloroplastic−like [MW = 16.21 kDa]

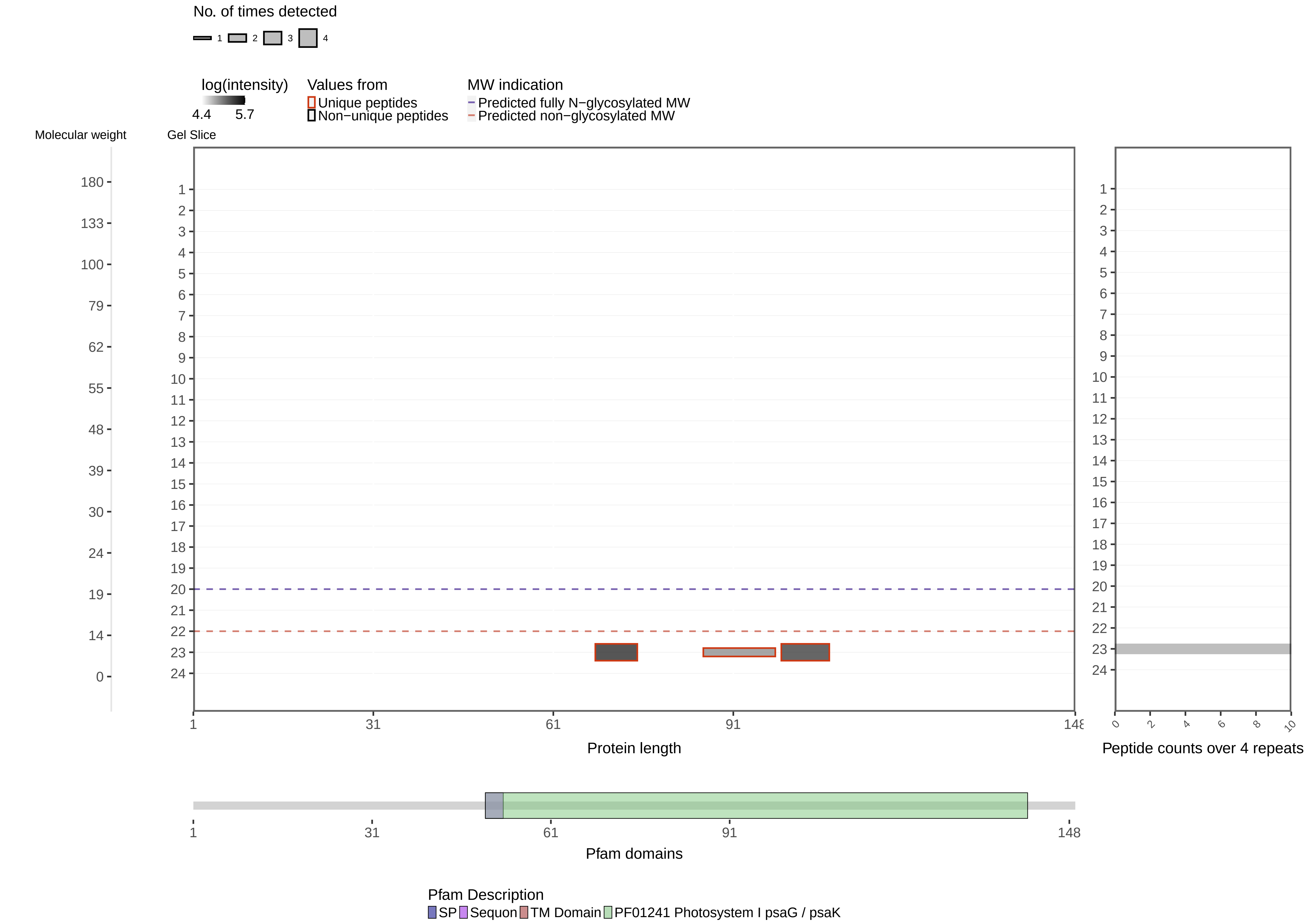


Protein sequence:

LSLFLGRFVFFNFQRENVAKQGLPKQNGISHFEAGDSRAKEYVSLLKSNDPVGFNIVDVLAWGSIGHIVAYYILATSSNGYDPKFFG*

NbL08g07810.1 DNA_damage−inducible_protein_1 [MW = 44.75 kDa]

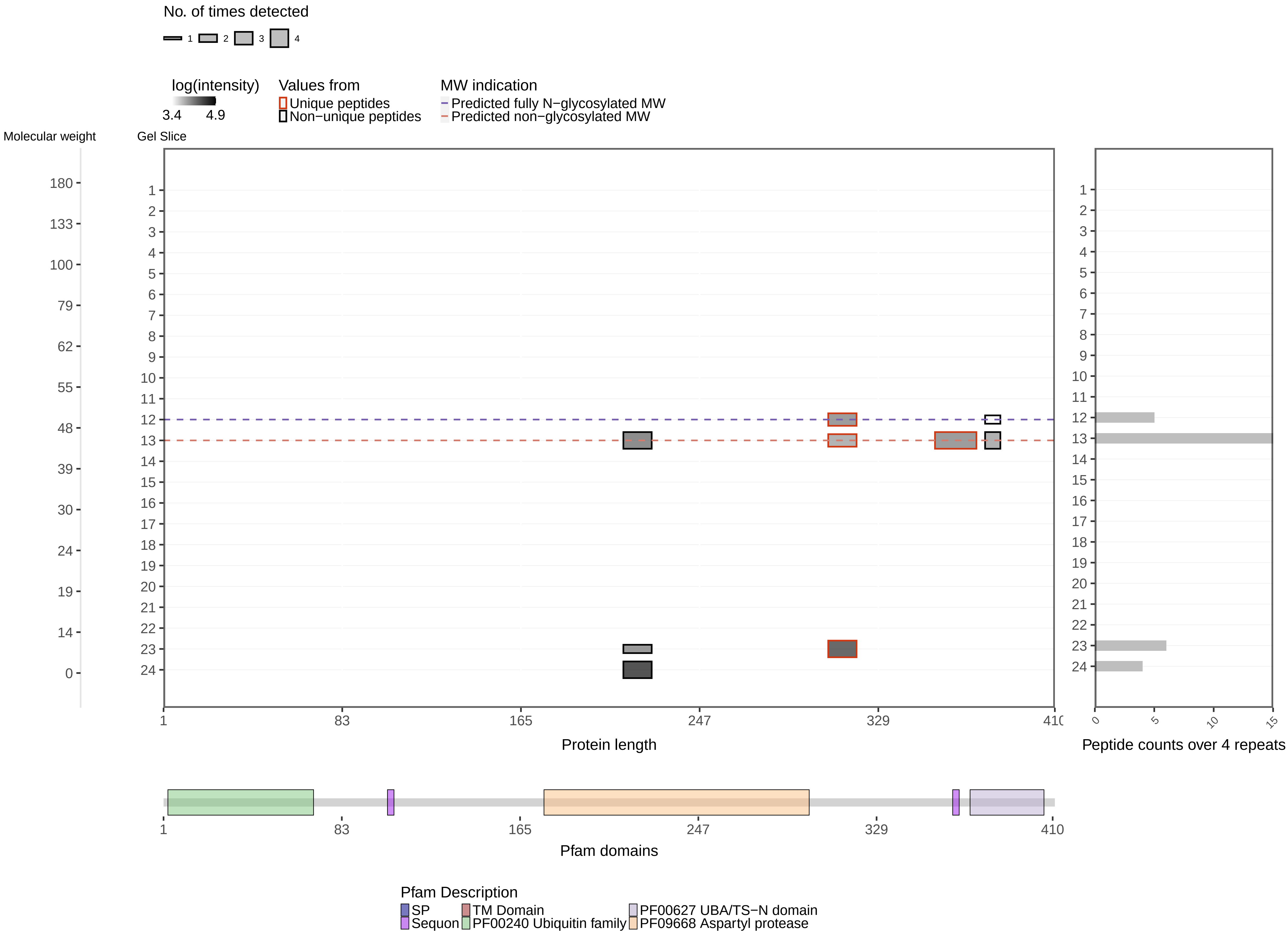


Protein sequence:

EAFARVVMLYVDMEVNGHPLKAFVDSGAQSTIISKSCAERCGLLRLLDTRYKGIAHGVGQSEILGRIHVAPIKIGQIFYPCSFVVLDSPNMEFLFGLDMLRKHQCMIDLKENVLRVGGGEVAVPFLQEKDIPGHFLDEERFAKEASSSGAQATSGAKEKTGPAKGGPSGGAPGNSTQGTEFEAKVAKLVELGFGRDAVIQALKFFDGNEEQAAGYLFGG*

### NbL08g07780.1 heat_shock_cognate_protein_80−like [MW = 80.18 kDa]

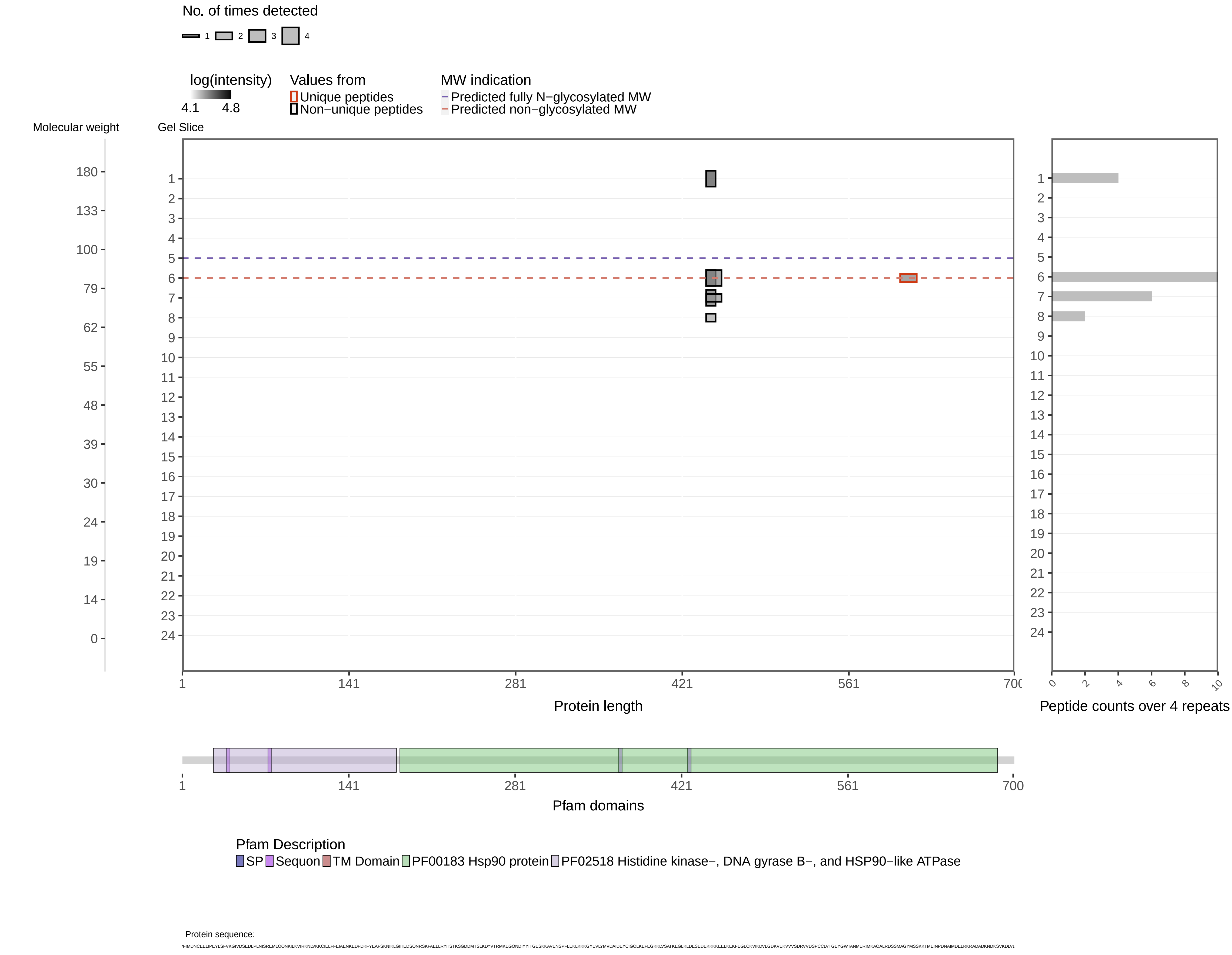

### NbL08g07740.1 protein_CWC15_homolog [MW = 26.86 kDa]

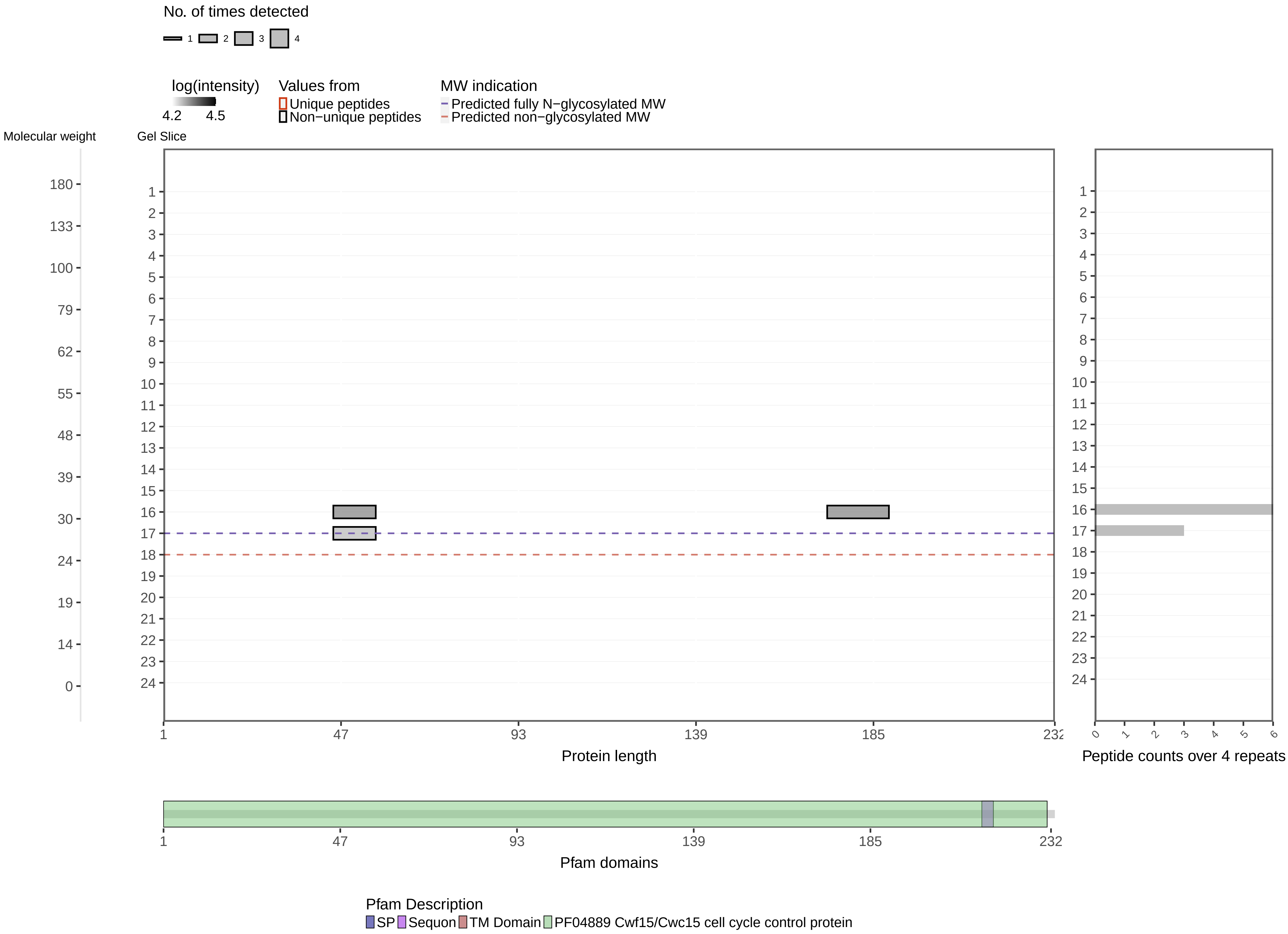


Protein sequence:

PRSADADDADVDVKSGDESDDDDDDDEDDTEALLAELEQIKKEKAEEKLRKERQEQEEELKAKEAELLRGNPLLNHQAQPTSFSVKRRWDDDVVFKNQARGEMKAAKRFINDTIRNDFHRKFLQKYMK*

### NbL08g07710.1 nudix_hydrolase_2−like [MW = 31.93 kDa]

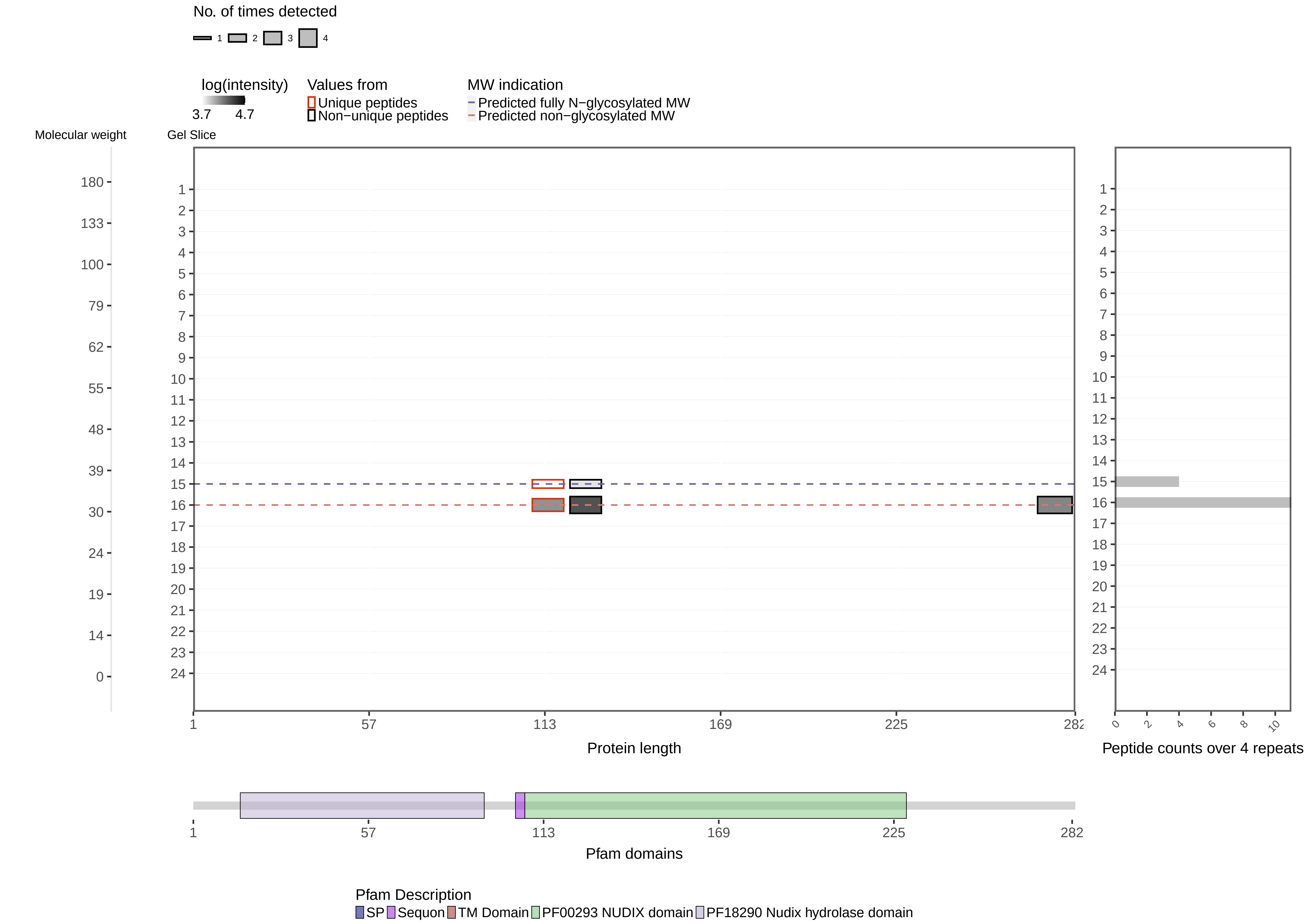


Protein sequence:

ENSGRLKGTAVWKIPTGILEEGEDIFEGAIREVKEETGIDTEFLEVLAFROTOKALFGKSDLFFICMMHPLSFDIOKODLEIEAAOWMPLEEYAAOPFVOKHGIFKYTKDLCLVKAERNYPGFTPVPITSFFDGSTSFLYVNKDGLDOESSAIHL*

### NbL08g07460.1 autophagy−related_protein_8C−like [MW = 13.74 kDa]

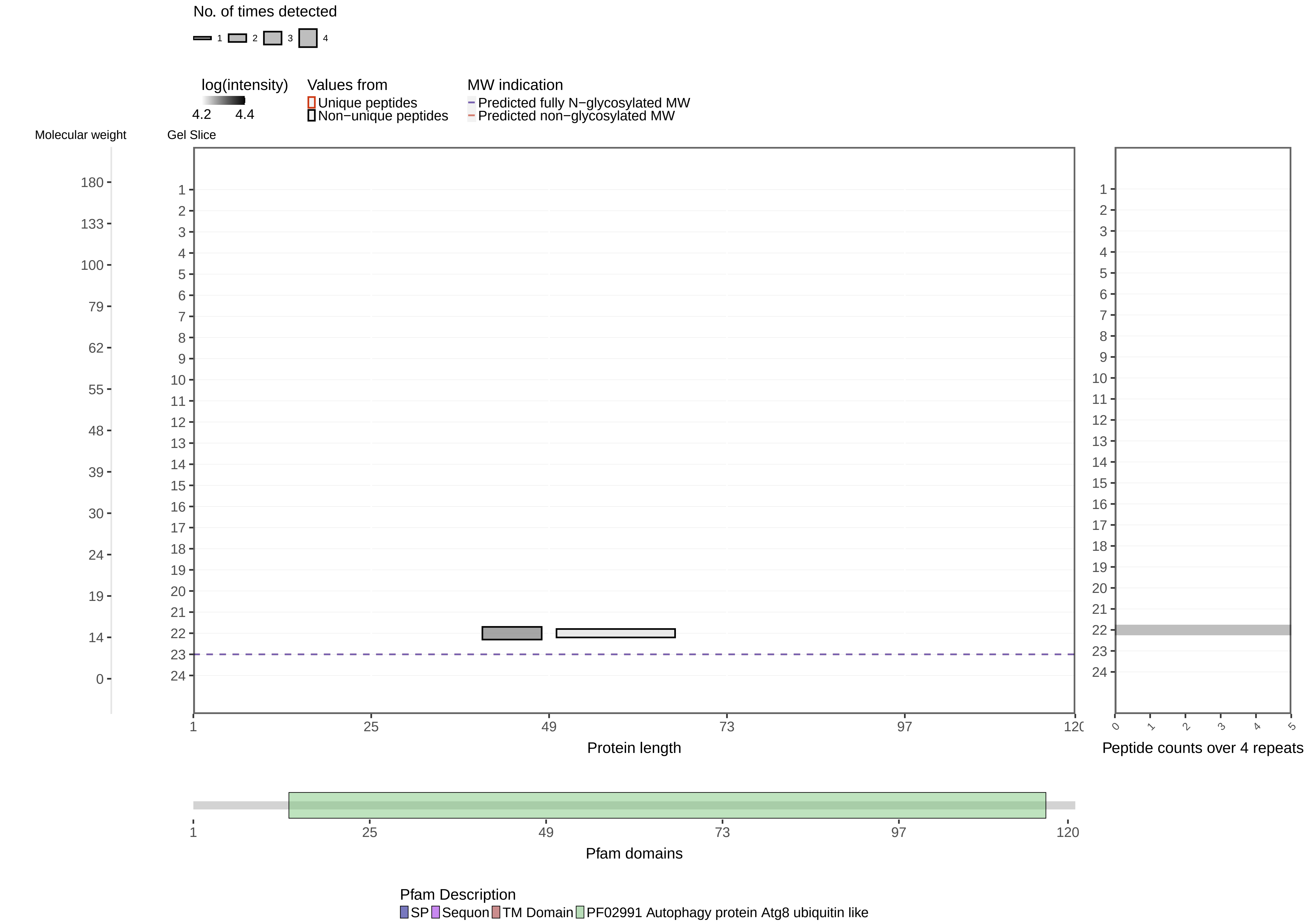


Protein sequence:

KKKYLVPADLTVGQFVYVVRKRIKLSAEKAIFIFVKNILPPTAAMMSAIYEEHKDEDGFLYMTYSGENTFGSF*

NbL08g07370.1 tryptophan_synthase_beta_chain_1 [MW = 51.19 kDa]

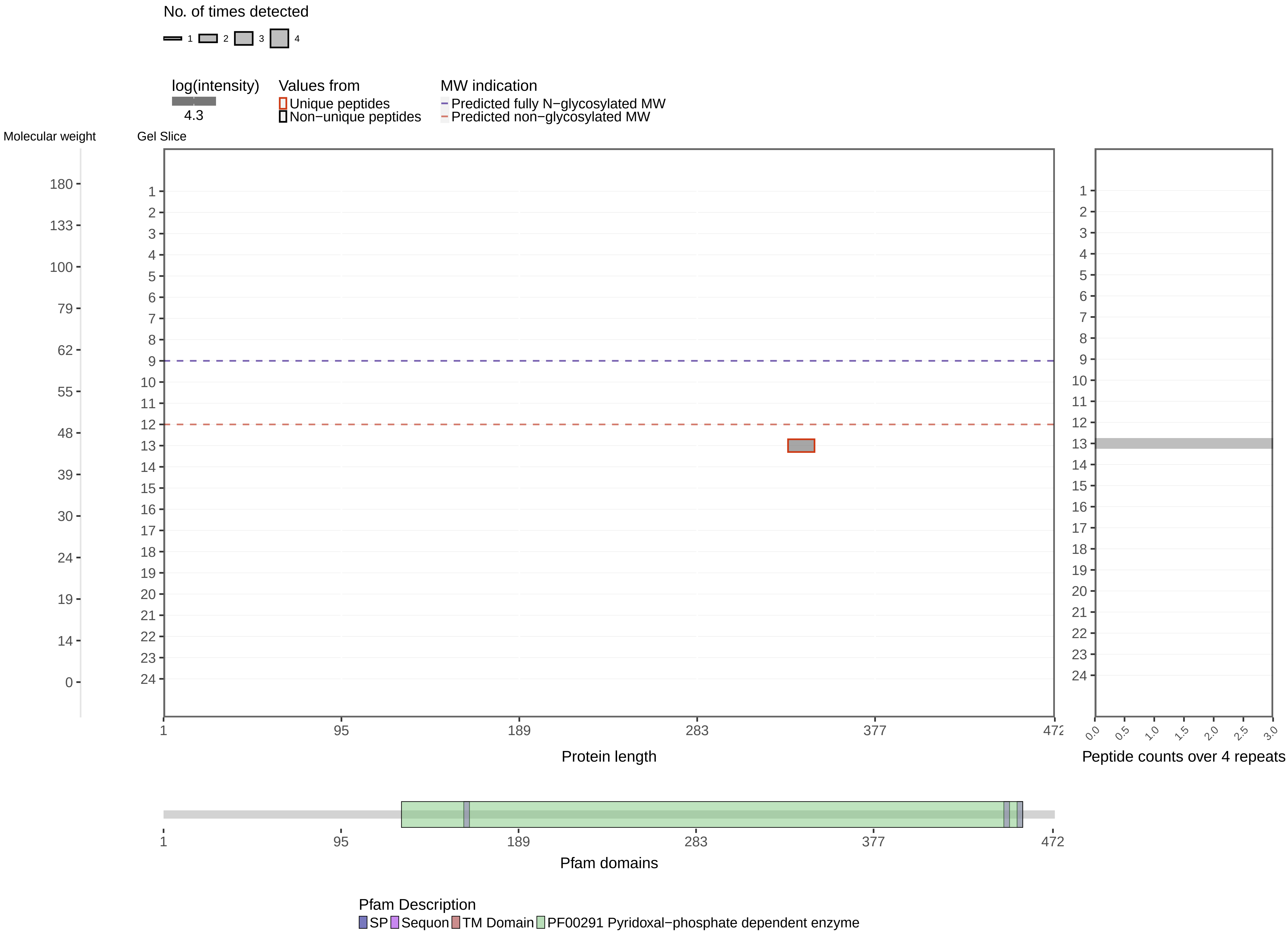


Protein sequence:

VFRMRLLGAEVRGVHSGTATLKDATSEAIRDWVTNVETTHYILGSVAGPHPYPMMVREFHAVIGKETRKQALEKWGGKPDVLVACVGGGSNAMGLFHEFVDDEDVRLVGVEAAGFGIDSGKHAATLTKGEVGVLHGAMSYLLQDEDGQIVEPHSISAGLDYPGVGPEHSFLKDLGRAEYYSITDEEALEAFKRLSRLEGIIPALETSHALAYLEKLCPTLPNGTKVVLNCSGRGDKDVHTAIKYLKV*

### NbL08g07290.1 trafficking_protein_particle_complex_subunit_3−like [MW = 20.89 kDa]

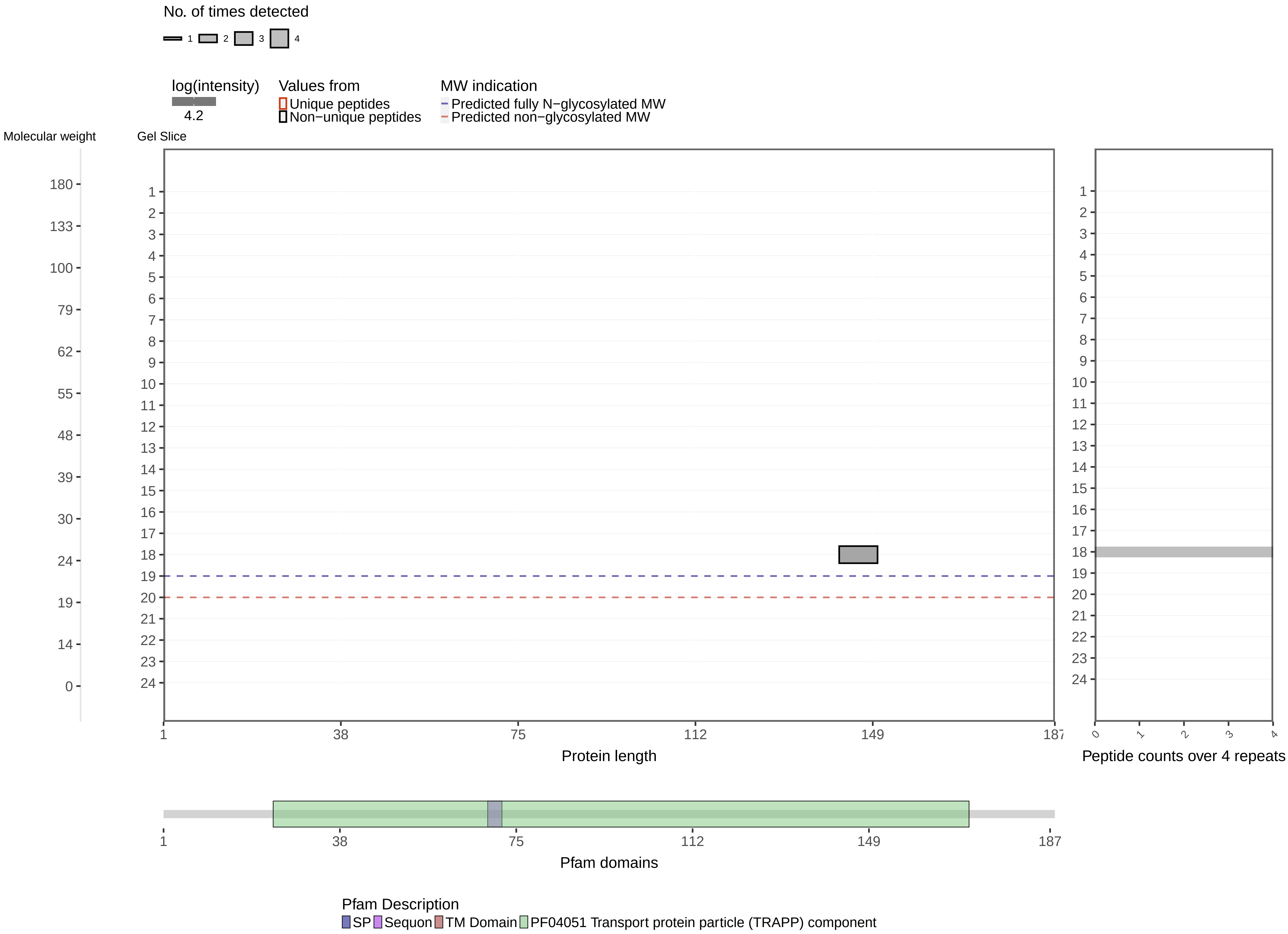


Protein sequence:

VIAKVGLKMFLGVTATVGNWDAEGTTCSLILEDNPLVDFVELPDTCQGLYYCNILSGVIRGALEMVSMKTEVTWIRDMLRGDDAFELQLKLLRQVPEEYPYKDDE*

NbL08g07230.1 50S_ribosomal_protein_L24_chloroplastic−like [MW = 21.15 kDa]

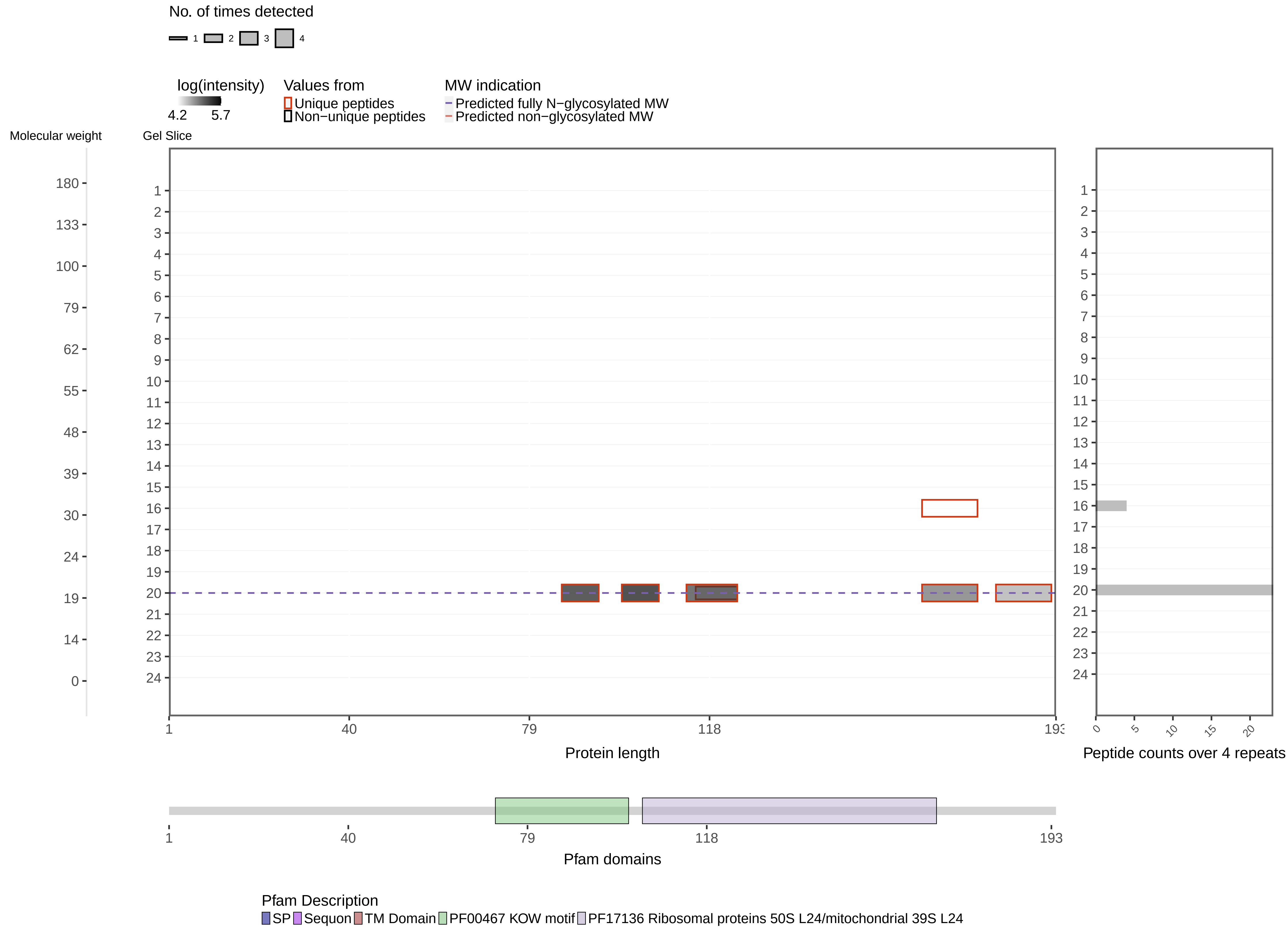


Protein sequence:

DKGKIGEITEIIKHNSKVVVSEINLKTKHVKSRDENEPGQIIKIEAPIHSSNVMLYSKEQKVASRVGHKTLDNGKRVRYLVKTGEIIDSAENWKRAVKETEKTTEAVAATS*

NbL08g07210.1 60S_ribosomal_protein_L8 [MW = 34.62 kDa]

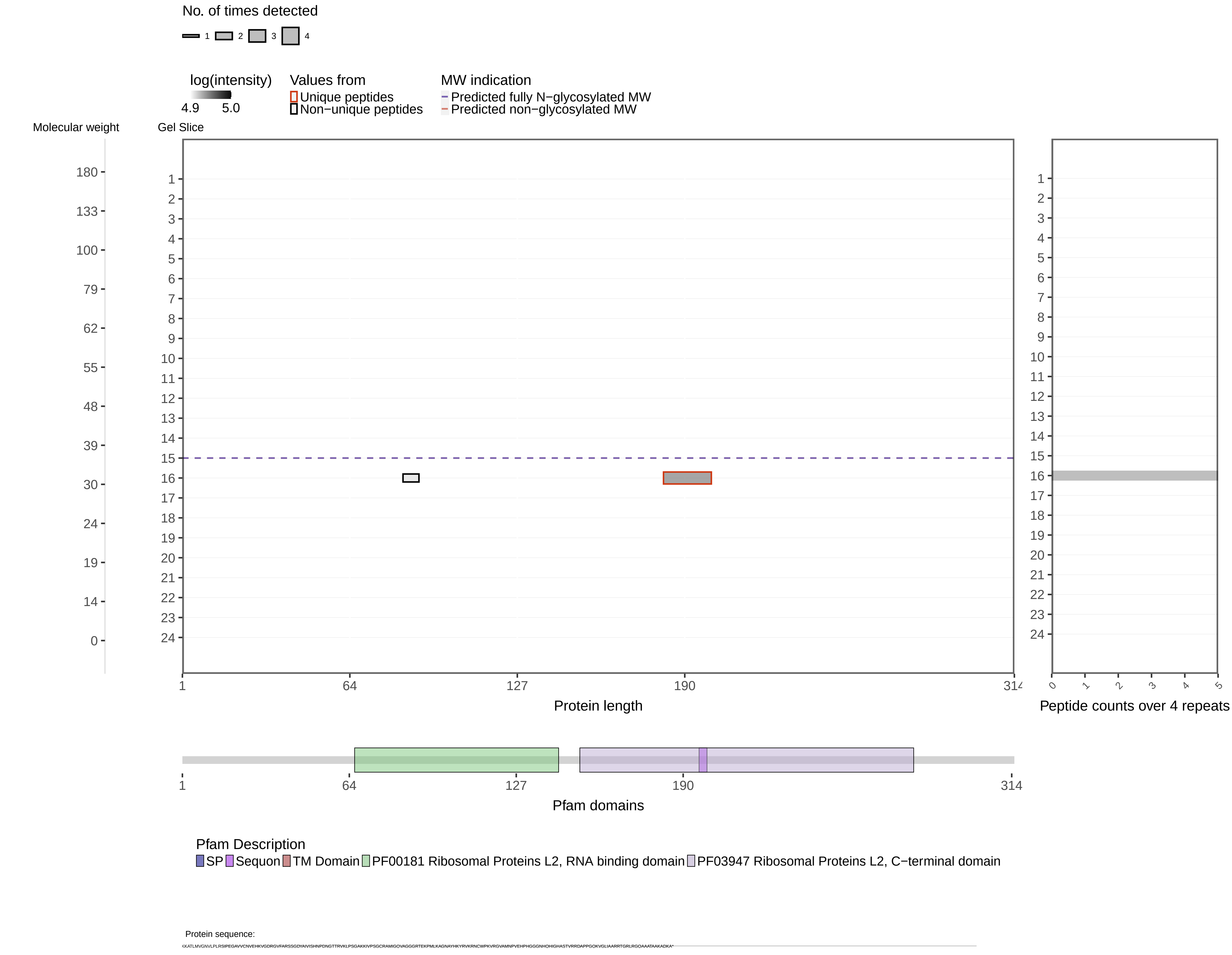

### NbL08g07080.1 uncharacterized_LOC107808755 [MW = 27.53 kDa]

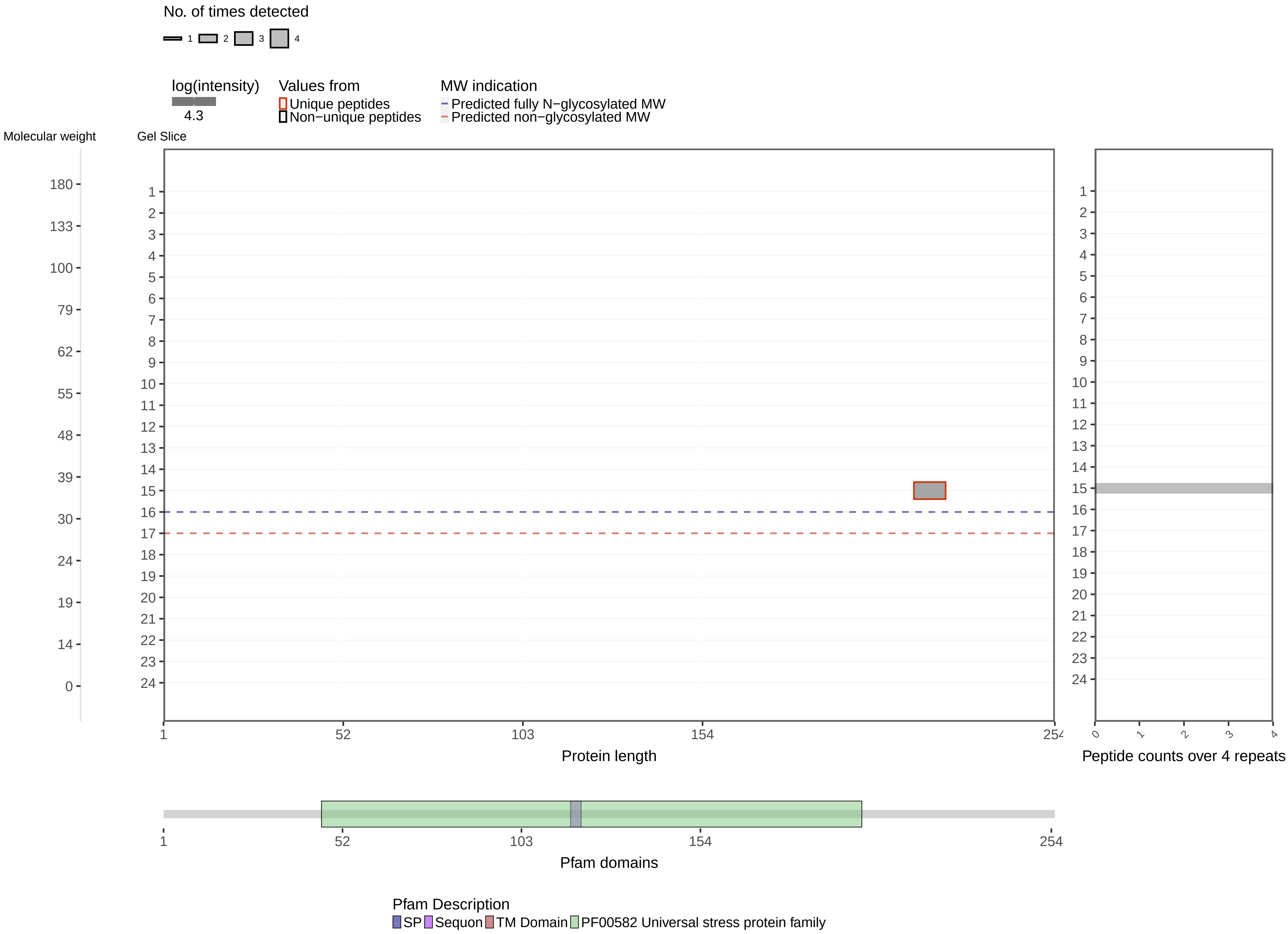


Protein sequence:

ONFTATKSSALAQPLVDAQIPYKIHIVKDHDMKERLCLEVERLGLSAVIMGSRGFGATKRGNDGRLGSVSDYCVRHCVCPVVVVRYPDEKDGGNAARKSVGSVASAAKEYEEETEYHDASDDRQGSQPAIGKGSKTDN*

### NbL08g06850.1 thioredoxin−like [MW = 19.92 kDa]

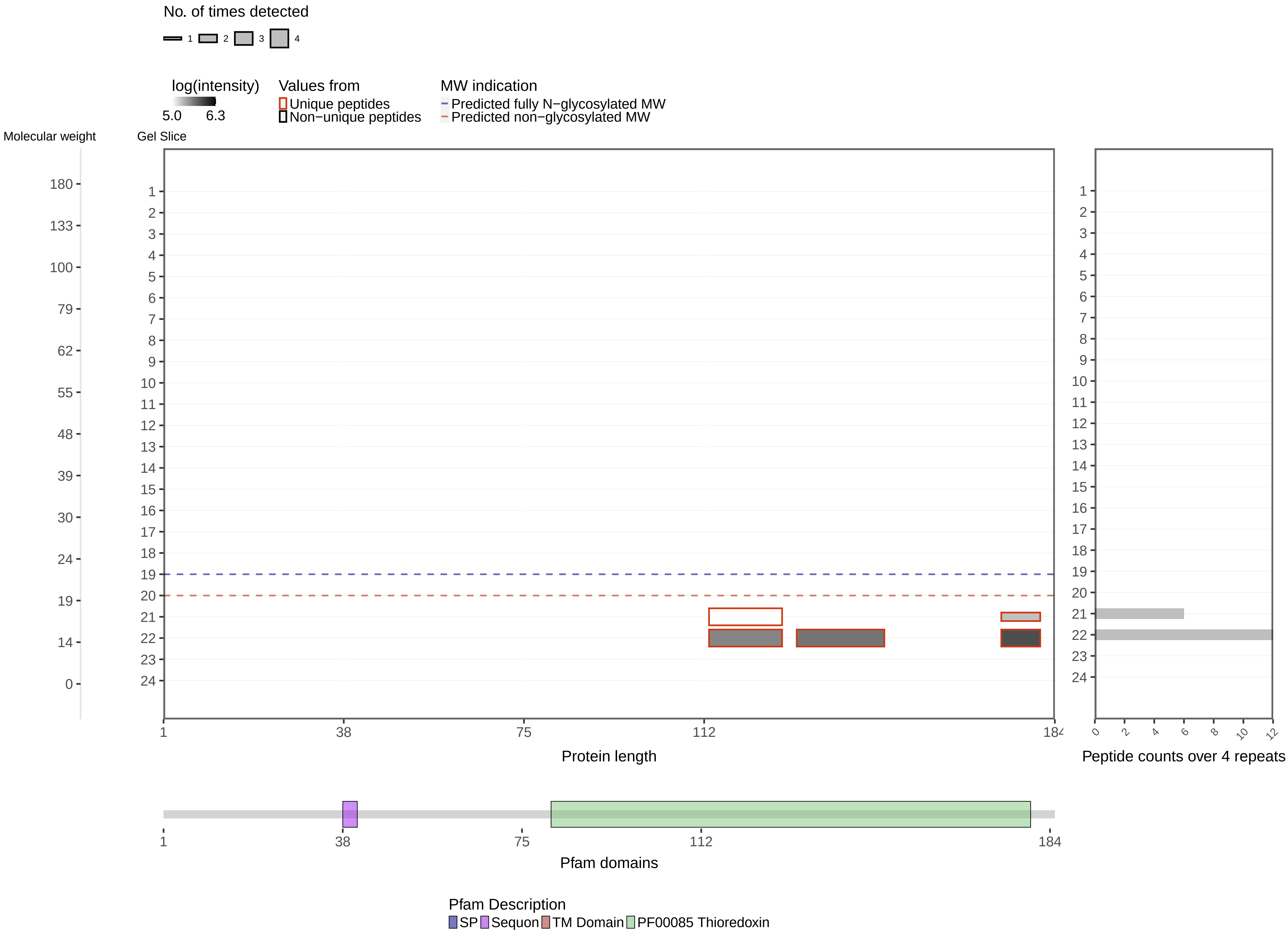


Protein sequence:

TAVQVPAVTDKTWKSLVIESDLPVLVEFWAPWCGPCRMIHPVIGELSQEYAGKFKFFQLNTDESPSTASEYGIRSIPTVMIFKGGEKKDAVIGAVPKTTLTTSIEKFL*

### NbL08g06660.1 malate_dehydrogenase_mitochondrial [MW = 36.23 kDa]

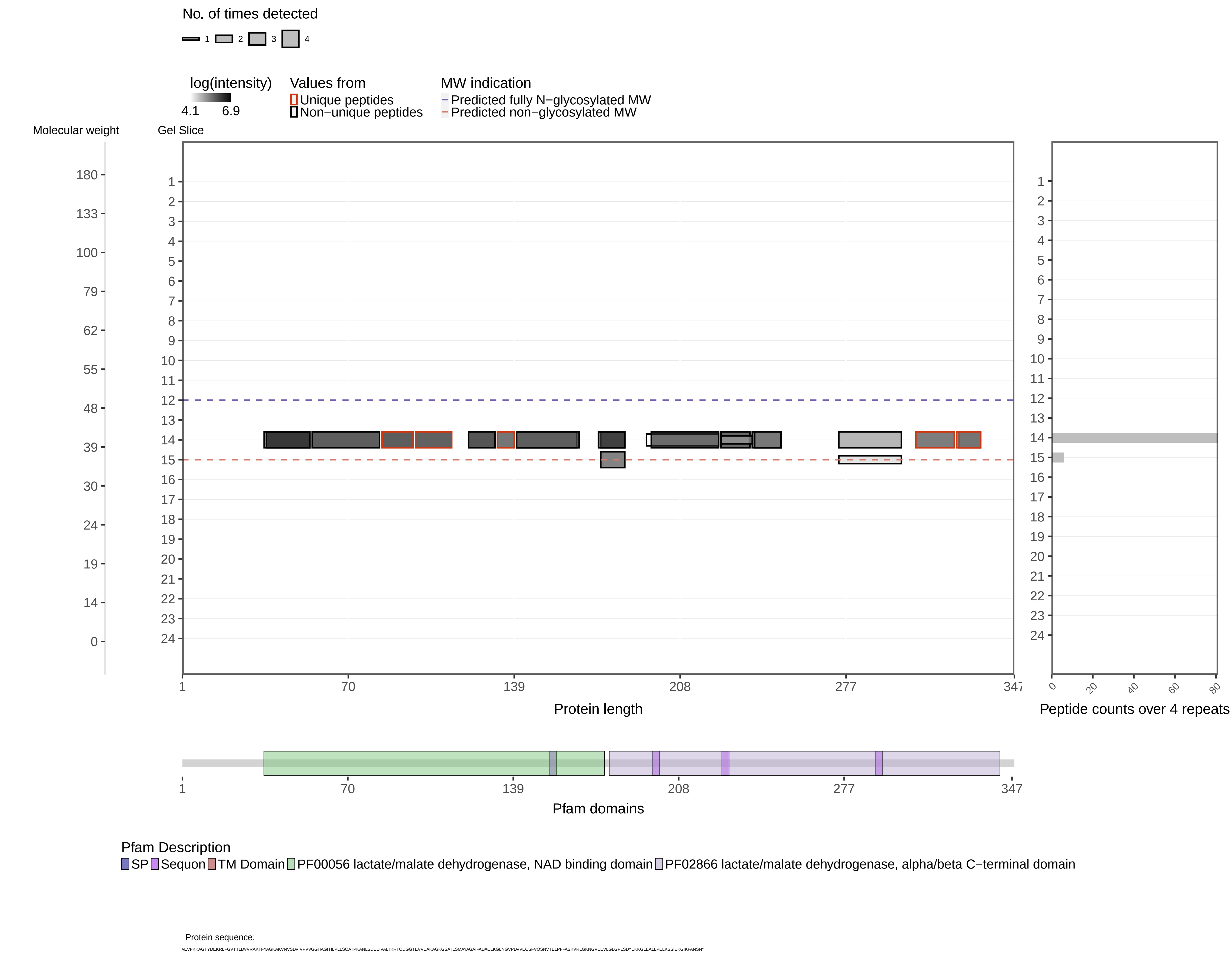

### NbL08g06580.1 uroporphyrinogen_decarboxylase_1_chloroplastic [MW = 46.32 kDa]

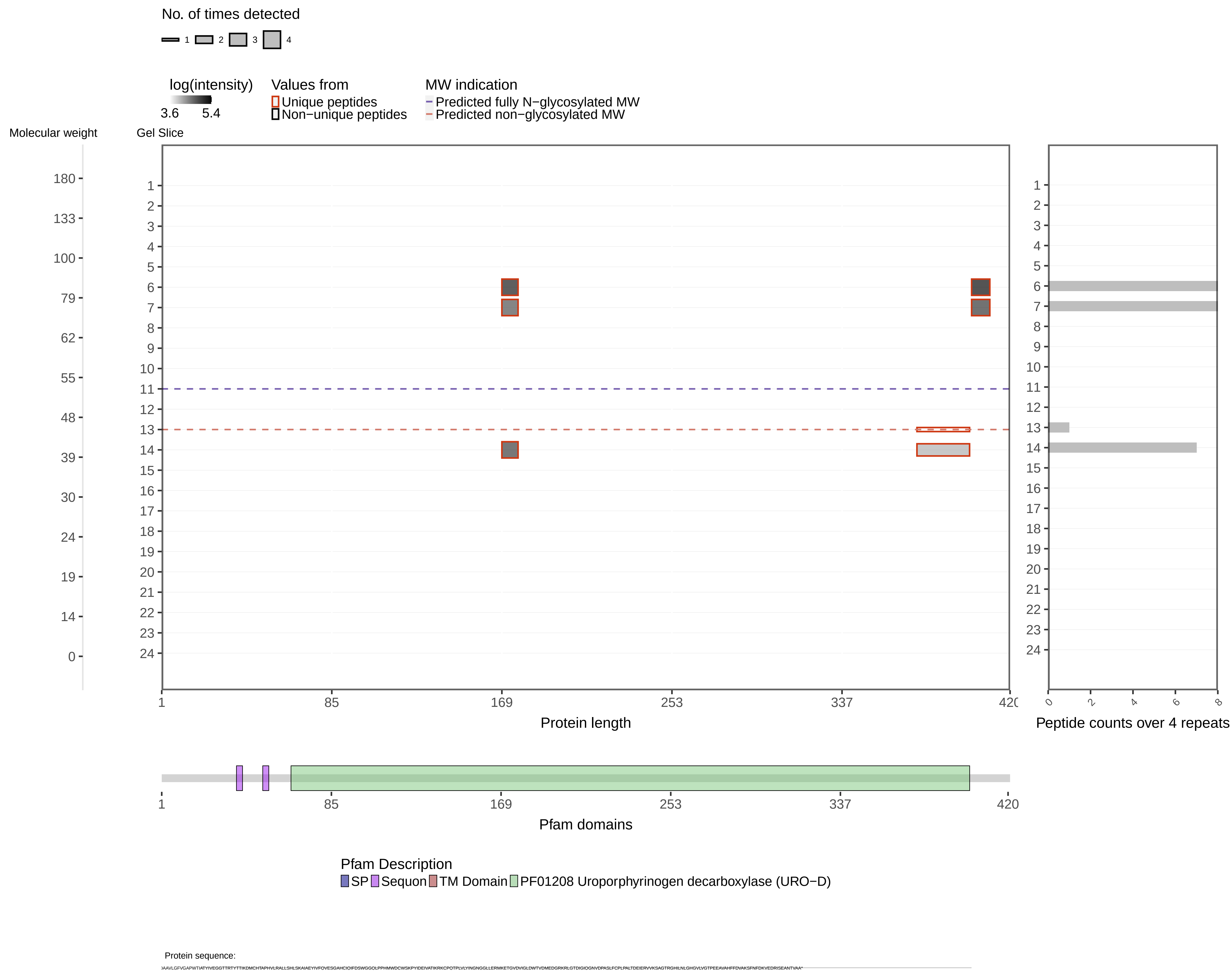

### NbL08g06560.1 multiprotein−bridging_factor_1b [MW = 15.25 kDa]

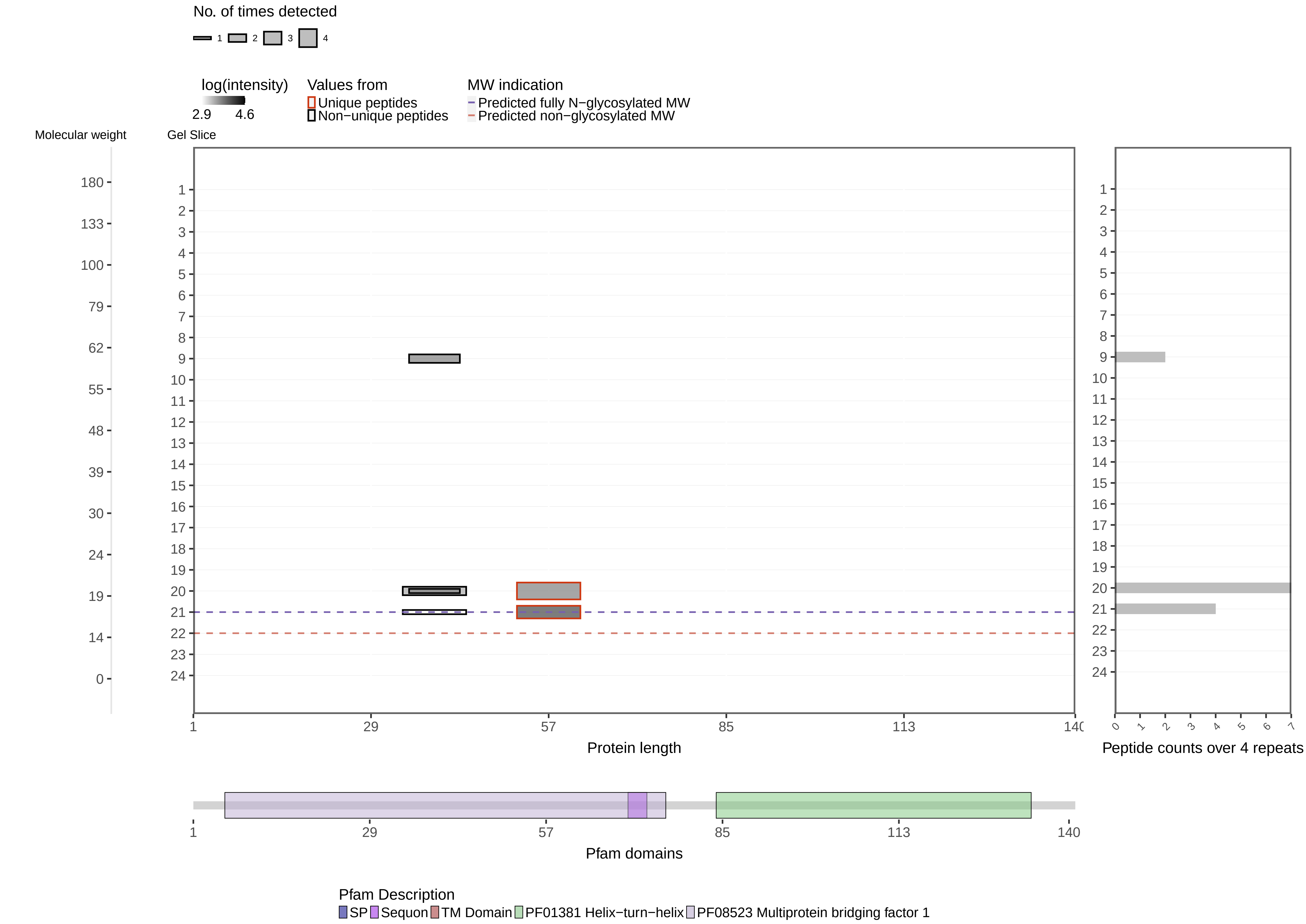


Protein sequence:

STSLNTRTLDEDTENLSHEKVPTELKKAIMQARQDKKLTQAQLAQLINEKPQIIQEYESGKAIPNQQIISKMERALGAKLRGKK*

NbL08g06510.1 gamma−glutamyl_hydrolase_2−like [MW − SP = 35.7 kDa]

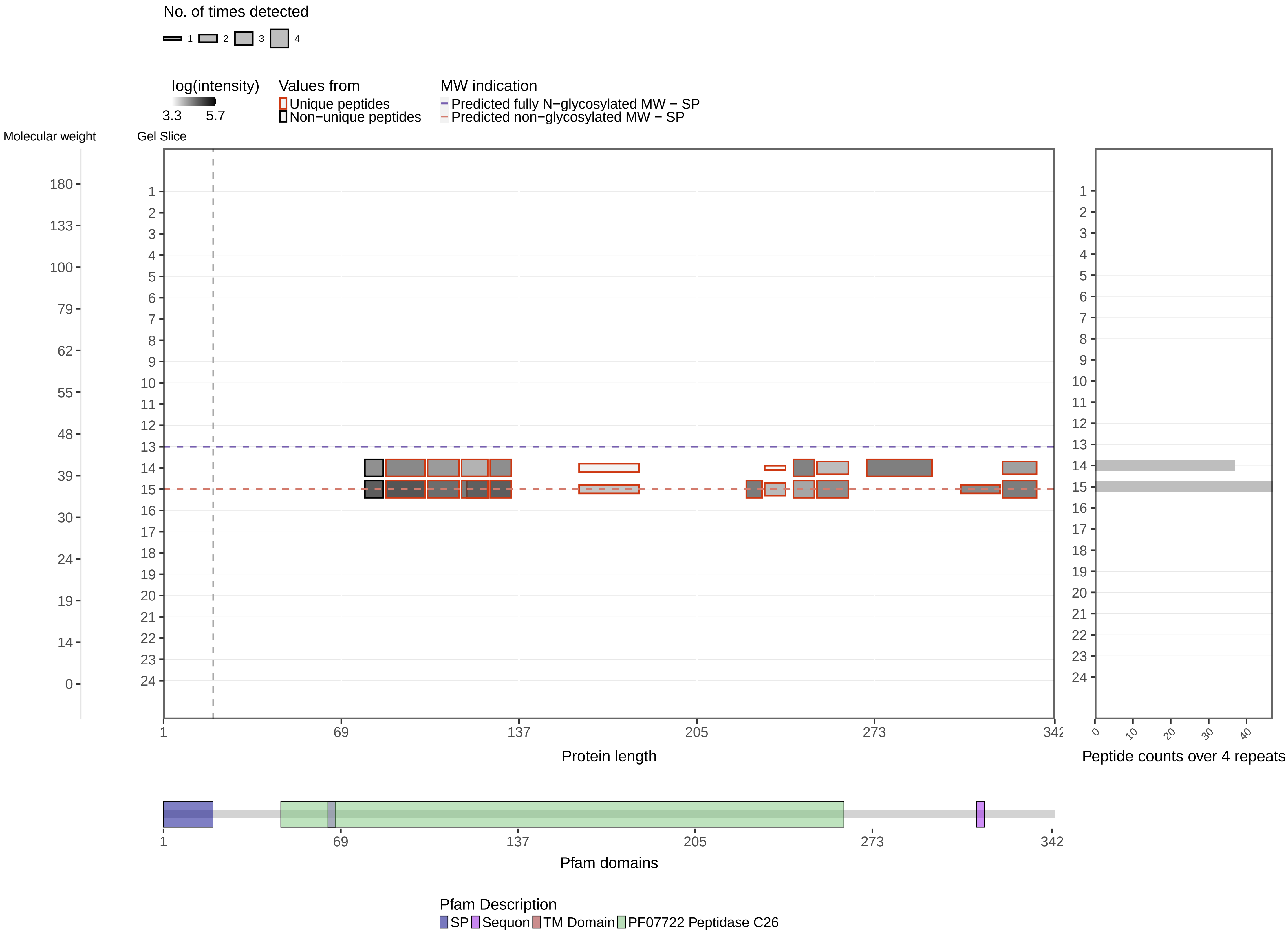


Protein sequence:

NNILEEFSAANQASTVQFVENVKLERTVFGRFPPVLIKKMSTHCLVMQNHHFGISPERFQANKDLCSFFRILTTSTDKKNKVYVSTAQARRYPVTALQWHPEKNVFEWRSSQIPHTEDAIQVTQHVANYFVSEGRKSSNKPVPSKLLDNLIYNYSPSFAGKAGGSFDEVYLFSPRPALSSL*

NbL08g06470.1
trifunctional_UDP−glucose_46−dehydratase/UDP−4−keto−6−deoxy−D−glucose_35−epimerase/UDP−4−keto−L−rhamnose−reductase_RHM1
[MW = 76 kDa]

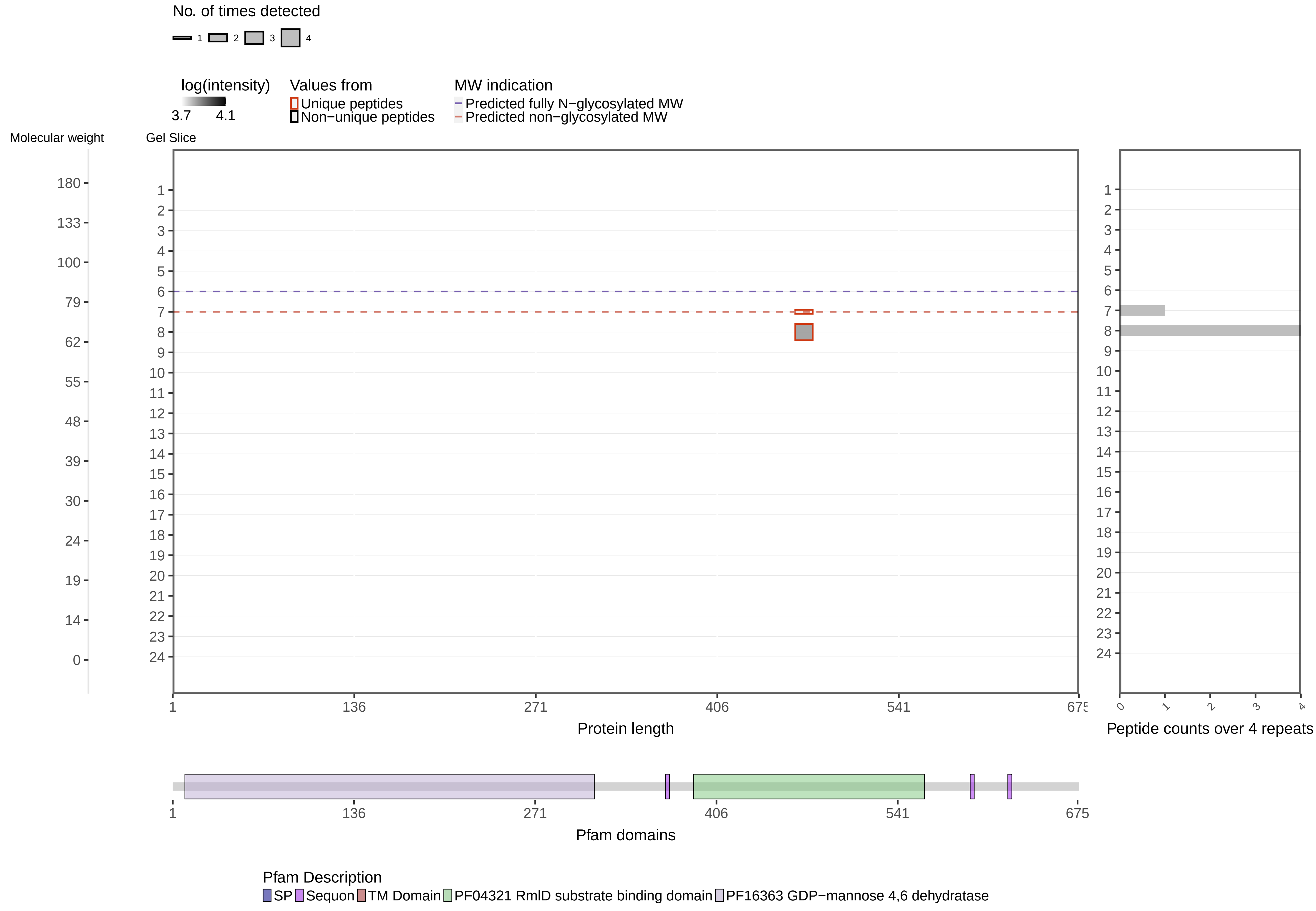


Protein sequence:
[illegible]

### NbL08g06350.1 peroxisomal_(S)−2−hydroxy−acid_oxidase [MW = 40.66 kDa]

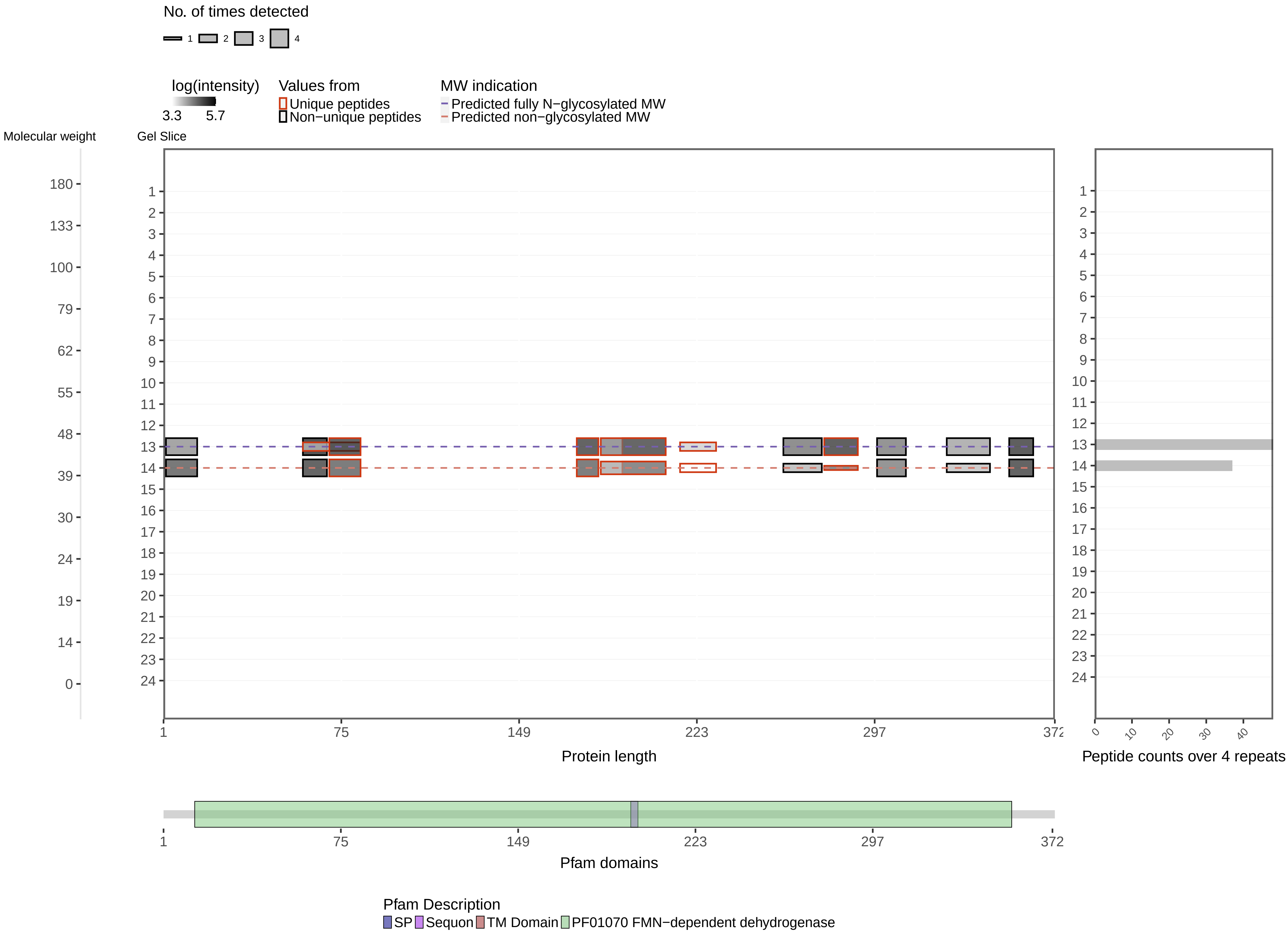


Protein sequence:

TLQPFLTLKNFEGLDLGKMDEANDSGLGSYVAGQIDRSLSWKDIQWLQTITSLPILVKGVLTAEDARIAVQAGAAGIIVSNHGARQLDYVPATIMALEEVVKAAQGHVPVFLDGGVRRGTDVFKALALGASGVFIGRPVVFSLAAEGEAGIRKVLQMMRDEFELTMALSGCRSLKEITRNHIVADWDAPRAALPTPRL*

### NbL08g06280.1 ultraviolet−B_receptor_UVR8 [MW = 46.97 kDa]

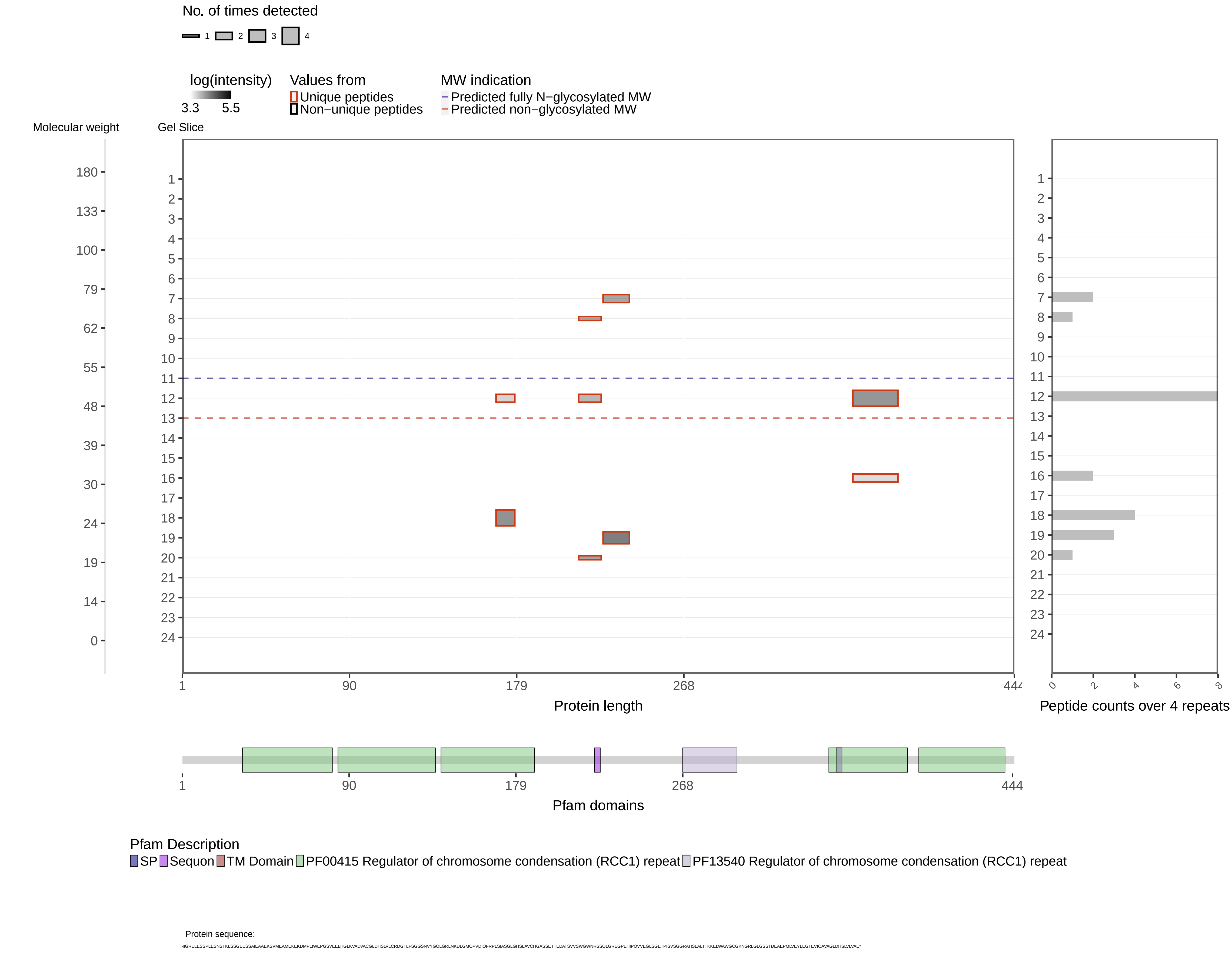

NbL08g06270.1 chlorophyll_a−b_binding_protein_8_chloroplastic [MW = 29.34 kDa]

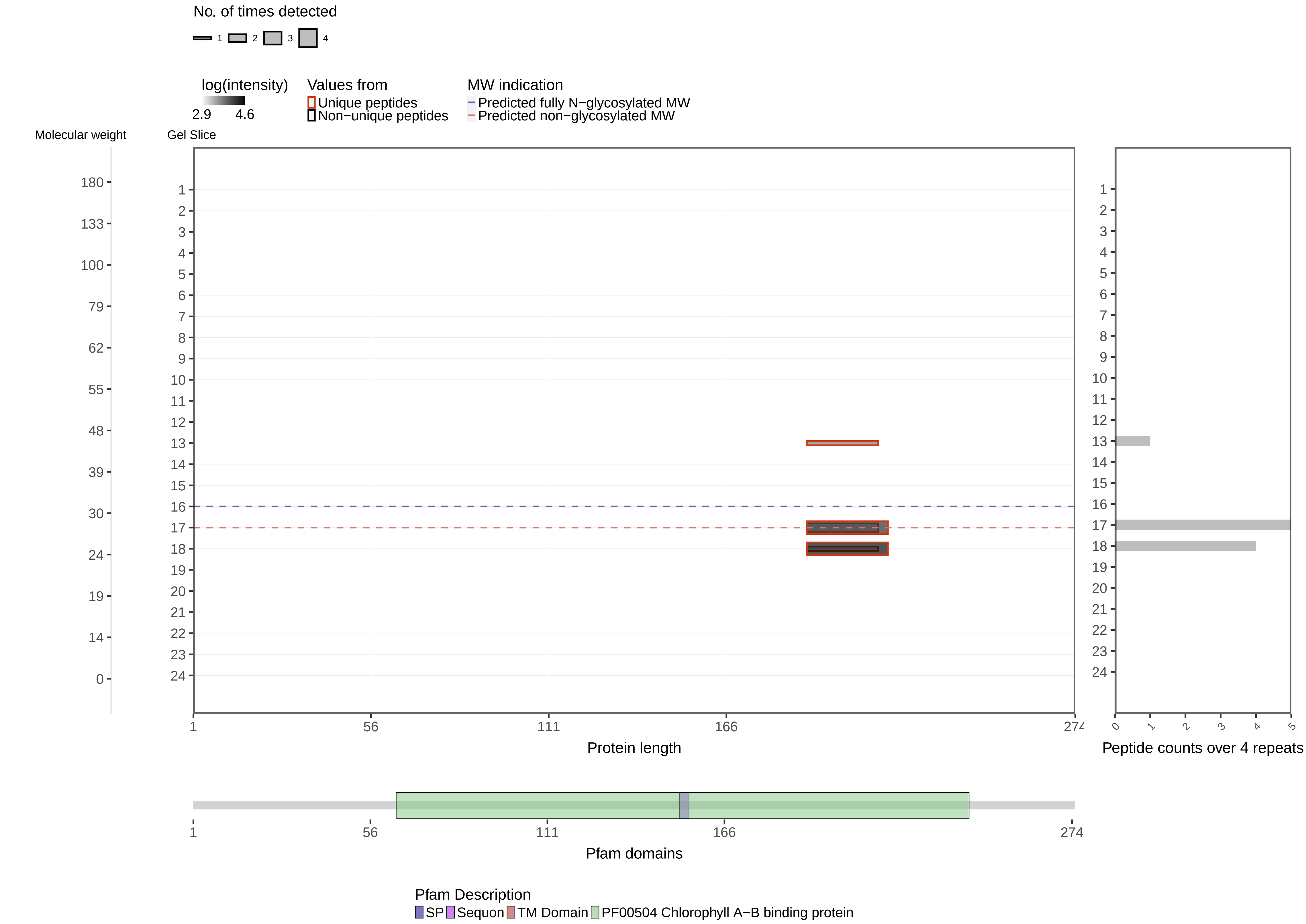


Protein sequence:

LIPPETALPWFQTGVIPPAGTYNYWADNYTLFVLEMALMGFAEHRRFQDWAKPGSMGKQYFLGLEKGLGGSGEPAYPGGPFFNPLGFGKDEKSMKDLKLKEVKNGRLAMLAILGYFIQGLVTGVGPYQNLLDHLADPVNNNILTNFKFH*

### NbL08g06260.1 ras−related_protein_RABB1c−like [MW = 23.15 kDa]

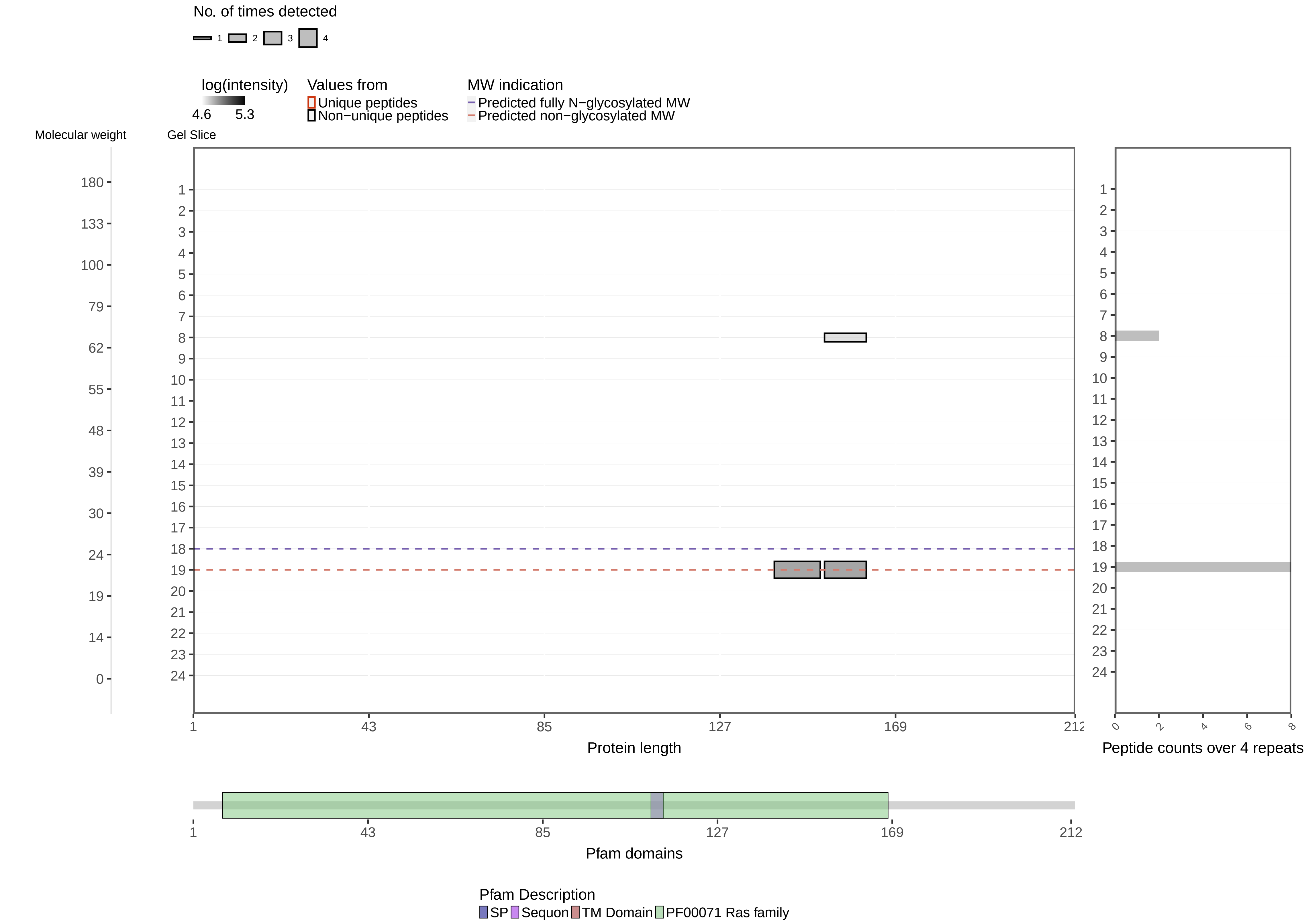


Protein sequence:

HLASWLEDARQHANANMTIMLIGNKCDLAHRRAVSTEEGEQFAKEHGLIFMEASAKTAQNVEEAFIKTASTIYKKIQDGVFDVSNEANGIKFGYGGLPGPSGGRDGAASQGGGCCS*

### NbL08g06160.1 ribose−phosphate_pyrophosphokinase_4−like [MW = 35.93 kDa]

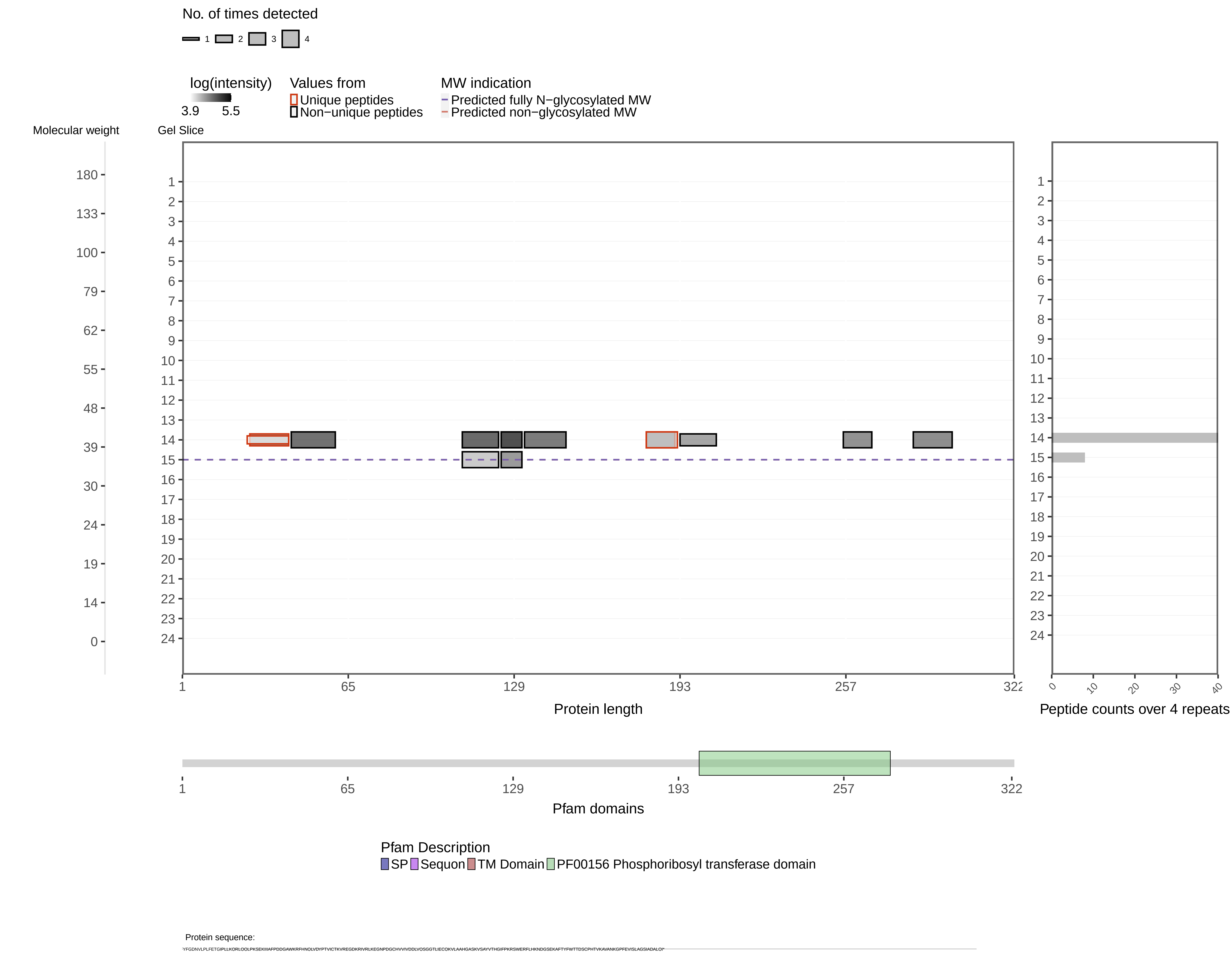

### NbL08g06070.1 glutamyl−tRNA(Gln)_amidotransferase_subunit_A_chloroplastic/mitochondrial [MW = 60.72 kDa]

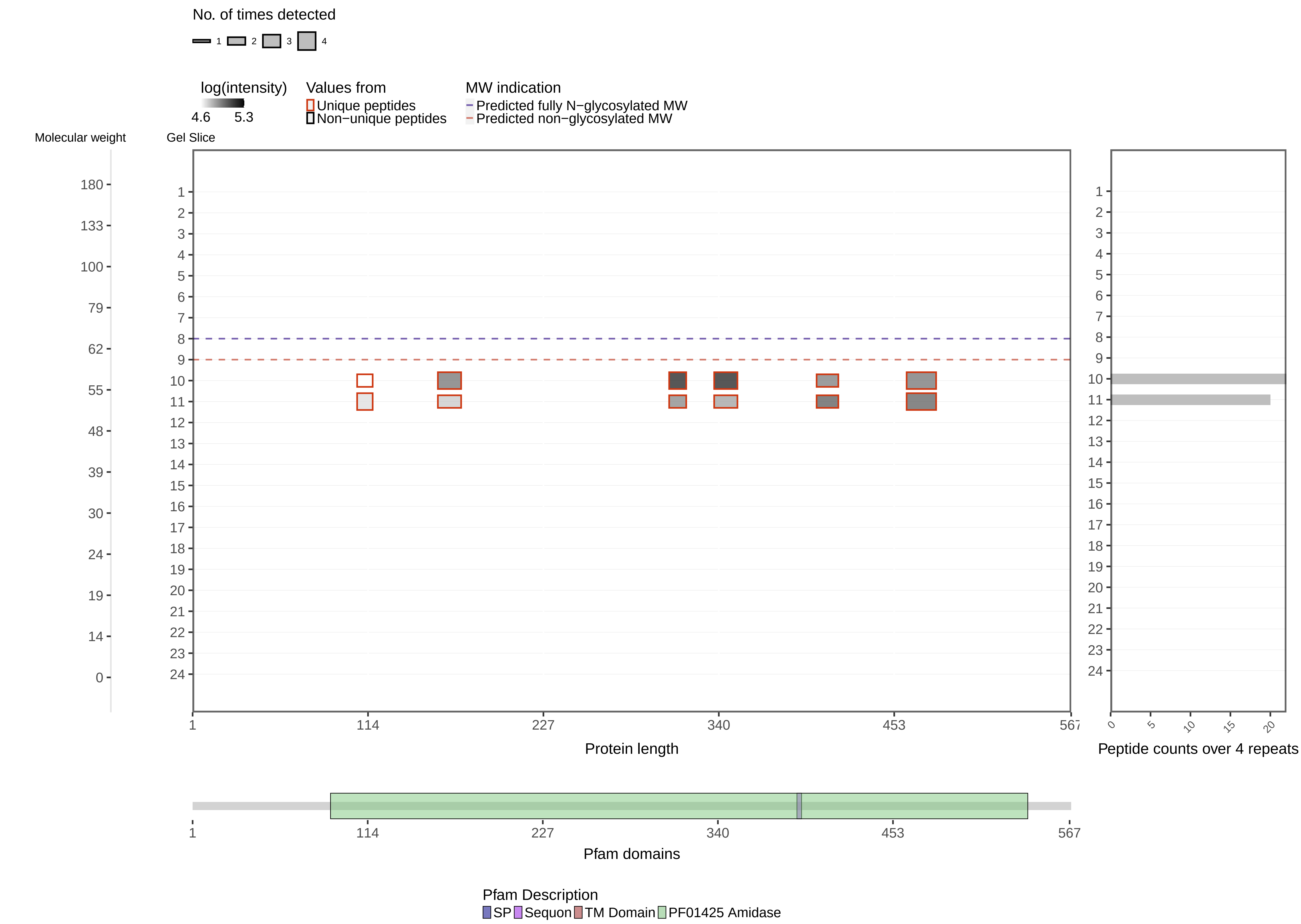


Protein sequence:

VAYASSLDVIGCFGSSVADAGILLHAISGHDKFDATSSKREVPDFASQFIARDHLDSKPLKGLRVGLIRETIEDGVDPEIITSIRGAASHLEELGCTVTEVSLPSFSLGLPAYYILASSESSSNLSRYDGIRYGKQVVADELNSLYGESRAGGFGSEVKMRILMGTYALSAGYYDAYYKRAQQVRTLVRENFRAALEENDILISPAAPSAAYRIGEKKNDPLSMYAGDIMTVNVNLAGLPALVLPCGFVDSSSVALPVGIQMIGAAFEEEKLLKVGHIFEQTLQGCSFIPPIVADELAC*

### NbL08g05820.1 serine/threonine−protein_phosphatase_PP2A_catalytic_subunit−like [MW = 35.47 kDa]

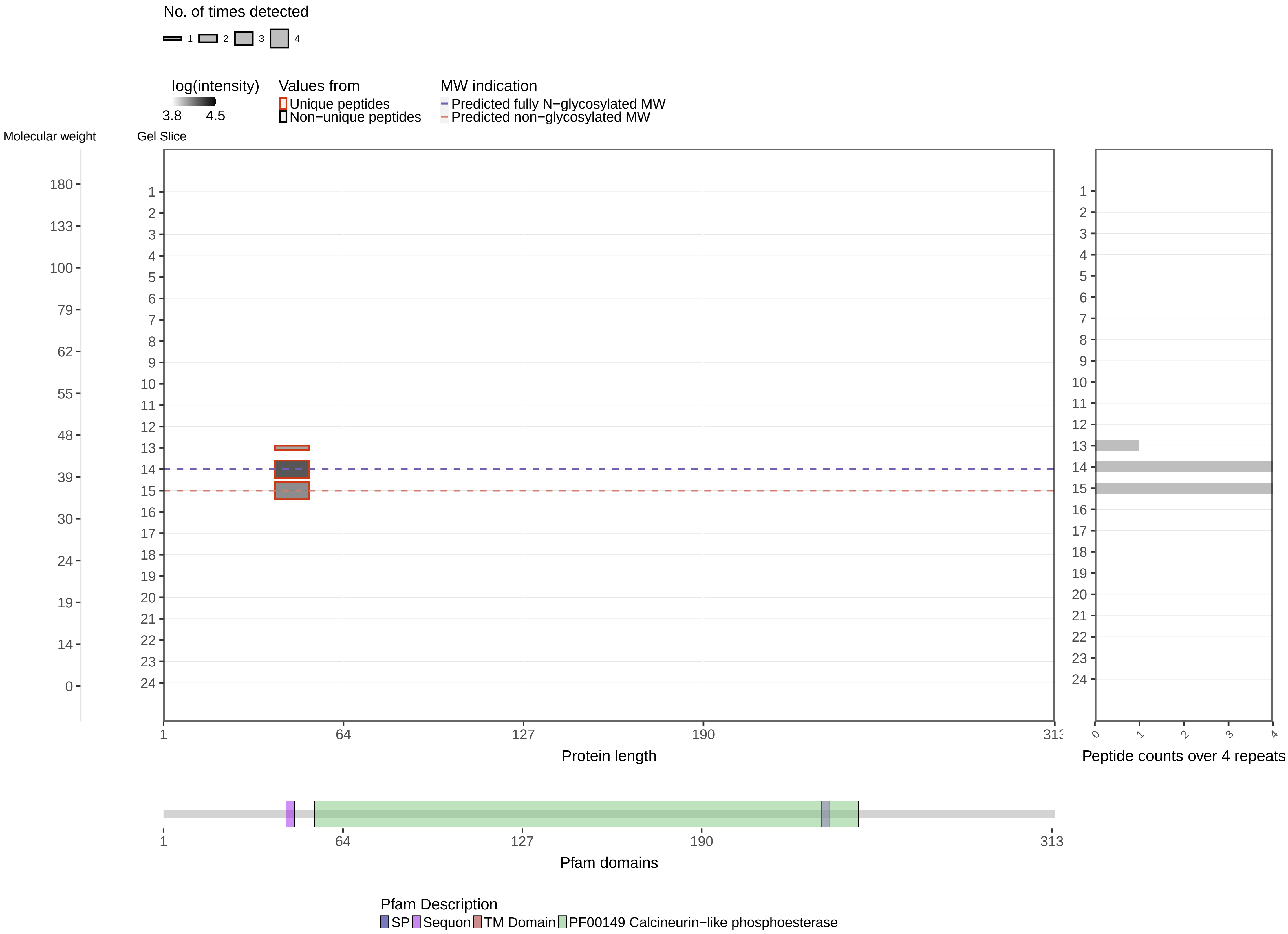


Protein sequence:

NVWKTFTDLFDYFPLTALVESEIFCLHGGLSPSIETLDNIRNFDRVQEVPHEGAMCDLLWSDPDDRCGWGISPRGAGYTFGQDISEQFNHTNNLKLIARAHQLVMEGFNWAHDQKVVTIFSAPNYCYRCGNMASILEVDDCNGHTFIQFEPAPRRGEPDVTRRTPDYFL*

### NbL08g05740.1 protein_YLS3−like [MW = 25.74 kDa]

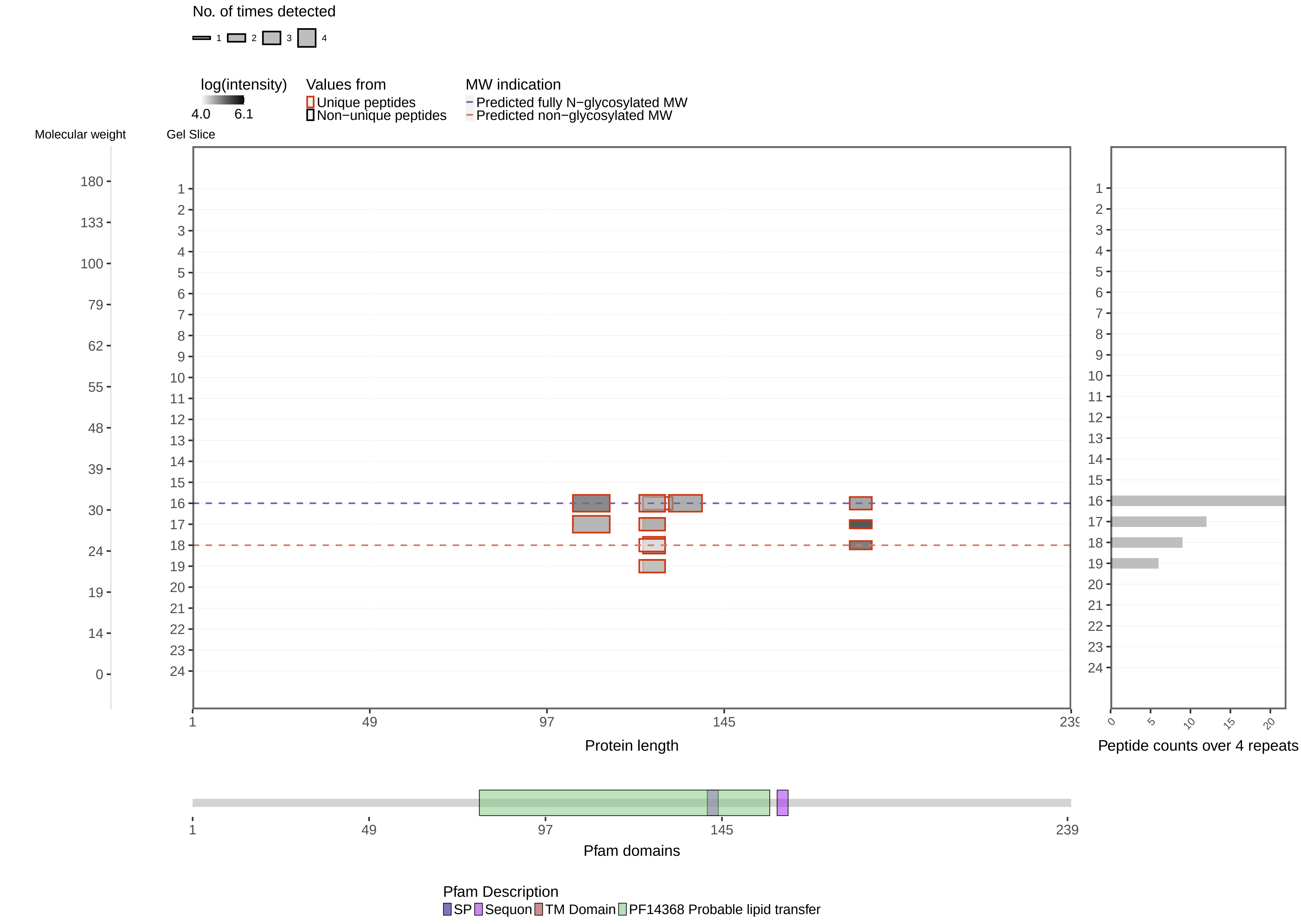


Protein sequence:

CCTGLKEVLDKSKKCLCVLVKDRNDPSLGLKINATLALSLPTLCHAPVSATNVSMCPELLHLPPNSPDAKVFEDFAKSAKGSSTASSAPVSGSSNGKTATTANEKNDGGDRRKWMGIVEMTMGFLFIMVL*

NbL08g05350.1 peptidyl−prolyl_cis−trans_isomerase_CYP28_chloroplastic−like [MW = 35.57 kDa]

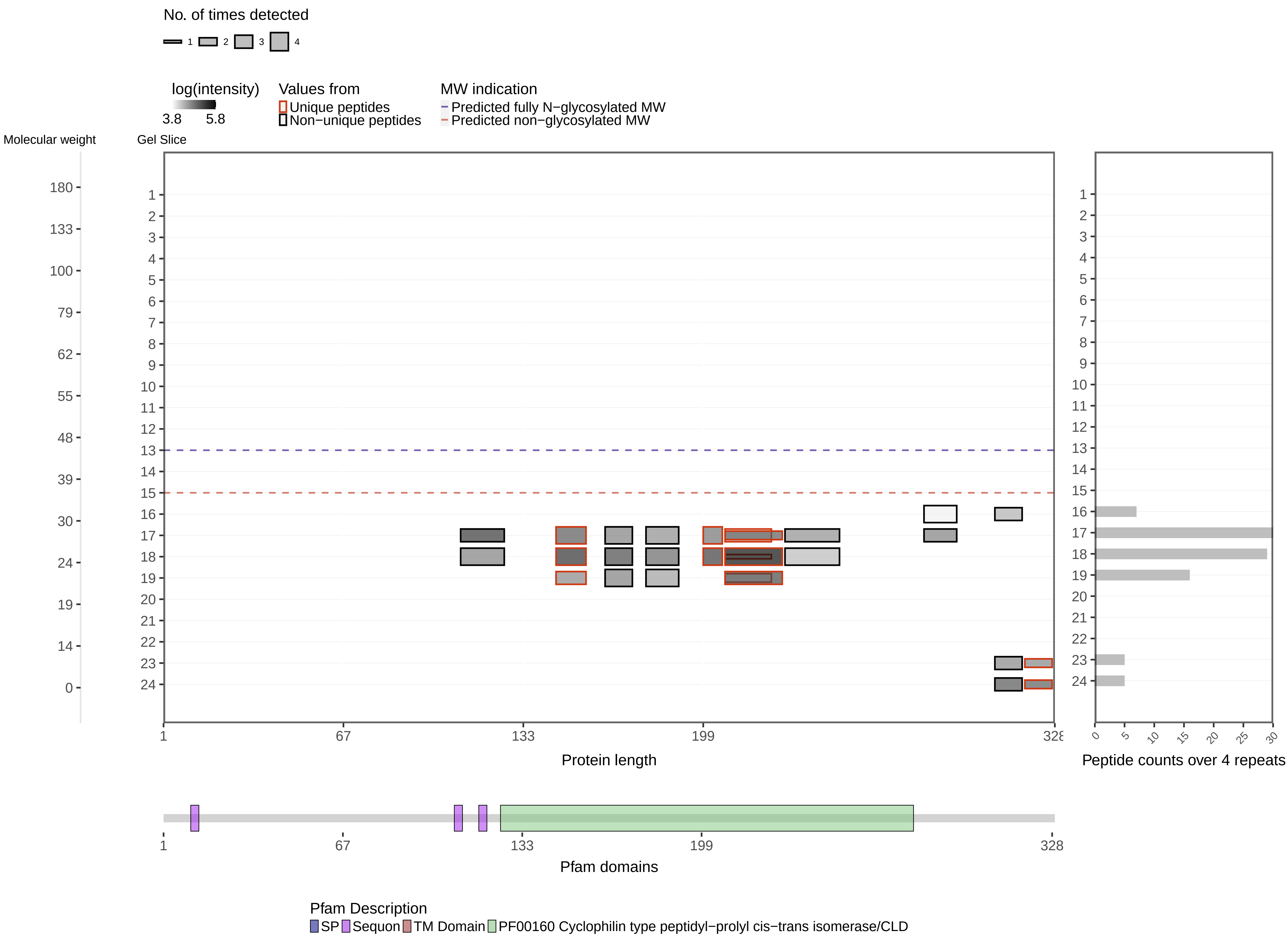


Protein sequence:

STYKGTLVQKIFPGQFFMAGRQGRRDKGEVKPPLELIRNTETIESKAFILEHNRGGIVSLPLSENDDDDDLKLNPKYHNVEFLITTGPGPVPQLDSKNIVFGTVLEGMDIVTSIAAIPTYKPSENIRQYNGFAELLGDGRAKTARAIWNKPQKTVYISDCGELKVAKPTLSPSLP*

### NbL08g05220.1 4−coumarate−−CoA_ligase_2−like [MW = 61.82 kDa]

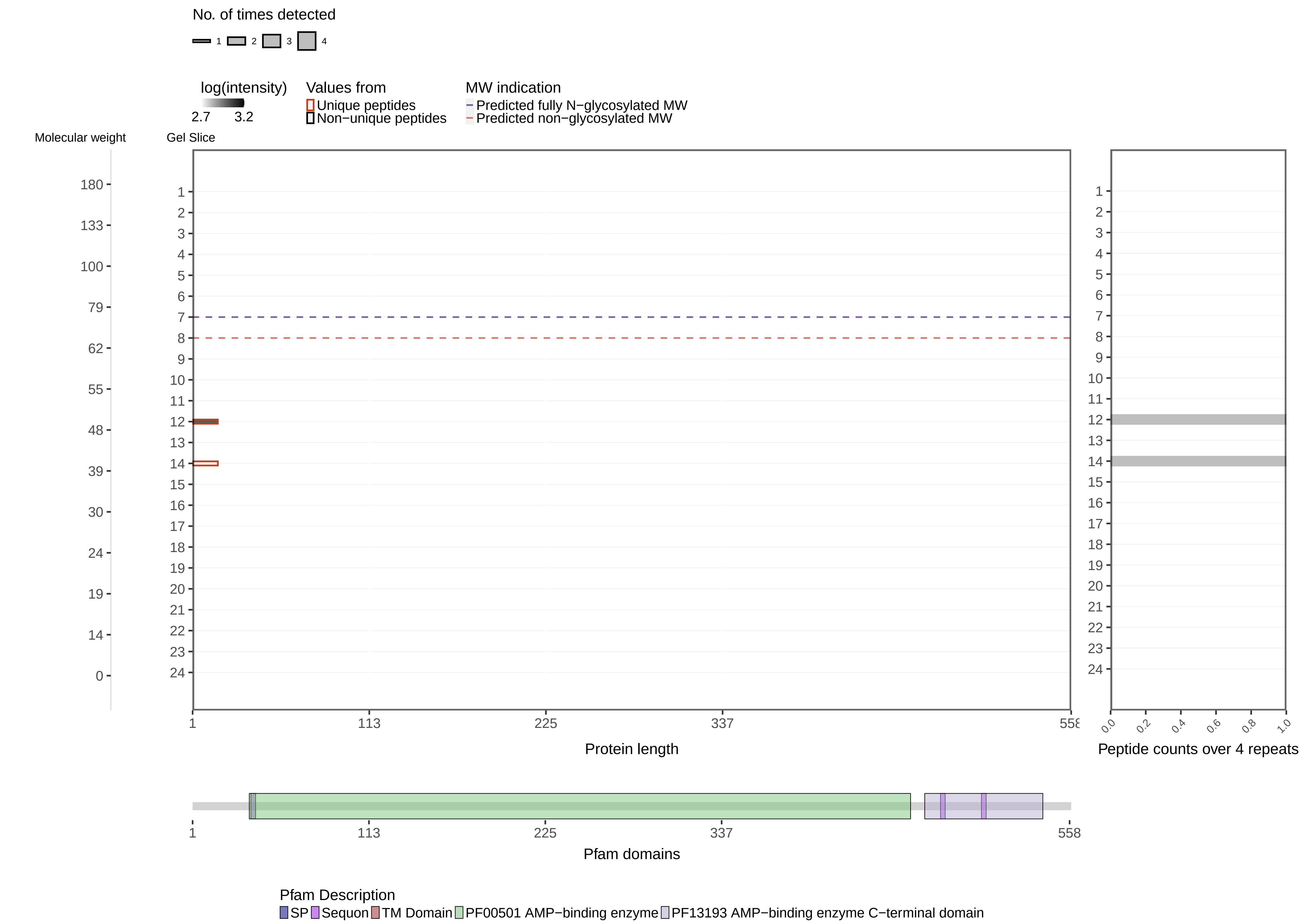


Protein sequence:

GAAILIMQKFEIKGLMELVEKYKVTIAPFVPPIVLAIAKCPLVHKYDLSSIRMIMSGAAPMGKELEDTVRAKLPNAILGQGYGMTEAGPVLSMCLAFAKQPFEVKSGSCGTVVRNAEMKIVDIDTGASLPLNHAGEICIRGDQIMKGYLNNPEATEETIDKEGWLHSGDIGYIDEDDQVFIVDRLKELIKYKGYQVAPAELEALLISNPNITDAAVVPMEDEVAGEVPVAFVVRANGSKISEEEIKKYISEQVIFYKRINRVFFTEAIPKAPSGKILRKDLRAKLAAGLAC*

NbL08g05090.1 bifunctional_dethiobiotin_synthetase/78−diamino−pelargonic_acid_aminotransferase_mitochondrial [MW = 95.8 kDa]

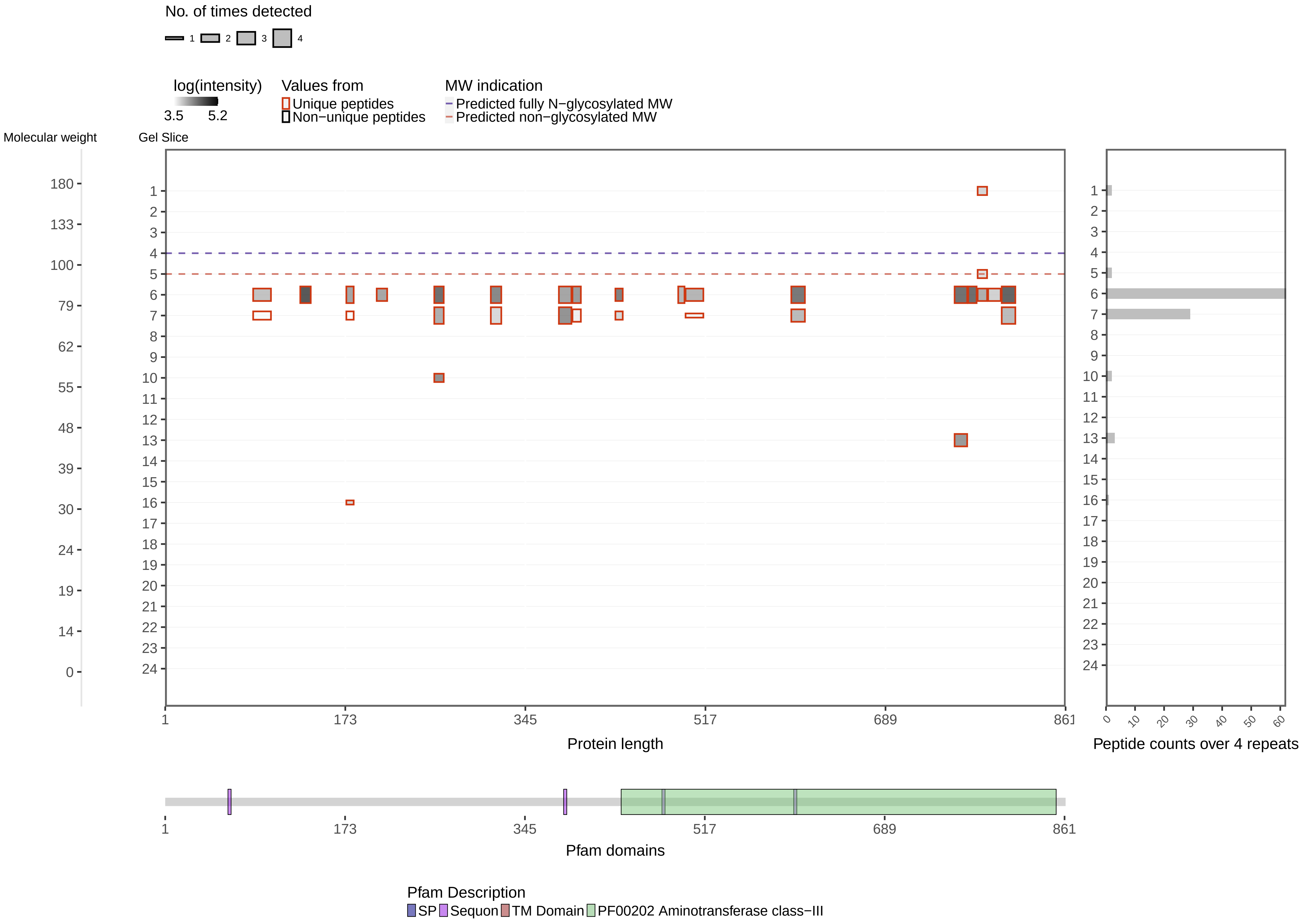


Protein sequence:

[illegible]

NbL08g05050.1 FGGY_carbohydrate_kinase_domain−containing_protein−like [MW = 69.08 kDa]

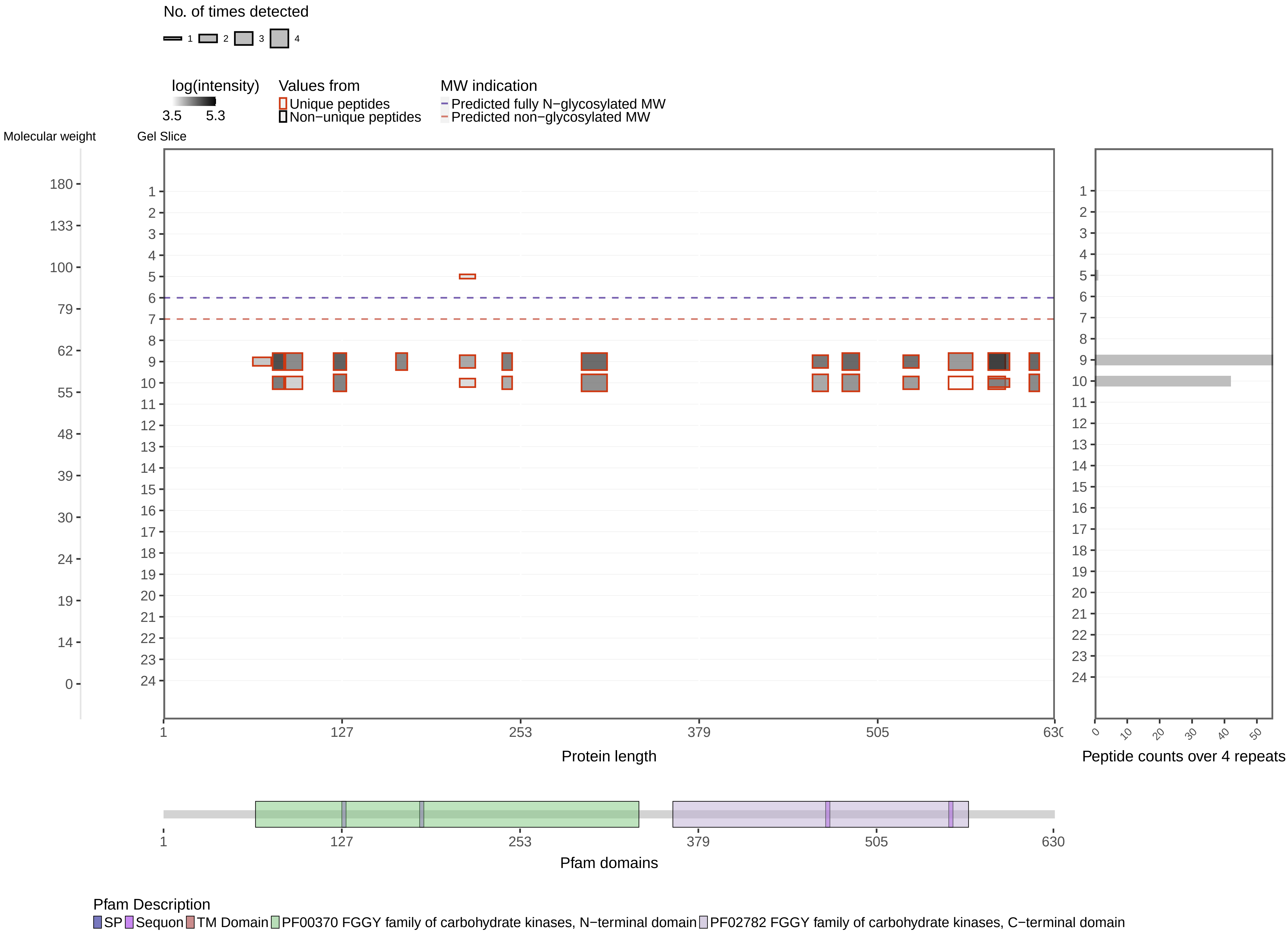


Protein sequence:

VAFPGHALGSGLTLDAAKELGLMAGTPVGTALIDAHAGGVGVMESVPASDSESIVDEDAISRRMVLVCGTSTCHMAVSRTKLFIPGVWGPFWSAMVPEYWLTEGGQSATGALLDYIIENHVASPHLANRAASRRISIFDLLNELLESMKQEEGSQFISALTKDLHILPDFHGNRSPIADPKSKGMISGLTLDTSEKQLALLYLATVQGIAYGTRHIVEHCNANGHKIDTLLACGGLAKNHLFVQEHVDIIGYPIILPRENESVLLGASILGAVASKKYPTVREAMKAMNAAGQVVYP[illegible]

### NbL08g04960.1 xylose_isomerase−like [MW − SP = 50.79 kDa]

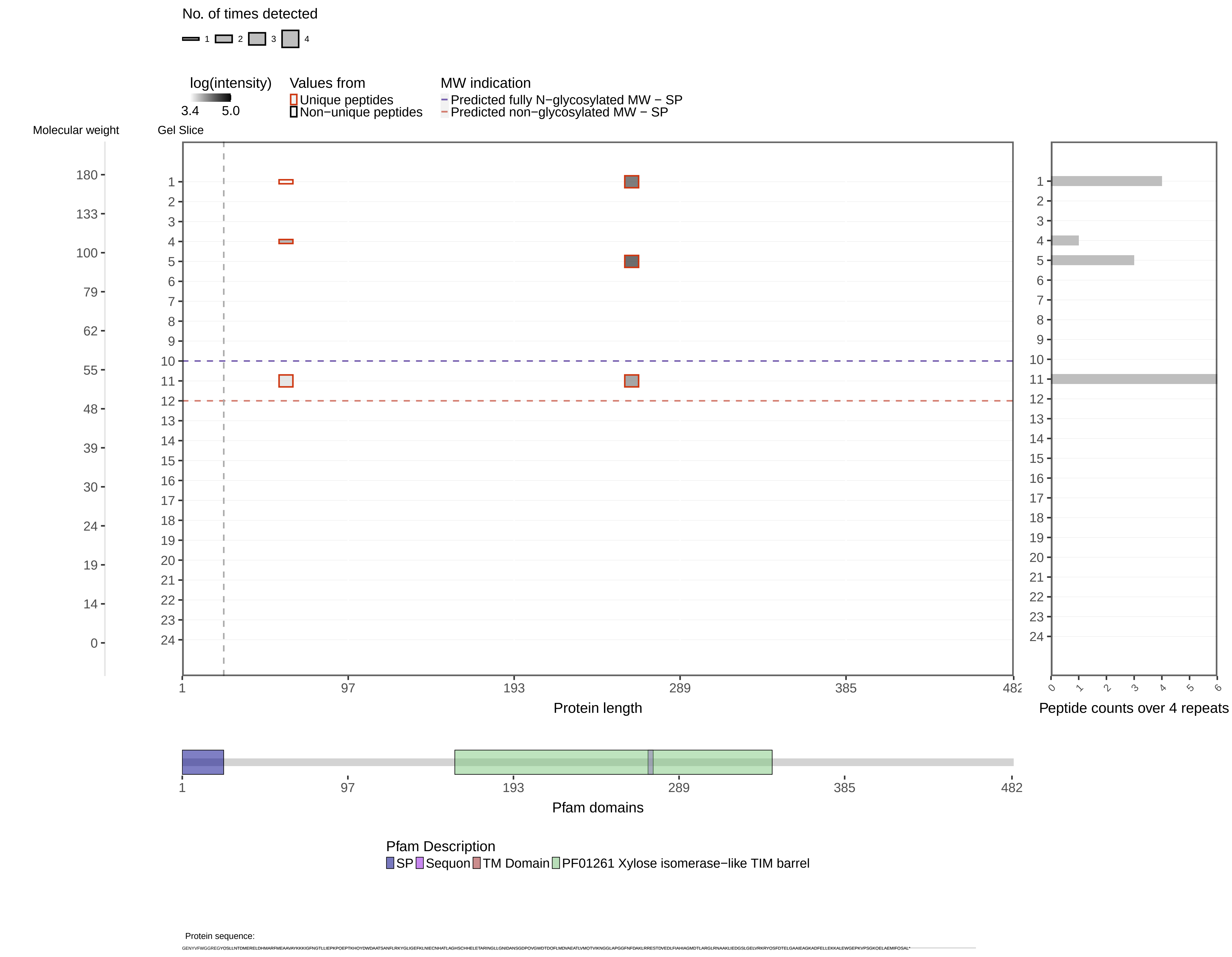

### NbL08g04950.1 ADP−ribosylation_factor_1 [MW = 20.31 kDa]

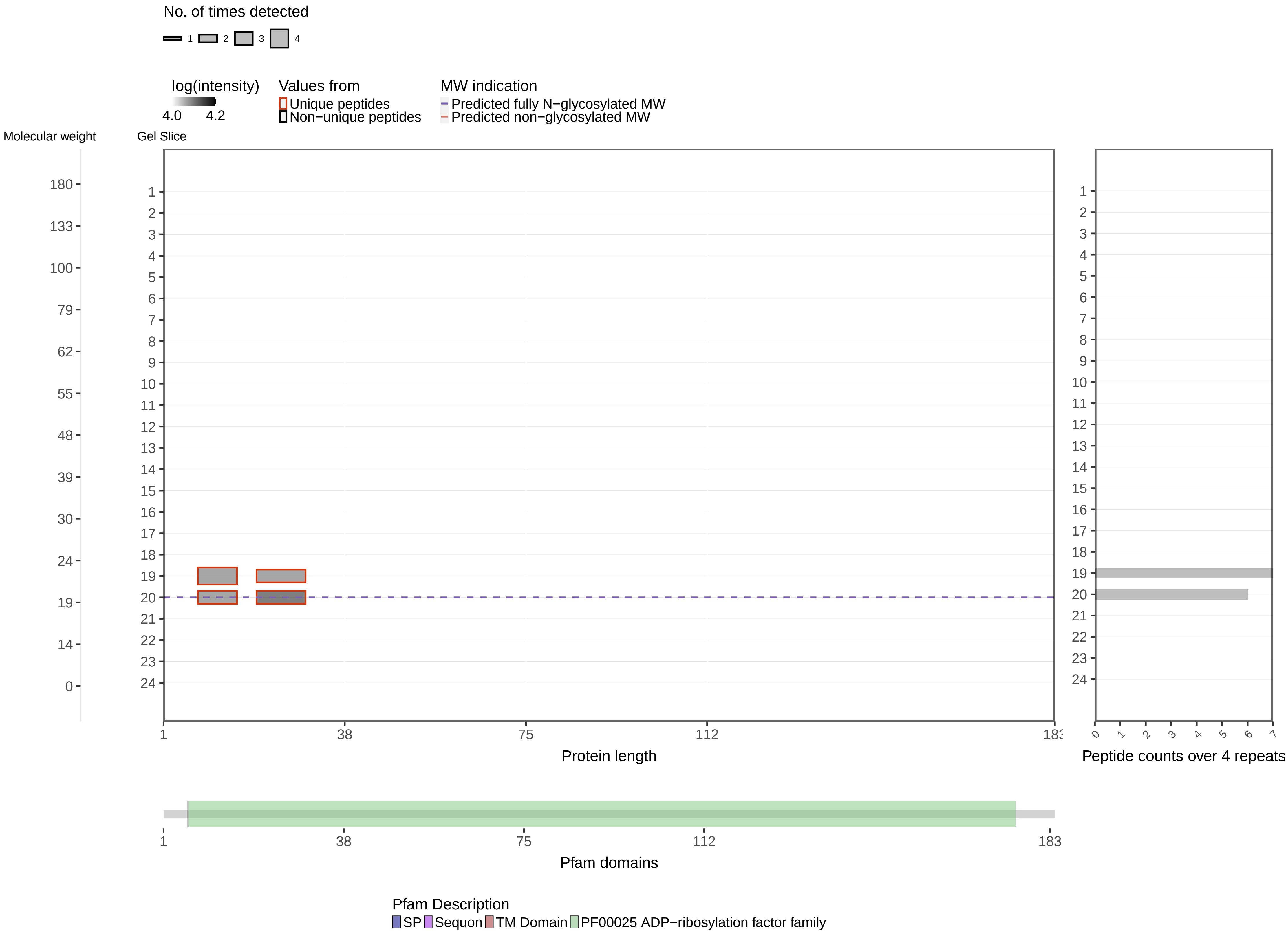


Protein sequence:

/RCYFPNTQAIIYVVDSSDTDRLVVAKEEFHAILEEEELKGAVVLVYANKQDLPGALDDAAVTEALELHKIKNRQWAIFKTSAIKGEGLFEGLDWLSNTLKSGGS*

### NbL08g04810.1 adenylosuccinate_lyase−like [MW = 60.93 kDa]

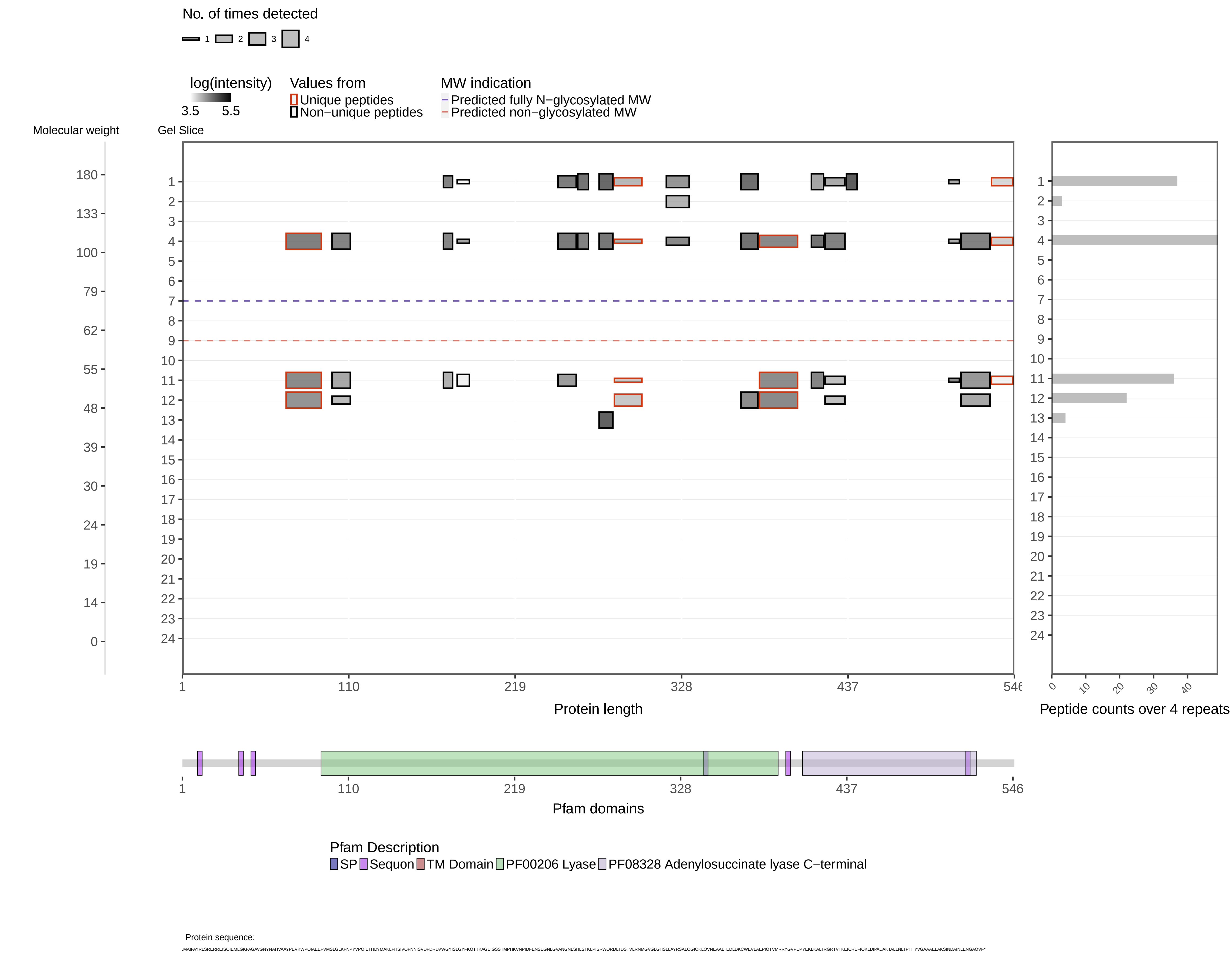

NbL08g04560.1 tetraspanin−8−like [MW = 30.12 kDa]

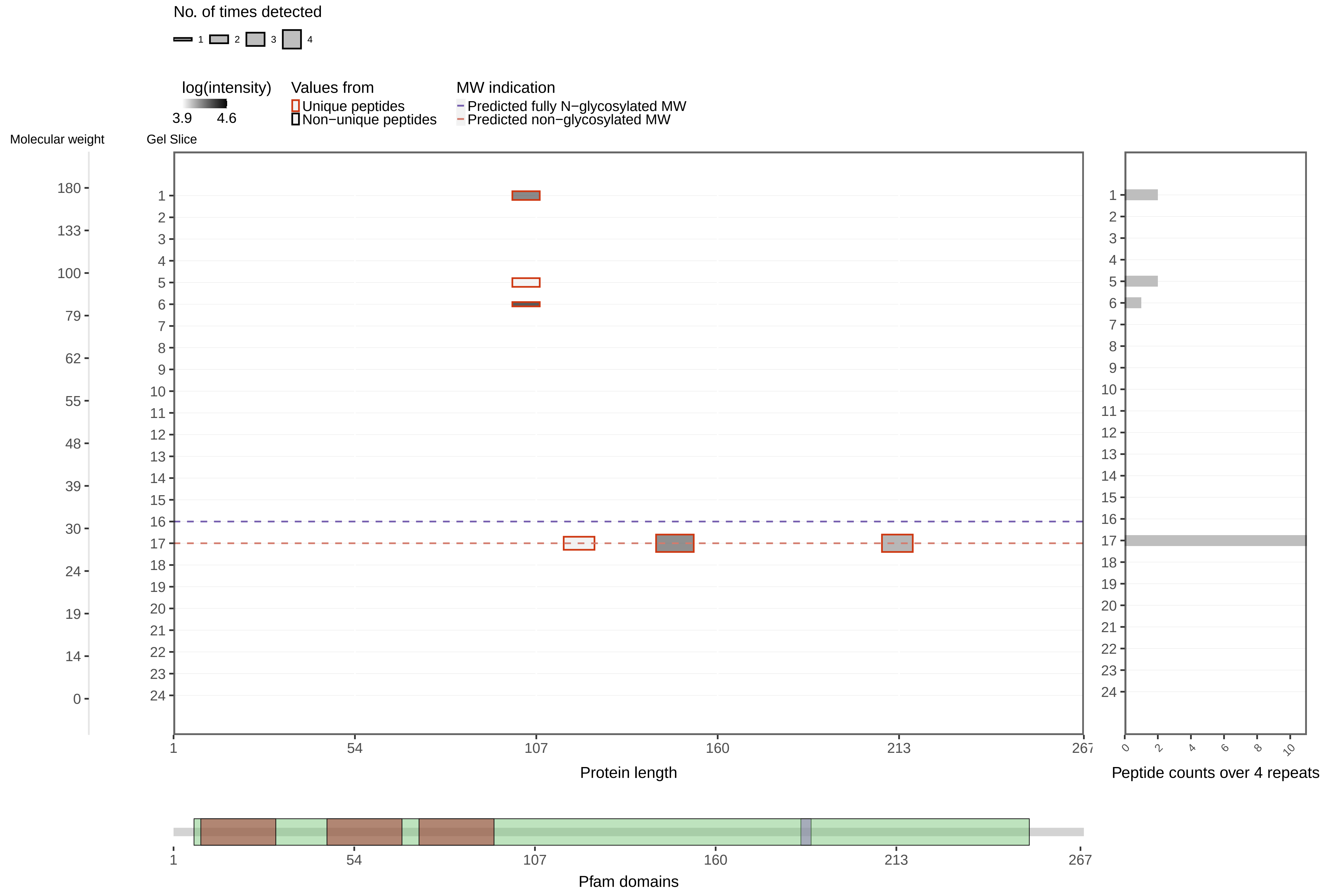


Protein sequence:

RVDNHWSRIHSCLQDSKICQSLIEGSNTKADDFFKKHLSALQSGCCKPSNDCNFEYVSPTNWTRTPSSSLDNPDCTAWSNDPKILCYGCQSCKAGLLDNLKSDWKRVAVLNIIFLVFLIVVYSIGCCAFRNNREDNAWKRYP*

NbL08g04470.1 tetratricopeptide_repeat_protein_1−like [MW = 31.06 kDa]

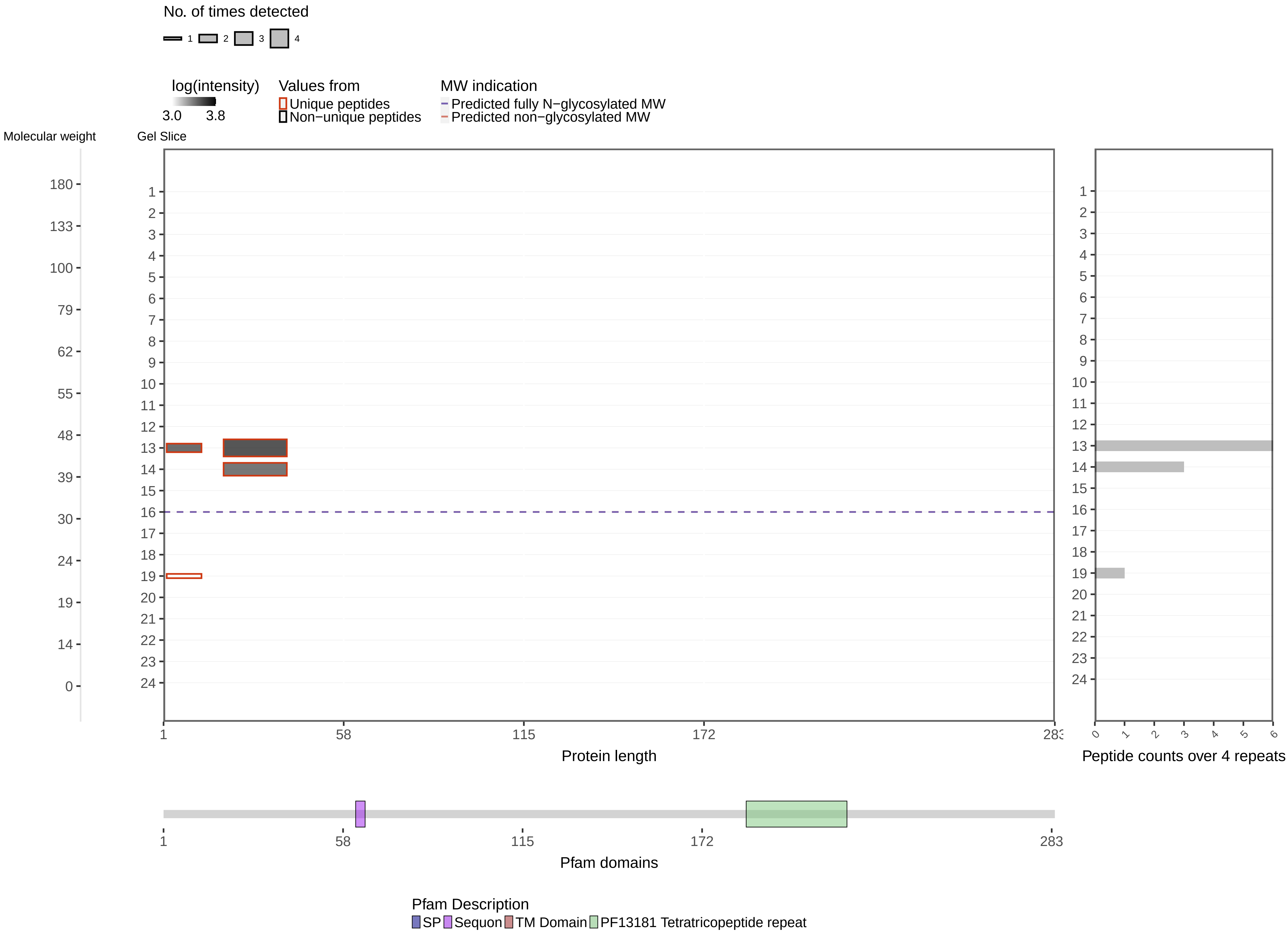


Protein sequence:

:ALSKYELALQVAADIPSSTEIRSICHANRAACFSKLGKHGETIKECTKALELNPTYMKALVRRAEAHEKLEHYDEAVTDMTKIMELEPSNDQARRSVMRLKPLADEKREKMKEEMIGKLKEMGNSILGRFGMSVDNFKAVKDPNTGSYSVSFQK*

### NbL08g04360.1 putative_GDP−L−fucose_synthase_2 [MW = 34.78 kDa]

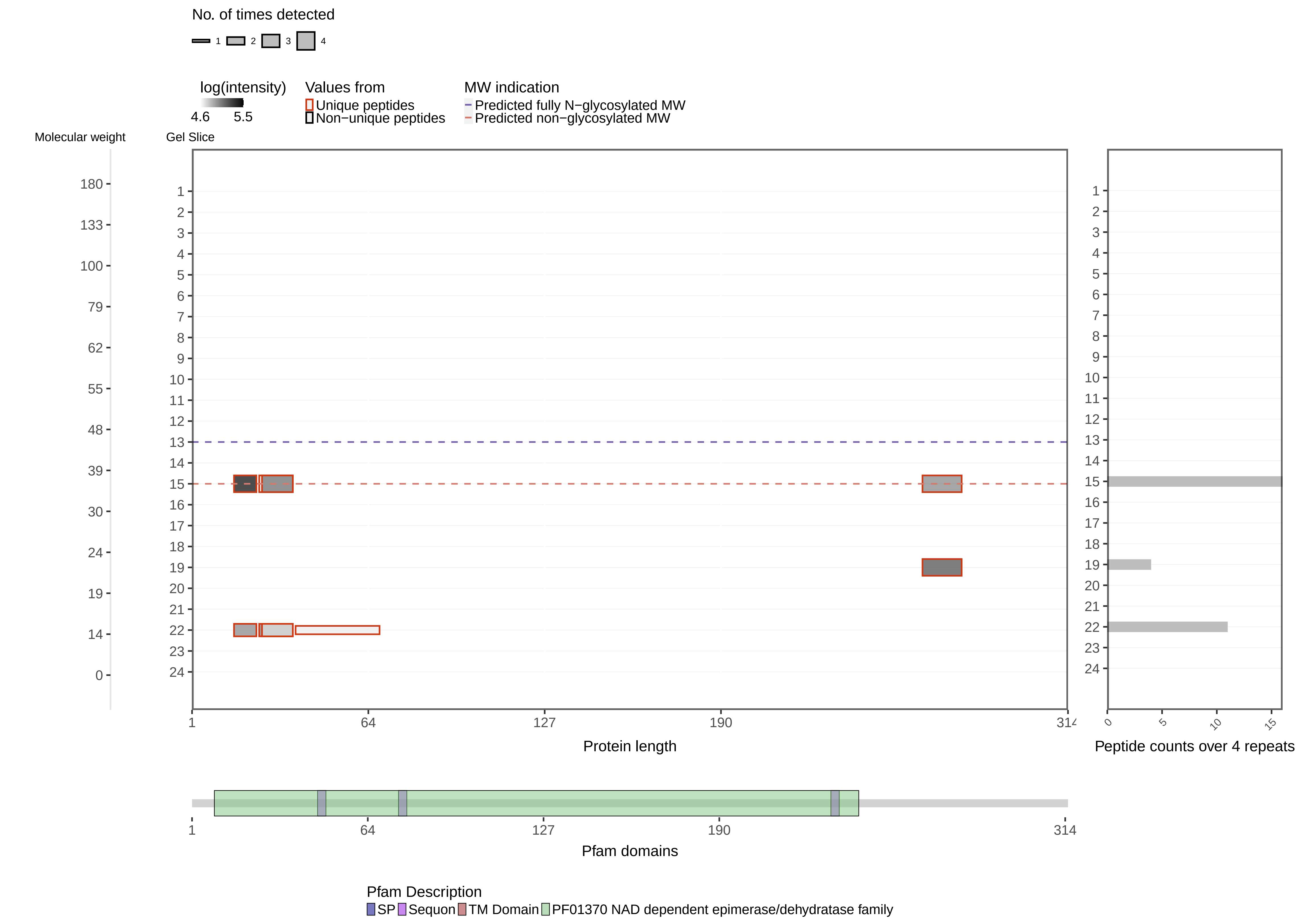


Protein sequence:

IKMCOAYRLOHNFDAISAMPTNLYGTNDNFHPENSHVLPALLRRFHEAKINNLDKVVVWGTGSPLREFLHVDDLADAVIFLLENYSDLEHVNVGSGSEVTIKELAELVKEVVGFKGELVWDSTKPDGTPRKLMHTSKLVGLGWTPKISLRDGLVDTYKWYLENYGKO*

NbL08g04330.1 clustered_mitochondria_protein [MW = 202.26 kDa]

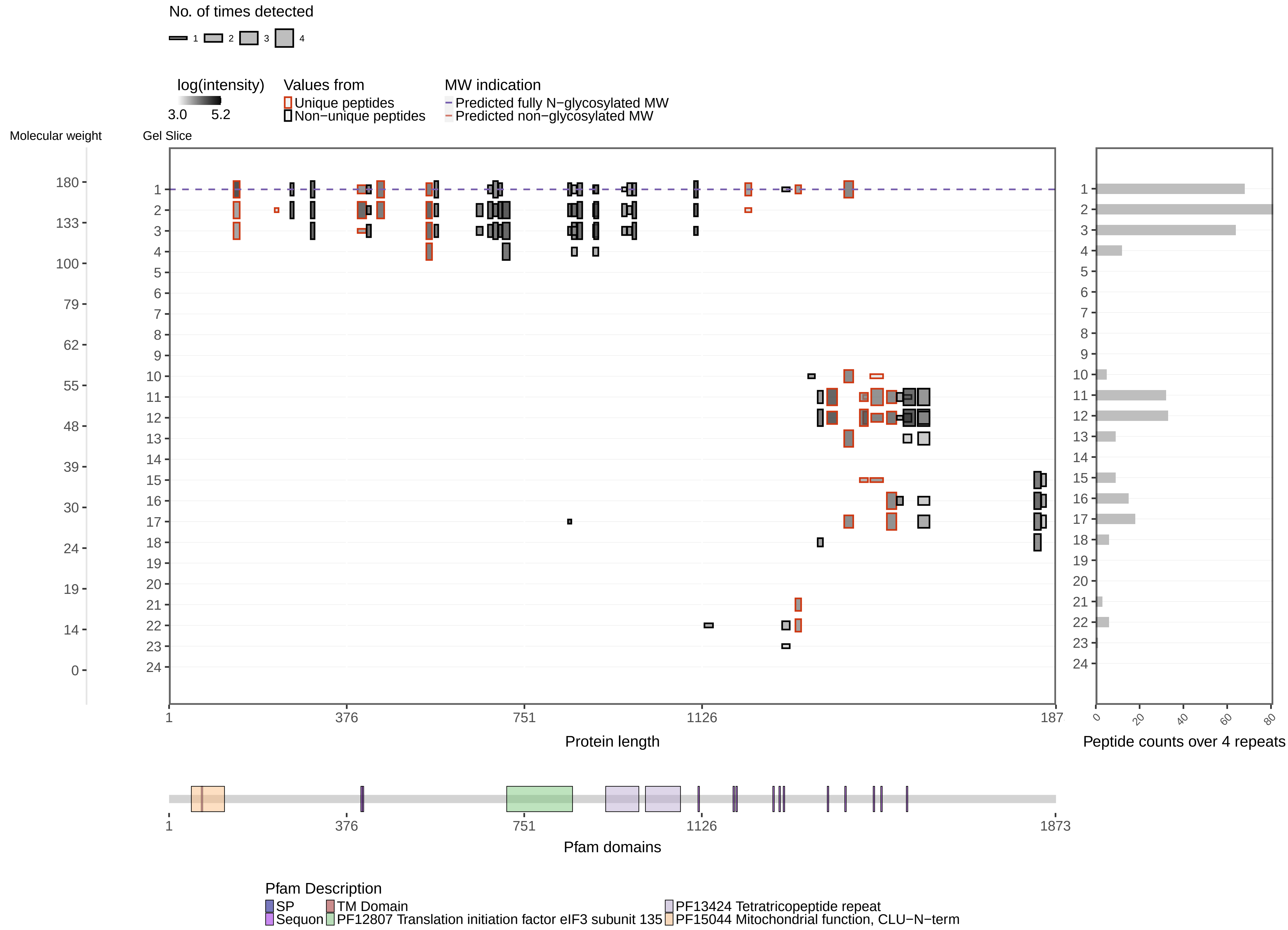


Protein sequence:

YSLLAVVLYHTGDFNQATIYQQKALDINERELGLDHPDTMKSYGDLAVFYYRLQHTELALKYVNRALYLLHLTCGPSHPNTAATYINVAMMEEGLGNVHVALRYLHEALKCNQRLLGADHIQTAASYHAIAIALSLMEAYSLSVQHEQTTLQILQAKLGPDDLRTQNFLQDAAAWLEYFESKALEQQEAARNGTPKPDASISSKGHLSVSDLLDYIAPDAEMKAREAQKKQARAKVKGKAGQNGGIATDEFEKDELLSPNSPVVENSSDKENKSELDNKLELQIADSTAKQSDHI

NbL08g04250.1 V−type_proton_ATPase_subunit_H−like [MW = 51.53 kDa]

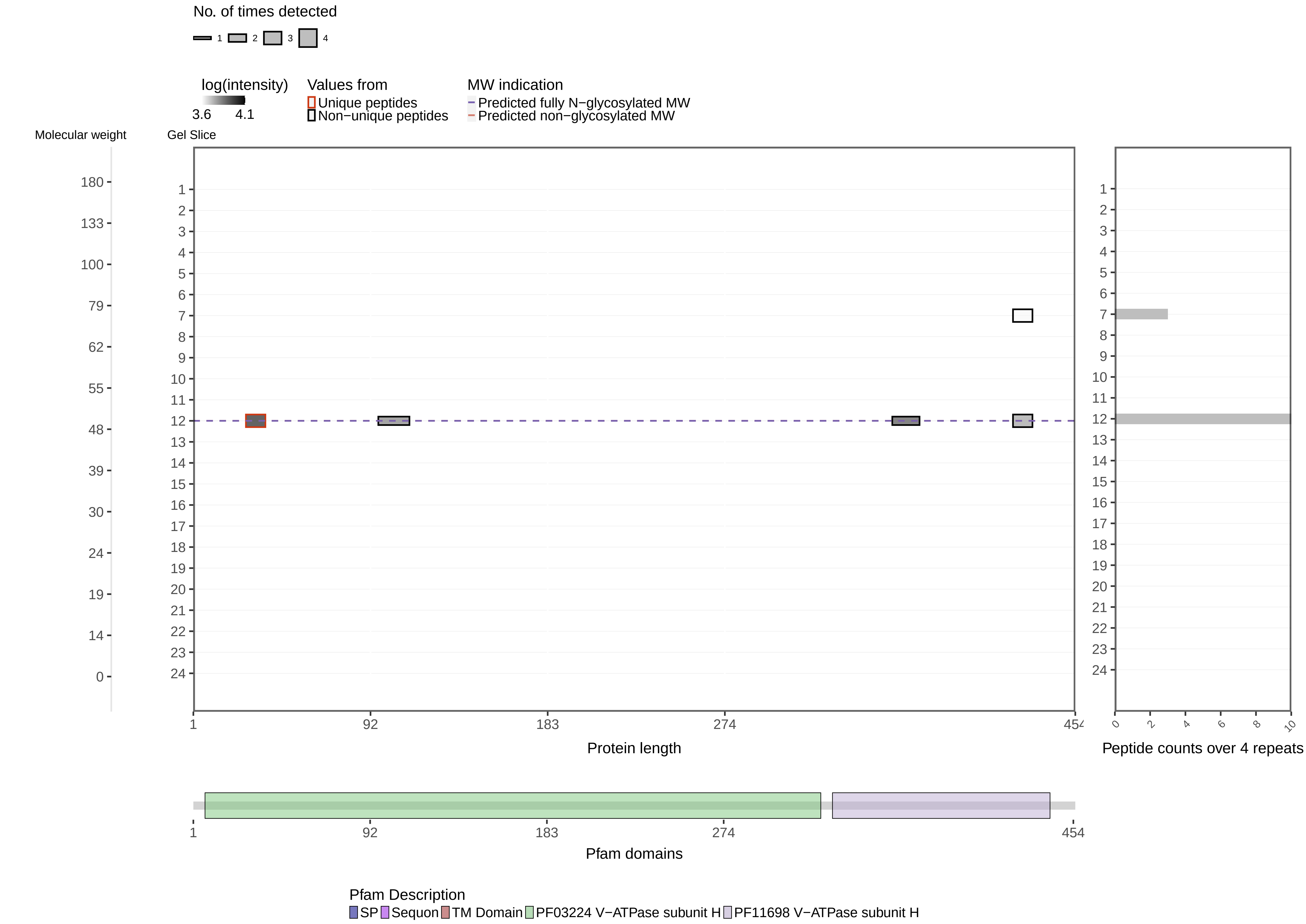


Protein sequence:

LVPLISPASTQQSIQLLYETCLCVWLLSYYEPAIEYLATSRALPRLIEVVKGSTKEKVVRVVILTLRNLLSKGSFSILMVDLGVLQIVQSLKAQAWSDEDLLDALNQLEEGLKDNIKKLSSFDKYKQEVLLGHLDWSPMHKDPLFWRENINSFEENDFQILRVLITILDTSSDPRTLAVACYDLSQFIQCHSAGRIIVNDLKAKERVMKLLNHENAEVTKNALLCIQRLFLGAKYASFLQV*

### NbL08g04080.1 DNA−directed_RNA_polymerases_II_IV_and_V_subunit_11−like [MW = 13.62 kDa]

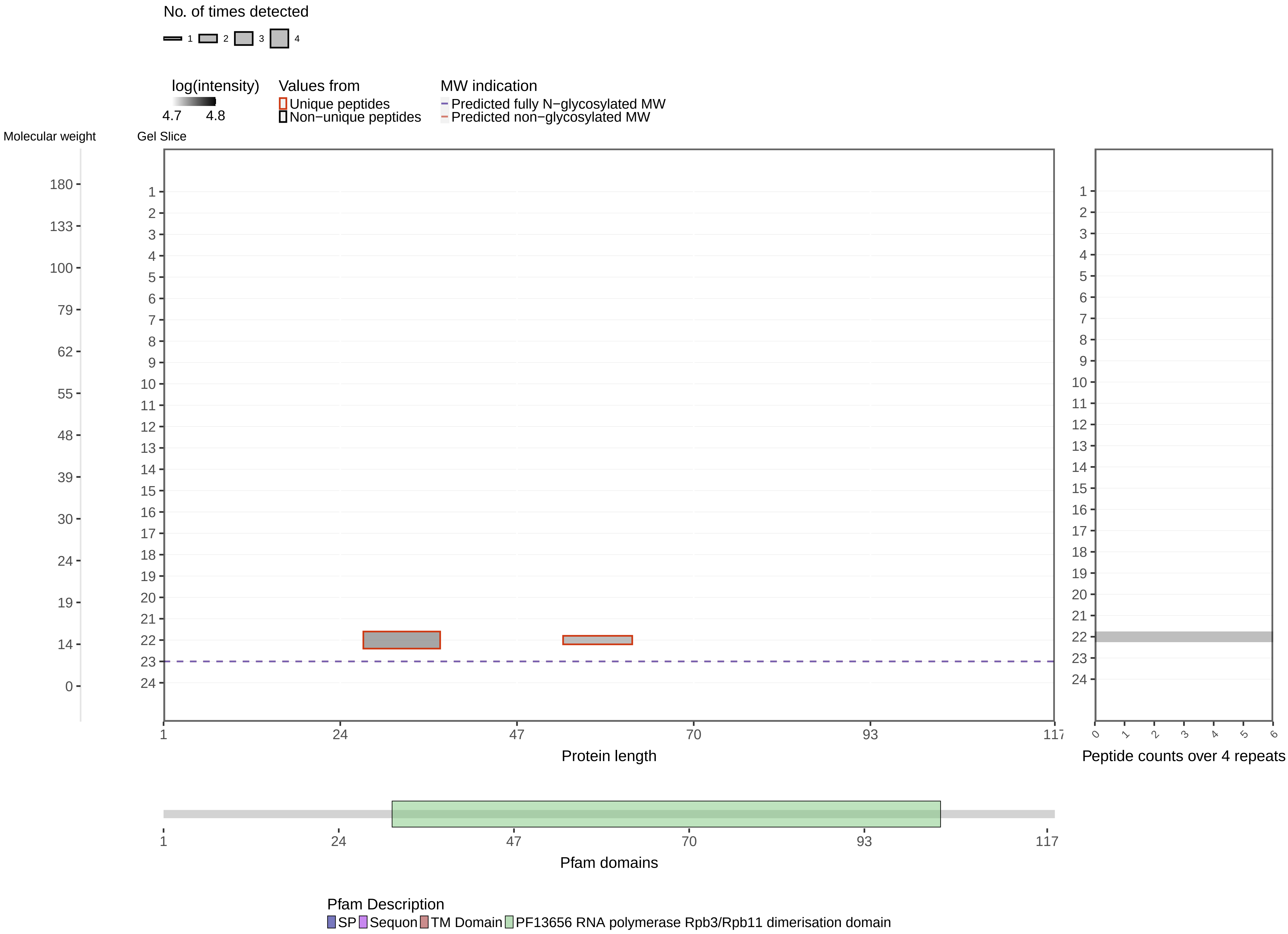


Protein sequence:

.RMQLHRDDNVLFAGYKLPHPLQYKILLRIQTTSQSSPMQAYNQAINDLDKELDHLKNQFEGELAKHSREY*

NbL08g03960.1 ruBisCO_large_subunit−binding_protein_subunit_alpha−like [MW = 62.11 kDa]

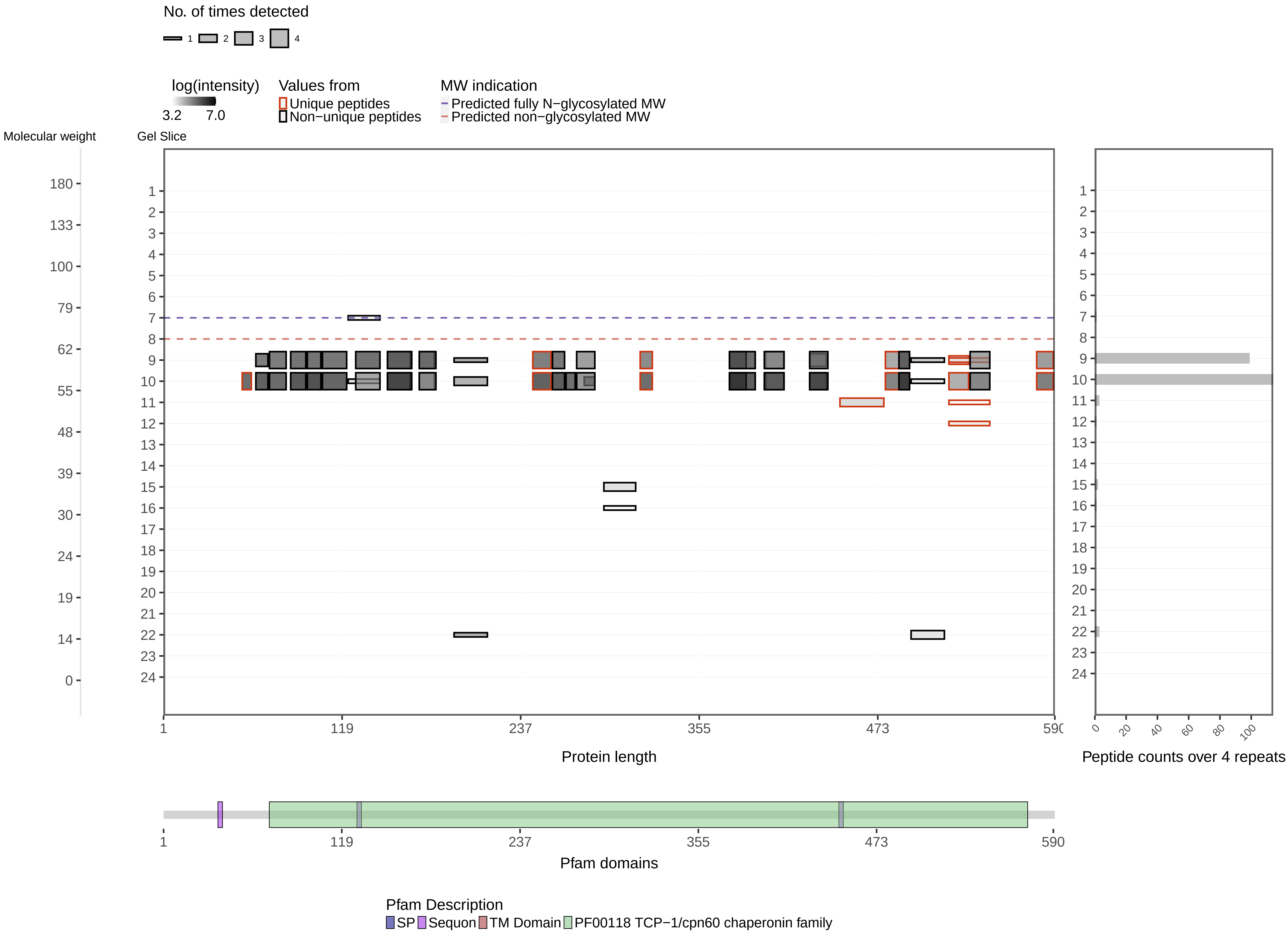


Protein sequence:

NbL08g03760.1 cell_division_cycle_protein_48_homolog [MW = 89.57 kDa]

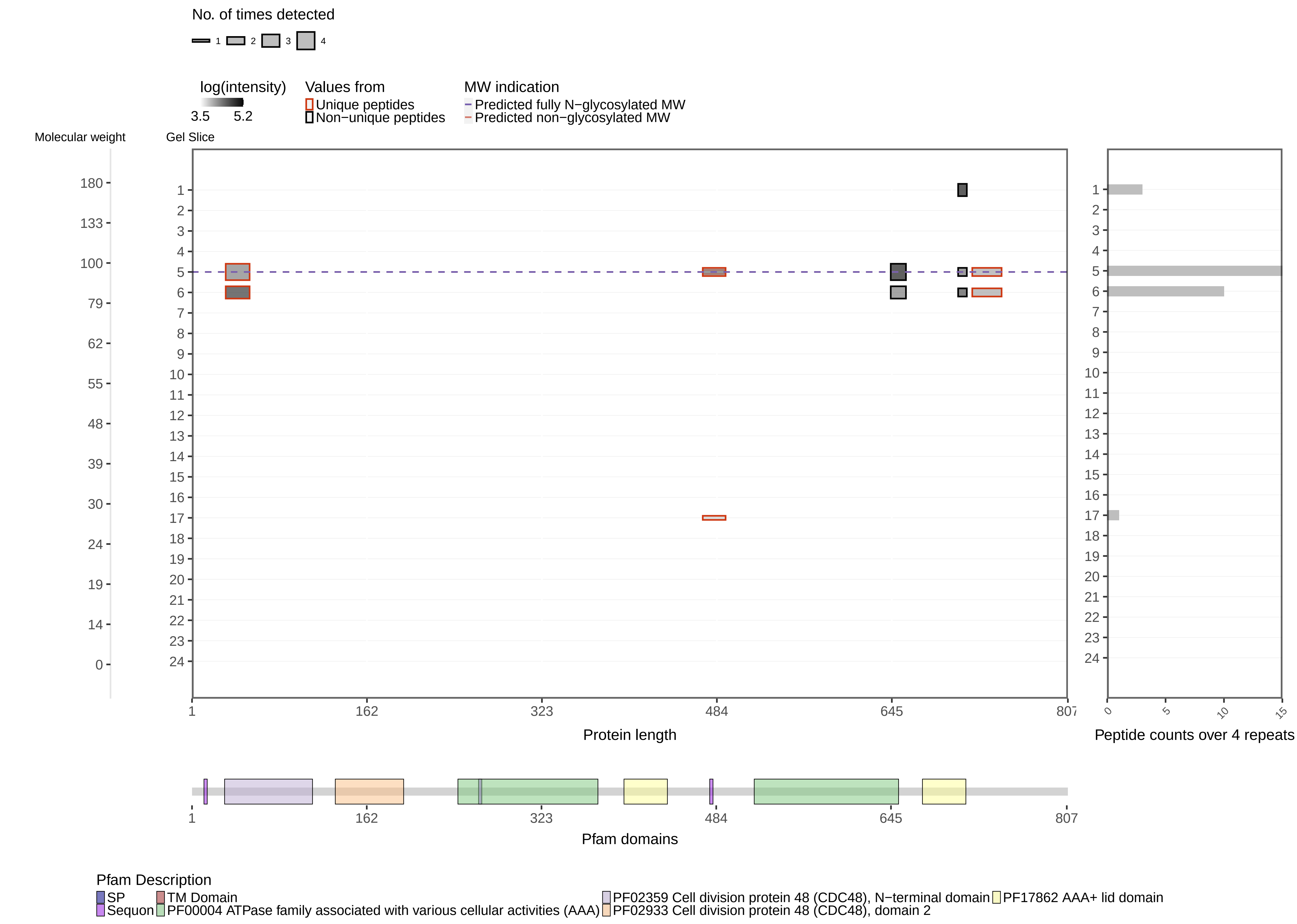


Protein sequence:

### NbL08g03740.1 elongation_factor_1−alpha [MW = 49.34 kDa]

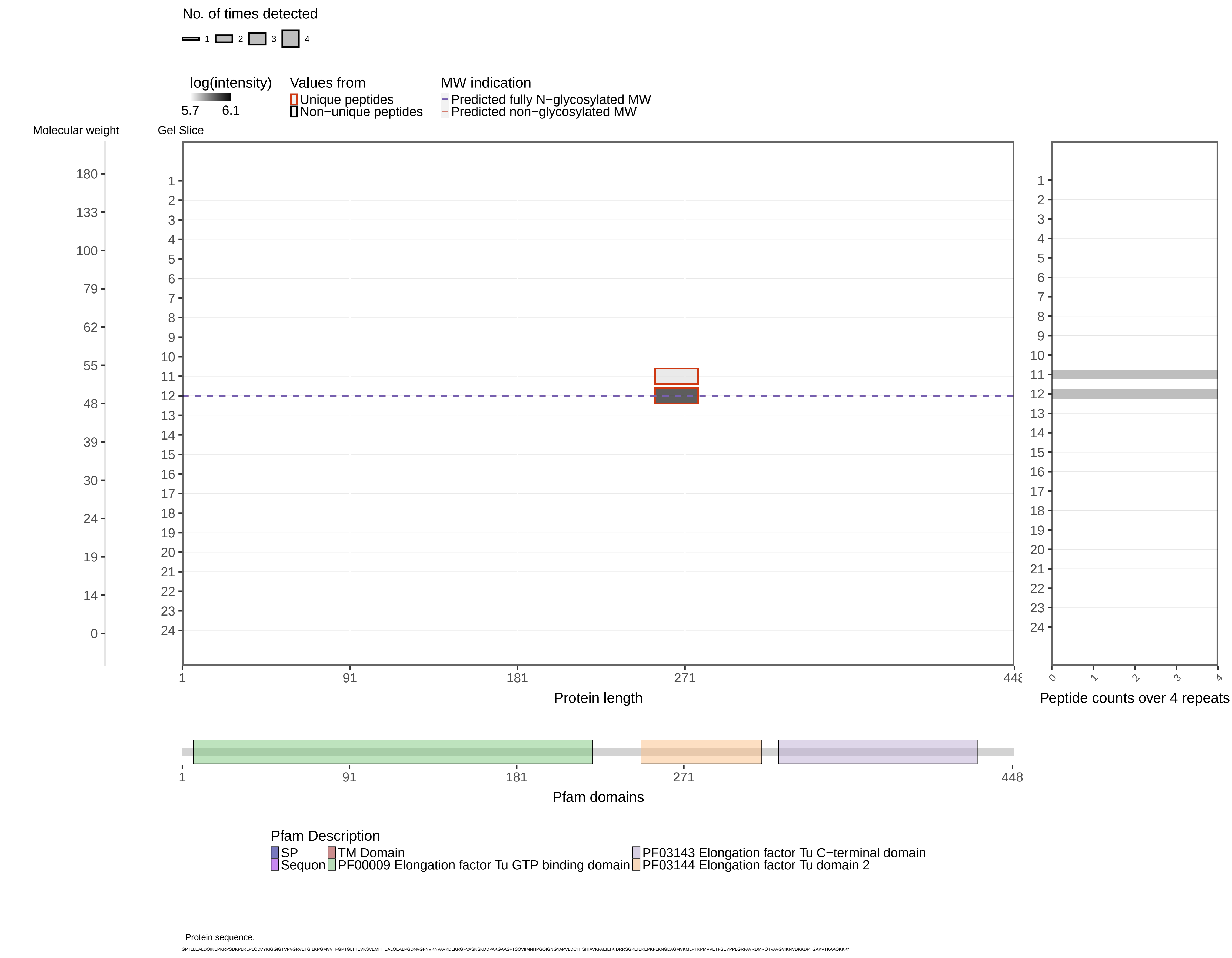

### NbL08g03640.1 uncharacterized_LOC109233996 [MW = 130.24 kDa]

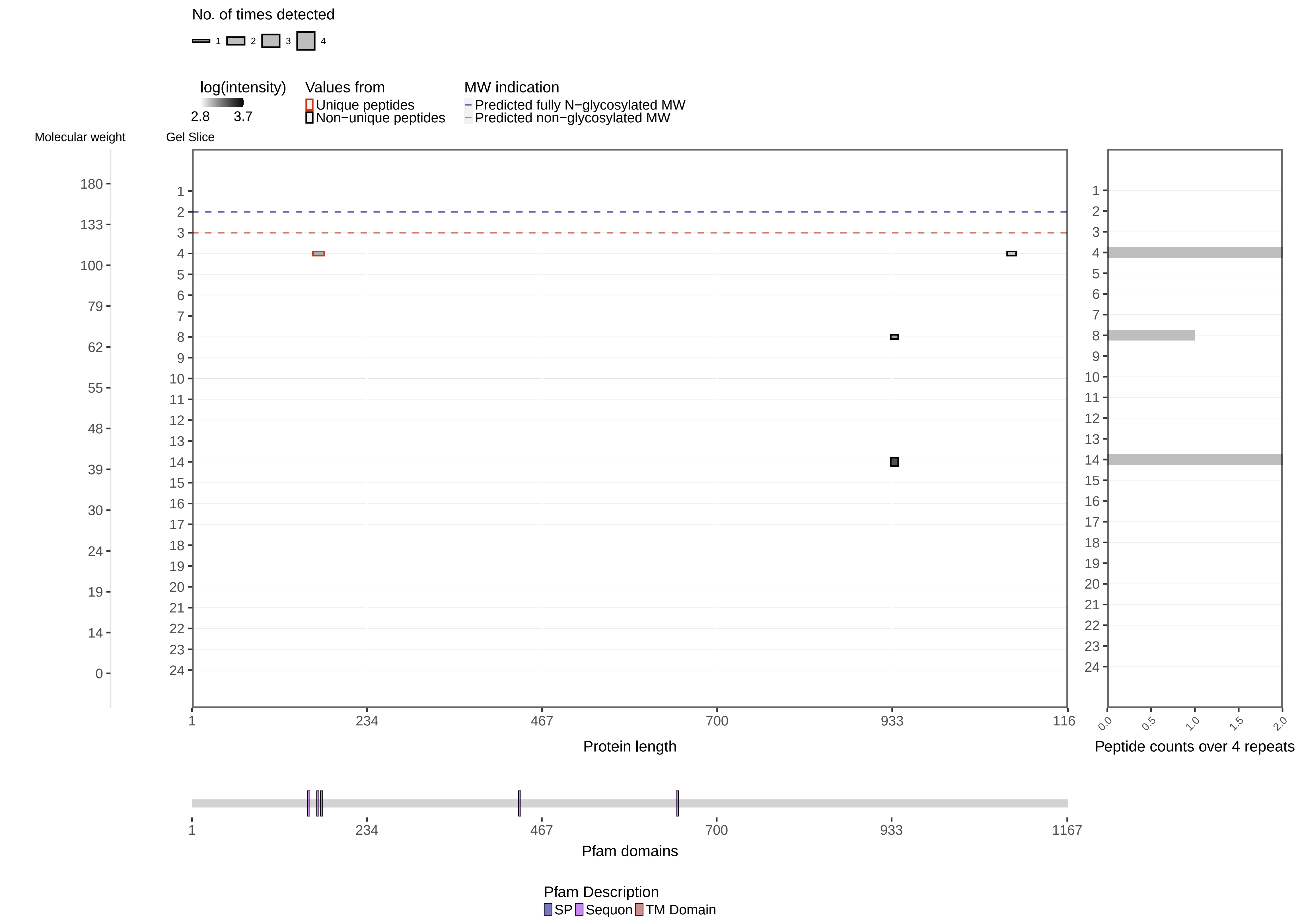


Protein sequence:

[illegible]

NbL08g03530.1 interactor_of_constitutive_active_ROPs_3−like [MW = 70.85 kDa]

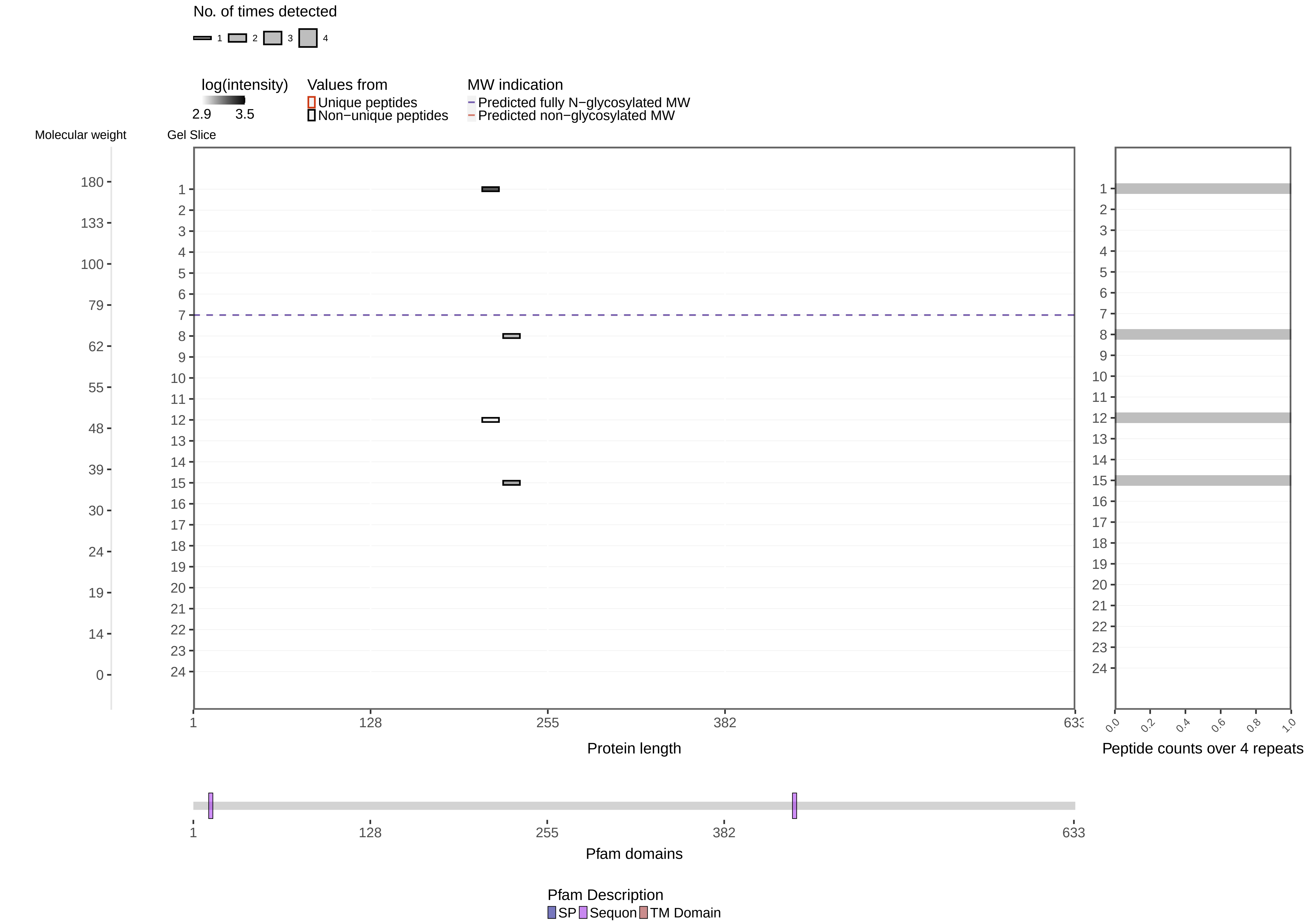


Protein sequence:

[illegible]

NbL08g03500.1 arogenate_dehydratase/prephenate_dehydratase_2_chloroplastic [MW = 43.21 kDa]

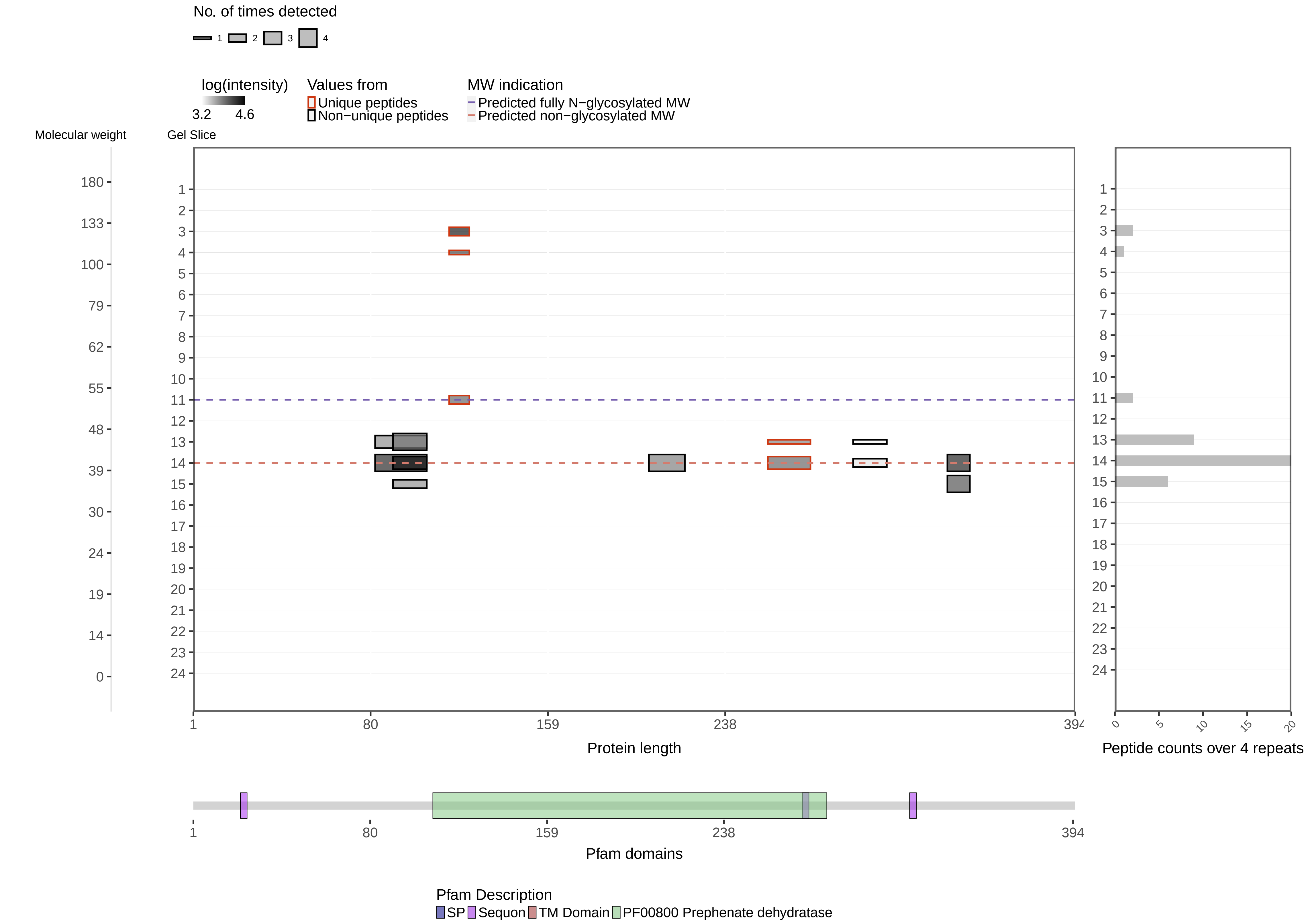


Protein sequence:

LAIRHCLMANNGVKIEDLKRVLSHPQALAQCENTLTKLGLVREAVDDTAGAAKYIAFQKLKDAGAVASLAAARIYGLNVLAQDIQDDSDNVTRFLMLAREPIIPGTDKPFKTSVVFSLDEGPGVLFKALAVFAMRNINLTKIESRPLQKQALRVLDDSTDGFPKYFPYLFYVDFEASMADQRAQNALGHLKEFATFLRVLGSYPSDSSVA*

NbL08g03250.1 2−dehydro−3−deoxyphosphooctonate_aldolase_1 [MW = 31.9 kDa]

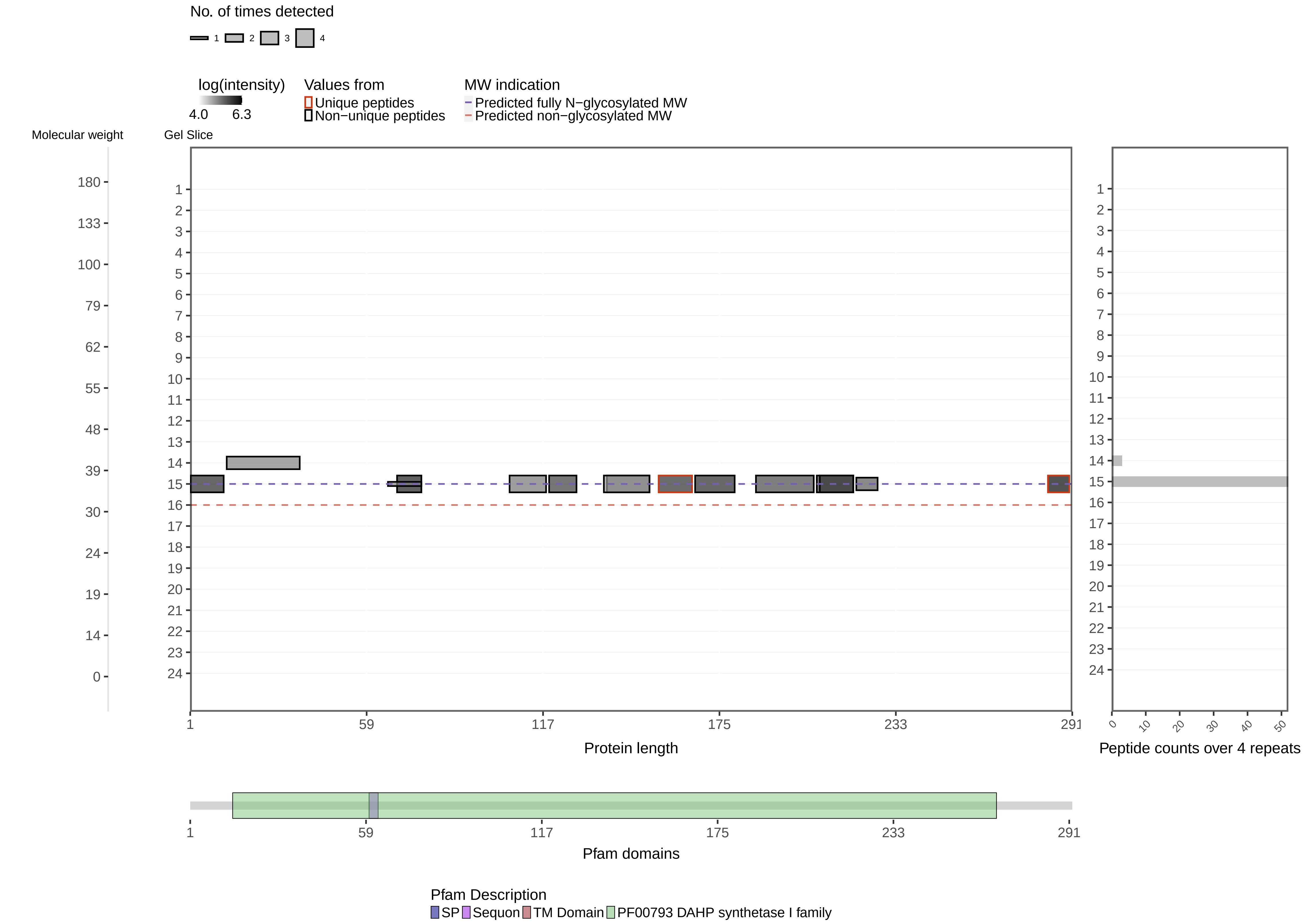


Protein sequence:

KGQFCAPSVMANSAEKVRLAGNHNVMVCERGTMFGYNDLIVDPRNFEWMREANCPVVADITHSLQQPAGKKLEGGGVASGGLRELIPCIARTAVAVGIDGIFMEVHNDPLSAPVDGPTQWPLRHLEELLEELVTLGRVSKGKQQFKIDLTPFRD*

### NbL08g03010.1 receptor−like_protein_kinase_HERK_1 [MW − SP = 89.5 kDa]

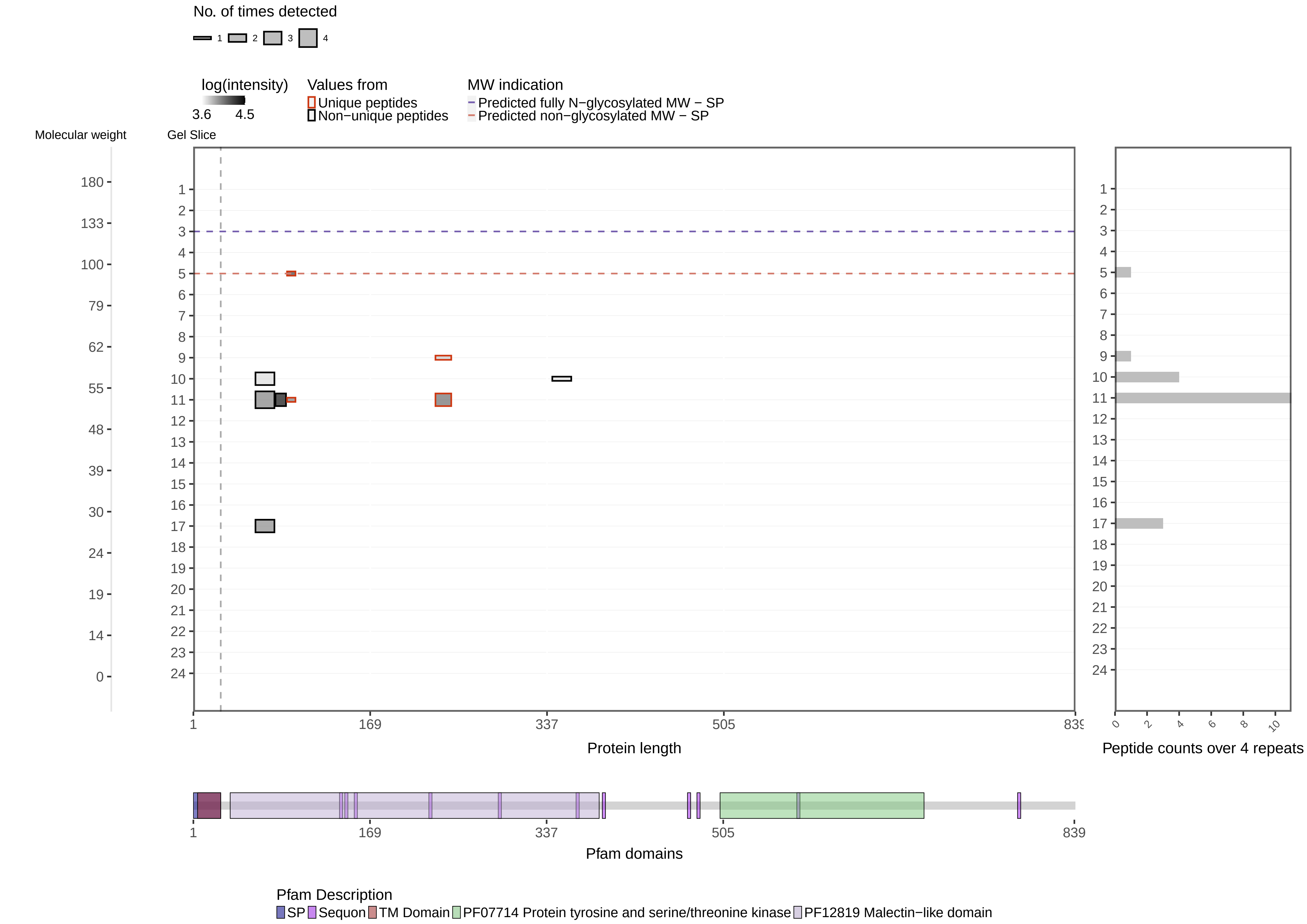


Protein sequence:

SRAKKNVGVIVGVSVGVPLILLIVGILFCMHRRKKQELLAQSKIWIPLSVNGGNSHTMGSKYSNGTTISAASNLSYRVPFPALLEATSNFDESLVIGIGGFGKVYRGVLCDGTKVAVKRGNPKSQQGLAEFRTEIEMLSQFRHRHLVSLMGYCDEKNEMILVYEYMENGTLKSHLYGSDLPSMSWKQRLEICIGSARGLHYLHTGYAKAVIHRDVKSANILLDESFMAKVADFGLSKTGPELDQTHVSTAVKGSFGYLDPEYFRRQQLTEKSDVYSFGVVLFEVLCARPVIDPSL

NbL08g02920.1 60S_ribosomal_protein_L35−like [MW = 14.19 kDa]

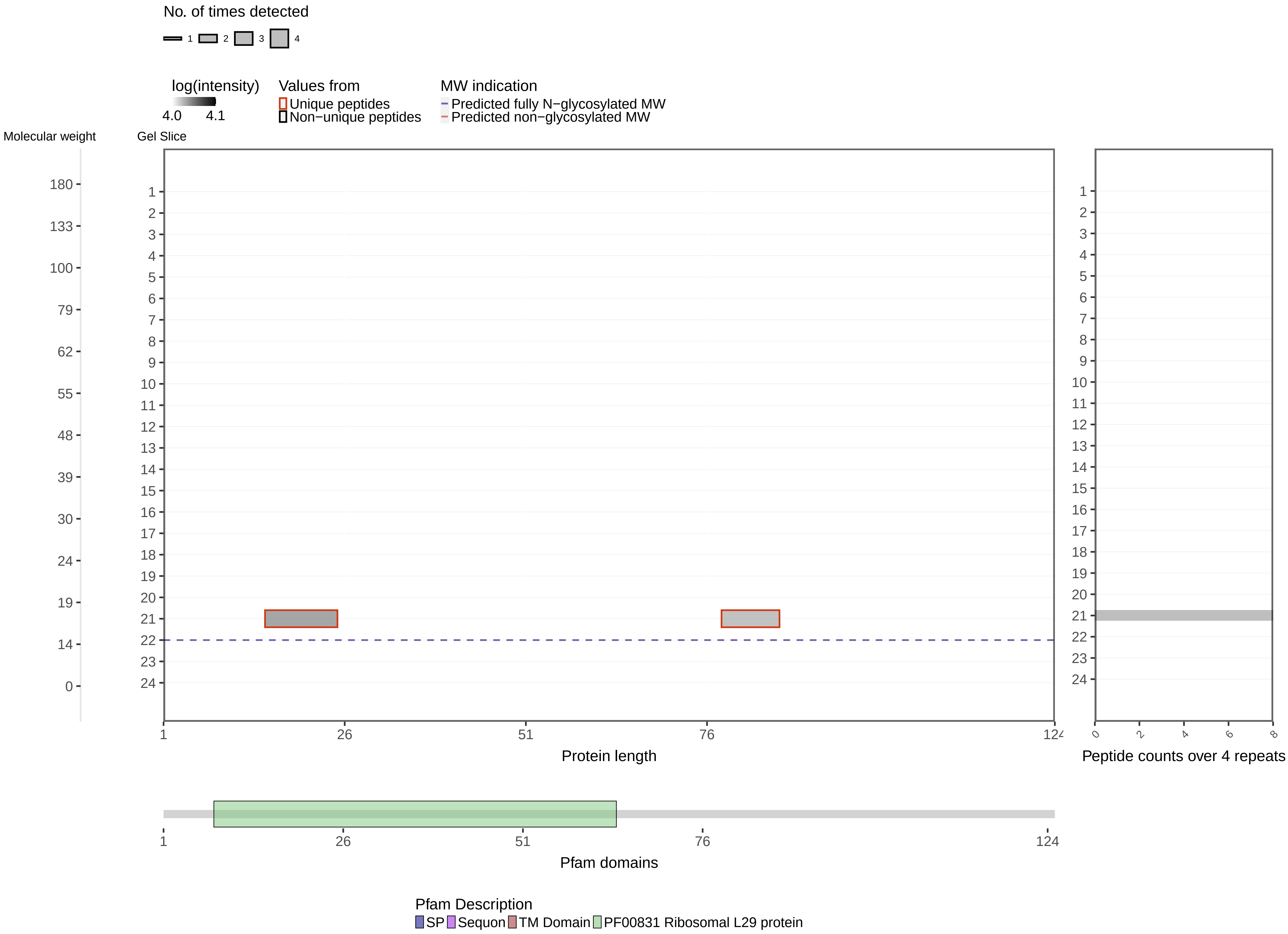


Protein sequence:

VVRLSIAQVLTVISQKQKTALREAYKNKKYLPLDLRPRKTRAIRKRLTKHQASLKTEREKKKEMYFPIRKYAIKV*

### NbL08g02790.1 monofunctional_riboflavin_biosynthesis_protein_RIBA_3_chloroplastic−like [MW = 57.86 kDa]

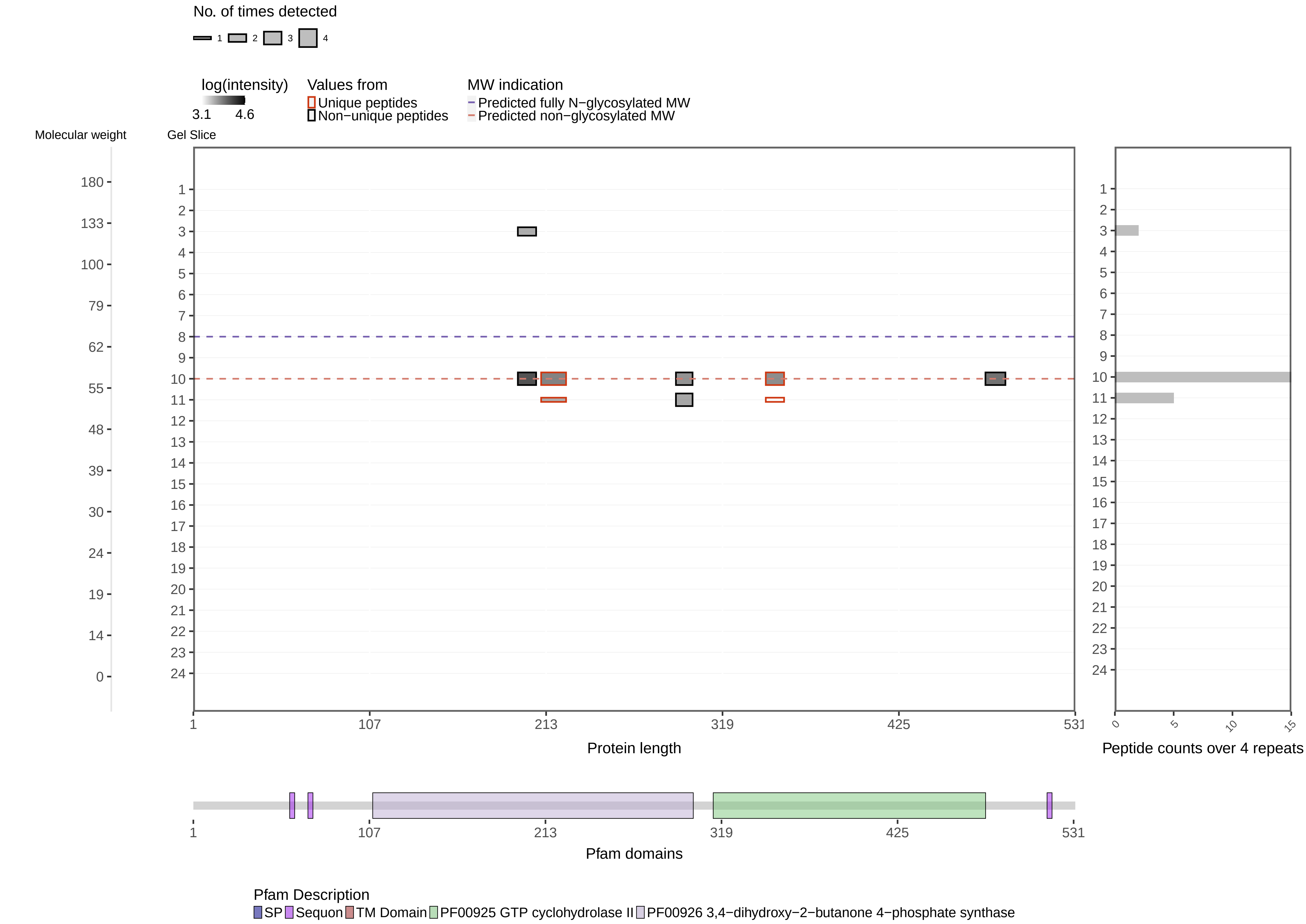


Protein sequence:

DLVMSAGLKPVSVLSAIVDKNDGSIASMLILKNLALENKIPIVSITDLVRYRRKRENLVERTAVSRLPTKWGLFEAYCYRSKLDGTEHIAVVKGDIGNGQDALVRVHSECLTGDIFGSRRCDCGNQLDLAMQLIEEAGRGVVIYLRGHEGRGIGLSNKLQAYNLQDQGHDTVEANLELGFAADAREYGIGAQMLRDIGVRTMRLMTNNPAKFTGLKGYGLAVIGRVPVLTPFTEENRKYLETKRTKMSHIYGSDVQGPIKASINRSLKKQDPSQERKES*

### NbL08g02600.1 actin−depolymerizing_factor_2−like [MW = 15.94 kDa]

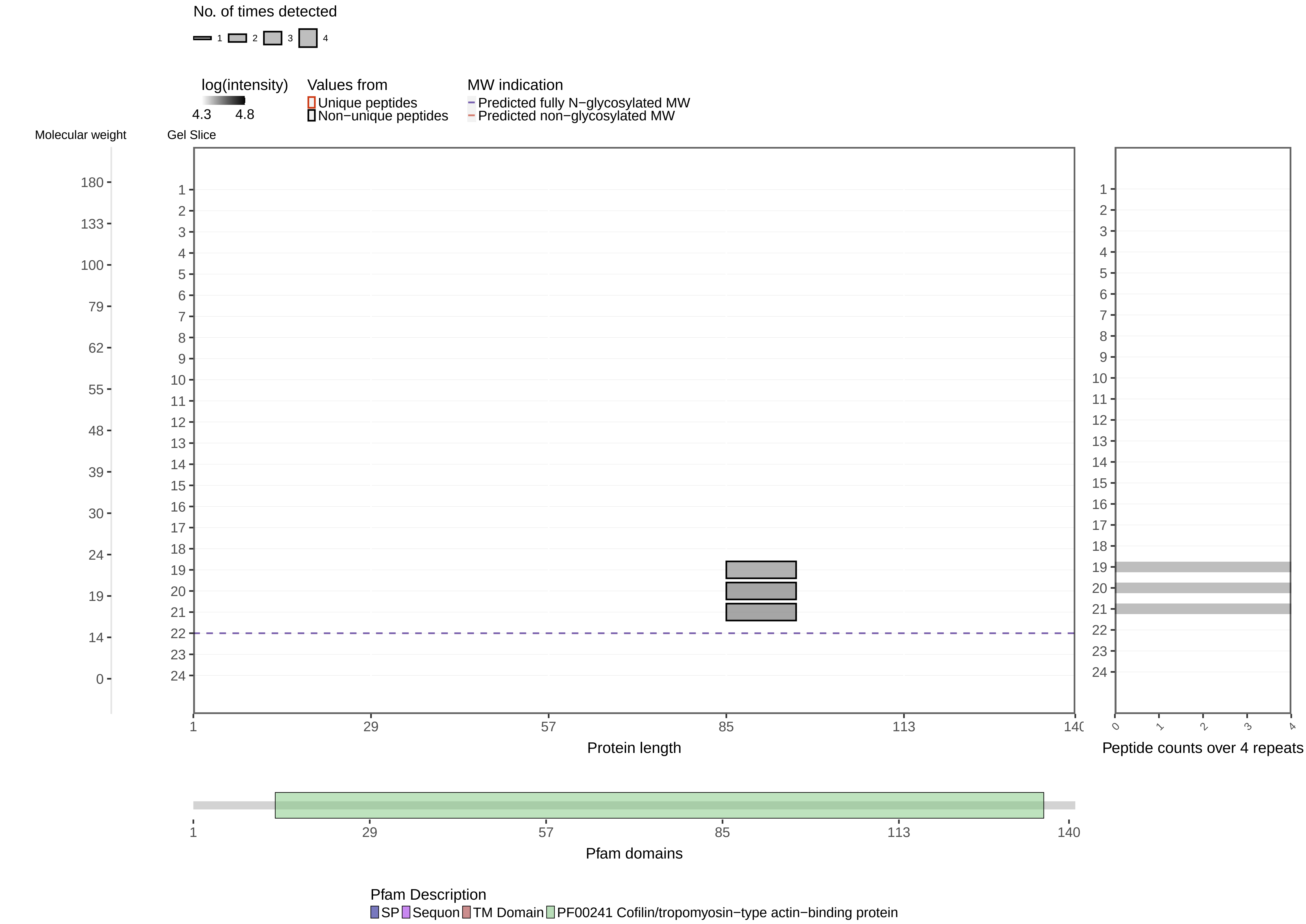


Protein sequence:

AASLPADECRYTVFDFDFVTEEGCQKSKIFFIAWSPDTAKVRSKMIYASSKDRFKRELDGIQVELQATDPTEMGLDVFKSRTT*

### NbL08g02560.1 UDP−glucuronic_acid_decarboxylase_6 [MW = 39.13 kDa]

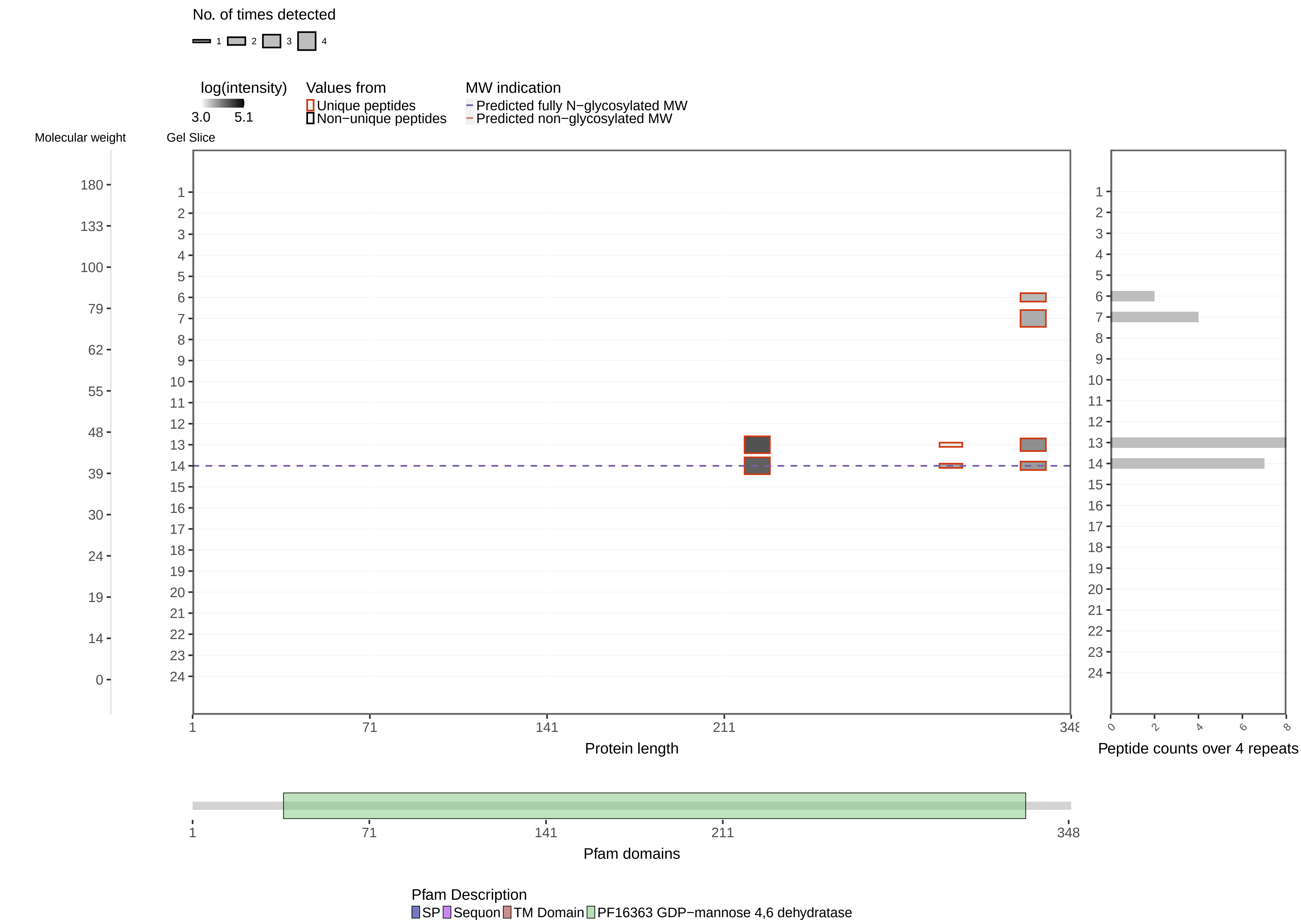


Protein sequence:

EEYWGNVNPIGVRSCYDEGKRVAETLMFDYHRQHGIEIRIARIFNTYGPRMNIDDGRVVSNFIAQALRDEPLTVQAPGTQTRSFCYVSDMVNGLIQLMEGEKTGPMNIGNPGEFTMIELAELVKELINPKVEIKSVENTPDDPRQRKPDITKAKELLGWEPKVELRDGLPLMEEDFRLRLGVSKKI*

### NbL08g02520.1 osmotin−like_protein [MW − SP = 24.46 kDa]

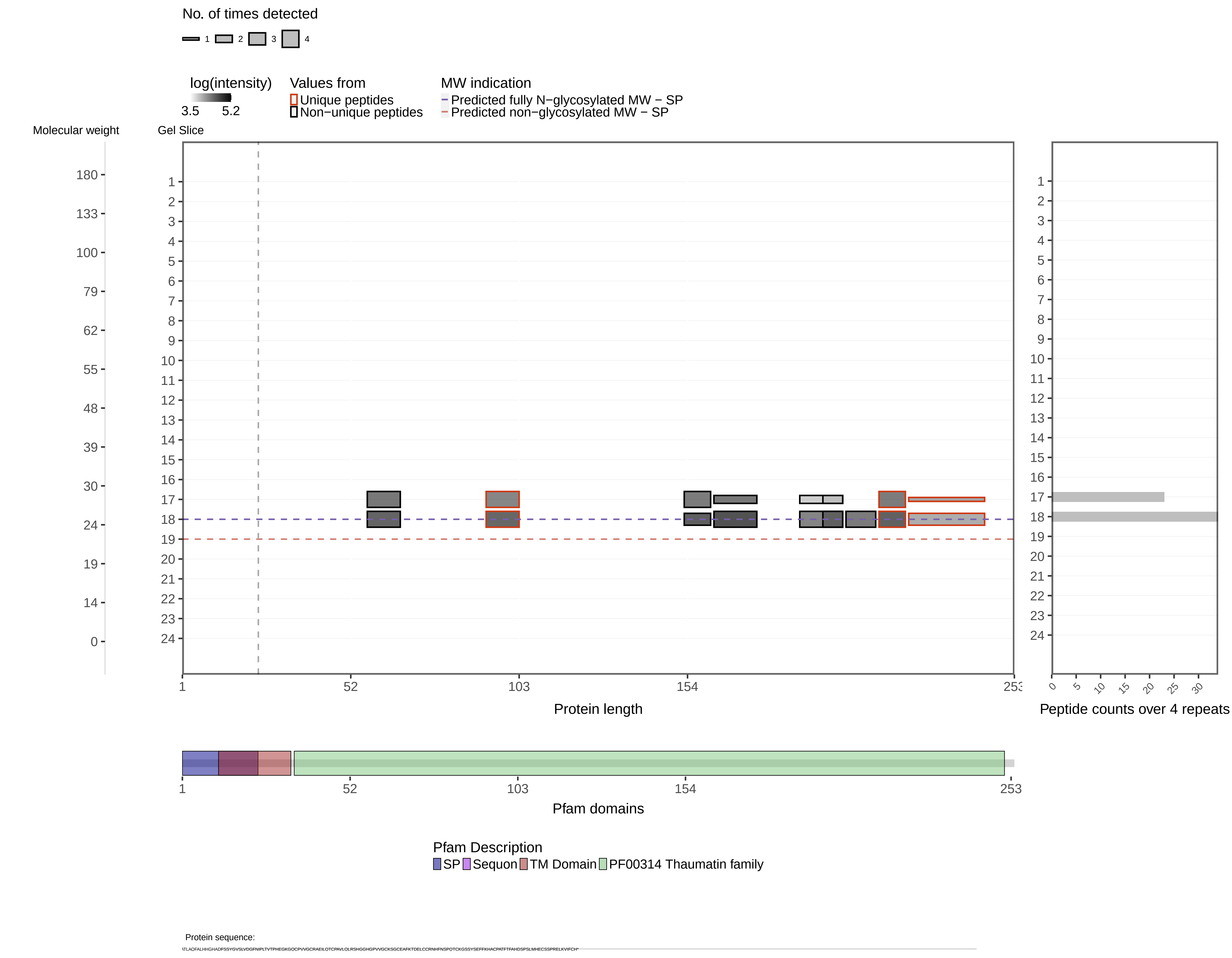

### NbL08g02380.1 actin [MW = 41.71 kDa]

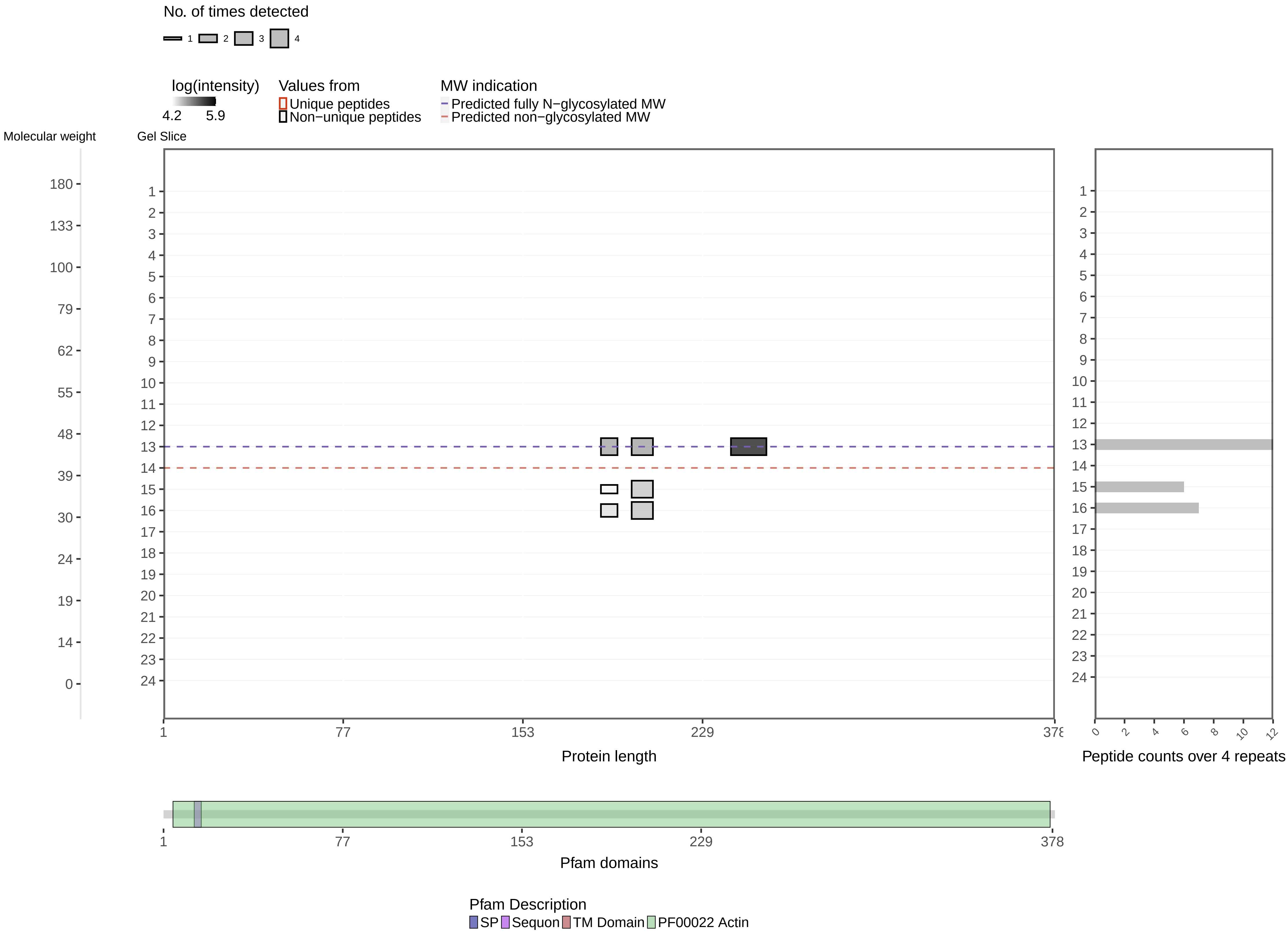


Protein sequence:

HAILRLDLAGRDLTDHLMKILTERGYSFTTSAEREIVRDVKEKLSYIALDFEQEMDTSKTSSSVEKSYELPDGQVITIGAERFRCPEVLFQPSMIGMEAAGIHETTYNSIMKCDVDIRKDLYGNIVLSGGSTMFPGIADRMSKEITALAPSSMKIKVVAPPERKYSVWIGGSILASLSTFQQMWIAKAEYDESGPSIVHRKCF*

### NbL08g02290.1 dicarboxylate_transporter_1_chloroplastic−like [MW = 62.78 kDa]

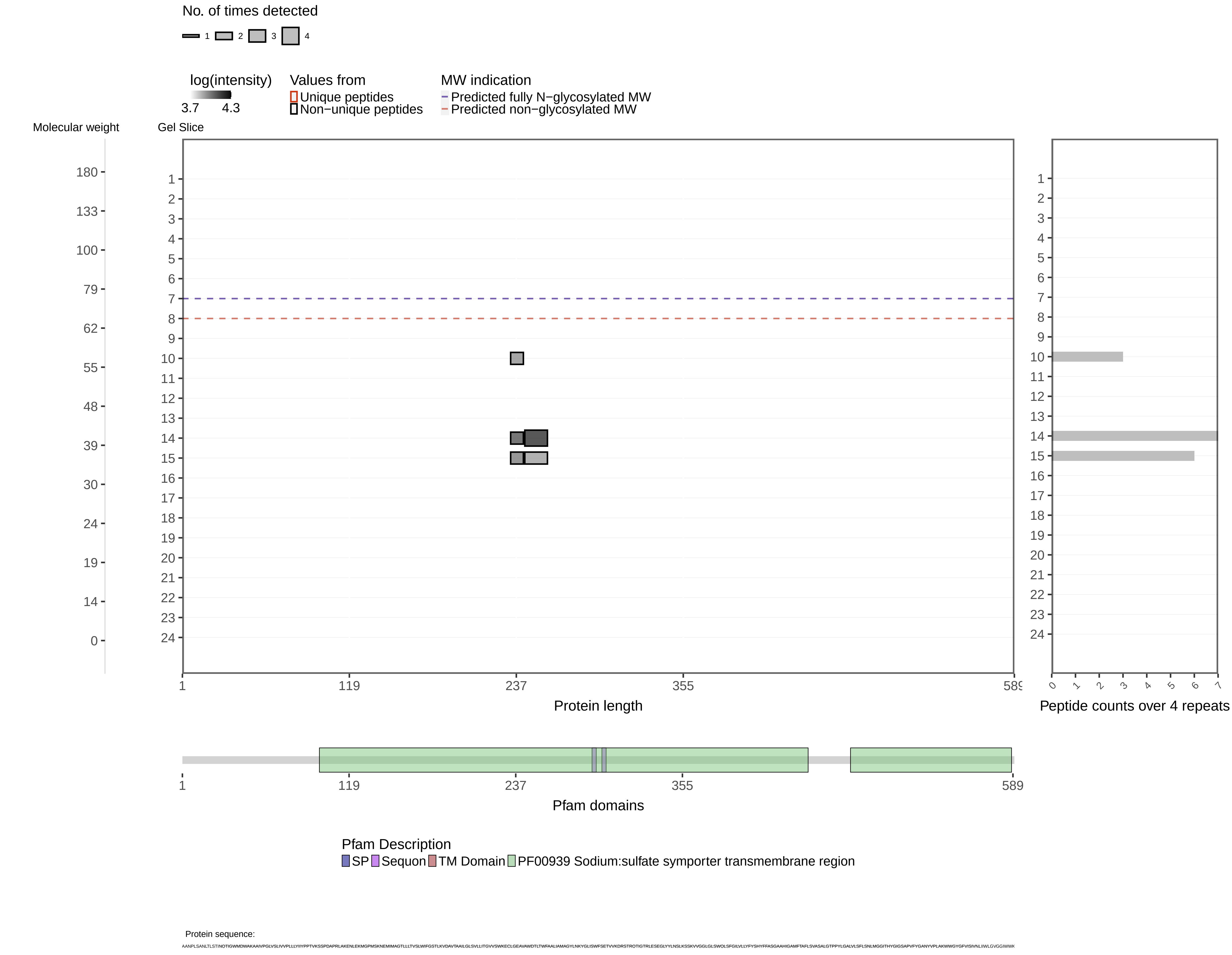

NbL08g02260.1 uncharacterized_LOC107821402 [MW = 25.51 kDa]

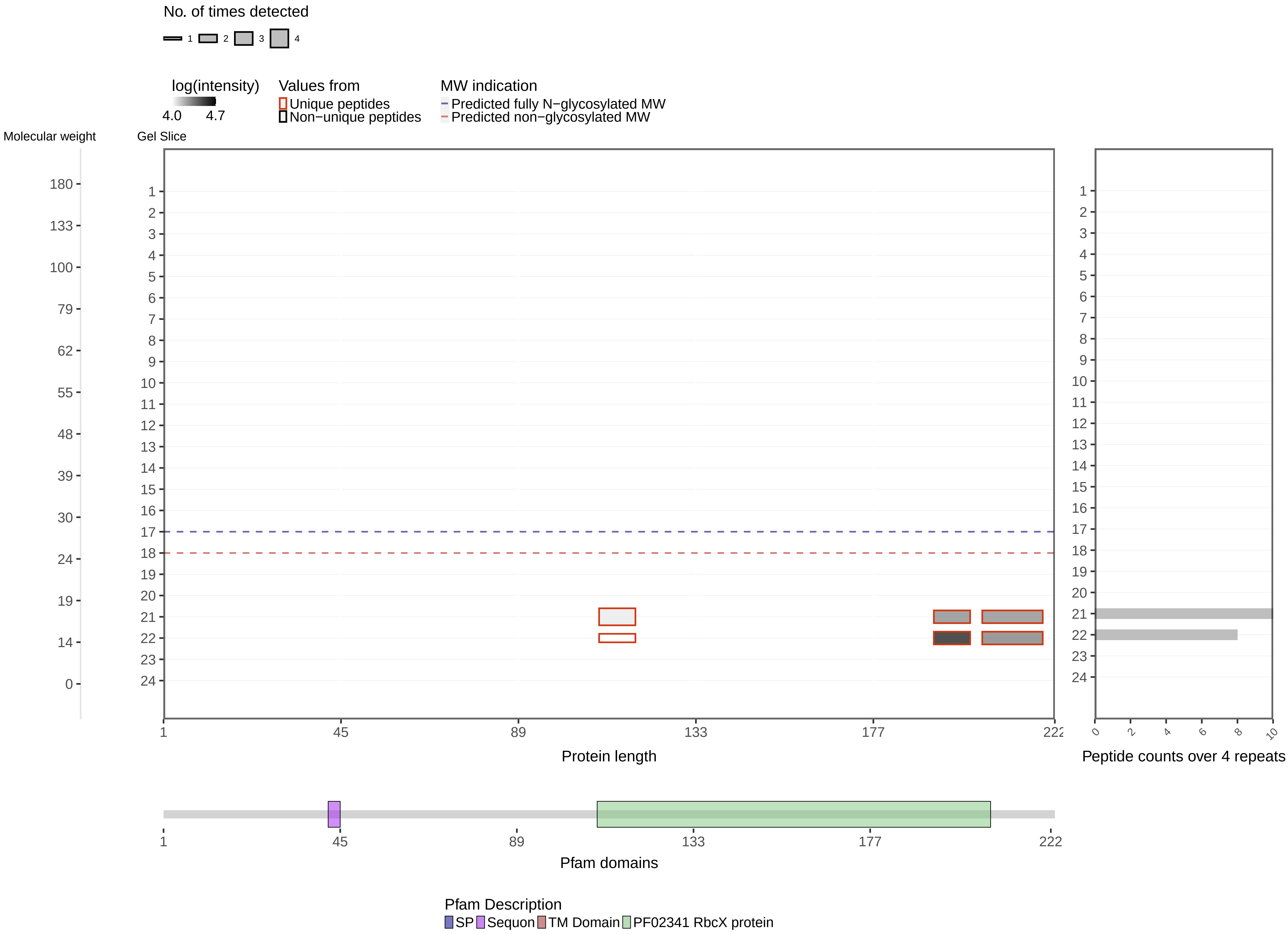


Protein sequence:

EQYEYGFDDIKTQILNYFTFKAVRTILNQLNEMDSPQYQWFNDYVTVNKLRDGKRFIRNLAKEKQELAEKVMVTRLSLYAKWIKKCDHEEIYNRISDQNVEVMRERLMETVIWPSDDTNSERVG*

### NbL08g02170.1 sulfite_reductase_1_[ferredoxin]_chloroplastic [MW = 82.25 kDa]

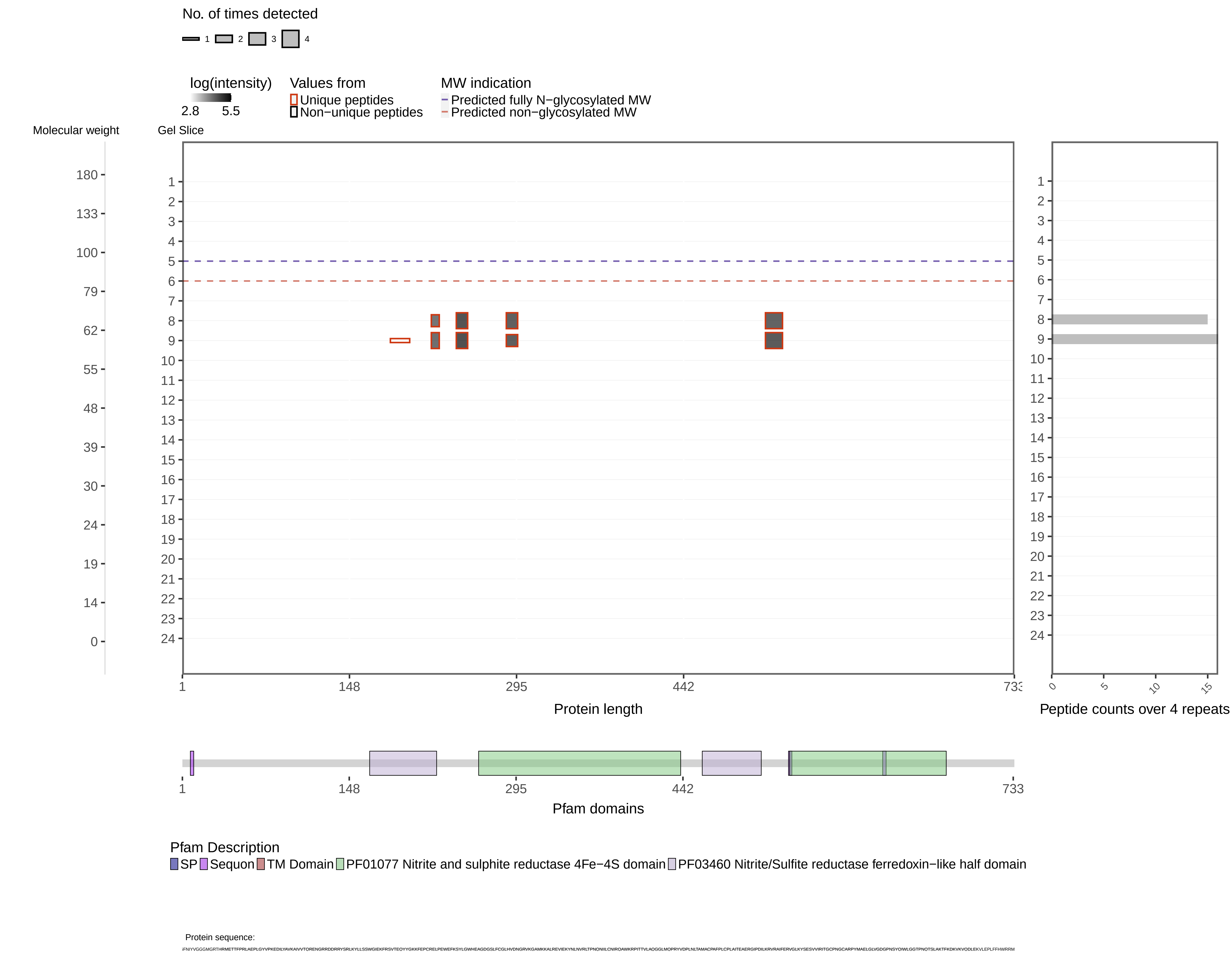

### NbL08g02150.1 glucosidase_2_subunit_beta−like [MW − SP = 65.13 kDa]

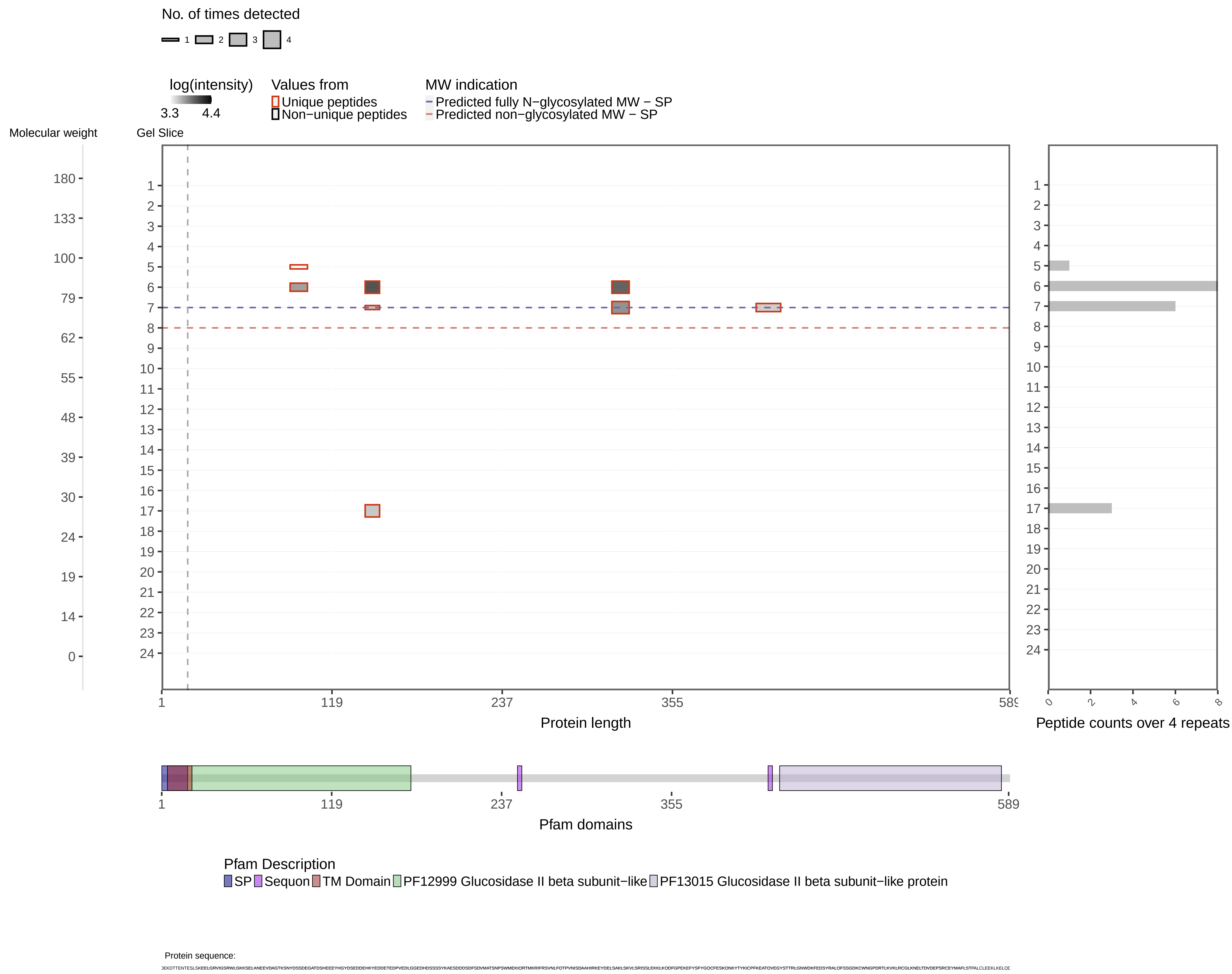

NbL08g02070.1 glucan_endo−13−beta−glucosidase_4 [MW − SP = 50.3 kDa]

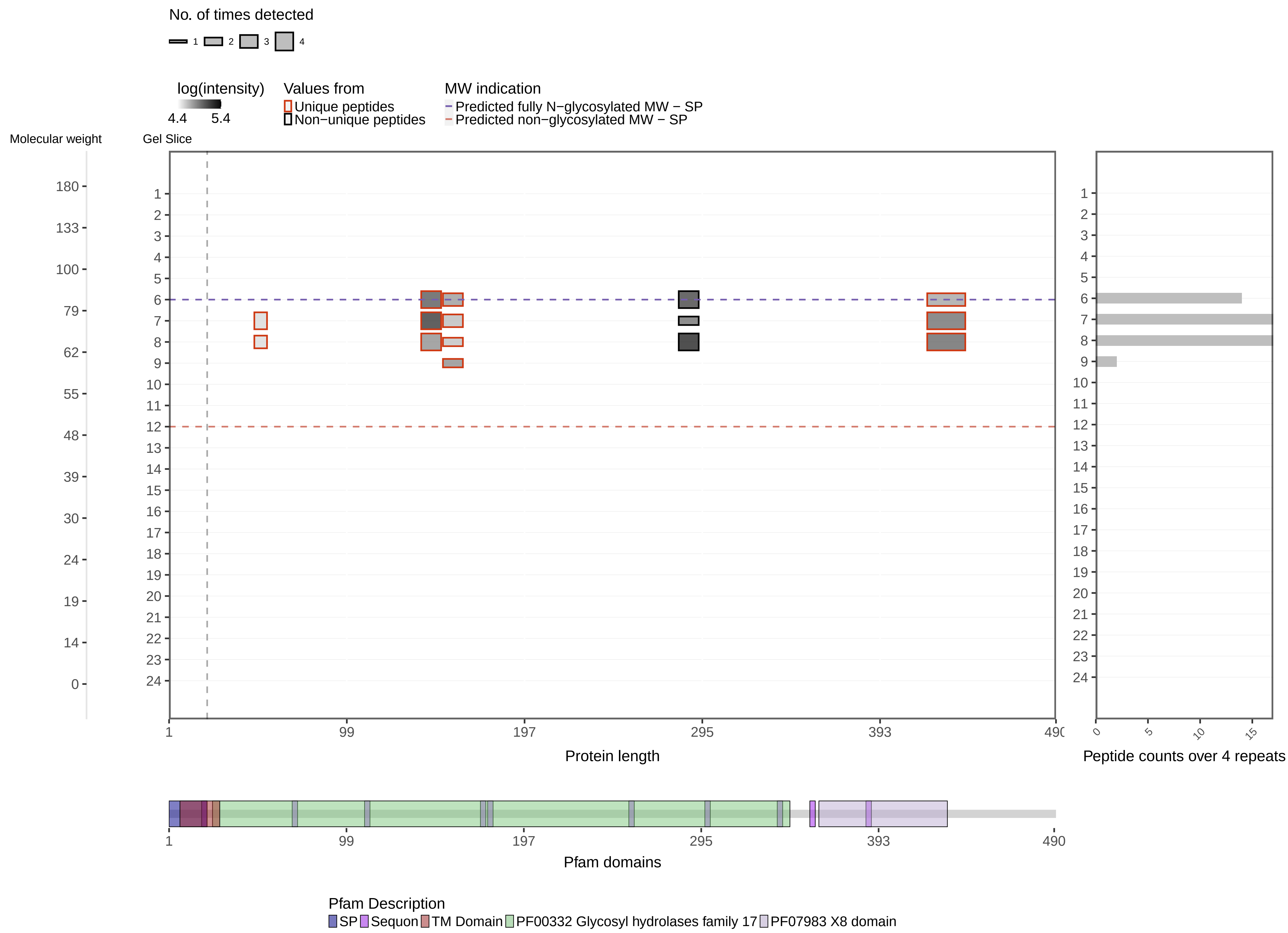


Protein sequence:

YVSMFDALVDAAYNSIAALNFSDIPIVVTETGWPWDGGVNEPDATKENAETFNNNLIRRVSNDTGPPSQPKNPINTFIYEMFNEDKRPGPISERNWGIFSTNGSEVYSLSLGSSAGISDNSTAVFCIAKKGADENKLQDGINWACGQGHANCSAIQSGQPCYFPDTLQNHASYAYNDYYQRMRSVGGTCDFDGTATTTTQDPSSKTCKFTGSSTPGGLFPPAAFGPISPTSQSSTIRASVIVYTLAVFFCFVDA*

### NbL08g01770.1 CDT1−like_protein_a_chloroplastic [MW = 66.1 kDa]

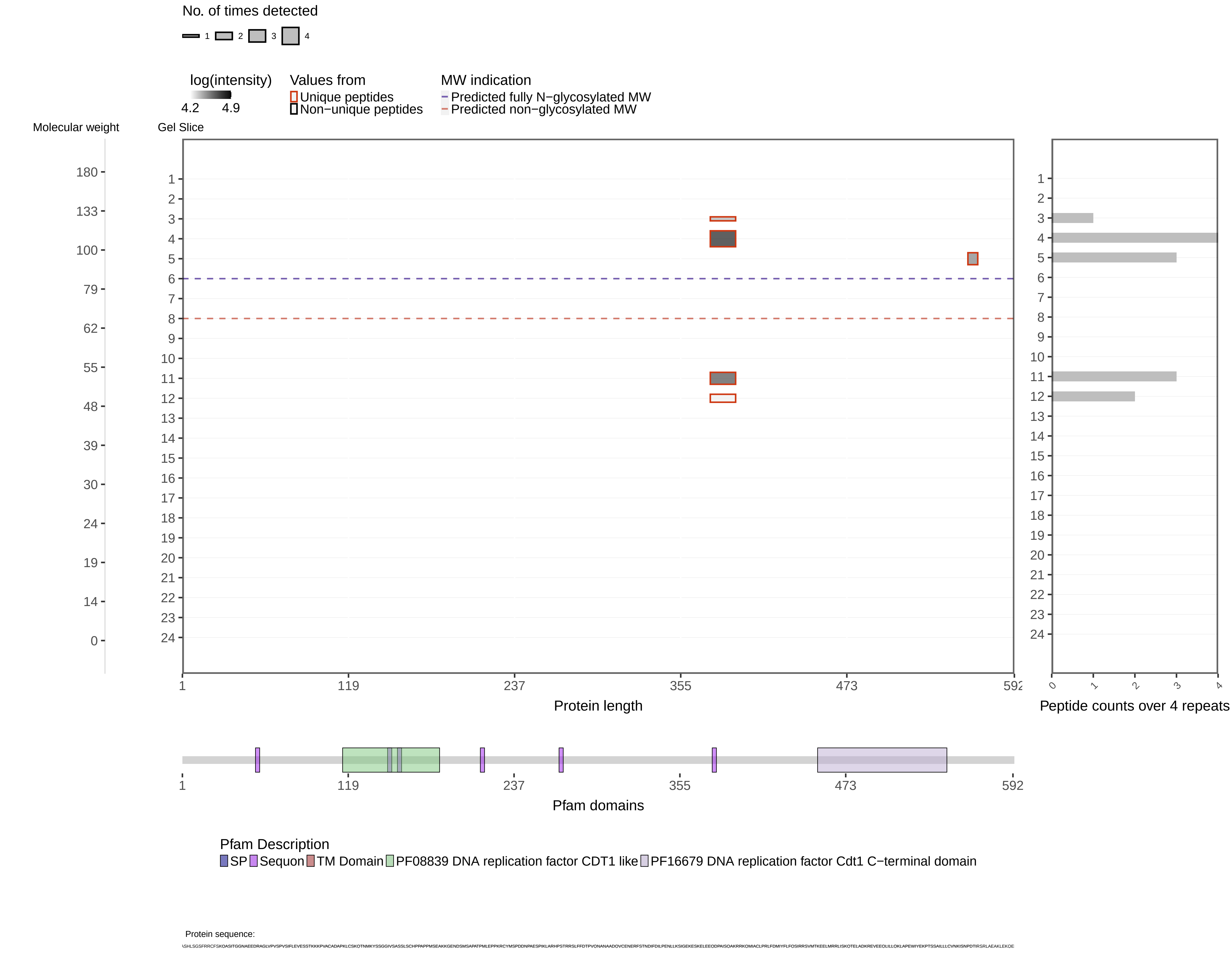

### NbL08g01650.1 formate−−tetrahydrofolate_ligase [MW = 67.33 kDa]

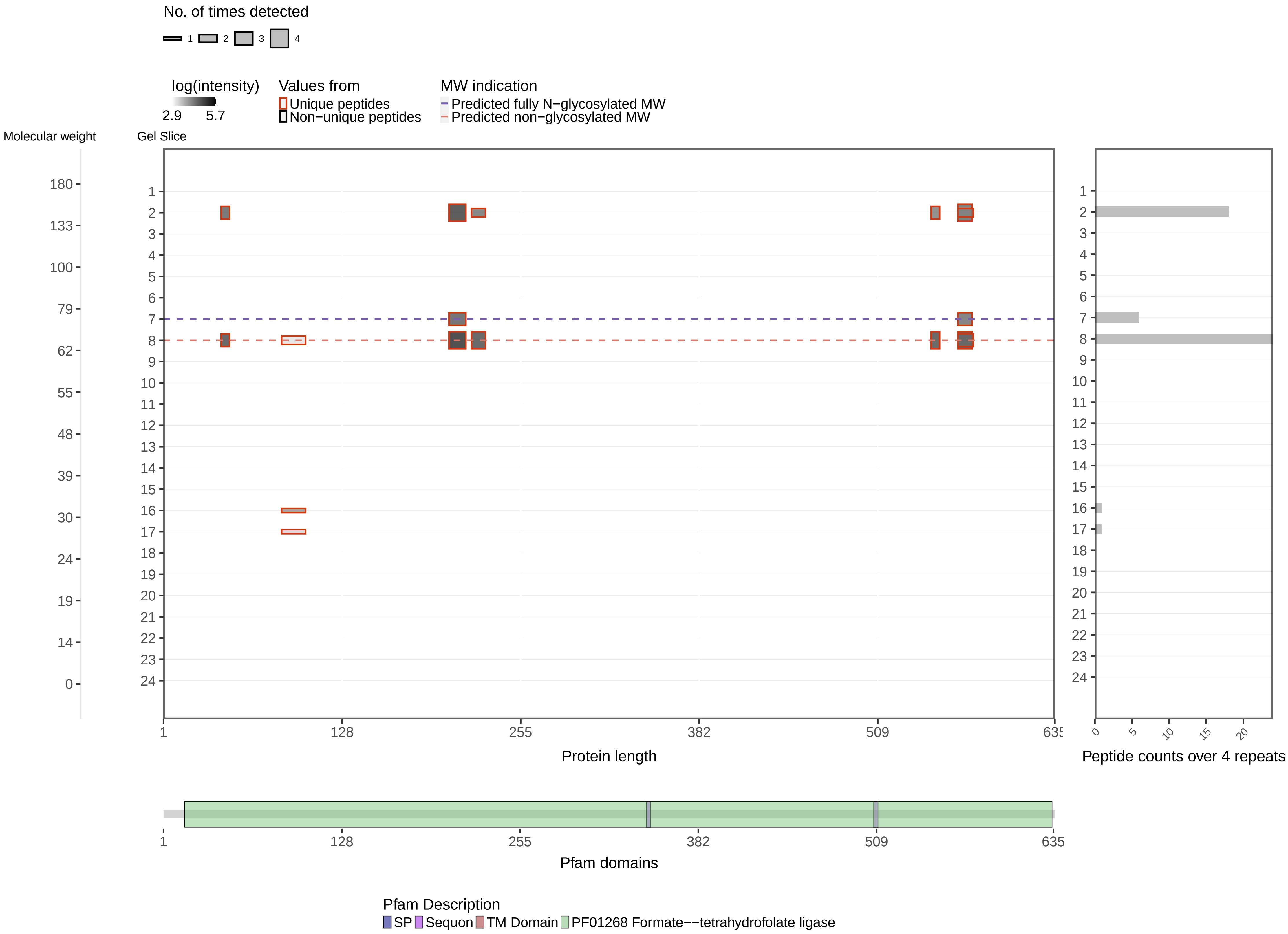


Protein sequence:

DLGVGGALTVLLKDAINPTLMQTLEGTPVLVHAGPFANIAHGNSSIVADKIALKLVGPGGFVVTEAGFGSDIGTEKFMNIKCRYSGLKPQCAVIVATVRALKMHGGGPDVTAGRPLDRAYVTENVALVEAGCVNLARHISNTKAYGANVVVAVNAFATDTEAELNAVKVAAFAAGAFDAVICTHHAHGGKGAVDLGIAVQKACENATQPLRFLYPLDIGIKDKIAAIAKSYGADGVEYSEQAEKQIEMYSKQGFSNLPICMAKTQYSFSHEAAKKGAPSGFILPIRDVRGSIGAGFIYP

### NbL08g01610.1 peroxidase_N−like [MW = 18.03 kDa]

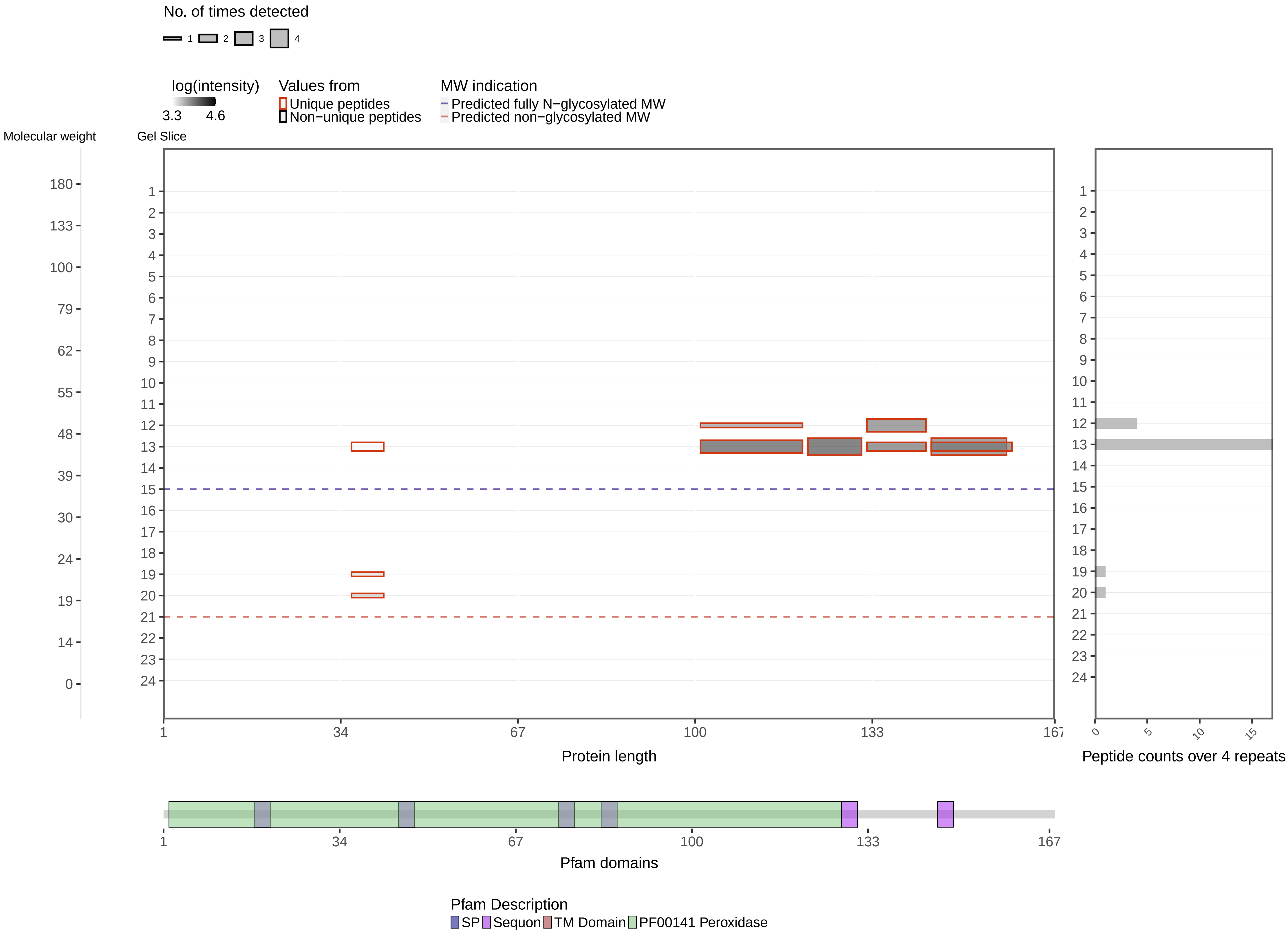


Protein sequence:

TSDGNNTAPLDRNSTDLFDNHYFKNLINGRGLLESDQVLFSSDDAVQTTKILVETYSNSSKFFFSDFVNSMIKMGNISPLVGSDGEIRKNCRVVNS*

### NbL08g01590.1 probable_plastid−lipid−associated_protein_12_chloroplastic [MW = 44.66 kDa]

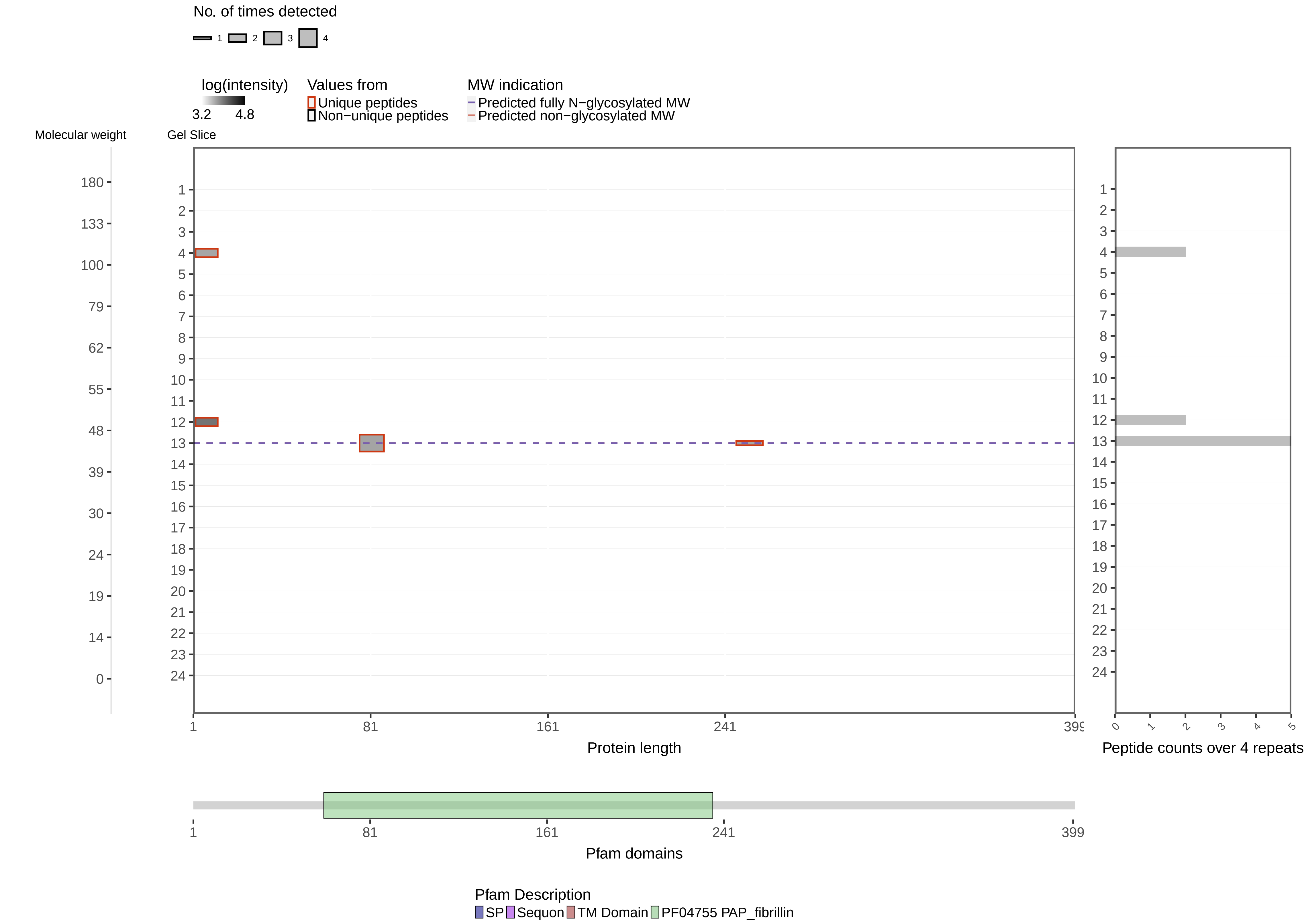


Protein sequence:

SFKFLPVKVPYPVPFRLLGDEAKGWLDTTYLSPSGNLRISRGNKGTTFVLQKEADPRQKLLSSISTGTSVEKAIDEFISLNQNVANPELEFLEGEWQMIWSSQVETDSWVENAGNGLMGMQIVKPNGQLKFLVEIFFGIRFSMTGKYEKSGSNTYDVIMDDGAFVAGVYGIPVEMESKFTIEILYTDDKIRISRGYNKILFVHVREDGSKKK*

NbL08g01470.1 enoyl−[acyl−carrier−protein]_reductase_[NADH]_chloroplastic−like [MW = 41.8 kDa]

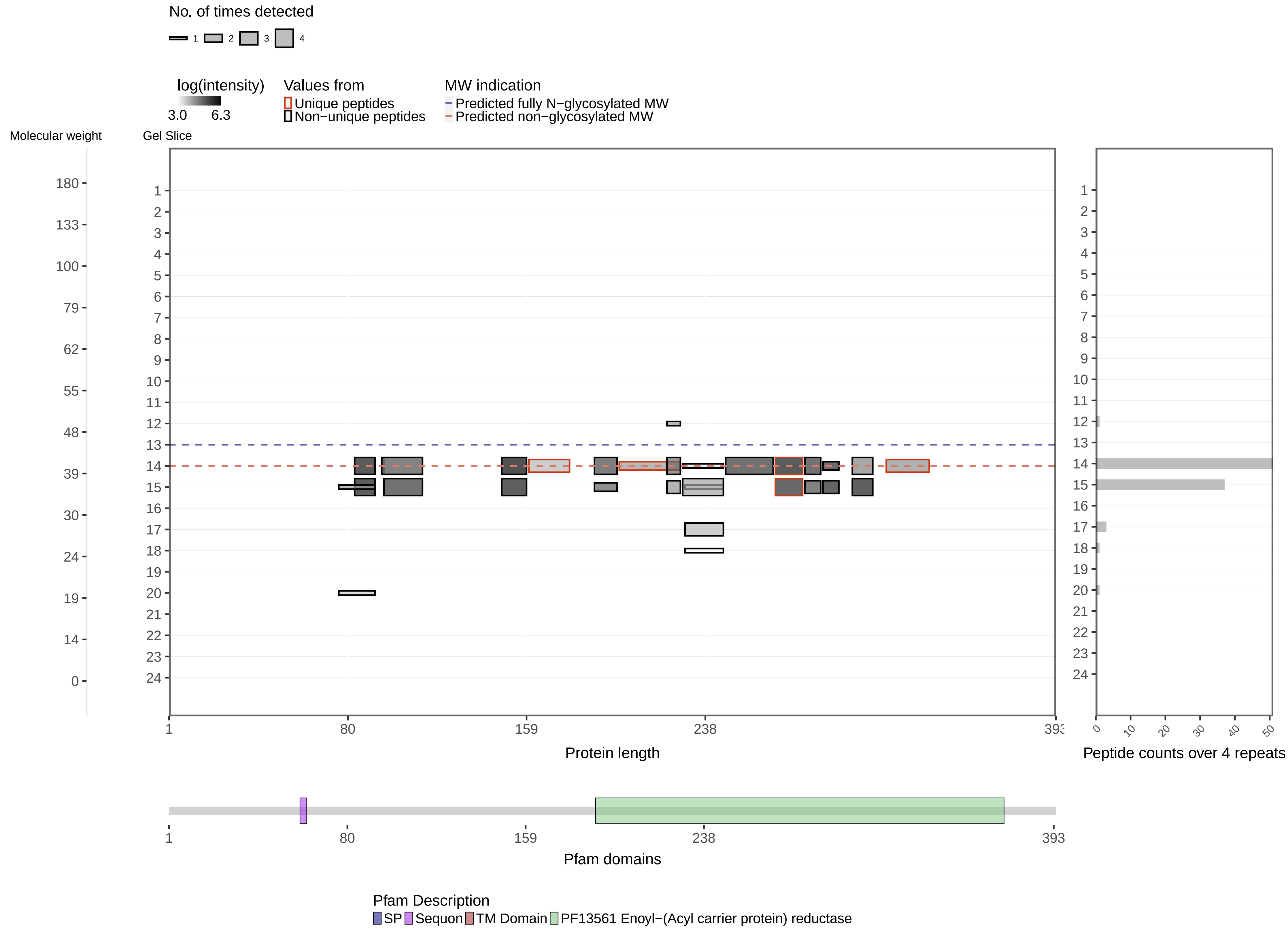


Protein sequence:

YAGSSKWTVSEVAESVKEDFGSVDILVHSLANGPEVTKPLLETSRKGYLAAISASSYSYVSLLKHFLPIMNPGGSSISLTYIASERVIPGYGGGMSSAKAALESDTRVLAFEAGRKKKVRVNTISAGPLRSRAAKAIGFIDMMIDYSIANAPLOKELSADEVGNTAAFLASPLASAITGSVIYVDNGLNAMGVGIDSPLFKELNIPKSAE*

### NbL08g01460.1 non−structural_maintenance_of_chromosomes_element_4_homolog_A−like [MW = 41.72 kDa]

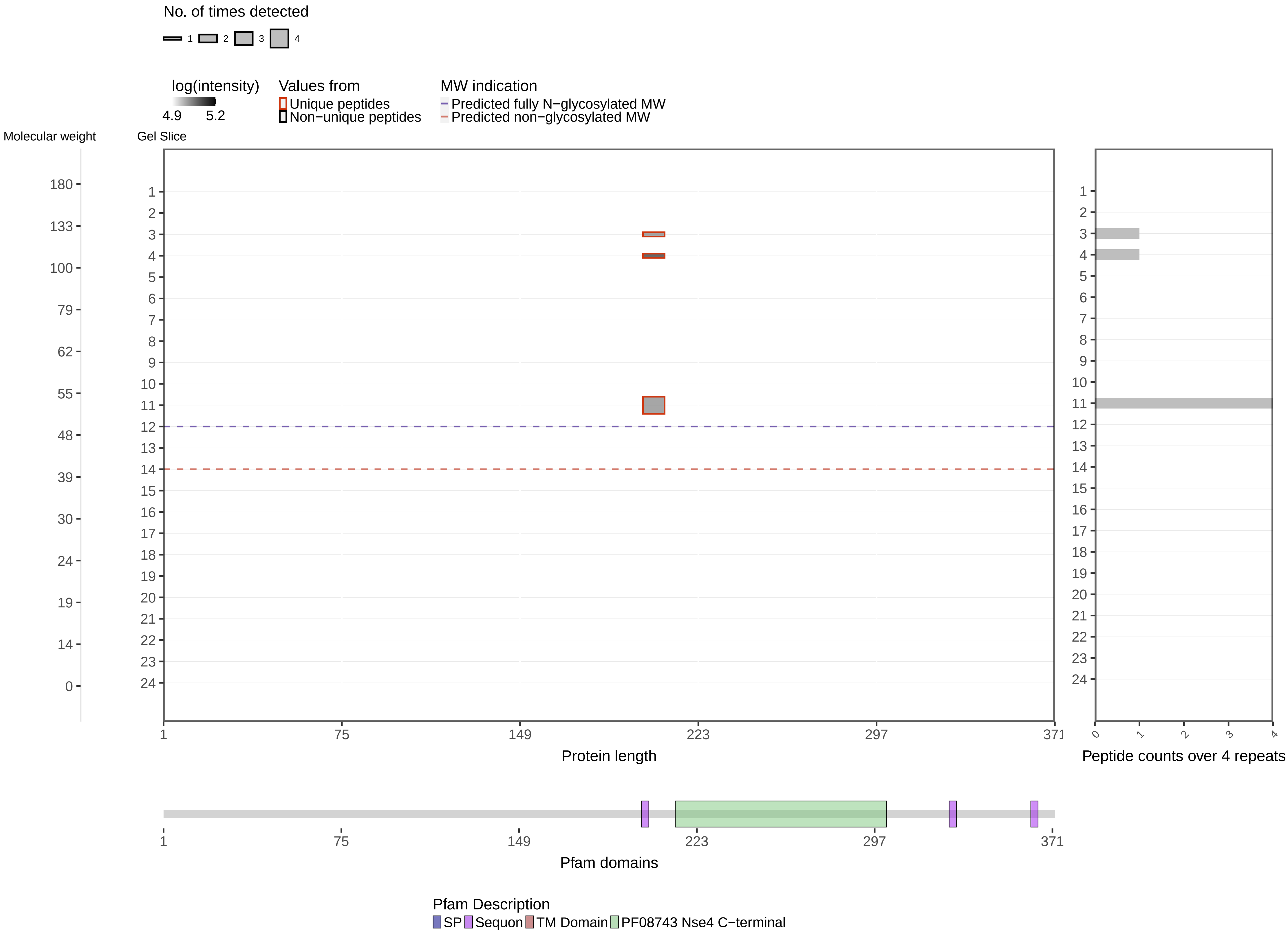


Protein sequence:

RPRTKVHSRACPNELEVNTEEKTDTDKNISTMFQILRKKKRVKLESLIMNRKSFAQSIENLFTLSFLVKDGRVFVDVDESGSHFVSPRNGPSAAAVKSGEVKYSHFVCRLDFADWELMKNVVPEGEELMPNRDILMNPGITKAEPVPITDSRPILNVTWVKKFSRNRGKVLRKASEVDDSPEIGDANICNGSLKRKLL*

NbL08g01420.1 UPF0061_protein_azo1574−like [MW = 70.97 kDa]

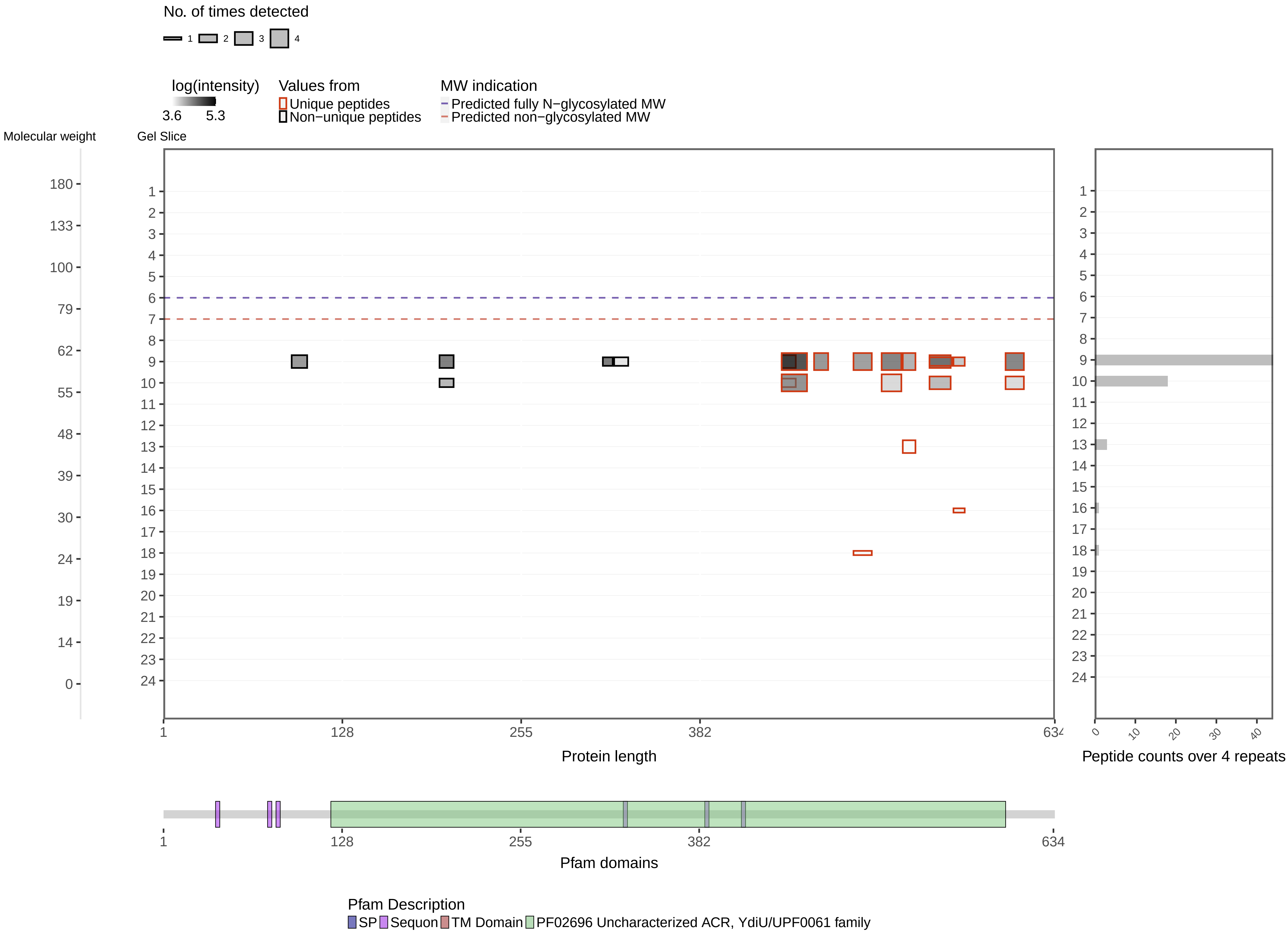


Protein sequence:

GKKDLEIVRTLADYAIRHHFPHLENMSKSESISFSTGEEDDSAVDLTSNKYAAWAVEVAERTASMIARWQGVGFTHGVMNTDNMSVLGLTIDYGPFGFLDAFDPSFTPNTTDLPGRRYCFANQPDIGLWNVAQFAKTLSTAQLLNDKESNYAMERYGIKFMDDYEAIMTKKLGLAKYNKQMIGDLLKNMAVDKVDYTNFFRLLSNVKADLTIPADQLLIPLKAALLDIGMERKEAWTSWVKSYIQELSTSGVSDEERKASMNSVNPKYVLRNYLCQSAIDATEQGDFGEVRRLL

NbL08g01380.1 G−type_lectin_S−receptor−like_serine/threonine−protein_kinase_RLK1 [MW − SP = 88.54 kDa]

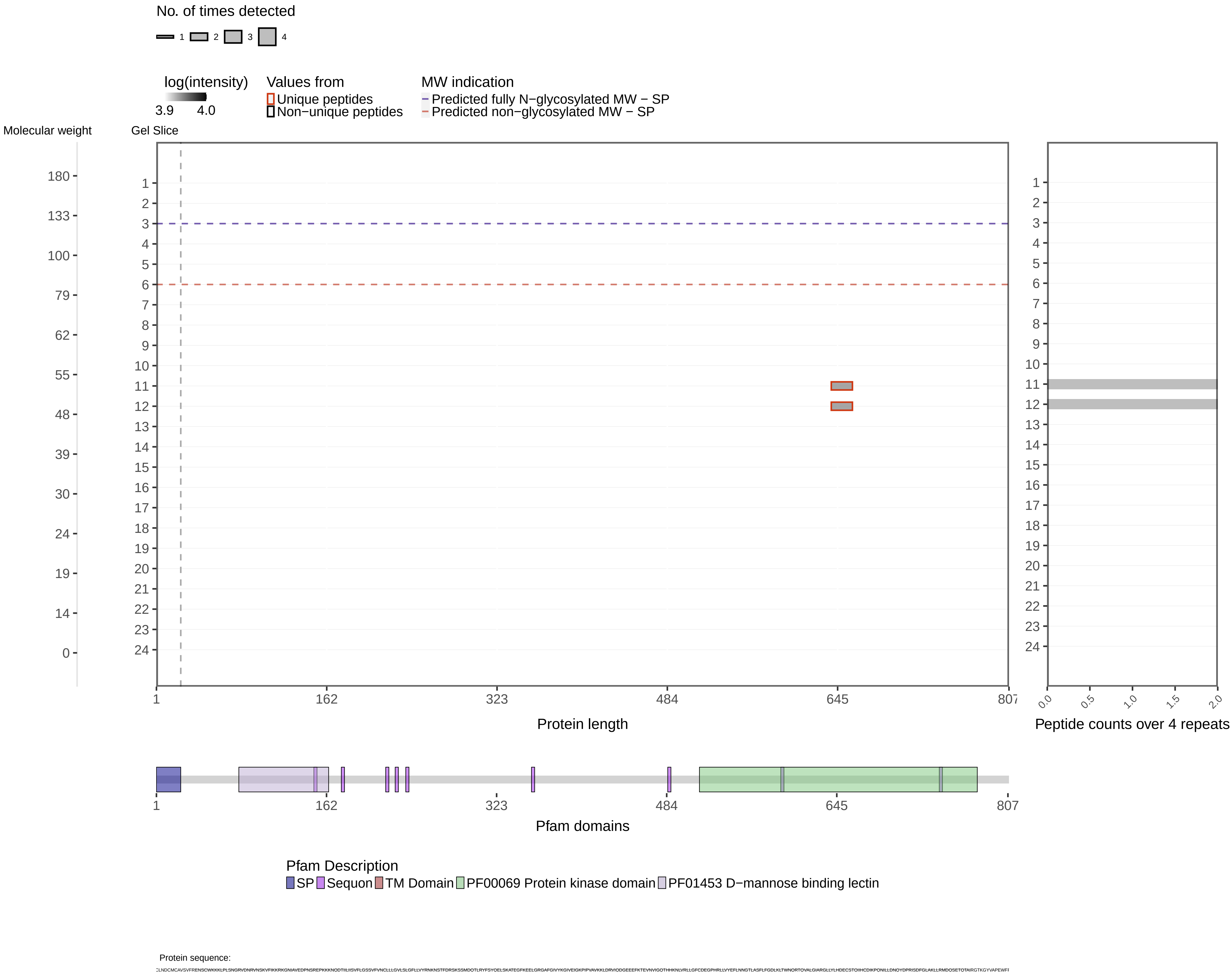

### NbL08g01370.1 sorbitol_dehydrogenase [MW = 38.56 kDa]

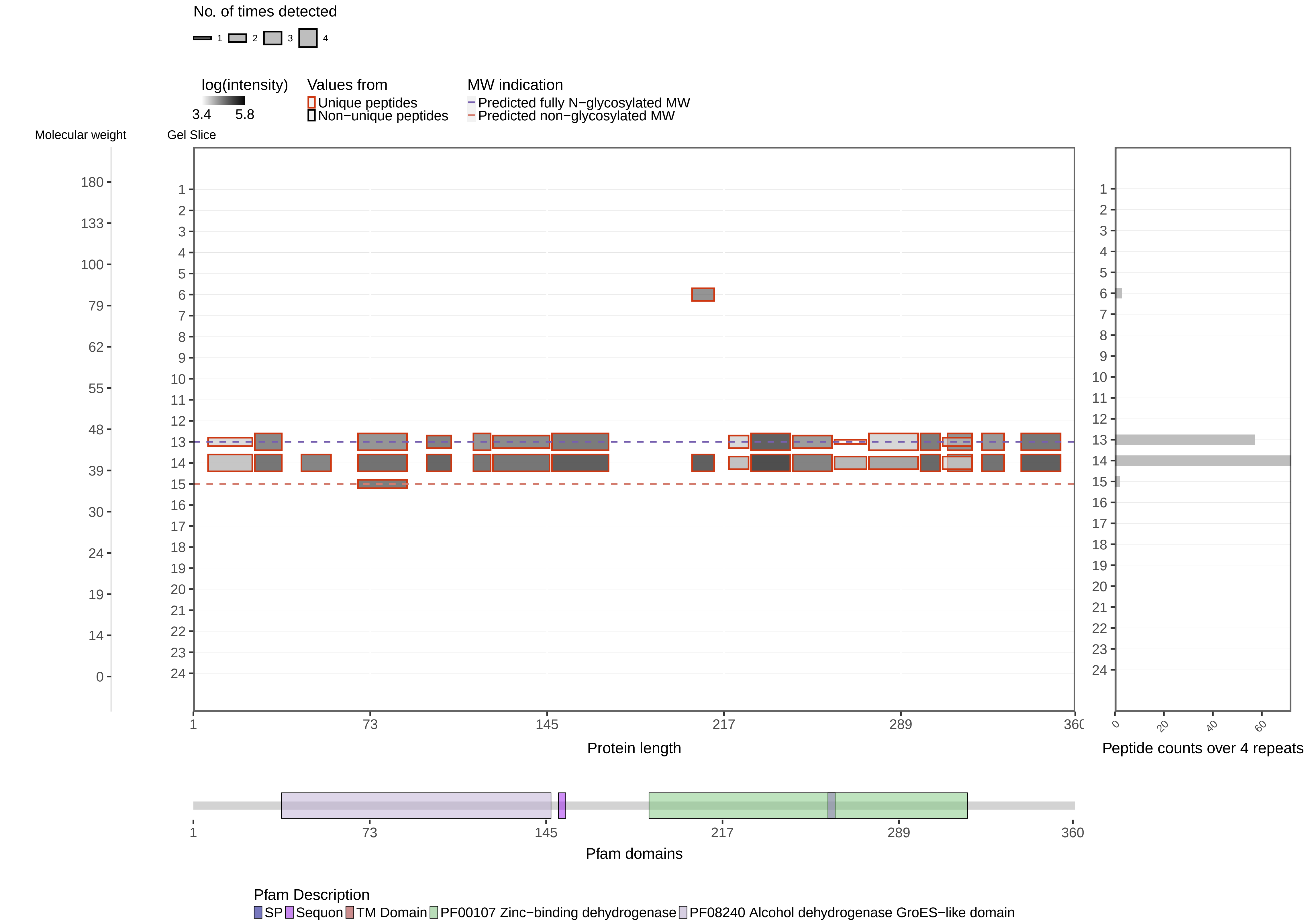


Protein sequence:

GVHACRRANVGPETNILVLGAGPIGLVTLLAARAFGAPRIVIVDVDDYRLSVAKKLGADETVKVSTNIQDVAADIENIQKAMGAGIDVSFDCAGFNKTMSTALGATCPGGKVCLVGMGHHEMTVPLTPAAAREVDVIGIFRYKNTWPLCLEFLRSGKIDVKPLITHRFGFSQKEVEEAFETSAGGGDAIKVMFNL*

NbL08g01350.1 linoleate_13S−lipoxygenase_2−1_chloroplastic−like [MW = 102.16 kDa]

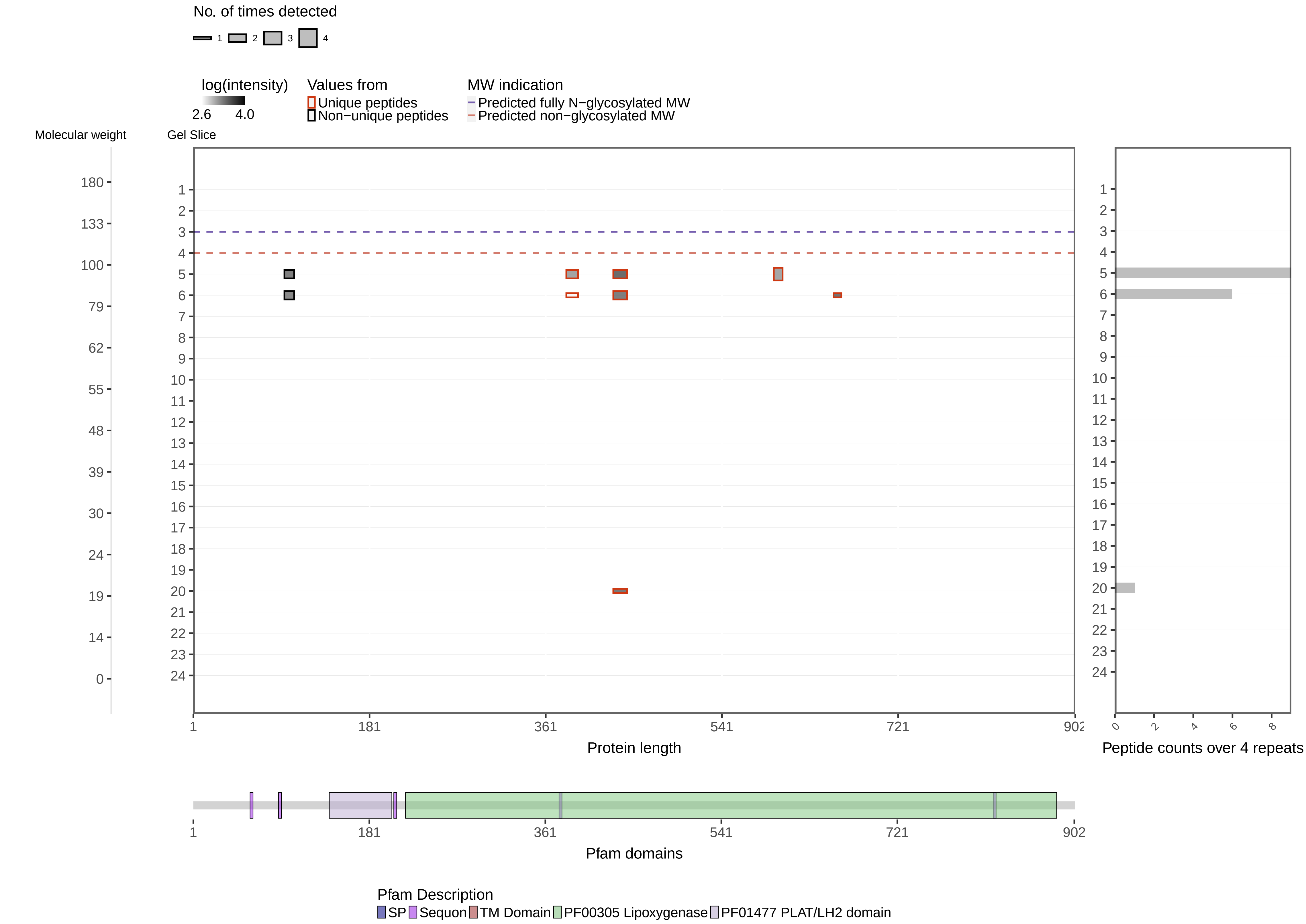


Protein sequence:

SAITKELIELEIAGFMTVEEAVKQKKLFILDYHDLLLPYVNKVNELKGRVLYGSRTLFFLTPDGTLRPLAIELTRPPVYDKPQWKEVYCPTWHATGSWLWKLAKAHVLAQDSGYHQLVSHWLRTHCATEPYIIATNRQLSAMHPIYRLLLPHFRYTMEINSLARDALINANGIIENSFFPGKYSMELSSVAYDLEWRFDRQALPEDLISRGMAVKDSNAPYGLKLTIEDYPFANDGLVLWDILKQWVTDYVNHYYTETELIEFDTELQAWWSEIKNVGHGDKKDAPWWPELKTPDD

### NbL08g01330.1 uncharacterized_LOC107827241 [MW = 39.62 kDa]

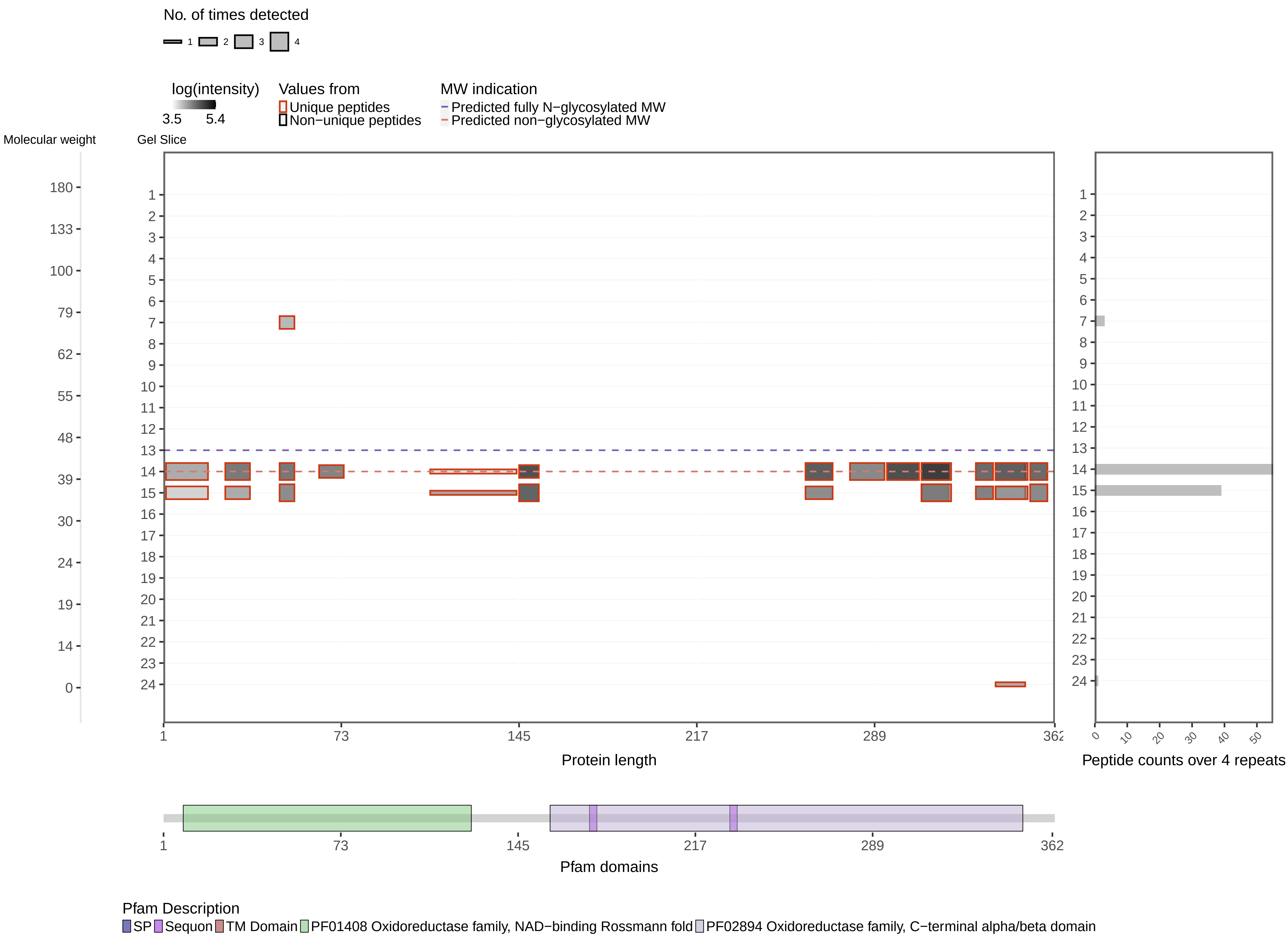


Protein sequence:

MNNSNPYFSSSWRRQFVGGFILDMGVHFVAGLRMLVGCEIASLSAMNSHVDKTLPPPDNISSMFQLENGCSGVFVMVVSSKSPKIIWRVVGLNGVLQVERGNKDGKHGYSIHLHMADGQSKSFFYPFSGVTDELKTFLSDVSTANLKKDHSSHIDPRLSFAEGARDVAVLDAMLESGKRQGALVQVKKF*

### NbL08g00990.1 chromatin_structure−remodeling_complex_protein_SYD−like [MW = 354.49 kDa]

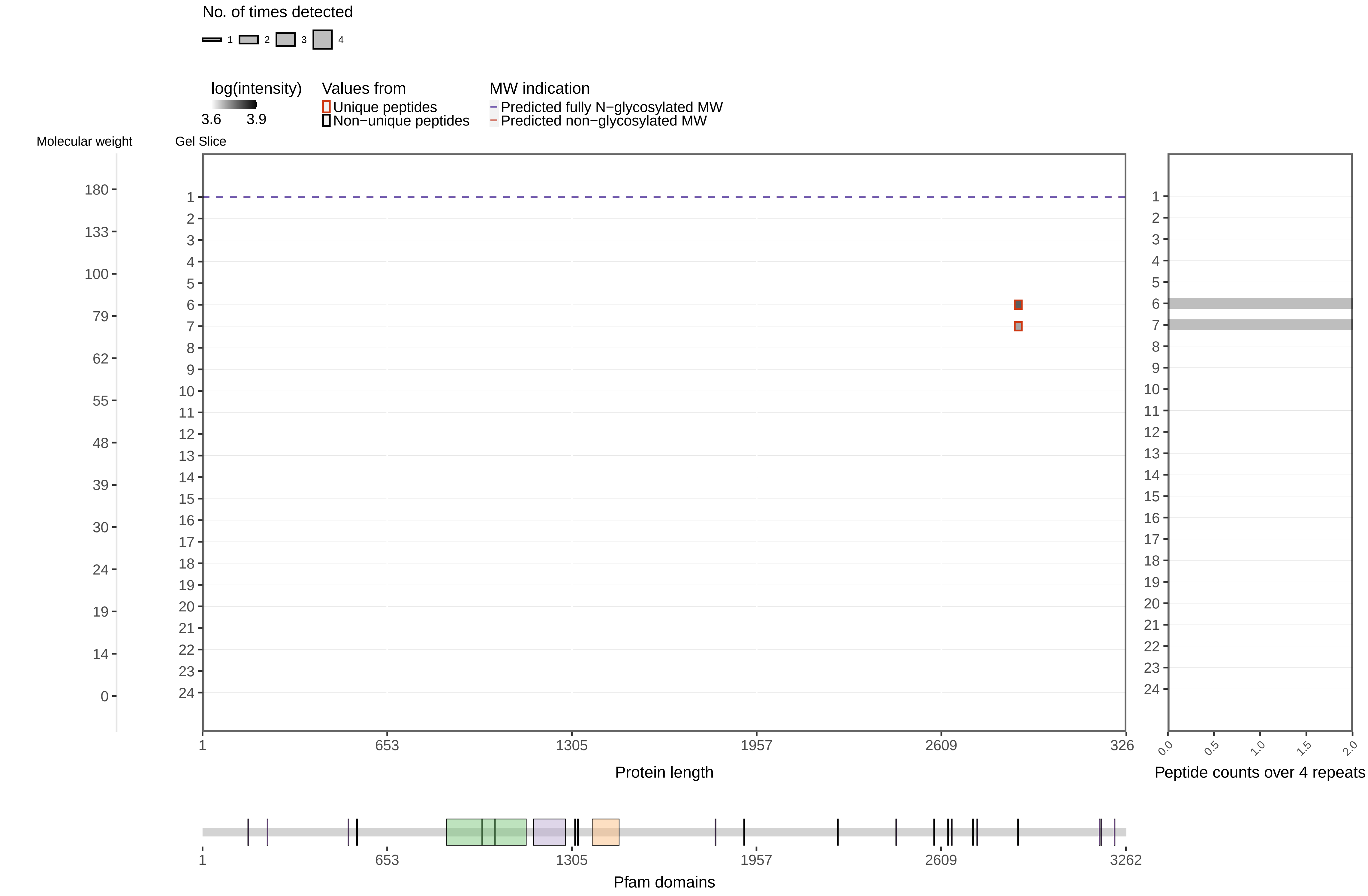


Protein sequence:

/ESTSPTILOSATAVAPCHOPVVPSVASLSDPPCPPTIGOGRGRGRGRGRGRGRGRGRKVENGGEAPGRRGKROSVTTEALLPPPTOAISEPVSAAOGVNVLSSSSHHMPPTPPSIGKPDLVPLVVAGLGSKDLGHAPASMRDANKELNSVSMMPLASSFTGKEVISVSTVPVIPSSSSSODPSSISPAVHSSSKNHSEDHRSFSAAOTEAIPOADAISVVPHTSPSADTEPSSATPGPLLTLTSKDSGSILPAPVPASSSACMELTSVCPVPVVASSCSSMGPASSAPAPLLLA

NbL08g00760.1 OTU_domain−containing_protein_6B−like [MW = 35.42 kDa]

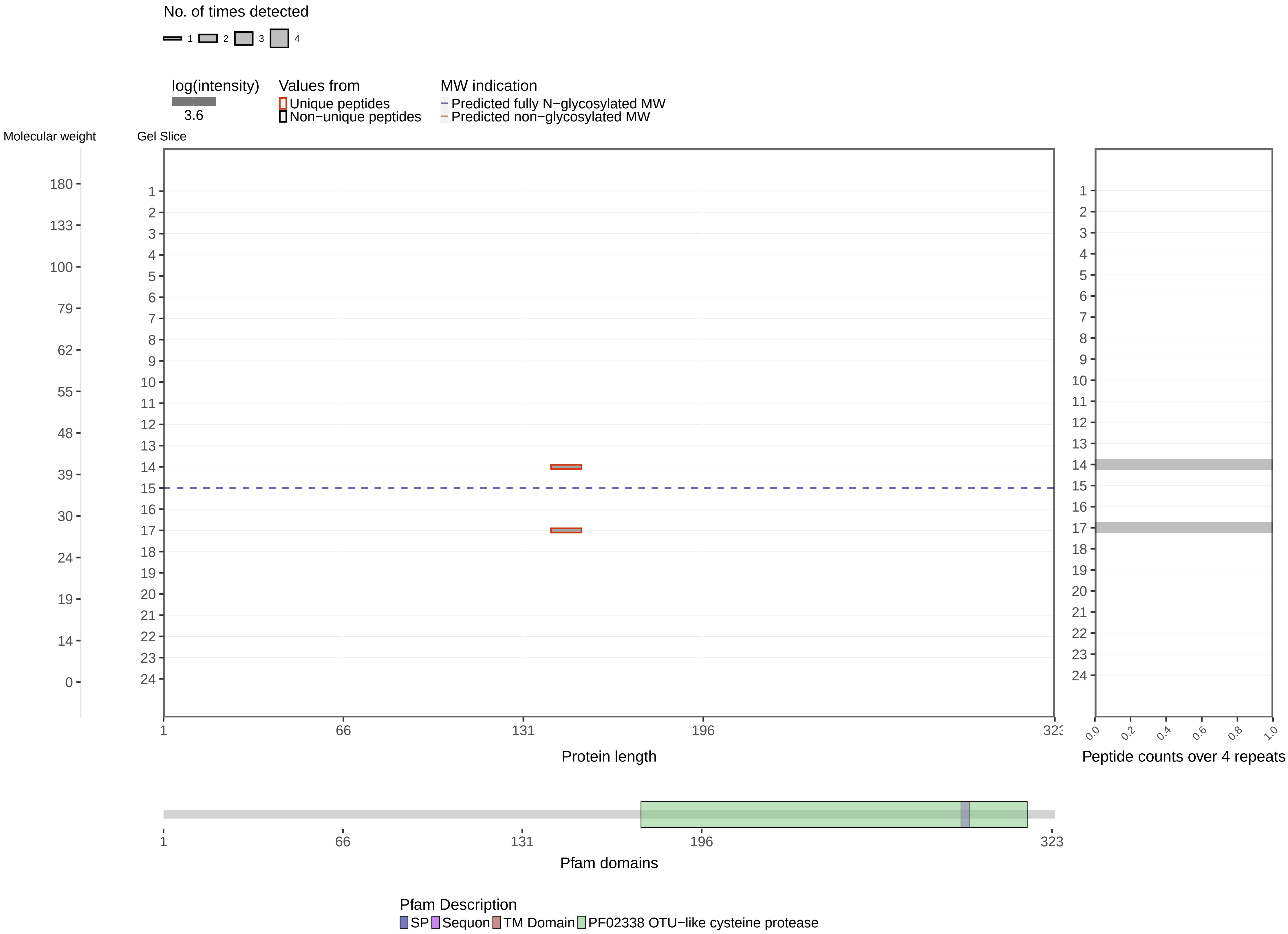


Protein sequence:

IIVSDRVIENEKLDSKLEPLGLMVNEIKPDGHCLYRAIENQLAVGSGGSSPYTYLELRQMVAAYMRKHATDFLPFFLSENAEGGESDDSLVERFENYCREVESTAAWGGQLELGALTHILKKHIMIFSGSFPDVEMGKEYKSDNGSGSSTSSIMLSYHMHAFGLGEHYNSVVPSSA*

### NbL08g00710.1 L−ascorbate_oxidase [MW − SP = 61.76 kDa]

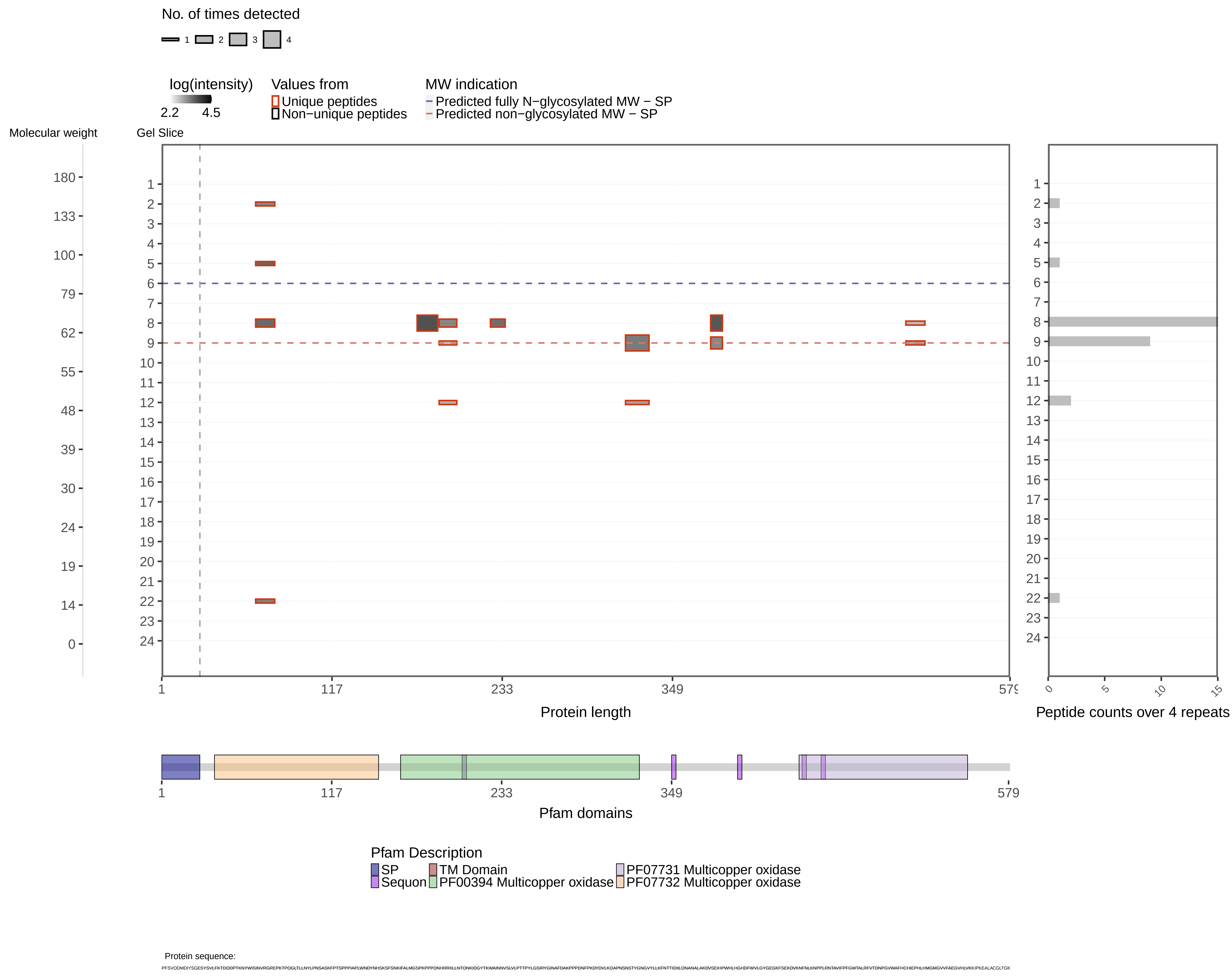

### NbL08g00620.1 GDP−mannose_46_dehydratase_1 [MW = 41.73 kDa]

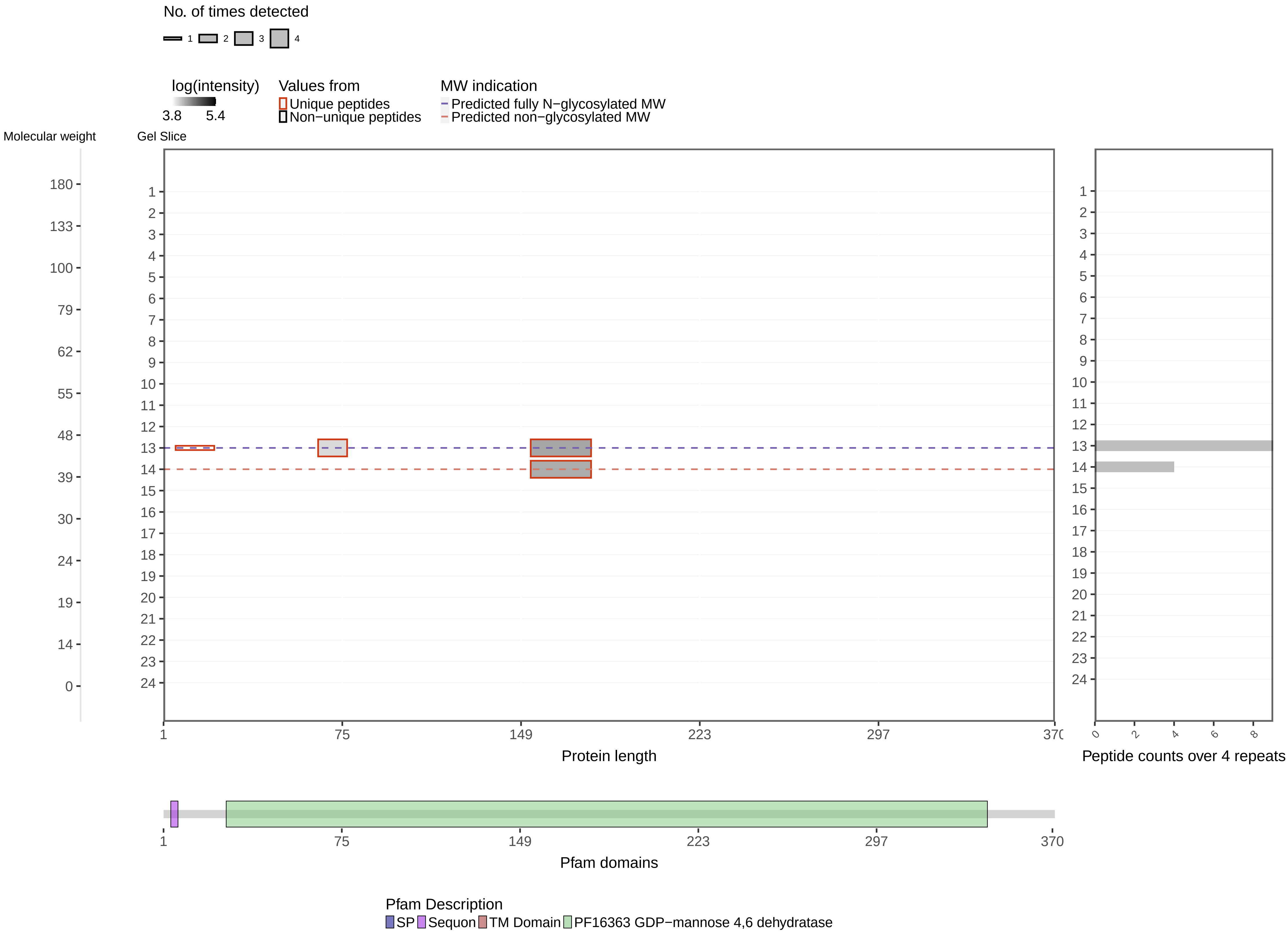


Protein sequence:

FHPRSPYAVSKCAAHWYTVNYREAYGIFACNGILFNHESPRRGENFVTRKITRAVGRIKIGLQSKLFLGNLQASRDWGFAGDYVEAMWMMLQQEKPDDYVVATEEAHTVEEFLEVAFRYVGLNWKDHVVIDKRYFRPTEVDNLKGDSSKARKVLGWKPRVGFEQLVKMMVDEDIELAKREKVLVDAGYMDAQQQP*

### NbL08g00470.1 THUMP_domain−containing_protein_1−like [MW = 44.15 kDa]

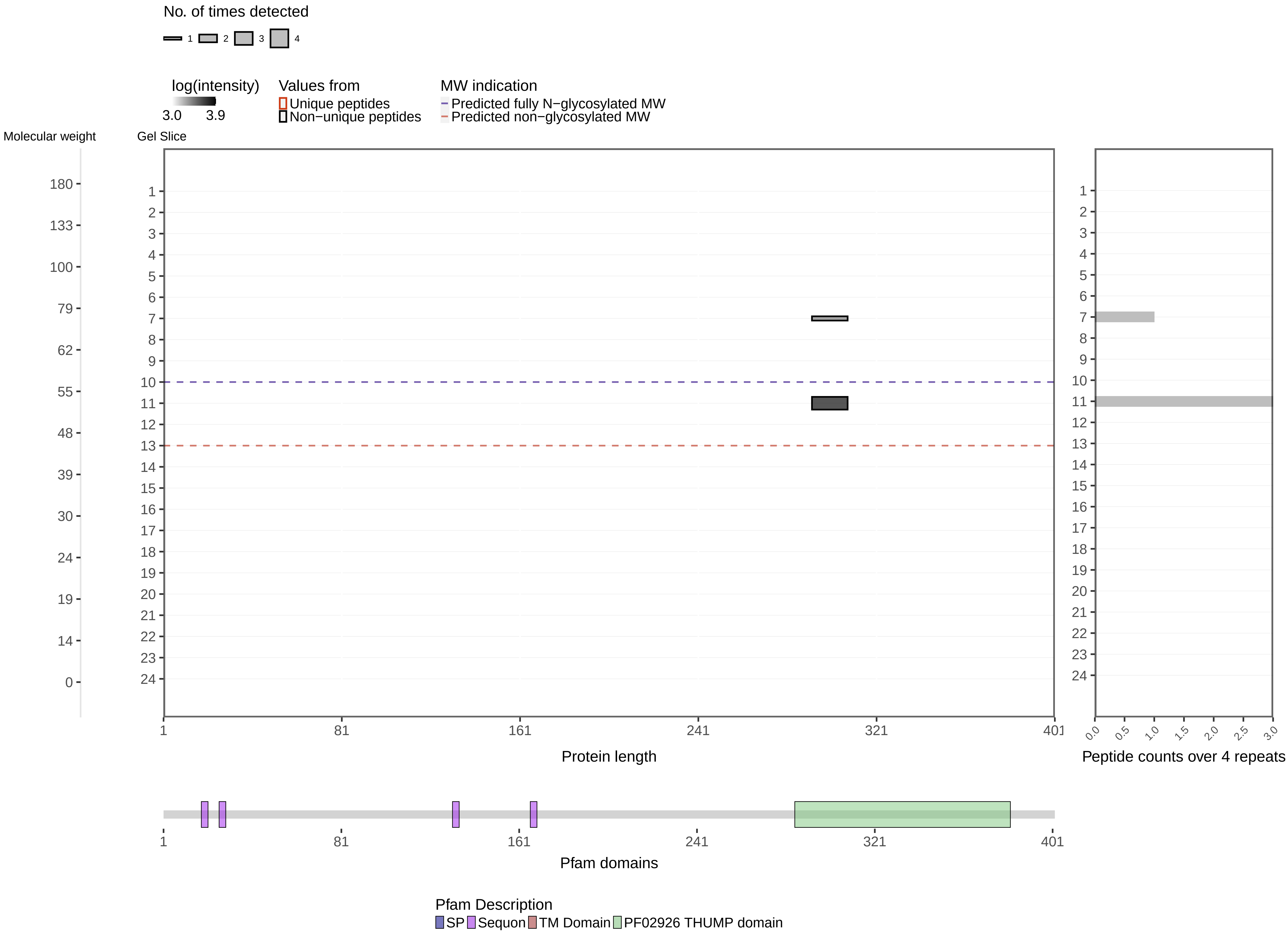


Protein sequence:

TTHEQKDGEAGEPPAKKQCVEREATKTGNNVVGKTEEKSVDKLIEAELAELGDKSKRRFTYLDSGCNGVVFVQMRKKDGDPNPKEIIQHMMSSLASTKRHVSRFILRVLPVEVTCYASEEEIGRAIKPLVEQYLPVETDTPKKFAVLYEARANTGINKMKIIDTVAKSVPSPHKVDLSNPEINIVVQIVKTVCLLGVVENYKELSKYNLRQLTSKN*

### NbL08g00200.1 importin−5−like [MW = 46.74 kDa]

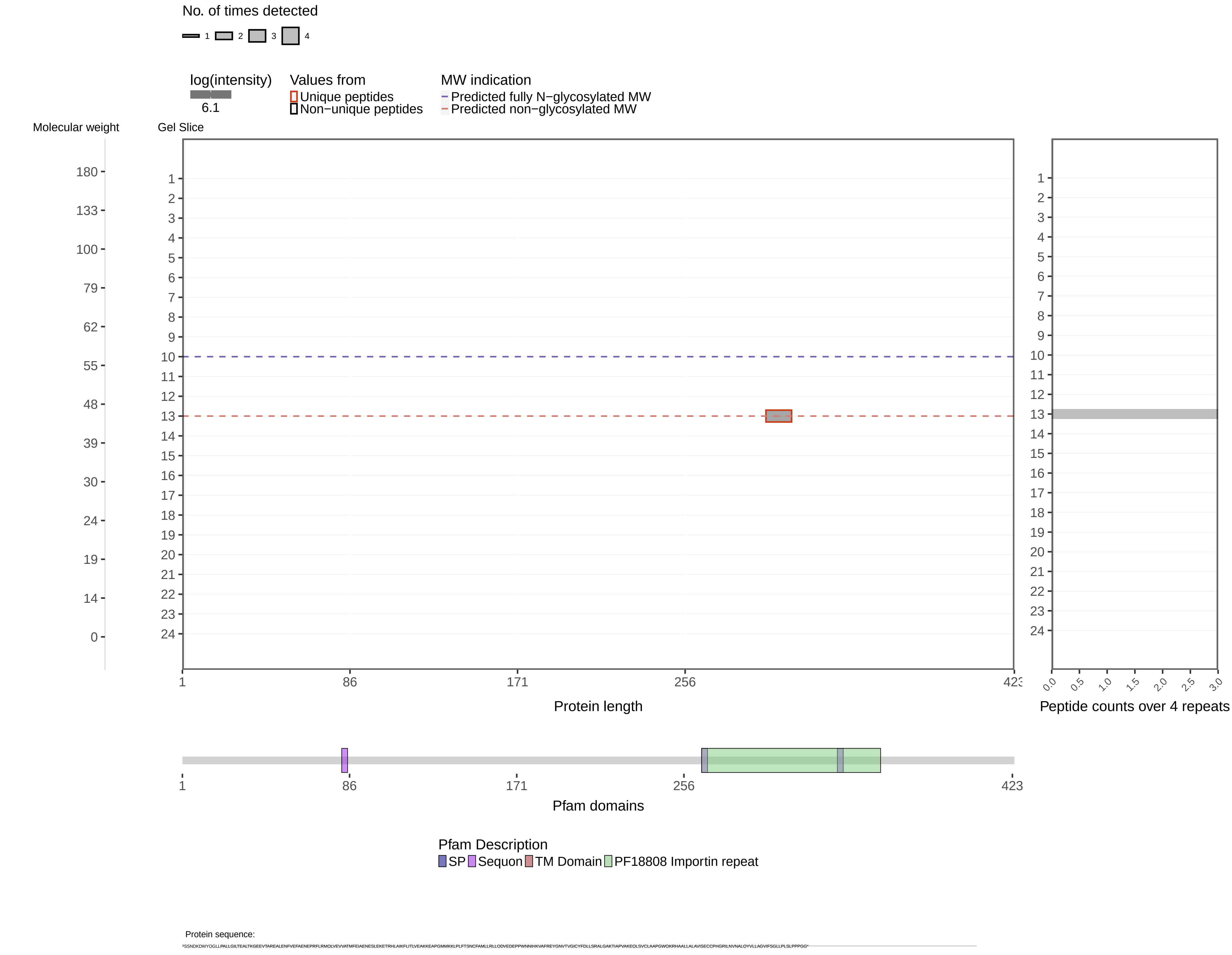

### NbL08g00130.1 mitochondrial_dicarboxylate/tricarboxylate_transporter_DTC−like [MW = 31.67 kDa]

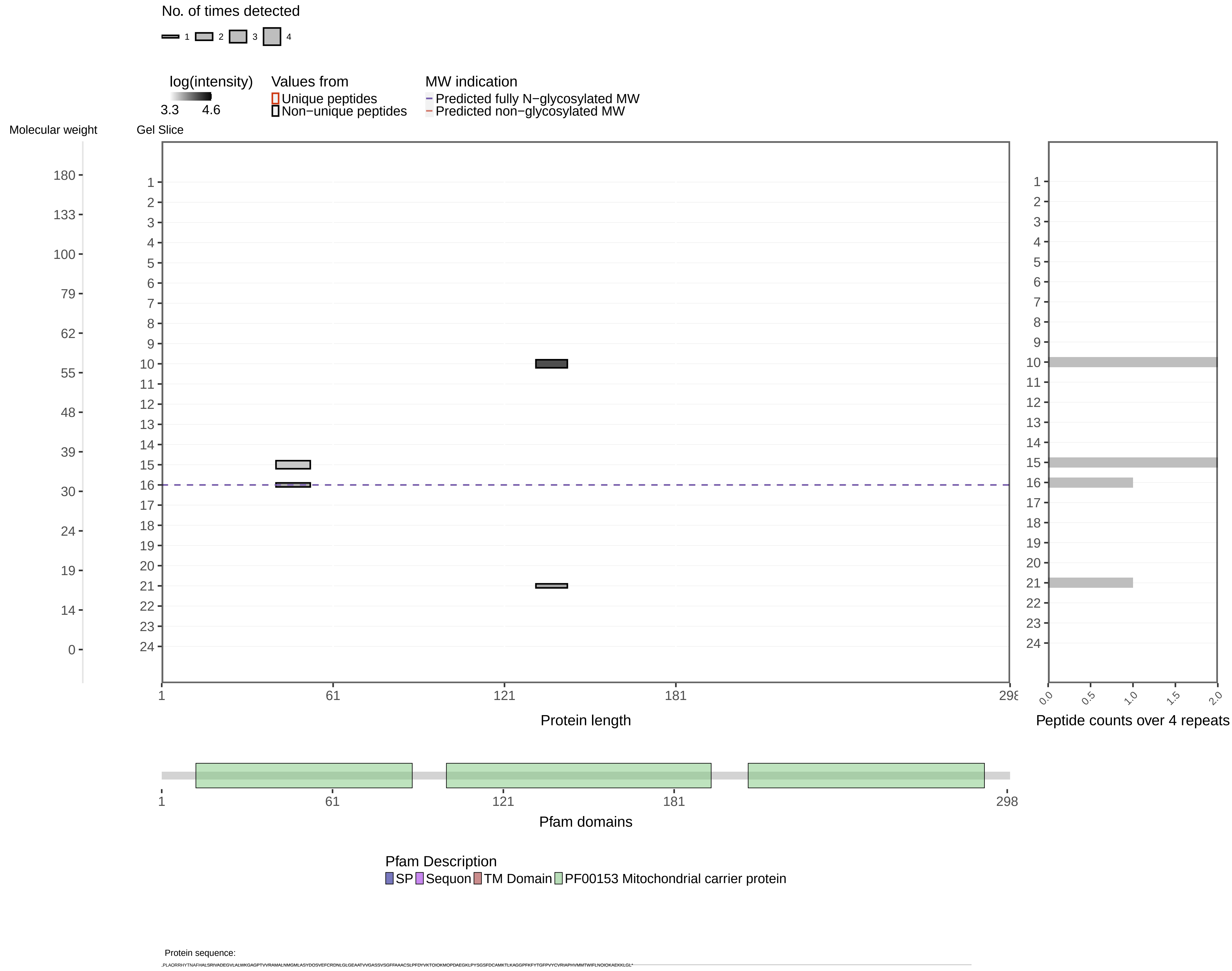

### NbL08g00070.1 isocitrate_dehydrogenase_[NADP] [MW = 46.84 kDa]

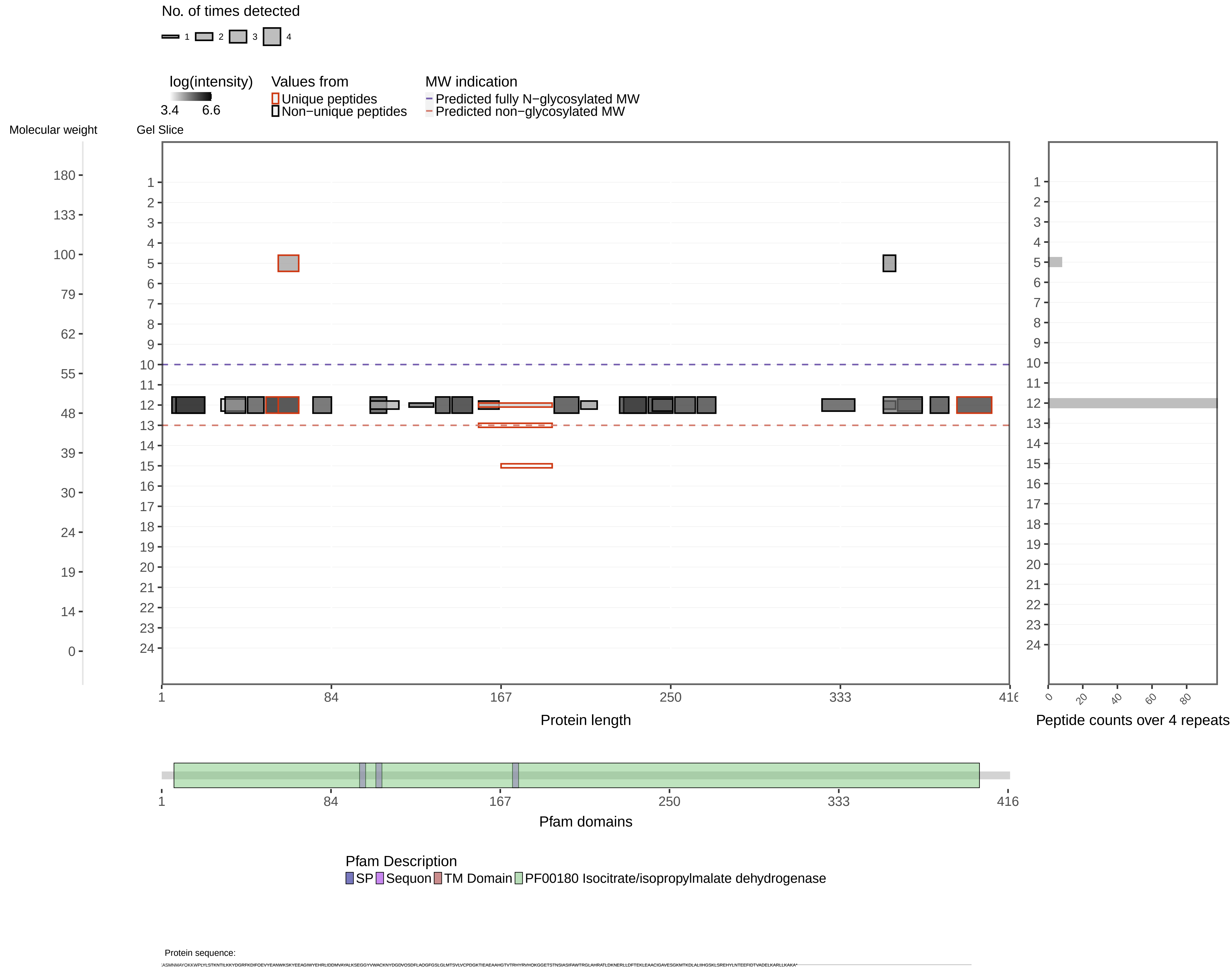

### NbL08g00030.1 lysine−−tRNA_ligase_chloroplastic/mitochondrial [MW = 27.41 kDa]

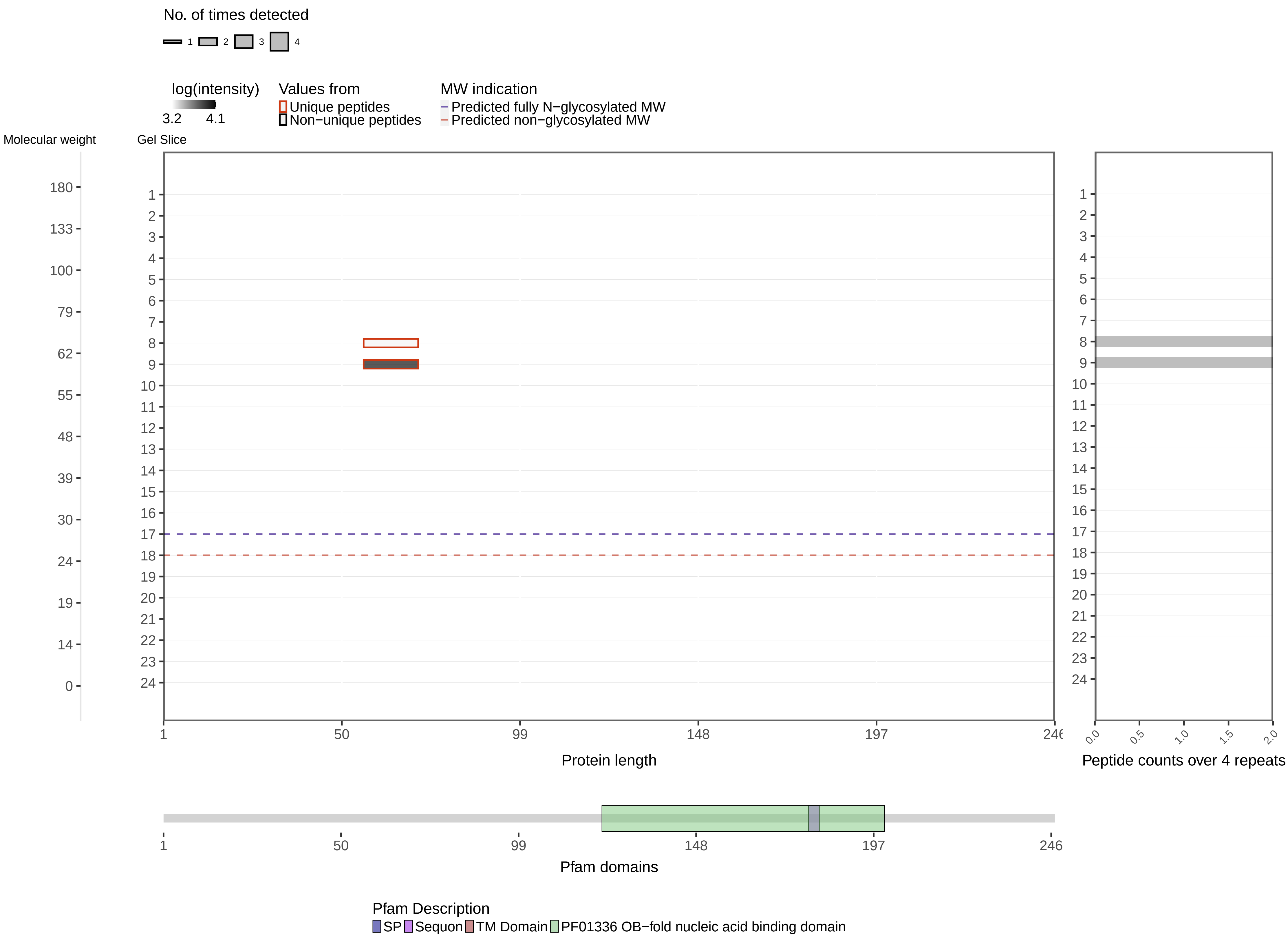


Protein sequence:

GNGEESNSENDYVSVSGRIVARRAFGKLAFLTLRDDSGMIQLYCEKERLVDDQFEQLKTLVDVGDILGANGSIKRTEKGELSVCVNSFSILTKSLLPLPDKYHGLTDVDKRYRQRYVDMIANPEVADTFRKRAKVG*

NbL08g00020.1 lysine−−tRNA_ligase_chloroplastic/mitochondrial−like [MW = 32.79 kDa]

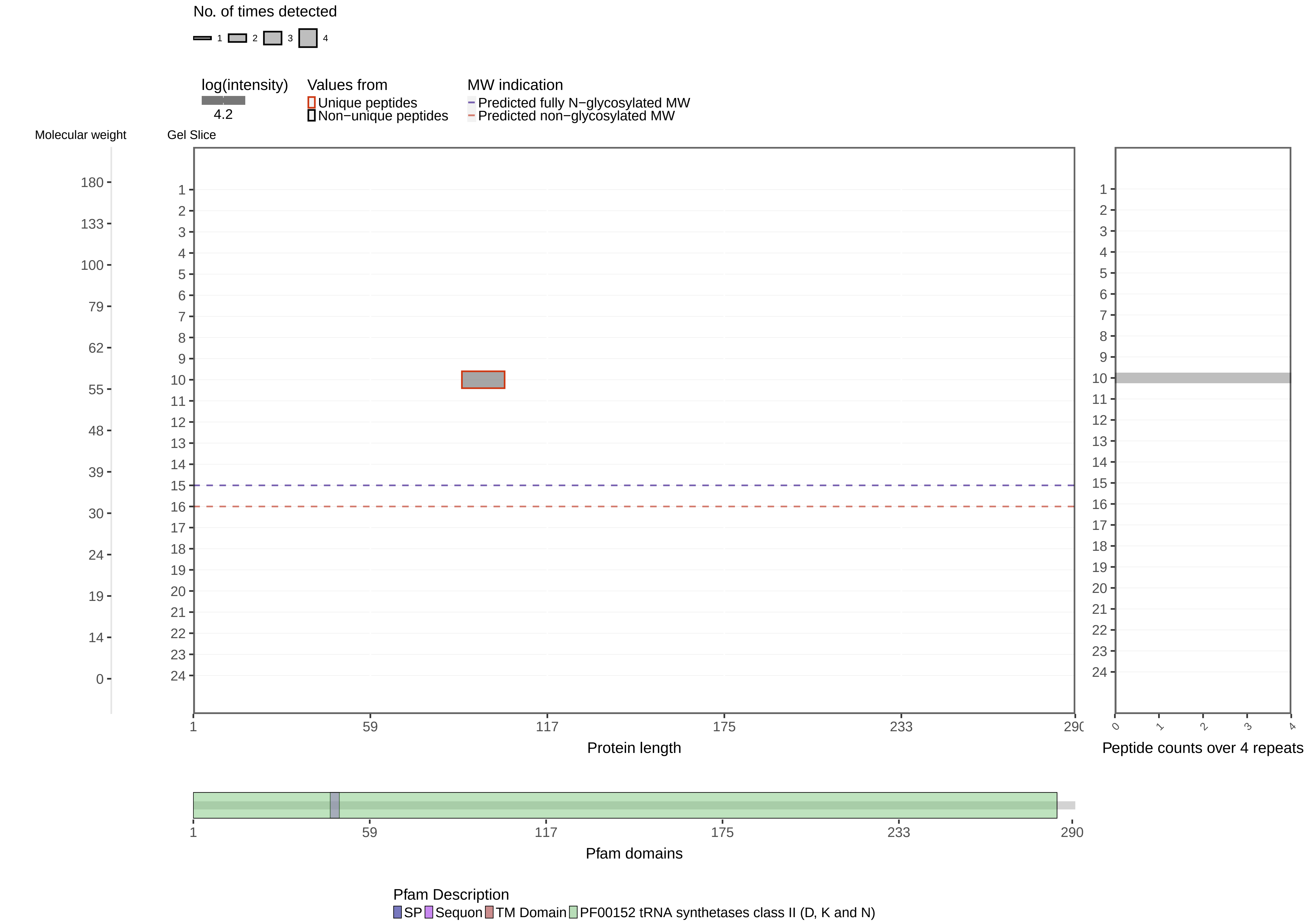


Protein sequence:

GHLVNEVFEMIVEPKLLQPTFVLDYPIEISPLAKPHRRHSGLTERFELFICGRELANAFSELTDPLDQRRRLEEQVKQHNEKKVASGLNMAGAEGQENKDDDDDSYKVTLDEDFLTALEYGMPPASGMGLGIDRLVMLLTNSASIRDVIAFPVLKIQQ*

### NbL07g17970.1 ATP_synthase_subunit_alpha_mitochondrial [MW = 55.25 kDa]

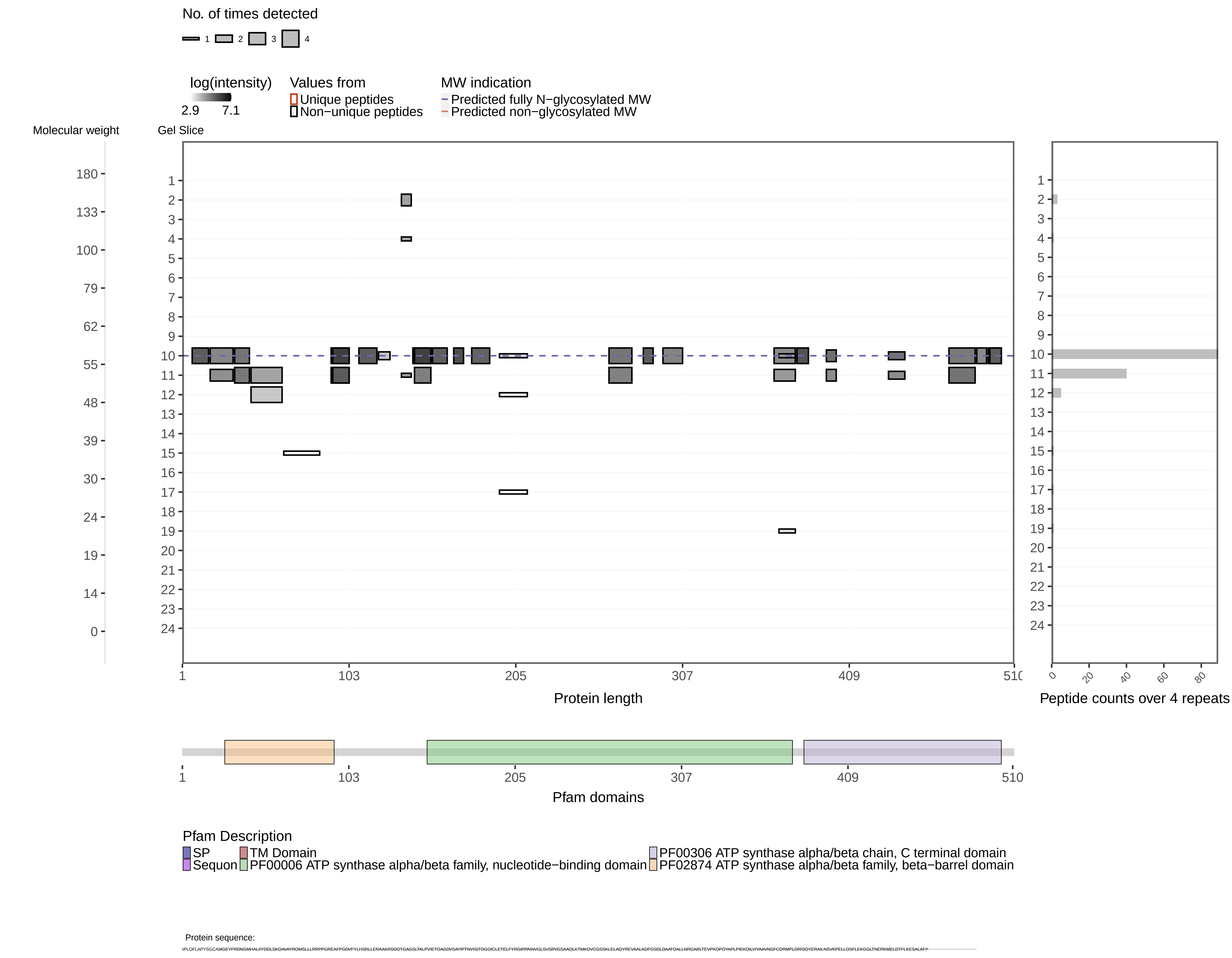

### NbL07g17960.1 ribosomal_protein_S3_mitochondrial−like [MW = 145.57 kDa]

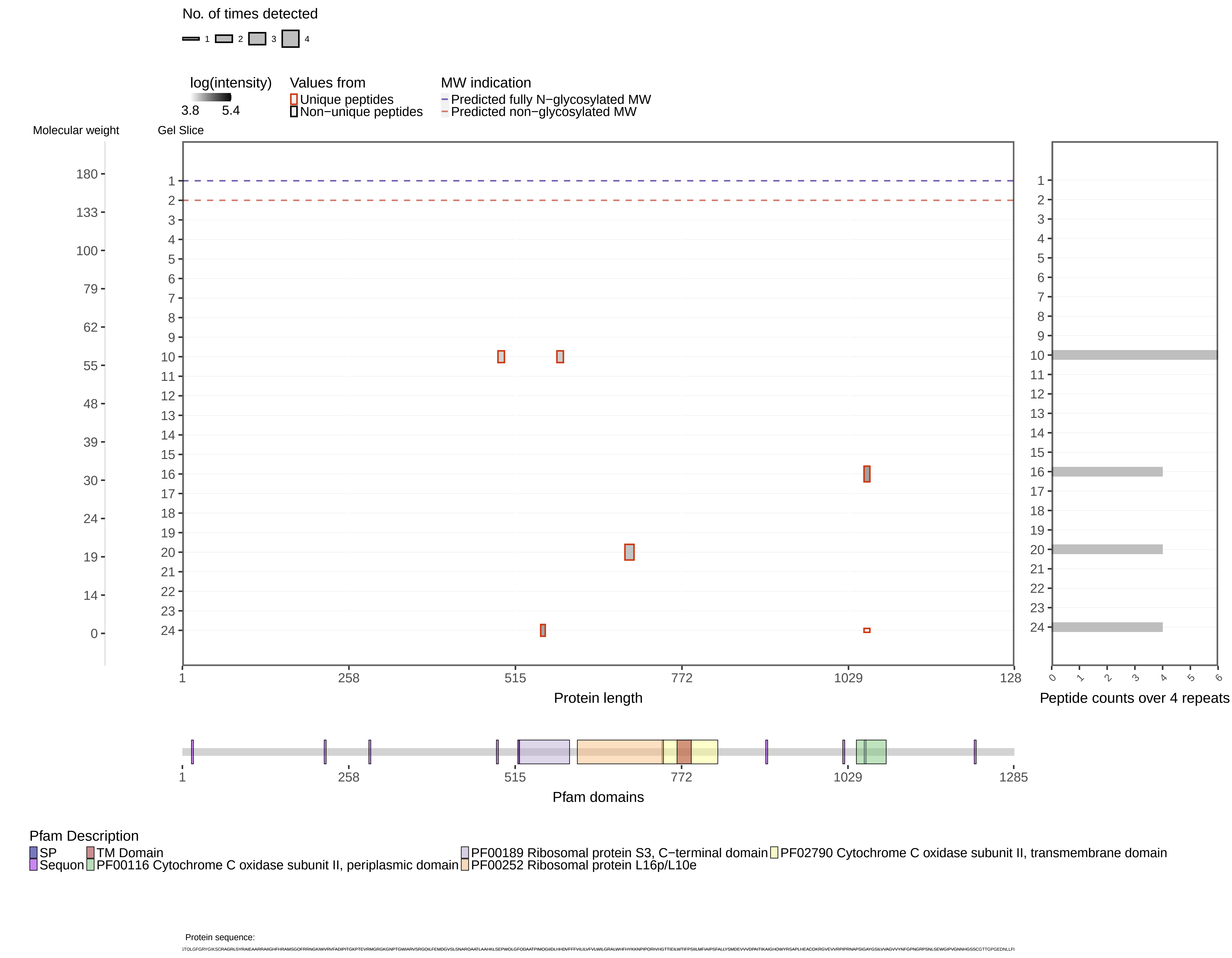

NbL07g17940.1 NADH_dehydrogenase_[ubiquinone]_iron−sulfur_protein_3 [MW = 22.86 kDa]

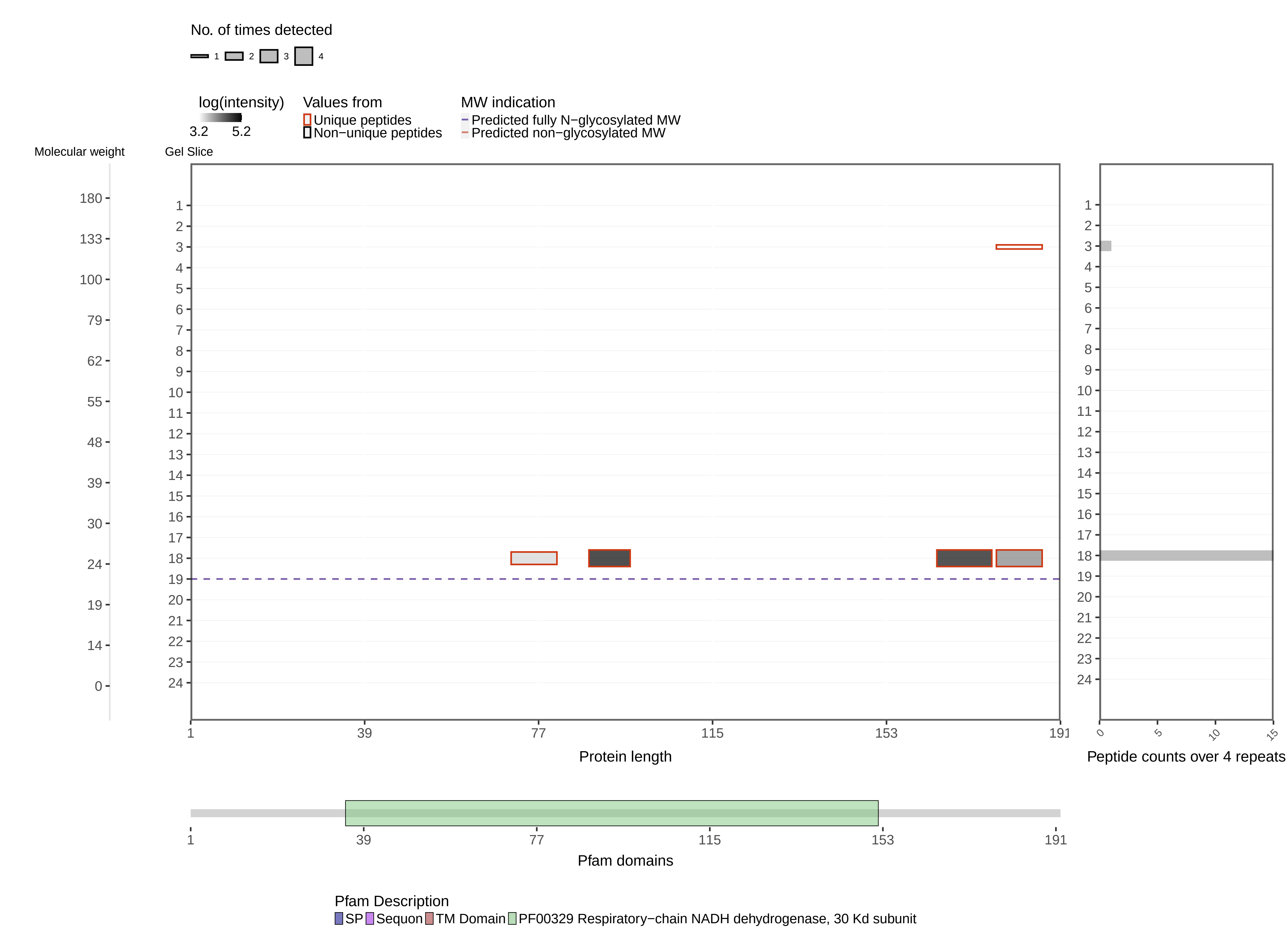


Protein sequence:

ISRIRVQTSADEVTRISSVVSLFPSAGWWEREVWDMFGVSFINHPDLRRILTDYGFEGHPLRKDFPLSGYVEVRYDDPEKRVVSEPIEMTQEFRYFDFASPWEQRSDG*

### NbL07g17930.1 ribosomal_protein_S4_mitochondrial [MW = 32.36 kDa]

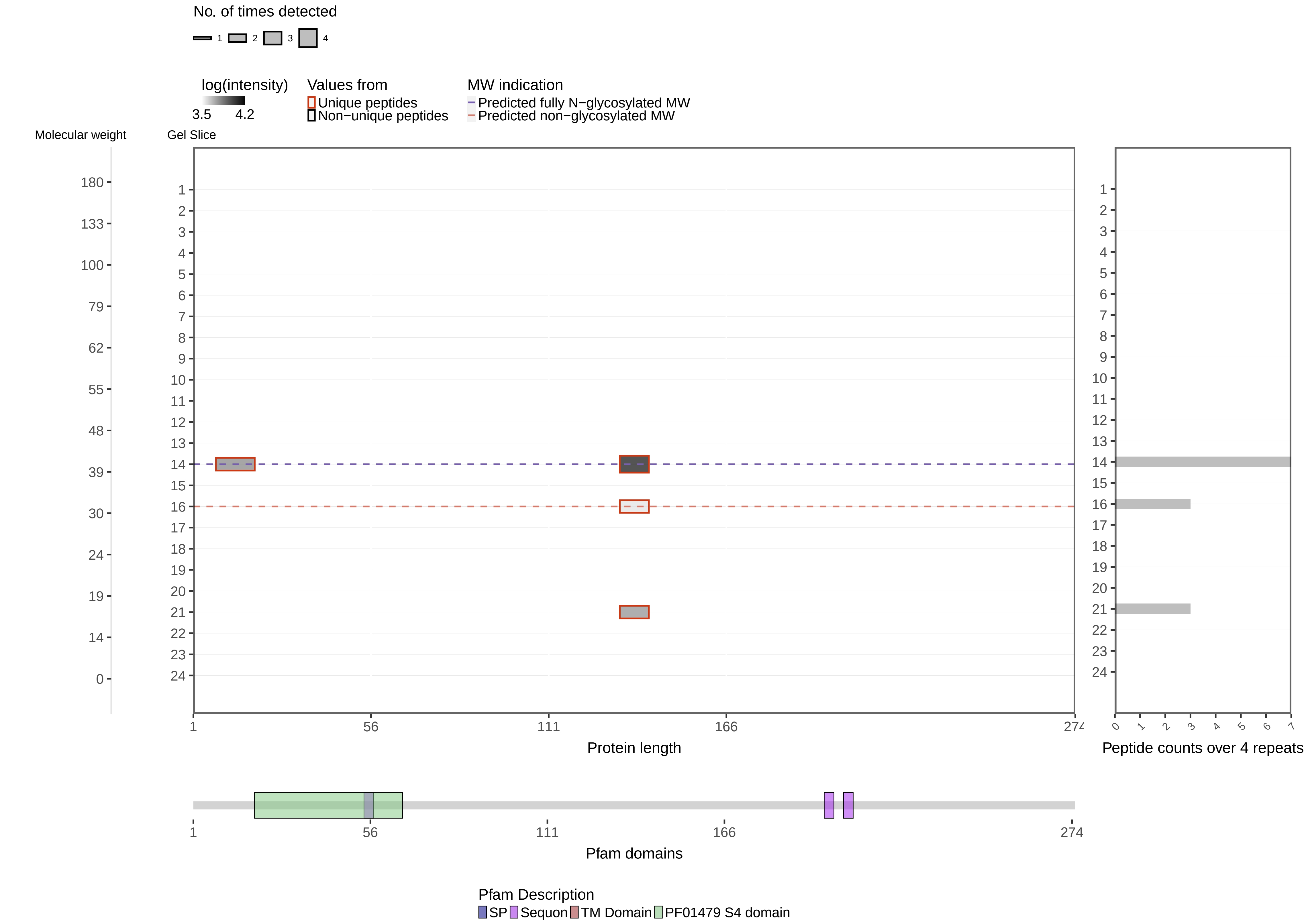


Protein sequence:

SRFLQQLRSSMQEEDLERTKKFGSEKVCLGSSFAEHNRMKRNLYHFKSLFLSKRRNEKNRNIPTRTRSPIVYNSSLYSNSTYCSASPHQFTKKIKIKRIELPTHYLEVNYRTPKAVVSYGPNIGHIPHDIRLKDLNLLLWSGKGRGQNI*

### NbL07g17900.1 photosystem_I_P700_chlorophyll_a_apoprotein_A1 [MW = 83.02 kDa]

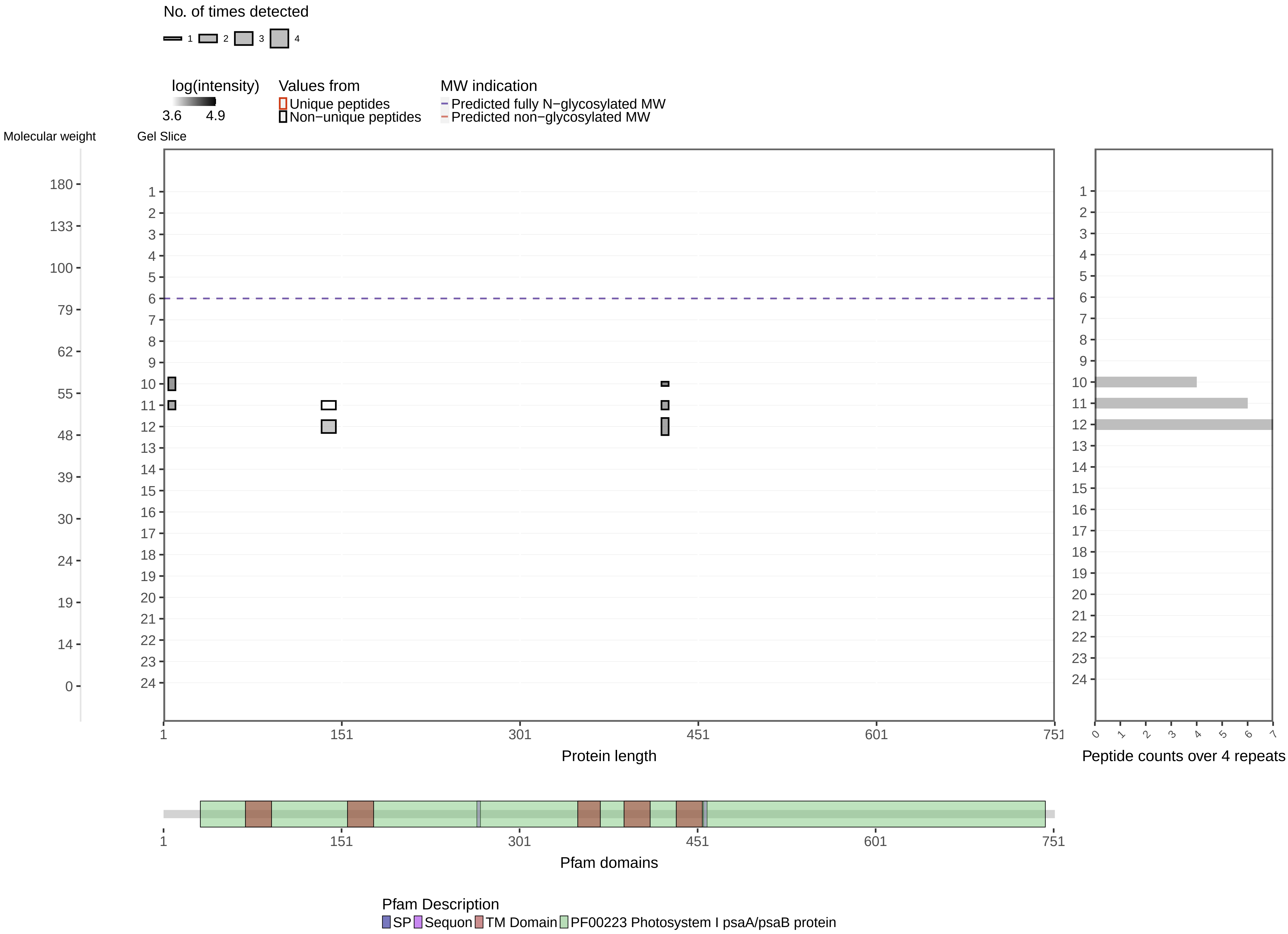


Protein sequence:

/VAHHMYSMPPYPYLATDYGTQLSLFTHHMWIGGFLIVGAAAHAAIFMVRDYDPTTRYNDLLDRVLRHRDAIISHLNWACIFLGFHSFGLYIHNDTMSALGRPQDMFSDTAIQLQPVFAQWIQNTHALAPGATAPGATASTSLTWGGGDLVAVGGKVALLPIPLGTADFLVHHIHAFTIHVTVLILLKGVLFARSSRLIPDKANLGFRFPCDGPGRGGTCQVSAWDHVFLGLFWMYNAISVVIFHFSWKMQSDVWGSVSDQGVVTHITGGNFAQSSITINGWLRDFLWAQASQVI

NbL07g17890.1 acetyl−coenzyme_A_carboxylase_carboxyl_transferase_subunit_beta_chloroplastic [MW = 58.28 kDa]

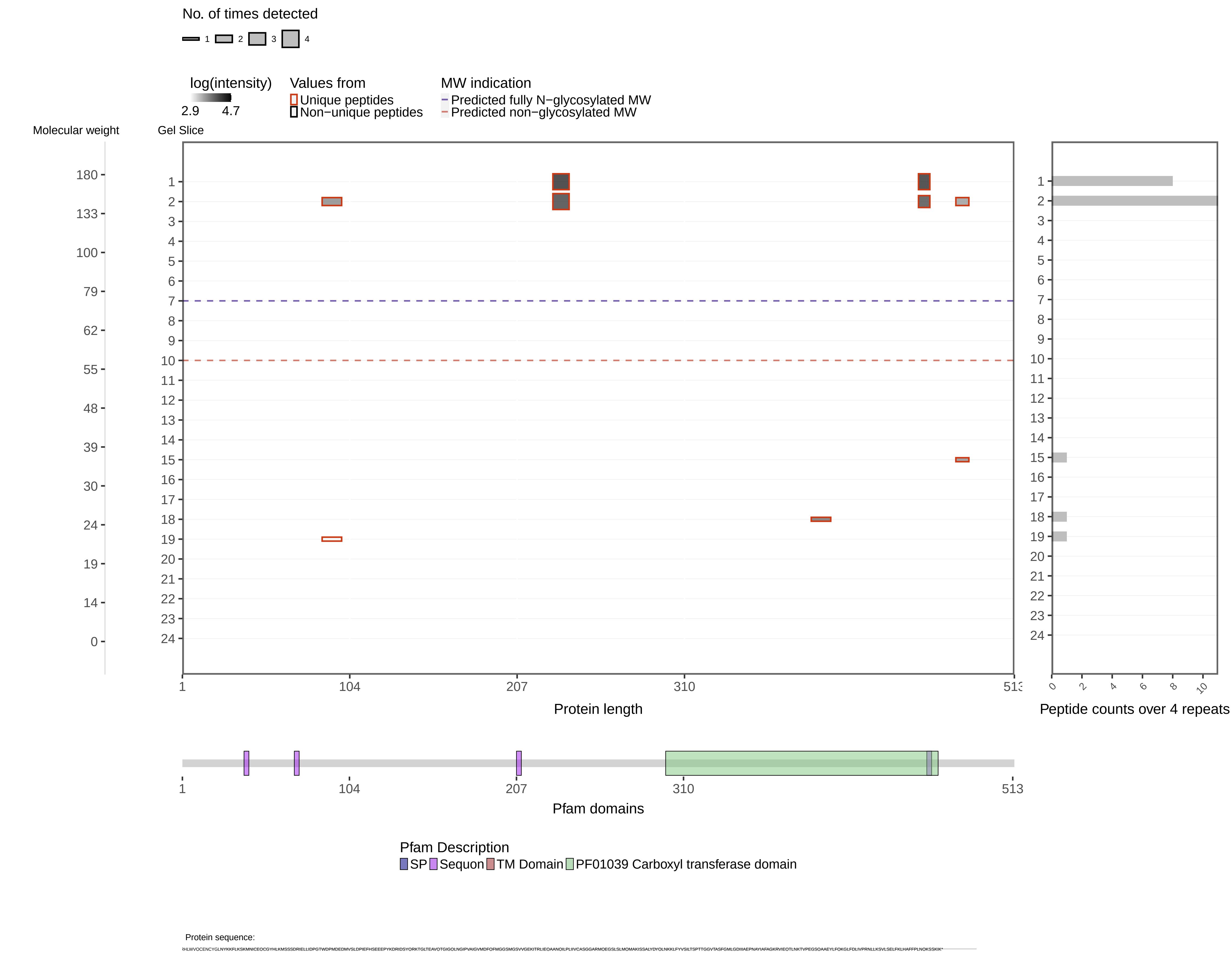

### NbL07g17750.1 stromal_processing_peptidase_chloroplastic−like [MW = 140.54 kDa]

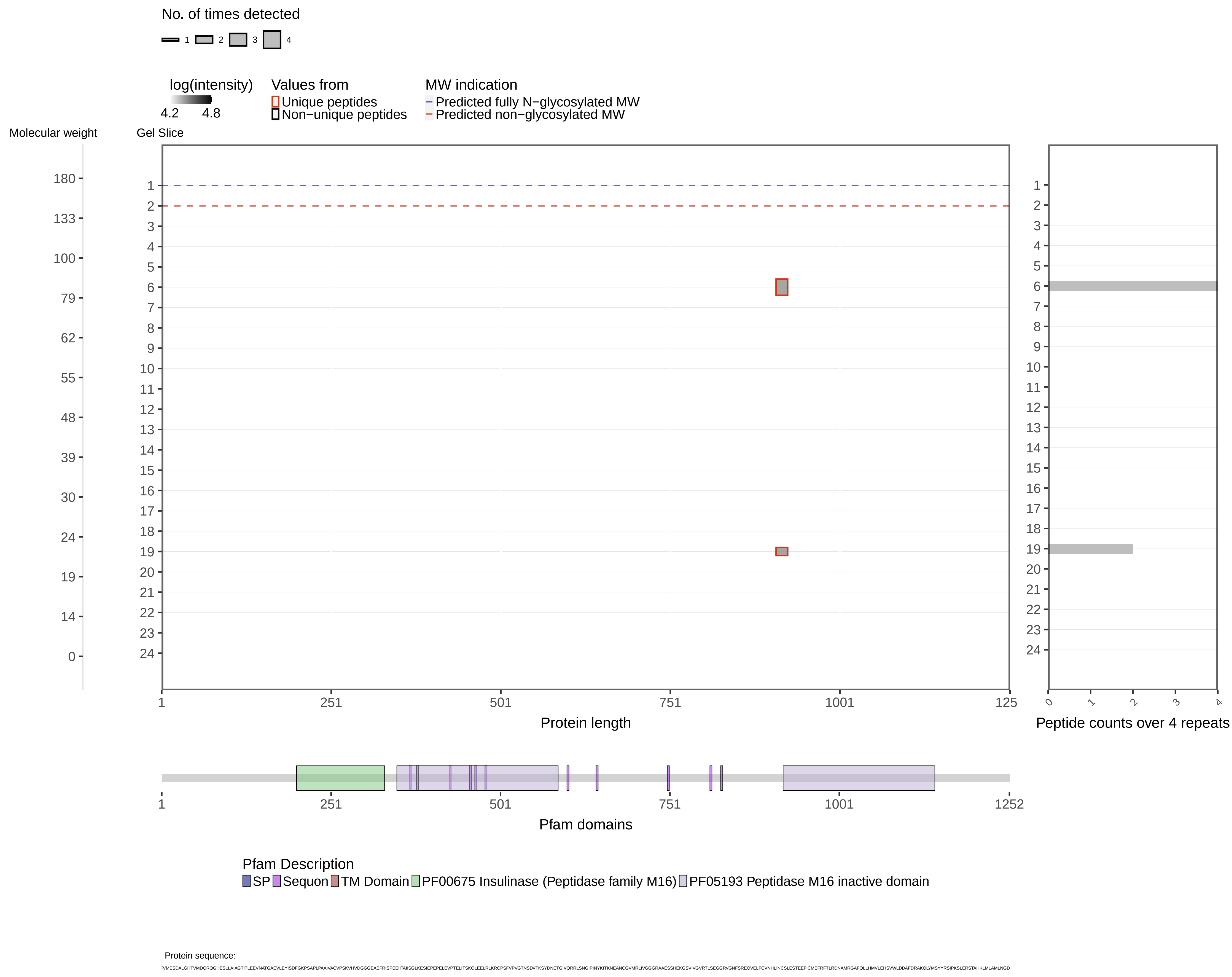

### NbL07g17730.1 acyl_carrier_protein_1_chloroplastic−like [MW = 14.99 kDa]

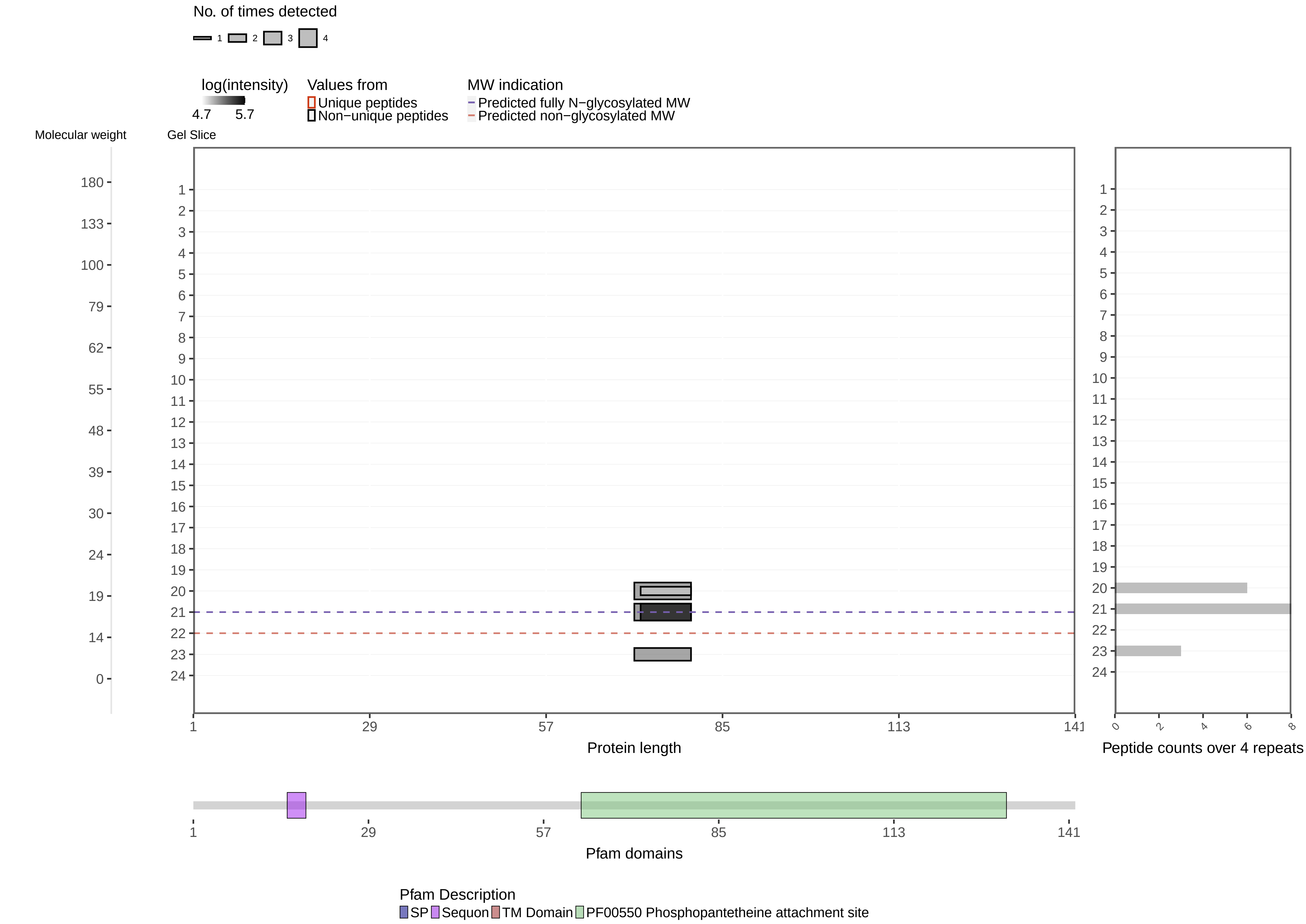


Protein sequence:

CAKPETVEKVISIVRKQLALPADTKVNAESTFNTDLGADSLDTVEIVMALEEEFGIAVEEENSETIVTVQDAADLIEKLVEKKAA*

### NbL07g17710.1 serine_carboxypeptidase−like_45 [MW = 56.08 kDa]

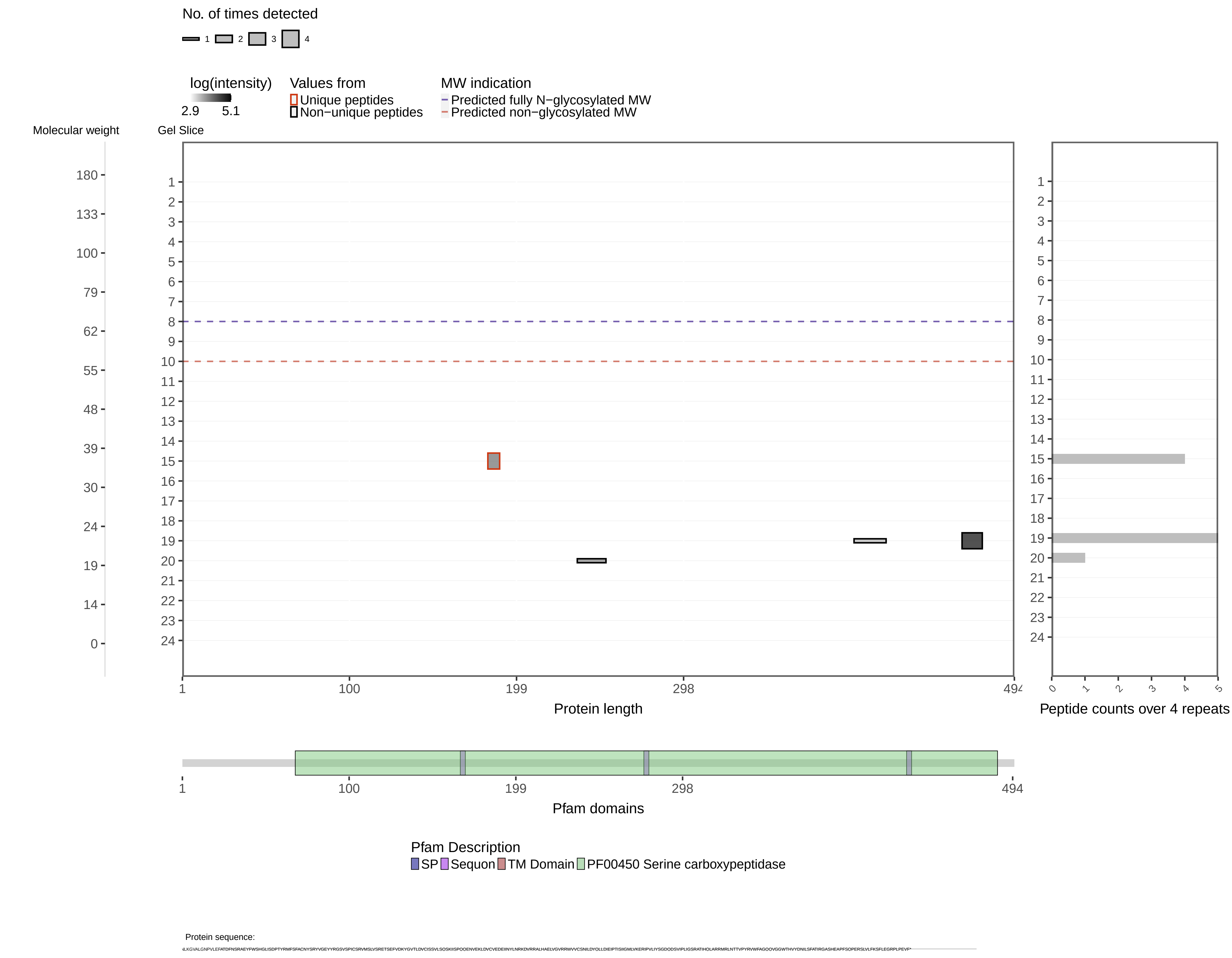

### NbL07g17690.1 ATP−dependent_Clp_protease_ATP−binding_subunit_clpA_homolog_CD4B_chloroplastic−like [MW = 102.37 kDa]

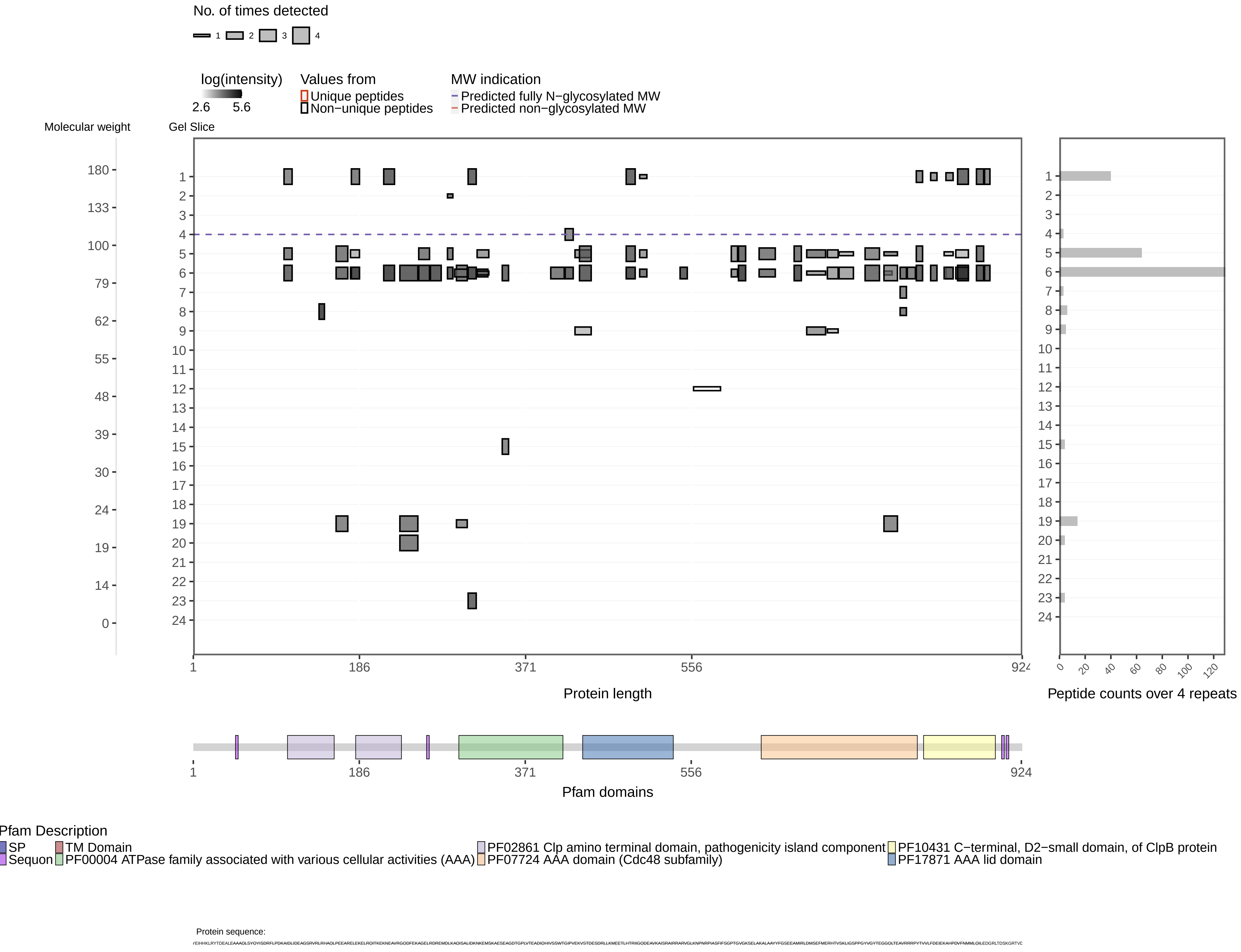

NbL07g17640.1 magnesium_protoporphyrin_IX_methyltransferase_chloroplastic−like [MW = 35.24 kDa]

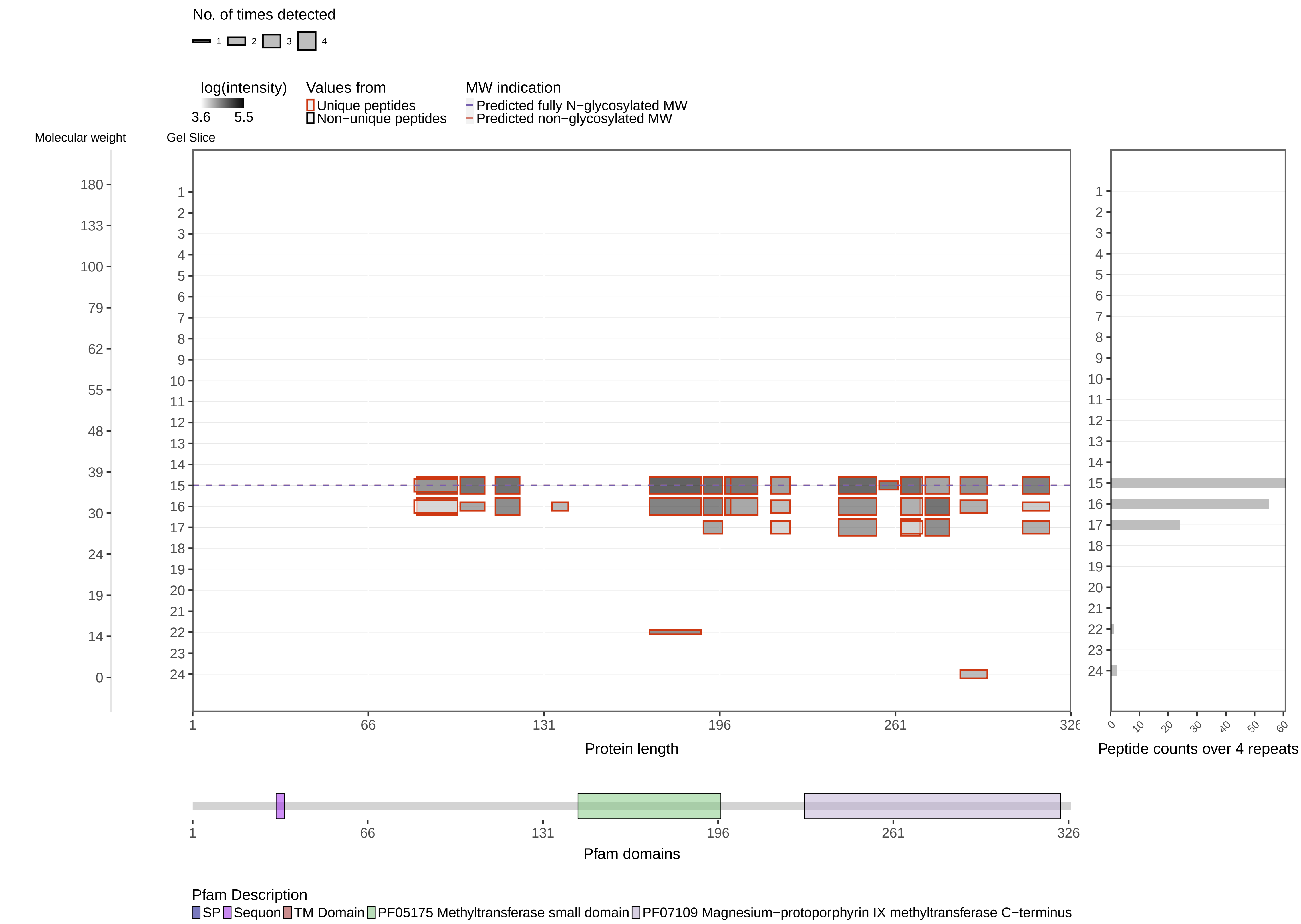


Protein sequence:

OGVTVCDAGCGTGCLSIPLAKEGAVVSASDISASMVAEAOKOAOEELFKAKDDOSPAPVMPKFEVKDLESLDGKYDTVVCLDVMIHYPOSKADGMIAHLASLAENRLILSFAPKTFYYDLLKRIGELFPGPSKATRAYLHAEADVERALOKAGWRIRKRGLIATOFYFAKLIEAVPA*

NbL07g17590.1 glycine−−tRNA_ligase_1_mitochondrial−like [MW = 70.13 kDa]

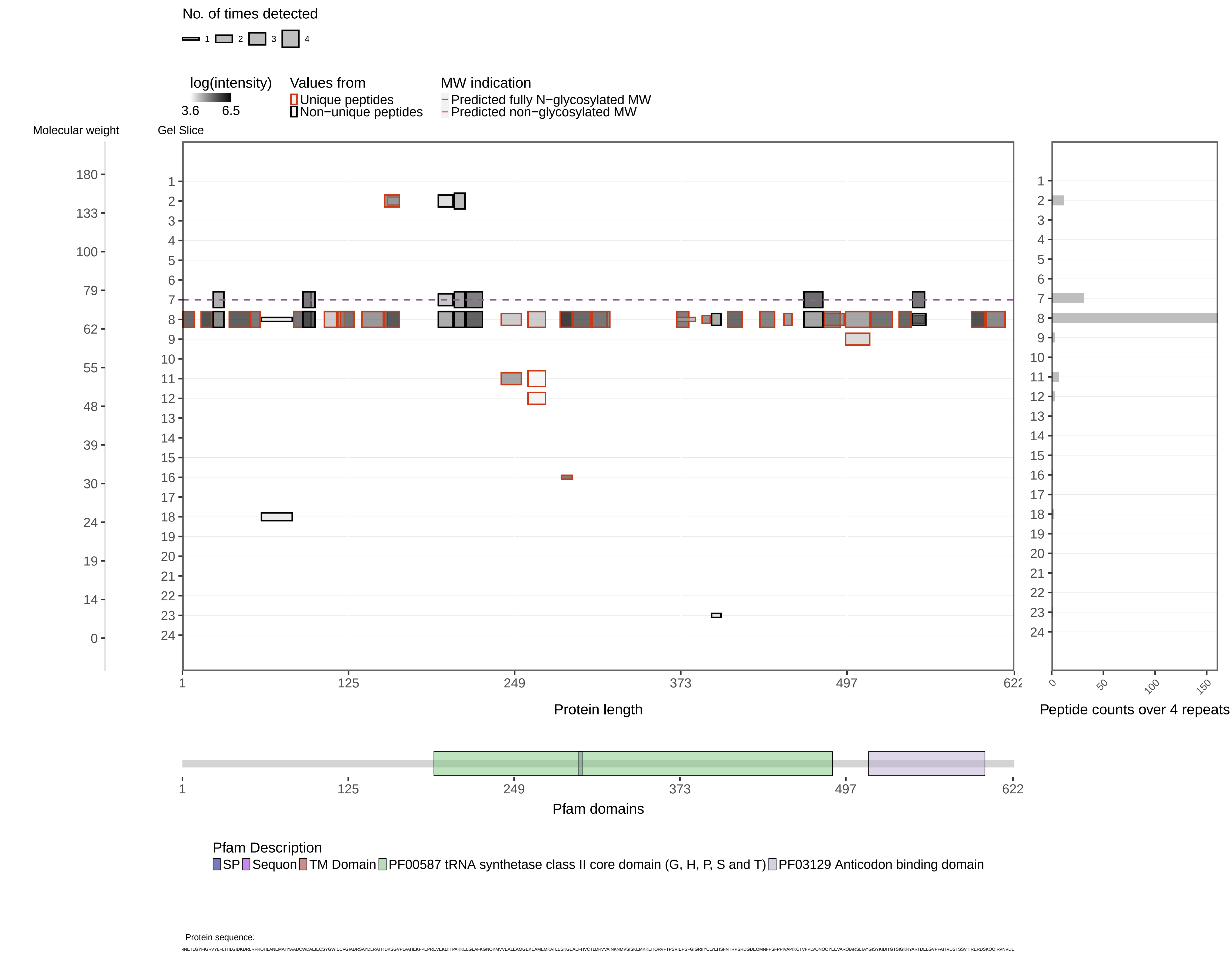

### NbL07g17560.1 uncharacterized_LOC107786132 [MW = 17.64 kDa]

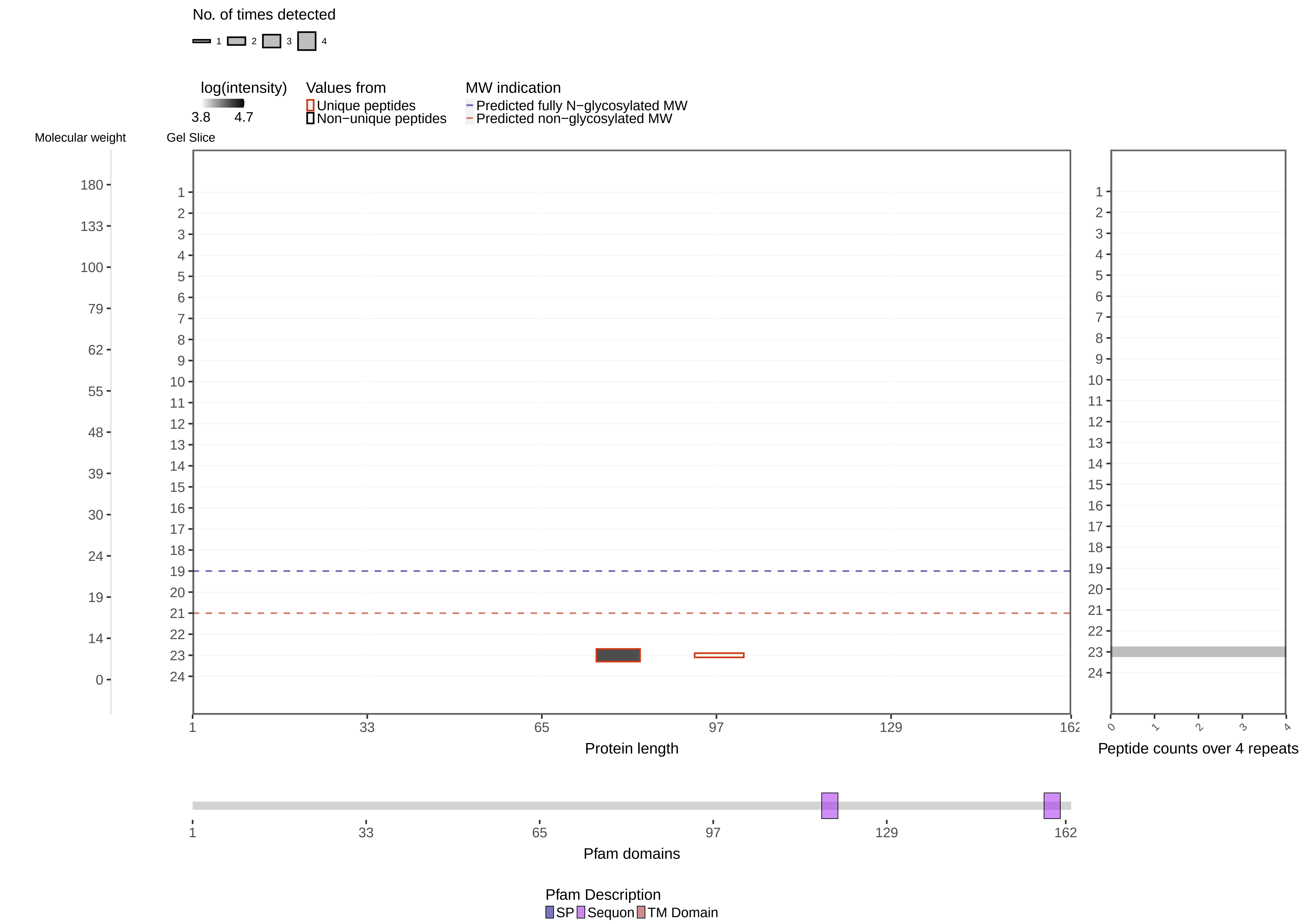


Protein sequence:

KQAYICRDCGYIYNDRTPFEKOPDKYFCPVCGAPKRRFRSYEPKVTKDANSTEVRKARKEQLKRDEAIGNALPIAIGVGVVALAALYFYVNNSS*

NbL07g17510.1 staphylococcal_nuclease_domain−containing_protein_1 [MW = 107.79 kDa]

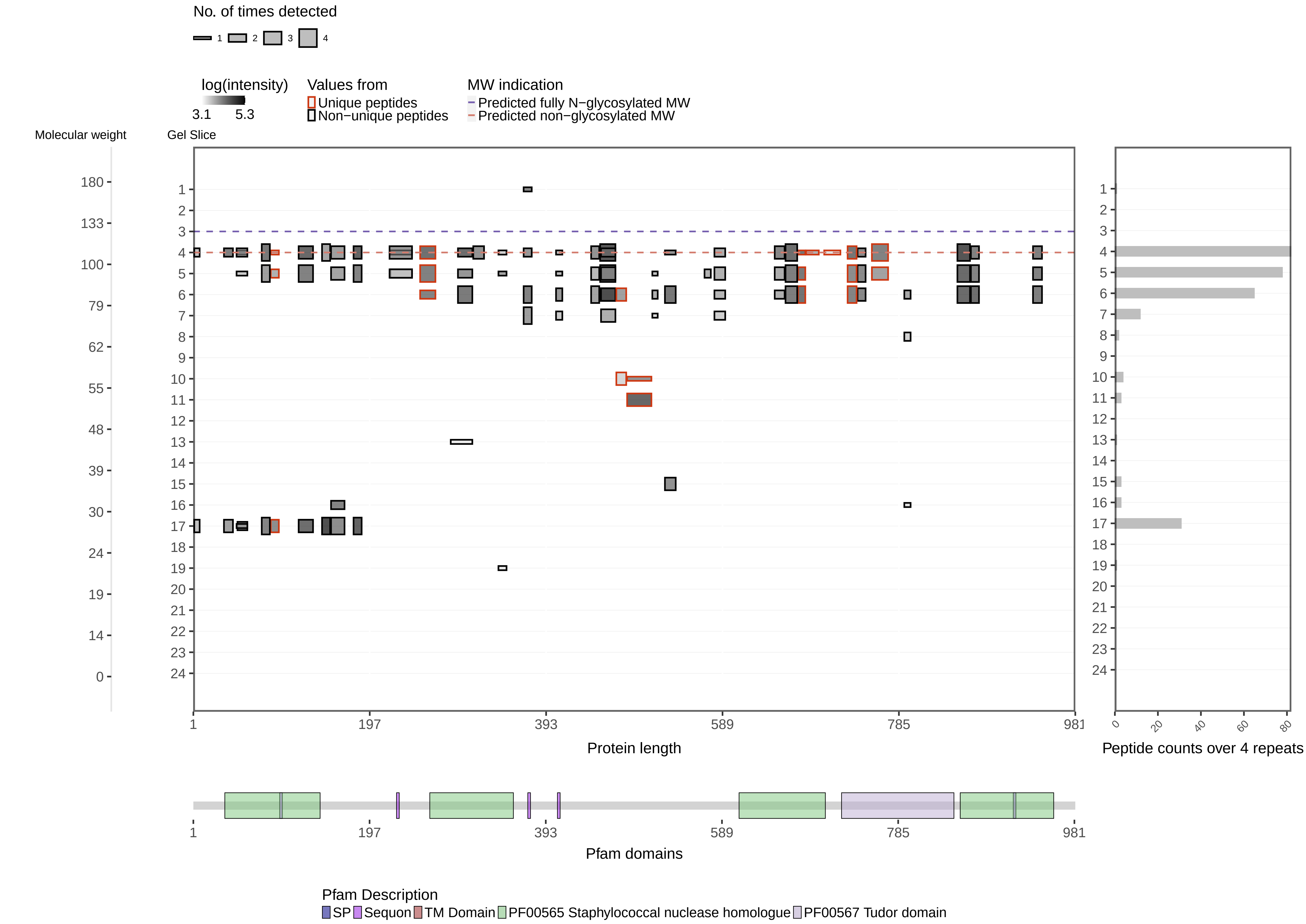


Protein sequence:

### NbL07g17500.1 glutamyl−tRNA(Gln)_amidotransferase_subunit_A−like [MW = 69.08 kDa]

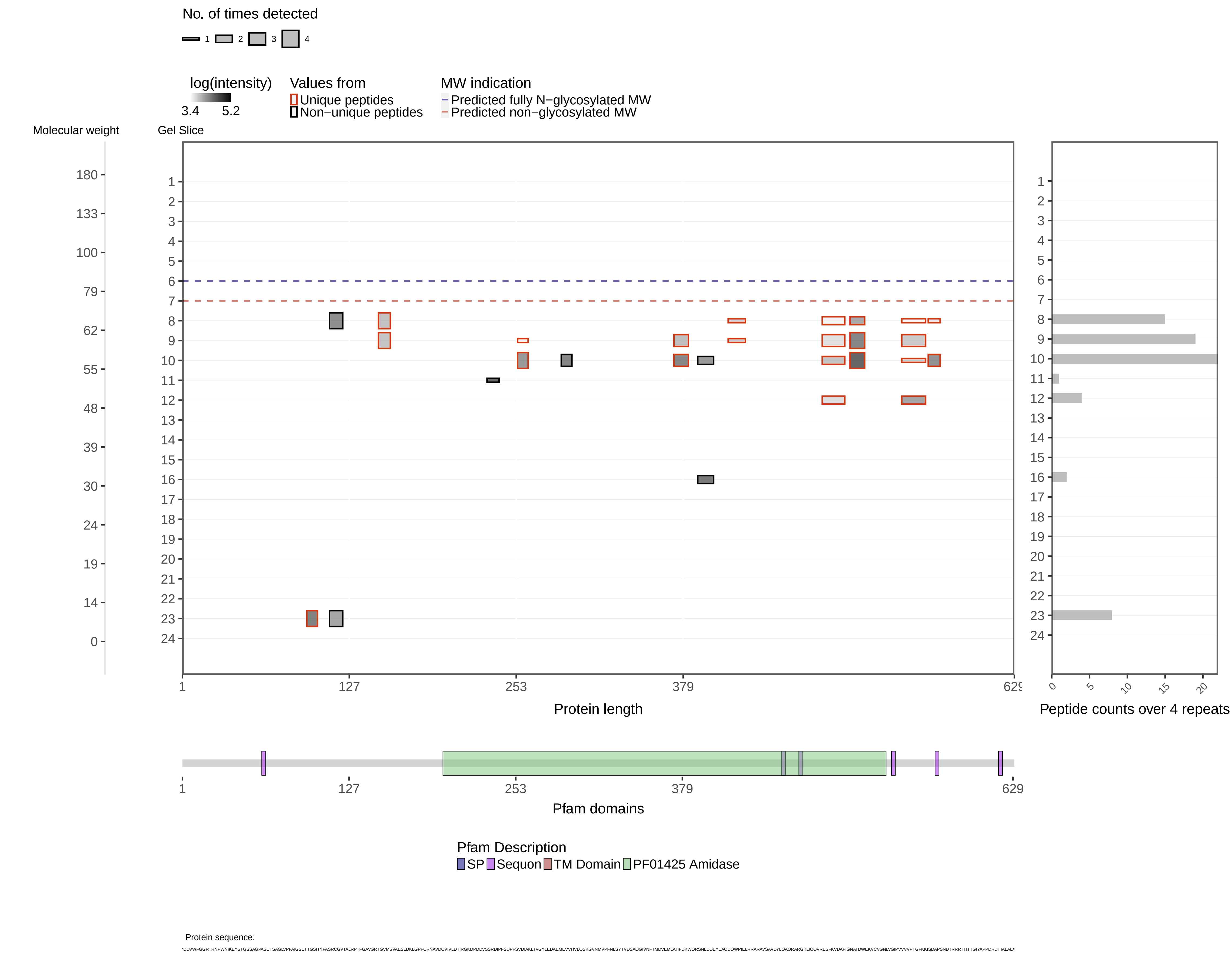

NbL07g17460.1 inositol_polyphosphate_multikinase_alpha−like [MW = 33.53 kDa]

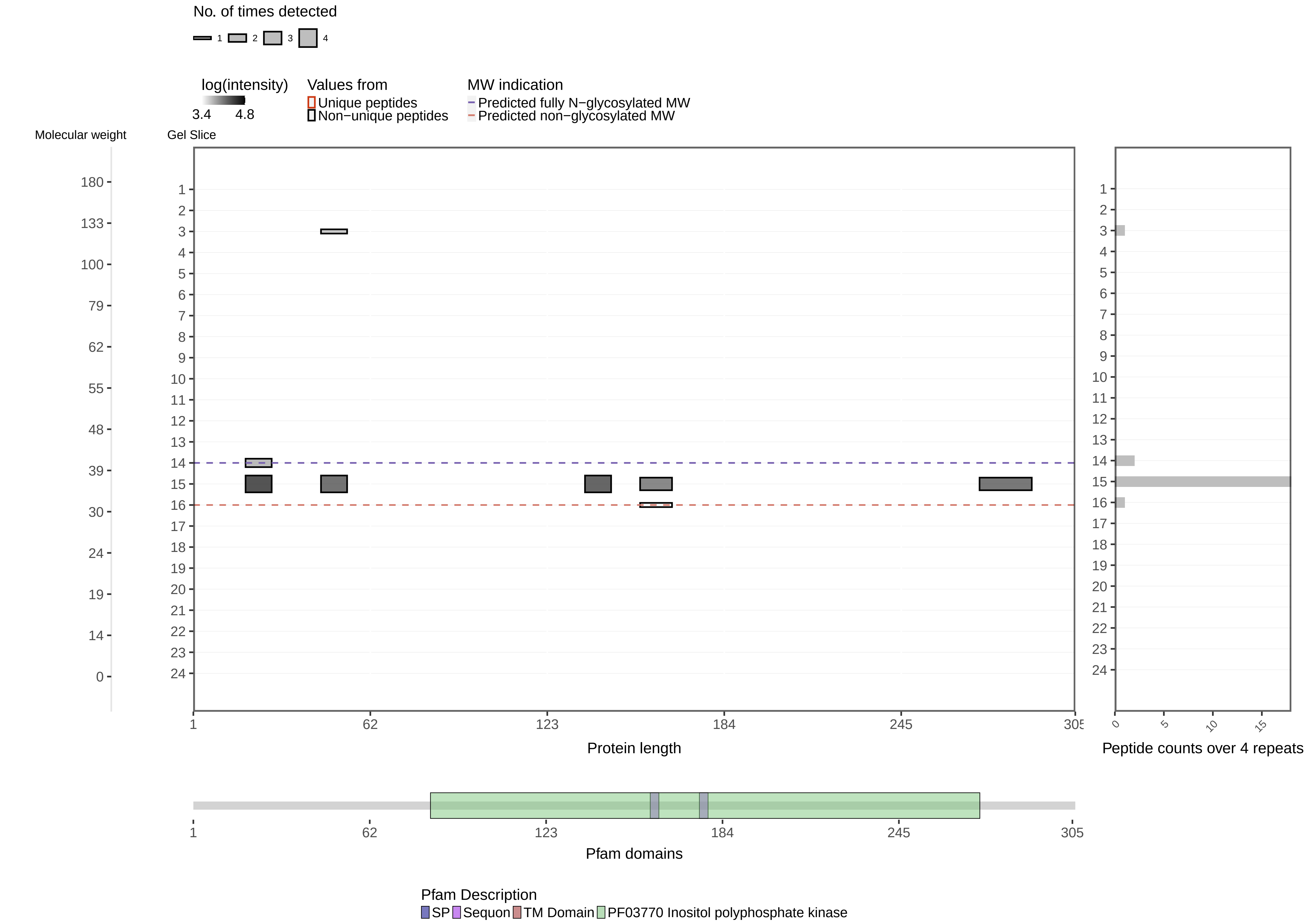


Protein sequence:

GIQIYGNKESGYWKPERTSIQNLSAEEVKLILKKFVSSNASTDSDLEPDCAFAATVYGGSTGILSQLLELKAWFEDQTMYHFYSCSILITFEKELALEGQNPGAQIKLIDFAHVSEGRGVIDHNFLGGLCSLIKFISEILTAPNEHTTEVSSKPDHQNLISSDNSVN*

### NbL07g17430.1 multiple_organellar_RNA_editing_factor_1_mitochondrial−like [MW = 48.38 kDa]

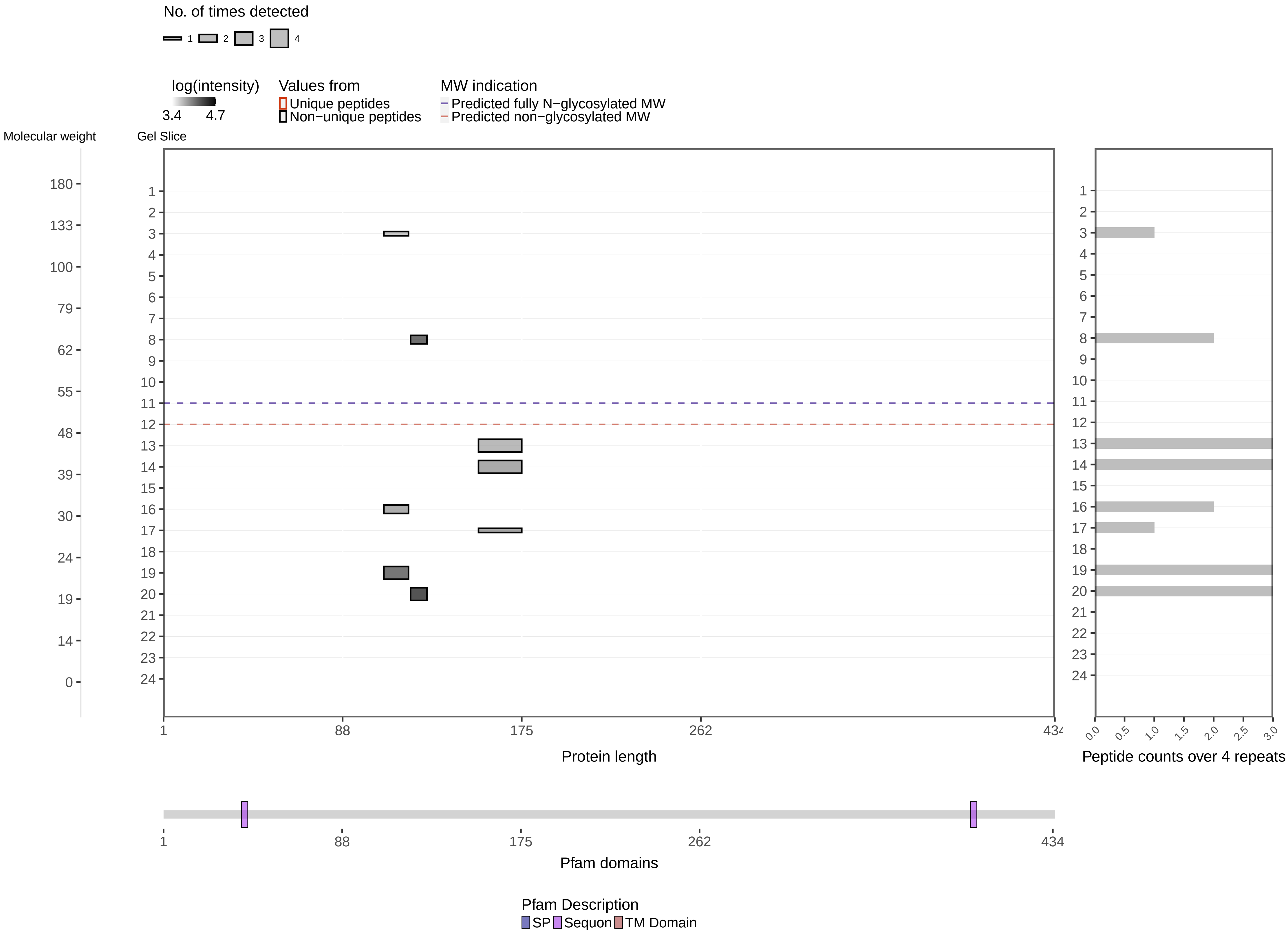


Protein sequence:

[illegible]GNYGPPQSPPQQNFGPPQGAPAQLNFGPPQGAPAQQNYGAAQGPPPQQNYAAAQGPPPQQNYGQPRYPPPQQNYGSQQYPPPPQQIYSAPQSPPQKNYGRPPNPLPQQSYSAPQNVPSQWNHGPQQSFSPQQNYGPQGAGDPWHQTPLGSSPGGWDNSQGGRQDSRPQYQVNYNQGEQGNYSPQRDHRGYAPPEERGFRGDNQHYAPPEQRGFRGDNQR*

### NbL07g17340.1 ACT_domain−containing_protein_DS12_chloroplastic−like [MW = 30.99 kDa]

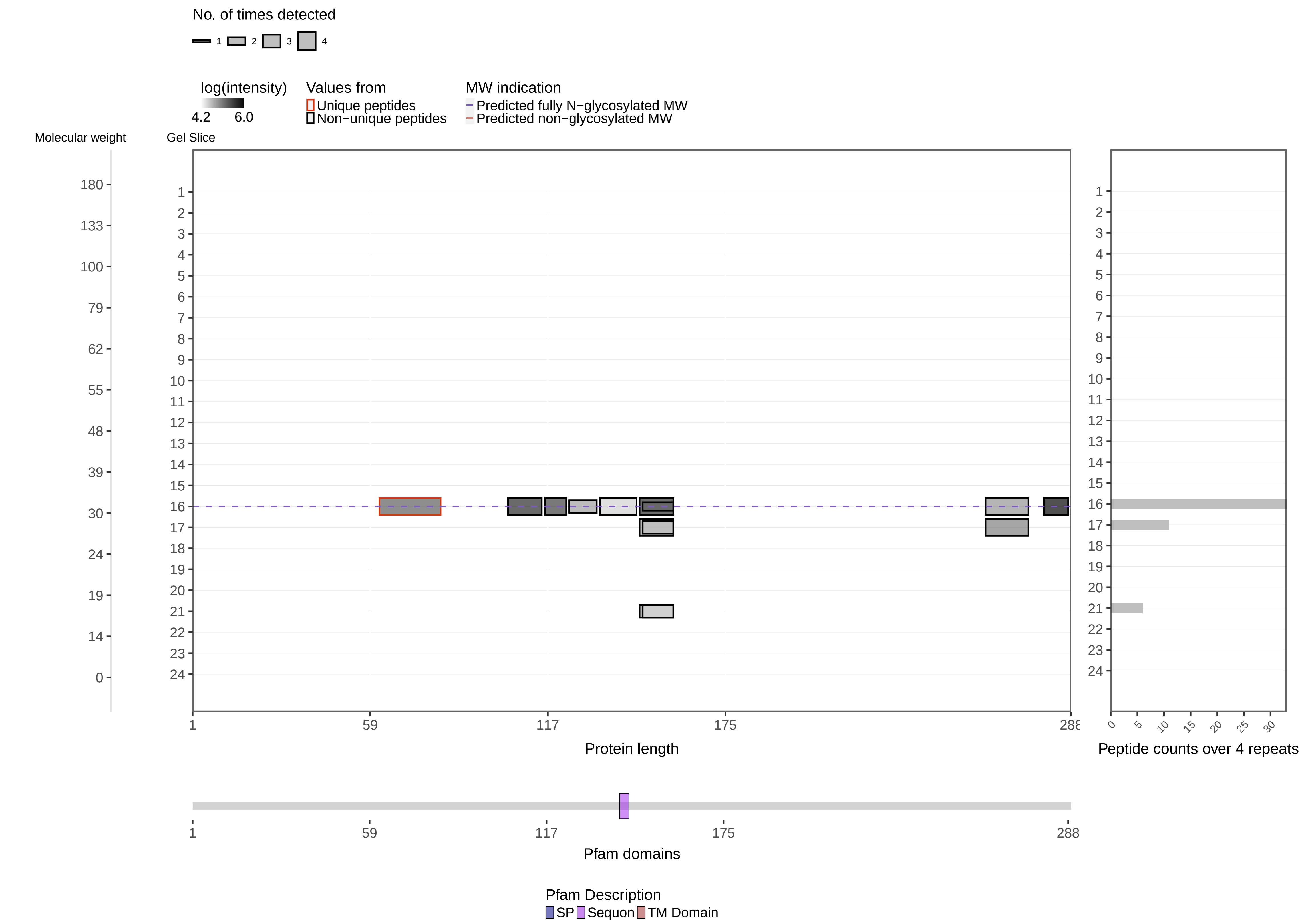


Protein sequence:

SGKHTTLCITNASTGRKVDDPEQLEAIRLTIINNMIEFHPESSALLAMGEVFGAYQPNEKLDVDIATHIHVYDDGPDRSLLYVEAADRPGLIVDLVKIITEINIDVESGEFDTEGLLAKAKFHVSYKNKALIKPLQQVLANSLRYYLRRPTTEESSF*

NbL07g17330.1 4−coumarate−−CoA_ligase_1 [MW = 59.83 kDa]

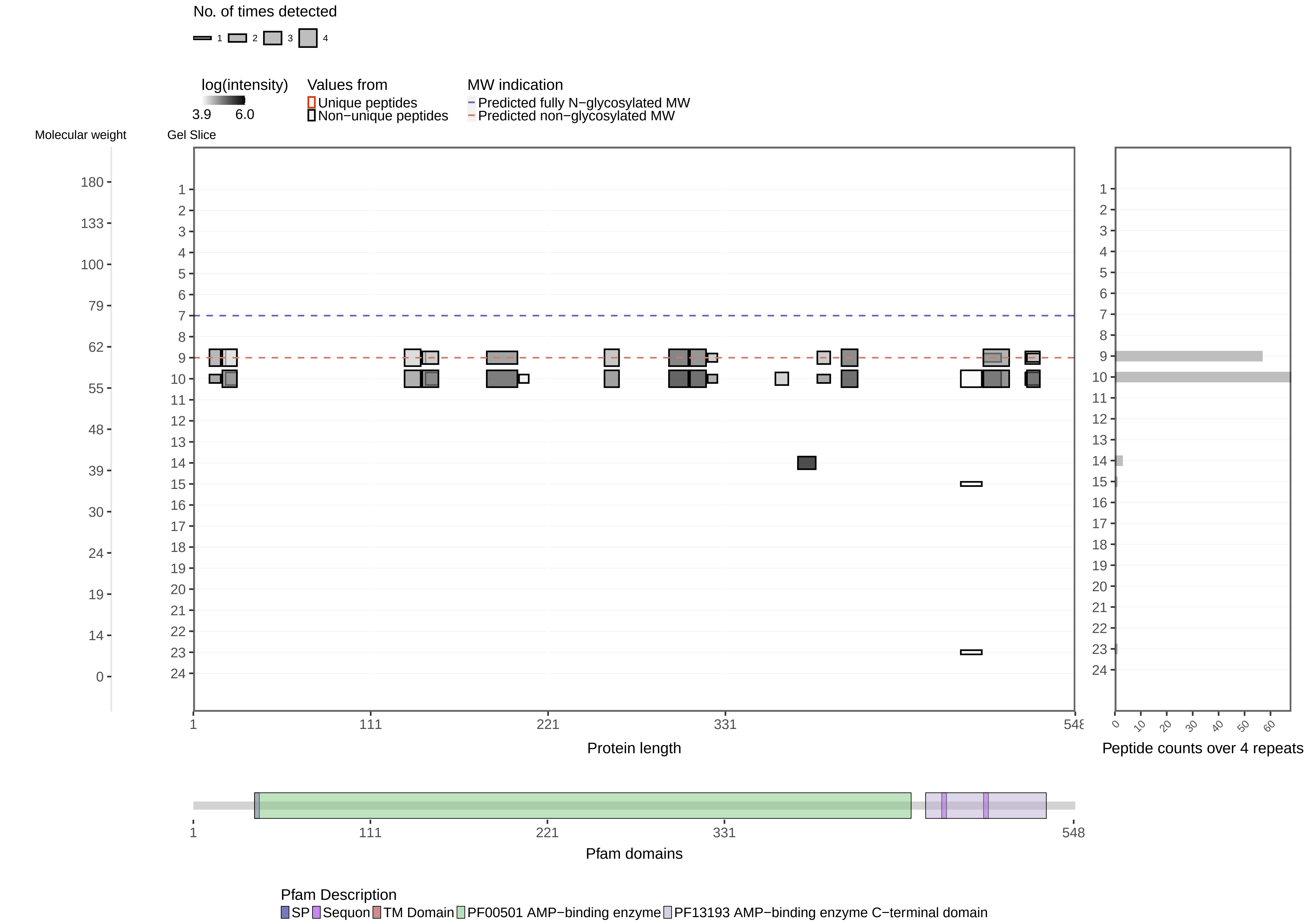


Protein sequence:

MQKFDIAPFLELIQKYKVSIGPFVPPIVLAIAKSPIVDSYDLSSVRTVMSGAAPLGKELEDAVRTKFPNAKLGQGYGMTEAGPVLAMCLAFAKEPFDIKSGACGTVVRNAEMKIVDPDTGCSLPRNQPGEICIRGDQIMKGYLNDPEATTRTIDKEGWLHTGDIGFIDEDDELFIVDRLKELIKYKGFQVAPAEIEALLLNHPNISDAAVVPMKDEQAGEVPVAFVVRSNGSAITEDEVKDFVSKQVIFYKRVKRVFFVETVPKSPSGKILRKDLRARLAAGVPN*

NbL07g17240.1 uncharacterized_LOC107804369 [MW = 33.72 kDa]

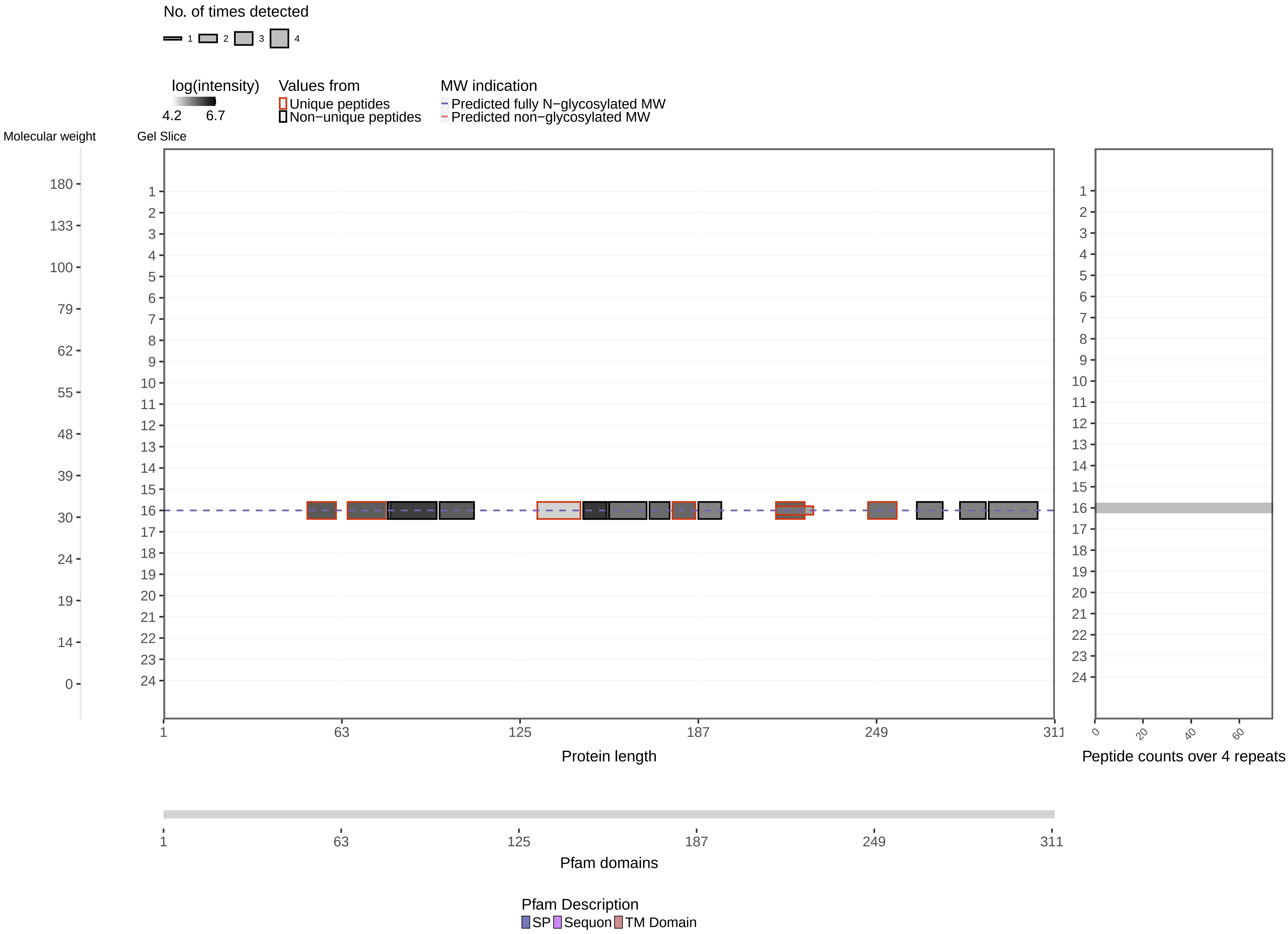


Protein sequence:

VSKLGEPPTDGKTDTNPEGLTAAYGKWASAVAARLQTGGLSCKVLEKEPFQKQMLEKLIWICAFMLVGARHPGATVGAVEKEYRSEVSQLIAELAAAAAAEKGLVFEDAMEDRLCAYSRAVAHFPTAVKEFKWRNGWFYSLSEKAIAEGKPDPCPLHTAWLKELKIV*

NbL07g17110.1 lamin−like_protein [MW − SP = 15.84 kDa]

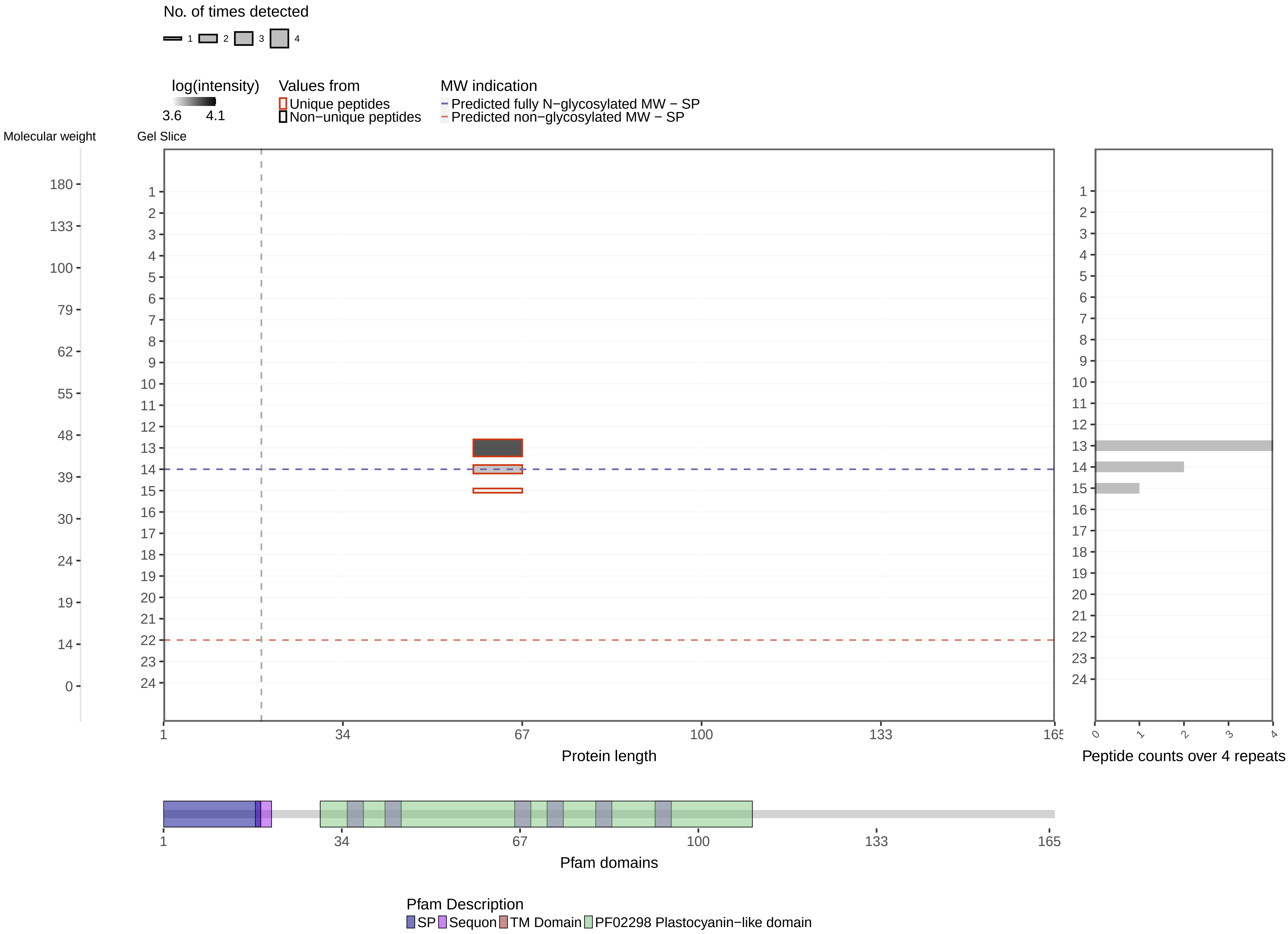


Protein sequence:

YDNCTTEGALGNWTGGKDFIMLNKTQRYYFICGTGGCLNGMKVSVVVHPLPSPPKSAISAEHSSKKSAAPVAARGFGSLLAMSFALFGLTFSVWI*

### NbL07g16970.1 COBW_domain−containing_protein_1−like [MW = 50.16 kDa]

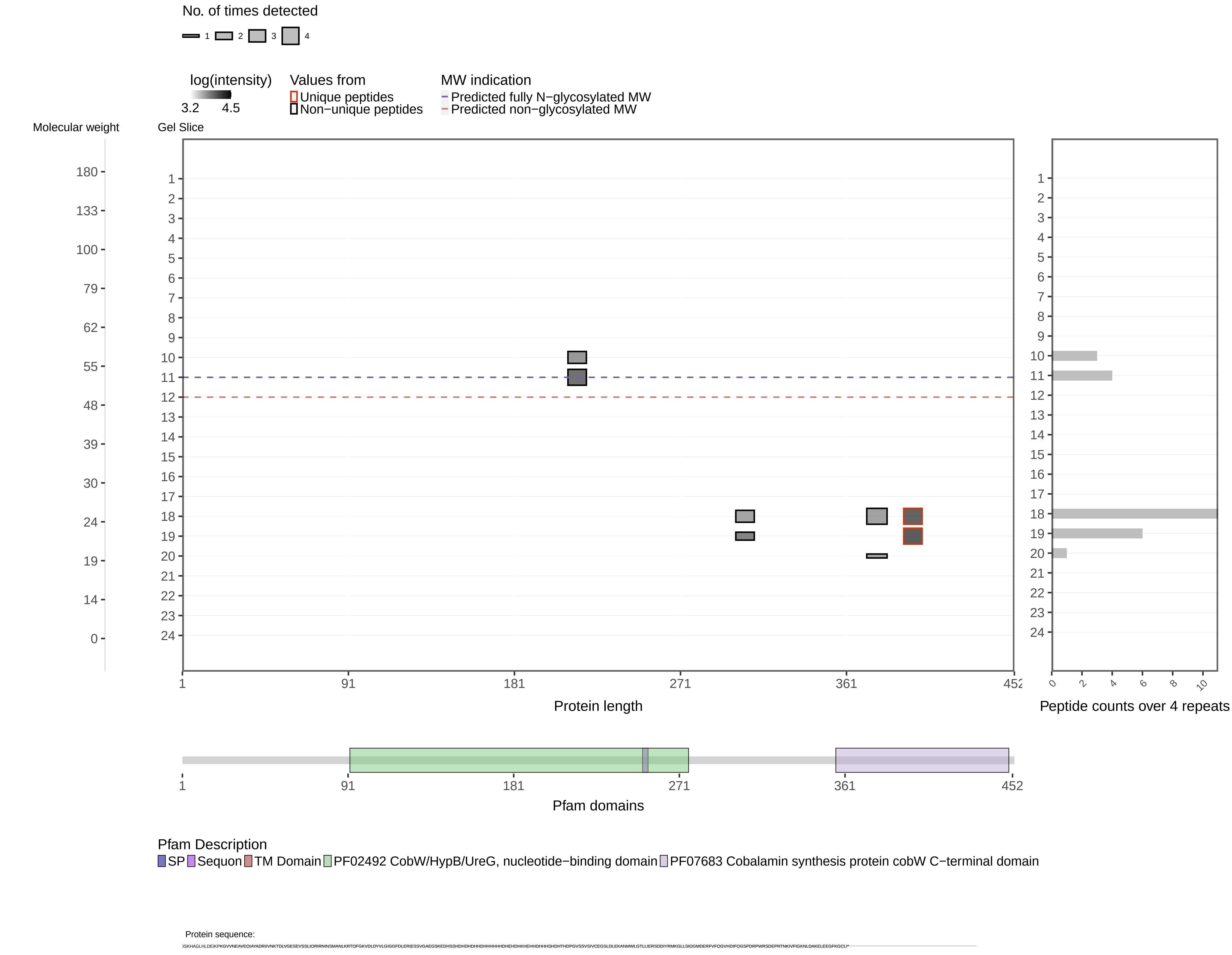

NbL07g16940.1 proline−−tRNA_ligase_cytoplasmic−like [MW = 57.17 kDa]

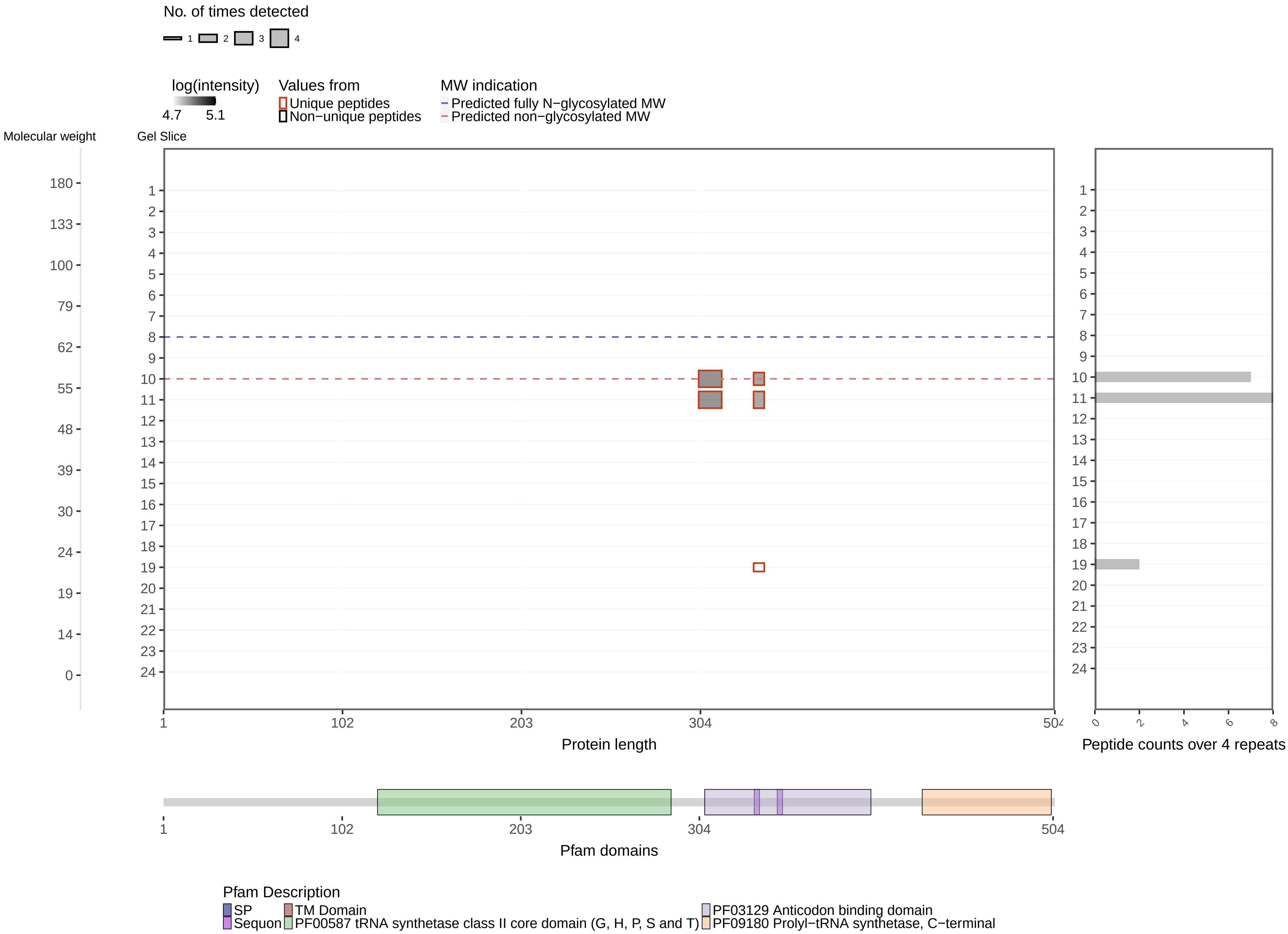


Protein sequence:

NbL07g16930.1 50S_ribosomal_protein_L3−2_mitochondrial−like [MW = 32.61 kDa]

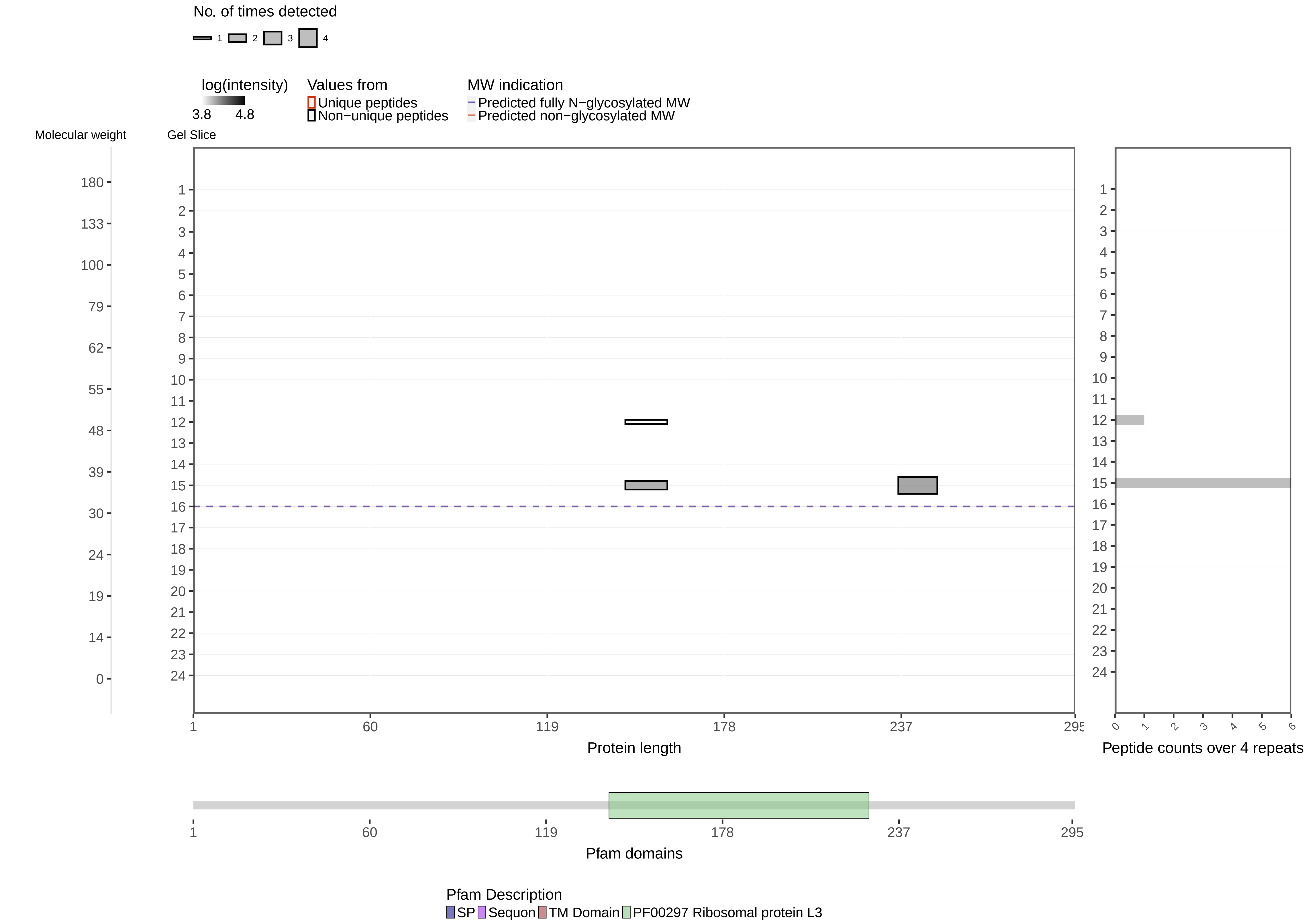


Protein sequence:

GTSISVRHFVPGQYVDVTGITRGKGFQGGMKRWGFKGMPASHGASLSHRSIGSTGQRDAPGKVFKGKKMPGRMGGVQRTVKNVWVYKIDPARNLMWVKGQVPGAEGNFVFIKDSVYKKPDISMLPFPTYFAPEDEDEDDLEPLVADLGGTDPFMAAD*

### NbL07g16880.1 uncharacterized_LOC109223891 [MW − SP = 55.3 kDa]

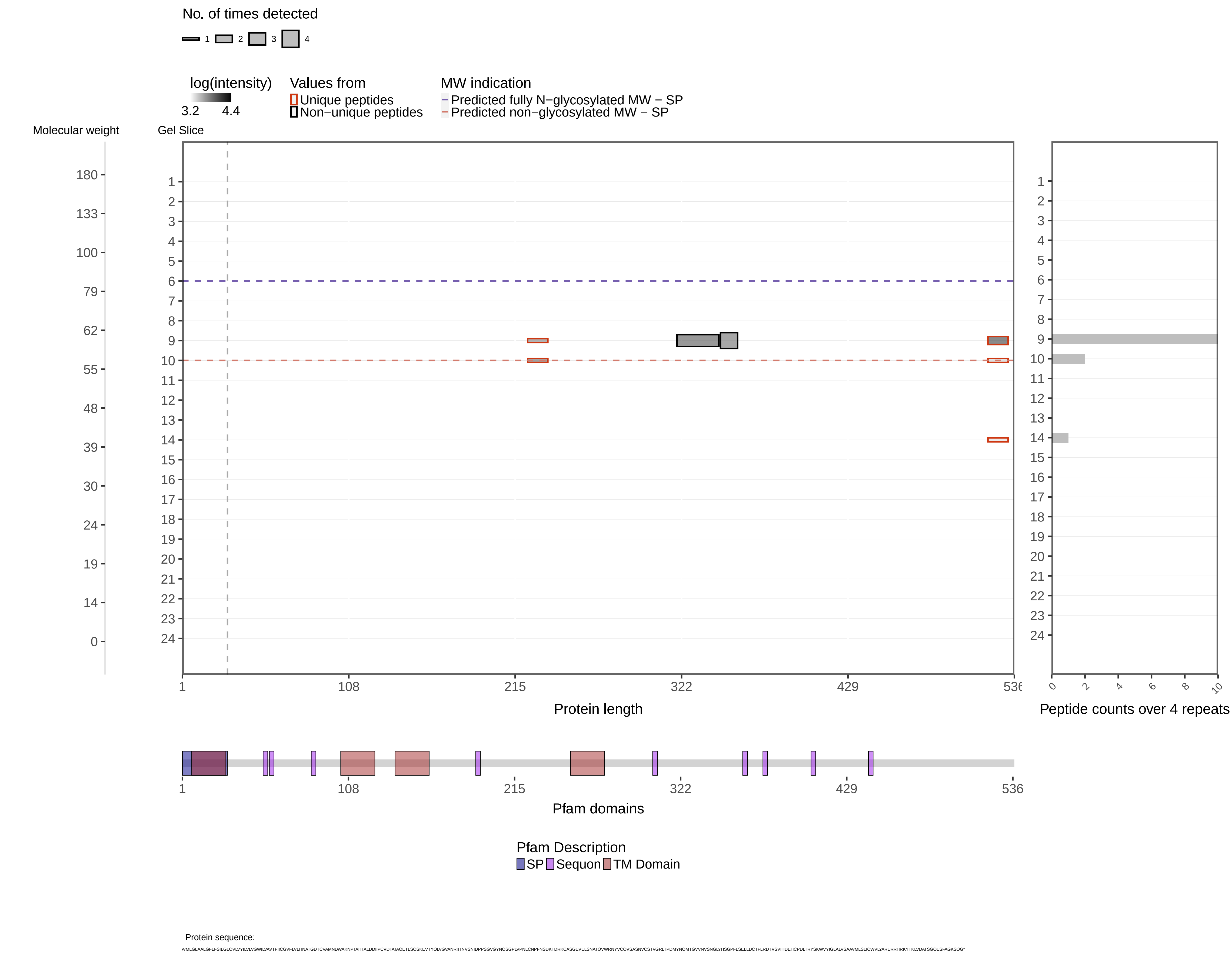

NbL07g16480.1 uncharacterized_LOC104218509 [MW = 31.46 kDa]

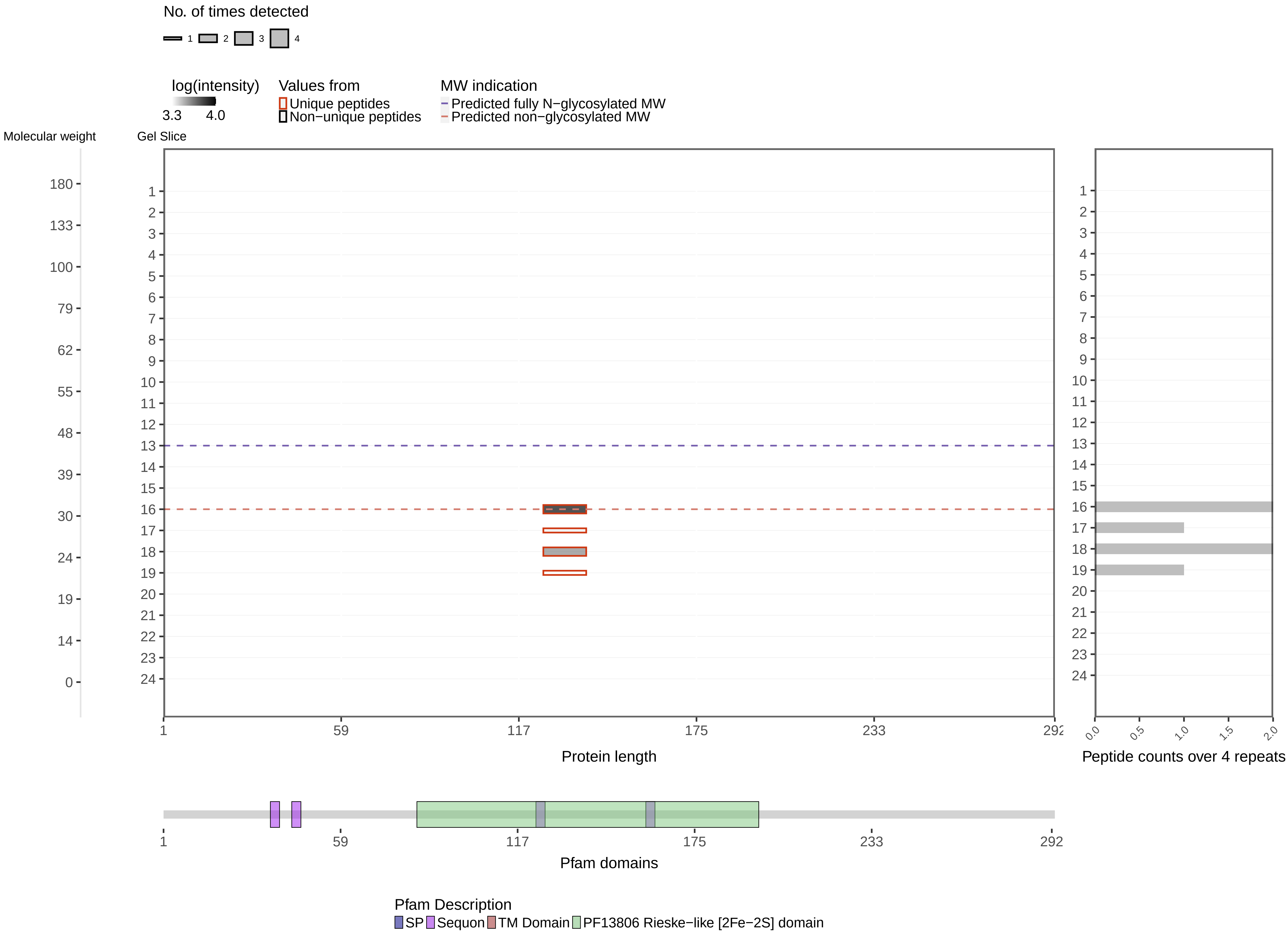


Protein sequence:

EGLLNAKLTODGCIVCPTTDSTFYLONGSIKEWYPNNPVLRVLTPALRNLFVYPVKTDDENIYISMRGGGKVDASAEIVFSGKAOPGVTASDVNVDEVRMVIDEGOEAFGFTGRNELINGKAAVIGFLLLLDFELLTGKGLLKGTGLLDFIYSVSDAFK*

### NbL07g16410.1 40S_ribosomal_protein_S5 [MW = 31.93 kDa]

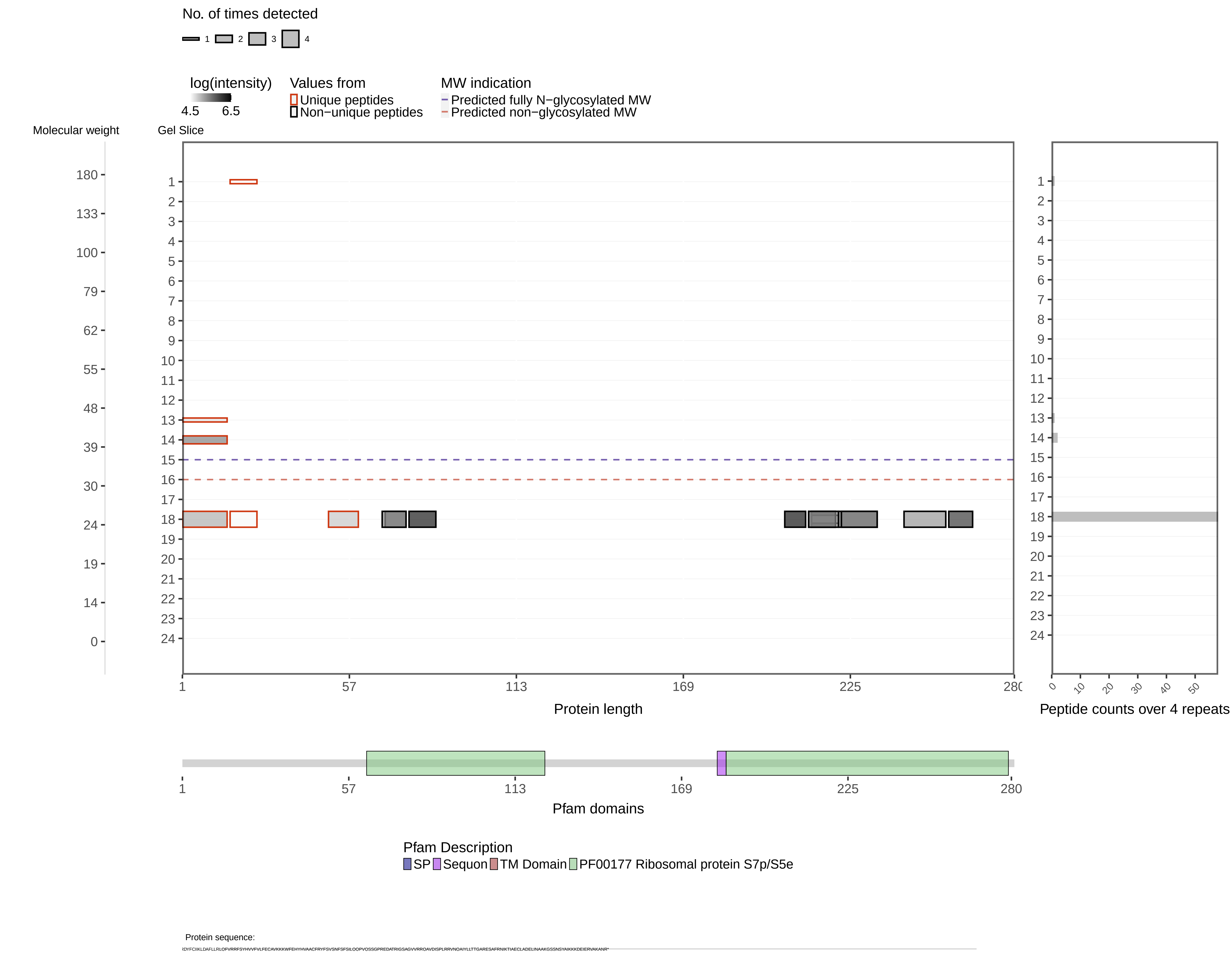

### NbL07g16360.1 cytochrome_c [MW = 12.12 kDa]

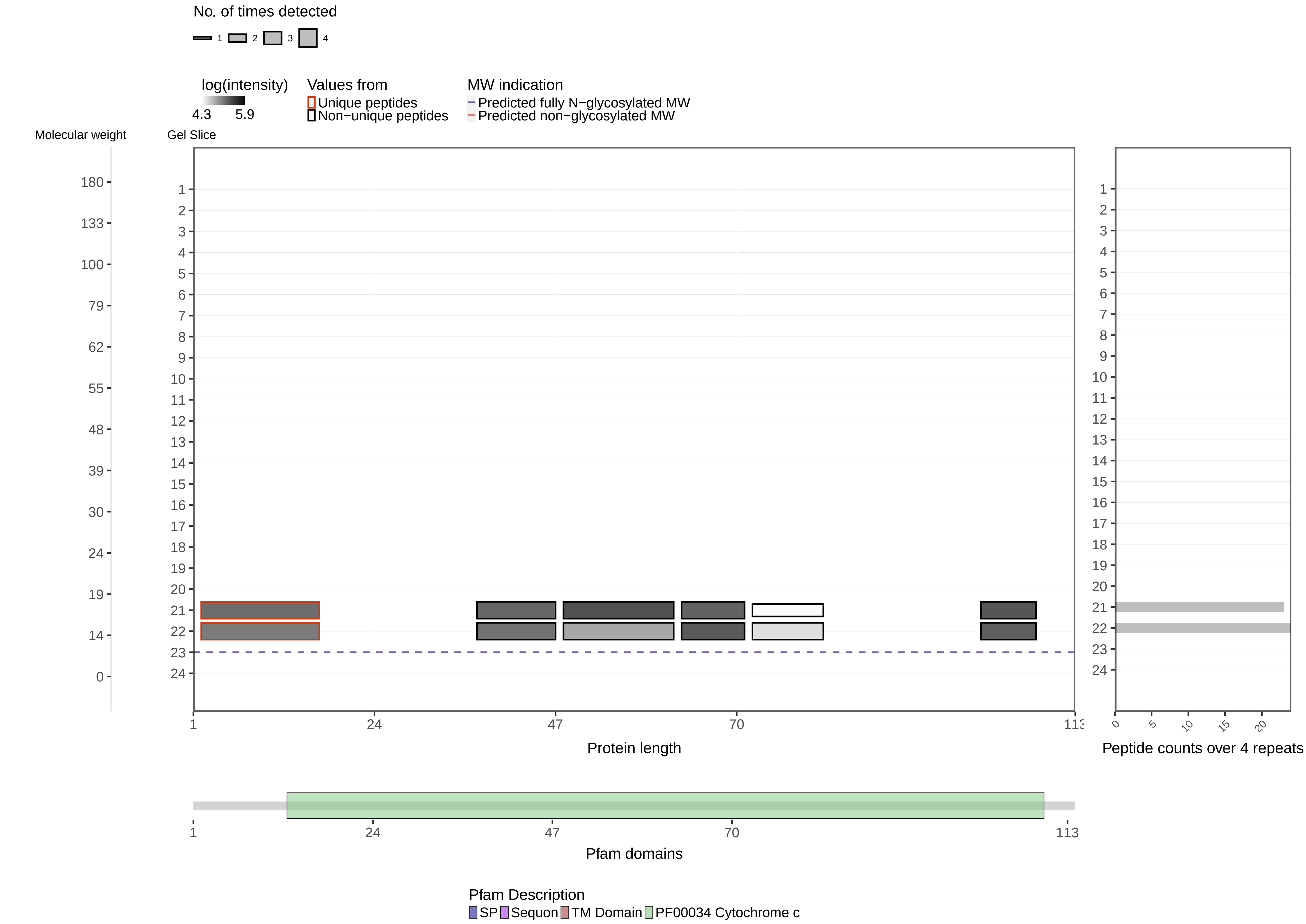


Protein sequence:

GLFGRQSGTTAGYSYSAANKNMAVMWEEKTLYDYLLNPKKYIPGTKMVFPGLKKPQERADLIAYLKSATA*

### NbL07g16260.1 bifunctional_aspartokinase/homoserine_dehydrogenase_chloroplastic−like [MW = 100.18 kDa]

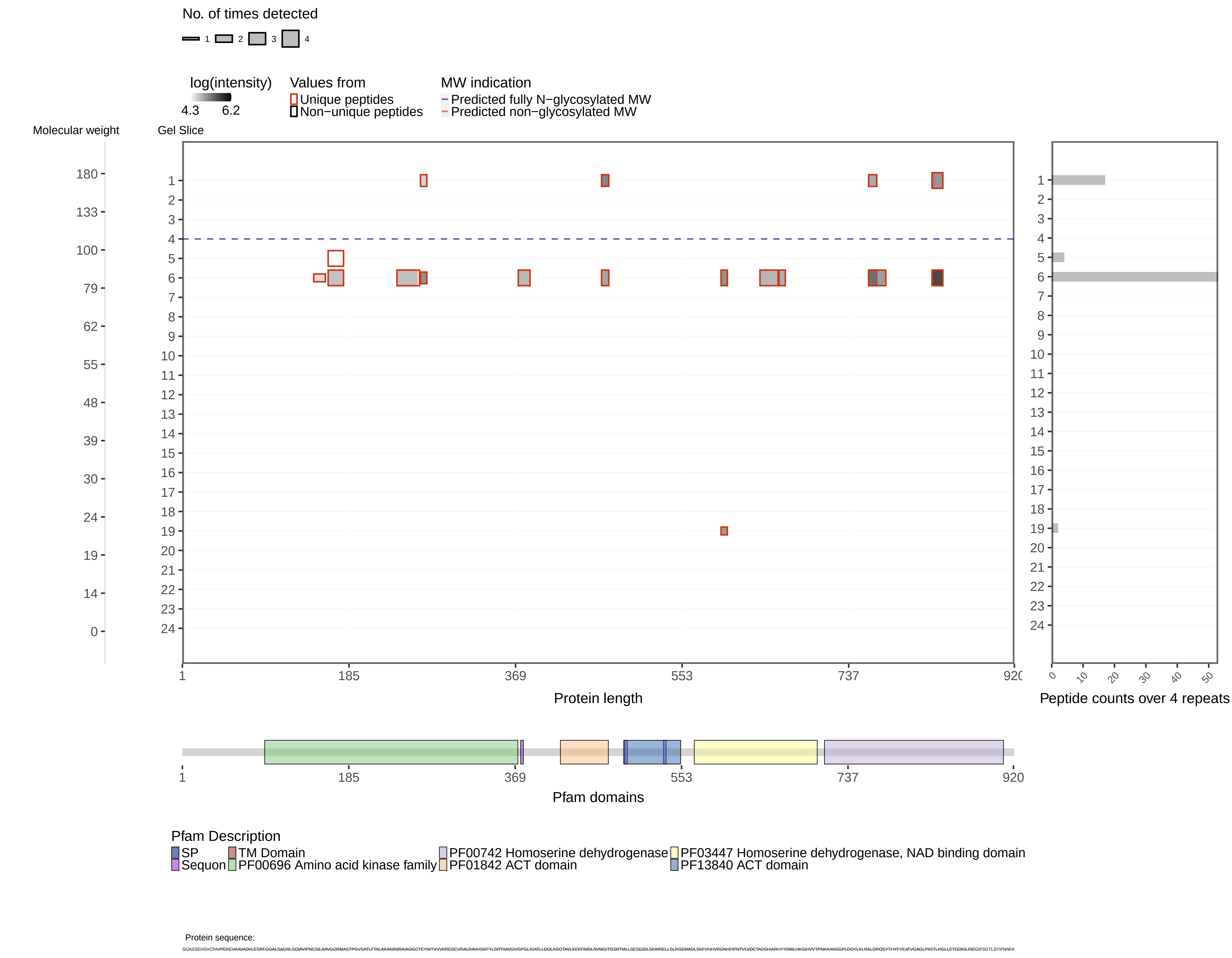

### NbL07g16190.1 far_upstream_element−binding_protein_1−like [MW = 84.18 kDa]

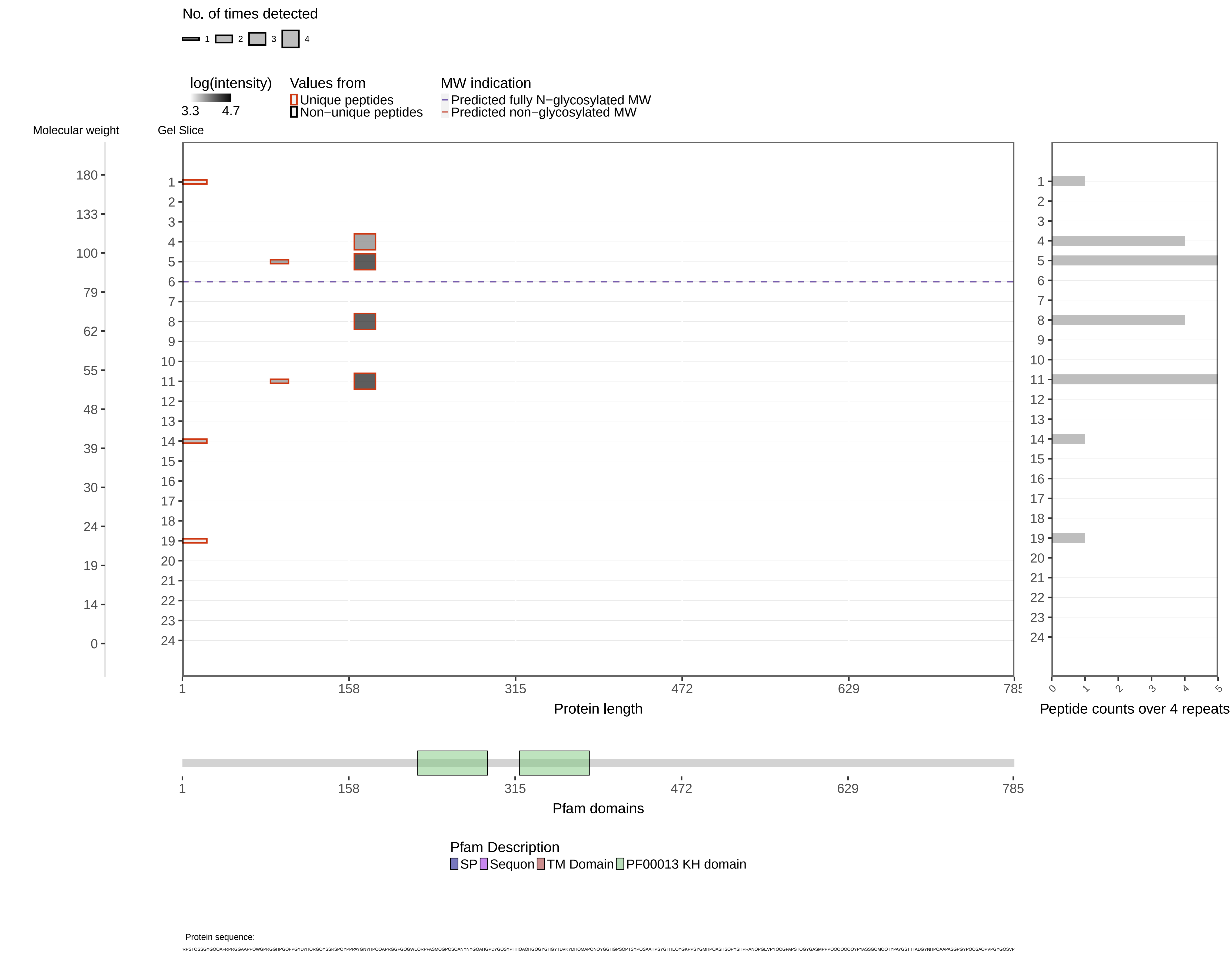

NbL07g16110.1 receptor−like_protein_12 [MW − SP = 49.4 kDa]

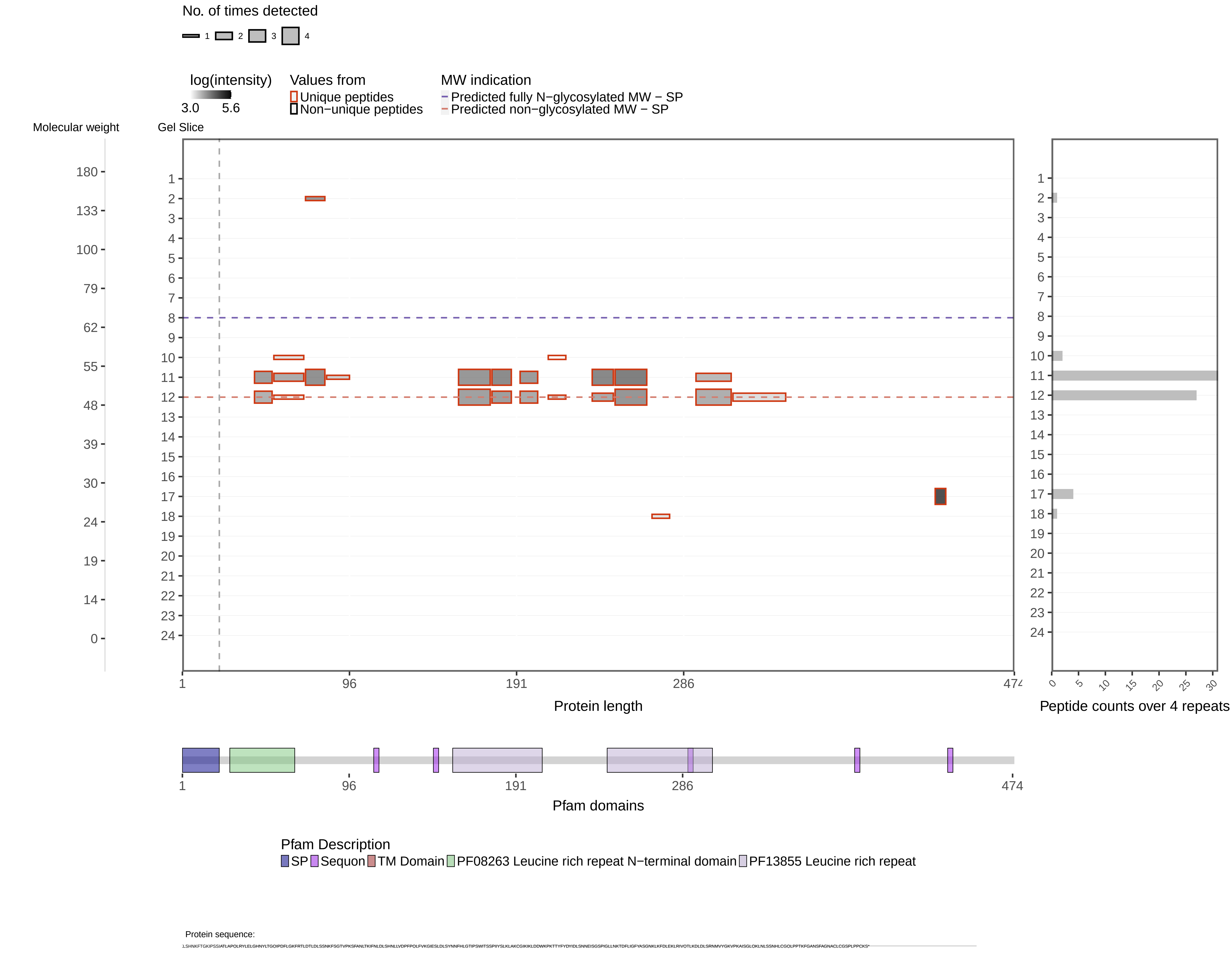

NbL07g16010.1 probable_sucrose−phosphate_synthase_4 [MW = 118.64 kDa]

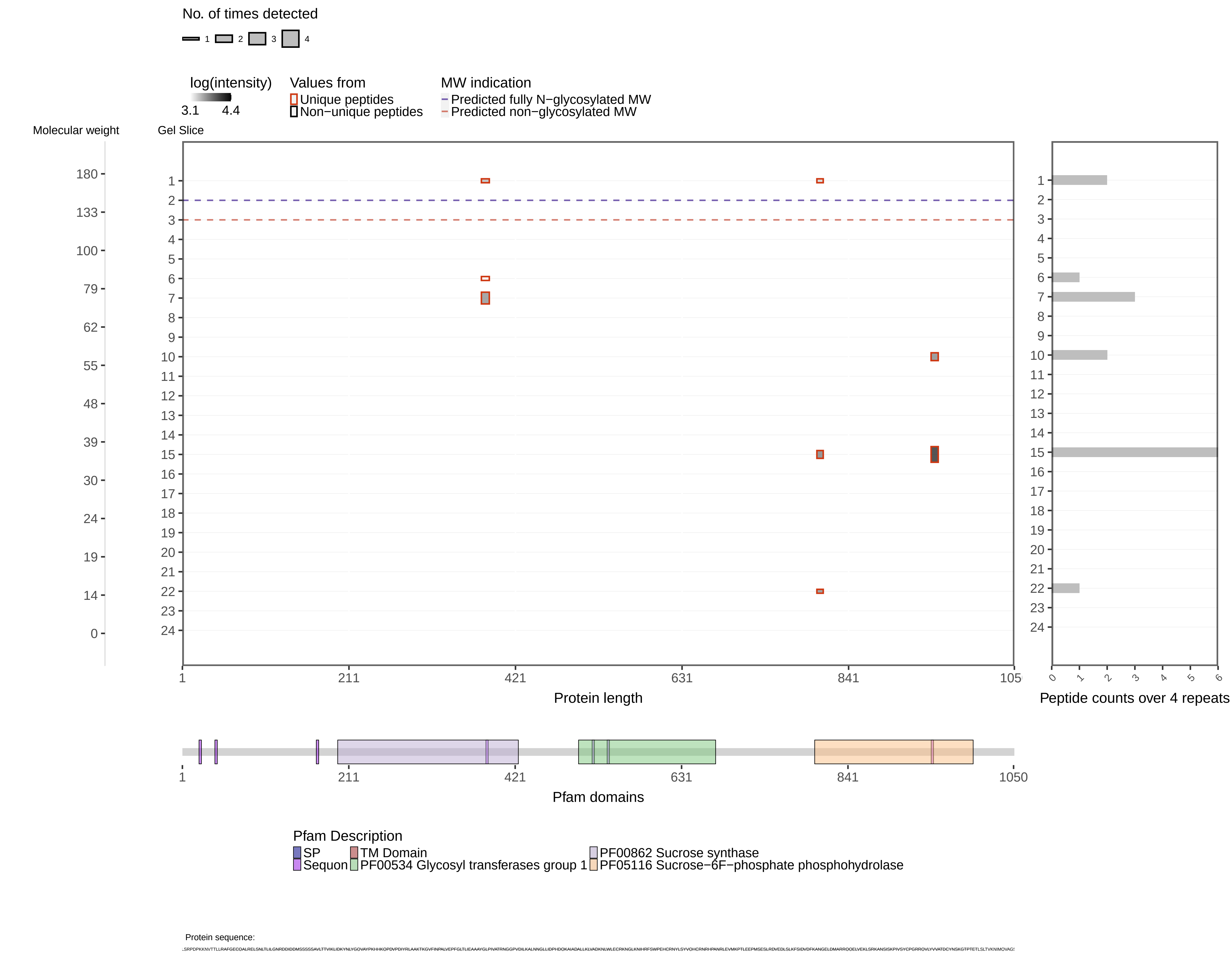

NbL07g15950.1 single−stranded_DNA−binding_protein_WHY2_mitochondrial [MW = 26.67 kDa]

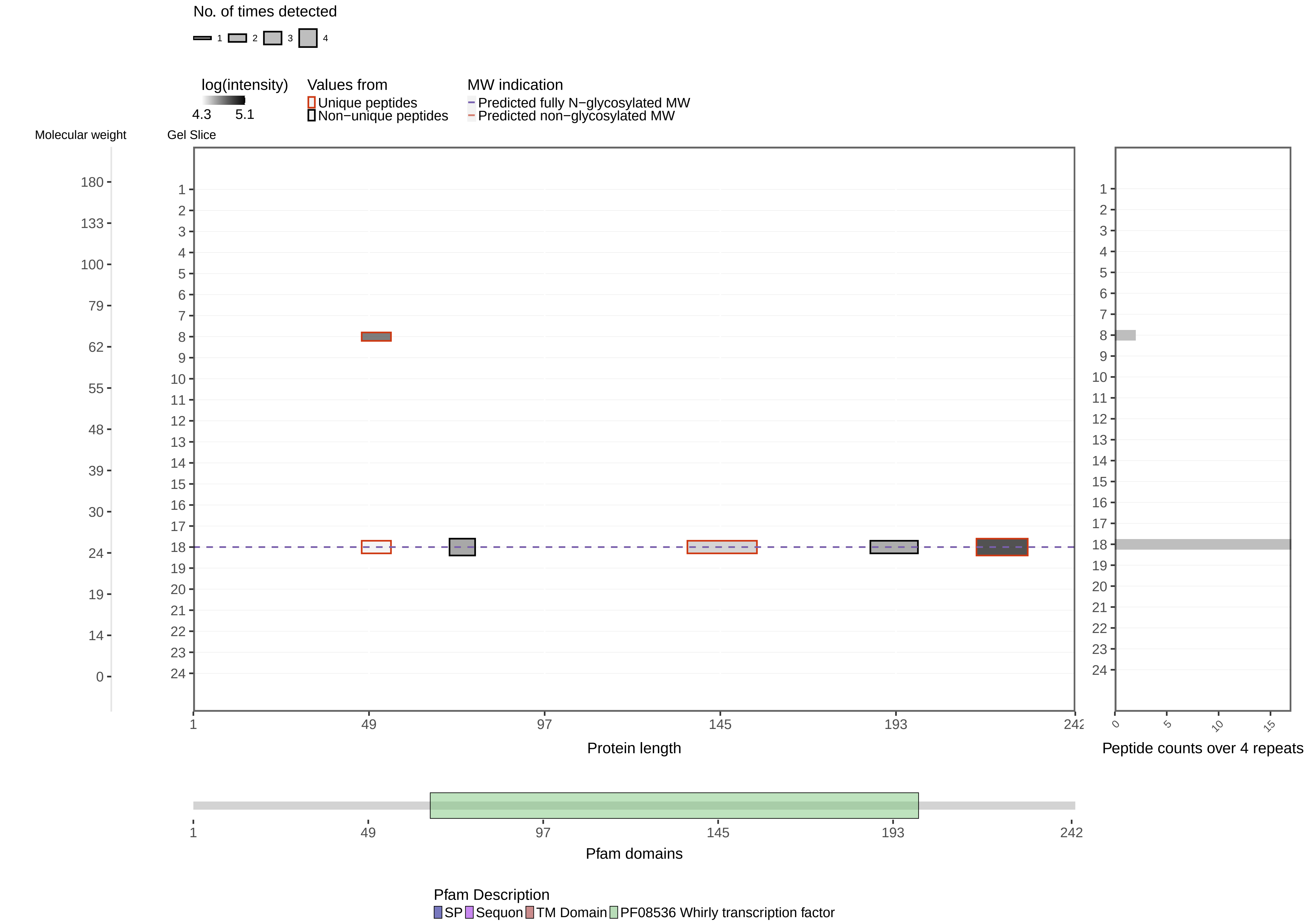


Protein sequence:

RKYDWEKRQLFALSATEVGSLISIGTRDTCEFFHDPSMLSSNAGQVRKSLSFKPNADGSGYFVSLSVVNNNLKTNDRFTVPVTTAEFAVMRTAFSFALPHIMGWDRFINQPLESISQSSSKVVPQLMETEWDR*

NbL07g15820.1 endochitinase_3−like [MW − SP = 33.6 kDa]

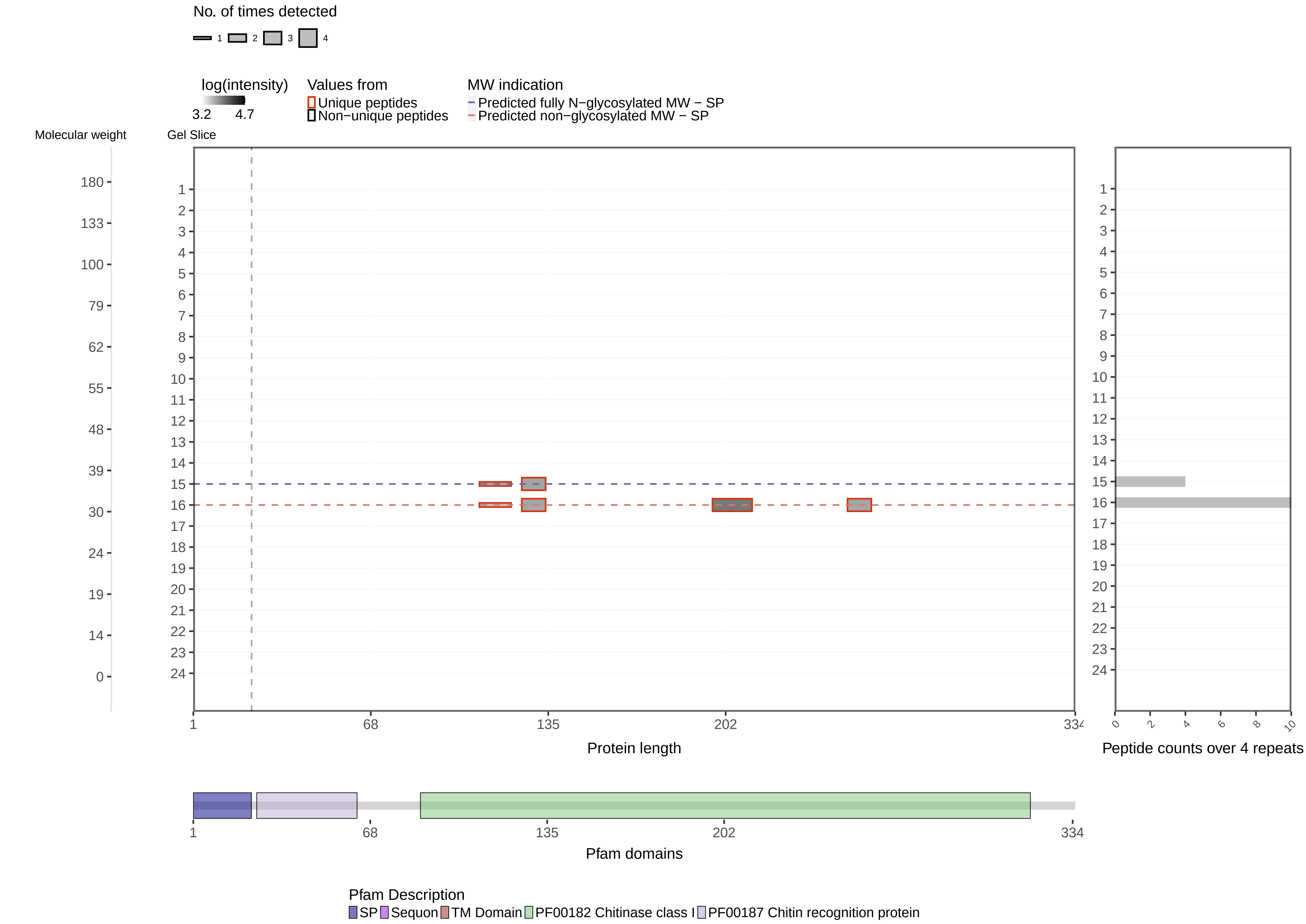


Protein sequence:
NDTAPDGRYAWGYCYLREOGNPPSYCVOSSOWPCAPGOKYYGRGPIOISYNYNYGPCGRAIGONLLNNPDLVATNAVVSFKSAIWFWMTAOSPKPSCHDVITGRWTPSAADRAANRLPGYGVITNIINGGLECGHGSDARVODRIGFYRRYCSILGVSPGDNIDCGNORPFNSGLLLETM*

### NbL07g15700.1 SKP1−like_protein_1A [MW = 17.51 kDa]

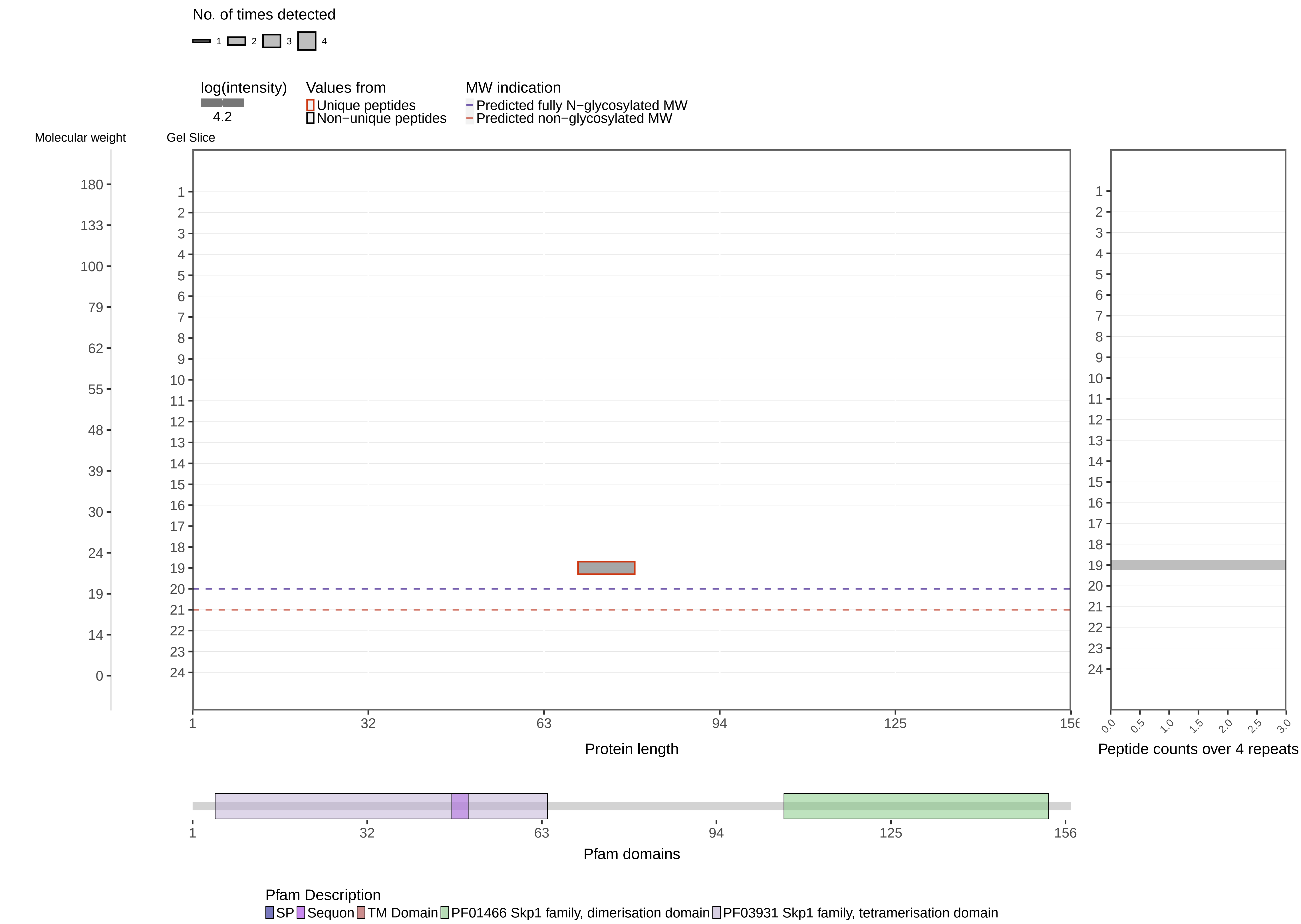


Protein sequence:

DATKAEDKASEDELKAFDSDFVKVDQATLFDLILAANYLNIKSLLDLTCQTVADMIKGKTPEEIRKTFNIKNDFTPEEEEEVRRENAWAFE*

NbL07g15500.1 ubiquitin_recognition_factor_in_ER−associated_degradation_protein_1−like [MW = 36.25 kDa]

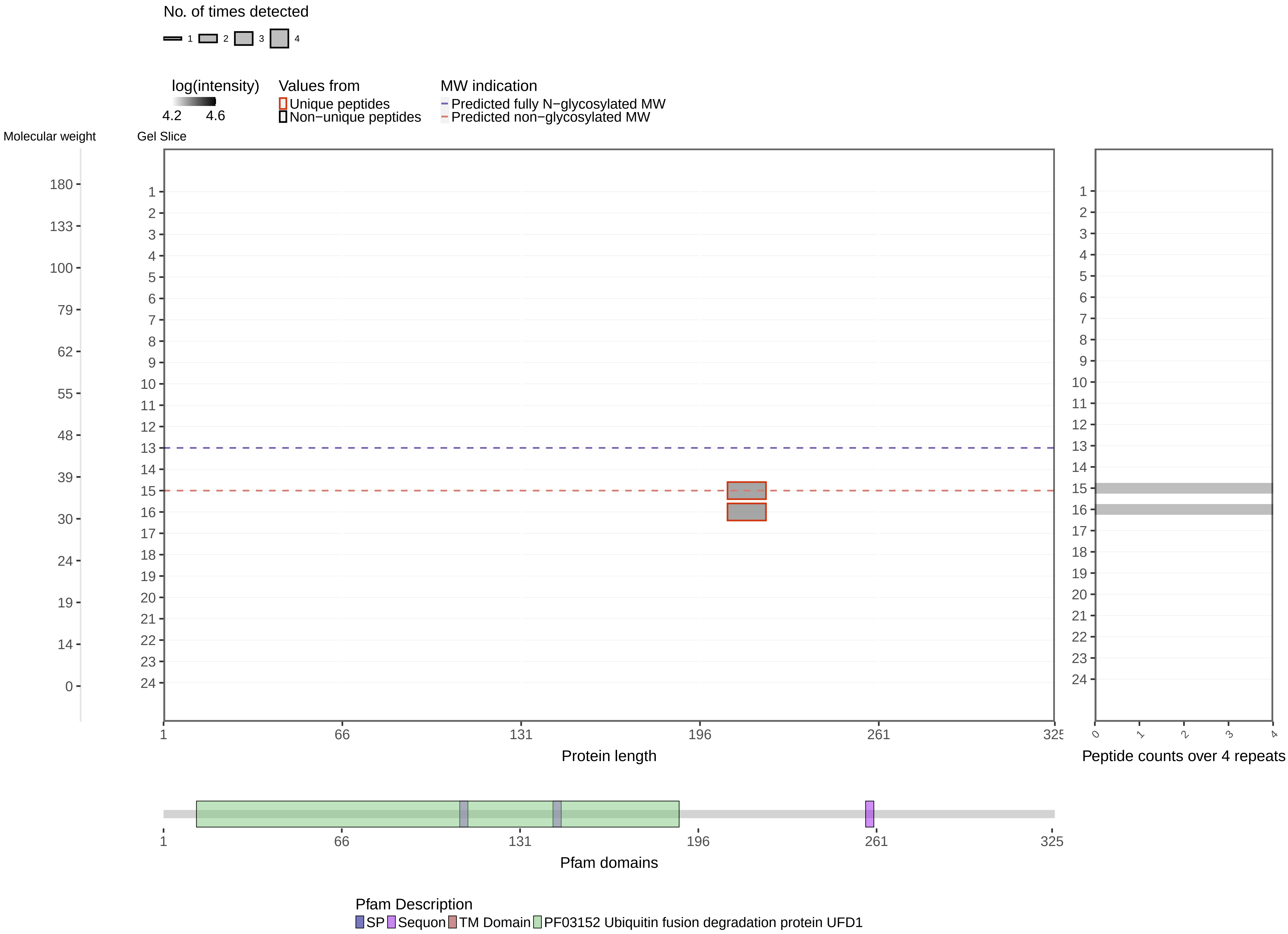


Protein sequence:

SIMVAYNNKKYYIDIVESKPSNAISIIETDCEVDFAPPLDYKEPERPVPSLPSKNPAEGEEAAAEEEPKFNPFTGSARRLDGKPLKQQPPPVSTSGSKDKRPDMANATGQSATGSSSQSSNRQTQGKLVFGSNANRNPDKQKEPVKEPVKEEPQKKEEPKFQAFTGKKYSLRG*

### NbL07g15490.1 epimerase_family_protein_SDR39U1_homolog_chloroplastic [MW = 38.97 kDa]

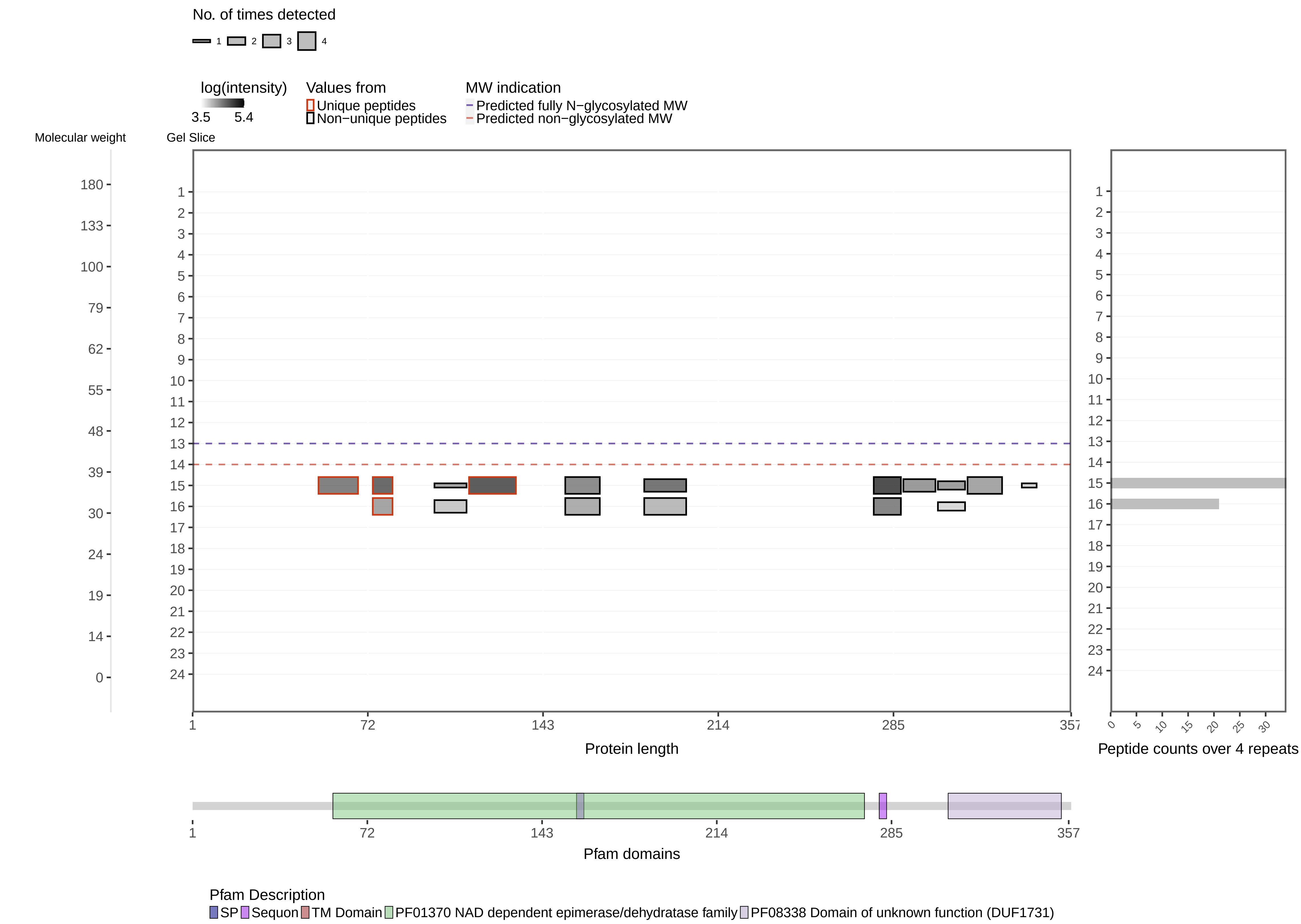


Protein sequence:

‹VLISASAVGYYGTSETRVFDEOSPSGNDYLAEVCREWEGAALKVNKDVRLALIRIGVVLGKDGGALAKMIPIFMMFAGGPLGSGNQWFSWIHLDDLVDLIYEALSNPSYKGVINGTAPNPVRLAEMCEOLGSVLGRPSWLPVPEIALKAVLGEGASVVLDGQKVVPTRVKELGFPFKYQYIKDALKAIMS*

### NbL07g15380.1 uncharacterized_LOC109223501 [MW = 23.51 kDa]

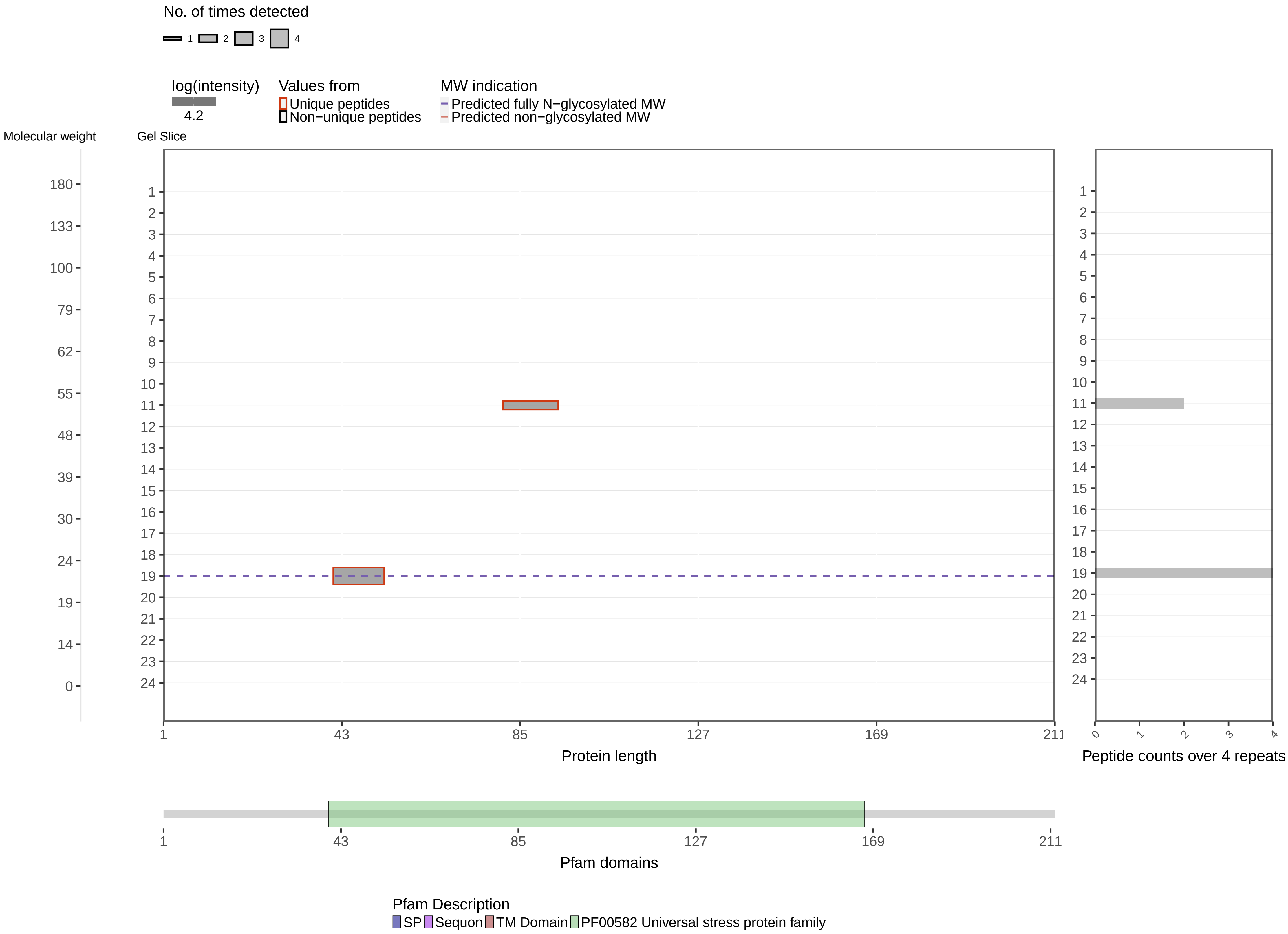


Protein sequence:

MEKLAIEAFEVAMVKTVARIVEGDAGKVICKEAERLKPAAVVMGTRGRSLIKSVLQGSVSEHCFHQCKTAPIIIVPGKGTLLHLFCFSKFFFWVVCEQNVILLSRSRGGICNLMGTCQV*

NbL07g15370.1 glycine−rich_RNA−binding_protein−like [MW = 17 kDa]

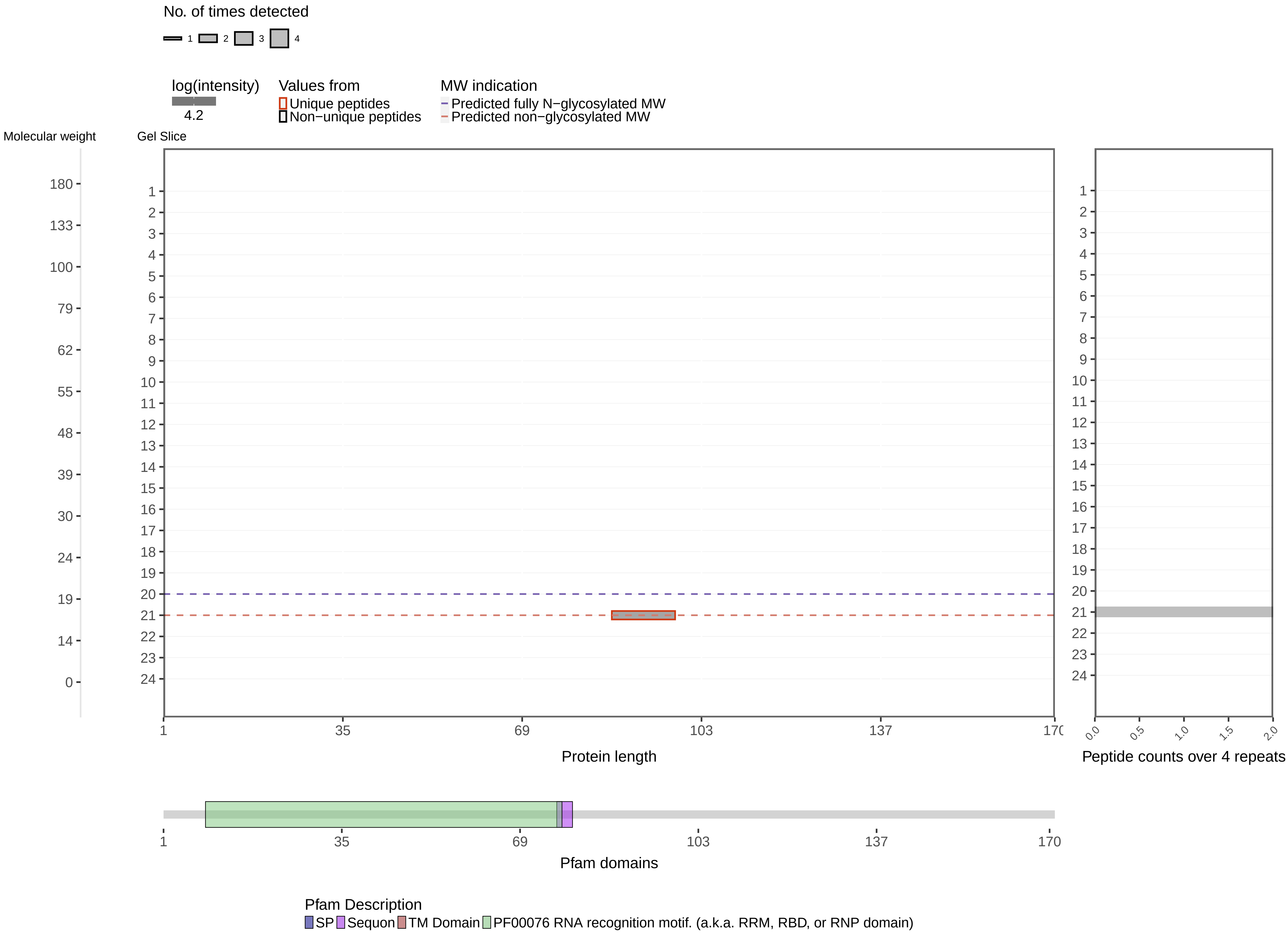


Protein sequence:

ITVNEAQSRGSGGSGGGGGGFRGGRREGGGGYGGGGYGGGRREGGGGGYGGGGGGYGGGRDRGYGGGGDRGYGGDGGSRYSRGGGDSDGNWRN*

NbL07g15350.1 coatomer_subunit_gamma [MW = 98.44 kDa]

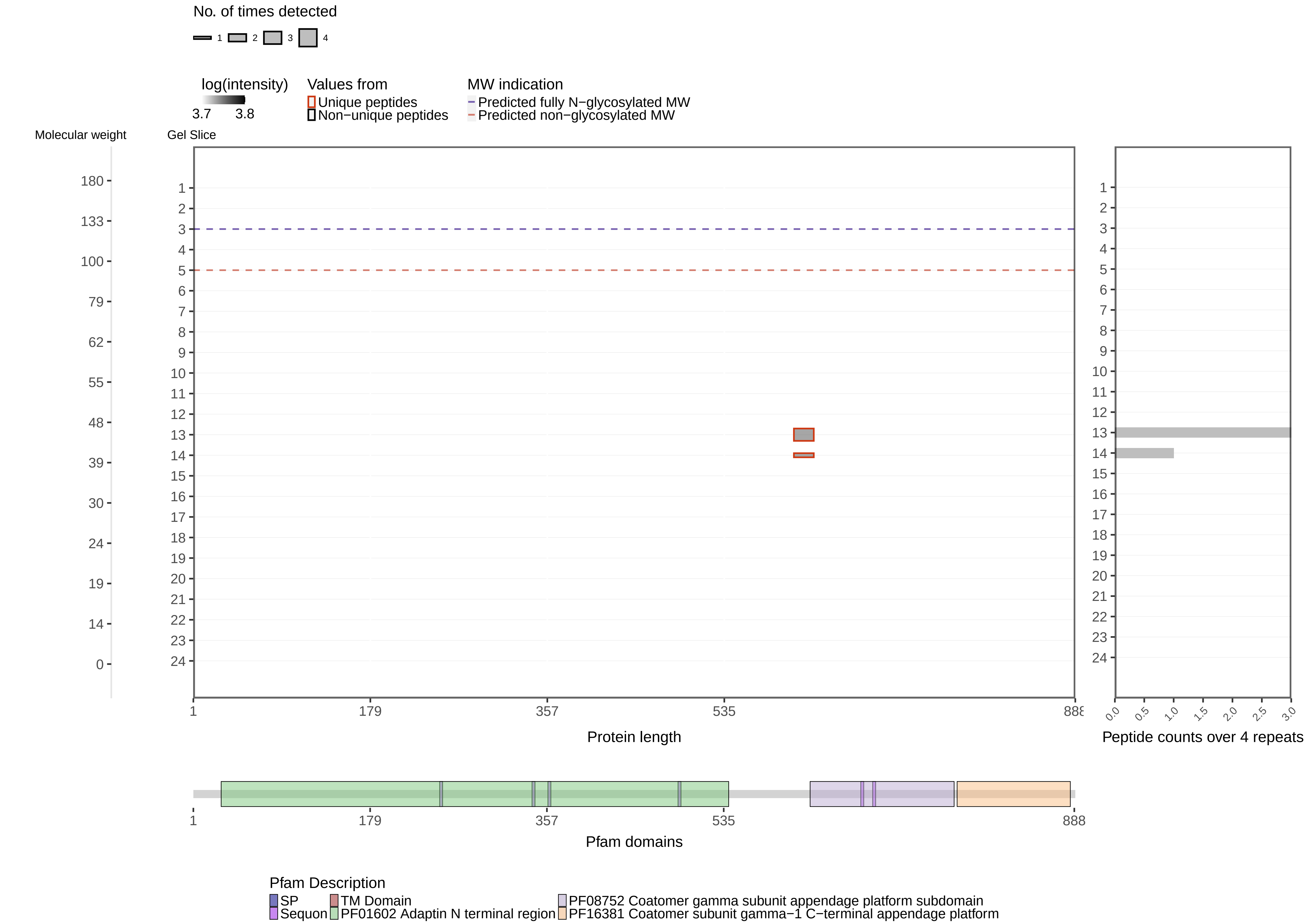


Protein sequence:

PDAKESGLLHLCEFIEDCEFTYLSTQILHFLGNEGPKTSDPSKYIRYIYNRVILENATVRASAVSTLAKFGALVDSLKPRIFVLLNRCLFDSDDEVRDRATLYLNTLGGDGAVVETDDEVKEFLFGSLSVPLINLETSLKNYEPSEEAFDIFSVPKEVKSQPLAEKKAPGKKPTGLGAPPVGPTSTVDSYERLLSSIPEFASYGKLFKSSAPVELTEAETEYAVNVVKHIFDSHVVFQYNCTNTIPEQLLENVSVIVDPSEAEEFSEIASKPLKSLPYDTPGQTFVAFEKLEGVRAVGK[illegible]

NbL07g15060.1 probable_cinnamyl_alcohol_dehydrogenase_1 [MW = 38.99 kDa]

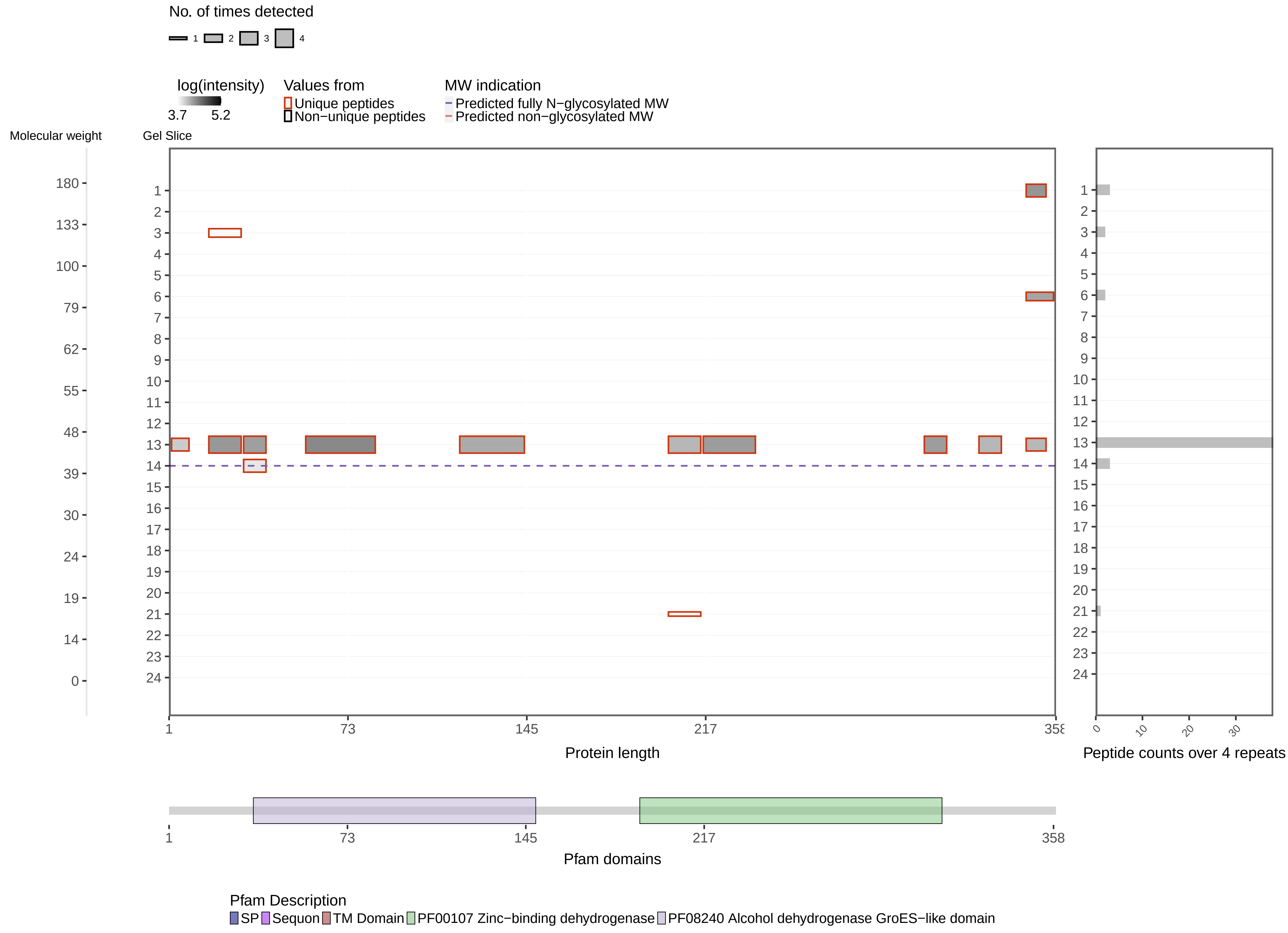


Protein sequence:

LCAGLTVYSPLNHFGLKOSGLRGGILGLGGVGHMGVKIAKAMGHHITVISSSDRKKEEALEHLGADEYLVSSDTTRMOEAADSLDYIIDTIPVFHPLEPYISLLKLDGKLILMGVINTPMOFITPMVMLGRKTITGSFIGSMKETEEVLEFCKEKGLTSOIEVVKMDYVNTAMERLEKNDVRYRFVIDVAGSKLDO*

NbL07g14960.1 beta_carbonic_anhydrase_5_chloroplastic−like [MW = 33.54 kDa]

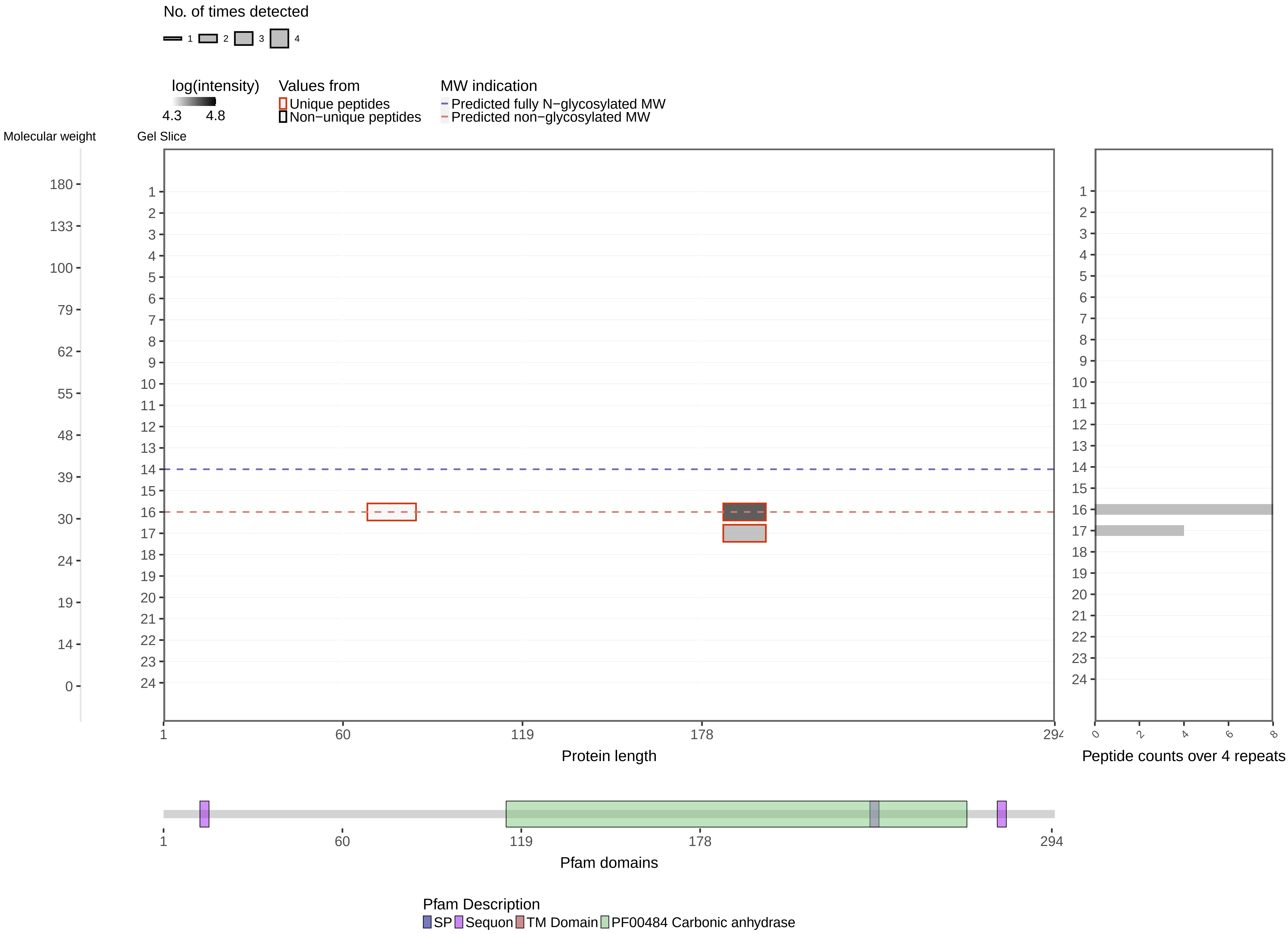


Protein sequence:

EAFVVRNVANLVPPYENGPSETNAALEFAVNTLKVQNILVIGHSRCGGIRALMSMDSETPSSFIRSWVVVGKSARARTKAAASNLCFDQQCKHCEKESVNCSLMNLFTYPWIKDKVNKGELLIHGGYYDFVDCSFEKWTLENNSRVDDQVSIRNREFWC*

### NbL07g14910.1 proline_iminopeptidase−like [MW = 48.03 kDa]

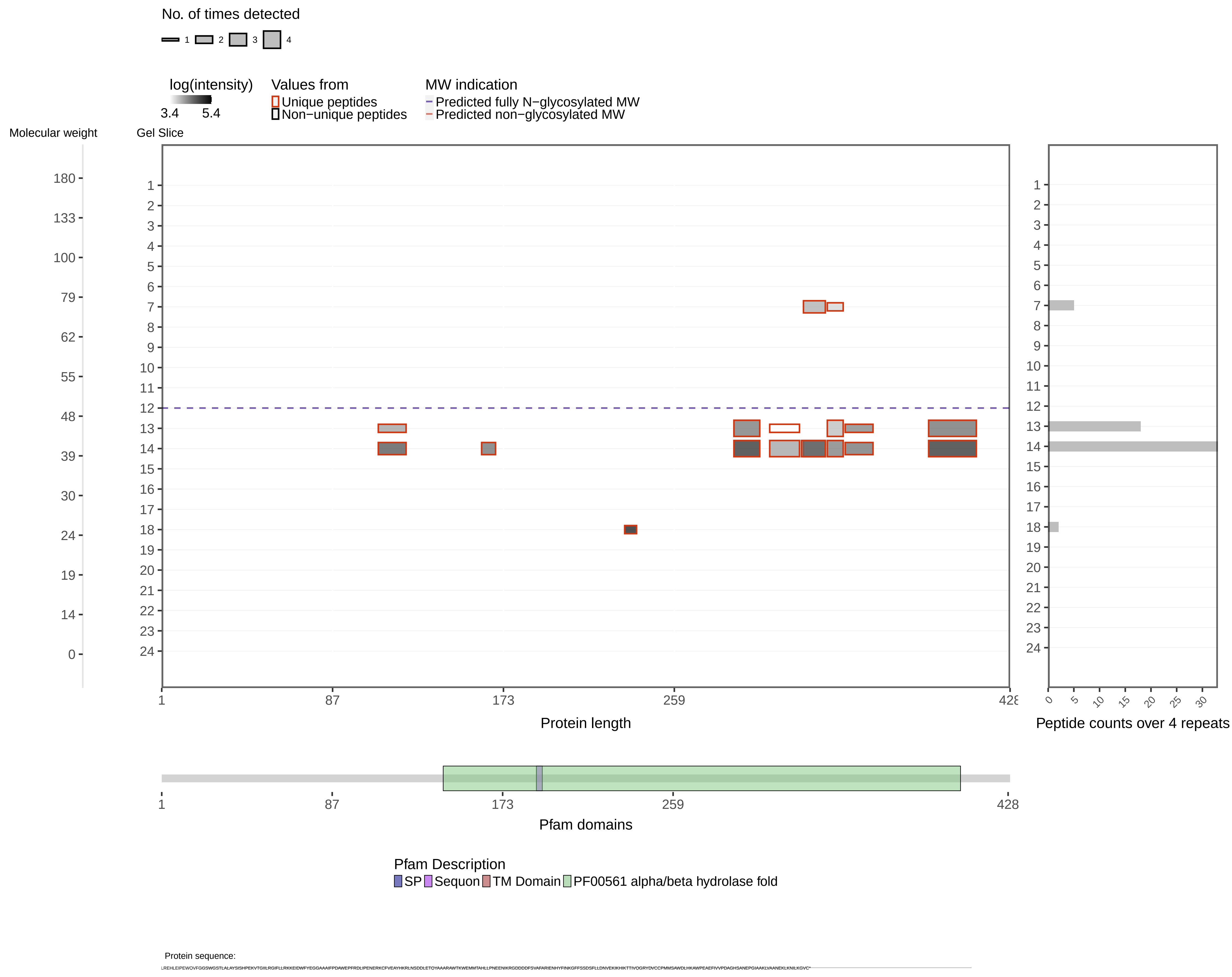

NbL07g14860.1 glucan_endo−13−beta−glucosidase_8−like [MW − SP = 49.94 kDa]

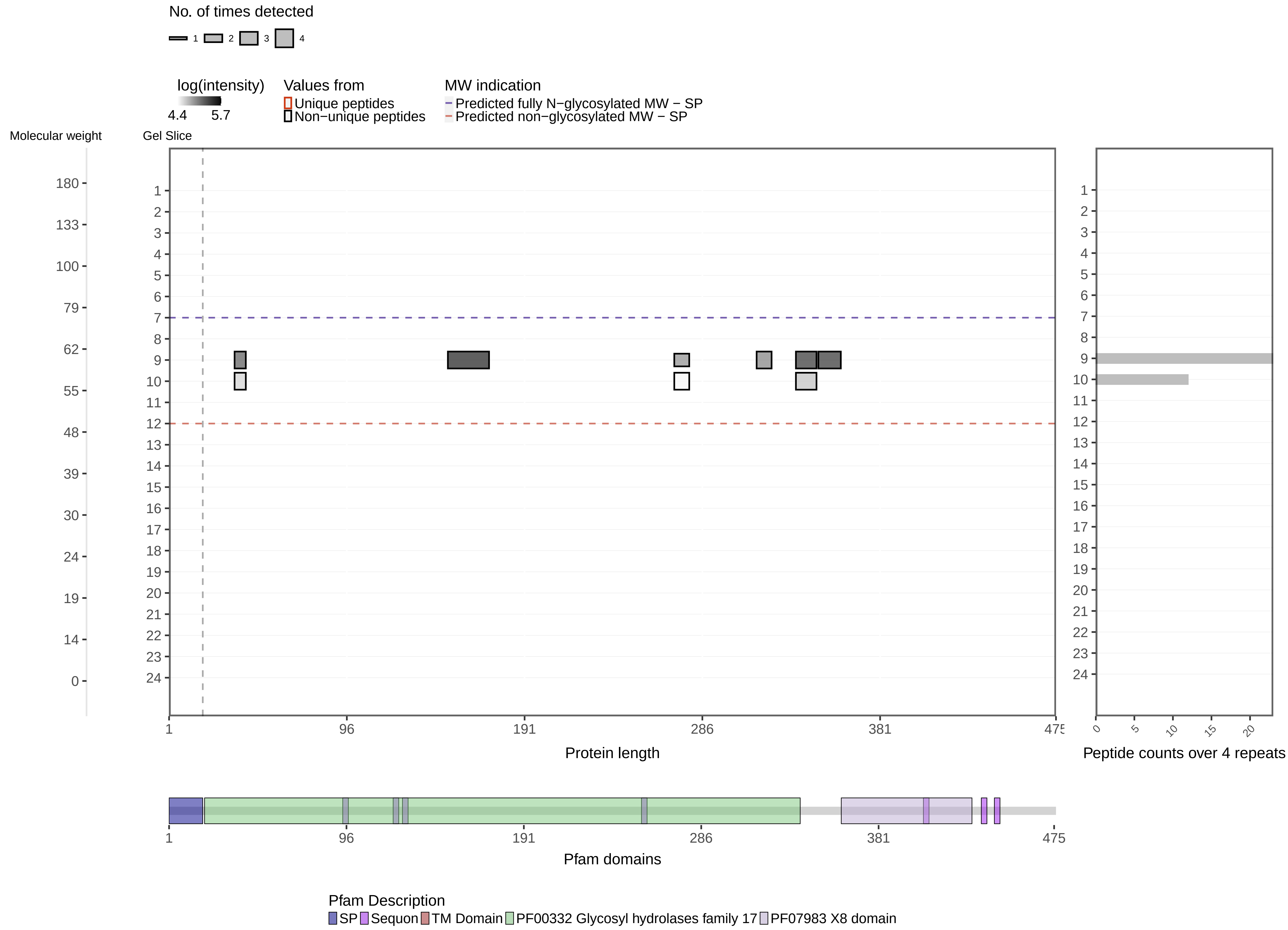


Protein sequence:

LDKGVTYTNVFDANFDTLVSALRAAGVGNMTILVGEVGWPTDGDKNANVNLAYRFFKGLLSRLAANTGTPLRPGYIEVYLFGLIDEDGKSIAPGNFERHWGIFRYDGQPKFAMDLTGQGQDKFLVGAQNVEYLPKKWCMLNPNAKDLSKLADNIDYACTFSDCTALGYGSSCNGLDANGNASYAFNMYYQAQNQGDFSCGFQGLAMVTDQNISQANCNFTIQIAYSFSQKVLPGPGVLAFLTAFTFILL*

### NbL07g14810.1 asparagine−−tRNA_ligase_cytoplasmic_1−like [MW = 64.8 kDa]

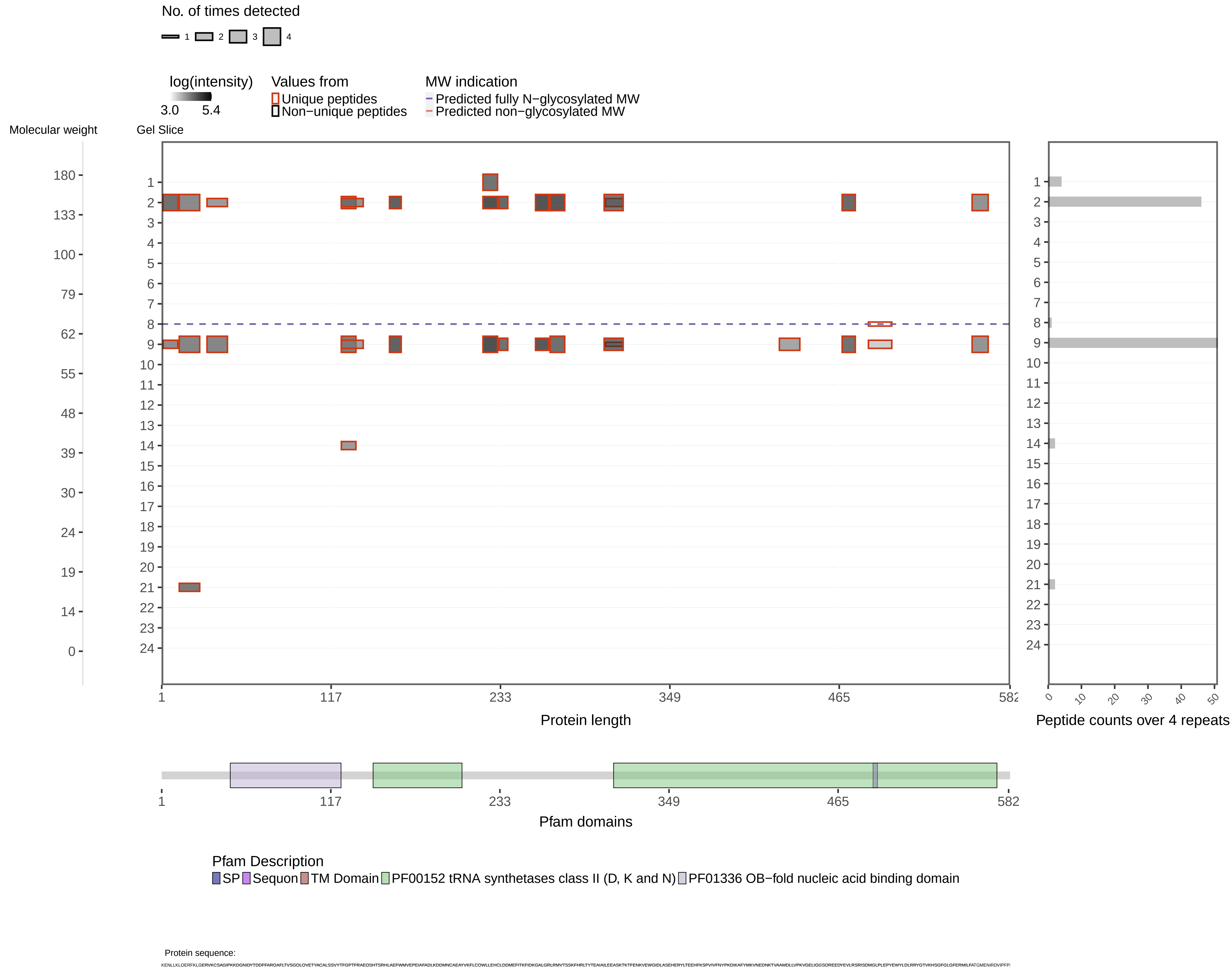

### NbL07g14780.1 costars_family_protein [MW = 12.13 kDa]

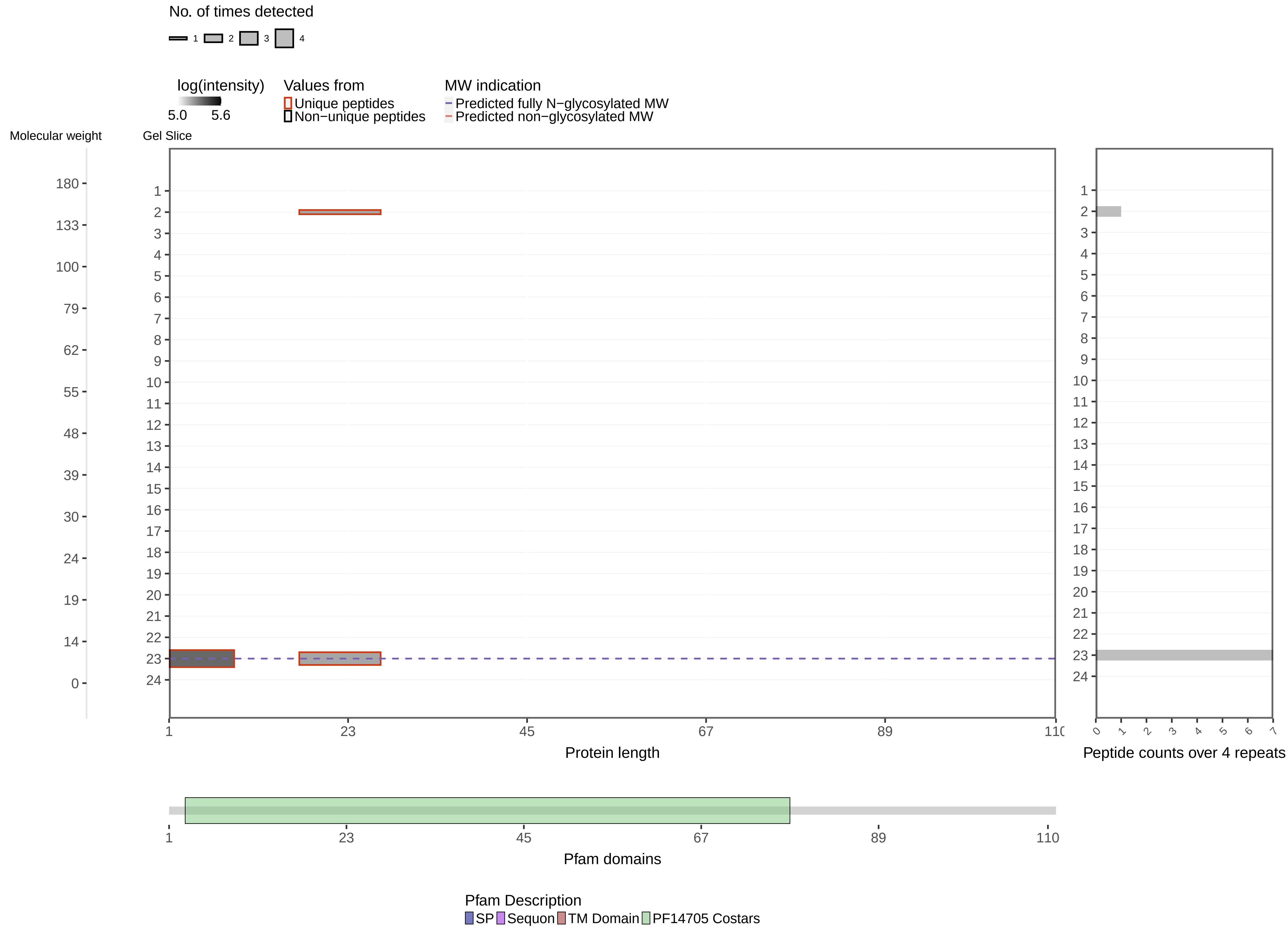


Protein sequence:

NIFEALVGTLRAAKKRKILAYDGELLLQGVHDNVEIILKPPSVTSGASALFILSYAPRVAWKQPLCPFG*

NbL07g14700.1 polypyrimidine_tract−binding_protein_homolog_3−like [MW = 49.15 kDa]

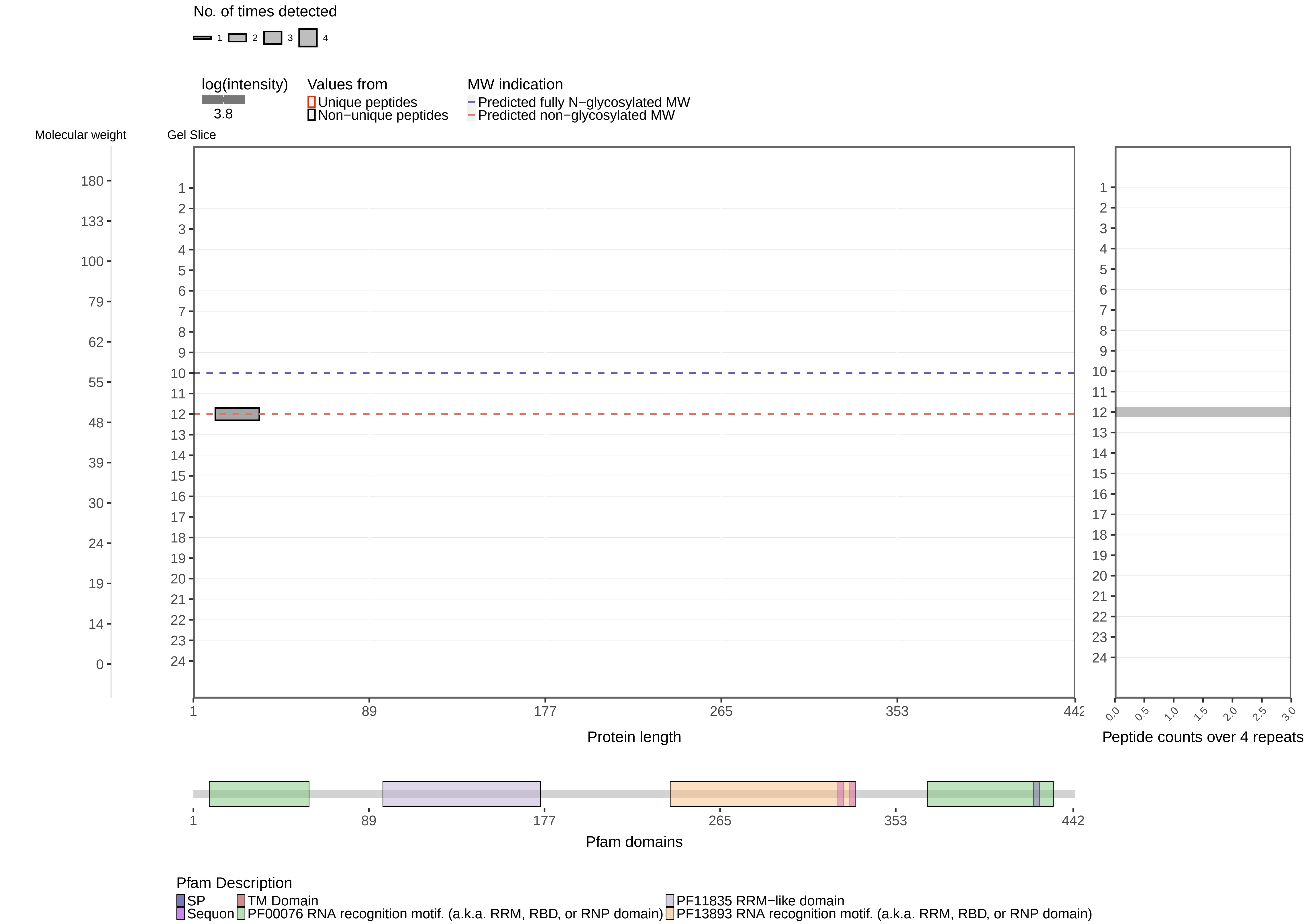


Protein sequence:

SQQGYGDMYSFQGSGAHAGGFPQMGNAEAIAAAFAGGLPPGISGTNDRCTILVSNLDSDRIDEDKLFNLFSLYGNIVKIKILRNKPDHALVQMGDGFQAELAVHFLKGAILFDKRLEVNFSKHPNITTGPDTHDYSNSNLNRFNRNAAKNYRYCCSPTKMIHLSSLPQDATEEEIVAHLEEHGSIINSKLFEMNGKQQALVLFEKEEQATEALVCKNATSLGGSTIRISFSQLQSI*

### NbL07g14600.1 adenylyl−sulfate_kinase_3−like [MW = 23.08 kDa]

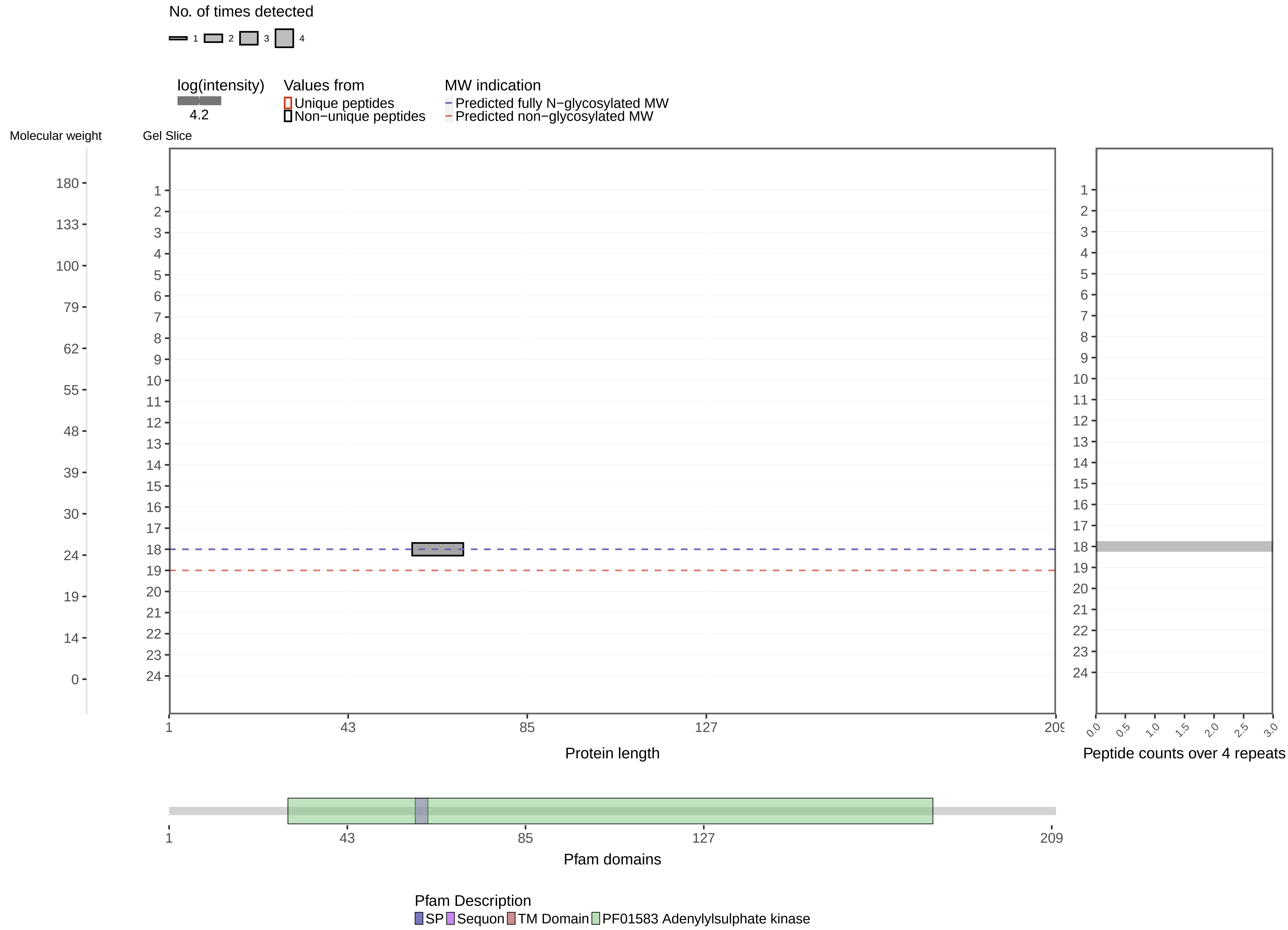


Protein sequence:

RVGEVSKLFADAGLICIASLISPYRRDRDACRALFPDKSFIEVFMNMPLELCEGRDPKGLYKLARAGKIKGFTGIDDPYEPPLNCEIEIQEKDGVVPTPHEMAGQVVTYMEDRGFLKAQ*

### NbL07g14450.1 photosynthetic_NDH_subunit_of_subcomplex_B_2_chloroplastic−like [MW = 38.82 kDa]

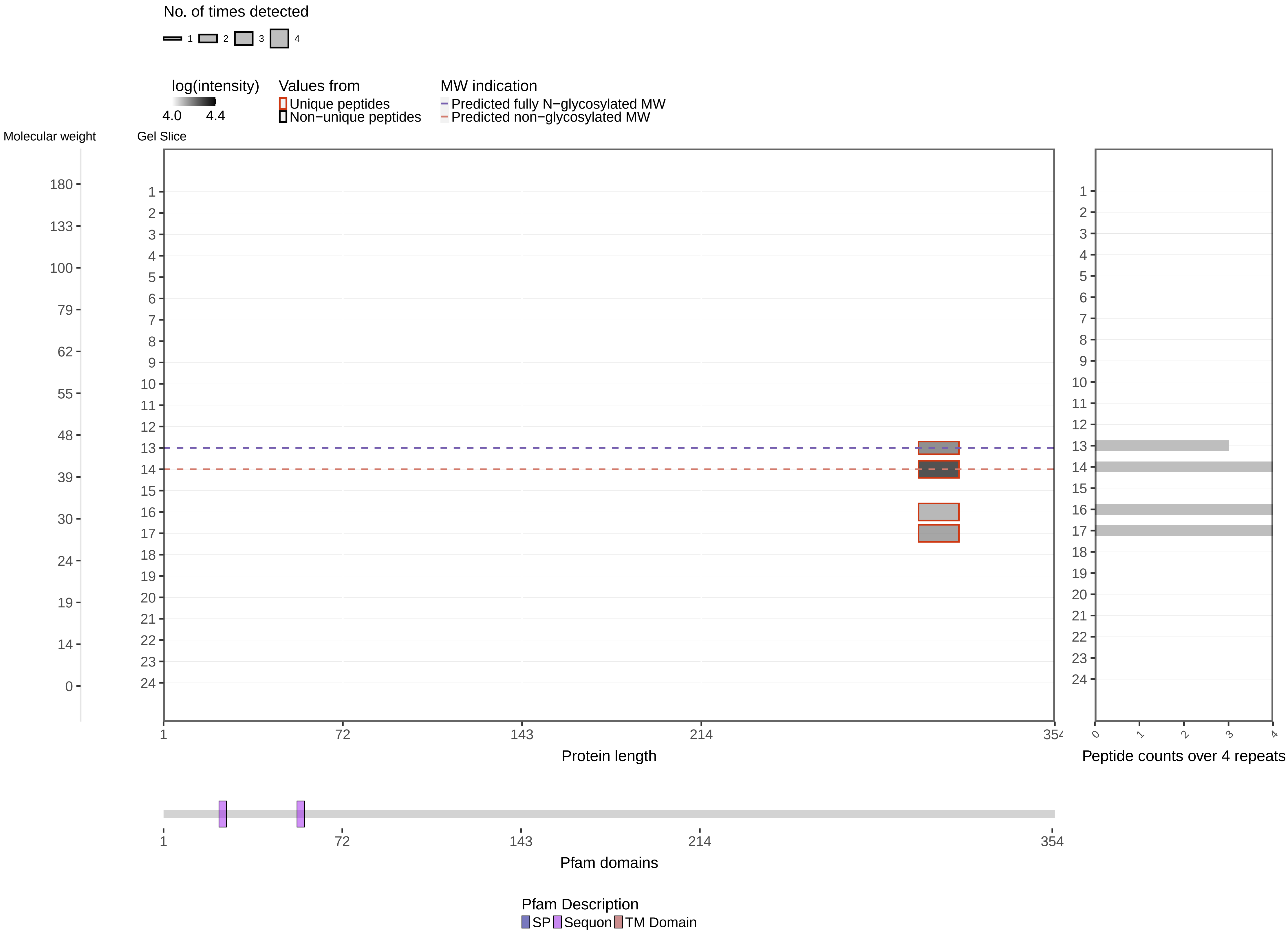


Protein sequence:

MATAVIVKNNGRKPVKLMSAILSHLKFKTRGGTGIQGLRSCTYCAHPPLSSPFEILSPGEAMKTEEPGLFSFGWEPELKPGIWSAQDVPITVLKHKLSRLYSVPPEERAKEFYNSIPSKYETIDQGRELFFRIIRMGFEDIFLSSPGSFSDKYGKEYFICTGPASMLVPLVVNPGEEWRGAQVIEHDNL*

NbL07g14400.1 probable_phosphatase_SPAC5H10.03 [MW = 31.48 kDa]

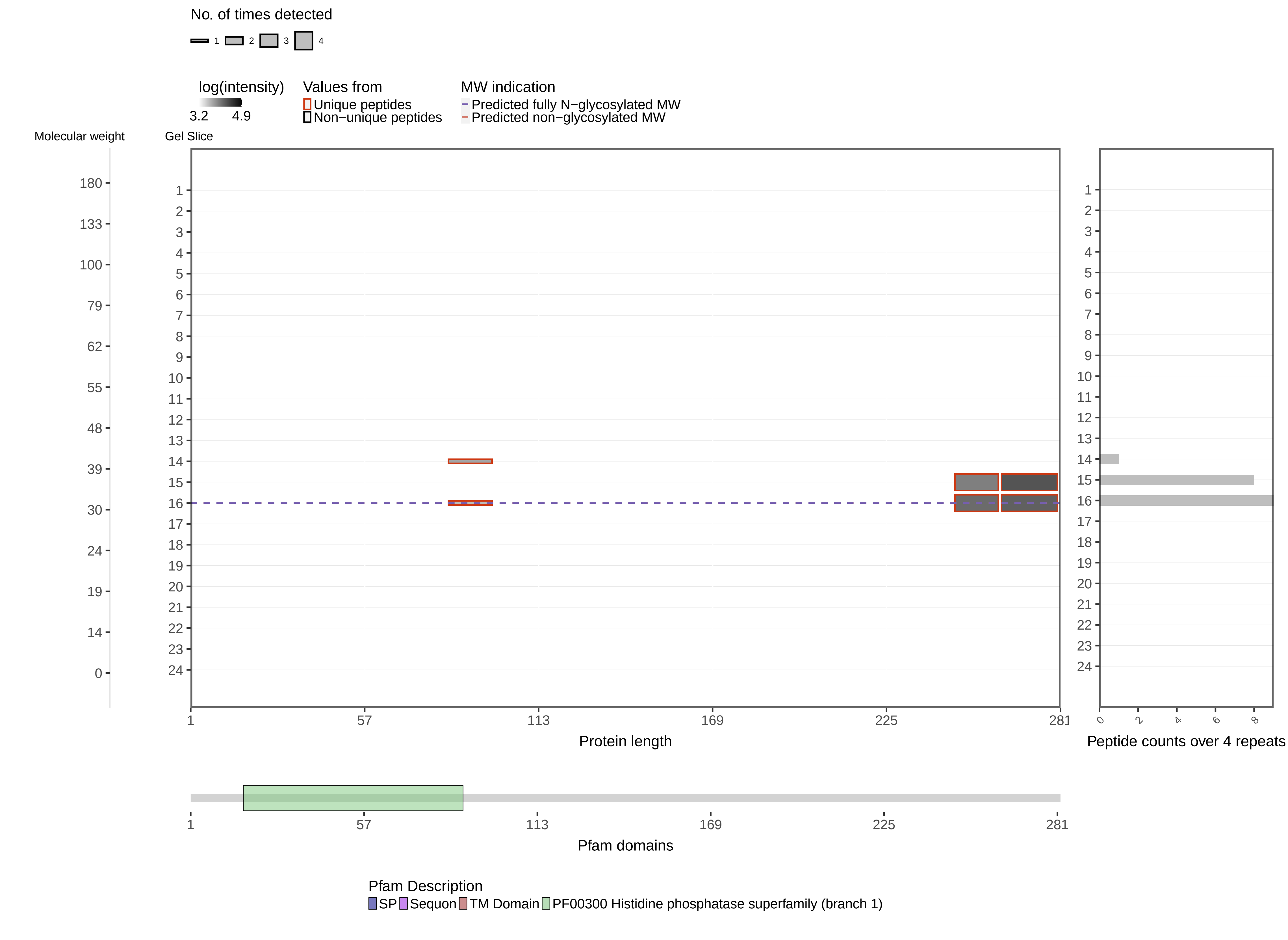


Protein sequence:

PIIAHELCREHLGVHPCDSRRSVREYQCLFPAIDLSLIESDEDTLWKANVRETKEEVMARGIKFLNWLLTRKEKEIAVVSHNGLLFHTLNAFGTDCHPFVKKEICKRFDNCELRSMVIFDRSMIGSDSSITYYPGKIPSAAILPSEAADESSGER*

### NbL07g14170.1 probable_receptor−like_protein_kinase_At5g24010 [MW − SP = 89.02 kDa]

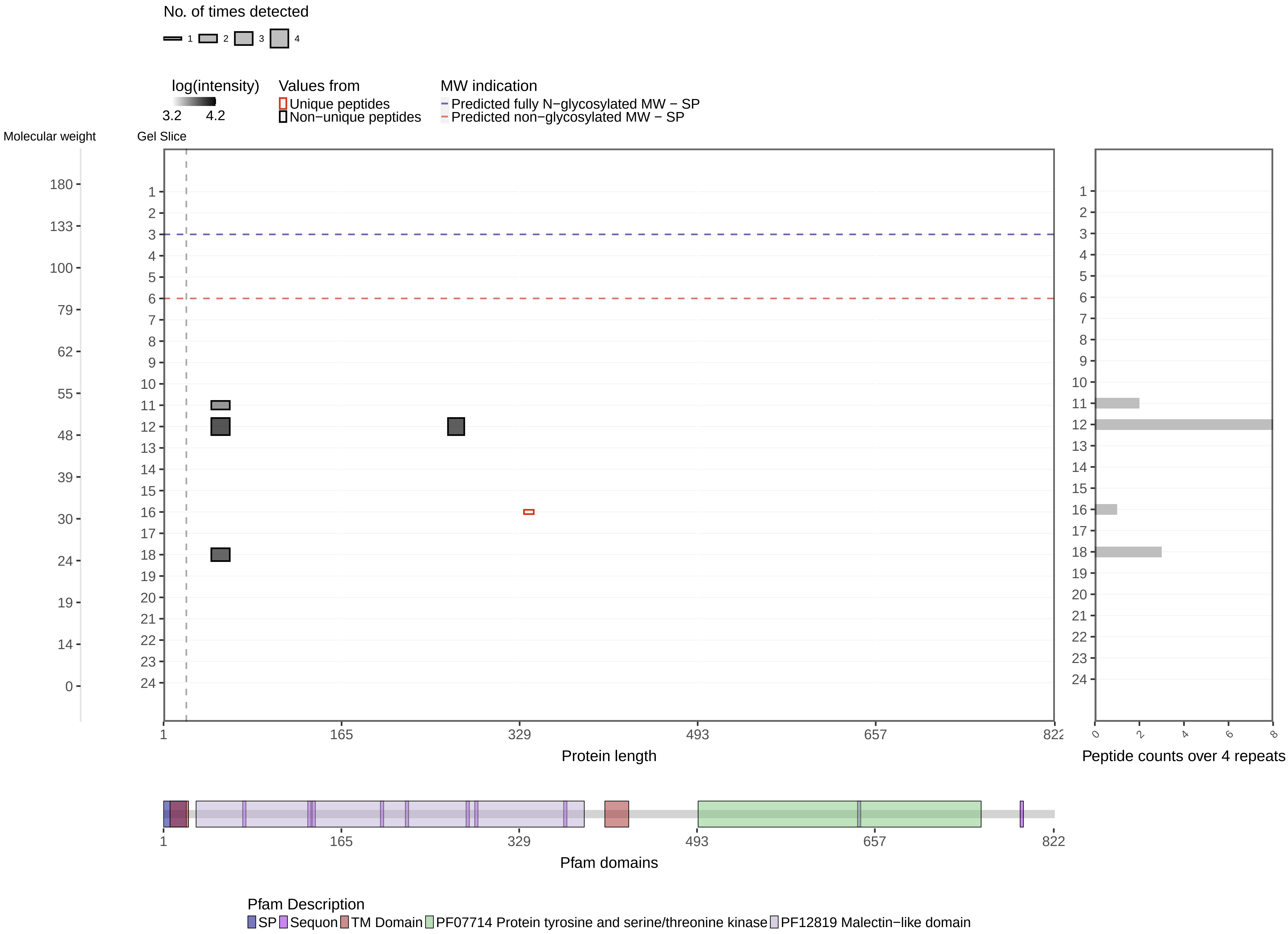


Protein sequence:

KKRNVWVIVSSVLGSFILLSVVILVALLVFVCRKKKLKPKRSESAGWTPLRQYGSTSHGTLSDGTSVGPNGFLGQRIPFAEIQSATNNFDKSLIVGSGGFGMVYKAVLGDNRKVAIKRGVPGSRQGLPEFHTEITVLSKIRHHHLVSLVGYCEEQSEMILVYEYMEKGPLKRHLYGPGITPLSWKQRVEICIGAARGLHYLHTGFAQGIIHRDIKSTNILLDENYVAKVADFGLSRTGPCLNETHVSTGVKGSFGYLDPEYFRRQQLTDKSDVYSFGVVLFEVLCARPAIDPLLSRE

### NbL07g14110.1 60S_ribosomal_protein_L21−1 [MW = 18.64 kDa]

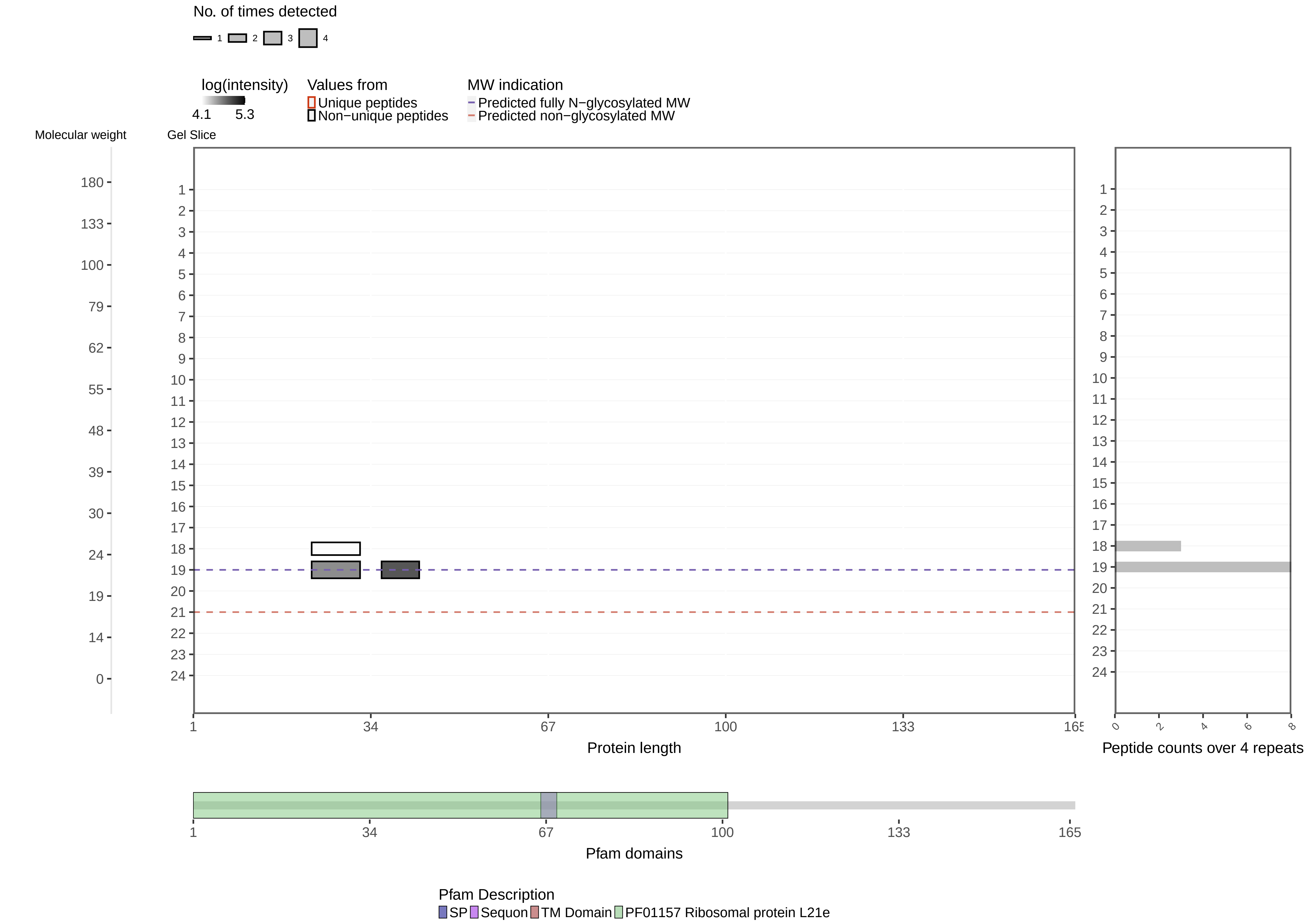


Protein sequence:

KRAVGVEVNKQVRNKILRKRIHVRIEHVQLSRCTEEVKERIKKNDQLKAEAKARGEVISTKRQPQGPKPGFMVEGATLETVTPIPYDVVNDLKGGY*

### NbL07g14080.1 UDP−glycosyltransferase_74F2−like [MW = 51.4 kDa]

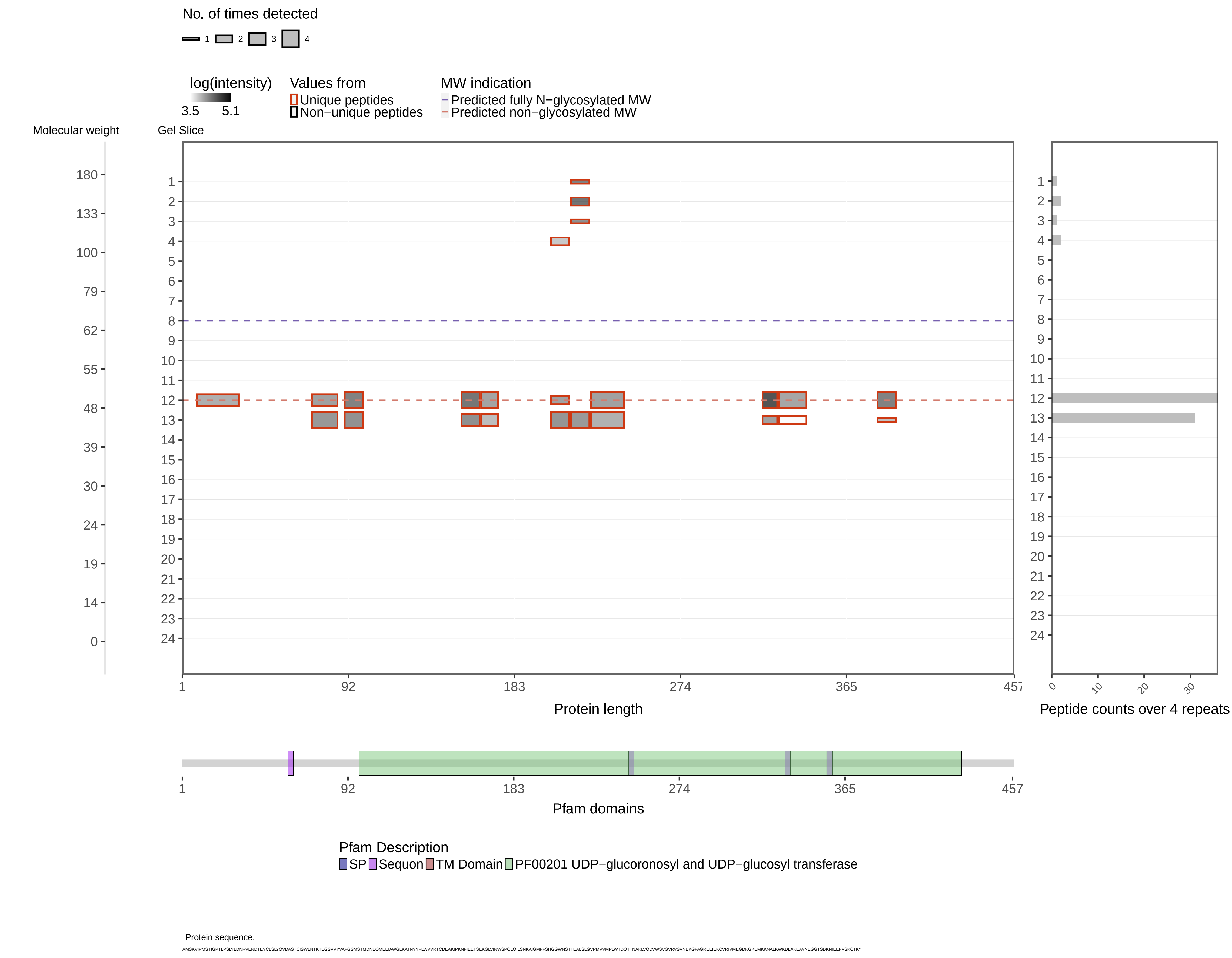

### NbL07g14010.1 uncharacterized_LOC107782443 [MW = 42.27 kDa]

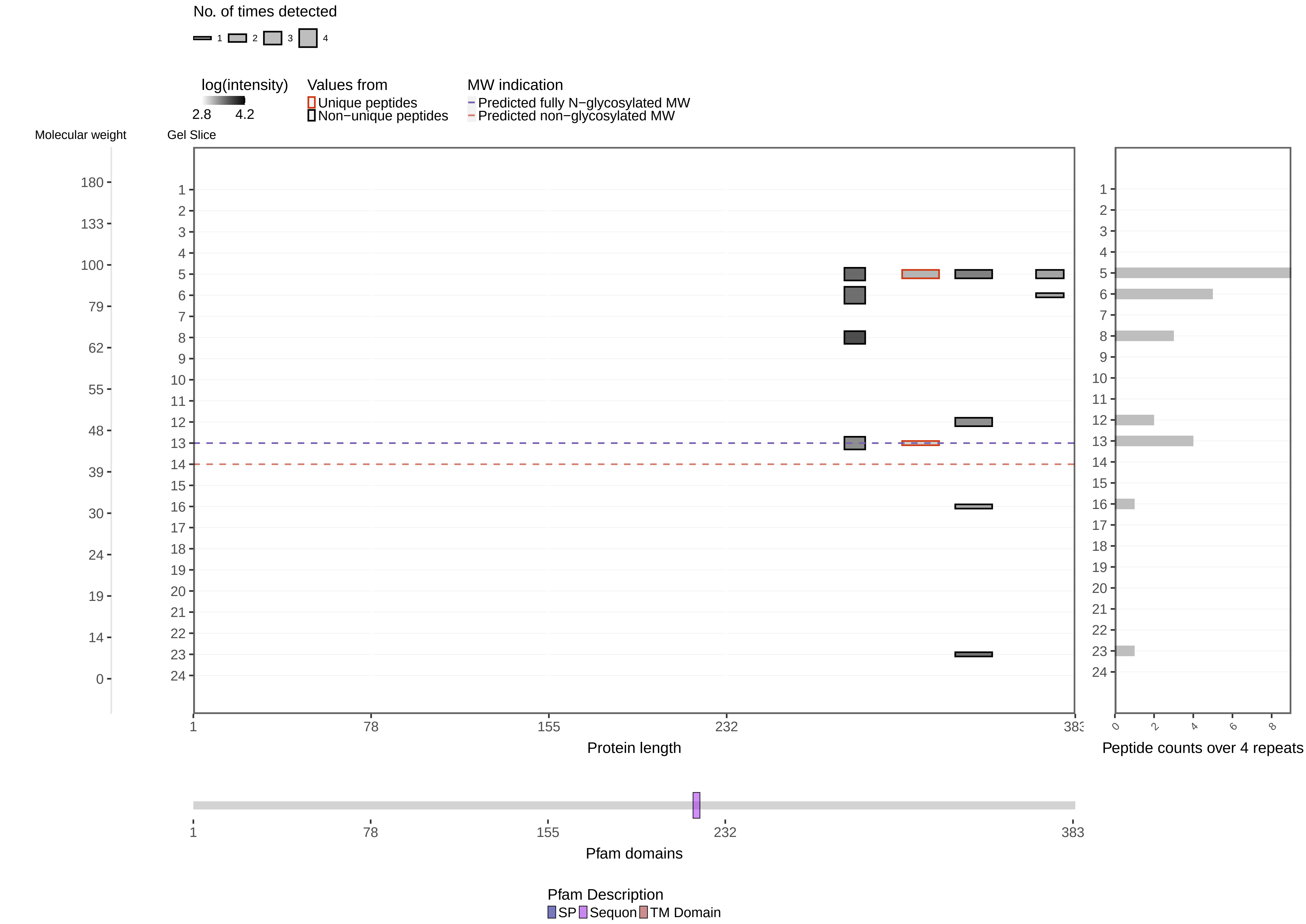


Protein sequence:

ERGAFLVQQAMRAFRANNIDSAKSRLTMCADDIRGQLERLGNTSELCSQLGVVLGMLGDCCRATGDAASAVTYFEESVNFLVKVPKDDLEITHTLSVSLNKIGDLKYYDDDLEAARSHYIKALDVRRNAIKQQSAPSQIIDIATSLAKVADVDRNLGNEDAAIDGFEEAIKMLESLELNPEEASLEQRRLSVLQFLNSQMEKKQPA*

### NbL07g13950.1 nudix_hydrolase_23_chloroplastic [MW = 31.96 kDa]

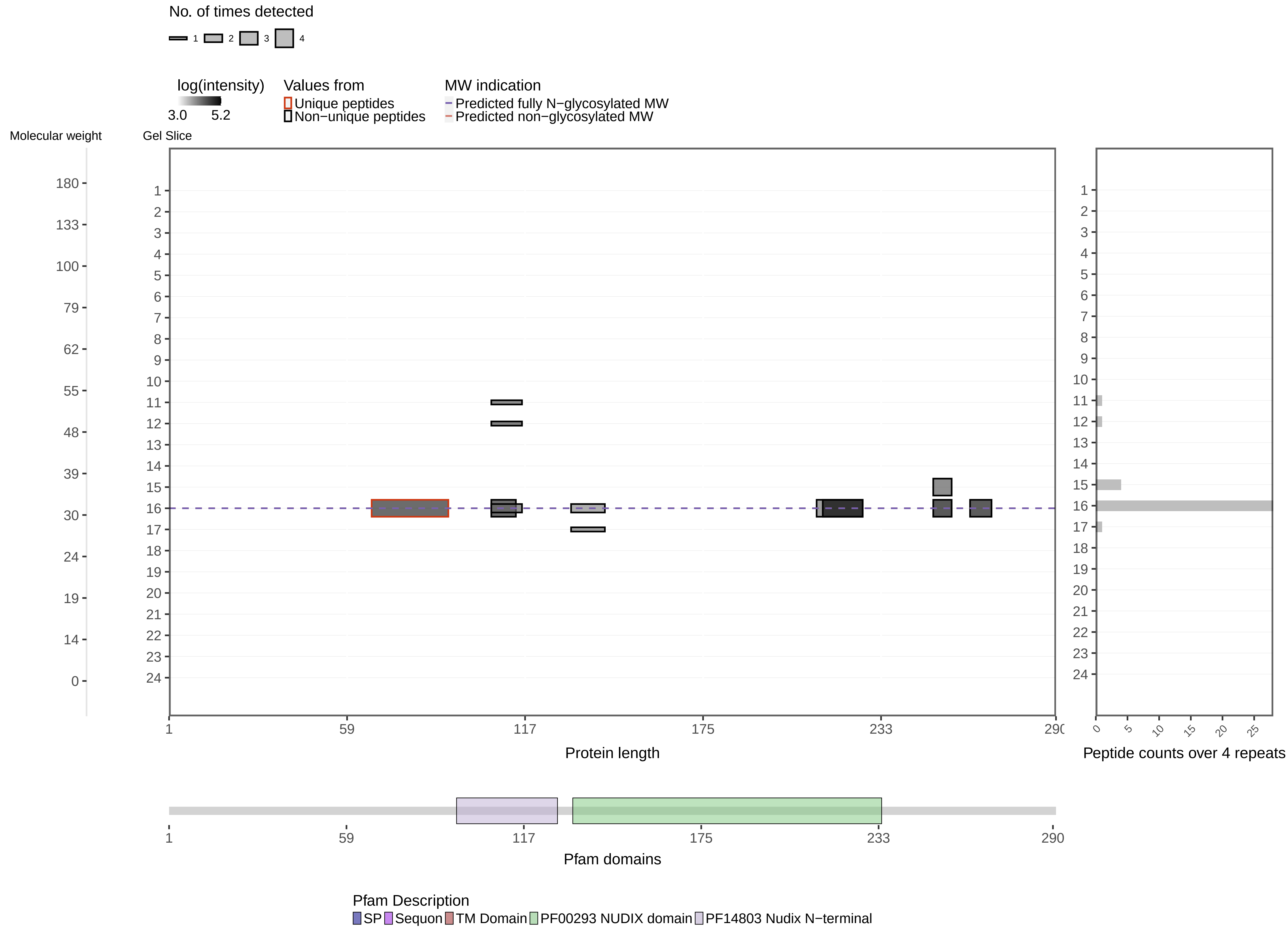


Protein sequence:

KMVVGCLIEHDKKILMCRRKIQPSYGLWTLPAGYMEIGESAAEGAIRETLEEANAEVEVQSPFAQLDIPLIGQTYMIFLAKLKTPHFSPGPESSECRLYELDDIPFDSLAFSSMLVTLKLYIEDIKVGRPKFHYGIINKRPGTSPSEIHAYTLDYHMQS*

### NbL07g13940.1 probable_plastid−lipid−associated_protein_13_chloroplastic [MW = 32.82 kDa]

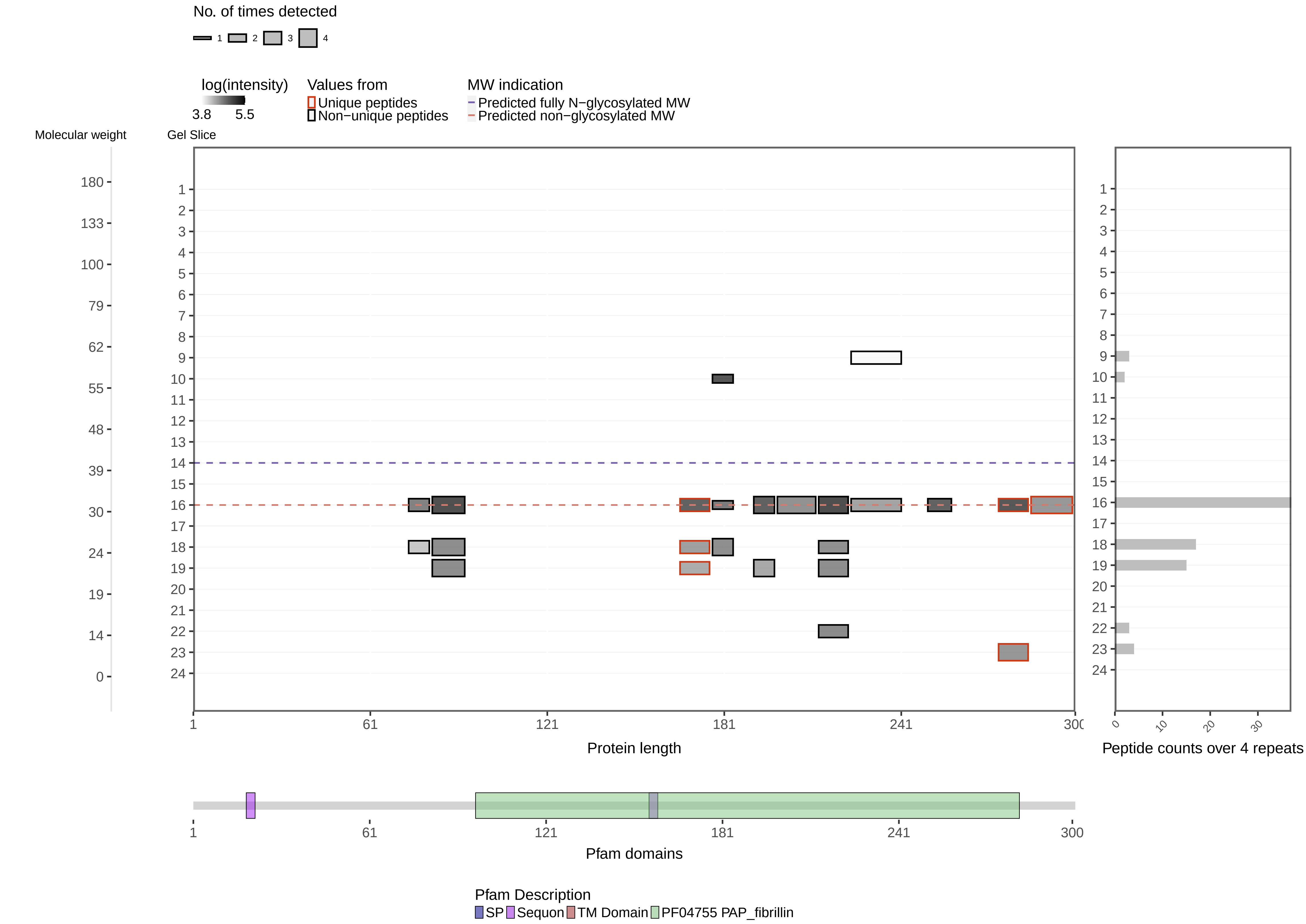


Protein sequence:

AGSPVLIFAKFLFGRIPPTLVNLSKLDVLIKDGYGTATAQVKLLNSIENKFIISTKYSVEGPLRMKEEYVEGTFESPKVNEEAVPEQLRGAFGQAFNTVQQLPVPIRDAVSSGVKVPLGGTFQRLIMISYLDDEILIVRNAAGEPEVLTRLEAPPETEPITEYES*

NbL07g13890.1 zinc_finger_CCCH_domain−containing_protein_14−like [MW = 30.67 kDa]

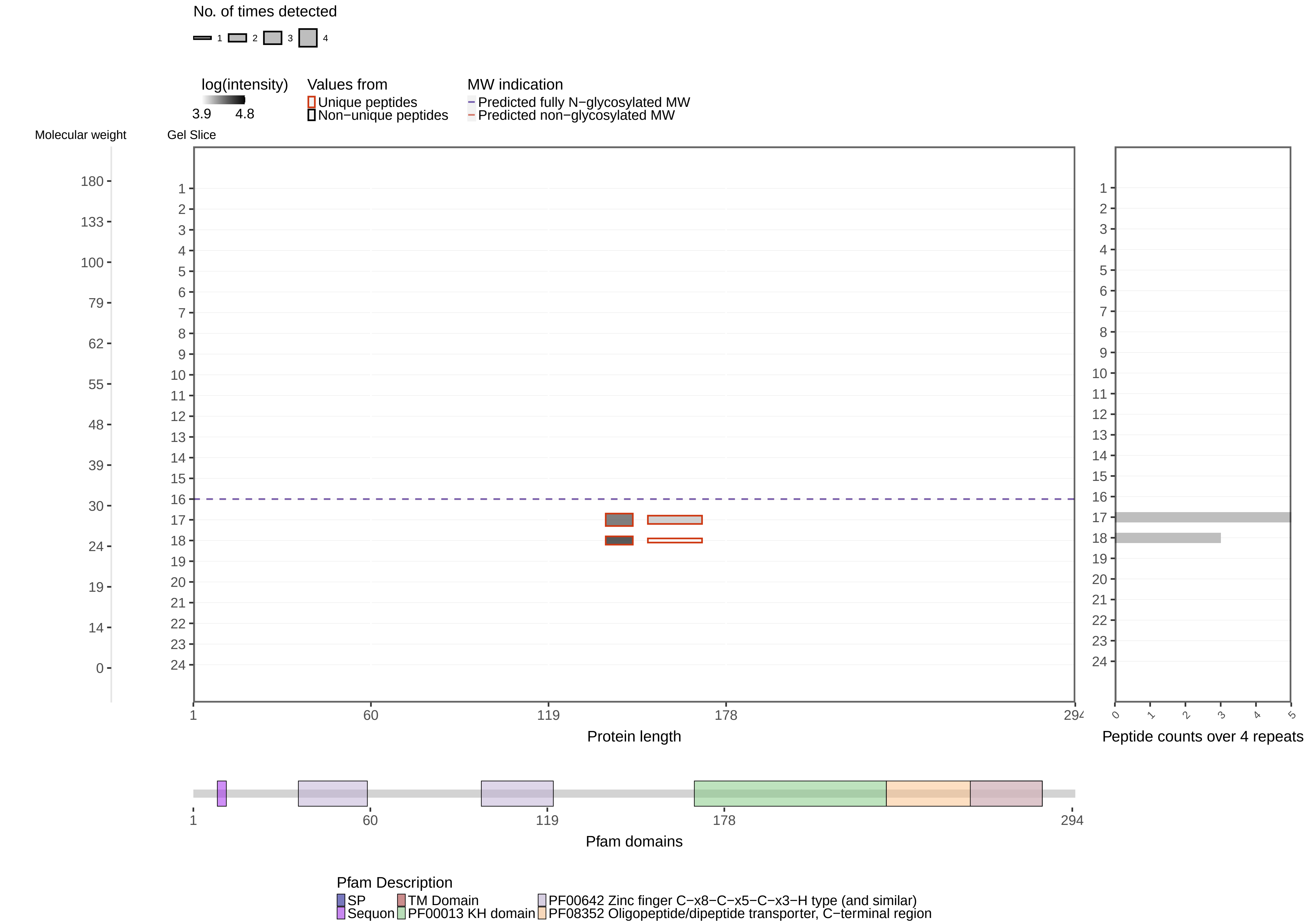


Protein sequence:

QEDPRAMGGGPVPGRFGGRMEPPISGPAASFGTSATAKISVDASLAGAIIGKGGVNSKQICRQTGAKLAIREHETDKNLRNIELEGTFEQISQASAMVRELISSLGSVGGPGRAPAVQGGPPPQMSNYKTKLCENFAKGSCTFGERCHFAHGAAELRKTGV*

NbL07g13880.1 enoyl−[acyl−carrier−protein]_reductase_[NADH]_2_chloroplastic−like [MW = 41.35 kDa]

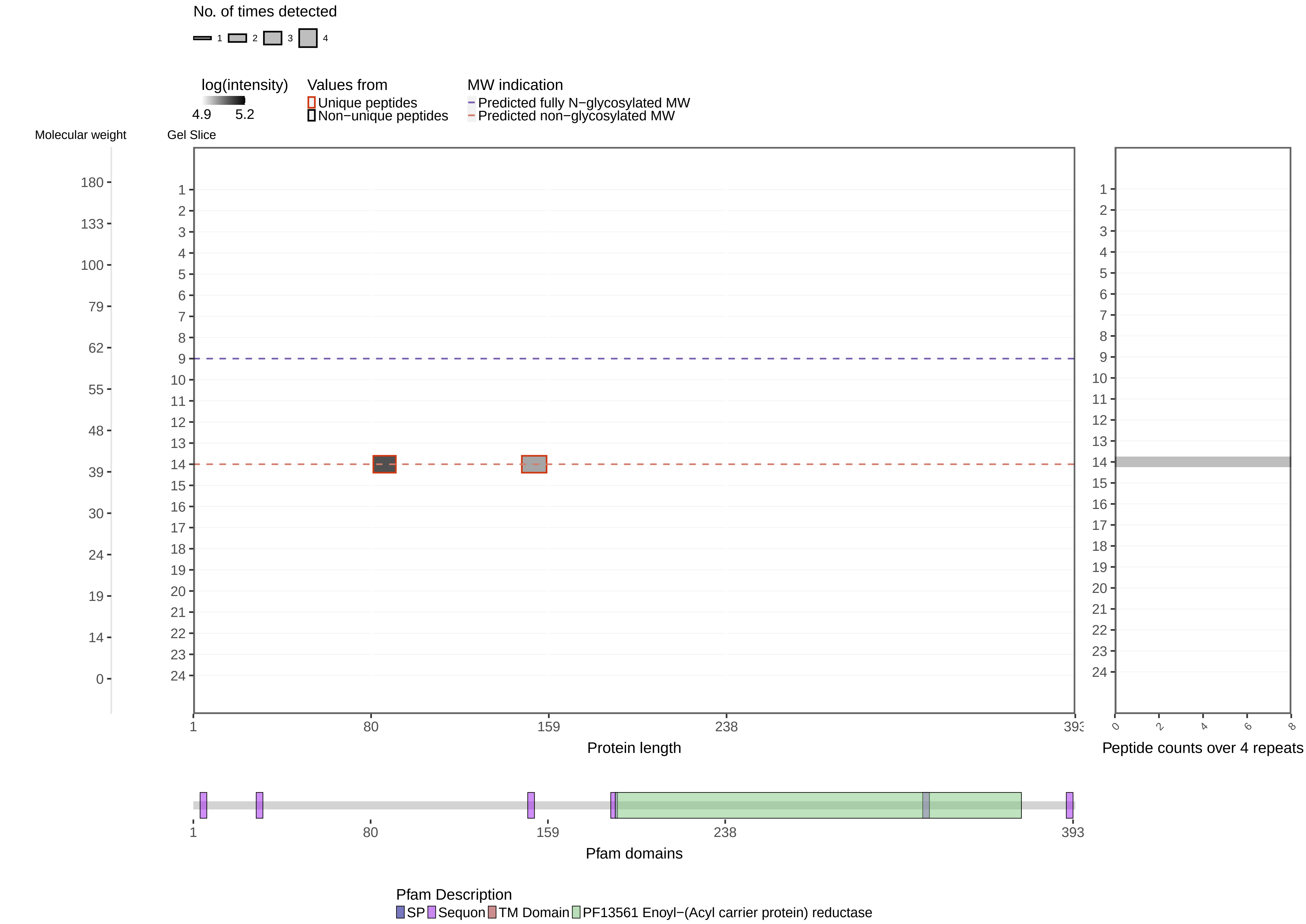


Protein sequence:

YAGSSNWTVKEAVESVKQDFGTIDILVHSLANGPEVSKPLLETSRKGYLAAVSASSYSFVSLLRHFLPIINPGGASISLTYIASERIIPGYGGGMSSAKAALESDTKVLAFEAGRKNRVRVNTISAGPLGSRAAKAIGFIDMMINYSLENAPLQKELYAEEVGNAAAFLVSPLASAITGATIYVDNGLNAMGVGVDSPAFEGLDIPKDNKS*

### NbL07g13690.1 phosphoenolpyruvate_carboxykinase_[ATP]−like [MW = 73.46 kDa]

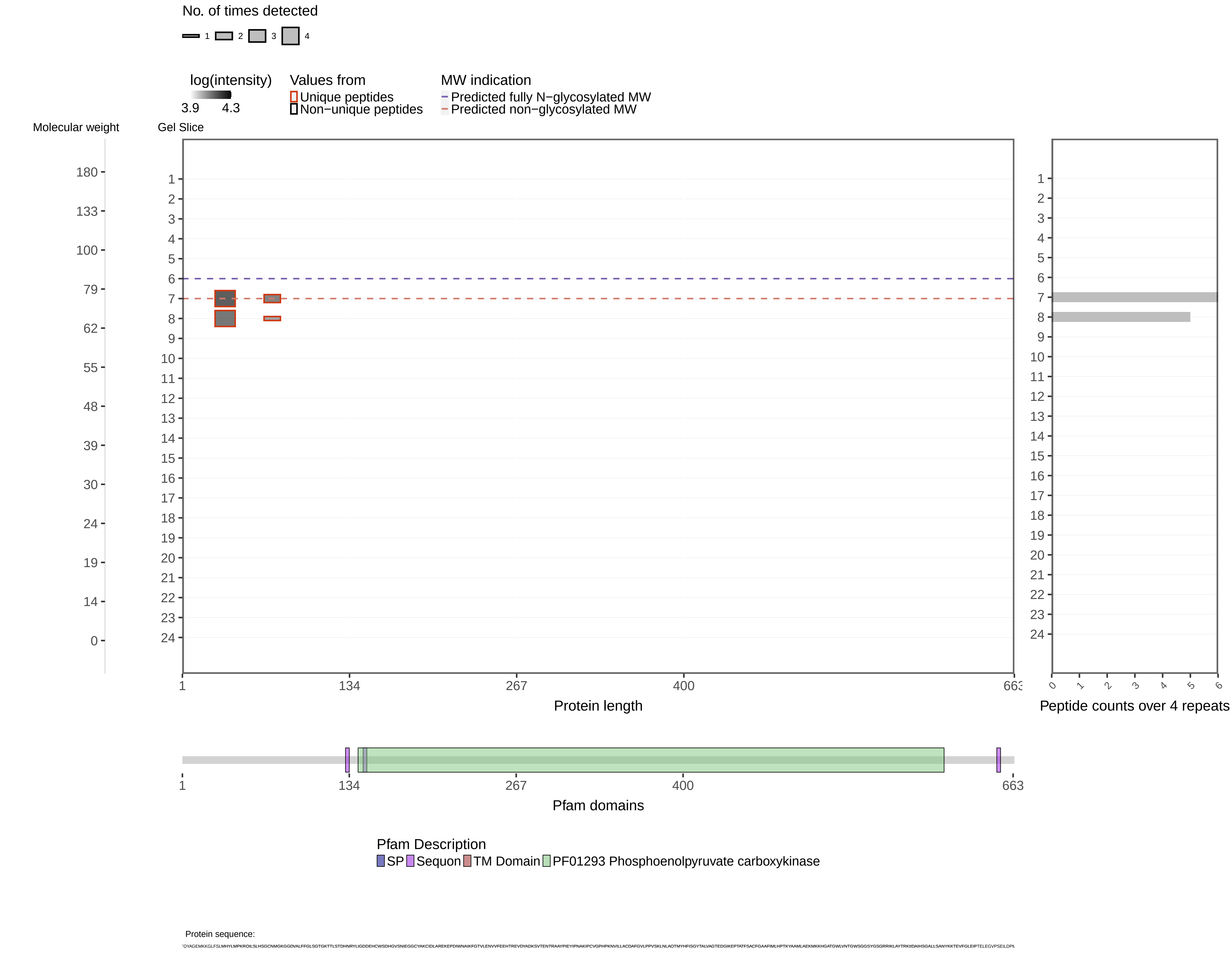

NbL07g13660.1 ATP−dependent_Clp_protease_proteolytic_subunit_4_chloroplastic [MW = 37.88 kDa]

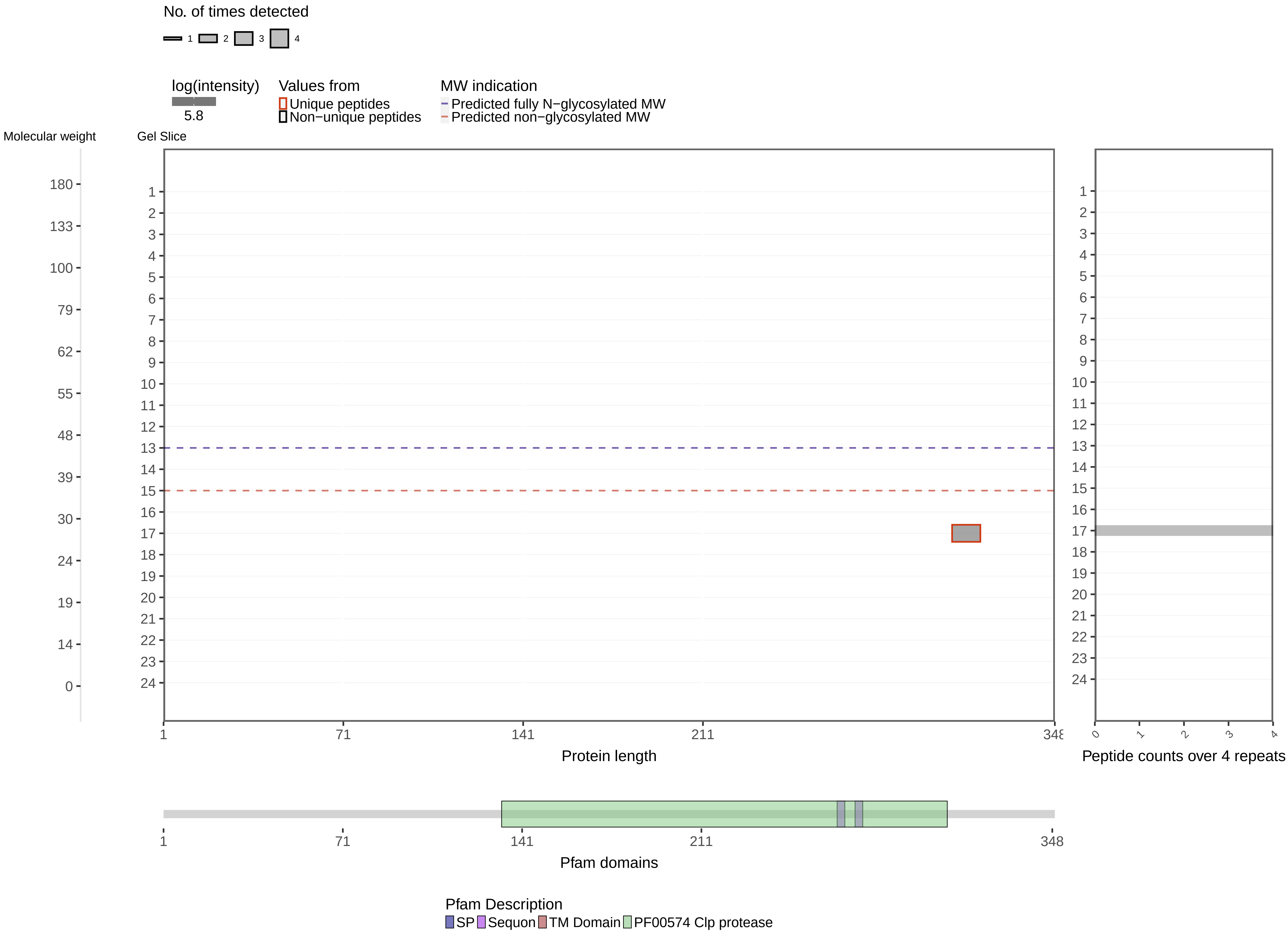


Protein sequence:

IISQLLLLDAMDSSKDIRLFINSPGGSLSATMAIFDVVKLVRADVSTVALGIAASTASIILGGGTKGKRYAMPNARIMIHQPLGGASGQAIDVEIQAREIMHNKDNVTKIISNCTGRSYEQVQKDIDRDRYMSPIEAVEYGLIDGVIDRHSIIPLEPVPERVKSTLNYQEISKDPKKFLNPEIPDDEIY*

NbL07g13430.1 phosphoribulokinase_chloroplastic−like [MW = 45.31 kDa]

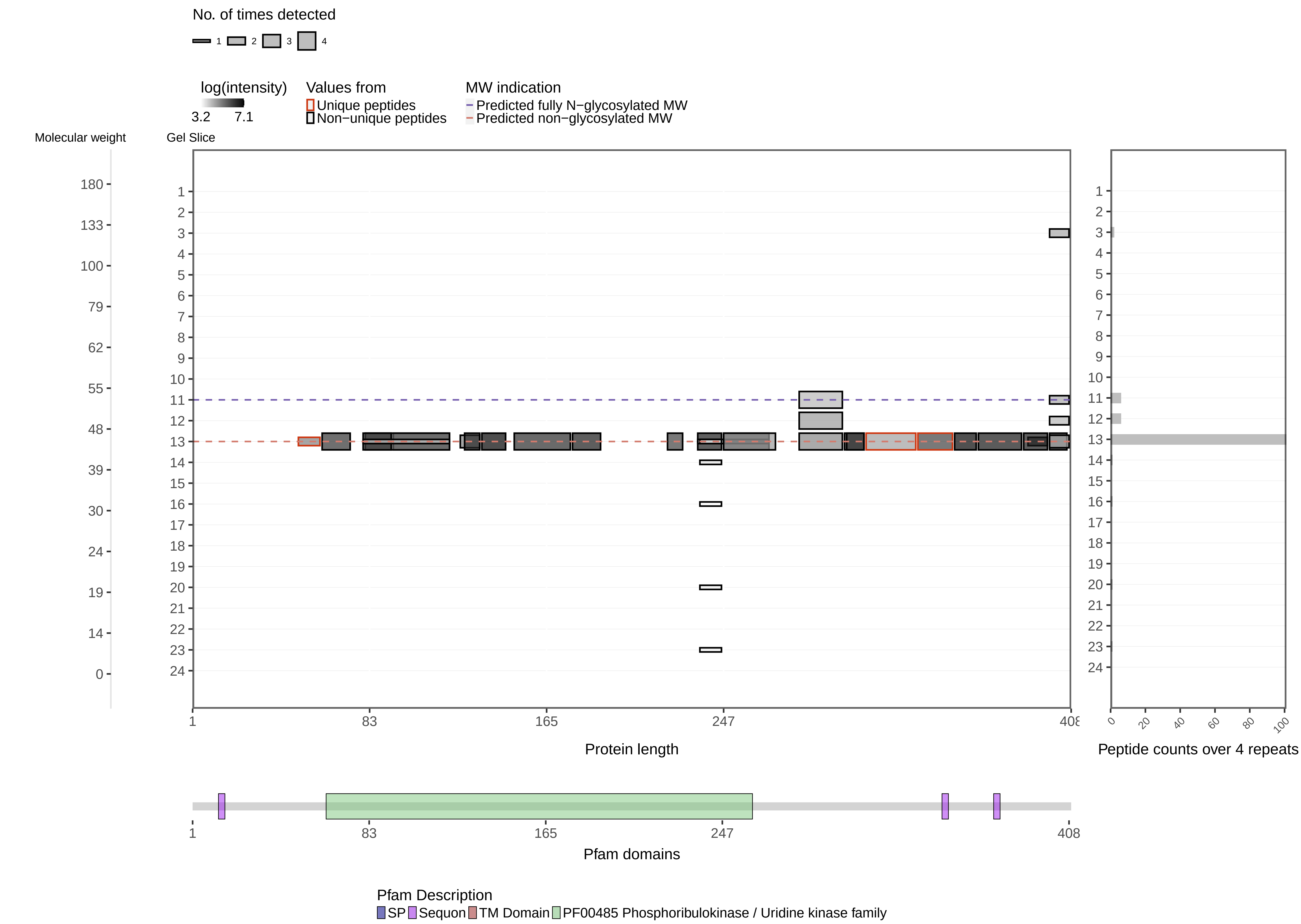


Protein sequence:

VRELLDFSIYLDISNEVKFAWKIQRDMAERGHSLESIKASIEARKPDFDAYIDPQKQYADAVIEVLPTQLIPDDNEGKVLRVRLIMKEGVKNFNPVYLFDEGSTISWIPCGRKLTCSYPGIKFTYGPDTYFDNEVTVLEMDGQFDRLDELIYVESHLSNLSSKFYGEVTQQMLKHADFPGSNNGTGFFQTIVGLKIRDLYEQIVASRAAAPVQAAKA*

NbL07g13350.1 uncharacterized_LOC109217894 [MW = 34.88 kDa]

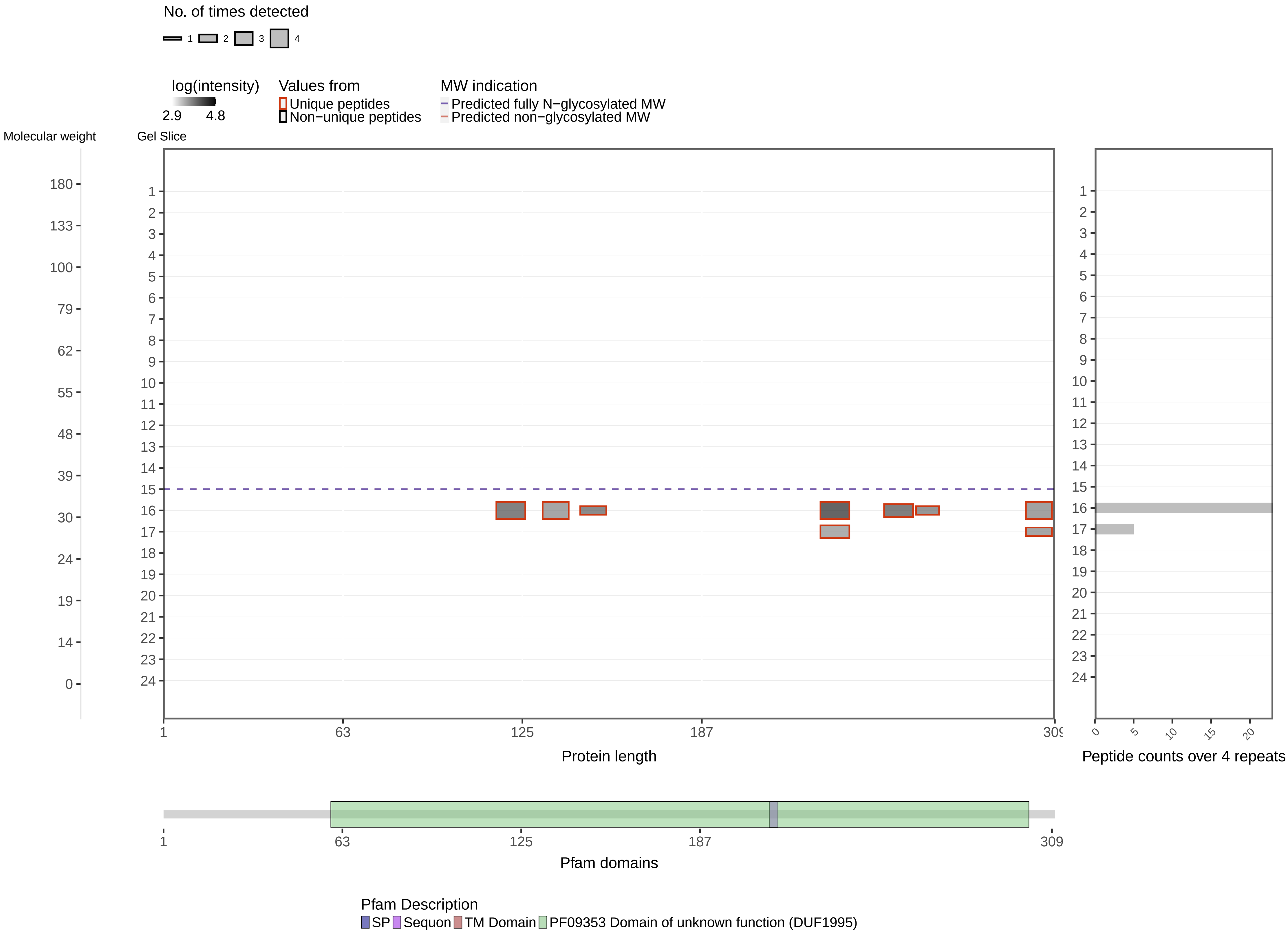


Protein sequence:

KFAROSIFGGASFKLDYLTKPSFFEDFGFTEKVKMADRVKPEDELFIVAYPYFNVNEMLVVEELYKAAVSNTSRKLIIFNGELDRIRSGYYPSFFYPKLAALSKTLFPKMETVYYIHNFKGPNGGVLFRCYPGPWKVLRRVRNAYICLHOOESMPSLKEVALDILPLA*

### NbL07g13180.1 ornithine_aminotransferase_mitochondrial−like [MW = 52.16 kDa]

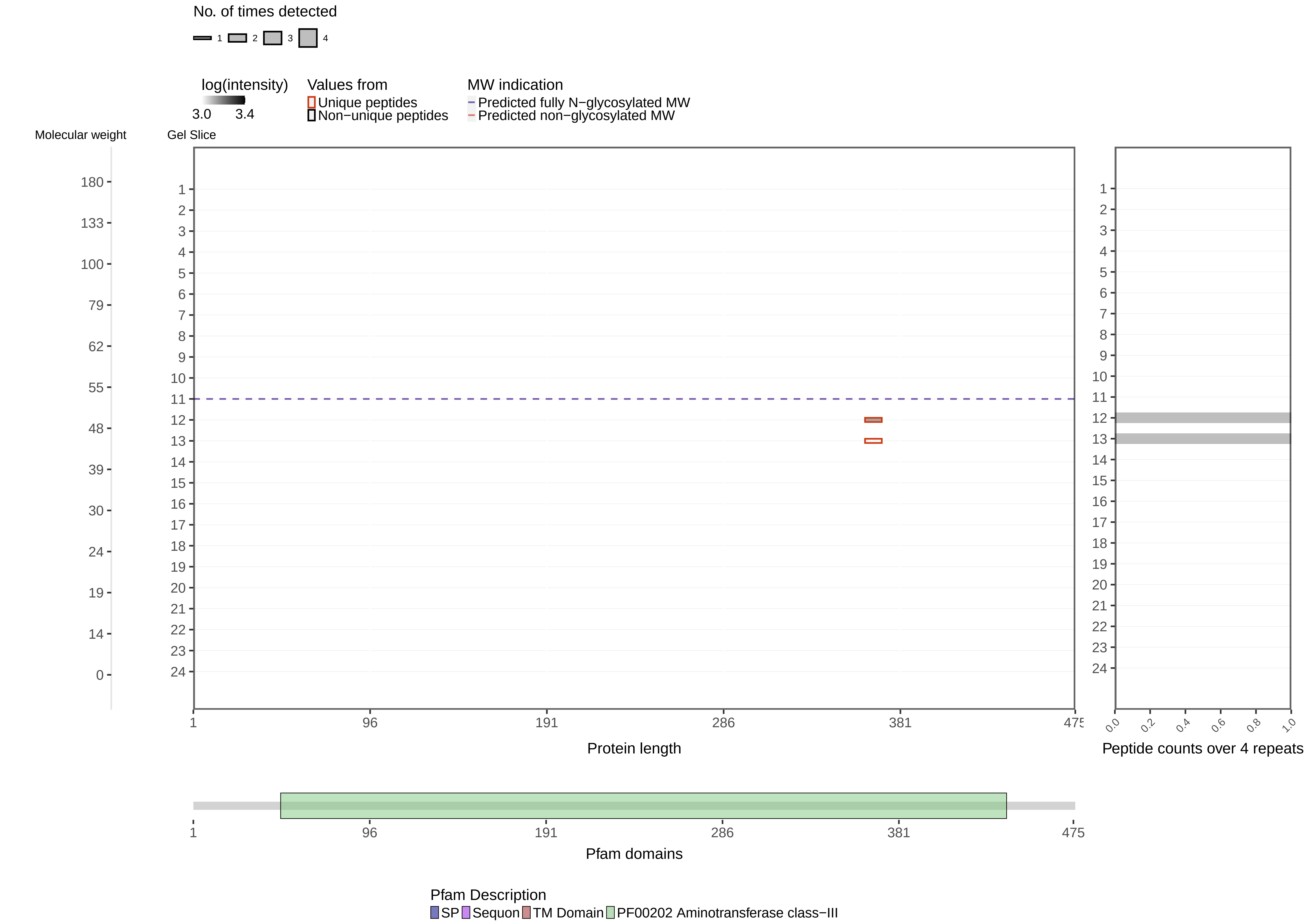


Protein sequence:

[illegible]IAGFLFEPIQGEAGVIIPPEGYLKSVRDLCSKYNILMIADEIQSGLARSGRLLACDWEEVRPDVVILGKALGGGVLPVSAVLADKDVMVCIQAGEHGSTFGGNPLASAVAIASLDVIREEGLAERSAQMGEQLRHQLMKVQEQFPNFIKEVRGKGLFNAVELNSKSLLPVTAYDICMKLKERGILAKPTHDSIIRLTPPLSMSLDELQEGSKVLHDVFELDLPKMQKEKPATVSHAASSVCDRCGRNLYGSS*

NbL07g13170.1 KH_domain−containing_protein_At4g18375−like [MW = 66.49 kDa]

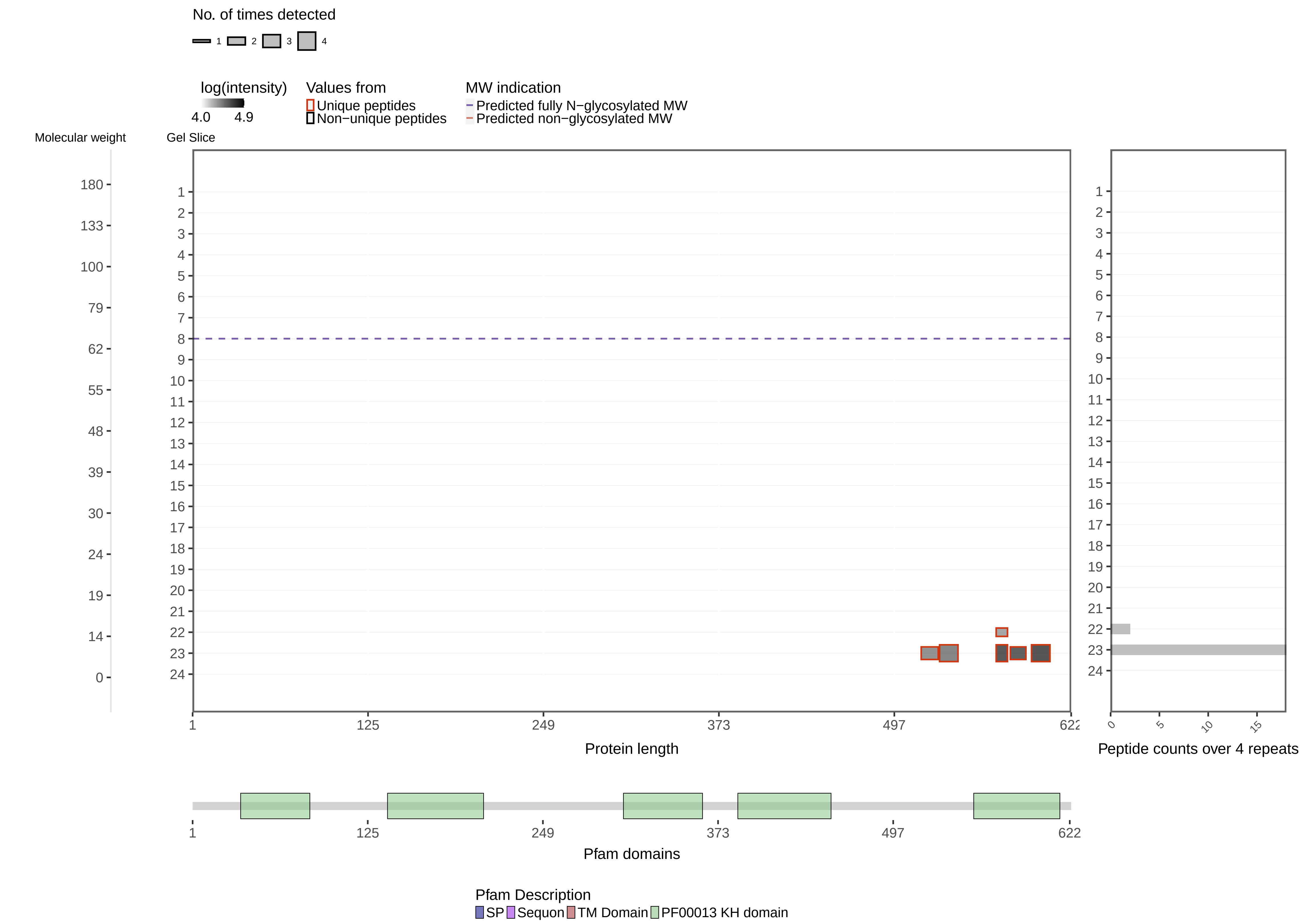


Protein sequence:

GASQTEELTIRVLCPTNNIGRVIGKGGASIKSVRQASGARIEVGDAKADRDQCIITVISTESVDDLKSMAVEAVLLLQGKINDDDEDSVTFRLLVPSKIIGCIIGKGGSIVNEIRKRTRADVRISKGQKPKCADSNDELVEVAGEISSVRDALIQIVLRLRDDVLKVRESNFNPSAGADSLRAGGTGHSLAPVLPNVPPTASLSYEHRGETGNGLGMRSSGSLYGHESFSRGDDGYGAYSSYSSKIYGGLPPPTALEMVIPGHAAGKVIGKGGSNIDNIRKISGASVDIIDSKSSRGD

NbL07g13130.1 pyruvate_dehydrogenase_E1_component_subunit_beta−3_chloroplastic−like [MW = 44.58 kDa]

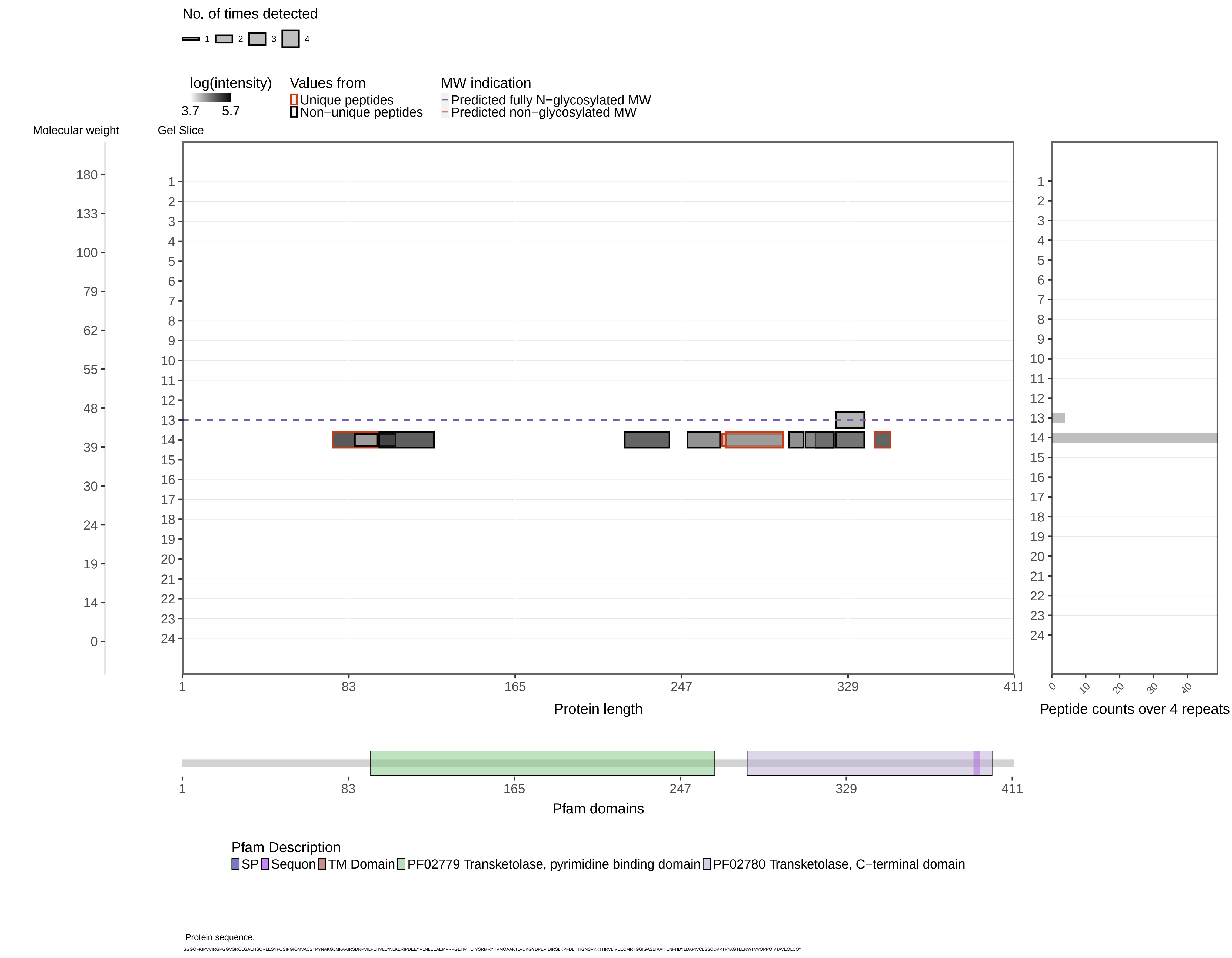

NbL07g13090.1 calcyclin−binding_protein−like [MW = 24.07 kDa]

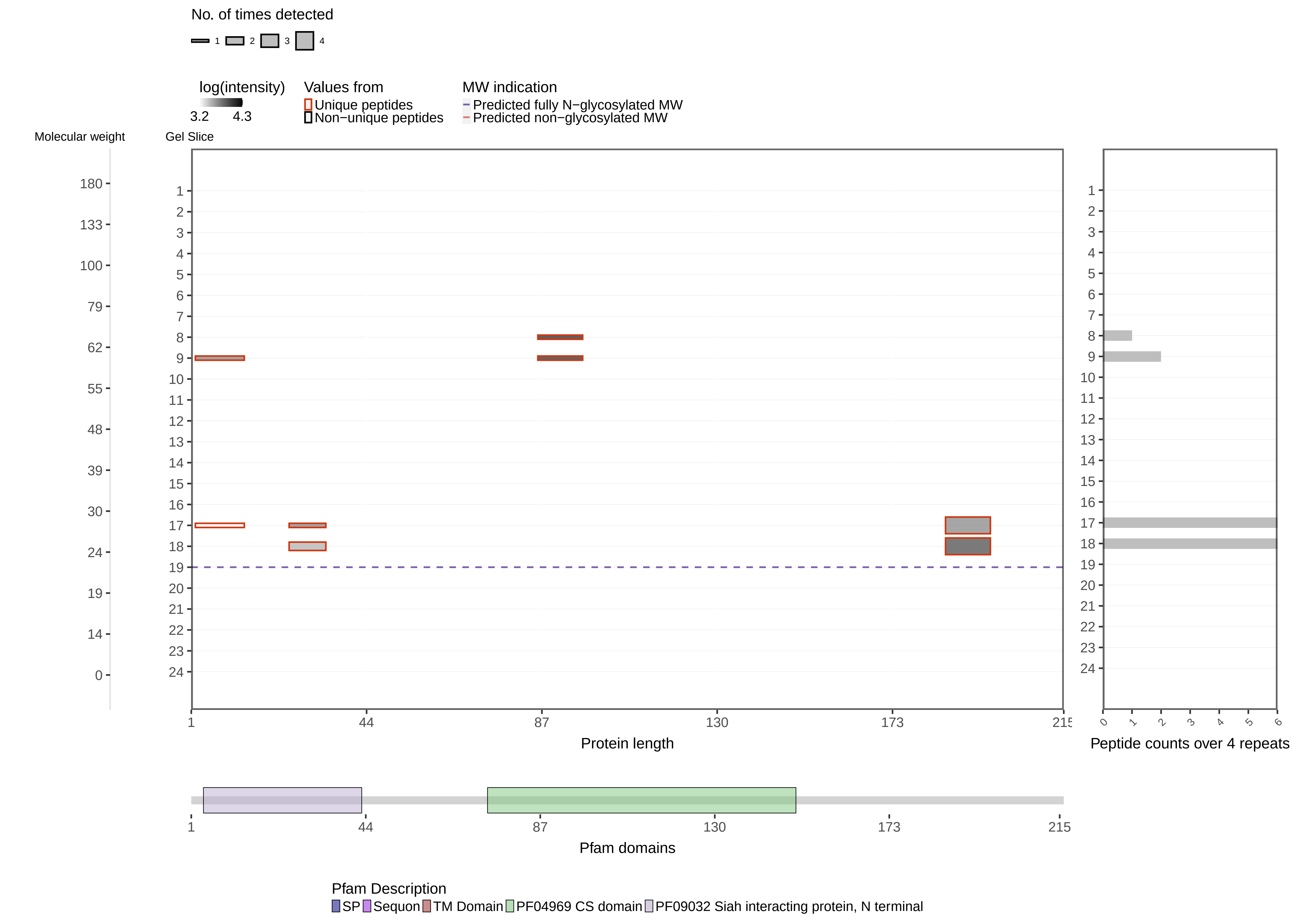


Protein sequence:

EKAETEFKPMSFDAKFHDVHGKNFRFSLPKLNKEIVPEKCKVLVKPTRVVITLIKASKGNWLDLHYKEDKFKPNLDKEKDPMAGIMDLMKNMYEDGDEEMKRTIAKAWTDARSGKAADR*

### NbL07g13030.1 prefoldin_subunit_6−like [MW = 14.77 kDa]

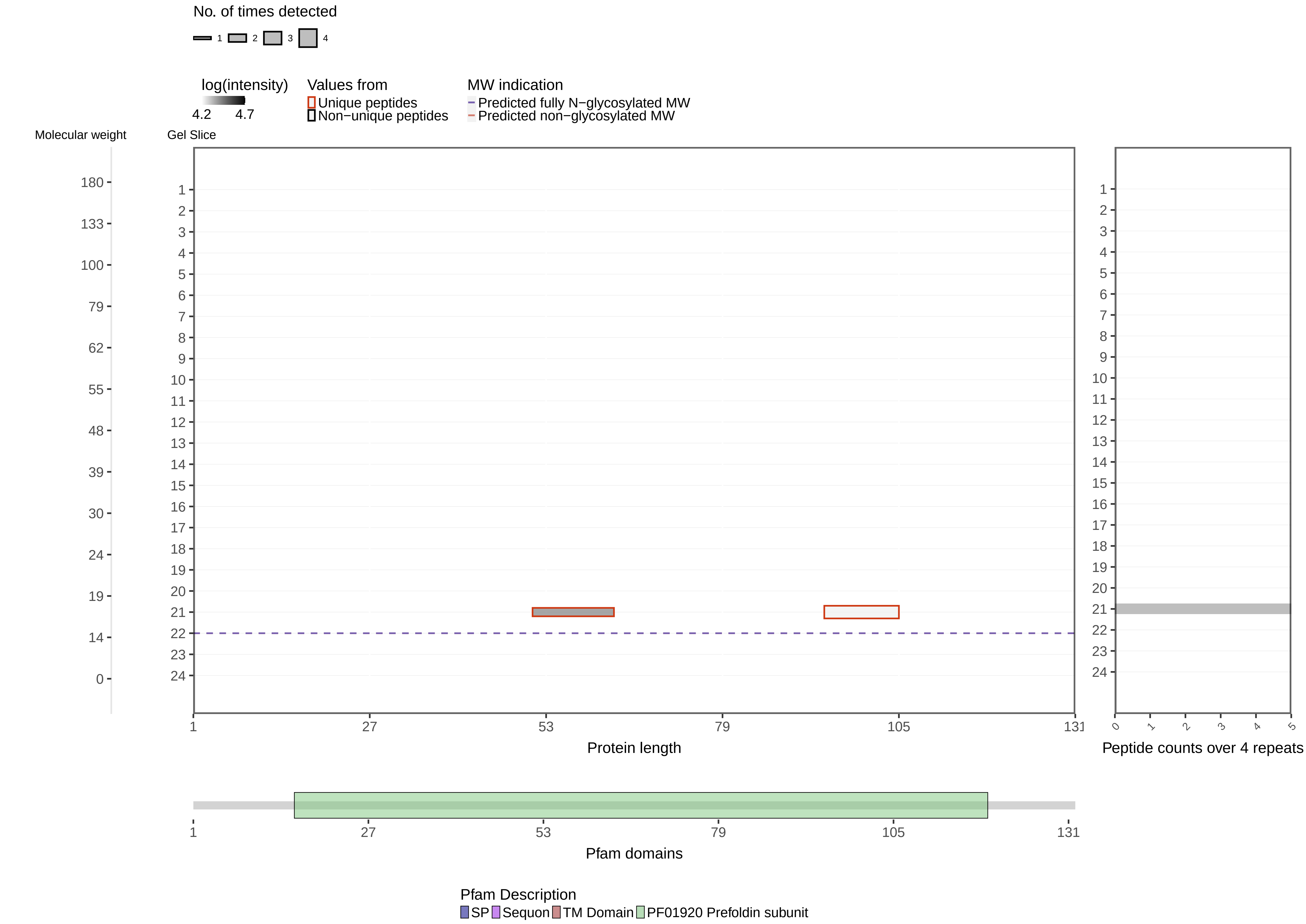


Protein sequence:
LDPLNEDANVYKLIGPVLVKQDLAEARANVKKRIEYISAELKRLDSTLQDLEEKQNSKKETILKLQQKIQSLQAGKGKA*

### NbL07g13010.1 3−oxoacyl−[acyl−carrier−protein]_synthase_I_chloroplastic−like [MW = 49.93 kDa]

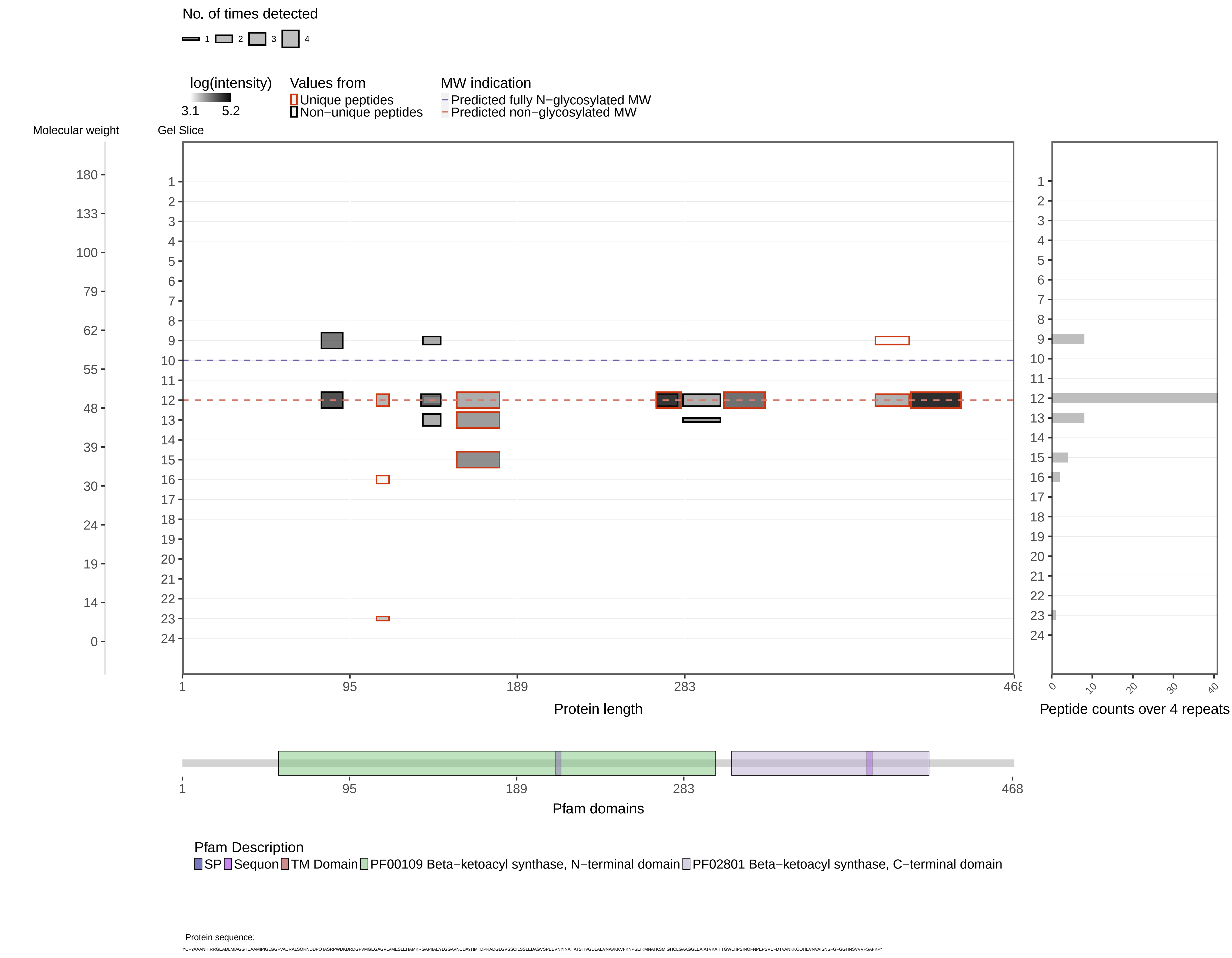

NbL07g12900.1 phosphomannomutase/phosphoglucomutase [MW = 67.34 kDa]

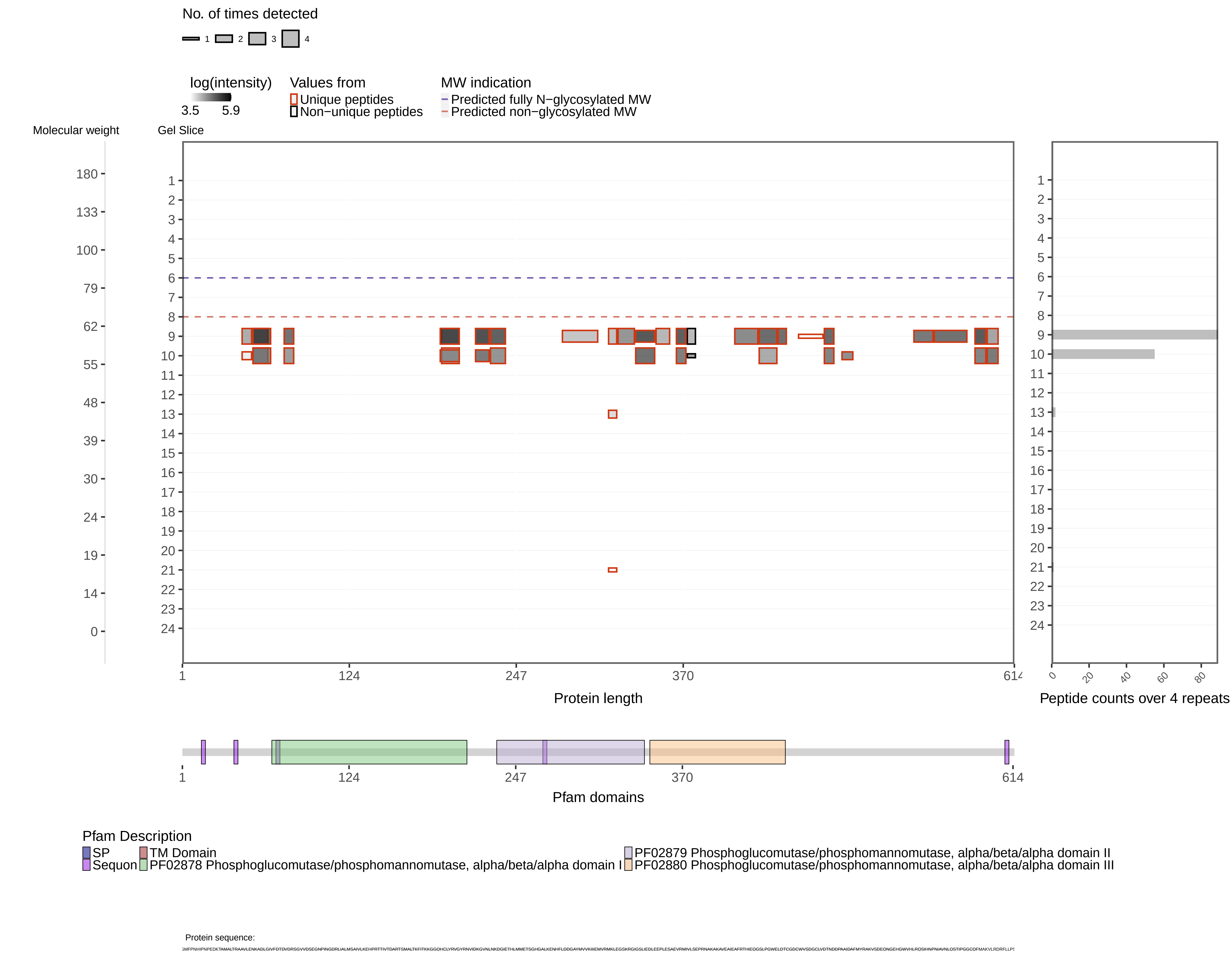

NbL07g12800.1 caspase_8_associated_protein_2 [MW = 53.19 kDa]

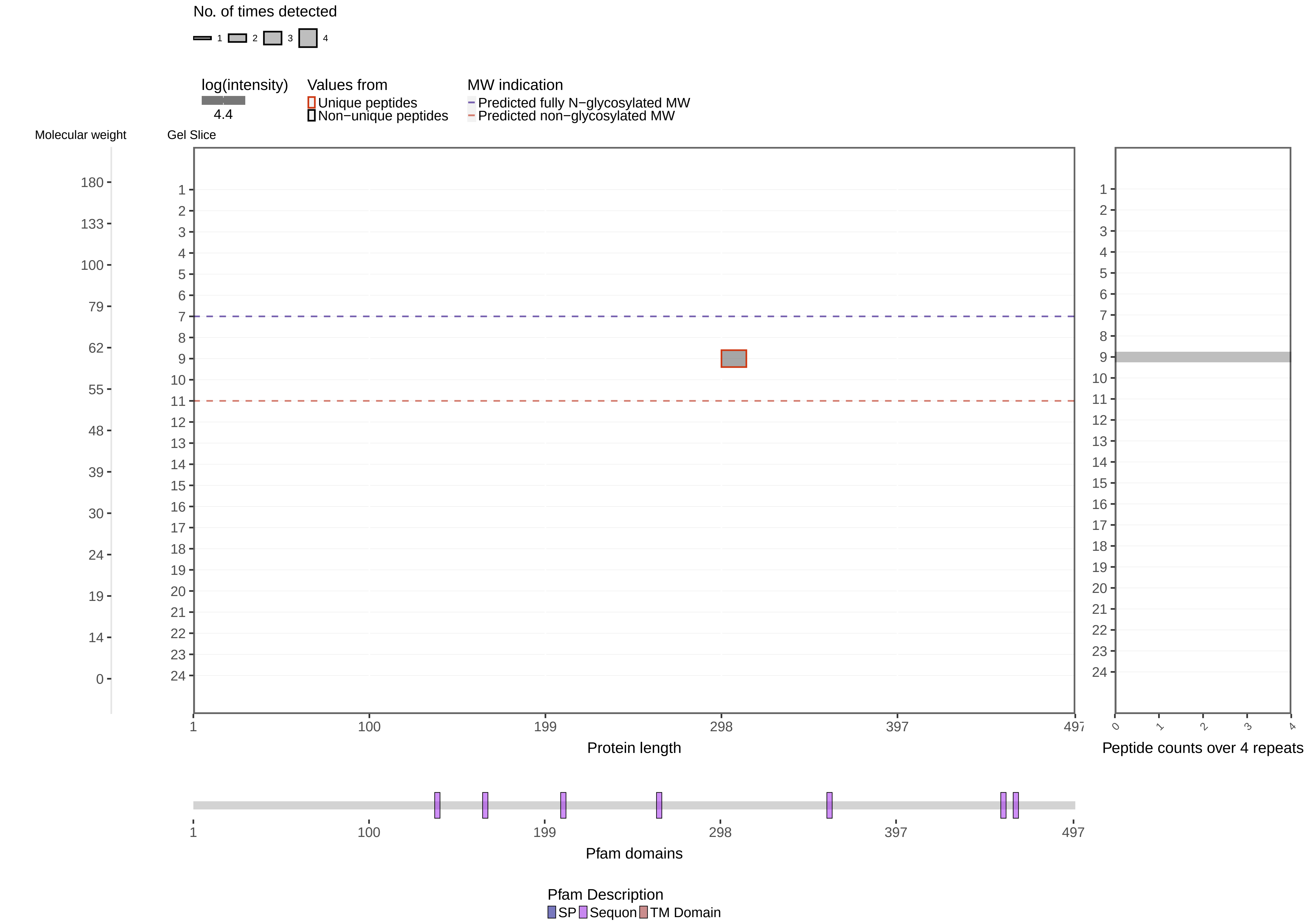


Protein sequence:

QFLIQSNSPVICTEPVEKPEIPFHSNDSLKECVRNFFNLGGDTTTTVLAGQPDDTTVQSDKMASILNPQSDITGVKAKSLAKTNGDTAQTSQQAVTATNLETTTNPHAAVMEKGKSDHQRLNSSTVVDSDSASTTATVPTFVTPISHNAQQYVMPSSISVRQPLNAFSGNDILACQHSPLVDKTADTKKESSDPTQQKETTPEVSILSKSPMPNAAEGNNISNRLCNETVPAVLSRPEIGSKLQSDANRPINVRDDIEQK*

### NbL07g12780.1 acetylornithine_deacetylase [MW = 52.99 kDa]

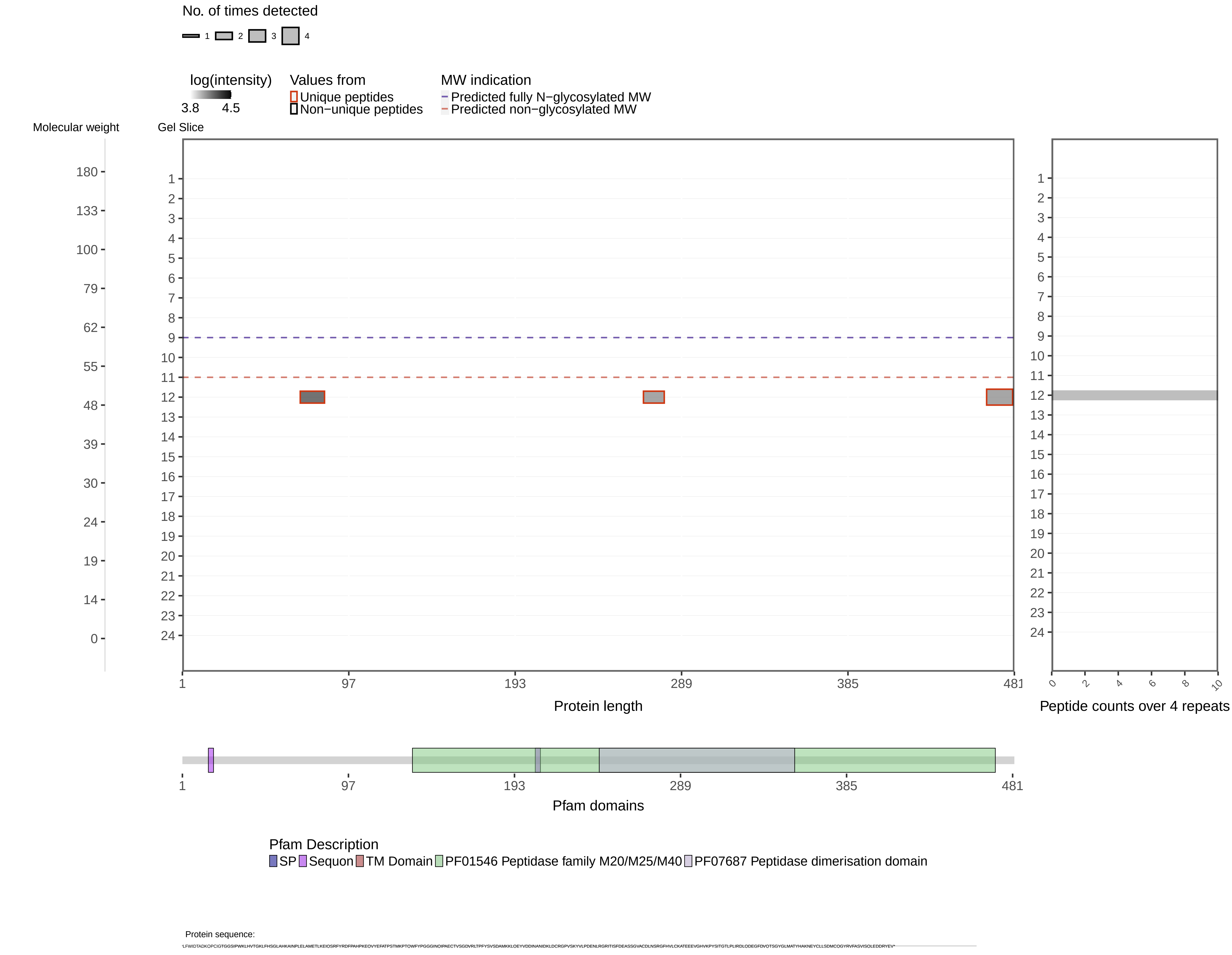

NbL07g12700.1 28_kDa_ribonucleoprotein_chloroplastic [MW = 31.67 kDa]

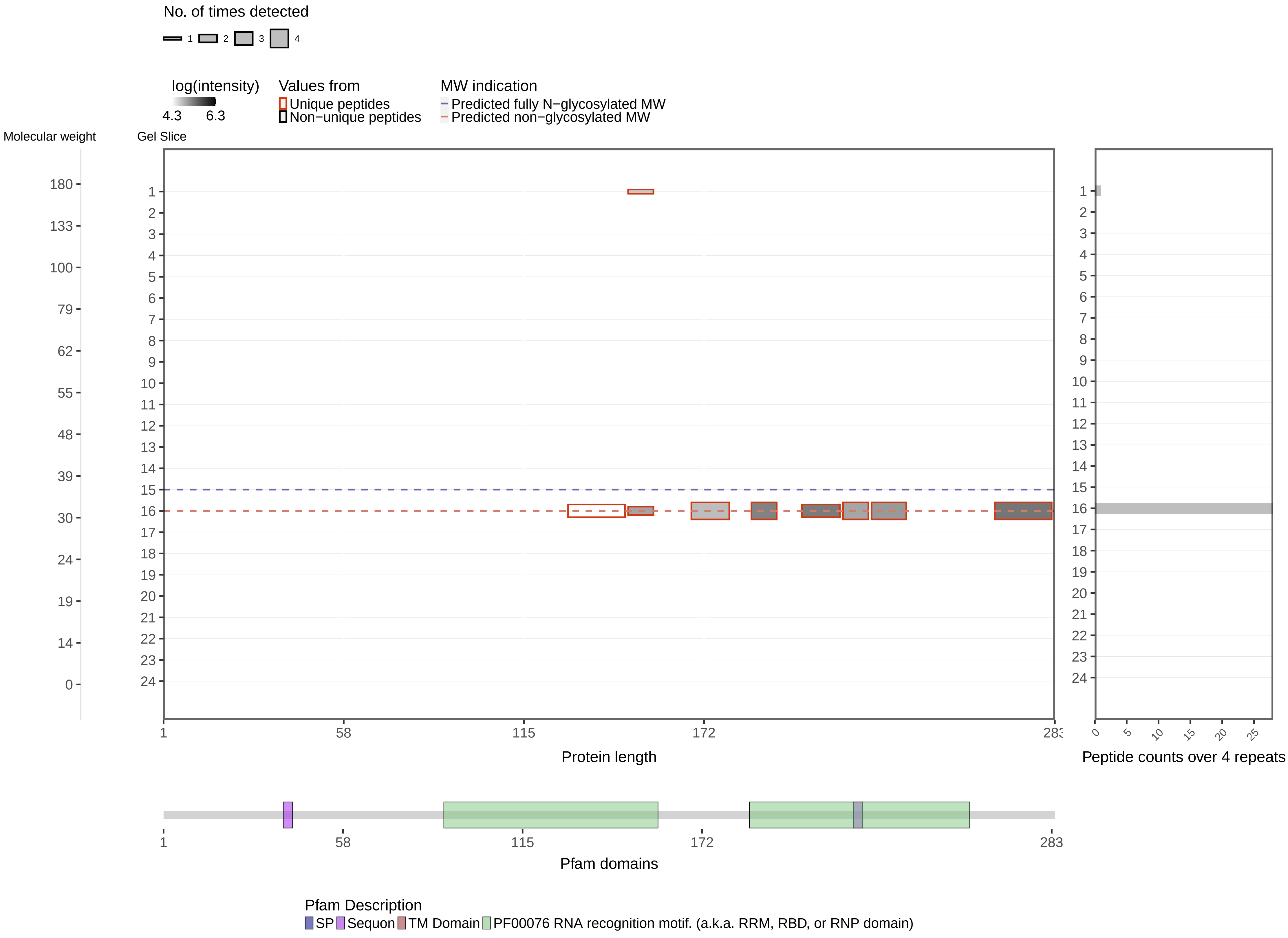


Protein sequence:

AFVTMSSGEEAQAAIDKLDSYTLLGRIIRVEFAKRFKKPAGAPPPPTPPRGETRHKLYVSNLAWKVRSTHLRDFFSTNHNPVSVRVVFDNASGRAAGYGFVSFDTKEEAEAALSAFDGKELMGRPIRLKFSERDADGSENNEEPIQESPEEP*

NbL07g12610.1 plastoglobulin−1_chloroplastic−like [MW = 37.21 kDa]

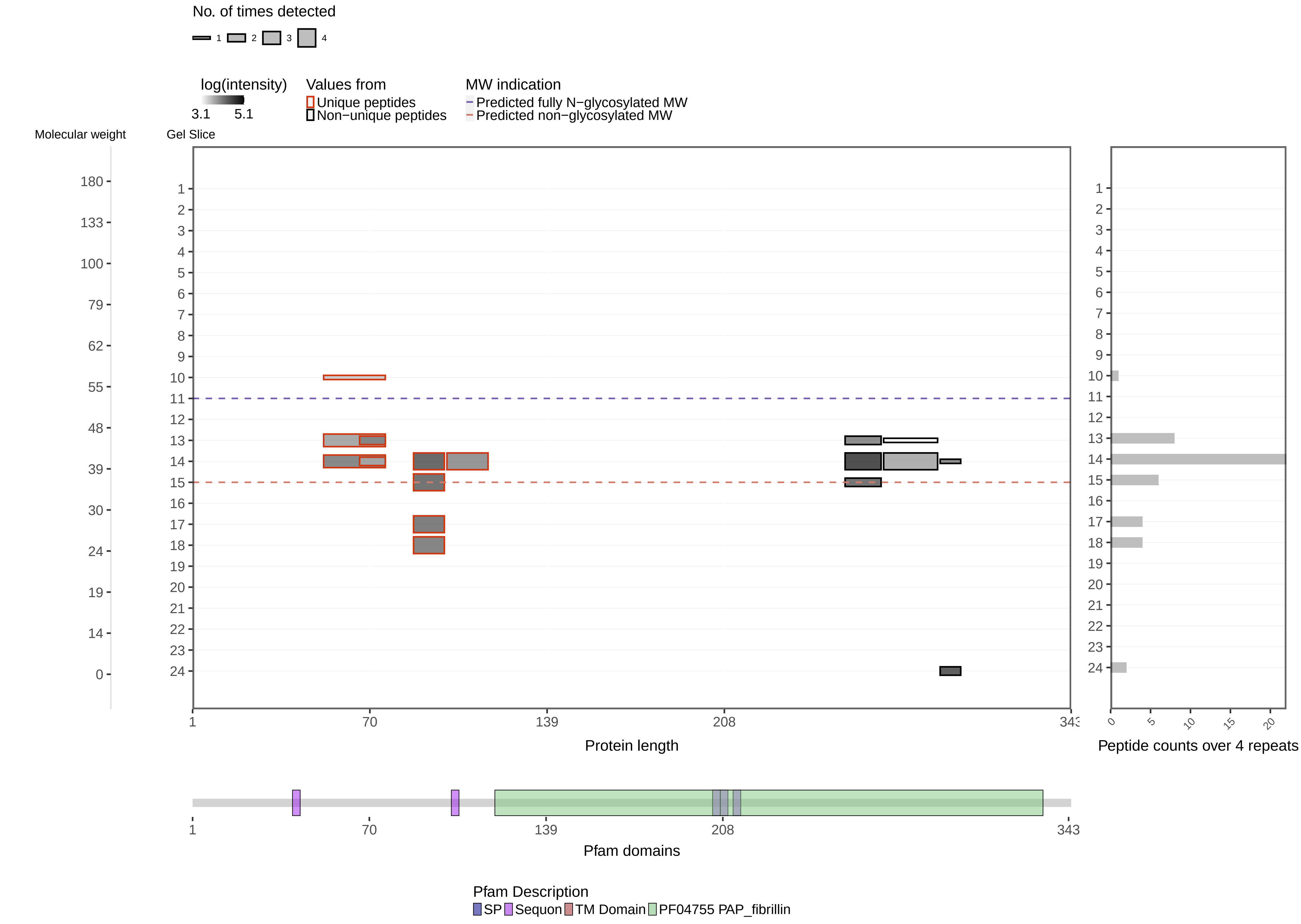


Protein sequence:

ANPTPAPTECPELLDGNWILVFTAFSELLPLLAAGSIPLVKVEKISQGINTSNFTIENSTTLSSPVATLSFSAAATFEVRSPSRIQVEFKEGNFKPPEIKSNIELPENVDVFGQKISLSPVQQSLGPLENAVAGIARTISGQPPLKIPIPGERTKSWLLTTYLDKDLRISKGDGGIFVLVKEGSSLLY*

NbL07g12530.1 nascent_polypeptide−associated_complex_subunit_beta−like [MW = 17.57 kDa]

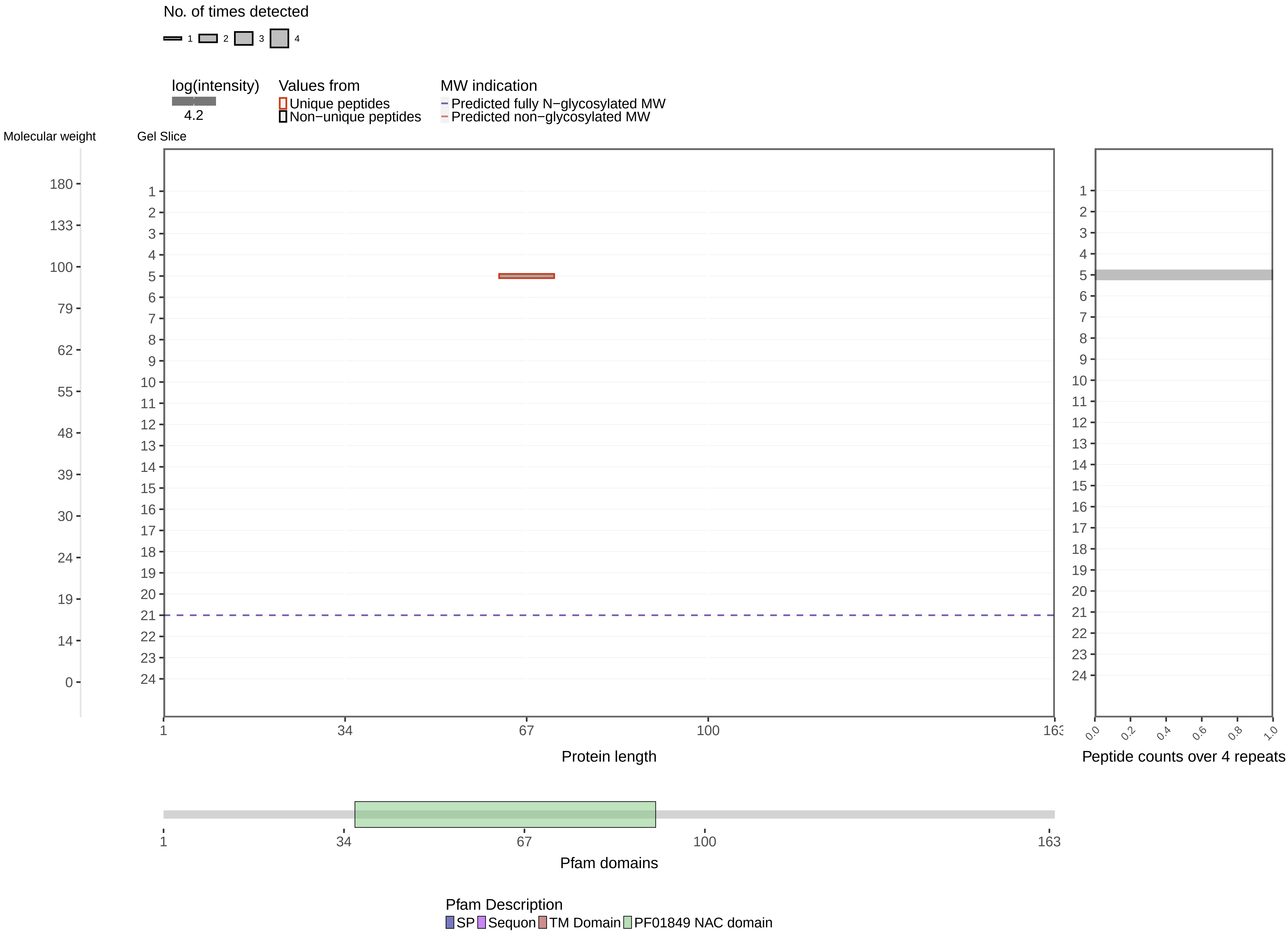


Protein sequence:

NPKVQASIAANTWVVSGSPQTKKLQDILPQIIHQLGPDNLENLKKLAEQFQKQAPTGAPEGGAAAQEDDDDEVPEPVAGQTFEAAAAEEGHTS*

NbL07g12520.1 villin−4−like [MW = 109.25 kDa]

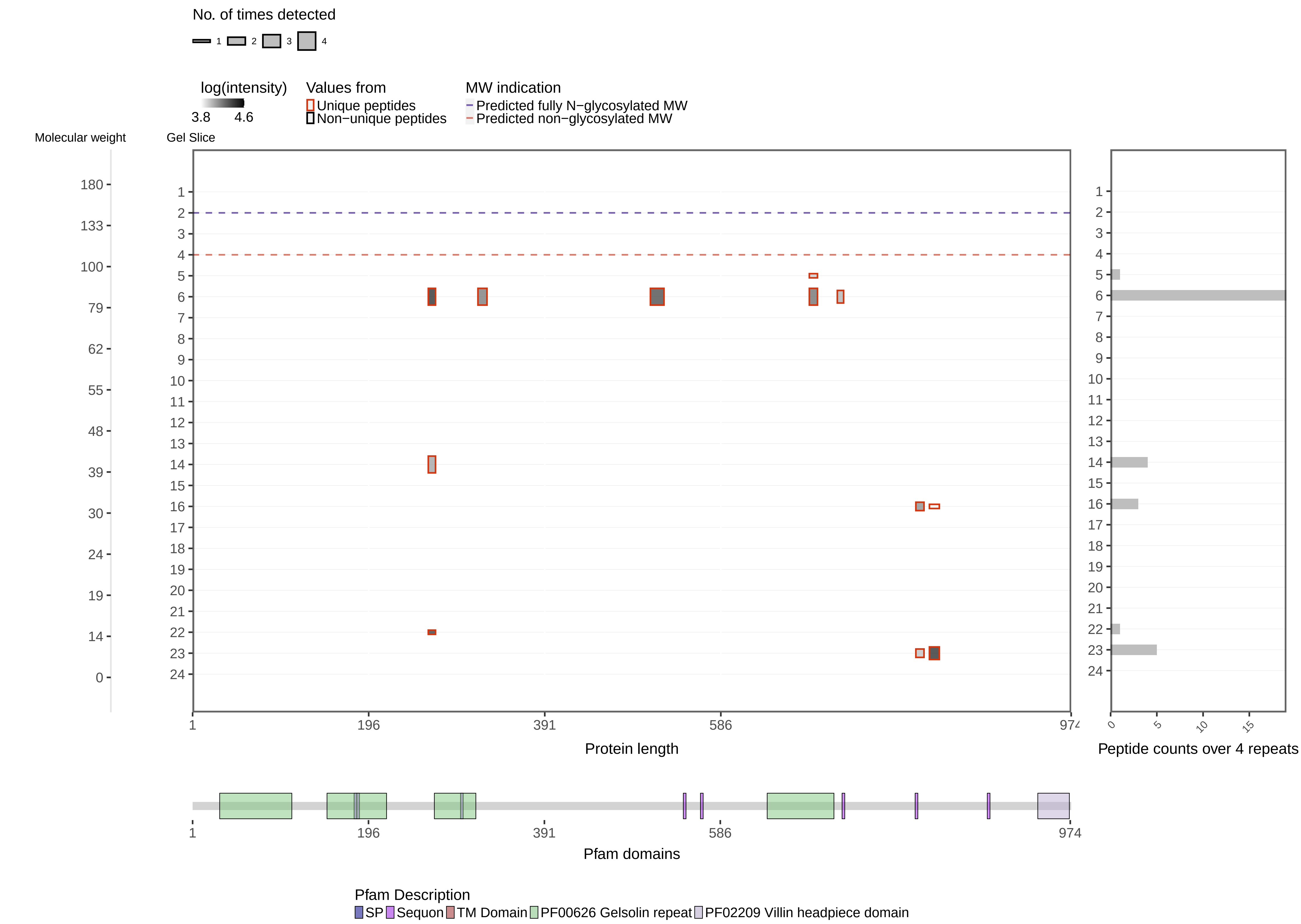


Protein sequence:

RIYEGYEPLQFFVIFQSFIVFKGGLSEGYKKHLVEKELADDTYKEDGIALFRIQGTGPDNMQSIQVEPVASSLNSSYCYILHSGSSVFTWTGNLTTSEDQELVERQLDLIKPDMQSKLQKEGAESEQFWDILSGKSEYPSEKIGRDAESDPHLFSCTFSKGDLKVTEIYNFNQDDLMTEDIFILDCHSDIYVWVGQLVEYKNKMQALSIGEKFLEYDFLMEKLSHQAPIYIVMEGSEPPFFTRHFSWDSNKSAMHGNSFQRKLTLVKNGGPPPMDKPKRRTPVSYGGRSAAPE

### NbL07g12480.1 uncharacterized_LOC109237233 [MW = 39.83 kDa]

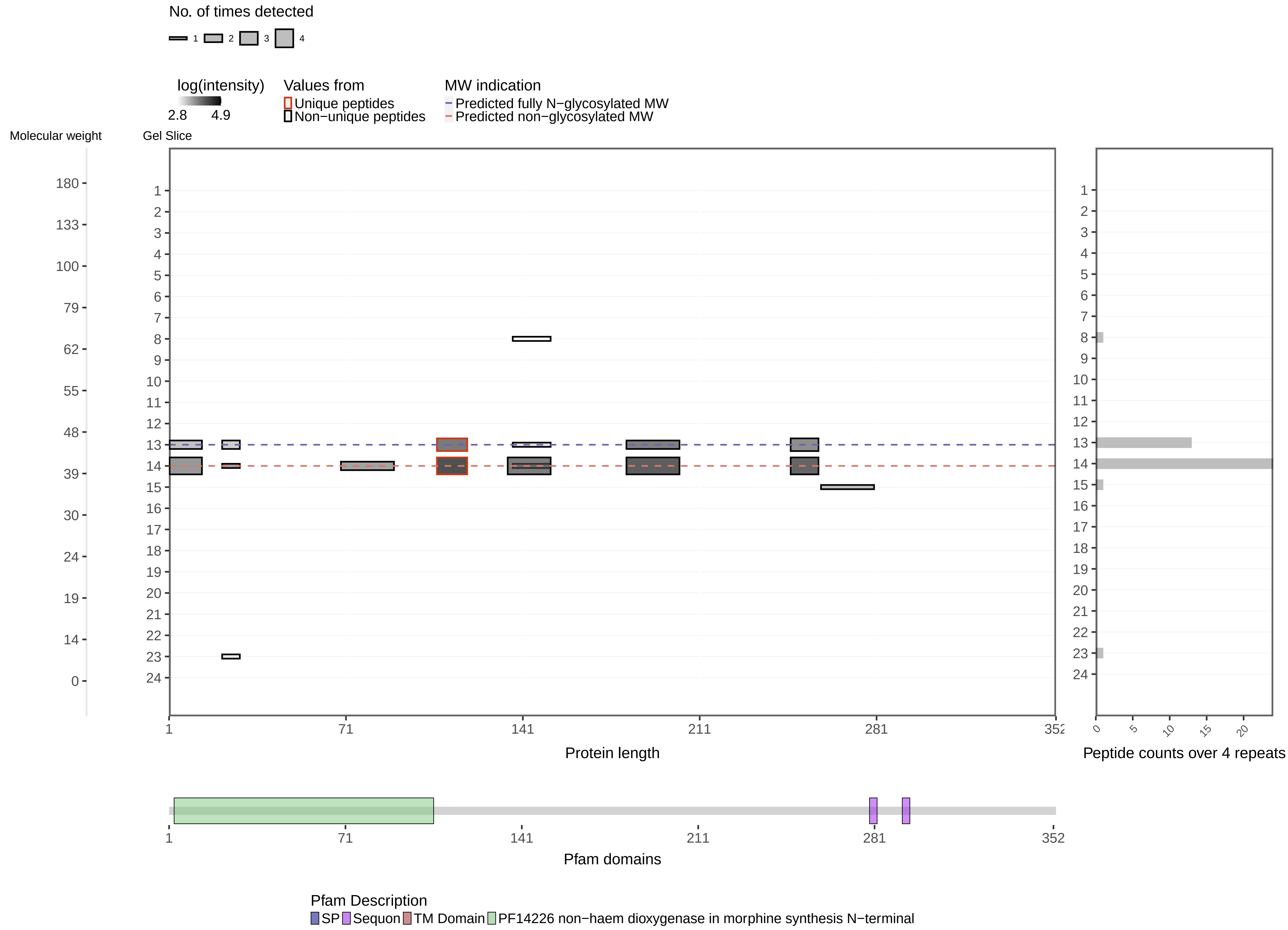


Protein sequence:

KMIAAIECVAEMVAIGFGLSKDAFTSLMQQGPHLLAPTGSDLRRYAKEGTVFAGYHYDLNFLTIHGRSRFPGLSIWLRNGQKMEVKVPVGCLLIQTGKQIEWLTAGECMAGMHEVVVTNRTTDAIRRALEQNRSLWRVSSTLFAHIASDAVLKPLGHFSDTPLASKYPPICAGEFVEQELAIINLKGNNGA*

### NbL07g12460.1 kirola−like [MW = 16.54 kDa]

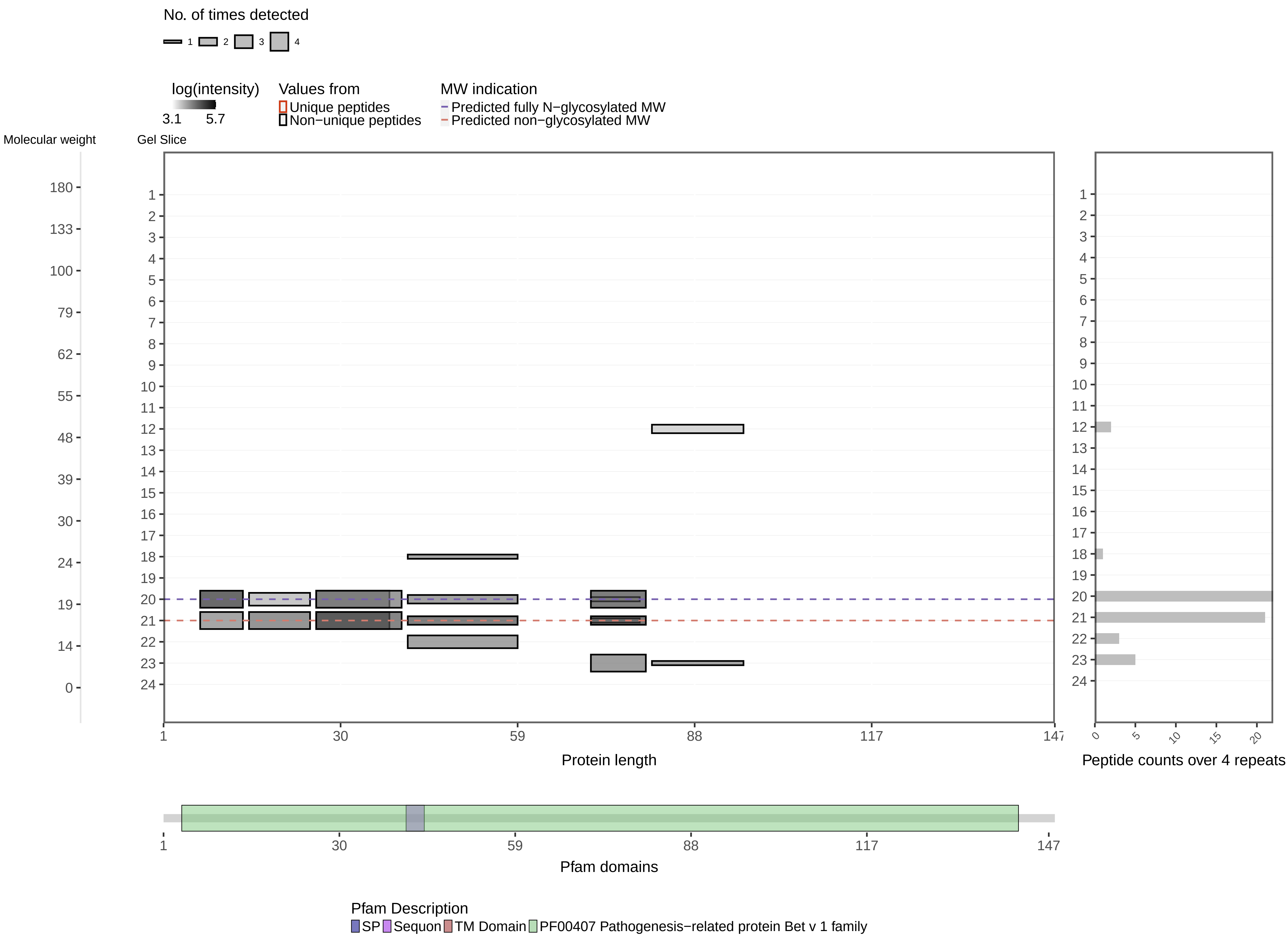


Protein sequence:

YVLGGKERHAKQVLHVDDAKKSITFNFVEGYMNELYKSITATLTAEGNWMTWTLVYEKLNENTPEPLDLLEFAICLLKDIEPHHVGK*

### NbL07g12170.1 14−3−3_protein_9−like [MW = 30.04 kDa]

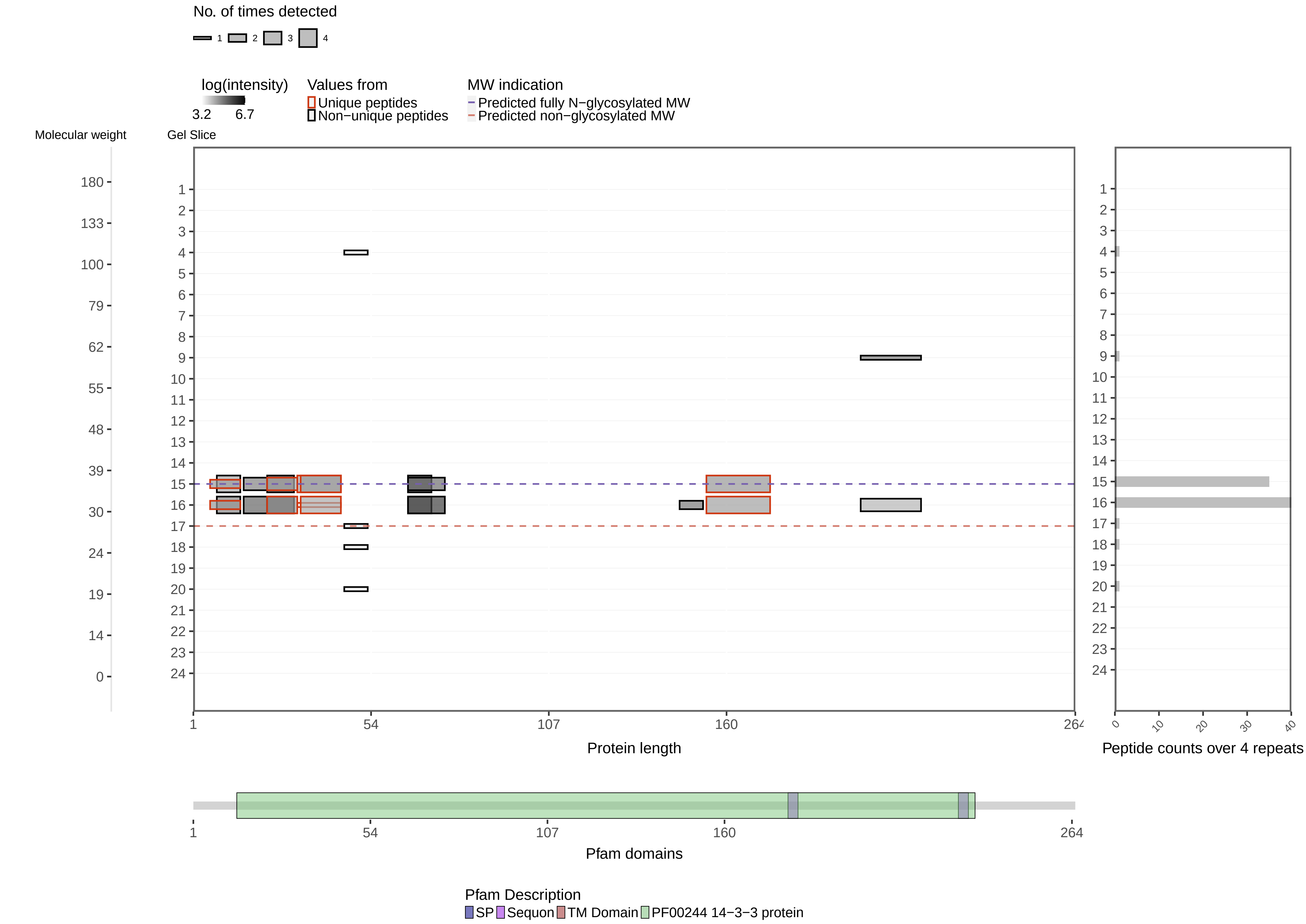


Protein sequence:

GESSVFYYKMKGDYYRYLAEFKTGTDKKEVSDLSLKAYQSATTTAEAELPPTHPIRLGLALNFSVFYYEILNSPERACHLAKQAFDEAISELDSLNEDSYKDSTLIMQLLRDNLTLWTSDIPEDSGIREVIFSICLDVFVYSTTCG*

NbL07g12160.1 uncharacterized_LOC109215312 [MW = 41.09 kDa]

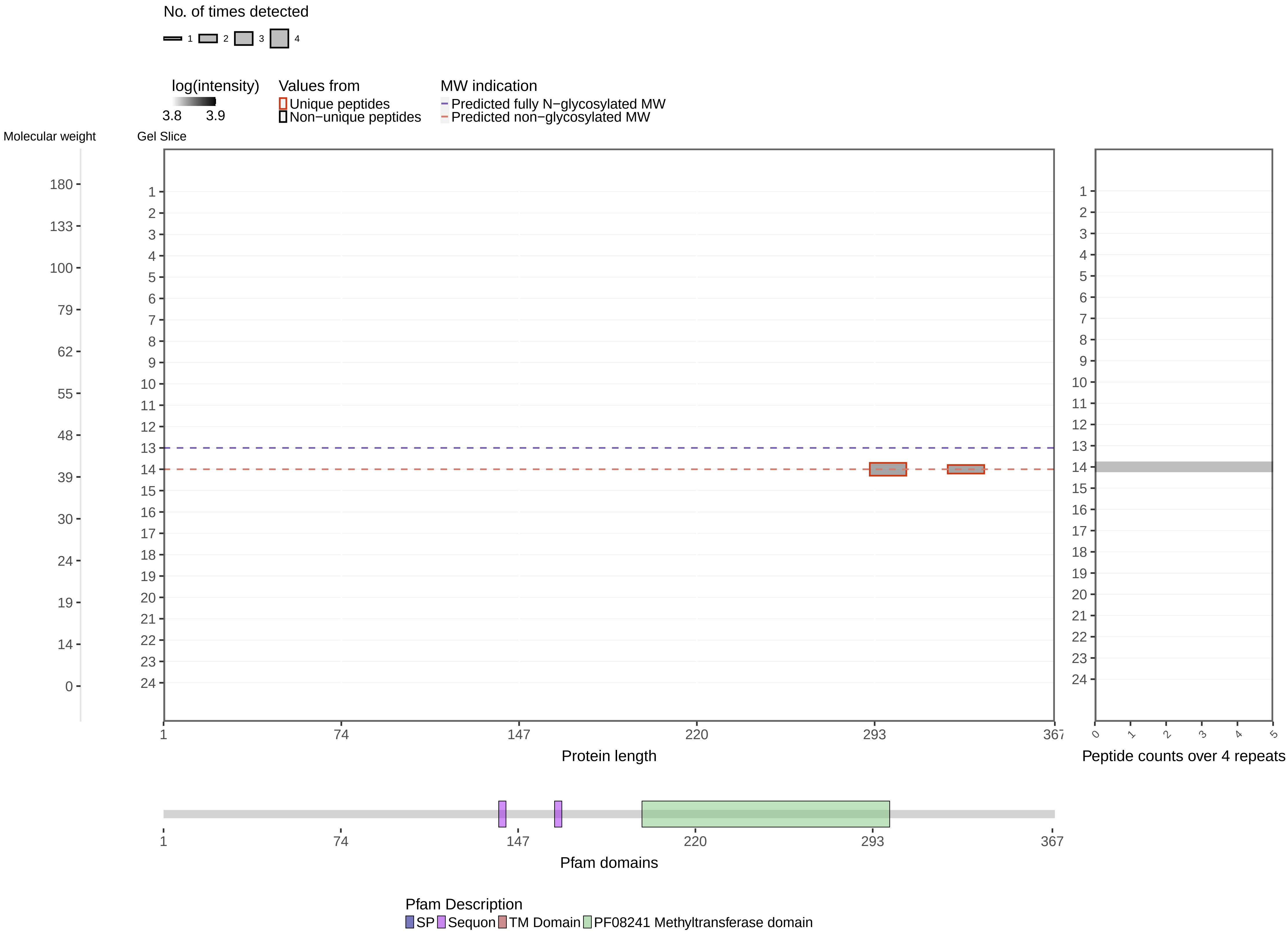


Protein sequence:

IQIVRGNWLDAIEKHHQQYSGDFSIRDILDIGCSVGVSTGYLADRYPSAKVTGLDLSPYFLAVAQYKEKKRNQRKNPISWIHANGENTGLPSKSFDLVSIAYVFHECPERAIRNLVKESFRILRPGGTVAITDNSPKSKILQELPPVLFTLMKSTEPFLDEYYLTDLERVMKETGFANVQTVLTDPRHRTVTATVPY*

### NbL07g12040.1 cytochrome_b−c1_complex_subunit_9−like [MW = 11.68 kDa]

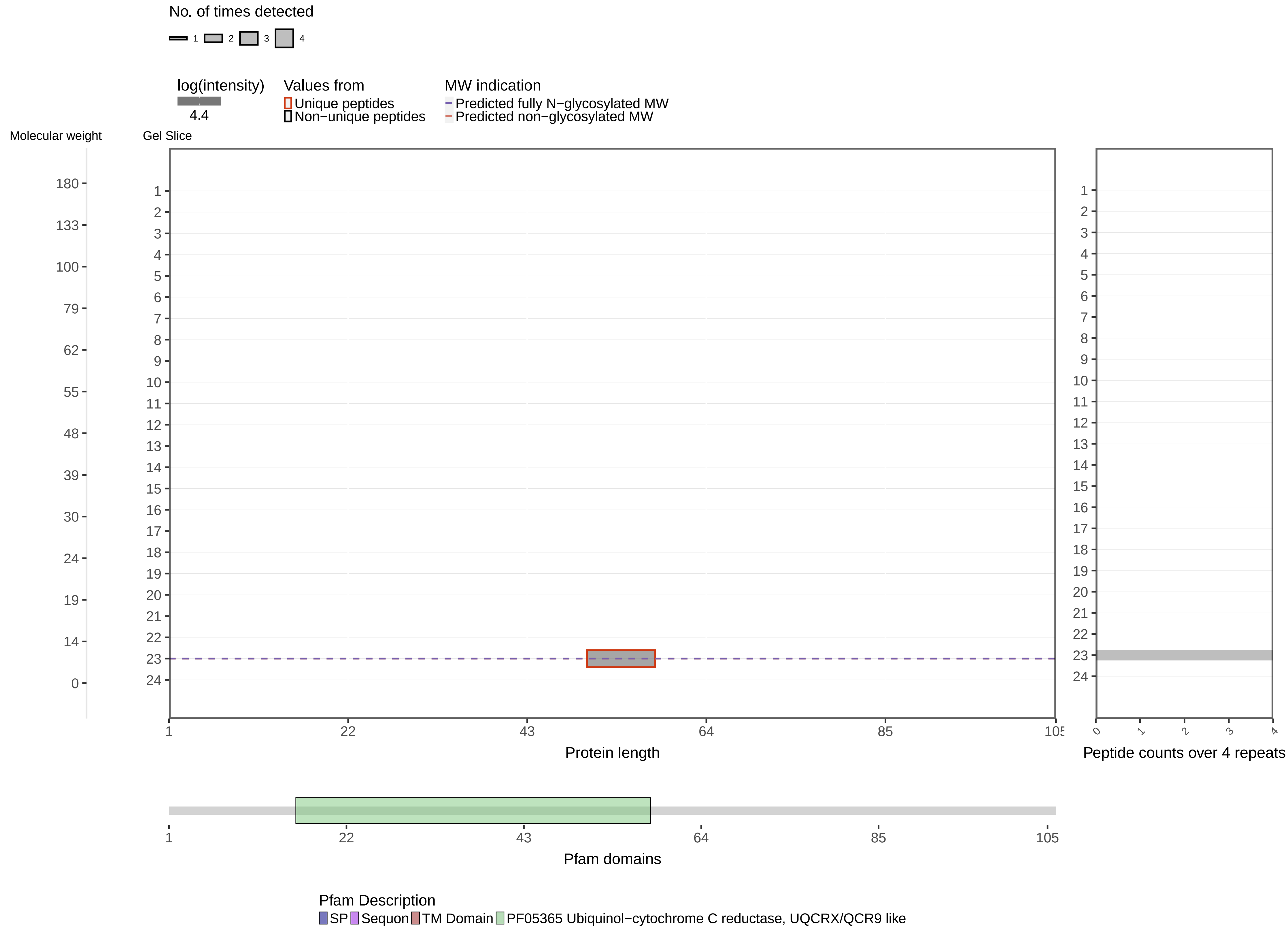


Protein sequence:

GERAVDYGVHRLWEMNNVGKTGKGSLQLVLPYVSPFSKSFWHLSSISLDYLPLLLYKLWGFCLILA*

### NbL07g12010.1 glucan_endo−13−beta−glucosidase_14−like [MW − SP = 39.55 kDa]

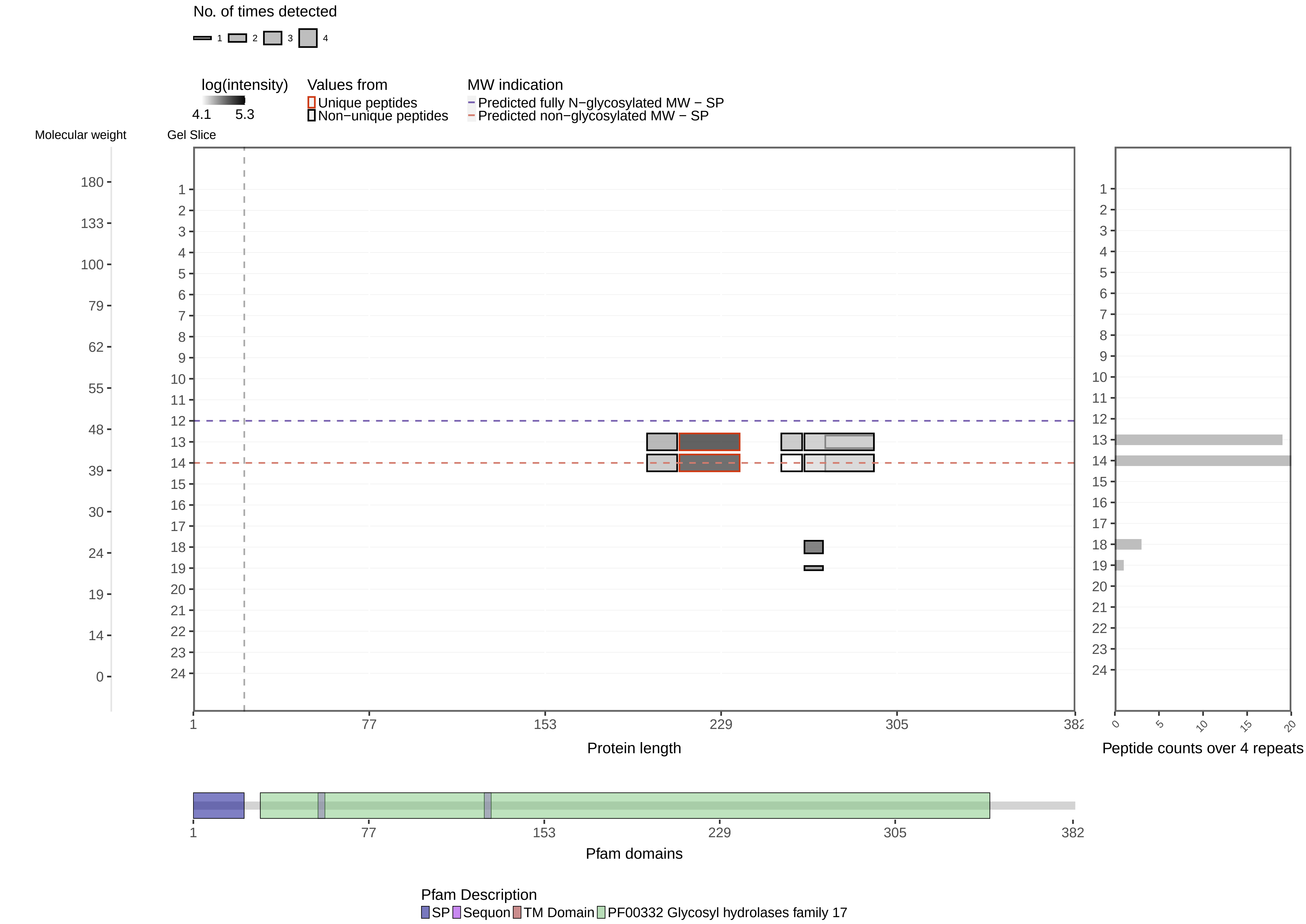


Protein sequence:

RQDLAEYIHGILNFHAQTKSPFLINAYPFFAYKDSPDQVPLDYVLFRPNAGTLDPITNLKYDNMLYAQIDAVHSAIKAMGYSDIQVKVSETGWPSKGDPNEFGATPENAAQYNGNLFRRIQQNQGTPAKPSEPIDIYVFALFNENLKPGPASERNYGLYYPDGTPVYNIGLQGFLPRMDYSAAKKNALSISAILLLILVFLIQF*

### NbL07g11910.1 N−carbamoylputrescine_amidase [MW = 33.53 kDa]

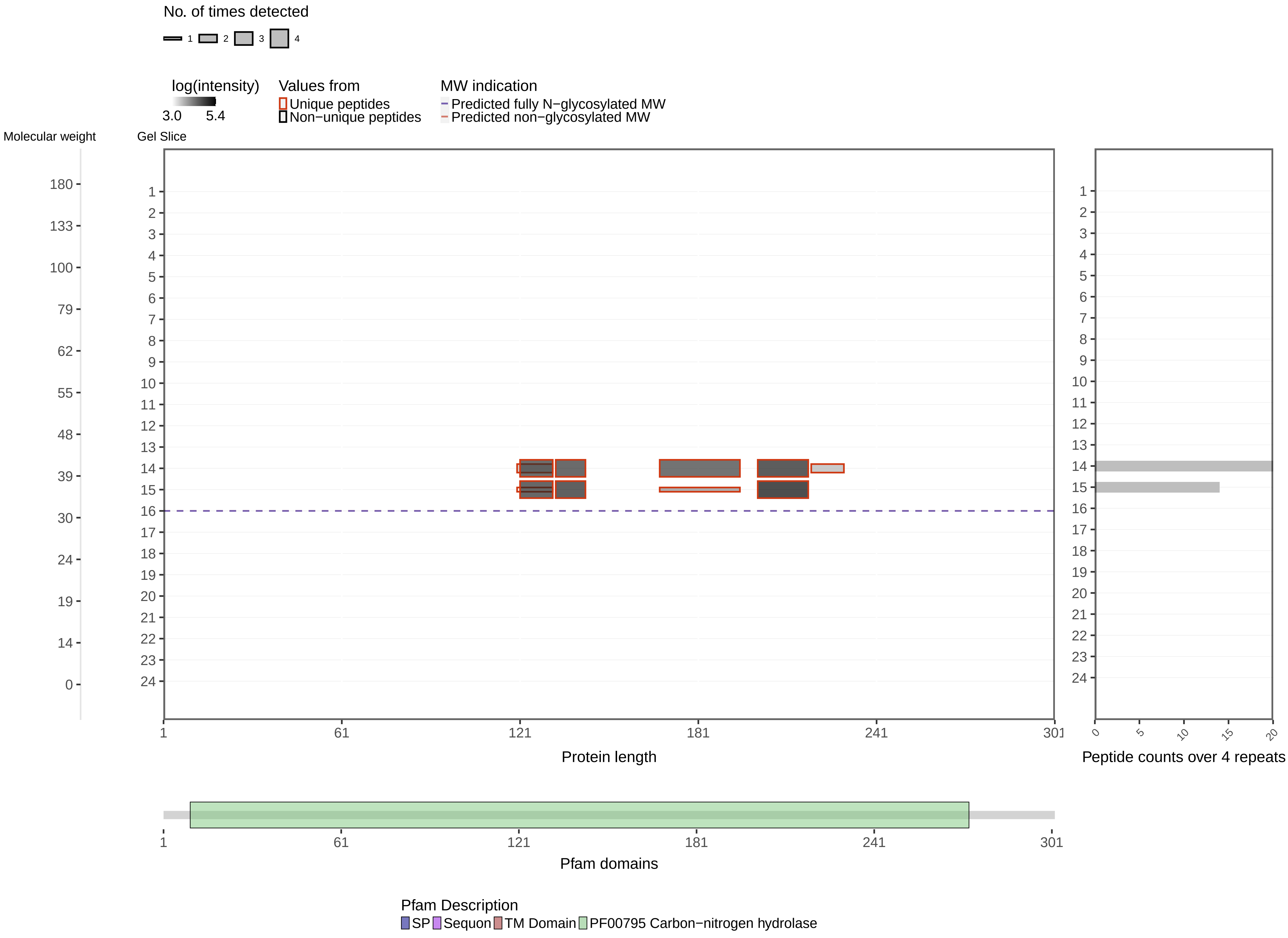


Protein sequence:

[illegible]TGFKVFQTKYAKIGVAICWDQWFPEAARAMALQGAEVLFYPTAIGSEPQDDGLDSRDHWRRVMQGHAGANVVPLVASNRIGKETIETEHGKSAITFYGYSFIAGPTGELVAAADDKEEVVLVAQFDLDKIKSKRQSWGVFRDRRPDLYKVLMTLDGSNPVN*

### NbL07g11830.1 long_chain_acyl−CoA_synthetase_4 [MW = 51.08 kDa]

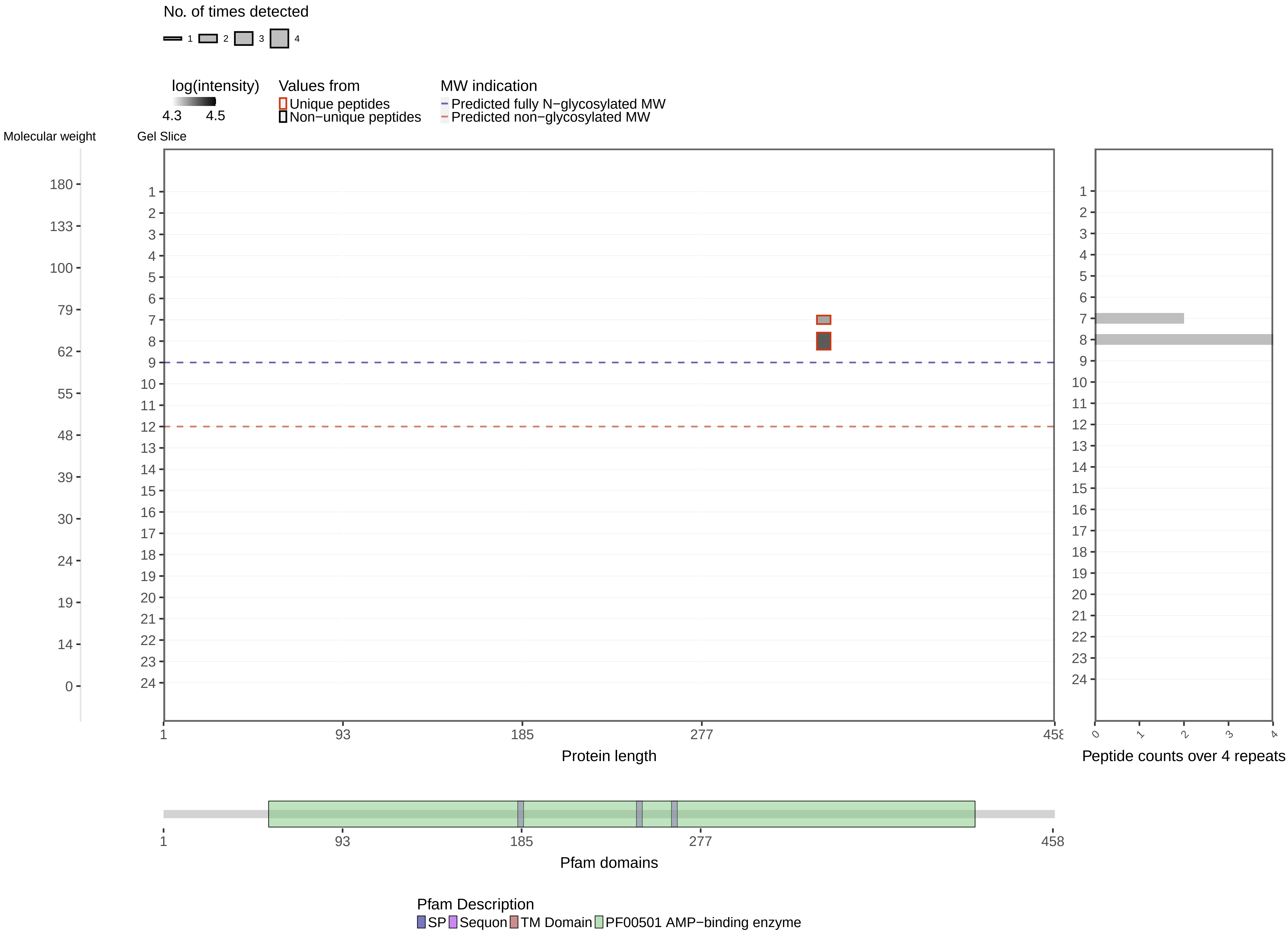


Protein sequence:

OFDLPVKKKEDICTIMYTSGTTGDPKGVLISNNSIVTLIAGVORLLGSVNESLTVDDVYLSYLPLAHIFDRVIEECFINHGASIGFWRGDVKLLTEDIGELKPTIFCAVPRVLDRIYSGLOOKISSGGLLKSTLFNLAYAYKHHNLKKGRKHFEASPLSDKVVFSKVKEGLGGKVRLILSGAAPLAAHVEAFLRVVACCHVLOGYGIPFFONIMSLAEIWMKKFLAFSFYIKHRWYNVKPSRCRT*

### NbL07g11700.1 nitrilase/nitrile_hydratase_NIT4B [MW = 37.67 kDa]

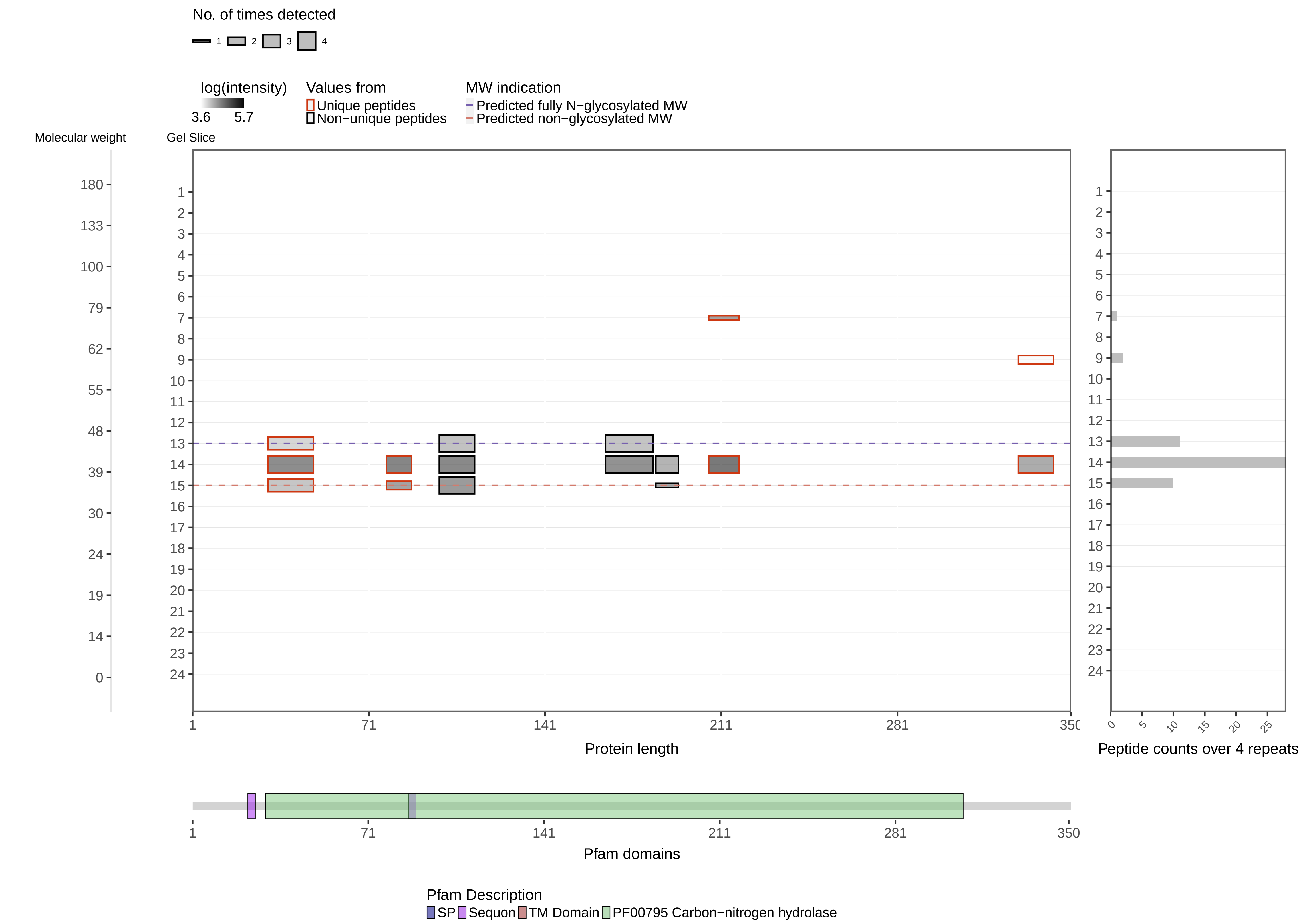


Protein sequence:

ALERIIWGFGDGSTIPVYDTPLGKIGAAICWENRMPLLRTAMYAKGIEIYCAPTADARDVWOASMTHIALEGGCFVLSANOFCRRKDYPPPPEYVFSGTEEDLTPDSIVCAGGSVIISPSGAVLAGPNYEGEALISADLDLGEIARAKFDFDVVGHYARPEVLSLIVRDHVVSPVSFTSTSSKAESSPK*

### NbL07g11610.1 uridine_5'−monophosphate_synthase [MW = 51.56 kDa]

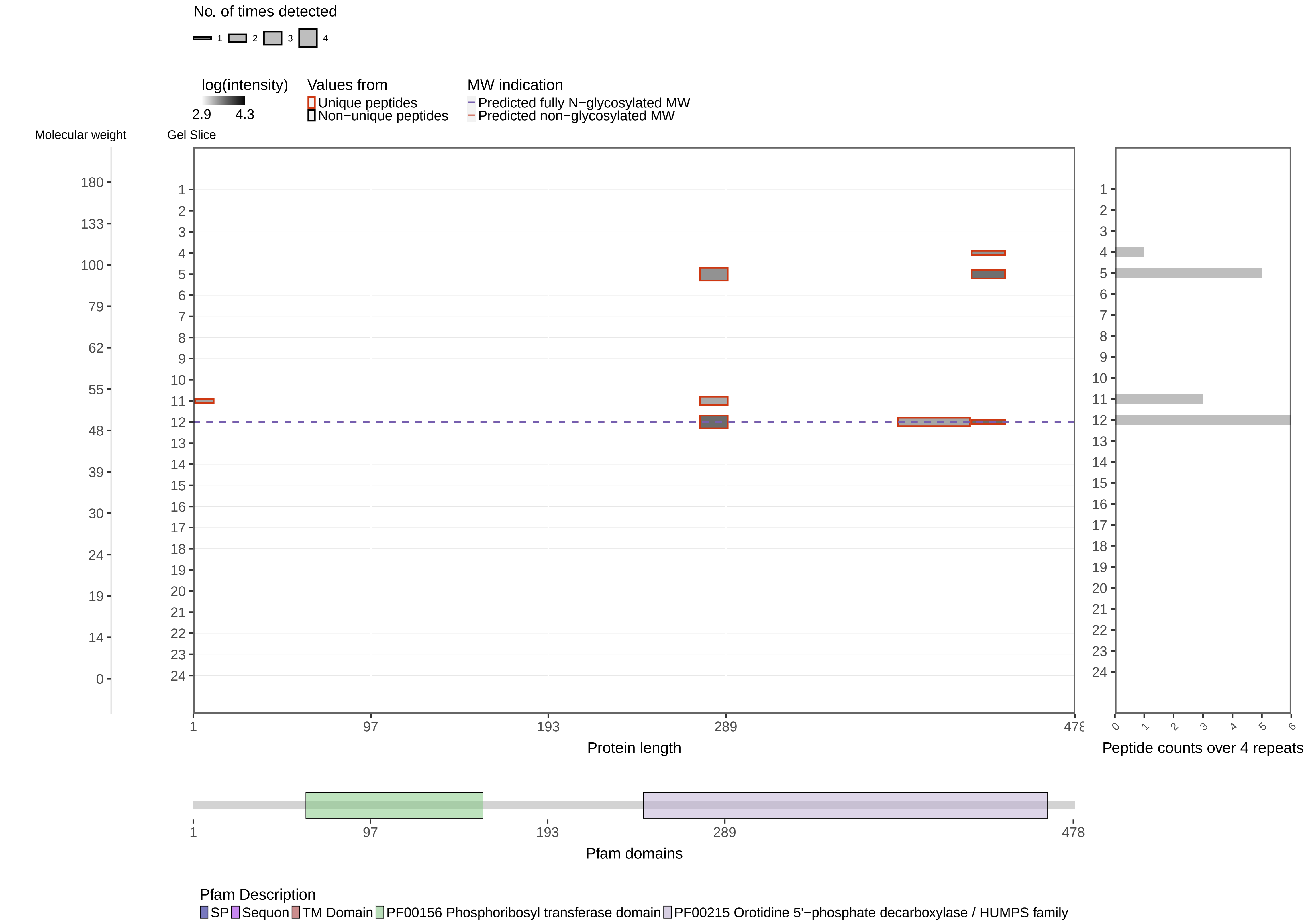


Protein sequence:

KIAKNPTGKKLFEIMVQKESNLCLSADVATAAELLDIADKVGPEICMLKTHVDILPDFAPDFGSKLRSIADKHNFLIFEDRKFADIGNTVTMQYEGGIFRILDWADITNAHIISGPGIVDGLKLKGLSRGRGLLLLAEMSSAGNLATGAYTAAAVKIAEDHSDFVIGFISVDPASWPNGPGNPSLIHATPGVQLVKGGDALGQHYNTPSAVIADRGSDIIIVGRGIIKAANPMEAAREYRLQGWDAYLVNCK*

### NbL07g11020.1 beta−glucosidase_BoGH3B−like [MW = 26.94 kDa]

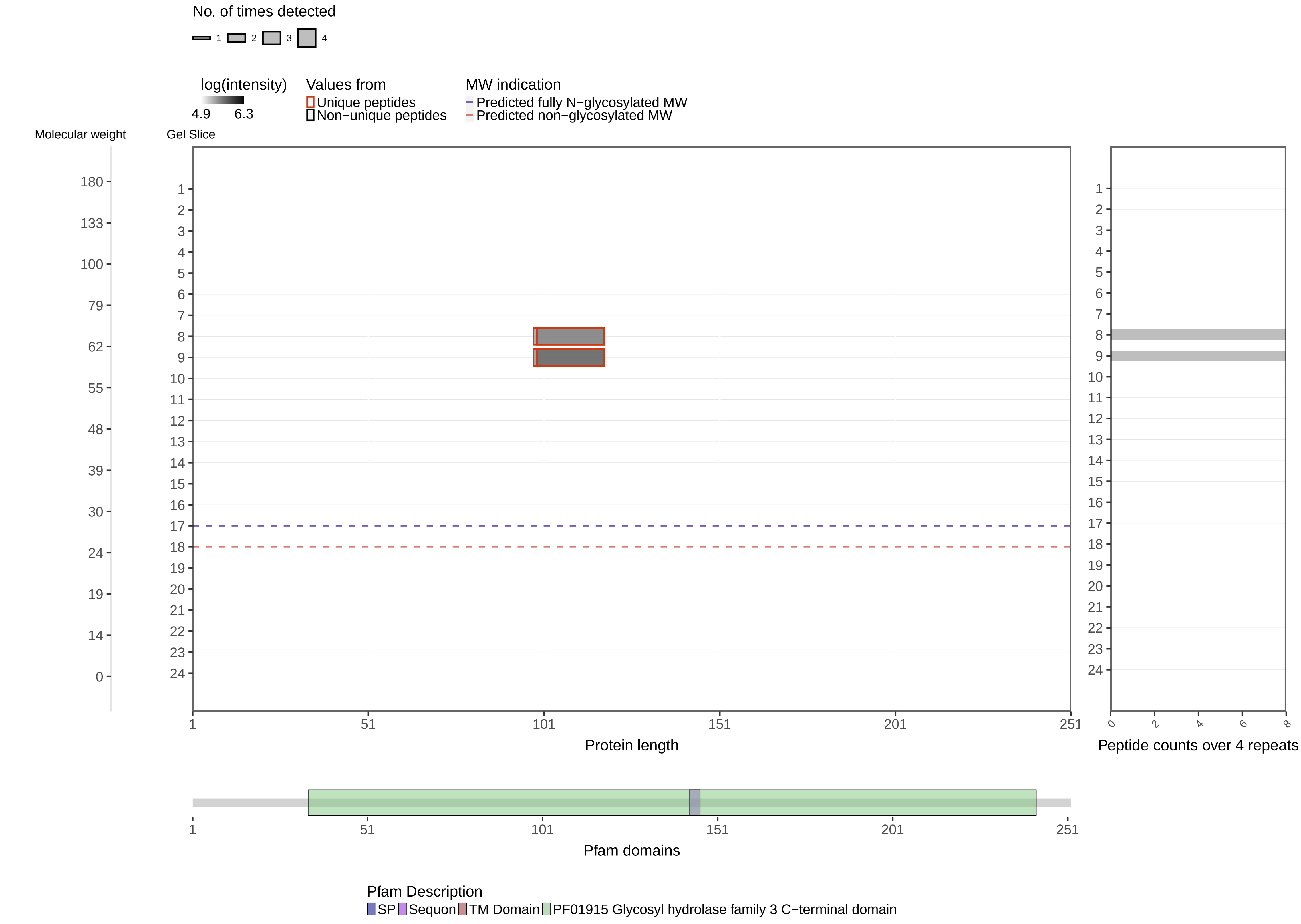


Protein sequence:

DENFVKSNKFSYAIVVVGEVPYAEMFGDSSNLTIAEPGPSTISHVCGAVKCVVVVVSGRPVVLEPYVSKMDALVAAWLPGTEGQGVADALFGDYGFTGKLARTWFKRVDQLPMSFDDAHVDPLFPFGFGITTKPVKGY*

### NbL07g10990.1 abscisic−aldehyde_oxidase−like [MW = 148.32 kDa]

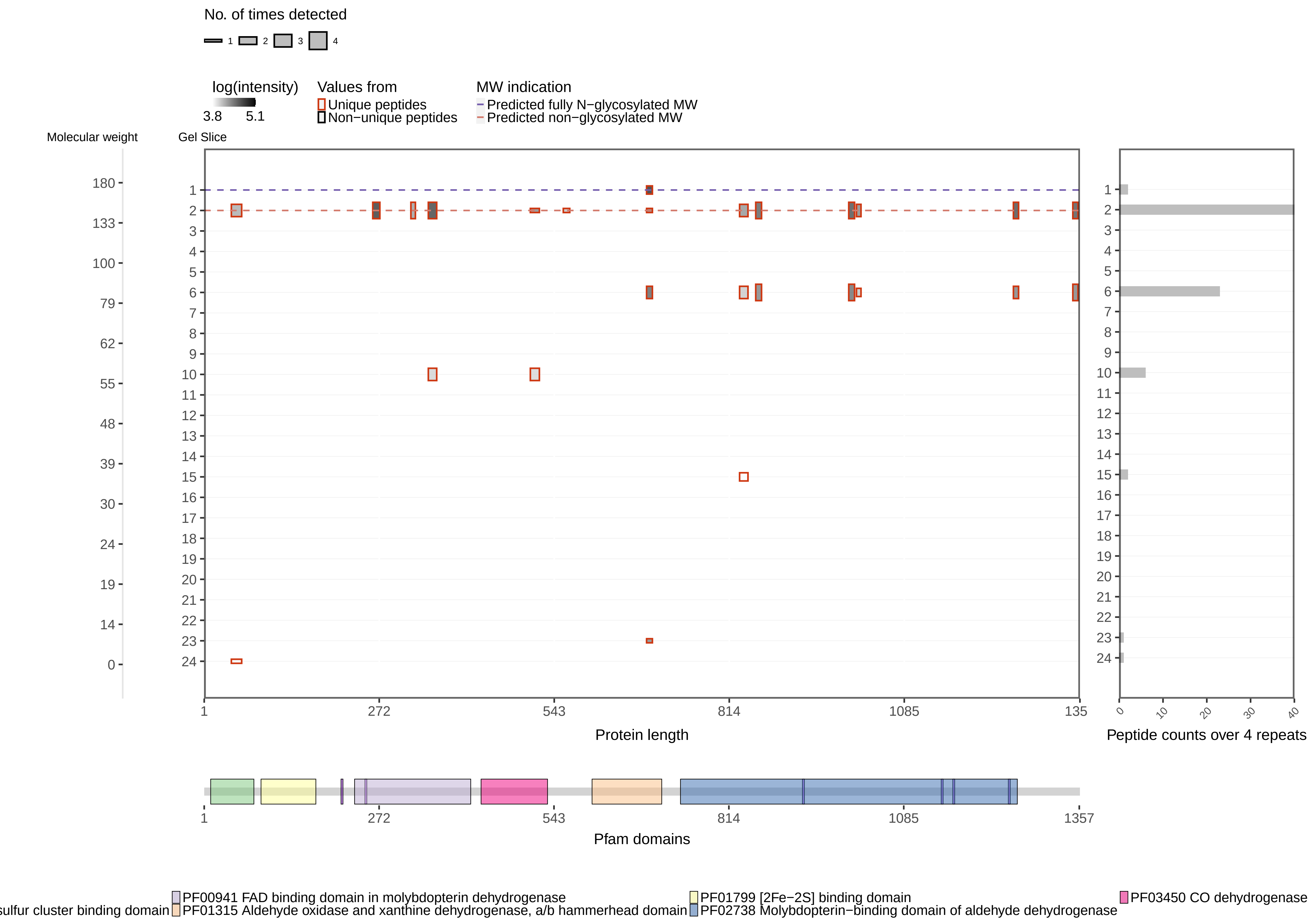


Protein sequence:

NbL07g10910.1 3−methyl−2−oxobutanoate_hydroxymethyltransferase_1_mitochondrial−like [MW = 37.84 kDa]

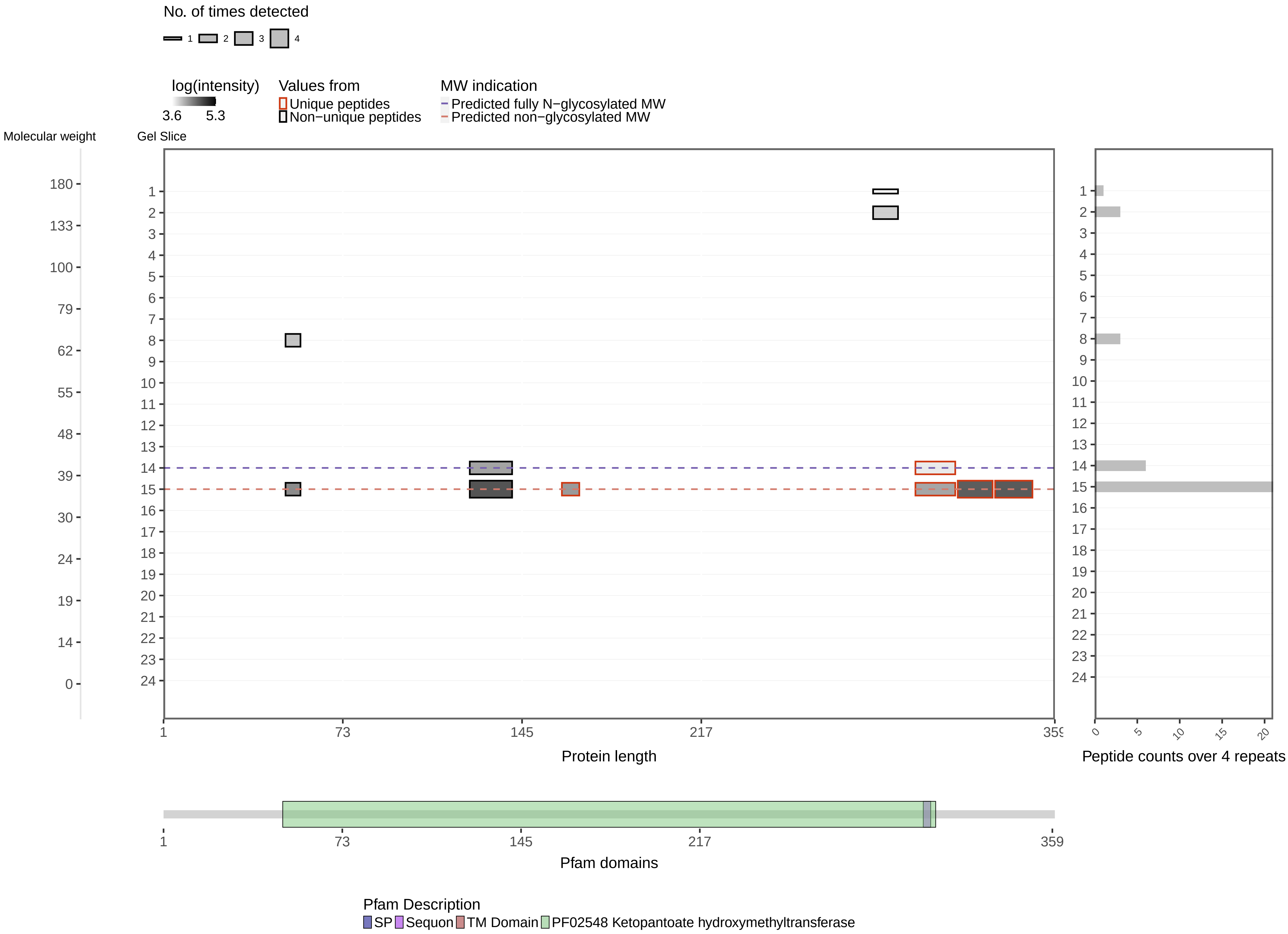


Protein sequence:

SRITAAKAIVEAGIAVIGHVGLTPOAISVLGGFRPOGRNVDSAVKVVETAMALOEVGCFSVVLECVPAPVAAAATSALOIPTIGIGAGPFCSGOVLVYHDLLGMMOHPHHAKVTPKFCKOFGOVGDVINKALVOYKEEVTNGSFPSSAHSPYKIGAADMDGFFSELOKLGFSDAASAAAAAAEKIOTANSPA*

NbL07g10830.1 60S_ribosomal_protein_L30−like [MW = 12.43 kDa]

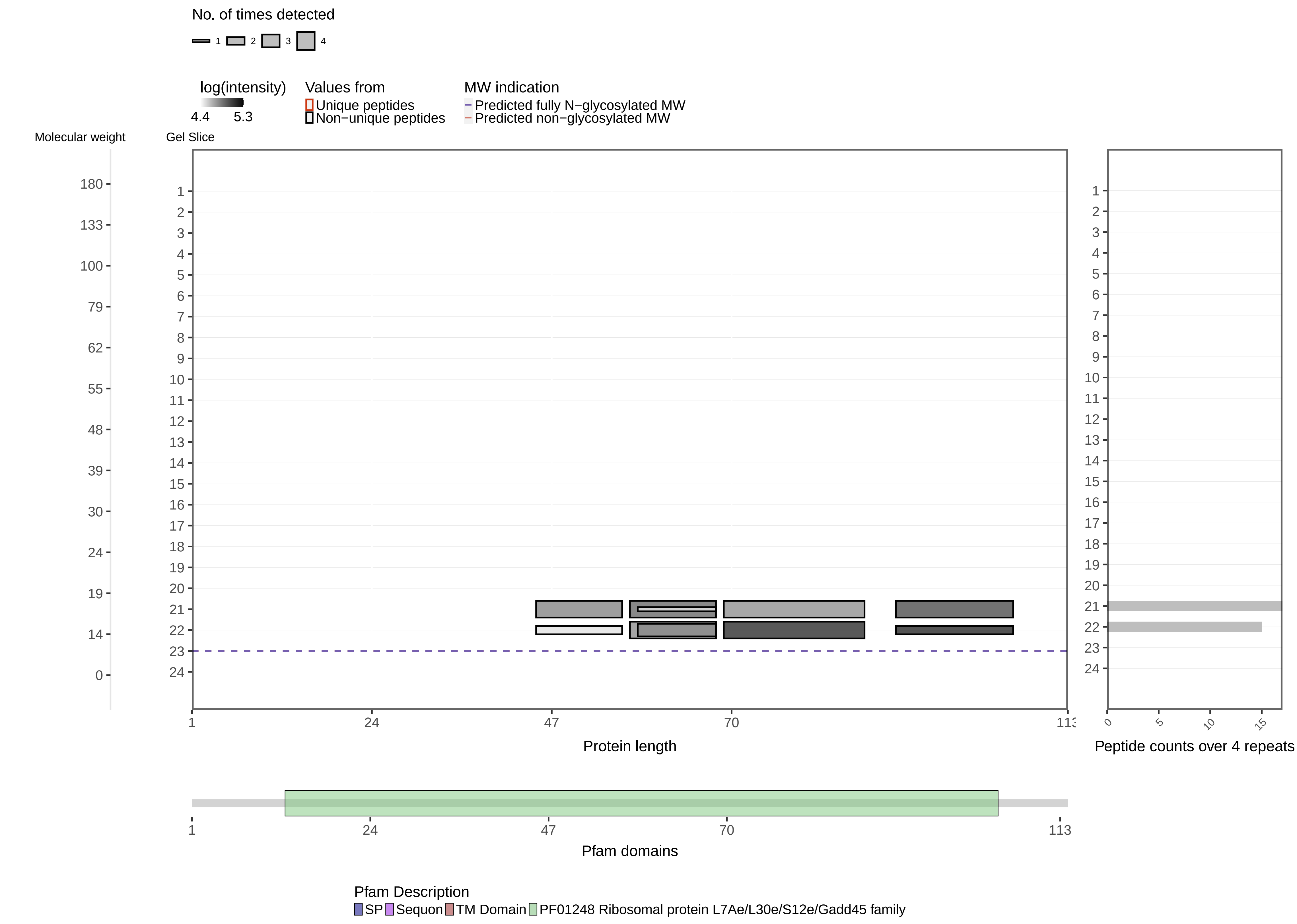


Protein sequence:

GKLIIIANNCPPLRKSEIEYYAMLAKVGVHHYNGNNVDLGTACGKYFRVCCLSIIDPGDSDIIKSMPDDK*

### NbL07g10810.1 agmatine_coumaroyltransferase−2−like [MW = 23.01 kDa]

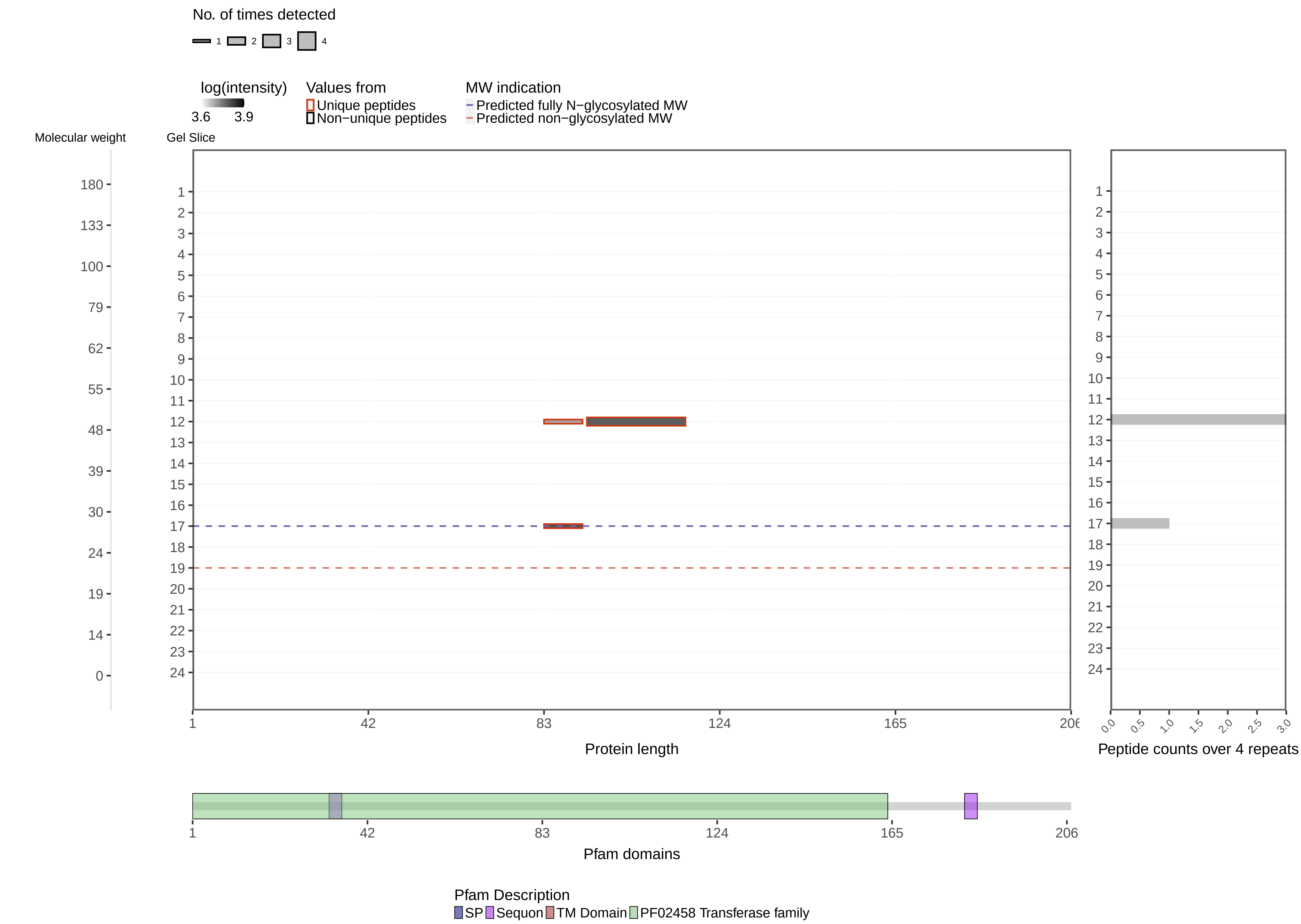


Protein sequence:
/RFIEASVDATLDEVLPLKPSPSLLKLHPSMKDVVELIQVQVTRFTCGSVVVGFTGHHMIADGHAASNFCYRLSVFKATRNWKPVRRRDTKNRTSGGTSNAYDFDELKFGNEITL*

NbL07g10780.1 homoserine_dehydrogenase [MW = 44.38 kDa]

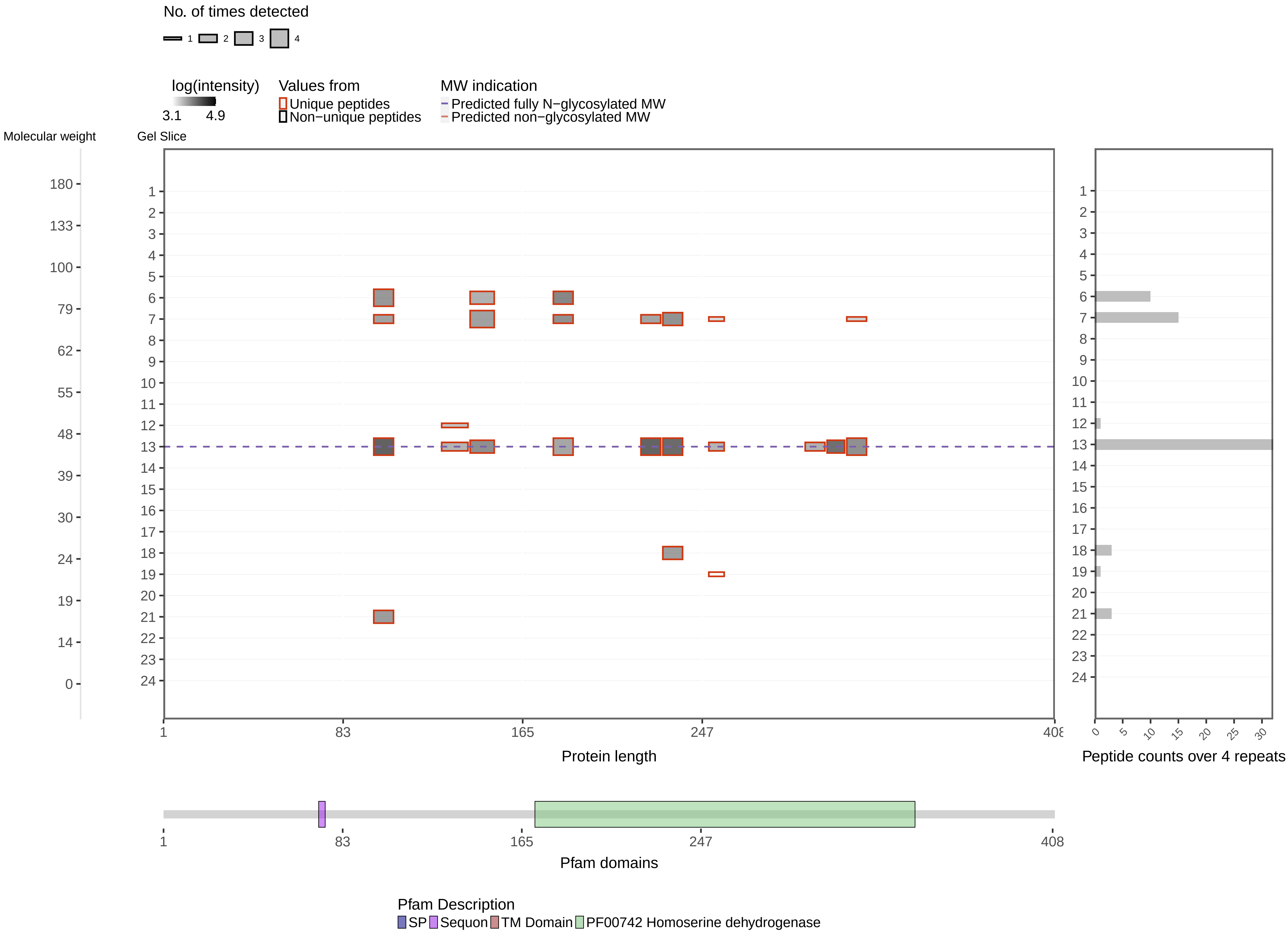


Protein sequence:

SLSGTLGYVMSEVEDGKPFSOVVSAAKSLGFTEPDPRDDLGGMDVARKALILARLLGRHINLDDMKIESLYPEEMGPDVMPLEDFLANGLPLLDKNIEDRIKOASANGNVLRYVCLIDDTRCEVGIOEVPKDSALGRLRGSDNVVTWVSOHAESLFEOLLISVLRLKPIFVITHHLLSPCKTLCSLRWRYTAVVTRNNRWLSKVLEOEMIPROLVS*

### NbL07g10690.1 feruloyl_CoA_ortho−hydroxylase_2−like [MW = 40.28 kDa]

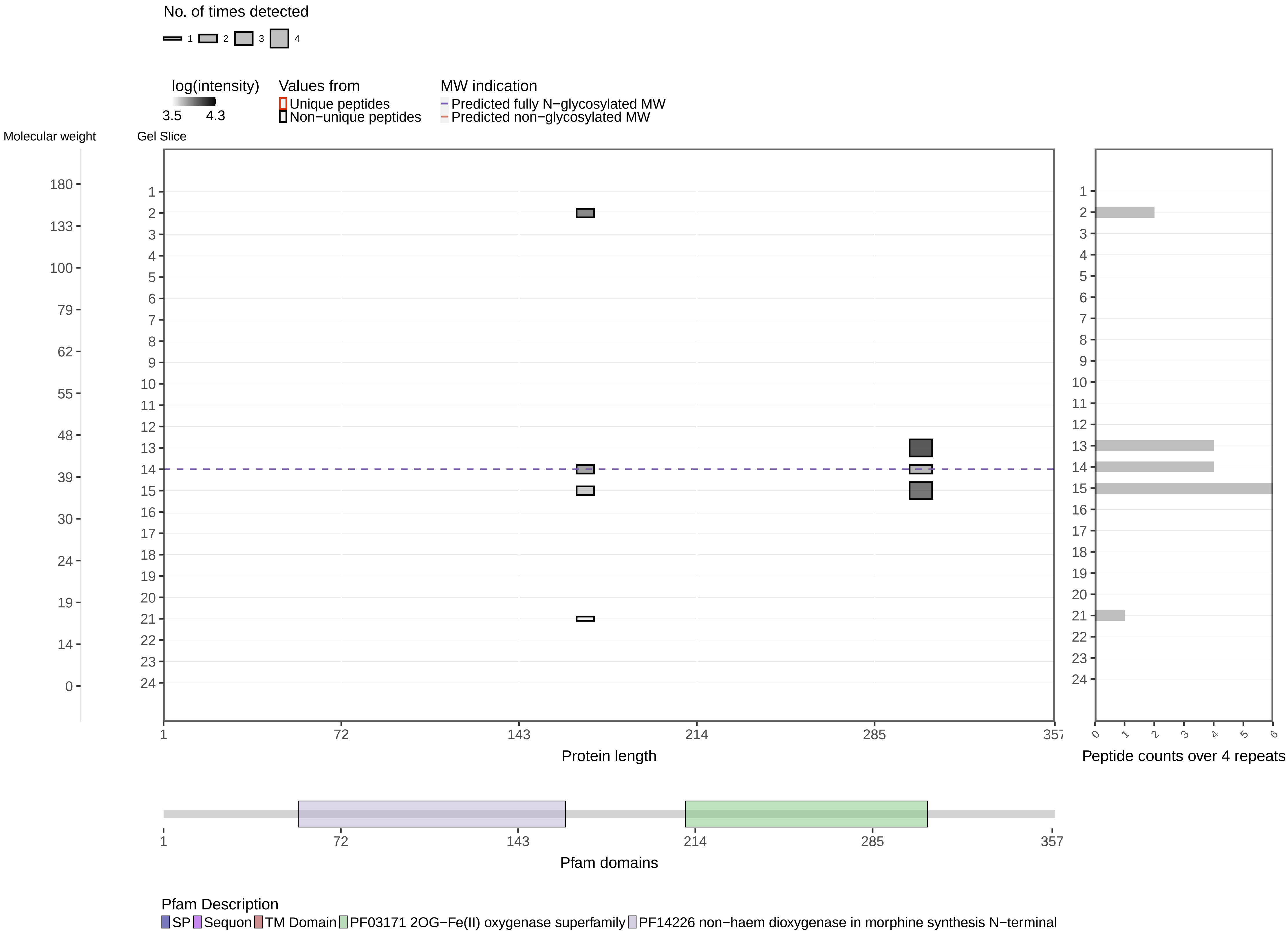


Protein sequence:

RDQVLEYQKWAIPLAKKLLEILLKGLNVNENIDESLKPLLMGTMAINMNYYPPCPNPSLTIGCRRHCDVSCITILLQDDTGGLFVRGTKDNNWIHVTPVKGALAVNIGDSLQIMSNDRYKSVEHCAAVDSSKARISIPFFVNPRLDTVIGPFPQVLENGEKPVYKKVLFSDYWEYFFSKRPSGKASLDFAKI*

NbL07g10600.1 elongation_factor_1−beta−like [MW = 24.81 kDa]

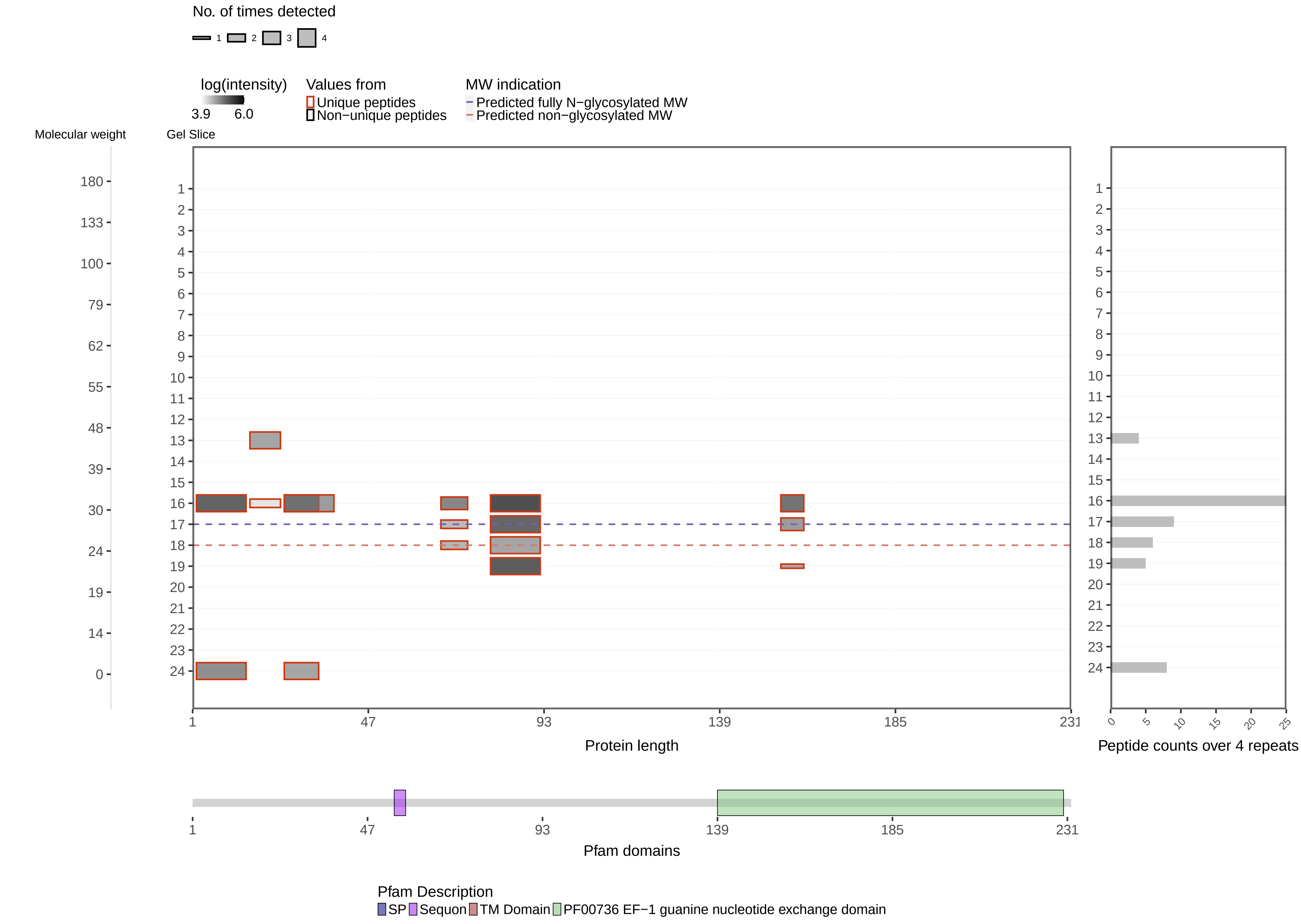


Protein sequence:
DDDDDIDLFGEETEEEKKAAEAREATKASTKKKESGKSSILLDVKPWDDETDMKLLEEAVRSVQQEGLTWGAFSLTAKLVPVGYGIKKLQIMLTIVDDLVSVDTLVEEKLTVEPINEYVQSCDIVAFNKI*

### NbL07g10590.1 calmodulin−7−like [MW = 20.77 kDa]

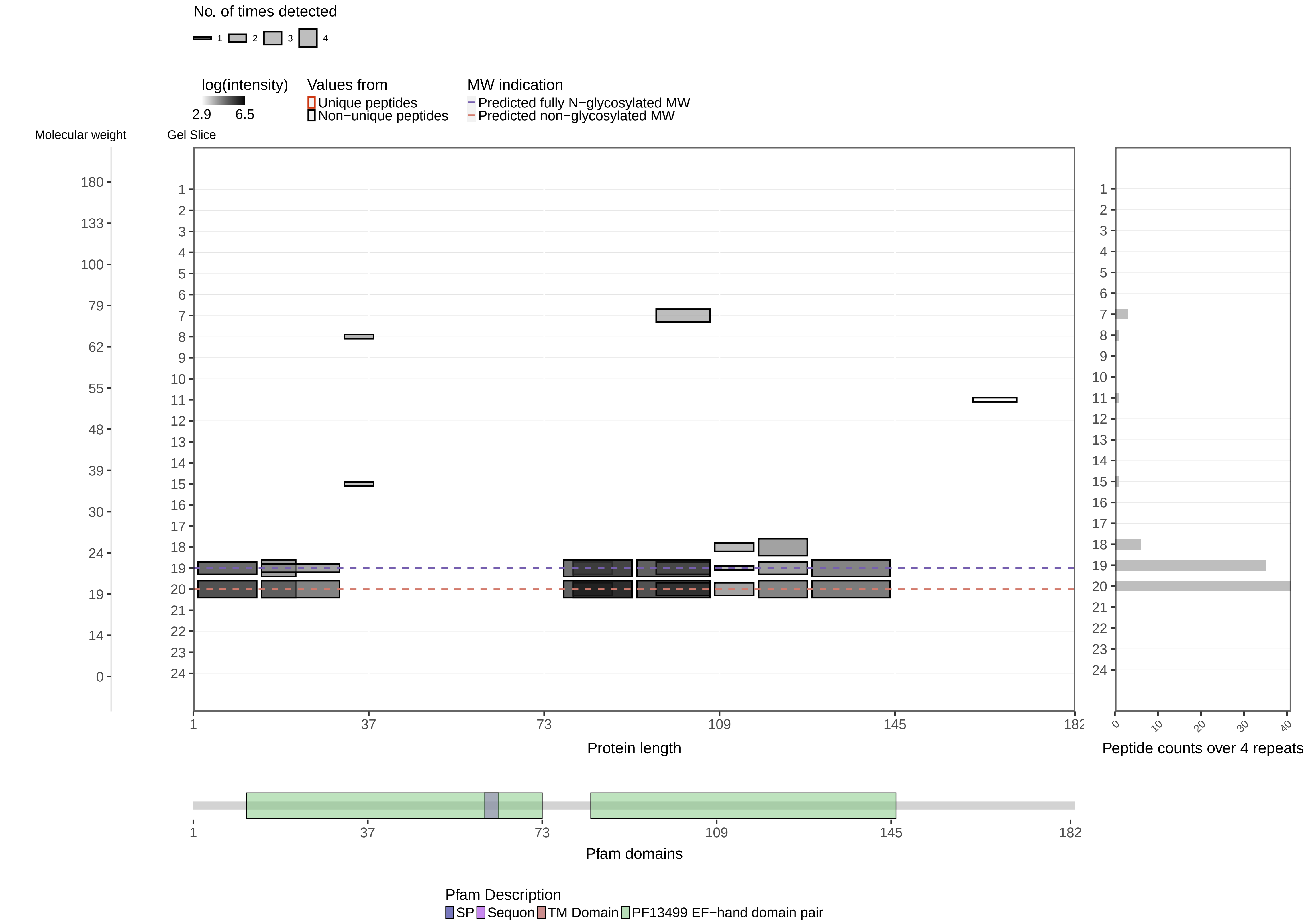


Protein sequence:

KDTDSEEELKEAFRVFDKDQNGFISAAELRHVMTNLGEKLTDEEVDEMIREADVDGDGQINYEEFVKVMMANRRRRRIEESKRGYSISQSNGRKSRKRDRCTIL*

### NbL07g10570.1 probable_calcium−binding_protein_CML13 [MW = 16.48 kDa]

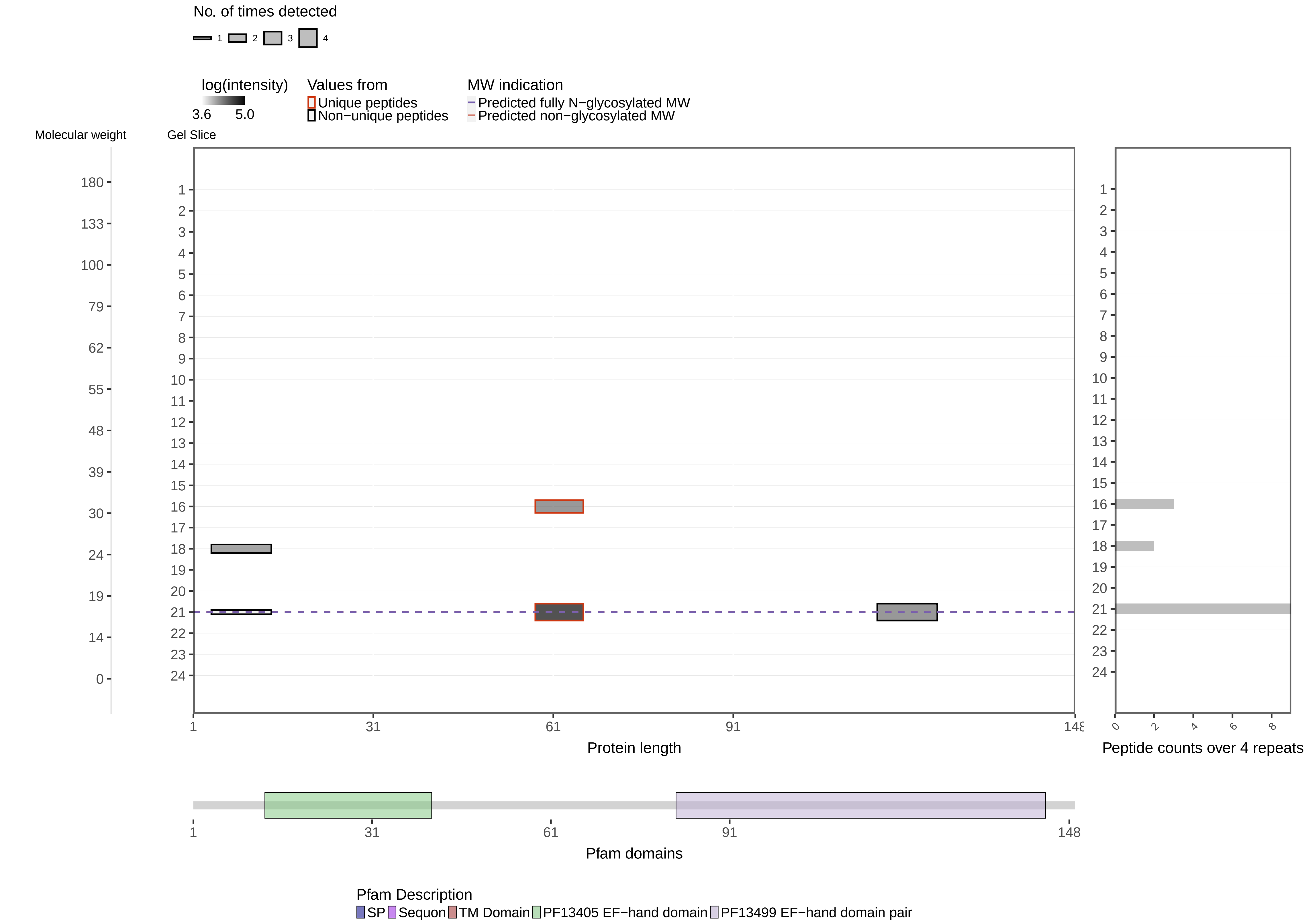


Protein sequence:

ᴾFDFNRFLDLMSKHLKPEPFDRQLRGAFKVLDKDGTGFVVVSDLKHILTSIGEKLEPAEFDEWIREVDVGSDGKIRYEDFIARMVAK*

### NbL07g10260.1 somatic_embryogenesis_receptor_kinase_2−like [MW − SP = 15.43 kDa]

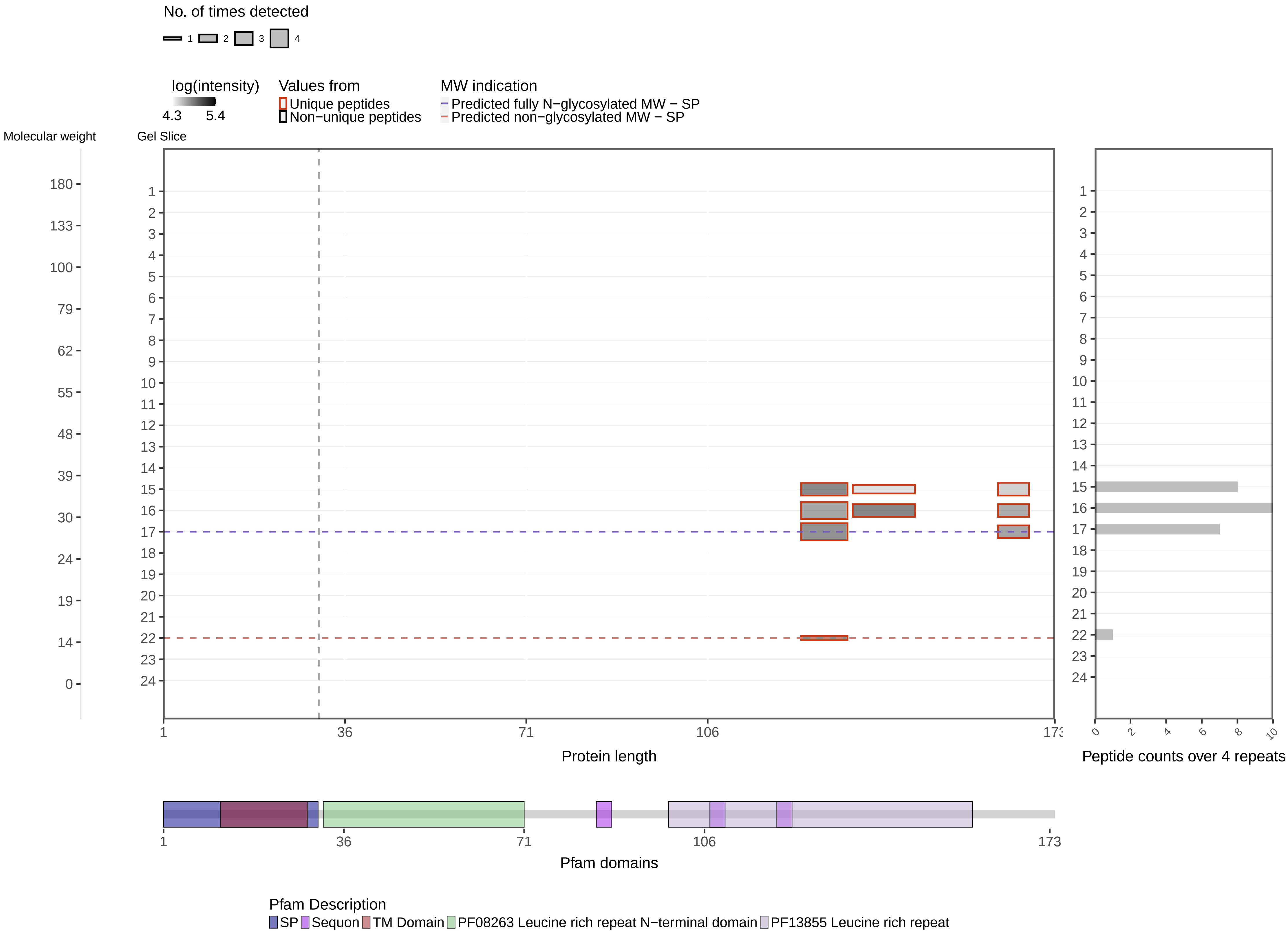


Protein sequence:

NSVVRVDLGAANLTGTLVPELGMLANLQYLQVENNSFSGAIPSQLGNLTRLLSLGLENNKLSGSIPYSLGNLKSLRWMRLNGNKLSGKIPISVLKLIYWG

NbL07g10160.1 photosynthetic_NDH_subunit_of_lumenal_location_5_chloroplastic [MW = 27.38 kDa]

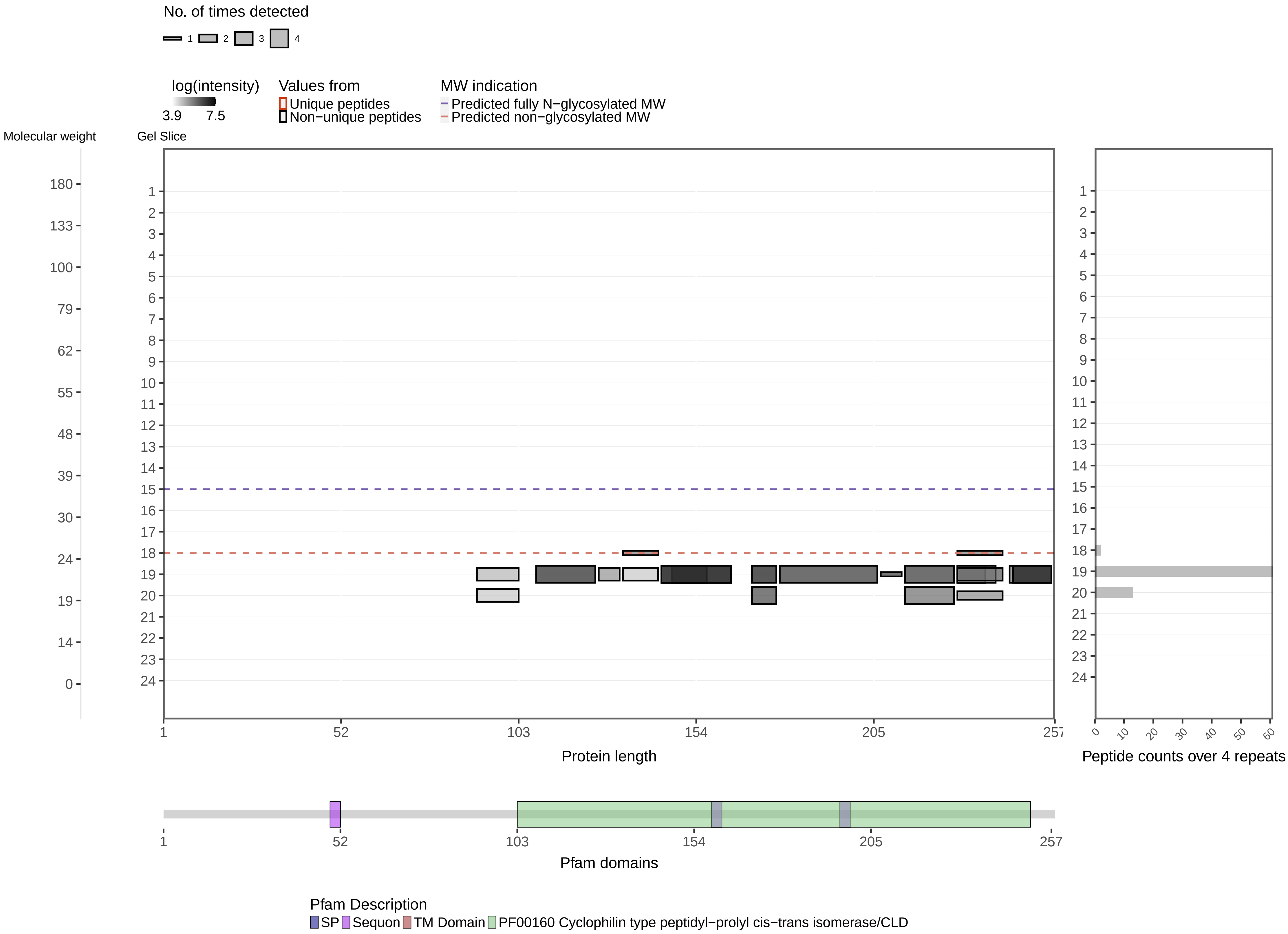


Protein sequence:

PQTAENFRALCTGEKGFGYKDSAFHRVIKDFMIQGGDFDKGNGTGGKSIYGRTFKDENFKLTHTGPGVVSMANAGPNTNGSQFFICTVKTPWLDERHVVFGQVLEGMDIVKLIESQETDRGDRPRKRVVISDCGELPVA*

NbL07g10090.1 heat_shock_protein_90−1−like [MW = 94.64 kDa]

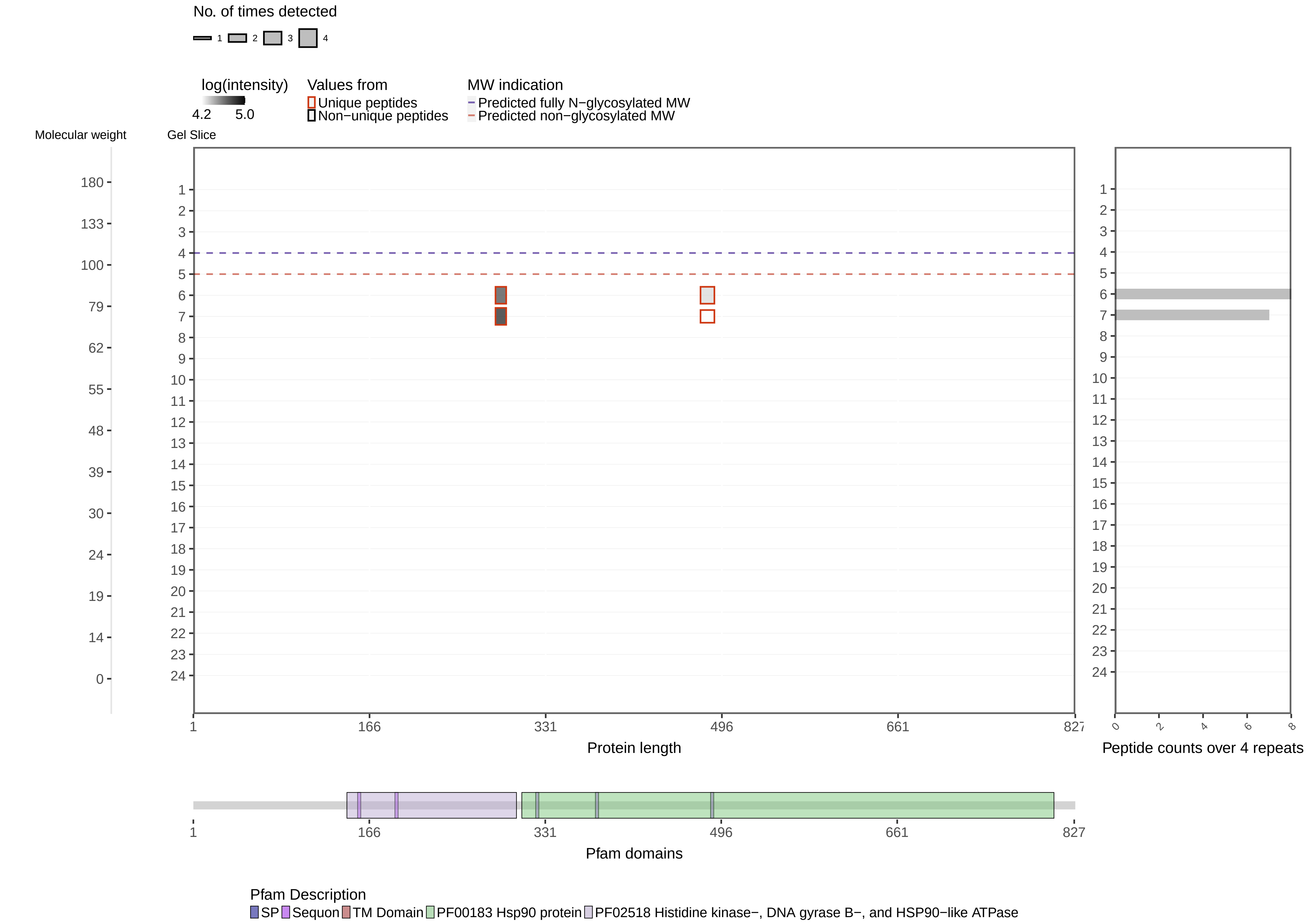


Protein sequence:

NbL07g09960.1 alpha−14−glucan−protein_synthase_[UDP−forming]_2−like [MW = 41.03 kDa]

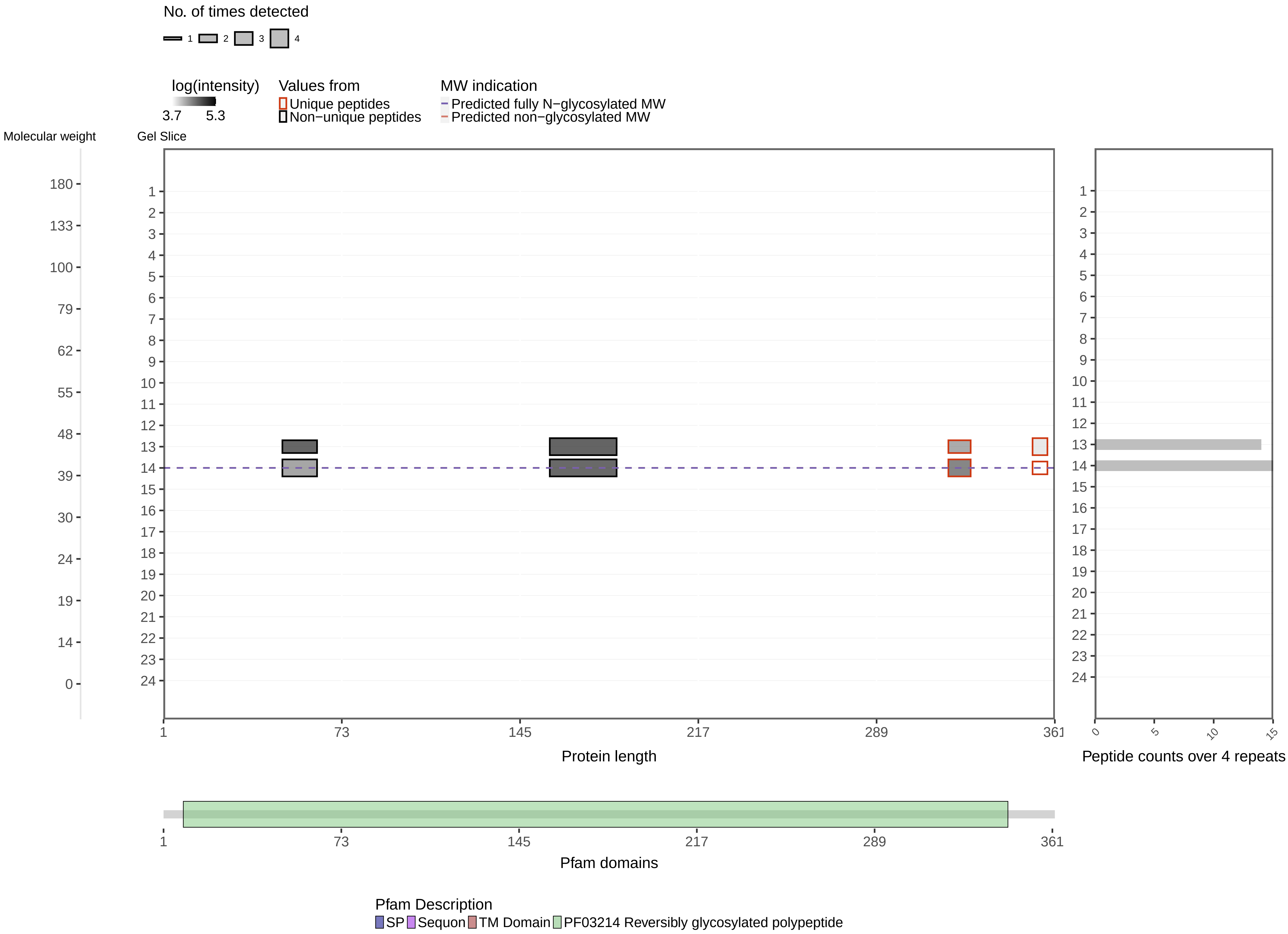


Protein sequence:

LNIPDYDAPTQLVKPRERNTRYVDAVMTIPKGTLFPMCGMNLAFNRELIGPAMYFGLMGDGQPIGRYDDMWAGWCTKVICDHLGLGIKTGLPYIWHSKASNPFVNLKKEYKGIYWQEEIIPFFQAVTLPKECTTVQQCYIELAKQVKEKLSNIDPYFTKLAEAMATWIEAWDELNPTGENLAKLSISDGKAK*

NbL07g09620.1 haloacid_dehalogenase−like_hydrolase_domain−containing_protein_3 [MW = 28.41 kDa]

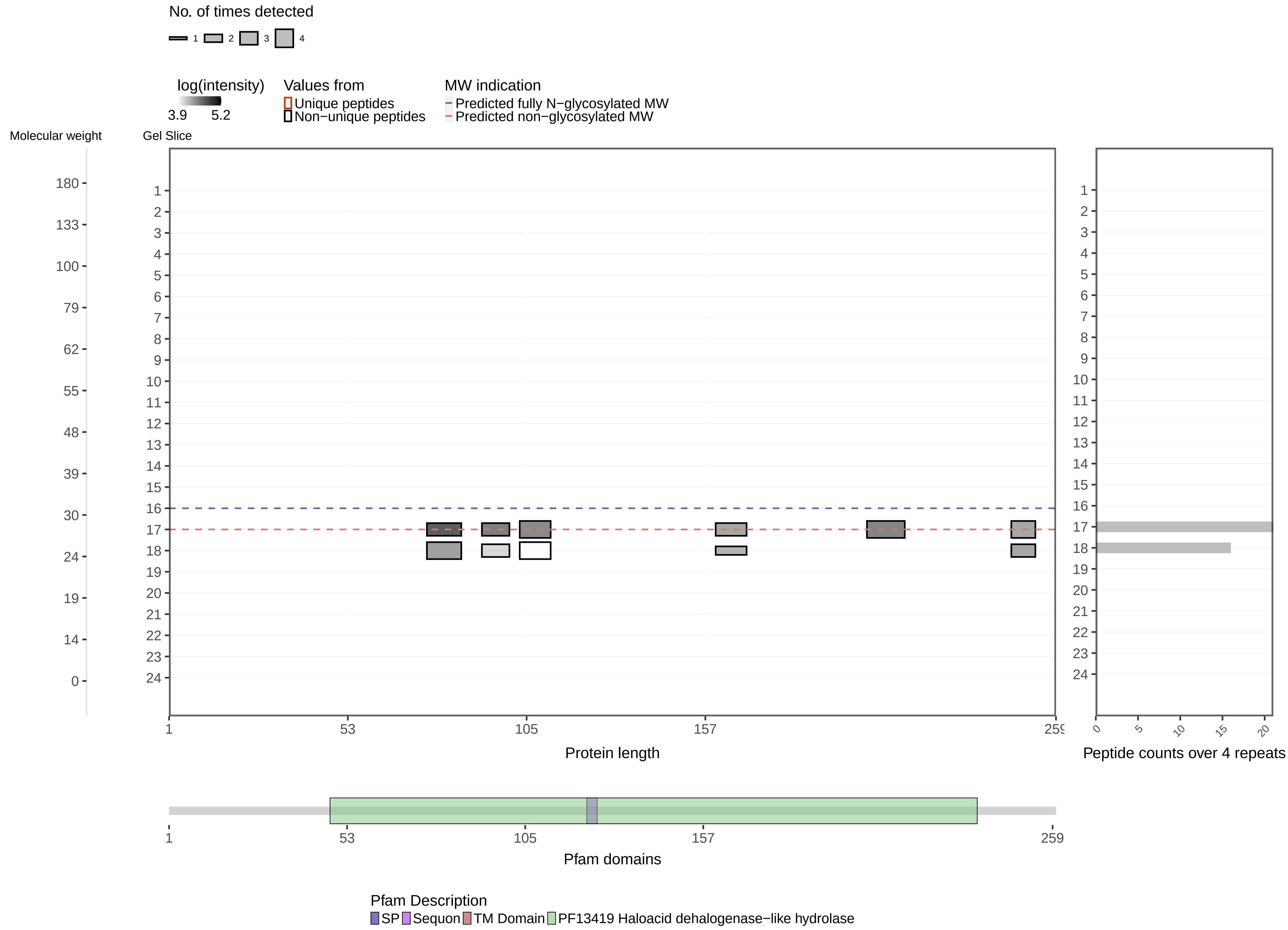


Protein sequence:

VAEATGCSNNTYFEEVYQHYASGGAWHLPAGAYDTISILKHSGVKLAVVSNFDTRLTRILKELKVLDLFDAVIISSEVGYEKPDIRIFRAALDQICVEAIKTVHVGDDEKADKGGANAAGIDCWLWGVDVKSFADIQNRLLTTA*

### NbL07g09410.1 26S_proteasome_non−ATPase_regulatory_subunit_7_homolog_A [MW = 34.95 kDa]

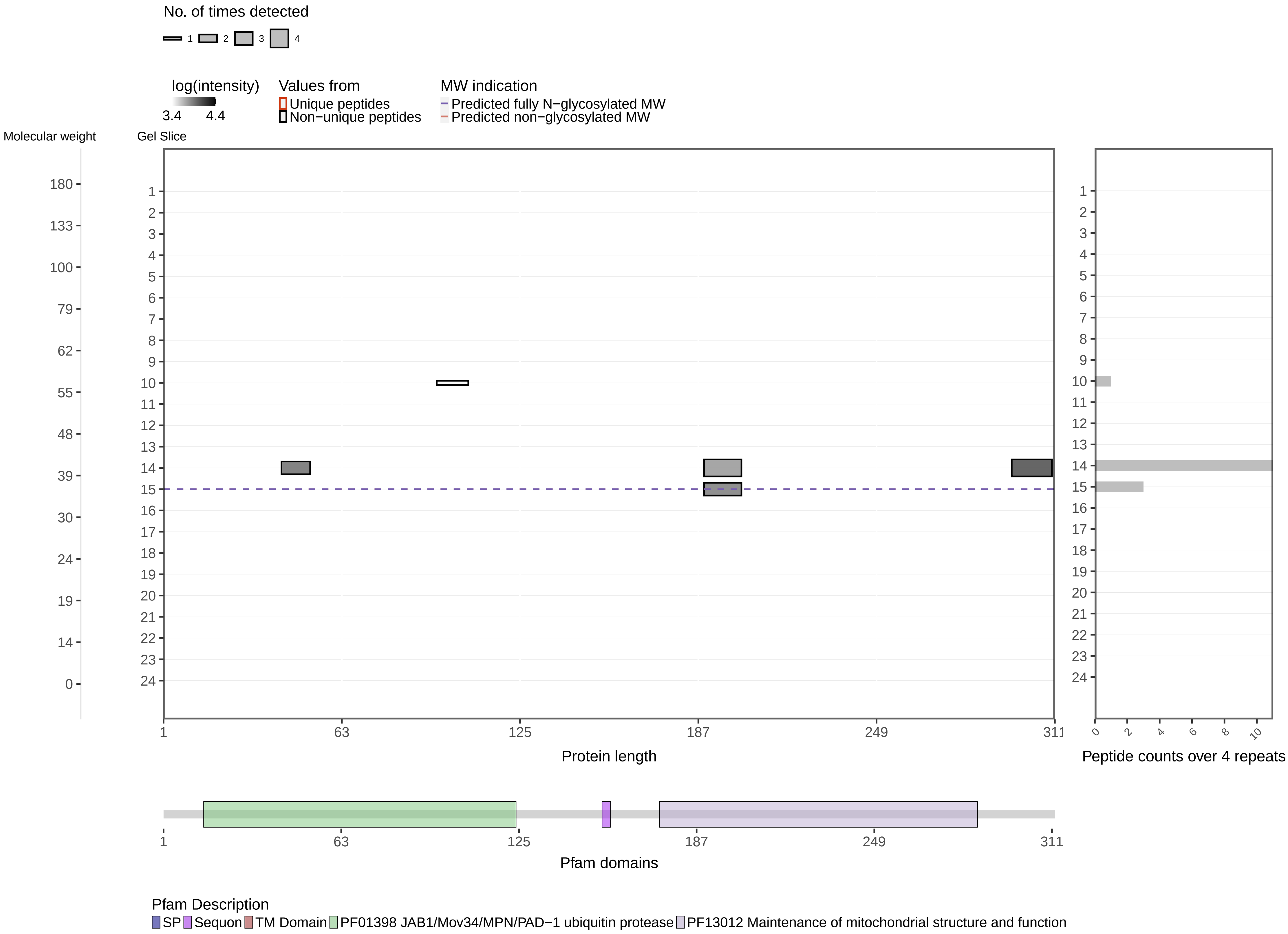


Protein sequence:

PTKAYYAVEEVKENATQKSQKVFVHVPSEIAAHEVEEIGVEHLLRDVKDTTISTLATEVTGKLAALKGLDARLQEIRSYLDLVIDGKLPLNHEILYHLQDVFNLLPNLNVAELVKAFSVKTNDMMLVIYLSSLIRSVIALHNLINNKMLNKEHEKAEDSKPLAVPTAAGS*

### NbL07g09290.1 biotin_carboxylase_1_chloroplastic [MW = 58.44 kDa]

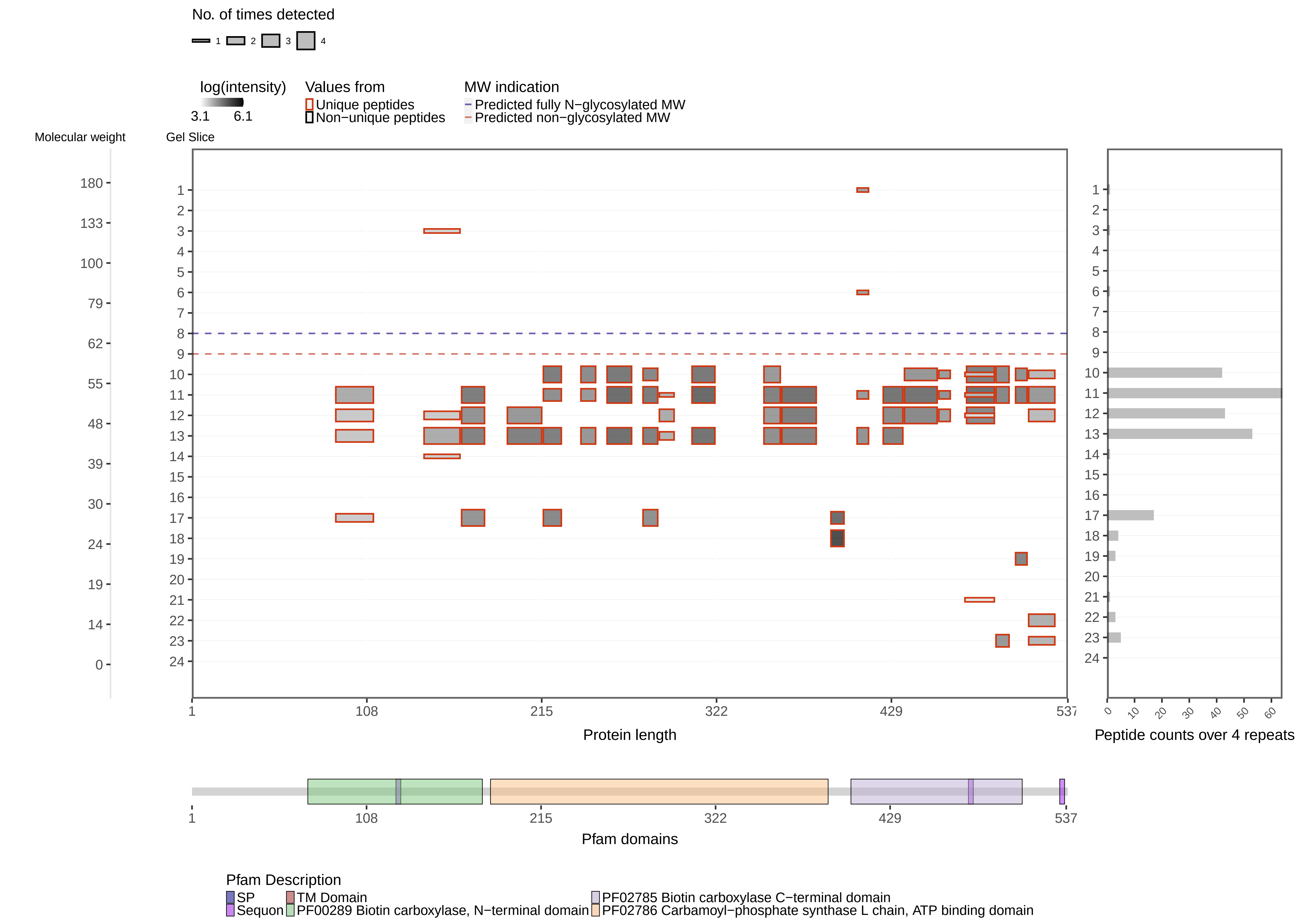


Protein sequence:

### NbL07g09270.1 thioredoxin_X_chloroplastic [MW = 24.31 kDa]

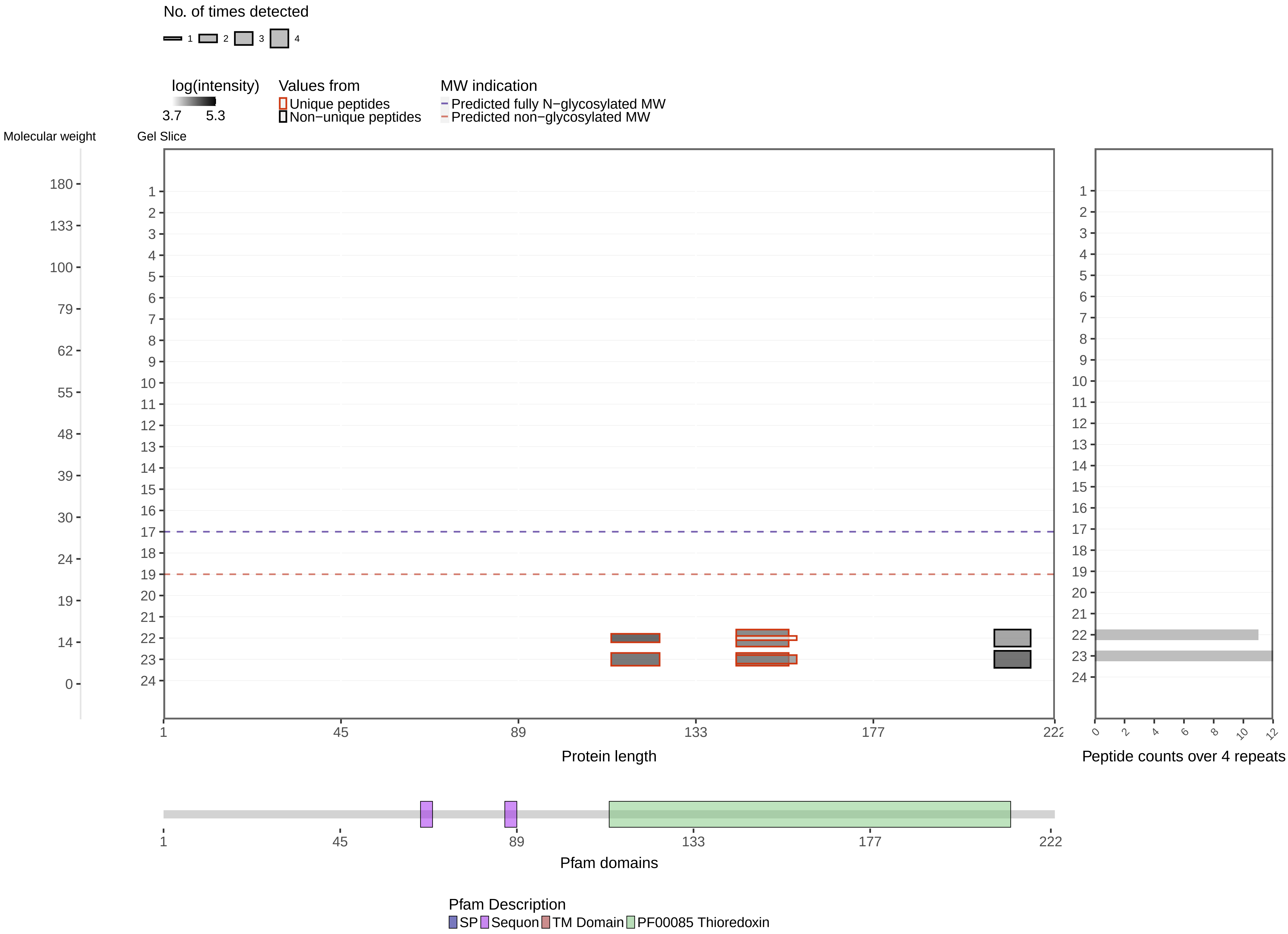


Protein sequence:

SSSSTTFRIRSGGLREIKEDEFSDIVLKSDRPVLVEFVATWCGPCRLIGPAMESVAEEYKEKITVVKIDHDANPKLIEEFKVYGLPTLILFKDGKEVPDSKREGAITKVKLKEYLDGLLKTVSVA*

NbL07g09170.1 glutamate−−glyoxylate_aminotransferase_2−like [MW = 22.58 kDa]

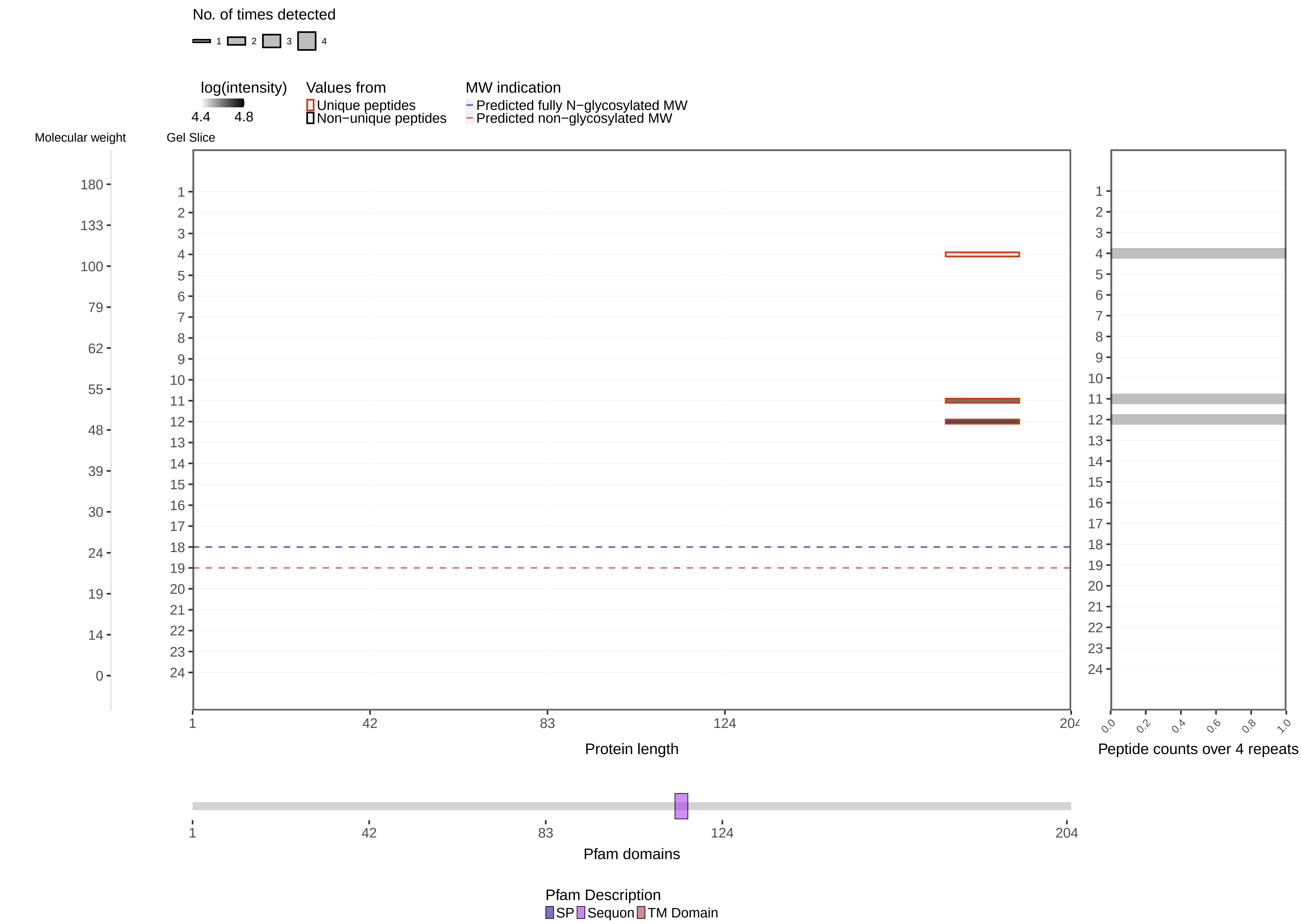


Protein sequence:

ILESLRRRAHIMTDGFNSCRNVVCNFTKGAMYSFPQIQLPCGAIDAAKKLGKAPNVLYCLKLLEATGISTVPGSGFGQKEGVFHLRTTILPAEEDMPSIMSSFKKFNDDFMKQYE*

### NbL07g09080.1 uncharacterized_LOC107808191 [MW = 48.21 kDa]

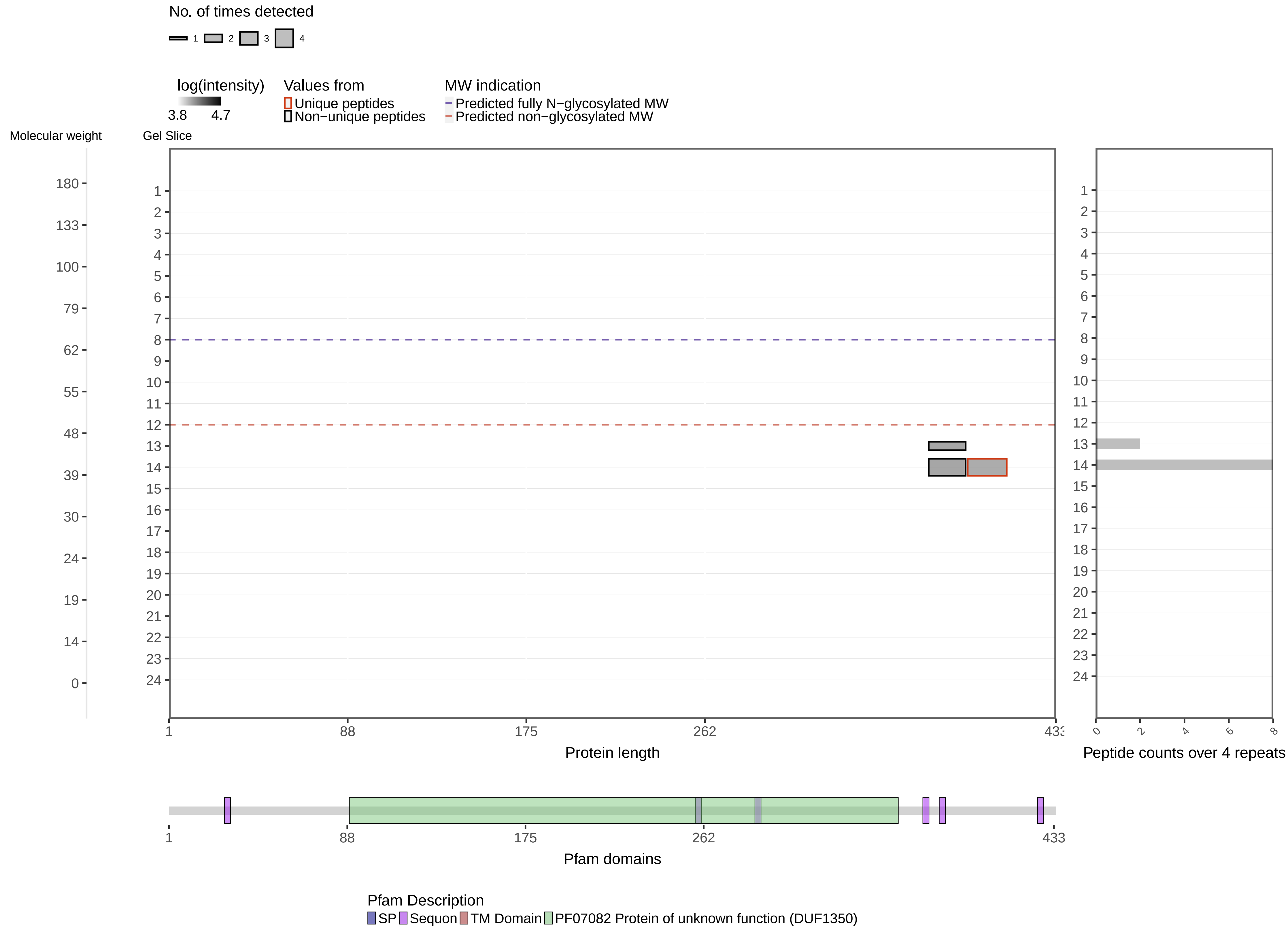


Protein sequence:

NGNVLMAFNNKDASLAVPFFSSVLAPMMQNIGPVLSDIASSPTIRLGAEMTMKQIENLSPSIVKQVFPLVEQLPPLYMDLIKGKDNFTPRPEETQRLIRSYYGISRNLLIKFKDDTIDETSTLAQVLSSGSAISSMLDMSIRSLPGDHALPLQQALPDVPPGMTDAVNRSGELLANLTAGTPWESVAKEVGNTLGNVDSILLSSGNSKDLDLLVETITSWMLSNSSPRLLQ*

### NbL07g09040.1 protein_SLOW_GREEN_1_chloroplastic [MW = 40.19 kDa]

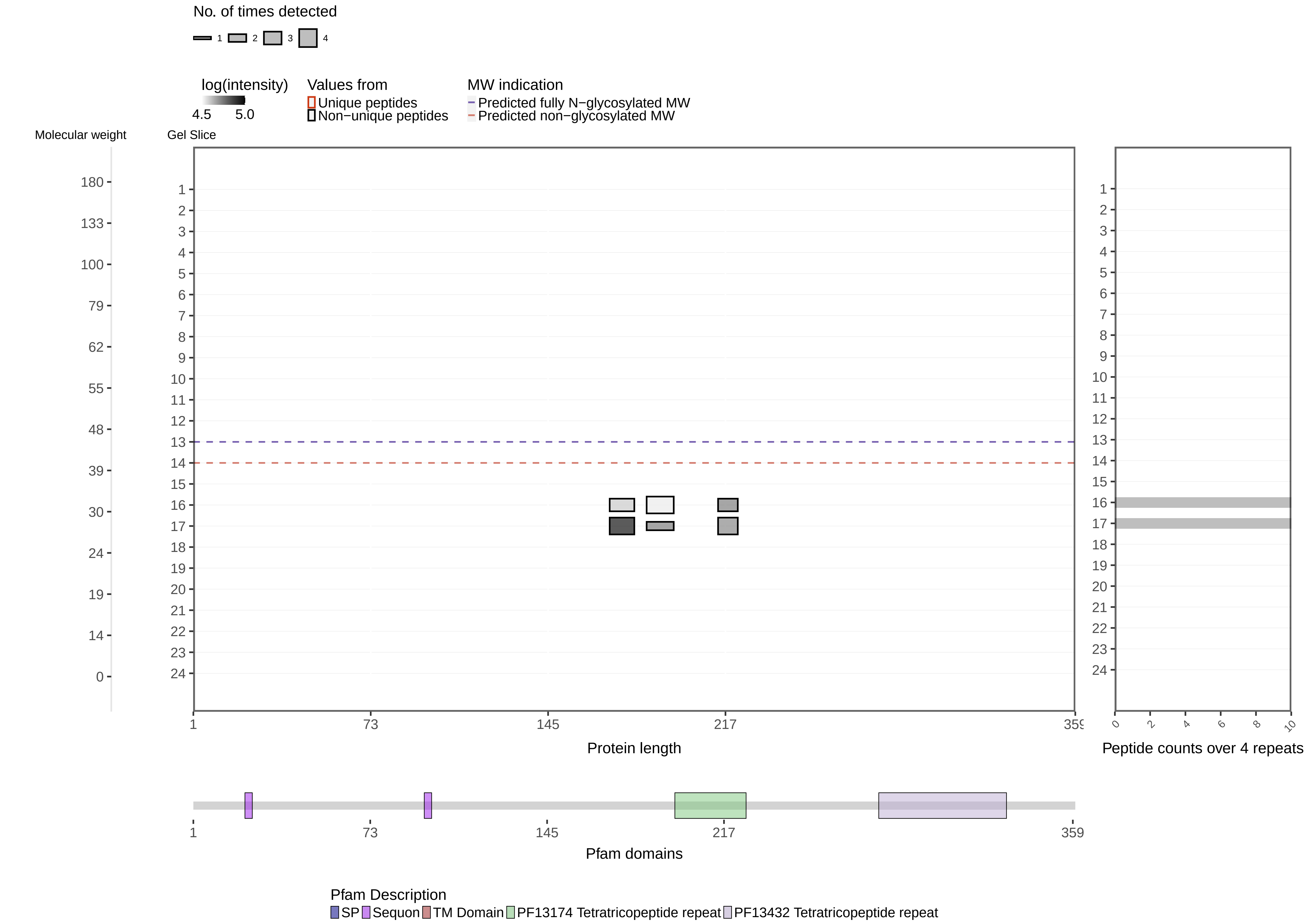


Protein sequence:

EKLEAGEDEESLKILKKLSSAQPENTEWKFLTARLLNEMGKVQEAREVFEEILSKNPLSFEALFENALLMDRSGEGVKVVQRLEEALKIAEEENKVKEGRDVRFIMAQVQFLQKNVEDALRSYDELEKEDPKDFRPYFCKGMIYSLLDRNKEAREQFAKYRELSPKKFEVEGYLQTPLSRMKLFGTDKKER*

### NbL07g08920.1 glutamate_decarboxylase_4 [MW = 55.85 kDa]

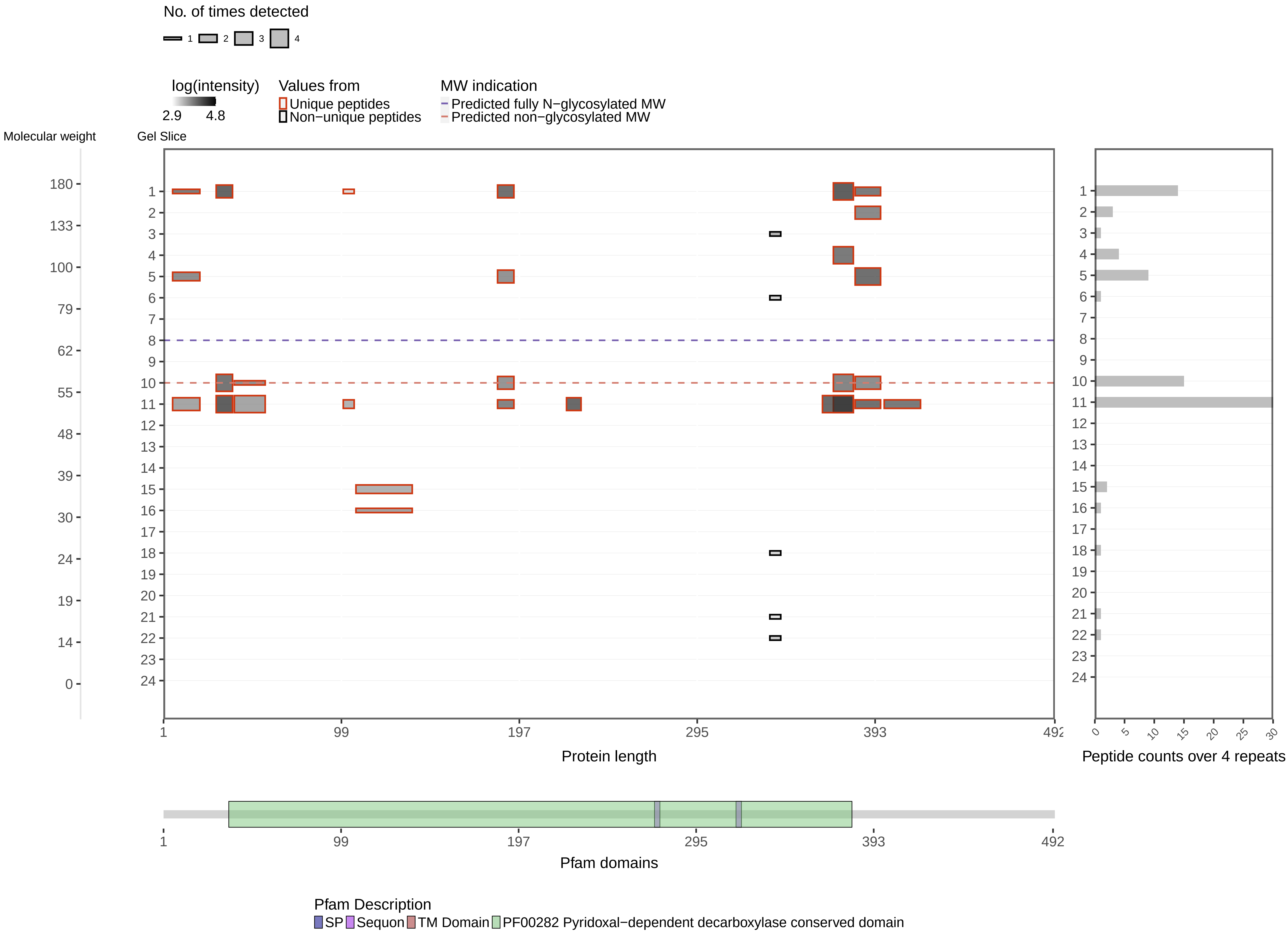


Protein sequence:

NKLTGWNTPIHVDAASGGFIAPFLYPELEWDFRLPLVKSINVSGHKYGLVYAGIGWVIWRTKQDLPEELIFHINYLGADQPTFTLNFSKGSSQVIAQYYQLIRLGYEGYRNVMENCRENAIVLREGLEKTGRFNIVSKDEGVPLVAFSLKDNSHHNEFEVSETLRRFGWIVPAYTMPADAQHVTVLRVVIREDFSRTLAERLVLDIVKVLHELDTLPARLSAKLEEVKLVKNGKKSEVEVQREVTNYWKKFVLARKAPVC*

NbL07g08620.1 probable_S−adenosylmethionine−dependent_methyltransferase_At5g38100 [MW = 41.61 kDa]

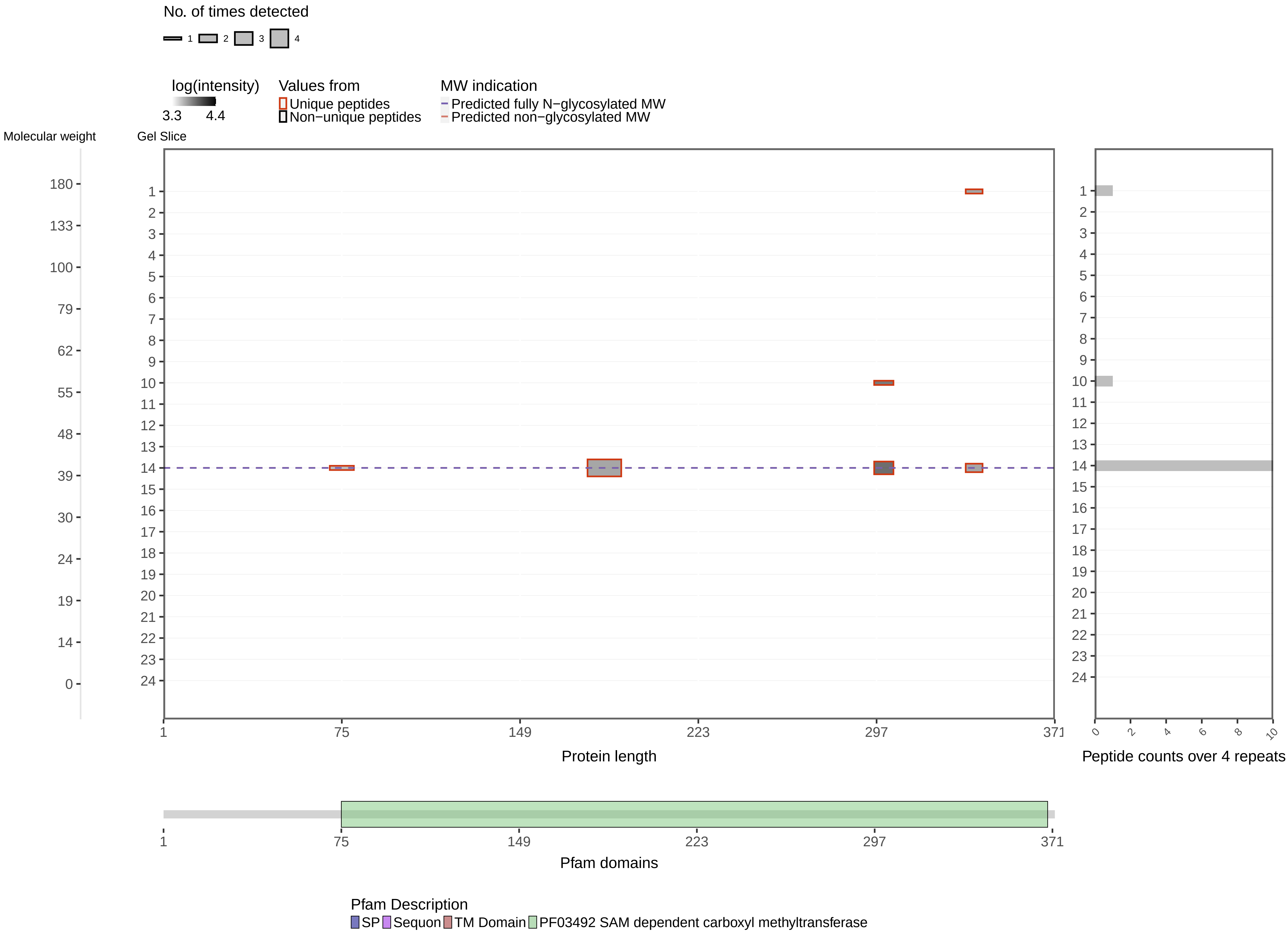


Protein sequence:

WLSKSPEELLDENSPAWNKGLIHYVDAPNIAVVNAYVSQFDKDMEVFLNARAEEIVPGGIMVLVSPFSGYRLLKFFGSSLVDMVNEGMIDESLVDAFNVPVYFPSPEDMTKVVERNGCFSIERIELAYPQSKLVNEADAKSLMTNLRAVLEGVFINHFGRKIADEAFSRTILKSEDISAWMKANYEKPCQLFVALKRK*

### NbL07g08610.1 aspartokinase_2_chloroplastic−like [MW = 61.31 kDa]

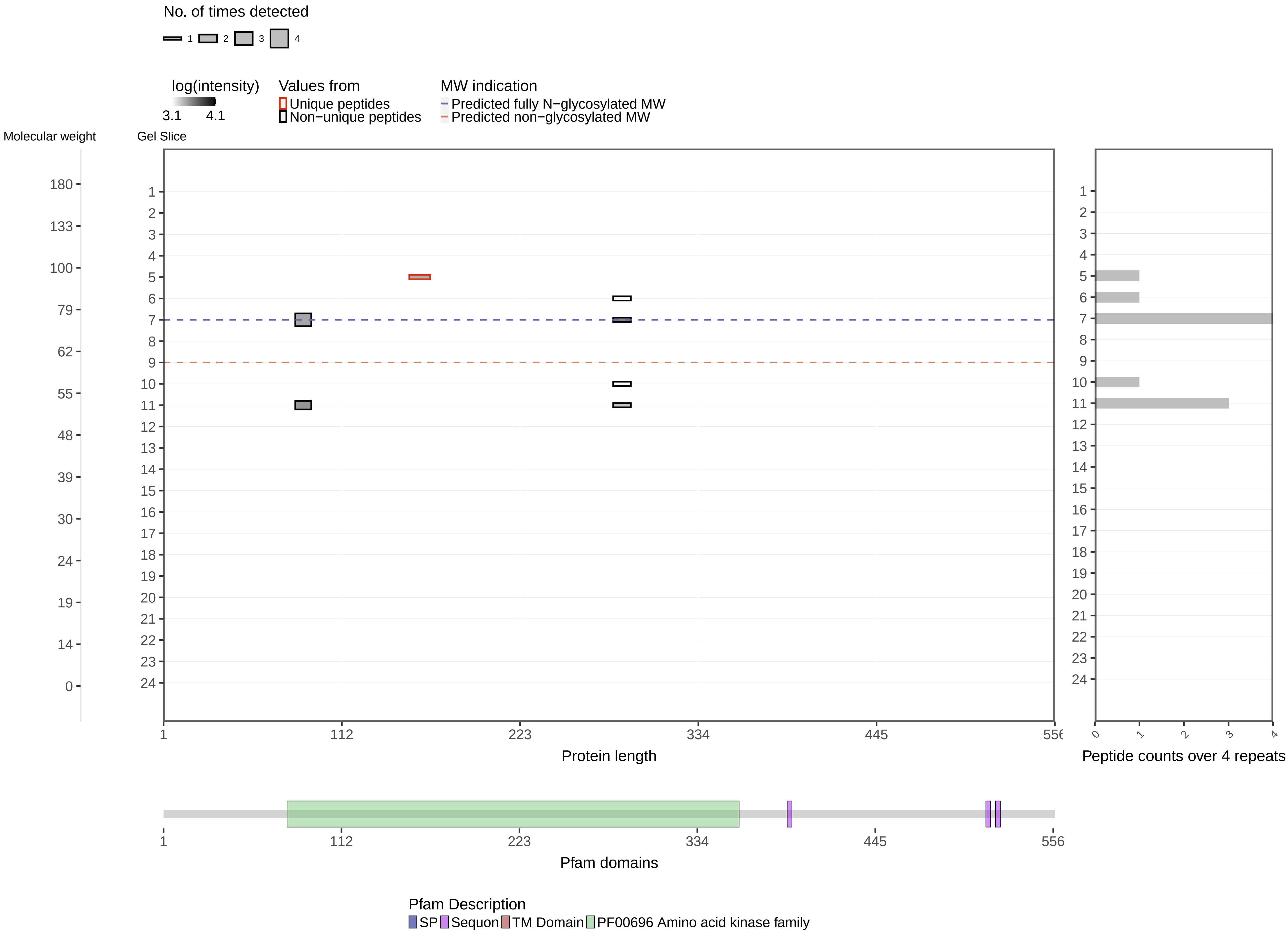


Protein sequence:

GFLGKGWRTCAVTTLGRGGSDLTATTIGKALGLREIQVWKDVDGVLTCDPNIYPRAQPVPYLTFDEAAELAYFGAQVLHPQSMRPAREGDIPVRVKNSYNPKAPGTLICRARDMSKAVLTSIVLKRNVTMLDIVSTRMLGQFGFLAKLLQVFSIFEDLGISVDVVATSEVSISLTLDPSKLWSRELIQQELDHVVEELEKIAIVNLLQHRSIISLIGNVQRSSLILEKAFHVLRTNGINVQMISQGASKVNISLIVNDSEAEQCVRALHKTFFESDLSELVWENGHVTPLST*

### NbL07g08590.1 aspartate−semialdehyde_dehydrogenase−like [MW = 44.53 kDa]

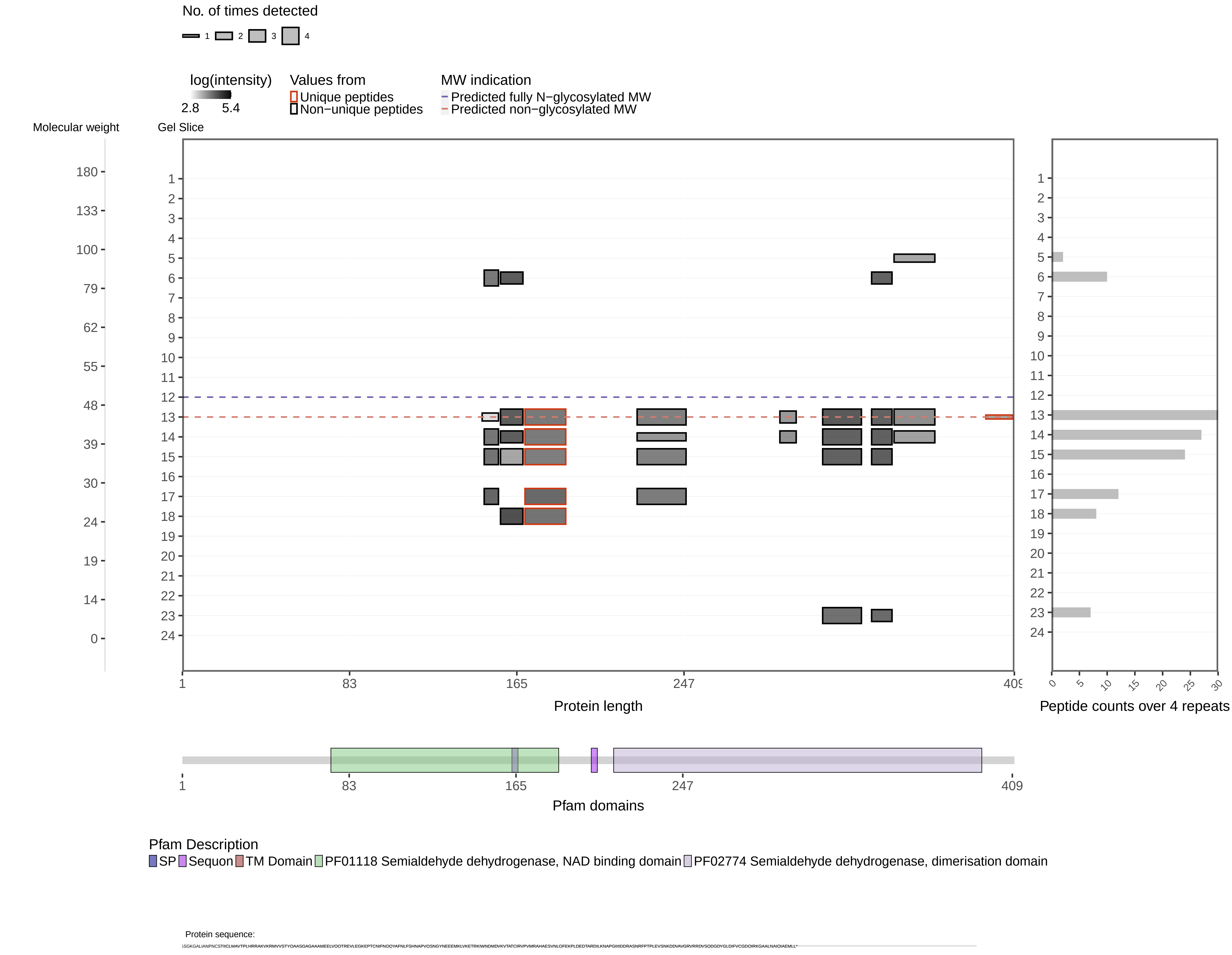

NbL07g08370.1 secoisolariciresinol_dehydrogenase−like [MW = 29.58 kDa]

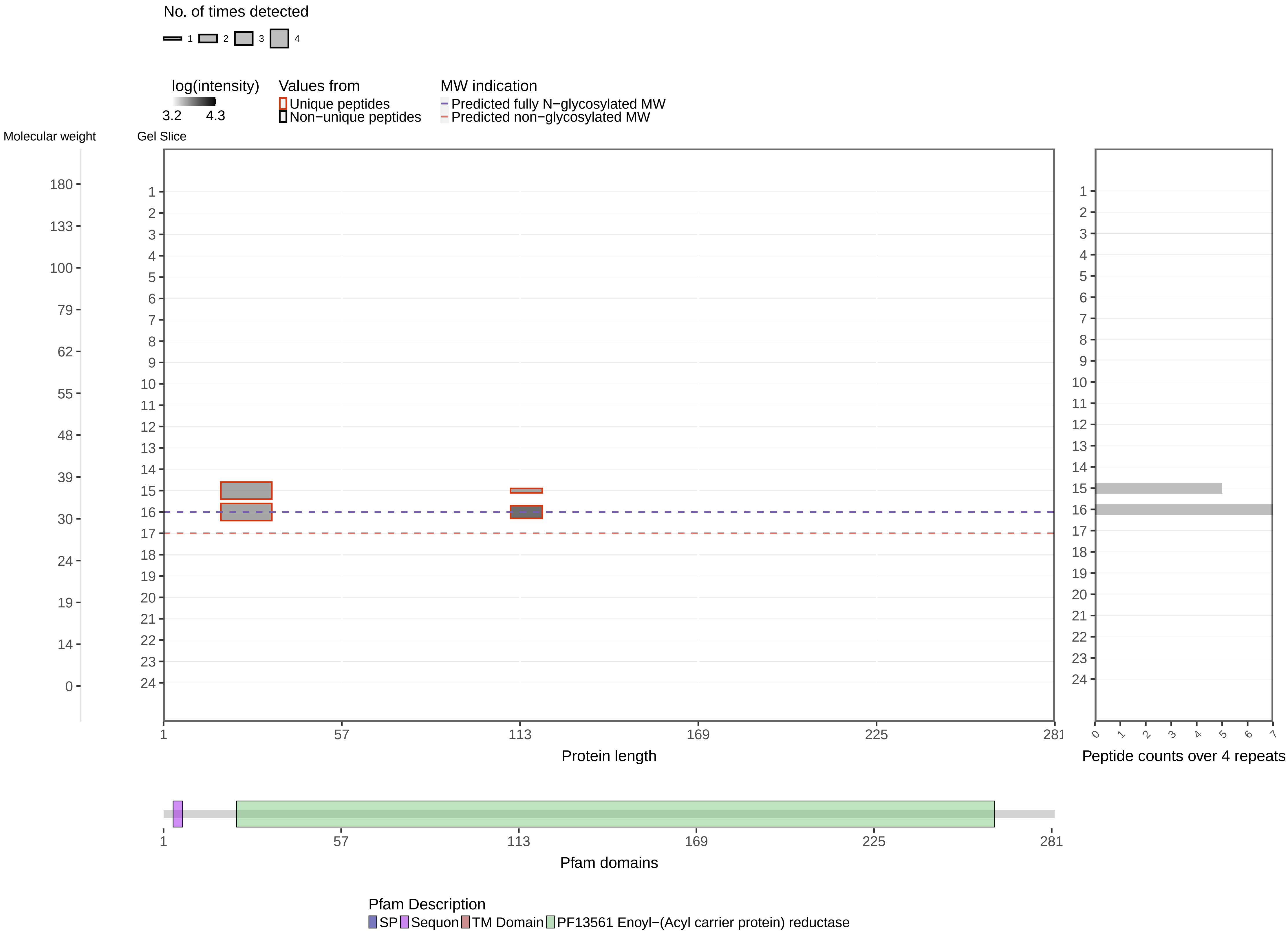


Protein sequence:

AFLGAKHAARVMIPAKTGCILFTASVTAVVAGMSPHPYASSKHAVVGLANNLCAELGQYGIRVNCITPSVVATPMVSKSVWGIDQSKVDEAVMKSANLKEAKLEVEDVAAAAVYLASDESKYVSGLNLLIDGGYSKTNPLFAMCIREILSDF*

### NbL07g08340.1 PHD_finger_protein_Alfin1−like [MW = 28.16 kDa]

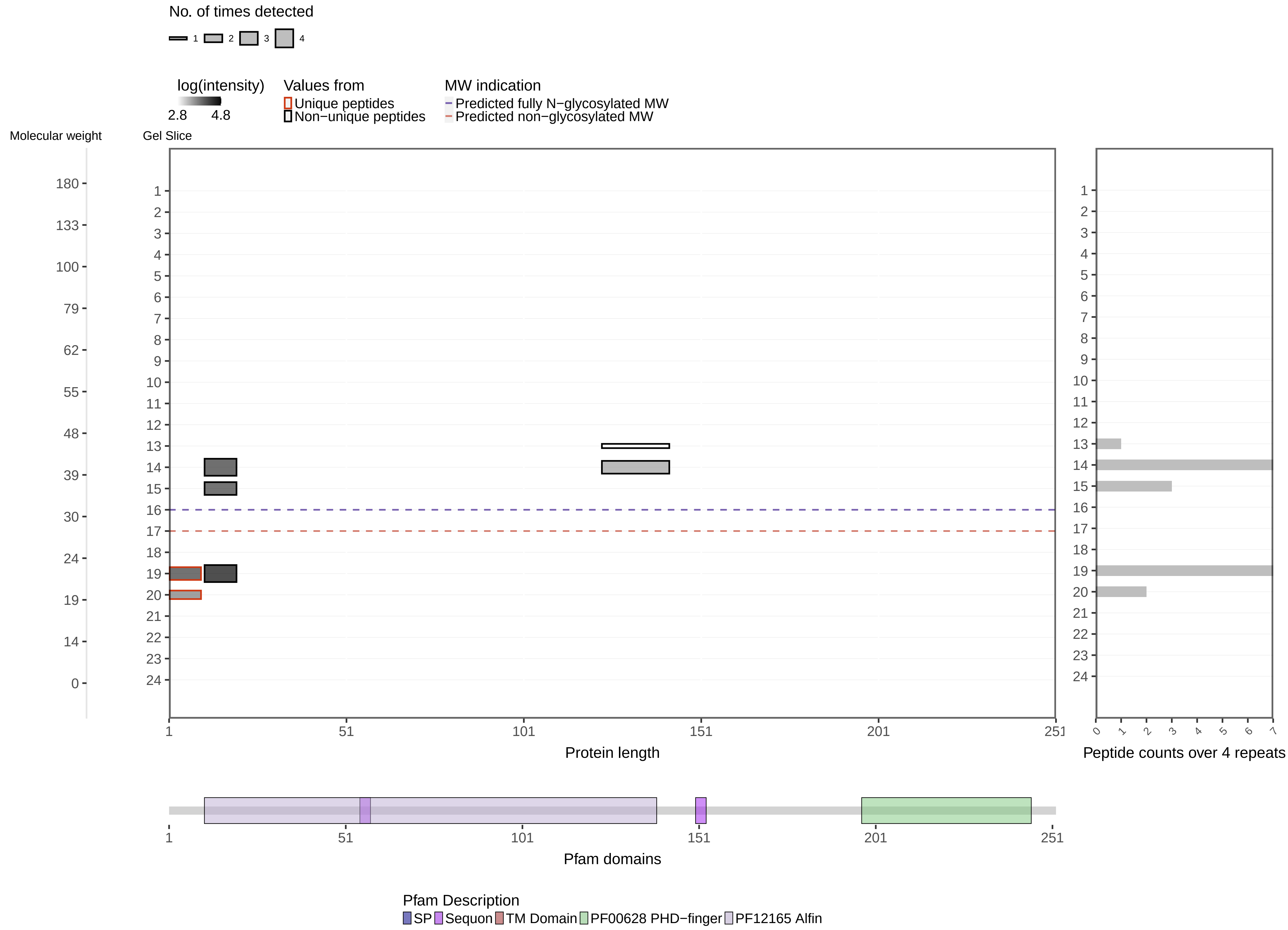


Protein sequence:

FGKSERKRLFQMINDLPTVFEVVTGAAKQARDAAHNNSSKSKSSGKFRQSEPQHKGVKVSPPKEEDESGEEVEEEDEQGATLCGACGDNYGQDEFWICCDVCEKWFHGKCVKITPAKAEHIKQYKCPSCSNKRARV*

NbL07g08200.1 pectin_acetylesterase_10−like [MW − SP = 43.79 kDa]

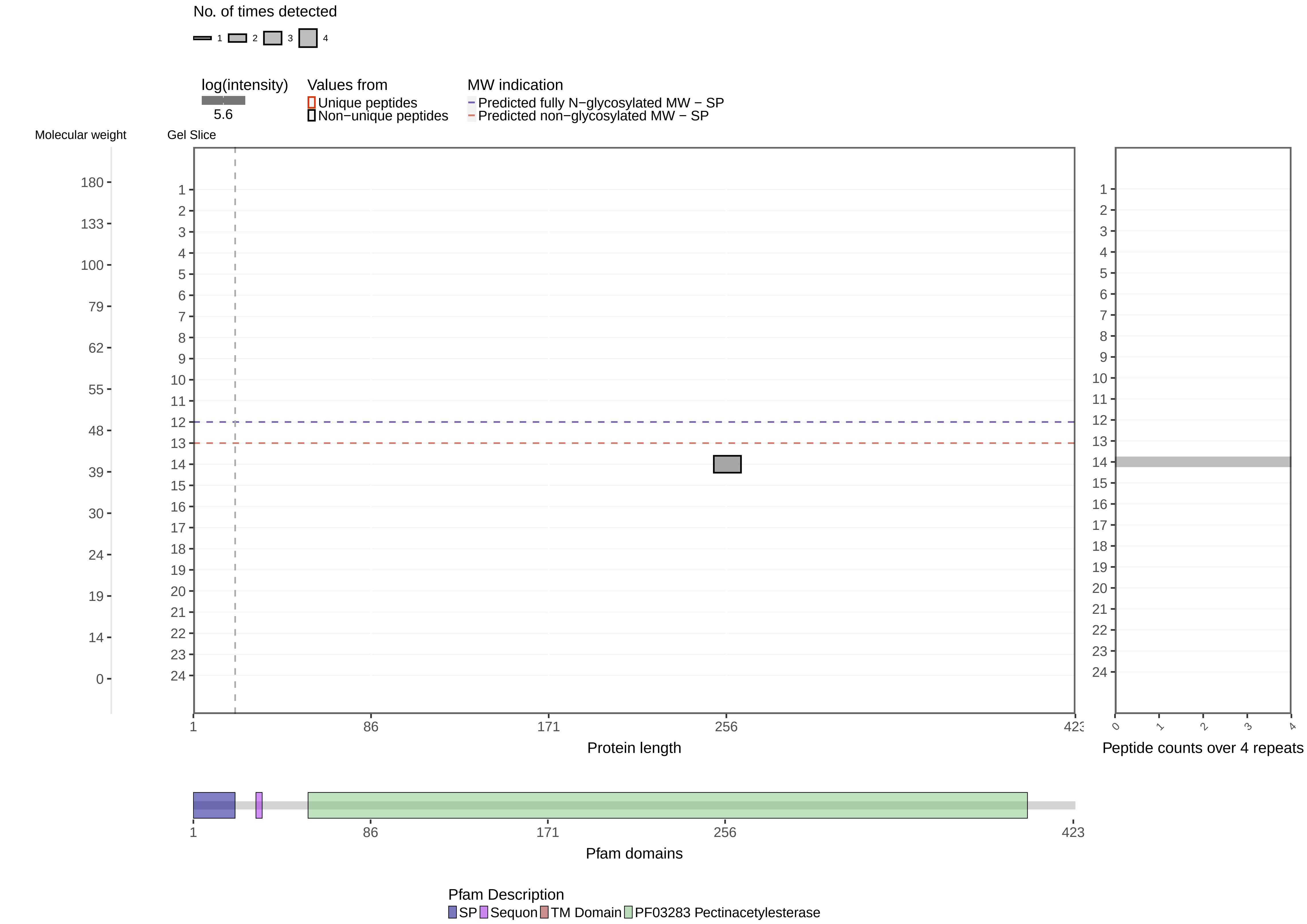


Protein sequence:

ALLSGCSAGGLASILHCDEFRTLFPSSTKVKCLSDAGLFMDATDVSGGHALRNLYQGVVSVQGLQKTLPSTCTNKLDPTSCFFPQNLIGNIKTPLFLLNAAYDSWQVQSSLAPPVADPKGLWHDCKLNNEQCSASQIQFLQGFRNDMLNAVKGFAASTQNGLFINSCFAHCQSERQDTWFADDSPLINNKPVALAVGDWYFDRAGVKSIDCAYPCDKTCHNLVFK*

NbL07g08180.1 eukaryotic_translation_initiation_factor_5B−like [MW = 47.16 kDa]

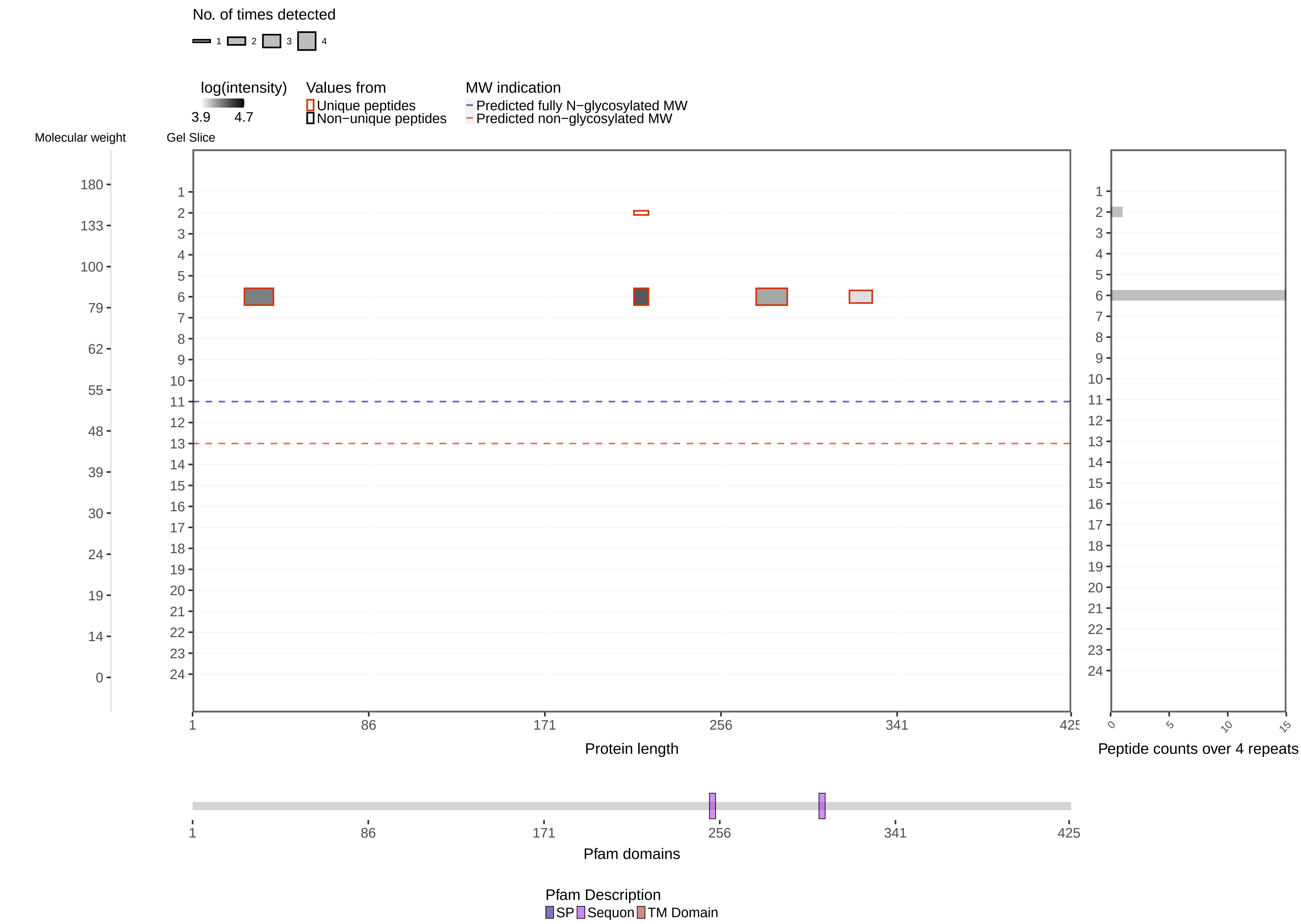


Protein sequence:

AEQNKETIVANVEDKLAEQPEVRKKIEQEPAETKPENTEAATEVPLKEEVEENDTKDAKTNQEDERKSVVTEAKEIETPEKAPEVEIASRSIELIAENGNNKDEINVTTEANEATEAEKKVDEATATVTEKSGEESQKPETETKVEEETAGEVEPKLEQEAKADAAKDGGERKTLEEAFKEEVPAKPKQSNNLISKMKQSLVKAKKAIMGKSPSSKSETKNEVTAK*

### NbL07g08090.1 ATP−dependent_Clp_protease_proteolytic_subunit−related_protein_1_chloroplastic−like [MW = 42.52 kDa]

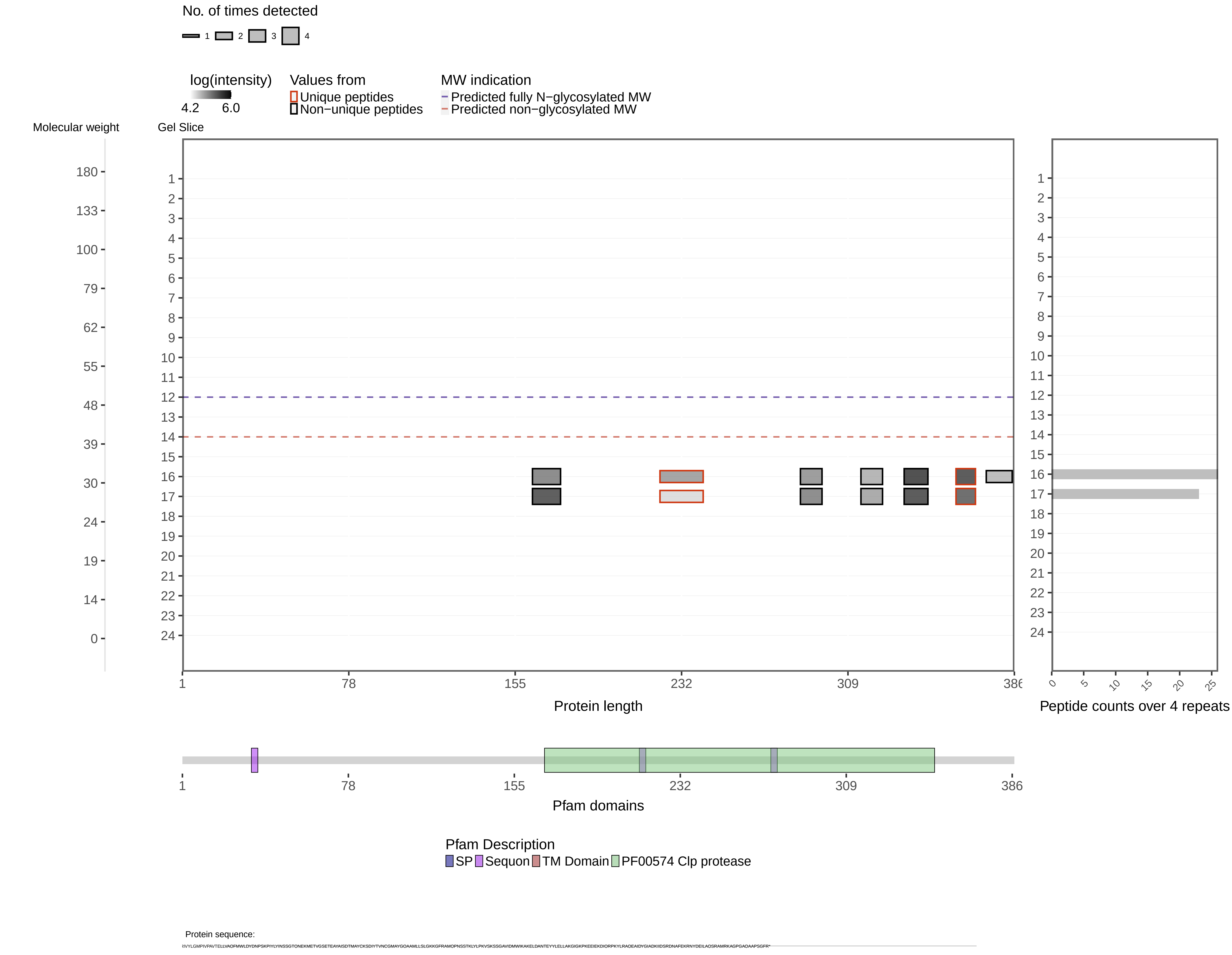

### NbL07g08030.1 3−oxo−Delta(45)−steroid_5−beta−reductase−like [MW = 43.69 kDa]

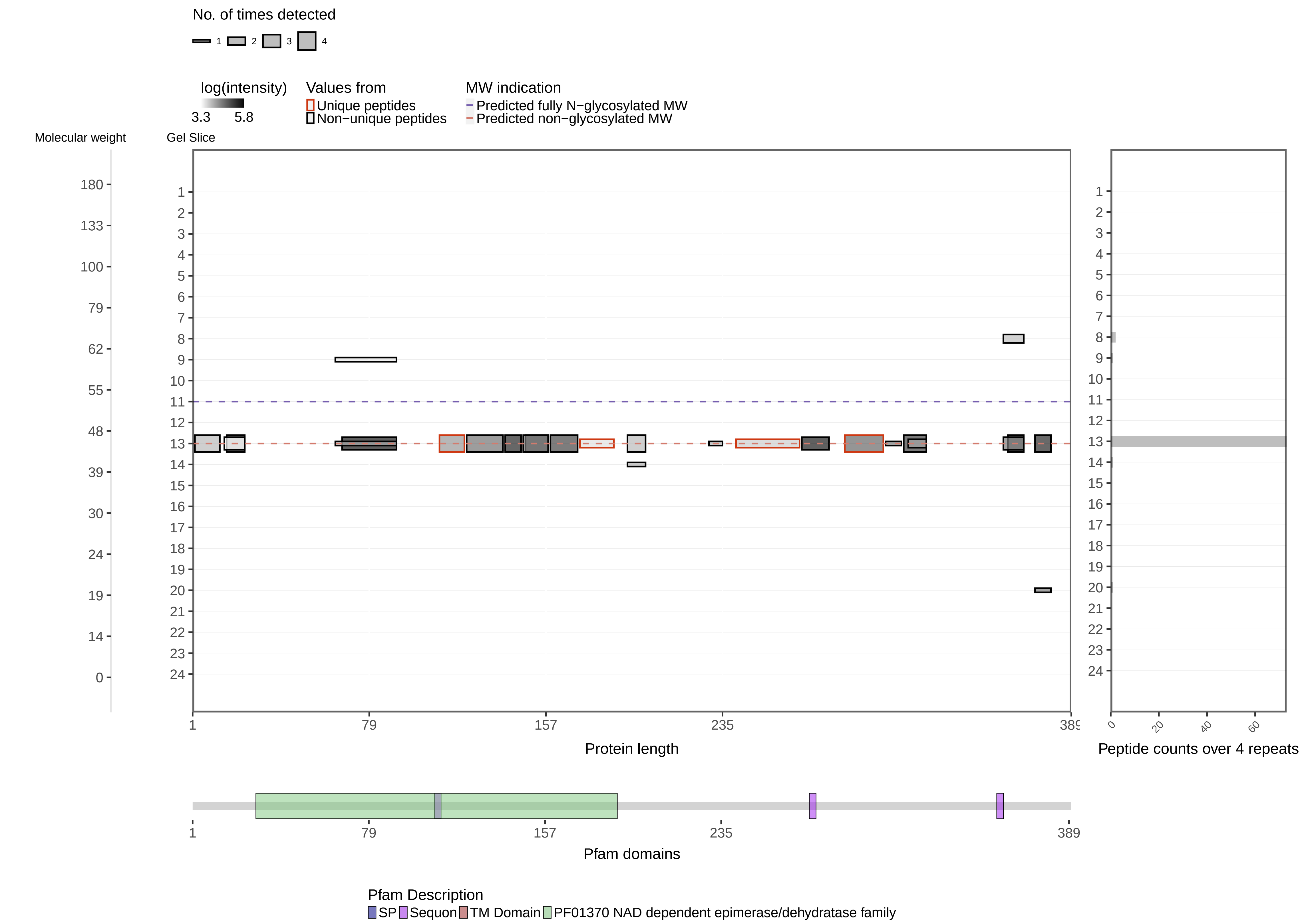


Protein sequence:

VLFKEIEKKEGLTWSVHRPGTIFGFSPYSLMNIVGTLCVYAAICKHEGLPLKFPGVKAAWDGYSDCSDADLIAEHQIWAAVDPYAKNEAFNVSNGDVFKWKHFWKVLAEQFGVGAAEFDEVNRCTLAEMMKDKGPVWDEIVKENGLTPTKLEDVGVWWFVDLILAGDCPLDTMNKSKEHGFLGFRNSQKAFISWIDKVKAYKVVP*

### NbL07g08010.1 omega−amidasechloroplastic−like [MW = 31.66 kDa]

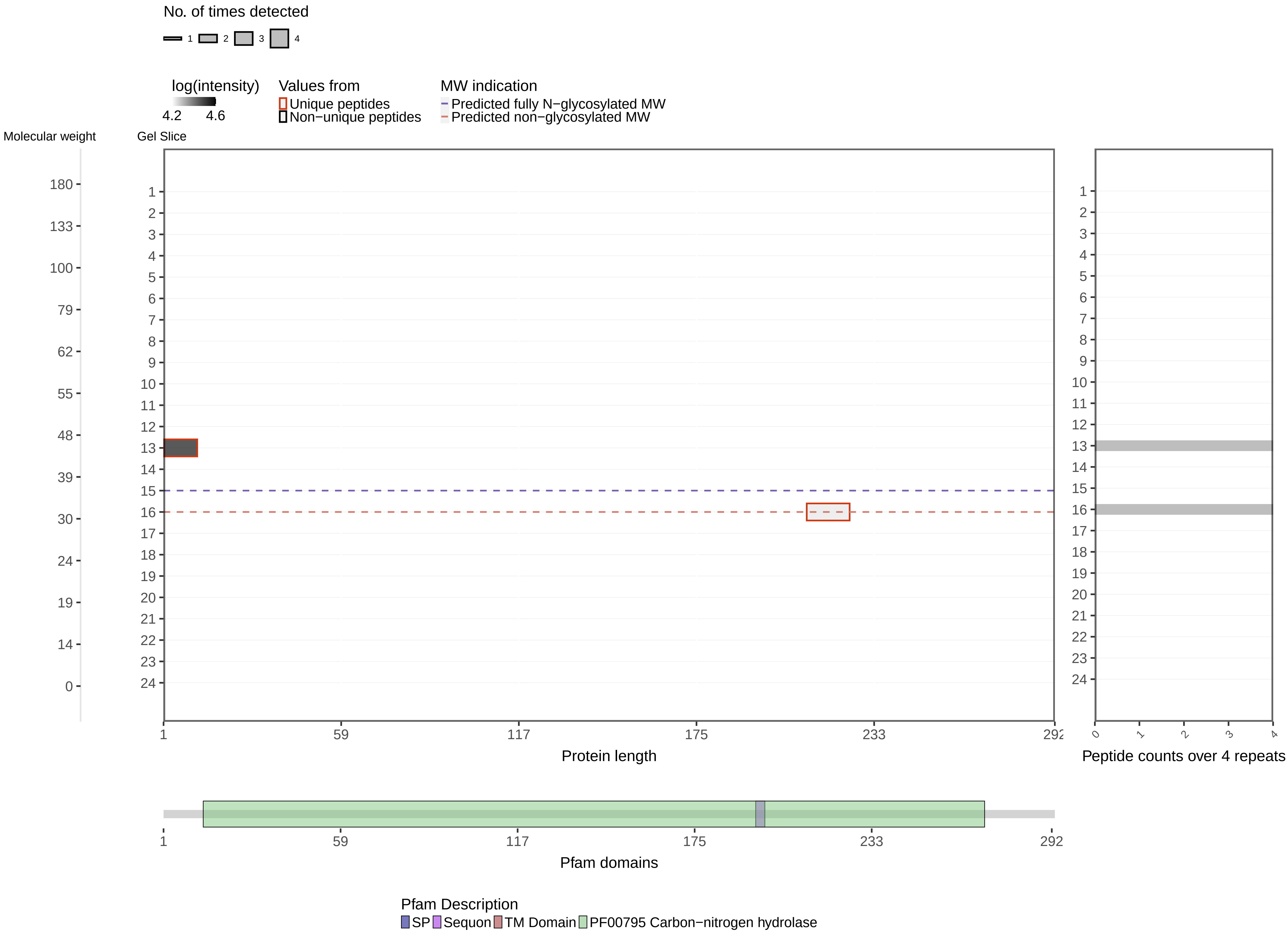


Protein sequence:

KPLPGEVOFKESDSISAGDKPTVVDTDFGCIGIGICHDIRFAELAMSYSARGAHLICYPGAFNMSTGAALWELEORTRAVDNOLYVASCSPSRDSASSYIIWGHSTVVGPMGEIIATTGHEEAVLIADIDYAAIOLARESFPLKSORRDDIYOFVDLLG*

NbL07g08000.1 upstream_activation_factor_subunit_spp27−like [MW = 20.71 kDa]

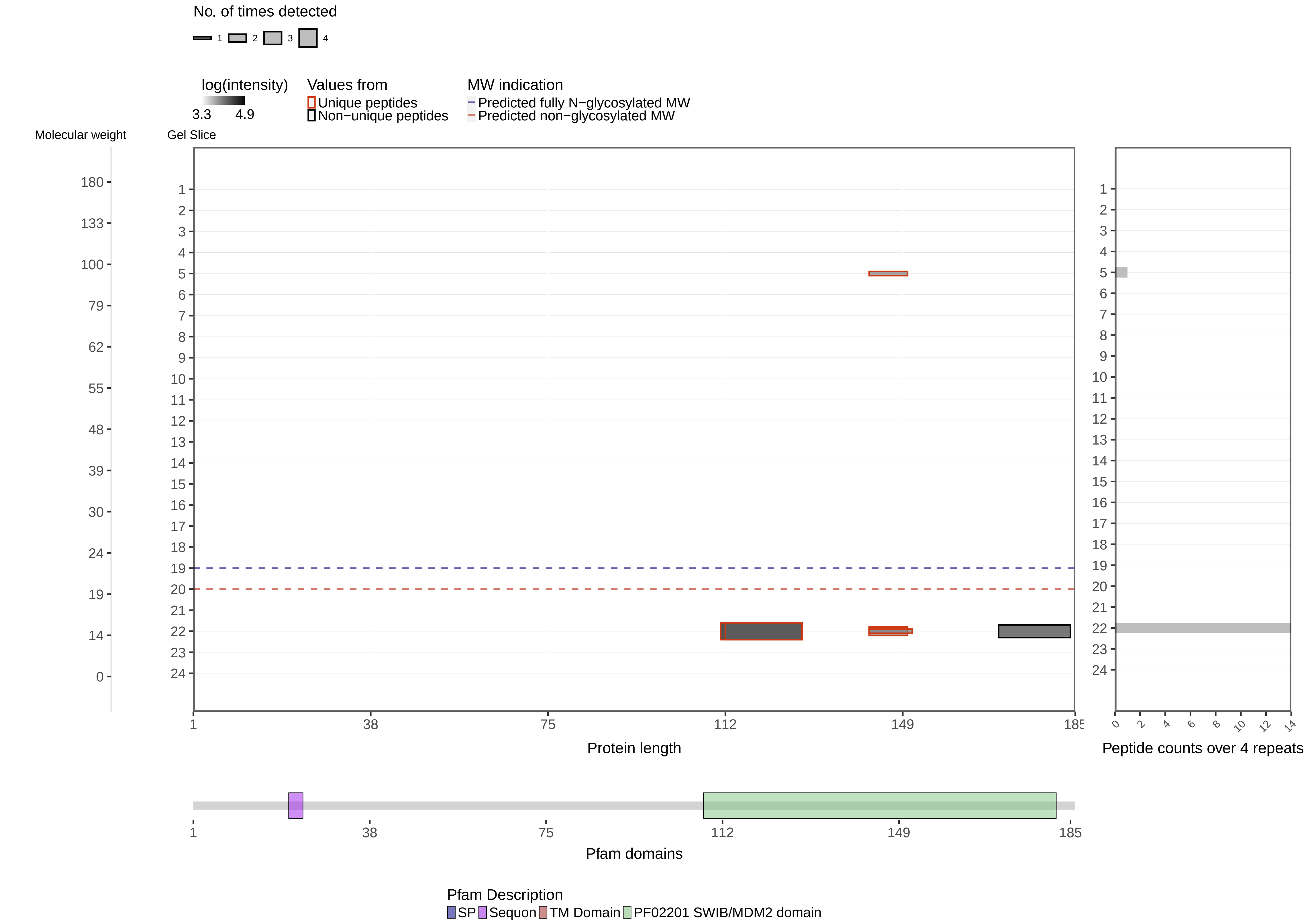


Protein sequence:

LPRVRMVTFAASSKPATTEPNKREPRGIMKPRRVTPEMQAFLGGLPEIPRTQALKFIWAHIKKHNLQDPEDKKVIVCDEKLKNIFGGRDRVGFLEIAGLISPHFLK*

### NbL07g07850.1 beta−ureidopropionase [MW = 46.63 kDa]

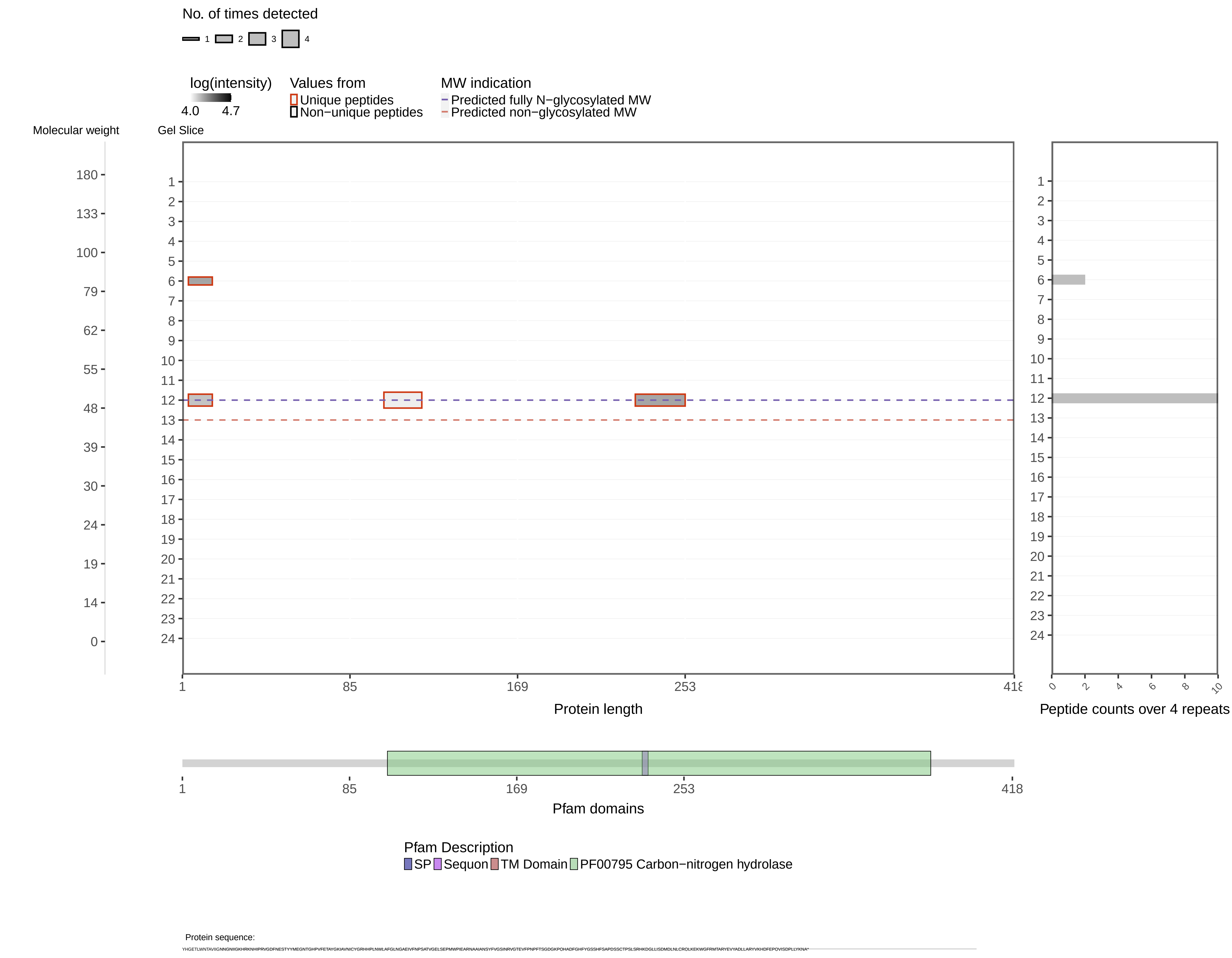

NbL07g07720.1 porphobilinogen_deaminase_chloroplastic−like [MW = 39.35 kDa]

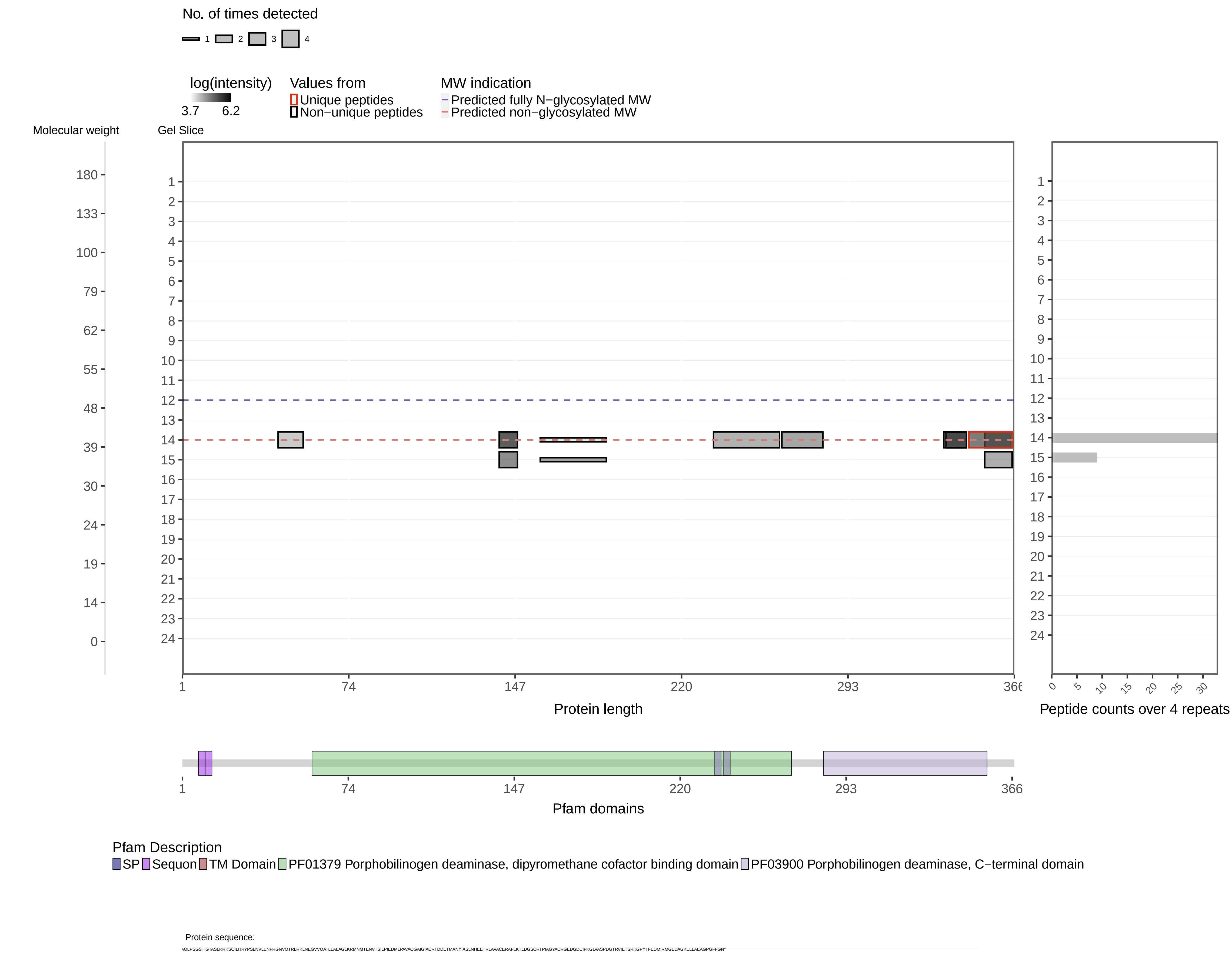

### NbL07g07590.1 phosphoglycerate_kinase_cytosolic [MW = 42.32 kDa]

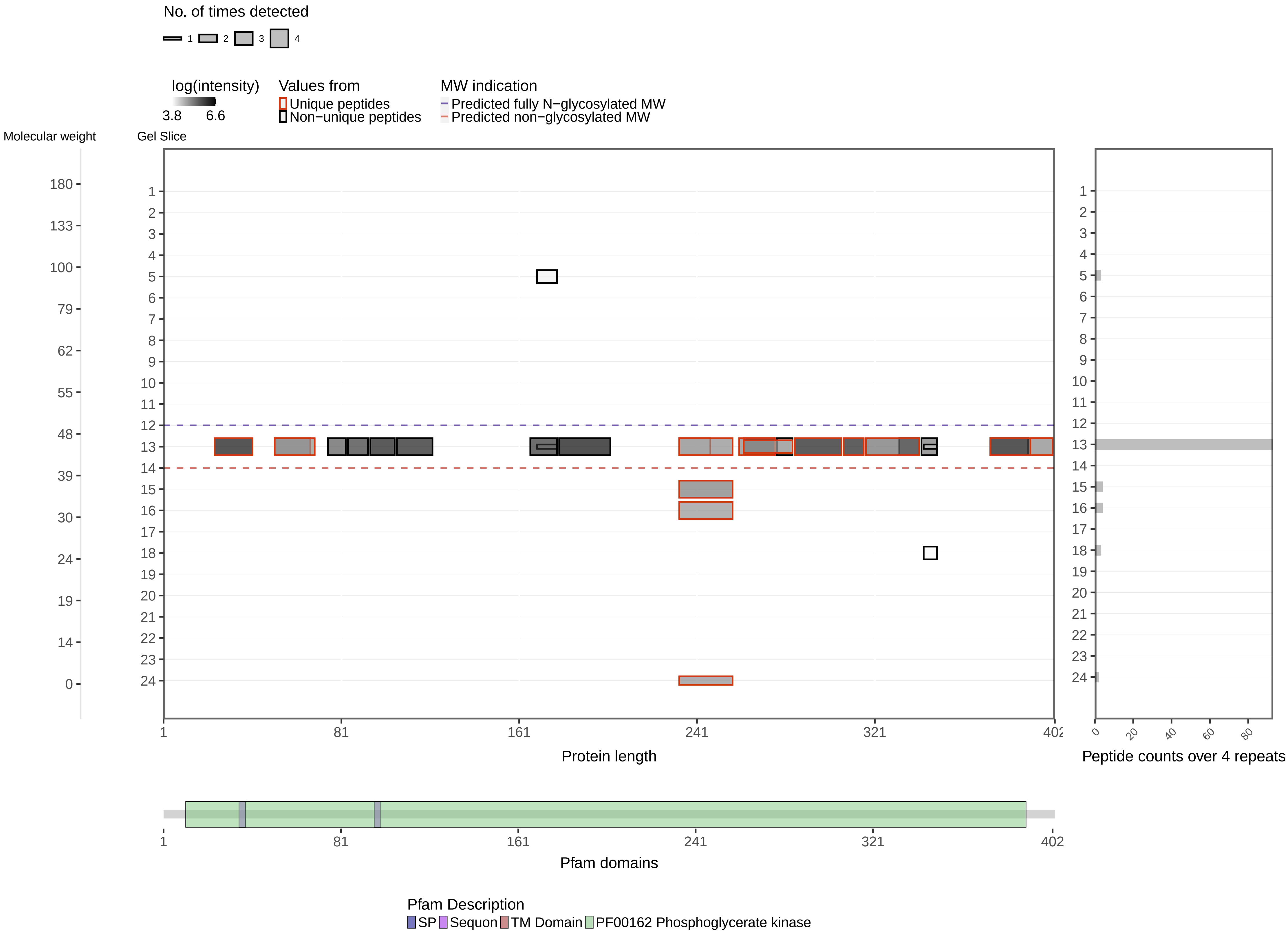


Protein sequence:

ANPOKPFAAIVGGSKVSSKIGVIESLLEKVDVLLLGGGMIFTFYKAOGYAVGSSLVEEDKLDLATSLMEKAKAKGVSLLLPTDVVIADKFAADANSKVVPASEIPDGWMGLDIGPDAIKSFGSALDTTKTVIWNGPMGVFEFDKFAAGTEAIAKKLAELSGKGVTTIIGGGDSVAAVEKVGLAEKMSHISTGGGASLELLEGKPLPGVLALDDA*

### NbL07g07580.1 phosphoglycerate_kinase_chloroplastic [MW = 51.12 kDa]

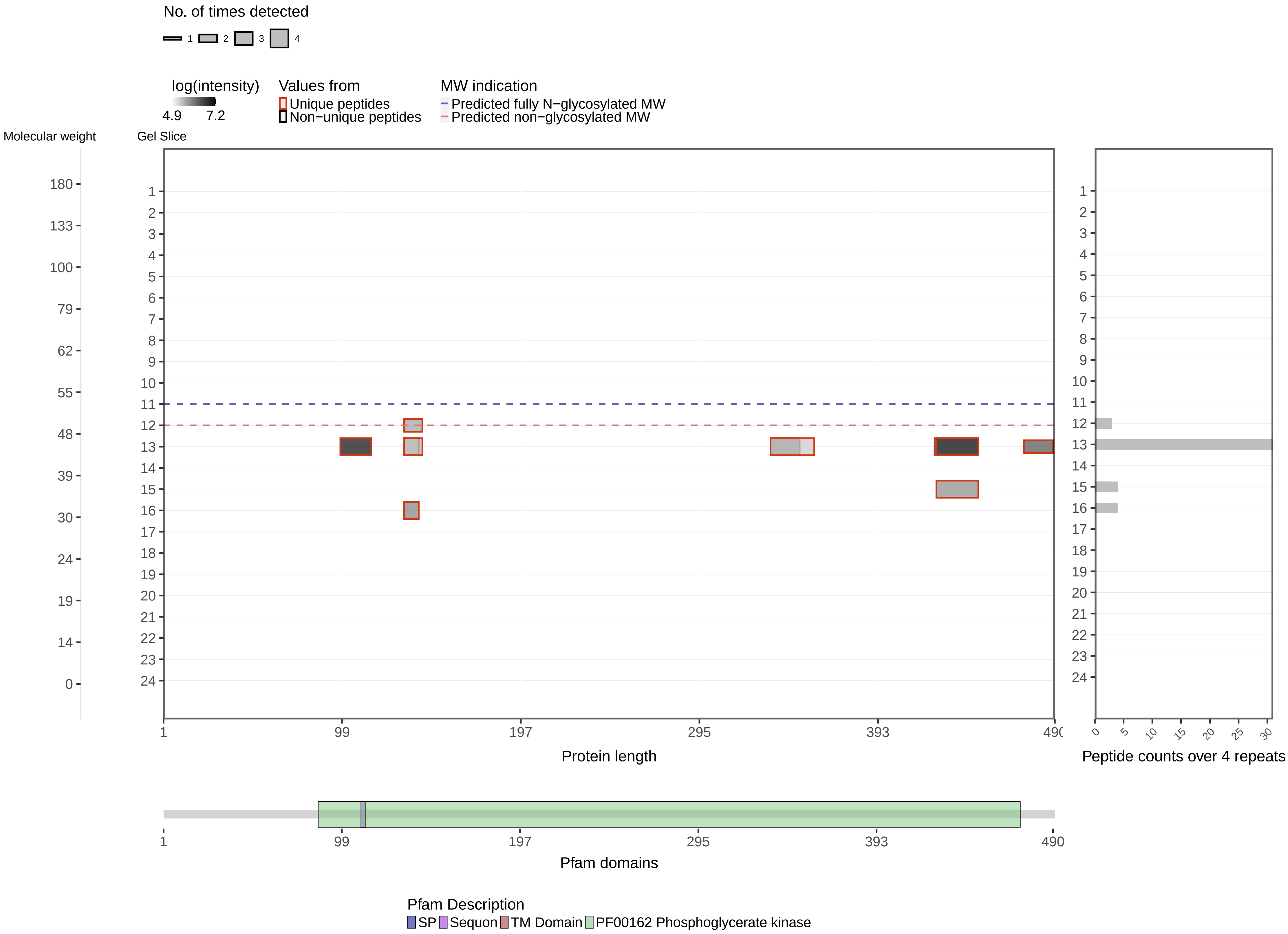


Protein sequence:

ASTEGVTKFLKPSVAGFLLQKELDYLVGAVSNPKRPFAAIVGGSKVSSKIGVIESLLEKCDILLLGGGMIFTFYKAQGLSVGSSLVEEDKLELATSLLEKAKEKGVSLLLPSDVVIADKFAPDANSKVCMLSFSHIVPSSAIPDGWMGLDIGPDSVKTFNDALDTTKTVIWNGPMGVFEFDKFAVGTEAIAKKLADLSGEGVTTIIGGGDSVAAVEKVGVASVMSHISTGGGASLELLEGKVLPGVIALDEADAPVAV*

NbL07g07200.1 glucose−1−phosphate_adenylyltransferase_large_subunit_2_chloroplastic/amyloplastic−like [MW = 61.7 kDa]

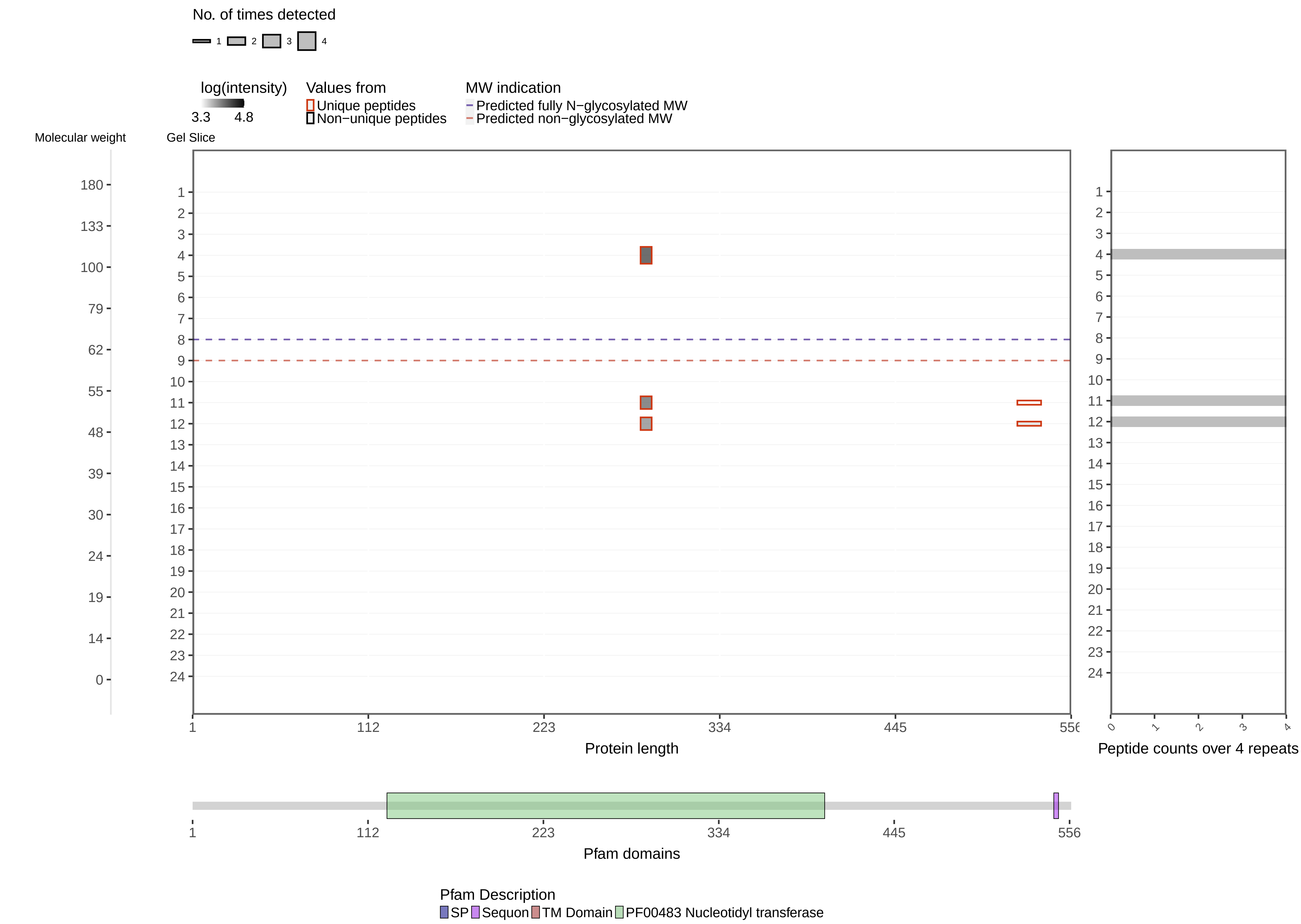


Protein sequence:

### NbL07g06740.1 isoamylase_3_chloroplastic [MW = 86.88 kDa]

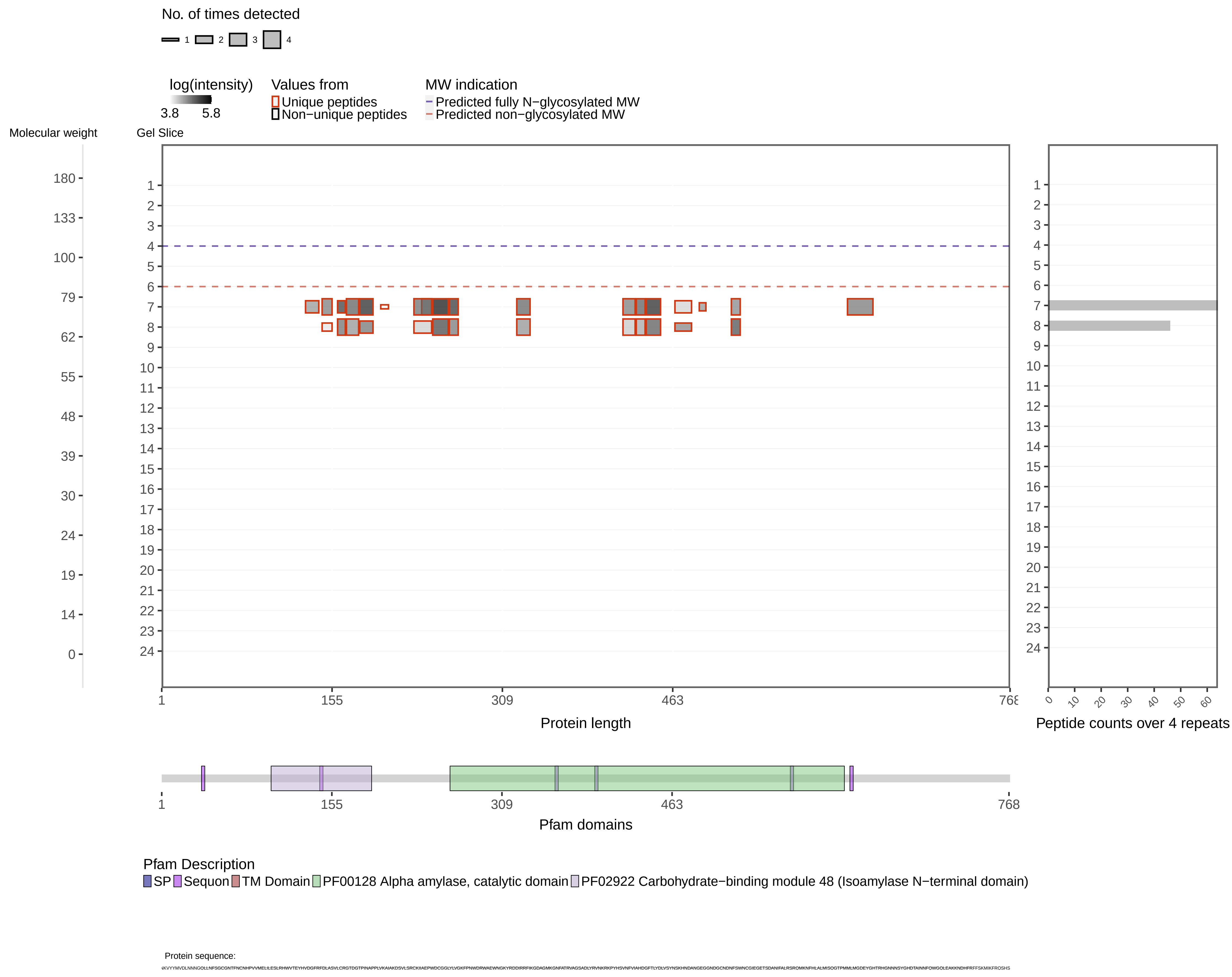

### NbL07g06650.1 proline−rich_receptor−like_protein_kinase_PERK1 [MW = 69.69 kDa]

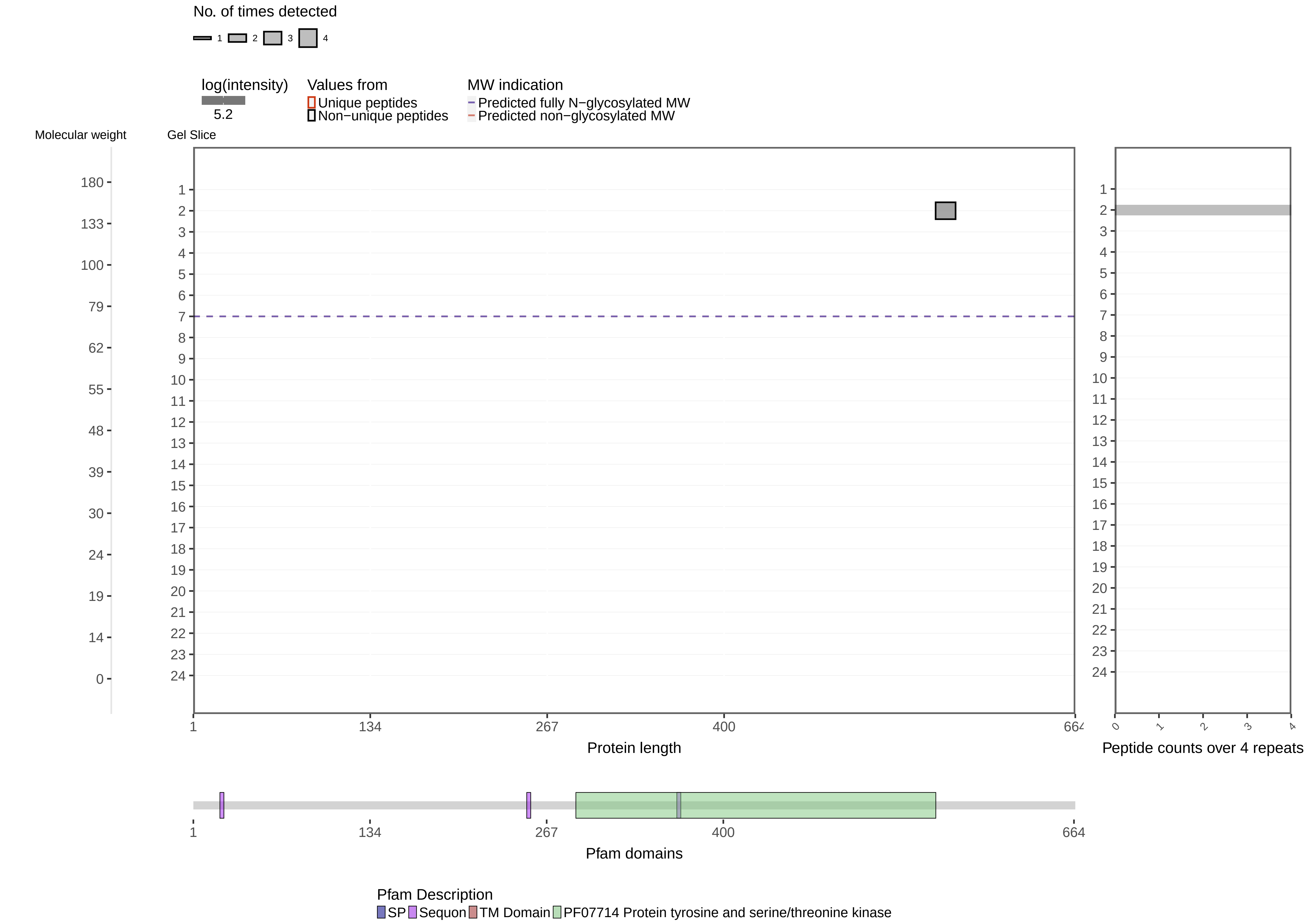


Protein sequence:

AGSGQGEREFQAEVEIISRVHHKHLVSLVGYCITGSQRLLVYEFVPNNTLEFHLHGKGRPPLDWPIRLKIALGSAKGLAYLHEDCQPKIIHRDIKAANILIDFNFEAKVADFGLAKLTSDVNTHVSTRVMGTFGYLAPEYASSGKLTEKSDVFSYGIMLLELITGRRPVDSSQTYMDDSLVDWARPQLTRALEDEKFDSLIDPRLGNDYNHNEVARMVACAAACVRHSARRRPRMSQVVRALEGDVSLSDLNEGIRPGHSTVYSSHGSSDYDASQYNEDMKKFRKTALGSQEYG

### NbL07g06540.1 GEM−like_protein_1 [MW = 32.31 kDa]

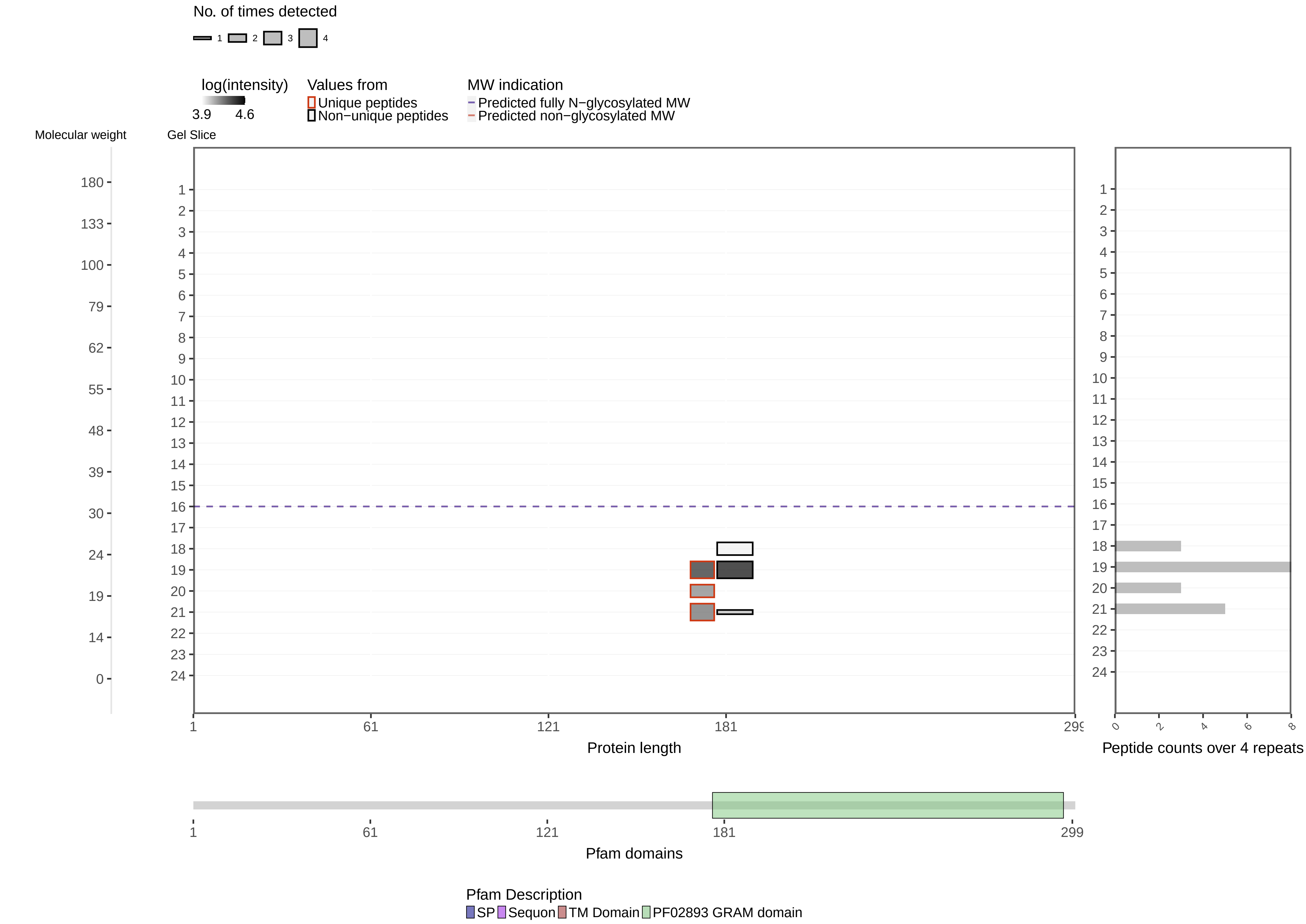


Protein sequence:

TKKAEDLAGNMWQHLKTSPSLADAAMGRIAQGTKVLAEGGYEKVFQQTFETVPEEKLFKSYACYLSTSAGPVMGVLYLSTAKLAFCSDNPLSYKVGDRTEYSYYKVVLPVDQLKAISPSTSKINPAEKYIQIISVDNHEFWFMGFVNYDSAVKNLQGALLGHHS*

NbL07g06500.1 3−hydroxyisobutyryl−CoA_hydrolase−like_protein_3_mitochondrial [MW = 41.64 kDa]

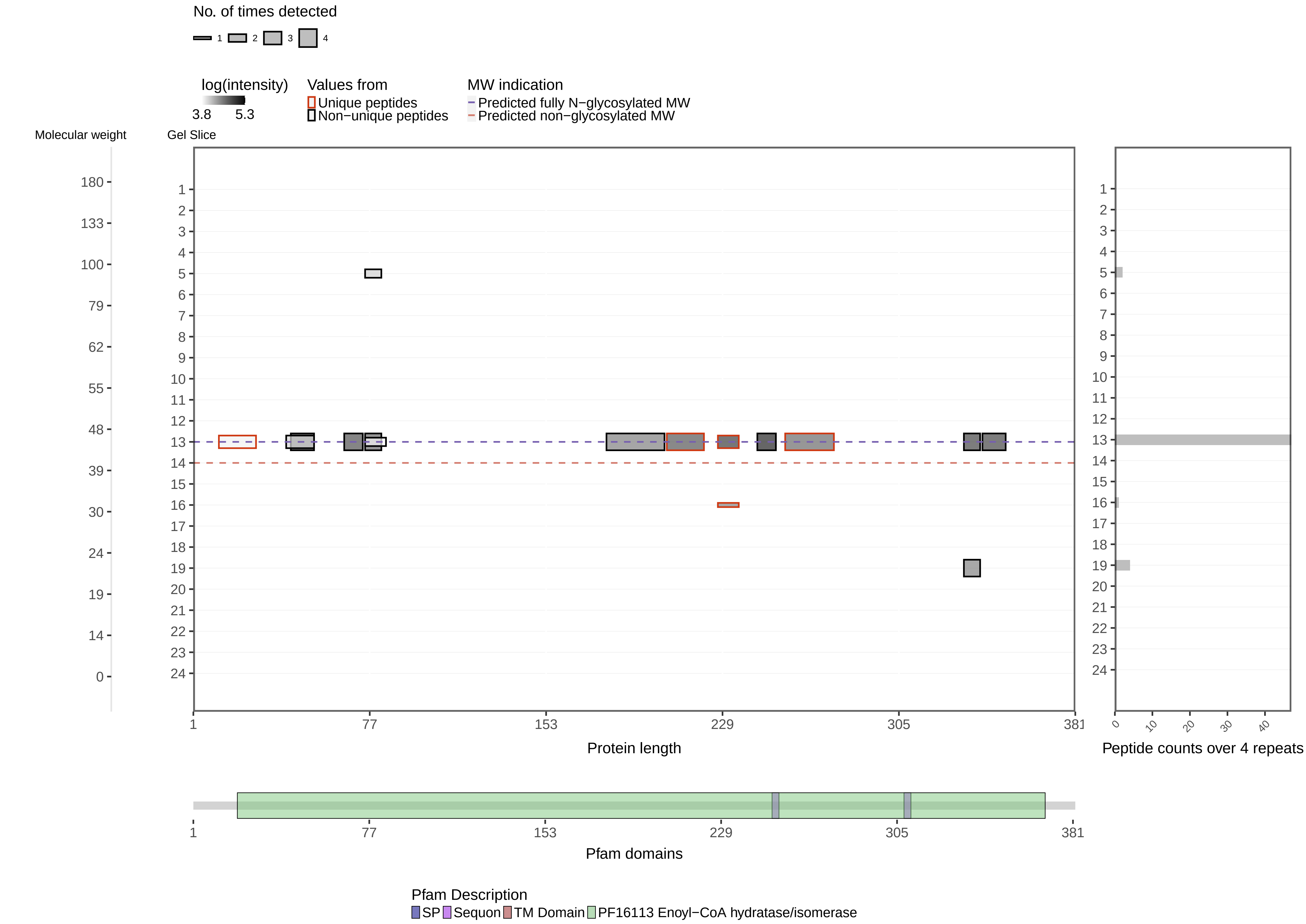


Protein sequence:

SRISTPADALYVGLGSHYVPSANLDTLKEAFLAATFSQDPDEEIKTLLEKYSIDPDSESRLKSLLPRIISTFGGNKSVKNIIEELDNHQQSADTLVAEWAKDALQGLRKGAPFSLCLTQQYFSRVASACGKNESTLSKLPVVMRTEYRIAIRSALRNDFAEGVRAILVDKDQNPRWNPSSLEEVNLNEVEVLFEPLSPEVELNV*

NbL07g06420.1 peroxisomal_fatty_acid_beta−oxidation_multifunctional_protein_AIM1−like [MW = 80.69 kDa]

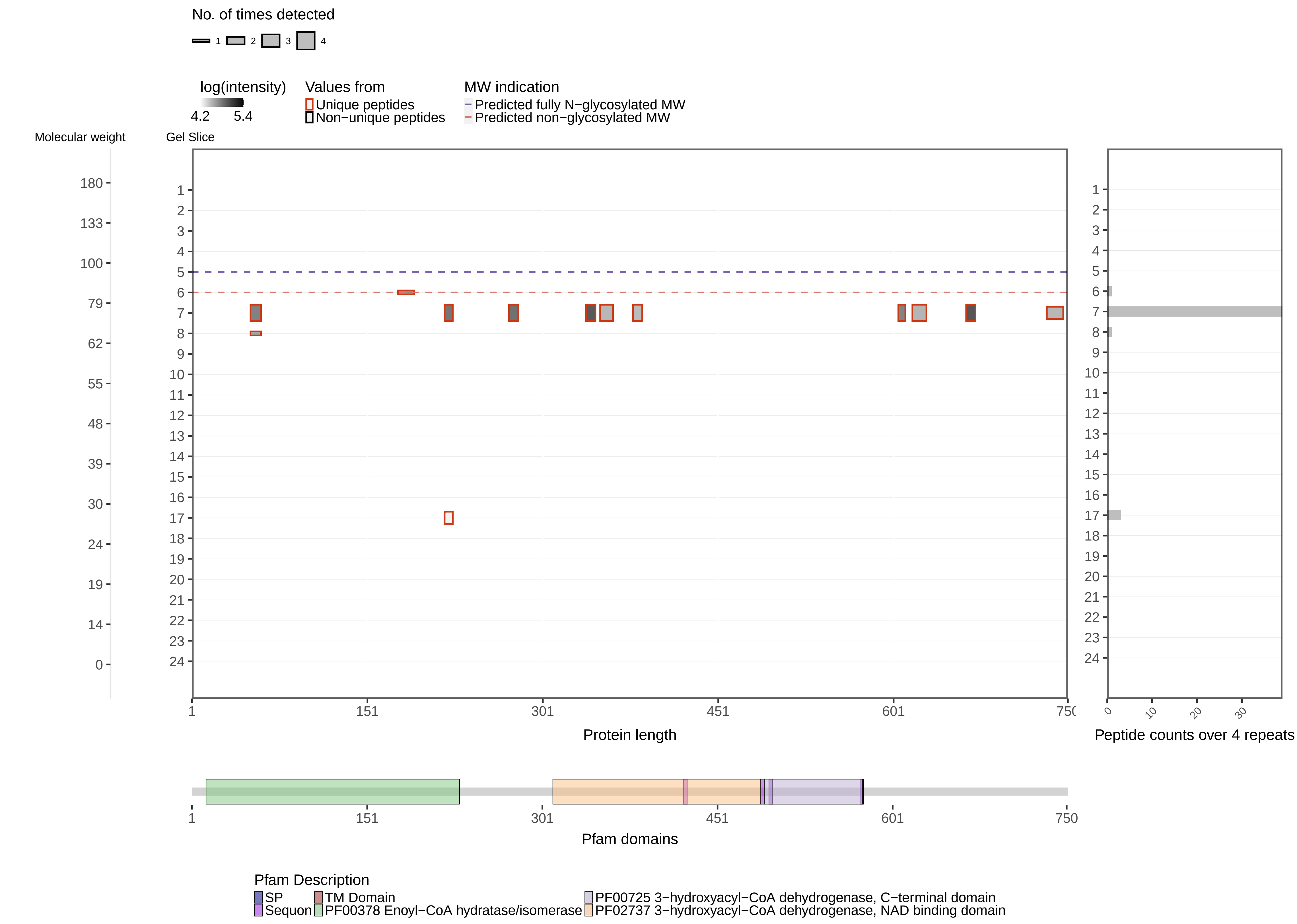


Protein sequence:

[illegible]

NbL07g06380.1 peroxiredoxin−2B−like [MW = 19.05 kDa]

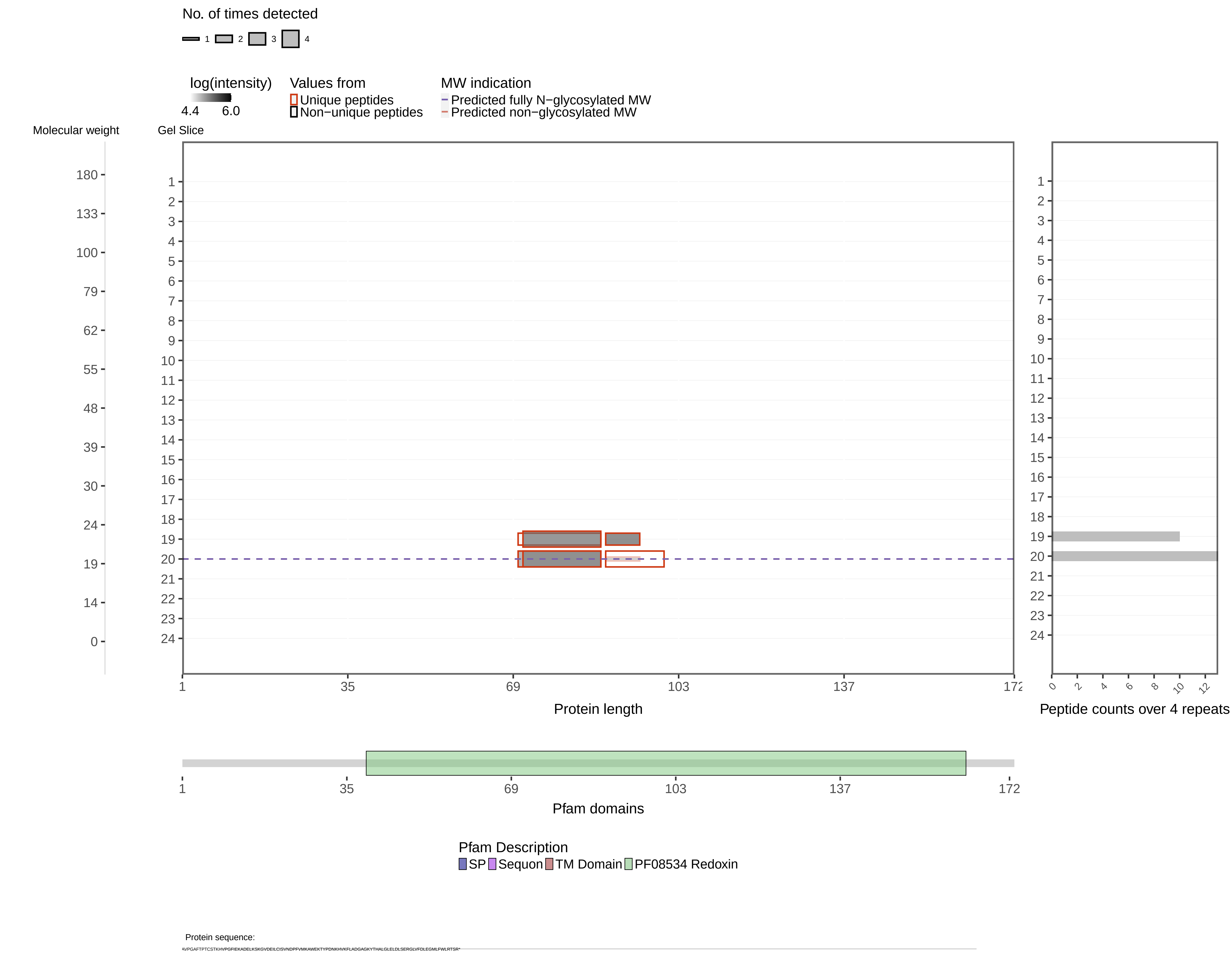

NbL07g06340.1 uncharacterized_LOC104245598 [MW = 113.69 kDa]

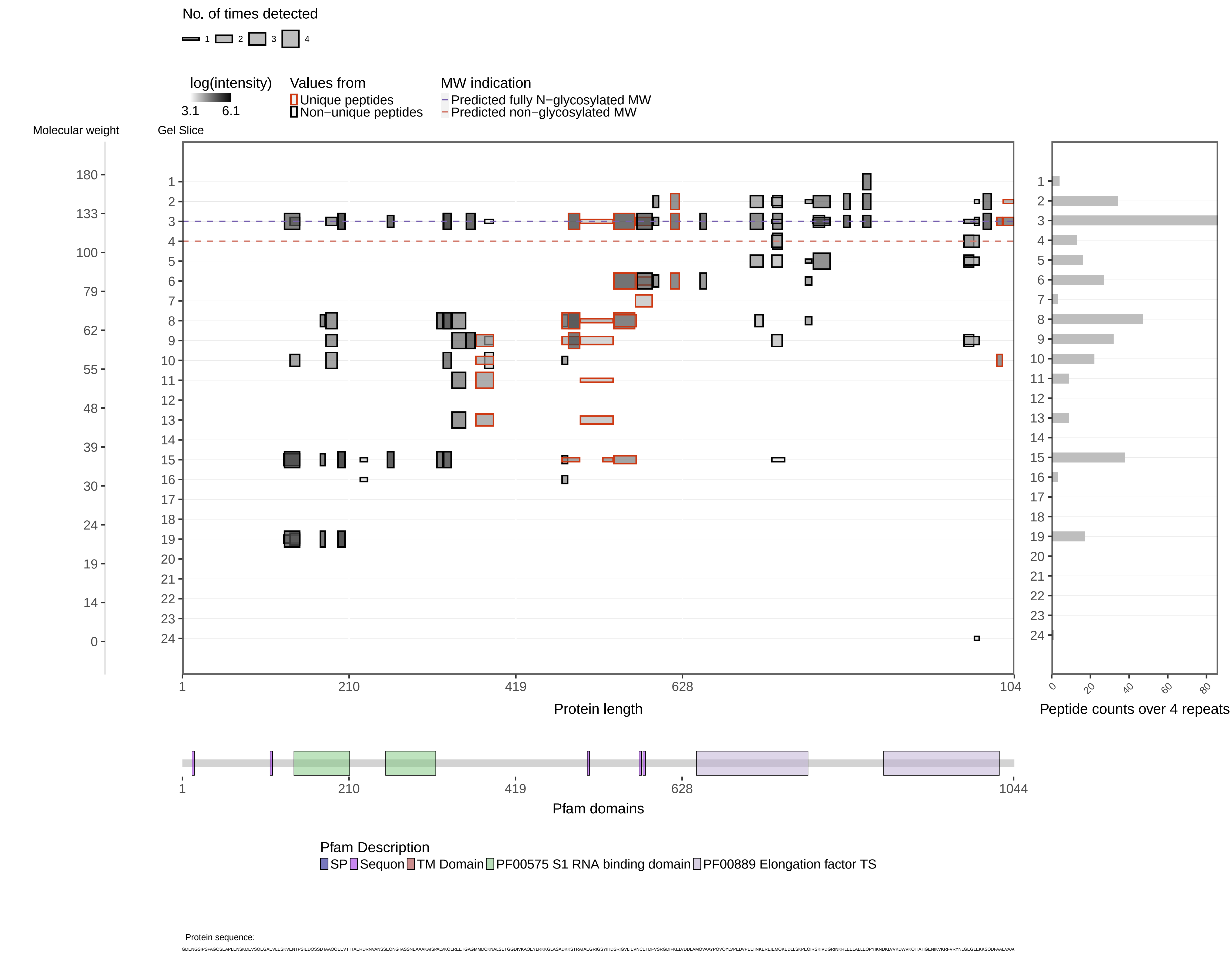

### NbL07g06070.1 NADPH−−cytochrome_P450_reductase−like [MW = 78.44 kDa]

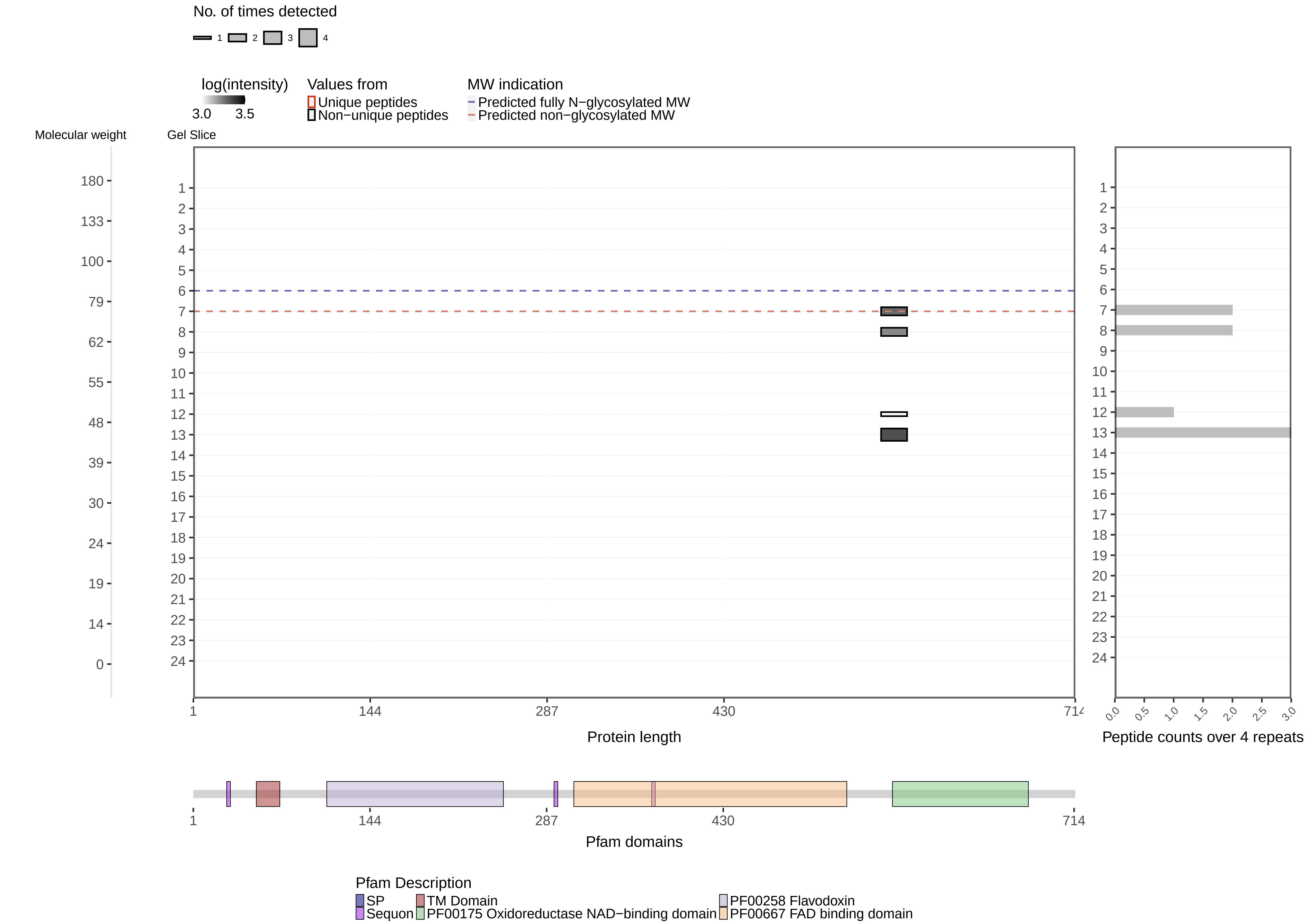


Protein sequence:

### NbL07g06060.1 (DL)−glycerol−3−phosphatase_2 [MW = 27.26 kDa]

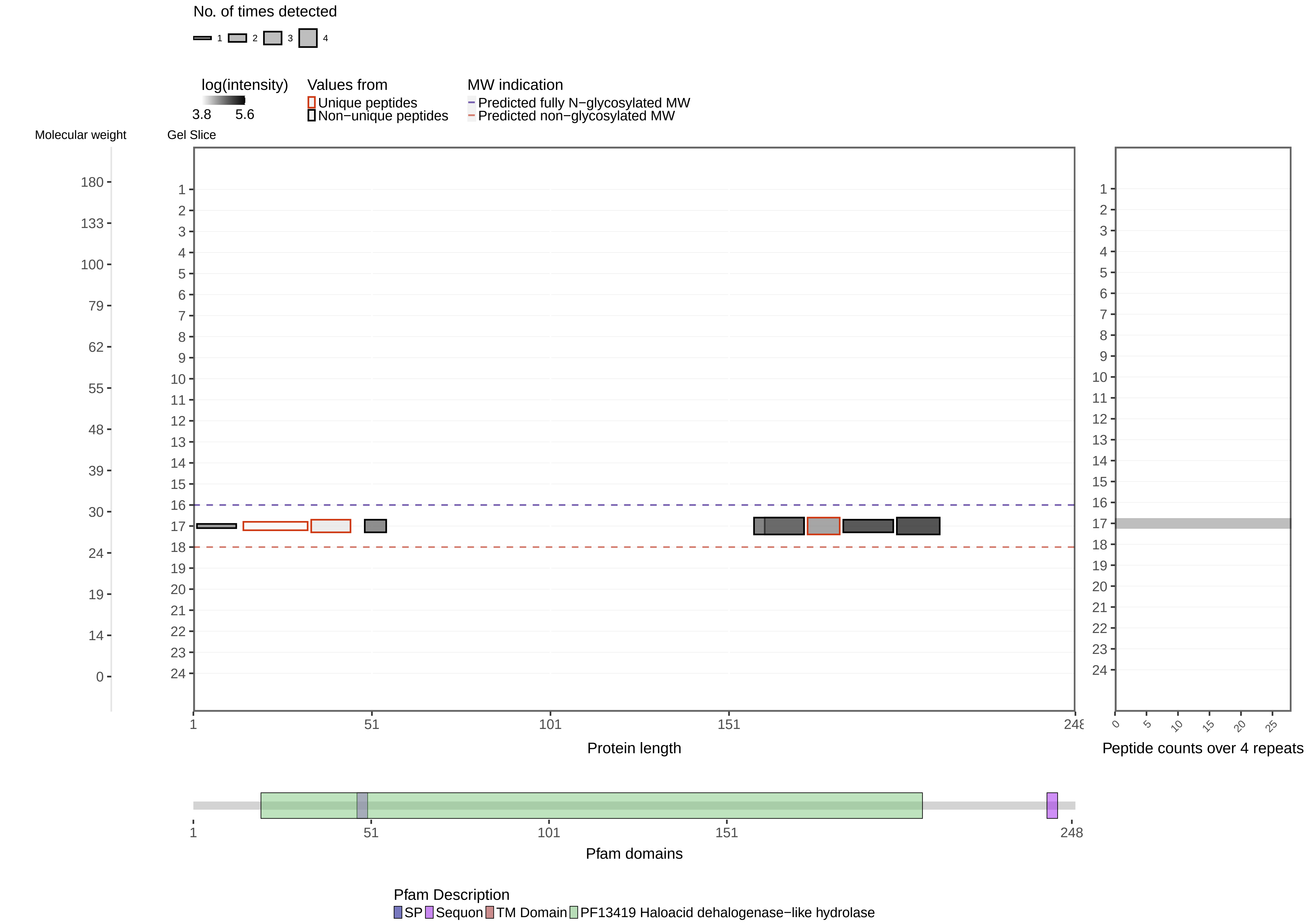


Protein sequence:

HLHANGIPICVATGTHTRHFDLKTQRHGELFSLMHHIVRGDDPEVKQGKPSPDIFLAAANRFEGGPVDPQRILVFEDAPSGVLAAKNAGMSVVMVPDARLDSSYHEVADQVLSSLLDFNPADWGLPPFENSSTKLD*

NbL07g06050.1 uncharacterized_LOC109234215 [MW = 68.88 kDa]

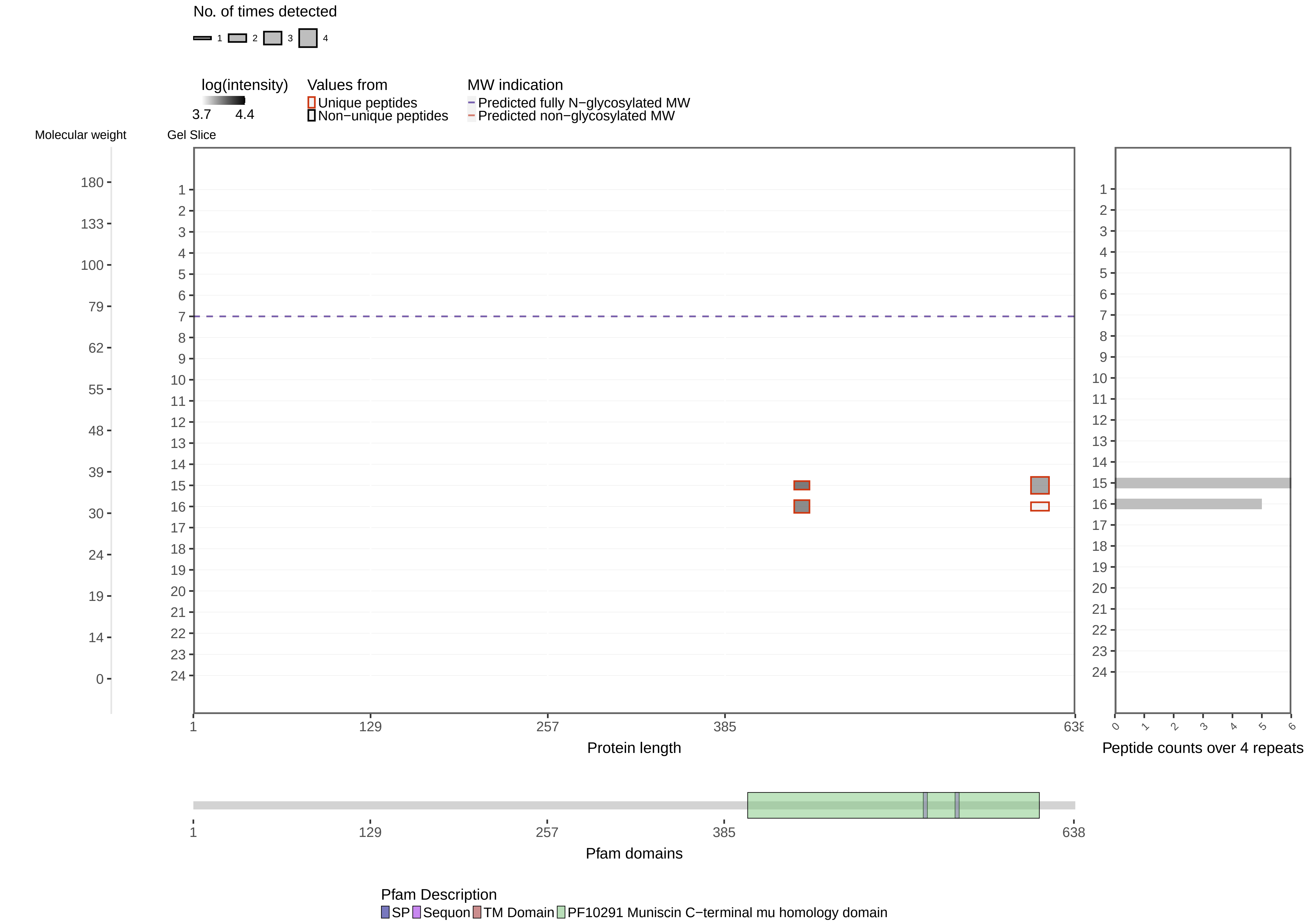


Protein sequence:

HIGVEGFEGDYGGIEFSNDGSTLQQDFEGINDAWGGGLDASEYVGTKKVKKDQGLGGLELLETSEPPKAAAGAATDGGAGKKLEDILVKKMNGPEMFITEEISAEFRESLLARVGLMGVVFLRTLPPKSSDDTETEFSFKVEGTAGVKRFVMQNSHVSSLGNGLFHVKTAPSNEPIPIIKYSLLPRLTPLPLRIRLVKRLSGTLLSVMLQYVANPDLPVPLTNVTFVLKLPVDPTLLKVTPKAVLNRSERELKWHVDEIPLKGHPGKLRARFPVDINDDDGGEELVLFGYVKFS

NbL07g06000.1 ribulose_bisphosphate_carboxylase_small_chain_clone_512−like [MW = 19.34 kDa]

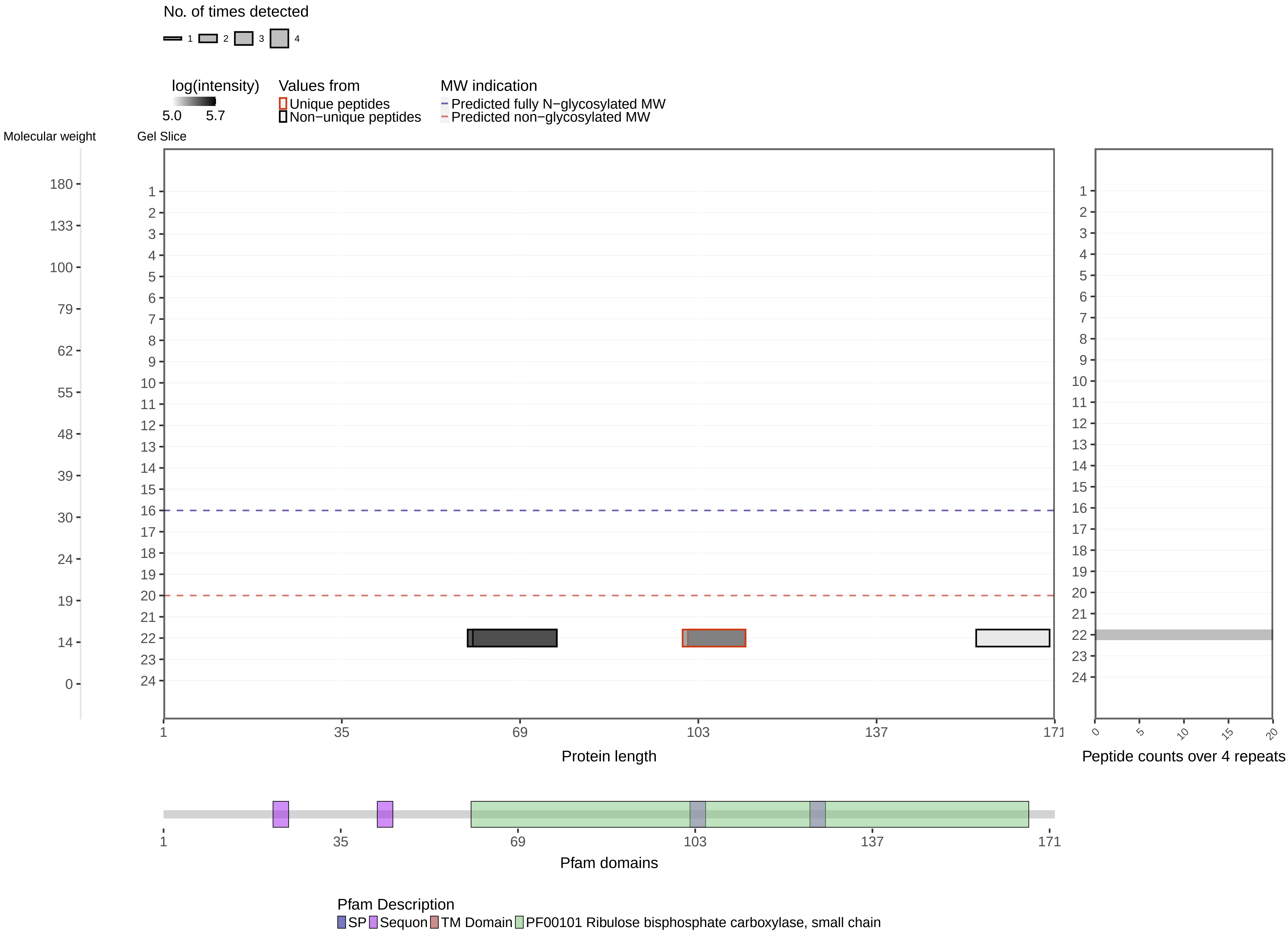


Protein sequence:
SIAKEVDYMIQNGWIPCLEFDQVGYVRRENSSMPGYYDGRYWTLWKLPMFGCNDSSQVLNEIQECKKAYPNAFIRCLAFDNVKQVQCMAFLIQKPAAA*

### NbL07g05970.1 putative_glucose−6−phosphate_1−epimerase [MW = 35.48 kDa]

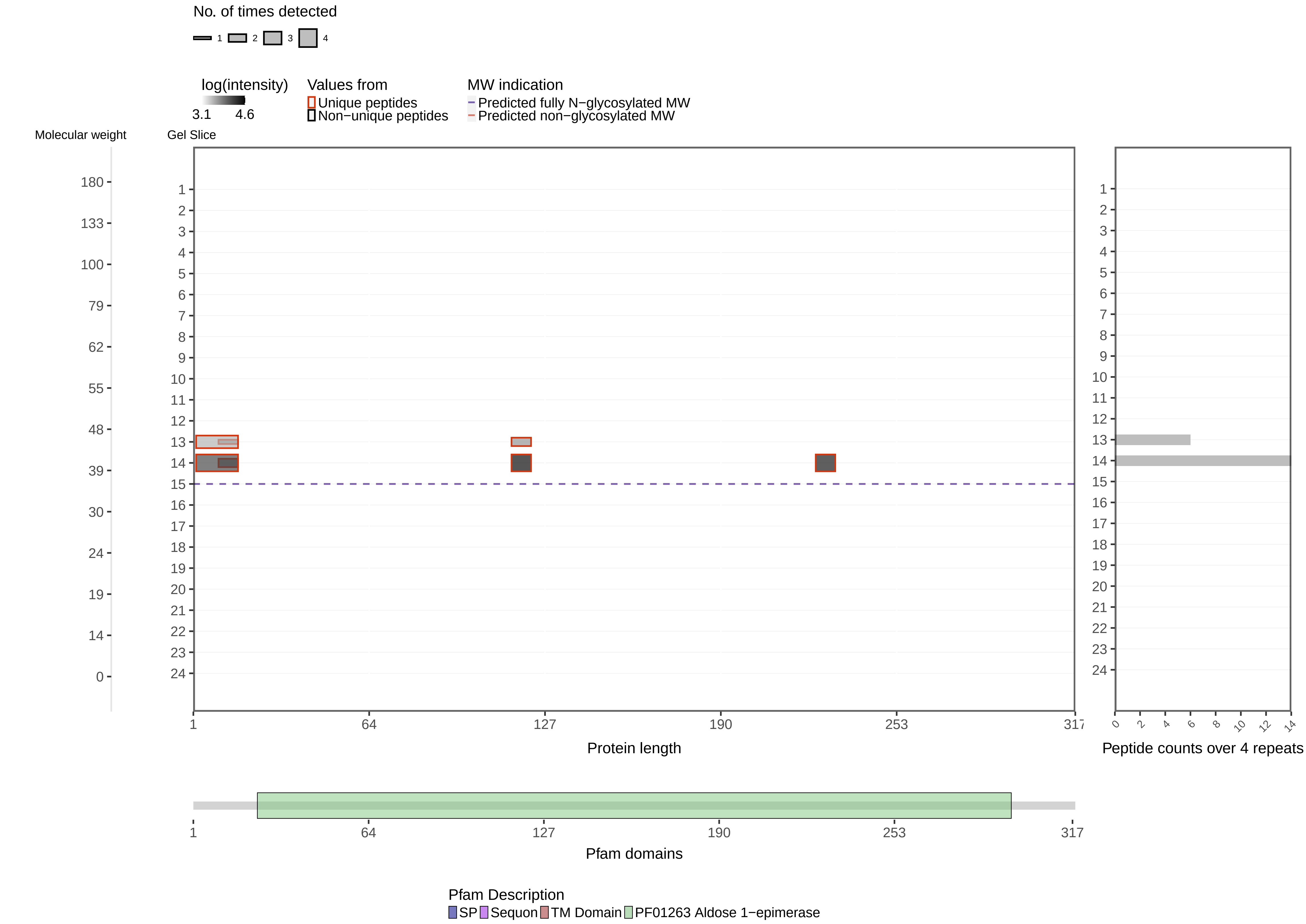


Protein sequence:

GDLMLTSRIRNTNTDGKPFSFTFAYHTYFSVSDISEVRVEGLETLDYLENKERFTEOGDAITFESEVNKIYLSTPTKVAILDHEKKRTFVLRKDGLPDAVVWNPWDKKAKAMADFGDEEYKHMLCVEAAAVERPITLKPGEEWRGROELSAVPSSYCSGOLDPKKVLLGS*

### NbL07g05920.1 acetyl−coenzyme_A_synthetase_chloroplastic/glyoxysomal−like [MW = 75.03 kDa]

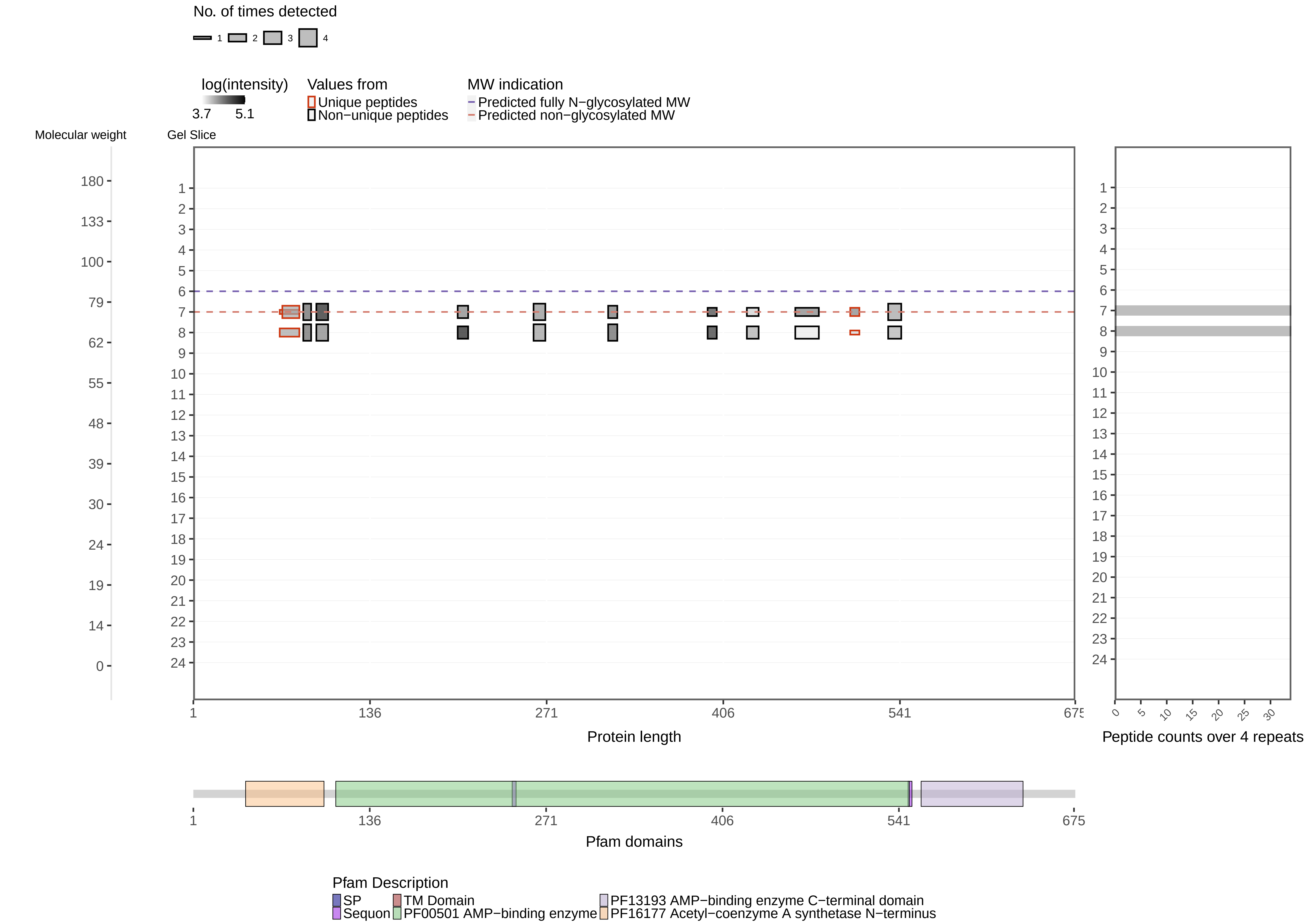


Protein sequence:

[illegible]

### NbL07g05900.1 plasma_membrane_ATPase_4 [MW = 105.31 kDa]

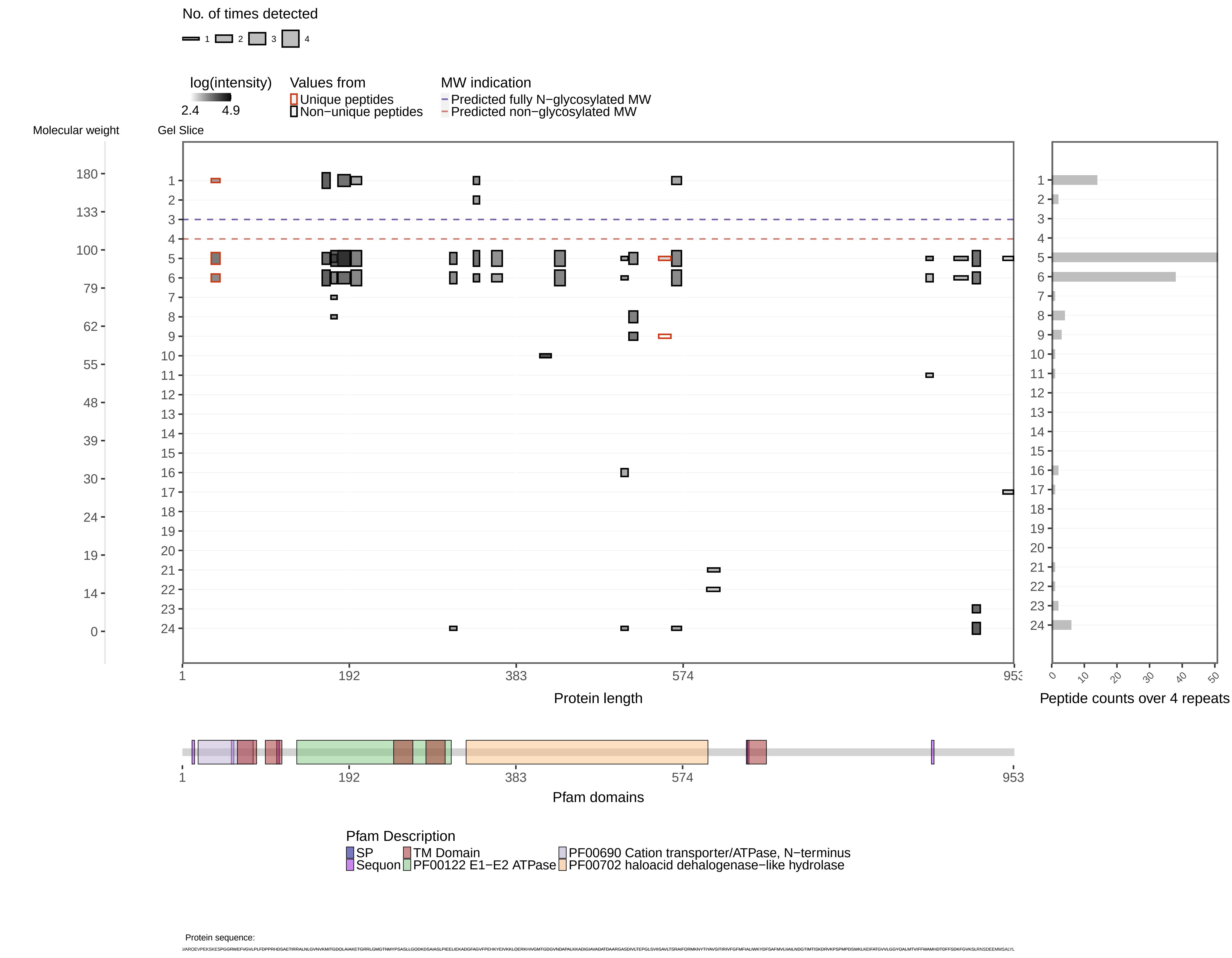

NbL07g05850.1 glucan_endo−13−beta−glucosidase_5−like [MW − SP = 52.51 kDa]

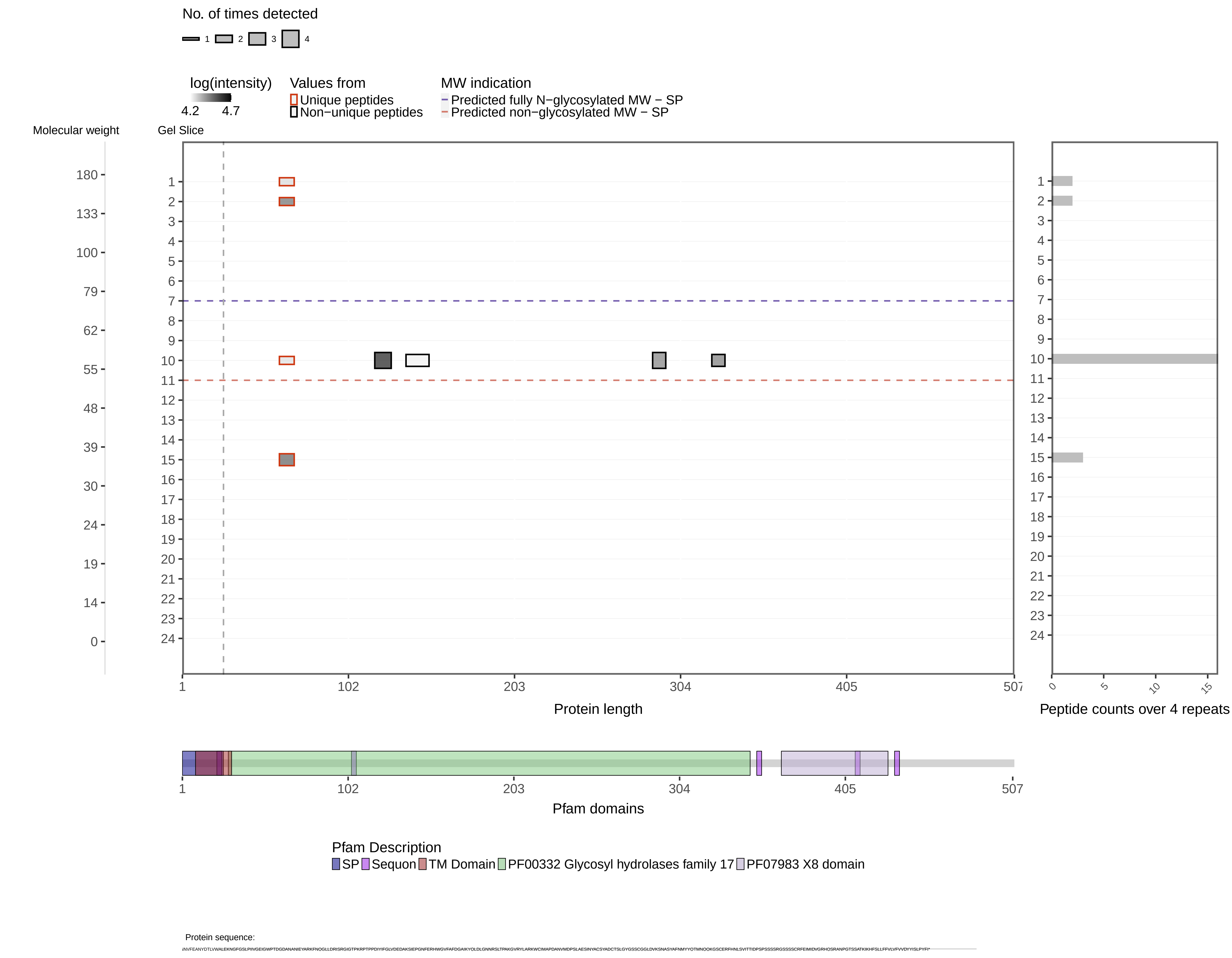

NbL07g05830.1 pectinesterase/pectinesterase_inhibitor_PPE8B−like [MW − SP = 56.17 kDa]

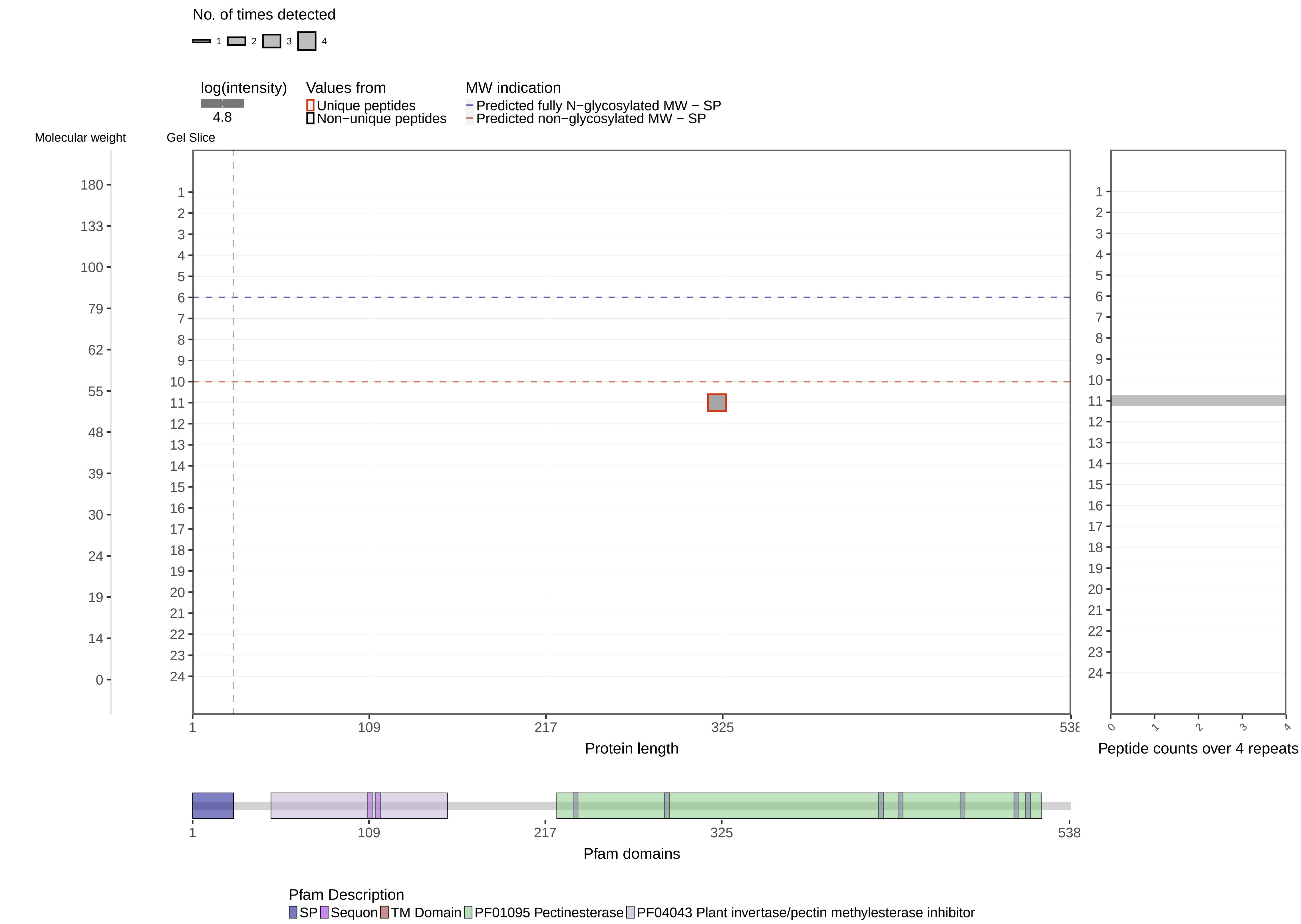


Protein sequence:

IKKGIYNEYVEISKKKWNILMIGDGIDVTVISGNRSFIDGWTTYRSATFAVKGQGFIARDMTFENTAGPQKHQAVAFRSDSDLSVLFRCALRGYQDTLYAHSMRQFYRNCIITGTVDFIFGDGTVVFQNCQILARKGLPEQKNTITAQGRKEAAETTGFSIQFSNISGEPDLLASLNSTQTFLGRPWKAYSRTVIMQSYMSNVIKPQGWLEWNGNISLDTLYYAEYQNYGPGAGLGARVNWPGYHLLNDSSQANNFTVAQFILGNSWLPSTGVKYTAGLAV*

### NbL07g05700.1 lysine−−tRNA_ligase−like [MW = 71.04 kDa]

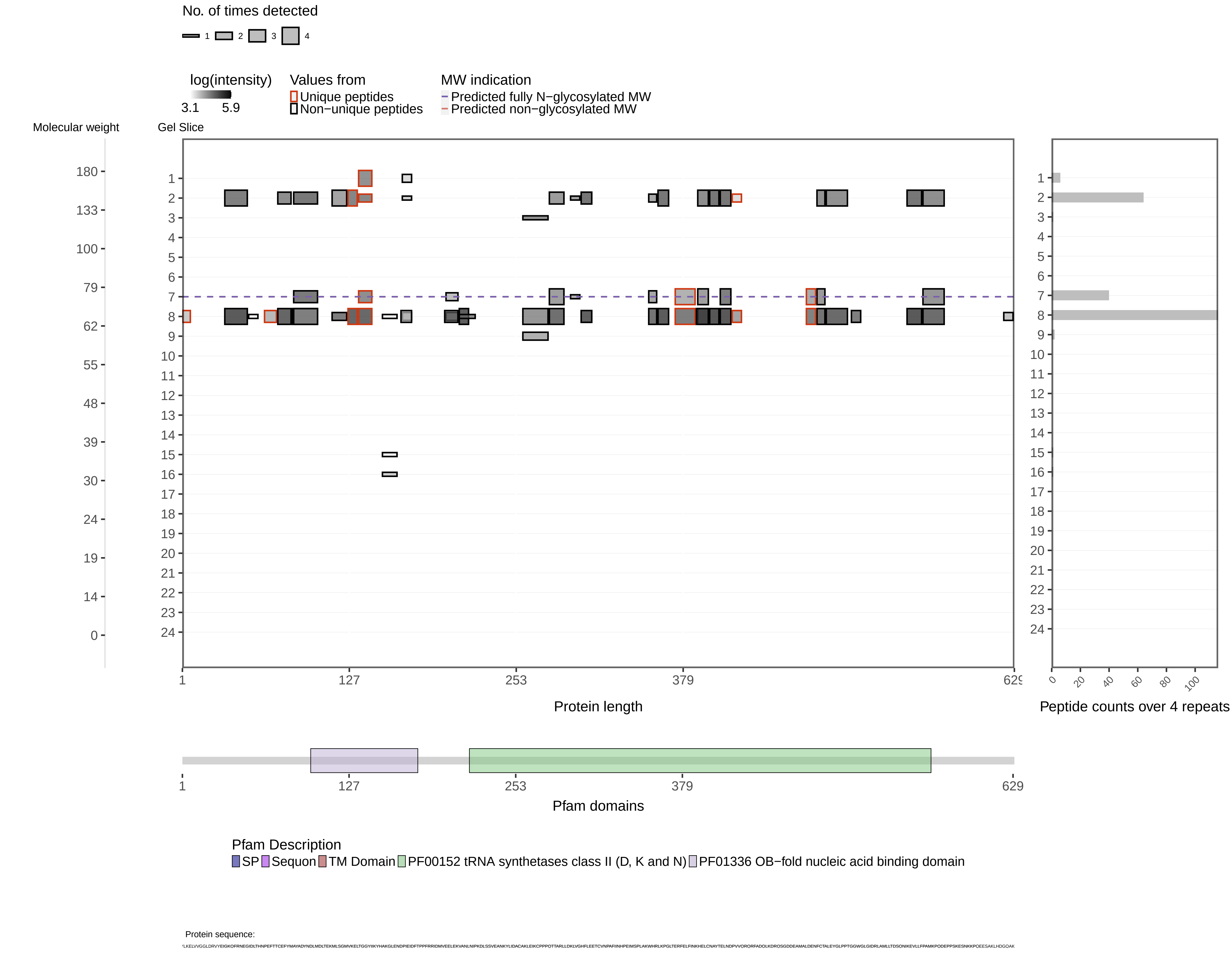

NbL07g05520.1 peptide_deformylase_1A_chloroplastic [MW = 30.84 kDa]

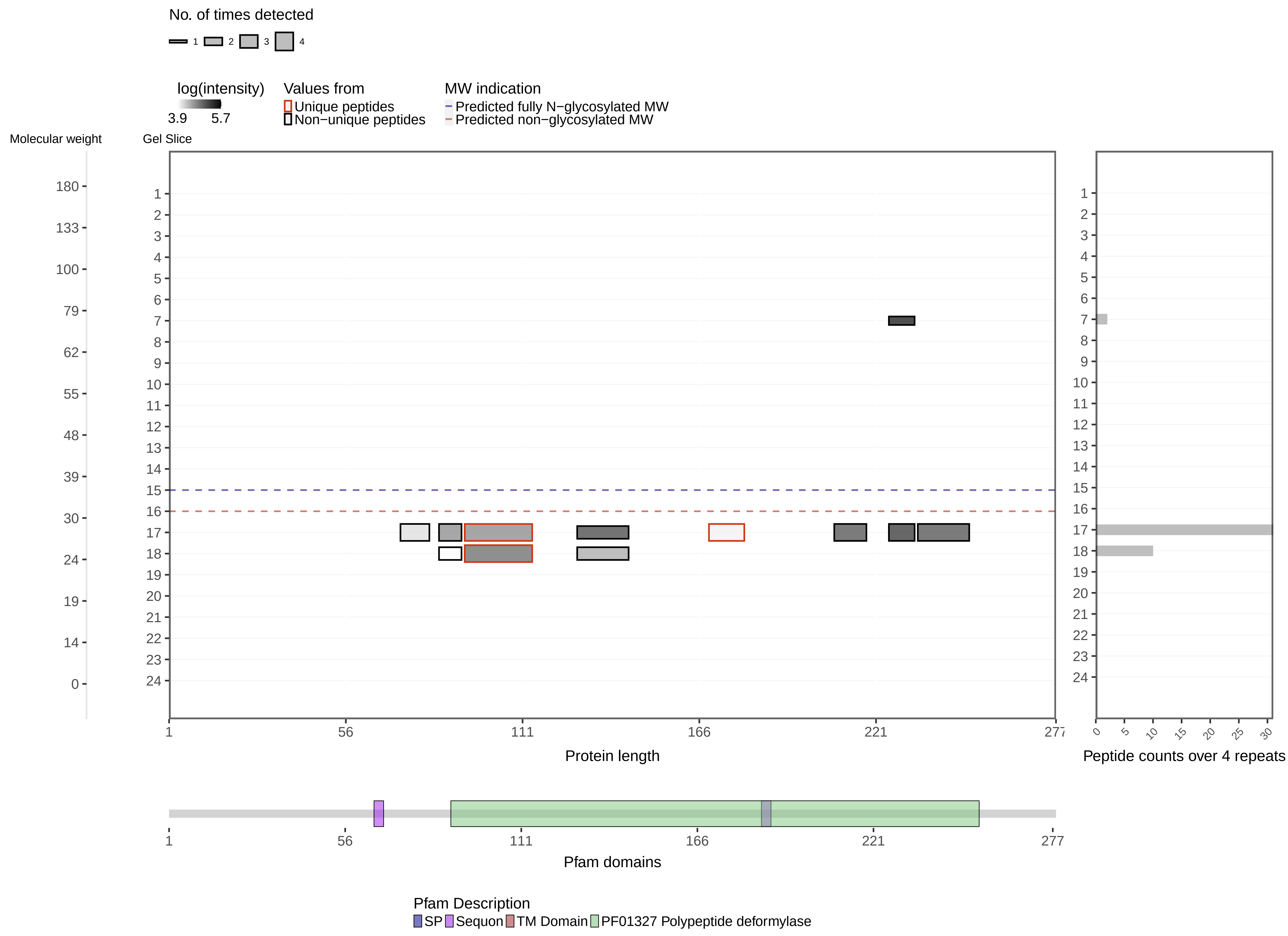


Protein sequence:

VMRNAPGVGLAAPQIGIPLKIIVLEDTNEYISYAPKDETKAQDRRPFDLLVIINPKLKKKGNKTALFFEGCLSIDGFRAVVERHLEVEVGGFDRNGKAIKVDASGWQARILQHEYDHLDGTLYVDKMVPRTFRTIENLDLPLAAGCPKLGVC*

NbL07g05420.1 ADP−ribosylation_factor−like_protein_2 [MW = 27.1 kDa]

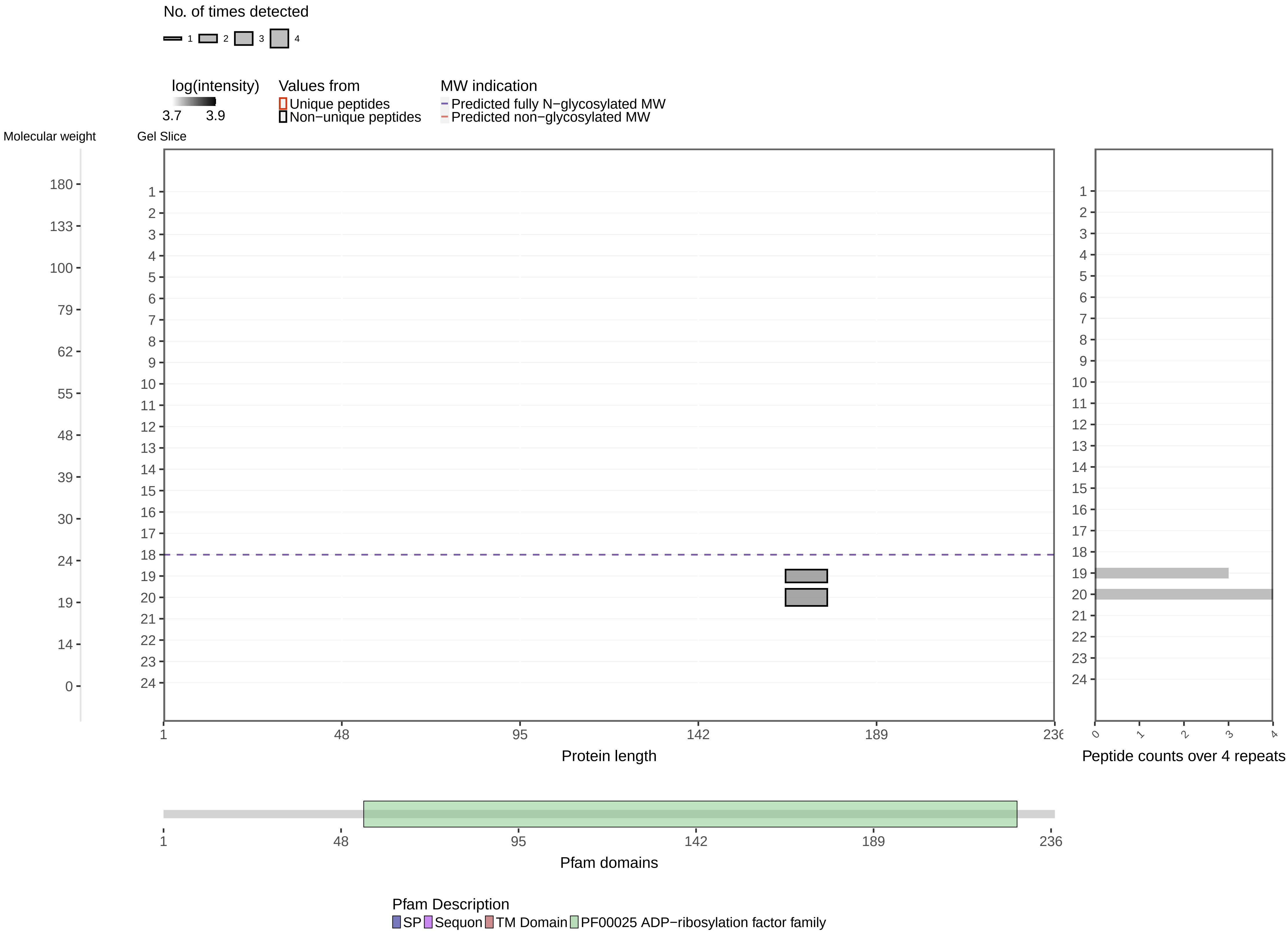


Protein sequence:

YOKYTLNIWDVGGOKTIRSYWRNYFEOTDGLVWVVDSSDLRRLDDCRYELHNLLKEERLSGASLLIFANKODIQGALSPDEIAKVLNLEAMDKSRHWRIVGCSAYTGDGLLDGFDWLVODIASRIYMLD*

NbL07g05400.1 isoamylase_1_chloroplastic [MW = 89.22 kDa]

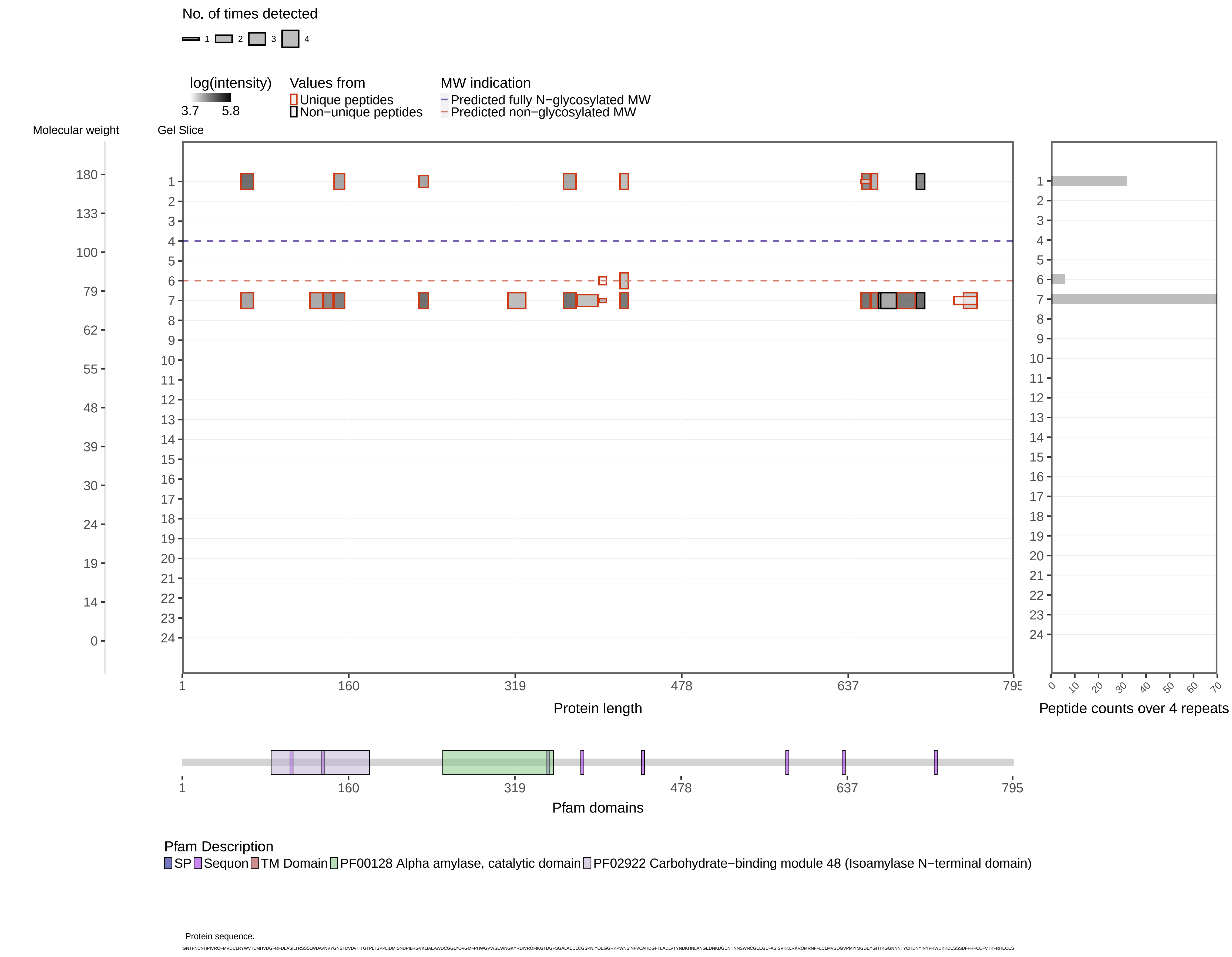

### NbL07g05300.1 60S_acidic_ribosomal_protein_P3−like [MW = 12 kDa]

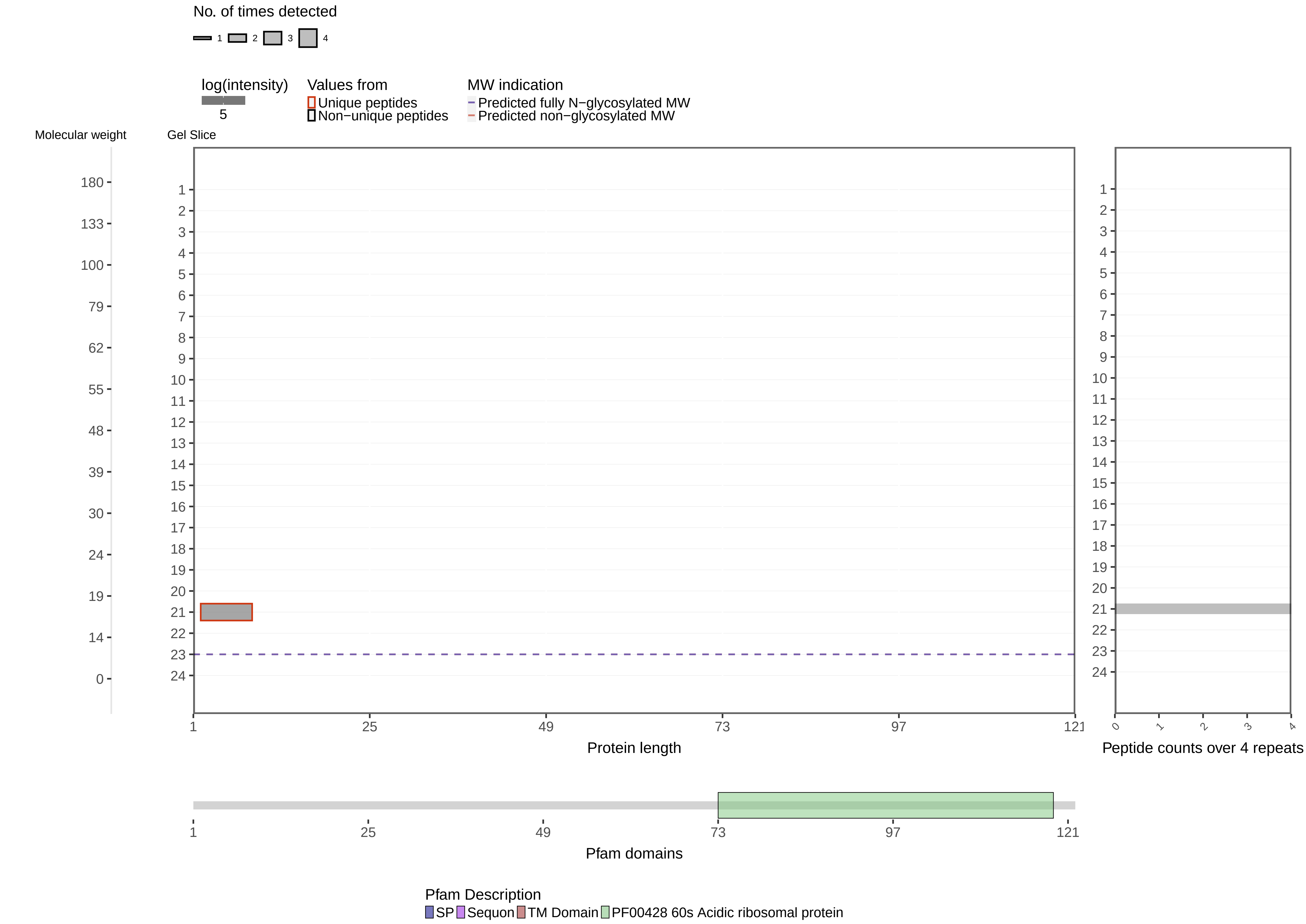


Protein sequence:

SGGVQSSFSYVTPSSAVFQVIIGGGGGGGFIGGGAAAAAPSGGAAAAEAPAAEEKKEEKEESEDDDMGFSLFD*

NbL07g05130.1 26S_proteasome_non−ATPase_regulatory_subunit_6_homolog [MW = 44.29 kDa]

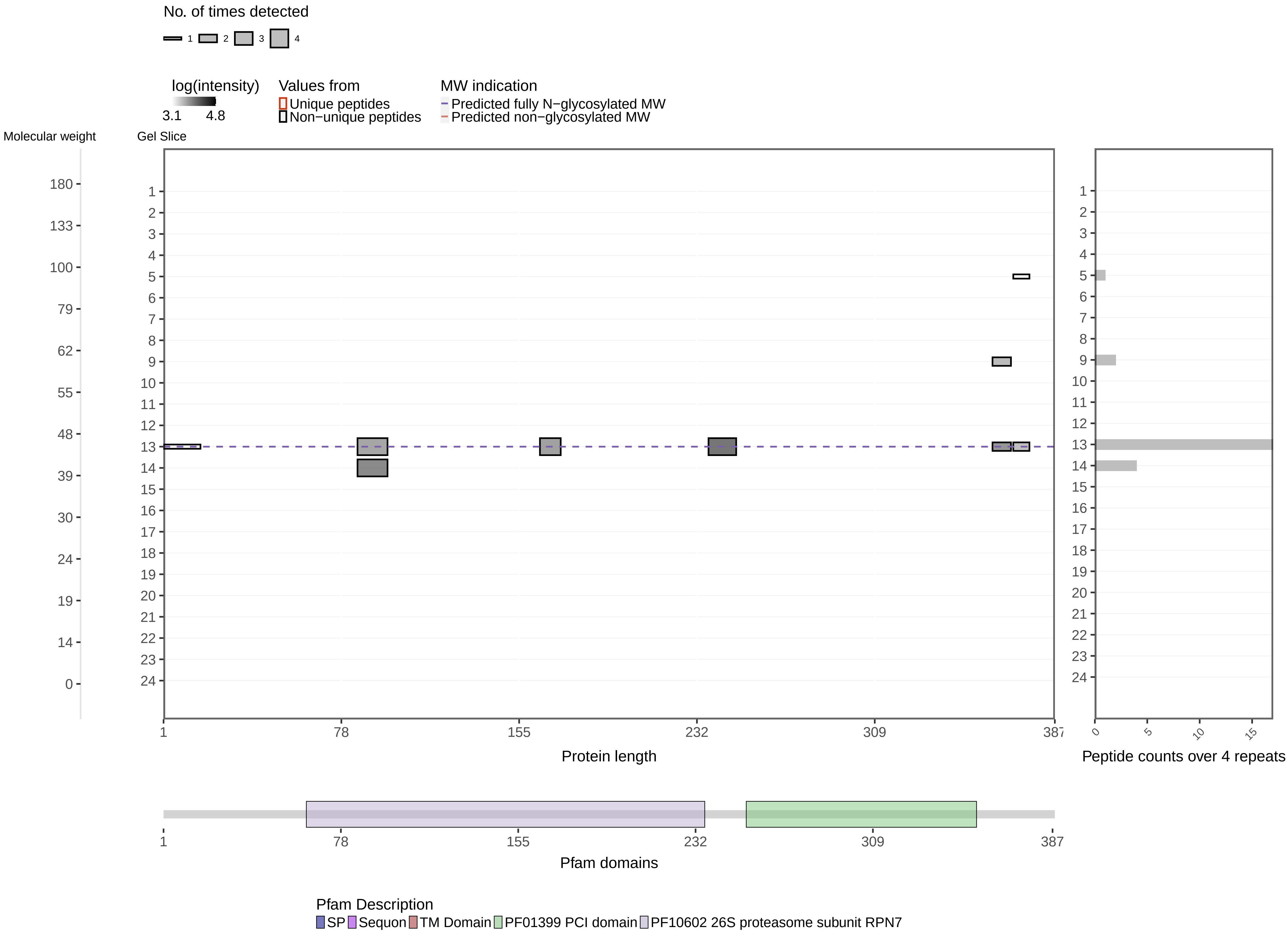


Protein sequence:

LKVYEGLFCMSTRNFKKAADLFLDSISTFTTYEIFPYDTFIFYTVLTSIITLDRVSLKQKVVDAPEILTVIGKIPYLSEFMNSLYECQYKSFFSAFAGLTEHIKLDRYLQPHFRYFMREVRIVVYSQFLESYKSVTIEAMAKAFGVSEDFIDLELSRFIAAGKLHCKIDKVAGVLETNRPDAKNALYQATIKQGDFLLNRIQKLSRVIDL*

### NbL07g05110.1 ER_membrane_protein_complex_subunit_8/9_homolog [MW = 23.09 kDa]

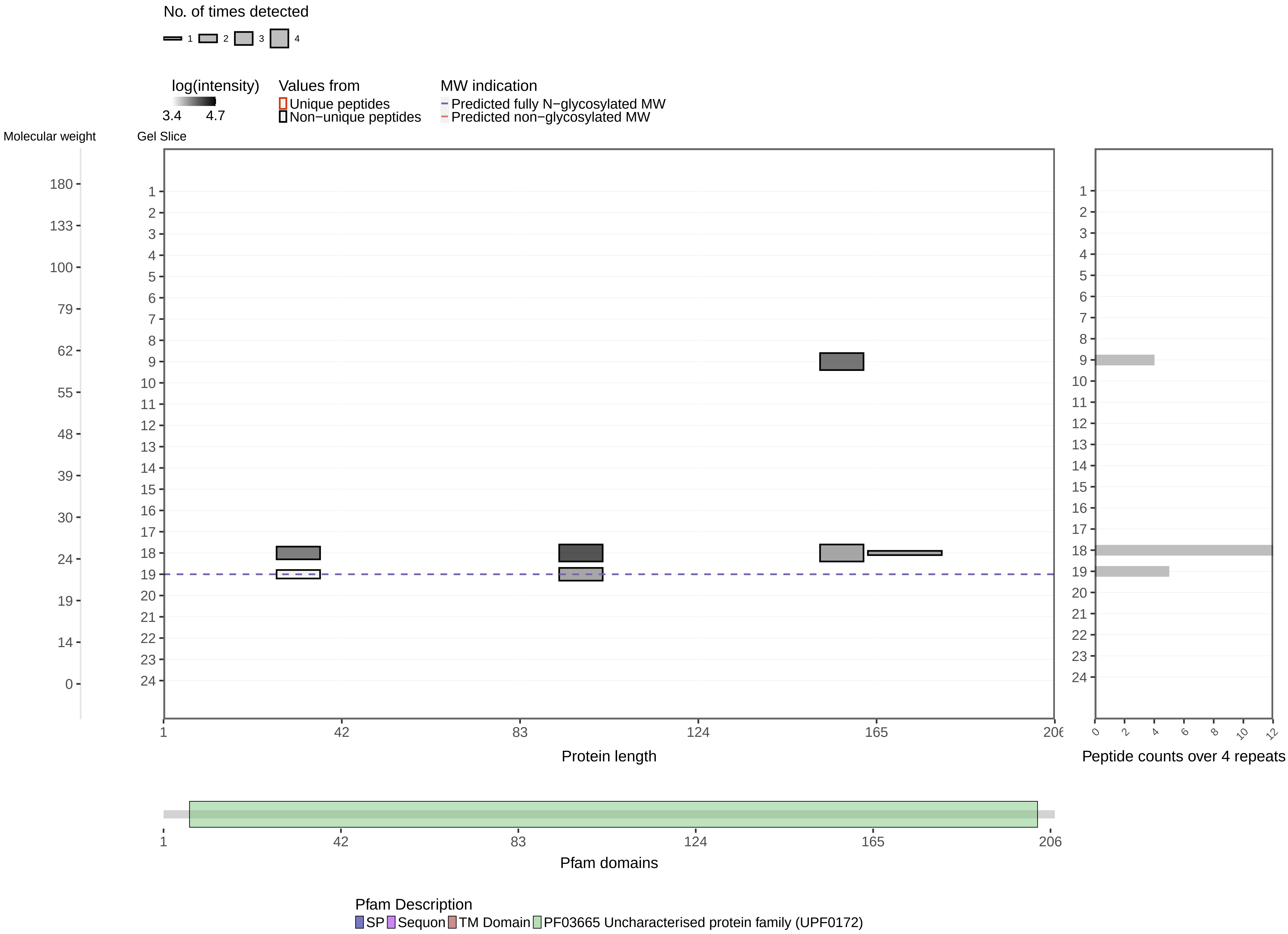


Protein sequence:

RFDDFEVGNVAKNIADHIFKYFPQAALLLLDNKKLEALLKEGKDRSPVMQLFTKEVSRSWKLVGTDASSPLKIKEPSANIVLLDYISSGKWKDIVDFDDHLDDISKDWLNPELFK*

### NbL07g05030.1 subtilisin−like_protease_SBT2.5 [MW − SP = 85.14 kDa]

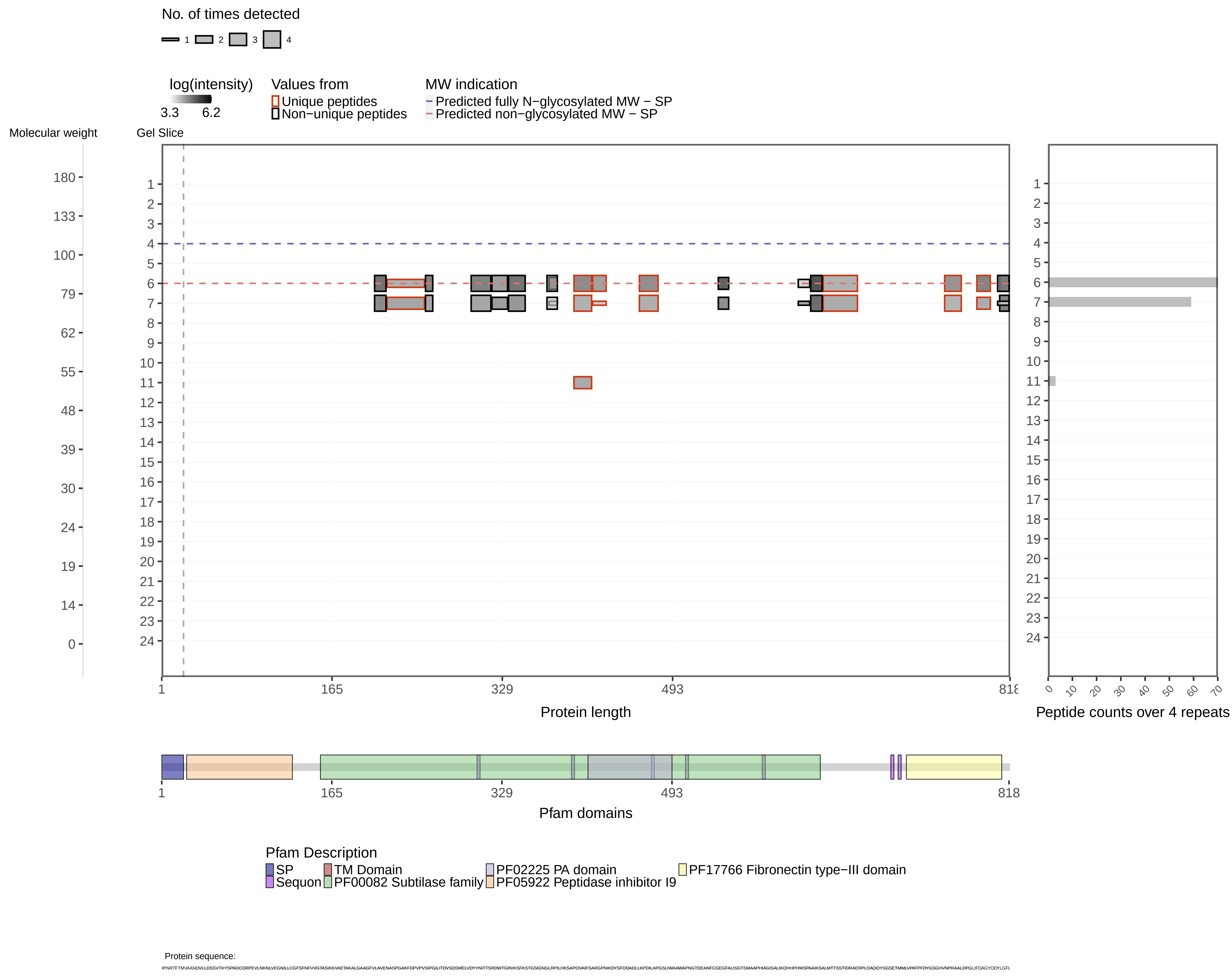

### NbL07g04970.1 inosine−5'−monophosphate_dehydrogenase_2−like [MW = 53.27 kDa]

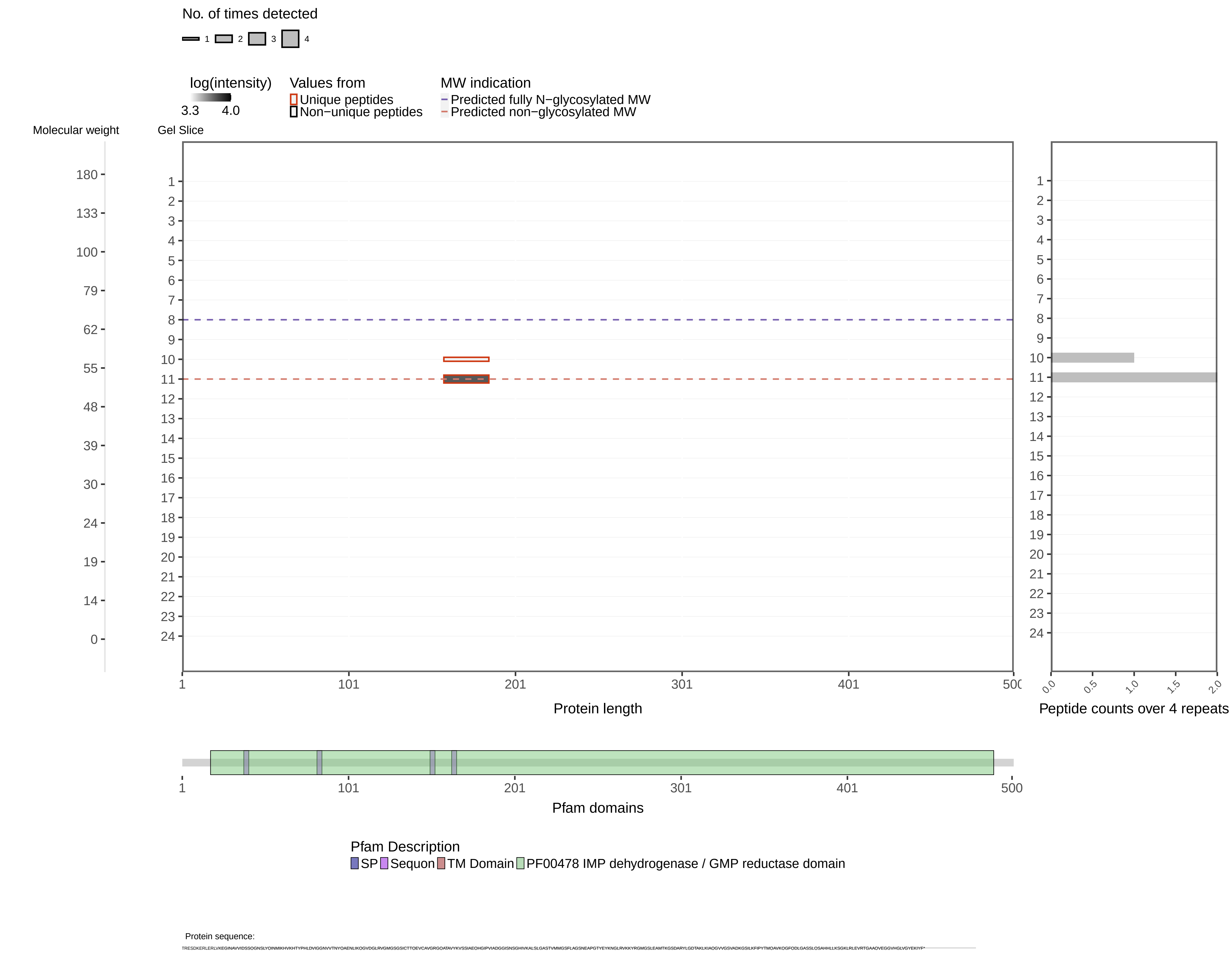

NbL07g04880.1 transcription_factor_BTF3_homolog_4−like [MW = 17.34 kDa]

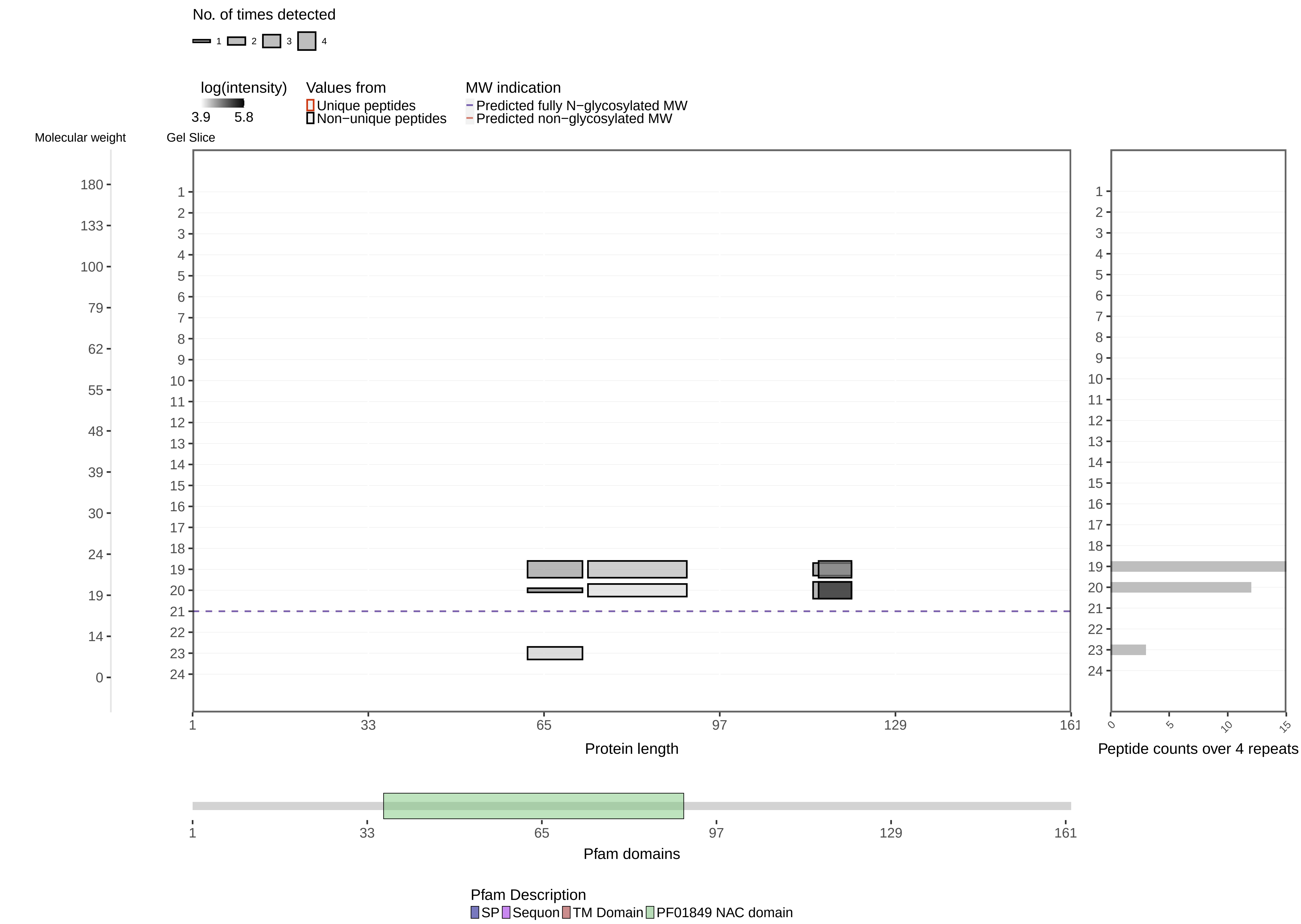


Protein sequence:

-INPKVQASIAANTWVVSGSPQTKKLQDILPQIIHQLGPDNLENLKKLAEQFQKQGAAAGTGEAAAQEDDDDEVPDLVAGETFEGAAEEGHTS*

### NbL07g04800.1 ABC_transporter_F_family_member_4−like [MW = 84.86 kDa]

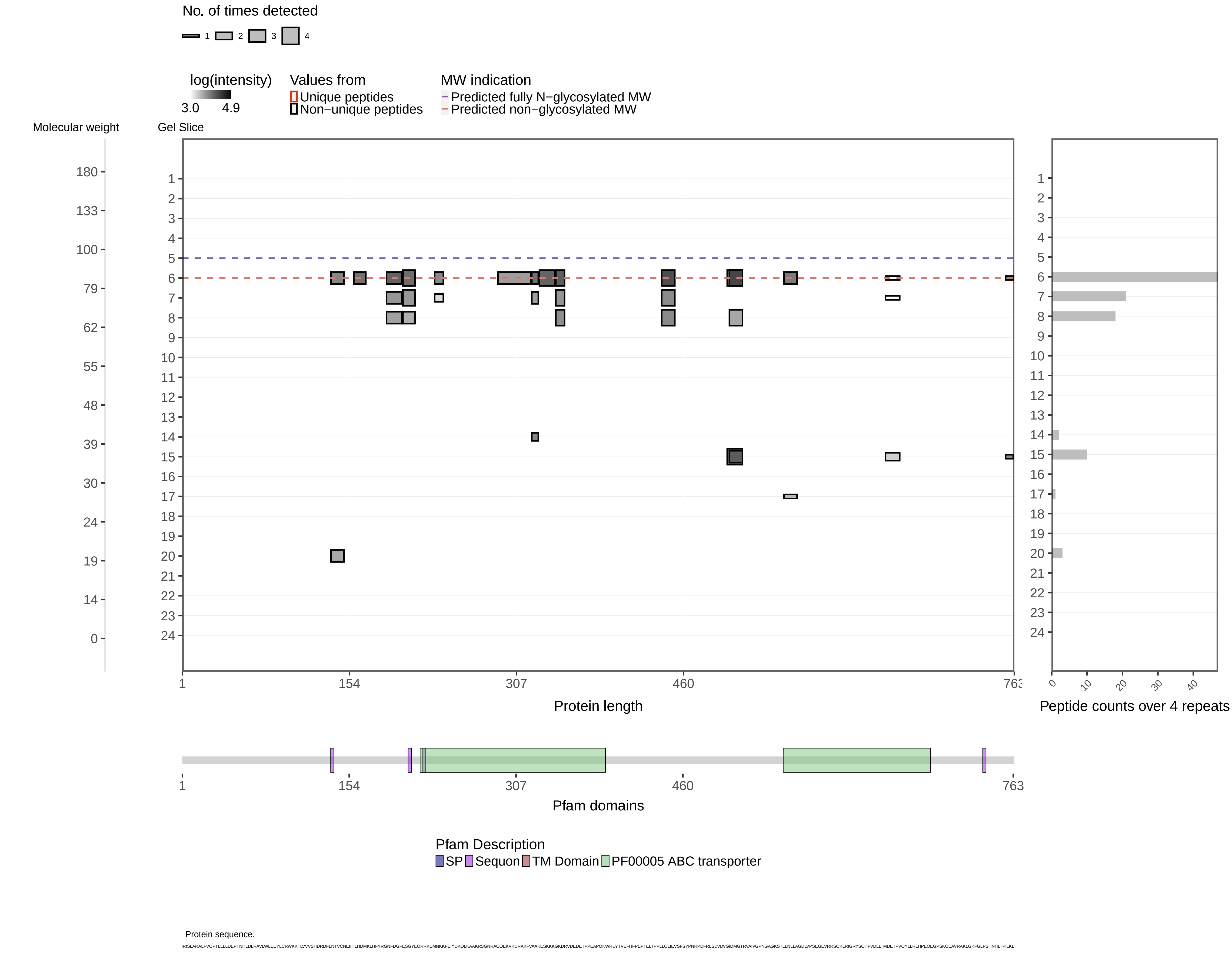

### NbL07g04570.1 macrophage_migration_inhibitory_factor_homolog [MW = 14.34 kDa]

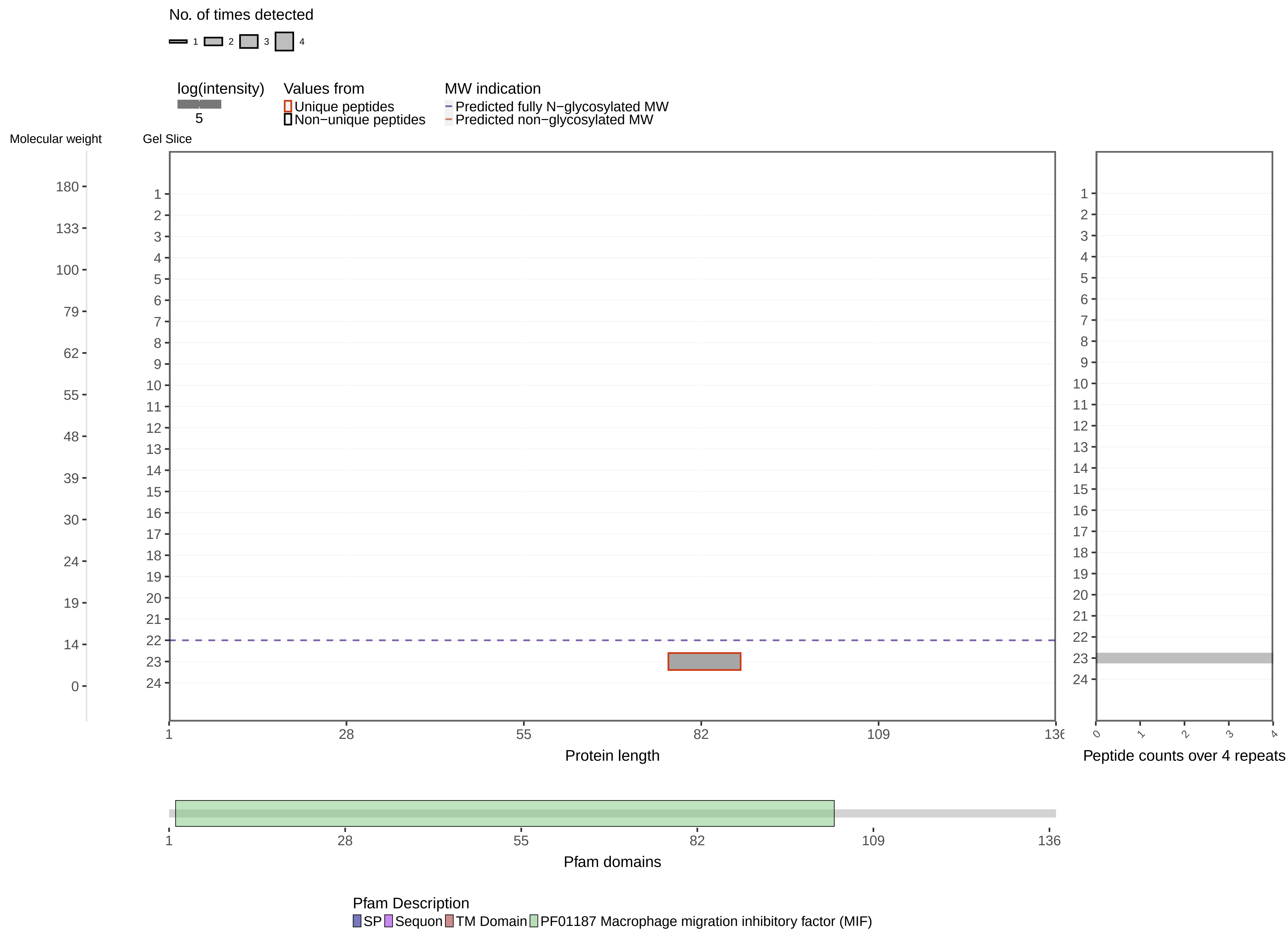


Protein sequence:

EAPAAYGELISIGDLGPGVNGKLSSTIAEILQTKLSIDSNRFYIKFYDSPVSLYKREMISLSSLTPTPPWATTGDRQVGLN*

NbL07g04500.1 60S_ribosomal_protein_L7−4−like [MW = 28.09 kDa]

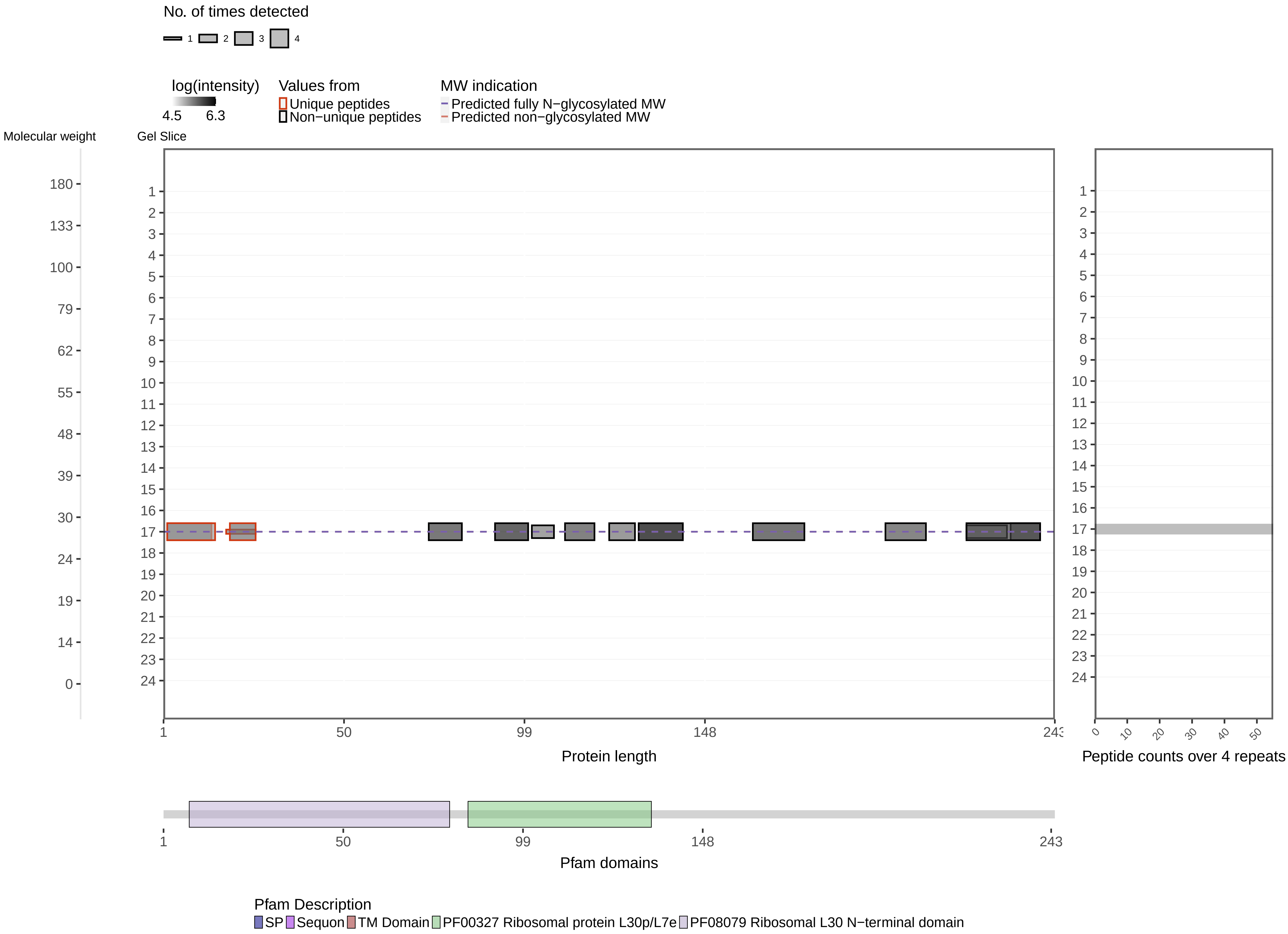


Protein sequence:

LRQIFNGVFLKVNKATVNMLHRVEPYVTYGYPNLKSIRELIYKRGYGKVDKQRIALTDNSVIEQVLGKHGIICIEDLVHEIITVGPHFKEANNFLWPFQLKAPLGGLKKKRNHYVEGGDAGNRENYINELIRRMN*

### NbL07g04460.1 calcium−transporting_ATPase_10_plasma_membrane−type [MW = 122.03 kDa]

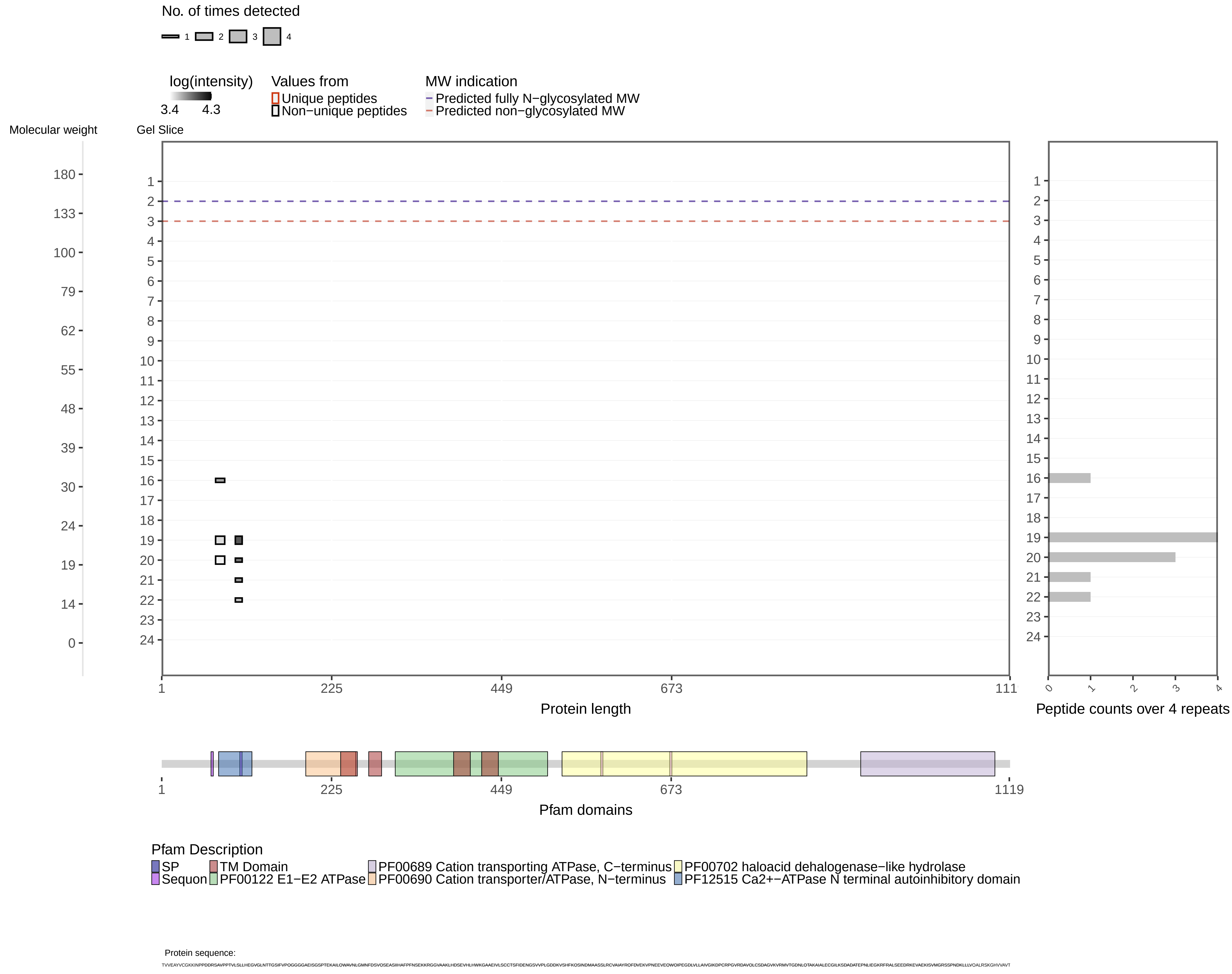

### NbL07g04430.1 uncharacterized_LOC107797264 [MW = 11.23 kDa]

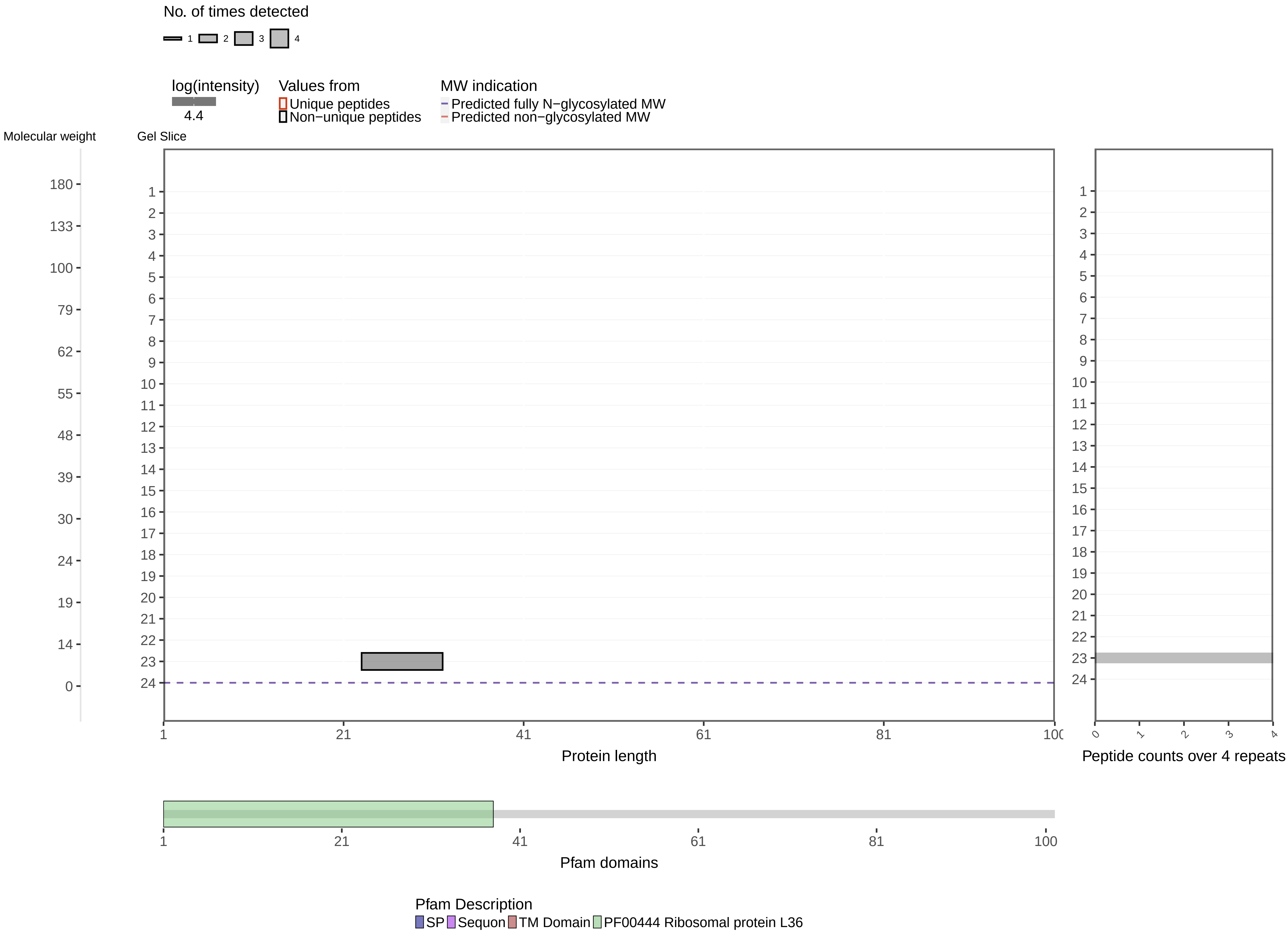


Protein sequence:

RQGYSTFASEGLISTTTPVMTRVTQDTSVTEGIRTGLTYLVPVKTEPTVTPWWKRGLVSVLFNSK*

NbL07g04350.1 methyl−CpG−binding_domain−containing_protein_10−like [MW = 56.84 kDa]

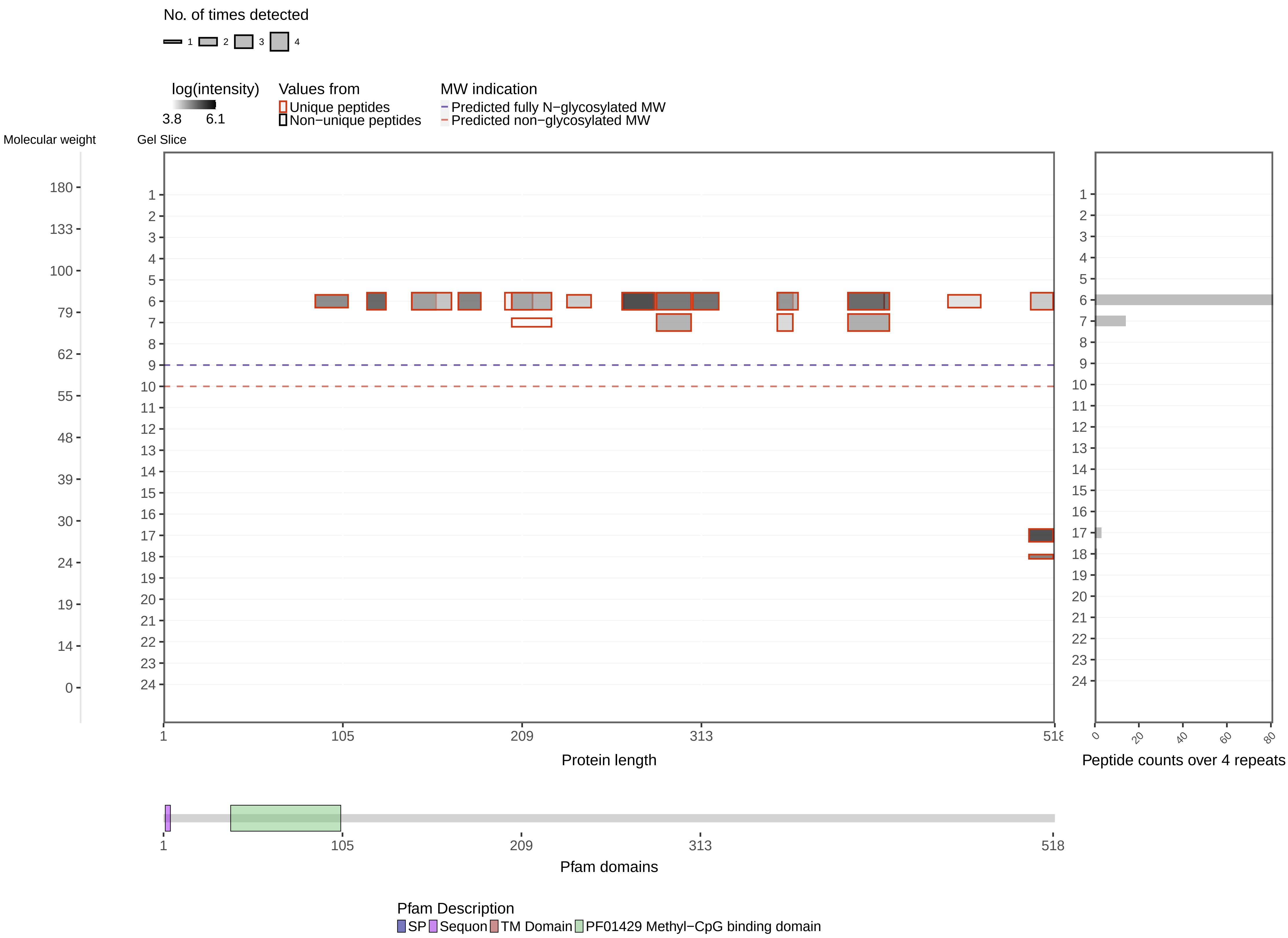


Protein sequence:

SKVADVEKDVKMAGNAAEKKDEMHSSNVDAVEENQADRKSDGQTEDGKAEEAAEGDKDIKMADNVQHPVGEASVVKEAPLEKEPEATKDEEGKDVQVQESVEQKLTETKKEDPSAAVEKDVKMADNAAEKKDEMHSSNVDAVKENQADKKSEGQTEDGKAEDAPAEAAEGDKDVKMADNFQHPEDVEEASVVKEAPLGKEADVPEATRDEEGKDVQVQESVEQKPAEKKKEDLSAAEGKKHQPEQEQRDENGAARKSTGDGDALMNSDIN*

NbL07g04300.1 blue_copper_protein−like [MW − SP = 32.65 kDa]

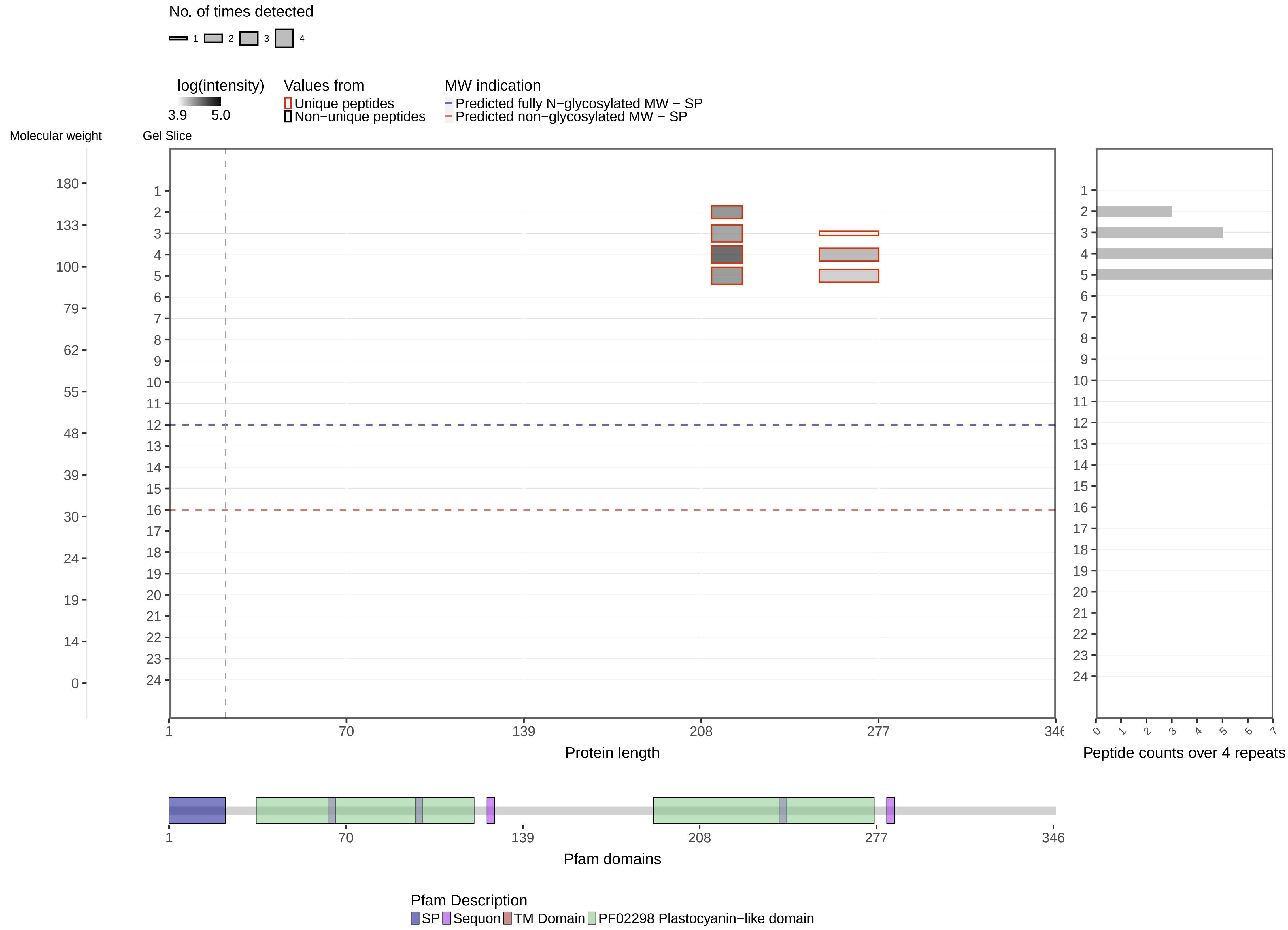


Protein sequence:

APSLAATPASAPAPSVAAQTFIVGGNMGWNVPTTGGPNAYQTWANGKSFKVGDTLVFNFVDGRHNVAMVNKAAYDSCNTTSPINTISTGPARITLTNSGENYYMCTFPNHCSLGQKLAINVTGTSAAAPMPSIAATPSGSTVPSVPSGDSPVTSPPPPSASAPSLVLAALPVTFLSLAFARLLN*

### NbL07g04180.1 uncharacterized_LOC109234414 [MW = 15.69 kDa]

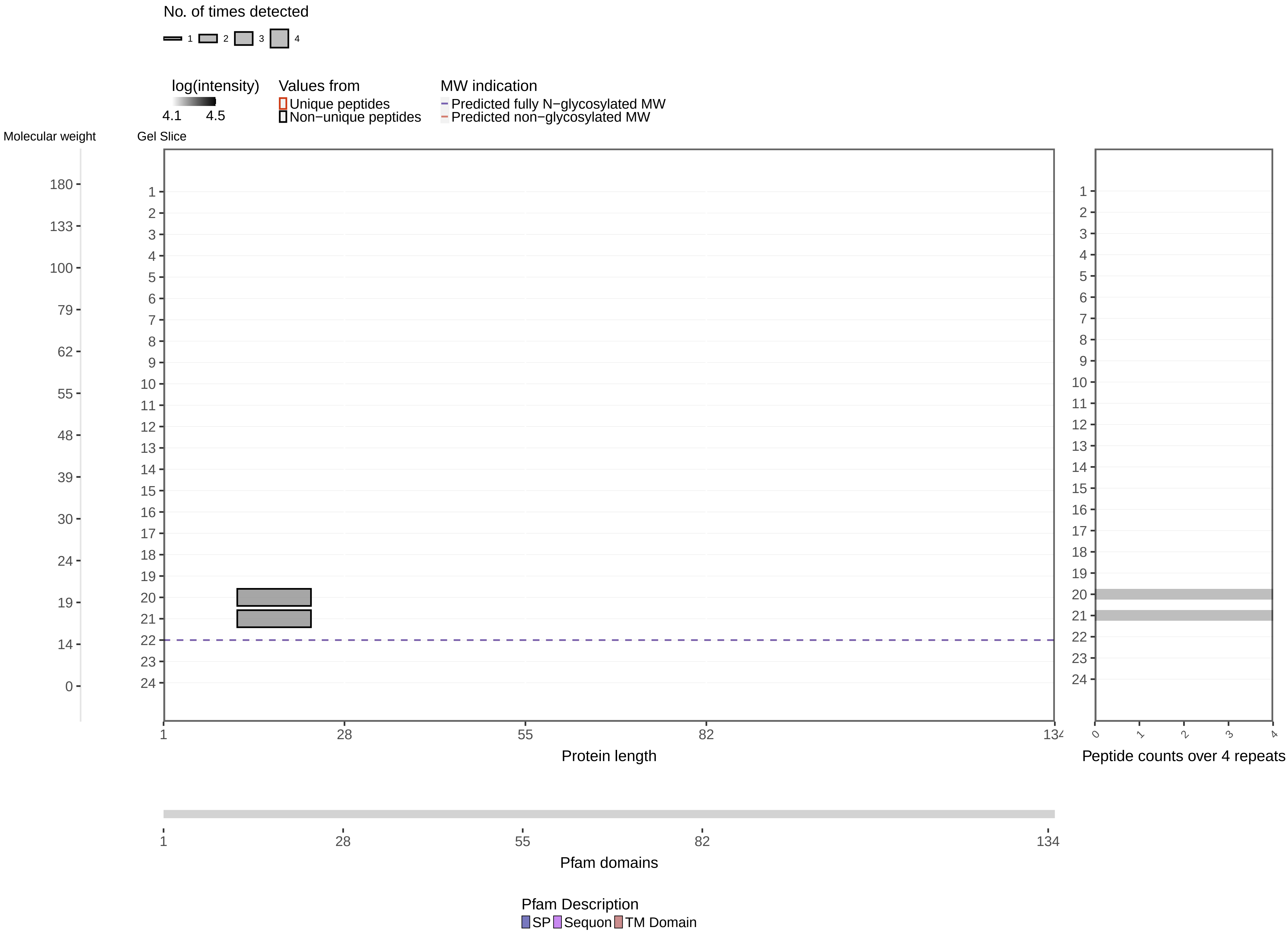


Protein sequence:

ARLRKEVLLAGQDWPYDPERKEMRTKQKGHKCDRISAEKREKTAELMQKMPDMLADYRKRRWERKMKAEEDAARRAVQE*

NbL07g04100.1 probable_galactinol−−sucrose_galactosyltransferase_6 [MW = 100.53 kDa]

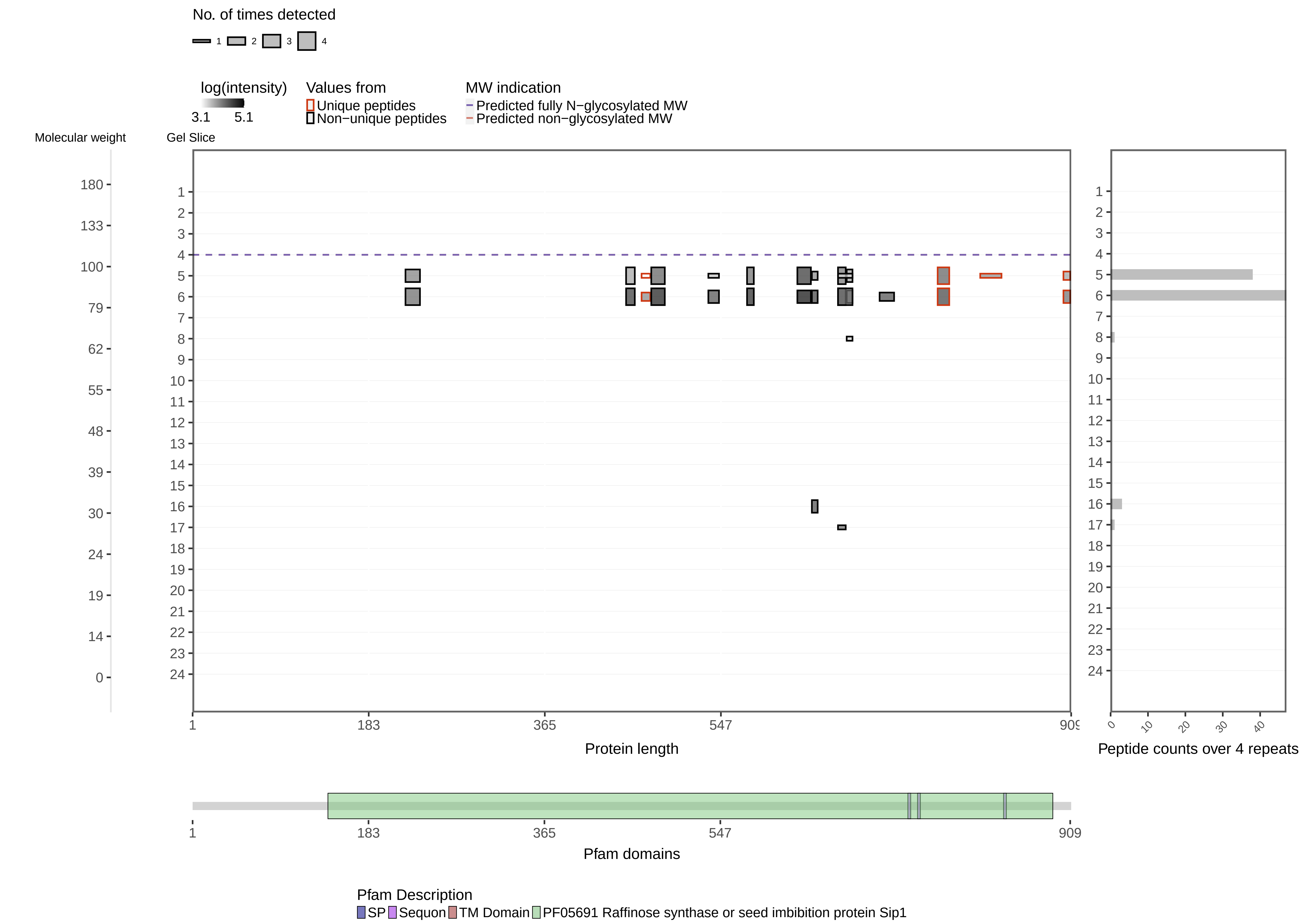


Protein sequence:

SVKGMEEYGSVMKYPEITKGVMENEPGWRTDAIAVQGLGLVNPKSAYKFYNEMHSYLASAGVDGLKVDVQCILETLGDGLGGRVELTKQYHQALDASVSKNFPDNGCIACMSHSTDALYCSKQTAVVRASDDFYPRDPVSHTIHIACVAYNSVFLGEIMVPDWDMFHSLHPAAEYHGSARALSGGPVYVSDAPGKHNFDVLRKLVLRDGSVLRARLPGRPTKDSLFSDPSRDGVSLLKIWNMNKYTGVLGVYNCQGAAWNTVERKNTFHQTNSEAITGYVRGRDVHFIS

### NbL07g04060.1 12−oxophytodienoate_reductase_3 [MW = 43.55 kDa]

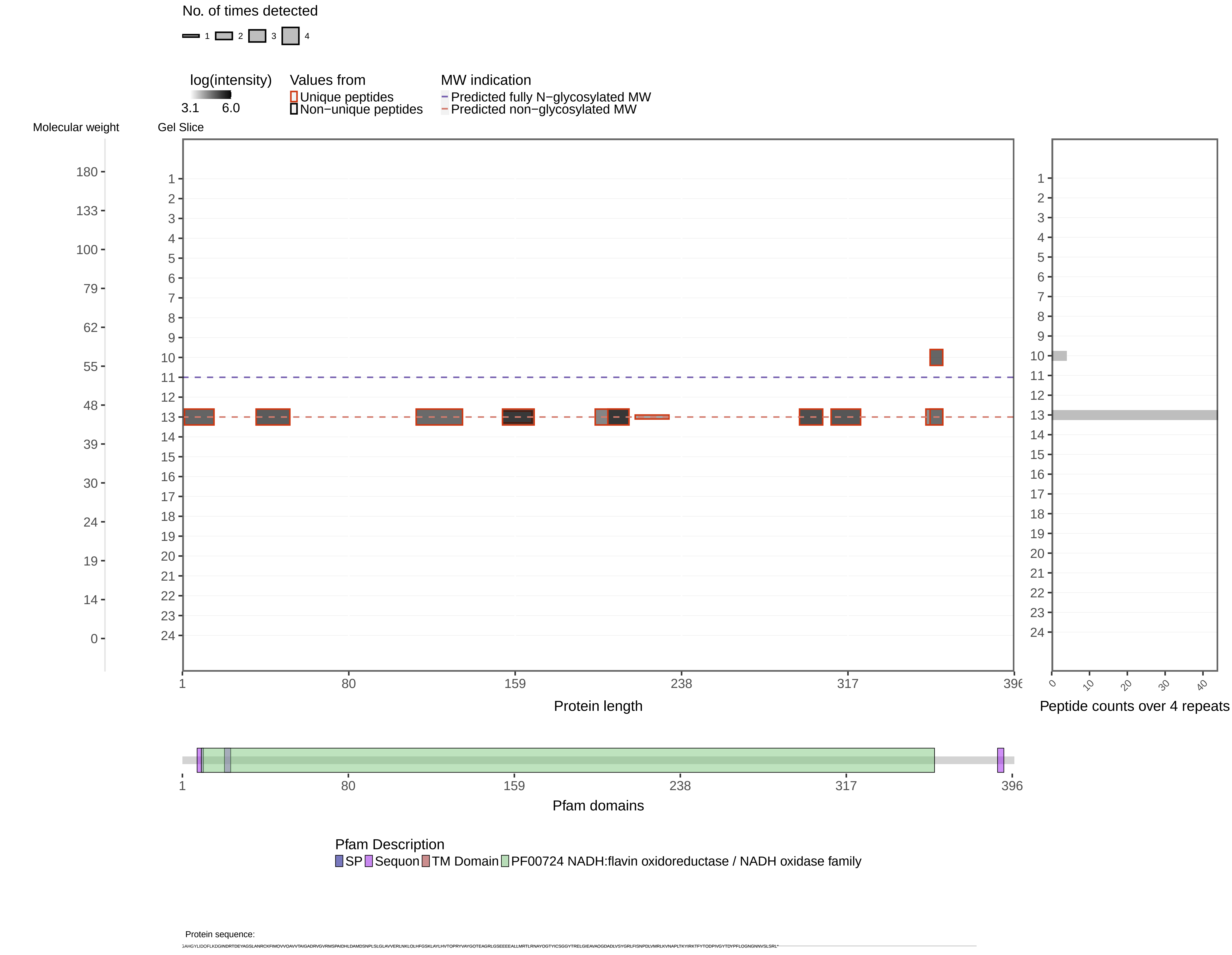

NbL07g03960.1 probable_sucrose−phosphate_synthase [MW = 118.81 kDa]

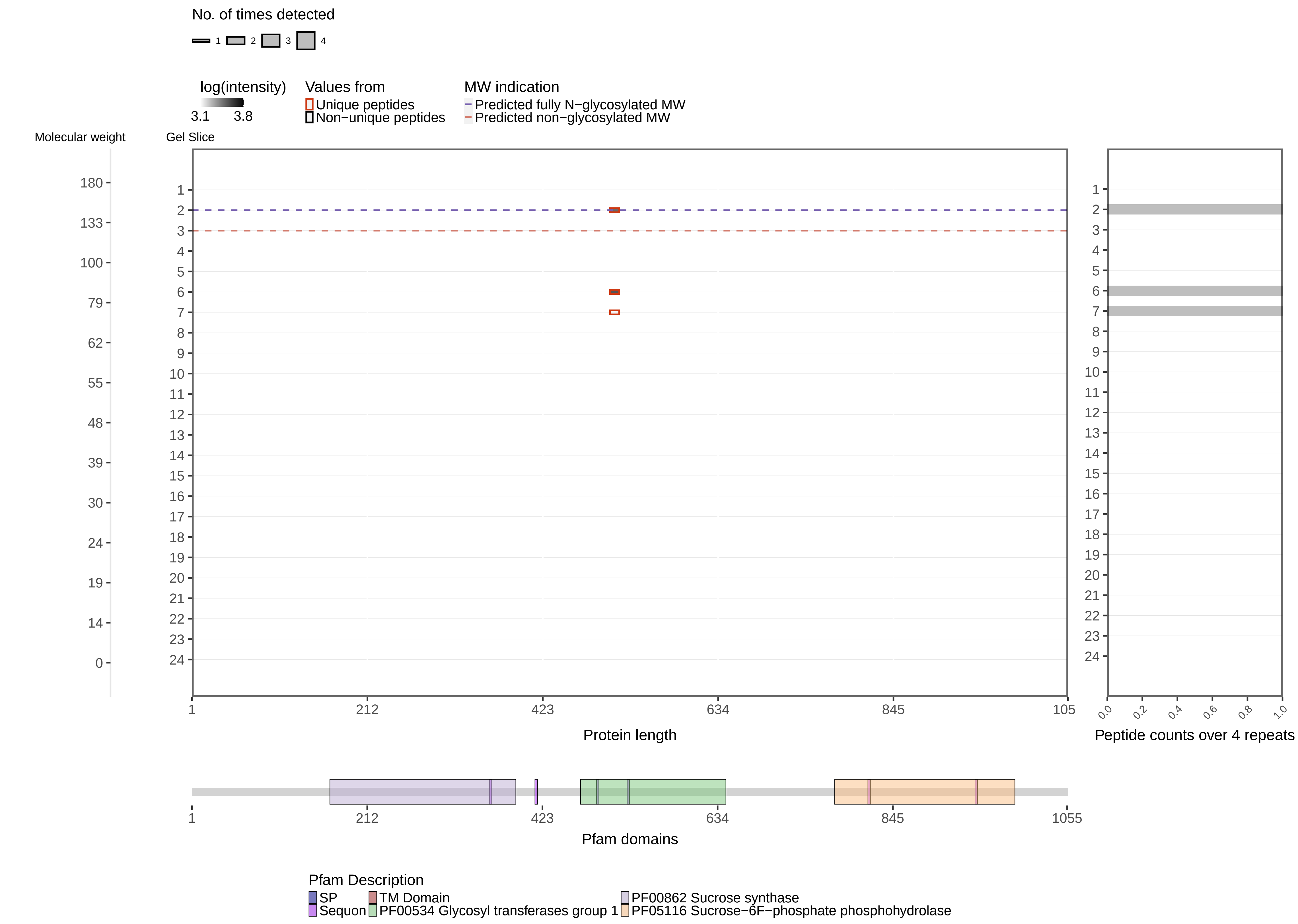


Protein sequence:

[illegible]

### NbL07g03880.1 cytokinin_dehydrogenase_7 [MW = 57.94 kDa]

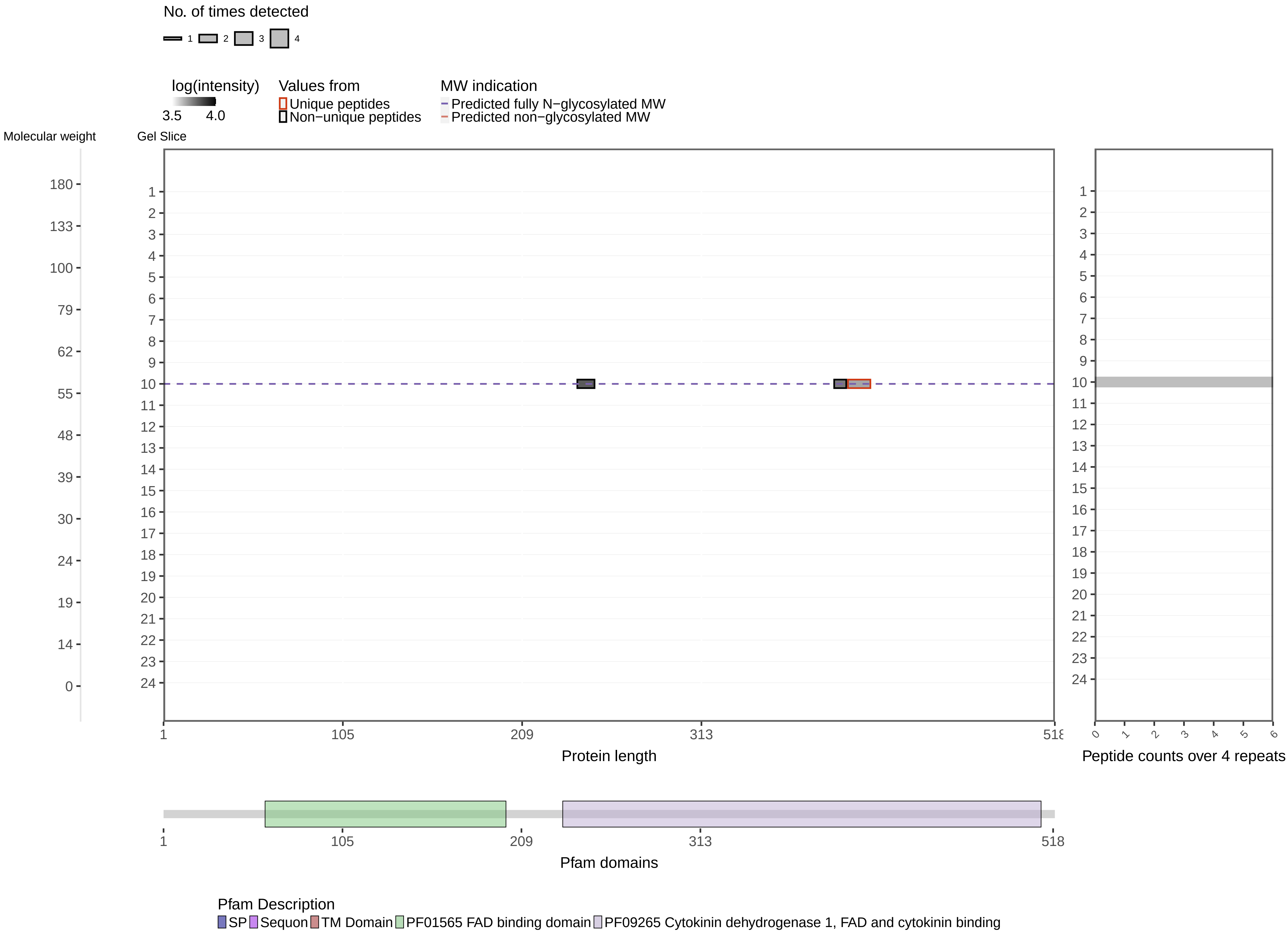


Protein sequence:

DEFTRDAELLVMSPESFDYVEGFVFVNSDDPVNGWPSVPLDSNHPFDPTHLPTNTGPVLYCLEVALHYHNHDHPSTVNMMVEKLLGRLRFIEPLMFEVDLTYMDFLLRVKRVEQMARDNGIWDAPHPWLNMFVSKRDIAEFNRIVFQNILKDGINGPILTYPLIRSKWNSRSSVVLPKGEIFYLVALLRFSHAHPKESEINEMVAQNQEILQTCTNNGFDFKLYLPHYKSTEEWKKHFGDQWGRFVERKSQFDPKAVLAPGQKIFTRNYQR*

### NbL07g03860.1 uncharacterized_LOC109237049 [MW = 26.65 kDa]

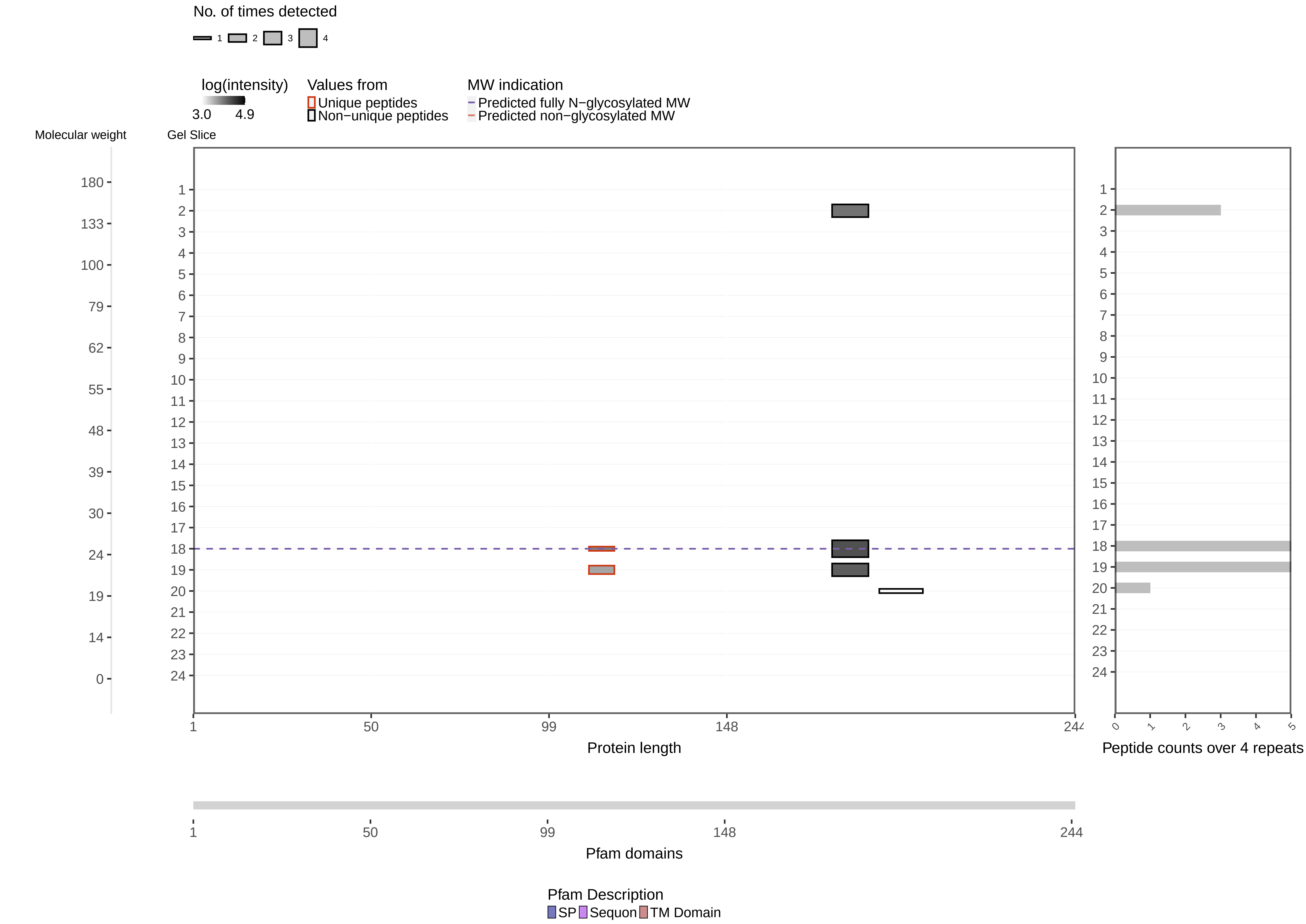


Protein sequence:

SLEVDQPKYGDCRNILRSGPASSLRVNIRAVAQYASDDGNGESASNDVDQCLSALEGLDSLLLRASRNDPGASIDSMKAKITIALNALDSLLKTVPADMLDKGKIVADSYFFADEEDVTPEKLDPNLKQLESIL*

### NbL07g03630.1 DEAD−box_ATP−dependent_RNA_helicase_56−like [MW = 48.44 kDa]

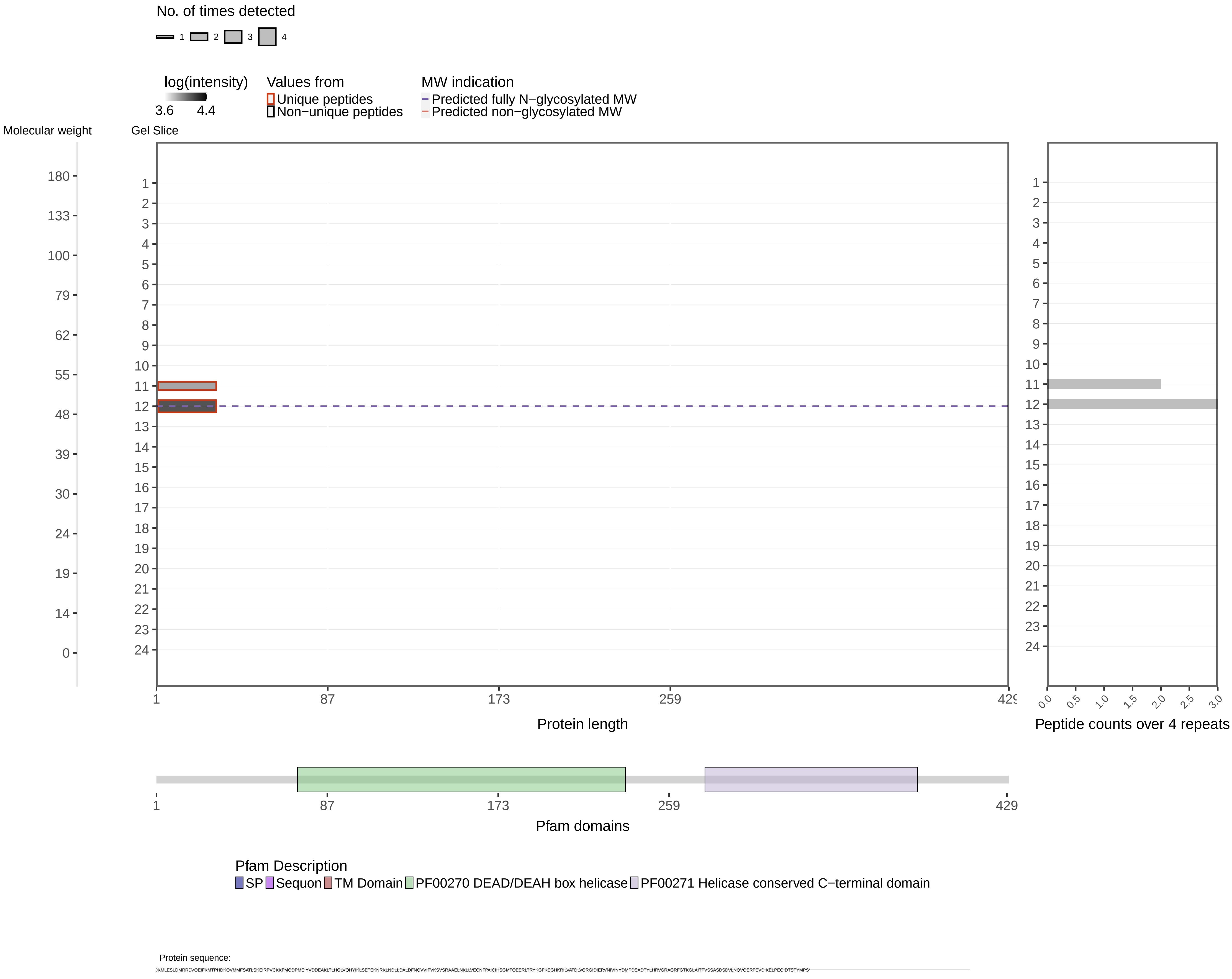

NbL07g03500.1 protein_WVD2−like_5 [MW = 48.15 kDa]

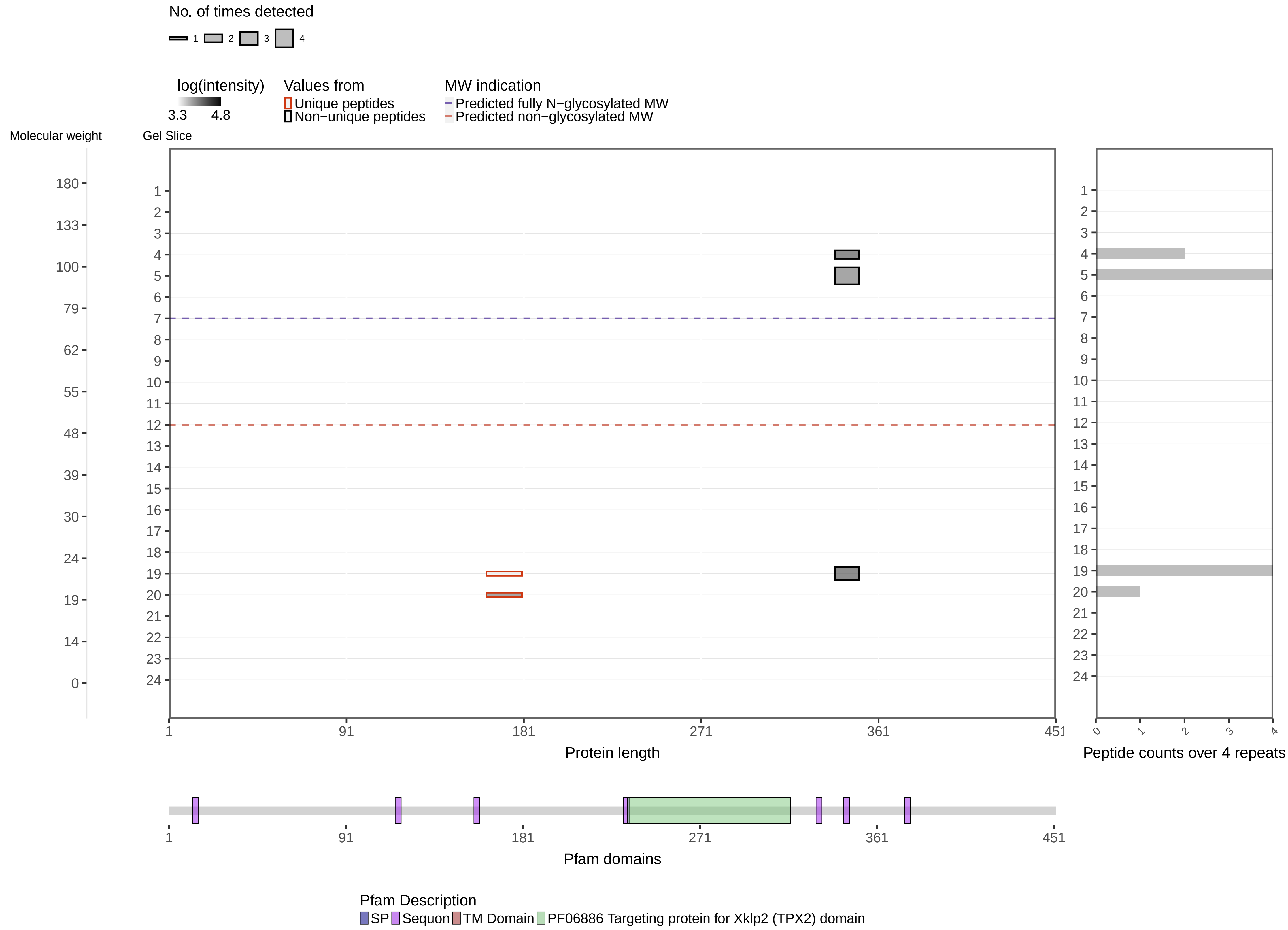


Protein sequence:

EESPSPAADAKPRKVGALPTYNMSFKCDERAEKRKEFYSKLEEKIHAKEVEKNNLQALSKACQEAEIKMLRKSLKFKATPMPSFYQEPPPPKVELKKIPTTRAKSPKLGRKKSSPTRVVNDTAMHTSRLSLDENATQNPAKVHAPDIVKKPSRKSLPKLPSQKVNLSINSKKPSSLKTSISKETNEAANNNMNEVASQPNNVLEQTNEIEATSAVETSQSKTVVEAQAETNVVQQSIAVE*

### NbL07g03330.1 ATP_synthase_subunit_b'_chloroplastic−like [MW = 24.05 kDa]

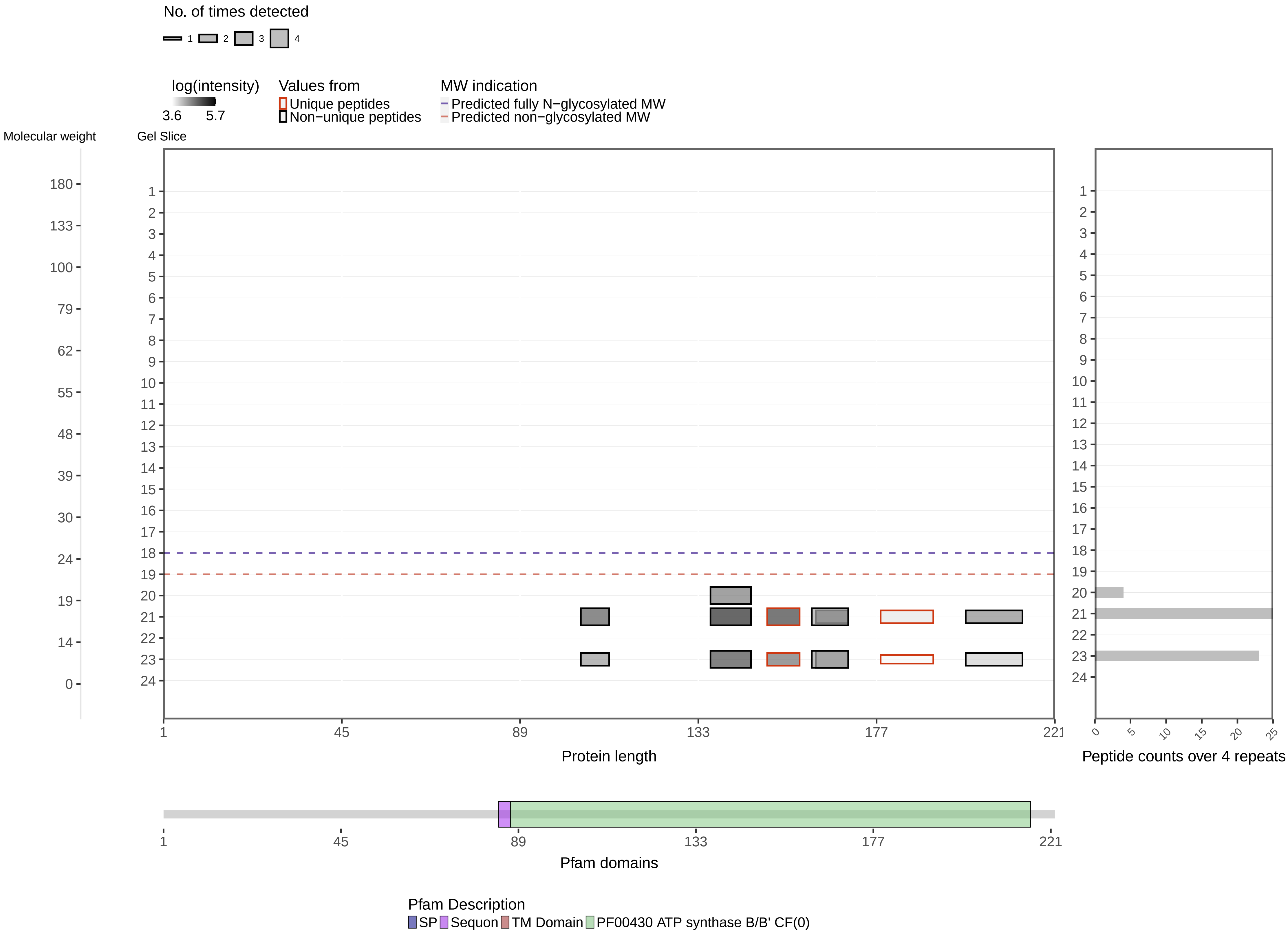


Protein sequence:

LDKLYFSPLGKFMDERDSAIKEKLNSVKDTSAEVKQLEDQAAAIMKAARAEVSAALNKMKKETQLEVEQKIAEGRKKVEVELQDALASLDKQKEETIKSLDSQIAALSDEIVKKVLPVSN*

NbL07g03080.1 protein_MEMO1−like [MW = 19.57 kDa]

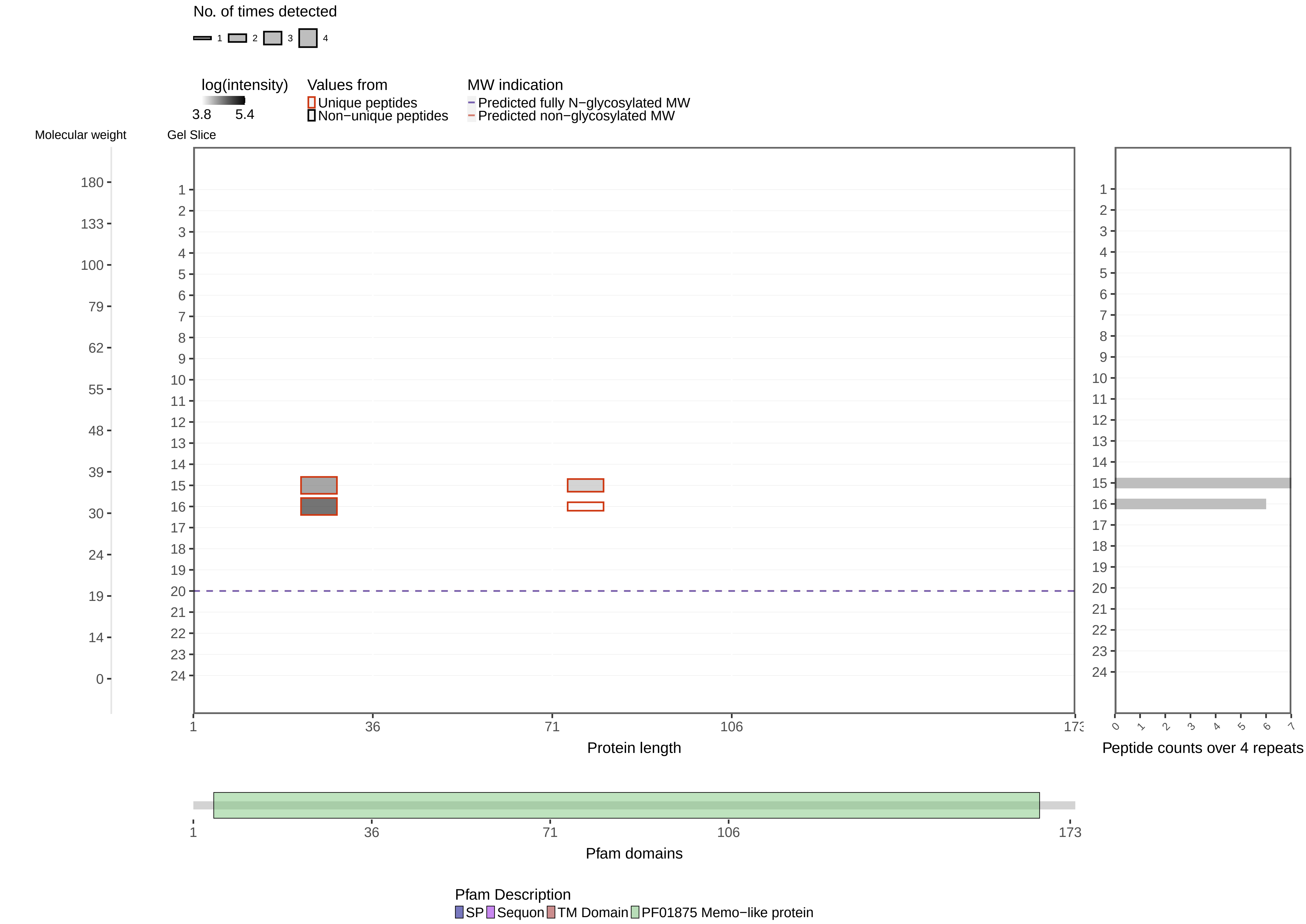


Protein sequence:

RFNFMHYDKSHGAIYKSIEALDKMGMDIIETGDPDAFKOYLLETDNTICGRHPISVFLHMLKHSSTKIKIRFLRYEOSSOCKSMRDSSVSYASAVGKIDA*

### NbL07g03000.1 40S_ribosomal_protein_S11 [MW = 17.7 kDa]

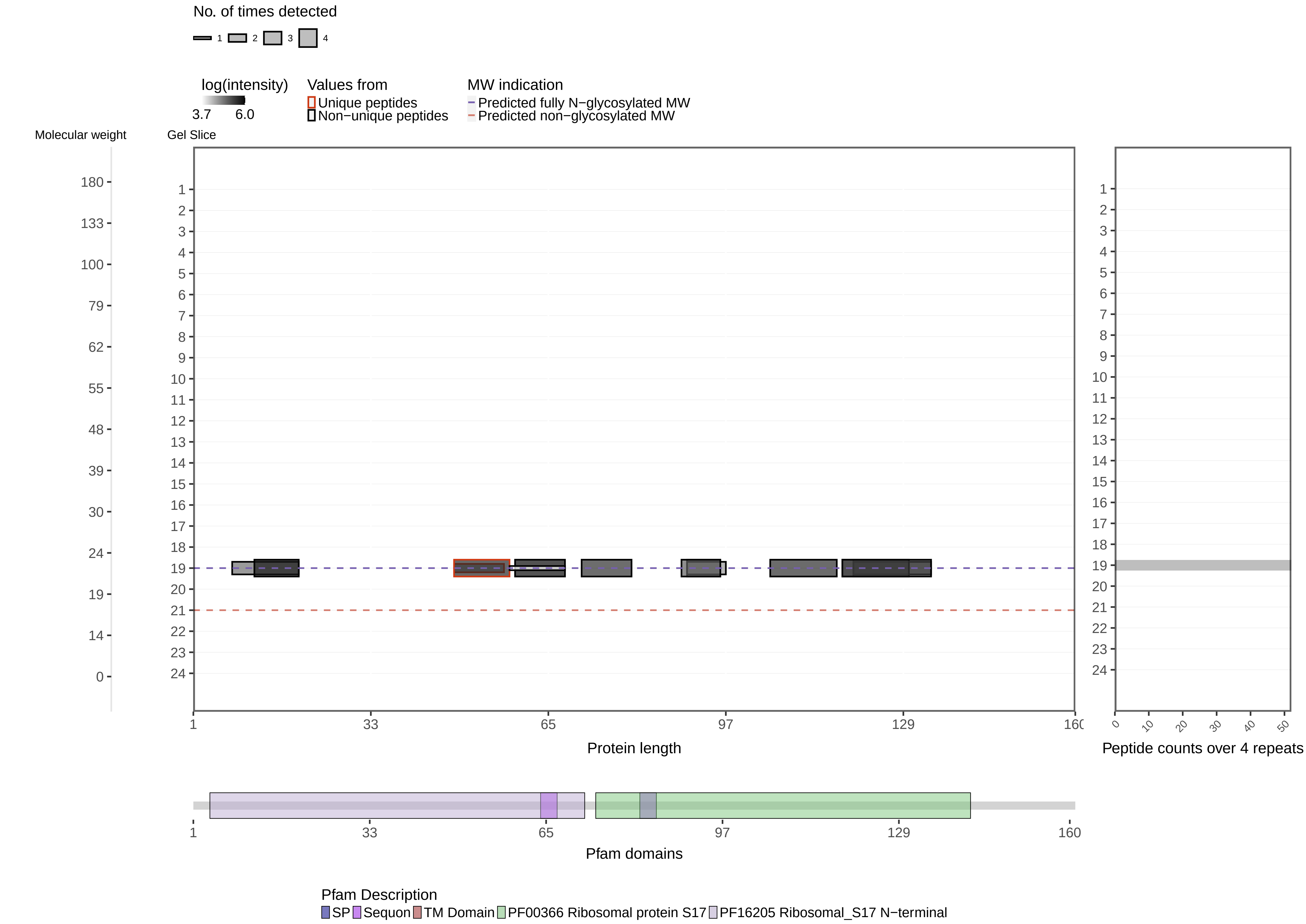


Protein sequence:

SIRGRILAGTCHSAKMNRTIIVRRNYLHYVKKYQRYEKRHSNIPAHISPCFRVKEGDHVIIGQCRPLSKTVRFNVLKVIPAGSAGGGKKAFTGM*

NbL07g02870.1 lignin−forming_anionic_peroxidase [MW − SP = 32.48 kDa]

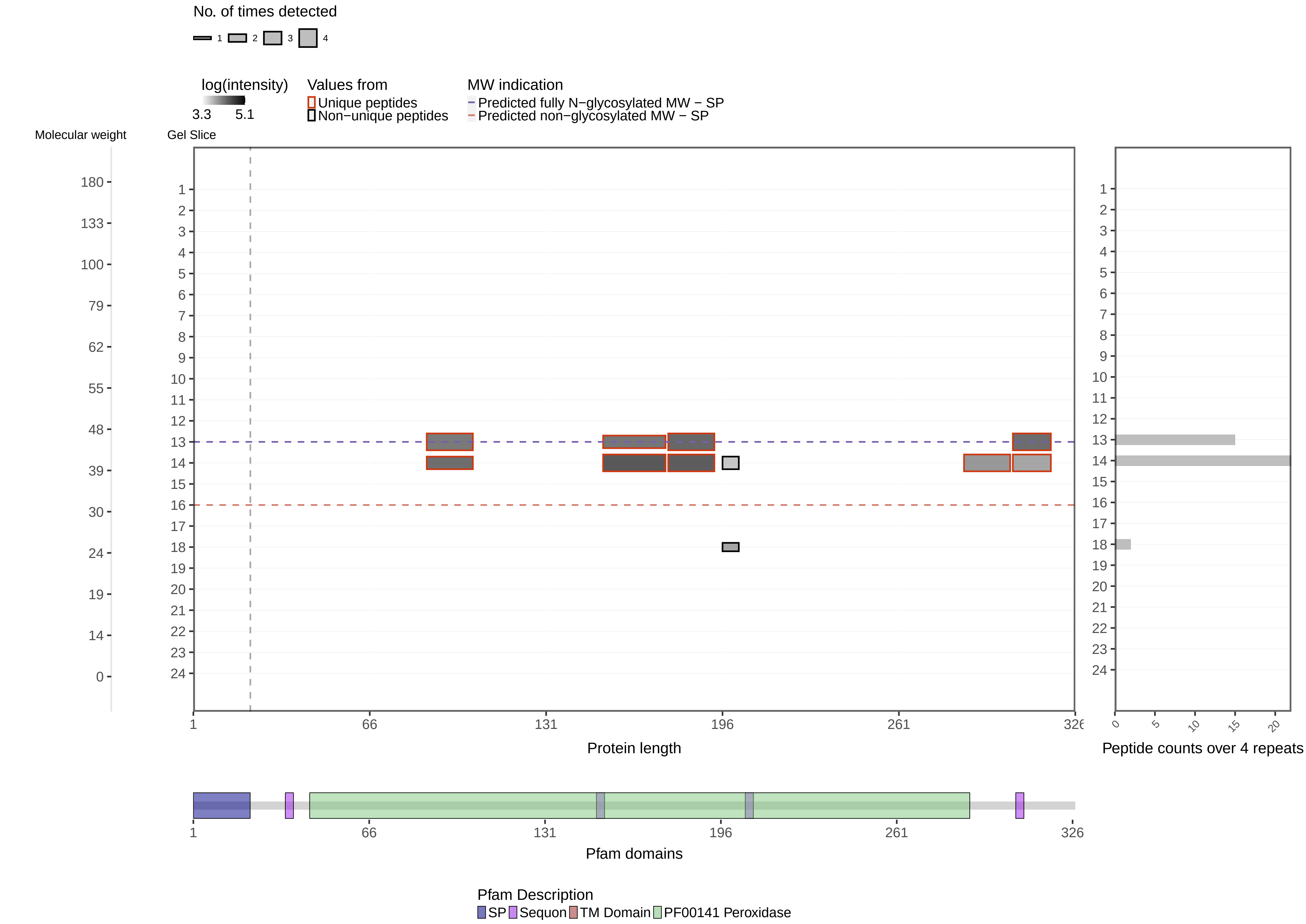


Protein sequence:

RSGAESDIPSPFETLAVMTPQFTNKGMDLTDLVALSGAHTFGRARCGTFEQRLFNFSGSSNPDPTVDATFLQTLQGICPQGGNNGNTFTNLDISTPNDFDNDYFTNLQNNQGLLQTDQELFSTTGSATIAIVNRYASSQTQFFEDFISSMIKLGNISPLTGTNGEIRTDCKRVNS*

### NbL07g02850.1 formate−−tetrahydrofolate_ligase [MW = 67.34 kDa]

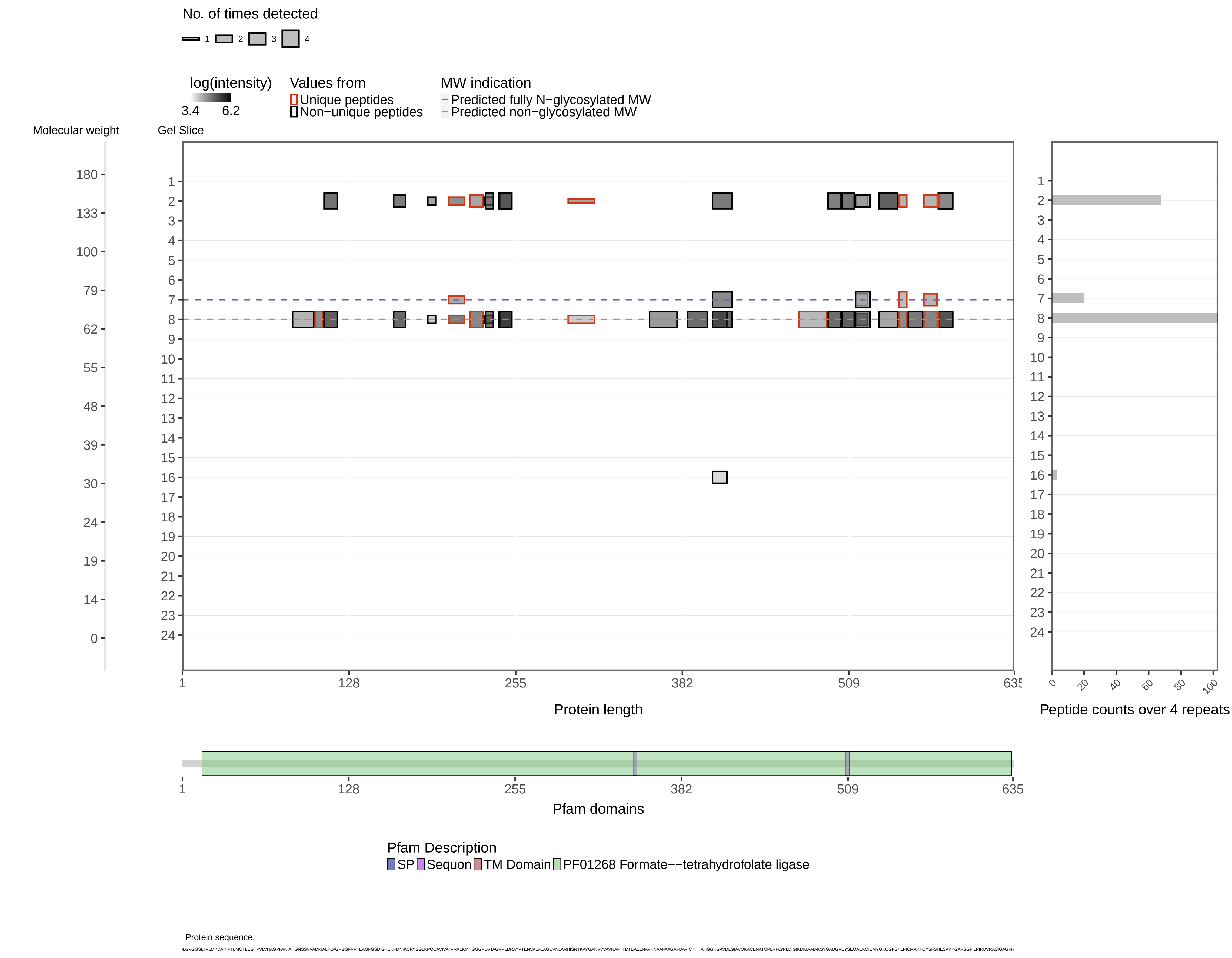

### NbL07g02630.1 arogenate_dehydrogenase_2_chloroplastic−like [MW = 42.77 kDa]

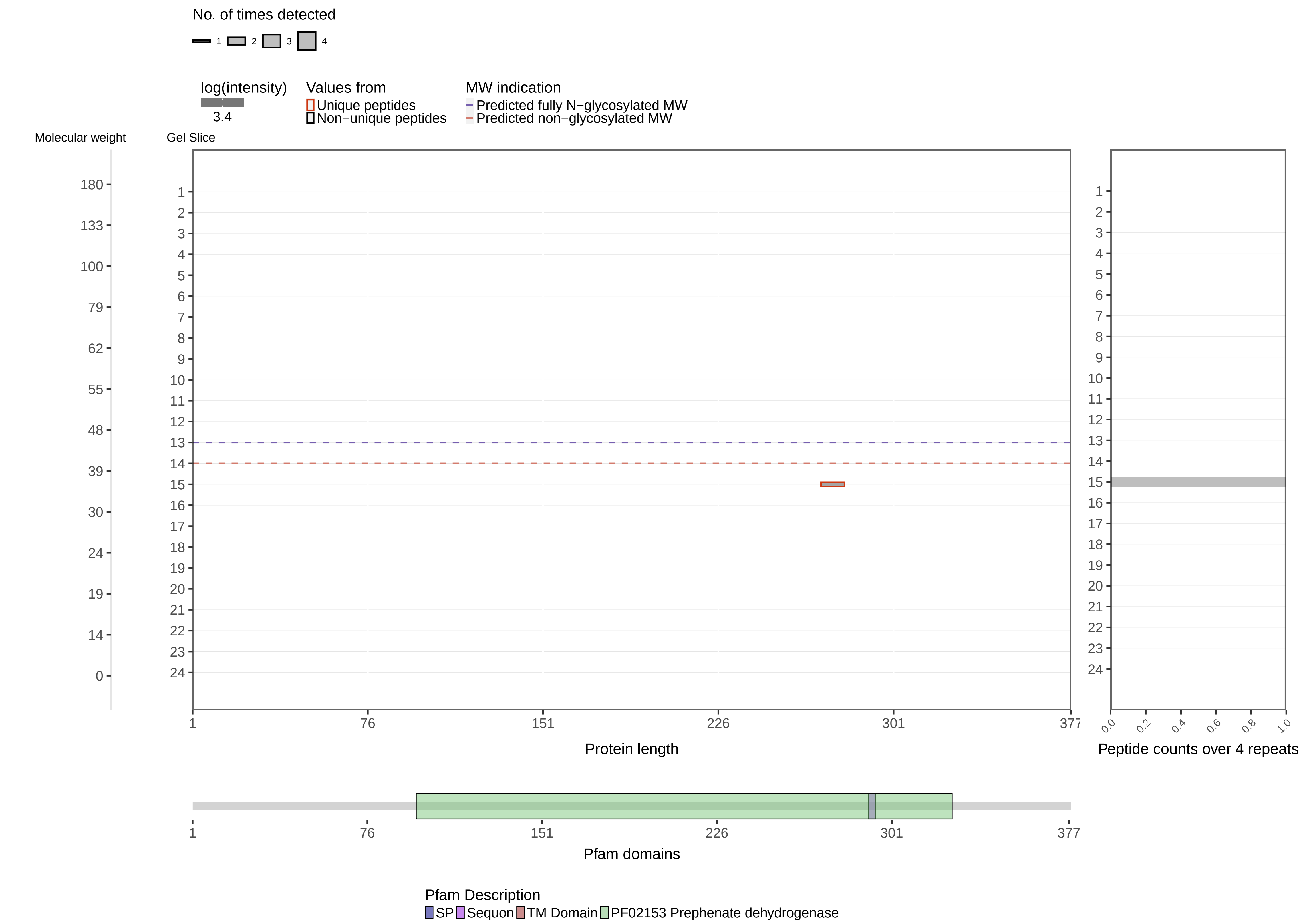


Protein sequence:

NIFLQVLPSHFDILCTHPMFGPESGKDSWKDLAFVFDKVRIGEGDSRKGRVDRFLDIFEKEGCRMVPMTCAEHDRHAAGSQFITHTMGRVLEKLDLETTPINTKGYETLLNLVENTSSDFDLYYGLFMYNRNAMEQLERLDLAFEALKKELFGHLHEVLRKQLFGKAEEAGQRRILTKLPKNGYALPAPSSEAVKSENN*

NbL07g02590.1 actin−related_protein_2/3_complex_subunit_3 [MW = 19.55 kDa]

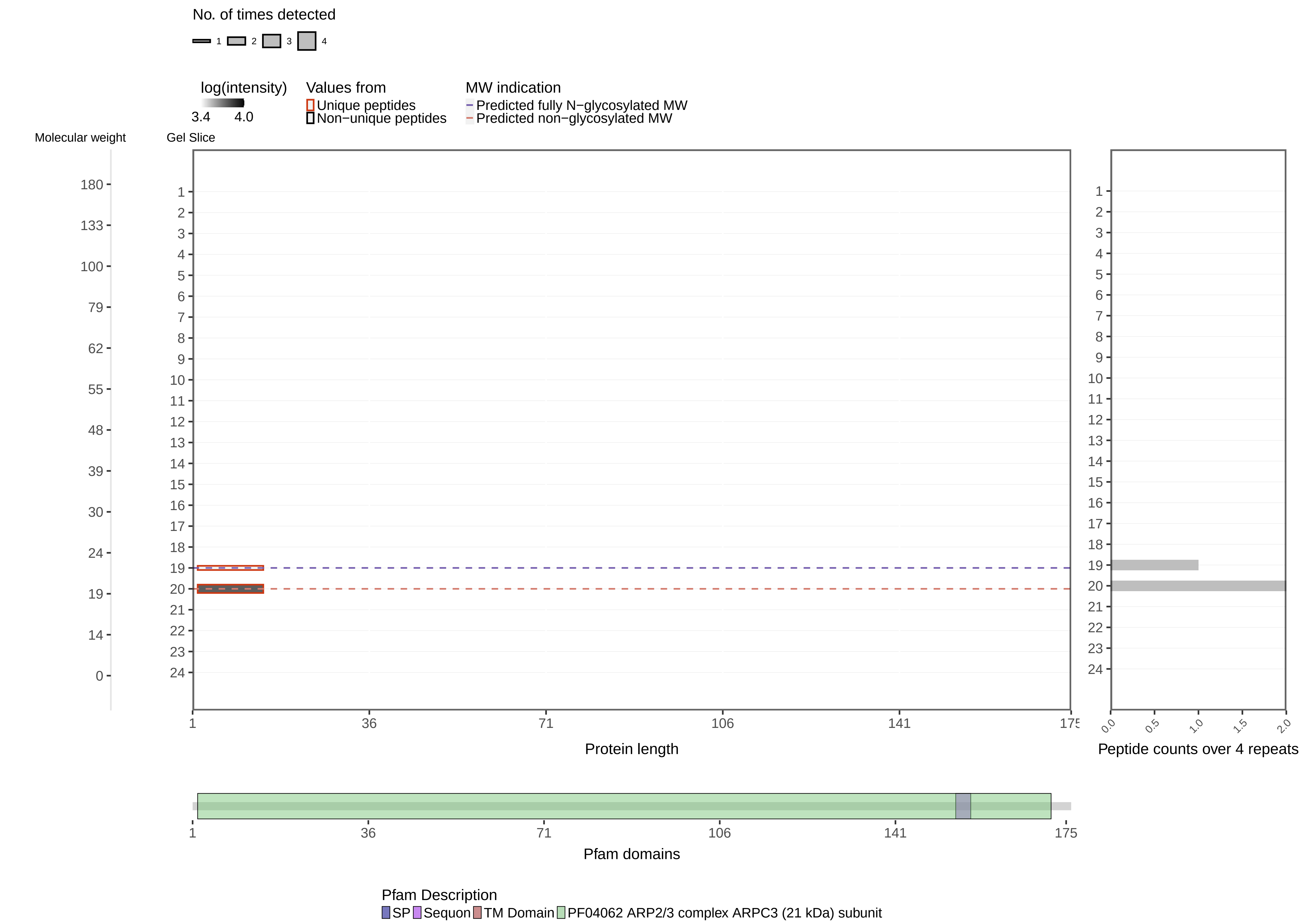


Protein sequence:

INIALKRLEGCRTLAEGTKAIINLGLETVPVPGESGFPFPGLFTAAQSKKEAELFRNYLRQIREETSGRLLSVAYRHNGTPNKWWLAFAKRKFMNMVLP*

### NbL07g02470.1 uncharacterized_LOC104229534 [MW = 27.08 kDa]

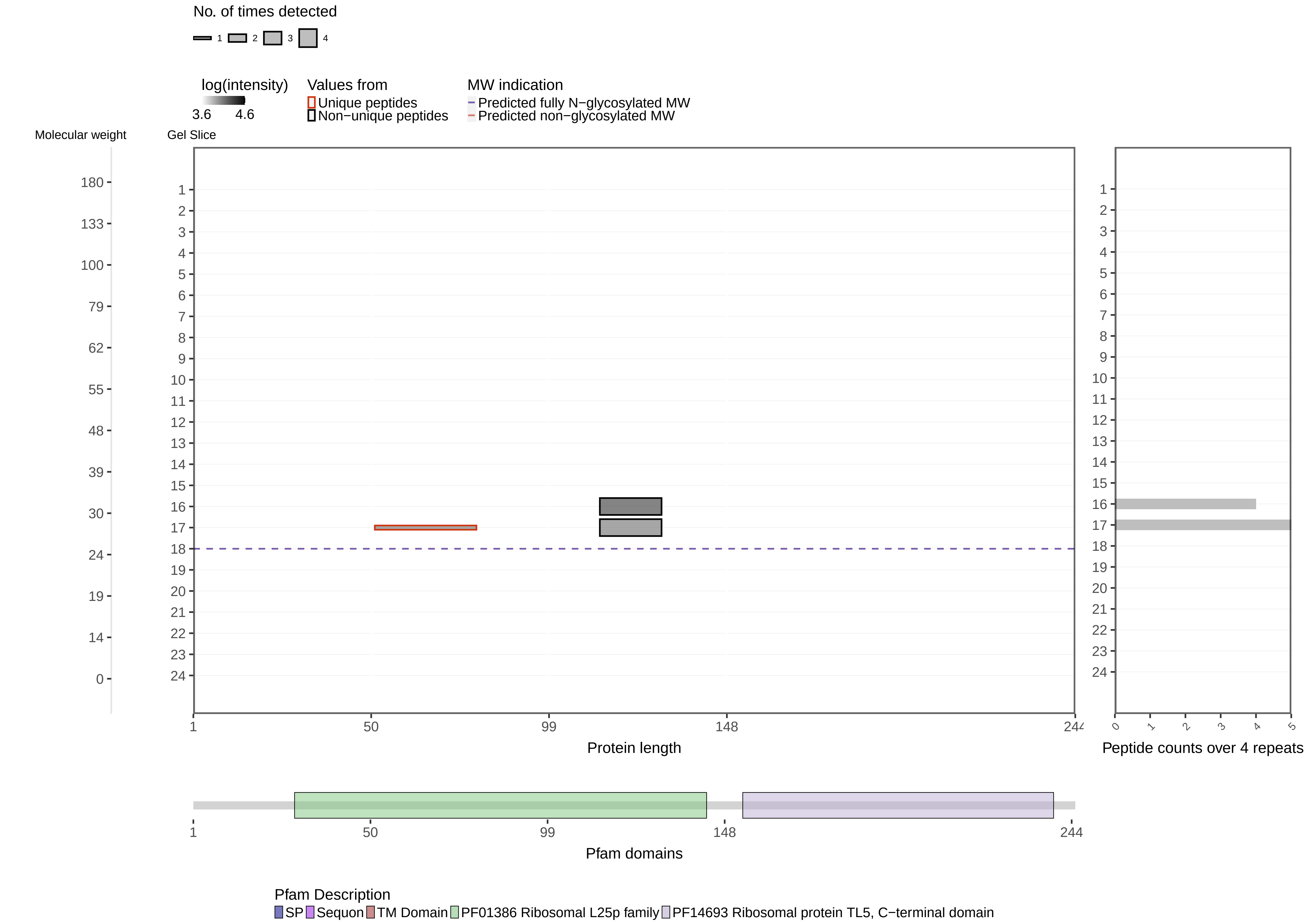


Protein sequence:

PLQIRAGSGSSTLLESGNVLPIKIHRDEETGKILNLVFVWAEEGTKLKVDVPVVFKGEEECPGLKKAGGSLNKIRPTLRFLCPAEHIPQKIEVDASQLDVEDKVFLHDIDLHPAWKLLSKNEAIPVCKIKATSIDS*

NbL07g02370.1 flagellar_radial_spoke_protein_5−like [MW = 49.44 kDa]

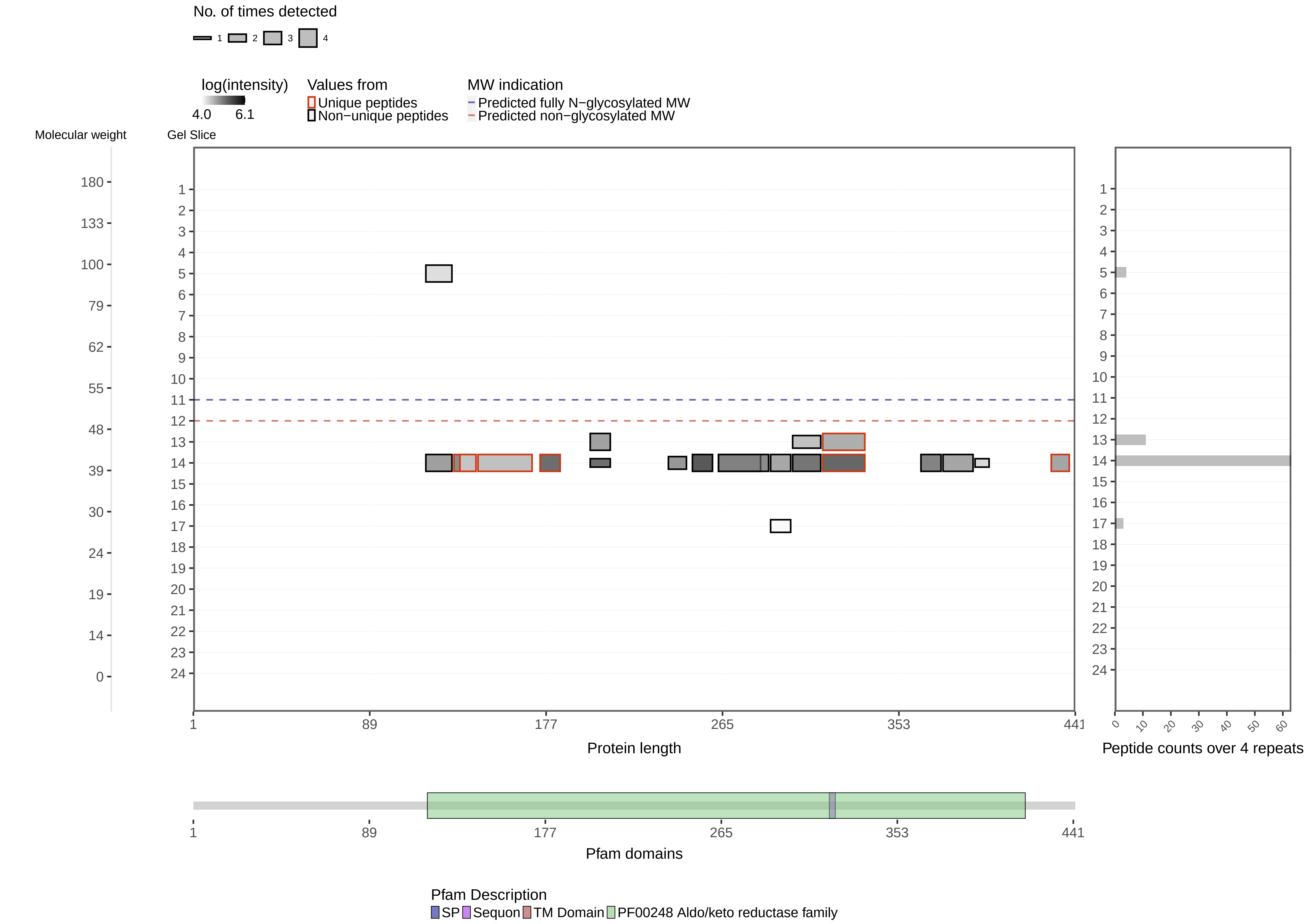


Protein sequence:

/SRKRMDVASLDMLQFHWWDYANSGYLDALKHLTDLKEEGKIKTIALTNFDTERLRIILENGIPVVSNQVQHSLVDMRPQQKMAELCQLTGVKLITYGTVMGGLLSEKFLDTNLSIPFAAPPLNTPSLQKYKRMVDAWGGWSLFQELLRILKTVANKHGVSIPTVAVKYILDQPTVAGSMVGVRLGLSEHIKDVNAVFSLVLDEEDISSILQVTNKGKDLQKVIGDCGDEYRRA*

### NbL07g02360.1 uncharacterized_LOC104119615 [MW = 20.29 kDa]

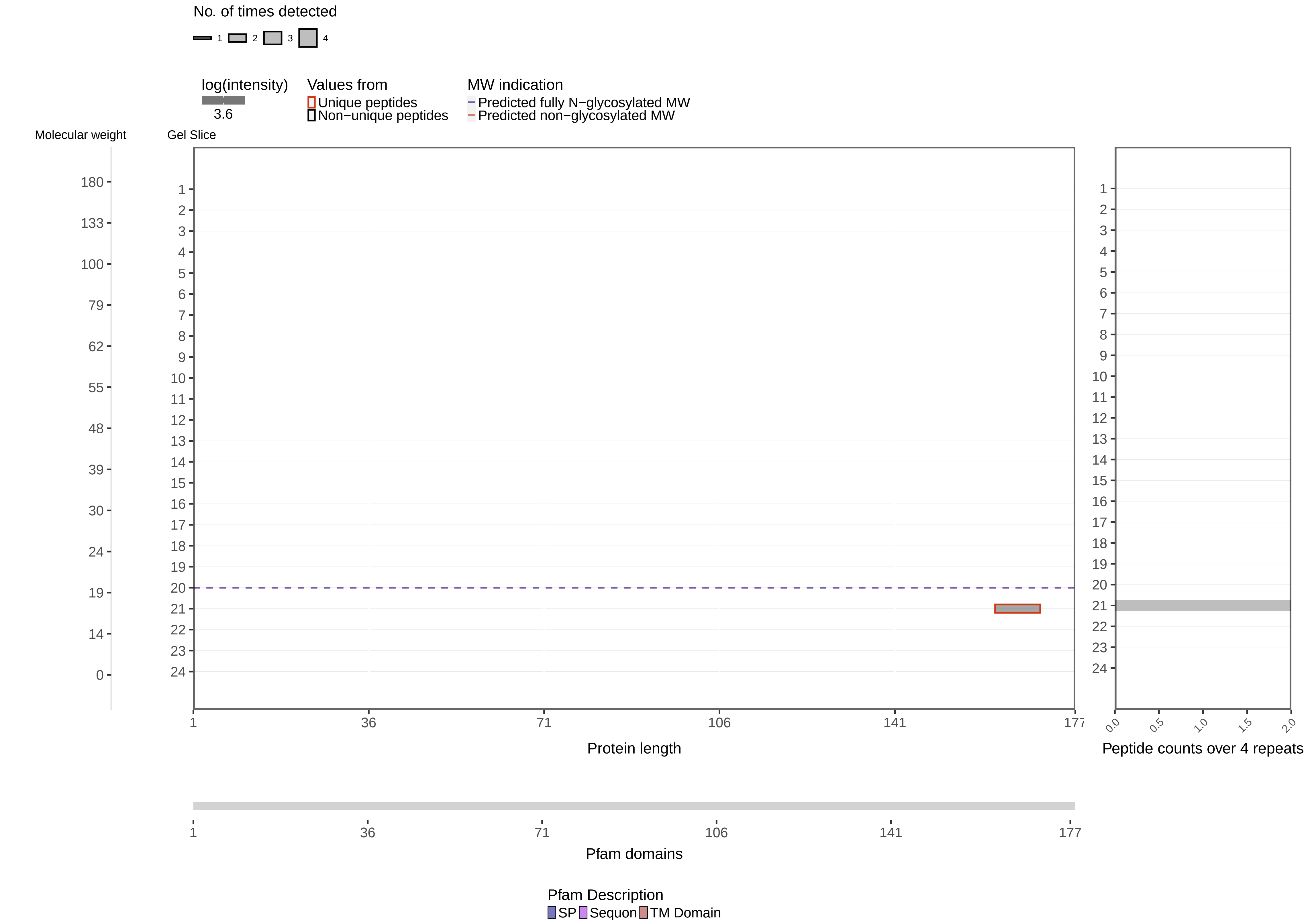


Protein sequence:

DKELRQRLQDSEQKLGLSMPLDQAKERATQLESEVTLLERRLILASGAEGMDGFRQRWSLHGRLTDTKKRLEALKGGMEKRKKDEPAESEPTKKRWFFW*

### NbL07g02330.1 60S_acidic_ribosomal_protein_P2−1−like [MW − SP = 9.75 kDa]

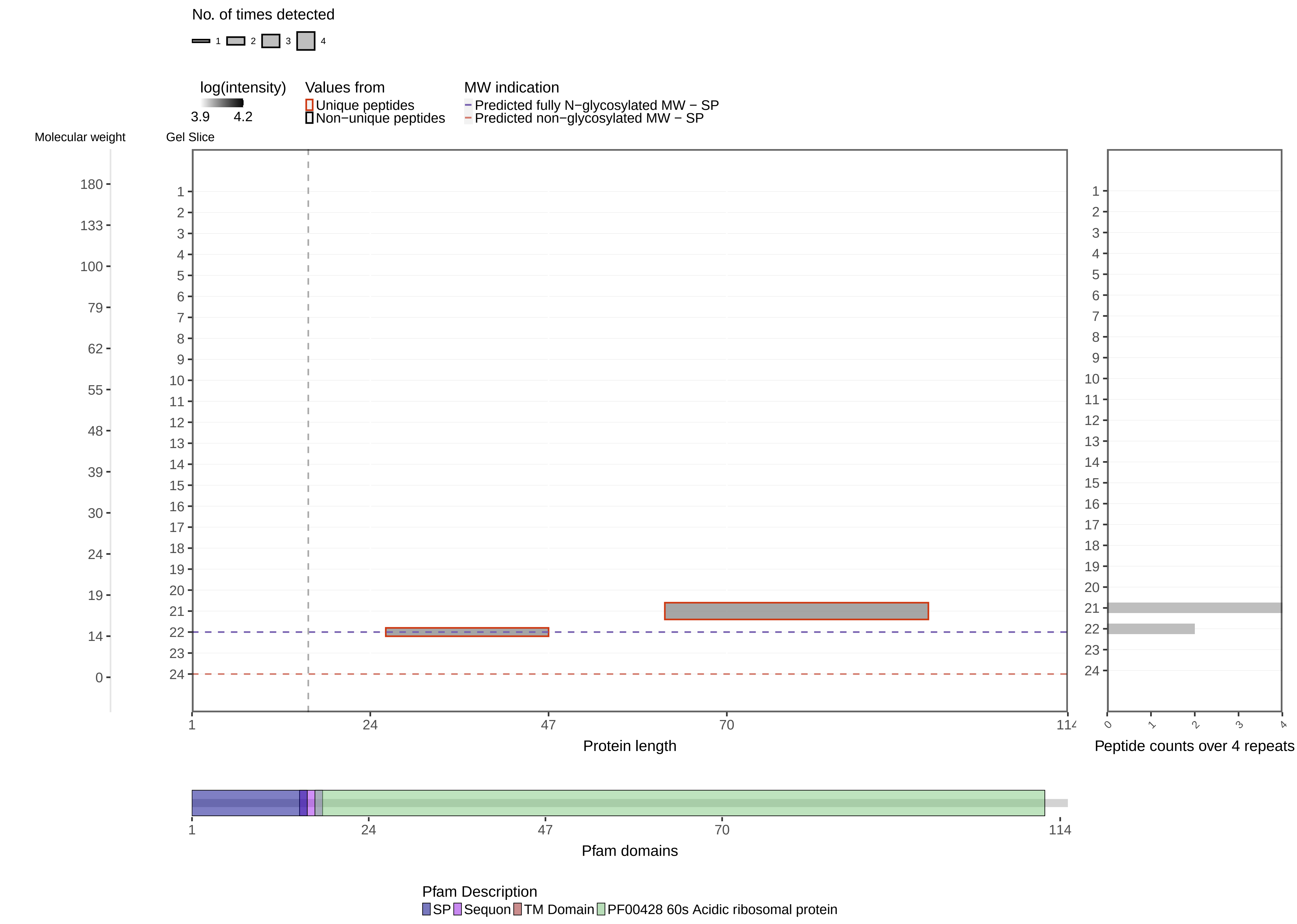


Protein sequence:

[illegible]VKGKDITELIAAGREKLASVPCGGGGGVAVAAPAGGSAAAAPAAEEKKEEKKEEKEESDDDMGFSLFD*

### NbL07g02260.1 putative_low_molecular_weight_protein−tyrosine−phosphatase_slr0328 [MW = 29.54 kDa]

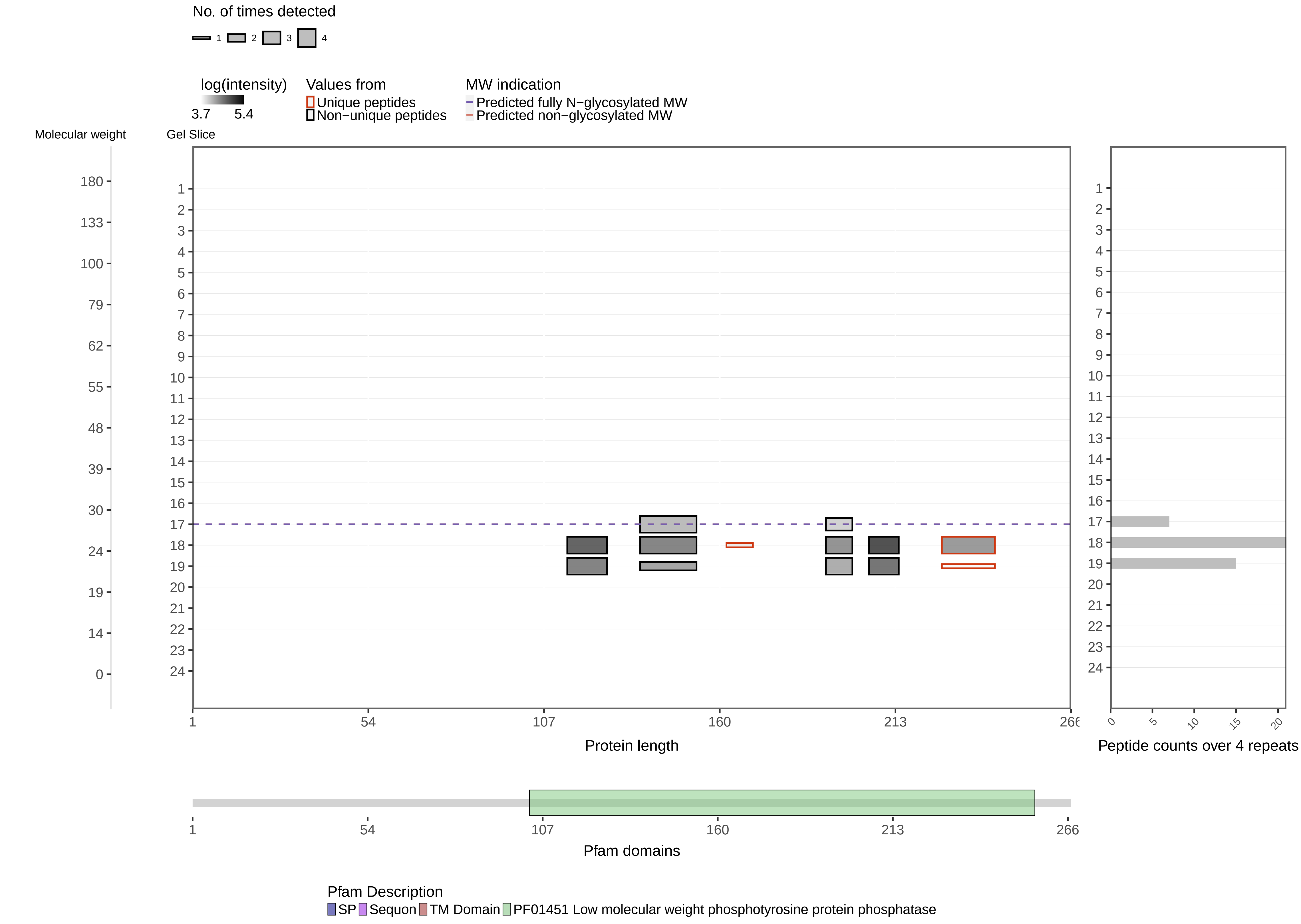


Protein sequence:

GVFTDLVKKRGLDSKFKIDSAGTINYHEGNEADPRMRAAAKRRGIAITSISRPIRPSDFKDFDLILAMDEQNKADILGALERWIHRETLPADAPEKVRLMCSYCKKHKETEVPDPYYGGAQGFEKVLDLLEDACDSLLESIVAEKKI*

NbL07g02230.1 photosystem_II_CP43_reaction_center_protein−like [MW = 46.54 kDa]

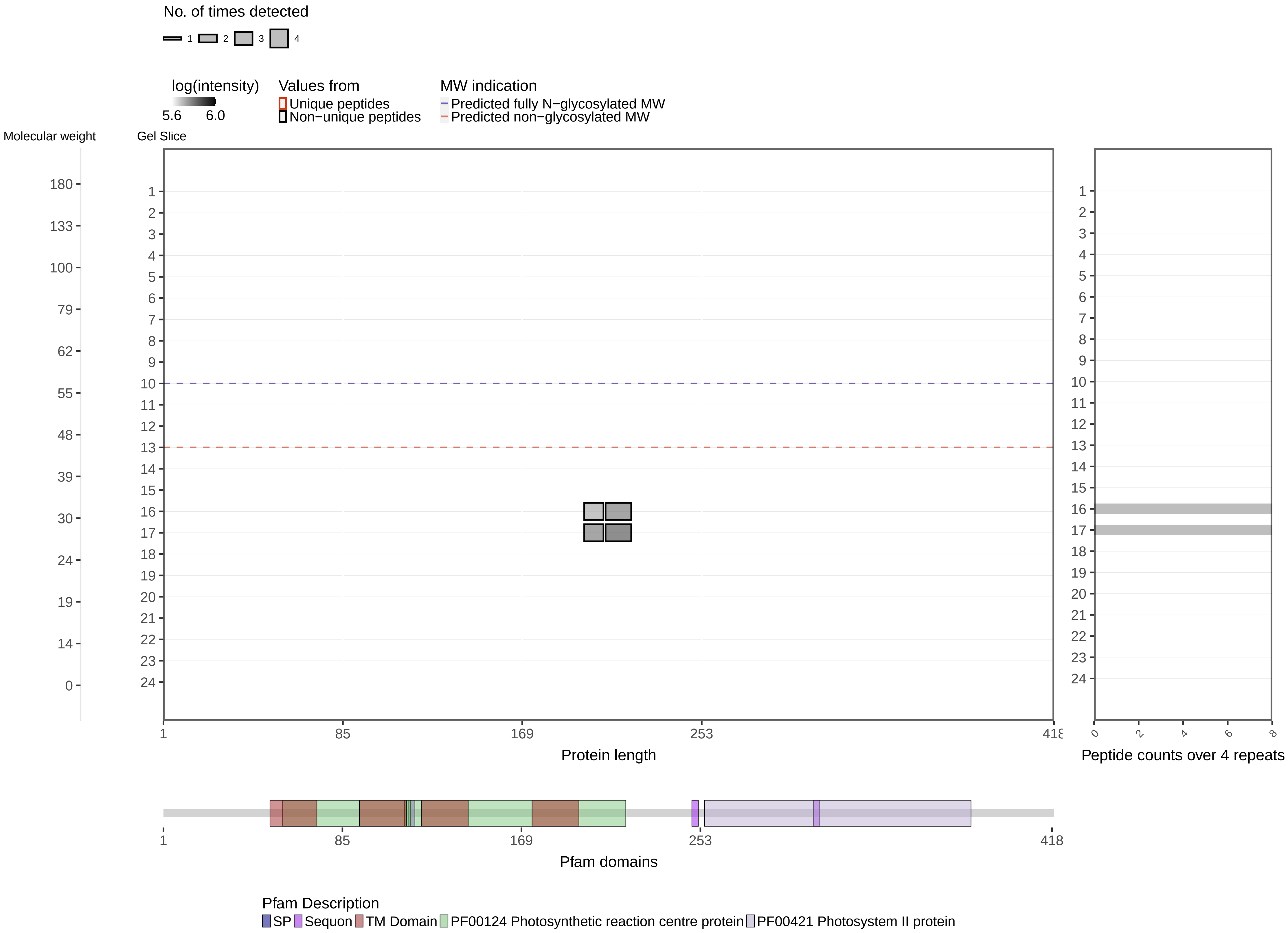


Protein sequence:

RAYDFVSQEIRAAEDPEFETFFYTKNILLNEALAGRDQETTGFAWWAGNARLINLSVLGFGGIYHALLGPETLEESFPFFGYVWKDRNKMTTILGALSVFGFIACCFVWFNNTAYPSEFYGPTGPEASQAQAFTFLVRDQRLGANVGSAQGPTGLGKYLMRSRPEKSFLEERNYAFWDLRAPWLEPLRGPNGLDLSRTTFEQKAFVMKPFQALPFCRSTTAI*

### NbL07g02170.1 bifunctional_protein_FolD_1_mitochondrial−like [MW = 42.17 kDa]

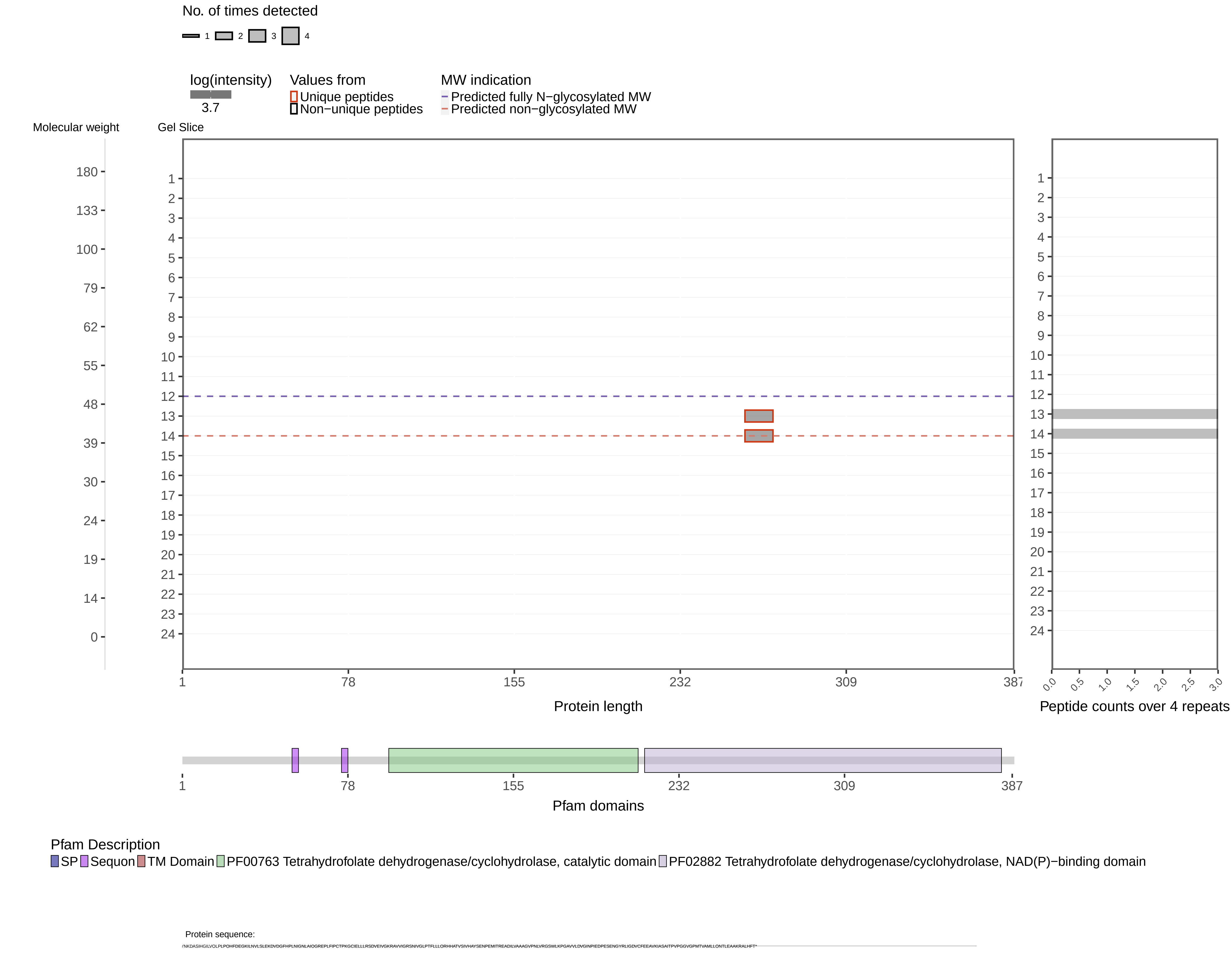

NbL07g02150.1 uncharacterized_protein_At5g22580−like [MW = 12.26 kDa]

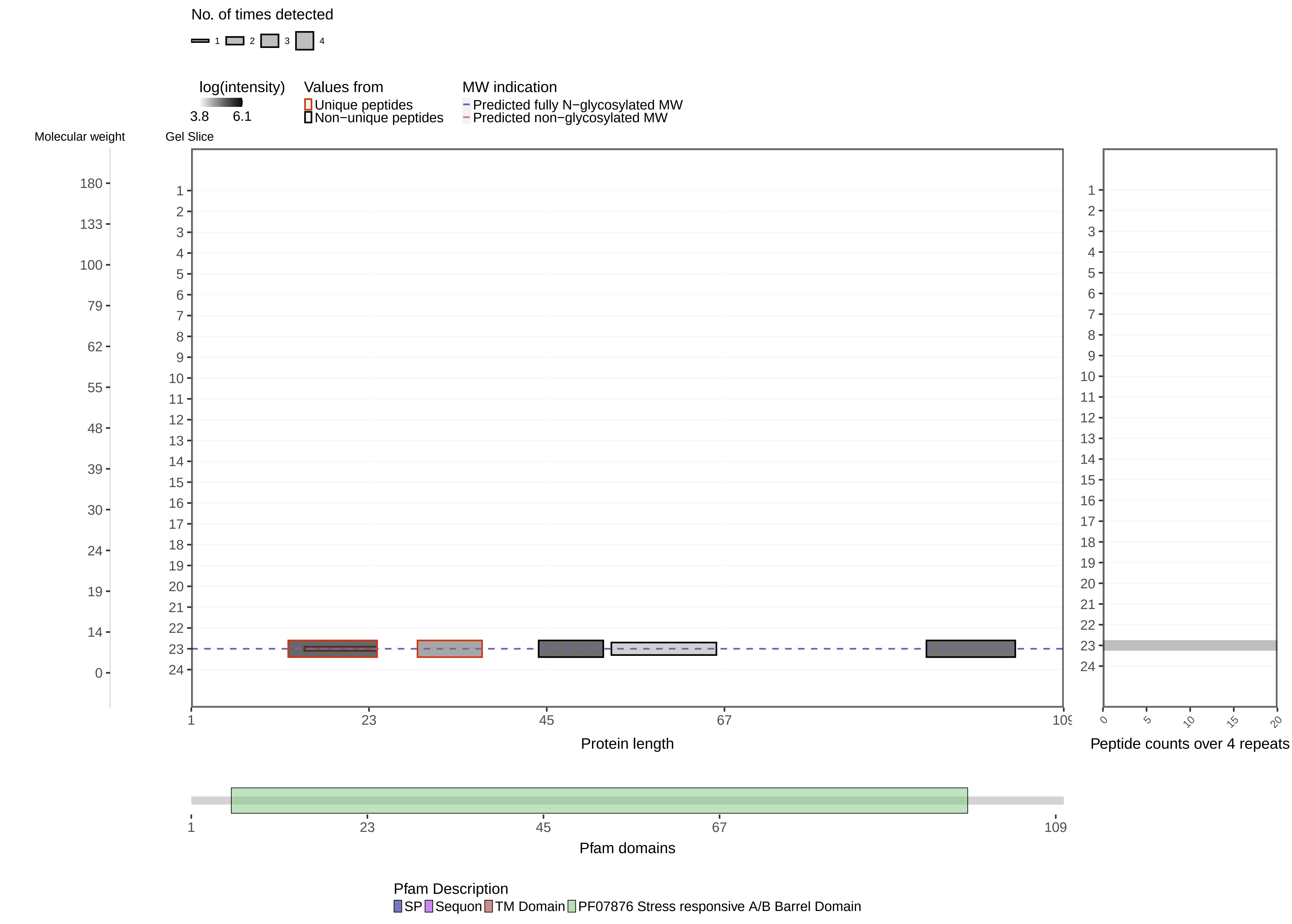


Protein sequence:

GKDAESHEMLRQGFTHAFLMTFNNKDDYTKFVSHPNHVEFSATFSTVIDKAVLLDFPAISVKAPAPA*

### NbL07g02110.1 arogenate_dehydratase/prephenate_dehydratase_6_chloroplastic−like [MW = 47.2 kDa]

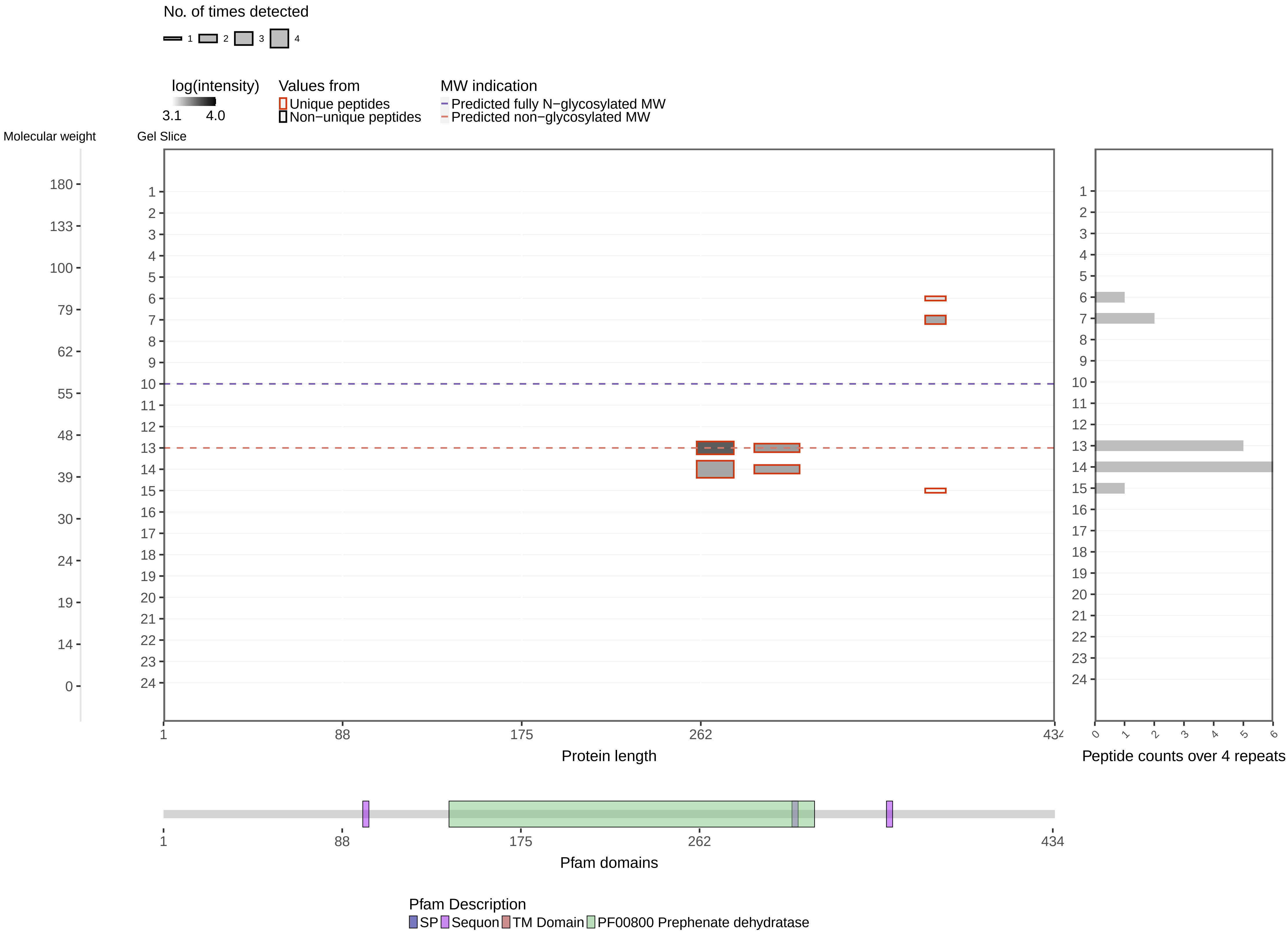


Protein sequence:

LLRHRLHIVGEVQLPVNHCLLALPGVRKEYLTRVISHPQALAQCELTLTKLGLNVAREAVDDTAGAAEYIAGNNLRDTAAIASARAADLYGLQILAEGIQDDSSNVTRFVMLAREPIIPRTDRPFKTSIVFAHHEGTSVLFKVLSAFAFRNISLTKIESRPHRNRPIRVVDDANVGTAKHFEYMFYVDFEASMADVRAQNALAEVQEFTSFLRVLGSYPMDMTPWCPSRED*

NbL07g01930.1 UDP−D−apiose/UDP−D−xylose_synthase_2−like [MW = 43.26 kDa]

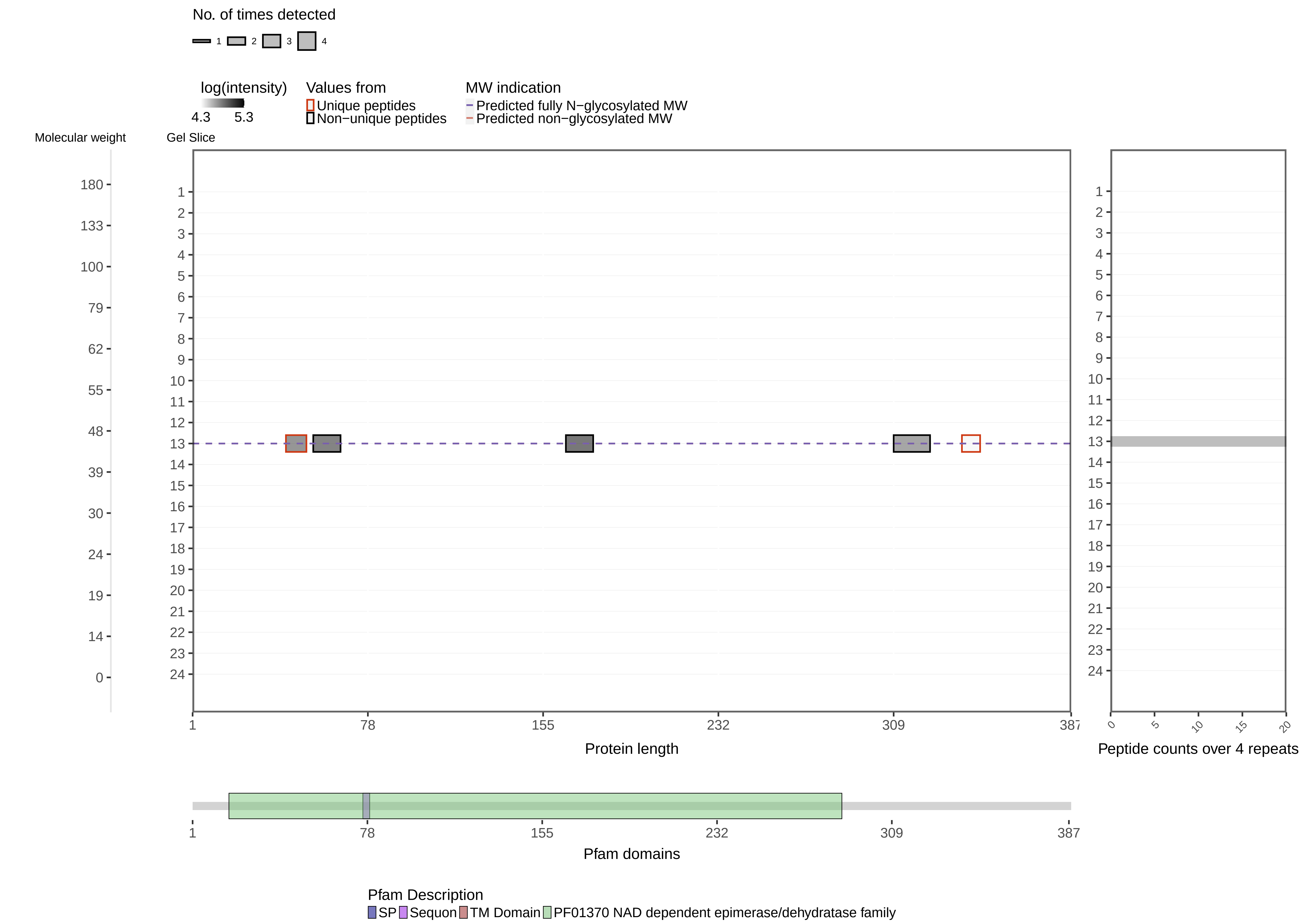


Protein sequence:

'ACAKQLIERLVYAEGAENGLEFTIVRPFNWIGPRMDFIPGIDGPSEGVPRVLACFSNNLLRREPLKLVDGGESQRTFIYIKDAIEAVLLMIENPARANGHIFNVGNPNNEVTVRQLAEMMTQVYSKVSGESSIETPTIDVSSKEFYGEGYDDSDKRIPDMTIISRQLGWNPKTSLWDLLESTLTYQHRTYAEAVKQAMSKTTAN*

NbL07g01840.1 VAMP−like_protein_YKT61 [MW = 22.61 kDa]

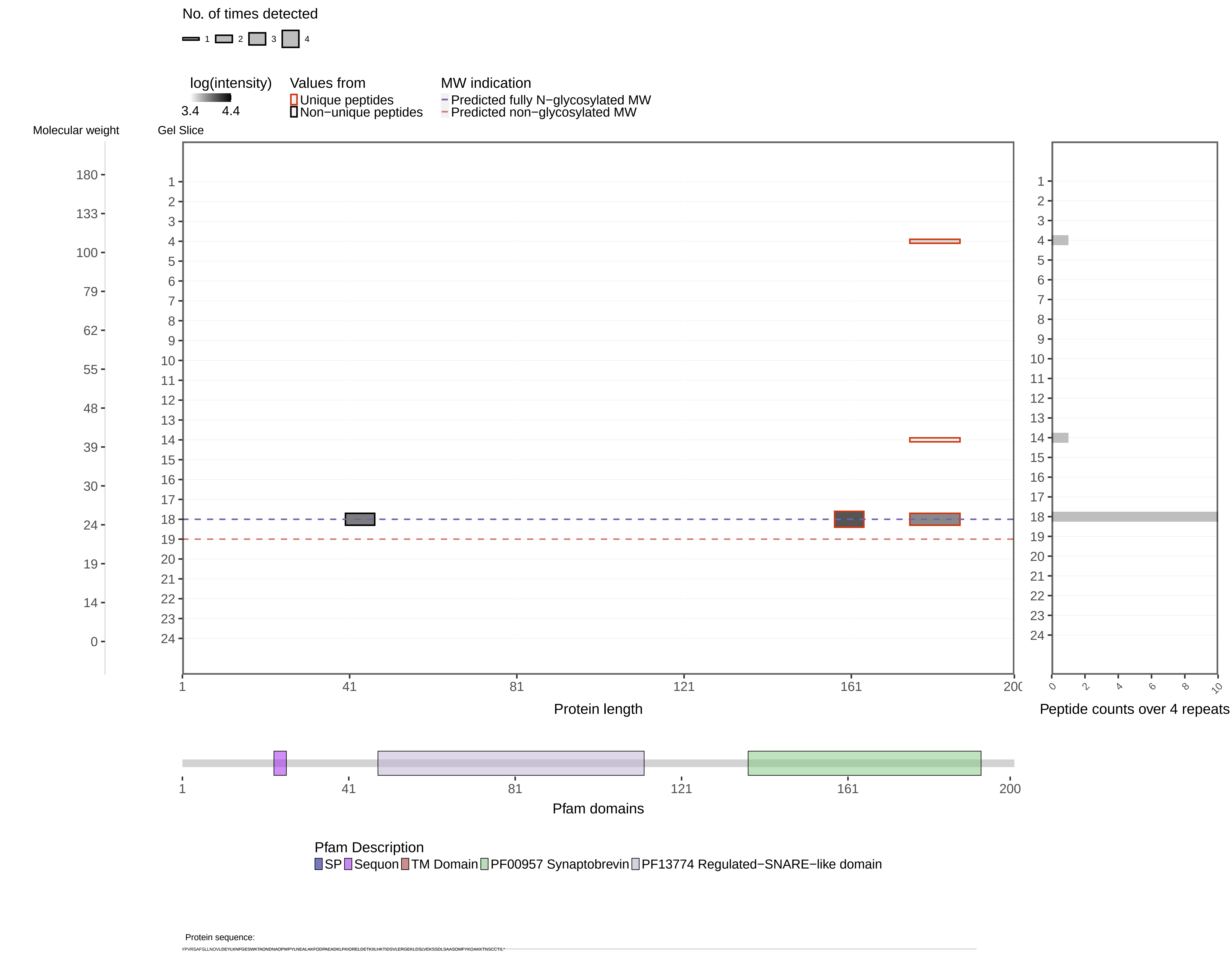

### NbL07g01830.1 temperature−induced_lipocalin−1−like [MW = 21.37 kDa]

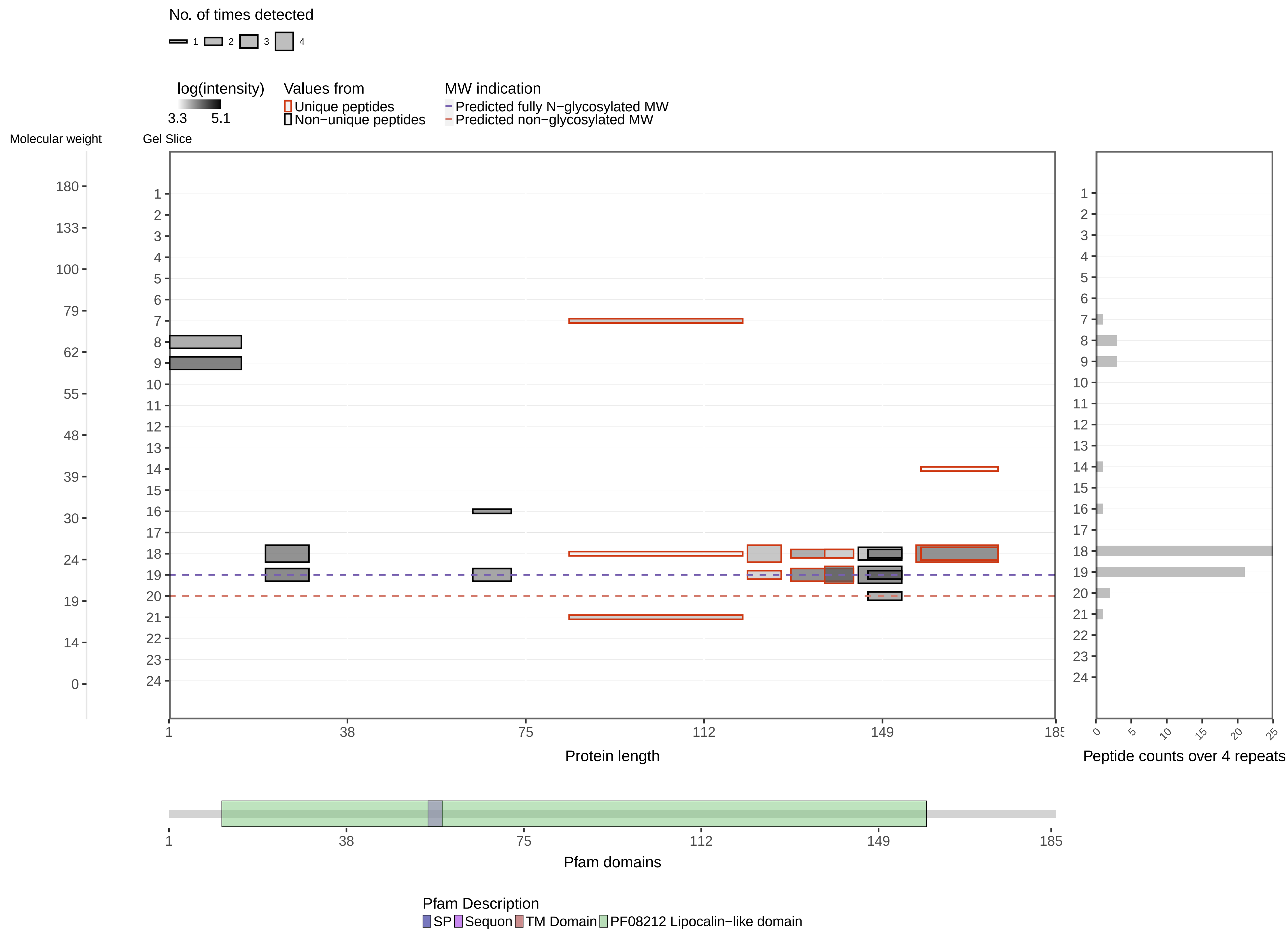


Protein sequence:

[illegible]DEAKLKVKFYLPPFLPIIPITGDYWVLYIDDDYKYALIGQPSRNYLWILCRKTHLDEEKYNMLIEKAKEVGYDVSKLRKTPQSDPPPEGELPQDTKGIWWLKSIFGR*

### NbL07g01690.1 S−norcoclaurine_synthase_1−like [MW = 17.01 kDa]

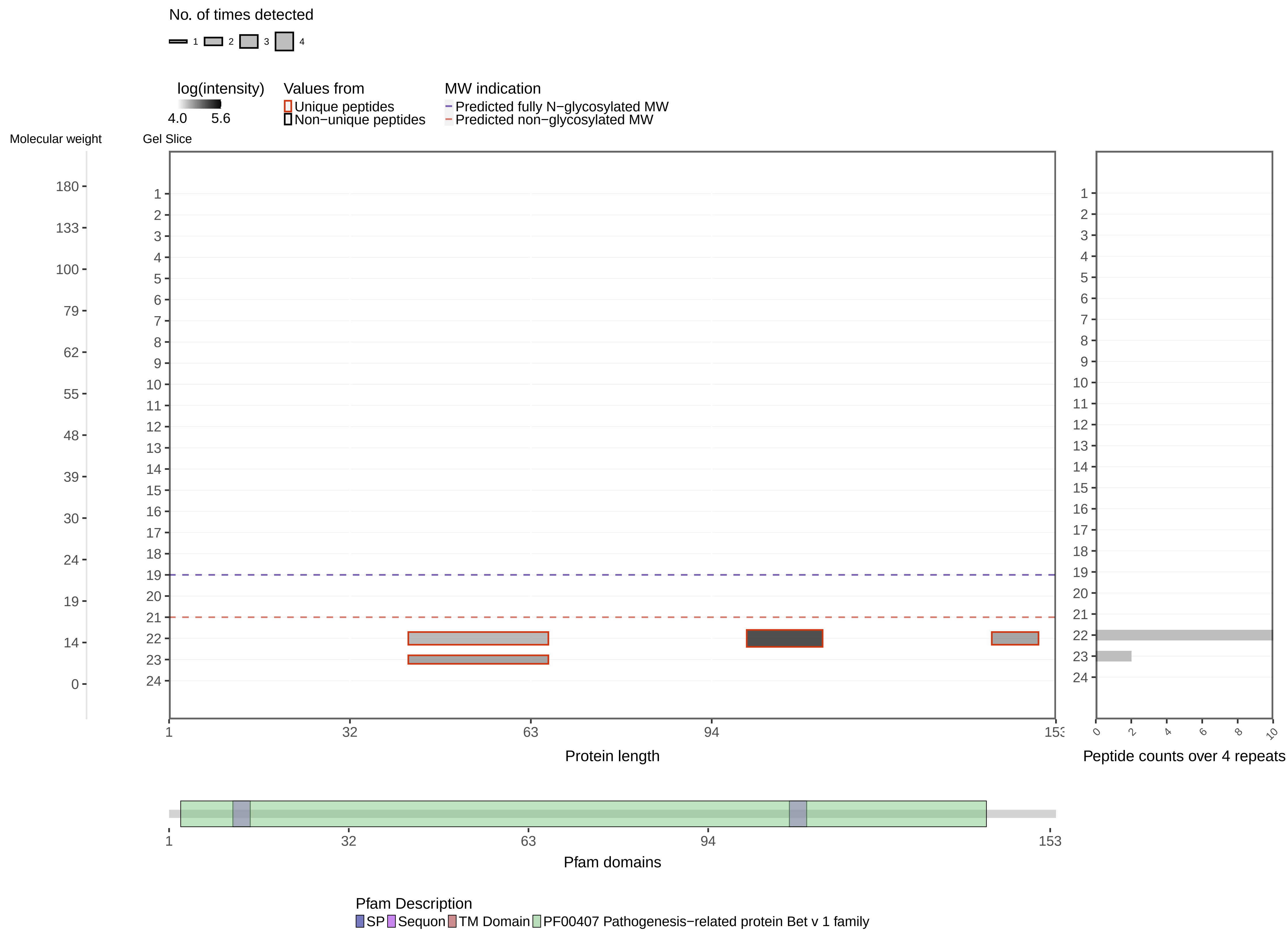


Protein sequence:

ꞩNAPYKEKFVSIDDEKRVKEVEIVEGGYLDLGFSFYGIKFEVIEKDENLSIVKITIDFETKDAENIHLTIGNIQAFVAIMKATVEYFSQKSK*

NbL07g01670.1 NADP−dependent_glyceraldehyde−3−phosphate_dehydrogenase [MW = 25.97 kDa]

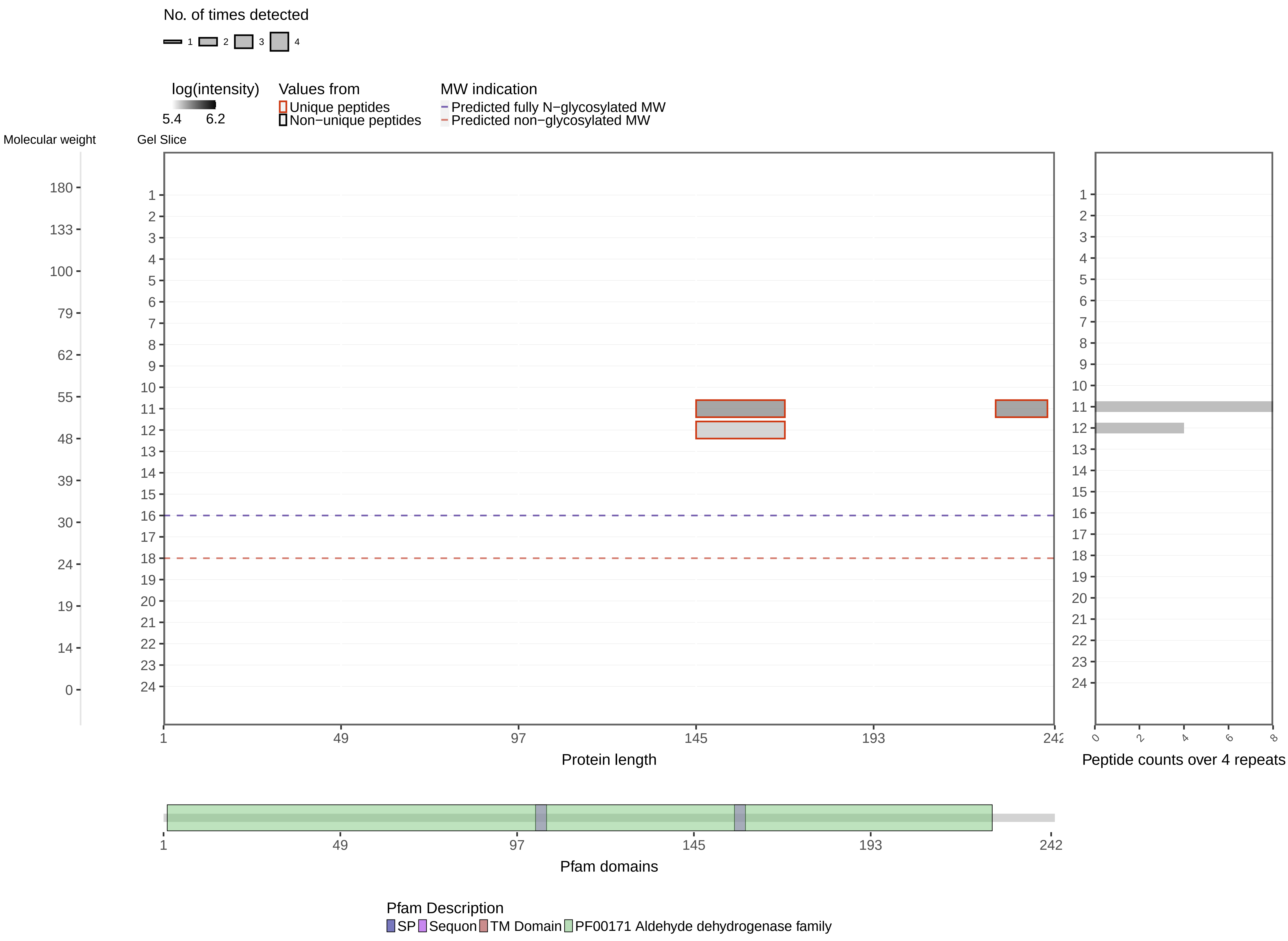


Protein sequence:

QYKREGNLIWPLLLDNVRPDMRIAWEEPFGPVLPVIRINSVEEGIHHCNASNFGLQGCVFTRDINKAILISDAMETGTVQINSAPARGPDHFPFQGIKDSGIGSQGITNSINMMTKVKTTVINLPTPSYTMGKL*

NbL07g01570.1 eukaryotic_translation_initiation_factor_5A−3 [MW = 17.54 kDa]

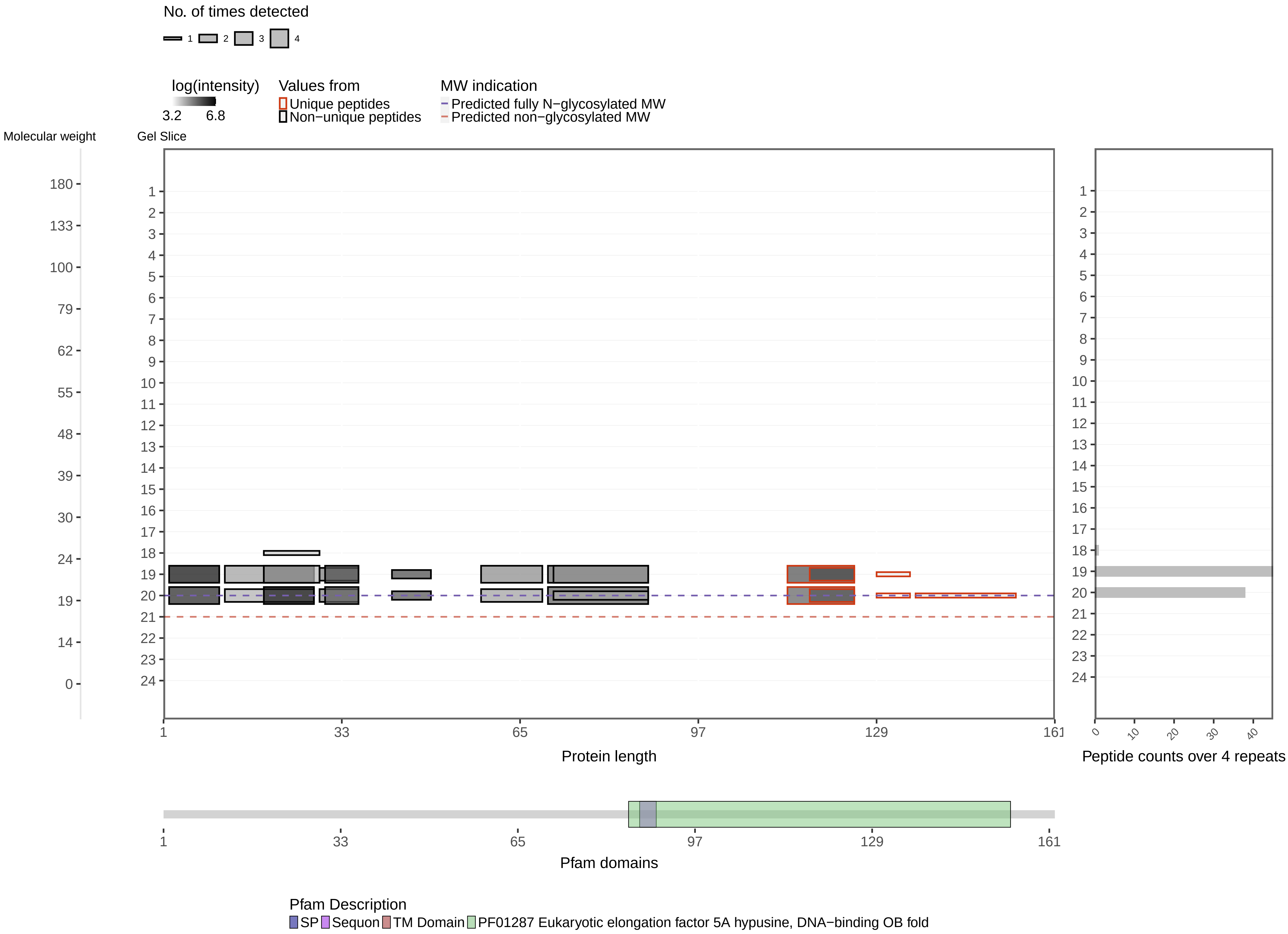


Protein sequence:

GKKLEDIVPSSHNCDVPHVNRTDYQLIDISEDGFVSLLTESGNTKDDLRLPTDEALLKQVKDGFQEGKDLVVSVMSAMGEEQINAVKDIGTKN*

### NbL07g01330.1 shikimate_O−hydroxycinnamoyltransferase−like [MW = 48.45 kDa]

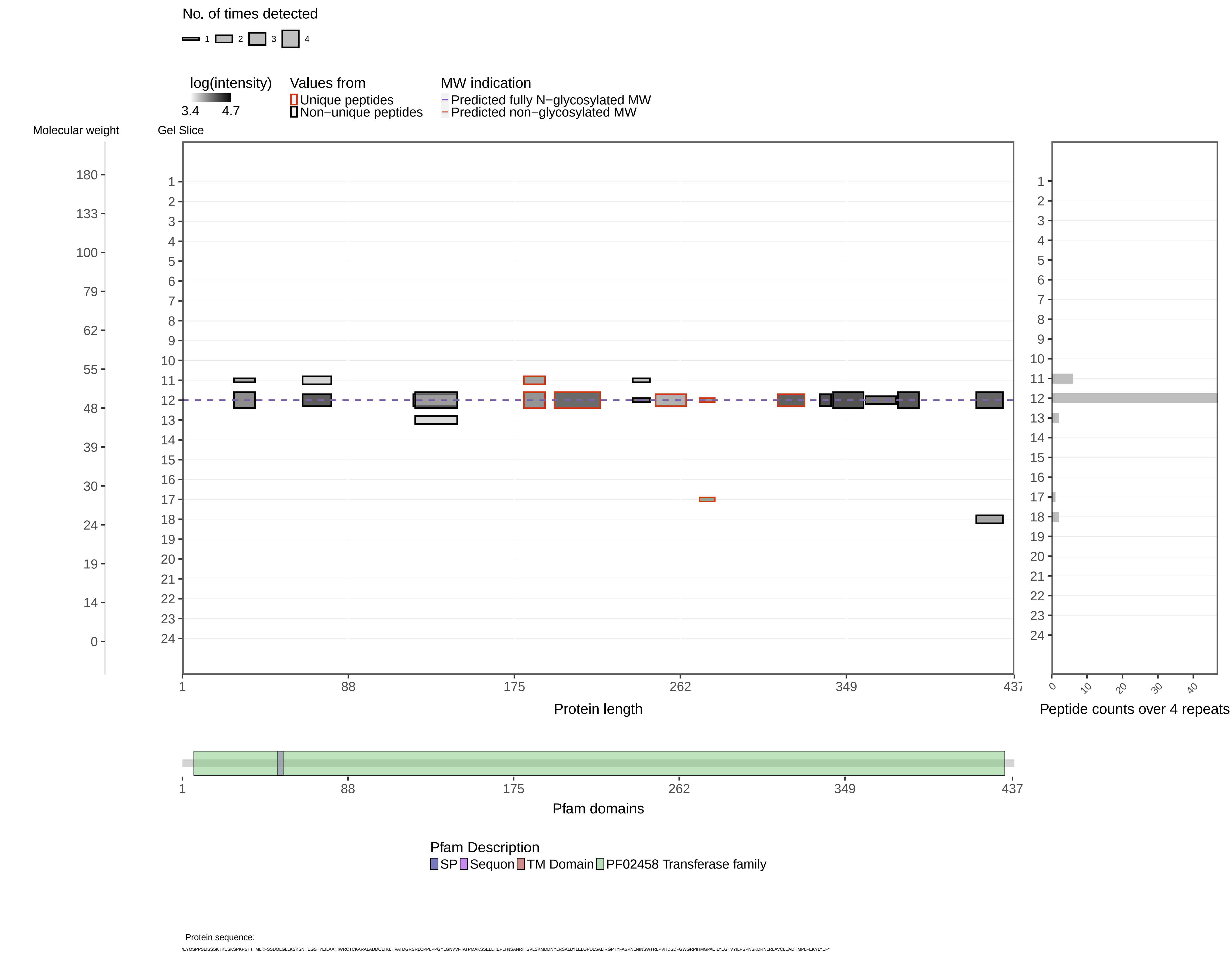

### NbL07g01050.1 protein_disulfide_isomerase−like_1−4 [MW − SP = 62.13 kDa]

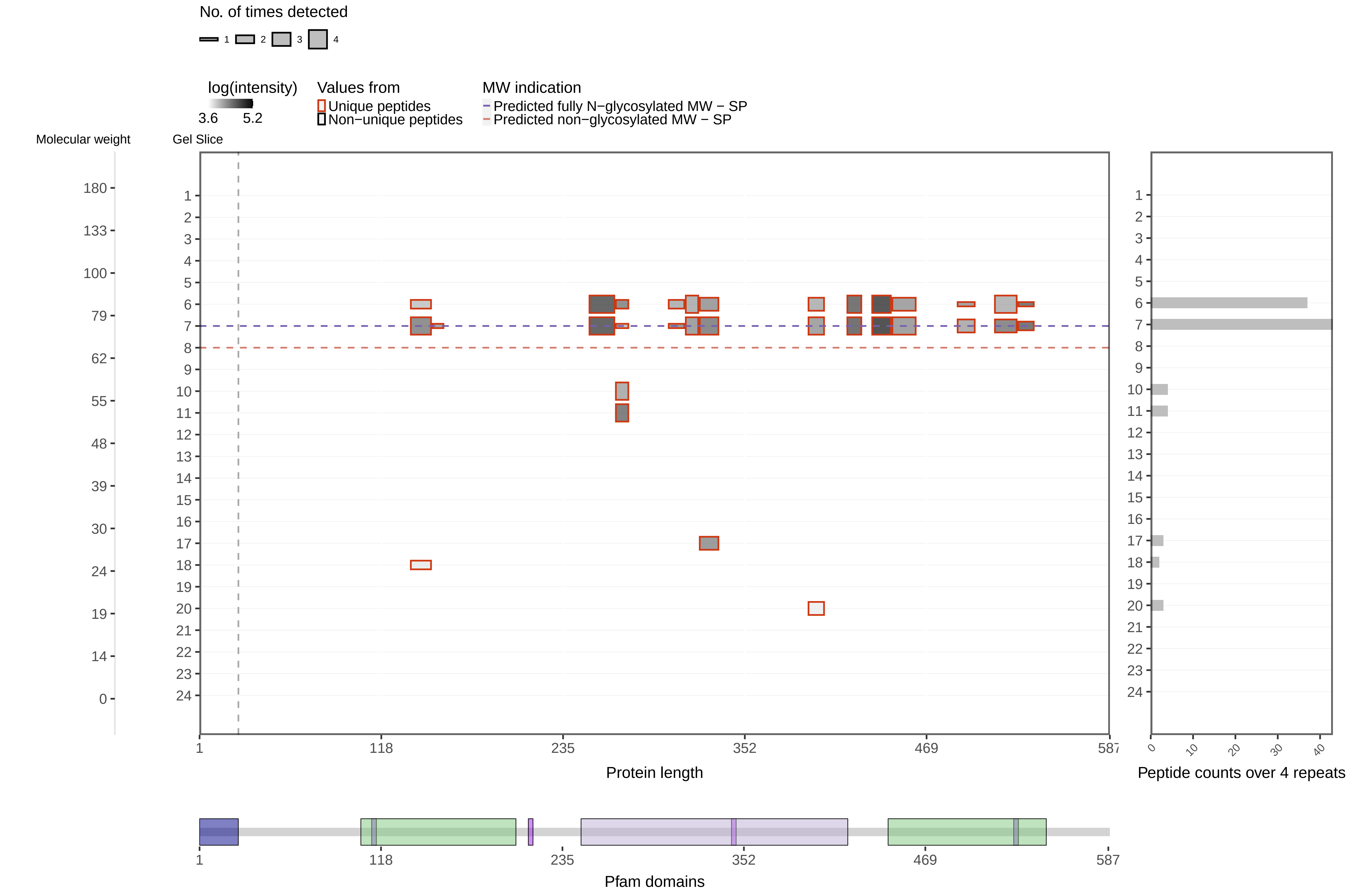


Protein sequence:

NbL07g01040.1 4−hydroxy−3−methylbut−2−en−1−yl_diphosphate_synthase_(ferredoxin)_chloroplastic [MW = 82.2 kDa]

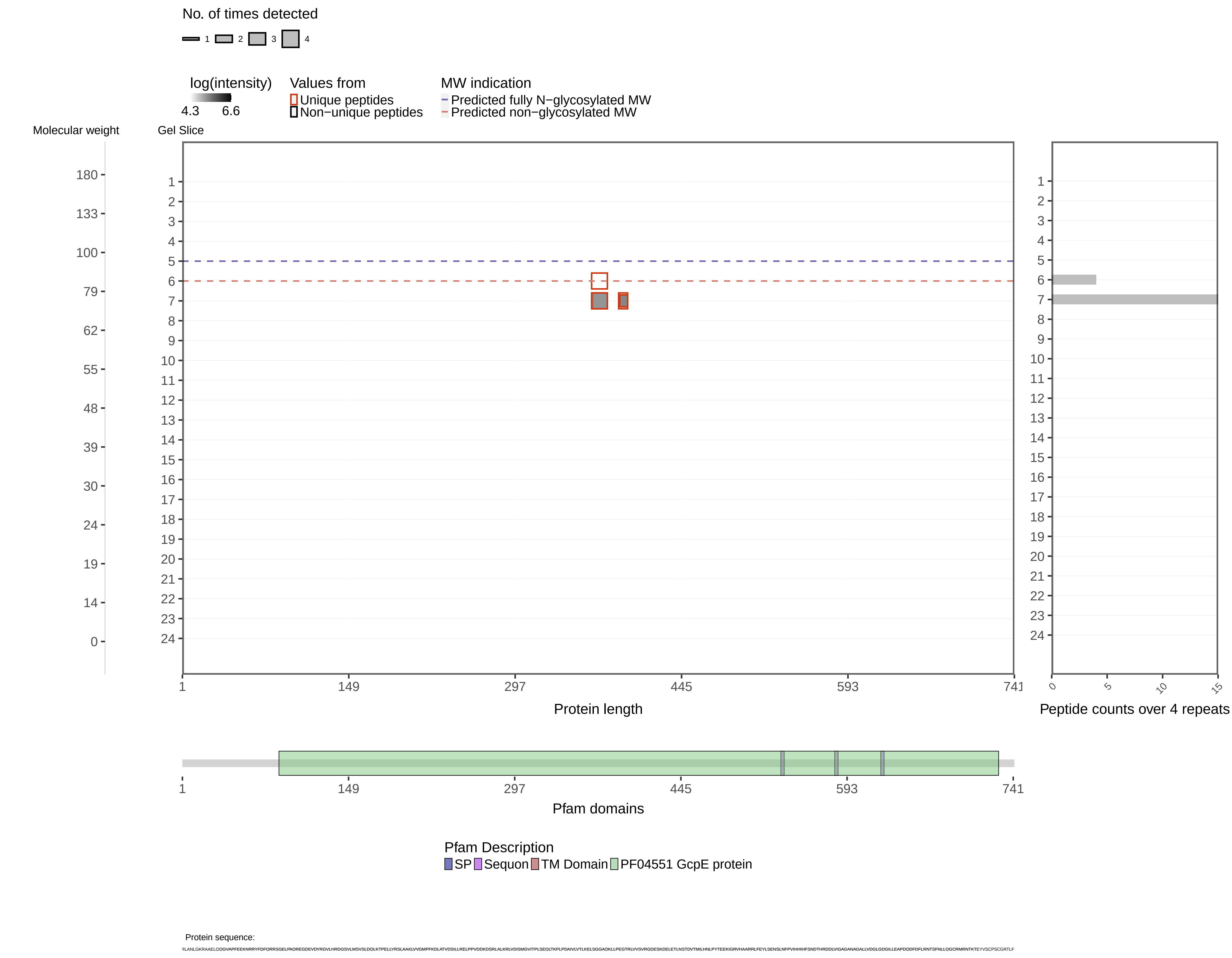

NbL07g00850.1 beta−galactosidase_8 [MW − SP = 89.69 kDa]

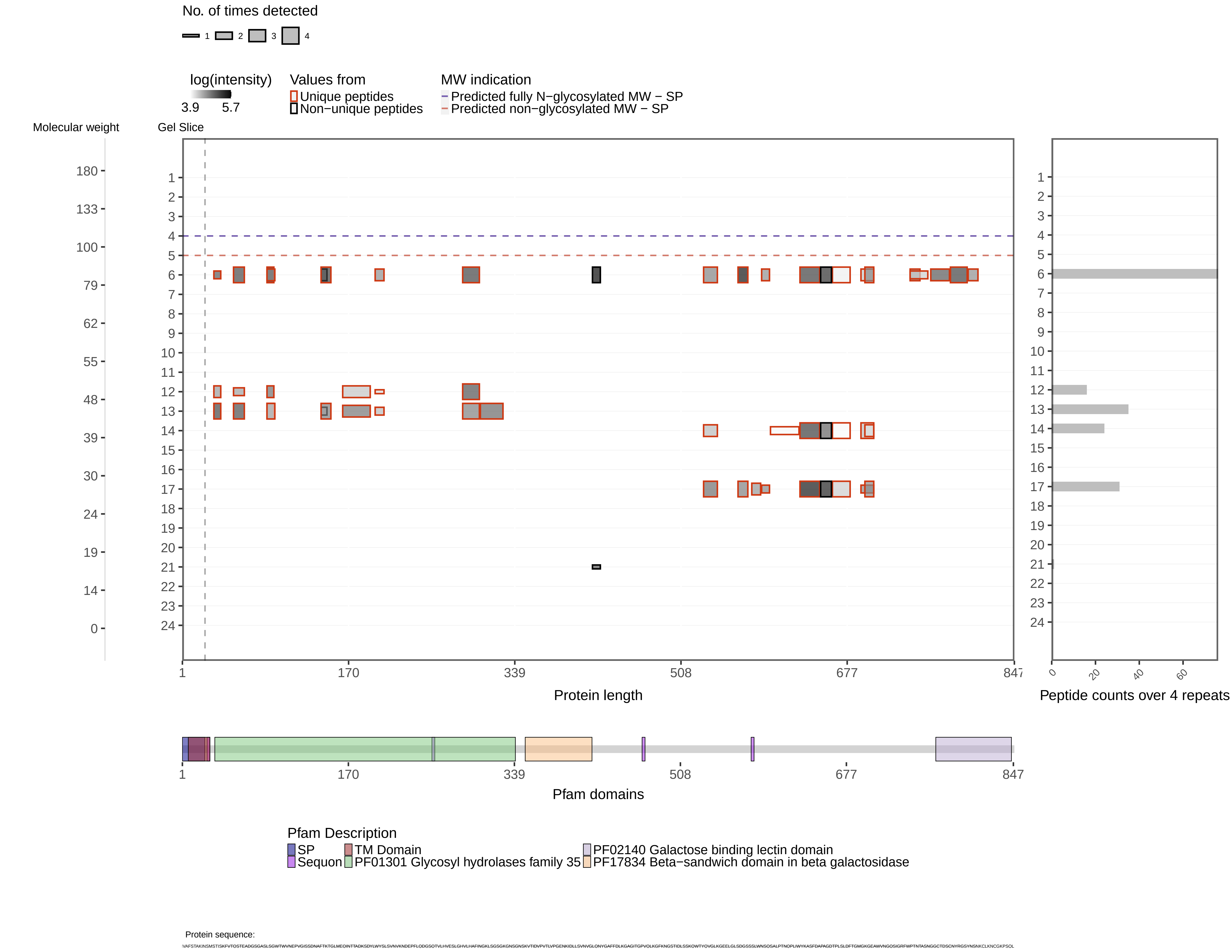

### NbL07g00810.1 proteasome_subunit_beta_type−4−like [MW = 27.61 kDa]

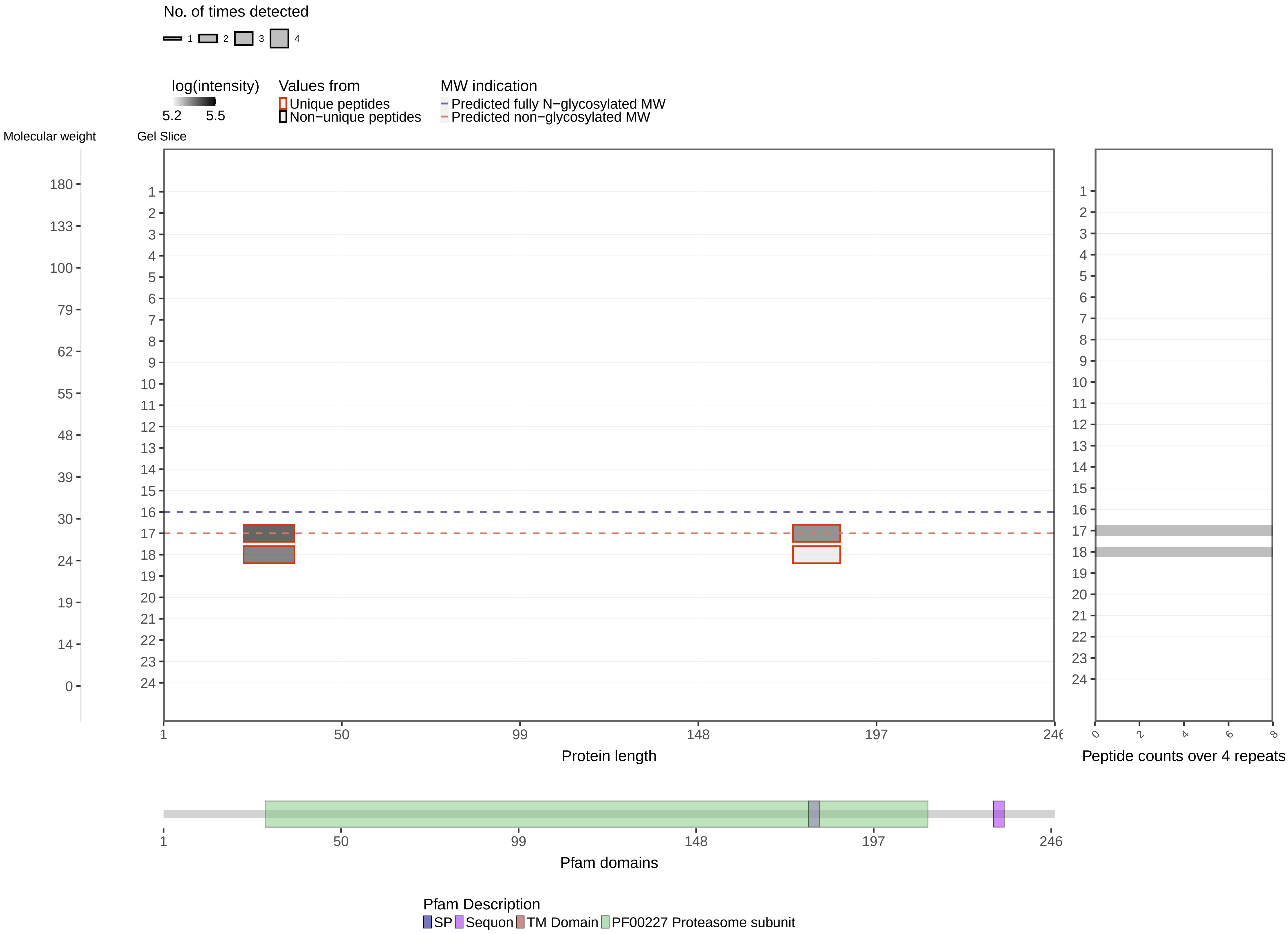


Protein sequence:

NYLTRMMYNRRNKFNPLWNSLVLGGVKNGQKYLGMVSMIGVHYEDDHVATGFGNHLARPILRDEWHENLSYEEGVKLLEKCMRVLLYRDRSAVNKLQIAKITEEGVTVSQPYSLKTFWNFSAFQNPTVGAEGSW*

### NbL07g00760.1 4−coumarate−−CoA_ligase−like_7 [MW = 59.73 kDa]

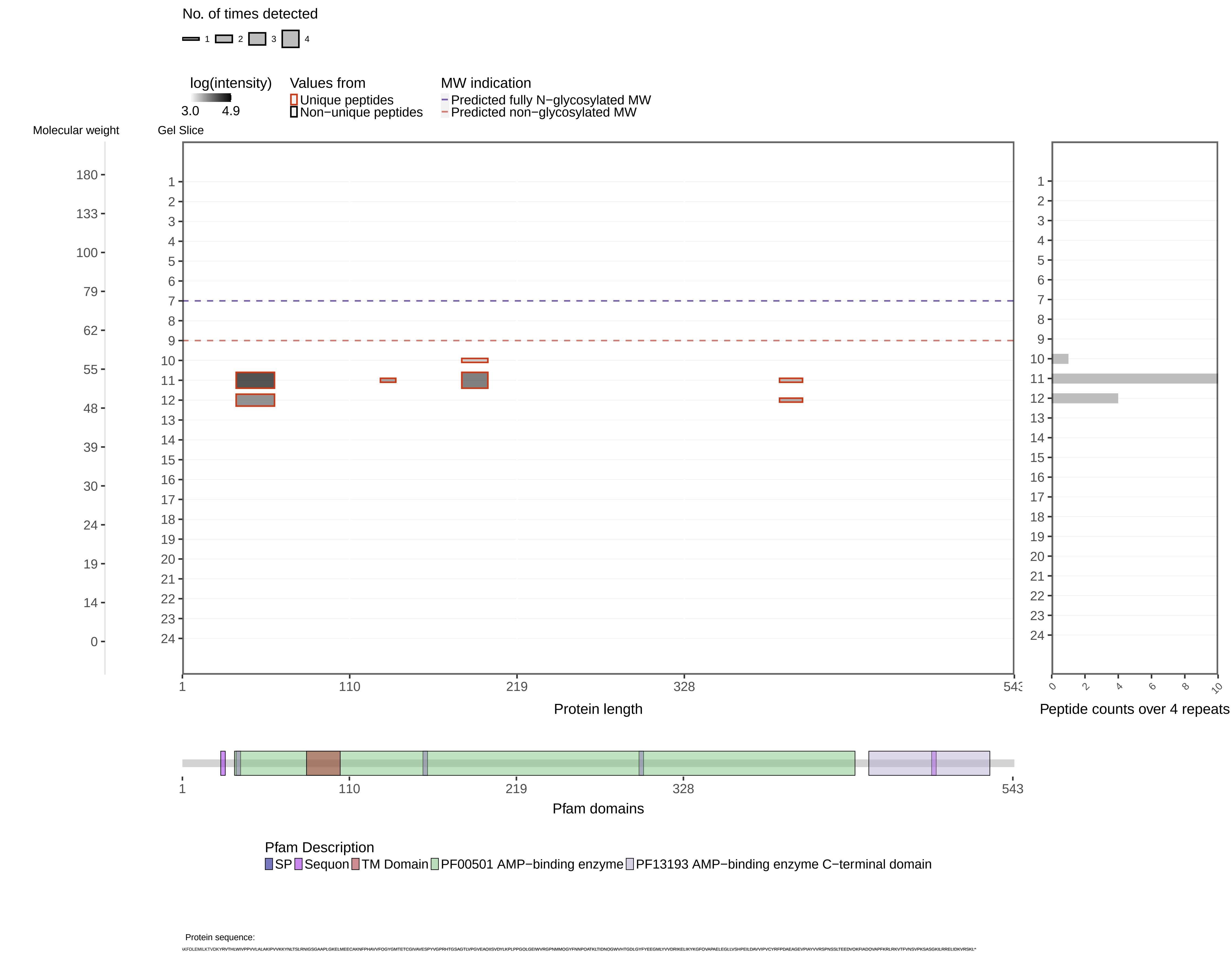

NbL07g00730.1 probable_inactive_purple_acid_phosphatase_29 [MW − SP = 40.34 kDa]

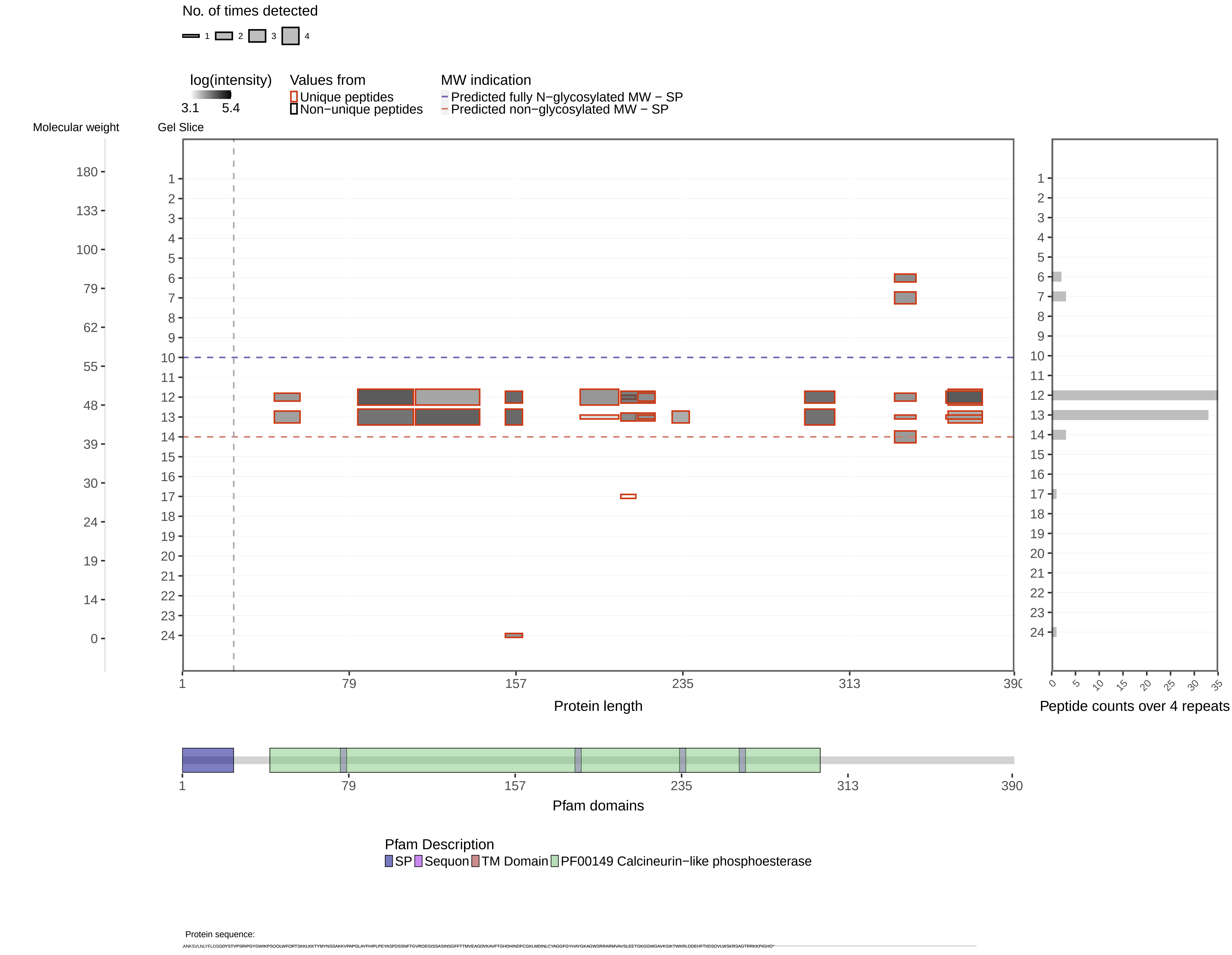

### NbL07g00710.1 xyloglucan_endotransglucosylase/hydrolase_protein_31−like [MW − SP = 31.42 kDa]

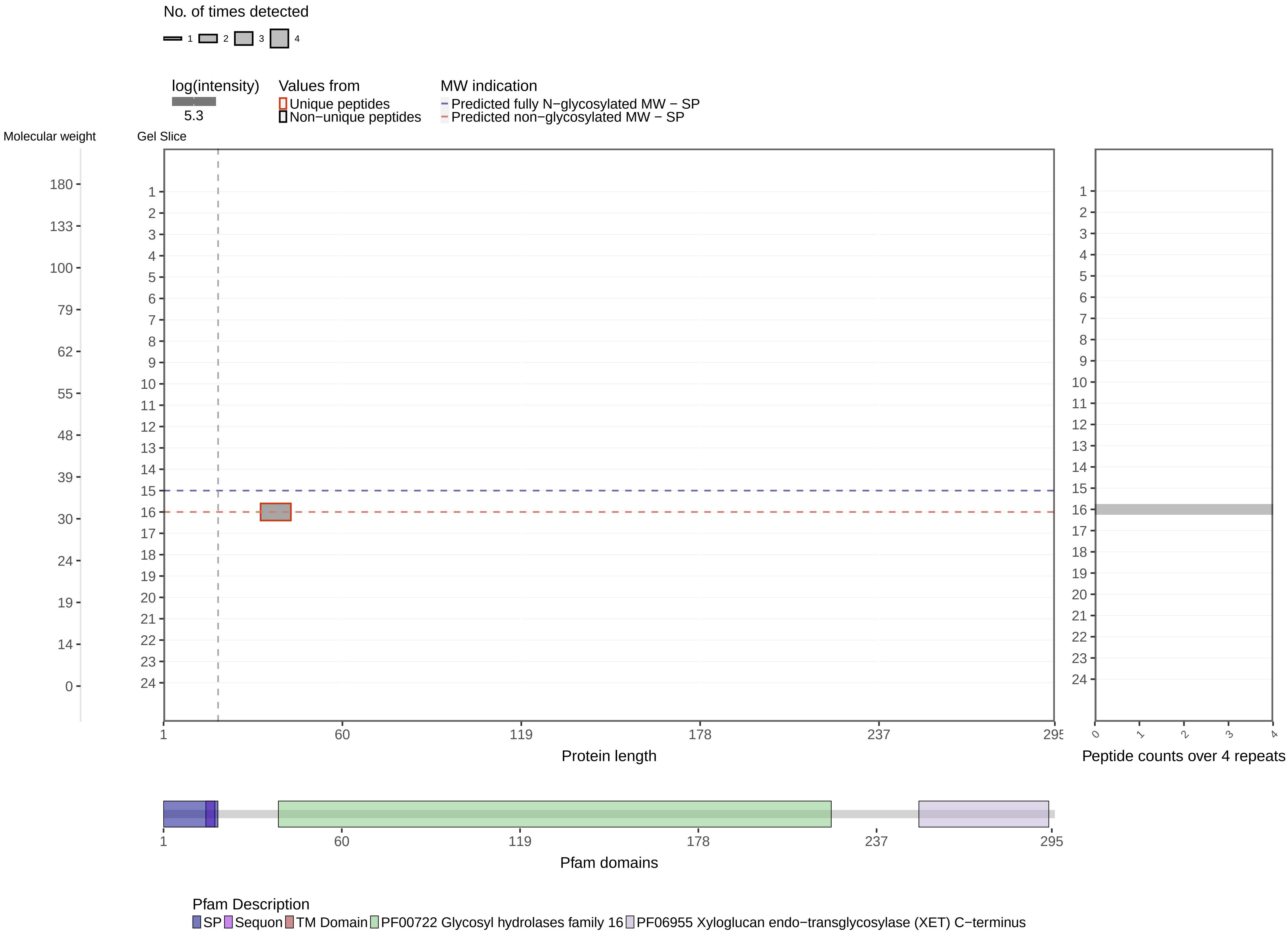


Protein sequence:

IRGSGDGNIIGREMKFHLWFDPTQAYHNYAILWDPNEIIFFVDDVPIRRYPRKNDATFPQRPMYVYGSIWDASSWATEEGRIKADYRYQPFVGKYNNFKIAGCTANENPWCGRSPSSSPSRAGGLSRQQIAAMLWVQRNYKVYDYCRDPRRDHTHTPEC*

### NbL07g00700.1 vacuolar_protein_sorting−associated_protein_24_homolog_1−like [MW = 25.08 kDa]

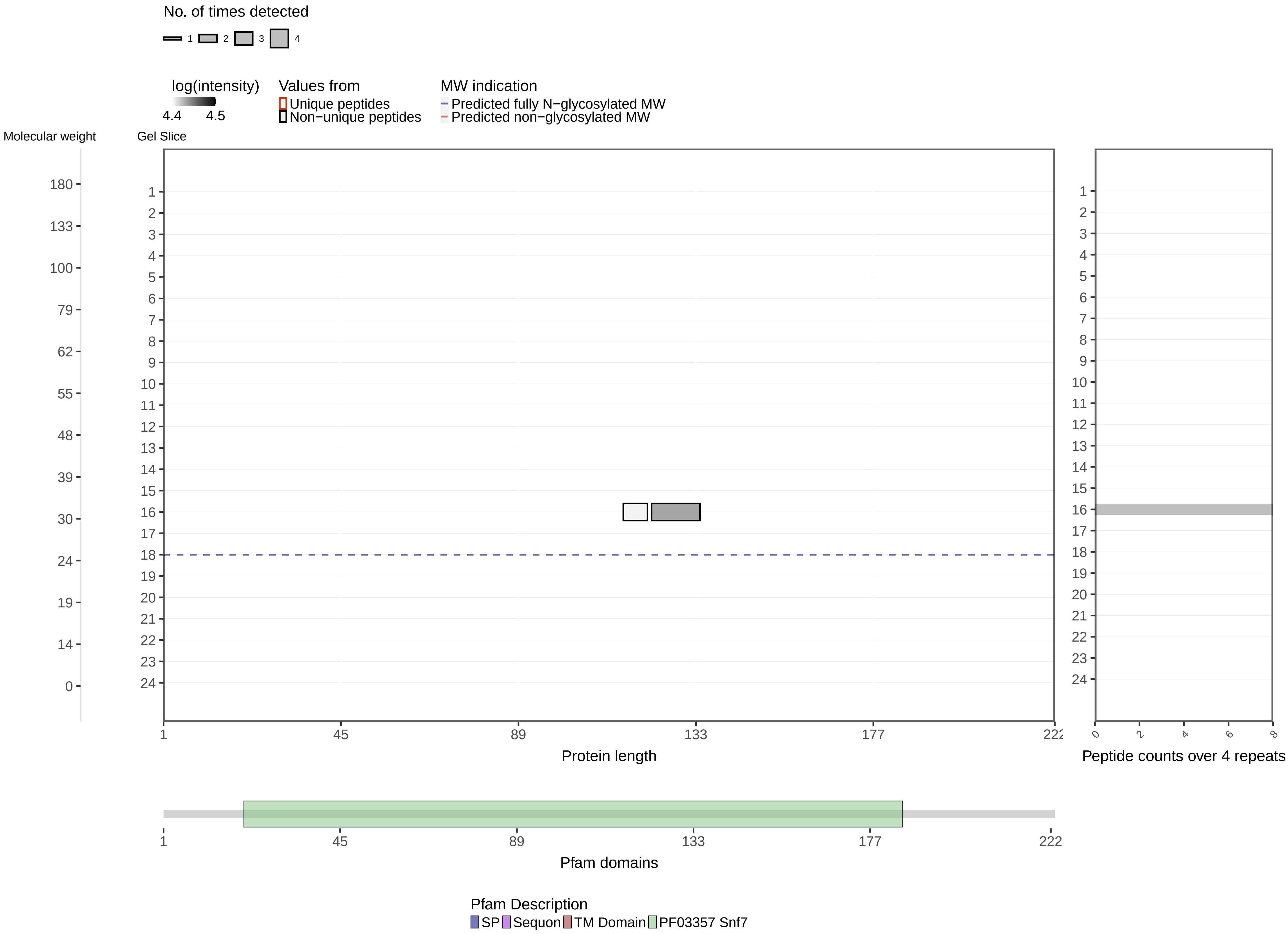


Protein sequence:

AIARTVGHLSKSAEVMKLVNNLMKAPEVAMTMQEFSKEMTKAGVMEEMVNDAVDSALDSEDIEDEIEEEVDKVLTAIAGETTALPEAVRKQKLKQPAQAVEDAEGADEEEDLEELRARLAKVRS*

NbL07g00690.1 serine_carboxypeptidase−like [MW − SP = 53.94 kDa]

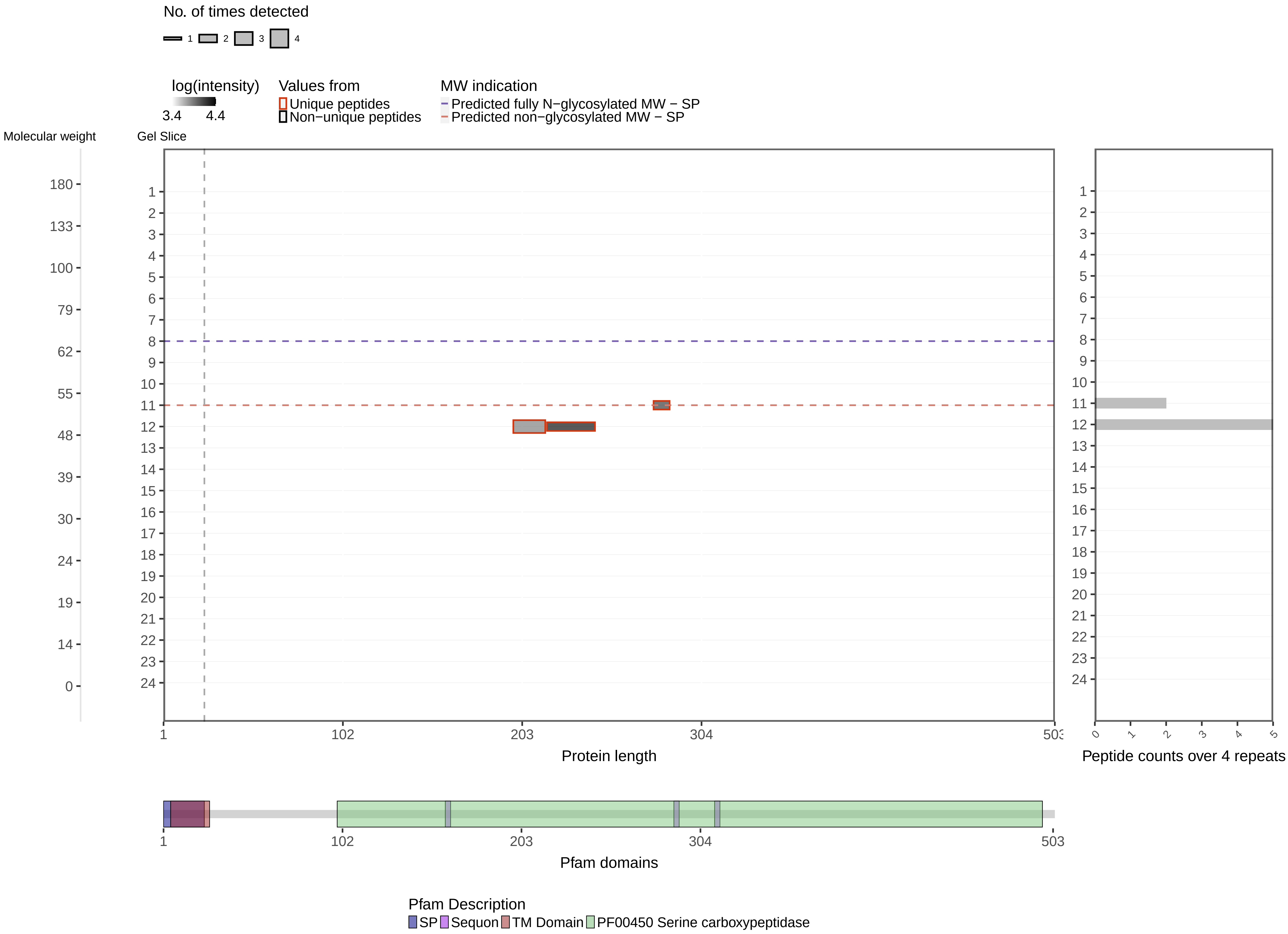


Protein sequence:

FASRVNQGNKNKEGININLKGFAIGNGLTNPEIQYKAYTDYALDMKLINQTDYDAINKLYPQCQQEITLCGNGSEDACLDAFGRCTNIFSSIMDVVSDKNYYDIRKKCEGNLCYDFSGMETYLNDNQVKRALGVPNSIDFVSCSSTVYQAMETDWMKNLEVGIPSLLEDGINLLIYAGEYDLICNWLGNSNWVHAVEWSGQNDFVAAPSVSFTVDGEEKGIQKNYGPLTFLKVHDAGHMVPMDQPKAALEMLQRWTTQGKLS*

### NbL07g00660.1 T−complex_protein_1_subunit_beta [MW = 57.01 kDa]

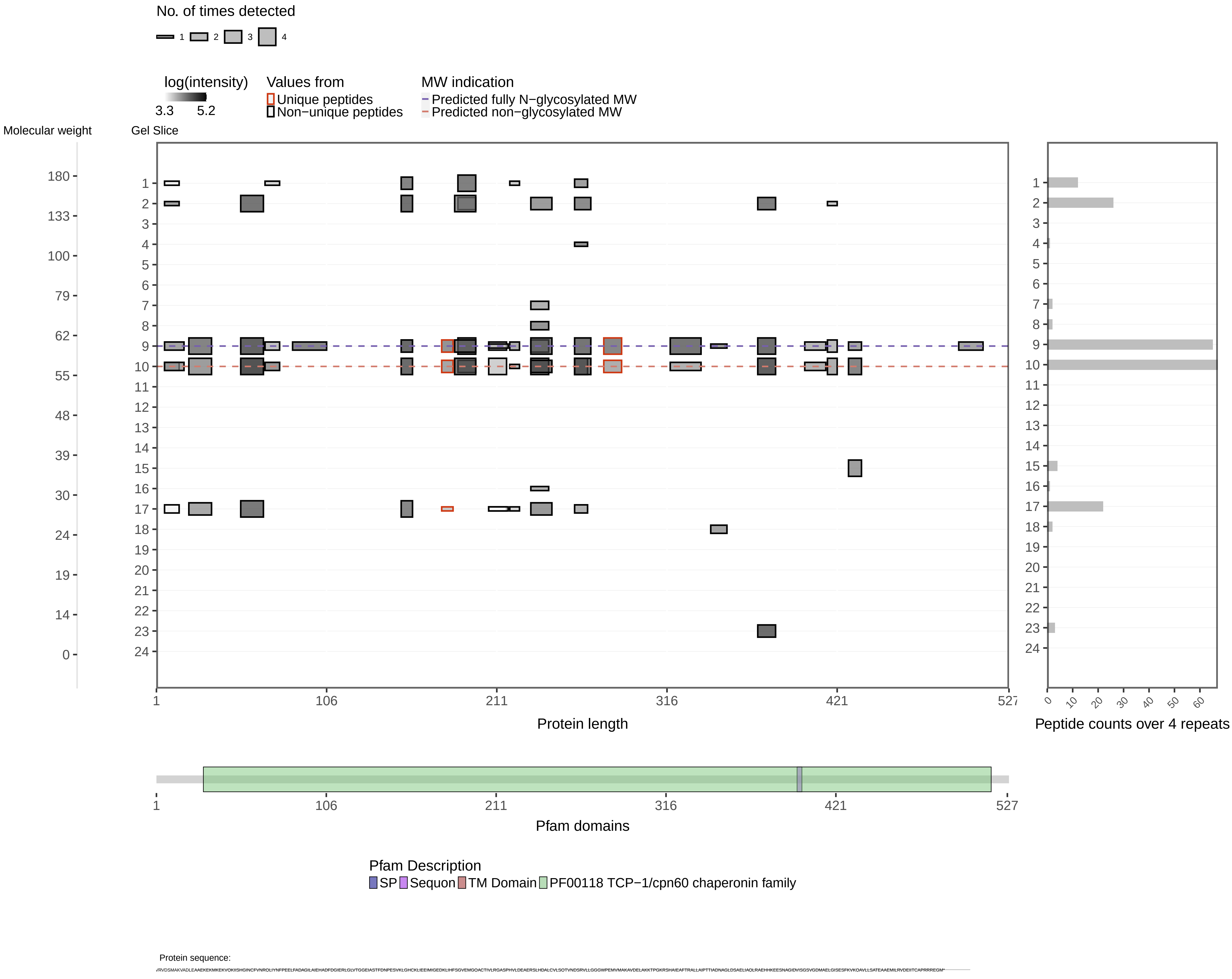

NbL07g00650.1 alpha−L−fucosidase_1 [MW − SP = 55.91 kDa]

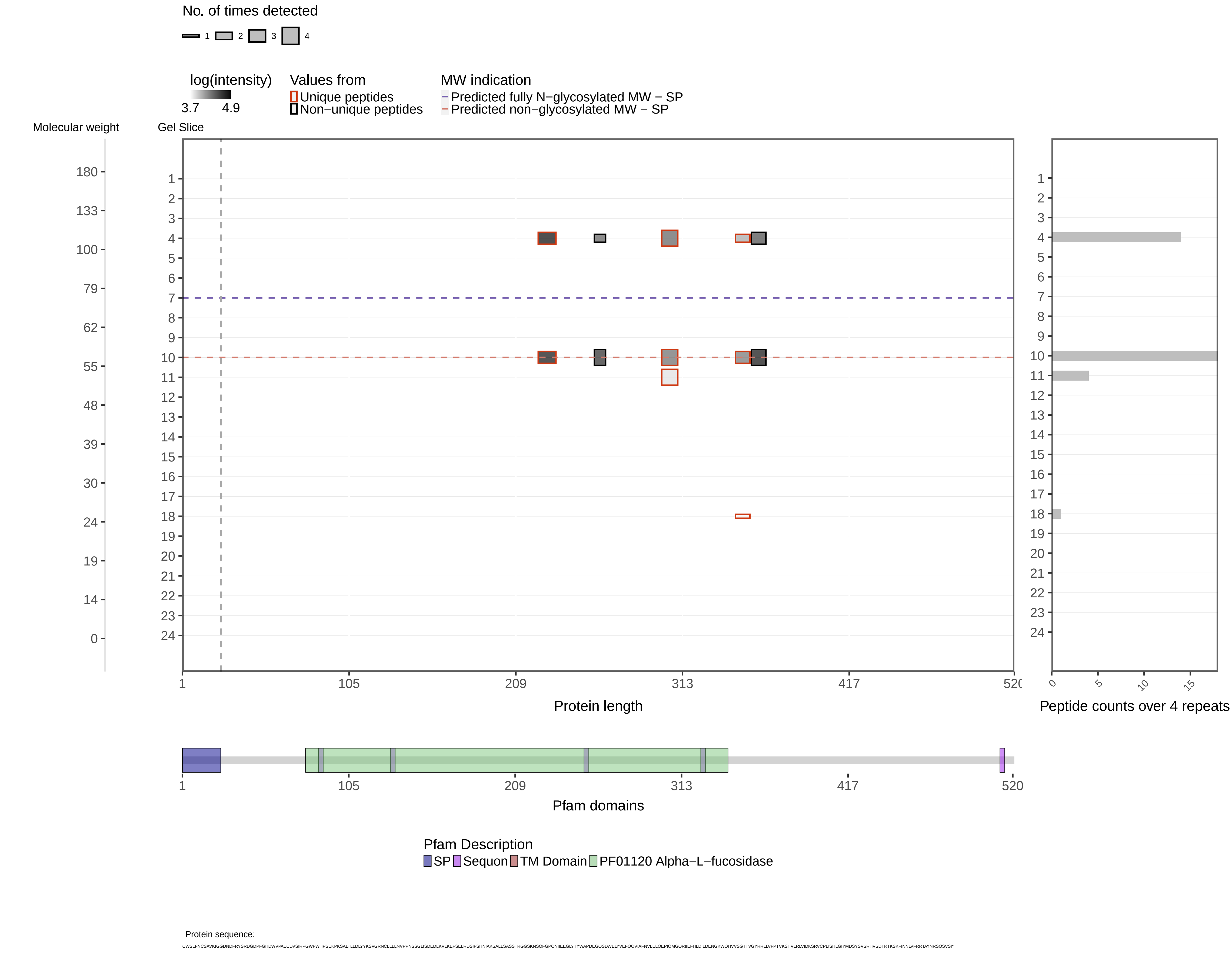

### NbL07g00630.1 lactoylglutathione_lyase [MW = 26.03 kDa]

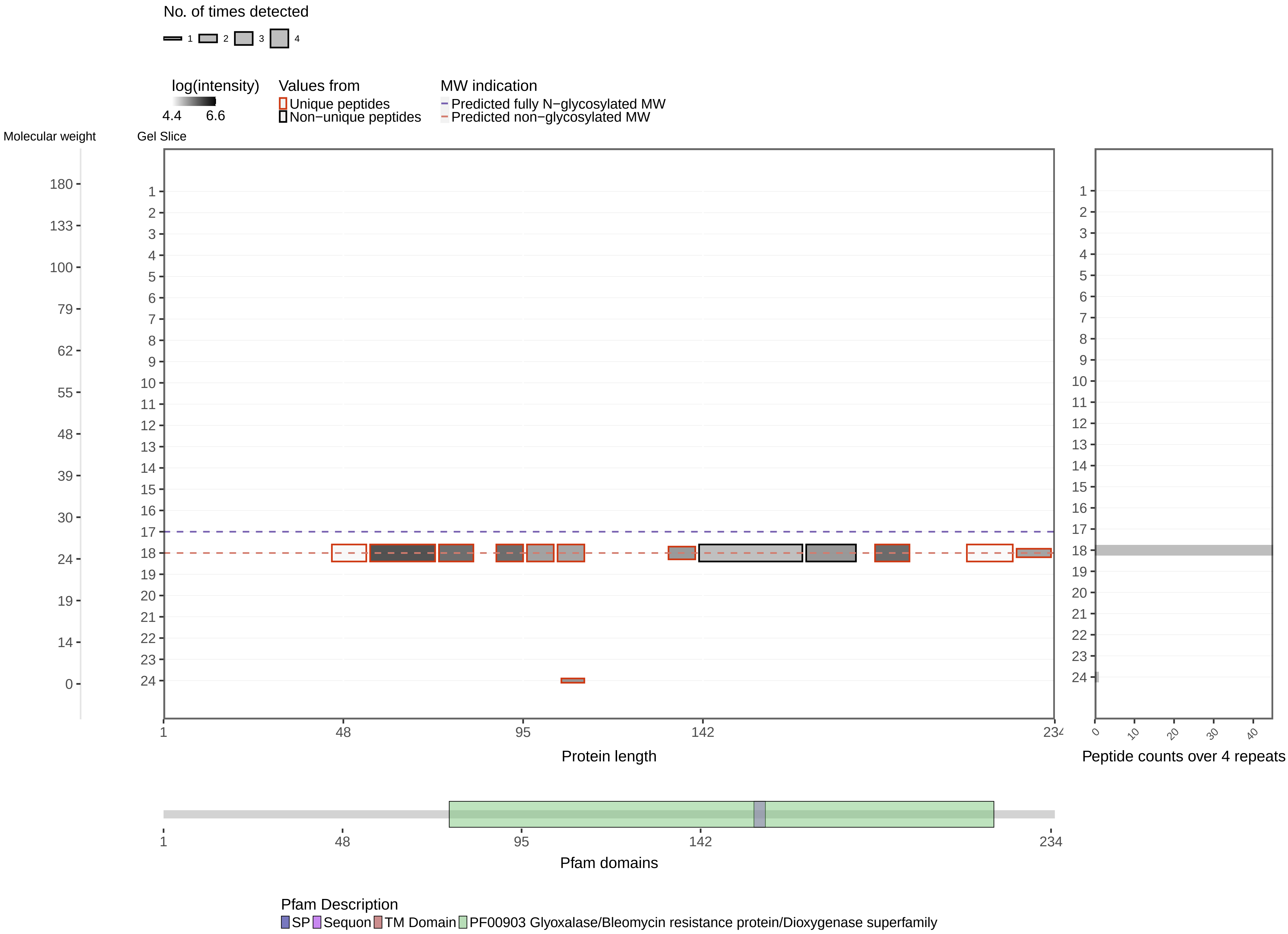


Protein sequence:

DFPEMKFSLYFMGYEDTSSAPSDPAERTAWTFSQKATLELTHNWGTESDPNFTGYHNGNSEPRGFGHIGVTVDDVYKACERFEQLGVEFVKKPFDGKMKGIAFVKDPDGYWIEIFDTKTIKEVAGAAS*

### NbL07g00510.1 superoxide_dismutase_[Cu−Zn]_chloroplastic [MW = 22.58 kDa]

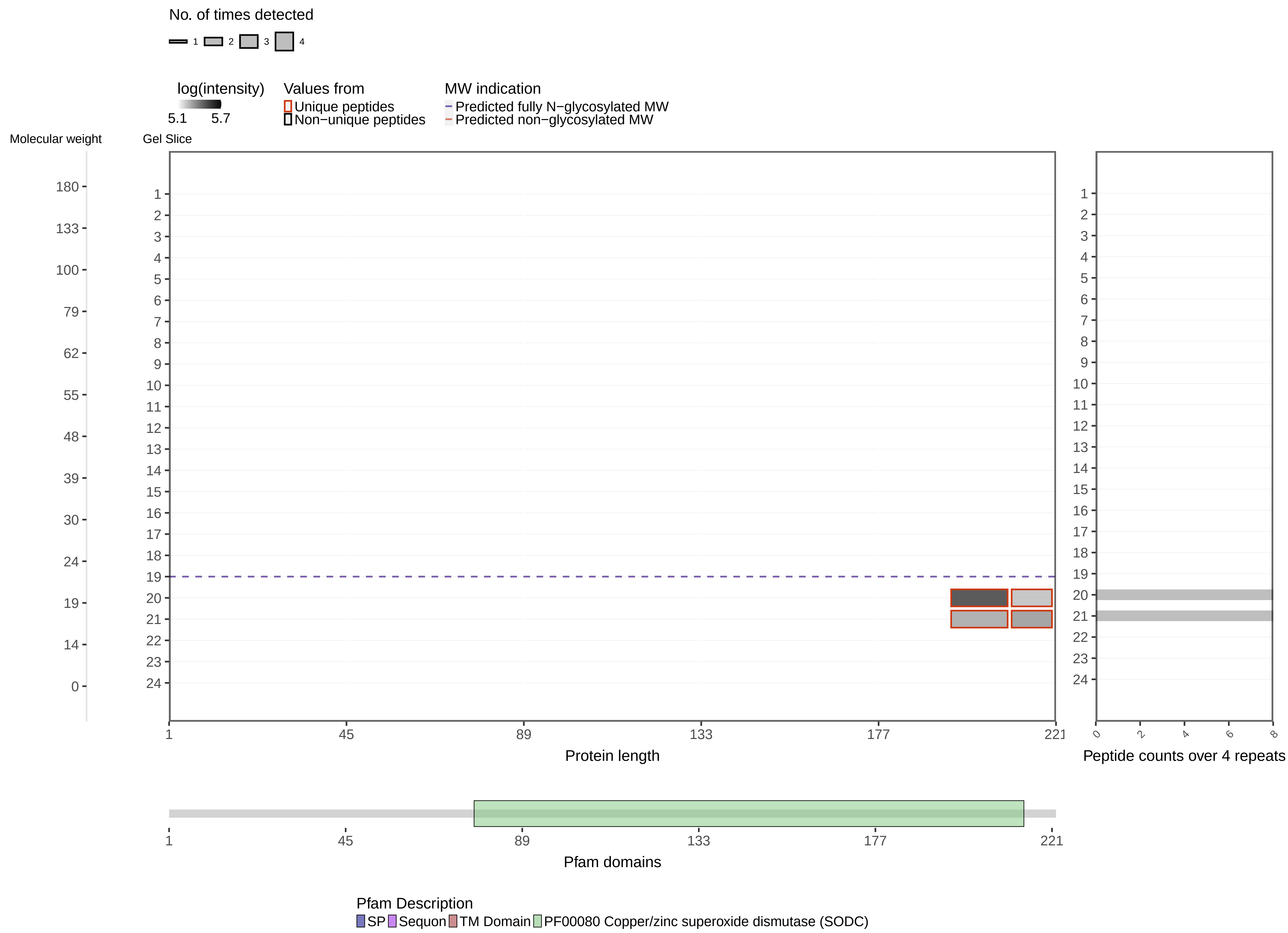


Protein sequence:

KVRITGLTPGLHGFHLHEFGDTTNGCMSTGPHFNPDGKTHGAPEDEIRHAGDLGNIVANADGVAEATIIDNQIPLTGPNSVIGRALVVHELEDDLGKGGHELSLTTGNAGGRLACGVLGLTPI*

NbL07g00490.1 50S_ribosomal_protein_L9_chloroplastic−like [MW = 19.77 kDa]

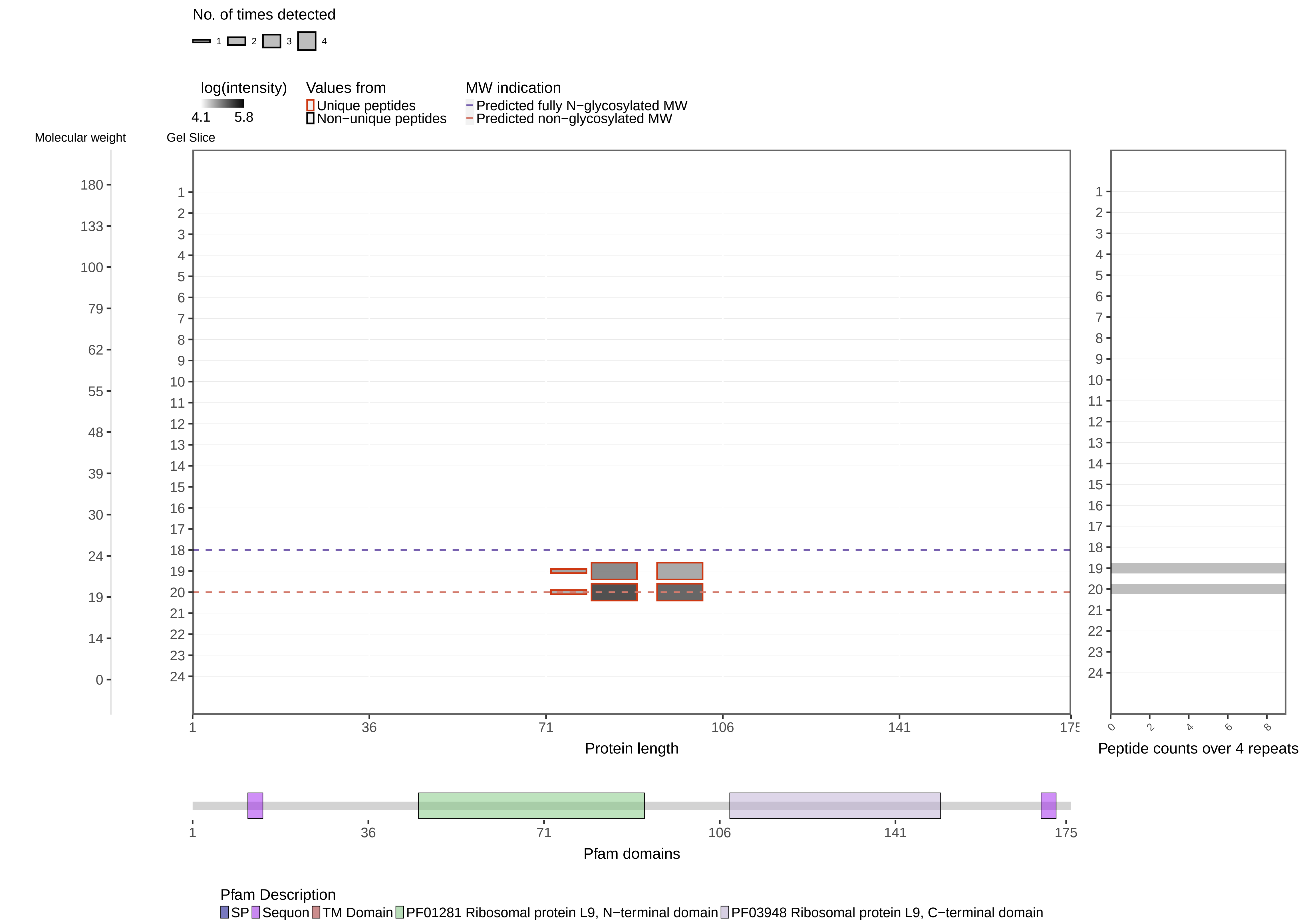


Protein sequence:

PMGKAQIVTPTLIKEMKIEDERIEAEKQRVKEEALQLARIFETVGAFKVKRKGGKGKQIFGSVTAQDLVDIIKHSYRGRWTRKLCLCLRFEKLENMSLN*

### NbL07g00400.1 serine_carboxypeptidase−like_51 [MW − SP = 47.24 kDa]

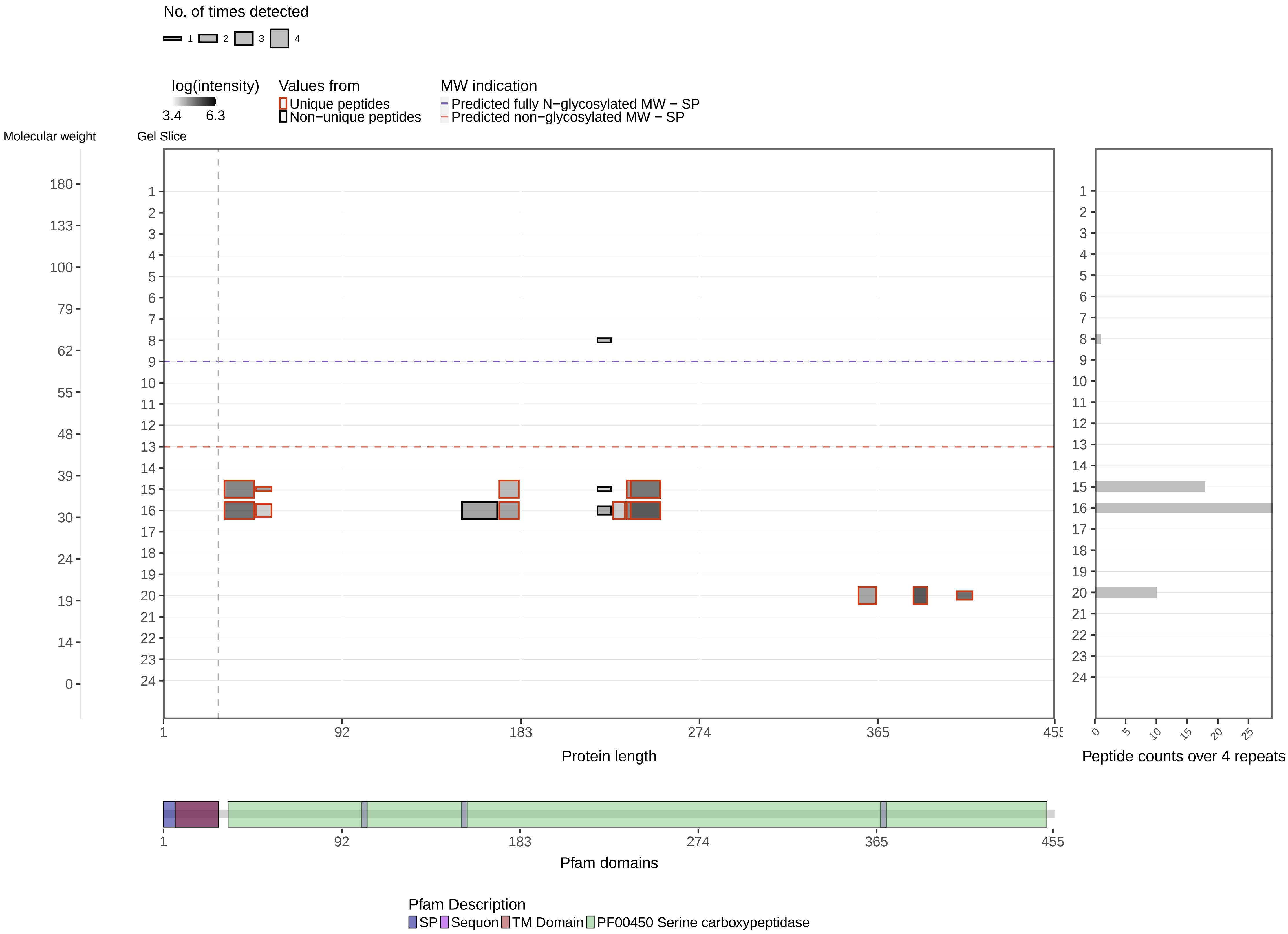


Protein sequence:

.LKDVSRIDGNGLQKSNSVALKIKQQINAGQYEAATNSWRDLENIISDNSNSVDFYNFMLDSGMDPLSLTASELSQGISMKRYSRYLQSSRITPGSDVDLDTLMNGAVKKKLKIIPQNVQWGGQSDSVFNALFGDFMKPRIDEVDELLAKGVNVTVYNGQLDLICATKGTEAWVEKLKWEGLKTFLKMDRSAIYCGGDKSTKAFTKSYKNLHFYWILGAGHFVPVDQPCVALDMVASITK*

NbL07g00230.1 ATP_synthase_subunit_beta_chloroplastic−like [MW = 55.16 kDa]

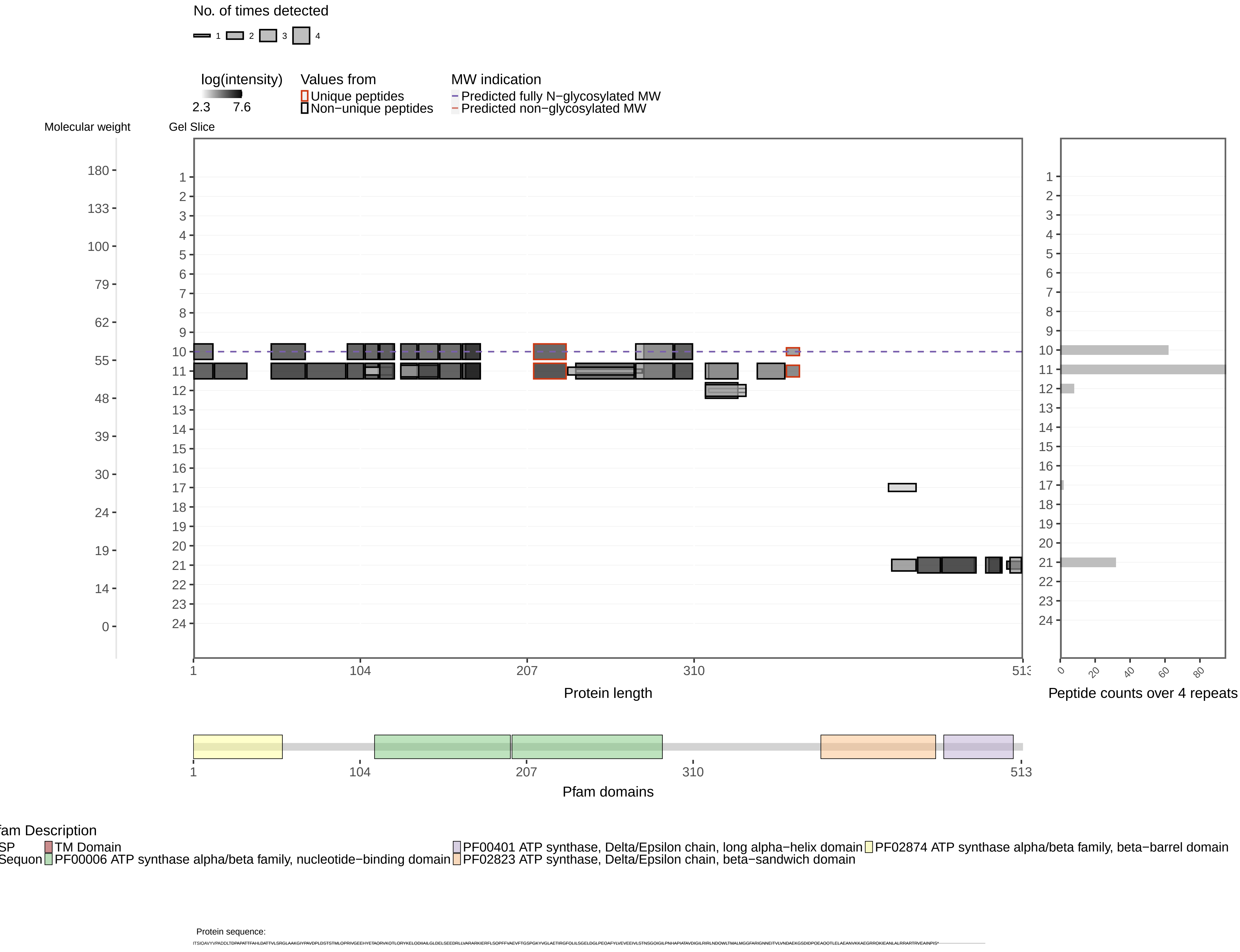

NbL07g00220.1 chitinase−3−like_protein_1 [MW − SP = 39.81 kDa]

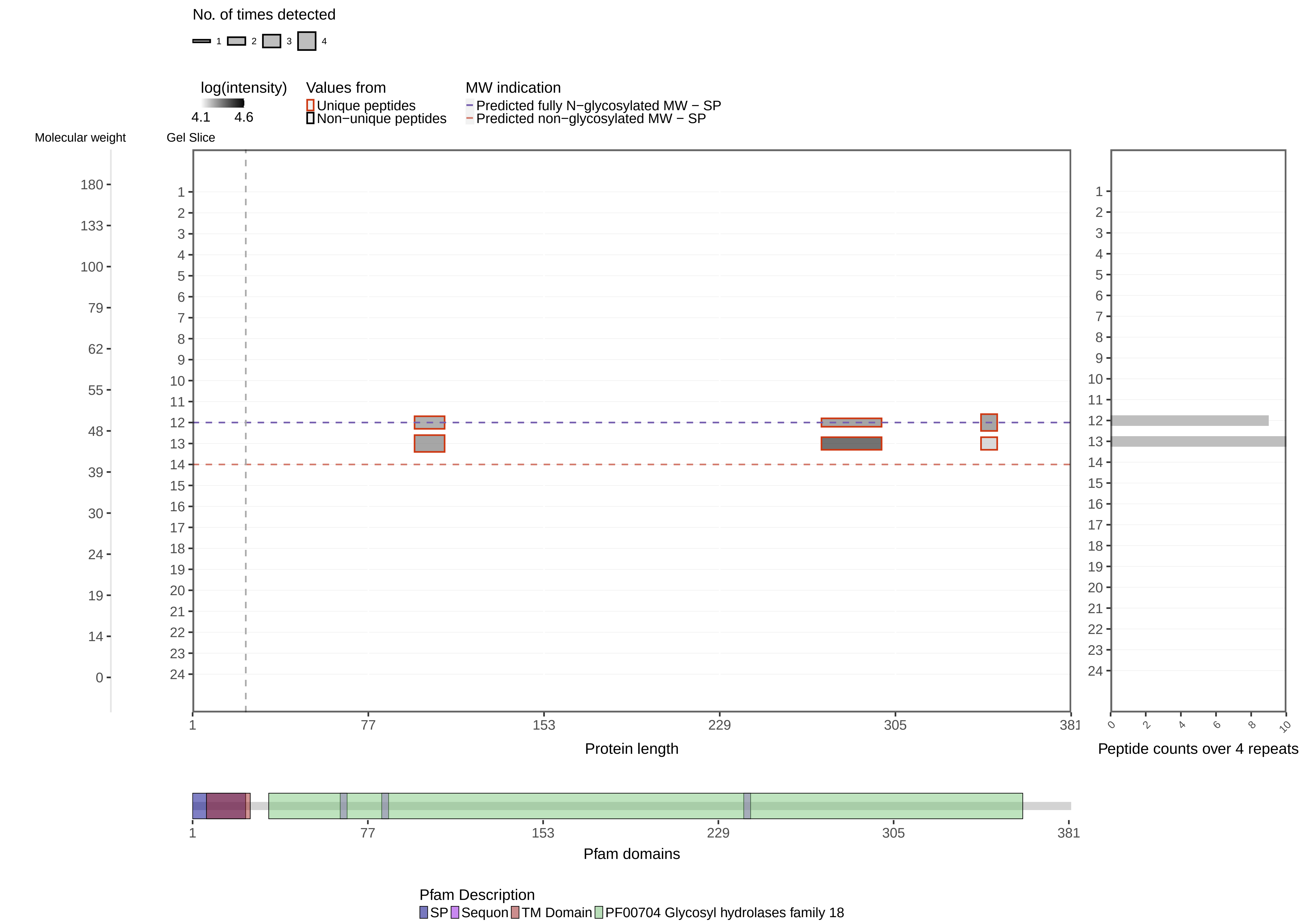


Protein sequence:

PLLITAAVYFSVDFFLSDEFRSYPVPSINKNLDWINLMCYDYRGSWDTSATGAPAALFDTKSNISTSYGVRSWIKAGAIRSKLIMGLPLYGRTWKLKDPNVHGIGAPGIGVGPGDEGTMTYREVEKFNEENNAKVVFDIATVSTYSVAGNSWIGYDDSRSVAMKIGYAQAQRLRGYFFWAVAGDLDWKISRTGPTPTAGAPIL*

### NbL06g21090.1 alcohol_dehydrogenase−like [MW = 41.25 kDa]

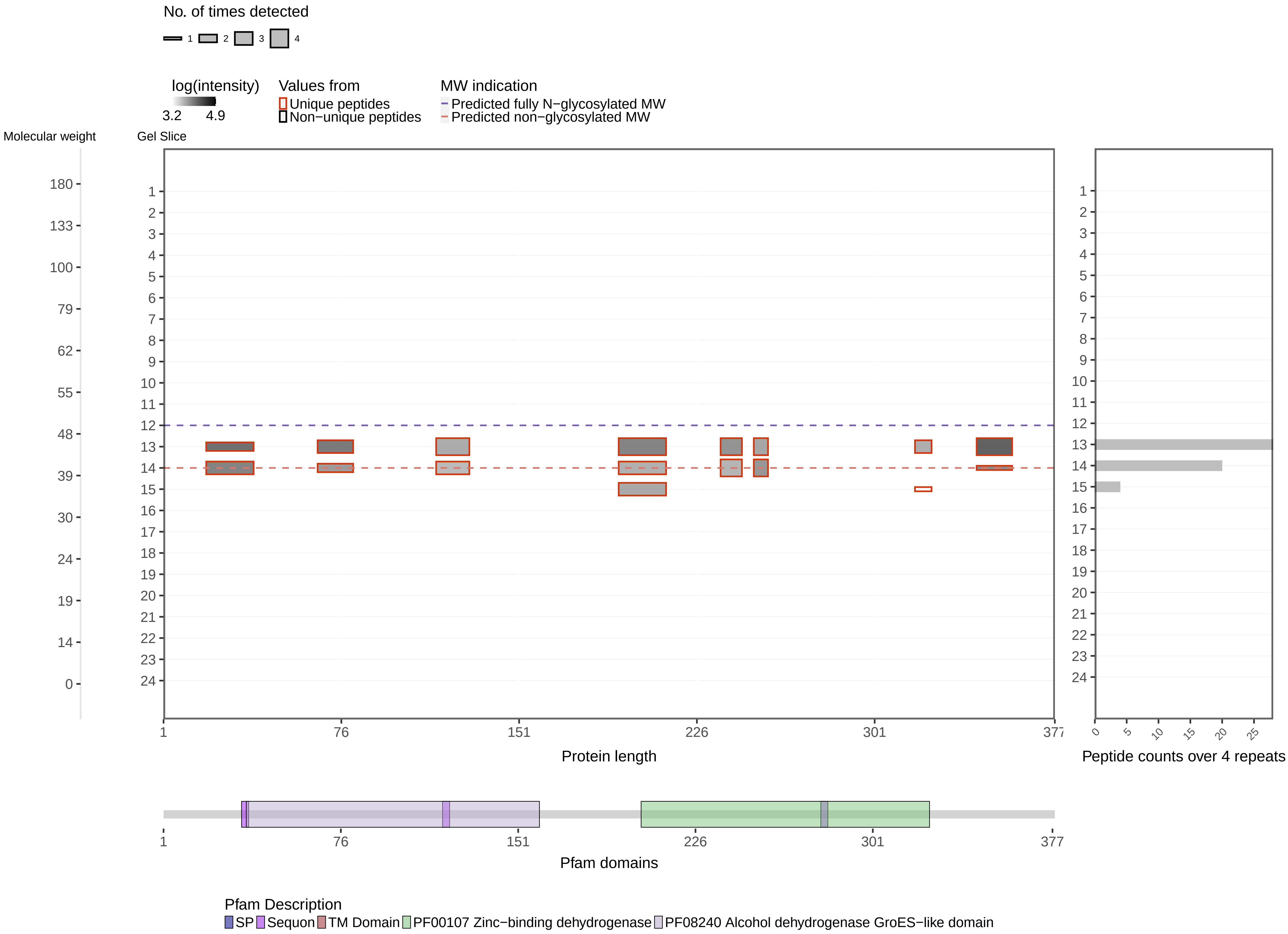


Protein sequence:

CCAFTTGFGATFKDVNIEKGSTVAVLGLGGVGLGVVEGAKQKEAAKIIGIDIHEMKSEKAKKFGVTDFINIKISEKSISELVKEATGGLGVDYFFECTNVPHLMINETIQSTRMGYGTLILLGAGLELDWQMSYVPLMLGRTLKGSIYGGIRTRTDLPSIFDKCINKEIKLDELLTHEVSLNDVNKAFEYLKEPNCVKVLIKF*

### NbL06g21020.1 D−amino−acid_transaminase_chloroplastic−like [MW − SP = 44.73 kDa]

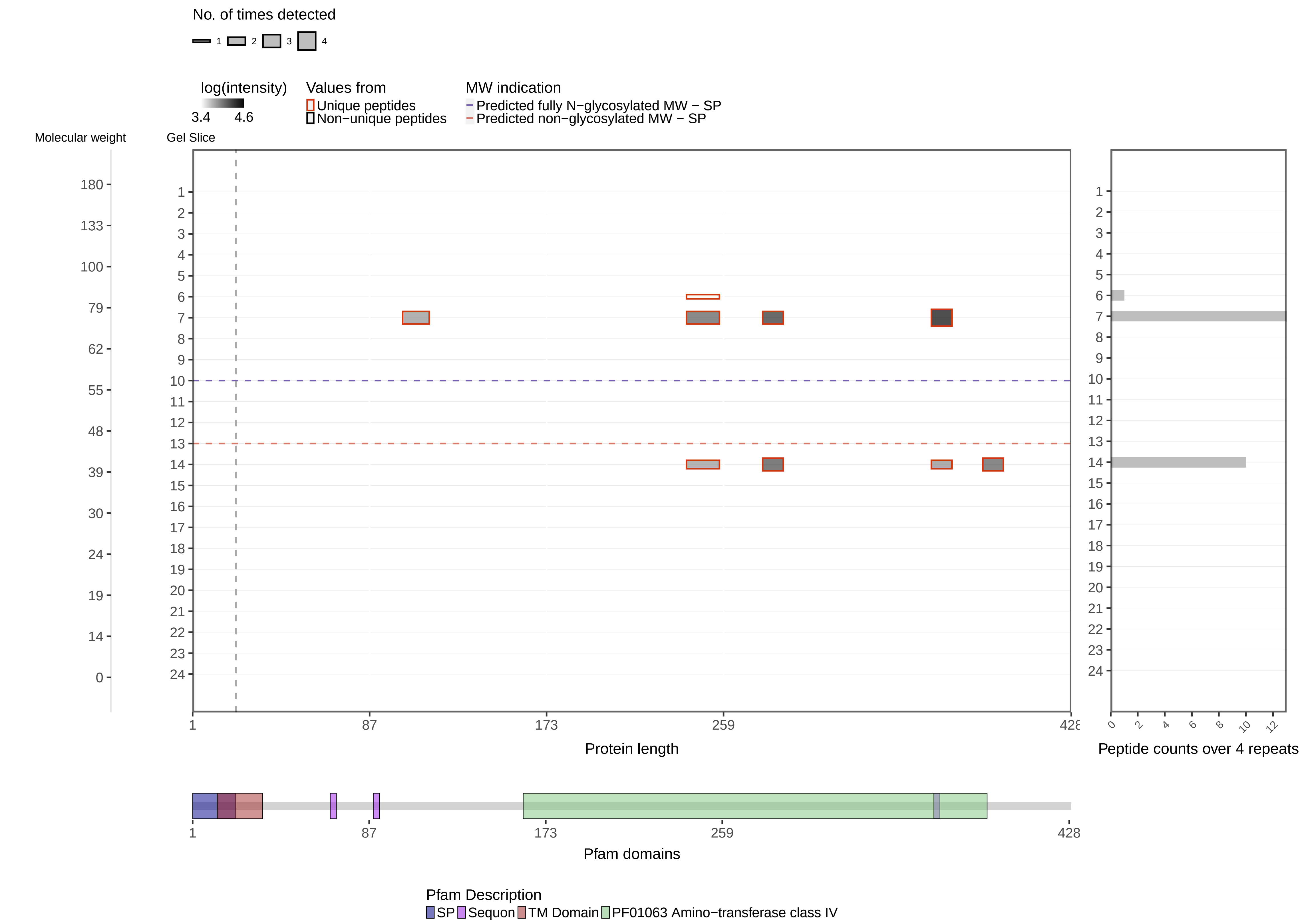


Protein sequence:

RILIRTVSVSOCRKGSLRYWLSAGPGDFOLSPSGCPRAALYAIVIODOSPPDHRGIRVVTSSIPIKPPOFAVMKSVNYLPNALSKMEAEENDAYAAIWLDADGFVAEGPNMNVAFVTKKKELLMPSFDKILSGCTAKRVLVLAAKLVKEGKLRGVKIDNVSVEDGKRADEMMLIGSGVLVRSVVOWDEEIIRDGREGPVTOALLNLLLEDMKSGPASVRVPVPY*

### NbL06g20910.1 pectinesterase_31−like [MW = 35.58 kDa]

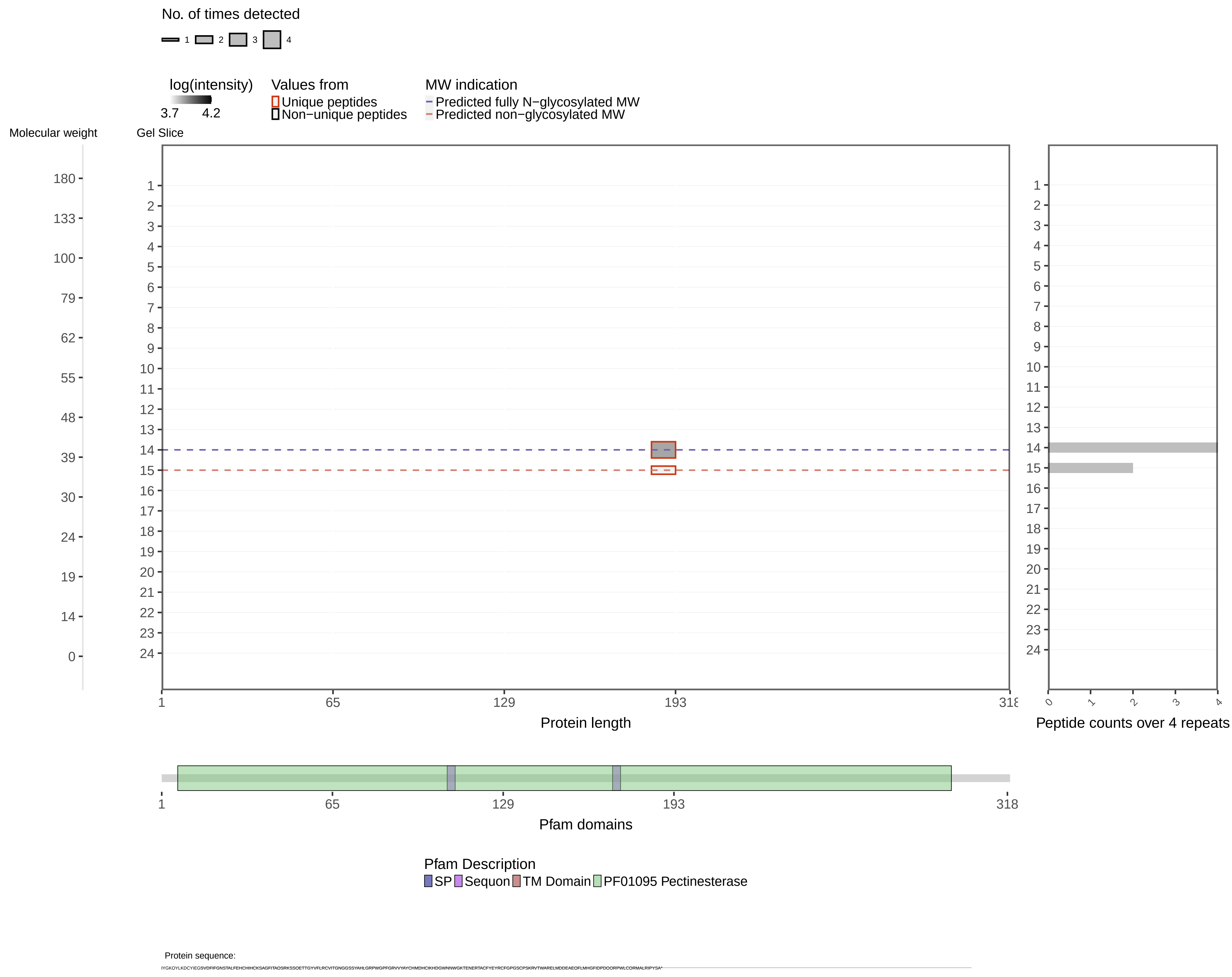

### NbL06g20870.1 clathrin_interactor_EPSIN_1−like [MW = 59.7 kDa]

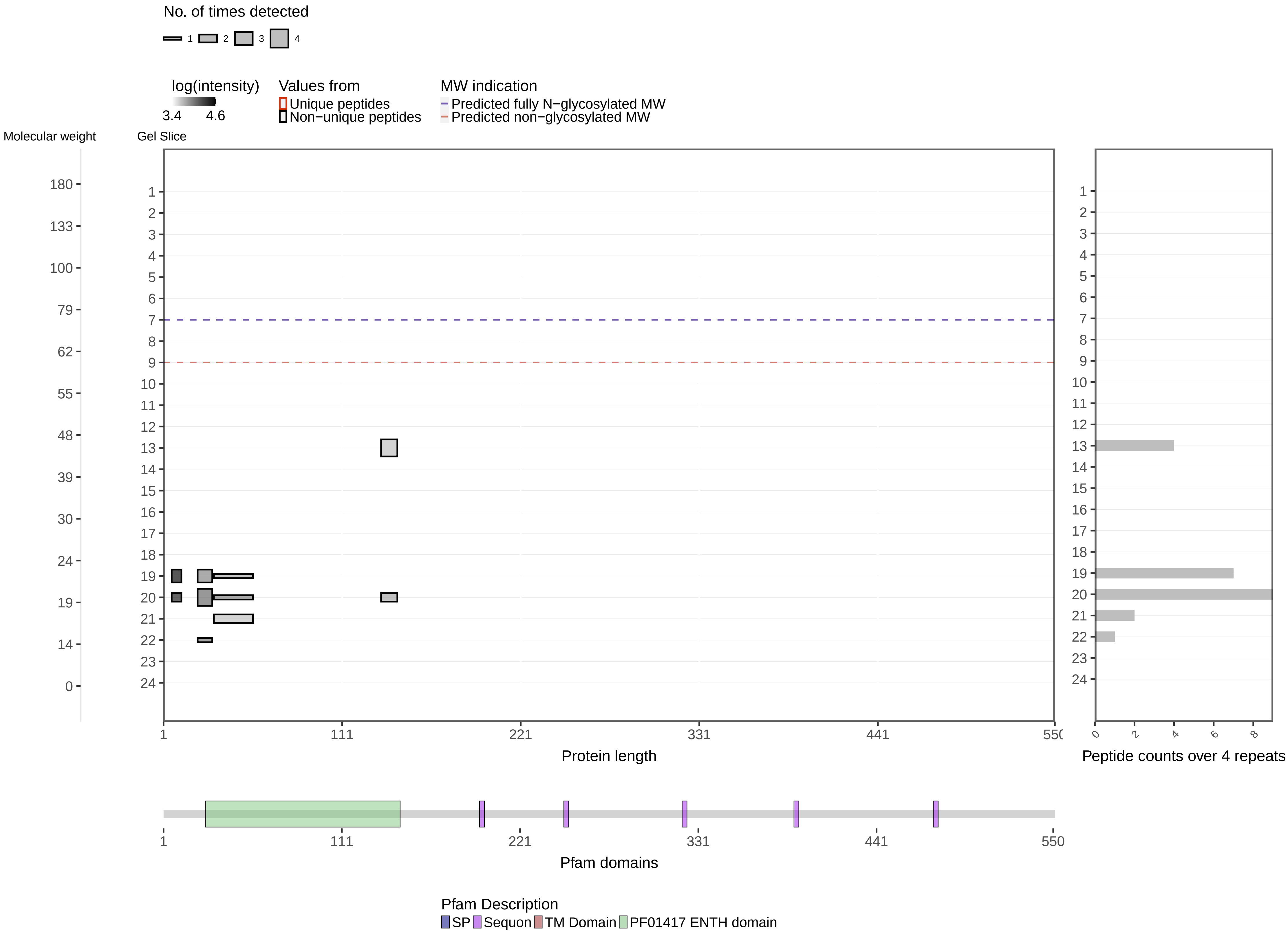


Protein sequence:

NPPHGAAVSSSKYEEDFDDFDPRGTSSTKPSTKKSDQVDLFGQNLIGDLLDVPTPVPADNSTVTSHPSEIDLFAKANFVSVKPQSEINVDLFASQPASSFAASSTIDFFSAPDPVVQSDIRSSKSDKINTTTVDPFAAVPLNTFDSSDPFGKFNSHADPVSVASENANRGGNQEETPSKLDKSPVEAKPAPKKGDFQVSSGIWADSLSRGLIDLNISAPKKVNLADVGIVGGLTDGSDVKEKGPTTFYMGRAMGQGTGLGQSGFTSTSIGGDDFFSNHQNYQFGSFQK*

### NbL06g20850.1 UPF0587_protein_C1orf123_homolog [MW = 18.51 kDa]

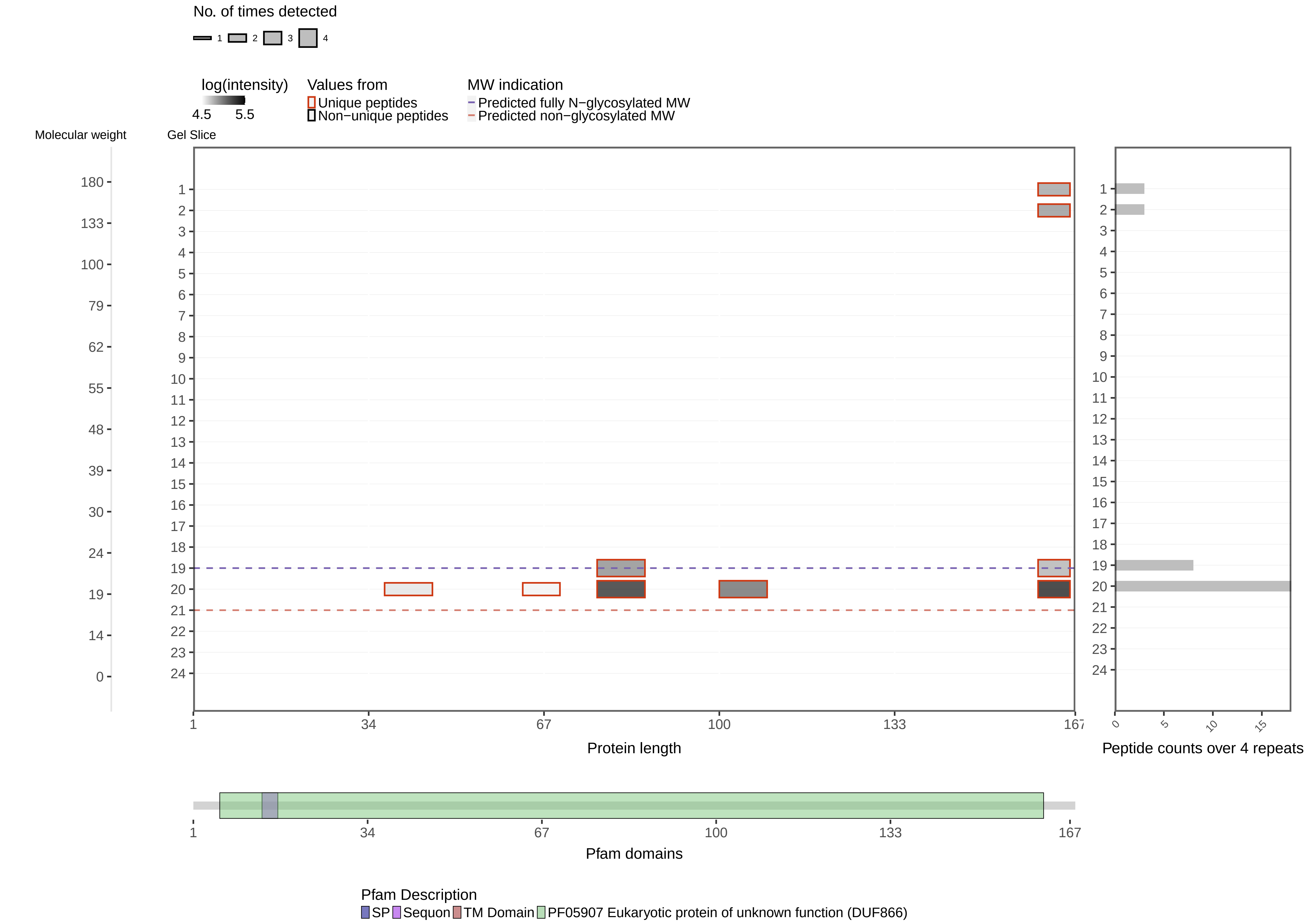


Protein sequence:

KFCSRDGTVTMITGRGRPLTHGDSQAGKYAPLMLFECRGFEPLDFAFLGEWQAESLEGTKFENIYLSGDEFADYDEKGECPVMISNLRATFNVVK*

### NbL06g20840.1 uncharacterized_LOC107809971 [MW = 35.11 kDa]

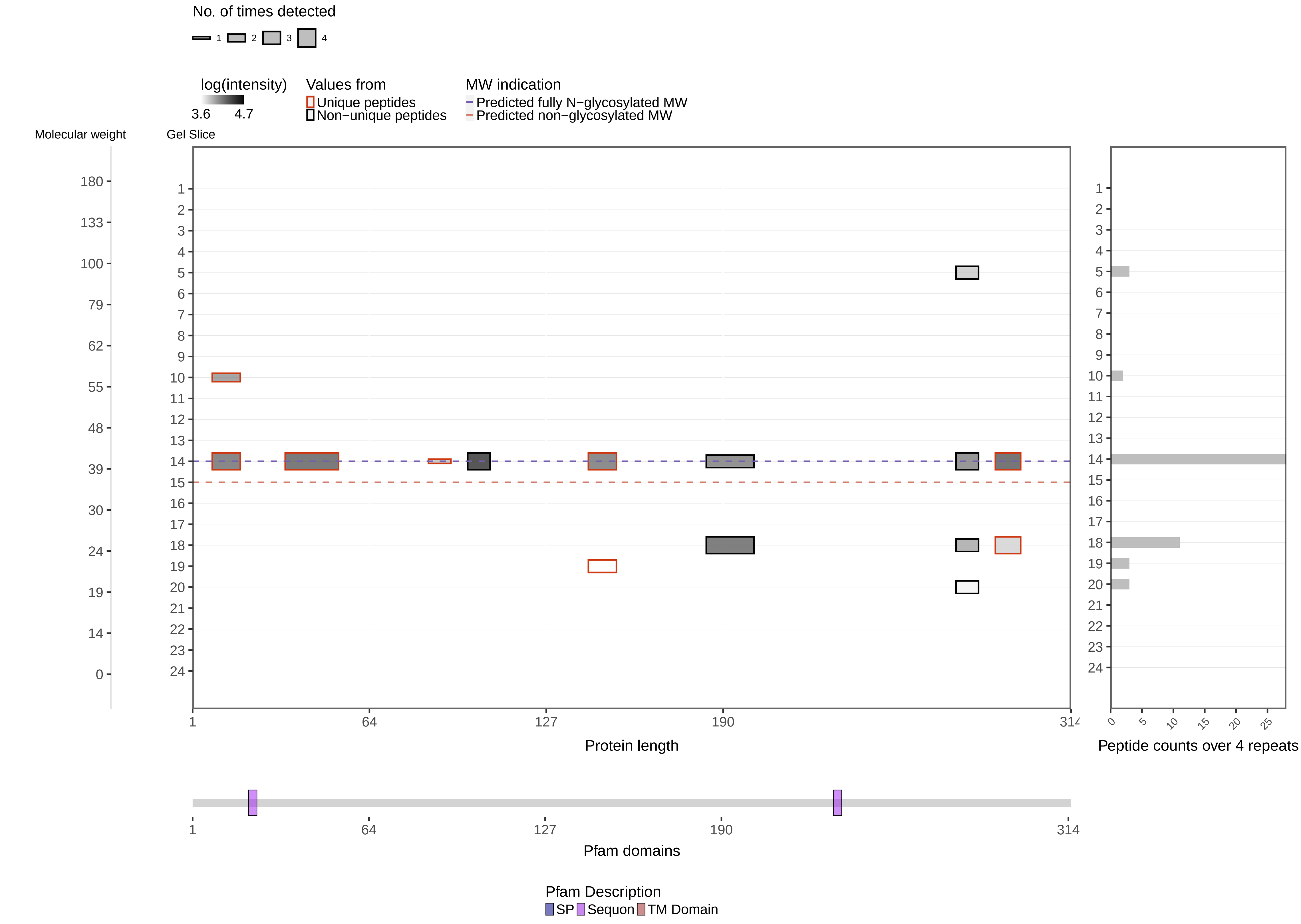


Protein sequence:

SFQEGKTLESSQFDLVILHVGAGQRTNGLKDLDHVNRLVGDLMQMAHPGTEVGSRLHMSVVLSYGVVREDDDSMYSITDTRPENNSKLSRIFPRQSYTIKEGKPRPNVRQFCPMLVAQWQNAVTRKDMVEAYSYKNFKERGANLVIPADRFLHEVAFKMWKAPKYGA*

NbL06g20820.1 probable_leucine−rich_repeat_receptor−like_protein_kinase_At1g68400 [MW = 73.08 kDa]

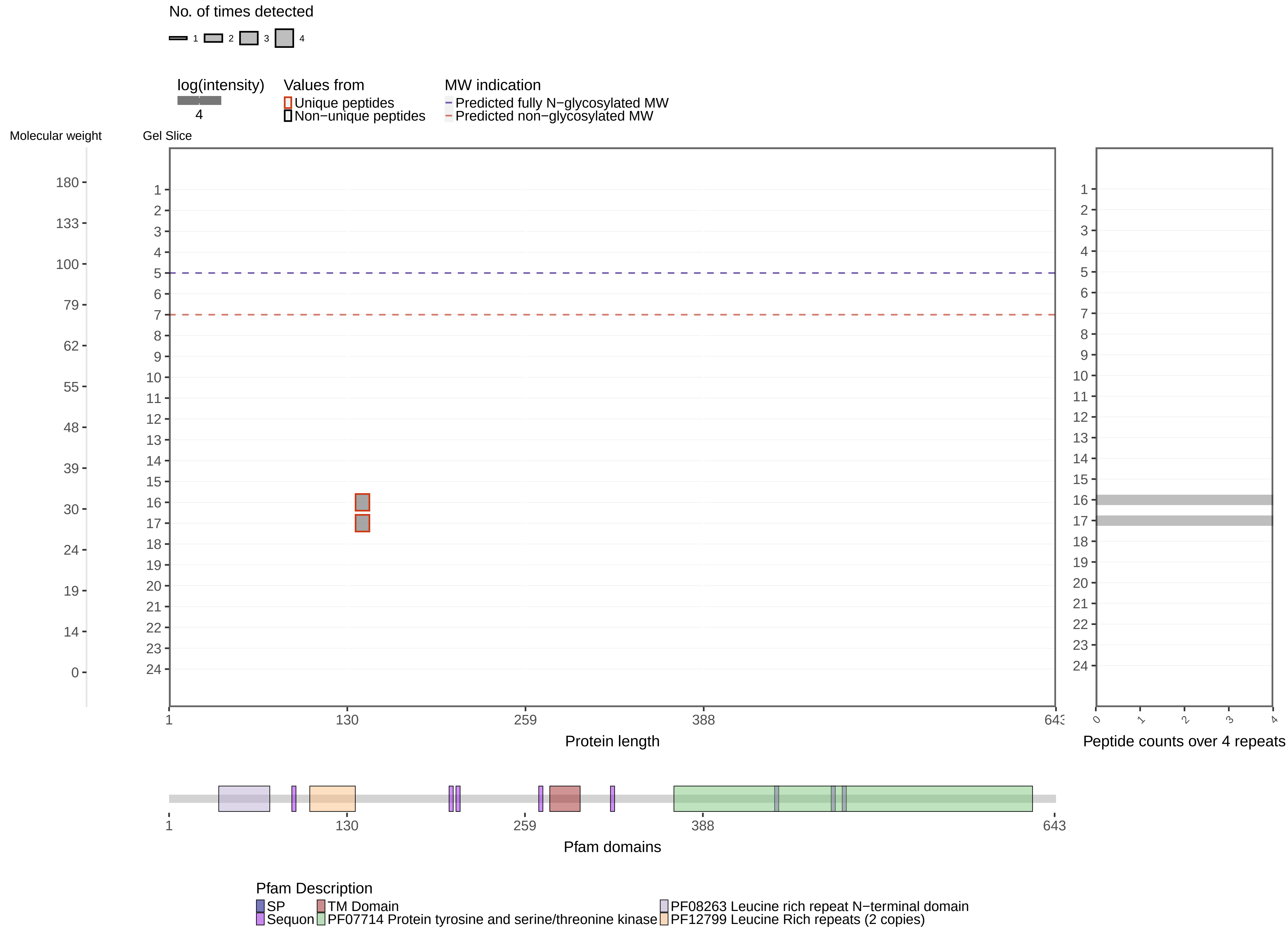


Protein sequence:

NKVKKYHDISNSSSLYHSFEYGHILKDKKSDQFATMVCFEGCKGFNKVEDLLKASAEMLGKGSVGTSYKVAMDCKDVVVVKRVIEKLKKMKDLDGFLRLIGGLRHQNVVSLRAYYSSKEELLLVYDFLPNGSLHNLLHGNRGPGRTPLDWSTRLKYALGAAKGLAFLHSYNKTKICHGNFTSSNILIDHFGNACISDICLHLLLQMPISSNNGYKAQELLTQNNMNTNQNPRKFSQKCDVYSFGVVLLEILTGKIATNEGEISLAKWVQRVVNKEWTWDVFDFELARYKEMED

### NbL06g20780.1 UDP−D−apiose/UDP−D−xylose_synthase_2−like [MW = 43.34 kDa]

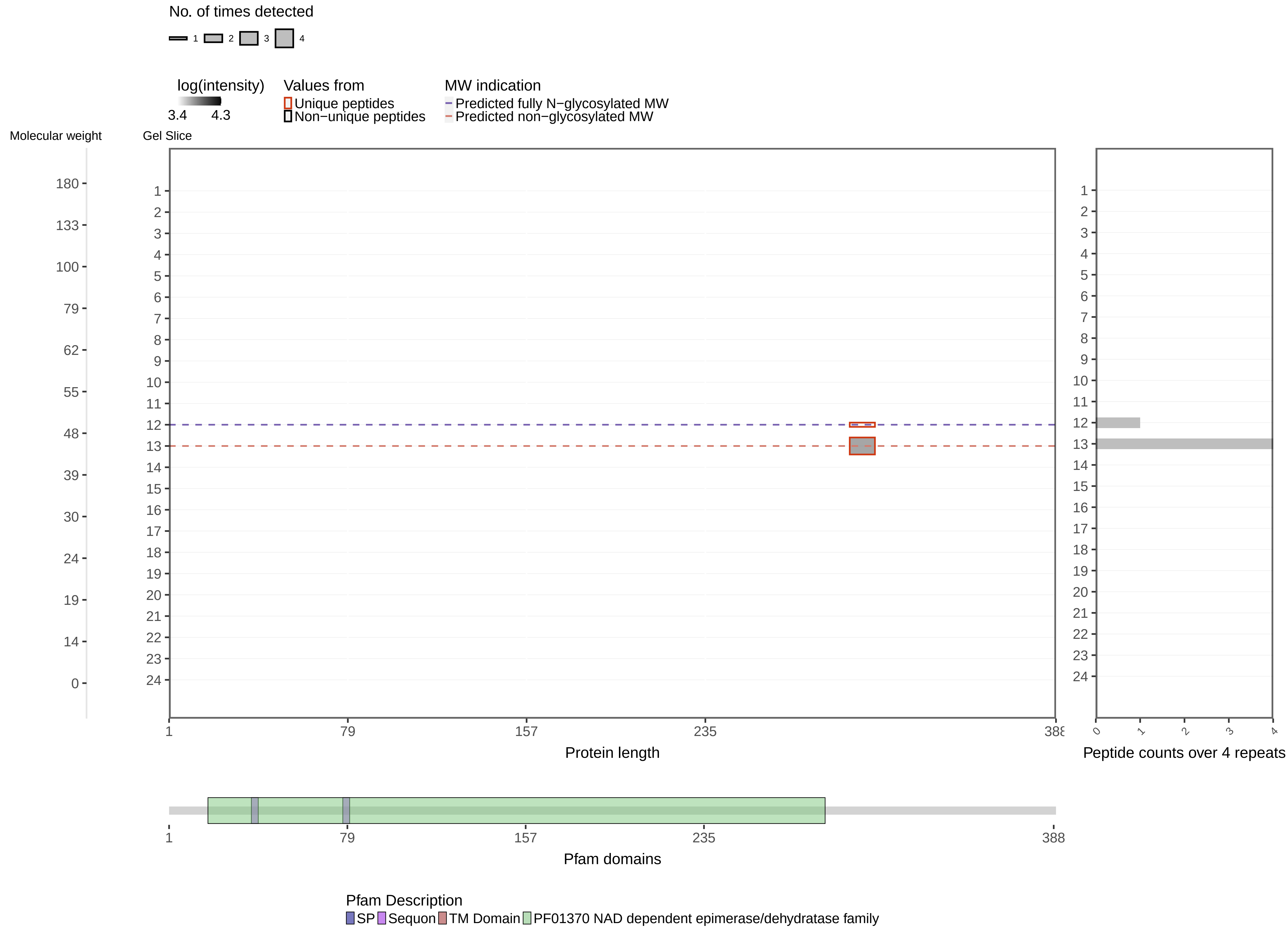


Protein sequence:

YACAKOLIERLIYAEGAENGLEFTIVRPFNWIGPRMDFIPGIDGPSEGVPRVLACFSNNLLRHEPLKLVDGGHSORTFIYIKDAIEAVLLMIENPTRANGOIFNVGNPNNEVTVROLAEMMTOMYSKVSGESPPETPTIDVSSKEFYGEGYDDSDKRIPDMTIINROLGWNPKTSLWDLLESTLTYOHRTYAEAVKOAMSKTTGN*

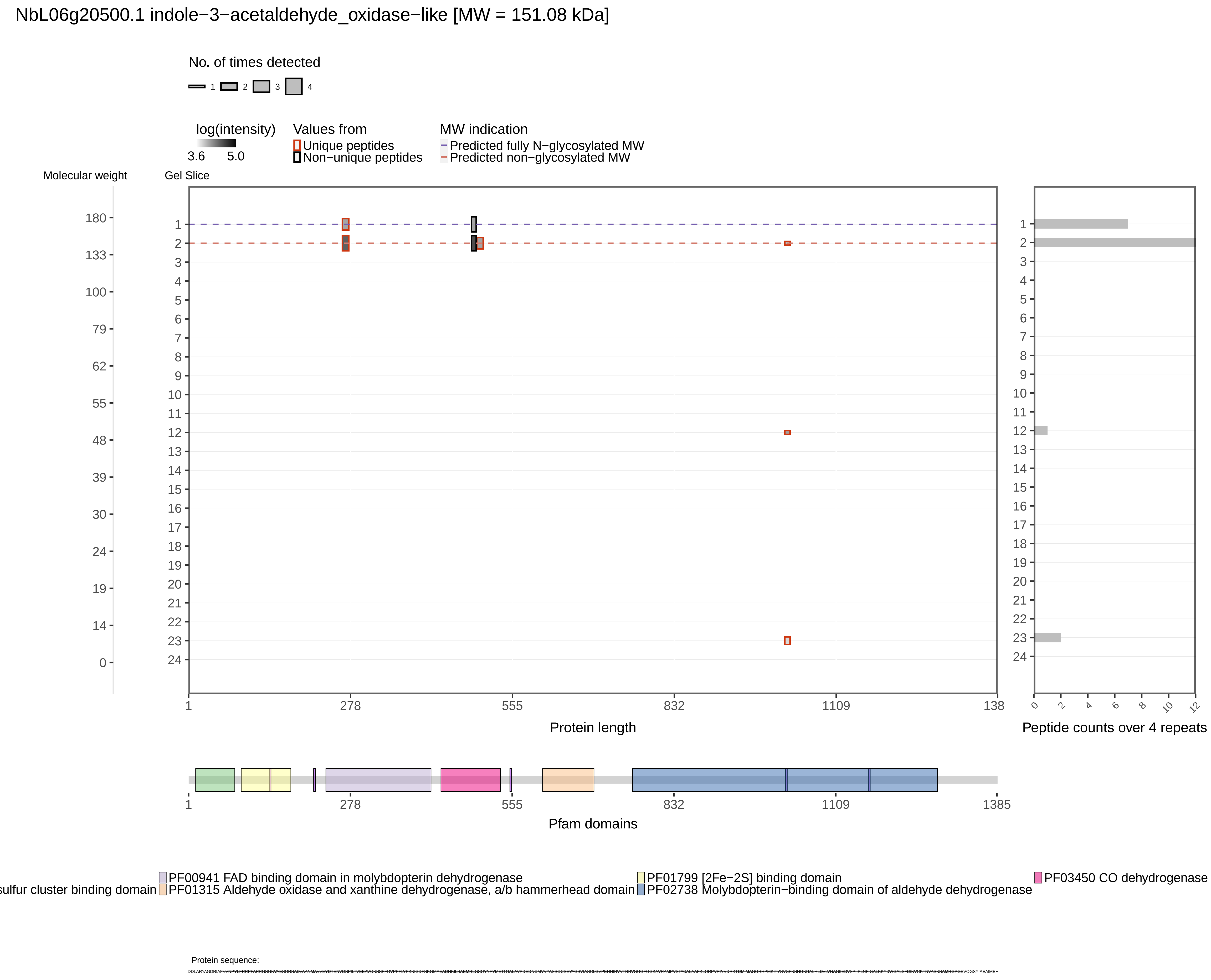

NbL06g20500.1 indole−3−acetaldehyde_oxidase−like [MW = 151.08 kDa]
No. of times detected
1 2 3 4
log(intensity)
3.6 5.0
Values from
Unique peptides
Non−unique peptides
MW indication
Predicted fully N−glycosylated MW
Predicted non−glycosylated MW
Molecular weight
Gel Slice
Protein length
Peptide counts over 4 repeats
Pfam domains
PF00941 FAD binding domain in molybdopterin dehydrogenase
PF01799 [2Fe−2S] binding domain
PF03450 CO dehydrogenase
sulfur cluster binding domain
PF01315 Aldehyde oxidase and xanthine dehydrogenase, a/b hammerhead domain
PF02738 Molybdopterin−binding domain of aldehyde dehydrogenase
Protein sequence:

NbL06g20430.1 5−methyltetrahydropteroyltriglutamate−−homocysteine_methyltransferase−like [MW = 91.8 kDa]

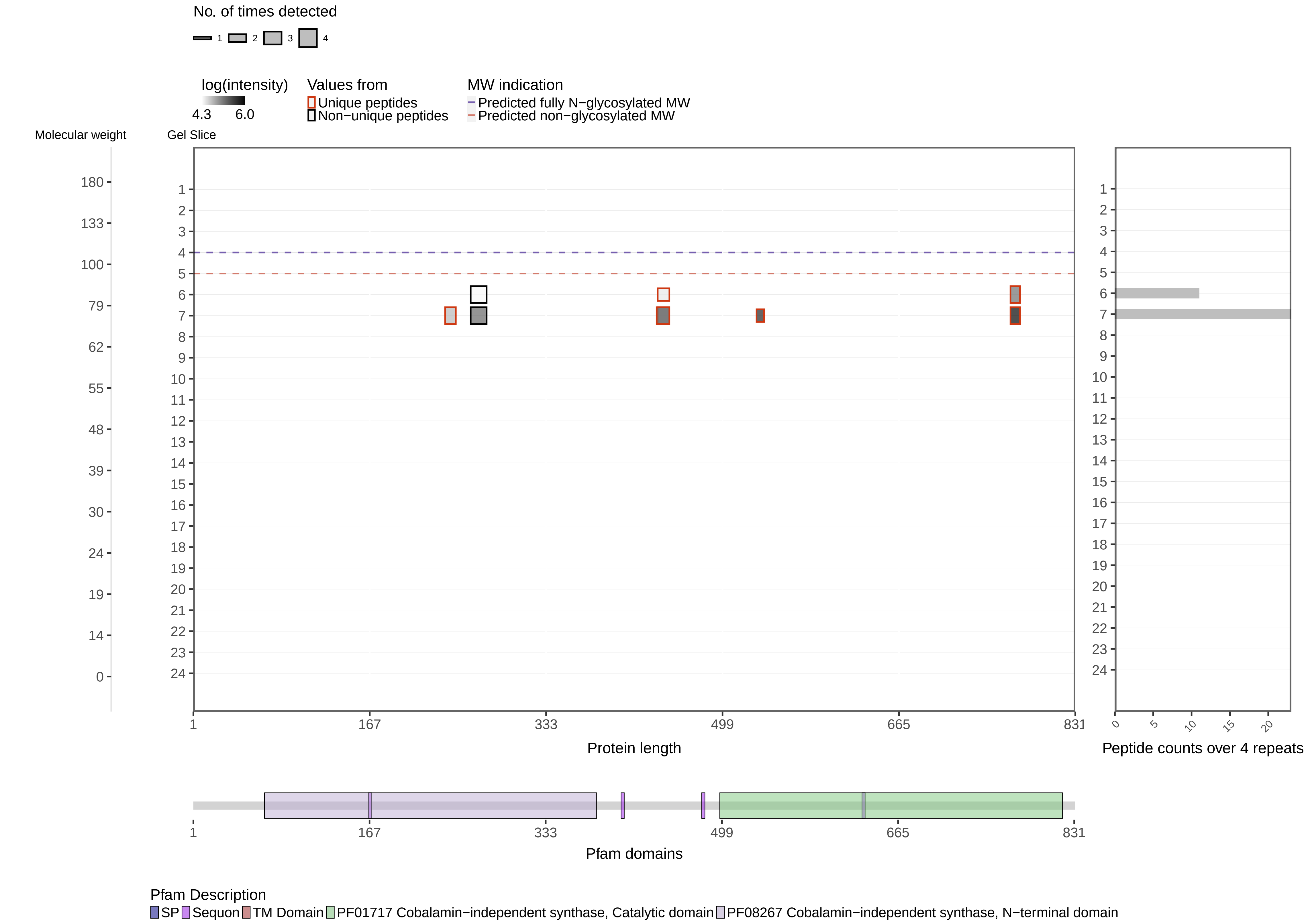


Protein sequence:

[illegible]

NbL06g20350.1 uncharacterized_LOC104214226 [MW = 22.22 kDa]

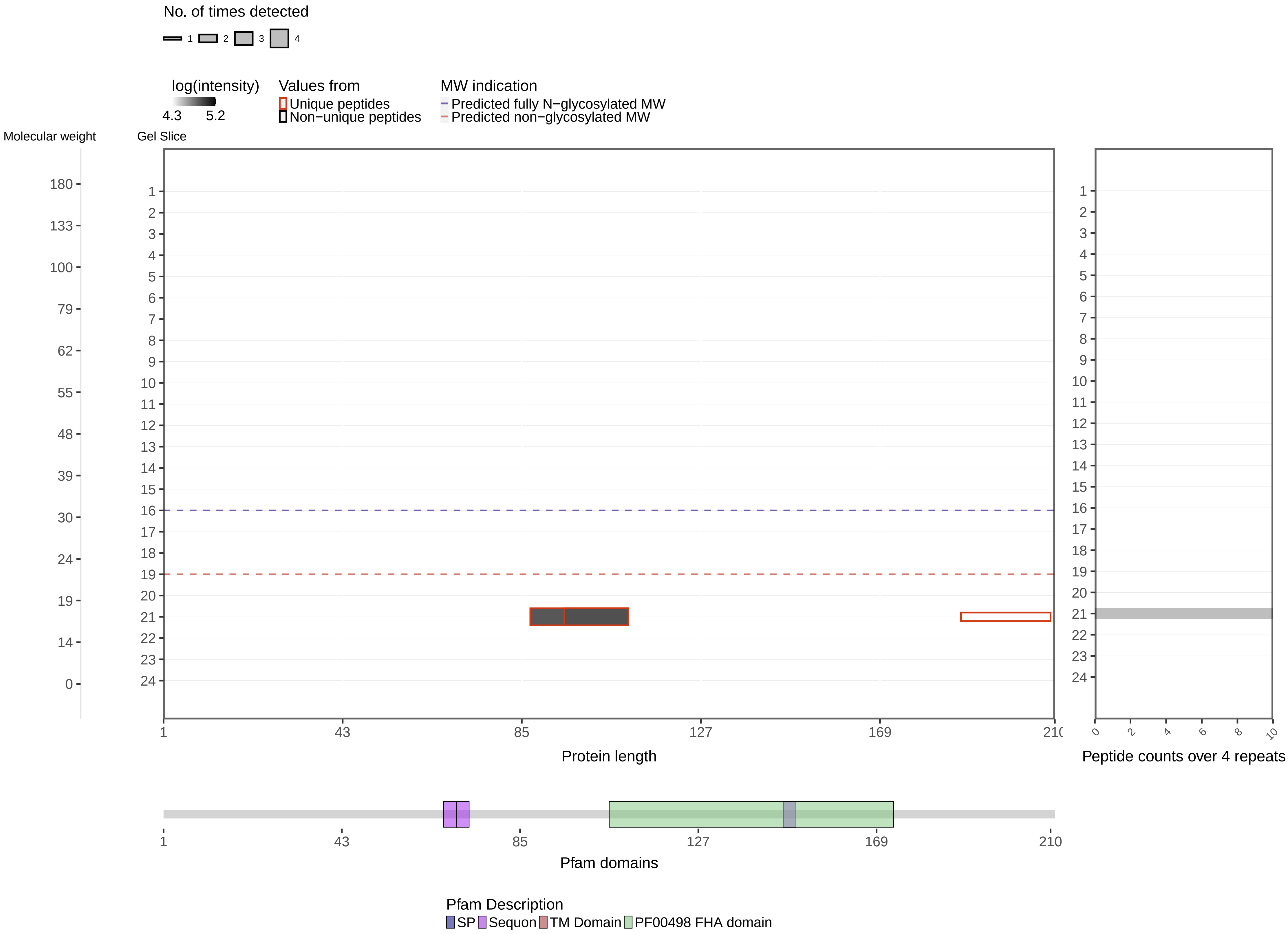


Protein sequence:

TAKPGSLEIPSAGAVTVGRVAEKADMVIPVPTVSAVHALLQSMEEYLVVTDLDSTNGTFIDEKRLRPGVAAAALPGSLLTFGDTNLAIFRVAKLEKMETSSEPEEAEANTEEEEPSSS*

NbL06g20340.1 molybdopterin_biosynthesis_protein_CNX1−like [MW = 70.39 kDa]

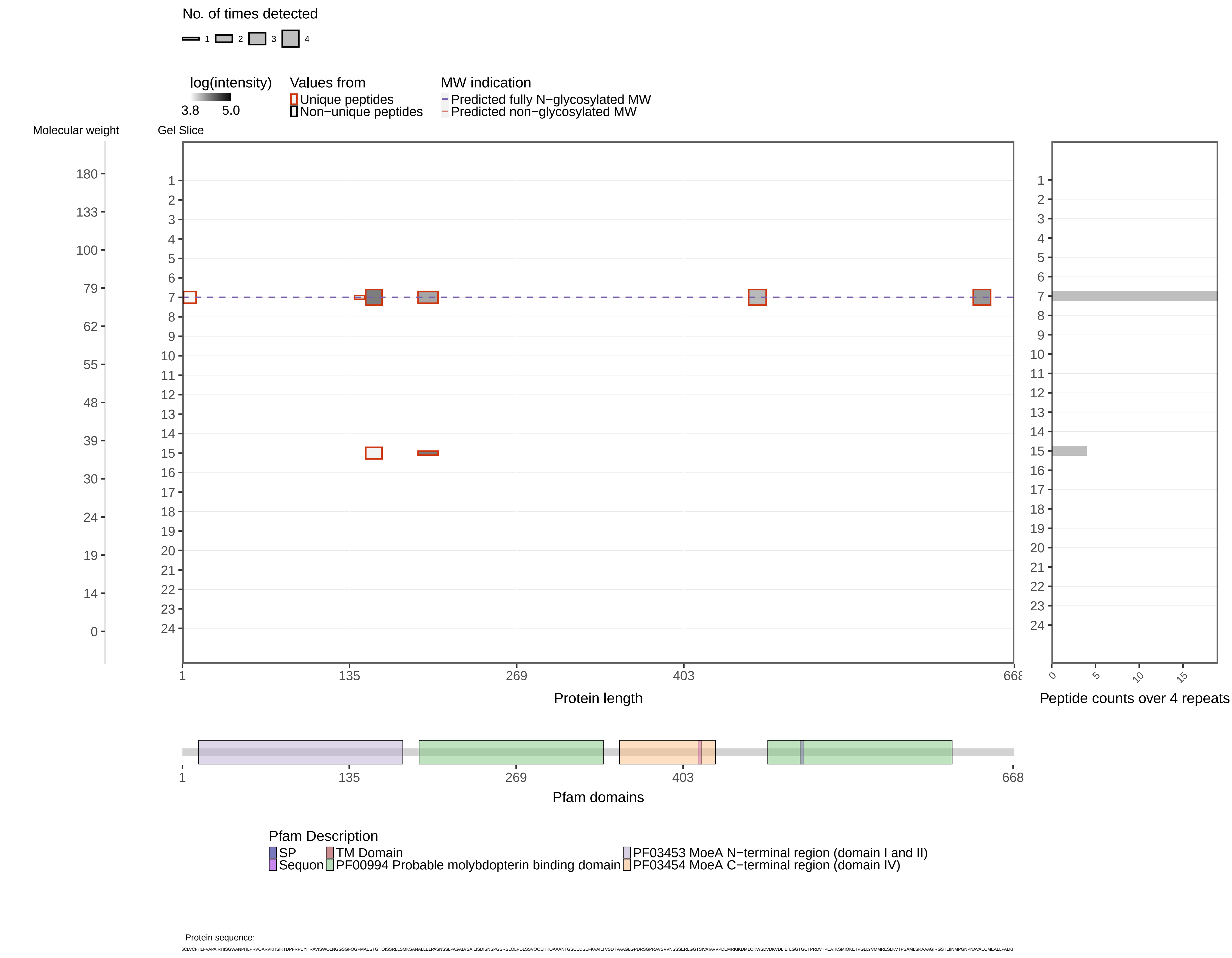

NbL06g20300.1 4−diphosphocytidyl−2−C−methyl−D−erythritol_kinase_chloroplastic/chromoplastic−like [MW = 45.21 kDa]

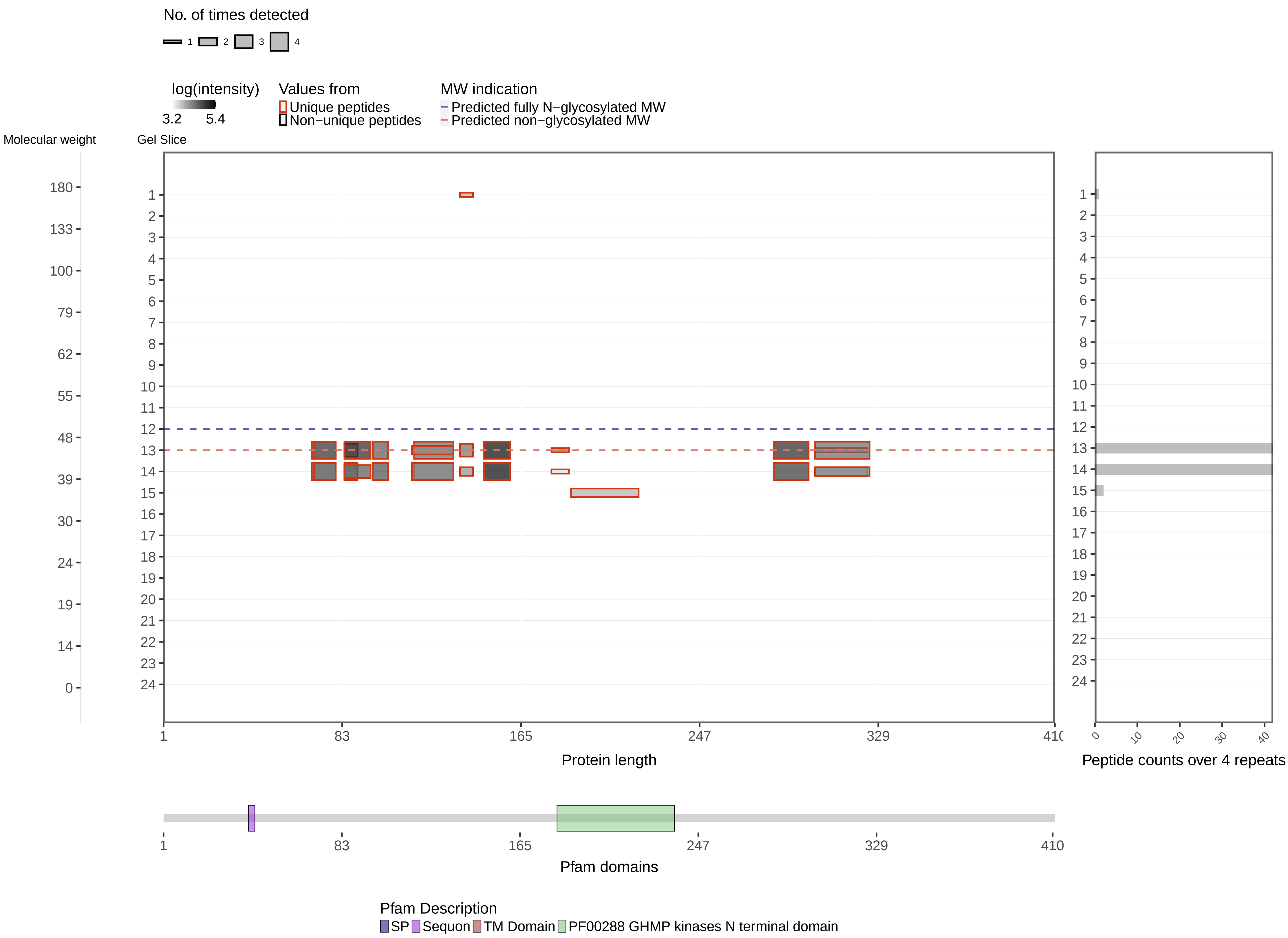


Protein sequence:
GLGGGSSNAATALWAANQFSGCLATEKELQEWSGEIGSDISFFFSHGAAYCTGRGEVVQDIPSPIPFDIPMVLIKPQQECSTAEVYKRFLLDQTSNVDPLSLLEKIKTSGISQDVCVNDLEPPAFEVLPSLKRLKQRVIAAGRGQYDAVFMSGSGSTIVGVGSPDPPQFVYDDEEYKDVFLSEASFITRAANQWYEEPLSAGNYADQAEFSGSFDNS*

### NbL06g20260.1 UBP1−associated_protein_2B−like [MW = 52.41 kDa]

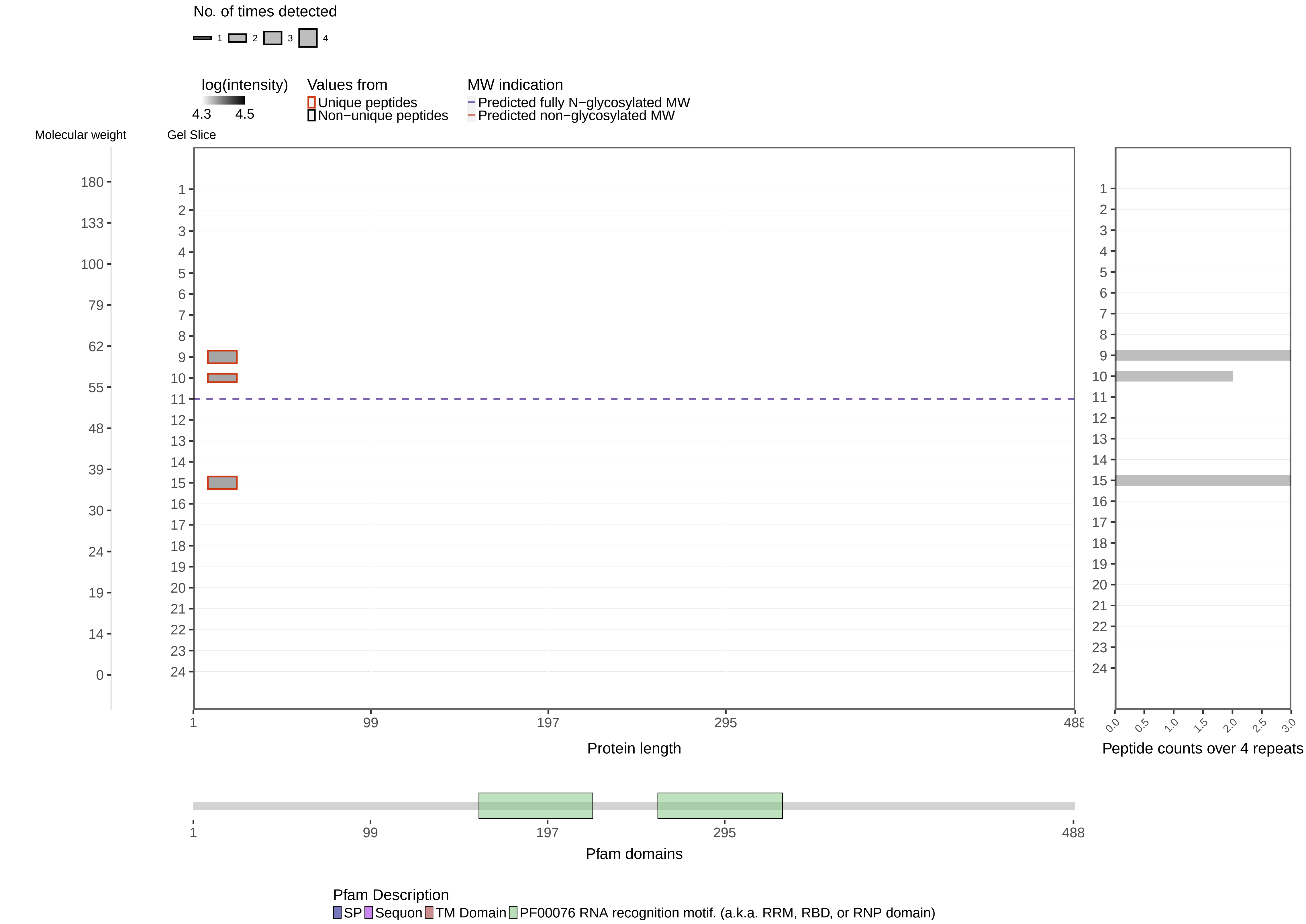


Protein sequence:

GPVPAPPPTAAAVPPVSEYTQRKIFVSNVAADLDPQKLLEFFSKFGEVEEGPLGLDKQTGKPKGFCLFVYKSVESARKALEEPHKSFEGHTLHCQKAVDGPKHNKNYYQQHQQPQQHYPQHQQHQQYYQHPAKKGKYSGGSGSGASASASAGHLMAPSGPAPVGFNPAVAPALGQALTALLATQGAGLGIGNLLGGFSGPVNPQGVPPVMNNAGGYGAQGAAGGYGAQPMQYQNPQIQGGARPQGGAPYPGYGGH*

### NbL06g20250.1 protein_argonaute_4−like [MW = 101.42 kDa]

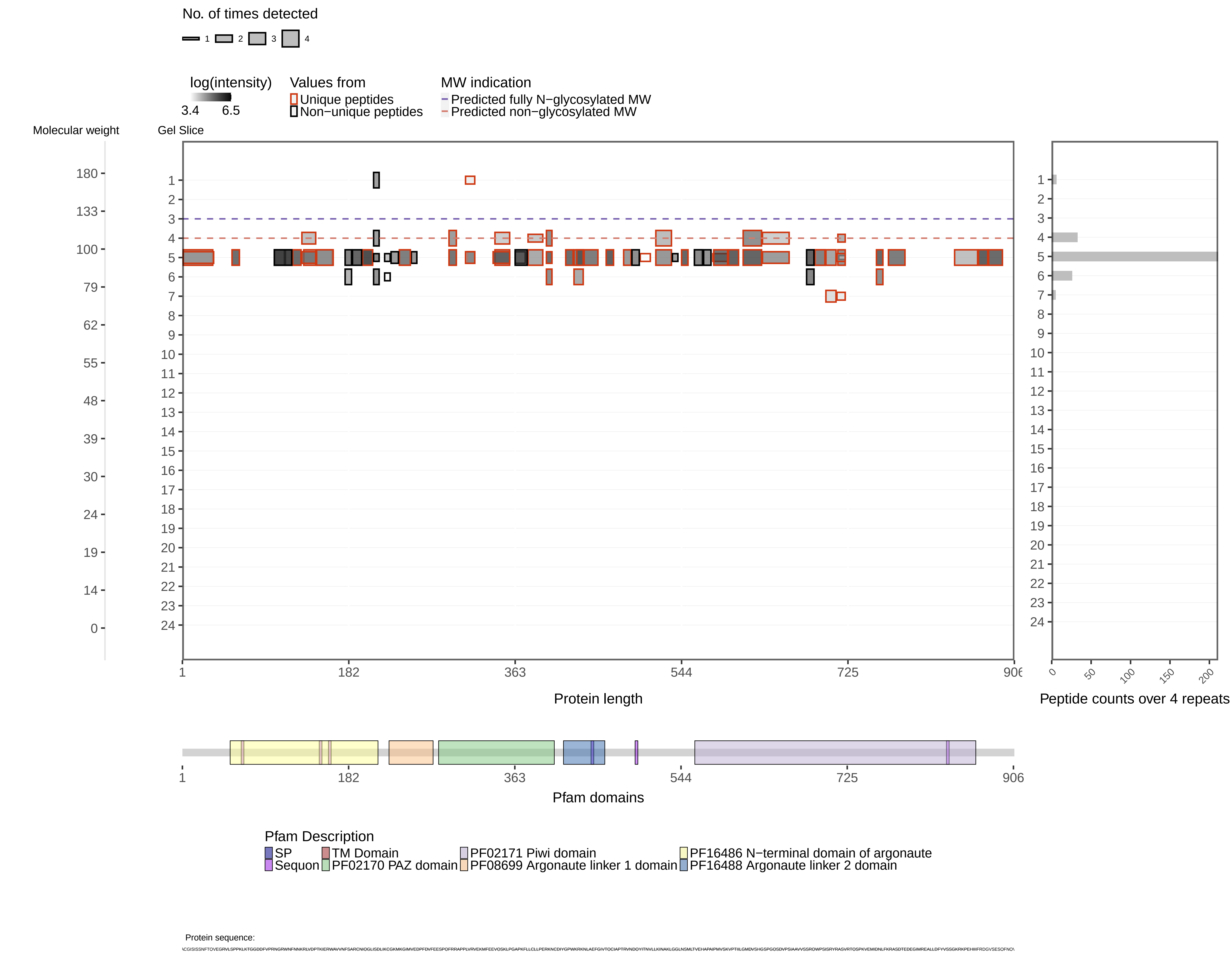

NbL06g20210.1 phytochromobilin:ferredoxin_oxidoreductase_chloroplastic−like [MW = 39.86 kDa]

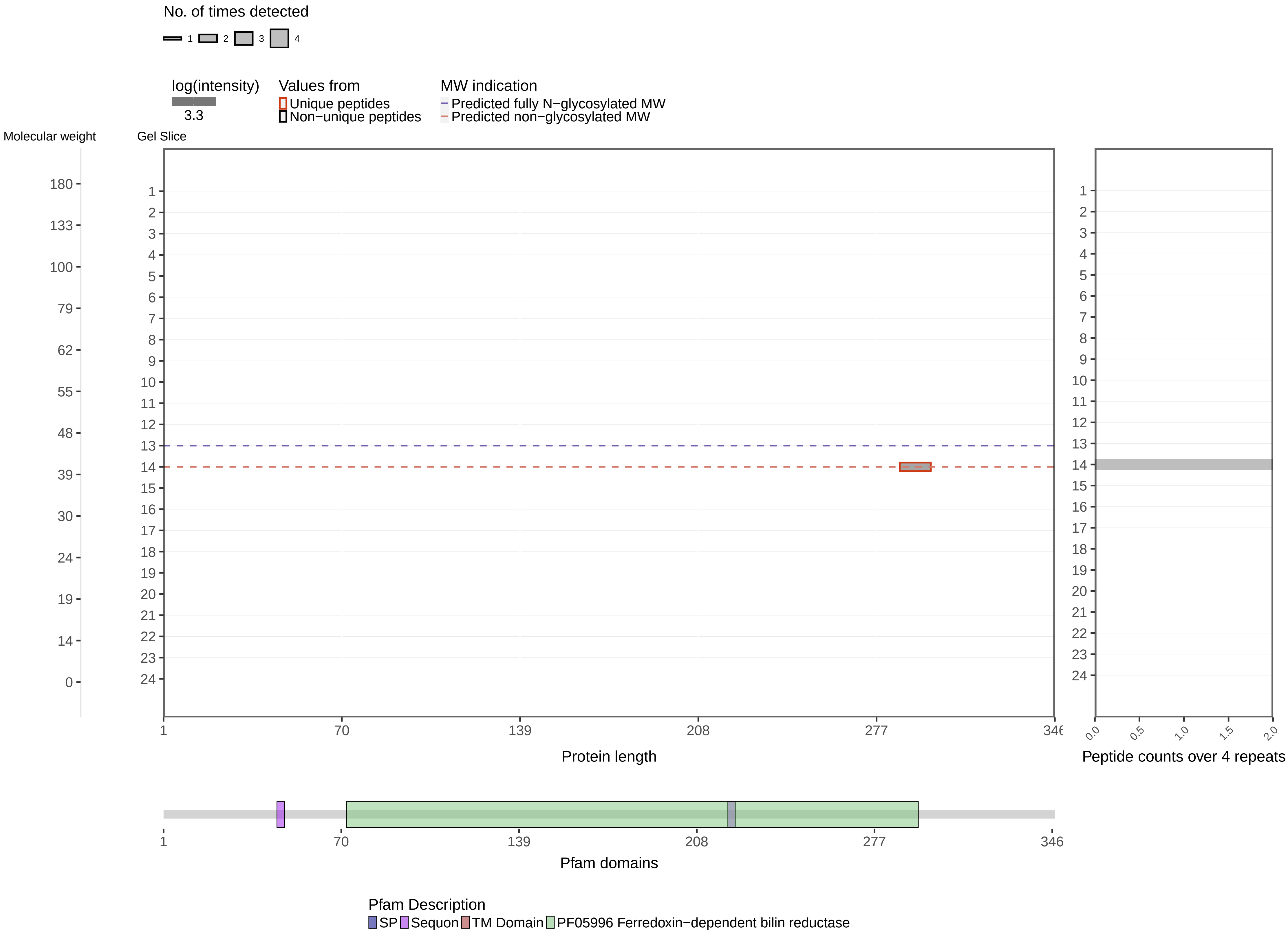


Protein sequence:

NPLHDVMDQEDYKEKYYKDLIPLGLKYSDLLPWGGKLTSESLKFFSPIVIWTRFSSSPYNHSVLFSAFKDYYKAWLELTDKSEEETDAFOVARNREAOHRYLTWRAEKDPGHGVLKRLVGEALAKDVISEFLFNGVNKLGSKTFLDYFPEYRCEDGRVNEKRSMIGKSFENRPWNARGEFIGNII*

NbL06g20200.1 uncharacterized_protein_At2g37660_chloroplastic−like [MW = 36.5 kDa]

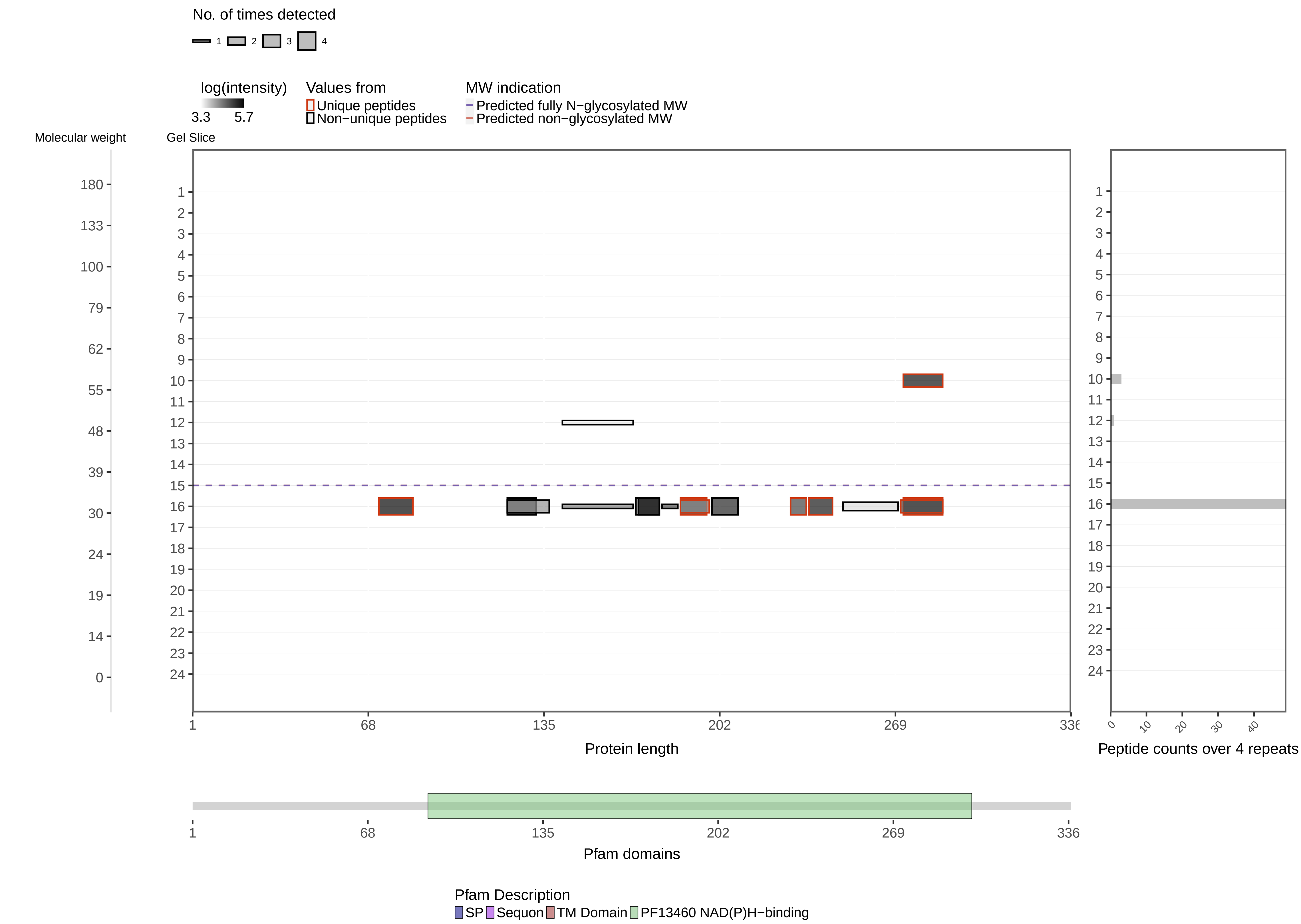


Protein sequence:

THVICCTGTTAFPSRRWDGDNTPERVDWEGVRNLVSSLPQSLKRIVLVSSIGVTKFNELPWSIMNLFGVLKYKKMGEDFVRNSGLPFTIIRAGRLTDGPYTSYDLNTLLQATAGERRAVLIGQGDTLVGEVSRLVVAEACIQALDIDFTEGQTYEINSIPGDGPGSDTLKWQELFRDSHNN*

NbL06g19970.1 glucan_endo−13−beta−glucosidase_acidic_isoform_GL161 [MW − SP = 35.98 kDa]

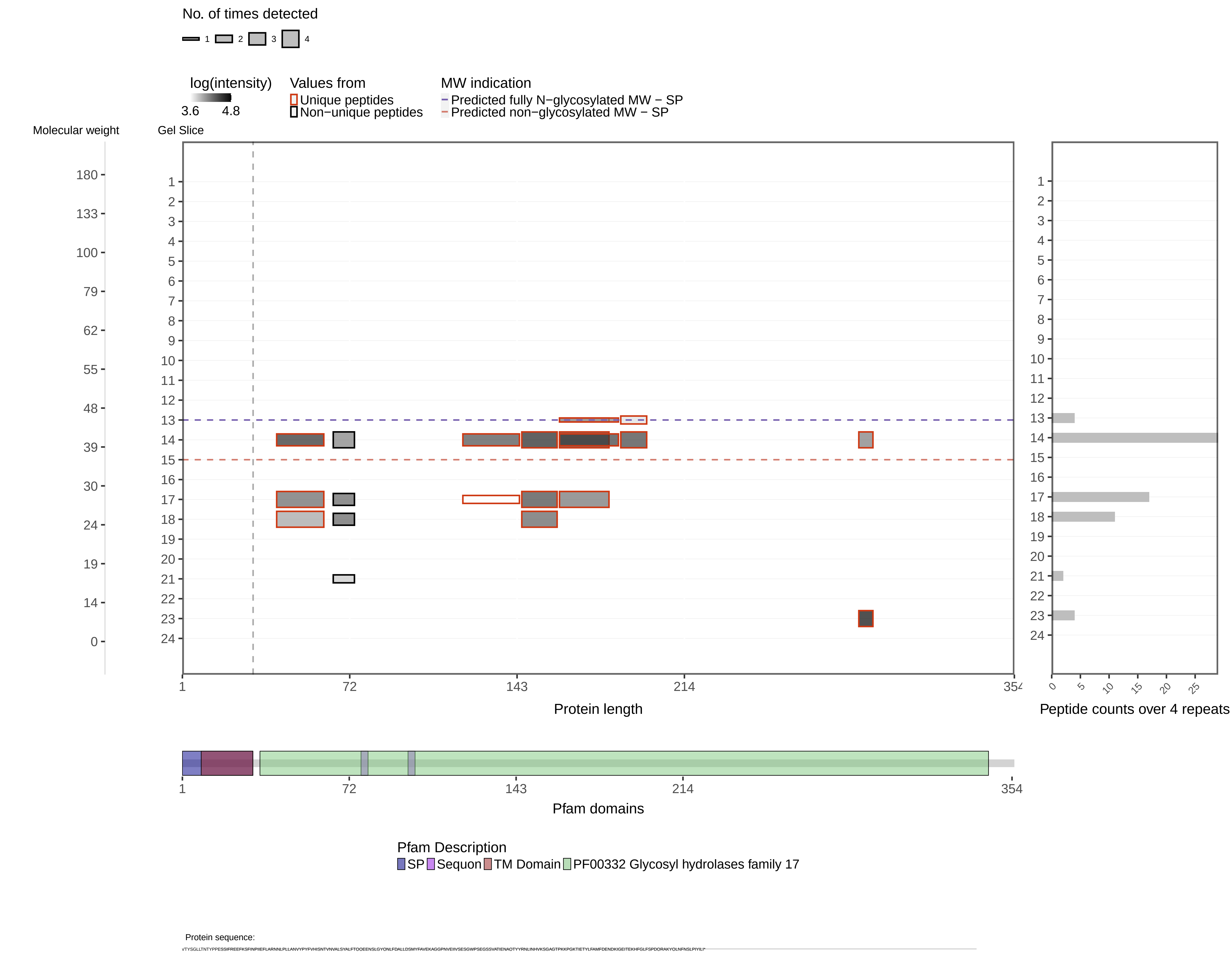

NbL06g19940.1 cinnamoyl−CoA_reductase_1−like [MW = 35.7 kDa]

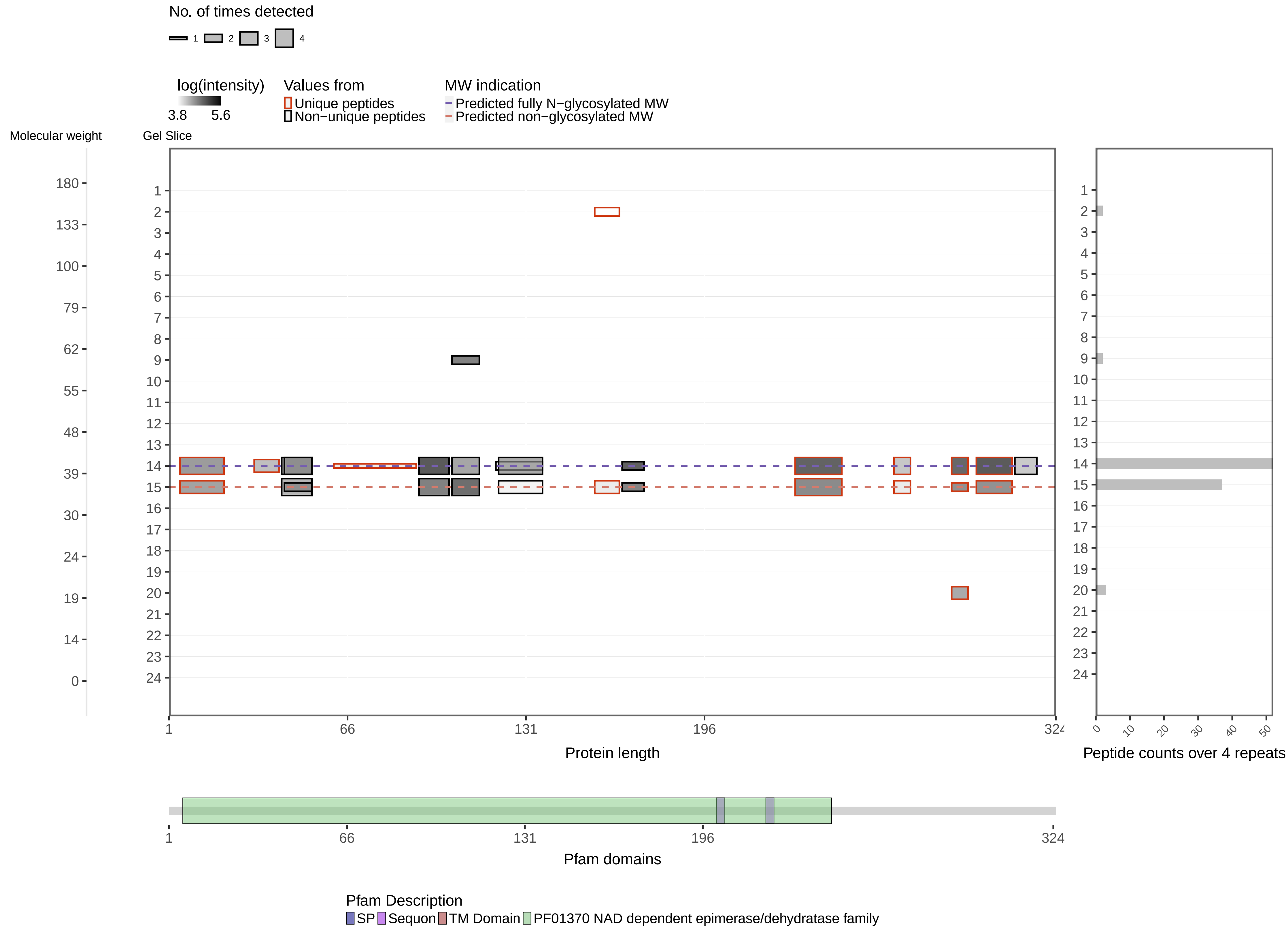


Protein sequence:

/SDPHLCRESQMWYVLSKTLAEDAAWKFVKEKAIDMVTINPAMVIGSLLQPTLNTSCAAVLQLINGAVTYPNATLGWVNVKDVALAHILAFENPSANGRYLMVEAVAHYSELVKILREHYPTMKLPEKCVDDKPFPPKYQVNIERAKQLGVEFTPLAESIKETVESLKEKKFYSDA*

NbL06g19920.1 cinnamoyl−CoA_reductase_1−like [MW = 43.22 kDa]

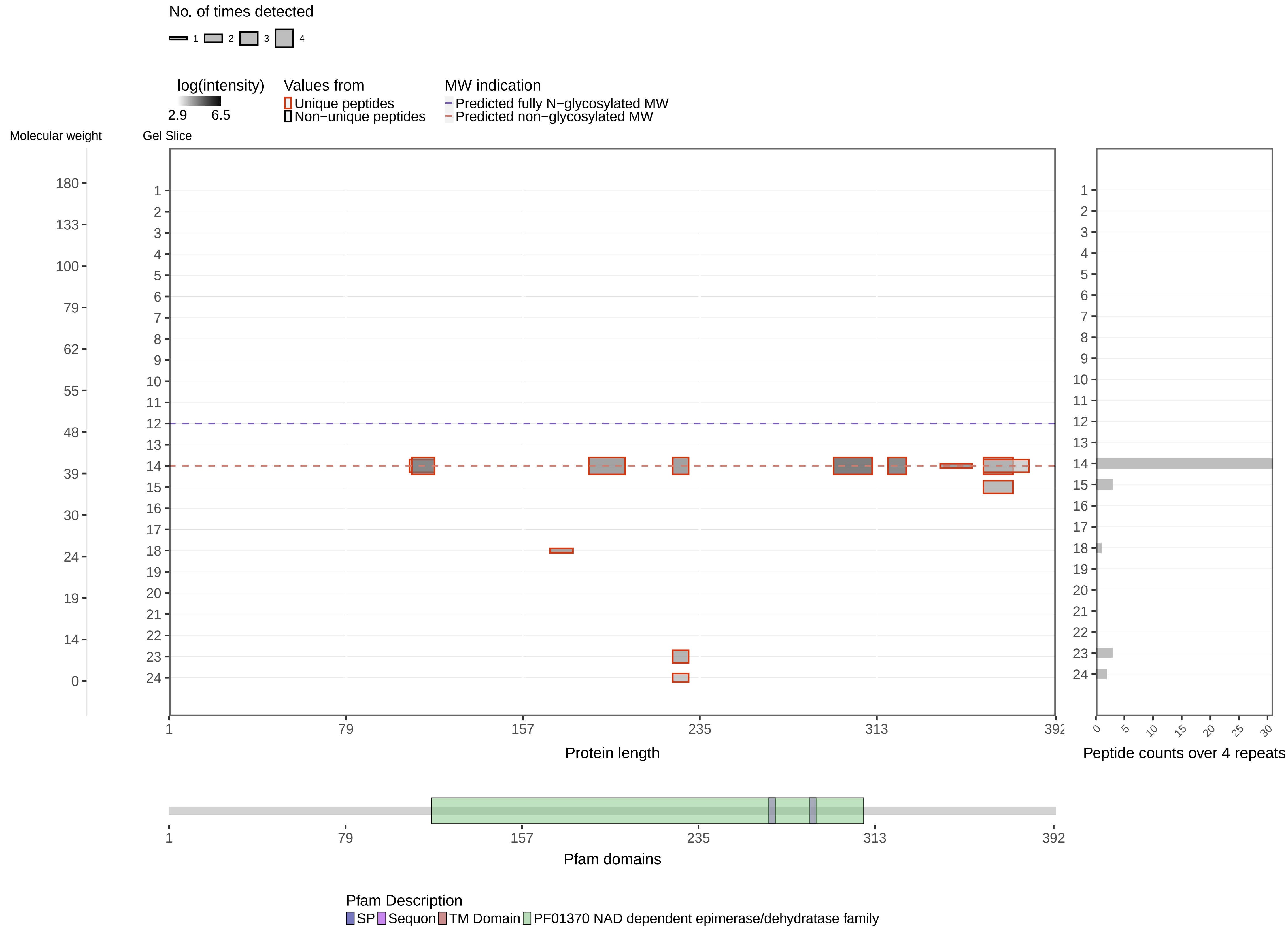


Protein sequence:

VVLTSSIAAVAYSGQPRTPEVVVDESWWTSPDYCREKQLWYVLSKTLAEDAAWKFVKEKGIDMVAINPAMVIGPLLQPTLNTSSGAVLNLVNGAETYPNSTFGWVNVKDVANAHILAFENPSANGRYLMVERVAHYSDILKILRDLYPTMRLPEKCADDNPLMQNYQVSKERAKSLGVEFTPLEESIKETVESLKEKRFFGGSSAM*

### NbL06g19700.1 ADPATP_carrier_protein_mitochondrial [MW = 41.85 kDa]

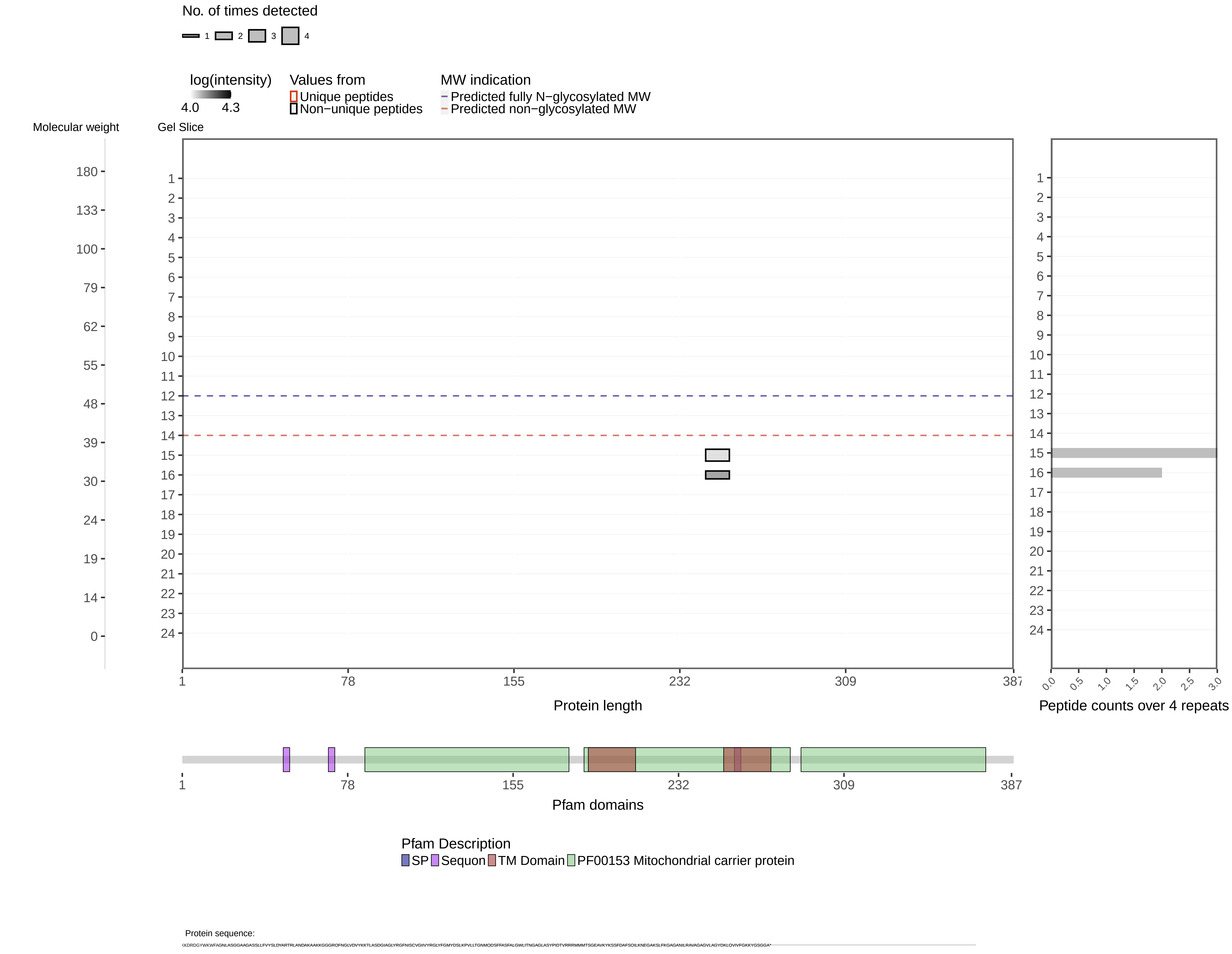

### NbL06g19560.1 zinc_finger_CCCH_domain−containing_protein_14−like [MW = 35.68 kDa]

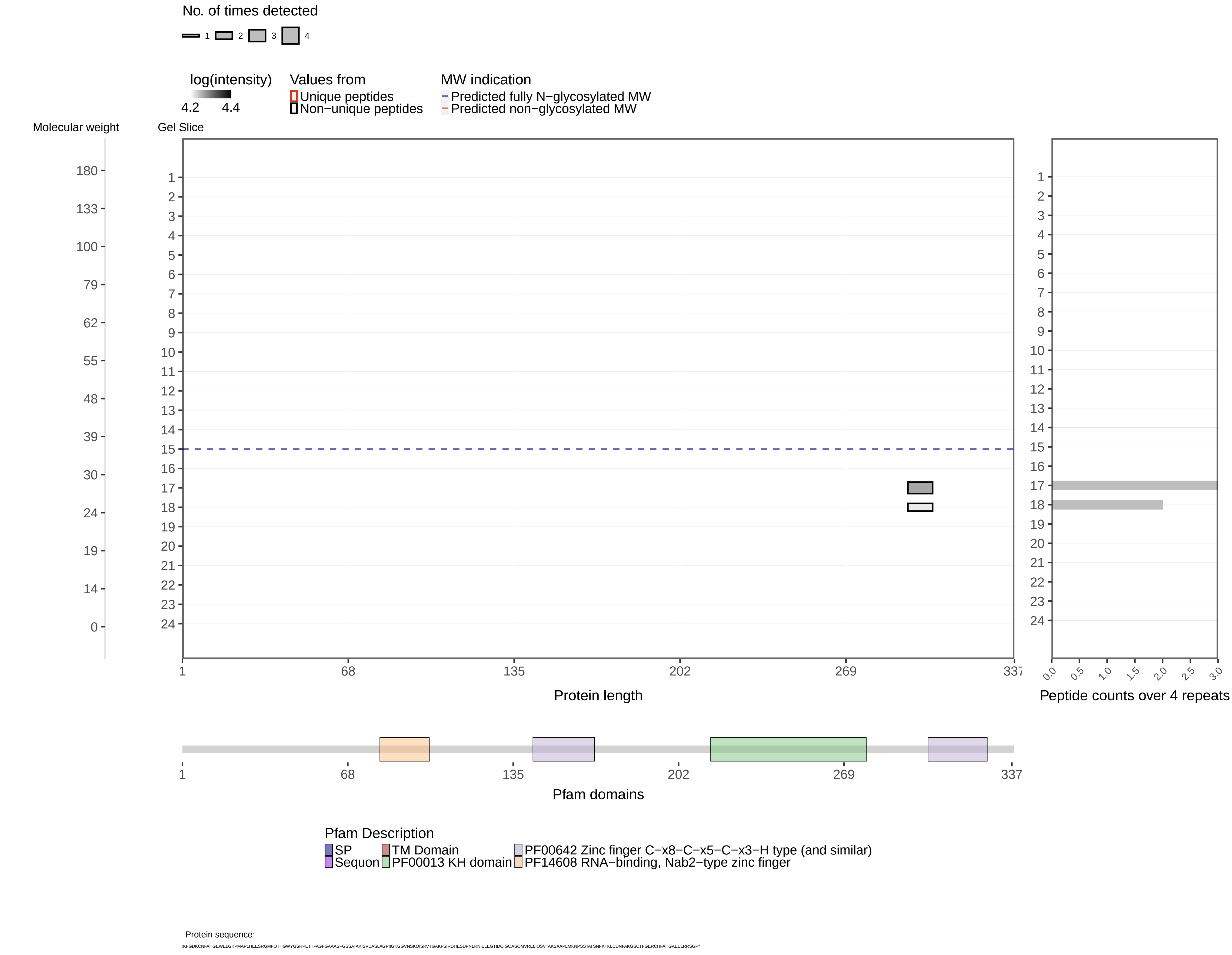

### NbL06g19430.1 vinorine_synthase−like [MW = 50.09 kDa]

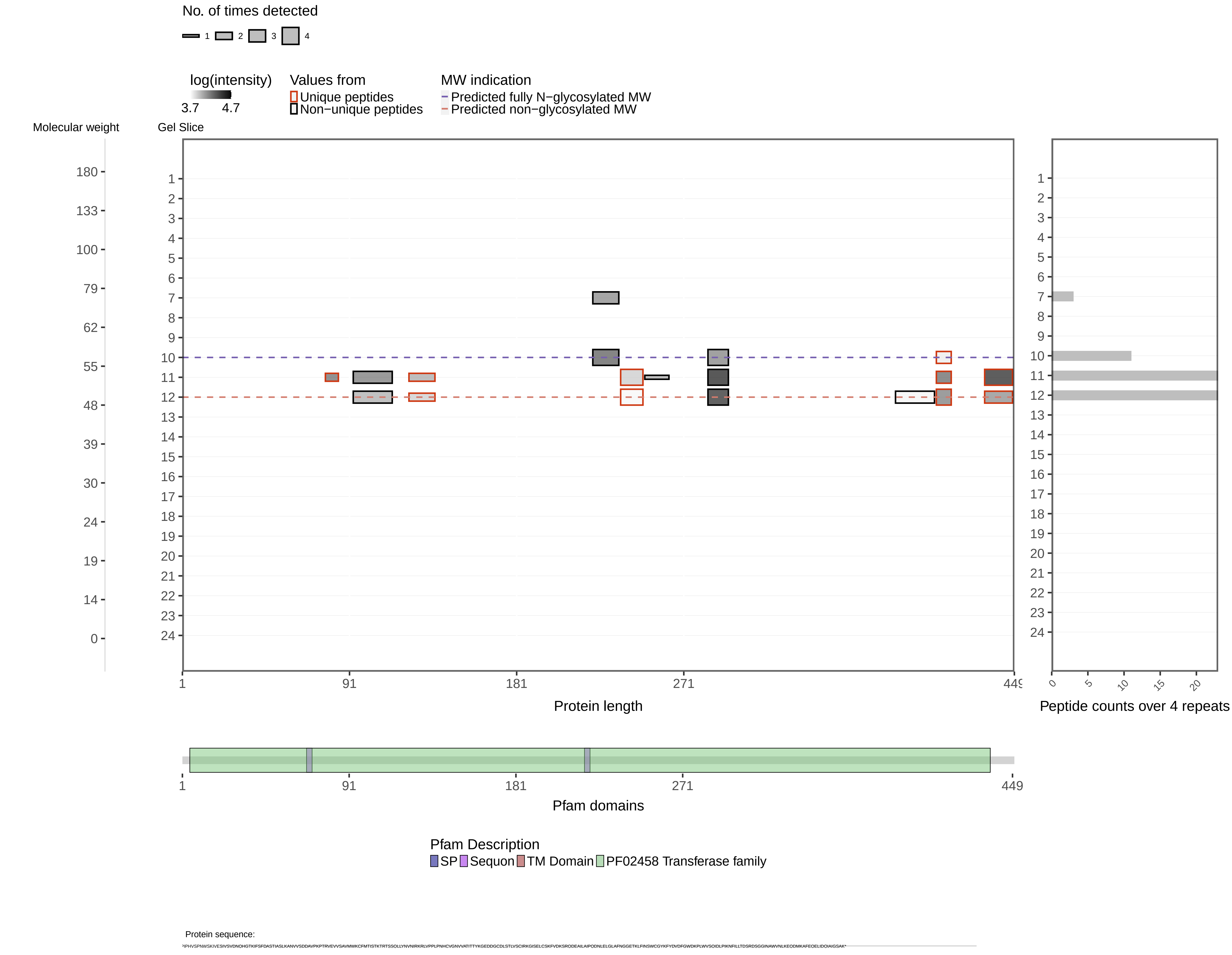

### NbL06g19370.1 uncharacterized_LOC104215593 [MW − SP = 16.75 kDa]

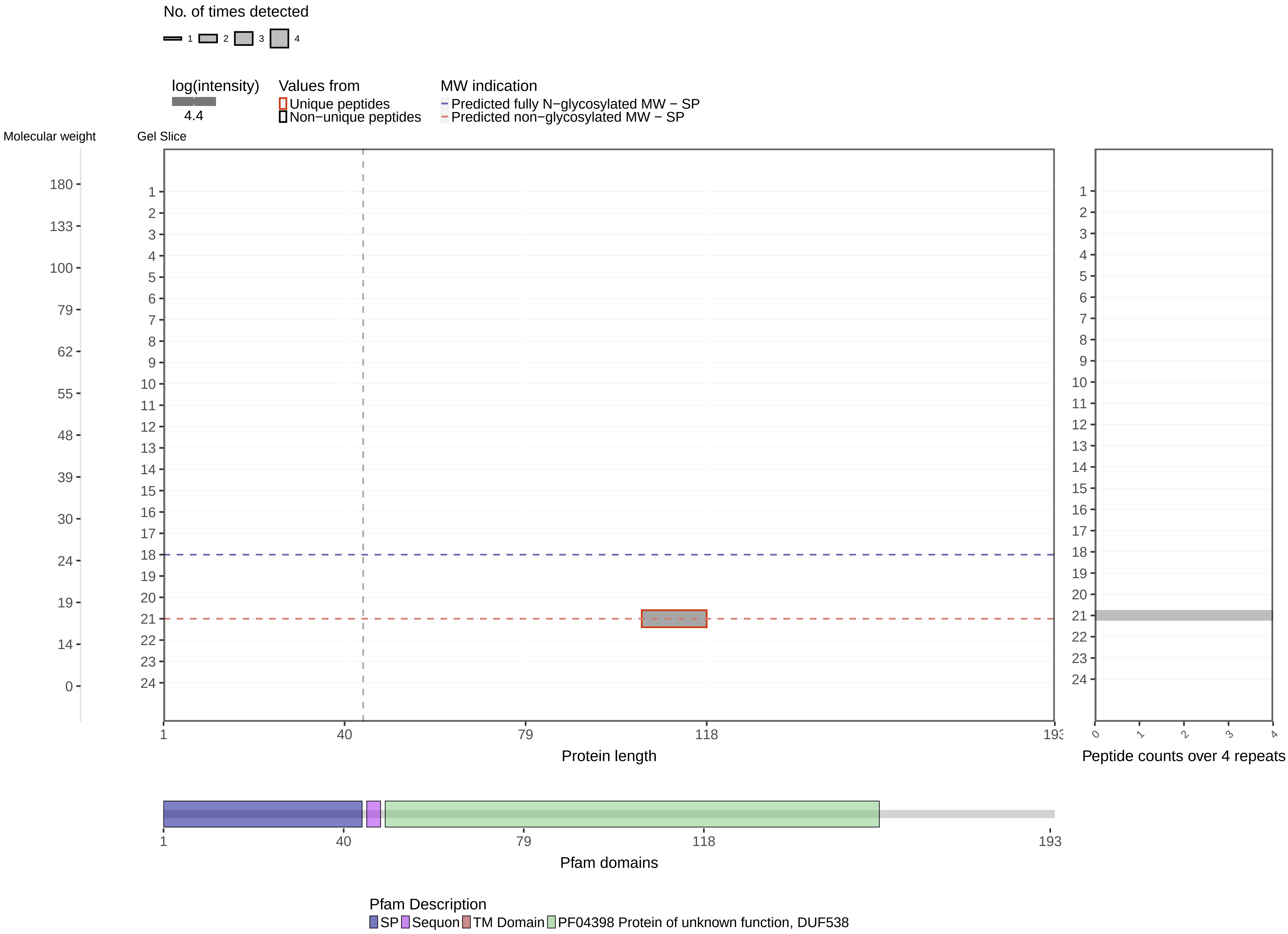


Protein sequence:

KPCYVEFEYLVYYAEKITGKLSIGSITDLDGIQVKRFFLWLNVNEIRVDLPSSGSIYFQVGIINKKLDMNQFETVHSCRDNALALCGGSLRQVLQLPDPVNDMPMLITE*

### NbL06g19340.1 serine/threonine−protein_phosphatase_PP2A_catalytic_subunit [MW = 35.64 kDa]

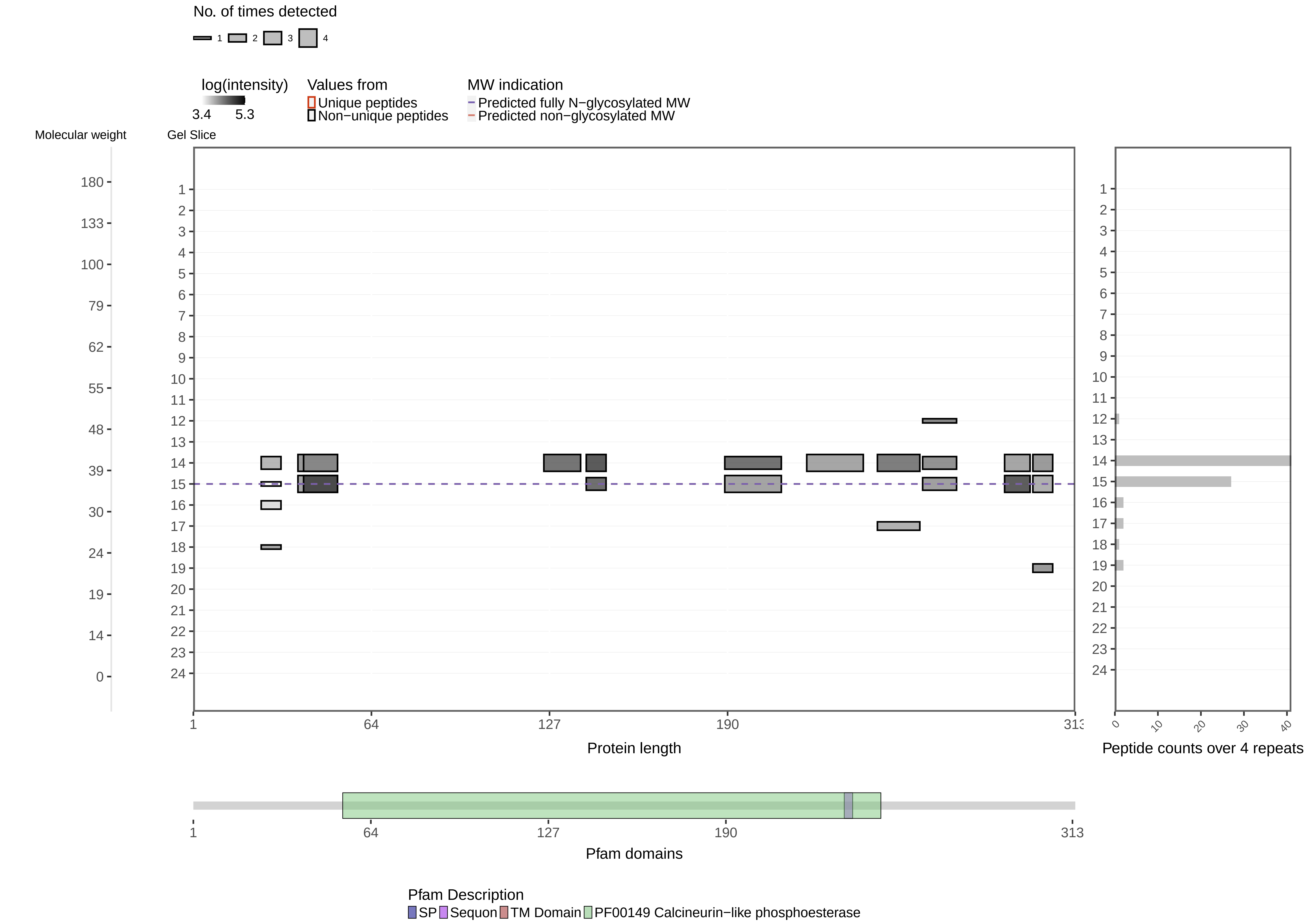


Protein sequence:

NVWKTFTDLFDYFPLTALVESEIFCLHGGLSPSIETLDNIRNFDRVQEVPHEGAMCDLLWSDPDDRCGWGISPRGAGYTFGQDISEQFNHTNNLKLIARAHQLVMEGFNWAHDQKVVTIFSAPNYCYRCGNMASILEVDDSRERTFIQFEPAPRRGEPDVTRRTPDYFL*

### NbL06g19240.1 importin−5−like [MW = 123.2 kDa]

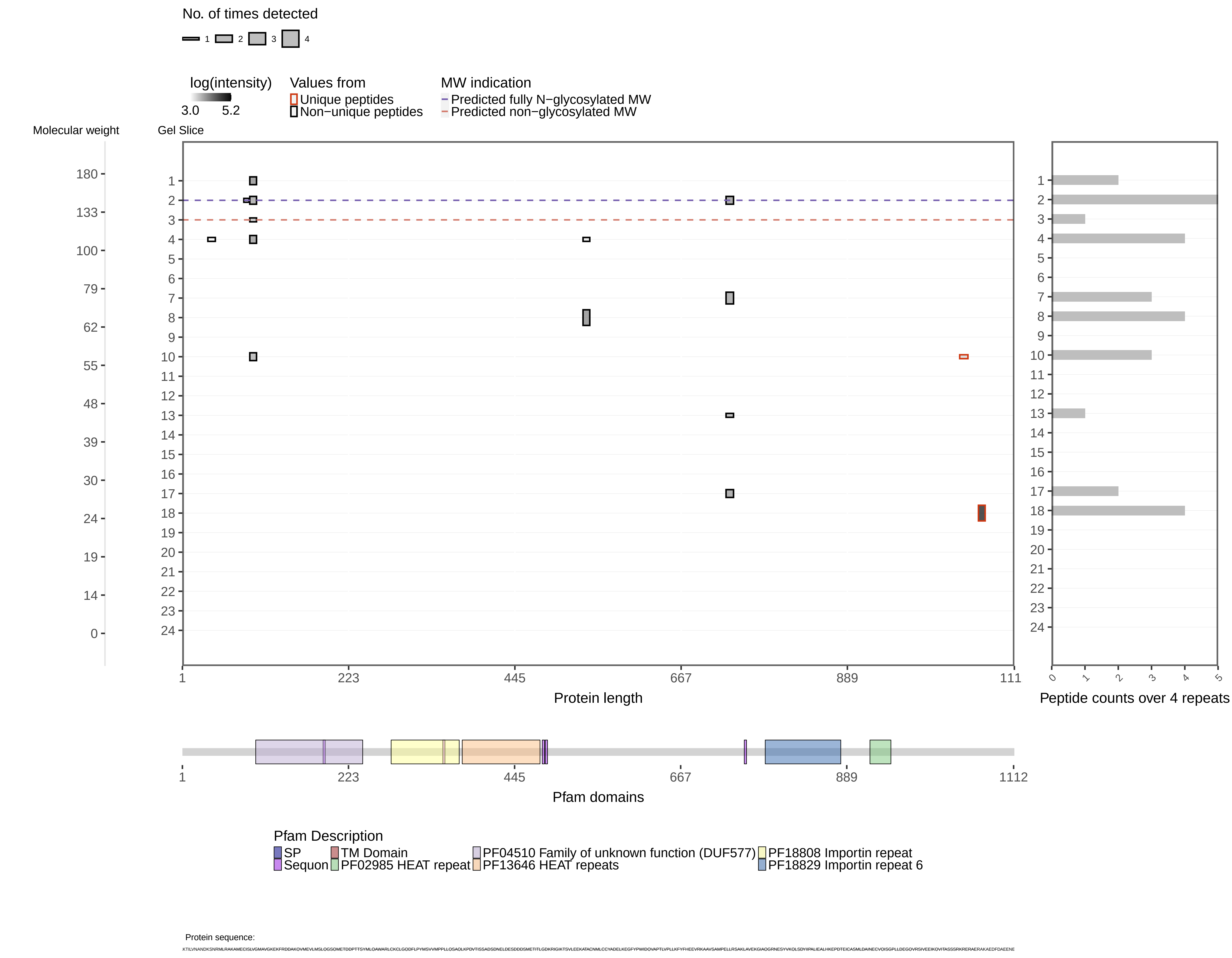

NbL06g19230.1 uncharacterized_LOC104209961 [MW = 46.08 kDa]

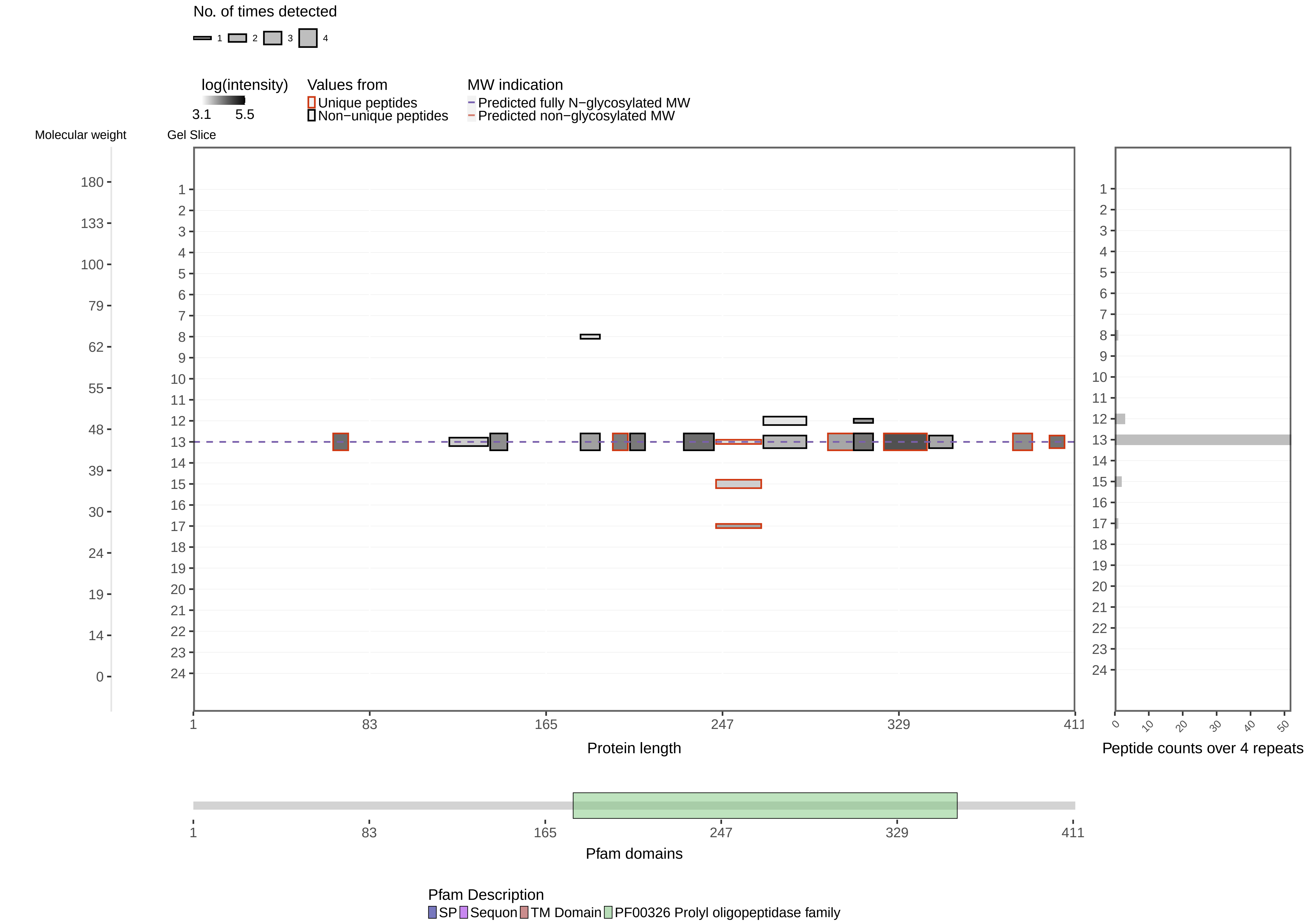


Protein sequence:

HGERATSMTTYRDALISSWKKGDTMPFIFDTVWDLIKLADYLTEREDIDSSRIGITGESLGGMHAWFAAFADSRYAVVVPIIGVQGFQWAIEHDRWQARVDSIKAVFEEACADLGKNEIDKEVVQKVWDRIAPGLASQLDSPYTVPVIAPRPLLILNGAEDPRCPIAGLDIPKSRTHKACENAGCPLNFKVIAEGGIGHKMTPLMVKEASNWFDRFLKV*

### NbL06g19210.1 mitochondrial_amidoxime_reducing_component_2−like [MW = 35.42 kDa]

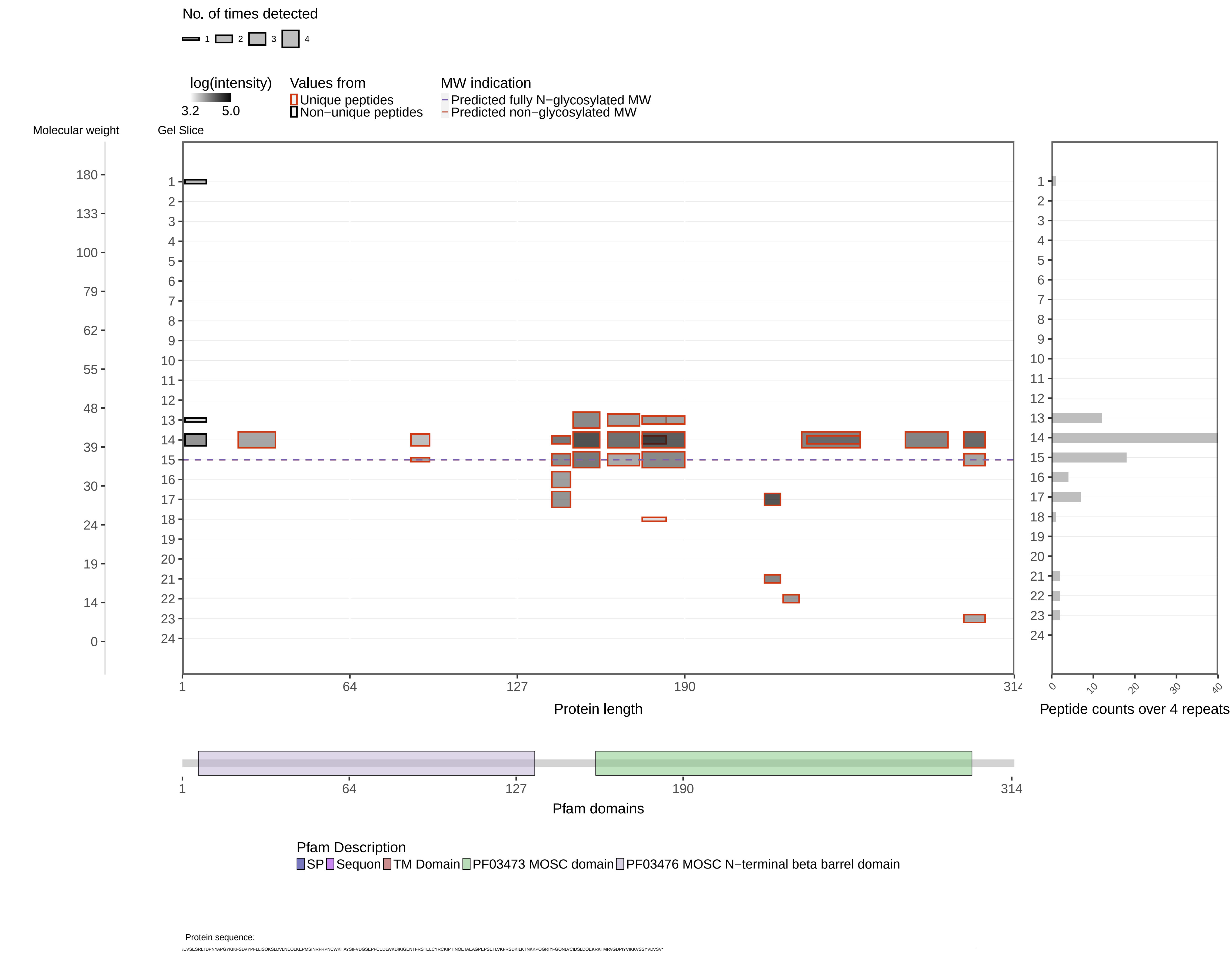

### NbL06g19120.1 chaperonin_60_subunit_beta_2_chloroplastic [MW = 65.26 kDa]

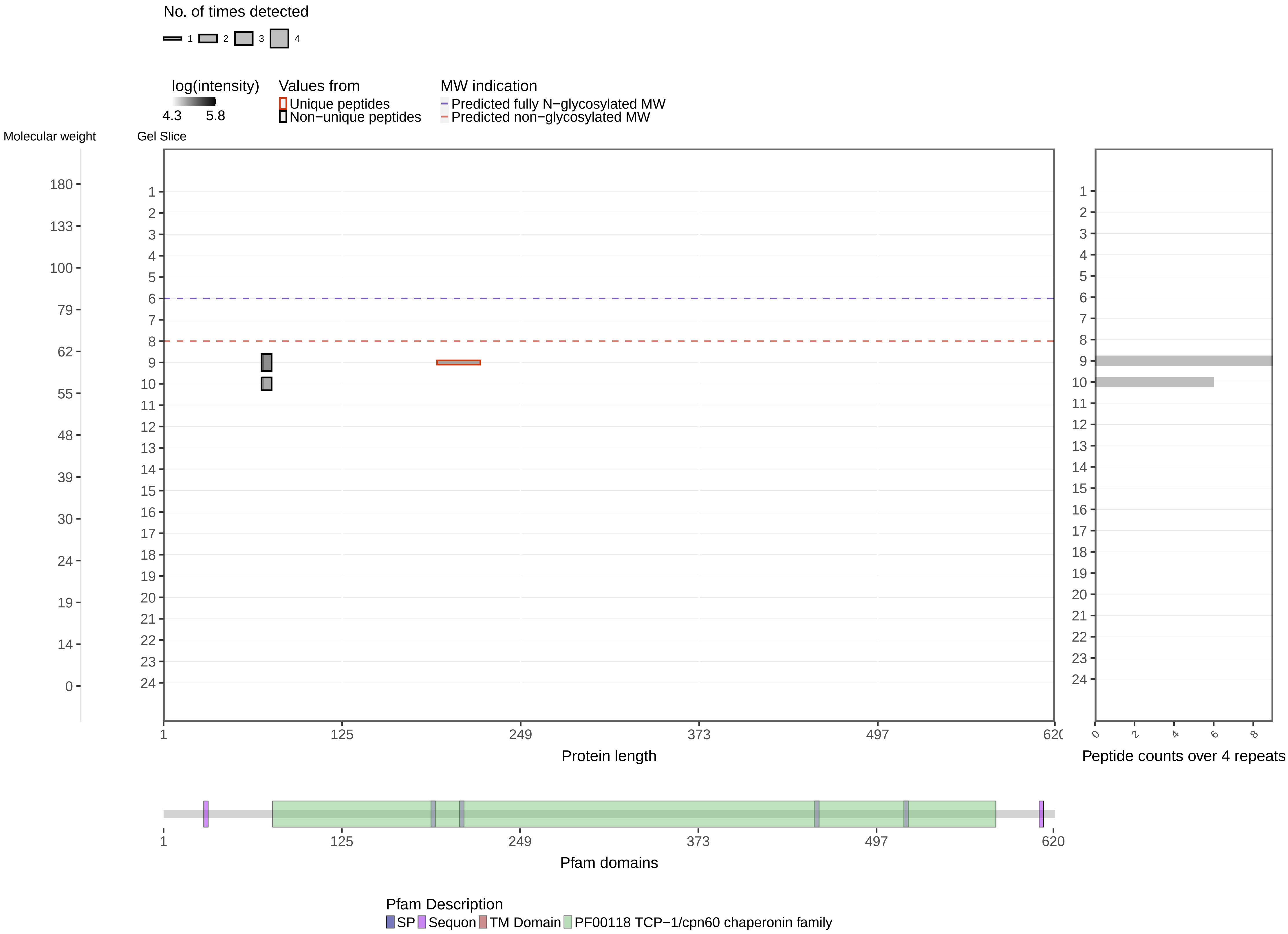


Protein sequence:

IRNGYPILIIAEDIEQEALATLVVNKLRGALKVAALKAPGFGERKSQYLDDIATLTGGTVIRDELGLTLDKADKEVLGHASKVVLTKDATTIVGDGSTQDAVNKRVAQIKNLIEAADQDYEKEKLNERIAKLSGGVAVIQVGAQTETELKEKKLRVEDALNATKAAVEEGIVVGGGCTLLRLASKVDAIKETLENDEEKVGADIVKRALSYPLKLIAKNAGVNGSVVSEKVLSSDNPKFGYNAATGNYEDLMAAGIIDPTKVVRCCLEHASSVAKTFLMSDCVVVEIKEPEPAVAGSPMD

### NbL06g19080.1 uncharacterized_LOC104245894 [MW = 76.08 kDa]

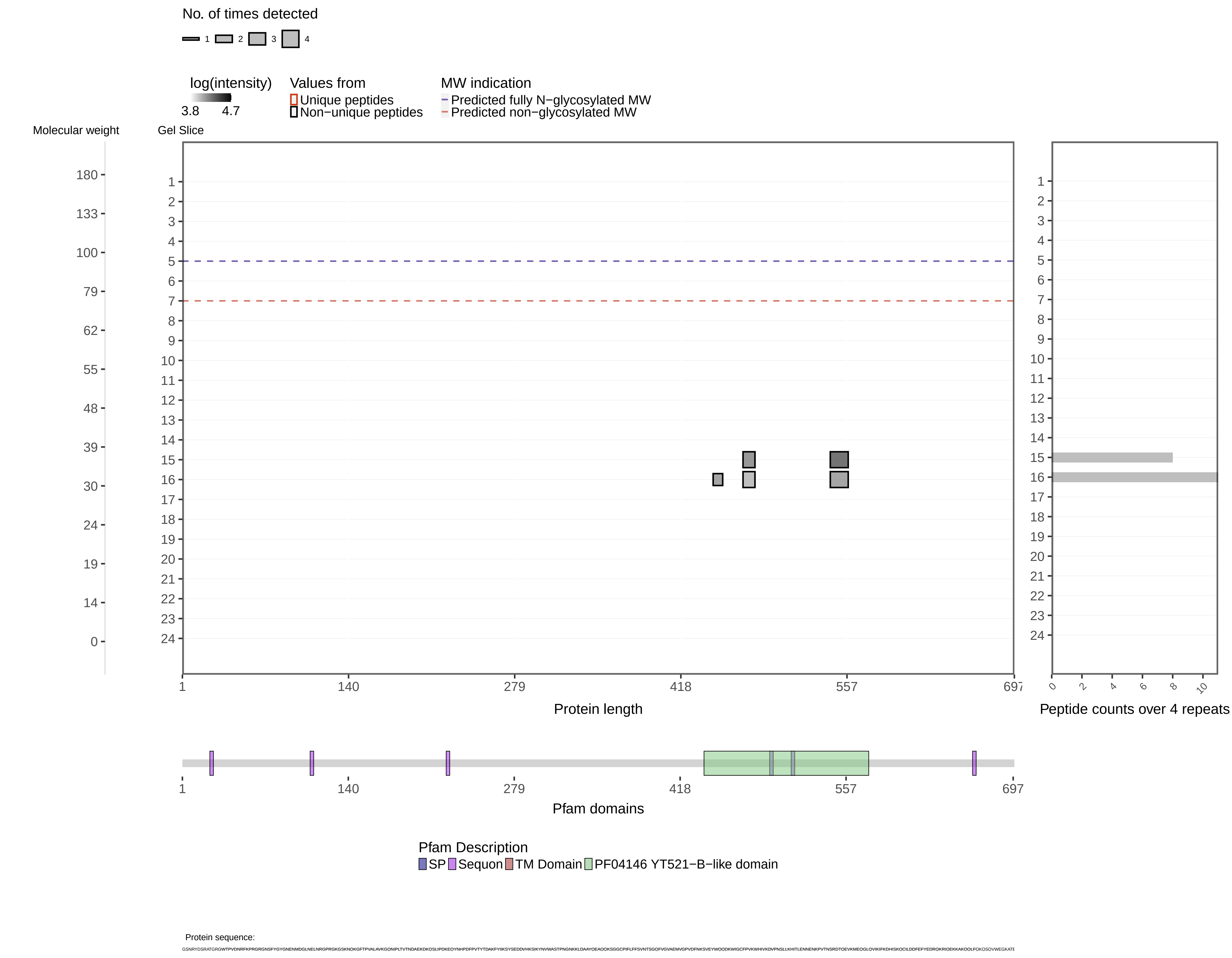

### NbL06g19060.1 2−oxoisovalerate_dehydrogenase_subunit_beta_1_mitochondrial−like [MW = 42.76 kDa]

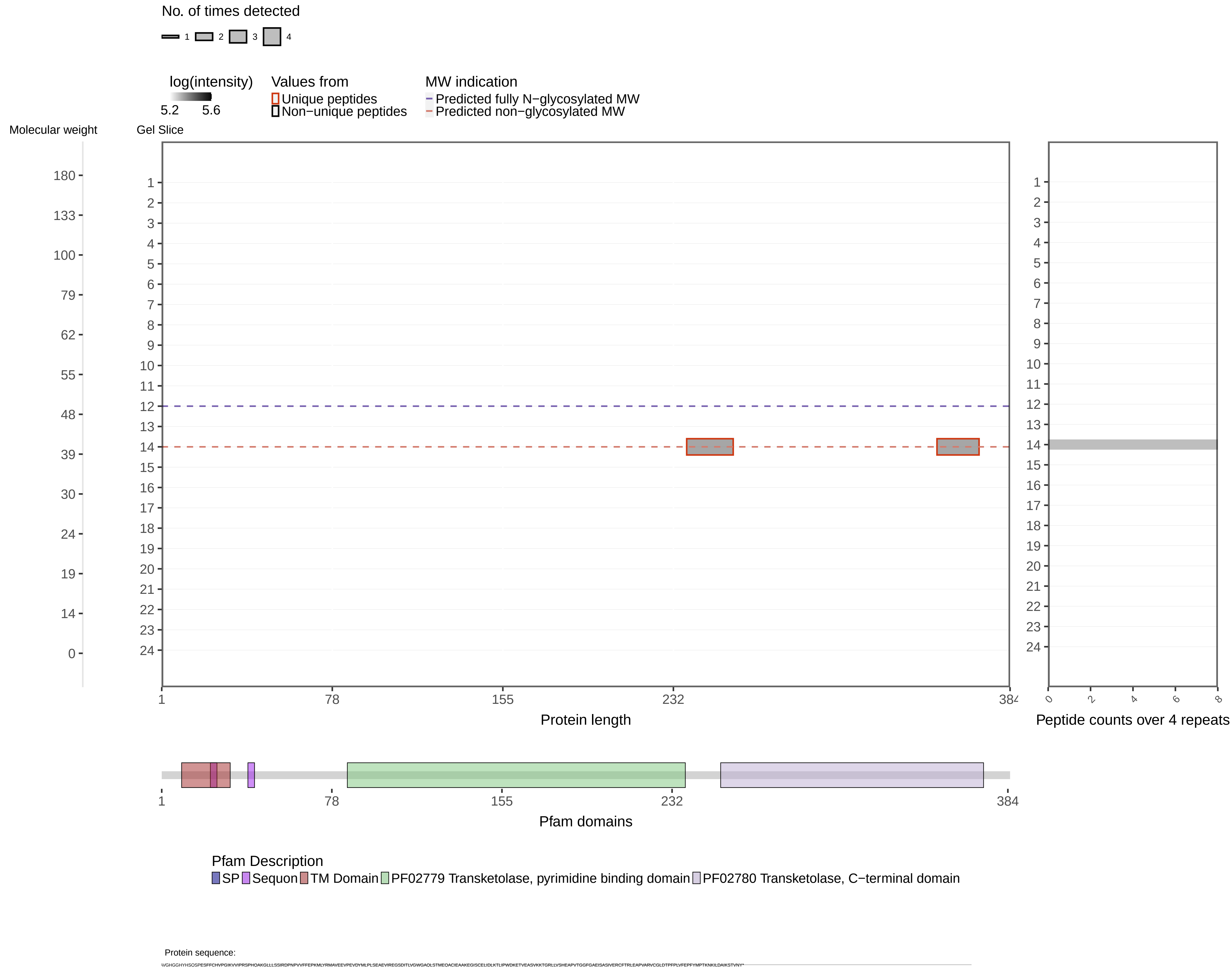

NbL06g19040.1 sm−like_protein_LSM7 [MW = 10.71 kDa]

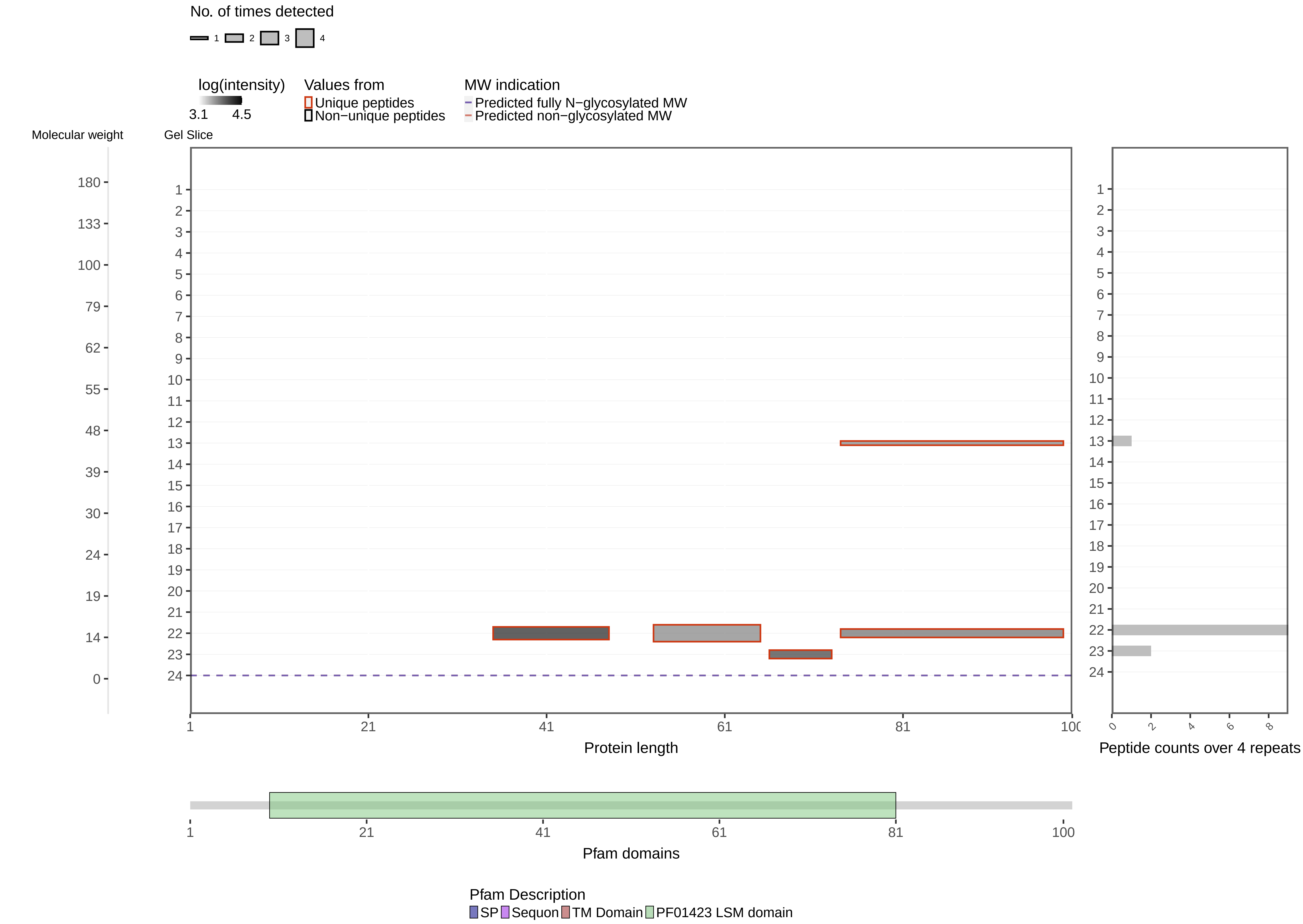


Protein sequence:

YDQLLNLVLDEAKEYLRDADDPLKTTEQTRALGLIVCRGTAVMLVAPTDGTDEISNPFVQPDGA*

NbL06g18990.1 pyruvate_kinase_cytosolic_isozyme [MW = 54.95 kDa]

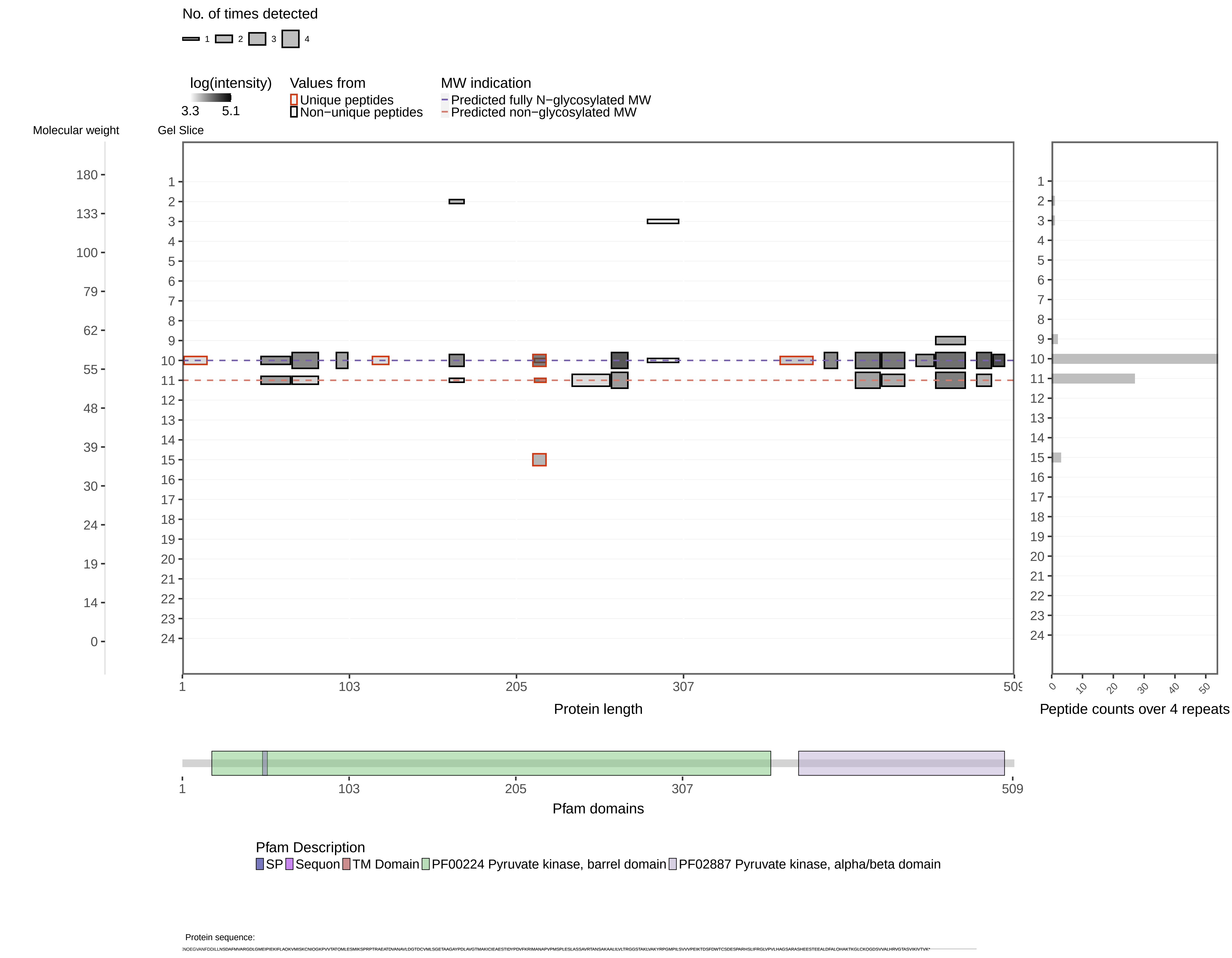

NbL06g18830.1 oxygen−dependent_coproporphyrinogen−III_oxidase_chloroplastic [MW = 44.85 kDa]

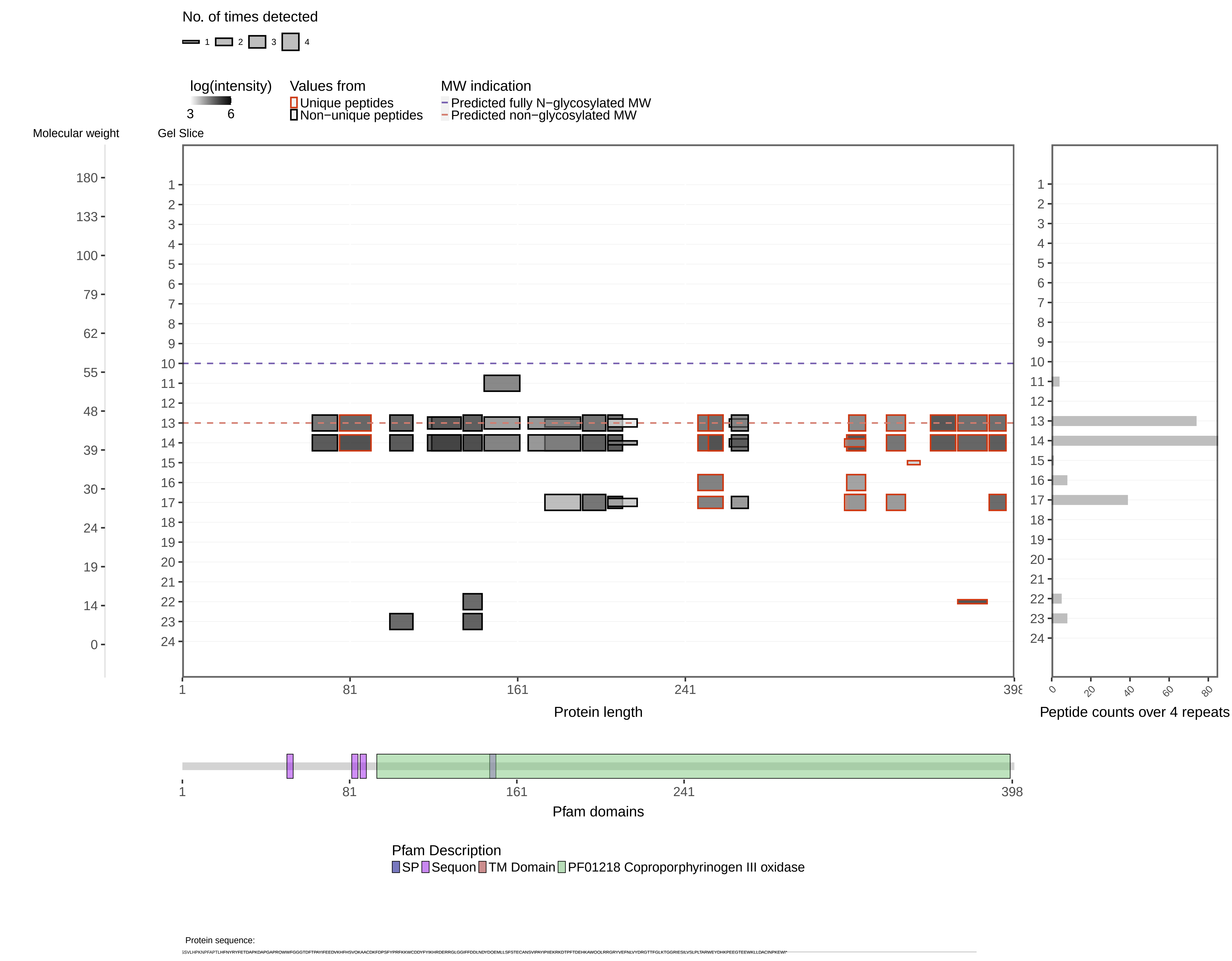

NbL06g18700.1 4−alpha−glucanotransferase_DPE2 [MW = 82.38 kDa]

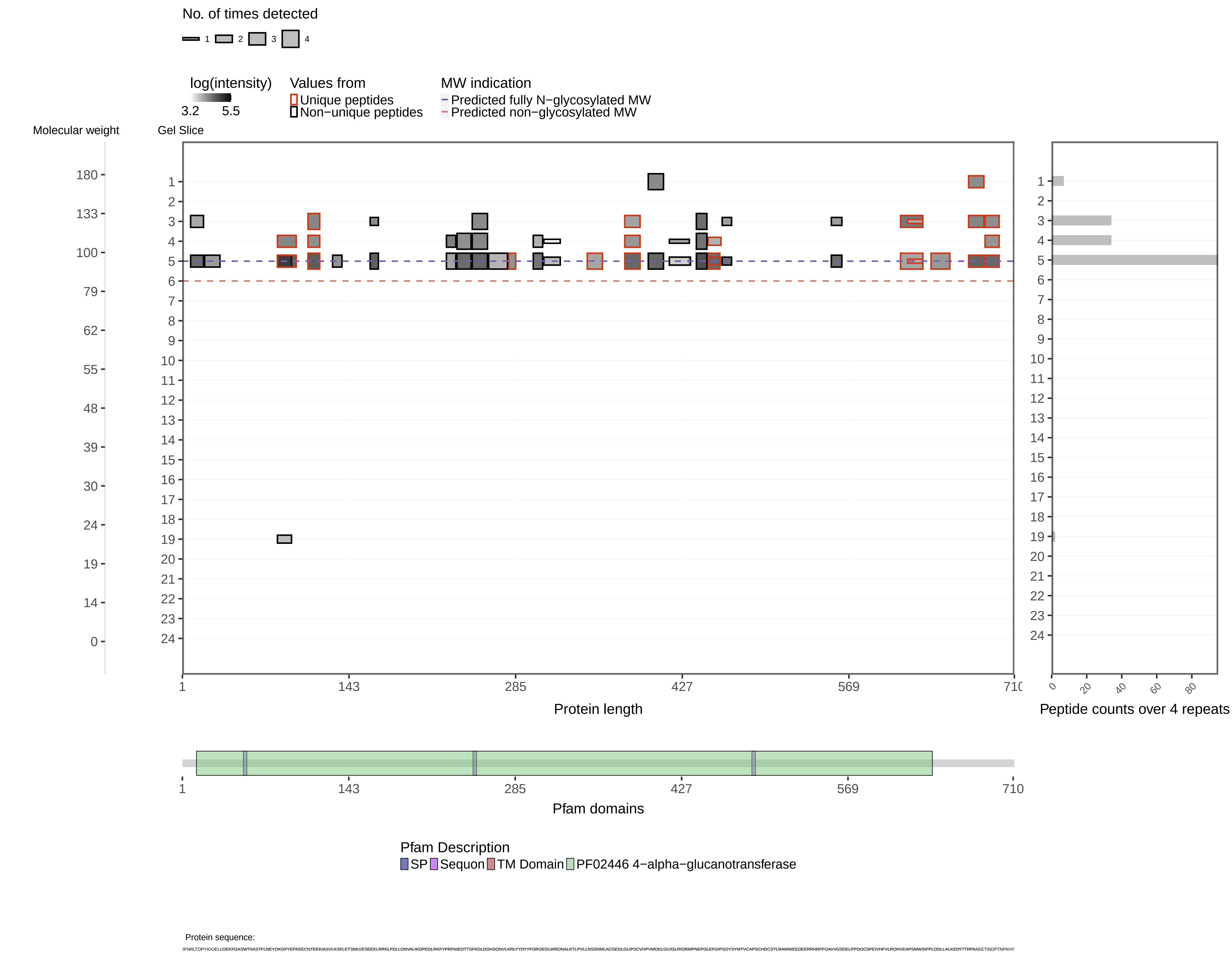

NbL06g18610.1 probable_polyribonucleotide_nucleotidyltransferase_1_chloroplastic [MW = 104.22 kDa]

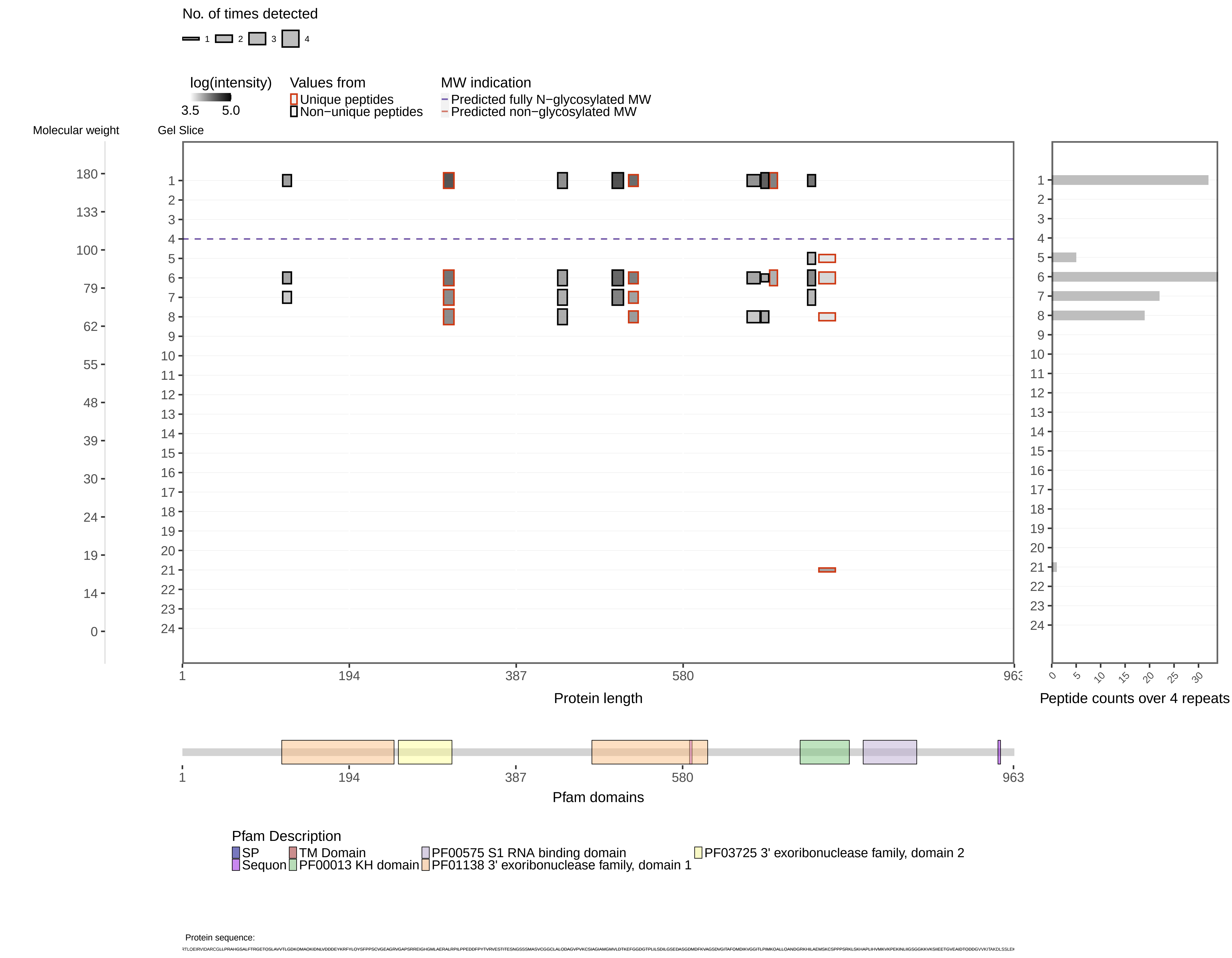

### NbL06g18570.1 39S_ribosomal_protein_L46_mitochondrial−like [MW = 26.98 kDa]

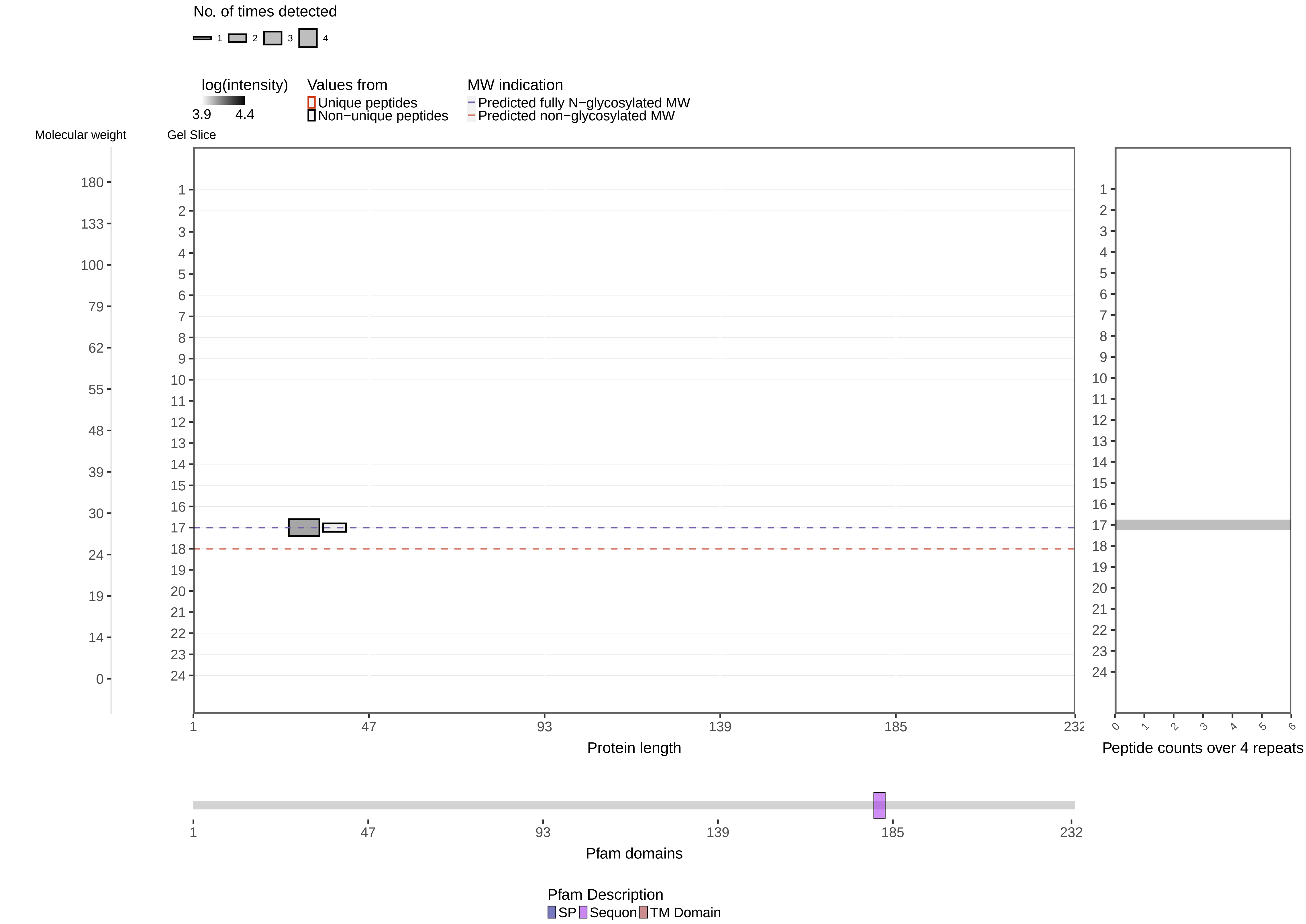


Protein sequence:

QRALDKRLYLLVHGTTHGSPSEKPVWHFPEKVYESEETLRKCAESALESVIGDLSHTYFVGNAPMGHIVVQPTEAKENTSFKRFFFKSQVIATNKFDIRKCDDFVWVTKDELLEYFPEQAEFLSKMIIS*

NbL06g18550.1 IAA−amino_acid_hydrolase_ILR1−like_3 [MW − SP = 45.5 kDa]

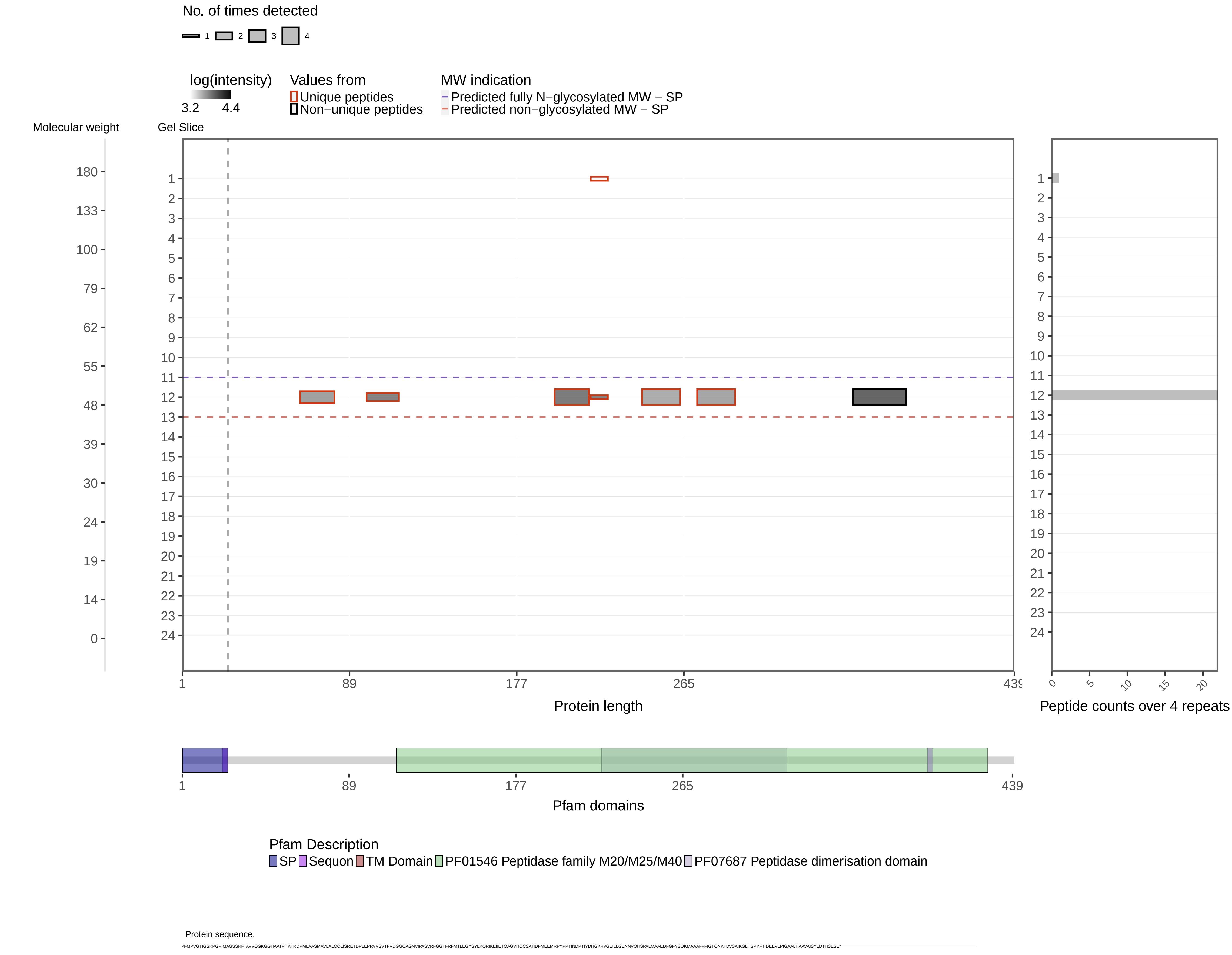

### NbL06g18400.1 DNA_ligase_1−like [MW = 41.94 kDa]

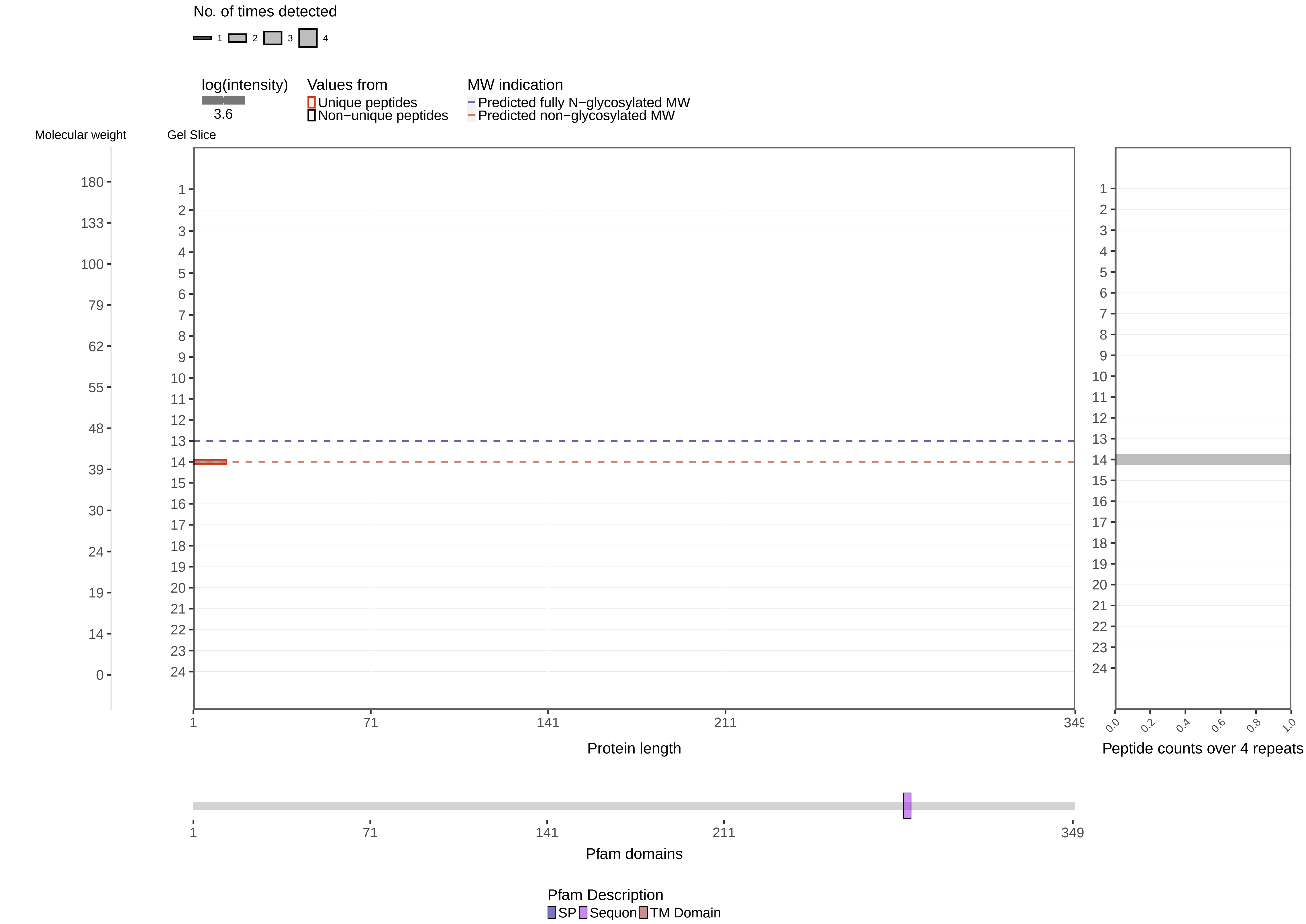


Protein sequence:

KDKESKEKEDKSKQDLEVEGEEEKDDKNDKKADEKKDKKGKKKDKDEEKKEKRKEESLEETERKKDKEKKHKKEKKDKDKEKKDKSGVELEEEMEEEKDDGKEKKKHKEKKHKEETEKGNKTKNREELEADLEAHGRKEKETKKDKEKKCKEDKDKVDEKGRETEVTTREIKIGGENRMPDEEAEH*

NbL06g18340.1 ubiquitin_carboxyl−terminal_hydrolase_12−like [MW = 134.63 kDa]

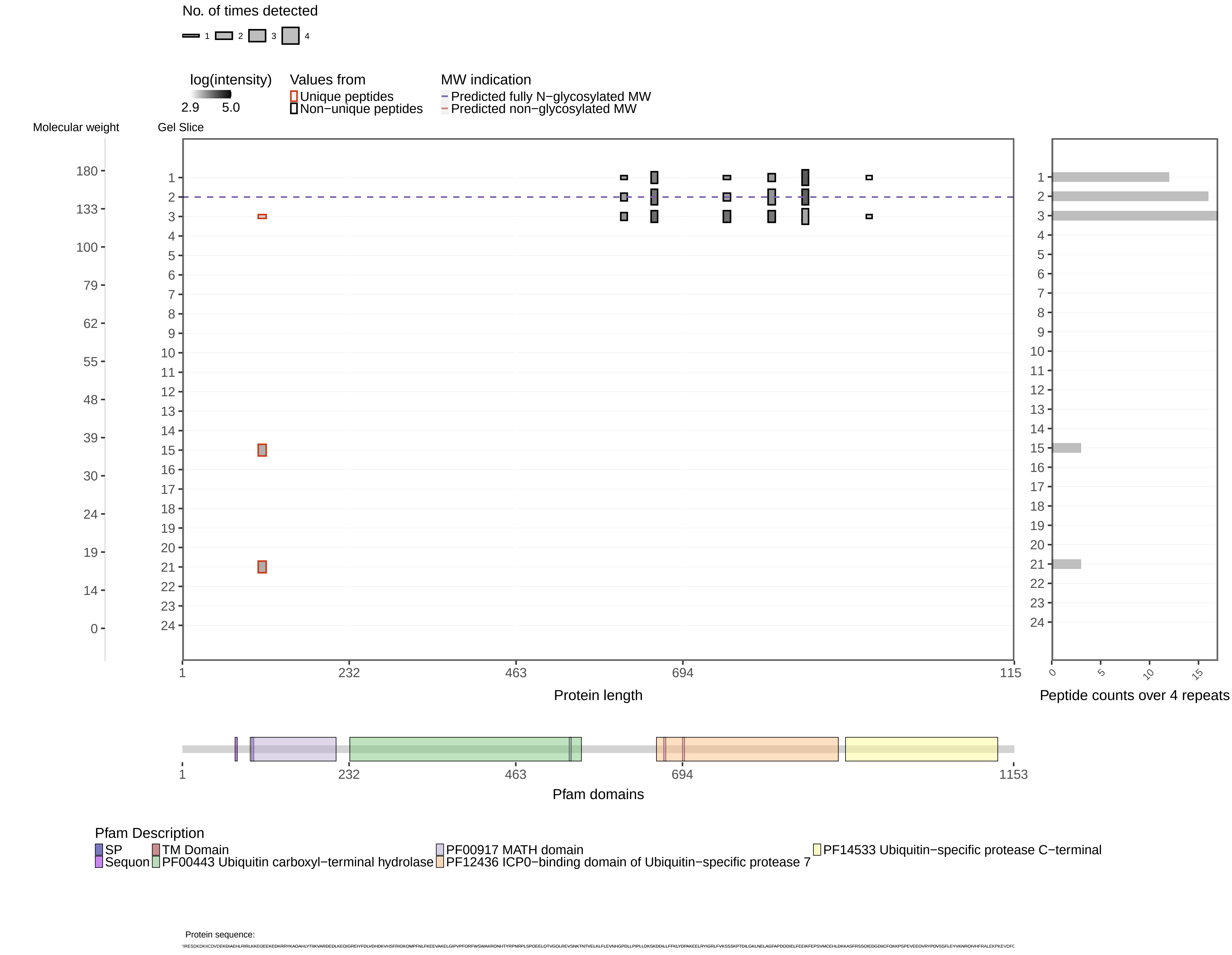

### NbL06g17860.1 probable_60S_ribosomal_protein_L14 [MW = 15.27 kDa]

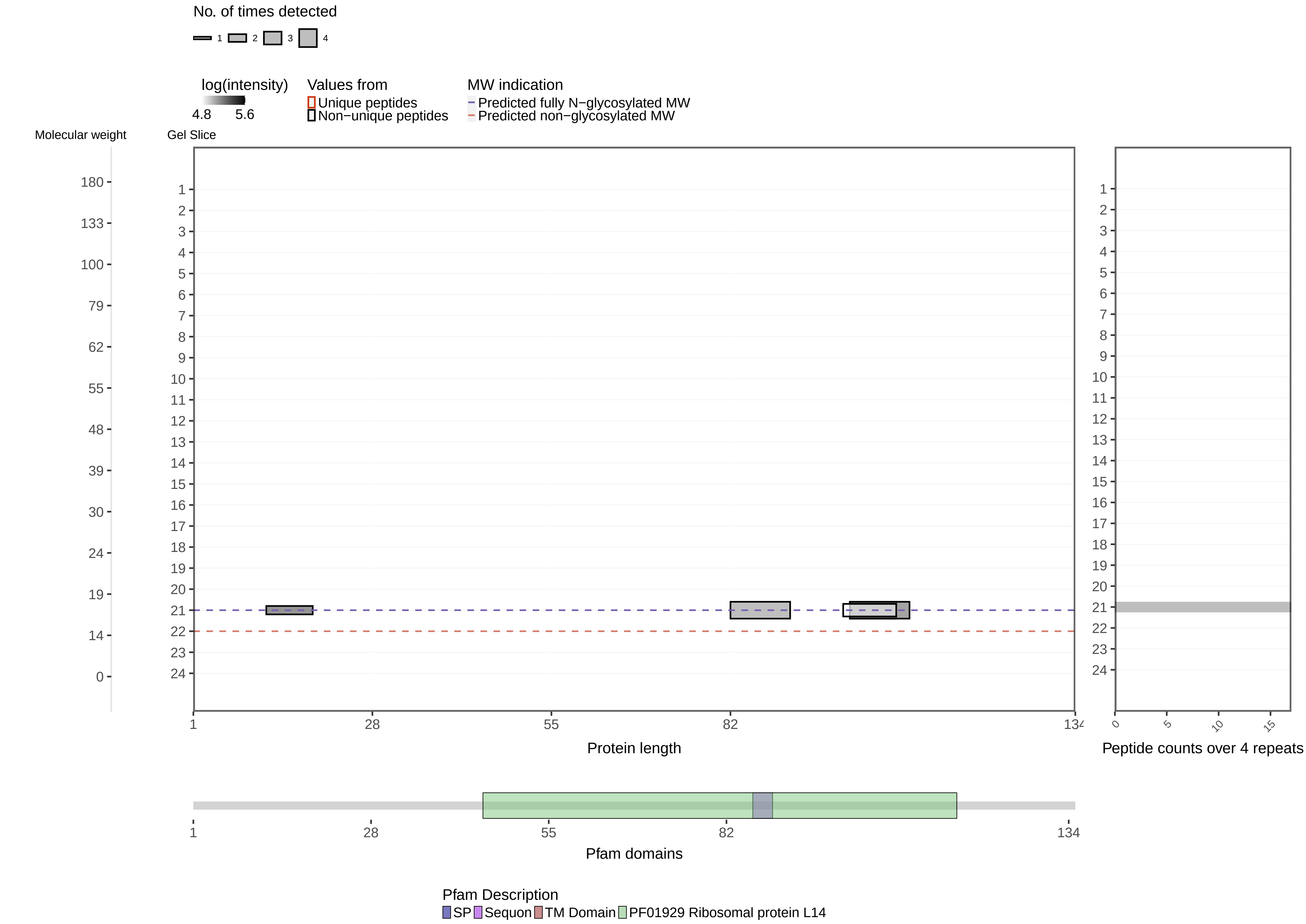


Protein sequence:

SLTDIKIDIKRIPKKKTLVAAMEAADVKTKWENSSWGRKLIVQKRRAALNDFDRFKLMLAKIKRAGVVRQELAKLKKTAA*

### NbL06g17650.1 cysteine−rich_repeat_secretory_protein_3−like [MW − SP = 28.85 kDa]

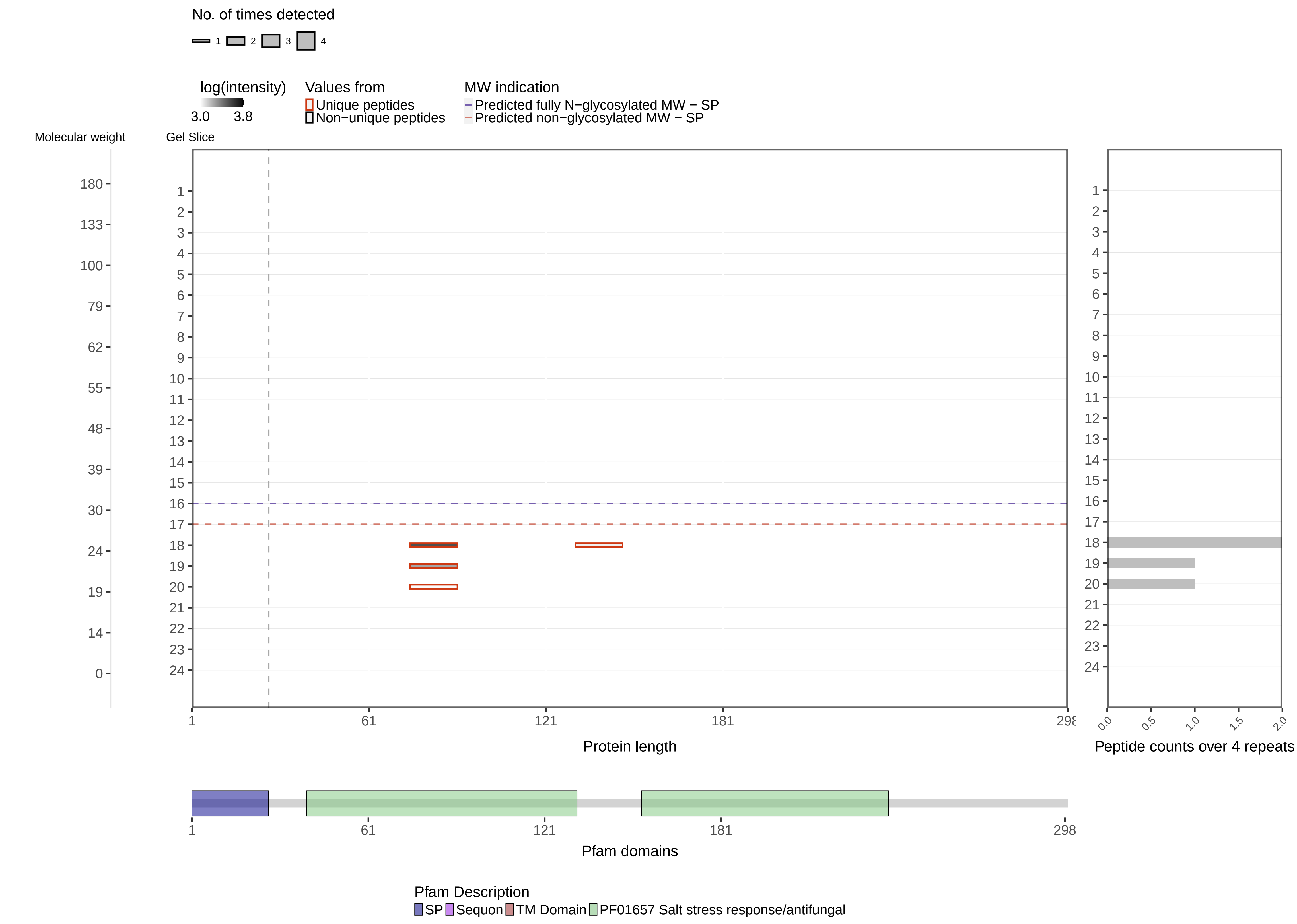


Protein sequence:

TVGPTELLYKVCASTQASGDGFLDRLETALGKIAKGVFSSGNKGFYTGGYQSVYVLGQCEGDLESGDCVNCVKNAAEEAKNQCGSSISAQIYLQQCYIRYTYYPSGVPTKSFSPSSGIRKSTQKTVAIVLGGLVGVGLVLACLLFTKSAFKKKSYVKYGG*

NbL06g17590.1 photosystem_II_reaction_center_Psb28_protein−like [MW = 19.96 kDa]

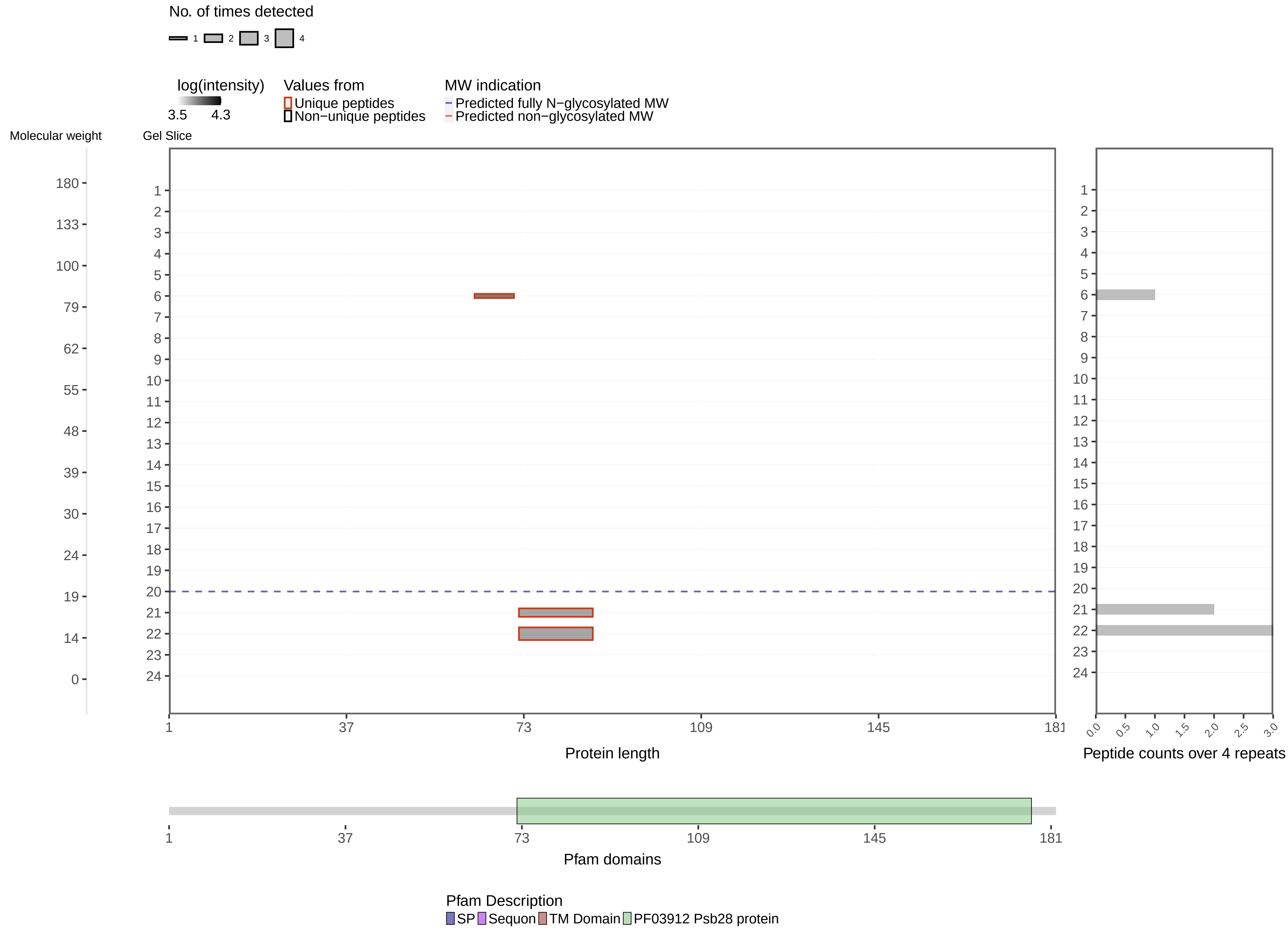


Protein sequence:

DEQTIPDVKLTKSRDGTNGMAIFRFDNPSVFDSSGEVGDITGFYMIDEEGVLSSVDVNAKFVNGKPAGVEAKYIMRTPREWDRFMRFMERYANANGLQFVKS*

### NbL06g17540.1 tyrosine_decarboxylase_1−like [MW = 40.62 kDa]

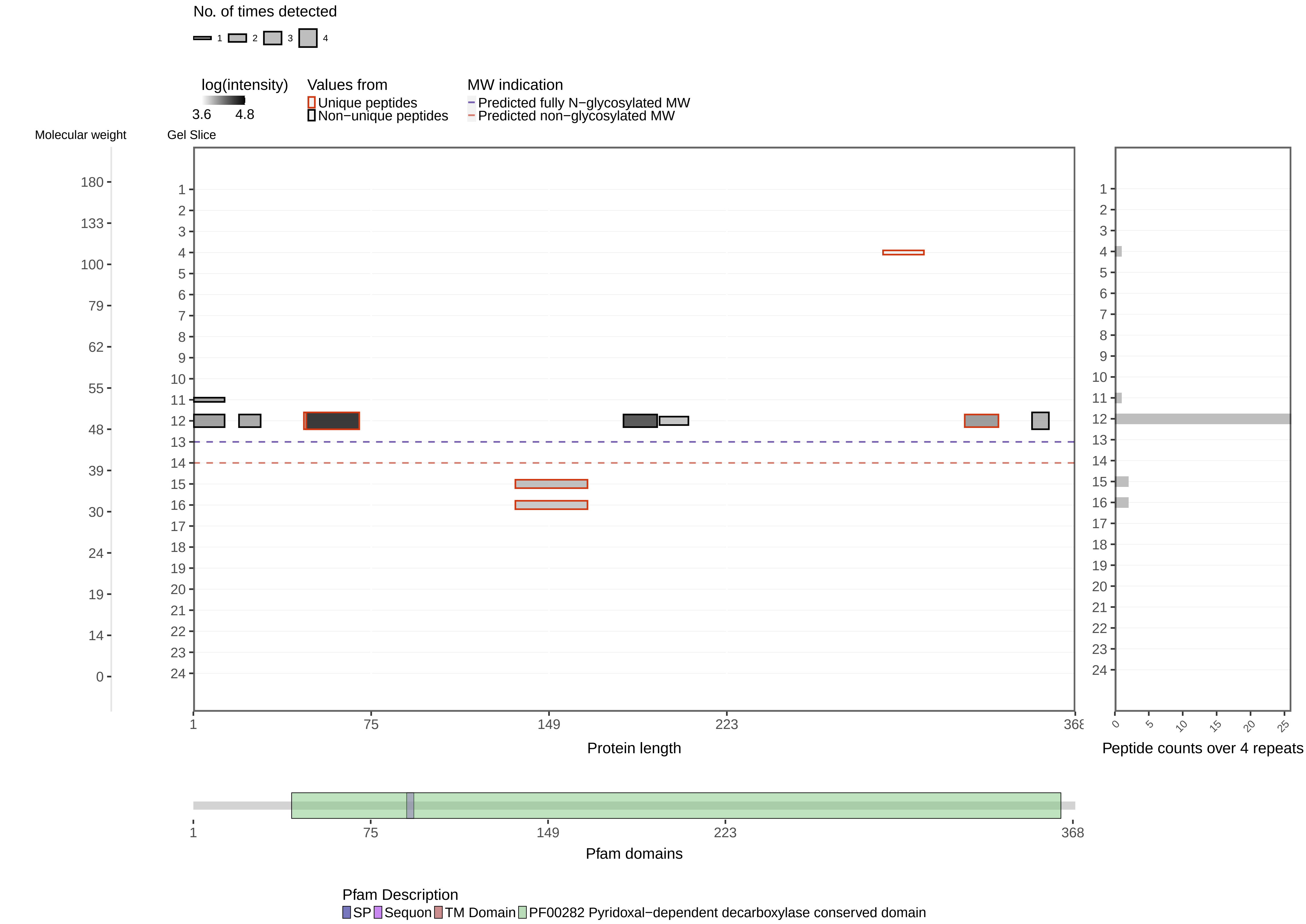


Protein sequence:

RVGKDAIGKLVVYCSDQTHSALQKACQIGGIHPENFRMLKAVSCRDYALSPDALSEAVSHDLATGLIPFFFCATIGTTSSTAVDPLLALGKITKSNSMWFHVDAAYAGSACICPEFRHYIDGVEEADSFNMNAHKWFLTNFDCSALWVKDRSALIQSLSTNPEFLKNKASQGNLVVDYKDWQIPLGRRFRSLKRGWY*

### NbL06g17510.1 alcohol_dehydrogenase_class−3−like [MW = 40.71 kDa]

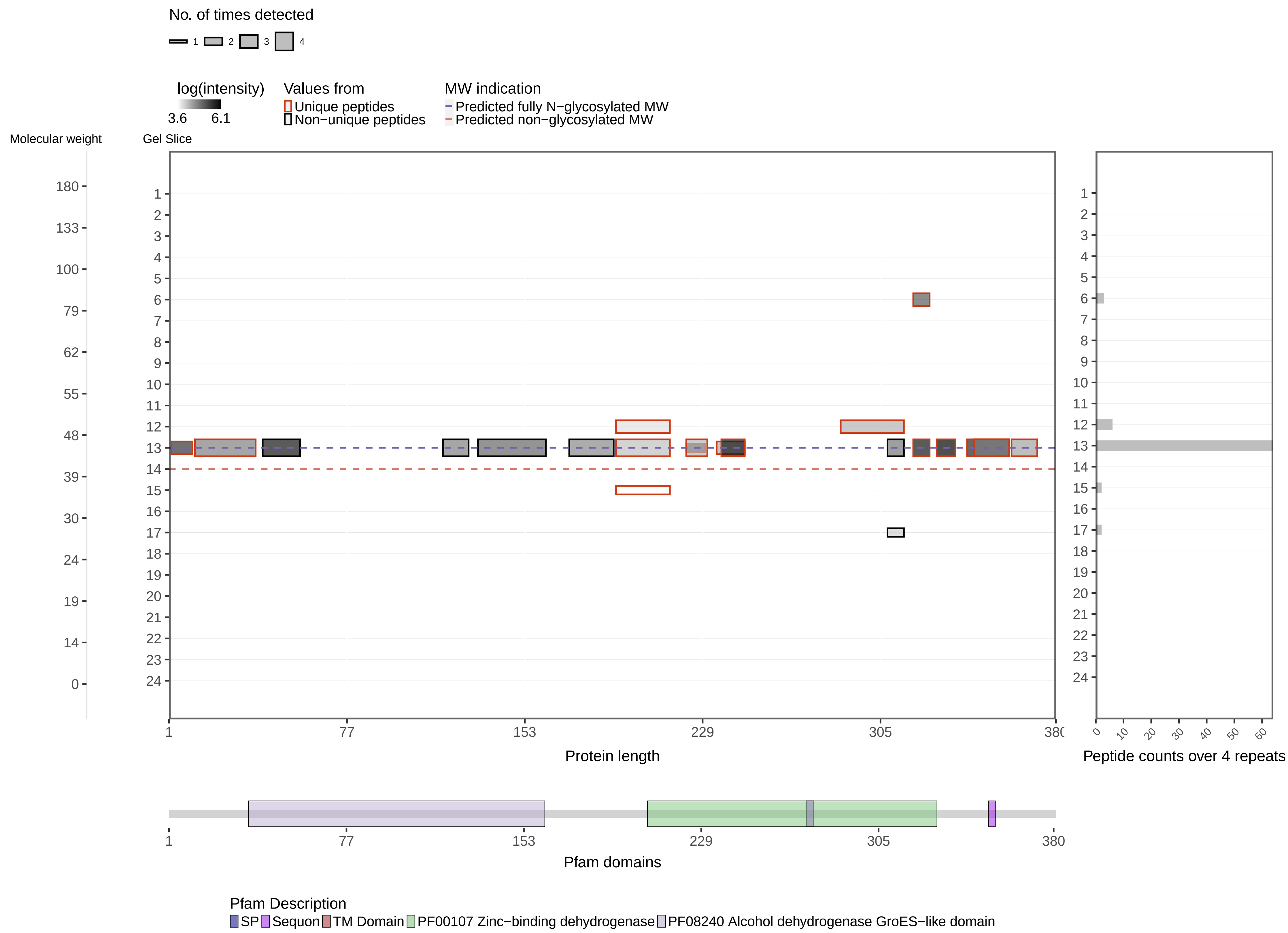


Protein sequence:

CGVPTGLGAVWNTAKVEPGSIVAVFGLGTVGLAVAEGAKAAGASRVIGIDIDSKKFDRAKNFGVTEFINPKEHEKPIQQVIVDLTDGGVDYSFECIGNVSVMRAALECCHKGWGTSVIVGVAASGQEISTRPFQLVTGRVWKGTAFGGFKSRSQVPWLVDKYLKKEIKVDEYITHNMTLADINKAFDLMHDGSCLRVVLDMFV*

### NbL06g17450.1 ribokinase−like [MW = 43.7 kDa]

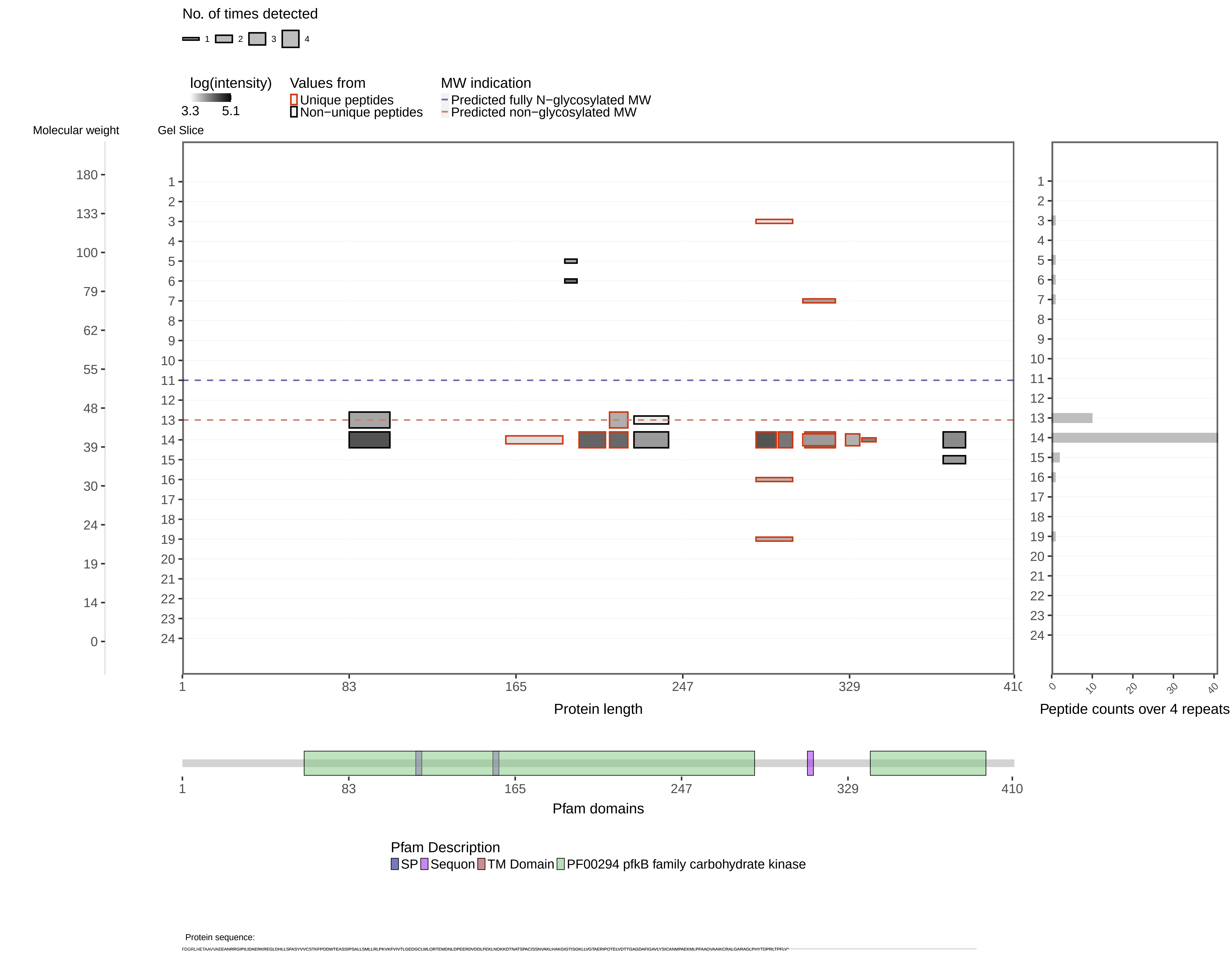

### NbL06g17350.1 GDSL_esterase/lipase_At4g28780−like [MW − SP = 37.22 kDa]

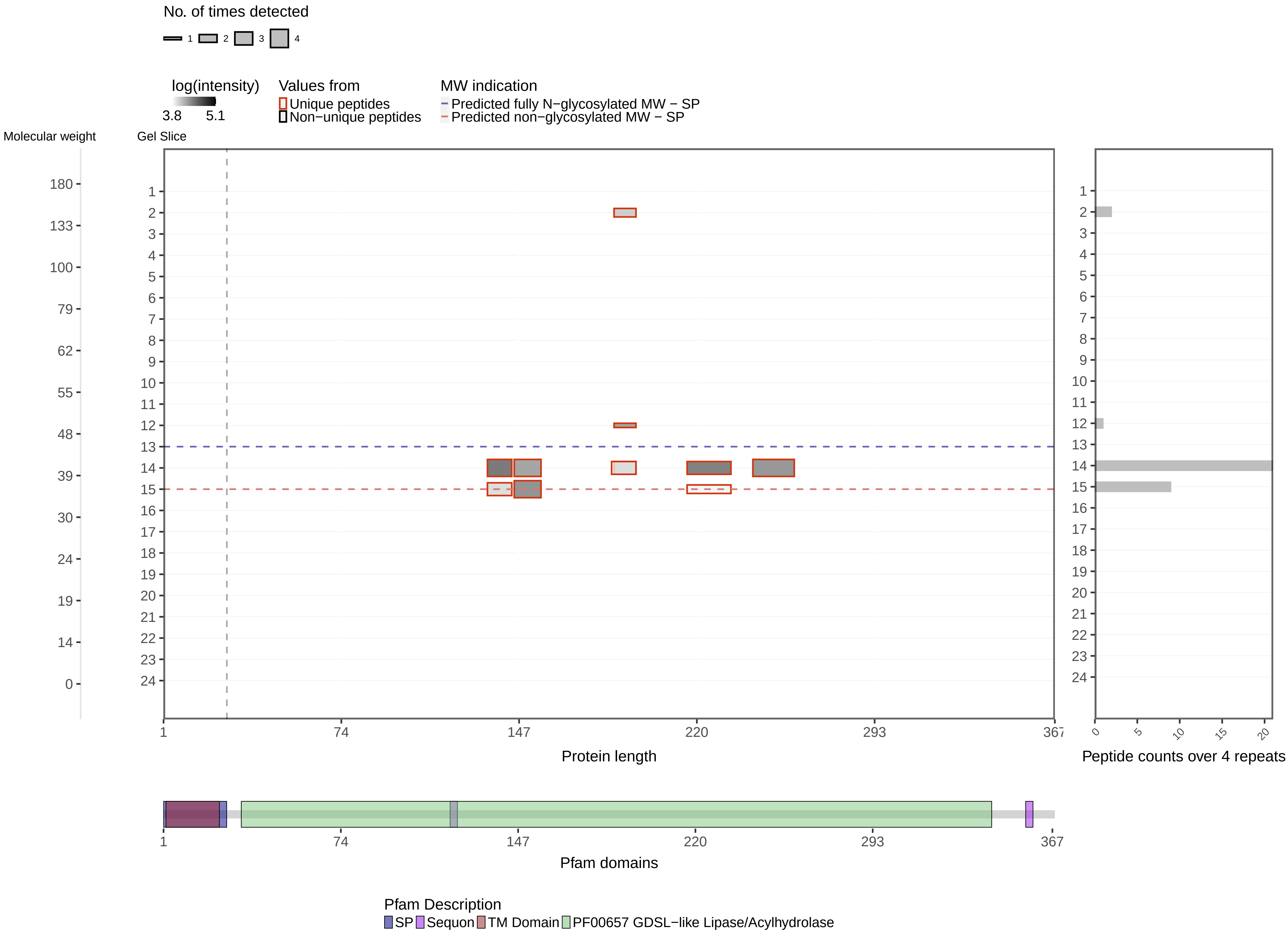


Protein sequence:

YVNNYFLTPVSARRIQYNIQDYSRYLVTEYRKILKRLHDLGGRRVLVTGTGPLGCVPAELATRSSNGECAKEPQQAAAAFNPLLVQMIKRLNQELGSDIFVAVNAMEMQNDFINNPKAFGFVTSKIACCGQGPYNGIGLCTAASNLCPNRDEYAFWDPFHPSEKANKIIVKTIYSGSDKYITPMNLSTIMAVESV*

### NbL06g17180.1 3−ketoacyl−CoA_thiolase_2_peroxisomal−like [MW = 49.08 kDa]

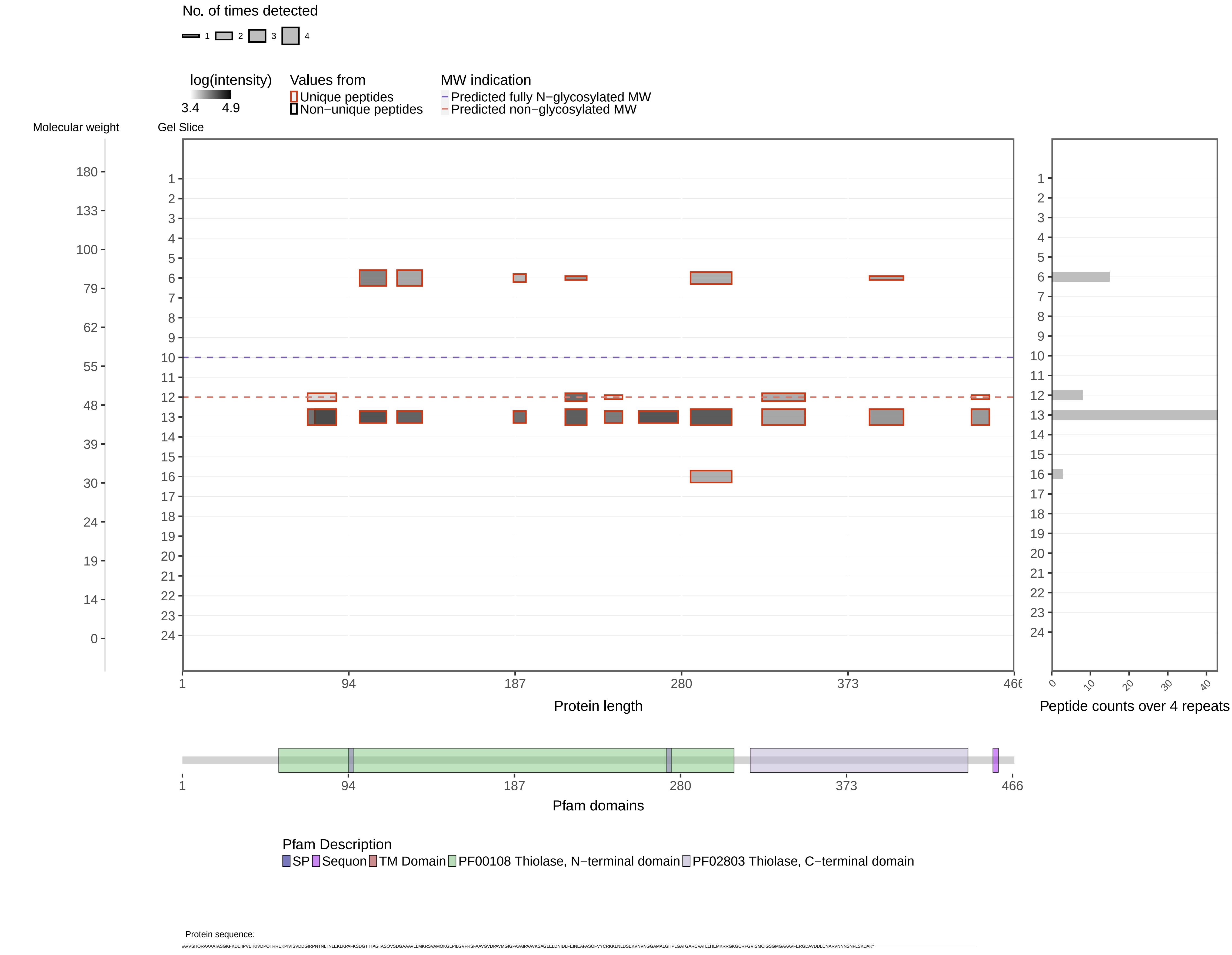

### NbL06g17110.1 pantoate−−beta−alanine_ligase [MW = 34.92 kDa]

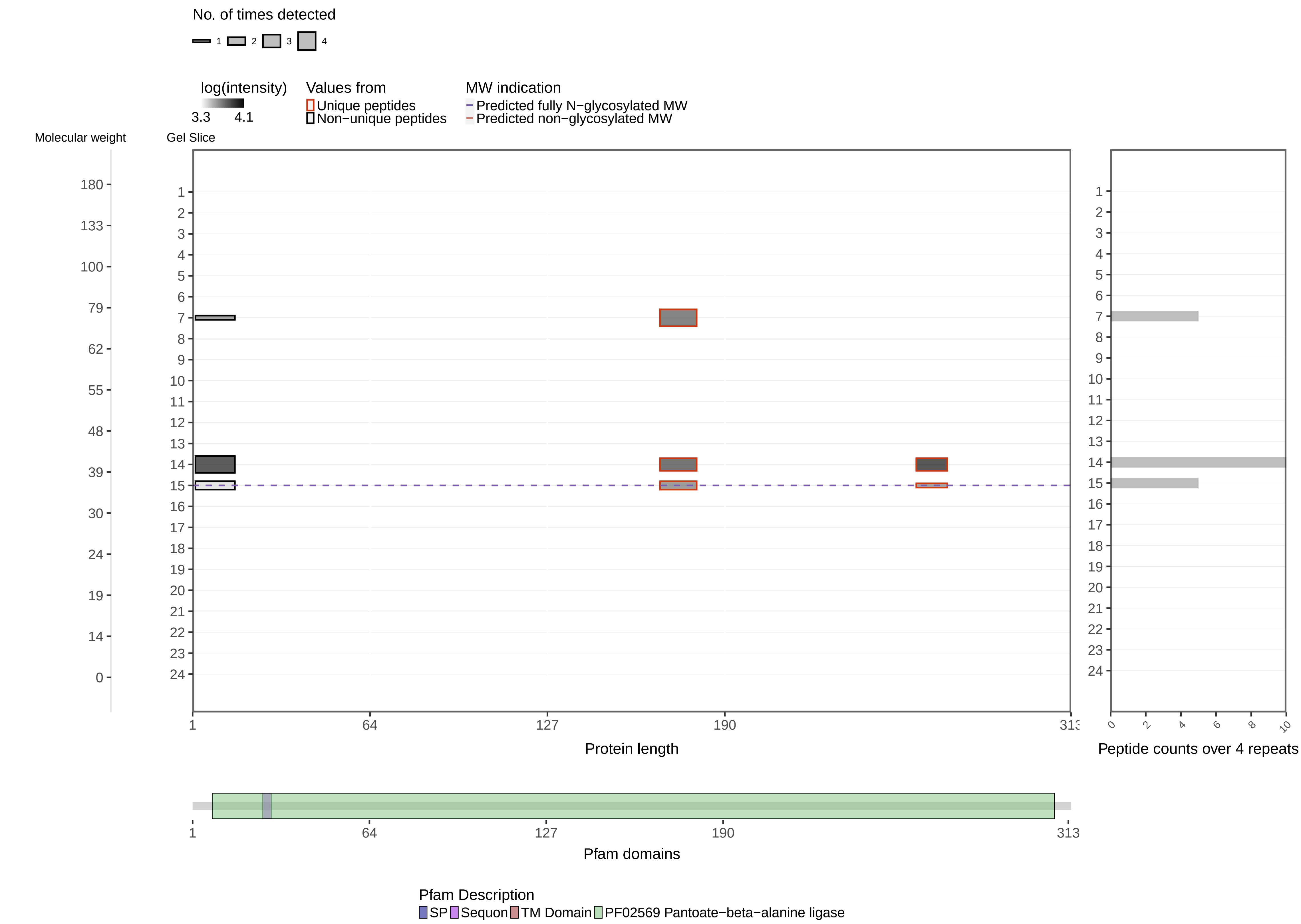


Protein sequence:

VENLEKGLCGKSRPVFFRGVATIVTKLFNIVEPDVAVFGKKDYOOWRIIORMVRDLDFGIKVIGSDIVRGHDGLAMSSRNVKLSPEDROKALSISRALSKAKVEAEKGOVNCGELIKSAIOTIDEAGGKVDYAEIVEOESLEPVETIKRPVVFCVAAWFGKVRLVDNMEINI*

NbL06g17090.1 uncharacterized_protein_At4g13200_chloroplastic−like [MW = 24.09 kDa]

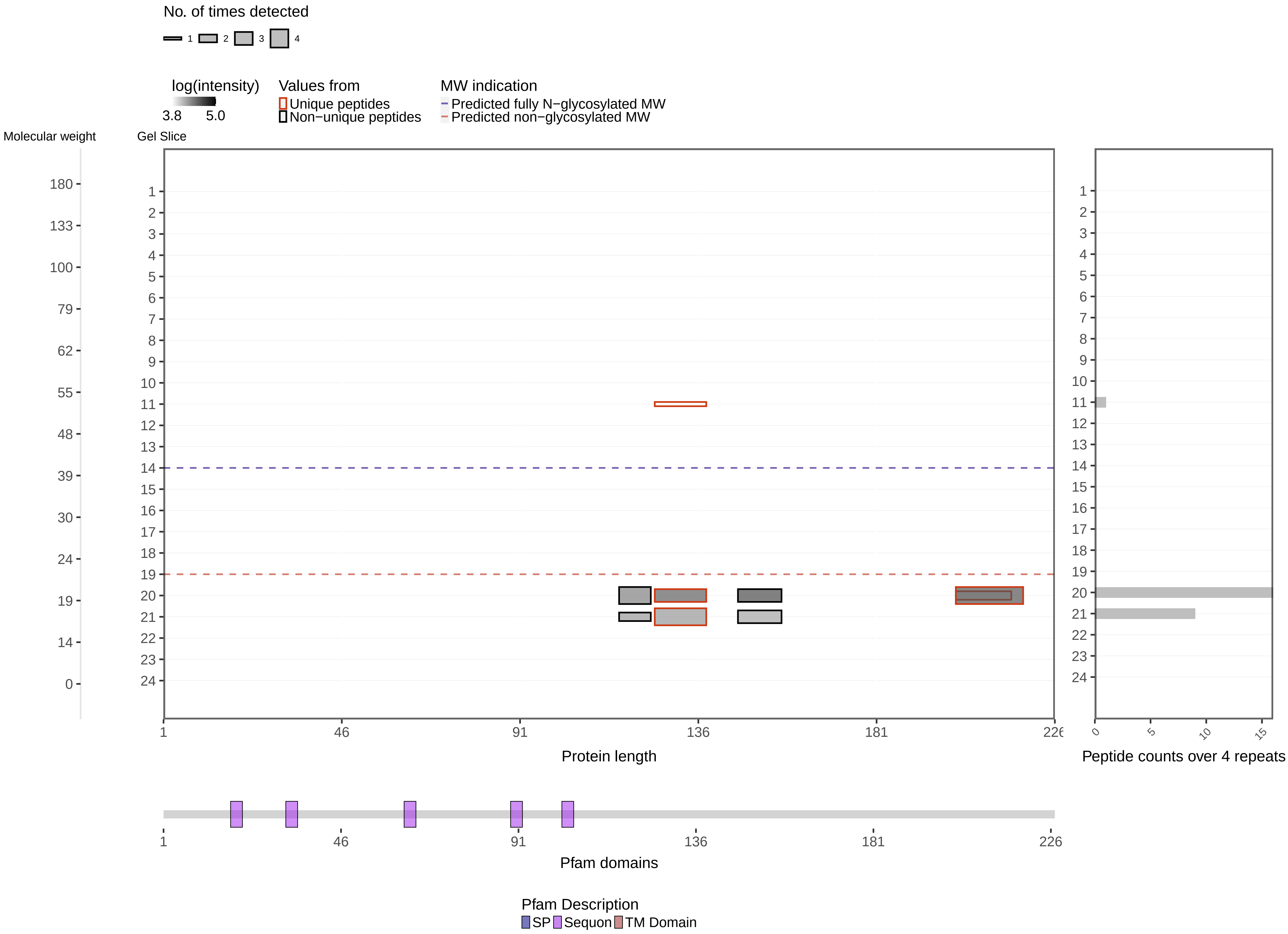


Protein sequence:

ENESKNILDAFFLGKALAEAVTERIESTVGEFLGTVGRLQAEQQKQVQDFQEEVLERAKQAKEKAARETMDTQGLIPNSFTADTSTAGTGVTSATSQQTRINSQPAMIDSEDGSKDSKLGLSNGD*

### NbL06g16870.1 small_ubiquitin−related_modifier_1−like [MW = 11.39 kDa]

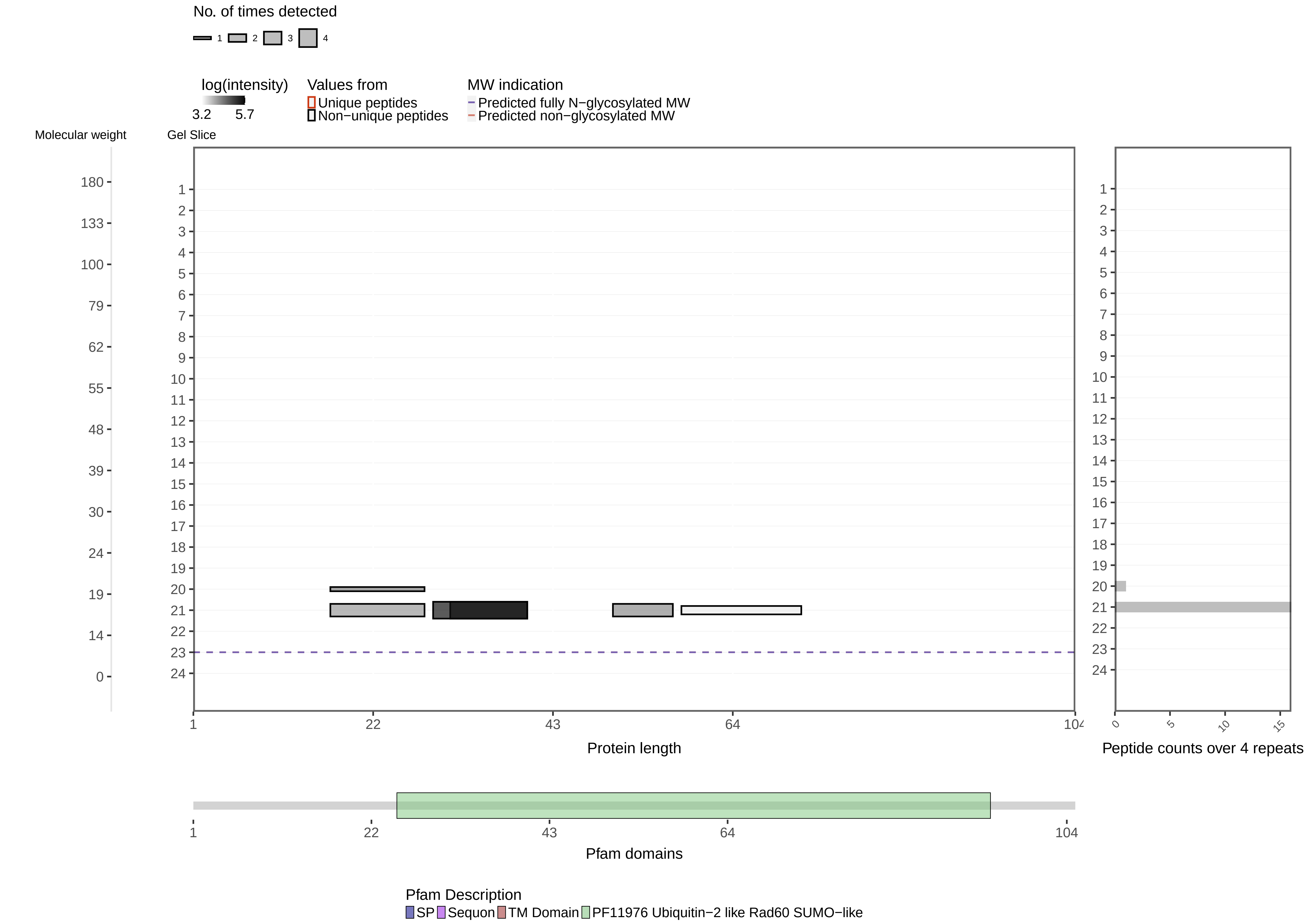


Protein sequence:

FRIKRNTQLRKLMNAYCDRQSVDMNSIAFLFDGRRLKGEQTPDELEMEEGDEIDAMLHQTGGNDA*

### NbL06g16860.1 uncharacterized_LOC109228095 [MW = 65.48 kDa]

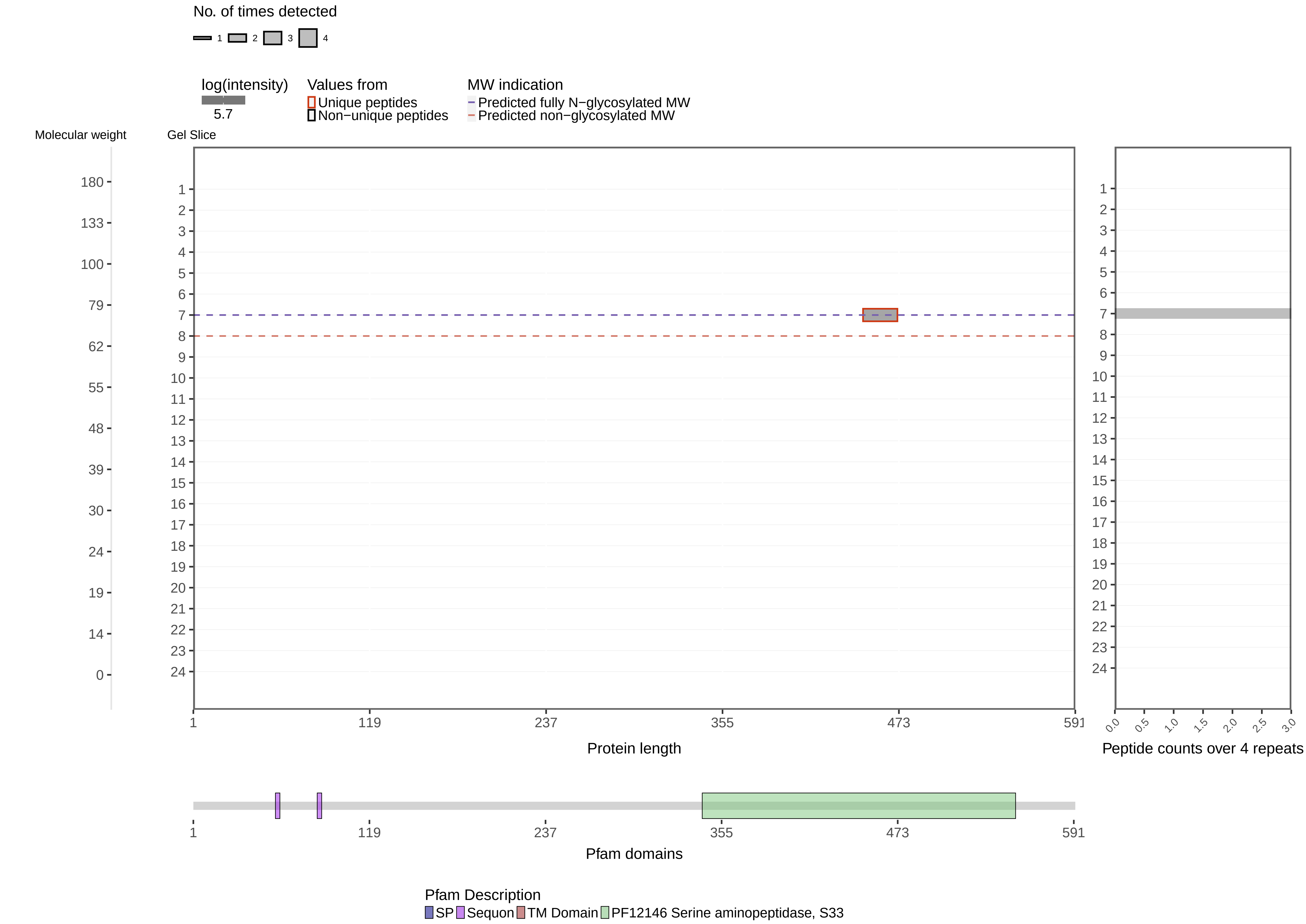


Protein sequence:

TREKSAVLTSQAGELSKKLVNIFEEGQRSVSSPLFDLQERLTTRIGDFQEQLDLILKYDWDFDHYLEEDVPAAMEYIKVQTRPKDDKLLAIGHSMGGILLYARLSWCGLEGREPGLAAVITLASSLDYTSSKSALKLLLPLADPAQVLNVPVIPLRTLLAAAYPLSSRAPYVLAWLNEMISATNMMHPDSLKKLVQNSFCNIPAKLLLQLATAFQEQGLRDRSGKIYYKDHLRKSSVPVMALAGDLDQICPPEAVYETVKLIPENLVTYKVFGDADGPHYAHYDLVGGHMATEQLYPC

### NbL06g16850.1 SEC23−interacting_protein−like [MW = 24.62 kDa]

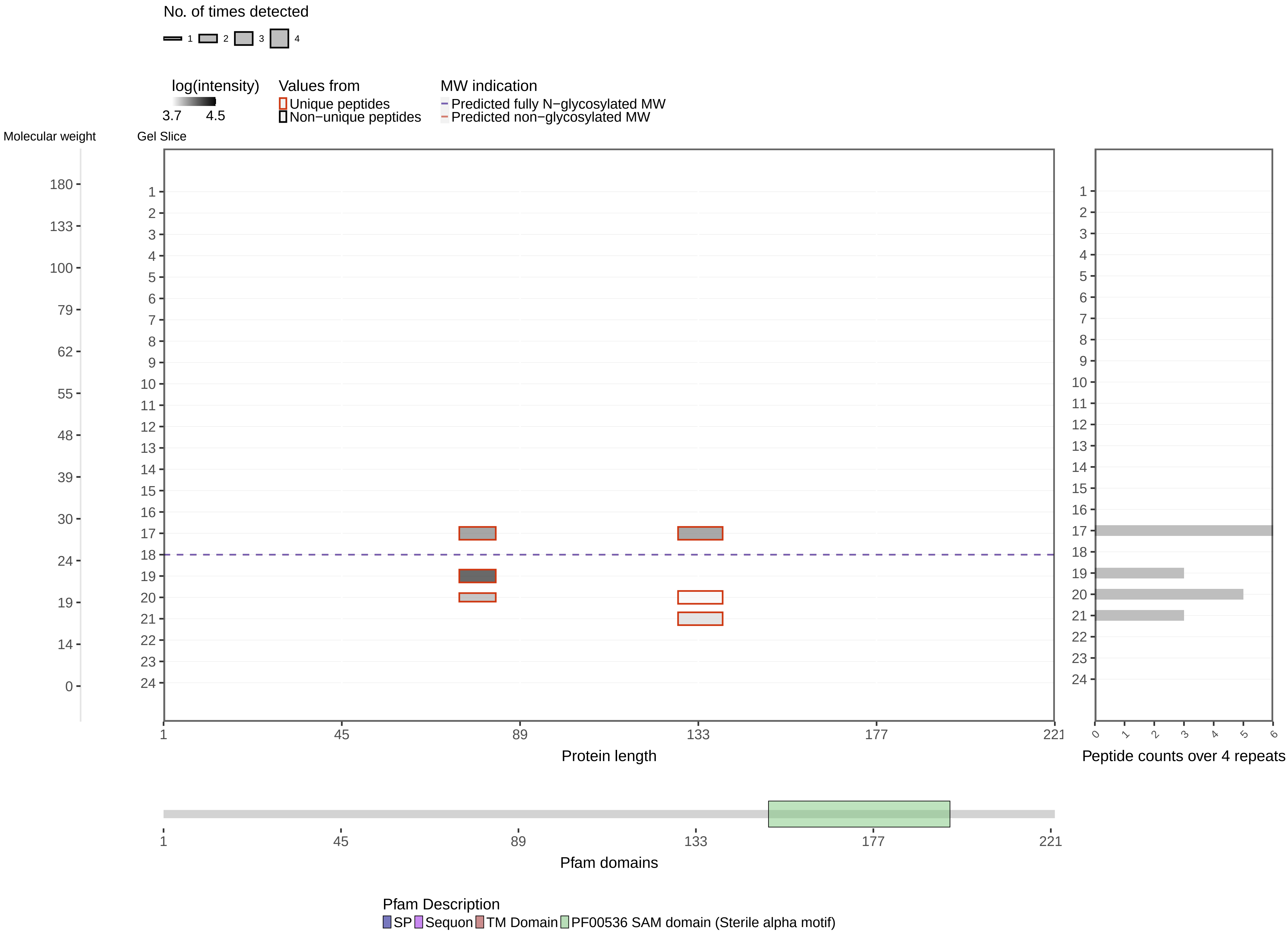


Protein sequence:

KLSGTVYSQTVENVPPKAKLKVAPEISKPVRRSVIAEAPAMETKKVASTVSKKKSQQKVQSVDSFLQSLGLEKYAITFQAEEVDMAALVHMTDEDLKAMGIPMVSIVMLYENSFEIVTSCFSFAS*

### NbL06g16510.1 peptidyl−prolyl_cis−trans_isomerase_FKBP15−1−like [MW = 23.15 kDa]

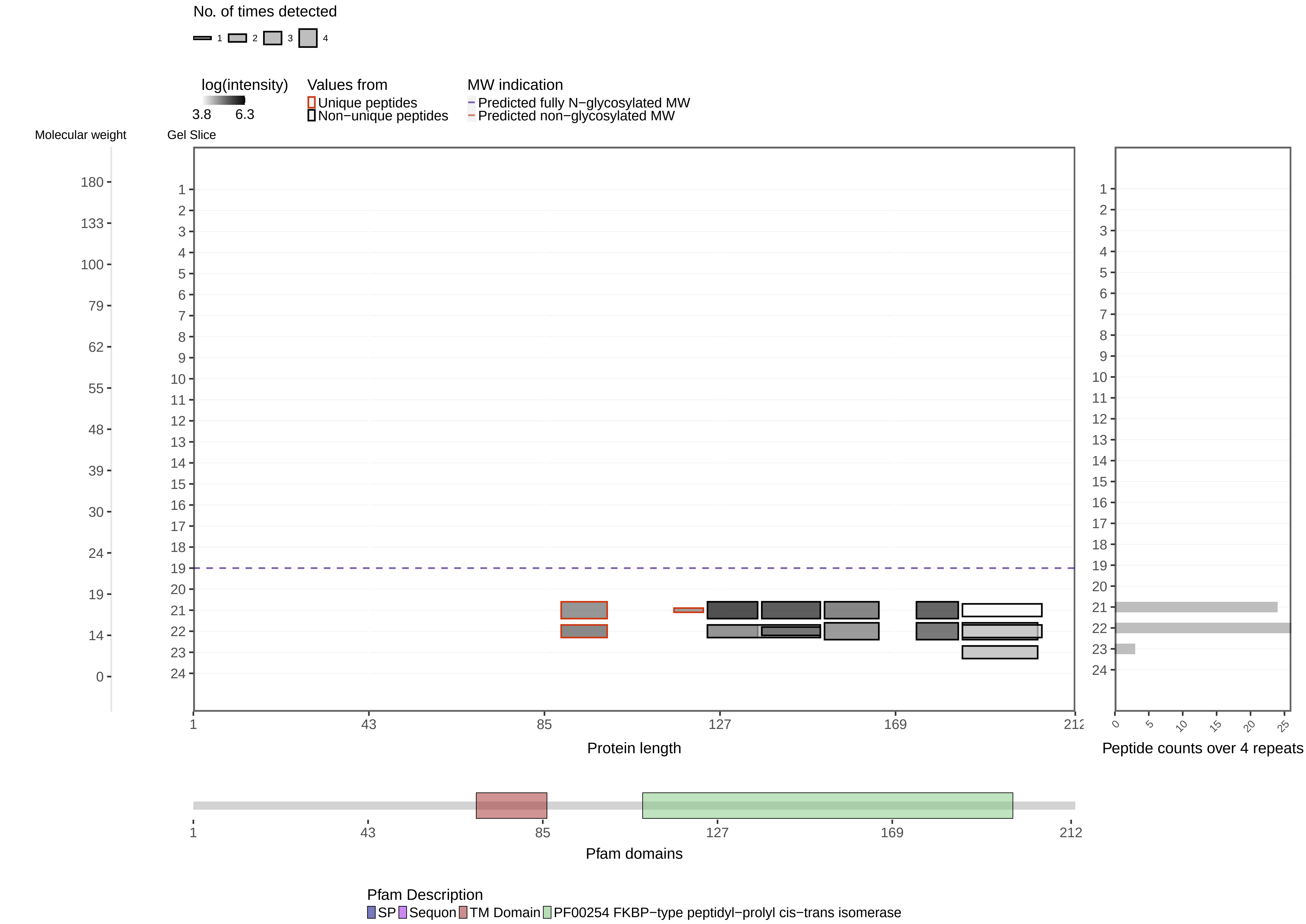


Protein sequence:

ELQIGVKFKPKSCELKAHKGDRVSVQYSGKLTDGTVFDSSYERNDPIEFELGSGQVIKGWDQGLLGMCVGEKRKLKIPAKLGYGESGSPPKIPGGATLVFDTELVAVNGKKSAADSEL*

NbL06g16490.1 40S_ribosomal_protein_S8 [MW = 25.11 kDa]

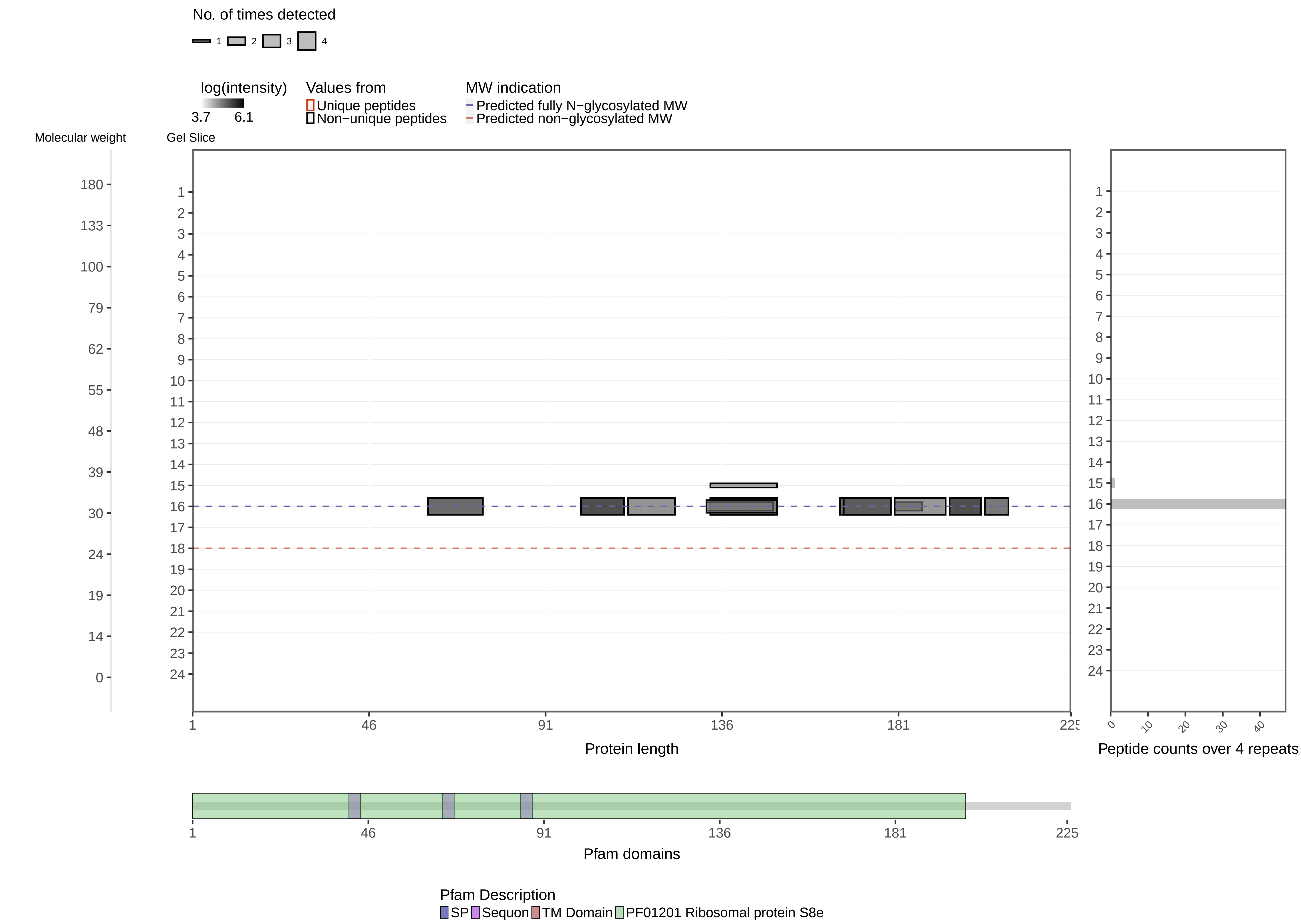


Protein sequence:

SAIVQVDAAPFKQWYLQHYGVDIGRKKKGPAKKETTEEGEGAAAAAEETKKSNHVLRKIETRQKDRKLDPHIEEQFGGGRLLACISSRPGQCGRADGYILEGKELEFYMKKLQKKKGKAGAGGAA*

NbL06g16380.1 cytochrome_b561_and_DOMON_domain−containing_protein_At3g25290−like [MW − SP = 23.16 kDa]

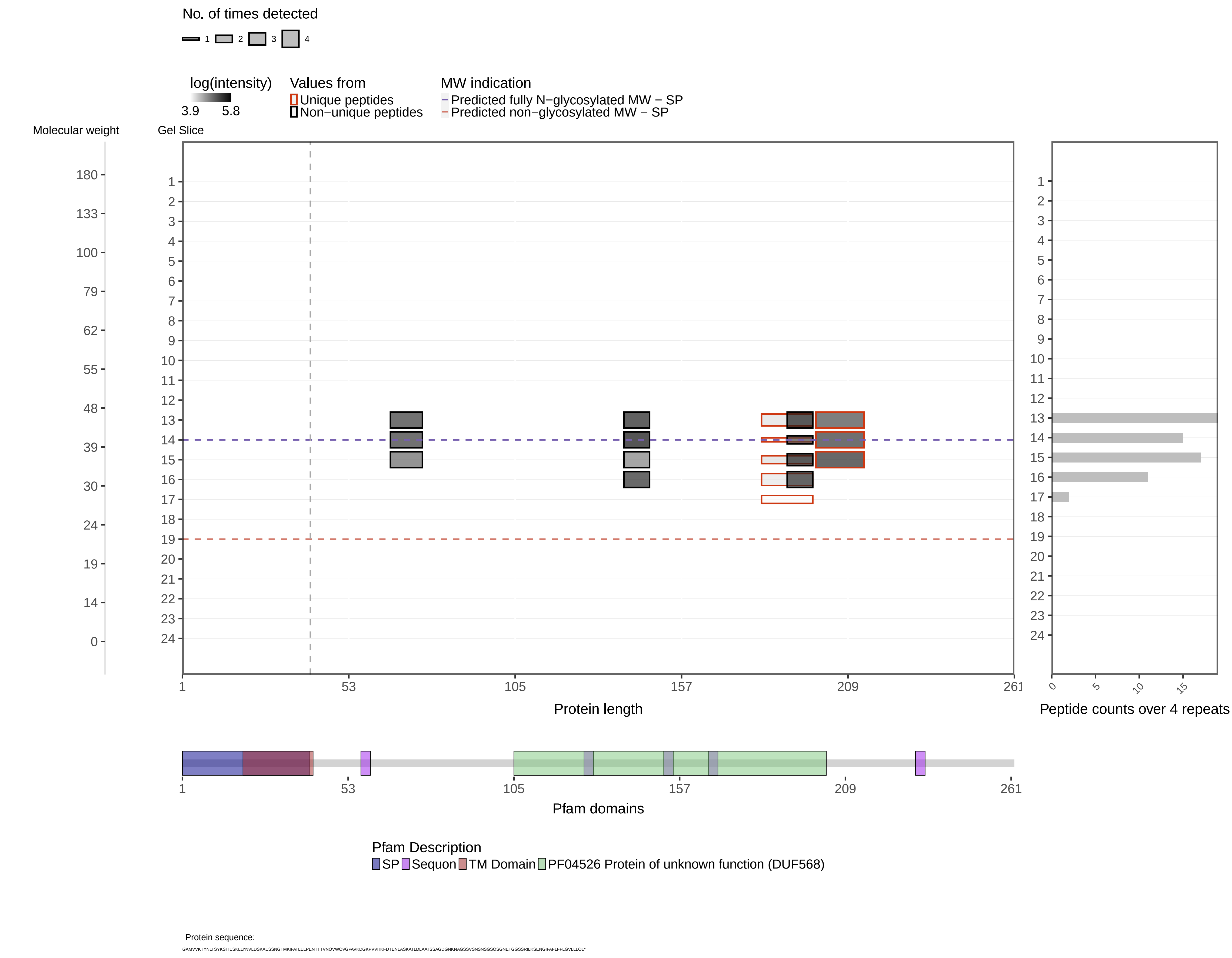

NbL06g16360.1 cytochrome_b561_and_DOMON_domain−containing_protein_At3g25290−like [MW − SP = 39.53 kDa]

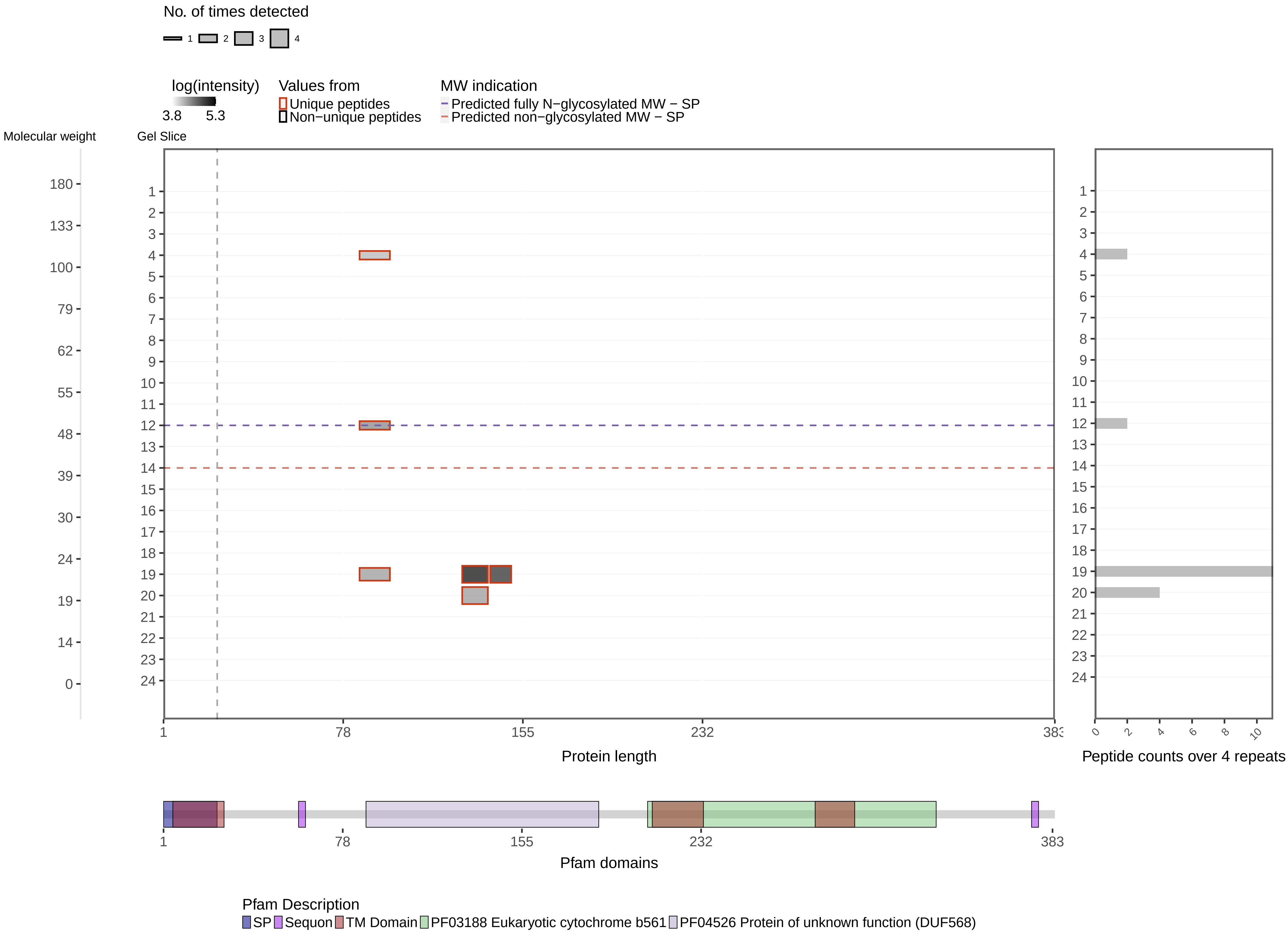


Protein sequence:

QPANLNSKGTLDLLSGESKSGASGDSKVKRKNIHGILNAISWGILFPVGIIIARYLRTFKSADPAWFYLHVSCQCSAYIIGVAGWATGLKLGSESKGITYDTHRNIGIALFSLATLQVFALLLRPKKDNKYRFYWNIYHHGVGYTMLVLSLINVFKGLDILDPATKWRSAYIGILVVLGAIALLLEVITWIVVLKRKSNKSTKPYDG*

### NbL06g16230.1 uncharacterized_protein_At5g48480−like [MW = 16.86 kDa]

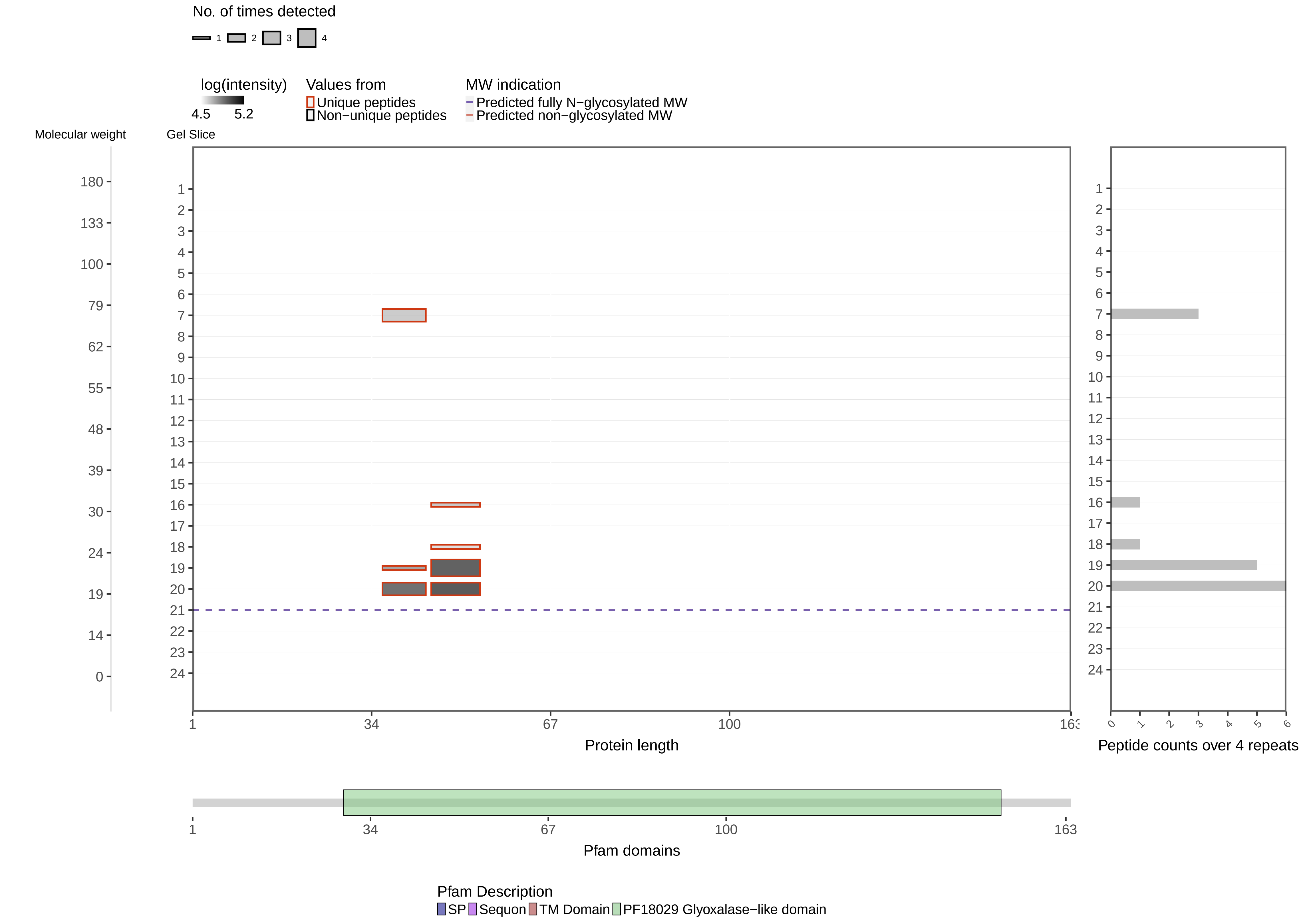


Protein sequence:

LILSAELKLGSTTFLVSDSDLTDDSSAPVKTAGGYVFCLETEDVEGAVAMAVTAGAVSEGEIAEGDGACCGGRVGKLKDPYGNIWLICSPSKKCE*

### NbL06g16190.1 LOC107783758 [MW − SP = 19.43 kDa]

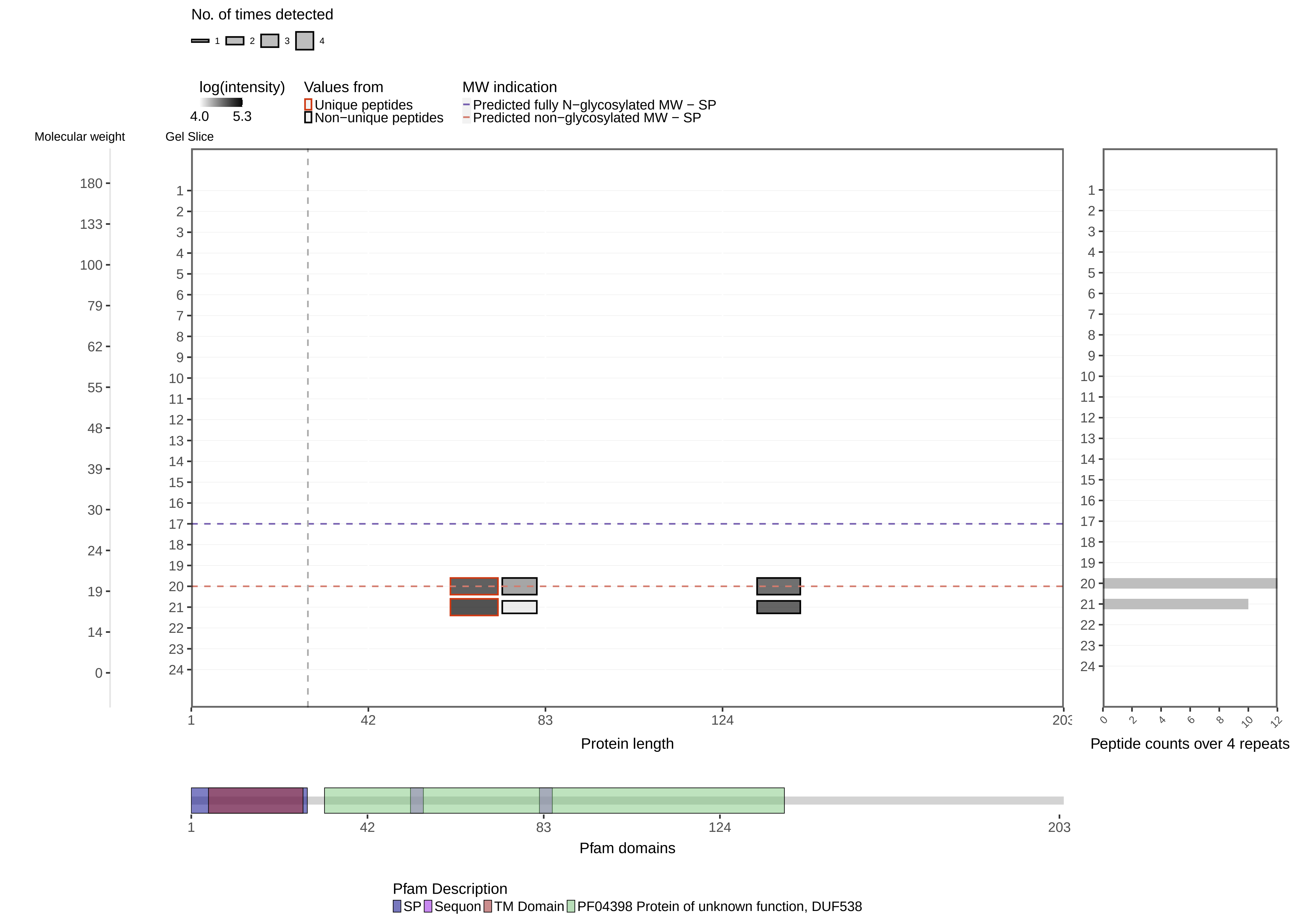


Protein sequence:

YGQIHALSGIEAQDLFLWFPVKDIRVDIPSSGLIYFNVGVVSKQFSLSSFETPRDCTAVQLMDLQDDMHIADAASKTVRLHLYVPCNSGTLAKSQSEELRYKFDQGNMGRKAIL*

NbL06g16180.1 monocopper_oxidase−like_protein_SKU5 [MW − SP = 63.38 kDa]

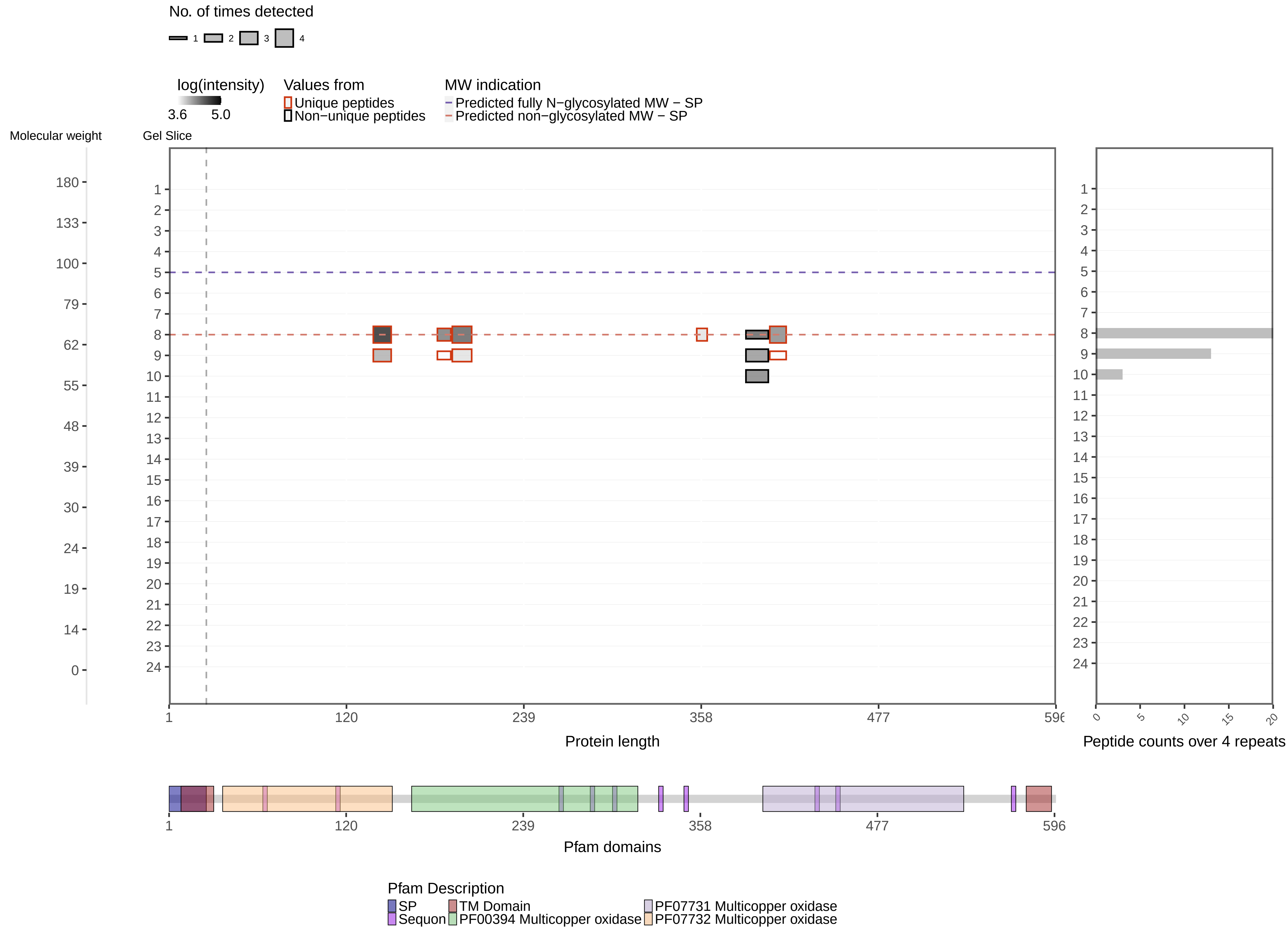


Protein sequence:

### NbL06g16060.1 ribulose_bisphosphate_carboxylase_large_chain [MW = 29.58 kDa]

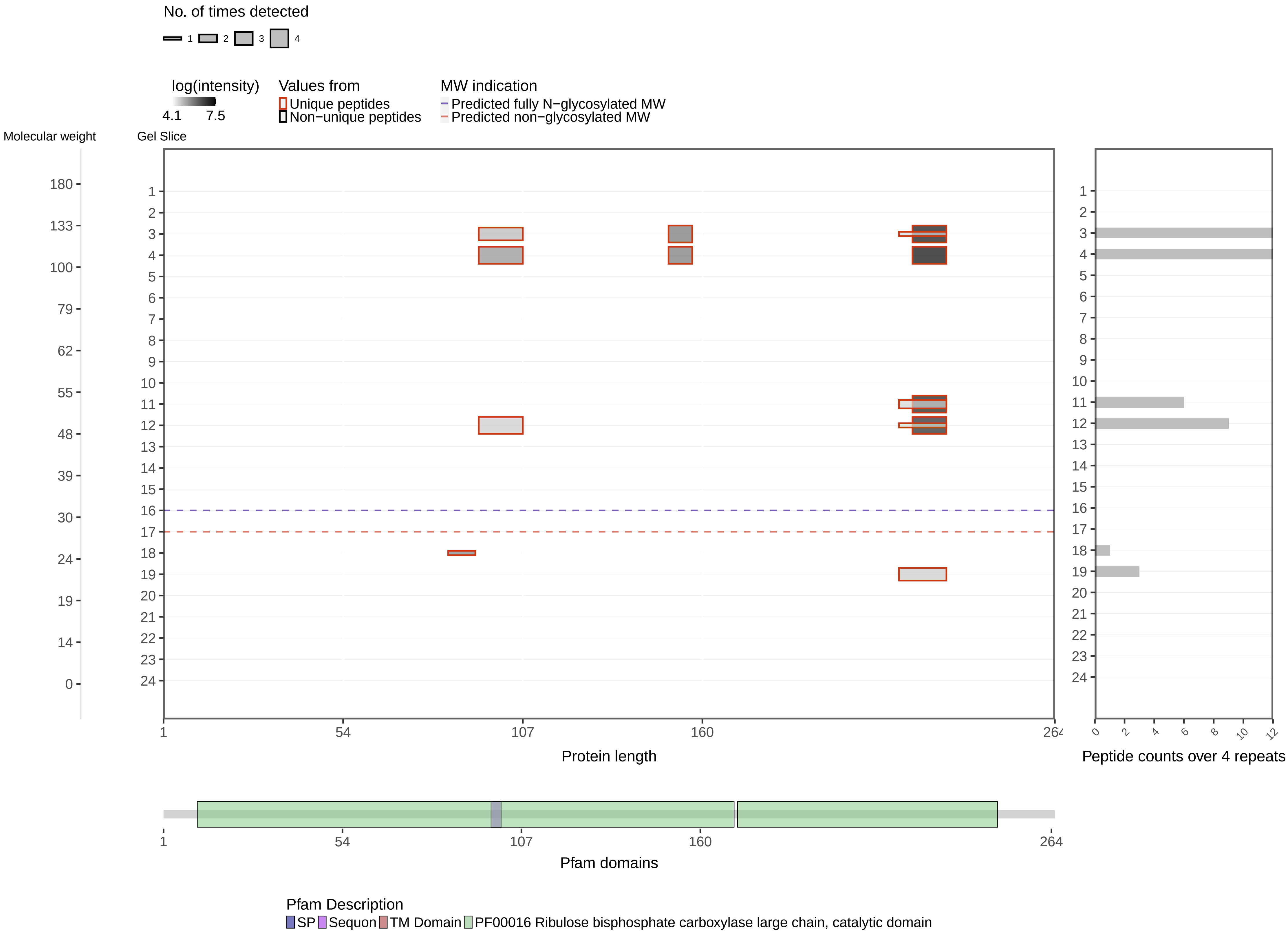


Protein sequence:

/MHDYLTGEFTANTGLAHYFQDNGLLFHIHRAMHEVIDRQKNHGIHFREVSRLALPALTEIFGDDSVLHFGGGTLGHPWGNAPGAVANRVALEACVKARNEGRDLAQEGNEIIREDSEWSPELDDACEVWKEIVFNFAASDVLDK*

NbL06g15950.1 33_kDa_ribonucleoprotein_chloroplastic [MW = 35.34 kDa]

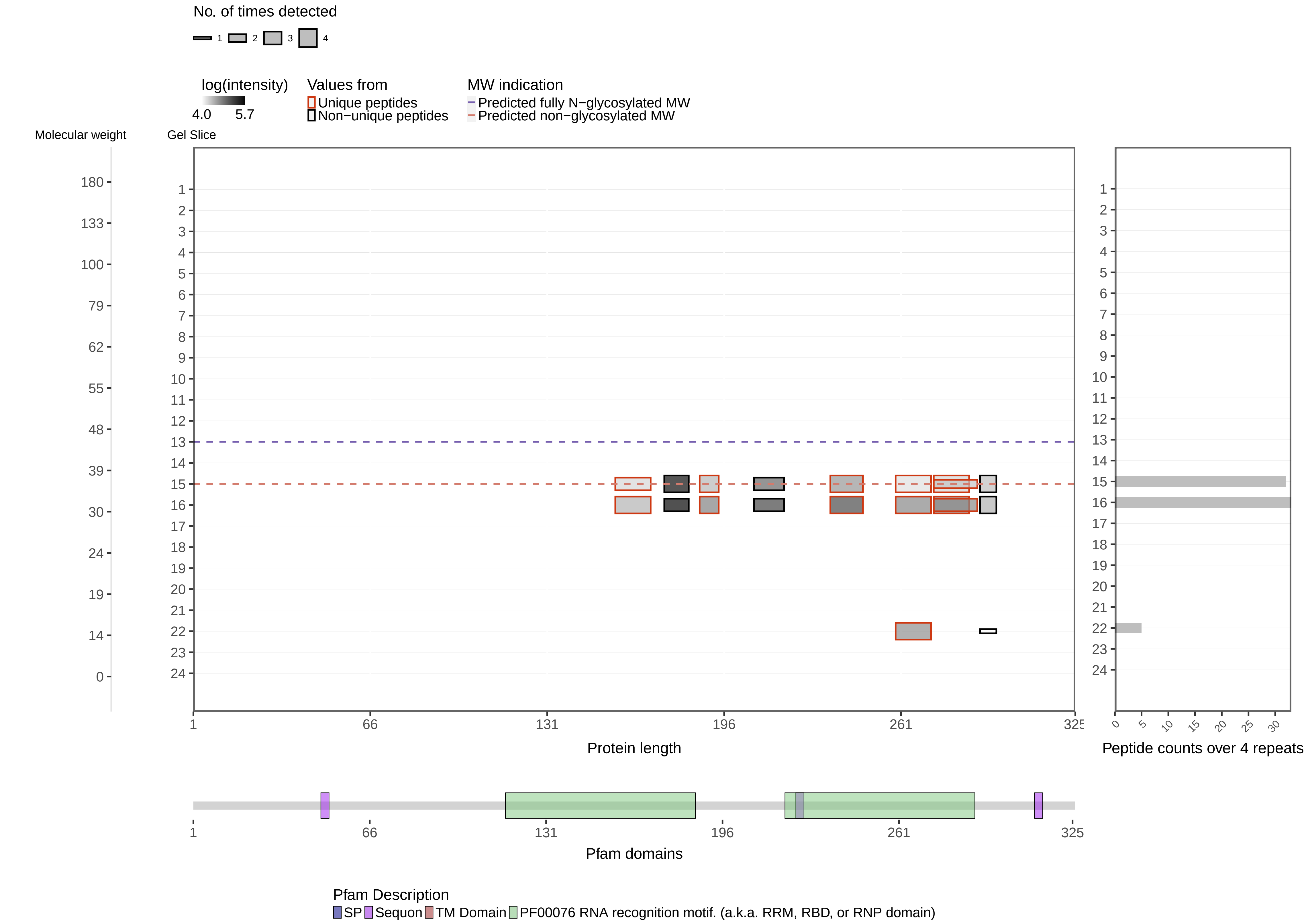


Protein sequence:
RSRGFAFVTMGSVEEAKEAIRLFDGSQVGGRTVKVNFPEVPKGGEREVMSAKIRSTYQGFVDSPHKLYVANLSWALTSQGLRDAFVDQPGFMSAKVIYDRSSGRSRGFGFITFSSAEAMKSALDTMNEVELEGRPLRLNVAGQKASLSSPPLVETSPENDSENSELLSSLSS*

NbL06g15910.1 putative_phosphatidylglycerol/phosphatidylinositol_transfer_protein_DDB_G0282179 [MW − SP = 14.46 kDa]

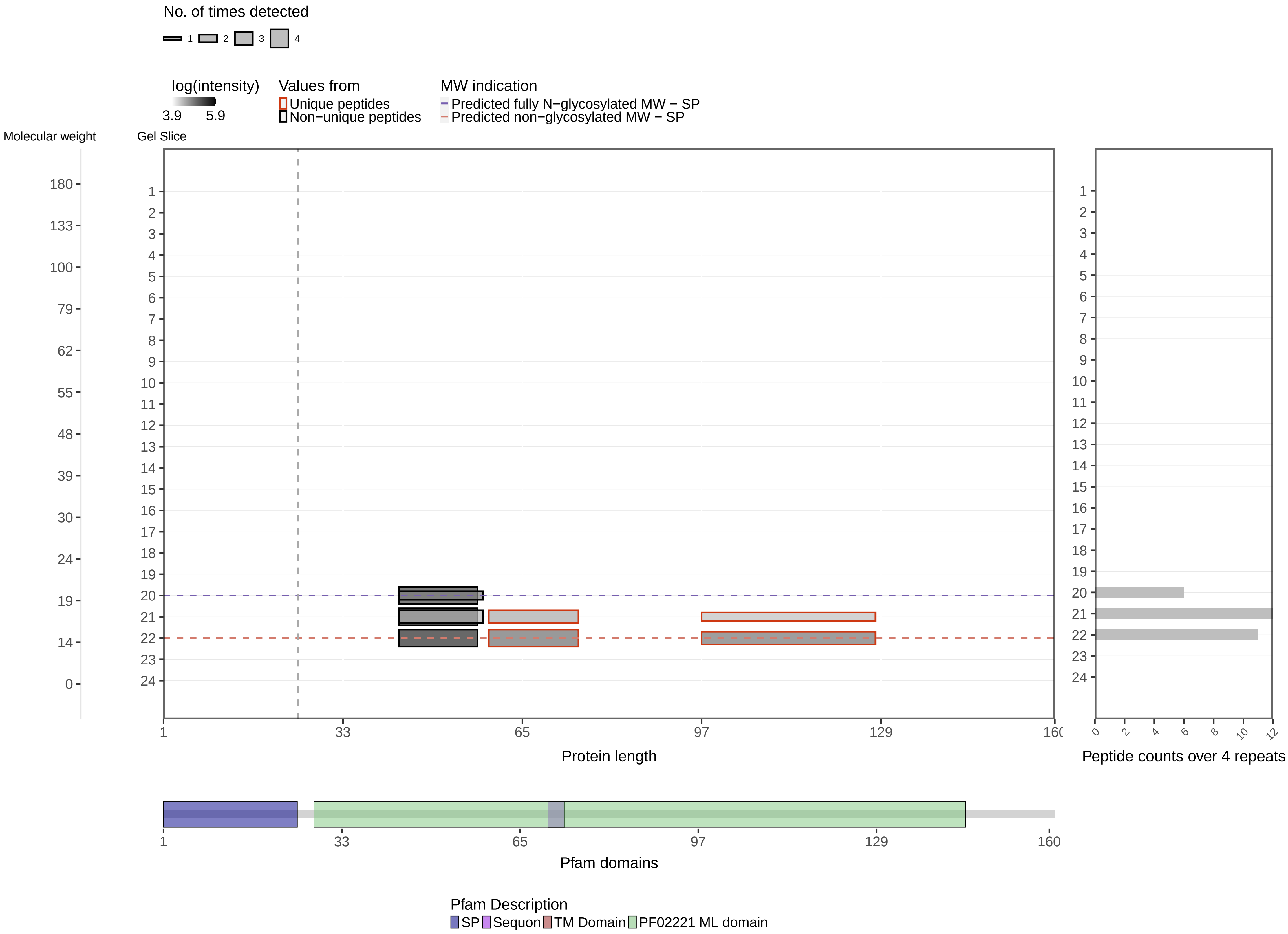


Protein sequence:

GQNLTGGKLVIDVNYLFFHVHHEAIDLCKETSCPASGDFVISHSQELPGFTPPGSYTLTMKMVDDKNHQLSCISFGFSIGFIAESQALAADS*

NbL06g15810.1 fasciclin−like_arabinogalactan_protein_17 [MW − SP = 46.83 kDa]

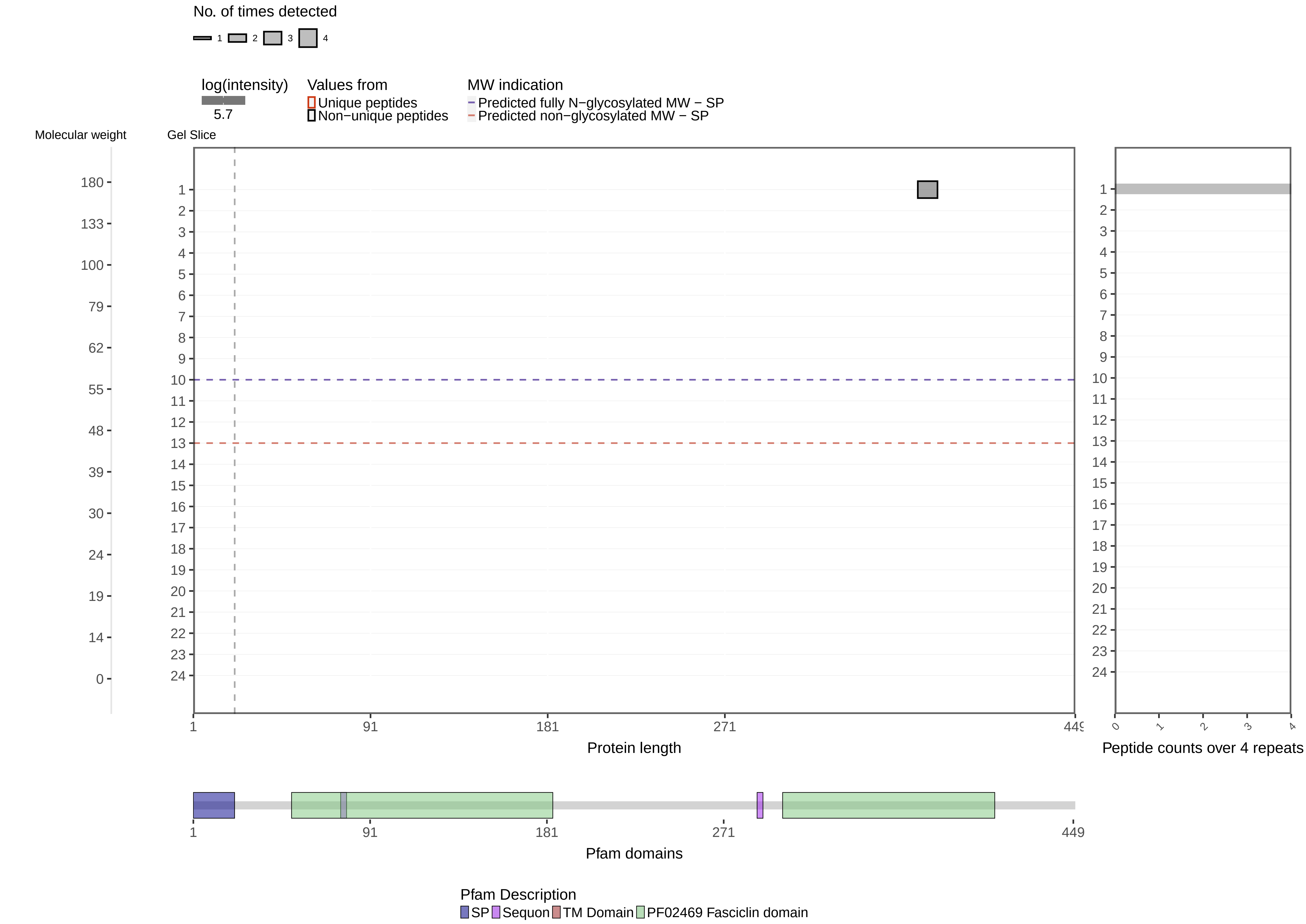


Protein sequence:

TNRLKKKPTPVPAGAPPVLPIYSAMAPGPSLAPAPAPGPGGPHHHFDGESQVKDFIQTLLHYGGYNELADILVNLTSLATEMGRLVSEGYVLTVLAPNDEAMAKLTTDQLSEPGAPEQIMYYHLIPEYQTEESMYNAVRRFGKVKYDTLRLPHKVVAEEADGSVKFGVAEGSAYLFDPDIYTDGRISVQGIDGVLFPVEEIKAAPVAAPVAKVVAKPRRGKLMEVACSVMGSFC*

NbL06g15640.1 histone_H1−like [MW = 31.58 kDa]

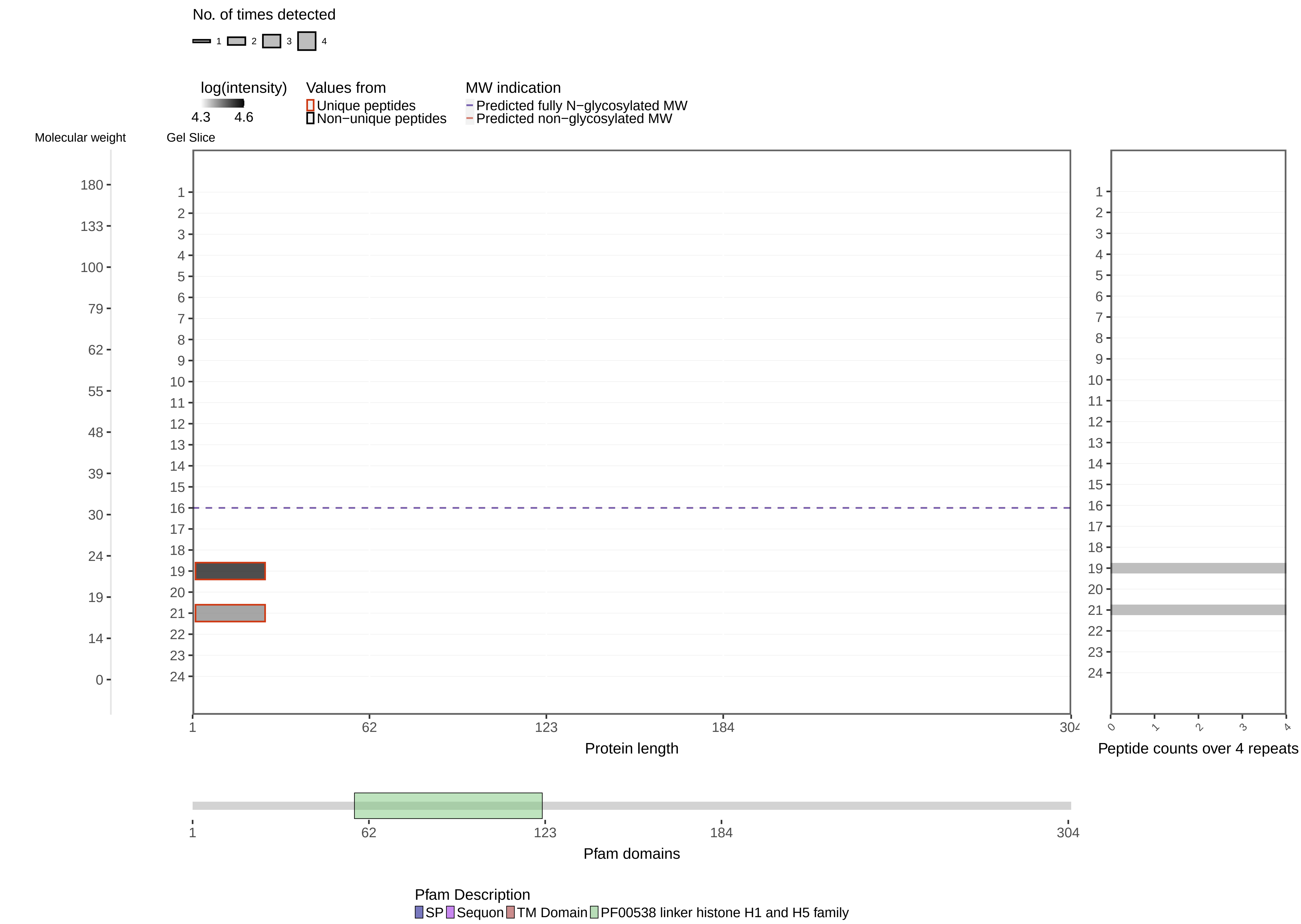


Protein sequence:

'AAKPKAAKAKPAGKPKAKPAGKPKAKPAAKPKAKPAAKPKAKPAAKAKPAAKAKPAAKAKPAAKAKPAAKPKPAAKAKPAAKPKPKAAVAPKPKATPVKAKAAPKPKAKETKPAKVARTTTRTTPSRKAAPKATPAKKEPAKKTPARSAKSPAKKASTRRGGRK*

### NbL06g15610.1 glutathione_reductase_chloroplastic [MW = 60.28 kDa]

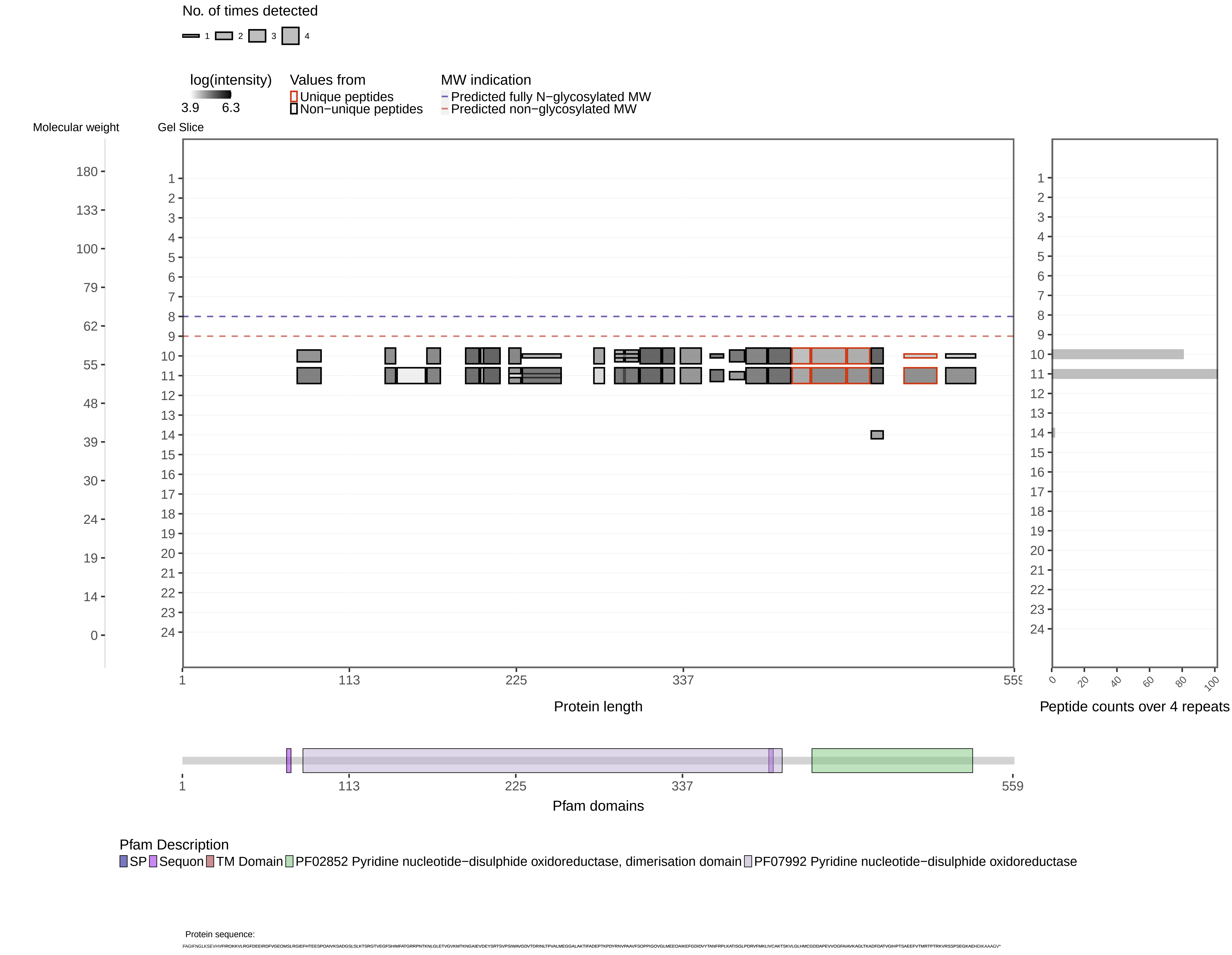

### NbL06g15110.1 acyl−CoA_dehydrogenase_family_member_10 [MW = 92.74 kDa]

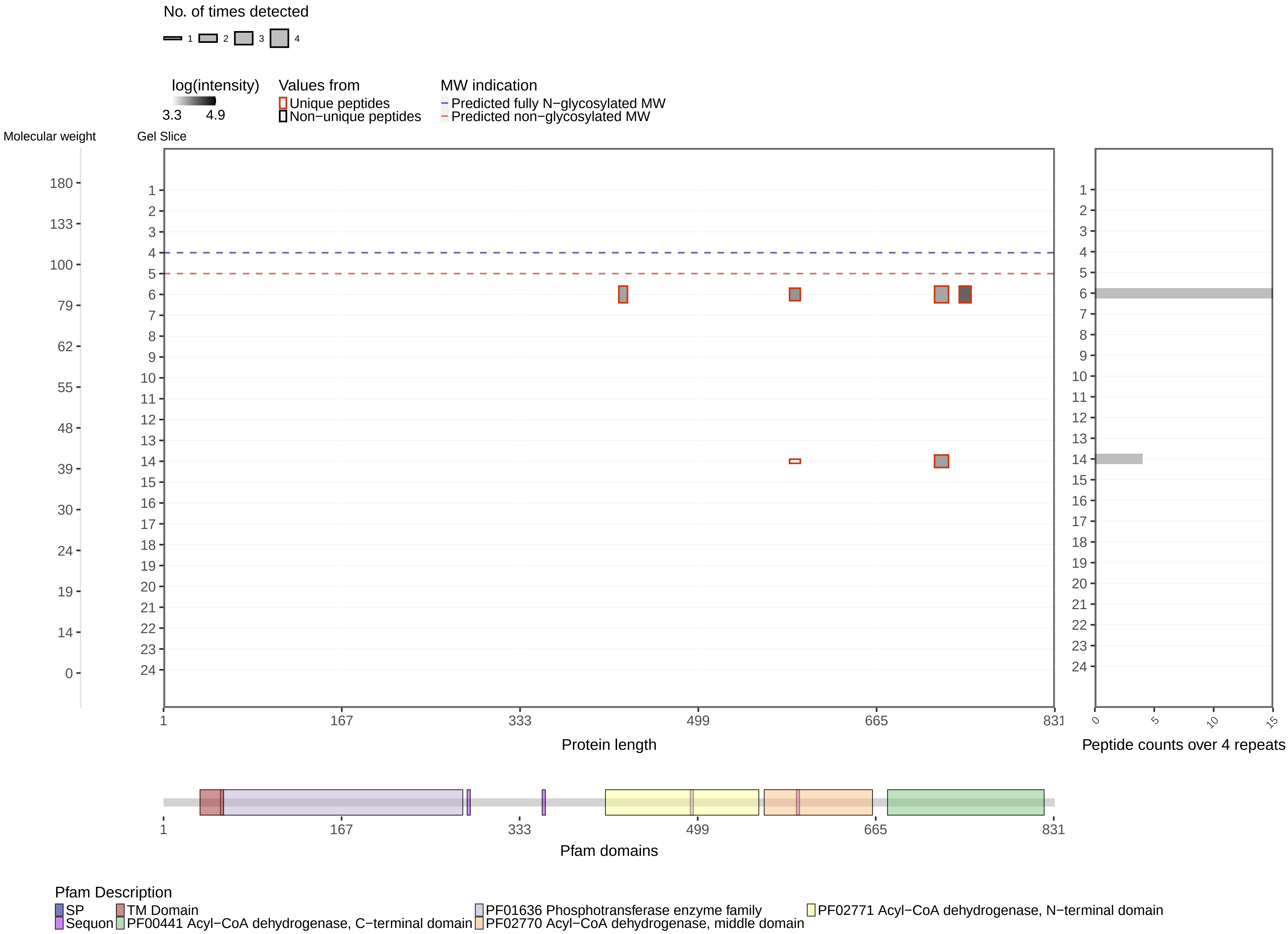


Protein sequence:

[illegible]

NbL06g15030.1
lipoamide_acyltransferase_component_of_branched−chain_alpha−keto_acid_dehydrogenase_complex_mitochondrial−like [MW = 55.26 kDa]

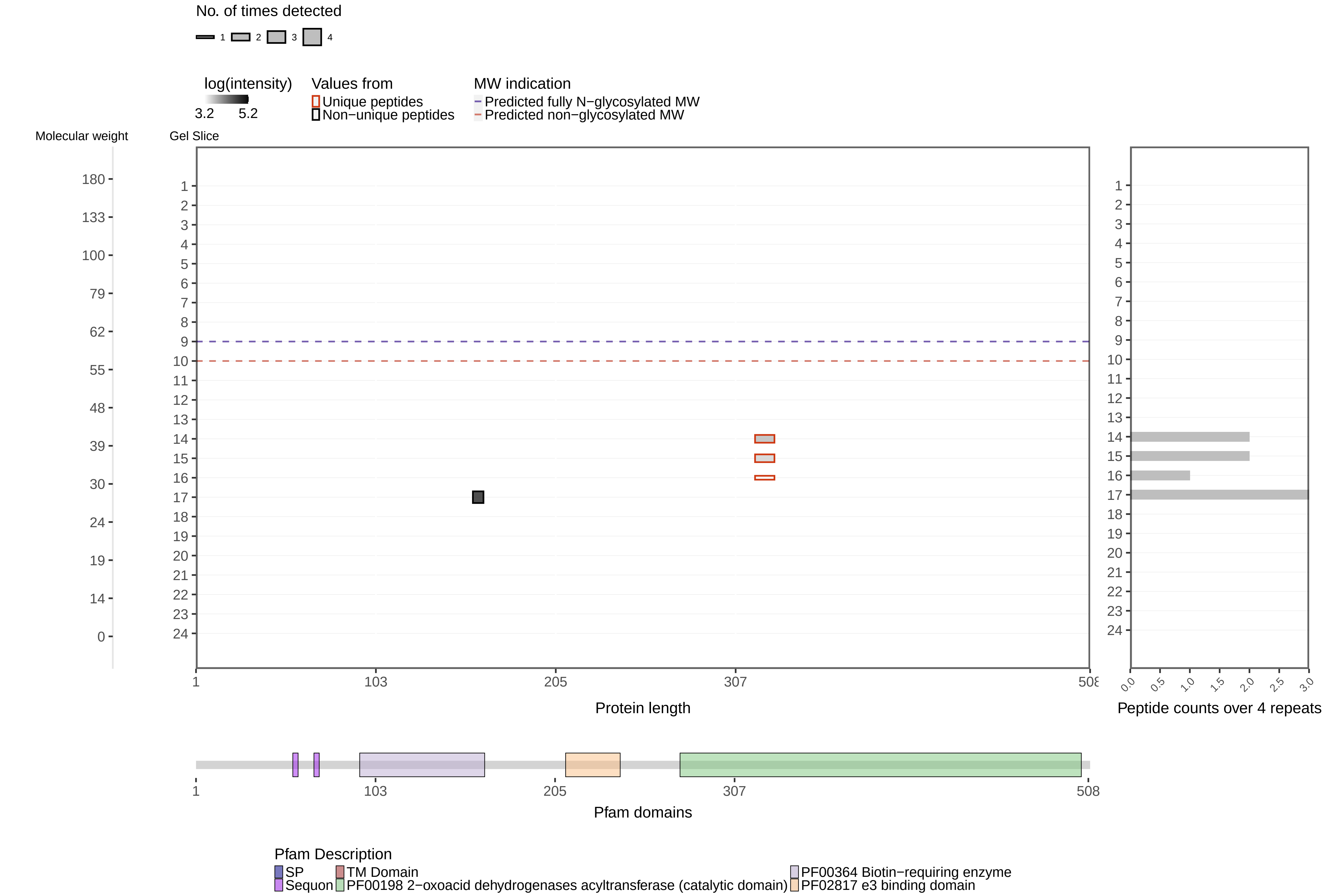


Protein sequence:

EDVTNYAMOKGLINEALECAOOKHSEVSSLIGGGYEDKTLOLRGYORAMVKSMTLAAKIPHFYFIEEMNCNALVELKKSFONENSDPEIKHTFLPVLIKSLSMALTTHPMLNSRFSEESYEVILKGSHNIGIAMATPNGLVVPNIKNVOSLSILEITKELTRLOKLAKINKLSPDDISGSTITLSNIGGIGGKFGSPLINSPEVAIIGLGRIOKIAOFGEDGDVYPASVMTINIGADHRVLDGATVARFCNDWKKFVEKPDLLLLHTR*

### NbL06g15000.1 dynein_light_chain_1_cytoplasmic−like [MW = 24.65 kDa]

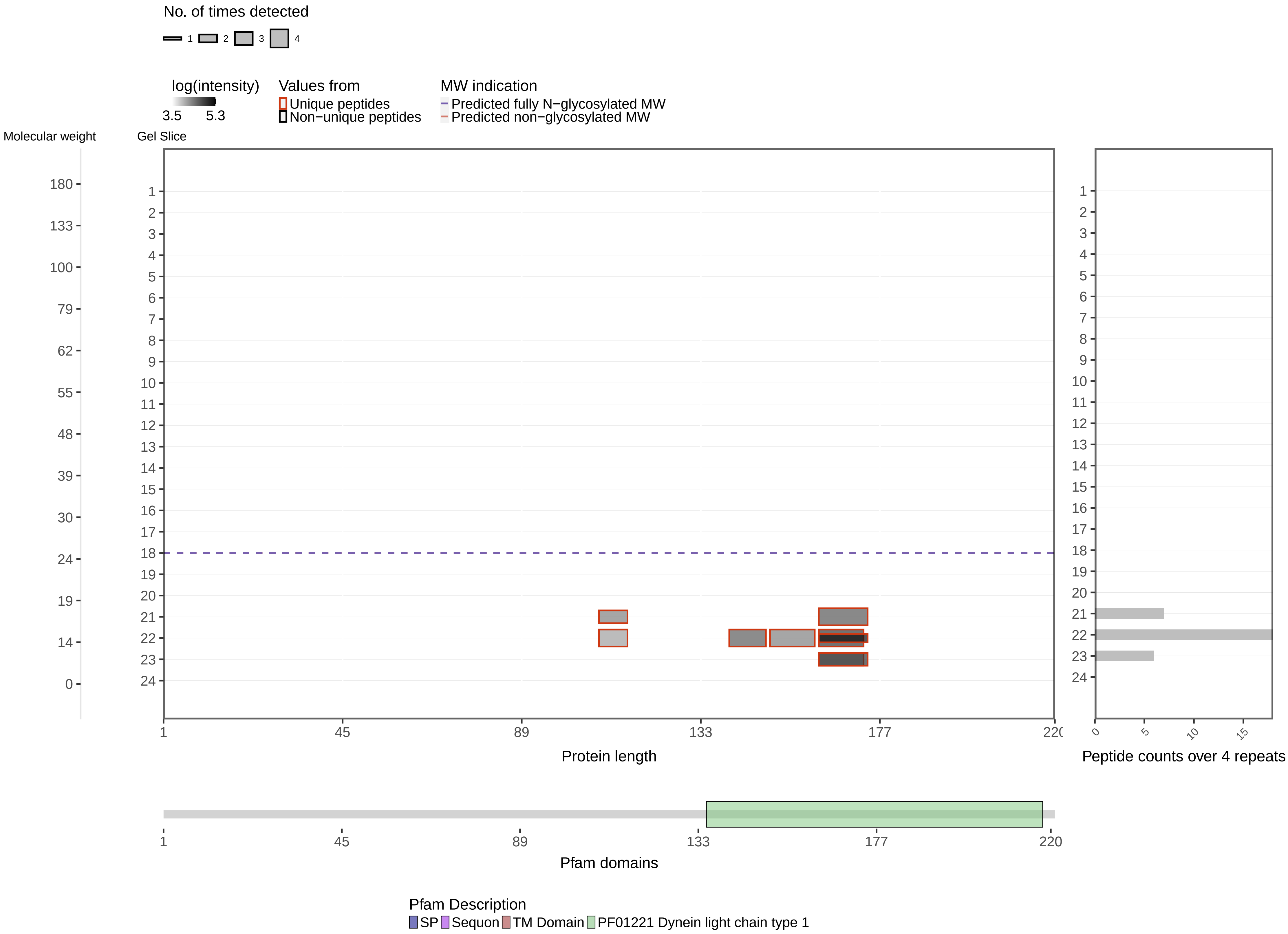


Protein sequence:

IFEDMSEDAKRTGGVPVAKTTSDDRRTSSGSVPSPIGKKITIKSADMKPDVQKEAVDIAIAAFEKYNVEKDVAEQIKKEFDKKYGPTWHCIVGKNFGSYVTHETNHFVYFYLDSKAVLLFKSG*

### NbL06g14840.1 CAP−Gly_domain−containing_linker_protein_1−like [MW = 16.47 kDa]

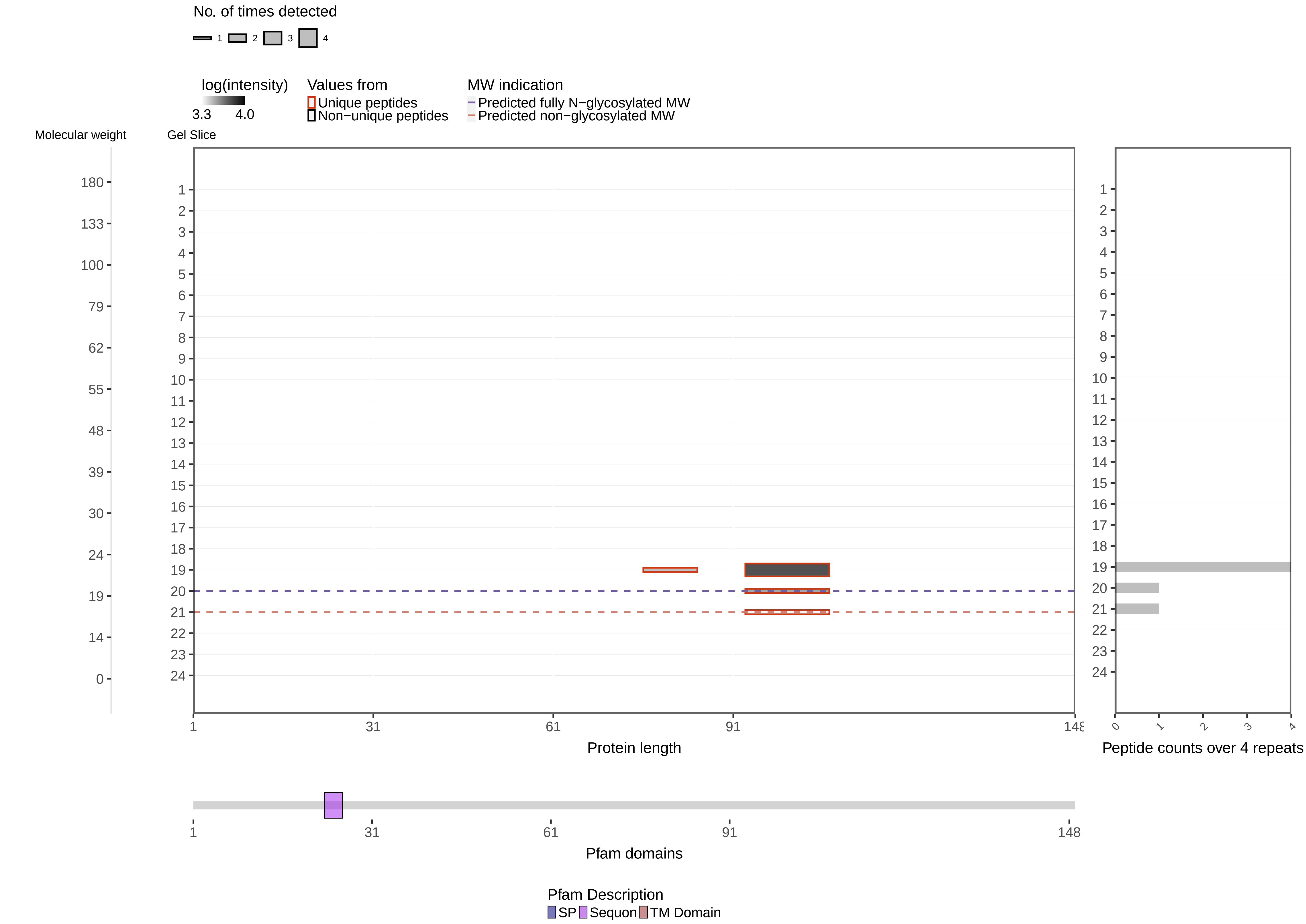


Protein sequence:

VKVYRDELSELKKSLNVEVDQLRSEFQELKTTLQQQQEDVTASLRNLGLQDVSEETKEAEDAKPNNDEEKSQEELPKDSGKDTDQ*

### NbL06g14640.1 branched−chain−amino−acid_aminotransferase_3_chloroplastic−like [MW = 45.22 kDa]

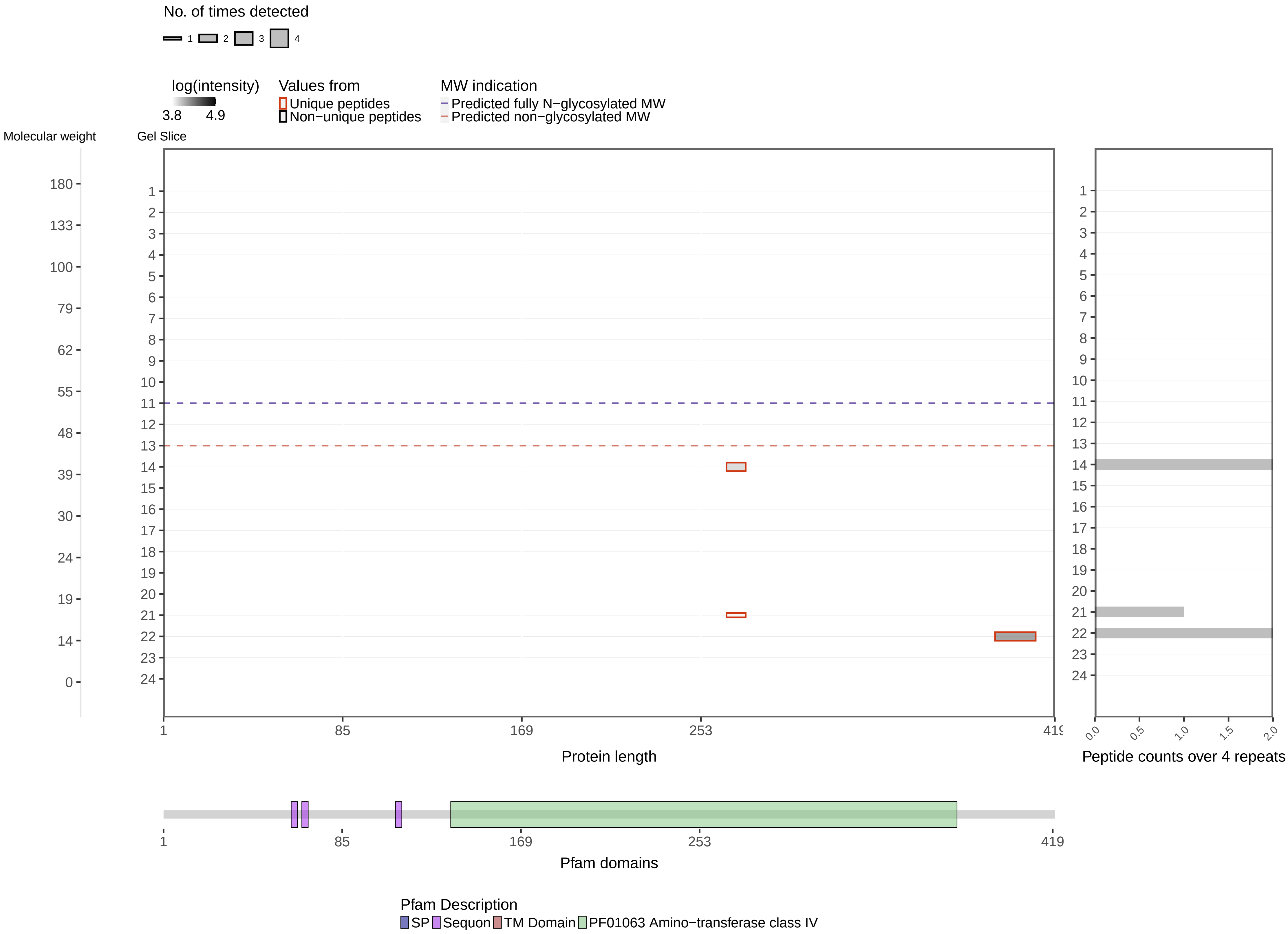


Protein sequence:

WIPPPGKGSLYIRPLLMGSGAVLGVAPAPEYTFLIYVSPVGNYFKEGMAPINLVIETEMHRATPGGTGGVKTIGNYAAVLKAQSAAKAKGYSDVLYLDSVNKRYLEEVSSCNVFIVKGNLITTPAIKGTILPGITRKSIIDVAVSQGFQVEERHVSVDELLGADEVFCTGTAVVVSPVGSITHMGKRITYGSDGVGRVSQQLYSALTSLQMGLSEDKMGWIVELK*

### NbL06g14540.1 60S_ribosomal_protein_L26−1−like [MW = 16.77 kDa]

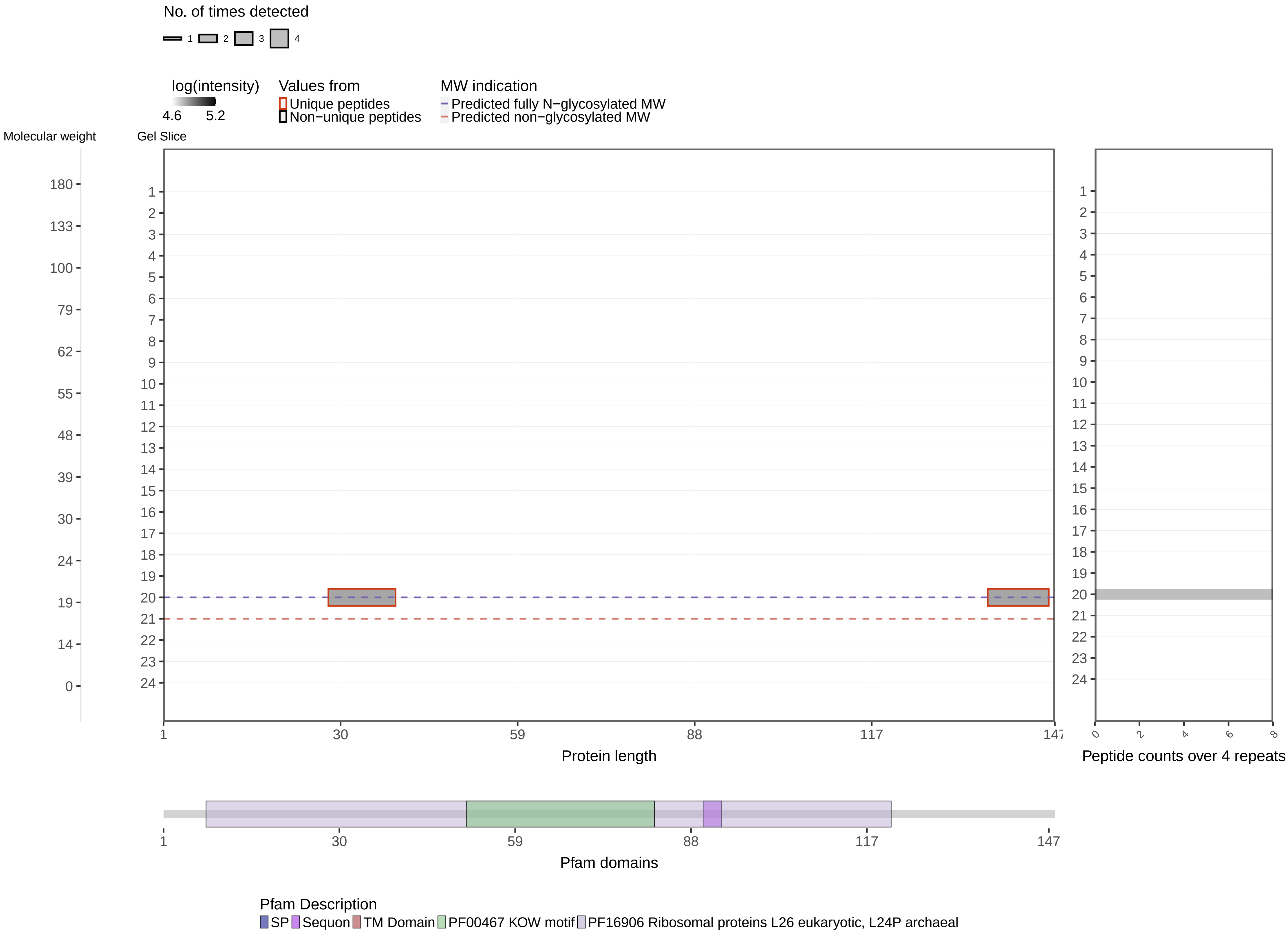


Protein sequence:

GTYKGREGKVVQVYRKKWVIHIERITREKVNGSTVNVGIHPSKTVITKLRLDKDRKSLIDRKAKGRAAADKDKGTKFTTEDIMQTVD*

NbL06g14490.1 subtilisin−like_protease [MW − SP = 82.64 kDa]

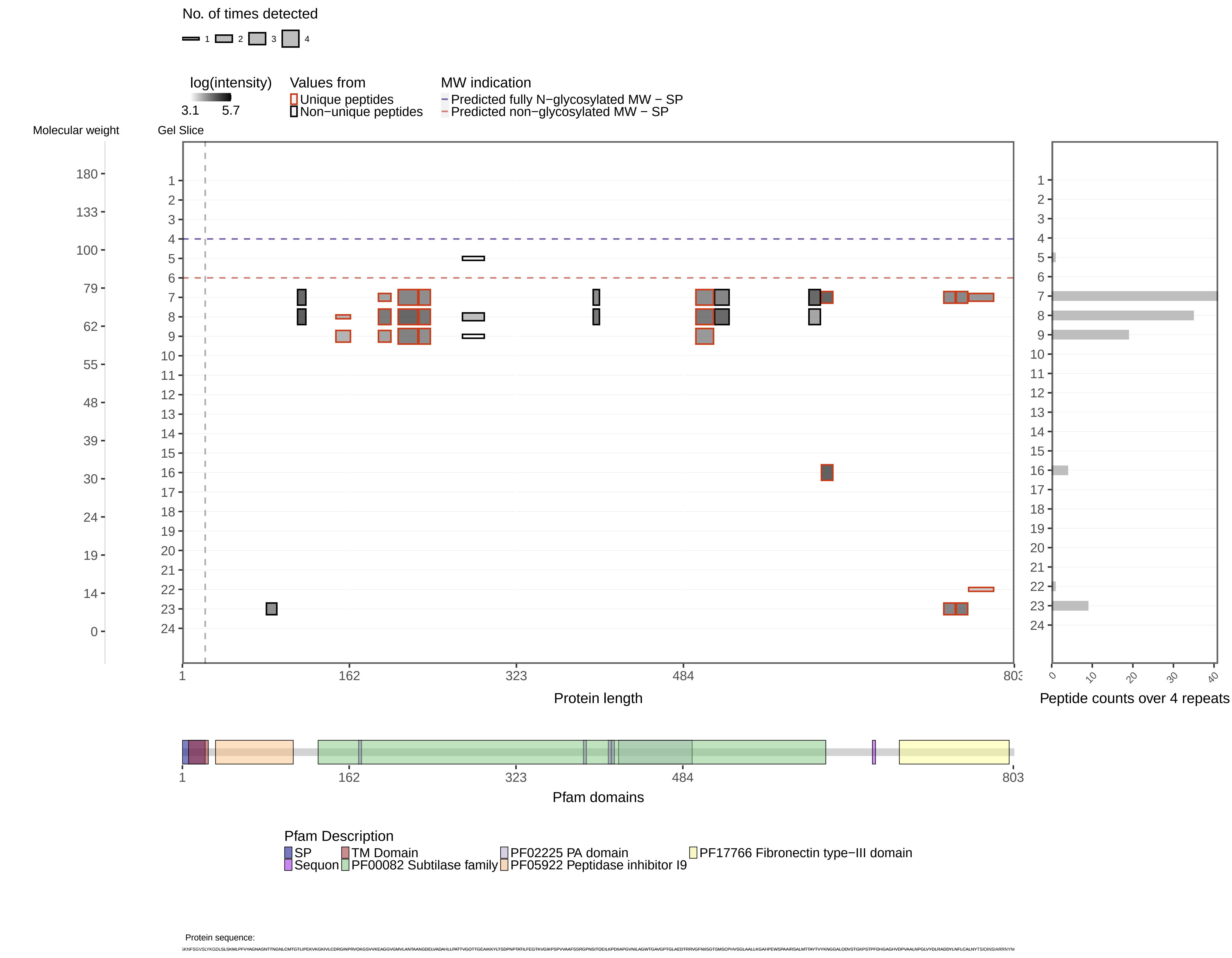

### NbL06g14470.1 tRNA−dihydrouridine(16/17)_synthase_[NAD(P)(+)]−like [MW = 49 kDa]

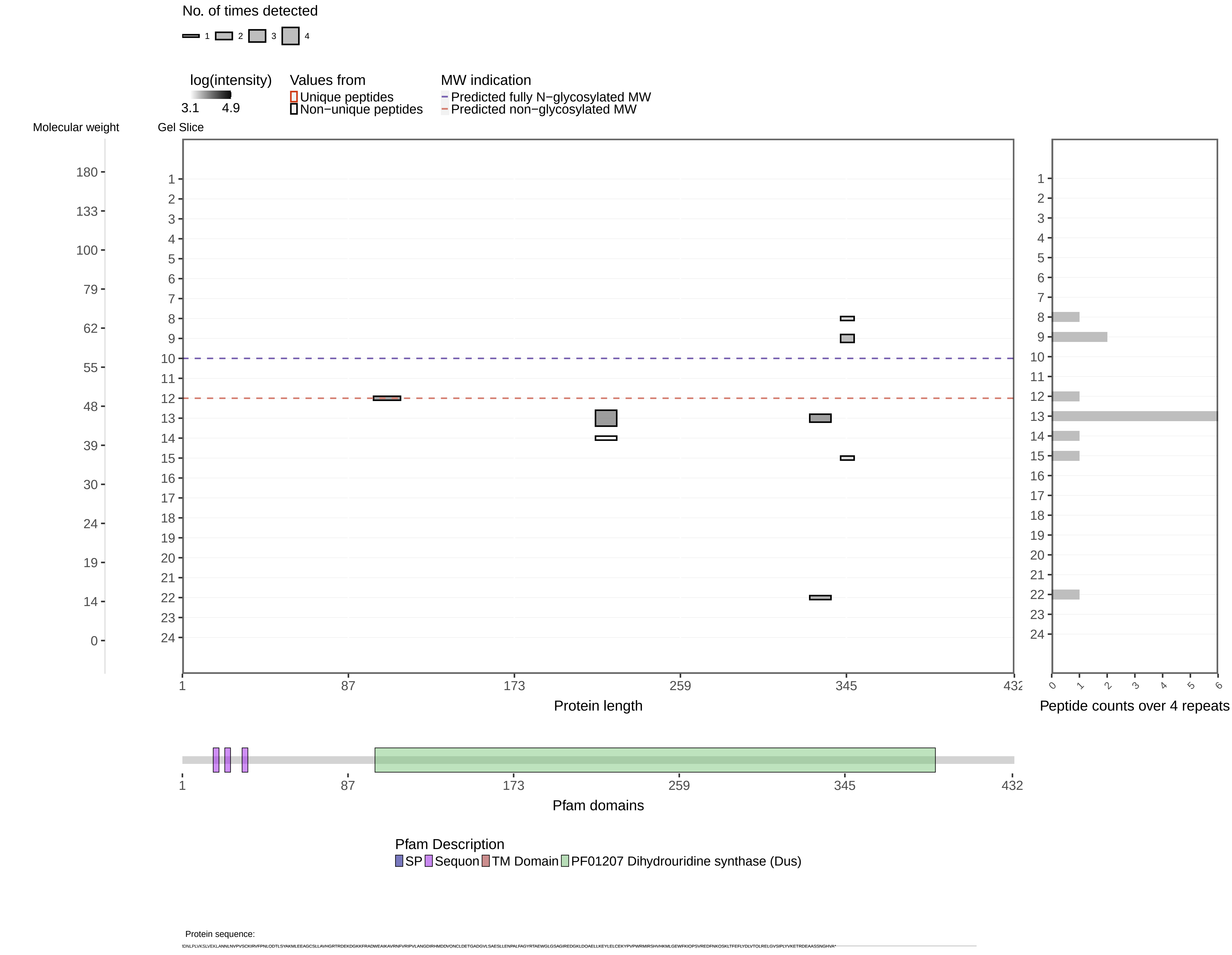

NbL06g14360.1 probable_phosphoribosylformylglycinamidine_synthase_chloroplastic/mitochondrial [MW = 154.1 kDa]

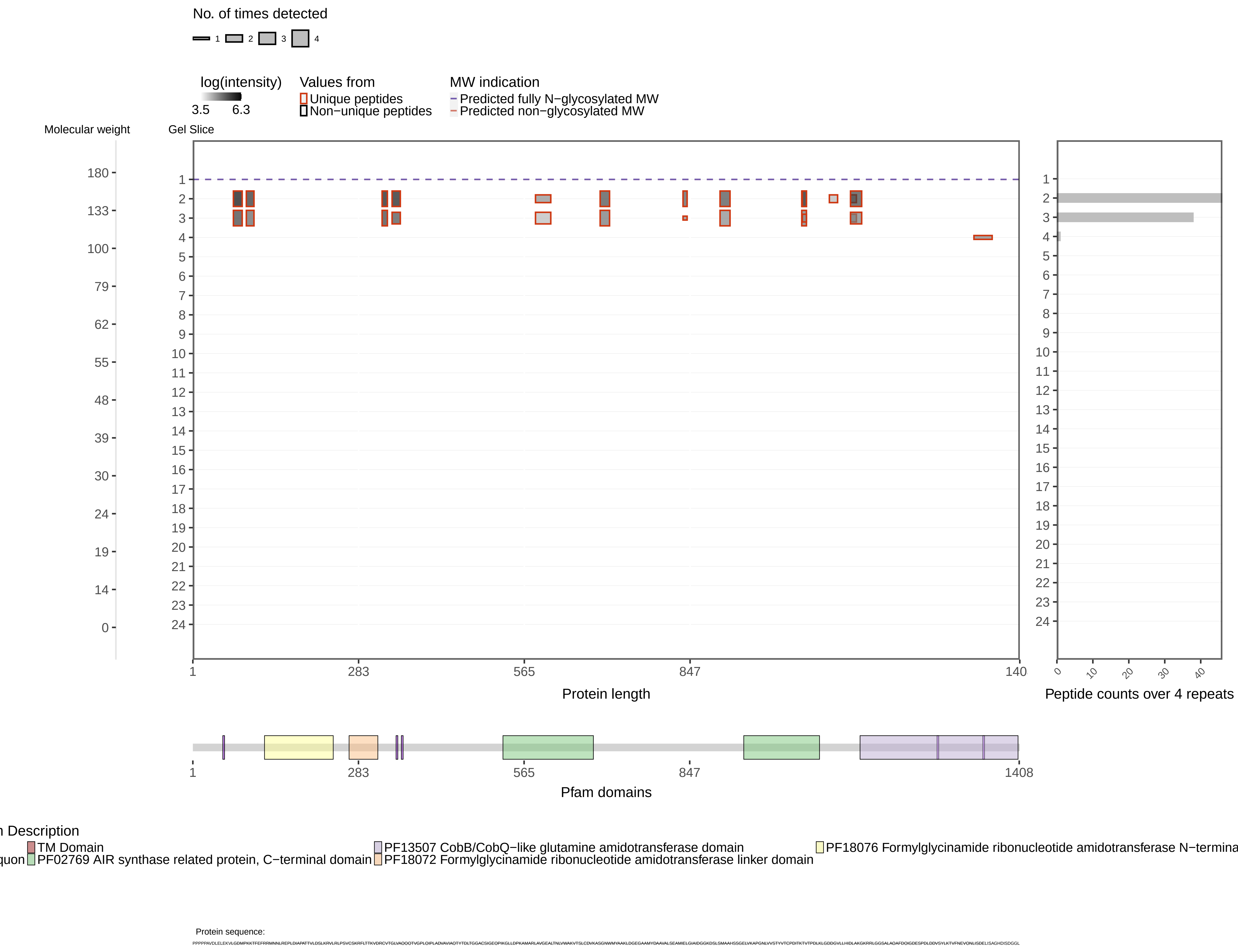

### NbL06g14330.1 red_chlorophyll_catabolite_reductase_chloroplastic [MW = 36.47 kDa]

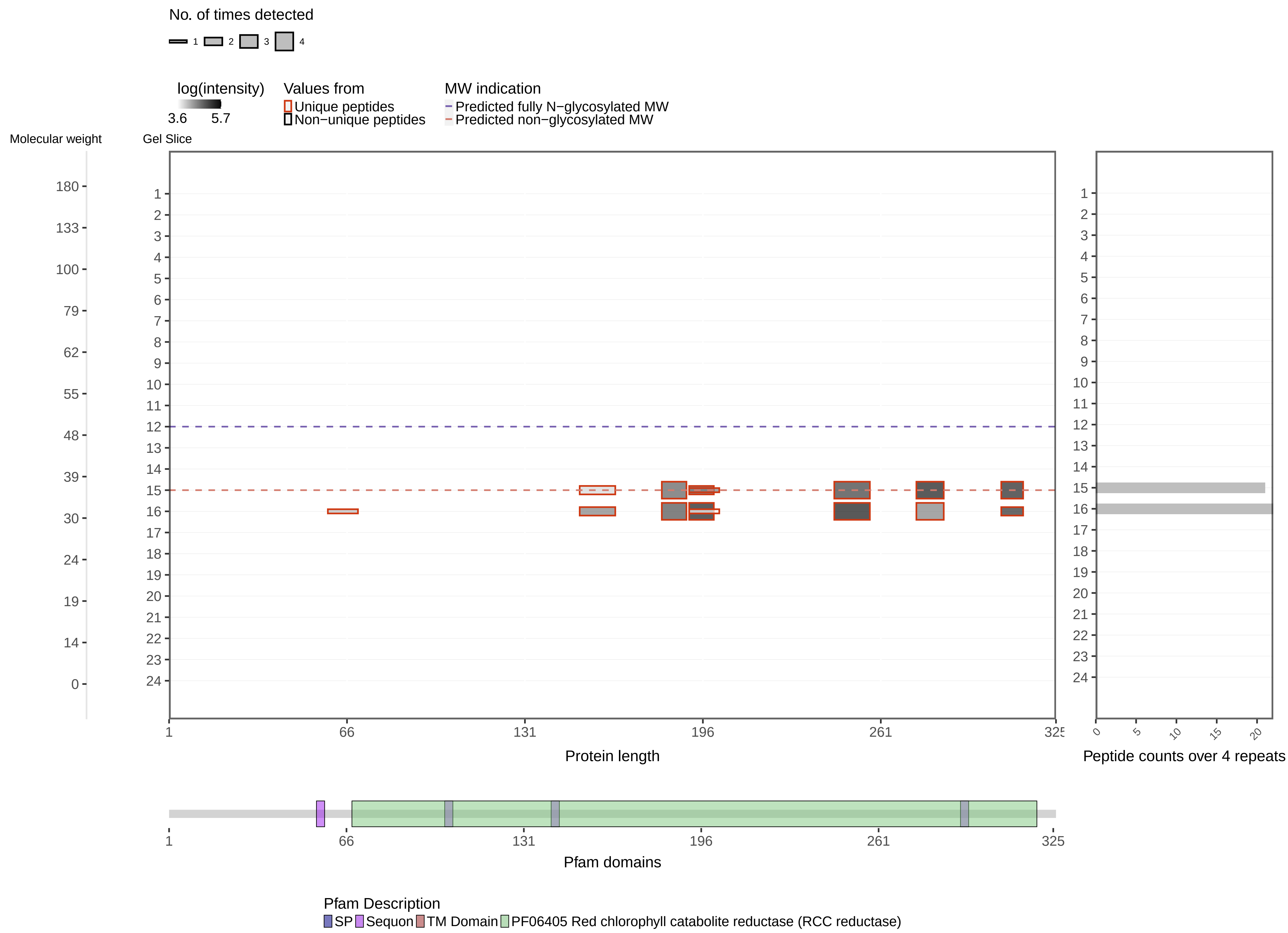


Protein sequence:

SAYLRPSTDAPNFLMEVIRSSPTSLILILDLPPRKDLVQHPDYLKNFYEETQLDKQRQLLEKLPEVKPYFSSSLYIRAVVSPSAILVSIETEASQAVRIDEIIQDHISPVAKVMLETWLDLCACPERRLTEDESTALAKRDKIFKNKTIEIDLESSFPRLFGQEVANKVLGVLREIYNA*

### NbL06g14260.1 patatin−like_protein_3 [MW − SP = 42.86 kDa]

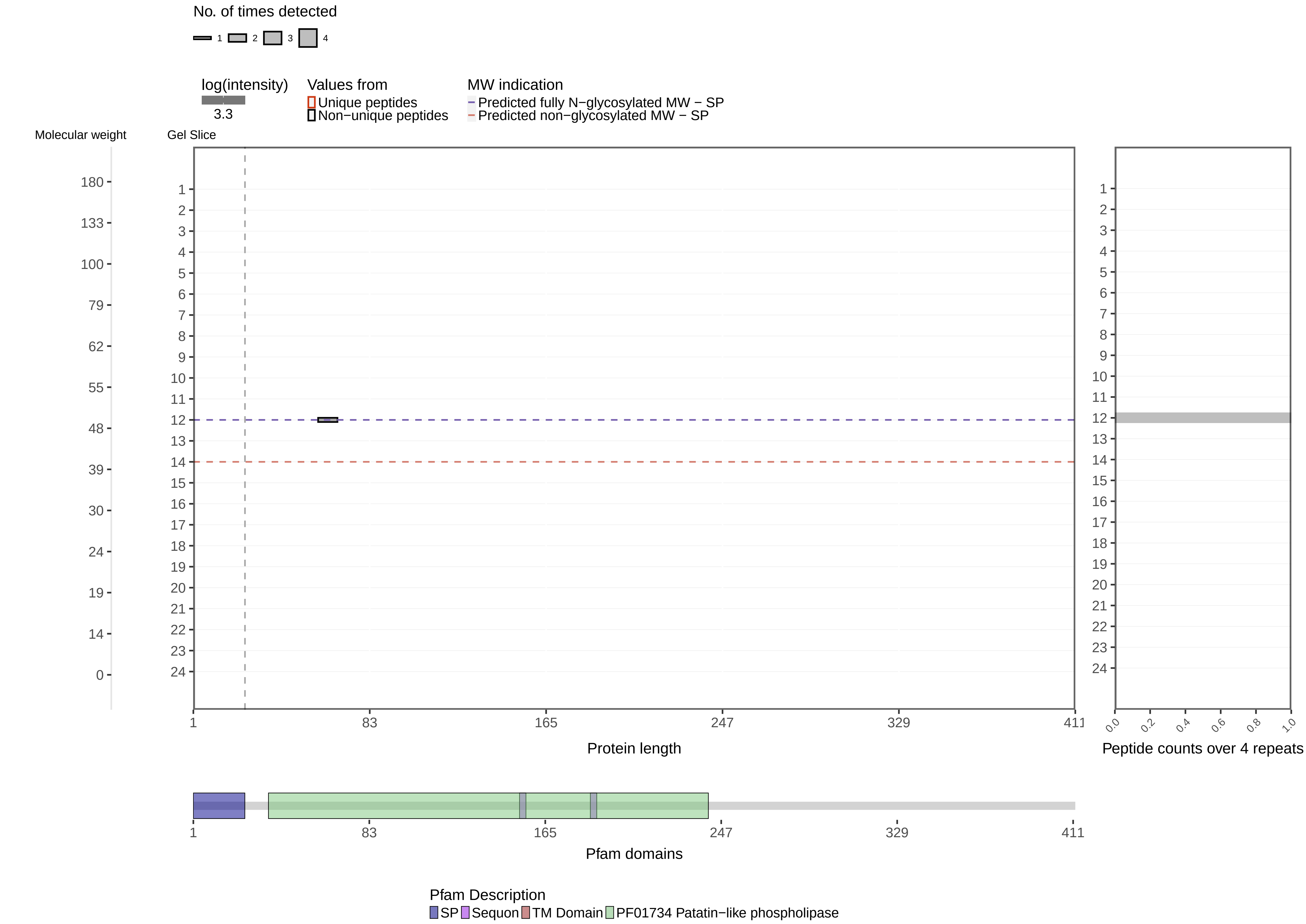


Protein sequence:

.STSAAPTFFPVHYFESRDTOGRIRTFDMIDGGVAANNPTOMAITHISKEIMRGKFEYEEMERMESKKMLVVSLGTGIGKHOGYNAASASKWGLLGWVYNNGOTPILDVYNDASADMVDIHVSTMFOTLRTEKNYLRIOEDNLIGDAASMDIATTKNMOKLVEIGNNLLKKAVSRVNLETGOYEPVOGEGTNEEALVHFAKLLSOEKKLRYNN*

NbL06g13790.1 alpha−14_glucan_phosphorylase_L−1_isozyme_chloroplastic/amyloplastic [MW = 110.32 kDa]

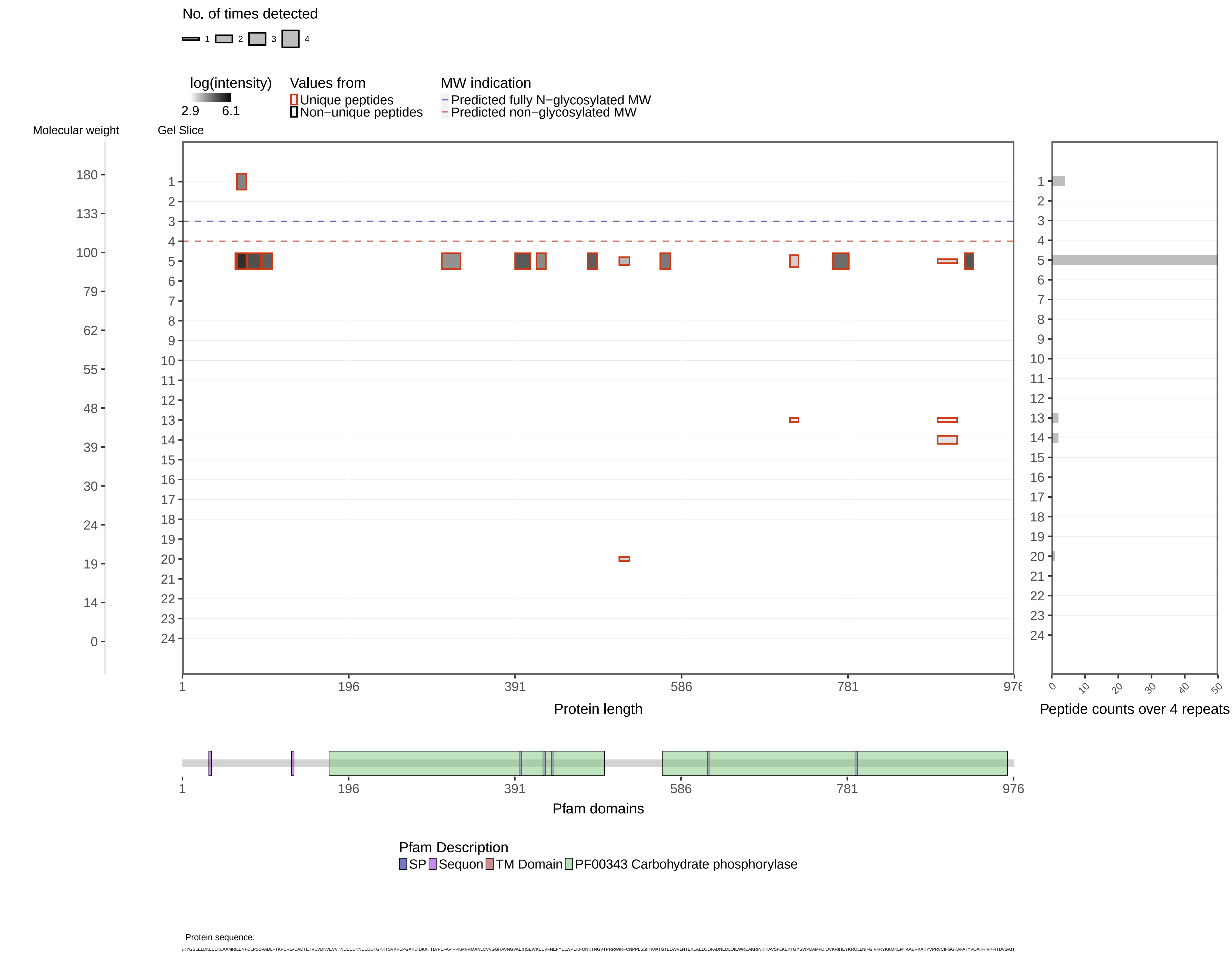

### NbL06g13670.1 cystathionine_gamma−synthase_1_chloroplastic−like [MW = 59 kDa]

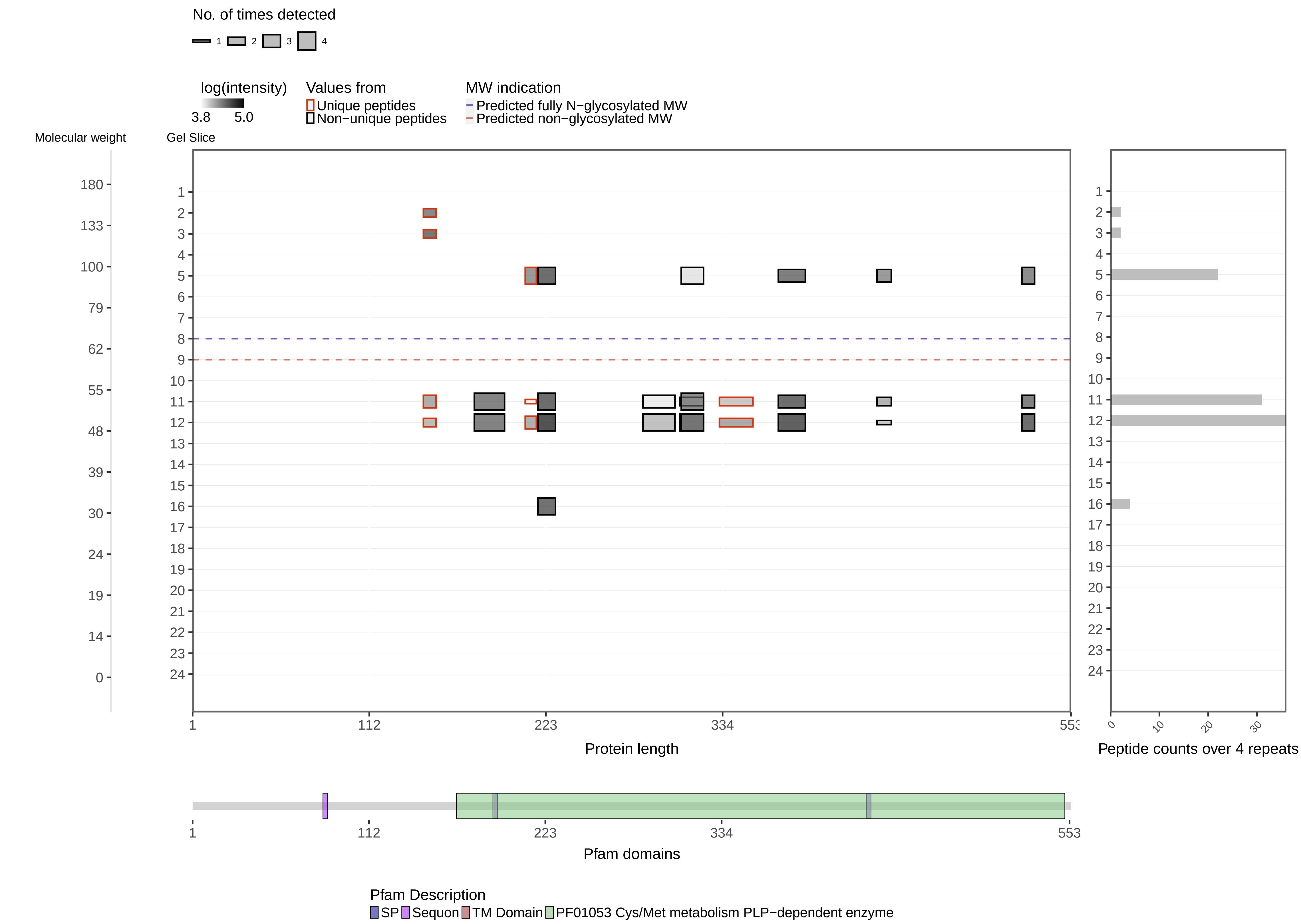


Protein sequence:

[illegible]

NbL06g13570.1 transcription_factor_PRE1−like [MW = 10.16 kDa]

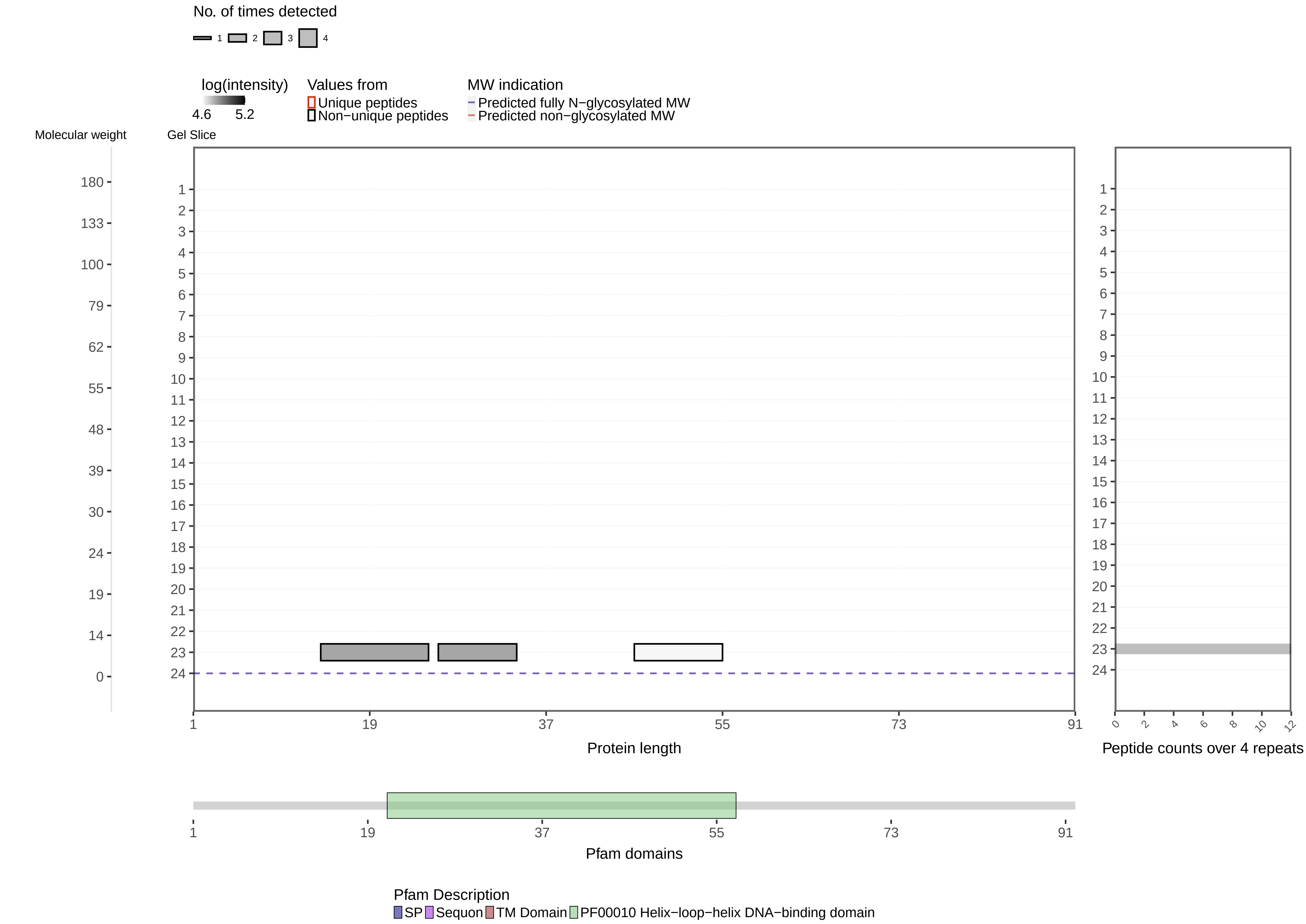


Protein sequence:

EIRTRRSSKASASKVLQETCNYIRSLNREMDDLSDRLSQLLSTVDADSPEAAIIRSLII*

NbL06g13480.1 hypersensitive−induced_response_protein_2 [MW = 15.96 kDa]

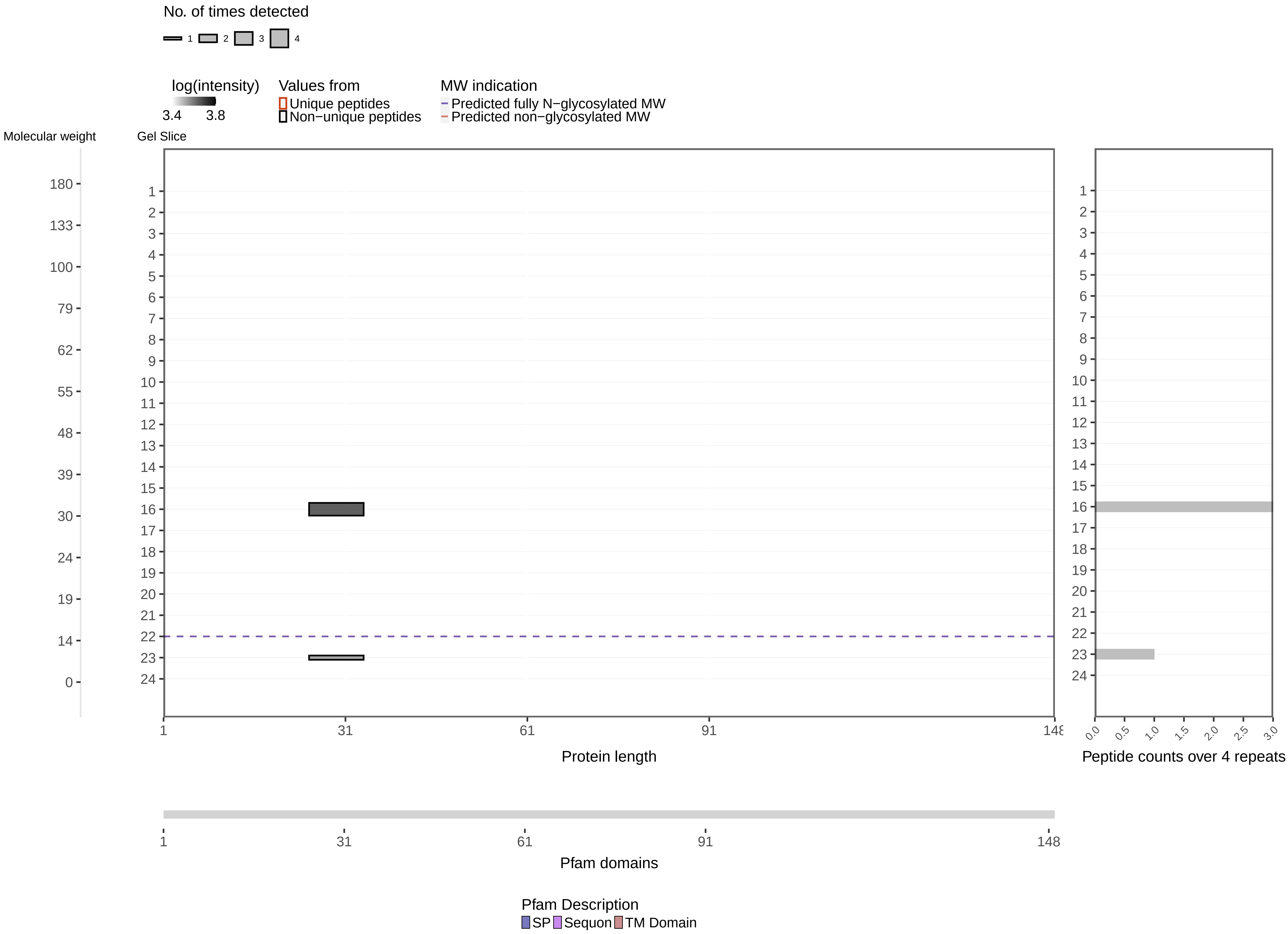


Protein sequence:
KYSSGLGIARQRQAIVDGLRDSVLAFSENVPGTTAKDMVLDTMKEIGATSKSYAVFIPHGPGAVKDVVRVFSRLKVFSISLTKTKDS*

### NbL06g13430.1 protein_transport_protein_SEC13_homolog_B−like [MW = 32.65 kDa]

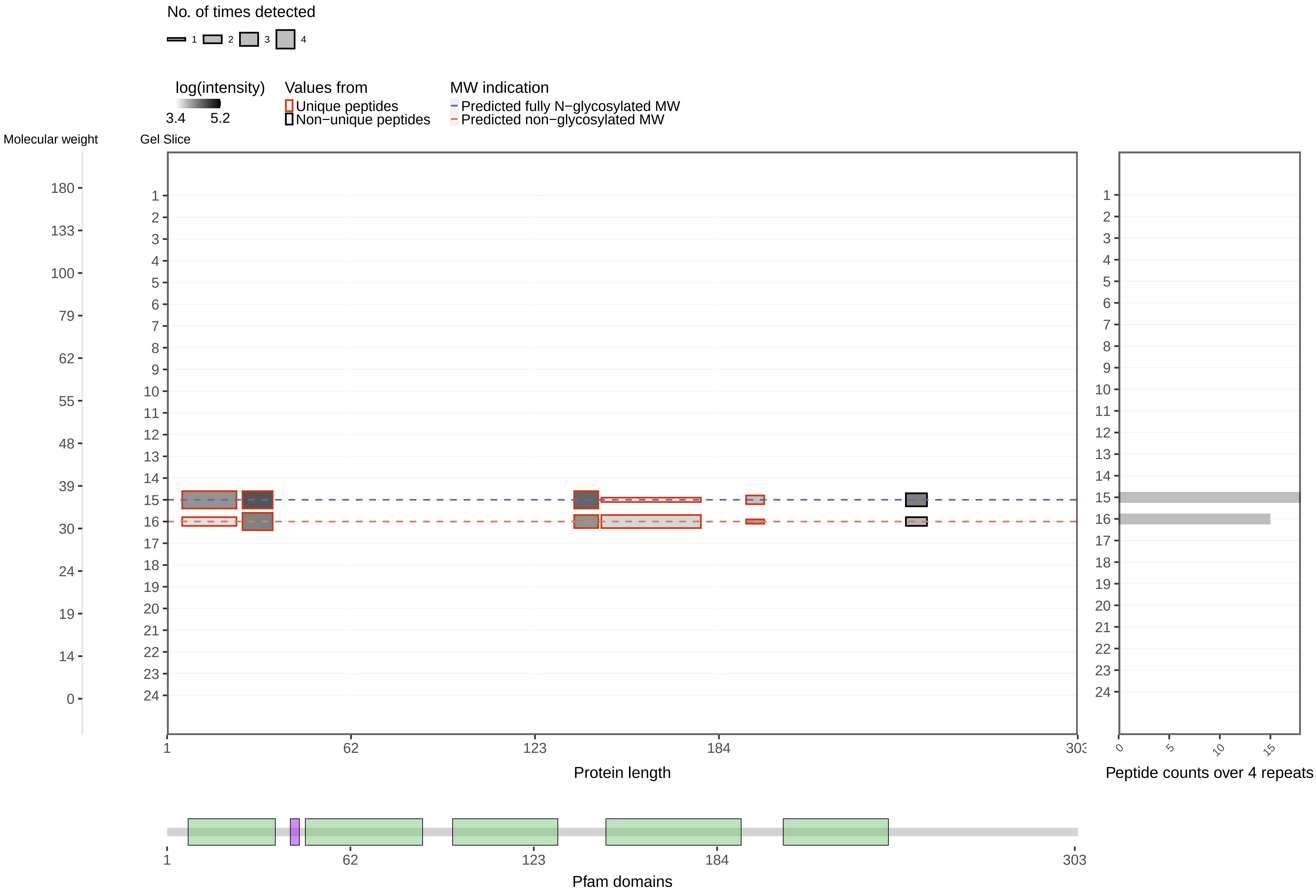


Protein sequence:

DTTKIDOAHPVGVTAVSWAPSMAPGALVGSGVLEPVOKLASGGCDNTVKVWKLYNGVWKMDCFPALOKHTNWVRDVAWAPNLGLPKSTIASASEDGTVVIWTVAKEGDOWEGKVLKDFKTPVWRVSWSLTGNLLAVAAGDNDVTLWKEAVDGEWOOVTSVDO*

### NbL06g13360.1 phosphoglucan_phosphatase_LSF1_chloroplastic−like [MW = 66.9 kDa]

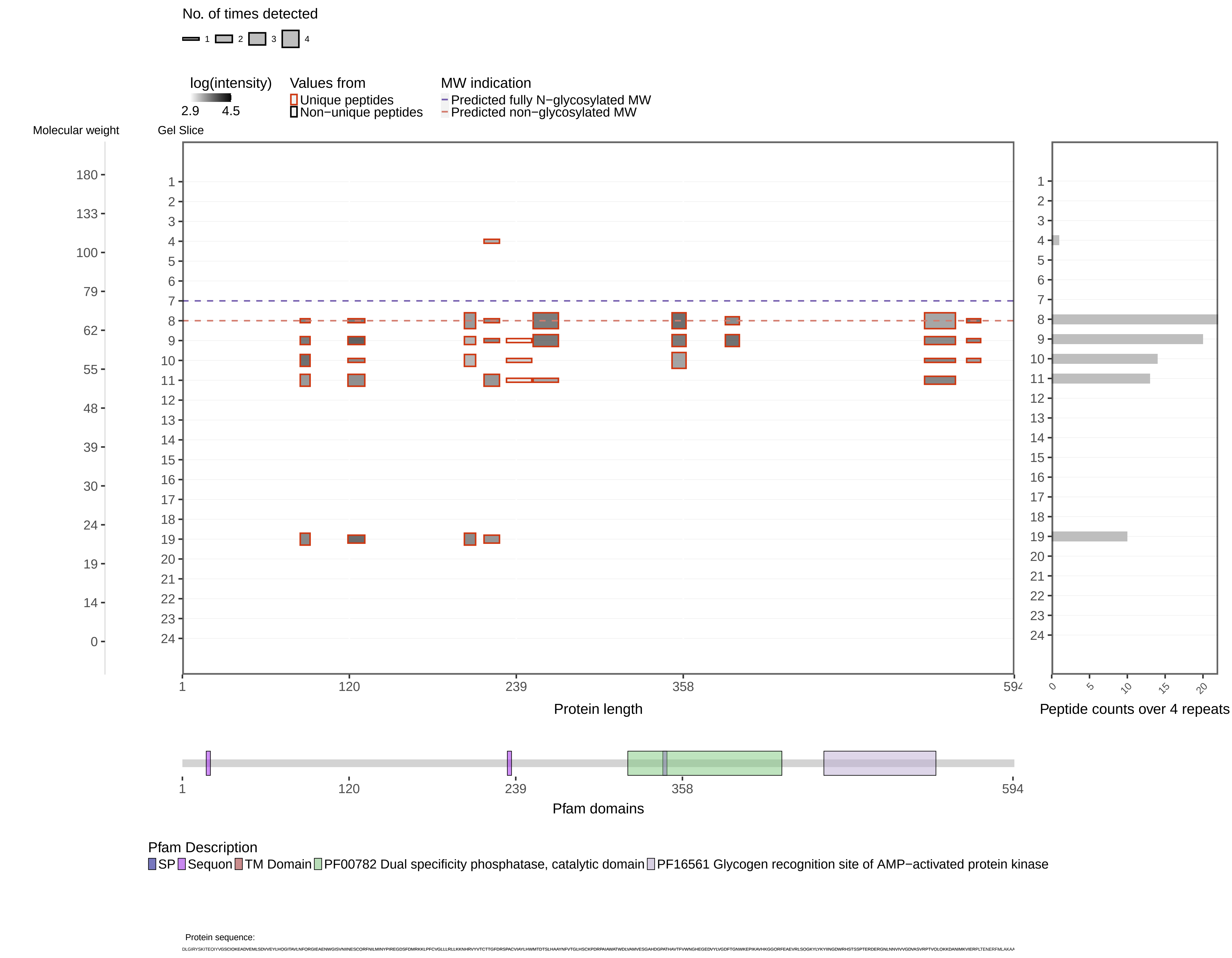

NbL06g13350.1 50S_ribosomal_protein_L12_chloroplastic−like [MW = 19.66 kDa]

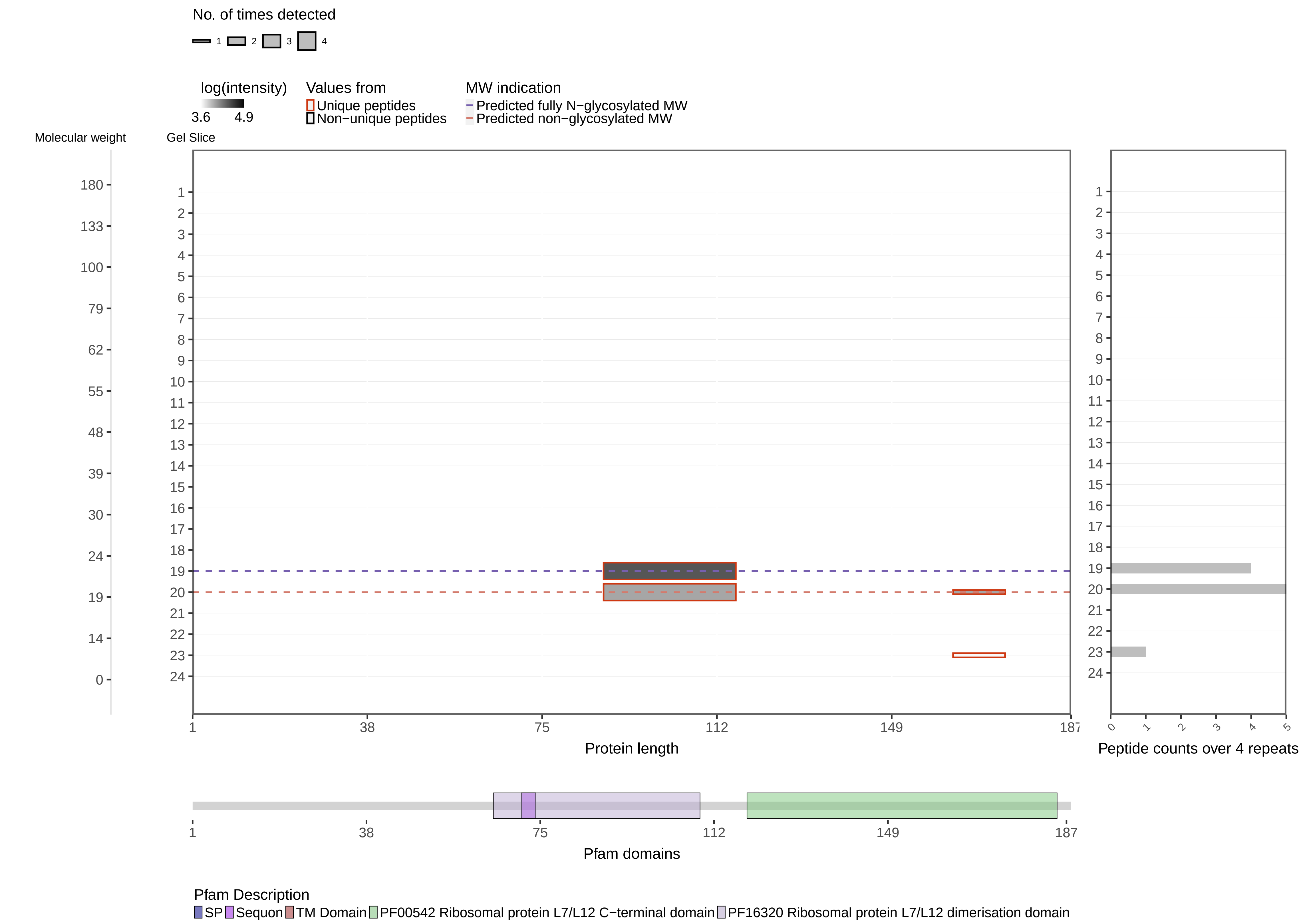


Protein sequence:

LVEYLQDKLGVTAASFAPAAVSAAPAAAEAPAVVEEKTEFDVIIEEVPSNARIAAIKAVRALTNLALKEAKELIEGLPKKFKEGISKDEAEEAKKQLEEAGAKVSIA*

NbL06g13340.1 universal_stress_protein_A−like_protein [MW = 19.79 kDa]

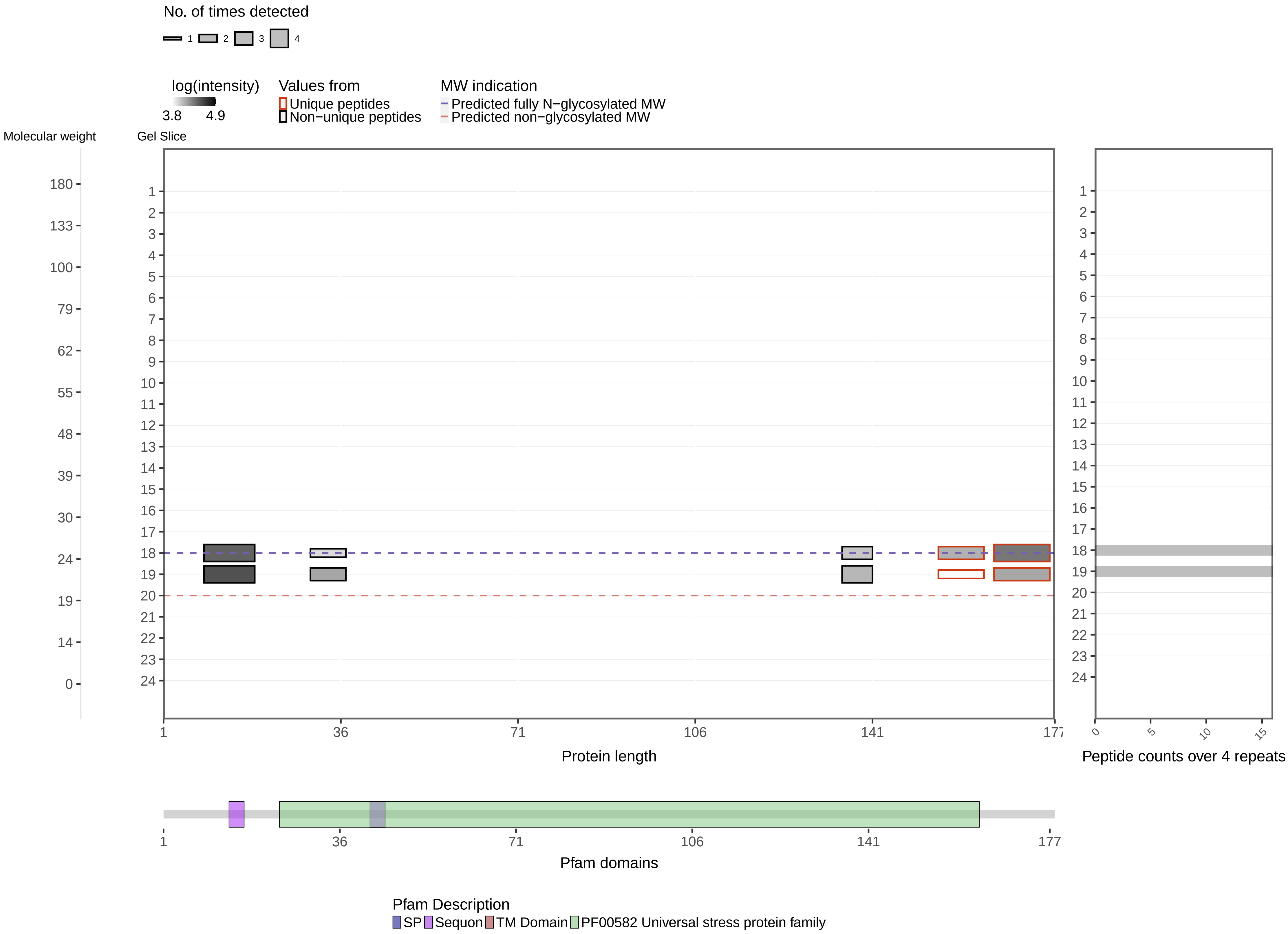


Protein sequence:

VLKHRDKIRGLHLLEYFVNRCHEIGVPCEAWIKQGDPKEVVCHEVKRVQPDLLVVGCRGLGPFQRVFVGTVSEFCVKHAECPVVTIKRSAAETPQDPIDD*

NbL06g13100.1
dihydrolipoyllysine−residue_acetyltransferase_component_4_of_pyruvate_dehydrogenase_complex_chloroplastic−like [MW = 49.25 kDa]

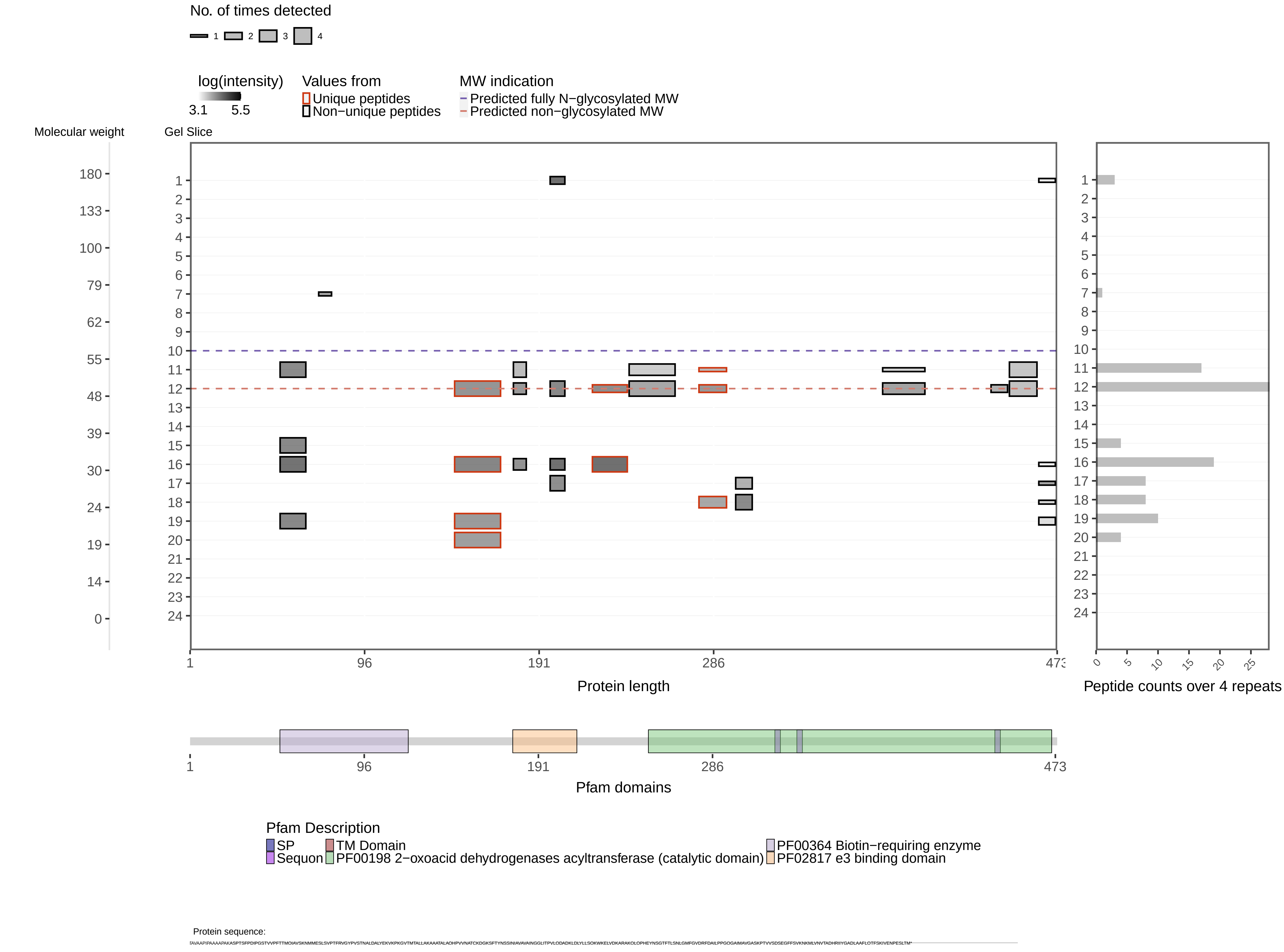

NbL06g13030.1 actin−like [MW = 41.68 kDa]

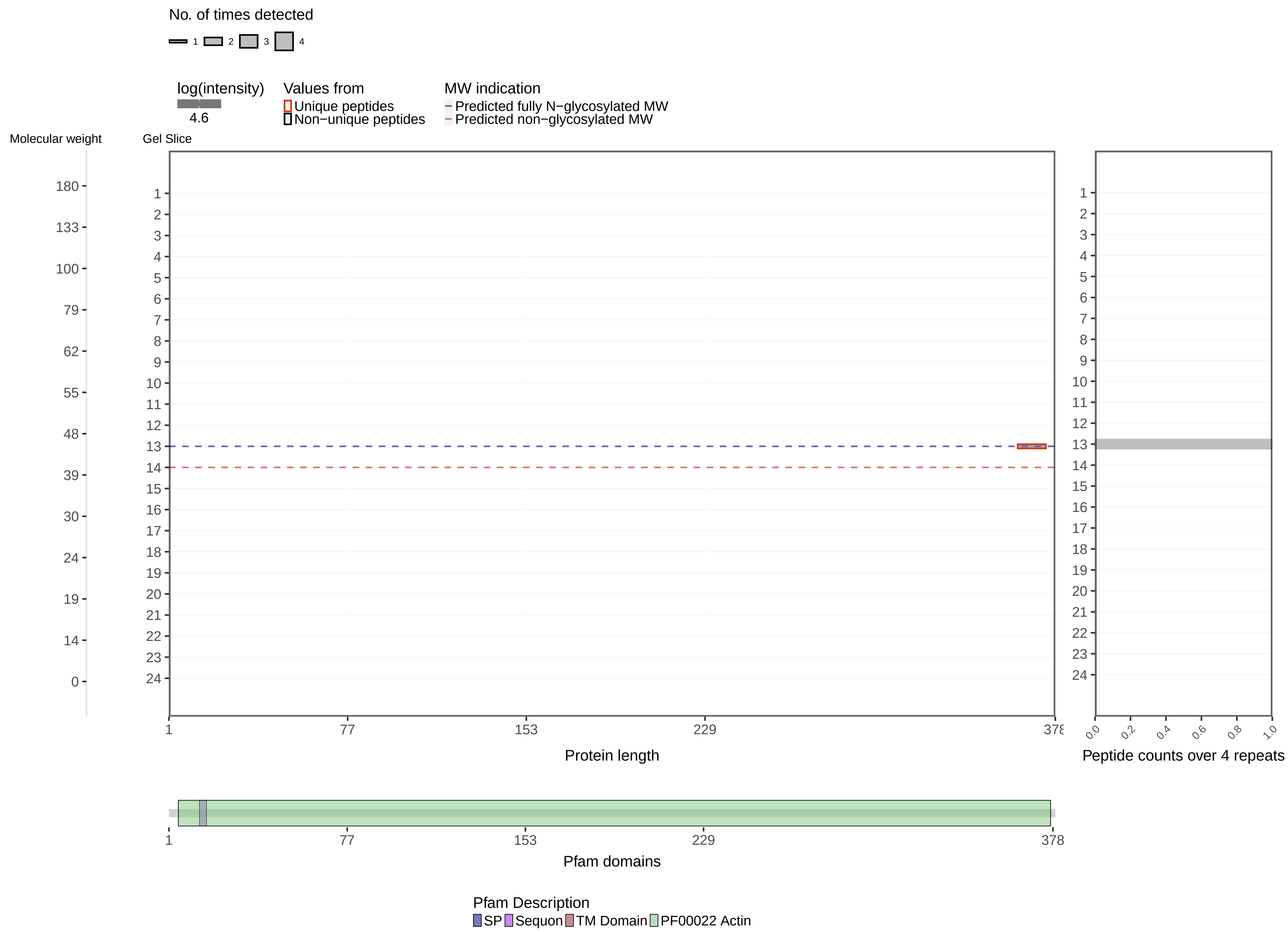


Protein sequence:

PHAILRLDLAGRDLTDNLMKILTERGYSFTTTAEREIVRDVKEKLSYIALDYEQELETSKSGSSVEKTYELPDGQVITIGAERFRCPEVLFQPSMIGMEAAGIHETTYNSIMKCDVDIRKDLYGNIVLSGGSTMFPGIADRMSKEITALAPNSMKIKVVAPPERKYSVWIGGSILASLSTFQQMWIAKAEYDEAGPSIVHRKCF*

### NbL06g13010.1 photosynthetic_NDH_subunit_of_lumenal_location_1_chloroplastic [MW = 25.66 kDa]

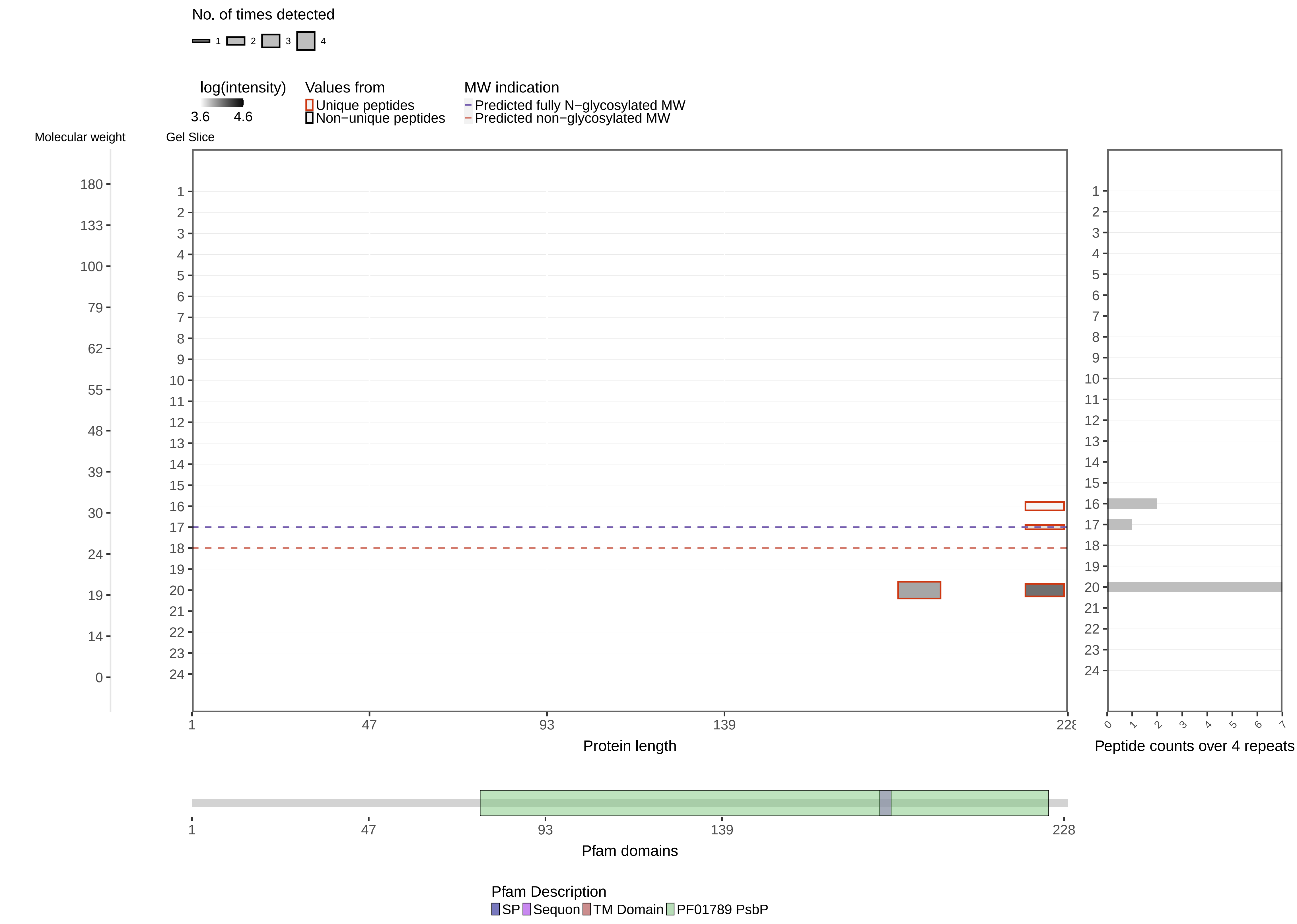


Protein sequence:

RGHDSAFKDRYLQLQNVRLSFIPTDKSDIHDLGPMEEVVANLVKHVYSAPNQVANIMEMRERSTDGKNYCTFEYVLTSPNFSRAAFASIAIGNGRYYTLIVGANERRWRKFRNKLKVVADSFQVLDI*

NbL06g12960.1 transcriptional_corepressor_LEUNIG−like [MW = 76 kDa]

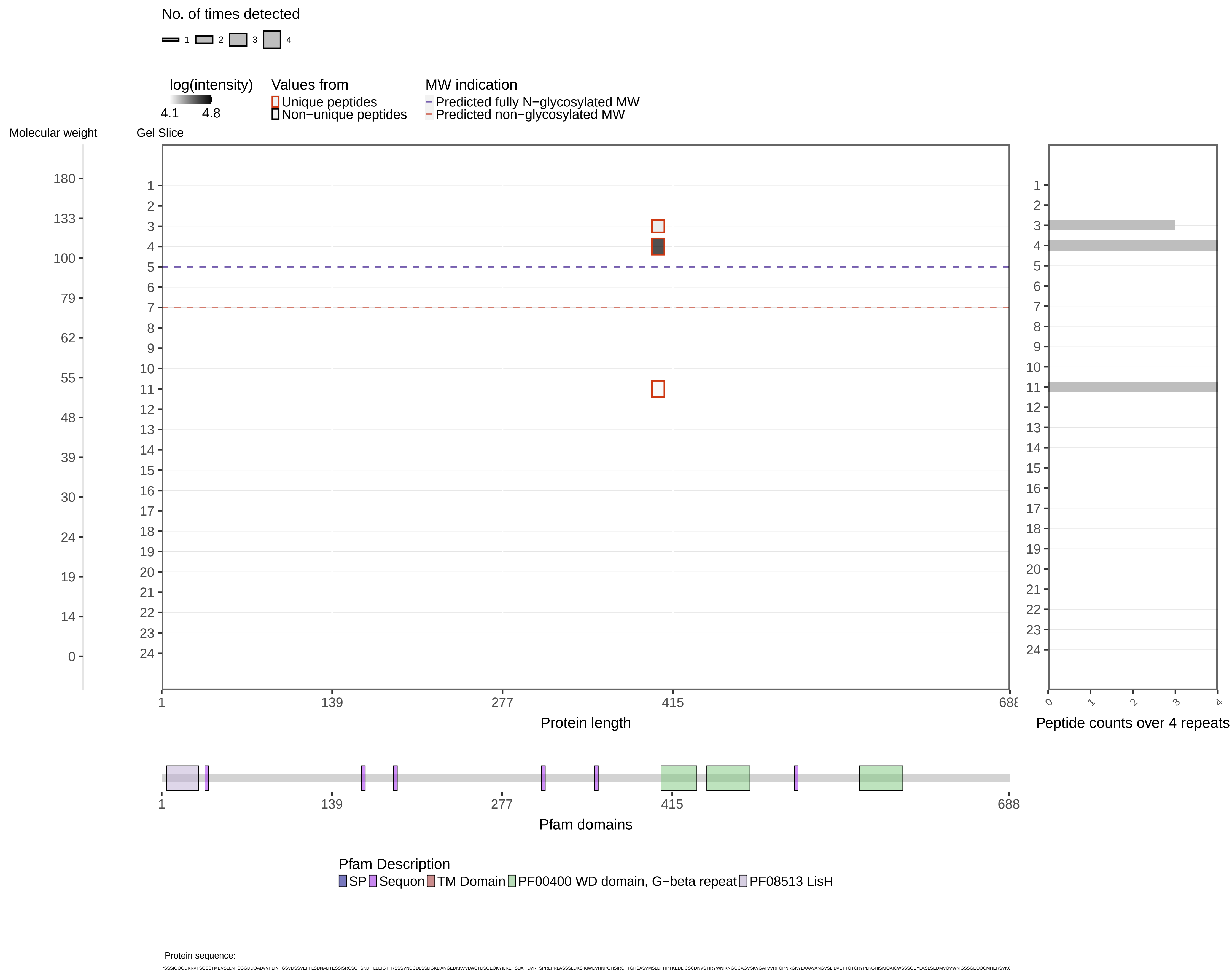

### NbL06g12900.1 beta−D−xylosidase_1 [MW − SP = 81.3 kDa]

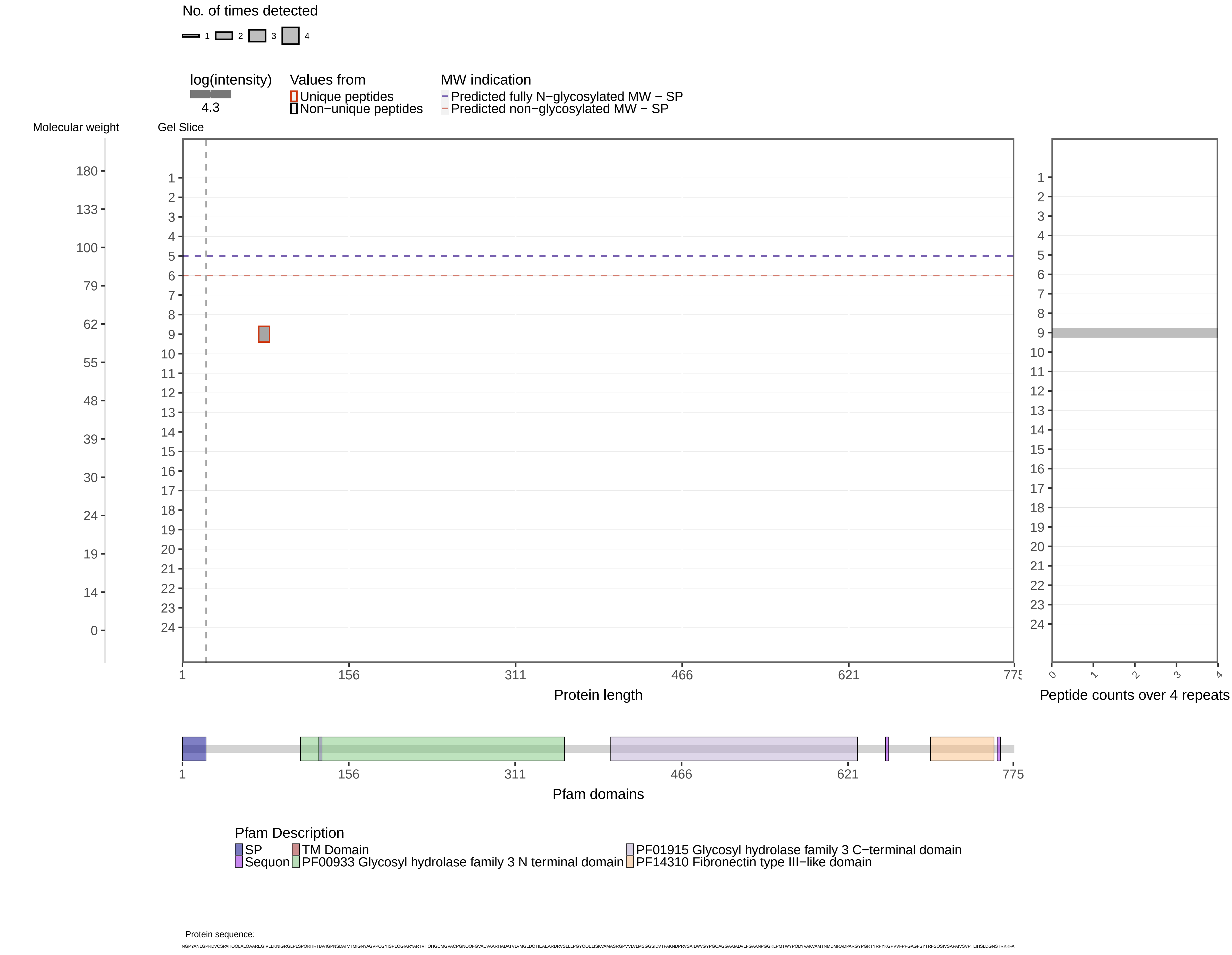

NbL06g12840.1 uncharacterized_LOC109232177 [MW = 389.76 kDa]

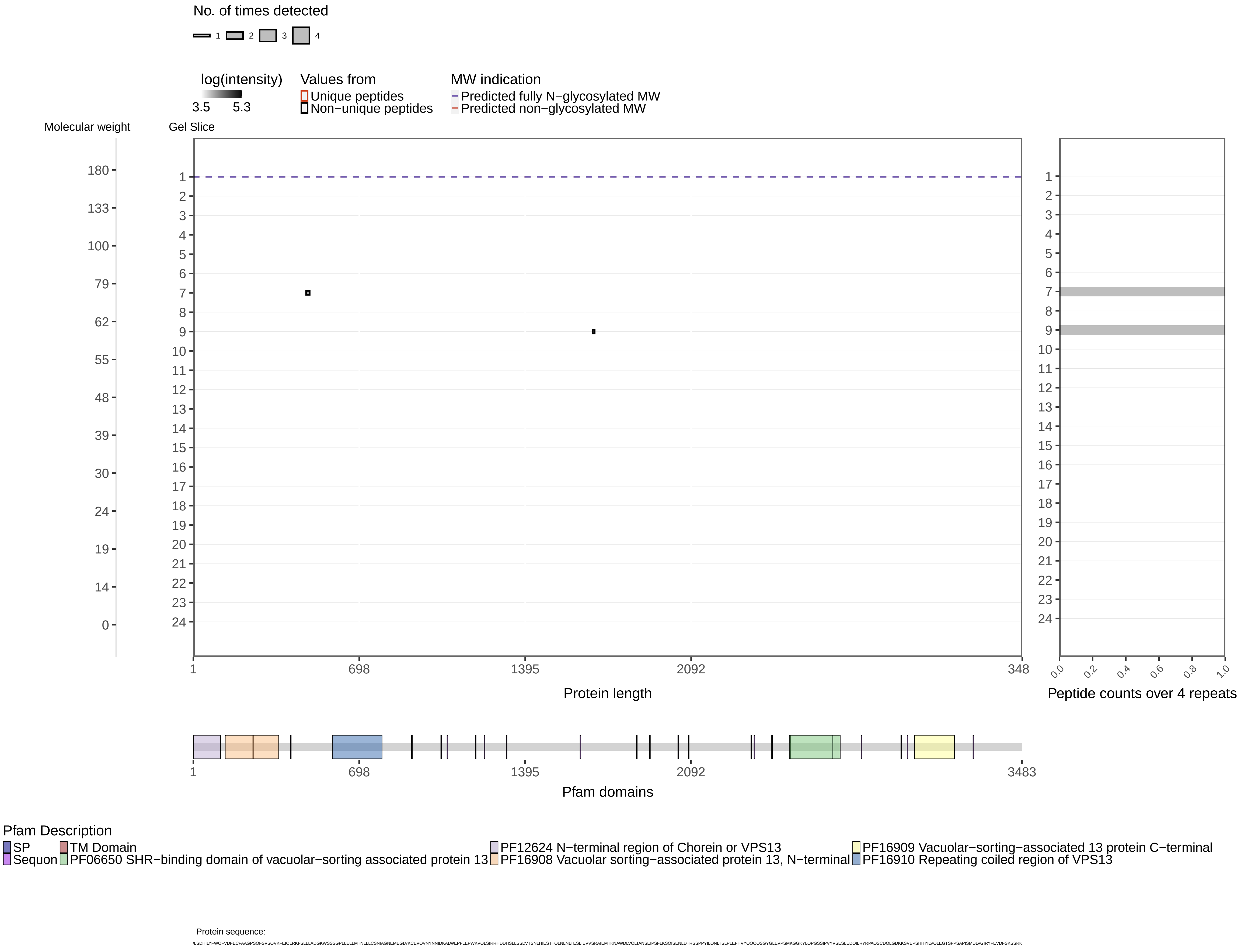

### NbL06g12820.1 pyruvate_dehydrogenase_(acetyl−transferring)_kinase_mitochondrial−like [MW = 42.22 kDa]

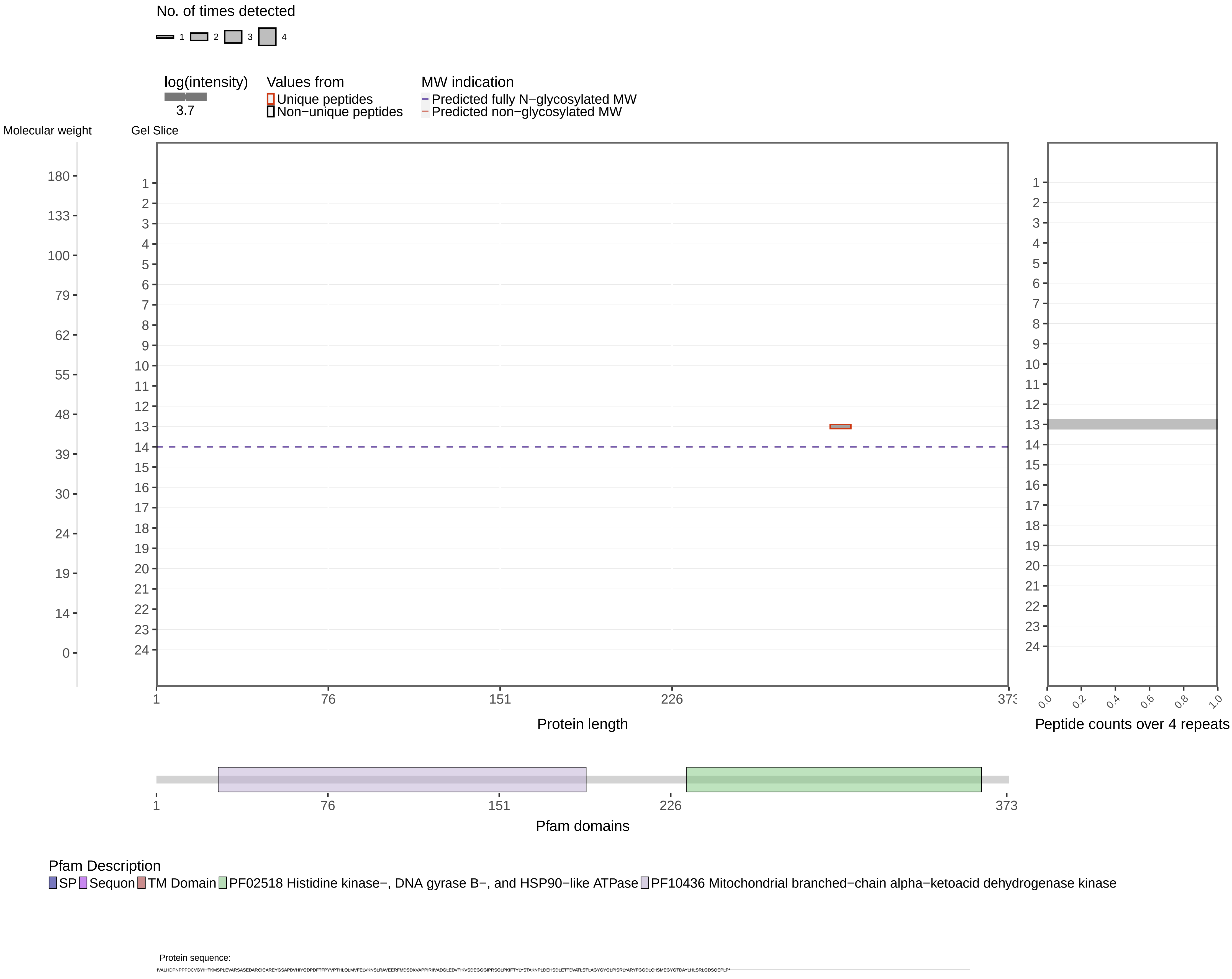

### NbL06g12690.1 galactokinase [MW = 54.62 kDa]

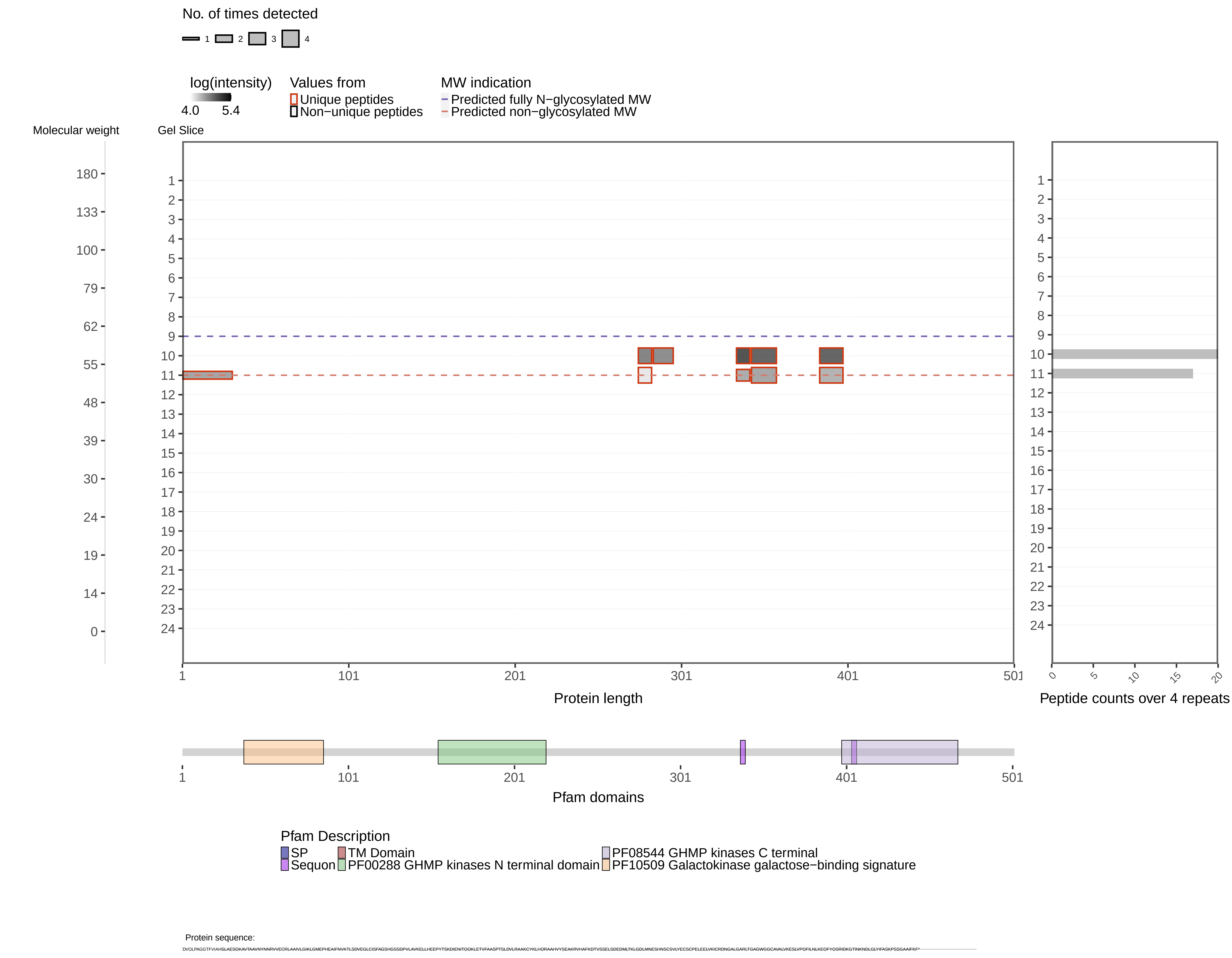

NbL06g12520.1 uncharacterized_LOC104235511 [MW = 35.45 kDa]

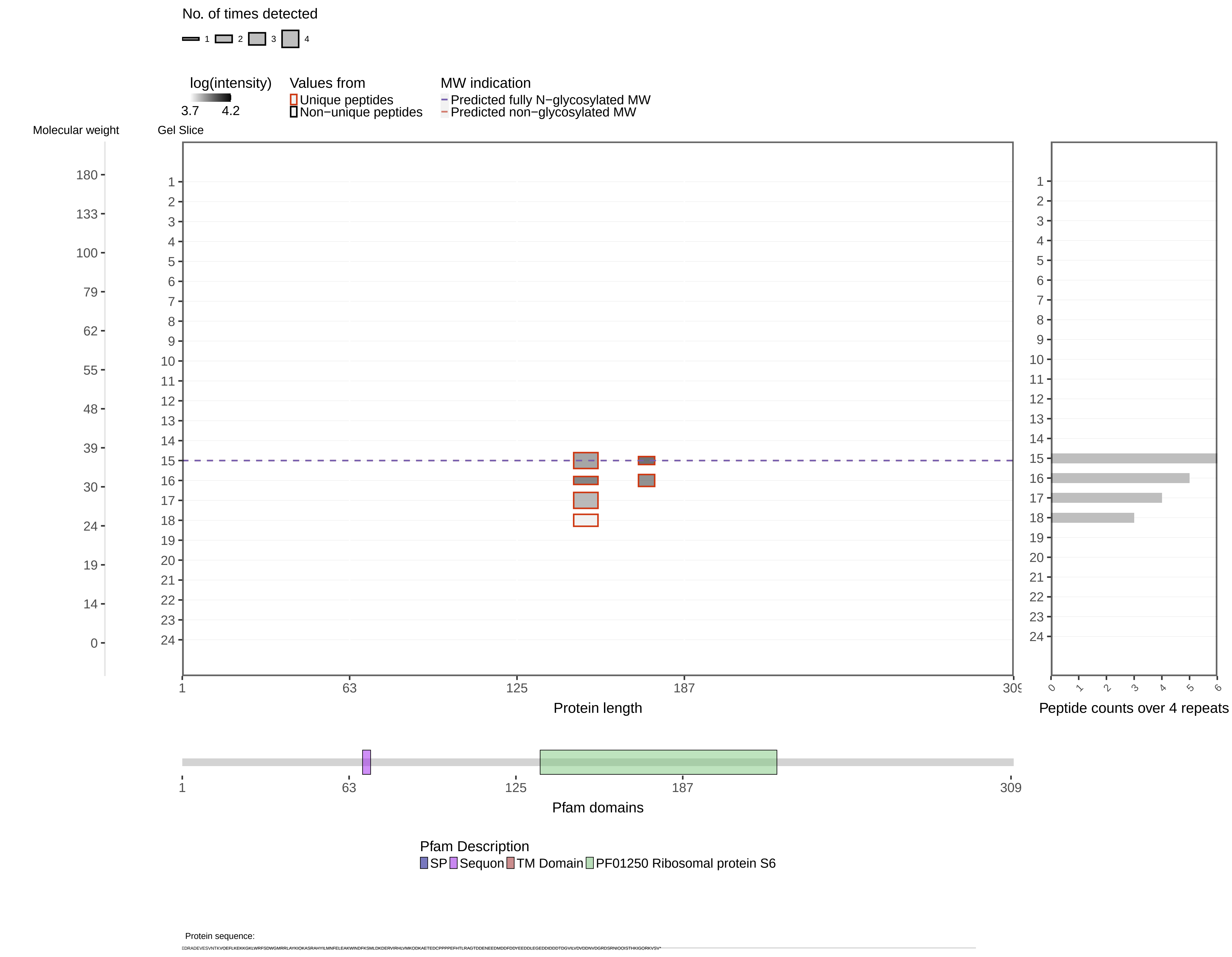

### NbL06g12480.1 probable_enoyl−CoA_hydratase_1_peroxisomal [MW = 29.26 kDa]

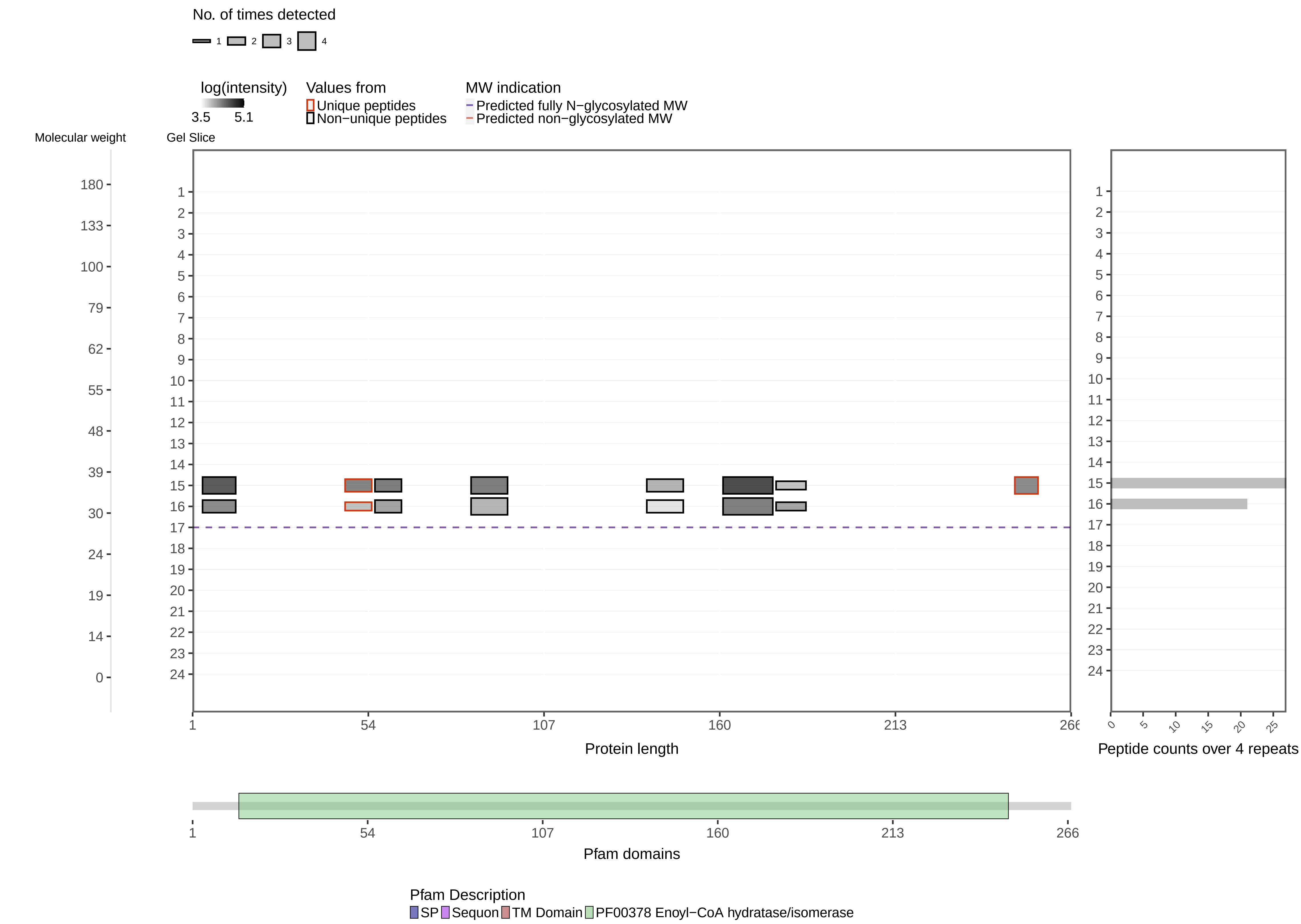


Protein sequence:

DLLVASNDAKFLDTHARFGIFPSWGLSQKLSRVIGPSRAREASLTAMPISAEQAERWGLVNYVVDRSELLKKARQIAEAIIKNNHDLVLKYKAVINDGFKQDLARALALEKERAHVYYDGMSKDHFKKLQEFIADRSSSKPSSKL*

NbL06g12430.1 ATP−citrate_synthase_beta_chain_protein_2 [MW = 65.58 kDa]

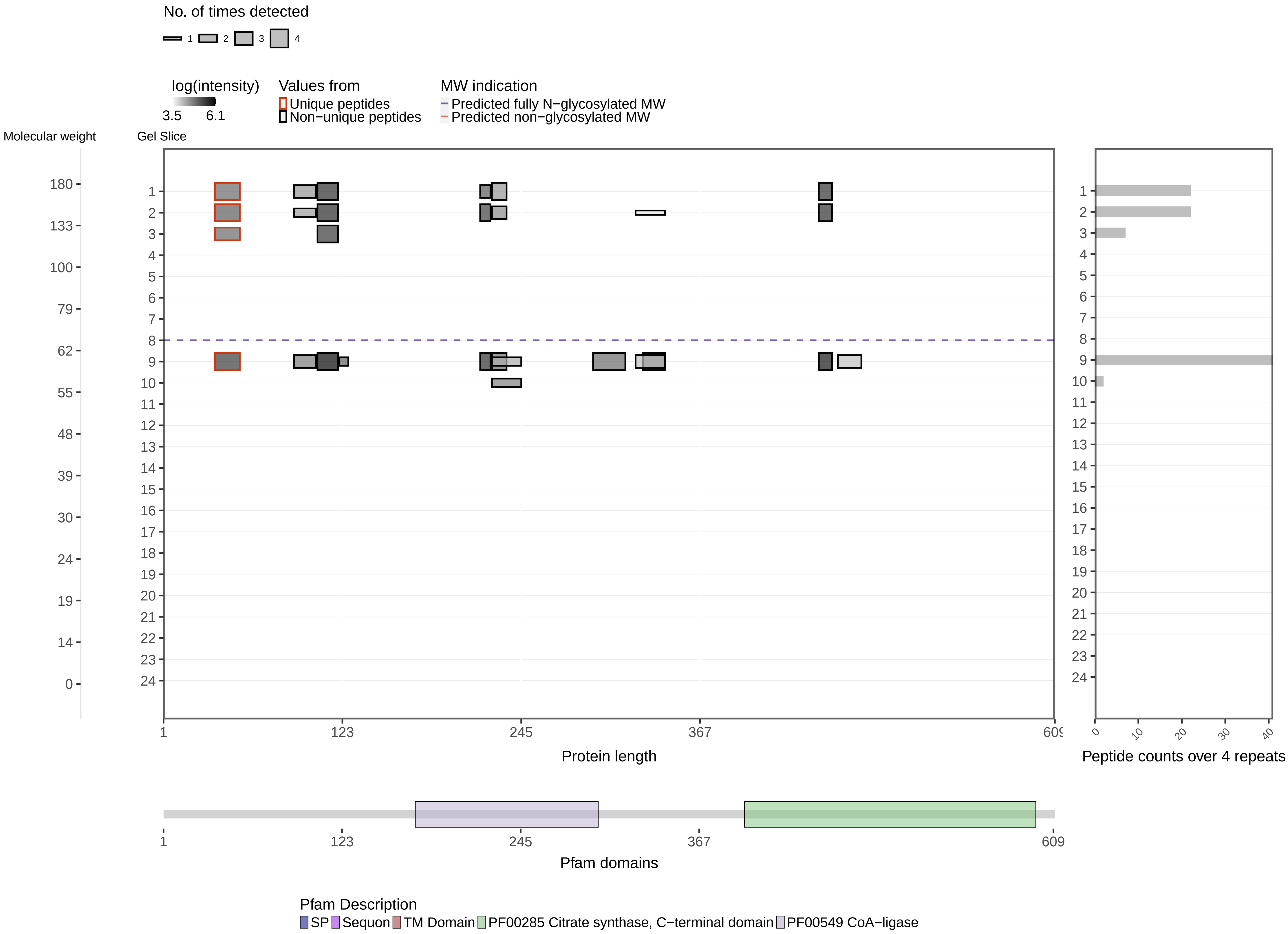


Protein sequence:

ALRDAGAVVPTSYEAFEGAIKDTFEKLVEAGKTTPVKEVSPPQIPEDLSTAIKCGKVRAPTHIISTISDDRGEEPCYAGVPMSSIVEQGLGVGDVISLLWFKRSLPRYCTRFIEICVMLCADHGPCVSGAHNSIVTARAGKDLVSCLVSGLLTIGPRFGGAIDDAARYFKDAYDKGLTPYEFVESMKKKGIRVPGIGHRIKRGDNRDKRVELLQLYARENFPSVKYMEYAVQVETYTLSKANNLVLNVDGAIGSLFLDLLAGSGMFTKPEIDEIVEIGYLNGLFVLARSIGLIGHTFDQK

### NbL06g12380.1 glucan_endo−13−beta−glucosidase_basic_vacuolar [MW − SP = 37.23 kDa]

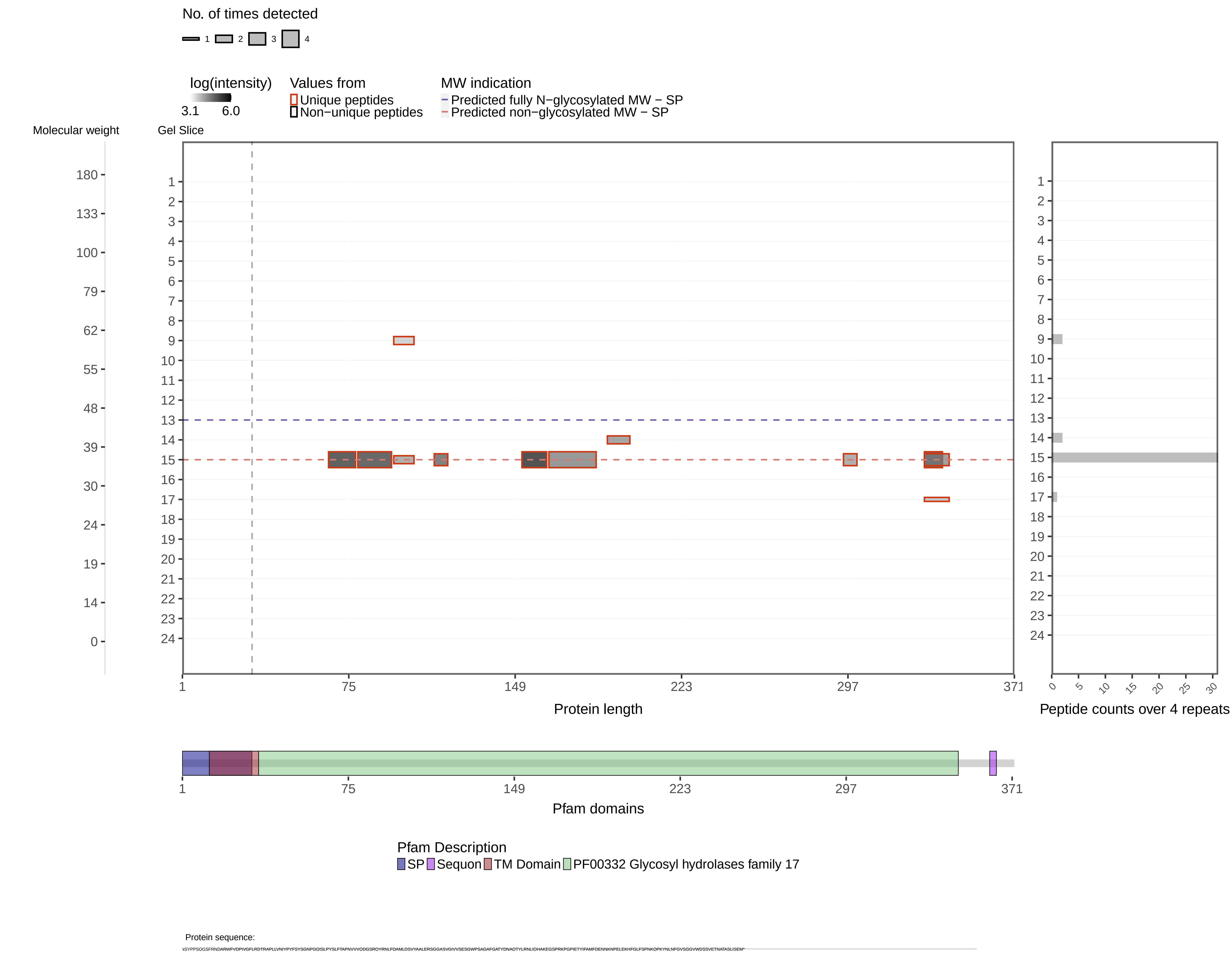

### NbL06g12330.1 endoplasmin_homolog [MW = 101.65 kDa]

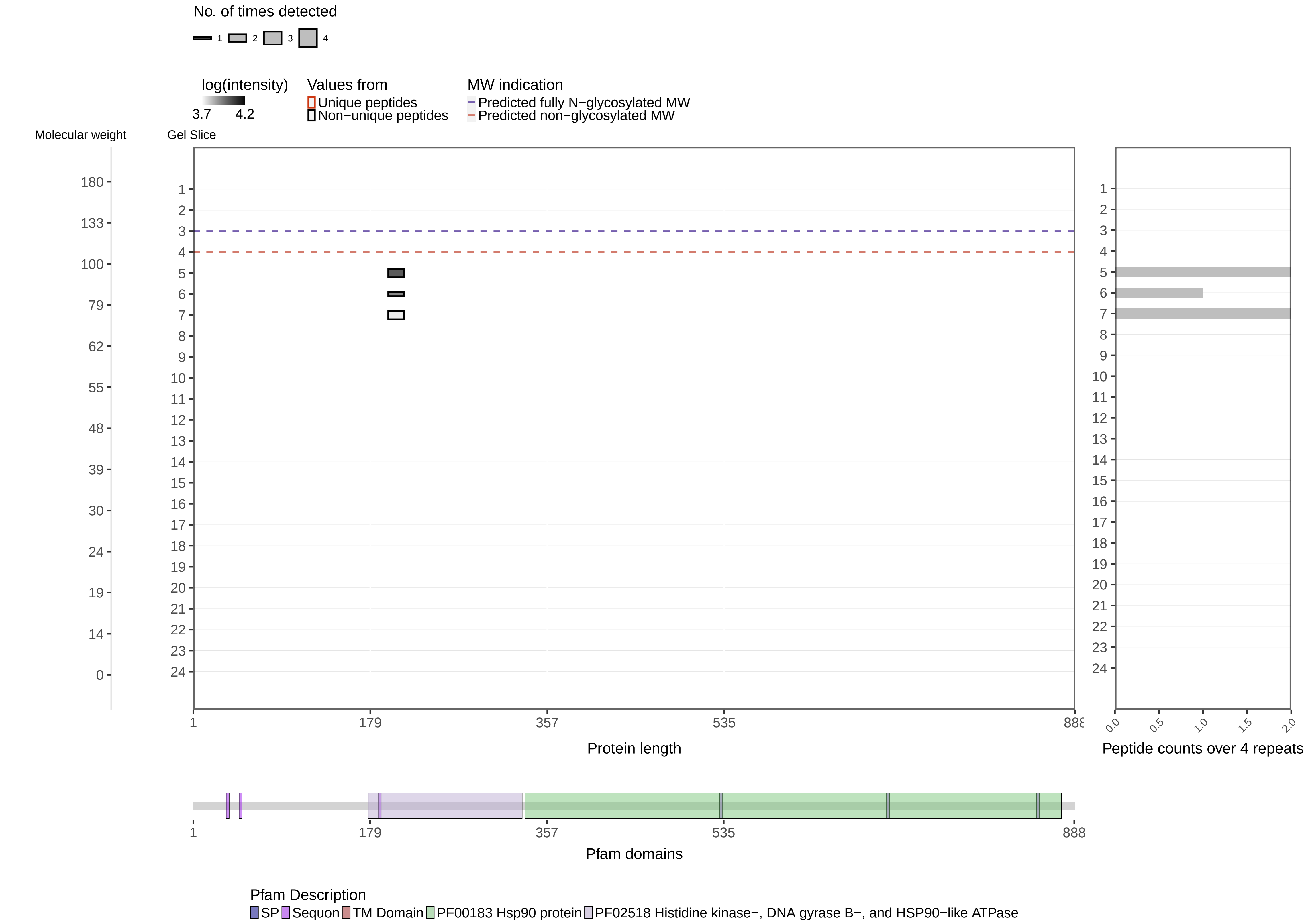


Protein sequence:

VTEEEYTKFYHSLAKDFSDEKPLAWSHFNAEGDVEFKAVLFVPPKAPHDLYESYYNSKKSNLKLYVRRVFISDEFDELLPKYLSFLMGLVDSDTLPLNVSREMLQQHSSLKTIKKKLIRKALDMIRKIADEDPDESNDKDKKDIEESSADNEKKGQYAKFWNEFGKSVKLGIIEDATNRNRLAKLLRFETSKSDGKLTSLDQYISRMKAGQKDIFYITGASKEQLEQSPFLERLTKKNYEVIFFTDPVDEYLMQYLMDYEDHKFQNVSKEGLKLGKDSKAKELKESFKELTKWW

NbL06g12200.1 12−dihydroxy−3−keto−5−methylthiopentene_dioxygenase_1 [MW = 22.26 kDa]

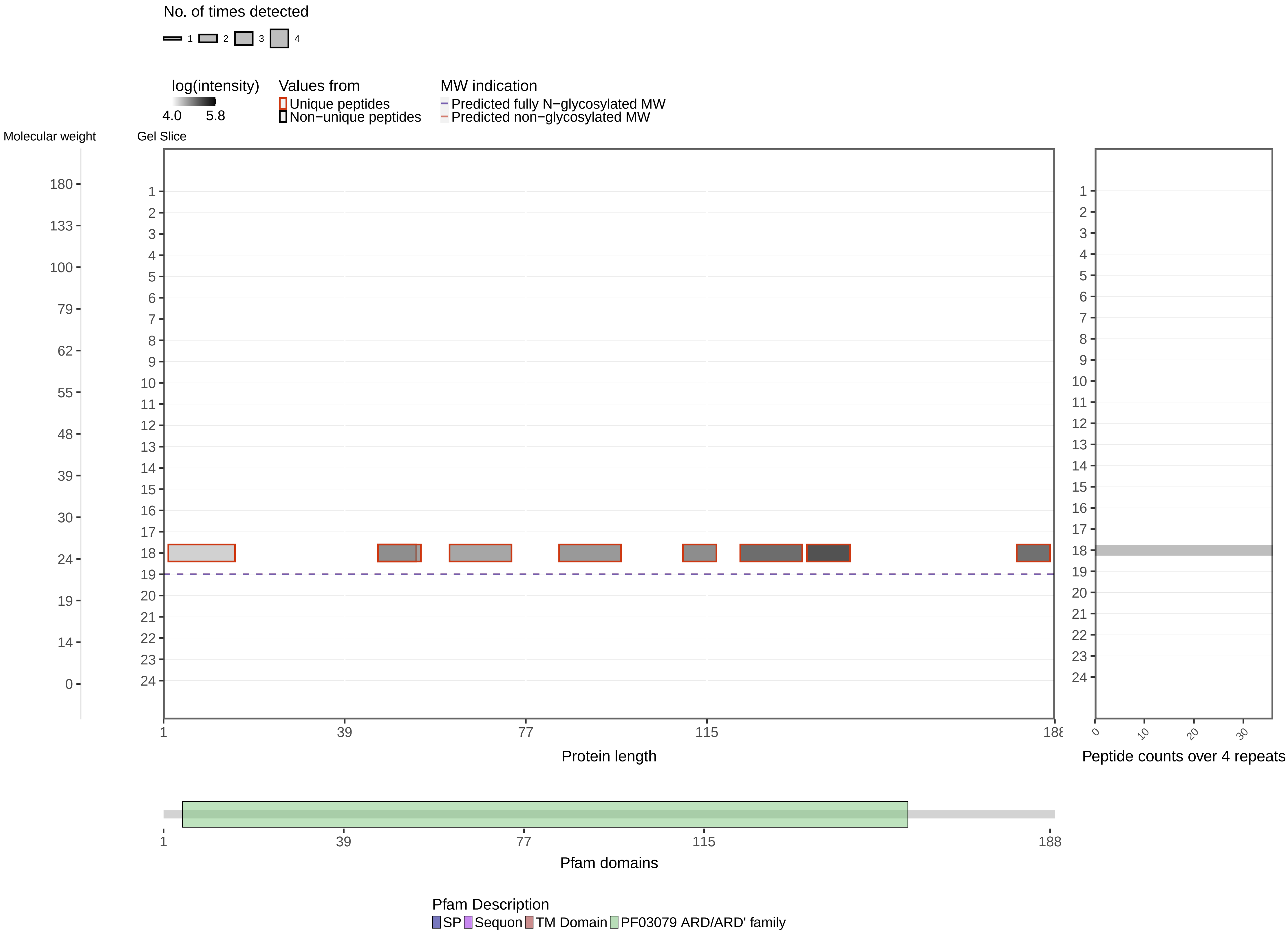


Protein sequence:

QKLKNFYTEHIHADEEIRYCLEGSGYFDVRDNEDHWIRIWIKAGDLIILPAGIYHRFTLDTGNYVKLMRLFVGEPVWTAHNRPQEDHPARREYIKSVTERVGVPLAAH*

### NbL06g12160.1 probable_histone_H2AXb [MW = 14.65 kDa]

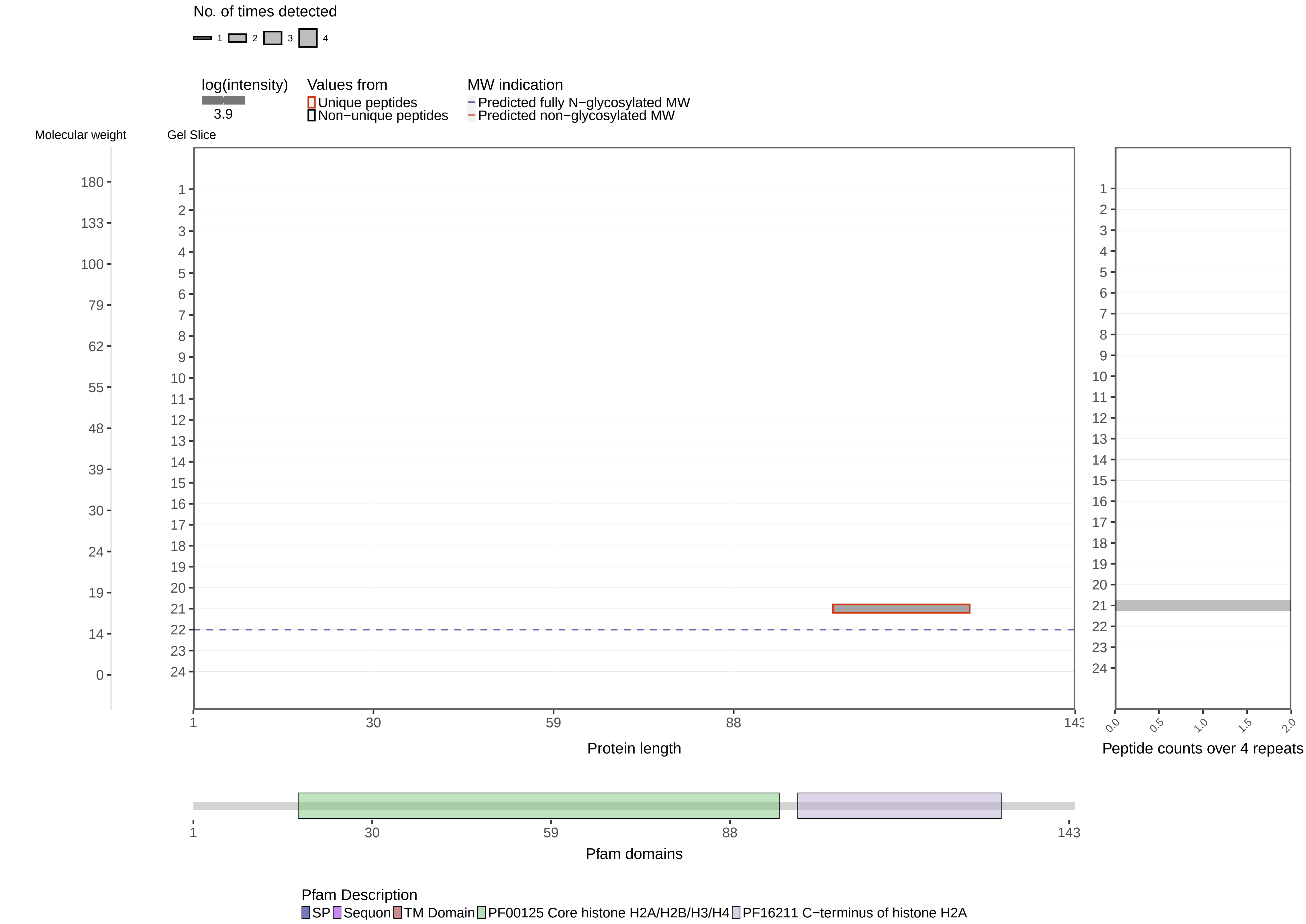


Protein sequence:

/YLSAVLEYLAAEVLELAGNAARDNKKNRIVPRHIQLAVRNDEELSKLLGQVTIANGGVLPNIHQNLLPKKAGSGKGDIGSASQEF*

### NbL06g12030.1 peroxidase_2−like [MW = 32.36 kDa]

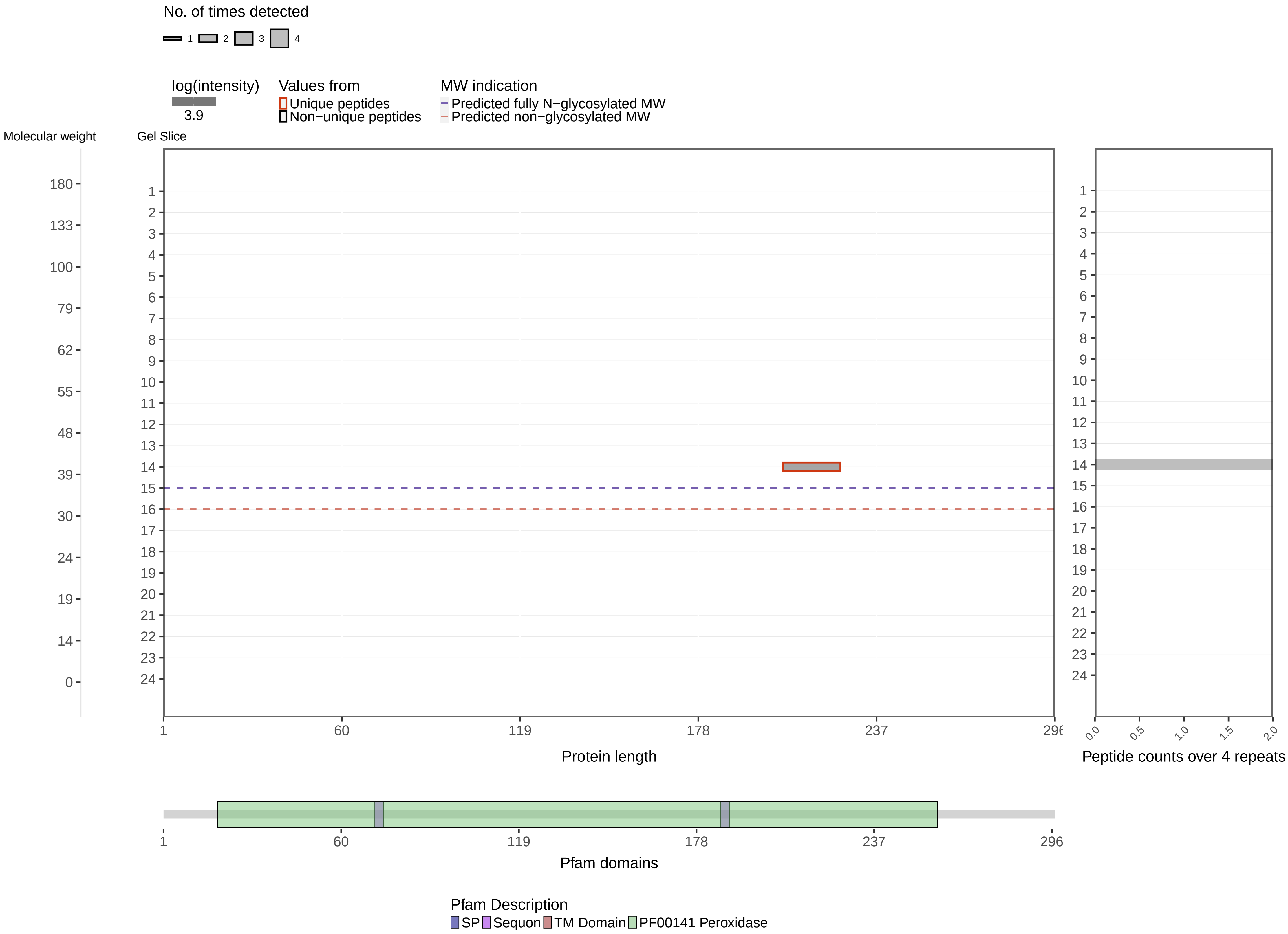


Protein sequence:

ANNDIPSPFMDLPALINSFKNQGLDKEDLVALSGGHTLGFAQCFTFRNRIYNETNNIDSTFASQRQANCPRSGGDSNLASLDPTAALFDSKYFSNLVSKKGLLHSDQALFSGGETDDLVKTYSTNLGTFSKDFAESMIKMGNIKPLTGNQGQIRVNCRKVN*

### NbL06g11920.1 COBRA−like_protein_7 [MW − SP = 68.86 kDa]

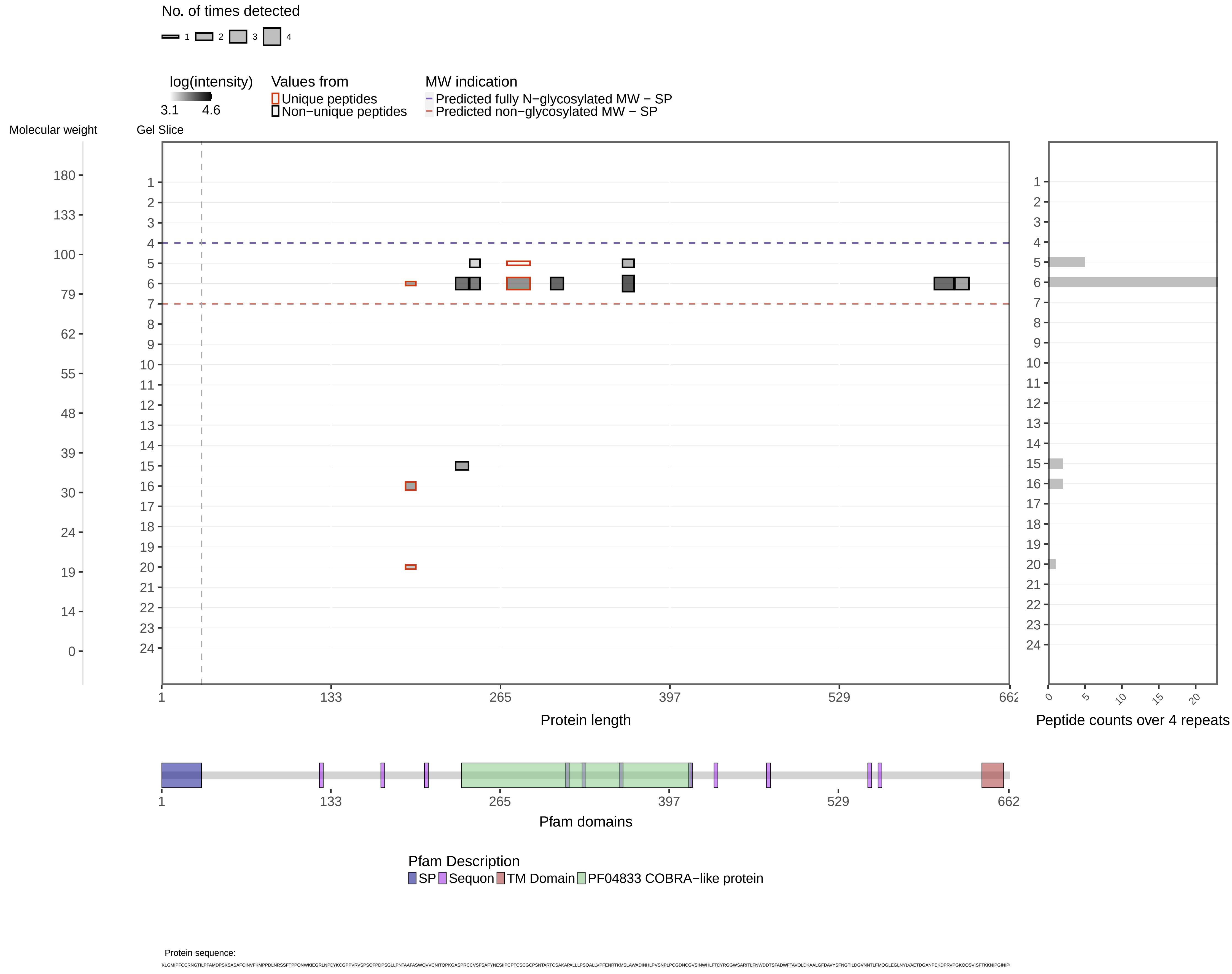

### NbL06g11810.1 acetate/butyrate−−CoA_ligase_AAE7_peroxisomal−like [MW = 62.55 kDa]

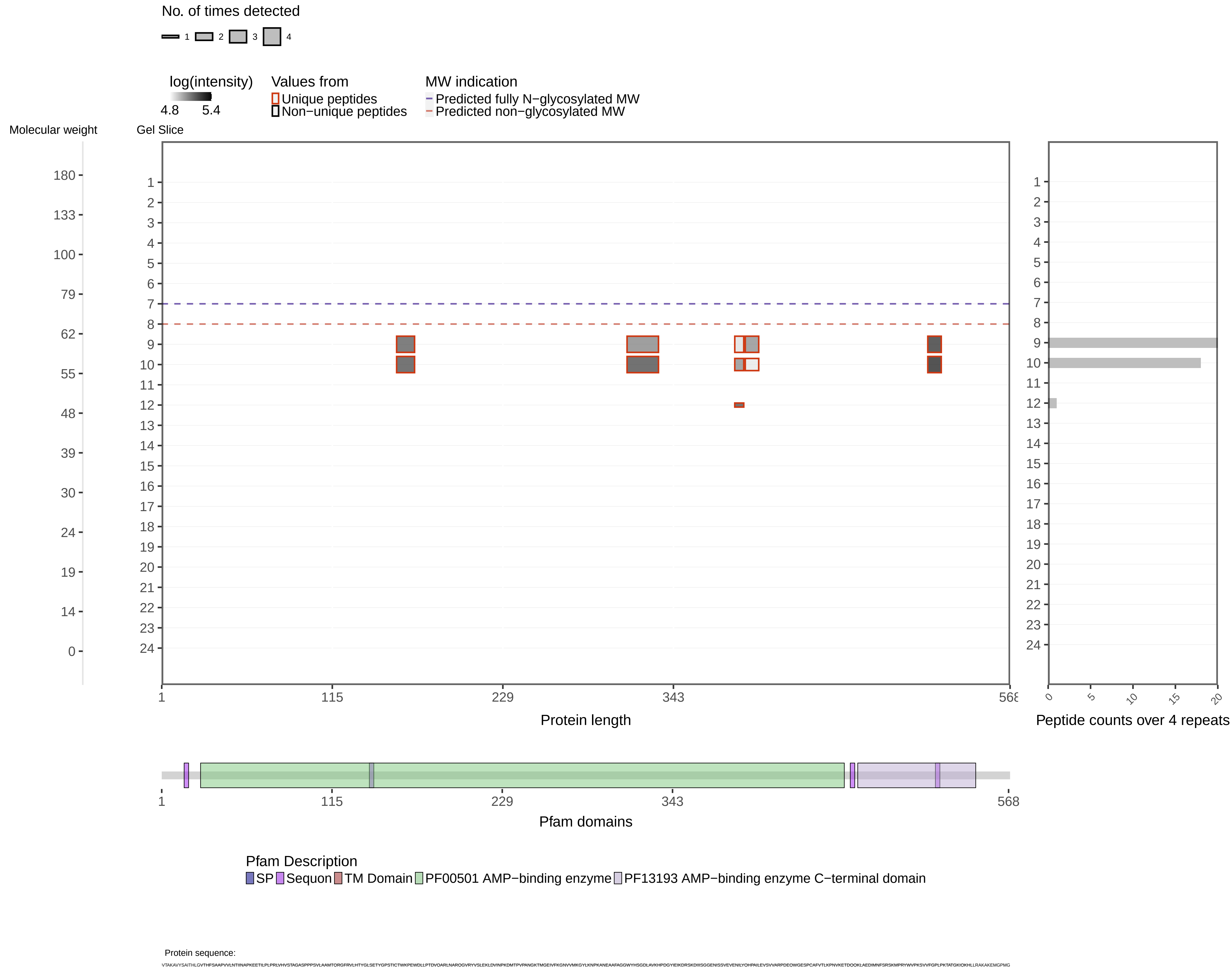

### NbL06g11790.1 GTP−binding_protein_SAR2 [MW = 21.96 kDa]

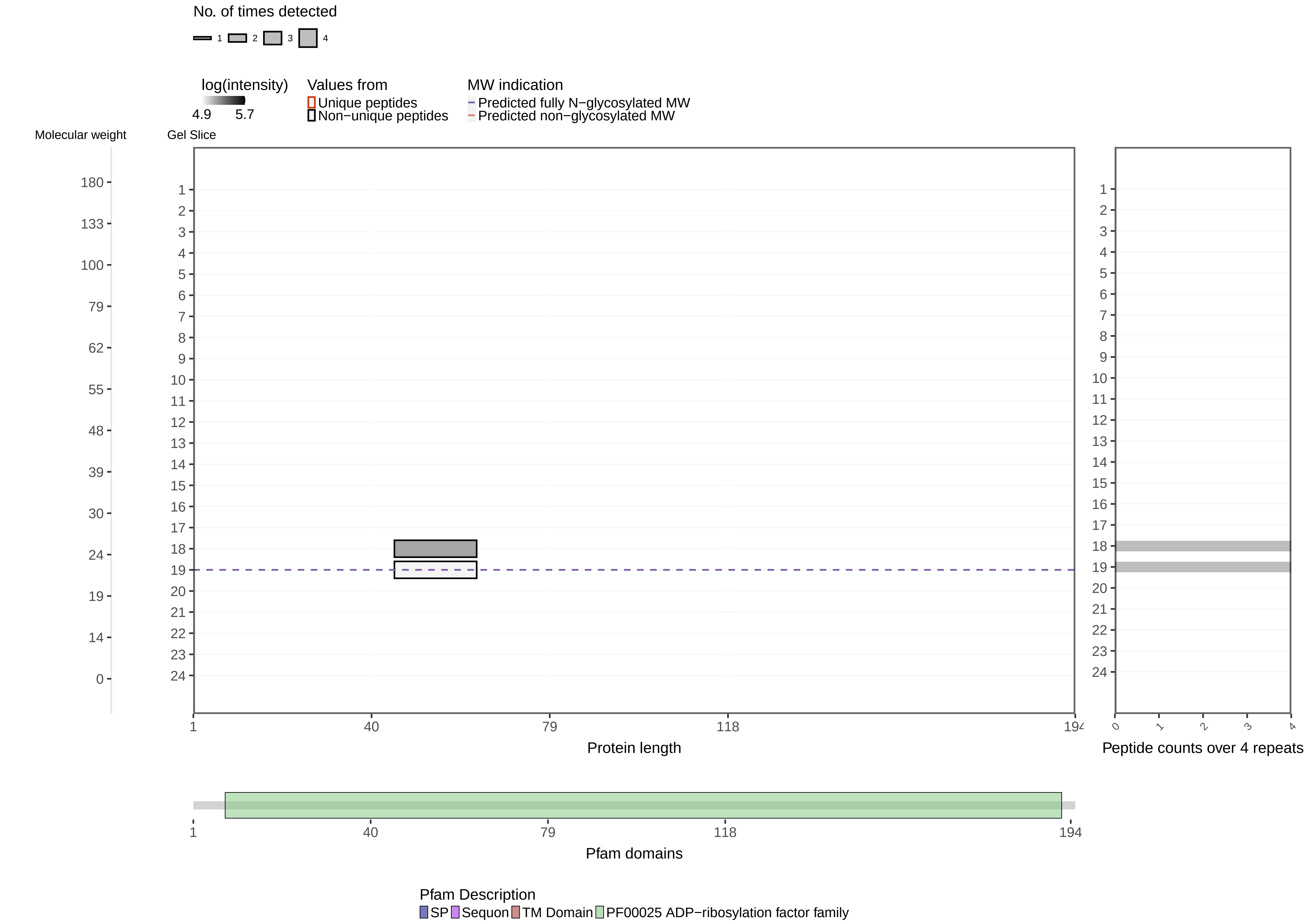


Protein sequence:

YYAKVDAVVYLVDAYDRERFPESKKELDGLLSDESLANVPFLILGNKIDIPYAASEDELRYHLGLTGVTTGKGNVNLAGTNVRPIEVFMCSIVRKMGYGEGFKWMSQYIK*

### NbL06g11660.1 proteasome_subunit_beta_type−2−A [MW = 22.59 kDa]

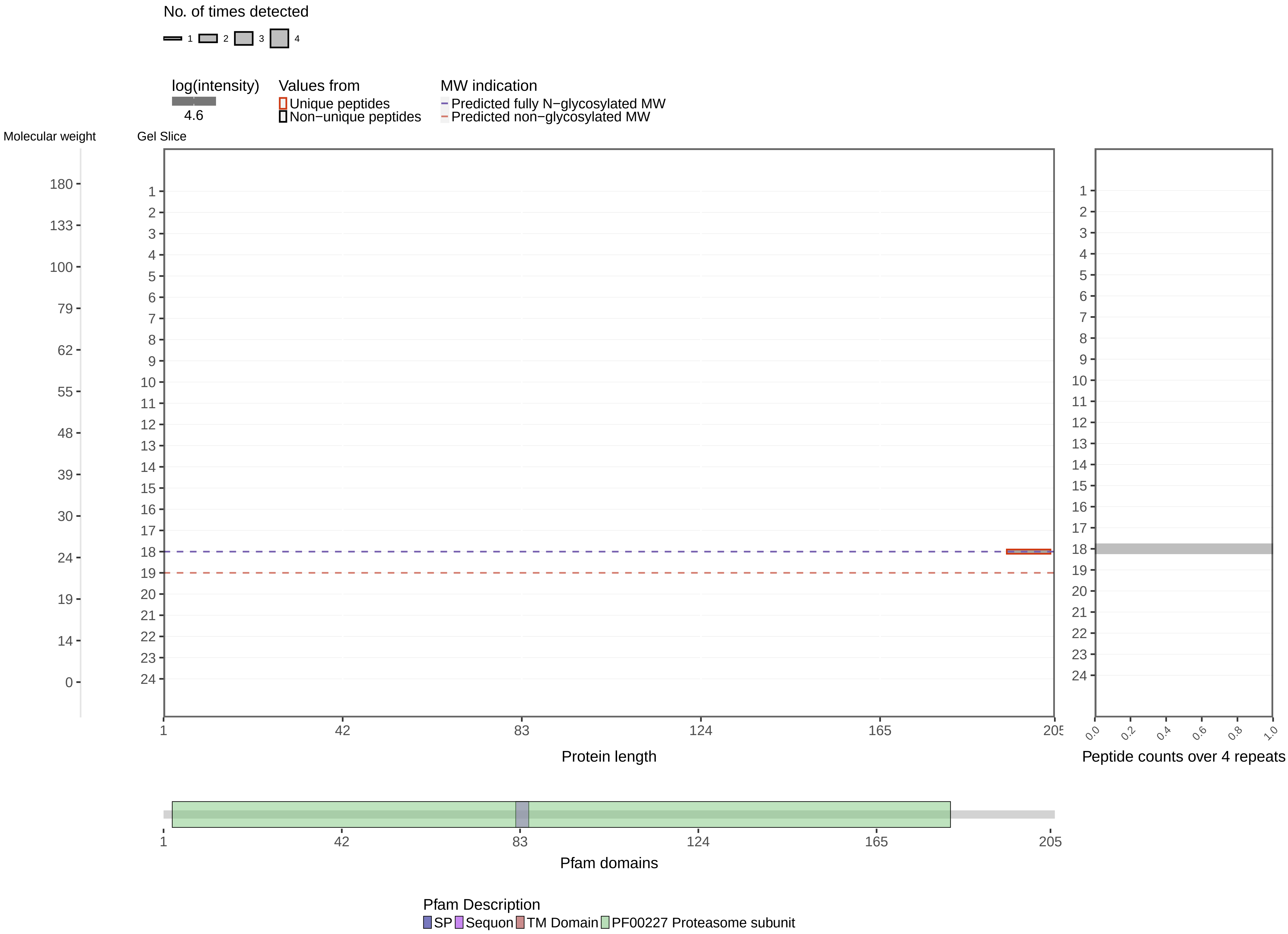


Protein sequence:

ATALRKNPYMVNIILAGYDKETGPSLYYVDYIATLHKVDKAAFGYGSYFSLAMMDRHYRKDMTVEEAVDLADKCILEIRSRLVVAPPNFVIKIVDKDGAREYAWRQSVKDAPVSDS*

### NbL06g11420.1 synthase [MW = 34.22 kDa]

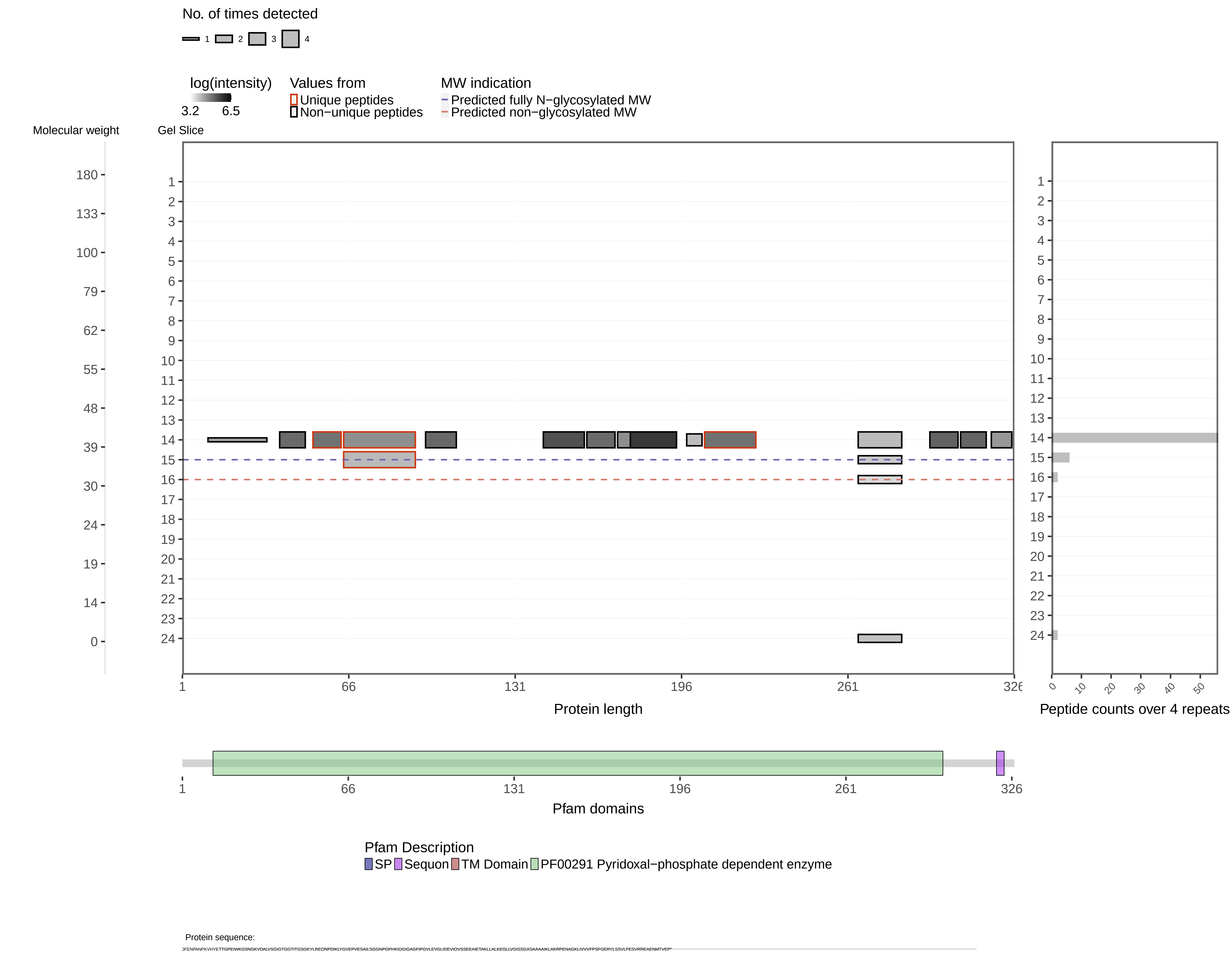

### NbL06g11280.1 heat_shock_70_kDa_protein_8 [MW = 62.51 kDa]

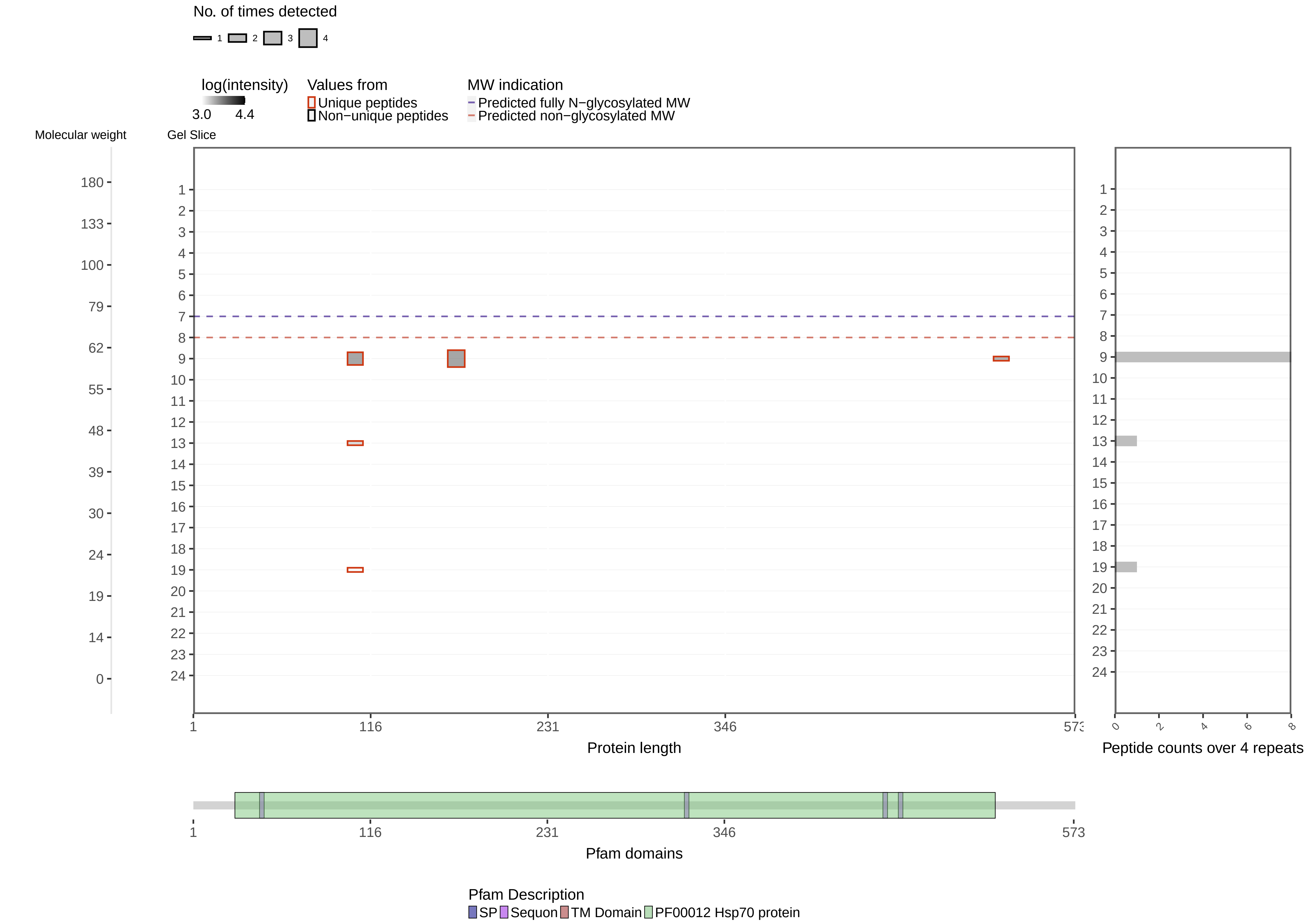


Protein sequence:

HLLPSMDHIFSSHGIEEIKRMGLLRVATQDAIHKLSSEMSVPIDVVLGNGTKISKFLERAEFEEVNKEVFQKCETLIERCLRDAKLEAEDVHDVIIVGGCSYVPKIRNIVMSICKREELYSGMNPLEAAVRGAALEGAVASGISDPFGSLDLLTIQATPLSIGIQADGSNFVPIIYQNTTTPVRKEQNFTTVHDNQAEALIFVYEGDEKTVKDNHLLGYFKITGIPPVPKGVPEINVCMDIDASNVLRVFAGVIIPGAKCPPFMEVRMPTVDDGHAWSAQTLHRTYGSTLDLVTVKKKGC

### NbL06g11160.1 mavicyanin [MW − SP = 16.4 kDa]

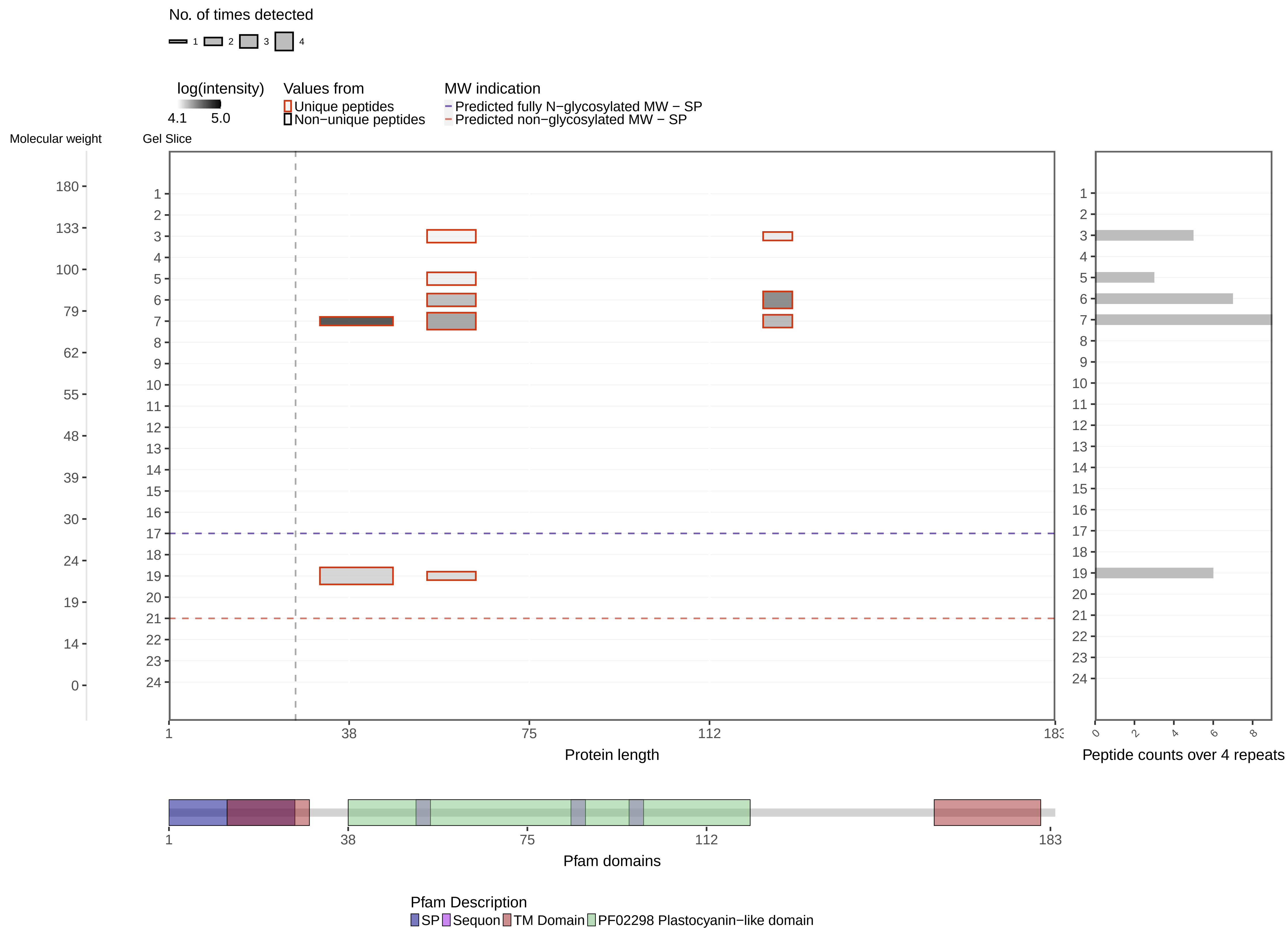


Protein sequence:

### NbL06g11080.1 RAN_GTPase−activating_protein_2−like [MW = 61.24 kDa]

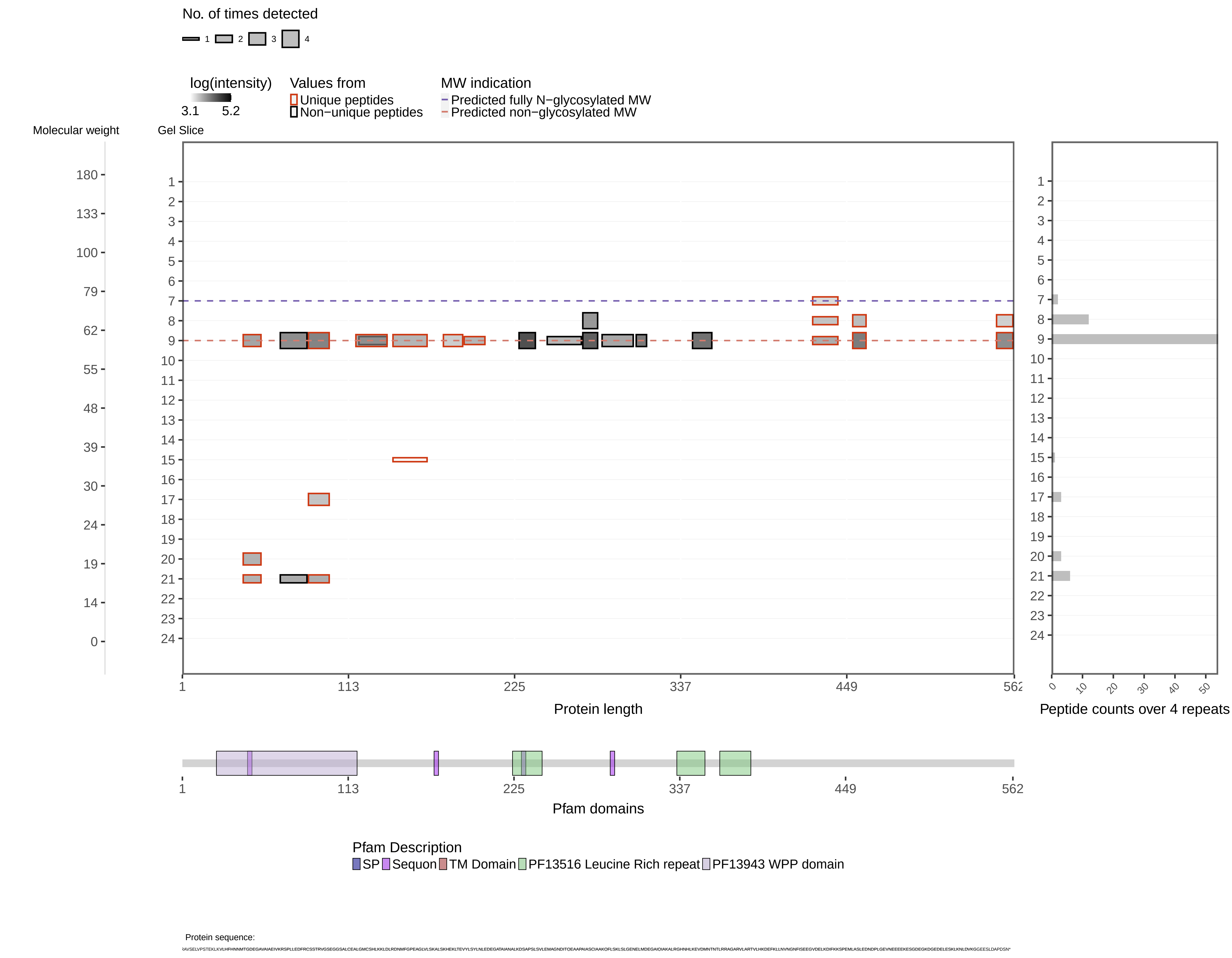

NbL06g11000.1 glucose−1−phosphate_adenylyltransferase_large_subunit_3_chloroplastic/amyloplastic−like [MW = 57.28 kDa]

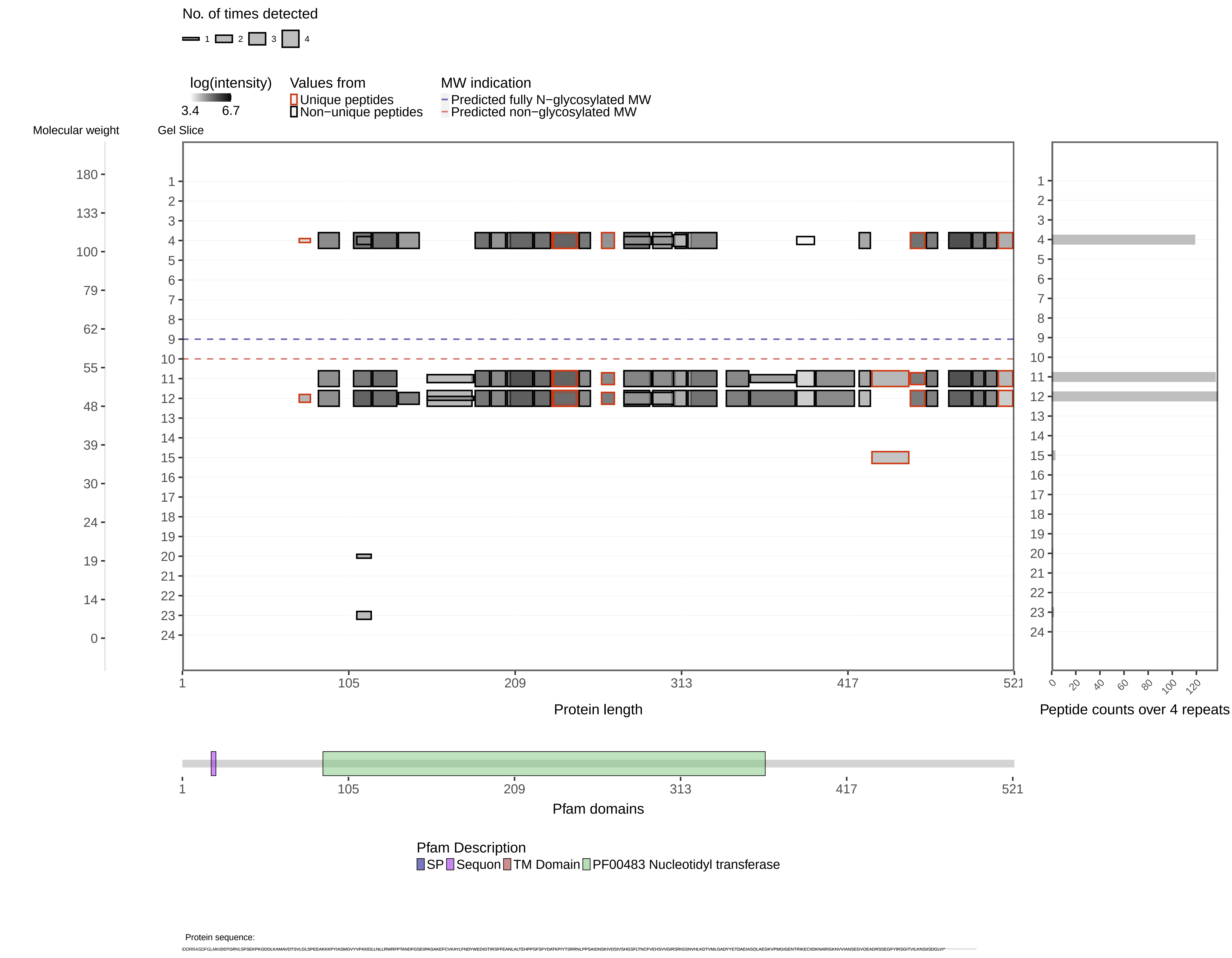

### NbL06g10970.1 peroxiredoxin−2F_mitochondrial−like [MW = 20.96 kDa]

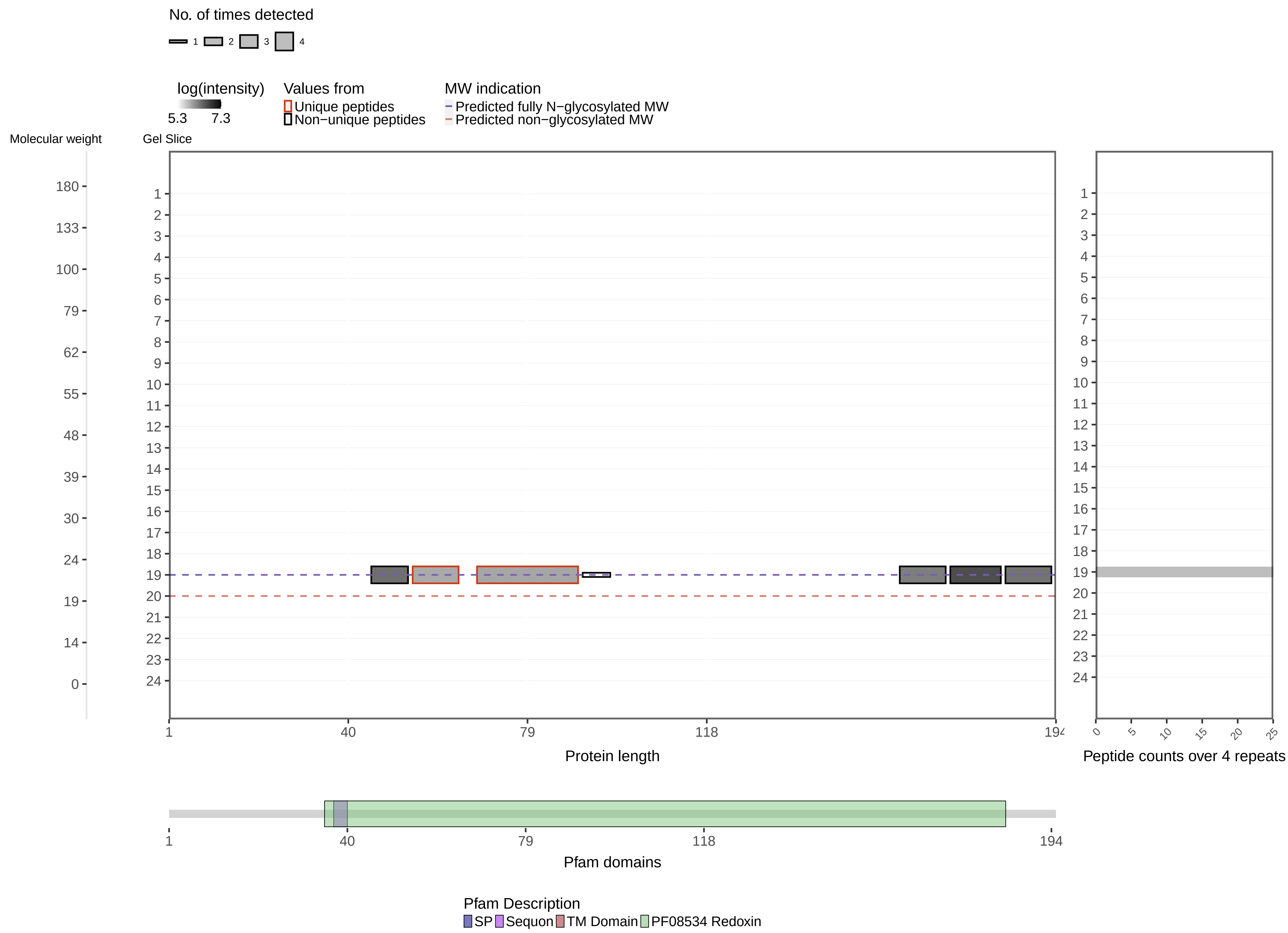


Protein sequence:

QHVPSYKNNIDKFKAKGIDSVICVAVNDPYVMNGWAEKLQAKEAIEFYGDFDGSFHKSLDLTIDLCAALLGPRSHRWSAYVDDGNVKVLNVEEAPSDFKVSGGDVILGQI*

### NbL06g10940.1 NEDD8−activating_enzyme_E1_catalytic_subunit [MW = 49.57 kDa]

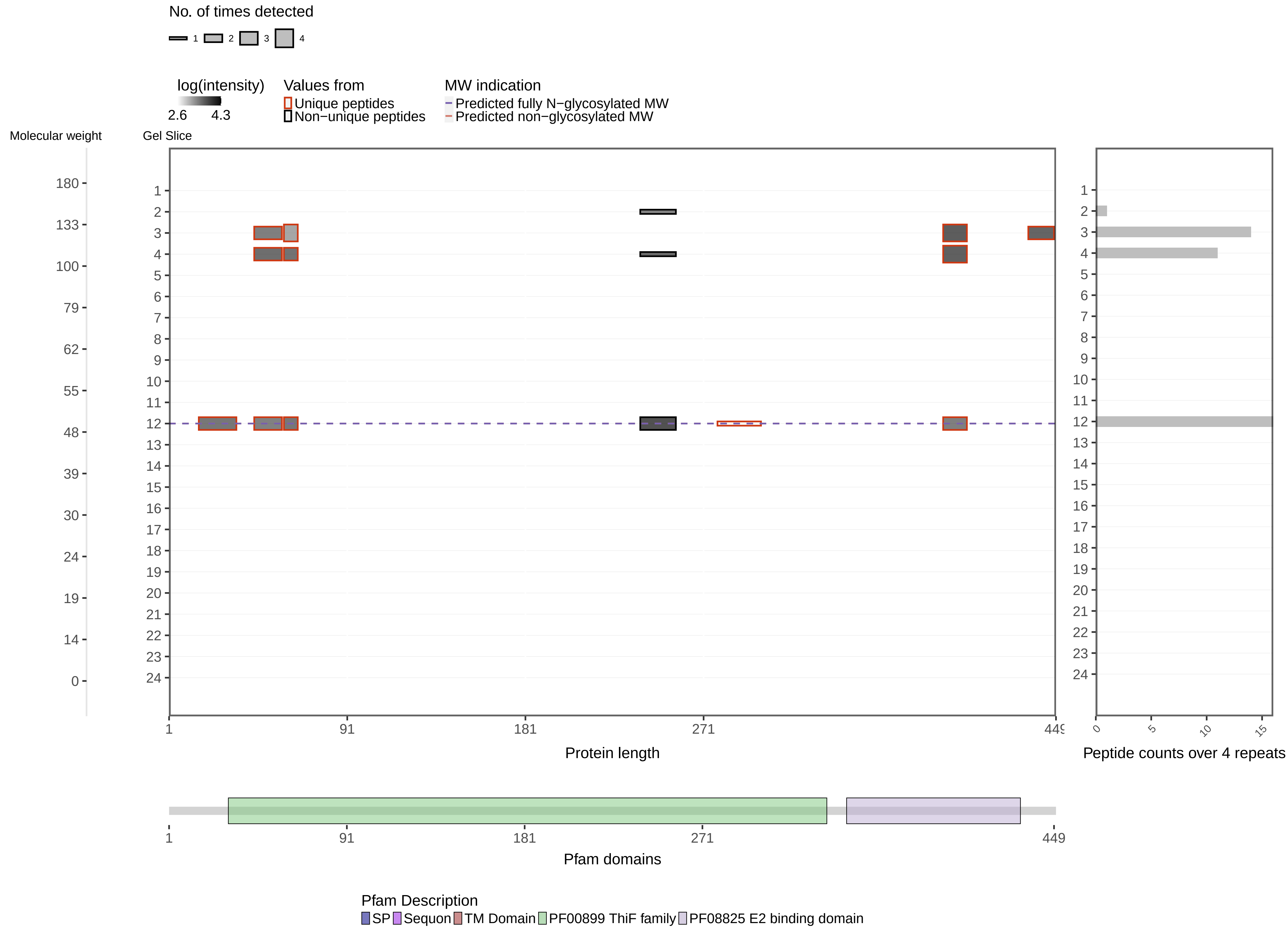


Protein sequence:
TLAETPRTAAHCIEYAHLIKWDEVHSGKTFDPDDPEHMQWVYSEAVKRAELFGILGVTYSLTQGVVKNIIPAIASTNAIISAACALETLKLVSGCSKTLSNYLTYNGVEGLHTKVTEFVRDKECLVCSPGILIELEASVTLKKFIGLLEDHPKLLLTRVSVTYRGKNLYMQAPPVLEEMTRSNLDLPLFELMSKTPRDIVHVTGTAGKGDTKQSCSRKLRVVFKGIDGVTDMDMAGGA*

NbL06g10920.1 ATP−dependent_(S)−NAD(P)H−hydrate_dehydratase [MW = 22.38 kDa]

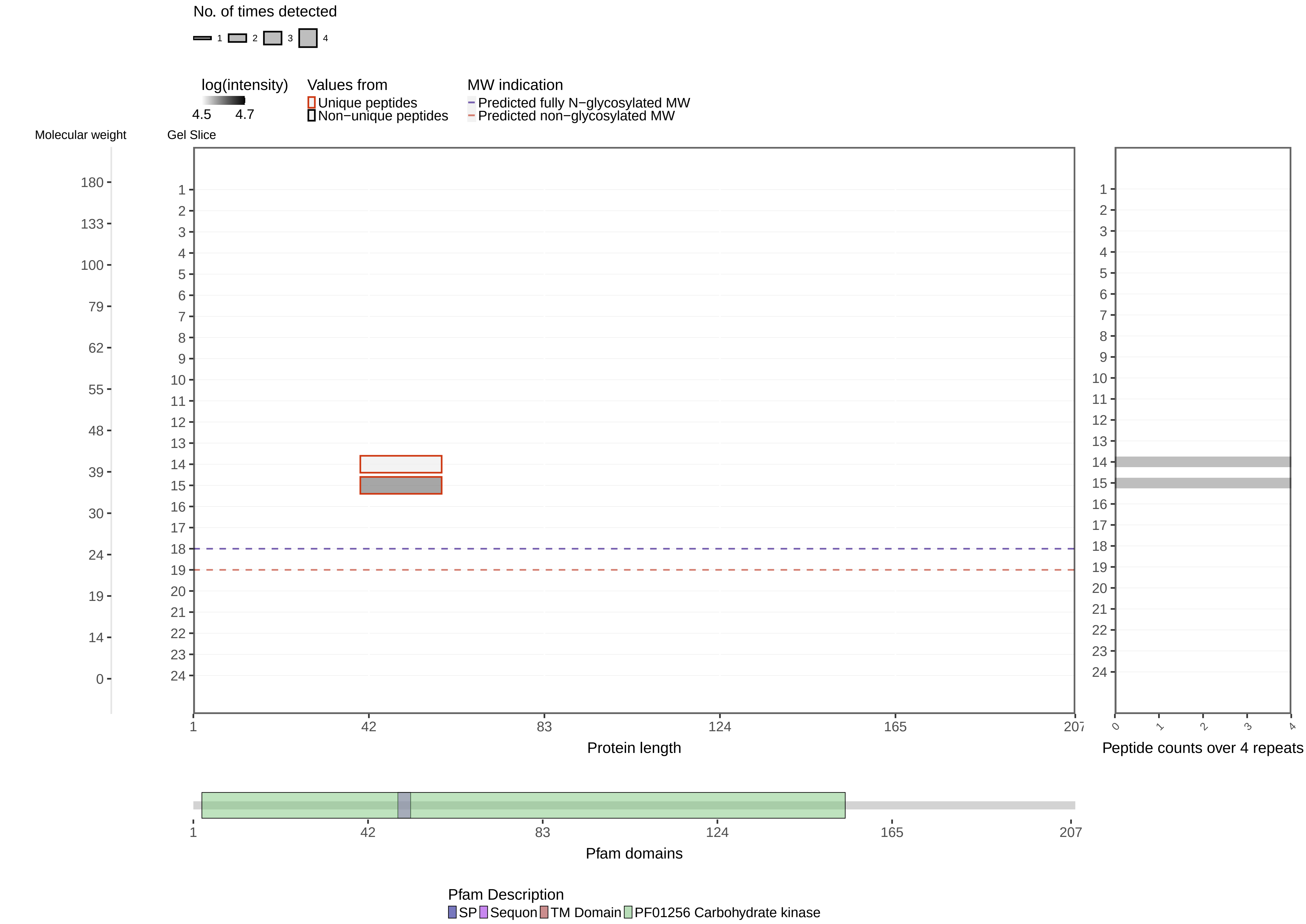


Protein sequence:

RCGGQGDILSGSIAVFLSWARQYAAKGEVSMNPTILGCVAGSALLRRAASLAFETKKRSTLTGVIIECLGRRKYAQSNRNFMEGFMGAQHMYSKLEEAFLWQRAKGKQRRKNNGK*

### NbL06g10900.1 stem−specific_protein_TSJT1−like [MW = 30.04 kDa]

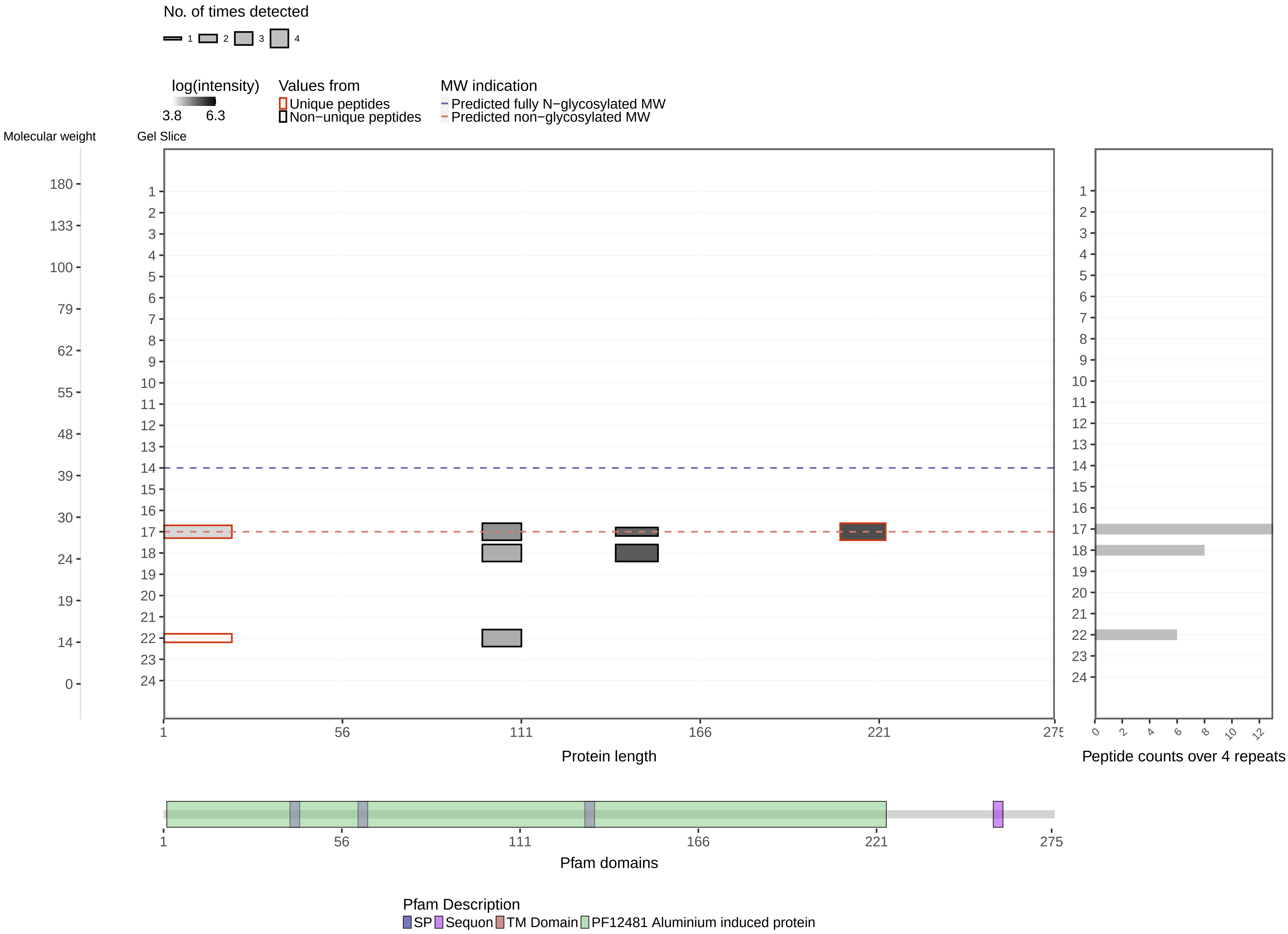


Protein sequence:

'GHLEGNFTFIVFDKSTSTLFVATDOVGKVPLYWGITADGYVAFANDADLLKGACGKSLASFPOGCFYSTSVGELRSYENPKNKITAVPATEEEIWGTKYMDPVEISGGRFGRCSCSHKIEDLVFLSELCWKNESREWOKMOGRALVGAM*

### NbL06g10770.1 UPF0613_protein_PB24D3.06c [MW = 37.06 kDa]

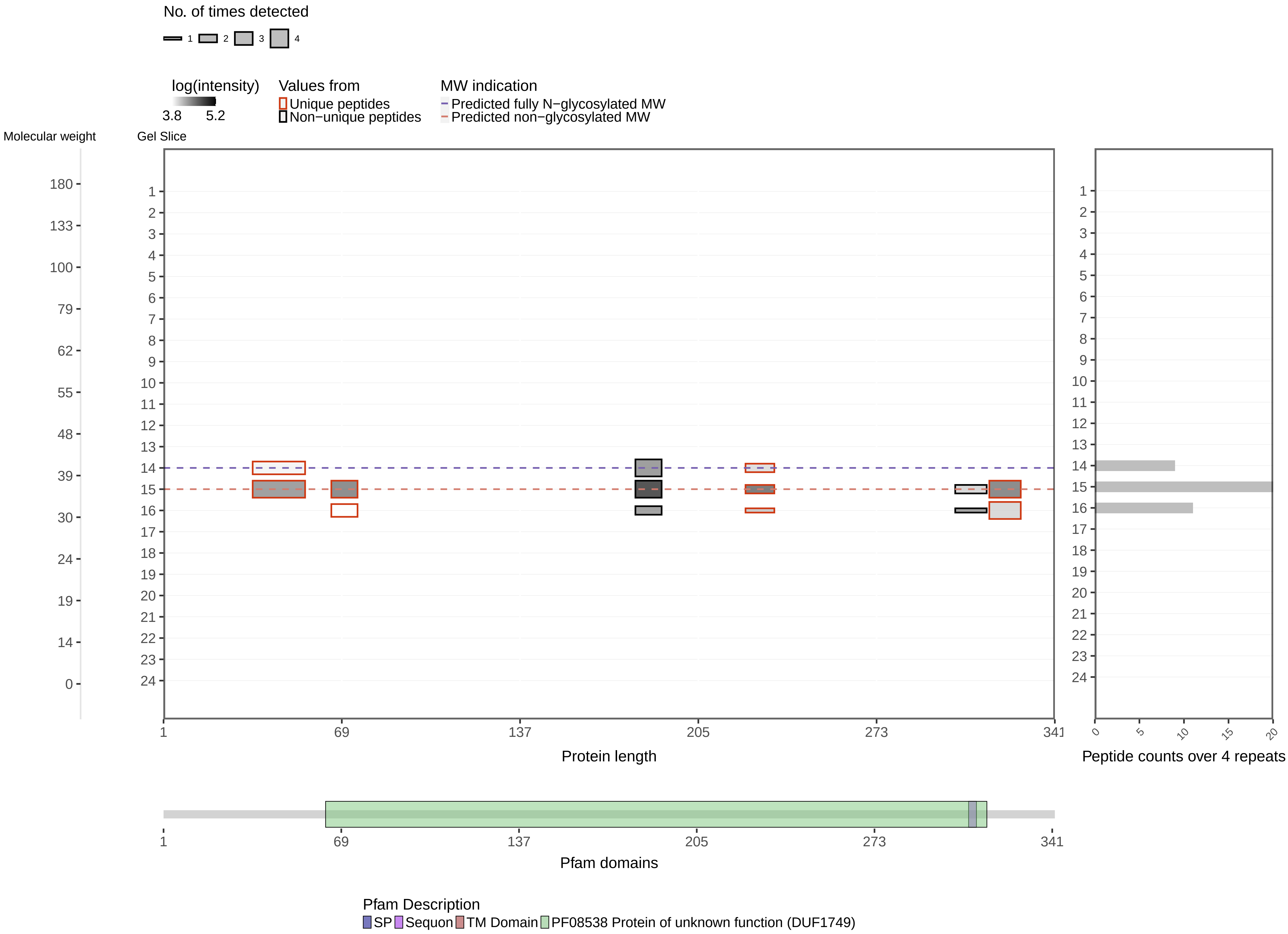


Protein sequence:

STGCQDIVYYMRTNAACSRAVRAAILQAPVSDREYKATLPDTASMIDLASNMISEGRESELMPREANPDAPITAFRYHSLCAYNGEDDLFSSDFSDDQLKQRLGHMSNTPCQVIFSMGDEYVPEYVDKKALVDRLCRAMGGAEKVEIEHGNHSLSNRVAEAVESIMSFVKREGPSGWDDPWN*

### NbL06g10700.1 NA [MW = 29.94 kDa]

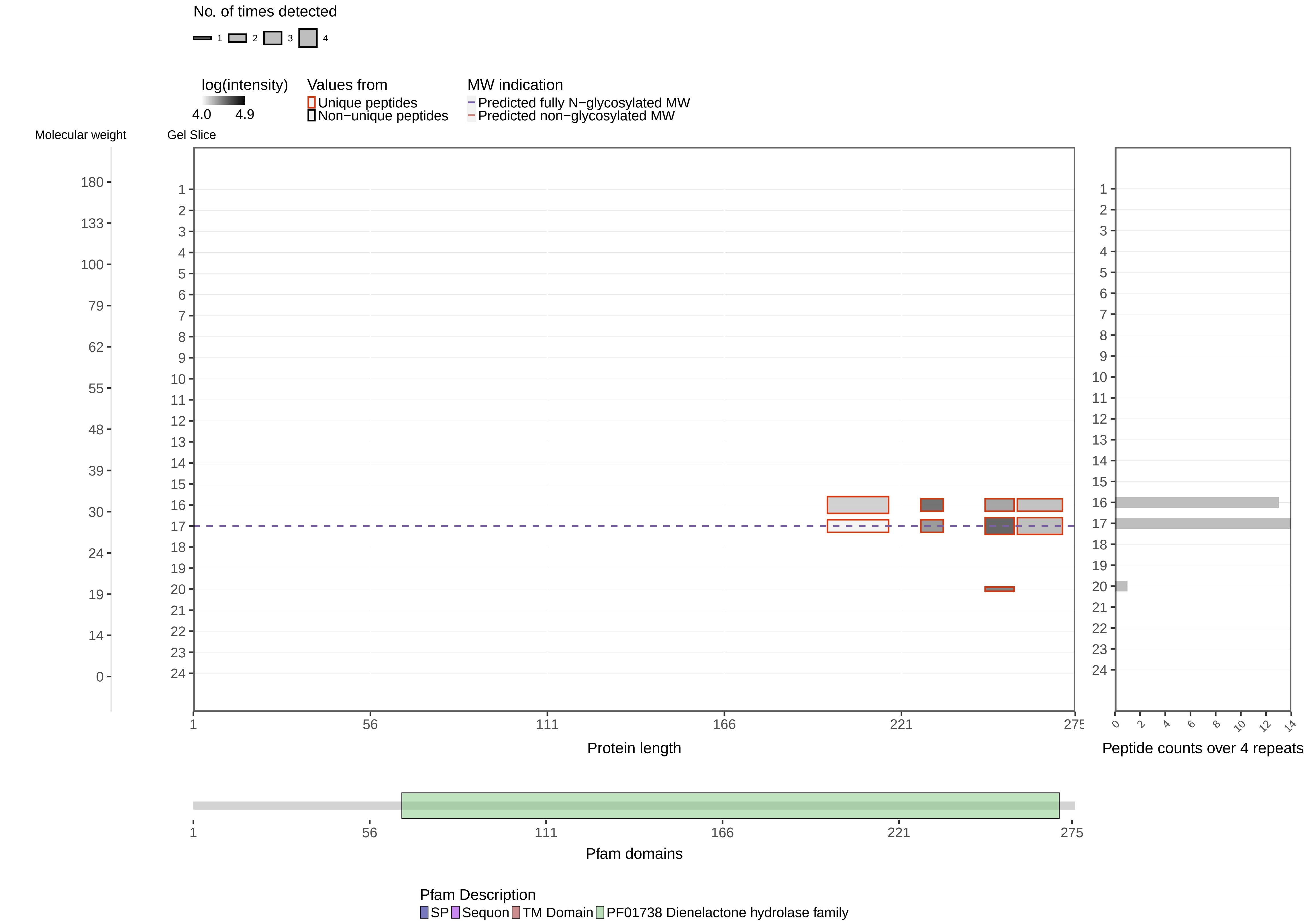


Protein sequence:

QTWLIDHGVDKGCEDAKAVVESLKSKGVSAVGAAGFCWGGMVVSKLAKYETSIGAAVILHPGPITVDEINEVKVPTAILAAEHDHIFLPDQAKQLGDALSAKSEIESFVKIFPGVQHGWTVRYNVEDESAVKAAEESHSDMLNWFIKFIN*

NbL06g10590.1 glutamine_synthetase_chloroplastic−like [MW = 47.53 kDa]

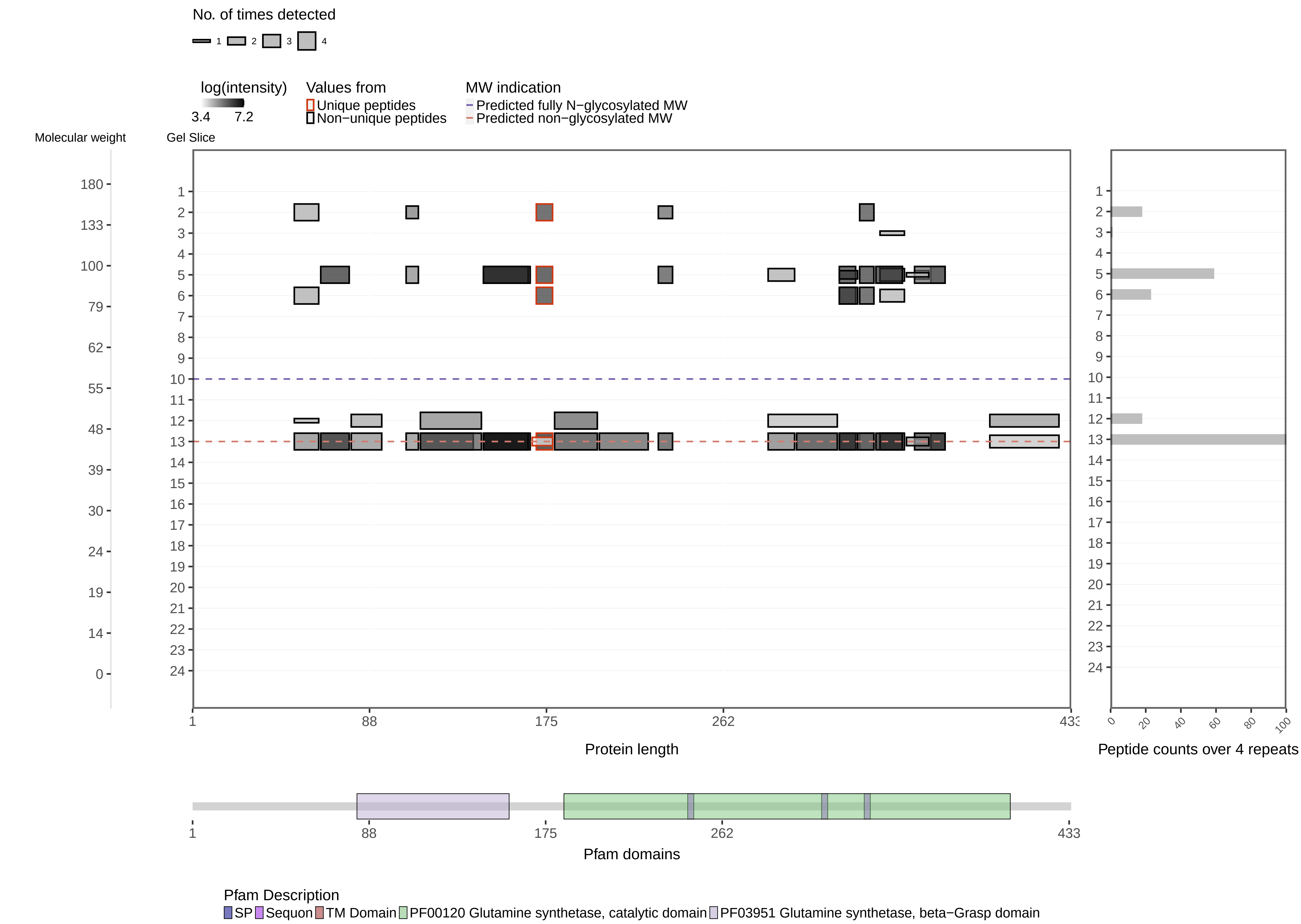


Protein sequence:

WPVGGYPGPQGPYYCGAGADKSFGRDISDAHYKACLYAGINISGTNGEVMPGQWEFQVGPSVGIEAGDHIWCARYILERITEQAGVVLSLDPKPIEGDWNGAGCHTNYSTLSMREEGGFEVIKKAILNLSLRHKEHISAYGEGNERRLTGKHETASIDKFSWGVANRGASIRVGRDTEKQGKGYLEDRRPASNMDPYVVTGLLAETTILWEPTLEAEALAAQKLALNV*

### NbL06g10410.1 pyruvate_phosphate_dikinase_chloroplastic [MW = 105.82 kDa]

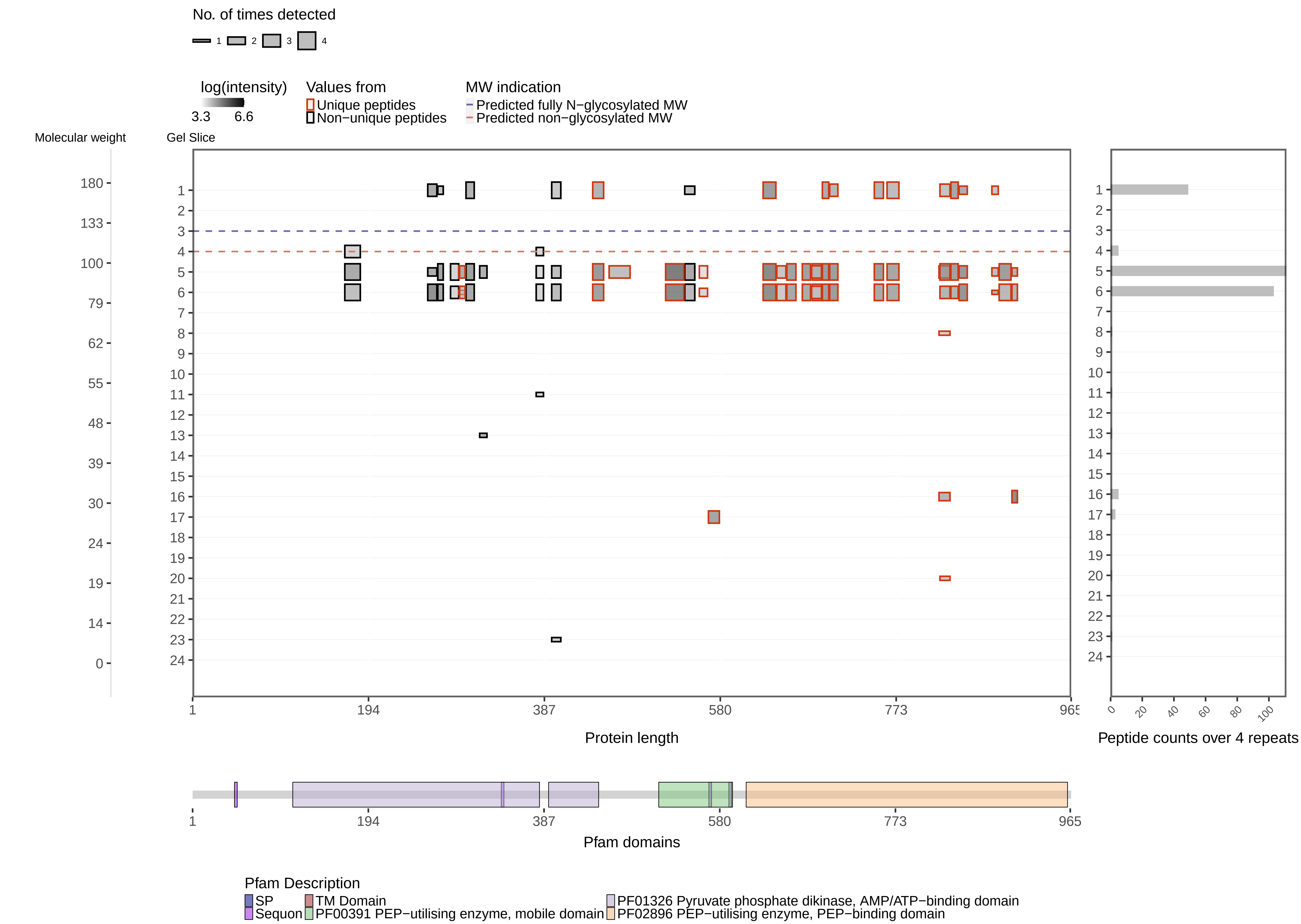


Protein sequence:

NbL06g10370.1 uncharacterized_protein_At4g15545−like [MW = 36.06 kDa]

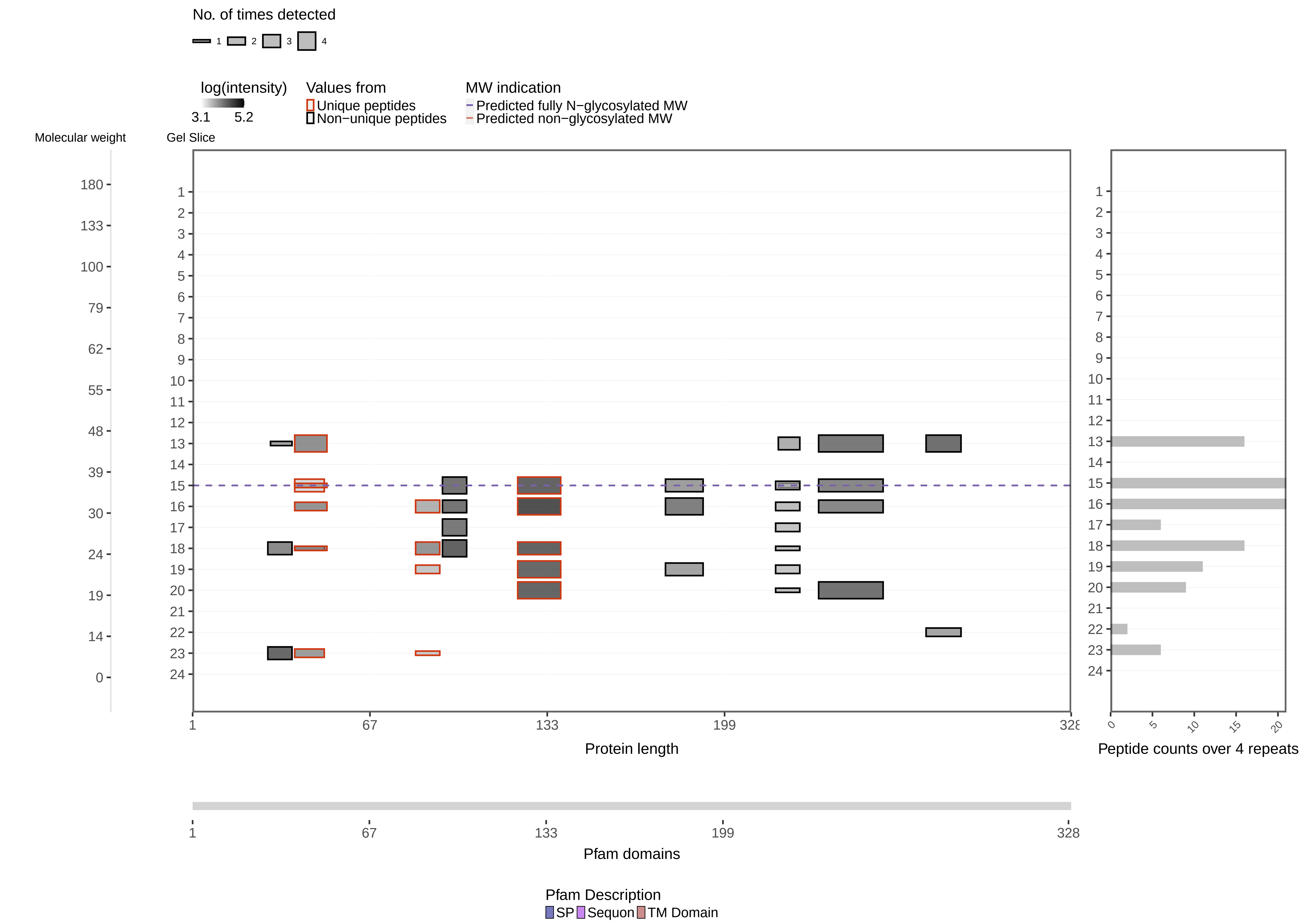


Protein sequence:

AQTQSTESGHSYSEDAESETSRPRISPGLLLASQTSTPRLTPPGSPPSLSASVSPSRSPRPLSPRRHSISFSTTRGMFDDRSSVYSSTSSQYGSMSGLETGSQTARARVDGKEFFRQVRSRLSYEQFGAFLTNVKELNSHKQTKEDTLRKAAEIFSDENKDLYTIFEGLITRNAQ*

### NbL06g10350.1 probable_prolyl_4−hydroxylase_4 [MW − SP = 29.96 kDa]

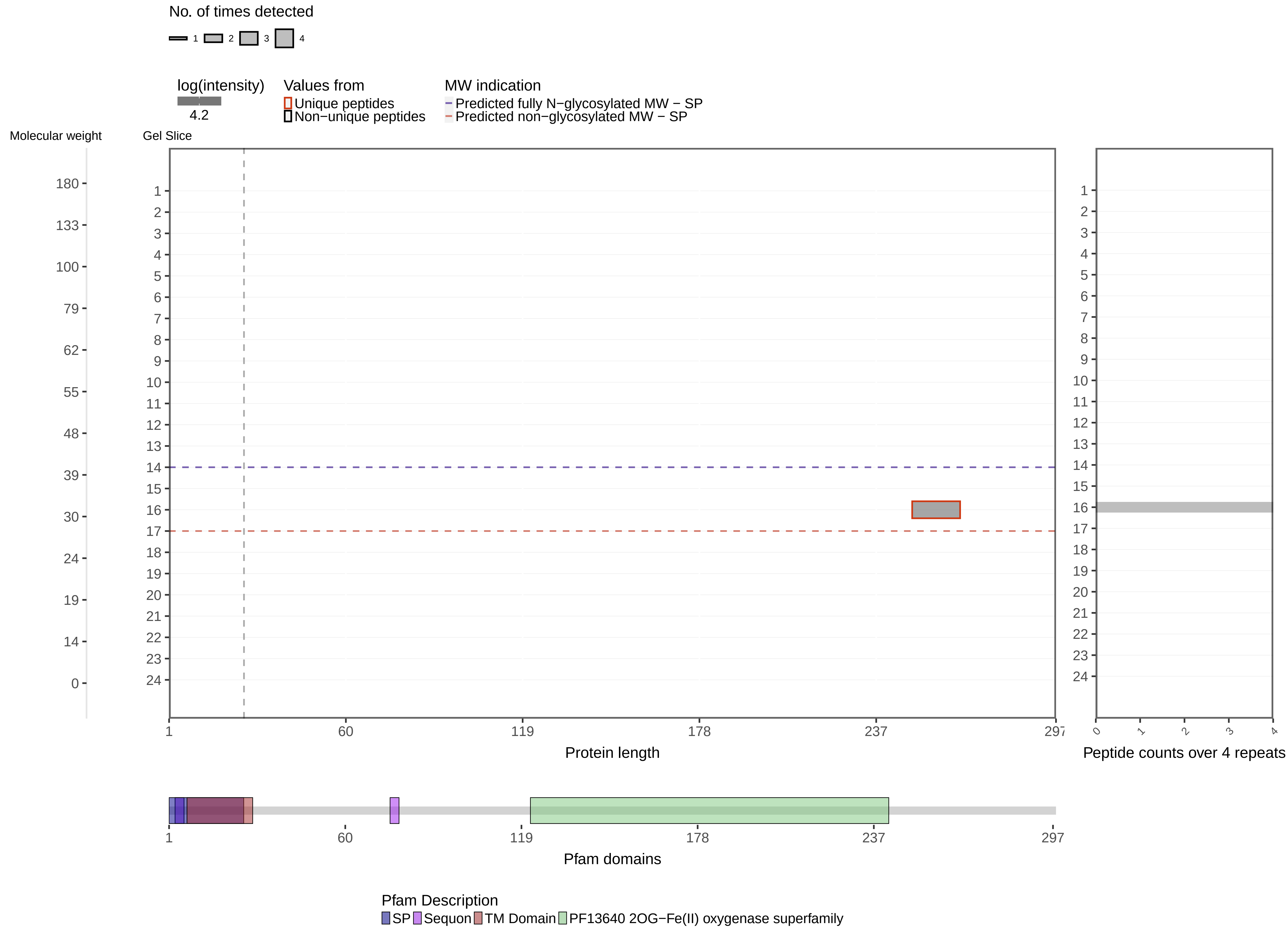


Protein sequence:

FSDKVNIVRGGHRLATVLMYLSDVEKGGETVFPEAEESTRRRSMAADNSLSECARKGIAVKPRKGDALLFFSLHPNAVPDPMSLHGGCPVIEGEKWSATKWIHVDSFDKIVGSEESCADQNENCERWAALGECTKNPEYMVGSSDLPGSCRKSCKAC*

NbL06g10330.1 uncharacterized_LOC109225403 [MW = 100.08 kDa]

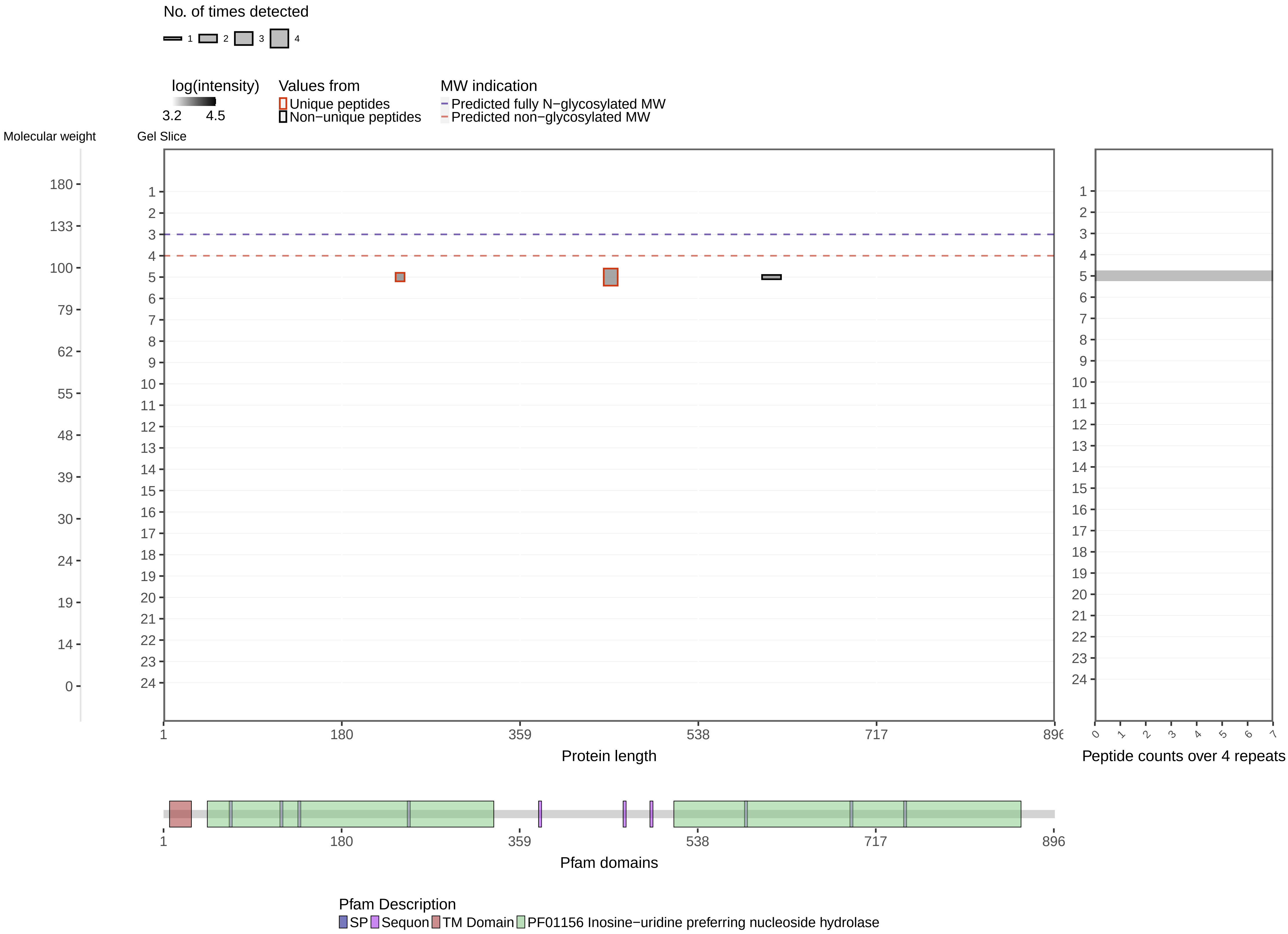


Protein sequence:

RGRCQDGYTKEVVGSGGVPVLVAVRAKPNRNASSVLDREFYVSFLDVLNQRKNSGKFNFSTQFPYYKEKLHKPDFRGKHLGKNVVFDMDMSAGDFLALFYLLKLPVEEINLKAIIVSPTGWANAATIDSVYDLLHMMGRDDIPVGLGDMFAMNQSDPVFSAVGECKYNKVIPQGSGGFLDSDTLYGLSRSLPRSPRRYTAENSVKFGAPRDTDHPELRQPLALEVWESVVKSLDSGSKVTILTNGPLTNIAKILLAGKNYTKAIQGILIVGGHINYDNTEKGNVINVPSNRFG

NbL06g10190.1 isoleucine−−tRNA_ligase_chloroplastic/mitochondrial [MW = 123.75 kDa]

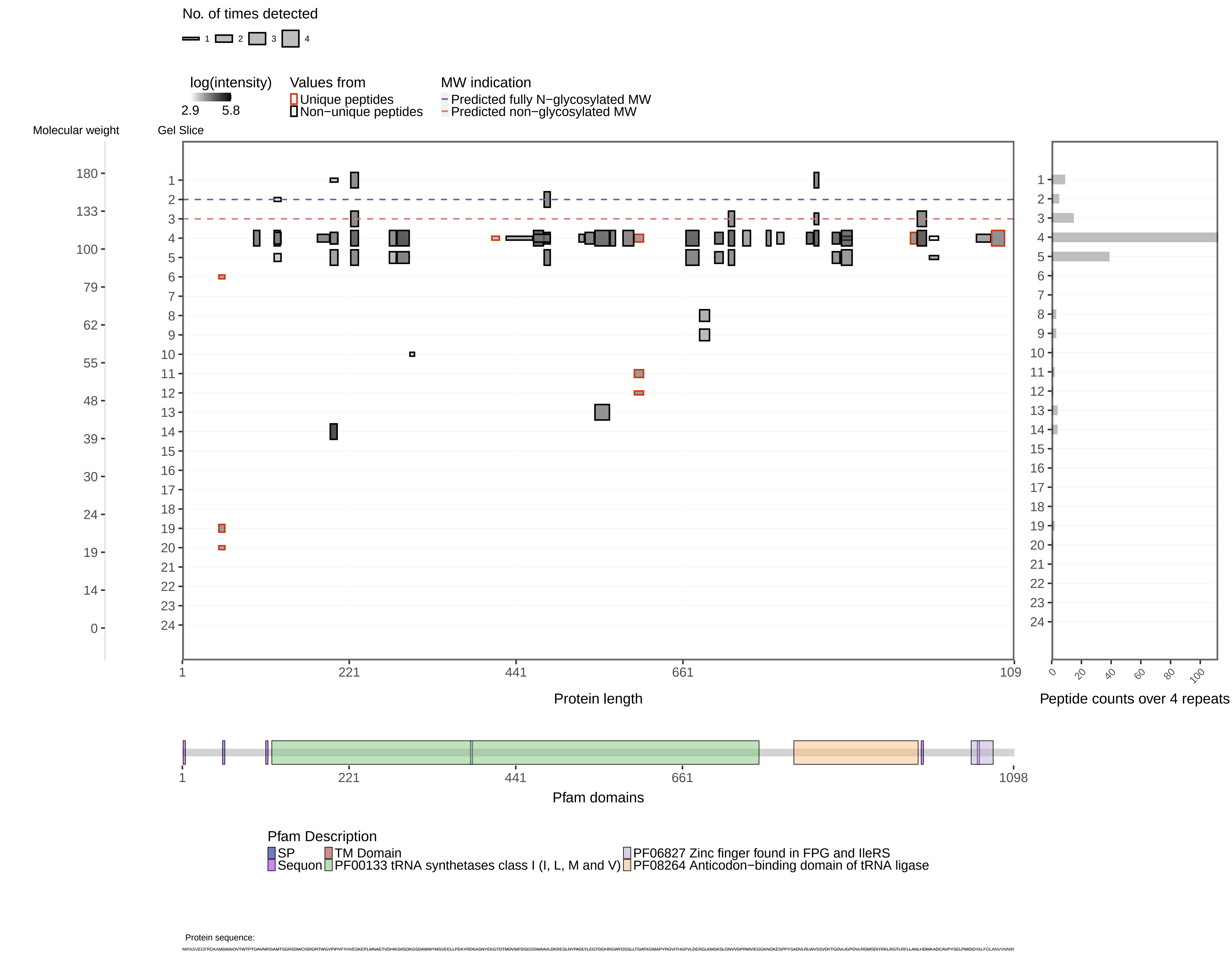

### NbL06g10160.1 heat_shock_factor−binding_protein_1−like [MW − SP = 10.83 kDa]

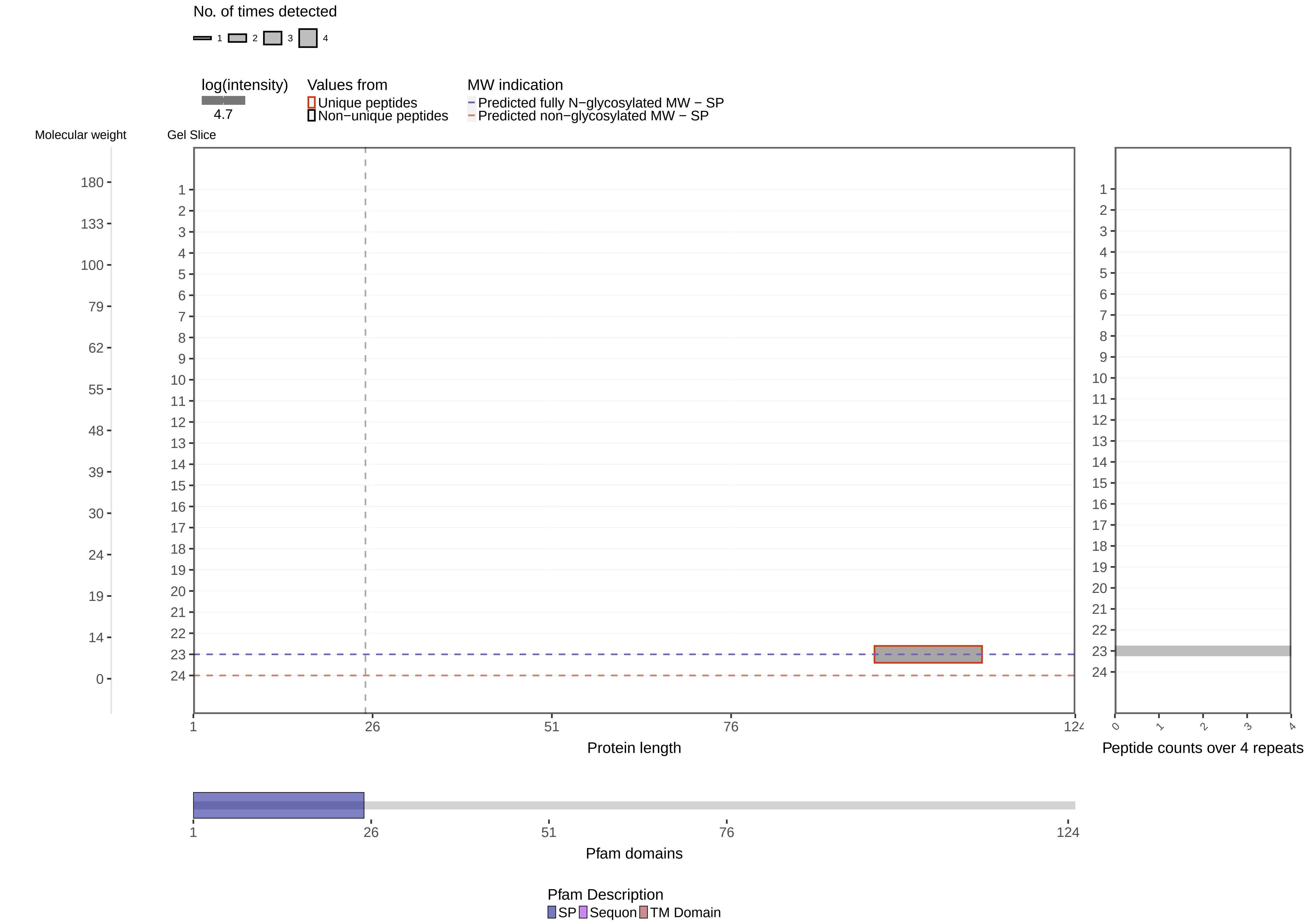


Protein sequence:

QMQTSILYIASIFCEINLEKVCLVLDYAALDEMGNRIDELEQSINDLRTEMGQEGSPSPSAALKSKDEPKSAEDST*

### NbL06g10050.1 acyl_carrier_protein_1_chloroplastic−like [MW = 14.05 kDa]

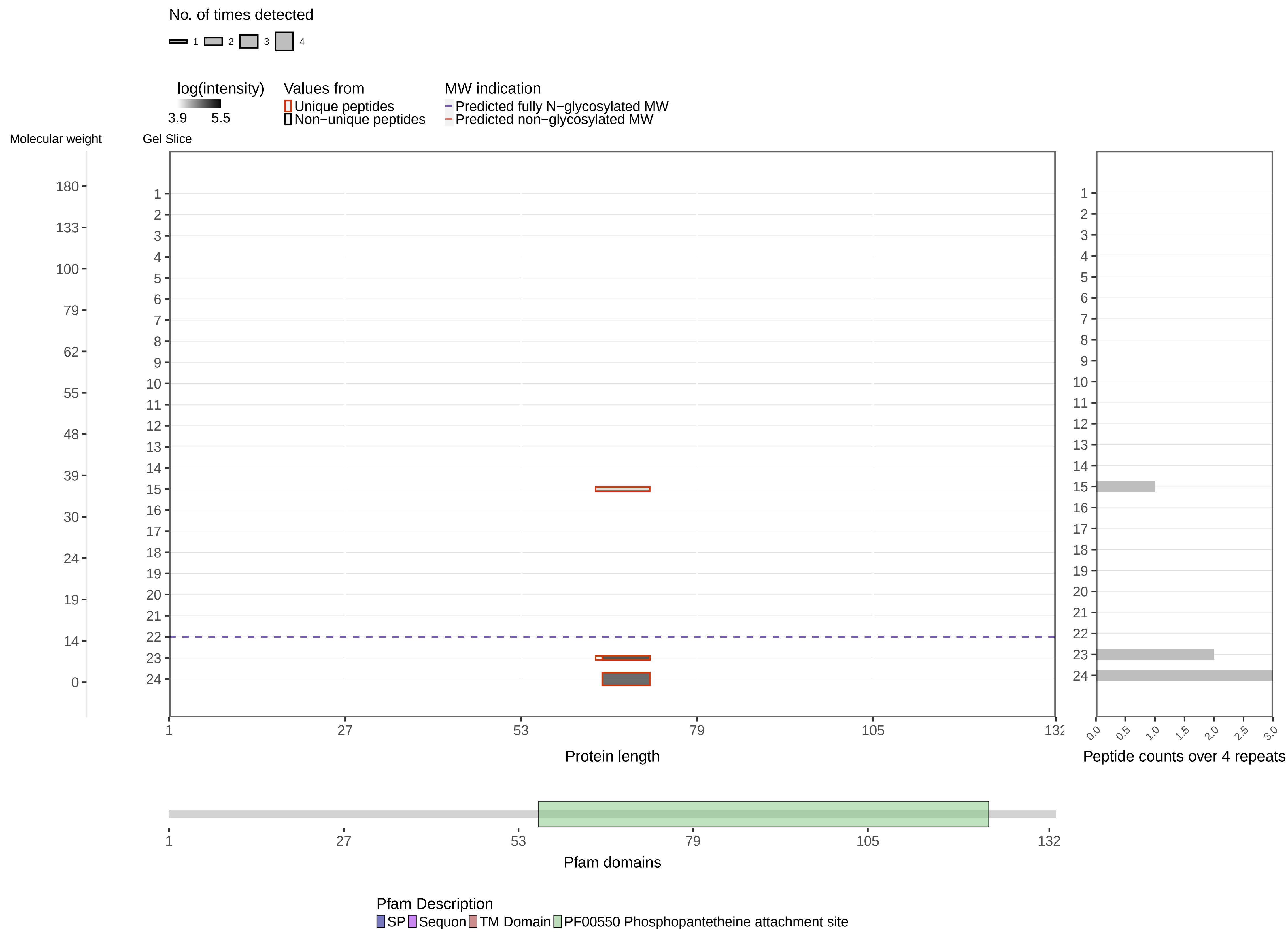


Protein sequence:

PETVDKVCEIVKKQLALTDEKVVNGESKFTALGADSLDTVEIVMGLEEAFGITVEEENAQTIATVQDAADLIEDLISKKA*

### NbL06g10040.1 bifunctional_3−dehydroquinate_dehydratase/shikimate_dehydrogenase_chloroplastic−like [MW = 57.51 kDa]

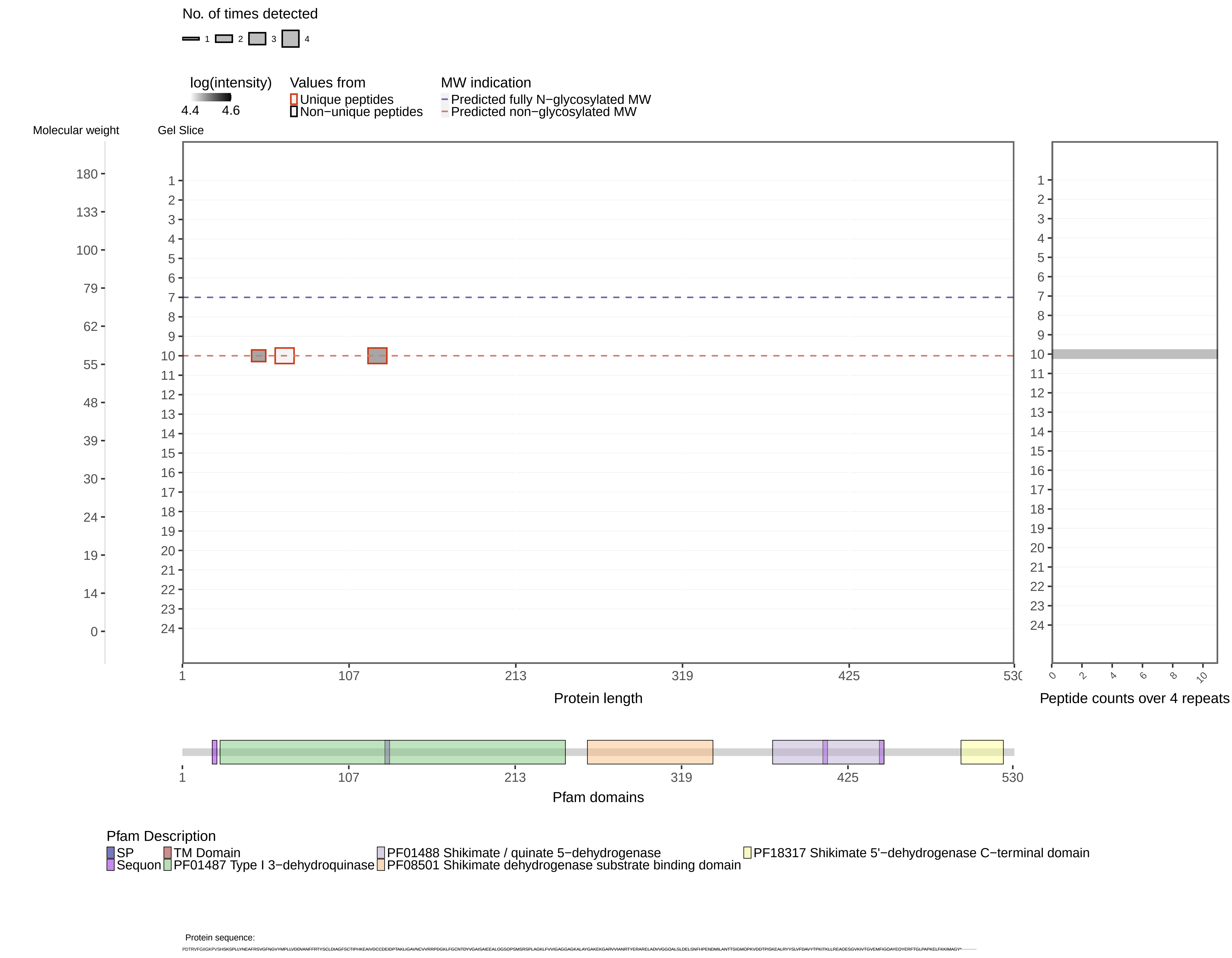

NbL06g09910.1 probable_xyloglucan_endotransglucosylase/hydrolase_protein_8 [MW − SP = 32.04 kDa]

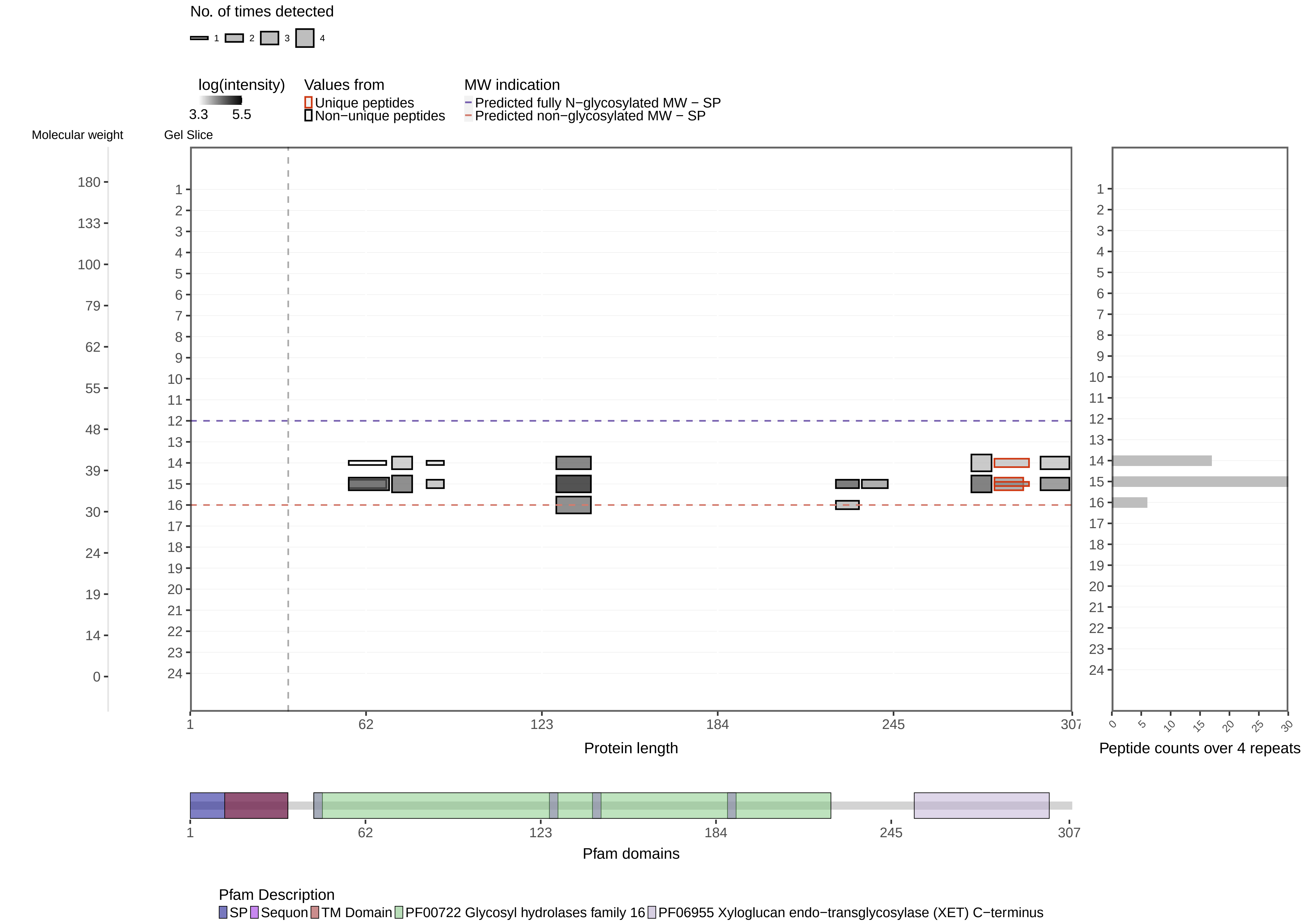


Protein sequence:

GNREMRHNLWFDPTEDFHTYSILWNTHQIVFFVDRVPIRVYKNANYTNNFFPNEKPMYLFSSIWNADDWATRGGLEKTNWKNQPFVSSYKDFSVDGCQWKDPFPACVSTTTKNWWDQYNSWHLSNDQKMDYAWVQRNLVTYDYCQDKERFPKKPEECWLNPWE*

### NbL06g09730.1 probable_acyl−activating_enzyme_6 [MW = 68.46 kDa]

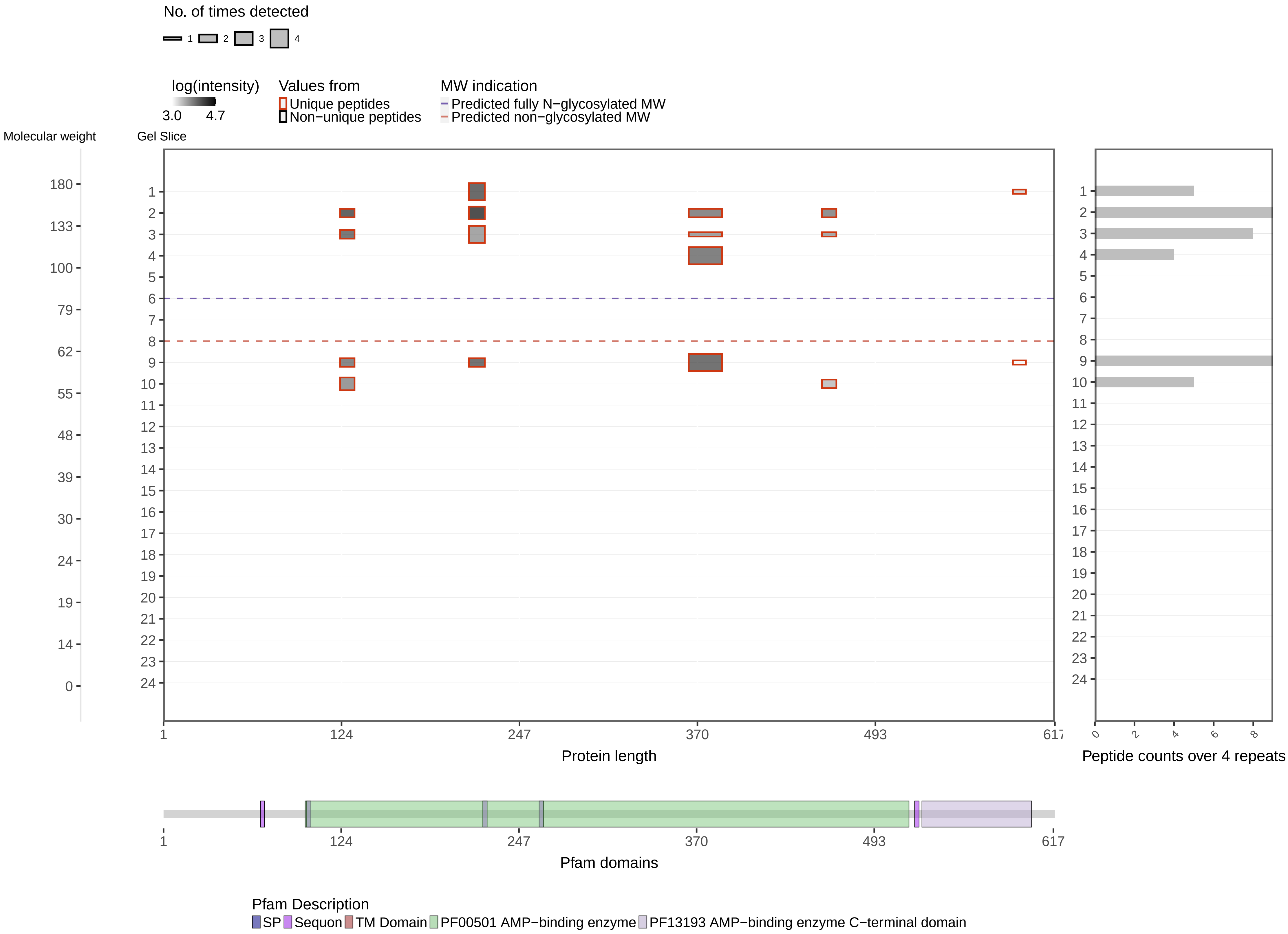


Protein sequence:

LWTLPIFHANGWGFCWGMAAVGGTNVCLRRGHDASSIYEAIHQHKVTHMCGAPVVLNMLSTYPKRQPLDSPVYIYTAGAPPPPAVLDRTESLGFVISHGYGLTETGGPAVTCAWKQEWDHLPQSERAQLKSRQGVRMVGFTEVDVVDFETGISIERNGIAIGEIVLRGGSVMLGYLKDPEGTSKCMKNDGWFYTGDVGVMHSDGYLEVKDRLKDVIICGGENISSVEVESILYKHPAVNEAAVVAKPDNYWGETPCAFISLKTDAEKMTEKELRDFCKKNLPLYMVPRTV

### NbL06g09640.1 thioredoxin_reductase_2 [MW = 35.16 kDa]

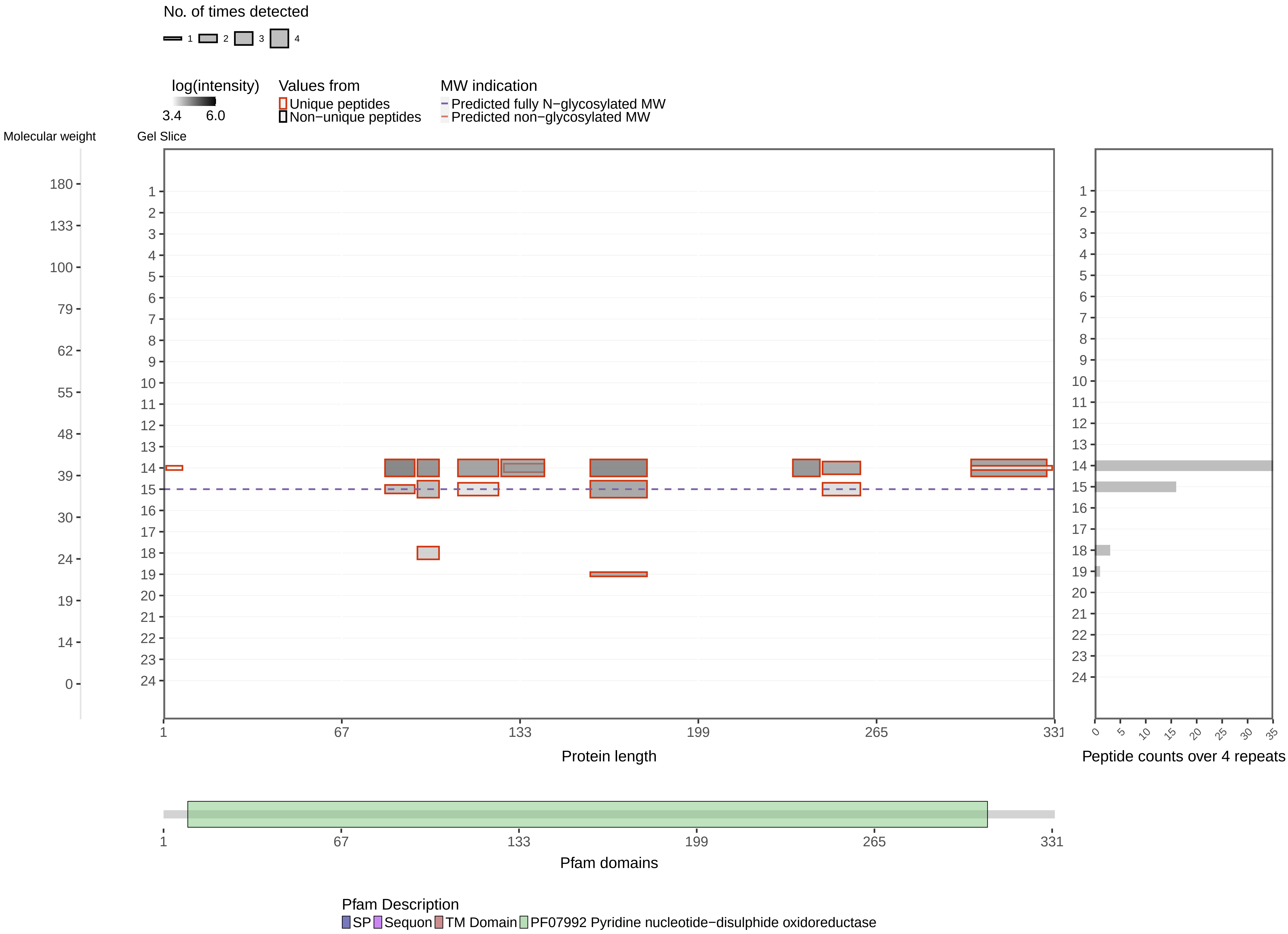


Protein sequence:

AAPIFRNKPLAVIGGGDSAMEEATFLTKYGSKVYIIHRRDEFRASKIMRNRALSNPKIEVIWNSAVVEAYGEKVLGGLKVKNVVNGEVSDLKVSGLFFAIGHEPATRFLDGQLELDSDGYVVTEPGTTKTSVRGVFAAGDVQDKKYRQAITAAGSGCMAALDAEHYLQEIGAQEGKSD*

### NbL06g09550.1 nucleoside−triphosphatase_THEP1 [MW = 21.87 kDa]

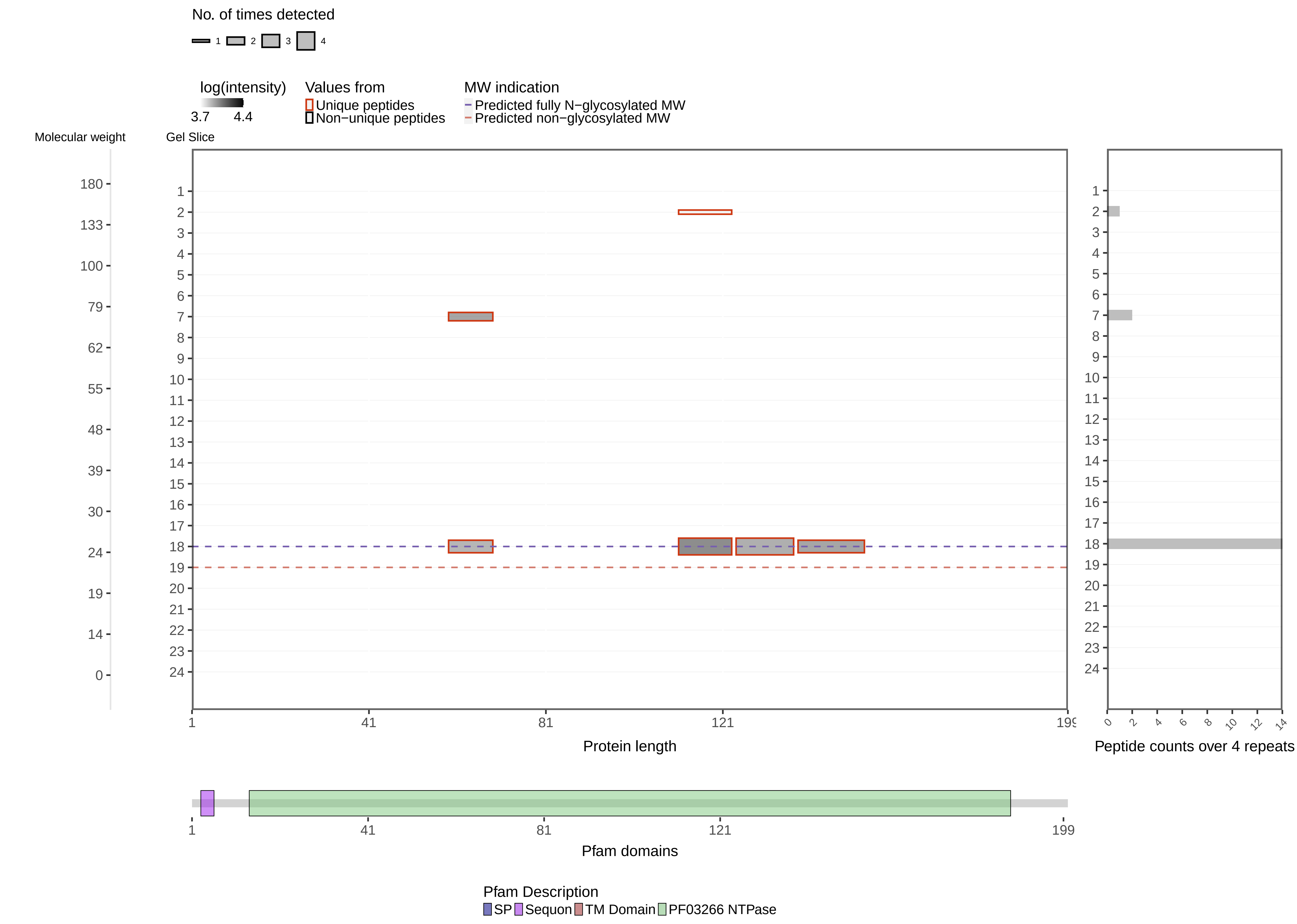


Protein sequence:

WPTVGRYRVDVASFESLALPELQVREDTDLFIIDEVGKMELYSSSFFPAVLKVLQSNIPLLASIPIPKAGRDIPGVARMRNHPGASVFTLNANNRDSMREQICSTLADMLQKP*

### NbL06g09510.1 ubiquitin_domain−containing_protein_DSK2b−like [MW = 59.16 kDa]

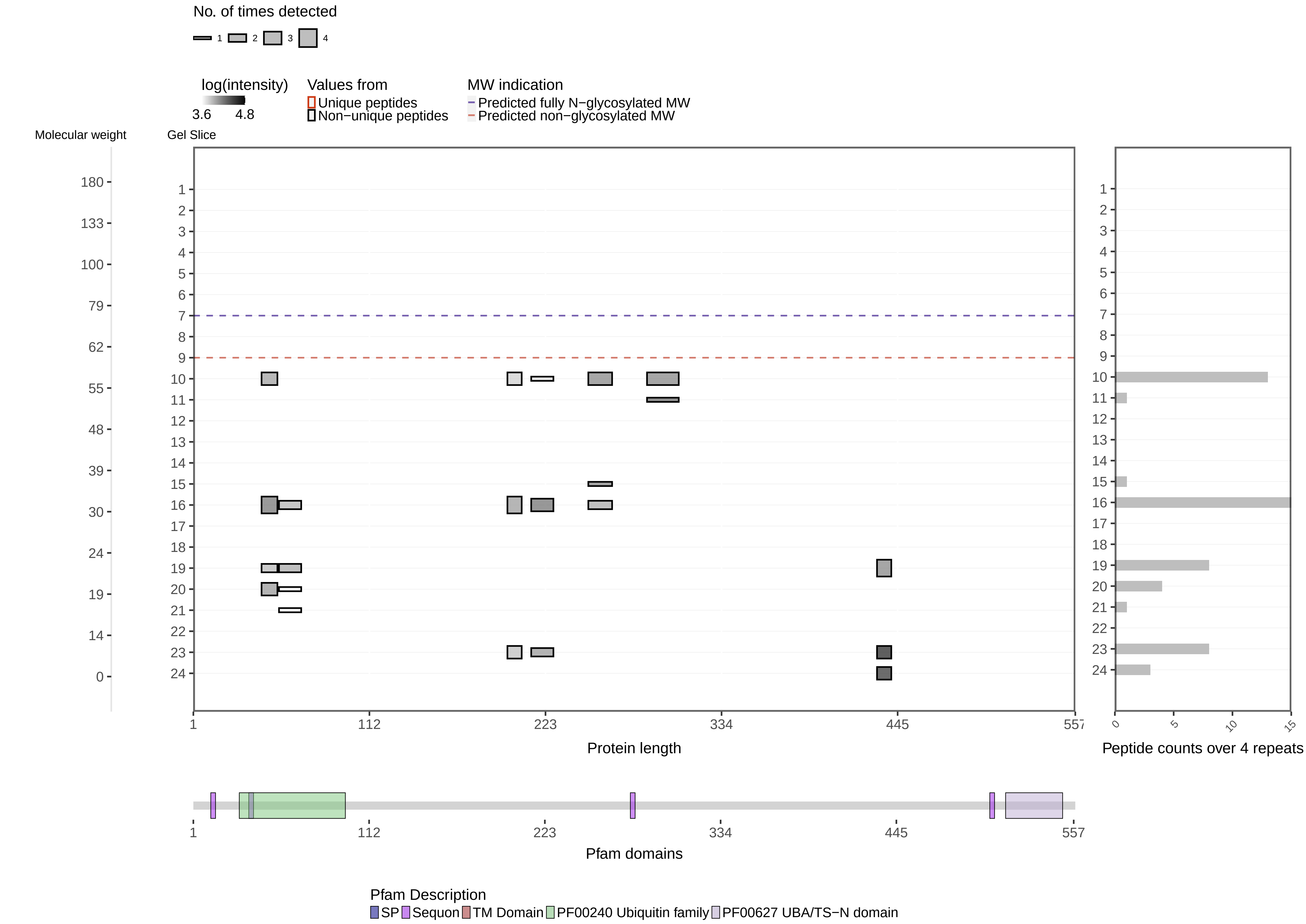


Protein sequence:

MYENVQEPLLNATTLSGDTRNDVGSNPFAALLGAQGGGQGRHQSNNPPTSGSETTDNPPAPNTNPLPNPWASAGTGAAQANTPARSNTAADTRAAPLGGLGGLASPGLEQMLSGMPDTTSLNQMMQNPAISQMMQSLLSNPQYMNQVLGLNPQLRSMLDSNSHLREMMQNPEFIRQLTSPETMQQLMTFQHGLLSQAWRQTNQQVGQNAGGAAGDNGMEMLMNMFGGLETGGLGVPNRSNVPPEELYATQLTQLQEMGFFDTQENIRALIATAGNVHAAVER

NbL06g09140.1 probable_serine_protease_EDA2 [MW − SP = 52.01 kDa]

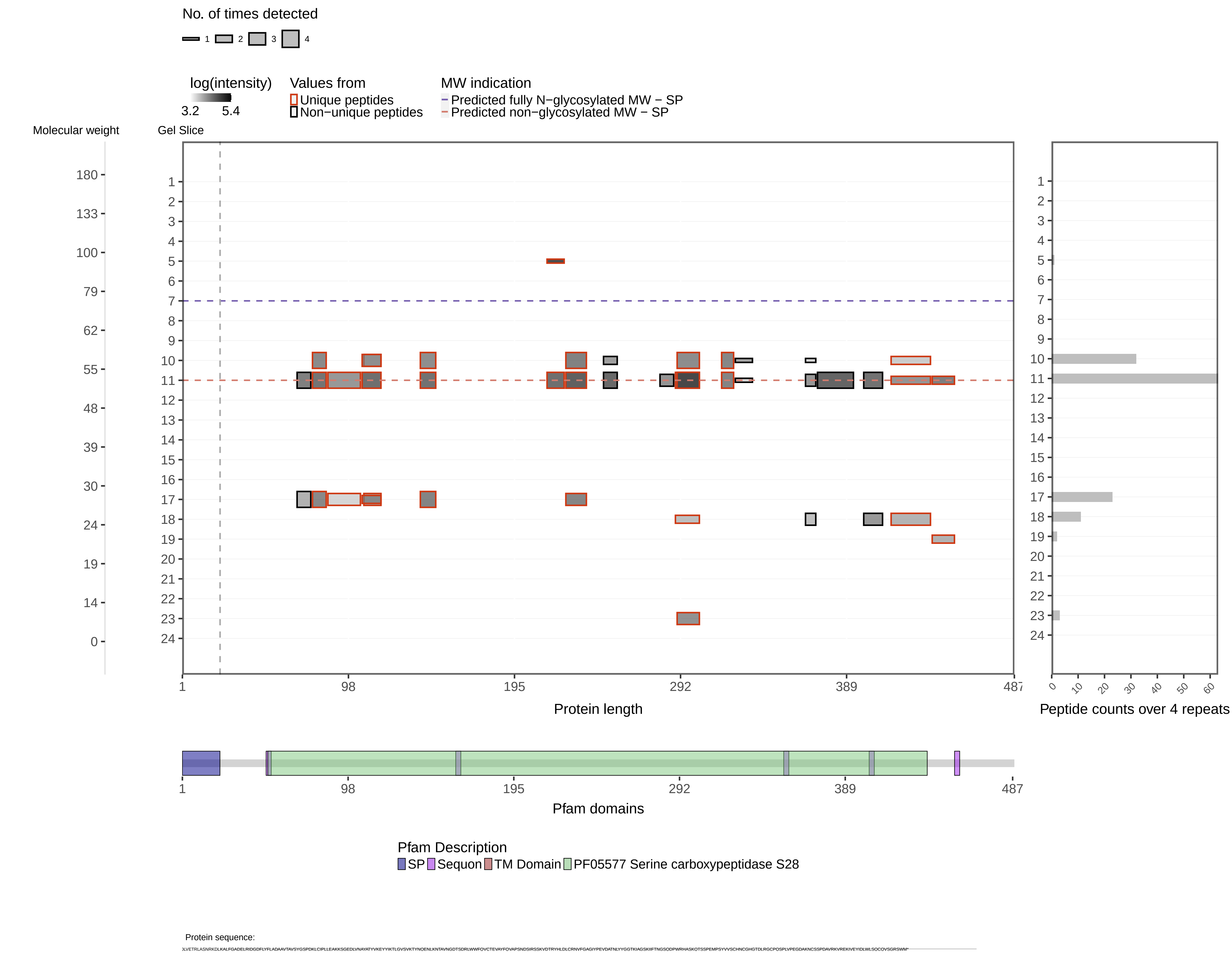

### NbL06g09080.1 thioredoxin_H−type [MW = 15.27 kDa]

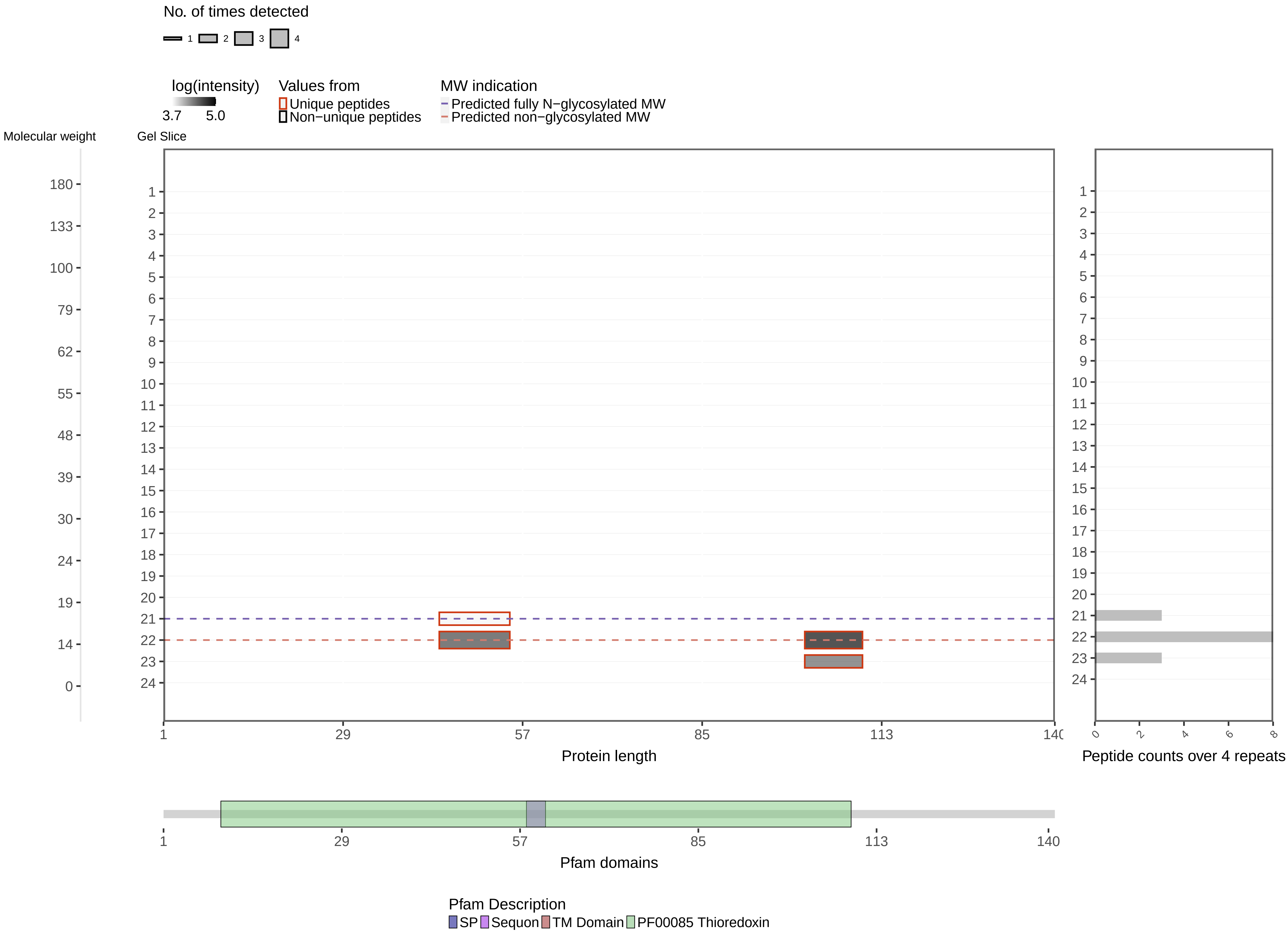


Protein sequence:

PNVTFLKVDVDELKSVATDWAVEAMPTFMFLKEGNIVDKVVGAKKEELQQAIAKHMEYSFGLKNICGCIKGAELAVGSTTAFD*

### NbL06g08750.1 thylakoid_membrane_protein_slr0575−like [MW = 31.08 kDa]

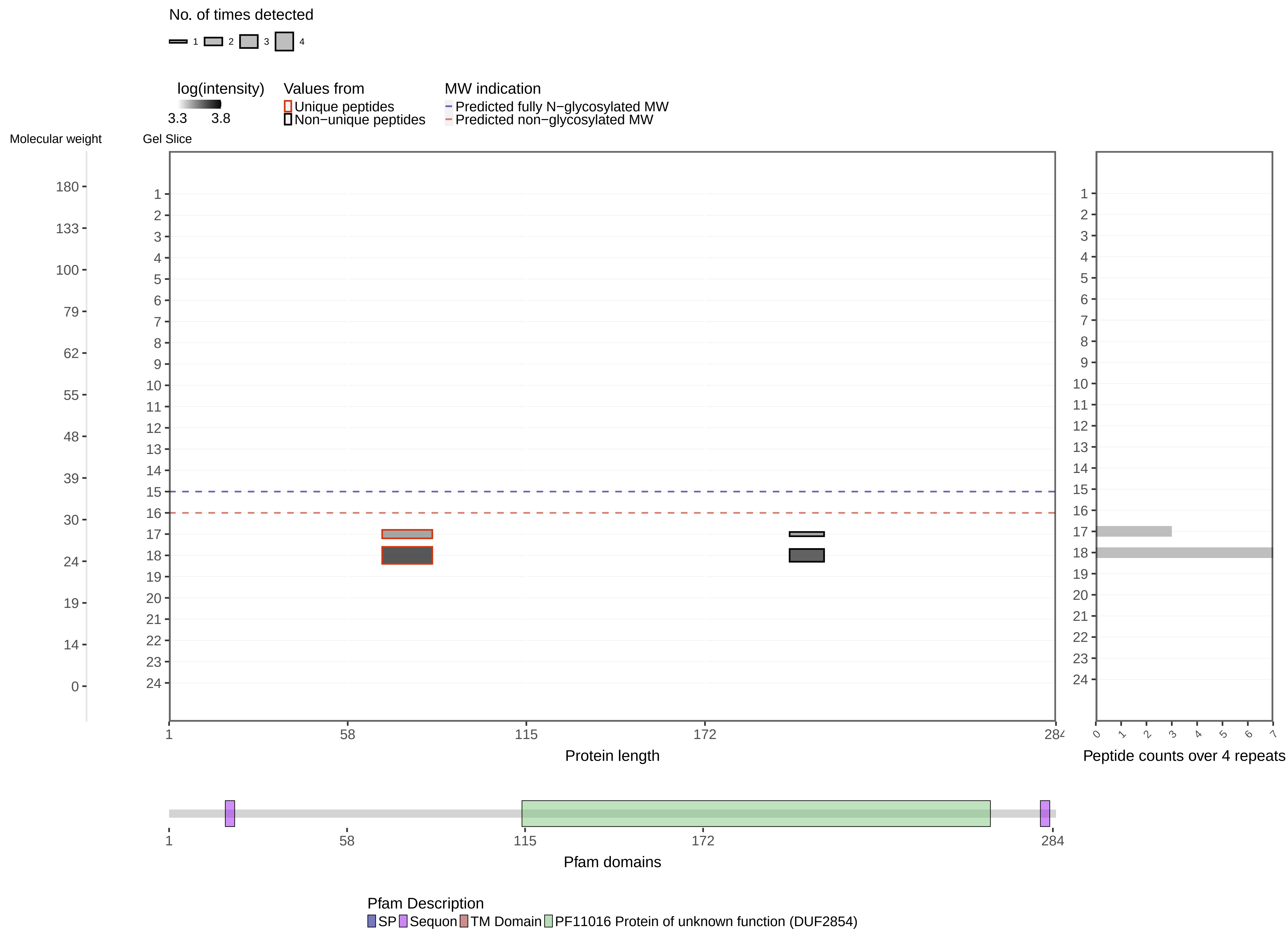


Protein sequence:

NSALMLTYGFPLTIIGMALKYAELKPVPCITYSDAOLLREKCATPILKOVRSDVIRYRYGDEOHLDESLKRIFOFGLGGGIPRRSAPVLOMIREEVTEDGKYCLVLVFEAKALOLSDLEKROAKFASFFGPGISAEIAKGEKDALYEVRLISNTTL*

### NbL06g08740.1 14−3−3_protein_4−like [MW = 29.38 kDa]

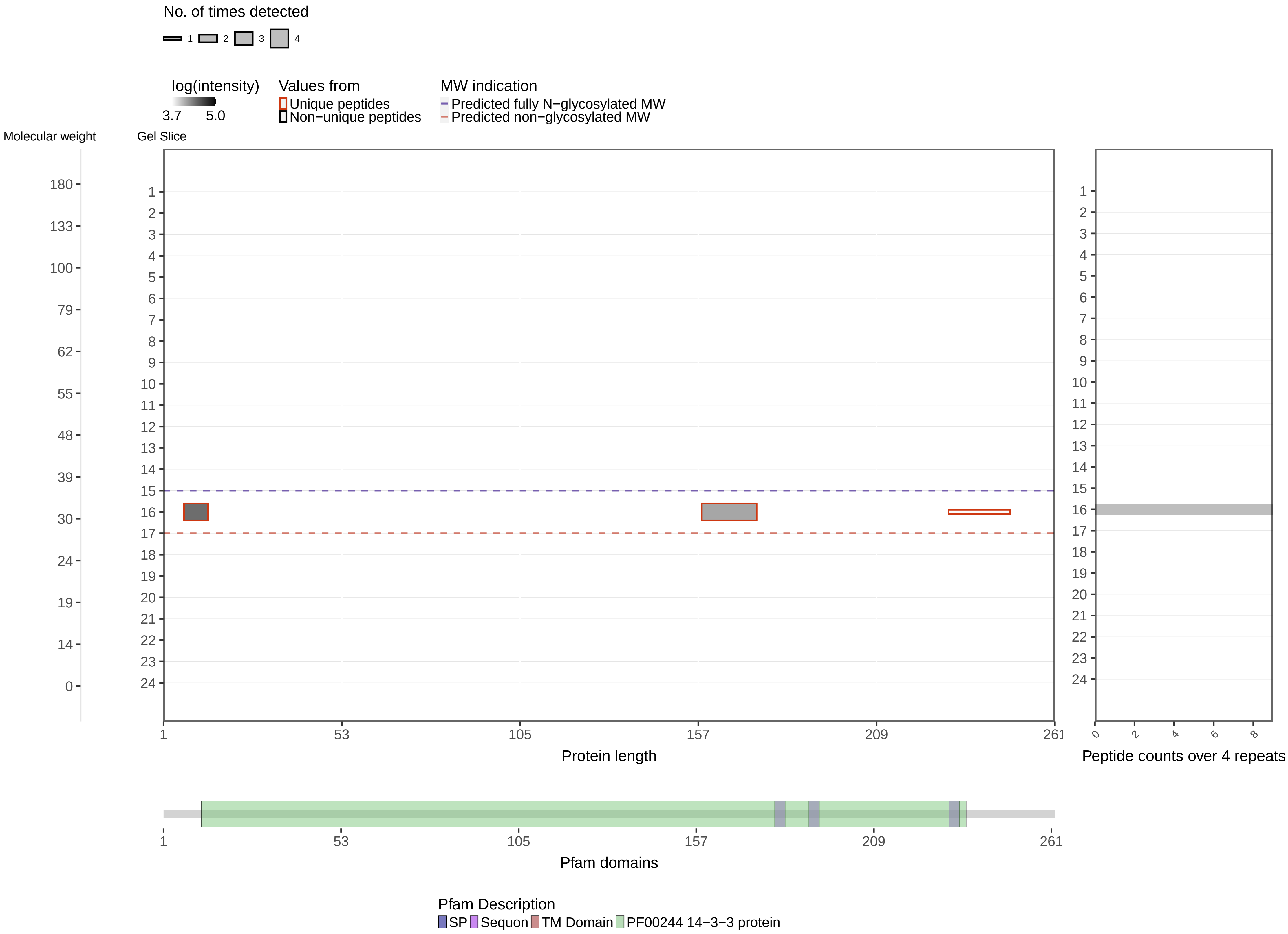


Protein sequence:

TAESKVFYLKMKGDYHRYLAEFKTGAERKEAAENTLLAYKSAQDIALDELAPTHPIRLGLALNFSVFYYEILNSSDRACNLAKQAFDDAIAELDTLGEESYKDSTLIMQLLRDNLTLWTSDTTDDAGDEIKEASKRESGDGEQ*

NbL06g08680.1 ribulose_bisphosphate_carboxylase_small_chain_chloroplastic [MW = 20.34 kDa]

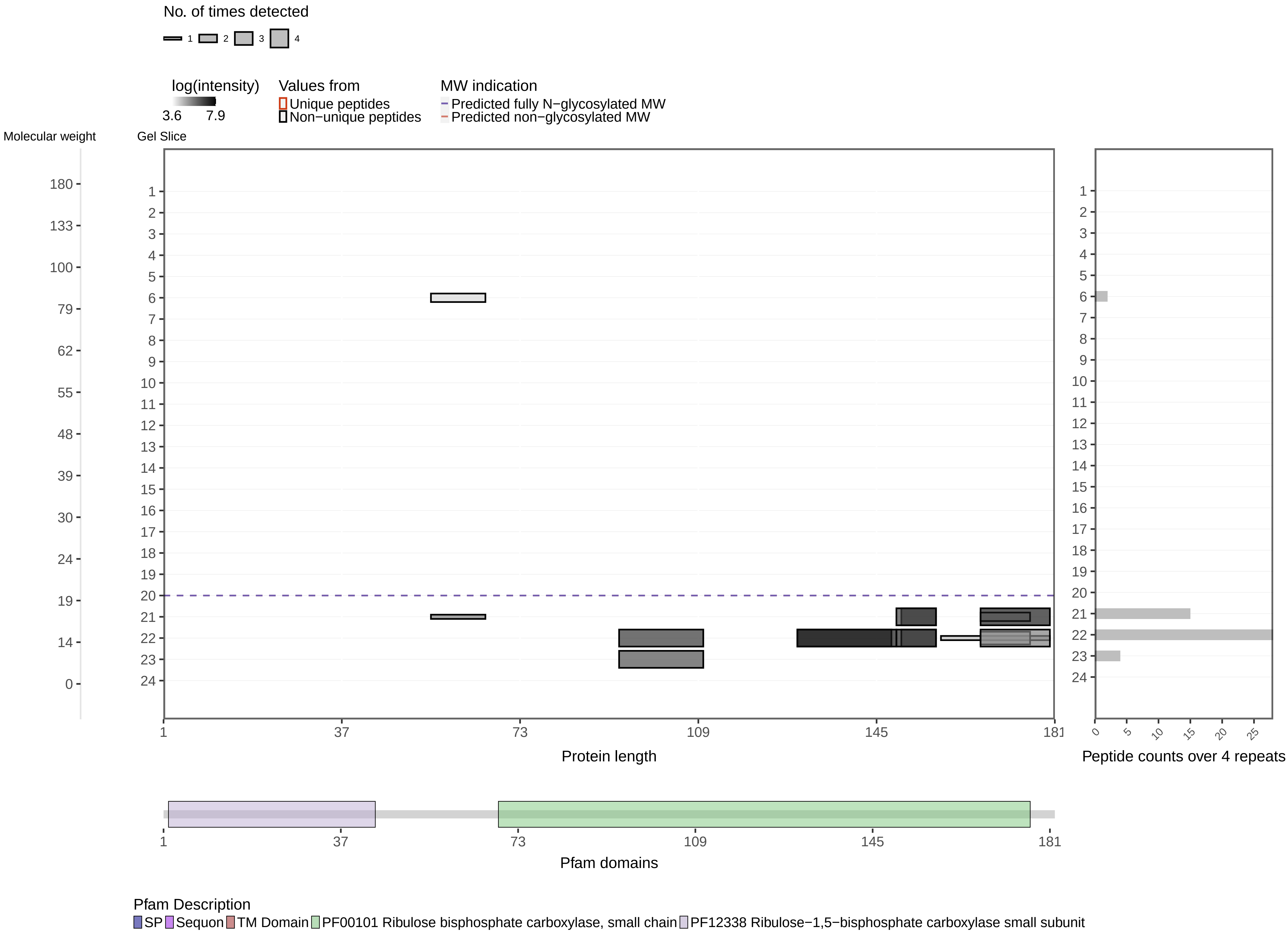


Protein sequence:

LSVEQLLSEIEYLLKNGWVPCLEFETEHGFVYREHHKSPGYYDGRYWTMWKLPMFGCTDATQVLAEVEEAKKAYPQAWIRIIGFDNVRQVQCISFIAYKPEGY*

### NbL06g08590.1 non−specific_lipid−transfer_protein_2−like [MW = 16.55 kDa]

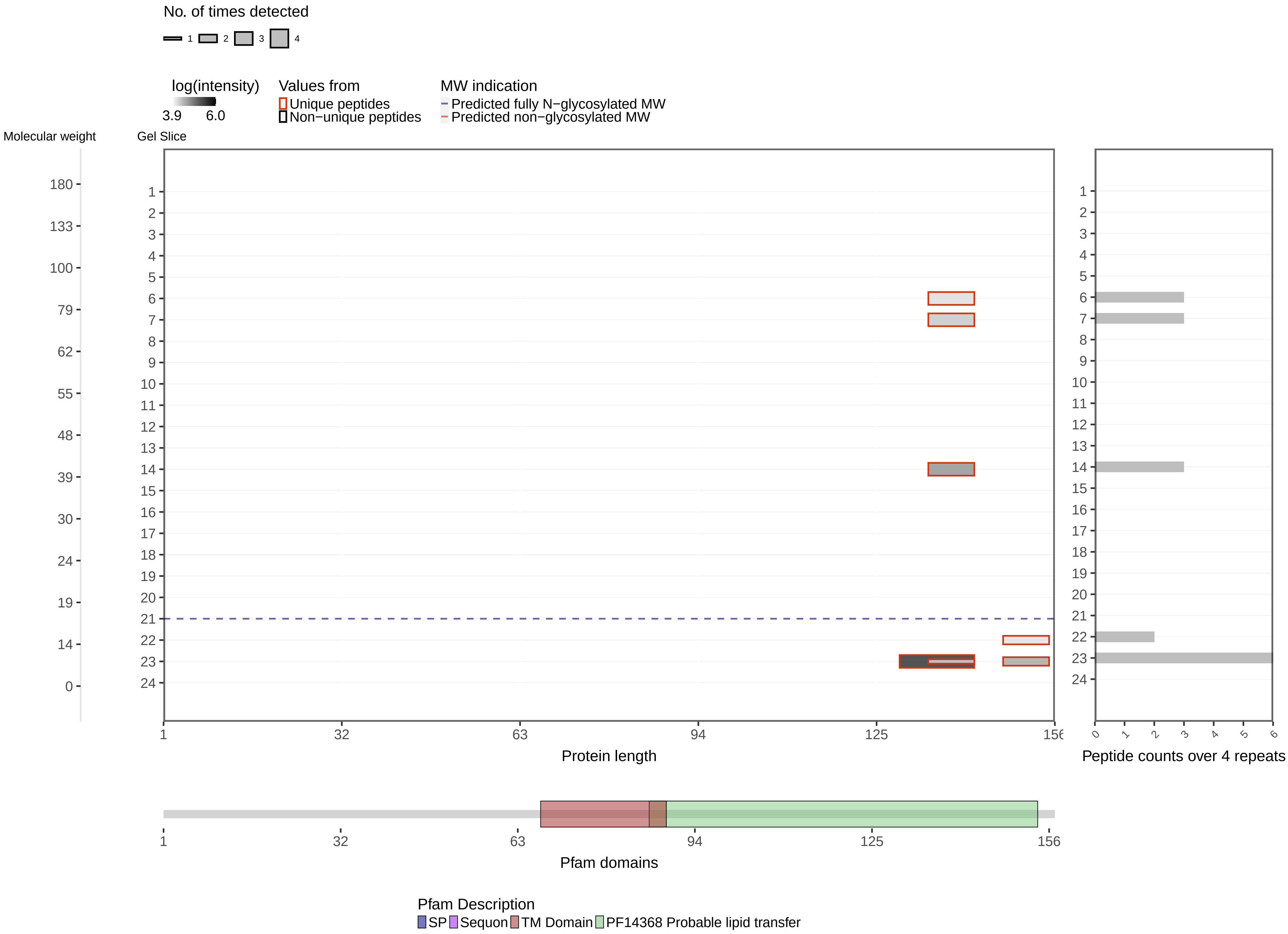


Protein sequence:

SCAAIFLVVTLILFLGEFLVTEAVTCSAVELSPCAGAITSPRPPSSECCAKLREQKPCLCGYLKDPNLRPYVNSPNAKRVAKSCGVPTPSC*

NbL06g08370.1 GDSL_esterase/lipase_At1g29670−like [MW − SP = 37.24 kDa]

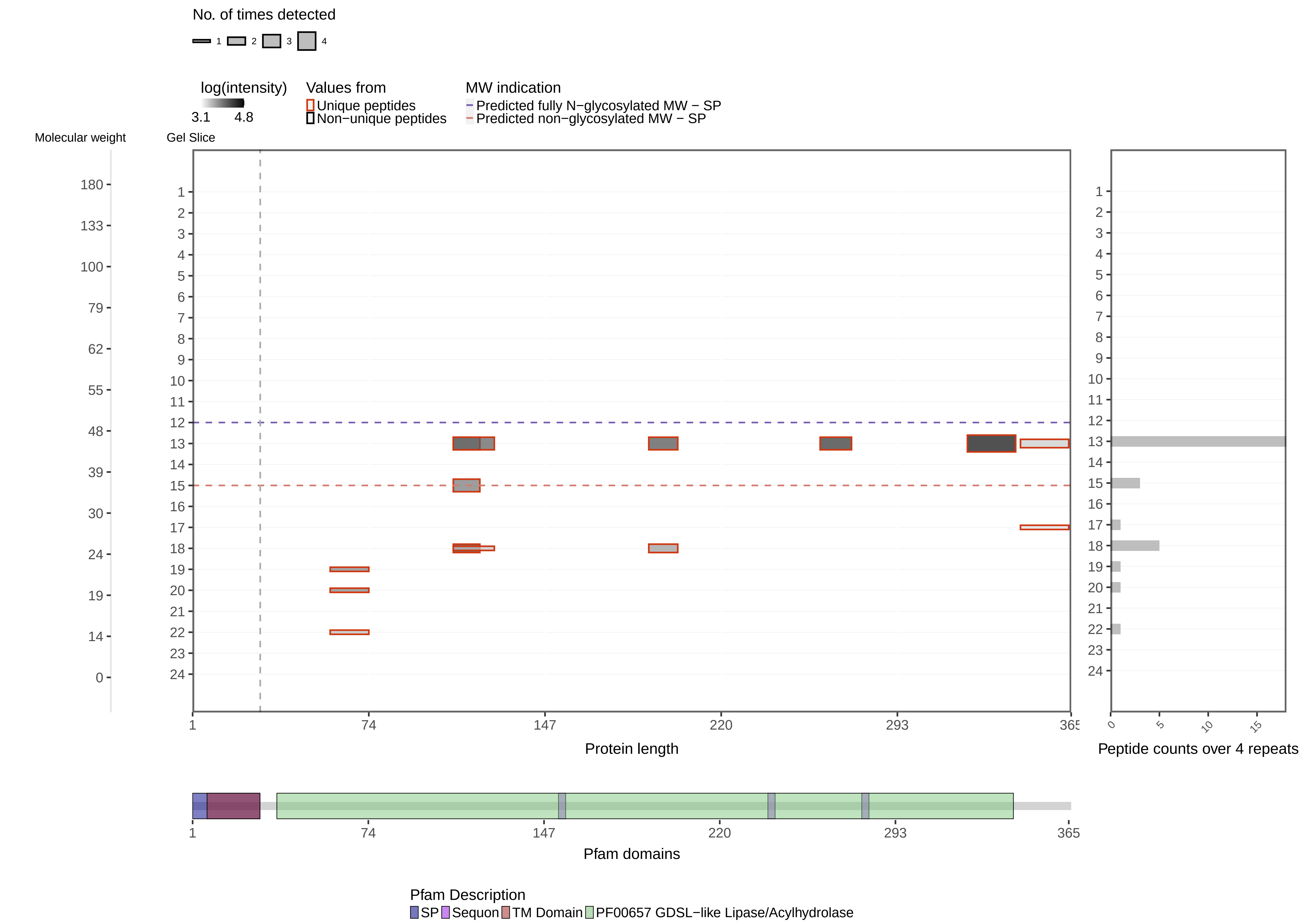


Protein sequence:

MGNNDYLNNYLKPQFYLSSRLYTPDQFATVLVRQFSEQLRTLYSYGARKVAVSNIGLLGCLPEELEEFGGNASGCVDFINNYAKLFNDKLKLLIDDLNTNLPNARFIYINQTSISSGGPTPVGFTVDSSCCITSETIAEGQCRIGEIPCSNRDQYIFFDNYHPTEVANMATARRSFNALLPSDAYPTDISQLVQT*

### NbL06g08190.1 GDSL_esterase/lipase_At5g45910 [MW − SP = 38.42 kDa]

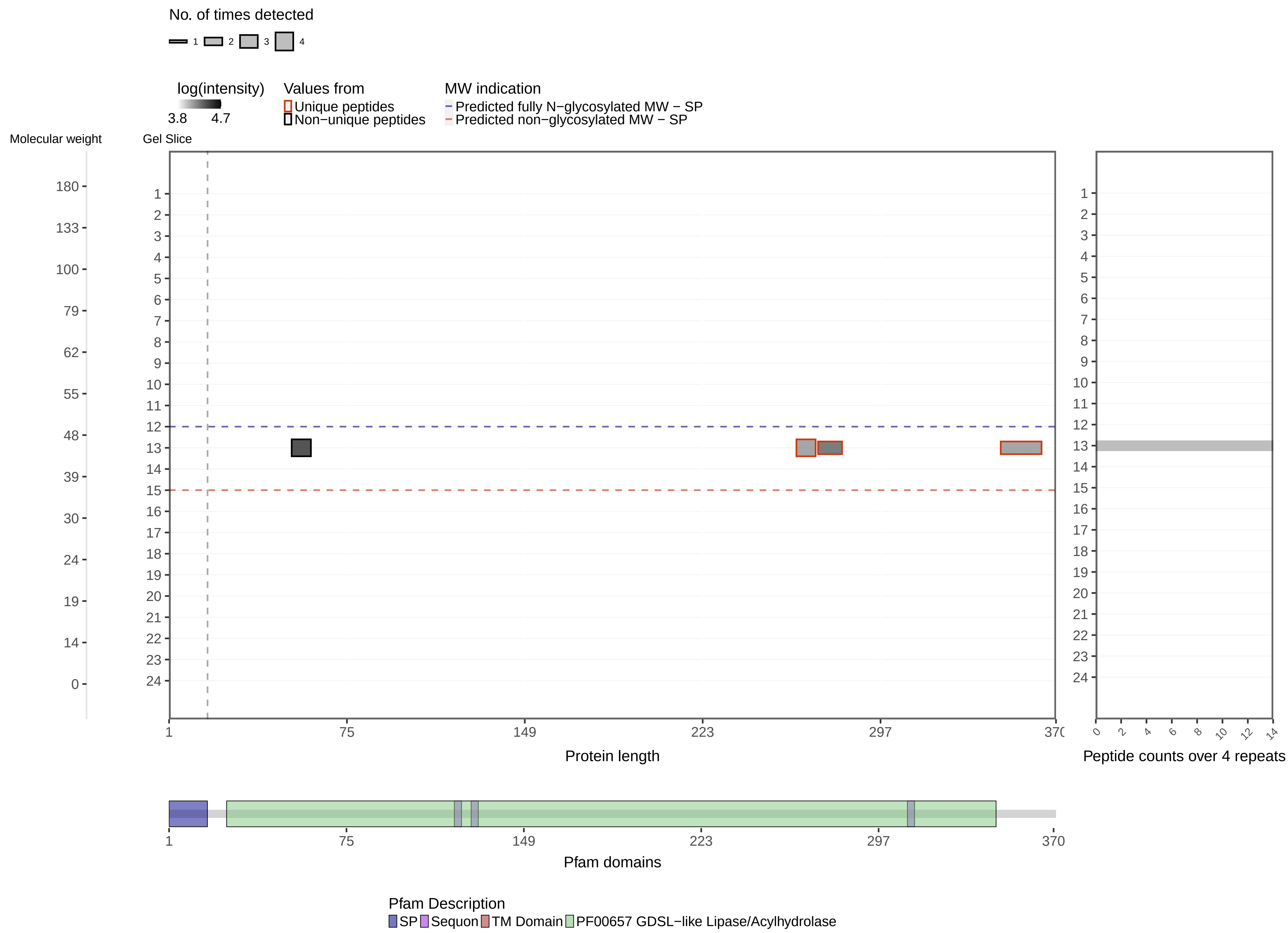


Protein sequence:

YPYFVGGSIKKLKALVPAVVETIIEATSVLIEEGAVELIVPGNLPIGCSAVYLTLFGTQNKAAYDKNGCLKTYNAFSKYHNAQLKLSLEKLREEYPHAKIIYADYYGAAKRFFHAPKHYGFSNTLVACCGGGGPYNFNNSARCGHIGSKACLDTSKLANWDGIHLTEAAYHHIAKGLLNGPFTSPPLTFSPIKQIKLT*

### NbL06g07940.1 epoxide_hydrolase_A−like [MW = 35.77 kDa]

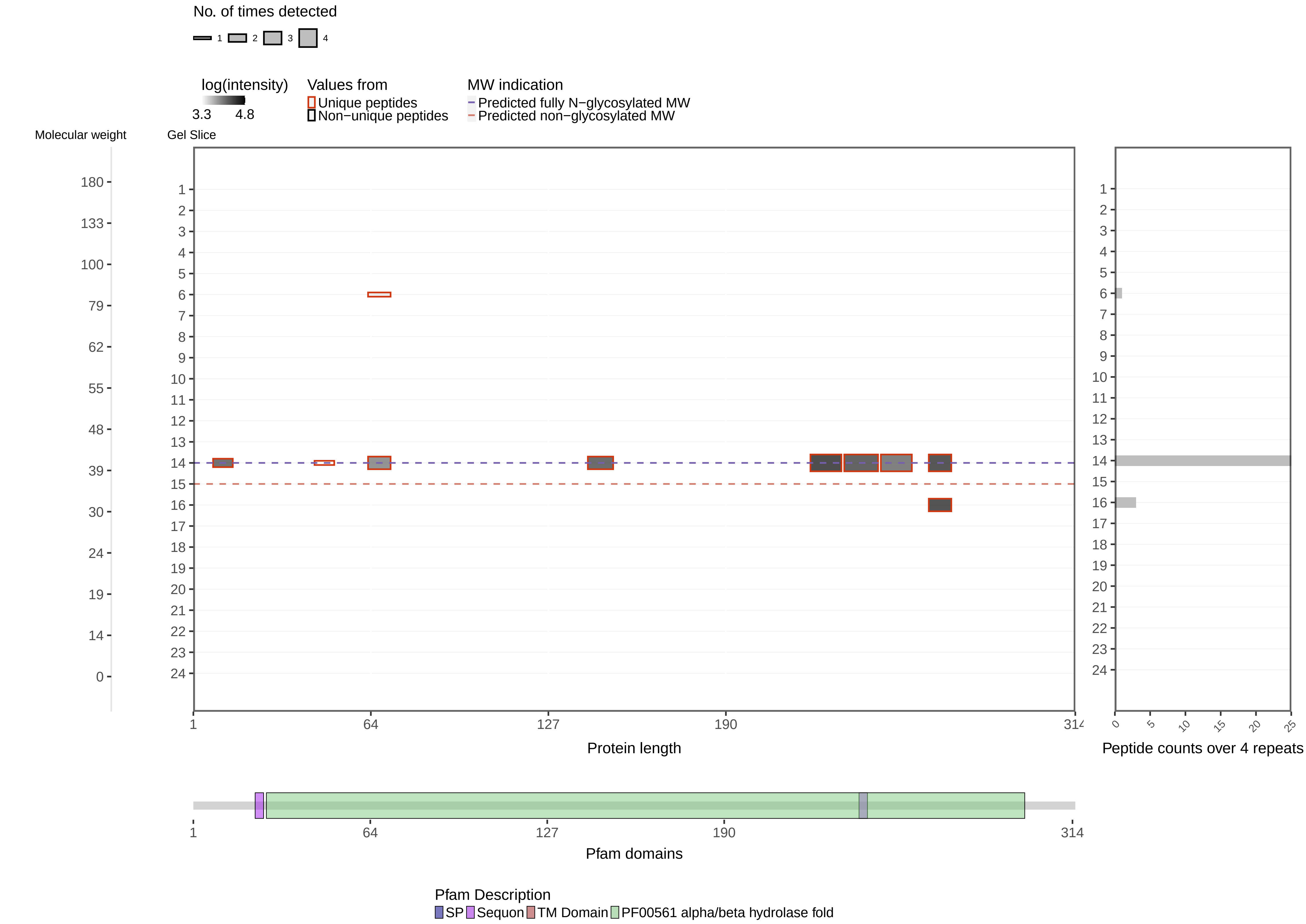


Protein sequence:

EGFYISRWREPGRAEADFGRFDTKTVVRKIYILFSRSEIPIADENQEILDLVEPSTPLPFWFSEKDLAFYGALYEKSGFLTALQVPYRAMGEQLNLTADPKIQVPALLLMGEKDYVLKFPGIEGYTRSEKLKSFIPNVEIIFMPEGTHFVQEQSPDEVNQLLLNFLHSNS*

NbL06g07810.1 acylsugar_acyltransferase_3−like [MW = 36.87 kDa]

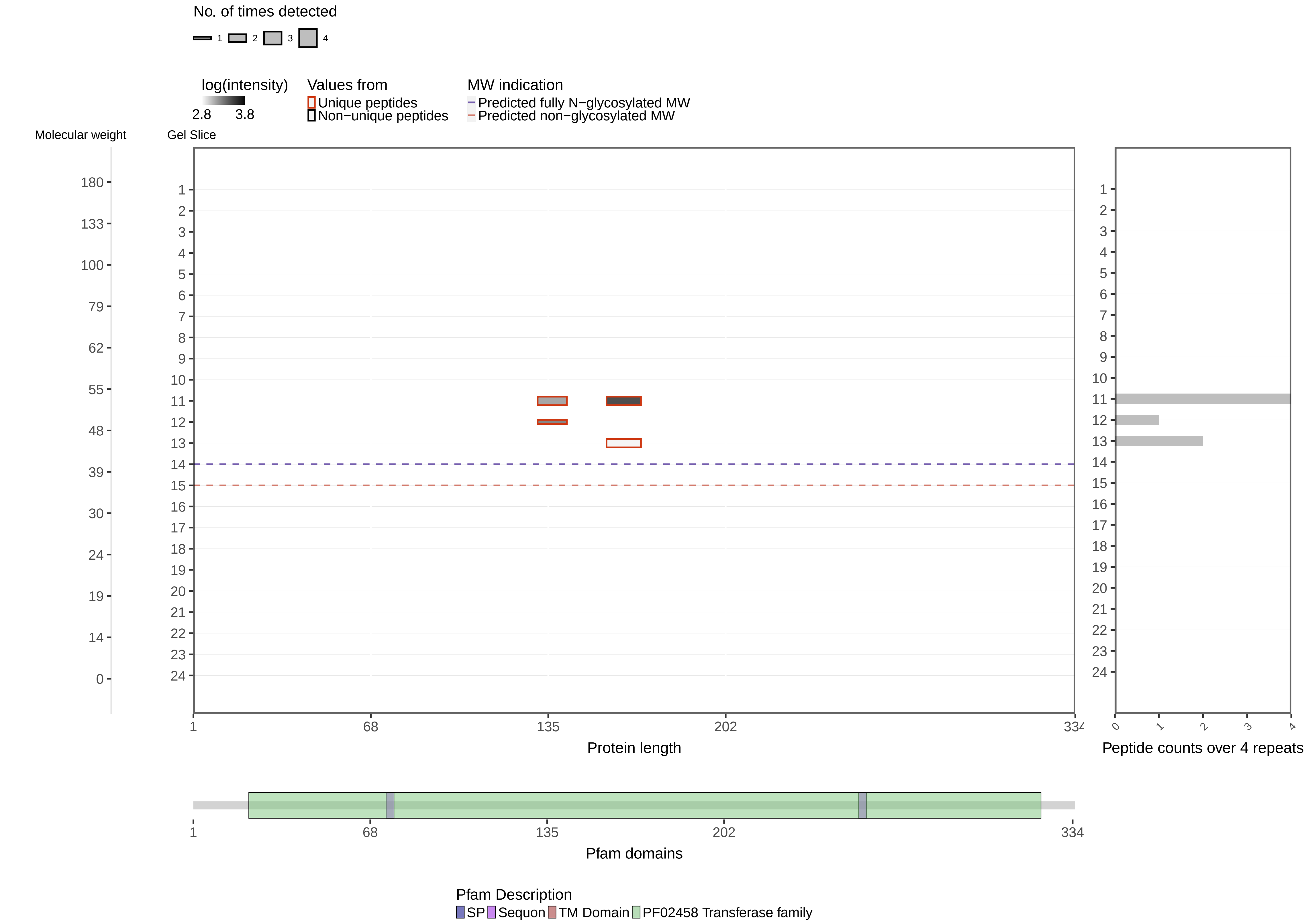


Protein sequence:

KAAAASSSANIQPSKLVHFLNVRTMTKPRLRRSAIGNLVSTFTTEATODIELPRLVONLRKEVEVAYKKDOVNENELVLETVDSMKKGKLPFEEKAENSTMYFCSNVCOFPLYSVDFGWGKPERVCLANGPSKNYFFLKDYKSGOGVEARVMLEKOHMSAFECDEELLOFTSSSVPSL*

NbL06g07480.1 uncharacterized_GPI−anchored_protein_At1g61900−like [MW − SP = 45.73 kDa]

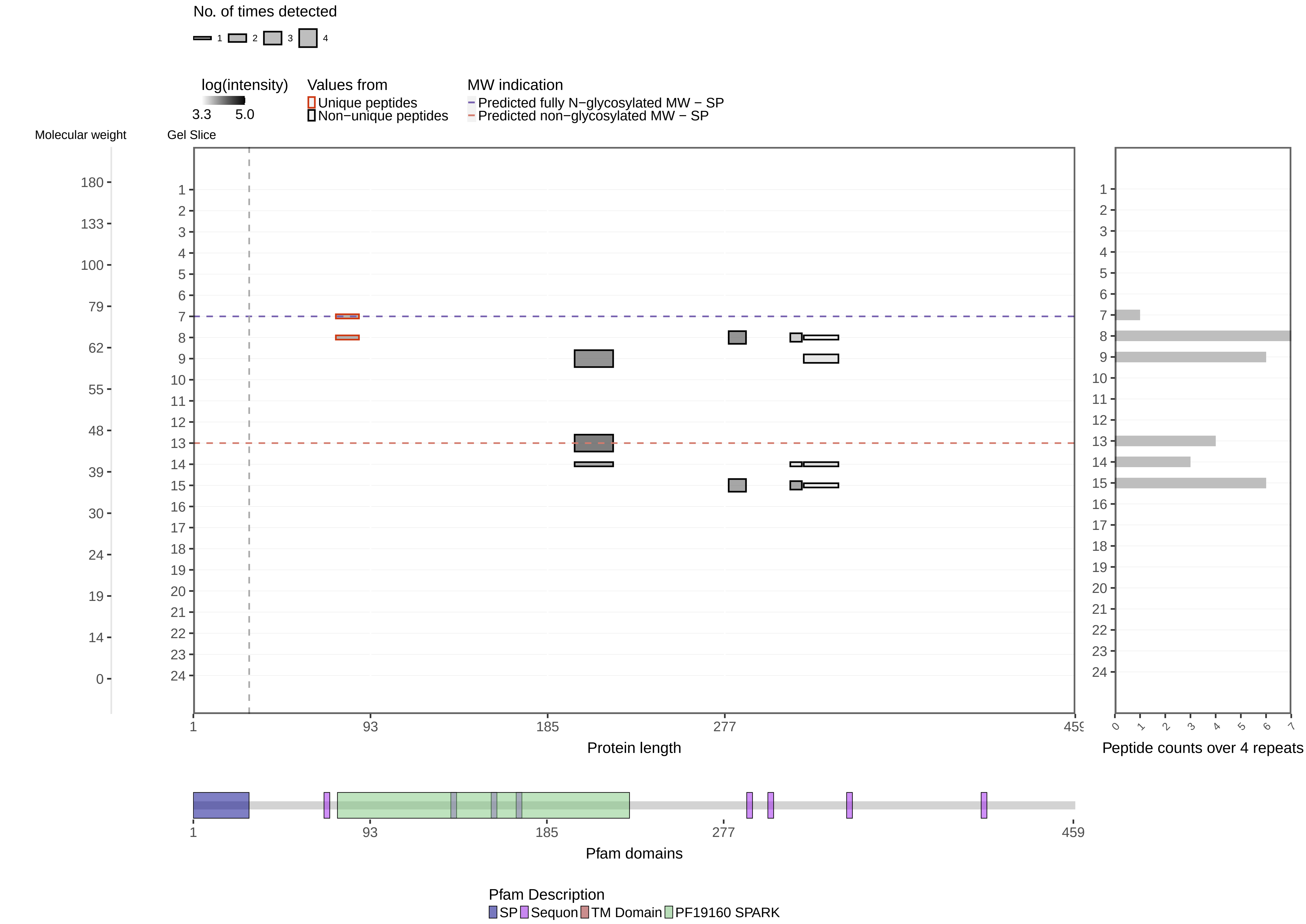


Protein sequence:

KKLALKAYDLLSMDGSHALADHTTRVNDCKSIVHRWLASKLDPAVAKDVLRGLSNCKNNKVCPLAFPGMKNITKACGNGMNNQTTCCNTVERYVSHLQRQSFVTNLQALDCAASLGLKLQKANVSKNVYNLCHISLKDFSVQVAPEVSGCLLPSLPSDAILDQSTGISFVCDLNDNIPAPWPSMSQLPASSCNKSVRIPALPAAASGQISKGLNMWSHMLLMASMILGICCISNAANLAY*

### NbL06g07470.1 UPF0051_protein_ABCI8_chloroplastic−like [MW = 61.18 kDa]

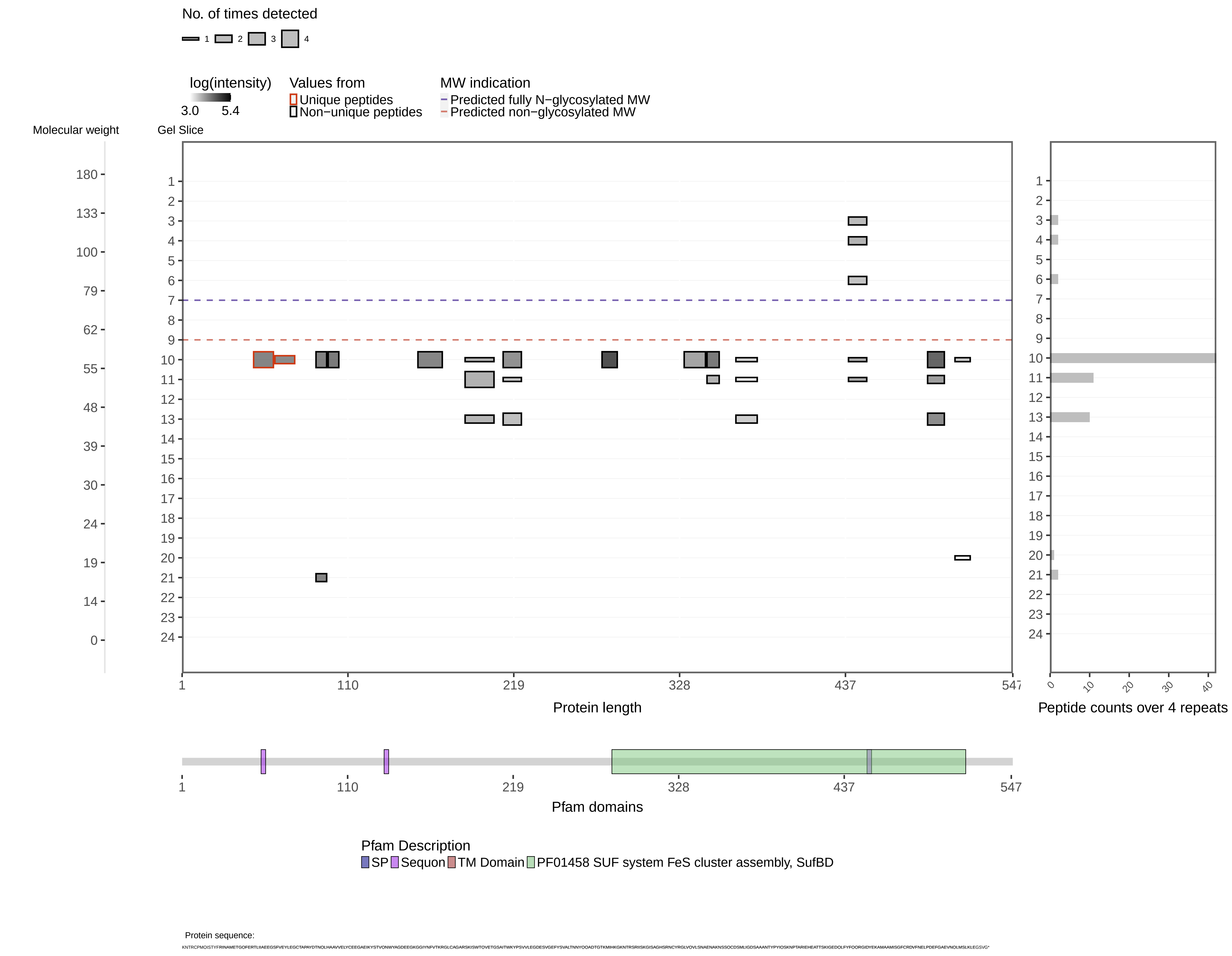

NbL06g07450.1 polyadenylate−binding_protein_RBP45−like [MW = 46.33 kDa]

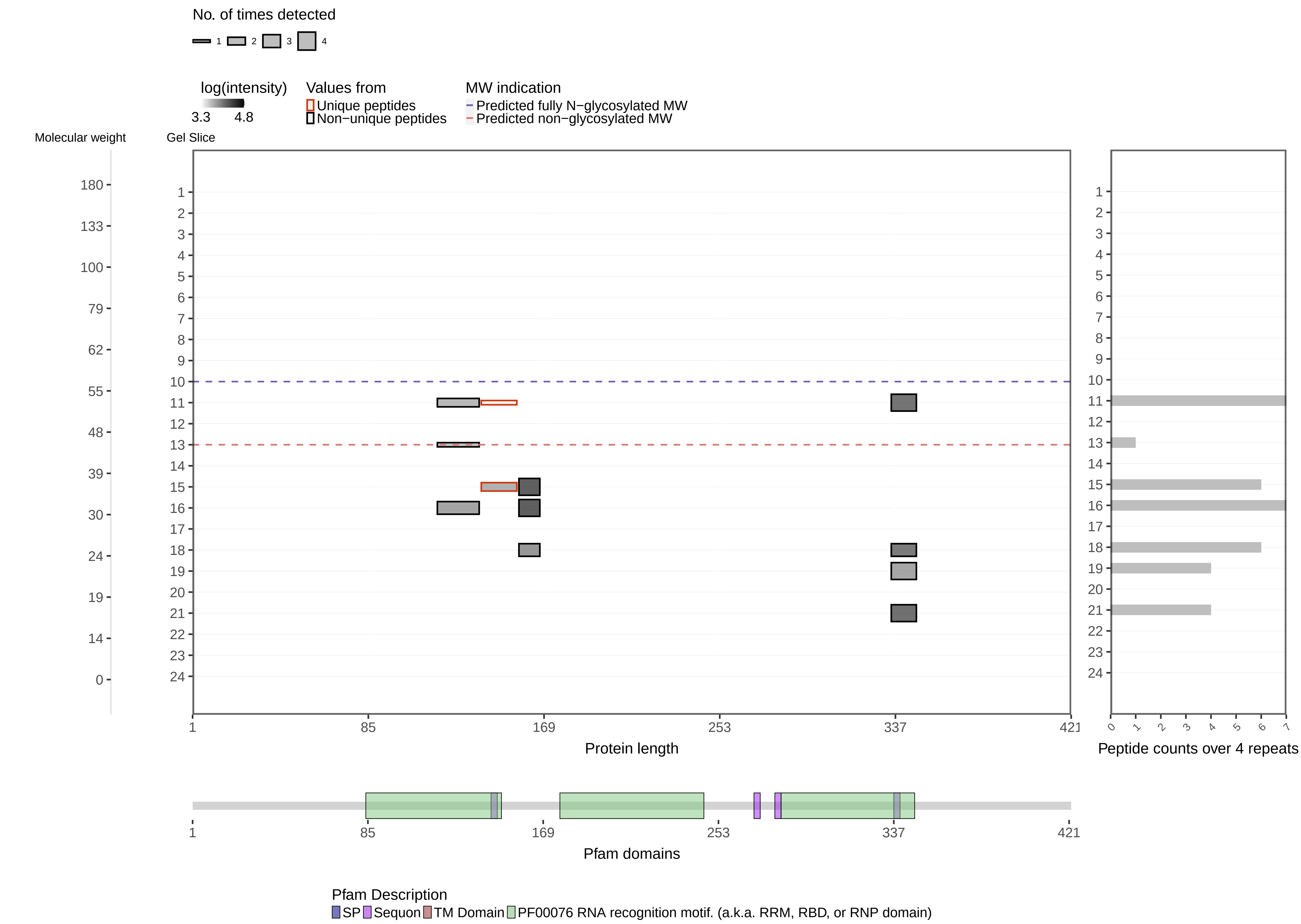


Protein sequence:
SYPSVKGAKVVTDRVTGRTKGYGFVKFGDESEQLRAMTEMNGKFCSTRPMRIGPAANKNSVAGQVQASYQSTNGTQNEDDPSNTTIFVGNLDSDVTDDHLRQVFGHYGQLLHVKIPVGKRCGFVQFADRSCAEEALRVLNGTQLGGQSIRLSWGRSPSNKQVVQPQADPNQWNGGGYYGYTAGYDAAAAAAYGYAQPAQDPNLYYAGYAGYGNYPPQQQQQPQ*

NbL06g07410.1 vacuolar_protein_sorting−associated_protein_28_homolog_2−like [MW = 23.36 kDa]

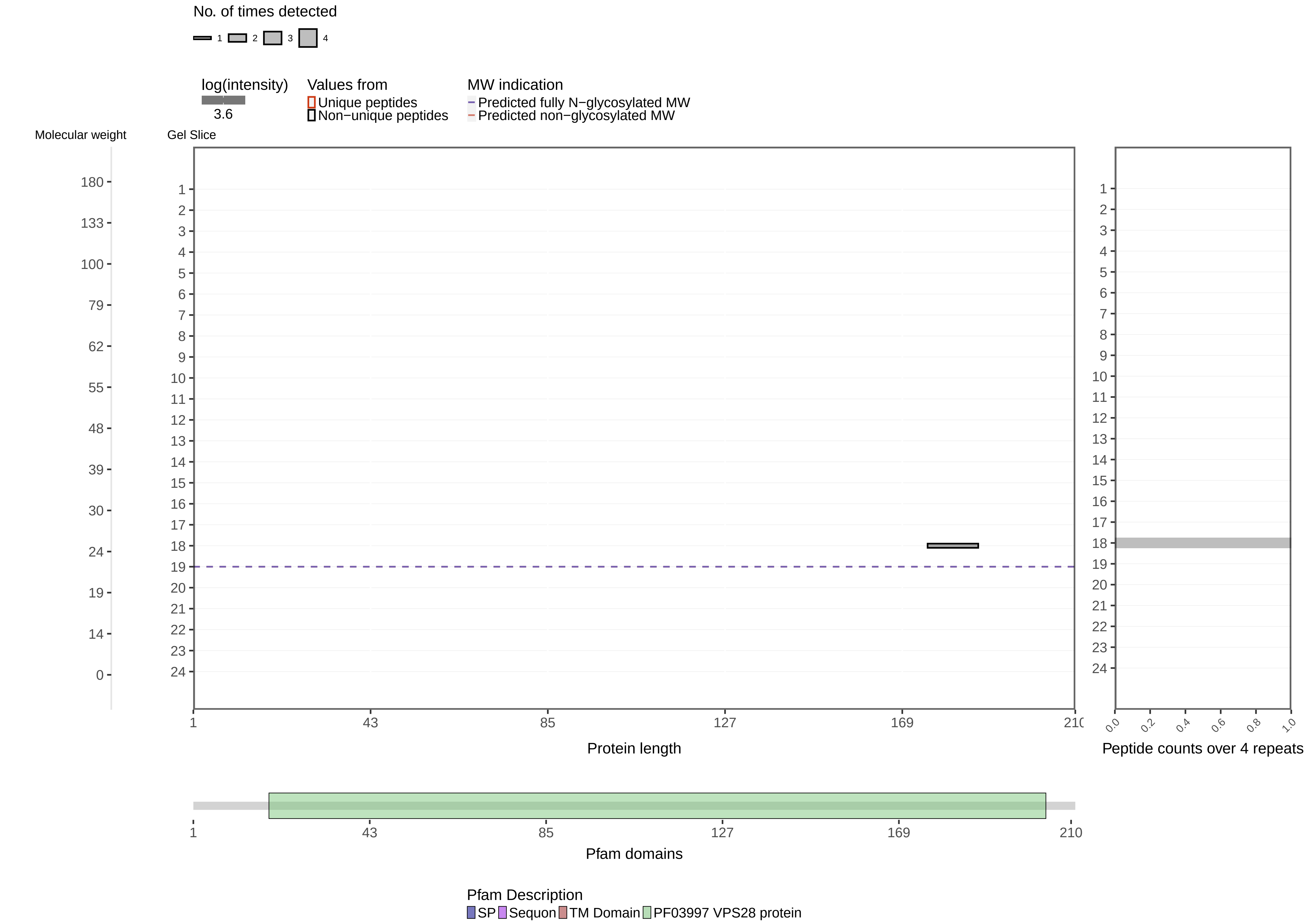


Protein sequence:

/PATVEHRAAASMSAITSAAVVAECVQNFITAMDSLKLNMVAVDQVHPLLSELSSSLNKLSILPVDFEGKTKMREWLSRLSKMGAADELTEQQARQLHFDLESSYNSFMAALPTDGT*

### NbL06g07300.1 eukaryotic_translation_initiation_factor_4E−1−like [MW = 24.79 kDa]

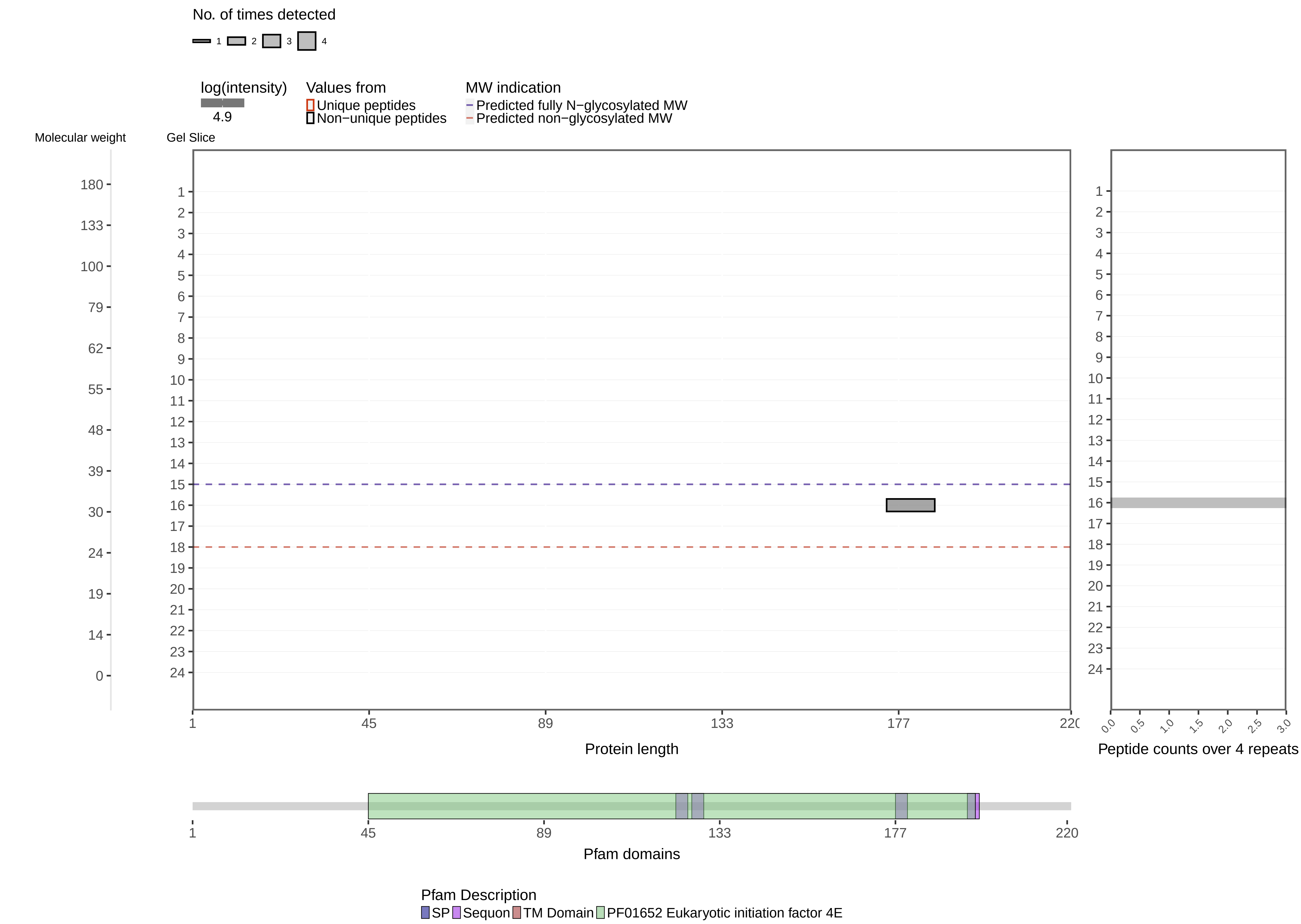


Protein sequence:

GADFHCFKHTIEPKWEDPVCADGGNWTMNFSKGKSDTSWLYTLLAMIGHQFDHEEEICGAVVSVRNKGDKIALWTKNAANETAQVSIGKQWKEFLDYNSSIGFTFHVDSMRLGRGAKNSYTV*

NbL06g06640.1 polyphenol_oxidase_E_chloroplastic−like [MW = 67.04 kDa]

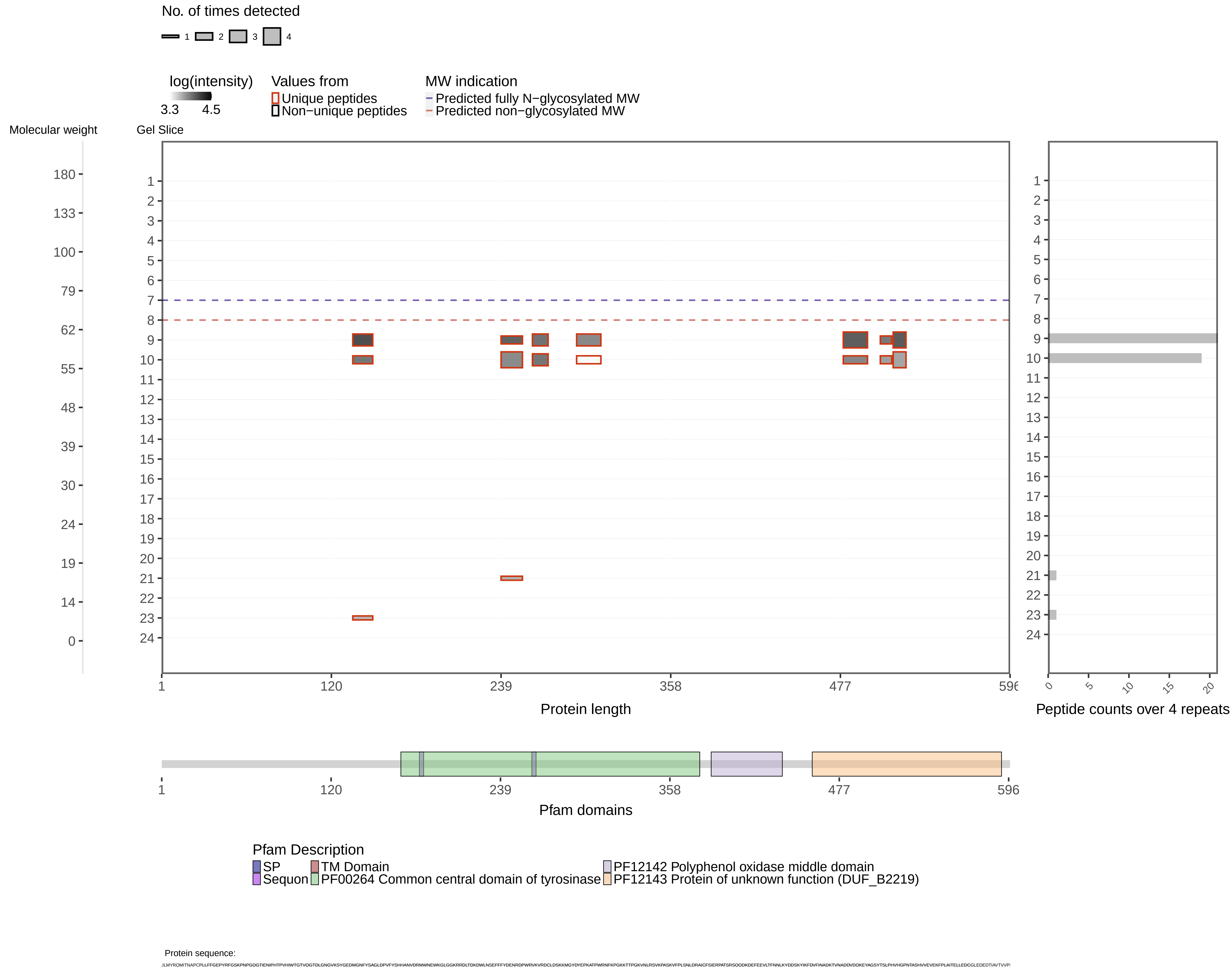

NbL06g06540.1 branched−chain−amino−acid_aminotransferase_2_chloroplastic−like [MW = 35.11 kDa]

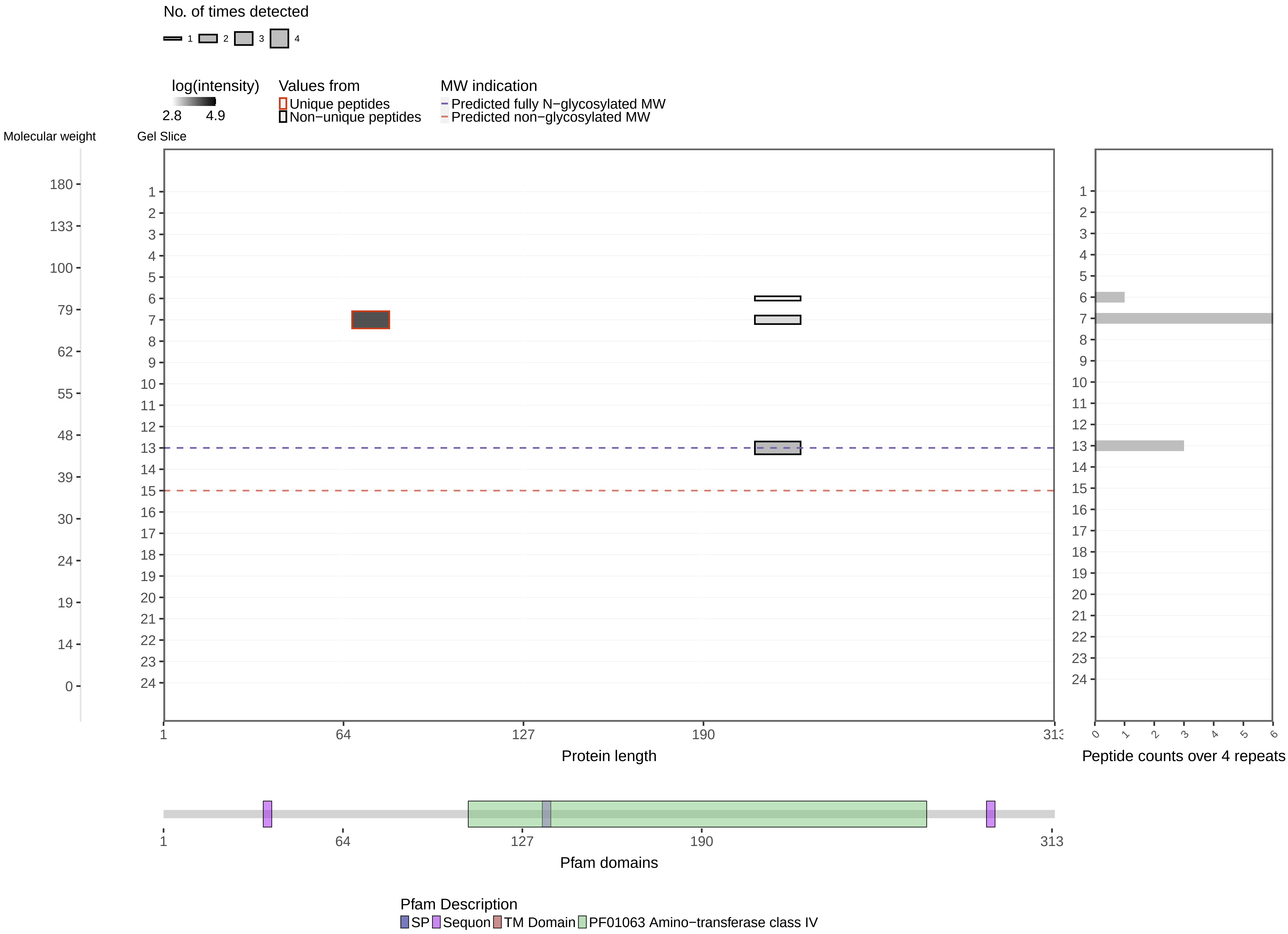


Protein sequence:

PLTLYVEEEHHRASQEGAGGVKSITNYGPVLKAIKRAKERGYSDVLYLDSVNNRYIEEVSAANIFLVKGRVIATPIPSGTILEGITRKSDIDIAHDLGYQVEERLIEAEELITADEVFCTGTALGVAPIGSITCKNKRIEYKVSSELISEQVNTTLVAIQKGIIEDKRDWVLEIK*

### NbL06g06470.1 tubulin_alpha−3_chain [MW = 49.74 kDa]

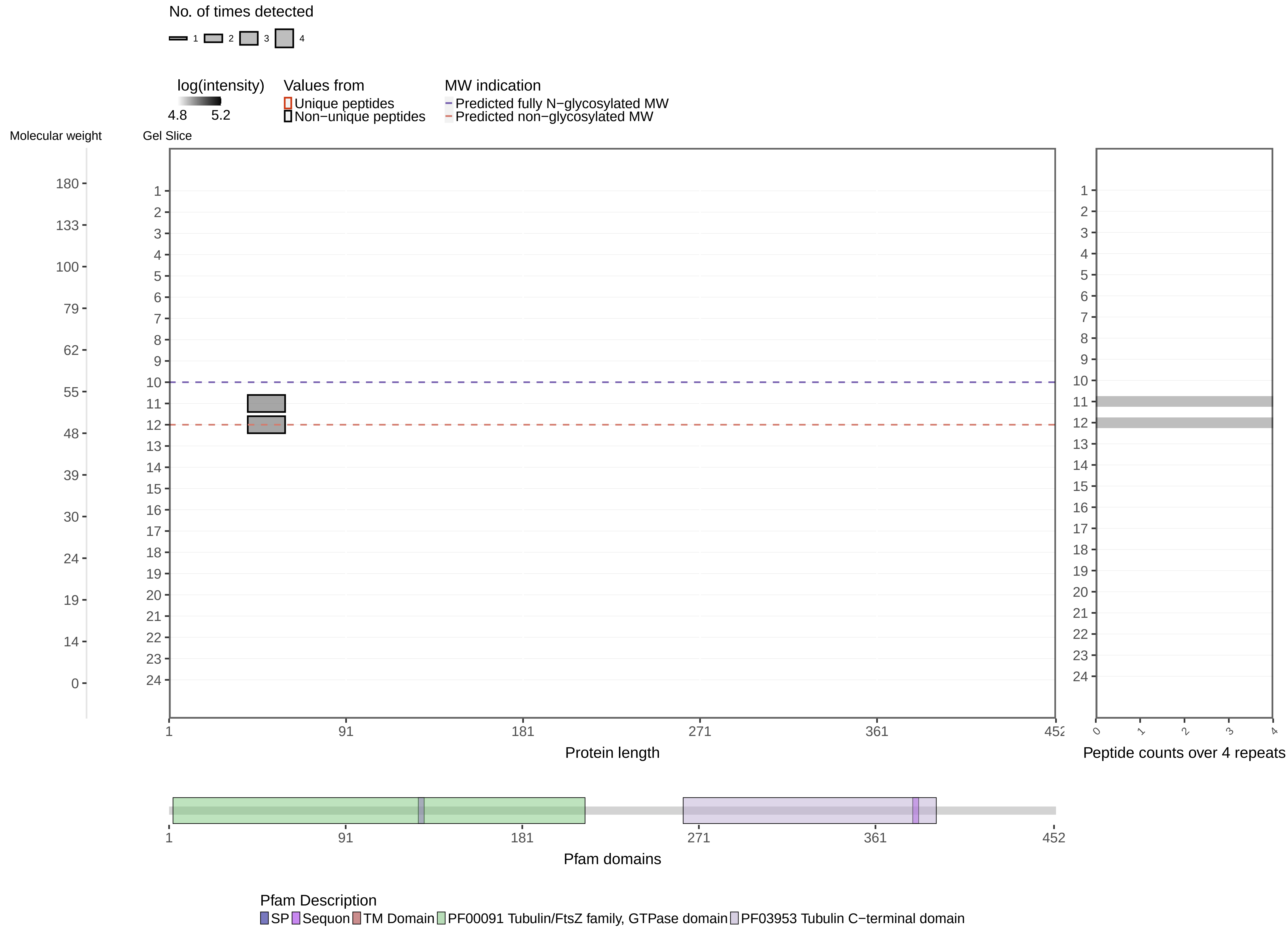


Protein sequence:

RSLDIERPTYTNLNRLISQVISSLTASLRFDGALNVDVNEFQTNLVPYPRIHFMLSSYAPVISAEKAYHEQLSVAEITNSAFEPSSMMVKCDPRHGKYMACCLMFRGDVVPKDVNAAVATIKTKRTIQFVDWCPTGFKCGINYQPPTVVPGGDLAKVQRAVCMISNSTSVAEVFSRIDHKFDLMYAKRAFVHWYVGEGMEEGEFSEAREDLAALEKDYEEVGAEVDDEEGGDEGDEY*

### NbL06g06400.1 transcription_initiation_factor_IIA_subunit_2 [MW = 24.11 kDa]

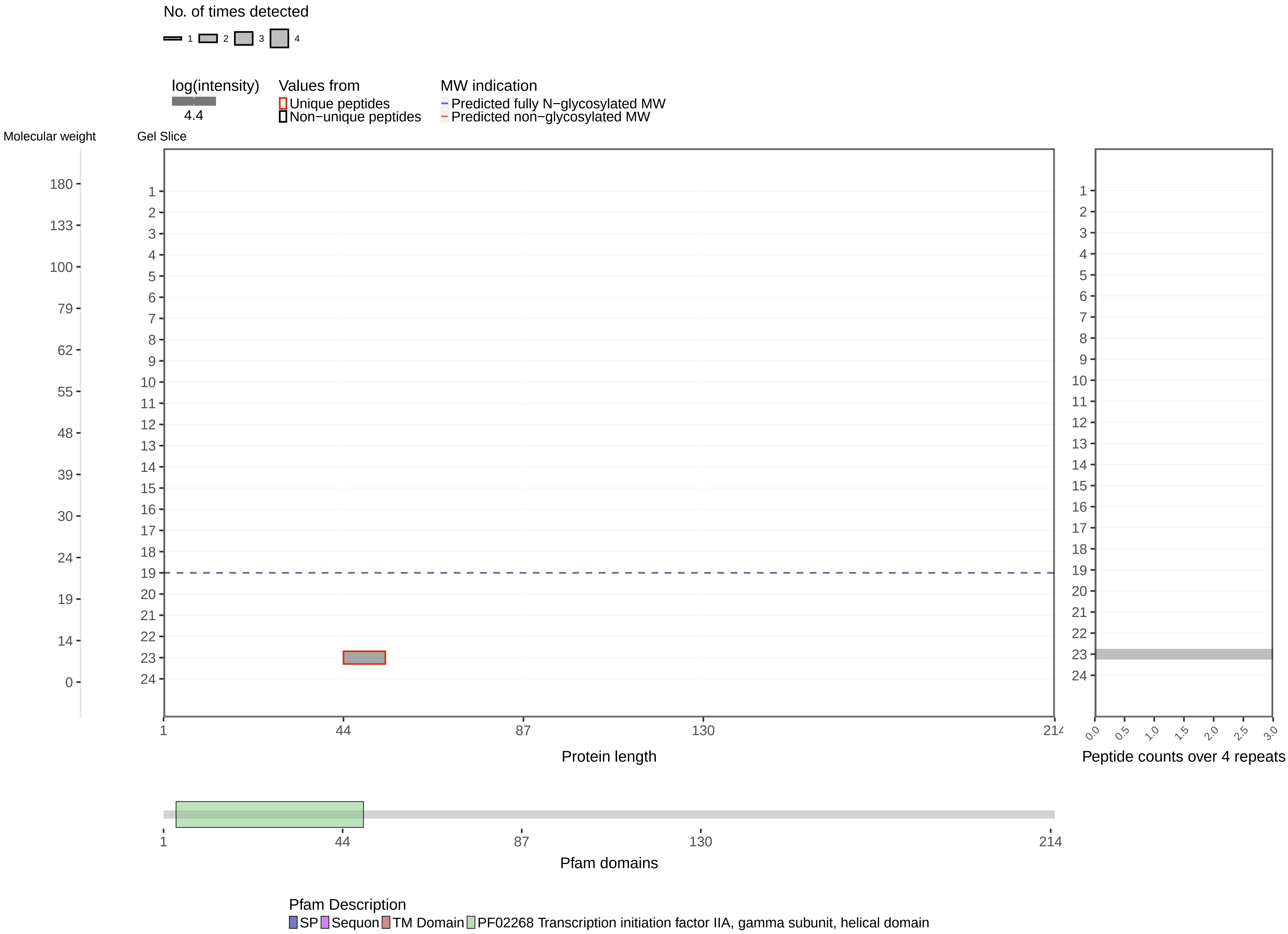


Protein sequence:

MSVRDIFIPTDSVTMCGPLSYKMLCSRAKNARRLLAALRLWHVTQSCSHSEKAADGGNFLLQICISIFNIEIDVREIWEQRMQGKCDYIHSGYISRQPFTFIFVQTSKHLCLSVVKLLYL*

### NbL06g06340.1 aspartyl_protease_AED3−like [MW − SP = 44.55 kDa]

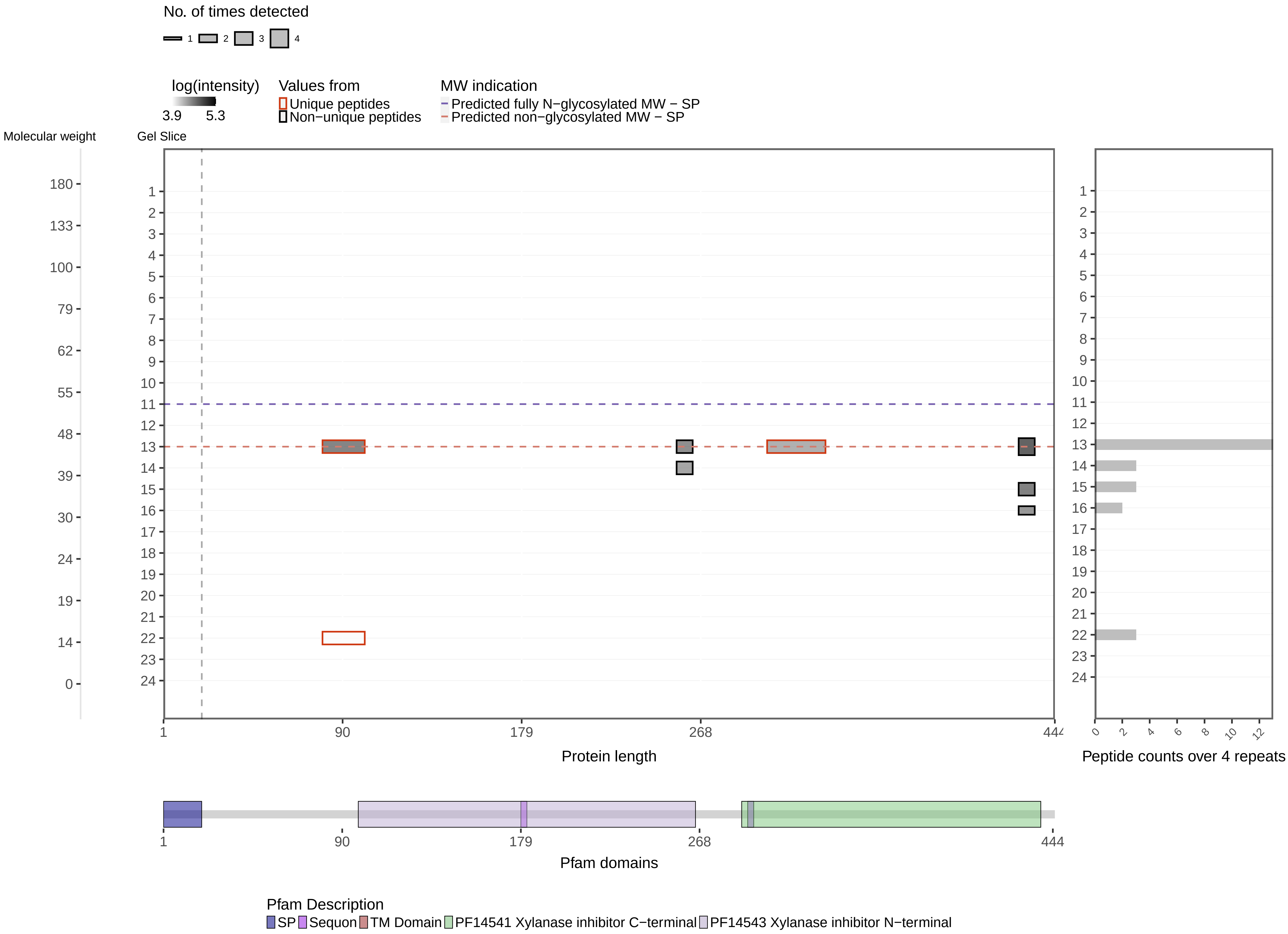


Protein sequence:

AFGCISAVSGNAIPPQGLLGLGRGTMSLMSQSGSLYSGVFSYCLPSFKSYYFSGSLKLGPLGQPKNIKTTPLLKNPRRPSLYYVNLTSVSVGRILVPIAPEQLGFDPNTGAGTIIDSGTVITRFVPPAYNAIRDEFRKQVKGPFSSLGAFDTCFASPNEAGAPSITLHFEGMDLKLPMENTLIHSSATPLACLAMAAAPSNVNSVLNVIANLQQQNLRILFDTANSRVGIAREVCN*

### NbL06g06090.1 14−3−3−like_protein_B [MW = 30.88 kDa]

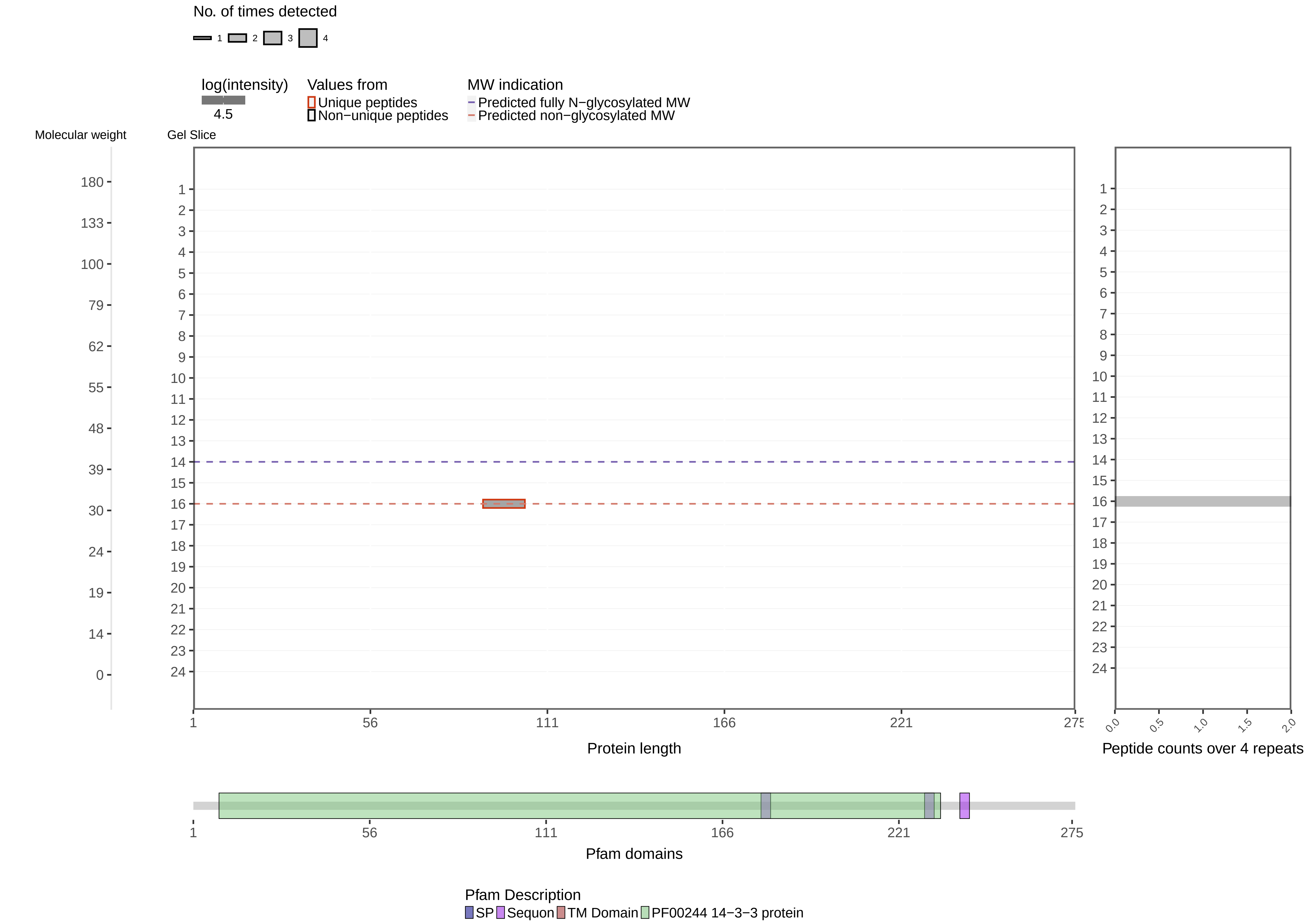


Protein sequence:

KMKGDYHRYLAEFKTGAERKEAAESTLSAYKAAQDIANAELAPTHPIRLGLALNFSVFYYEILNSPDRACNLAKQAFDEAIAELDTLGEESYKDSTLIMQLLRDNLTLWTSDMQVNNSATFFVFVCFLSVRCIARVLAFESKGNNNNNPV*

### NbL06g05820.1 fructose−16−bisphosphatase_cytosolic−like [MW = 34.22 kDa]

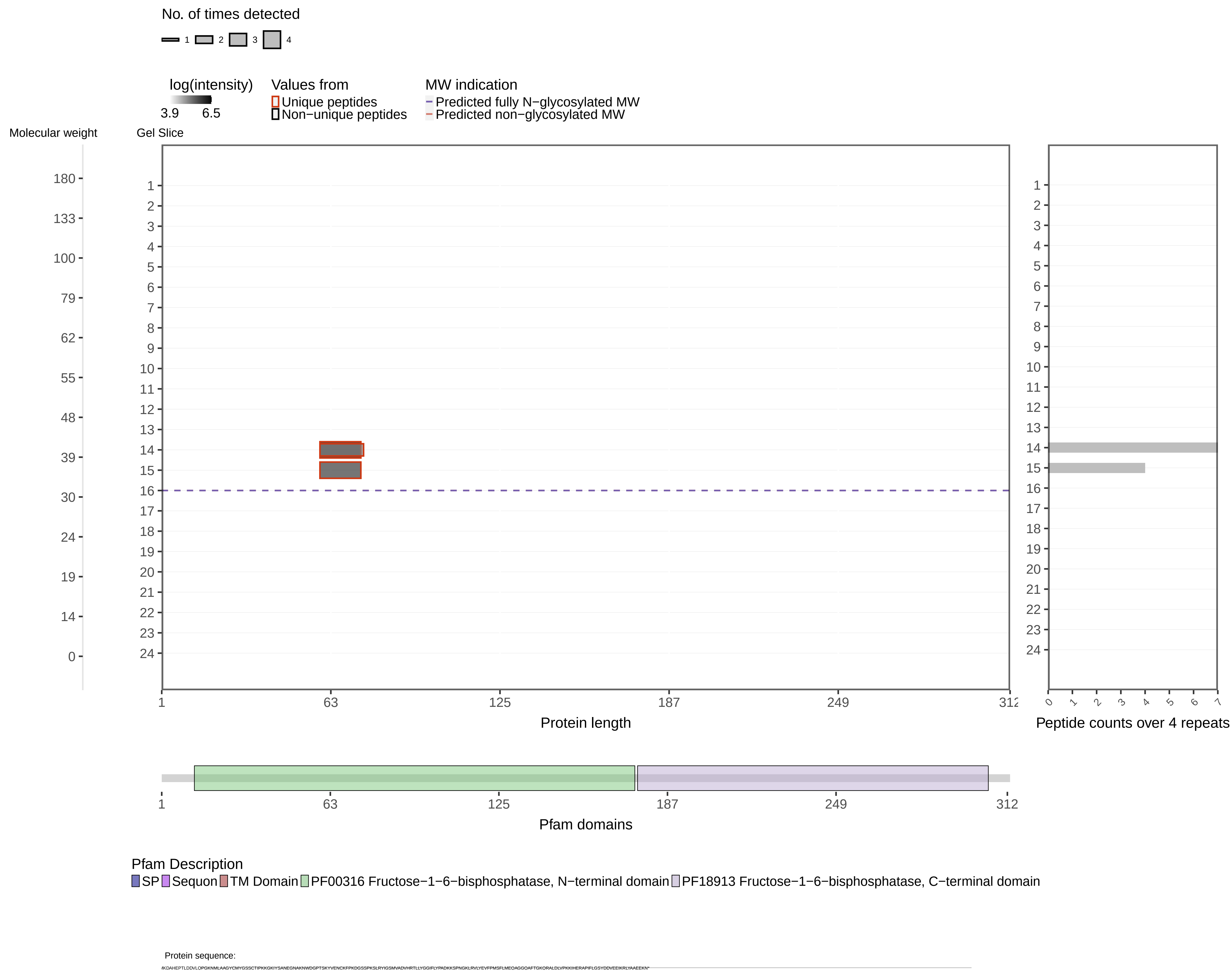

### NbL06g05710.1 secoisolariciresinol_dehydrogenase−like [MW = 29.62 kDa]

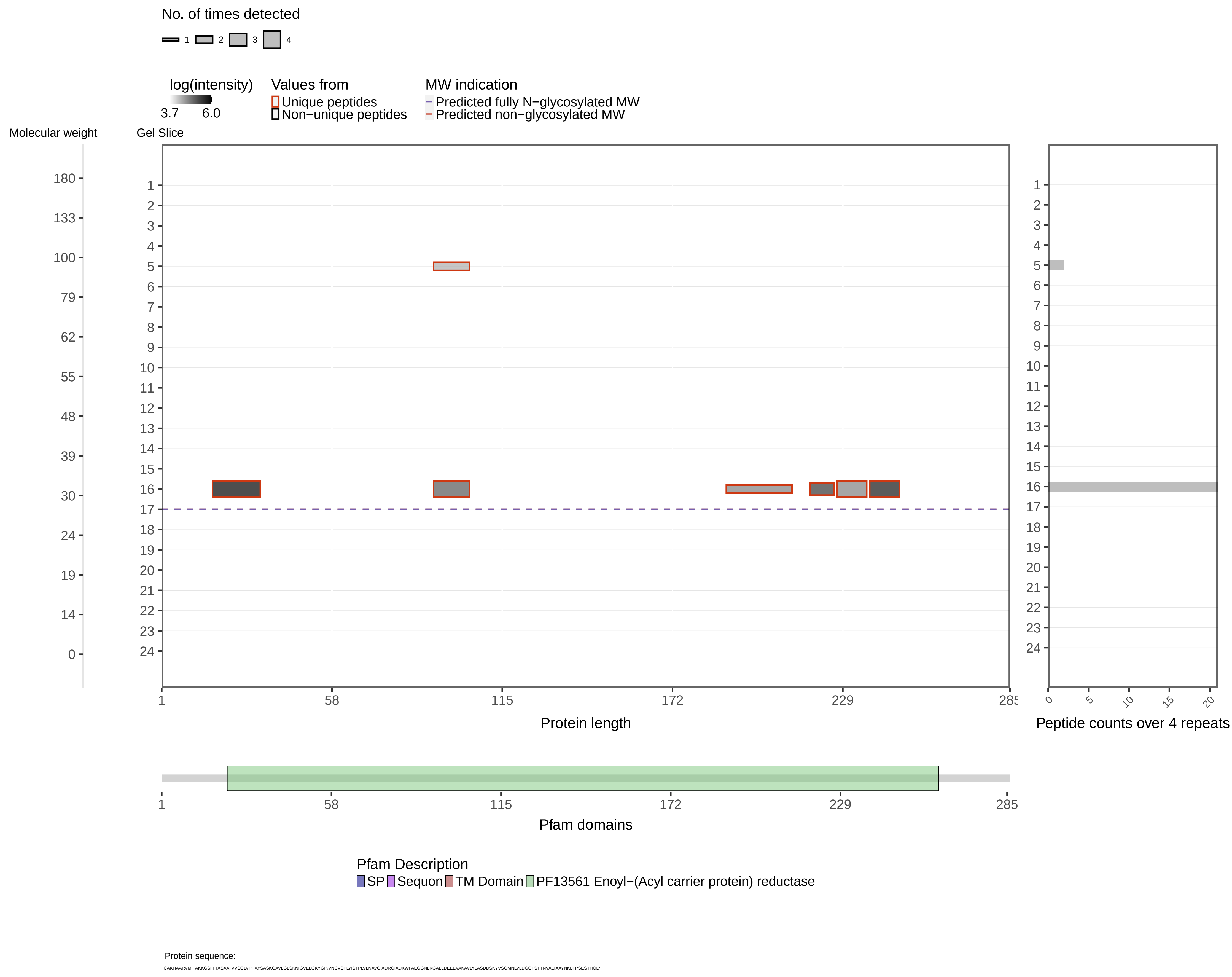

### NbL06g05620.1 secoisolariciresinol_dehydrogenase−like [MW = 29.75 kDa]

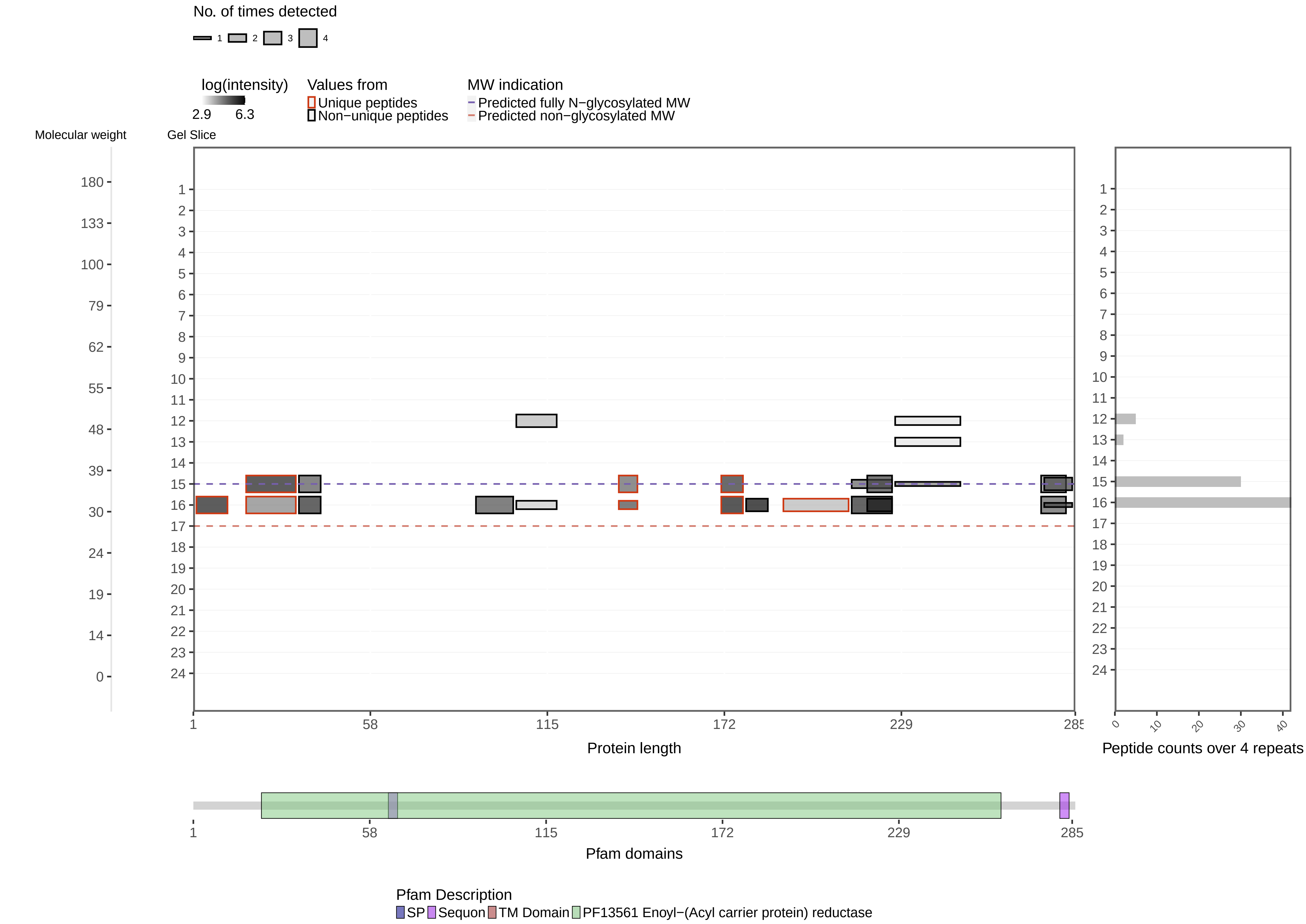


Protein sequence:

CAKHAARVMIPAMKGVIIFTASASTEVYGTAIPHTTLASKNALVGLSKNVGVELGKYGIRVNCVSPYYISTPPILNGFGIDKQTADKWFAEAGNLKGALLGVEDVANAVLYLASDDSKYASGMNLVLDGGFSTTNVAITEAYKKLFPSTTNKTE*

### NbL06g05610.1 secoisolariciresinol_dehydrogenase−like [MW = 30 kDa]

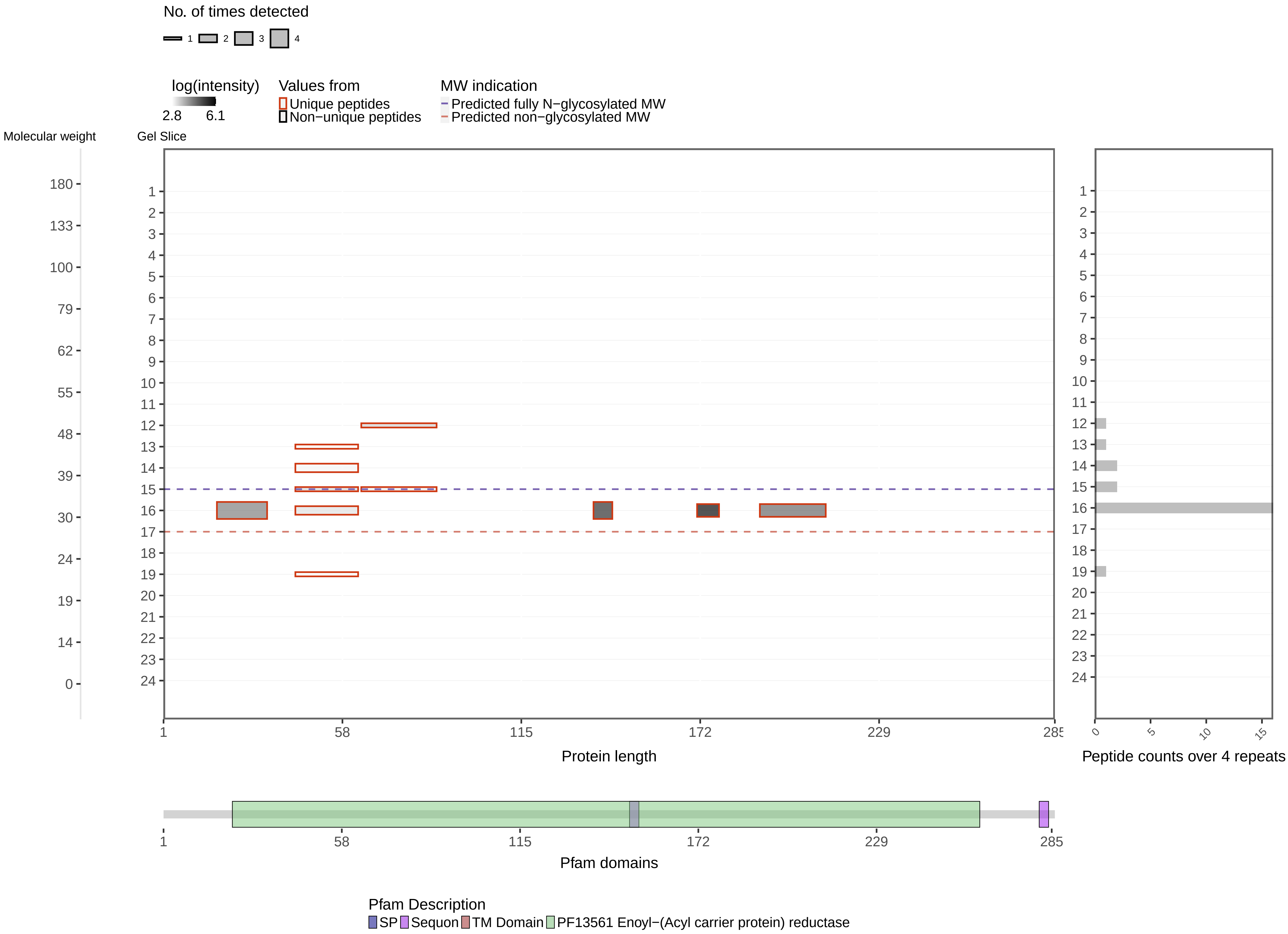


Protein sequence:

AKHAARVMIPFNKGSIIFNASASTEVCVSAIMHTSLASKNAVVGLSKNVGVELGKYGIRVNCVSPYYISTPLILNGYGIDKOTADKWFAEAGNLKGALLGVEDVANAVLYLASDDSKYASGMNLVLDGGFSTTNVAITEAYKKLFPSTTNKTE*

NbL06g05510.1 uncharacterized_oxidoreductase_At4g09670 [MW = 39.72 kDa]

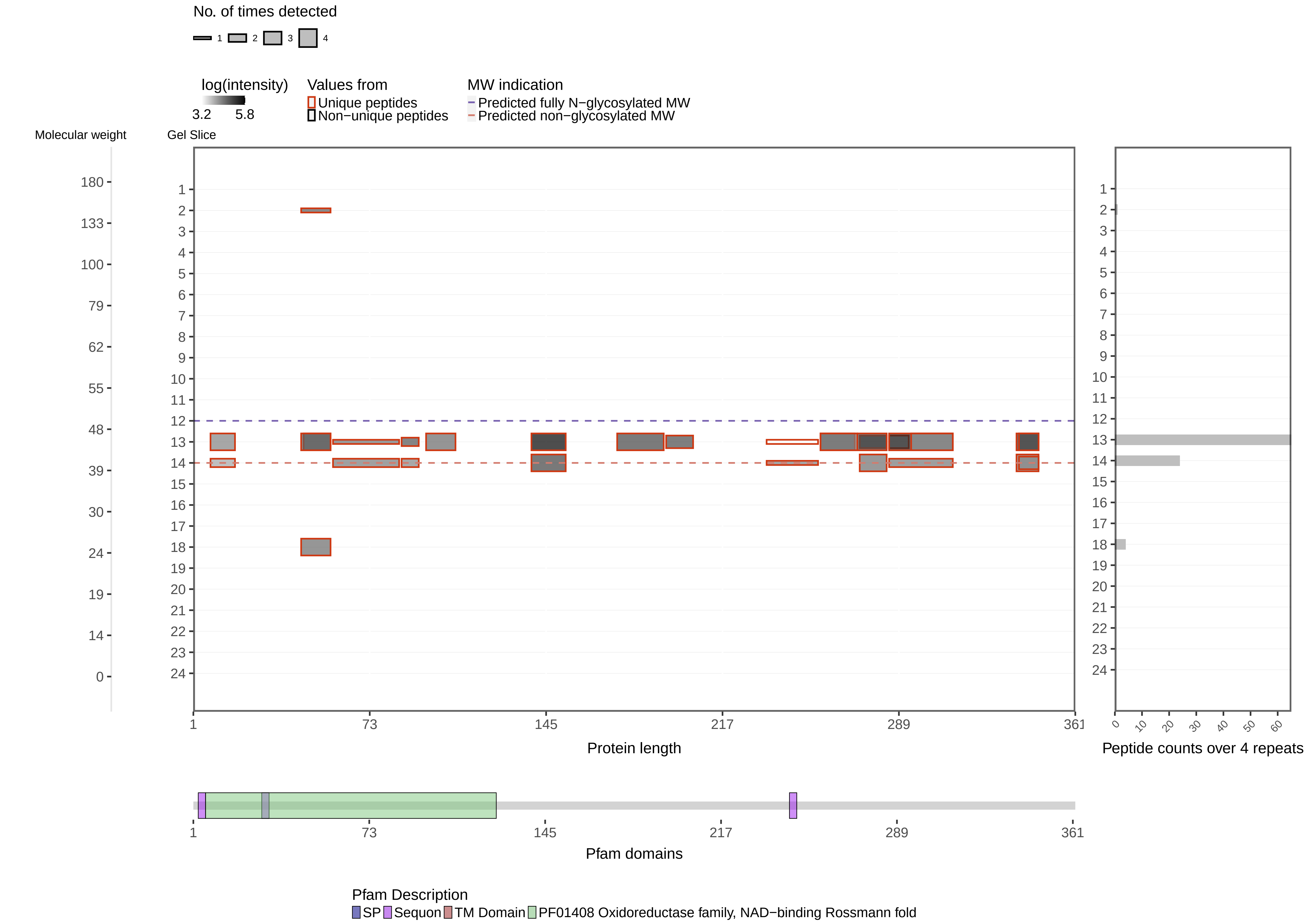


Protein sequence:

IFLENDIRVKPDLDALGALGDVGWYCIRGILWAADFELPKSVIALRNPVFNKAGVVISCSASLSWEDGKVASFHCSFLSNLTMDLTAVGTKGTLHIHDFVIPFEEKKASFVSAVESGFKELVTGWEPKPSEHTVMTDFPOEALMVKEFSRLVASIKNEGSKPEKKWPTLSRKTTLVLDAVKASIEKGFEAVEIIS*

NbL06g05470.1 ATP_synthase_delta_chain_chloroplastic−like [MW = 26.76 kDa]

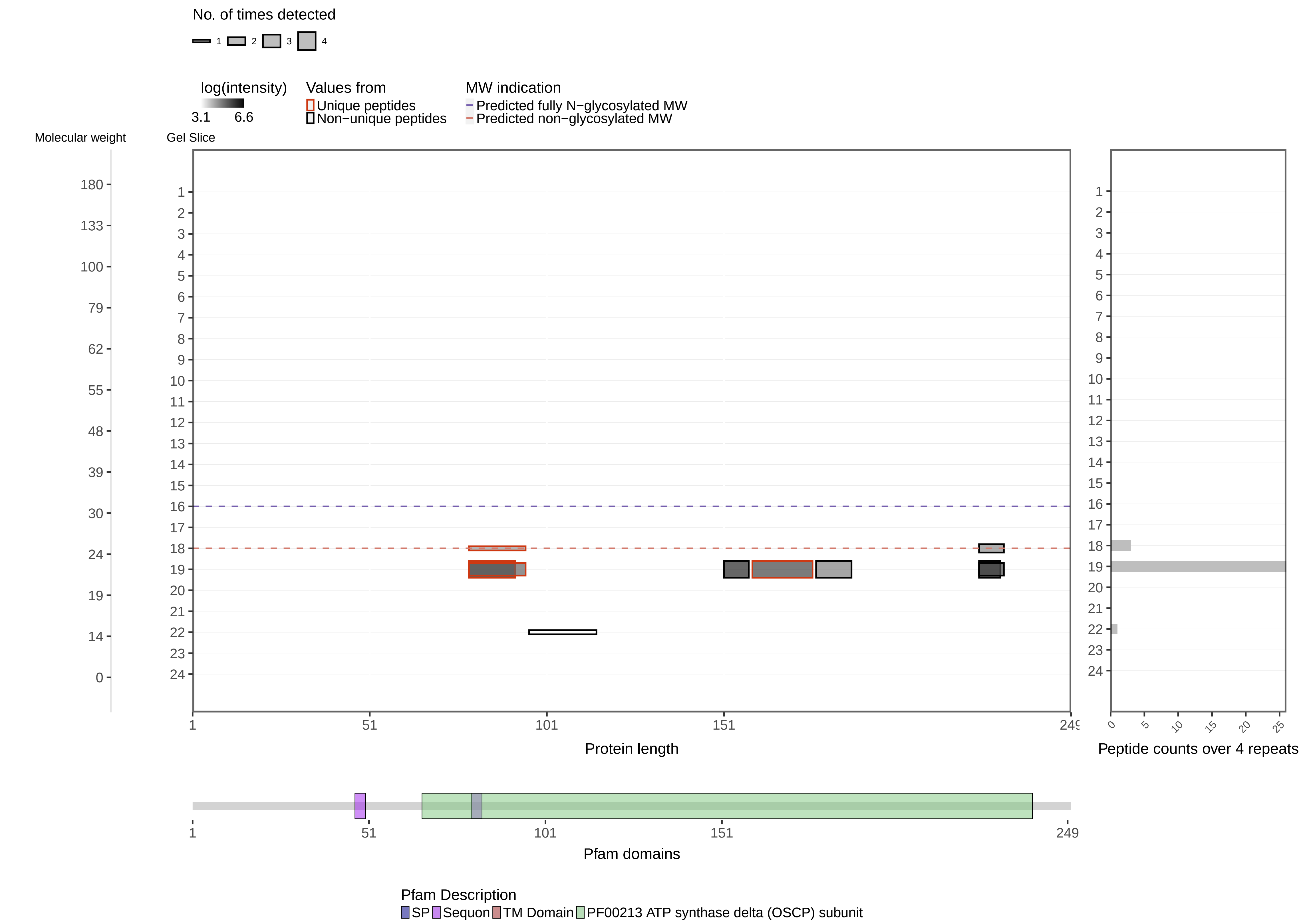


Protein sequence:

PIVGEEKKRELVDEIVSSSSIQPHVANFLNILVDMKRVELIKEIVKEFEKVYNTLTDTELAVVTSVVKLESQHLAQIAKGVQRLTGSKNVRIKTAIDESLVAGFTIRYGNSGSKLIDMSVKKQLEDIAAQLEIGDIQLAV*

### NbL06g05160.1 glutathione_S−transferase_T1−like [MW = 29.72 kDa]

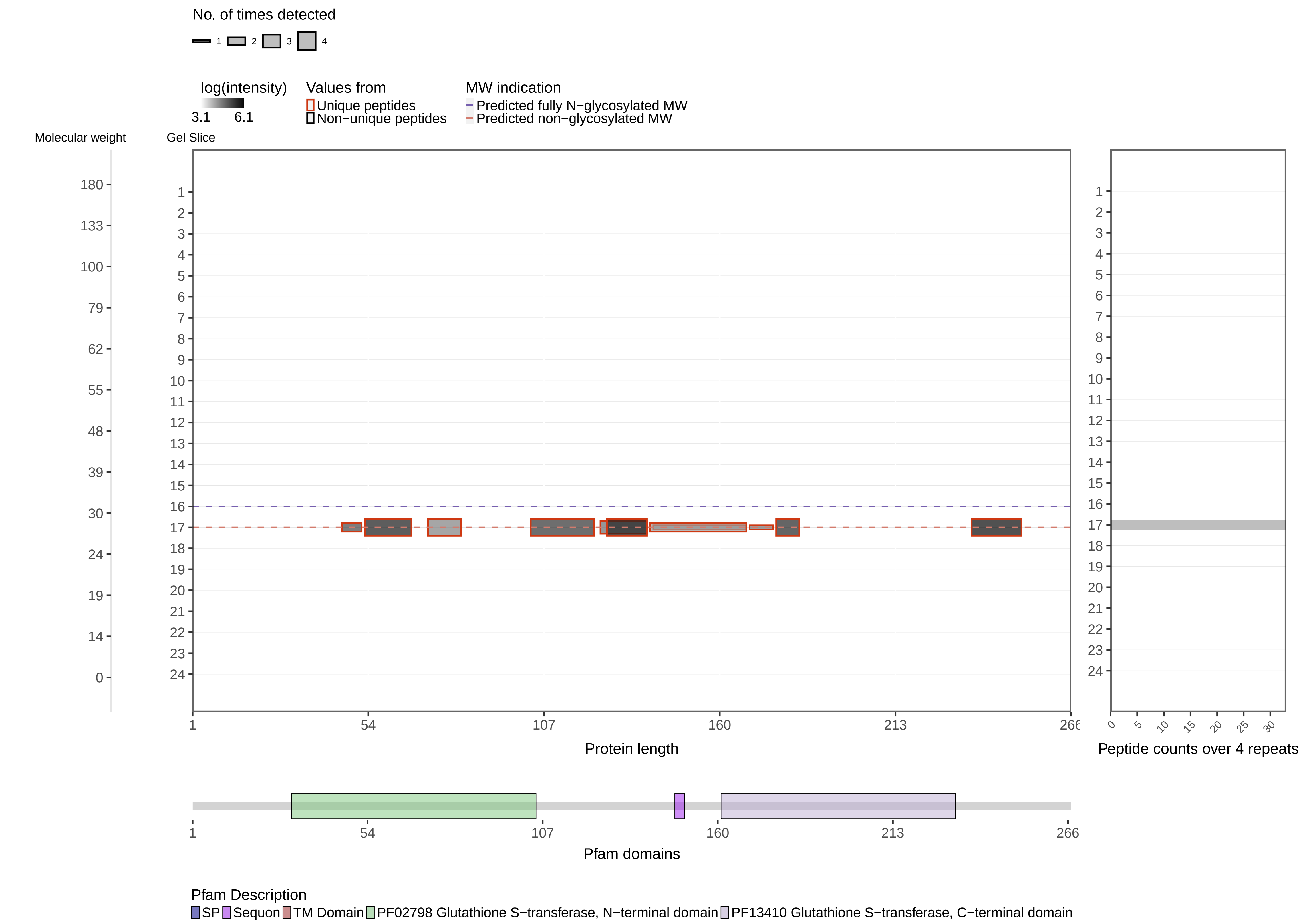


Protein sequence:

QKRAKTESVLDWHHSNLRRGSVGYVLNSTLAPALGLPLNPQAAAEGEKVLSASLAKIDTYWLQKDGSFLLGNSQPSIADLNLVCEIMQLEVVDEKDRHRILSAHKNVLKWIDDVKSATASHFDEIHATLFKVKEIFQKQRSAGAS*

### NbL06g05150.1 probable_glutathione_peroxidase_8 [MW = 19.4 kDa]

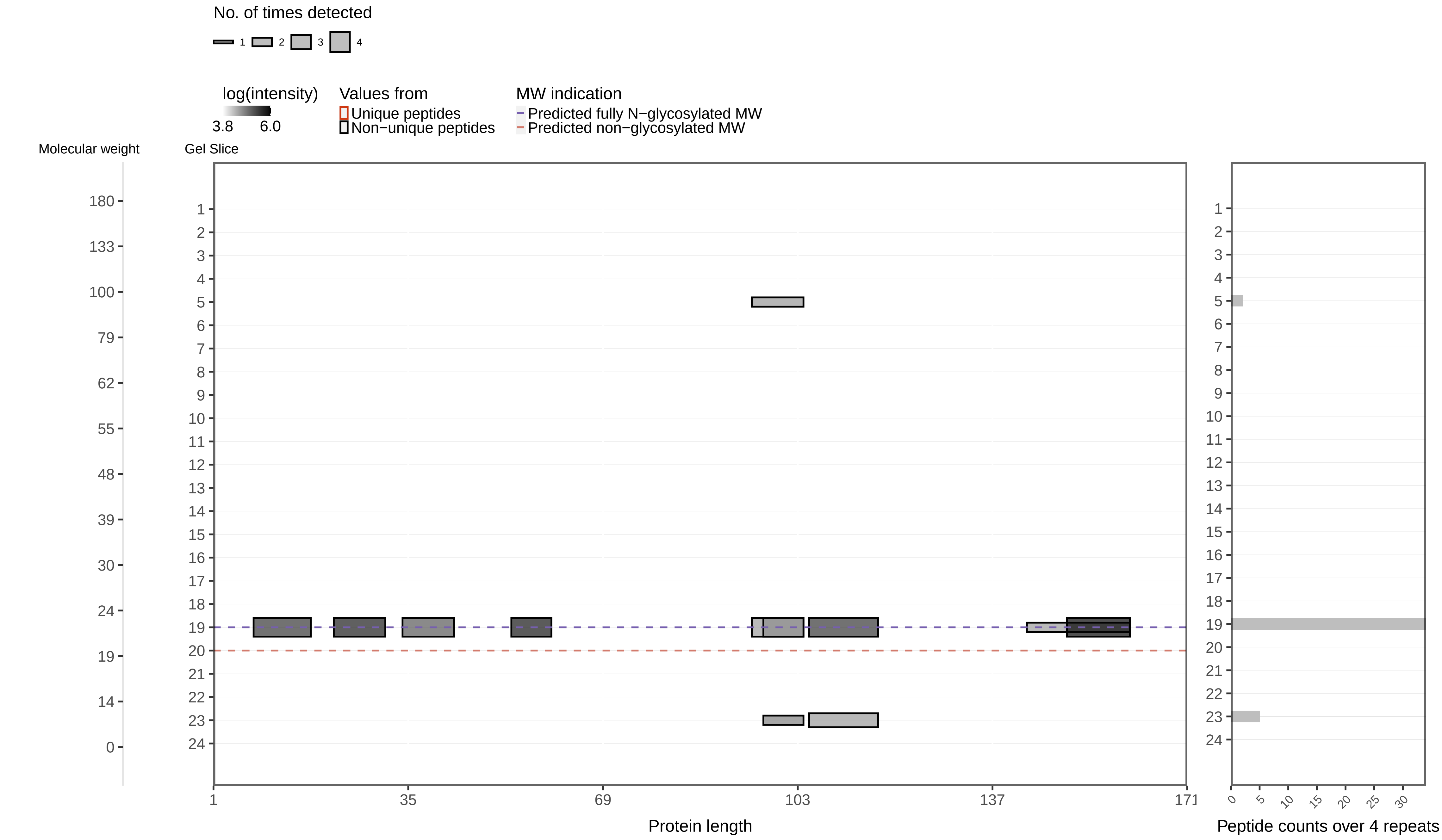


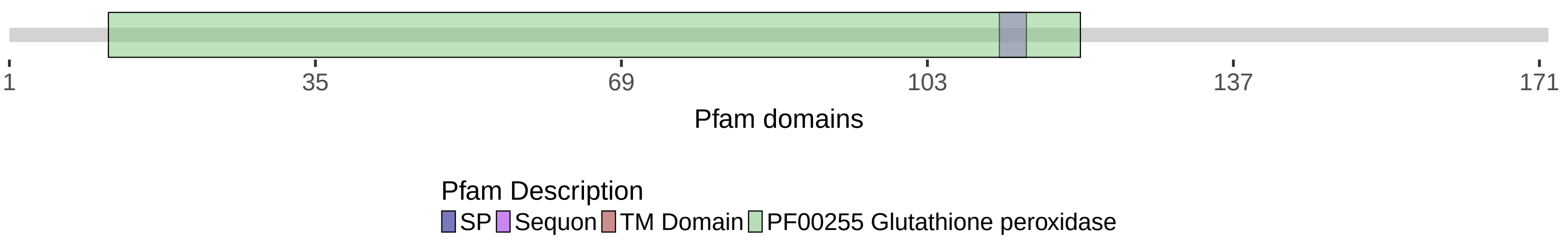


Protein sequence:

PCNQFGEEEPGSNDQILDFVCTRFKSEFPIFDKIEVNGENASPLYKLLKSGKWGIFGDDIQWNFAKFLVDKNGQAADRYYPTTSPLTIERDIKKLLEIV*

### NbL06g05030.1 V−type_proton_ATPase_subunit_E−like [MW = 32.94 kDa]

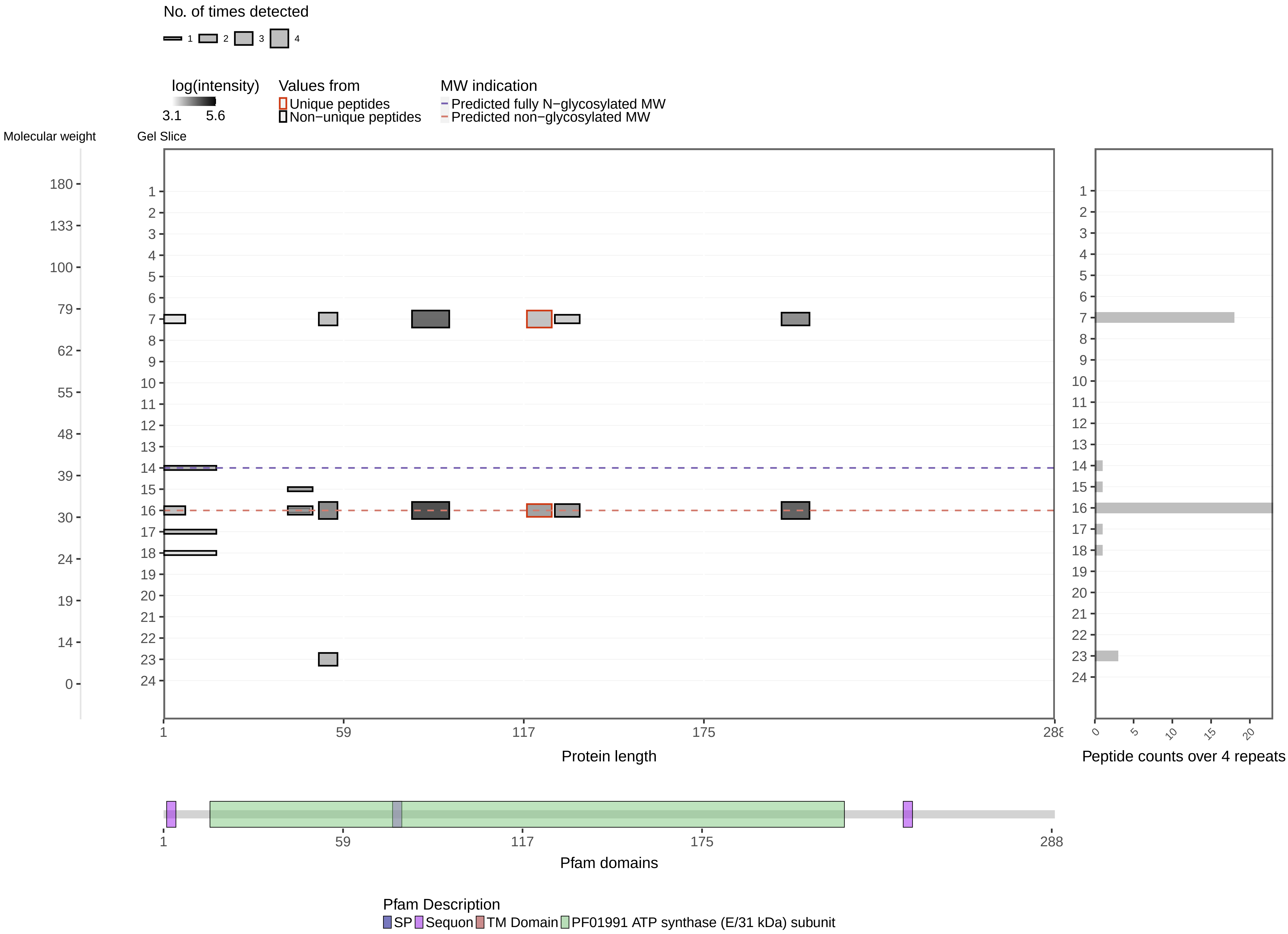


Protein sequence:

PSVLLRCRKDDVHLVEDVLDAAKEEYAEKAMVHSPEIIVDHIHLPPAPSHHNAHGPSCSGGVVLASRDGKIVCENTLDARLEVVFRKKLPEVLALVLISVLLVILCSCLANFSSFPSTSIYTFVVYLPSYICLLQMHPMLFFRCHVDELLSILHDHYL*

### NbL06g05000.1 6−phosphogluconate_dehydrogenase_decarboxylating_2_chloroplastic−like [MW = 54.17 kDa]

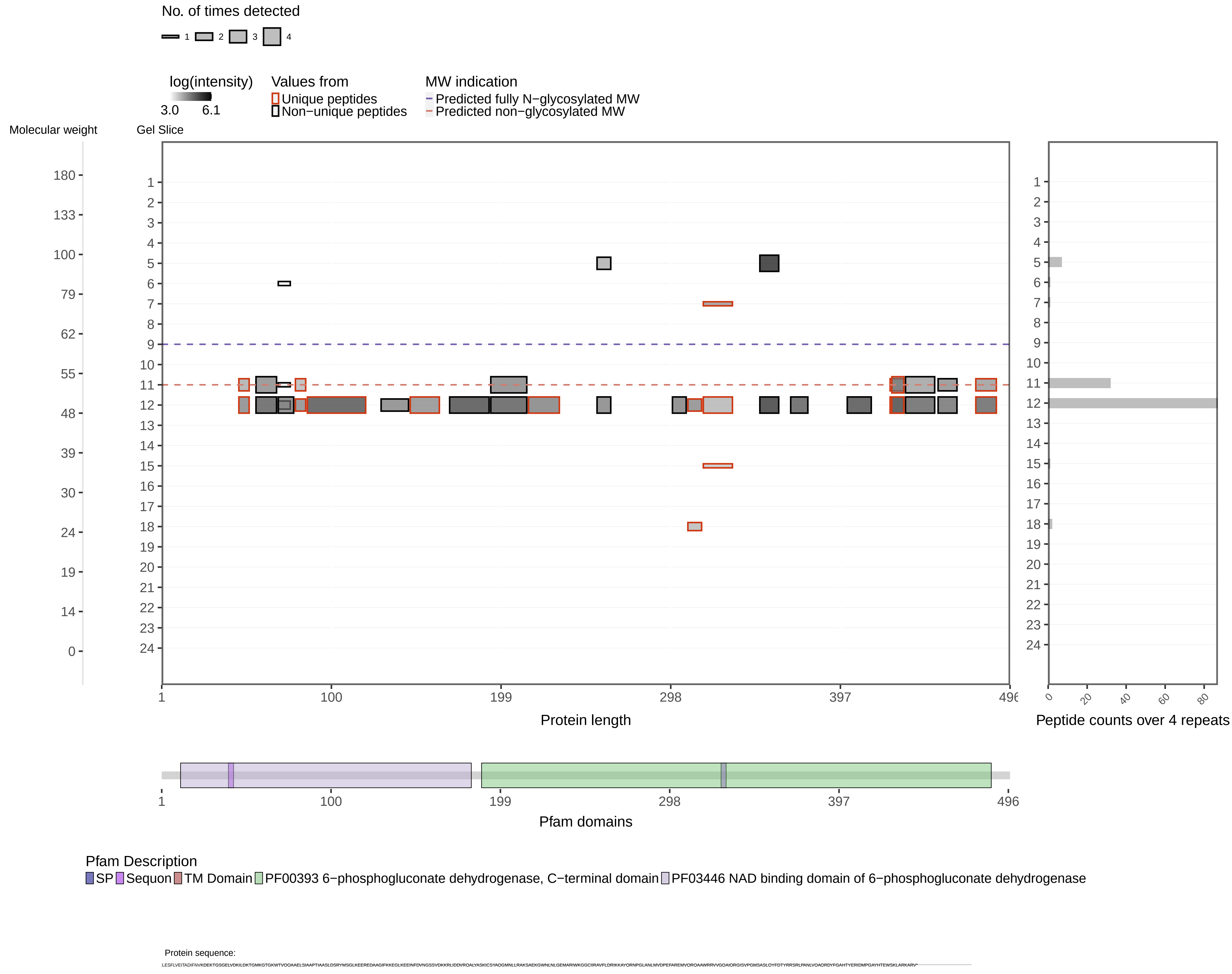

NbL06g04800.1 30S_ribosomal_protein_S6_alpha_chloroplastic−like [MW = 24.76 kDa]

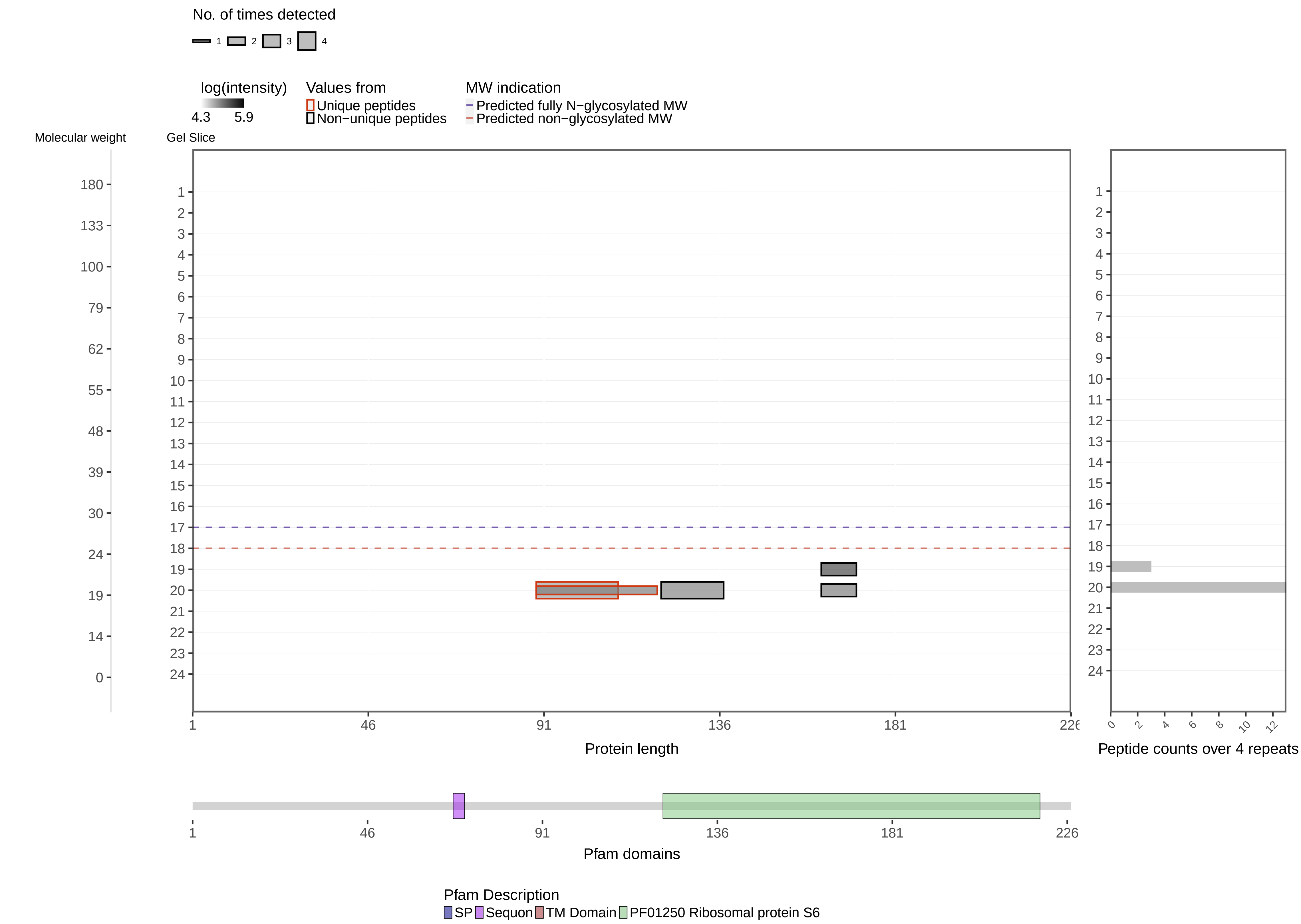


Protein sequence:

[illegible]GSGIAAVEDKEEPQCPPGMRPYETMAVLRPDMSEDERLALTQKYEELLVAGGGMYVEVFNRGVIPLAYSIRRKNKAGETNTYLDGIYLLFTYFTKPQSMTALEATLNTDDDVIRSSSFKIRKRKY*

NbL06g04750.1 glucan_endo−13−beta−glucosidase_11−like [MW − SP = 43.02 kDa]

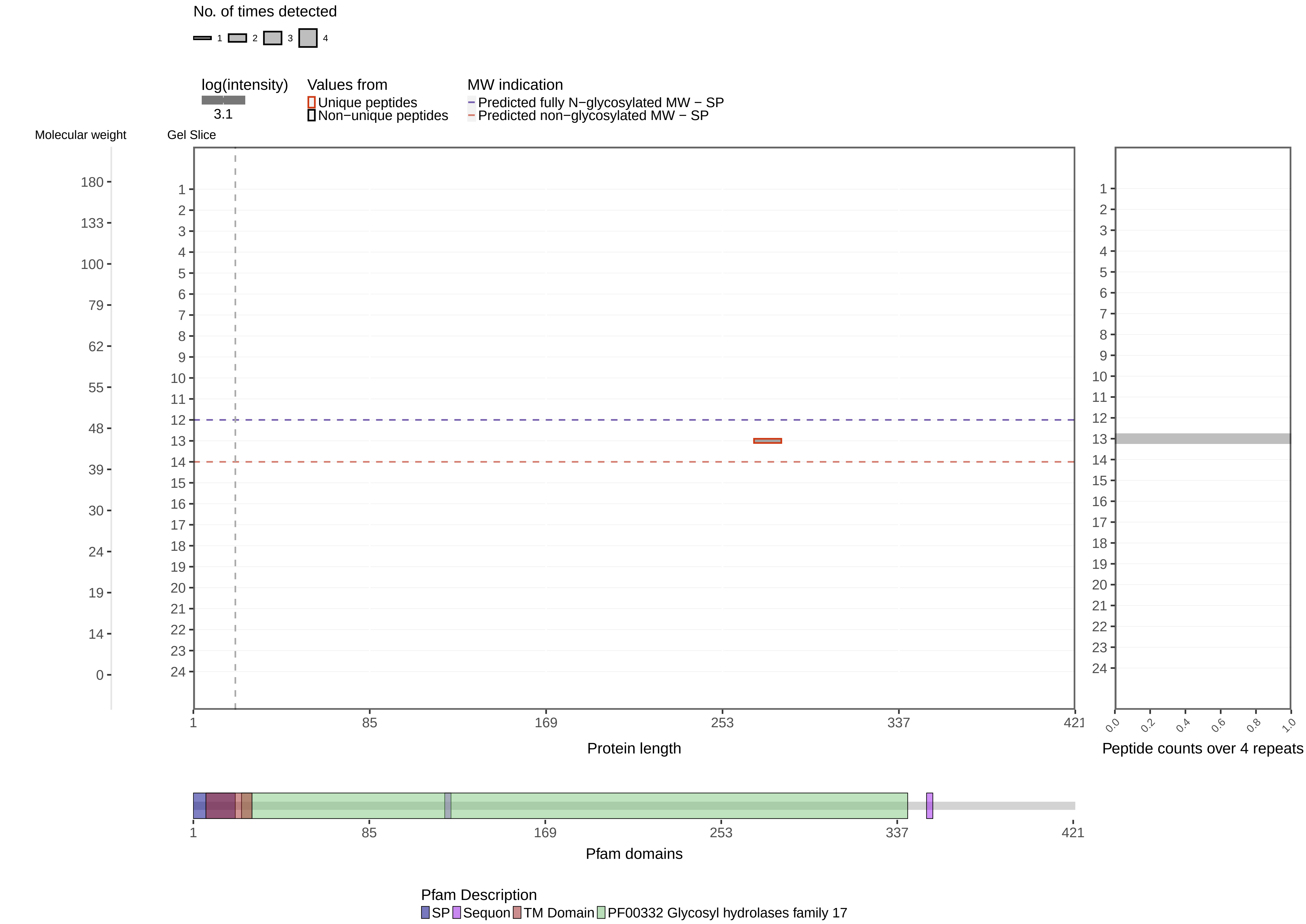


Protein sequence:

FAYKANPKHVPLDYVLFQPNPGNVDPVTNLHYDNMLYAQIDAVHSALASIGYKNVCVQISETGWPSKGDGDEVGATPDNARKYNCNLIKLVSQKKGTPLRPNTNLNIYVFALFNENLKPGPTSERNYGLFKPDGSPSYPLGFAGINAVGTNSSSSSSPATGTGSSSSTSSSTPAWNSPDGYLSITSDSGRVQFCWISAILGLISLCLFSTARNCMSCHLR*

### NbL06g04740.1 lysosomal_beta_glucosidase−like [MW − SP = 65.91 kDa]

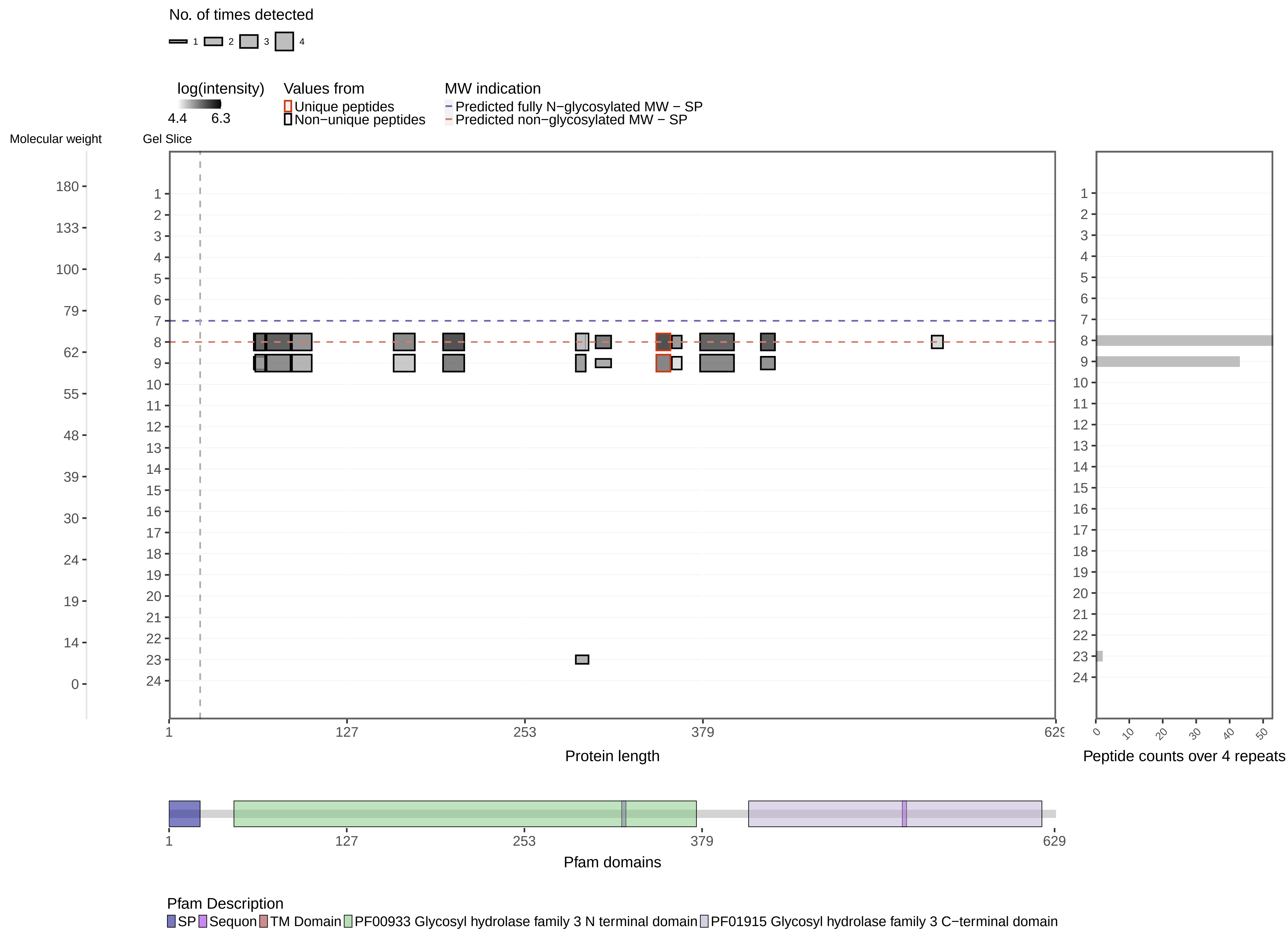


Protein sequence:

### NbL06g04730.1 soluble_inorganic_pyrophosphatase_PPA1 [MW = 24.27 kDa]

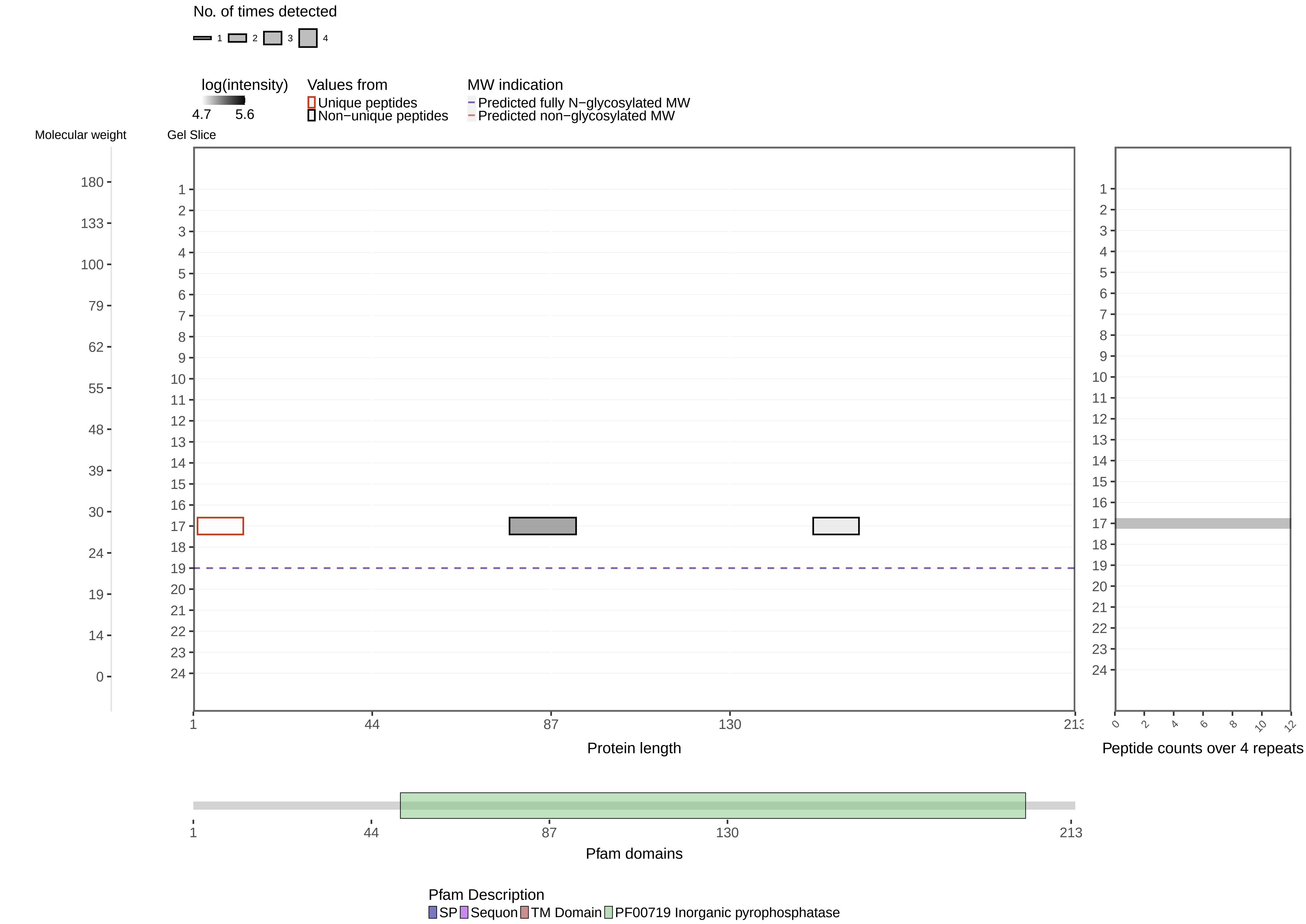


Protein sequence:

RTLCEDNDPMDVLVLMQEPVLPGCFLRARAIGLMPMIDQGEKDDKIIAVCADDPEYRHYTNINQLPPHRLAEIRRFFEDYKKNENKEVAVDEFLPPSTAVDAIQYSMDLYAEYILHTLRK*

### NbL06g04700.1 probable_cinnamyl_alcohol_dehydrogenase_1 [MW = 38.58 kDa]

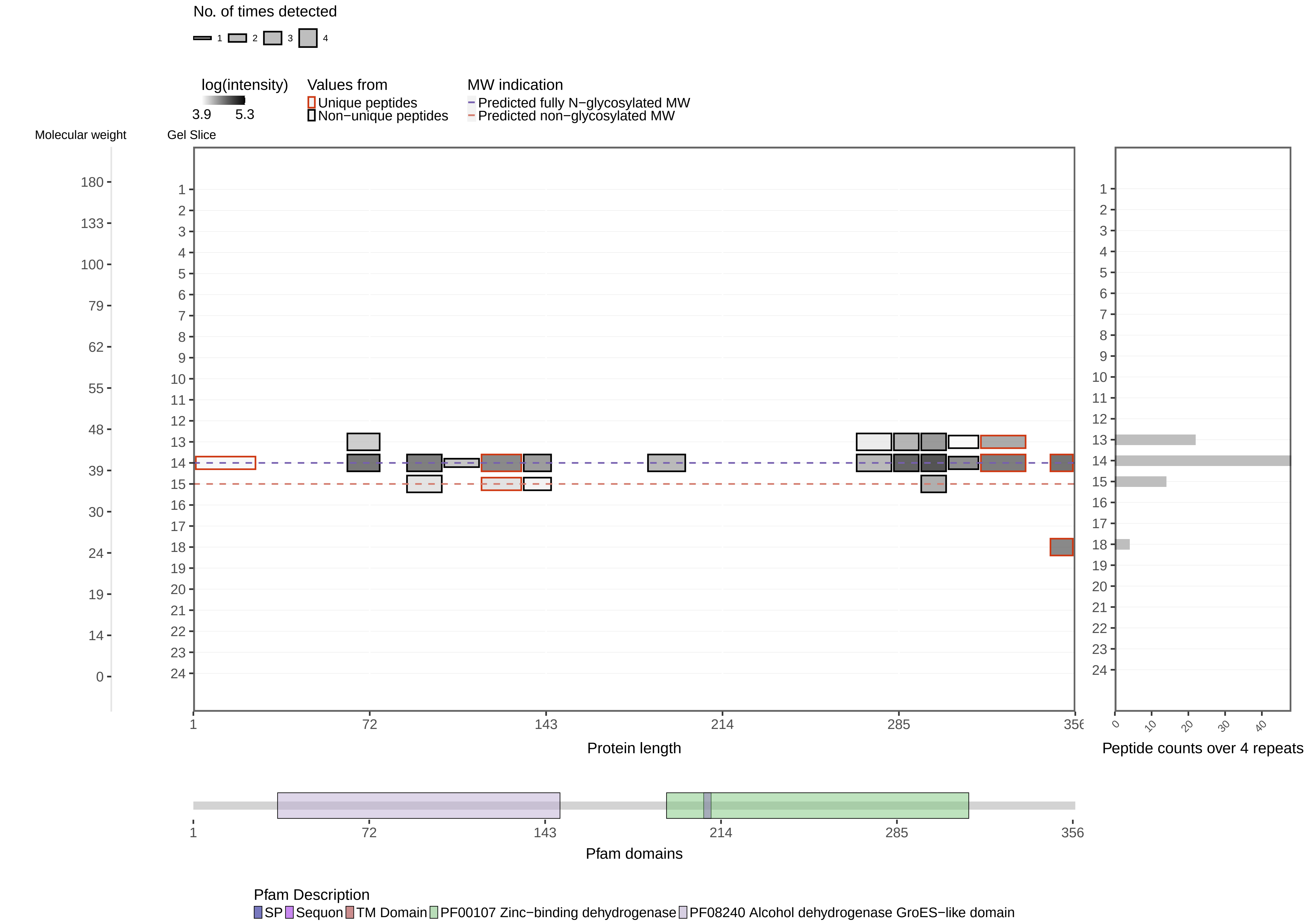


Protein sequence:

LCAGITVYTPMMRHNMNQPGKSLGVIGLGGLGHLAVKFGKAFGLNVTVFSTSISKREEALKLLGADNFVVSSDEQQMTALGMSFDFMINTASGDIPFDPYLSLLKTAGILVLVGFPTEVKFSPGNLVLGMKSIVGSVTGGTKQTQEMLDFCASHKIYPEIEIVPIQYANEALERLIKKDVNYRFVIDVGNSLK*

NbL06g04280.1 uncharacterized_LOC107769601 [MW = 33.74 kDa]

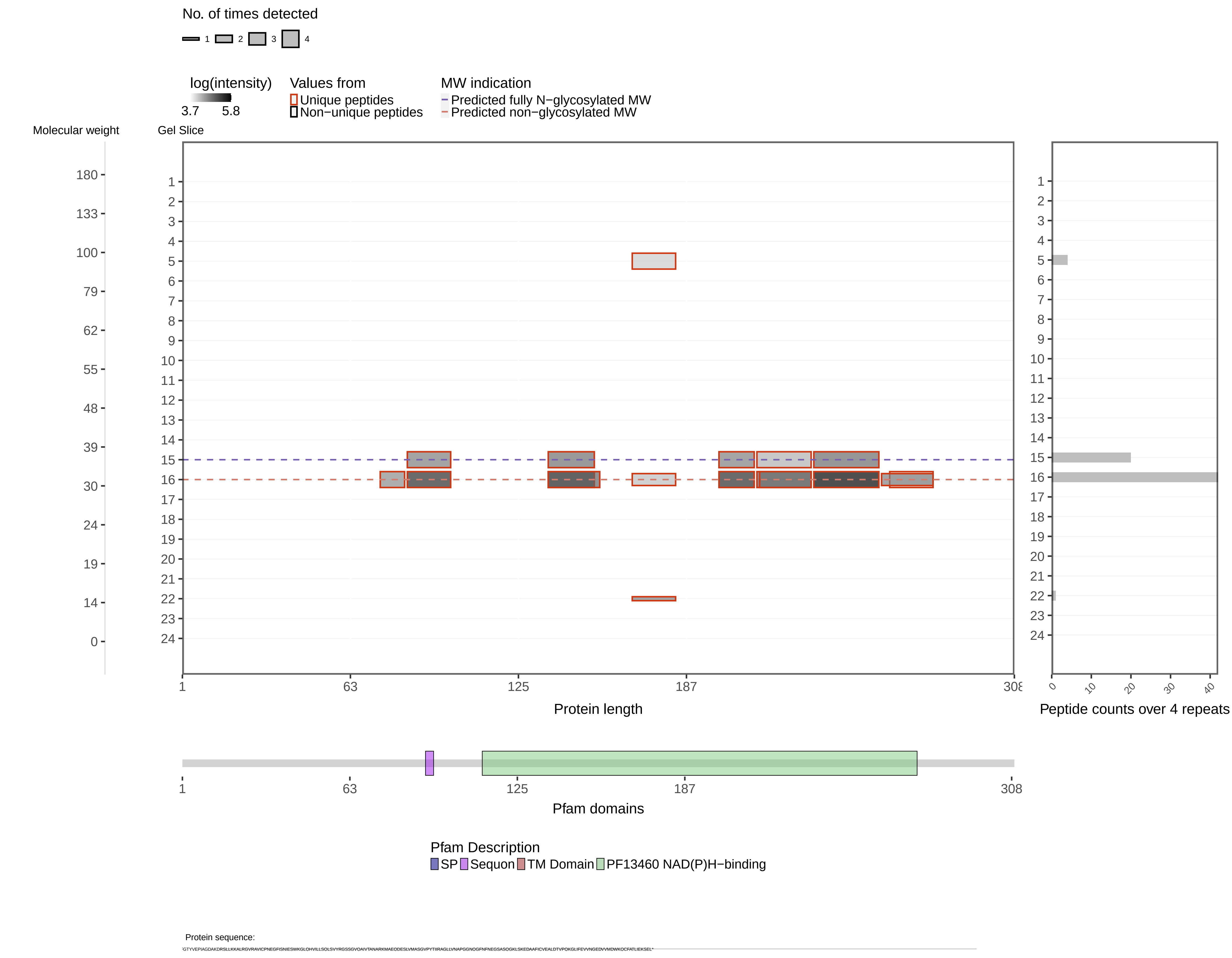

### NbL06g04270.1 uncharacterized_LOC104210585 [MW = 89.55 kDa]

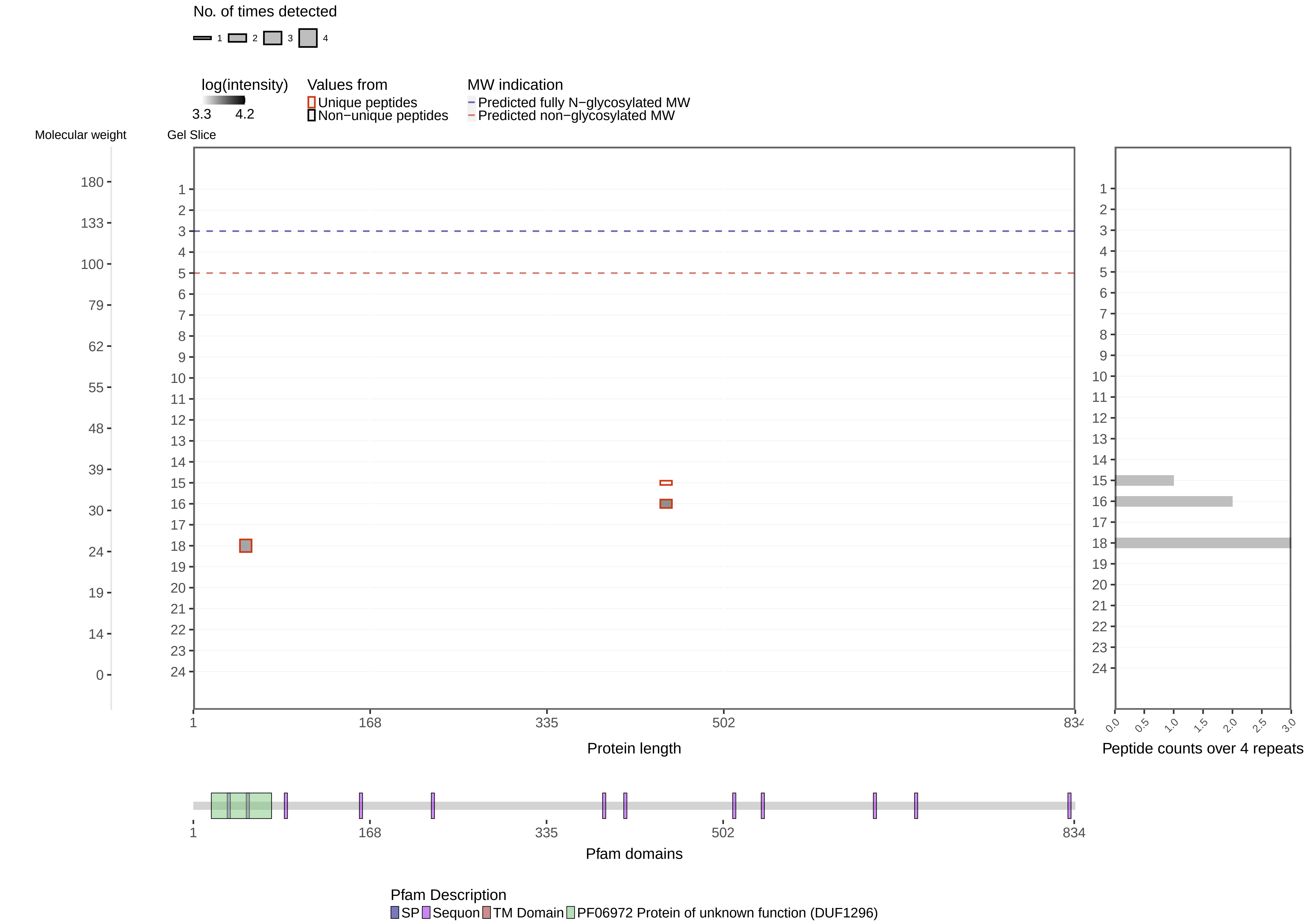


Protein sequence:

SRLNISDNVIIAEHIRVSETDRCRLTFGSFGAELKPKDLDEESQTESSRLSVLVSEPSTDEPVGSKQLDLADDRVQNPGLTSPGSGVSLDQKLADNRESSSPEDLGNYTDVGLVQDNRASYTPPESQQQQDASNLSSFSTYDPQTGYDIPYFRPAVDEALHGQGAQEALSSHAANSMPASTIPMVQQVQQHQPIAQMYPQVHVSHFANLMPYRHIFSPVYVPPMAMPGYSSNPAYPHPPNGSNYLLMPGGGSHLSANGLKYGIQQFKPVPTGSASGFGNFTSPTGYAI

### NbL06g04210.1 beta−galactosidase [MW = 94.38 kDa]

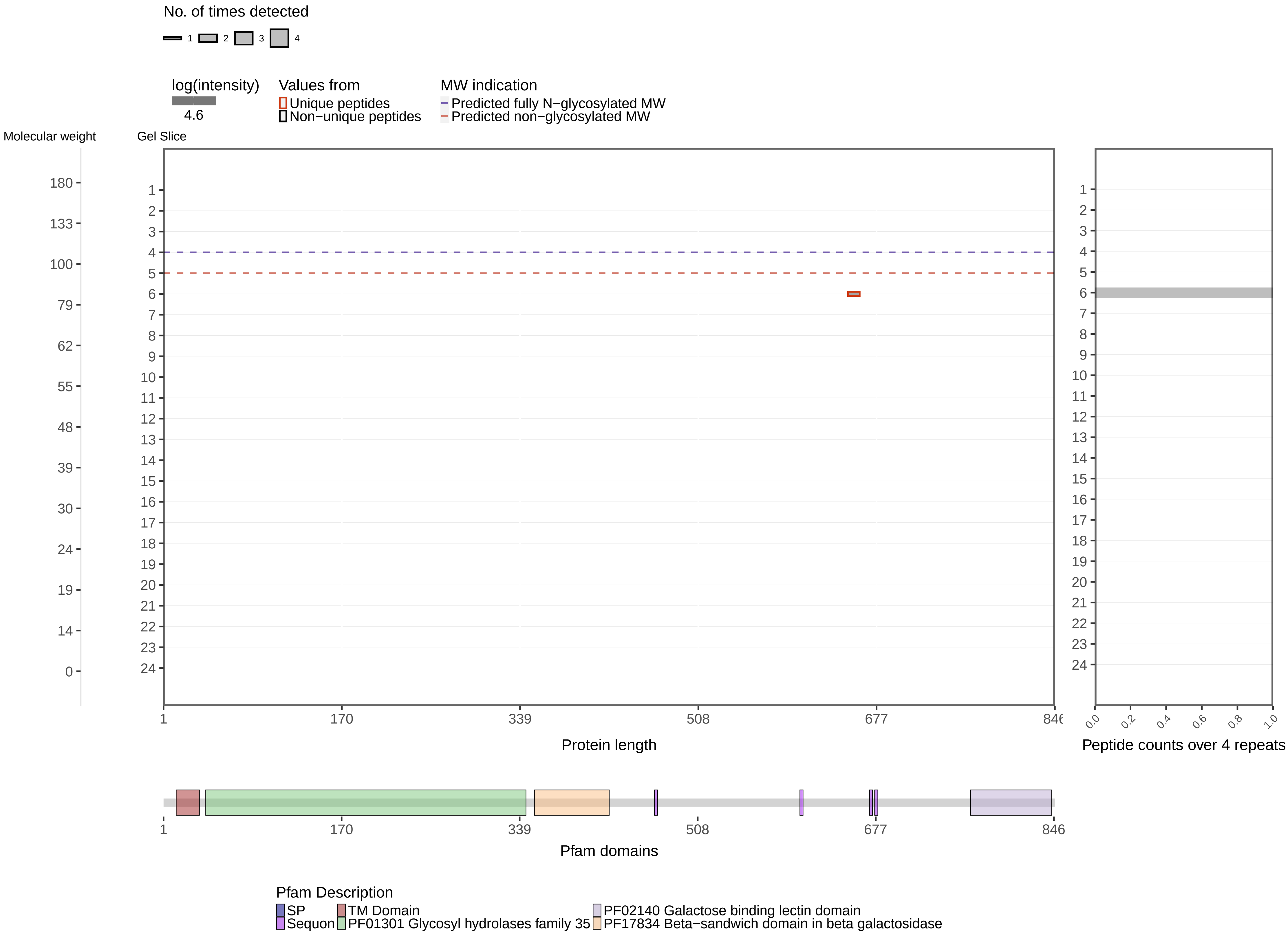


Protein sequence:

[illegible]

NbL06g04170.1 ubiquitin_carboxyl−terminal_hydrolase_6−like [MW = 54.66 kDa]

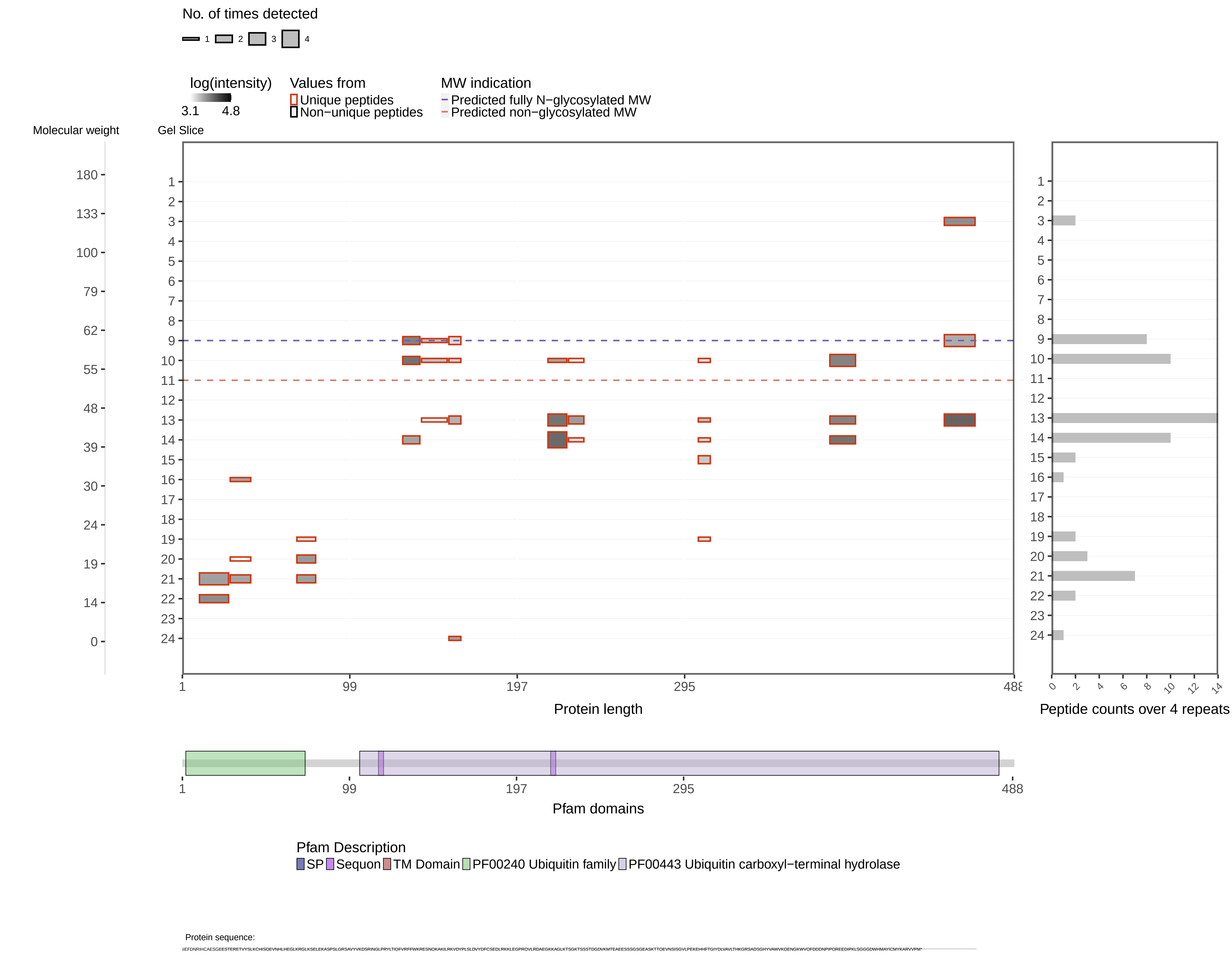

NbL06g04130.1 60S_ribosomal_protein_L28−1−like [MW = 15.84 kDa]

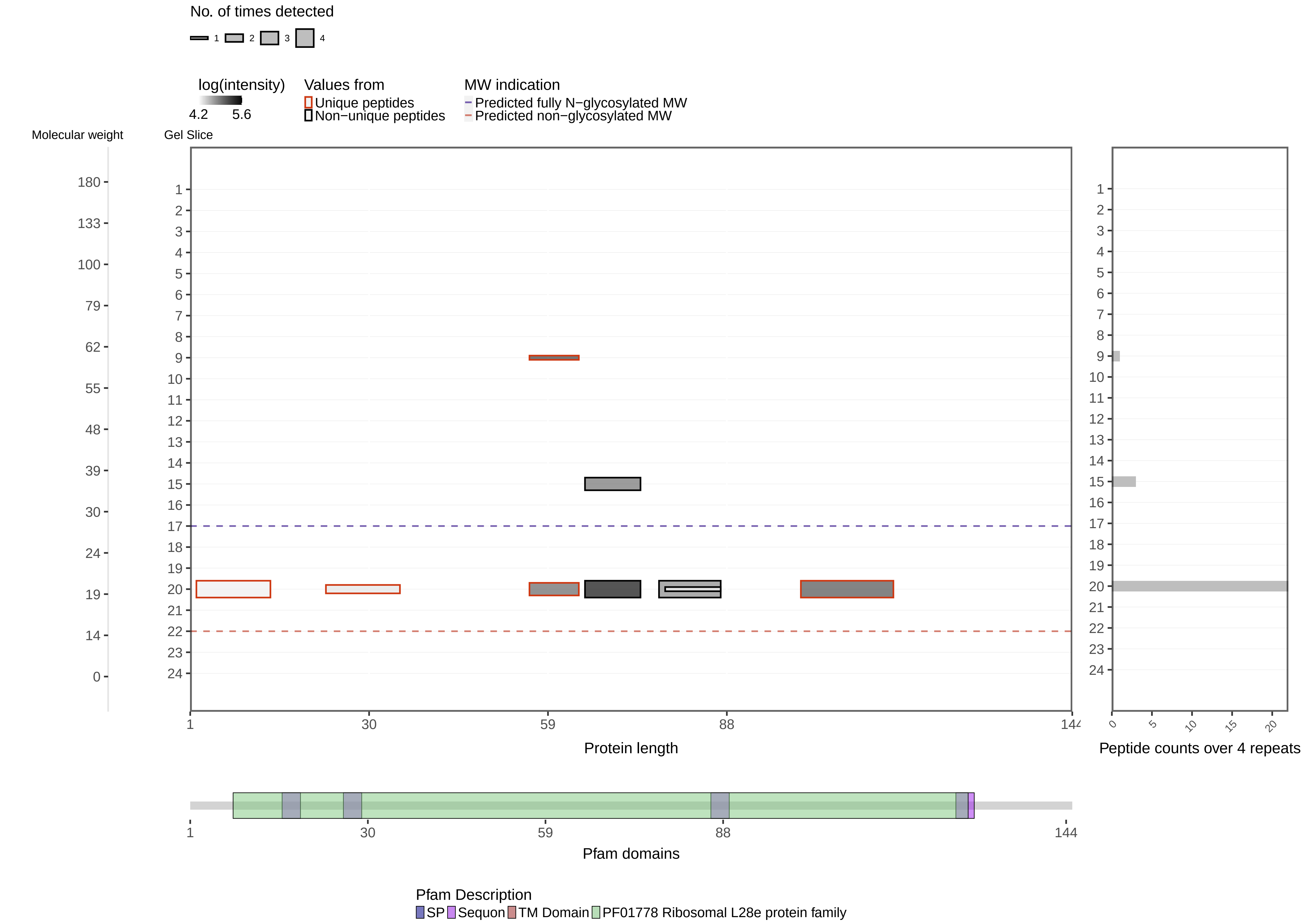


Protein sequence:

QAVGKDQSVLLATTKTKKQNKPSSLLNKSAMKKEFRRMAKAVSNQVADNYYRPDLKKAALARLSAVNRSLKVAKSGVKKRNRQA*

### NbL06g04050.1 NADP−dependent_malic_enzyme_chloroplastic [MW = 70.38 kDa]

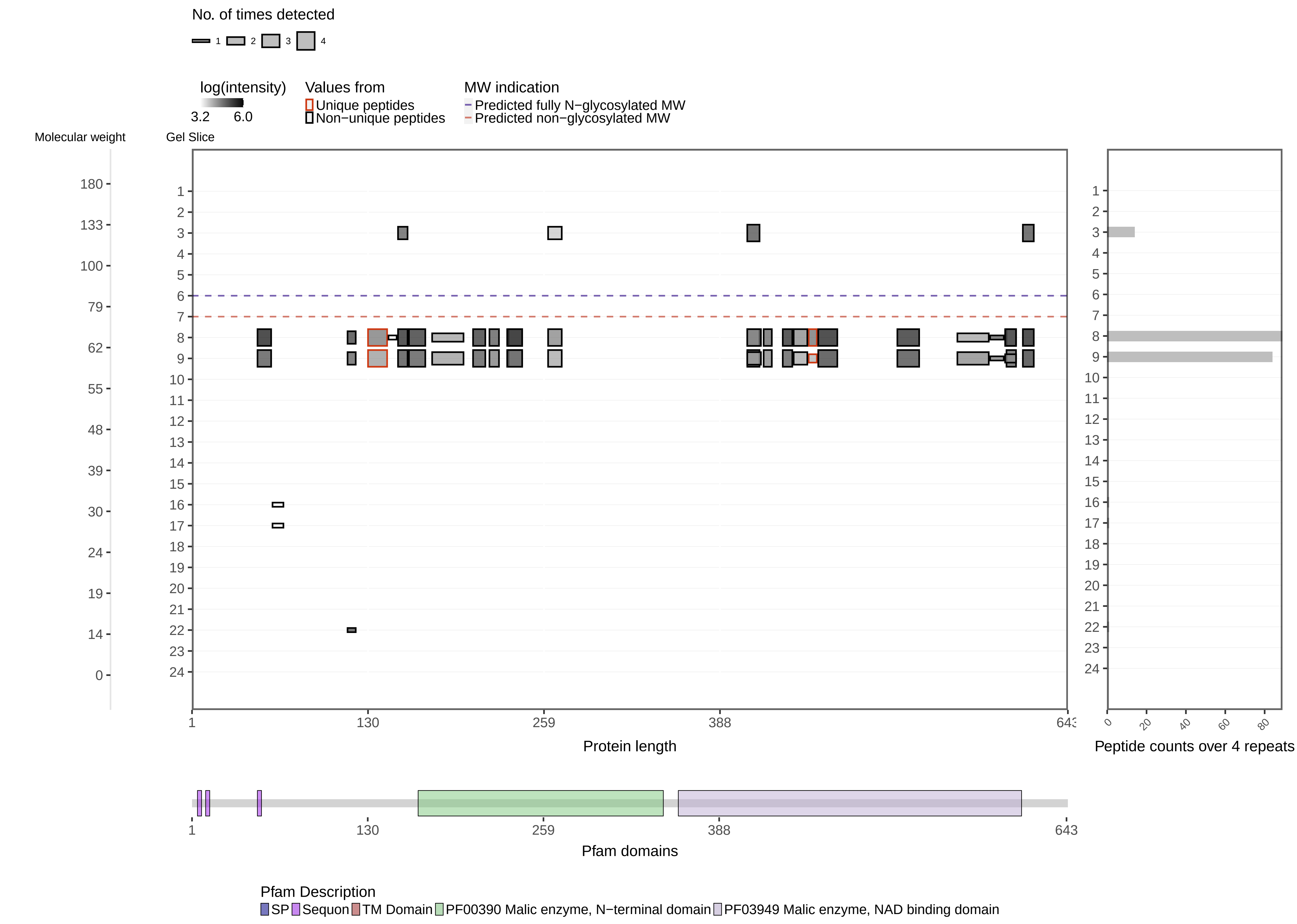


Protein sequence:

### NbL06g03900.1 cytochrome_b6 [MW = 24.15 kDa]

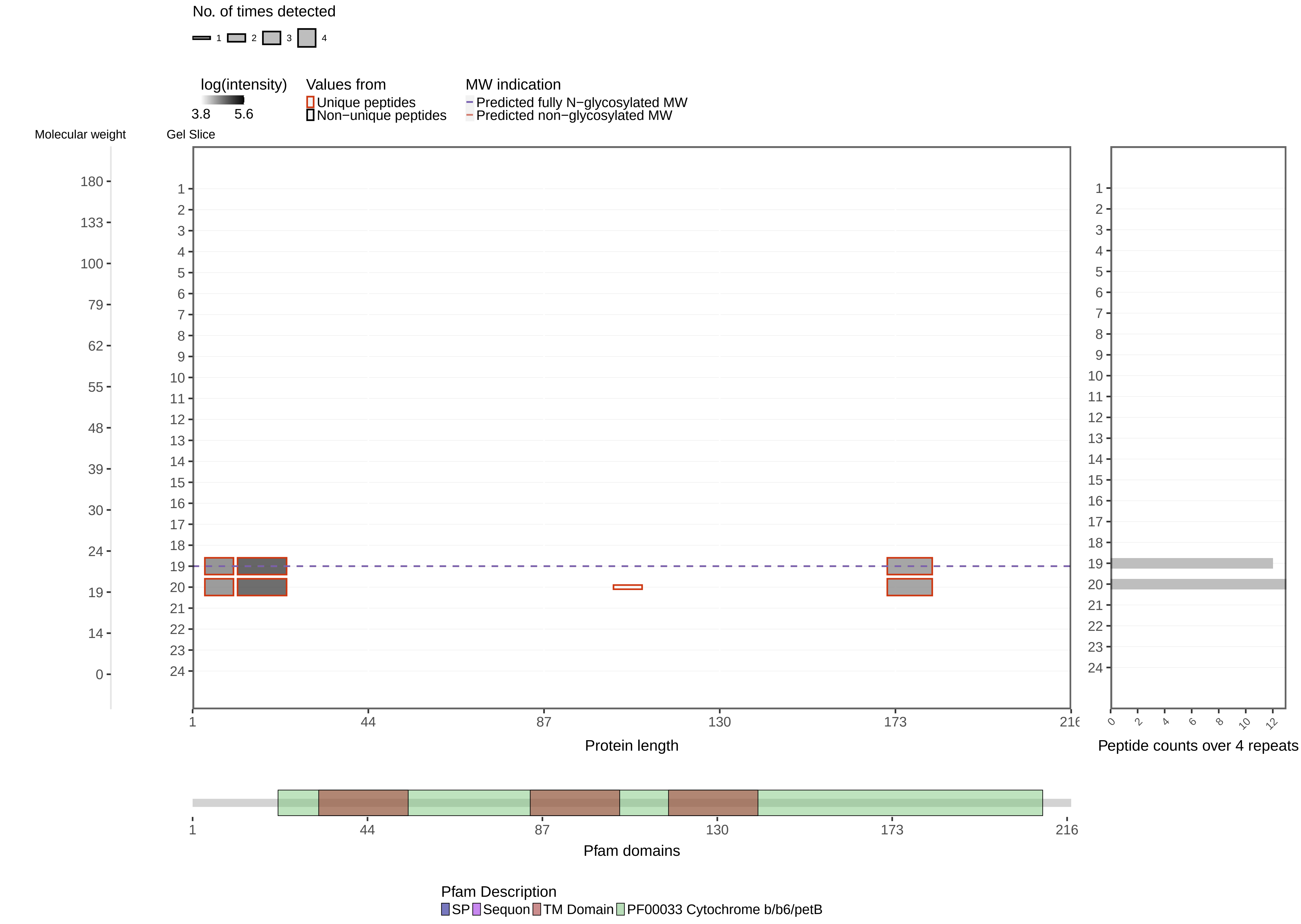


Protein sequence:

'LMMILHVFRVYLTGGFKKPRELTWVTGVVLAVLTASFGVTGYSLPWDQVGYWAVKIVTGVPDAIPVIGSPLVELLRGSASVGQSTLTRFYSLHTFVLPLLTAVFMLMHFLMIRKQGISGPL*

NbL06g03810.1 heat_shock_70_kDa_protein_15−like [MW = 94.07 kDa]

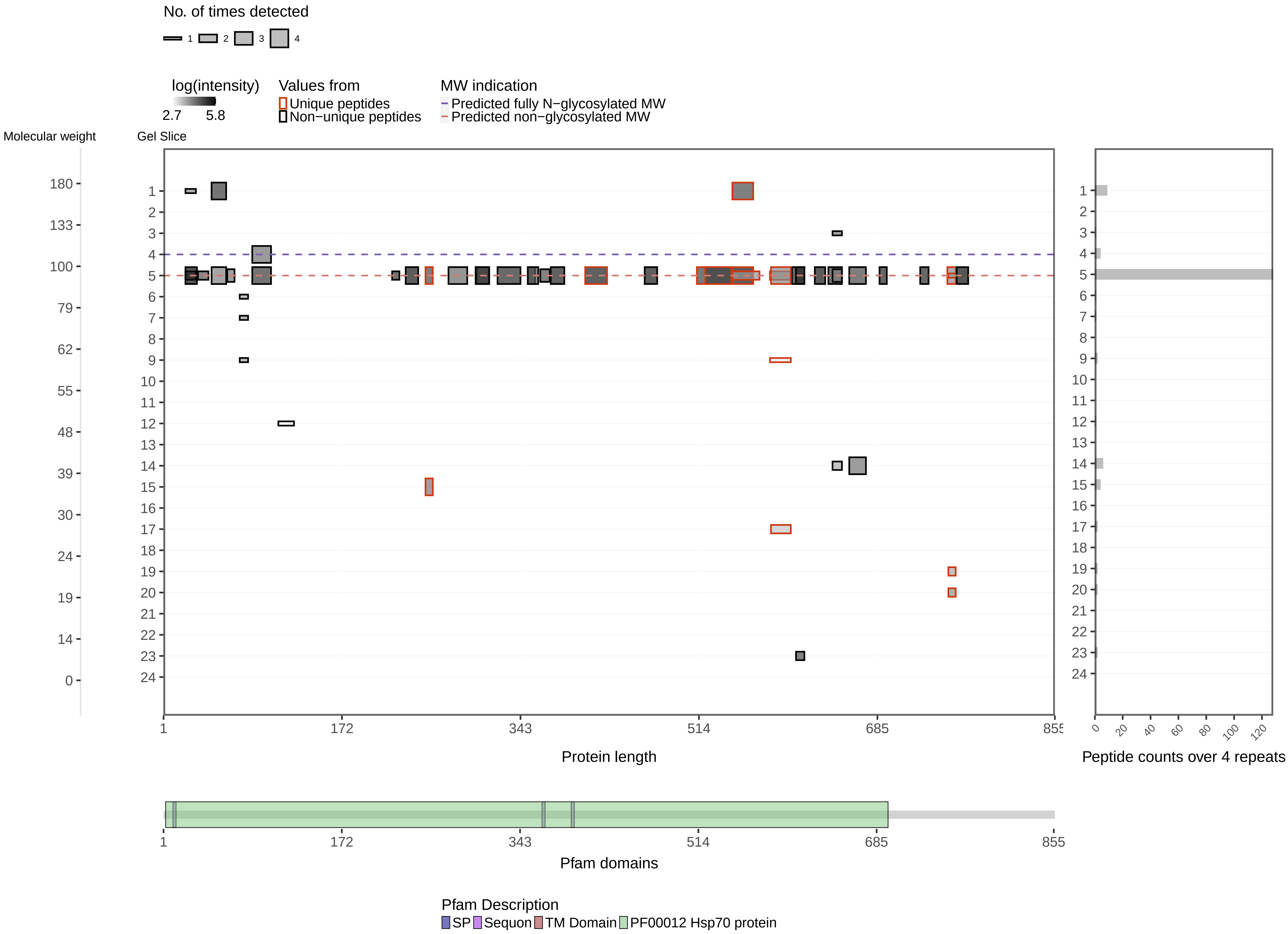


Protein sequence:

NbL06g03640.1 pyrophosphate−energized_vacuolar_membrane_proton_pump−like [MW = 80.23 kDa]

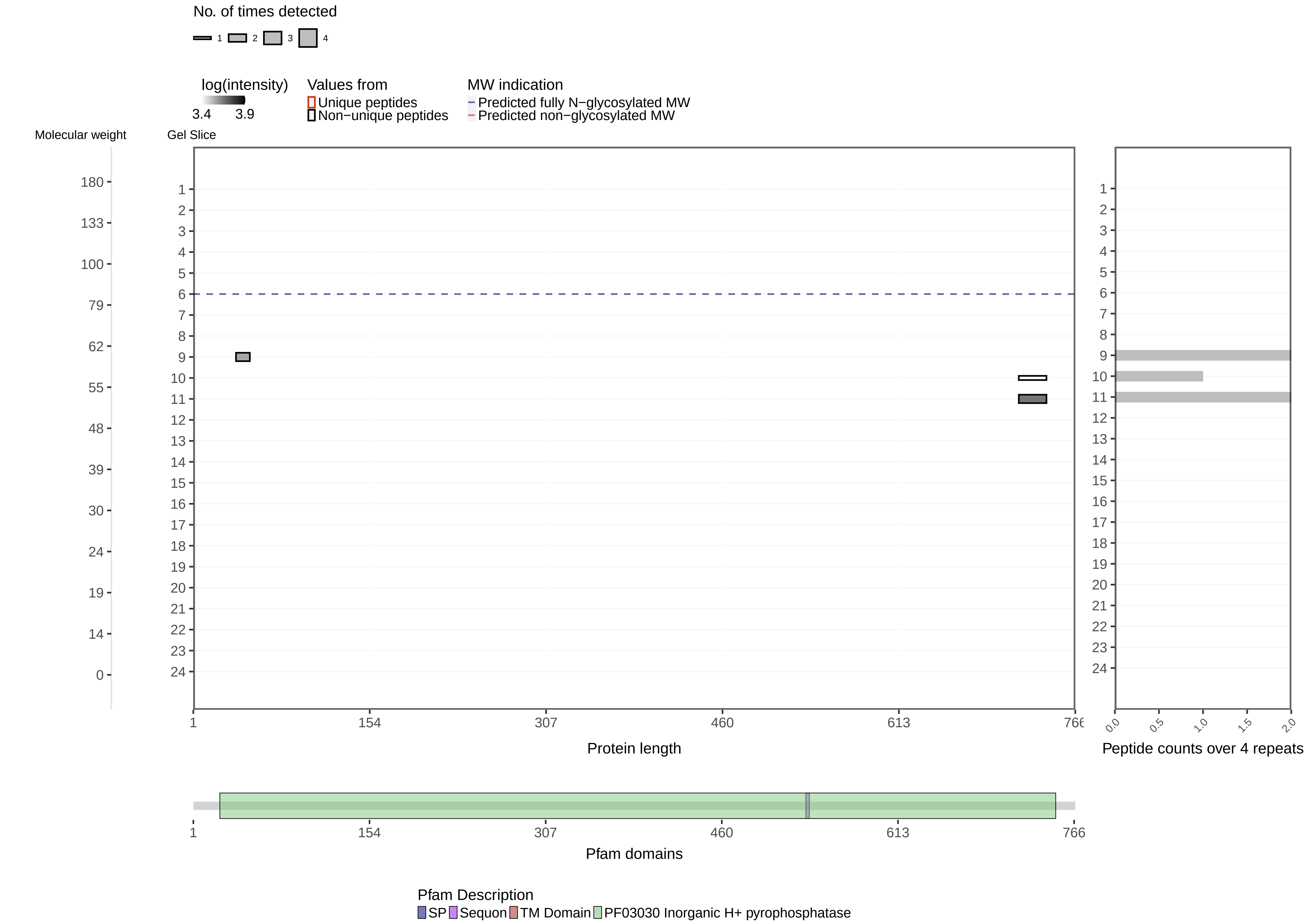


Protein sequence:

MTVGIAIVTWTCLPSSFTIFNFGAQKVVKNWQLFLCVAVGLWAGLIIGFVTEYYTSNAYSPVQDVADSCRTGAATNVIFGLALGYKSVIIPIFAIAIAIFVSFSFAAMYGIAVAALGMLSTIATGLAIDAYGPISDNAGGIAEMAGMSHRIRERTDALDAAGNTTAAIGKGFAIGSAALVSLALFGAFVSRAAITTVDVLTPQVFIGLIVGAMLPYWFSAMTMKSVGSAALKMVEEVRRQFNTIPGLMEGTAKPDYATCVKISTDASIKEMIPPGALVMLTPLIVGIFFGVETLSGVLAGALVSG

NbL06g03590.1 uncharacterized_LOC107791394 [MW = 29.7 kDa]

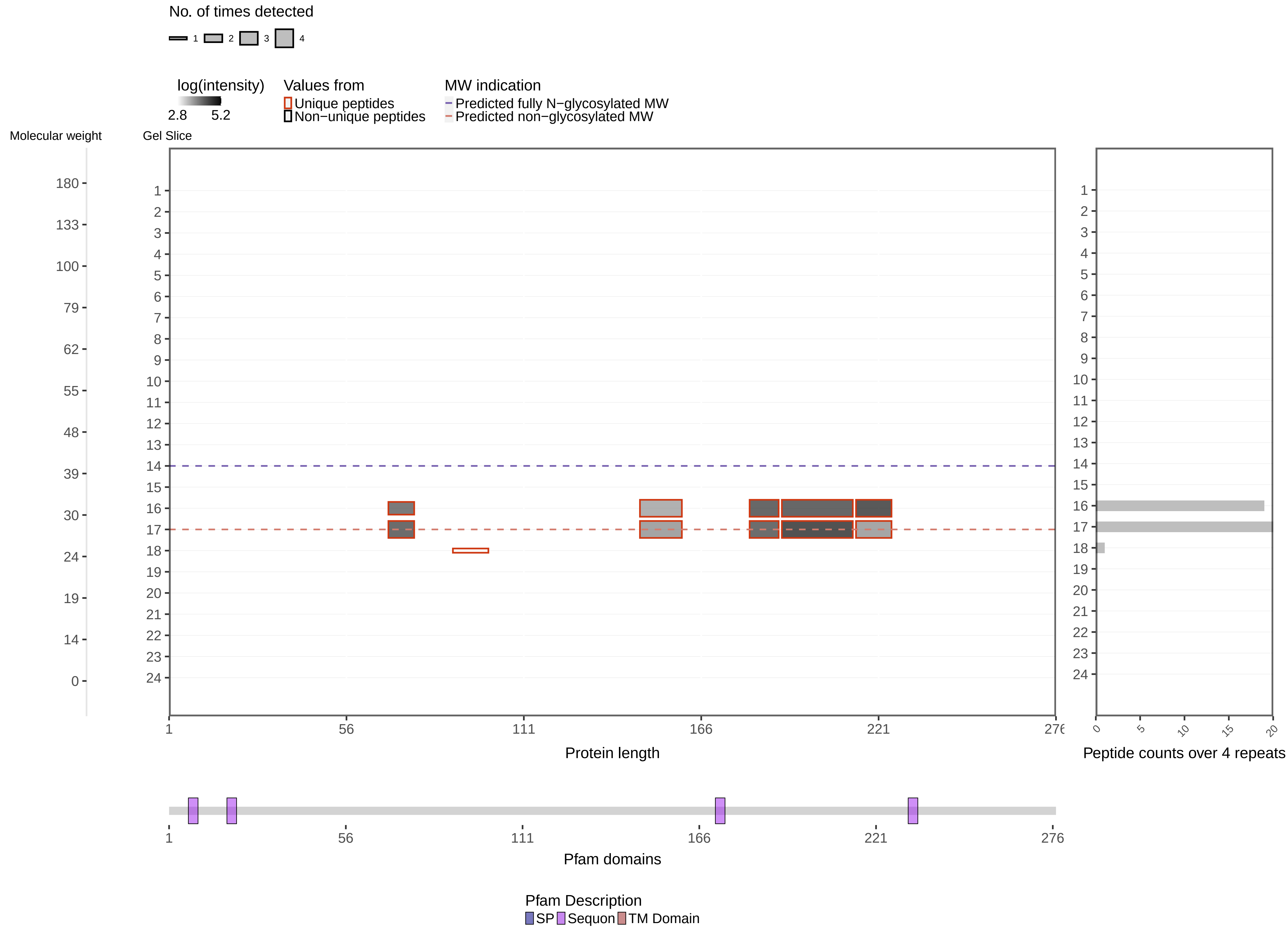


Protein sequence:

FNQPIMCGGEPRAMLMKTRGKADPPIYTIQICIPKHALSLIFSFTNGTEWDGPYRLQFQVPNAWRNKPTEFFTEGLAGELSQEGACDKAIFPDTNIVITRCAMVGNLSVEGGDRCDLDIVPGCIDPSSPHFNPLANVDDGSCPPYSDAED*

### NbL06g03340.1 guanine_nucleotide−binding_protein_subunit_beta−like_protein [MW = 37.69 kDa]

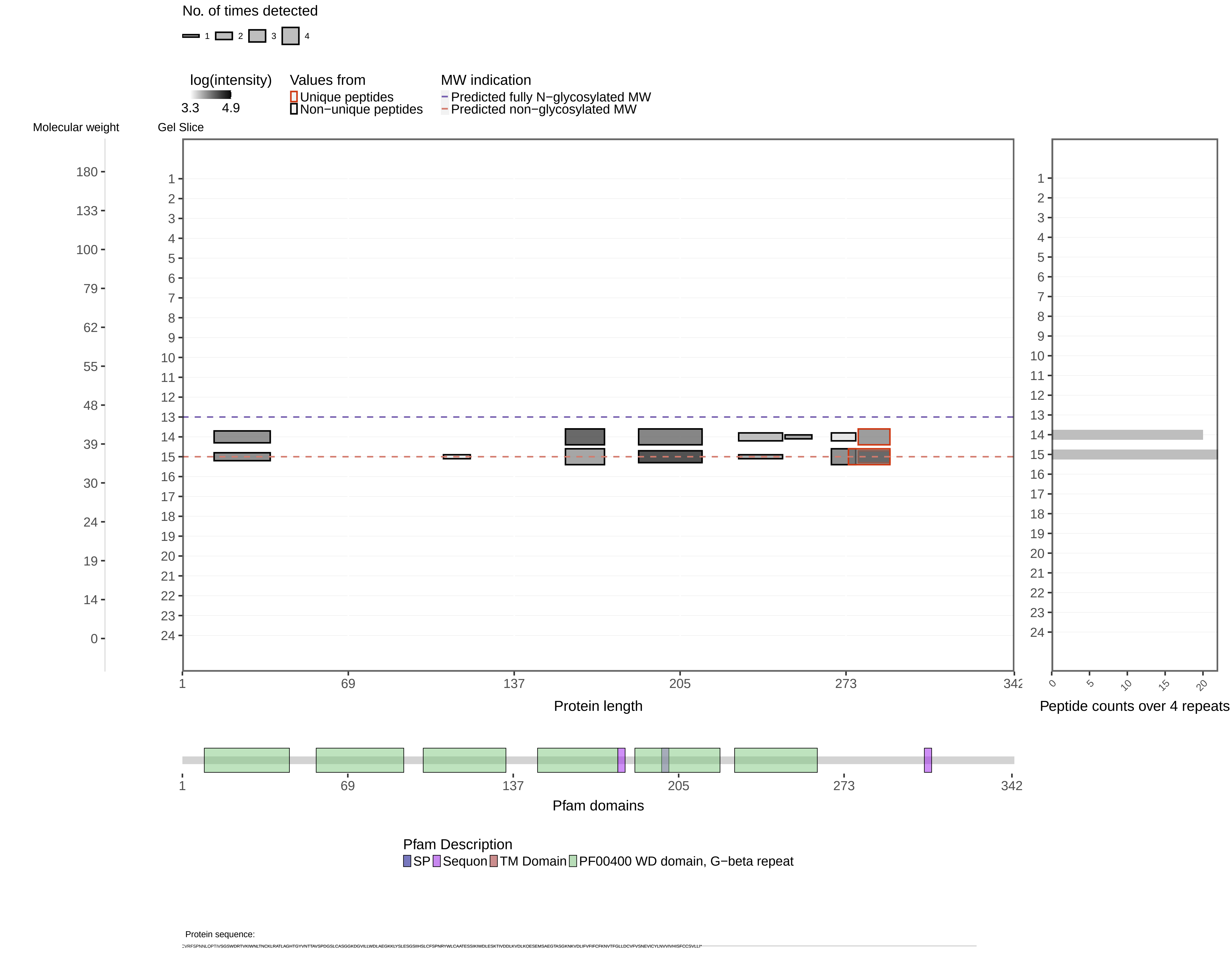

NbL06g03220.1 psbP−like_protein_1_chloroplastic [MW = 26.71 kDa]

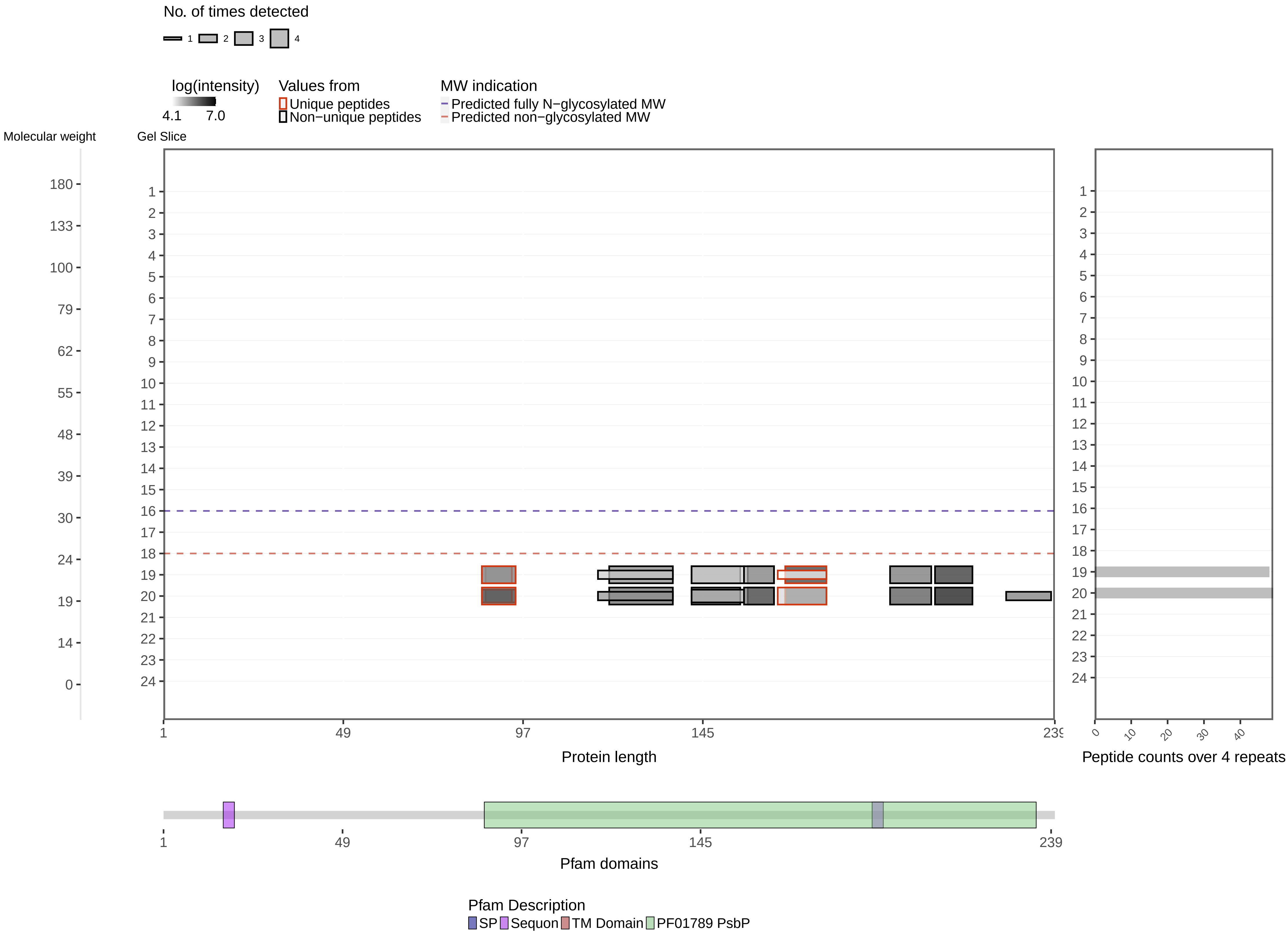


Protein sequence:

GWQEVVVEGQDKVFKDVIEPLESVSVNVIPTSKQDIRDFGSPQEVAETLIKKFLSSPSQKTKLIEASQYDVEGKAYYTFEFVAQAPNFTRHGLTAVCIGNGKFYTLTTGANERRWEKMKDRLHTVVDSFQIFNV*

### NbL06g03050.1 serine/threonine−protein_kinase_16−like [MW = 38.69 kDa]

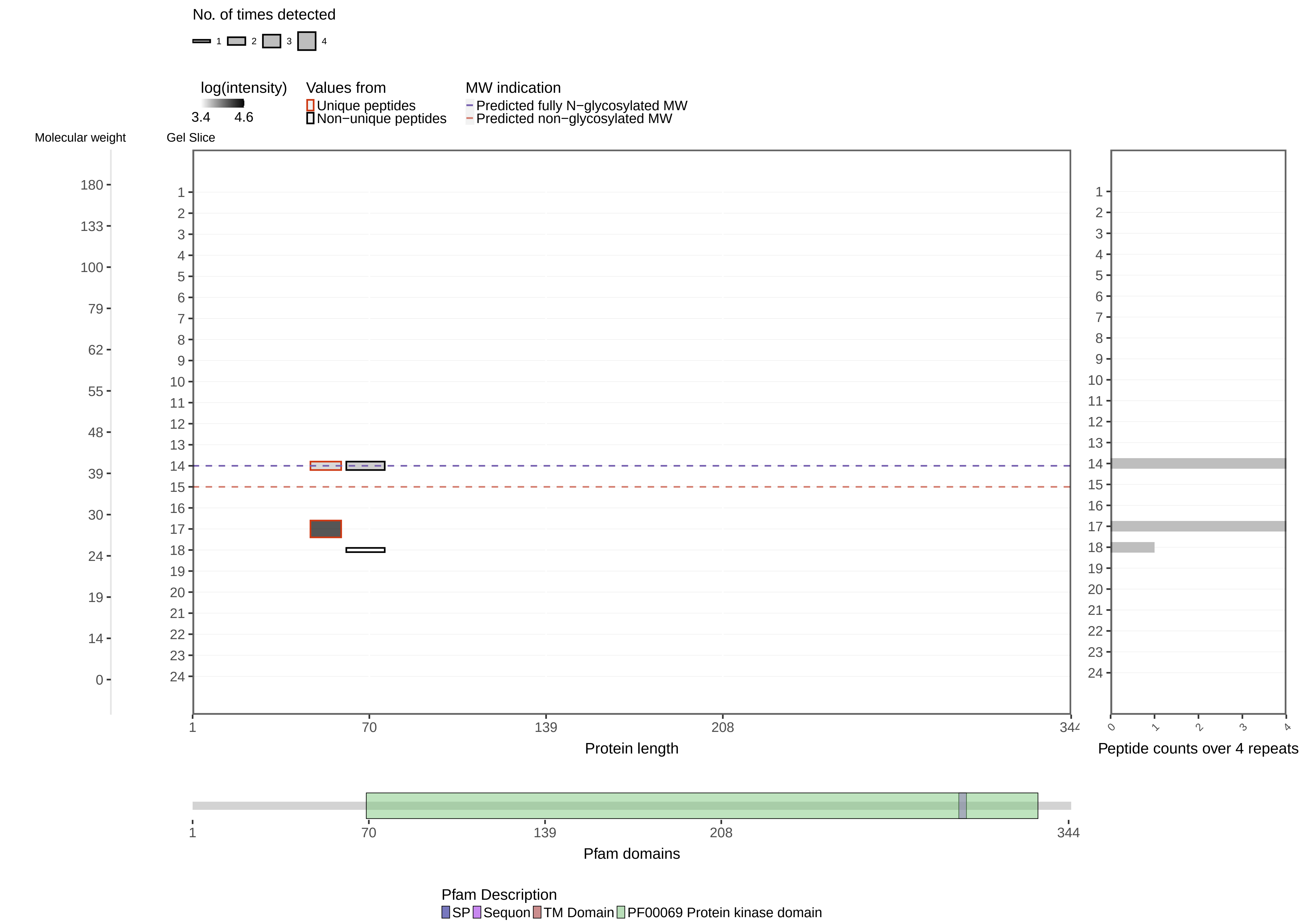


Protein sequence:

QIFRQLCAGLKHMHNFDPPYAHNDVKPANVLLTHKKGQTPLTKLMDFGNARPARRQIRSPSEALQLQEWASEHVAAPFRAPELWDCPSQCDIDERTDIWSLGCTLFAIMYGVSPFEYAFEESDESPQLAIVNAQIKWPTEPNTSYPDALHQFVTWMLQPQATVRPRIDDIVIHVDKLISKYSTN*

### NbL06g03040.1 agmatine_deiminase [MW = 41.91 kDa]

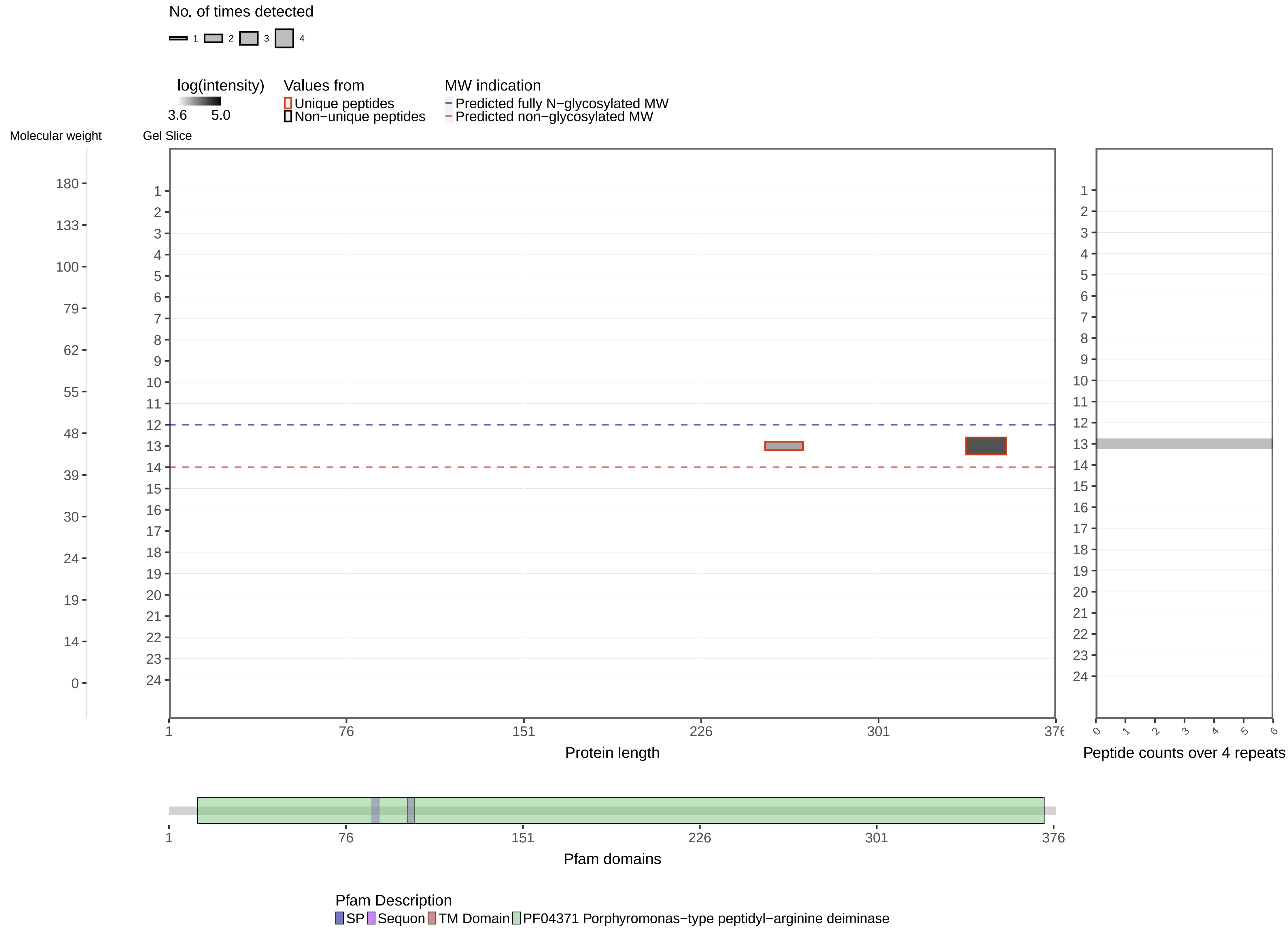


Protein sequence:

GTCLTTEECLLNKNRNPHLSKEQIENELKAYLGVRKIIWLPRGLFGDDDTNGHIDNMCCFVKPGVVLLSWTDDQSDPQYERSIEALSILETTFDANGRKLQVIKLHIPGPLCMTDEEADGIFQDGDAKPRTSGTRLAASYVNFYIANGAIITPQFGDKKWDEEAIRVLSLAFPDHEVVGIEGAREIVLGGGNIHCITQQQPSGP*

### NbL06g03020.1 H/ACA_ribonucleoprotein_complex_subunit_2−like_protein [MW = 16.97 kDa]

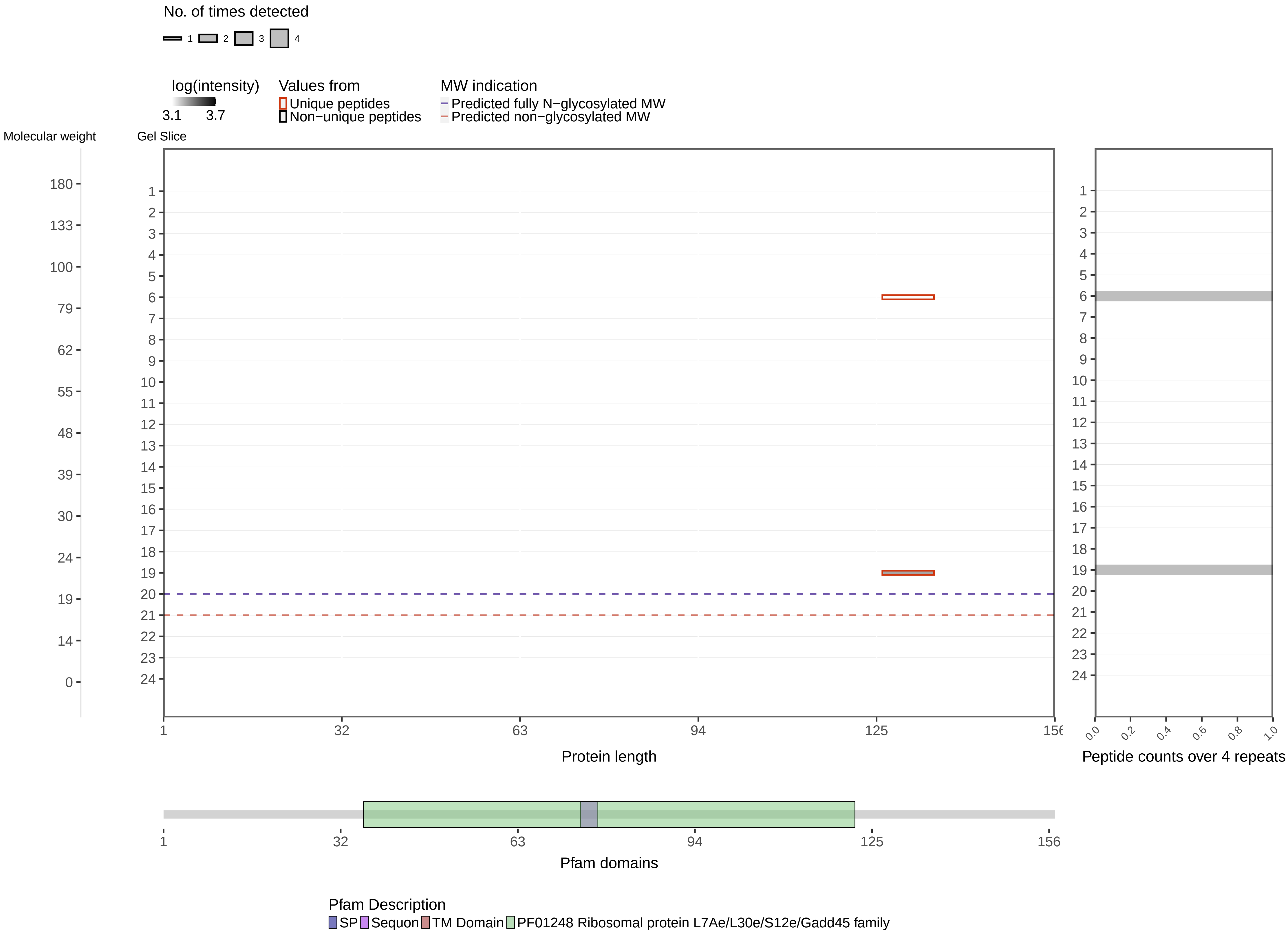


Protein sequence:

QKGVCVIAGNISPIDVITHVPILCEEAEIPYVYVSSKEDLANAGATKRPTCCVLVQTKPAKGELSQEDQEKLKADYDQVVSEVGEMASSMF*

NbL06g02670.1 ATP_synthase_subunit_gamma_mitochondrial−like [MW = 35.28 kDa]

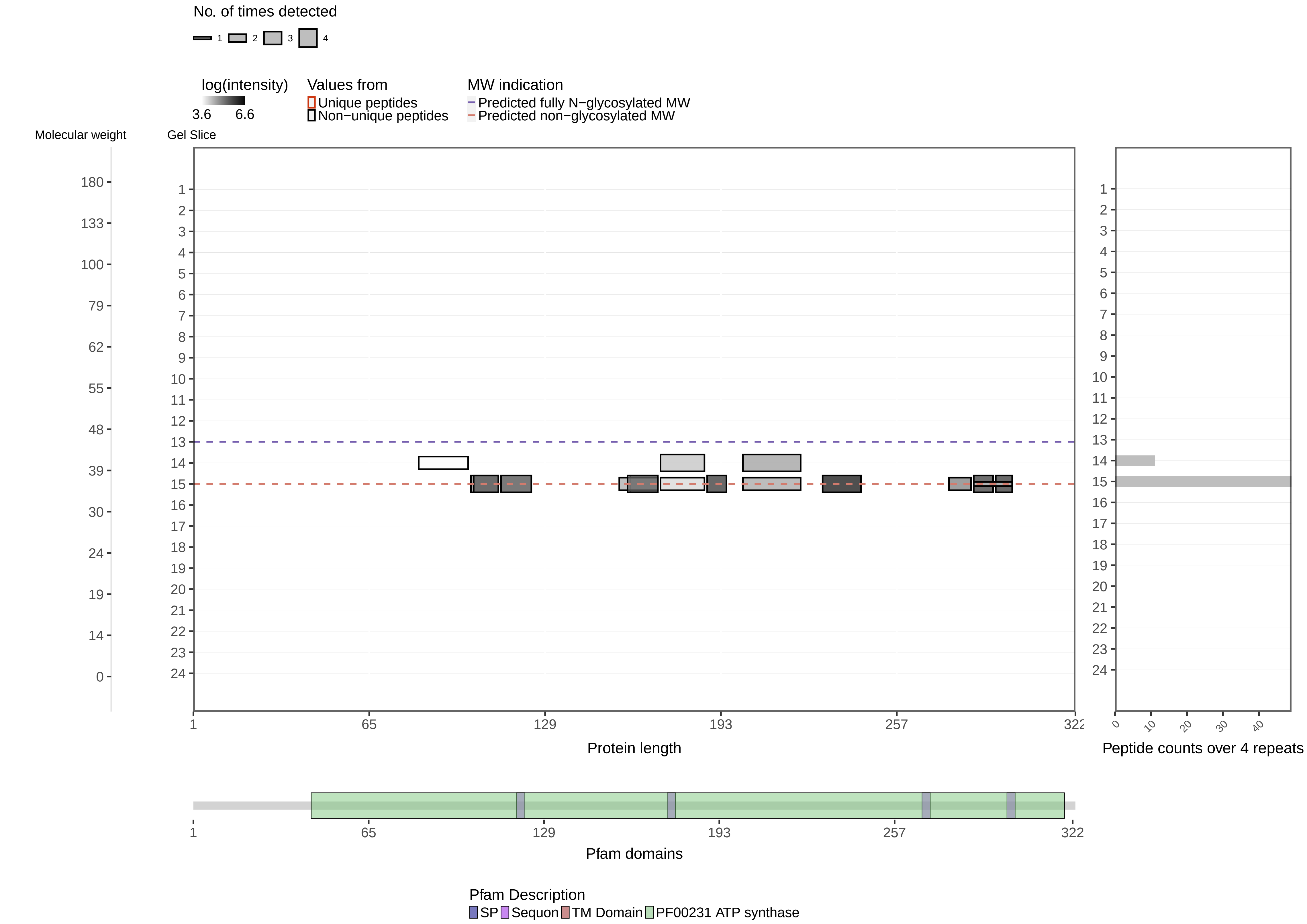


Protein sequence:

KAKAQLVRDSKKDIELIMTELQKNPLNYTQVSVIADDILKNVEFDALRIIYNKFQSVVAFLPTMSTVLSPEVVERESESGGKLGELDSYEVEGAETKSEVLQNLAEFQFSCVLFNALLENATSEQGARMSAMDSSSRNAGDMLDRLTLTYNRTRQASITTELIEIISGASALEG*

### NbL06g02600.1 40S_ribosomal_protein_S19−3−like [MW = 16.04 kDa]

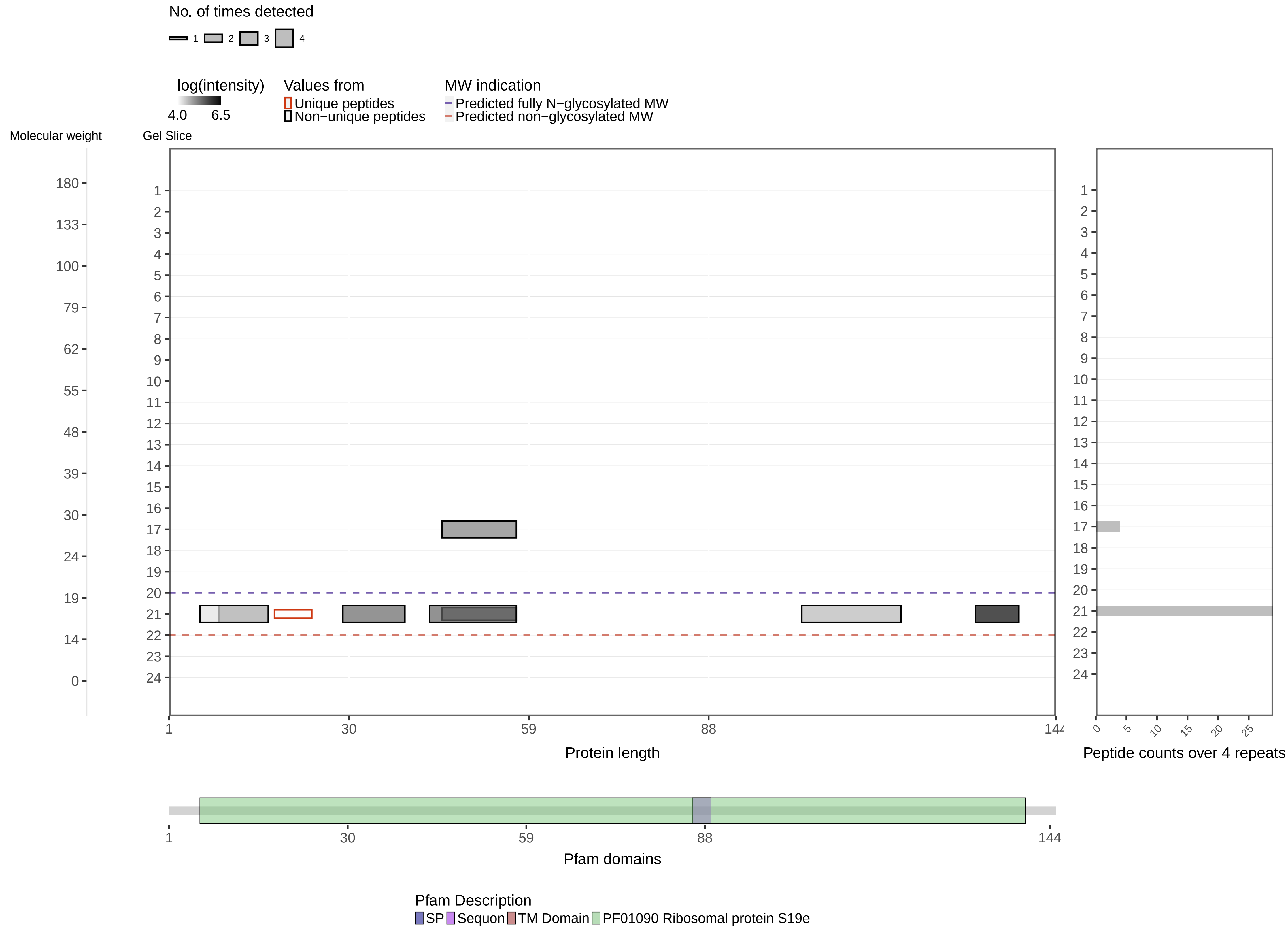


Protein sequence:

SMARKIYLRGGIGVGGFRRIYGGNQRNGSRPRHFCKSSGSVARNILQQLQNMNIVDFDPKGGRRITSNGQRDLDQVAGRIAAAL*

NbL06g02190.1 probable_voltage−gated_potassium_channel_subunit_beta [MW = 44.68 kDa]

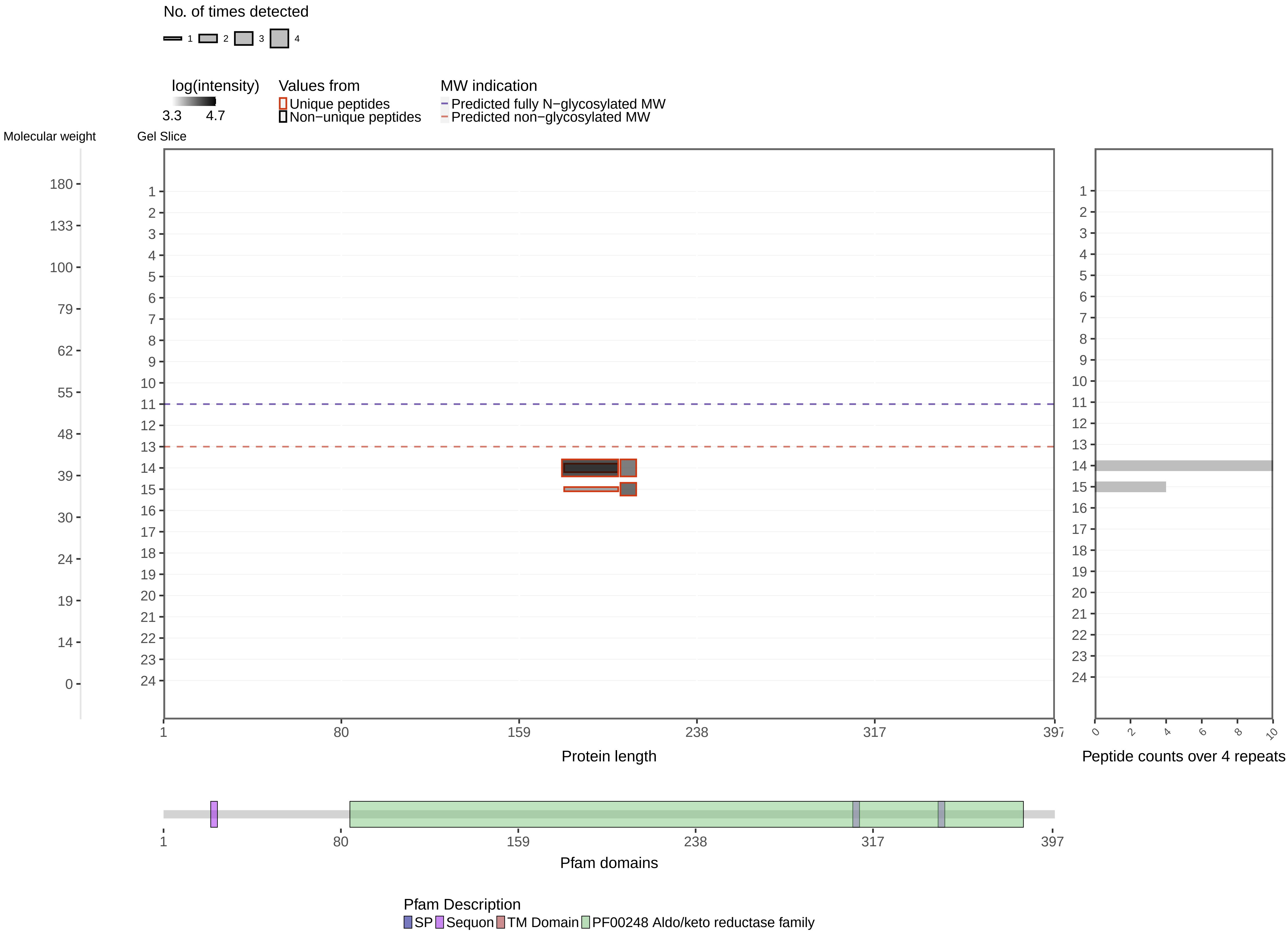


Protein sequence:

LIYCHRPDASTPIEETVRAMNYVIDKGWAFYWGTSEWSAQQITEAWGVAQRLDLVGPIVEQPEYNLLSRHKVESEYLPLYSNYGIGLTTWSPLASGVLTGKYNAGNIPADSRFALENYKNLANRSLVDDVLRKVDGLKPIAESLGVPLPQLAIAWCAANPNVSSVITGATKEYQIEENMKAINVIPMLTPAVMERIEAVVQSKPKRQDSYR*

### NbL06g02000.1 PLASMODESMATA_CALLOSE−BINDING_PROTEIN_3−like [MW − SP = 17.13 kDa]

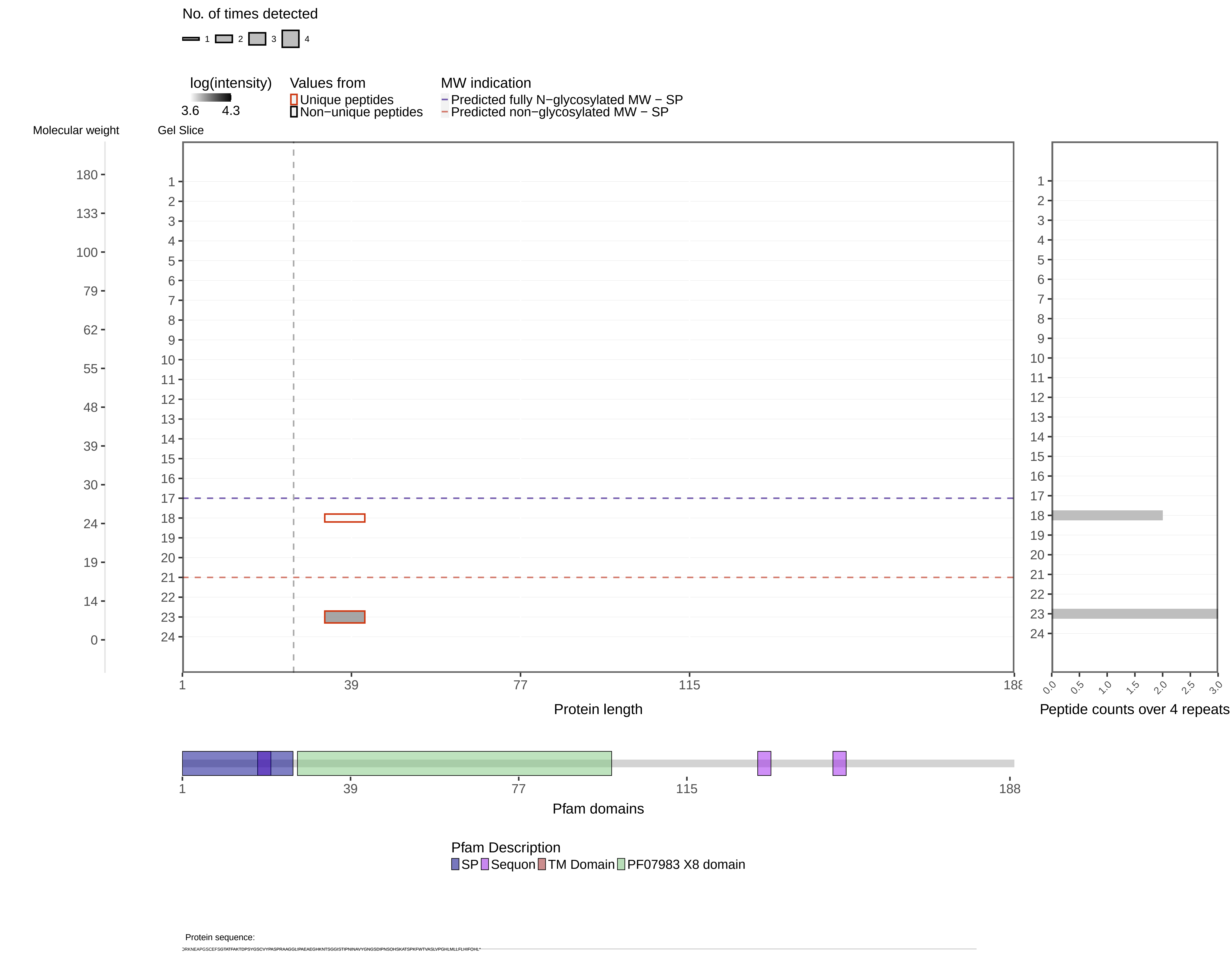

### NbL06g01990.1 stress_response_protein_NST1−like [MW = 46.51 kDa]

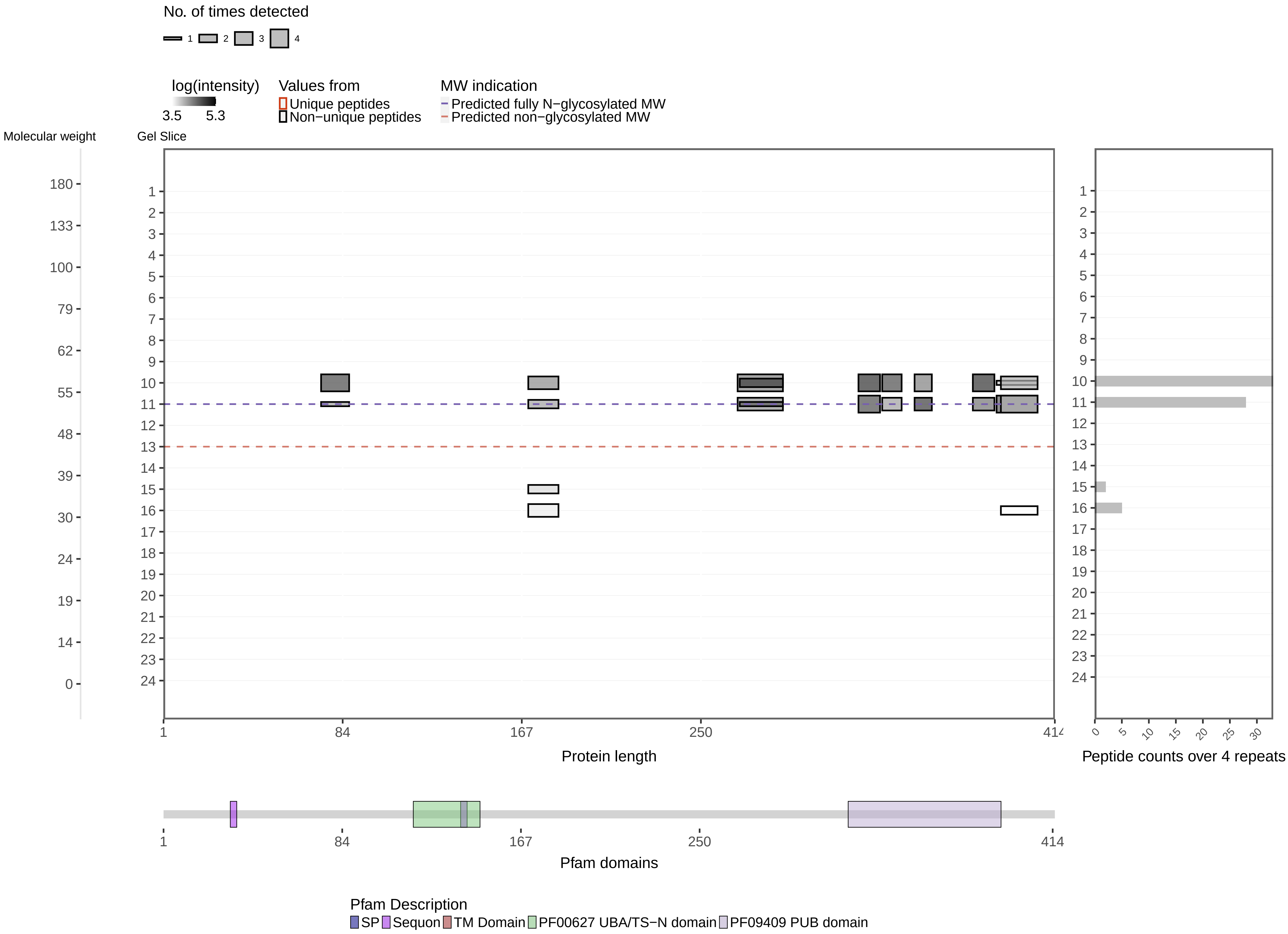


Protein sequence:

ERARKKKEEEERQREREREKERIRIGKELLEAKRMEEENERKRIIALRKAEKAEEQRAREKIRQKLEEDKAERRRKLGLPAEDPAPAKPSTPVVEEKKITLPVRPATKAERMRECLRTLKQTHKDDEAKVKTAFNTLLTYAKNVATNPNEEKYRKIRLSNAAFQDRIGKLQGGIEFLEHCGFEKIEGGEFLYLPREKVDMAVLNTAGTELNNAIKNPFFGVL*

### NbL06g01960.1 glutamic_acid−rich_protein−like [MW = 65.82 kDa]

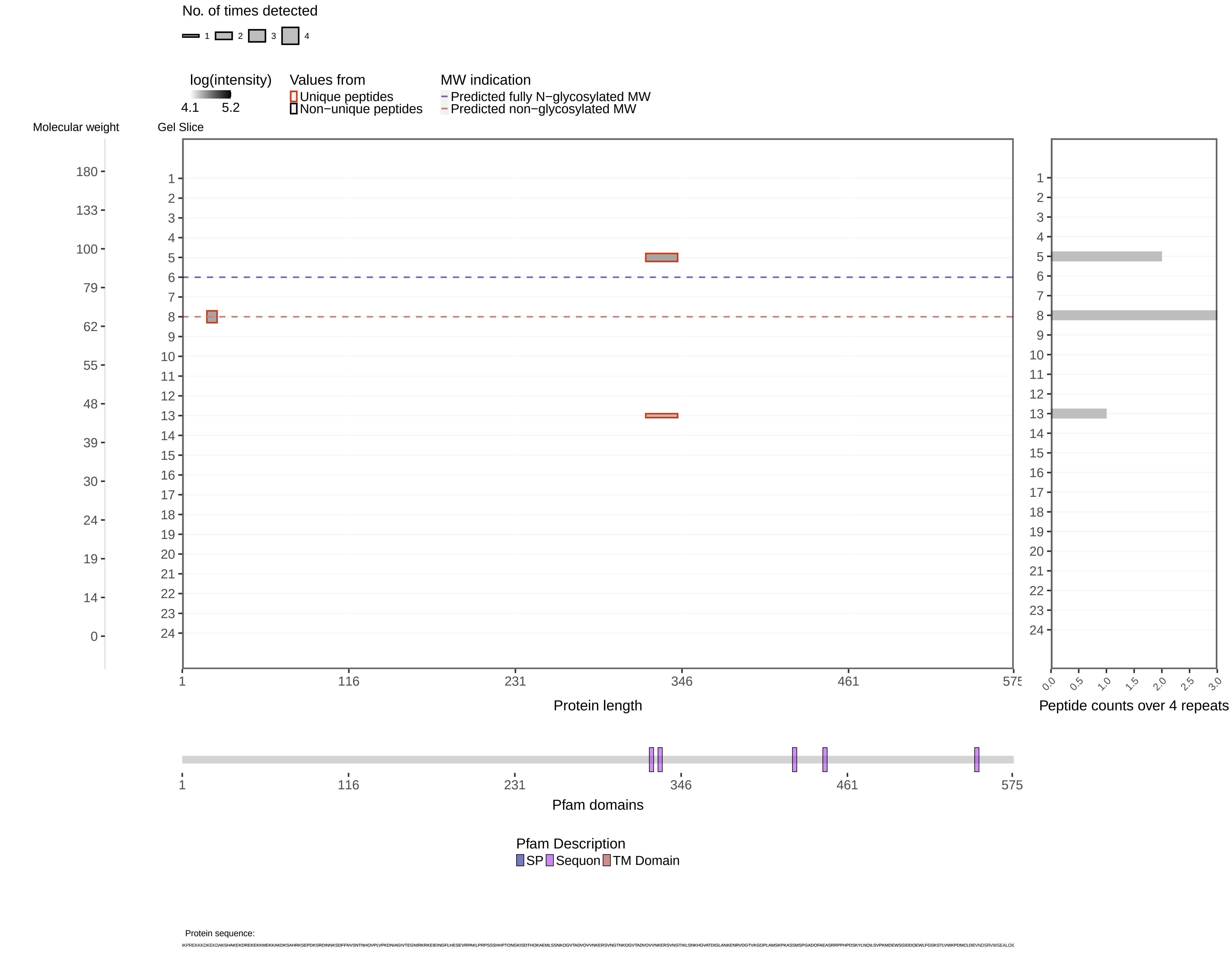

### NbL06g01850.1 probable_receptor_protein_kinase_TMK1 [MW − SP = 97.44 kDa]

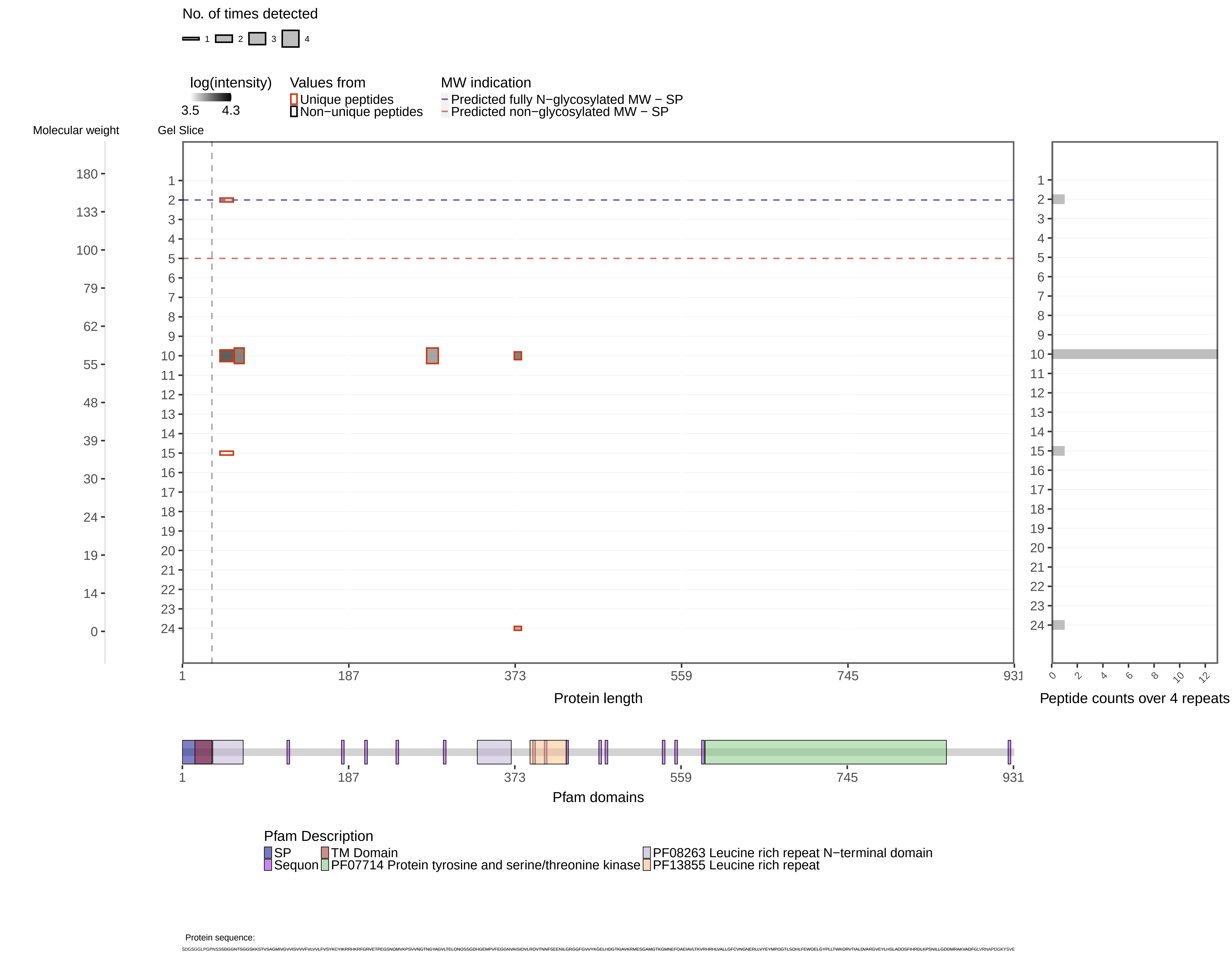

### NbL06g01770.1 serine/threonine−protein_phosphatase_4_regulatory_subunit_3 [MW = 17.87 kDa]

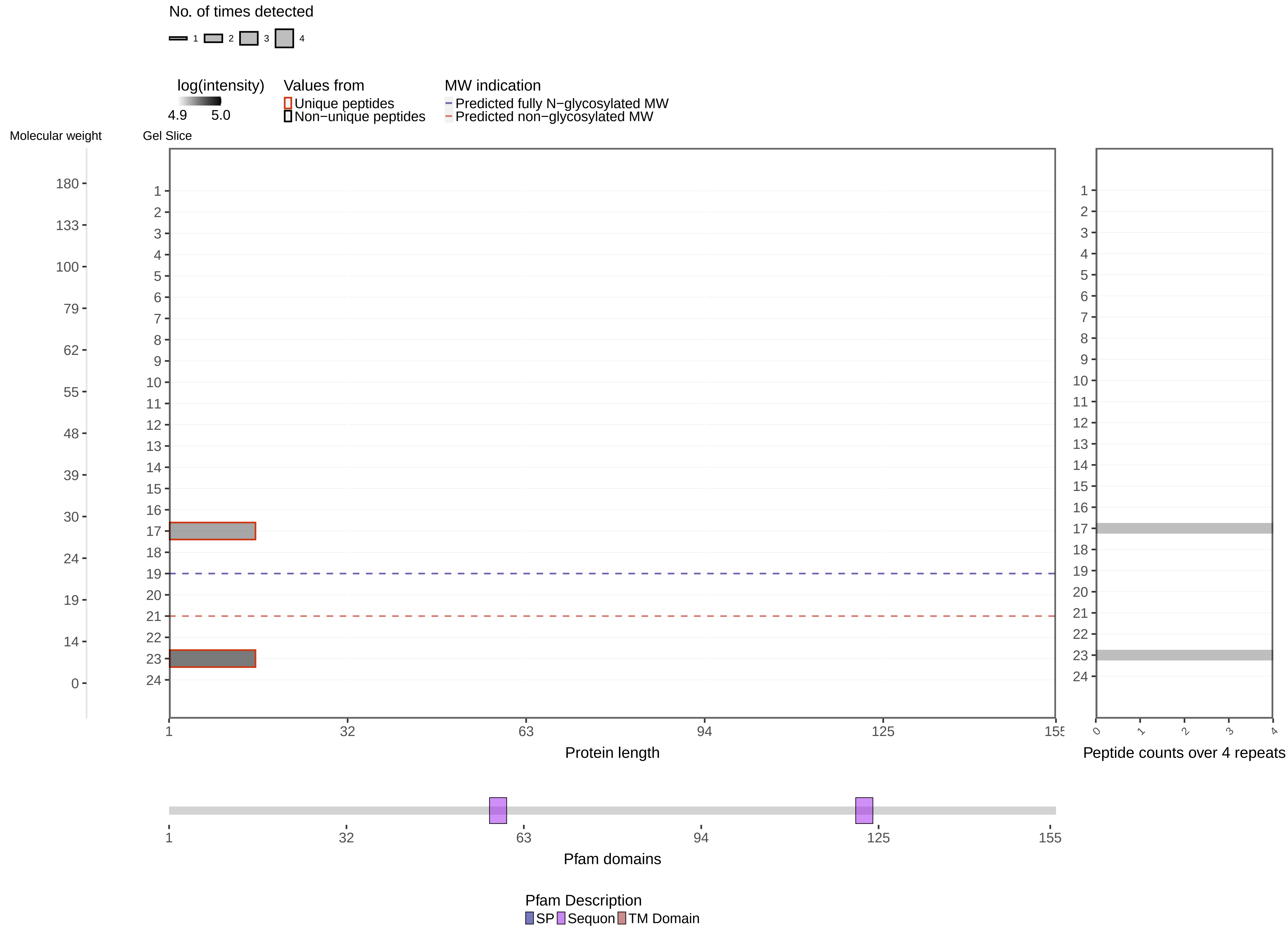


Protein sequence:

RISAEDIYRKQEDTIISWRDPEYSTELALSFQETTGCSYIWDNICNVQRNMHFSSLNNETFHSVNSDLRELPPVSFLRFPDTKDSCGEWHC*

NbL06g01720.1 major_allergen_Pru_ar_1−like [MW = 17.65 kDa]

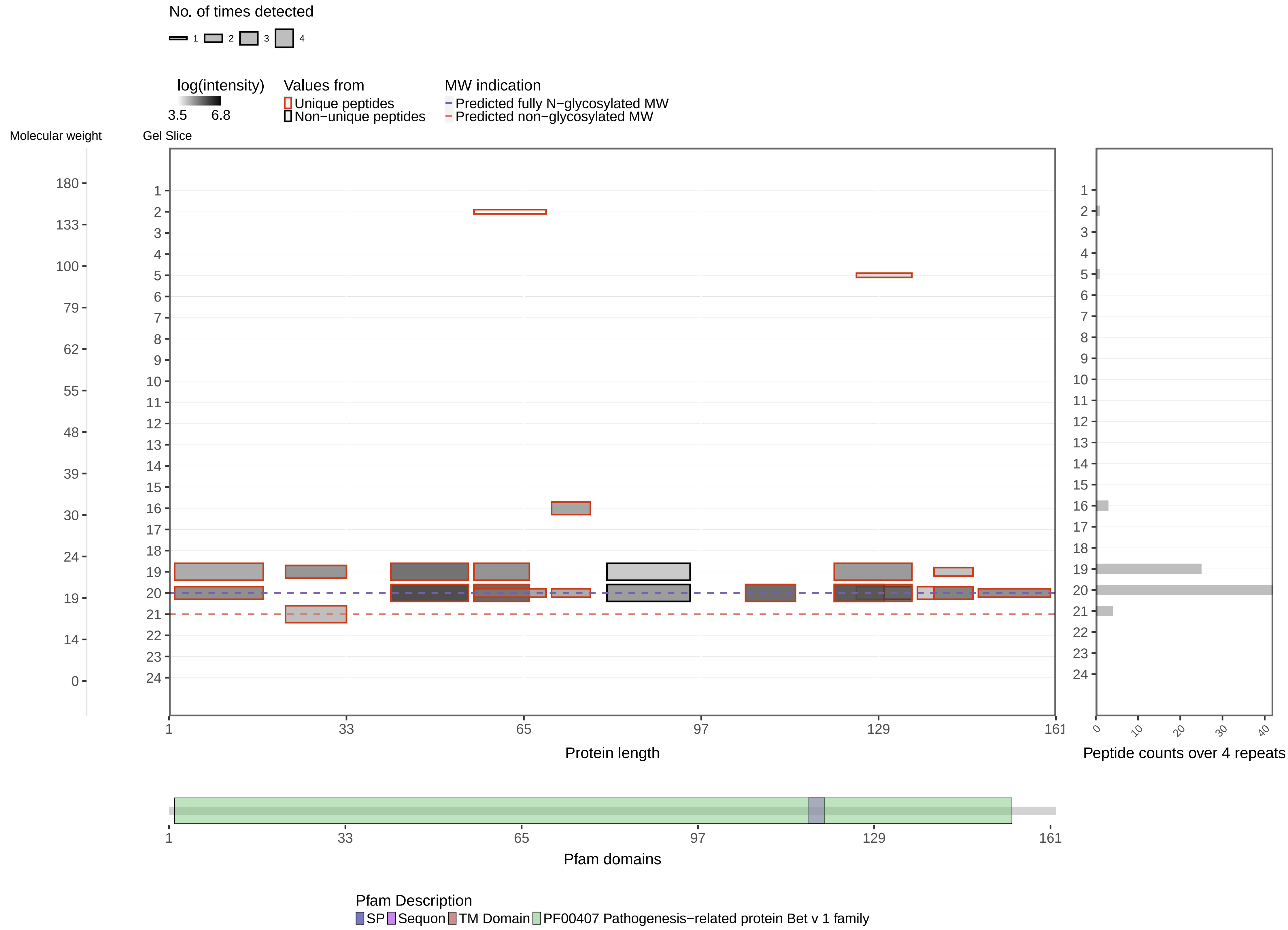


Protein sequence:

/KHRVDAIEKEKMTYAYTLIEGDTLMDKIDSISYEVKFESMPDGGCKGKNTSKYNTKPGVEIKEEEIKAGKEMAAGIVKAVEAYLVANPNEYV*

### NbL06g01640.1 glyoxysomal_fatty_acid_beta−oxidation_multifunctional_protein_MFP−a [MW = 79 kDa]

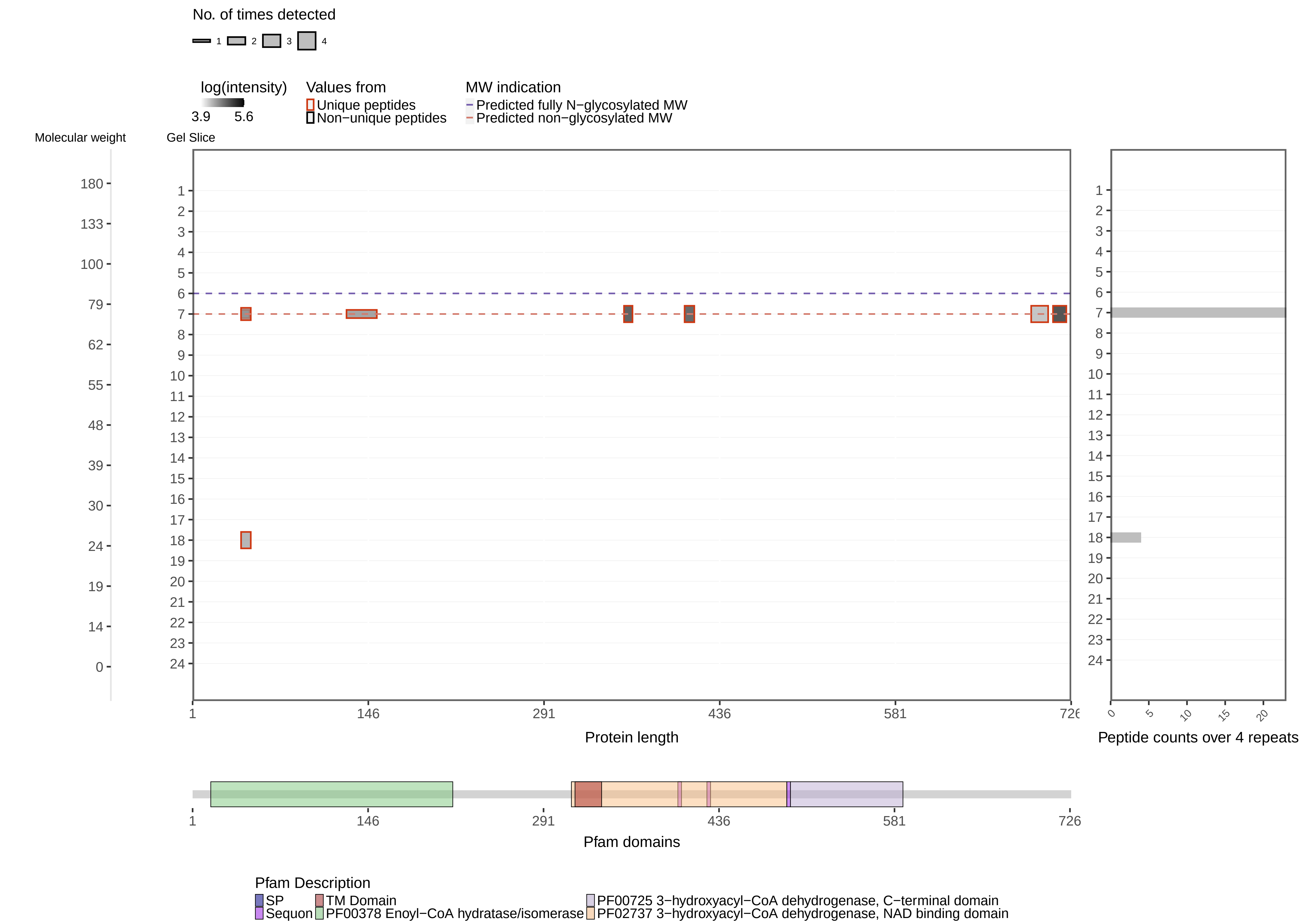


Protein sequence:

### NbL06g01620.1 5'−nucleotidase_domain−containing_protein_4−like [MW = 70.63 kDa]

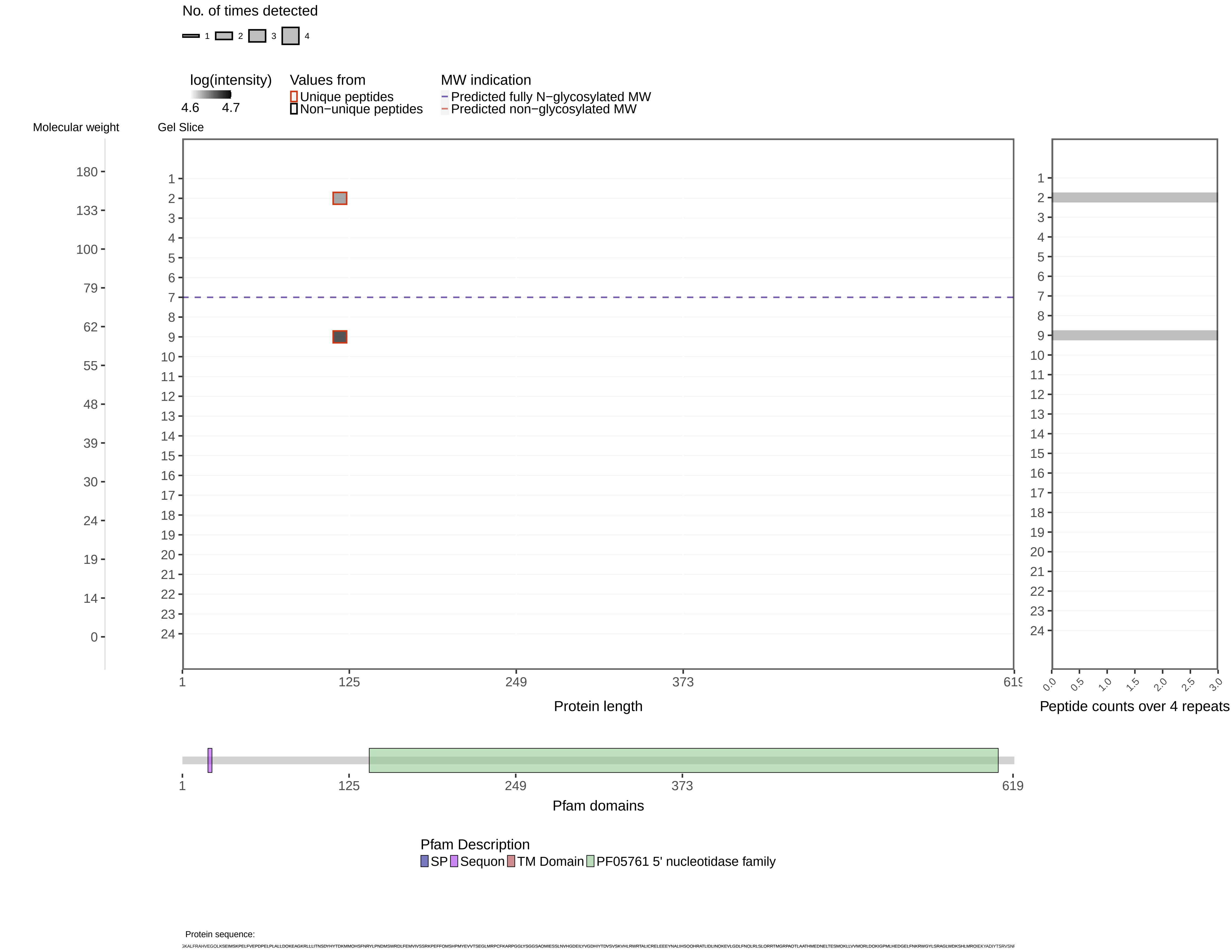

NbL06g01580.1 clathrin_interactor_EPSIN_2−like [MW = 97.44 kDa]

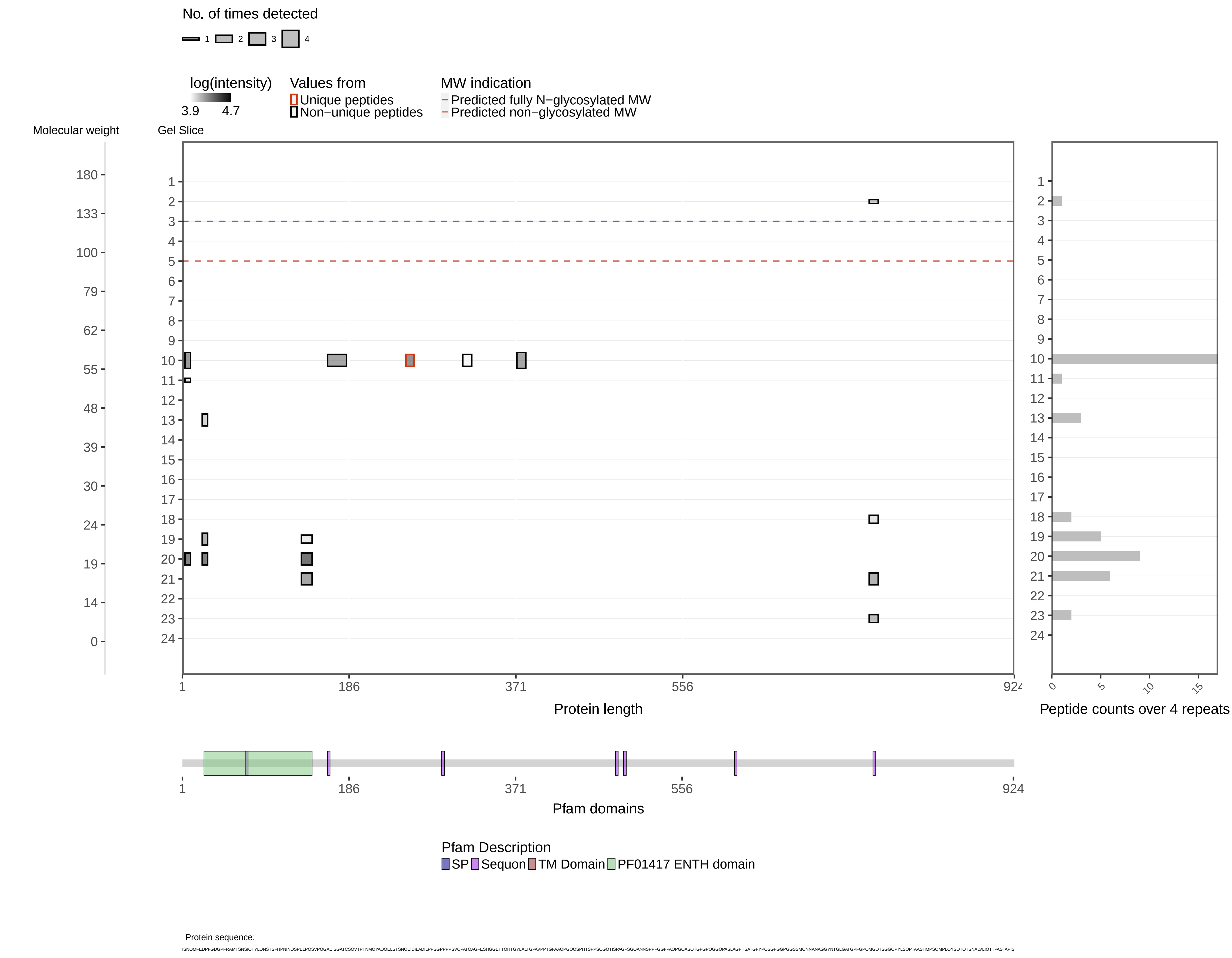

NbL06g01370.1 ATP−citrate_synthase_beta_chain_protein_2 [MW = 65.66 kDa]

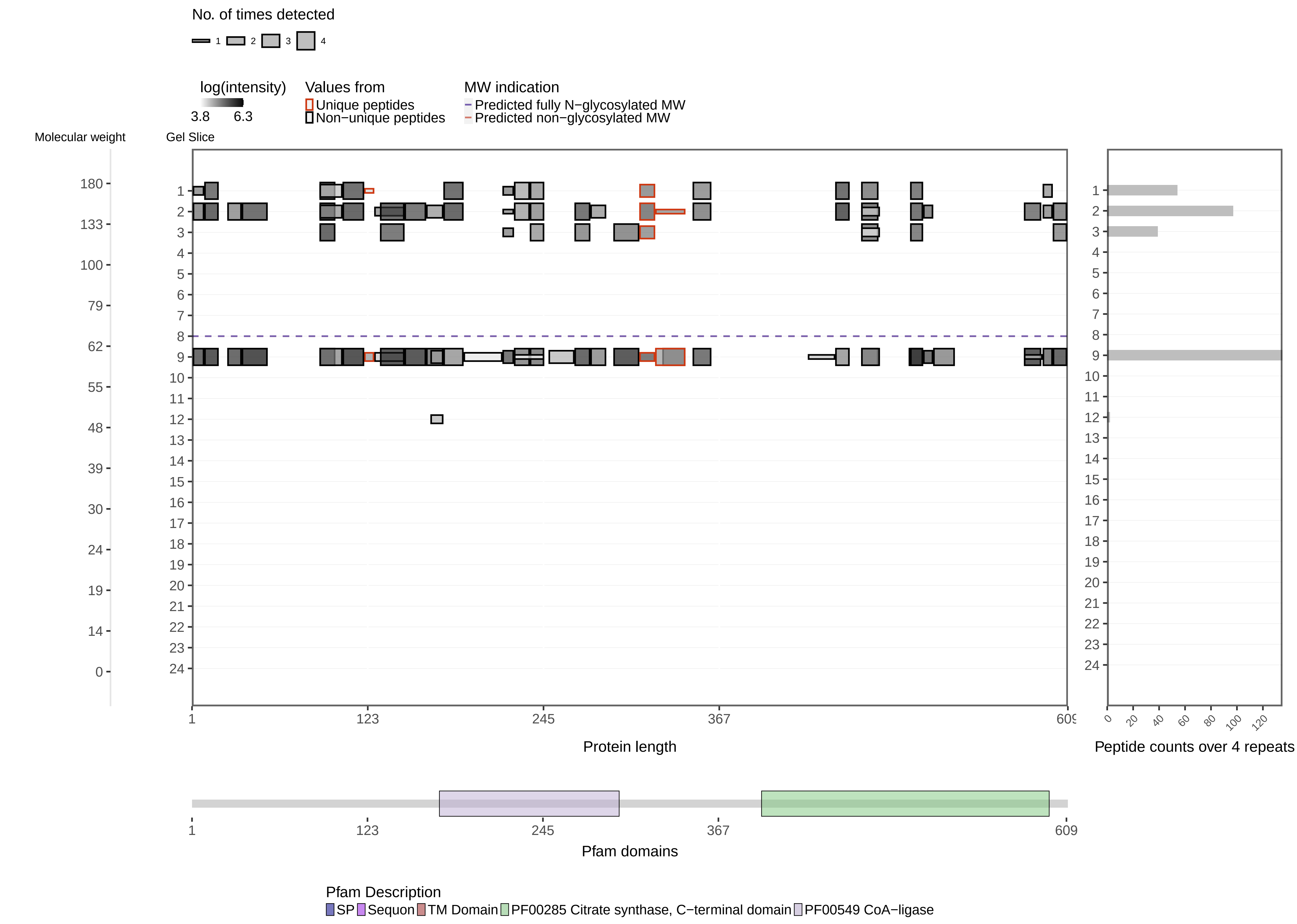


Protein sequence:

### NbL06g01320.1 huntingtin−interacting_protein_K−like [MW = 12.17 kDa]

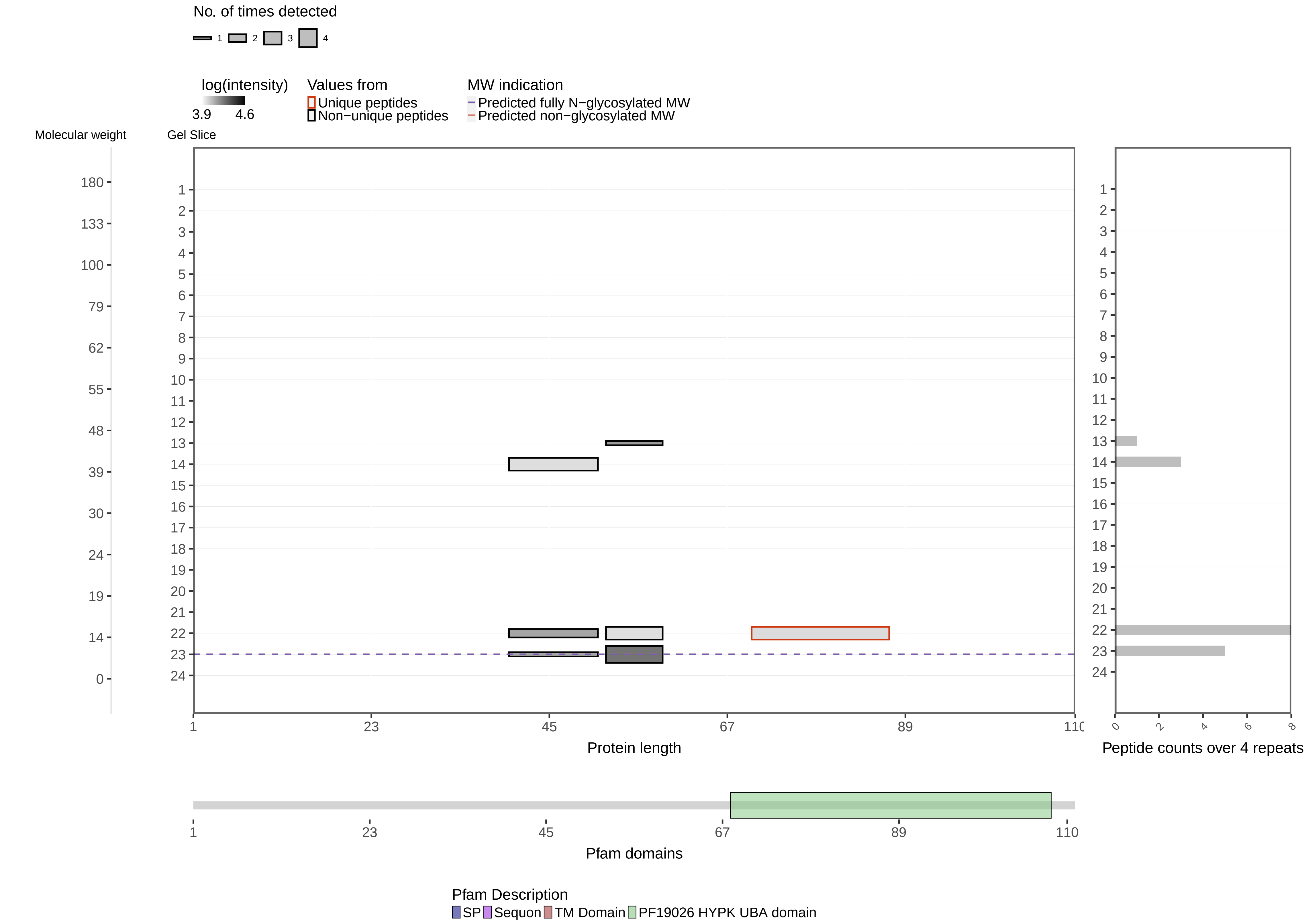


Protein sequence:

/QTAMASIYASKEADLQAMRMREKELAAVKINAADIDIIANELEVDKKVAERTLREHKGDAVAAIRHLLN*

NbL06g01290.1 cell_wall_/_vacuolar_inhibitor_of_fructosidase_1−like [MW − SP = 15.7 kDa]

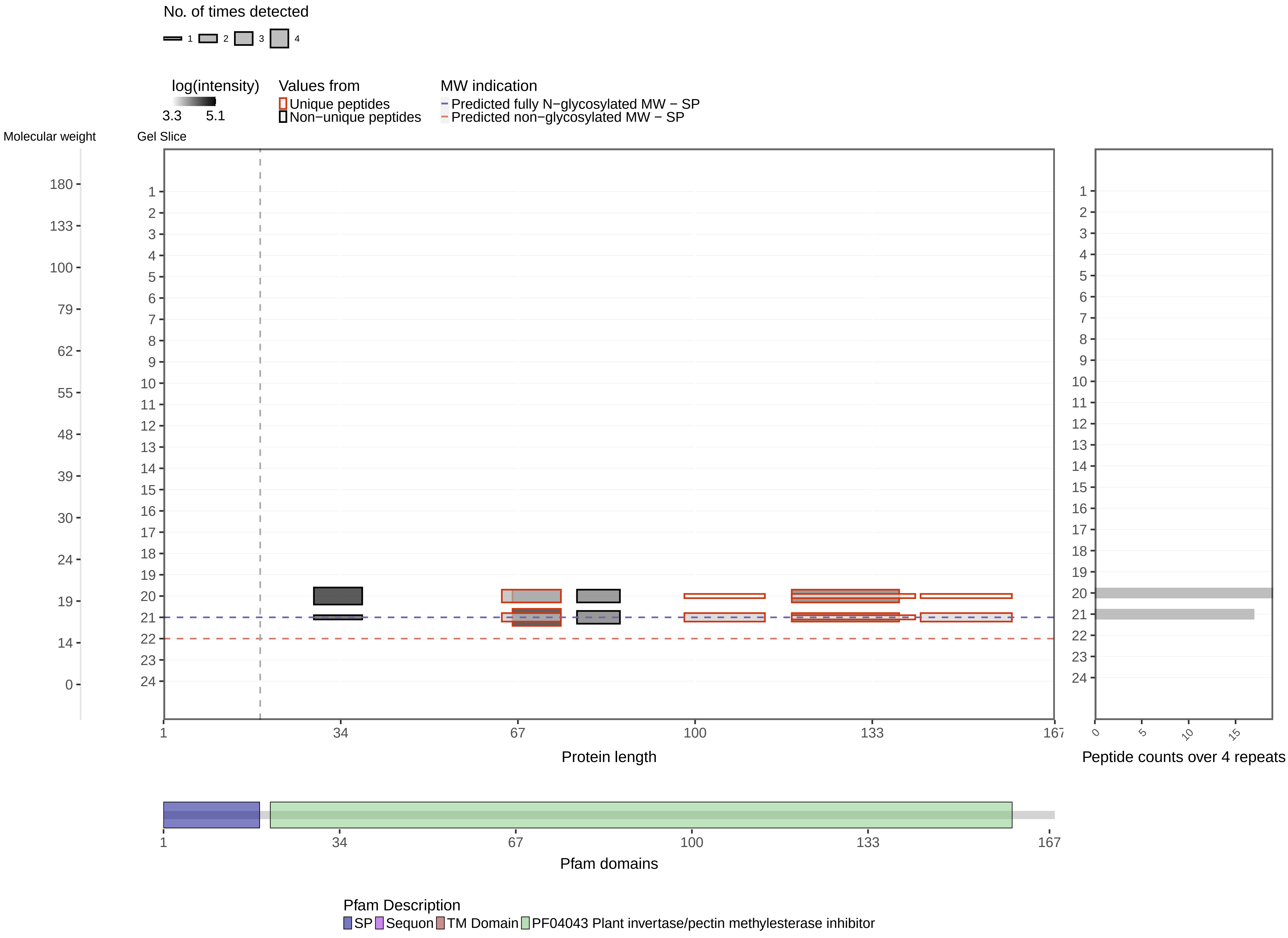


Protein sequence:

TISKLRHSNPPAAWKGPLKNCAFSYKVILTASLPEAIEALTKGDPKFAEDGMVGSSGDAQECEENFKGSKSPFSALNVAVHQLSDVGRAIVRNLL*

### NbL06g01240.1 serine_carboxypeptidase−like_33 [MW − SP = 51.62 kDa]

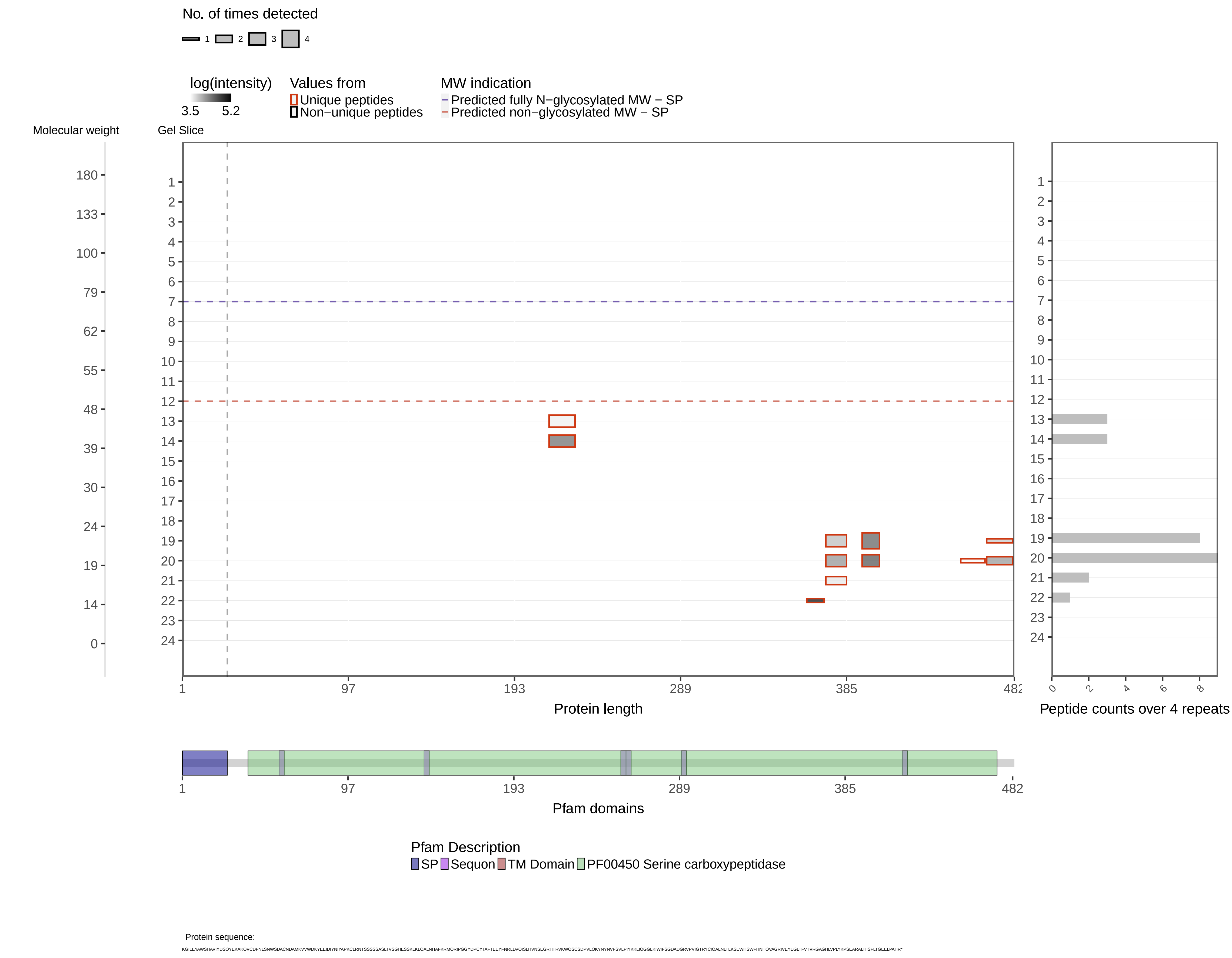

NbL06g01210.1 dihydrolipoyl_dehydrogenase_mitochondrial [MW = 53.49 kDa]

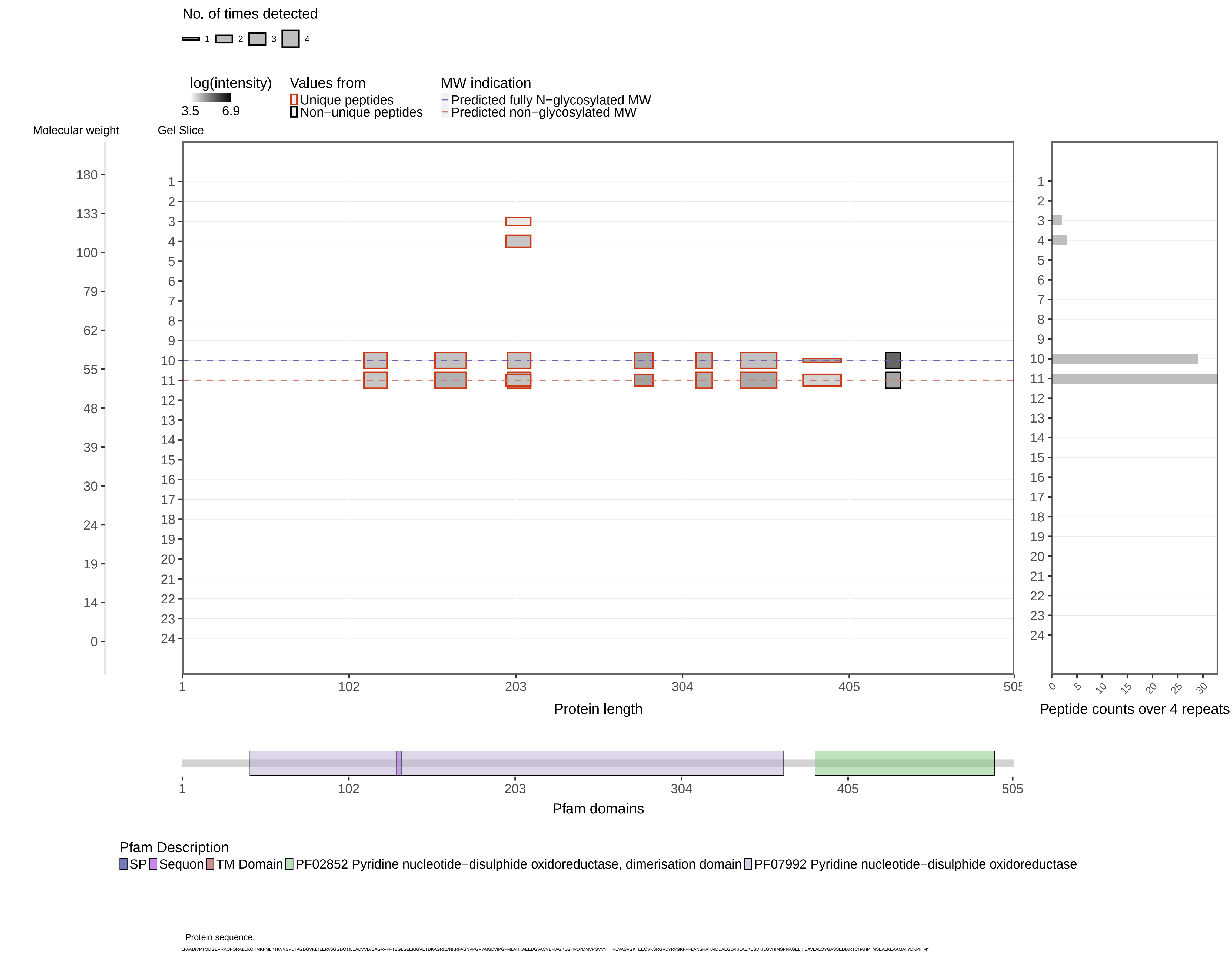

### NbL06g01180.1 ubiquitin−40S_ribosomal_protein_S27a−like [MW = 17.7 kDa]

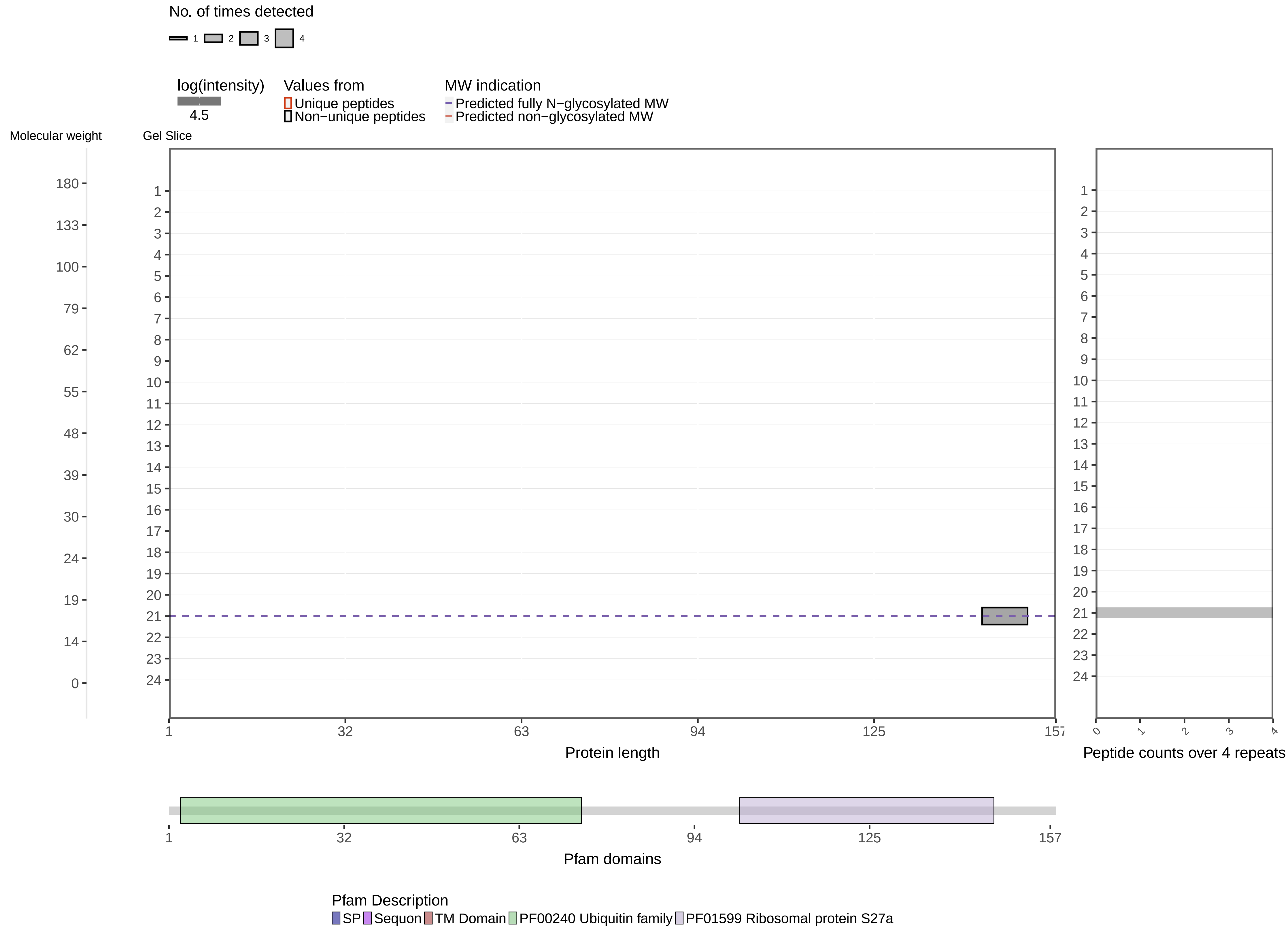


Protein sequence:

ILVLRLRGGAKKRKKKTYTKPKKIKHKKKKVKLAVLQFYKVDDSGKVQRLRKECPNAECGAGTFMANHFDRHYCGKCGLTYVYQKAGGD*

### NbL06g01150.1 S−adenosylmethionine_synthase_2 [MW = 48.13 kDa]

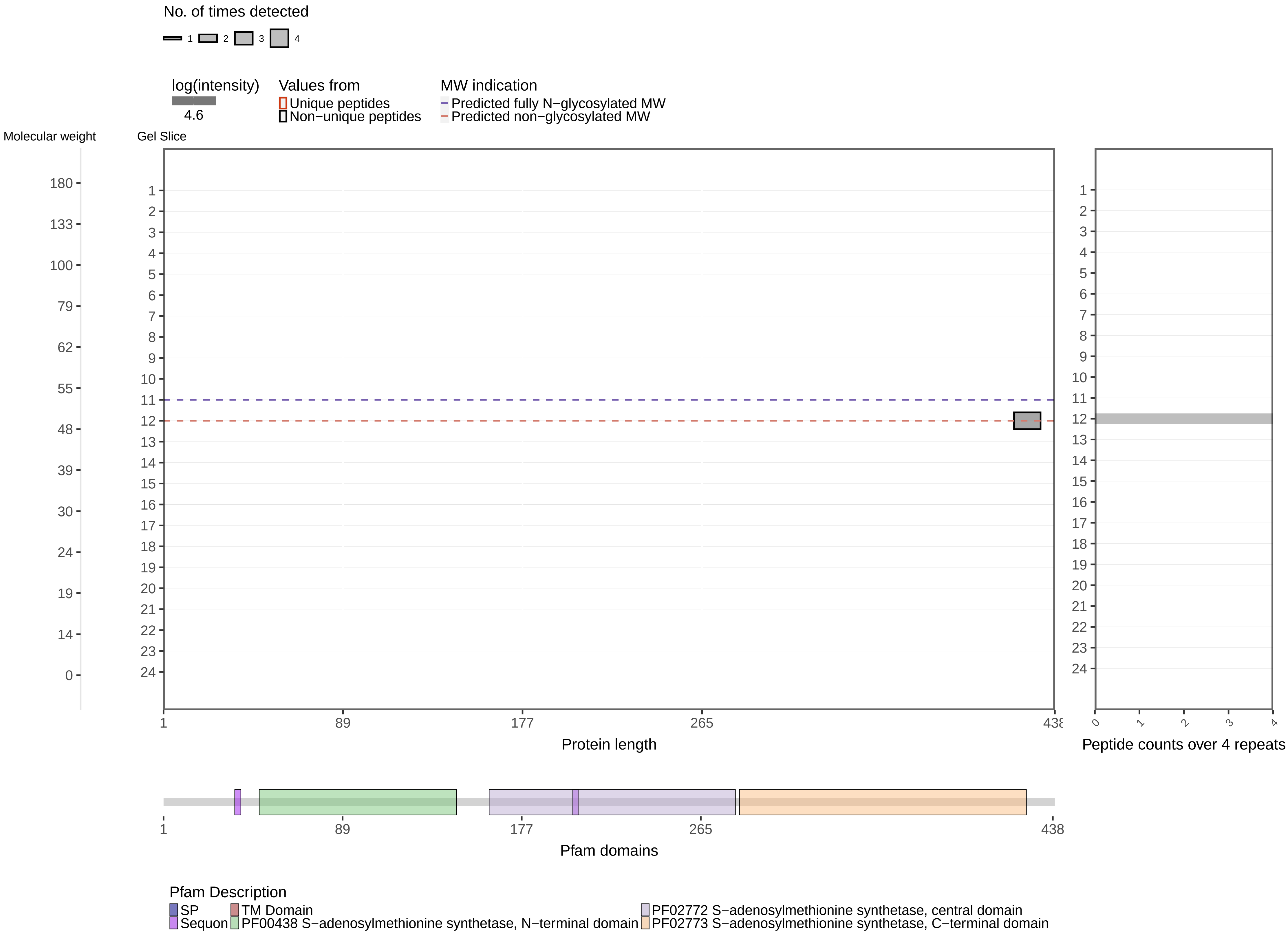


Protein sequence:

WLRPDGKTQVTVEYHNDNGAMVPLRVHTVLISTQHDETVTNDEIARDLKEHVIKPVIPEKYLDEKTIFHLNPSGRFVIGGPHGDAGLTGRKIIIDTYGGWGAHGGGAFSGKDPTKVDRSGAYIVRQAAKSIVANGLARRCIVQVSYAIGVPEPLSVFVDTYGTGKIPDKEILNIVKENFDFRPGMISINLDLLRGGNGRFLKTAAYGHFGRDDSDFTWEVVKPLKWEKPQA*

### NbL06g01090.1 [Pyruvate_dehydrogenase_(acetyl−transferring)]_kinase_mitochondrial−like [MW = 41.82 kDa]

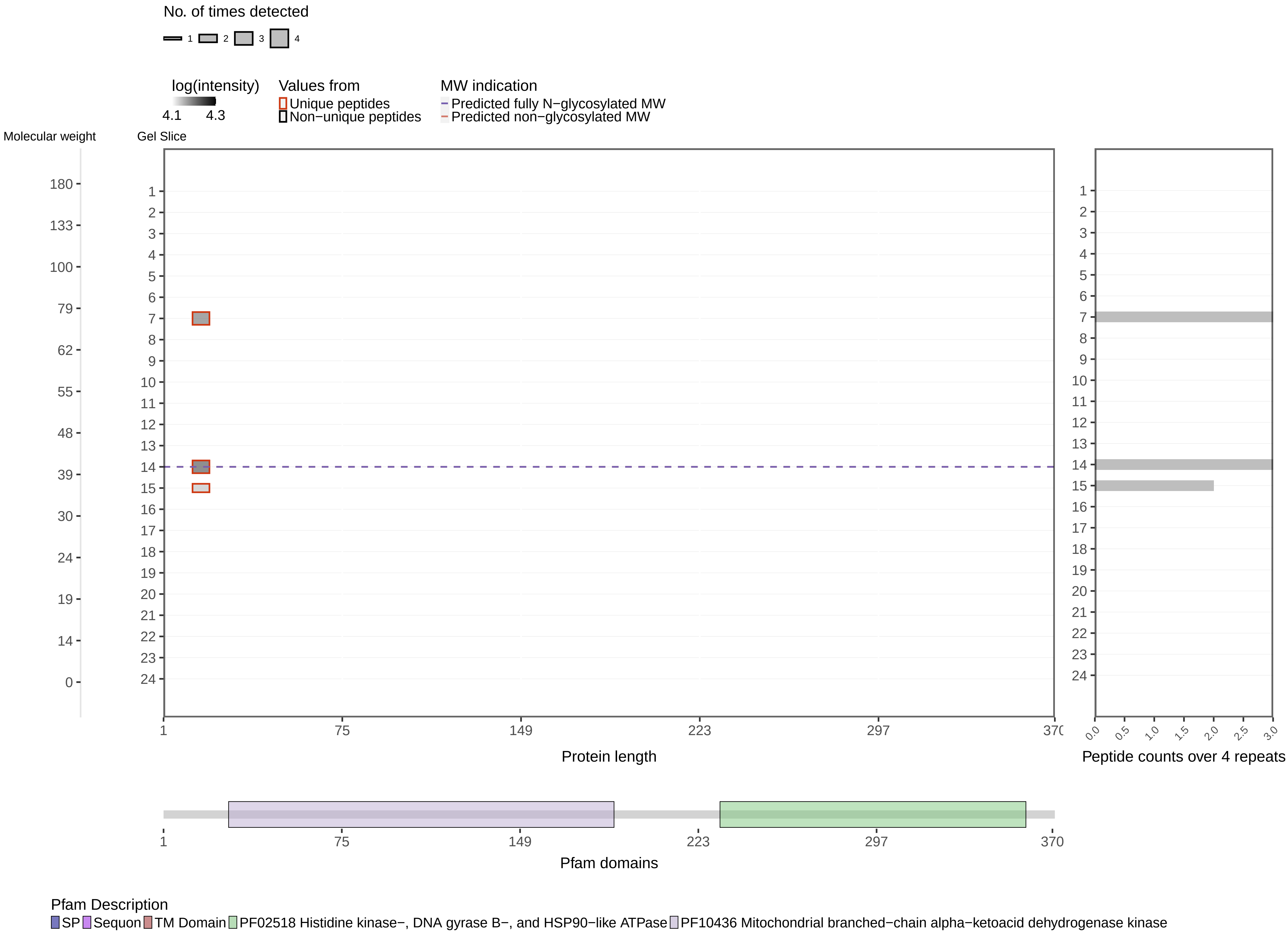


Protein sequence:

QHVALHDPNPPPNCVGYIHTKMSPLEVARTASEDARGICLREYGSAPDVNIYGDPDFTFPYVPTHLHMMVFELVKNSVRAVQERFMDSDKVAPPIRIIVADGLEDVTIKVSDEGGGIPRSGLPKIFTYLYSTARNPLNEHFTTTDVATSSTLAGYGYGIPISRLYARYFGGDLQIISMEGYGTDAYLHLSRLGDSQEPLS*

NbL06g01060.1 50S_ribosomal_protein_L18_chloroplastic−like [MW = 18.43 kDa]

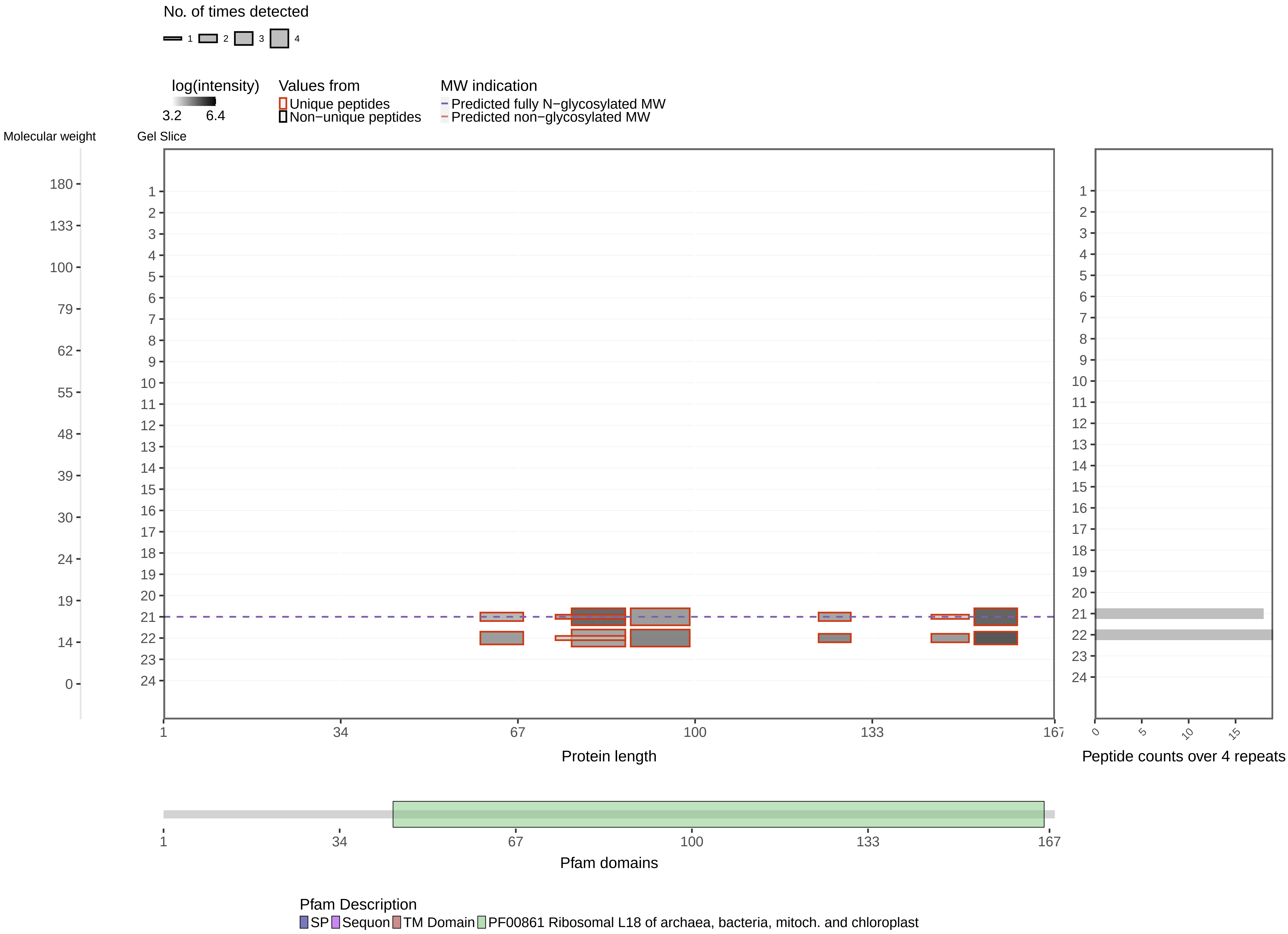


Protein sequence:

VFRSNKHIYVQVIDDSKMHTLASASTMQKPIAEEFDYSAGPTTVSSQDVAKKVGEVIAKACLEKGITKVAFDRGGYPYHGRIEALADAAREHGLQF*

NbL06g00970.1 protein_FAM32A [MW = 16.28 kDa]

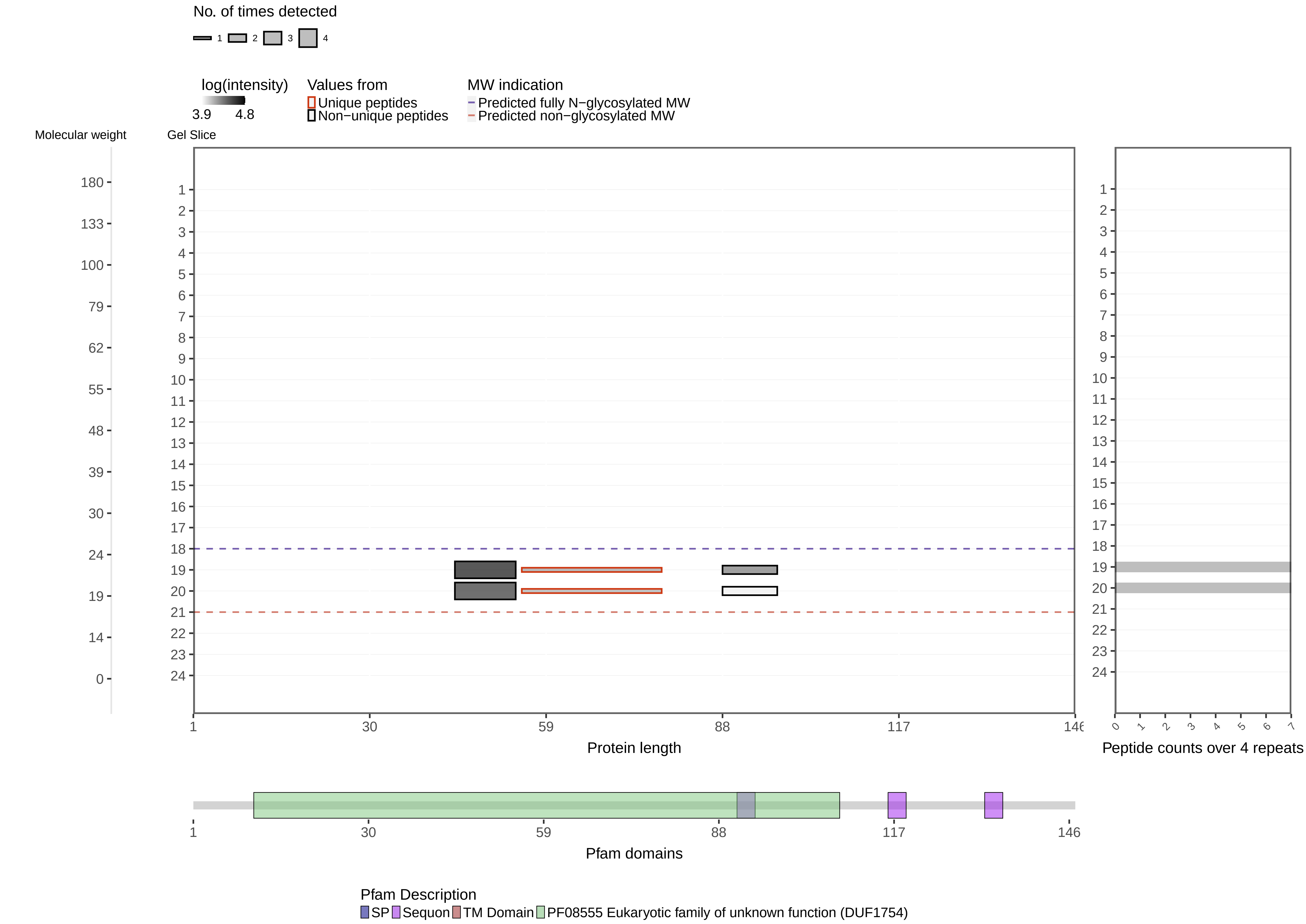


Protein sequence:

GSGSLDDPTKEESTDATKSVGEENAGRWDDNLTPAERRYIEQRERIDMHKMAKTANKSHRDRIEDFNQYLANMSEHYDIPKVGPG*

NbL06g00900.1 probable_histone−arginine_methyltransferase_1.3 [MW = 59.12 kDa]

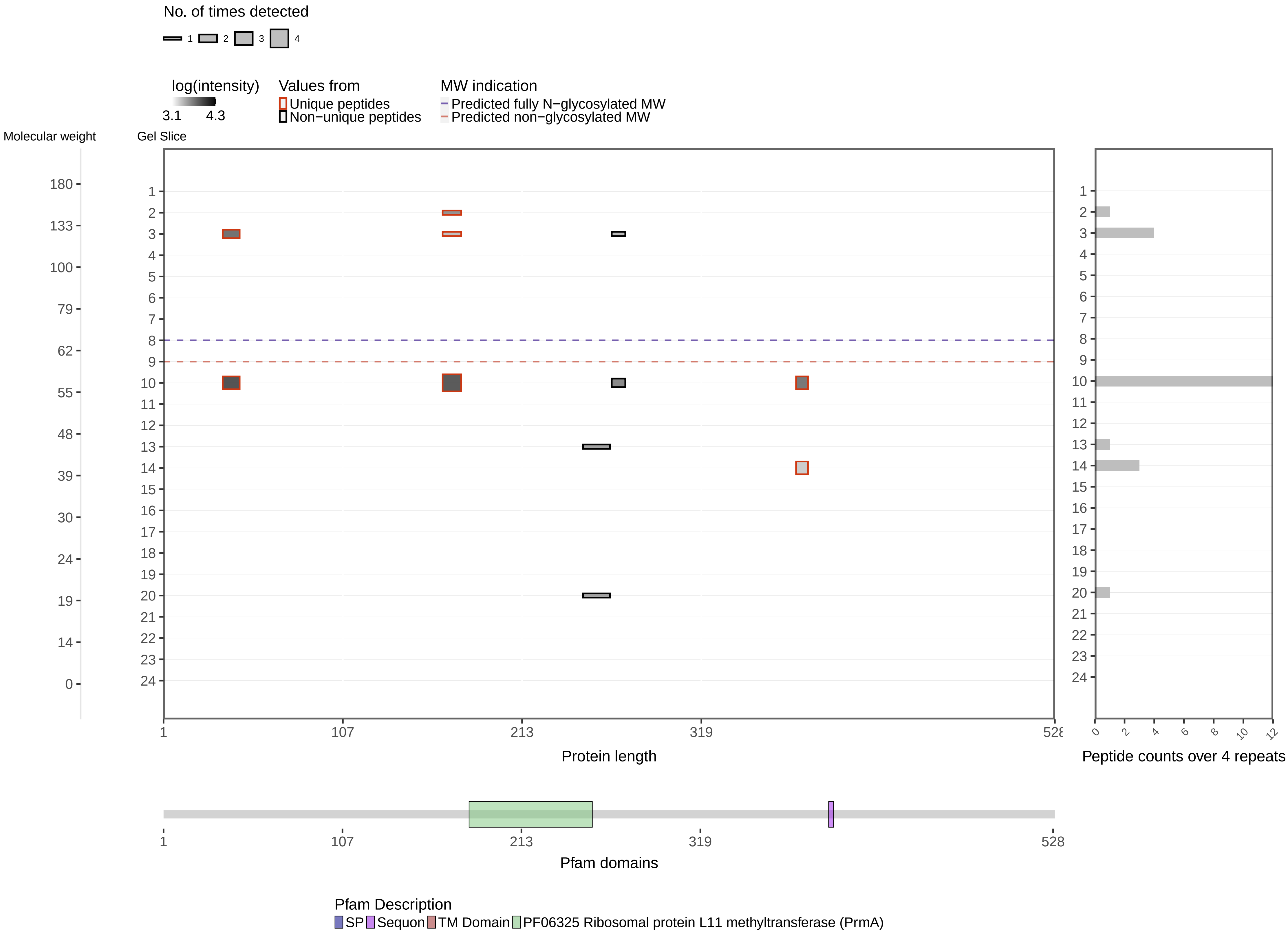


Protein sequence:

### NbL06g00820.1 glucosidase_2_subunit_beta−like [MW − SP = 65.35 kDa]

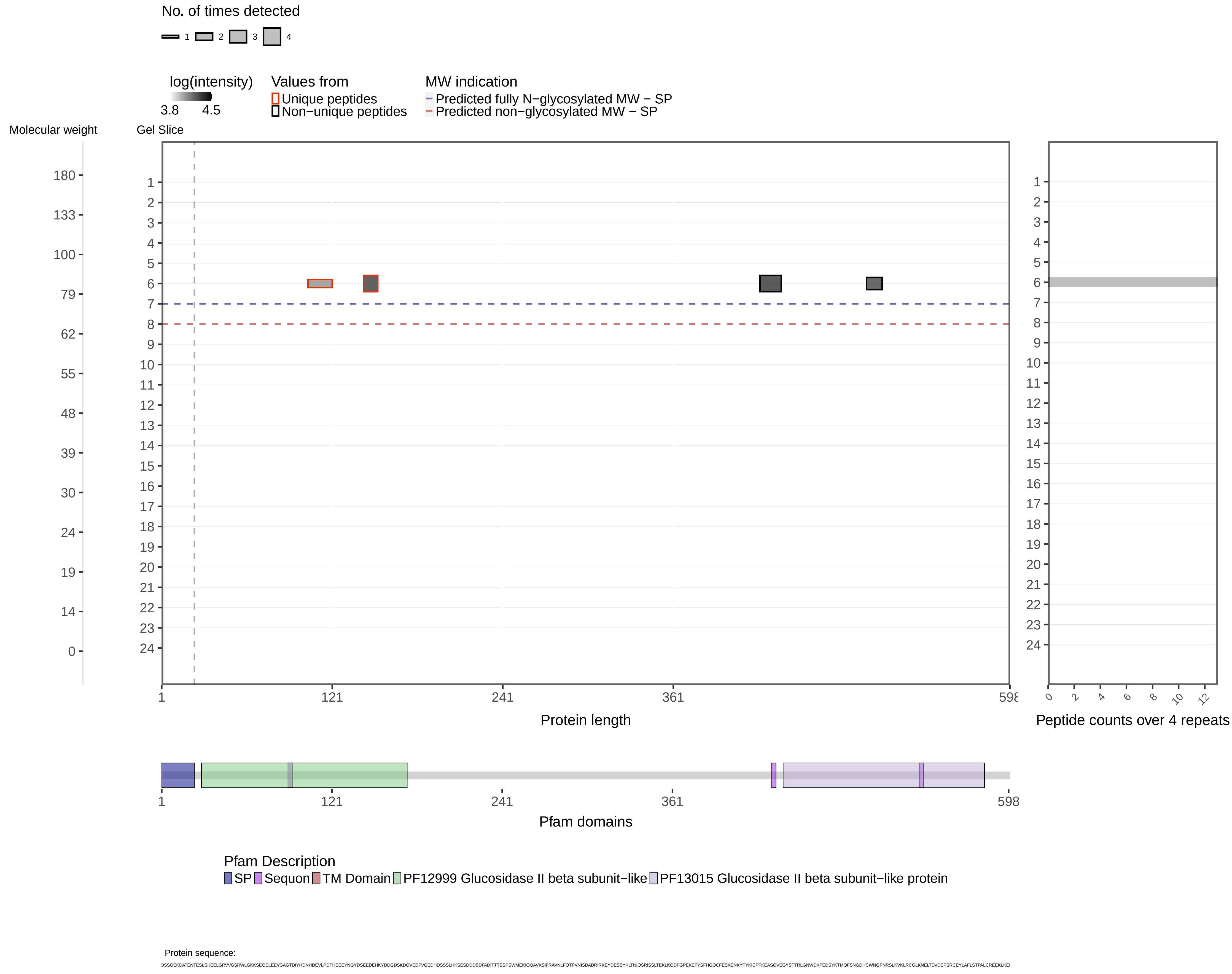

### NbL06g00690.1 rhodanese−like/PpiC_domain−containing_protein_12_chloroplastic [MW = 33.91 kDa]

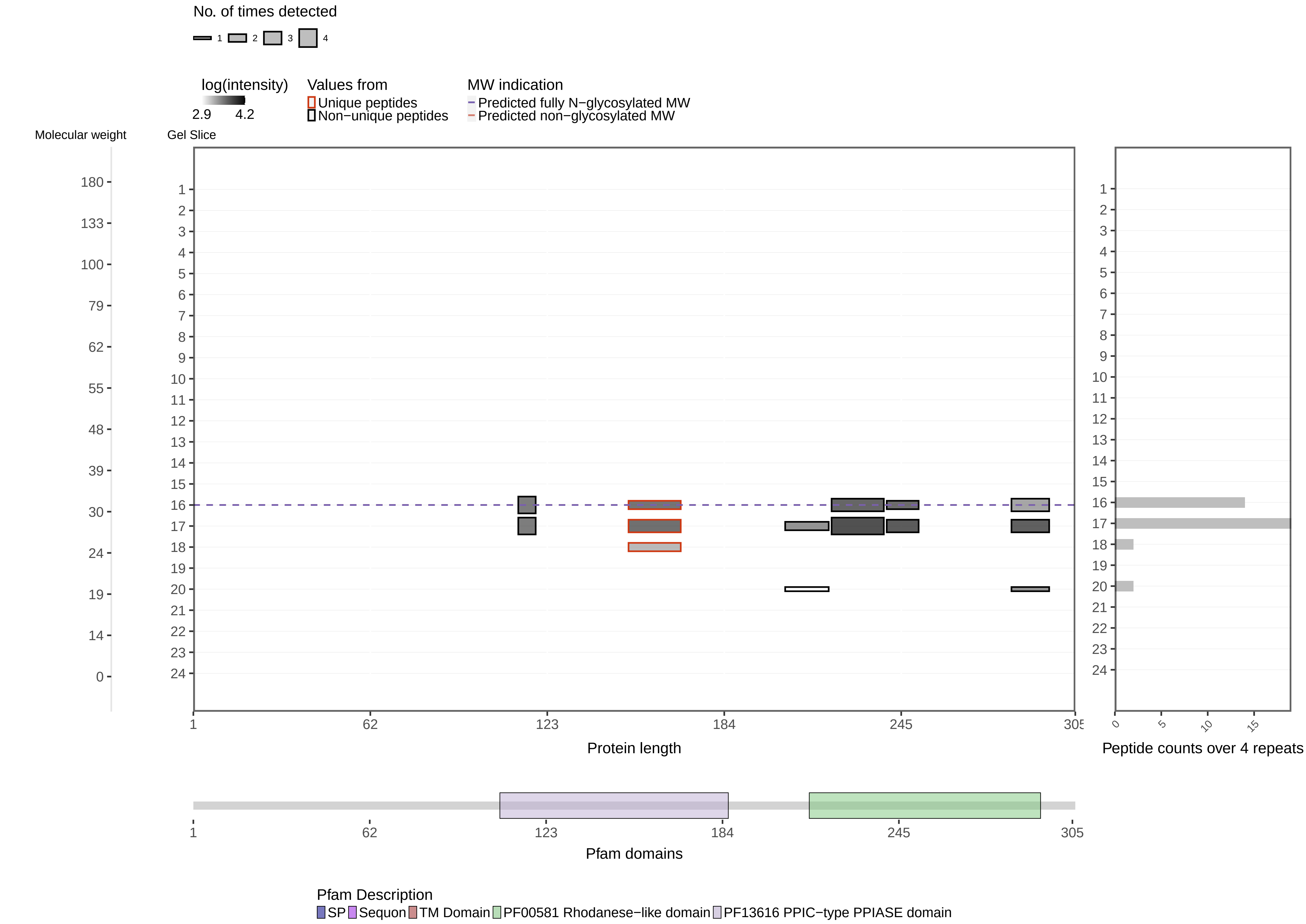


Protein sequence:

:GGMLGWVRKGQMVPEFEEAAFVAPLNKVVKCKTKFGWHLLQVLSEREESVLEHIQPNEFHVKLQDPSFLEEAQLIDVREPEEVAQASLPGFQVLPLRQFGSWGPEITTKFDPQKDTYVLCHHGMRSLQVAKWLQTQGFRKVFNIAGGINEYAVKVDPSIPTY*

### NbL06g00670.1 signal_recognition_particle_9_kDa_protein [MW = 17.72 kDa]

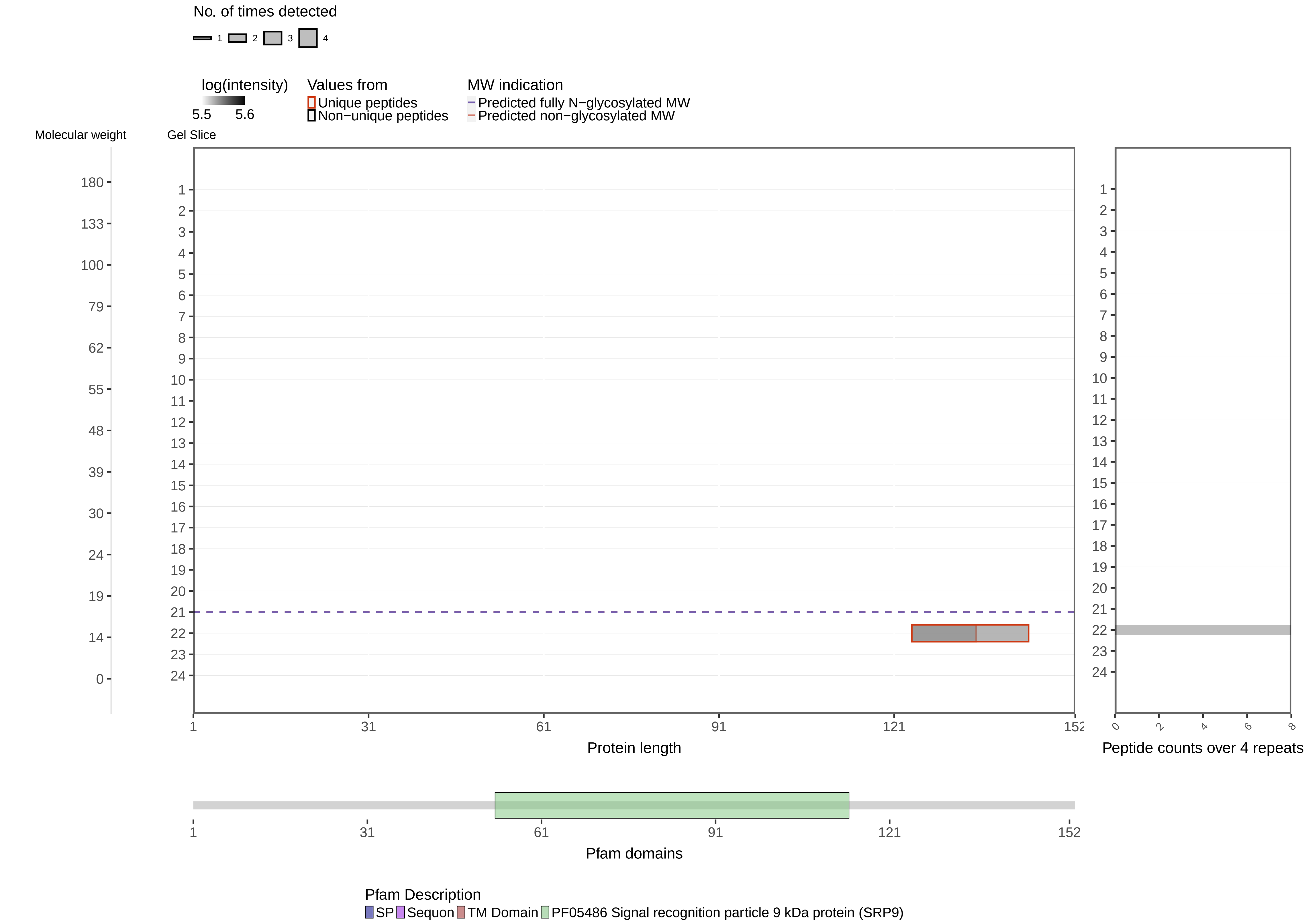


Protein sequence:

LFRADPEKTRYVMKYRHSEGKLVLKVTDDTECIKFRTDQAQDAKKMEKLNNIFFTLMARGPDADISEVGGKEQMEAQAPKKGRGRKQ*

NbL06g00500.1 serine−−glyoxylate_aminotransferase−like [MW = 43.92 kDa]

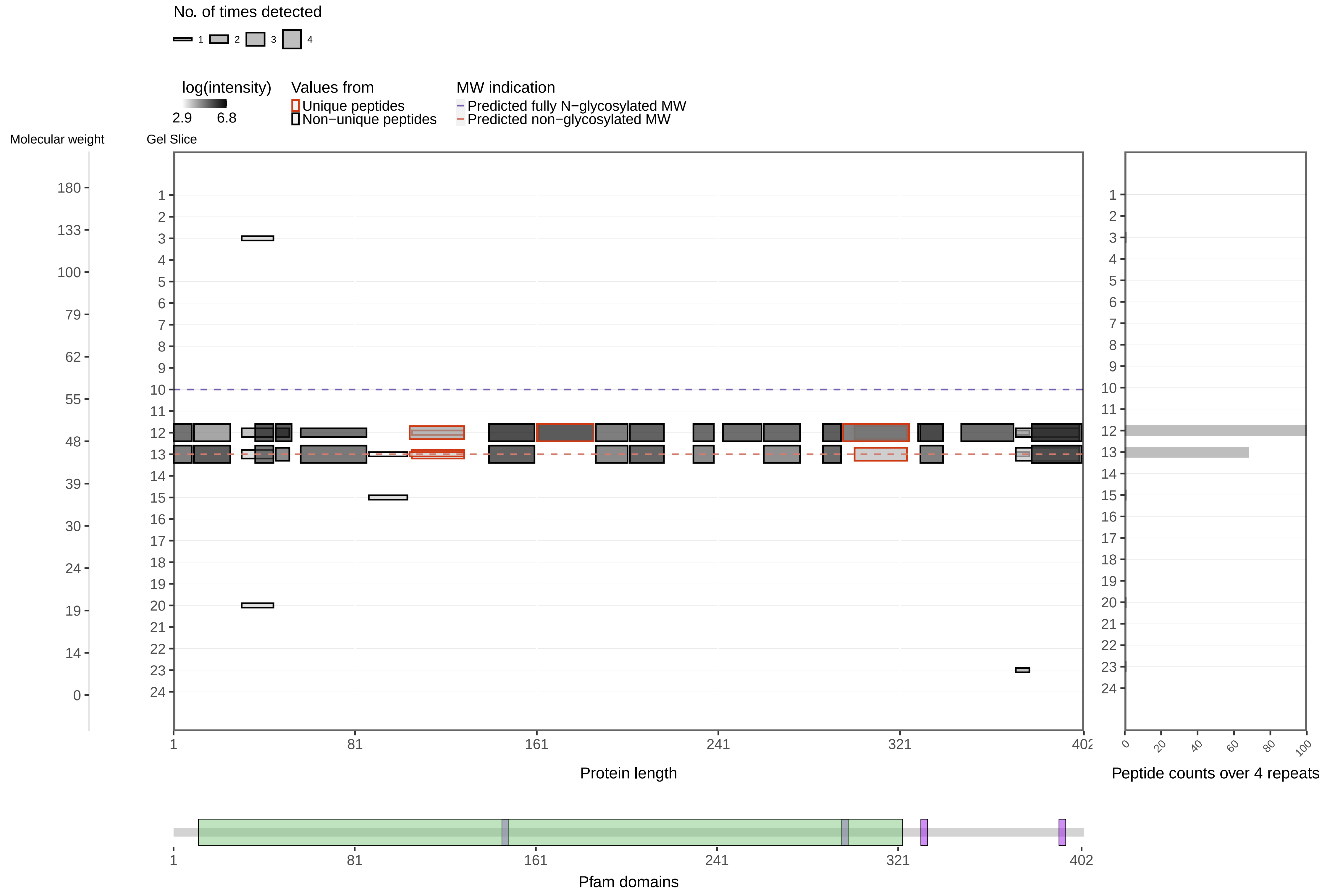


Protein sequence:

WGVDVALTGSQKALSLPTGLGIVCASPKALEAAKTAKSVRVFFDWSDYLKFYKMGTYWPYTPSIQLLYGLRAALDLLFEEGLENVIARHARLGKATRLAVEAWGLKNCTQKEEWFSDTVTAVVVPPYIDSAEVVKRAWKRYNLSLGLGLNKVAGKVFRIGHLGNLNELQLLGCLAGVEMVLKDVGYPVKFGSGVAAASTFLQNSTPLIPSRI*

### NbL06g00460.1 calmodulin−7 [MW = 22.62 kDa]

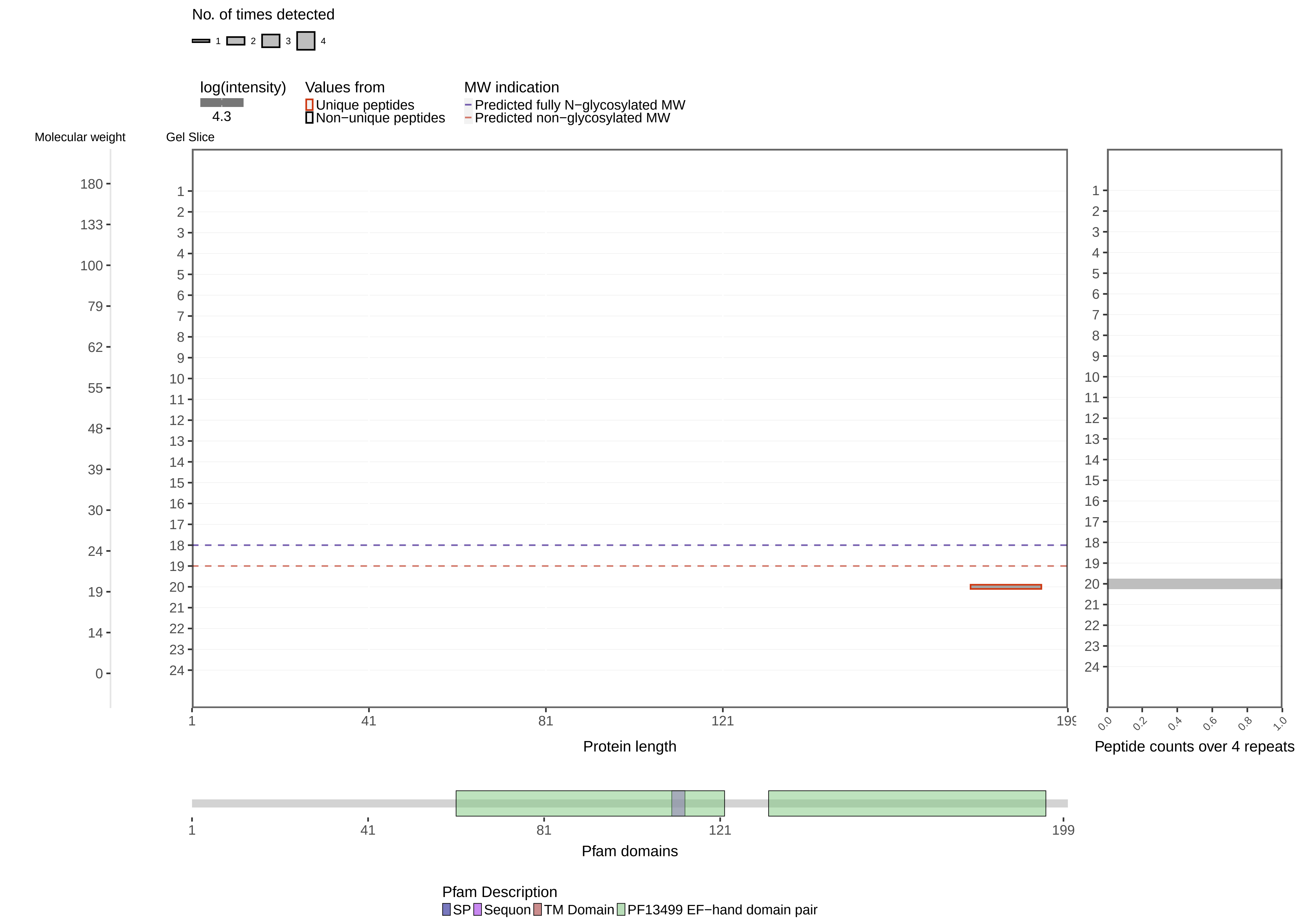


Protein sequence:

LGQNPTEAELQDMINEVDADGNGTIDFPEFLNLMARKMKDTDSEEELKEAFRVFDKDQNGFISAAELRHVMTNLGEKLTDEEVDEMIREADVDGDGHINYEEFVKVMMAK*

### NbL06g00340.1 uncharacterized_LOC107818673 [MW − SP = 94.01 kDa]

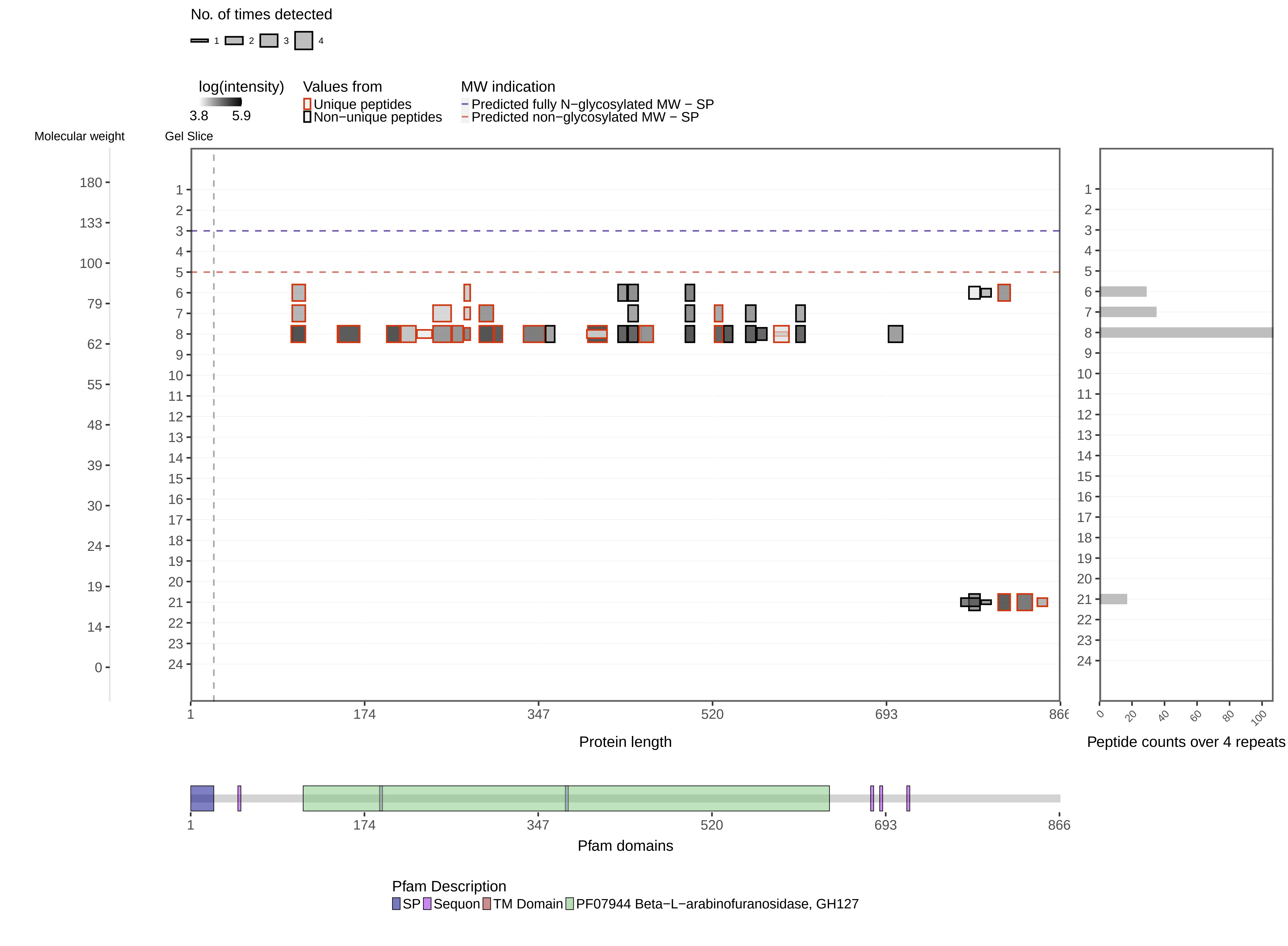


Protein sequence:

/SRHLFRWTKEMAYADYYERALTNGVLSIQRGRDPGVMIYMLPLLRGGSKAHSYHGWGTQFDSFWCCYGTGIESFSKLGDSIYFEEKGNTPSLYIIQYIPSTLDWKSGQVLVSQKIEPVVSWDNRLLVTITISSNGKATGVSSTLNLRIPSWTYSSGAKATLNGKDLSLPTPGNFLSITKKWIQGDKITLELPMSLRTEAIKDDRPEYASVQAILYGPYLLAGHSSGDWDIETKSSTDLSDVITYIPADYNSQLISLMQESGNATFVLTNSNQSIQMEKFPEAGTDAAVSATFRLISLN

NbL06g00290.1 50S_ribosomal_protein_L6_chloroplastic [MW = 25.15 kDa]

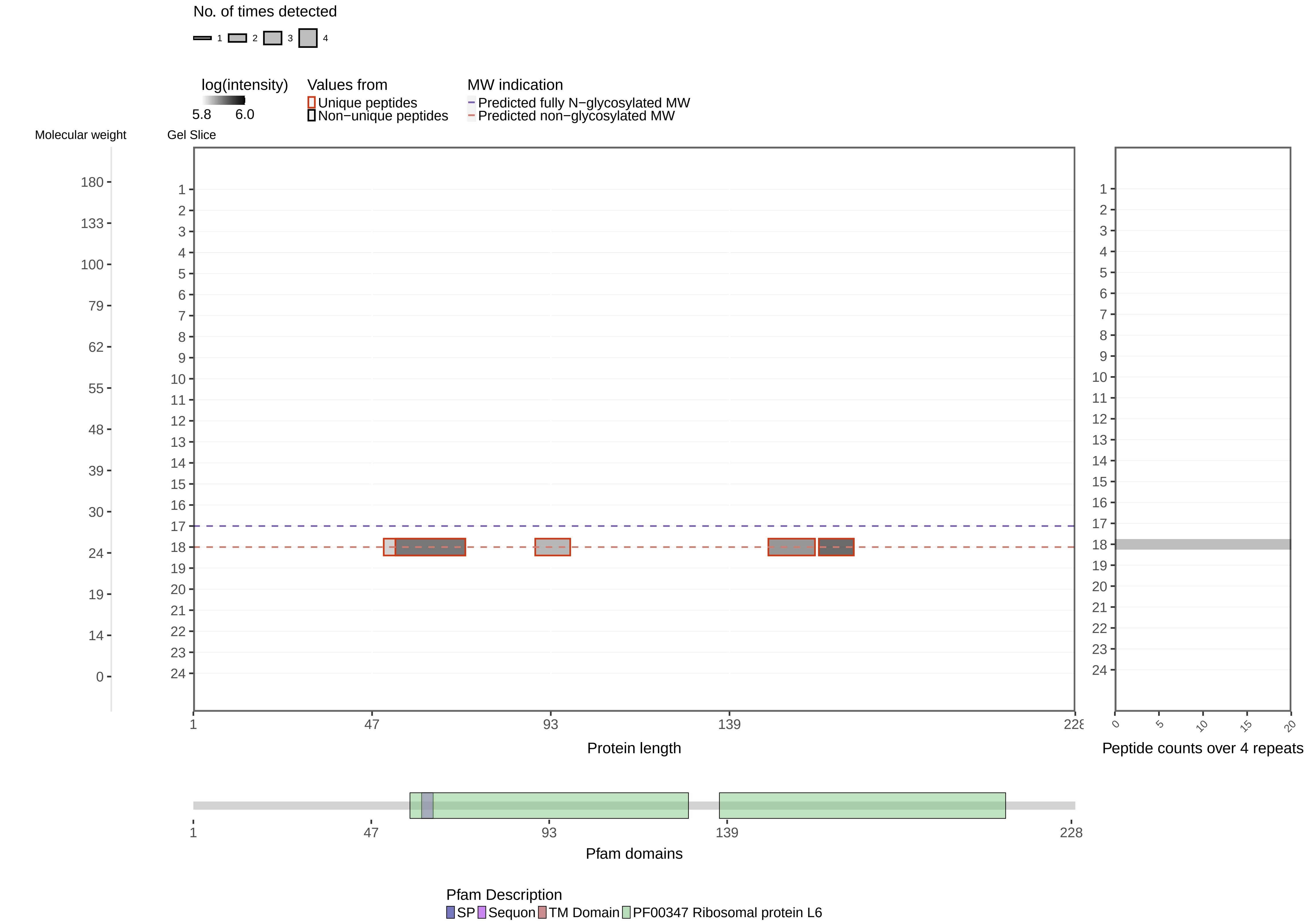


Protein sequence:
AVETRRANQMHGLFRTLTDNMVVGVSKGFEKKLQLVGVGYRATVEGKEIVLNLGFSHPVKMEIPNGLQVKVEENTRITVSGYDKSEIGQFAASIRKWRPPEPYKGKGVKYADEIVRRKEGKAGKKK*

### NbL06g00270.1 pleiotropic_drug_resistance_protein_1−like [MW = 162.13 kDa]

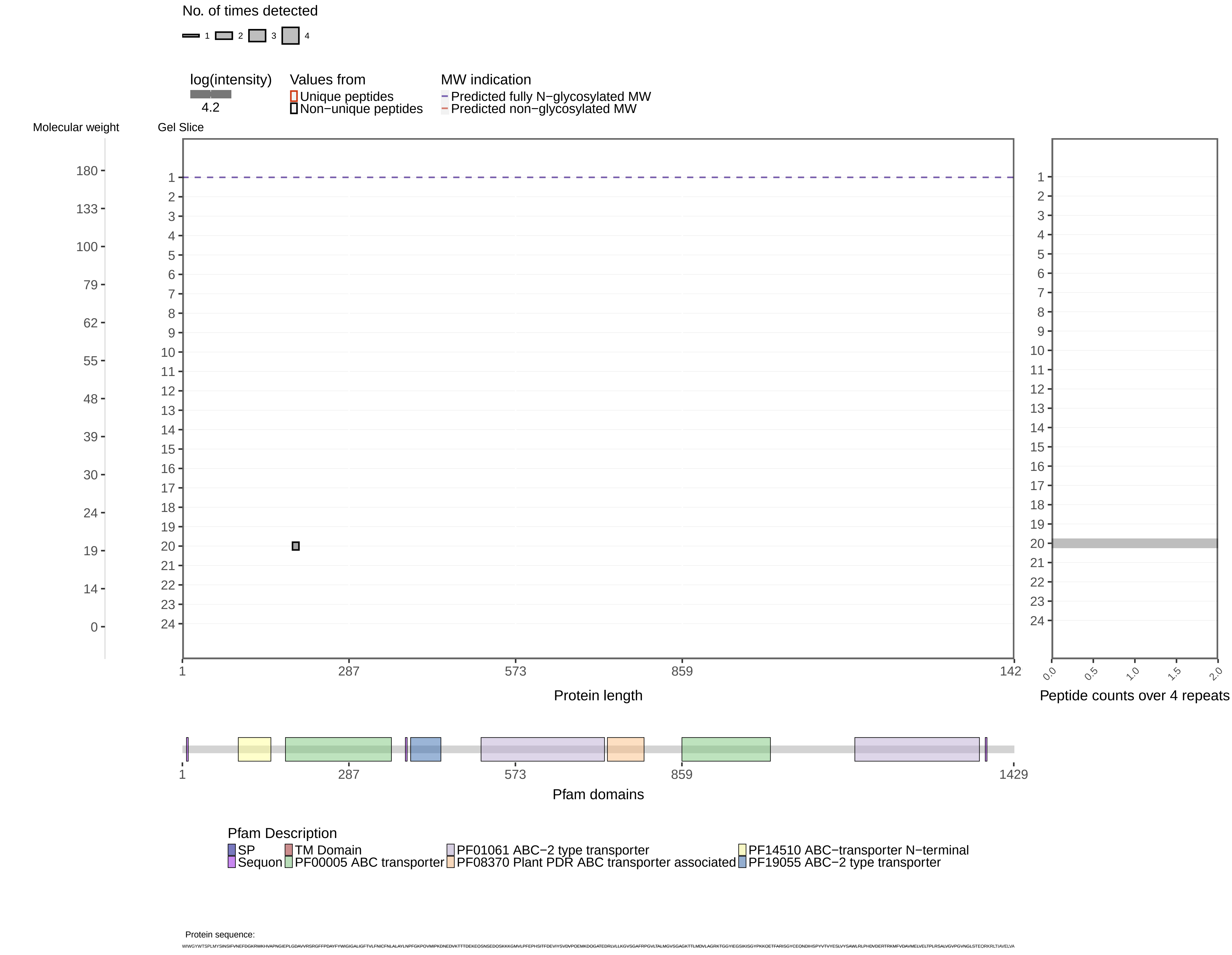

### NbL06g00260.1 selenoprotein_O−like [MW = 72.31 kDa]

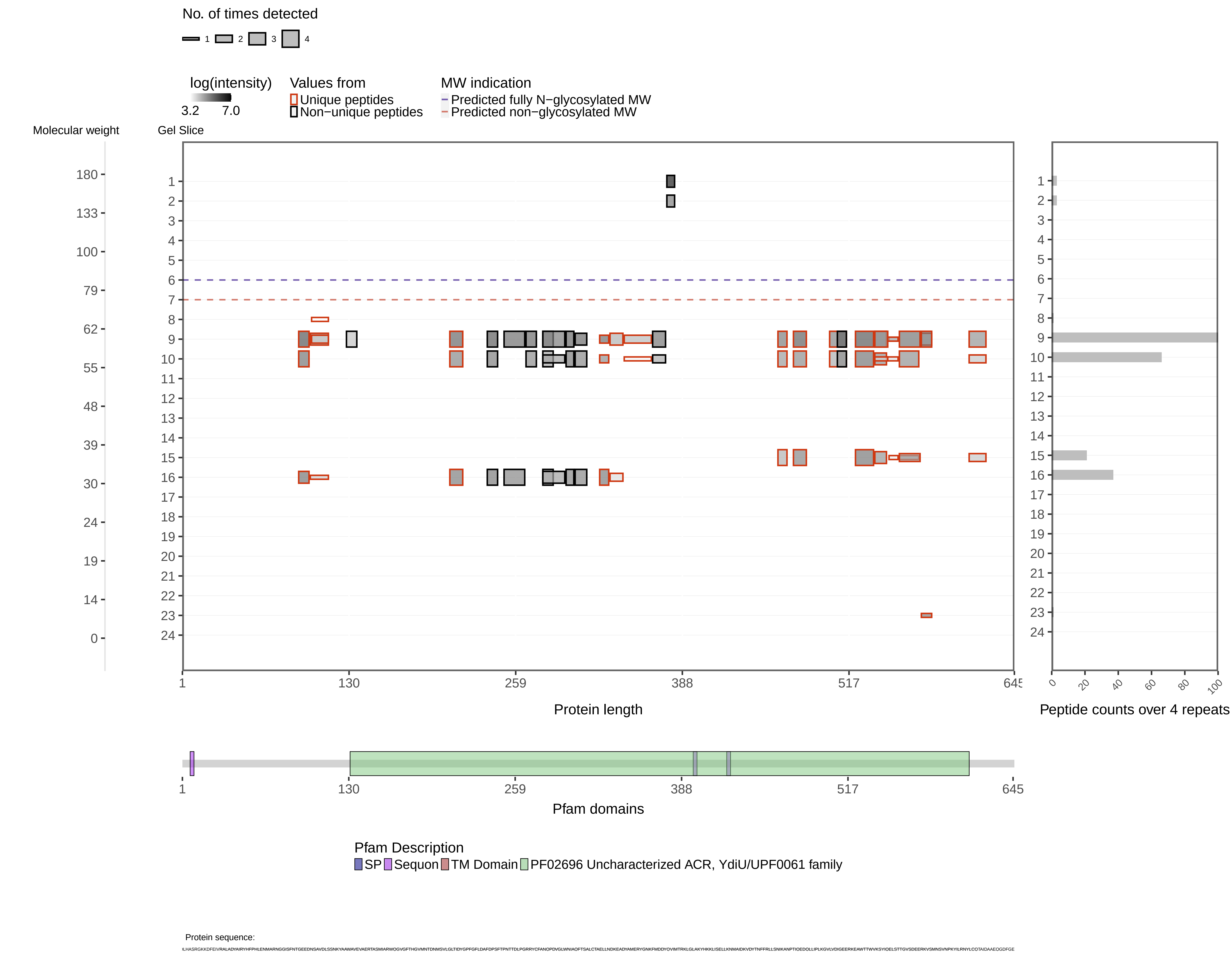

NbL06g00220.1 dehydrodolichyl_diphosphate_synthase_2−like [MW = 30.87 kDa]

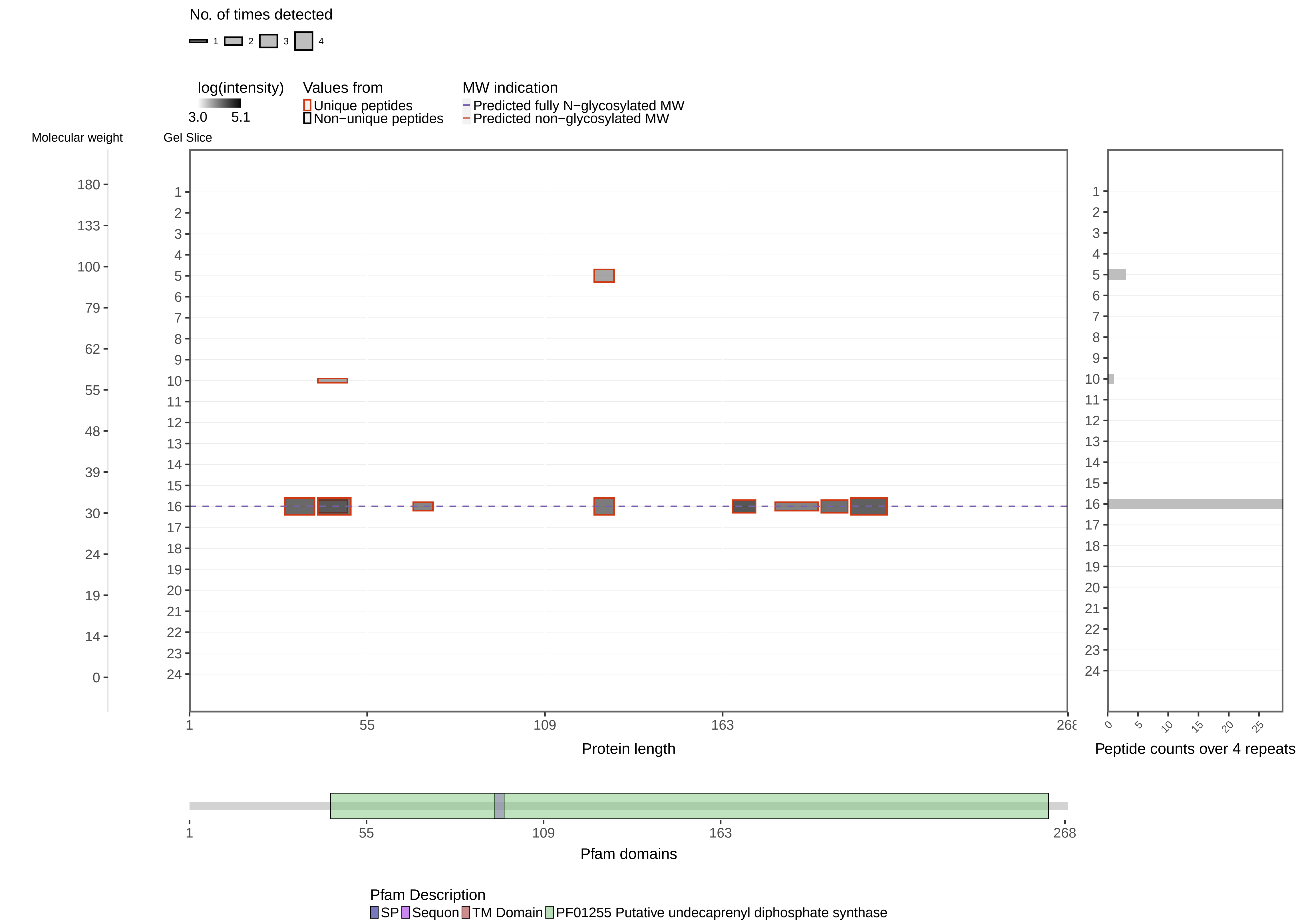


Protein sequence:

YGMKMSVIGDKSNLPNSLKEAINHAEETTKDNIGLHFVAALNYGGRYDITQATKNIATKVKDGVIQFEDINDKLIDQELETKNLEYPNPDLLIRTSGELRISNYMLWQLAYSEFCFVDKLFPDLDEADFVAALRSFQRRQRRYGGHKI*

NbL06g00150.1 DNA_(cytosine−5)−methyltransferase_CMT3−like [MW = 101.77 kDa]

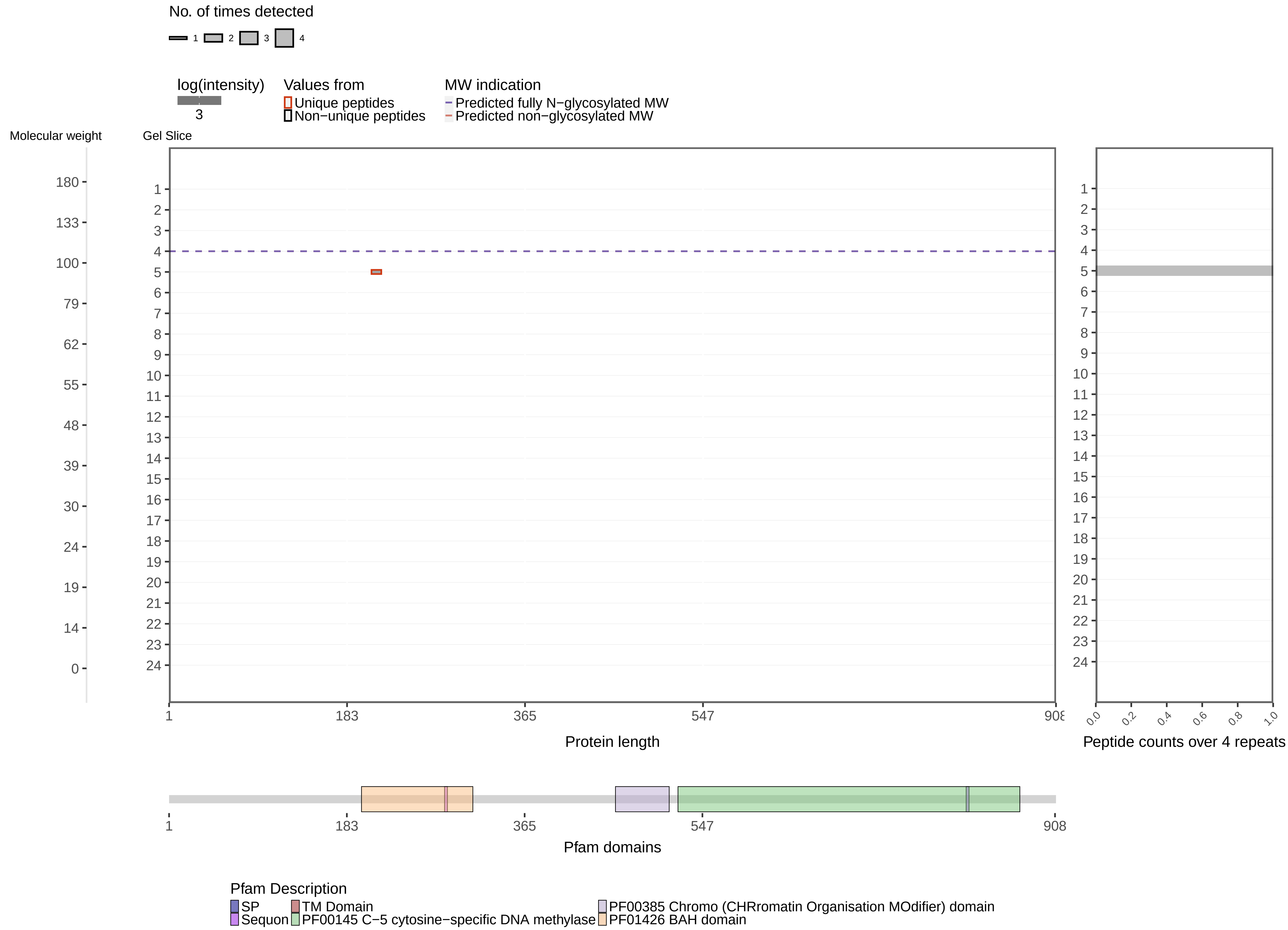


Protein sequence:

DDESEDSGDDKKGEIFEVEEILEVCYGDPNAVRKPGLYFKVRWQGYGPEEDTWEPIDGLSECPKKIREFVTKGFKANLLPLPGDVDVICGGPPCQGISGFNRFRNKENPLGDPKNQQLDVFMDIVDFLKPRFVLMENVVDLIKFSNGFLGRYALGRLVGMNYQARMGMMAAGAYGLPQFRMRVFMWGALSSEKLPQYPLPTHKVIVRGVIPTEFESNTVAFDEGMELELKKELFLGDALSDLPSVENNEQRDEMAYTNEPTSDFQHFIRLGRDGALGSVLYDHRPLQLN

### NbL06g00110.1 spermidine_coumaroyl−CoA_acyltransferase [MW = 49.93 kDa]

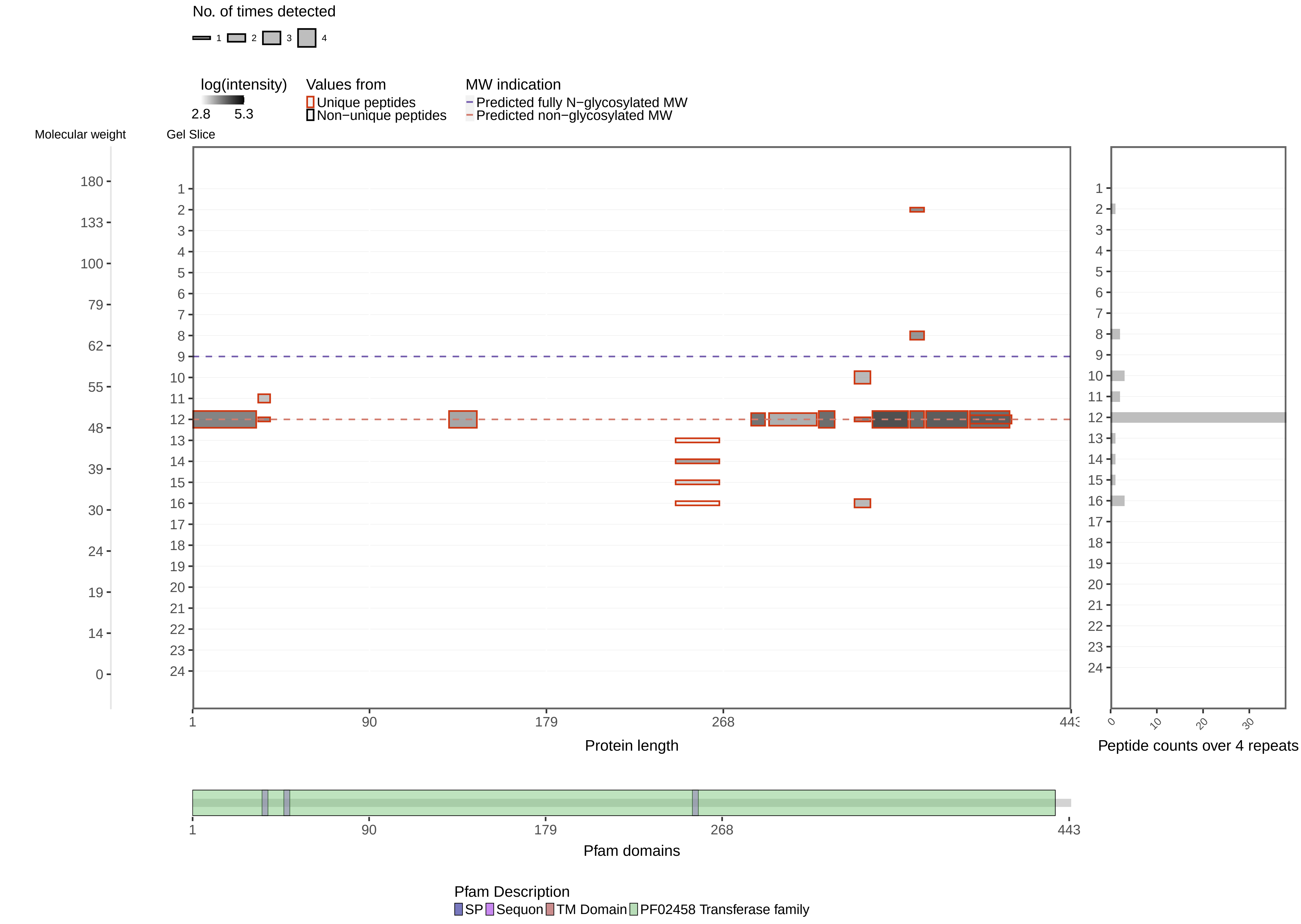


Protein sequence:

[illegible]EFFSLAKDSVPYLGSGKPAVRKCFEVKDEWLDRLKGFLHEQSGLNFTTFEALGAFIWRAKAKASKIPGDEMVKYAYLTNIRRTVKPALPAGYWGNGCVPIYVQLLAKDLINQPIWKTADAIKKSKSIITDEYVRSLIDFQELHYDEGITAGNRVSAFTDWRHVGHETVDFGWGGPVTVFPLSRHLVGSVEPCFFLPYSSATEGKKDGFRVLVCLQEEAMPVFKEEMKQLEHDLS*

### NbL05g22310.1 cysteine_proteinase_inhibitor_1−like [MW − SP = 9.97 kDa]

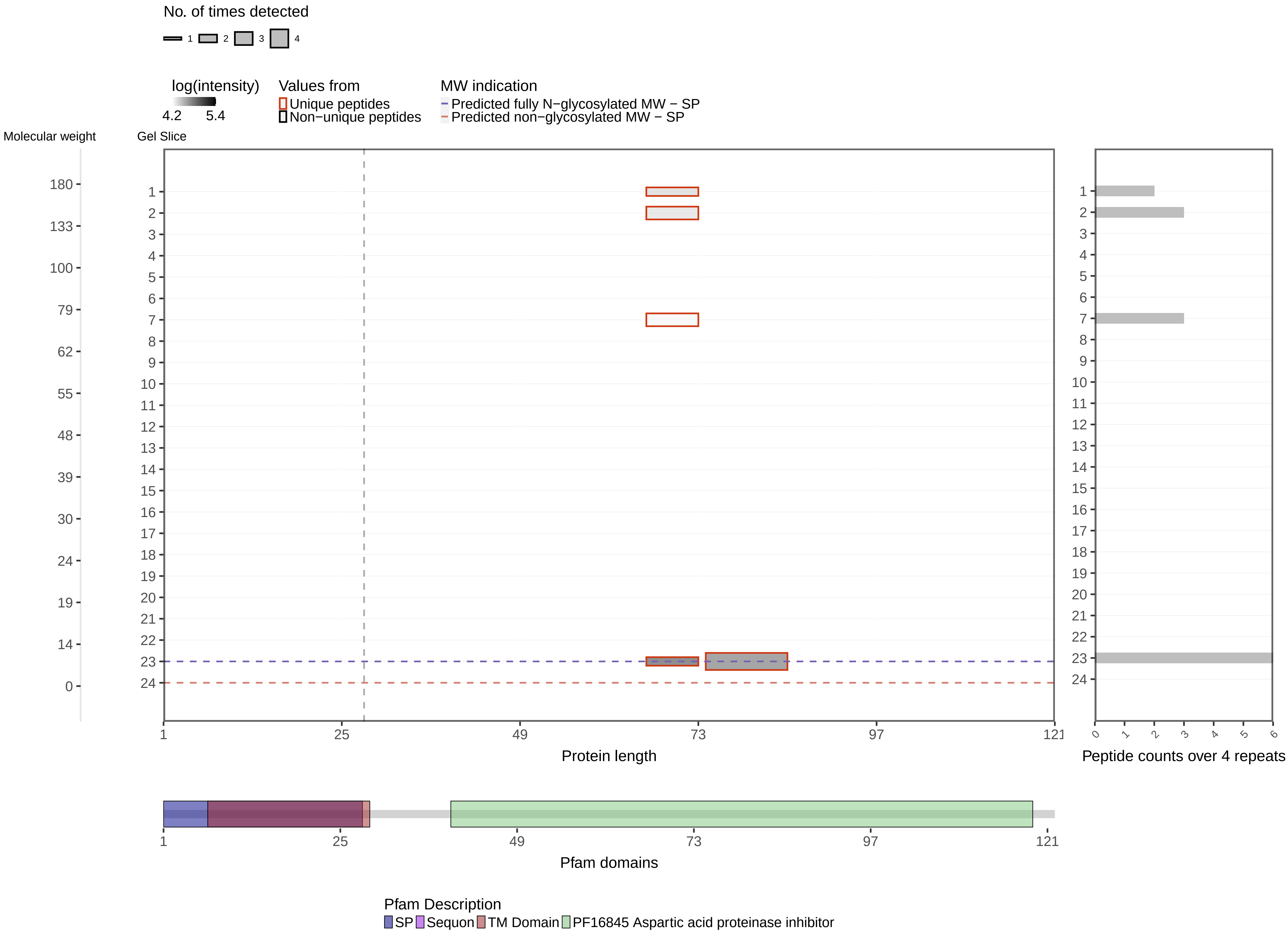


Protein sequence:

IGQFAVNEHNKAAGTKLEYQNIAKGETQVVAGTNYRLVINAKDGGSVHKYLAVVWDKPWEKVKKLTSFKKM*

NbL05g22270.1 induced_stolen_tip_protein_TUB8−like [MW = 16.18 kDa]

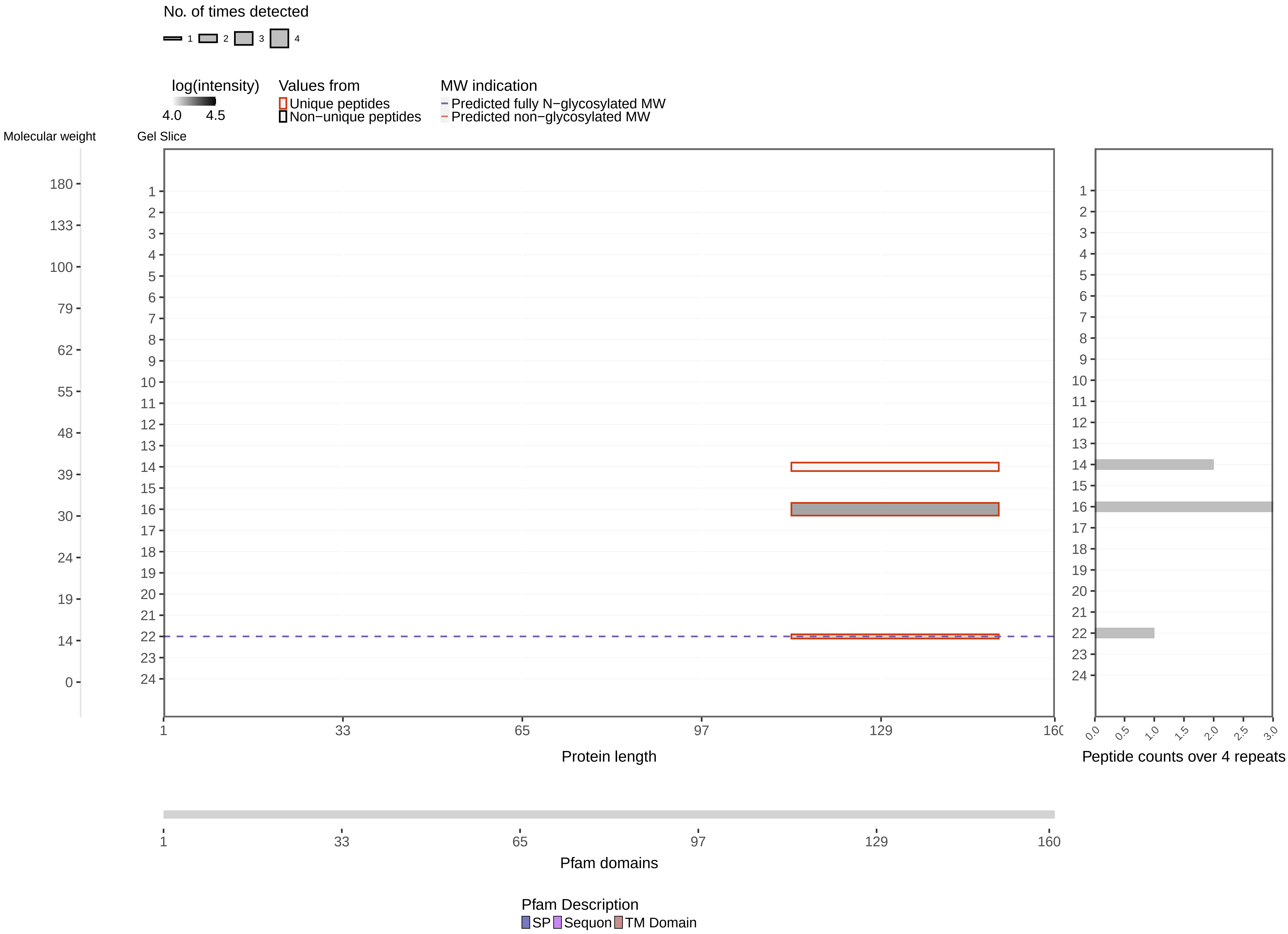


Protein sequence:

EPVAAAEPATTETQEPSTVGPVEEVAAPVDEVELEAAPVEEPETEKAVPVAEESAPVTEEAVAEETINAPAEEPVAAAESVEEKAVPVEKAEE*

### NbL05g22200.1 fumarylacetoacetase [MW = 50.04 kDa]

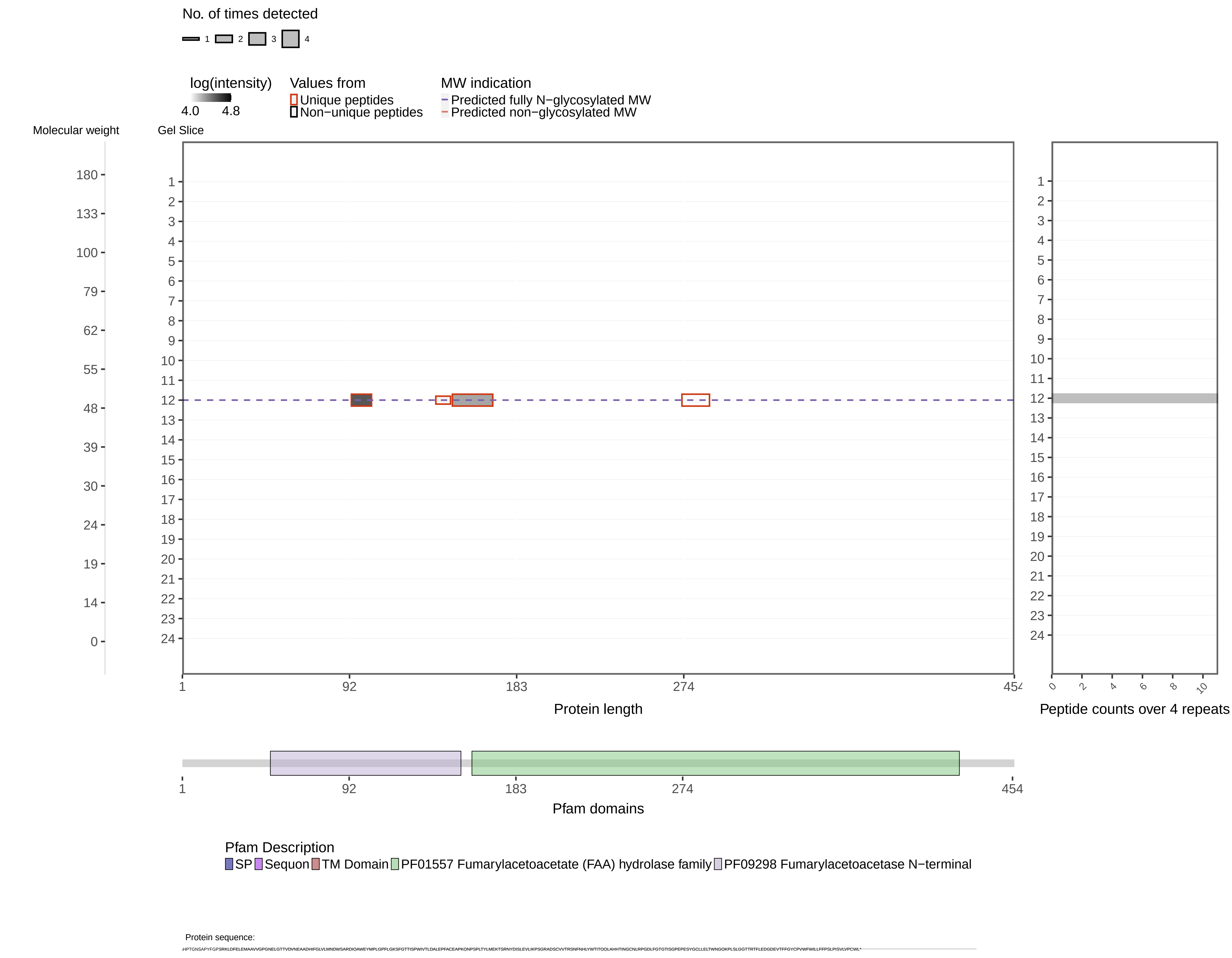

### NbL05g22170.1 malate_dehydrogenase_[NADP]_chloroplastic [MW = 48.4 kDa]

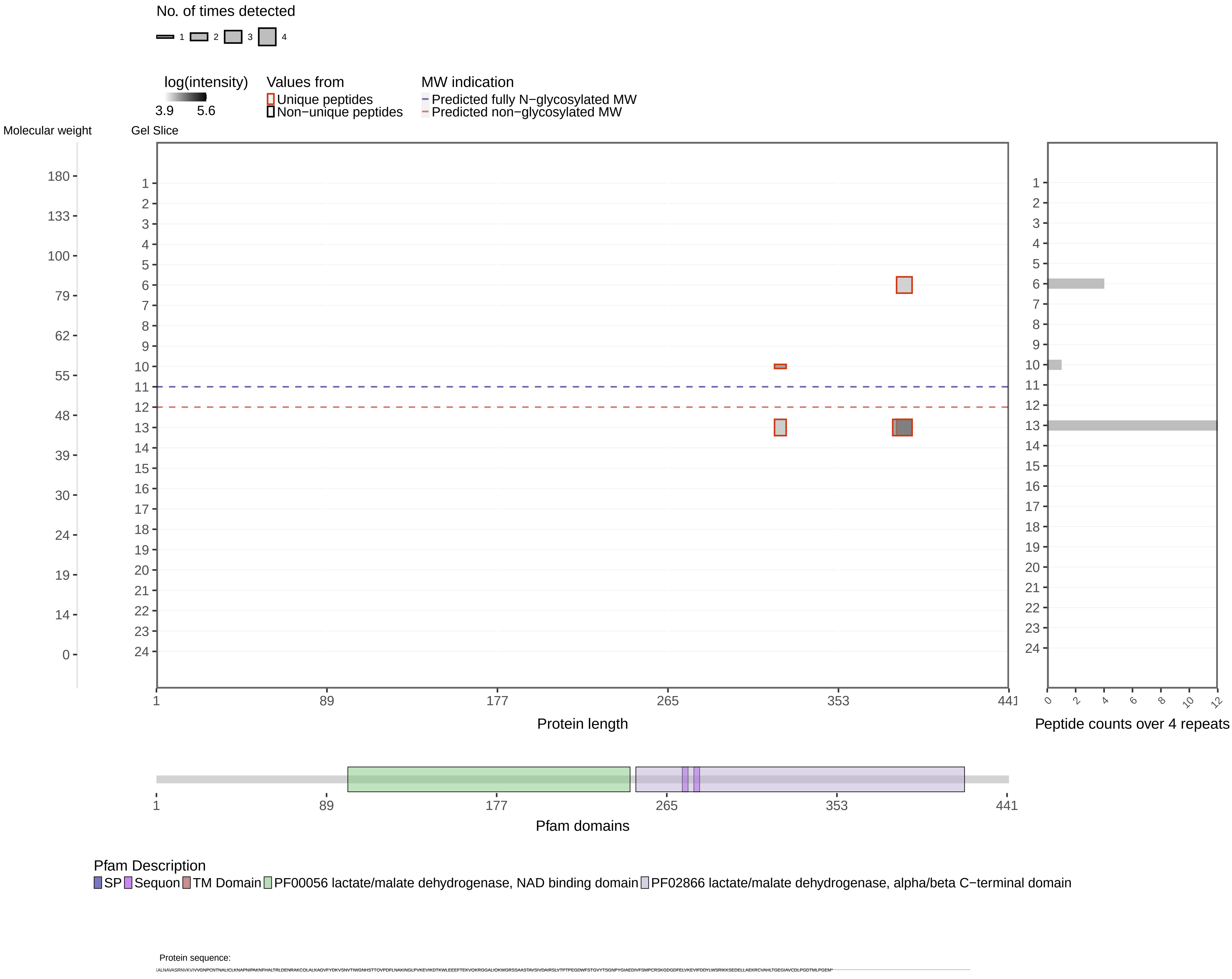

NbL05g22140.1 pullulanase_1_chloroplastic−like [MW = 105.99 kDa]

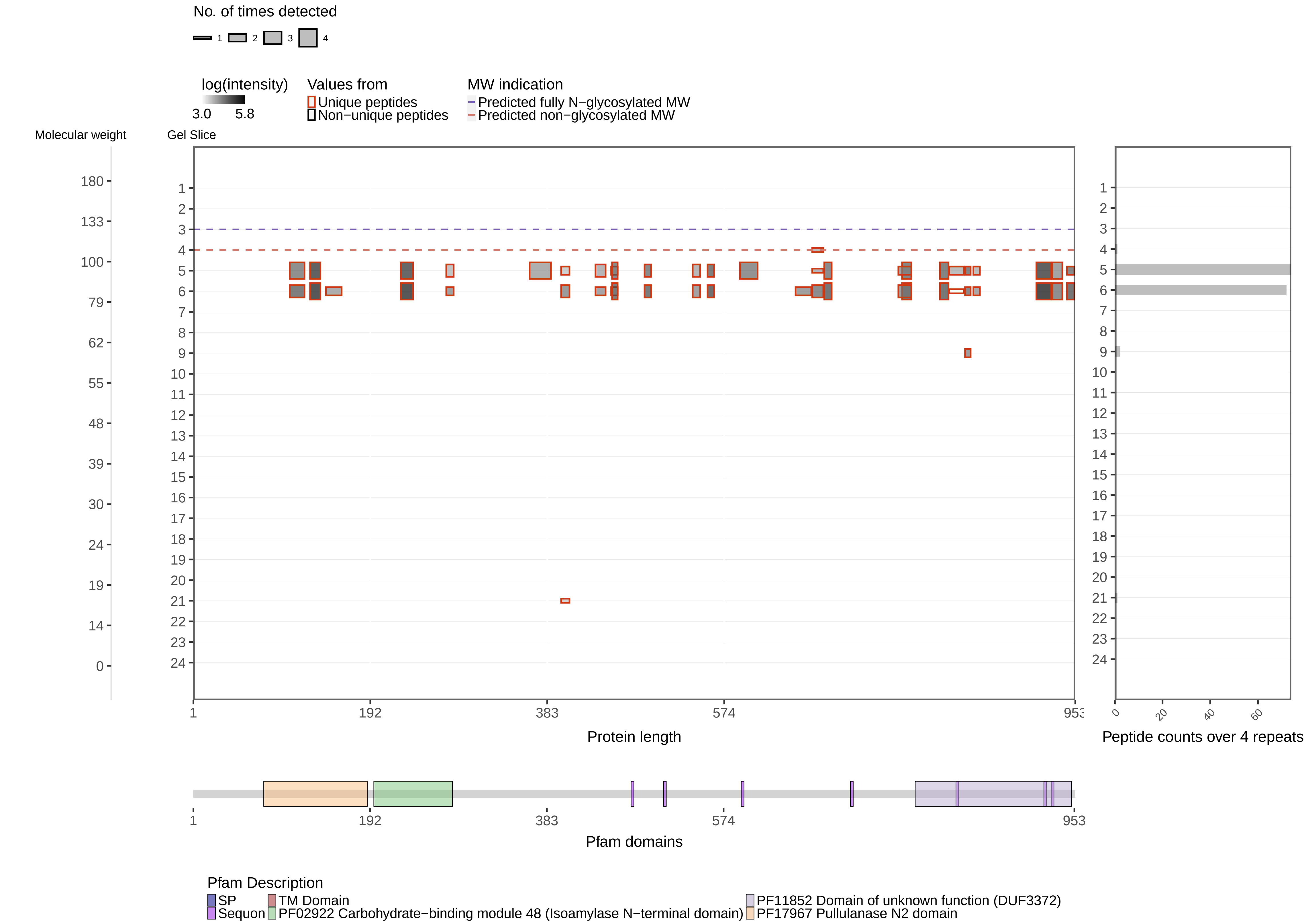


Protein sequence:

### NbL05g22050.1 serine_carboxypeptidase−like [MW − SP = 53.21 kDa]

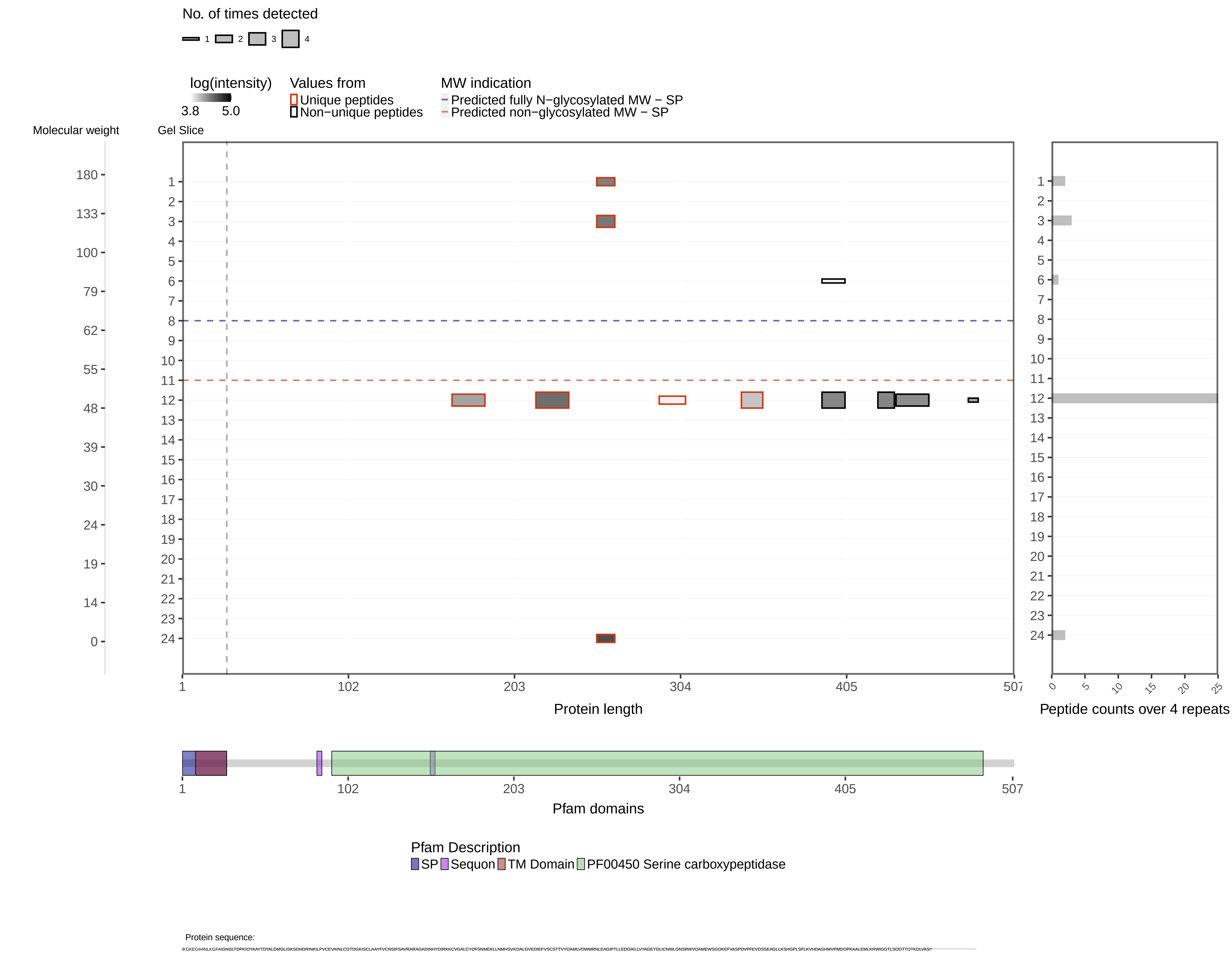

### NbL05g21990.1 kinesin−like_protein_KIN−UA [MW = 121.11 kDa]

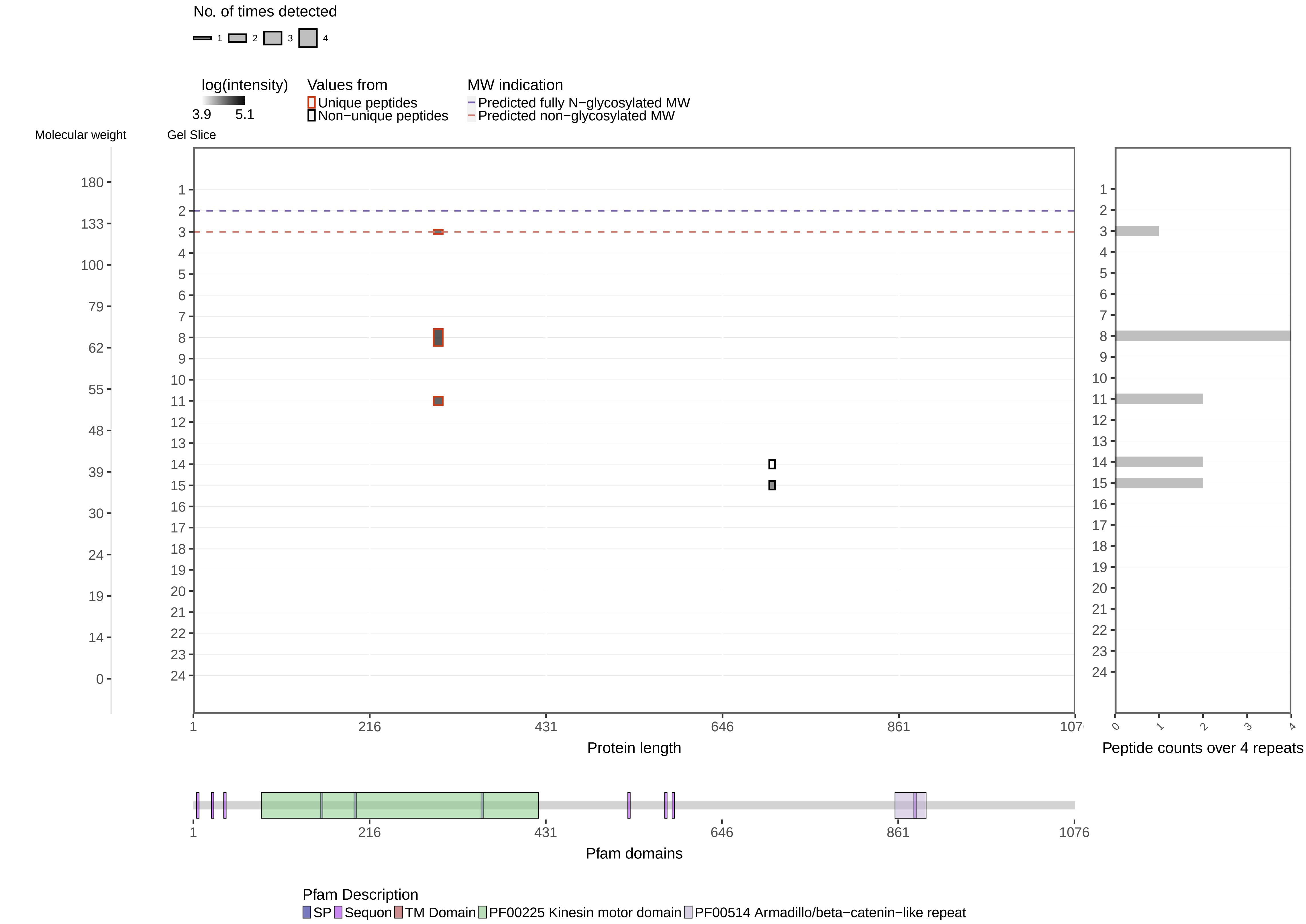


Protein sequence:

IVKLESDFNNSKLLEKENGCLKVELENVLKDFNREKIHNNLLQDEVVHLERSLNQSKQQQLENSSYQKILAENIQSYEKKIADLMKQLEDERAQSENAQQQLELMKEQLTGVEELMQHHQKQTSIYQKELADTTLVYEEKIAQLEQLLKDEHSHVENVKEQLHAIEEQFTDHETSVKIQREKEIDALRSRLEDMHQLYEPTVEELQTLKTEYQVLVSKKKALHDELHEVRQTLLIEEKQRKAAENELFNMKKLVPESEDGFEDKKSYMKQYTTSGSFNMHRSTESRERIFAHC

### NbL05g21970.1 hydroxymethylglutaryl−CoA_lyase_mitochondrial−like [MW = 50.25 kDa]

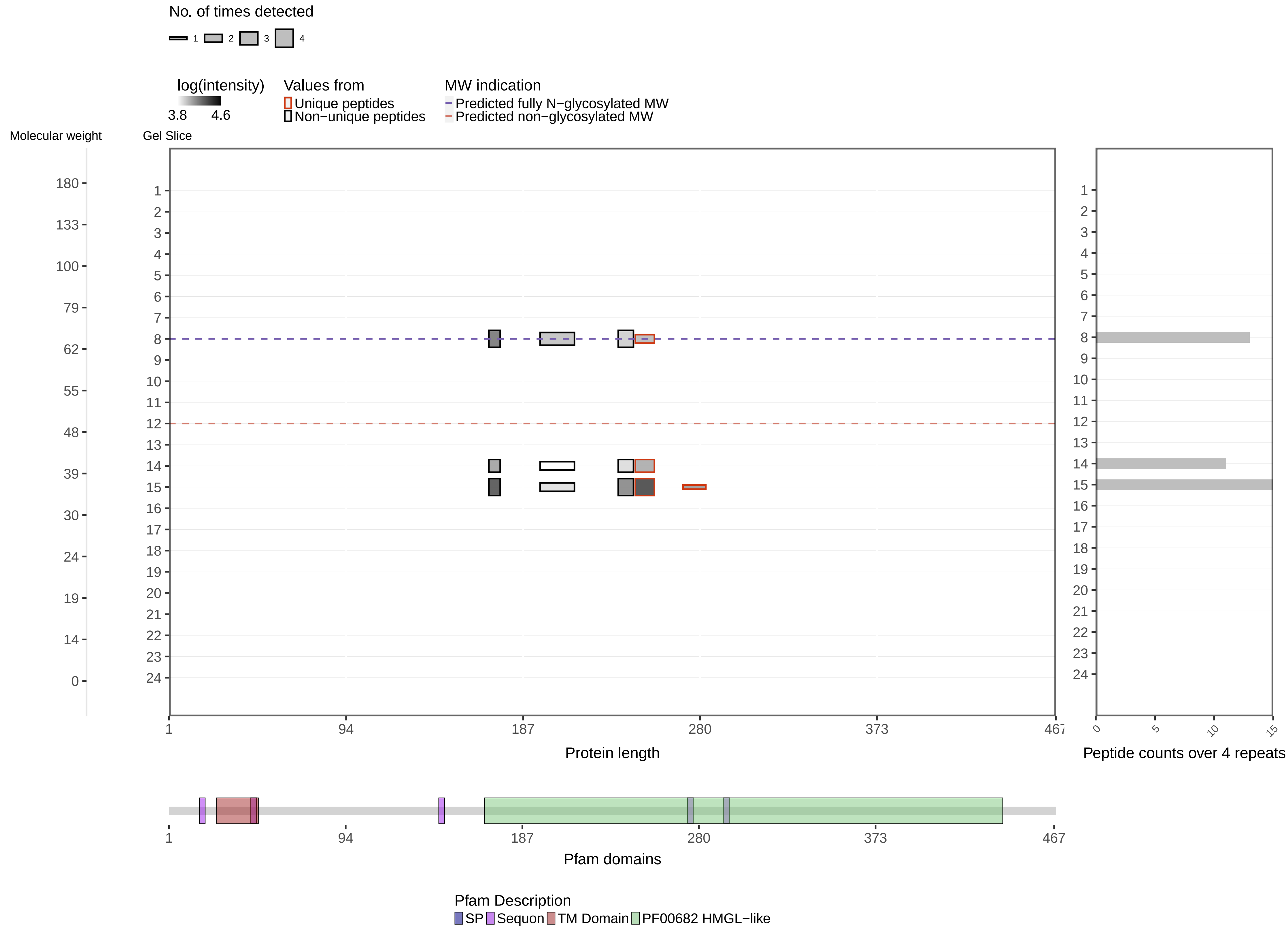


Protein sequence:
ADAKDVIEAVRSLTGVRLPVLTPNLKGFETAVAAGAKEVAIFAAASESFSRSNINCSIEDSLARYRDVALAAKNHSIPIRGYISCVVGCPVEGAVSPSKVAYVAKELIDMGCFEISLGDTIGVGTPGTVIPMLDAVTQVVPVERLAVHFHDTYGQALSNILVSLQMGISTVDSSVSGLGGCPYAKGASGNVATEDVVYMLNGLGVKTNVDMRKLLSAGDFICKHLGRPSGSKAAVALRRKTSTASKL*

### NbL05g21760.1 thioredoxin−like_3−2_chloroplastic [MW = 21.96 kDa]

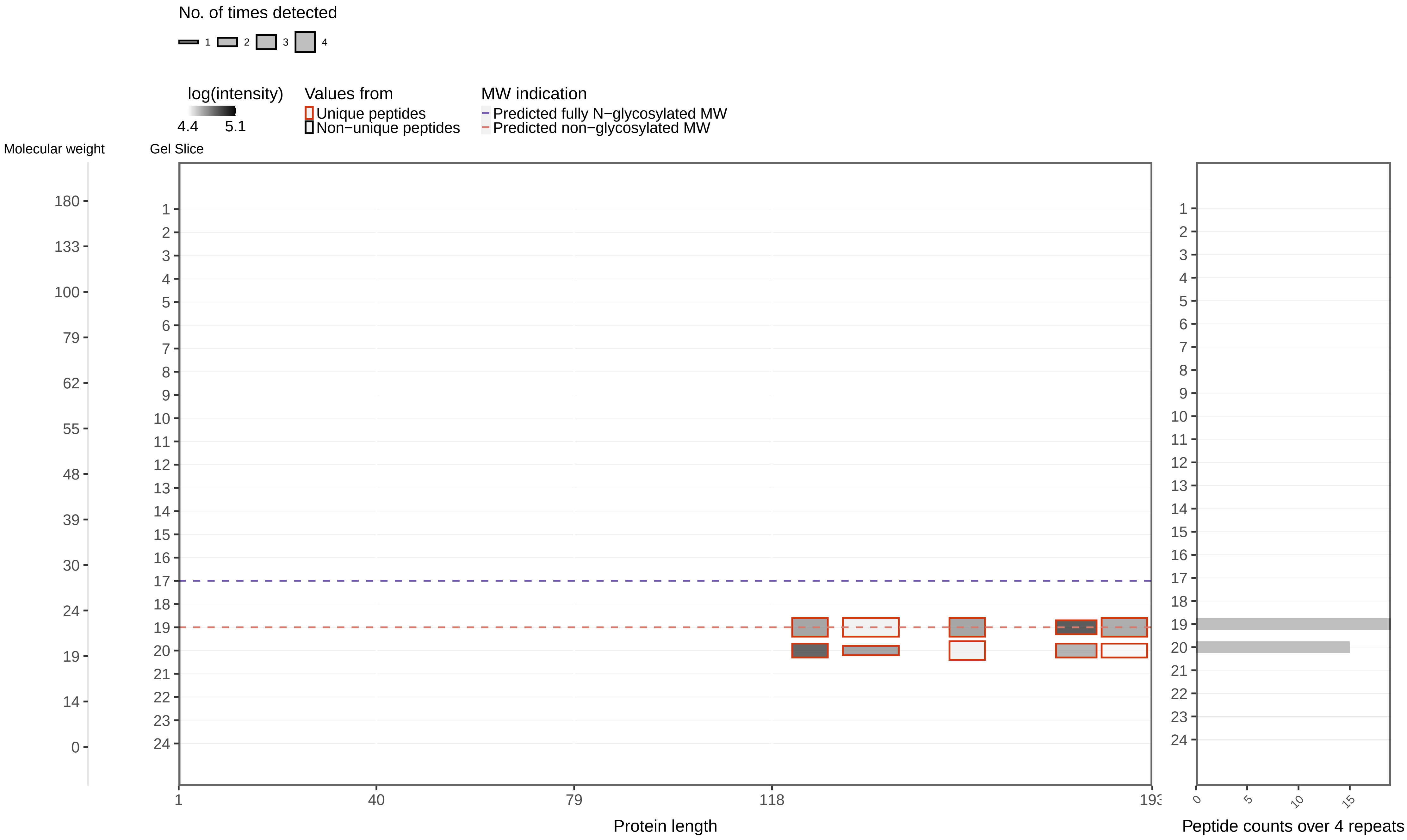


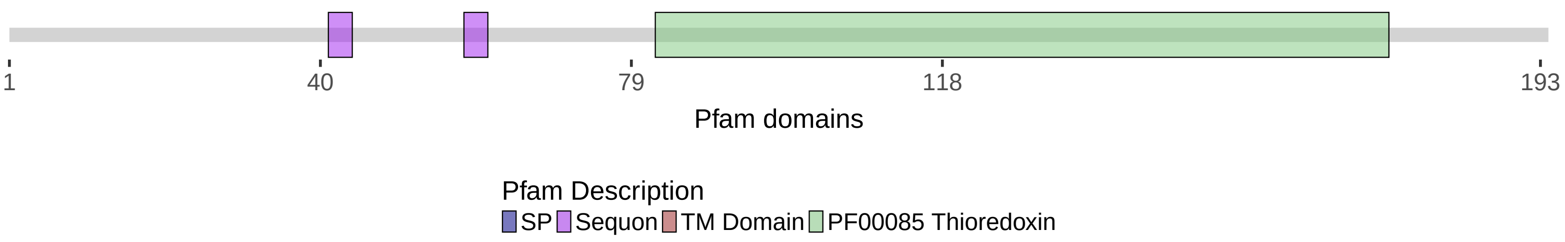


Protein sequence:

AQFDRVISEAQQIDESVVILWMATWCRKCIYLKPKLEKLAADYFPRTRFYCVDVNNVPHKLVVRAGITKMPTIQLWRDGNKQAEVIGGHKAYLVVTEVRDMIENEDNNL*

### NbL05g21730.1 actin−related_protein_7−like [MW = 39.68 kDa]

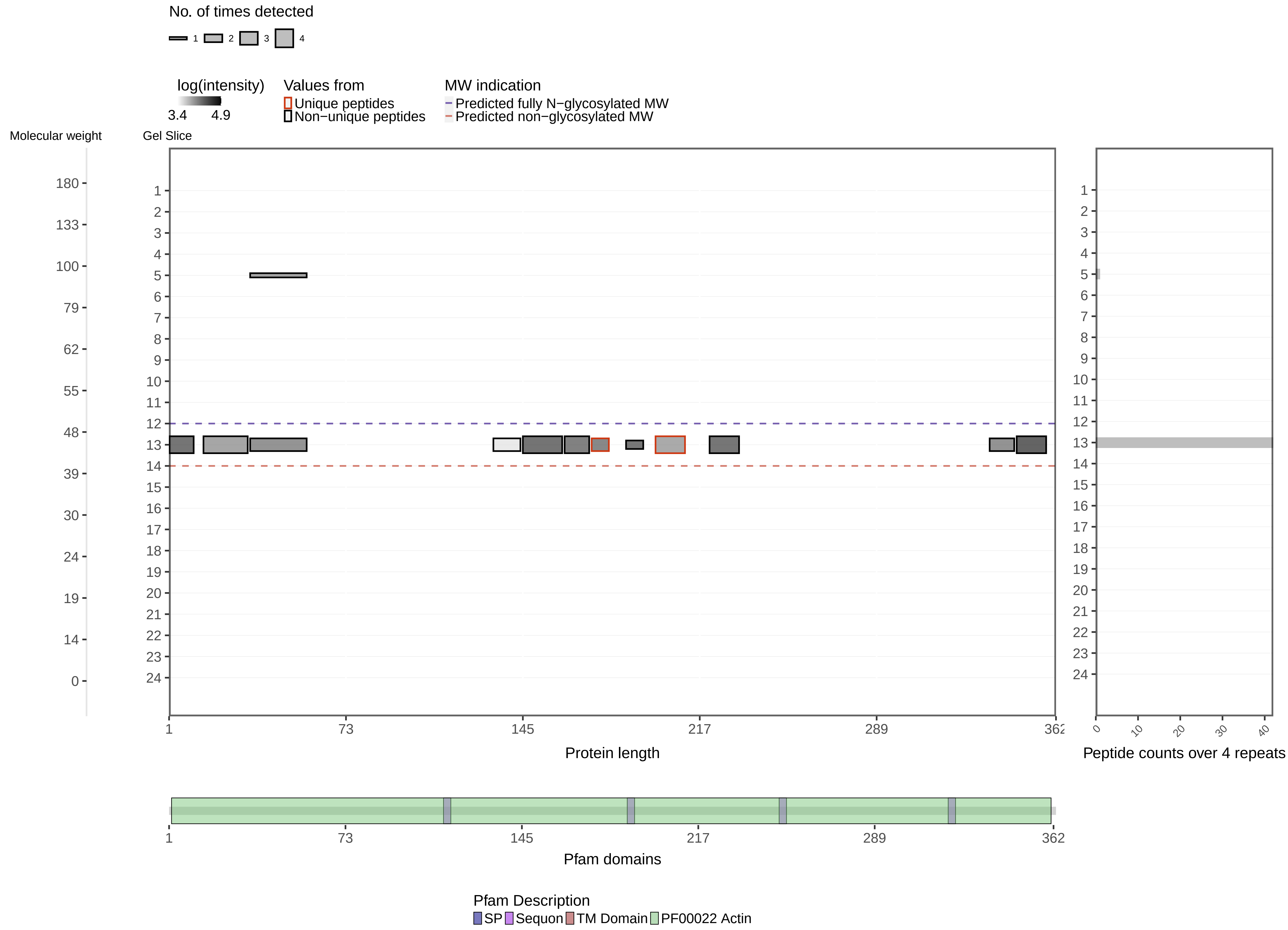


Protein sequence:

.DLTKLLADELTKSNPTVKLNISDVEKLKERYACCAADDIAYEKLOOSCLEEKHTLPDGOVITIGKERYTVGEALFOPSILGNDTHGIVEOLVHSISSVSSENHROLLENTVLCGGTATITGFEDRFOREAMLCSSAVRPSLVKPPEYMPENLTSYSAWIGGAILAKVVFPONOHITKADYDESGPSVVHRKCF*

### NbL05g21620.1 phosphoglycolate_phosphatase_1B_chloroplastic−like [MW = 40.92 kDa]

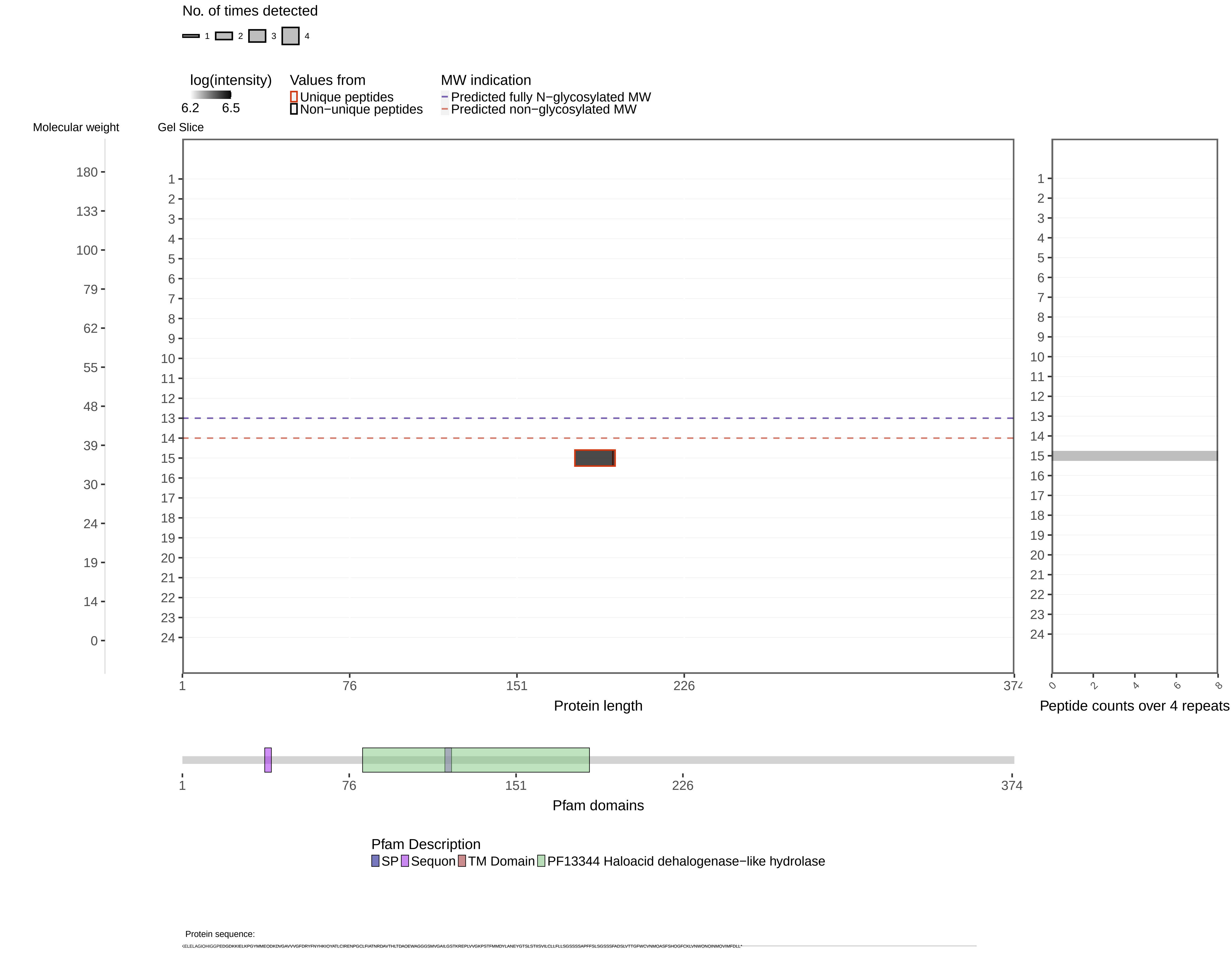

### NbL05g21550.1 protein_transport_protein_Sec24−like_At4g32640 [MW = 117.58 kDa]

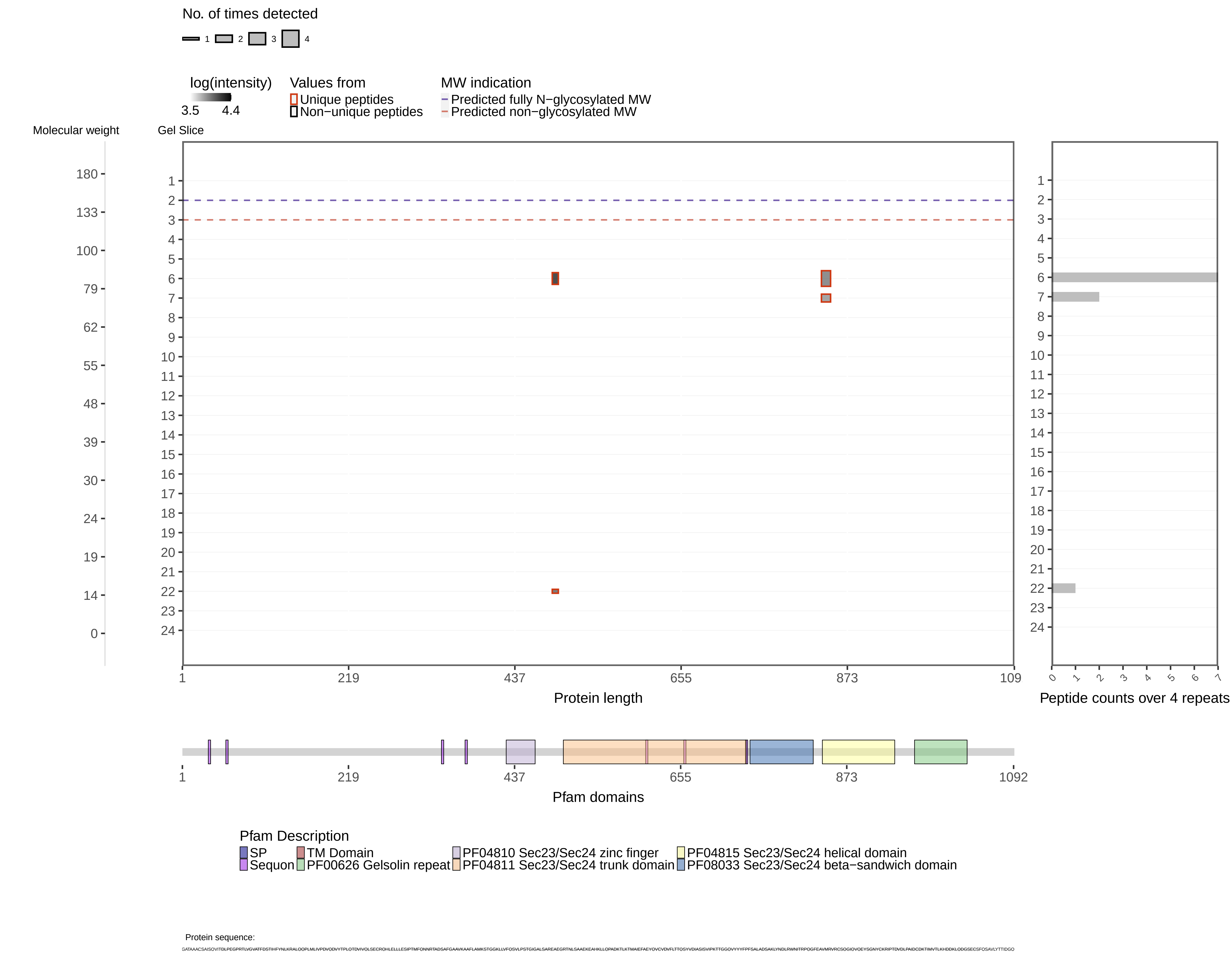

NbL05g21480.1 NADH_dehydrogenase_[ubiquinone]_iron−sulfur_protein_2−like [MW = 25.91 kDa]

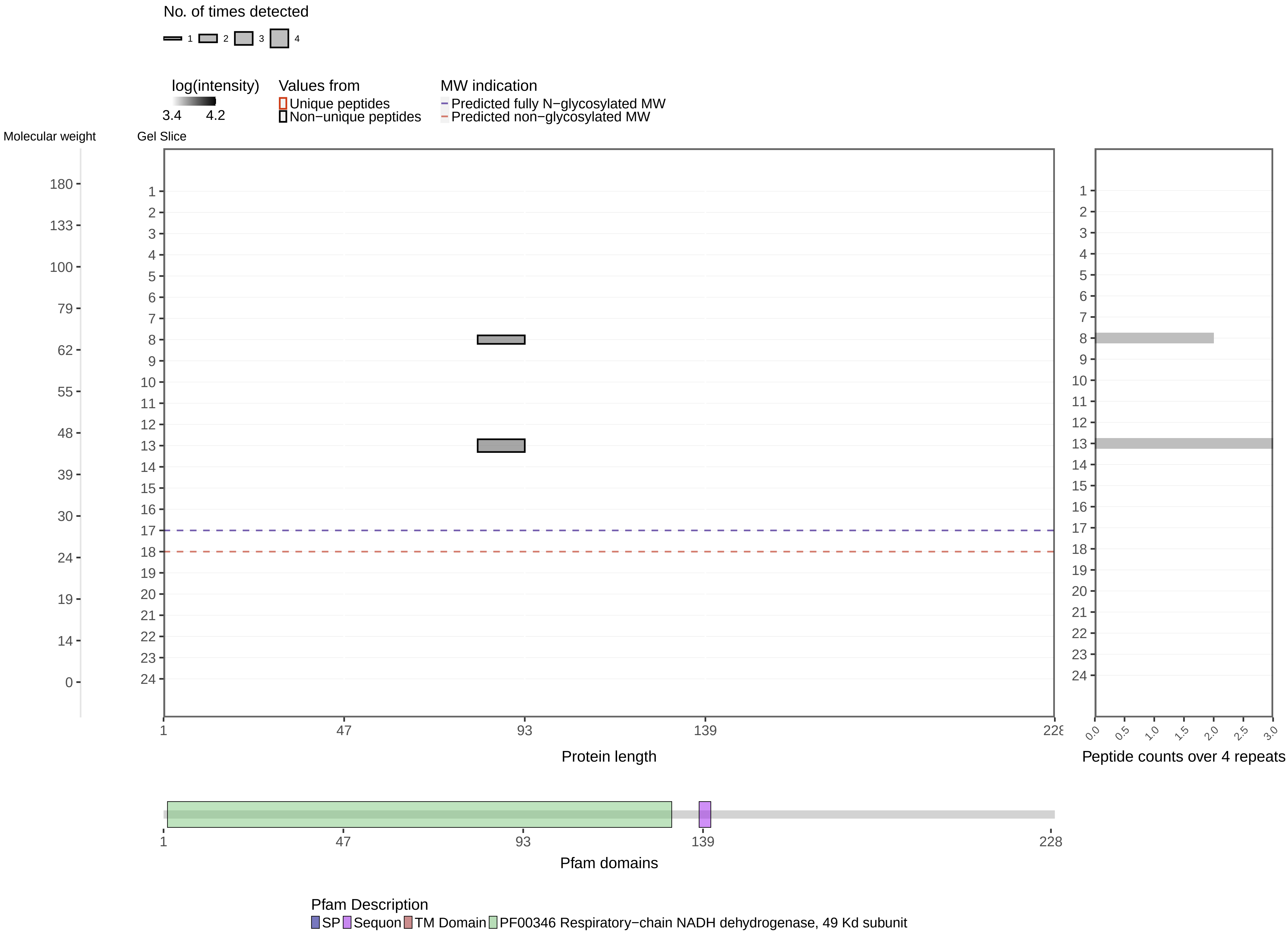


Protein sequence:

/MLKGSRICWDLRKAAPYDVHNQLDPDIPVAMSSITISNETIHGIVRRVKRISLPFLTETQVKLRLGTLWVGCWHNRVVPYRTAGGRAASFVVVLQDSNVLRPRSERAYRWTTAVPFLSELEHNHLIH*

NbL05g21230.1 acetolactate_synthase_small_subunit_1_chloroplastic [MW = 54.43 kDa]

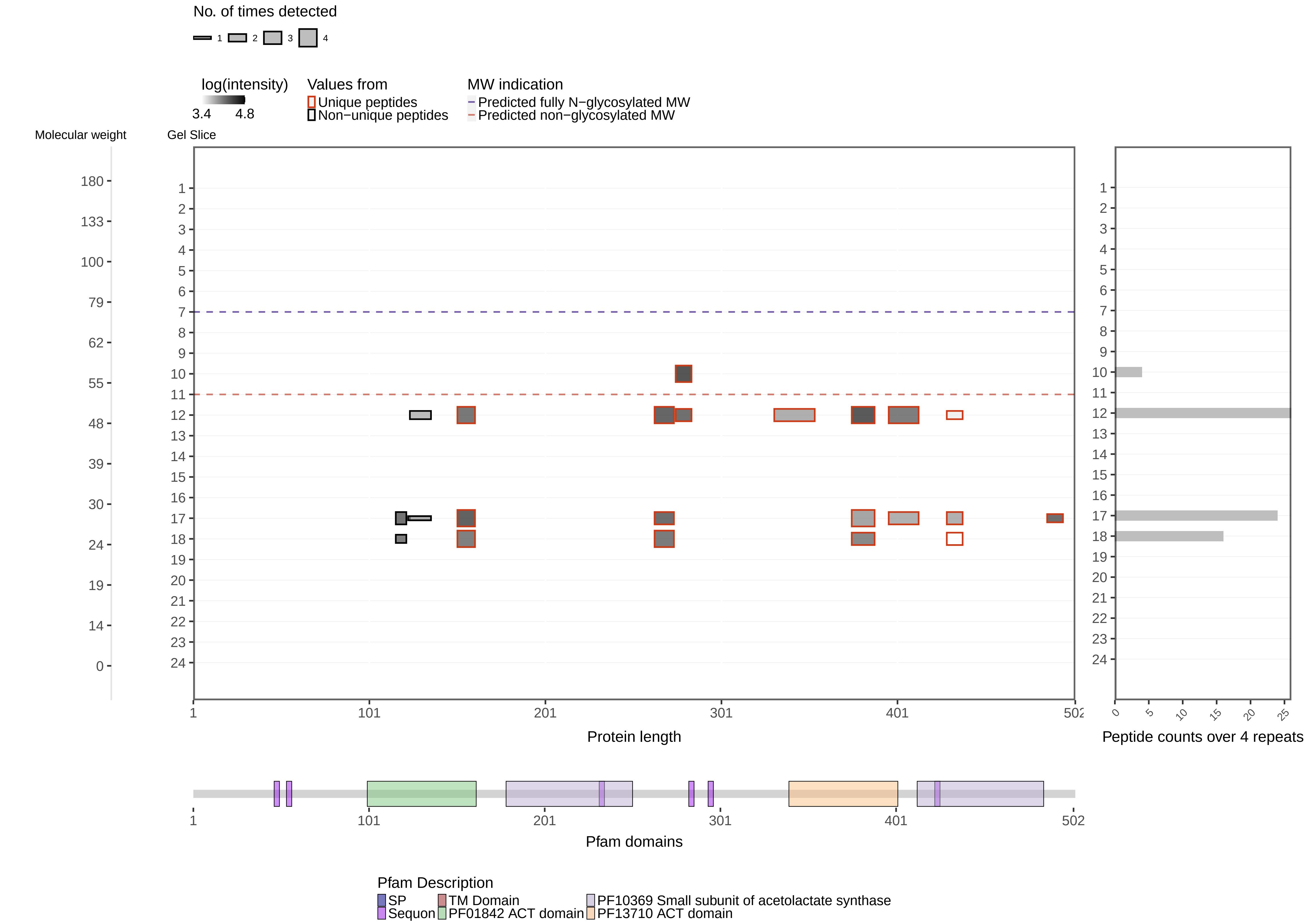


Protein sequence:

IKELARTGKIALRREKMGEDAPFWRFSAASYPDLEEKIPSVGILNNKSRSIPNEASNDSRGDVYPVESSDDISVQVLDAHWGYLYDGDLSGLRSHTISMLVNNAPGVLNLITGVISRRGYNVQSLAVGPAESKGLSRITTVVPGTDETIGKLVKQFYKLVDIHEVQDITHLPFAERELMLIKVAANATARRDVLDIASIFRAKPVDVSDHTVTLELTGDFNKLLALQRLLEPYGICEVARTGRVALVRESGVDSTSLRGYPLPL*

### NbL05g21060.1 membrane−associated_protein_VIPP1_chloroplastic [MW = 35.56 kDa]

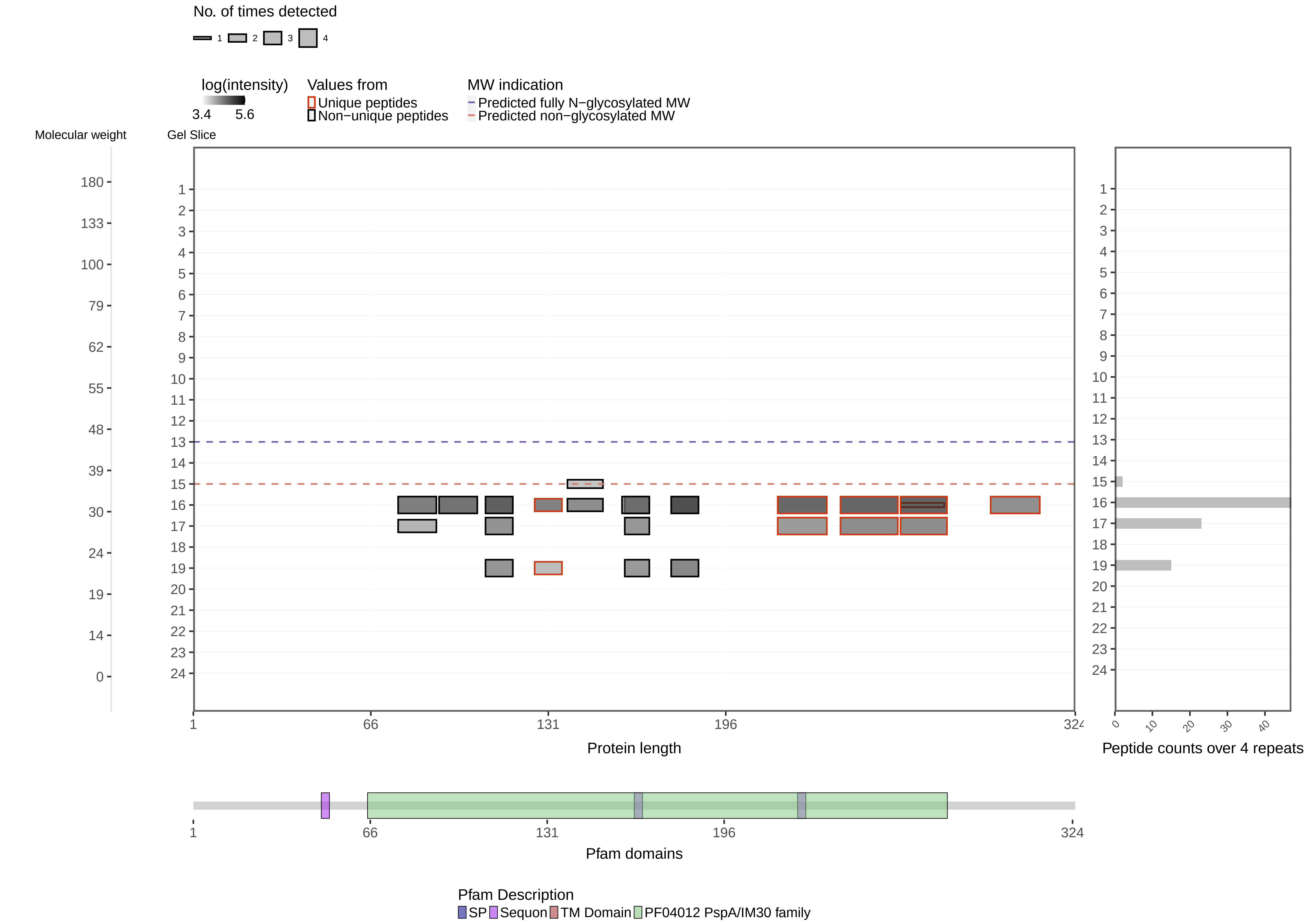


Protein sequence:

### NbL05g20990.1 haloacid_dehalogenase−like_hydrolase_domain−containing_protein_At4g39970 [MW = 34.41 kDa]

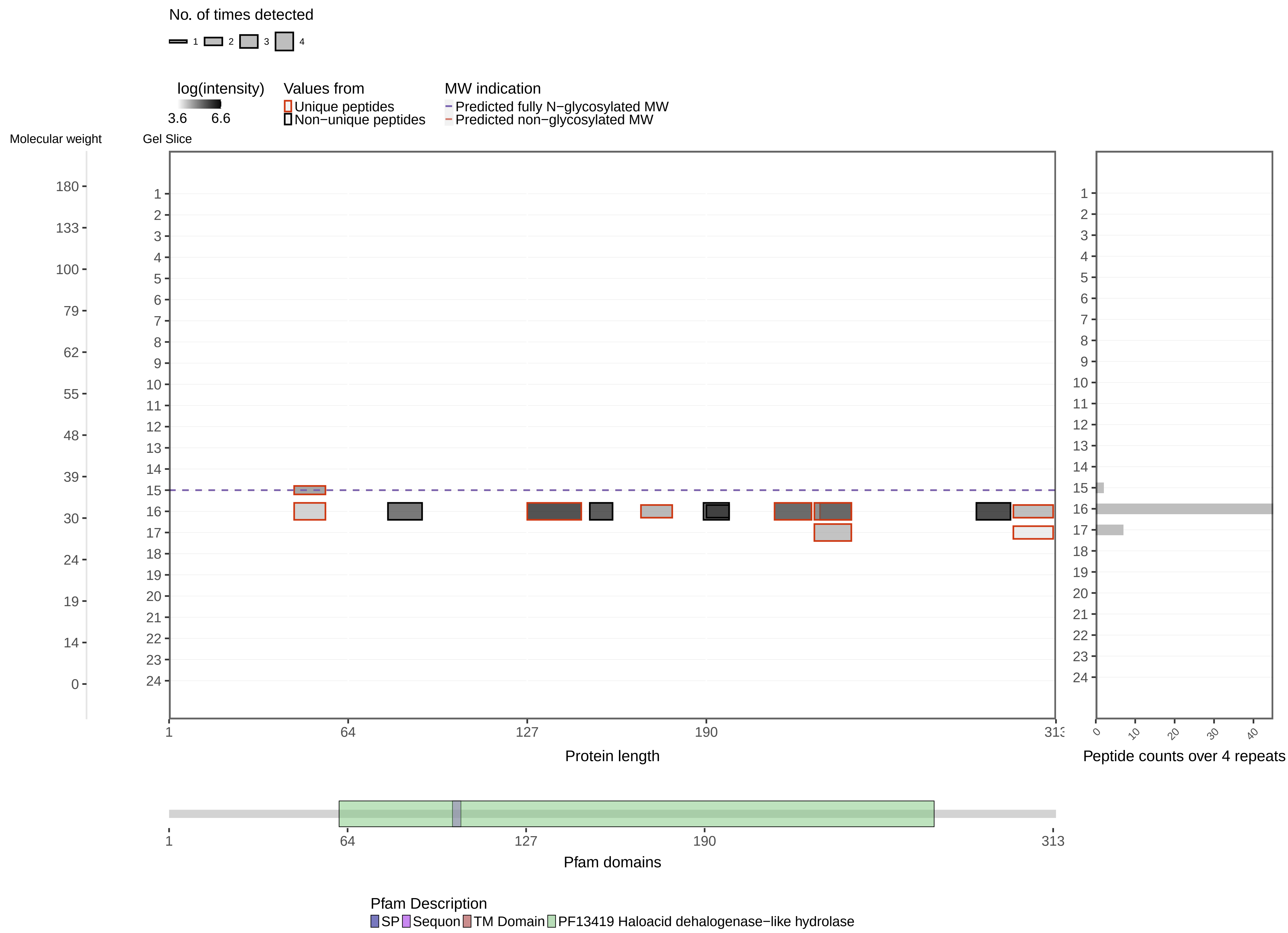


Protein sequence:

[illegible]DSNRAKLIDTLQDWKTERYKQIIKSGTVKPRPGVLRLMDEARAAGKKLAVCSAATKSSVIMCLENLLGLDRFHGLDCFLAGDDVKVKKPDPSIYITAAKKLGVSESNCLVVEDSVIGLQAATGAGMSCVITYTSSTADQDFKEAIAIYPDLSDVRFNDLESLLQDVVITK*

### NbL05g20920.1 phospho−2−dehydro−3−deoxyheptonate_aldolase_2_chloroplastic [MW = 57.49 kDa]

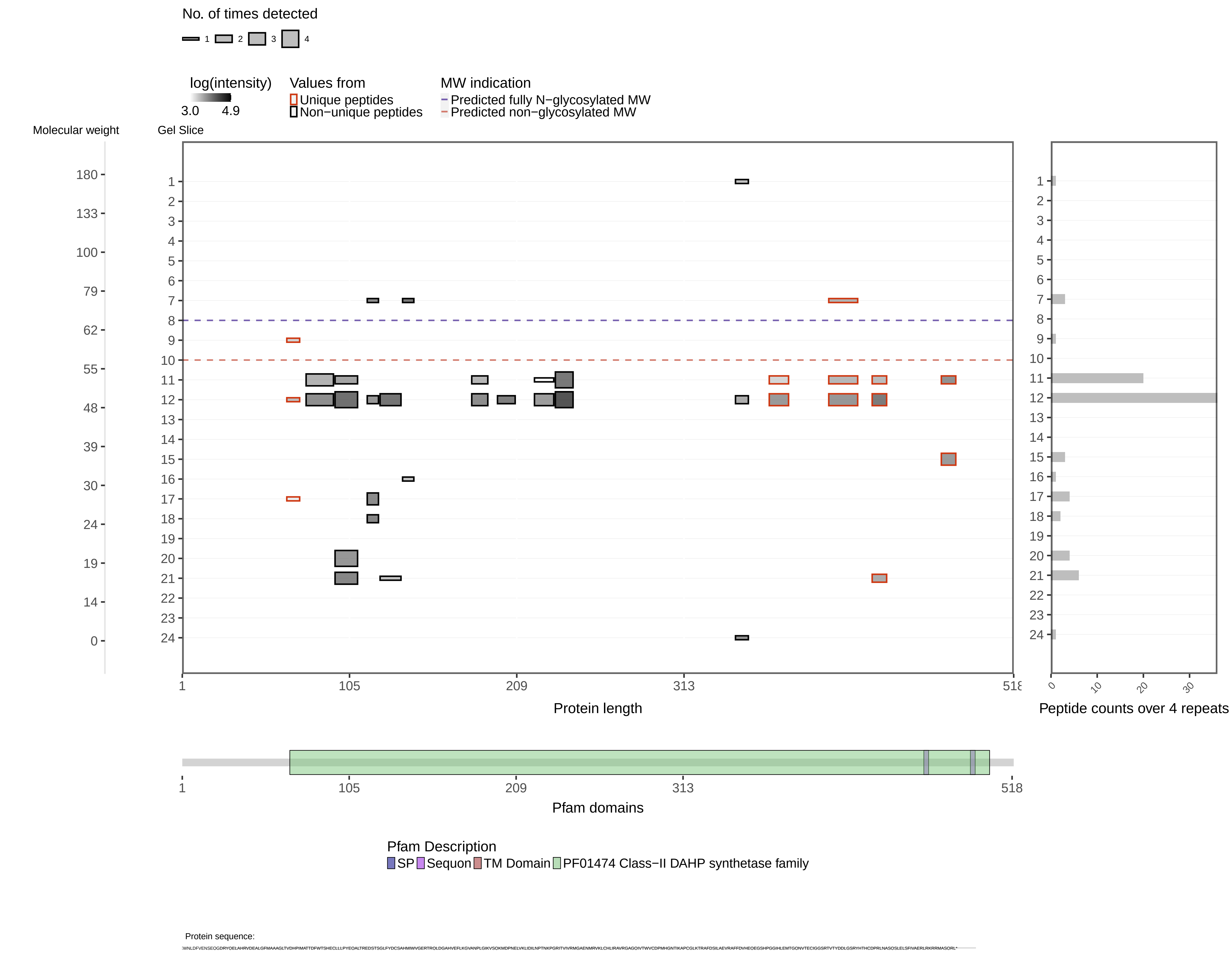

NbL05g20870.1 tRNA_(cytosine(34)−C(5))−methyltransferase−like [MW = 100.42 kDa]

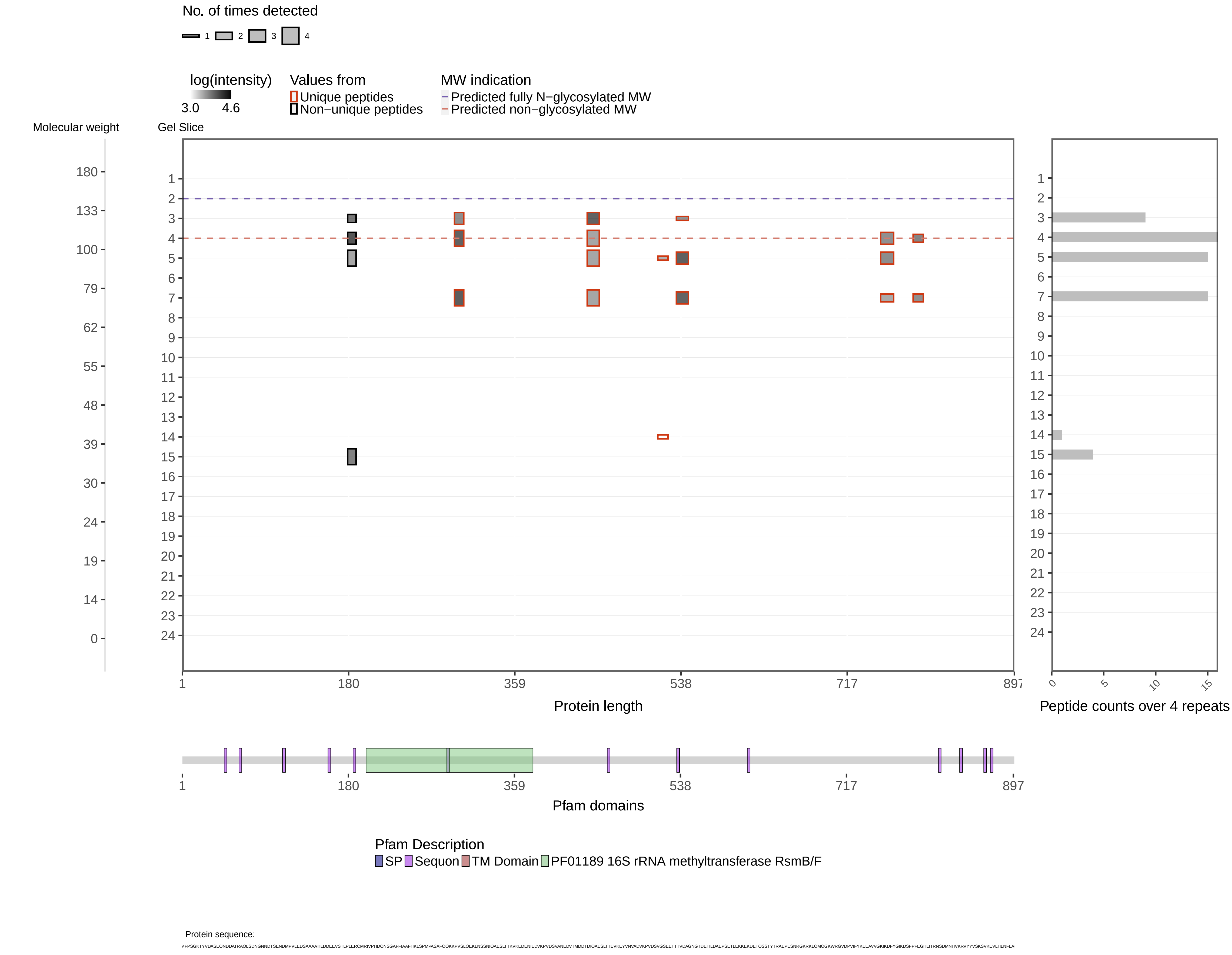

NbL05g20860.1 peroxidase_17 [MW − SP = 35.31 kDa]

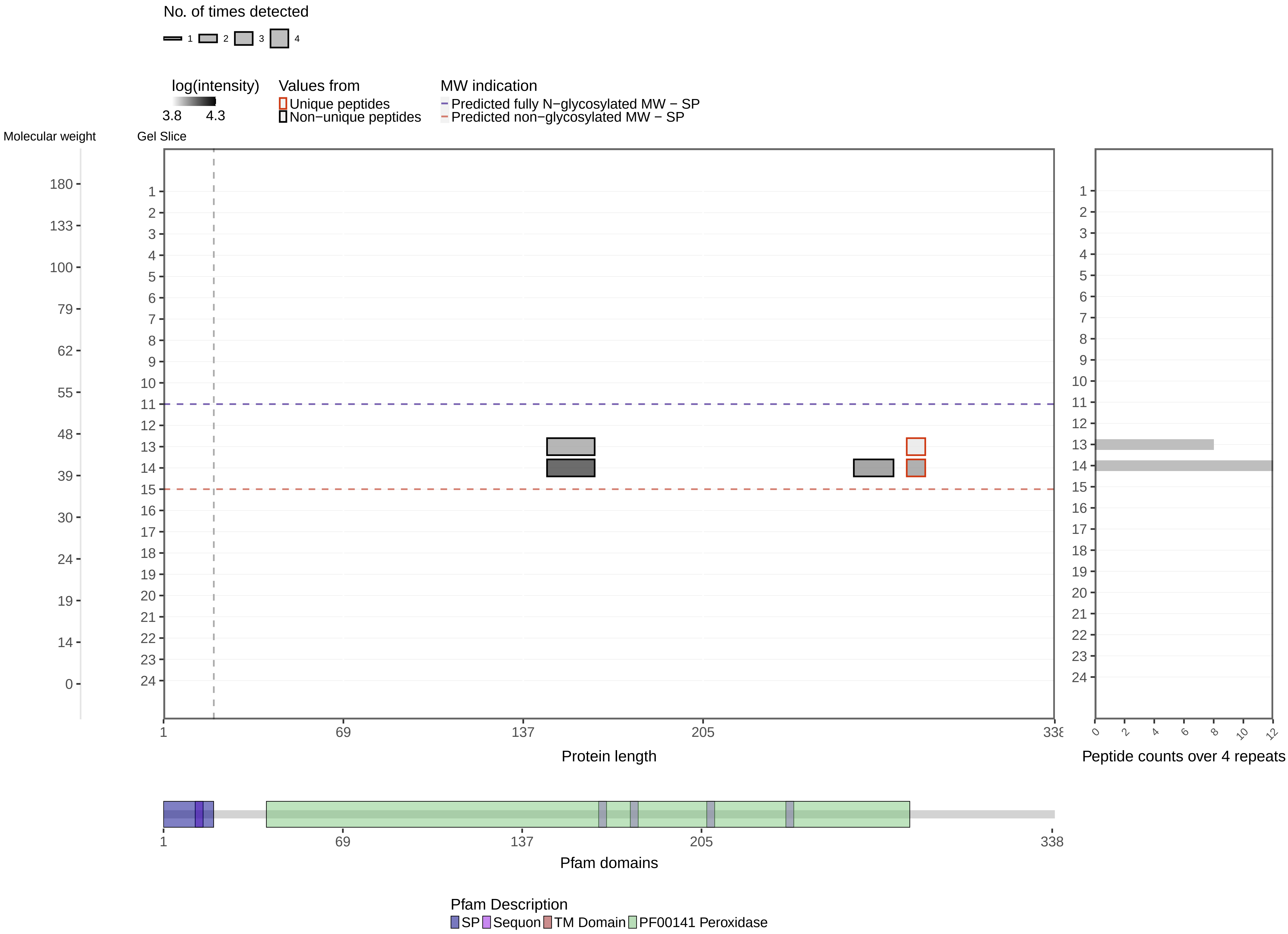


Protein sequence:

SNQIMPSPRSNATYLIDLFSRYNLSIKDLVALSGSHSIGKGRCFSIVFRLYNQSGSGRPDPTIEPRFREKLDKLCPLGGDGNVTGDLDATPEVFDNQYFKDLVNGRGFLNSDQTLFTNPLTREYVREFSVSQEKFFKAFVEGMITMGDLQSGRPGEIRRDCRVVNSHRPRVEVLFETSNGGE*

NbL05g20800.1 lanC−like_protein_GCL1 [MW = 50.23 kDa]

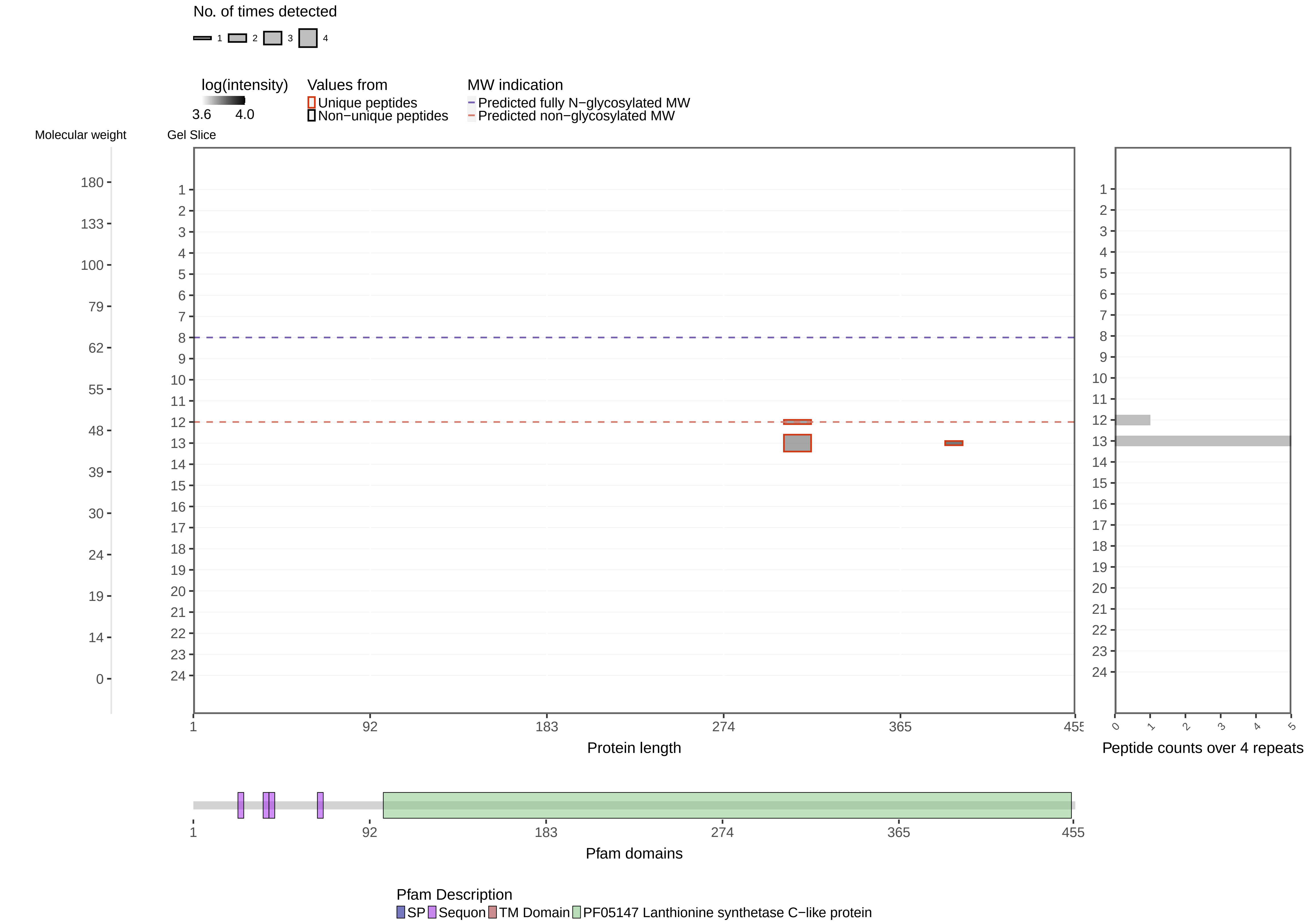


Protein sequence:

LWAALFIRKYIGAETVPDEYVMPVVEAILAGGRAGATDNPACPLMYRWHGTRYWGAAHGLAGILHVLLHFPLPQEDIKDVKETLRYMMSNRFLHSGNYPVSEGNPRDKLVQWSHGATGIAITMCKASEVLSNDREFRDAAVEGGEVVWKNGLVKKVGLADGASGNAYAFLSLFRLTGESIYEERARAFASCLYHNARTIVNARHHNEADHSYSLFQGLGGVACFLFDLLTPVNSRFPGFEL*

NbL05g20720.1 nicotinamidase_1−like [MW = 27.01 kDa]

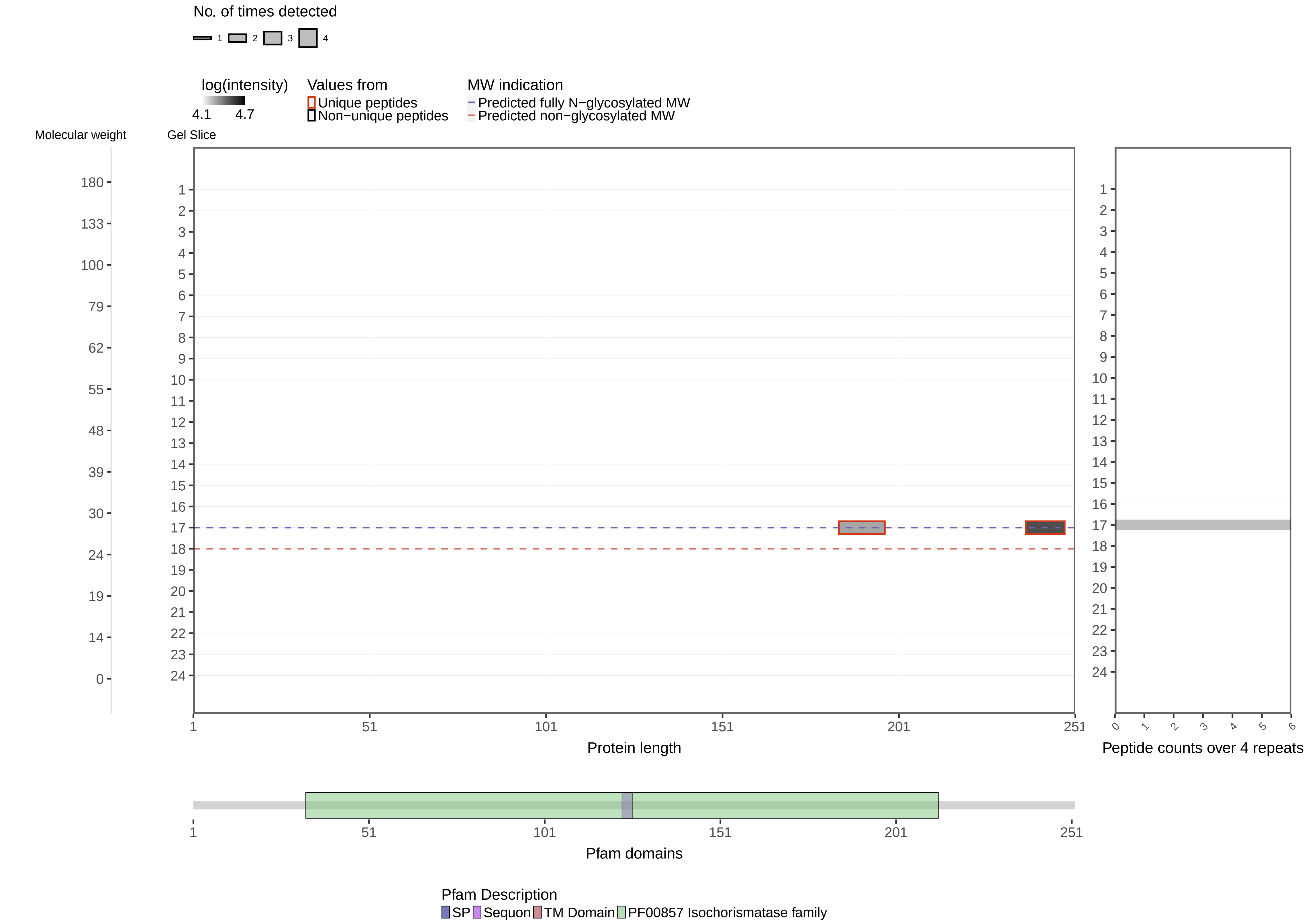


Protein sequence:

.VPALQWLESEPNVTLRCKDCIDGFLGSIEKDGSNVFVDWVKANEIKAILVVGICTDICVLDFVCSVLSARNRGFLSPLEDVIVYSRGCSTFDLPAQIANIKGALPHPQELMHHIGLYMAKGRGAKVVSEVSFDTSAKAT*

### NbL05g20680.1 uncharacterized_protein_At1g04910−like [MW − SP = 53.29 kDa]

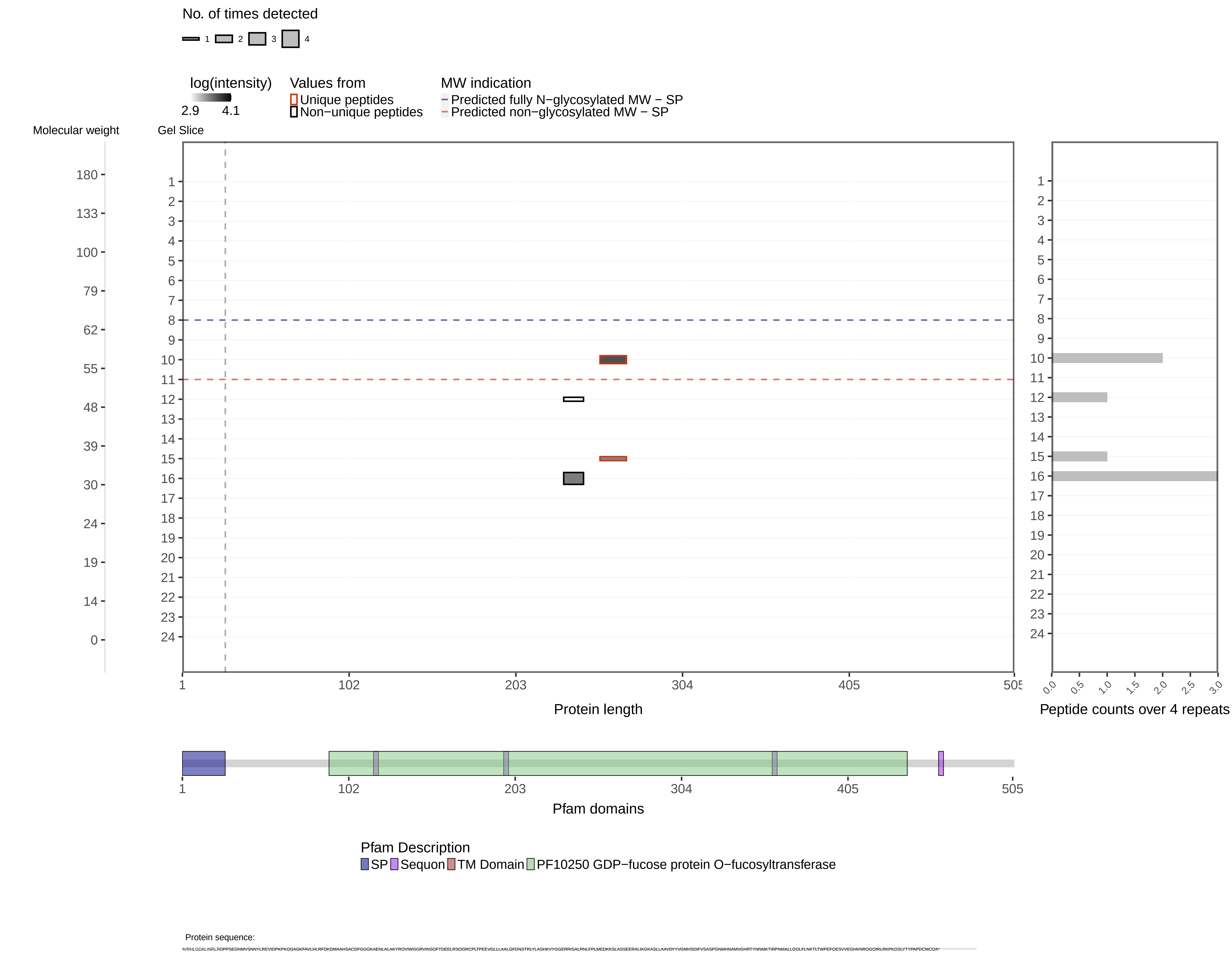

### NbL05g20560.1 14−3−3−like_protein_D [MW = 28.27 kDa]

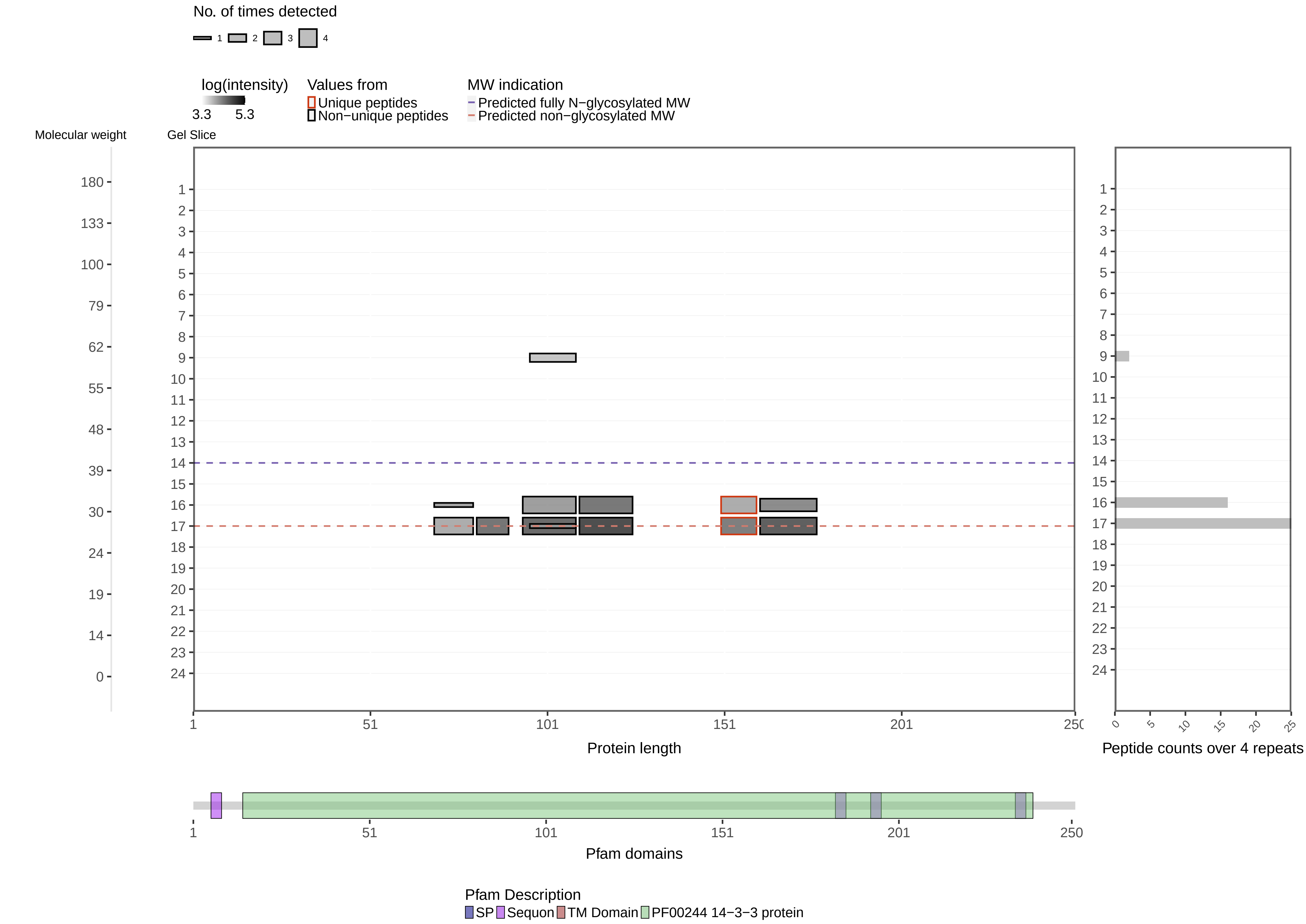


Protein sequence:

QYLIPSASAGESKVFYLKMKGDYYRYLAEFKVSNERKEAAEATMLAYKAAQDIALAELAPTHPIRLGLALNYSVFYYEILNASEKACSMAKQAFEEAIAELDTLGEESYKDSTLIMQLLRDNLTLWTSDMQEQMDEA*

### NbL05g20430.1 14−3−3−like_protein_16R [MW = 30.56 kDa]

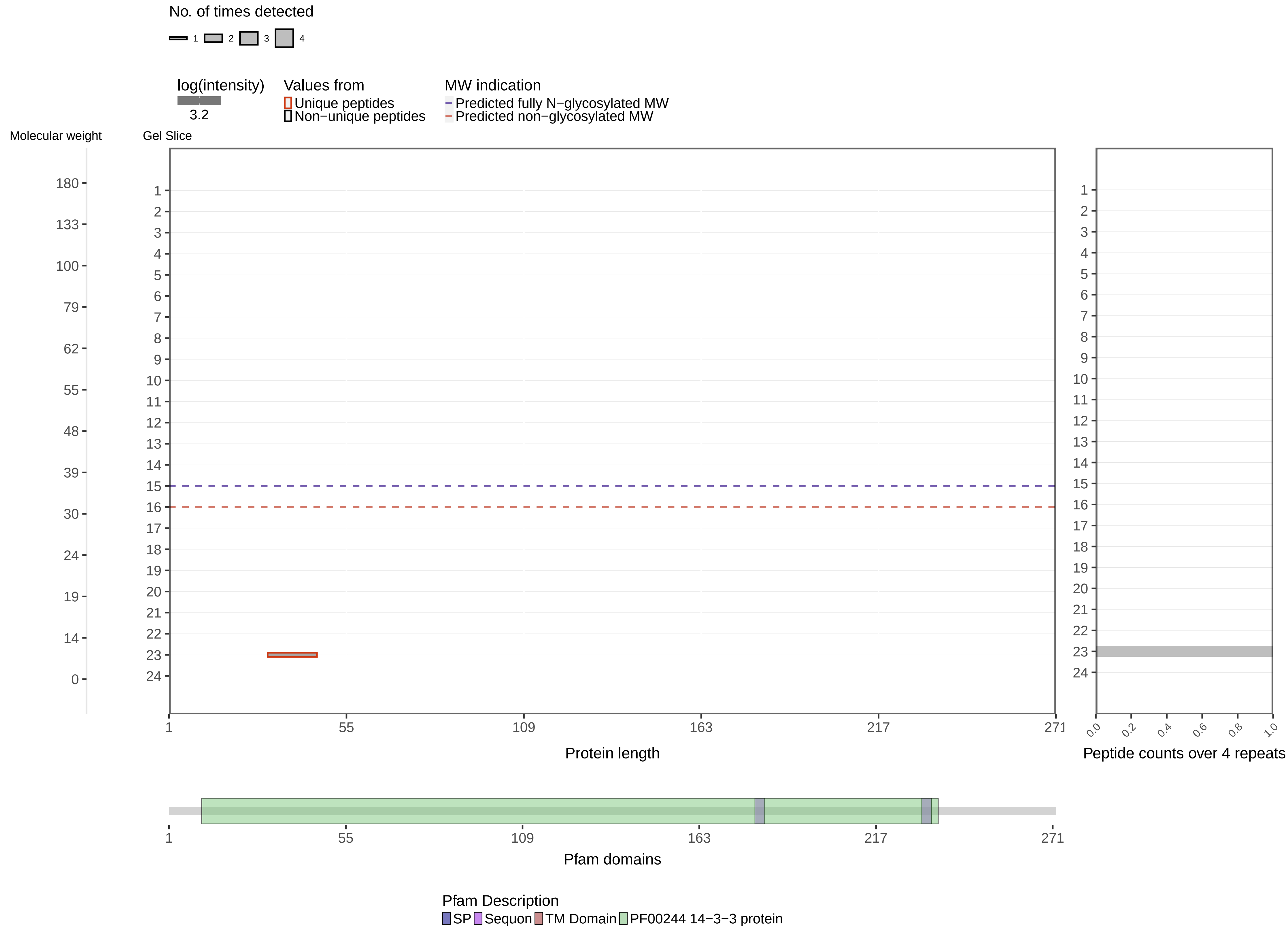


Protein sequence:

VFYLKMKGDYHRYLAEFKTGAERKEAAENTLSAYKSAQDIANTELAPTHPIRLGLALNFSVFYYEILNSPDRACNLAKQAFDEAIAELDTLGEESYKDSTLIMQLLRDNLTLWTSDMQVDISFLLFNKIVSSPFSYDFINLQVLLNLF*

### NbL05g20050.1 uncharacterized_LOC109208398 [MW = 12.05 kDa]

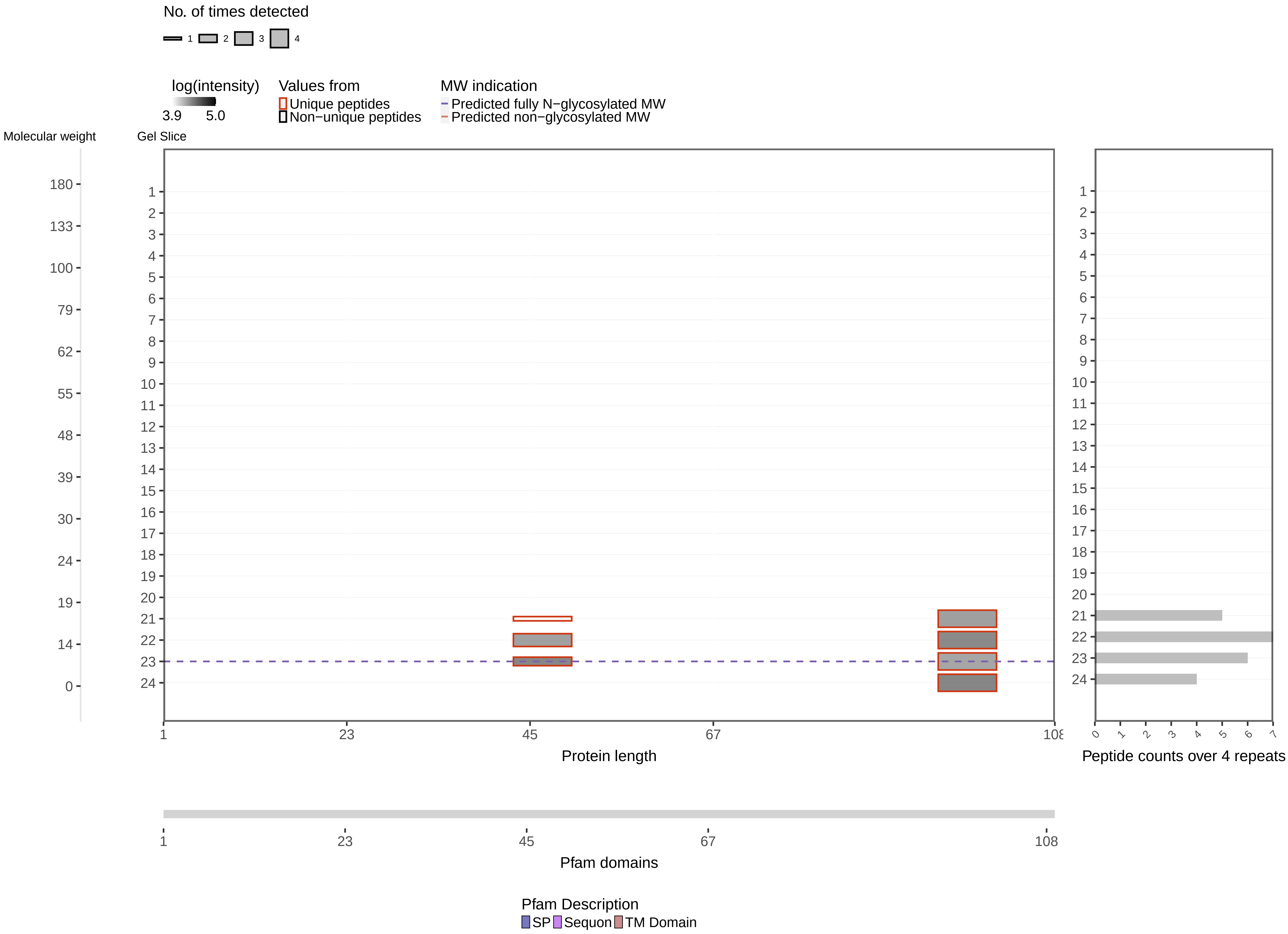


Protein sequence:

AKFKDSPLLKQLLEKSKQNKEKNKKSIEDKYCLRGAEWGIGDCSTAGMTPEERDSFIAMLKQRAVVE*

NbL05g19980.1 8−hydroxygeraniol_dehydrogenase−like [MW = 38.73 kDa]

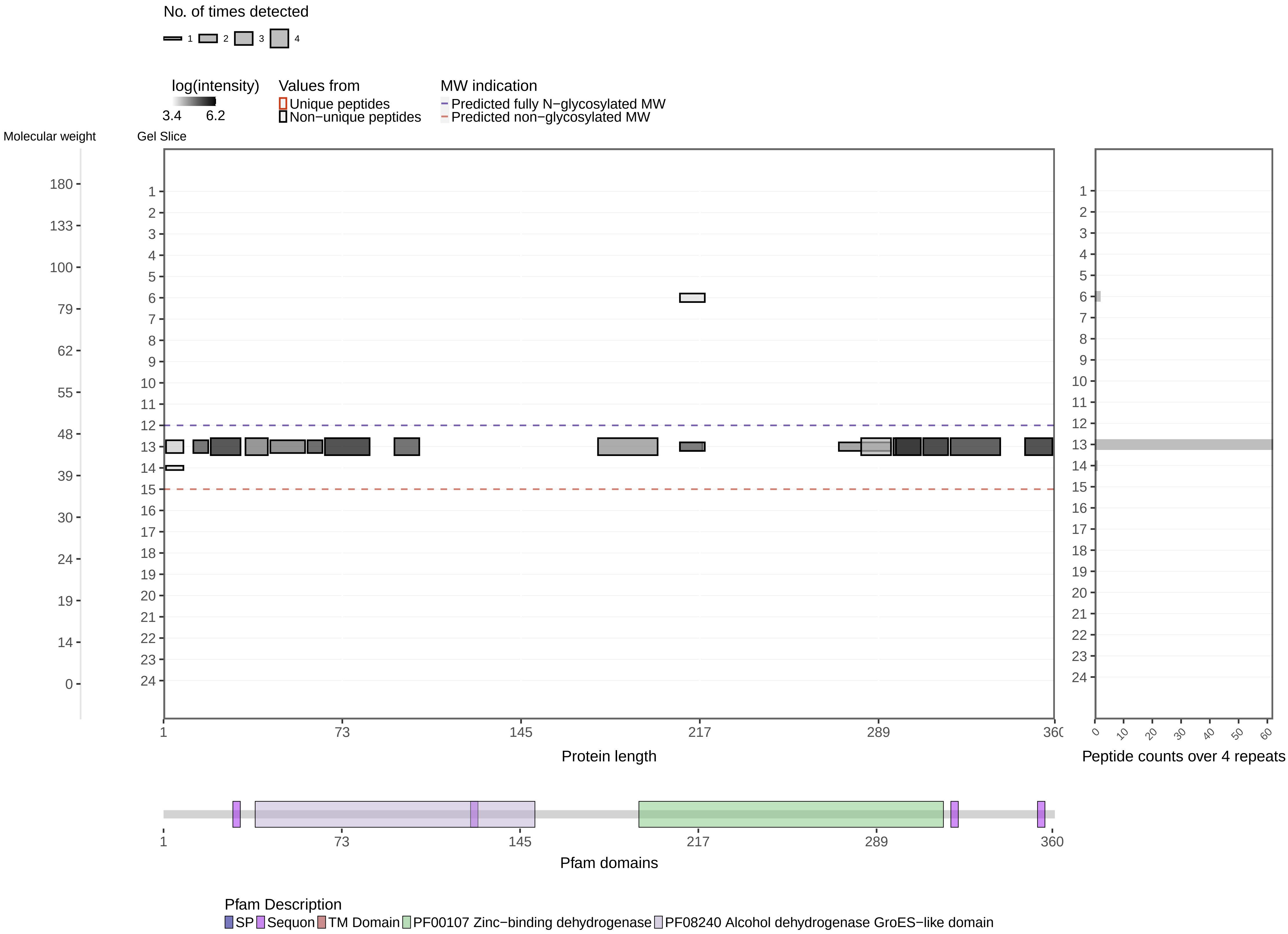


Protein sequence:

CAGITTYSPLKYFGLDKPGMHIGVVGLGGLGHMAVKFAKAFGAKVTVISTSIGKKDEALQHLGADLFVLSSQPQEIQGALNTMDGIIDTVSADHPLLPLLGMLKPHGKLVMVGAYVKPLQLPVFPLILGRKLVAGSAIGGVKETQEMLDFASKHNITPDVEVVPMDYVNTAVHRLLKSDVKYRFVLDIGNHSNSA*

### NbL05g19970.1 8−hydroxygeraniol_dehydrogenase−like [MW = 38.9 kDa]

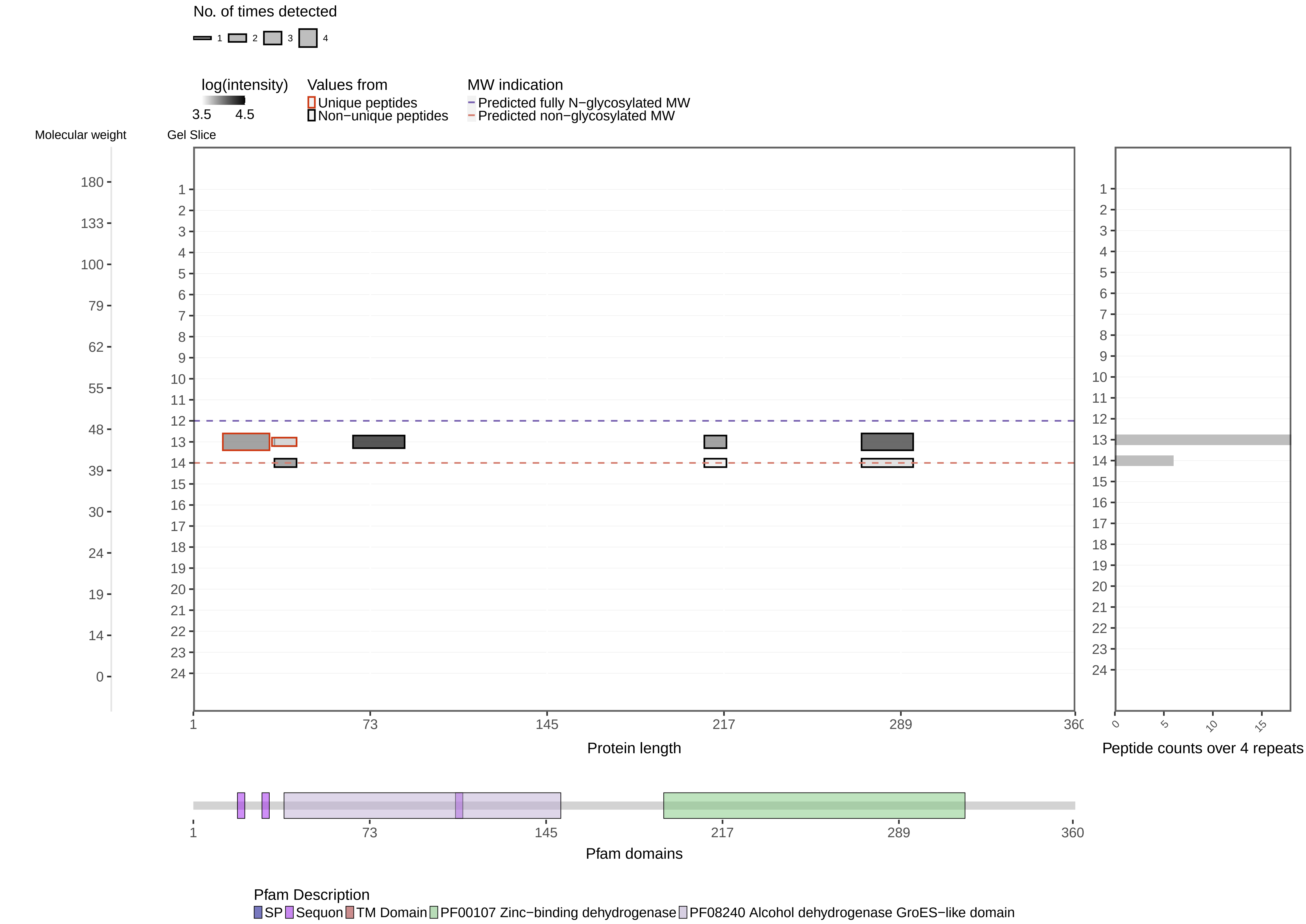


Protein sequence:

LCAGITTYSPLKYFEFDKPGLNIGVVGLGGLGHMAVKFAKAFGCKVTVISTSVNKKDEAIERLGADSFLLSGDPEOLOAASSTLDGIIDTVSADHPILPLLNILRPHGKLVVVGAPPKPLELPVFPIIMGRKIVGGSIIGGMKETOEMLDFAAKHTIIPDVEVVAMDYVNTALERLLKSDVKYRFVLDVGETLKAD*

NbL05g19960.1 8−hydroxygeraniol_dehydrogenase−like [MW = 27.54 kDa]

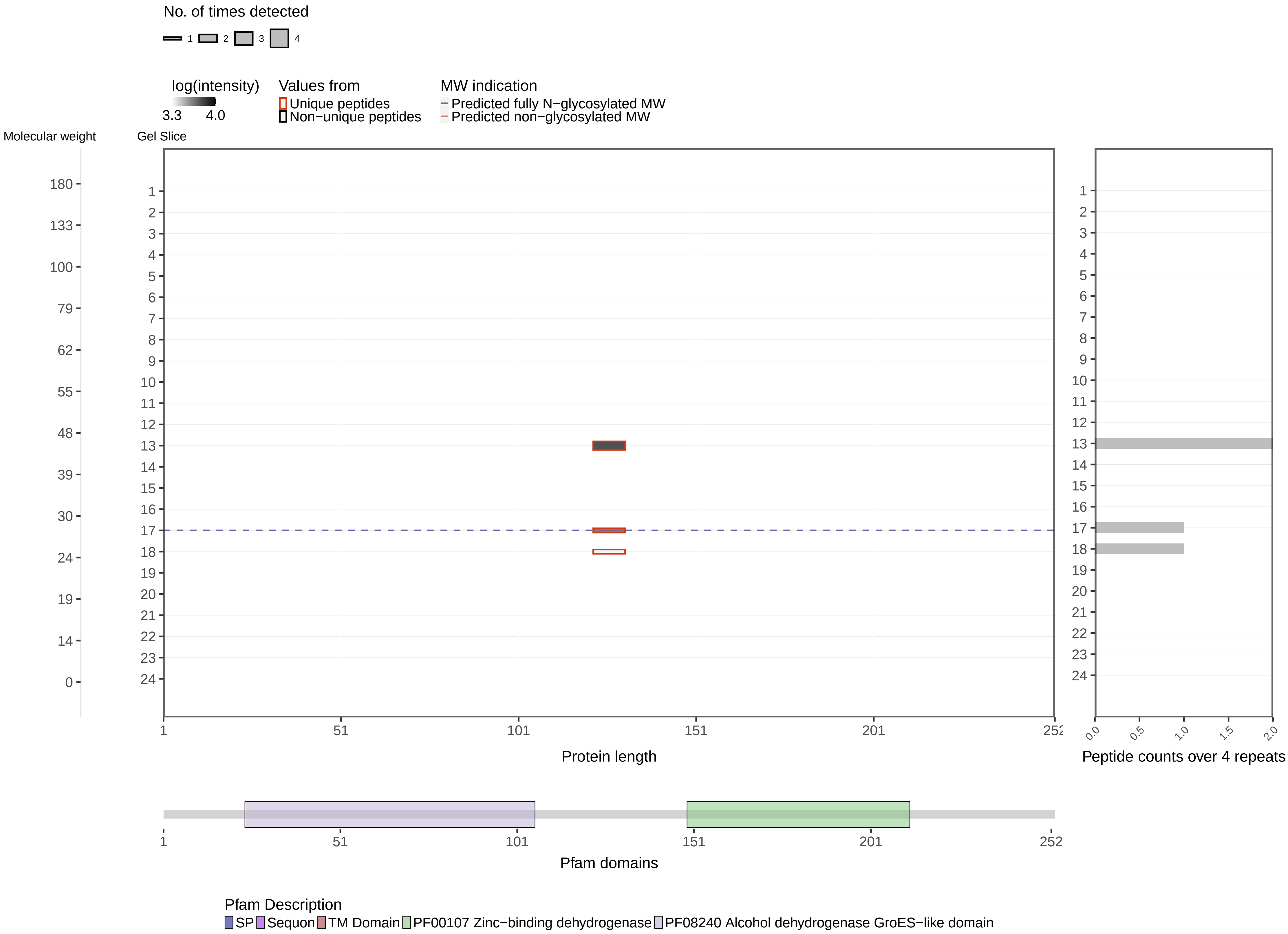


Protein sequence:

NLPMETAPLLRAGITTYSPLKYFGLDKPGMHIGVFCLGGLGHMAVKFAKAFGTKVTVISTSASKKQEAAERLGADAFLISRDTEQMQAAMSTLDGIIDTVSAVHPILPLLSLLNLMVSLLWLVHQKNQWSCQFFLYFWEGS*

### NbL05g19940.1 8−hydroxygeraniol_dehydrogenase−like [MW = 39.15 kDa]

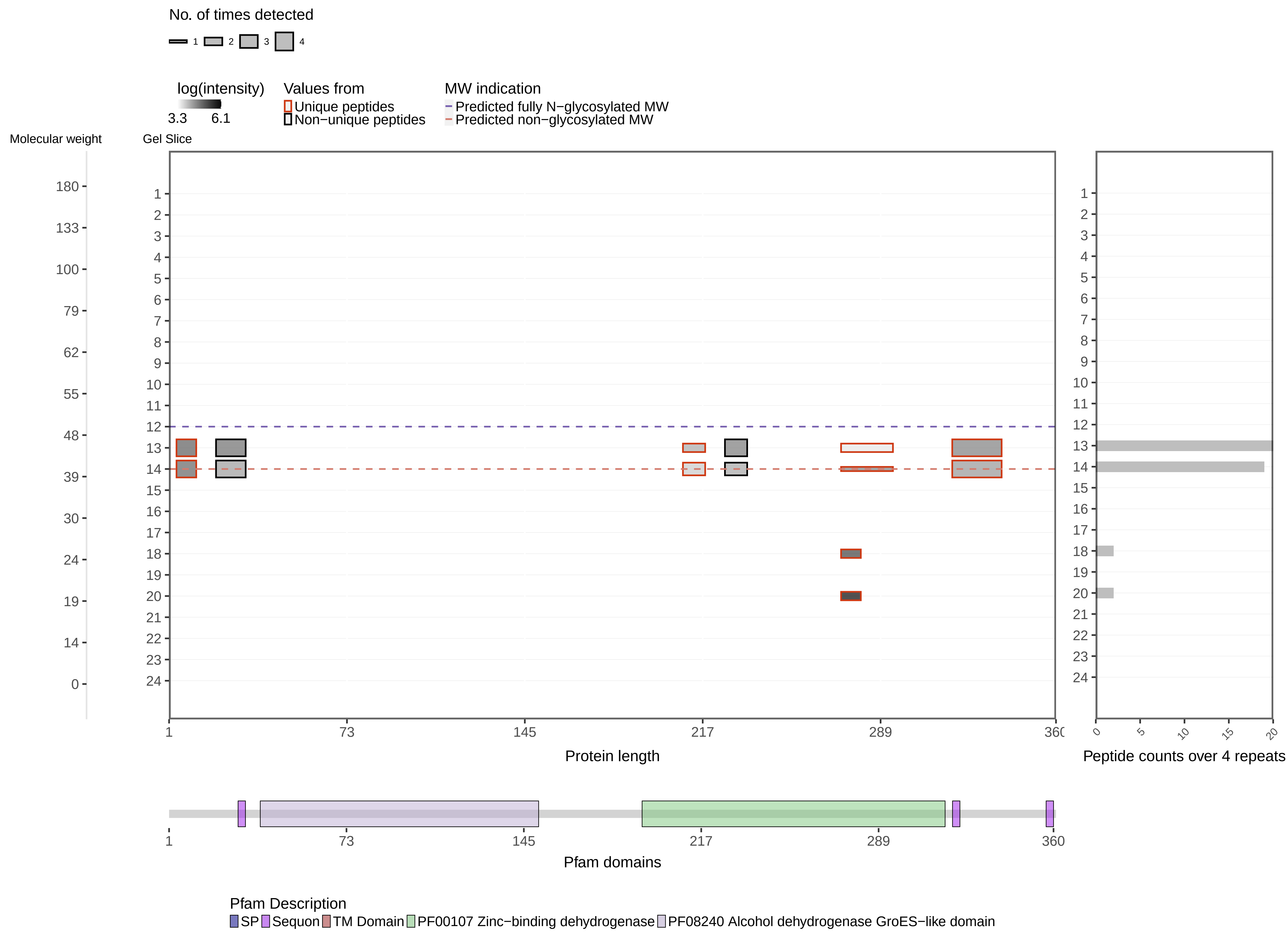


Protein sequence:

.LNAGITTYSPLKYFGLDKPGMHIGVVGLGGLGHMAVKFAKAFGTKVTVISTSVSKKQEAIERLGADSFLLSRDPDQMQAATGSLDGIIDTVSAVHPLLPLINLLKTHGKLVILGAPEKPLELPVFPLLLGRKLVAGSCIGGMKETQEMVDFAAKHNITPDVEVVPMDYVNTAIERLAKSDVKYRFVLDIGSTLNKS*

NbL05g19850.1 glutathione_S−transferase_DHAR3_chloroplastic [MW = 28.55 kDa]

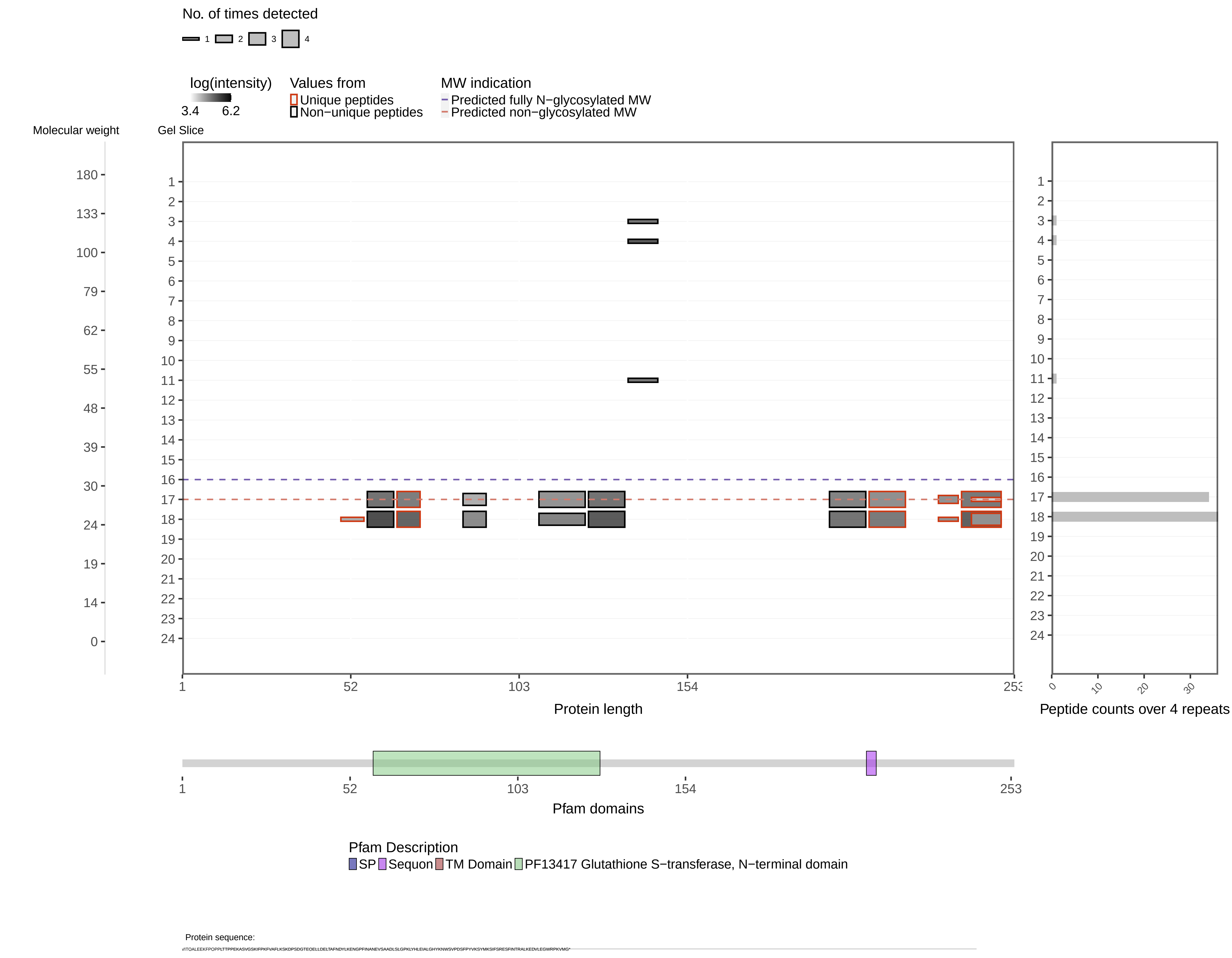

NbL05g19790.1 NADH_dehydrogenase_[ubiquinone]_iron−sulfur_protein_1_mitochondrial [MW = 84.82 kDa]

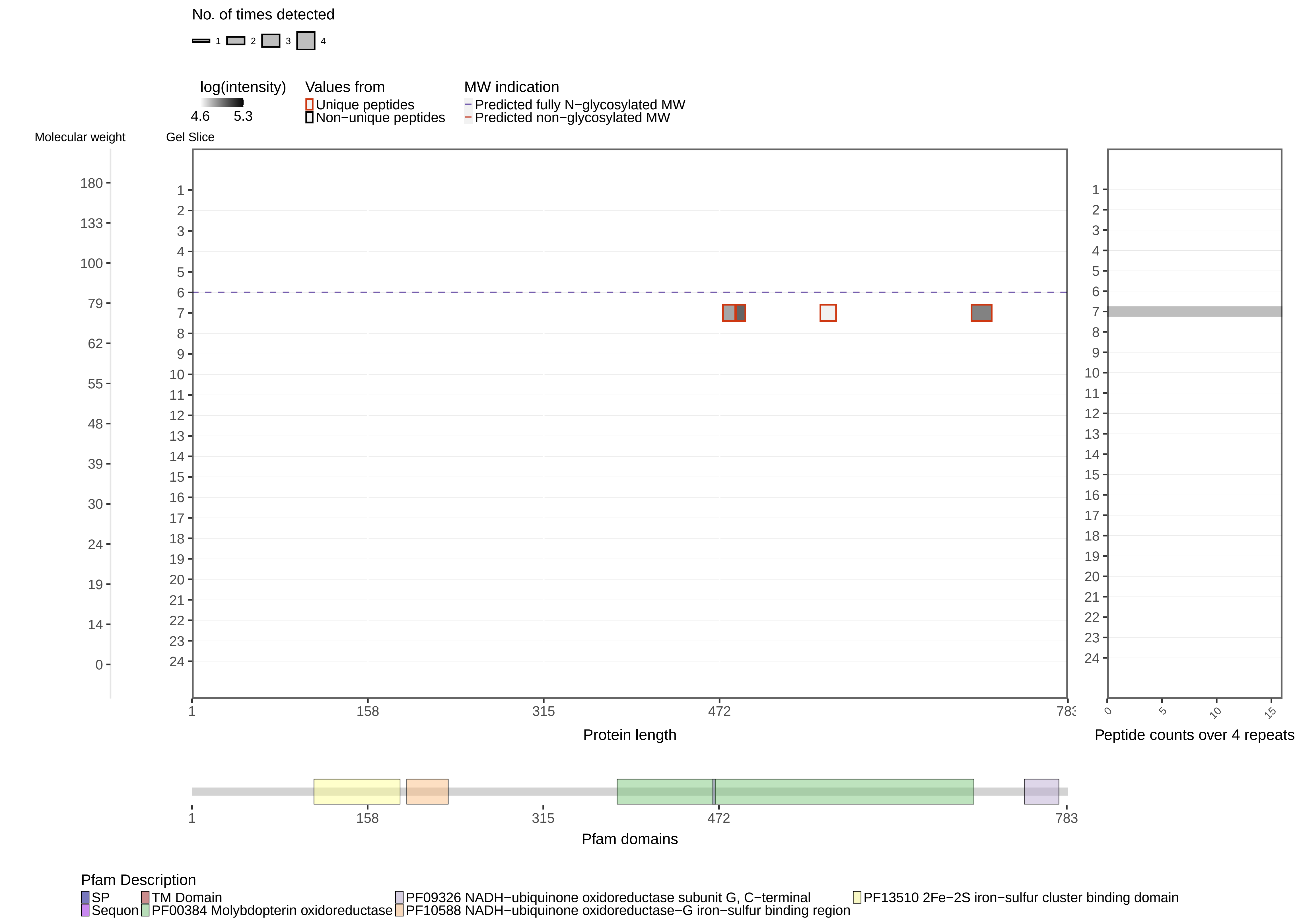


Protein sequence:

RQRLNDPMIRGADGRFKAVSWRDALAIVAEVMHQIKPEEIVGVAGKLSDAESMMALKDLLNKMGSNNIFCEGNGMQPNADLRSGYIMNTSISGLEKADAFLLVGTQPRVEAAMVNARIRKTVRANNAKVGYIGAAADFNYDHQHLGTDPQTLVEIAEGRHPFSSALKNAKNPVIIVGAGVFDREDKDAVFAAIDTIAKNNNVVRSDWNGLNVLLLNAAQAAALDLGLVPESEKCIDSAKFIYLMGADDVSLDKLPDDAFVVYQGHHGDRGVYRANVILPASAFTEKEGIYENT

### NbL05g19760.1 NADH−cytochrome_b5_reductase−like_protein [MW = 36.15 kDa]

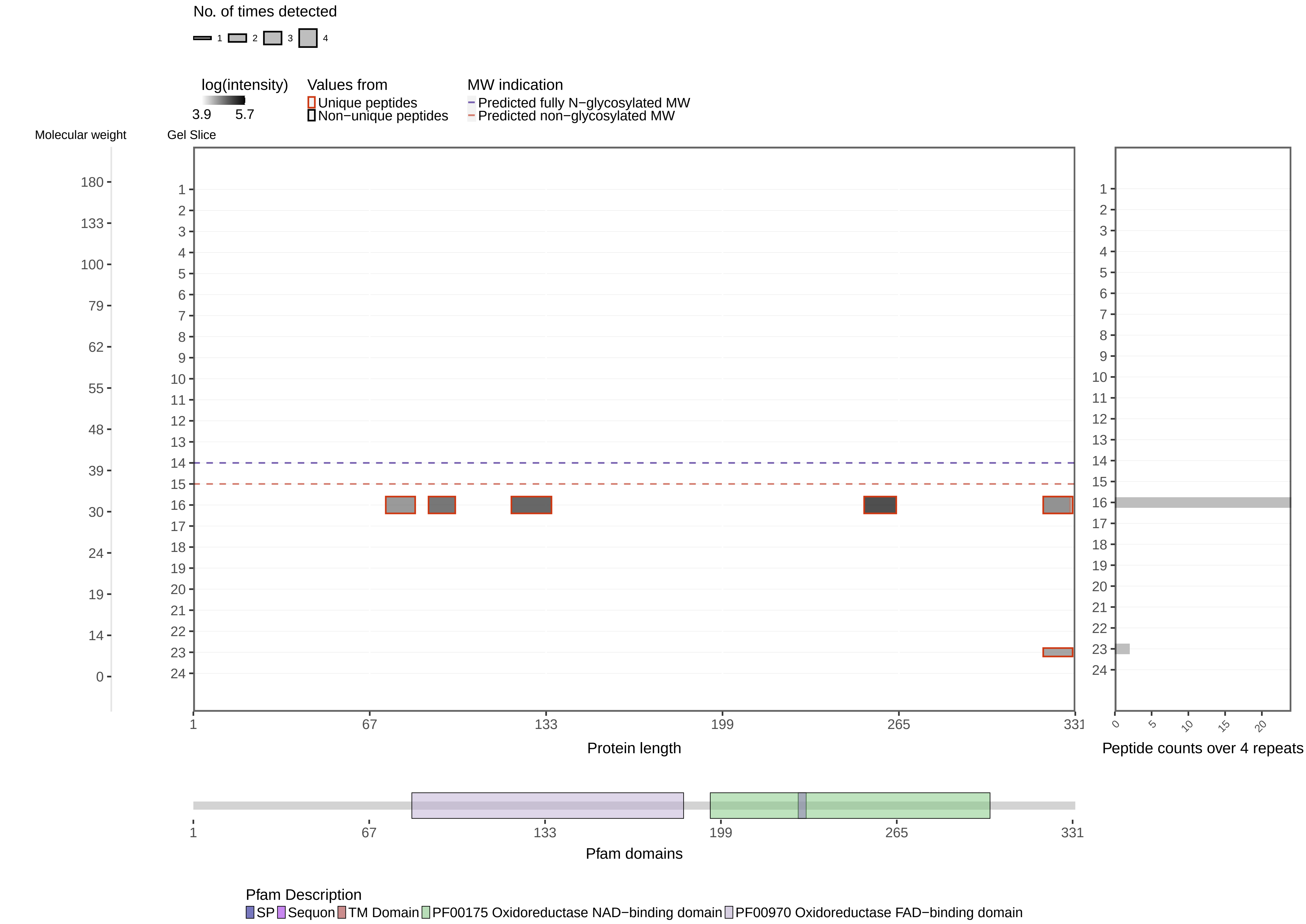


Protein sequence:

LIKVYPEGKMSQHFASLKPGDVLEVKGPIEKLRYSTNMKKHIGMIAGGSGITPMLQVIEAILKNPDDNTQVSLIYANVSPDDILLKKKLDVLAATHPNLKVFYTVDSPTNDWRGGIGYVSKDMIVKGLPPPSDDTLILVCGPPGMMKHLSGEKPKPREQGELTGLLKDVGFTENMVYKF*

### NbL05g19660.1 UPF0678_fatty_acid−binding_protein−like_protein_At1g79260 [MW = 18.46 kDa]

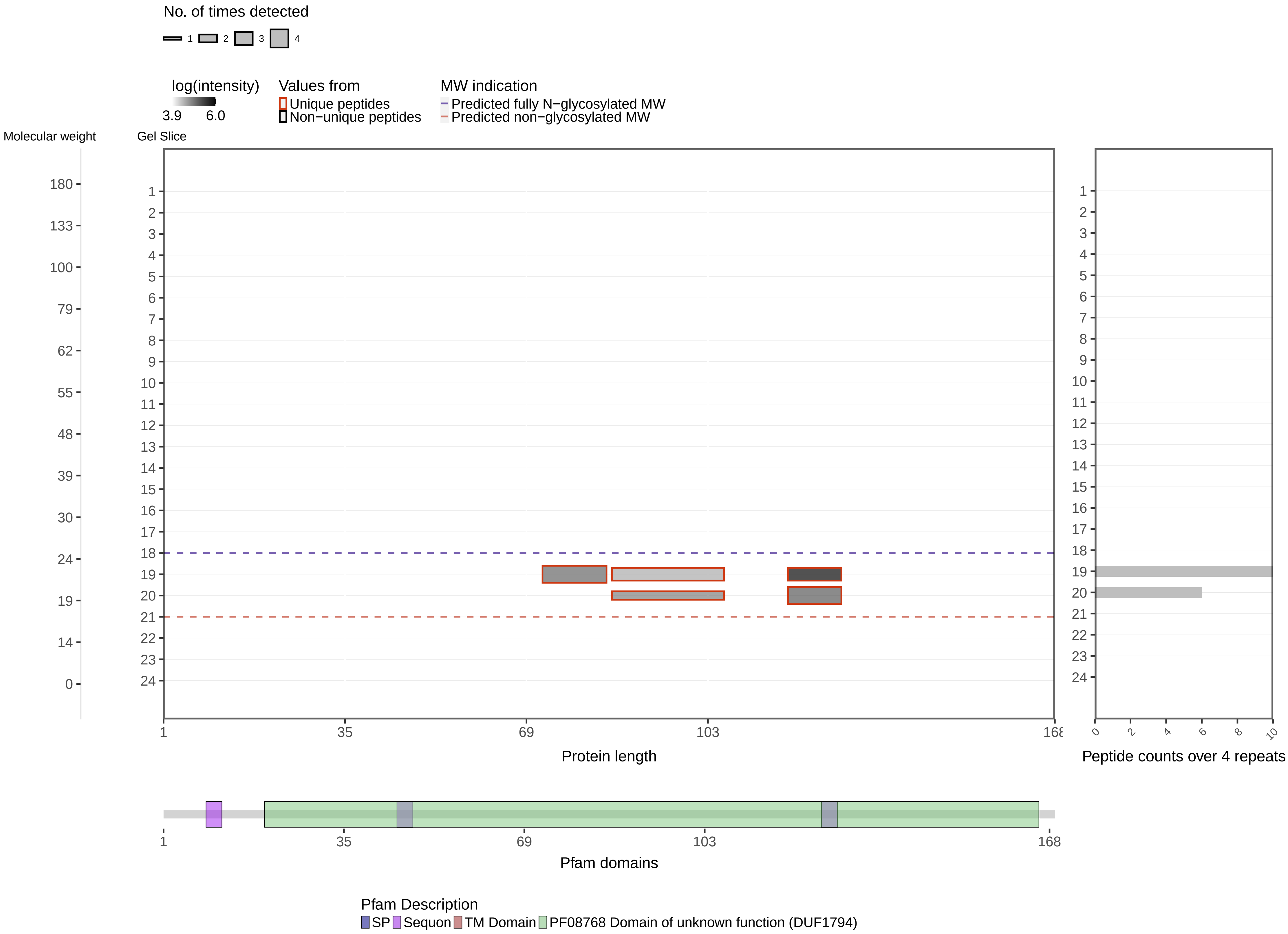


Protein sequence:

KSGQPMHSESGYWRPKPDGTIEAVIAQSTGLVEVQKGTYDMKEGVVKLSSELVGNASKVKEISRVFKVENCELSYVVEMATSLIGLQPHLKASLKKL*

NbL05g19610.1 glutamine_synthetase [MW = 39.08 kDa]

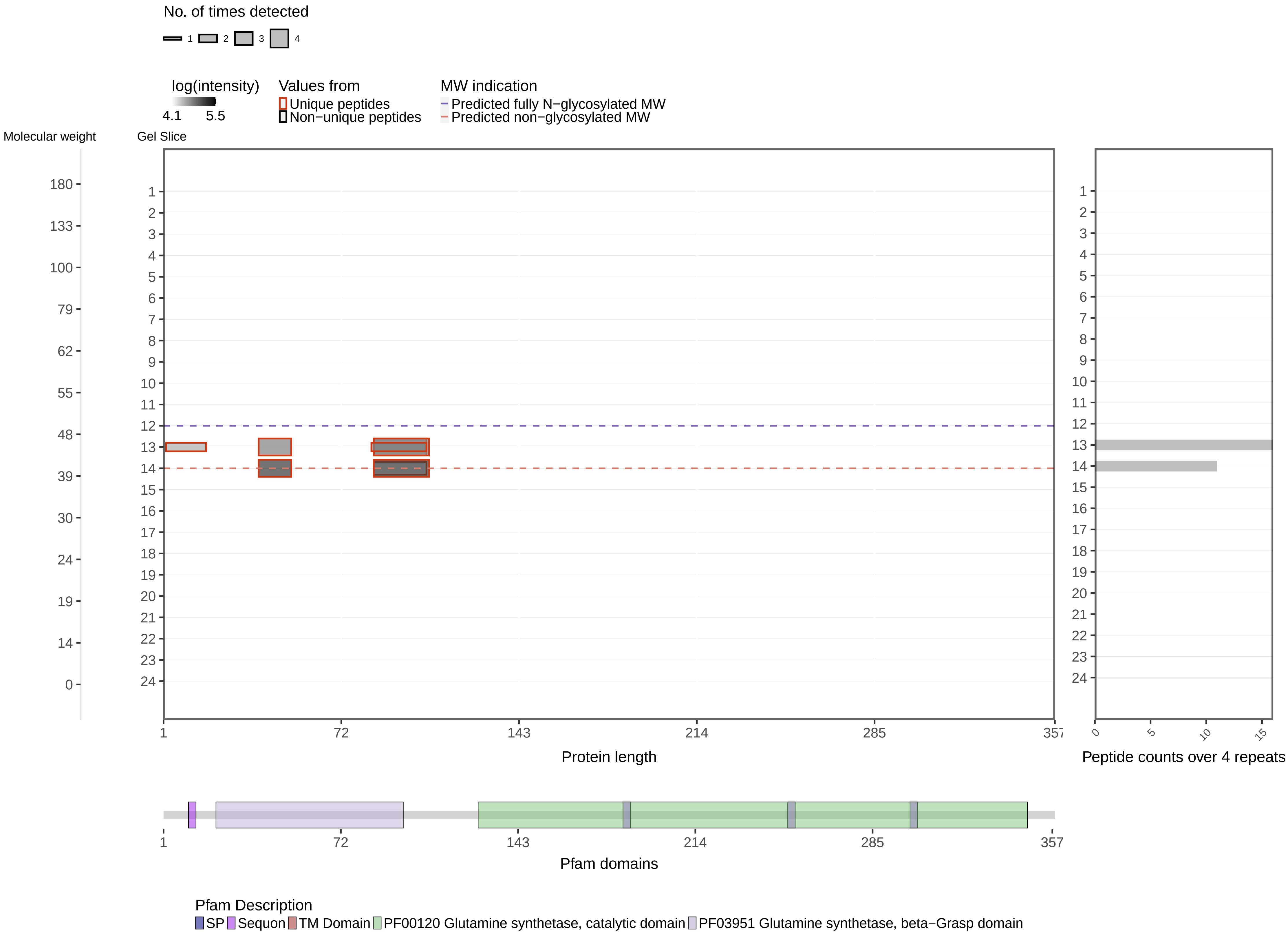


Protein sequence:

AFGRDIVDSHYKACLYAGINISGINGEVMPGQWEFQVGPSVGISAGDEVWVARYILERITEIAGVVVSFDPKPIPGDWNGAGAHTNYSTKSMREDGGYEVILKAIEKLGLKHKEHIAAYGEGNERRLTGKHETANISTFKWGVANRGASVRVGRDTEKAGKGYFEDRRPASNMDPYVVTAMIAETTIIWKP*

### NbL05g19390.1 peptidyl−prolyl_cis−trans_isomerase_CYP19−3−like [MW = 18.82 kDa]

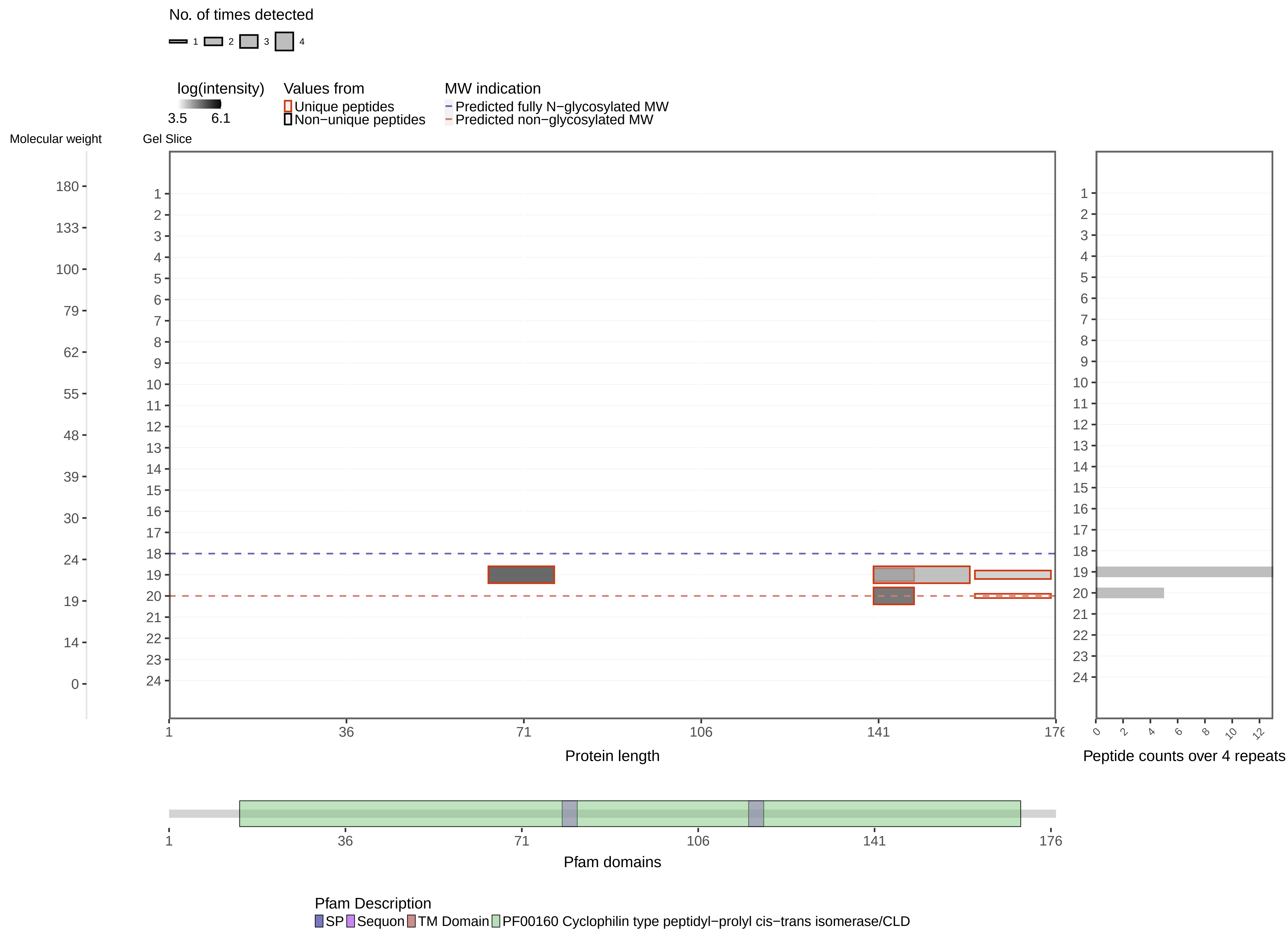


Protein sequence:

TKGNGTGGESIYGAKFADEDFSVIHTTPGLLSMANAGPNTNGSQFFITTVPTPWLDGKHVVFGKVVDGYSVVKEMENVGSDSGKTSCTVSIEDCGELKEN*

### NbL05g19120.1 probable_pectinesterase/pectinesterase_inhibitor_51 [MW = 63.84 kDa]

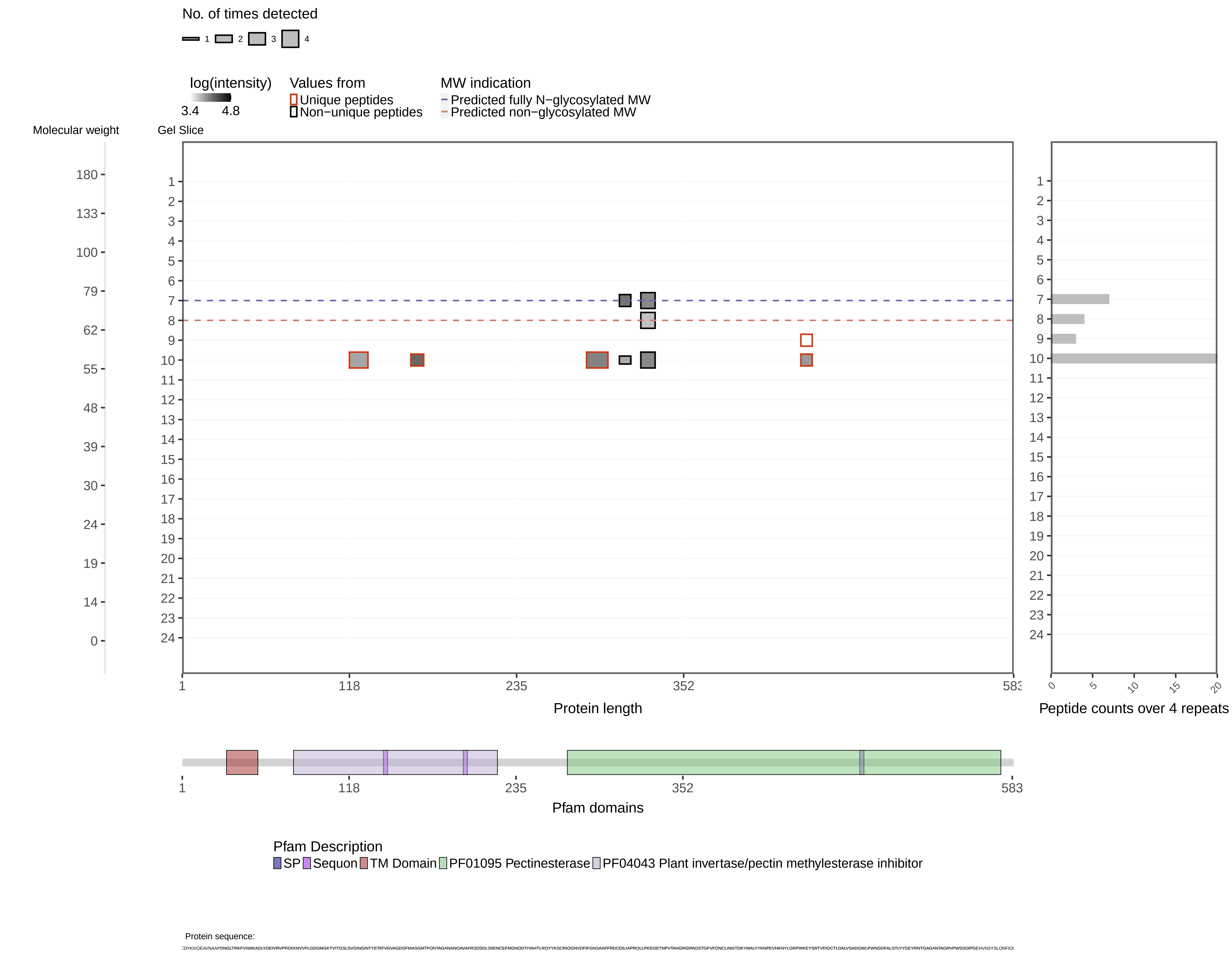

NbL05g19110.1 probable_pectinesterase/pectinesterase_inhibitor_51 [MW = 61.51 kDa]

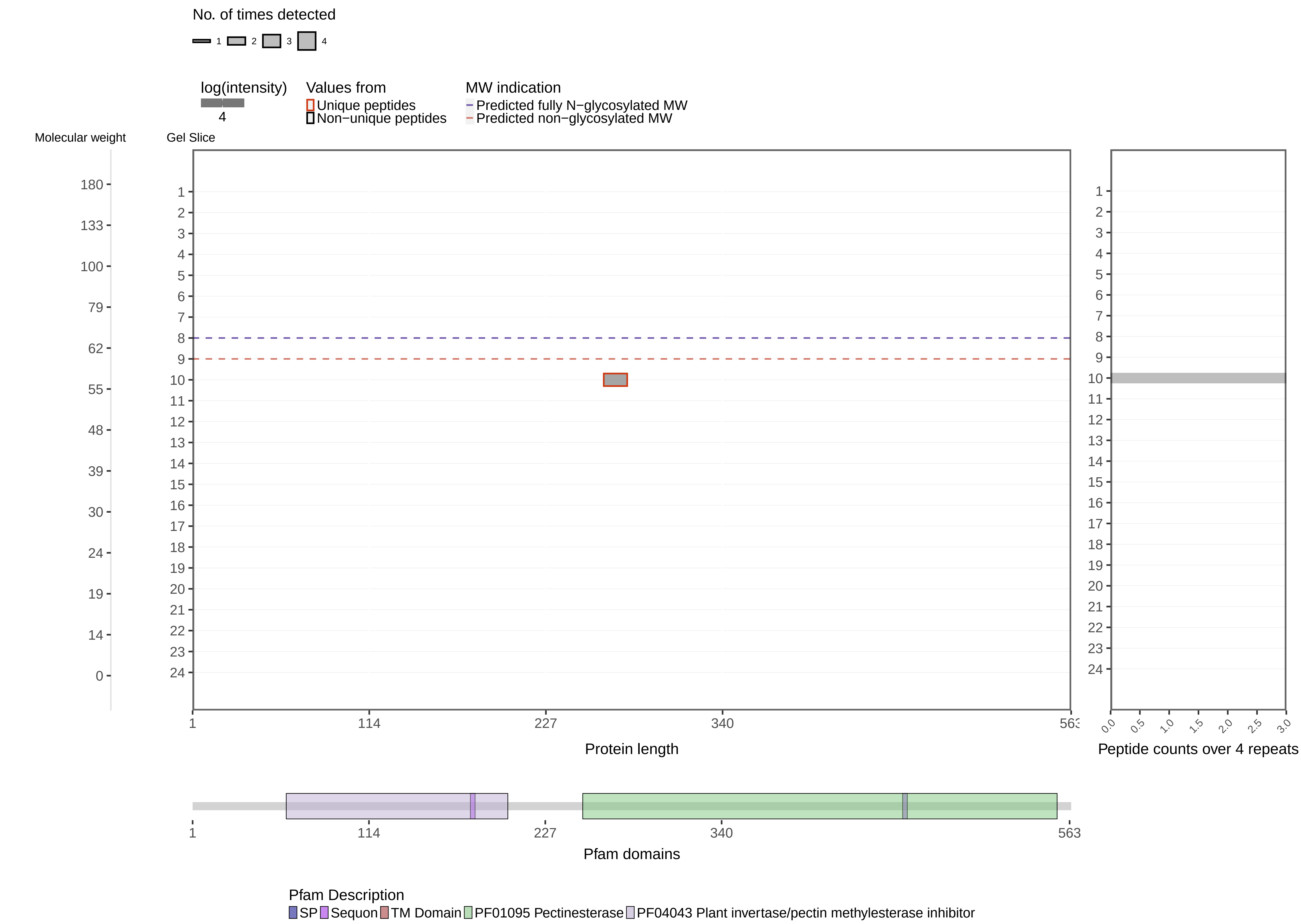


Protein sequence:

EAVNAAPDNQVTRKFVIWIKAGLYEEKVLVPLEKKNVVFLGDGMGKTVITGSLNVGMAGVTTYETATVGVVGDGFMASGITFQNTAGPVAHQAVAFRSDSDLSVIENCEFIGNQDTLYAHALRQYFKSCRIQGNVDFIFGNSAAFFQECDIFIAPRQVNPEKGEKNAVTAHGRIDPAQSTGFVFQNCLINGTEQYMALYYKNPKVHKNYLGRPWKEYSRTVFIECTLEALVLADGWLPWSGDFALKTLYYGEYRNTGAGANTAGRVQWSSQIPAEHVNSYSIQNFIQGDQWITT

### NbL05g19090.1 probable_pectinesterase/pectinesterase_inhibitor_51 [MW − SP = 55.98 kDa]

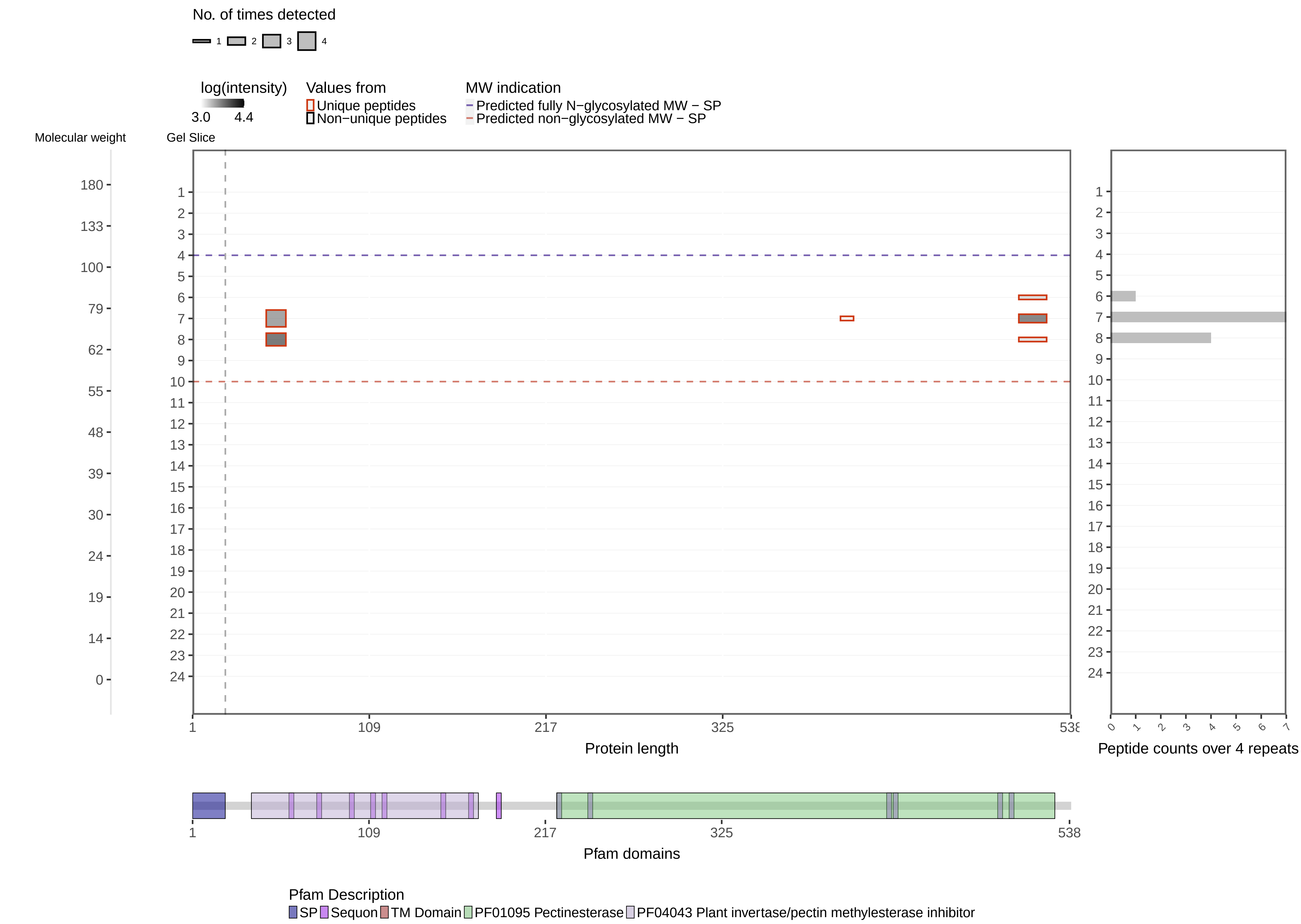


Protein sequence:

NbL05g18670.1 50S_ribosomal_protein_L15 [MW = 31.86 kDa]

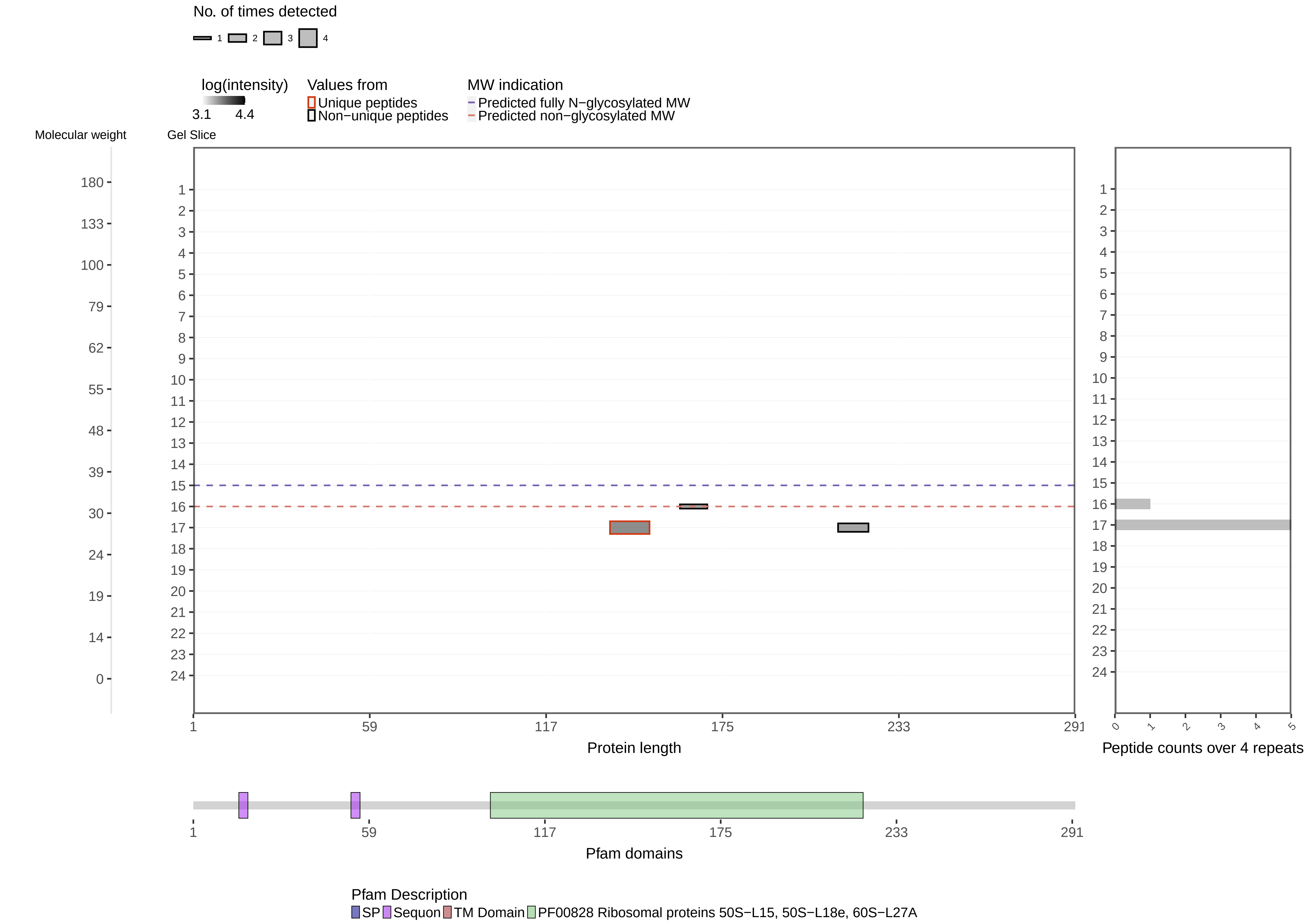


Protein sequence:

.PKRGFKNPFSLTFQPVGLGKIAKLINAGKIDSSELITMKTLKDSAAIGKQIGDGVRLMGRGAEHIQWPIHLEVSRVTVRAKAAVEAAGGSVRRVHYNKLGLRALLKPEWFEKKGRLLPRPARPPPKLKEKVDSIGRLPAPTKPIPFAAEETEAMSAAPA*

NbL05g18320.1 ycf3−interacting_protein_1_chloroplastic [MW = 33.92 kDa]

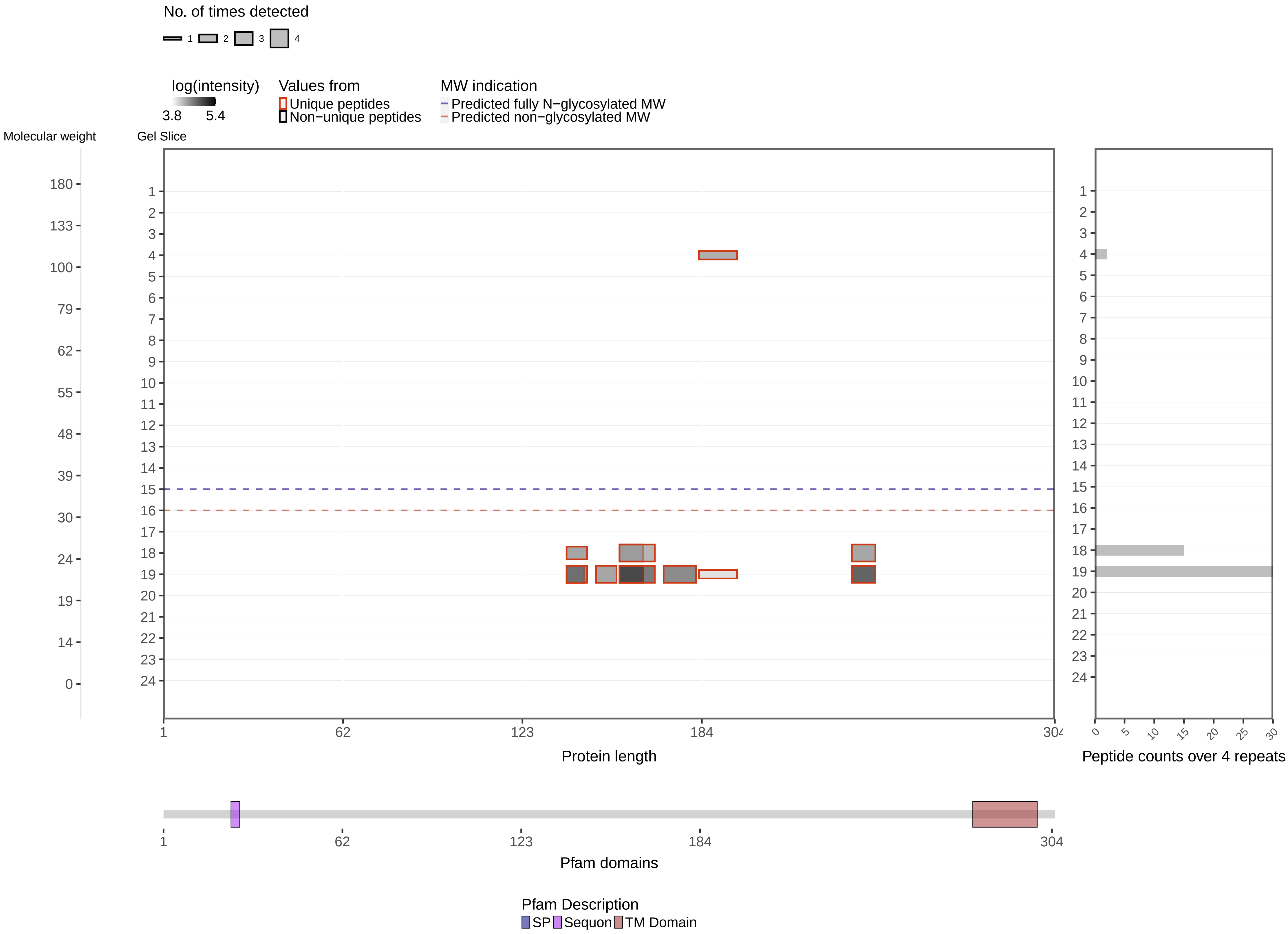


Protein sequence:

:SVLELLKRNRDMLFGEVKLTIMIEDPRDVERRRLLGIDDENAPTRDDLAAALEEINEGKVPKDFAALQMLAEEMNSWPNLEVEATKQNKPGRSLYAKATDTGIDPKEAAKRLKIDWDSAAEIDDSAQSDEPDVPPALGYGALYLVSAFPIIIGISVVLILFYNSLQ*

### NbL05g18270.1 plasma_membrane−associated_cation−binding_protein_1−like [MW = 28.57 kDa]

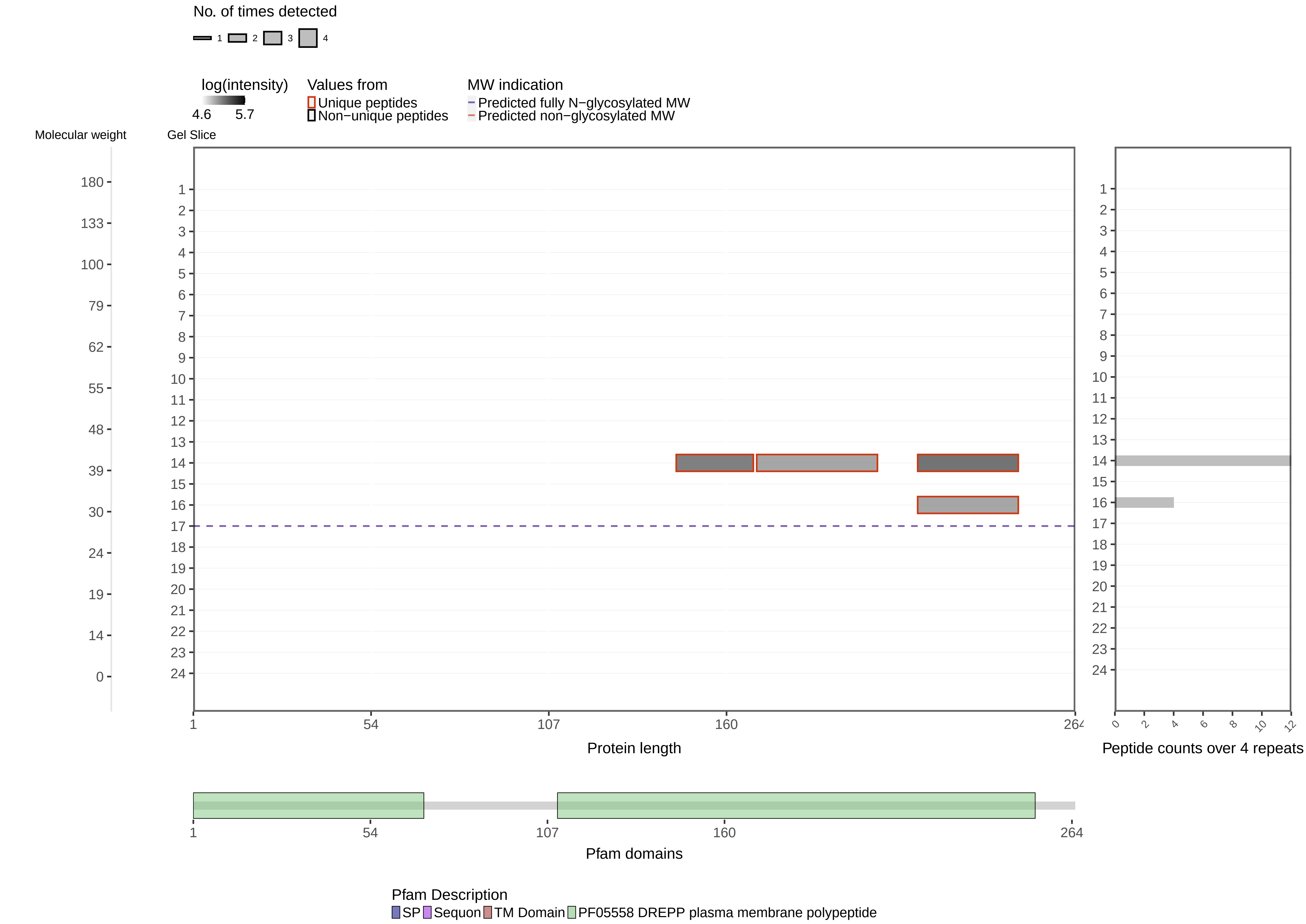


Protein sequence:

GLKKHSTQVQKFLDELVKIEFPGSKAVSEASSNFGPSYVSGPVFFVFEKVSTFIVTEDKKEEEAAAAPIAAASDAEATAVTPVEVKEKEIVVEAEAKKEEEPAPAPAPAPAAAPAAEVATAADVAPAKVEEAPAAVAAPEPPKA*

### NbL05g18240.1 polyadenylate−binding_protein_RBP45−like [MW = 46.09 kDa]

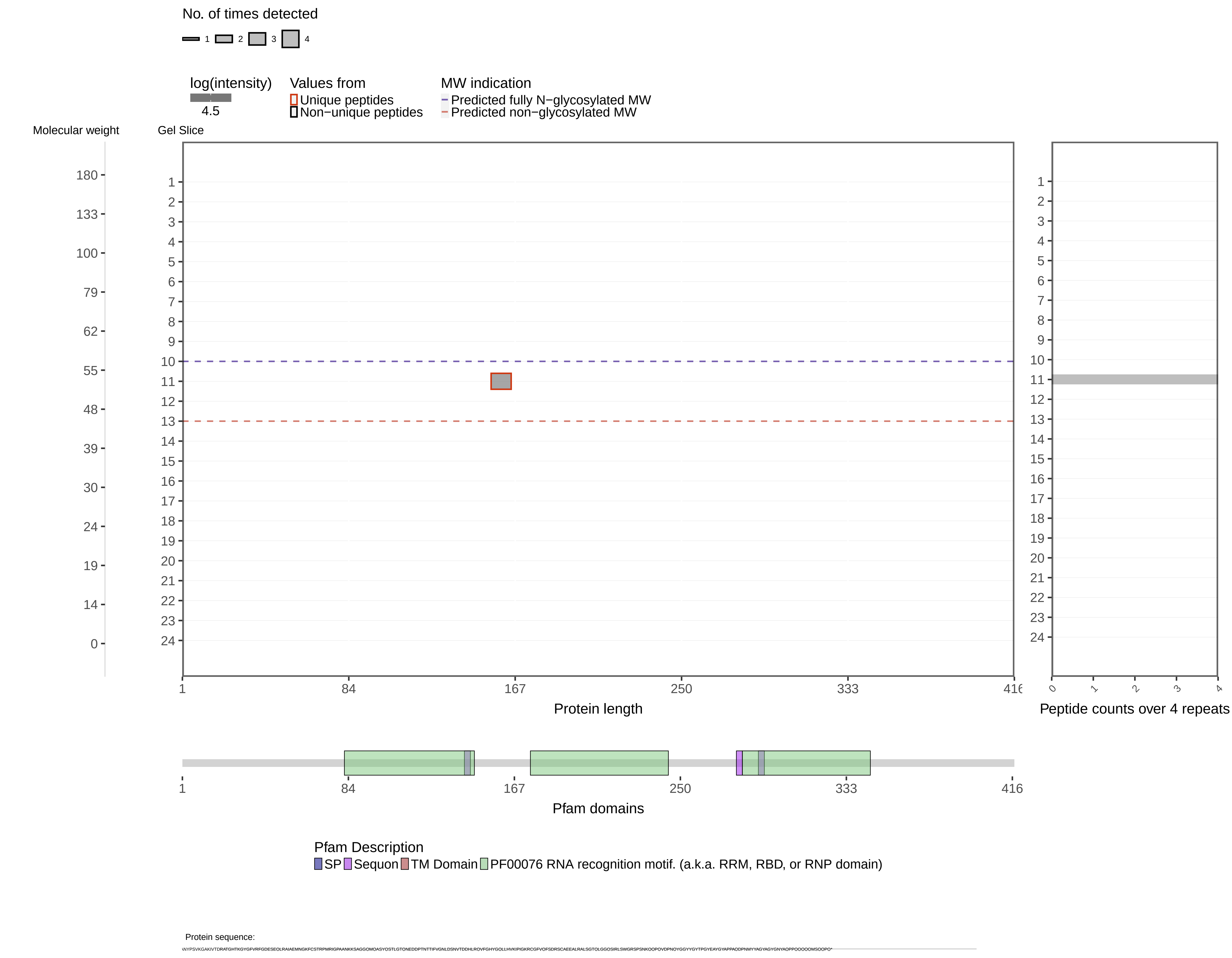

### NbL05g18210.1 adenosine_deaminase−like_protein [MW = 40.69 kDa]

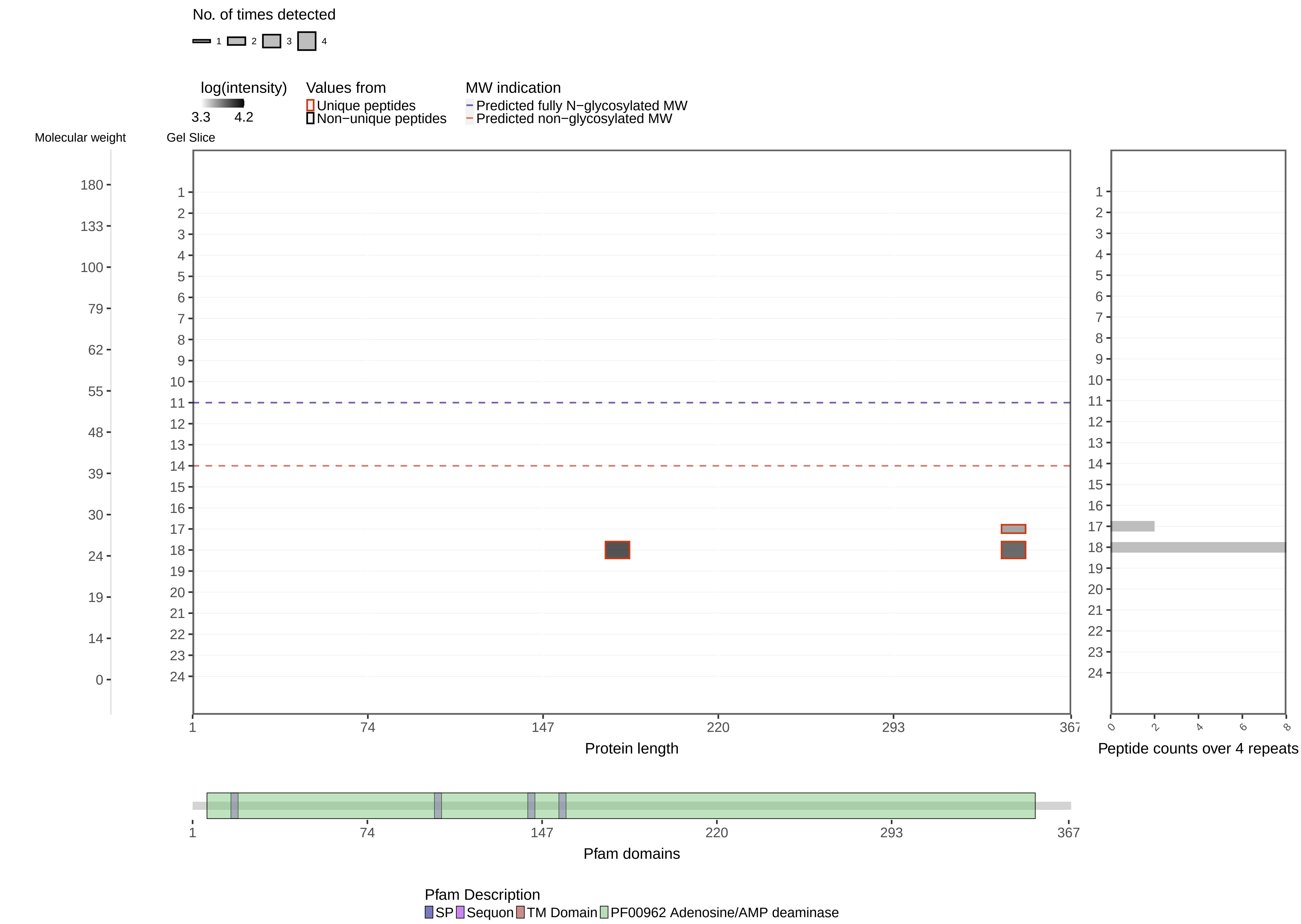


Protein sequence:

RRESTESAMETVKLALEMRHKGVVGIDLSGNPIIGEWLTFLPALEFAKEQGLLITLHCGEVPNPVEIHAMLSFLPARIGHACCFGEEEWAKLKSLKIPVEICLTSNIRTETISSLDIHHFADLYNSGHPIVLCTDDSGVFSTSVSGEYSLAASAFGIQKREMFQLARSGINFIFAGDKVKQELEQLFDLAAKGLEF*

### NbL05g18030.1 uncharacterized_LOC109207523 [MW = 15.6 kDa]

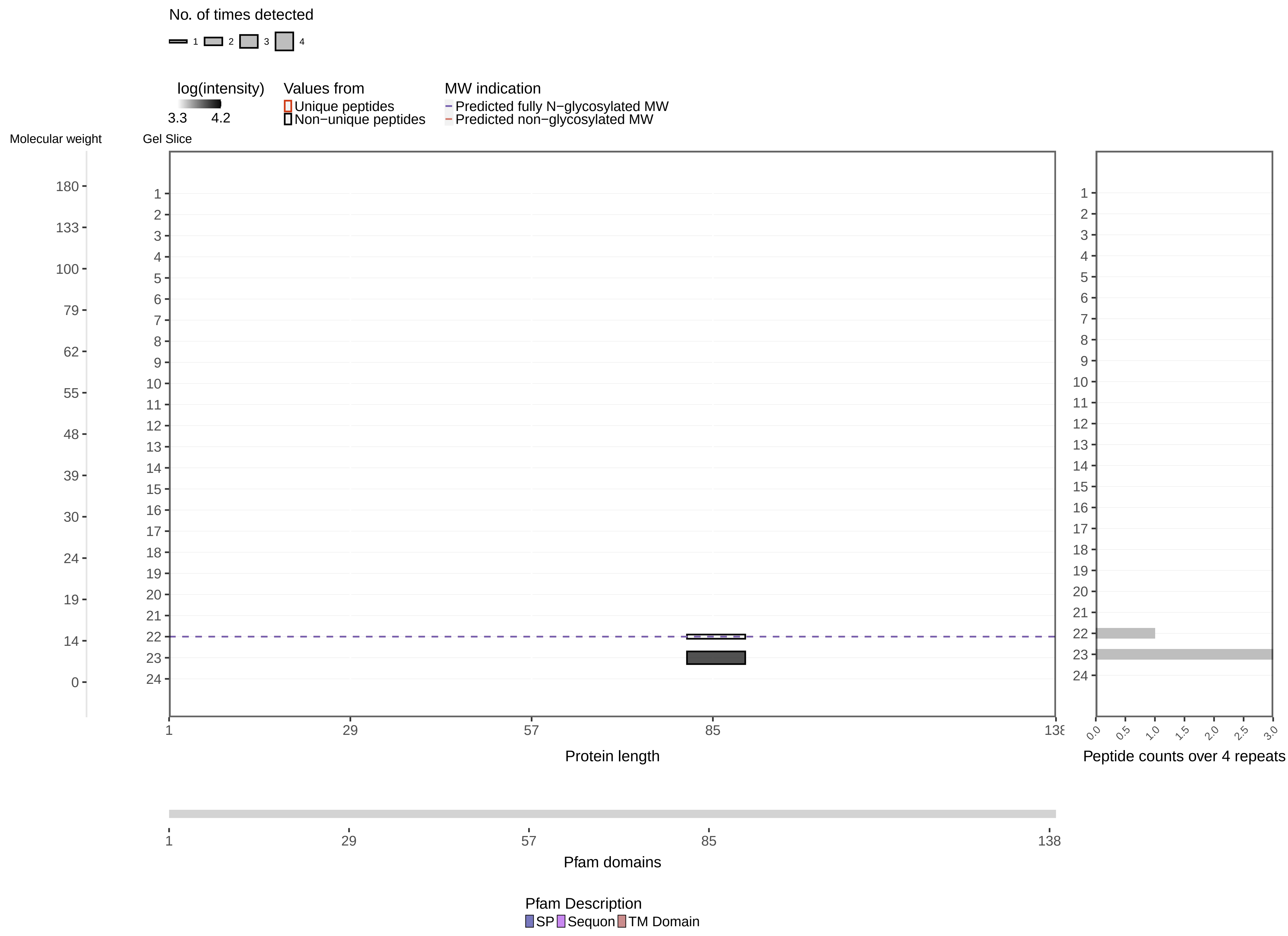


Protein sequence:

RSFLTLEEAGLVEVSGLSTHERFLCRLTISSLNLLRVIAEQEGCSIEELNAGRVCDWFLKDKLKREQNLESAVLQWDESDFLL*

### NbL05g17590.1 nudix_hydrolase_15_mitochondrial [MW = 33.98 kDa]

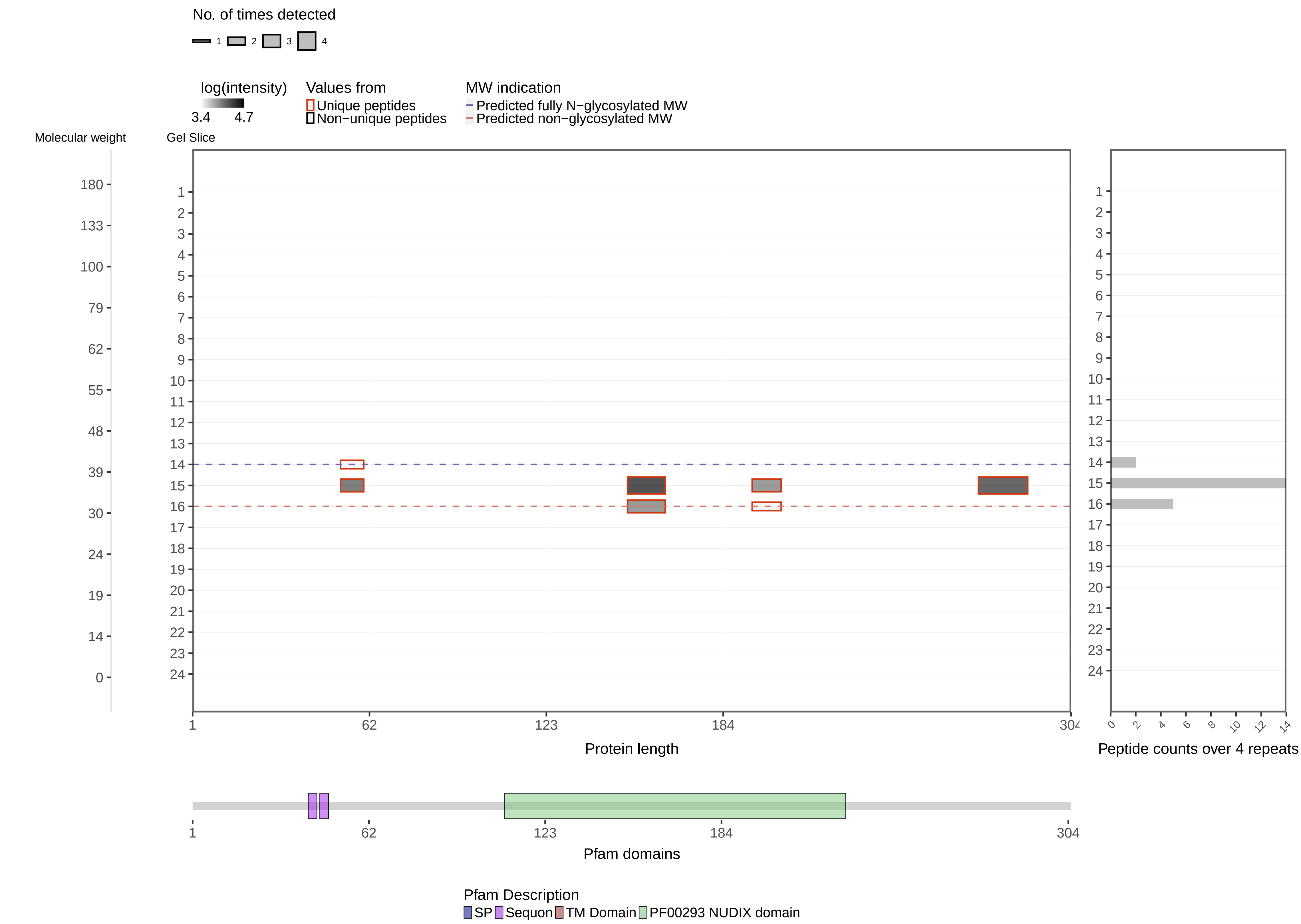


Protein sequence:

THSGEVALPGGKAEEGDASDADTATREAKEEIGLDPSLVNIVTCLEPFLSKHLLRVIPVIGILSNKKEFNPTPNIAEVEAVFDVPLAMFLKDENRRSEEREWMGEKYLIHLFDYELDNKKYLIWGLTAGILIRAASVVYQRPPAFLEQNPKFKLPKVVDKDTTMS*

### NbL05g17570.1 magnesium−chelatase_subunit_ChlI_chloroplastic [MW = 46.43 kDa]

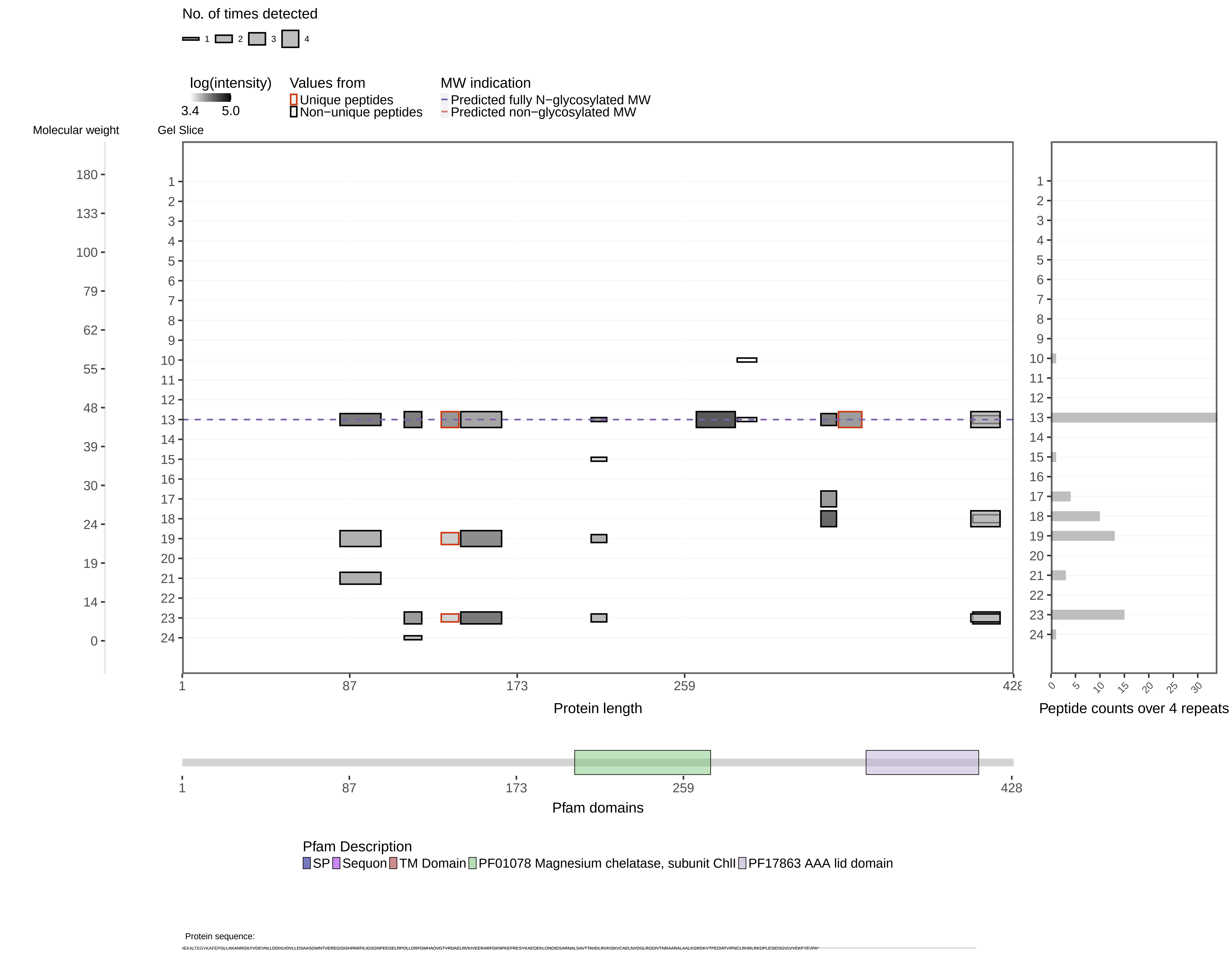

NbL05g17520.1 malonyl−coenzyme:anthocyanin_5−O−glucoside−6'''−O−malonyltransferase−like [MW = 50.68 kDa]

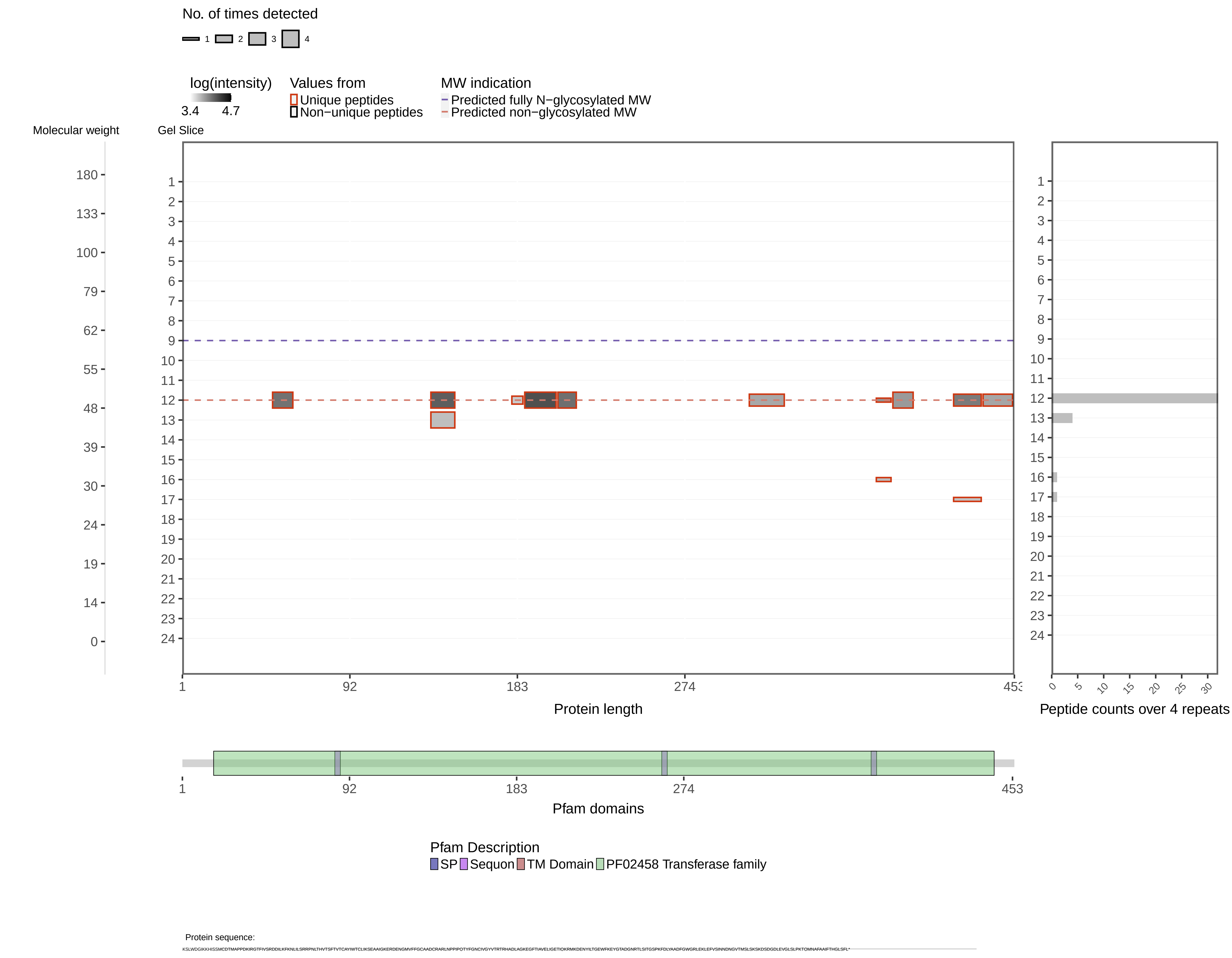

NbL05g17490.1 diacylglycerol_kinase_5−like [MW − SP = 53.96 kDa]

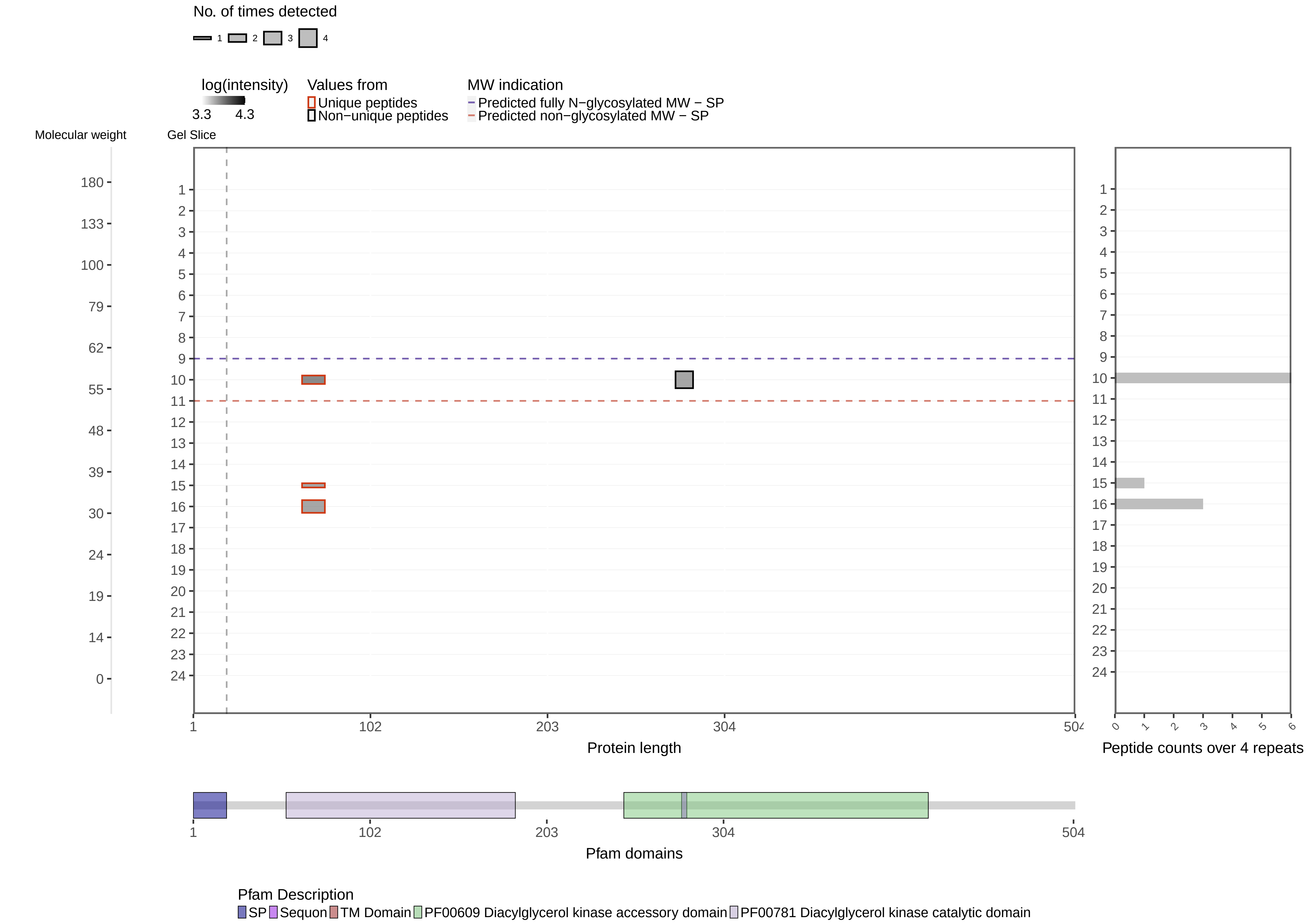


Protein sequence:

FRGGFWNYFSMGMDAQVSYAFHSERKMNPEKFKNQLVNQSTYARLGCTQGWFFASLFHPSSRNIAQLAKVKIMKRQGEWQDLHIPHSIRSIVCLNLPSFSGGLNPWGTPNSNKRRDRDLTPPFVDDGHLEVVGFRDAWHGMVLLAPNGHGTRLAQAHRIRFEFHKGAADHTFMRIDGEPWKQPLPVDDDTVVVEIAHLGQVKMLATHGCRSKSMHDPSSHVNHDGDDGDSGEEDSVVEEQRKFGAADTFKIPDEVDISHLS*

### NbL05g17420.1 malonyl−coenzyme:anthocyanin_5−O−glucoside−6'''−O−malonyltransferase−like [MW = 50.17 kDa]

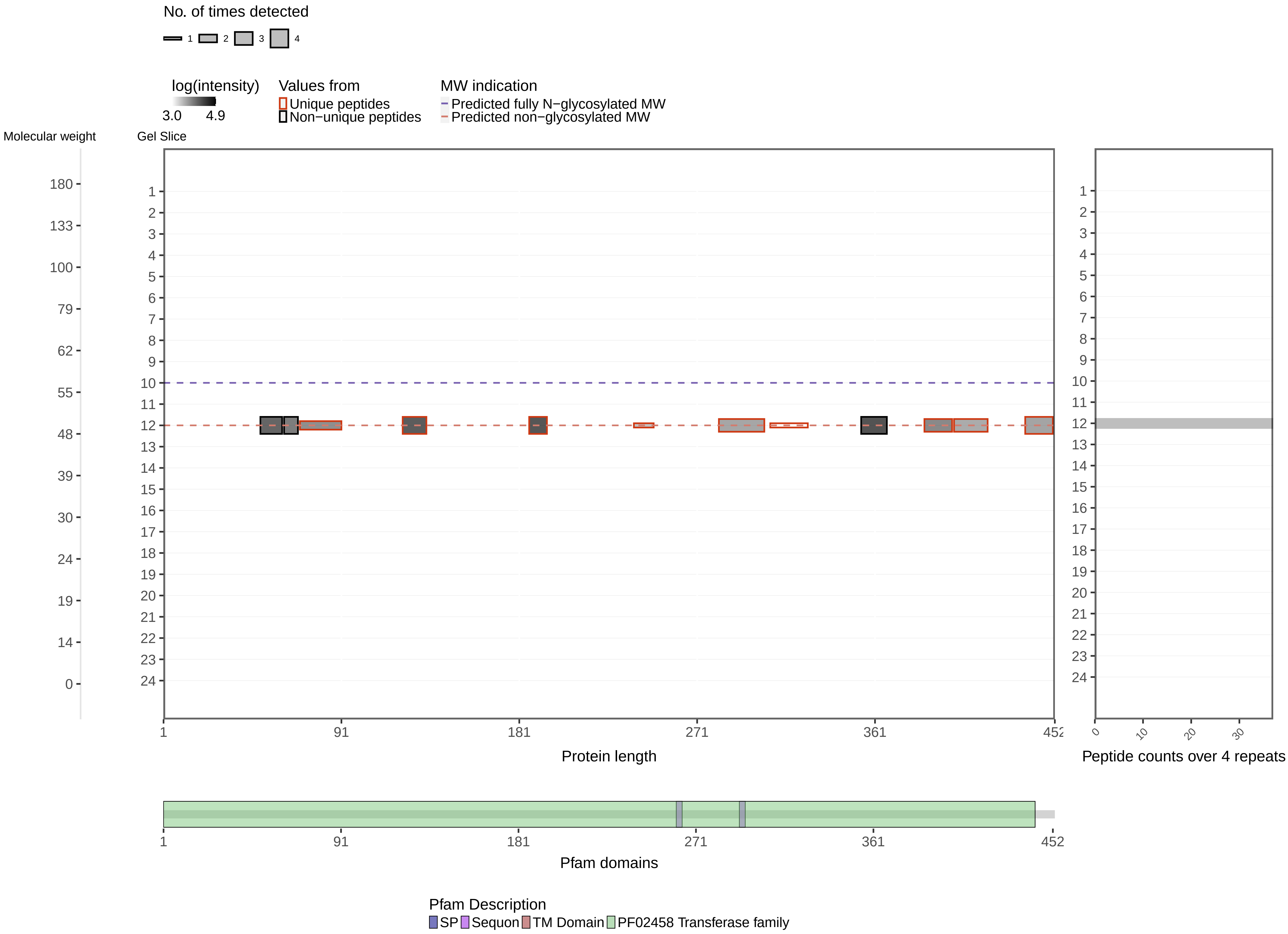


Protein sequence:

GMSIWDETKKNKLEMCDIVTPPDKVRCTFIVTQDVIRKLKNLMLSRRPNLTHVTSFTVTCAYVWTCLIKSEDAIREEIEENVTELFGCVADCRSRLNPPLPPSYFGNCLVTLIARASHVDLVGKEGFTIAVEVIGETIQKRMKDENYILTGEWFKEFCSGDIKRSLSIAGSPKLDLYAVDFGWEKPEKLEFVSIDNGDGVSMSLTKSKDSDGDLEIGLSLSKTRINAFAAMFTHGLSFL*

### NbL05g17320.1 jasmonic_acid−amido_synthetase_JAR1−like [MW = 68.18 kDa]

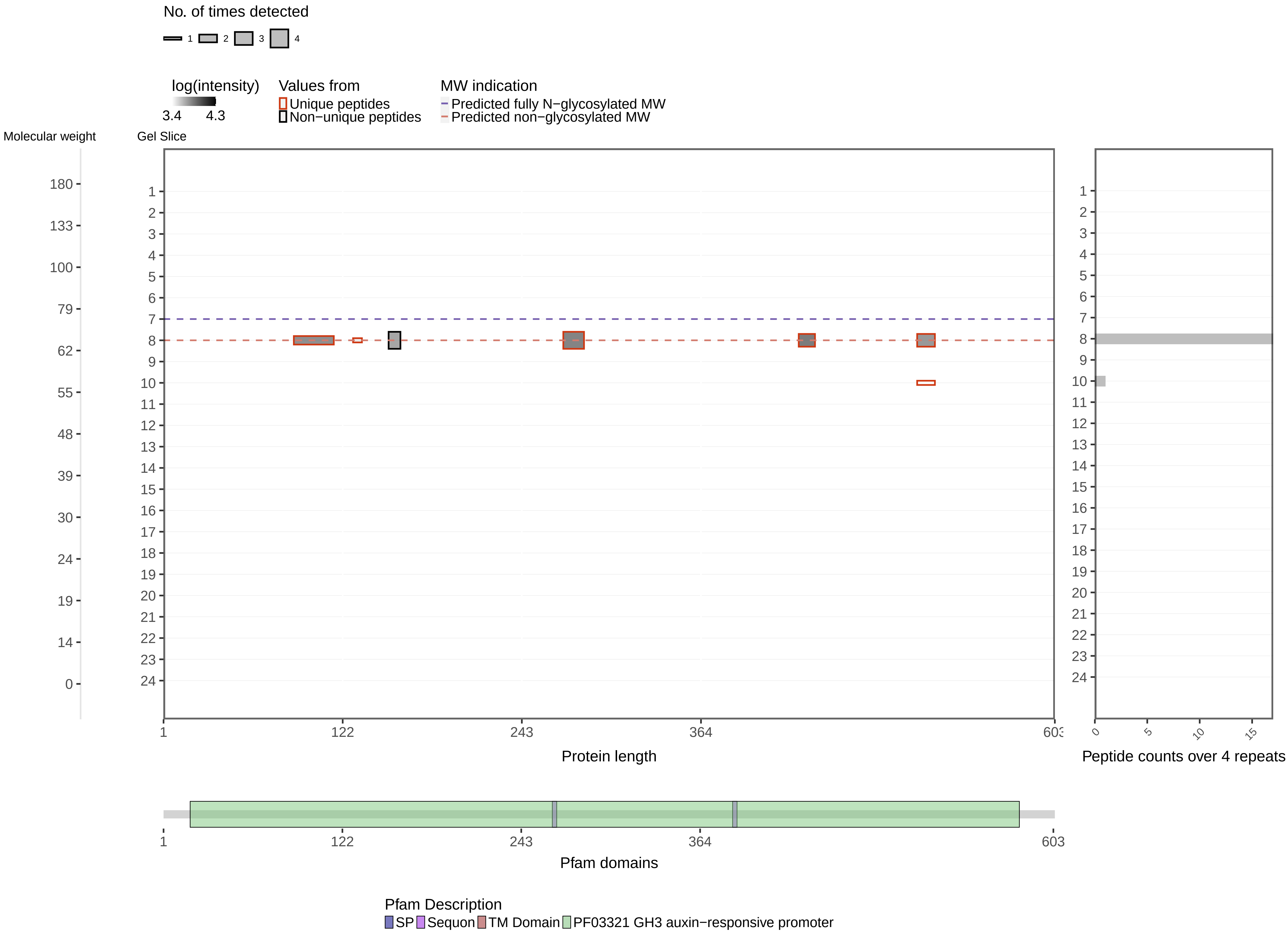


Protein sequence:

ICEEVEKEDWFSIIPRLWPNAKYVVYSIMTGSMQPYLTKLRHYAGKLPLVSADYGSTESWIGVNVEPSSPPEKVTFAVIPTFAYFEFIPLHRTKSQNCNNSQDANLATDDFIEDDPVPLSQVKIGQEYEIVLTTFTGLYRYRLGDVVEVAGFHKNTPKLNFICRRKLILTVNIDKNTEKDLQLVVERGSQILSKSRAELVDFTSHANVAKRPGHYVIYWEIKGEVEENVLGECCREMDASFVDHGYVVSRKTNSIGPLELCIVERGTFKKILEYFIGNGAALSQFKTPRCTSNQVLLTILN

### NbL05g17240.1 bifunctional_epoxide_hydrolase_2−like [MW = 35.39 kDa]

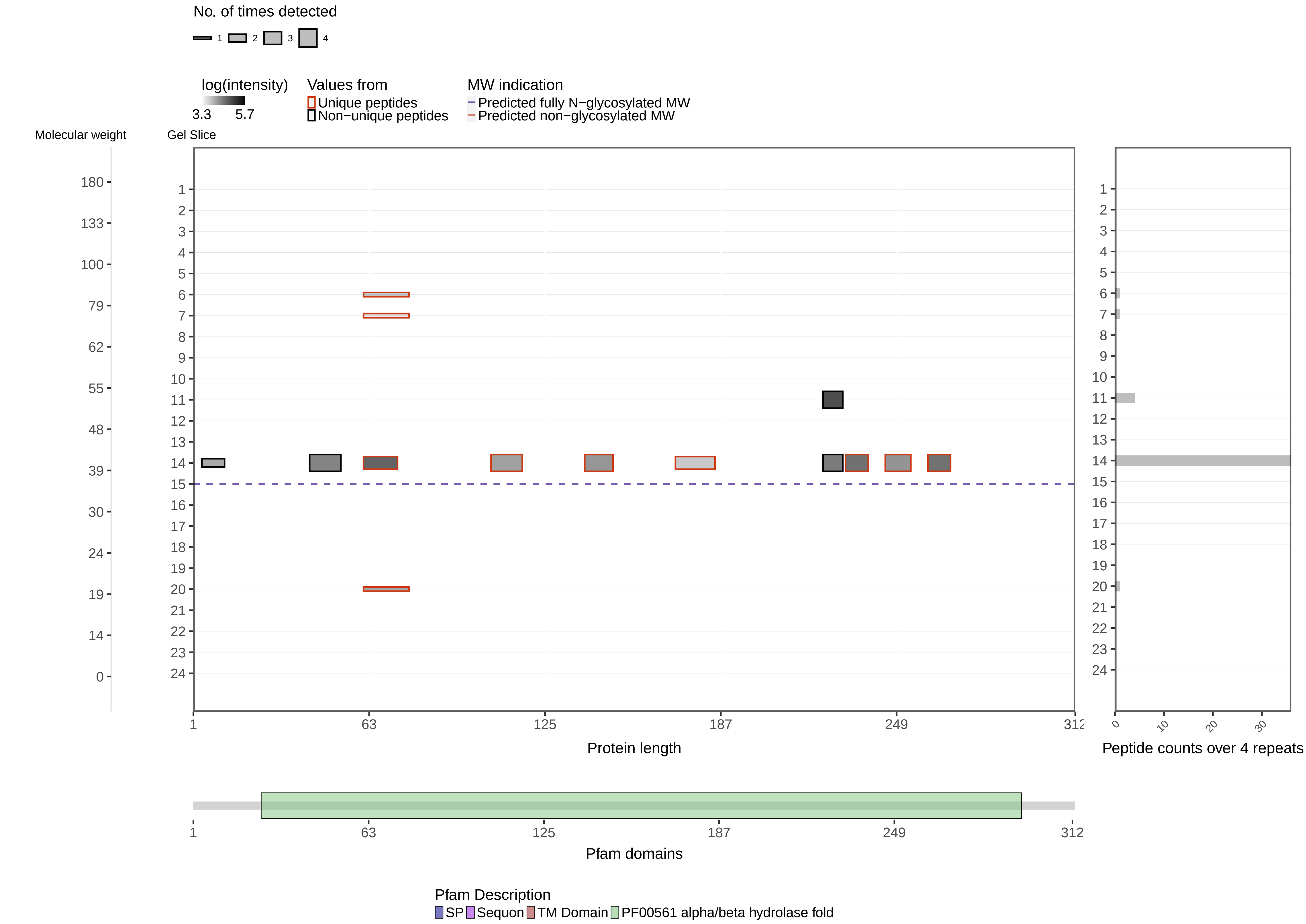


Protein sequence:

GFYMLRWOEAGRAEKDFGRFDTKTVVKNIYIMFSGSELPVAKDDEEIMDLVDPSAPLPDWFTEODLANYASLYEKSSFRTALOVPYRAWLEEYGVKDIKVKVPCLLVMGEKDYSLKFGGLEOYVKSGMVKEYVPNLETIFLPEGSHFVOEOFPEOVNOLIITFLKKLI*

NbL05g17230.1 lanC−like_protein_GCL2 [MW = 46.95 kDa]

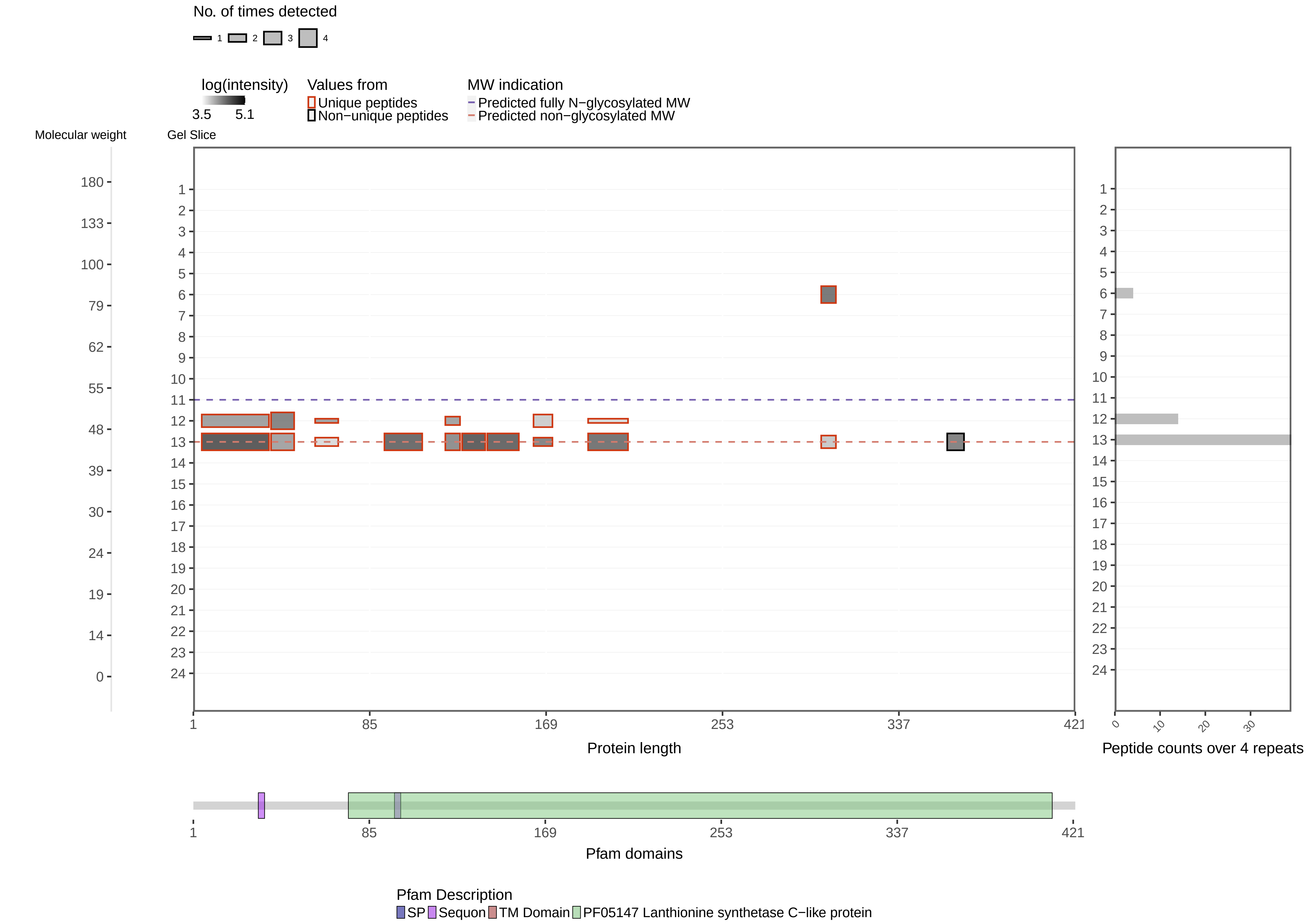


Protein sequence:

VNEIIENGRRLGGRGKSPLMYEWYGEMYWGAAHGLAGIMYVLMDFELKPDELEDVKETLRYMVKNRFPSGNYPTSEEDKRRDVLVHWCHGAPGIALTLVKAAELFGDREFLNAAVGAAEVVWNRGLLKHVGMCHGISGNAYVFLSLYQLTGNVEYLYRAKAFACFLLDRAHKLIAKGEMHGGDNPYSLFEGIGGMAYLFLDMTEPKEARFPAYGCEKVHAT*

### NbL05g17130.1 uroporphyrinogen_decarboxylase_1_chloroplastic [MW = 46.01 kDa]

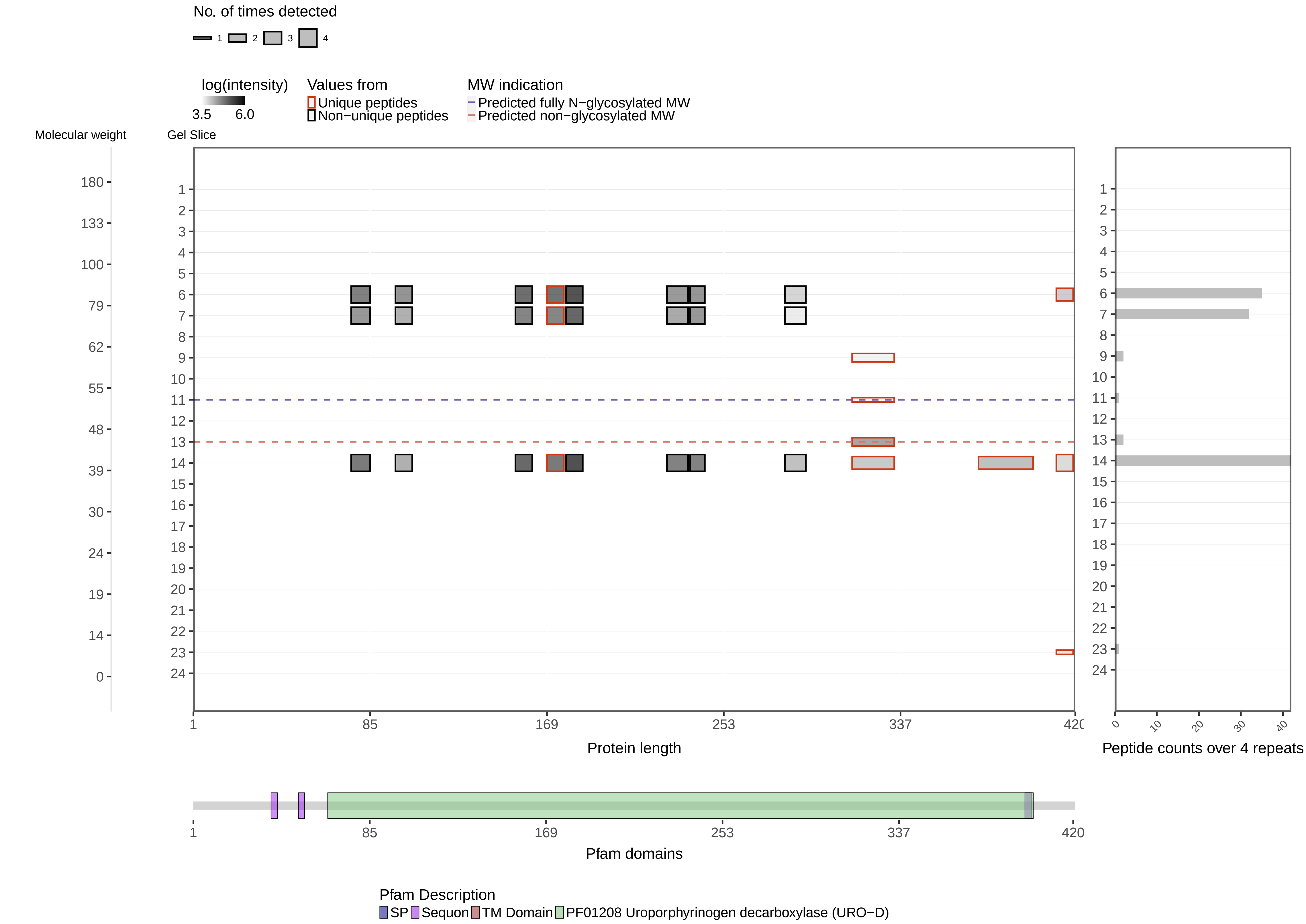


Protein sequence:

AAVLGFVGAPWTIATYIVEGGTTRTYTTIKDMCHTAPHVLRALLSHLSKAIAEYIVFQVESGAHCIQIFDSWGGQLPPHMWDCWSKPYIDEIVATIKRKCPQTPLVLYINGNGGLLERMKETGVDVIGLDWTVDMADGRKRLGTDIGIQGNVDPASLFCPLPALTDEIERVVKSAGTRGHILNLGHGVLVGTPEEAVAHFFNVSKSFNFDKVGDRLSEANTVAA*

NbL05g17110.1 phosphoenolpyruvate_carboxylase [MW = 110.41 kDa]

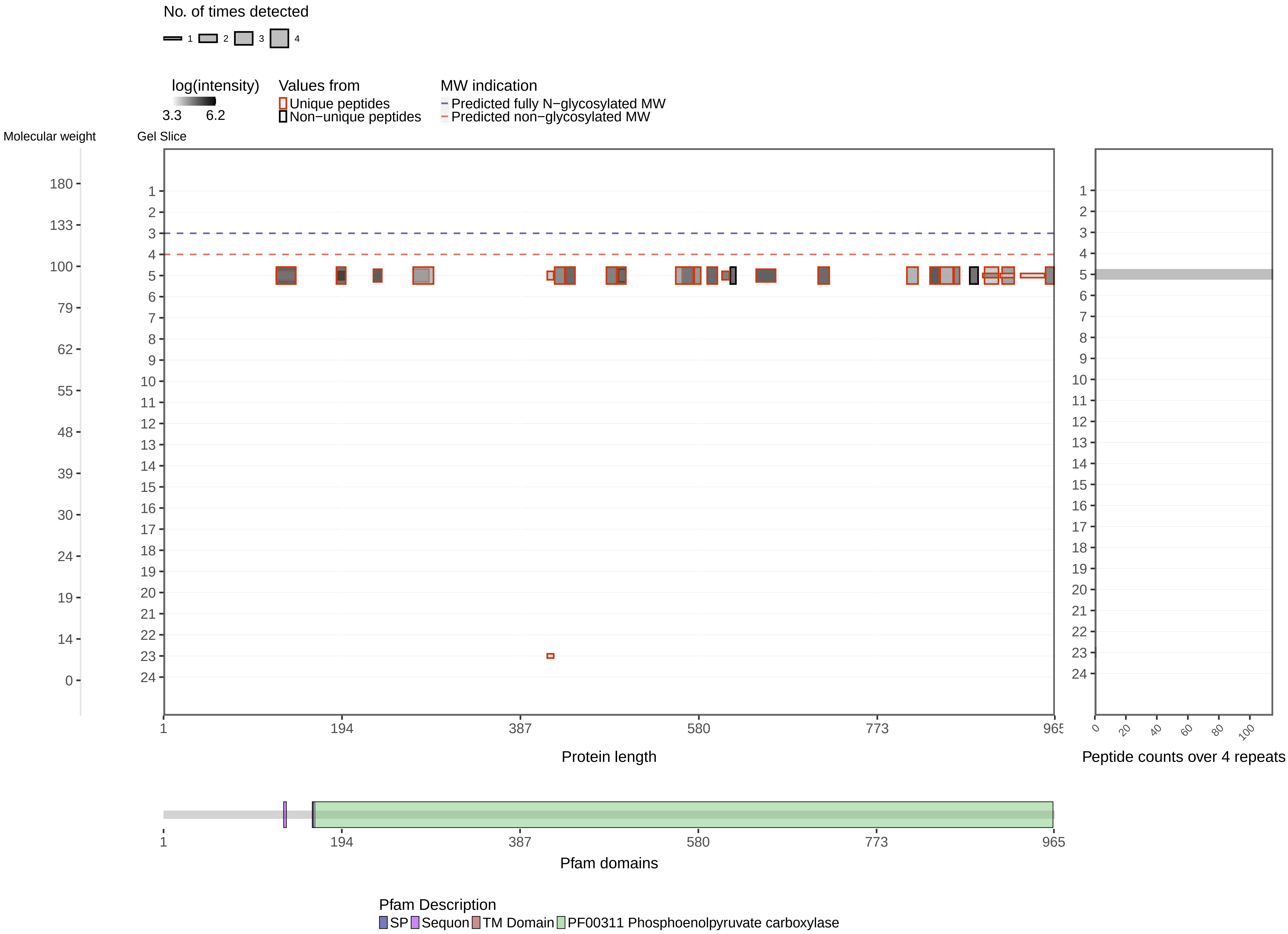


Protein sequence:

IGSYREWSEERRQEWLLSELSGKRPLFGPDLPRTEEIADVLDTLHVIAELPSDCFGAYIISMATAPSDVLAVELLQRECHVKQPLRVVPLFEKLDDLESAPAAVARLFSIEWYRNRINGKQEVMVGYSDSGKDAGRFSAAWQLYKAQEELIKVAKEHGVKLTMFHGRGGTVGRGGGPTHLAILSQPPDTIQGSLRVTVQGEVIEQSFGEEHLCFRTLQRFTAATLEHGMHPPVSPKPEWRALMDEIAVIATEKYRSIVFKEPRFVEYFRLATPELEYGRMNIGSRPSKRKPSG

### NbL05g17070.1 ubiquitin−conjugating_enzyme_E2_36 [MW = 17.22 kDa]

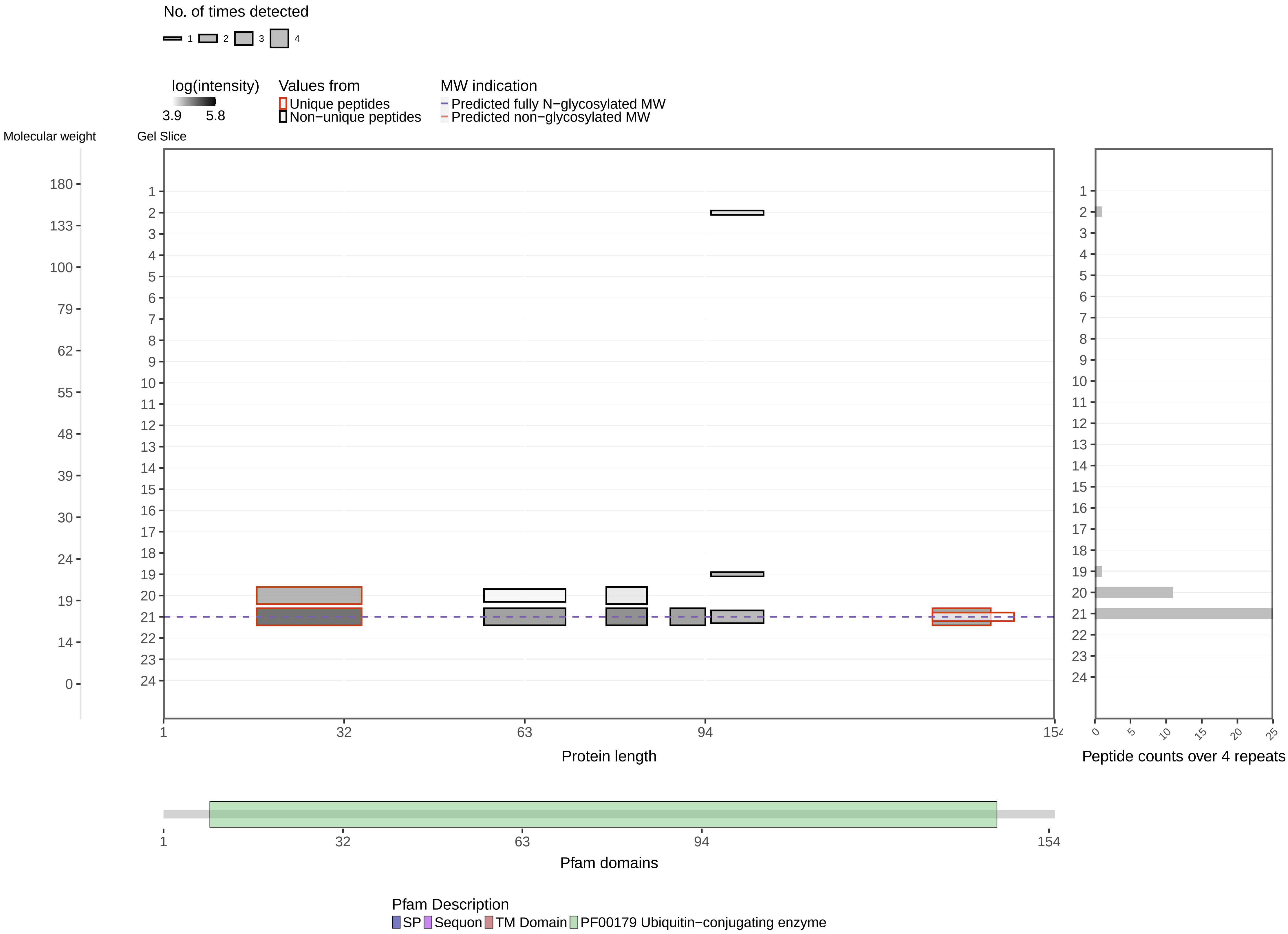


Protein sequence:

EYPMAAPKVRFLTKIYHPNIDKLGRICLDILKDKWSPALQIRTVLLSIQALLSAPNPDDPLSENIAKHWKSNEAEAVETAKEWTRLYASGA*

### NbL05g17020.1 malate_dehydrogenase_mitochondrial [MW = 41.53 kDa]

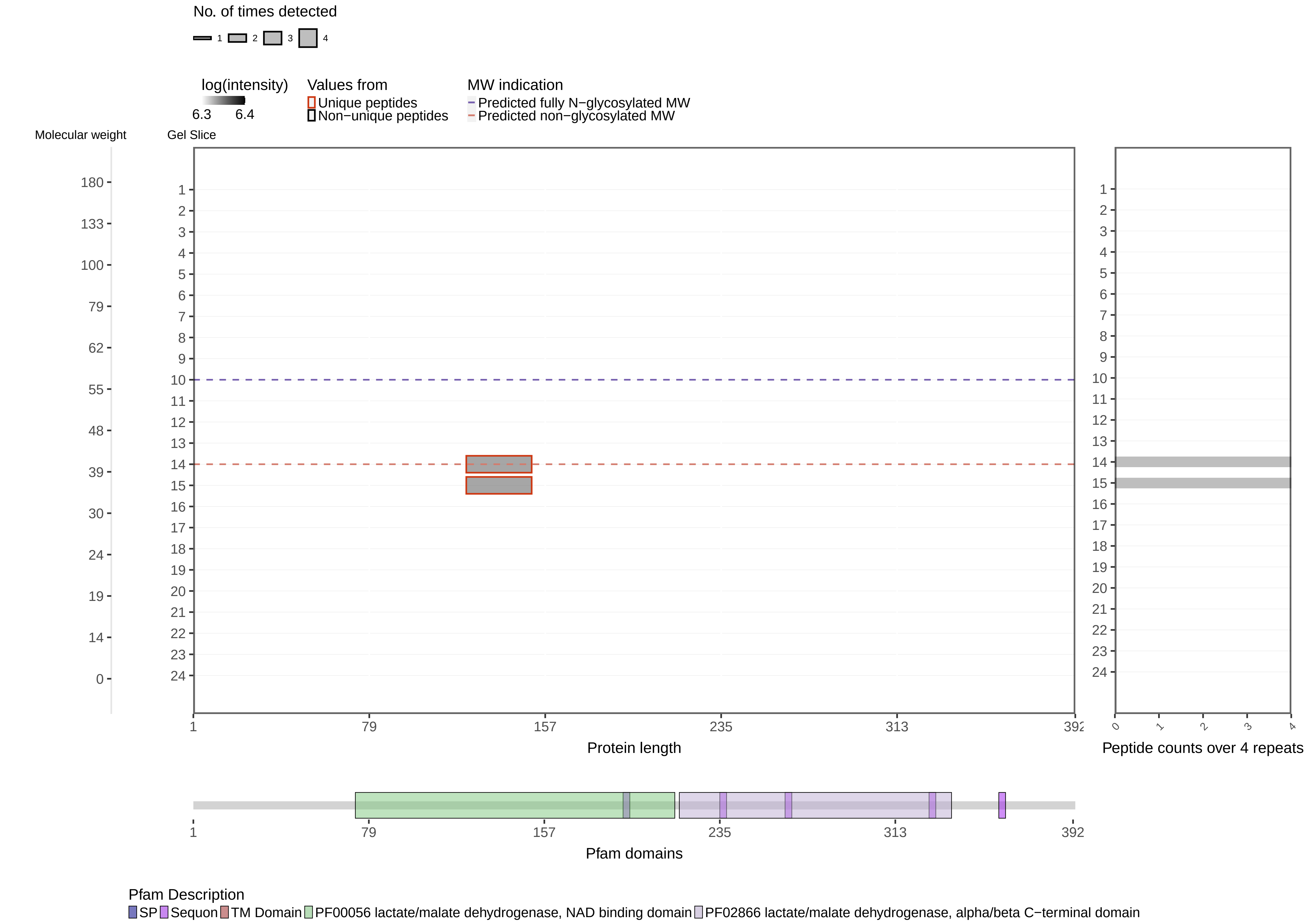


Protein sequence:

MISNPVNSTVPIAAEVFKKAGTYDEKRLFGVTTLDVVRAKTFYAGKAKVNVSDVIVPVVGGHAGITILPLLSQATPKANLSDEEIVALTKRTQDGGTEVVEAKAGKGSATLSMAYAGAIFADACLKGLNGVPDVVECSFVQSNVTELPFFASKYNTHIQALPNMPQSSSTPISNFSALKKNTELMNAKHGSSKIVQDKKLQRRQKE*

NbL05g16840.1 glutamyl−tRNA(Gln)_amidotransferase_subunit_A_chloroplastic/mitochondrial [MW = 57.18 kDa]

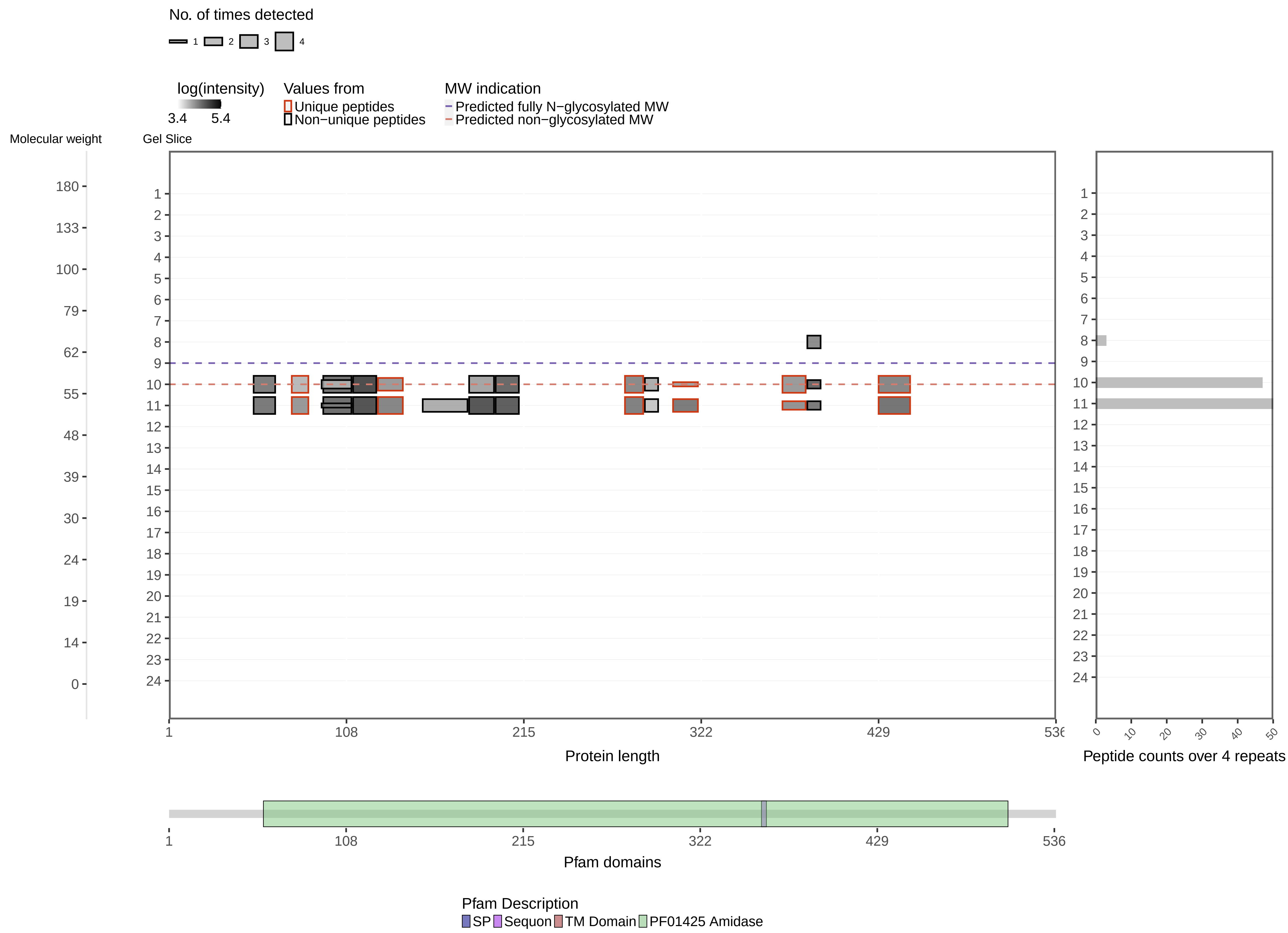


Protein sequence:

ADAGIILHAISGHDKYDATSSKREVPNFASQFIARDHLDSKPLKGLRVGLIRETIEDGVDPEVITSIRGAASHLEELGCTVTEVSLPSFSLGLPAYYILASSESSSNLSRYDGIRYGKQVFADELNSLYGESRAGGFGSEVKMRILMGTYALSAGYYDAYYKRAQQVRTLVRENFRAALEENDILISPAAPSAAYKIGEKKNDPLSMYAGDIMTVNVNLAGLPALVLPCGFVDSSSAALPVGIQMIGAAFEEEKLLKVGHIFEQTLQGCSFIPPIVVADELAC*

NbL05g16630.1 proteasome_subunit_alpha_type−2−A−like [MW = 25.65 kDa]

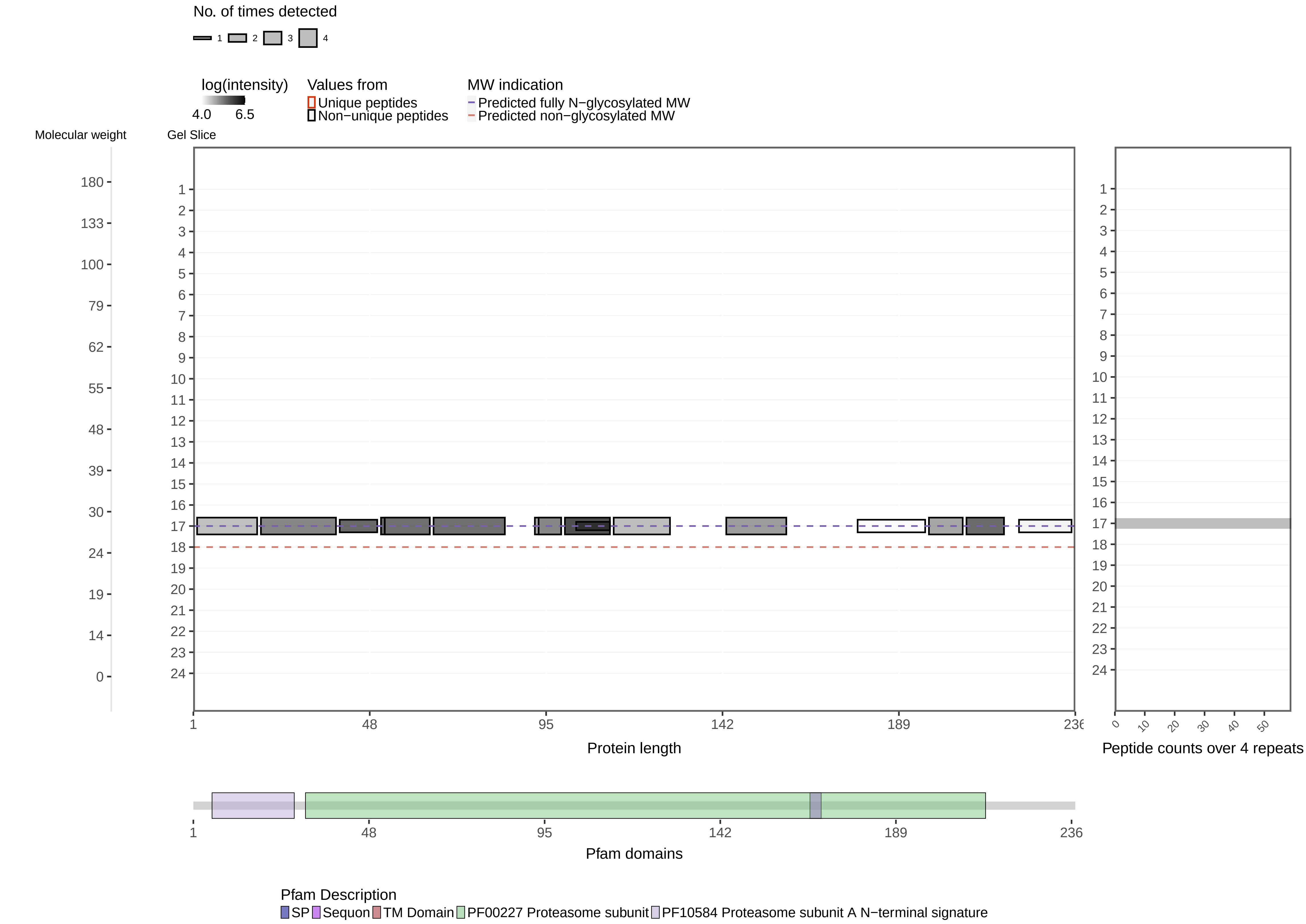


Protein sequence:

'IPVTQLVRETAAVMQEFTQSGGVRPFGVSLLVAGYDDKGPQLYQVDPSGSYFSWKASAMGKNVSNAKTFLEKRYTEDIELDDAVHTAILTLKEGFEGQISGKNIEIGIIGNDKVFKILTPTEIDDYLQEVE*

NbL05g16610.1 14−3−3_protein_9−like [MW = 21.44 kDa]

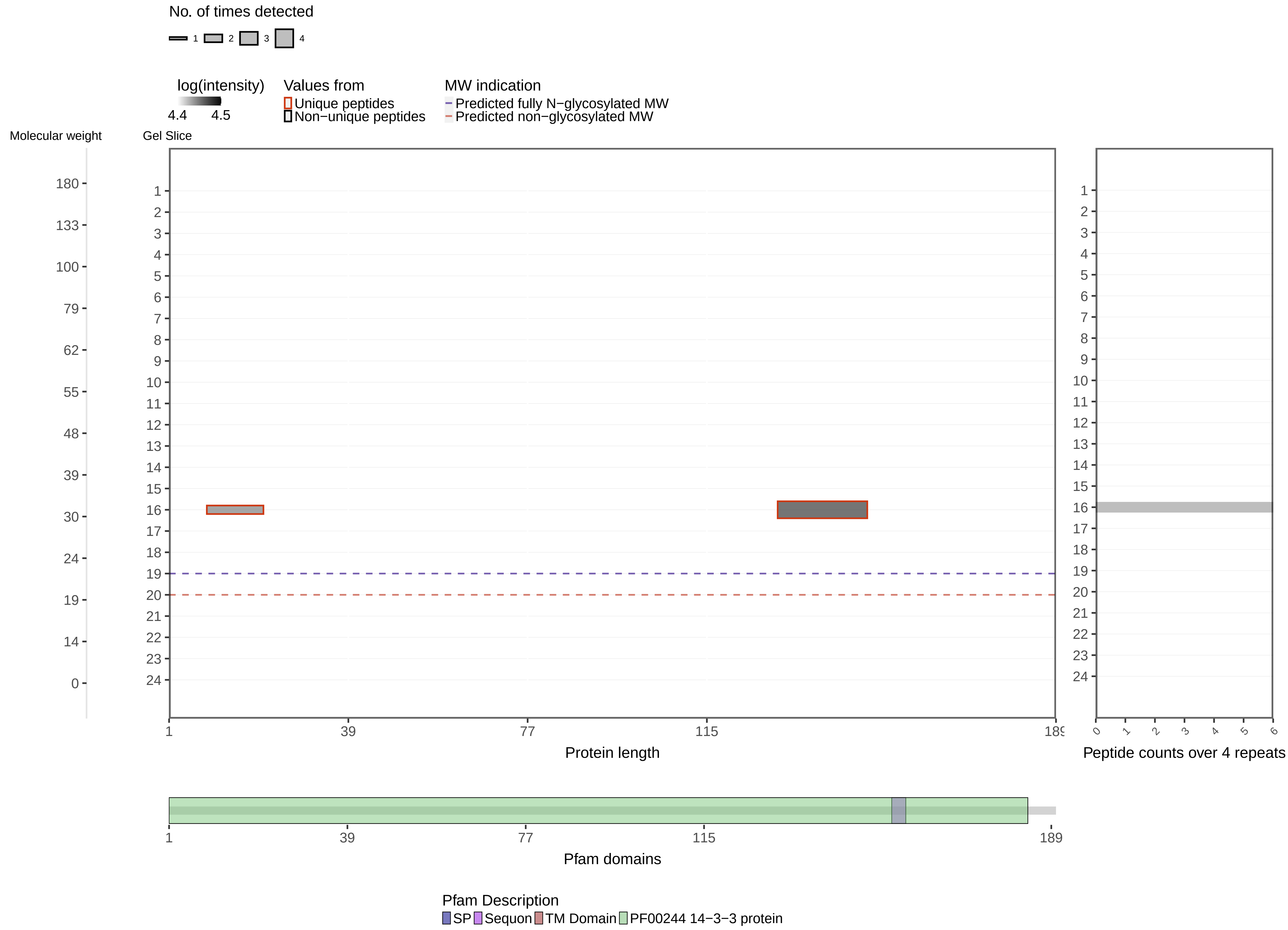


Protein sequence:

[illegible]TVIDEHLIPSCPAGESSVFYYKSEGDYYRYLAEFKTGTDKKEVSDLSLKAYQSATTTAETELPPTHPIRLGLALNFSVFYYEILNSPERACHLAKQAFDEAISEWIAE*

NbL05g16530.1 60S_ribosomal_protein_L19−3−like [MW = 24.82 kDa]

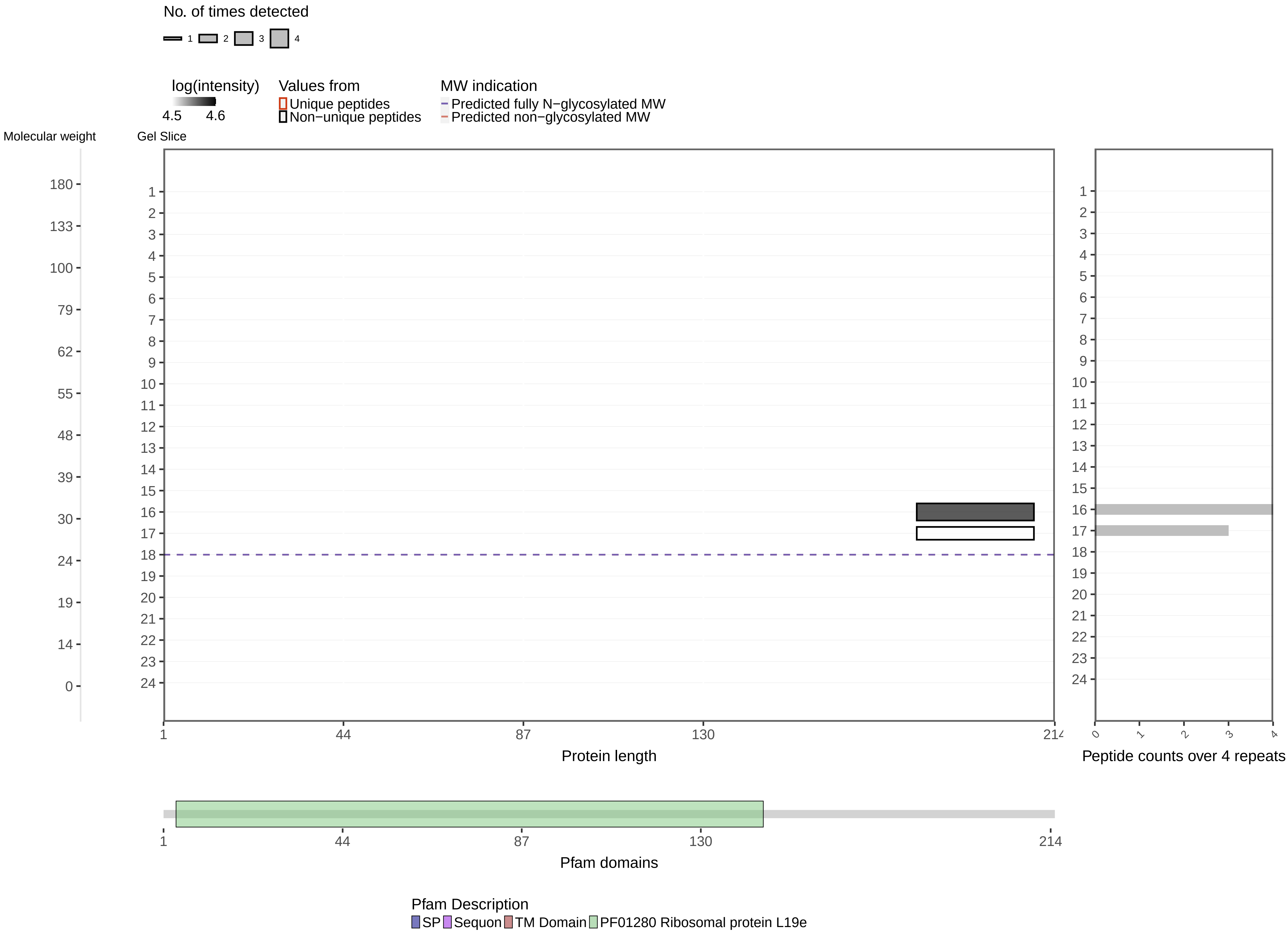


Protein sequence:

VMRRMRVLRRLLRKYRESKKIDRHMYHDMYMKVKGNVFKNKRVLMENIHKTKAEKAREKTLSDQFEARRAKNKASRERKFARREERLAQGPGERPVQPAAPAAAAPPAQPALGSKKSKK*

### NbL05g16470.1 importin_subunit_alpha [MW = 59.01 kDa]

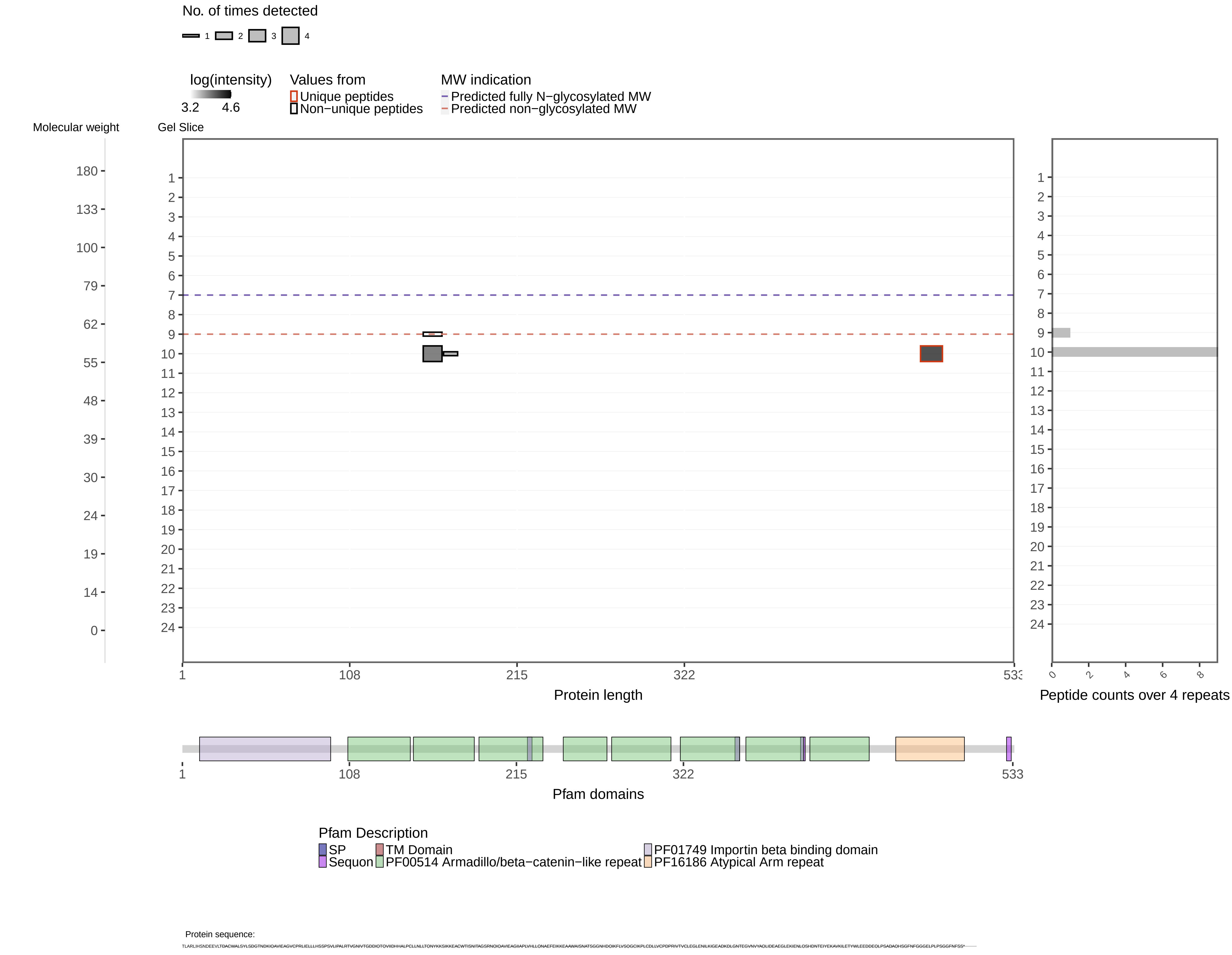

### NbL05g16350.1 calvin_cycle_protein_CP12−1_chloroplastic−like [MW = 14.35 kDa]

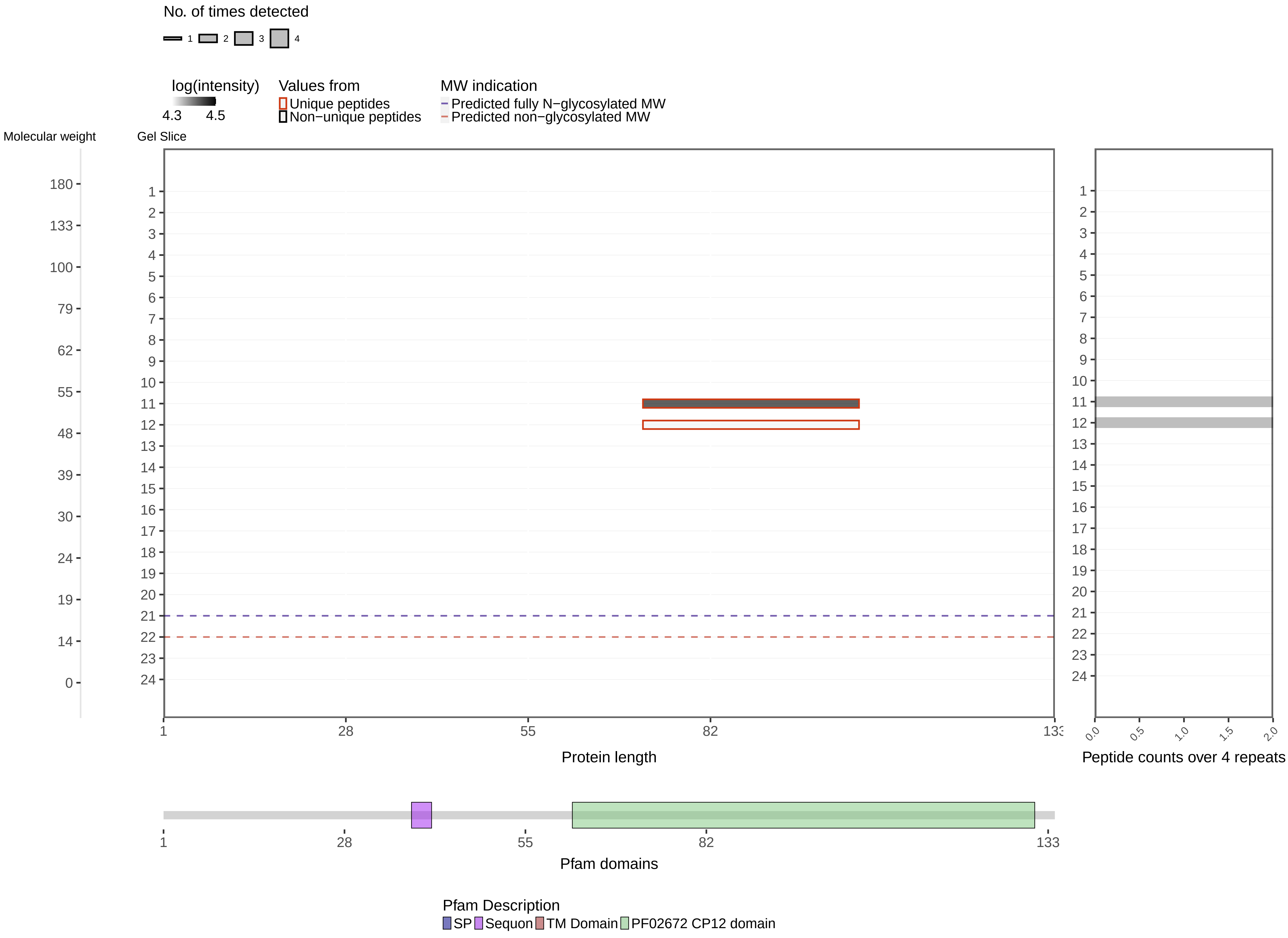


Protein sequence:

SATPDNKLSDLVAESVKEAEEACAENPVSGECAAAWDVVEETSAAASHARDKKKESSDPLENYCKDNPETDECRTYDN*

### NbL05g16240.1 V−type_proton_ATPase_16_kDa_proteolipid_subunit [MW = 11.42 kDa]

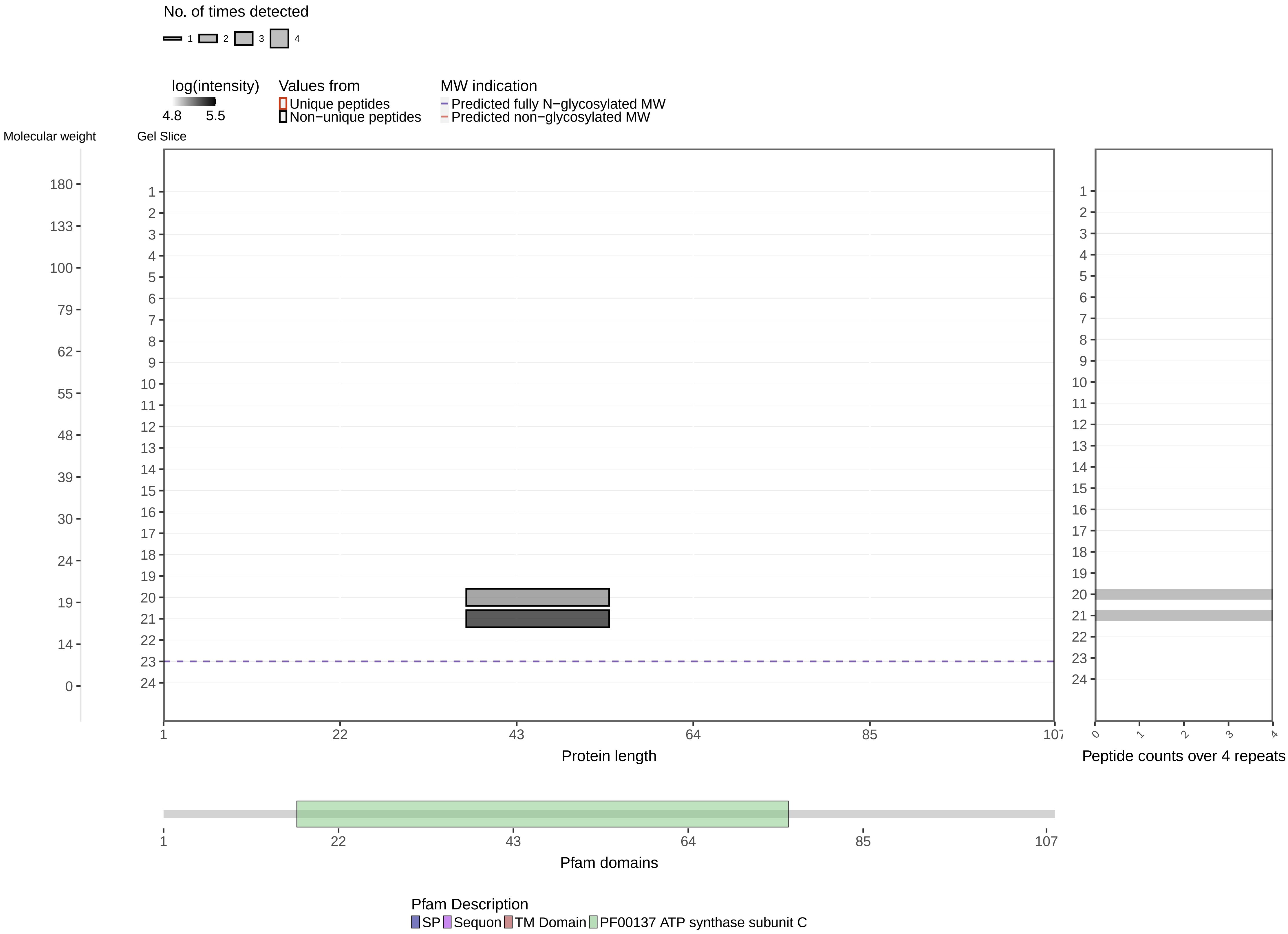


Protein sequence:

VGVASMGVMRPELVMKSIVLVVMPGVLGIYGMIIVVIISTGINPKTKLHVFWRLCLIFLTYIHFVLNV*

NbL05g16220.1 uncharacterized_LOC109215494 [MW = 79.01 kDa]

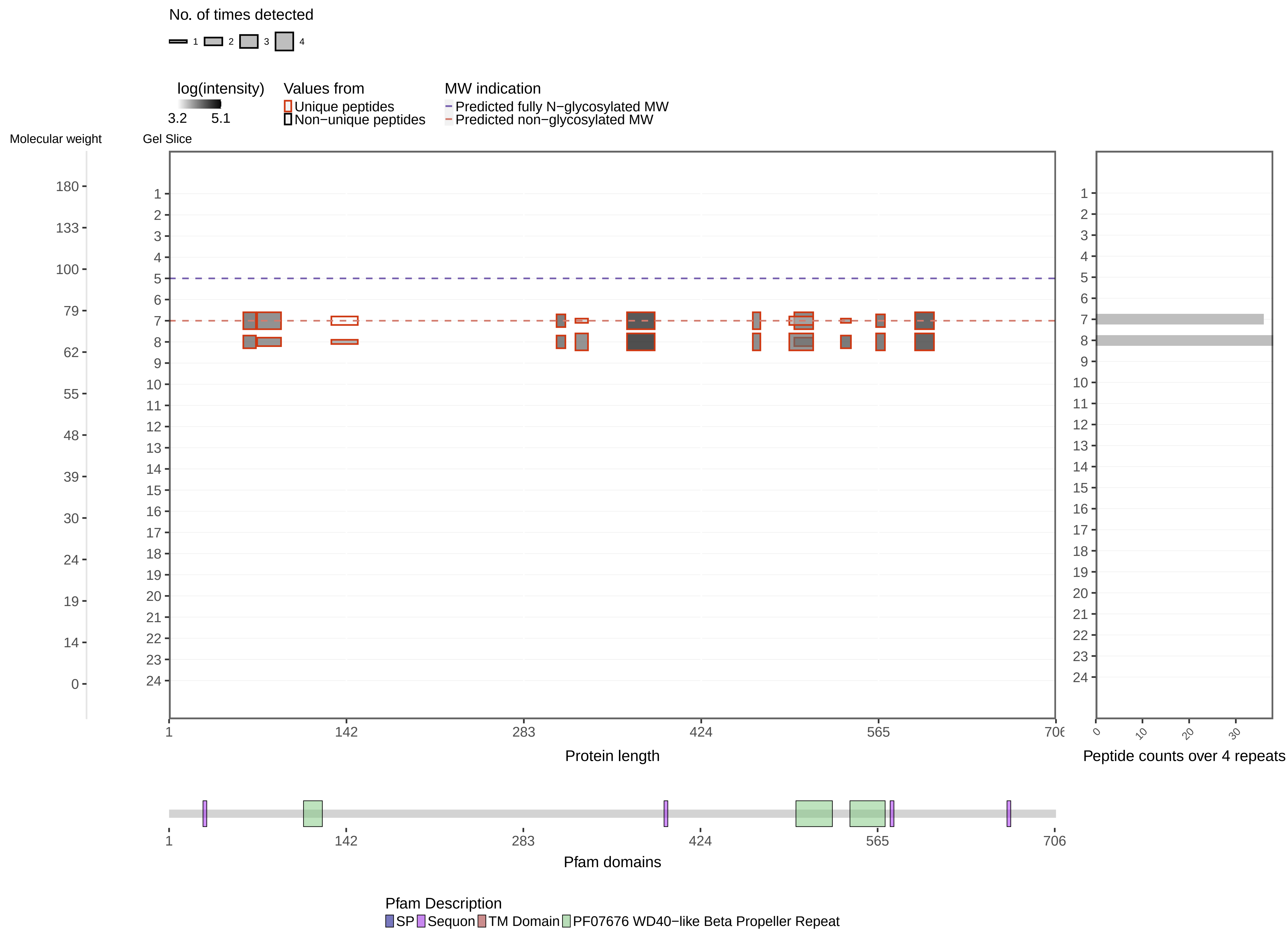


Protein sequence:

### NbL05g16150.1 nitrogen_regulatory_protein_P−II_homolog [MW = 23.6 kDa]

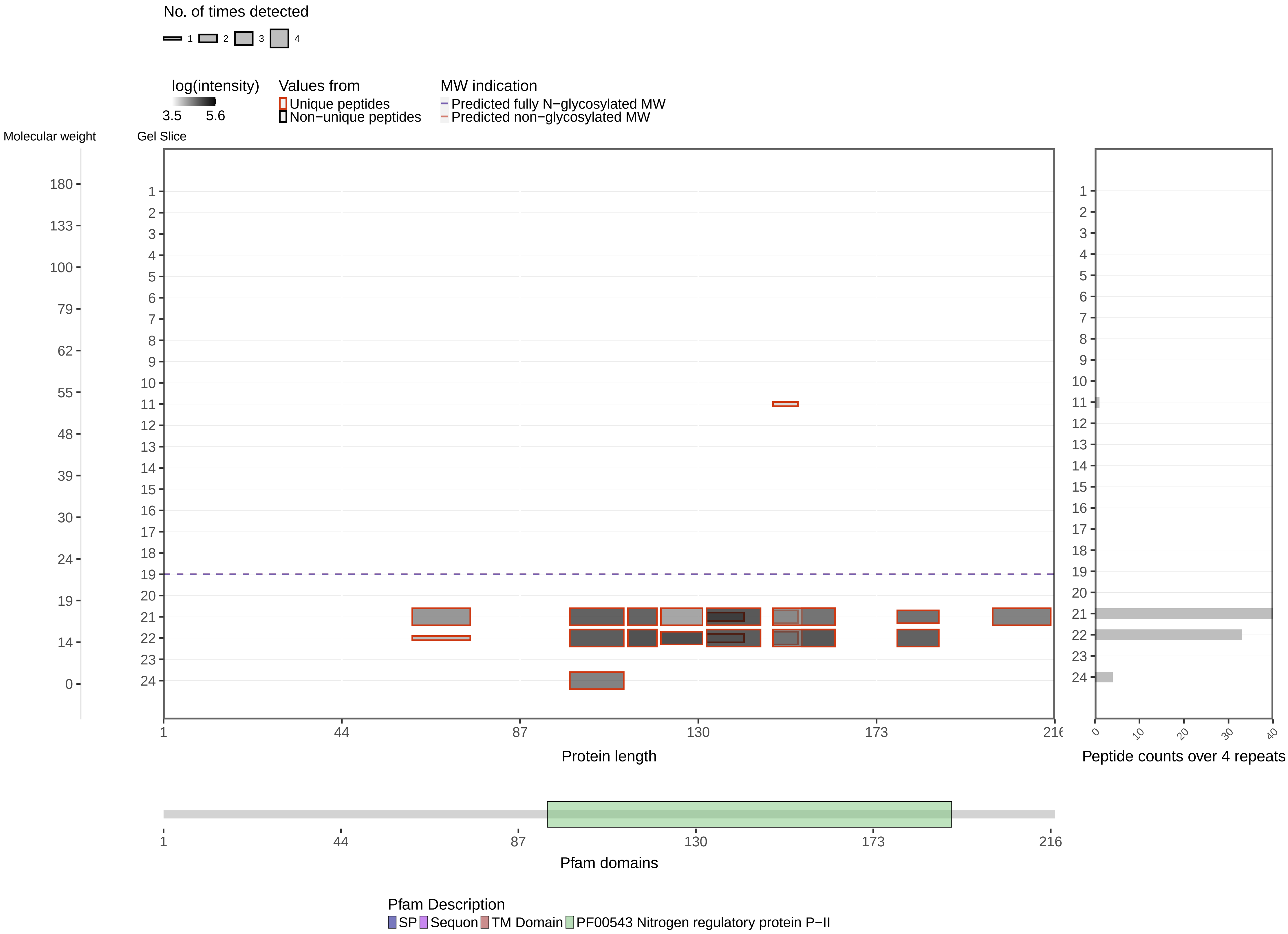


Protein sequence:

/RIQQVSSALLQMGIRGVTVSDVRGFGAQGGLTERQAGSEFSEDKFVAKVKMEIVVSKDQVEGVIAKIIEEARTGEIGDGKIFLIPISDVIRVRTGERGEKAERMVGGHADMSSALSTS*

NbL05g15890.1 GDSL_esterase/lipase_At2g23540−like [MW − SP = 38.27 kDa]

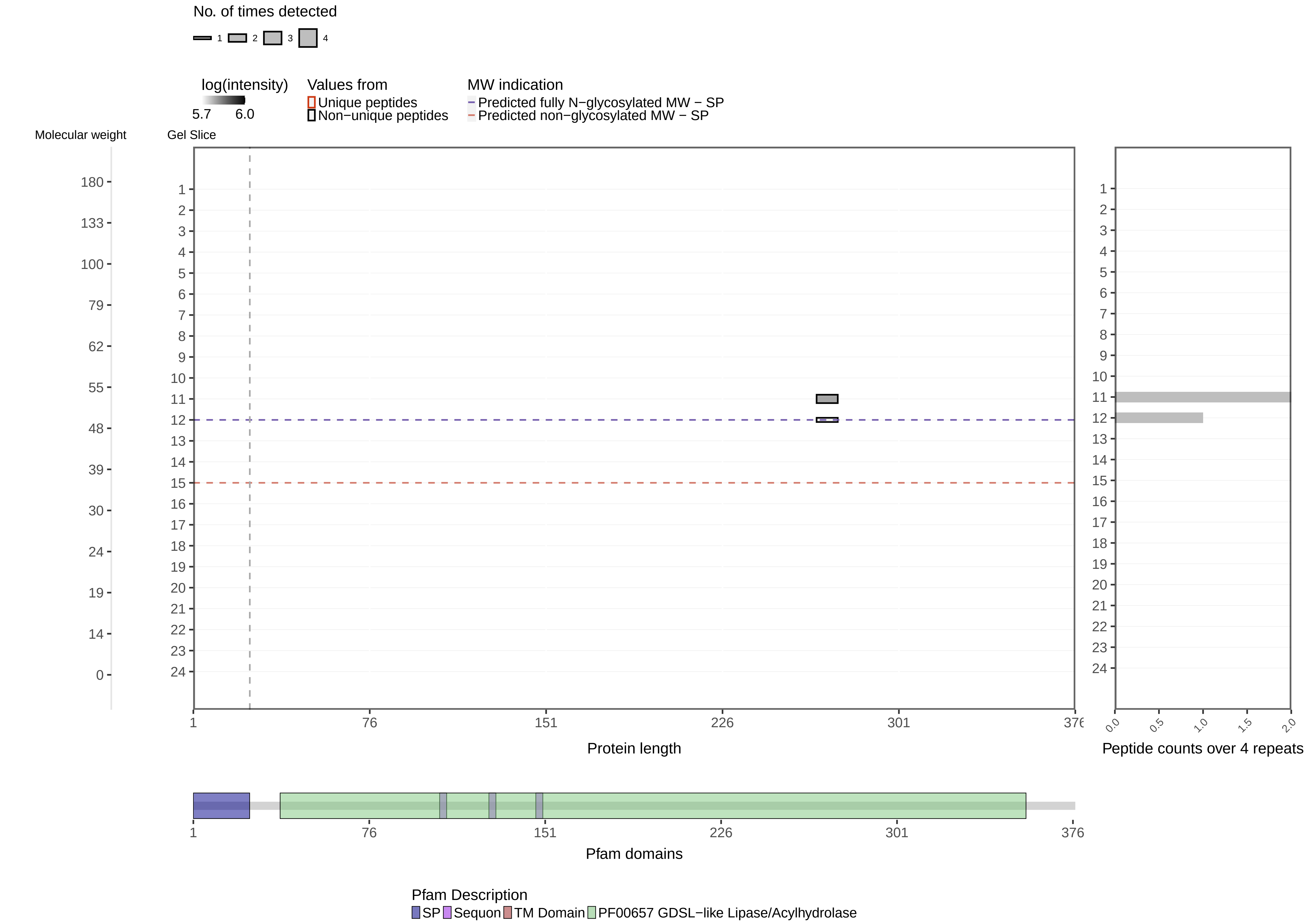


Protein sequence:

IGSNDFLNNYLLLVLSIGTRITESPDAFIDDLLSHLKAOLTRLYKLDARKFIIGNVGPSGCIPYOKTINOLKENECVELANKLALOYNARLKDMLSEMNKNLDGATFLHANVYDLVMELITNYDKYGFATATKACCENGGOFAGIVPCGPTSSMCSDGDKHVFWDPYHPTEAANLIIAKOLLEGDPTYISPMNLKOLRDL*

### NbL05g15840.1 40S_ribosomal_protein_S11−like [MW = 20.79 kDa]

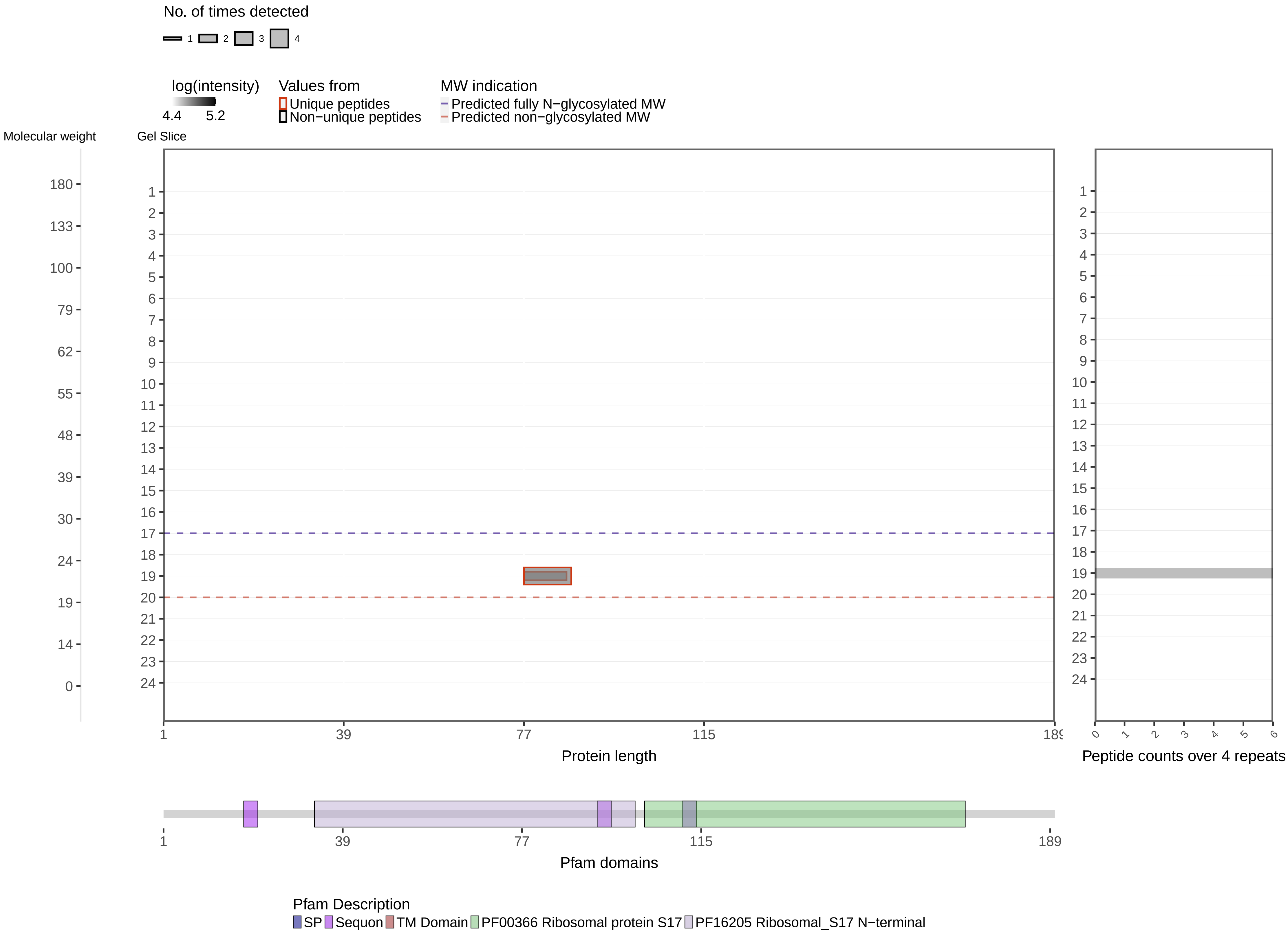


Protein sequence:

GFYIDKKCPFTGNVSIRGRILAGTCHSAKMNRTIIVRRNYLHYVKKYQRYEKRHSNIPAHISPCFRVKEGDHVIIGQCRPLSKTVRFNVLKVIPAGSAGGGKKAFTGM*

### NbL05g15650.1 60S_acidic_ribosomal_protein_P3−like [MW = 15.95 kDa]

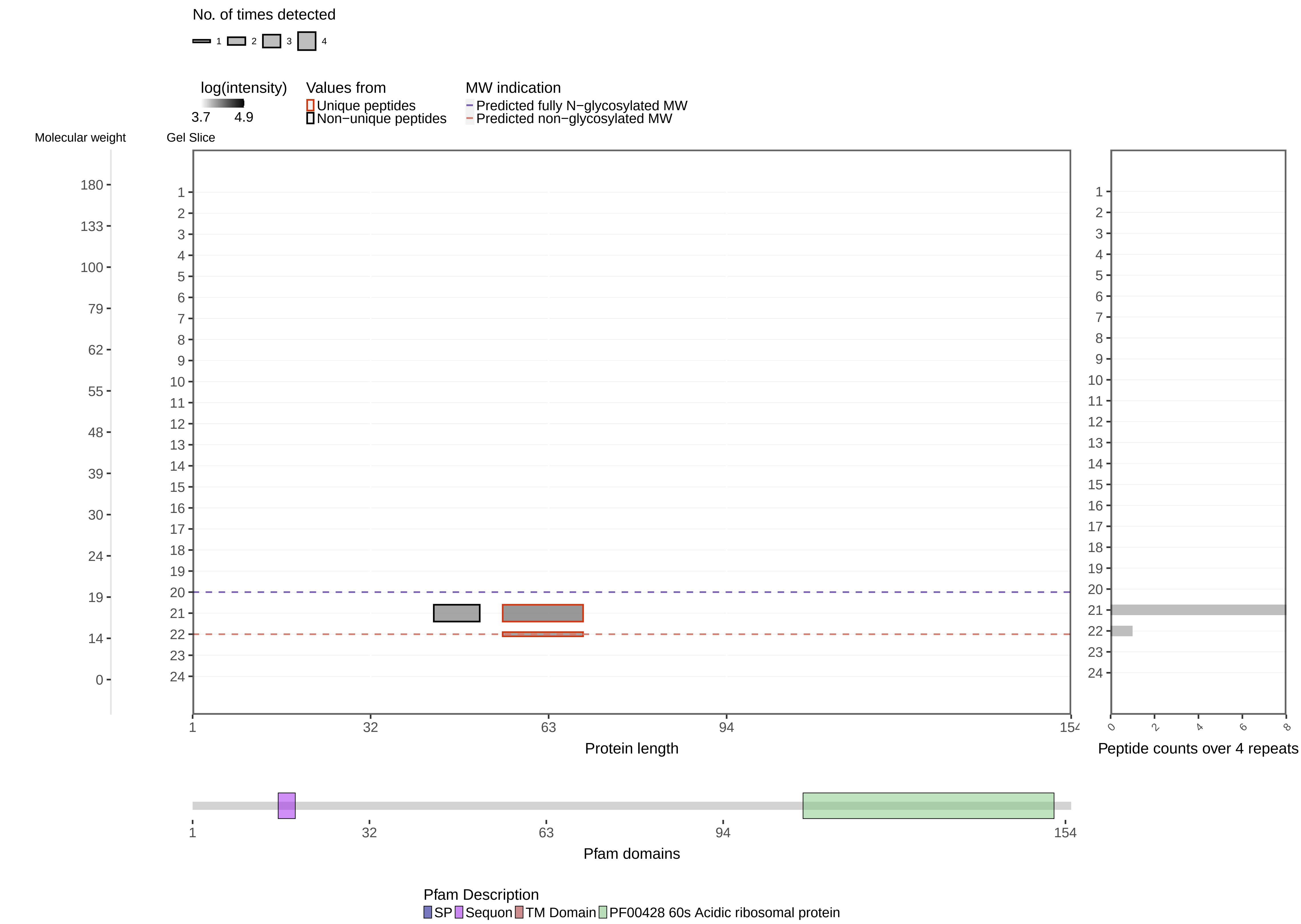


Protein sequence:

YDLQRKLVQAALSTDSSGGVQSSFSYVTPSSAVFQVIIGGGGGGGFIGGGAAAAAAPSGGAAAAEAPAAEEKKEEKEESDDDMGFSLFD*

NbL05g15520.1 phospholipase_A2−alpha [MW = 27.22 kDa]

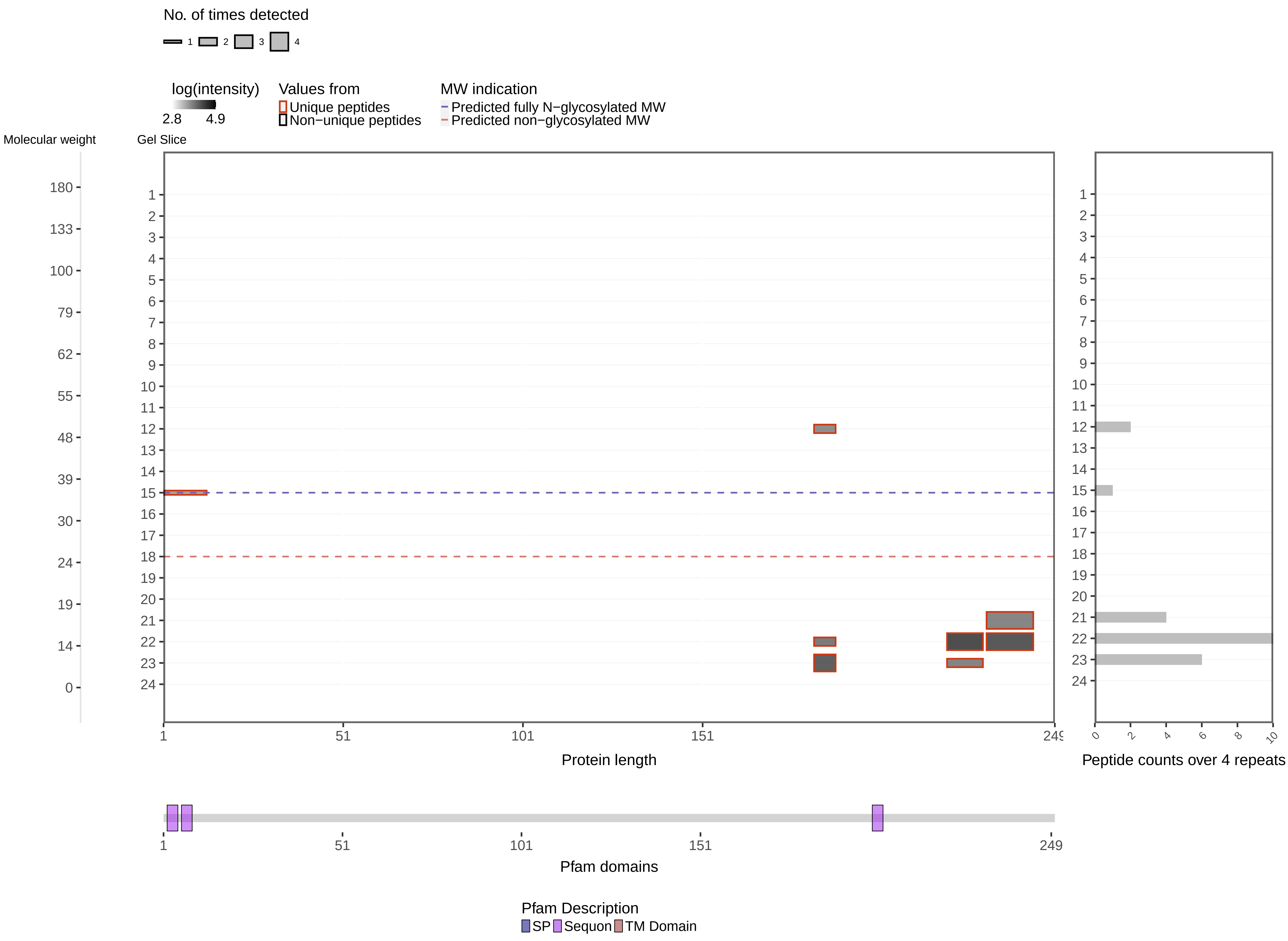


Protein sequence:

FSPISVHALNIGIETNAGISLEKECSRTCESKFCAVPPLLRYGKYCGVLYSGCPGEQPCDGLDTCCMKHDLCIQRKGNNYLNLECNQSFLNCVATFTKSGAPSFKGNSCSVGTVVRVITDVIDAAVVAGKIFKKP*

NbL05g15510.1 peptide_deformylase_1A_chloroplastic [MW = 30.83 kDa]

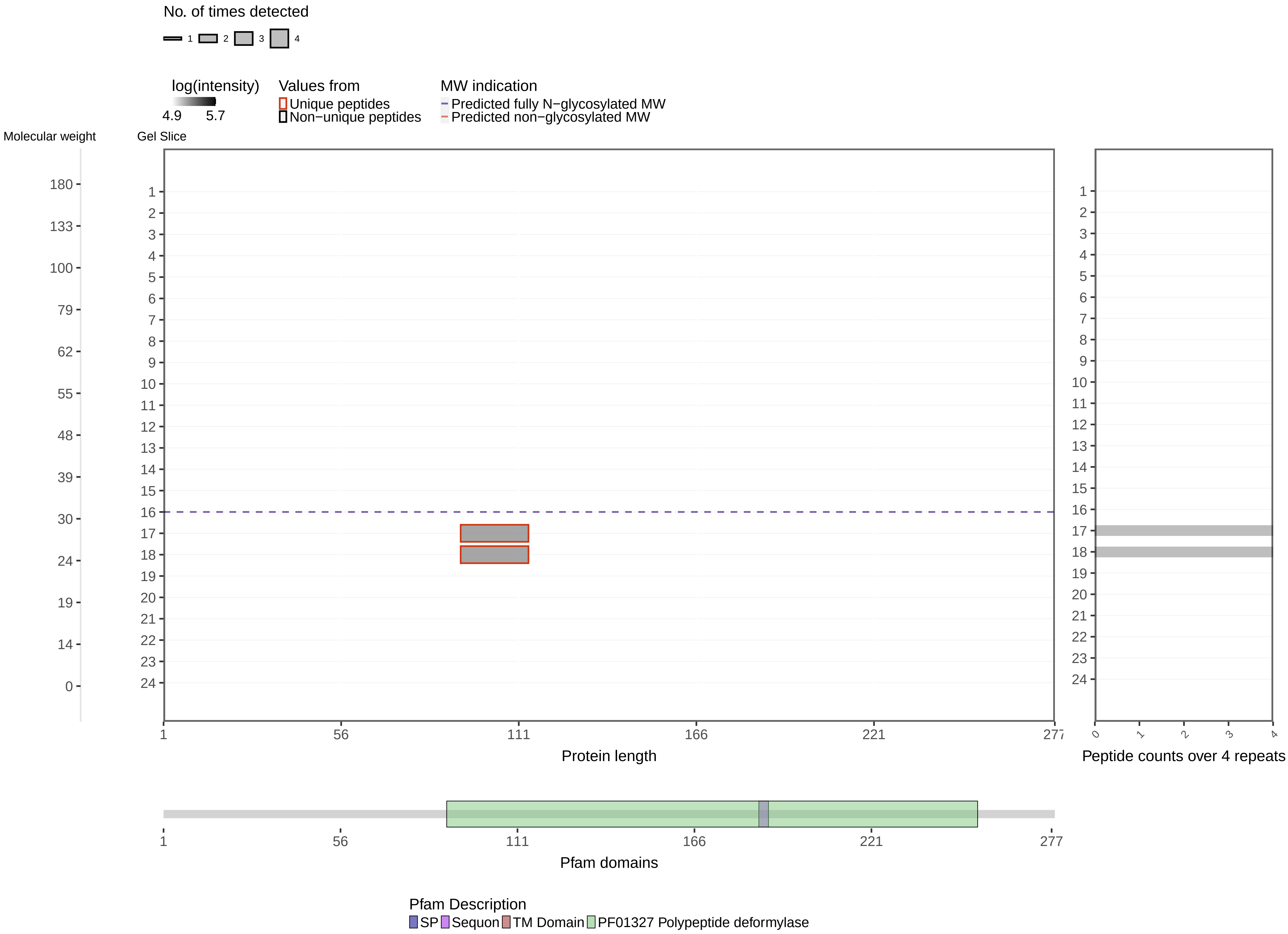


Protein sequence:

/MRNAPGVGLAAPQIGIPLKIIVLEDTNEYISYAPKDETKAQDRCPFDLLVIINPKLKKKGNKTALFFEGCLSVDGFRAVVERHLEVEVGGFDRNGKAIKVDASGWQARILQHEYDHLDGTLYVDKMVPRTFRSVENLDLPLAAGCPNLGVR*

### NbL05g15410.1 proteasome_subunit_beta_type−6−like [MW = 17.29 kDa]

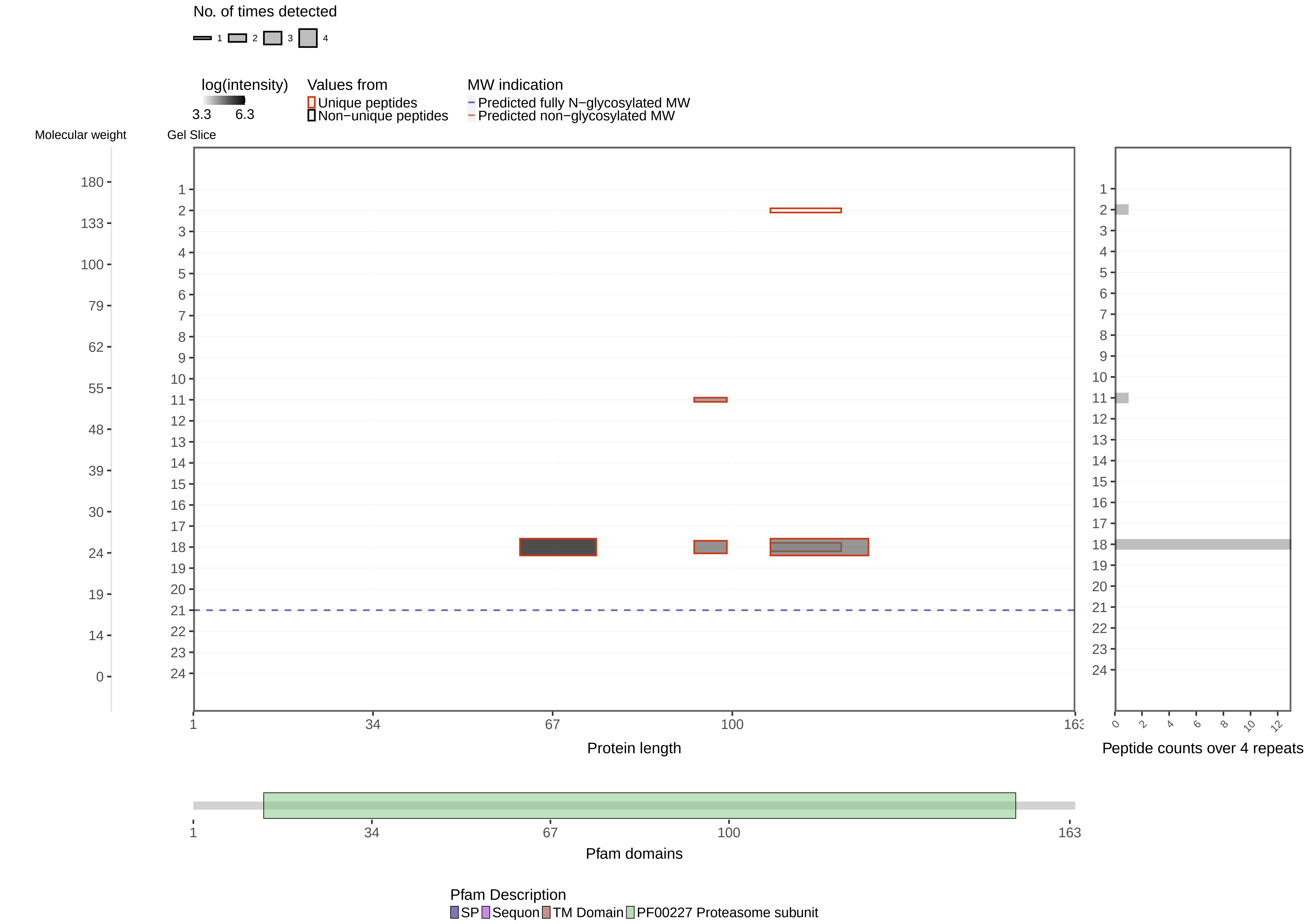


Protein sequence:

QVVSDYVRYFLHQHTIQLGQPATVKVAANLVRLLSYNNKAMLQTGMIIGGWDKYEGGKIYGVPLGGTLLEQPFAIGGSGSSYLYGFSSTKHGGKE*

### NbL05g15350.1 lysine−−tRNA_ligase−like [MW = 69.81 kDa]

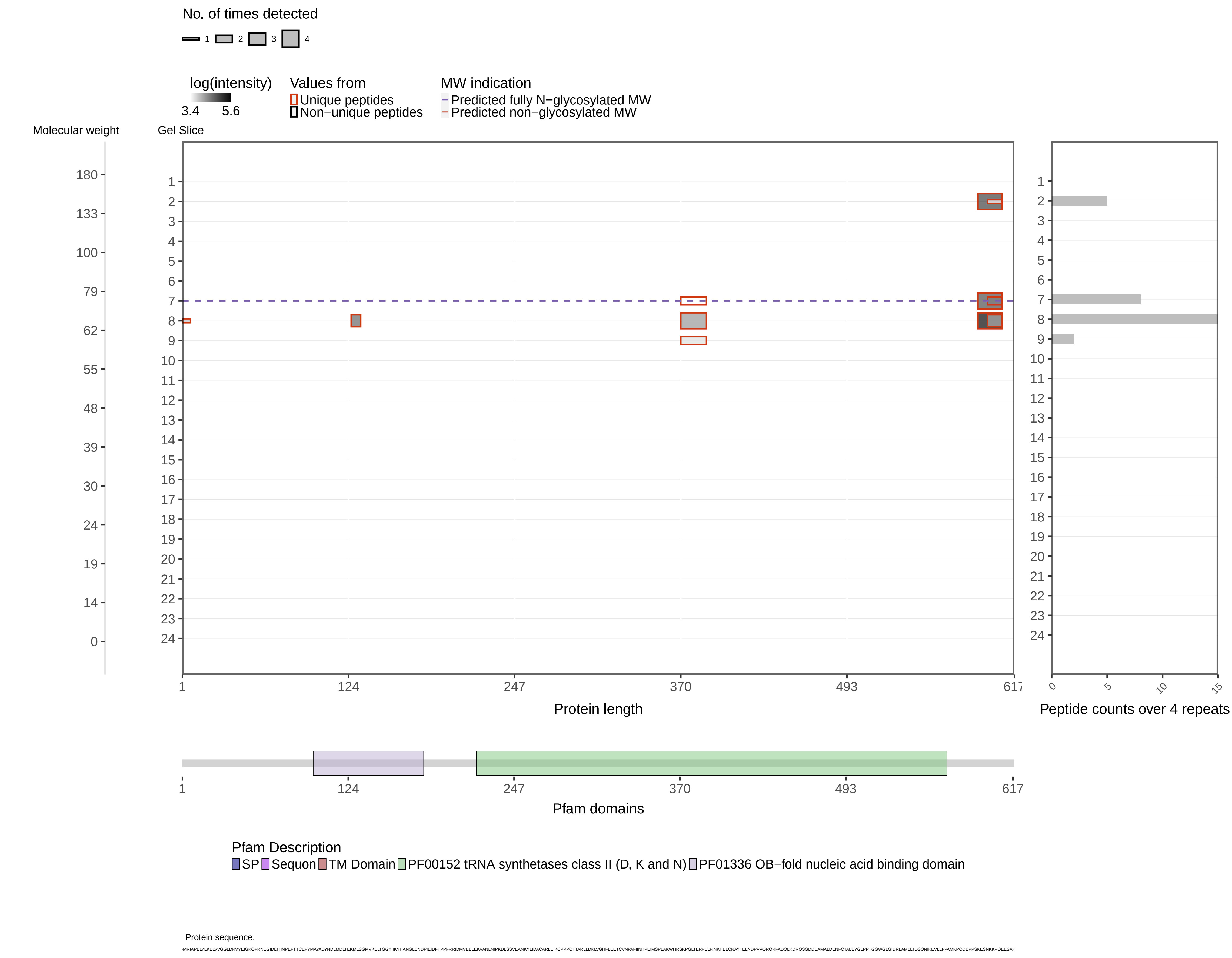

NbL05g15200.1 pectinesterase/pectinesterase_inhibitor_PPE8B−like [MW − SP = 56.68 kDa]

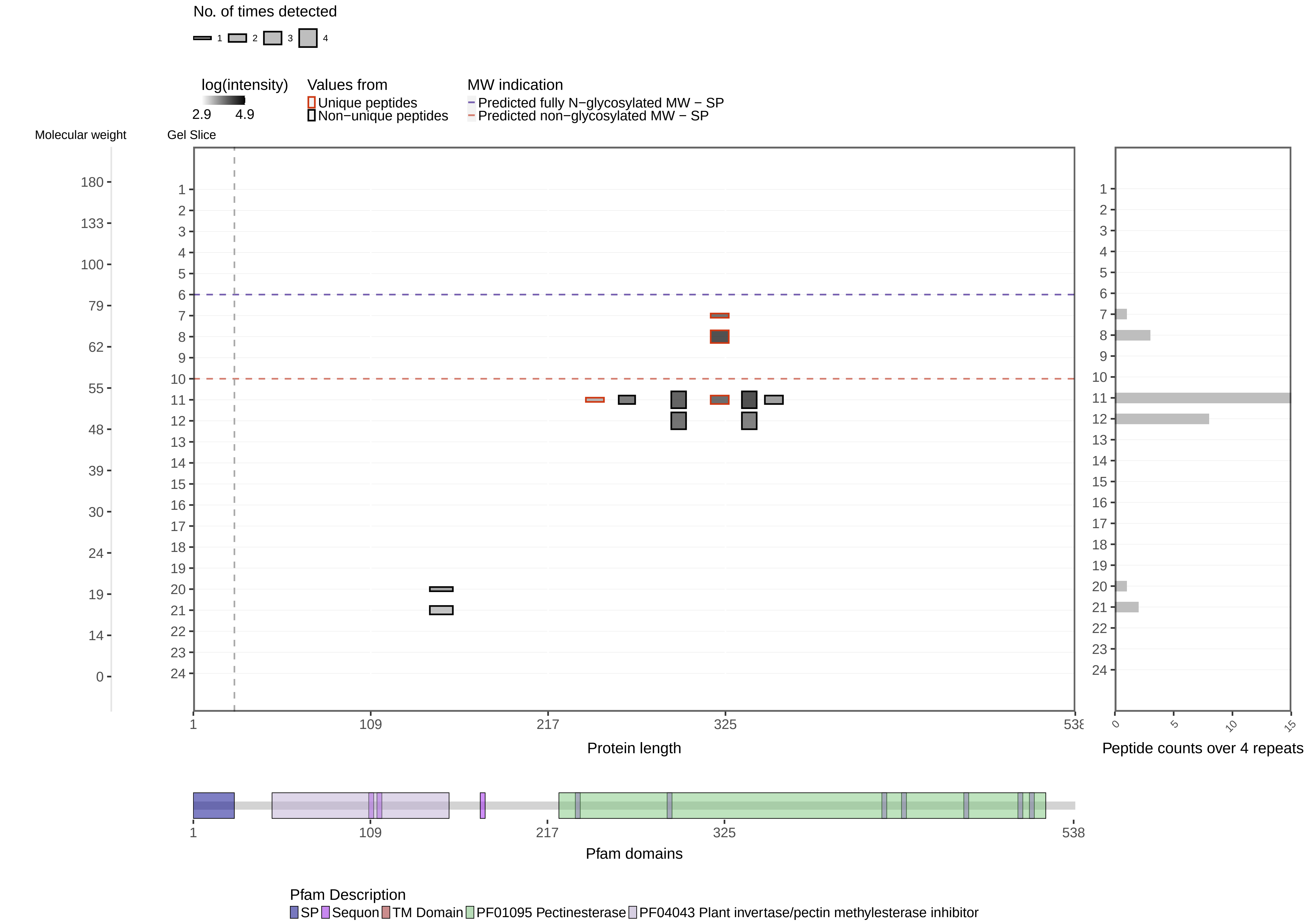


Protein sequence:

IIKKGIYNEYVEISKKKWNILMIGDGIDVTVISGNRSFIDGWTTYRSATFAVKGQGFIARDMTFENIAGPQKHQAVAFRSDSDLSVLFRCAFRGYQDTLYAHSMRQFYRNCVITGTVDFIFGDGTVVFQNCQILARKGLPEQKNTITAQGRKEAAETTGFSIQFSNISGEPDLLASLNSTQTFLGRPWKAYSRTVIMQSYMSNVIKPQGWLEWNGNISLDTLYYAEYQNYGPGAGLSARVNWPGYHLLNDSSQANNFTVAQFILGNSWLPSTGVKYTAGLAV*

### NbL05g15120.1 plasma_membrane_ATPase_4 [MW = 105.24 kDa]

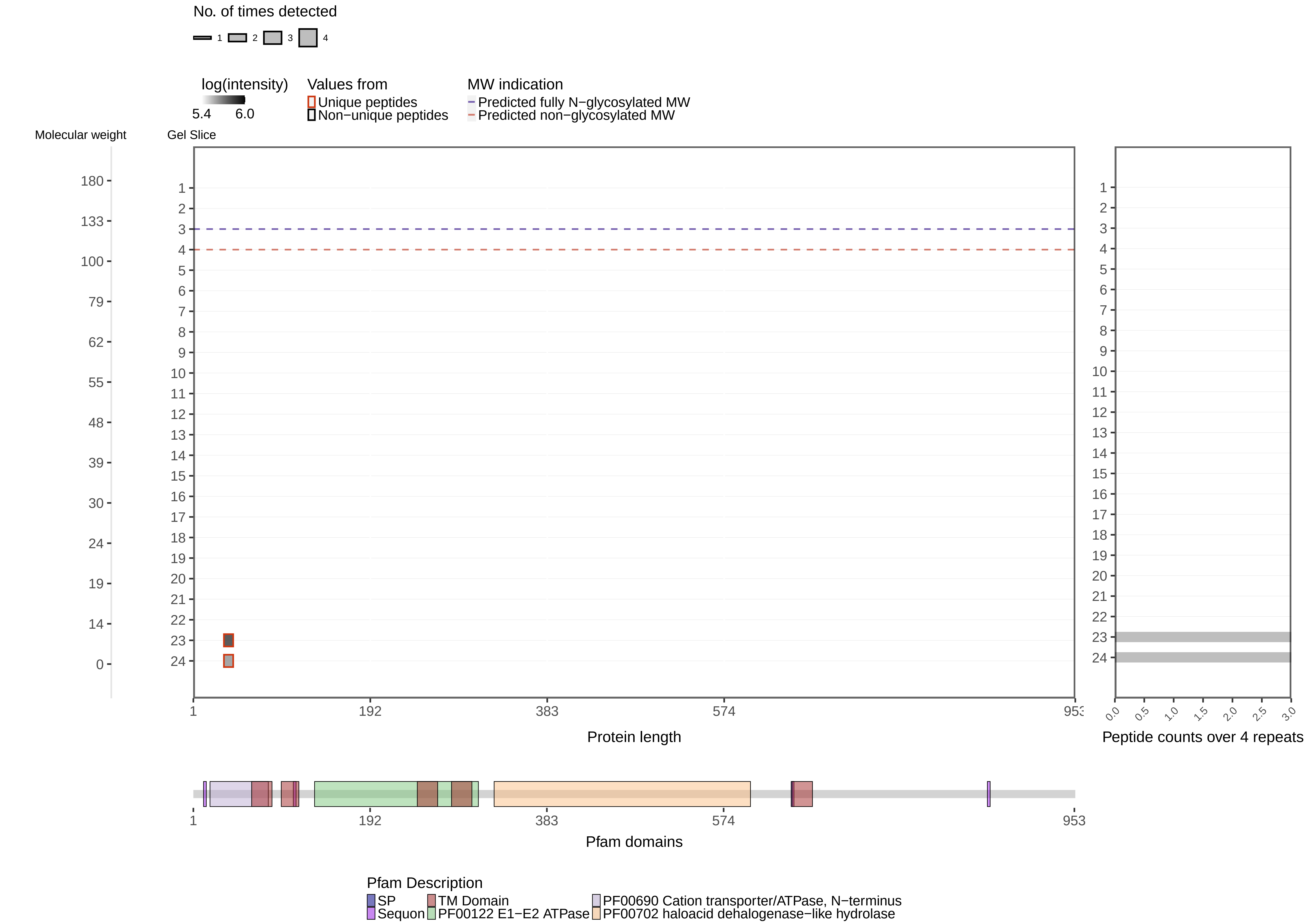


Protein sequence:

### NbL05g15070.1 putative_glucose−6−phosphate_1−epimerase [MW = 35.54 kDa]

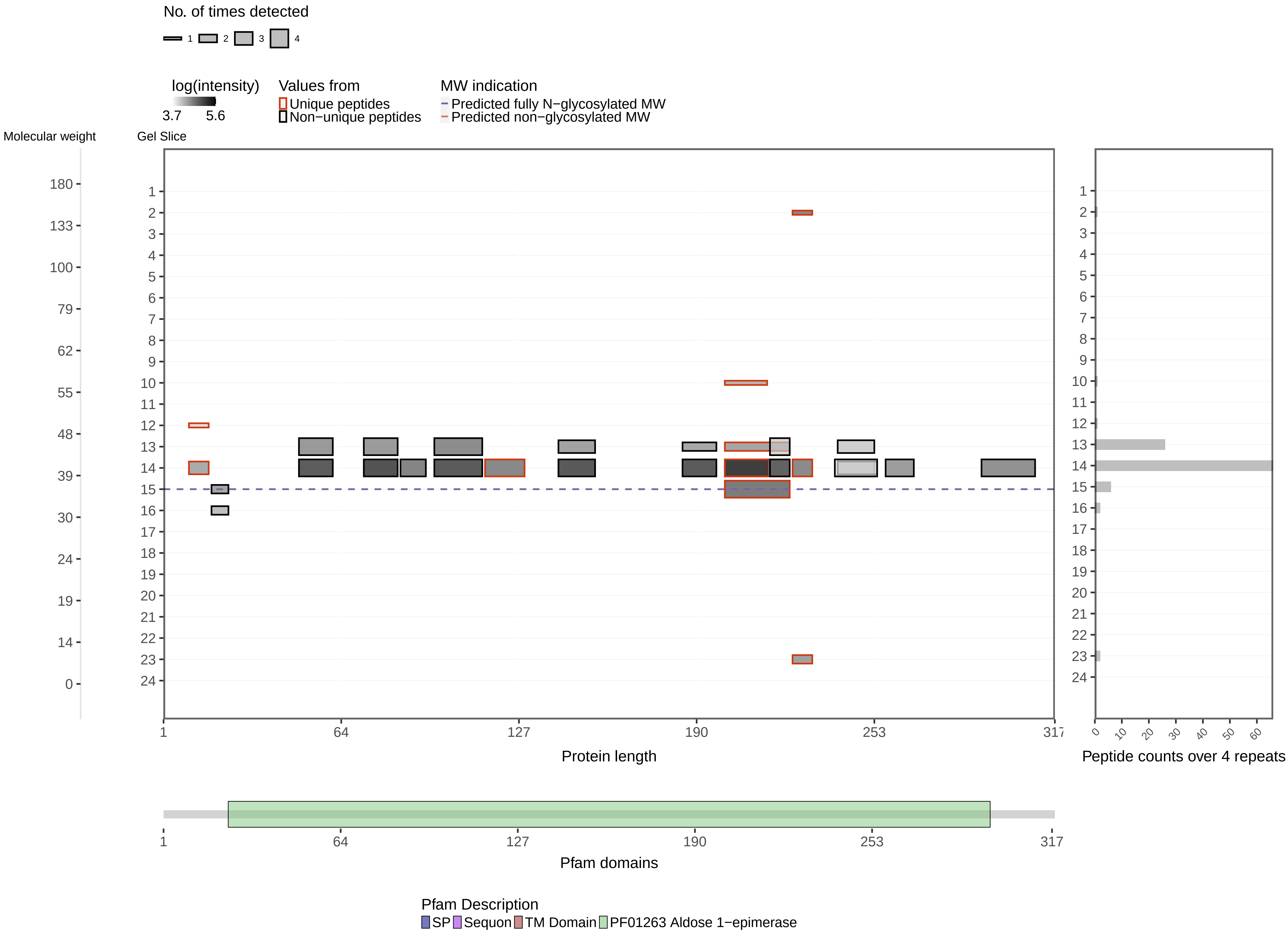


Protein sequence:

GDLMLTSRIRNTNTDGKPFSFTFAYHTYFSVSDISEVRVEGLETLDYLENKERFTEQGDAITFESEVDKIYLSTPTKIAILDHEKKRTFVLRKDGLPDAVVWNPWDKKAKAMADFGDEEYKHMLCVEAAAVEKPITLKPGEEWRGRQELSAVPSSYCSGQLDPKRVLLGS*

NbL05g14960.1 uncharacterized_LOC104245598 [MW = 115.33 kDa]

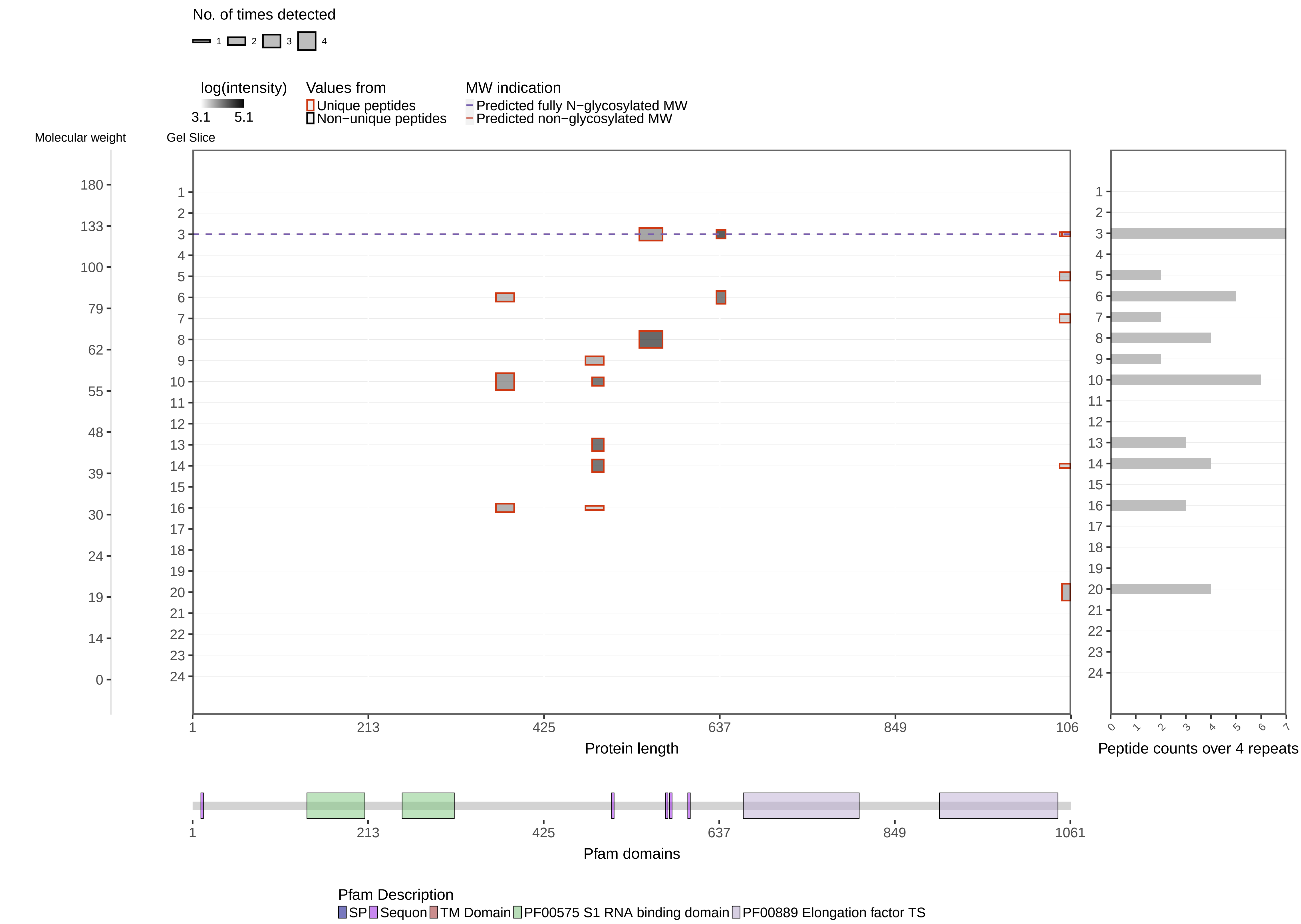


Protein sequence:

SQSEAPLENSKDEVSQEGAEVVESKVENTPSIEDQSSDTAAQQEEVATAAEQDRNVANSSEQNGTASSNEAAAKDRNVAKSSERNGTALSNEAAAKAISPALVKQLREETGAGMMDCKNALSETGGDIIKAQEYLRKKGLASADKKSSRATAEGRIGSYIHDSRIGVLIEVNCETDFVSRGDIFKELVDDLAMQVAAYPQVQYLVPEDVPEEIINKEREIEMQKEDLLSKPEQIRSKIVDGRINKRLEELALLEQPYIKNDKLVVKDWVKQTIATIGENIKVKRFVRYNLGEGLEKK

NbL05g14930.1 uncharacterized_LOC109234215 [MW = 71.67 kDa]

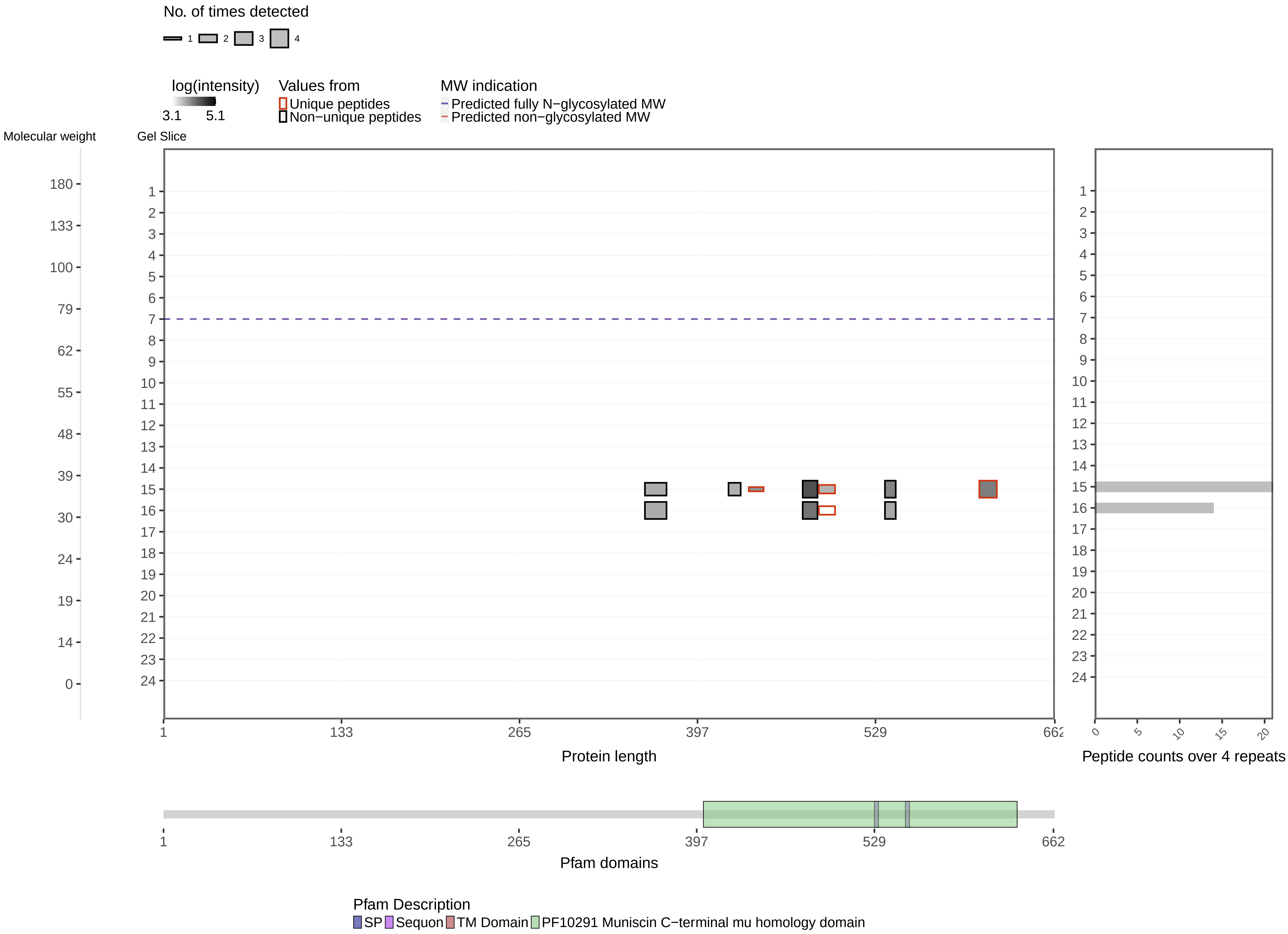


Protein sequence:

GIEFSNDGSTLQQDFEGINDAWGGGLDASEYVGTKKVKKDQGLGGLELLETSEPPKAAAGAATDGGAGKKLEDVLVKKMNGPEMFITEEINAEFRESLLARVGLMGVVFLRTLPSKSSDDKETEFSFKVEGTAGVKRFVMQNSHVSSLGNGLFHVKTAPSNEPIPIIKYSLLPHLTPLPLRIRLVKRLSGTLLSVMLQYVANPDLPVPLTNVTFVLKLPVDPTLLKVTPKAVLNRSERELKWHVDEIPLKGHPGKLRARFPVDINDDDGGEELVLFGYVKFSSQGPRLLSGISLC

### NbL05g14920.1 (DL)−glycerol−3−phosphatase_2 [MW = 27.28 kDa]

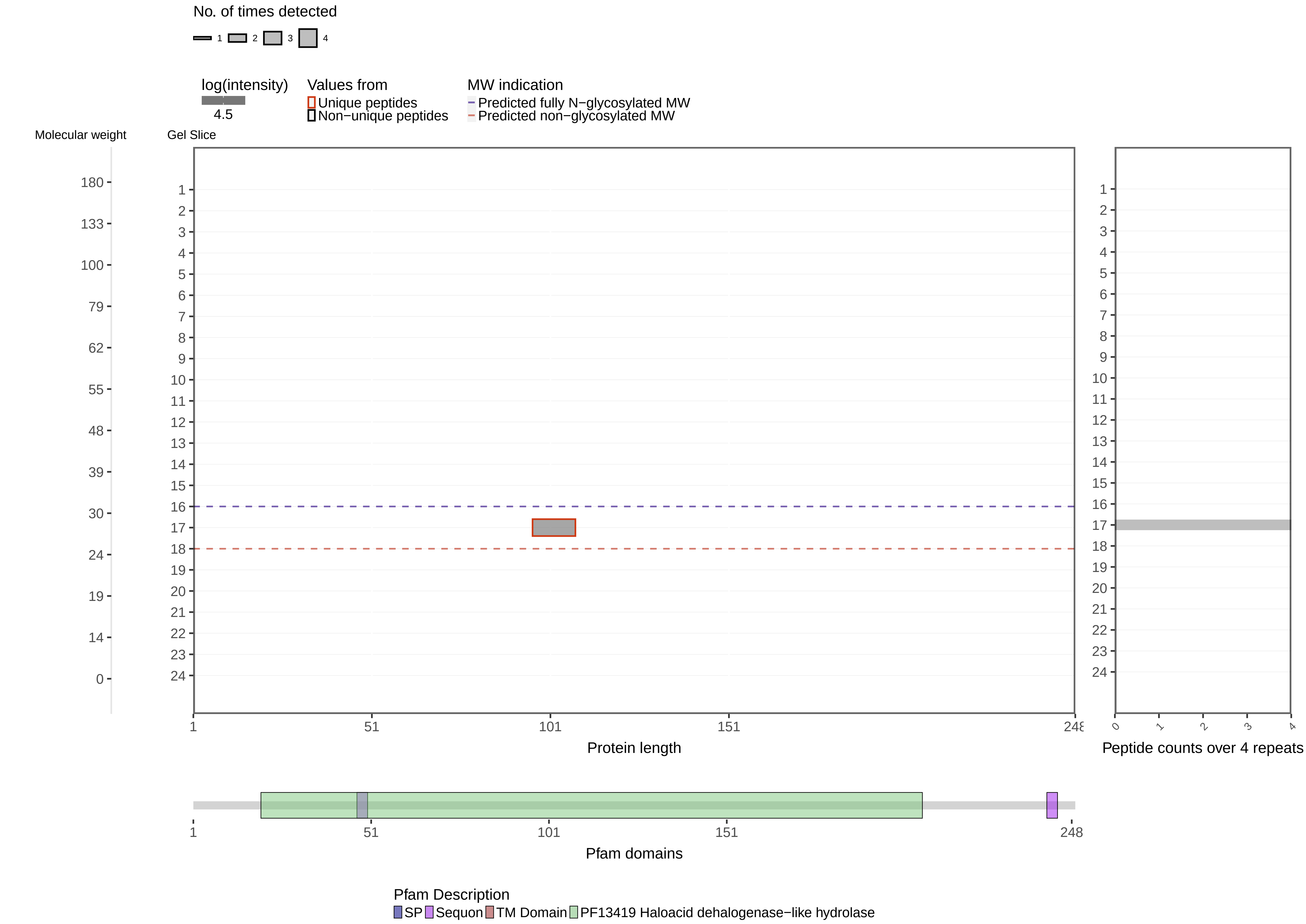


Protein sequence:

NbL05g14860.1 replication_protein_A_32_kDa_subunit_A−like [MW = 27.14 kDa]

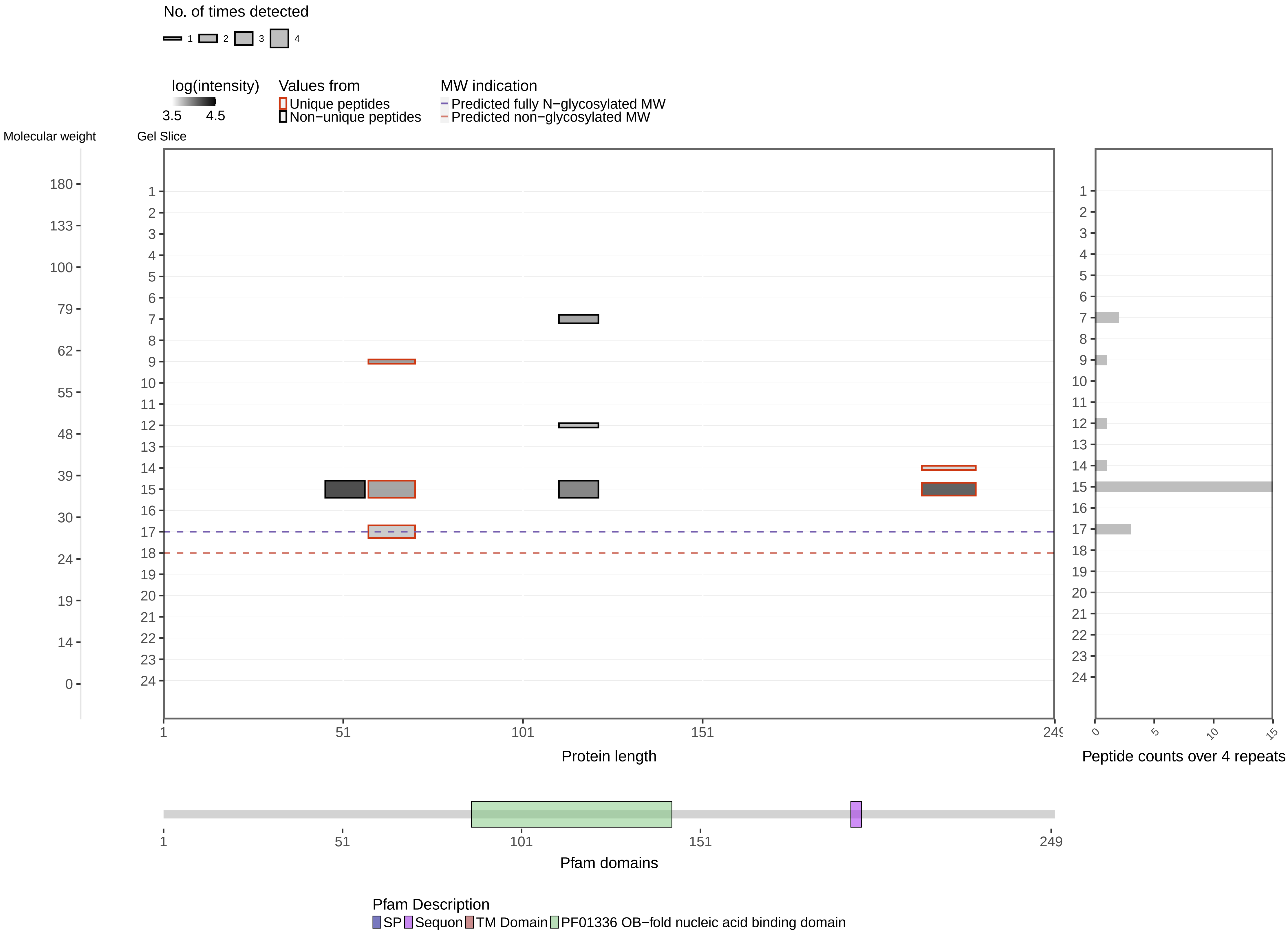


Protein sequence:

EVEEVSDGMYVRVHGHLKGFQGKTQLVIFAIRPITDYNEVATHFLECIYVHHCNRKPQSGISVSAPGQTDVSAAVSAPSSGFNSSQLSGHLSIDGLKGIEKAVIDYLEQPSSLSQEKGIHWNELLNNLKFLWRKSRKQ*

#### NbL05g14730.1 3−hydroxyisobutyryl−CoA_hydrolase−like_protein_3_mitochondrial [MW = 41.77 kDa]

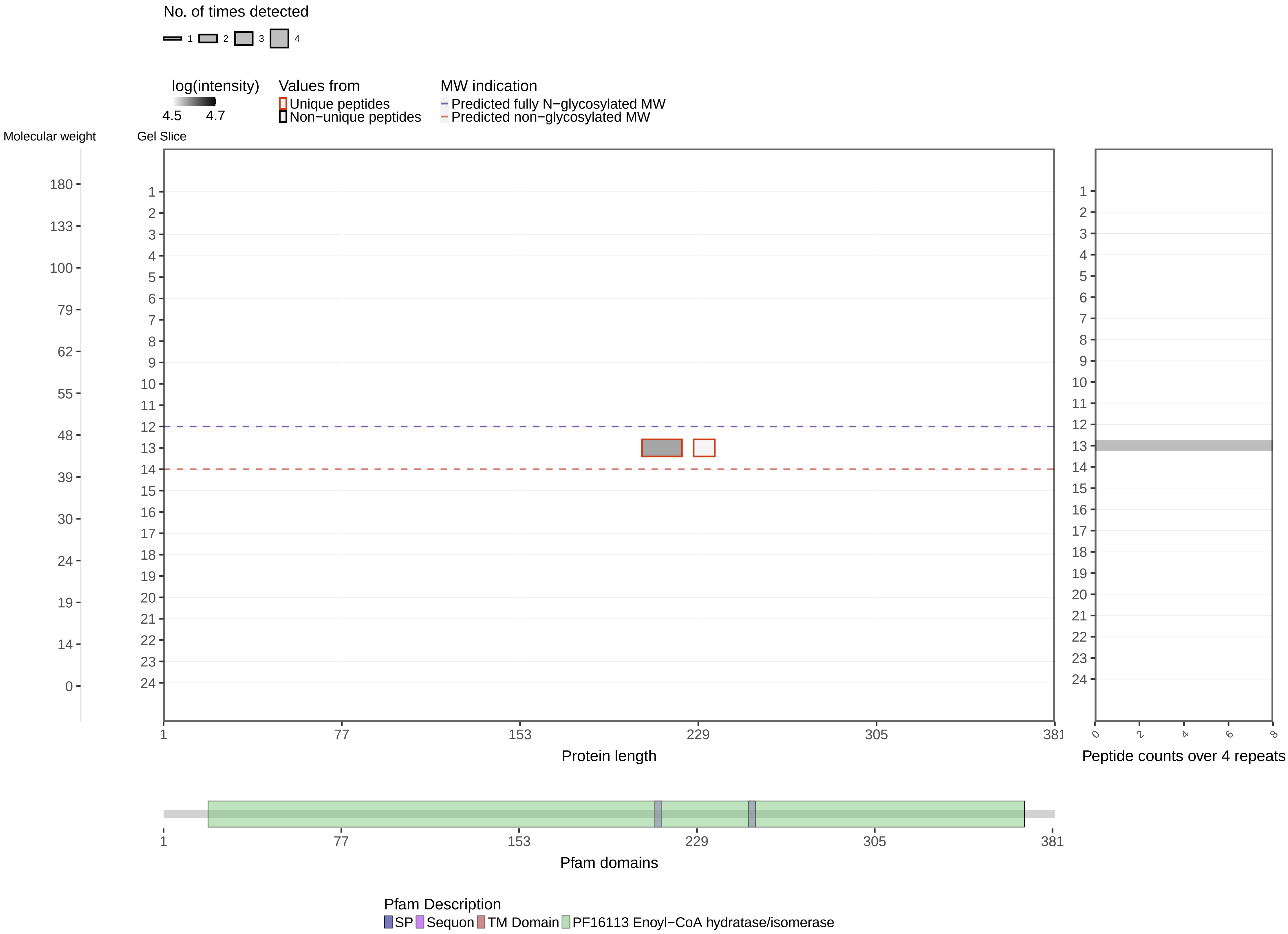


Protein sequence:

RISTPADALYVGLGSHYVPSANLDTLKEAFLAANFSQDPDEEIKRLLEKYSTDPDSESRLKSLLPRIISTFGGNKTVKNIIEELENHQQSADTLVAEWAKDALQGLRKGAPFSLCLTQHYFSRVASACGKNENTLSKLPGVMRTEYRIAIRSALRNDFAEGVRAILVDKDQNPRWNPSSLEDVNLNEVEALFEPLSPEVELNV*

### NbL05g14580.1 40S_ribosomal_protein_S14−2 [MW = 16.37 kDa]

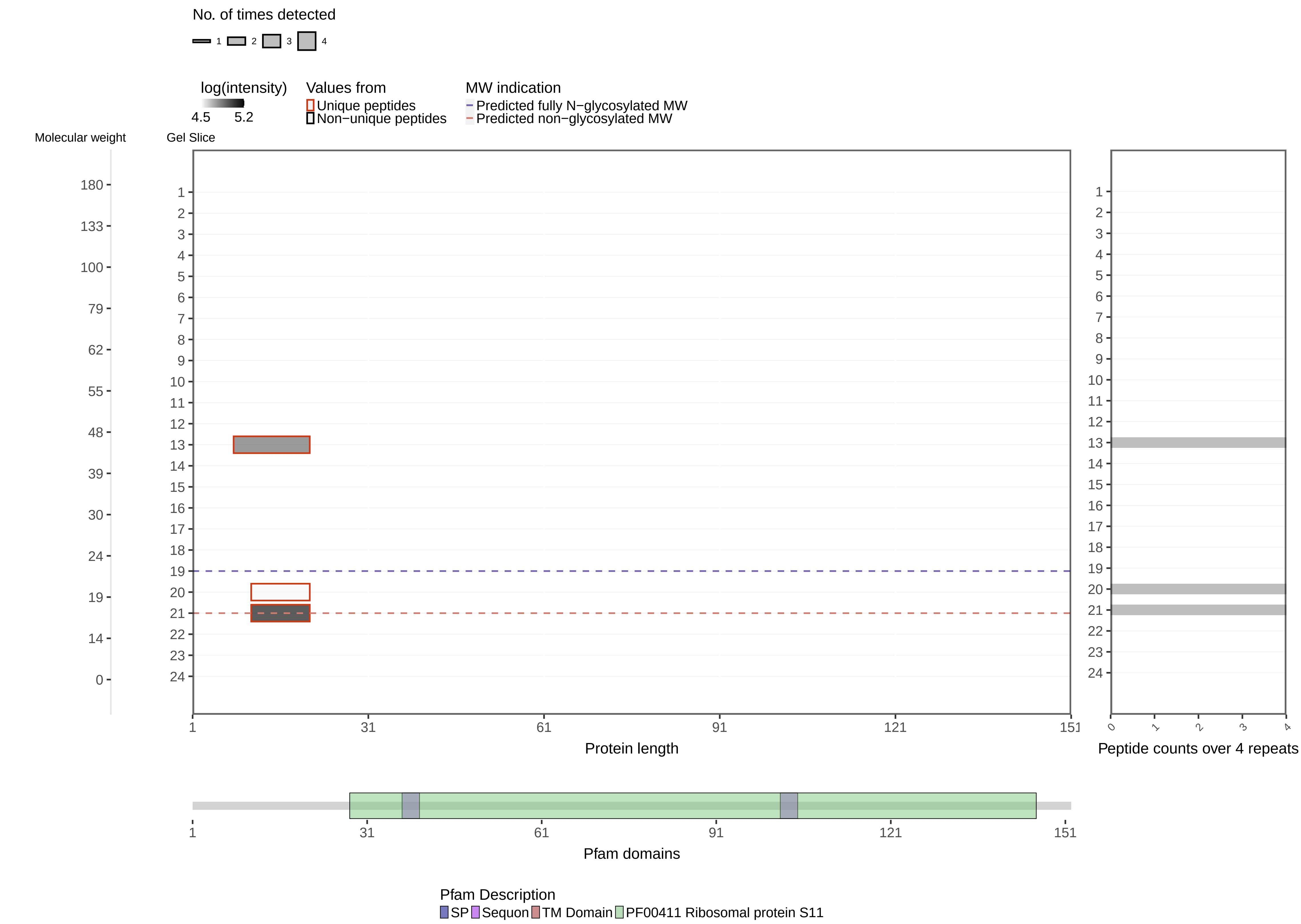


Protein sequence:

ADRDESSPYAAMLAAQDVSQRCKELGINALHIKLRATGGNKTKTPGPGAQSALRALARSGMKIGRIEDVTPIPTDSTRRKGGRRGRRL*

NbL05g14540.1 50S_ribosomal_protein_L7/L12−like [MW = 21.26 kDa]

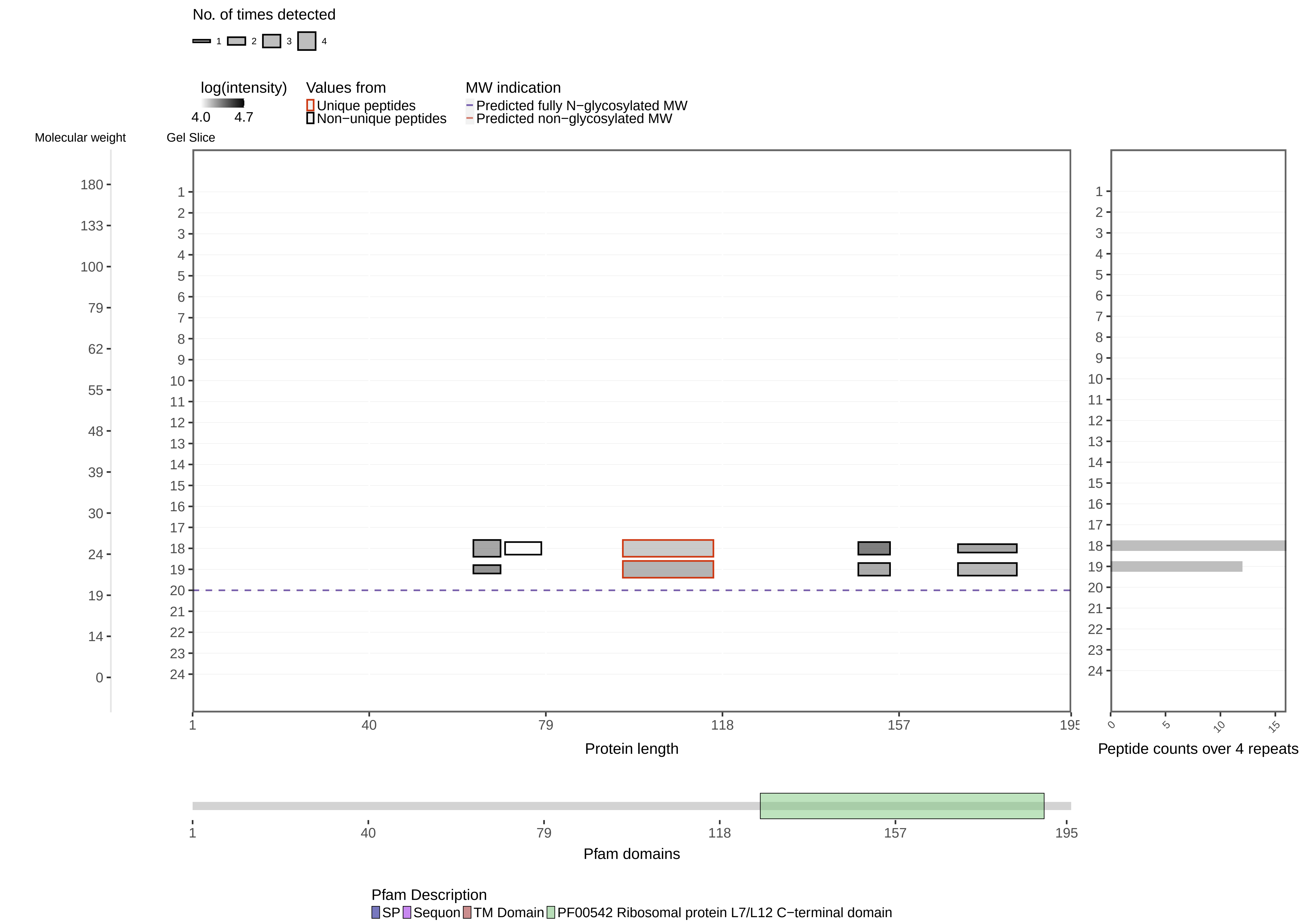


Protein sequence:

TLREILRHPNLKQISVEGIDLGAMGGAGGGPAKAEEKKAEKTSFDVKLEKFDAAAKIKVIKEVRSFTSLGLKEAKDLVEKVPAILKQGVTKEEANEIIEKIKAAGGVAVME*

### NbL05g14510.1 1−aminocyclopropane−1−carboxylate_oxidase_homolog_1−like [MW = 41.63 kDa]

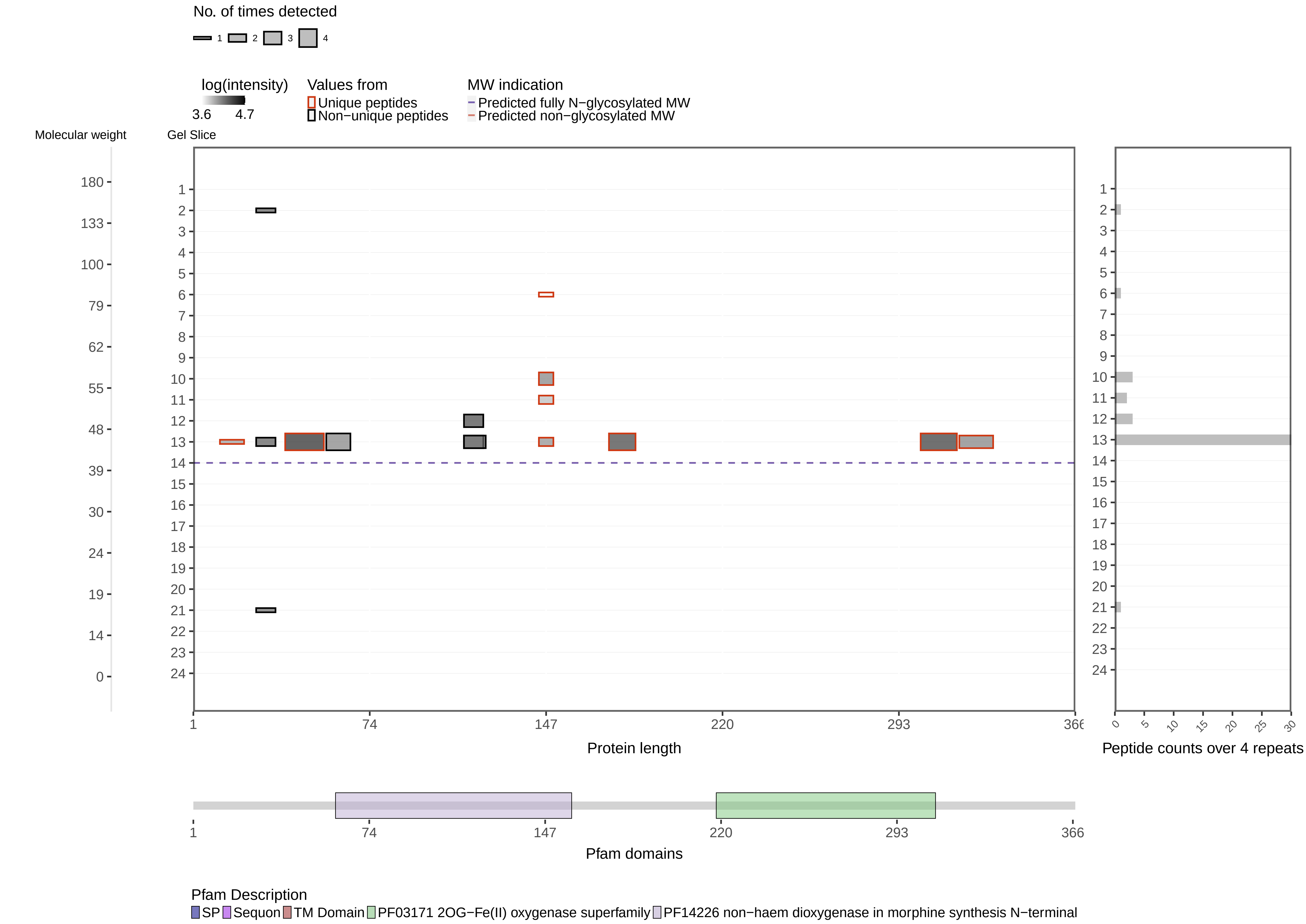


Protein sequence:

SVCRDILFEYTNQVMKLAYTIFELLSEALGLKPSHLNDMGCSEGLTALGHYYPACPQPELTLGAGKHADSNFLTVLLQDNIGGLQVLHQNHWVDVPPTPGALVVNIGDLLQLISNDKFKSVEHRVLANKVGPRISMACFCTTSFLPTERLYGPIQELISEHNPRKYKETTIKEYTTYYSNKGLDDTSALLHFRLKTK*

### NbL05g14450.1 heterogeneous_nuclear_ribonucleoprotein_R [MW = 53.64 kDa]

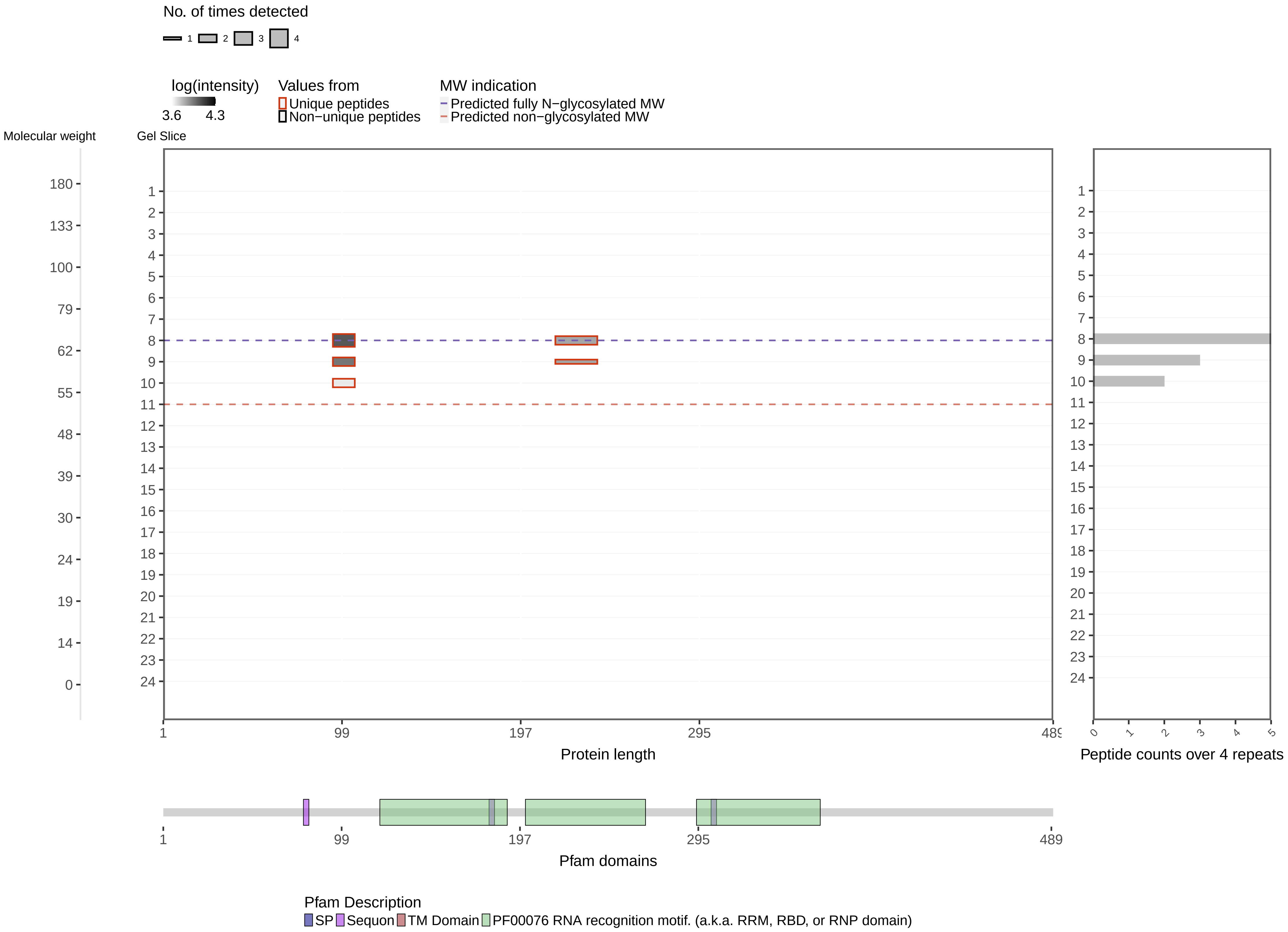


Protein sequence:

QNSGRNRGFAFIEYYNNACAEYSRQKMSNSDFKLDDNAPTVSWADPKNAETAASAQVKAVYVKNLPKNITQDKLKELFEHHGKIMKVVLPPAKPGQEQSRYGFVHFAERSSAMKALKNTERYEIDGKILECSLAKPQADQKSSGGSSSQKGPTLPTYPPRIGYGMVGAPYGALSAGYGAAGVGQPLVYGRGATPAGLAMMPMLLPDGRIGYVLQQPGVMHTPPPASRGGRSSGGGSSGGRRGDSGRGRSRYNPY*

### NbL05g14140.1 glucose−6−phosphate_1−dehydrogenase_chloroplastic [MW = 66.95 kDa]

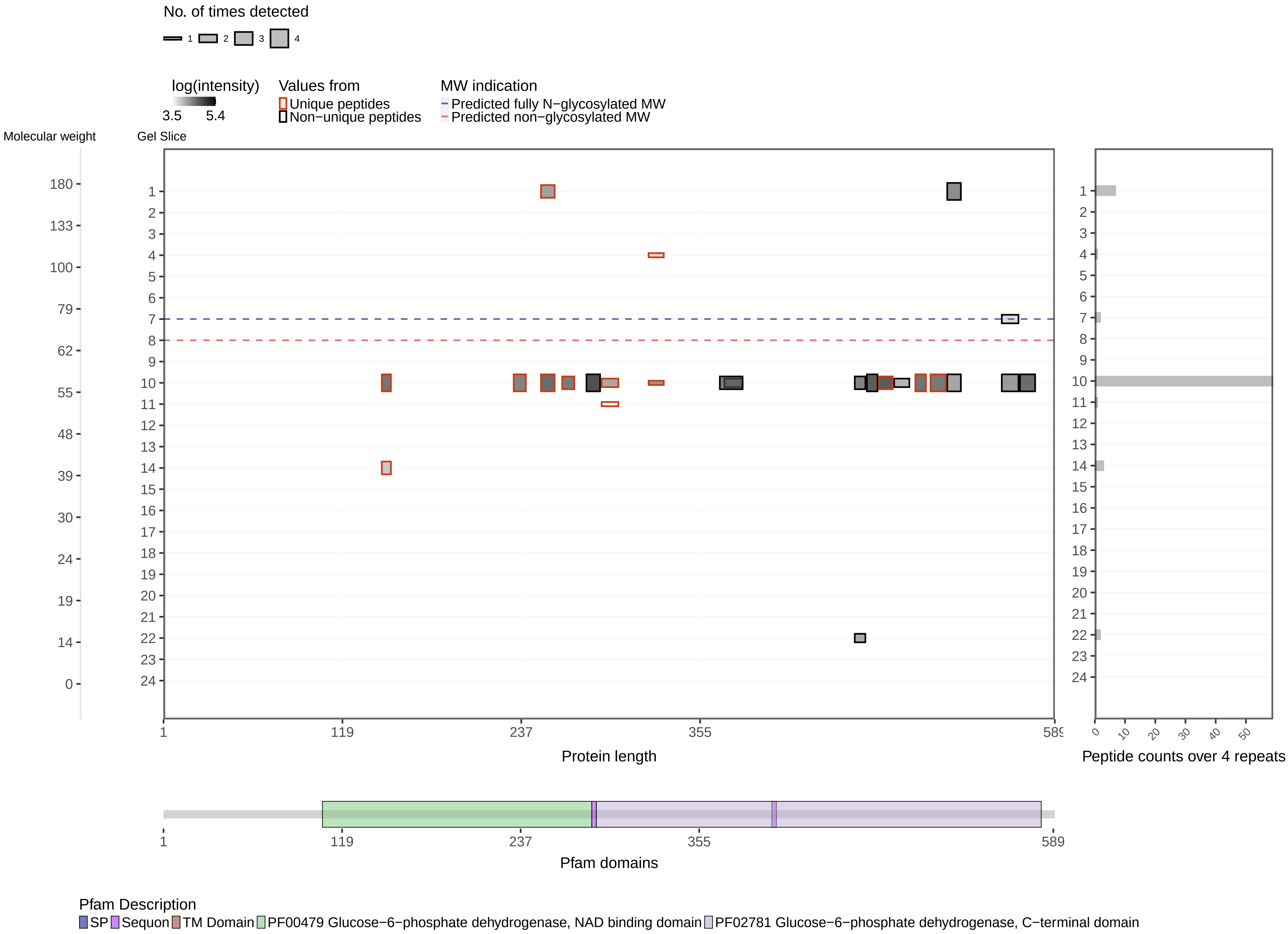


Protein sequence:

.VENLSVLRFSNLVFEPLWSRNYIRNVQFIFSEDFGTEGRGGYFDNYGIIRDIMQNHLLQILALFAMETPVSMDAEDIRNEKVKVLRSMRPLQLEDVVLGQYKGHSKGGKLYPAYTDDPTVPNGSITPTFSAAALFINNARWDGVPFLMKAGKALHTRRAEIRVQFRHVPGNLYKKNFGTDLDKATNELVLRLQPDEAIYLKINNKVPGLGMRLDRSDLNLLYKAKYRGEIPDAYERLLLDAIEGERRLFIRSDELDAAWALFTPLLKELEEKKIAPELYPYGSRGPVGAHYLAAKH[illegible]

### NbL05g14040.1 fasciclin−like_arabinogalactan_protein_2 [MW − SP = 41.23 kDa]

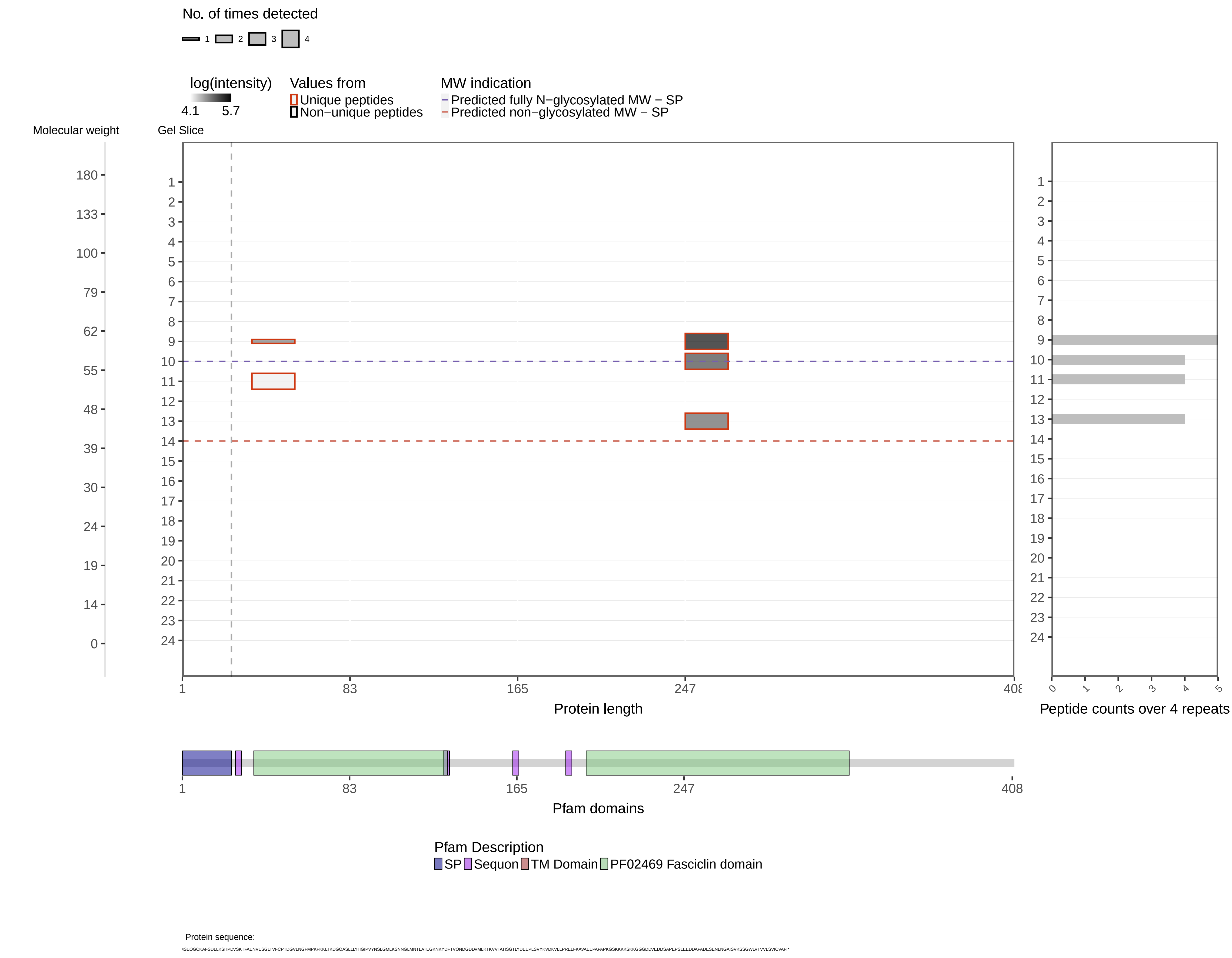

### NbL05g13960.1 acetyl−CoA_acetyltransferase_cytosolic_1−like [MW = 41.3 kDa]

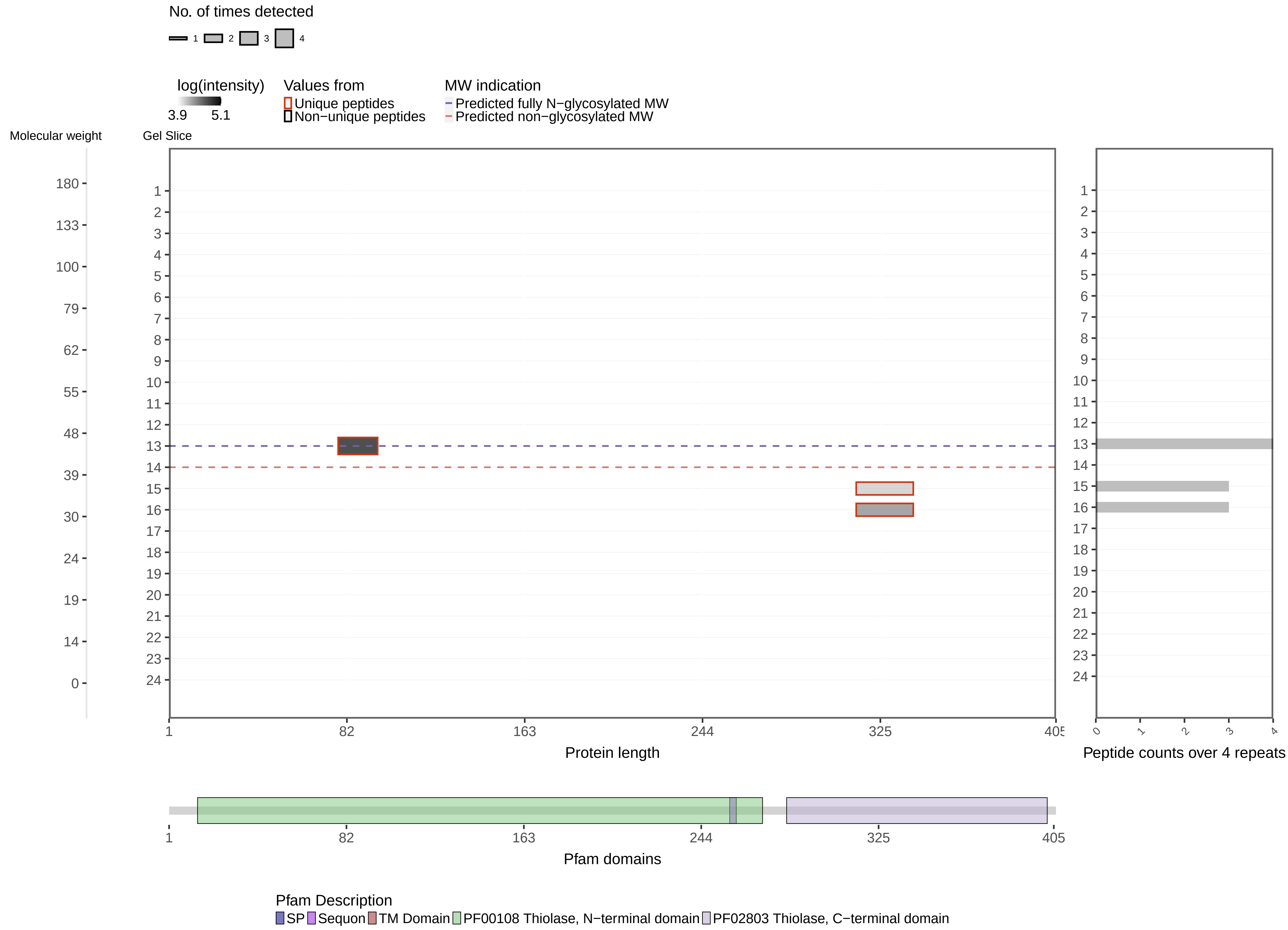


Protein sequence:

NbL05g13930.1 ribulose−15_bisphosphate_carboxylase/oxygenase_large_subunit_N−methyltransferase_chloroplastic−like [MW − SP = 57.77 kDa]

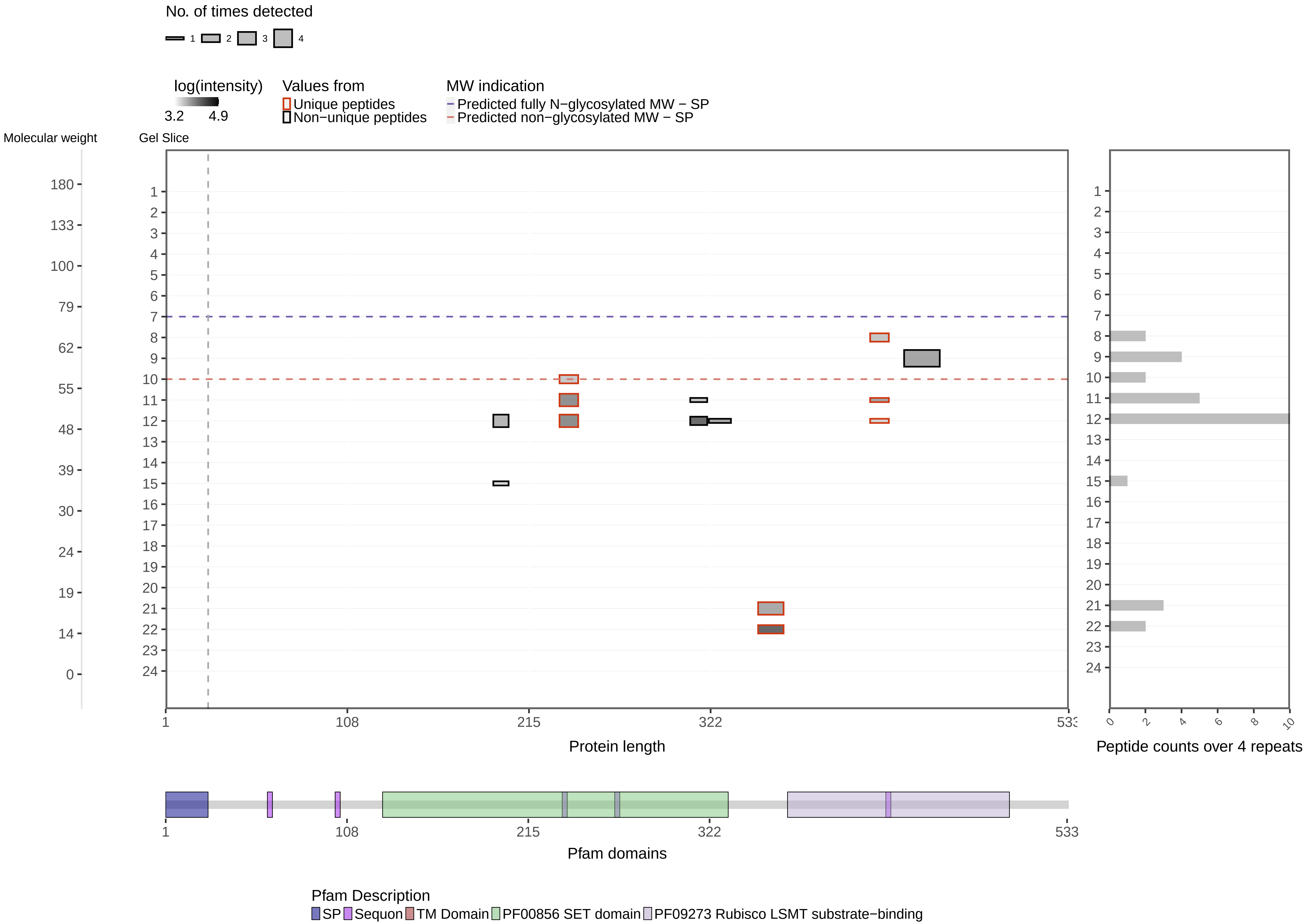


Protein sequence:

FPEEIFNIETFNWSFGILFSRLVRLPSMDGRVALVPWADMLNHNCEVETFLDYDKSSQGIVFTTDRAYQPGEQVFISYGRKSNGELLLSYGFVPREGTNPSDSIELSFSLKKYDKCYKEKVEALKKHGLSASECFPIQVTGWPLELMAFAYLVVSPPSMSRQFGEMAAAASNKSTSKKDMKYPEIEEEALQFILDSCESSISKYSKFLQASGEMDLDVTNPKQLNRRVFLKQLAVDLCTSERRILFRSQYILRRRLRDIRSGELKALKLFDGLRNLFK*

### NbL05g13540.1 23−bisphosphoglycerate−independent_phosphoglycerate_mutase [MW = 61.19 kDa]

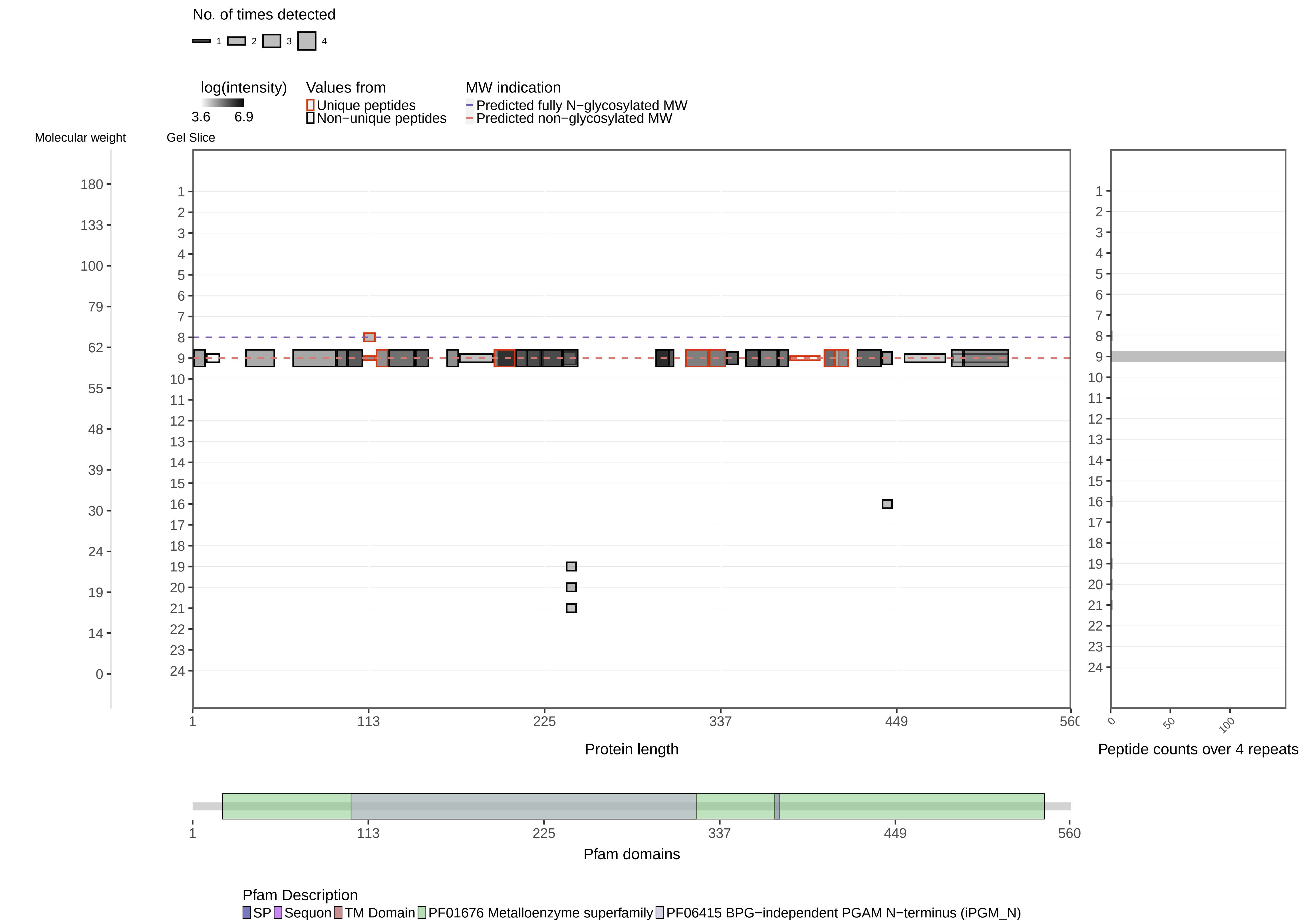


Protein sequence:

/DDNGKPVGPILDGDAVVTFNFRADRMVMLAKALEYENFDKFDRVRVPKIRYAGMLQYDGELQLPRHYLVSPPEIDRTSGEYLVRNGVRTFACSETVKFGHVTFFWNGNRSGYFNEKLEEYVEIPSDVGITFNVQPKMKALEIAERTRDAILSGKFDQVRVNLPNGDMVGHTGDIKATIEACKSADEAVKMILDAIEQVGGIYLVTADHGNAEDMVKRNKKGEPALDKNGNIQILTSHTCEPVPIAIGGPGLAPGVRFRQDLPTGGLANVAATFMNLHGYEAPSDYEPSLIEVVD

### NbL05g13420.1 probable_polyamine_oxidase_2 [MW = 54.5 kDa]

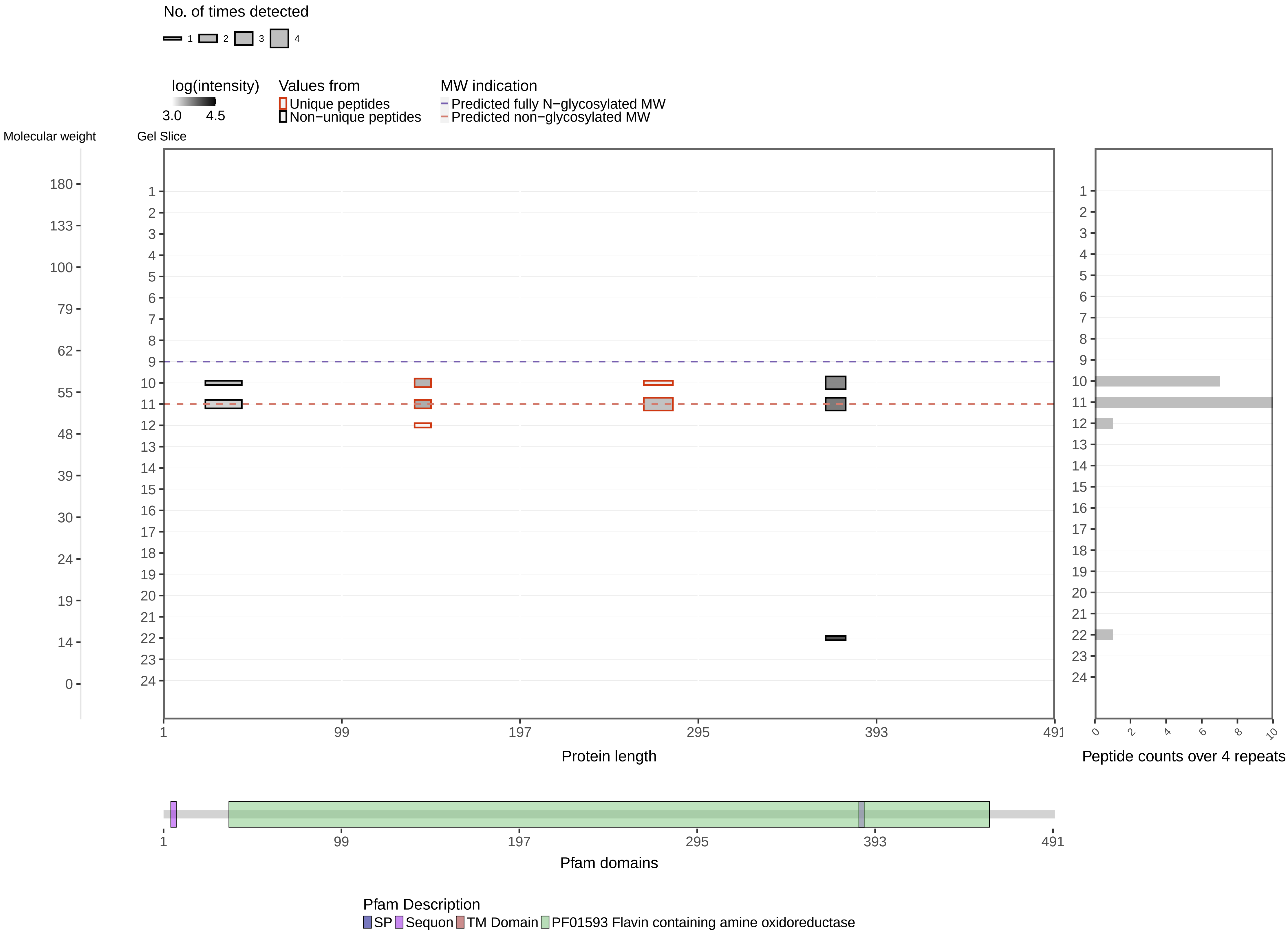


Protein sequence:

VINTLAKGLDIRLGHRVAQVVRRYNGVKVTVEDGRTFVADAAVVAVPLGVLKSKRIKFEPRLPEWKEAAINELGVGIENKIILHFEKVFWPNVEFLGIVAESSYECSYFLNLHKATGHPVLVYMPAGQLARDIEKLSDEDAANFAFKQLQRILPNSTTPIQYLVSHWGTDVNSLGSYSYDTVGKPHEFYERLRIPVDNLFFAGEATSMDYPGSVHGAYSTGLMAAEDCRMRVLERYGELDLFQPVMGEDTPVAVPLLISRM*

### NbL05g13400.1 uncharacterized_oxidoreductase_At1g06690_chloroplastic [MW = 39.81 kDa]

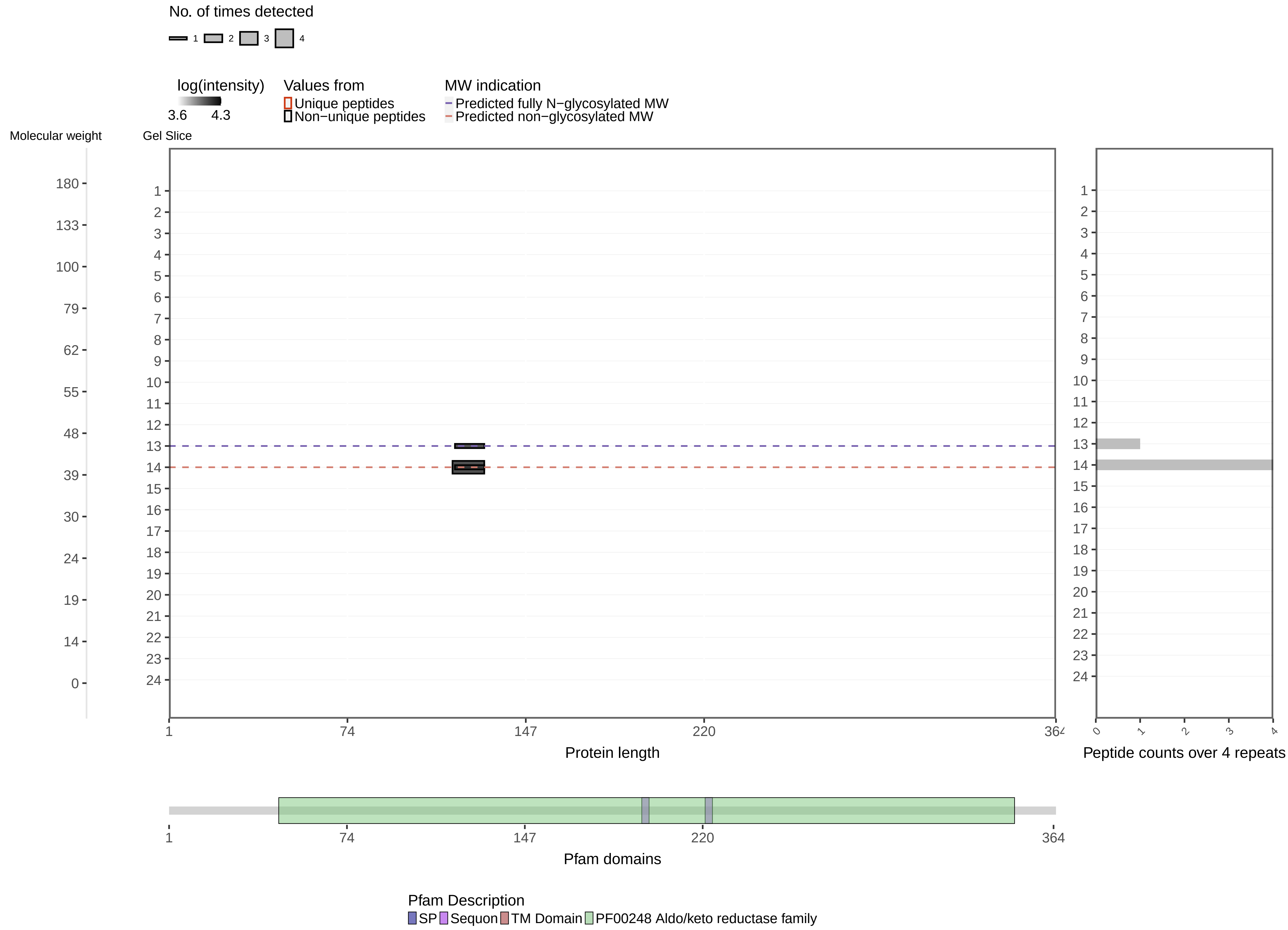


Protein sequence:

IWGNEGYIDGLGDAVEQGLVKAVGVSNYSEKRLRAAYEQLKRRGIPLASNQVNYSLIYRFPEQNGVKAACDELGVTLIAYSPIAQGALTGKYTPENPPSGPRGHIYTPEFLTKLQPLINRIKEIGESYSKTPTQVVLNWLIAQENVLPIPGAKNADQAKEFAGALGWRLTQDEVDELRSLALDVKPVTGFPVEKL*

### NbL05g13380.1 heat_shock_70_kDa_protein_17 [MW − SP = 97.08 kDa]

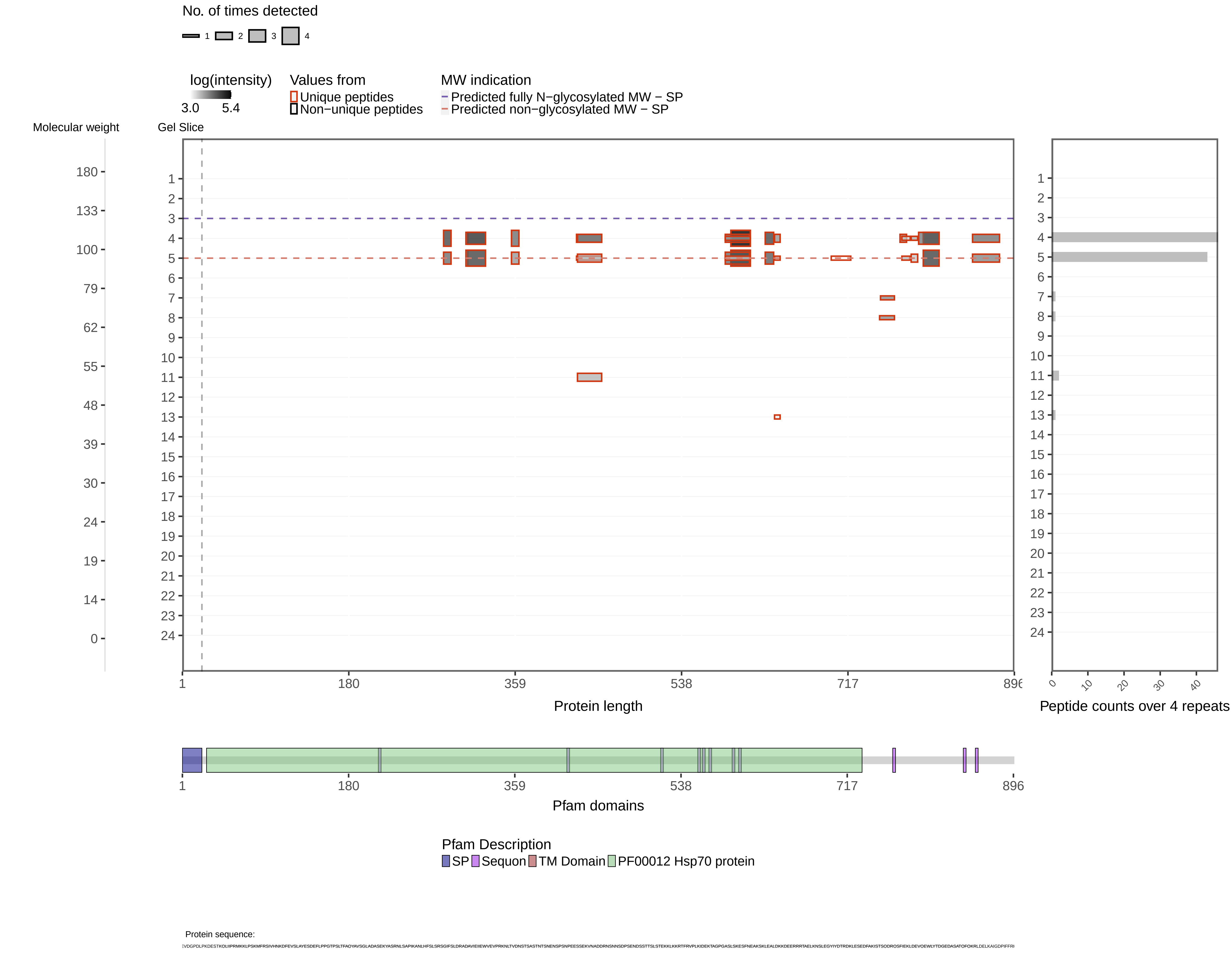

NbL05g13320.1 myosin_heavy_chain_cardiac_muscle_isoform−like [MW = 147.83 kDa]

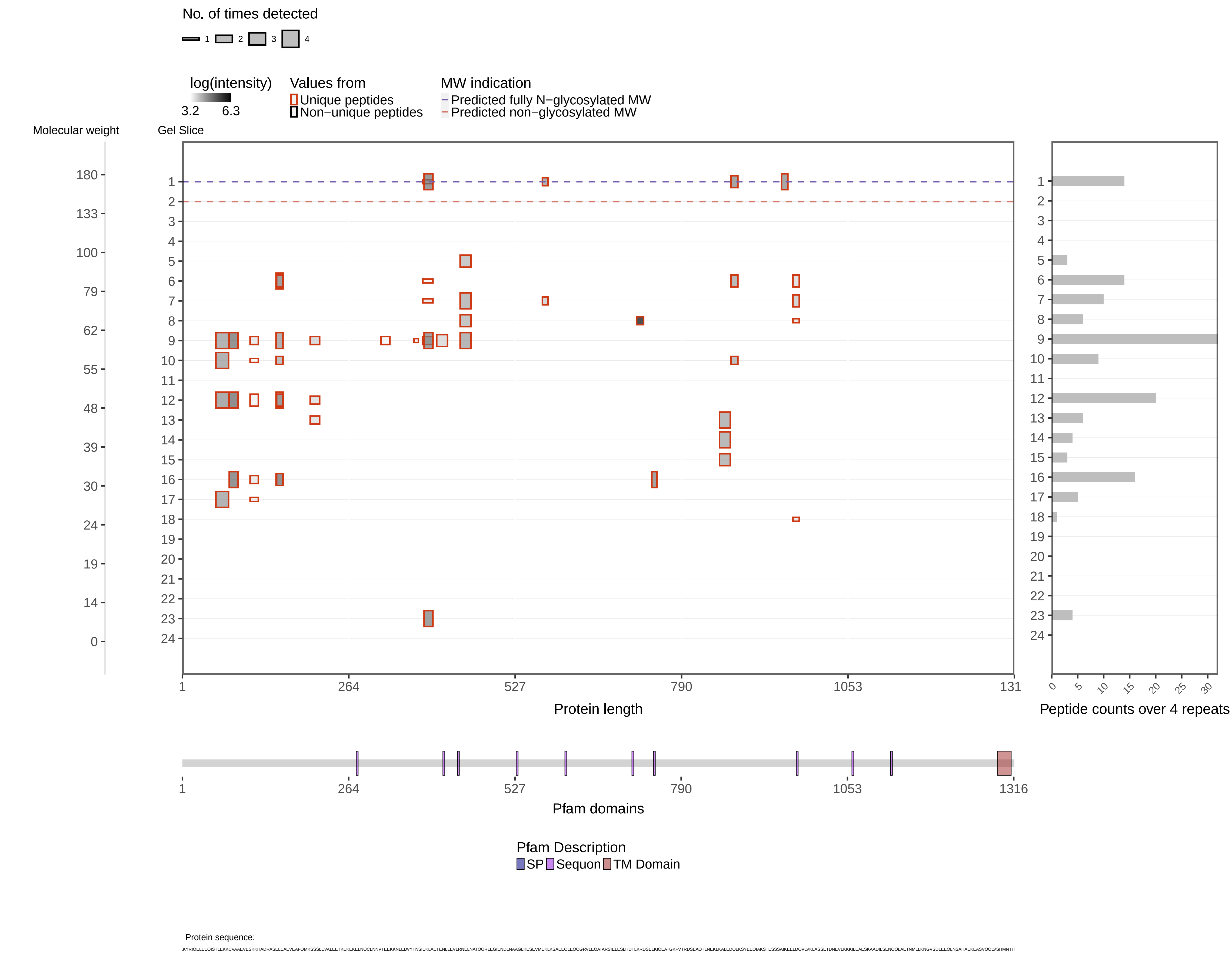

### NbL05g13310.1 gamma_aminobutyrate_transaminase_1_mitochondrial [MW = 56.9 kDa]

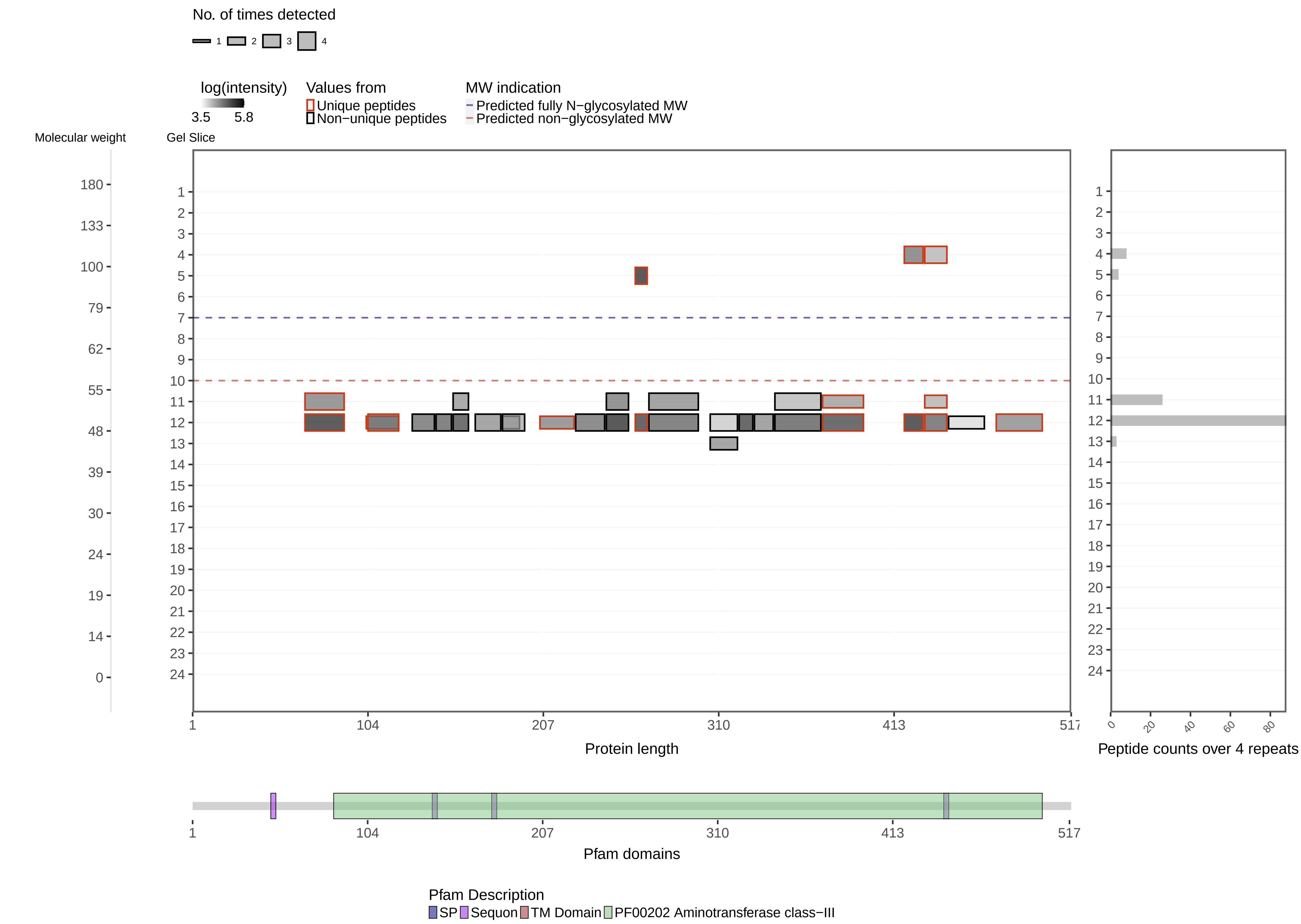


Protein sequence:

### NbL05g13250.1 UDP−glycosyltransferase_71E1−like [MW = 54.33 kDa]

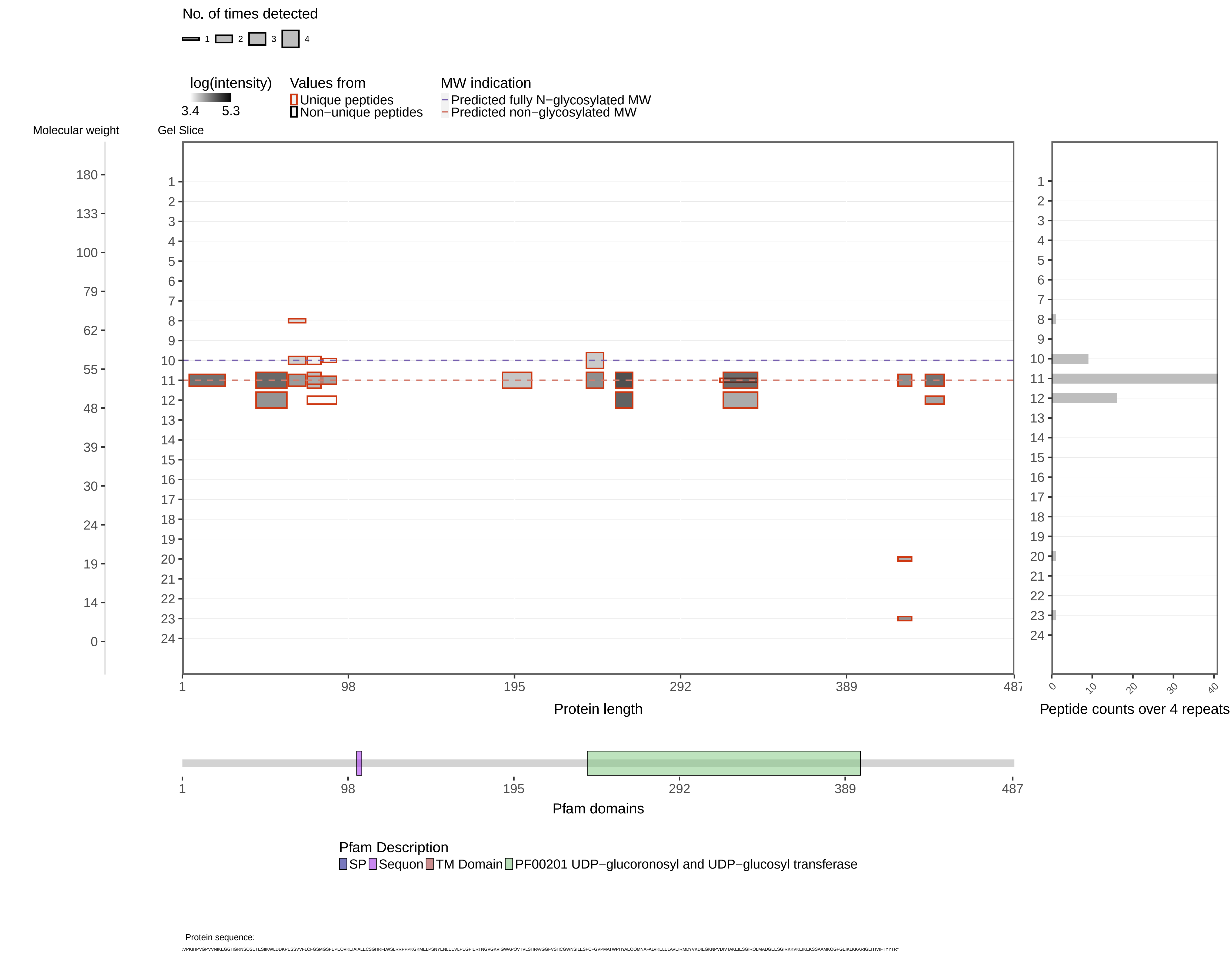

NbL05g13240.1 anthocyanidin_3−O−glucosyltransferase_6−like [MW = 53.58 kDa]

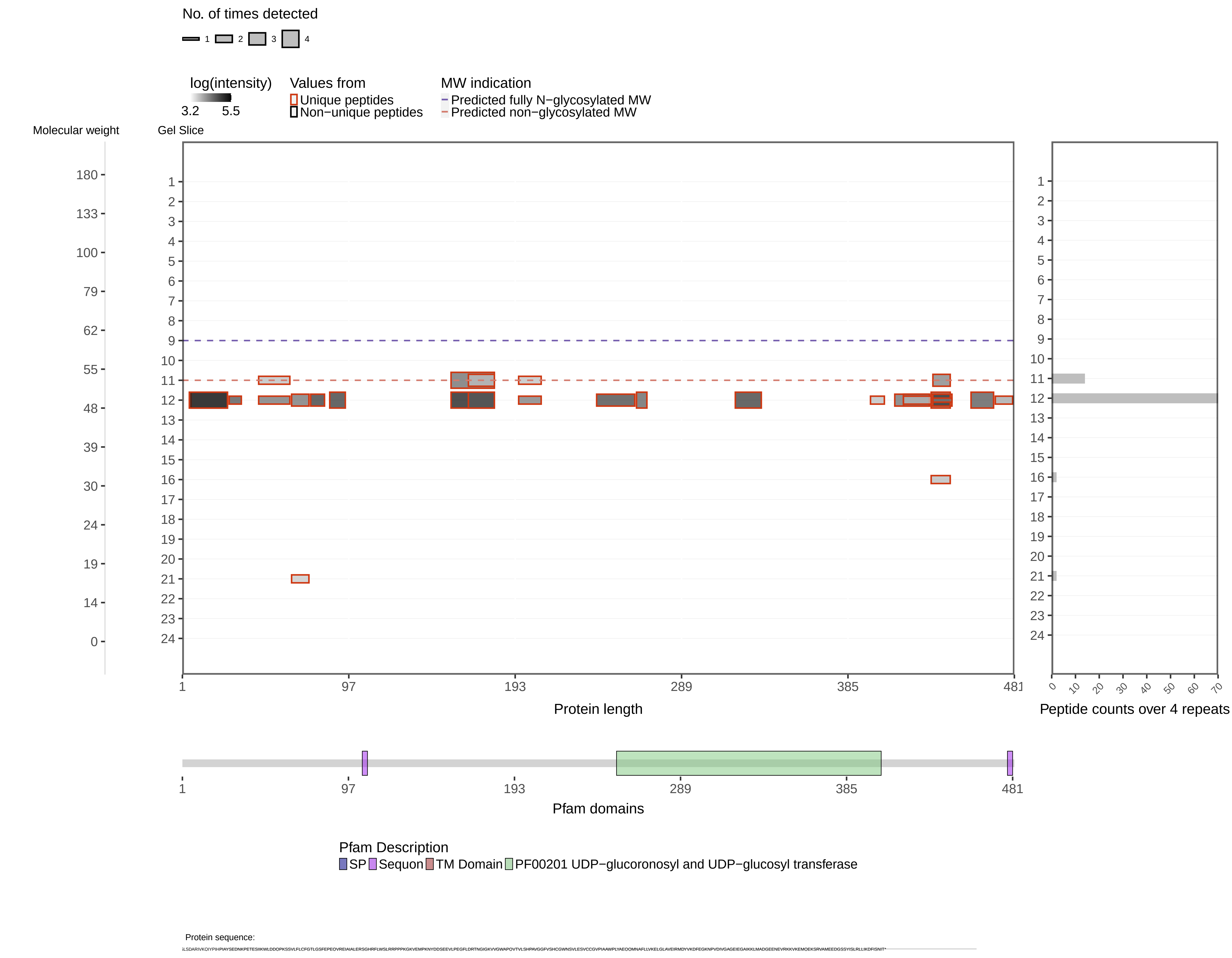

NbL05g13180.1 pyruvate_dehydrogenase_E1_component_subunit_alpha−3_chloroplastic−like [MW = 47.75 kDa]

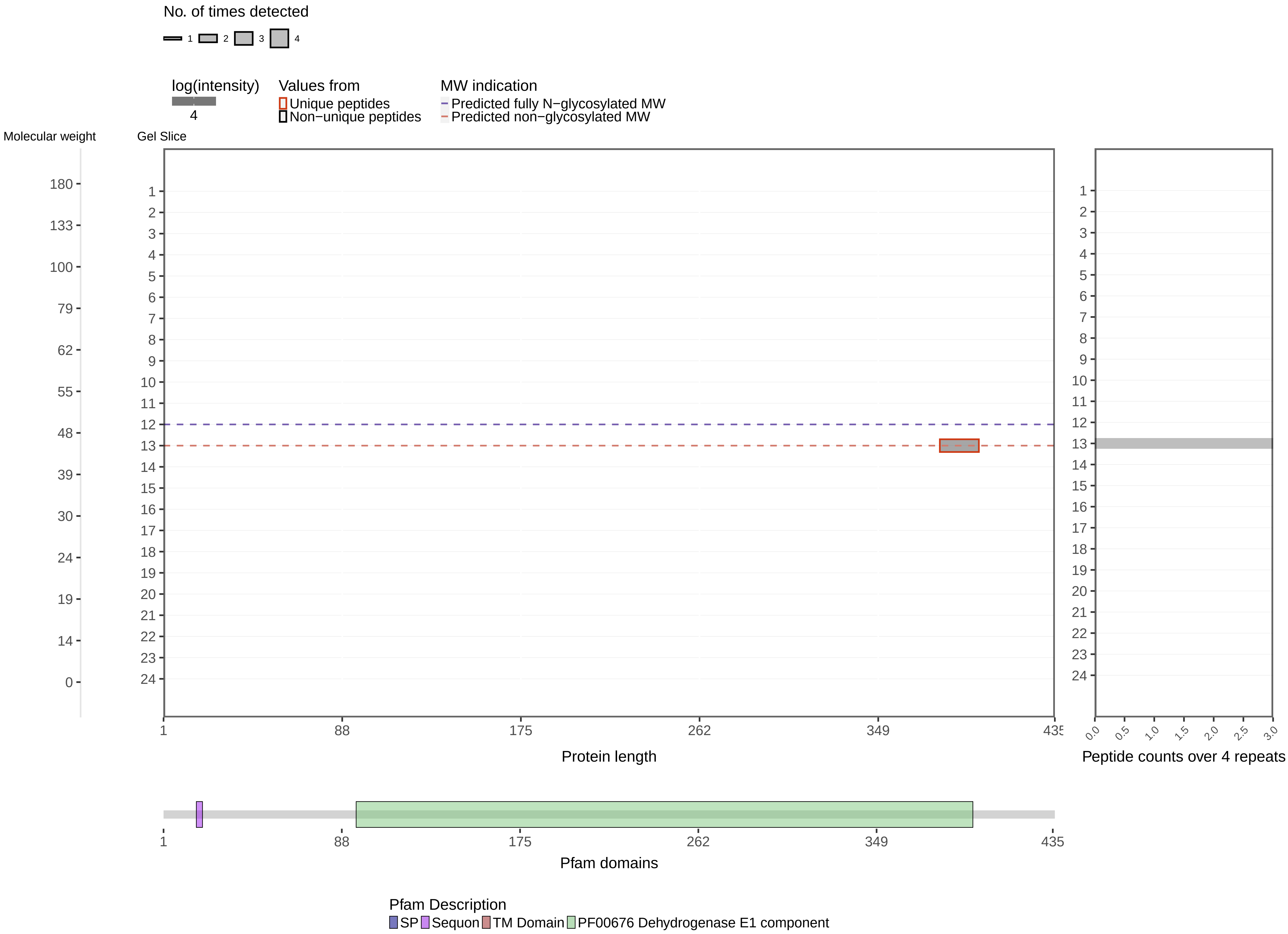


Protein sequence:

TGAAFTSKYRKEVLKGAECDHVTLAFFGDGTCNNGQFYECLNMAALWKLPIIFVVENNLWAIGMSHLRSTSDPEIWKKGPAFGMPGVHVDGMDVLKVREVAKEAIGRARRGEGPTLVECETYRFRGHSLADPDELRDPAEKNHYATRDPITALKKYMFENNLVSEAELKAIDKKIDEVVEESVDFADASPVPARSQLLENVFADPKGFGIGPDGRYRCEDPKFTEGTAHV*

### NbL05g13080.1 glycogen_synthase−like [MW = 61.76 kDa]

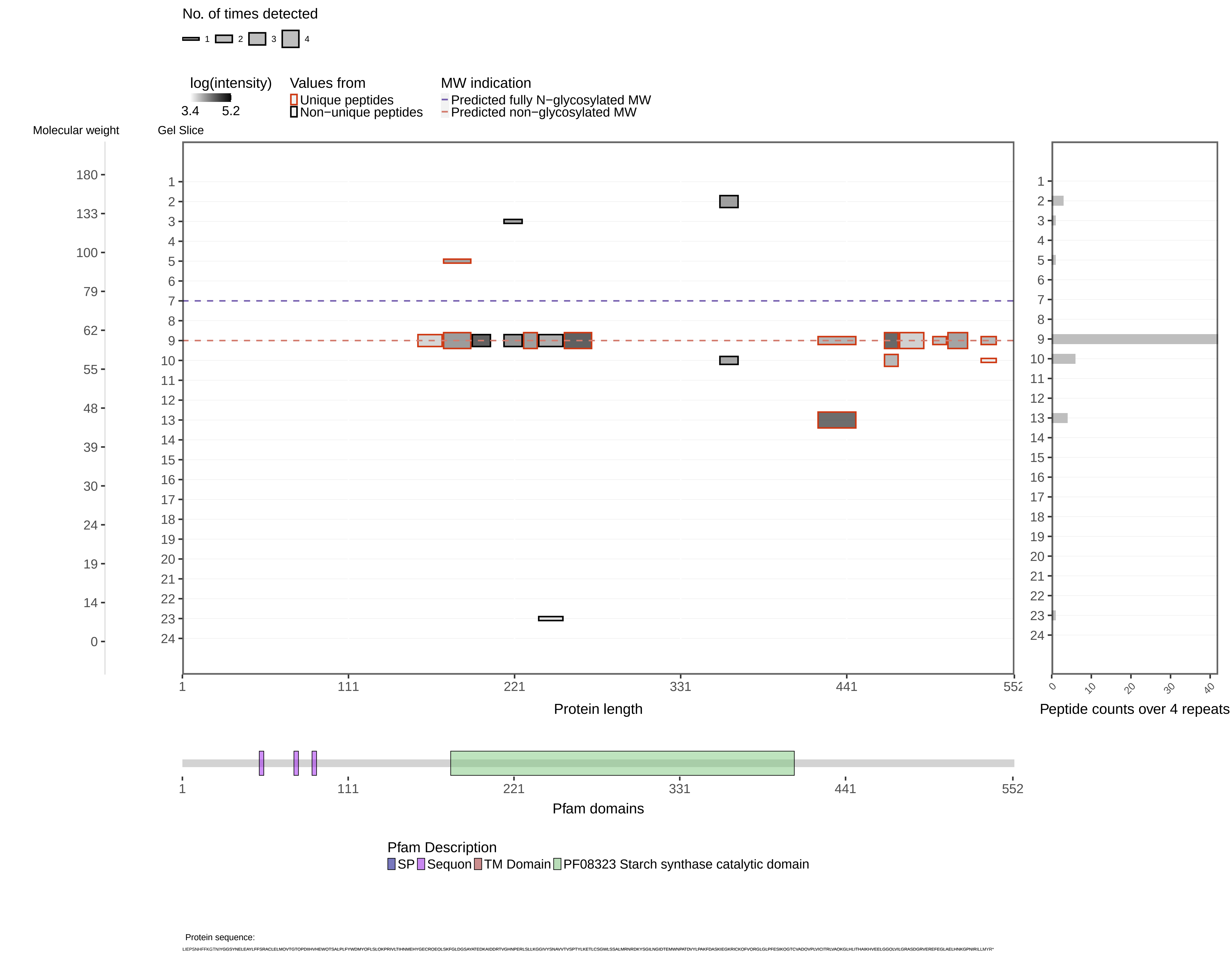

### NbL05g13000.1 eukaryotic_translation_initiation_factor_3_subunit_K−like [MW = 26.3 kDa]

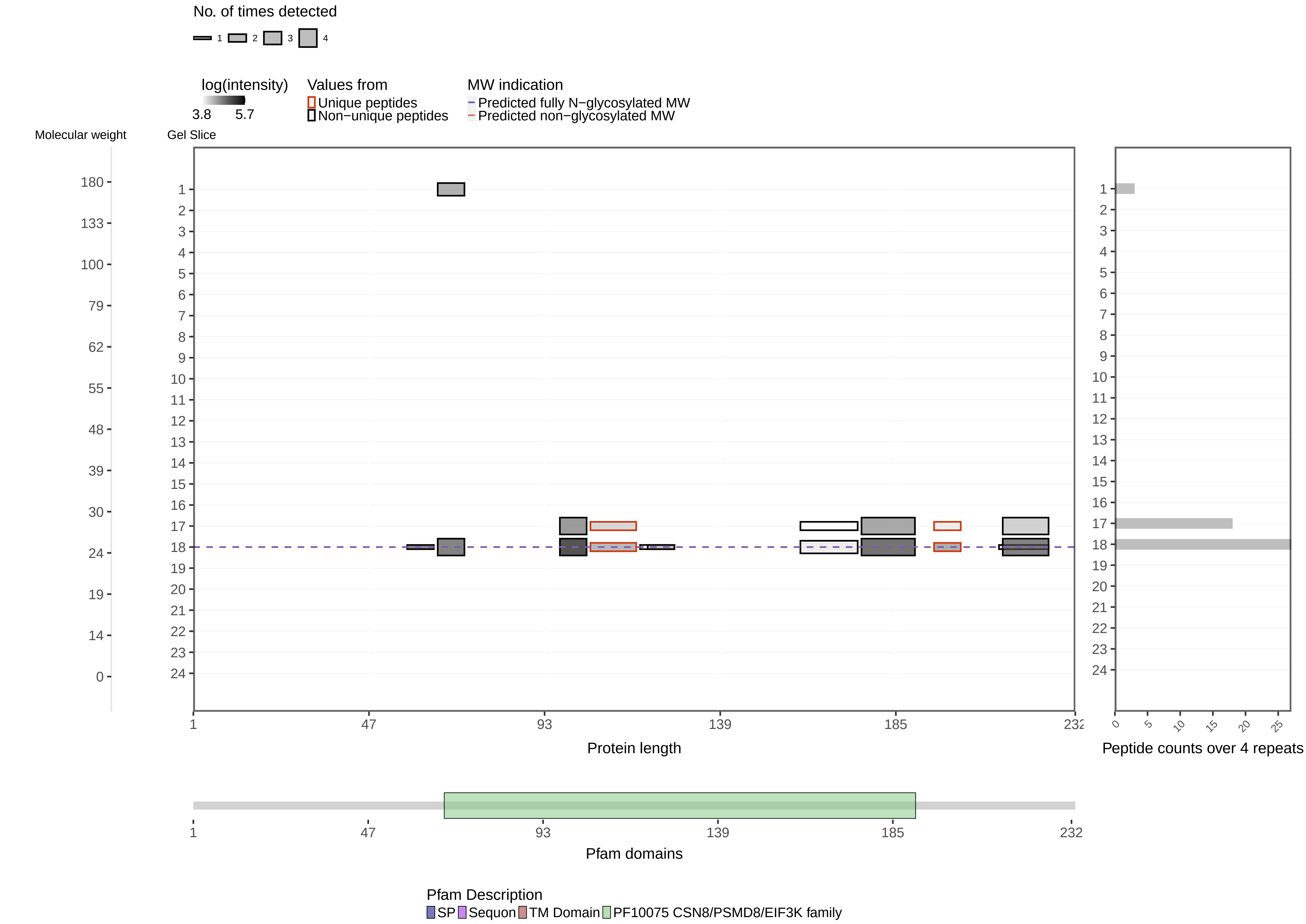


Protein sequence:

QFKTLILLSHYLETARFRQFWDEAAKSRHIVEVVPGFEQAIQAYAVHVLSITYQKVPRAIVAEAINIEGLSLDKFLEHQVSNNGWVIEKGQGKGQLITLPRTEFNHPELKKNTADSVPLEHVTRIFPILG*

NbL05g12990.1 sucrose_synthase [MW = 92.63 kDa]

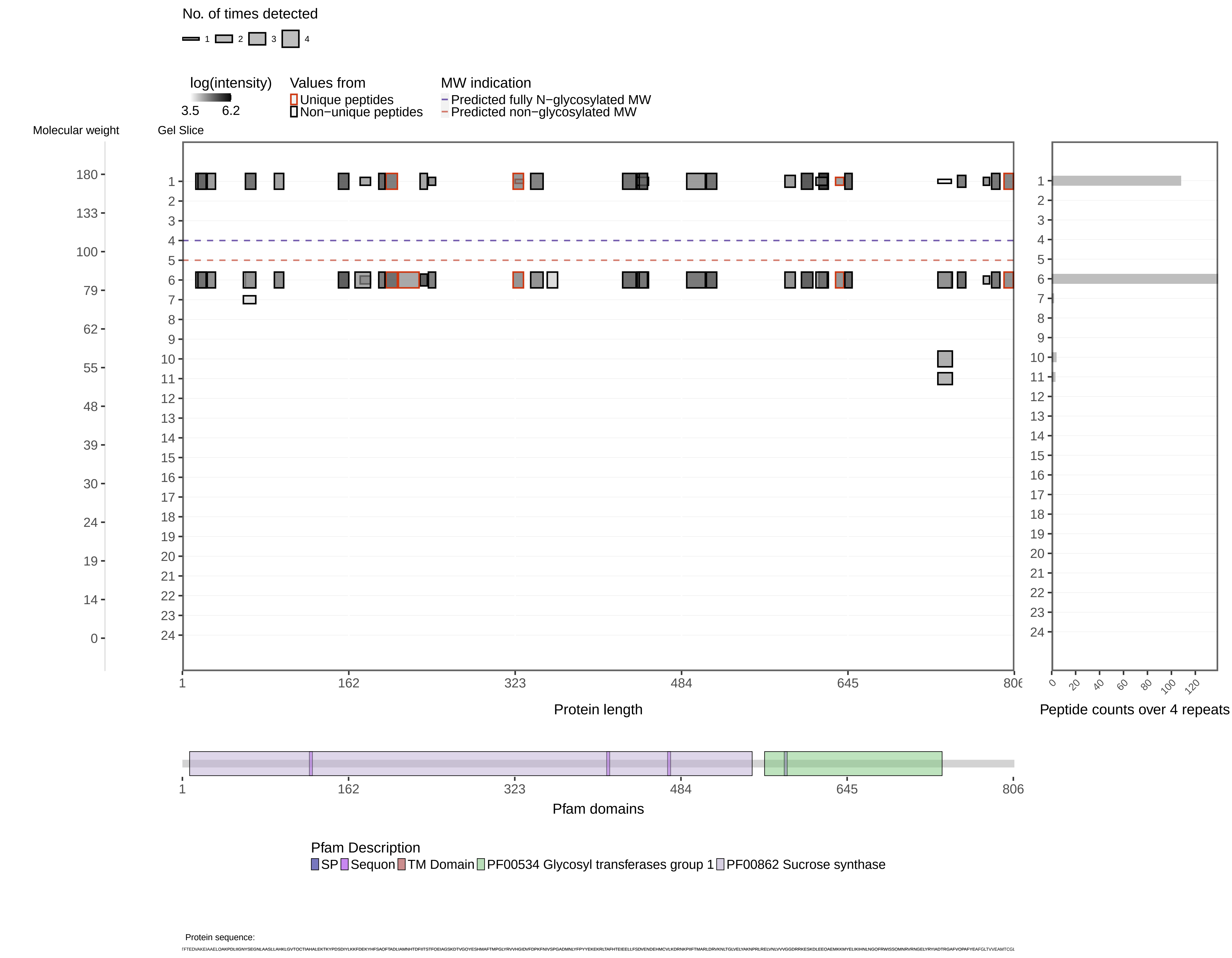

### NbL05g12970.1 sucrose_synthase−like [MW = 91.19 kDa]

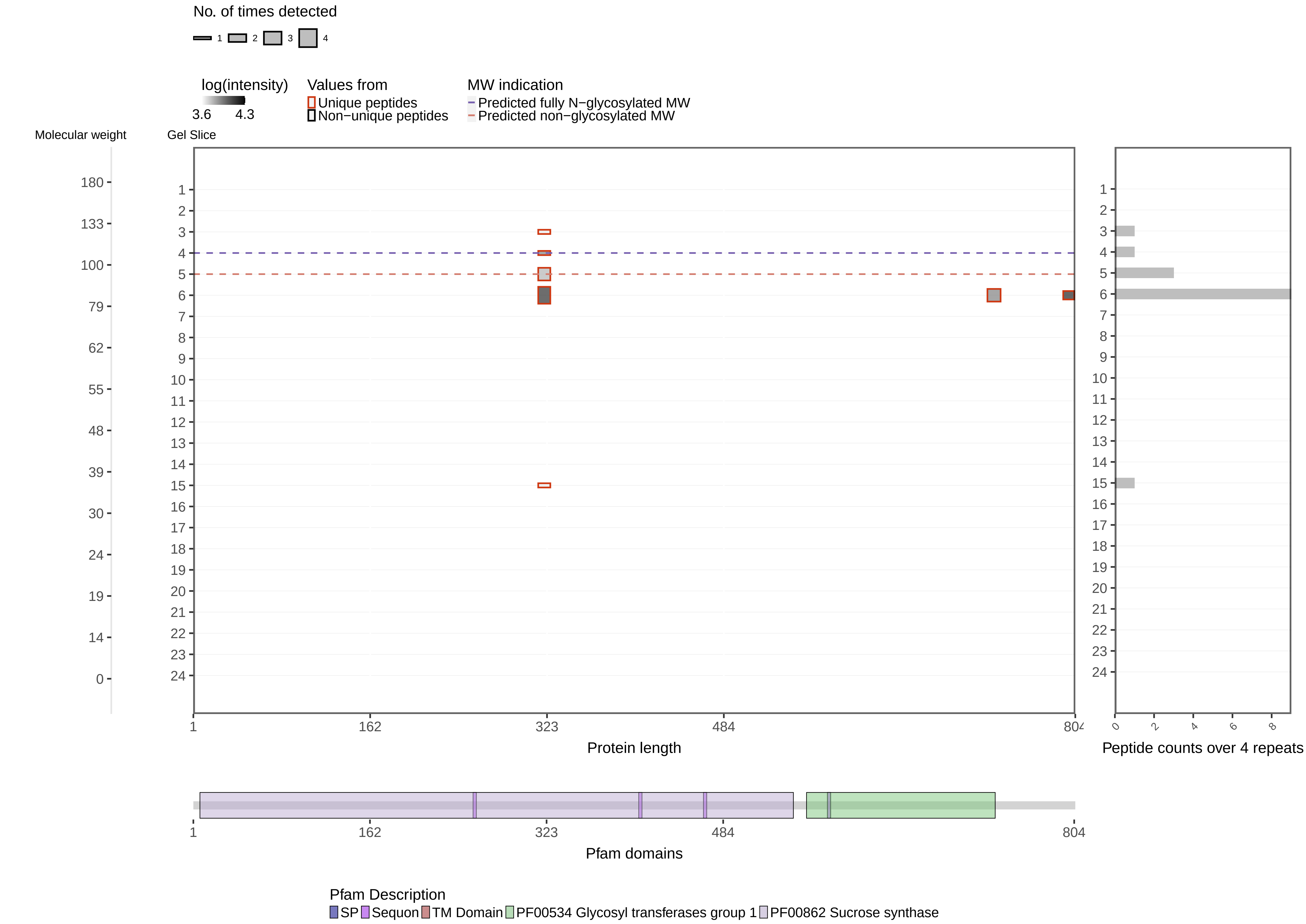


Protein sequence:

EELVKELQGKPDLIIGNYSEGNLAASLLAKKFGATQCTIAHALEKTKYSNSDLNWKKFDDKYHFSSQFTADLFAMNHTDFIITSTFQEIAGSKNTVGQYESHTAFTMPGLYRVVHGIDSFDPKFNIVSPGADMSIYFPYTEKEKRLTNFHPEIEELLYSPVENKDHLCVLKDRNKPILFTMARLDRVKNLTGLVEWYAKNARLRELVNLVVVGGDRRKESKDLEEQAEMKKMYDLIETYNLNGQFRWISSQMNRVRNGELYRYIADTRGAFVQPAFYEAFGLTVVESMTCGLPTFA

NbL05g12930.1 peroxiredoxin_Q_chloroplastic−like [MW = 23.7 kDa]

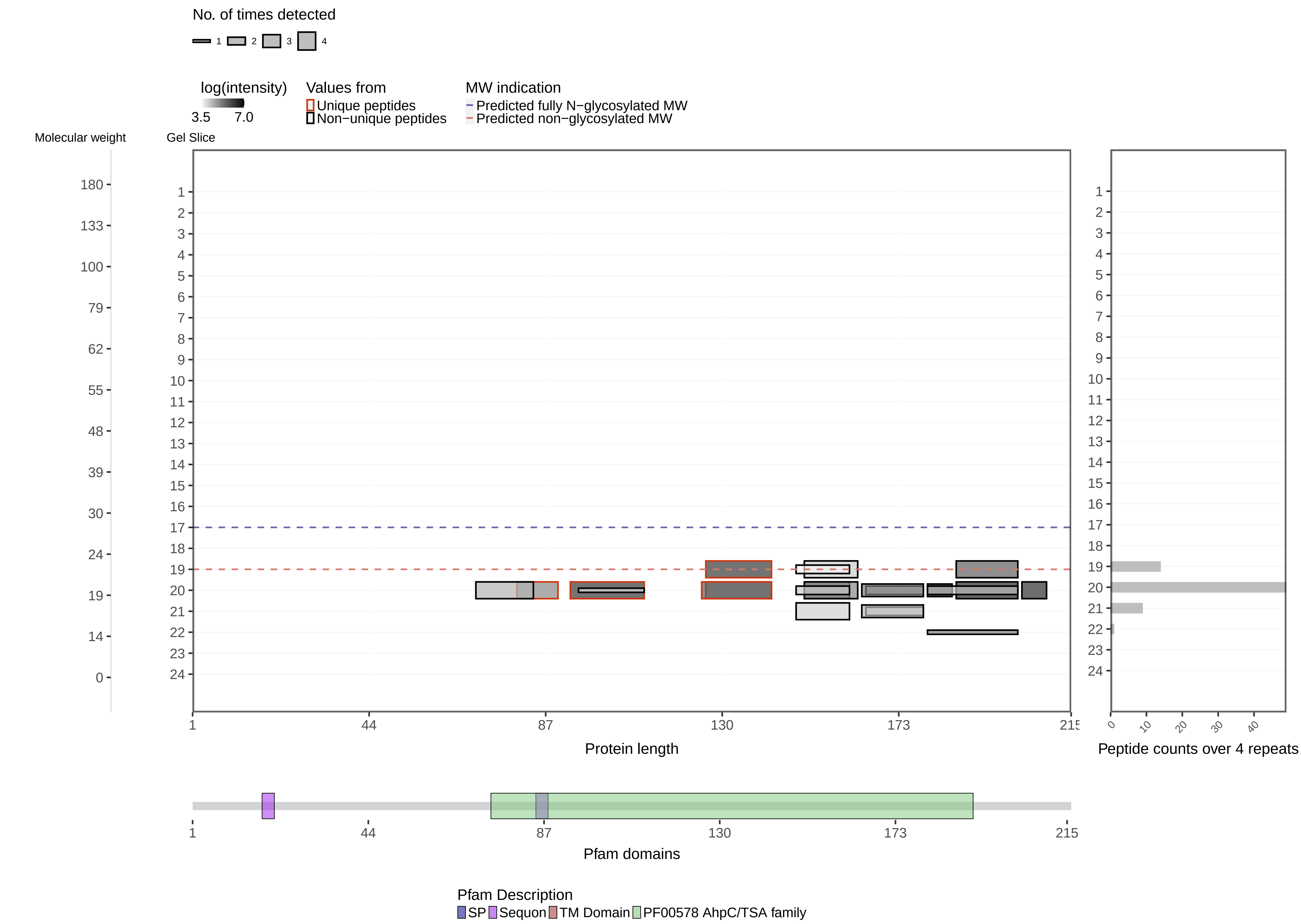


Protein sequence:

VVVYFYPADETPGCTKQACAFRDSYEKFKKAGAEVVGISGDDPASHKAFAKKYRLPFTLLCDEGNKVRKEWGIPGDLFGTLPGRQTYVLDKNGVVQLIYNNQFQPEKHIDETLKLLQSL*

### NbL05g12850.1 20_kDa_chaperonin_chloroplastic−like [MW = 26.47 kDa]

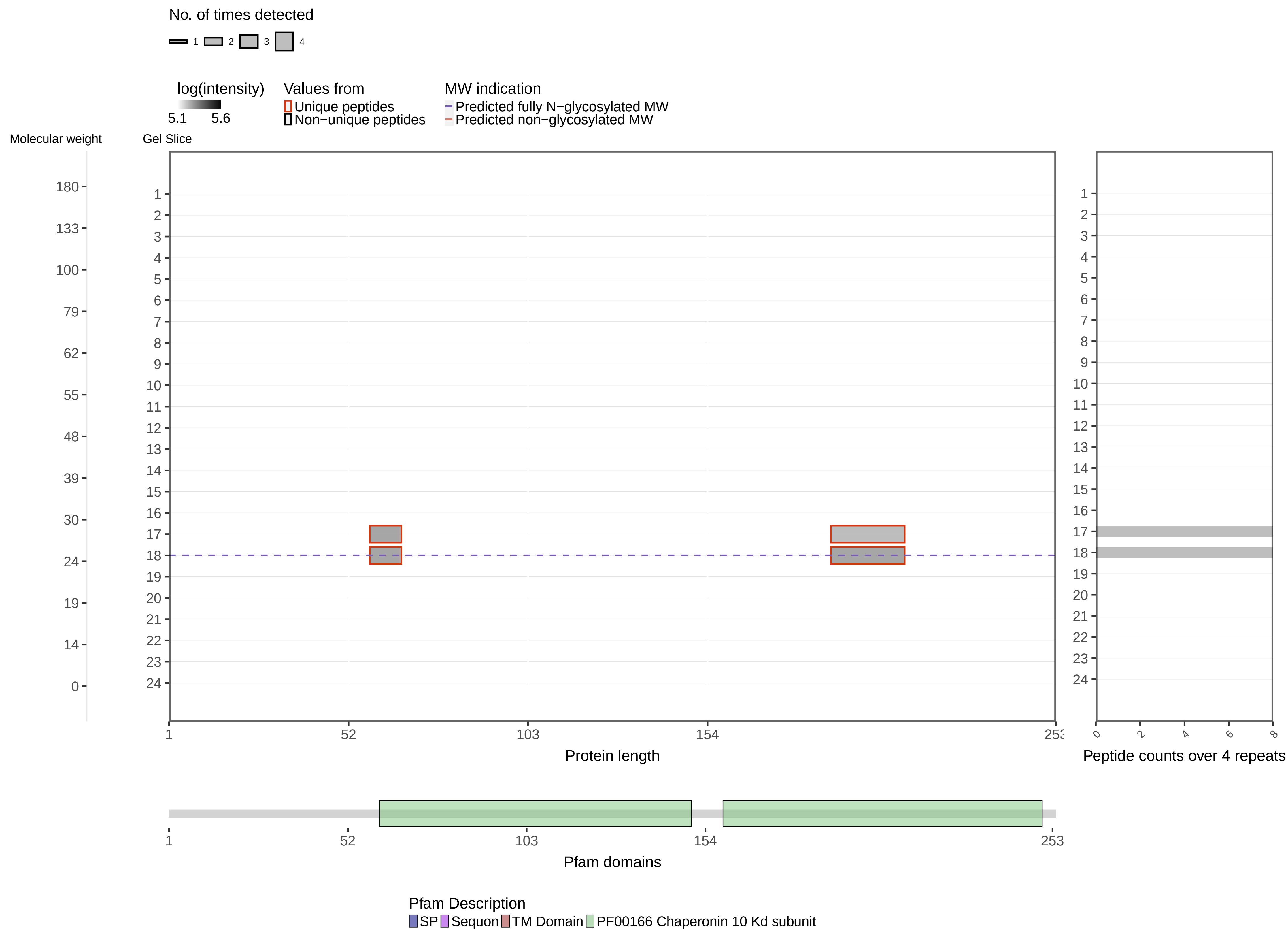


Protein sequence:

'DISVKTGAQVVYSKYAGTEVEFDGSKHLILKEDDIVGVLETDDIKDLQPLNDRVLIKVAEAEEKTAGGLLLTEAAKEKPSIGAVIAVGPGPLDEEGNRKPLSVSPGNTVLYSKYAGSEFKGADGSDYITLRASDVMAVLS*

### NbL05g12650.1 subtilisin−like_protease_SBT1.8 [MW − SP = 79.19 kDa]

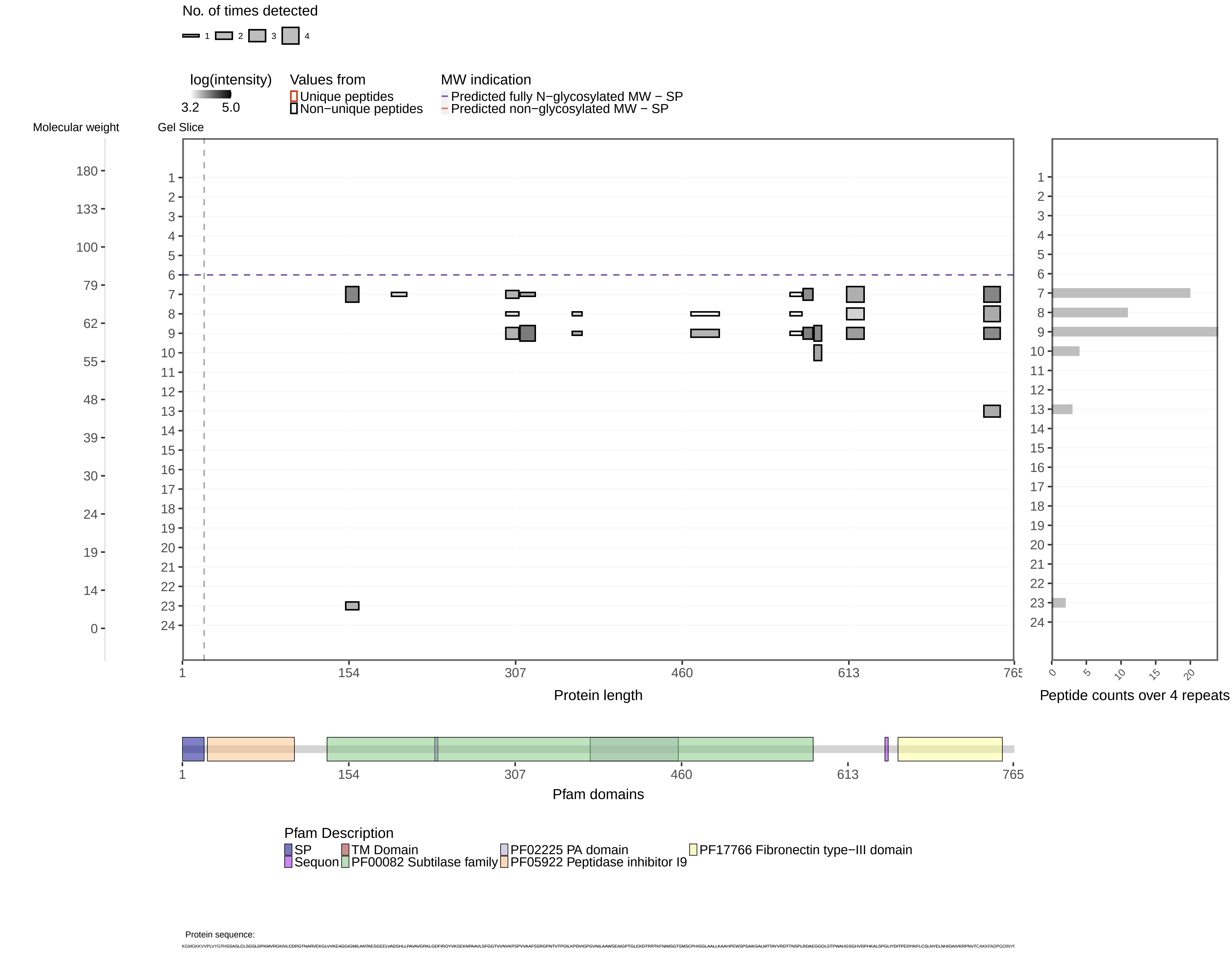

### NbL05g12610.1 cysteine_proteinase_3 [MW − SP = 37.02 kDa]

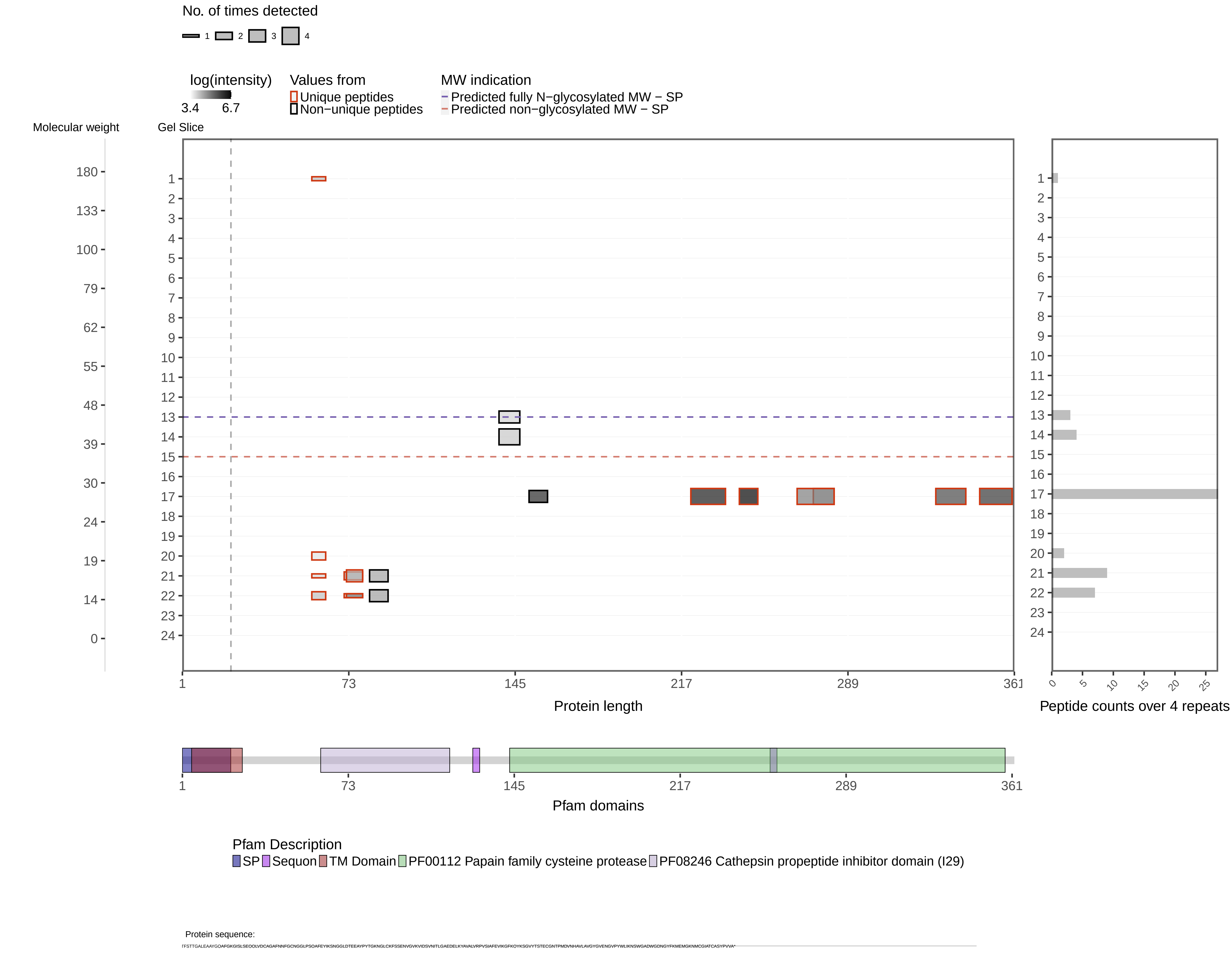

### NbL05g12600.1 tubulin−−tyrosine_ligase−like_protein_12 [MW = 100.24 kDa]

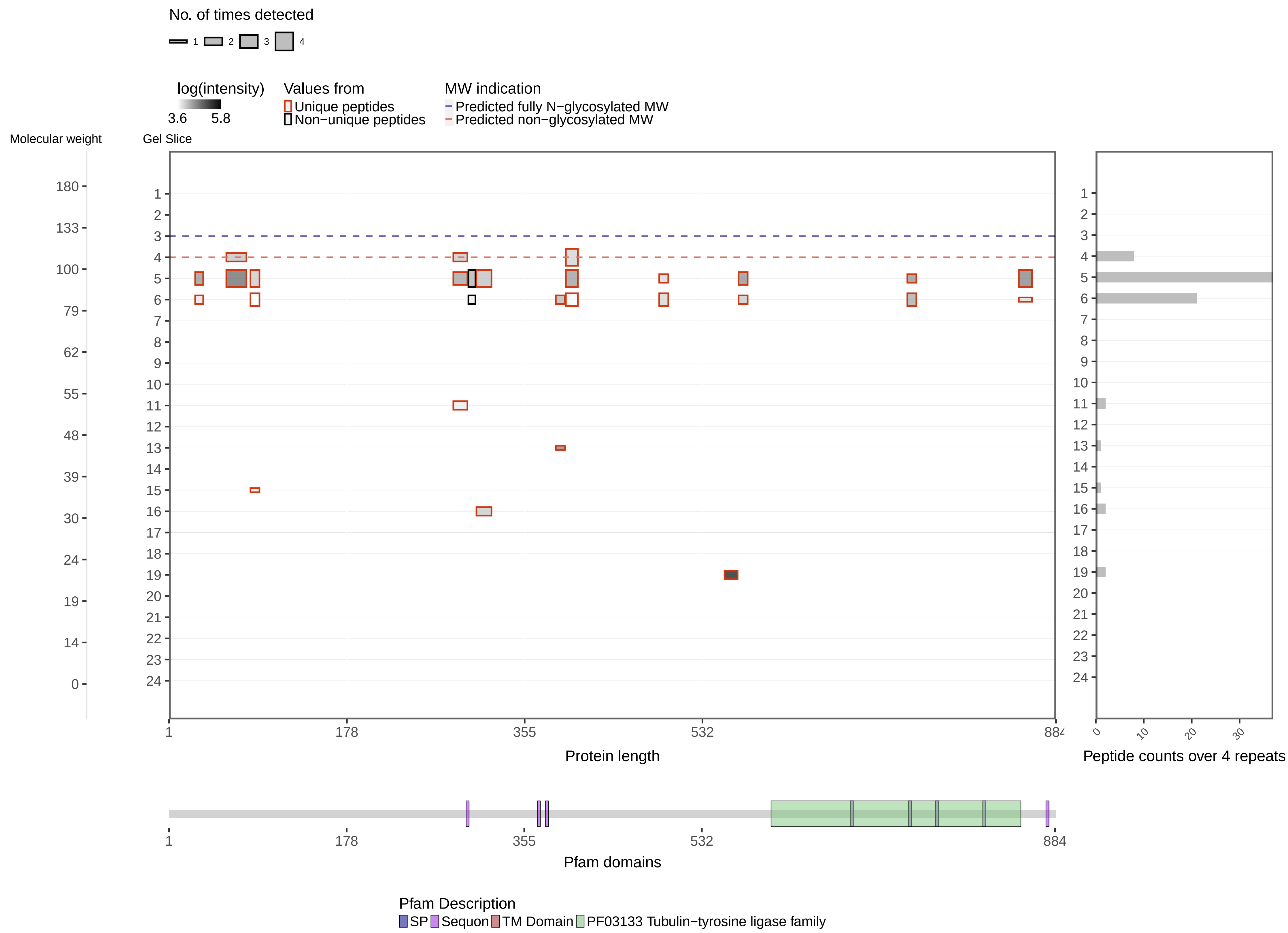


Protein sequence:

NbL05g12510.1 auxin−binding_protein_ABP19a−like [MW − SP = 19.89 kDa]

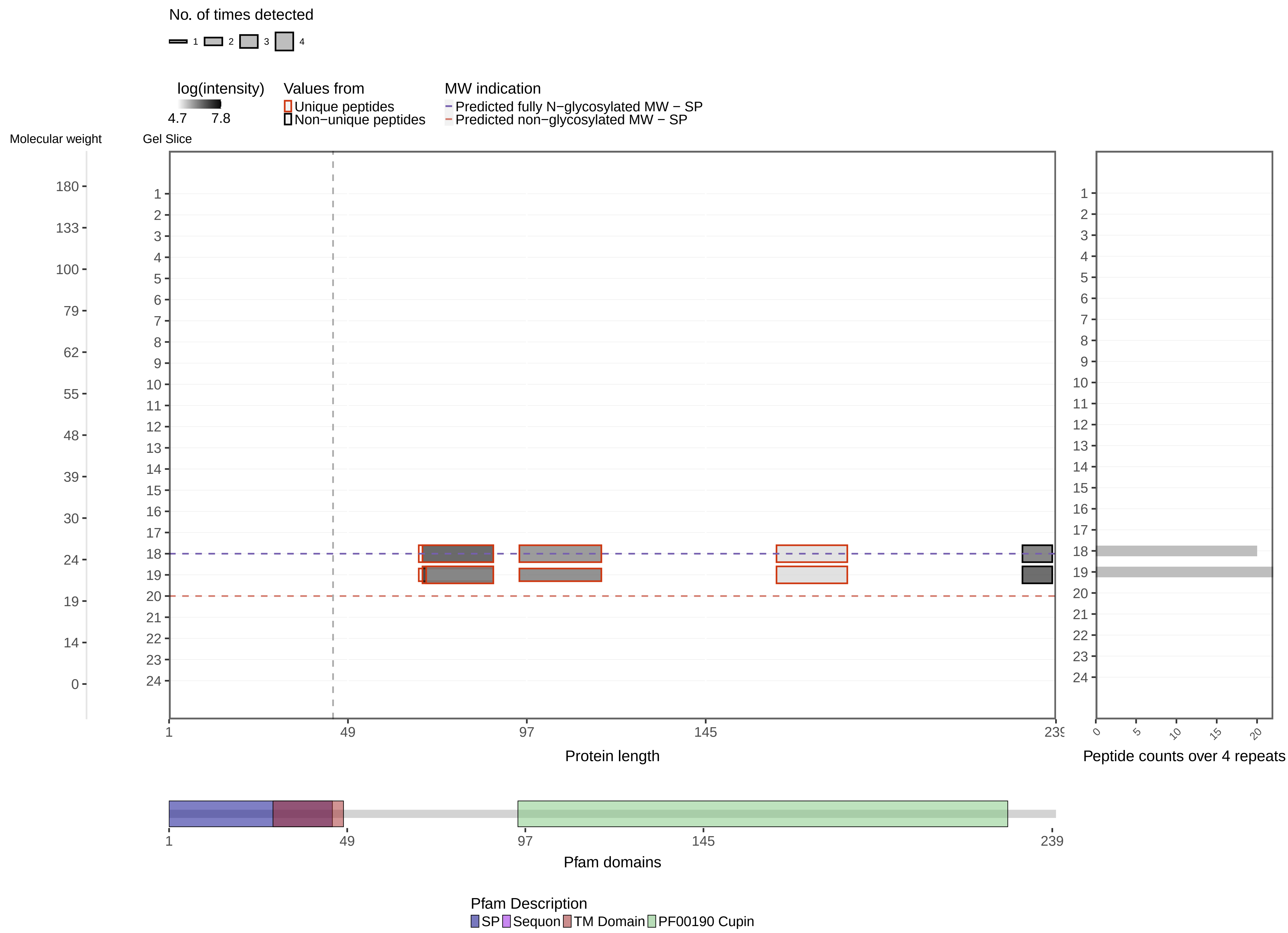


Protein sequence:

PGLNGLGISMARLDLAIGGVIPMHTHPGASEVLYVGSGEICAGFISSSDNKVFFKNLKQGDIMVFPQGLLHFQINSGKTAALAIVSFSSPTPGLQITDFALFANDLATEIVQATTFLDAATIKKLKGVLGGTN*

NbL05g12500.1 40S_ribosomal_protein_S27−2−like [MW = 12.23 kDa]

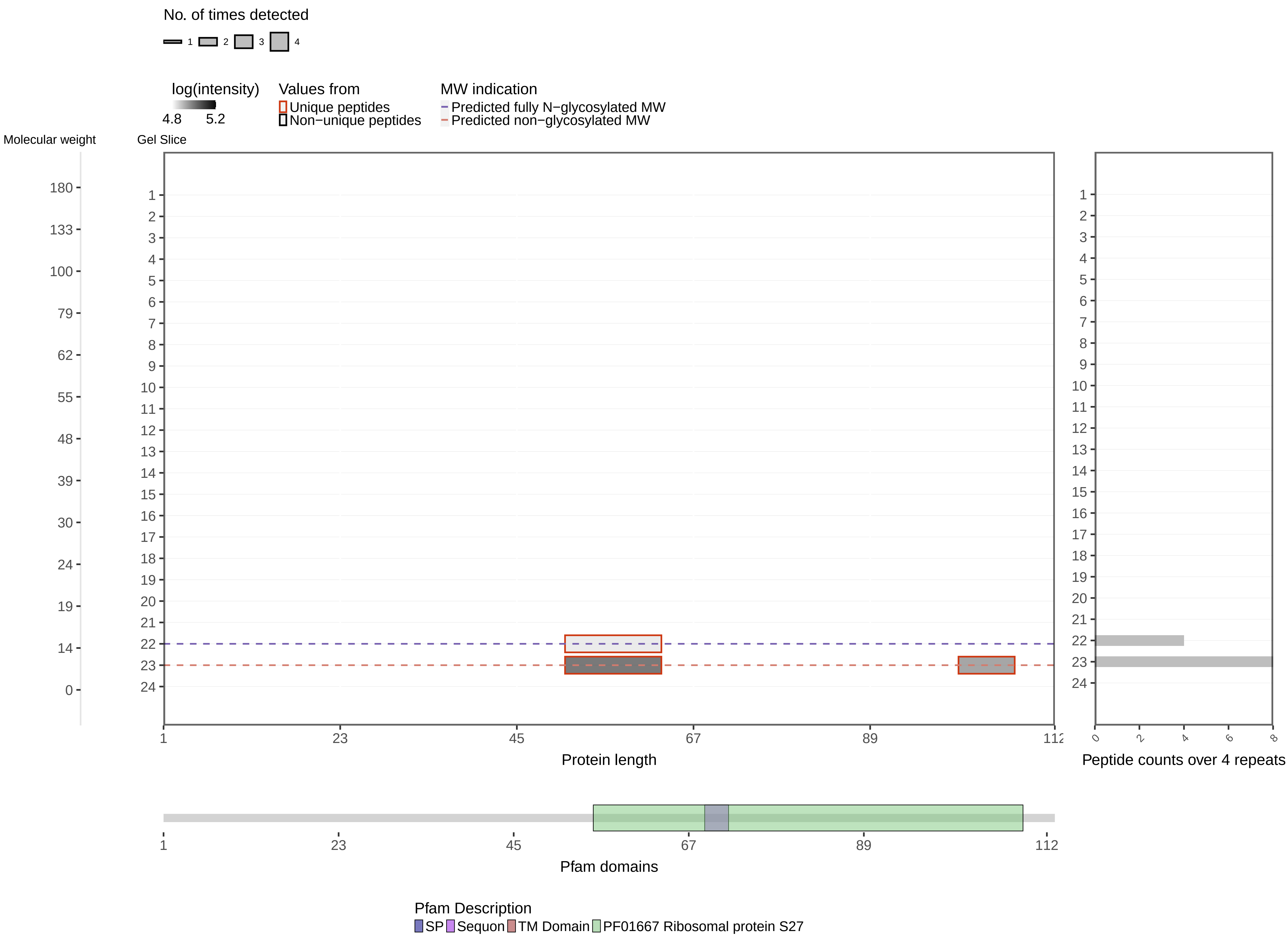


Protein sequence:

KHKLKRLVQSPNSFFMDVKCQGCFNITTVFSHSQTVVVCGNCQTVLCQPTGGRARLTEGCSFRRKGD*

### NbL05g12440.1 stress−response_A/B_barrel_domain−containing_protein_UP3 [MW = 33.43 kDa]

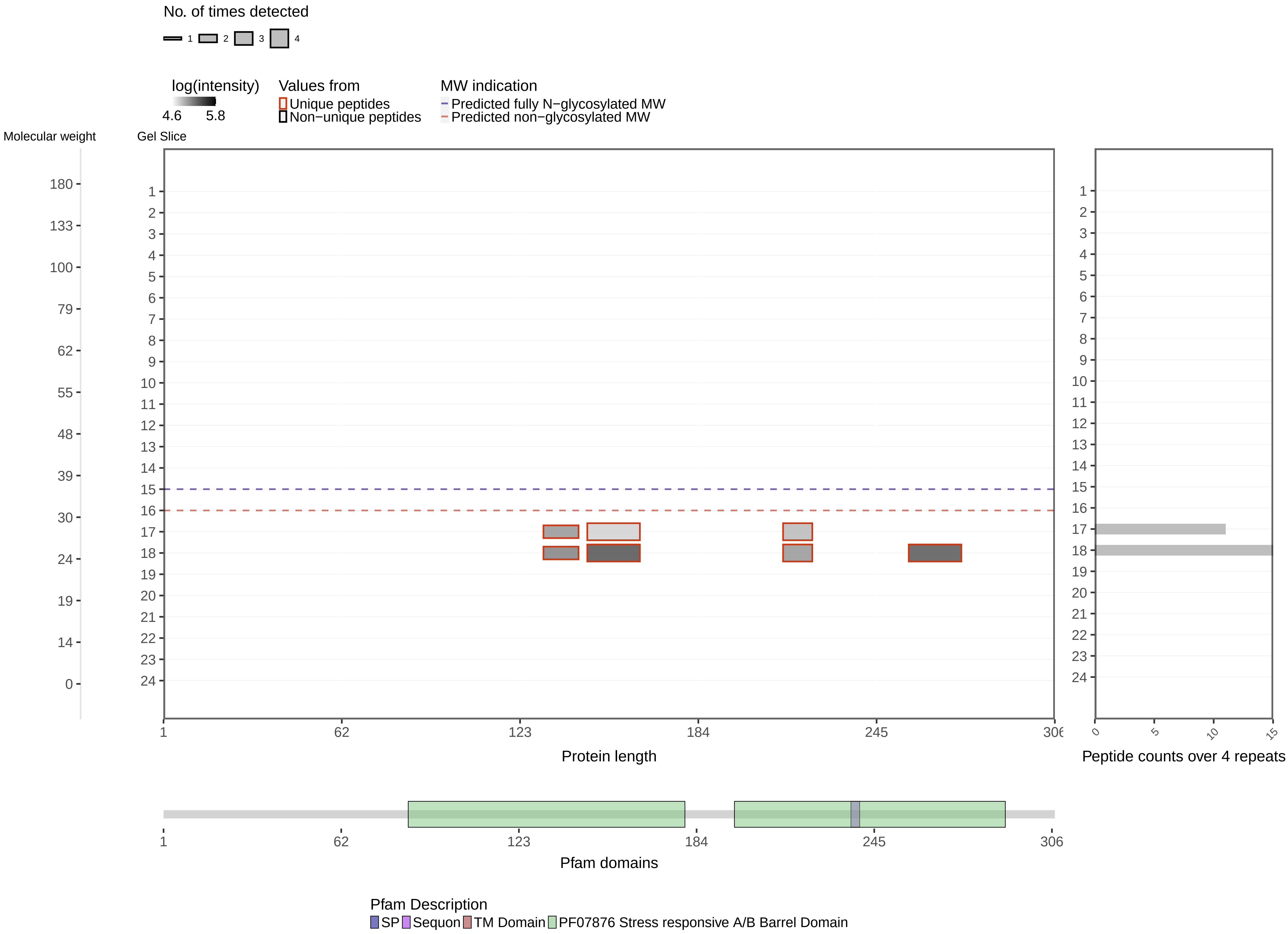


Protein sequence:

.HSRYKSISDLADYSAHPDHVSVVKEYVLPVVDDIMAVDWIPTDFSGPSEVPPGSALRVTFLKLKENLGENEKSQVLSAVGGIKEKFPGIEQLTVGENFSPGRAKGFSIASIAVFKGVNELEALESQTELANEQKEKVRDFLDGVVVVDYAVLPTQSPSLSVKISK*

NbL05g12300.1 glucose−1−phosphate_adenylyltransferase_small_subunit_chloroplastic/amyloplastic [MW = 57.24 kDa]

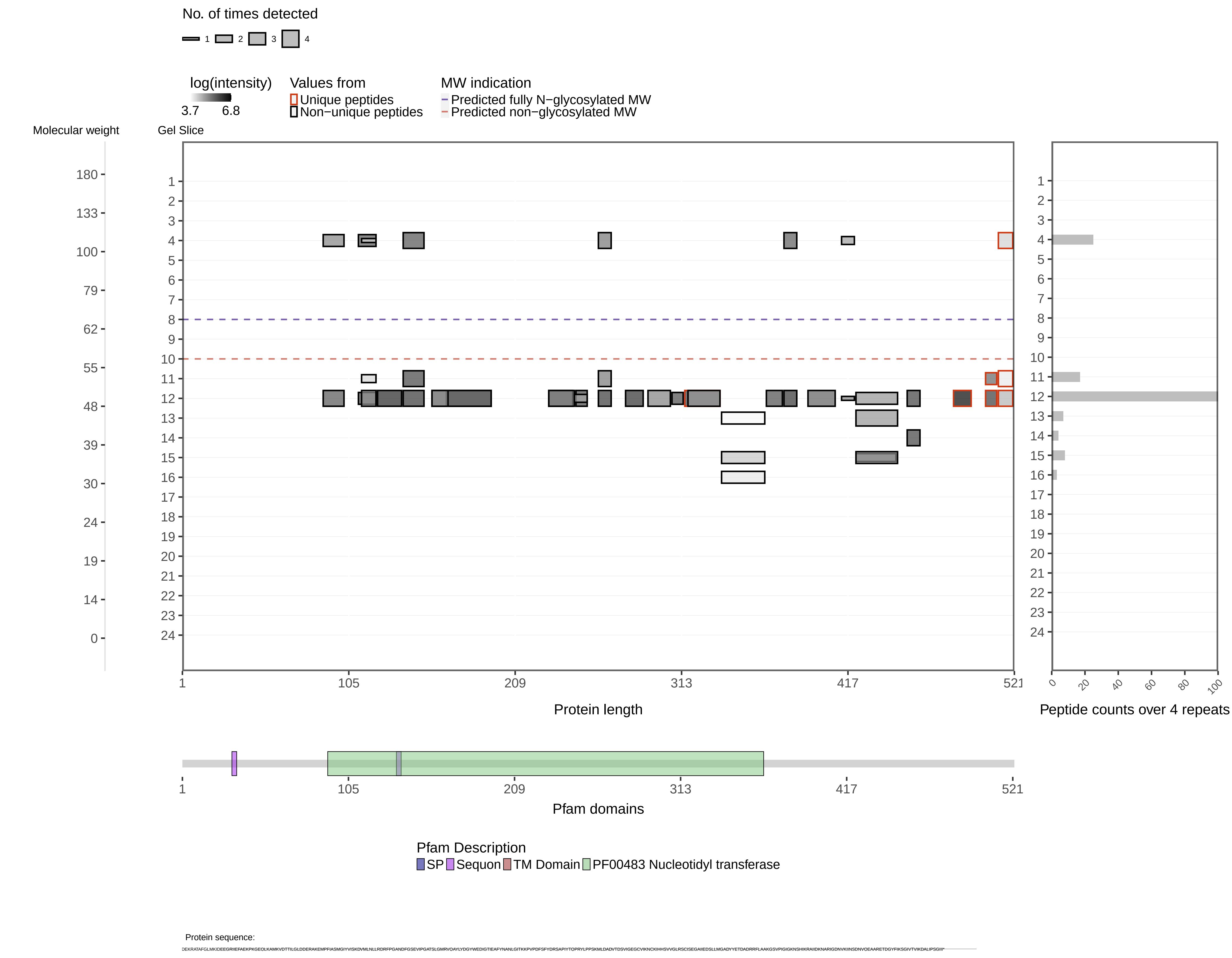

NbL05g12260.1 subtilisin−like_protease_SBT5.3 [MW − SP = 79.89 kDa]

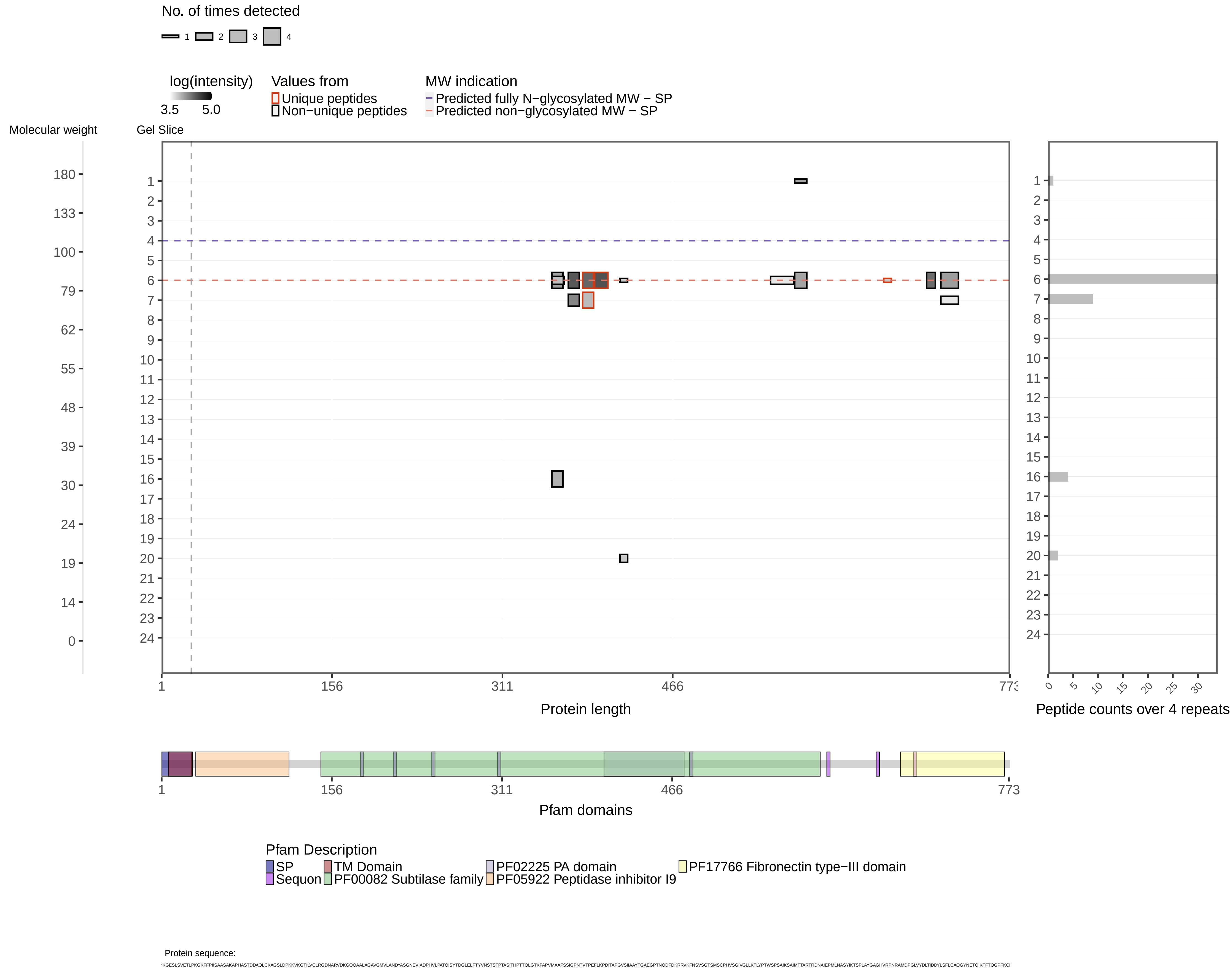

### NbL05g12070.1 peroxisomal_and_mitochondrial_division_factor_2−like [MW = 39.41 kDa]

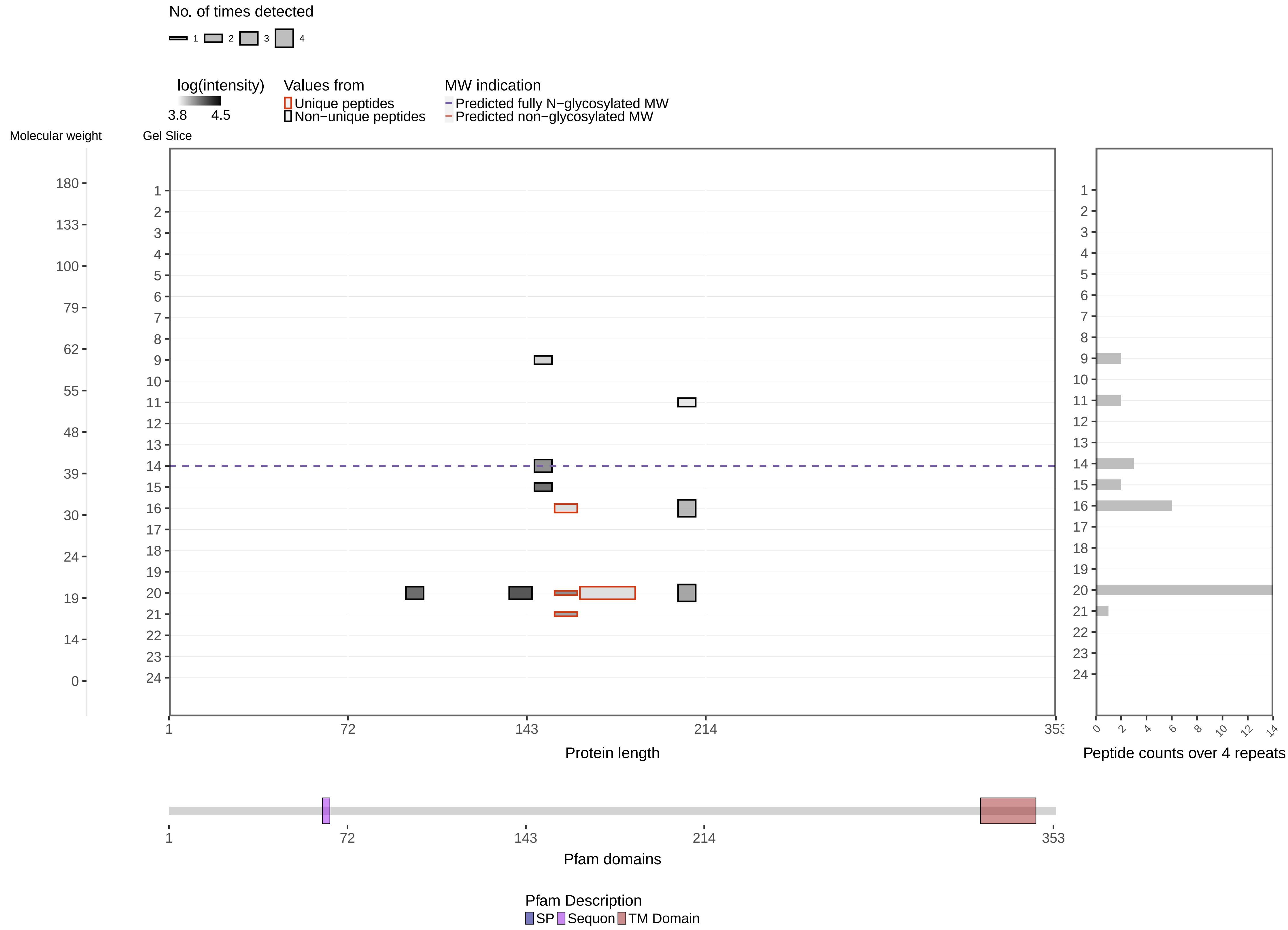


Protein sequence:

OHNLITAMNDLEGSNSELSGVKSTLEGLKSSENEMTVKLDAIESERNLLLSKLERLEASGNNORGEVEVKEEEIRGLKKHIEELEGTVVNNEEWEKEKTELHMVKEELEKRVKEMIARAAELEKKLEEKEKVITERLVASNVNGIPVGDYDKVGYFGAEVNLPVVAASSVVAVAVIGVVCYLKYGRKA*

NbL05g12030.1 ribose−phosphate_pyrophosphokinase_4 [MW = 35.74 kDa]

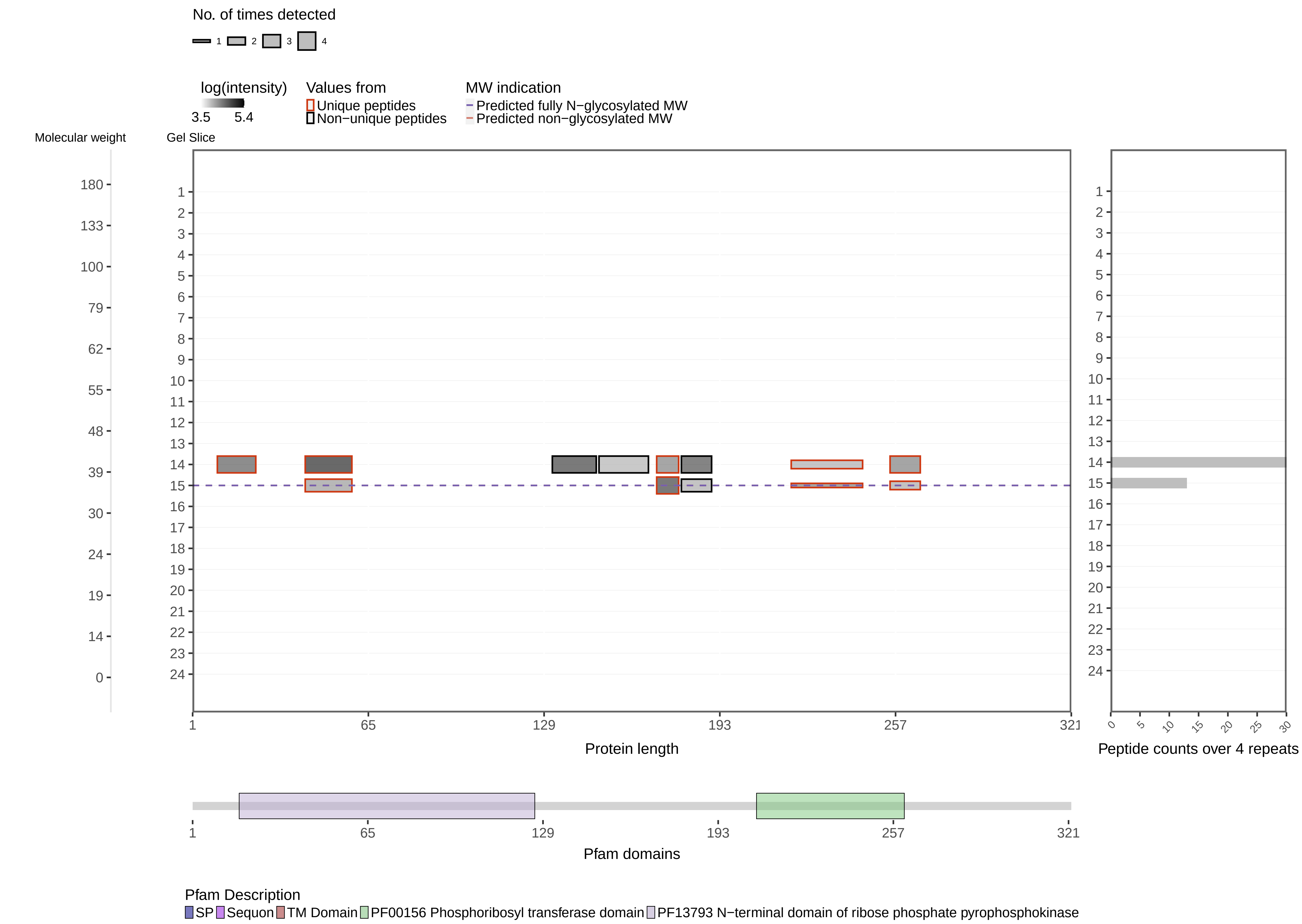


Protein sequence:

FYFGDHVLPLFETGIPLLKQRLQQLPQTEKIVIAFPDDGAWKRFYKQLGHYPAVICTKVREGDKRIVRLKEGNPAGCHVVIVDDLVQSGGTLIECQKVLAAHGAAKVSAYVTHGVFPKRSWERFIHKDDGSEKAFTYFWITDSCPLTVNAIADKAPFEVISLAGSIADALQT*

NbL05g11920.1 serine/threonine−protein_phosphatase_5 [MW = 61.32 kDa]

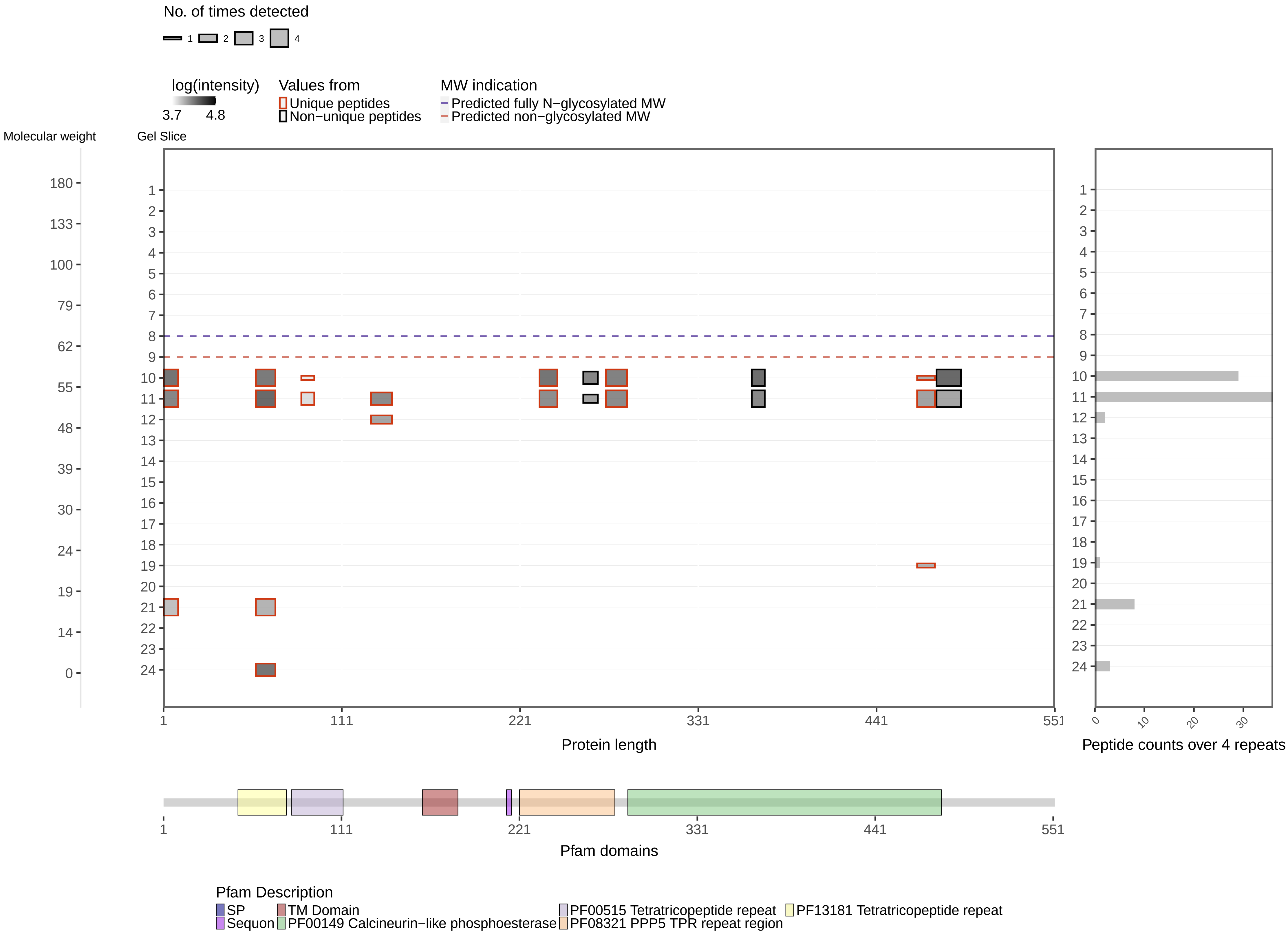


Protein sequence:

QVVLQTREMLRALPSLVDIIVPEGKHFTVCGDVHGQFYDLLNIFDLNGLPSEDNPYLFNGDFVDRGSFSLEVILTLFAFKCMCPSAIYLARGNHESKSMNKIYGFEGEVRSKLSETFVELFAEVFCCLPLAHVINGEVFVVHGGLFSVDGVKLSDIRAIDRFCEPPEEGLMCELLWSDPQPQPGRGPSKRGVGLSFGADITKRFLQENNLDLVVRSHEVKDEGYEIEHDGKLITVFSAPNYCDQMGNKGAFIRFEAPDMKPNIVTFSAVPHPDVKPMAYANNFLRMFS*

NbL05g11900.1 malonyl_CoA−acyl_carrier_protein_transacylase [MW = 39.78 kDa]

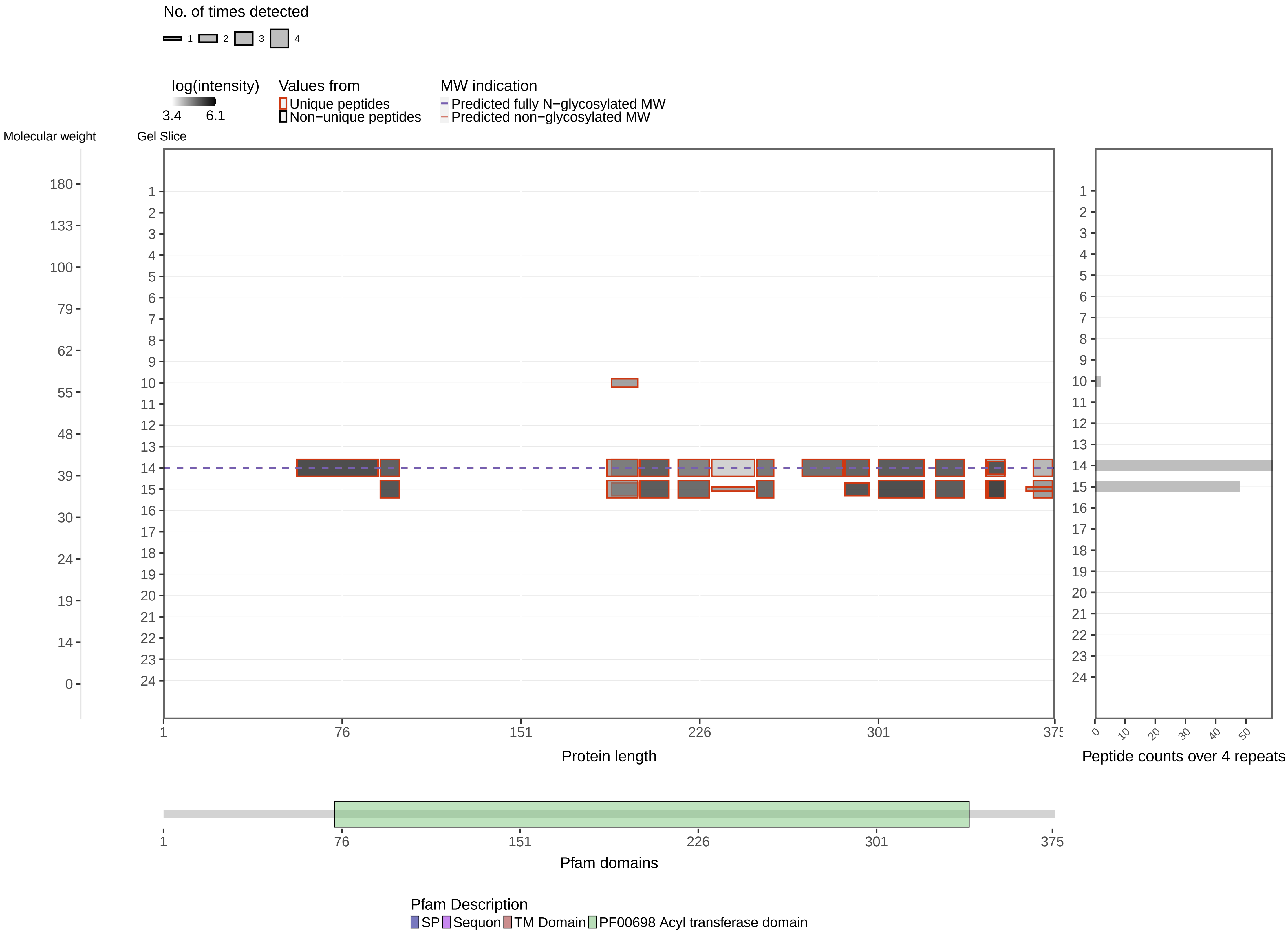


Protein sequence:

AFSFEDGLKLVKLRGEAMQDAADAAKSAMVSIIGLDSEKVQKLCDAANEEVDEANRVQIANFLCTGNYAVSGGLKGVEVVEAKAKSFKARMTVRLAVAGAFHTSFMNPAVSRLEDALGATEIRTPRIPVISNVDAQPHADPDTINKILARQVTSPVQWETTVKTLLTKGLKKSYELGPGKVIAGIVKRMDKGAVIENIGA*

NbL05g11840.1 protein_trichome_birefringence−like_43 [MW − SP = 37.79 kDa]

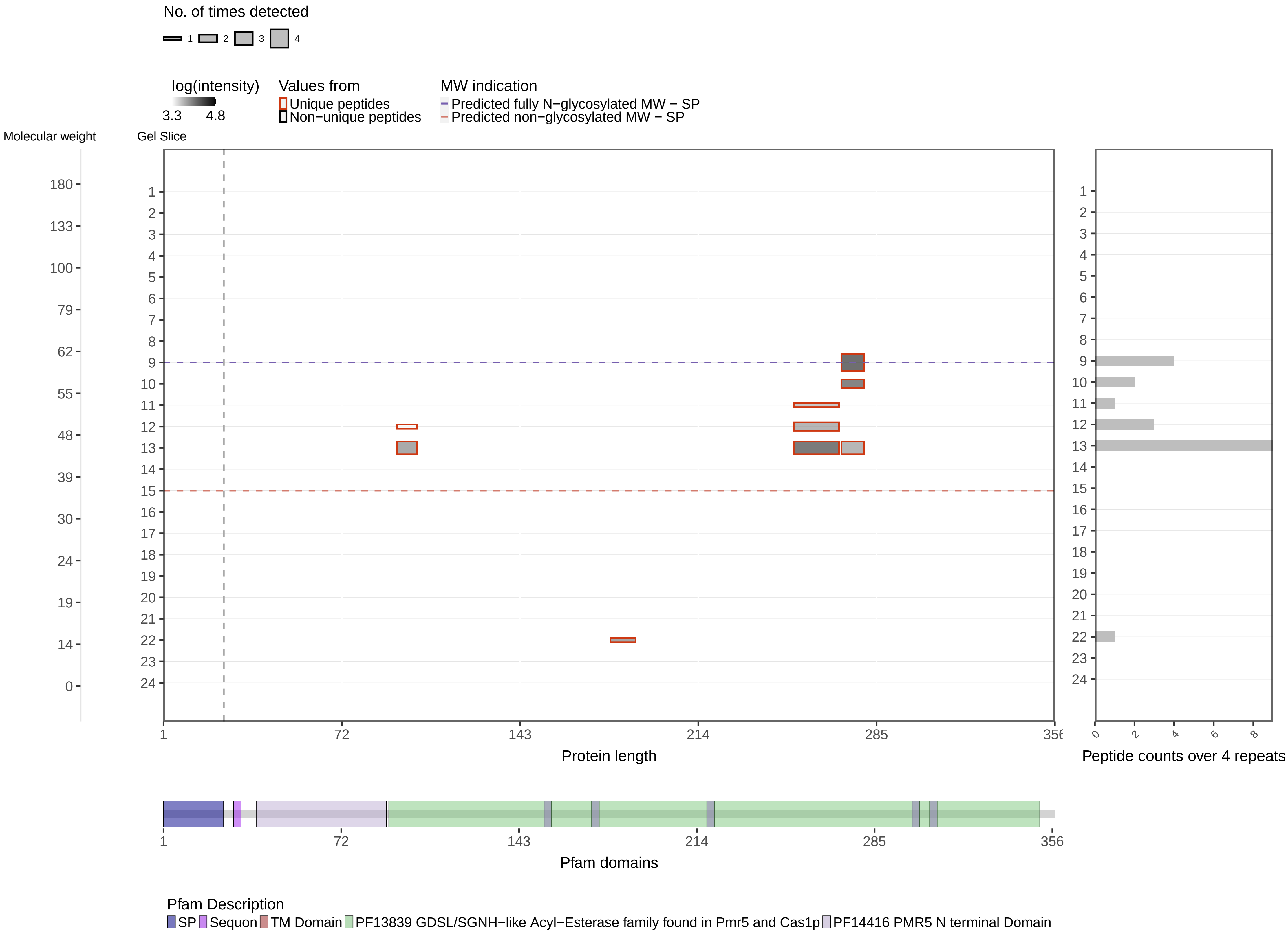


Protein sequence:

TENNGTRVLKLDSLSSAAQWKDMDVLIFDSWHWWLHTGRKQPWDLVQDGNSTYKDGPRLTLYEKALNTWAKWVDSEVDITKTKVFFQGVSPDHDNPTSSGSKTCEGVTQPLKSTEEVHQEELVLEKVLRGMNKSVYLLNITKLSQYRADGHPSVYGHGGHRDLDCTHWCLAGVVDTWNLFLNALLGL*

### NbL05g11690.1 probable_alphaalpha−trehalose−phosphate_synthase_[UDP−forming]_7 [MW = 97.34 kDa]

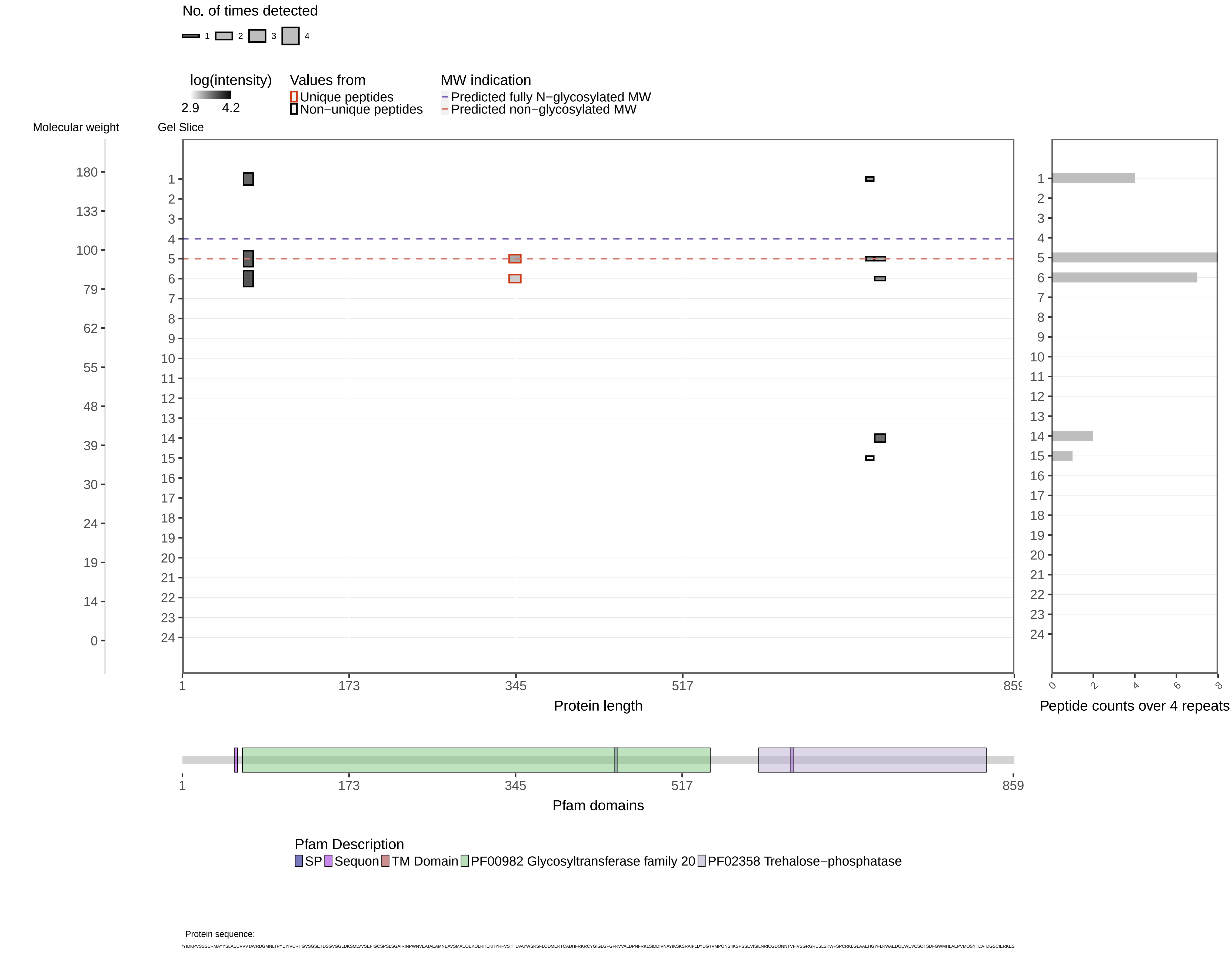

NbL05g11580.1 aspartate_aminotransferase_mitochondrial [MW = 46.91 kDa]

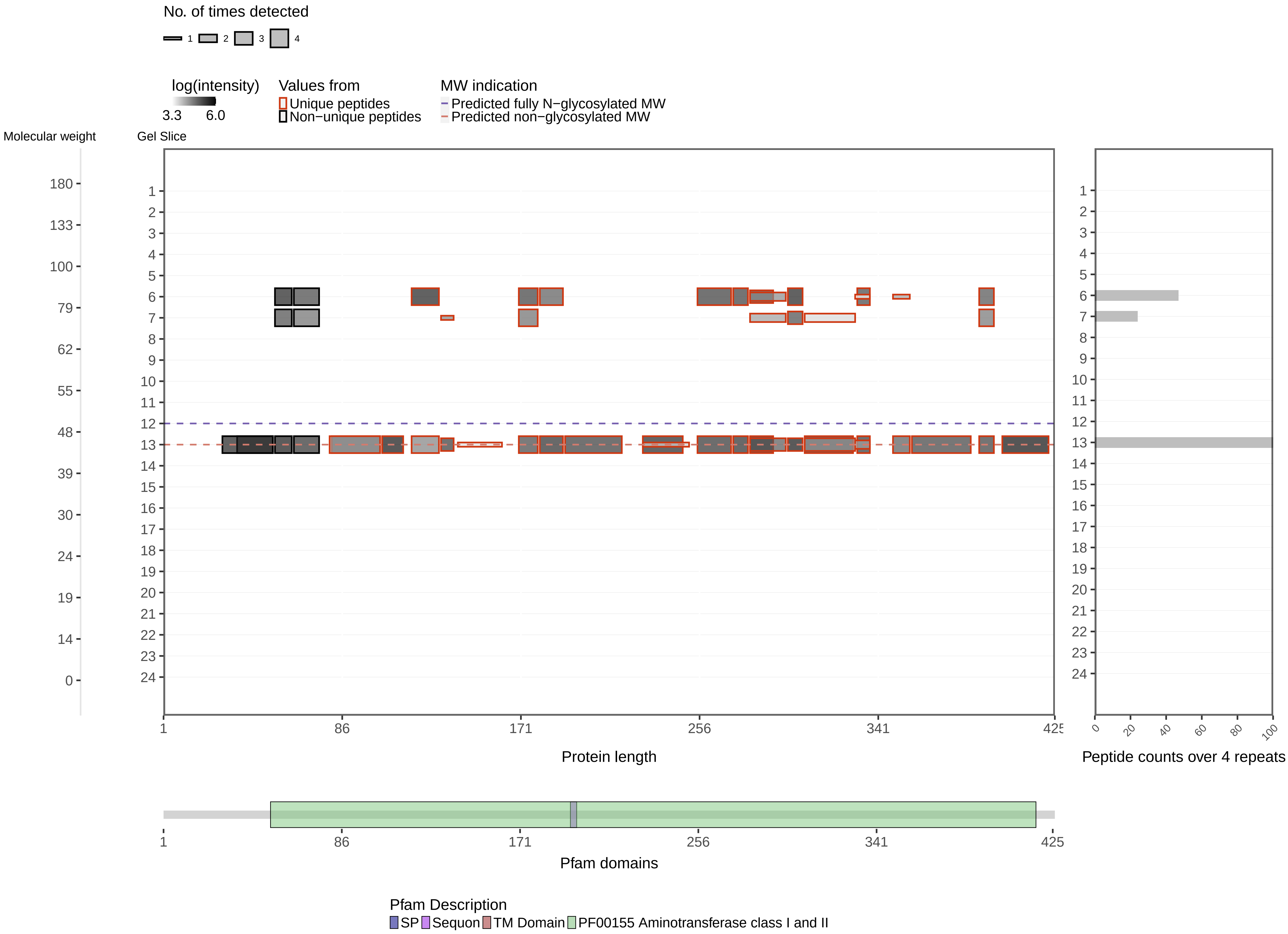


Protein sequence:

LHACAHNPTGVDPTEEQWREISHQFKVKGHFAFFDMAYQGFASGNPEKDAKSIRIFLEDGHPIGCAQSYAKNMGLYGQRVGCLSVVCEDEKQAVAVKSQLQQLARPMYSNPPVHGALVVATILGDPNLKKIWLGEVKGMADRIIGMRTALRENLEKLGSPLSWEHITNQIGMFCYSGMTPEQVDRLTKEYHIYMTRNGRISMAGVTTGNVGYLANAIHEVTKSA*

### NbL05g11560.1 peroxidase_P7−like [MW = 35.66 kDa]

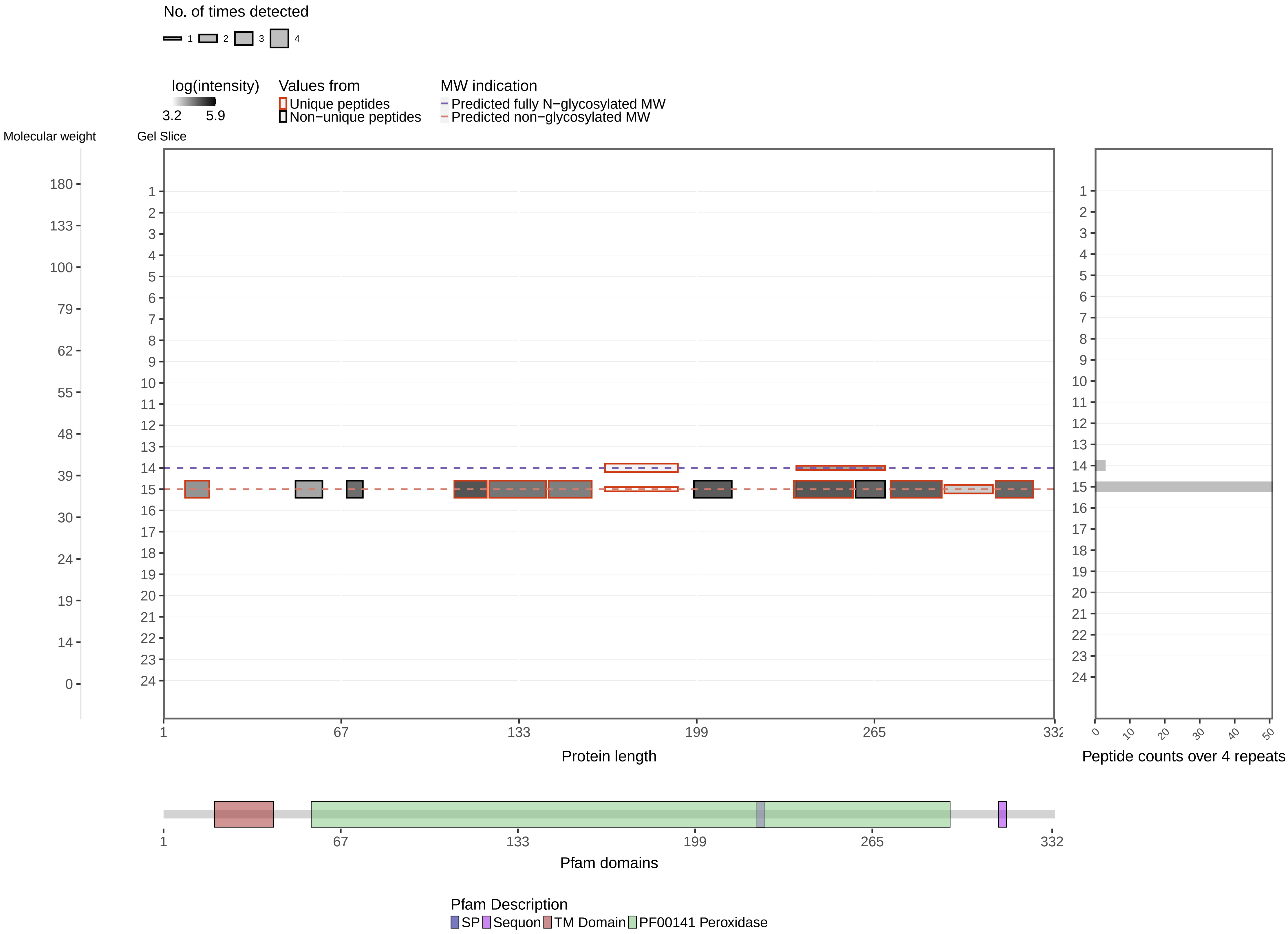


Protein sequence:

AVPLGRRDARTASOSAANAOIPAPTSSLSTLLSMFSAKGLNARDMTALSGGHTIGOARCTTFRTRIYNDTNIDAOFATTRRATCPSSGGDSNLAPLDIOTPNRFDNDYYONLVARRGLLHSDOELFNGGSODALVRSYSTNVAAFTSDFAAAMVRMGNISPLTGTNGEIRRNCRAIN*

NbL05g11230.1 uncharacterized_LOC107832338 [MW = 24.48 kDa]

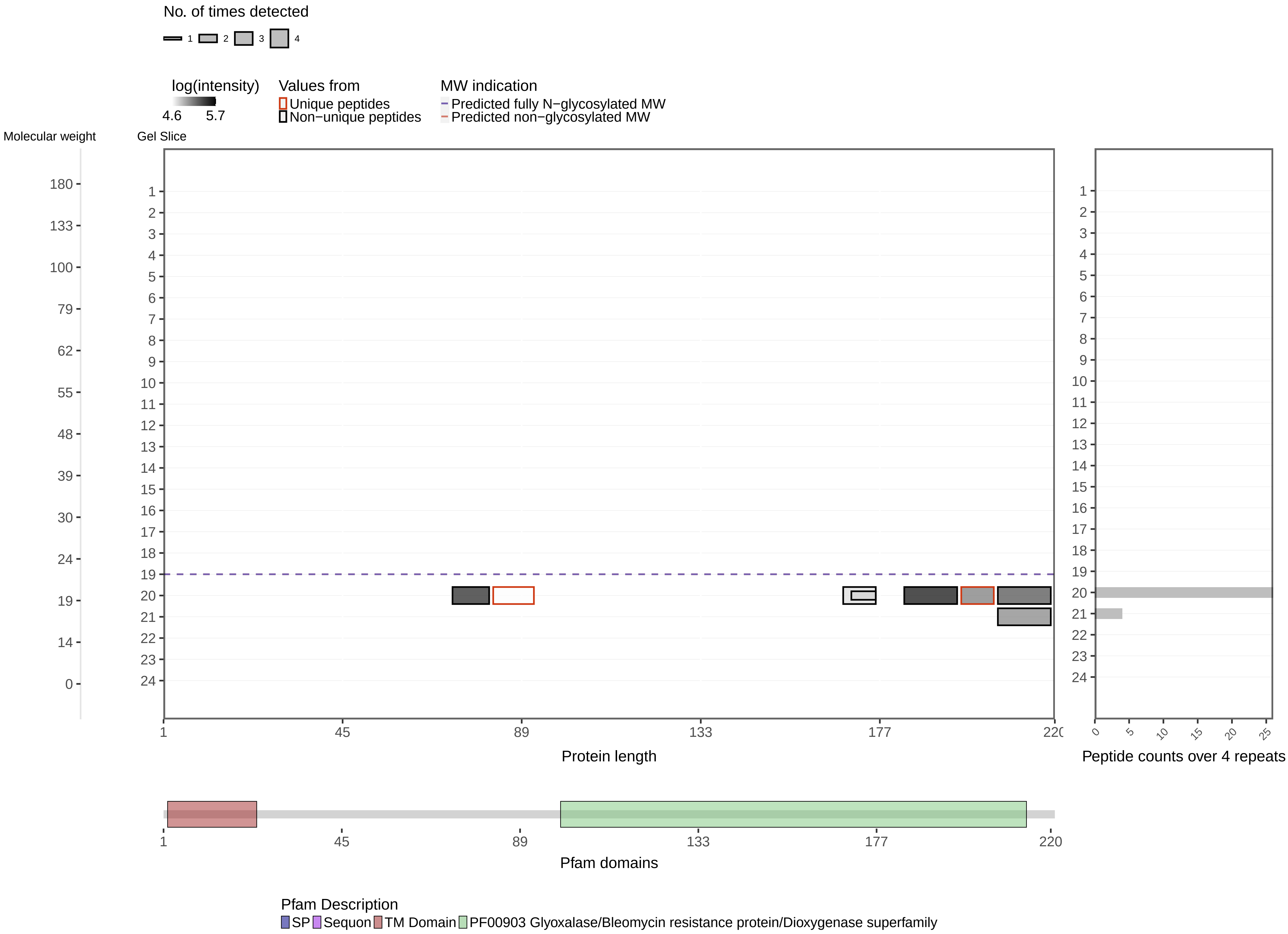


Protein sequence:

/SMHHVGILCENLERSLDFYQNILGLEINEARPHDKLPYRGAWLWVGSEMIHLMELPNPDPLTGRPEHGGRDRHTCIAIRDVSKLKVILDEAGIPYTLSRSGRPDIFTRDPDANALEFTQVDA*

NbL05g11050.1 translation_initiation_factor_IF−2_chloroplastic−like [MW = 108.29 kDa]

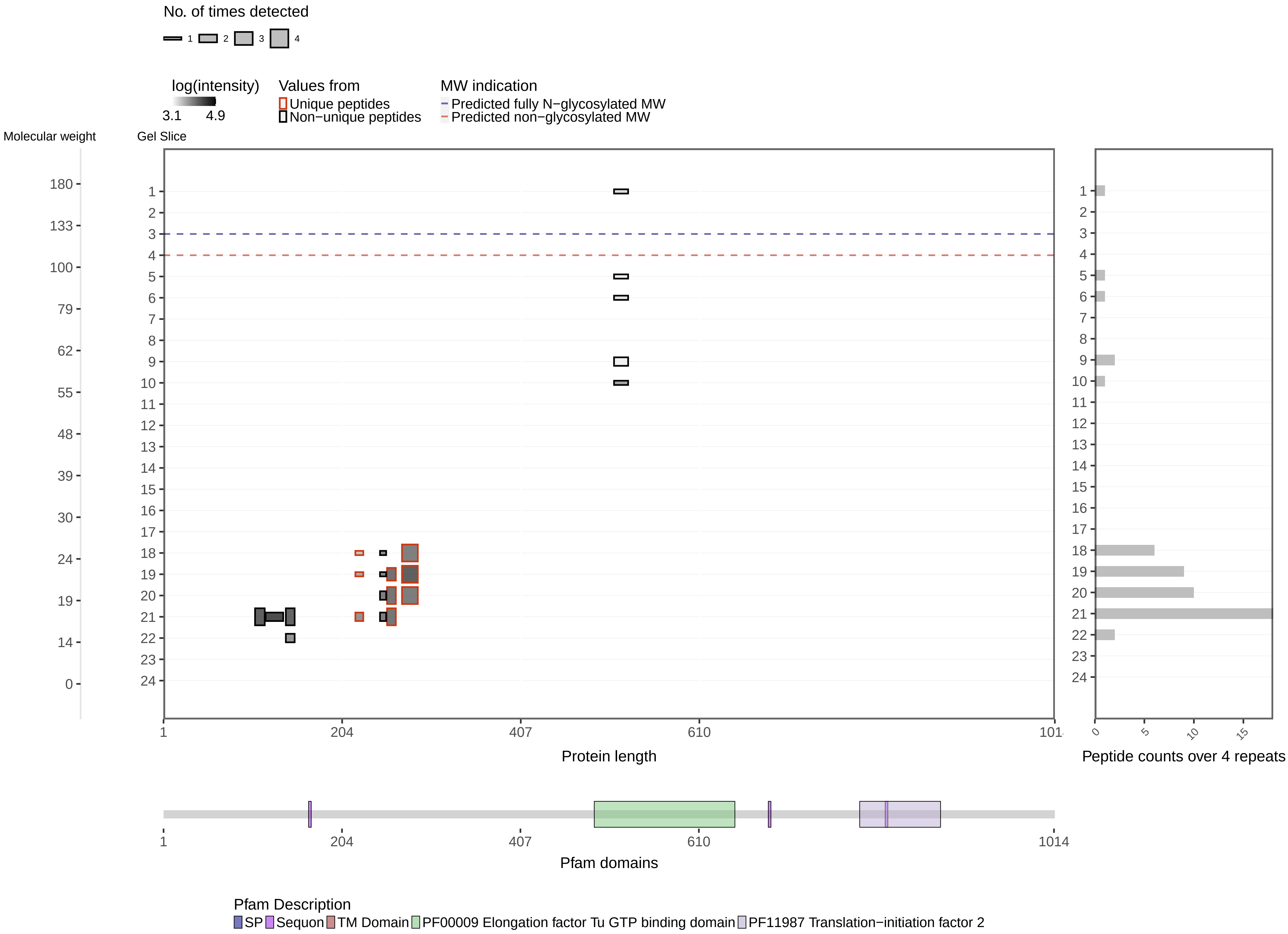


Protein sequence:

PVITIMGHVDHGKTTLLDHIRKTKVAASEAGGITQGIGAYKVQVPIDTKPQICVFLDTPGHEAFGAMRARGARVTDIAIIVVAADDGIRPQTNEAIAHAKAAGVPIVIAINKIDKDGANPERVMQELSSIGLMPEDWGGDVPMVQISALKGENIDDLLETVMLVAELQELKANPQRNAKGTVIEAGLDKSKGPVATFIVQNGTLKGGDVVVCGEAYGKVRALFDDKGKRVDEAGPSIPVQVIGLNNVPVAGDEFEVVGSLDVAREKAKTREESLRTERLSAKAGDGKVTLSSFASA

### NbL05g10980.1 probable_glutathione_S−transferase [MW = 25.71 kDa]

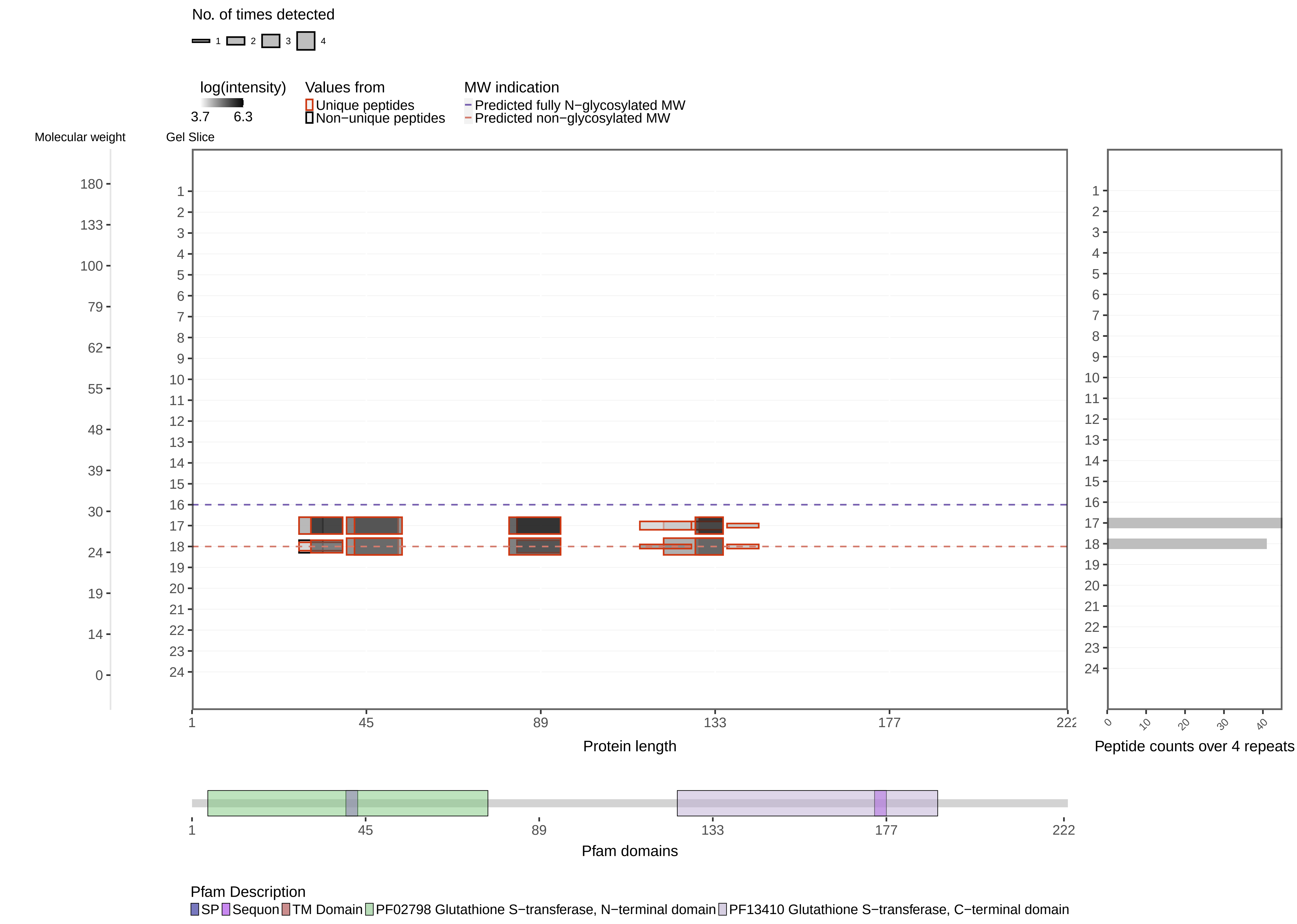


Protein sequence:

WADYIDKKLYDVGRKLWATKGEEQEAAKKDFIECLKVLEGALGDKPYFGGESFGFVDIALIGYYSWFYAYETFGNFSTEAECPKFVAWAKRCMQRESVAKSLPDQPKVLEFVKVLRQKFGLE*

NbL05g10970.1 probable_glutathione_S−transferase [MW = 16.3 kDa]

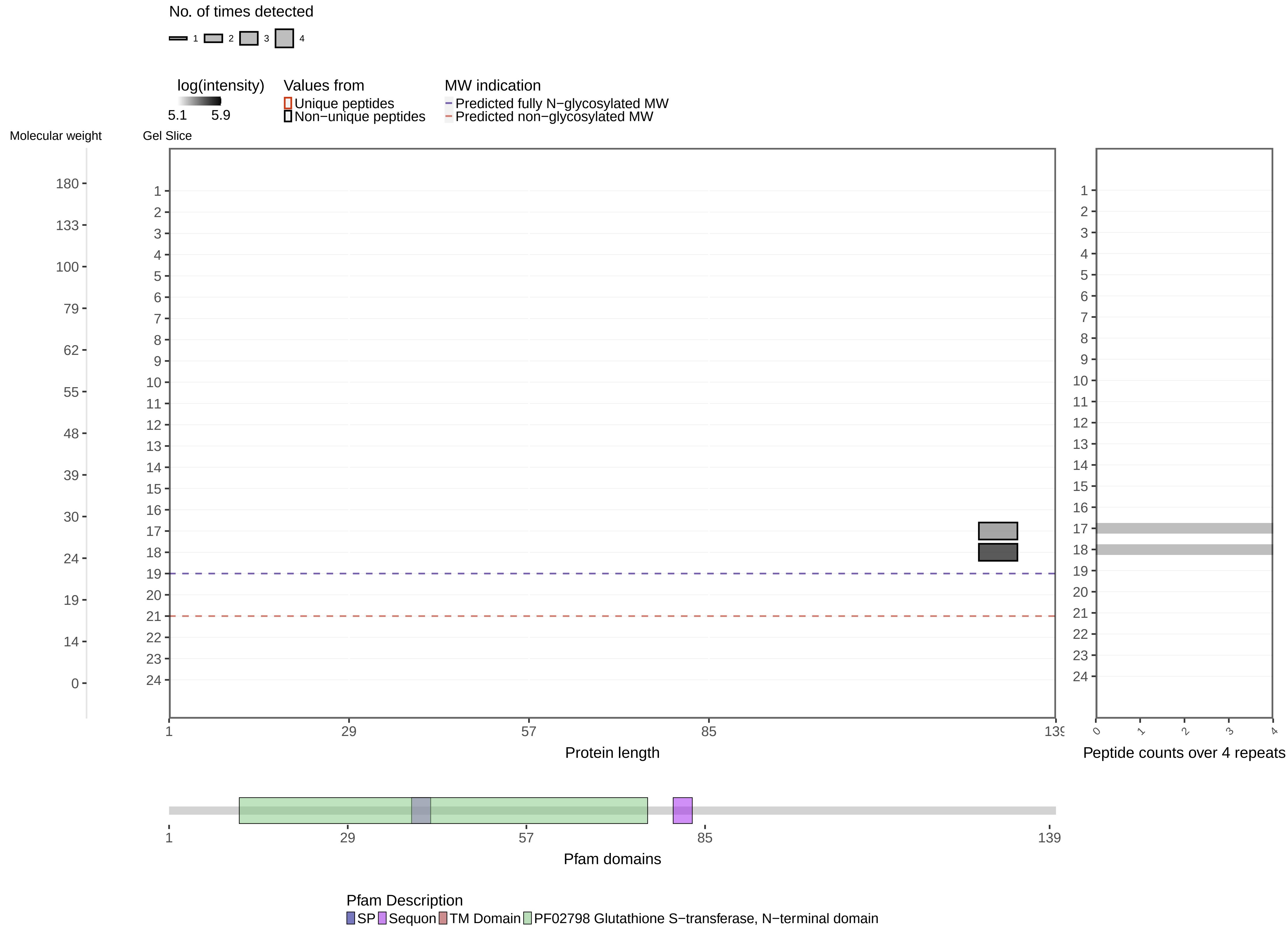


Protein sequence:
LIHNGKPICESLIIVEYIDEVWKNKTPFMPSDPYMRAQARFWADYIDKKIYESGKKMWTSKVEDQEAANKEFIECLKHWKGN*

NbL05g10960.1 probable_glutathione_S−transferase [MW = 29.88 kDa]

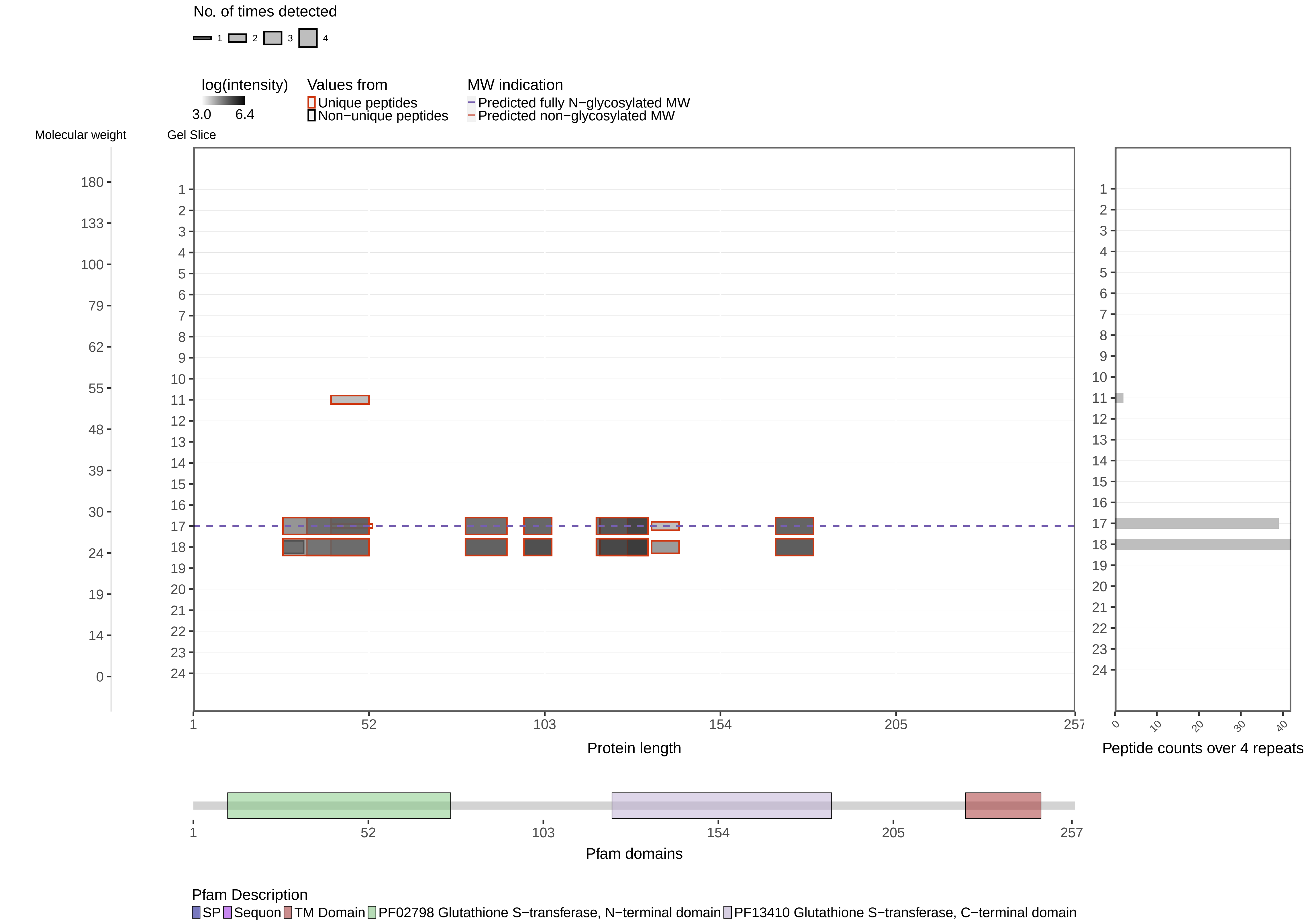


Protein sequence:

SKKEDQEAANKEFIECFKLLEEELGDKPYFGGESFGFVDMALIPYYTWFPAYEKFGNFNIEAECPKIVEWAKKCMQKESVSKSLADPDKVYNFIVMVCHYKHSPPIRIGQVPWMIGSRVMAMPFFLGILVSWSSKKQRIVA*

### NbL05g10930.1 peroxisomal_(S)−2−hydroxy−acid_oxidase−like [MW = 40.61 kDa]

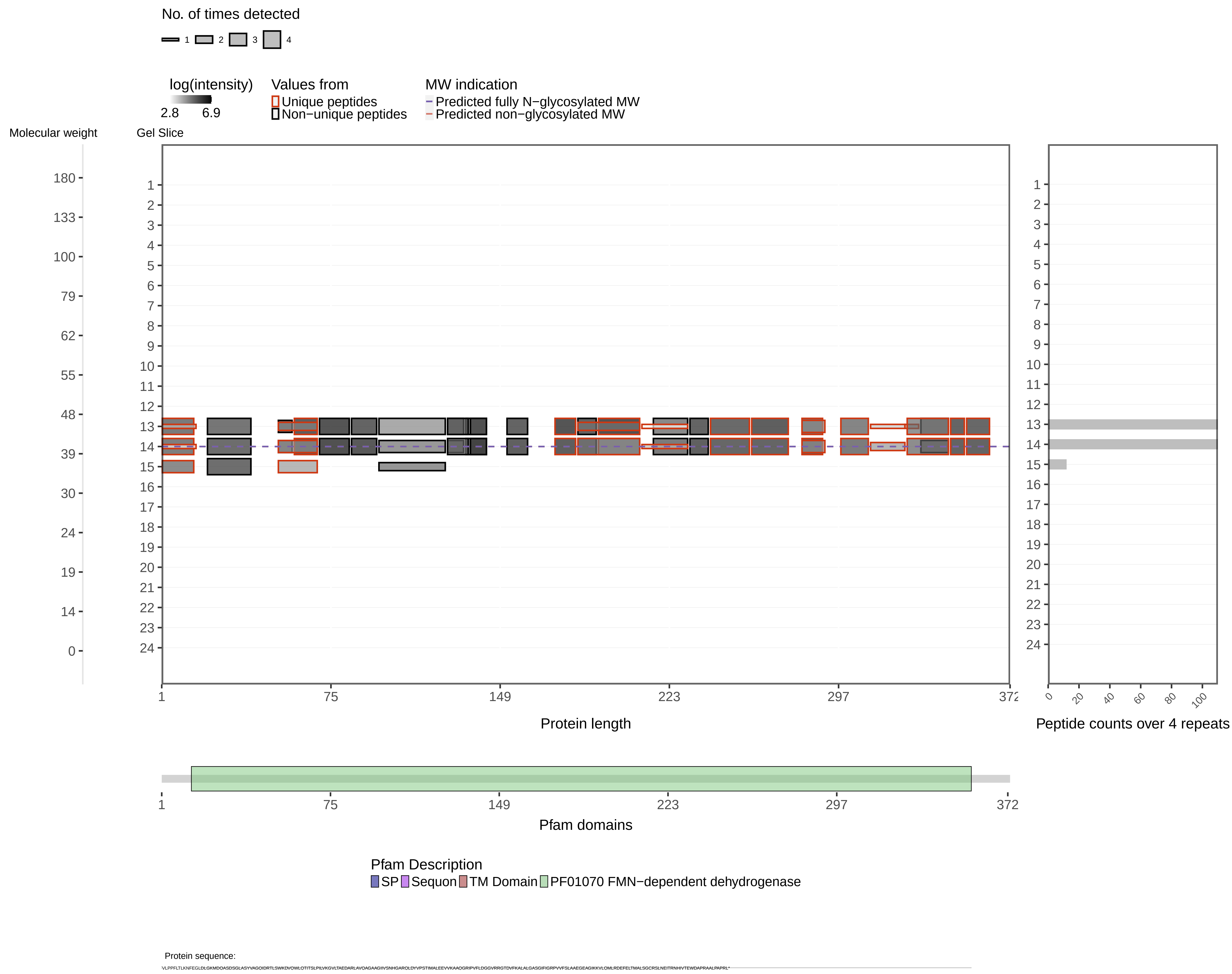

NbL05g10850.1 uncharacterized_LOC107809971 [MW = 35.03 kDa]

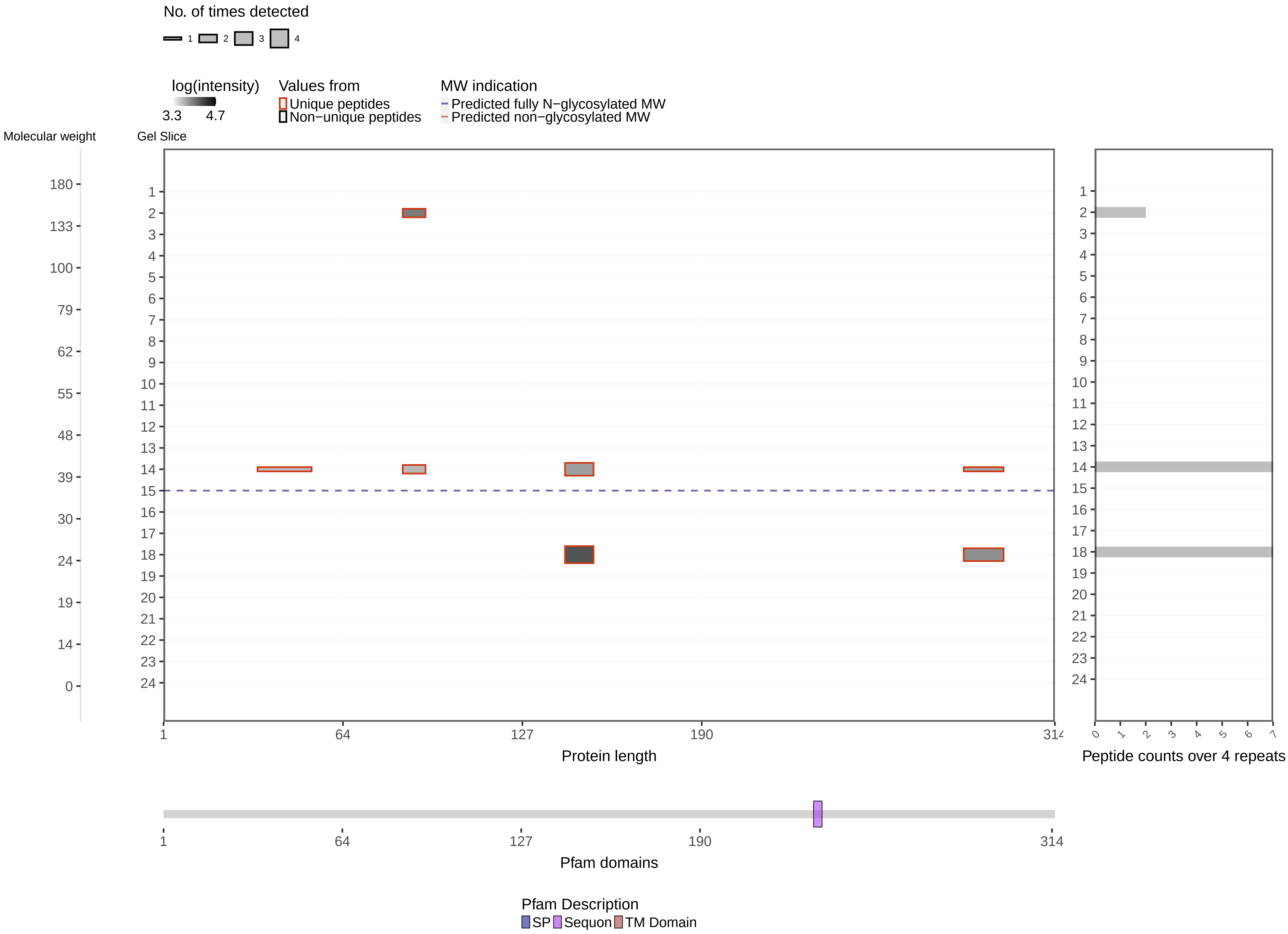


Protein sequence:

ƐFQQGKTLESSQFDLVILHVGAGQRTNVLKDLDHVNRLVGDLMQMAHPGTEVGSRLHMSVVFSYGVVREDDDSMYSIADTRPENNSKLSRIFPRQSYTIKEGKPRPNVRQFCPMLVAQWQNAVTRKDMVEAYSYKDFKEHGANLVIPADRLLHEVAFKMWKAPKYGA*

### NbL05g10830.1 vacuolar_protein_sorting−associated_protein_32_homolog_2−like [MW = 24.39 kDa]

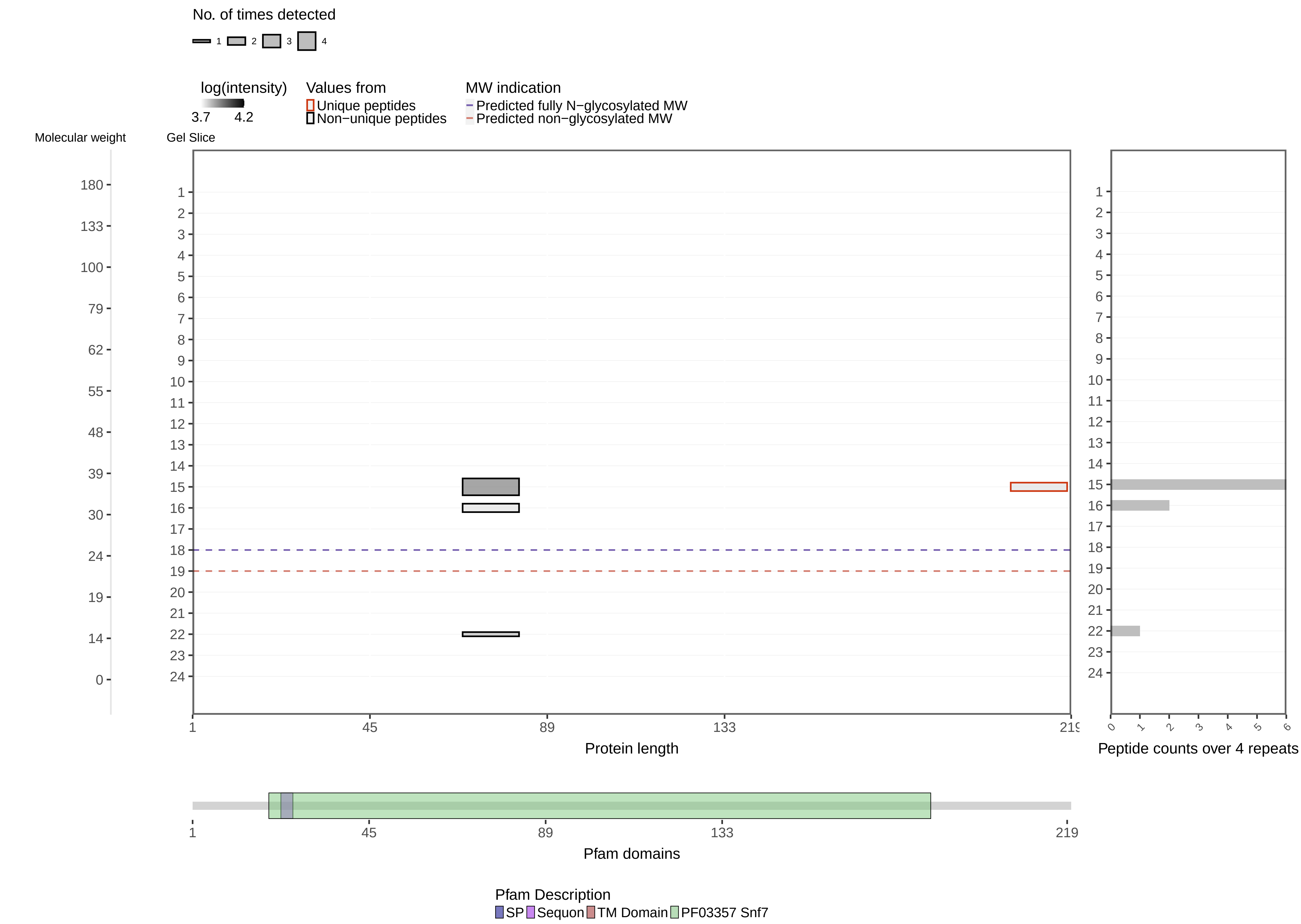


Protein sequence:

TETVDALRTGAAAMKAMQKATNIDDVDKTMDEINEQTENMKMIQEALSTPIGAAADFDEDELEAELEELEGAELEEQLLQPATTAPSAPIHLPTGRQQVRPTAQQNRAEEDELAALQAEMAL*

NbL05g10660.1 50S_ribosomal_protein_L24_chloroplastic [MW = 20.63 kDa]

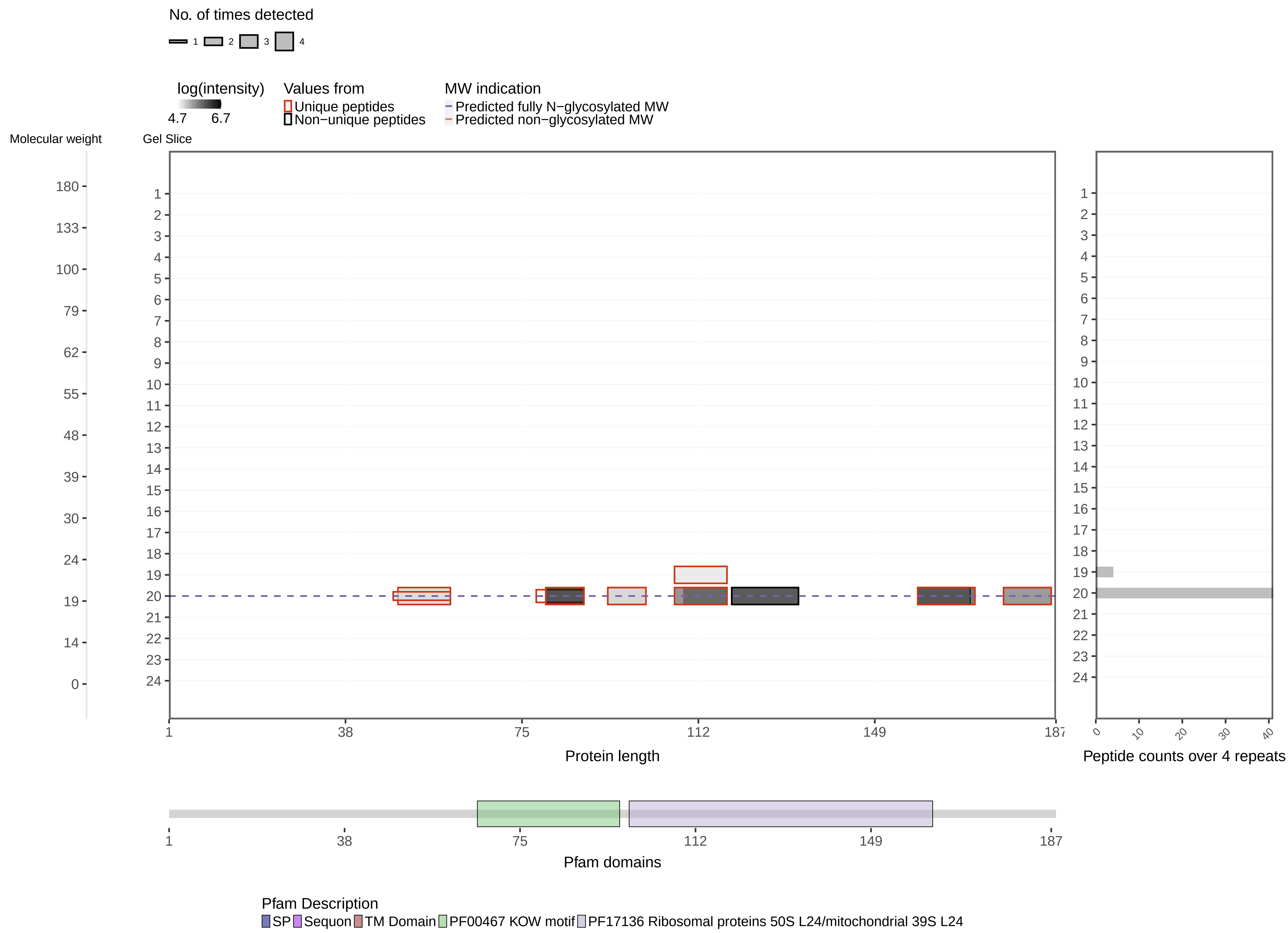


Protein sequence:

KVGEITEIIKHNSKVVVKDVNLKTKHVKSRSEDEPGQIVKIEAPIHSSNVMLYSKEQKVASRVGHKTLDNGKRVRYLIKTGEIIDSAENWKKAVKEKEKTTEAVAAAS*

### NbL05g10610.1 60S_ribosomal_protein_L8−like [MW = 30.94 kDa]

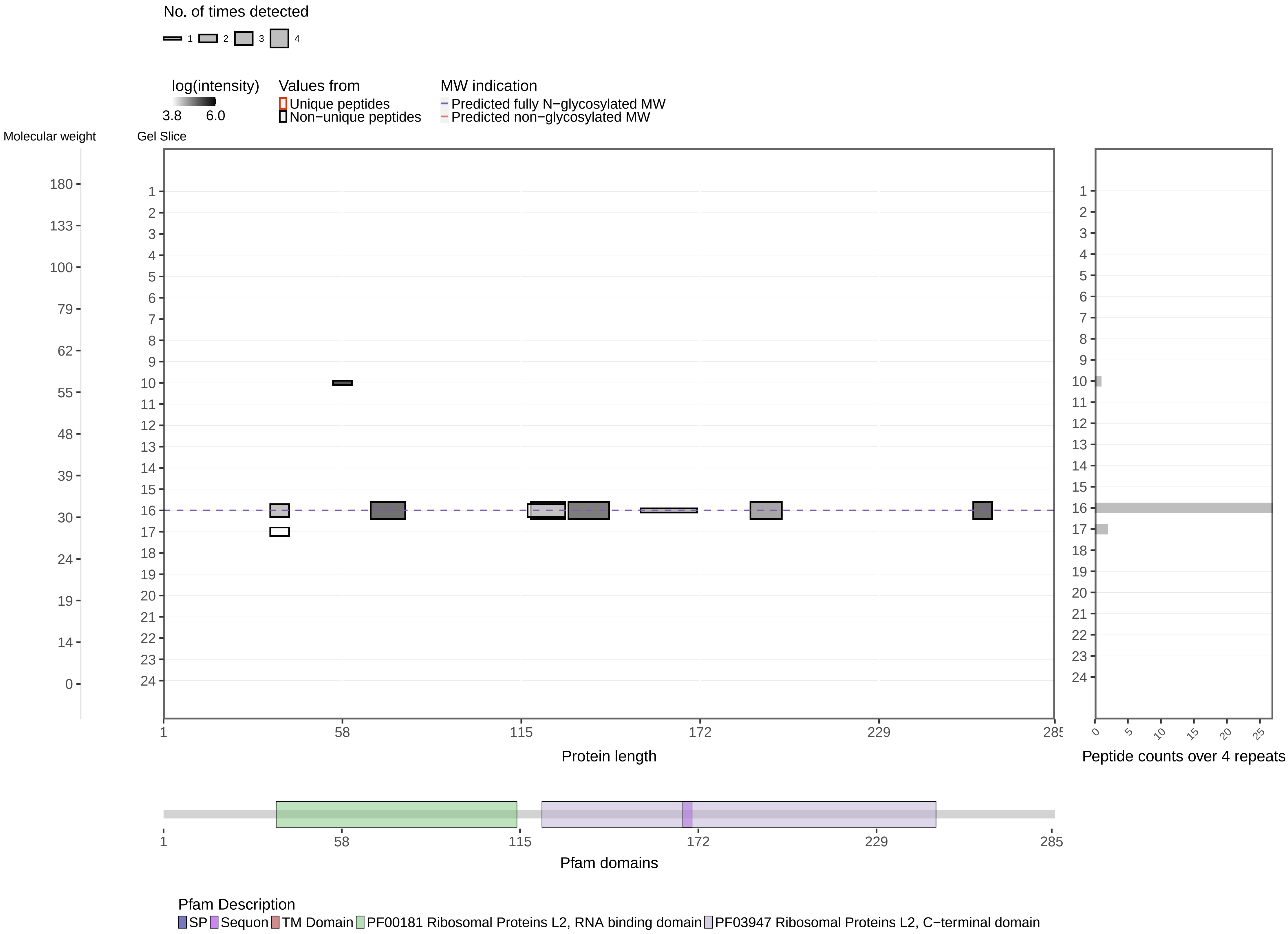


Protein sequence:

SIPEGAVVCNVEHKVGDRGVFARCSGDYAIVISHNPDNGTTRIKLPSGSKKIVPSGCRAMIGQVAGGGRTEKPMLKAGNAYHKYRVKRNCWPKVRGVAMNPVEHPHGGGNHQHIGHASTVRRDAPPGQKVGLIAARRTGRLRGQAAATAAKADKA*

NbL05g10490.1 uncharacterized_LOC107827684 [MW = 26.45 kDa]

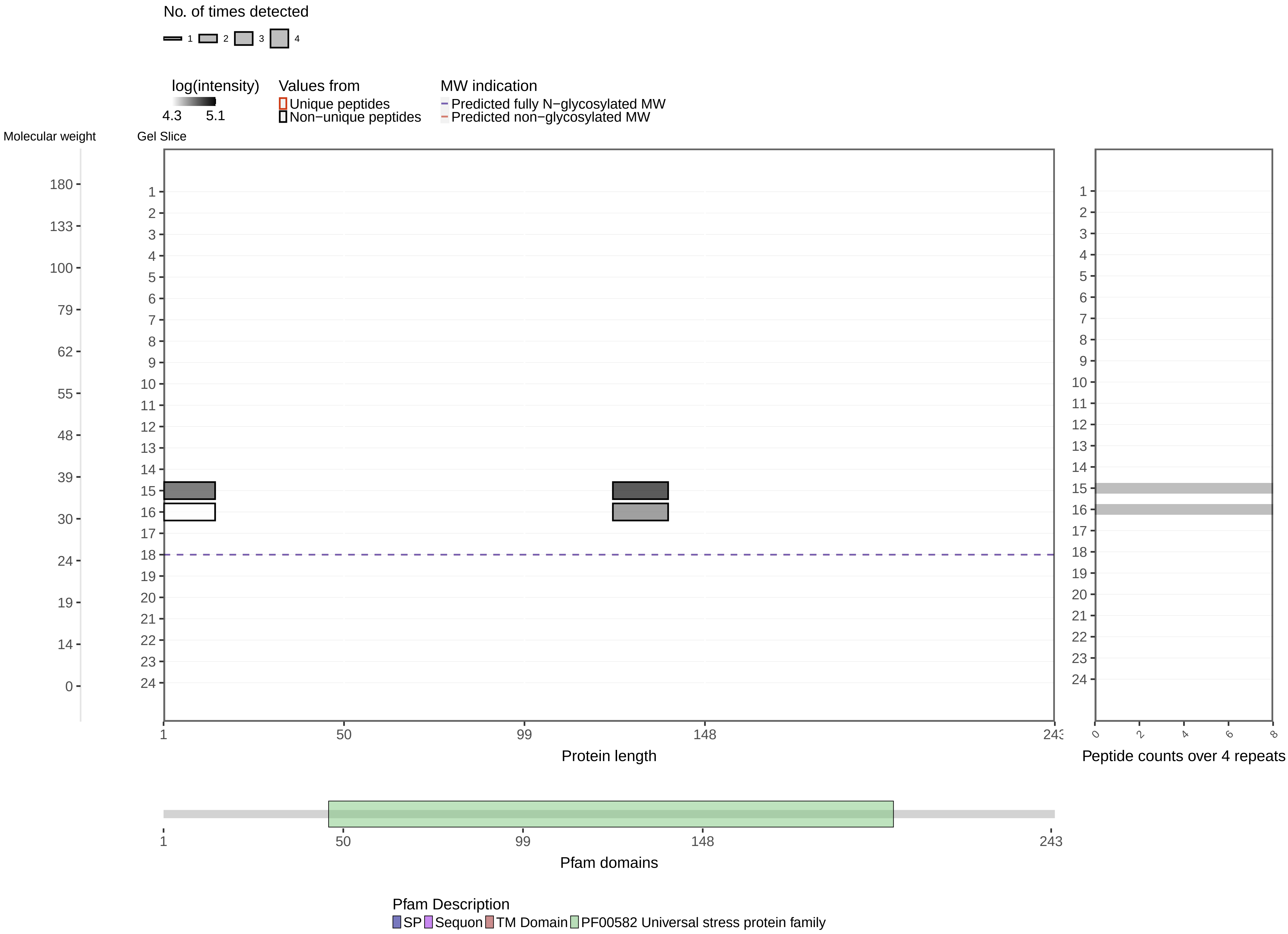


Protein sequence:

KLEDDFDTFTTTKASDISQPLVDAQIPFKIHIVKDHDMRERLCLEVERLGLSAVIMGSRGFGATKRGRDGRLGSVSDYCVRHCVCPVVVVRYPDDKDSGANAVAEPVVSVASAAEEDEEDAEYHDASDHRKDS*

### NbL05g10460.1 ubiquitin_carboxyl−terminal_hydrolase_14−like [MW = 88.88 kDa]

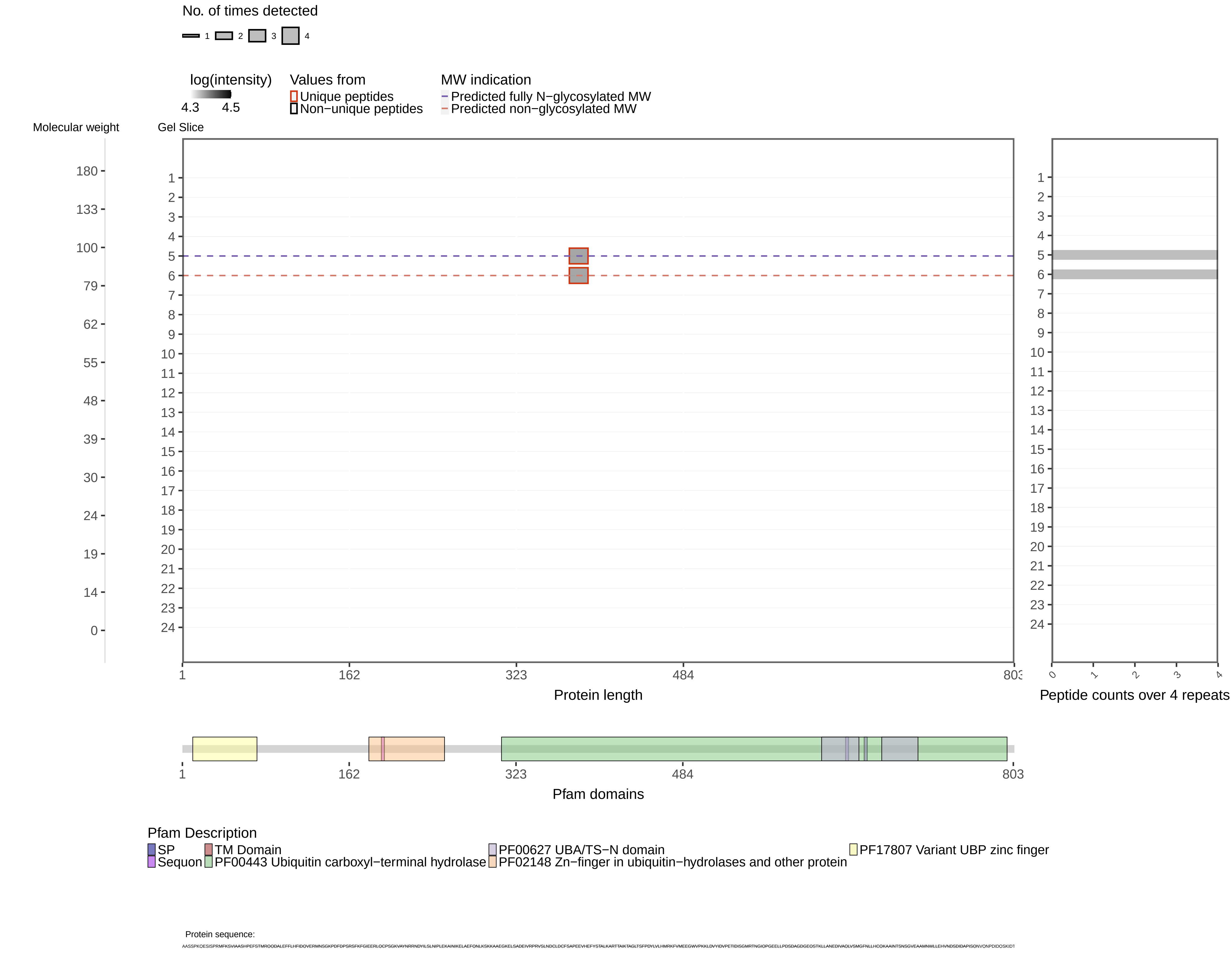

NbL05g10200.1 beta−glucosidase_18−like [MW − SP = 58.12 kDa]

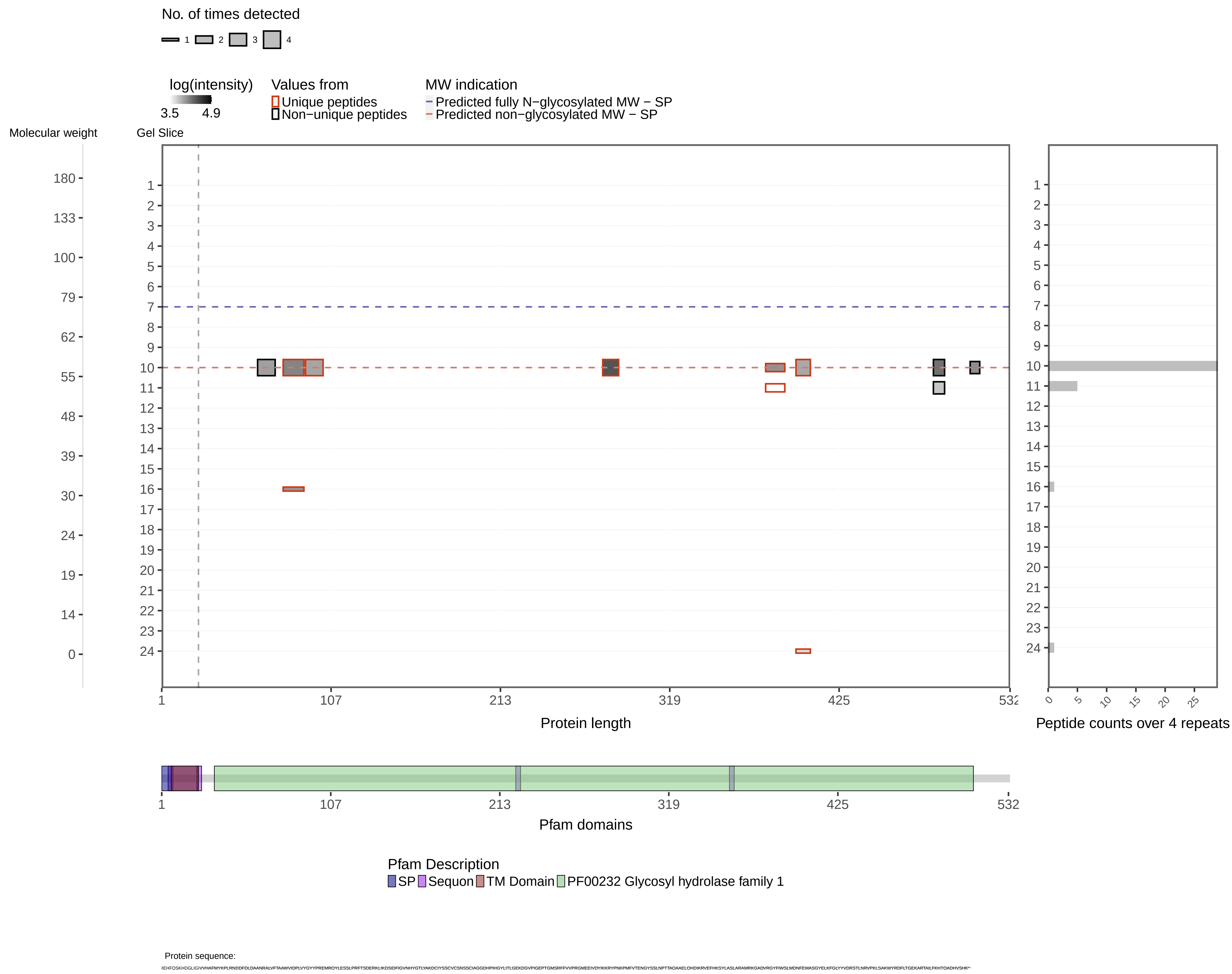

### NbL05g09870.1 probable_bifunctional_methylthioribulose−1−phosphate_dehydratase/enolase−phosphatase_E1_1 [MW = 56.55 kDa]

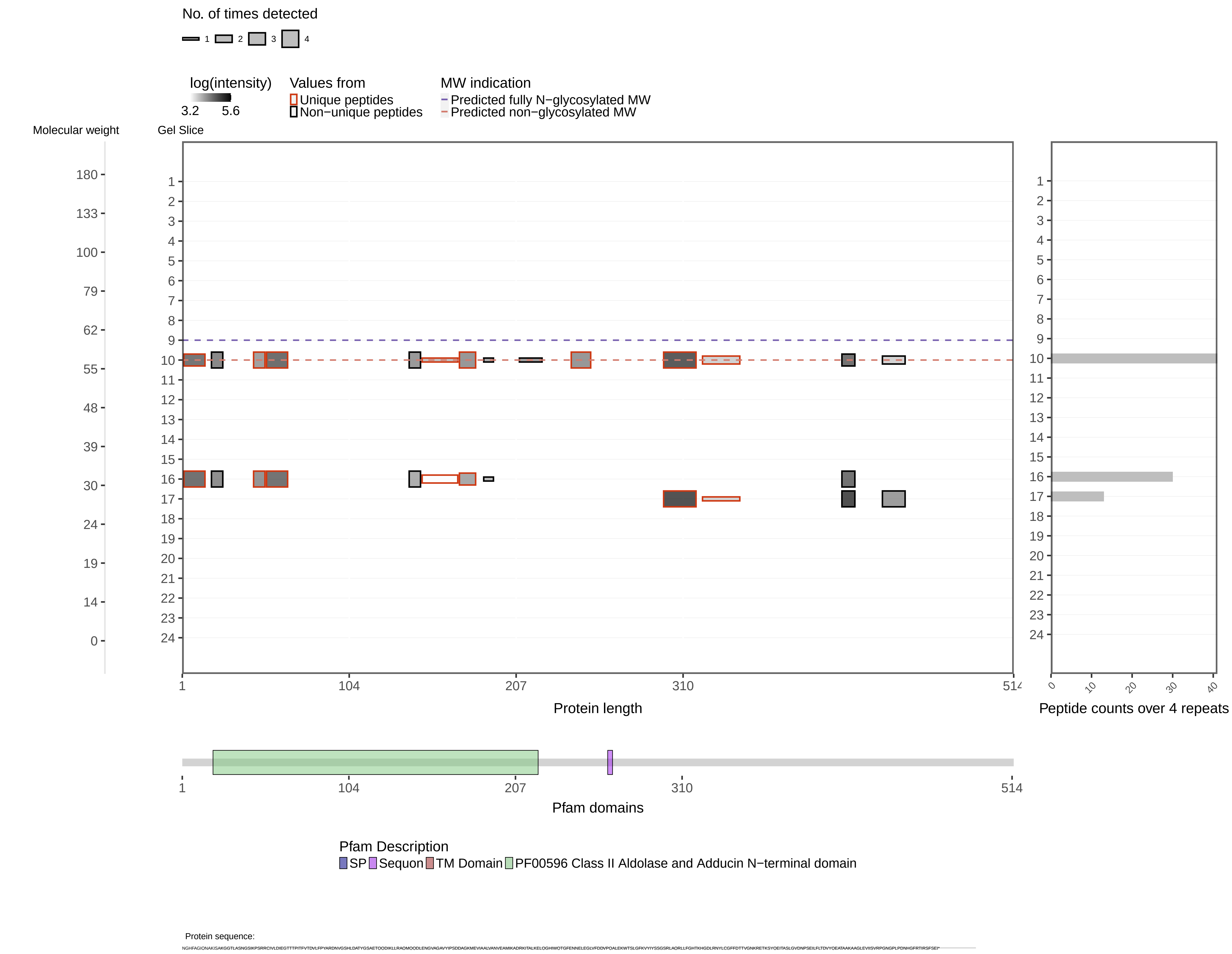

### NbL05g09830.1 3−hydroxybutyryl−CoA_dehydrogenase [MW = 32.1 kDa]

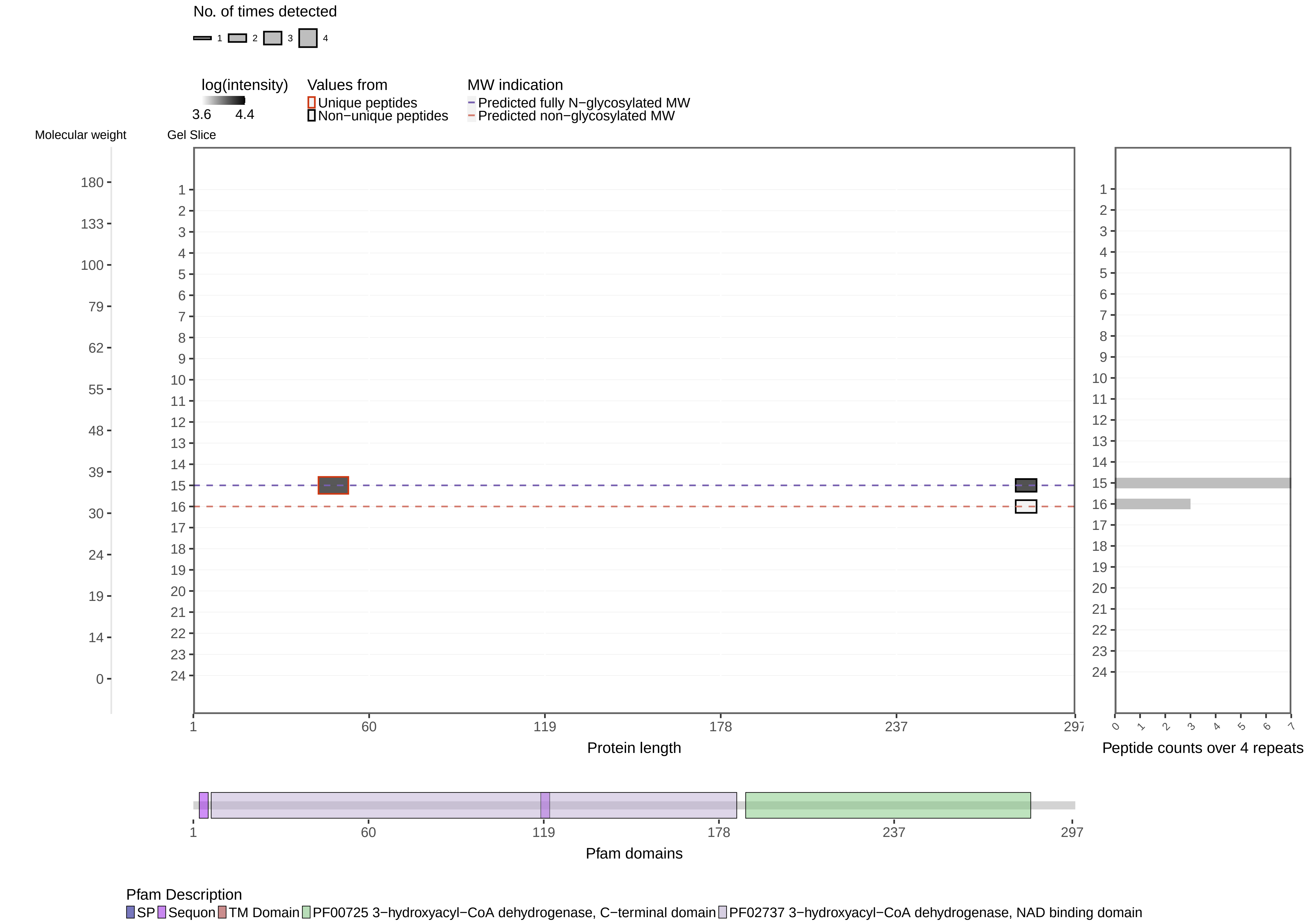


Protein sequence:

VIGMHFMNPPPIMKLVEIIRGADTSEDTYCATKCLAERFGKTVICSQDYPGFIVNRILMPMINEAFHTLYTGVATKEDIDTGMKLGTDHPMGPLELADFIGLDVCLSIMKVLHVGLGDDKYAPCPLLVQYVDGDRLGKKRGIGVYDYRSAAADTGKSPRL*

### NbL05g09540.1 protein_DJ−1_homolog_B−like [MW = 47.03 kDa]

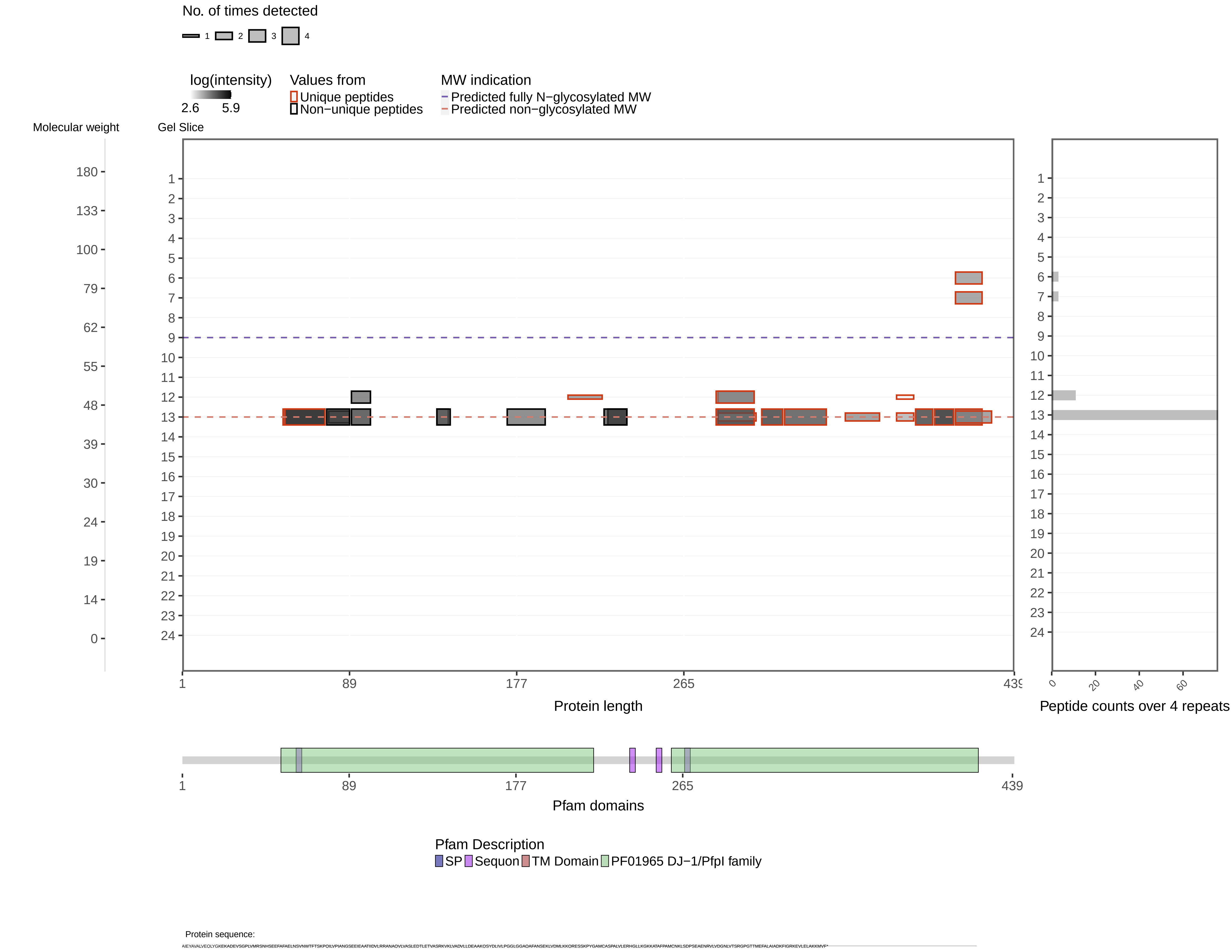

NbL05g09490.1 ubiquitin−conjugating_enzyme_E2_35 [MW = 19.29 kDa]

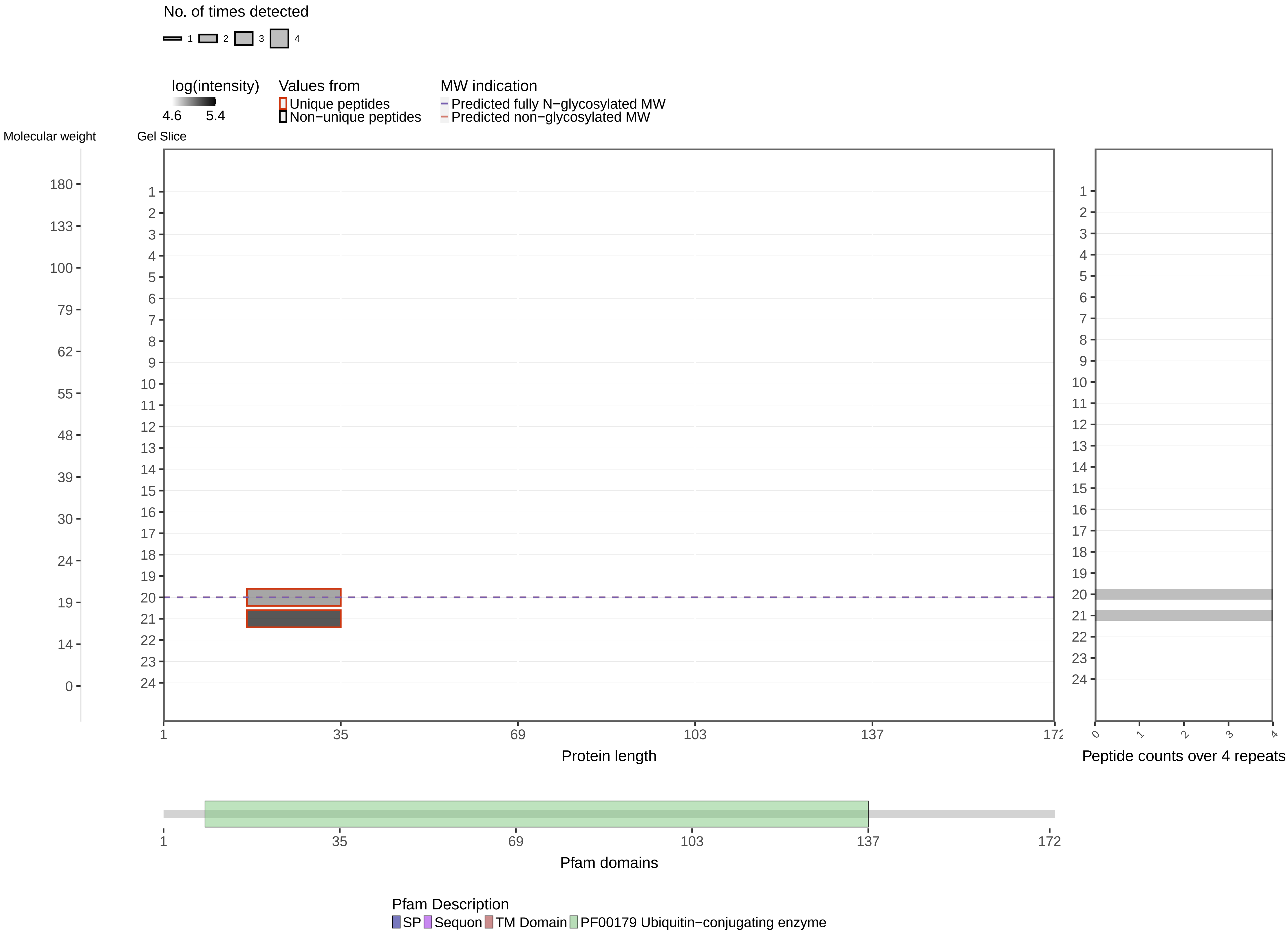


Protein sequence:

VRFLTKIYHPNIDKLGRICLDILKDKWSPALQIRTVLLSIQALLSAPNPDDPLSENIAKHWKSNEAEAVETGSLYSGLSIYCLQTSPDYANRYTFFFTPLI*

### NbL05g09440.1 phosphoenolpyruvate_carboxylase−like [MW = 36.72 kDa]

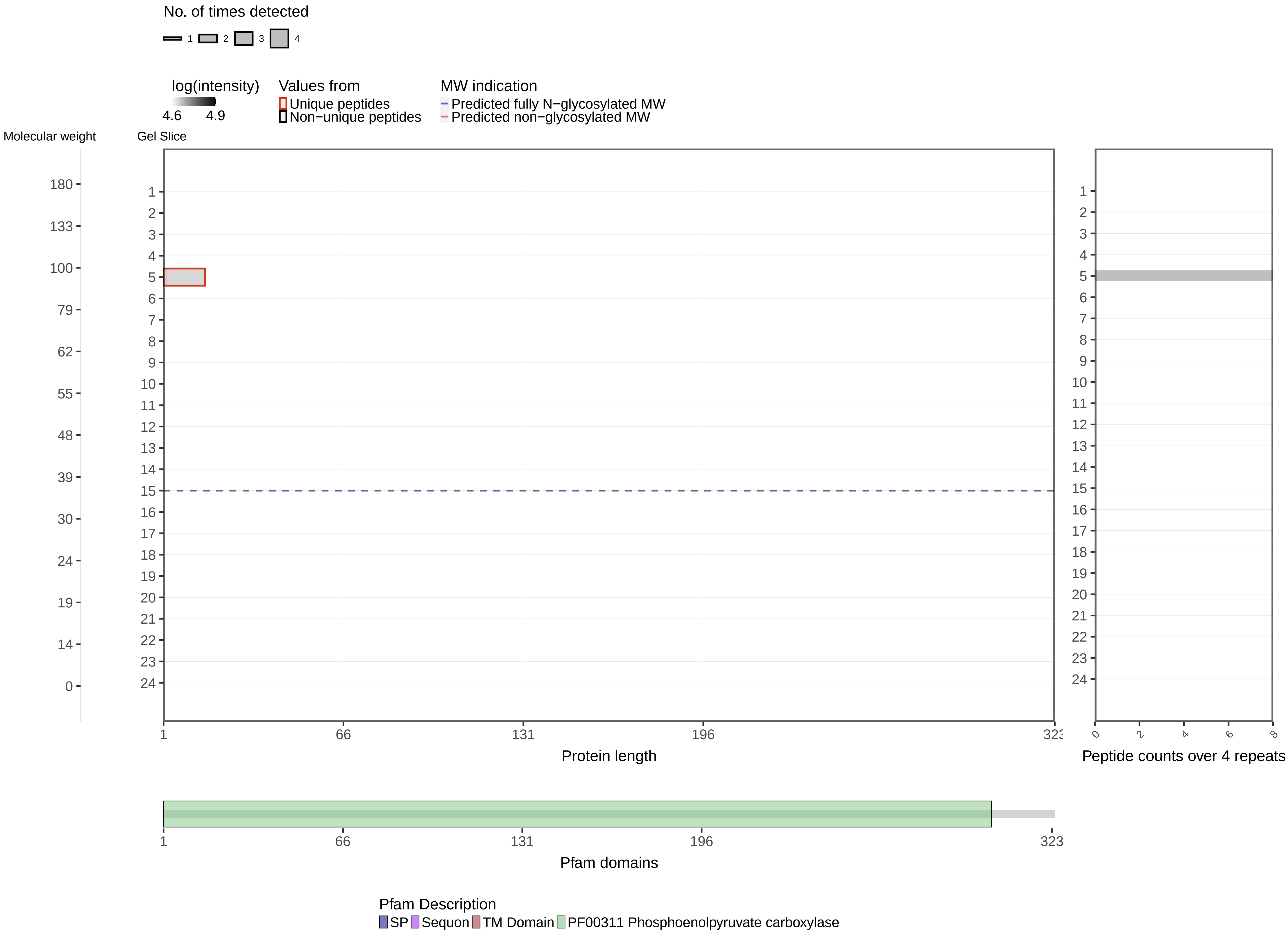


Protein sequence:

EEHLCFRTLQRFTSATLEHGMNPPLSPKPEWRALLDEIAVIATEKYRSIVFKEPPFVEYFRLATPELEYGRMNIGSRPSKRKPSGGIEPLRAIPWIFAWTQTRFHLPVWLGFGAAFKYALNKDVKNLHILQDMYNKWPFFRVTIDLVEMVFAKGRPRHCCFIRQASGFRRFVVLR*

NbL05g09420.1 epidermis−specific_secreted_glycoprotein_EP1−like [MW − SP = 26.05 kDa]

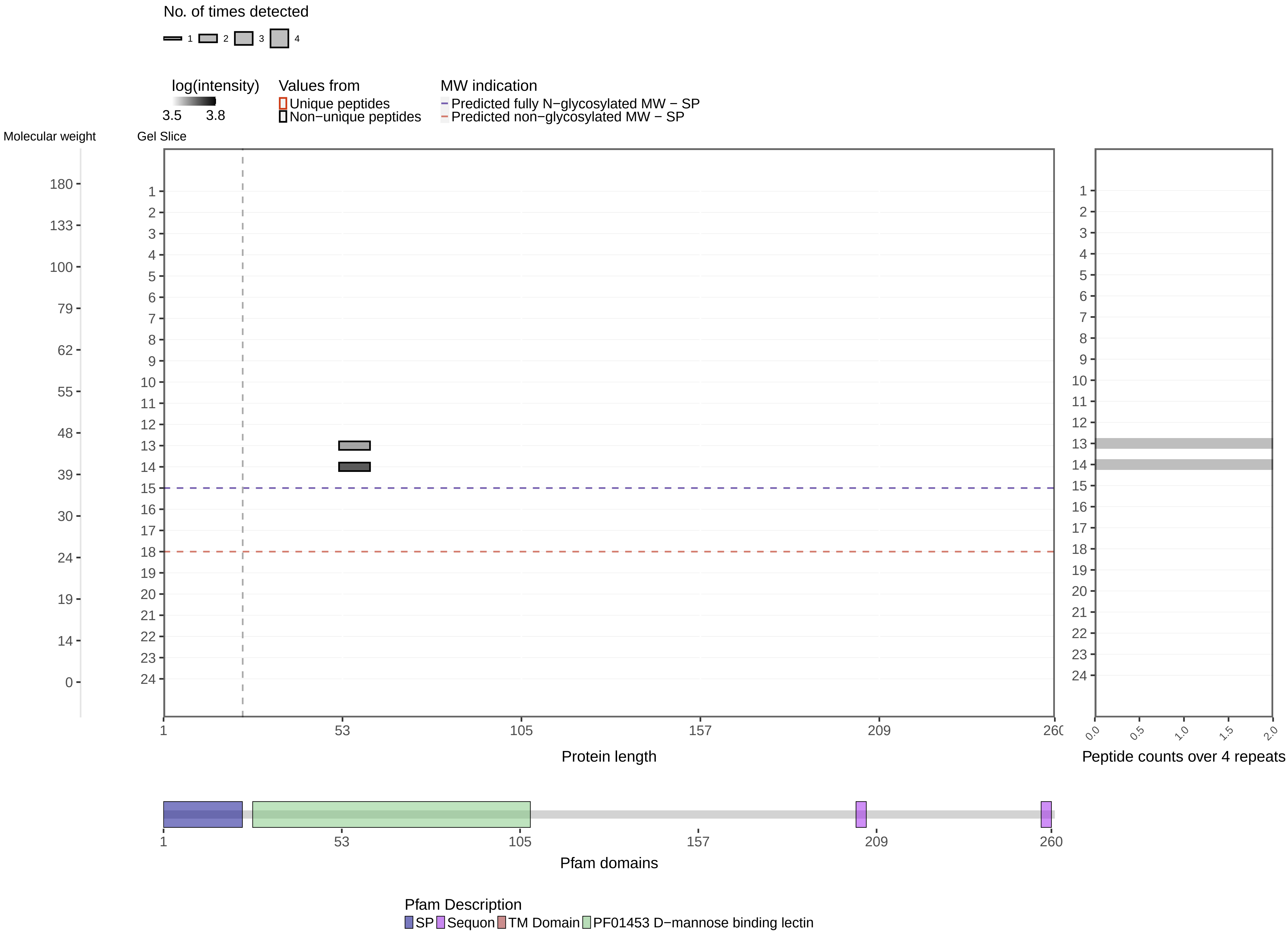


Protein sequence:

NANGPYSLEVKPKIFGCEFESSQEQGGKFLEGRMPGVYLETASLPKGRDKFVALEISGKNYMTWVLDAEIHLDAMGPGDAIKDRNNASTQDCAKALIFLRHRLDEVKDPLILWNGLKERYDNLKLVTLPQARDDENPKANSS*

### NbL05g09390.1 polynucleotide_3'−phosphatase_ZDP [MW = 46.52 kDa]

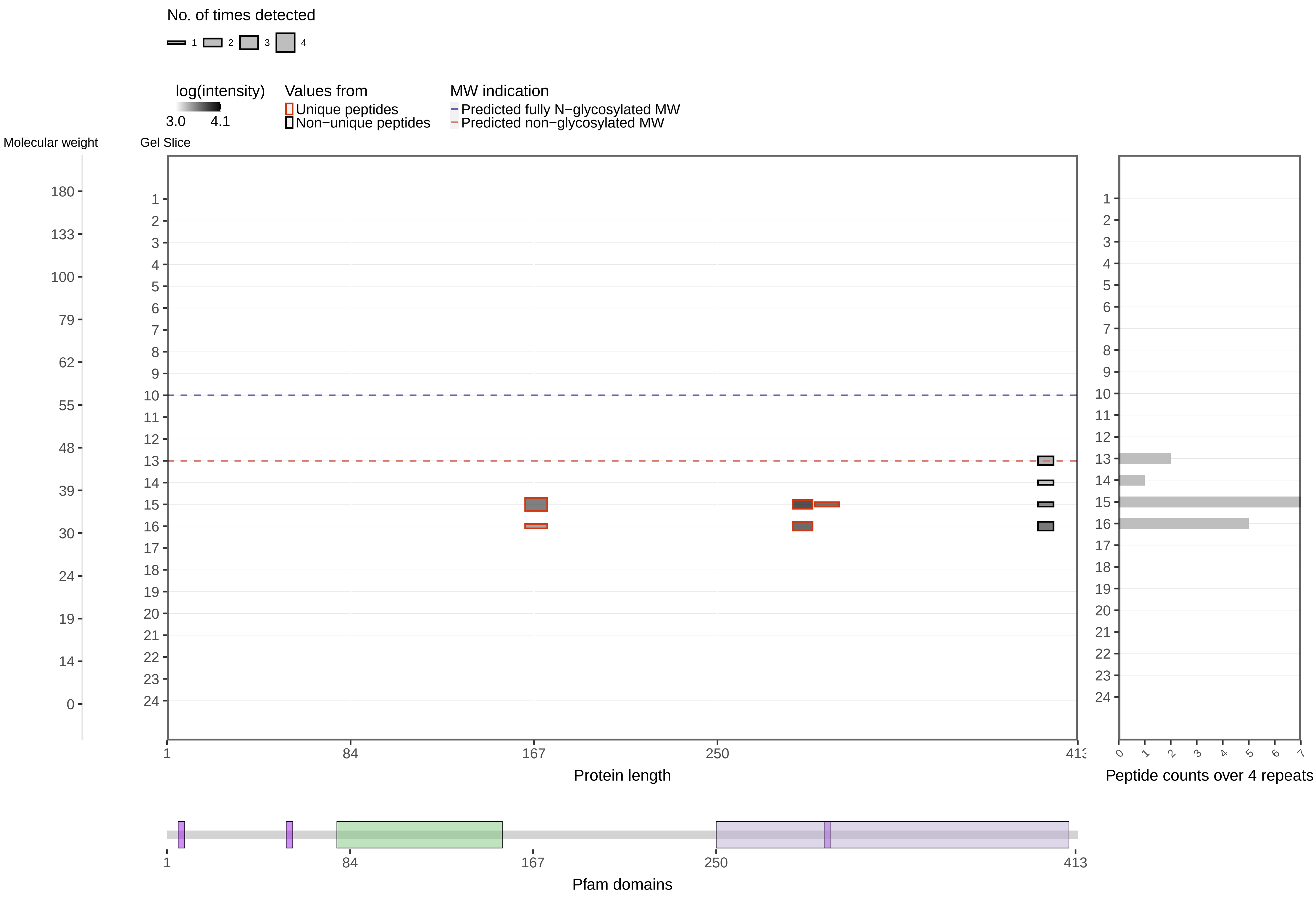


Protein sequence:

KQKRLKLSATDEGAELEIAFSTSDVKTTYKDAKLPPKWRAFQTIIYLEKDDVLHASHKIAAFDFDGCLAKTSVKRVGADAWSLMYPSIPEKMQSLYNDGYKLVIFTNESNIERWKKSRQAAVDSKIGRLEQFIKLAGVPFQVFIACGLSNSKPEDPFRKPKIGMWNIMKKQFNFGISFEMDKCFYVGDAAGRQGDHSDADIRFAQAIGLKFYVPEEFFGK*

NbL05g09370.1 multiprotein−bridging_factor_1b−like [MW = 15.33 kDa]

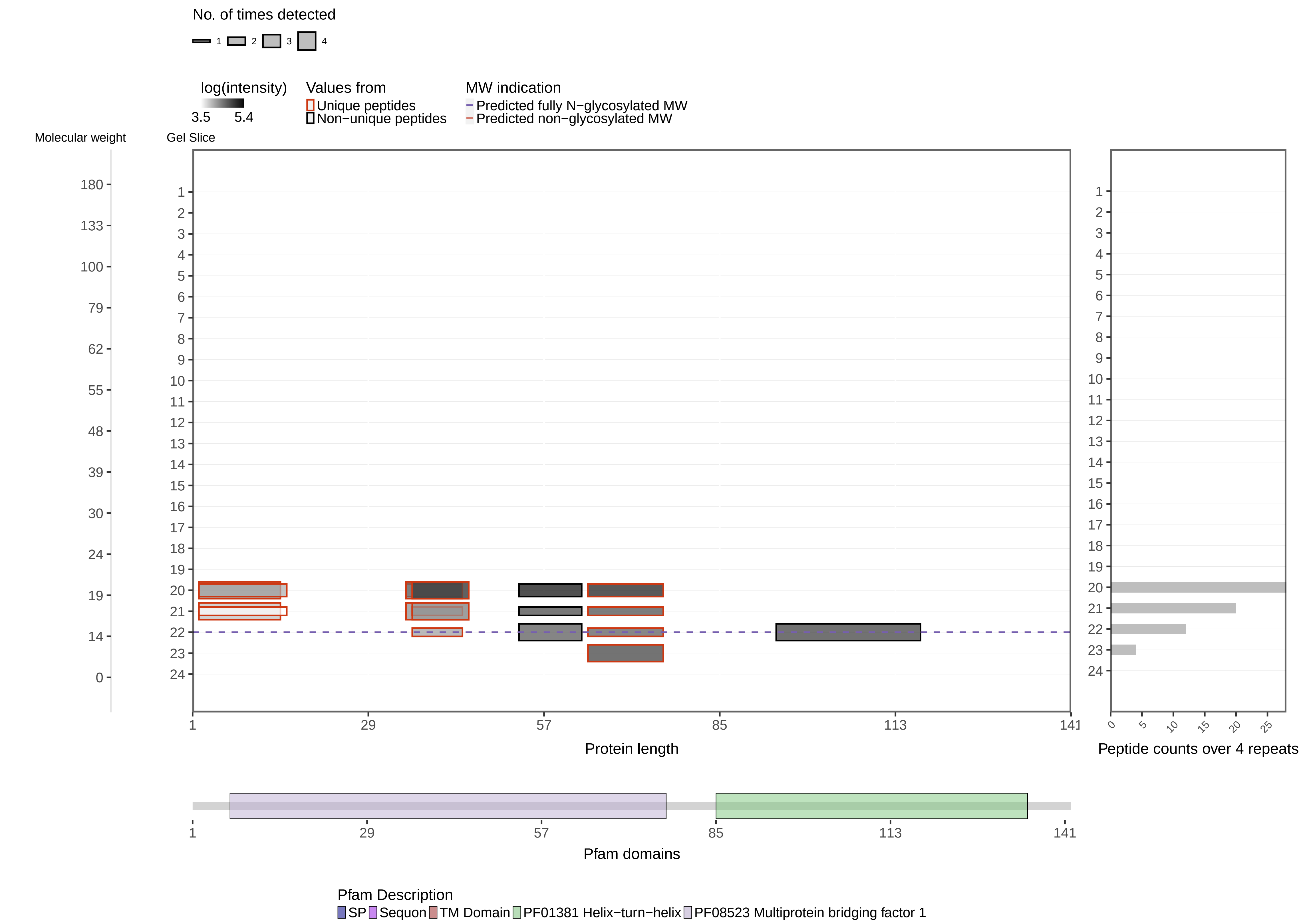


Protein sequence:

STTLNTRKLDEDTENLAHQKVPTELKKAIMQARQDKKLTQAQLAQLINEKPQIIQEYESGKAIPNQQIISKLERALGAKLRGKK*

NbL05g09160.1 peptide_methionine_sulfoxide_reductase_B1_chloroplastic−like [MW = 24.46 kDa]

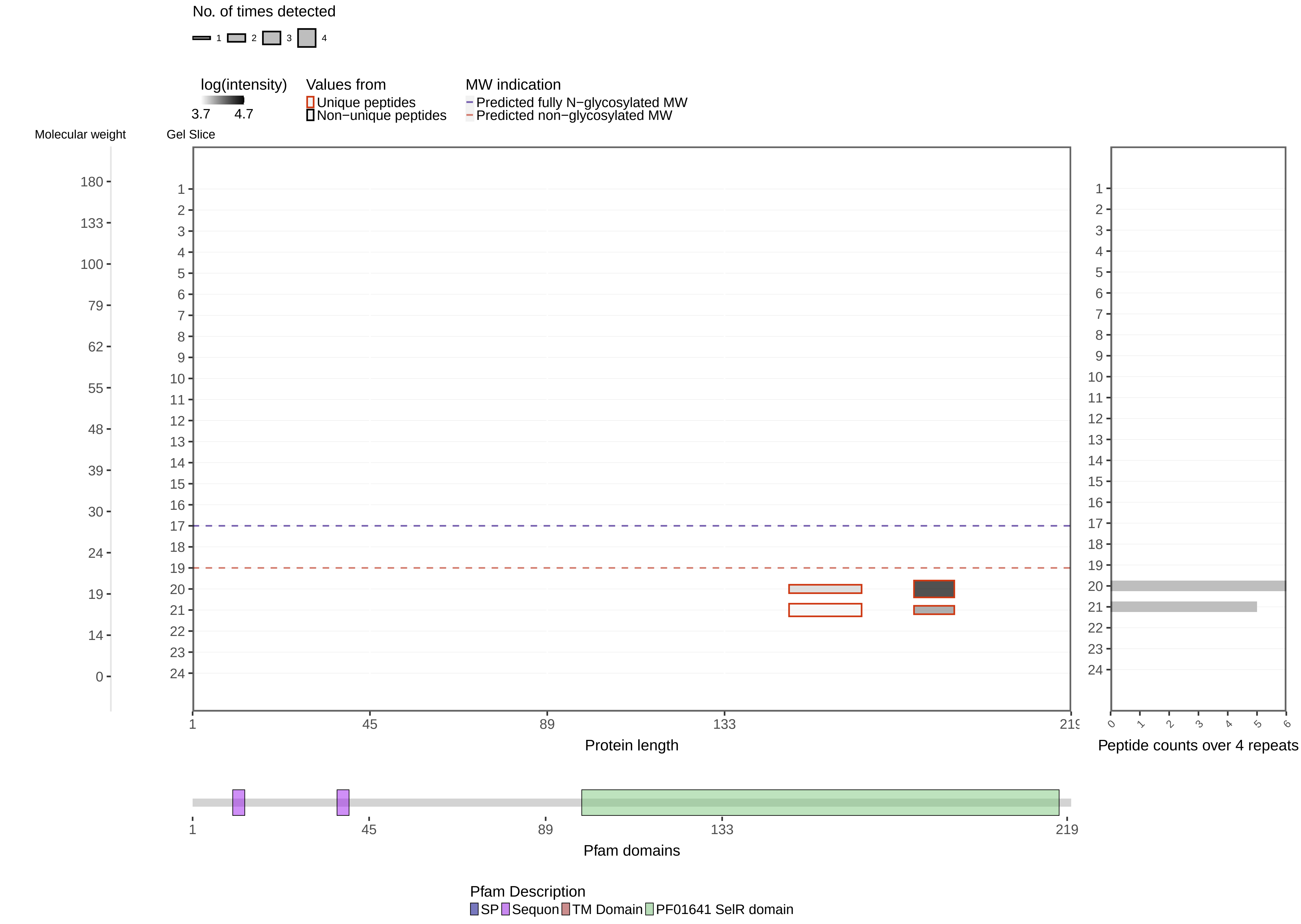


Protein sequence:

EWRKQLTNEQFYITRQKGTERAFTGEYWNTKIQGTYHCVCCDTPLFESSTKFDSGMGWPSYYQPVDNNVKSKMDLSIIFMPRQEVLCAVCDARLGHVFDDGPPPTGKRYCINSAALKLKPK*

### NbL05g09100.1 persulfide_dioxygenase_ETHE1_homolog_mitochondrial−like [MW = 32.71 kDa]

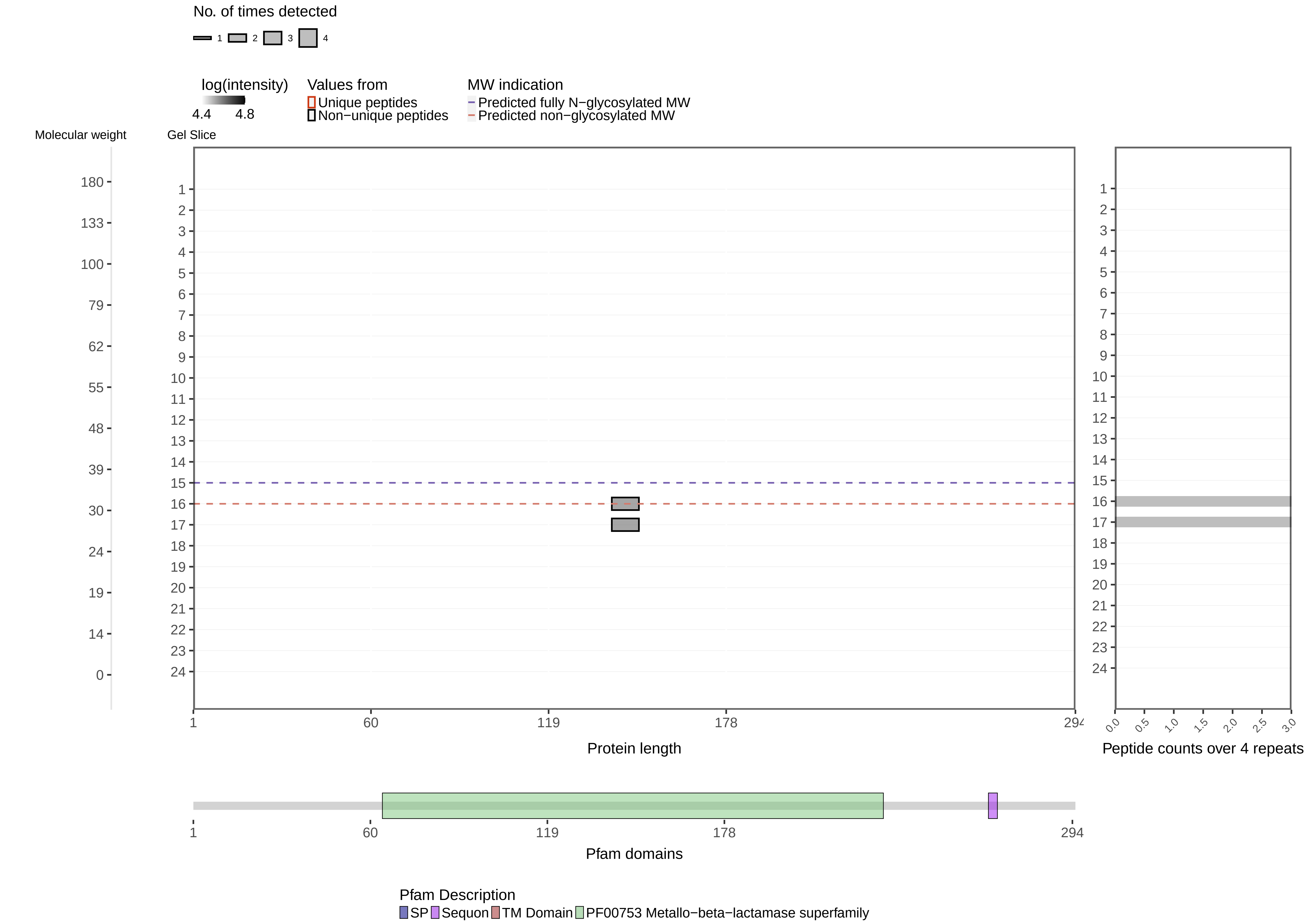


Protein sequence:

NAKADLFVEPGDKIHFGDIFLEFRATPGHTLGCVTYVTGEGPNQPQPRMAFTGDALLIRGCGRTDFQGGSSEKLYDSVYSQIFTLPKDTLVYPAHDYKGFTVSTVGEEMQYNPRLSKDKETFKNIMQNLNLSYPKMMDVAVPANMVCGLQETKSEAY*

NbL05g08970.1 heme−binding_protein_2−like [MW − SP = 23.45 kDa]

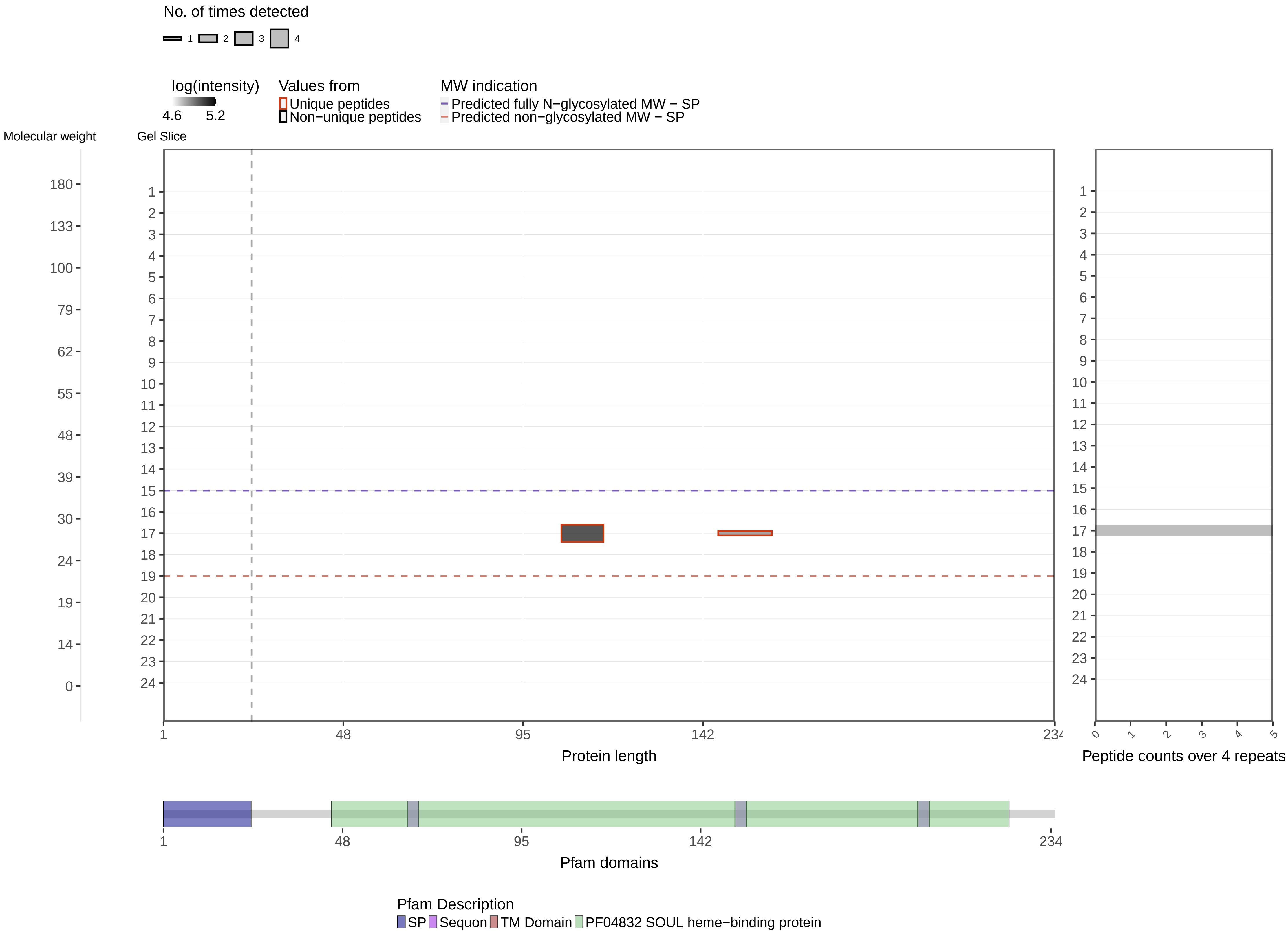


Protein sequence:

EMTAPVITQVKPSDGPFCASSFVVSFYVPKKNQPNPPPAKGLHVQNWSNTYVAVRQFGGFVADVDVAKEAAALSASIADTKWAAAVEKSHAADNTTMYTVAGYNSPFEFKDRVNEIWFTFDLDKASAI*

### NbL05g08850.1 thiamine_thiazole_synthase_chloroplastic [MW = 38.06 kDa]

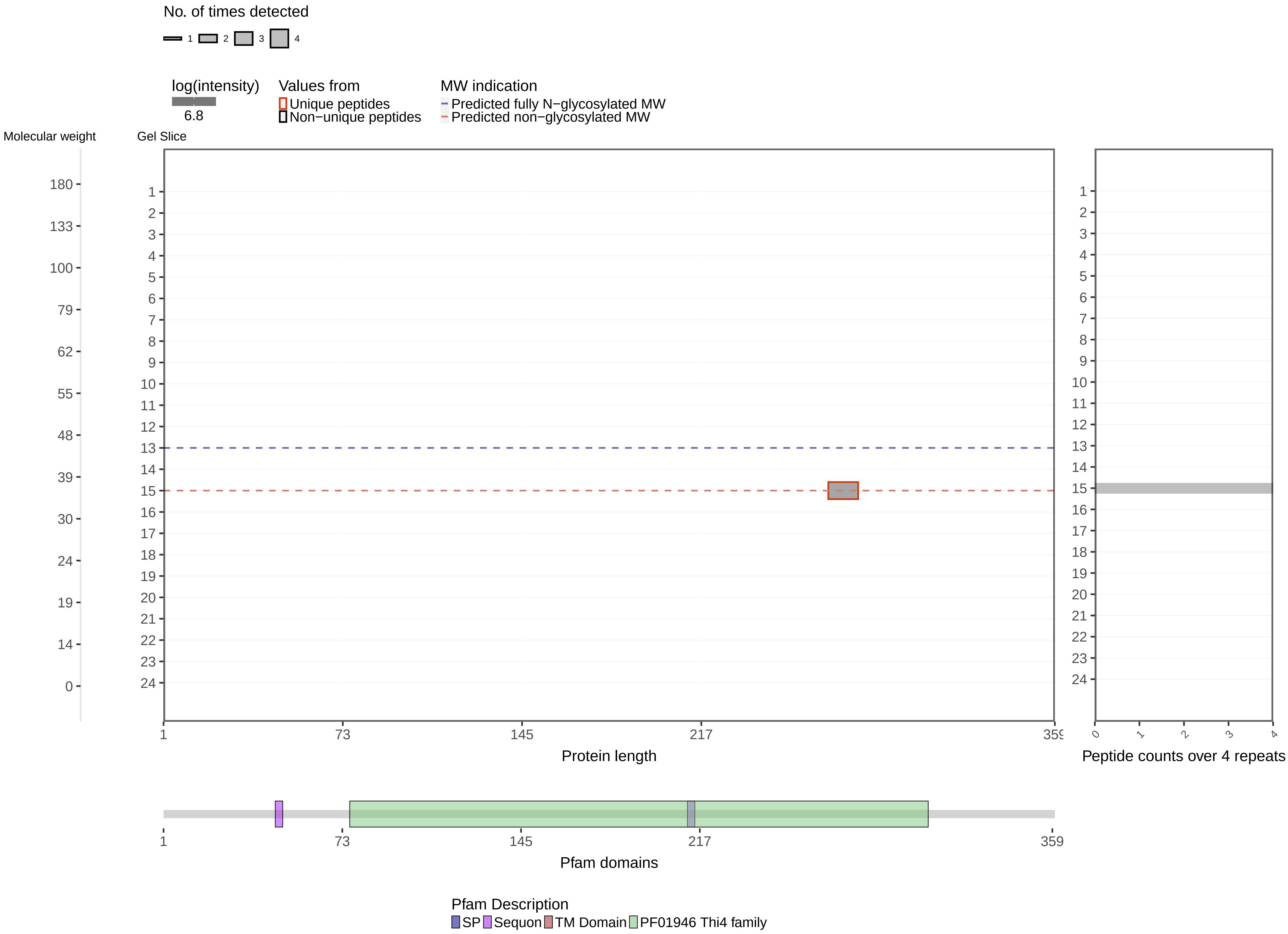


Protein sequence:

VIKHAALFTSTIMSKLLARPNVKLFNAVATEDLIVKNGRVGGVVTNWSLVSQNHDTQSCMDPNVMEAKIVVSSCGHDGPMGATGVKRLKSIGMINHVPGMKALDMNAAEDAIVRLTREVVPGMIVTGMEVAEIDGAPRMGPTFGAMMISGQKAAHLALRALGLPNALDGTAESTIHPELILAAADNAETADA*

### NbL05g08840.1 pectinesterase_2.1−like [MW = 61.88 kDa]

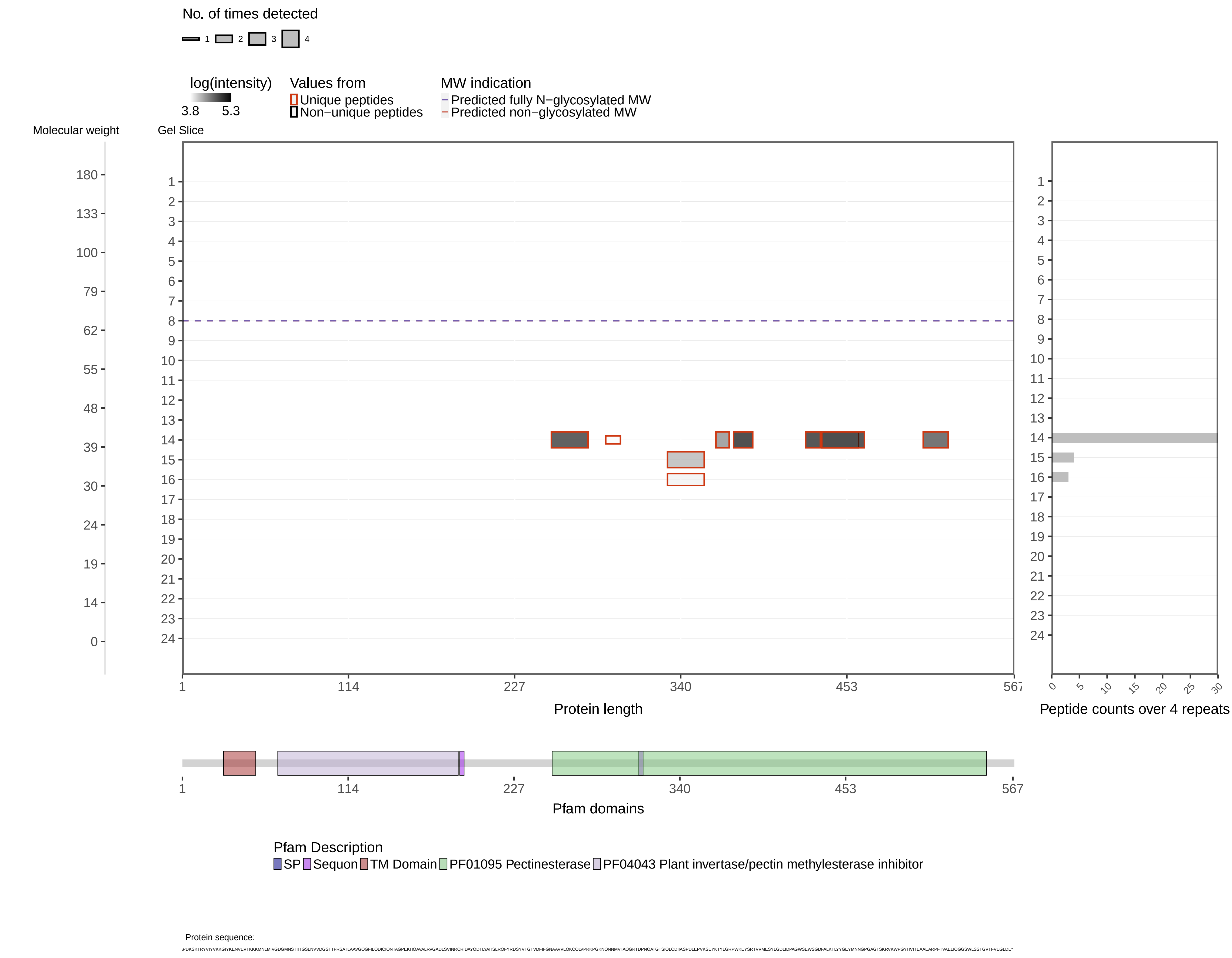

### NbL05g08740.1 probable_protein_phosphatase_2C_55 [MW = 40.07 kDa]

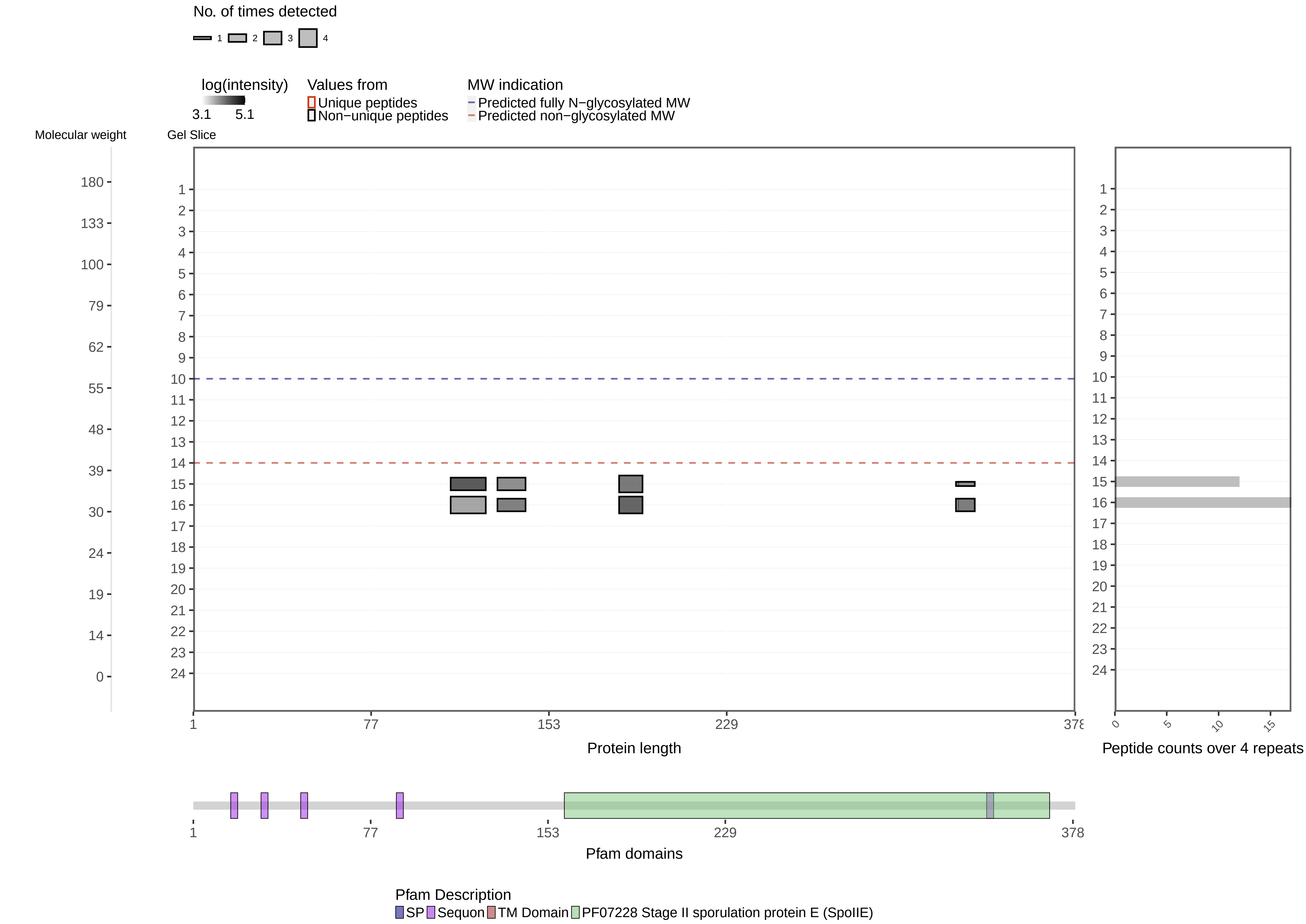


Protein sequence:

GIDAGLYARELMSNSLAAIREEPKGSIDLIRALDKAYMETKAKGSSTACIVALTEEGLYAVNLGDSGFMLVRHGYAAFKSPAOOHGFNFPYOLDCNNAGDSPSSAMVFKIAVAPGDVLIAGTDGLFDNLYDEDIKGVVLOSVEAGLGPOMTARRIAELAOORAKDPINSSPFSDGAOEAGFEHHGGKLDDITVVVSYIMNNENS*

### NbL05g08670.1 DNA_polymerase_I_B_chloroplastic/mitochondrial−like [MW = 91.9 kDa]

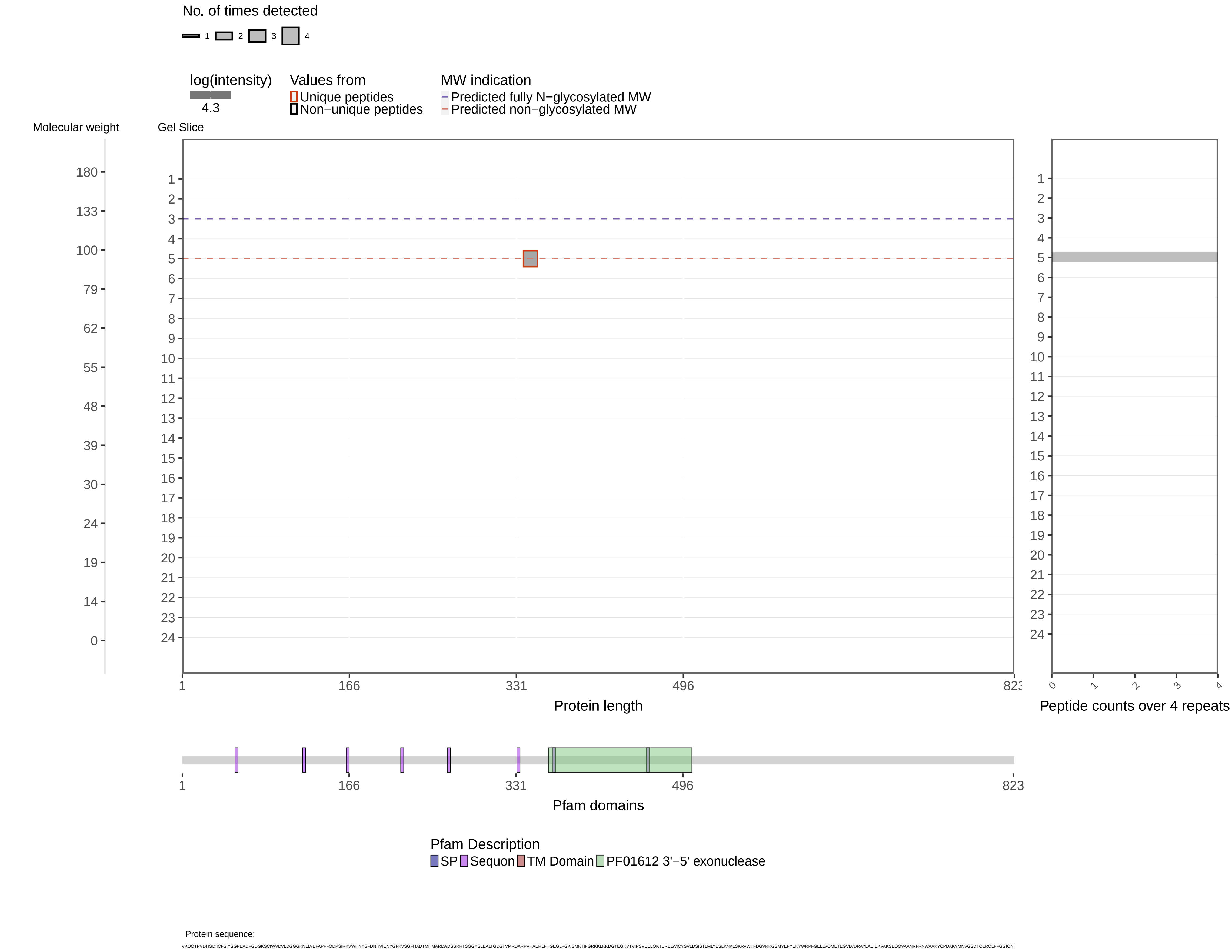

### NbL05g08540.1 polyadenylate−binding_protein_RBP45 [MW = 45.29 kDa]

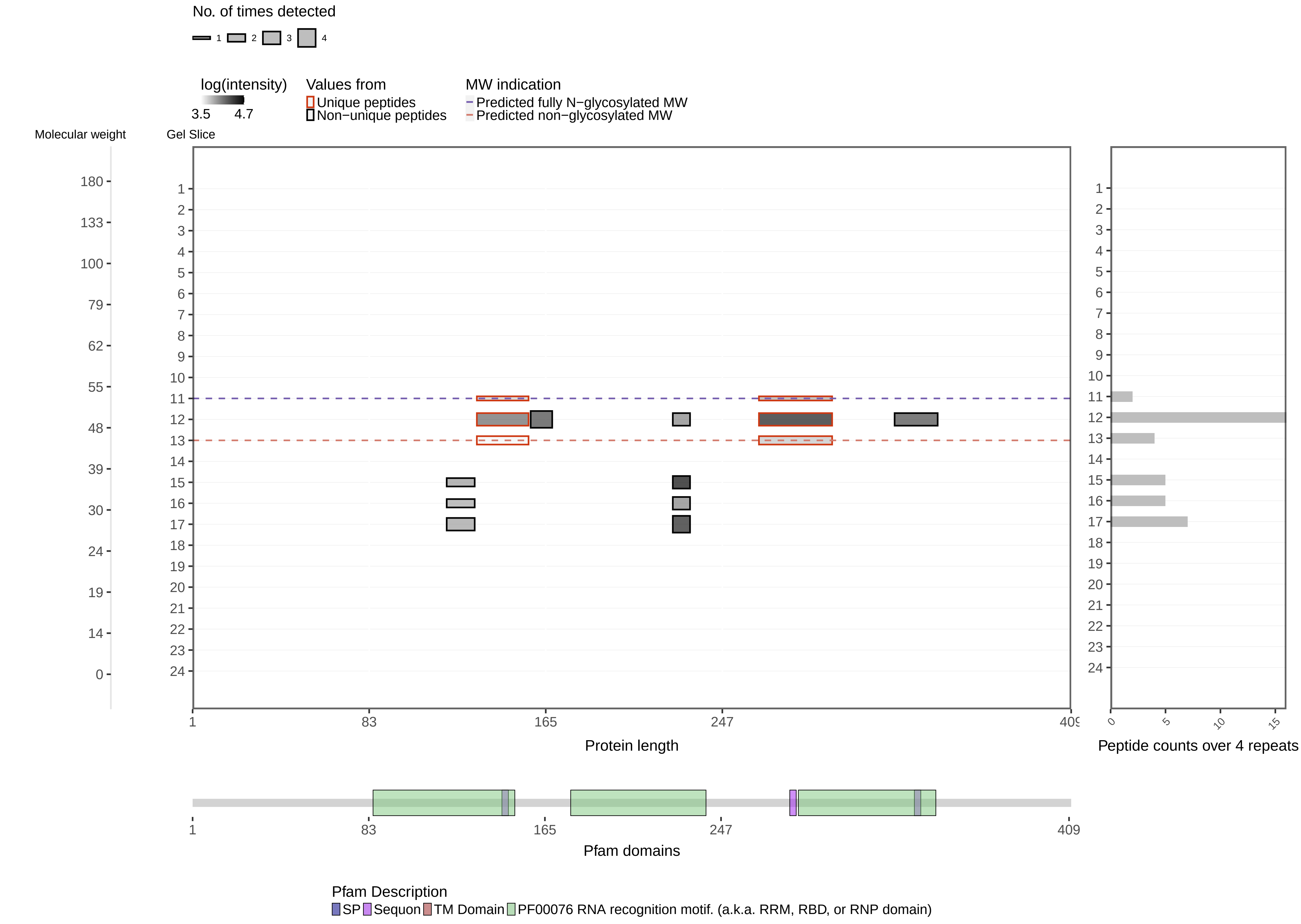


Protein sequence:

MLQETFKSVYSSVRGAKVVTDRITGRSKGYGFVKFADESEQLRAMTEMNGVLCSTRPMRIGPAANKKPVGTPQKVTYQNPQATQGESDPNNTTIFVGGLDPTVAEEHLRQVFSPYGELVHVKIVAGKRCGFVQFGTRASAEQALSSLNGTQLGGQSIRLSWGRSPSSKQTDQTQWGGSGGAYYGYGQGYEAYGYAPPPQDPNMYYGNYPGYTNYQQPQQ*

NbL05g08490.1 ubiquitin_thioesterase_OTU1 [MW = 23.32 kDa]

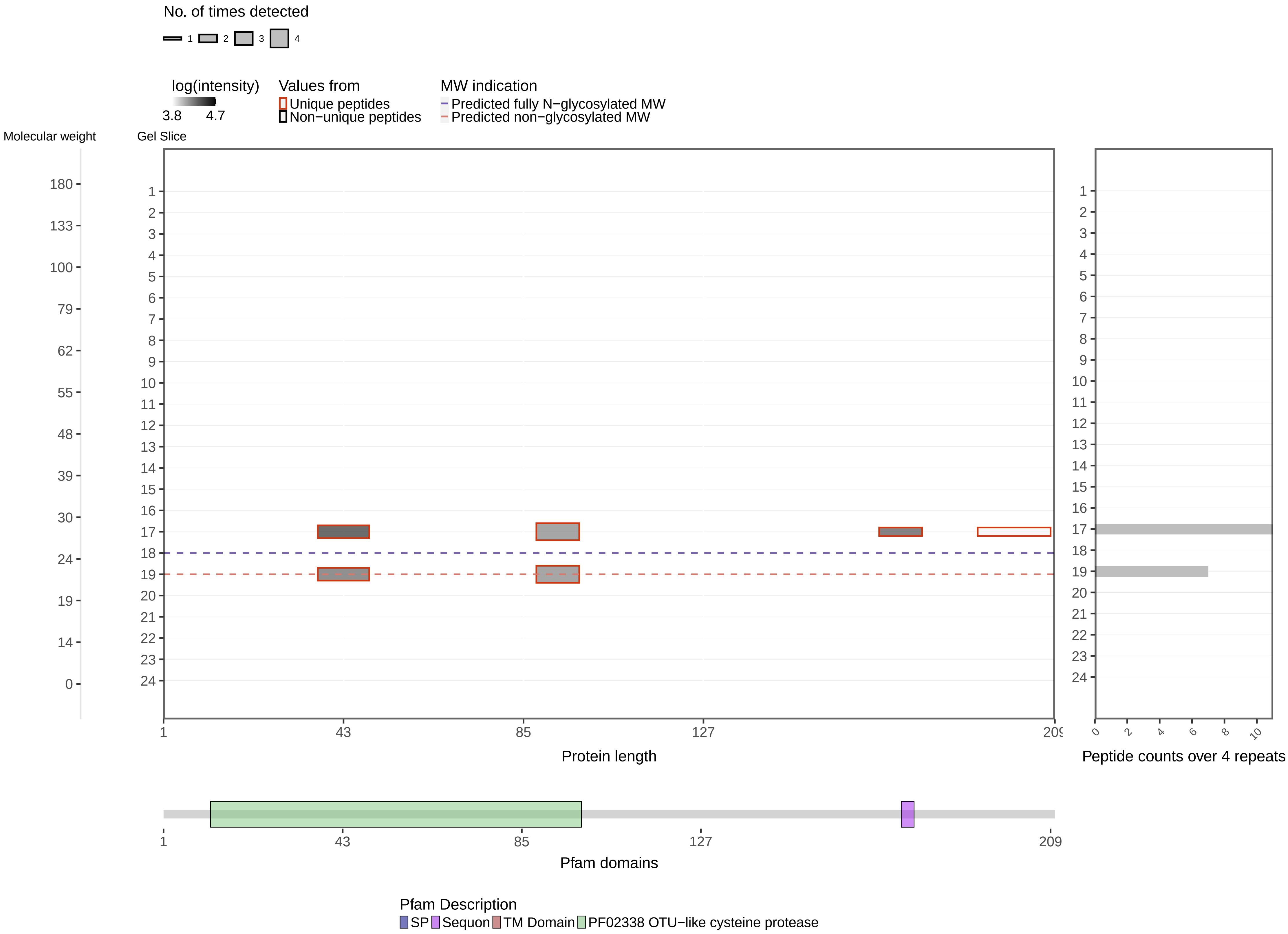


Protein sequence:

DIQTTRCDLYGQGKNYQERVMLIYDGLHYDALAMSPAEGAPEEFDQTIFTVQRDGTVGPAERLALDLVKEQQRKRSYTDTANFTLRCGVCQIGVVGQKEAVEHAQATGHVNFQEFK*

### NbL05g08430.1 aconitate_hydratase_cytoplasmic−like [MW = 51.01 kDa]

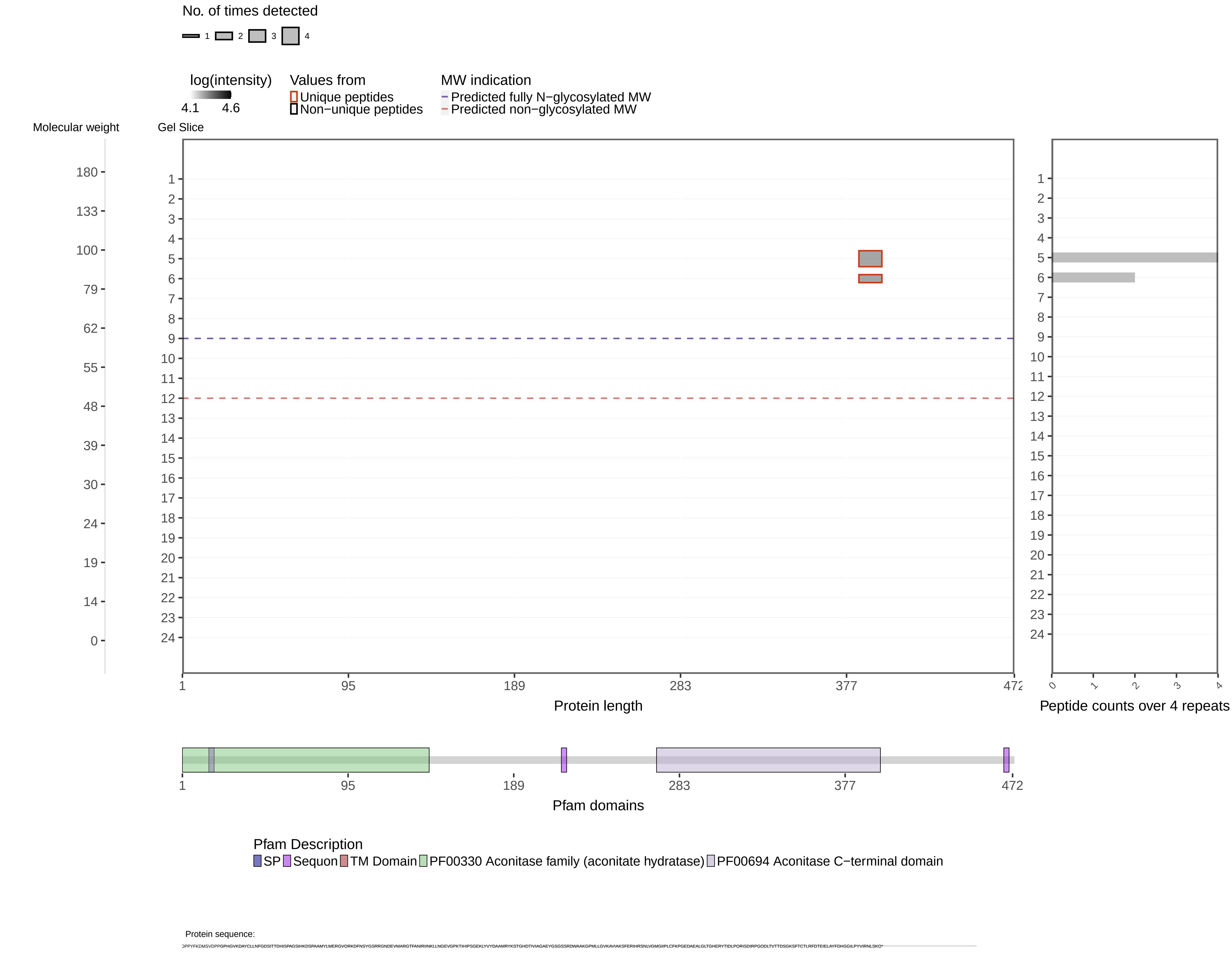

### NbL05g08410.1 mimitin_mitochondrial [MW = 21.49 kDa]

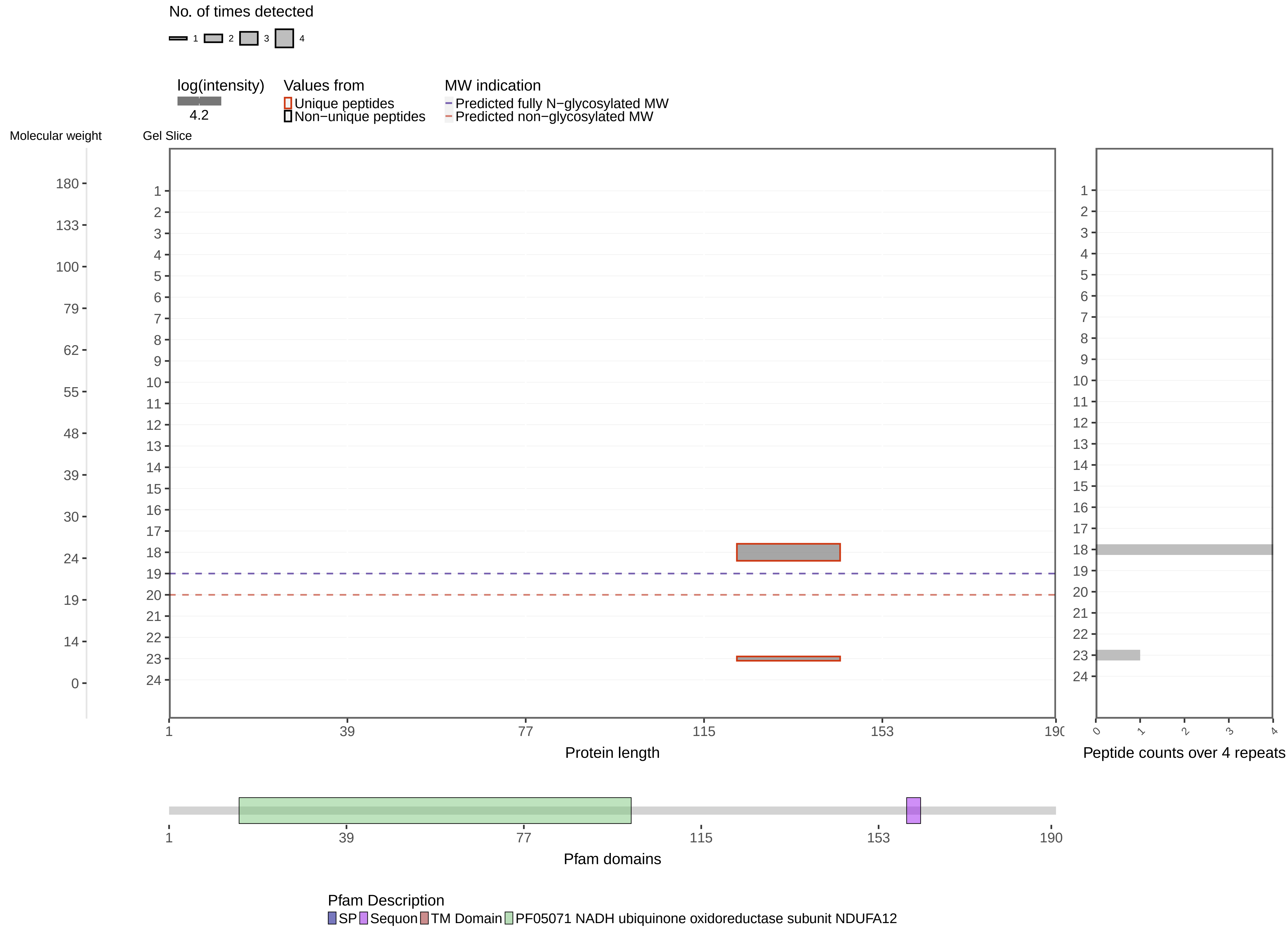


Protein sequence:

RREFVKLNIARLRKEEEERKAKEGKNKATSIGKADGPDLKSFVQQFPDASEGDIIPEASDTQKGVRHARSKETEAEKNMSYRLKQQPLSSEPTGSGESFRPGTWQPPS*

NbL05g08130.1 nardilysin [MW = 118.64 kDa]

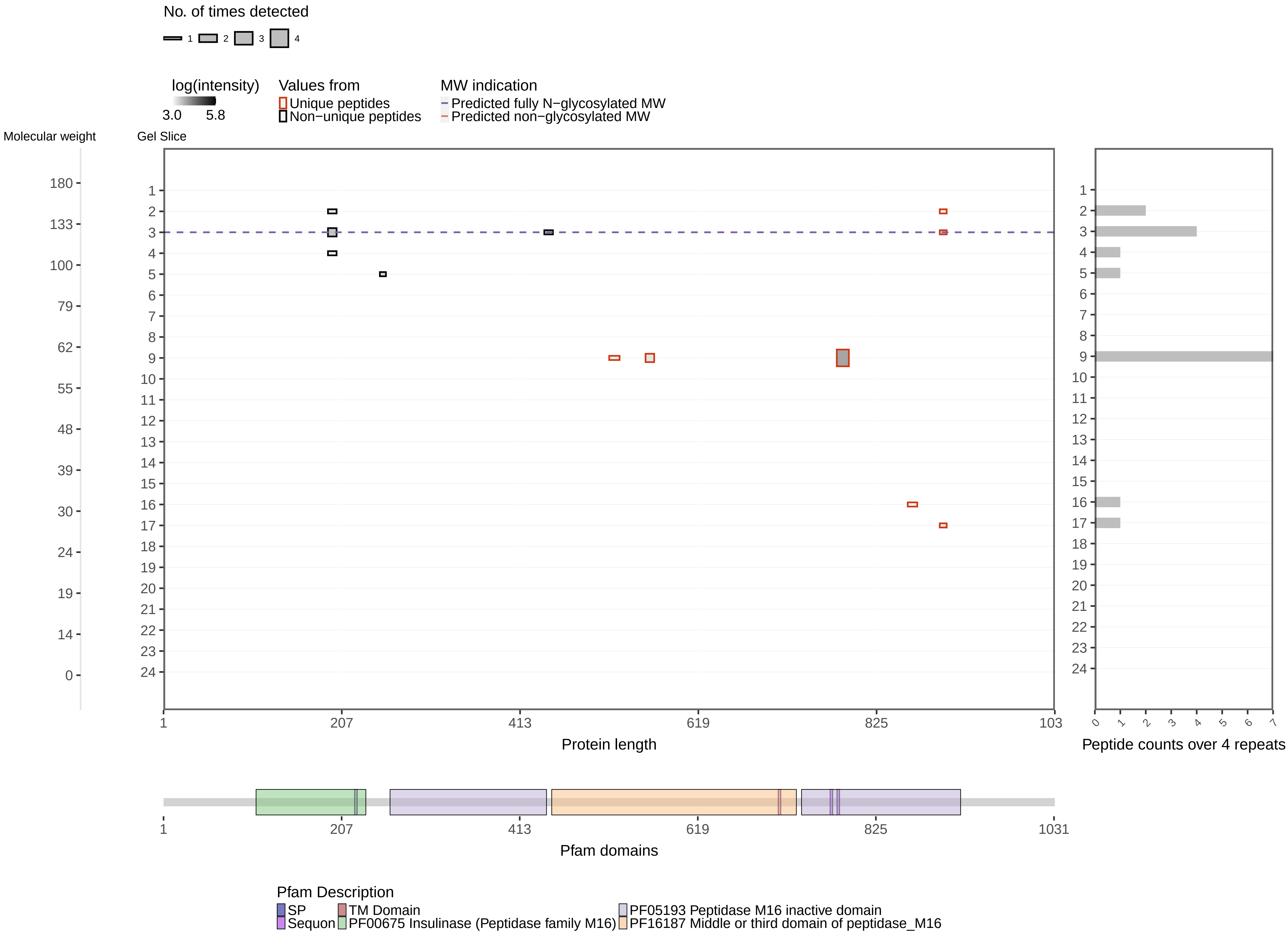


Protein sequence:

.DFFRPENMRVDIVTKSFQKSDDIQQEPWFGSRYVEEDIPSSLFELWKDPEEISVCLHLPAKNEFIPSDFSIRAEKVNCDSKNAKPRCILEEPLMKIWYKLDKTFKLPRANTYFRISLRGGYGNLKNALLTELFIHLLKDELNEIIYQASVAKLETSVSLYGDKLELKVYGFNDKLPVLLSKVLAMTKSFLPRDDRFMVIKEDMERTLKNTNMKPLNHSSYLRLQVLCQSFWNVEEKLLLLNDLTLAELKAFIPELLSQLYIEGLCHGNLLEEEALNISNIFRSNFSAQPLPSEMTHKEY

NbL05g07950.1 glucan_endo−13−beta−glucosidase_9 [MW − SP = 50.41 kDa]

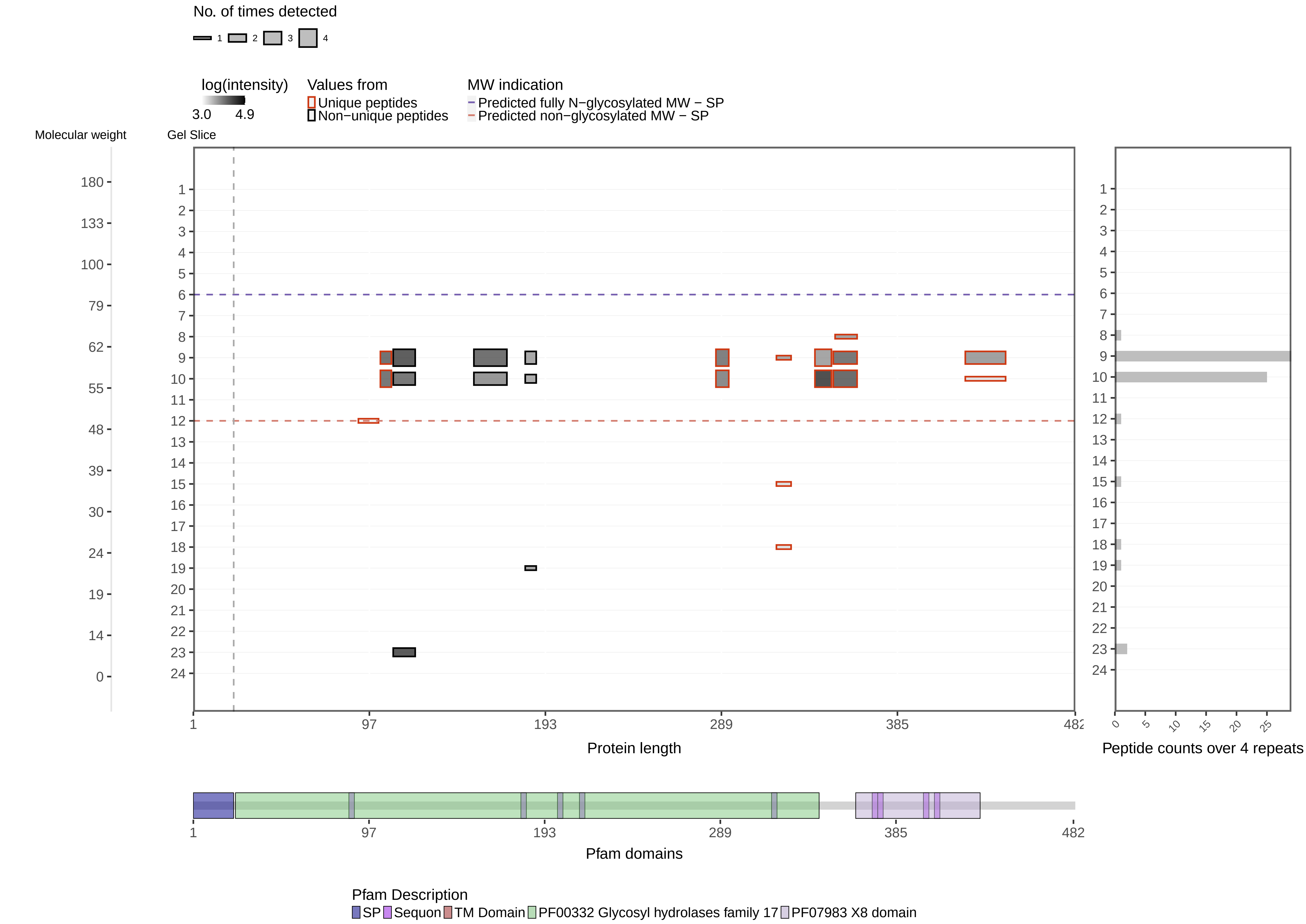


Protein sequence:

RRTYKNSFDLSYDTLVSALSSAGSDKMDIVIGQIGWPTDGAASATSSNAQVFMKKFLEHLHSRSGTPLRPKEPPIEAYILSLLDEDNRSIATGNFERYWGVFTFDGQAKYQLDLGQGSRKLVNAQNVQYLSSKWCVVNNNQNLSNATTRALEACSSADCSALSPGASCFNLSWPGNISYALNSYYQQHDQRADSCDFGGLGLITTVDPSVGTCRFIVQLRTSSSASFFISHWRITASTILLWLVGGGFGWT*

### NbL05g07930.1 40S_ribosomal_protein_S4−like [MW = 29.85 kDa]

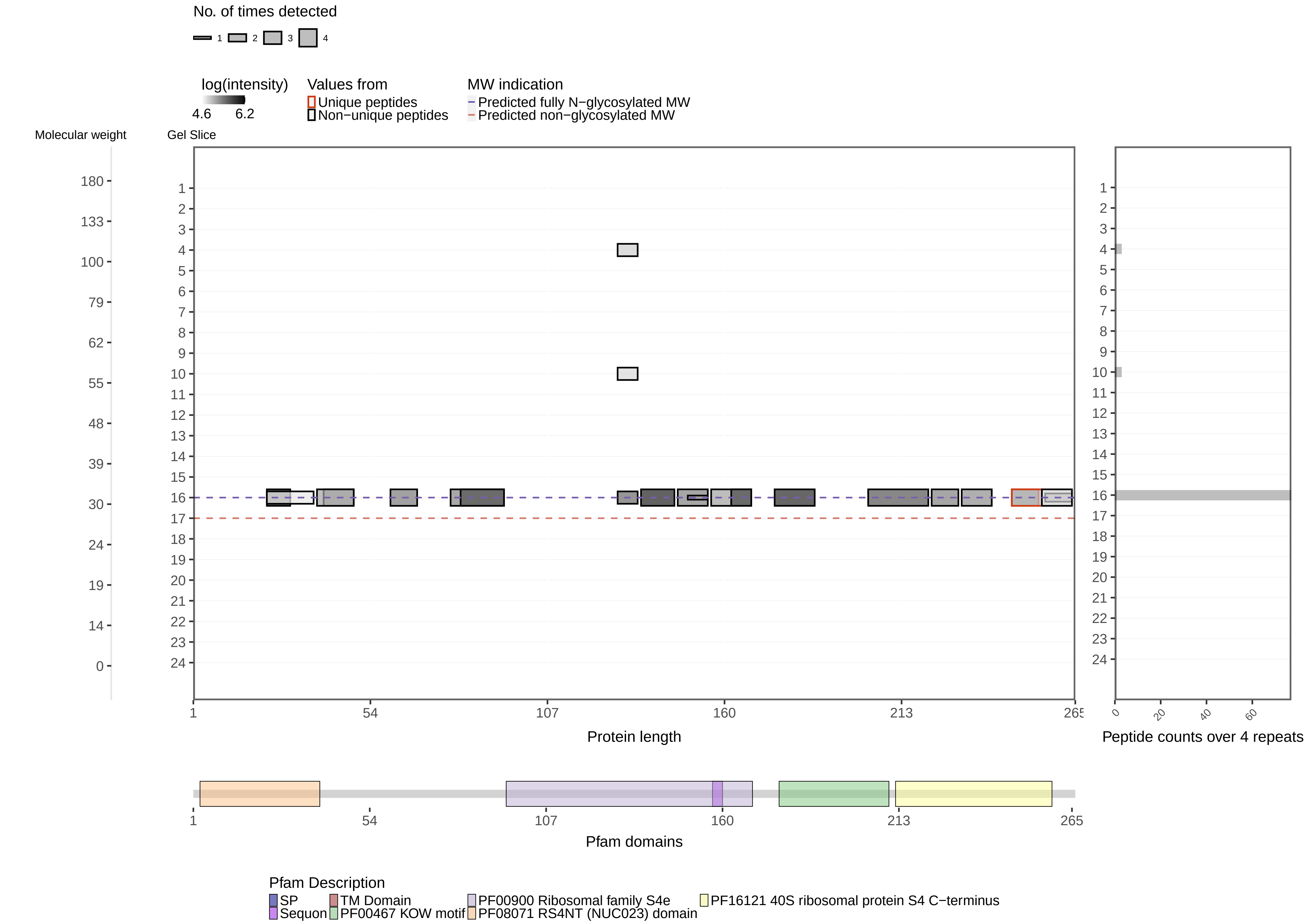


Protein sequence:

AKFKLCKVRSIQFGTKGIPYLNTYDGRTIRYPDPLIKANDTIKLDLESNKIVDFIKFDVGNVVMVTGGRNRGRVGVIKNREKHKGSFETVHIQDALGHEFATRLGNVFTLGKGTKPWVSLPKGKGIKLSIIEEAQKRLAAQSATTA*

### NbL05g07750.1 alpha−glucan_phosphorylase_H_isozyme [MW = 100.46 kDa]

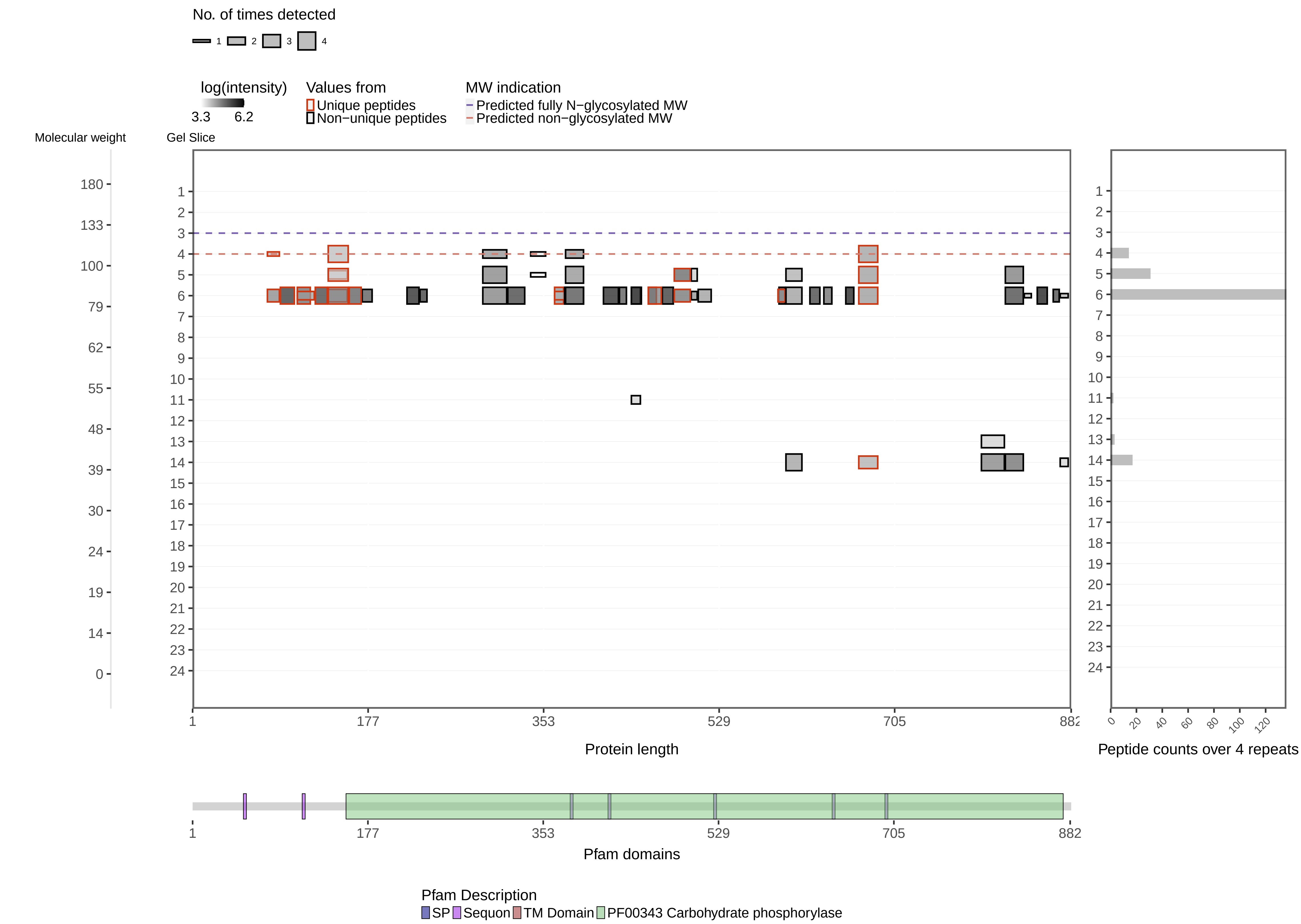


Protein sequence:

NbL05g07740.1 neutral_ceramidase−like [MW − SP = 82.2 kDa]

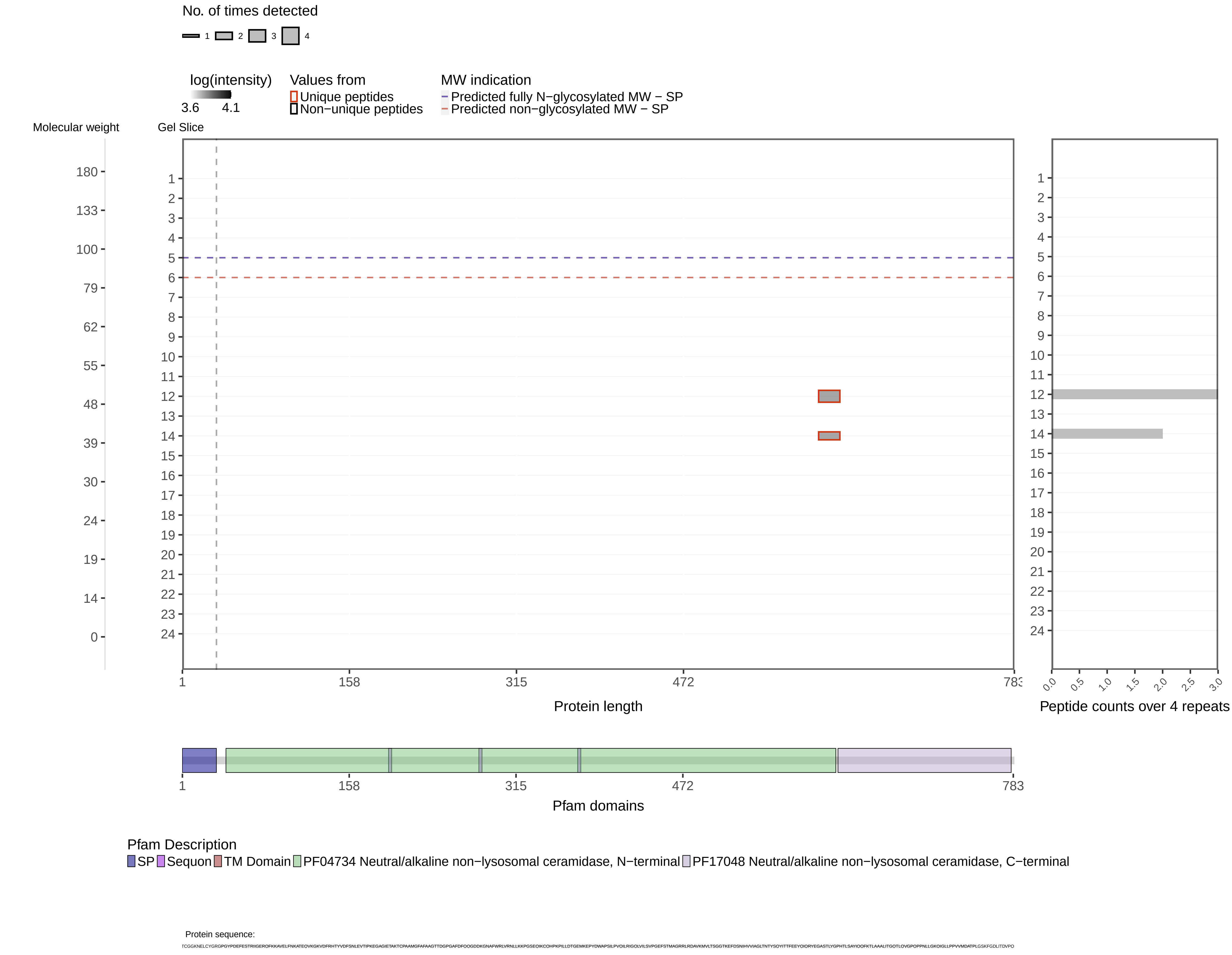

NbL05g07700.1 phosphoglucomutase_cytoplasmic−like [MW = 63.3 kDa]

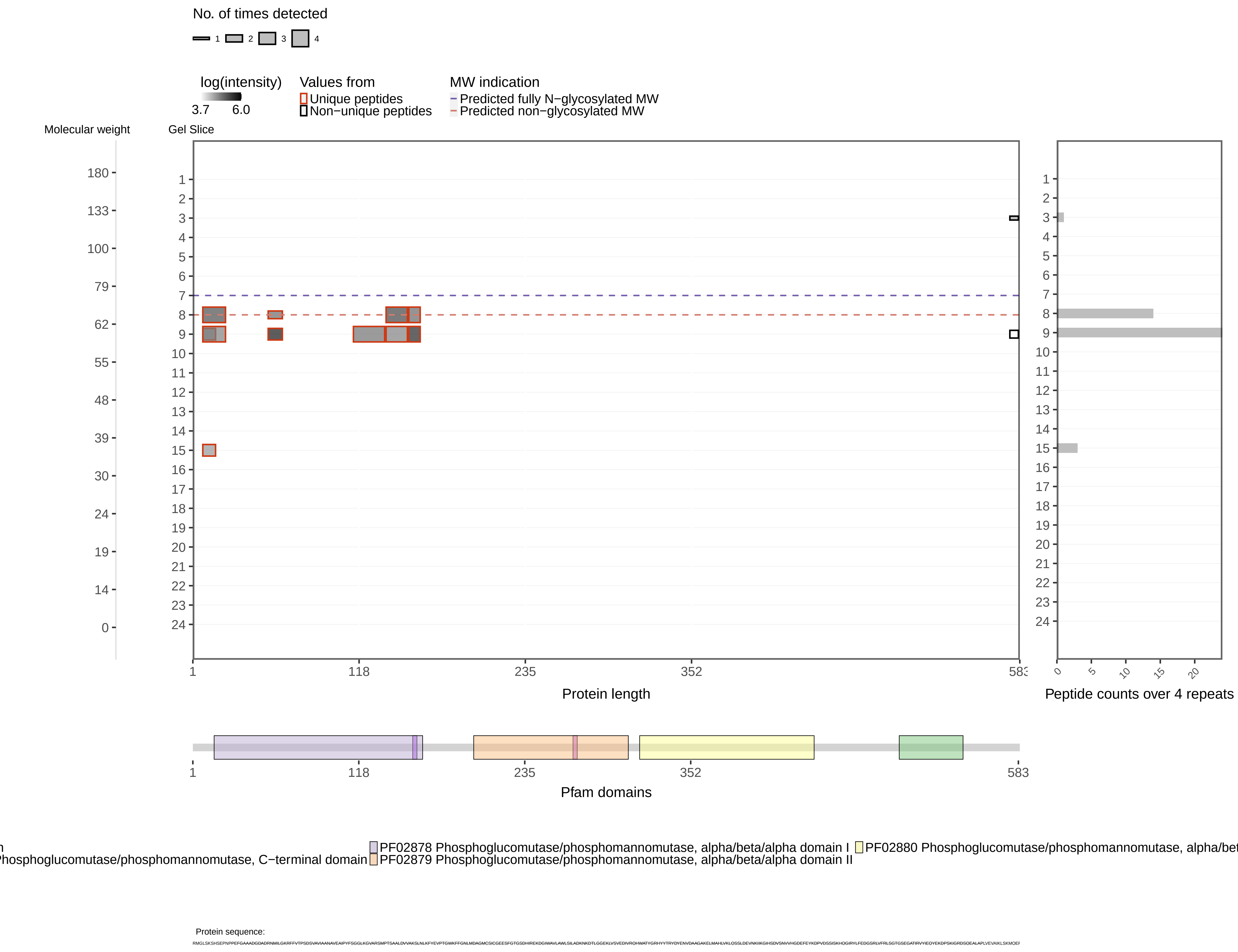

### NbL05g07580.1 probable_LRR_receptor−like_serine/threonine−protein_kinase_At1g06840 [MW − SP = 103.14 kDa]

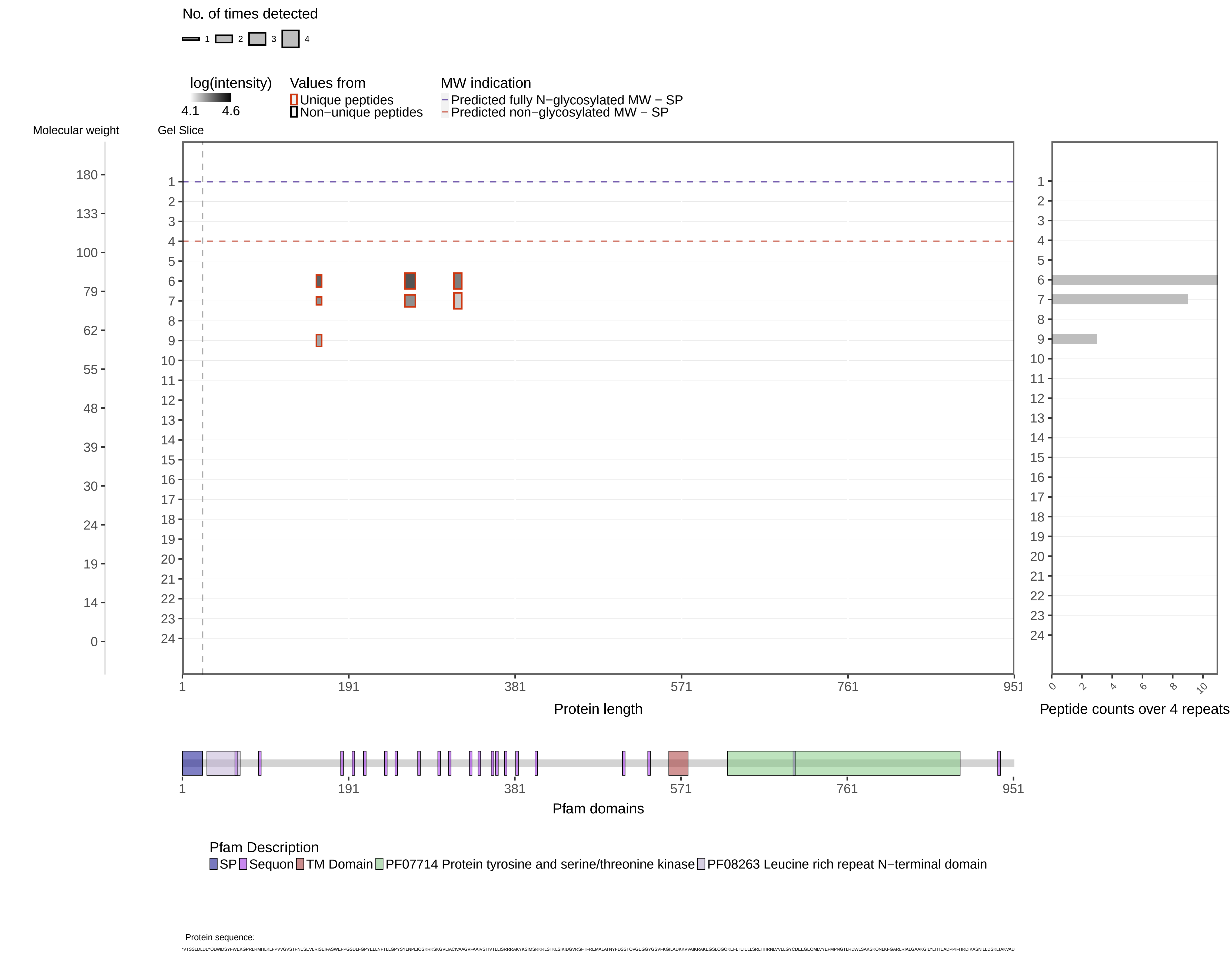

NbL05g07500.1 receptor−like_protein_kinase_FERONIA [MW − SP = 93.8 kDa]

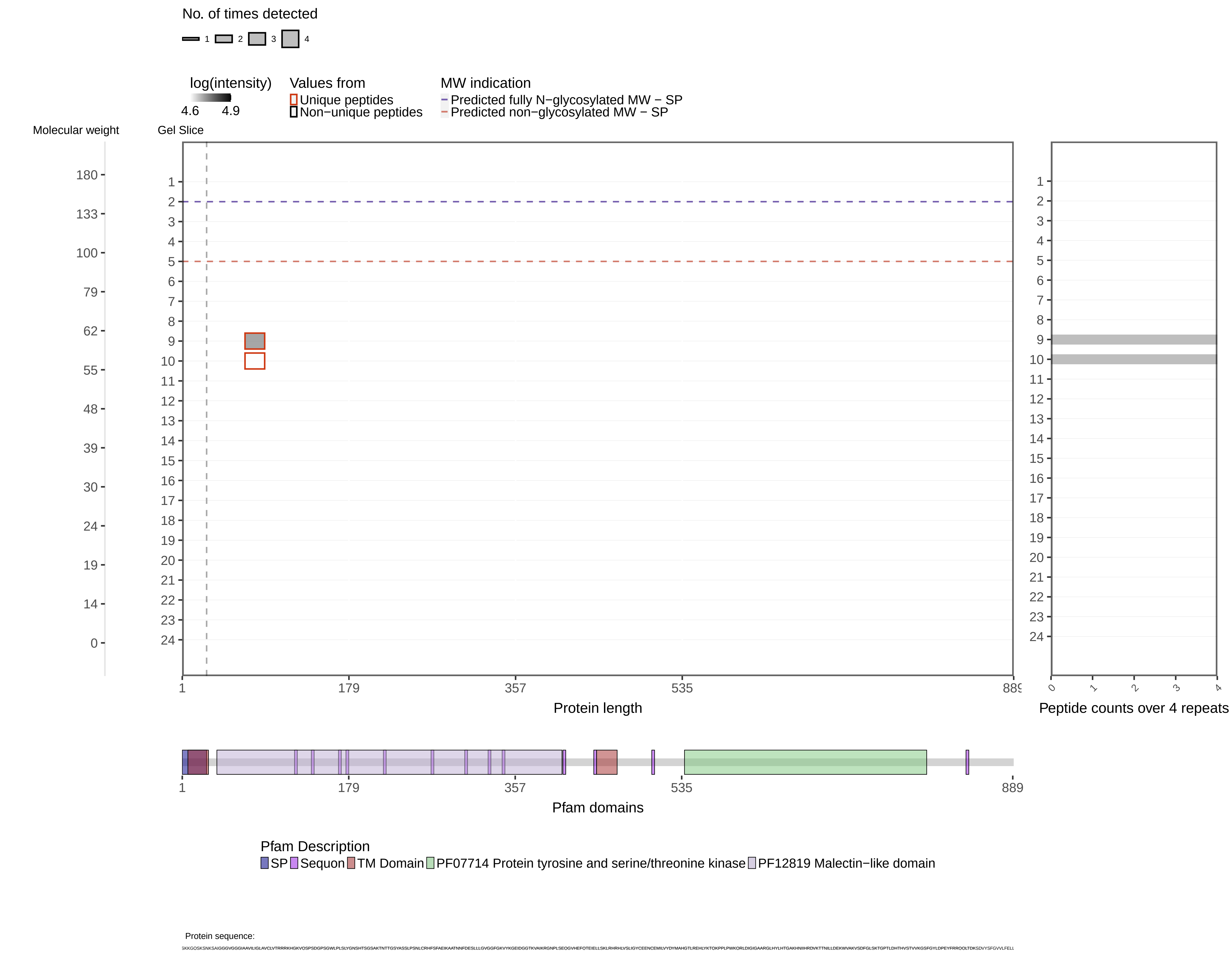

### NbL05g07470.1 vacuolar−sorting_receptor_1−like [MW − SP = 66.98 kDa]

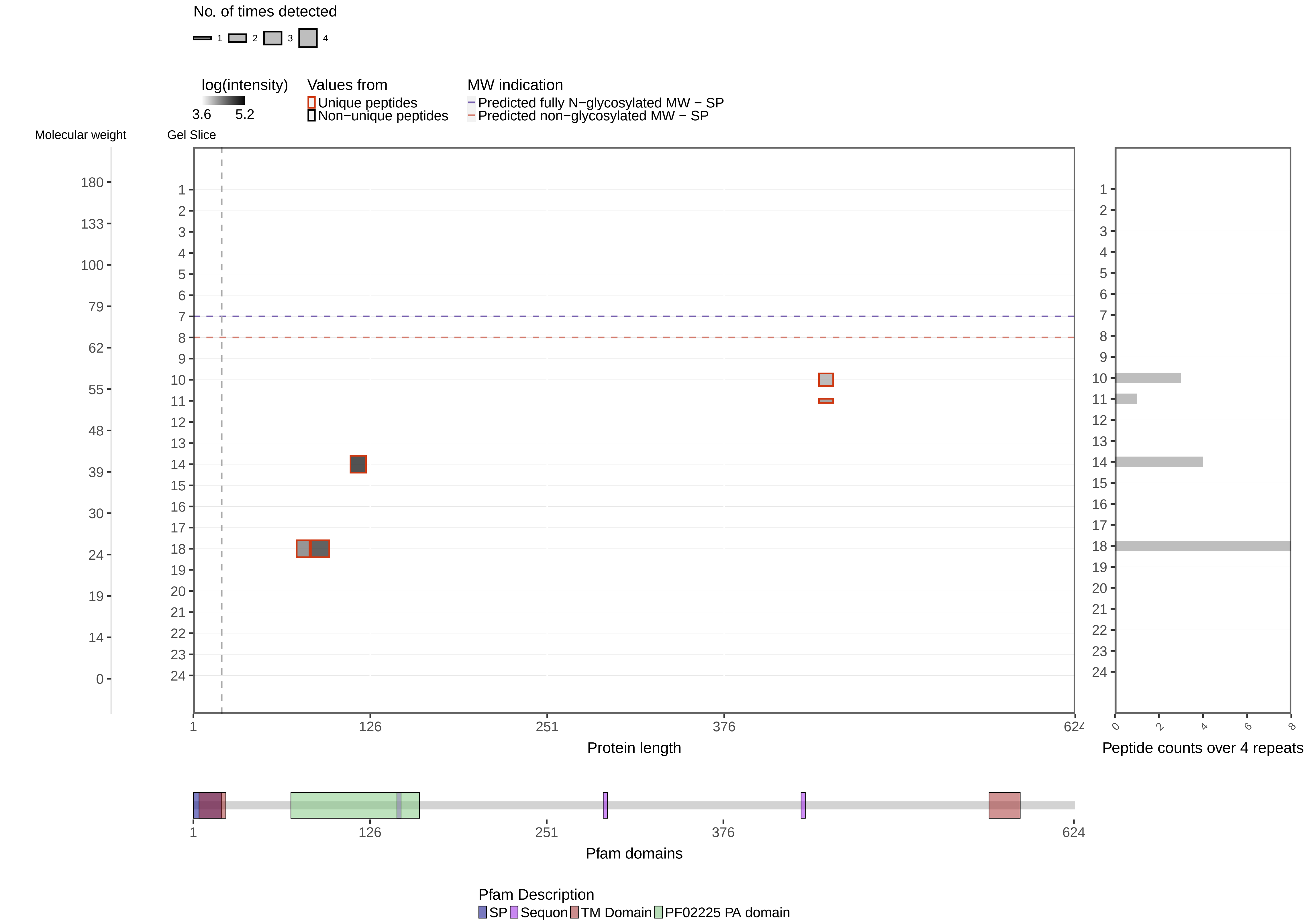


Protein sequence:

### NbL05g07380.1 non−specific_lipid−transfer_protein_1−like [MW − SP = 9.46 kDa]

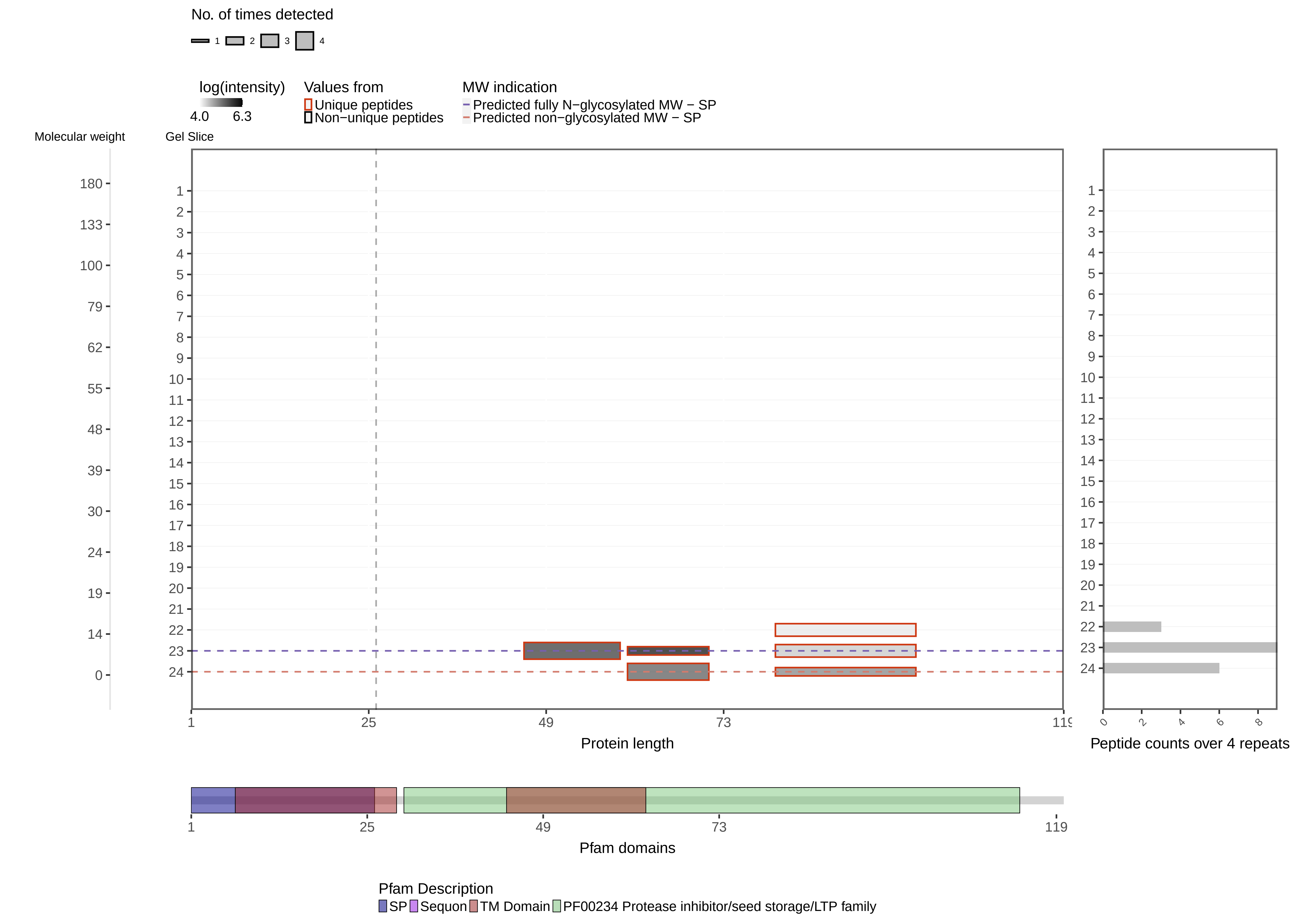


Protein sequence:

GGVIPTMCCNGIRSLFSAASTTODROTVCNCMKSAAGVISGVNLNLAAALPSKCGVNIPYKISPSIDCAKVO*

### NbL05g06990.1 ubiquitin−activating_enzyme_E1_1−like [MW − SP = 124.19 kDa]

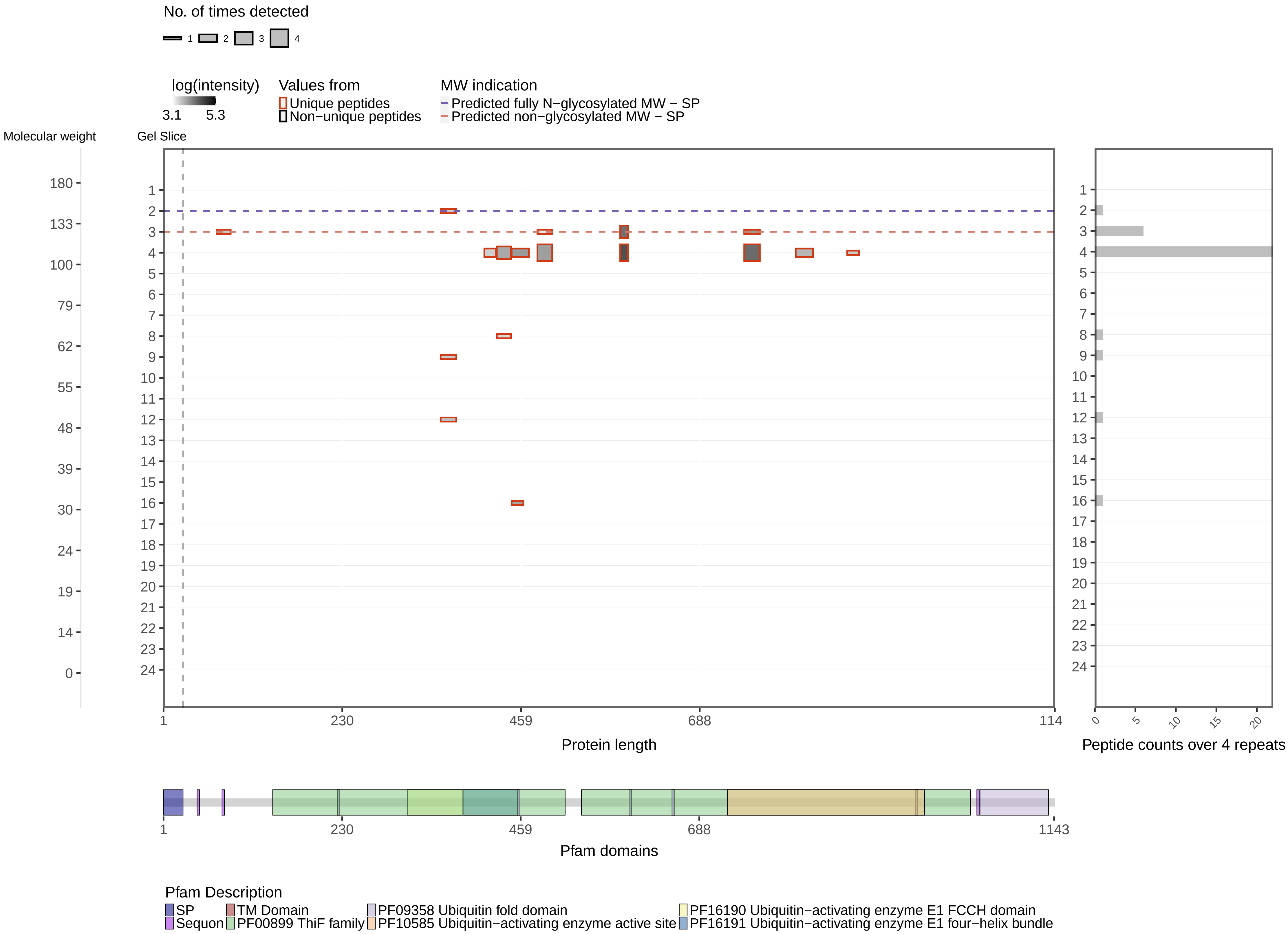


Protein sequence:

### NbL05g06630.1 GDSL_esterase/lipase_CPRD49−like [MW = 28.31 kDa]

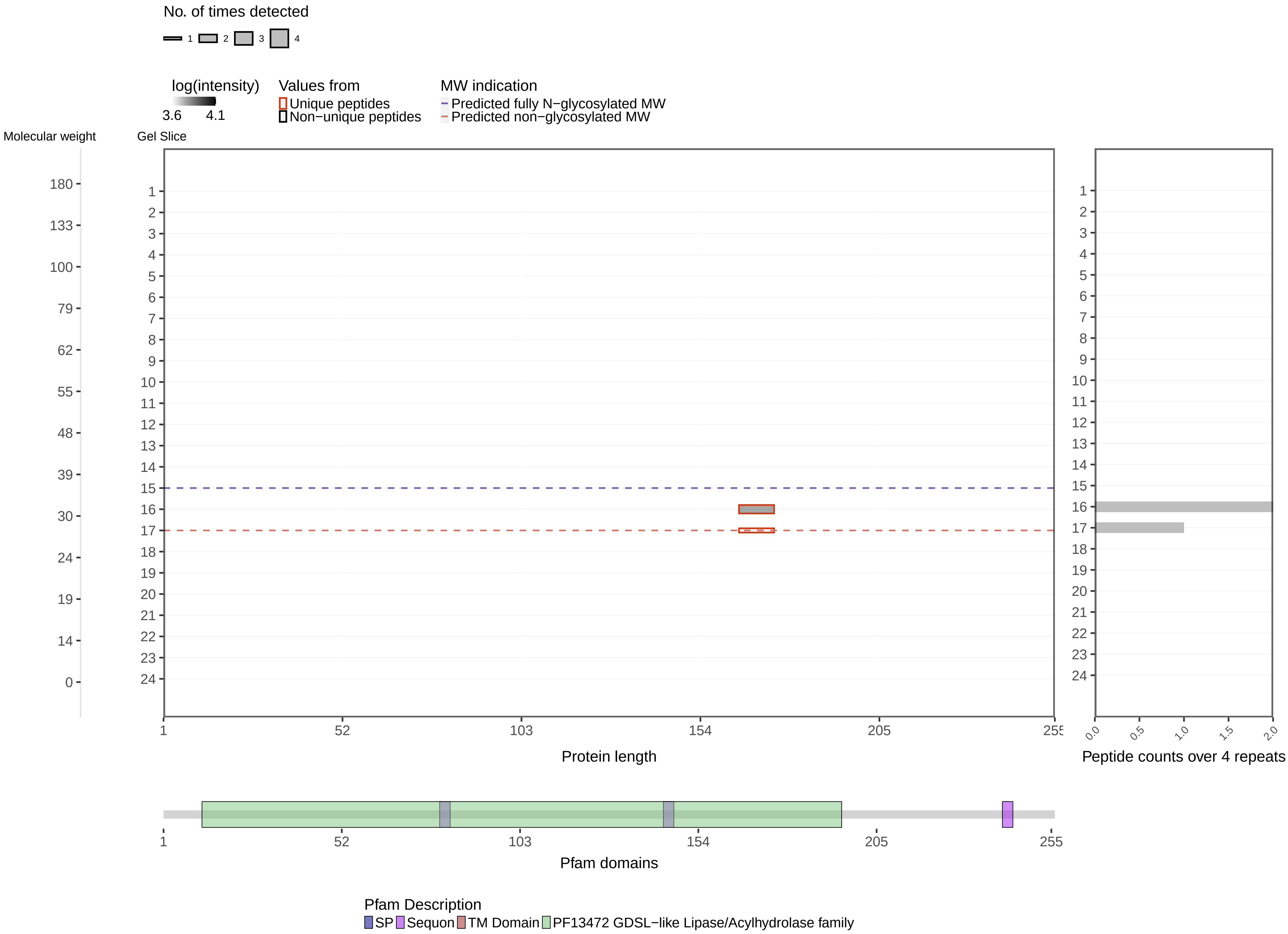


Protein sequence:

SDKTHIILLSAPAVNEAQITEVFGNGLGRSNGSCRKYSEACIQLGQELGVKVIDLWTTLQQRDDWLTACFTDGIHLSSEGSKLVAKEILKVLKEAEWEPNLHWRSLPIEFPEISPYIFVDPKGETAVNVSQLDLEWQTQWH*

### NbL05g06480.1 uncharacterized_LOC104227496 [MW = 53.13 kDa]

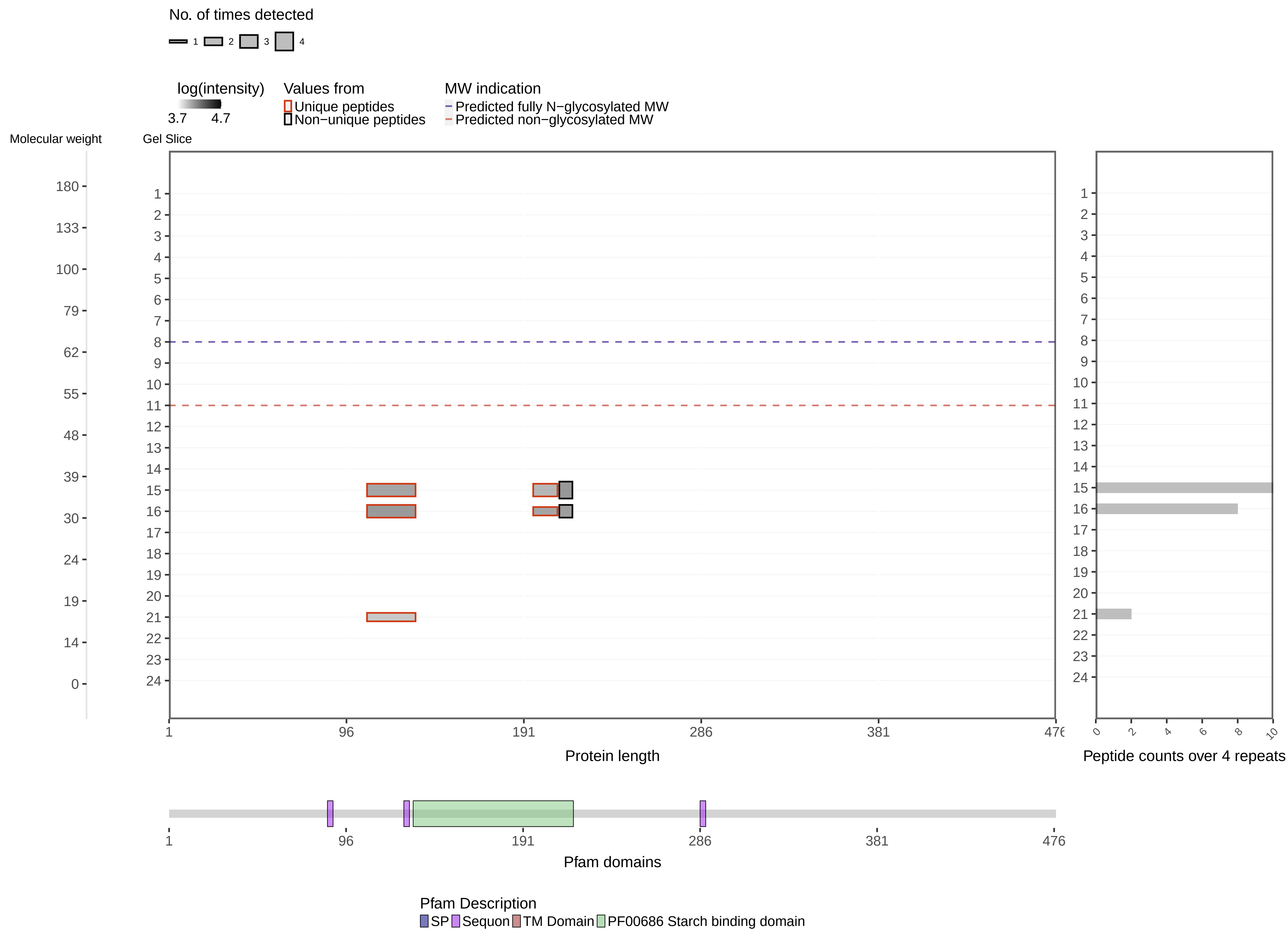


Protein sequence:

WDNAELQTIVEQEPAAEQSEESAVNPEILIAENLVPPTVVDVEDDINEKKSNEVLAMVAENITEVKEDVNTNLNEDTIIETKVIGKNGMVFNDEVLSSKKEPILVARETVPVLVPGLTQVLDMDKVIAEISVGSNGEELNVLLDKVQMNKVVTEGSLGLKSEEFNVPELSSNNEVATDLTHPRETPEMMLNVKQEVRGNEDIEKSEFVEGGDWPSGKPVEEIFDSDMKWGKRTLQKFFANLGFFKVELEG*

NbL05g06350.1 proteasome_subunit_alpha_type−4 [MW − SP = 27.04 kDa]

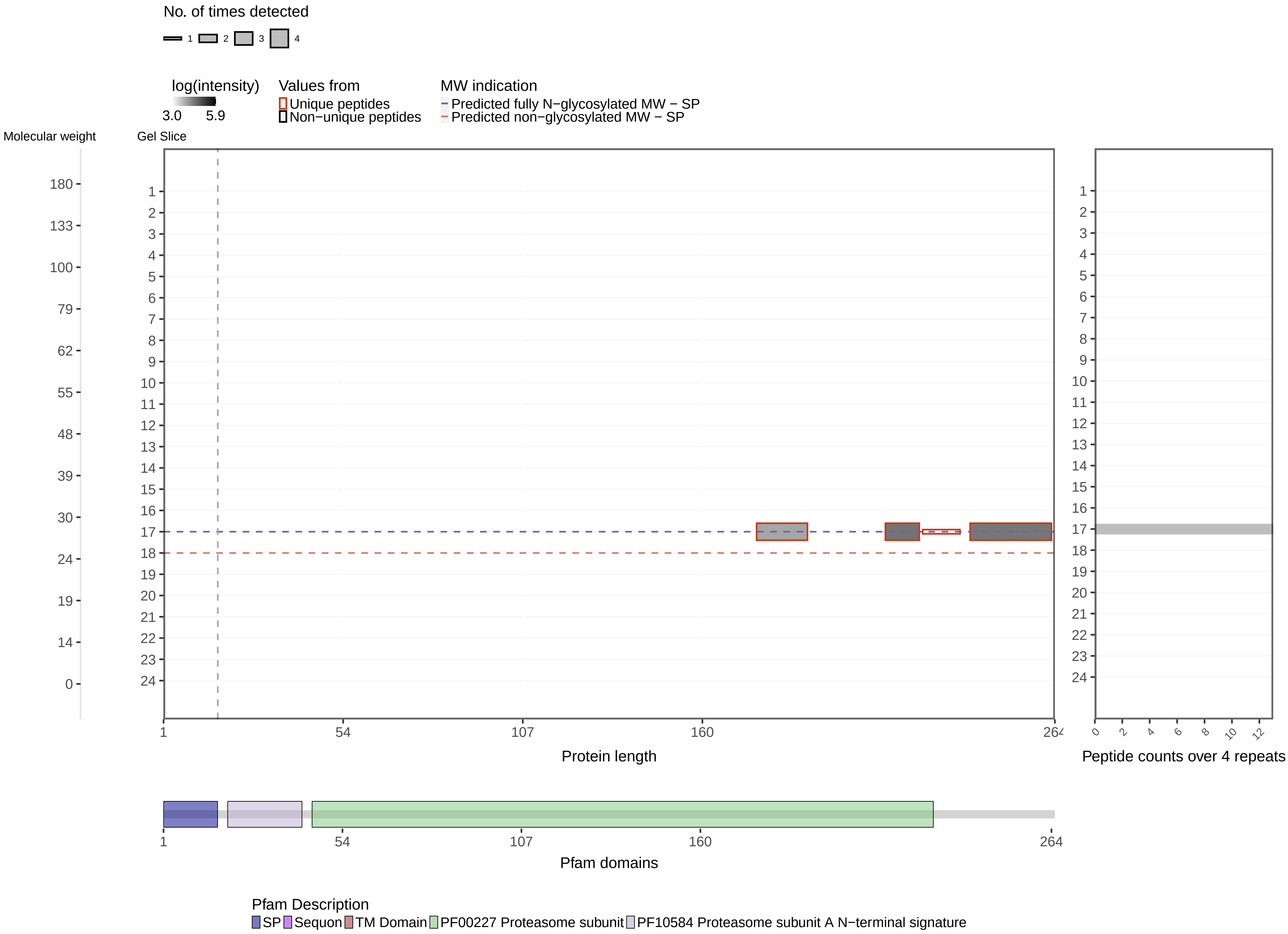


Protein sequence:

VEQLVQSLCDTKQGYTQFGGLRPFGVSFLFAGWDKNYGFQLYMSDPSGNYGGWKAAAIGANNQAAQSMLKQDYKDEITREEAVKLVLKVLSKTMDSTTLTSEKLELAEVFLFNGKVKYQVHSSESLNNLLVQSGLTQPAADS*

NbL05g06140.1 fructose−16−bisphosphatase_chloroplastic−like [MW = 44.84 kDa]

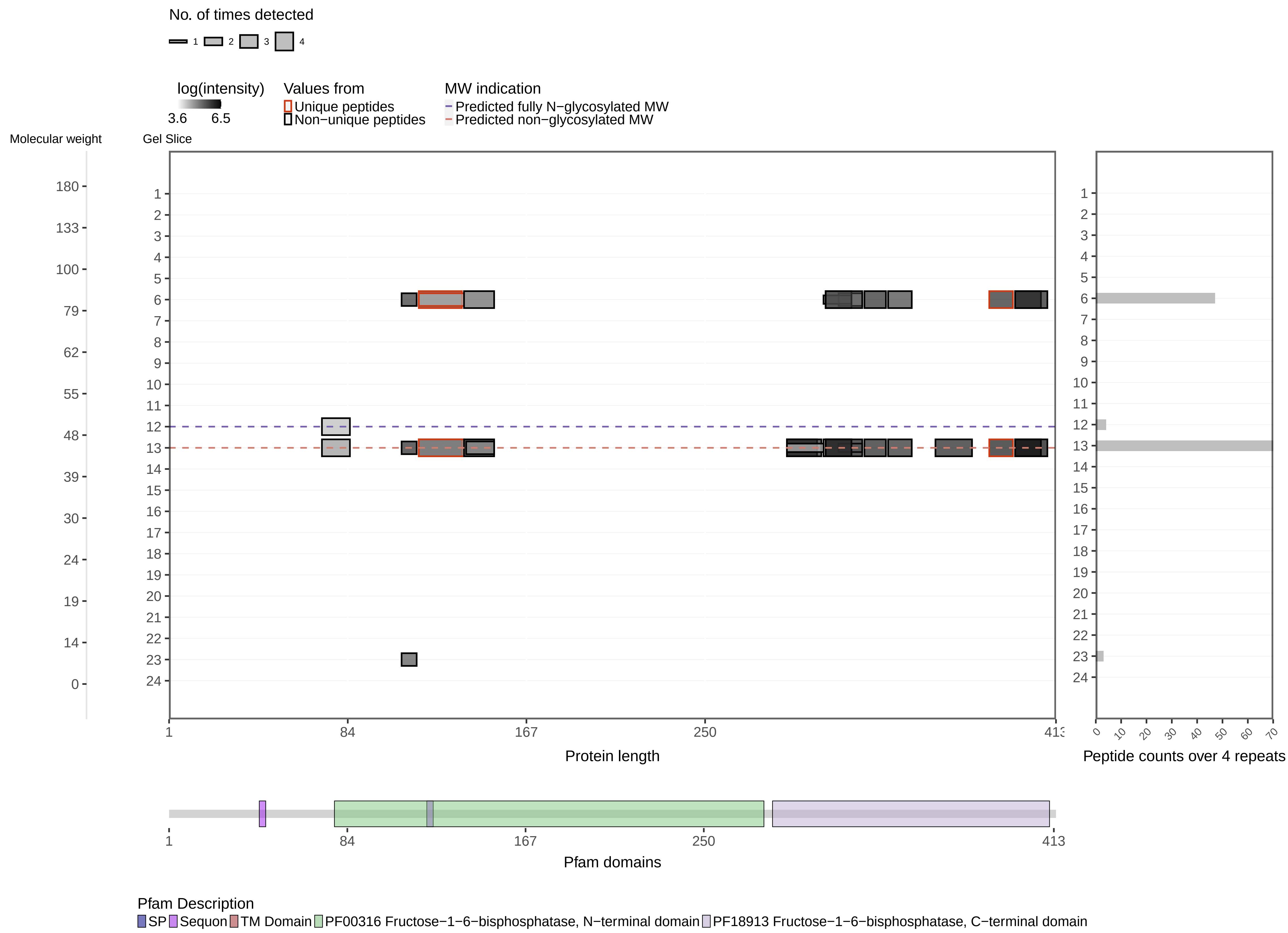


Protein sequence:

AAVSTGSIFGIYSPNDECLADLGDDSTLDNVEQRCIVNVCQPGSNLLAAGYCMYSSSVIFVVTLGNGVFAFNLDPMYGEFVLTQENIQIPKSGKIYSFNEGNYQLWDDKLKKYIDDLKDPGPSGKPYSARYIGSLVGDFHRTLLYGGIYGYPRDKKSKNGKLRLLYECAPMSFLVEQAGGKGSDGHQRVLDIQPTEIHQRVPLYIGSTEEVEKLEKYLS*

NbL05g06090.1 probable_glutathione_S−transferase [MW = 24.99 kDa]

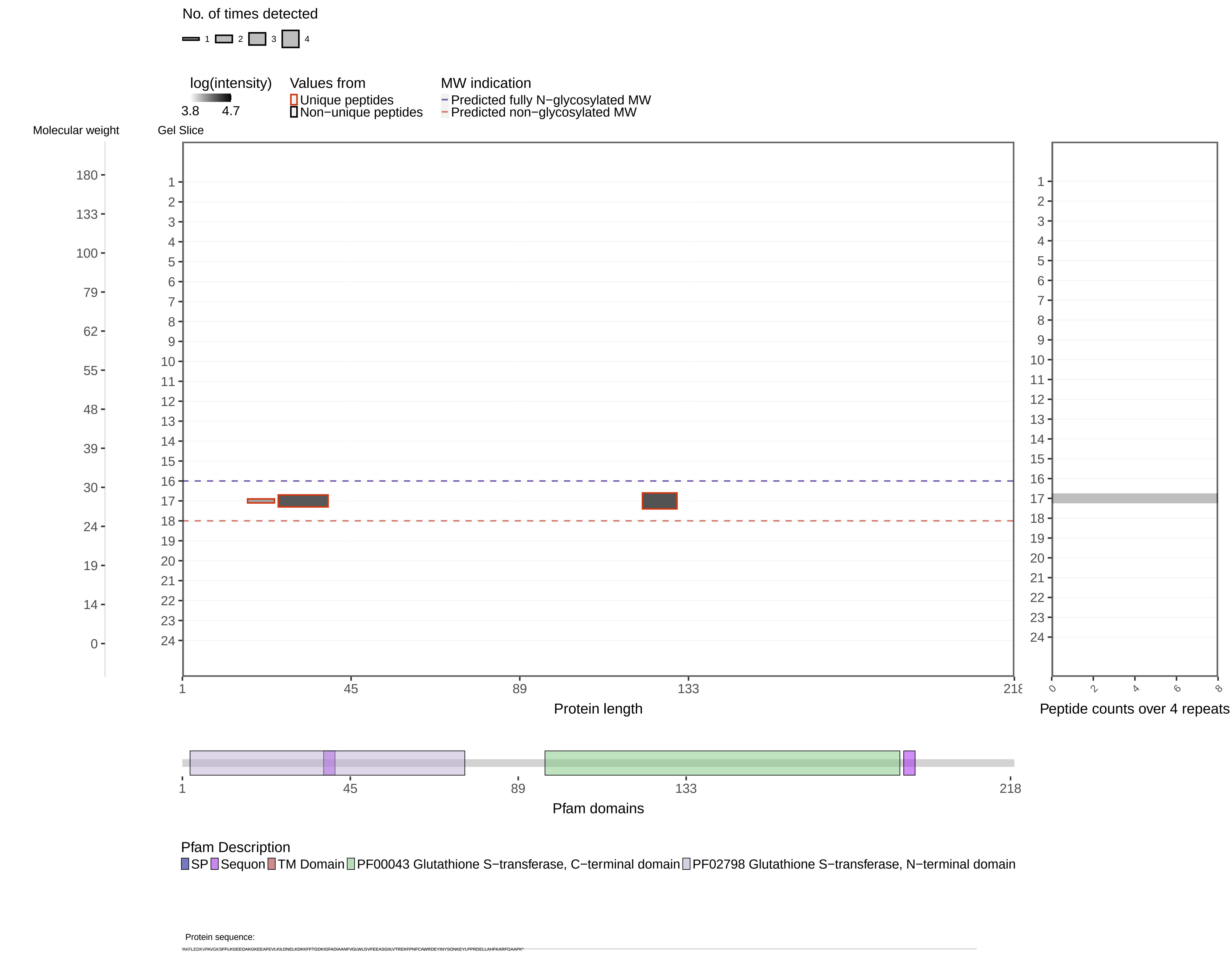

NbL05g05910.1 probable_proteasome_inhibitor [MW = 31.91 kDa]

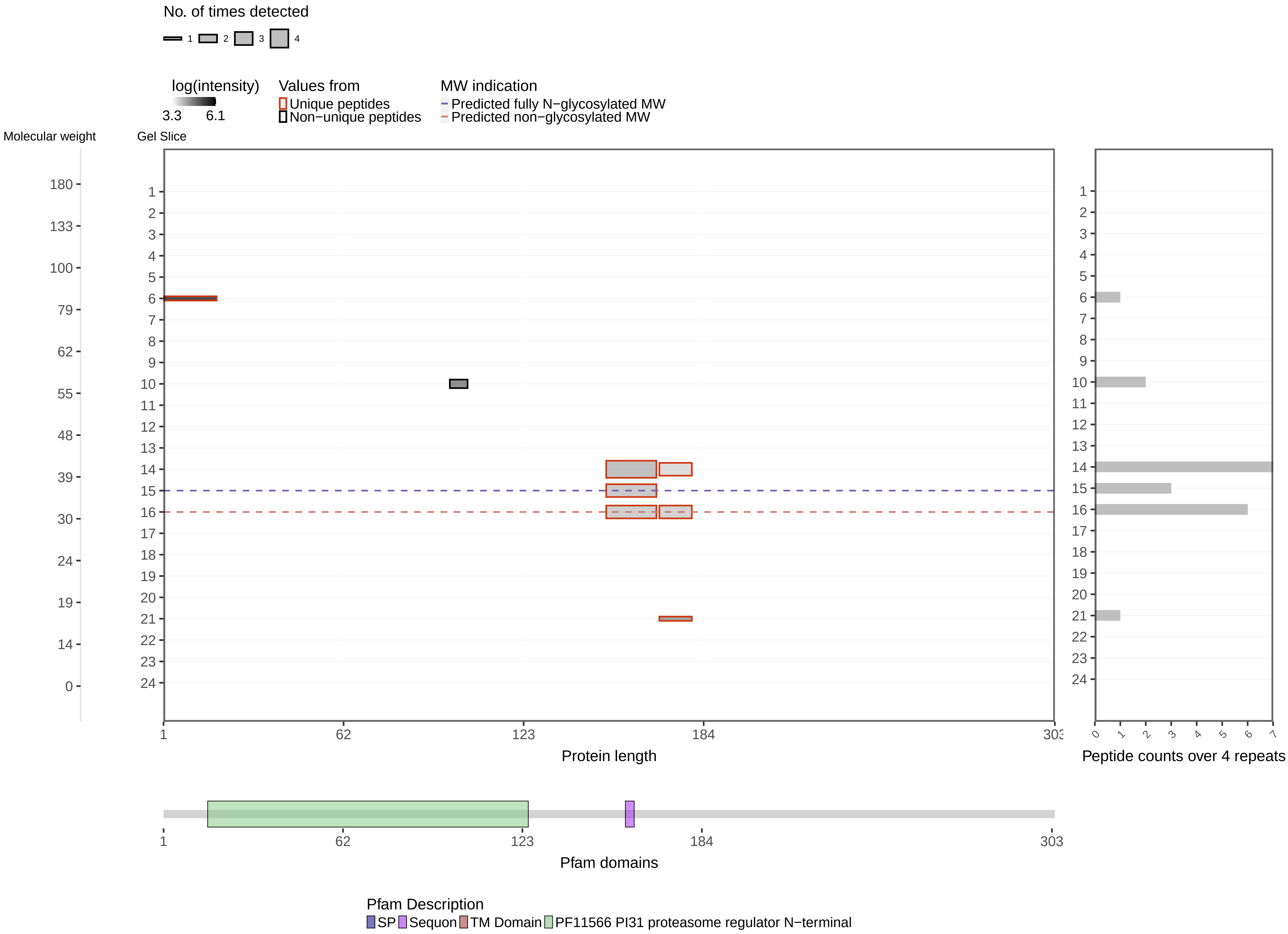


Protein sequence:

ELNKEVLSKYSGSSVANSSTQTSSFGKGGLEDQPSAELRSDYPYQPGSSFPSGYVVPPIPGSGGSDLFPGPGAGVYPTRGSPGGGSMLVGPNDPRFFGGMGGDPRLPGGLPGVLPGARFDPYGPSDVPGFEPGRFIRDPRRPGGGRPHPDLPHFGGDSDFI*

### NbL05g05840.1 uracil_phosphoribosyltransferase [MW = 30.85 kDa]

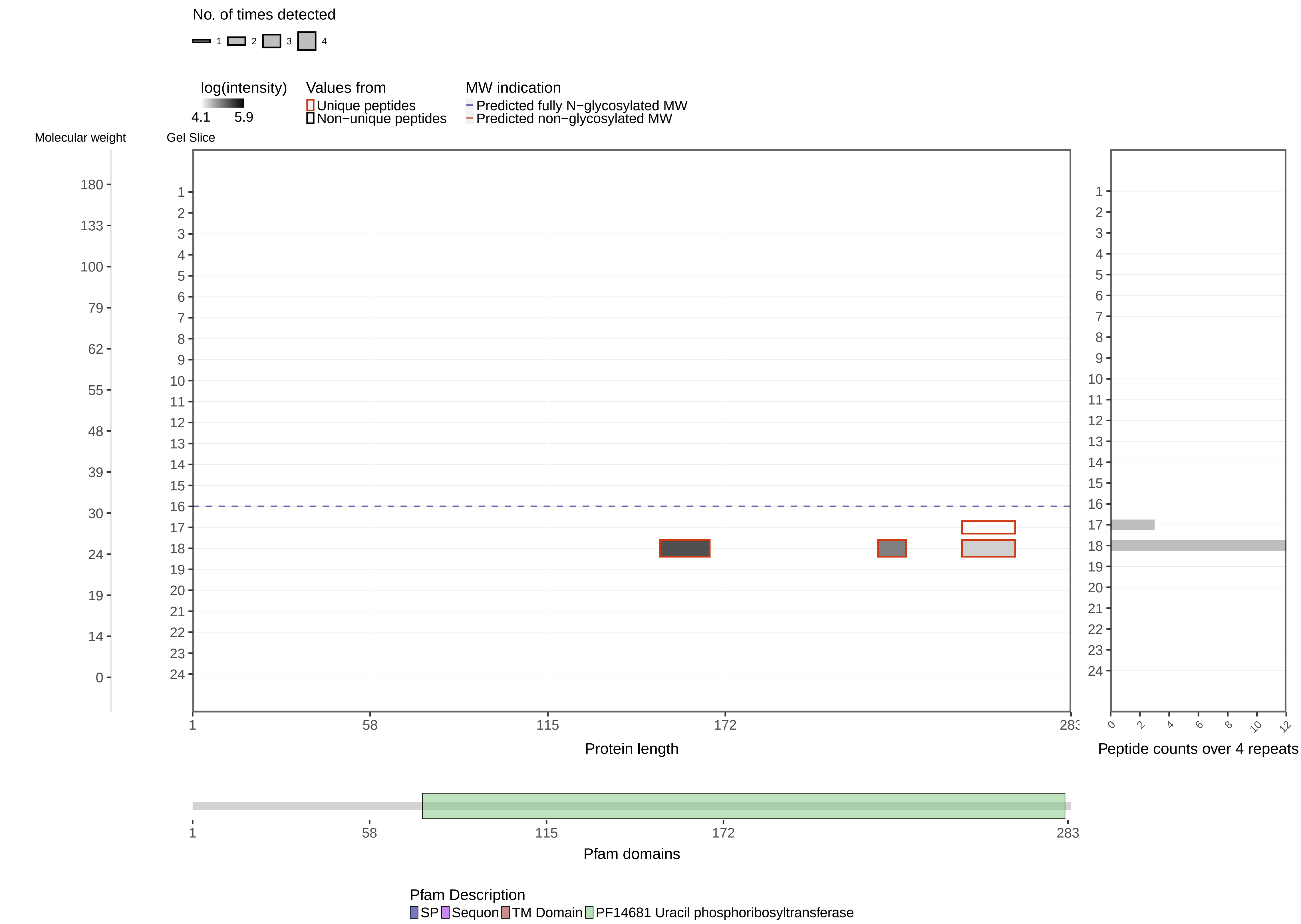


Protein sequence:

SPMGVASVEFVDPREPVAIVPILRAGLVLAEHASSILPATKTYHLGISRNEETLQPSVYLNKLPDKFPEGSRVFVVDPMLATGGTIVAAIDLIKEHGVDNSQIKVICAVGAPPALQKLSEKFPGLHVYAGILDPTVNDKGFIIPGLGDAGDRSFGT*

NbL05g05670.1 ribulose_bisphosphate_carboxylase/oxygenase_activase_1_chloroplastic [MW = 52.67 kDa]

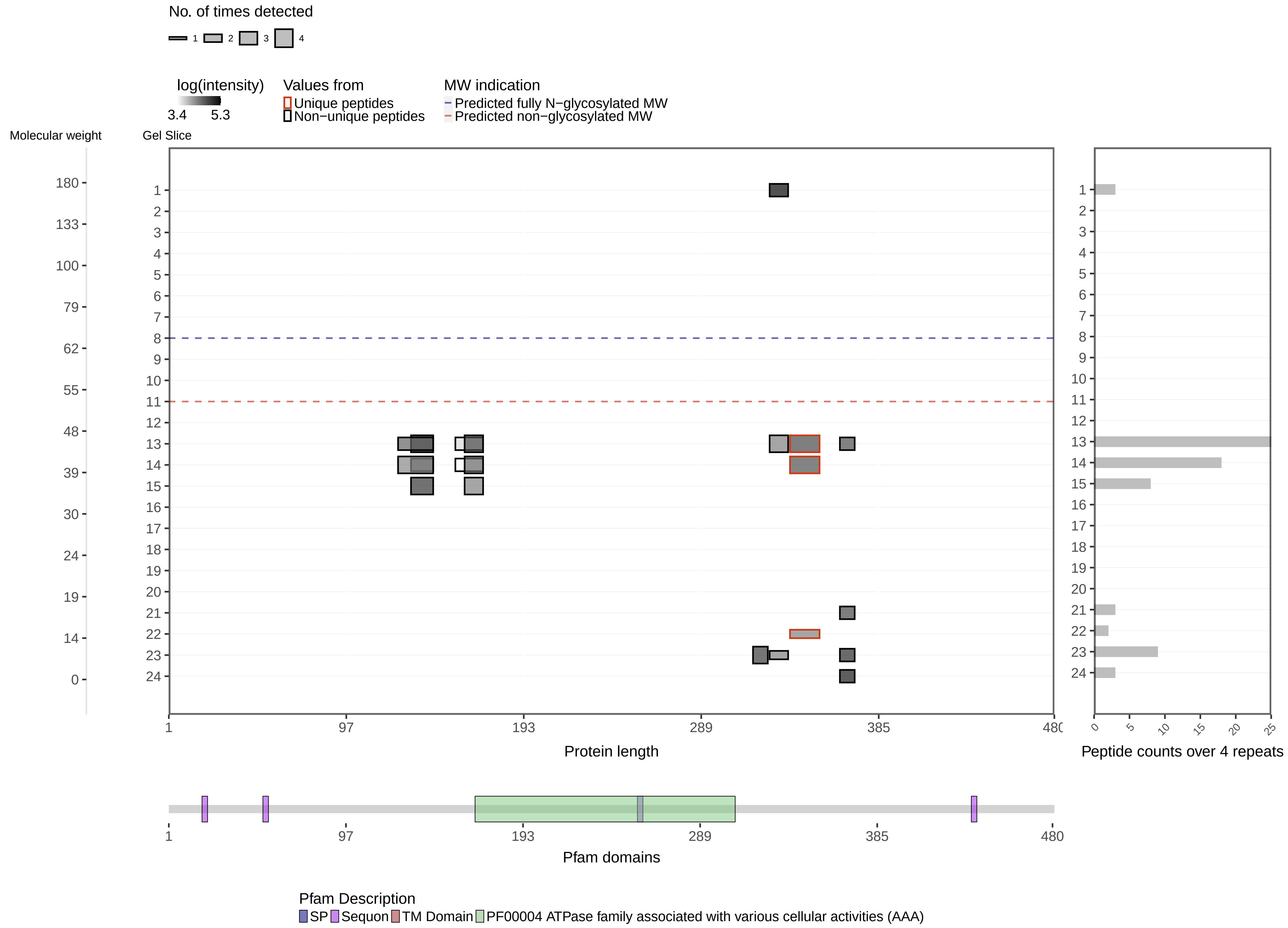


Protein sequence:
CLFINDLDAGAGRMGGTTQYTVNNQMVNATLMNIADNPTNVQLPGMYNKQENARVPIIVTGNDFSTLYAPLIRDGRMEKFYWAPTREDRIGVCTGIFRTDNVPAEDVVKIVDSFPGQSIDFFGALRARVYDDEVRKWVSGTGIEKIGDKLLNSFDGPPTFEQPKMTIDKLLEYGNMLVQEQENVKRVQLADKYLKEAALGDANADAINNGSFFTREEAQHADVPVGEGSTHSIATKLDSTSRAEDGSYLFKF*

NbL05g05510.1 uncharacterized_protein_At5g02240−like [MW = 32.25 kDa]

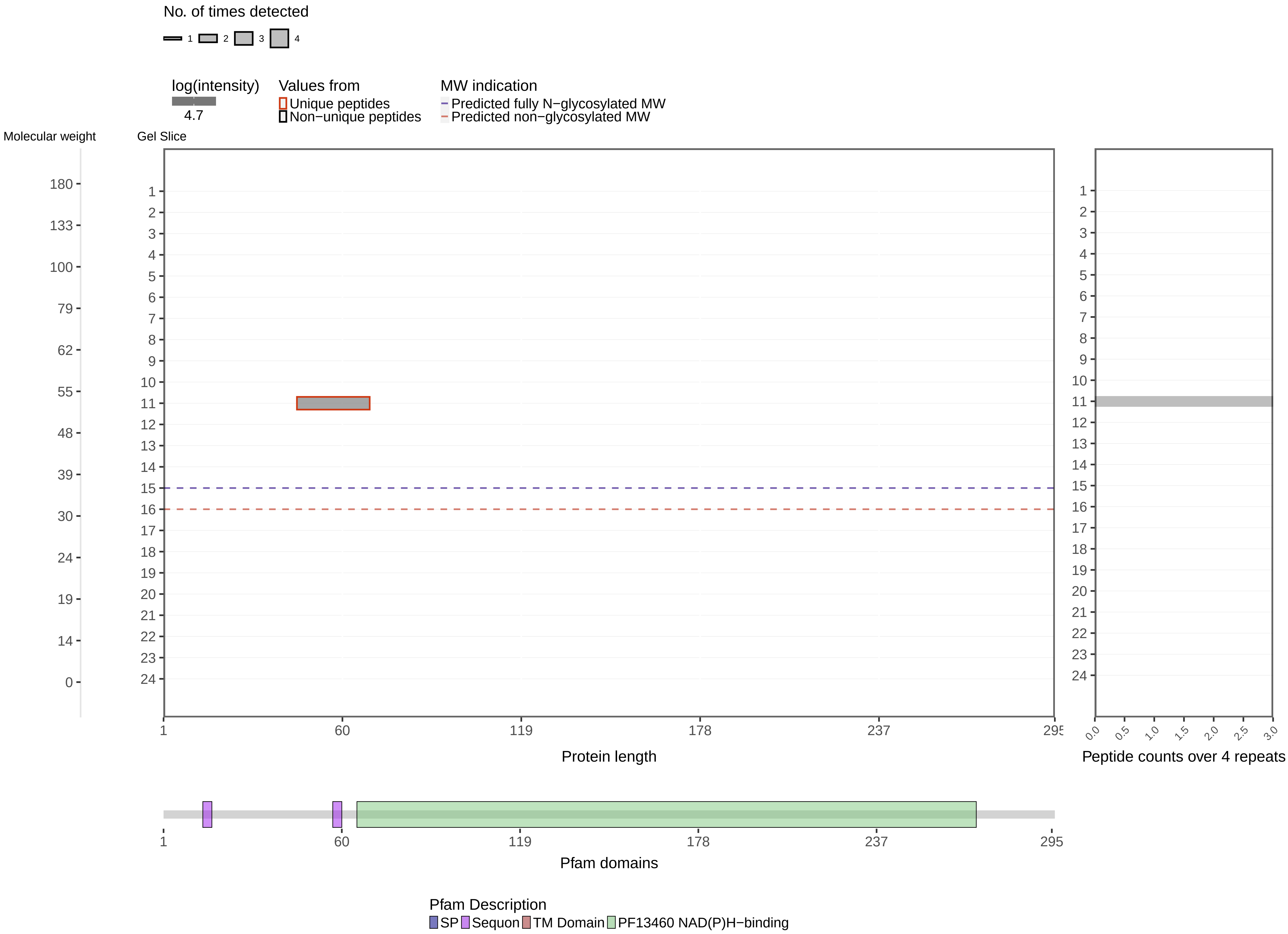


Protein sequence:

MKPGFDPTOGGRPEFYFEDGAYPEOVDWIGOKNOIDAAKAAGVKOIVLVGSMGGTNPNHPLNSIGNGNILVWKRKAEOYLADSGIPYTIIRAGGLODKEGGIRELLVGKDDELLOTETRTIARADVAEVCIOLLFVLYRDVCCFFDOLOCFTPTTHFL*

NbL05g05450.1 ribonuclease_II_chloroplastic/mitochondrial [MW = 92.73 kDa]

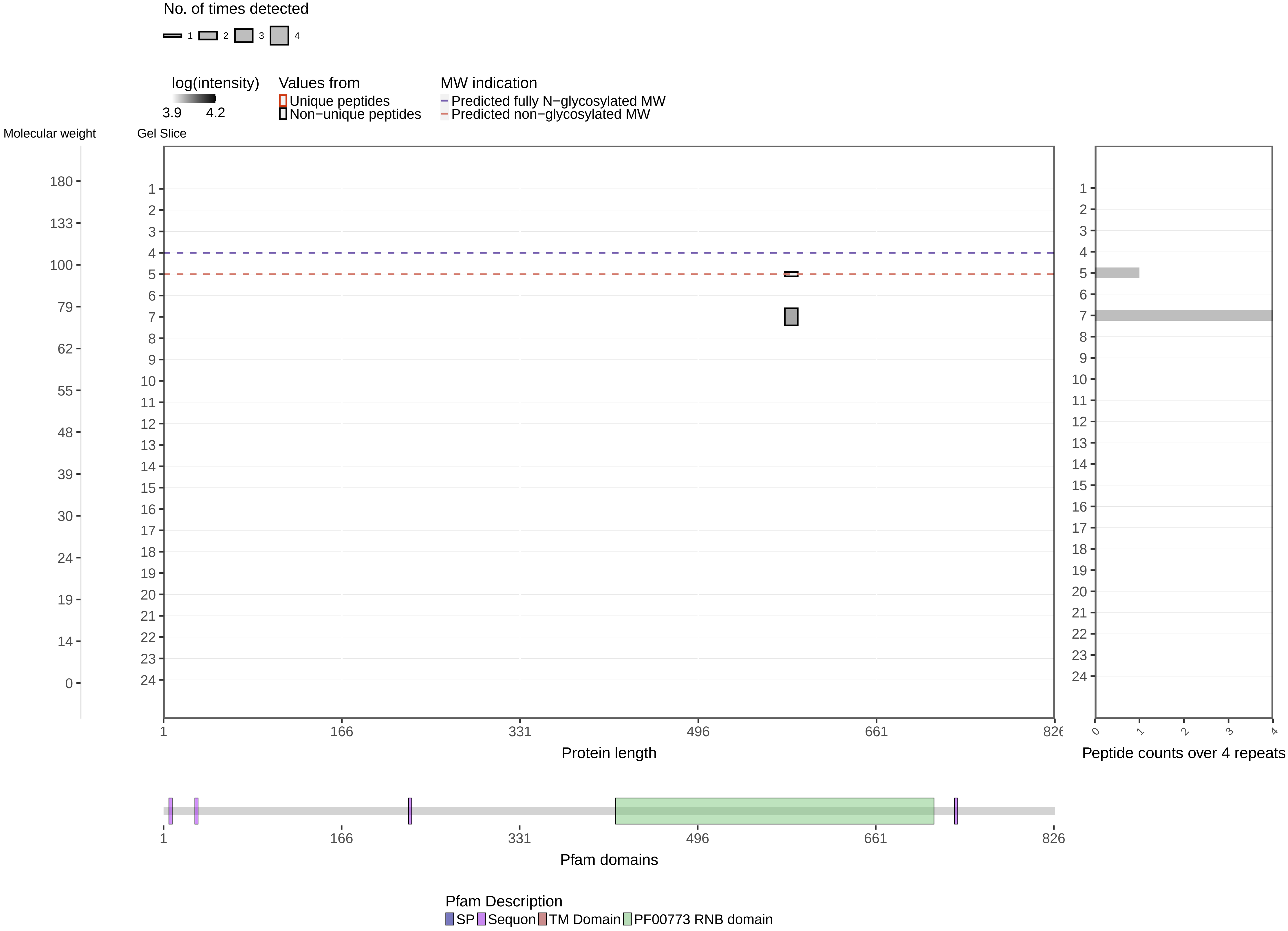


Protein sequence:

RDEILSVAESILSSPTDLDEADRVDLTDLKVYAIDVDEADELDDALSATRLQDGRIKVWIHVADPTSLLQPGSIIDKDARRRGTSIFLPTATYPMFPERLAMEGMSLKQGRLCNAVTVSVVLRSDGSIAEYSVENSIIKPTYMLTYESATELLHLNLEEEIELKVLSEAAALRLRWRREQGAIDTGTLETRIKVTNPDHPEPSIKLYVENQADAAMRLVSEMMVLCGEVIATYGSHNNIPLPYRGQPQSNIDTSAFEHLPEGPVRSSAIVRIMRAAEMDFRNPVRHGVLGLPGYVQFTS

### NbL05g05440.1 expansin−A3−like [MW − SP = 25.59 kDa]

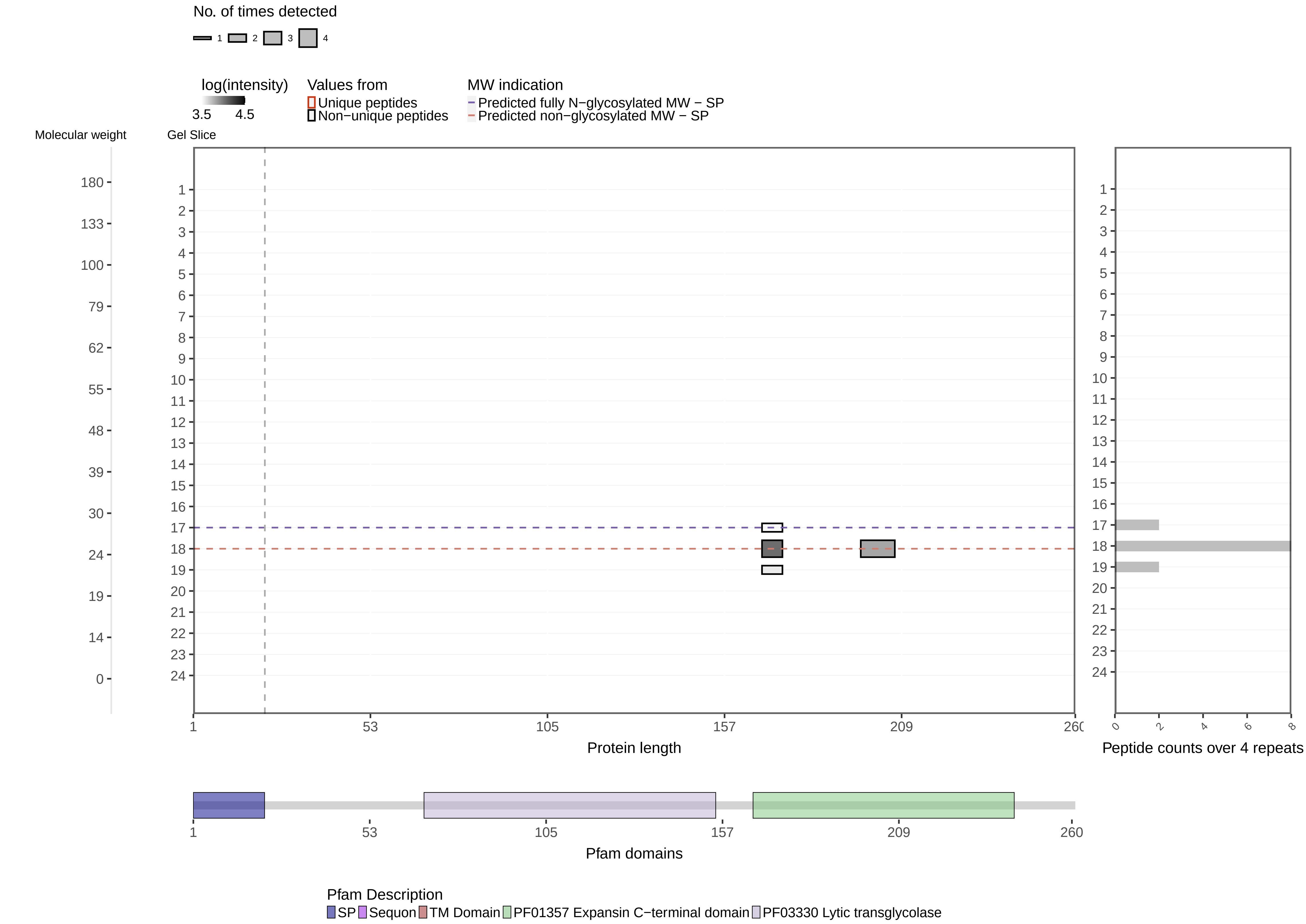


Protein sequence:

.PNDDGGWCNPPRPHFDLAMPMFLKIGQYRAGIVPVTYRRVPCRKAGGIRFTINGFRYFNLLLVTNVAGAGDIQKVLIKGTNTQWIAMSRNWGQNWQTNAVLVGQALSIRVKASDRPSVTSNNVAPPNWQFGQTFEGKNFRV*

### NbL05g05270.1 shock_cognate_70_kDa_protein_2−like [MW = 71.1 kDa]

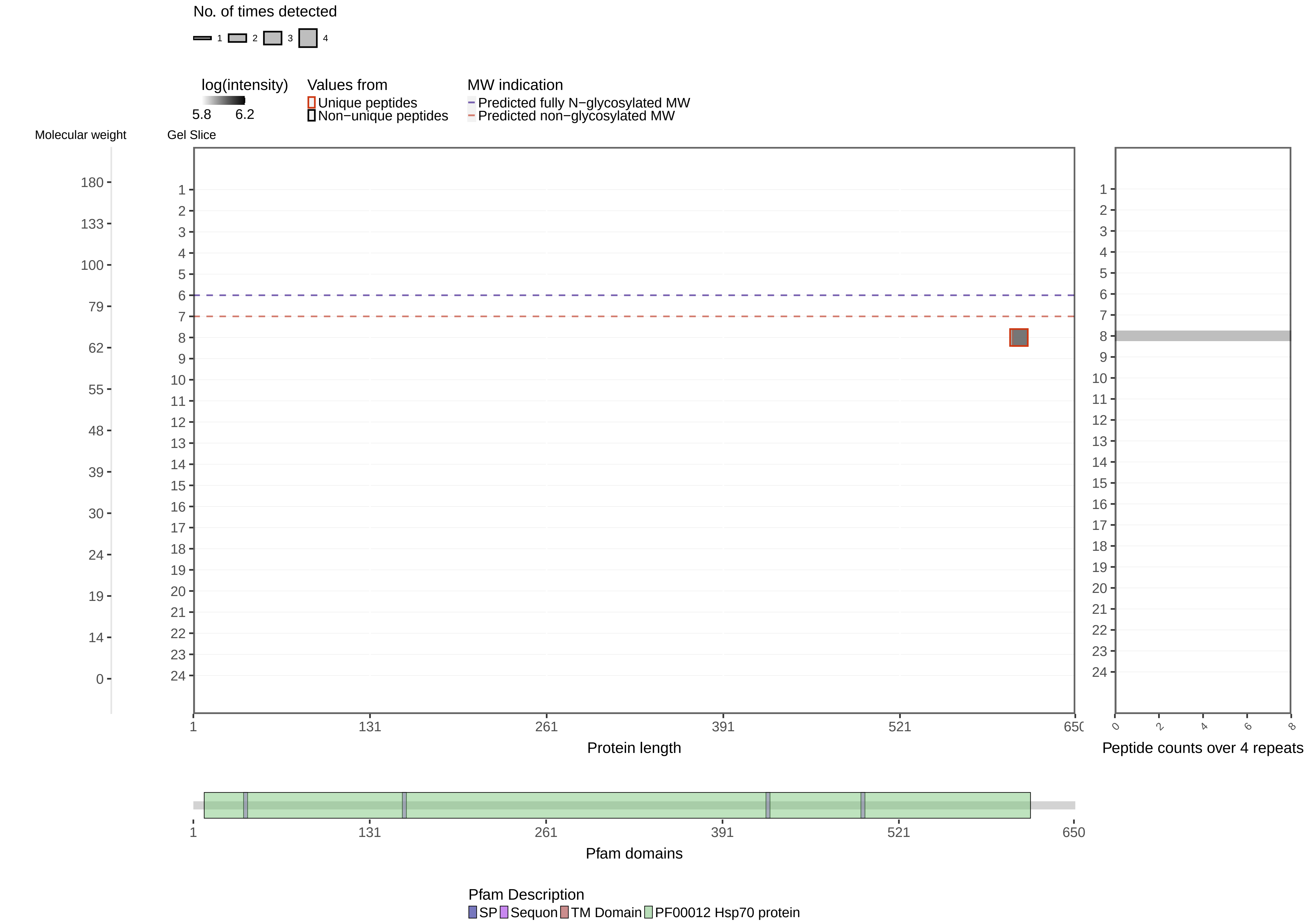


Protein sequence:

LFRKCMEPVEKCLRDAKMDKSTIHDVVLVGGSTRIPKVQQLLQDFFNGKELCKSINPDEAVAYGAAVQAAILSGEGNEKVQDLLLLDVTPLSLGLETAGGVMTVLIPRNTTIPTKKEQVFSTYSDNQPGVLIQVYEGERTRTRDNNLLGKFELSGIPPAPRGVPQITVCFDIDANGILNVSAEDKTTGQKNKITITNDKGRLSKEEIEKMVQEAEKYKSEDEEHKKKVEAKNALENYAYNMRNTVKDEKIASKLPEADKKKIEDAIEAAIQWLDANQLAESDEFEDKMKELESVCN

### NbL05g05150.1 5−oxoprolinase−like [MW = 137.99 kDa]

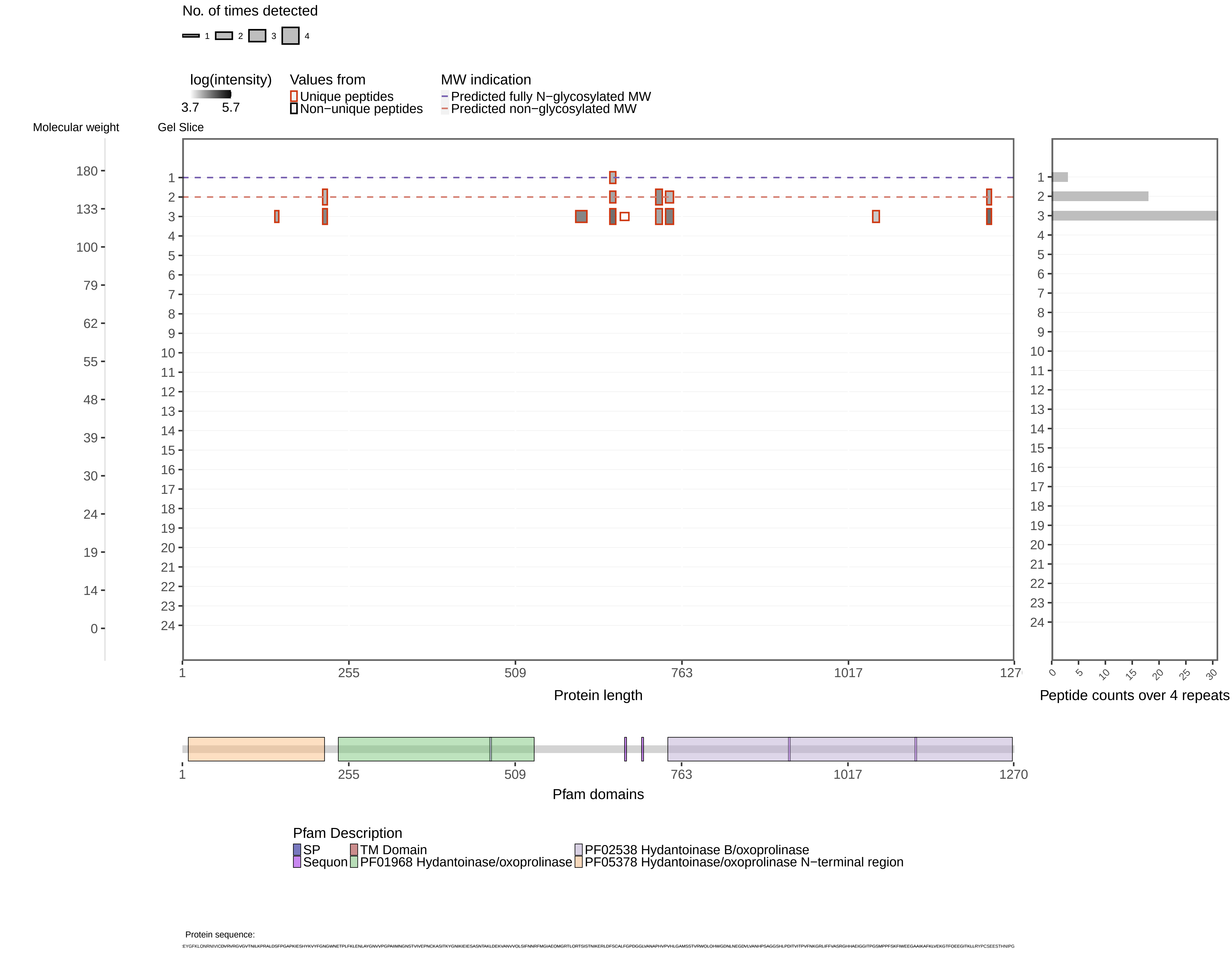

NbL05g05030.1 eukaryotic_translation_initiation_factor_3_subunit_A [MW = 111.63 kDa]

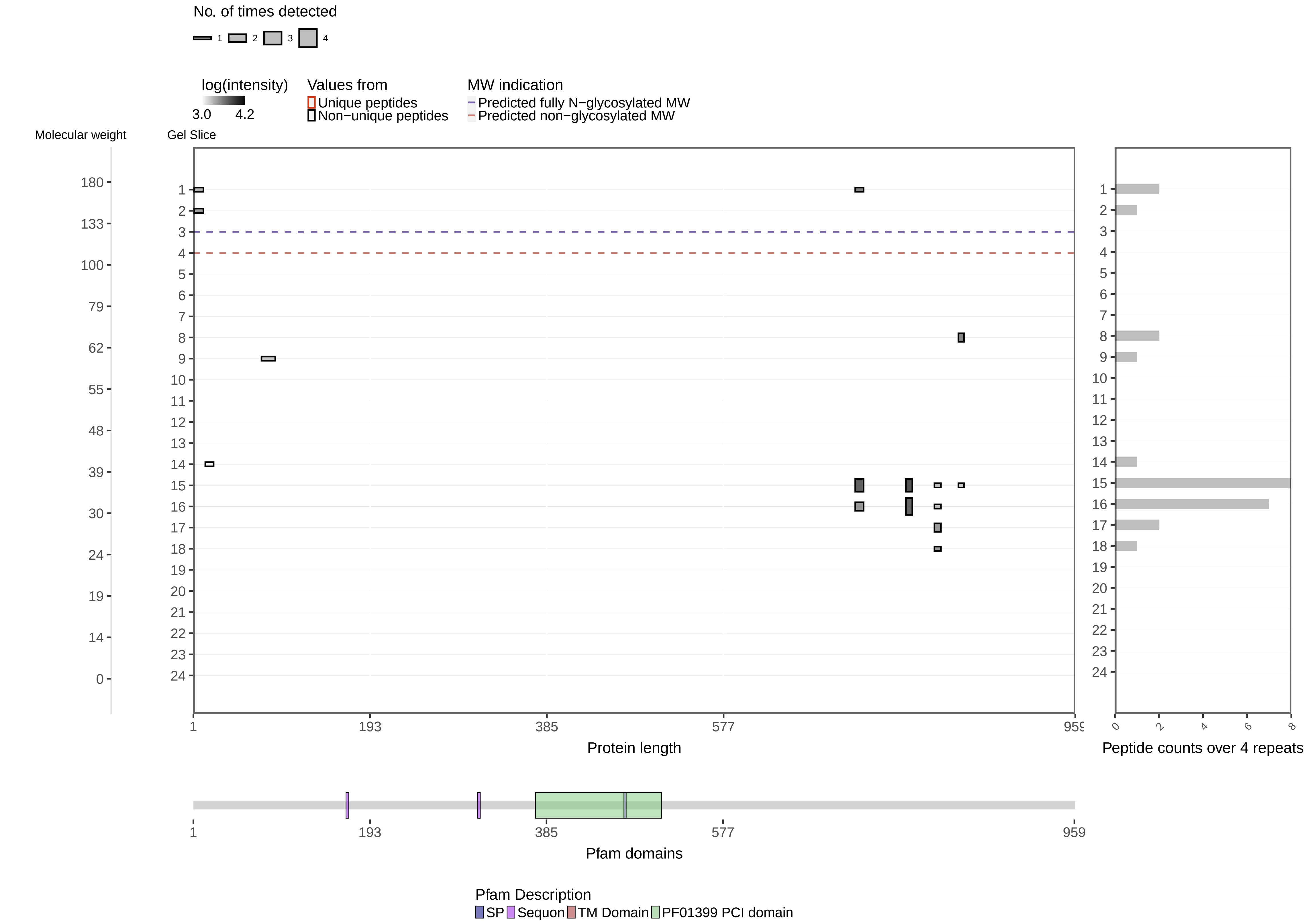


Protein sequence:

NISKMIPFFDFTVIEKISVDAVRRNFLAIKVDHMKGAVFFGKQSIEAEGLRDHLSLLAESLSKARTMIYPPAKKAAKLGEALSNLAEIVEKEHKRLLARKSIIEKRKEEQERLLLEMERVEETKRRELQKMTEEAEQKRIAVEYEQRRNQRILKEIEDRELEEAQALLHEAEKRSKRKKKPVLEGEKMTKQVIMELALNEQLRERQEMEKKLLKFAKTMDHLERAKREEAAPLIESAFKQRLAEEAALHEREQQQEIELSRQRHAGDLEEKRRLGRMLENKRILQEKVVSSREAE

### NbL05g05010.1 actin−depolymerizing_factor_1 [MW = 15.95 kDa]

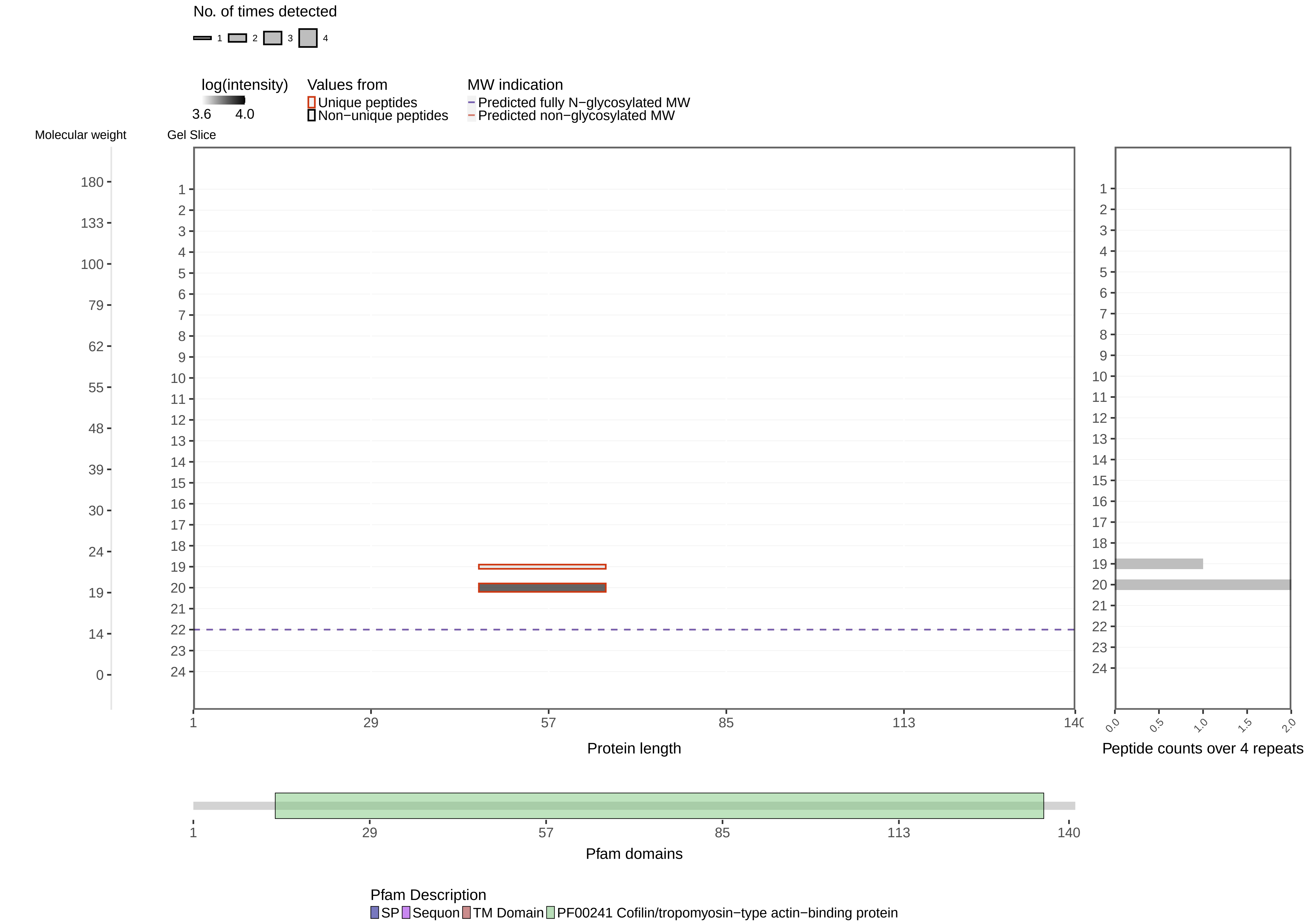


Protein sequence:

AASLPDNECRYAVYDFDFVTAENCQKSRIFFIAWSPDTSRVRSKMIYASSKDRFKRELDGIQVELQATDPTEMGLDVIKSRAN*

### NbL05g04990.1 arginine_biosynthesis_bifunctional_protein_ArgJ_chloroplastic [MW = 49.27 kDa]

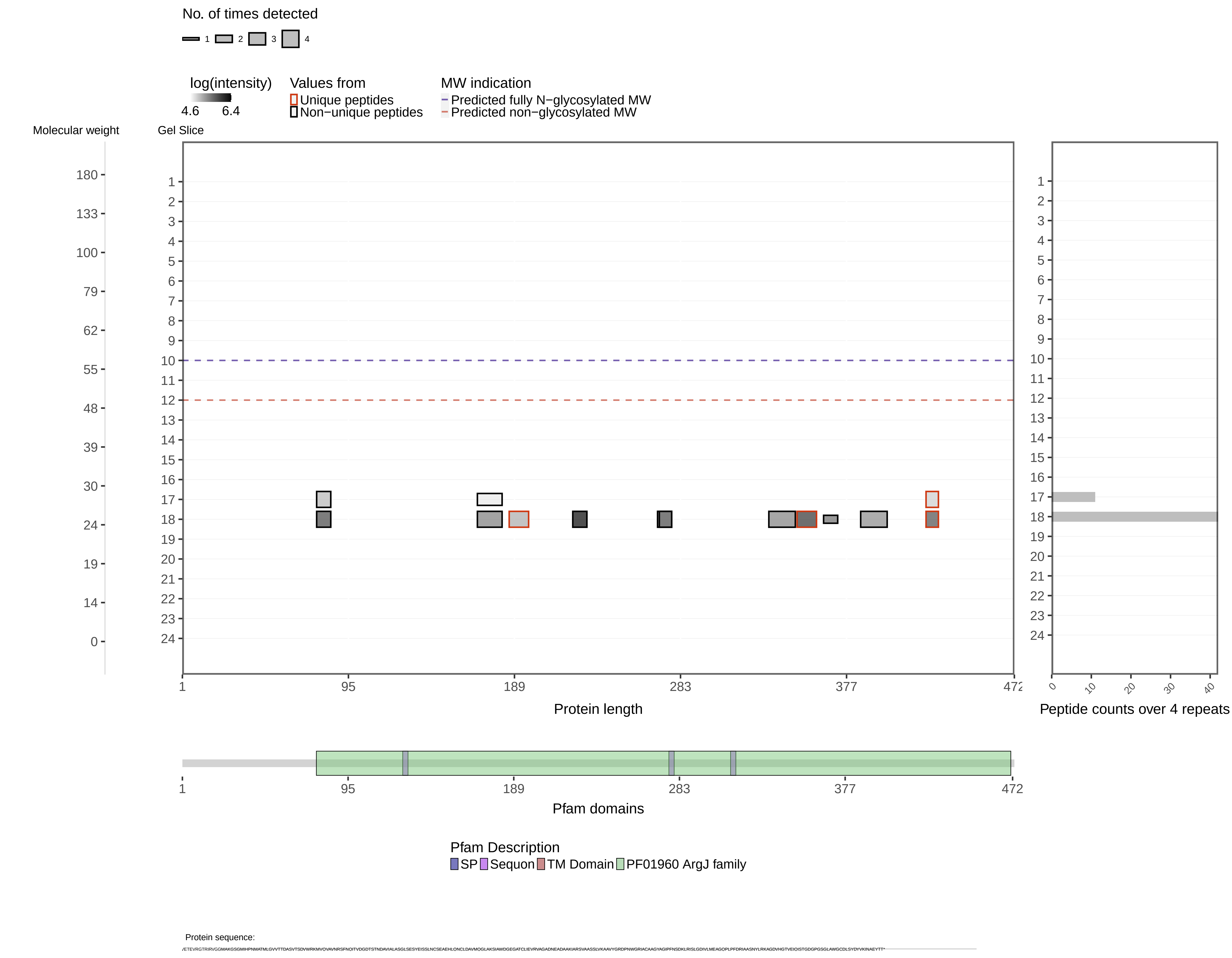

### NbL05g04950.1 soluble_inorganic_pyrophosphatase−like [MW = 24.46 kDa]

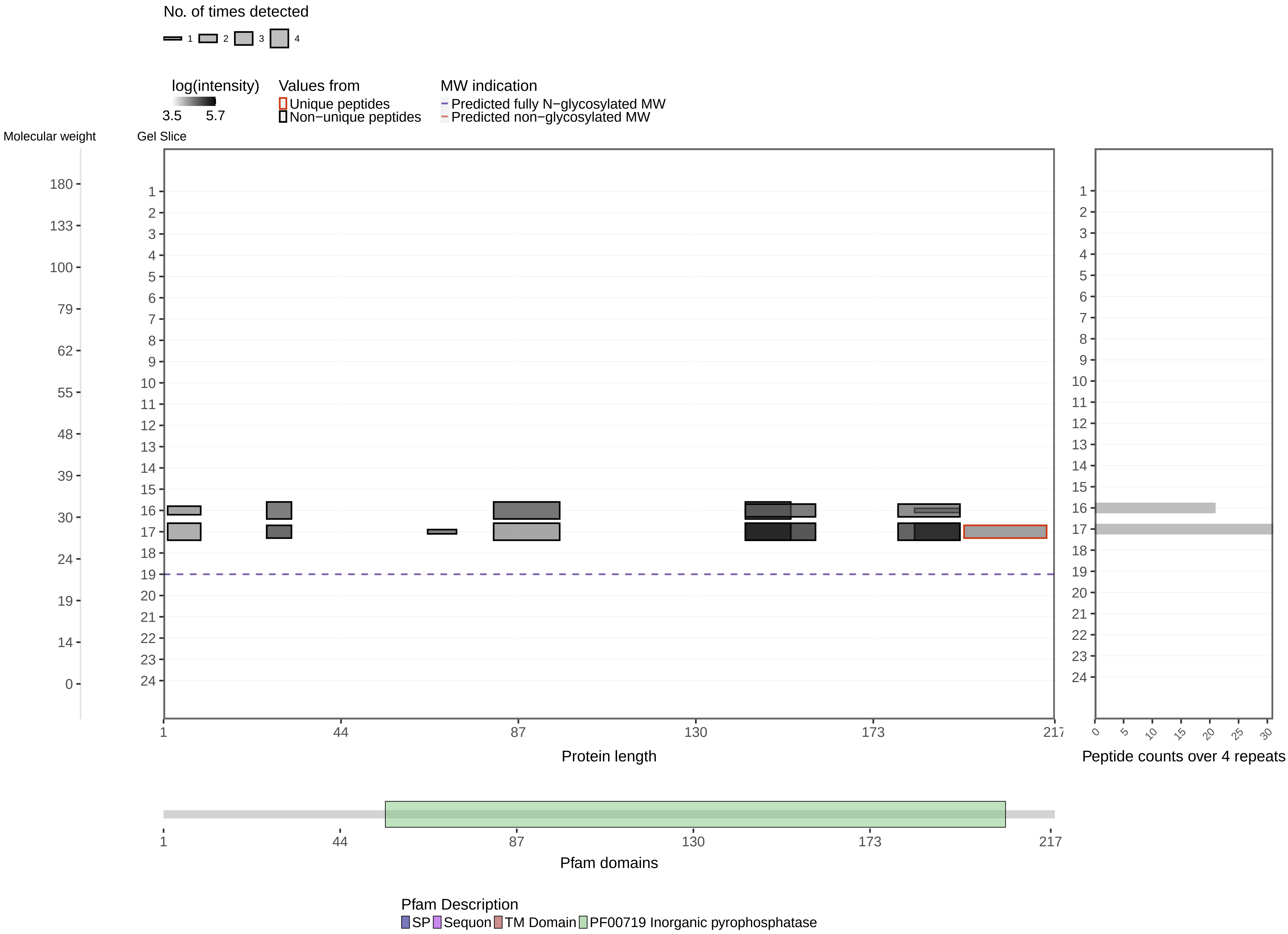


Protein sequence:

IPRTLCEDSDPLDVLVIMQEPVLPGCFLRAKAIGLMPMIDQGEKDDKIIAVCADDPEYKEYTDIKELPPHRLAEIRRFFEDYKKNENKEVAVNDFLPATKAFEAVQHSQDLYADYIVESLRR*

### NbL05g04920.1 ras−related_protein_RABE1c−like [MW = 23.99 kDa]

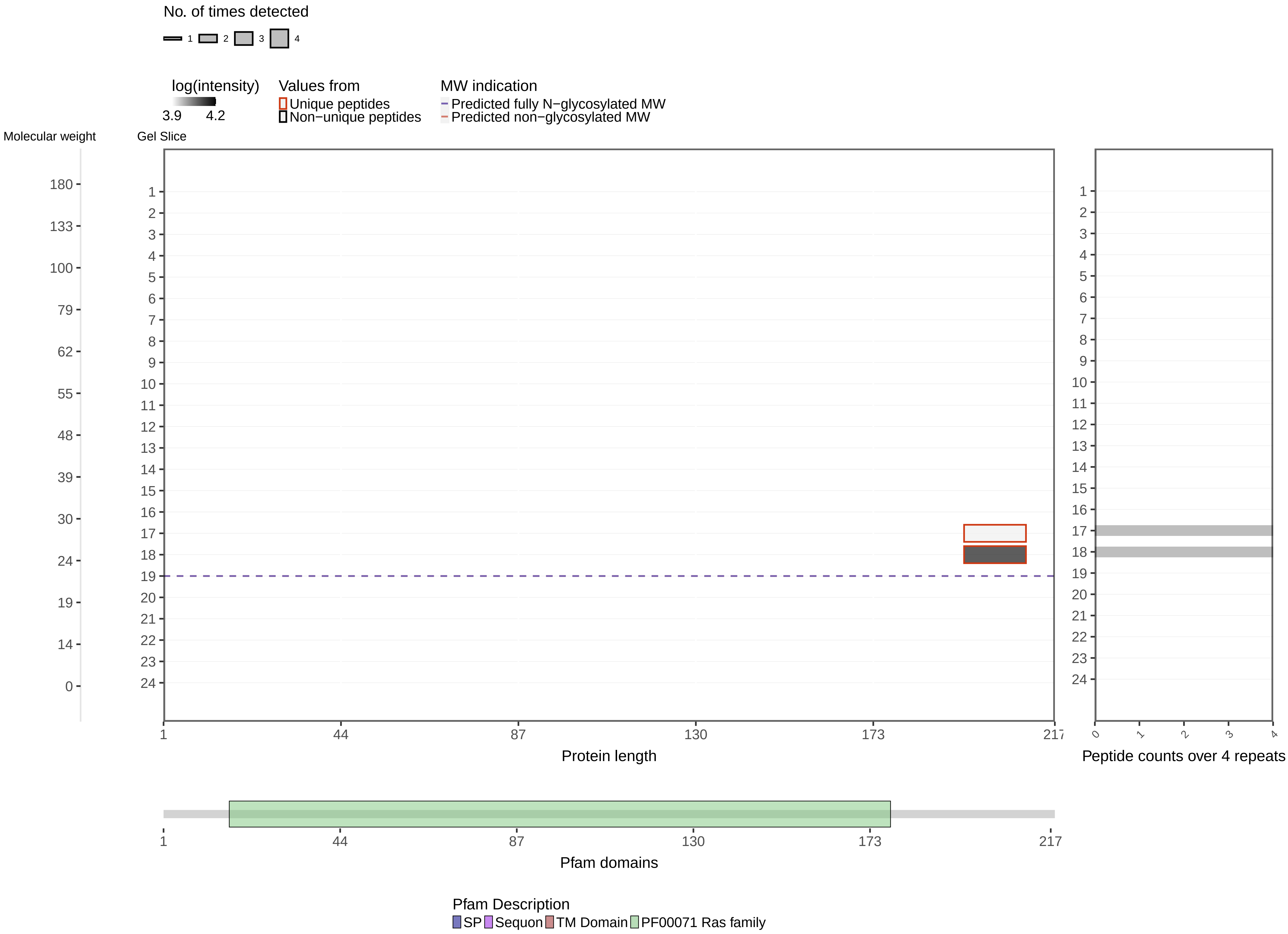


Protein sequence:

DESSFNNIRNWIRNIEQHASDNVNKILVGNKADMDESKRAVPTSKGQALADEYGIKFFETSAKTNMNVEEVFFSIARDIKQRLAESDSKAEPQTIRINQPDQAAGAAQSAQKSACCGS*

NbL05g04890.1 diaminopimelate_epimerase_chloroplastic−like [MW = 39.77 kDa]

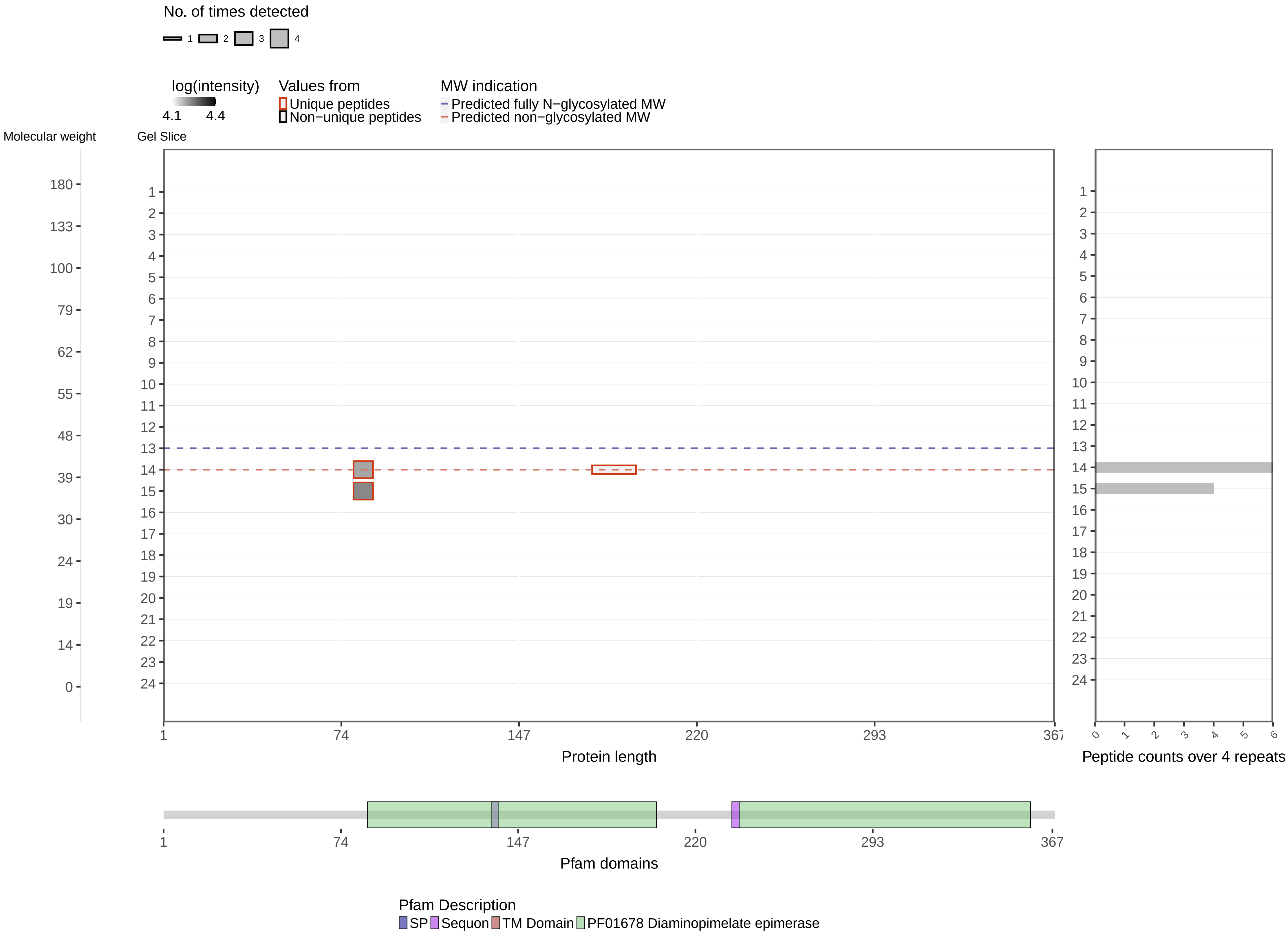


Protein sequence:

RGKRSFTIHTGGGLIVPEIQEDKVRVDMGEPILKASDVPTKLPPNRDQSVVKARLDVDGTAWNVTCVSMGNPHCVTFGTEQSQDLQVDELNLADIGPKFEHHVMFPARTNTEFVQVFSPTHLKMRVWERGAGVTLACGTGACAVVVAAVLEDRAARRCTVDLPGGPLDIEWSEKDNHIYMTGPAEVVFYGSVPL*

NbL05g04860.1 protein_SLOW_GREEN_1_chloroplastic−like [MW = 40.07 kDa]

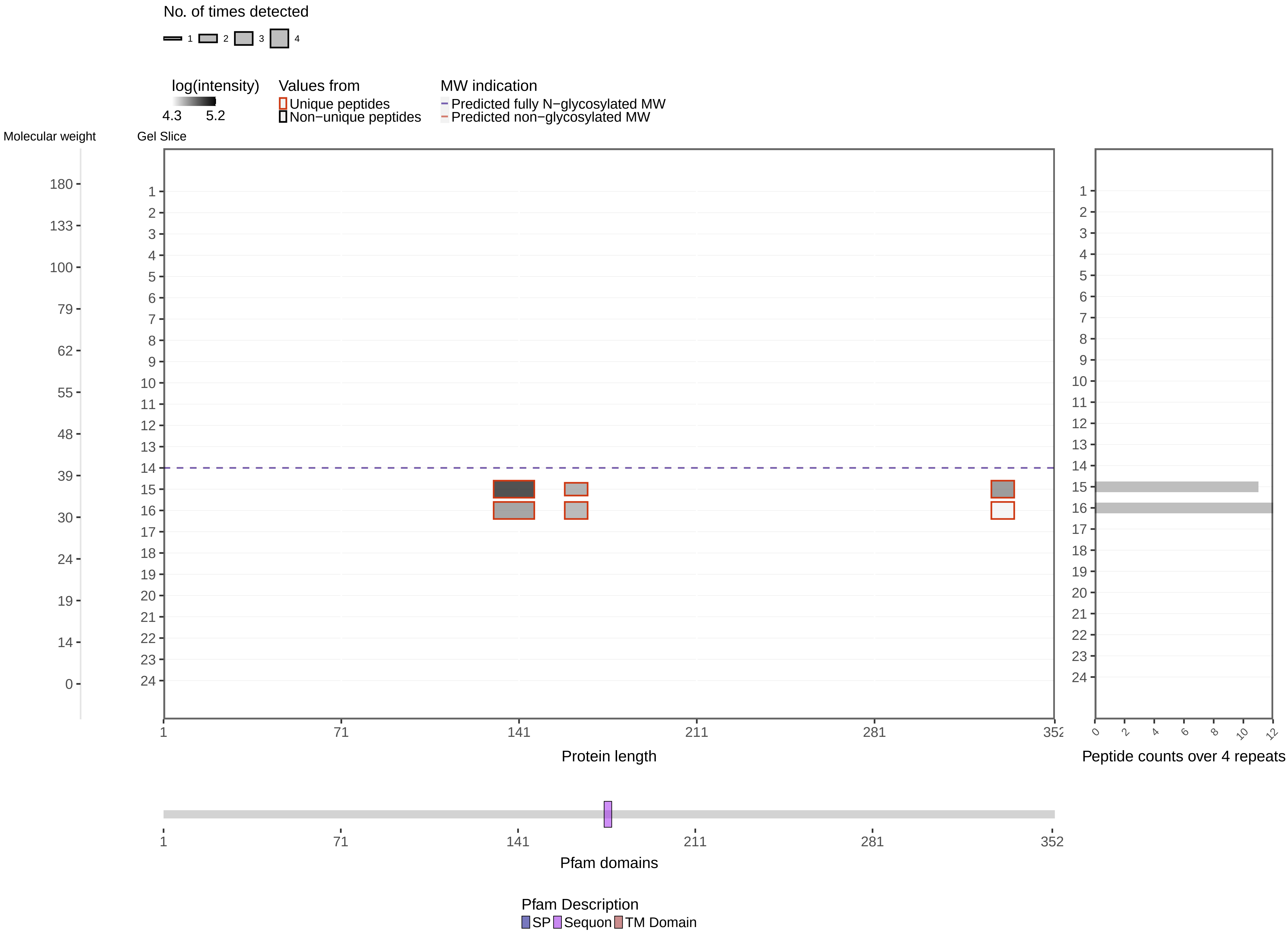


Protein sequence:
IIDKLIEVEPNESEWPLMKSHLYVNIGEVELAKVGFNEIIKKDPFRVEAYHGLVMAASQDESIEDLKGIEKMIEEGMKLCKKENKKTDLRDFKLLLAQIRVIEGKYEDALKVYGELVKEEPRDFRPYLCQGIIYTLLRKSSDAEKCFEKYRRLVPQGHPYARYFDENMIATKVFAQKVENERAGSKS*

### NbL05g04800.1 UDP−glucuronic_acid_decarboxylase_1−like [MW = 48.72 kDa]

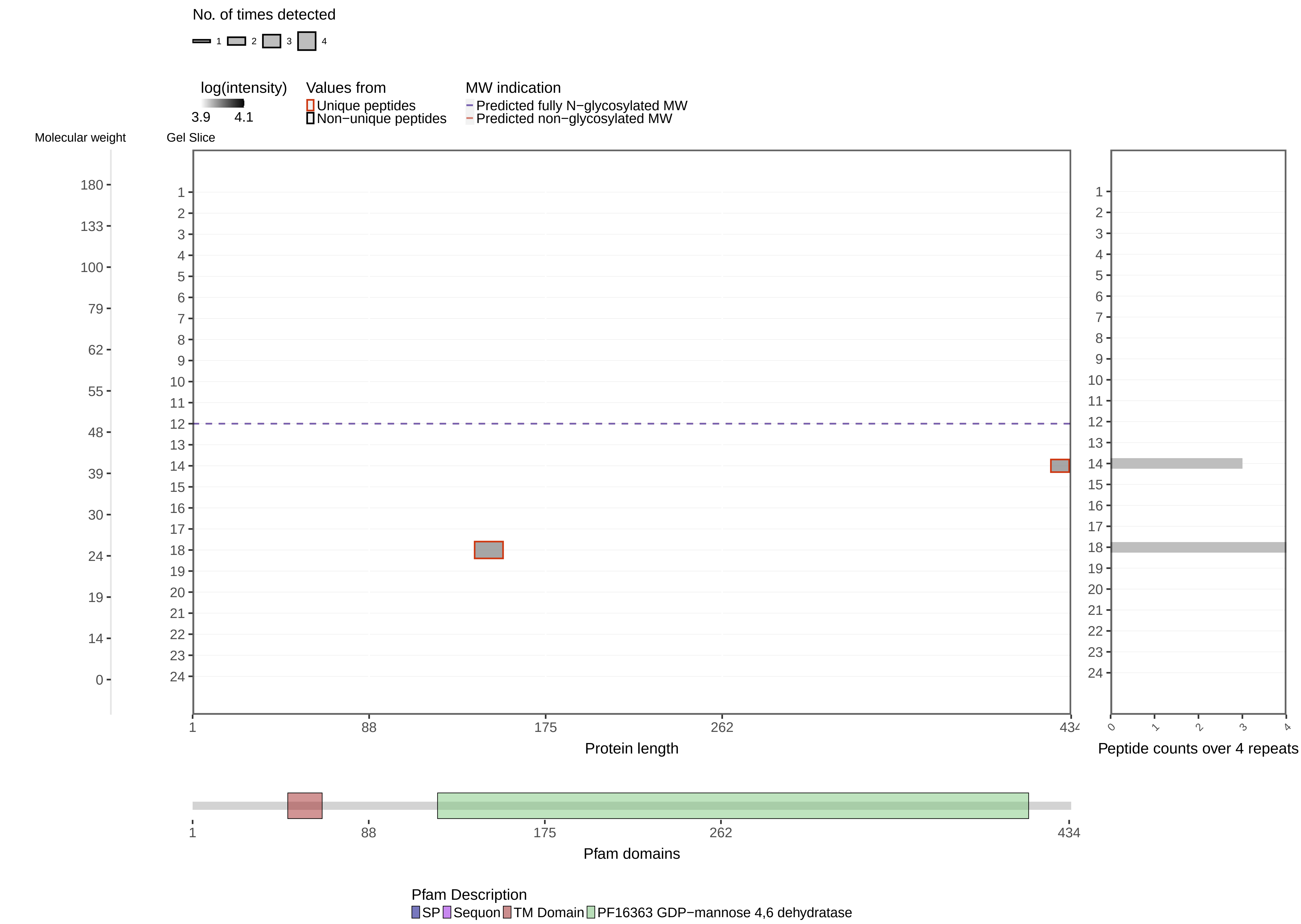


Protein sequence:

VMGTLNMLGLAKRIGARFLLTSTSEVYGDPLEHPQKETYWGHVNPIGVRSCYDEGKRTAETLTMDYHRGAGVEVRIARIFNTYGPRMCLDDGRVVSNFVAQAIRNQPMTVYGDGKQTRSFQYVSDLVDGLVALMEGEHIGPFNLGNPGEFTMLELAEVVKEVIDPSAKIEFRANTADDPHKRKPDISKAKELLNWEPKVPLREGLPLMVNDFRNRILNEDEGKEN*

NbL05g04580.1 protein_MODIFIER_OF_SNC1_11−like [MW = 21.42 kDa]

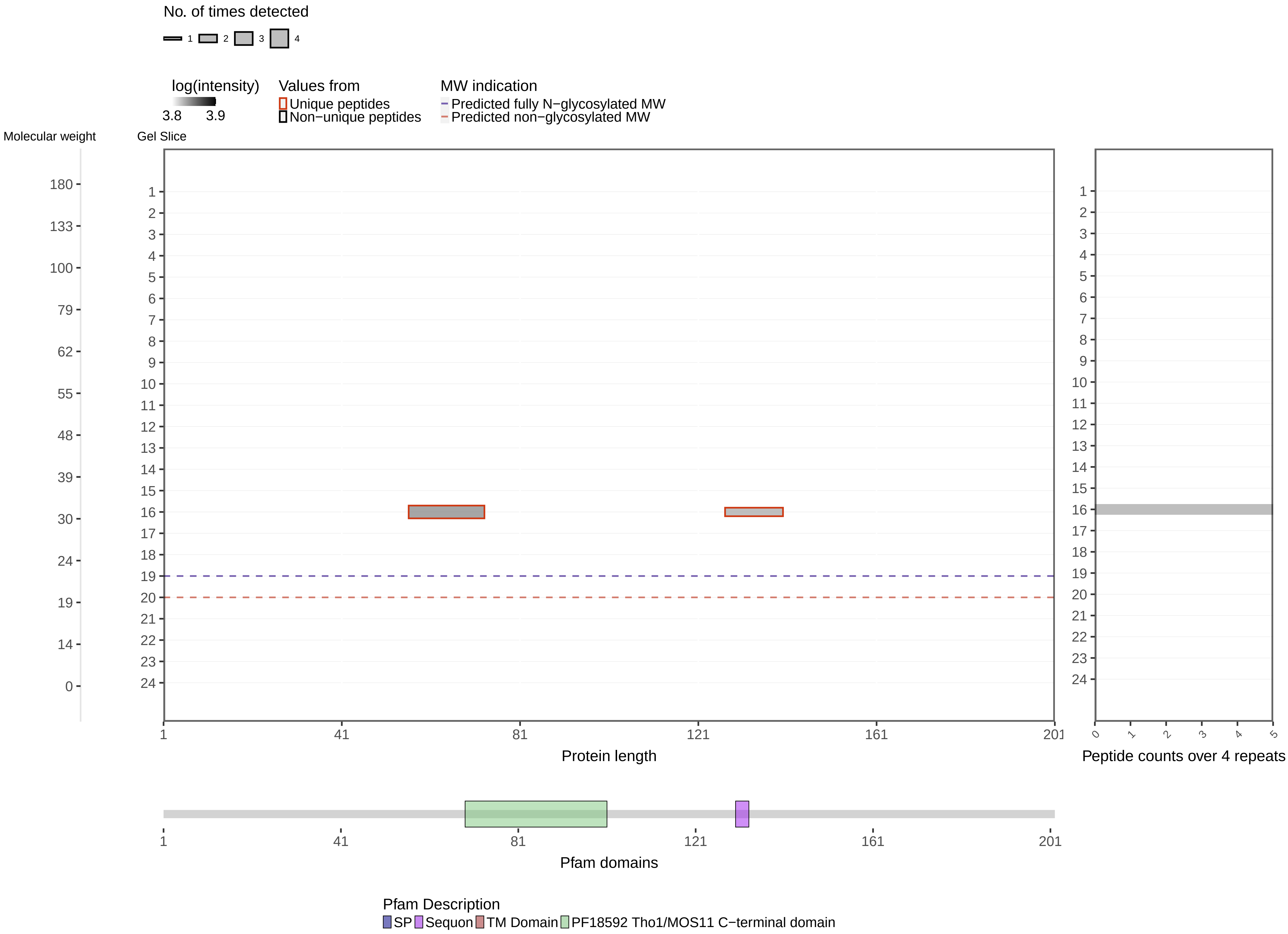


Protein sequence:

EQEKRNTRSERFGTASAAQGSDALKKSEEQKRKARAERFGINQSDSADEEAKKKARLTRFTSTVKADPVEEDKRKARALRFSQSQSGSQSQENGKGKIEQDTVVVEKAGGGT*

### NbL05g04470.1 60S_ribosomal_protein_L4−like [MW = 48.06 kDa]

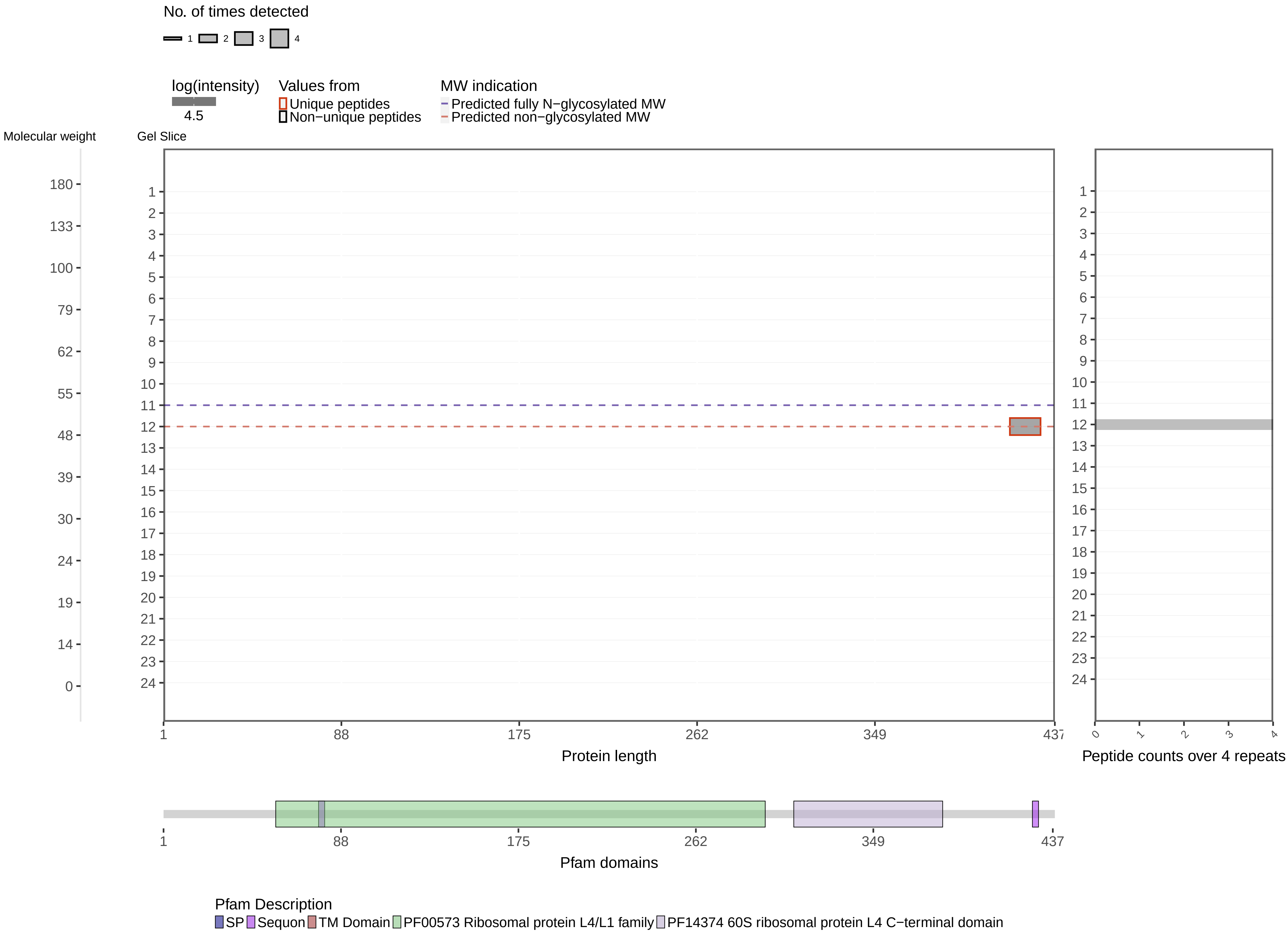


Protein sequence:

QVGAYPDAEKAKNSHAIRPGKGKMRNRRYISRKGPLIVYGTEGAKLVKAFRNIPGVEICHVDRLNLLKLAPGGHLGRFVIWTKSAYEKLDSIYGSFDKPSEKKKGYLLPRPKMVNADLARIINSDEVQSVVRPIKKDVNKRATLKKNPLKNLNVLLKLNPYAKTARRMSLLAEAQRVKAKKEKLDKKRHQITKEEASAIRSASHSWYKTMISDSDYTEFDNFSKWLGVSQ*

### NbL05g04450.1 L−ascorbate_peroxidase_2_cytosolic [MW = 27.6 kDa]

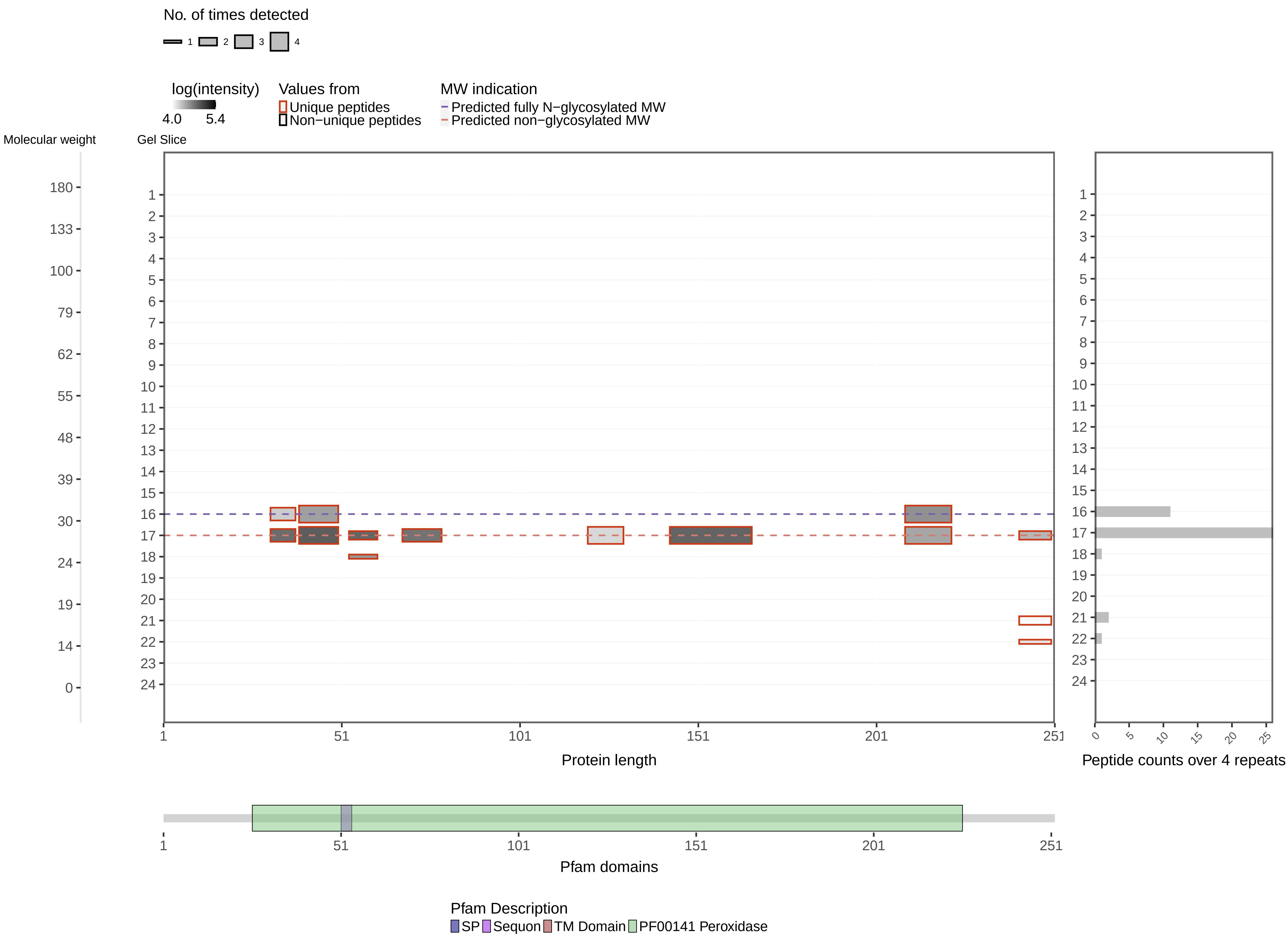


Protein sequence:

FHPGRQDKTEPPPEGRLPDATKGSDHLREVFGHMGLSDKDIVALSGGHTLGRCHKERSGFEGAWTNNPLIFDNSYFRELLSGEKEGLLQLPSDKALLEDPVFRPLVEKYAADEDAFFADYAEAHLKLSELGFADAE*

NbL05g04420.1 probable_LRR_receptor−like_serine/threonine−protein_kinase_At1g67720 [MW − SP = 100.33 kDa]

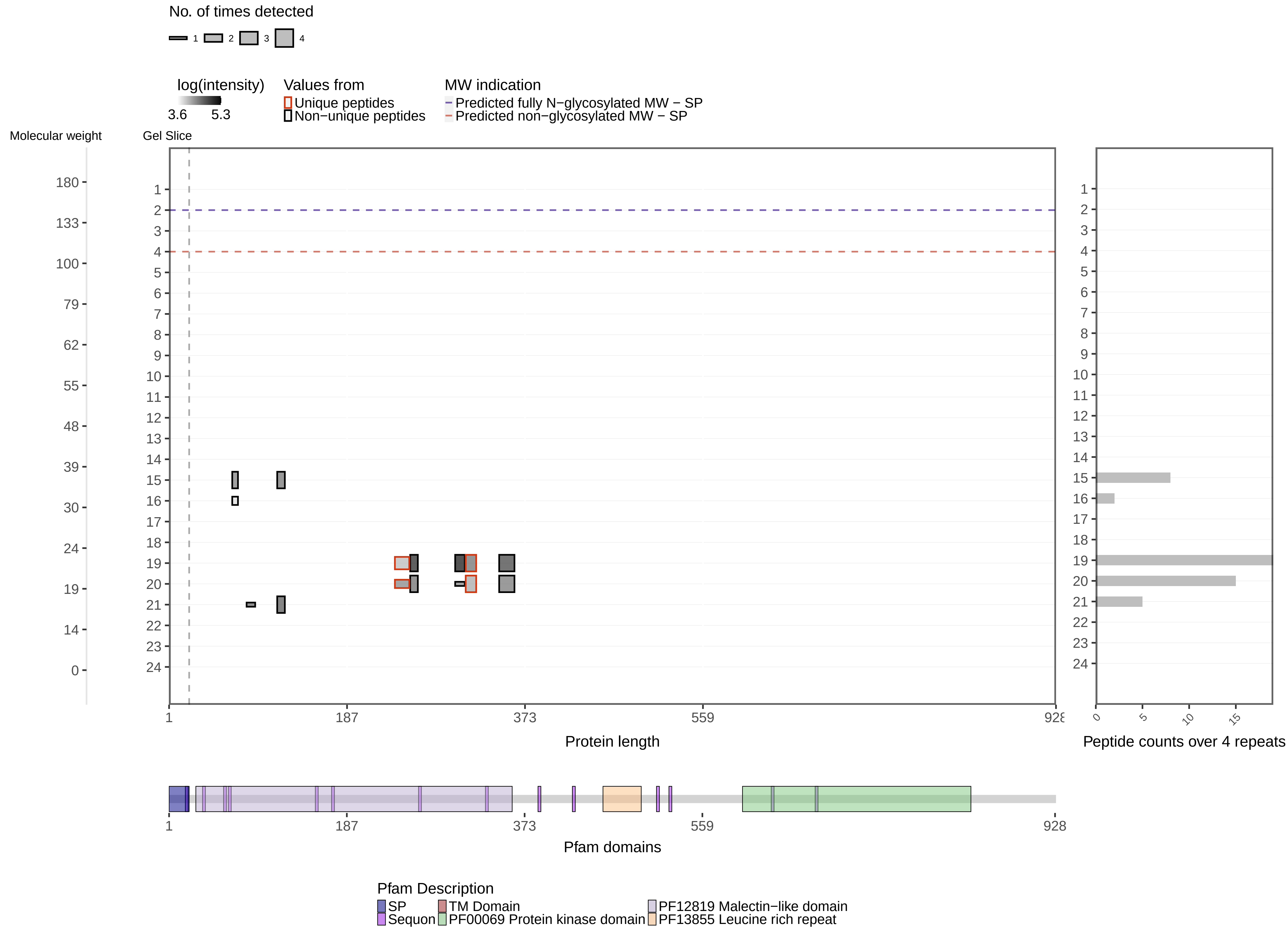


Protein sequence:

[illegible]

NbL05g04310.1 thioredoxin_domain−containing_protein_PLP3B−like [MW = 16.78 kDa]

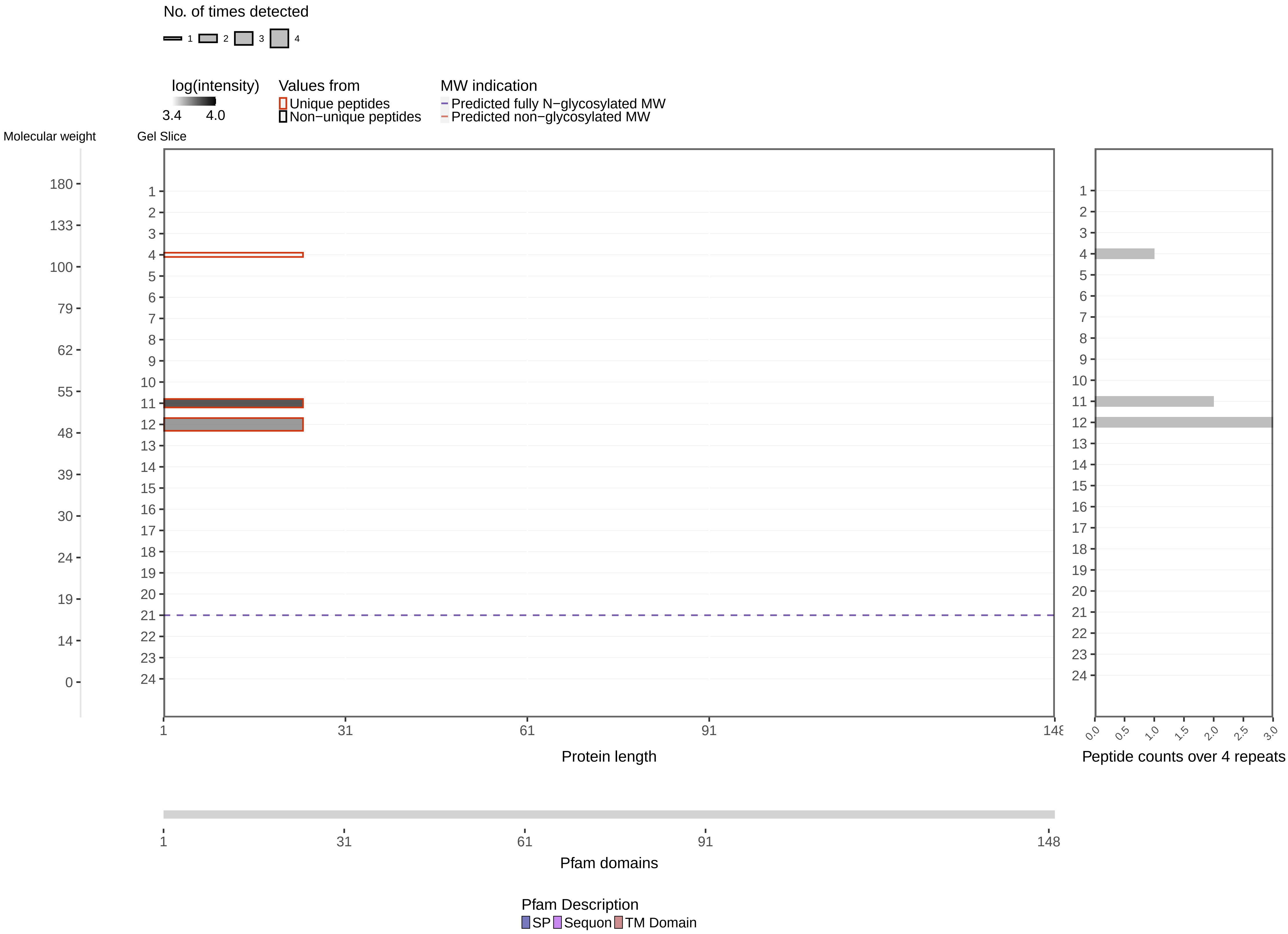


Protein sequence:

HADRIASLKKEAEKREALKRQGHGEYREIGEGDFLGEVTGCEKVIFHFYHREFYRCMIMDKHLKSLAPRHVDTKFLKLDVEEGNSFR*

NbL05g04150.1 valine−−tRNA_ligase_mitochondrial_1−like [MW = 128.48 kDa]

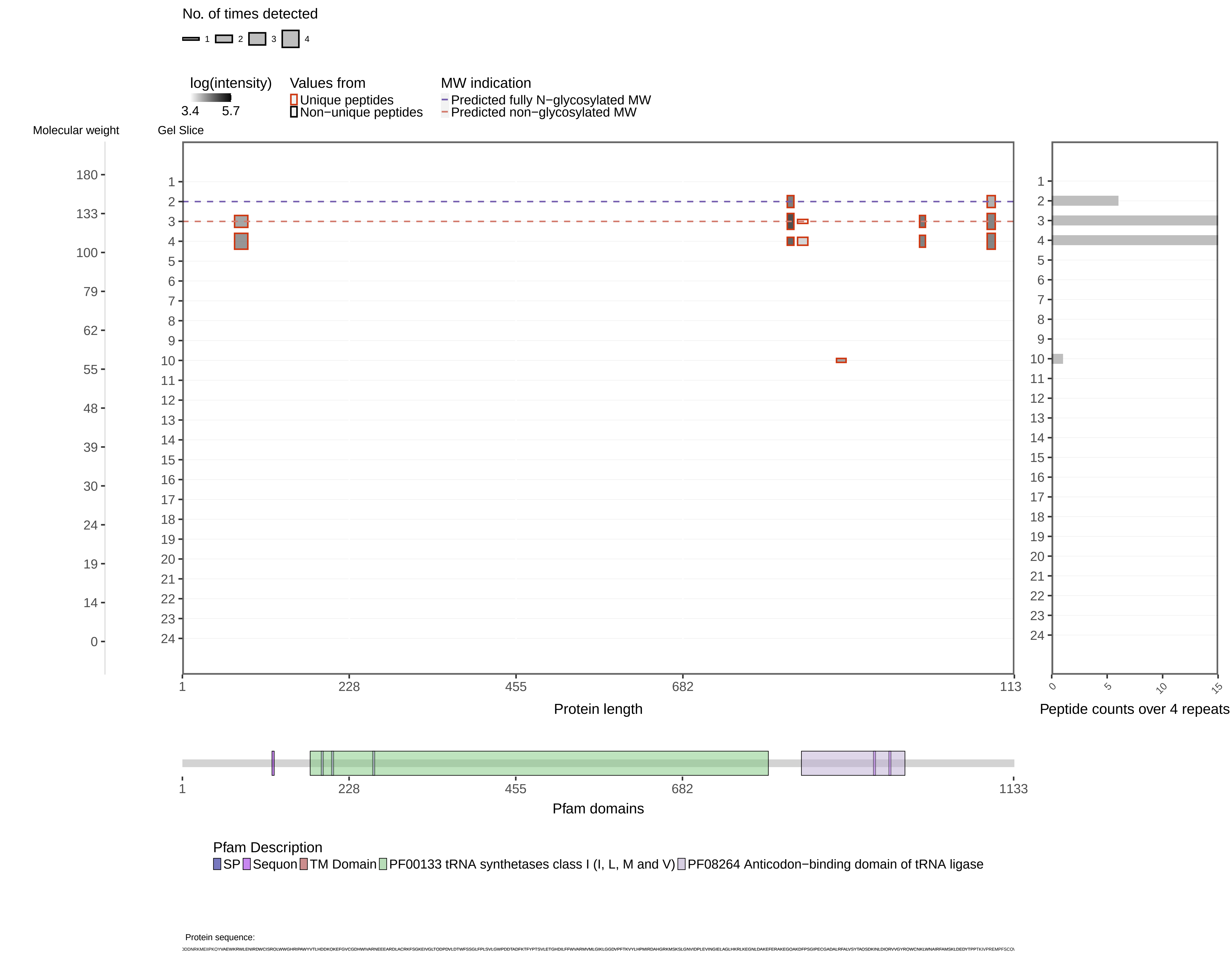

### NbL05g04020.1 PIP2−1−like [MW = 31.18 kDa]

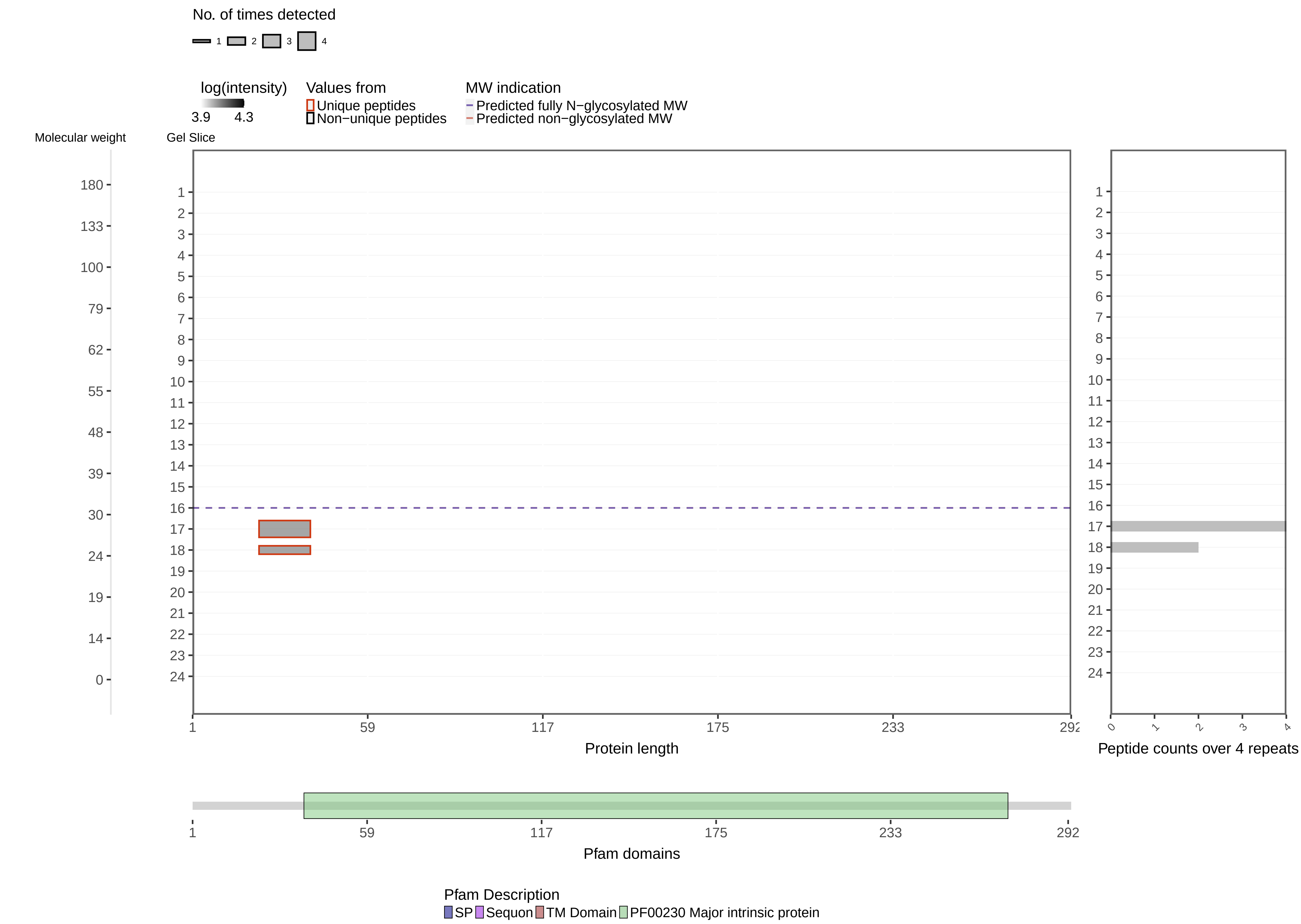


Protein sequence:

ILAQCLGAICGCGLVKAFQKAYYVKYGGGANTLNDGYSTGTGLGAEIIGTFVLVYTVFAATDPKRNARDSHVPVLAPLPIGFAVFMVHLATIPVTGTGINPARSFGAAVIYGKNKAWDDQWIFWVGPFIGAAIAAFYHQFILRAGAVKALGSFRSNA*

### NbL05g03930.1 29_kDa_ribonucleoprotein_A_chloroplastic [MW = 29.67 kDa]

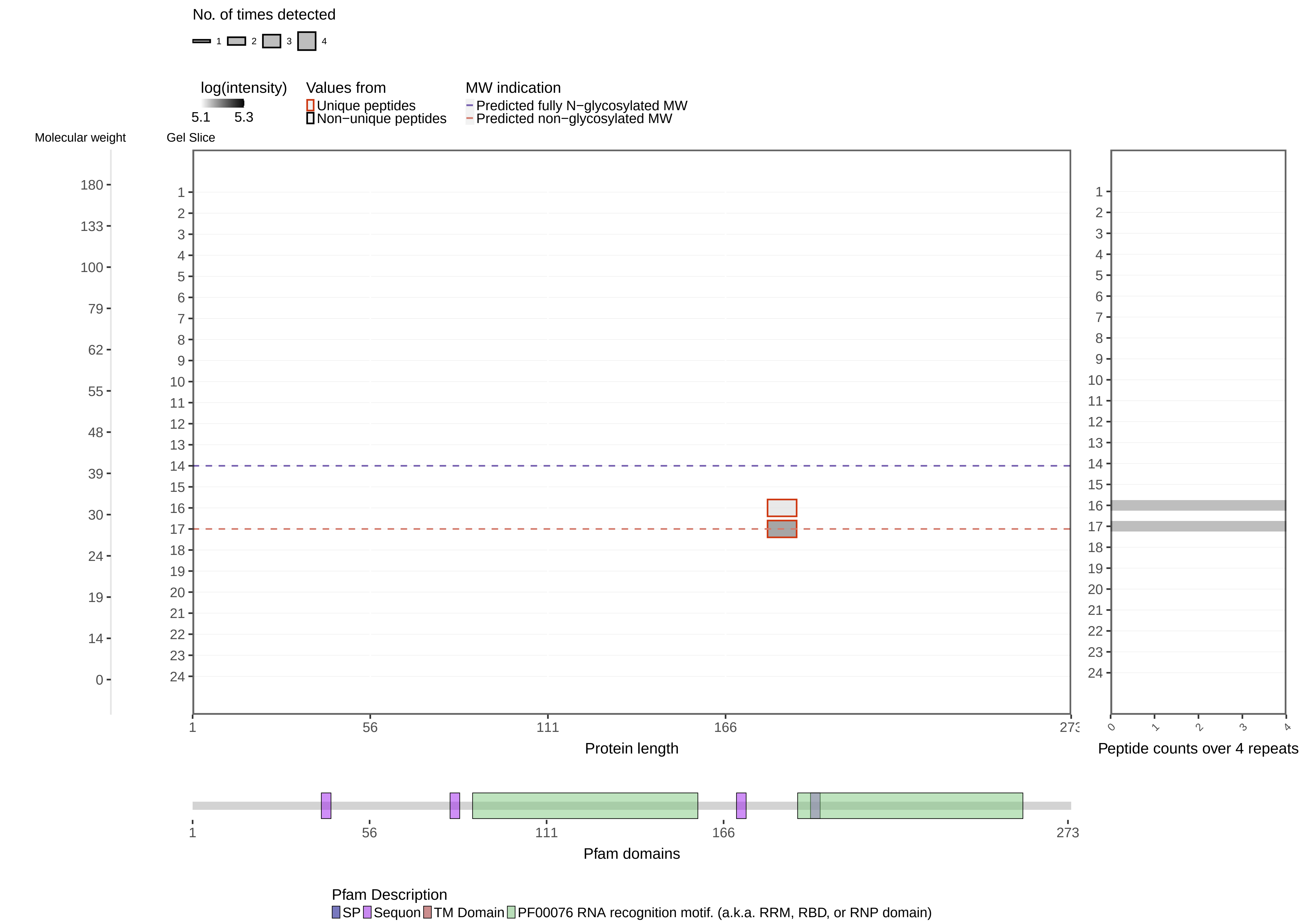


Protein sequence:

RGFGFVTMSSKEEVEAACQQFNGYELDGRELRVNSGPPPEKRENSSFRGGNRGGGSFDSSNRVYVGNLSWGVDQDALETLFSEQGKVVDAKVVYDRDSGRSRGFGFVTYSSAEEVNNAIESLDGVDLNGRAIRVSLAEARPPRRQF*

### NbL05g03910.1 ethylene−insensitive_protein_2 [MW = 143.76 kDa]

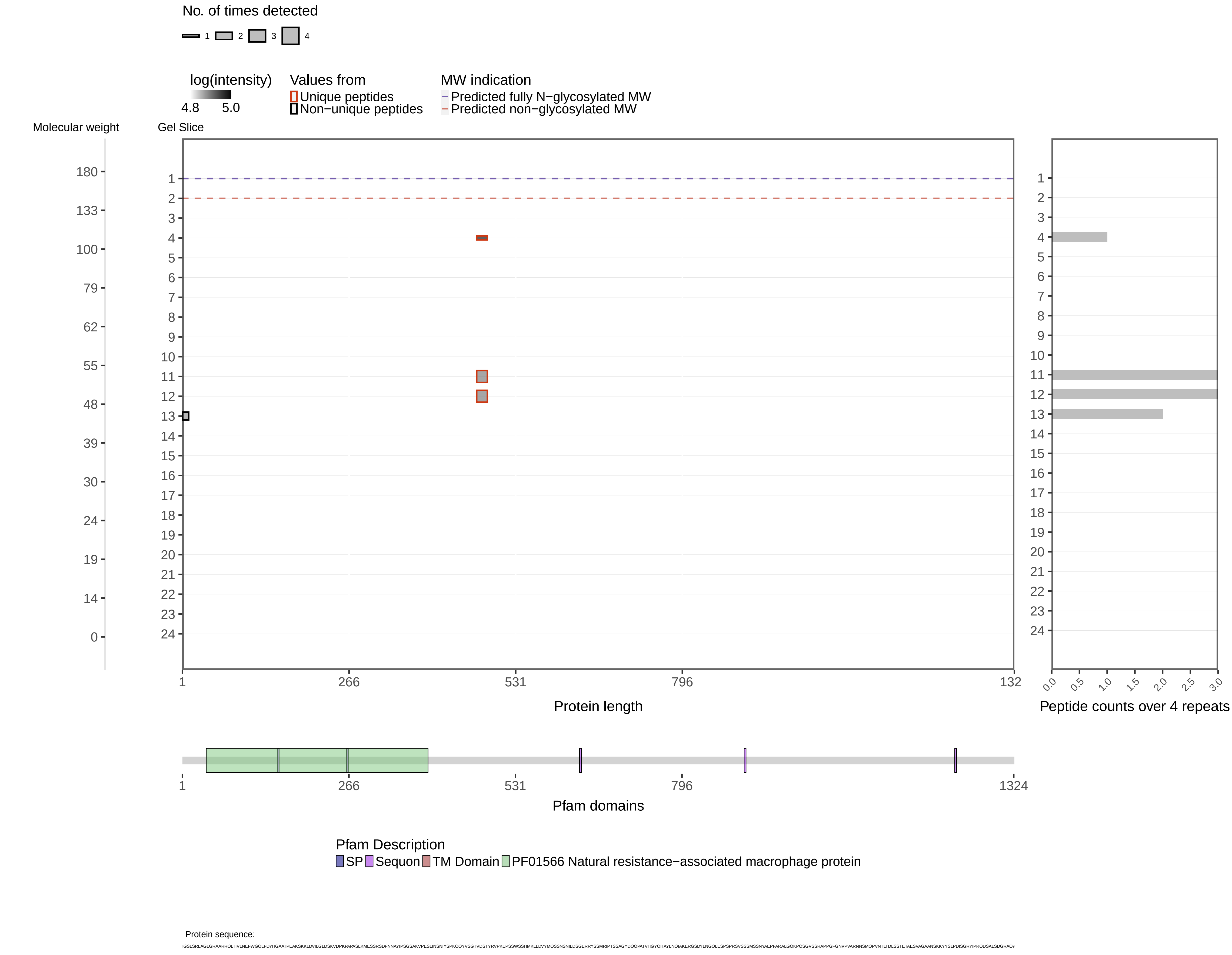

### NbL05g03880.1 ammonia−lyase−like [MW = 78.01 kDa]

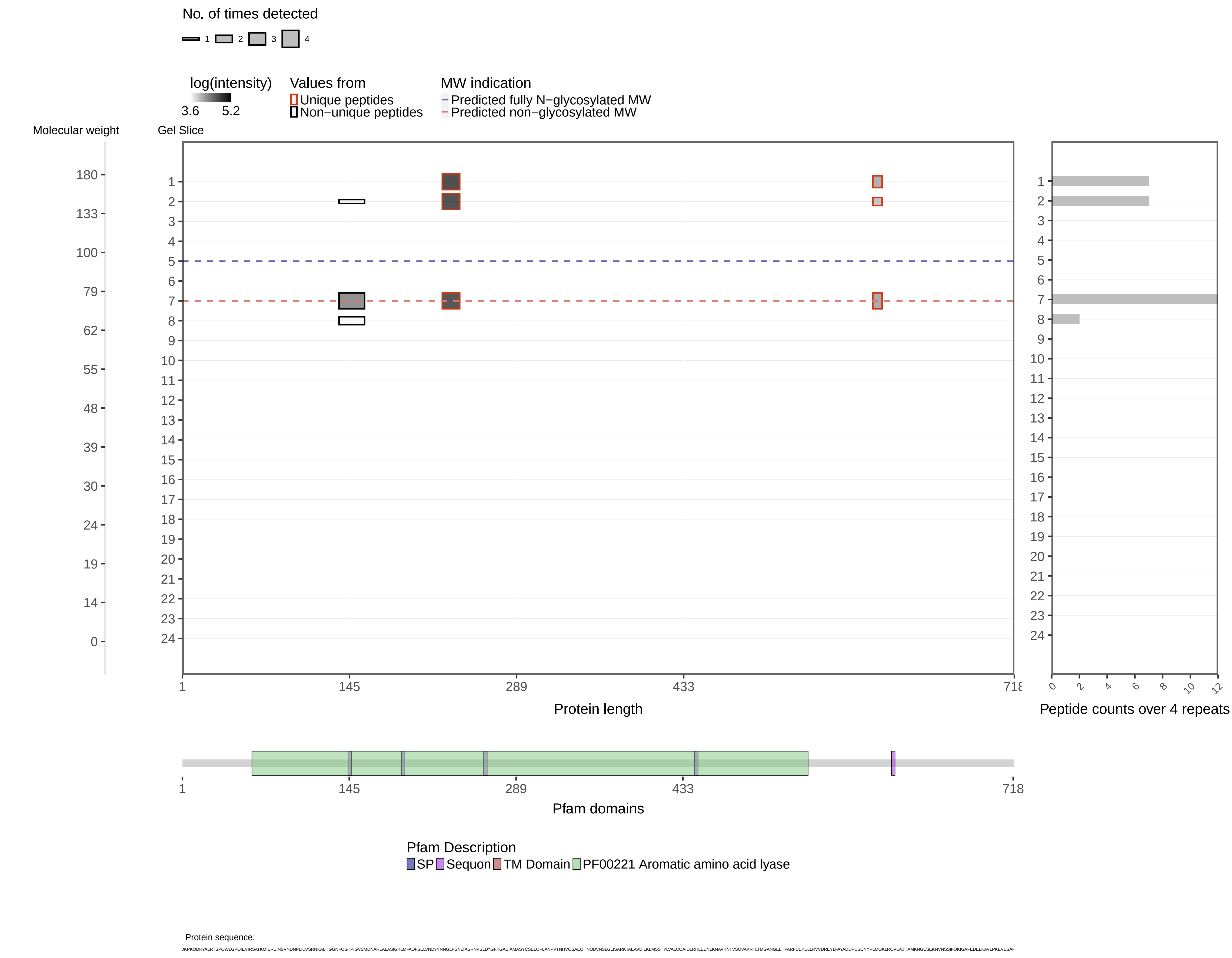

### NbL05g03850.1 kinase_2−like [MW = 37.55 kDa]

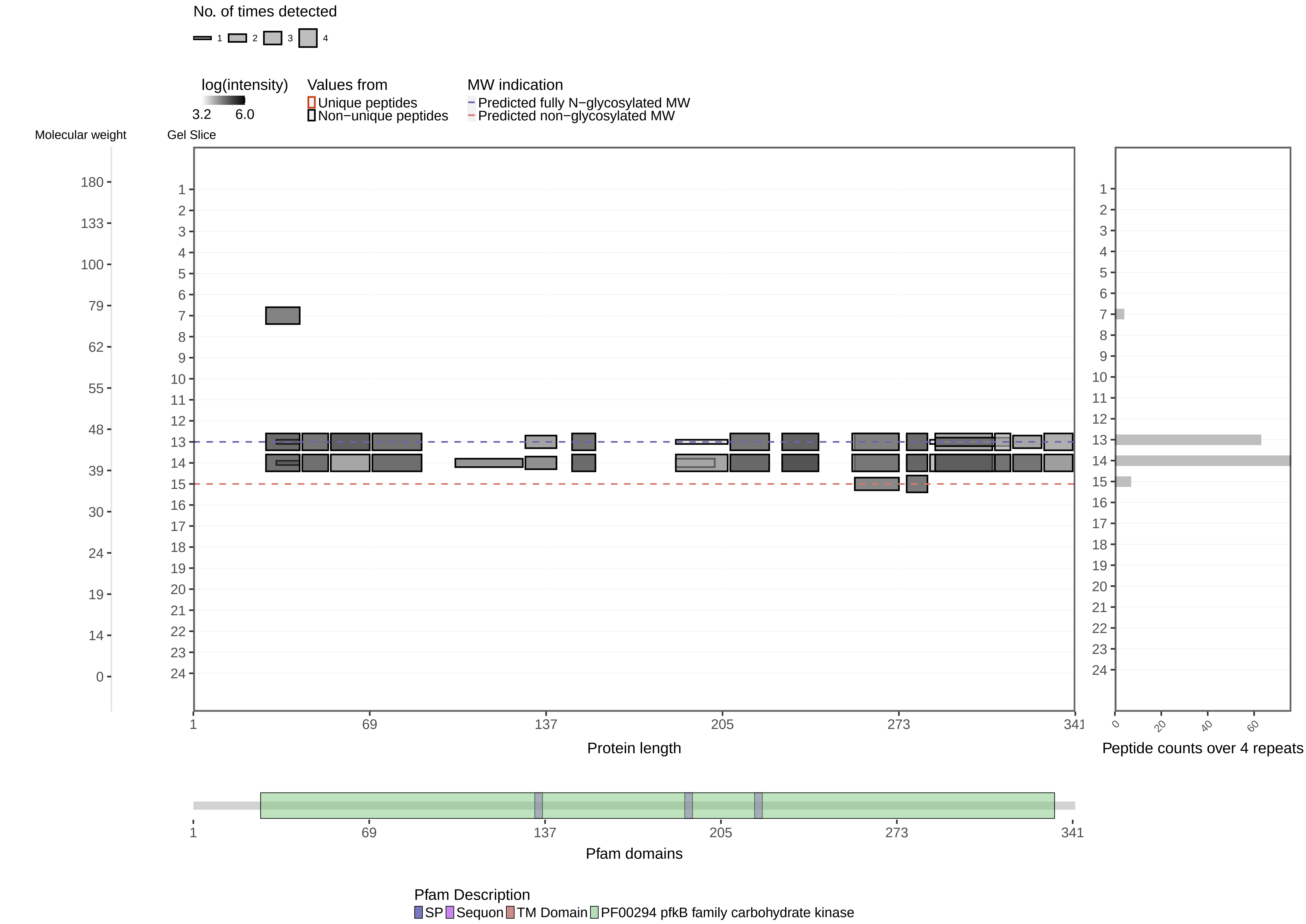


Protein sequence:

AKFYYIAGFFLTVSPESIQLVAEHAAANNKIFSMNLSAPFICEFFRDPQEKALPYMDFVFGNETEARTFSKVHGWETDNVEEIALKISEWPKASETHKRITVITQGADPVVVAENGKVKLFPVIPLPKEKLVDTNGAGDAFVGGFLSQLVQGKPVEDCVRAGCYASNVIIQRSGCTYPEKPDFA*

### NbL05g03730.1 cell_division_cycle_protein_48_homolog [MW = 89.69 kDa]

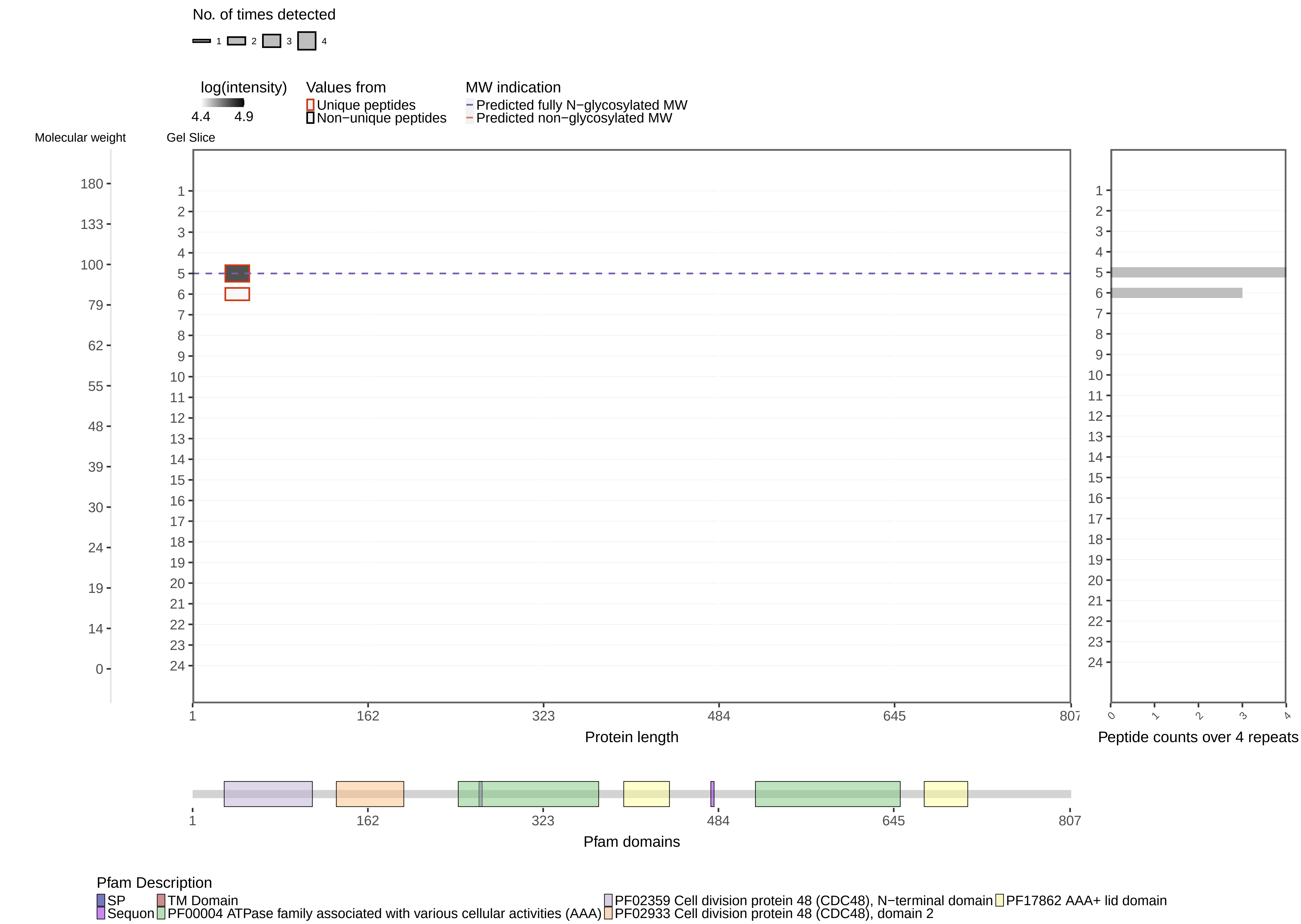


Protein sequence:

[illegible]

NbL05g03610.1
ABC_transporter_B_family_member_1_(LOC104237401)_transcript_variant_X1_mRNA_XM_009791552.1_PREDICTED:_Nicotiana_sylvestris_ABC
[MW = 146.58 kDa]

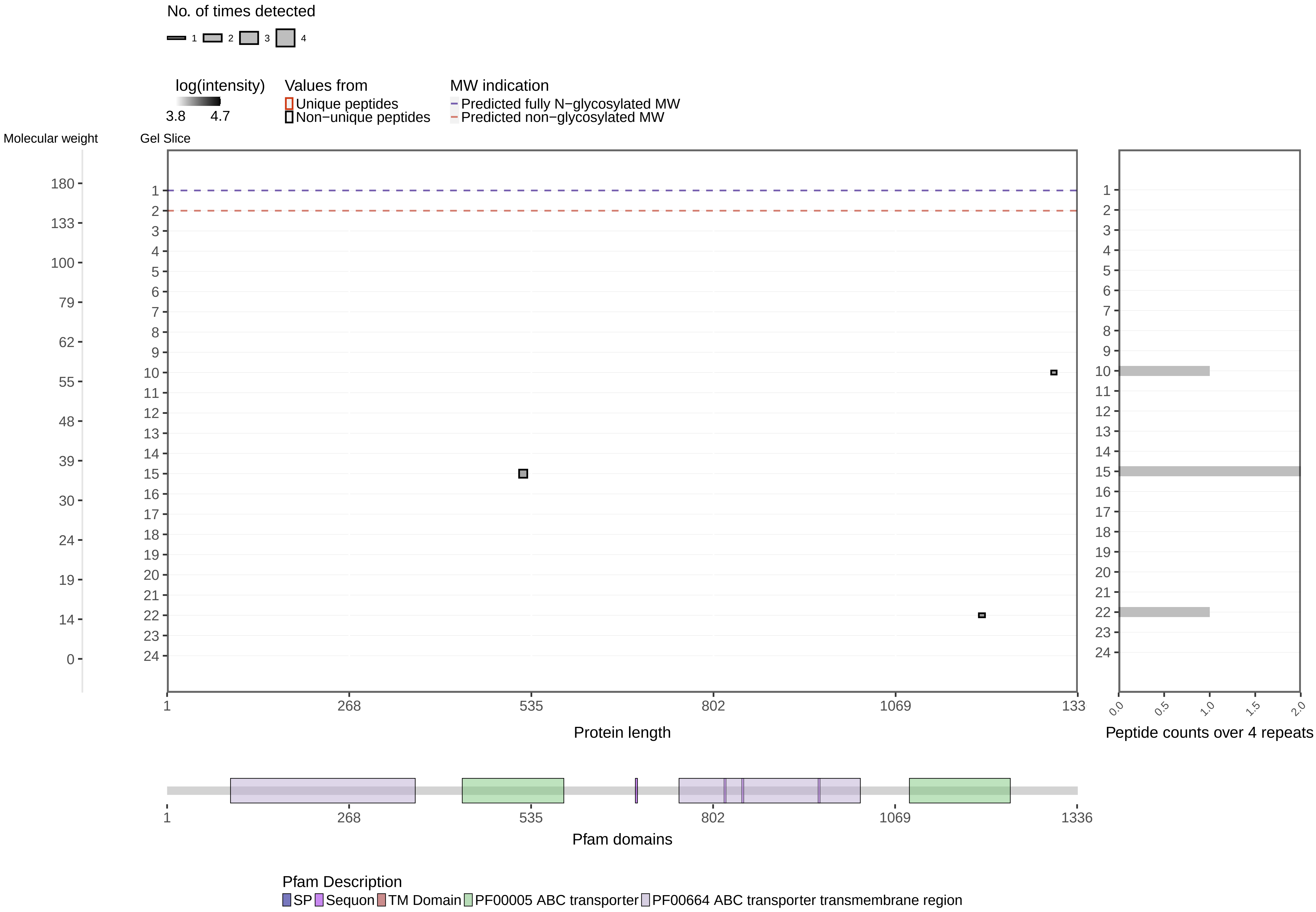


Protein sequence:
EAAHETALSNARKSSARPSSARNSVSSPIITRNSSYGRSPYSRRLSDFSTSDFSLSLDAAYSSYRHEKLAFKDQASSFCRLAKMNSPEWSYALVGSIGSVICGSLSAFFAYVLSAVLSVYYNPDHAYMSKQIAKYCYLLIGVSSAALIFNTLQHFFWDVVGENLTKRVREKMLDAVLKMEIAWFDQEENDSSRIAARLALDANNVRSAIGDRISVIMQNSALMLVACTAGFVLQWRLALVLIAVFPVVVAATVLQKMFMTGFSGDLEAAHARATQLAGEAVANVRTVTAFNSETKI

### NbL05g03540.1 S−adenosylmethionine_synthase_3 [MW = 42.62 kDa]

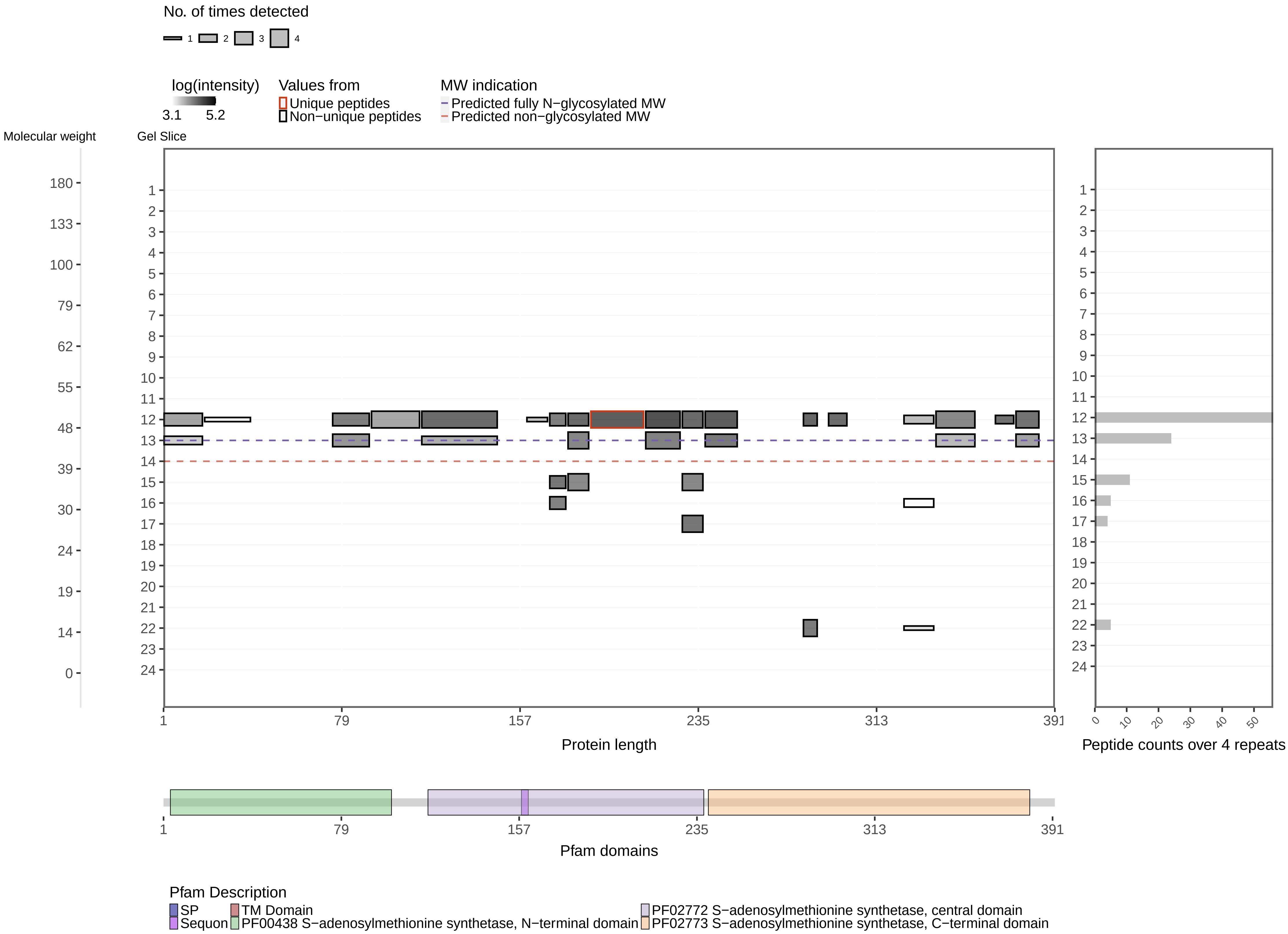


Protein sequence:

GAMVPIRVHTVLISTQHDETVTNDQIAQDLKEHVIKPVIPAQYLDEKTIFHLNPSGRFVIGGPHGDAGLTGRKIIIDTYGGWGAHGGGAFSGKDPTKVDRSGAYIVRQAAKSVVASGLARRCIVQVSYAIGVAEPLSVFVDTYKTGTIPDKDILALIKENFDFRPGMMSINLDLLRGGNFRYQKTAAYGHFGRDDPDFTWETVKVLKPKA*

NbL05g03090.1 threonine_dehydratase_biosynthetic−like [MW = 65.61 kDa]

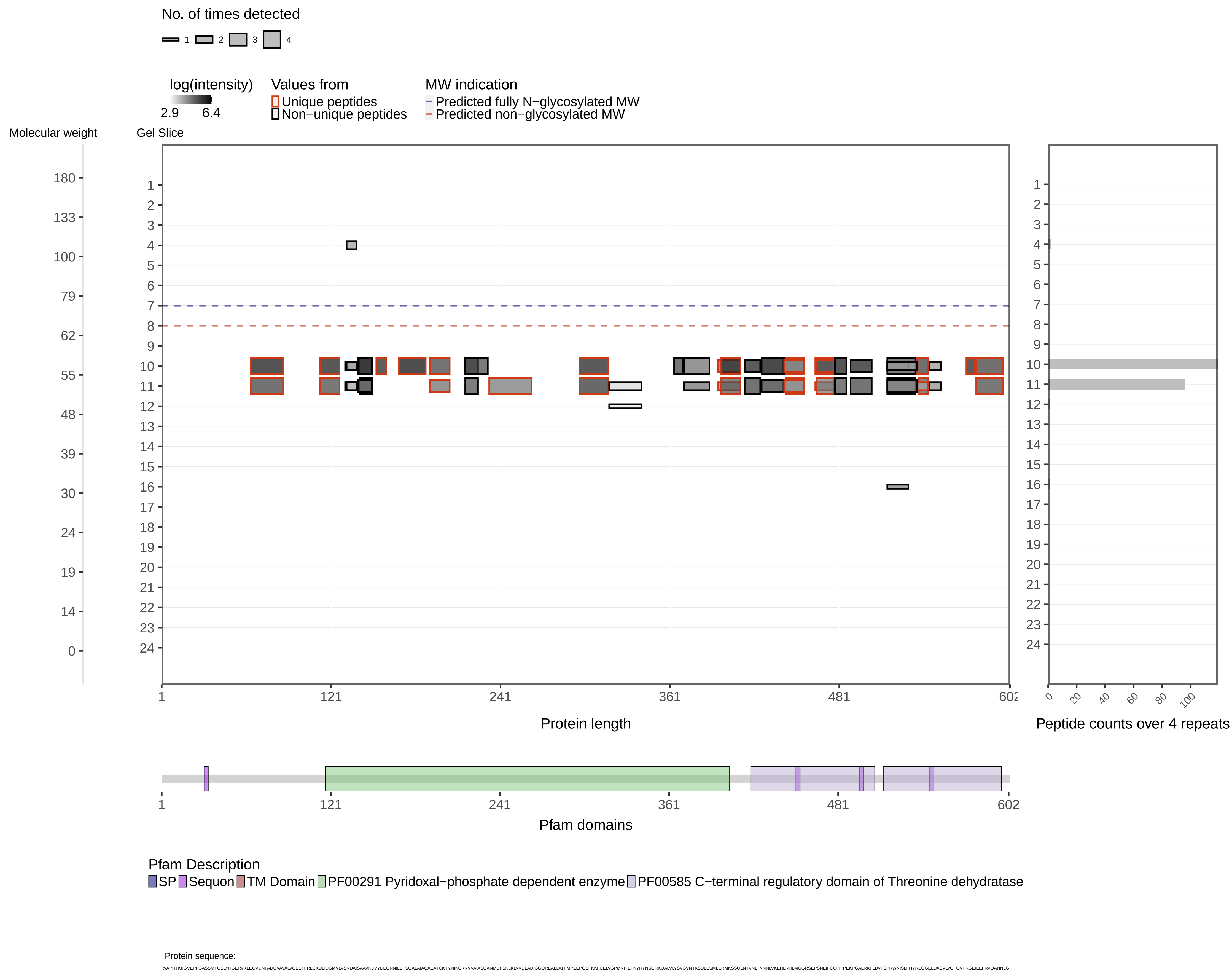

NbL05g02960.1 uncharacterized_protein_DDB_G0284311 [MW = 54.42 kDa]

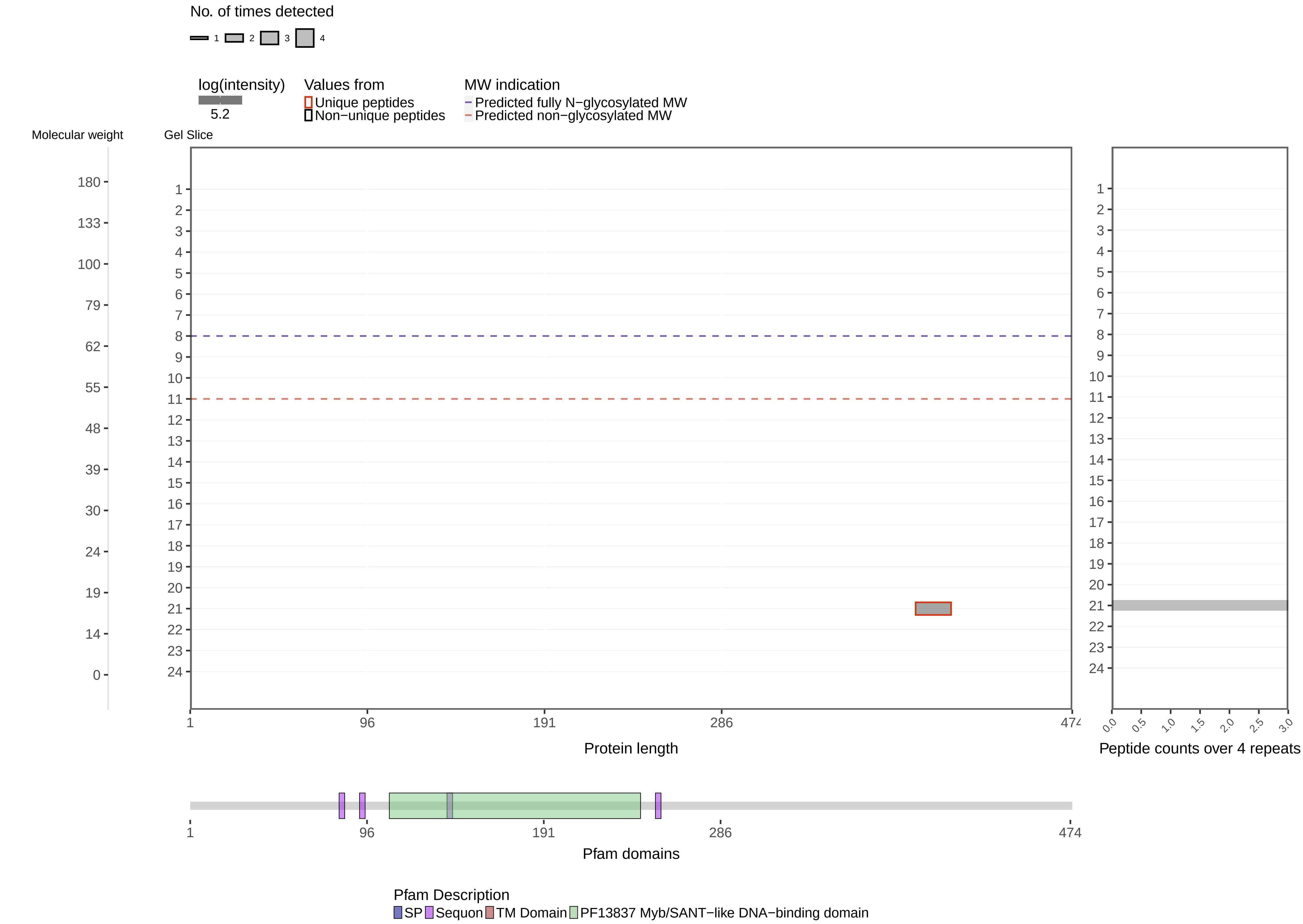


Protein sequence:

KEEVKKLLNSKHLFFREMCAYHNSCGHNNTSTGGGSGNVQLSTEVAIEVNNQAQKRCLHSSENARIEVHLEEGTKMAKGKSVDDDQEDDDEDEDDDEEMSGARGGHEHHHSHGNEYDDDMDERSRKRQRKSGSLSPMVQQLSSEVANVIQDGGRSSIEKRQWLKTRMVQLEEQHVGYLCQAFELQKQRLKWEKFTSKKEREMEKEKLINERKKLENERMVLLLRQKELELLDFHQHQNHNRNSDPSSVTG*

### NbL05g02910.1 geranylgeranyl_pyrophosphate_synthase_small_subunit_chloroplastic−like [MW = 36.27 kDa]

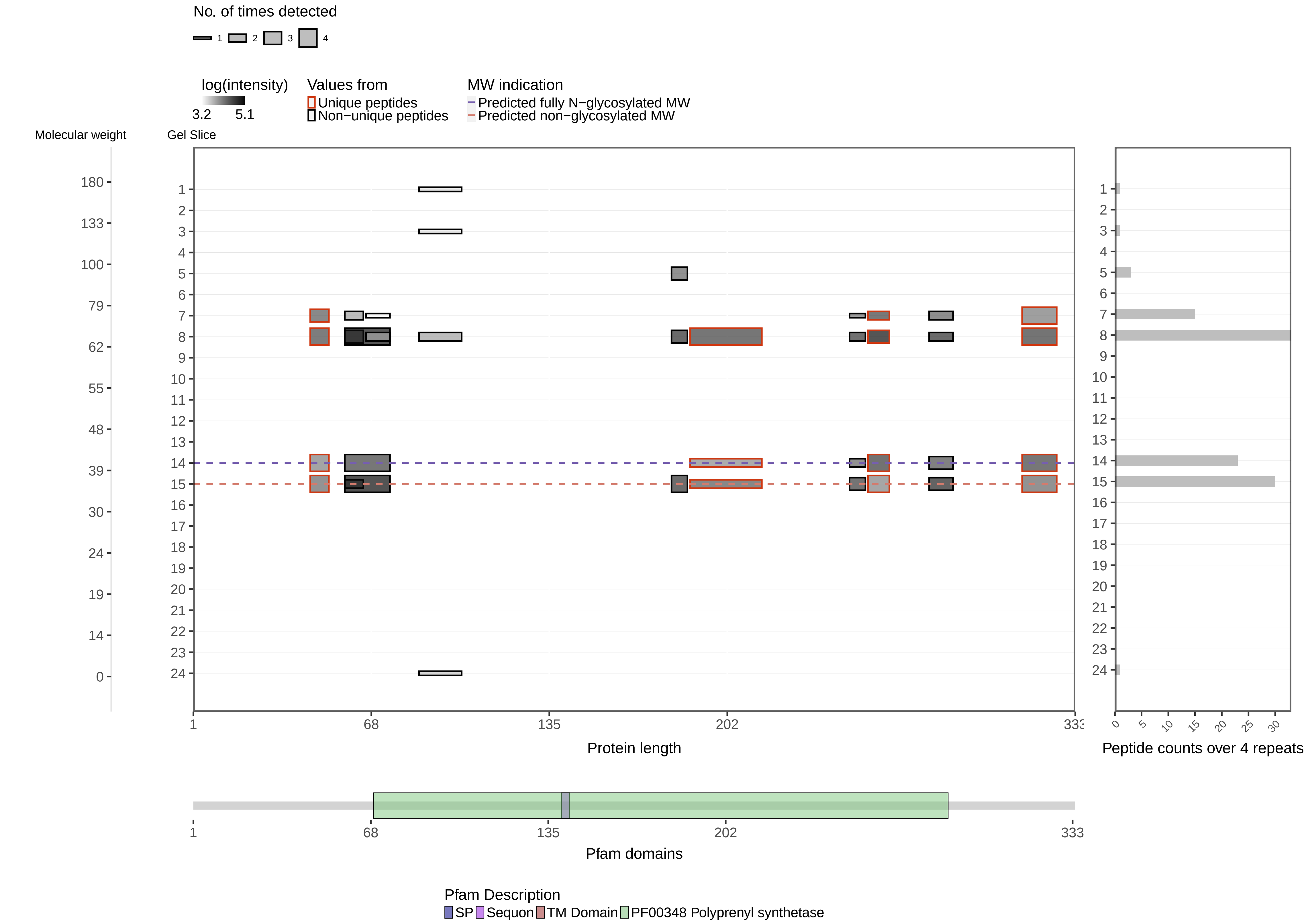


Protein sequence:

GDALFPLGFQHIVSHTPSDLVPEDRVLKVITEIARAVGSTGMAAGQFLDLEGGPNAVDSVQEKKYGEMGECSAVCGGLLAGASDEEIQHLRRYGRAVGVLYRVVDDILEAKETEDKGEGKKKKGKSYVSLYGIEKAKKVAEDLRAQAKRELDGLDKYGDKVIPLYSFVDYAADRGFSLDN*

NbL05g02840.1 peroxiredoxin−2E−2_chloroplastic [MW = 24.82 kDa]

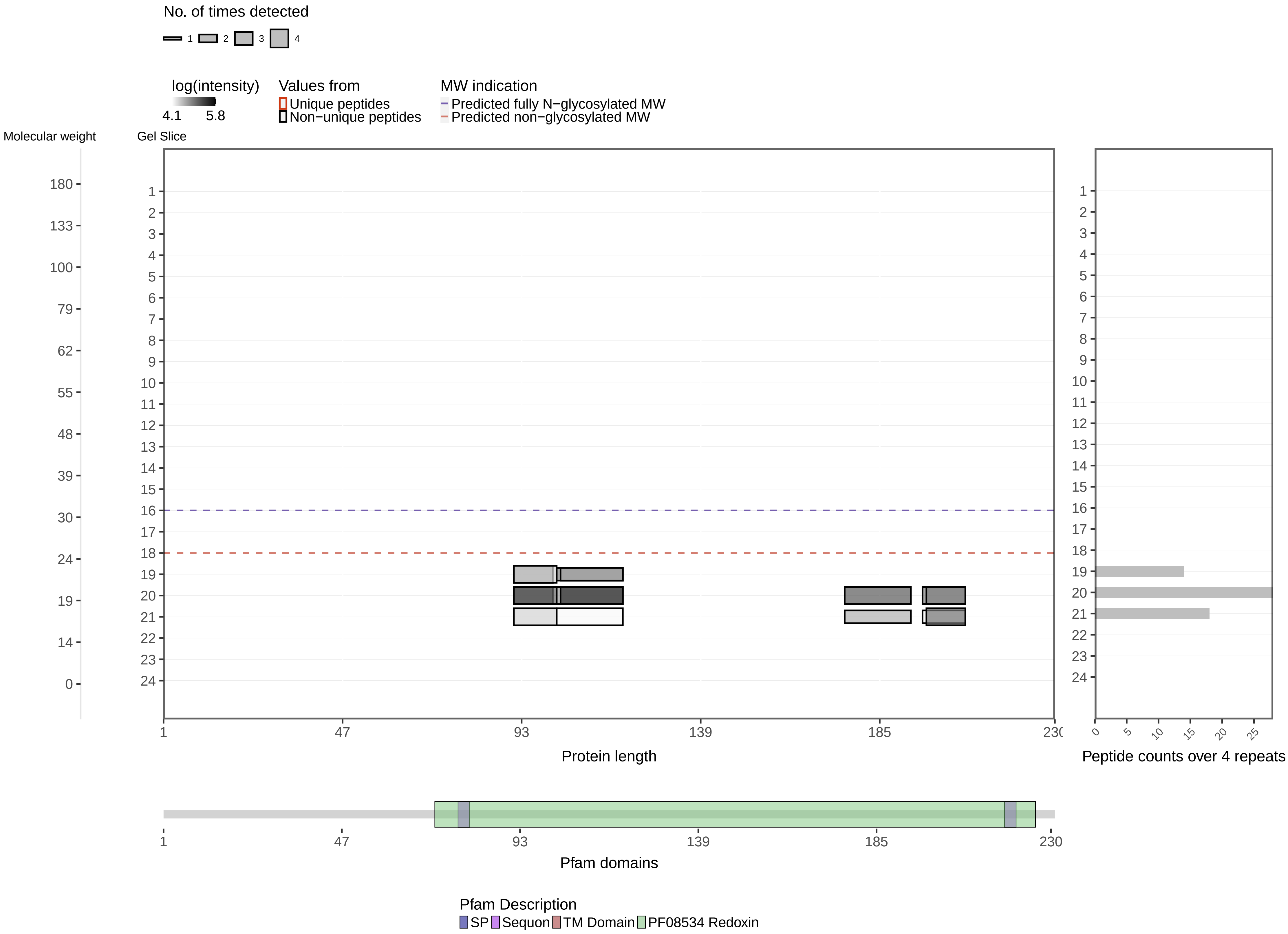


Protein sequence:

VLLAVPGAFTPTCSQKHLPGFVEKSKELKSKGVDTIACISVNDAFVMKYWKENLGINDEVMMLSDGNLEFTKAIGCELDLSDKPVGLGIRSRRYSMLVEDGVVKILNLEEGGAFNVSSADDILKAL*

NbL05g02830.1 enolase−like [MW = 58.33 kDa]

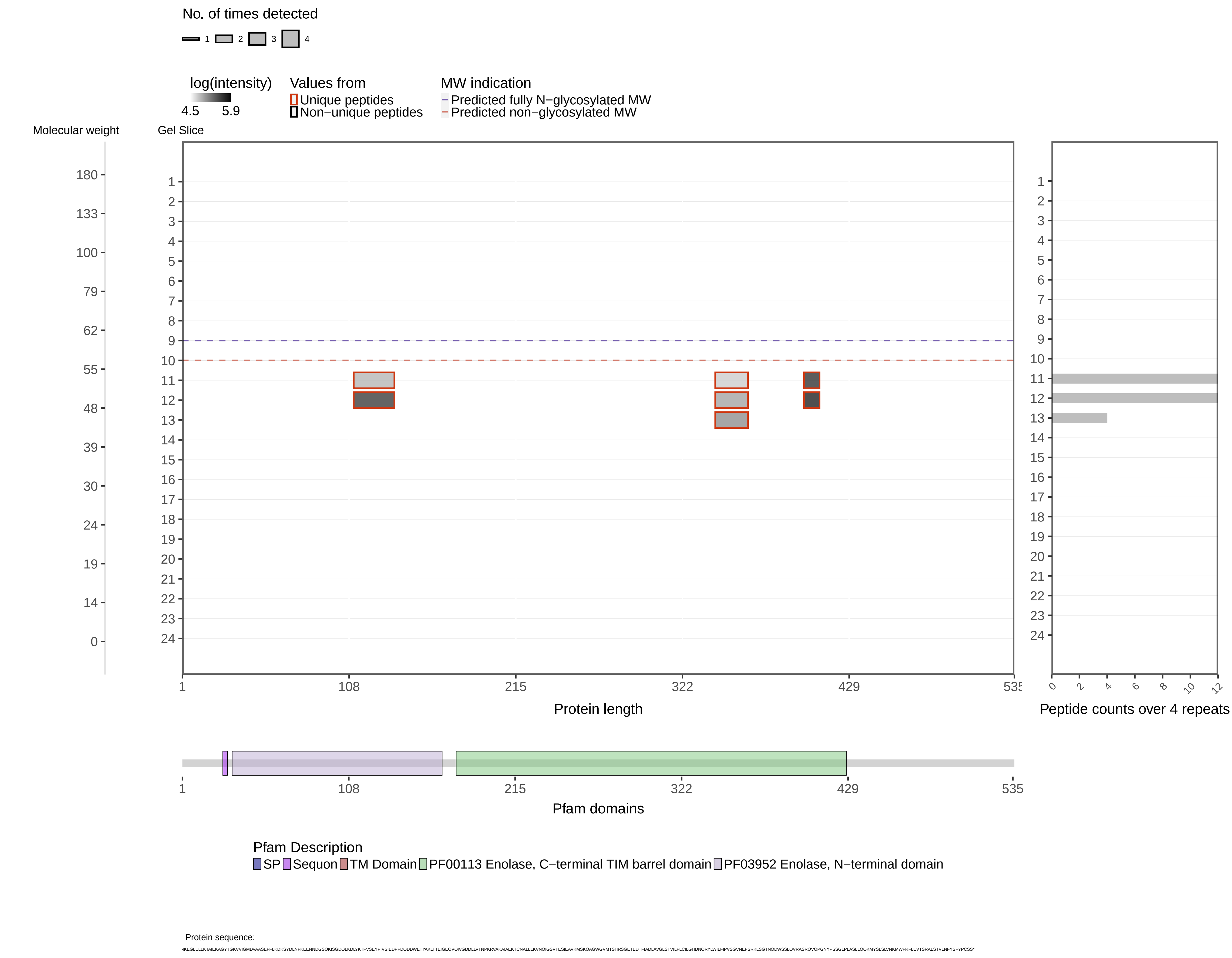

NbL05g02790.1 protein_Ycf2 [MW = 73.26 kDa]

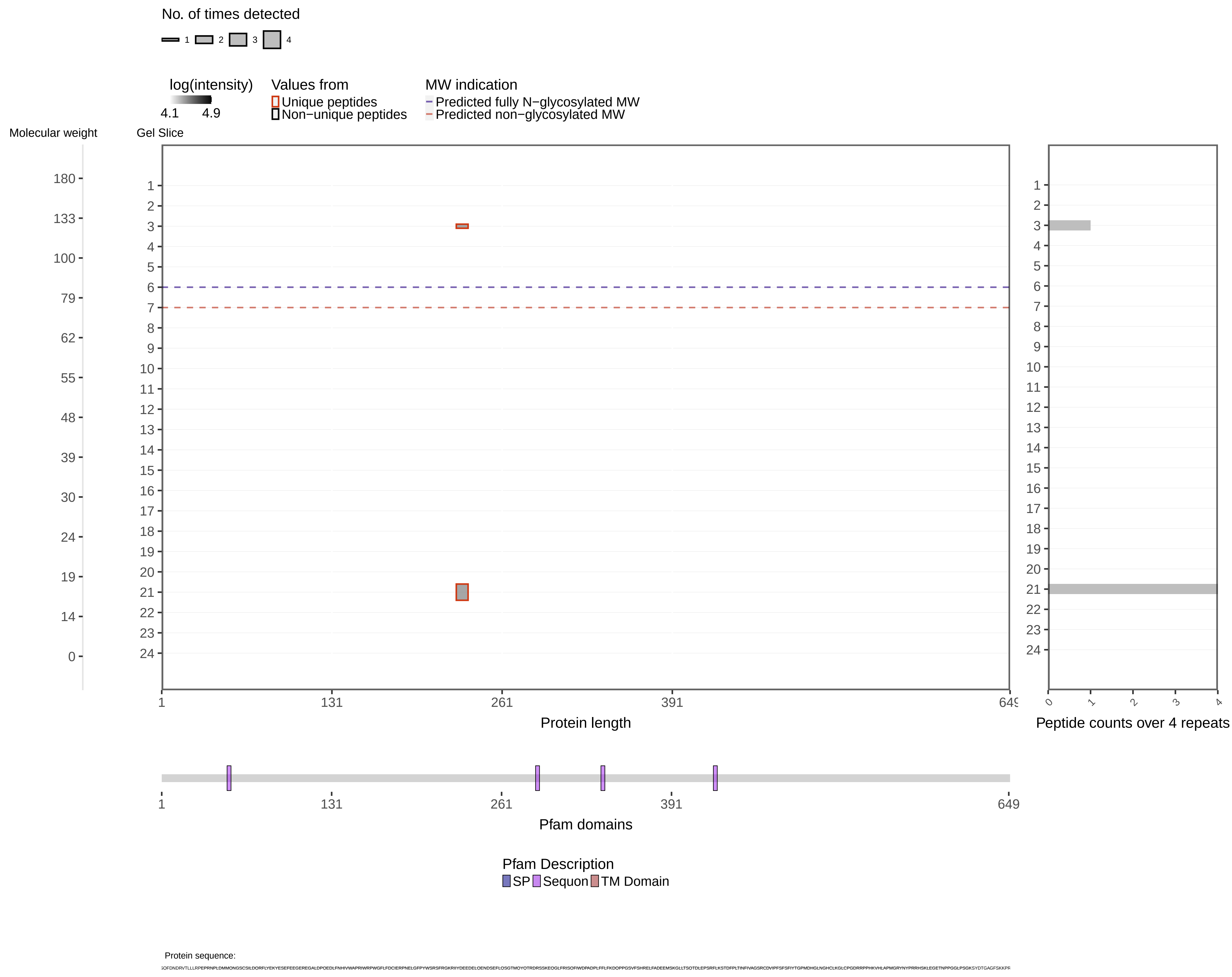

### NbL05g02730.1 photosystem_I_P700_chlorophyll_a_apoprotein_A2 [MW = 12.55 kDa]

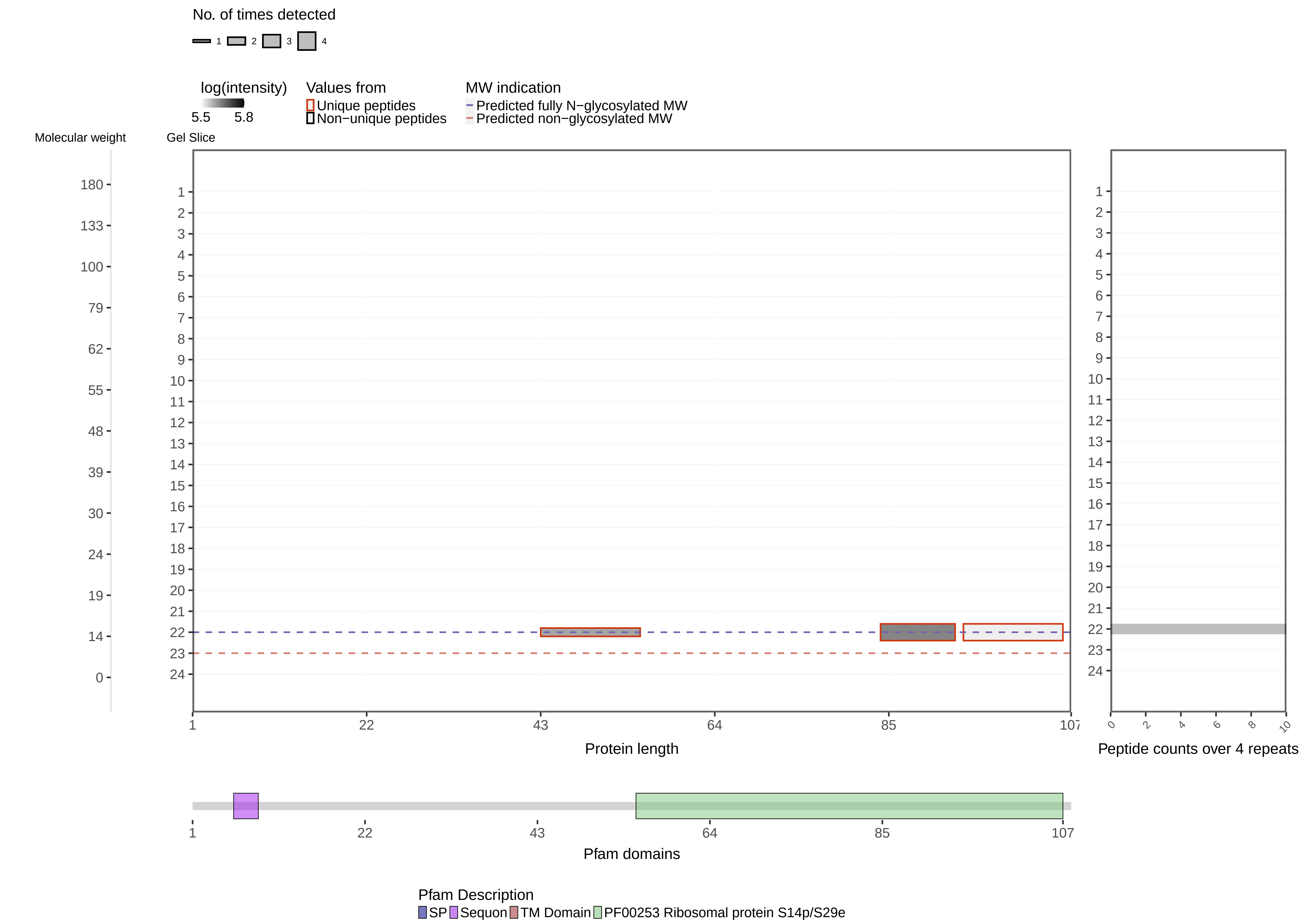


Protein sequence:

KVPSLSDKWEIYGKLQSLPRNSAPTRLHRRCFLTGRPRANYRDFGLSGHILREMVHACLLPGATRS

### NbL05g02710.1 30S_ribosomal_protein_S7_chloroplastic [MW = 17.37 kDa]

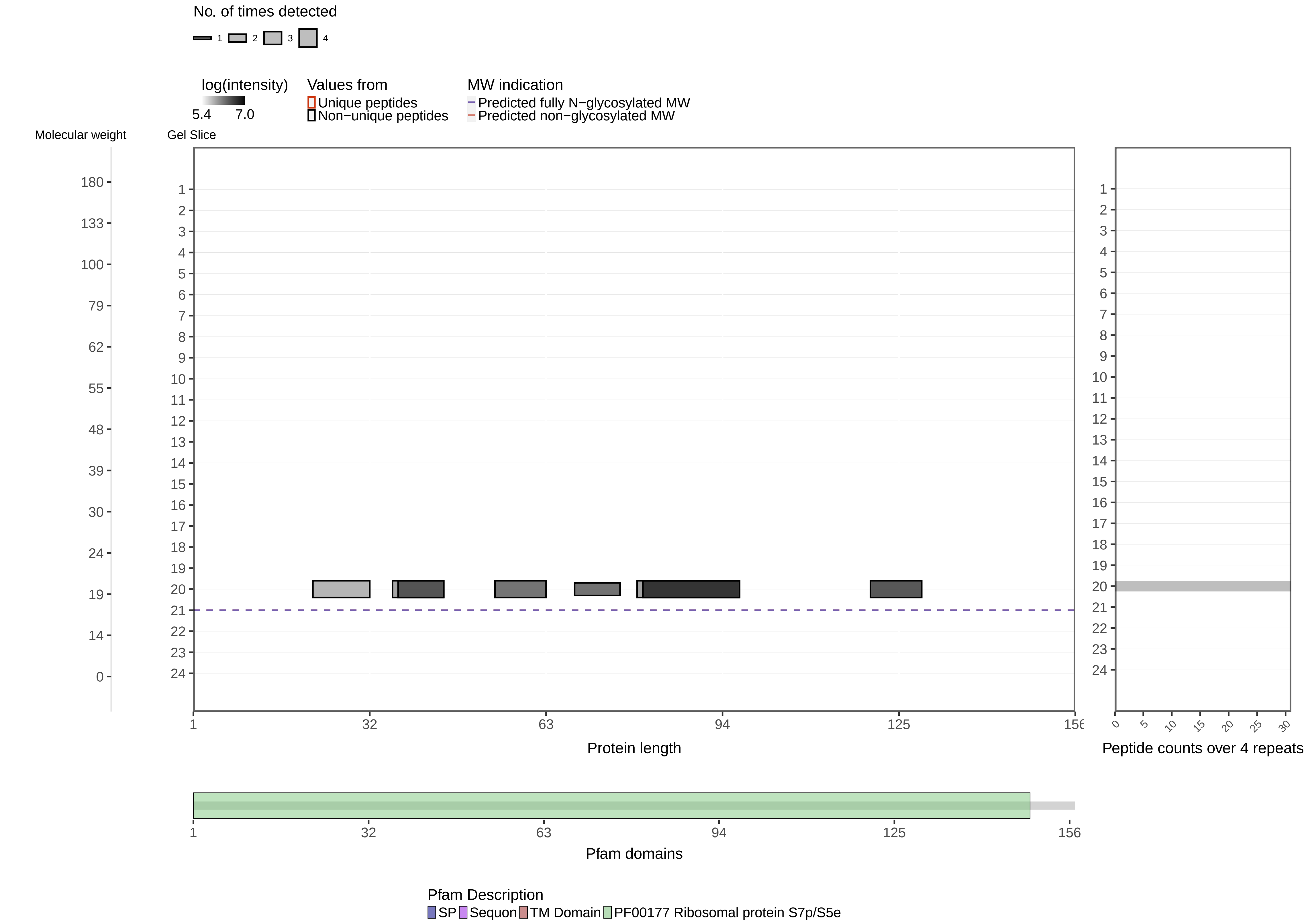


Protein sequence:

IRGVTPDITVKARRVGGSTHQVPIEIGSTQGKALAIRWLLAASRKRPGRNMAFKLSSELVDAAKGSGDAIRKKEETHRMAEANRAFAHFR*

### NbL05g02620.1 fructose−bisphosphate_aldolase_cytoplasmic_isozyme [MW = 41.49 kDa]

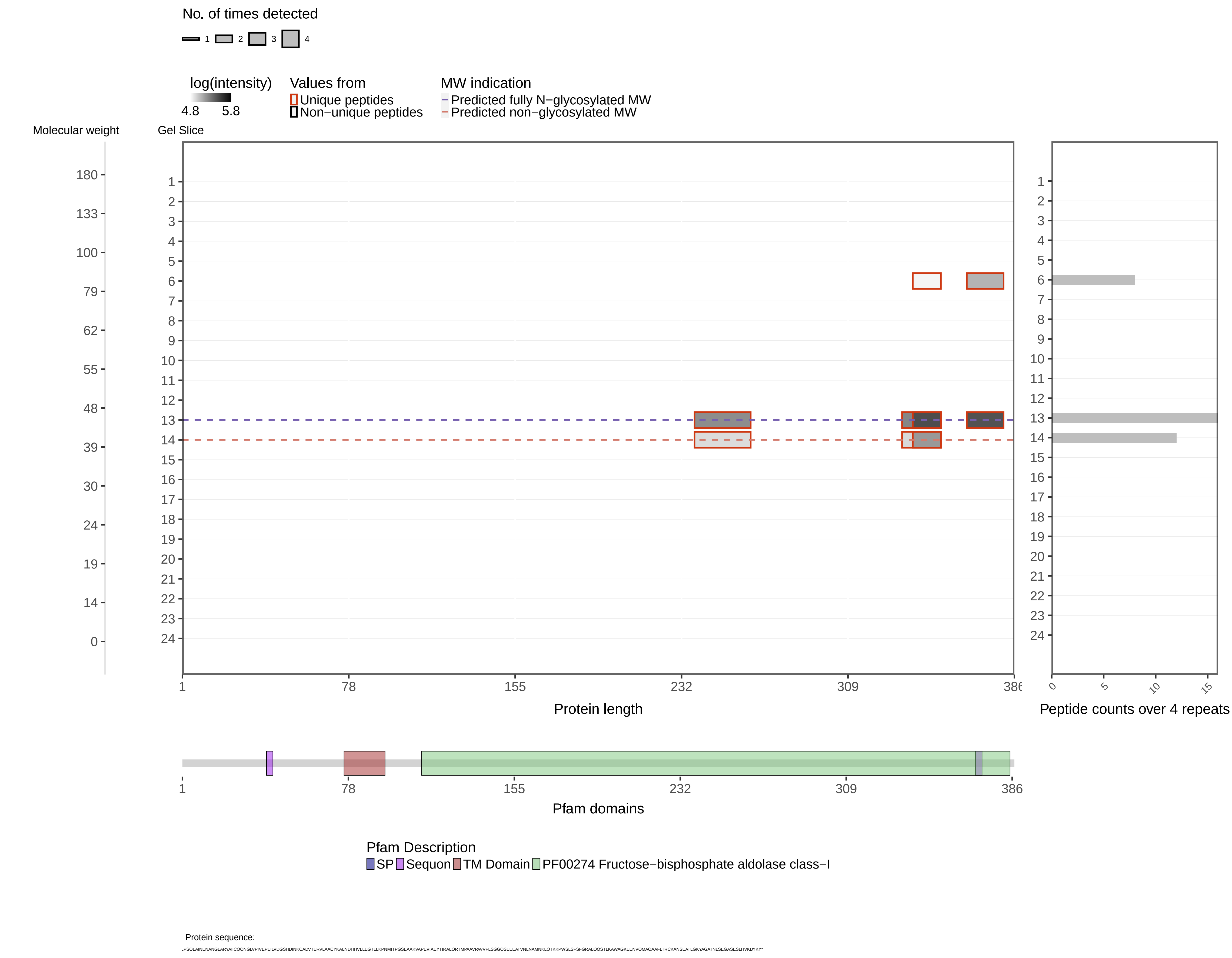

NbL05g02530.1 14−alpha−glucan−branching_enzyme_1_chloroplastic/amyloplastic−like [MW = 99.21 kDa]

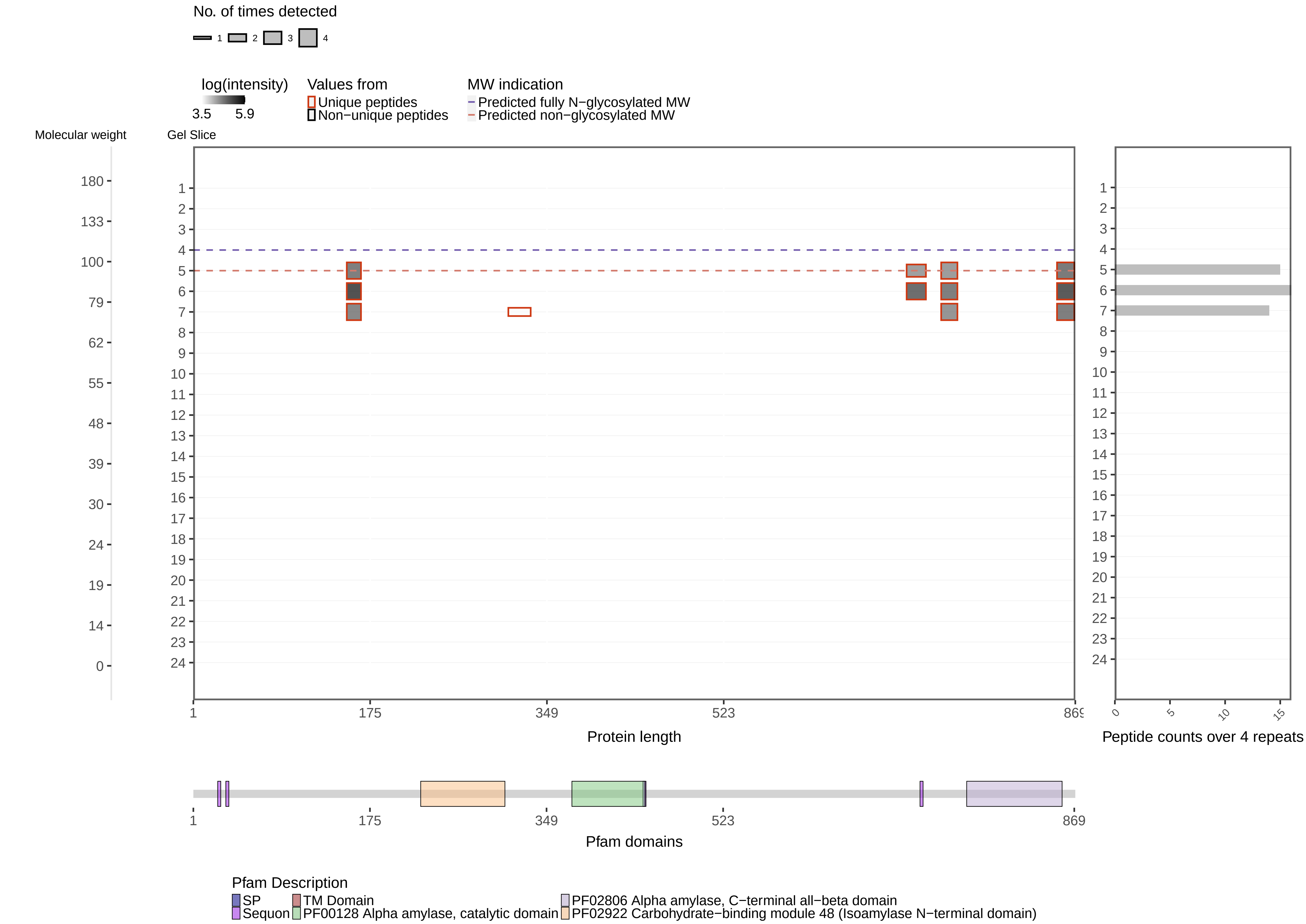


Protein sequence:

AHELGIVVLMDIVHSHASNNTLDGLNMFDGTDSCYFHSGSRGYHWMWDSRLFNYGHWEVLRYLLSNARWWLDEFKFDGFRFDGVTSMMYTHHGLSVTFTGNYNEYFGFATDVDAVVYLMLVNDLIHGLFPDAITIGEDVSGMPTFCIPVQDGGVGFDYRLHMAIADKWIELLKKRDEDWRVGDIVHTLTNRRWSEKCVSYAESHDQALVGDKTIAFWLMDKDMYDFMALDRPSTPLIDRGIALHKMIRLITMGLGGEGYLNFMGNEFGHPEWIDFPRAEQHLPDGQVI[illegible]

### NbL05g02520.1 uncharacterized_LOC109207925 [MW = 35.31 kDa]

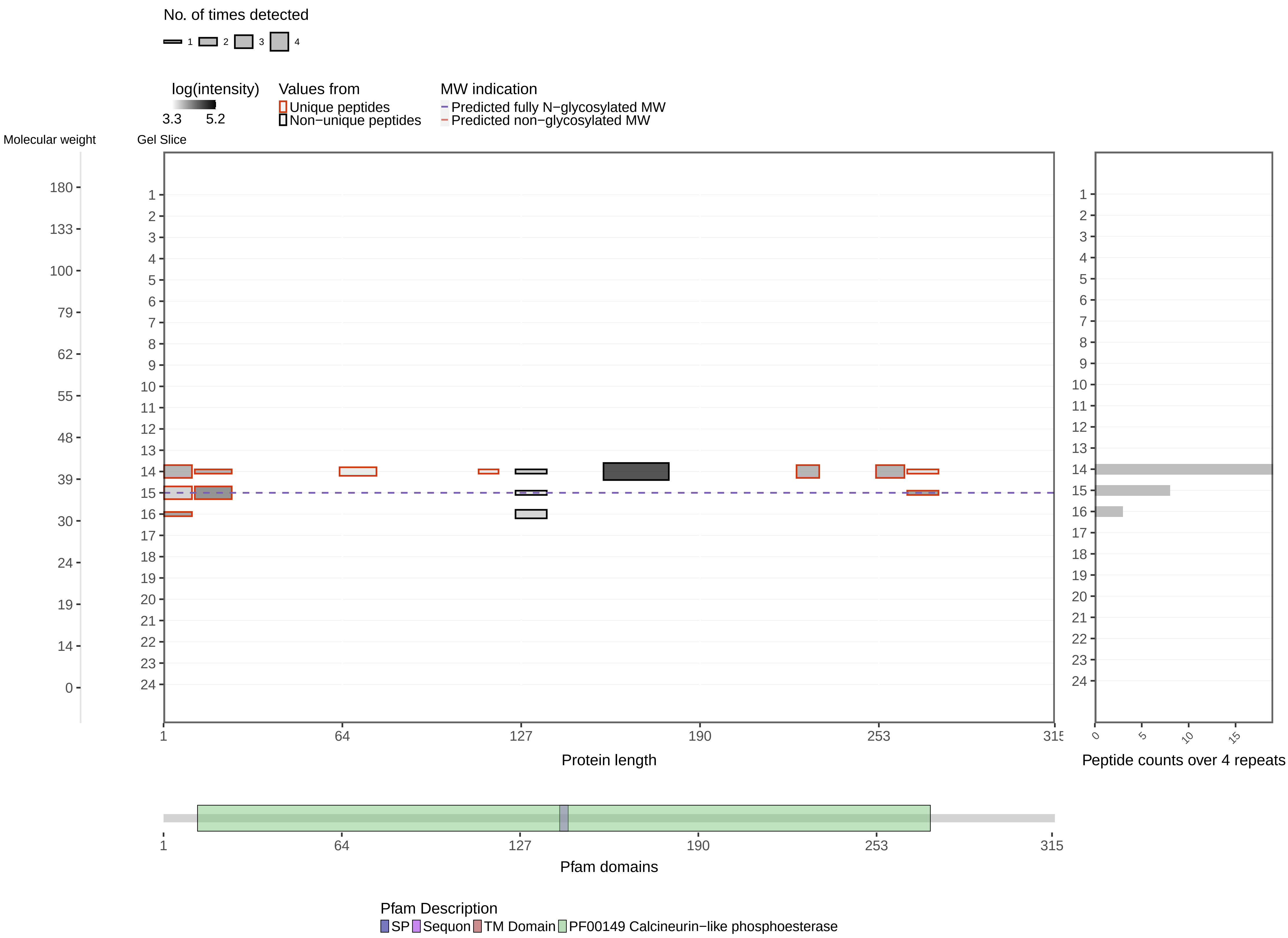


Protein sequence:

ITVKFNTVKGIEYKGSIYDAAPTFESYGVPHGSADLMKAVPDEHKKFLADLVWIHEEDDVCIKADEGIKSCKLIAVHAGLEKNKDVEEQIKTLKAKDTRISKVEALSGRKNVWEIPQELIRTPAIVVSGHHGKFHIEGLRLVIDEGGGLEGNPVAAVVLPSMEVVRDTDHLVK*

NbL05g02460.1 monodehydroascorbate_reductase [MW = 47.09 kDa]

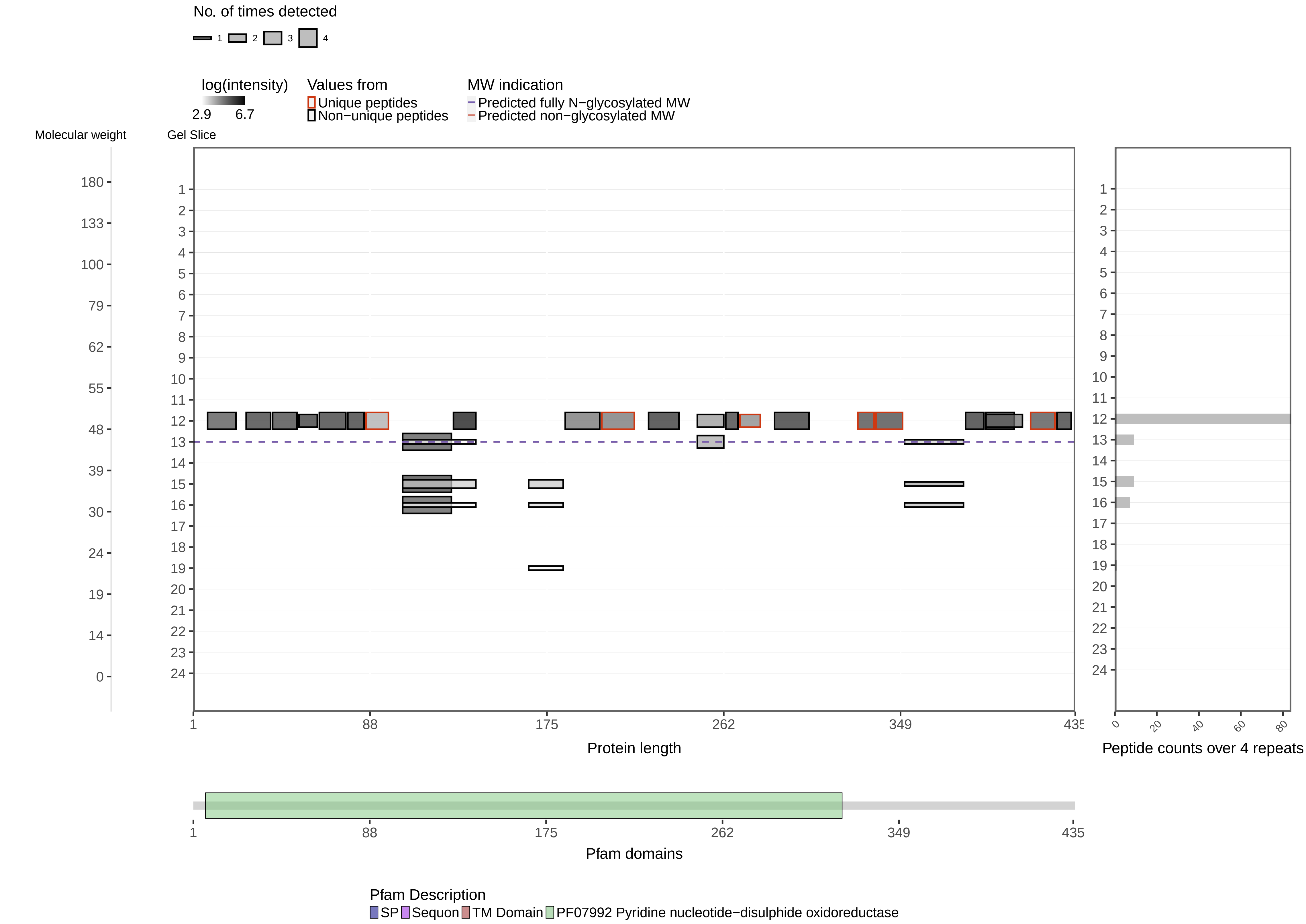


Protein sequence:

IAAFYEGYYQNKGVNIIKGTVAVGFDTHPNGEVKEVKLKDGRVLEADIVVVGVGARPLTTLFKGQVEEEKGGIKTDAFFKTSVPDVYAVGDVATFPLKMYNDIRRVEHVDHSRKSAEQAVKAIFASEQGKSVDEYDYLPYFYSRAFDLSWQFYGDNVGETVLFGDADPNSATHKFGQYWIQDGKVVGVFLESGTPEENKAIAKVAKVQPLVSSLDQLAKEGLSFASKI*

### NbL05g02360.1 DNA_ligase_1 [MW = 75.19 kDa]

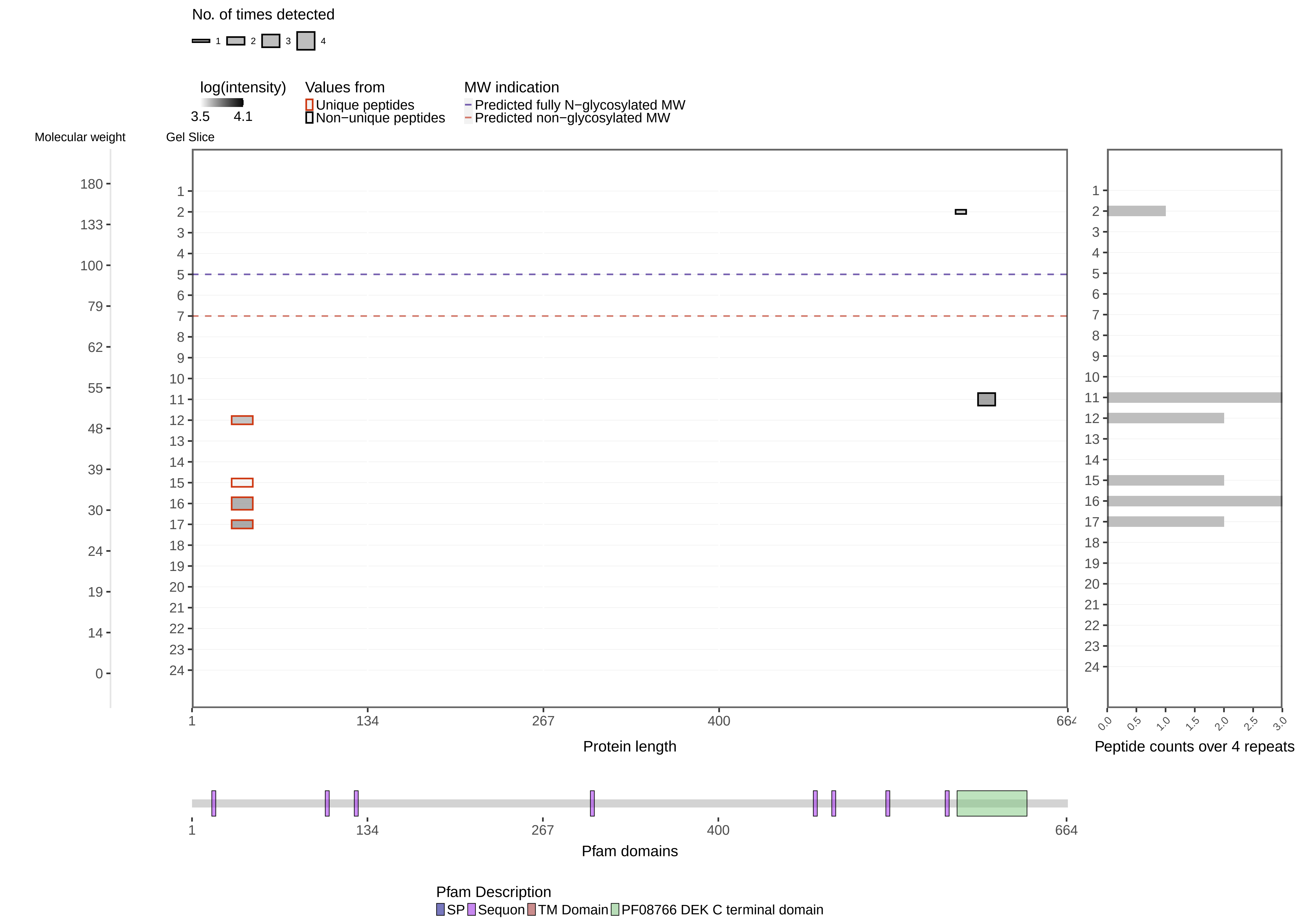


Protein sequence:

KQKMKVKEKLDKYVKEKLLEFCDVLDIPVSKATSRKEDIVTKLLDFLEAPHAMTSELLAEKEQSSKGKKRKRESKKSPSSTSVSSKGSTKSGKKTETAPQKADEKKTVHESEDESEQEDDHEEESEEEKANGVPEKSDDEMSEKGESEKESEPENESEEDRQKRKQSSNKSSTKKEPVRKAKTKKPAVAKLPSPAPKKTPSKSPKSSKHNASADASPKVVSAKKKKTEVIKEKSSIPKKSTSKENTGKKIVKGKNESKEDKLRPSDDELRNAICEILKEVDFNTATFTDILKQ

NbL05g02020.1 syntaxin−51−like [MW = 25.87 kDa]

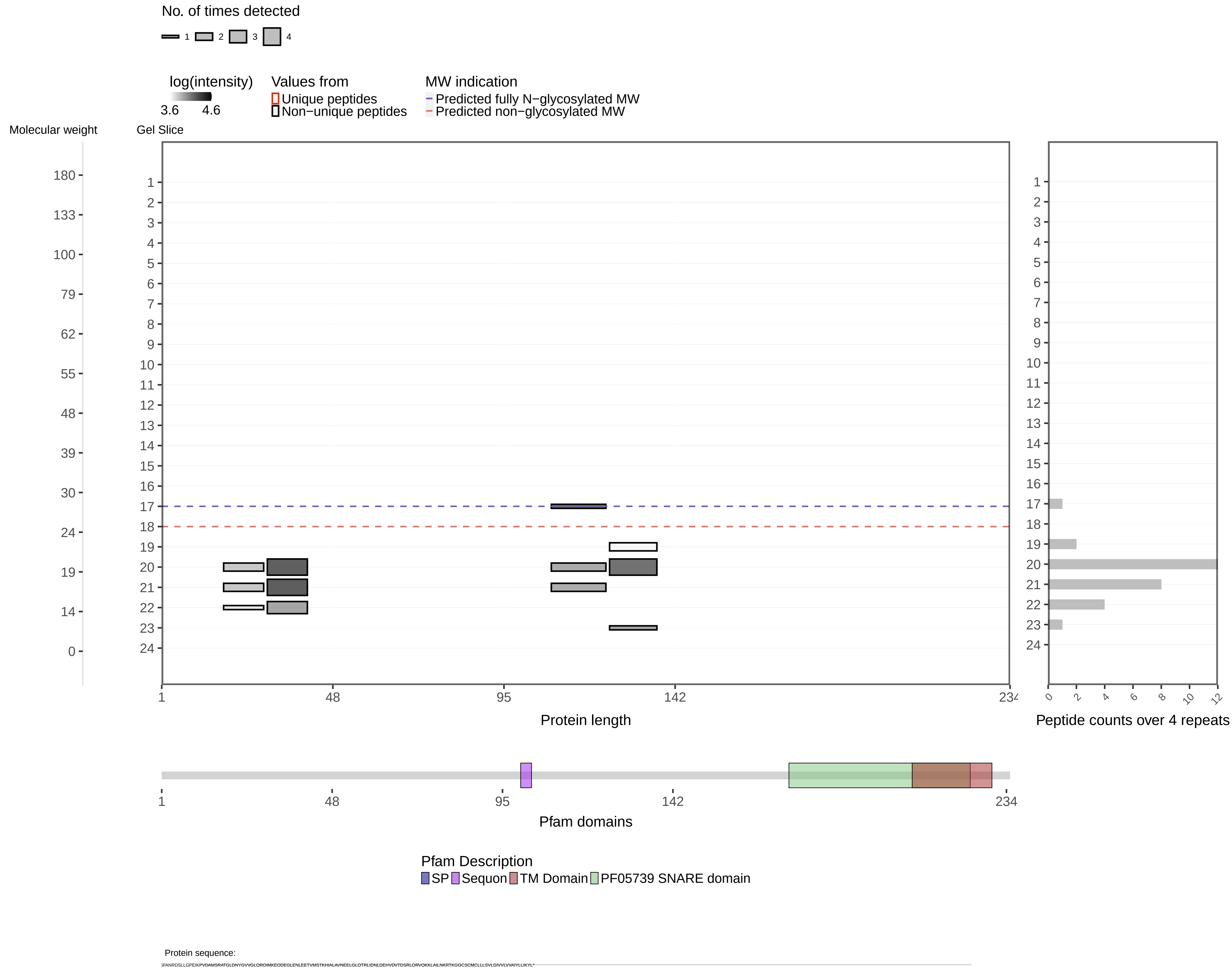

NbL05g02000.1 serine/threonine−protein_phosphatase_PP−X_isozyme_2 [MW = 35.72 kDa]

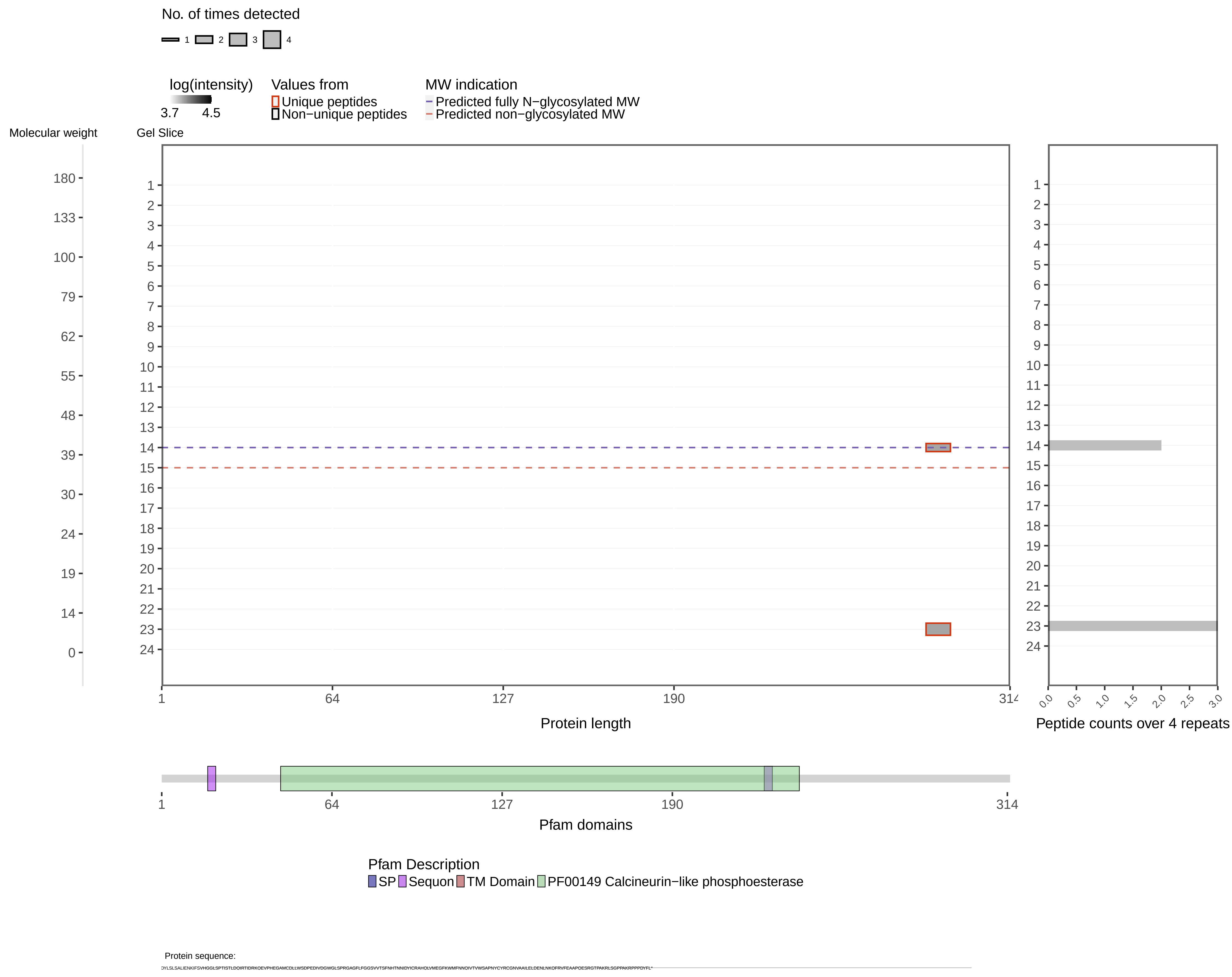

NbL05g01940.1 glycerophosphodiester_phosphodiesterase_GDPDL3−like [MW − SP = 79.67 kDa]

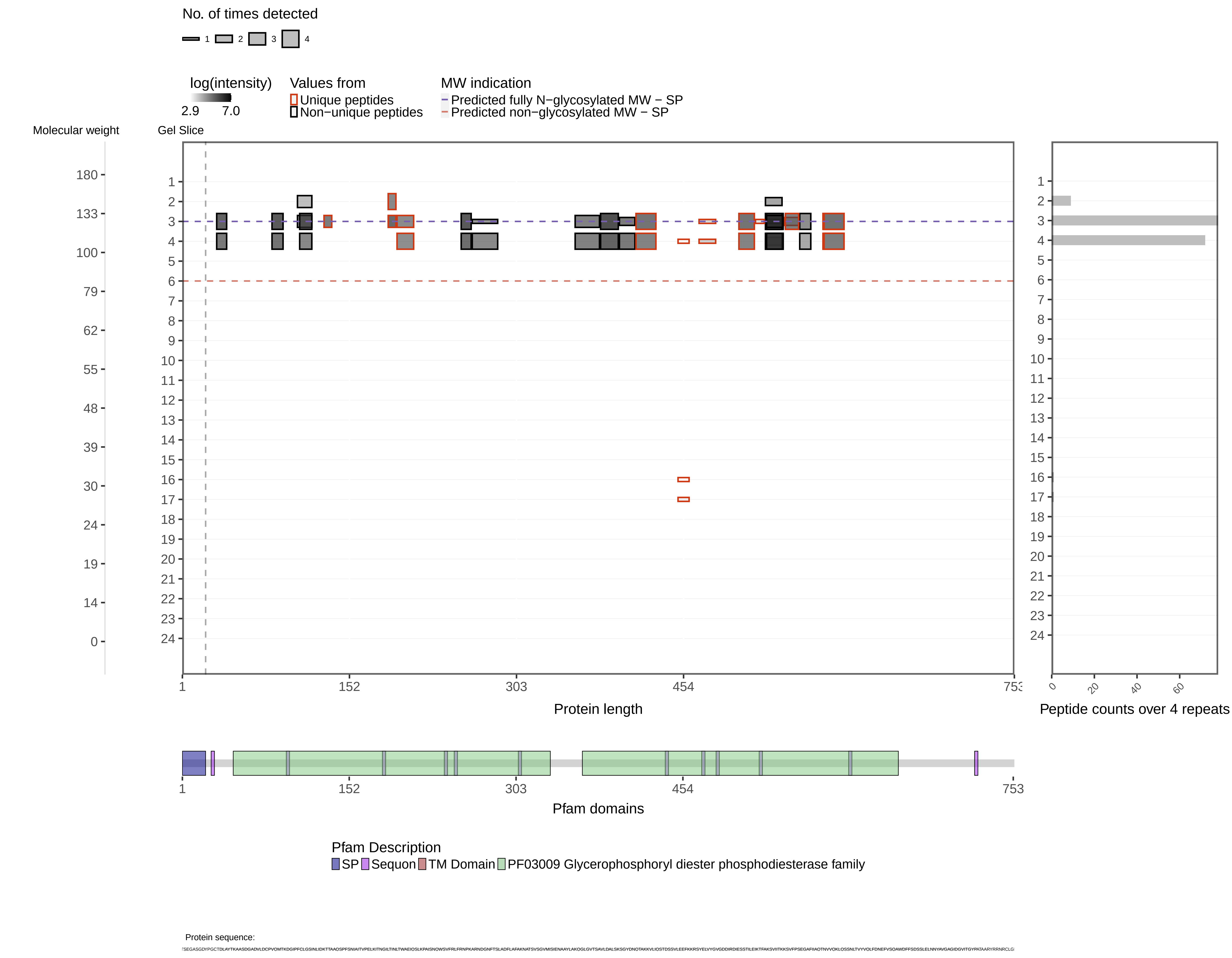

NbL05g01900.1 40S_ribosomal_protein_S8−like [MW = 28.87 kDa]

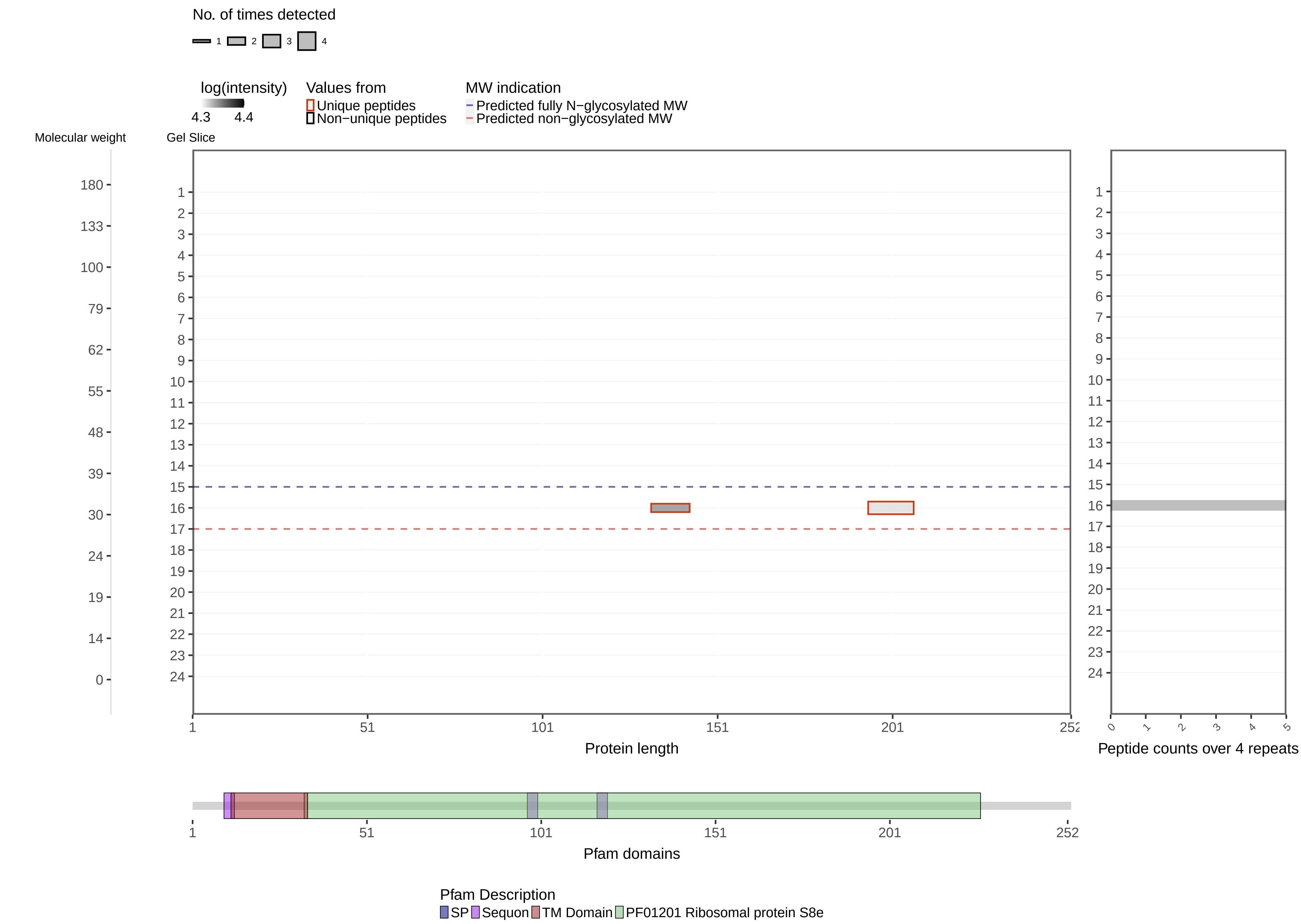


Protein sequence:

VVYNASNNELVRTQTLVKGAIVQVDAAPFKQWYLQHYGVDIGRKKKTAASKKEGEEAEAVPEEKKSNHVQRKLQKRQQDRKIDPHIEEQFSSGRLLAAISSRPGQCGRADGYILEGKELEFYVKKLQRKKGKGASGAA*

### NbL05g01890.1 uncharacterized_LOC107809878 [MW = 35.24 kDa]

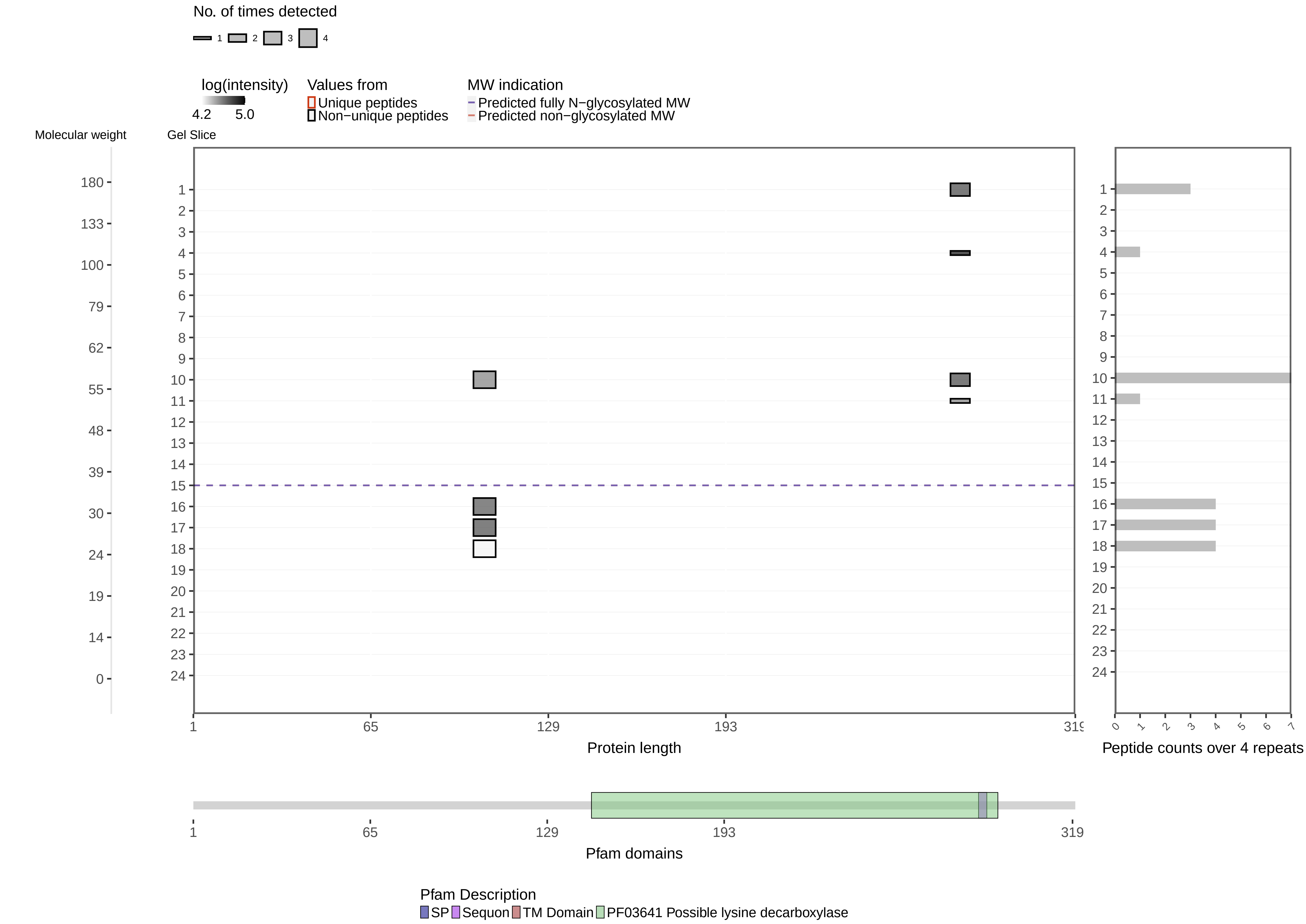


Protein sequence:

[illegible]AATQGALQAGKRVGGFKIGREAGEWTASNFHPYLPSESYLTCRFFSARKHGLVDAVVRCKSSERTAVVALPGGIGTLDEIFEVMALIQLERIGSQLPVPFLLMNYDSFYSNLLEFLNDCEKWGTVSKNEVASLWKVCNNNSEALAYLTEFYGLSPVETGRKDPESSECRSSP*

### NbL05g01760.1 imidazole_glycerol_phosphate_synthase_hisHF_chloroplastic [MW = 65.05 kDa]

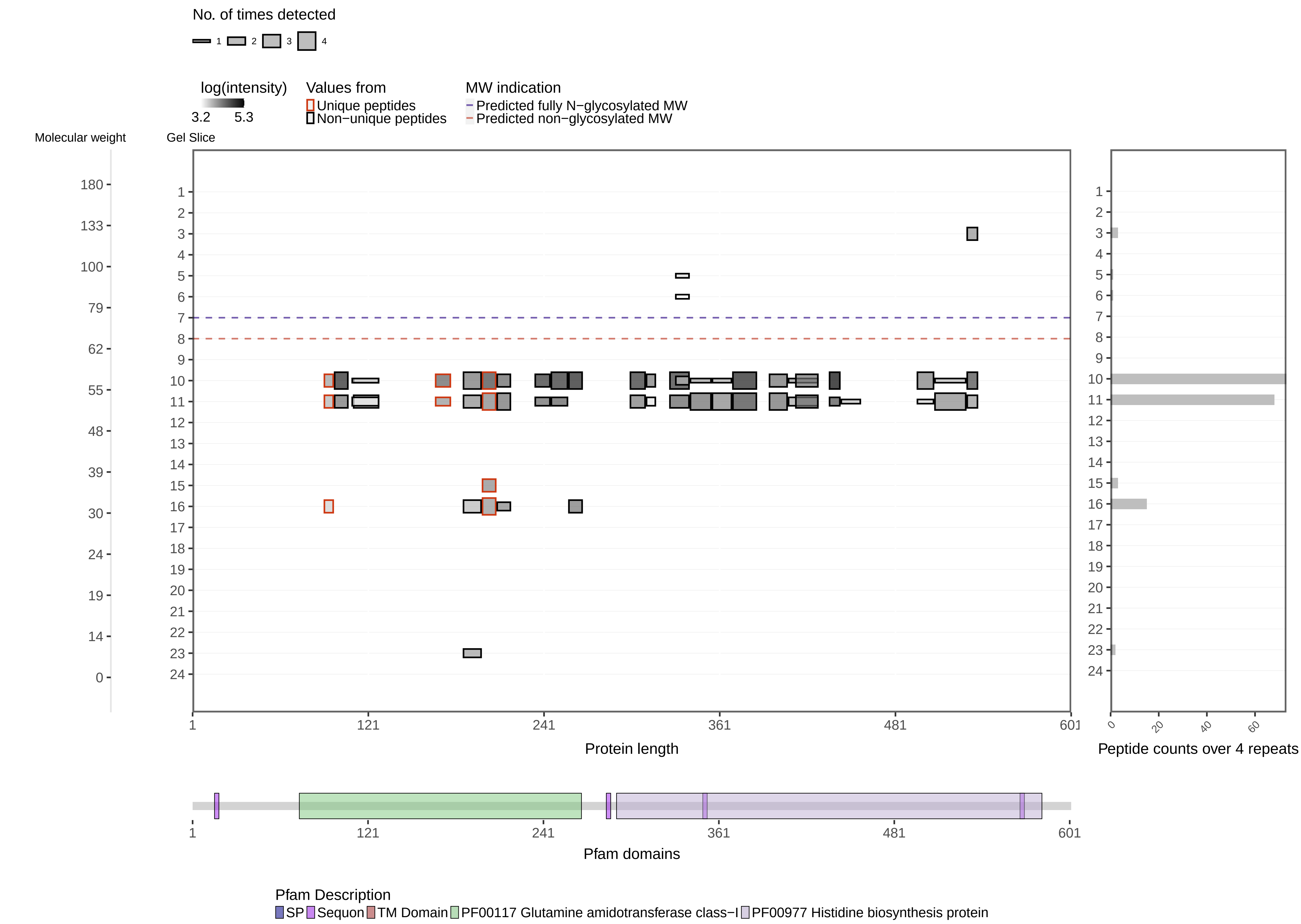


Protein sequence:

LAKRVIACLDVRTNDKGDLVVTKGDQYDVREHTKENEVRNLGKPVDLAGQYYKDGADEVSFLNITAFRDFPLGDLPMLQVLRYASENVFVPLTVGGGIRDFKDGNGRYYSSLEVASEYFRSGADKVSIGSDAVYAAEEYLKSGVKTGKSSLEQISRVYGNQAVVVSIDPRRVYVKDPKAVEFRTVKVRSPGPNGEEYAWYQCTVSGGREGRPIGAYELAKAVEELGAGEILLNCIDCDGQGKGFDIDLIKLISDAVSIPVIASSGAGTVEHFSEVFTETNASAALAAGIFHRKE

NbL05g01750.1 14−alpha−glucan−branching_enzyme_3_chloroplastic/amyloplastic [MW = 103.76 kDa]

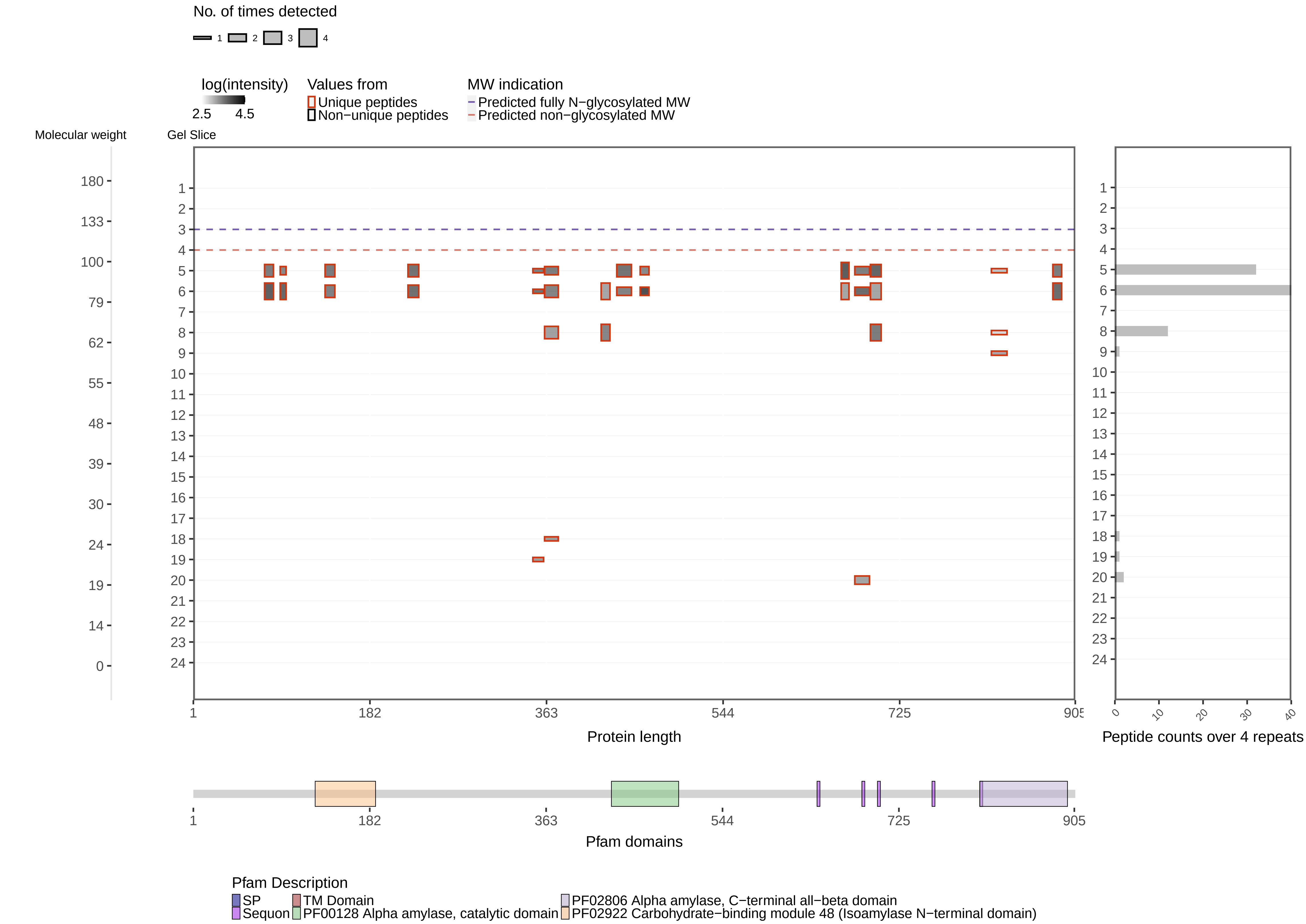


Protein sequence:

[illegible]

NbL05g01690.1 GDSL_esterase/lipase_At5g55050−like [MW − SP = 36.68 kDa]

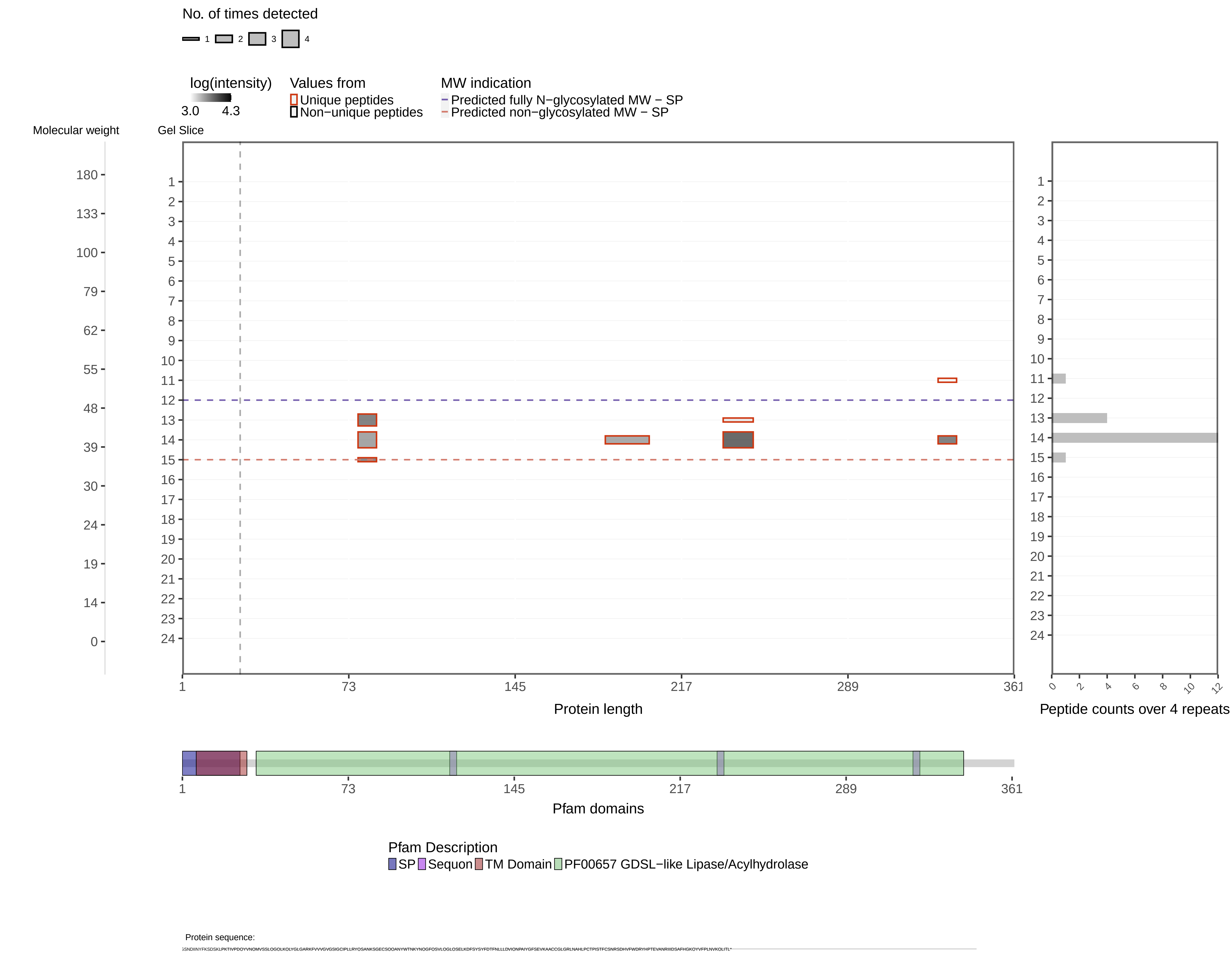

NbL05g01640.1 bifunctional_bis(5'−adenosyl)−triphosphatase/adenylylsulfatase_FHIT [MW = 24.35 kDa]

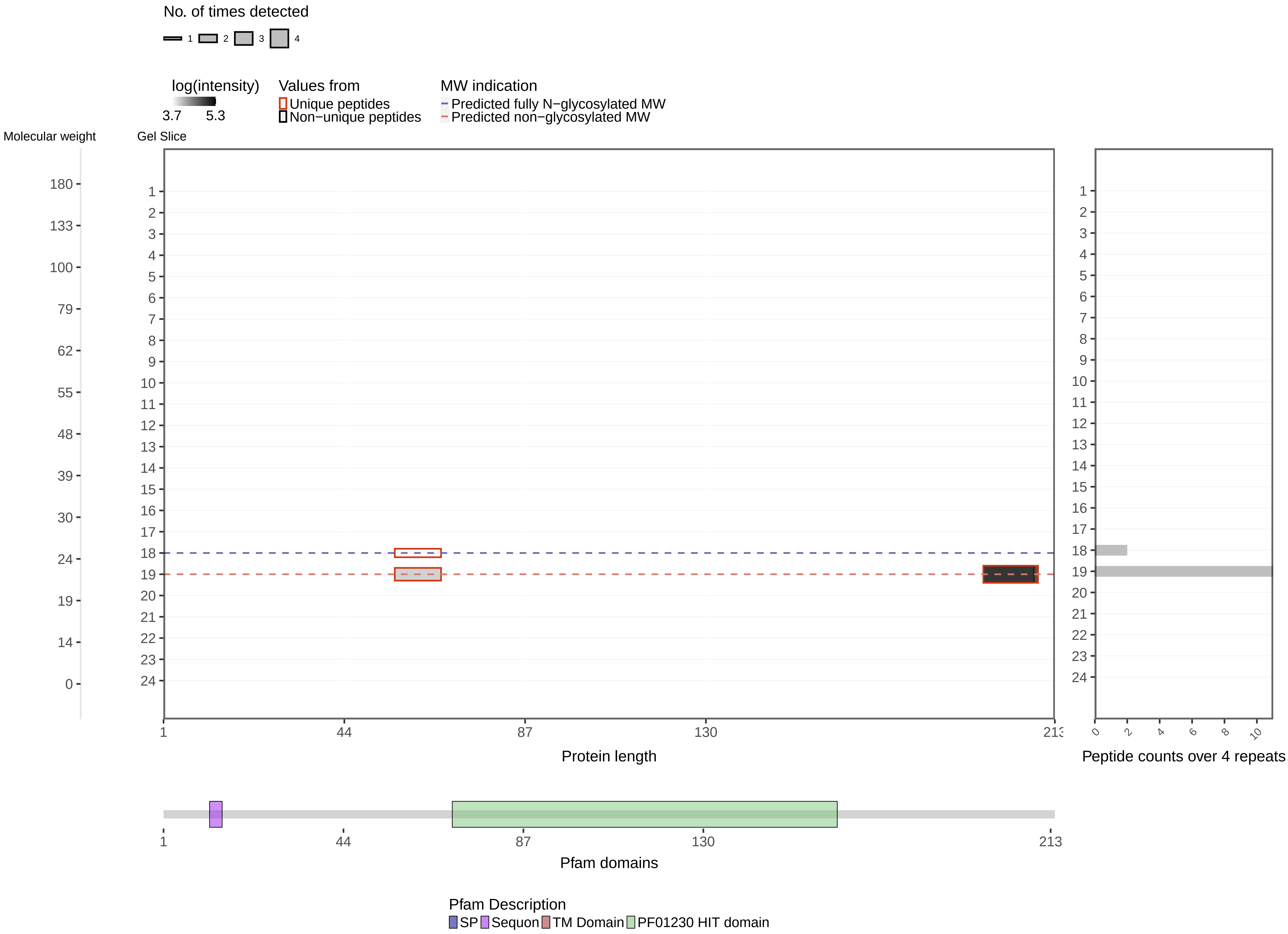


Protein sequence:
/LVCPRRQVKRFVDLTADETSDLWLTAQKVGKQLESYHKASSLTFAIQDGPQAGQTVPHVHIHIIPRKSGDFEKNDEIYDALDVKEKEMKQSLDLDKERKDRSIEEMAEEADEYRKLFQ*

### NbL05g01590.1 CBS_domain−containing_protein_CBSX3_mitochondrial [MW = 22.6 kDa]

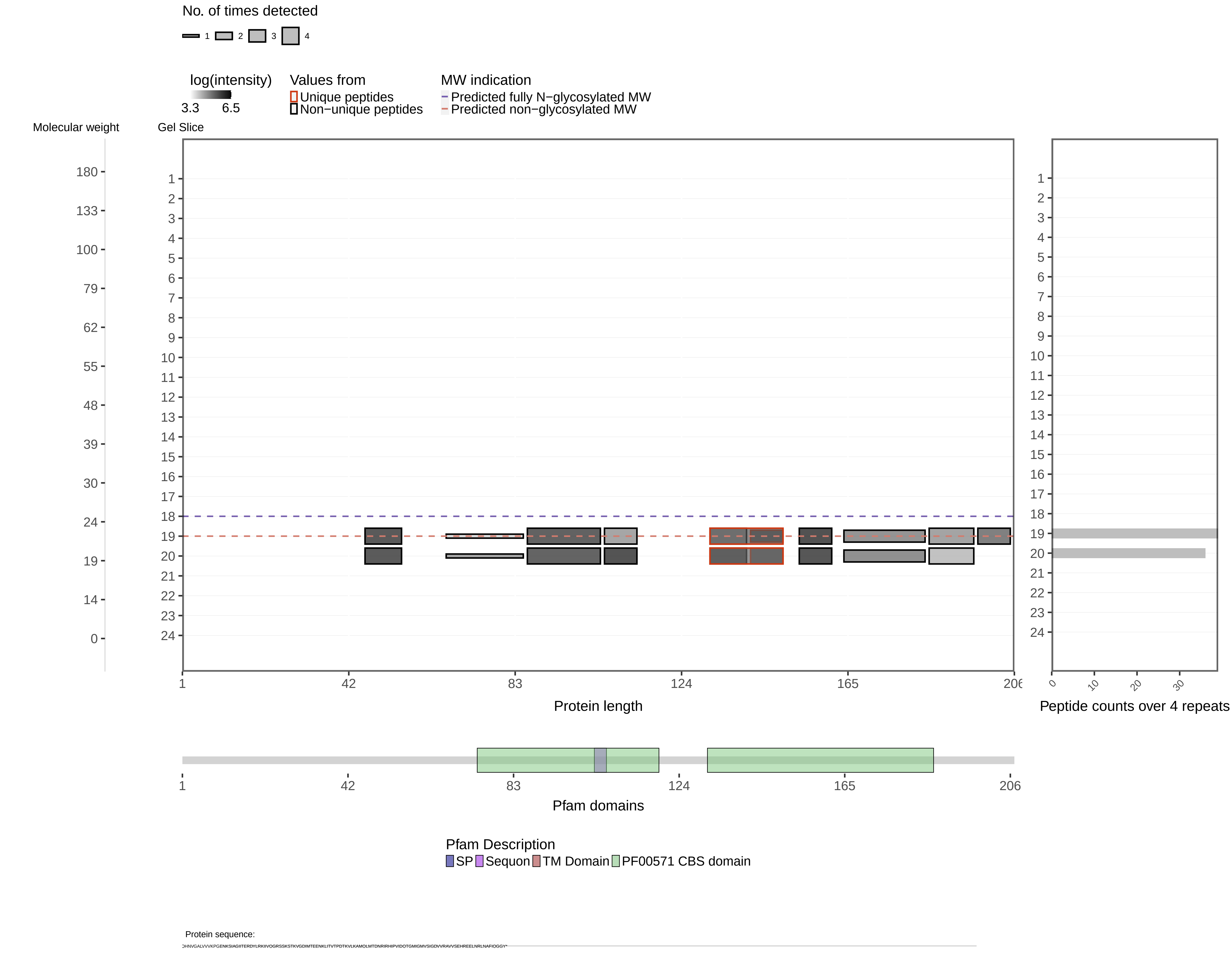

### NbL05g01570.1 oligoribonuclease−like [MW = 28.23 kDa]

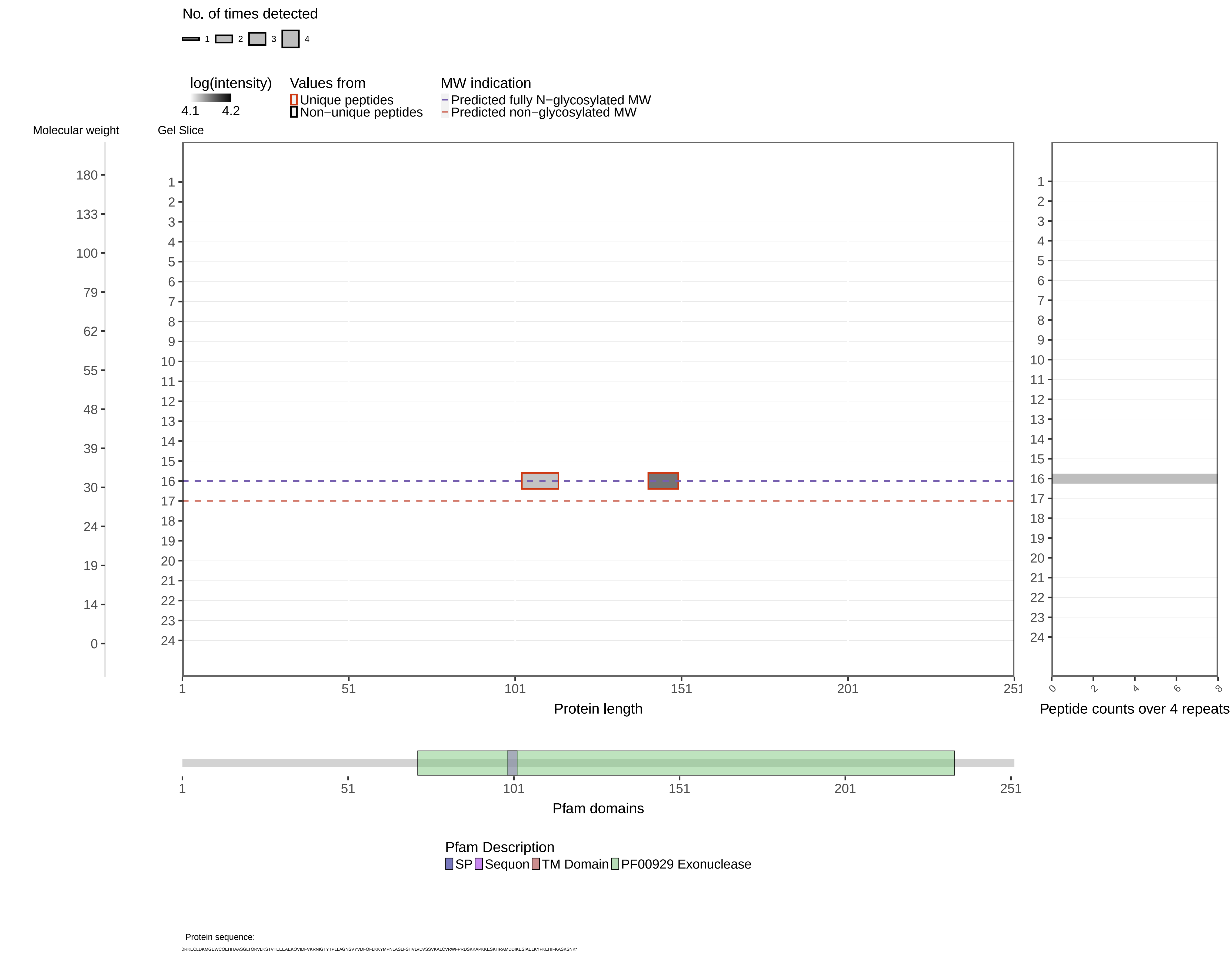

### NbL05g01530.1 uncharacterized_LOC107809774 [MW = 22.76 kDa]

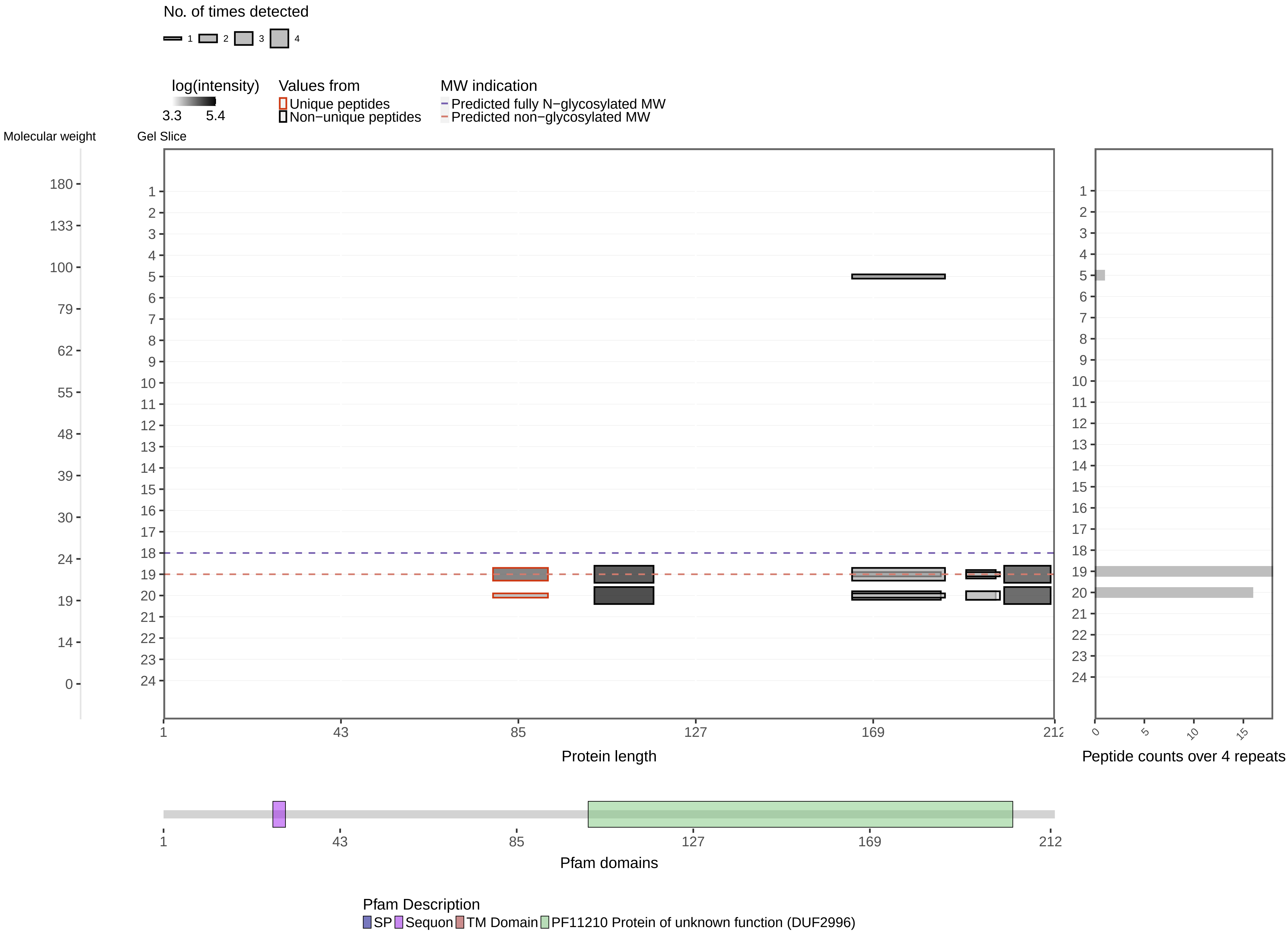


Protein sequence:

[illegible]PAAKAPAKSLPQMMEEDVIPSLKTILEAQDDISELELMFNDNKLEGSFSKKGNPYSFWAFFPDGLTGAKGFSLSSYGSGPSTVEPFLVDEKKITAKHVVFWVEKRLAAQGIIPVWNE*

### NbL05g01440.1 60S_ribosomal_protein_L5 [MW = 37.44 kDa]

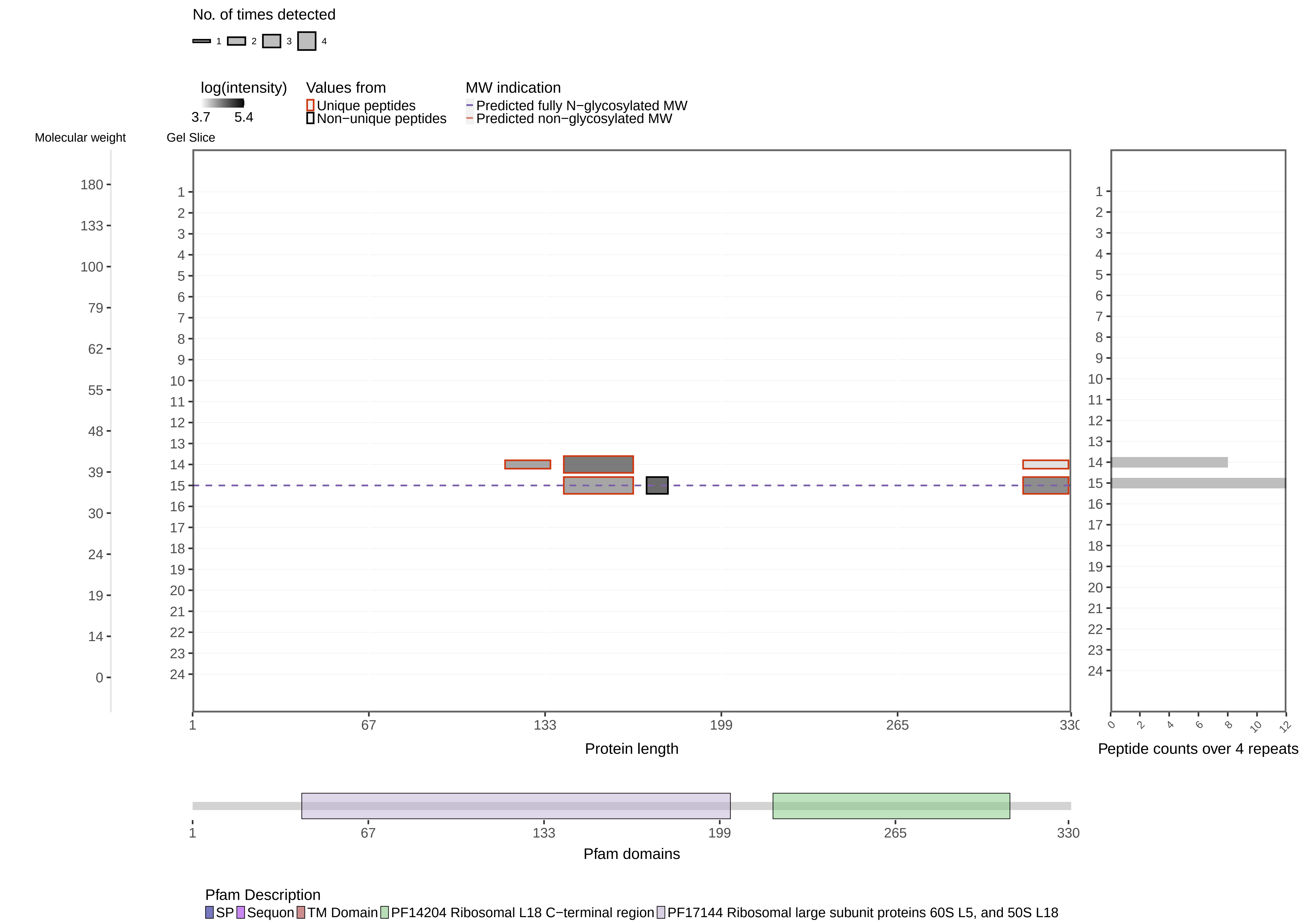


Protein sequence:

ATGEDYSVEPAESRRPFRALLDVGLVRTTTGNRVFGALKGALDGGLDIPHSEKRFAGFSKDSKOLDADVHRKYIYGGHVATYMKTLTEDEPEKFOSHFSEYIKOGLEADDLEEMYKKVHAAIRADPSPKKSEKOPPKOHKRYNLKKLTYEERKAKLIERLNALNSAAGNDEEEEEDD*

NbL05g01240.1 leucine−−tRNA_ligase_cytoplasmic−like [MW = 123 kDa]

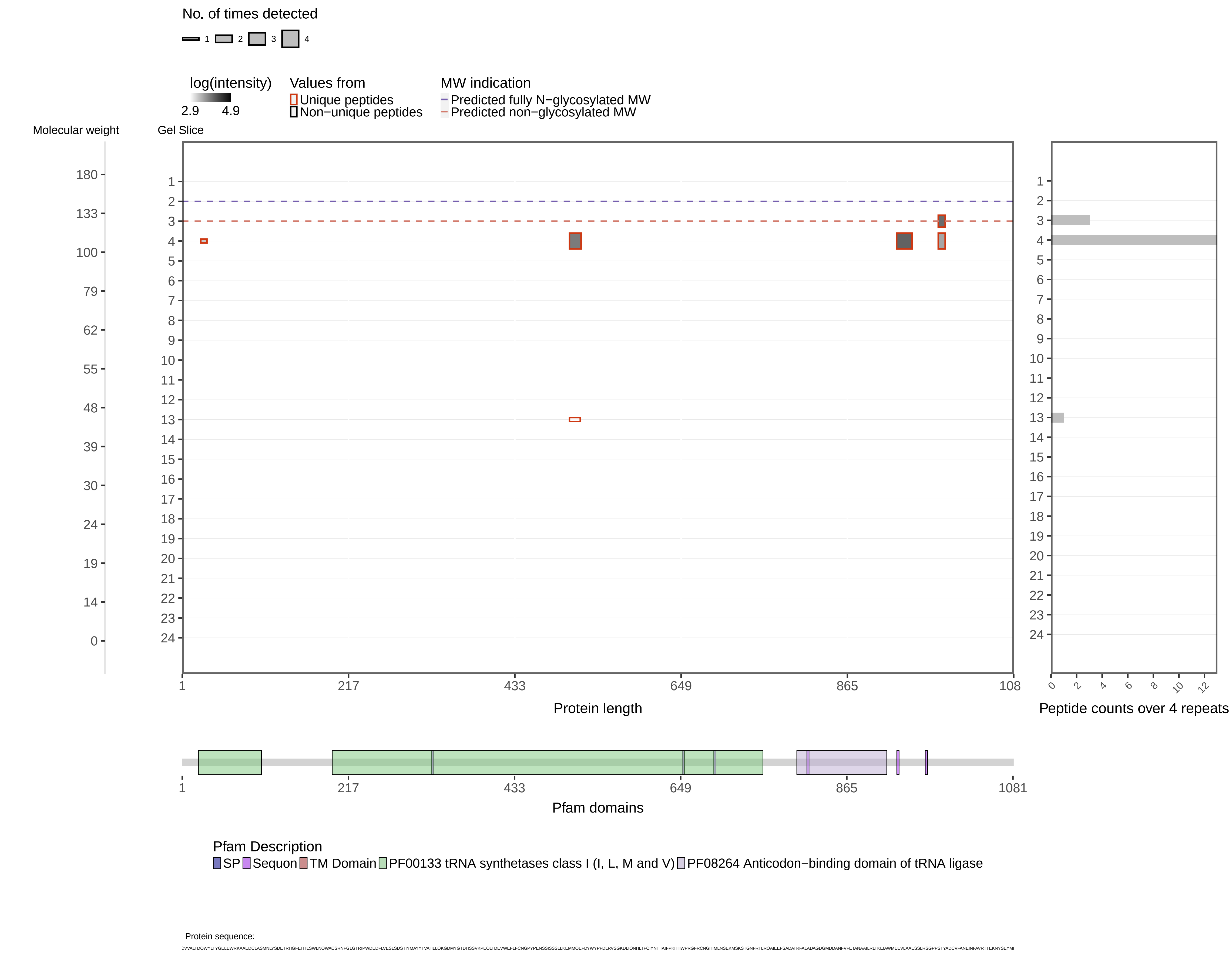

NbL05g01220.1 2−Cys_peroxiredoxin_BAS1_chloroplastic−like [MW = 34 kDa]

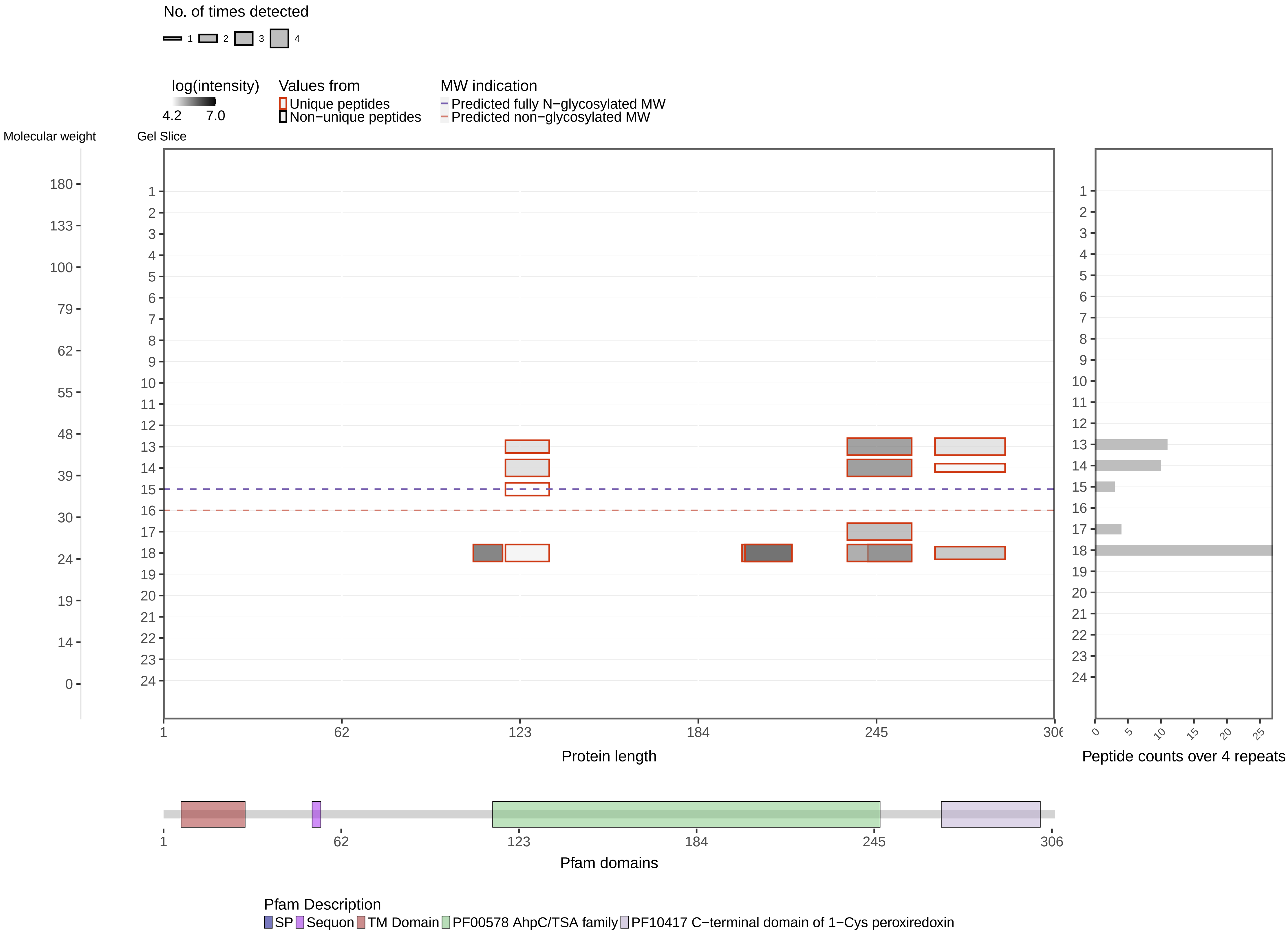


Protein sequence:

IGKKYVILFFYPLDFTFVCPTEITAFSDRYEEFQKVNTEILGVSVDSVFSHLAWVQTDRKSGGLGVLNYPLISDVTKSISKSYNVLIPDQGIALRGLFIIDKEGVIQHSTINNLAIGRSVDETLRTLQALQYVQENPDEVCPAGWKPGDKSMKPDPKGSKEYFASI*

NbL05g01120.1 ubiquitin−conjugating_enzyme_E2_variant_1D−like [MW = 16.71 kDa]

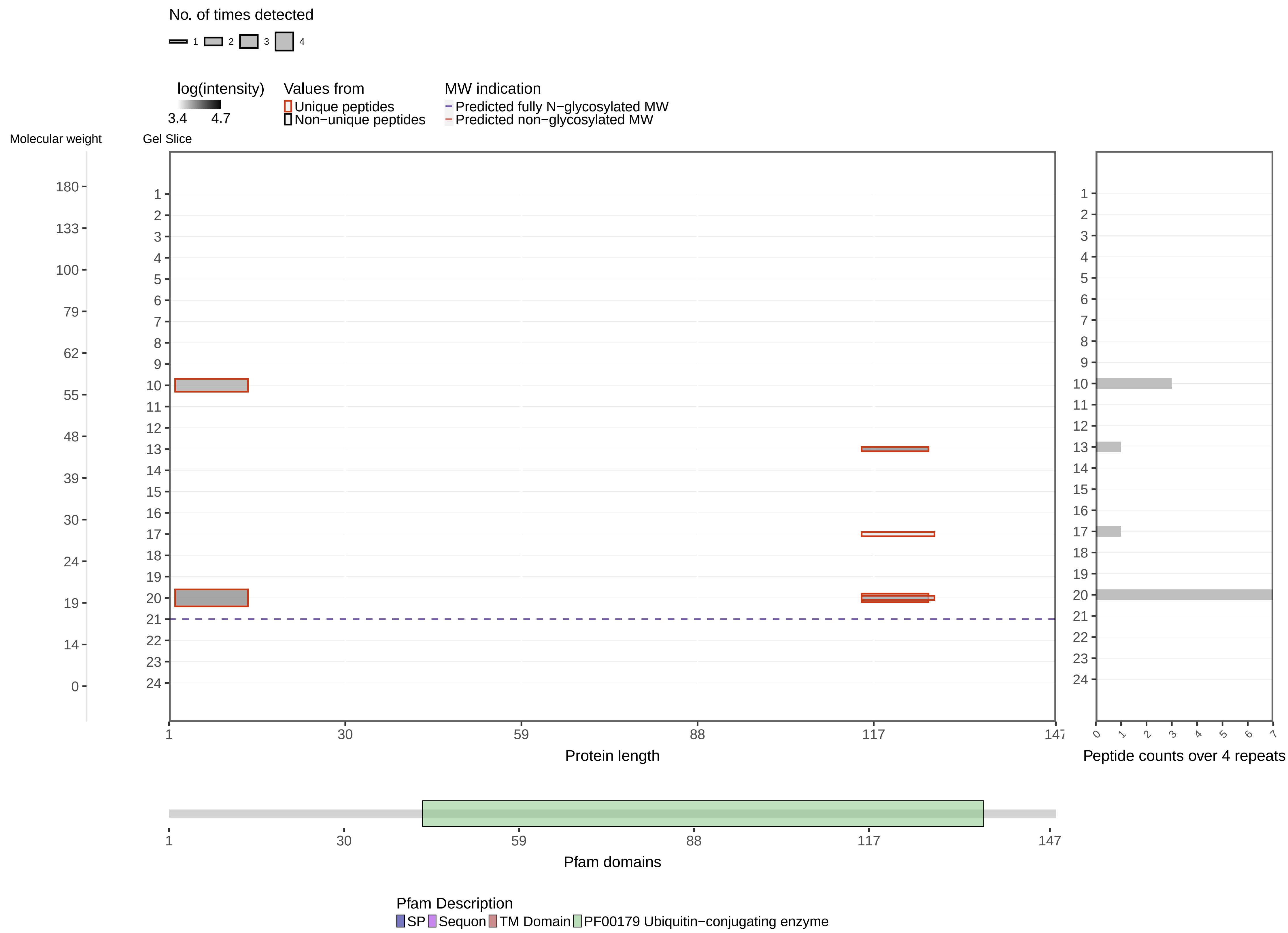


Protein sequence:

HEGRIYQLKLFCDKDYPEKPPSVRFHSRINMACVNHETGVVEPKKFDLLANWQREYTMEDILTQLKKEMASPHNRKLVQPPEGTYF*

### NbL05g00410.1 topless−related_protein_4−like [MW = 124.36 kDa]

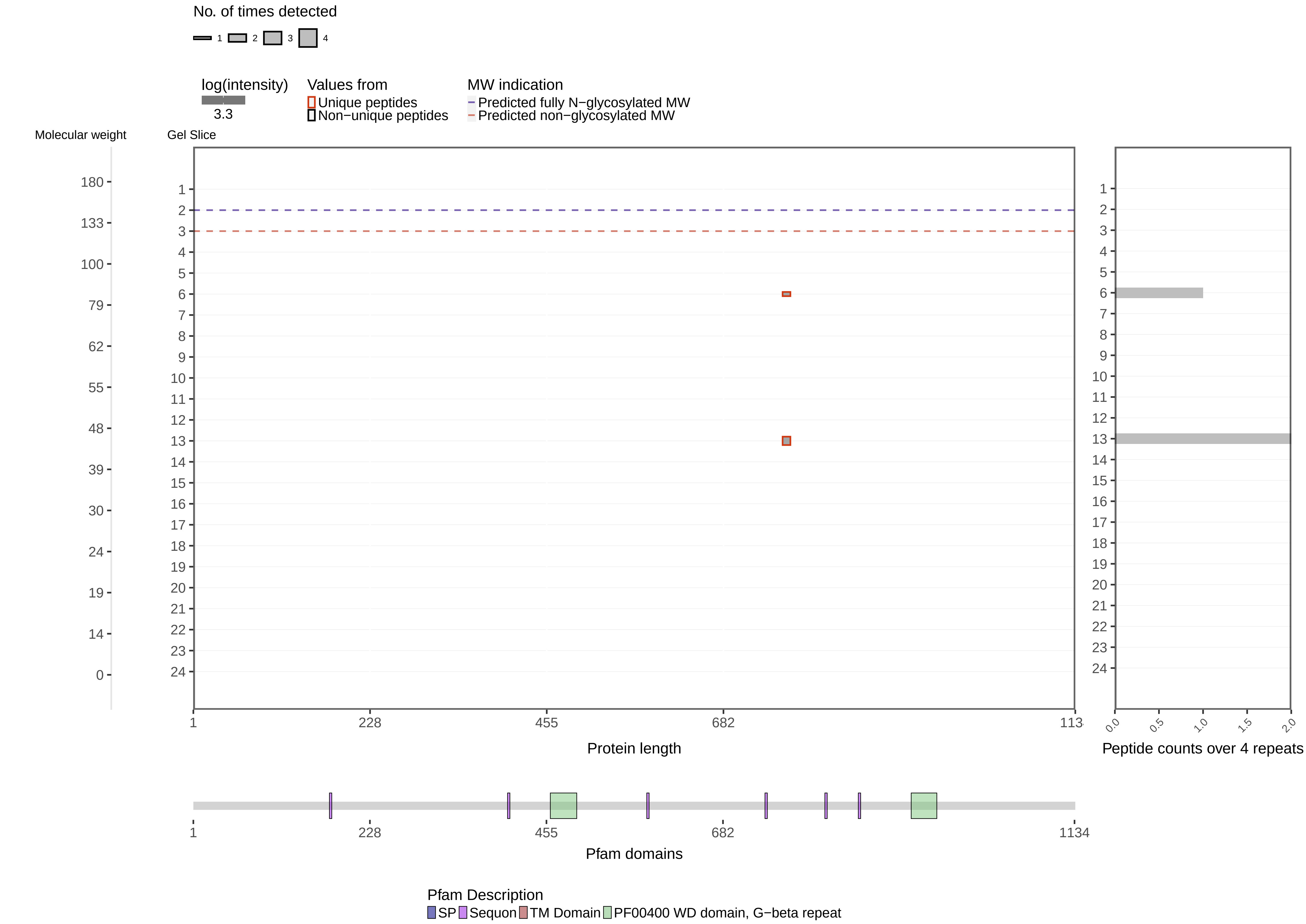


Protein sequence:

### NbL05g00330.1 malate_dehydrogenase_[NADP]_chloroplastic [MW − SP = 50.52 kDa]

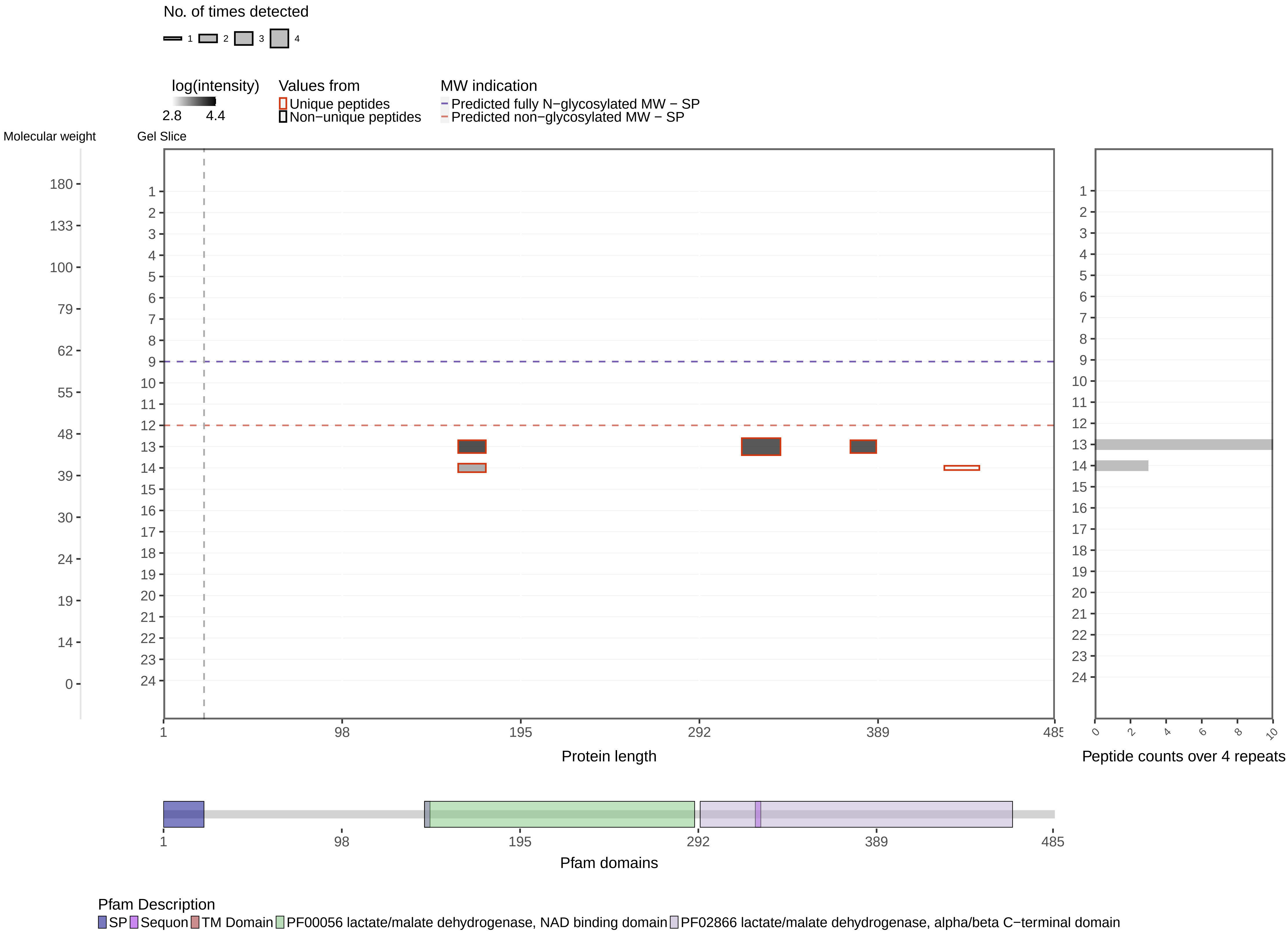


Protein sequence:

PGMERADLLDINGQIFAEQGKALNAVASRNVKVIVVGNPCNTNALICLKNAPGIPAKNFHALTRLDENRAKCQLALKAGVFYDKVSNVIIWGNHSTAQVPDYLNAKINGFPVKEVIRDTKWLEEEFTEKVQKRGGVLIQKWGRSSAASTAVSVVDAIRSLVTPTPEGDWFSTGVYTDGNPYGIAEGIVFSMPCRSSGDGDYELVKDVIIDDYLRSRIKKSEDELLAEKRCVAHLTGEGIAVCDLPGDTMLPGEM*

NbL05g00280.1 non−symbiotic_hemoglobin_2 [MW = 17.63 kDa]

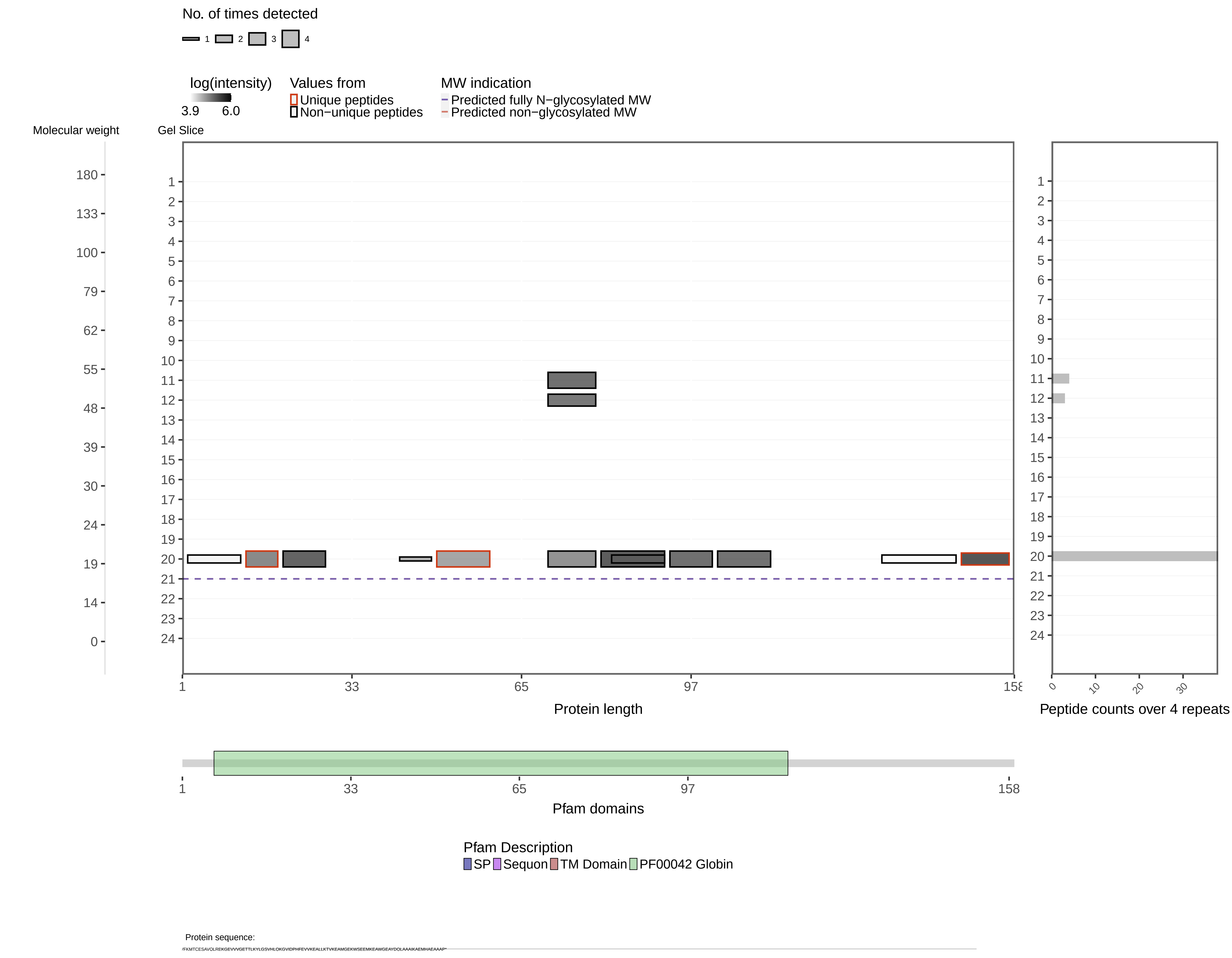

### NbL05g00230.1 wall−associated_receptor_kinase−like_20 [MW − SP = 65.73 kDa]

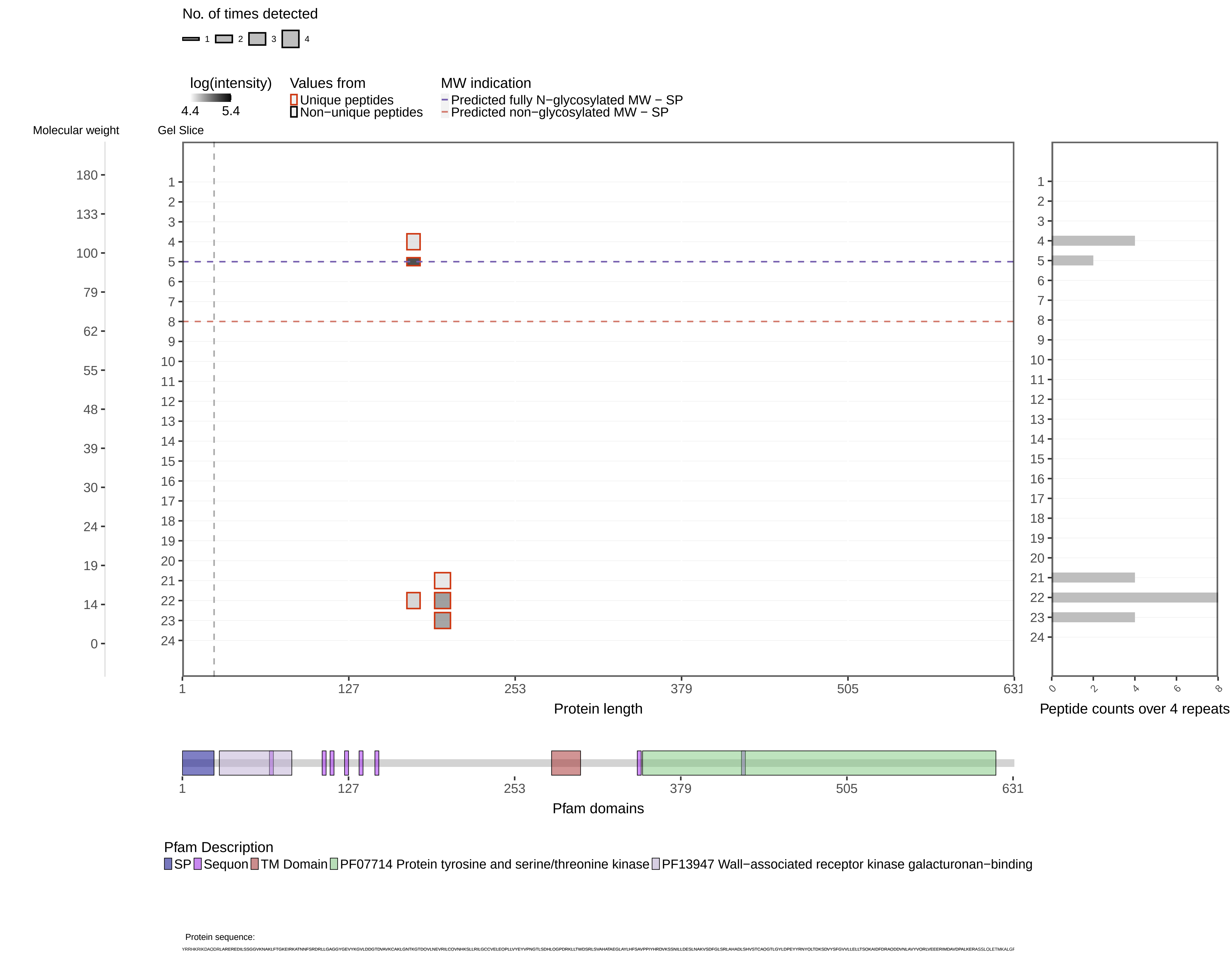

NbL05g00150.1 vacuolar_protein_sorting−associated_protein_27 [MW = 30.85 kDa]

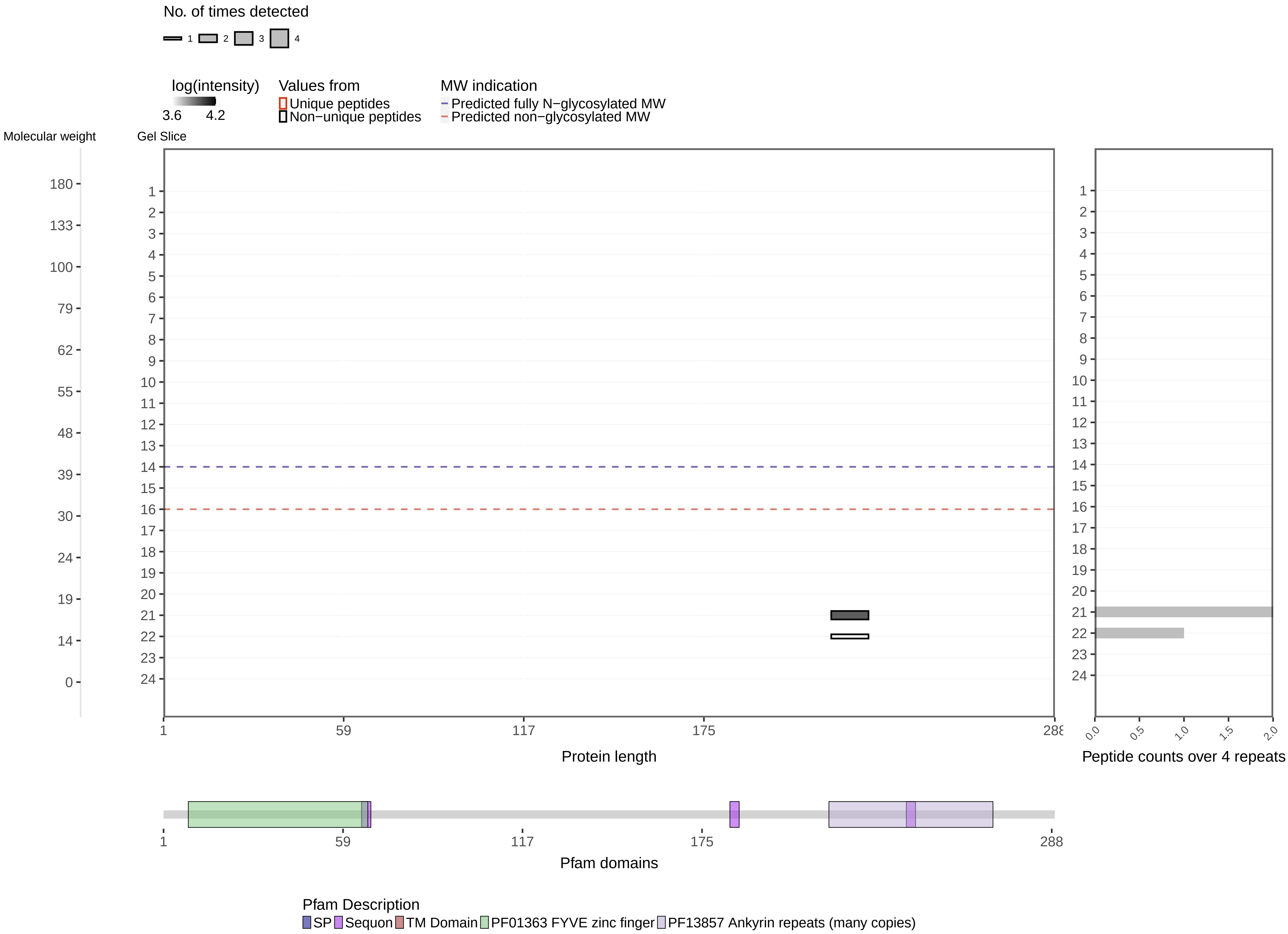


Protein sequence:

ATPTTSVTPQSKTMPNPIVNTNPKPKKENRIPTSRGSTSNNKQGTTFNTGPATNCSMETMSADYEVSGEGLREAIKNGDVSAAKKLLNQGVDANYLDKQGSSLLHLAAVFNRTEIAFALMESGASLYCKNSQGETPLDCAPATLQFKMKKKMEESGQ*

NbL05g00110.1 uncharacterized_LOC107765250 [MW − SP = 11.76 kDa]

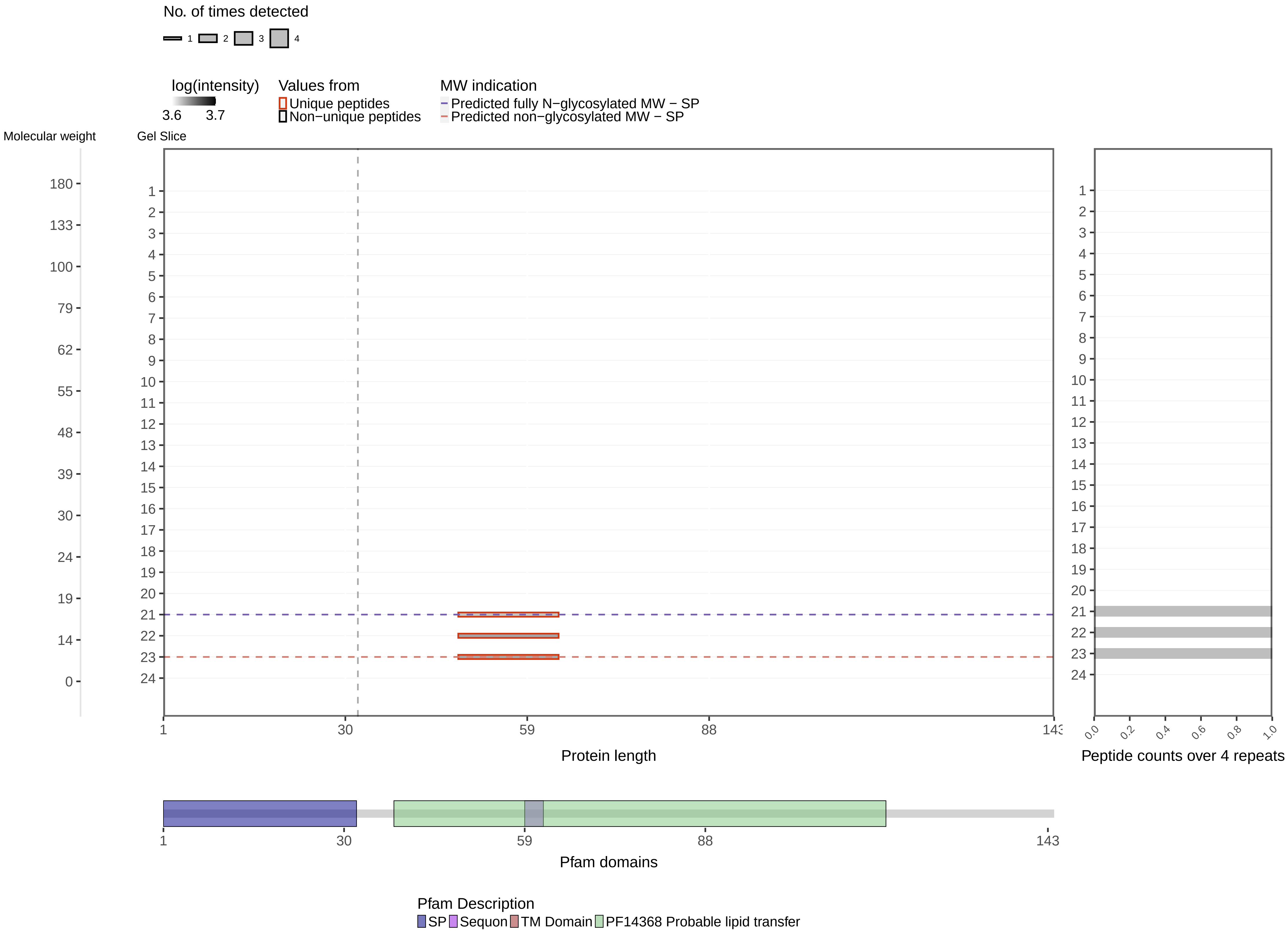


Protein sequence:

NASVSRSCCQQIQKLGQNPKCLCAVMLSNTAKDSGAKPEVAITIPKRCNLANRPVGYKCGRNLHVALKAAEDLDVFMDVMILLW*

NbL04g20090.1 uncharacterized_LOC109229657 [MW = 37.31 kDa]

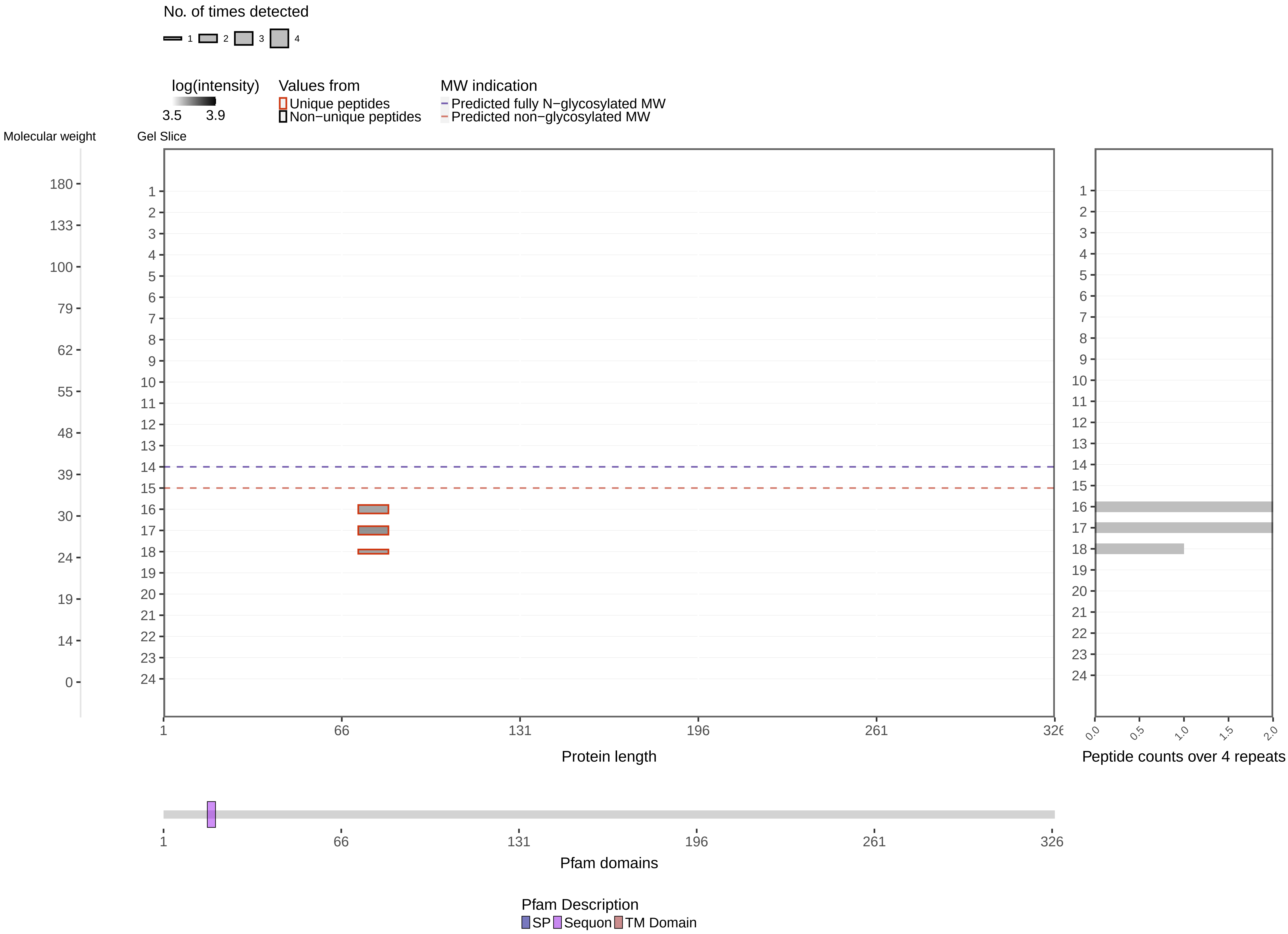


Protein sequence:

DTDVAEYLYKHNPDLDSLKSFLCKDLTKACSKASPPVPKNRAPGEPFVSKSSKEAEMEKLMRSMQDMPGAPGMQMYSREDLMNQNFGDEDADGDDDDDDEGGFPSKLGKVLKGKERKKNDWKQWIAKGIQDTSEVLKTHAYRVSHRMRNWWGARKVQWKKNSKTSKGEL*

### NbL04g19980.1 uncharacterized_LOC107790078 [MW = 34.77 kDa]

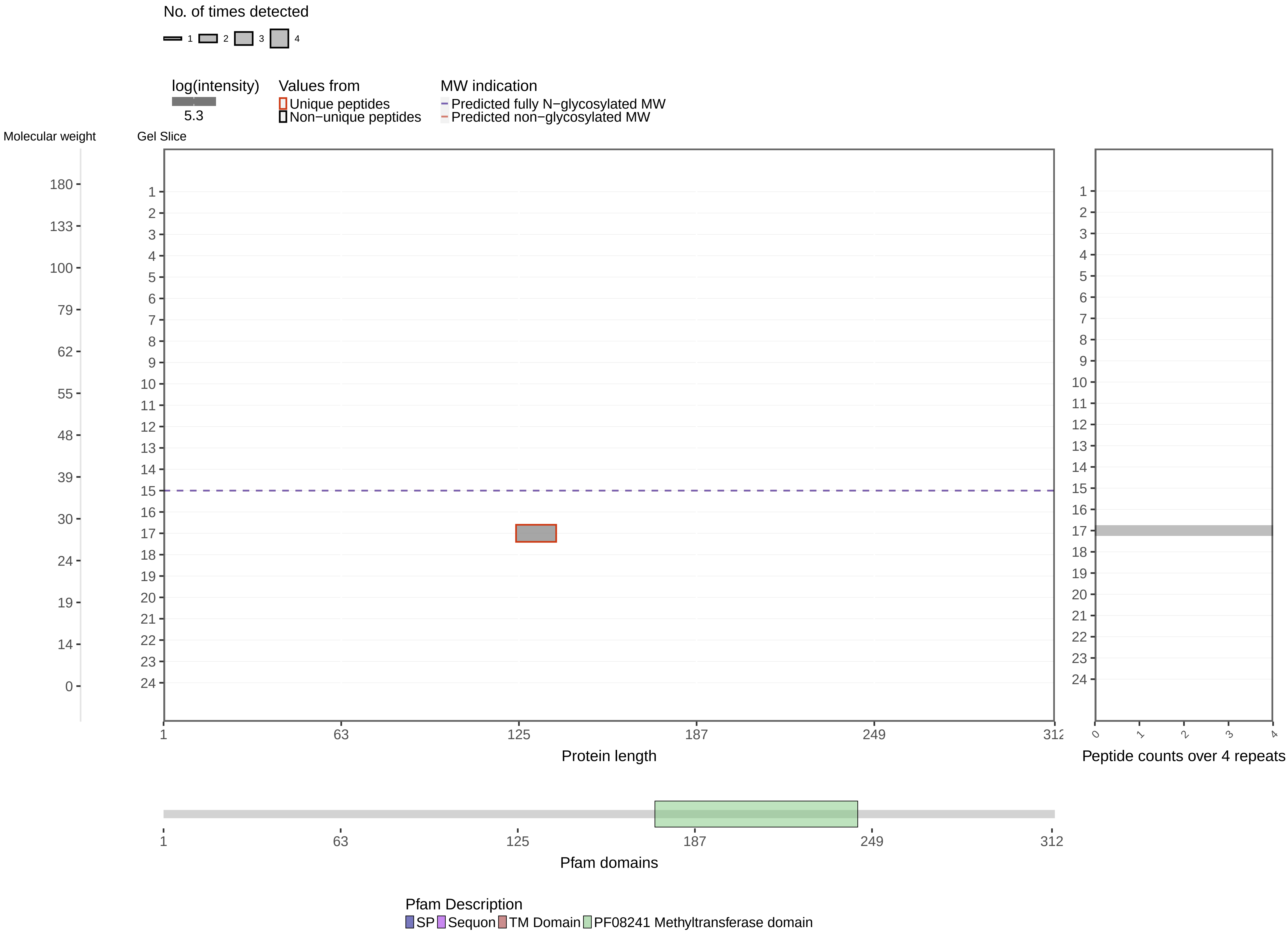


Protein sequence:

EVLPPSNAPGVAILDMCSSWVSHYPAGYKQERIVGMGLNEEELKRNPVLTEYVVQDLNSNPKLPFGDNTFDVITNVVSVDYLTKPLDVFKEMSRILKPGGLAIMSFSNRCFFTKAISIWTSTGDADHVMIVGSYFHYAGGFEPPQAVDISPNPGCSDPMYIVYSRKIATA*

### NbL04g19960.1 thioredoxin−like_2_chloroplastic [MW = 25.46 kDa]

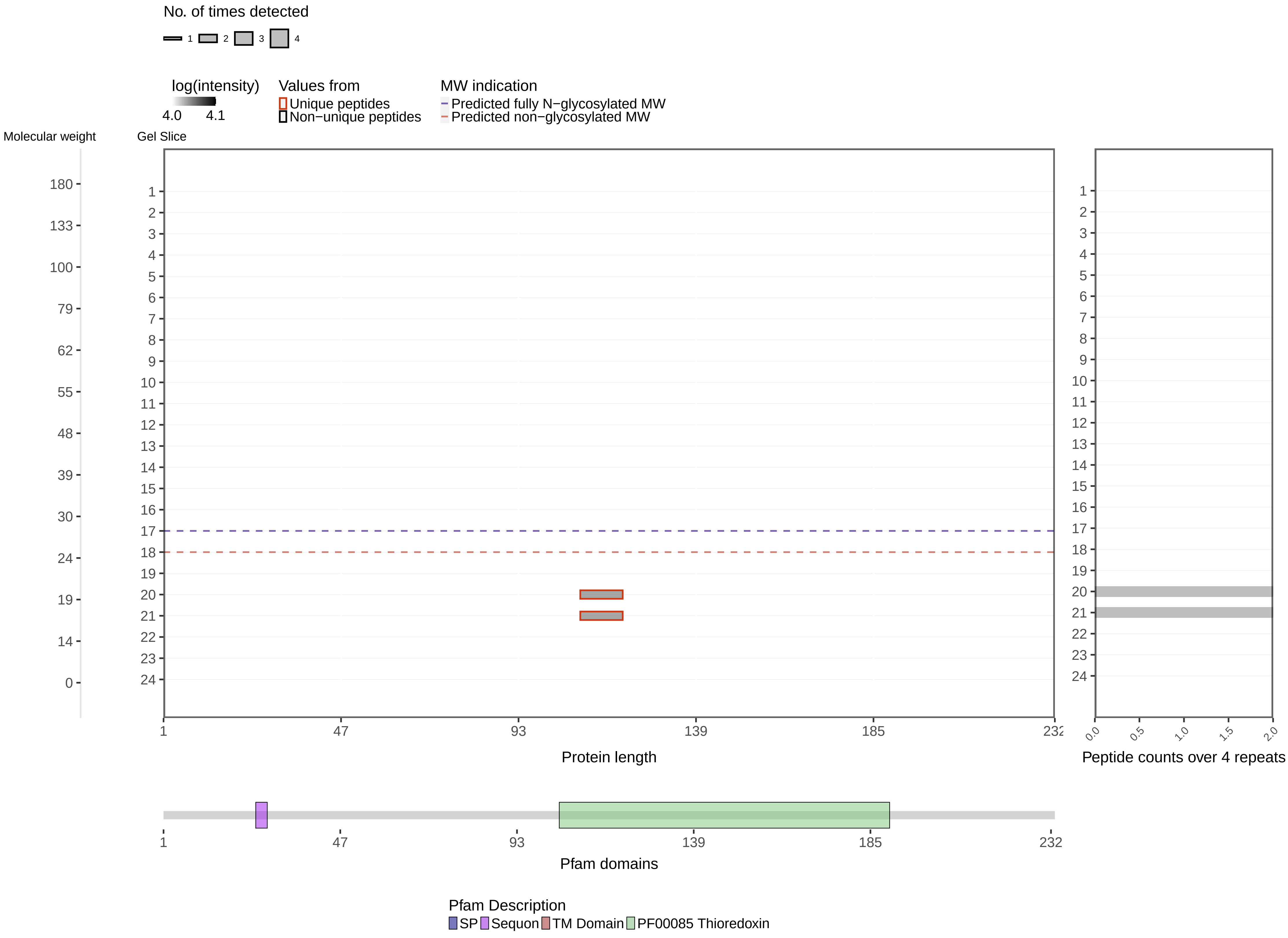


Protein sequence:

/DIHSTKEFLDALSQAGERLVIVEFYGTWCASCKALFPKLCRTAQENPEIMFLKVNFDANKPMCKALNVKVLPYFHFYRGADGQLESFSCSLAKFQKIKDAIQLHNTARCSIGPPIGVGDLTLELLSGTK*

### NbL04g19940.1 ectonucleotide_pyrophosphatase/phosphodiesterase_family_member_3−like [MW = 54.93 kDa]

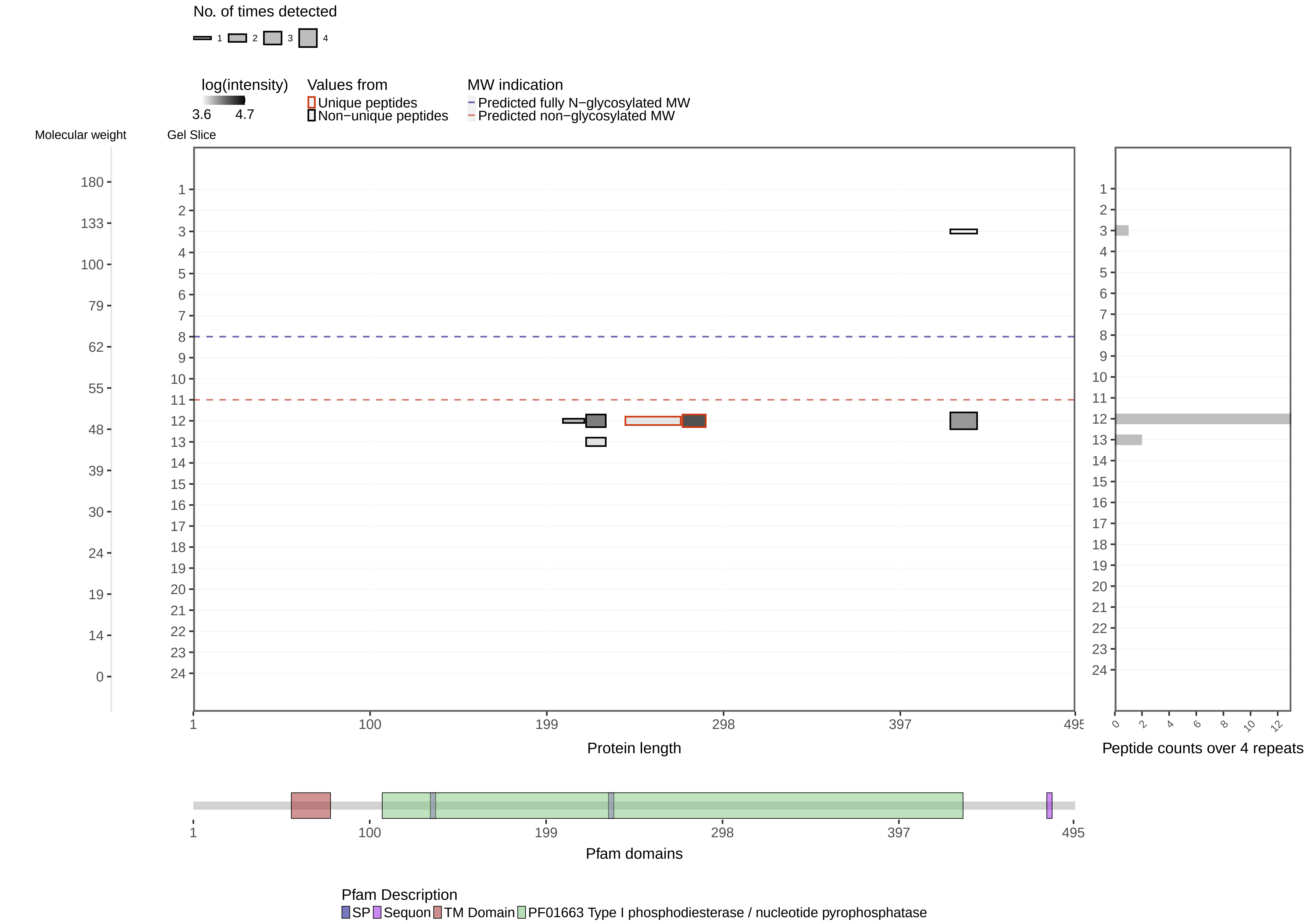


Protein sequence:

[illegible]

NbL04g19850.1 WD−40_repeat−containing_protein_MSI4−like [MW = 54.72 kDa]

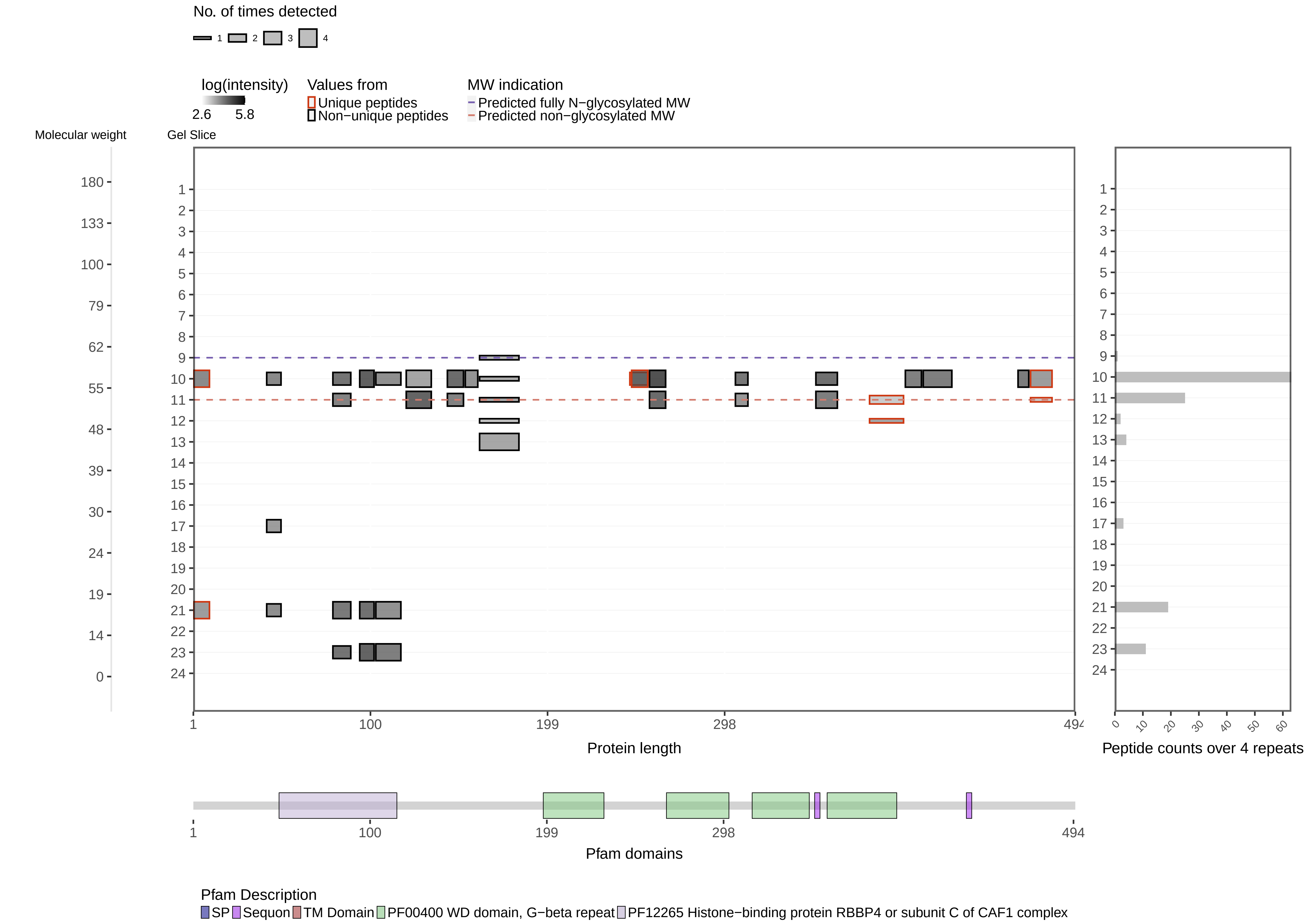


Protein sequence:

ISTLAADAKKLAGSAGSIIKSADNPSIGPRGIFQGHEDTVEDVQFCPSSSQEFCSVGDDSCLILWDARVGNNPVVKVEKAHDADLHCVDWNPHNDYLIITGSADNSVRLFDRRNLTSDGVGSPVHKFEHHTAAVLCVQWCPDRTSVFGSSAEDGLLNIWDYEKVGETTDTEAKSQDDPPPGLFFQHAGHRDKIVDFHWNASDPWTIVSVSDDLETSGGGGTLQIWRMSDLLYRQEEEVLAELQQFKDHVSNCAPKPQA*

### NbL04g19790.1 uncharacterized_LOC104097489 [MW = 29.08 kDa]

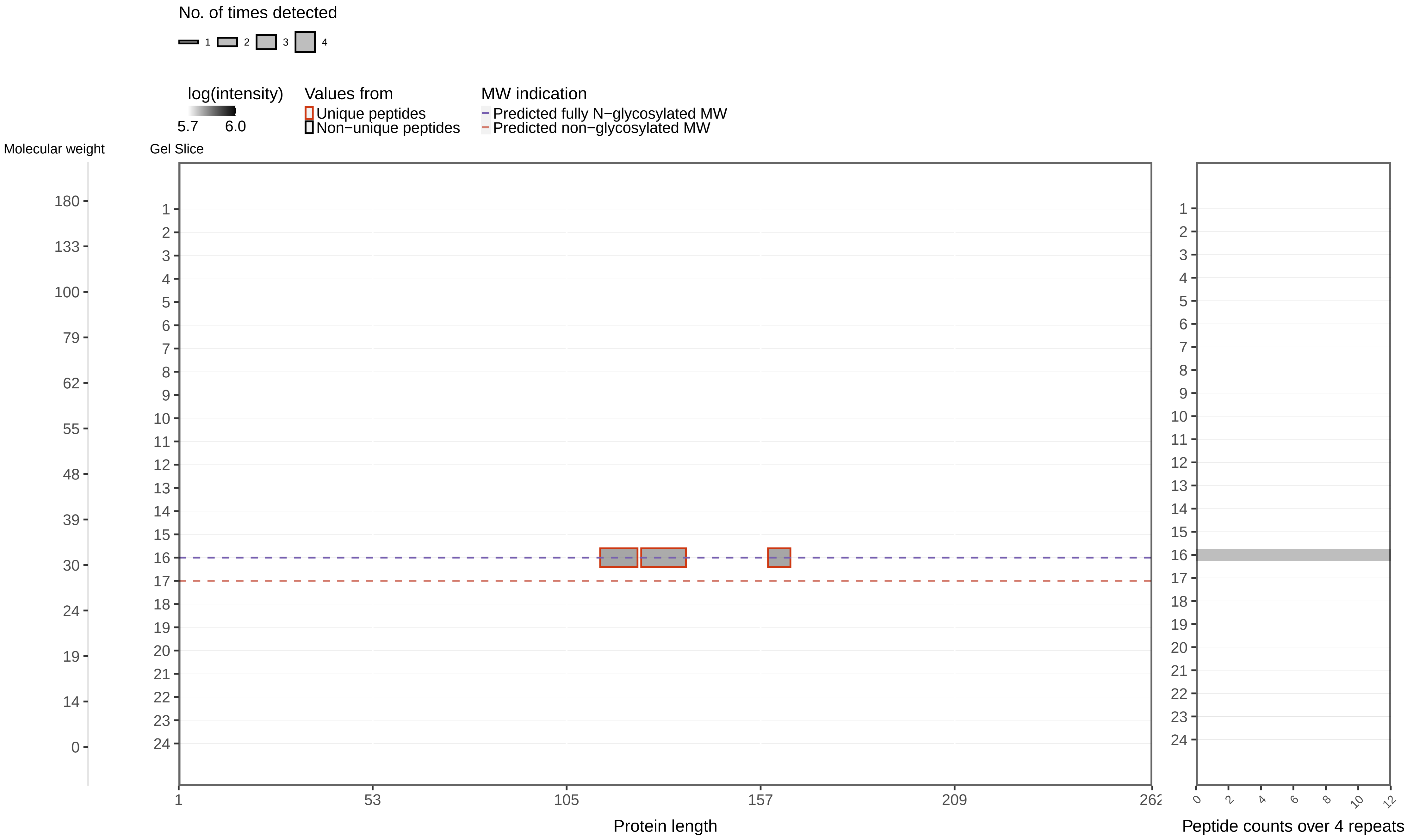


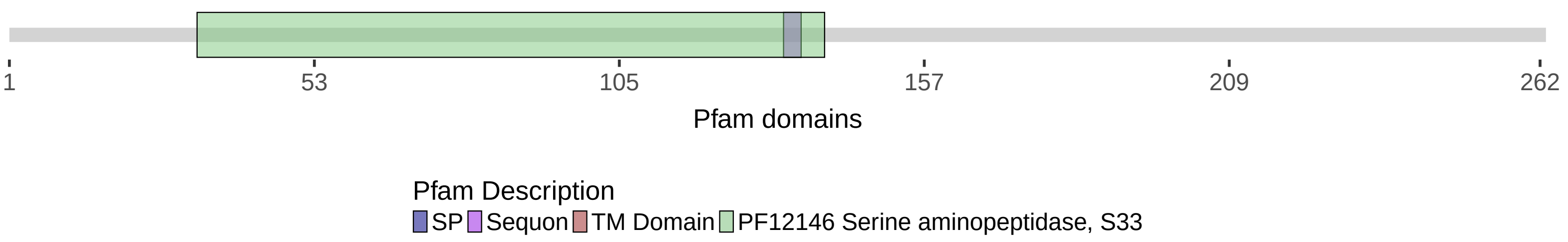


Protein sequence:

GDVVLLYASKYHDVHTVVNLSGRYNLEKGIAERLGEDFLEIIKKDGFIDVKNKAGEVDYRVSEESLMDRLHTNMHDACLQIDKGCRVLTVHGSADEIIRVEDALEFDKIIPNHKLLIIEGANHCYTSHLAELTPAILPFIKEGLQKN*

### NbL04g19770.1 proteasome_subunit_alpha_type−2−A [MW = 25.54 kDa]

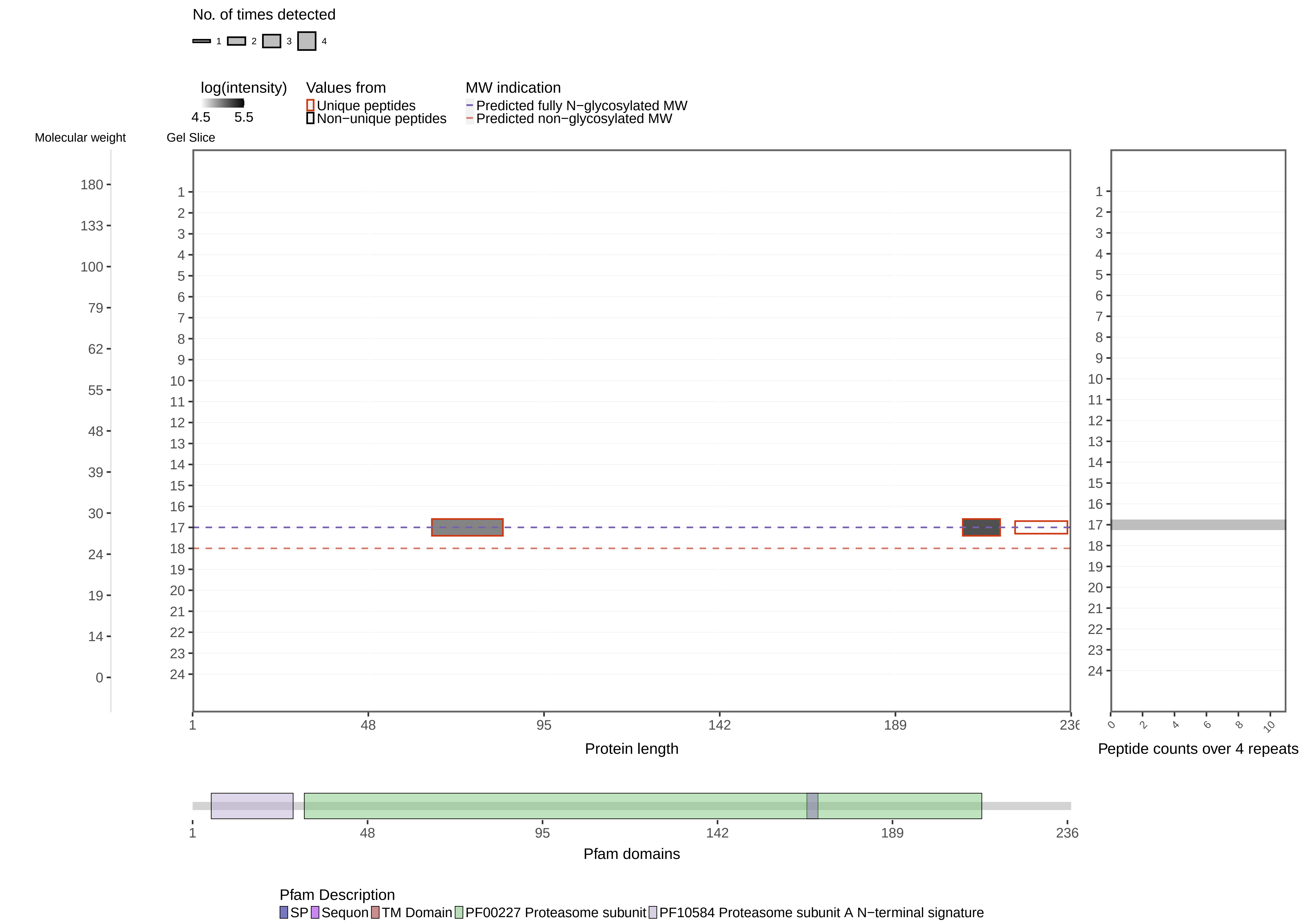


Protein sequence:

IPVTQLVRETAAVMQEFTQSGGVRPFGVSLLVAGFDDKGPQLYQVDPSGSYFSWKASAMGKNVSNAKTFLEKRYTEDIELDDAVHTAILTLKEGFEGQISGKNIEIGIVGTDKVFKVLTPAEIDDYLQEVE*

### NbL04g19730.1 probable_protein_phosphatase_2C_58 [MW = 32.19 kDa]

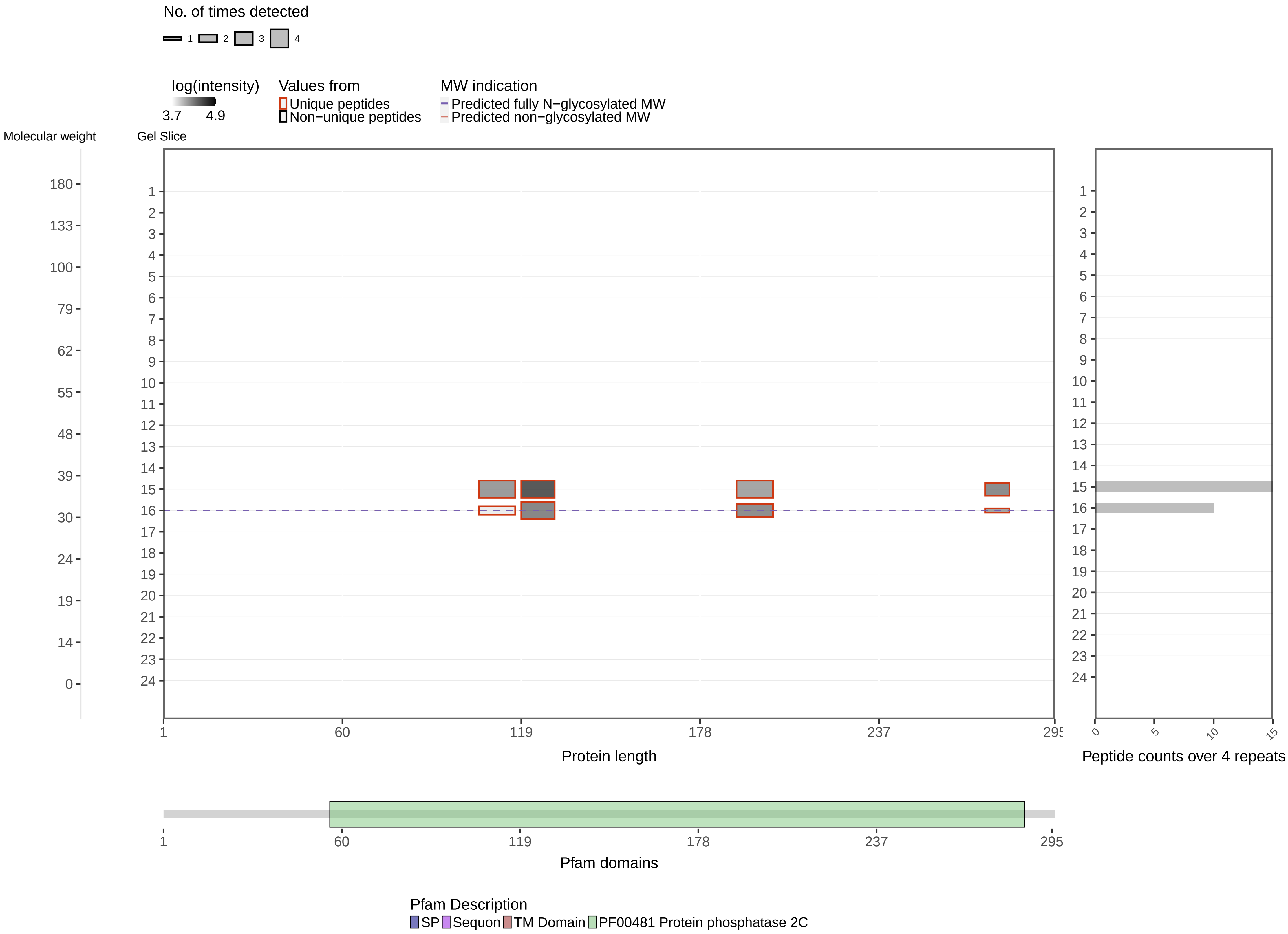


Protein sequence:

QLGKGGSTAVTAMLINGQMLVVANVGDSRAVISKKSVVKQLSVDHEPNREKEIIESKGGFVSNLPGDVPRVDGQLAVARAFGDKSLKKHLSSDPDVVIELIDNDVDLLILASDGLWKVMSNQEAVDCIKNMKDAKAAAKHLAELAISRKSKDDISCIVVKFQ*

NbL04g19690.1 uncharacterized_protein_At4g28440−like [MW = 16.18 kDa]

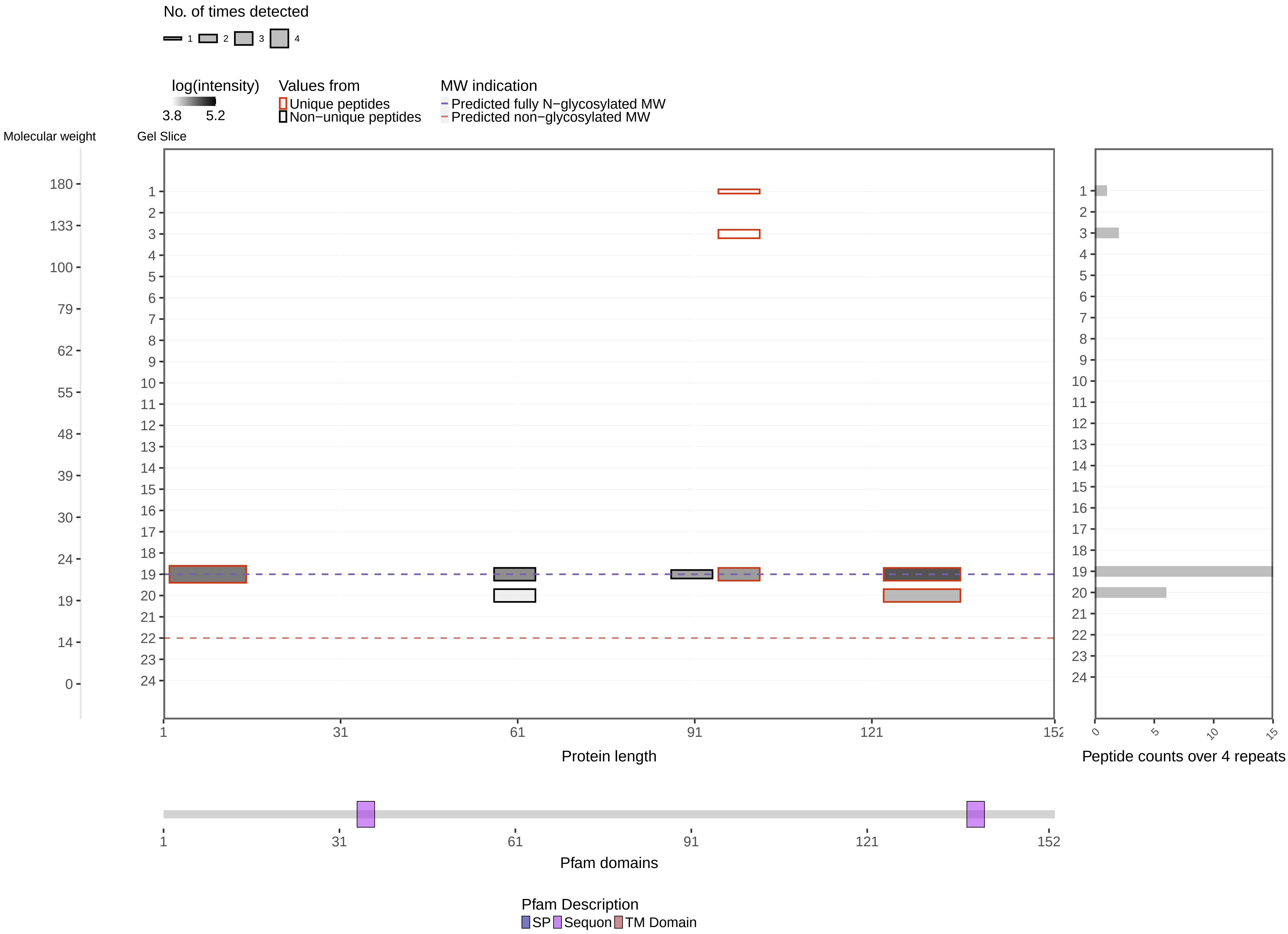


Protein sequence:

PRGPARLAECLVGDDTGSILFTARNEQVDMMKADAIVILRNAKIDMFKGSMRLAVDKWGRVEVTEPADFGVNEKNNLSLVEYELVNVEE*

### NbL04g19670.1 26S_proteasome_non−ATPase_regulatory_subunit_2_homolog_A−like [MW = 98.45 kDa]

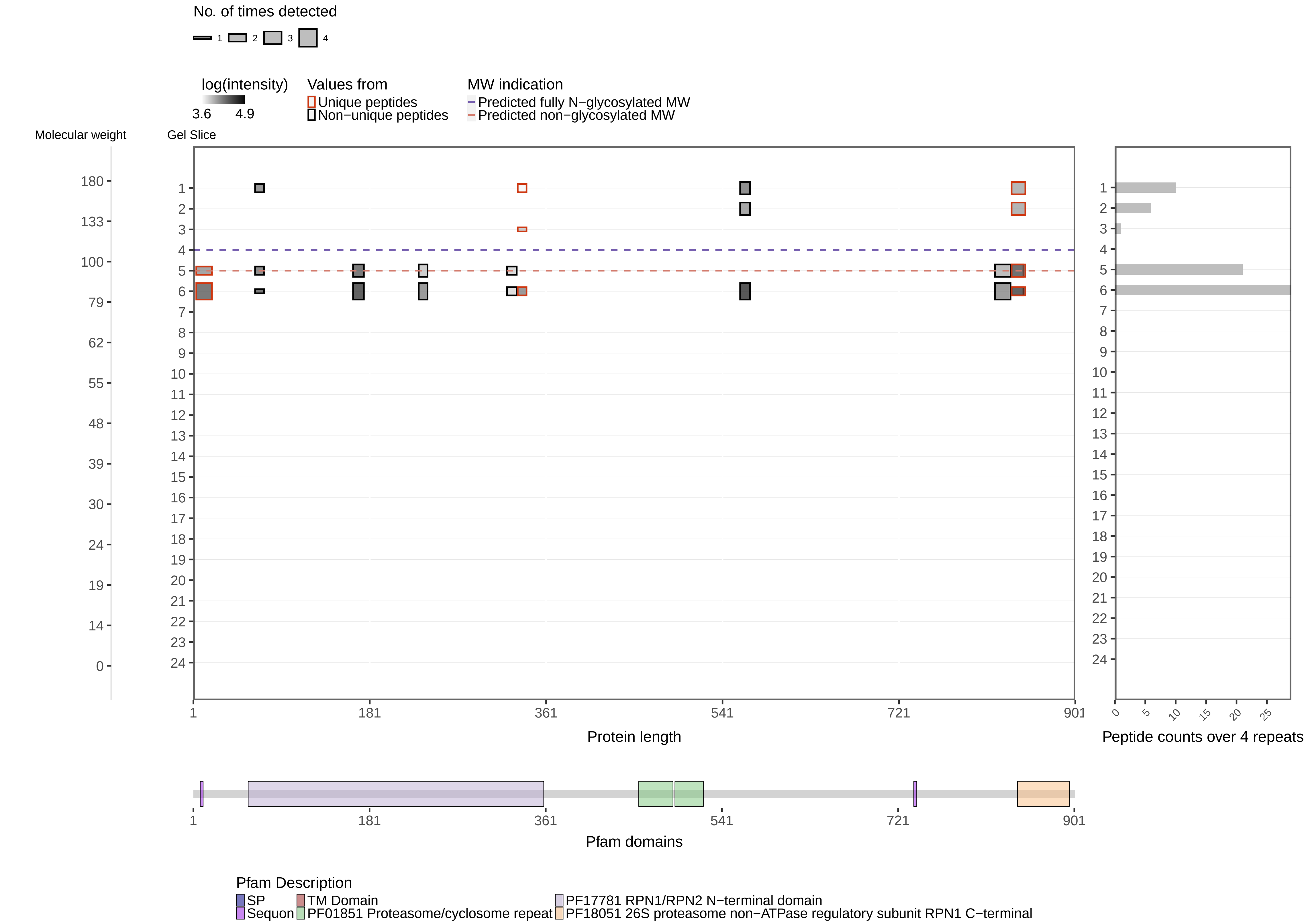


Protein sequence:

[illegible]

NbL04g19520.1 fasciclin−like_arabinogalactan_protein_9 [MW = 32.43 kDa]

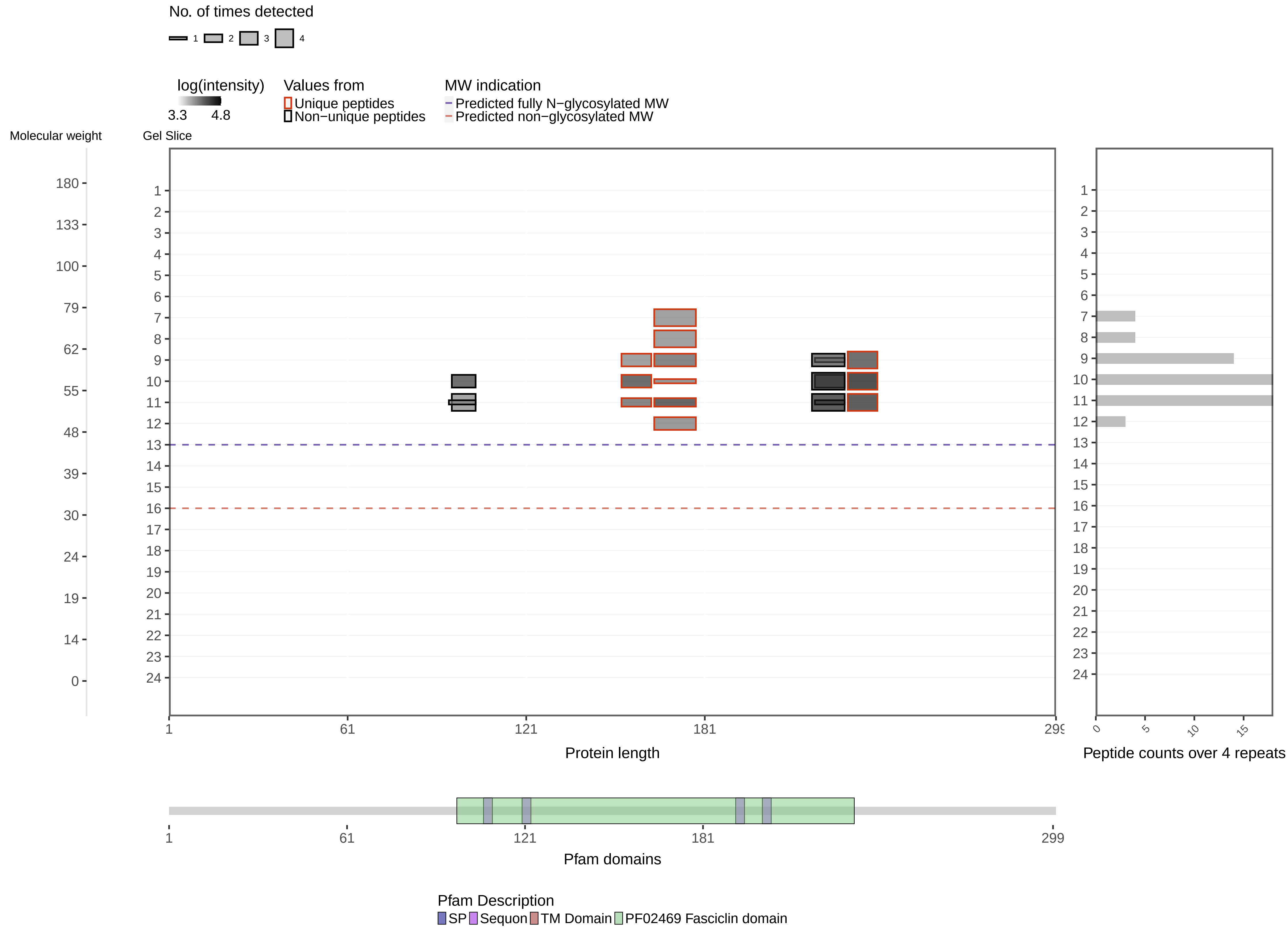


Protein sequence:
PGGTLNKLSDQQKVQLIQYHVVPKFYSFEDLQTVSNPVRTQATGQKGEAFGLNFTGQNNQVNVSSGAVETNIYNAIRKDPPLAVYQLDKVLIPLAFTDTKTPSSDDAPAPTDDSAKSGSTADKTKAKEPSPAPNGAKRINVGVFGLISGVFLFCMGALS*

### NbL04g19280.1 ketol−acid_reductoisomerase_chloroplastic [MW = 63.26 kDa]

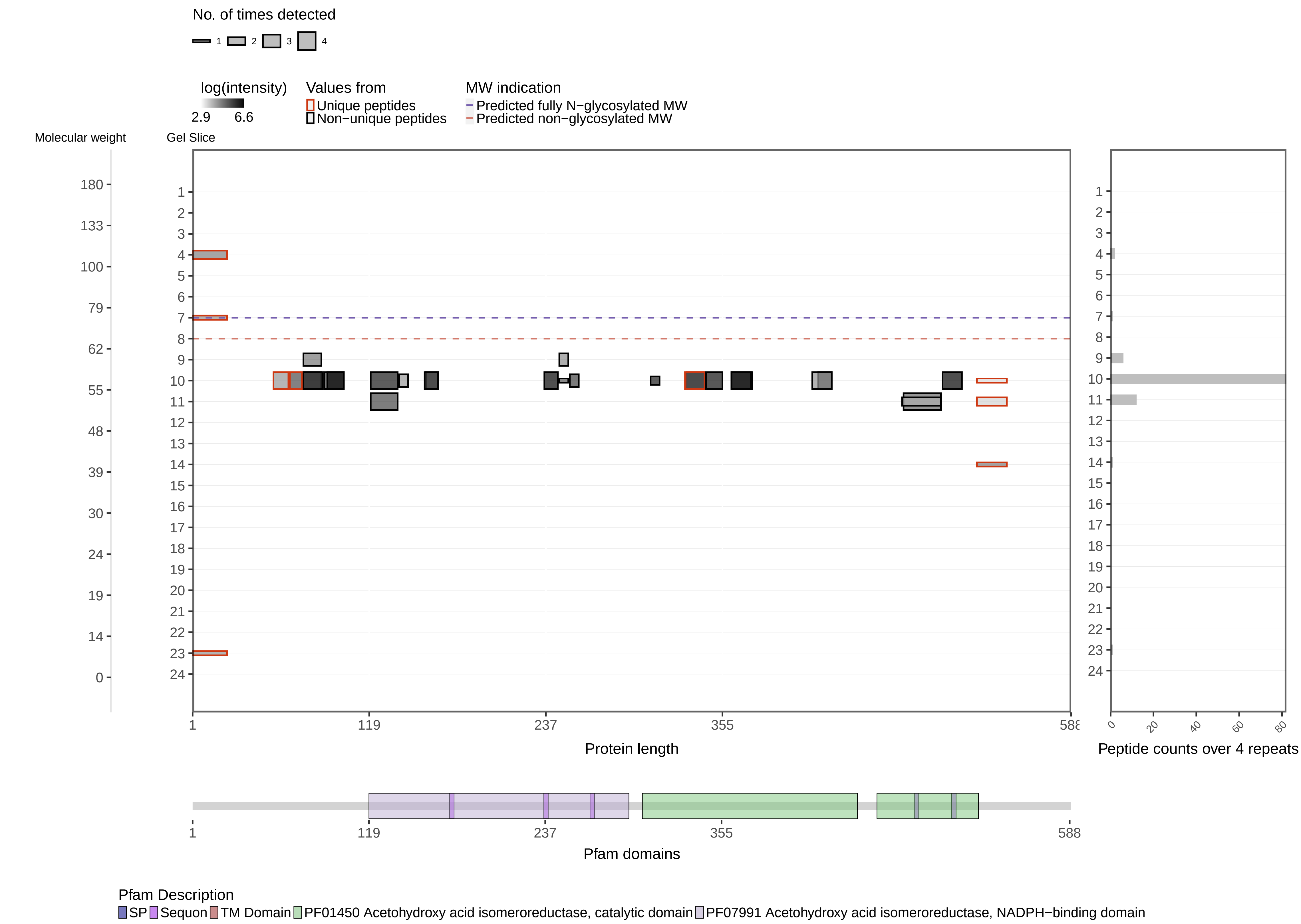


Protein sequence:

### NbL04g19260.1 14−3−3_protein_9 [MW = 29.54 kDa]

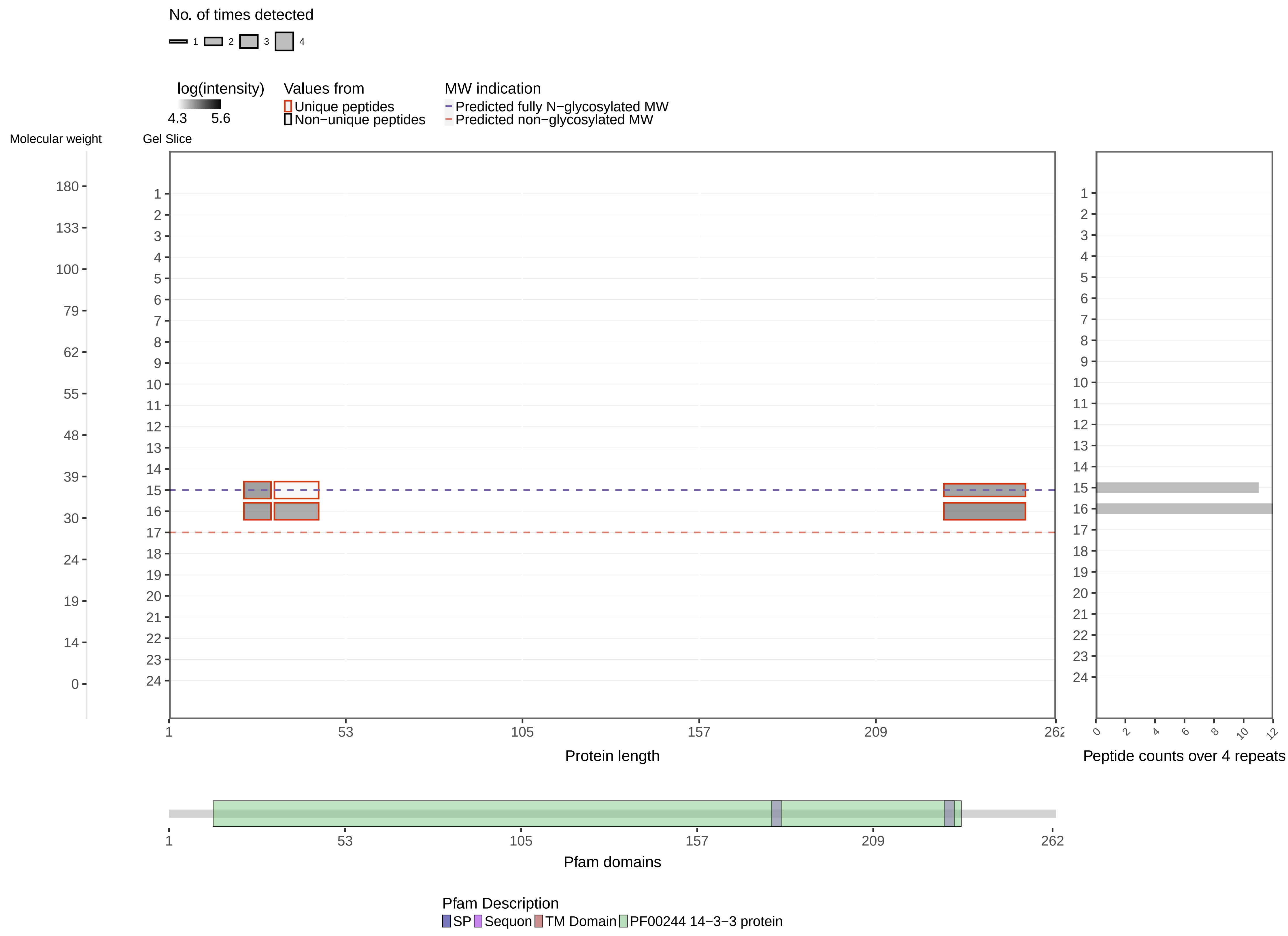


Protein sequence:

GESTVFYHKMKGDYYRYLAEFKSGNDKKEVAELSLKAYQSATTAAEAELPPTHPIRLGLALNFSVFYYEIMNSPERACHLAKQAFDEAISELDSLNEDSYKDSTLIMQLLRDNLTLWTSDLPEDAEDAQKGDSTNKAGGGEDAE*

### NbL04g19240.1 phosphoenolpyruvate_carboxylase−like [MW = 110.03 kDa]

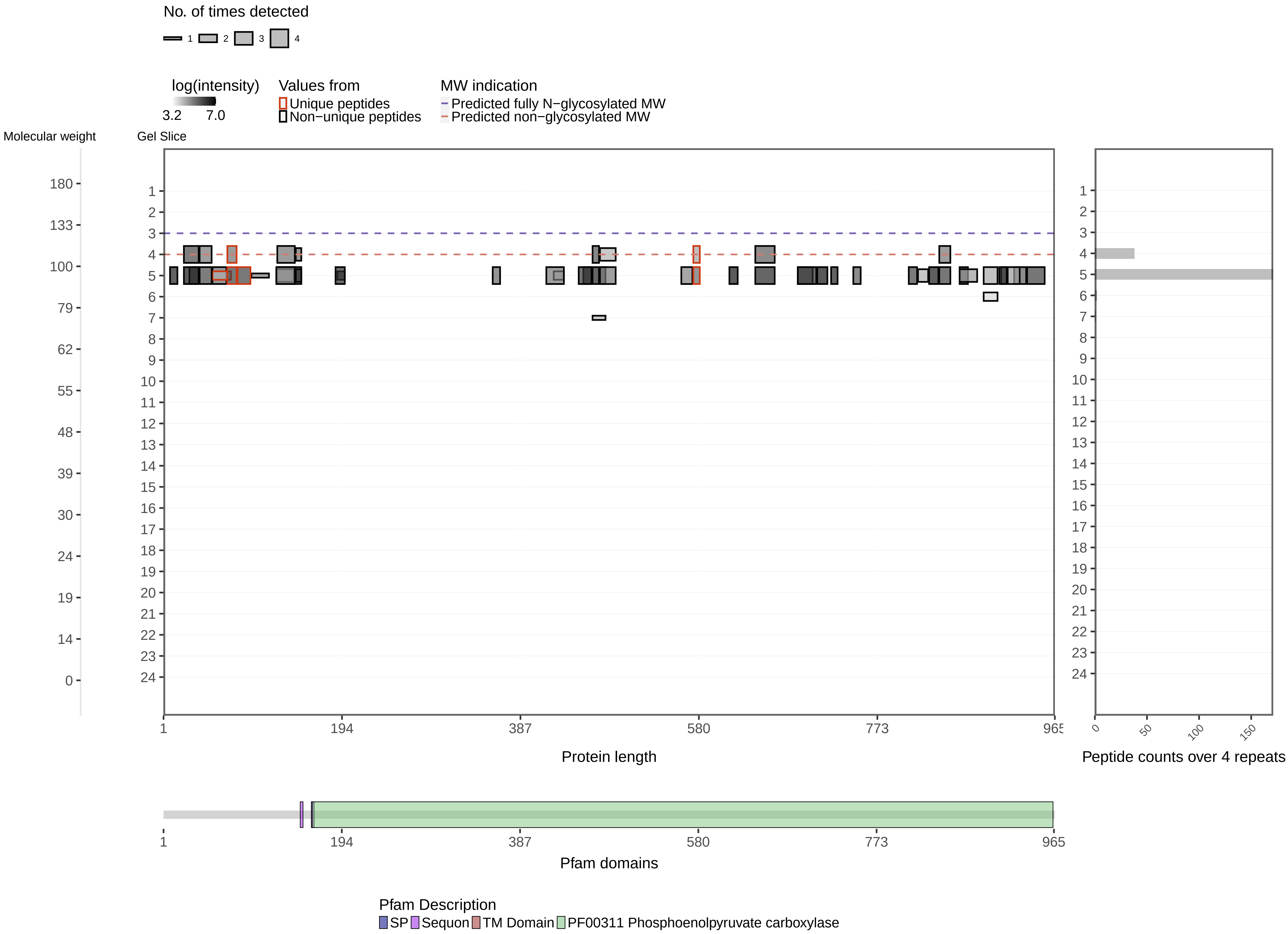


Protein sequence:

SYKEWSEEQRQEWLLSELSGKRPLFGPDLPKTDEIADVLDTFNVIAELPADNFGAYIISMATAPSDVLAVELLQRECHVKSPLRVVPLFEKLADLESAPASVARLFSIDWYRNRINGKQEVMIGYSDSGKDAGRLSAAWQLYKAQEELVNVAKEFGVKLTMFHGRGGTVGRGGGPTHLAILSQPPDTIHGSLRVTVQGEVIEQSFGEEHLCFRTLQRFTAATLEHGMNPPVAPKPEWRALLDEMSVVATKEYRSLVFQDPRFVEYFRLATPELEYGRMNIGSRPAKRKPSGGI[illegible]

### NbL04g18830.1 photosystem_II_repair_protein_PSB27−H1_chloroplastic−like [MW = 18.79 kDa]

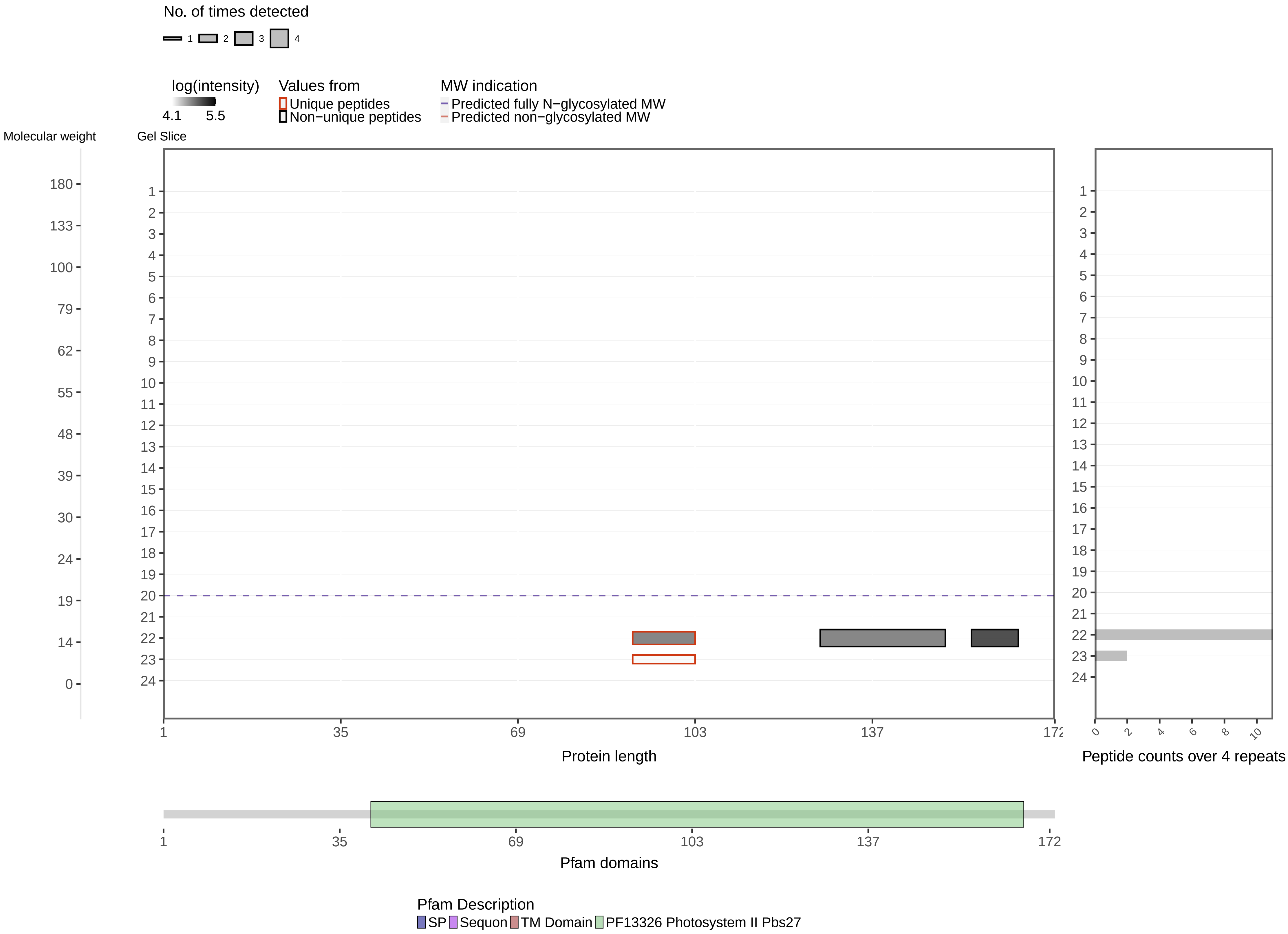


Protein sequence:

ASEVIKKVRSTLSMEKSDPNIADSVTELREASNYWVAKYRREKDLLGRASFRDIYSALNAVSGHYVSFGPTTPIPAKRKQRILEEMDTAEKALQRGR*

### NbL04g18670.1 lipoyl_synthase_2_mitochondrial−like [MW = 26.4 kDa]

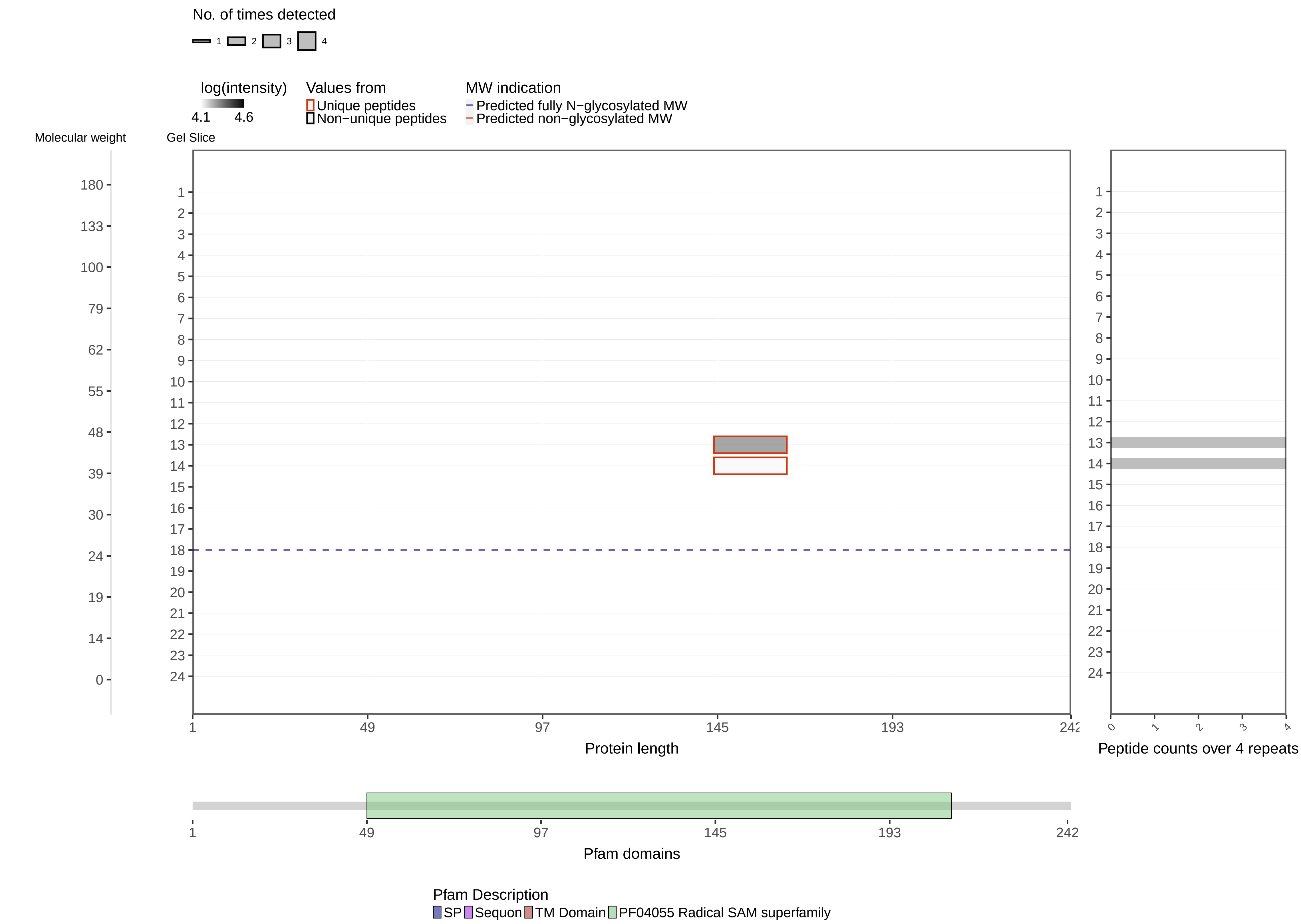


Protein sequence:

ETVQKLKALKPNMLIKALTPDFRGDPGCVEKVAKSGLDVFAHNIETVEELQSVVRDHRANFKQSMDVLKMAKGYAPANTLTKTSIMLGCGETPEQVCKTMEKVRAAGVDVMTFGQYMRPSKCHMPVSEYVTP*

NbL04g18290.1 GDSL_esterase/lipase_At4g26790−like [MW − SP = 36.27 kDa]

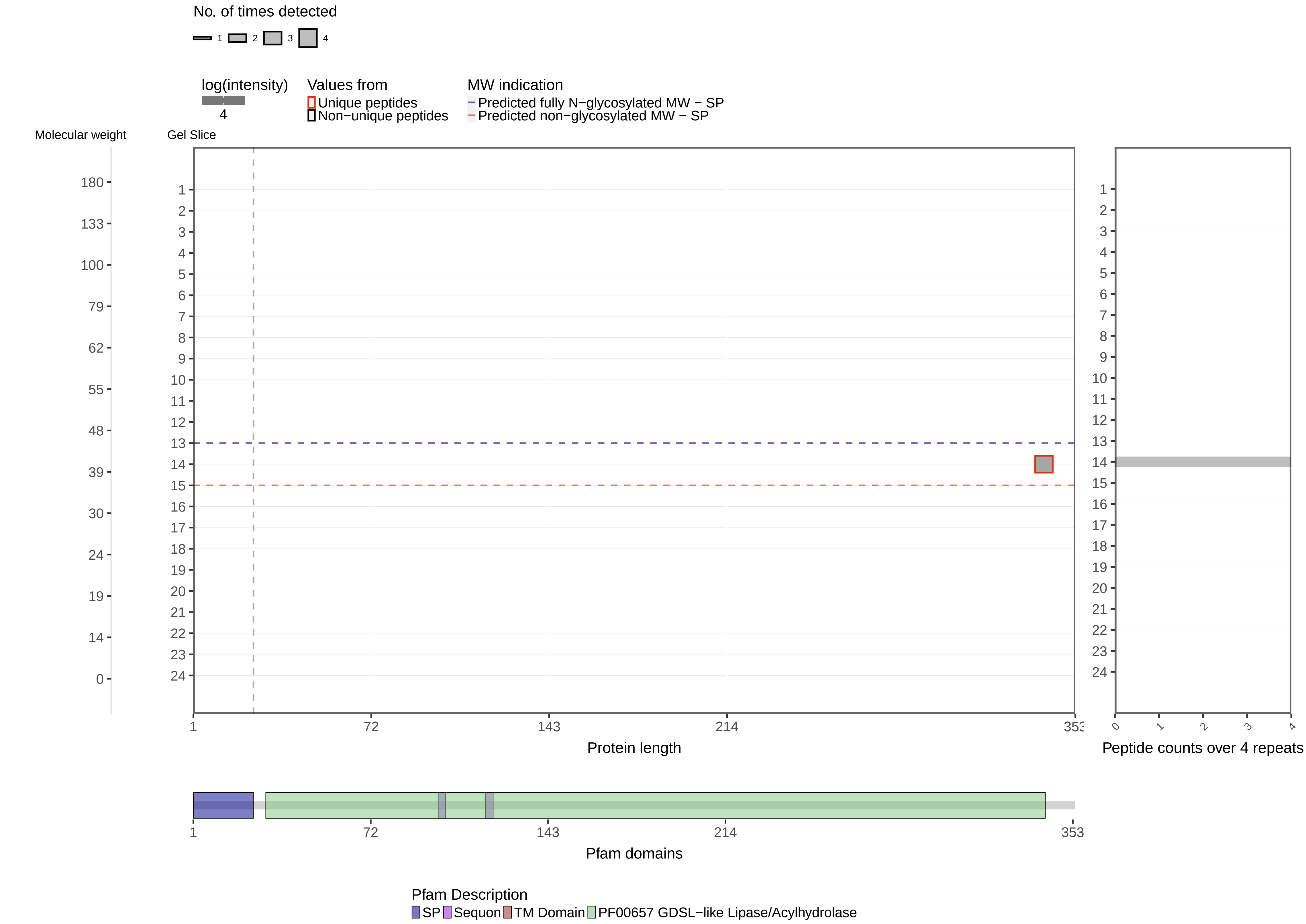


Protein sequence:

[illegible]TNDFLENYYTMPQRRSQYTVDQYQTFLIGIAKNFVTNLYNLGARKISLGGLPPMGCMPLERTSNMANGNECLESYNVVAMNFNDKLSGLVIELNKELSGIQVVLSNPYNVMLQMIKKPSLYGFEVASLACCATGLFEMGYACDPYNLFTCKDANKYIFWDAFHPTEKTDRIIADHVVKTALSKFLV*

NbL04g18230.1 L−ascorbate_oxidase_homolog [MW − SP = 57.45 kDa]

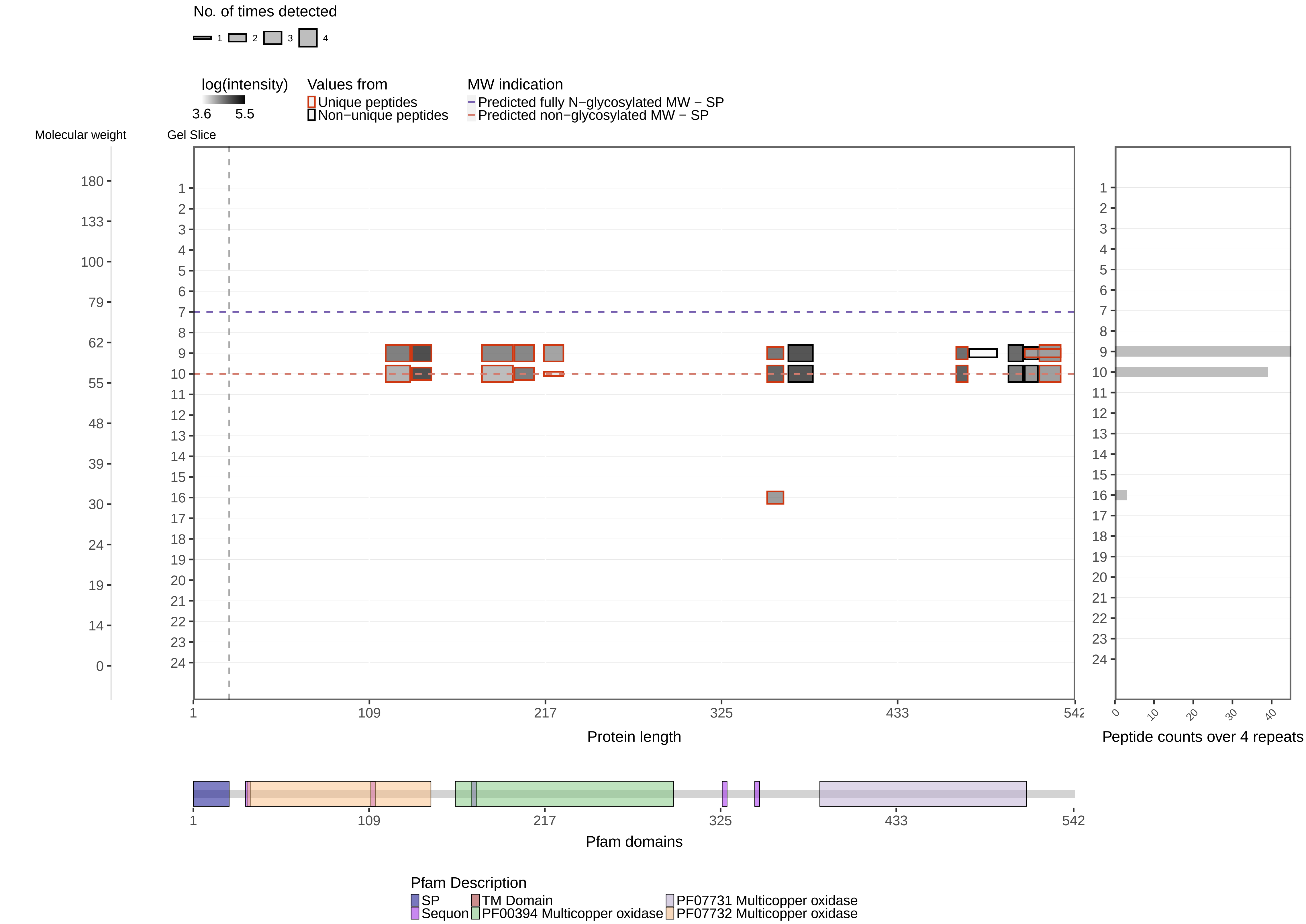


Protein sequence:

SYSVLLIADQPAKDYYIVVSTRFTSQVLTATSTLHYSNSAGSVSGPPPSGPTTEIDWSLNQARSLRRNLSASGPRPNPQGSYHYGLINTTRTIRLANSAPVINGKQRYAINSVSFVPADTPLKLADHFNIPGVFSLGSIPDSPTGGGAYLQTSVMAADFRAYTEVIFENLEVSVQSYHIDGHHFFVVGMGGGEWTPASRLTYNLRDTISRSTVQVYPKSWTALYMPLDNVGMWNIRSQNWARQYLGQQFYLRVYSPANSWRDEYPIPSNALLCGRASTRGILN*

NbL04g18220.1 L−ascorbate_oxidase_homolog [MW − SP = 57.85 kDa]

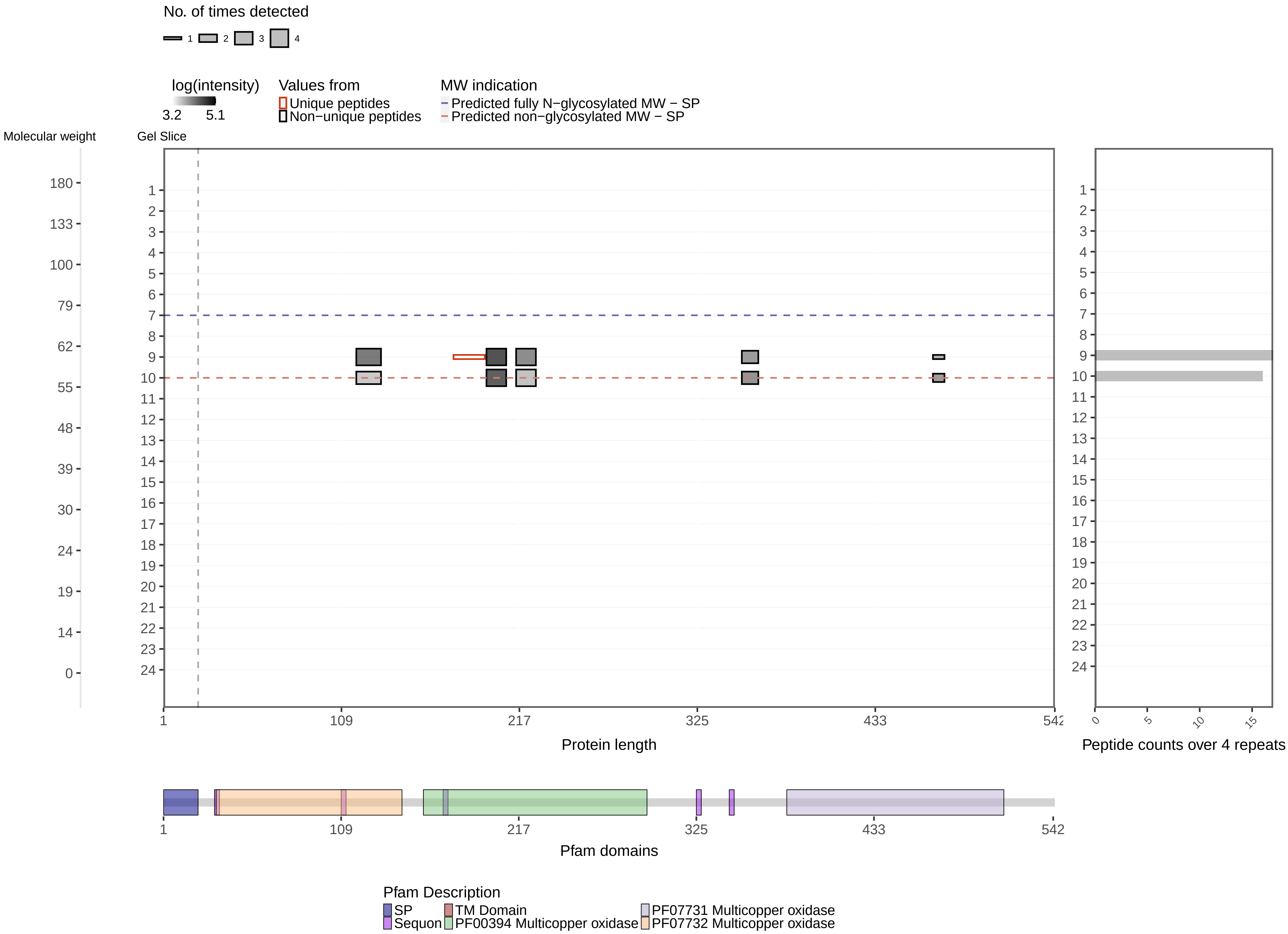


Protein sequence:

'SVLVTMDQPGQDYYIVASTRFTSQVLTATSILHYSNSAGSVSGPPPGGPTIEIDWSLNQARSIRQNLTASGPRPNPQGSYHYGLVNTTRTIRLANSAPMINGKQRYAVNSVSFIPADTPLKLADYFKIPGVFNLGSIQDNPTGGGGYLQTSVMAADFRAFVEVVFENPEDIVQSWHIDGHIFFVVGMDGGQWSPASRLNYNLRDGISRCTIQVYPKSWTALYMPLDNVGMWNIRSENWARQYLGQQFYLRVYSPANSWRDEYPIPKNALVCGRASGRRTRPL*

### NbL04g18140.1 aconitate_hydratase_cytoplasmic [MW = 108.63 kDa]

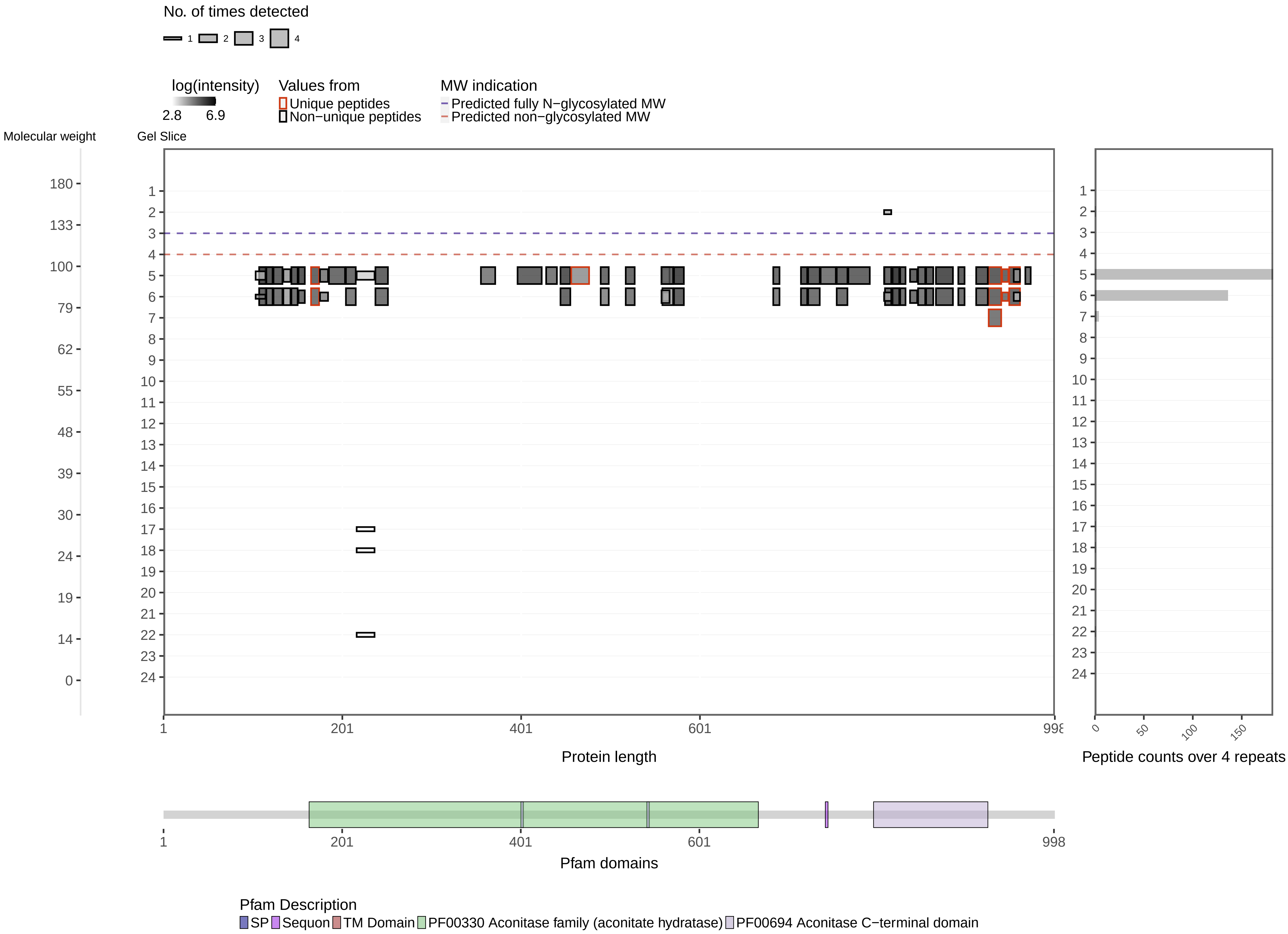


Protein sequence:

LKEMKSDWHACLDNKVGFKGFAVPKEVQDKVAKFDFHGQPAELKHGSVVIAAITSCTNTSNPSVMLGAALVAKKASELGLHVKPWVKTSLAPGSGVVTKYLLKSGLQKYLNQQGFNIVGYGCTTCIGNSGDLDESVASAISENDIVAAAVLSGNRNFEGRVHALTRANYLASPPLVVAYALAGTVDIDFEKDPIGVGKDGKDVYFRDIWPSTEEIAEVVQSSVLPDMFKSTYEAITKGNTMWNELSVPTAKLYSWDPNSTYIHEPPYFKGMTMDPPGPHGVKDAYCLLNFGD

### NbL04g17910.1 beta−amylase−like [MW = 65.45 kDa]

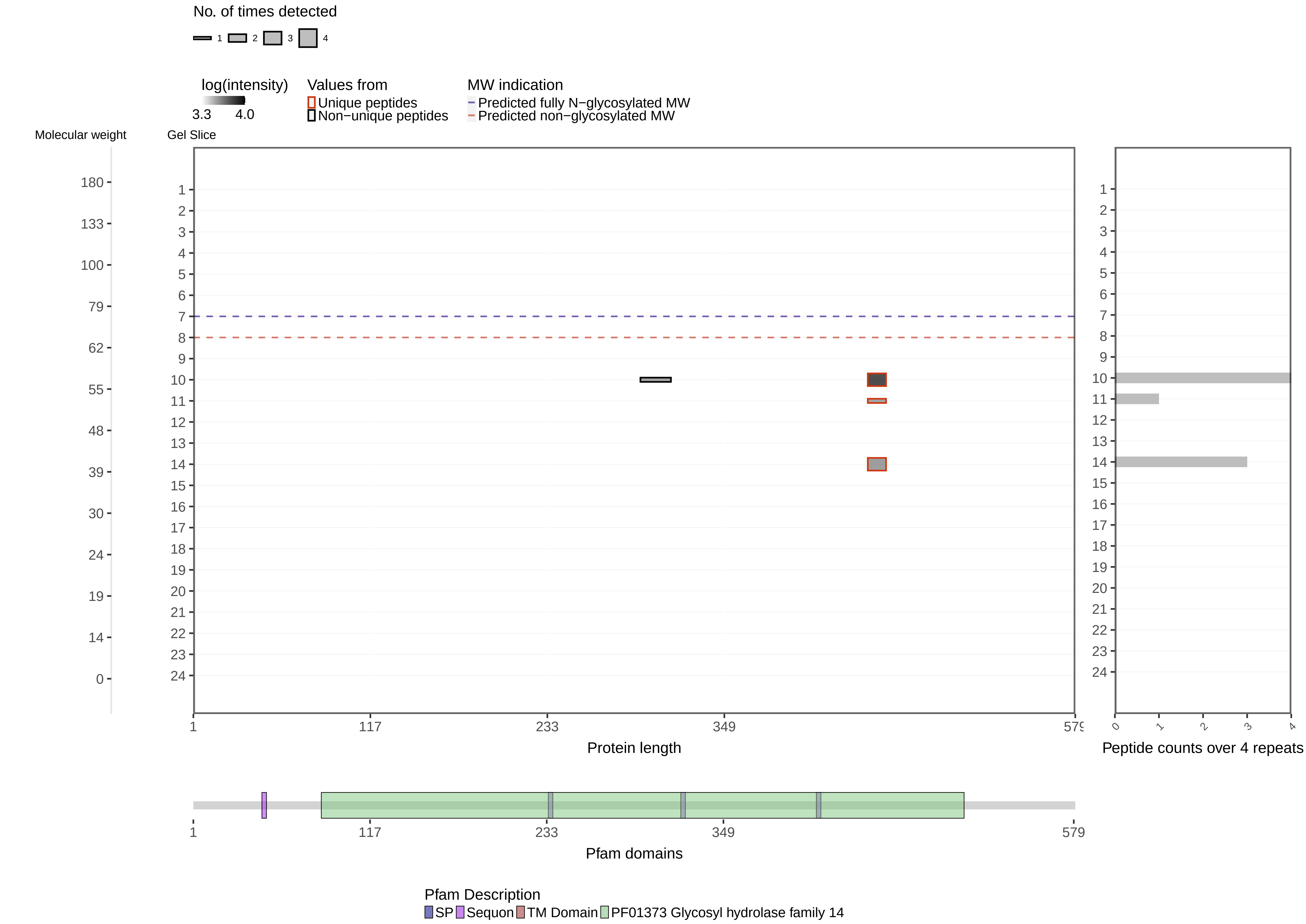


Protein sequence:

CYDKYMRTDFKEAVTKAGYPQWDLPDDAGTYNDIPVKTGFFGPNGTYLTEKGKFFLTWYSSKLLLHGDQILDEANKAFLGCKVKLSAKVAGIHWWYKDASHAAELTSGYFNLDNRDGYRPIARMLSRHHGTLNFTCLEMRNSEQPAYAKSGPQELVQQVLSVGWKENIEVAGENALSRYDGYAYNQILLNARPNGINKNGPPKQKMAGFTYLRLSEKLLQGRNFNTFKTFVKKMHADLDYCPEYEKLAPLSRSKMEISIDELVTATQRTKPFPWDEQTDARIGGILAEYWI

### NbL04g17790.1 xyloglucan_endotransglucosylase/hydrolase_protein_9 [MW − SP = 31.48 kDa]

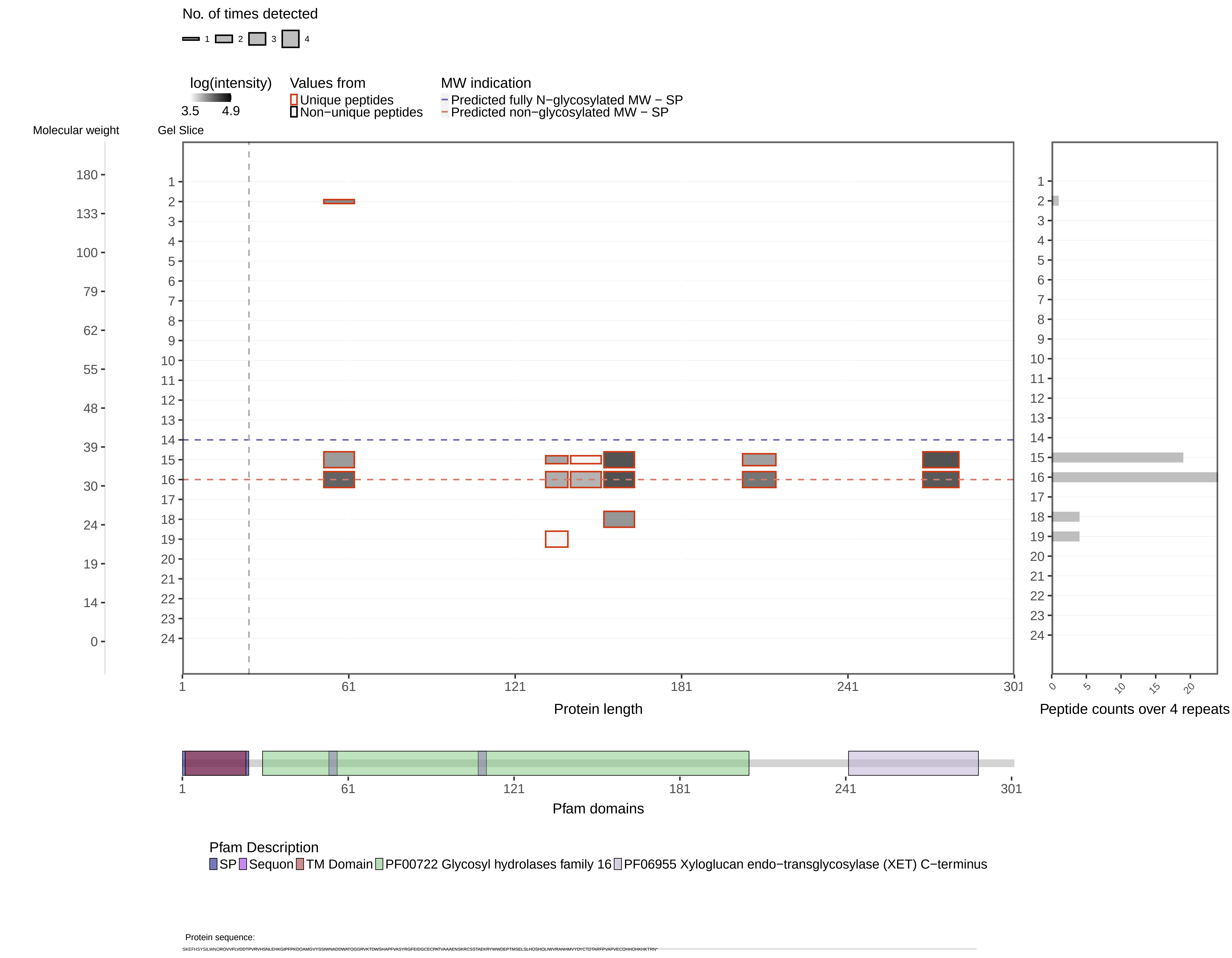

NbL04g17670.1 frataxin_mitochondrial [MW = 22.86 kDa]

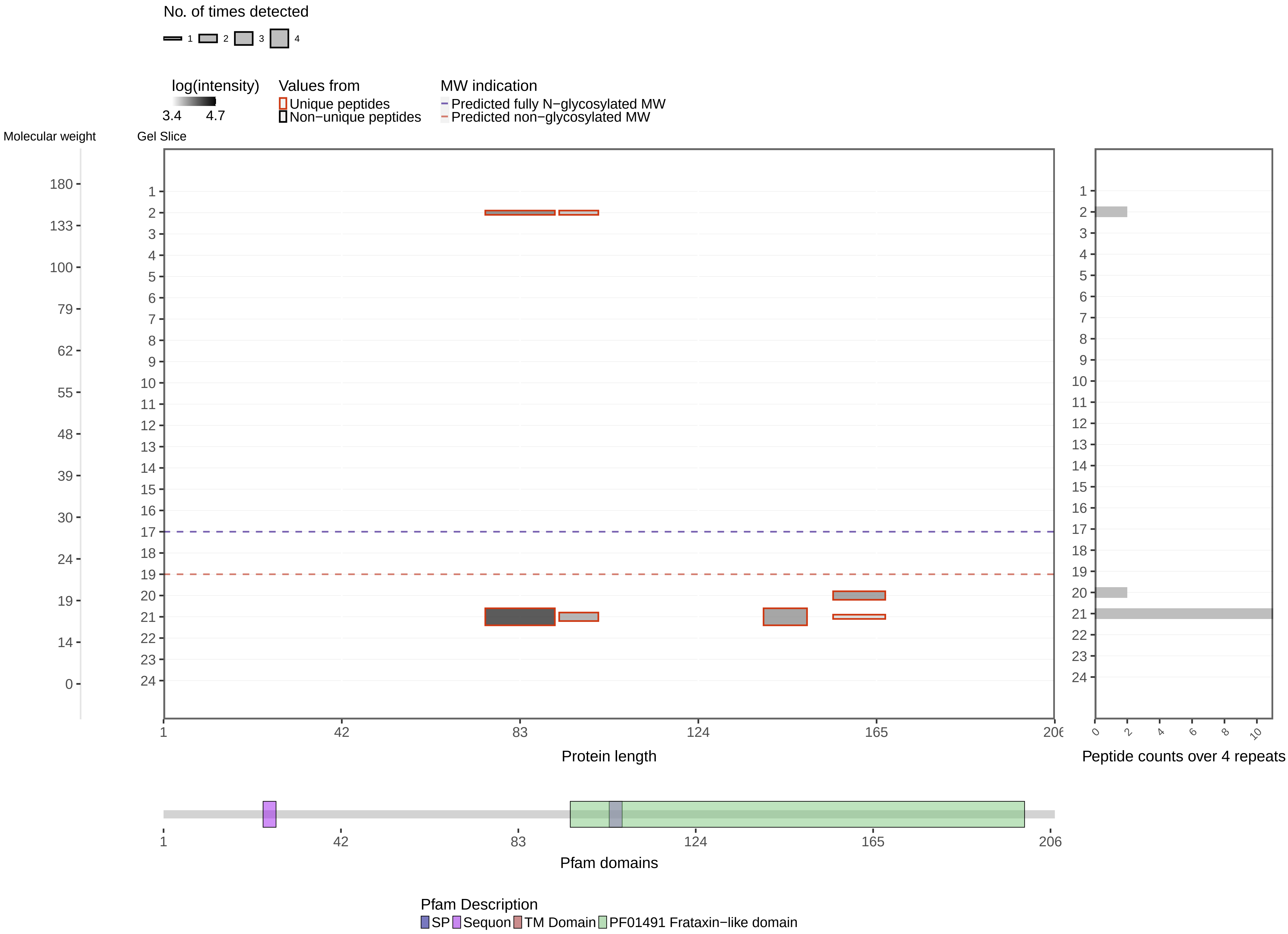


Protein sequence:

SLLQEDEYHRLANATIHDLLDKLEDYGDSVDIDGFDVDYGNEVLTLKLGDLGTYVINKQTPNRQIWMSSPVSGPSRFDWDQNSQGWIYRRTKANLRTVLEDELEKLCGSAITLS*

### NbL04g17520.1 uncharacterized_GPI−anchored_protein_At4g28100 [MW − SP = 31.99 kDa]

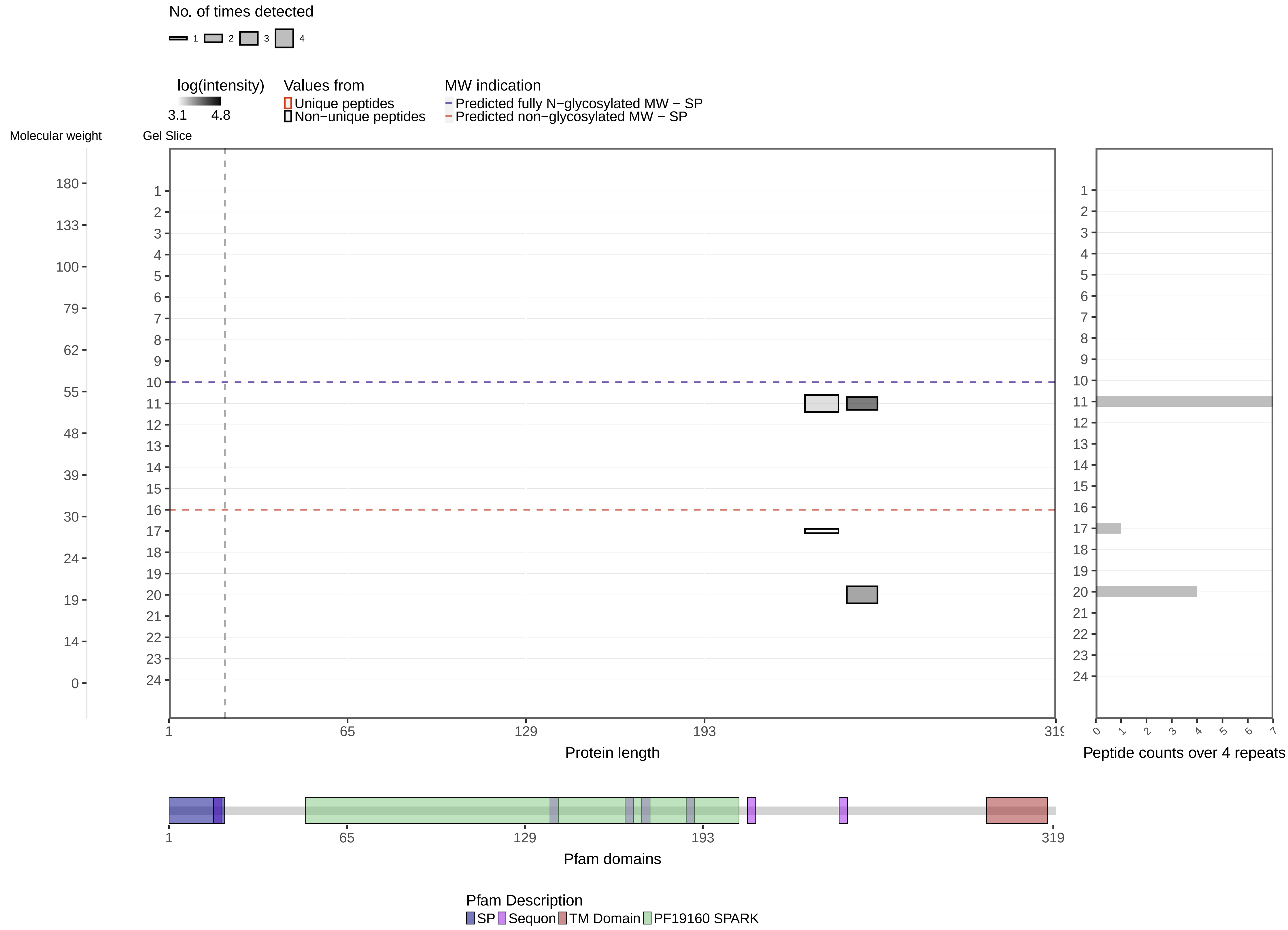


Protein sequence:

LCFCGIRLHQITSLSCPAAFNLTGSKNATPTAAVRNLERNCRNSSYSGCTRCLGALQKLNGDGKNRTHKMVDIGGDRVSKMLTRDCQLMGLTWLLARNKTAYIPTVSAVLRAIMYSSHPPHESKCSPDQENMPLAVDSLQFEKTDSSPPSIGVSRFAILFPFMPLIILLNYLFG*

NbL04g17500.1 protein_THYLAKOID_FORMATION1_chloroplastic [MW = 33.94 kDa]

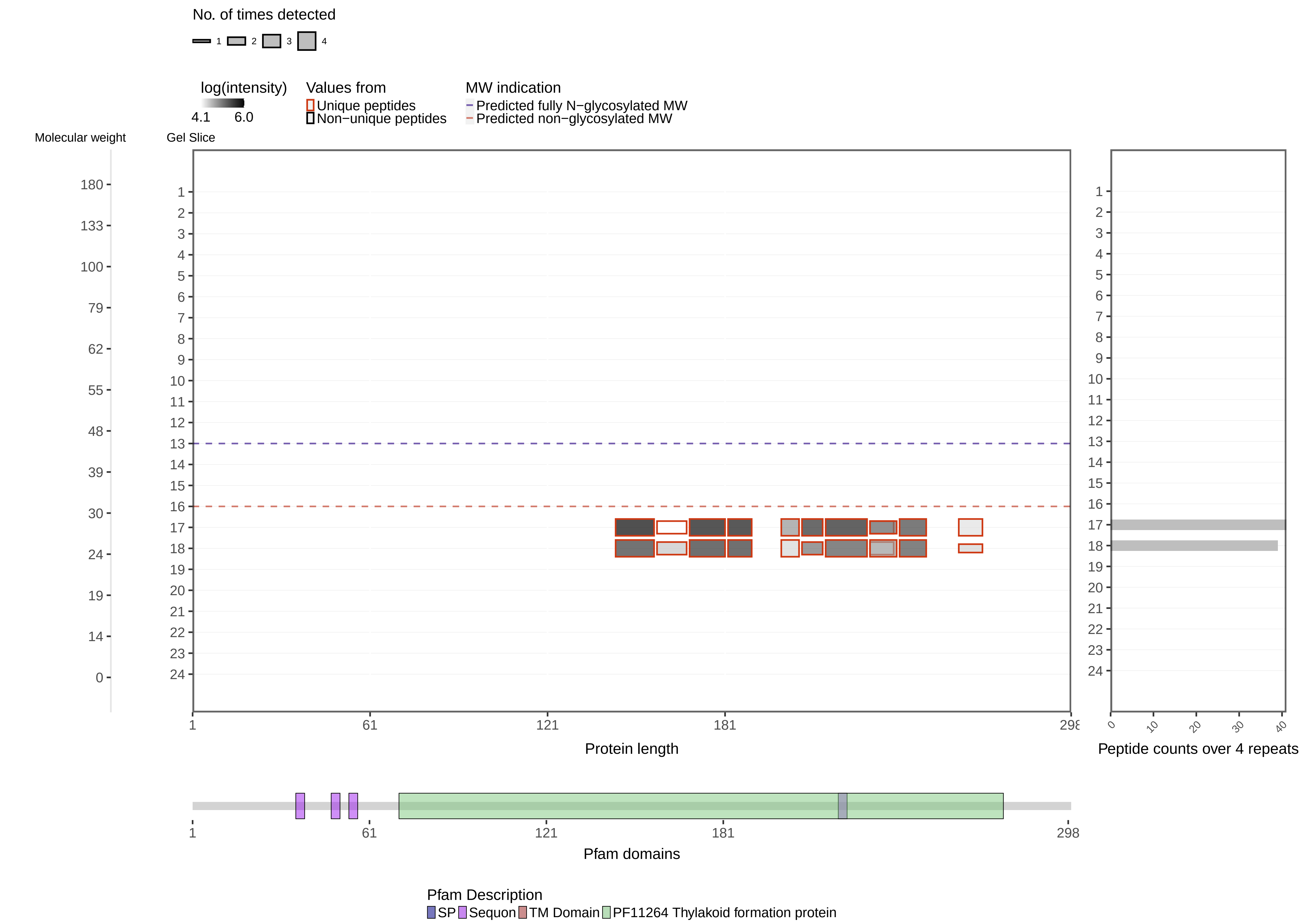


Protein sequence:

EDRDAIFKAYIEALNEDPVQYRADAQKFEEWARTQNANTLIDFSSRDGEVENILKDIAQRAGTKDSFCYSRLFAVGLFRLLELANVTDPTILEKLCASLNINKKSVDRDLDVYRNLLSKLVQAKELLKEYVEREKKKRGERESQKANEAVTKCLGDYQYAGR*

### NbL04g17440.1 fasciclin−like_arabinogalactan_protein_7 [MW − SP = 25.77 kDa]

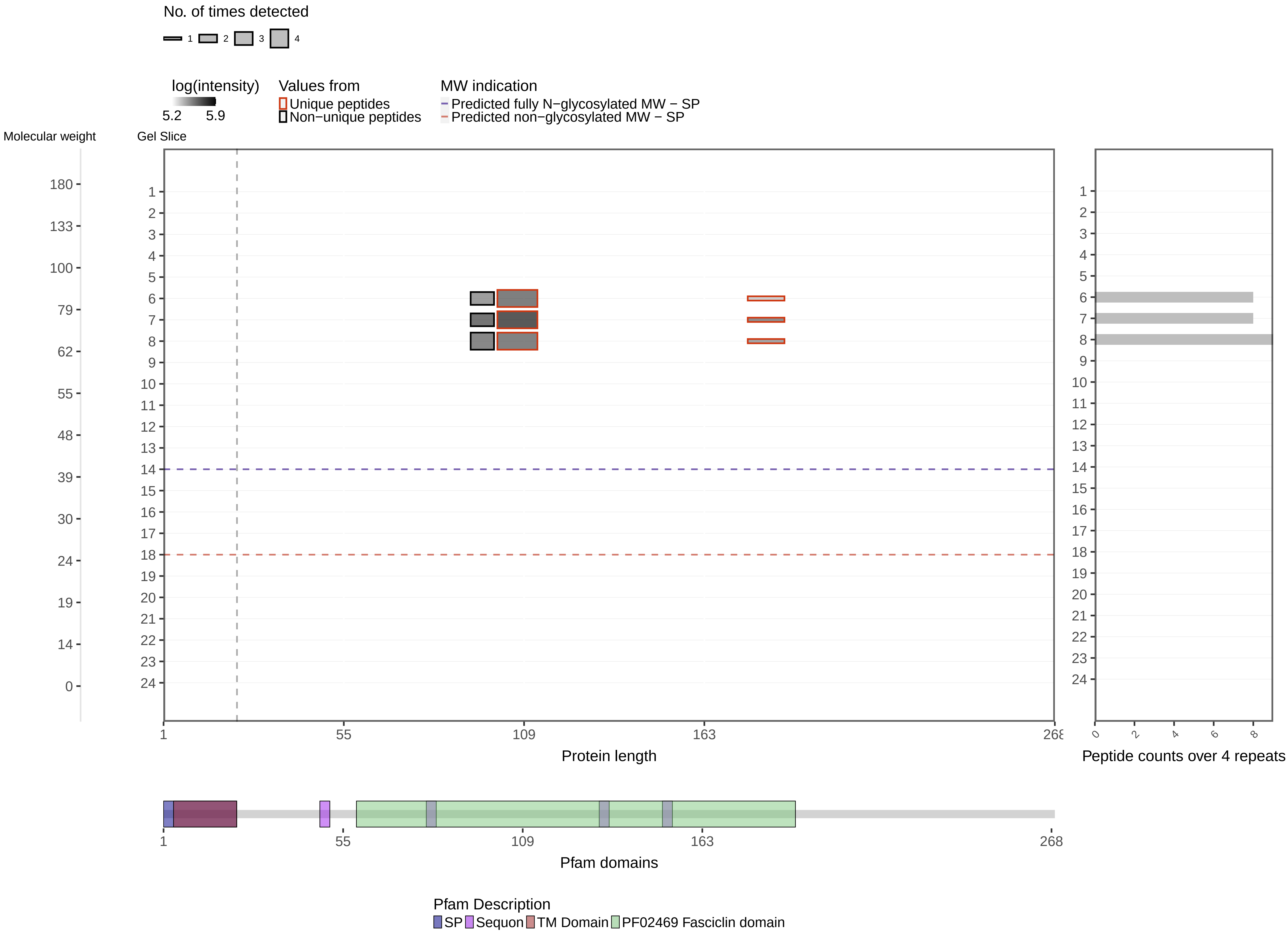


Protein sequence:

HYYSLADFKNLSDMSPINTFAGGNLYSLNFTYDSGTVHLNSGWSRTKVSSAVRATFPVAVYQVDKVLLPEAIFGTDLPPMPAPAPAPETDIAPTADSPAADQTGKARAASSPDSSSPSASHRMMSWGILNHLVLVIAGGFLMLLL*

### NbL04g17290.1 plant_UBX_domain−containing_protein_4−like [MW = 42.64 kDa]

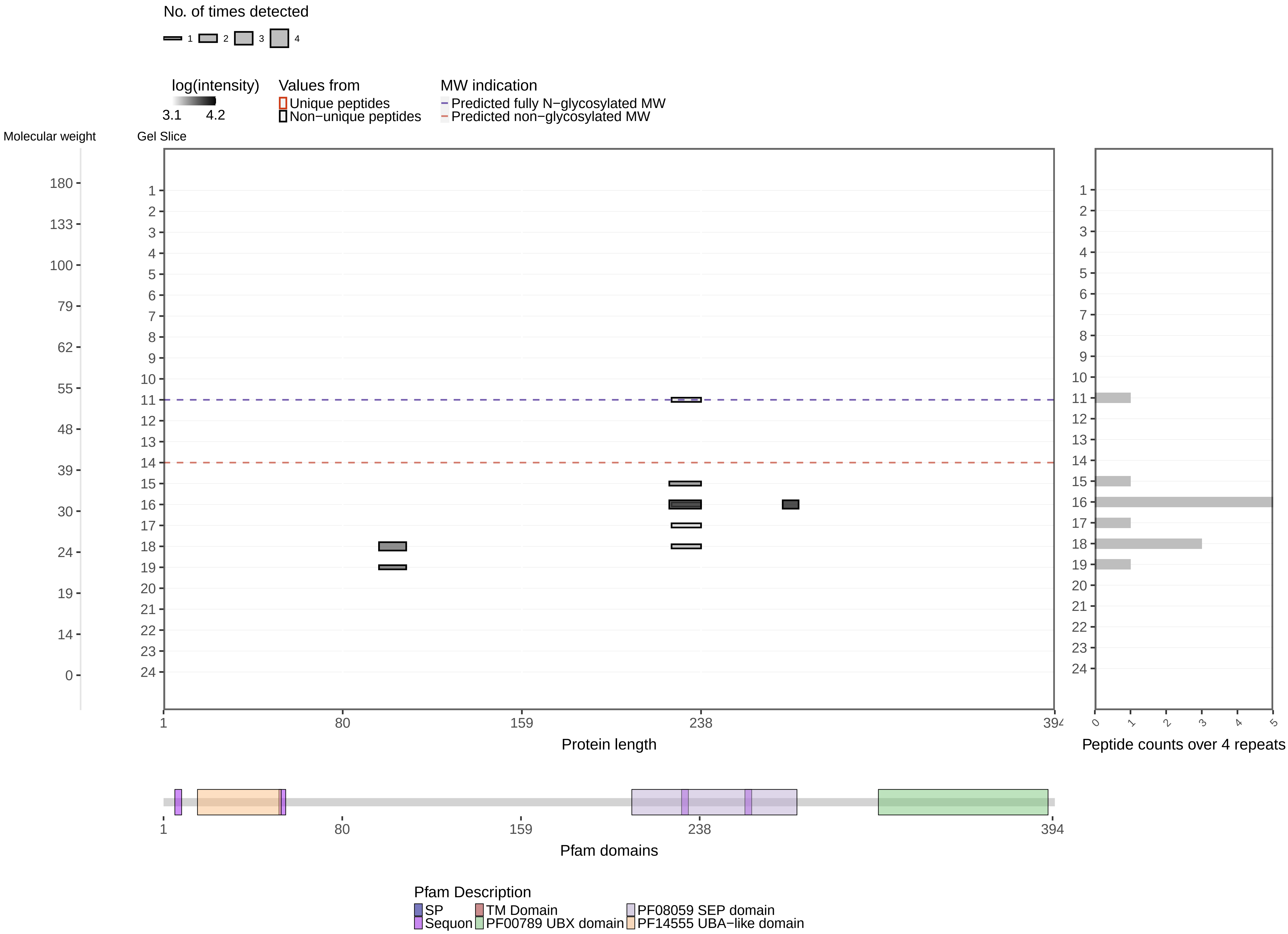


Protein sequence:

:SGTARRLTGETVSAAPOPPESITHVITFWTNGFTIDDGPLRRFDDPENASFLESIRKSECPKELEPEDRRTSVRVNLTRREEDCPVPEKRRATFOGVGRTLGSTSNAEOVDSIGAVSSFTAAPTPSVGLVVDOSOTSTSIOLRLADGTRMVSRFNFOHTIRDIRGFIDASRPGGPRSYOLOTVGFPPKOLSDLDOTIEOAGLANSVVIOKL*

NbL04g17270.1 pyrophosphate−−fructose_6−phosphate_1−phosphotransferase_subunit_beta−like [MW = 60.48 kDa]

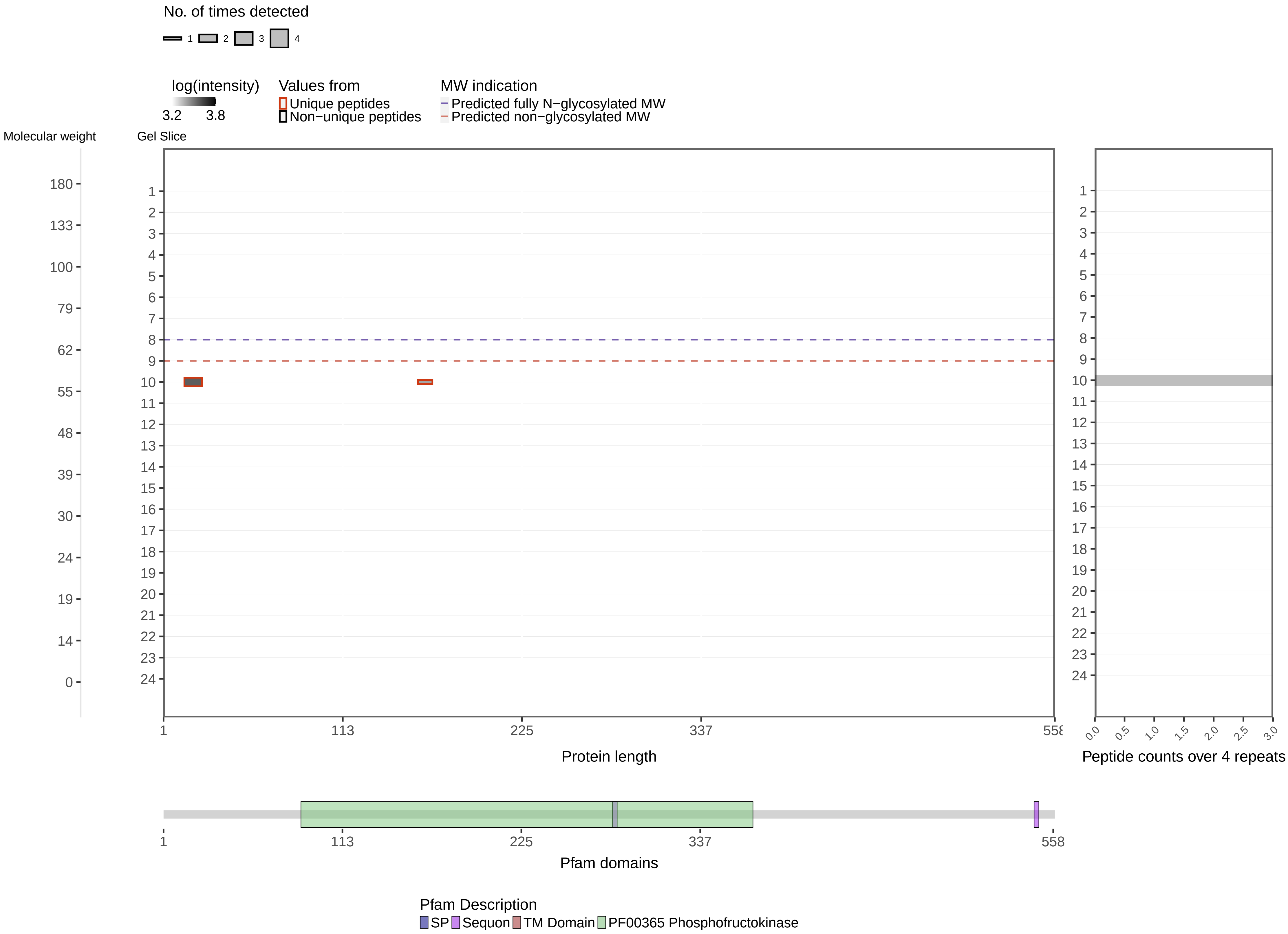


Protein sequence:

IGRAASHITLECALOTHPNITIIGEEVAAOKOTLKSVTDYIVGIICKRAKLGYNYGVILIPEGLIDFIPEVOOLIAELNEILANDVIDEAGNWKKKLRSOSHELFEFLPOAIOEOLLLERDPHGNVOVAKIETEKMLIOMVEAELENKKKEGSYTKHFKGOSHFFSYEGRCGLPSNFDSNYCYALGYGAGALLHSGKTGLISSVGNLRAPVEDWTVGGTALTSLMDVERRHGKFKPVIKKAMVELDGAPFKKFASLREEWALKNRYFSPGPLOFVGPTANEINHTLMLELGAKA*

NbL04g17180.1 heterogeneous_nuclear_ribonucleoprotein_1−like [MW = 49.76 kDa]

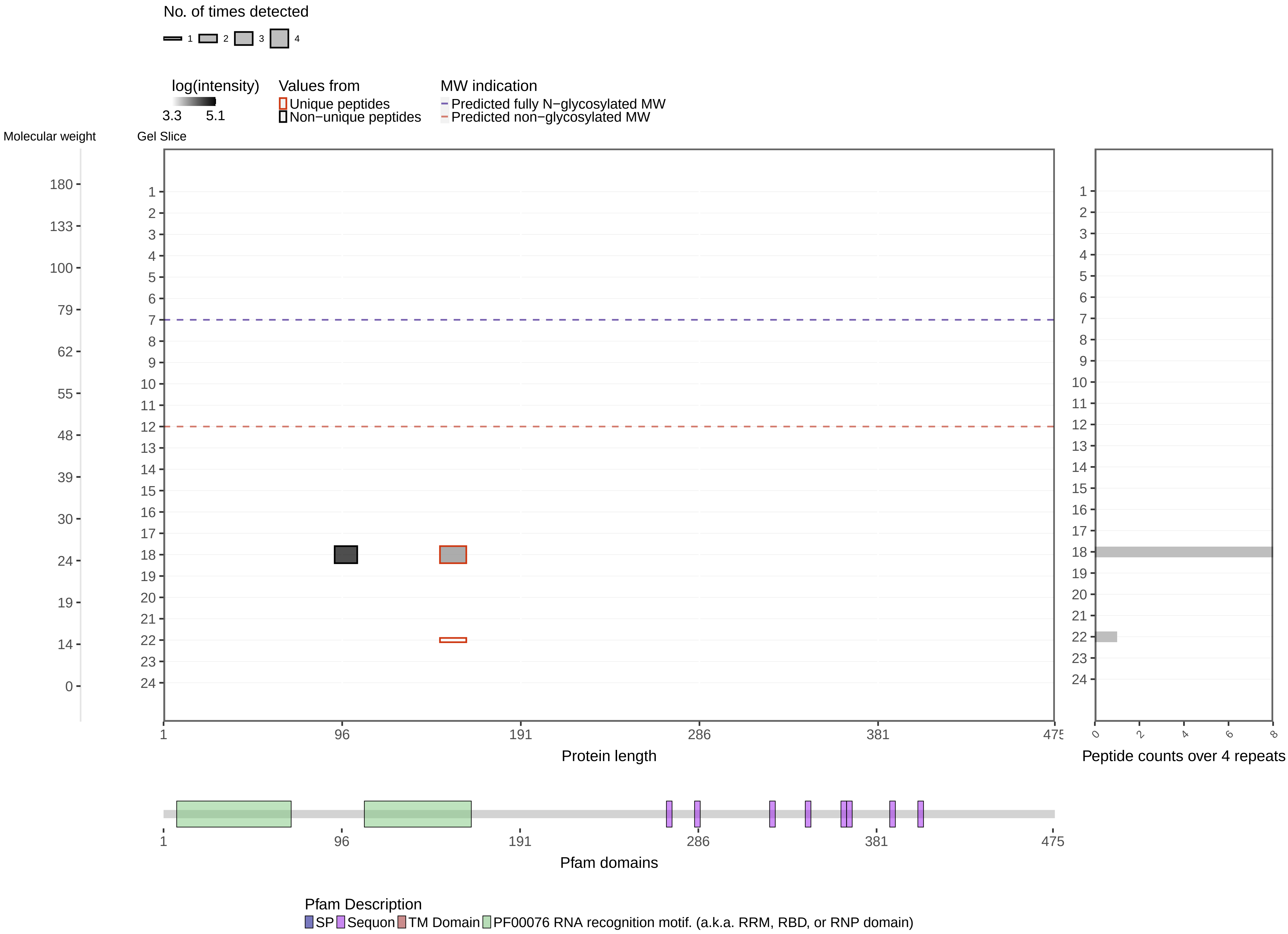


Protein sequence:

FSPIAAGDRSGFAQFGSGYGMGLNFEPGLSPGYGGNANFNSSLSYGRGMSPYYVNSSNRFGGPIGFDGGNGGSSSFFSSATRNLWGNGGLGYGTNSTSSSNFAVSGSGNMGAGNFSNTGVWGSSSISSQGGGNVSNQSGNLGYGGDDTLYGLSGGYGRNVTTSGAPASSYPASNGSYDGALADLYSGGLGYSDSDWRSANSERDGSGSIGYGLGTSPSDMPPKSSASYVGSYGVGKRQTNRGIGG*

### NbL04g17030.1 heat_shock_protein_83 [MW = 90.16 kDa]

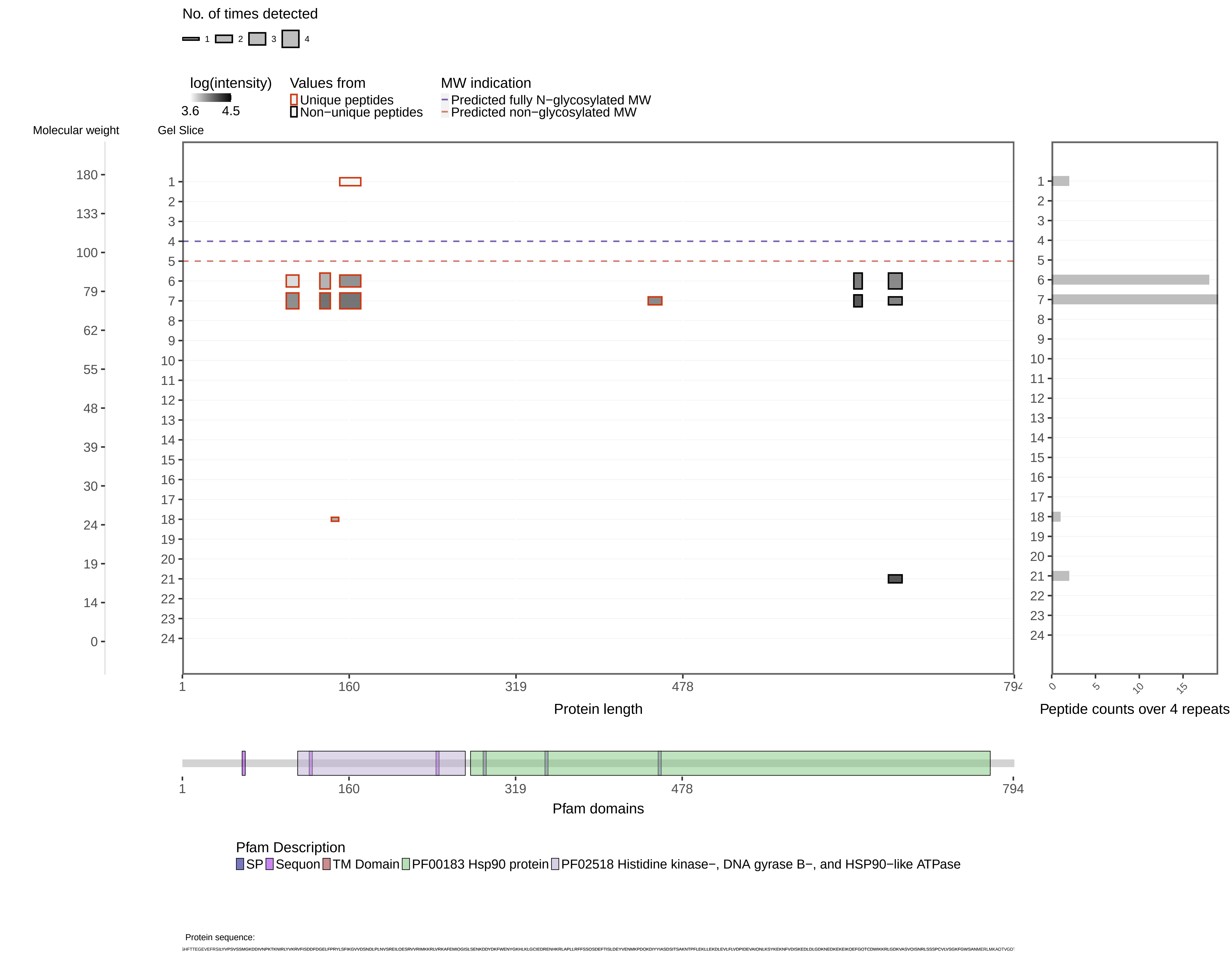

NbL04g16800.1 low−temperature−induced_cysteine_proteinase−like [MW − SP = 48.94 kDa]

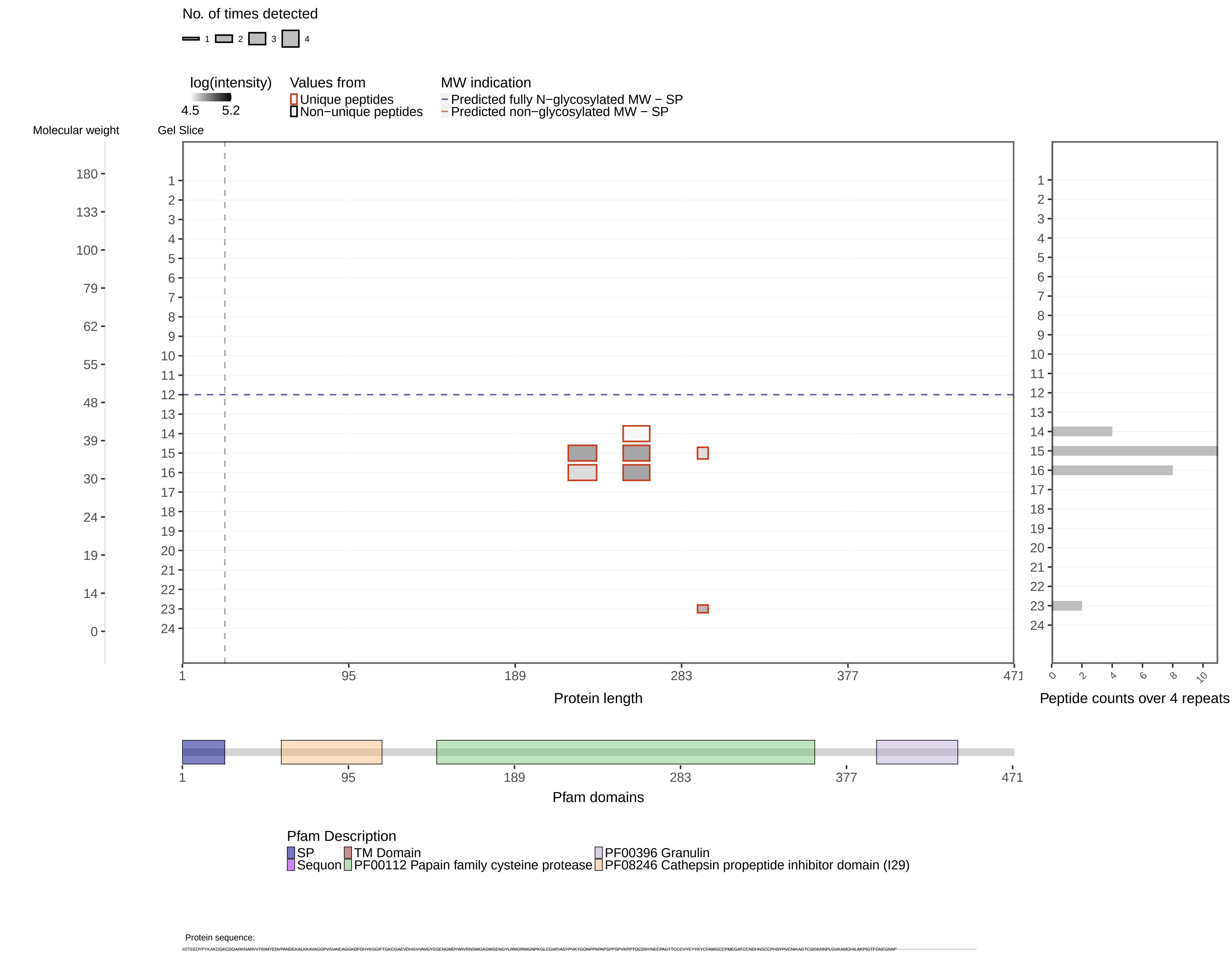

NbL04g16640.1 phosphoglycerate_mutase−like_protein_1 [MW = 31.4 kDa]

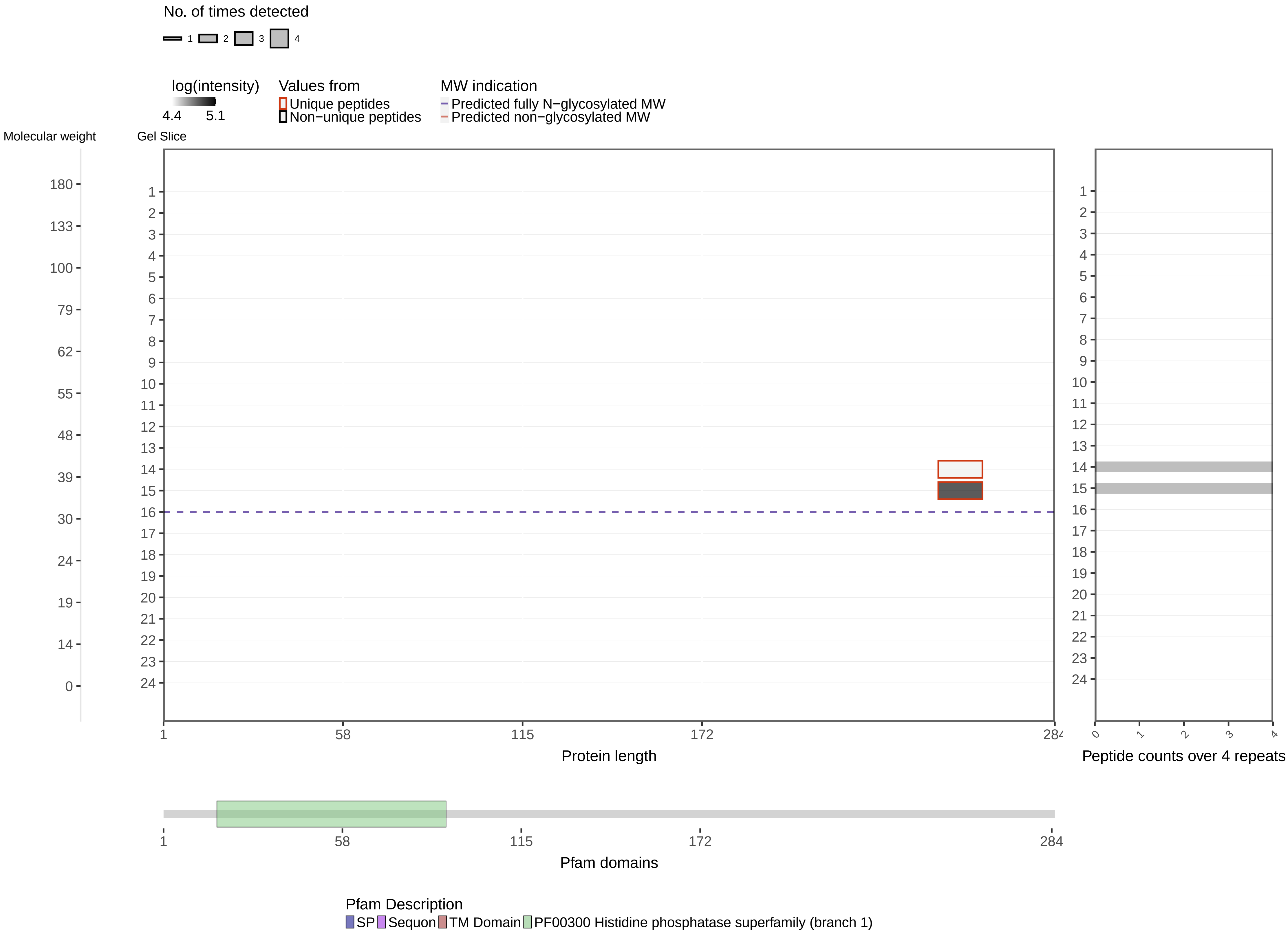


Protein sequence:

SVEGCREHLGVHWCDKRRSISEYKPLFPAIDFSLIECDDDVLWEPDIREPNEHLASRGMEFLKWLWTRKEKEIAVVTHSGFLIHTLSAFGNDCHPDVKNDICRPFKNCELRSMVIIDRGMIGSDSSTTDYPGKIPSGVDVPSDDTCINHPDGLLN*

### NbL04g16410.1 protein_DCL_chloroplastic [MW = 26.26 kDa]

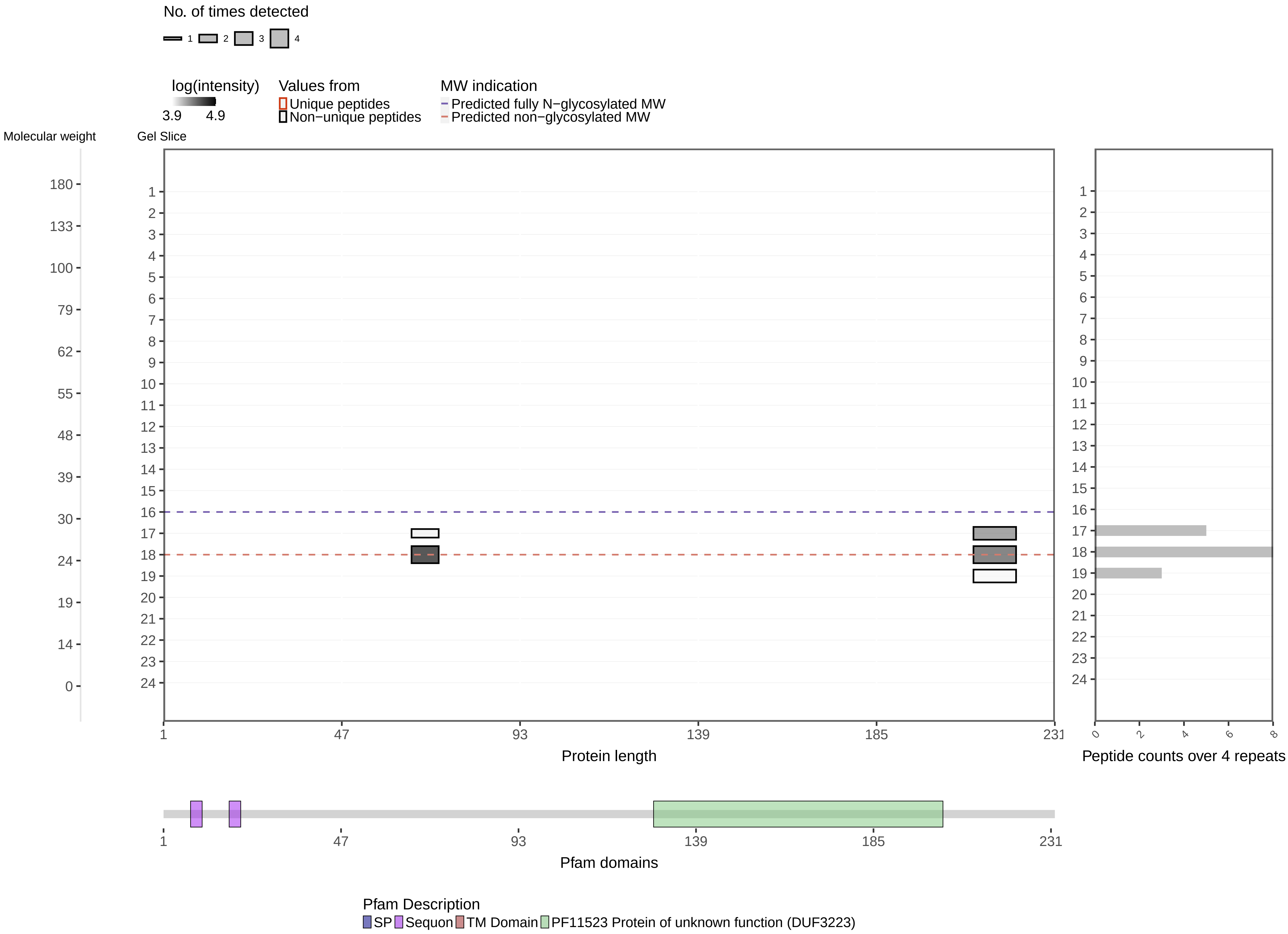


Protein sequence:

NGDEQNWIDWEDQILEDTVPLVGFVRMILHSGKYAVGDRLSPDHQRTILQRLLPYHPECEEKIGSGVDYITVGYHPDFESSKCLFIVRKDGELVDFSYWKCIKGLIRKNYPLYADSFILRHFRRRRRND*

### NbL04g16340.1 thioredoxin−like_4_chloroplastic [MW = 24.44 kDa]

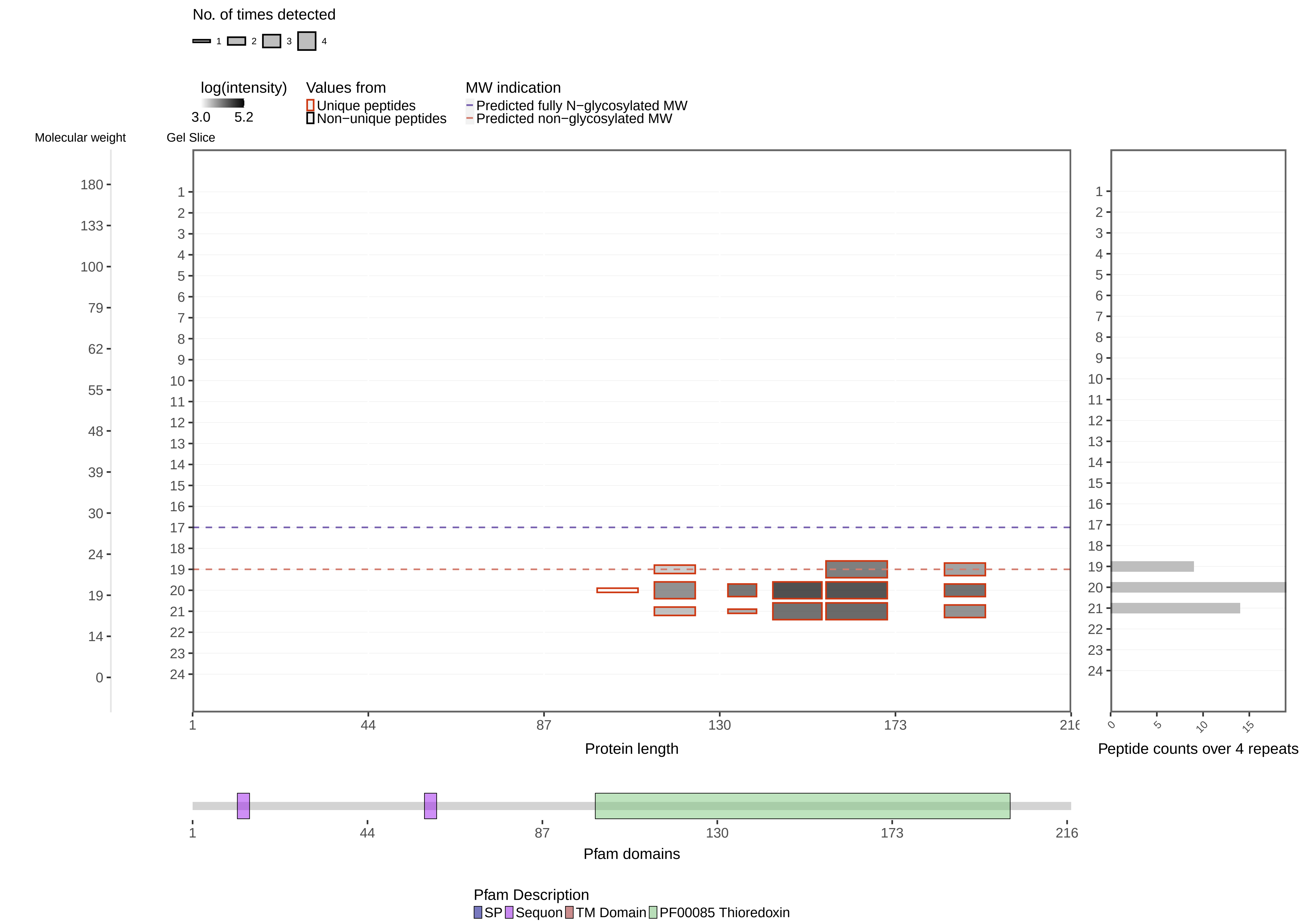


Protein sequence:

:CVREFKTDEEFFIILEKAKKAGSLVVVDFYRTACGSCKYIEQGFAKLCRGAGDEQAPVIFLKHNVIDEYDEQSEVAERLRIKTVPLFHFYKNGVLLEAFPTRDKERILAAILKYSAPASADA*

NbL04g16320.1 non−specific_lipid−transfer_protein−like [MW = 13.76 kDa]

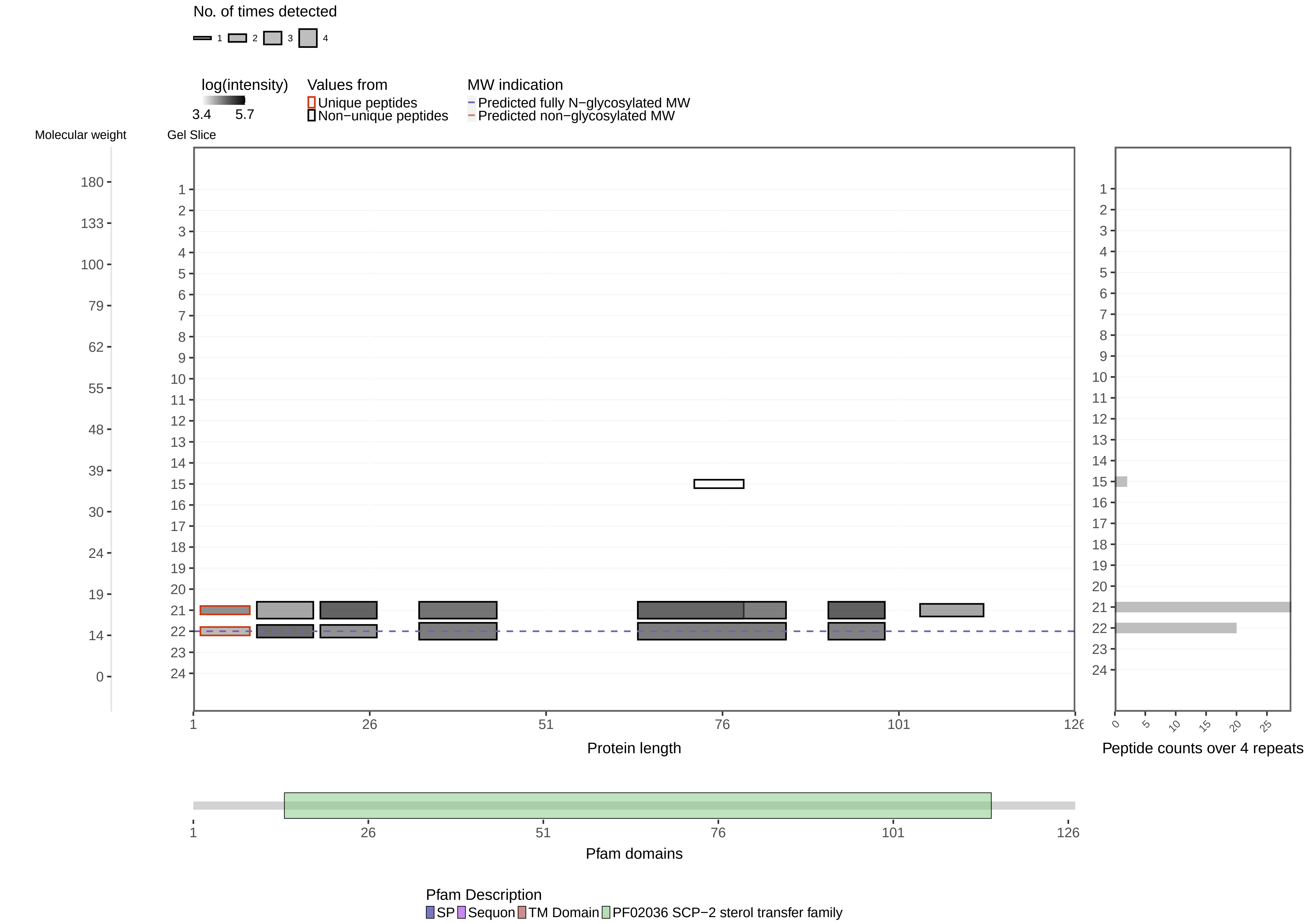


Protein sequence:

VYVVDLKKGETKEGPYEGGKPDATFSFKDDDFVKIATGKMNPQIAFMRGAMKIKGSISAAQKFTPDIFPKPSKM*

### NbL04g16240.1 protein_TIC_22_chloroplastic [MW = 31.49 kDa]

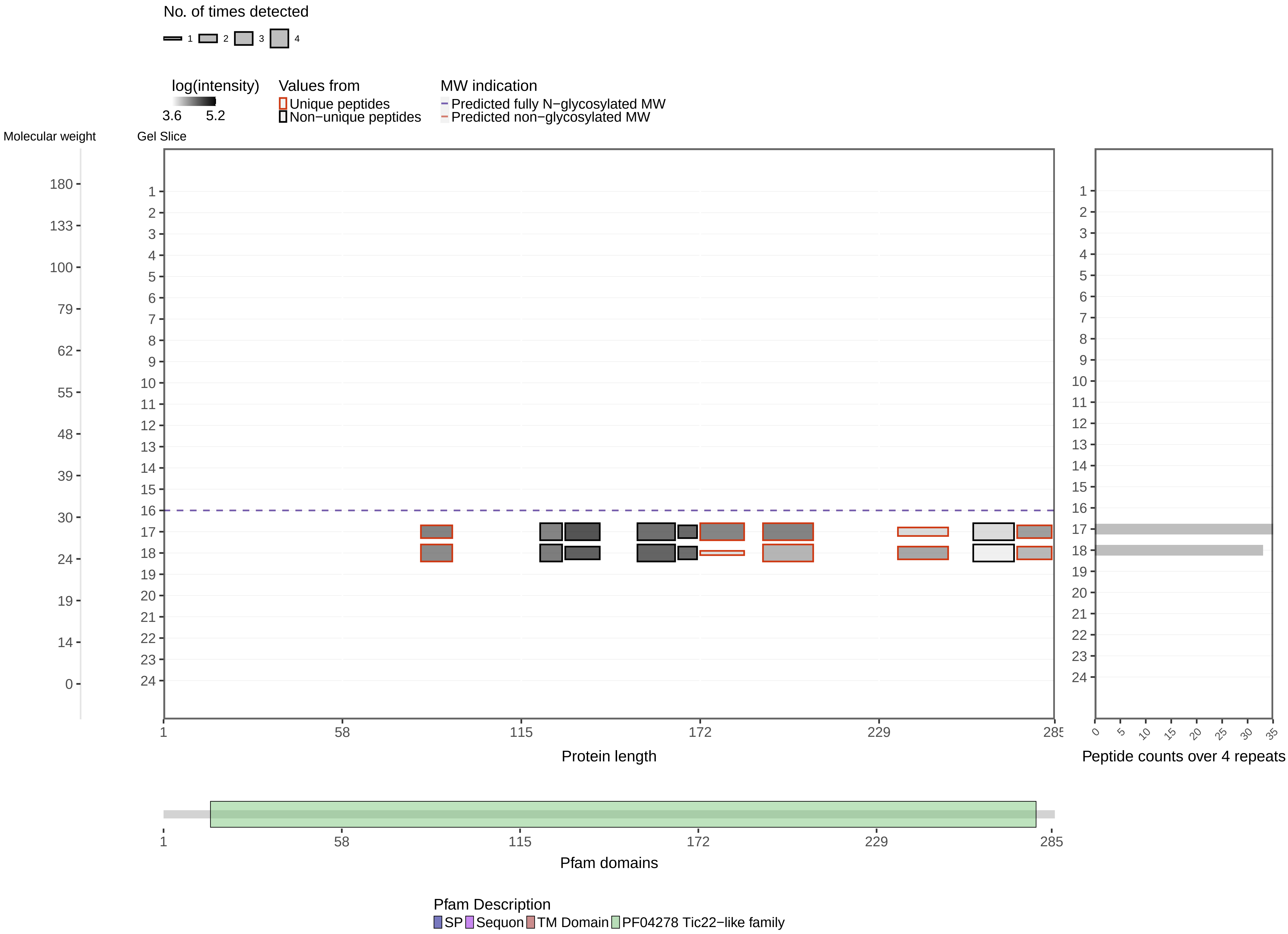


Protein sequence:

DAEAFLAQVRMRKGEVRGGAKVVPLTLDQVYMLKVEGIAFRFLPDPVQVQNAIELKASDVKTGFDGVPVFQSDLLLVKKRNRRYCPIYFRKEDIEKELSSRASRGGASQHIMVGSLEDVLKRMEISQRHSGWEDLIFIPPGKSHSQHIQDVTKV*

### NbL04g16180.1 uncharacterized_LOC107761413 [MW = 46.16 kDa]

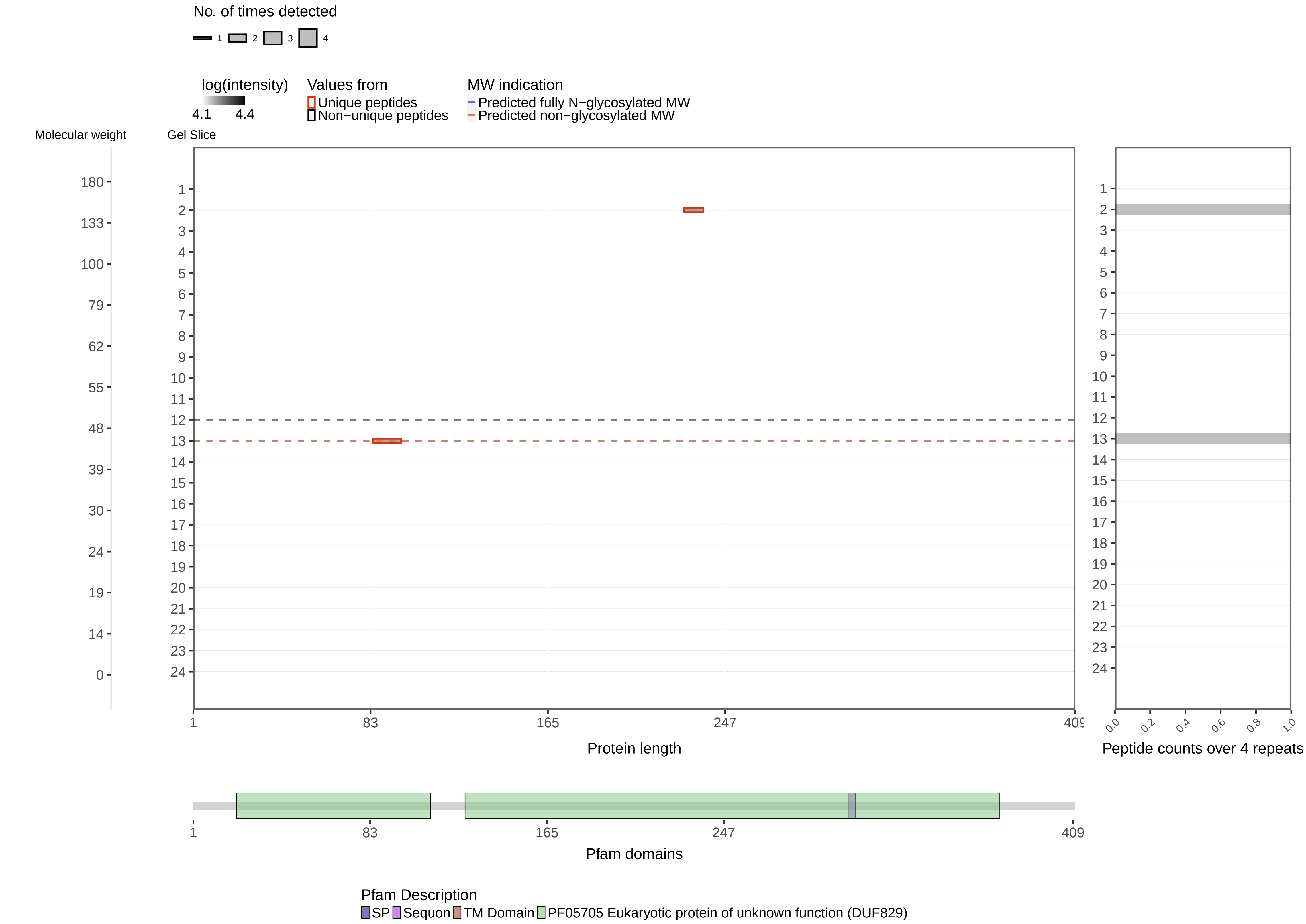


Protein sequence:

VQTLYASVSTGAPYLILCSEDDDLAPYQTICNFAQRLKDLGSDVKLLKWSISPHVGHYRYHQLEYKAAVTEVLGKAAMIYSQRIRQLEGEKMGLEGSHDEISDPLGNLRKAAATANQSFQRIALELNDRVFVPSSVEYHECSNVGSIQHEQKERYIPLSSPPRINAHGVLGQILFDVCVPKVVEDWDIRSSSSFSKASFPSTRRHSPFNPMKCIRRSRL*

NbL04g16150.1 uncharacterized_protein_At1g47420_mitochondrial−like [MW = 33.79 kDa]

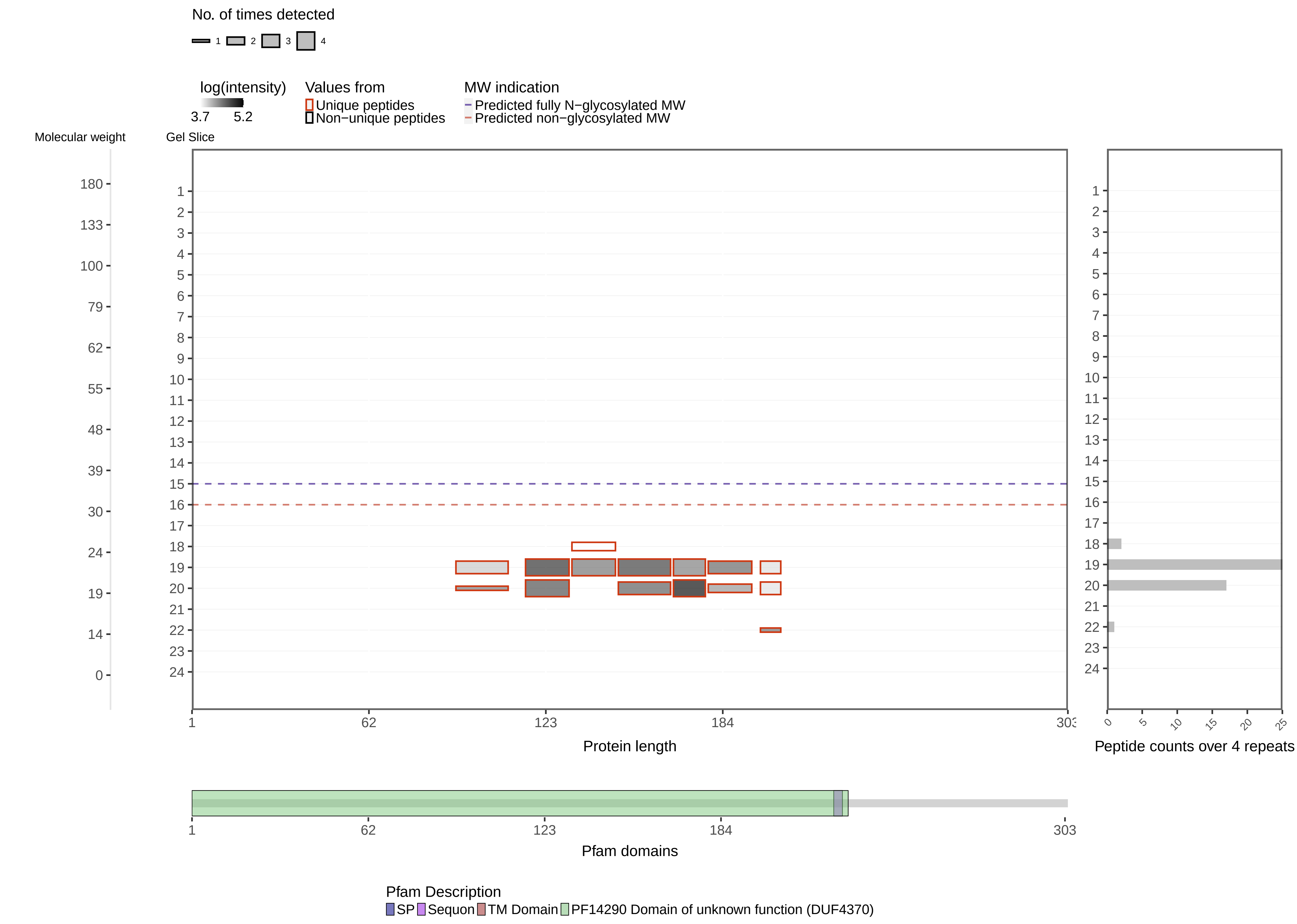


Protein sequence:

AEEFTGILTSLKMEIDDAIGLSGEDVKPLSKEVSDALQTVFQRYNAYMSAFGPEEGYLRKKVEMELGTRMIHLKMRCSGIDSEWGKVNLTEASWLHVFQFFCRLQLPTQASISAFSVVYNFLSNICITSYVSLLGDKKAVDTSCNLNSLVKLHSNSDFFFSMVPTKF*

### NbL04g16110.1 26S_proteasome_non−ATPase_regulatory_subunit_14_homolog [MW = 36.52 kDa]

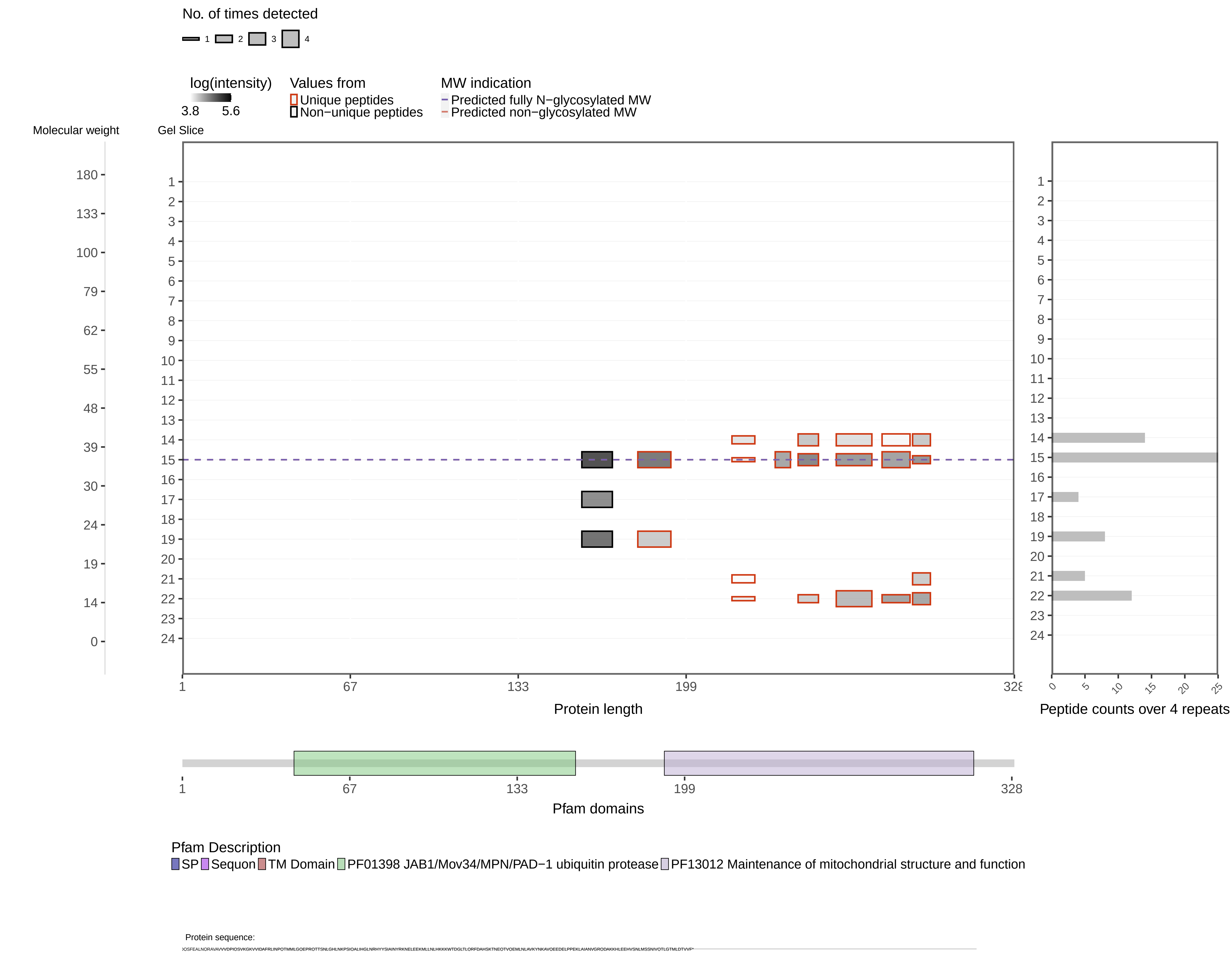

### NbL04g15990.1 uroporphyrinogen−III_synthase_chloroplastic [MW = 32 kDa]

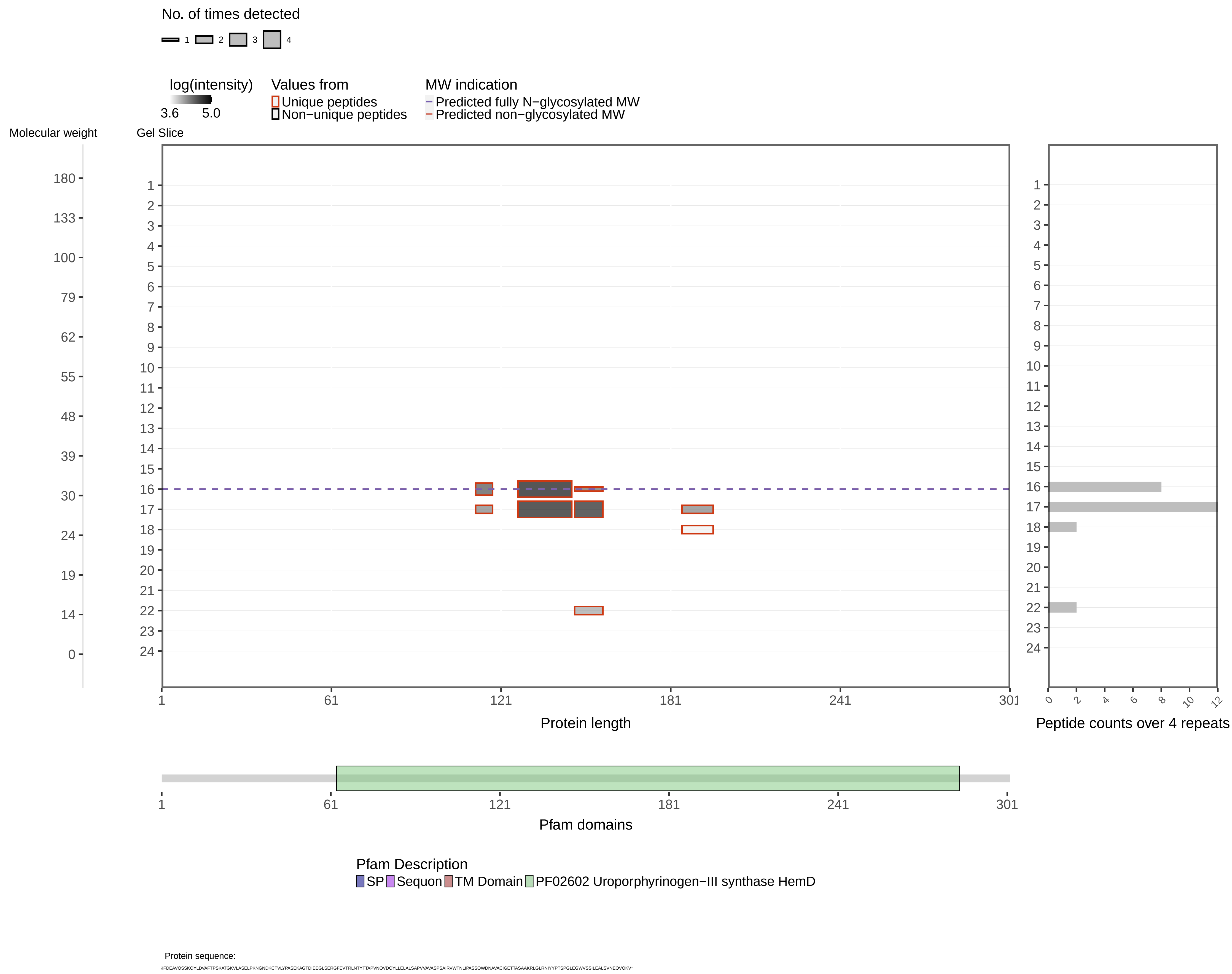

NbL04g15770.1 serpin−ZX [MW = 42.69 kDa]

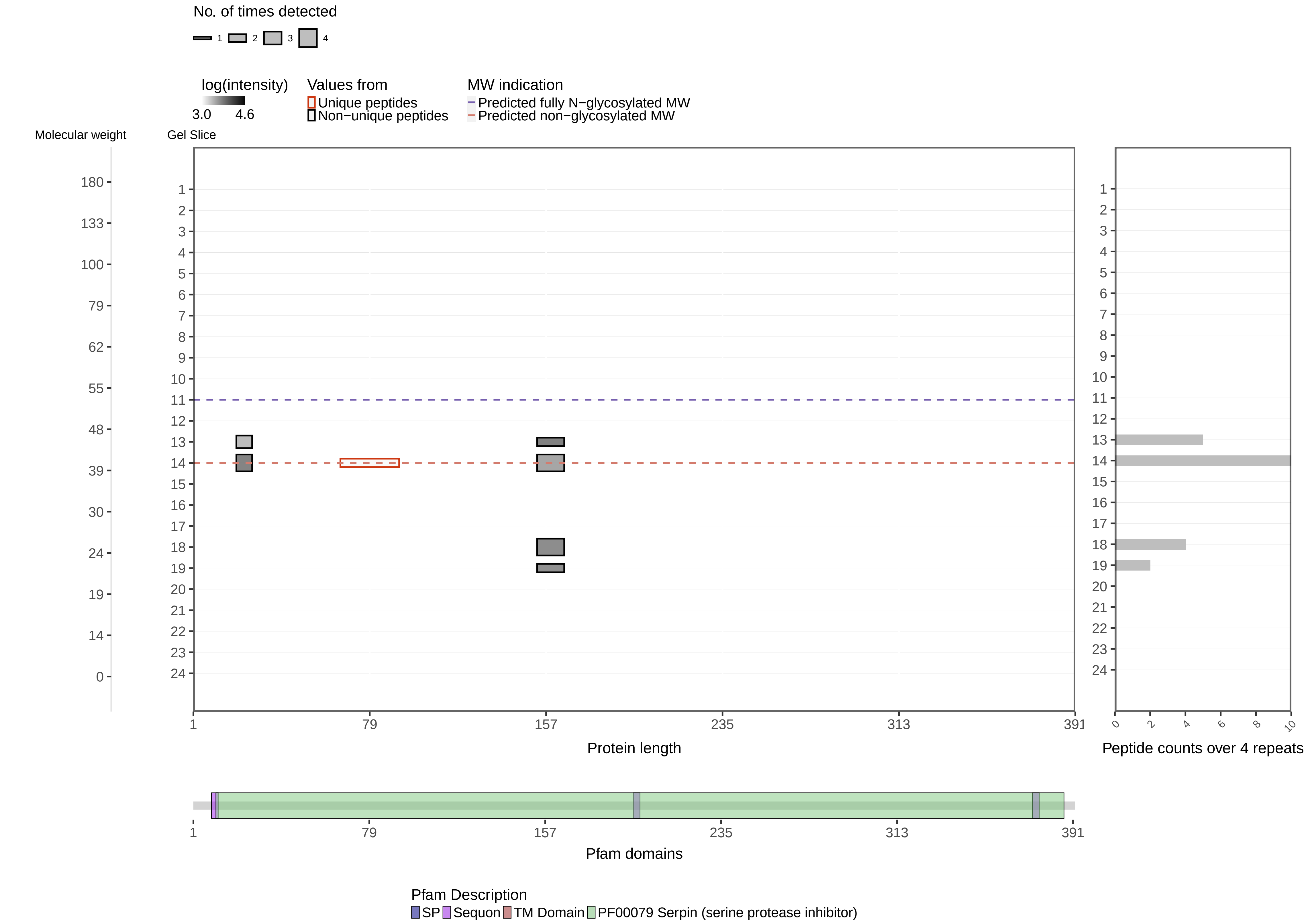


Protein sequence:

FDASVTKDNEFHLLNGTSVKVPFMTSKKKQYVKAFDGFKVLGLPYKHGEDRRRFTMYFFLPDANDGLPALVDKVSSESQFLESHLPYQKVEVGEFRIPKFKISFGFEASNILKGLGLVLPFSGDGLTEMVDSPVGSNLYVSSIFHKSFIEVNEEGTEAAAATAGVIKLRALMVEEKVDFVADHPYLFLIRENATGVVLFVGSVQNPLVD*

### NbL04g15650.1 plastidial_lipoyltransferase_2−like [MW = 33.52 kDa]

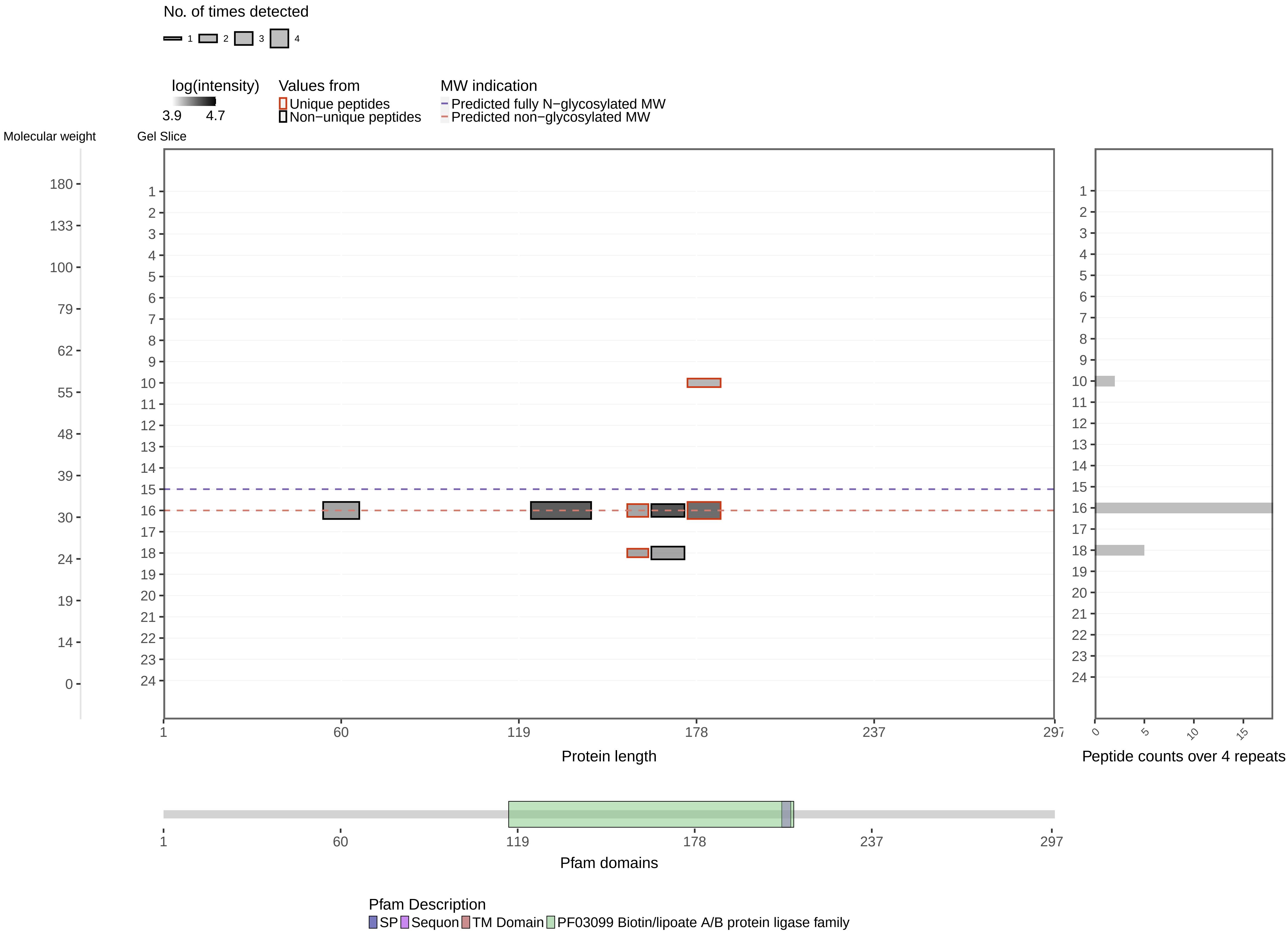


Protein sequence:

VMYPIINLRYHKMDLHWYLRSLEEVIIRVLSSSFSMEASRIDGLTGVWVEGRKLAAIGIRVSQWVTYHGLALNVTTDLTPFQQIVPCGIRDRQVGSIKGLLKGLPVSNGCGTENHQDADDVQLIDIAHKSLVKEFCEVFQVDIHFKPTPVLTSAEKPFSWLD*

### NbL04g15610.1 30S_ribosomal_protein_S9_chloroplastic [MW = 22.85 kDa]

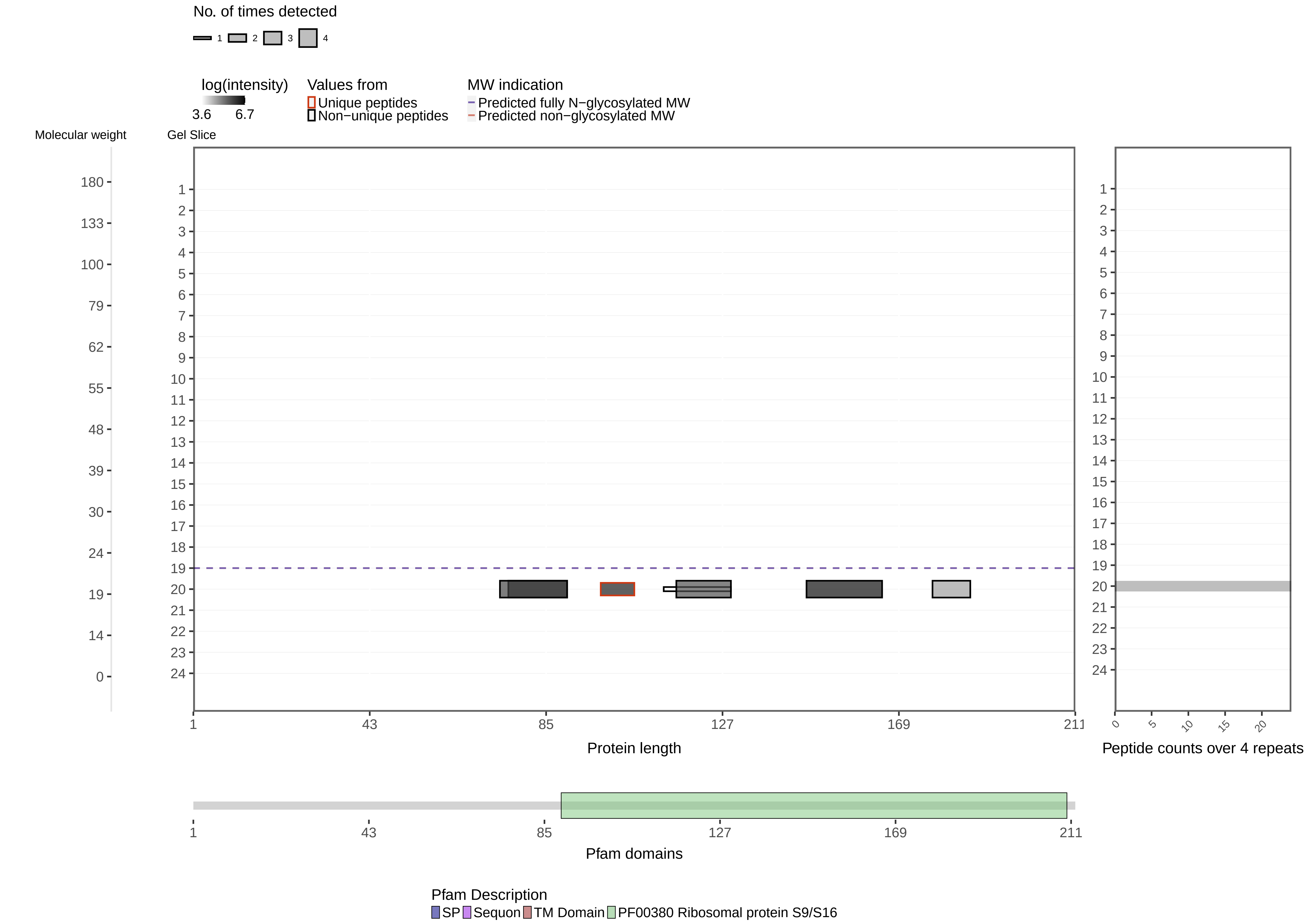


Protein sequence:

ASARVVLQEGSGKIIINYRDAKEYLQGNPLWIQYVKTPLATLGYEASYDIFVKVEGGGLSGQAQAISLGIARALLKVSESHRKPLREEGLLTRDARVVERKKVGLKKARKRPQFSKR*

NbL04g15140.1 uncharacterized_LOC109224904 [MW = 200.4 kDa]

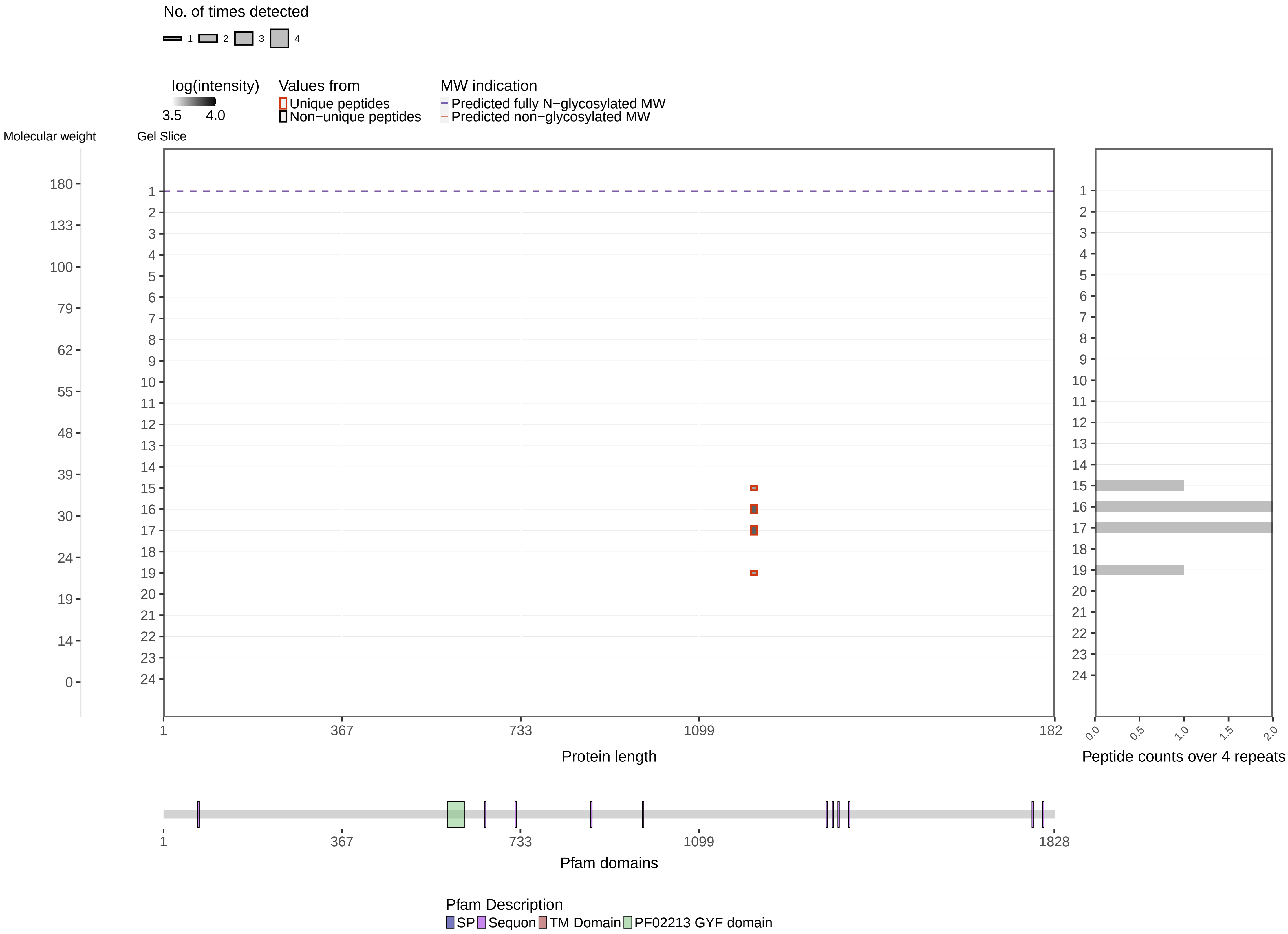


Protein sequence:

[illegible]

### NbL04g14860.1 maf−like_protein_DDB_G0281937 [MW = 25.31 kDa]

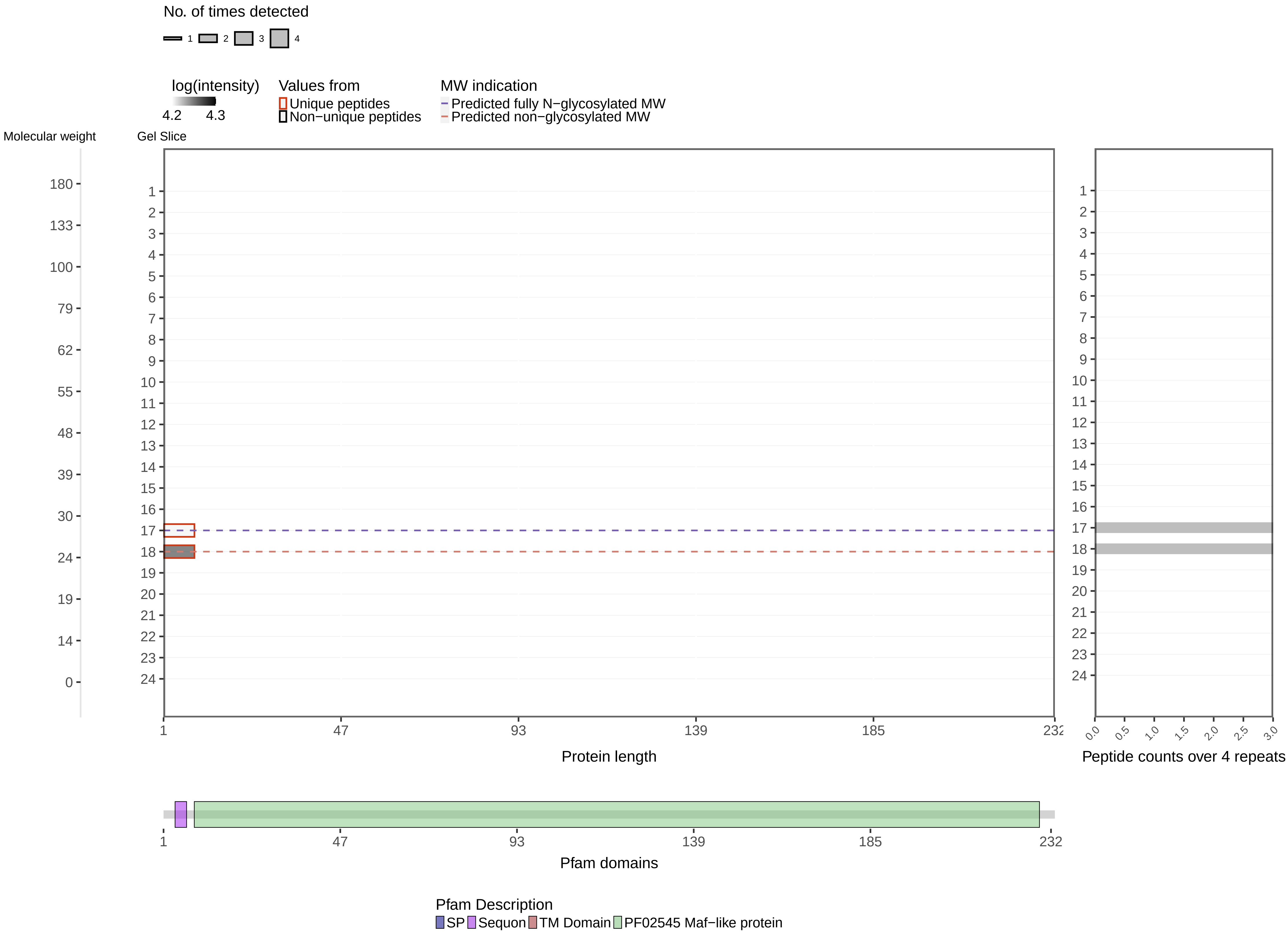


Protein sequence:

AEPTLLITCDQVVVYEGMVREKPSSEAEARQFMKDYANGHAATVSSVLVTNLVTGCRRGEWDKVEIYFHDIPDQVIDRLIEEGIVLYAAGGLIIEHPLILPCIKEVVGSTDSVMGLPKALTERLIKEVL*

### NbL04g14800.1 isoflavone_reductase−like_protein [MW = 34 kDa]

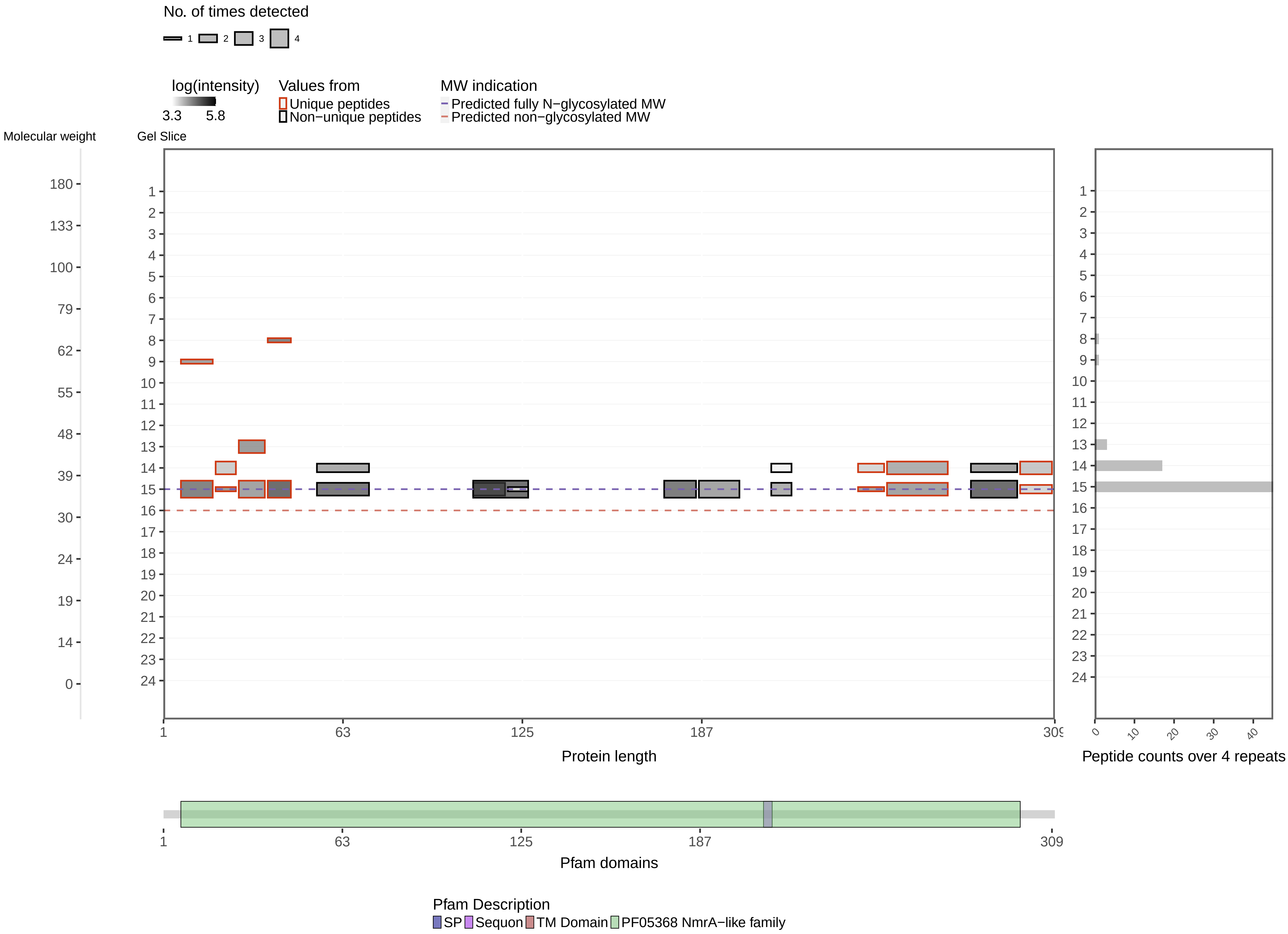


Protein sequence:

IEAEGIPYTYVSCNCFAGYFLPTMVQPGATAPPRDKVIIPGDGNVKAVFNEEHDIGTYTIKAVDDPRTLNKTLYIKPPKNTLSFNELVAVWEKMIGKTLEKIYLPEEQILKDIETSPMPLPVILAINHATFVKGDQTNFKIEPSFGVEASELYPDVKYTTVEEYLGHFI*

NbL04g14790.1 gamma_carbonic_anhydrase_2_mitochondrial−like [MW = 29.6 kDa]

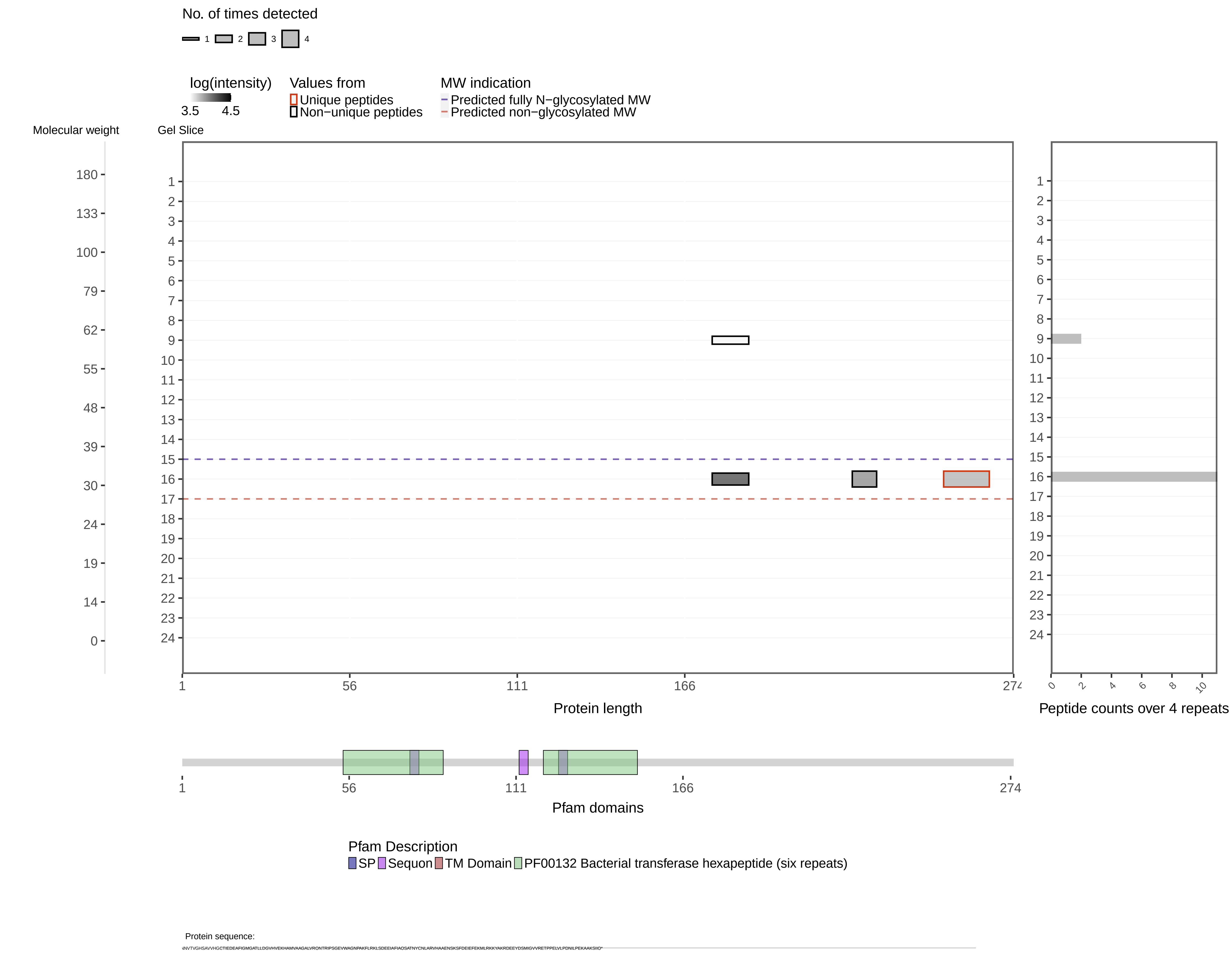

### NbL04g14770.1 proteasome_subunit_alpha_type−1−B−like [MW = 29.54 kDa]

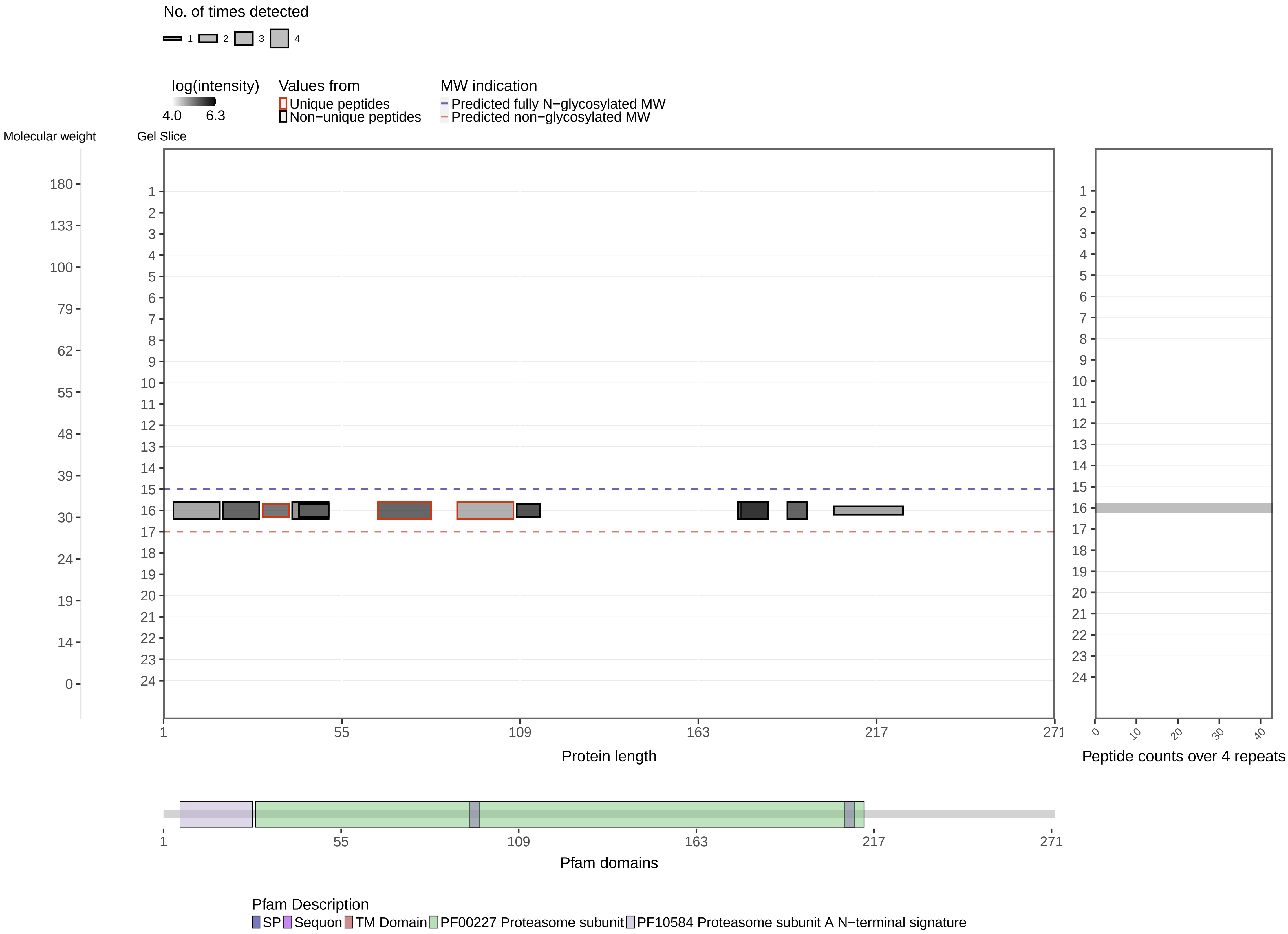


Protein sequence:

SWKRPYGVGLLVGGLDESGAHLYYNCPSGNYFEYQAFAIGSRSQAAKTYLERRFETFMDSSRESLLKDALFALRETLQGEKLTSTNCTVAVVGVGEAFHMLDKETIQALINEFEIAGEEAPAAAPDEAGSDEPAAAPPADQGAAPMDI*

### NbL04g14600.1 GTPase_HflX−like [MW = 62.93 kDa]

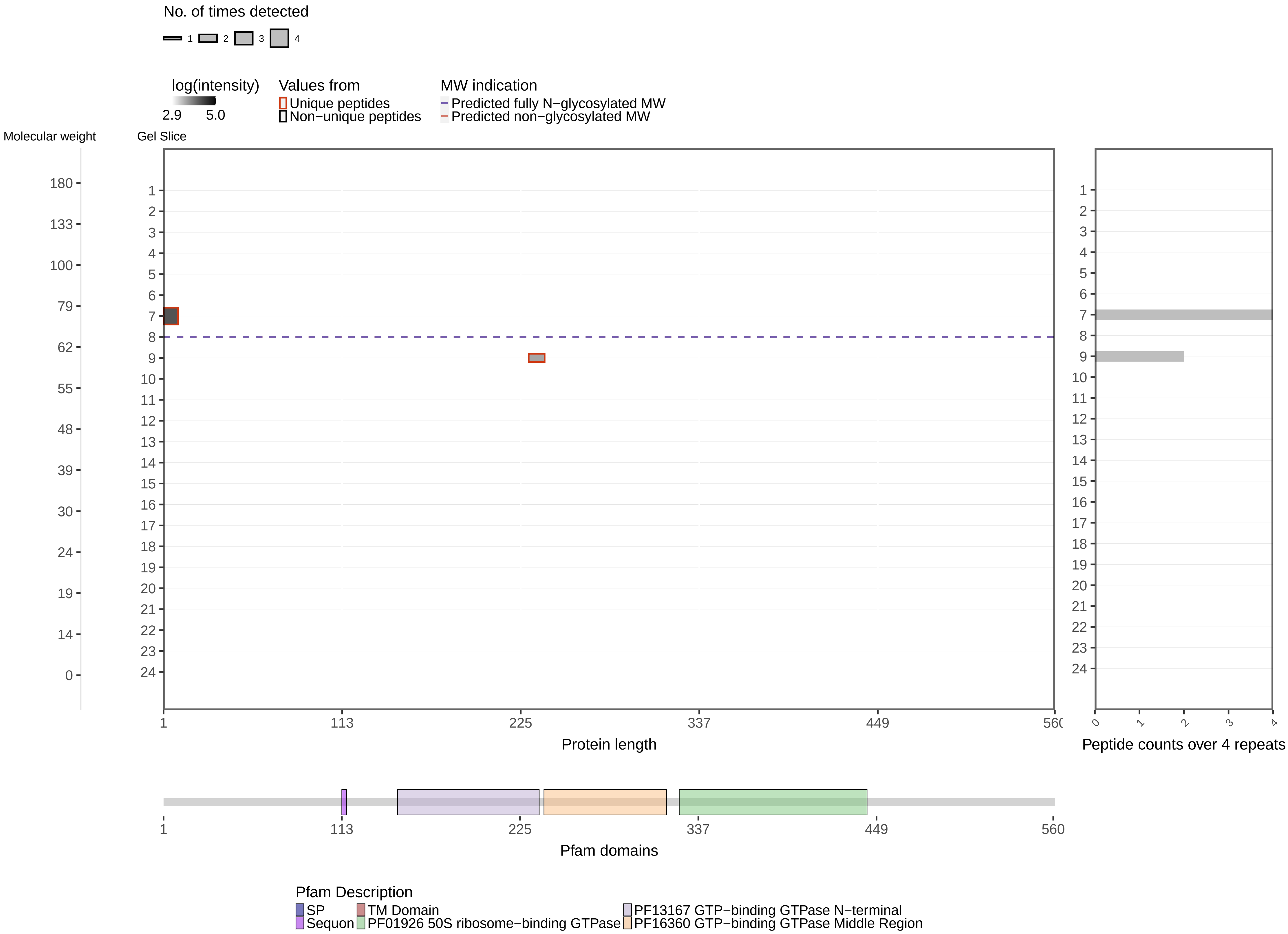


Protein sequence:

THLERQAGGQVKGMGEKQIEVDKRILRTQIGVLKKELESVRKHRKQYRNRRVSVPVPVVSLVGYTNAGKSTLLNRLTGADVLAEDRLFATLDPTTRRVQMKNGKEFLLTDTVGFIQKLPTTLVAAFRATLEEIAESSILVHVVDISHPLAELQIEAVEKVLSELDAVSIPKLMVWNKVDKARDPEKIKLEAKTKNDVVCVSALTGEGLDEFCNEIQNRLKDTMVWVEALIPFDKGELLSTIHHVGMVERTMHTACFFTSLNFKEYTEKGTLVRAHVPLRFARLLTPMRQMCVS*

### NbL04g14570.1 ribosomal_RNA_small_subunit_methyltransferase_I [MW = 41.51 kDa]

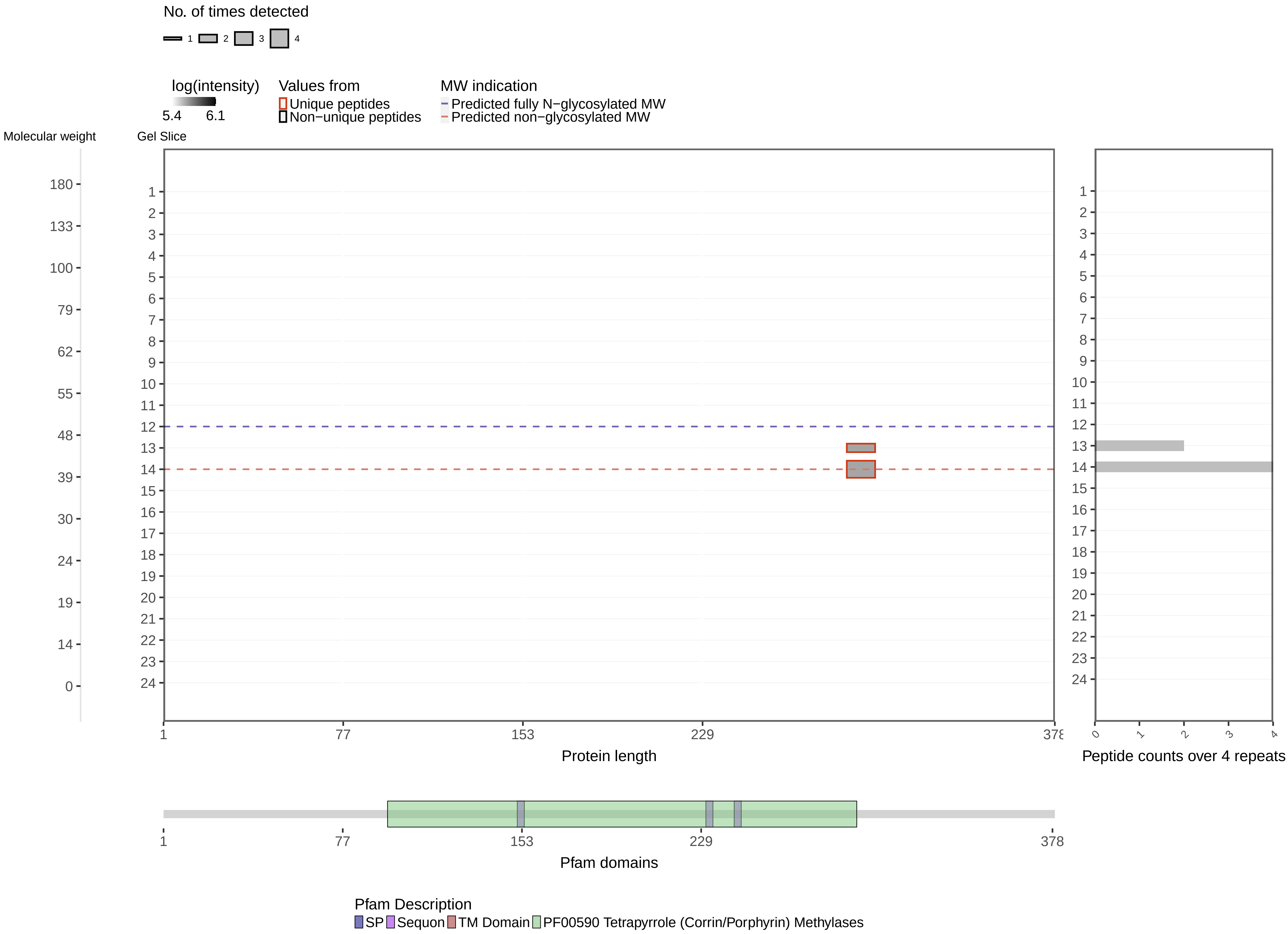


Protein sequence:

TPGISDPGMELAKLCVDKNVLVVPIPGPSAVVTALSASGLPTNEFTFVGFLPKHNSSRKEKLILSANESATQIFFIPPHRLSQFLEEAAAIFGESRQCVMAREMTKVHEEFWRGTIGEAKEAFLARQLKGEITFLIEGKSISIDESPSDSQLENELRELISEGHSLSMAVKLVASGKSVKRKAIYSLALRKFGGQLESEDE*

### NbL04g14540.1 thioredoxin_H−type_1−like [MW = 13.55 kDa]

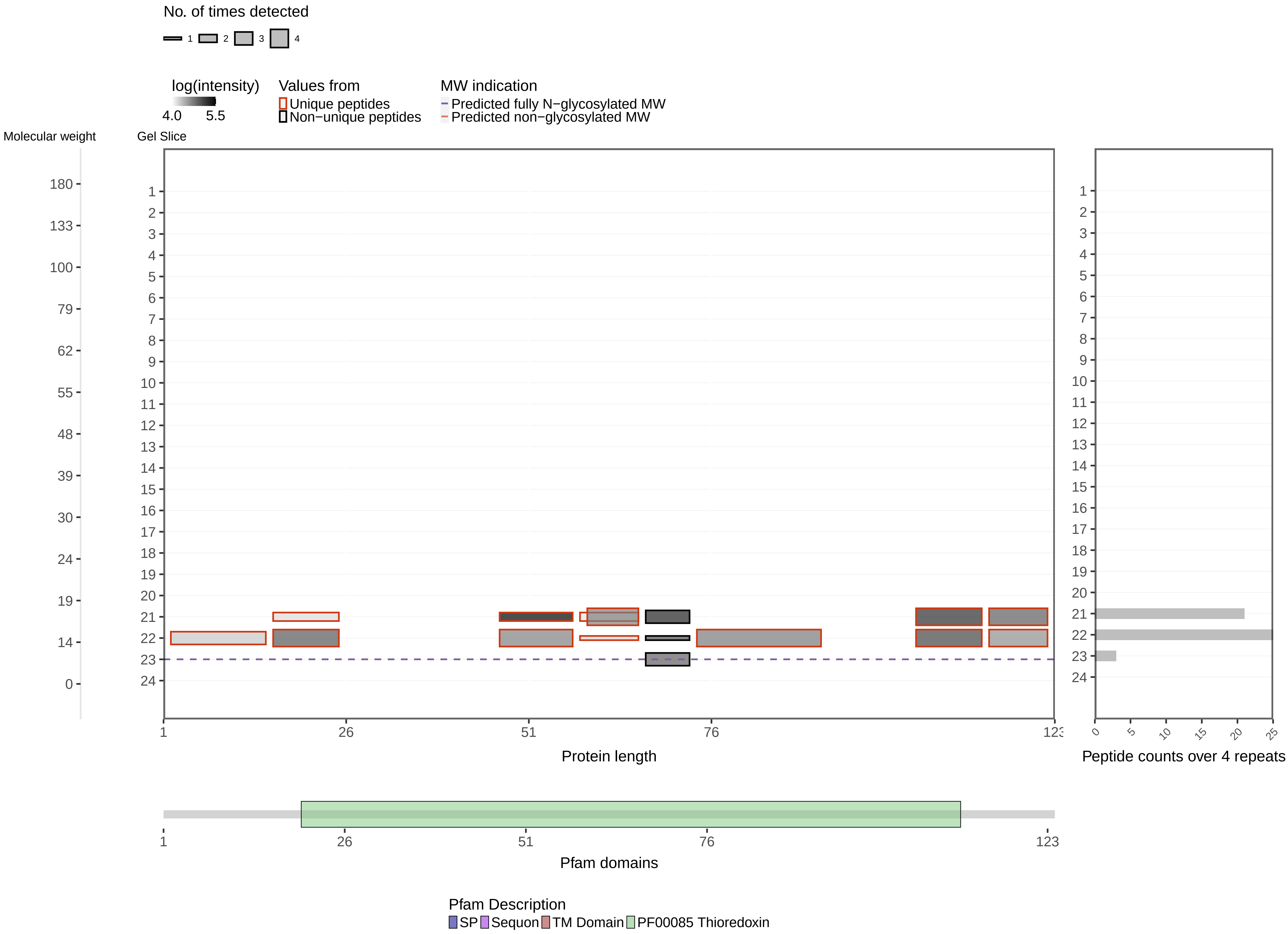


Protein sequence:

FIAPVLADIAKKMPHVIFLKVDVDELKTVAEEWSVEAMPTFVFLKDGKEVDRVVGAKKEELQQTIVKHAAPATVTA*

NbL04g14510.1 zingipain−2−like [MW − SP = 45.66 kDa]

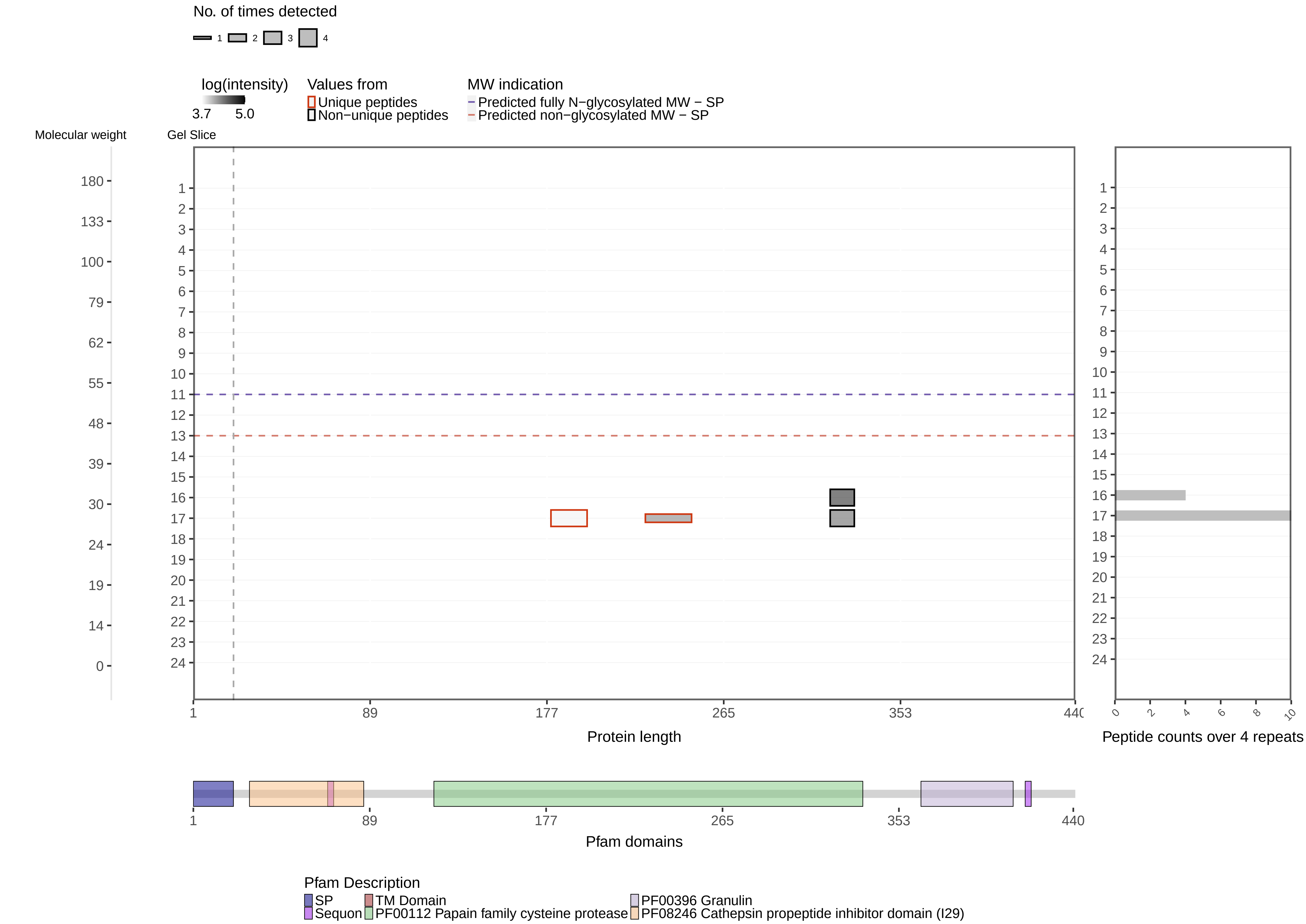


Protein sequence:

FNEREGTCNKNKLQRRVVTIDGYTDIPQNDEDKLLNAVAKQPVSVGICGSERAFQSYSKGIFTGPCSTALDHAVLIVGYGSENGVDYWIIKNSWGTSWGINGYMHMQRNSGNQEGICGINKLASYPTKSSPNPPSPPSPGPSKCSMFTSCGQGETCCCGWSLLGVCLSWKCCGLDSAVCCKDGRHCCPHDYPICDTNRNLCLKQMSNATIVQQPQKEAFSGKFGGLIYPF*

NbL04g14460.1 cysteine_proteinase_15A−like [MW − SP = 38.09 kDa]

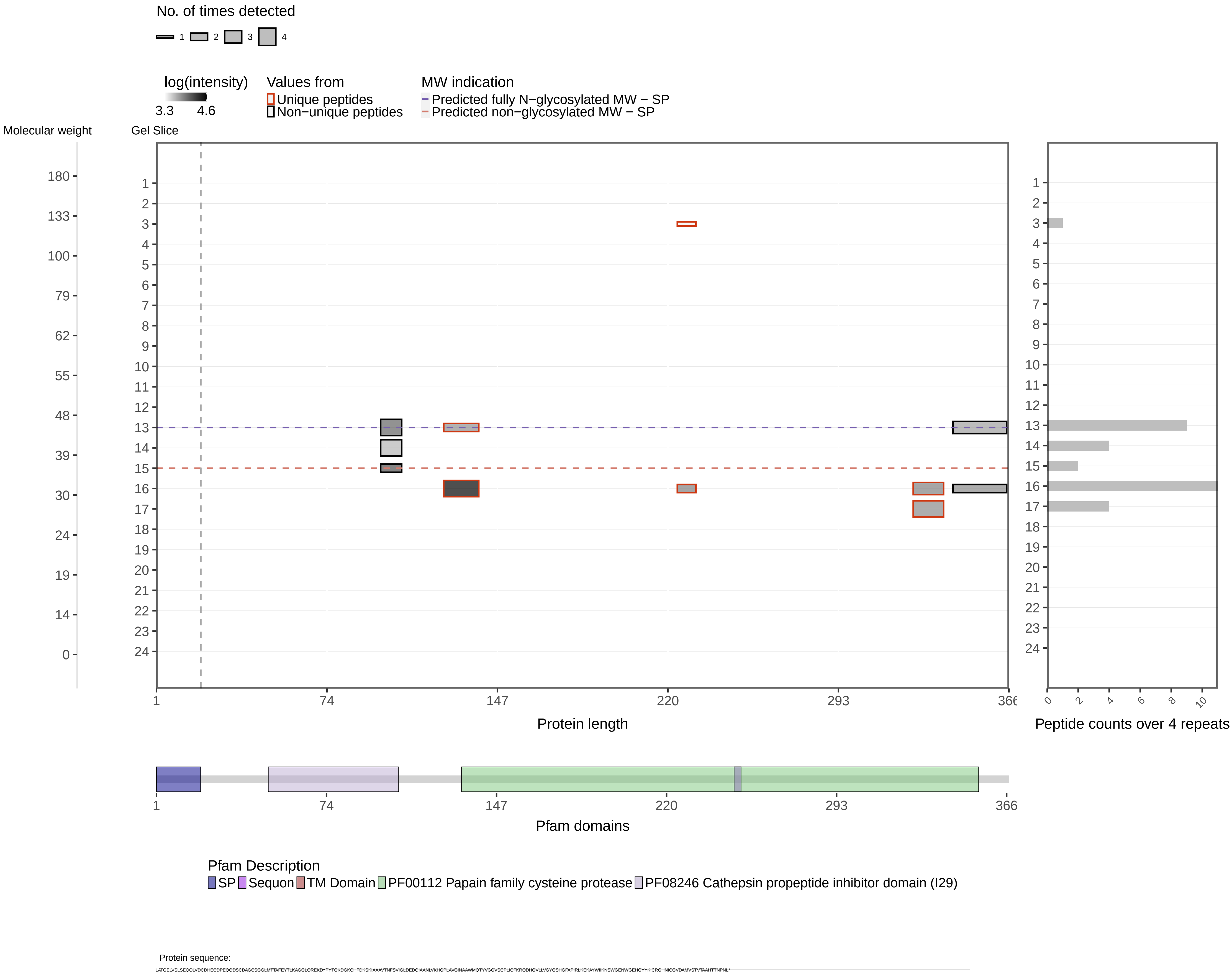

### NbL04g14430.1 coatomer_subunit_alpha−1−like [MW = 137.14 kDa]

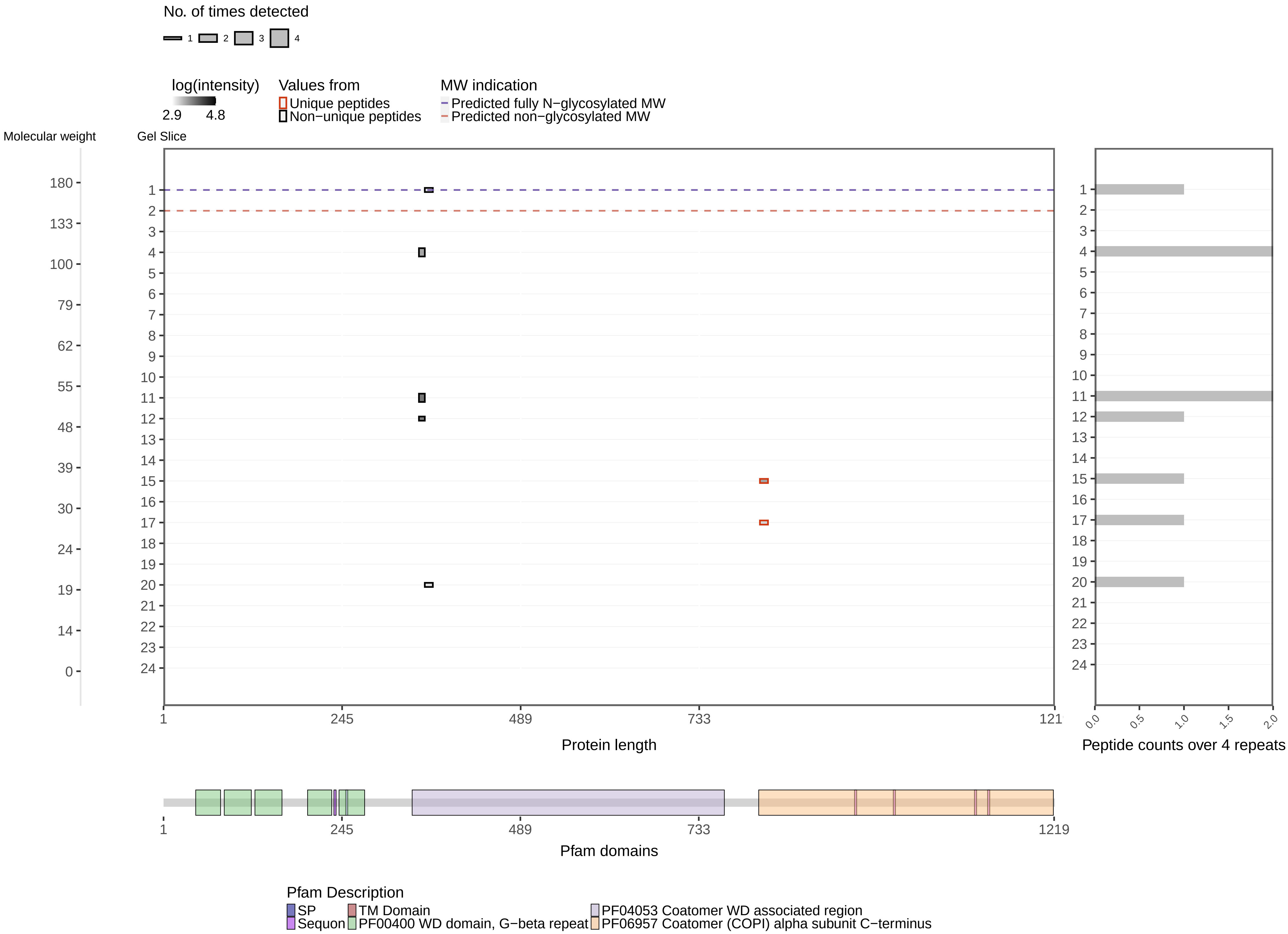


Protein sequence:

YIFKLALLKRRYDQVMSMIRNSELCGQAMIAYLQQKGFPEVALHFVKDELTRFNLALESGNIEIALESAKKLDQKDHWYRLGVEALRQGNAGIVEYAYQKTKNFERLSFLYLITGNLDKLSKMMKIADVKNEVMGQFHDALYLGDVRERVKILENAGHLPLAYITATTHGLKDIAERLGEELGENVPSLPKEKKASLLLPPTPILGGGDWPLLMVTKGIFEGGLDATVRGGHEEYEEAADANWGESLDIGEVENLQNGDISMVLEDEDGQEENDEEVGWDLEDLDLPPDTDTPK

NbL04g14070.1 endoglucanase_2−like [MW = 57.2 kDa]

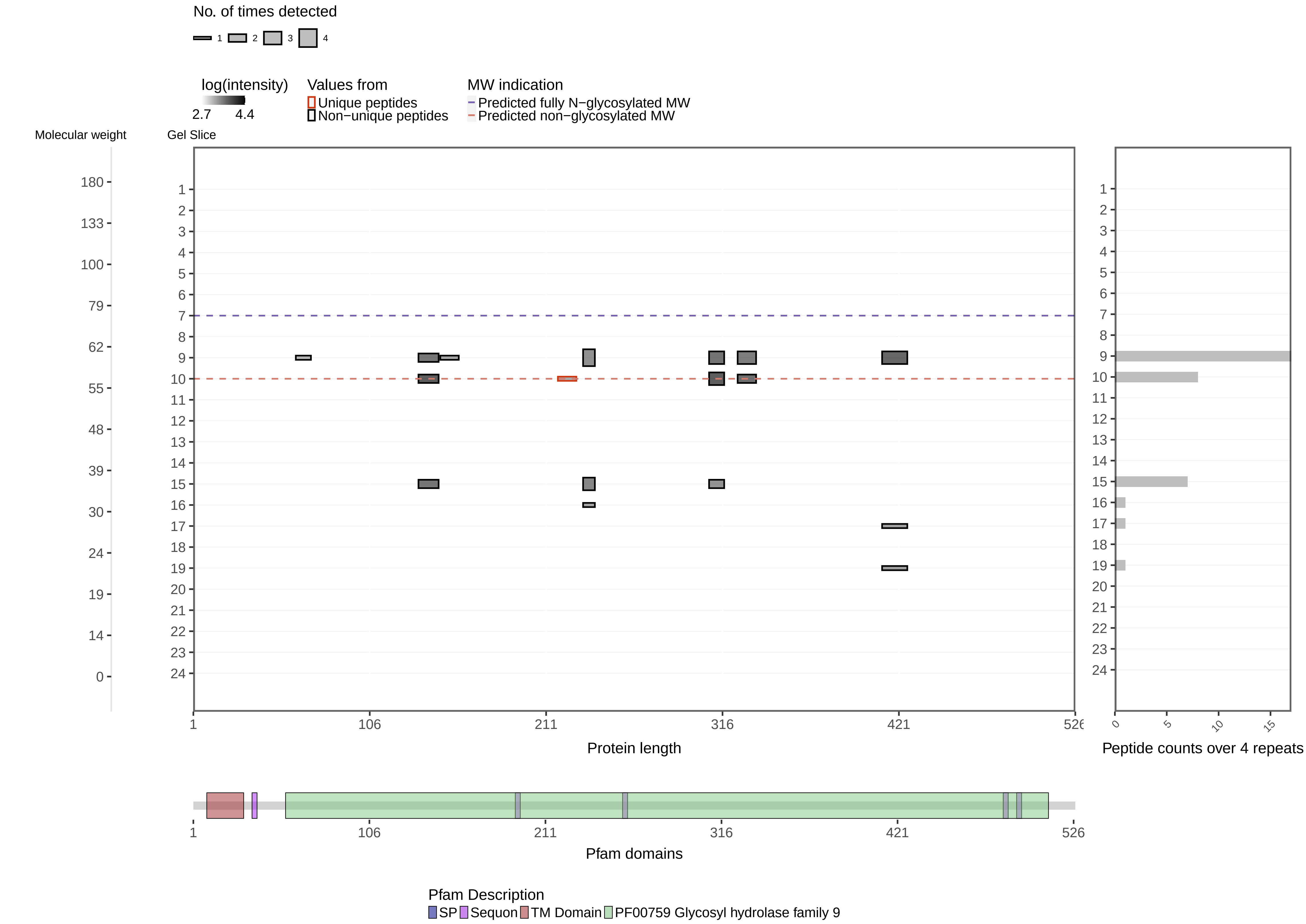


Protein sequence:

EVATYYNSTGYGDELLWAASWLYHATGDQSYYDYATGKNADSFGNFGNPTWFSWDNKLAGTQVLLSRVGFFDSKGTSNSETLQEYRKTAEAVMCGLLPKSPTATSSRTDSGLIWISEWNALQHPVASAFLAVLYSDYMLTSRTAKITCNGDAYTPSDLRKFAISQADYVLGDNPSKMSYLVGYGDNYPQYVHHRGASIPKDADTGCKDGWKWLDSTEPNPNVATGALVGGPFLNETYIDSRNNSMQAEPTTYNSAVIVGLLSGLVTTSSVVQSFT*

NbL04g14030.1 tRNA_(adenine(58)−N(1))−methyltransferase_non−catalytic_subunit_trm6−like [MW = 48.75 kDa]

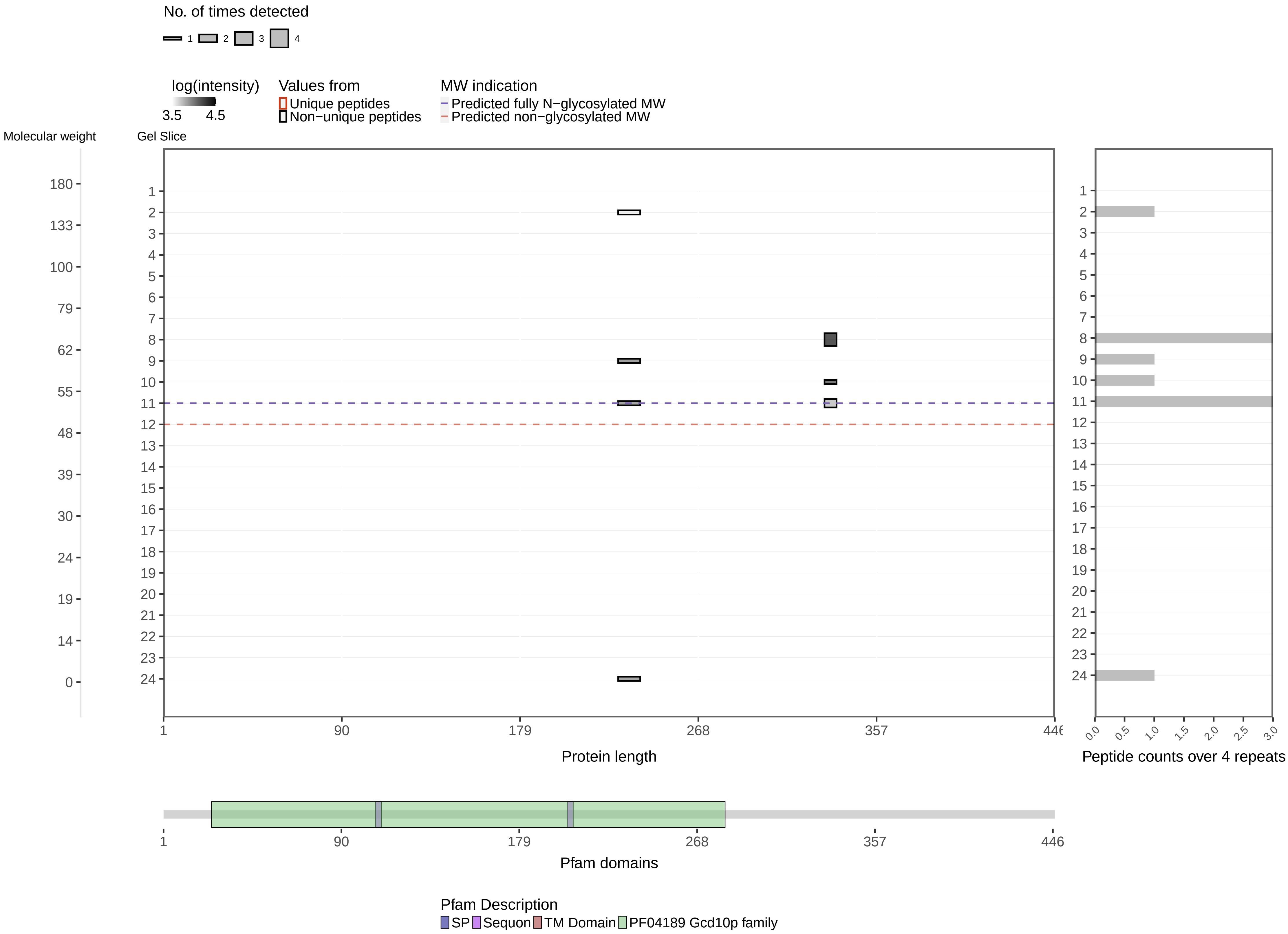


Protein sequence:

VLVVDTVGGILTGAIAERLGGTGYVCNTYRGSSPYPVDITRMFNLNDEICKRIVHASLSDLNSLKTGASVSESQEEVLPATHSGGLEEVGVSSEHSCMDITPESVSSPIIKPCKSIKPGQKAPLEAIKLWKENGFSSLIIAAPELDPWGIIKEVLPLLAFSAPFAVYHQYTQPLATCMHNLQIEKMAIGLQISEPWLREYQVLPSRTHPHMQMSAFGGYILSGTKISSSEPVTSSN*

### NbL04g13960.1 hexokinase−2_chloroplastic−like [MW = 54.04 kDa]

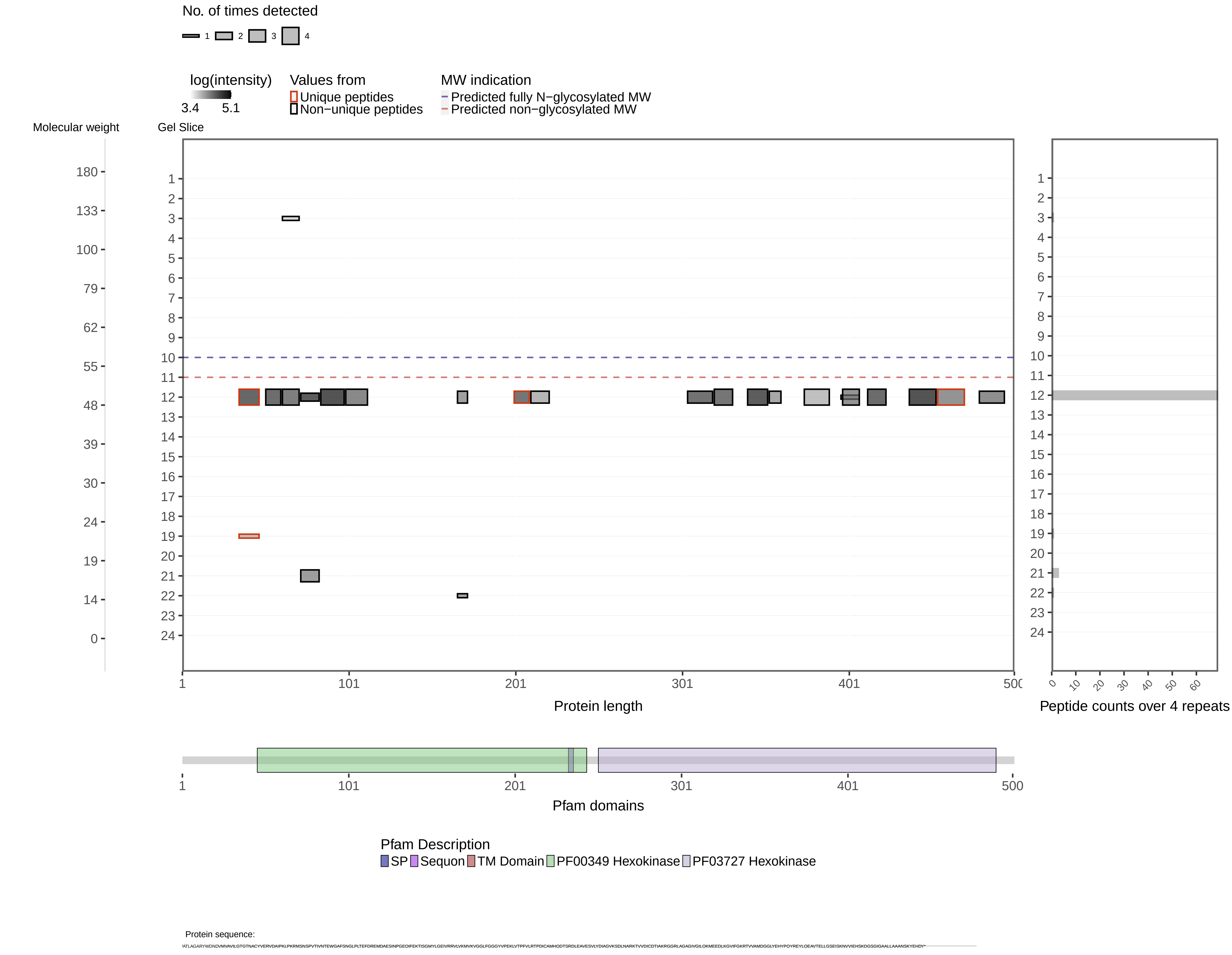

### NbL04g13880.1 tubulin_beta−1_chain [MW = 50.5 kDa]

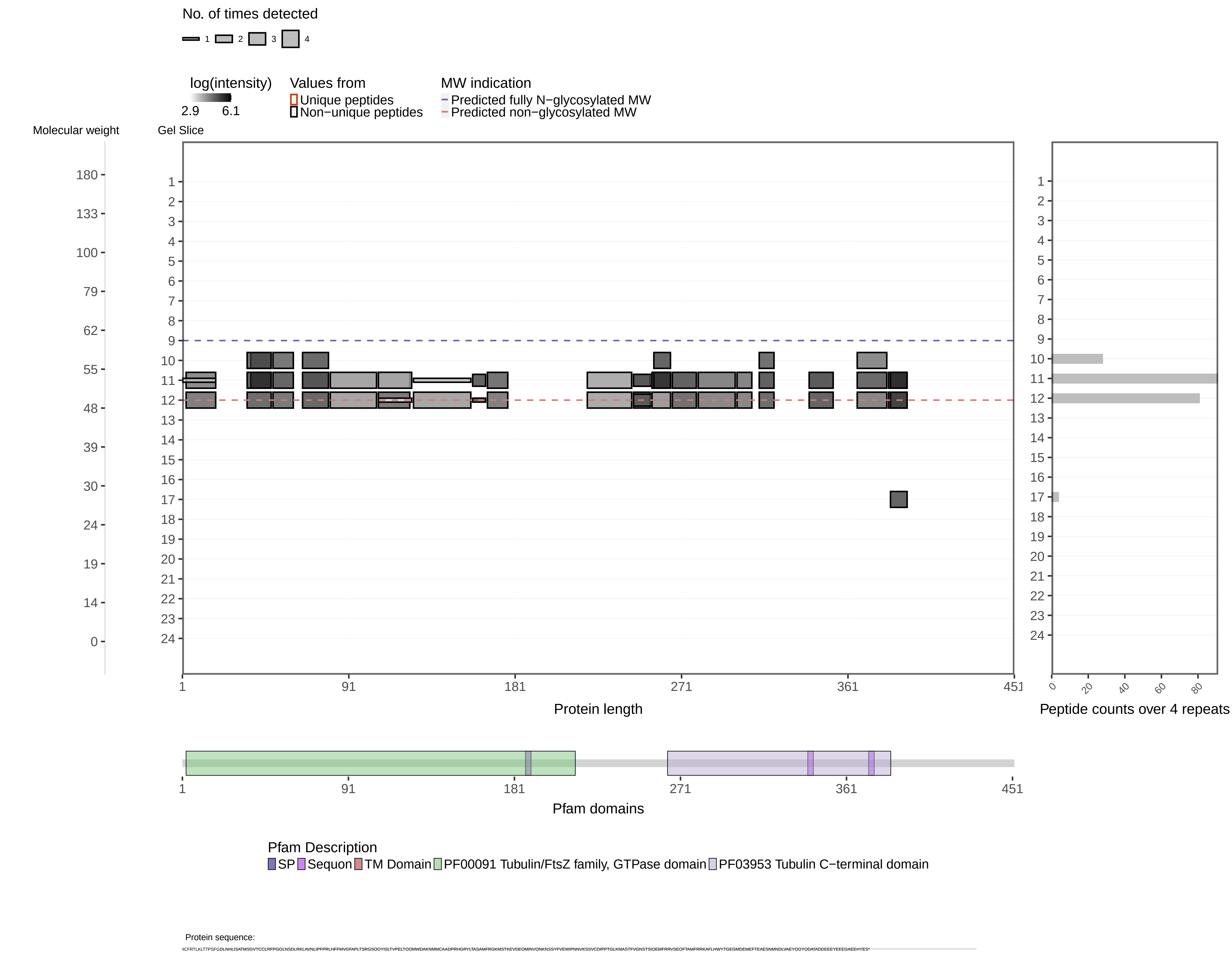

### NbL04g13830.1 thaumatin−like_protein_1b [MW = 39.48 kDa]

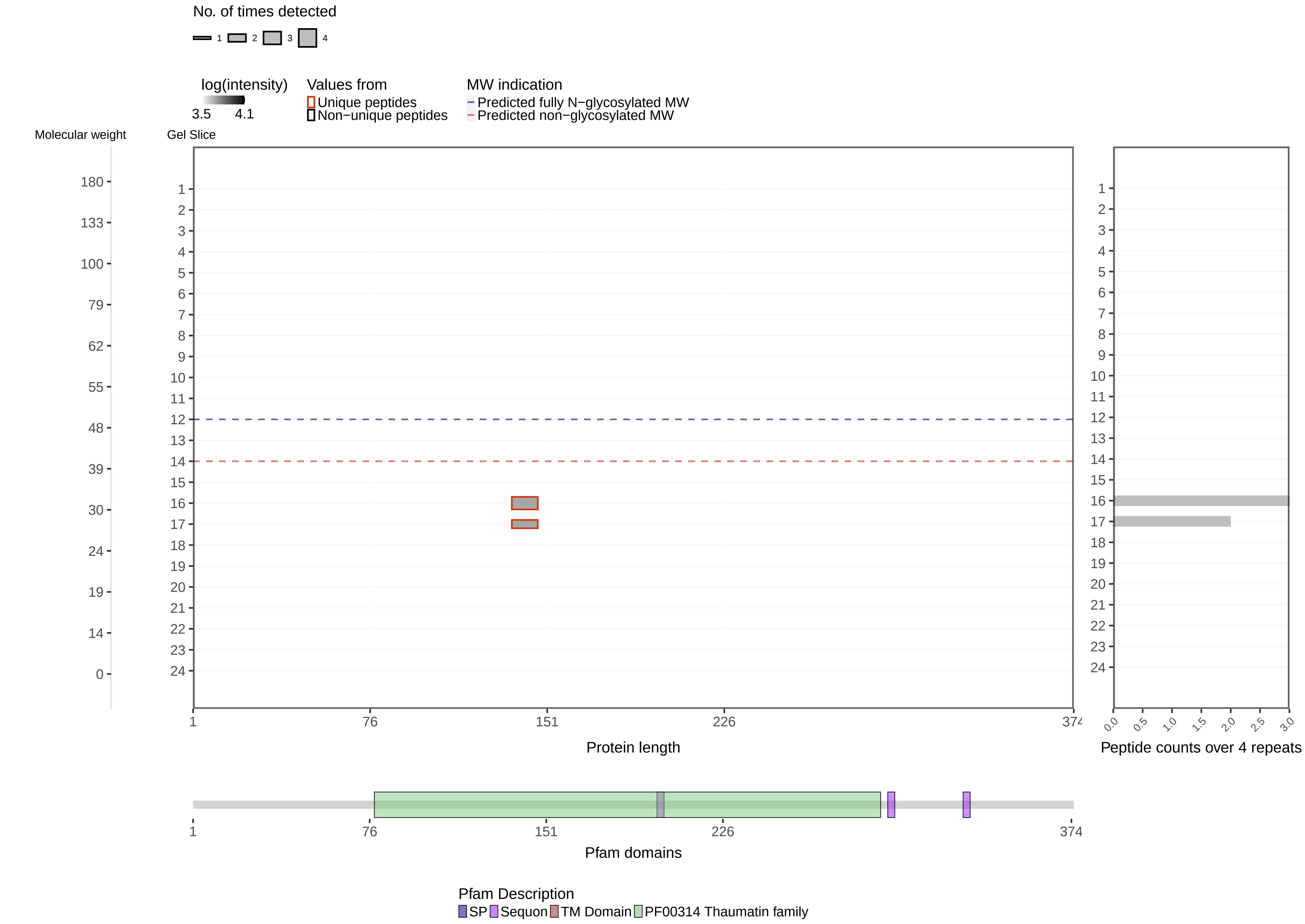


Protein sequence:

DVSLVDGYNLPMLVVPOGGSGNNCTSTGCVVDLNGACPSELKVTSVGGVNVACKSACEAFRODEYCCAGAYNSPSICKPSSYSMLFKKACPSAYSFAYDDKTSTFTCTGANYLITFCPSPNTSOKSSSTGGONRNGEPEDNSOSNNNVPSTINGSMVYDGAWDVSSASSTTCAHVLSSPAMAGSIAFLTAIWOLRHLF*

NbL04g13750.1 prostaglandin_E_synthase_2−like [MW = 35.55 kDa]

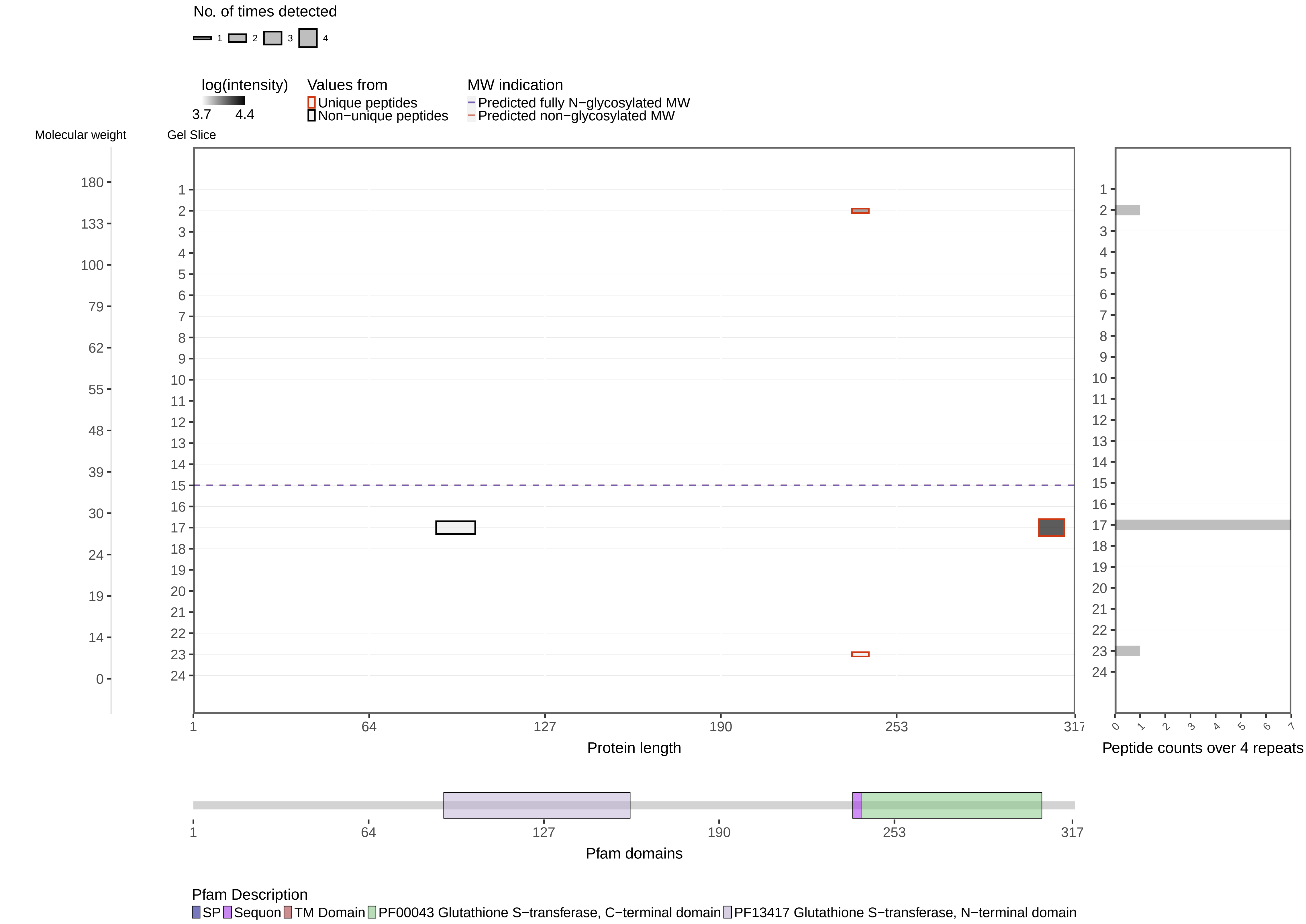


Protein sequence:

SSDIIDKLFEKVRSVDSTSNADEESKWRKWVDNHLVHVLAPNIYRTTSEALESFEYITSHGNFGFTERFTVKYGGAASMYFVSKRLKKKYNITDERAALYEAAETWVDALKGRDFLGGSKPNLADLAVYGVLRPIRYLKSGRDMVENTHIGDWYSRMESAVGLSARIQA*

### NbL04g13700.1 CO(2)−response_secreted_protease−like [MW − SP = 84.43 kDa]

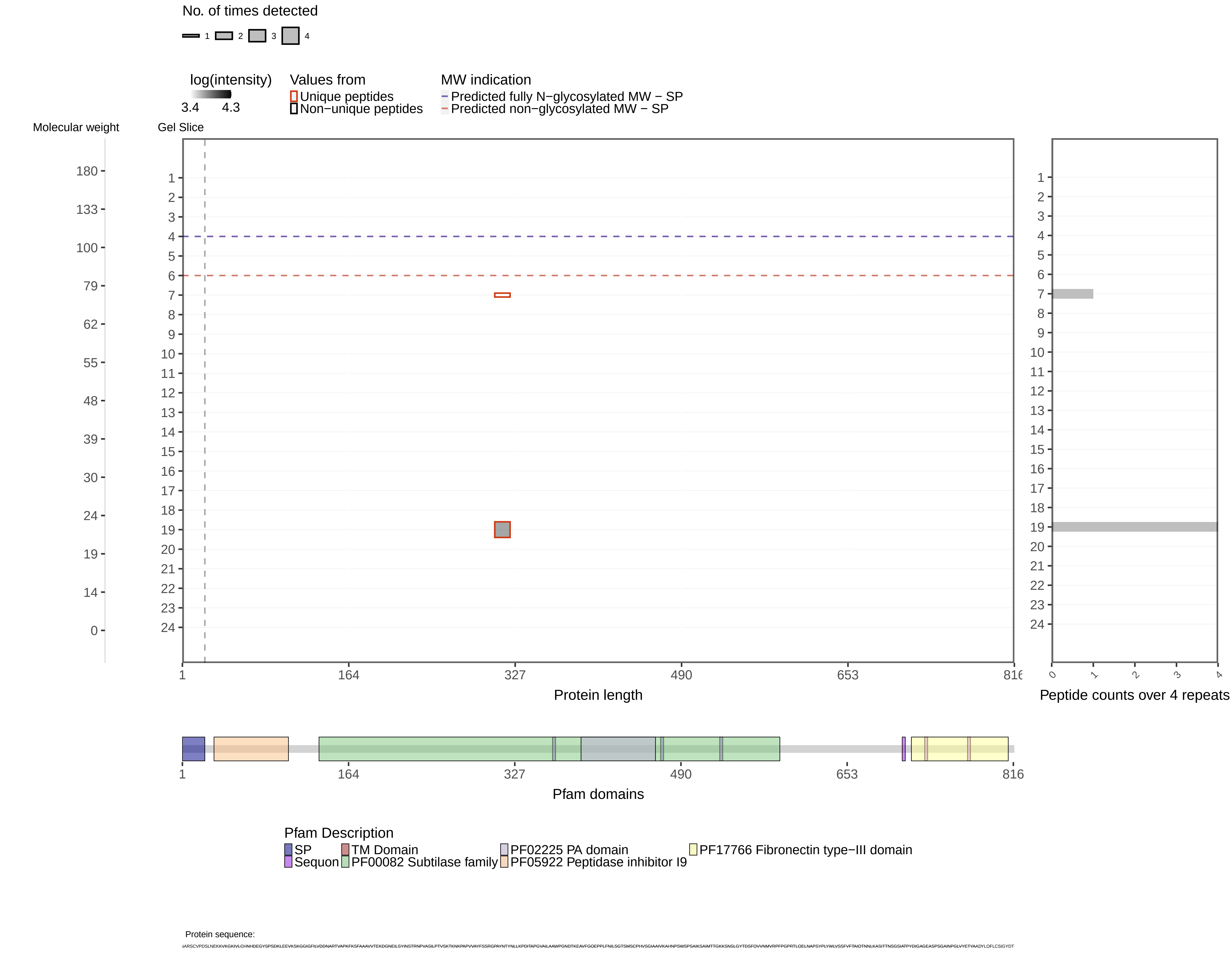

### NbL04g13640.1 expansin−A11−like [MW − SP = 25.23 kDa]

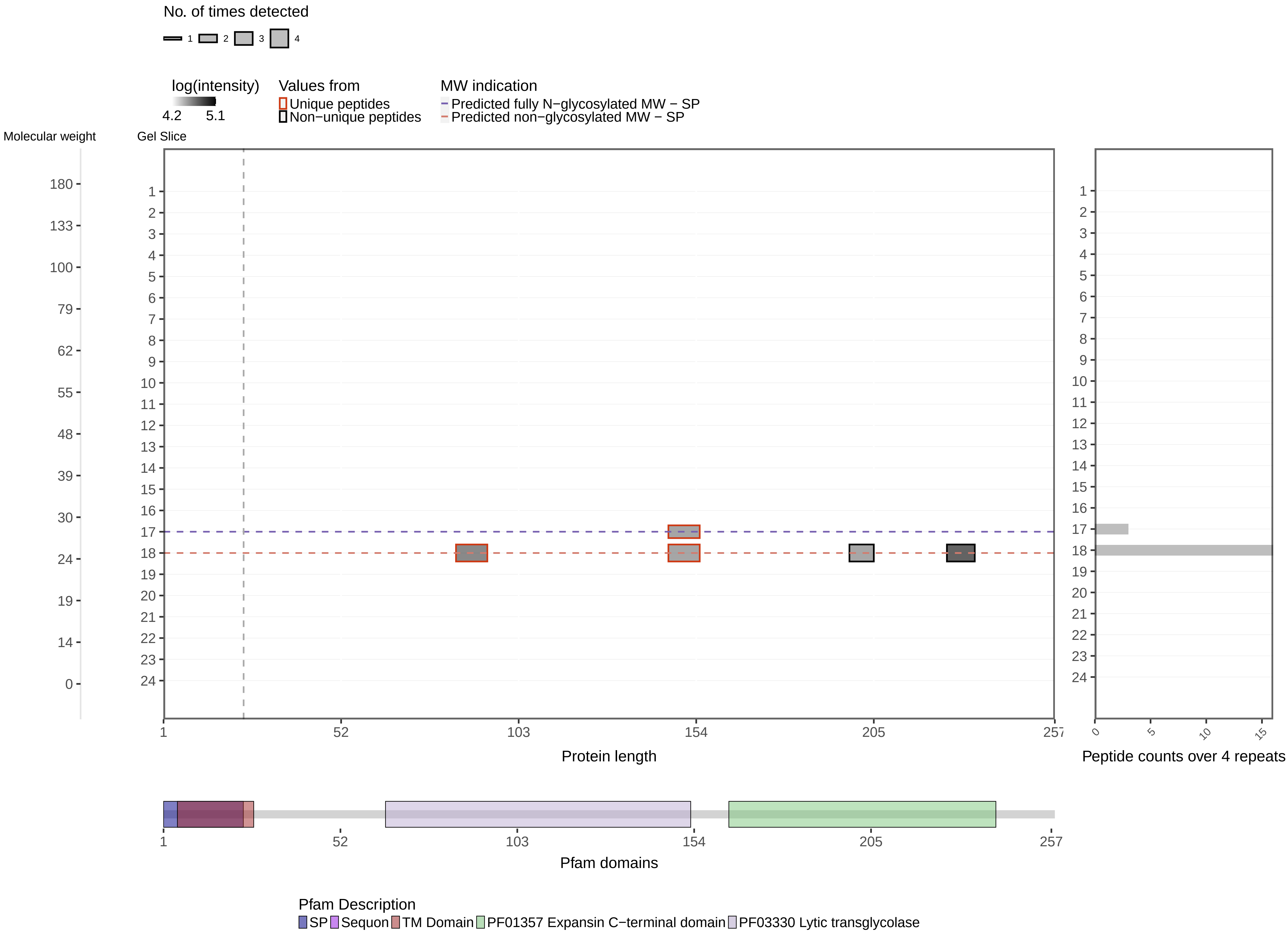


Protein sequence:

PSNNGGWCNPPRPHFDMAQPAWENIGIYRGGIVPVQYQRIPCEKKGGVRFTINGRNYFELLMISNVGGAGSVQSVQIKGSKTNWMTMSRNWGANWQSNAFLNGQPLSFKVTTTDGVTKTFLNAVPSNWHFGQTYSTSINF*

### NbL04g13630.1 ribonuclease_P_protein_subunit_p25−like_protein [MW = 28.06 kDa]

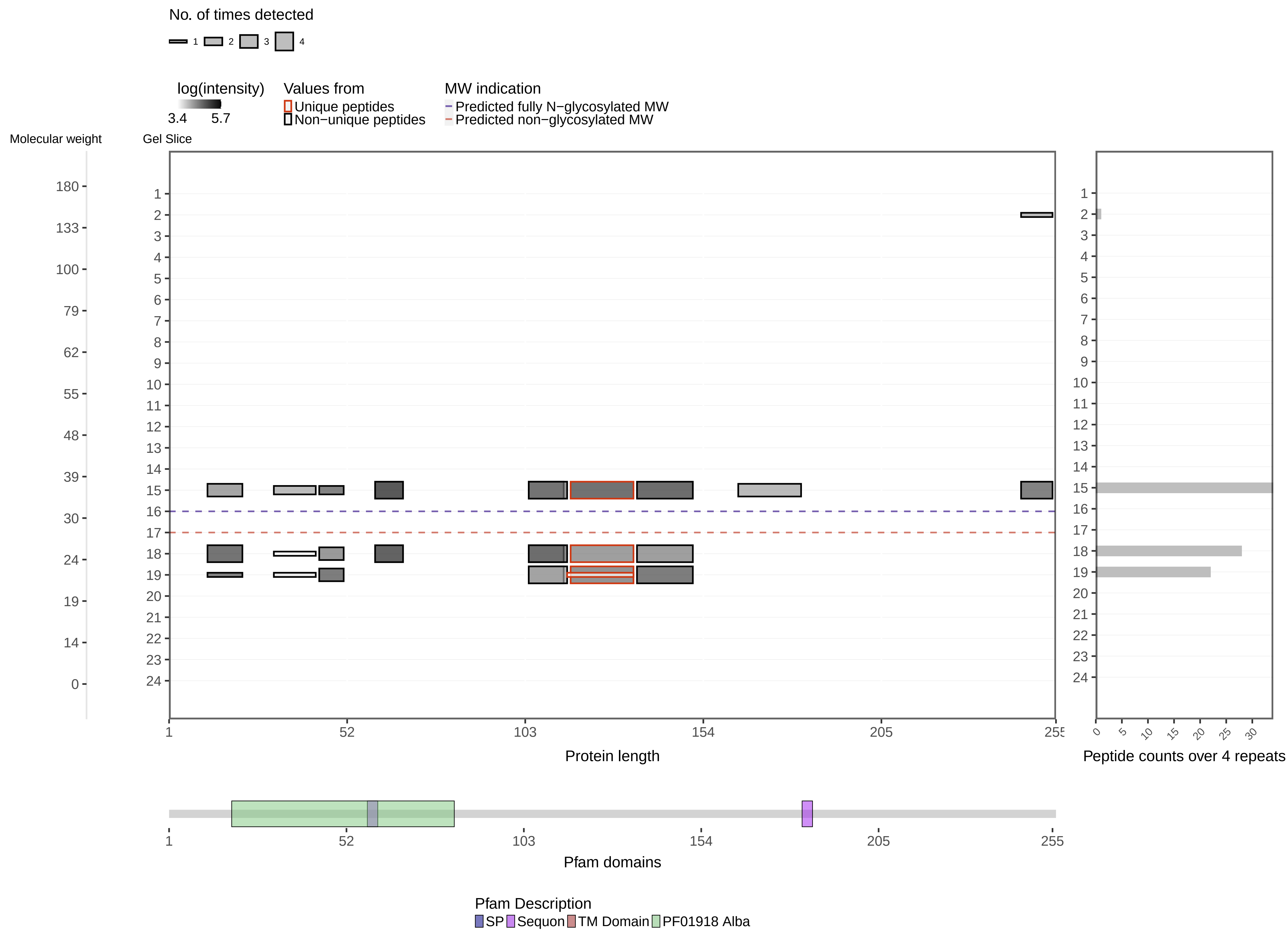


Protein sequence:

/GYQPPLPADQVKVLTDFDYDGEGSPSGGRRGRGGRGRGRSRGFAGNGFMSAEYDDGGYDRNRSYGRGRGRGRGRSFRGRGRGGYNGPQDTQQDAGFYNQEAPMQGRGRGRGRGTRGRGRGFRTTGPIHGGGA*

### NbL04g13360.1 enoyl−CoA_hydratase_2_peroxisomal−like [MW = 34.07 kDa]

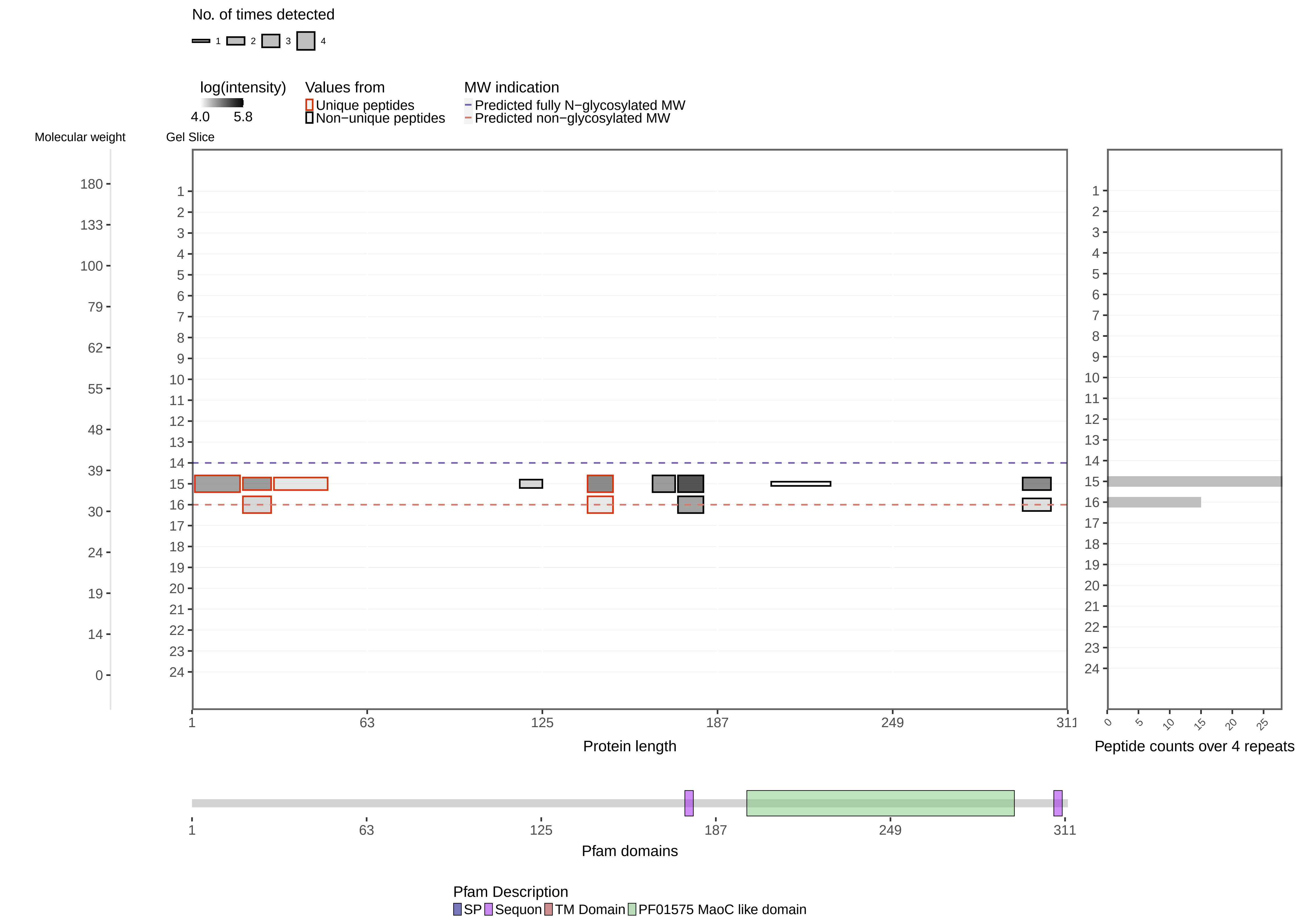


Protein sequence:

GEPLCMNRMSIYLRGSGGFSKSSRPYSYSRNPSNQTAFPKIPKSQPFAVFEGCTHASQALLYRLSGDYNPLHSDPLVASIAGFSRPILHGLCTLGFAIRAIIGRICRGDHTLIKSISGRFLLHVYPGETLITEMWPEASRVIYQVKVKERNRAVLLGFVDLDRINSSL*

NbL04g13340.1 L−ascorbate_oxidase_homolog [MW − SP = 57.82 kDa]

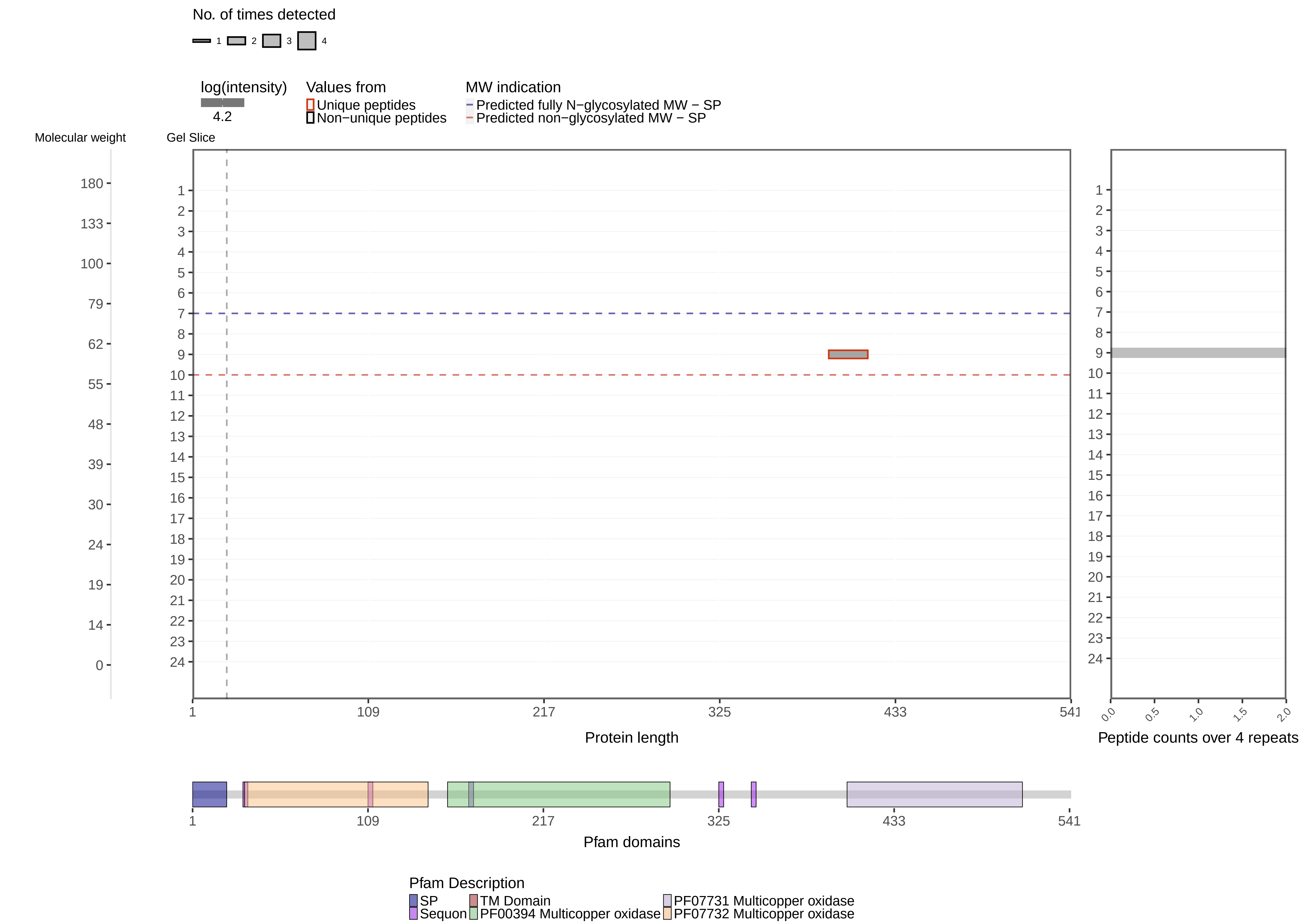


Protein sequence:

VLITADQEAKDHYIVVSSRFTSQVLTTTGVLHYSNSNTPVSGPPPGGPTIQIDWSLNQARSIRTNLSASGPRPNPQGSYHYGMINTTRTIRLASSAGQVNGKQRYAVNSVSFVPLDTPLKLLDYFKVGGFRVGSISDAPSGGGIFLDTSVLGVDYRQFIEIVFENNEDIVQSWQLNGYSFWVVGMDGGQWTQASRNGYNLRDAVARCTTQVYPKSWTAIYIALDNVGMWNLRTEFWARQYLGQQLYMRVYTTSTSLRDEYPIPRNARLCGKVAGRHTRPL*

### NbL04g13310.1 alcohol_dehydrogenase−like_7 [MW = 42.51 kDa]

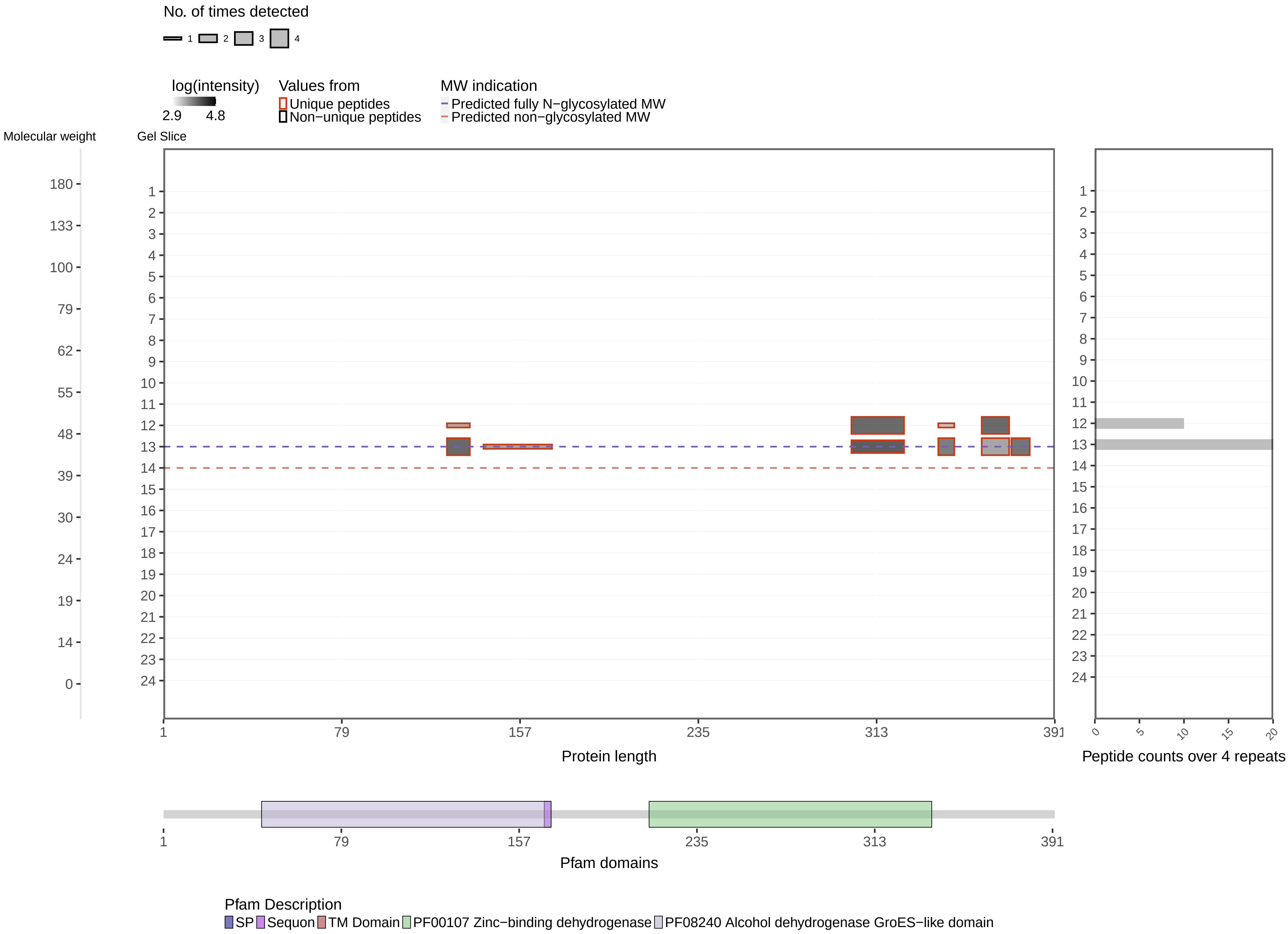


Protein sequence:

FSCGVSTGVGAAWKTANVEPGSTVVIFGLGAIGLAVAEGARLCDATRIIGVDINSDKFEIGKOFGVTDFINSNSCGDKPISOVIIEMTNGGADYCFECVGLGTLVOEAFACCRKGWGKTVVLGVDKPGAOLNLNSFEVLTSKKTLTGALFGGLKPKSDVPILVKRYLDKELOLDKFVTHEVNFEDINKAFDLLIOGKSLRCVIWMDK*

### NbL04g13290.1 phosphoprotein_ECPP44−like [MW = 23.61 kDa]

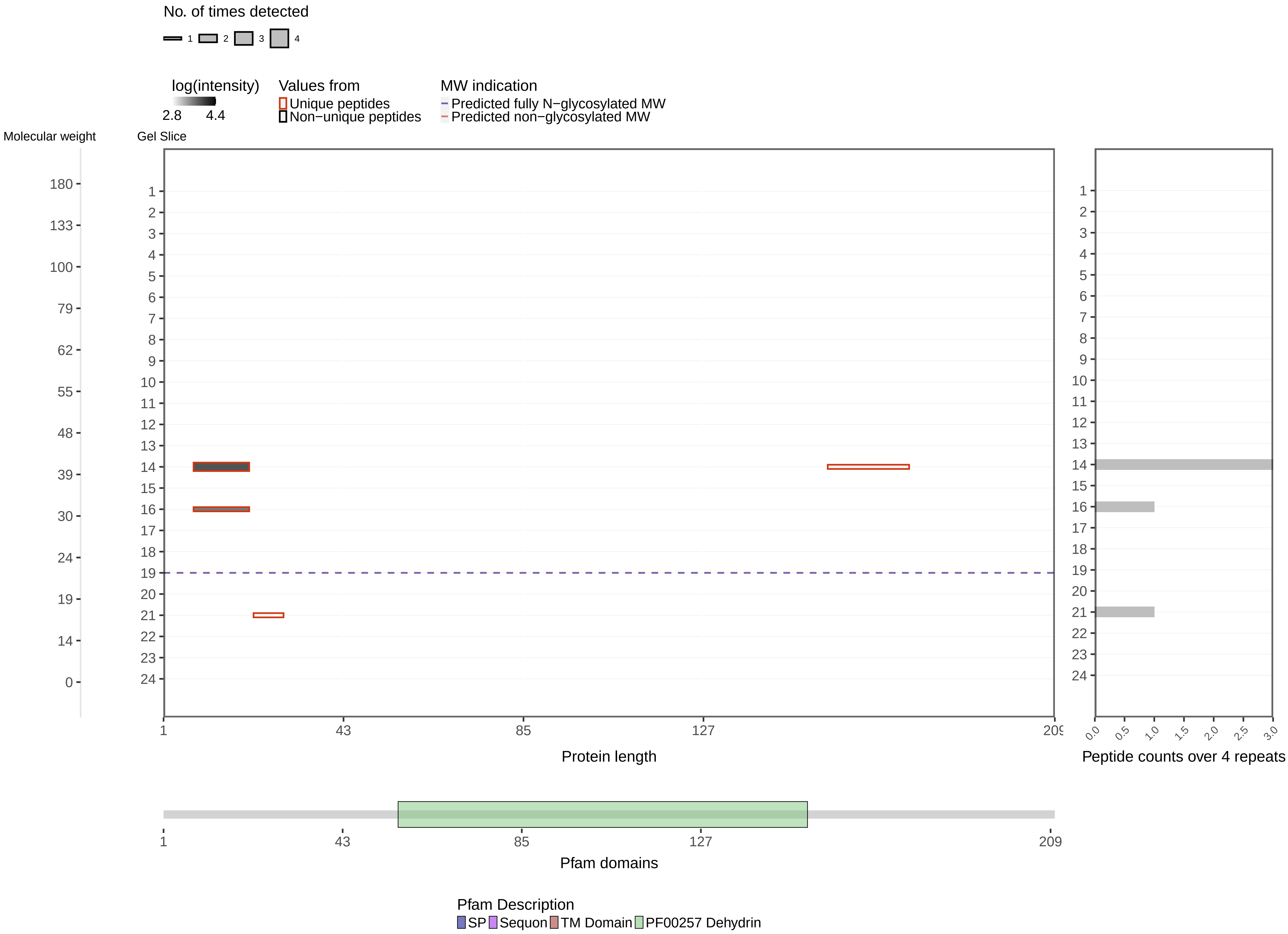


Protein sequence:

EDGQKIKKKKKKGLKDKIKEKISGEHKEEEKAGDTTVPVEKYEETEEKKGFLDKLKEKLPGGGQKKTEEVAPPPPAATEHEAEGKEKKGFLDKIKEKLPGYHSKTEEEKEKEKETASH*

### NbL04g13150.1 14−alpha−glucan−branching_enzyme−like [MW = 97.47 kDa]

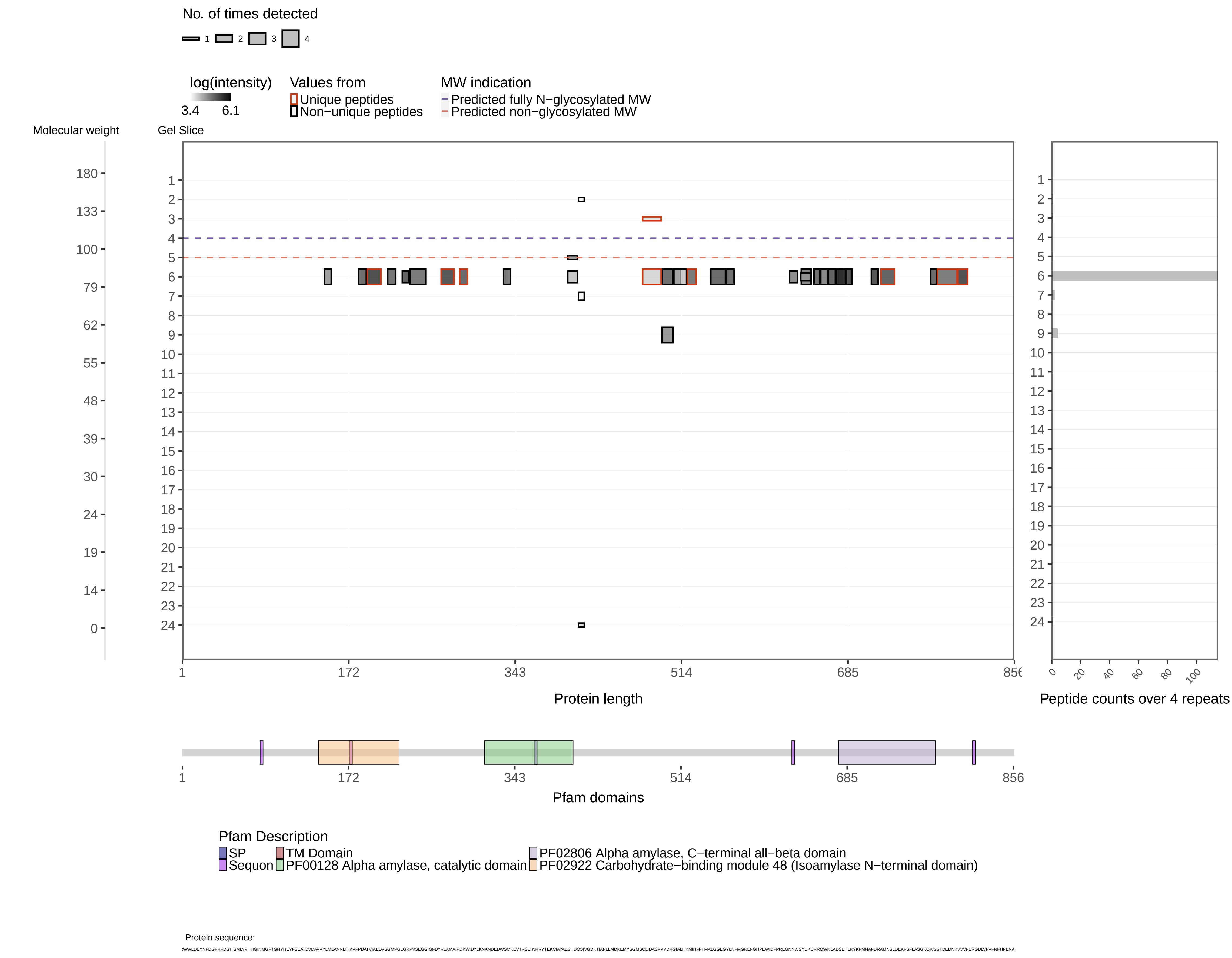

### NbL04g13100.1 isozyme_3_(LOC107771182)_mRNA_HF564633.1_Nicotiana_tabacum_mRNA_for_catalase_3 [MW = 57.02 kDa]

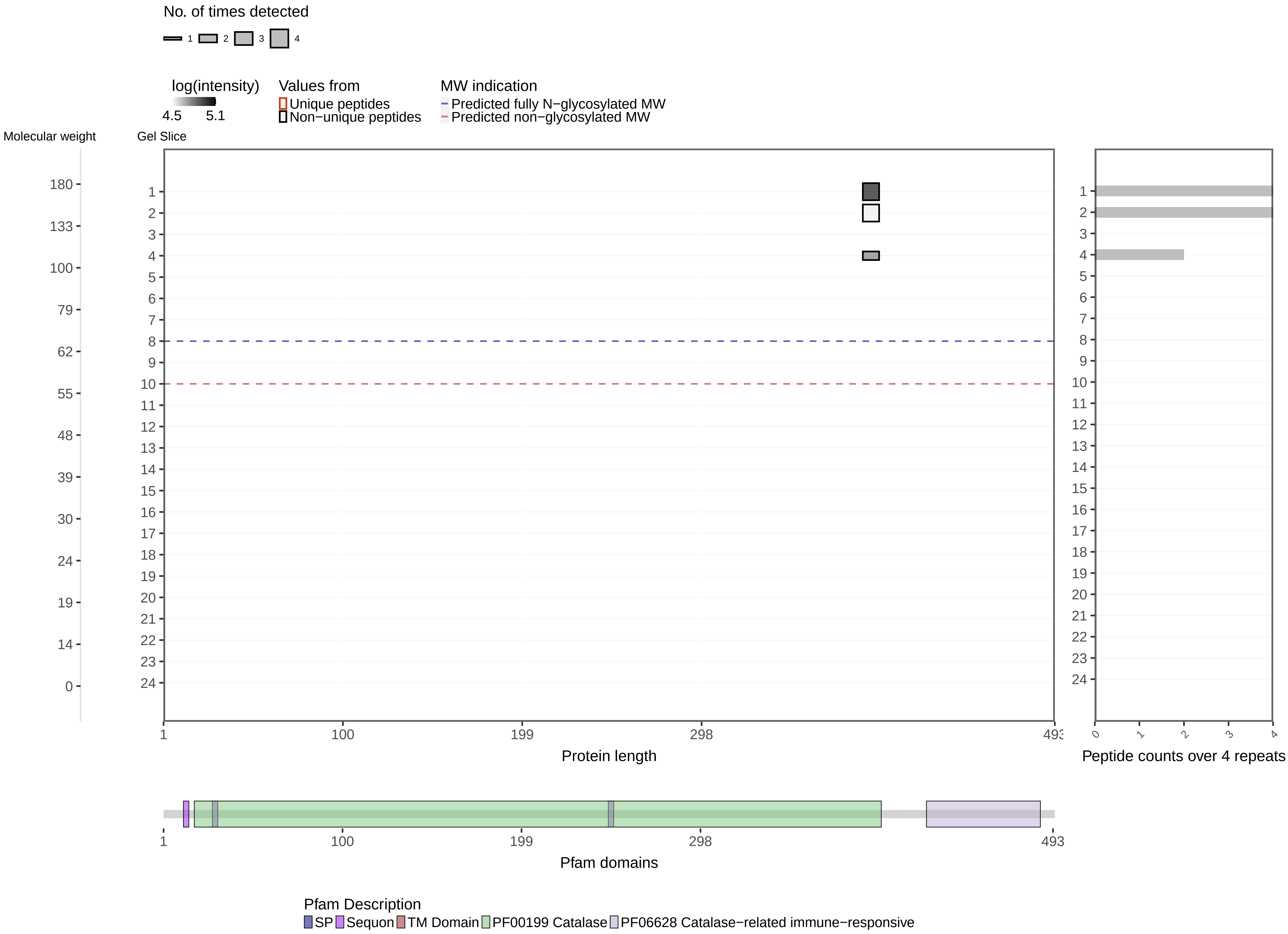


Protein sequence:

VKCLLEEEAIKVGGANHSHATKDLYDSIAAGNYPEWKLFIQIIDPDHEDRFDFDPLDVTKTWPEDILPLQPVGRLVLNKNIDNFFAENEQLAFCPAIVVPGVYYSDDKLLQTRIFSYSDTQRHRLGPNYLQLPANAPKCAHHNNHHEGFMNFMHRDEEVNYFPSRFDPCRHAEQYPIPPRVLTGKRDKCIIEKENNFKQPGERYRSWTLDRQERFICRWVDALSDPRVTHEIRSIWISYWSQADKTLGQKIASRLNVRPTM*

### NbL04g13010.1 epidermal_growth_factor_receptor_substrate_15−like [MW = 107.69 kDa]

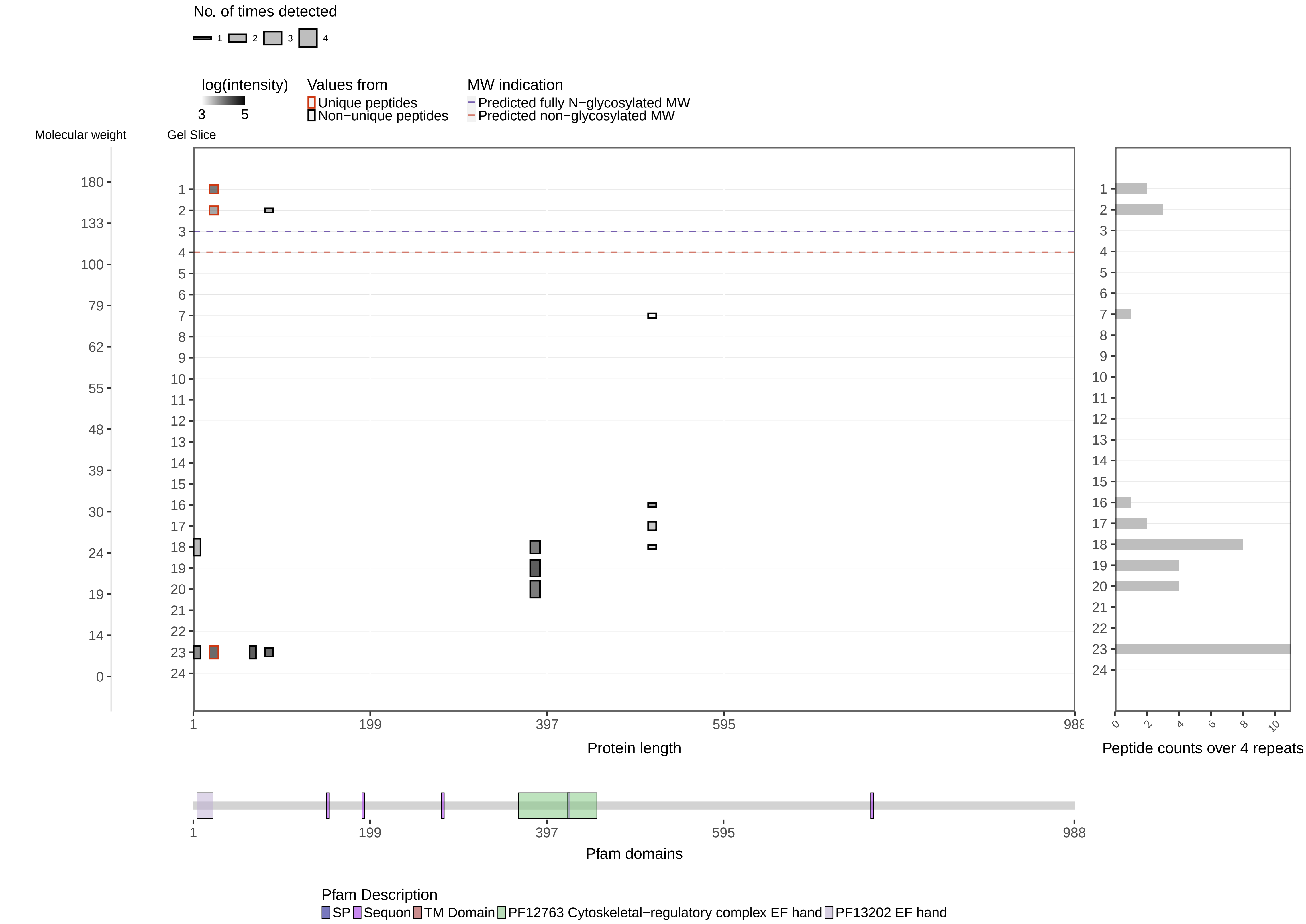


Protein sequence:

QGLPGVQPGAHPGLRPPTQGVFPQPDRSMQFNQQNARATSMDNSHMDQLSNGEQNKLEPKREDAVAGEEKDESKDNVLLDSREKLEFYRTKMQDLVLYKSRCDNRLNEITERALADKREAEMLAKKYEEKYKLVAEIASKLTIEEASFRDTQERKLELQQAIIKMEQGGSADGILQVRADRVQHDLEELLKALADRCKKHELNIKSTALIELPPGWQPGIPVISAVWDEDWDKFEDEGFSFDVAAPANSKSTSNQKEISPIHGDSPDSMSNADTKSENYSAKGNNNNFET

NbL04g12970.1 protein_canopy_homolog_1−like [MW − SP = 18.82 kDa]

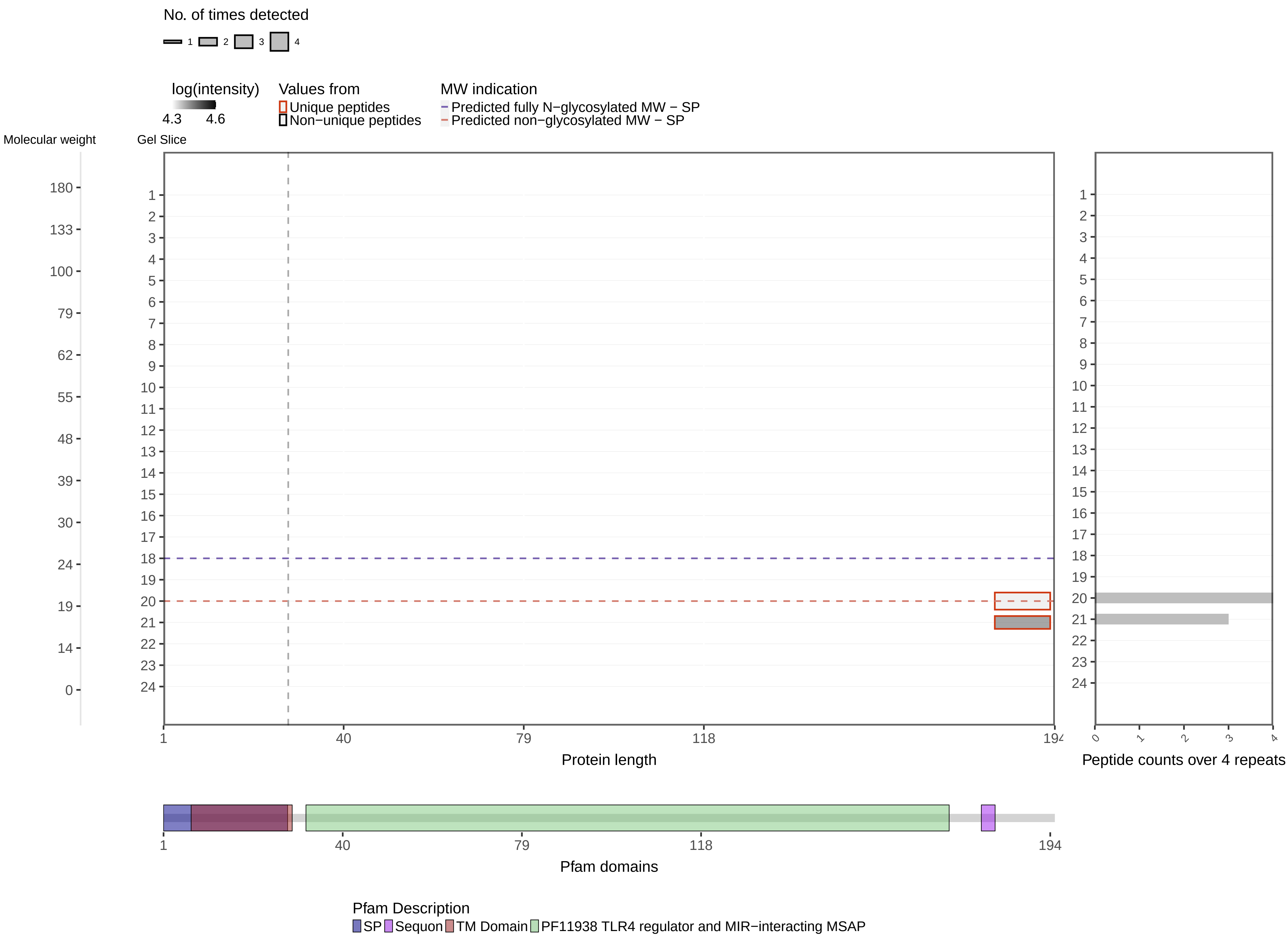


Protein sequence:

EDLCEKMQDYTLEKVDSSTKTWIKVNNWDLLKTNKQEARAHSKAISSFCGRLLEQTEDDFTELMKKGSVQVGDVSKLLCEDLSNYCNGKSSSNKTVGDENEDTDREL*

### NbL04g12940.1 glyceraldehyde−3−phosphate_dehydrogenase_B_chloroplastic−like [MW = 48 kDa]

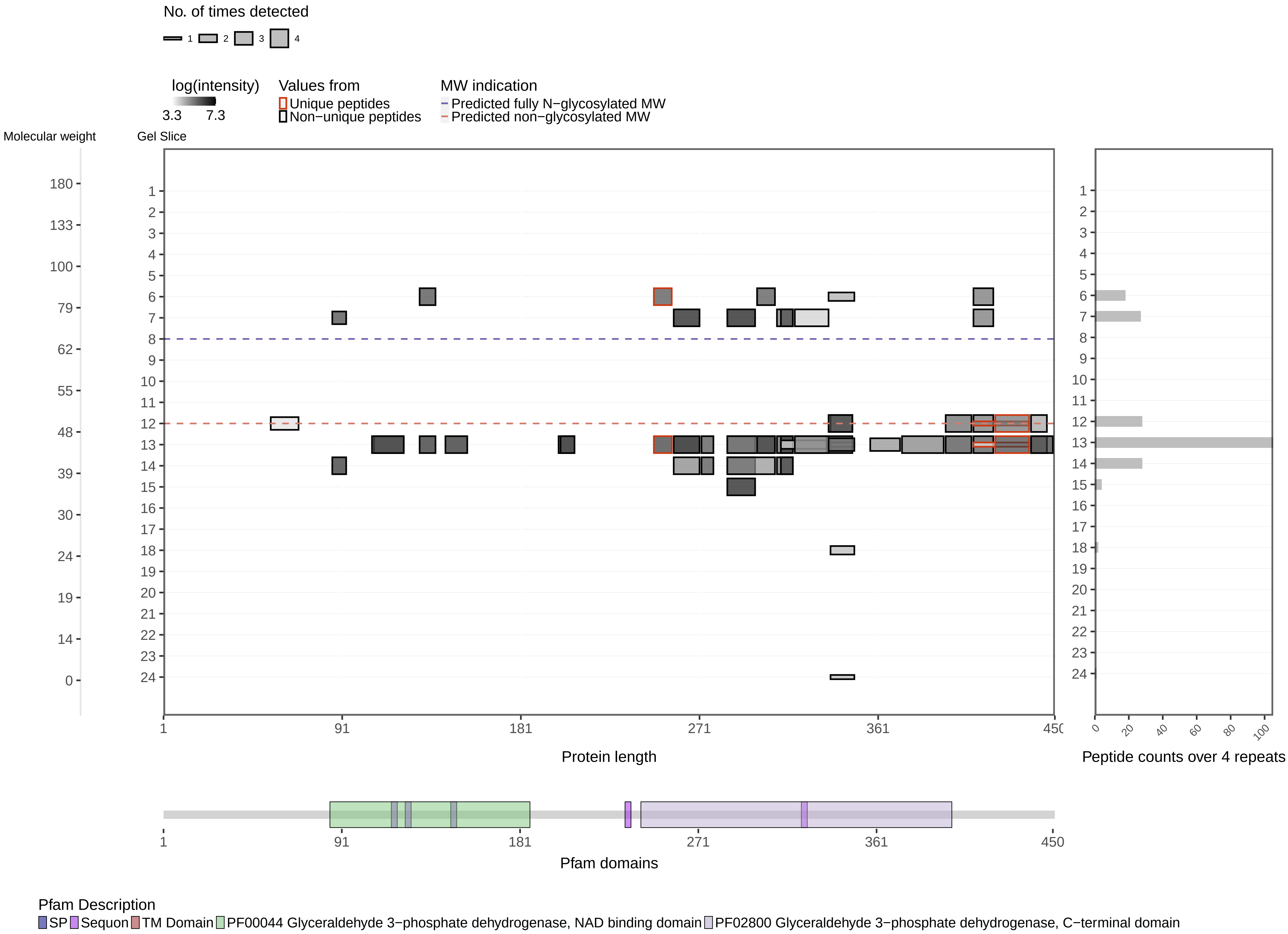


Protein sequence:

TYVVGVNEQDYSHEVADIISNASCTTNCLAPFVKVMDEELSIVKGTMTTTHSYTGDQRLLDASHRDLRRARAAALNIVPTSTGAAKAVSLVLPQLKGKLNGIALRVPTPNVSVVDLVVNVAKKGITAEDVNAAFRKAADGPLKGVLAVCDEPLVSVDFRCSDVSSTIDSSLTMVMGDDMVKVVAWYDNEWGYSQRVVDLAHLVANKWPGVAVAGSGDPLDEFCKTNPADEECKVYE*

NbL04g12870.1 thiamine_pyrophosphokinase_2−like [MW = 29.53 kDa]

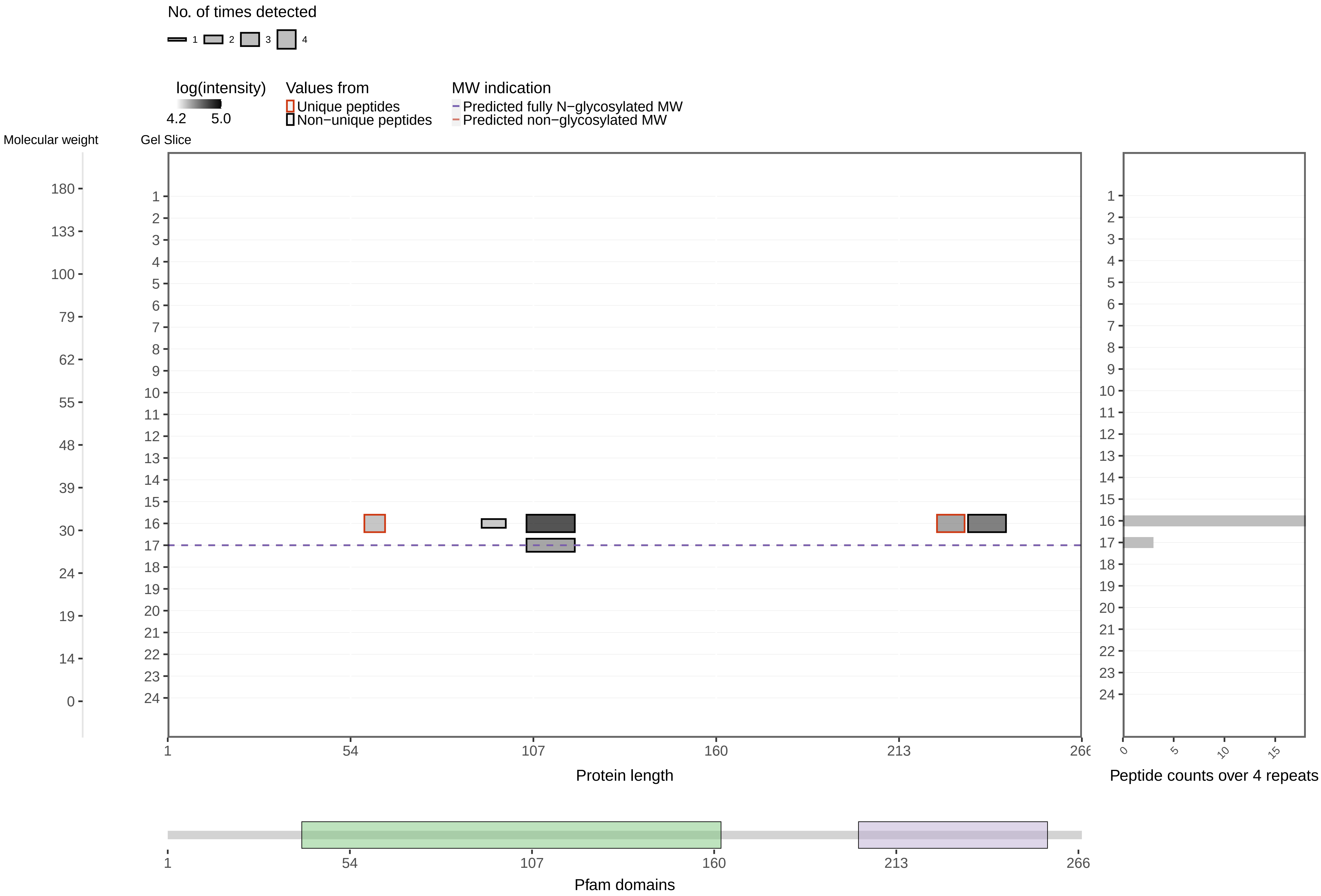


Pfam Description

SP Sequon TM Domain PF04263 Thiamin pyrophosphokinase, catalytic domain PF04265 Thiamin pyrophosphokinase, vitamin B1 binding domain

Protein sequence:

HKCVAYIRELLNLEHPNLCILVTGALGGRFDHEMGNINVICHFPSMPIILLSDDCLIQLLPSTHHHKIHIQSSVEGPHCGLIPIGTACGLKPIGTAAGRTTTTGLKWNLDETEMRFGGLVSTSNIVKENIVTVQSESDLLWTISIKKE*

NbL04g12720.1 cinnamoyl−CoA_reductase_2−like [MW = 43.43 kDa]

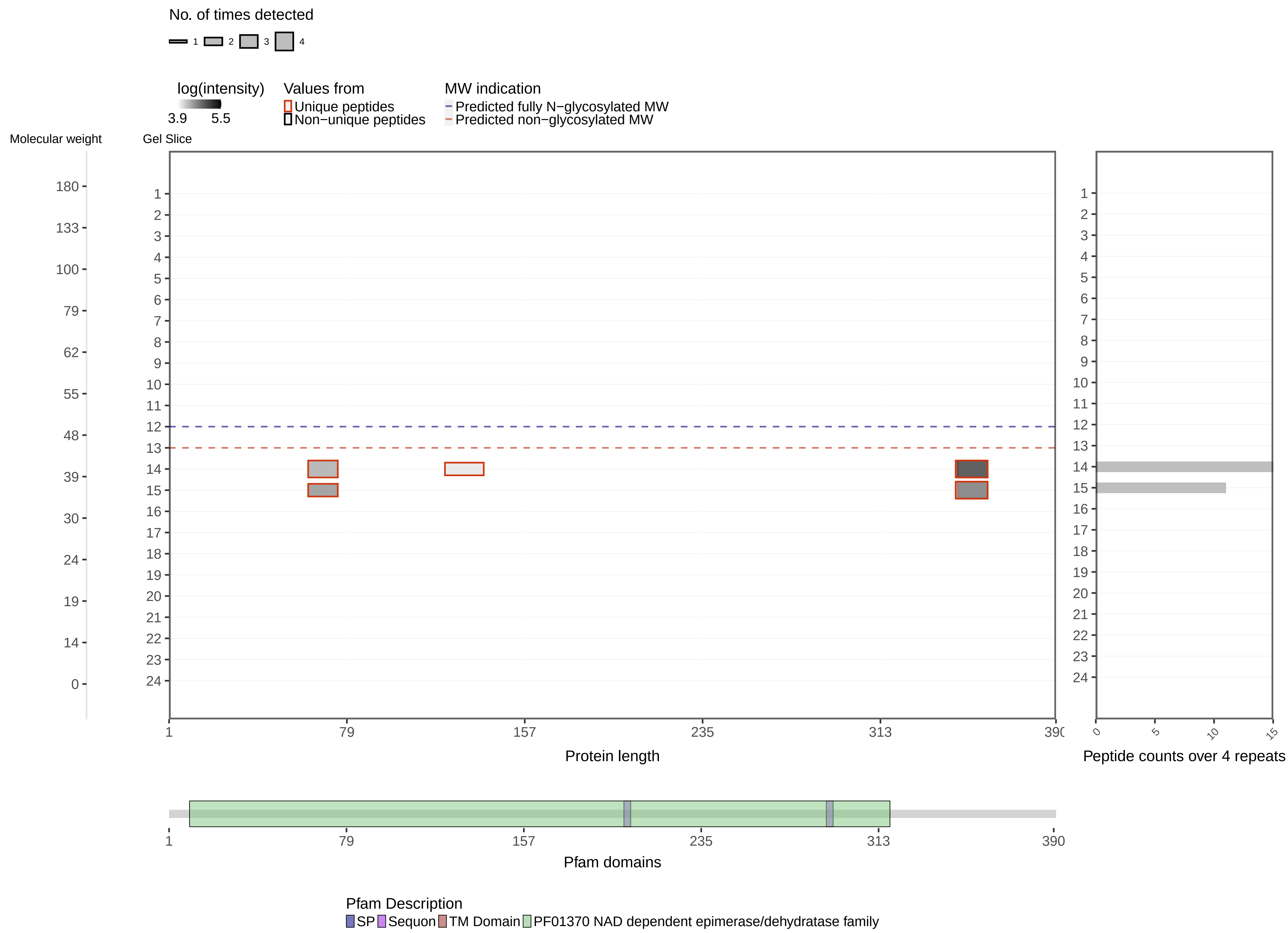


Protein sequence:

GLDLVTVLPTIVWGPMLQKTTNASSLILIKLLKSCPTFGSPKYFLLNYEWLAELLCAVAHLPQHGIYETLSRPVGHDIVLFTSIAWSMFSSFCSNGLSIALAMKRYEELENKTRFIIDVRDLAEALVLVYEKPEAEGRHICMAHKAKSQDLVDMLKRHYPNYNYPKRFTDVMEEGSYSTEKLQKLGLKCRPLEETLVDAVESYKQAGILD*

### NbL04g12620.1 pyrophosphate−−fructose_6−phosphate_1−phosphotransferase_subunit_alpha [MW = 67.33 kDa]

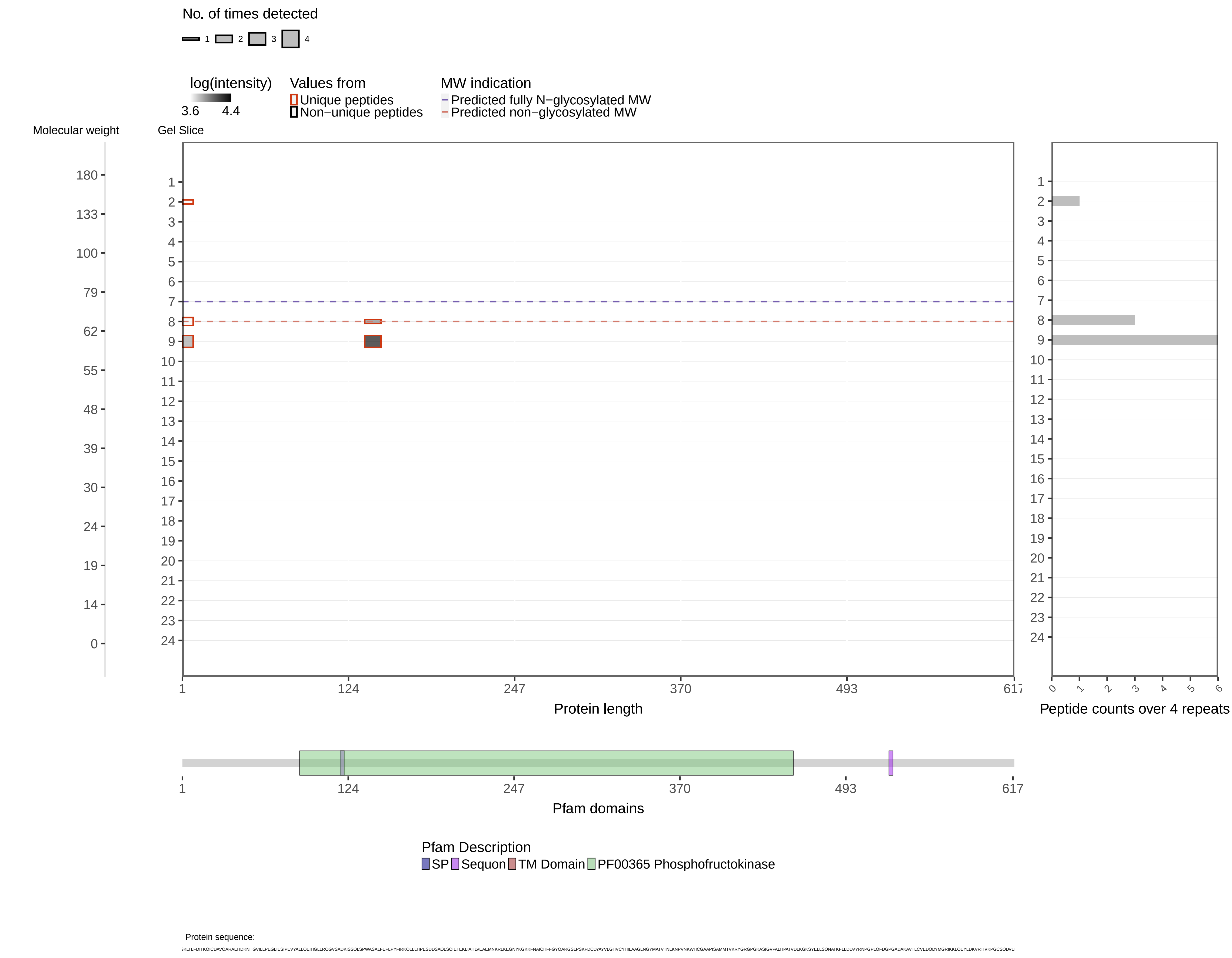

### NbL04g12580.1 calvin_cycle_protein_CP12−3_chloroplastic−like [MW = 13.91 kDa]

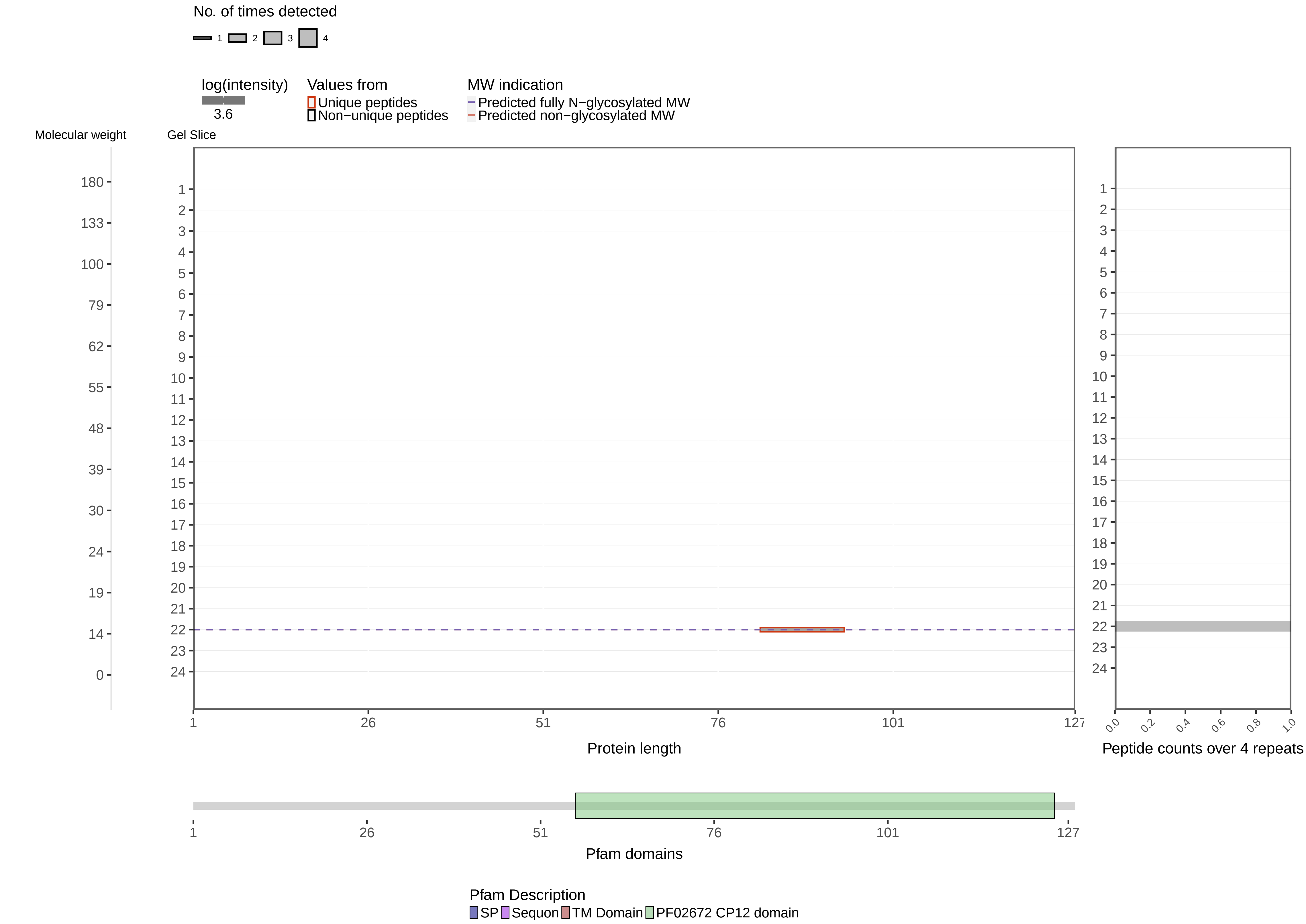


Protein sequence:

QMREKQLTEMIEKKVKEAKAVCGEDARSDECKVAWDEVEEVSQAKAHLRIKLQLNQDPLESFCQDNPETDECRIYED*

### NbL04g12210.1 50S_ribosomal_protein_6_chloroplastic [MW = 11.44 kDa]

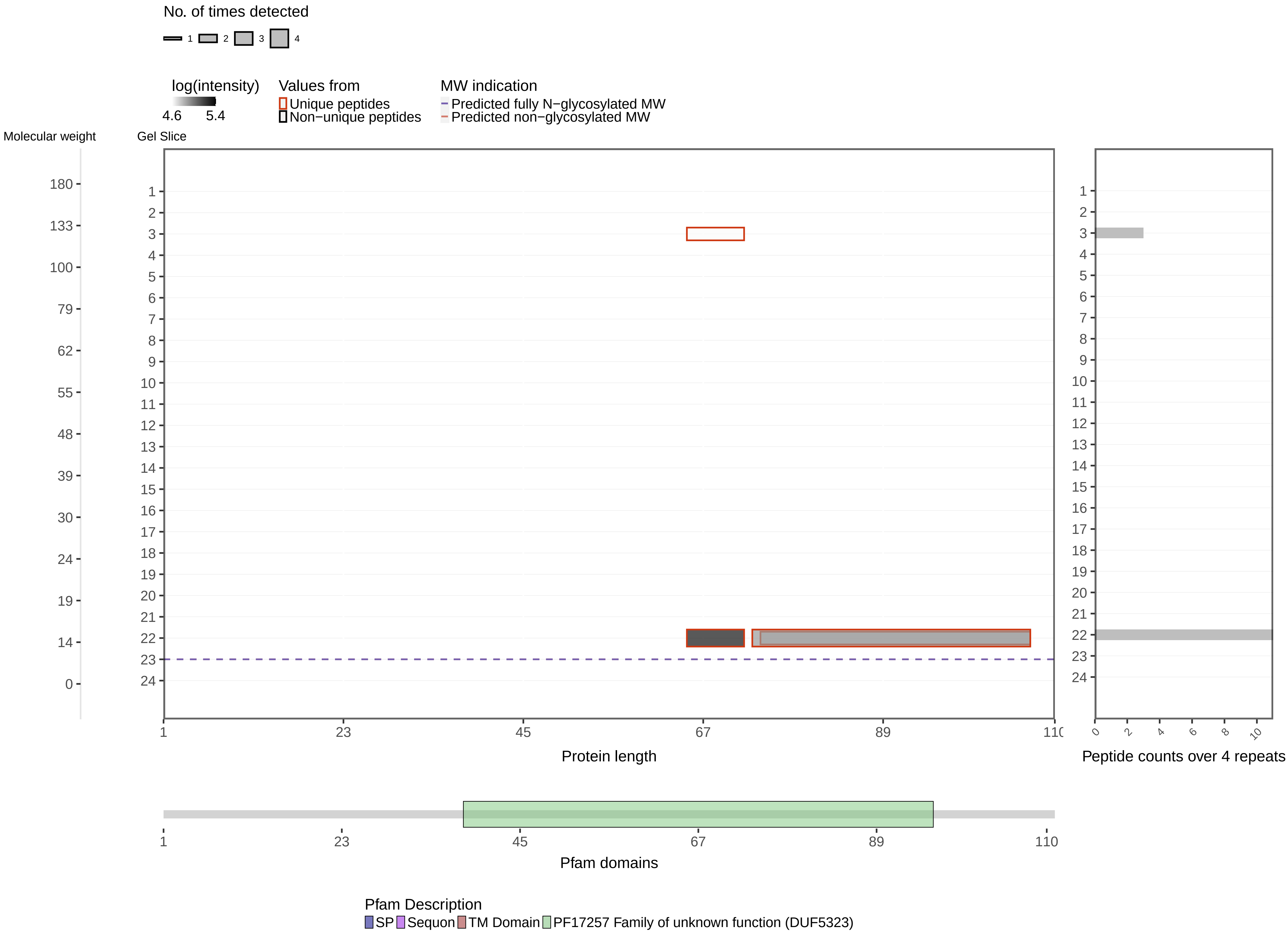


Protein sequence:

AIECSSRPQKKATAHHKKTRPRKTQAWDIRRGPAVYPPLPALPPEWSFATDESGQPASSGASEPKTE*

### NbL04g12160.1 uncharacterized_LOC104213280 [MW = 108.29 kDa]

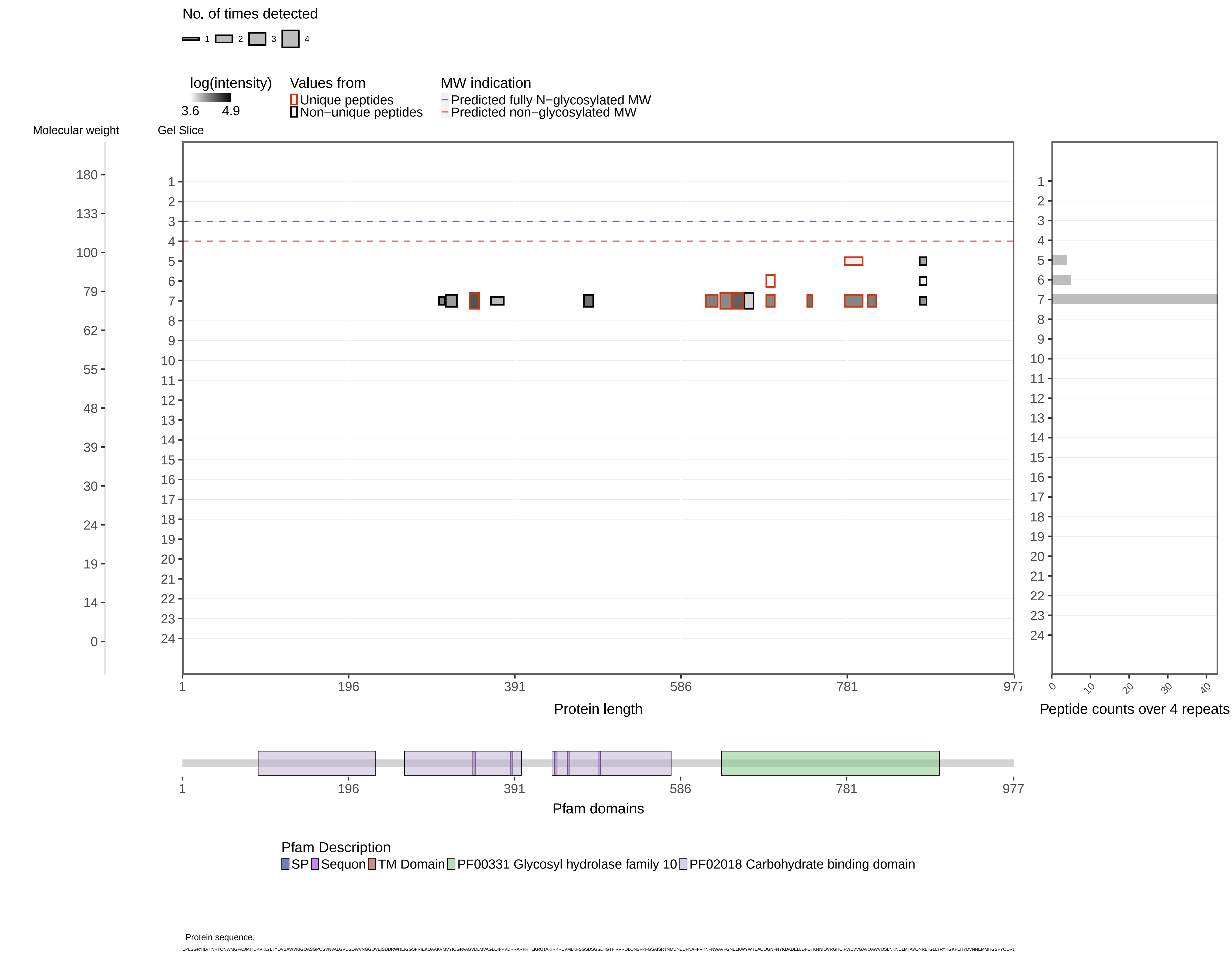

NbL04g11990.1 uncharacterized_LOC109240426 [MW = 23.78 kDa]

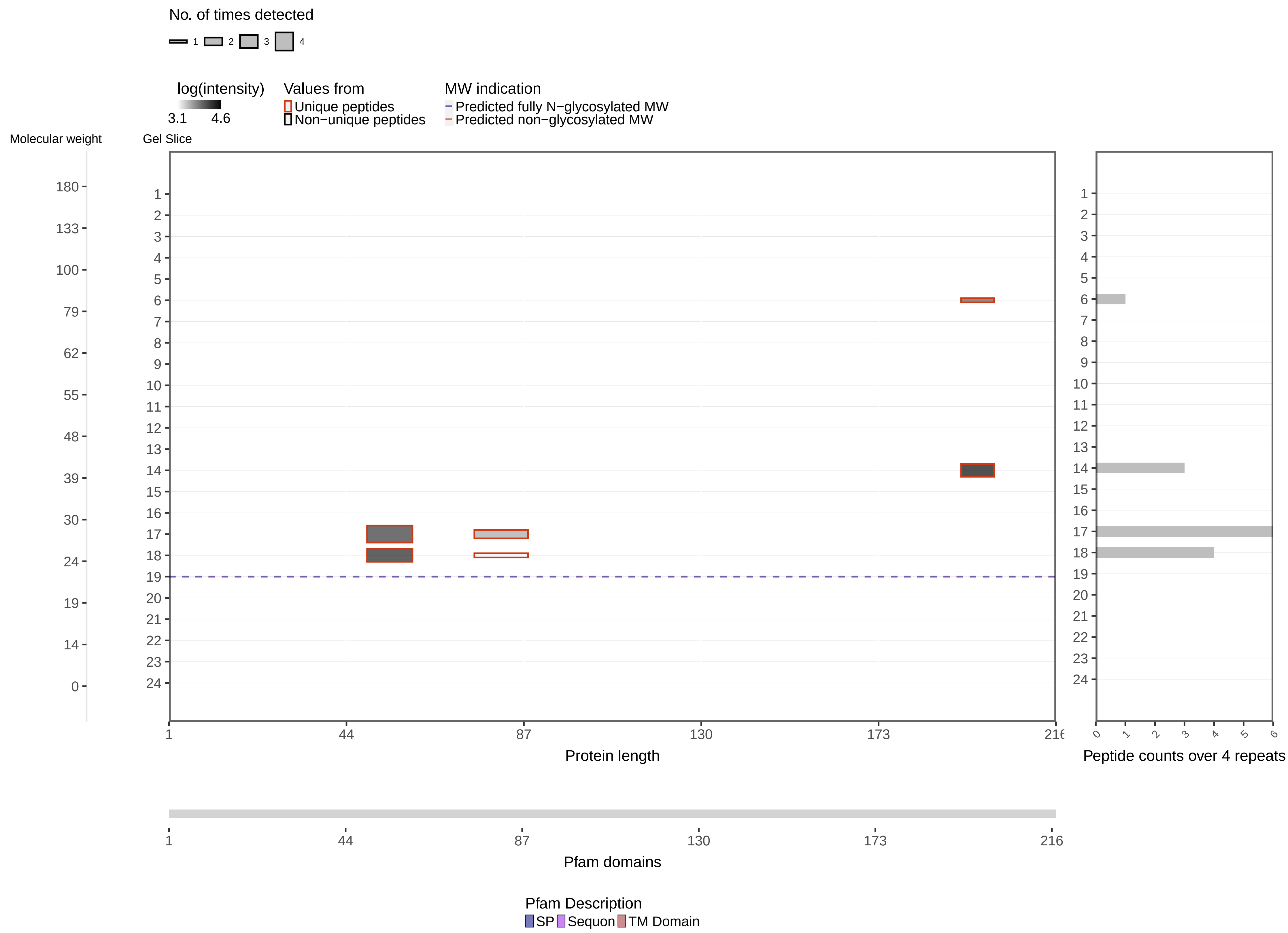


Protein sequence:

AFQSLPDISKAISELTVLKGVGPATASAVLAAFAPDIAPFMSDEAMVAANGNSKDYTLKQYLVLVDKLQAKSKELSAKGDCFTPSDVERALWSSAVGAKLDGLSQEPKEVSLKRQSKRKRVR*

### NbL04g11670.1 activating_signal_cointegrator_1_complex_subunit_2−like [MW = 93.59 kDa]

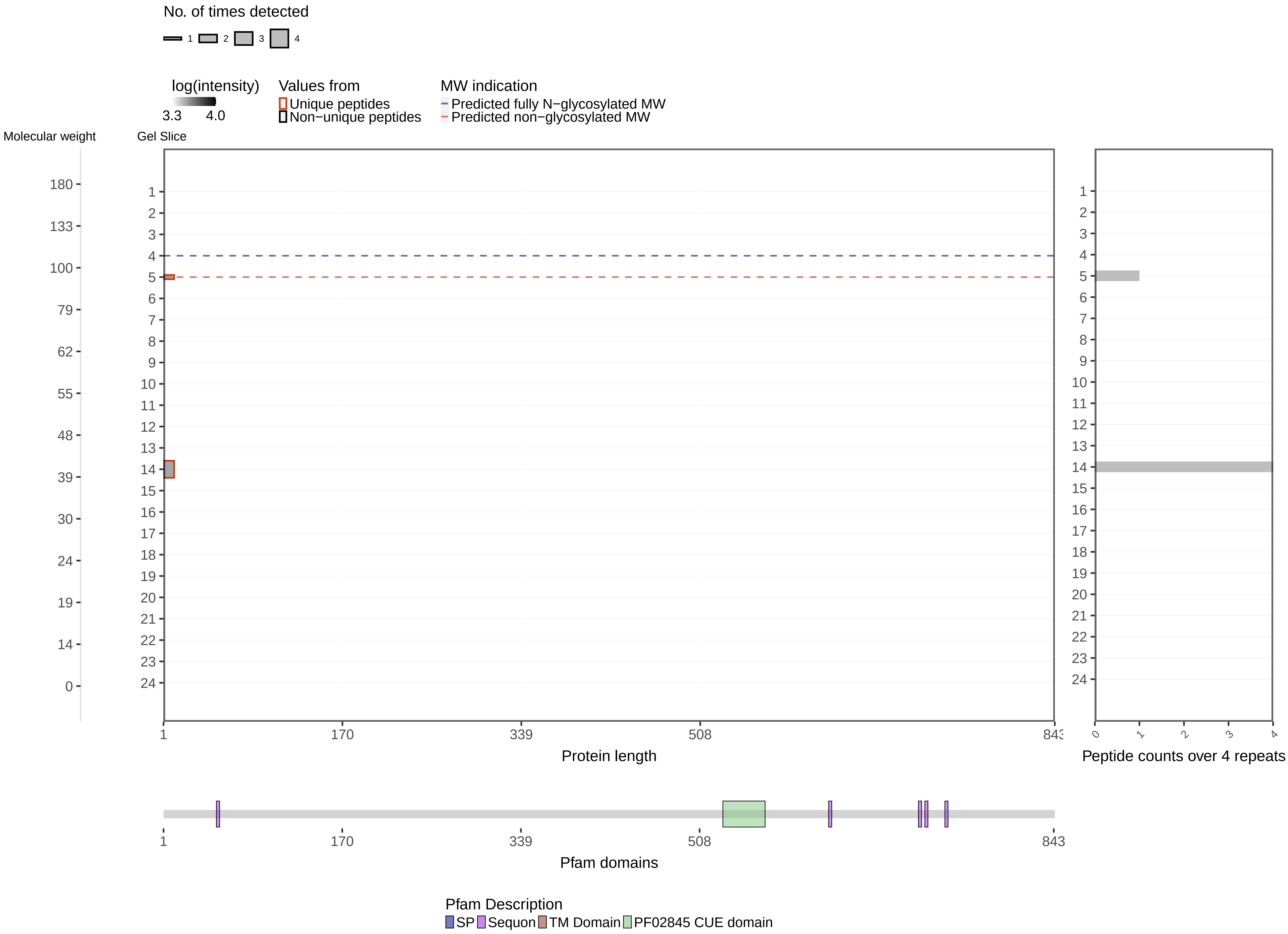


Protein sequence:

VESSPLPATMKMFPTNVEDPAIRADILVQSVRDINGDYSQALEGHSKGTFLQIIDKNHNIMSRIELLRNTGWISMDDDQFKFLSGIMVHPVEDNFGEAAHPAVSQKDNQPPVDEDNAIMESKISQIKDLFPDYGKGFLAACLEVYNLNPEEVIQRILEGTLHEDLQSLDISLEKISQPKSGVPSMTRNDKGKGKLVEPVPVPPRNIMPAATPYQAEGSSNSSTTSVGRFIRKTTTEEPASLTLDSREAKDLAKTIALSSQLEYEDEYDDSFDDLGLSIGDSAFEETESLQEKSNFGI

### NbL04g11430.1 subtilisin−like_protease_SBT1.7 [MW − SP = 78.45 kDa]

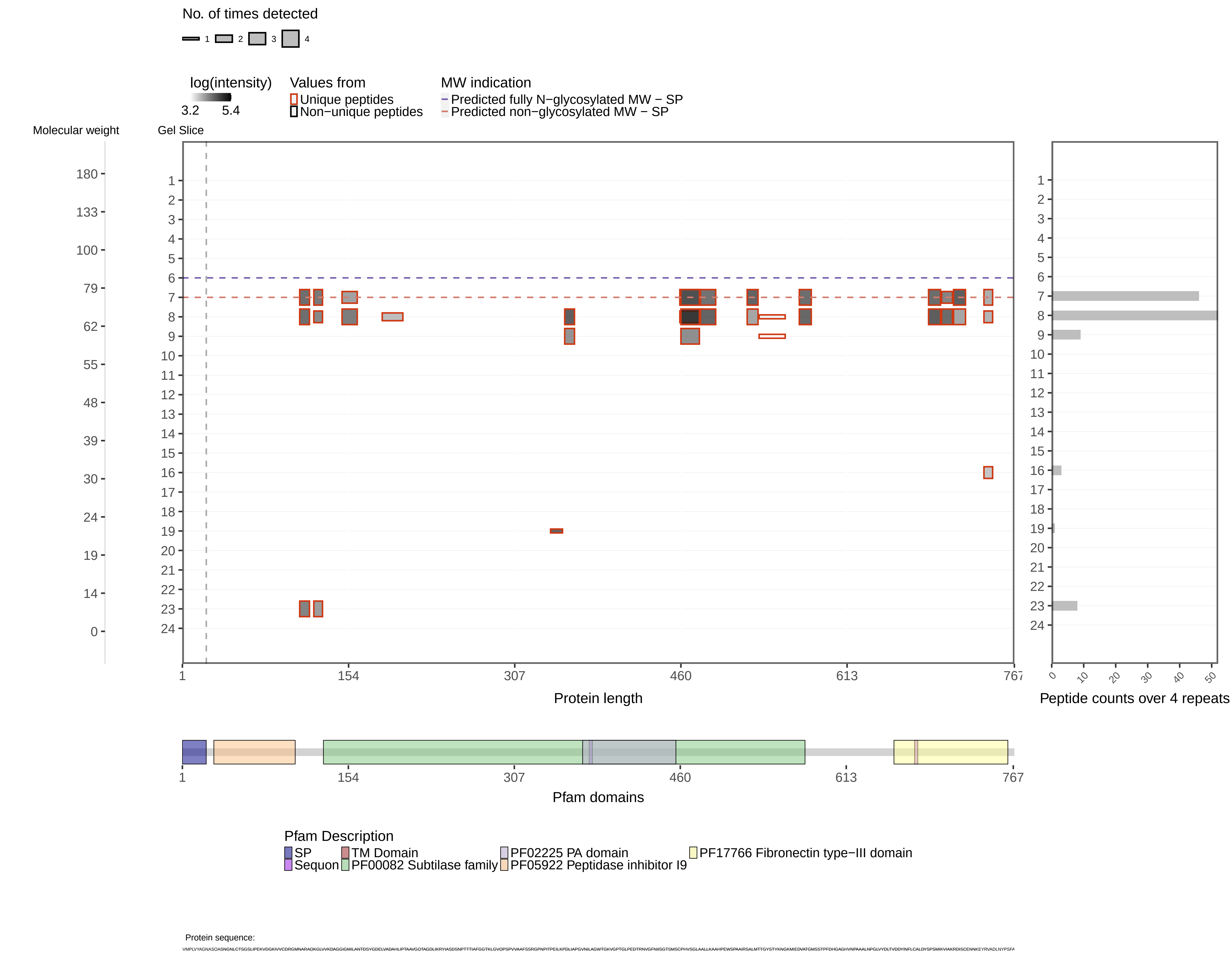

### NbL04g11170.1 geranylgeranyl_transferase_type−2_subunit_alpha_1 [MW = 81.01 kDa]

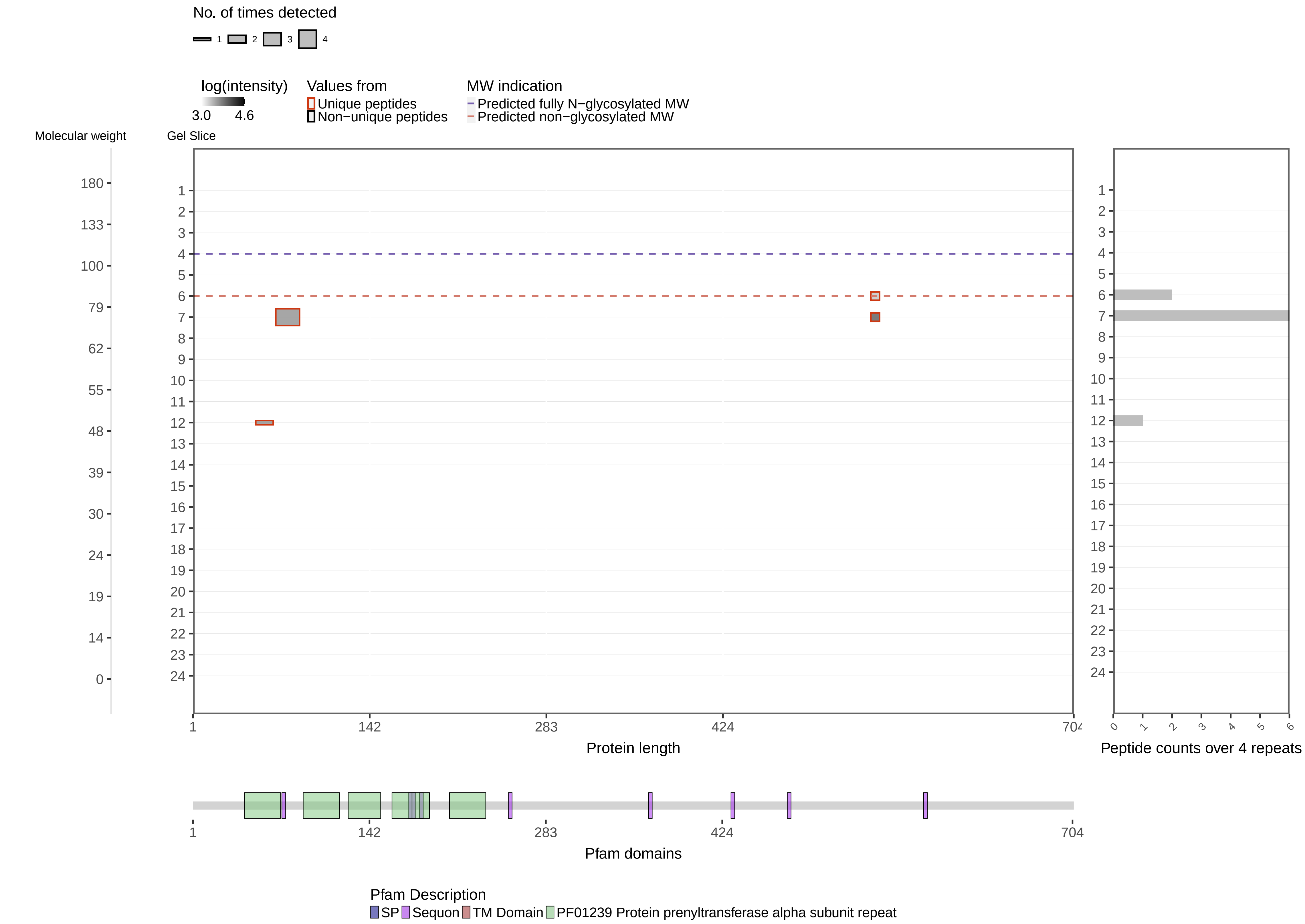


Protein sequence:

HANSLKSYQVKVNLARSQGIFSSTGAHHGNSSHIAFTVSISPYRSEHVDLKNEEKISWSEESFCPHEAQFLESVLANLFHIERTKVDEKVVDYQWNITTVNNEIAHYRELLATMNCKIGKLTLARLLMAHDTTVTGTSHRNVSHHTEVLQLYDDLKKMDPAHFHYYQDEYSLVLLKHIISNQELLLKHCCKYRDPSSPKINNFCLRLNDLSLSRIGSMEKLLWVQVLDLSNNQLQSLEGLEVMQLLTCLNVSHNKLCSFTALEPLRLLRSLKVLDISYNEIGAHSIDTRRYLCSSPL

NbL04g11060.1 argininosuccinate_lyase_chloroplastic [MW = 58.32 kDa]

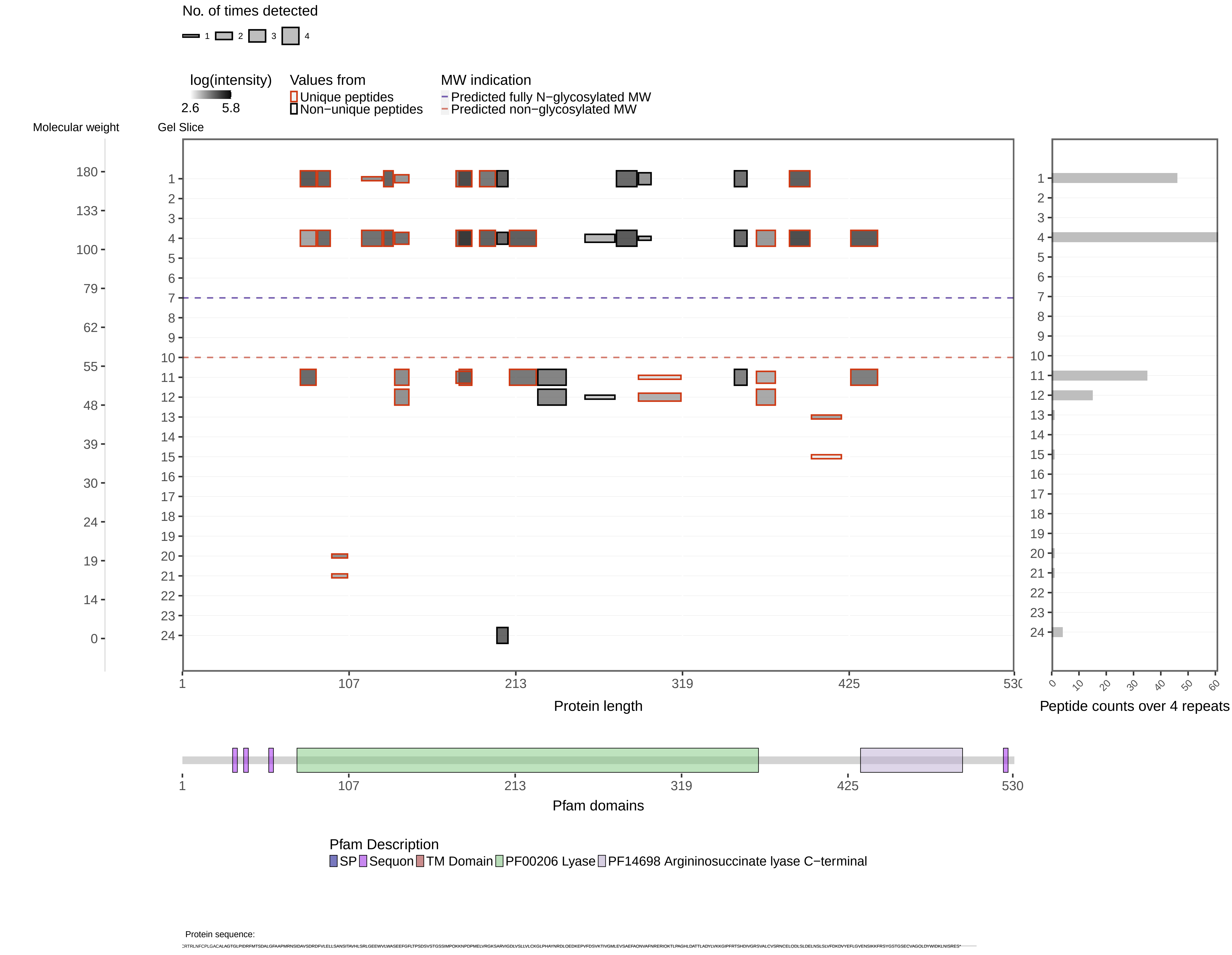

### NbL04g11000.1 chaperone_protein_DnaJ−like [MW = 46.42 kDa]

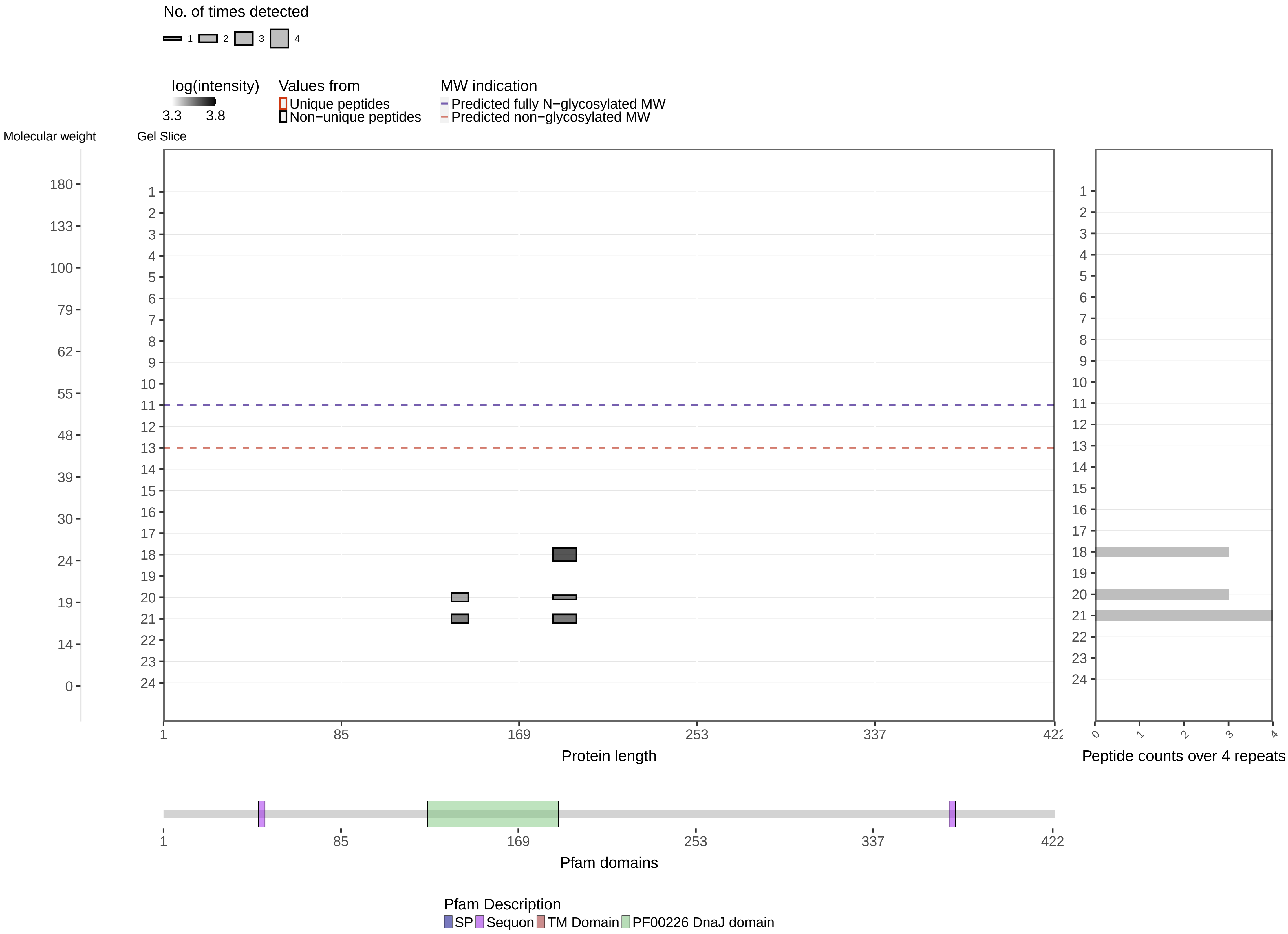


Protein sequence:

SRPKSYNKOOKRGDEKLPVRRSSRASGGSISGONONPSPVNFONRVPVOPOOPKTAATPVSGGMRGOSVVNLWTACPYCYNLFEYPRVYEGCCLRCEKCNRAFTATEILSMPPLVPGREAYYCIWGFFPMGFVNGDLENGKSHAAGDAGFPSWMPPMFSSEGNVNVANONATPTPAPAPAPAPAPVOAOAOTOAPASAPRPMSAPTGAKRRGRPRKYN*

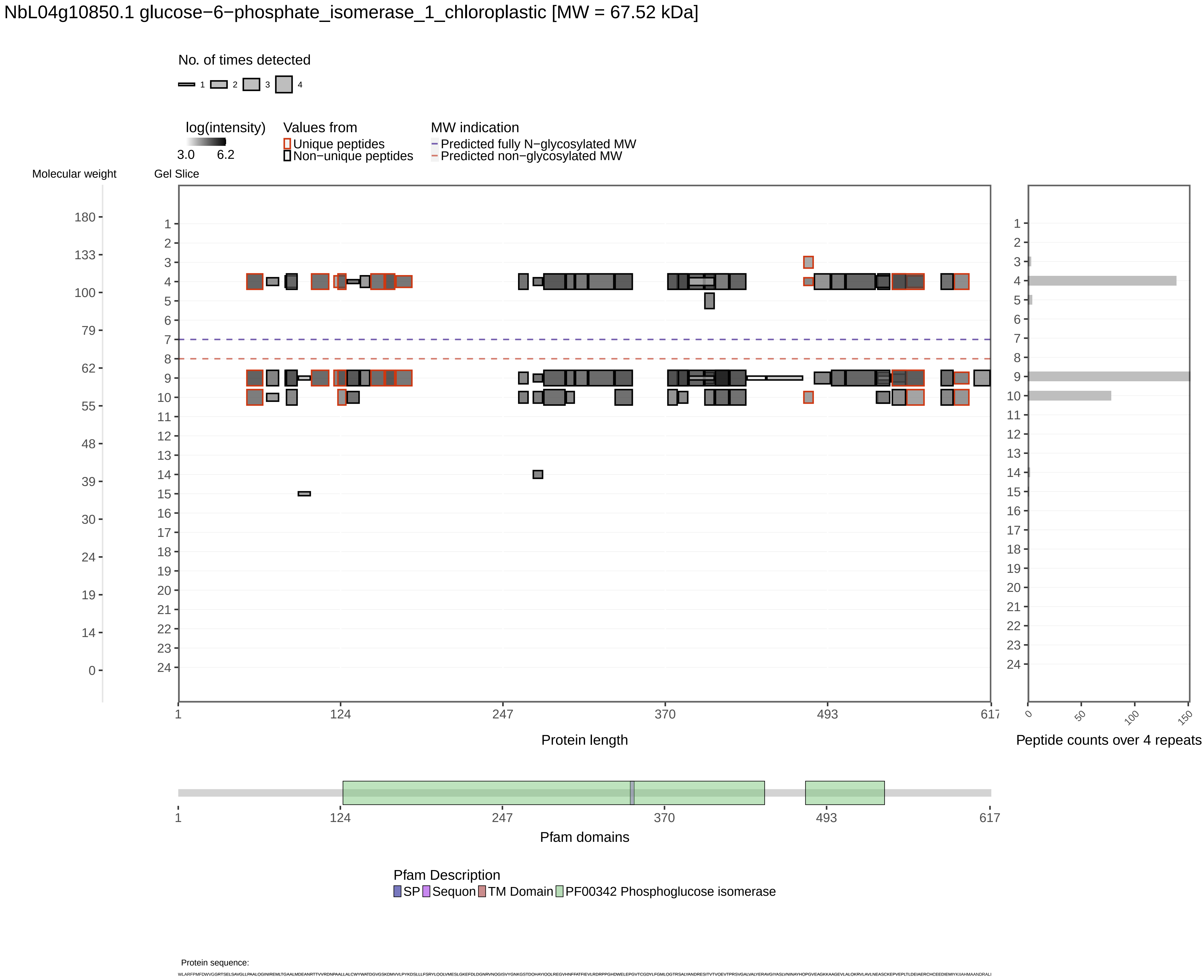
NbL04g10850.1 glucose−6−phosphate_isomerase_1_chloroplastic [MW = 67.52 kDa]
No. of times detected
1 2 3 4
log(intensity)
3.0 6.2
Values from
Unique peptides
Non−unique peptides
MW indication
Predicted fully N−glycosylated MW
Predicted non−glycosylated MW
Molecular weight
Gel Slice
180 133 100 79 62 55 48 39 30 24 19 14 0
1 2 3 4 5 6 7 8 9 10 11 12 13 14 15 16 17 18 19 20 21 22 23 24
1 124 247 370 493 617
Protein length
0 50 100 150
Peptide counts over 4 repeats
Pfam domains
Pfam Description
SP
Sequon
TM Domain
PF00342 Phosphoglucose isomerase
Protein sequence:

### NbL04g10760.1 14−3−3_protein_10 [MW = 30.01 kDa]

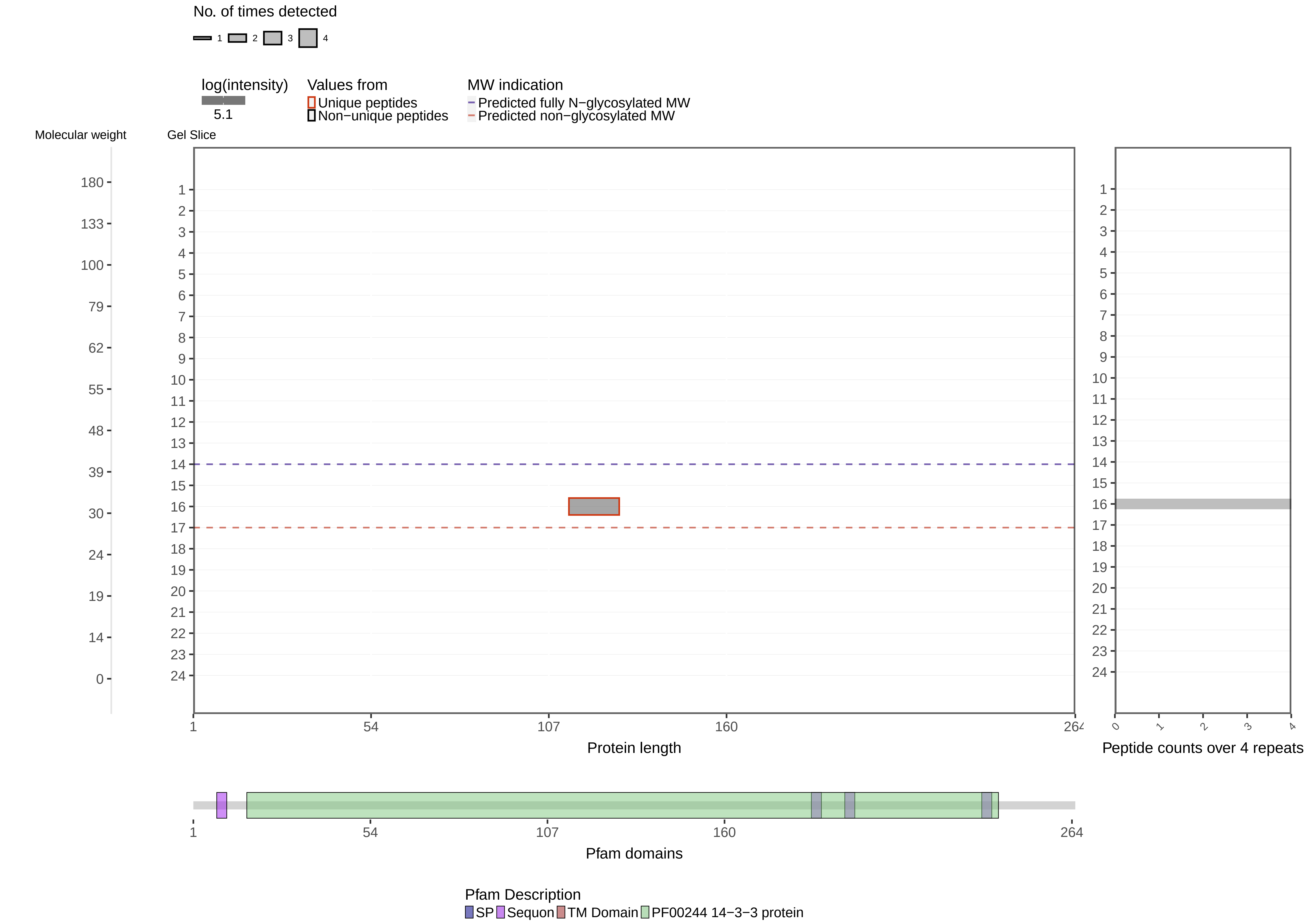


Protein sequence:

PSASTGESRVFYLKMKGDYYRYLAEFKVGDERKQAAEDTMNSYKAAQEIALADLPPTHPIRLGLALNFSVFYFEILNSSDKACSMAKQAFEEAIAELDTLGEESYKDSTLIMQLLRDNLTLWTSDAQVGTLLRAVITKHRHMQTR

NbL04g10720.1 esterase_AGAP003155−like [MW = 24.99 kDa]

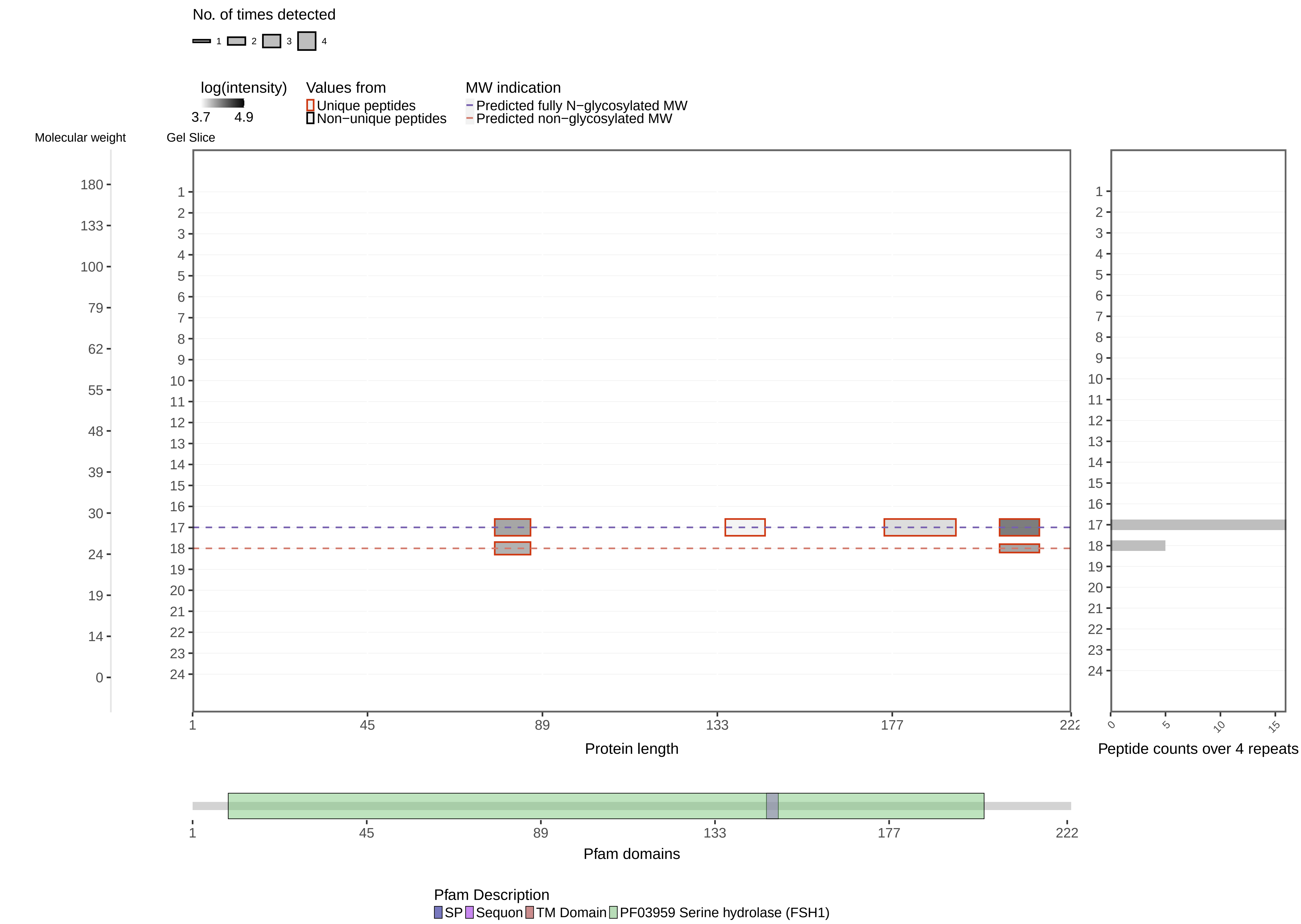


Protein sequence:

MIKHGPFDGLLGFSQGAILSGALSGLQEKGVALTKVPKIKYLIIIGGAQFKNKSVAEKAYSSSITCPSVHFLGEQDYLKKYGIELLESYVDPVVIHHPKGHTIPRFDEKGLESMLSFIEKVQAEISK*

NbL04g10700.1 purine_permease_3−like [MW = 14.4 kDa]

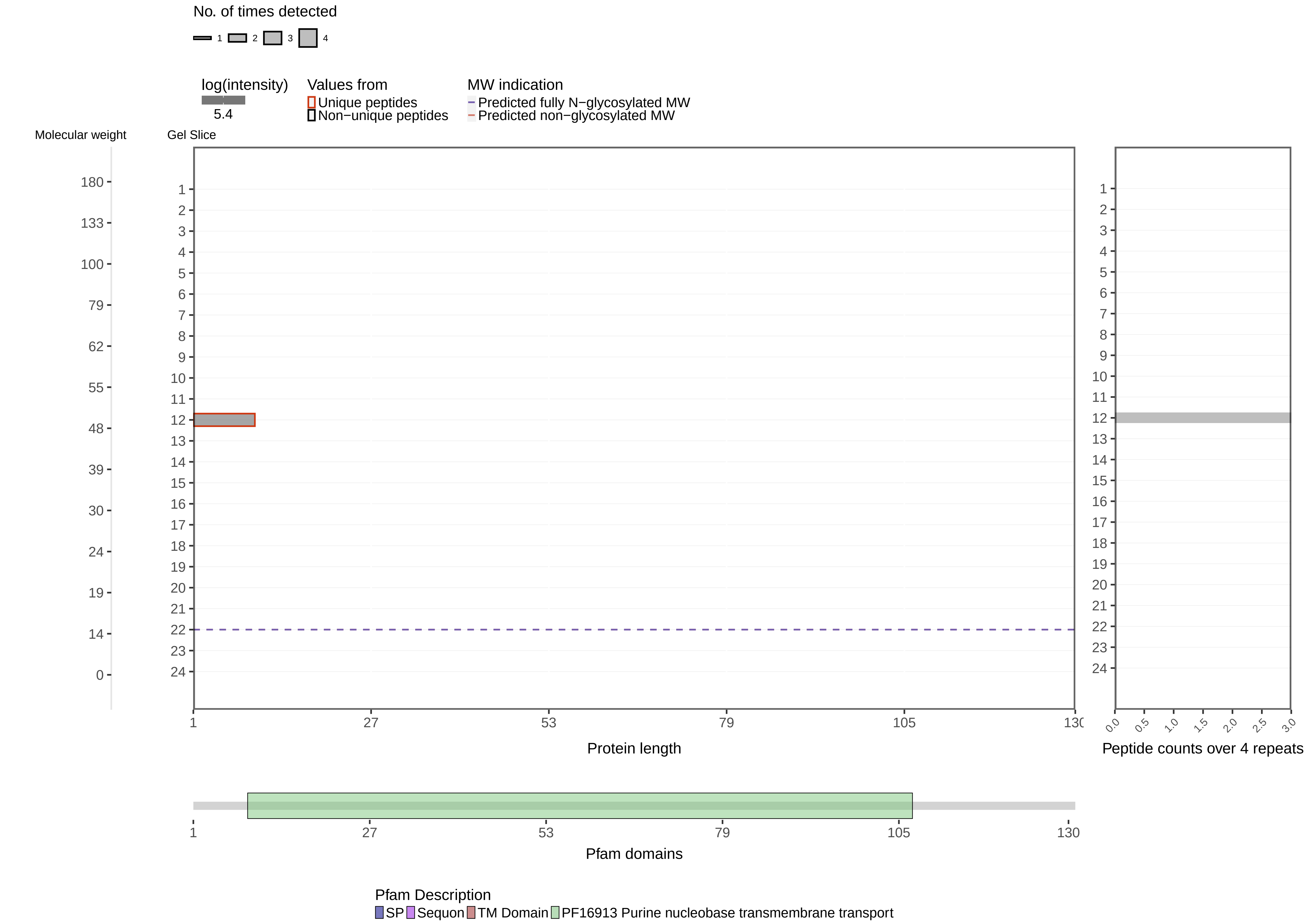


Protein sequence:

SSLLSGVIIAVLLPVTEVLAVIIFHEKFQTEKGVALFLSIWGFISYFYGEMKQHNKEKQTLELEMATTQSSNPPLESA*

### NbL04g10610.1 28_kDa_ribonucleoprotein_chloroplastic [MW = 32.84 kDa]

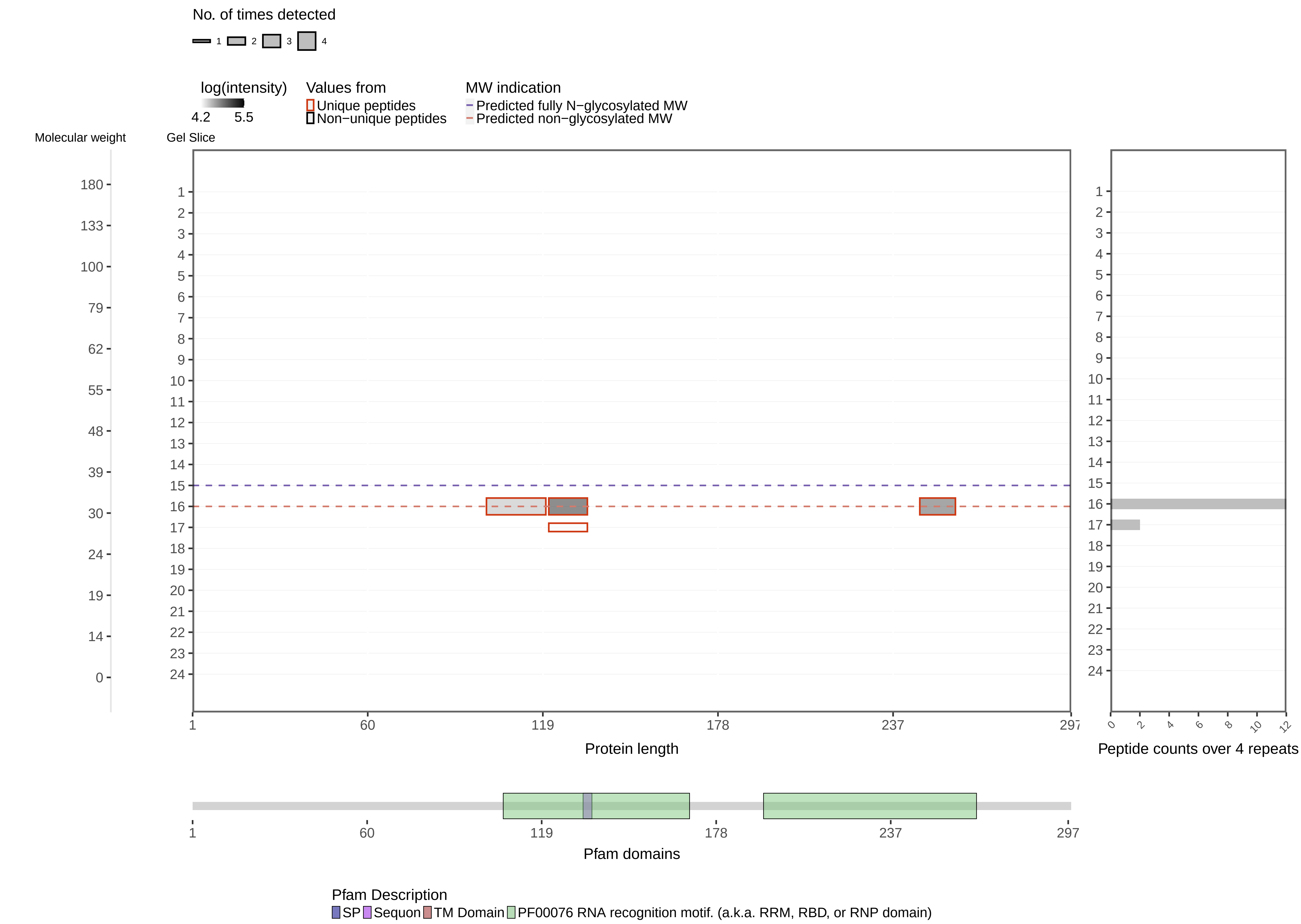


Protein sequence:

RGLVFVTMGSHEEAKAVLENLEAYELEGRPLRLAWAKPKTVKPSSPPPSKRLPVHNVFVDNLHFEARSKDLMEFFKANGADVVSAEVIFNSDNPRRSAGYGFVSFNTKVEADAALSSFDGKEFMGRPIRVAHSKRFMRGETKKILQSQEVSSELISVAE*

### NbL04g10500.1 14−3−3_protein_7 [MW = 29.34 kDa]

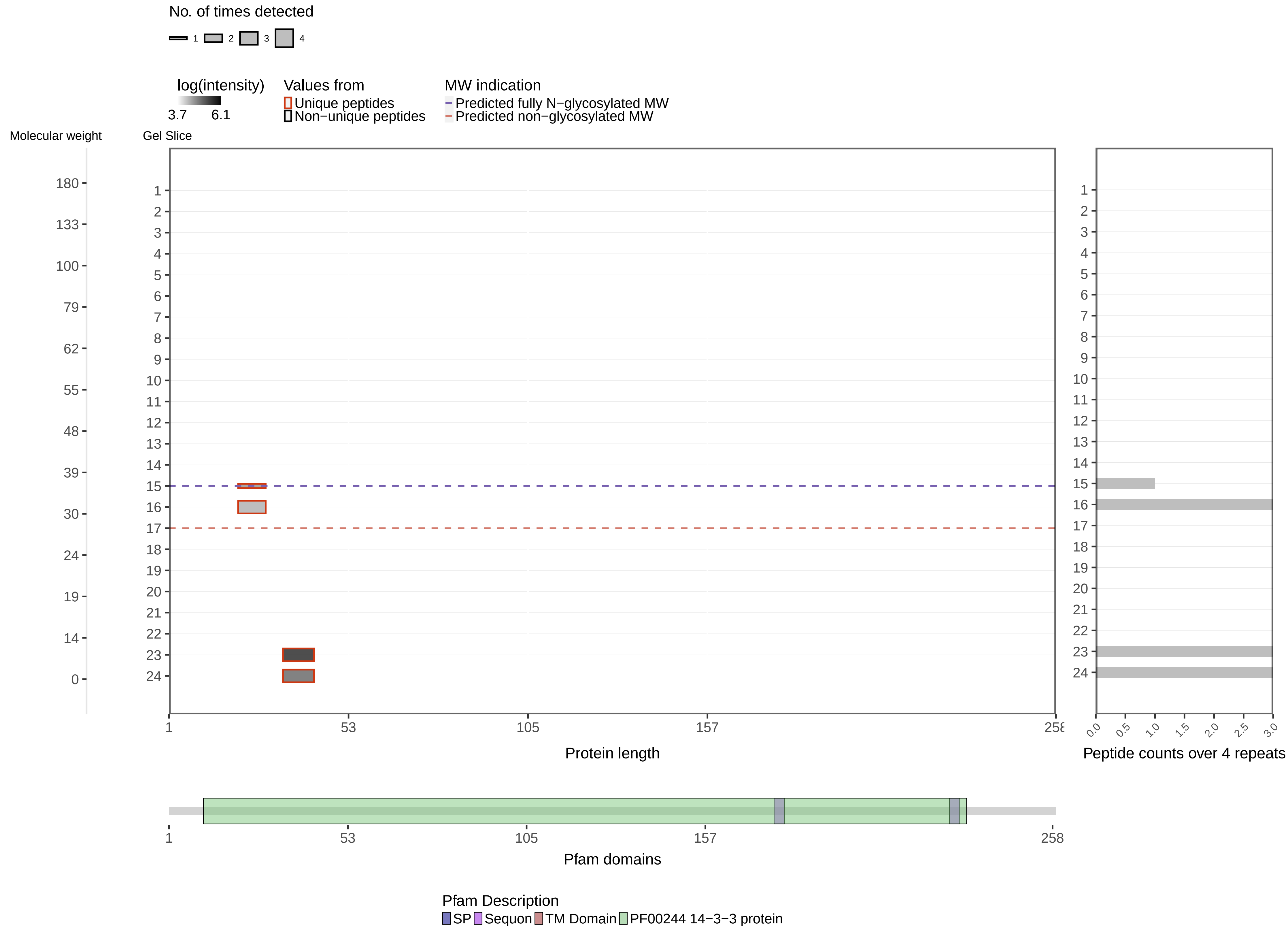


Protein sequence:

AGESTVFYYKMKGDYYRYLAEFKSGDDRKEAADQSLNAYEAATATASADLAPTHPIRLGLALNFSVFYYEILNSPERACHLAKQAFDEAIAELDSLSEESYKDSTLIMQLLRDNLTLWTSDLEEGGISASIISWLIIPFNMVR*

### NbL04g10330.1 aldolase_1_chloroplastic [MW = 59.94 kDa]

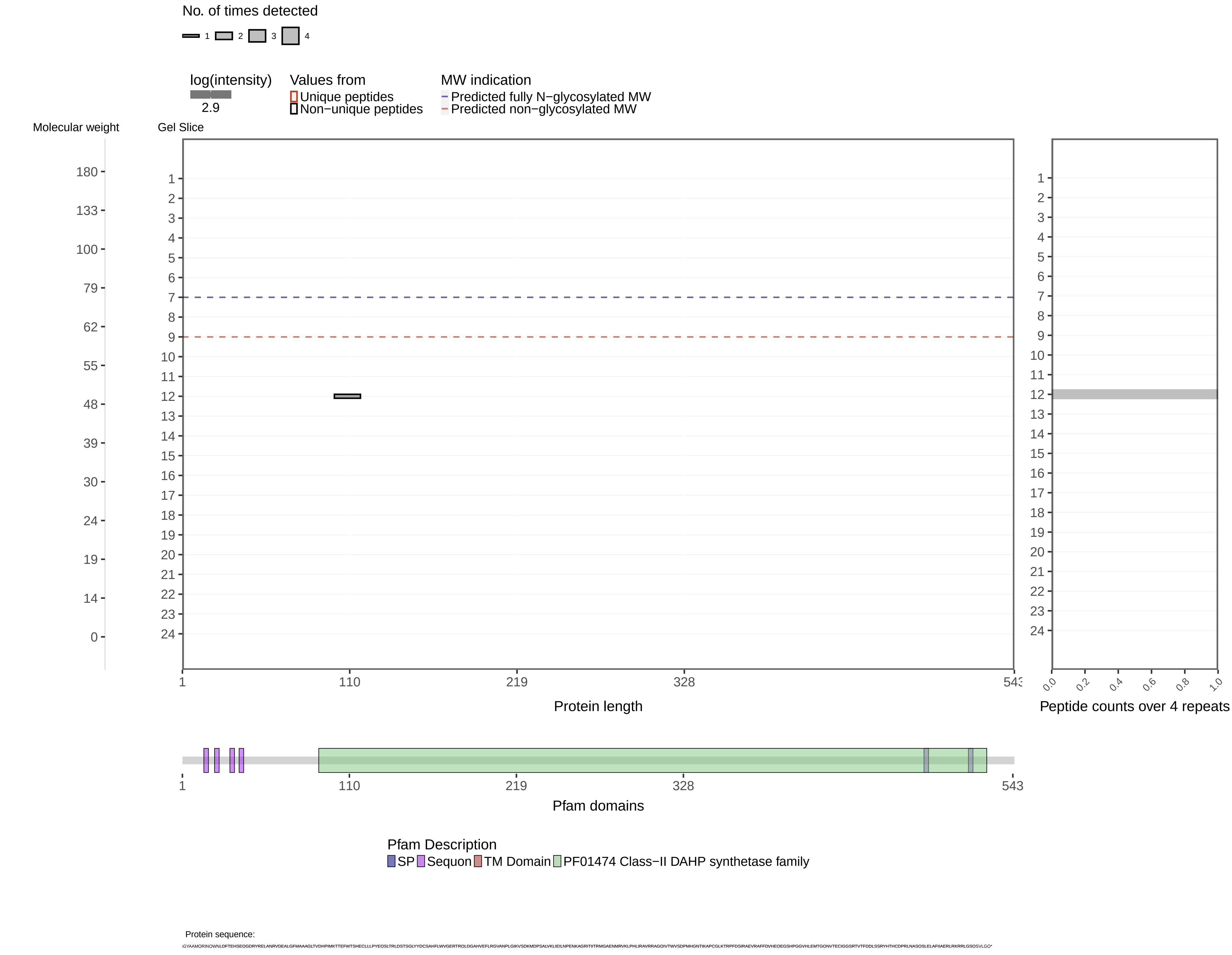

### NbL04g10260.1 thylakoid_lumenal_29_kDa_protein_chloroplastic−like [MW = 37.7 kDa]

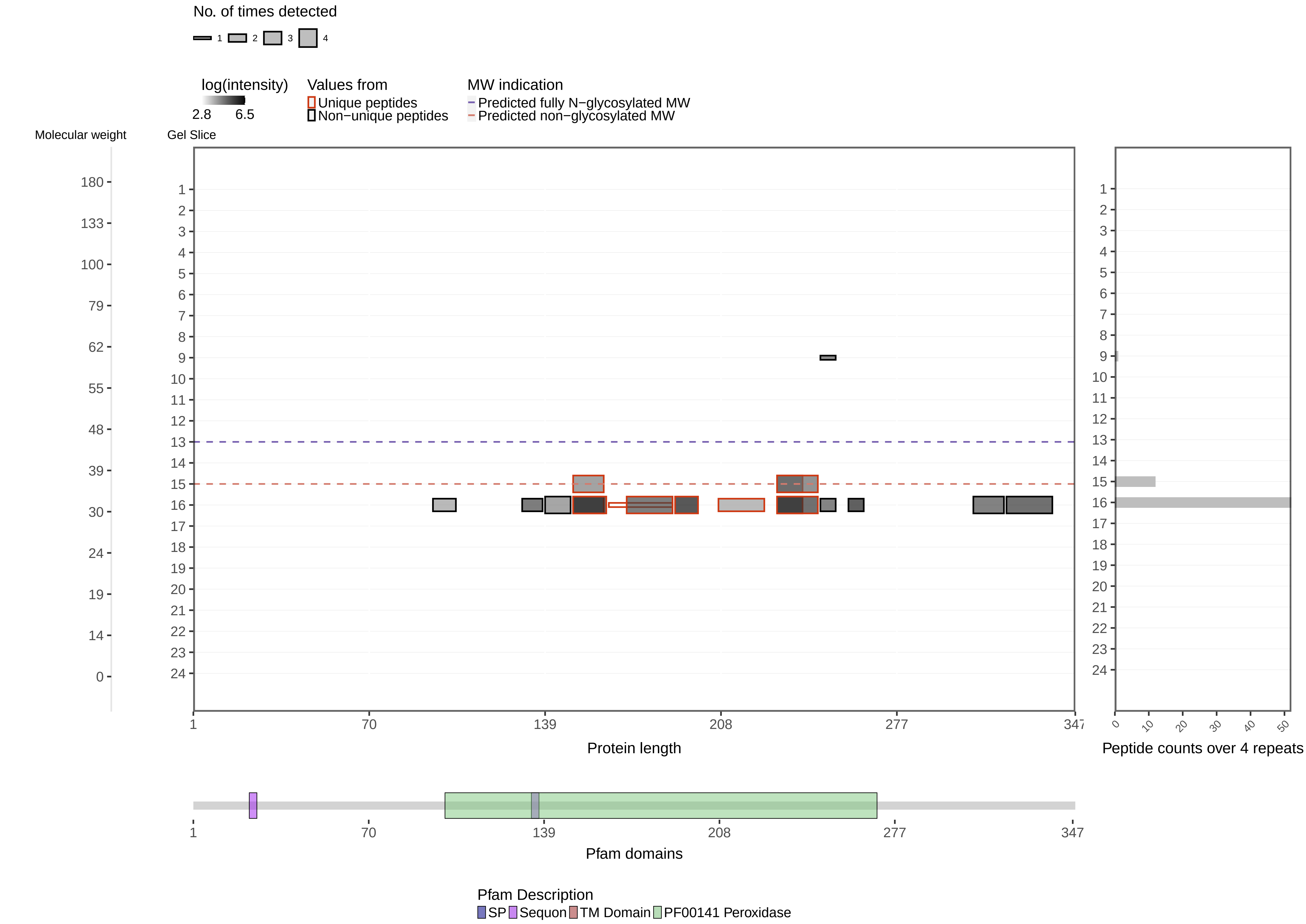


Protein sequence:
EIDADSKGGPISYADLIQLAAQSAVKSTFLASAISKCGGNVEKGSLLYSAYGSNGQWGLFDKQFGRSDAQEPDQEGRVPQWDKASVQEMKDKFKAVGFGPRQLAVMSVFIGPDQDATETLLASDPEVLPWVQKYQRSRETVSRTDYEVDLITTLTKLSCLGQQINYEAYSYPVQKIDVTKLKL*

NbL04g10250.1 serine−−tRNA_ligase_chloroplastic/mitochondrial [MW = 59.73 kDa]

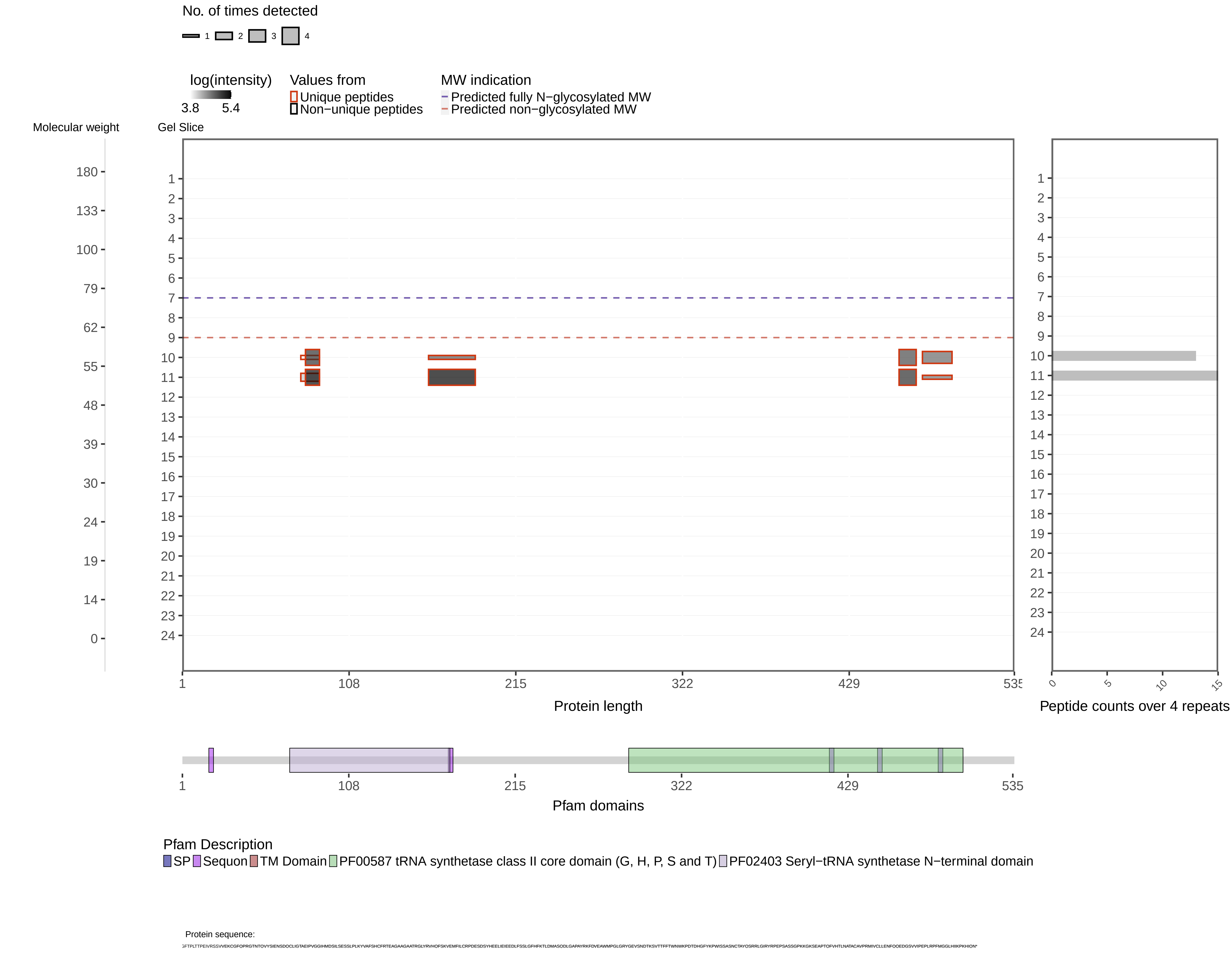

### NbL04g10240.1 cytochrome_c_oxidase_subunit_6b−1−like [MW = 20.02 kDa]

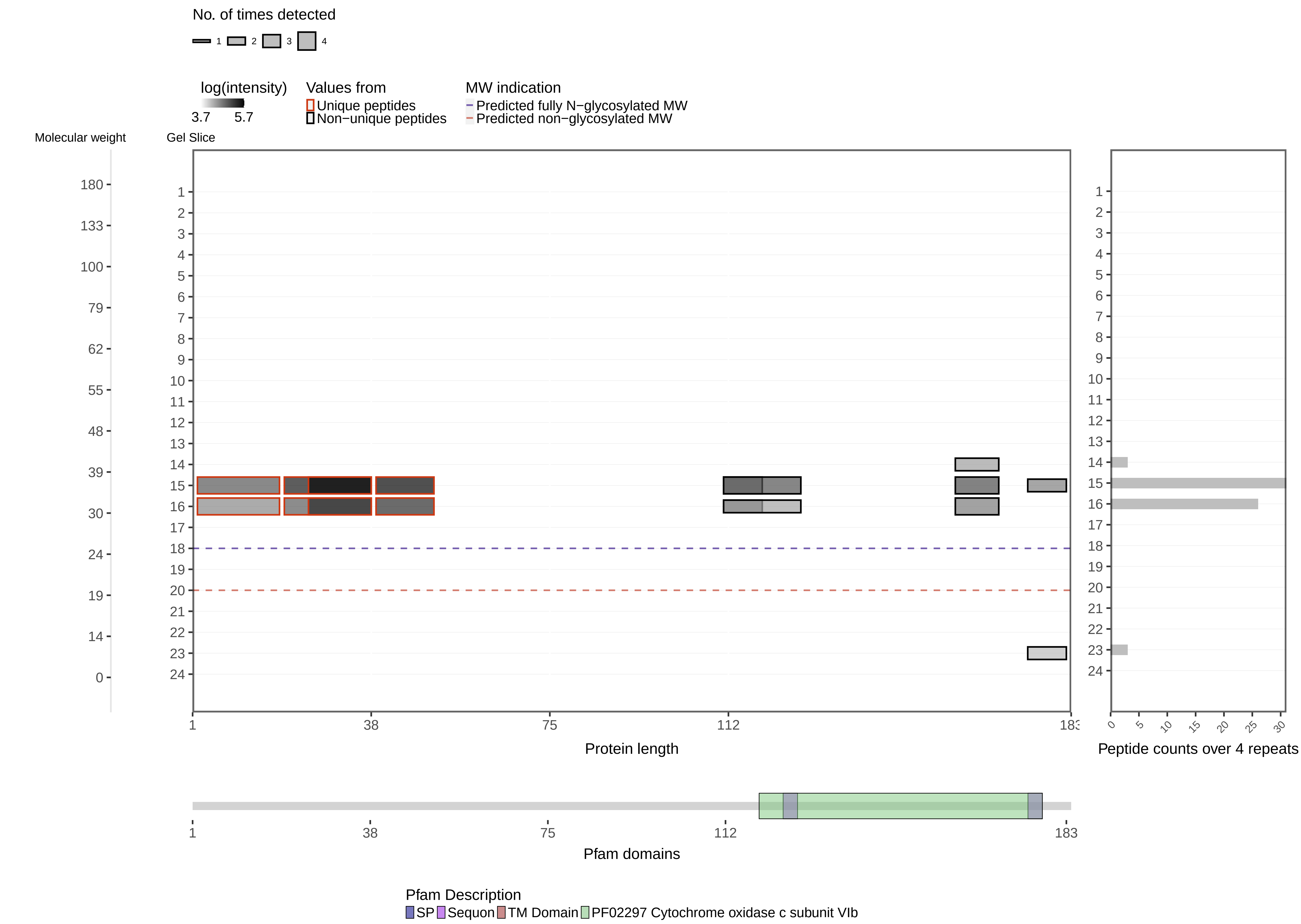


Protein sequence:

PVEASSDESSESTNDETSGNEESAVETPEIKLETAPADFRFPTTNQTRHCFTRYVEYHRCVAAKGEGAPECDKFAKYYRSLCPGEWIDRWNEQRENGTFPGPL*

### NbL04g10230.1 14−3−3−like_protein_C [MW = 29.36 kDa]

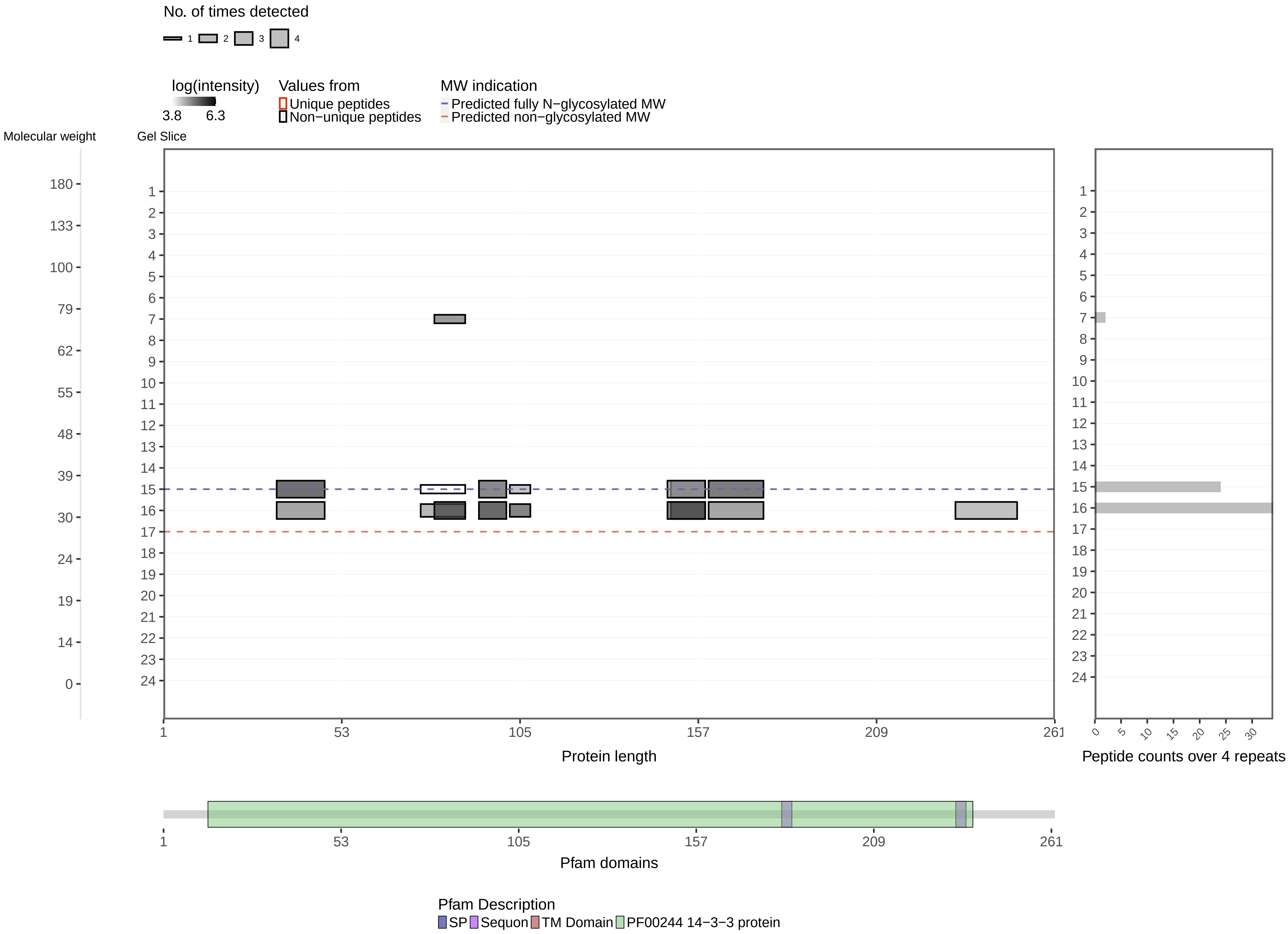


Protein sequence:

AASGDSKVFYLKMKGDYHRYLAEFKTGAERKEAAESTLTAYKAAQDIATTELAPTHPIRLGLALNFSVFYYEILNSPDRACNLAKQAFDEAIAELDTLGEESYKDSTLIMQLLRDNLTLWTSDMQDDGADEIKEDPKPDEAKN*

NbL04g10020.1 probable_beta−D−xylosidase_7 [MW − SP = 82.37 kDa]

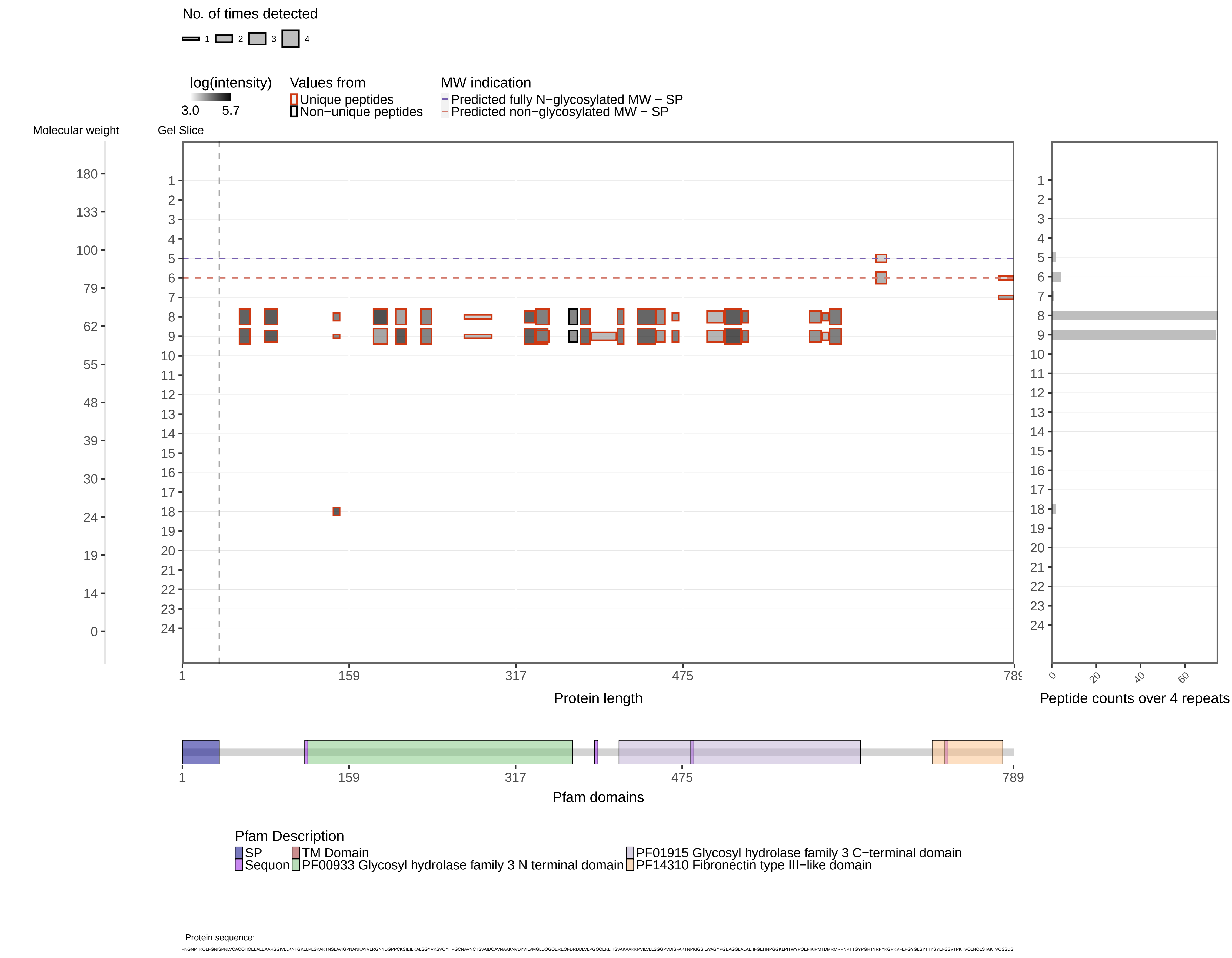

### NbL04g09850.1 uncharacterized_LOC107766695 [MW = 30.56 kDa]

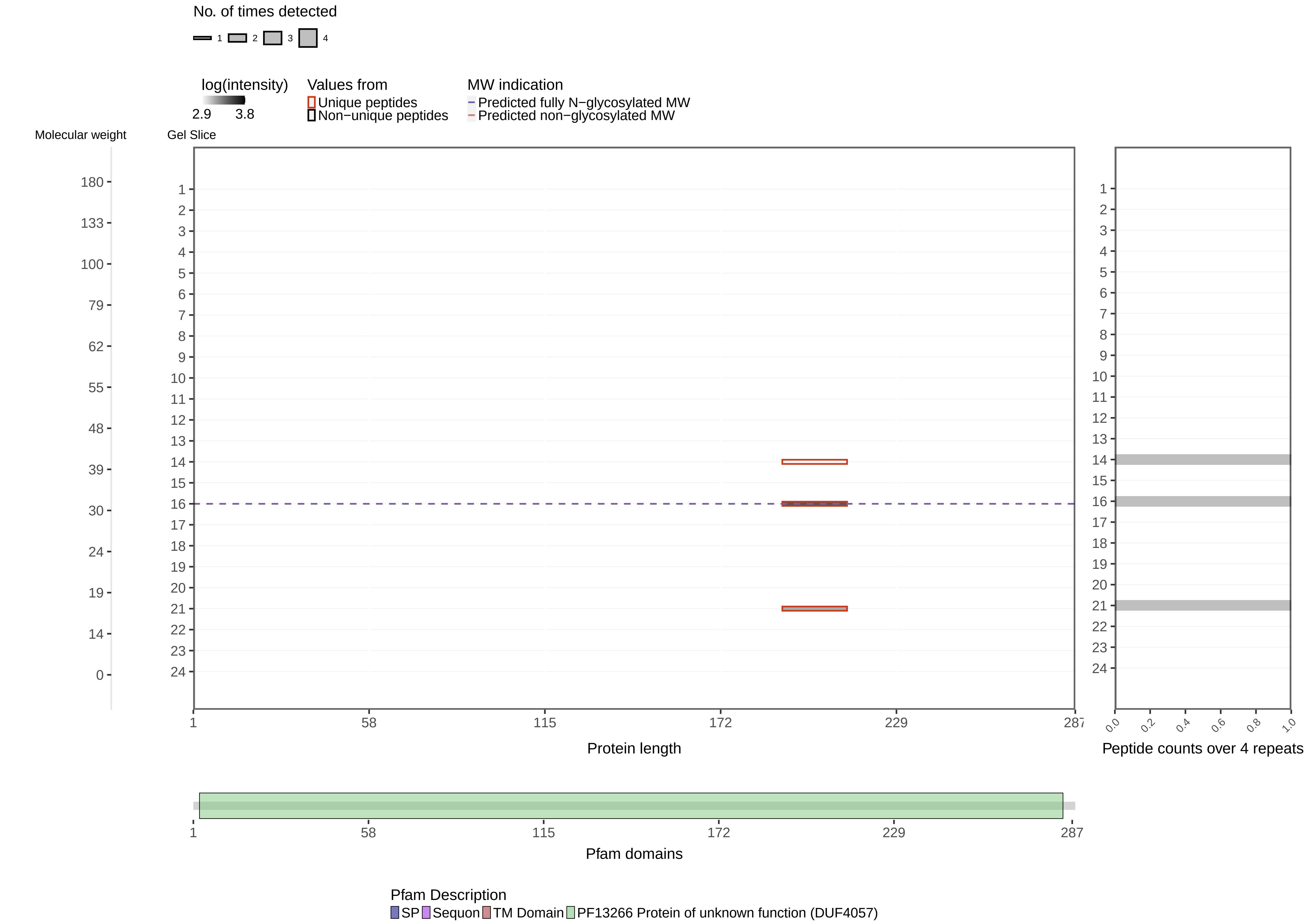


Protein sequence:

KPVTIPEVAKQRELSGTLESEAEMKLKKQLSDAKTKELVGHDIFAPPPEIQPRPLAARALALRESITIGEPTNNGDVSGDEQAVKTAKKIPEKKLAELTGNDIFKGDTPPASAEKHLSSAKLREMSGSNIFADGKVESRDFYGGVRKPPGGESSIALV*

### NbL04g09750.1 aspartic_proteinase−like_protein_1 [MW − SP = 54.41 kDa]

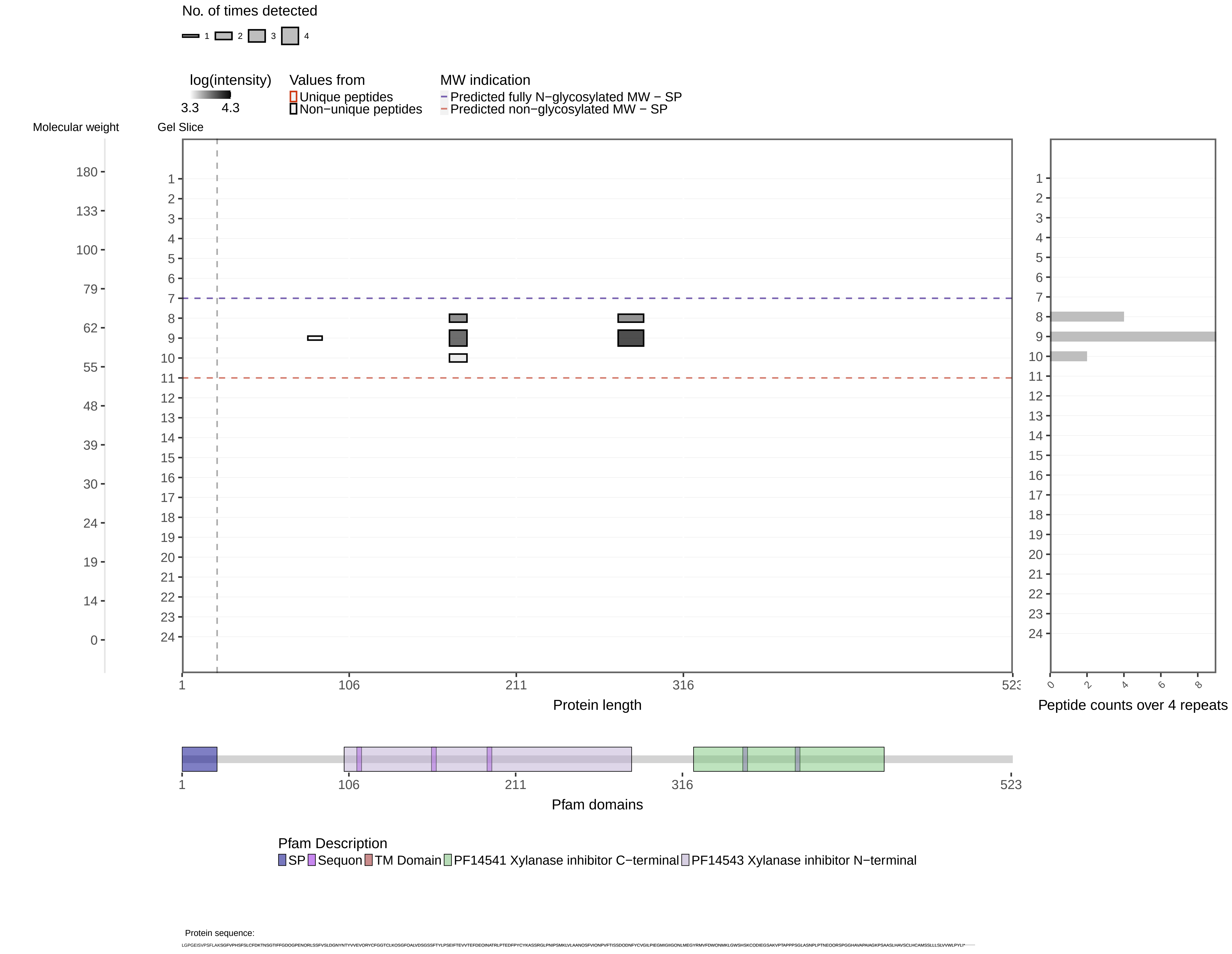

### NbL04g09740.1 60S_ribosomal_protein_L30−like [MW = 12.4 kDa]

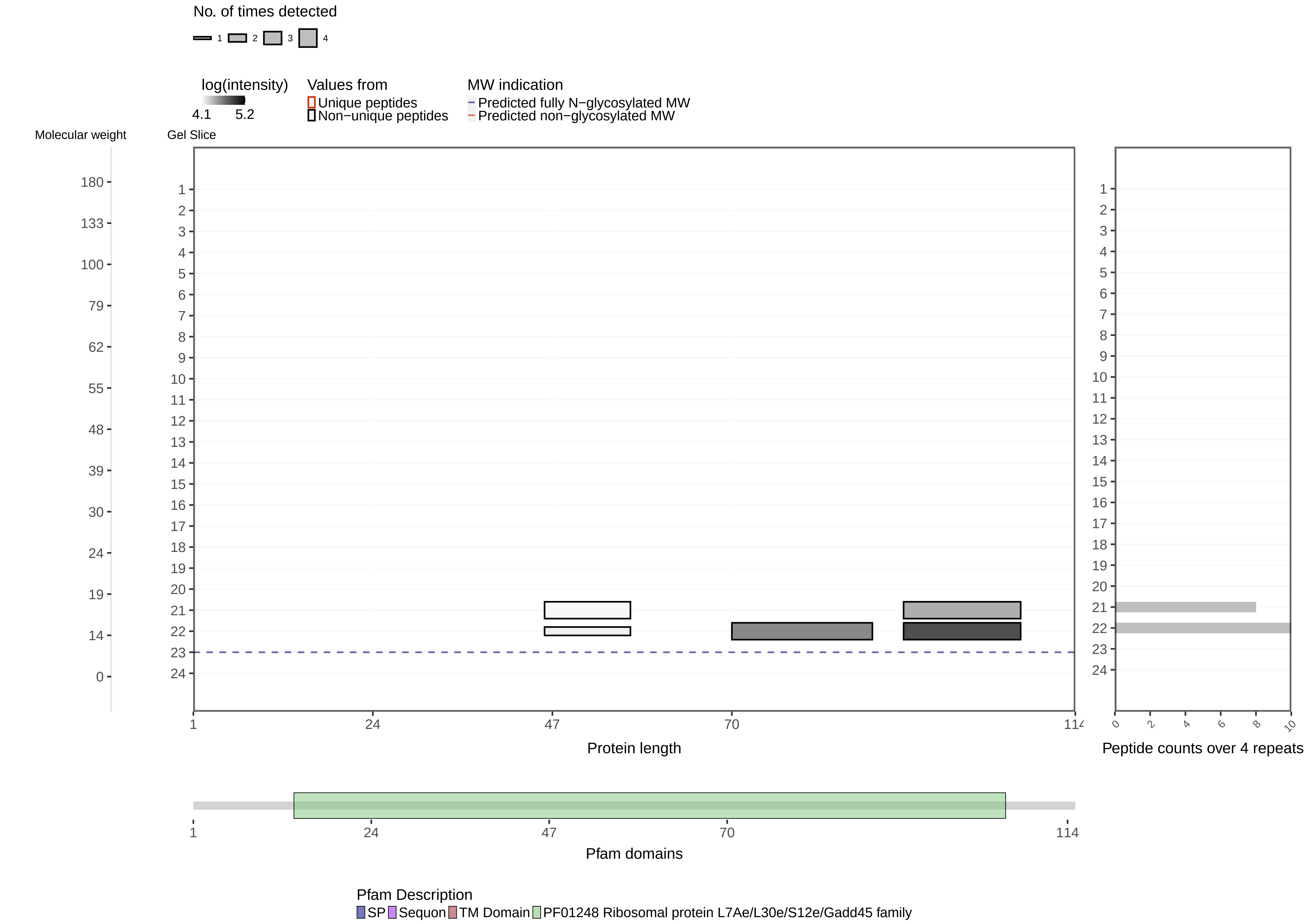


Protein sequence:

KGKLILISNNCPPLRKSEIEYYAMLAKVGVHHFNGNNVDLGTACGKYFRVSCLSIIDPGDSDIIKSLPGDH*

### NbL04g09530.1 uncharacterized_LOC107779703 [MW = 58.14 kDa]

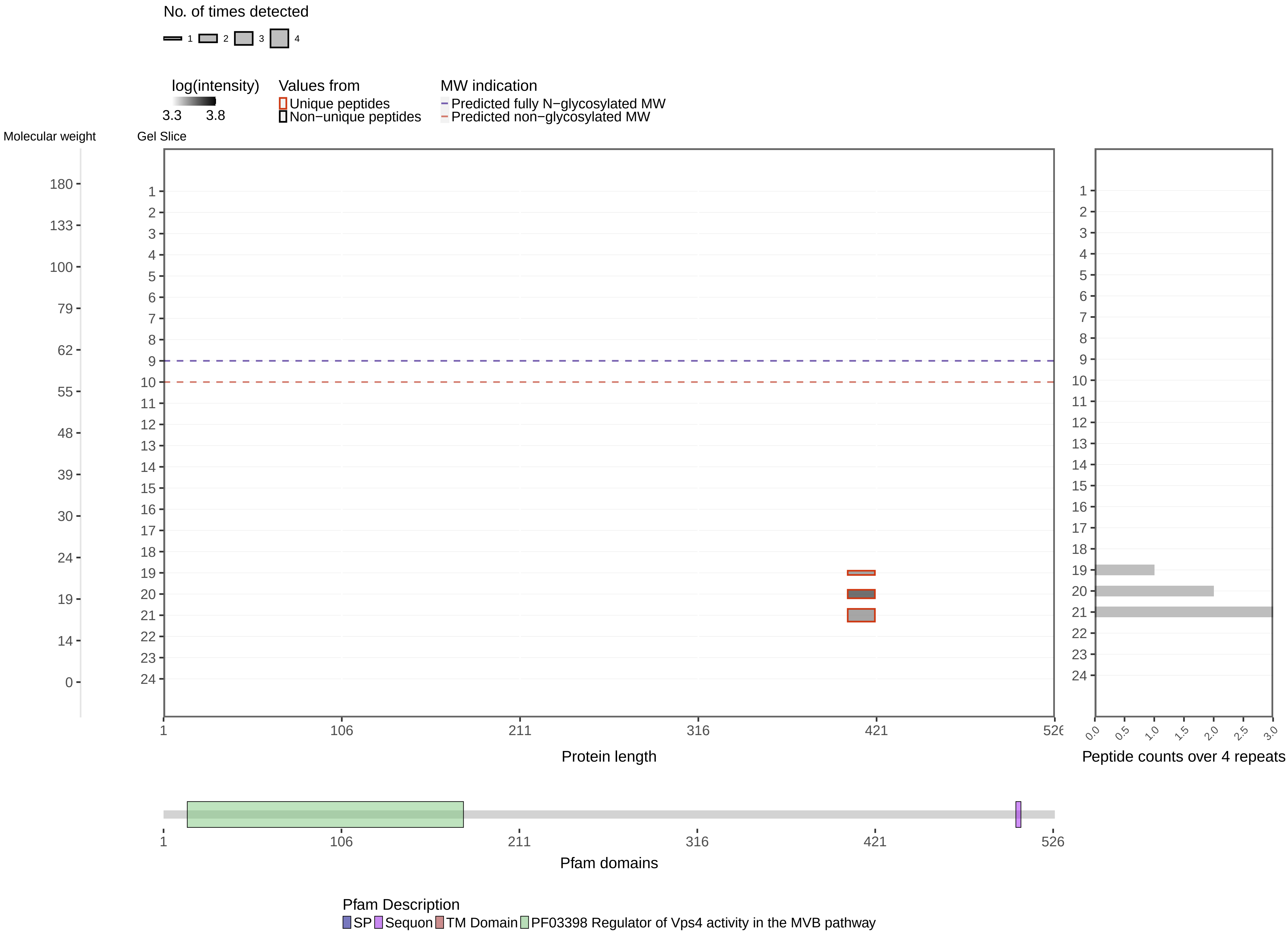


Protein sequence:

PSIGGVSAPEMLPFPASALSDSDEEIAKPSEDAELKSHKQNVERDEVLQEKVVSKDLGSPETLSAPEEEKQFLPFMVPPQRLSPFSSTQSTTTQSIQPPSITKTKAESDIDLQDVLAAANAAAESAERAAAAARSAANLAQFRISELTRKRSEEVPGSTSENSEQESRILEKAPFDLQHQLSNSDGIPSPLHGSPLEHQVSNVPSCDATVGFESFMSTNHHPAPGTALHQPQRLPSLDDETYYSYPNLFTTNGSNPSSRSQSFKDNIKSTHDK*

### NbL04g09390.1 uncharacterized_protein_OsI_027940−like [MW = 26.12 kDa]

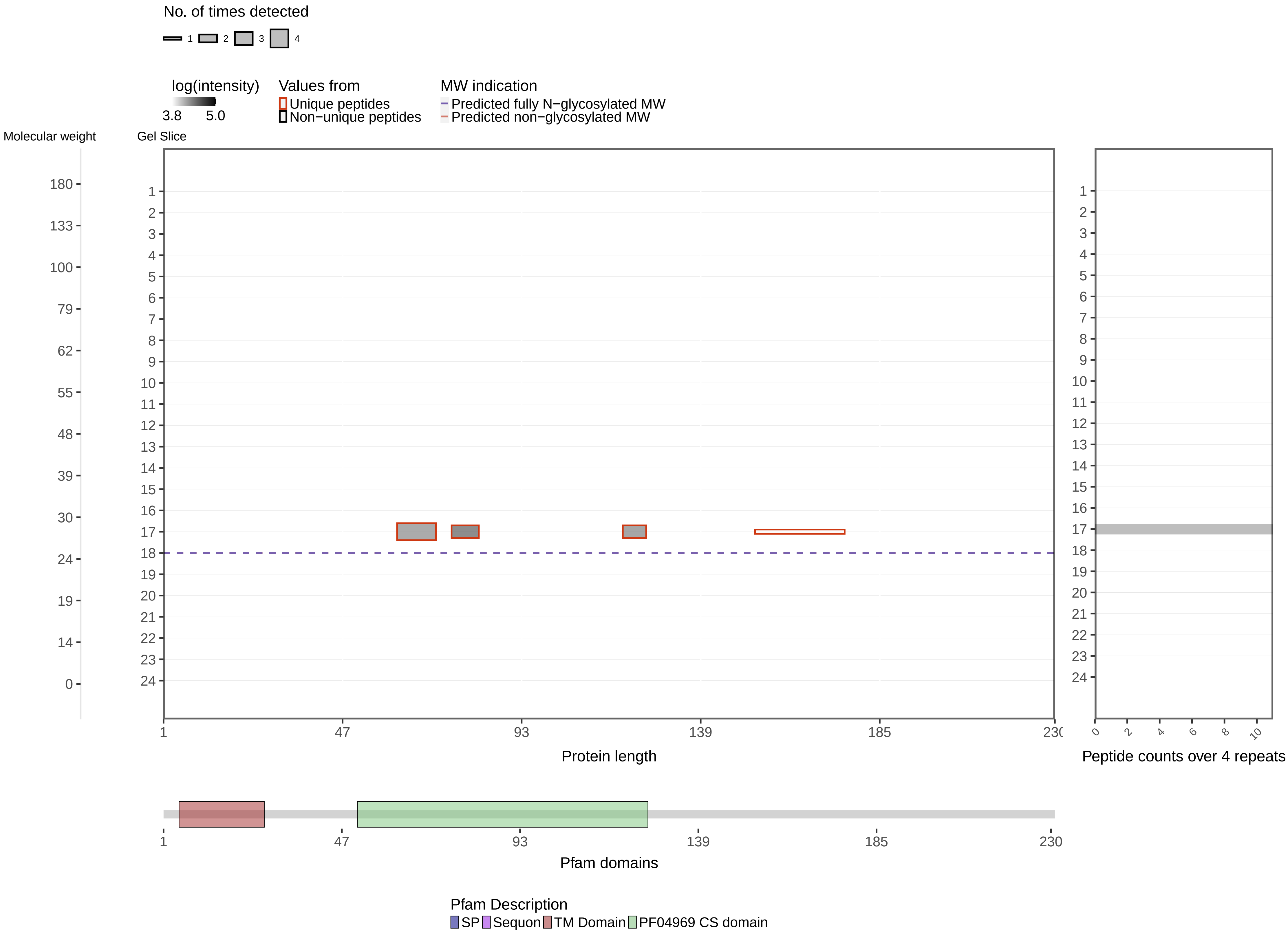


Protein sequence:

NVDESKTSTTSRSIVYLVKKAEDKWWSRLIKQEGVRPVFLKVDWDKWVDEDEEDTKPGTDMDFGDIDFSKLNMGGGPSDFDADVPEGDDDSDTEEEEVREEENSADNEPKAAVPPSTEPKAKA*

### NbL04g09320.1 probable_ATP_synthase_24_kDa_subunit_mitochondrial [MW = 27.47 kDa]

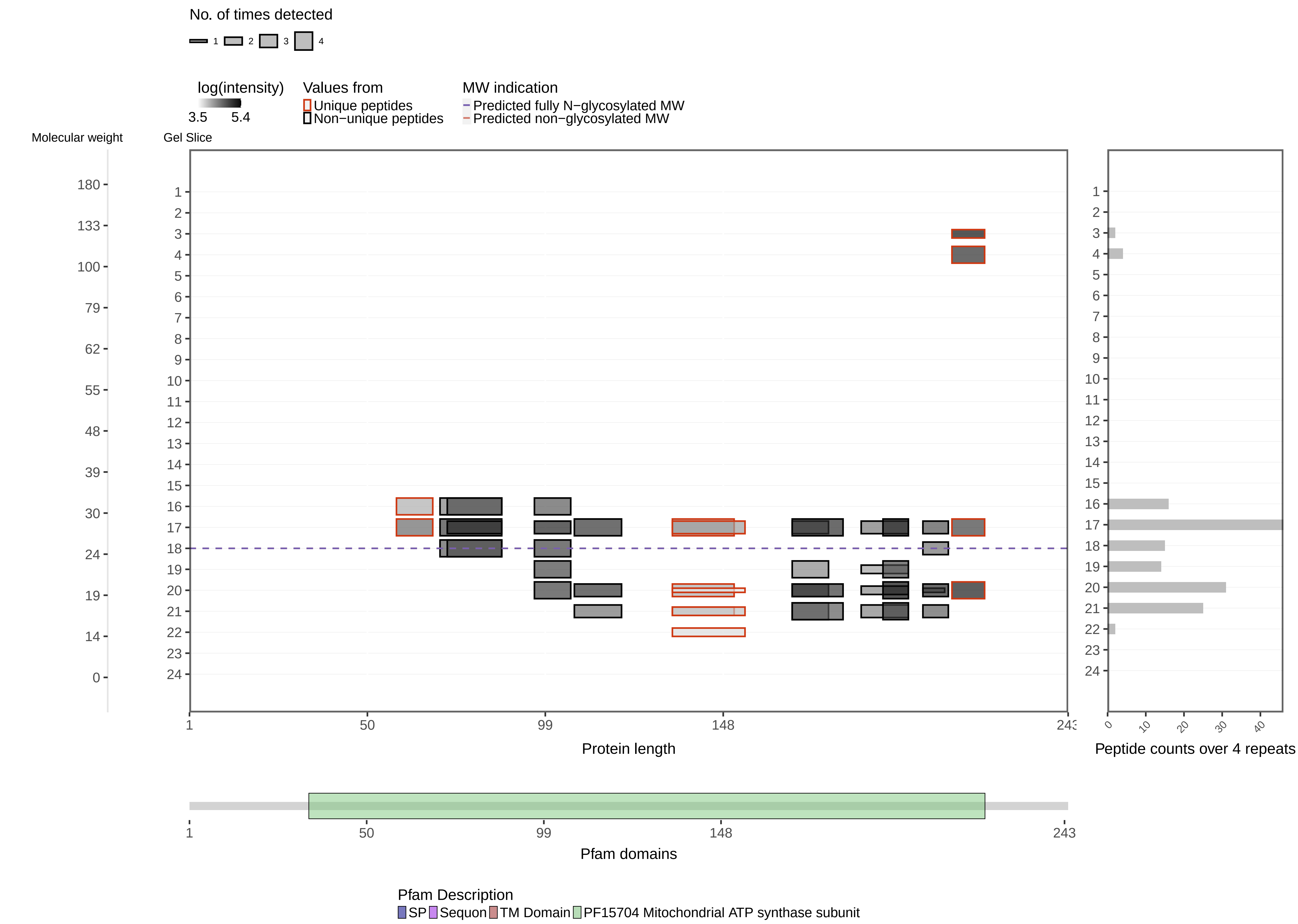


Protein sequence:

NIEVETHEIPDARTYLLKLKEMRVKRGLIDGQGIEDLQMAALDKVEKEIKKPLMRNDKKGIALLTAEFDKINQKLGIRKEDLPKYEEQLELKIAKAQLEELKKDALEAMEAQKKREEFKDEEMPSVKSLDIRNFI*

### NbL04g09270.1 hypoxanthine−guanine_phosphoribosyltransferase [MW = 24.52 kDa]

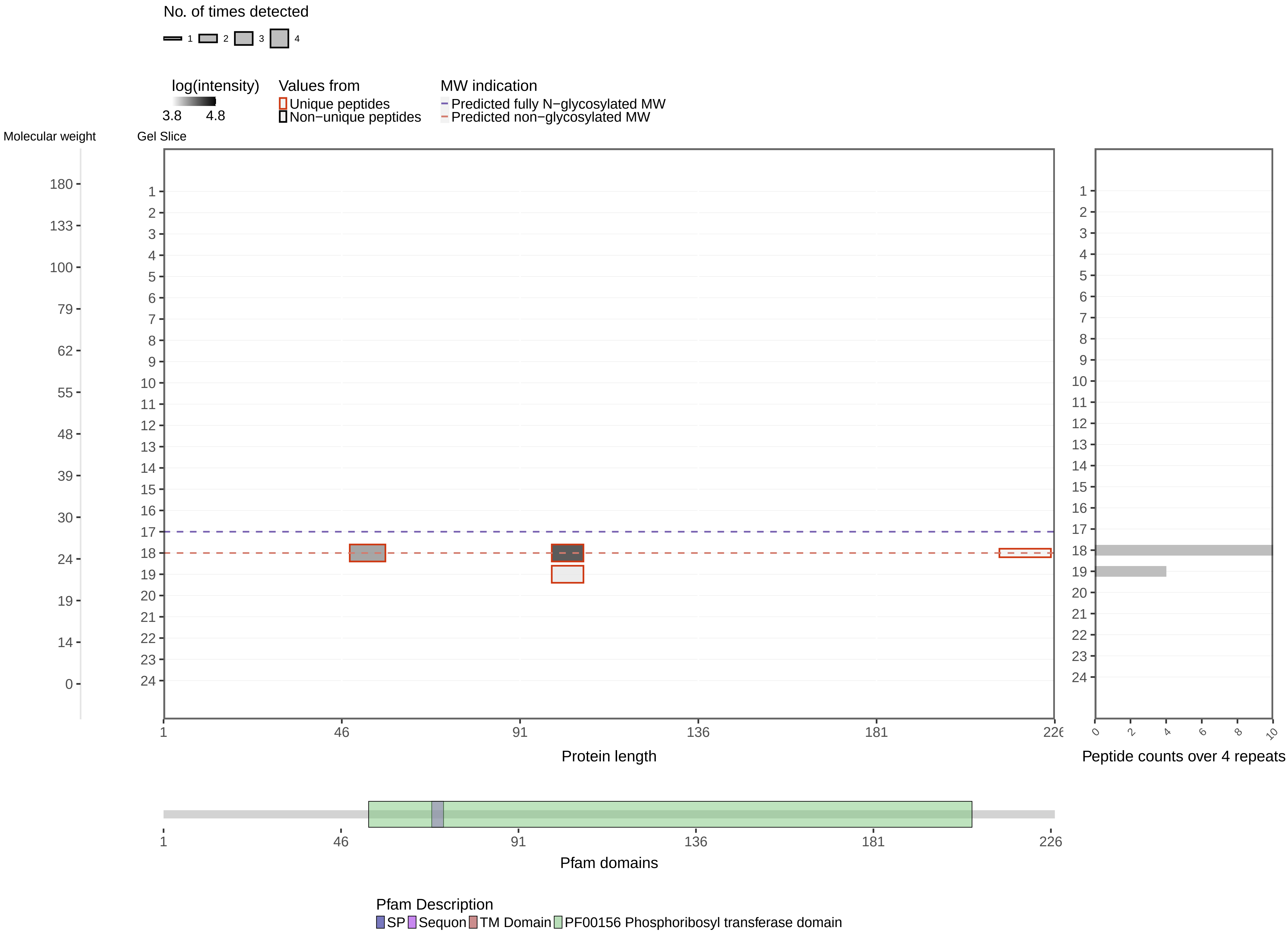


Protein sequence:

VDFVRVESYGSGTVSNGKPKISCDLKIDVTGKHVILVEDIVDTGNTLSCLIEHLKSKGASSISVCTLLDKPSKRKVNLELVGEGMFYRGFECPDYFVVGYGLDFAELYRNLPYVGVLKPEMYS*

### NbL04g09260.1 uncharacterized_LOC104248969 [MW = 20.43 kDa]

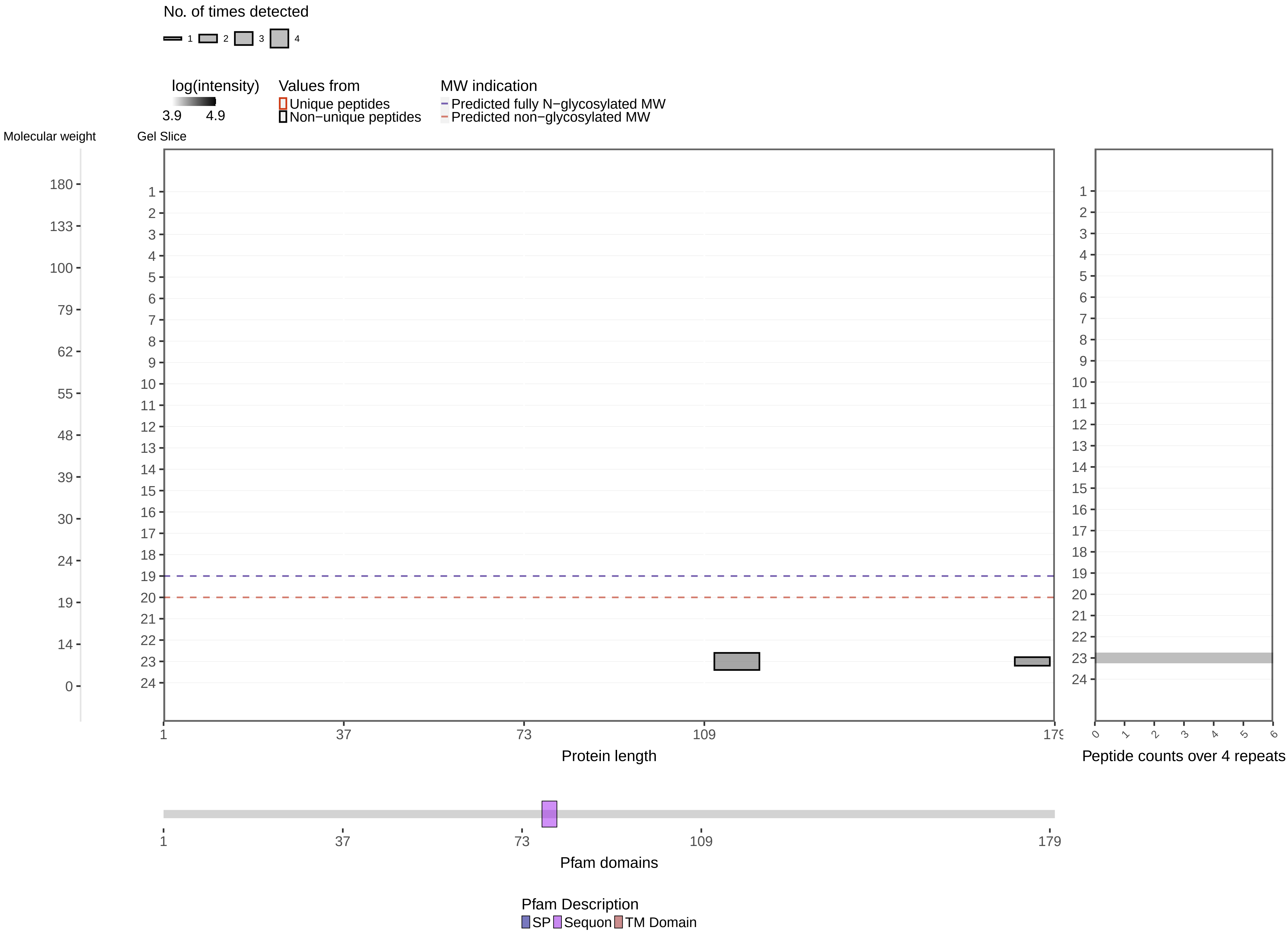


Protein sequence:

SSTQFIISQNVRKYCVDTDKEVDKINLKFAEAREEIESAMESKETVYFNEEAECARAAVKEVFDLYYGLLEKLSESEKGVIQRSMGLKIEQLKAELEQLNE*

NbL04g09150.1 xanthoxin_dehydrogenase−like [MW = 29.68 kDa]

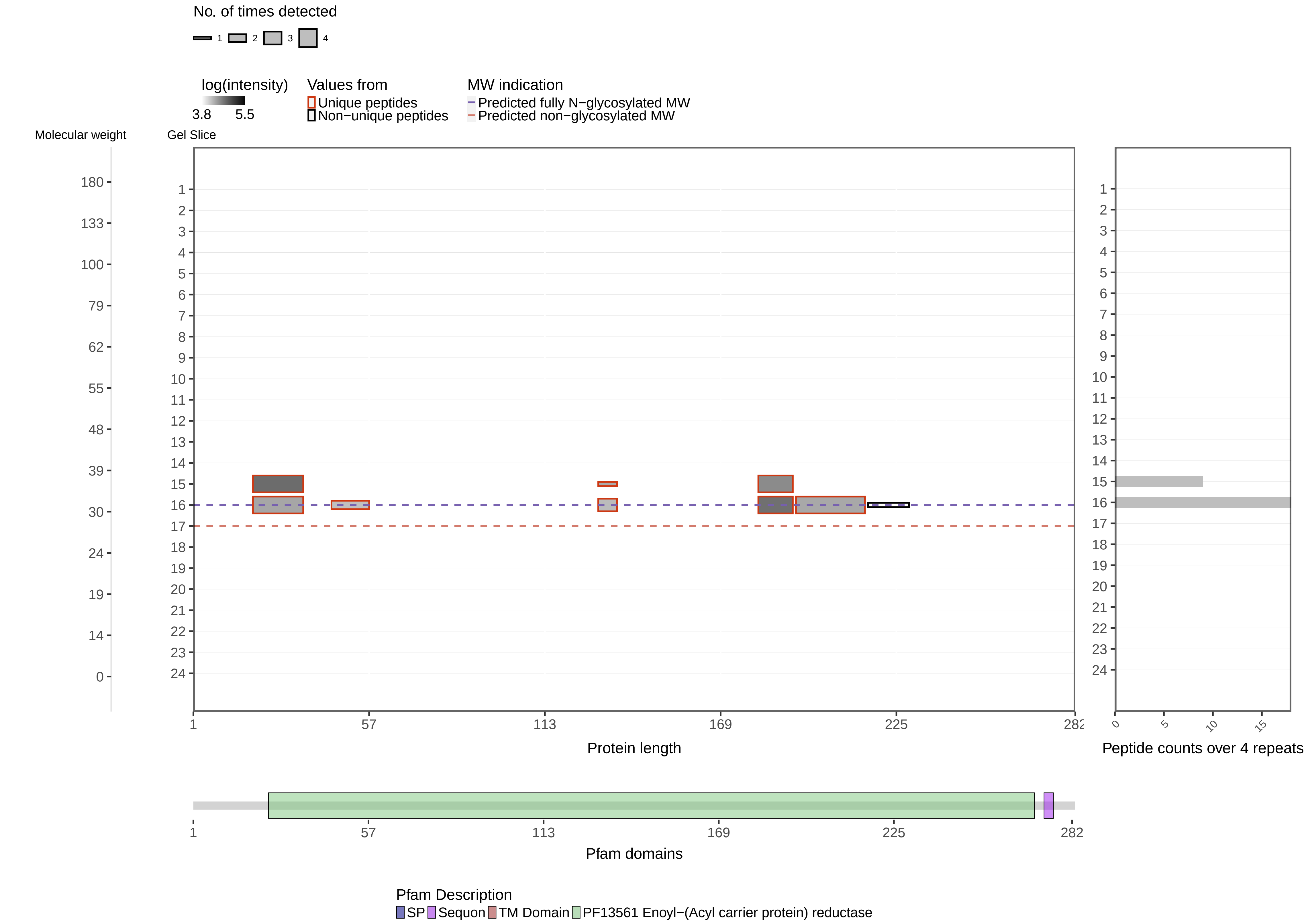


Protein sequence:

VKGVFLGMKHAARIMIPLKKGSIVSLSSAASAIAGLAPHAYTTSKHAISGLTKNVAAELGQHGIRVNCVSPYAVATGLSLPHLPEDQRNDDAIADFHAYFGKLANLQGVDLLVQDVANAVLFLASAEARYISGHNLMVDGGFSCVNHSLKVFRS*

### NbL04g09130.1 peroxidase_12−like [MW − SP = 36.38 kDa]

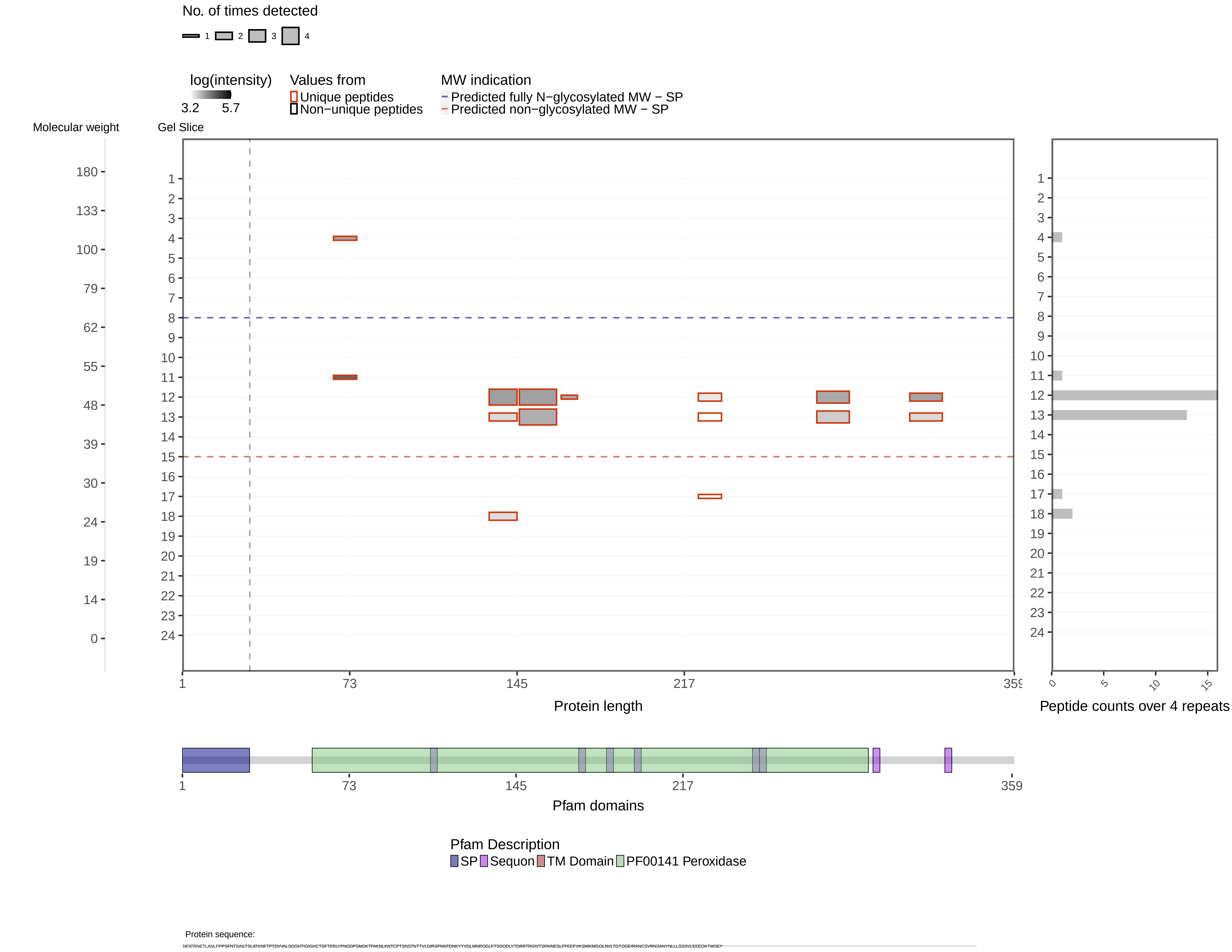

NbL04g09110.1 peroxidase_12−like [MW − SP = 35.87 kDa]

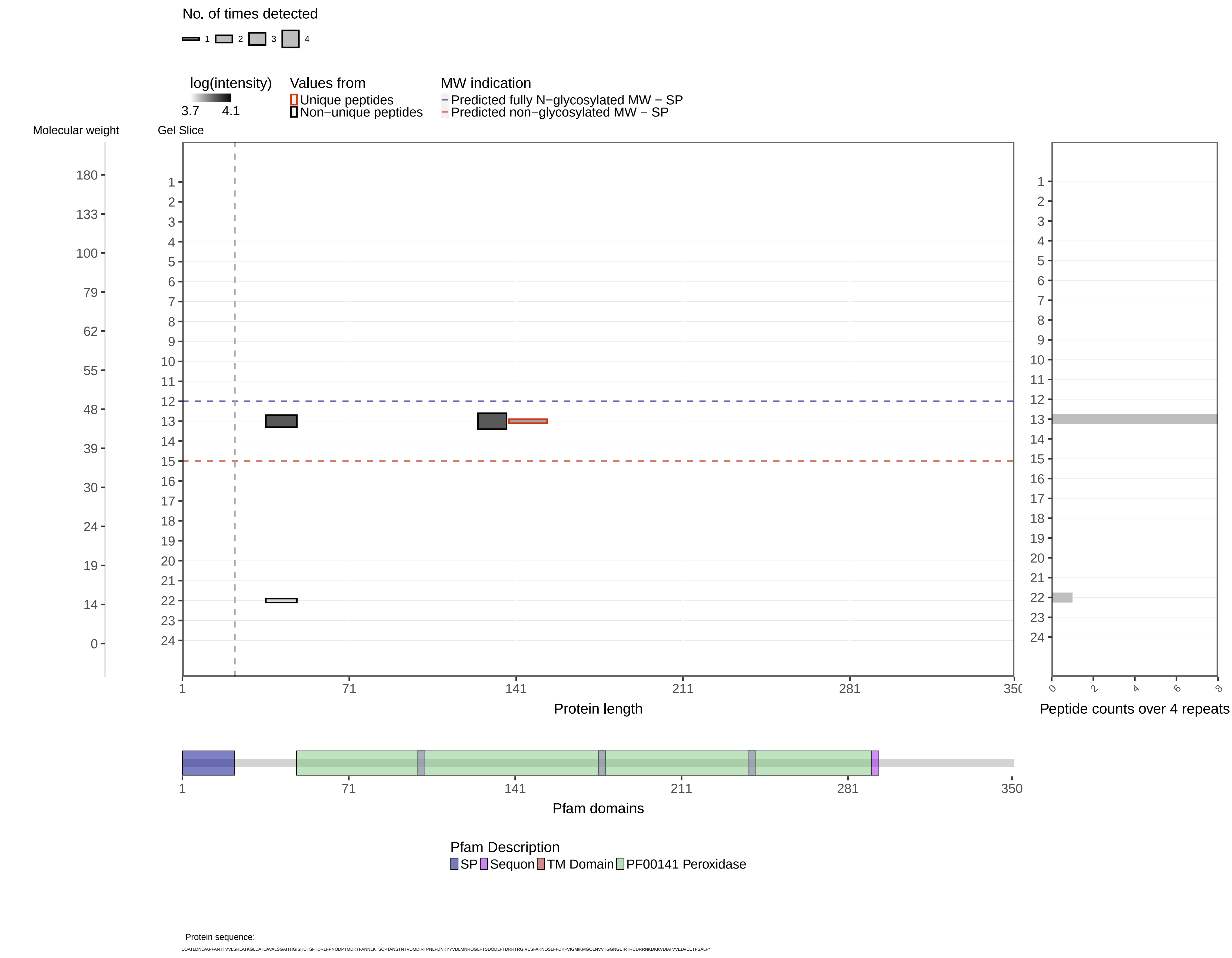

NbL04g09010.1 5−formyltetrahydrofolate_cyclo−ligase−like_protein_COG0212 [MW = 39.89 kDa]

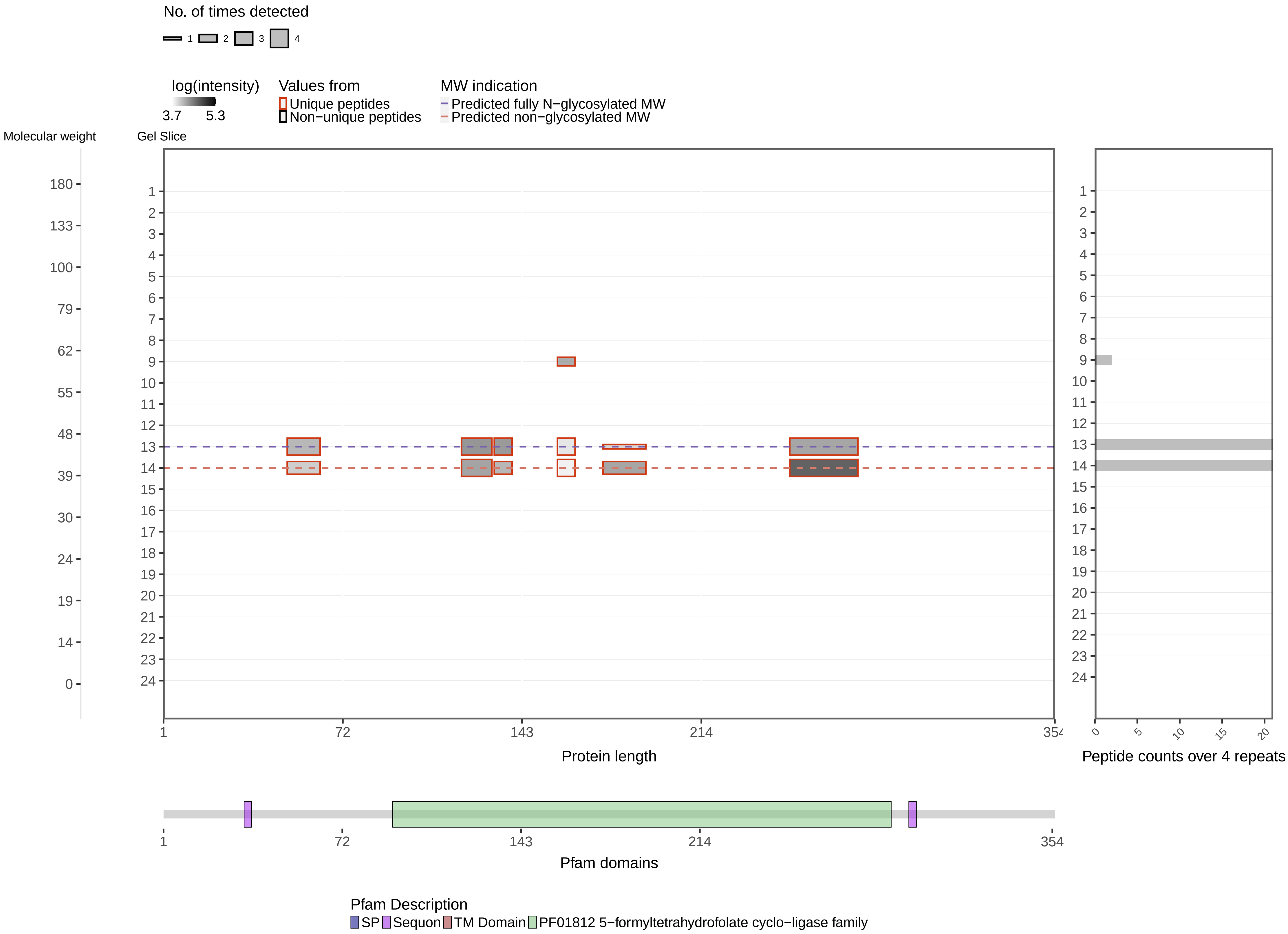


Protein sequence:

GGKKLLTPQPRLRTGFFSVLESSMLSPGTIKEACTSAGVAKYGRPIGLEEKIKVDLIVIGSVAVDPKTGARLGKGEGFAELEYGMLRYMGAIDDSTPVVTTVHDEQLVDDIPIDKLLIHDVPVDIICTPTQVIFTNTSIPKPQGIYWEKLYPEKLSQIRILRQLKSKIEQETGRKLPTGPSEKLPPTAQRRR*

NbL04g08930.1 aminoacylase−1 [MW − SP = 47.74 kDa]

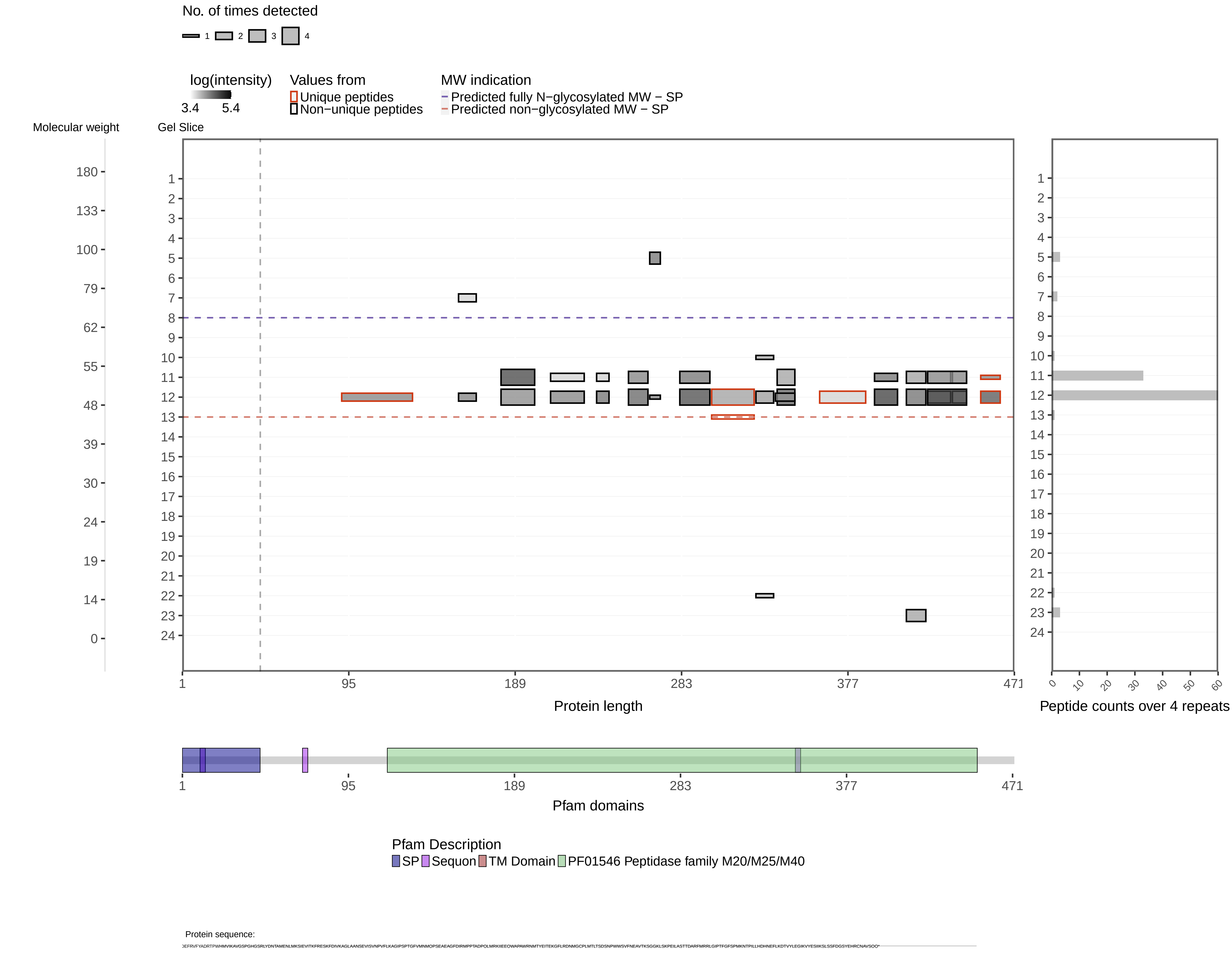

NbL04g08840.1 synthase/flavanone_3−hydroxylase−like [MW = 39.87 kDa]

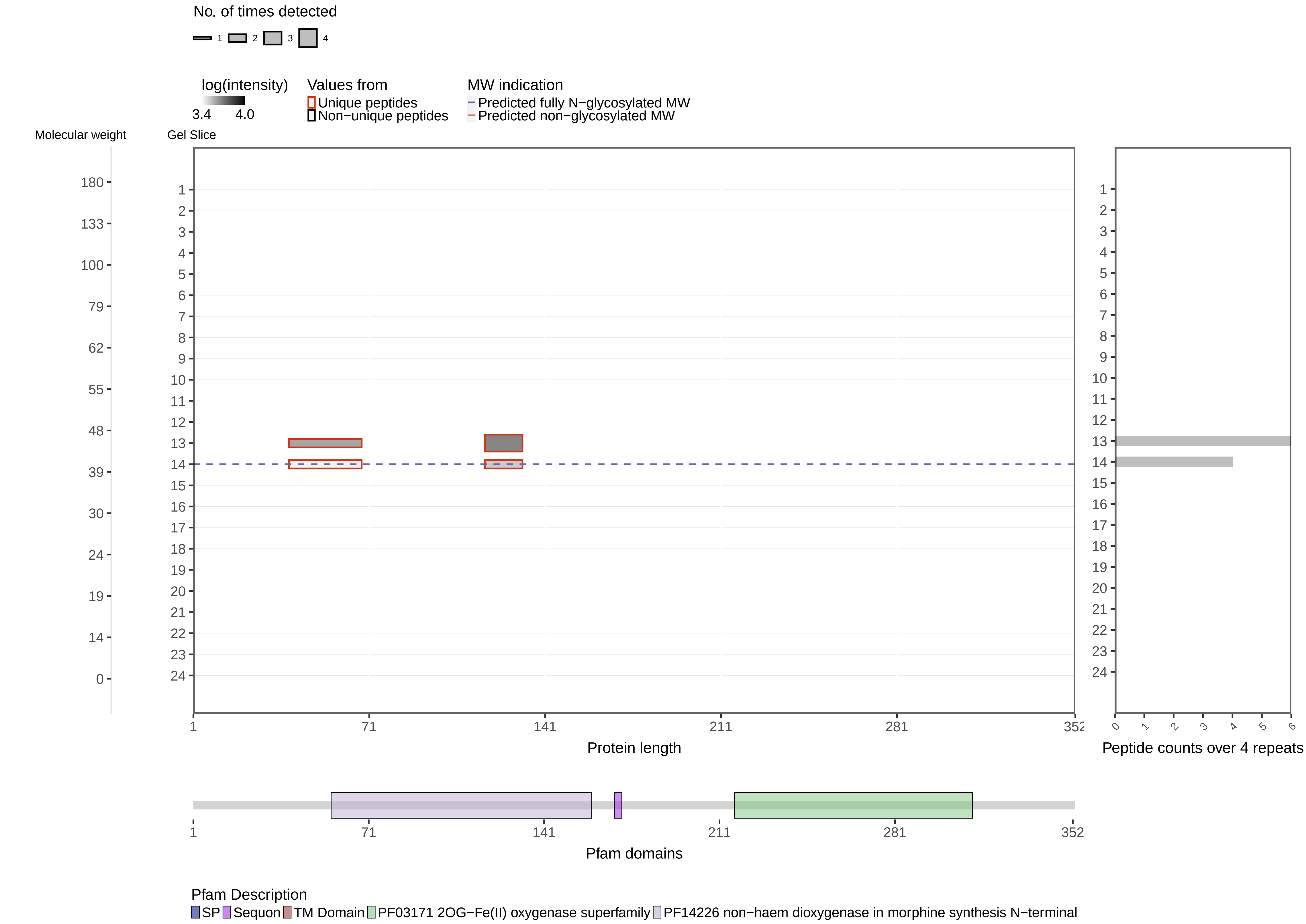


Protein sequence:

PSYRKNLSREANEEYAERMREVVEKIFKSLSLGLGLEAHEMMEAAGGEDIVYLLKINYYPPCPRPDLALGVVAHTDMSYITILVPNEVQGLQVFKDGYWYDVKYIPNALIVHIGDQVEILSNGKYKSVYHRTTVTKDKTRMSWPVFLEPPSEREVGPIPKLVNEANPPKFKTKKYKDYVYCKLNKLPQ*

### NbL04g08780.1 probable_N−succinyldiaminopimelate_aminotransferase_DapC [MW = 50.65 kDa]

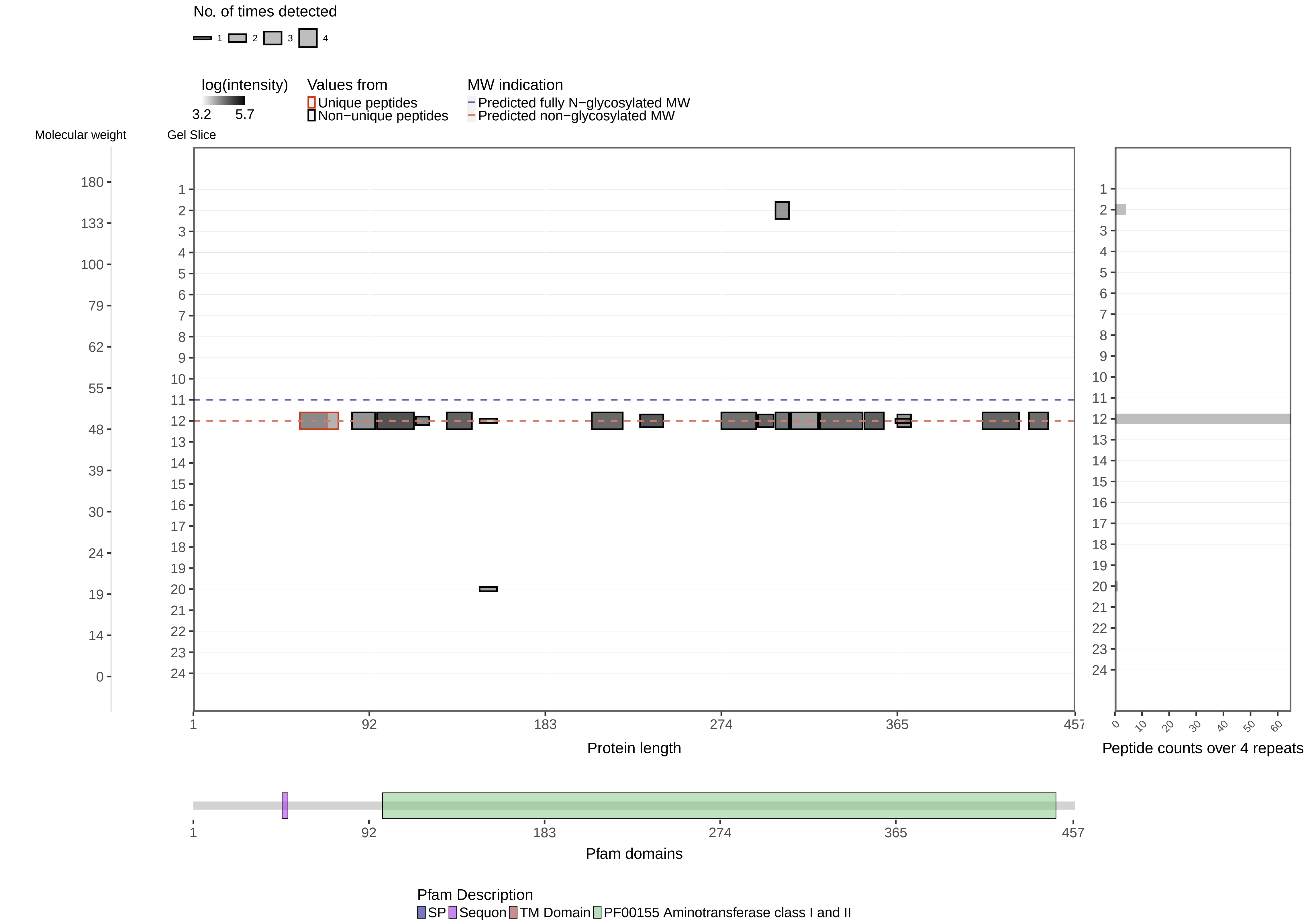


Protein sequence:

FSLPIDELKSAVSKNTRAILLNTPHNPTGKMFTREELNVIASLCIENDILVFADEVYDKLAFEMEHISIASLPGMYERTVTMNSLGKTFSLTGWKIGWAIAPPHLTWGVRQAHSYLTFATSTPMQYAAATALRAPDSYYEELKRDYSAKKAILVEGLKAAGFTVYPSSGTYFVVVDHTPFGLENDIAFCEYLIKEVGVVAIPTSVFYLNPEEGKNLVRFTFCKDDDTLKSAVQRMKEKLSKK*

### NbL04g08680.1 pathogenesis−related_protein_5−like [MW − SP = 27.88 kDa]

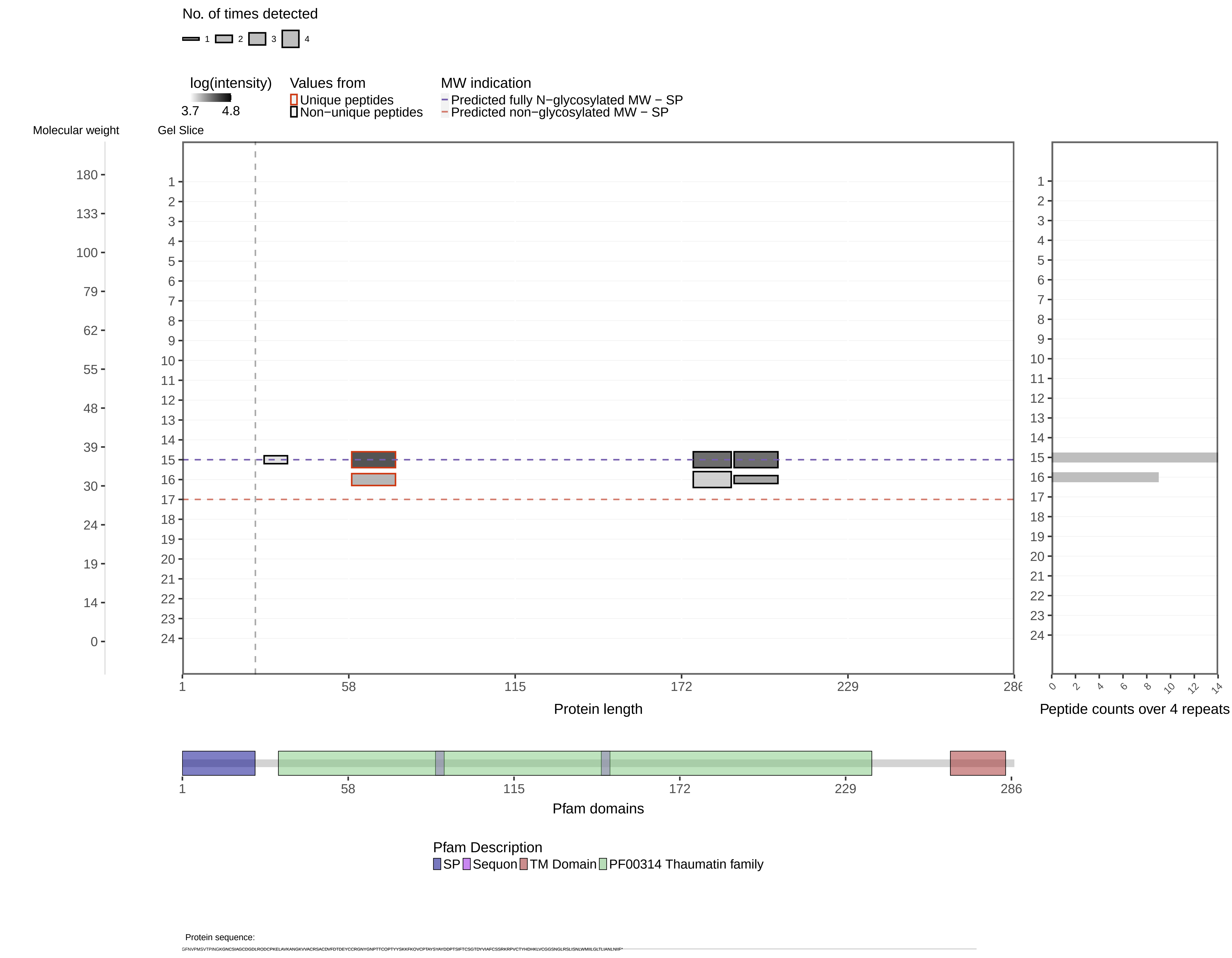

NbL04g08580.1 protein_GrpE−like [MW = 36.73 kDa]

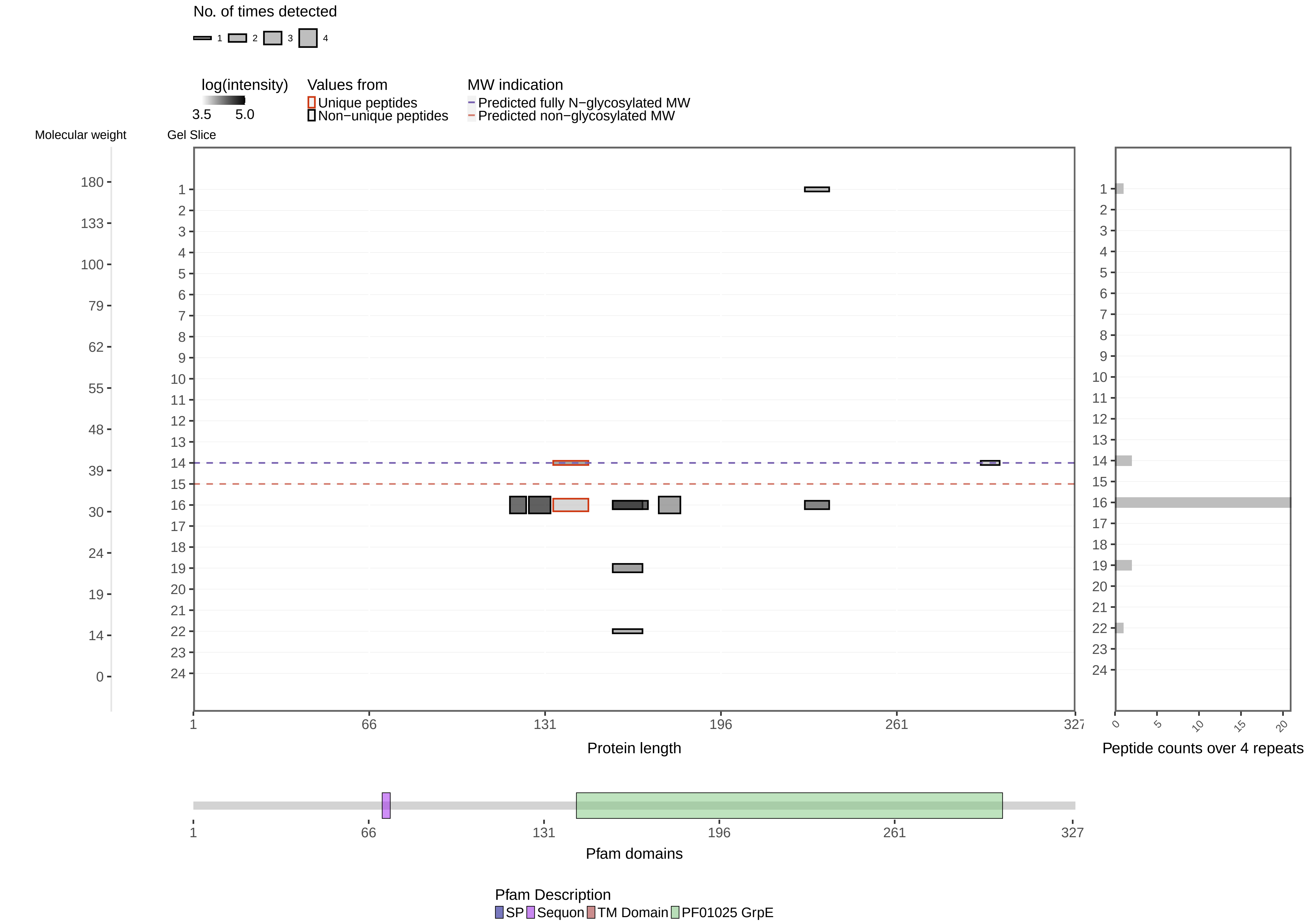


Protein sequence:

DDLAQKVSALSAEIDSGKEKYIRLQADFDNFRKRSEKERLTIRTITQGEIIESLLPMVDNFERAKRQIKLETETEKKIDASYQGIYKQFVEIMKSLRVAAVPTVGKPFDPALHEAVAREESQEFTDGIIIEEFRRGFLLGDRLLRPAMVKVSAGPGKRIPSSVAQKSAAATARVDDN*

### NbL04g08430.1 nitrate_reductase_[NADH]_2 [MW = 102.02 kDa]

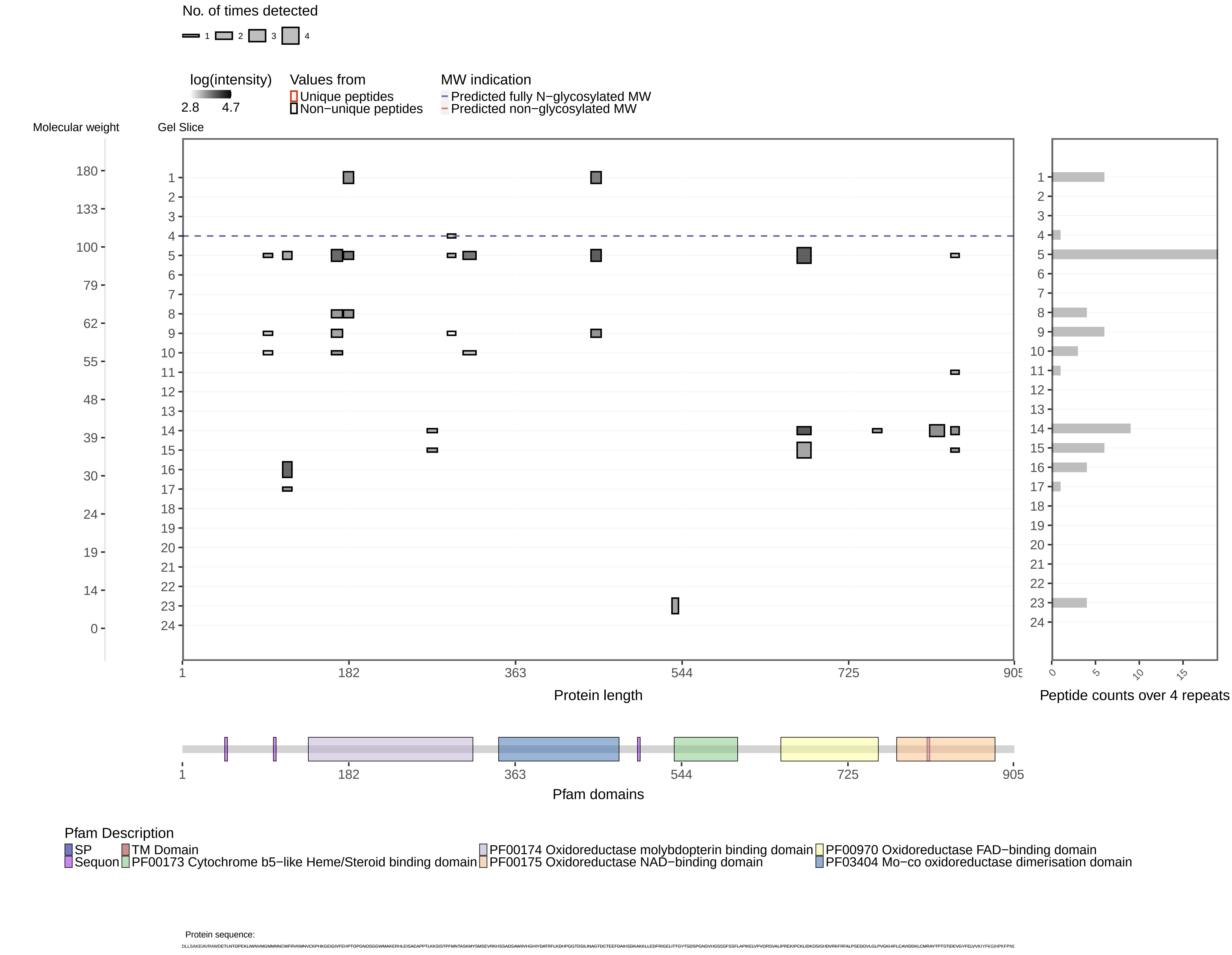

### NbL04g08230.1 eukaryotic_translation_initiation_factor_3_subunit_I−like [MW = 19.54 kDa]

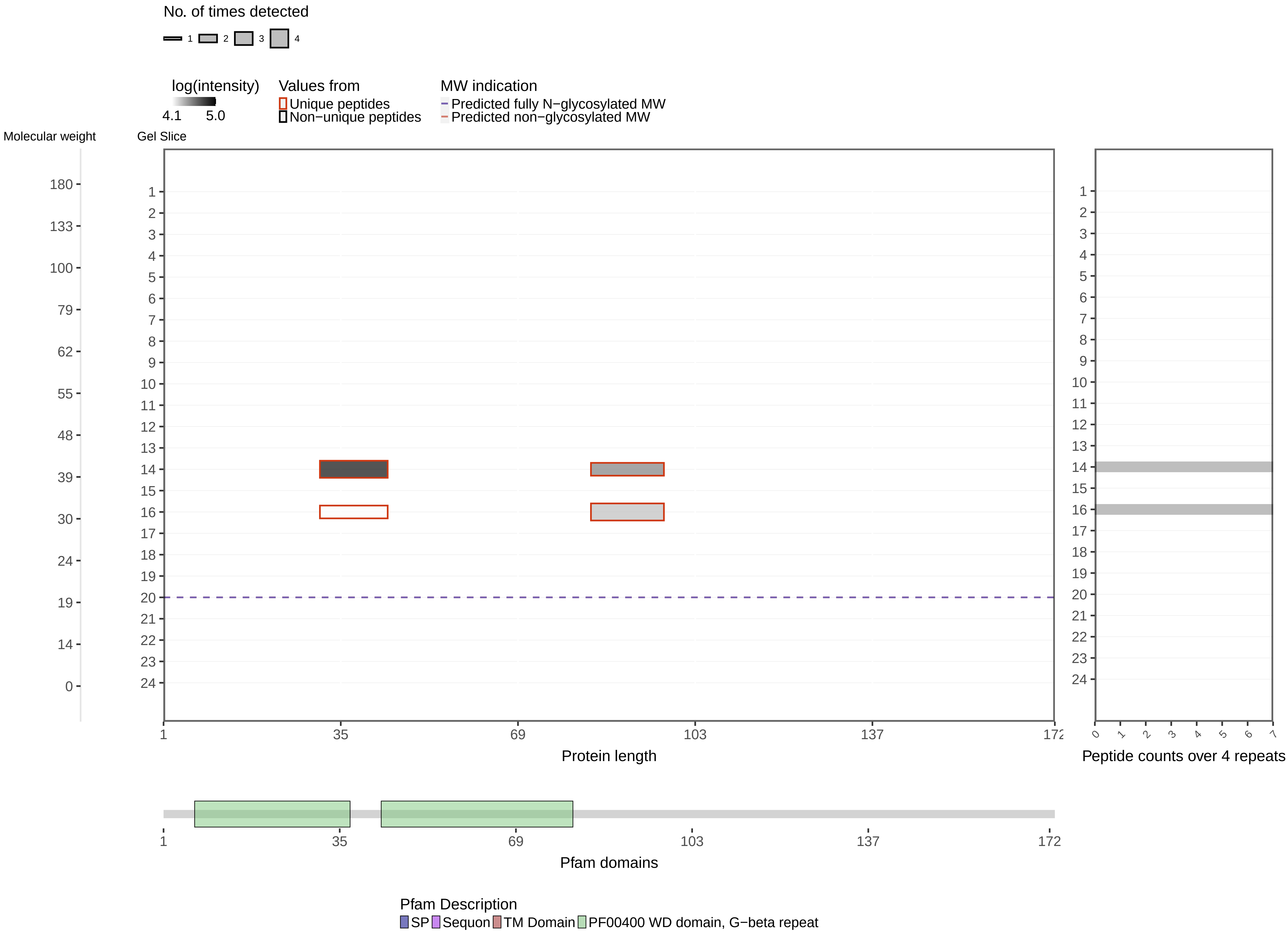


Protein sequence:

ADQSAKLWDVRTGTQLFTFNFDSPARSVDFSVGDKLAVITTDPFMGLPSAIHVKRISRDPSDRHPVHSLVSRLEYLYHWPLSPLRYYILVLEHDTFGYVK*

NbL04g08140.1 dihydrolipoyllysine−residue_acetyltransferase_component_5_of_pyruvate_dehydrogenase_complex_chloroplastic [MW = 49.49 kDa]

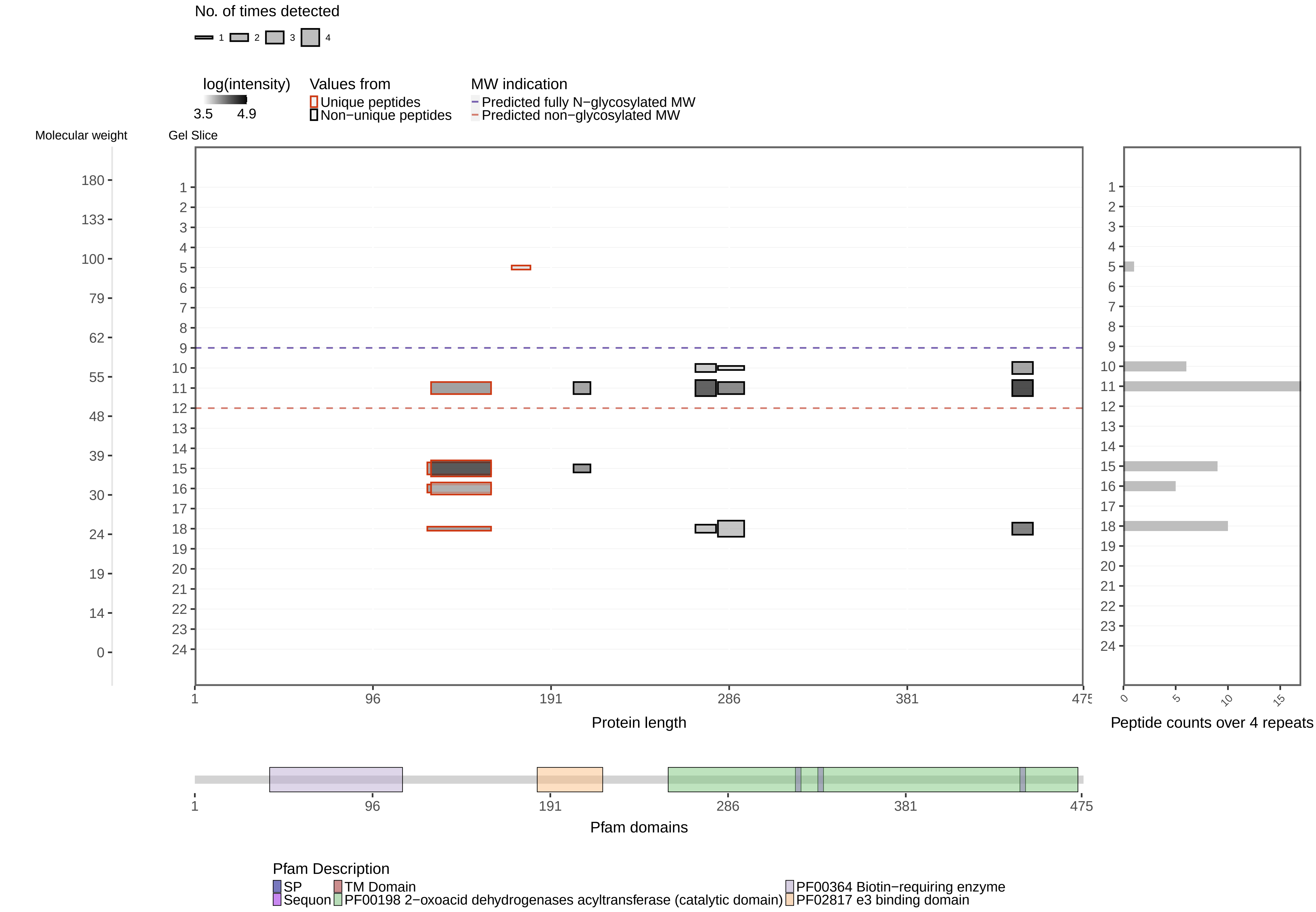


Protein sequence:

AASTPAPAVGTTVAQPSGGAPVAPAVELGTTVPFTTMQNAVSRNMVESLAVPTFRVGYTITTNALDALYKKIKSKGVTMTALLAKATALALAKHPVVNSSCRDGKSFTYNSSINIAVAVAIDGGLITPVLQDADKVDIYSLSRKWKELVDKARAKQLQPHEYTTGTFTLSNLGMFGVDLFDAILPPGTGAIMAVGASHPALVGTKDGRIGLKNQMQVNVTADHRVIYGADLASFLQTLAQIIEDPKDLTL*

NbL04g07980.1 uncharacterized_LOC109225373 [MW = 82.78 kDa]

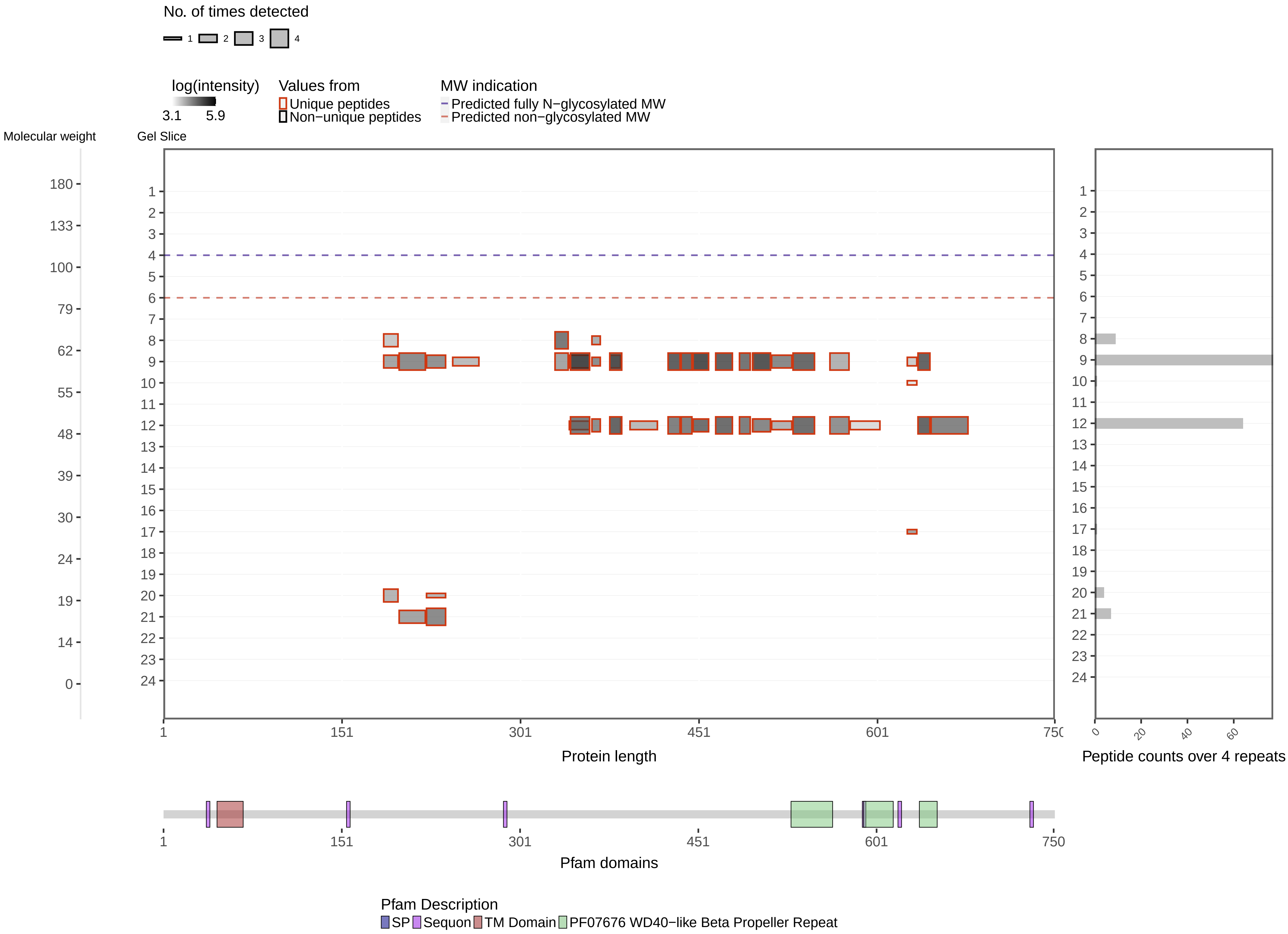


Protein sequence:

RASSEYRHIELFDVVLKKFIEITRPASPHAHHLNPFFSPDSSWVGYHKCRGTGNGKGSDTLLLENLKNPIPGISLFRVDGSFPSFSPSGDRIAYVKMPGLYVVNYDGSGLRQVSTRMAFSTAWDPKRKGVVYTSVGPTFASERTEVDVISINVDDKDLSYKQLTTGGQNNAFPSPSPDGKWIVFRSGRSGHKNLYVMDALEGETGGLYRLTEGSWTDTMCNWSPDDDWIAFASDRENPGSGSFDMFMVHPNGTGLRKVIQSGEGGRTNHPYFSPDGKYIVFTSDYAAVS

### NbL04g07770.1 NAD(P)H−quinone_oxidoreductase_subunit_M_chloroplastic [MW = 17.18 kDa]

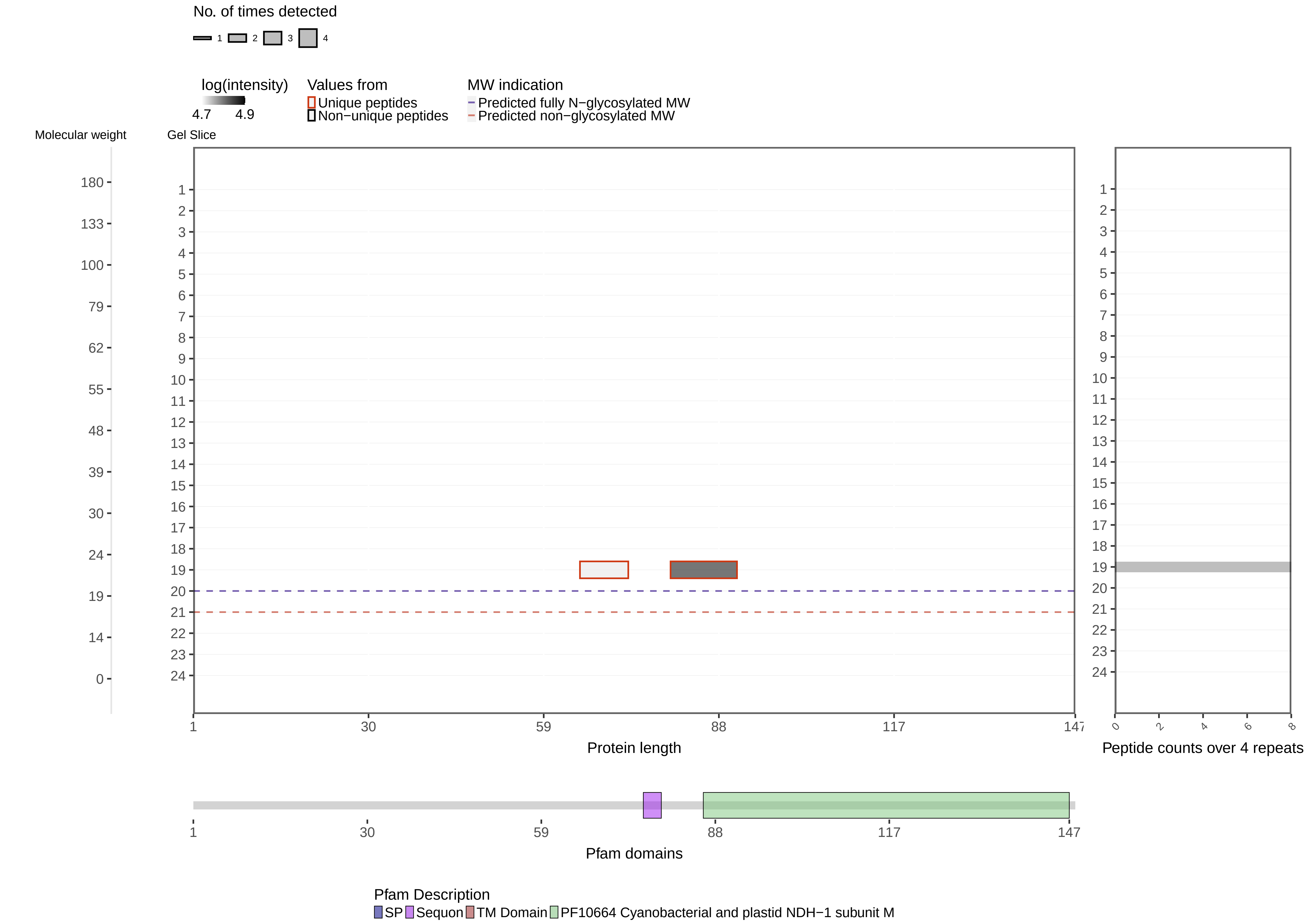


Protein sequence:

MKQPRPVEAQINVKSKNMSREYGGQWLSSATRHVRIYAAYIDPETFAFDQTQMDKLTLILDPTDEFVWTDDTCTKVYSYFQELVDHYE

### NbL04g07740.1 UDP−sugar_pyrophosphorylase [MW = 78.37 kDa]

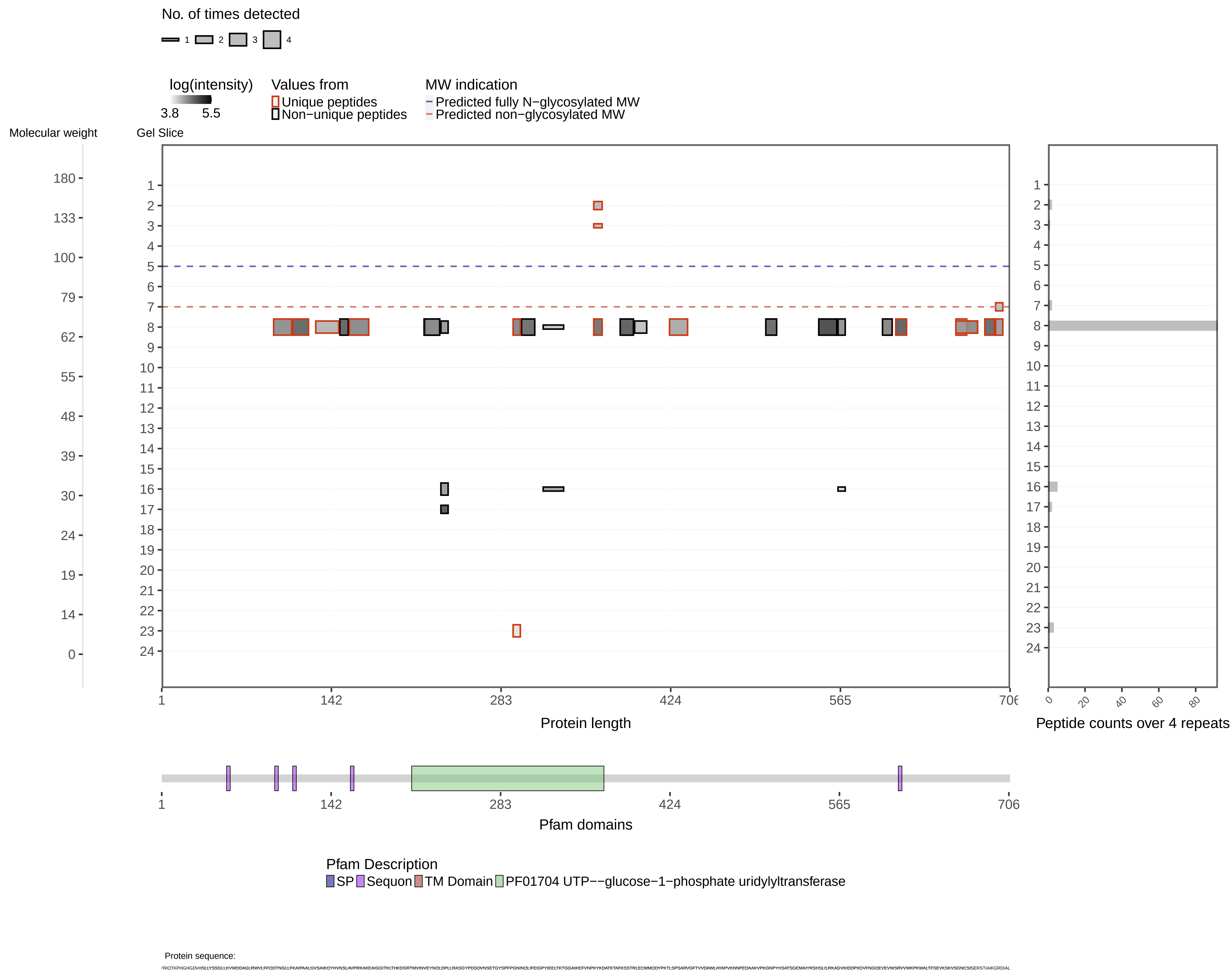

NbL04g07720.1 phosphoribosylformylglycinamidine_cyclo−ligase_chloroplastic−like [MW = 49.34 kDa]

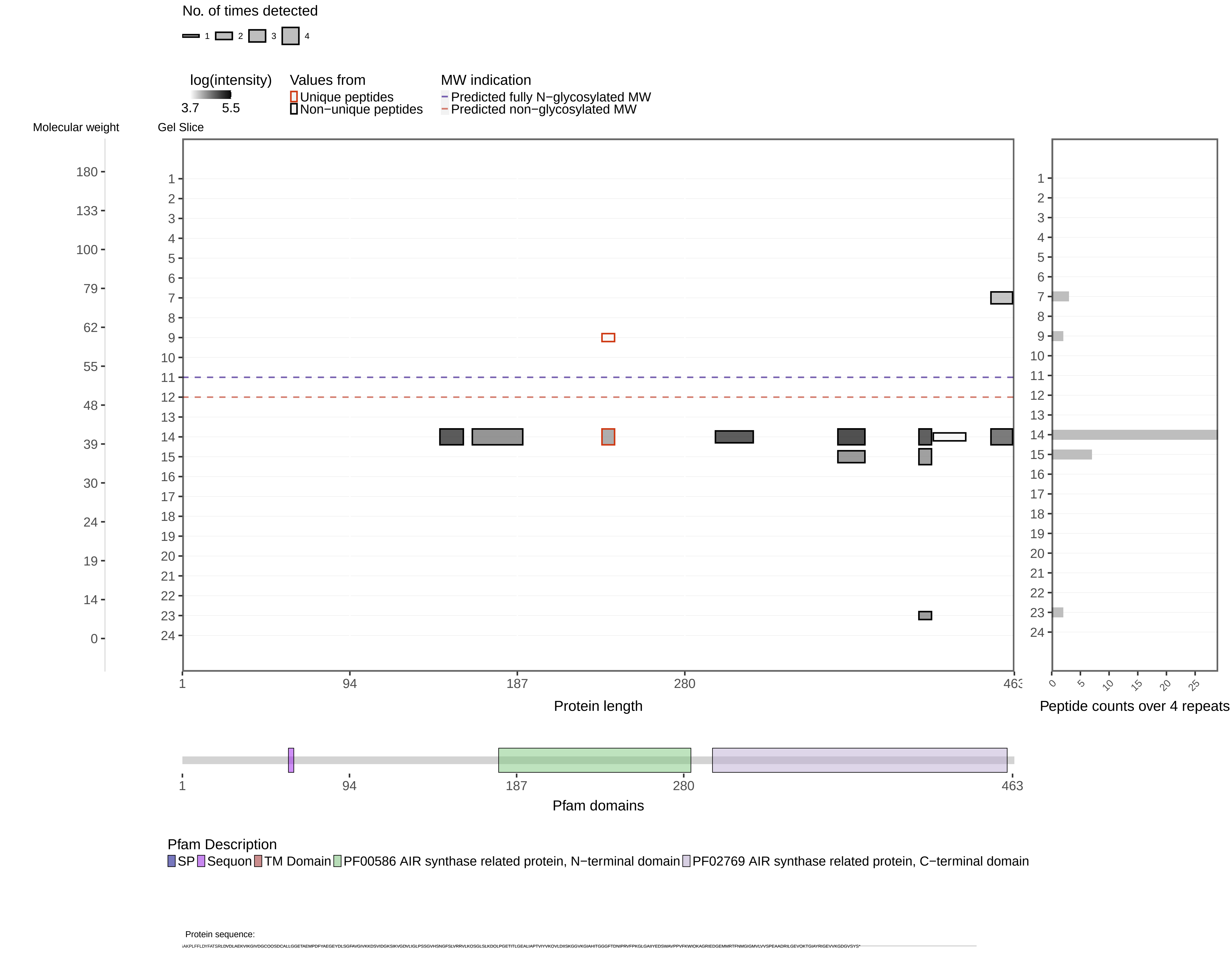

### NbL04g07640.1 probable_methyltransferase_PMT24 [MW = 85.49 kDa]

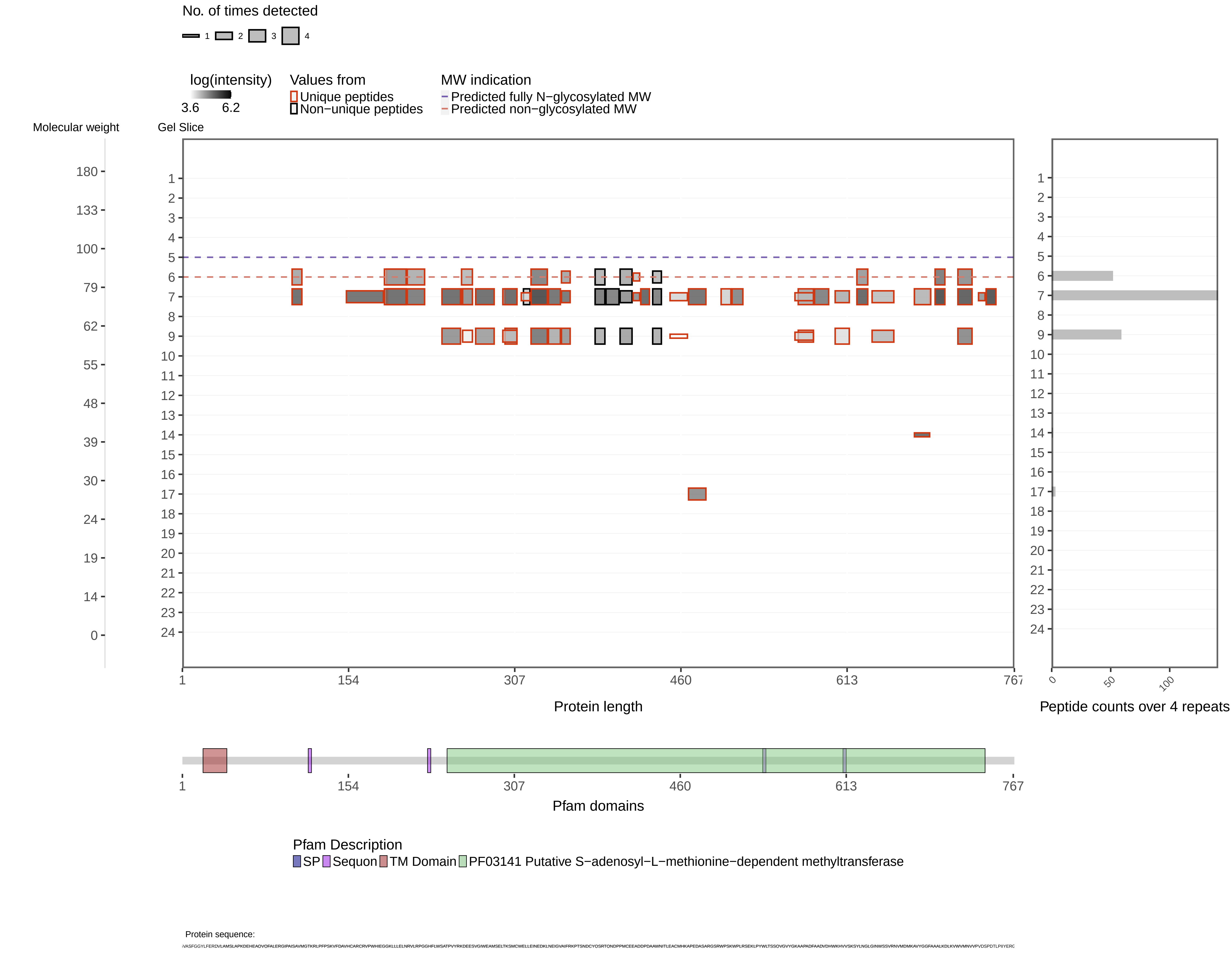

### NbL04g07620.1 TOM1−like_protein_2 [MW = 56.25 kDa]

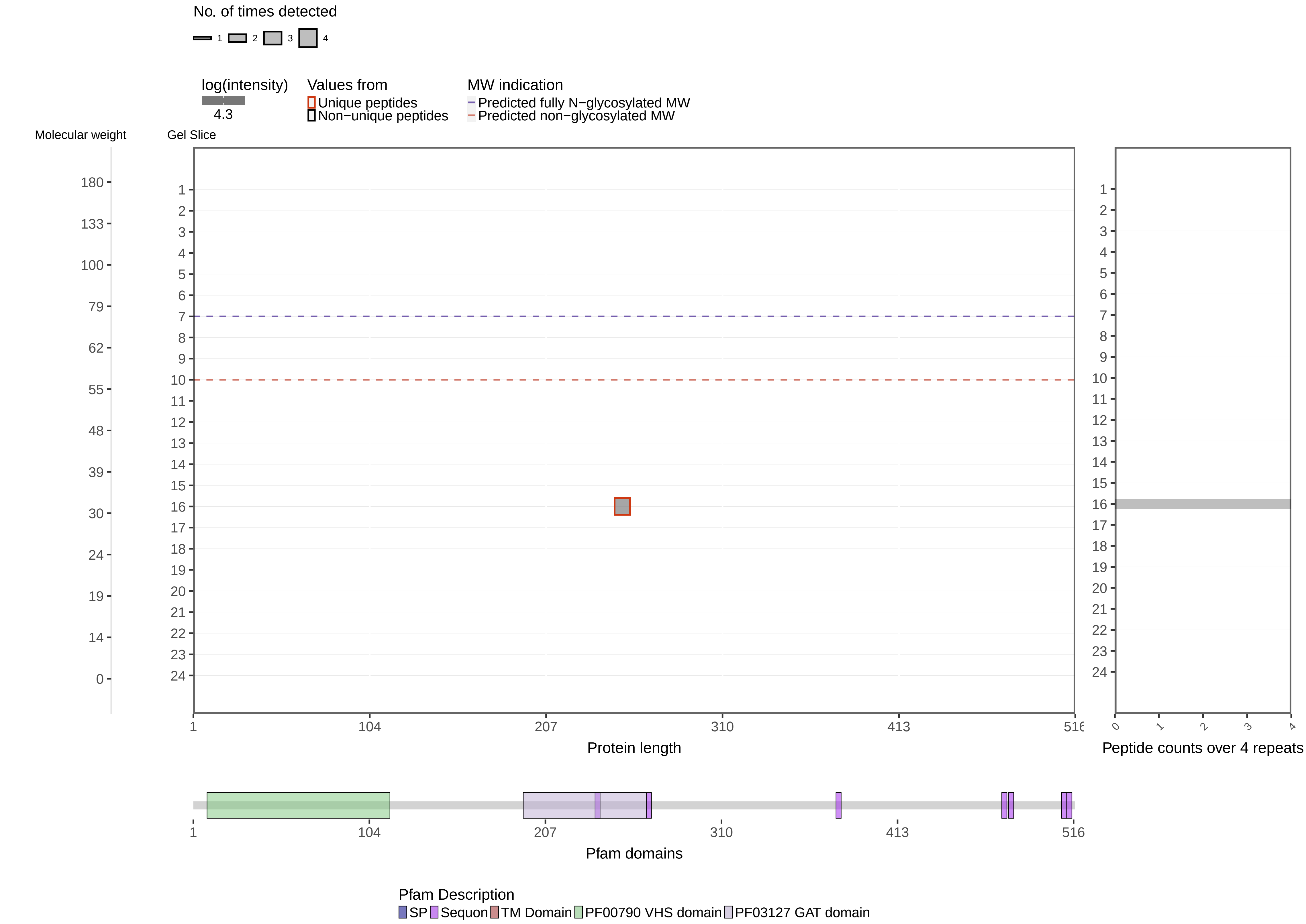


Protein sequence:

### NbL04g07590.1 aspartic_proteinase_Asp1−like [MW − SP = 45.2 kDa]

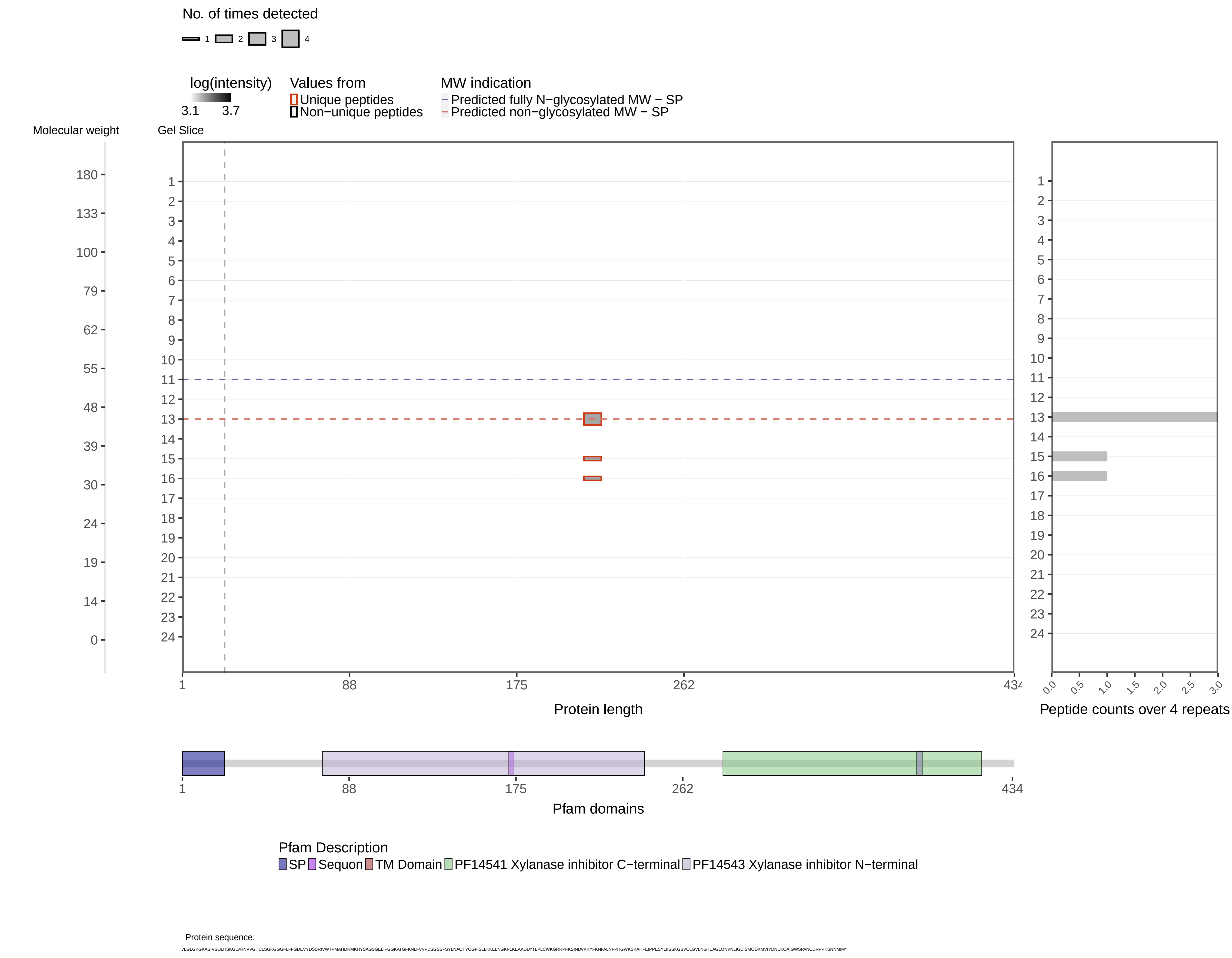

### NbL04g07580.1 2−oxoisovalerate_dehydrogenase_subunit_alpha_1_mitochondrial−like [MW = 51.33 kDa]

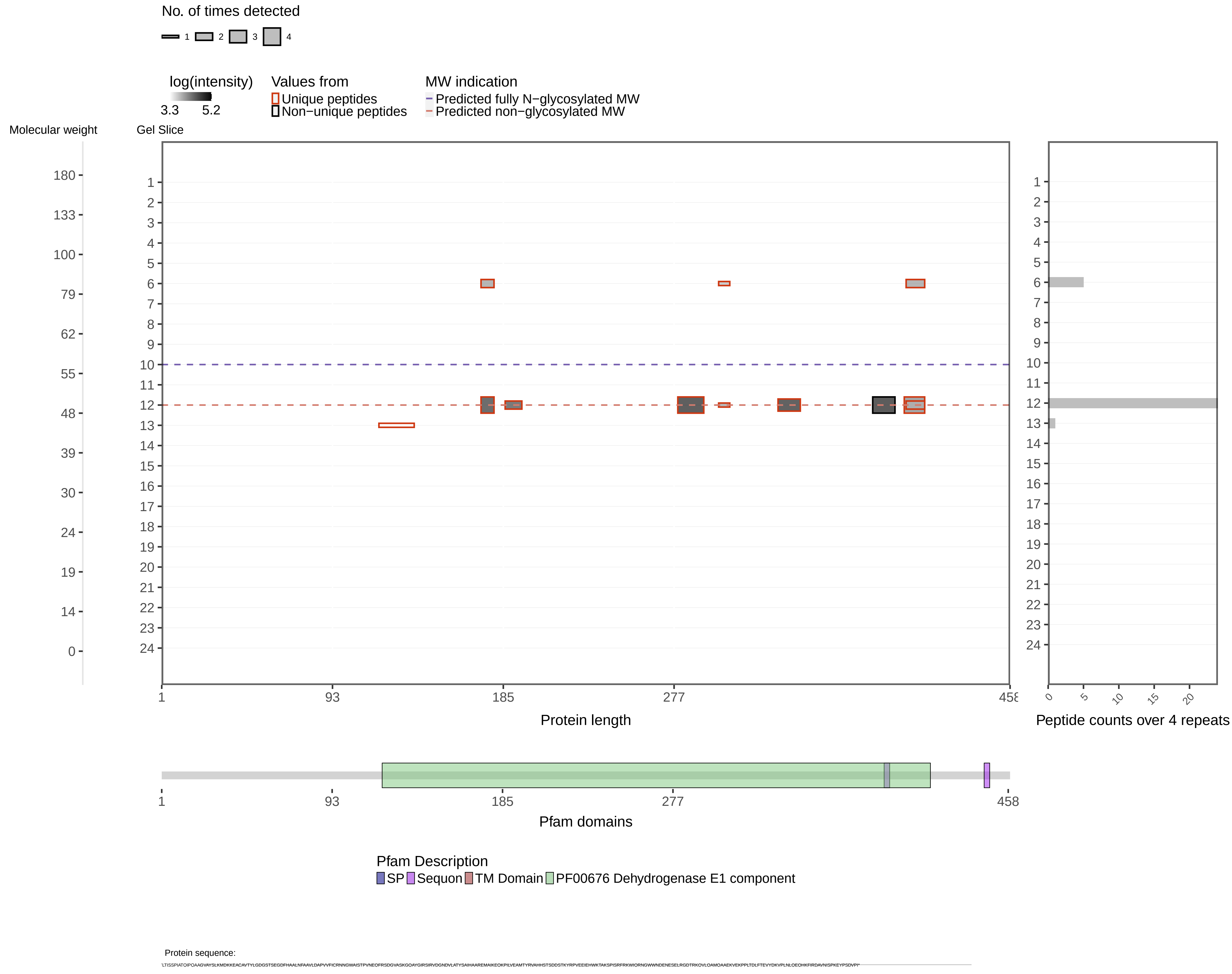

NbL04g07300.1 serine_carboxypeptidase−like_42 [MW − SP = 37.74 kDa]

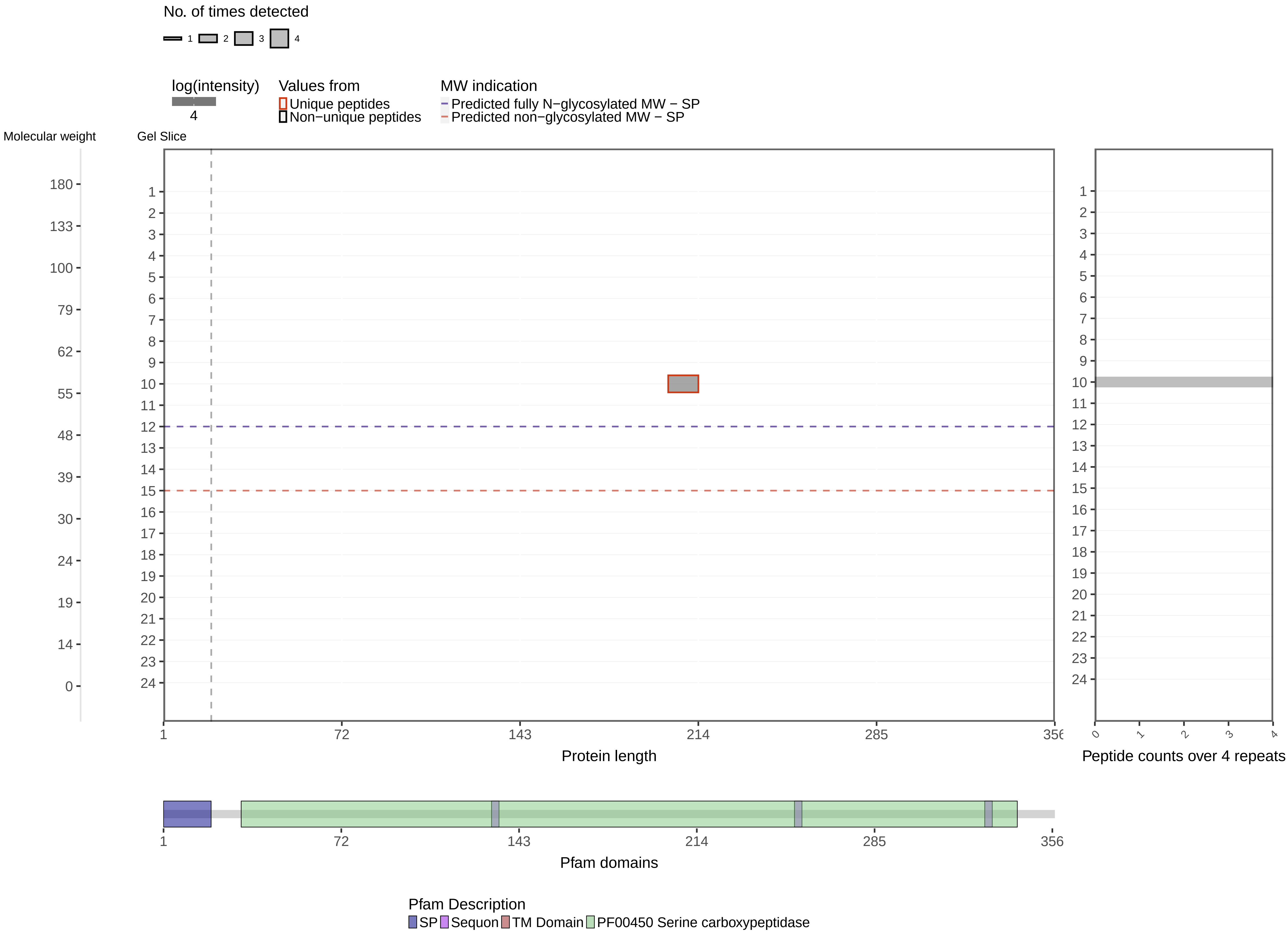


Protein sequence:

RSRELYLTGESYAGHYIPQLAVALLDHNEHSKEYKFNIKGVAIGNPLLNLDRDVPATYEYFWSHGMISDEVGLAIMNQCDFEDYTFGSPHNVSHECNNAISEANDIISKYINNYDVILDVCYPSIVEQELRLRKMATKMSVGVDVCMSYERRFYFNLLEVQKALHANRTNLAYSWSMCSGTGLLLLLQYSELQ*

### NbL04g07230.1 pathogen−related_protein−like [MW = 26.7 kDa]

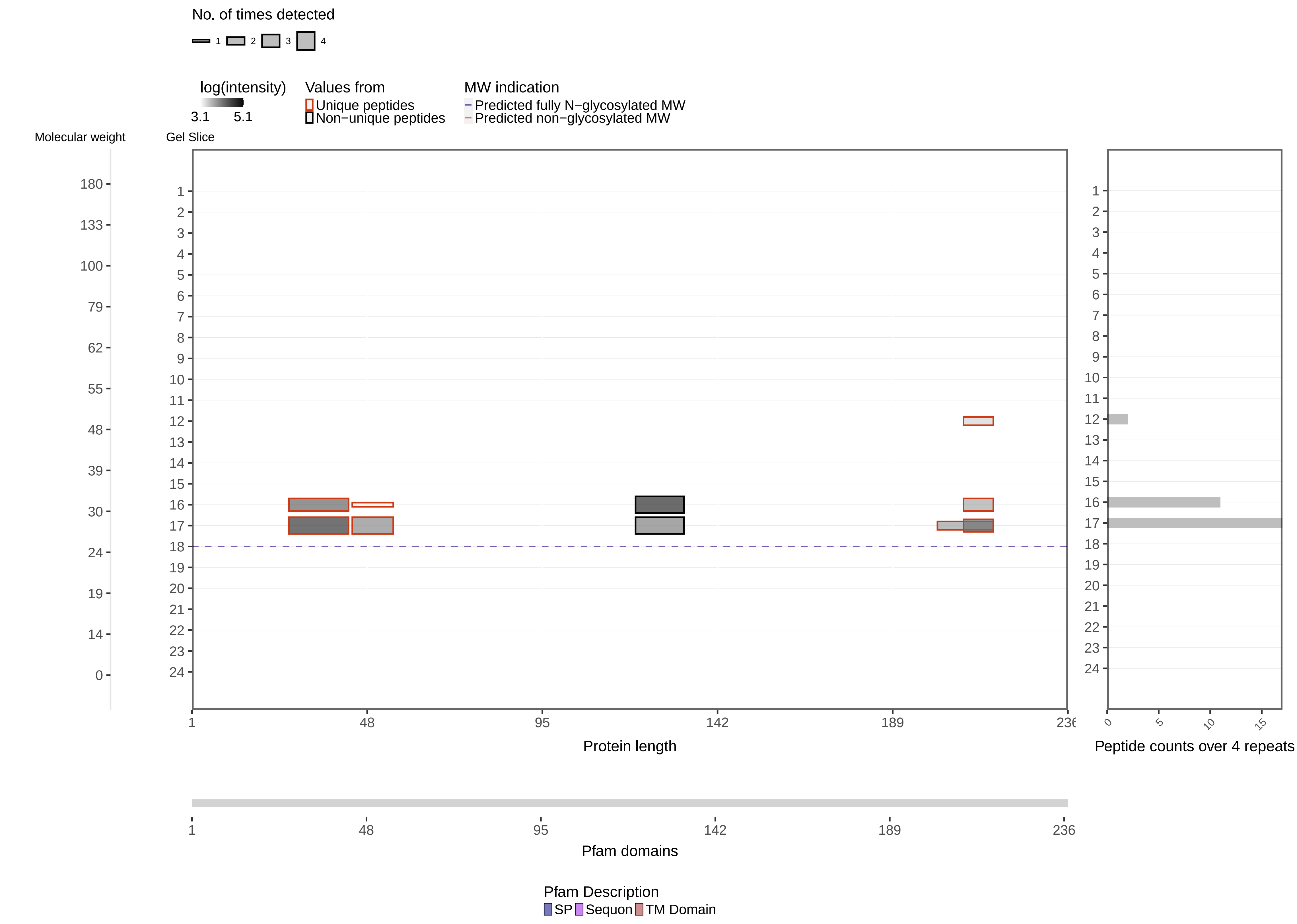


Protein sequence:

GGYNTFLQTSLPESVRVYNPDEETFESSQKVFRSIFLRGFAIEILHVYSGPPEIVYKFRHWGHMEGPFKGHAATGELVQLFGIGIFEVGKESNKIVKAEFFFDRGELLGPLVKGEKNGDSTNAMALACPFMK*

NbL04g07120.1 long_chain_acyl−CoA_synthetase_9_chloroplastic [MW = 76.01 kDa]

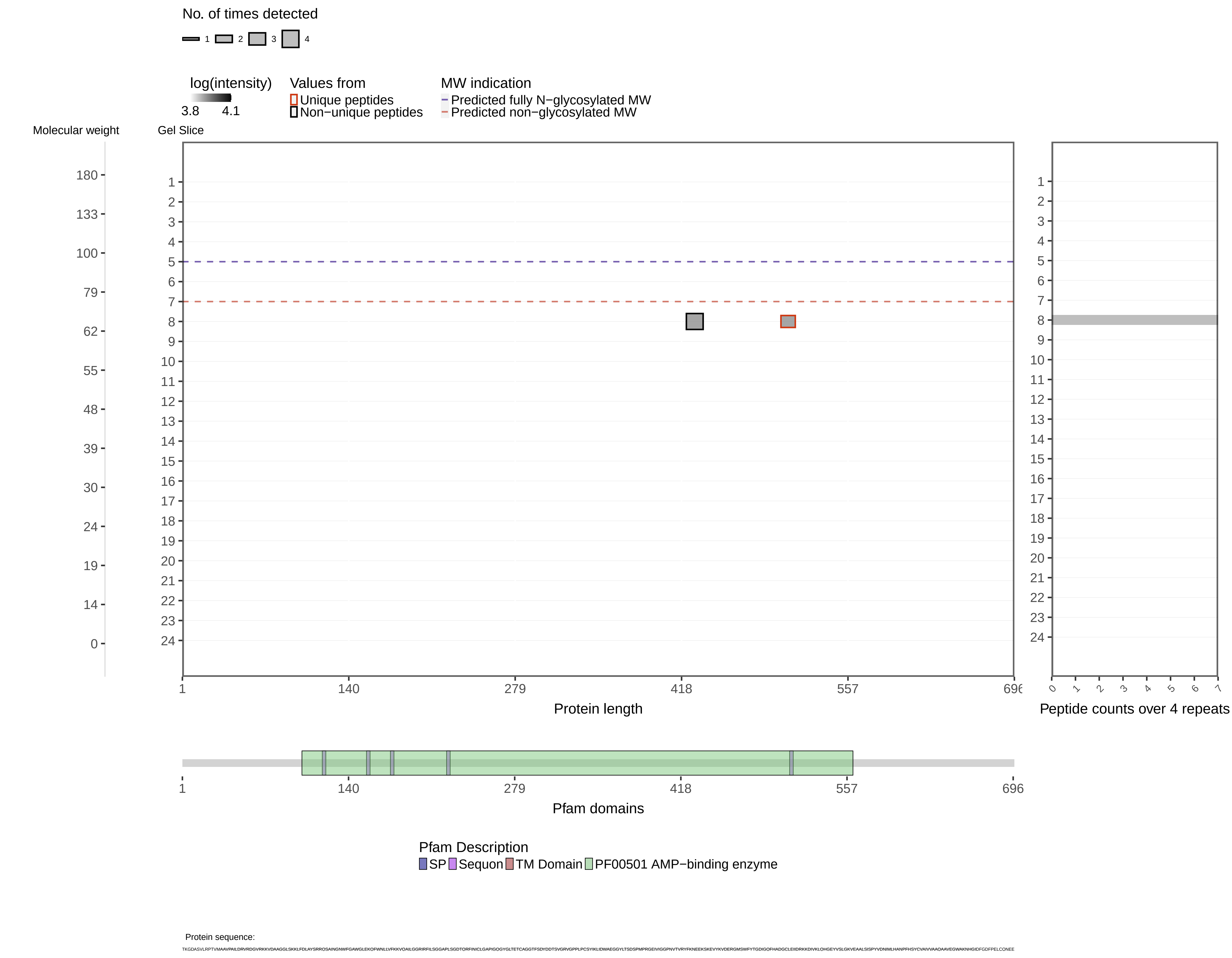

### NbL04g07090.1 50S_ribosomal_protein_L25−like [MW = 31.45 kDa]

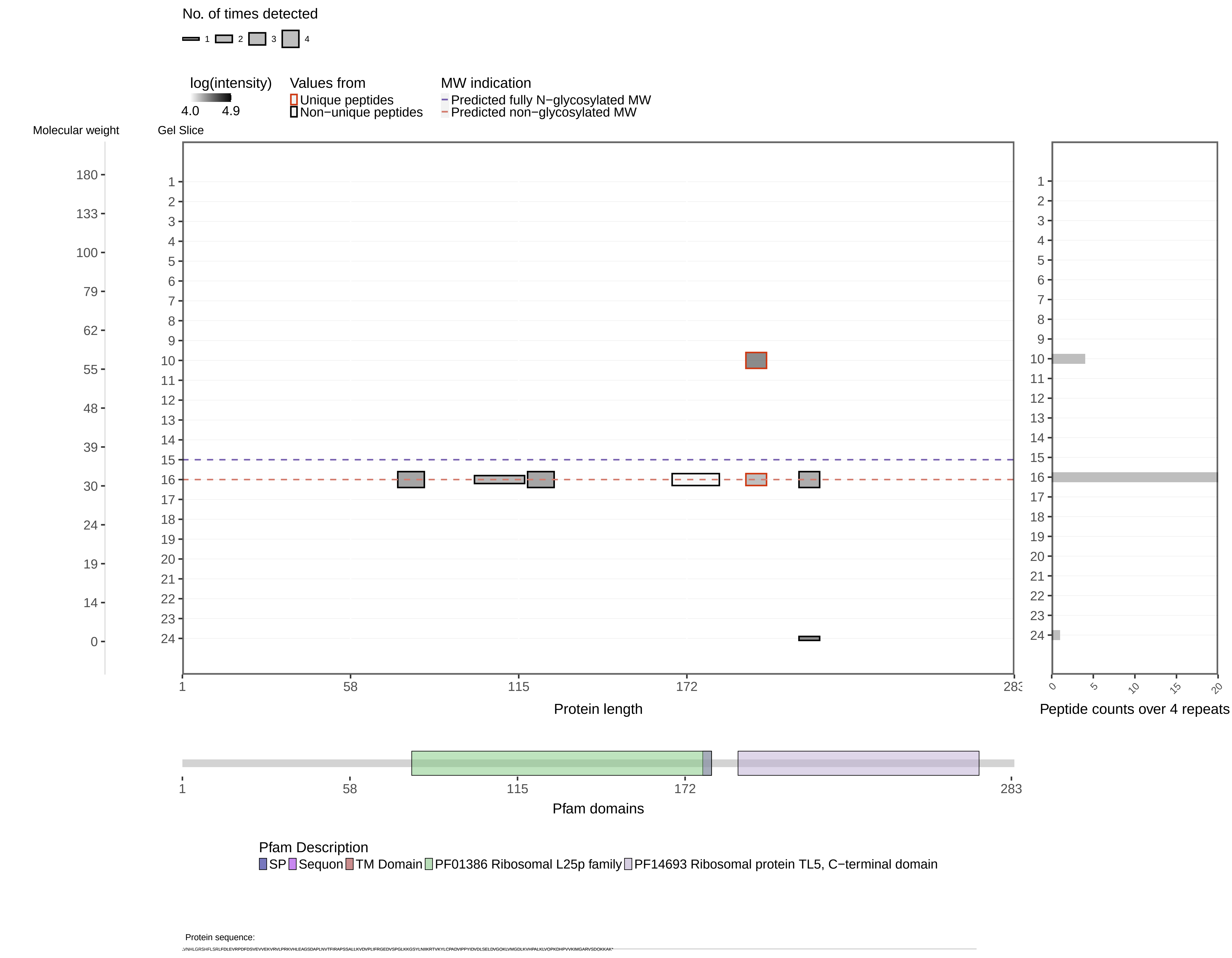

NbL04g07080.1 probable_L−ascorbate_peroxidase_6_chloroplastic [MW = 47.21 kDa]

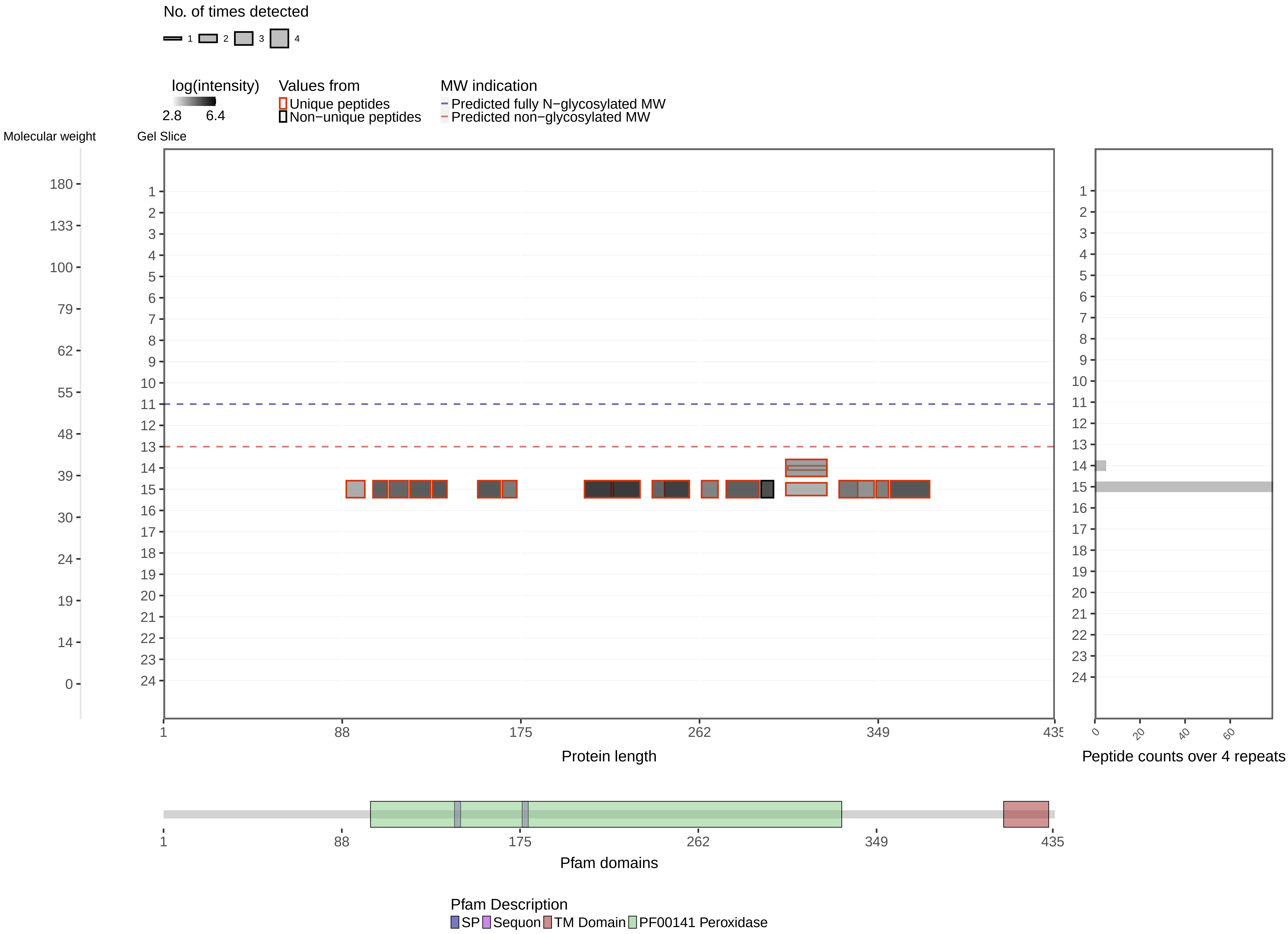


Protein sequence:

DVSVPEECPEEGRLPDAGPPSPASHLRDVFYRMGLNDKEIVALSGAHTLGRSRPERSGWGKPETKYTKDGPGNPGGQSWTVQWLKFDNSYFKDIKERRDEDLLVLPTDAALFEDSSFKEYAEKYAVNQDVFFKDYAEAHAKLSNLGAKFDPPEGFSIDNTPTQGQPEKFVAAKYSTGKRELSDAMKQKIRAEYEGFGGSADKPLPTNYFLNIMIVIGVLAIVTSLLGN*

### NbL04g06910.1 thioredoxin_Y1_chloroplastic−like [MW = 18.96 kDa]

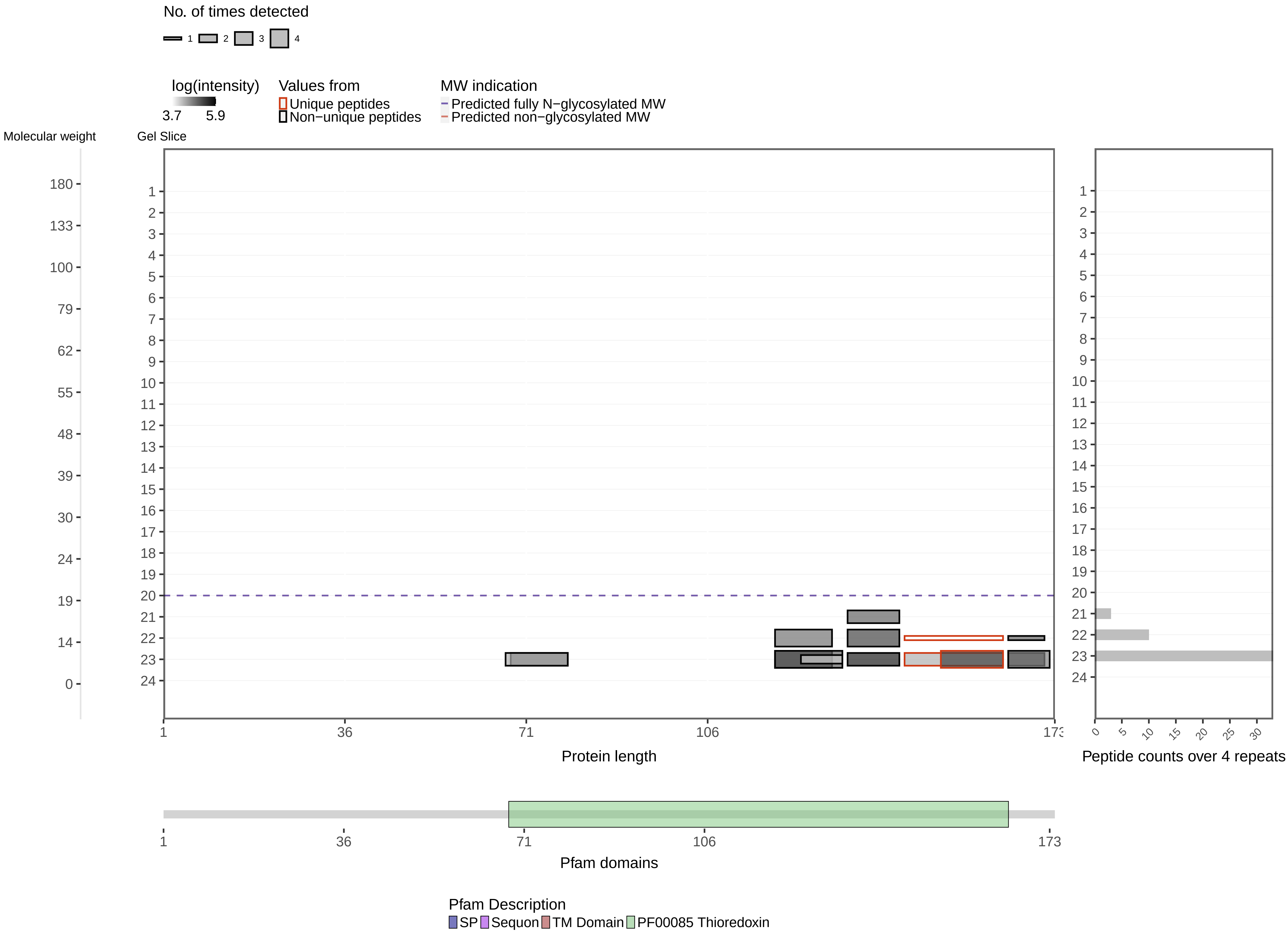


Protein sequence:

FEELLEKSEKPLLVDFYATWCGPCQFMVPILNEVGESMKDKIQVVKIDSEKYPALADKYKIQALPTFILFKDGEVCDRFEGALNAPTLIQRIESALEVKQ*

### NbL04g06790.1 trihelix_transcription_factor_GT−2−like [MW = 76.1 kDa]

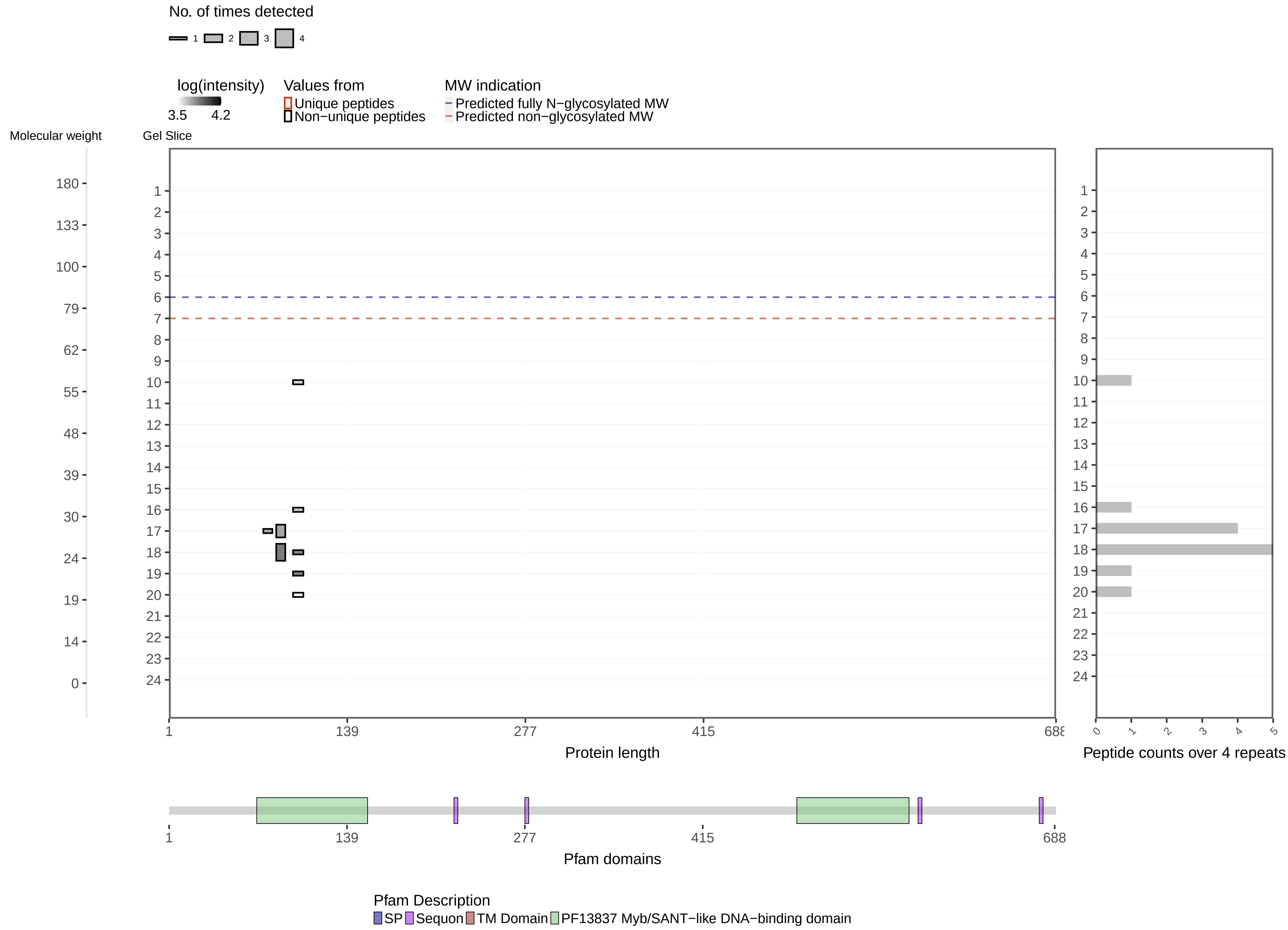


Protein sequence:

EKRERDRMVREEAWRVOEVARMNREHDLLVOERSTAAAKDAAVISFLOKITEOONIOIPSNINVAPPSAOVOIOLPENPPPPSETRSOVVHOOTOPTAVLVSPAPTOPSPATPAPISLPATIPAOSLPLTPPVPAKSVELTPKSDNGGEGYSPASSSRWPKAEVEALIMLRTOLDVKYOENGPKGPLWEEISAGMKKLGYNRNAKRCKEKWENINKYFKKVKESNKKRPEDSKTCPYFHOLDALYKEKAKNETSSSFSNPSASGFALNPENNPMMPIMARPEOOWPRPPHH

### NbL04g06760.1 fructose−16−bisphosphatase_cytosolic [MW = 37.19 kDa]

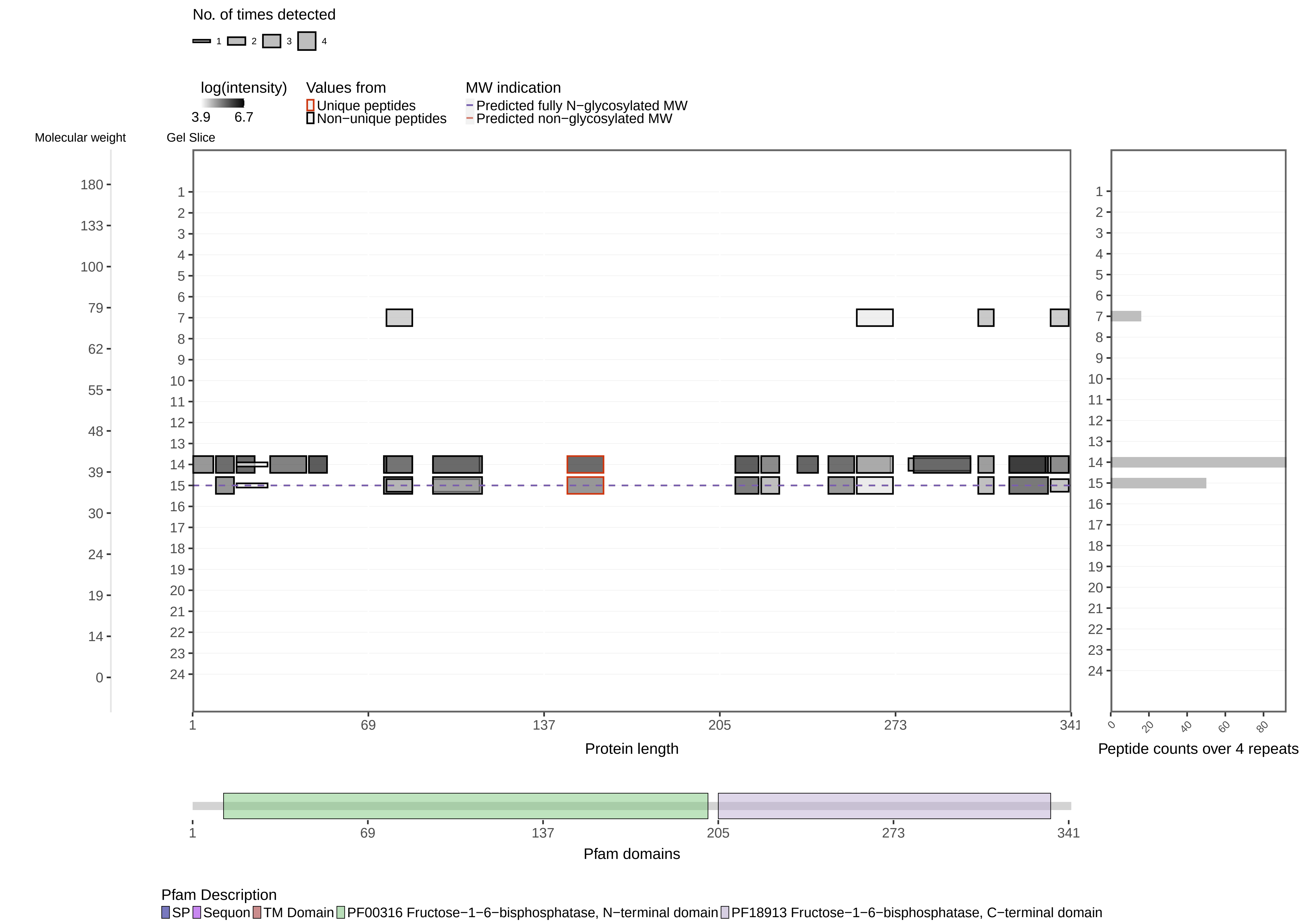


Protein sequence:

GKNMLAAGYCMYGSSCTLVLSTGSGVNGFTLDPSLGEFILTHPDIKIPKKGKIYSVNEGNAKNWDGPTSKYVQNCKYPTDGSSPKSLRYIGSMVADVHRTLLYGGIFLYPADKKSPNGKLRVLYEVFPMSFLMEQAGGQSFTGKQRALDLVPKKIHERAPIFLGSYDDVEEIKKLYAAEEQN*

NbL04g06740.1 uncharacterized_LOC107817613 [MW = 20.35 kDa]

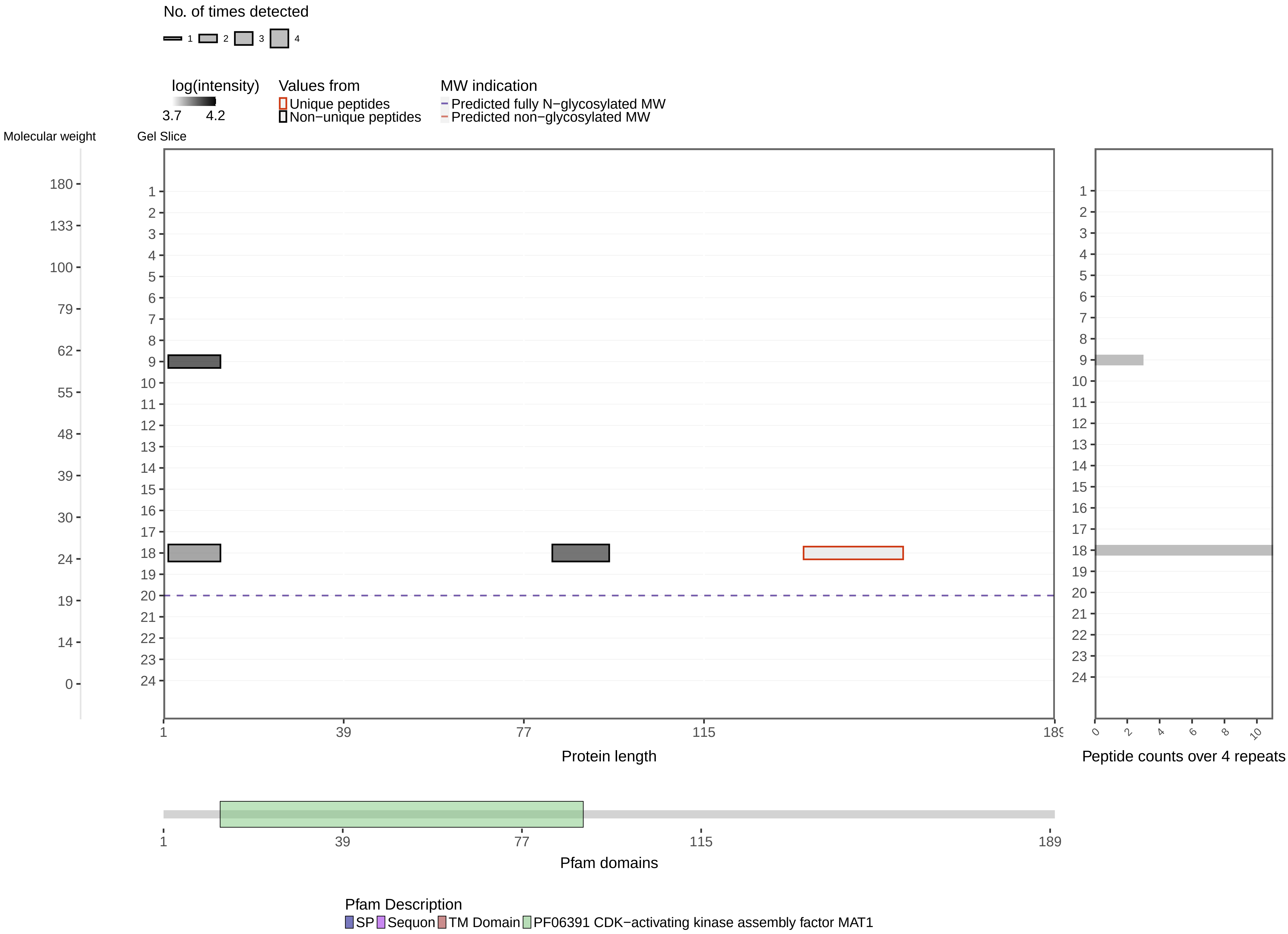


Protein sequence:

AEEYAAALAASKGQPTQPDGDAALGLGPQAGSSTGPQGHYAPAVAGGIAQPRPTQPLPVGSIPGLEYEDEETMKLKAEKAARAGGWSLEISRKRALEEAFGSLWV*

### NbL04g06600.1 RNA−binding_protein_2−like [MW = 27.37 kDa]

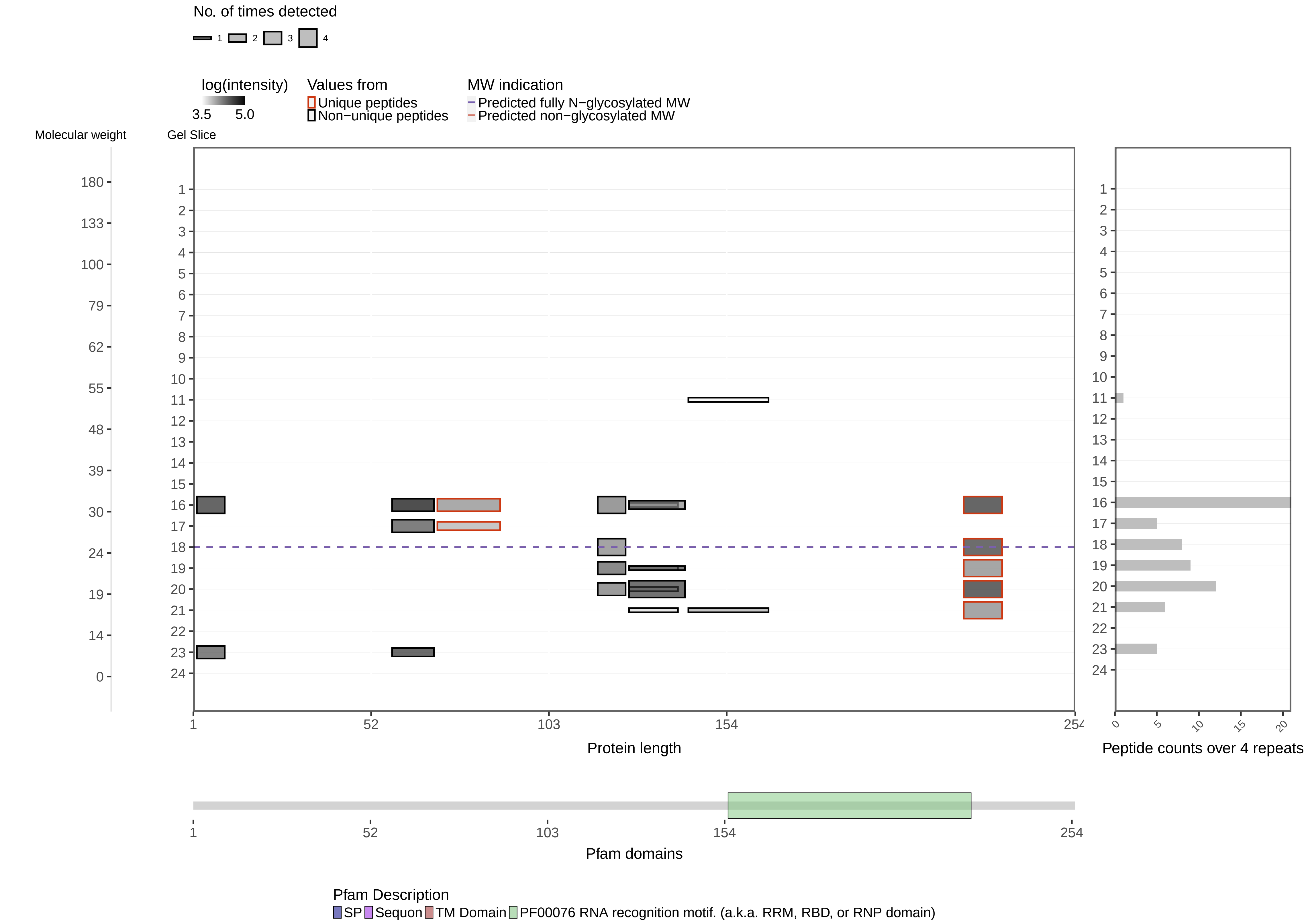


Protein sequence:

SARGPELAPNGRAMVLSGQMPVESLPRPRETLPLPPDASNTLYIEGLPADSSRREVAHIFRPFVGYKEVRLVRKESKHRGGDPLILCFVDFMDPACAATALSALQGYKMDEHDPDSAYLRLQFSKFPGPRSGGSGSRGKR*

NbL04g06540.1 cycloartenol_Synthase−like [MW = 86.02 kDa]

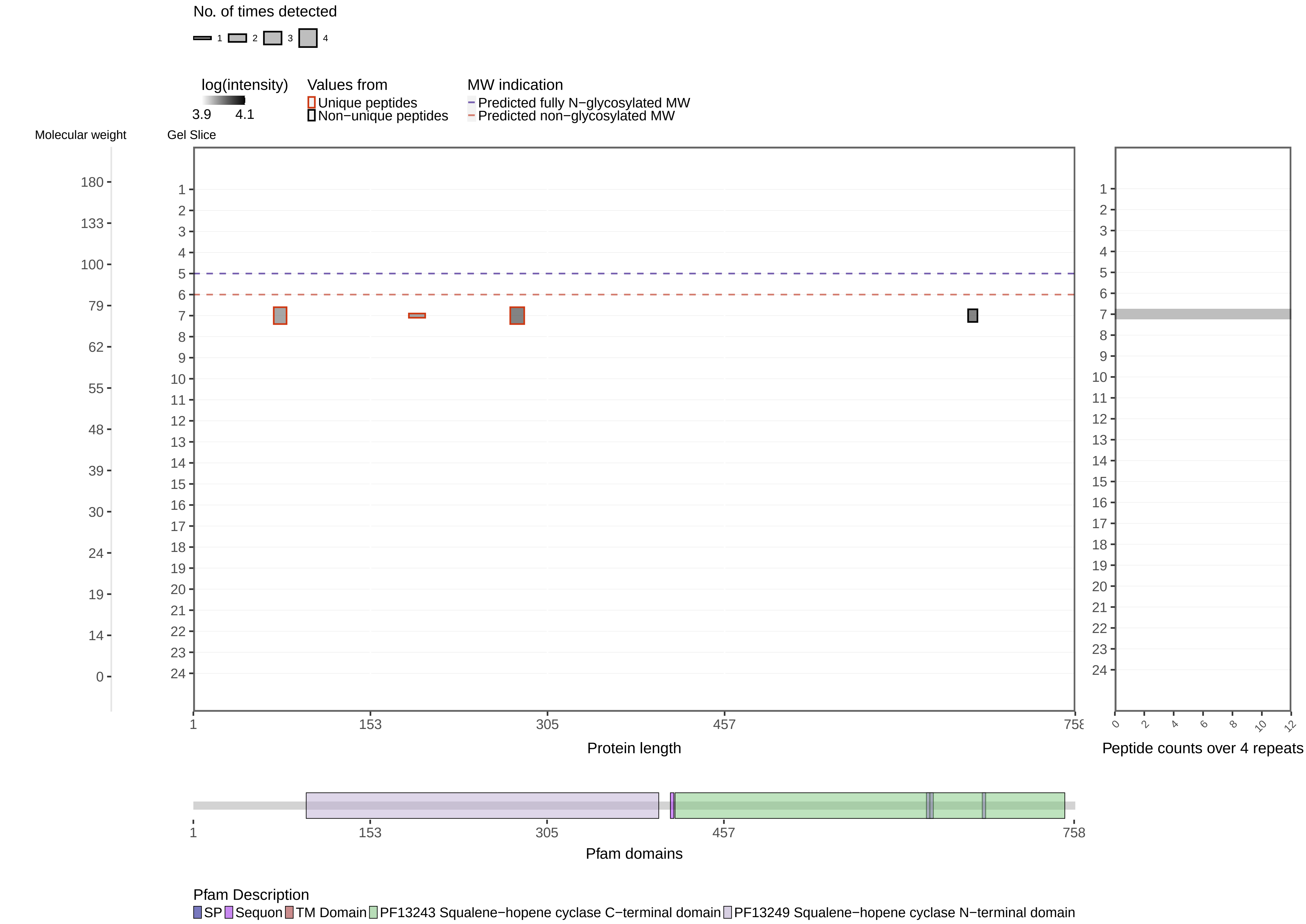


Protein sequence:

ICIGPVNKILNMLCCWVEDPHSEAFKLHLPRIHDYMWVAEDGMKMKGYNGSQSWDTSFAIQAIIATELGEEYGSALRKAHSFIKNTQVLDDCPGNLDFWYRHISKGAWPFSTADHGWPISDCTAEGLKASLLLSKLPSEIVGDPLEAKRFYDAVNVLLSLQNSGGGFATYELSRSYPWLEIINPAETFGDIVIDYPYVECSSAAIQALTAFKKLYPGHRKEEVERCIAKAAAFIEKIQEADGSWYGSWAVCFTYGTWFGVLGLLAAGRNYTNSSSIRKACDFLLSKQISSGGWGE

### NbL04g06400.1 uncharacterized_LOC109240968 [MW = 113.46 kDa]

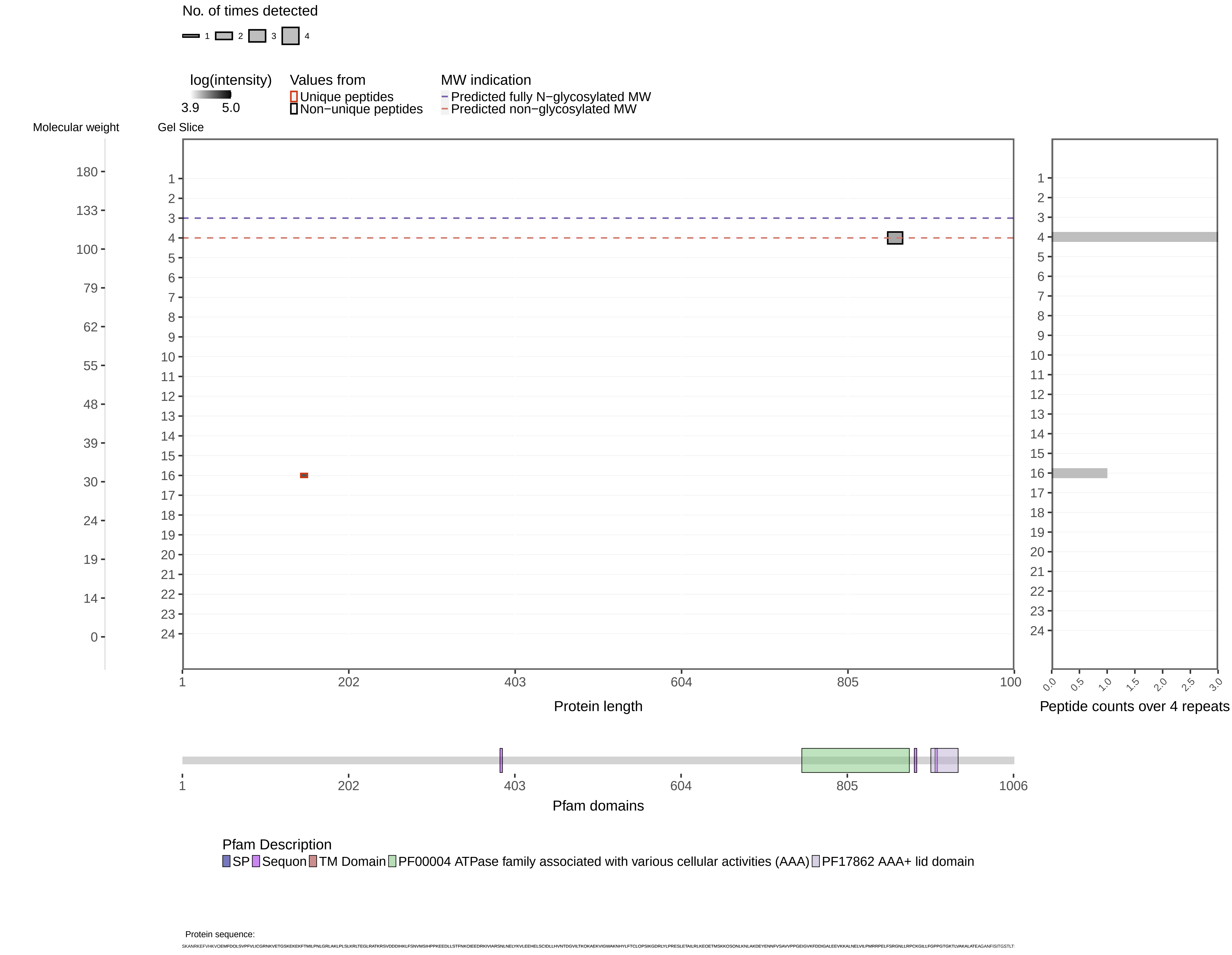

### NbL04g06390.1 nucleolin_1 [MW = 69.01 kDa]

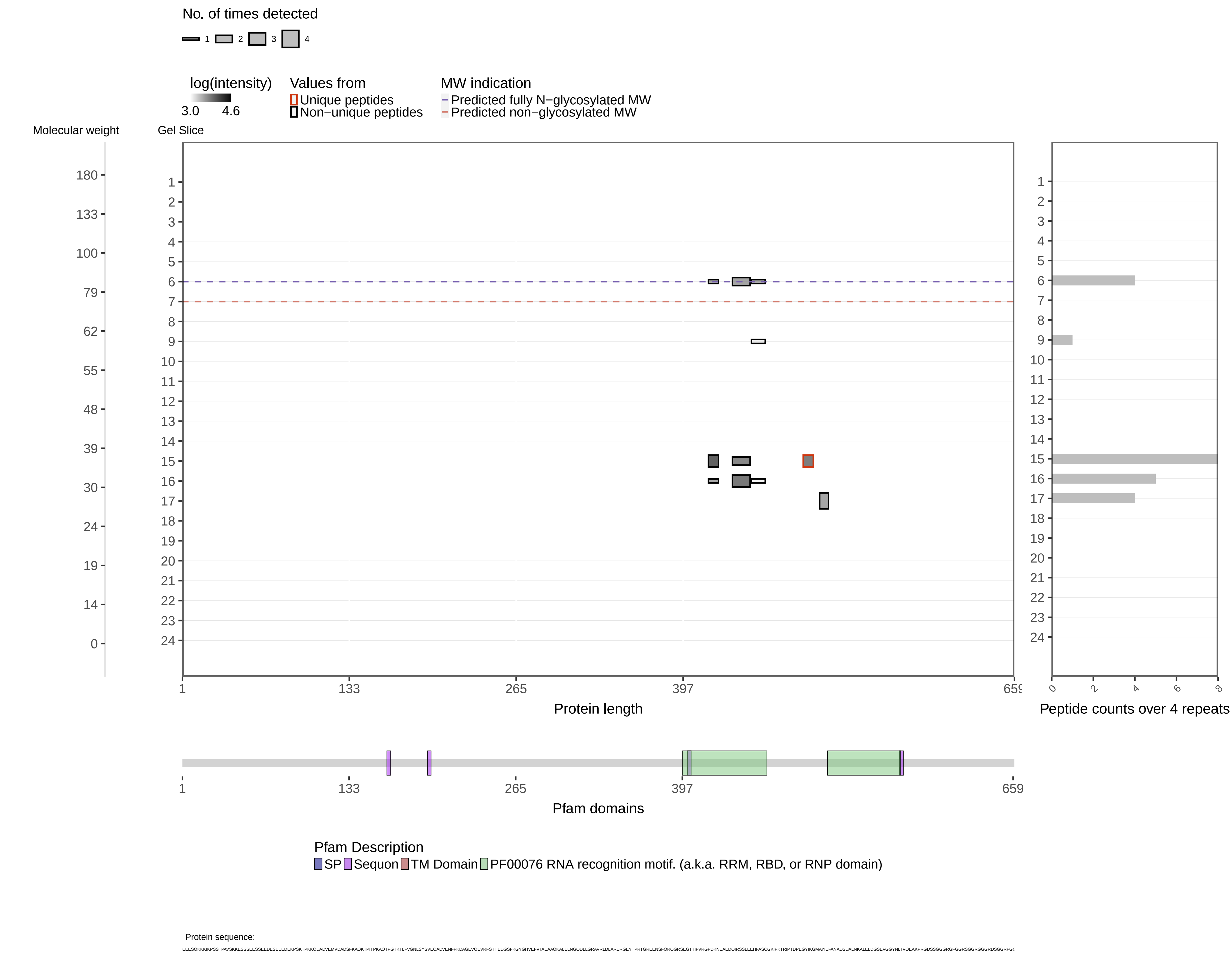

### NbL04g06300.1 succinate−−CoA_ligase_[ADP−forming]_subunit_alpha−2_mitochondrial [MW = 35.61 kDa]

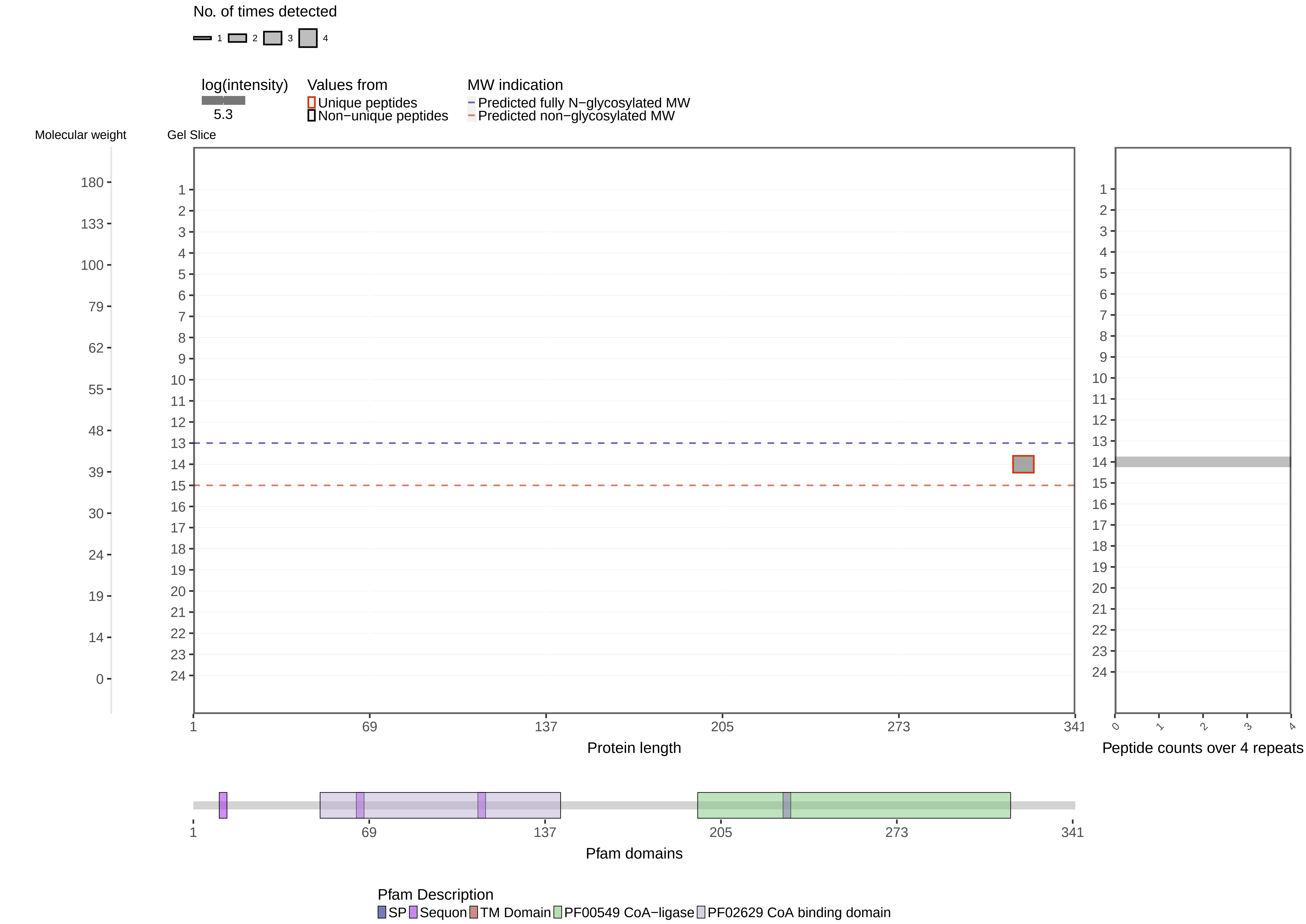


Protein sequence:

[illegible]LKKQSKTRLIGPNCPGIIKPGECKIGIMPGYIHKPGRIGIVSRSGTLTYEAVFQTTAVGLGQSTCVGIGGDPFNGTNFVDCLEKFIADPQTEGIVLIGEIGGTAEEDAAALIKESGTQKPVVAFIAGLTAPPGRRMGHAGAIVSGGKGTAQDKIKALKEAKVTVCESPAKIGVTMLDVFKQRGLA*

### NbL04g06220.1 photosystem_II_stability/assembly_factor_HCF136_chloroplastic [MW = 49.14 kDa]

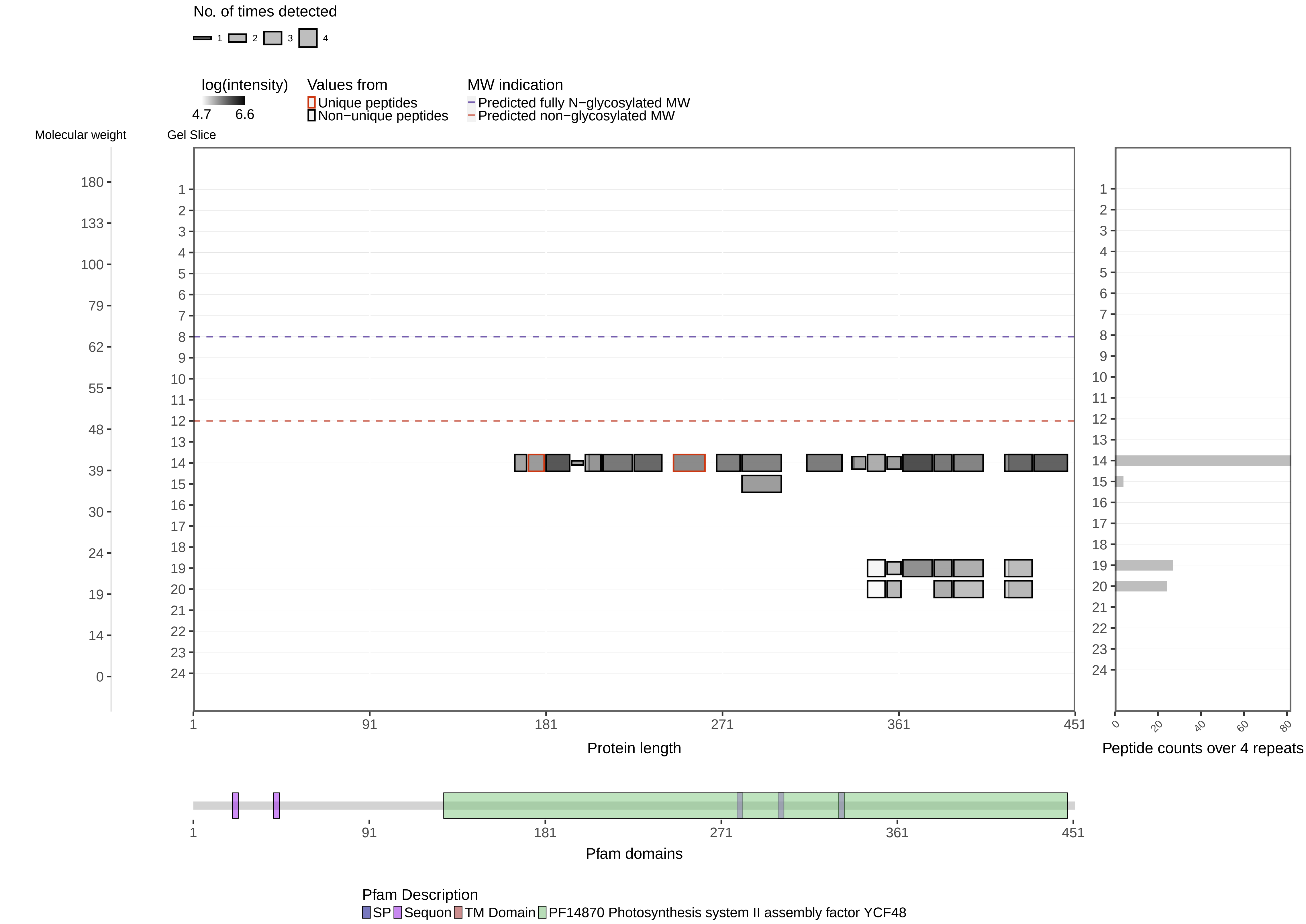


Protein sequence:

SDAGETWQRIPLSSQLPGDMVYIKATGEKSAEMVTDEGAIYVTSNKGYNWKAAIQETVSATLNRTVSSGISGASYYTGTFSTVNRSPDGRYVAVSSRGNFYLTWEPGQAYWQPHNRSVARRIQNMGWRADGGLWLLVRGGGLYLSKGTGLSEDFEEVSVQSRGFGILDVGYRSQDEAWAAGGSGILLKTKNGGKTWIRDKAADNIAANLYSVKFINDNQGFVLGNDGVLLKYLG*

### NbL04g06030.1 pyruvate_kinase_cytosolic_isozyme [MW = 54.99 kDa]

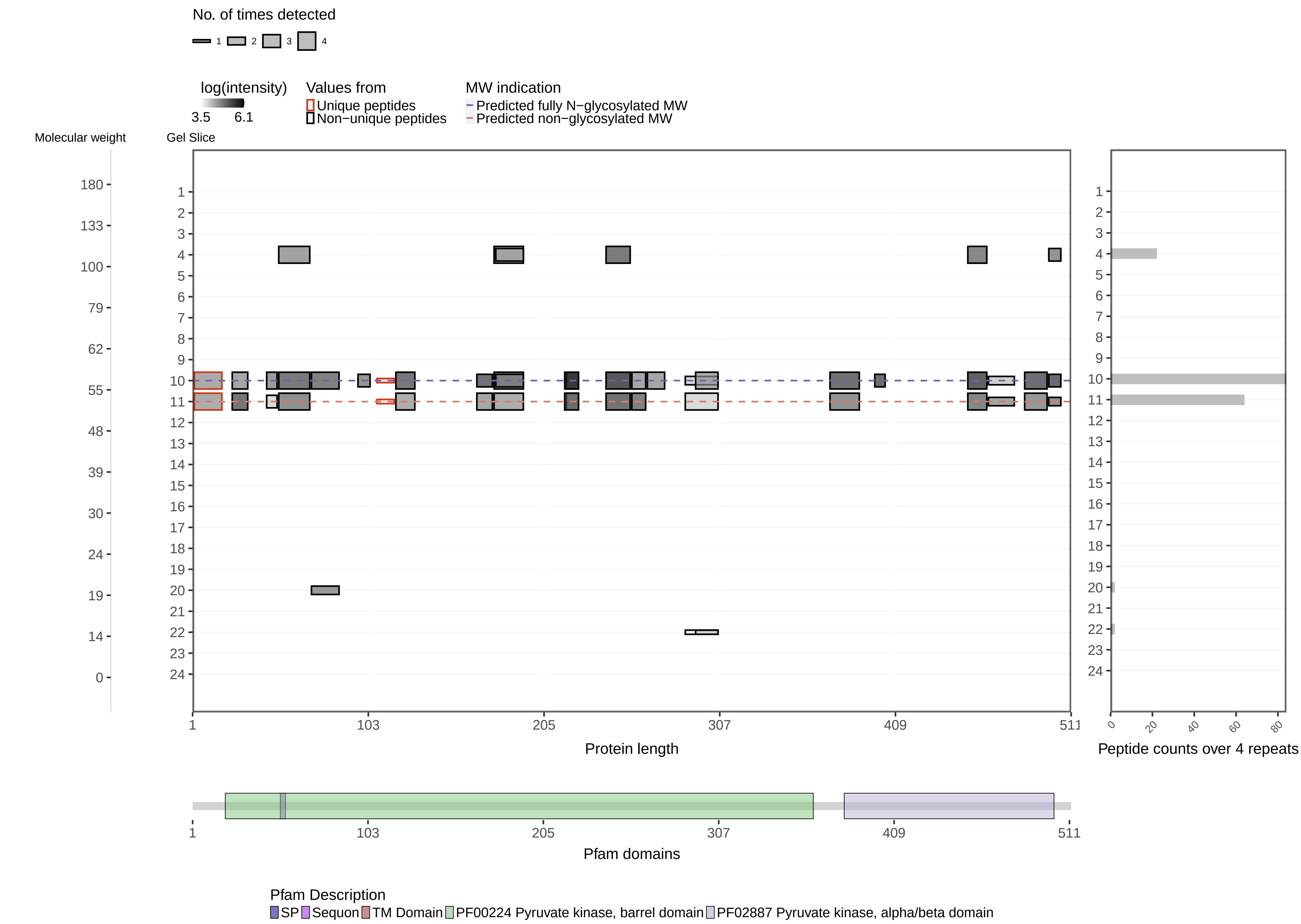


Protein sequence:

VENQEGVVNFDEILRETDSFMVARGDLGMEIPVEKIFLAQKMMIYKCNLVGKPVVTATQMLESMIKSPRPTRAEATDVANAVLDGTDCVMLSGESAAGAYPELAVKIMARICIEAESSLDYGAIFKEMIRSTPLPMSPLESLASSAVRAANKANAKLIVVLTRGGTTAKLVAKYRPAVPIVSVVVPVLTTDSFDWSITDETPARHSLIYRGLIPLLAEGSAKATDAESTEVILQASLKSATEKGLCTPGDSVVALHRIGAASVIKICLVK*

NbL04g05730.1 AP−1_complex_subunit_gamma−2−like [MW = 95.95 kDa]

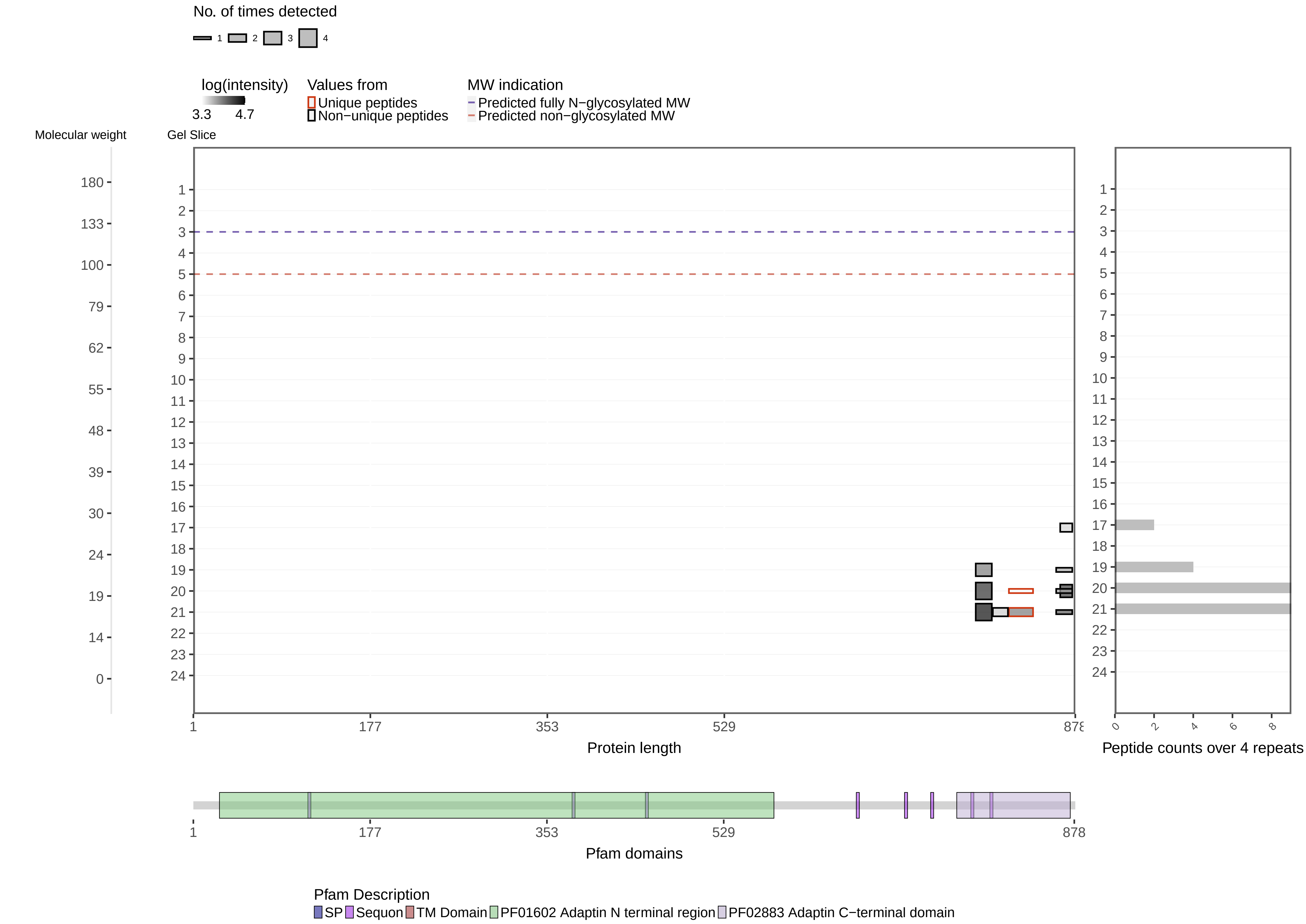


Protein sequence:

SEAGNYVKDEVWHALIVVITNTSDLHGYAVRSLYRAVQAAGEQETLVRVAVWCIGEYGEMLVSNLGRLDIEEHLTVTESDAVDVVETSIKSHSCDLTTRAMCLIALLKLSSRFPFCSQRINDIIGQYKGSFELELQQRAIEFNSIIERHQNMRSSLAERMPVLDEATFSGRRAGSVPAAVSTSKGVSVNLPNGAAKPSIAPVADLLDLSSDDAPAPSSSGGEFLQDLLGVNLTPVSLQSDTNQGQKSGTDALLDLLSIGTPPAQSSSSTAQMLSSNTENKSPLDILNRLSTPSAPSA

NbL04g05680.1 uncharacterized_LOC109225794 [MW = 39.85 kDa]

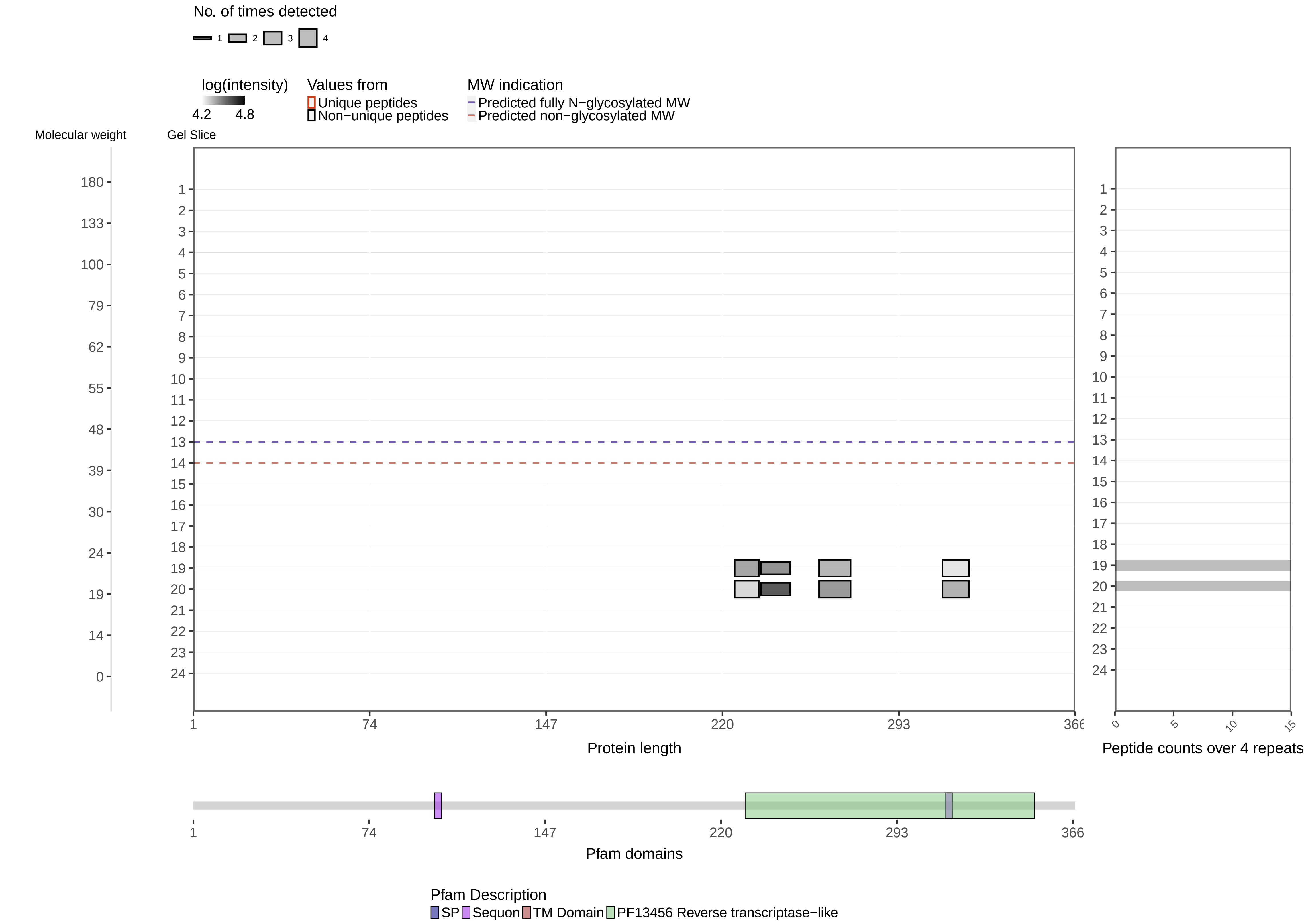


Protein sequence:

SRGGMPERMPKKRSQEVMLSECTDAVGSAVISNDFLRKHVKLDDHKVDQAIPSGRSCTLEFDGASKGNPGLAGAGAVLRADDGSFICRLREGLGVATNNAAEYRGIILGLKYALSKGFTNIRVQGDSKLVCMQIQGLWKVKNQNISTLYEQAKQLKDRFLSFRIIHVLRESNSDADVQANLGVELAEGQIQEEIDK*

### NbL04g05200.1 probable_inactive_purple_acid_phosphatase_2 [MW − SP = 70.79 kDa]

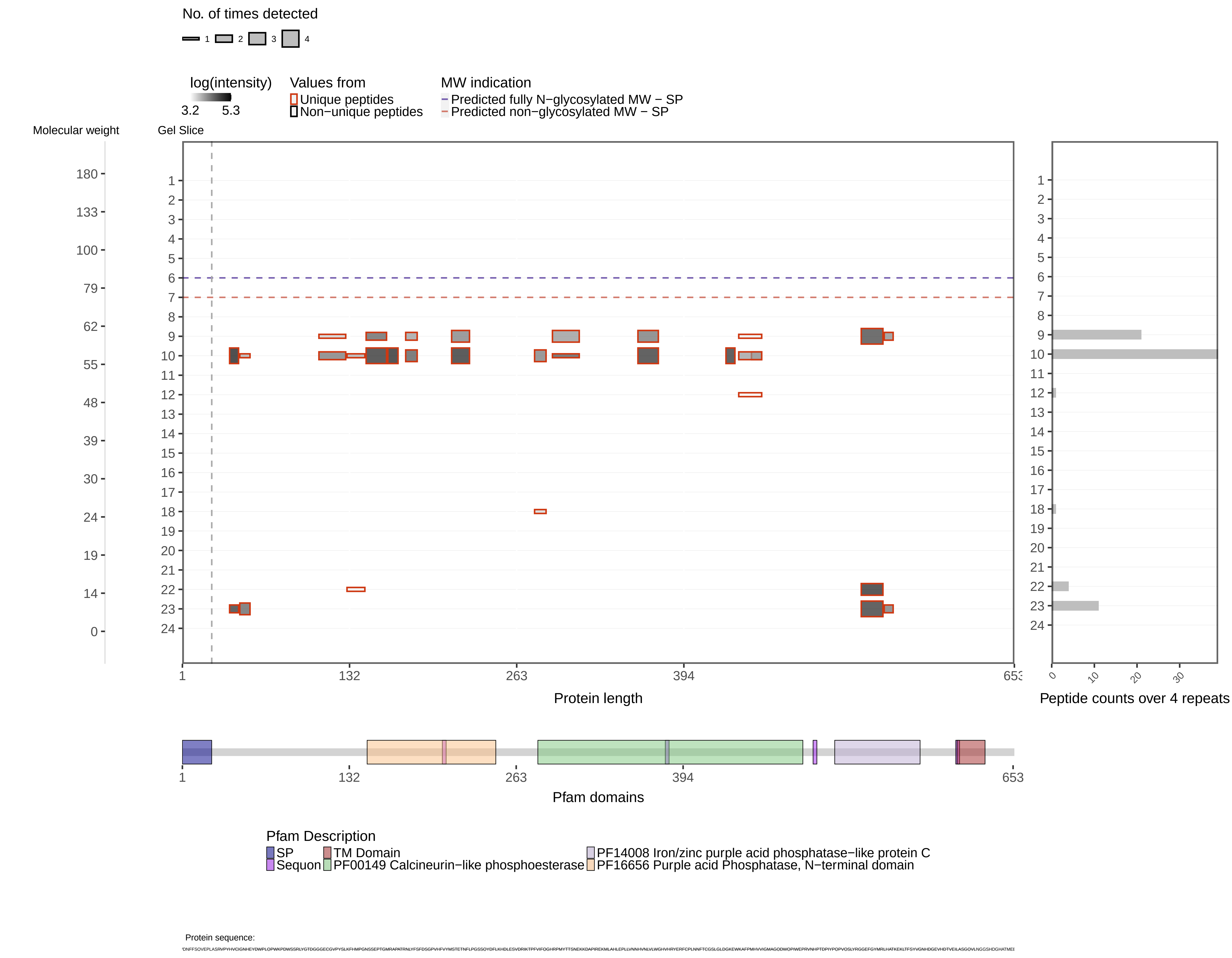

### NbL04g05180.1 PLASMODESMATA_CALLOSE−BINDING_PROTEIN_3−like [MW − SP = 27.13 kDa]

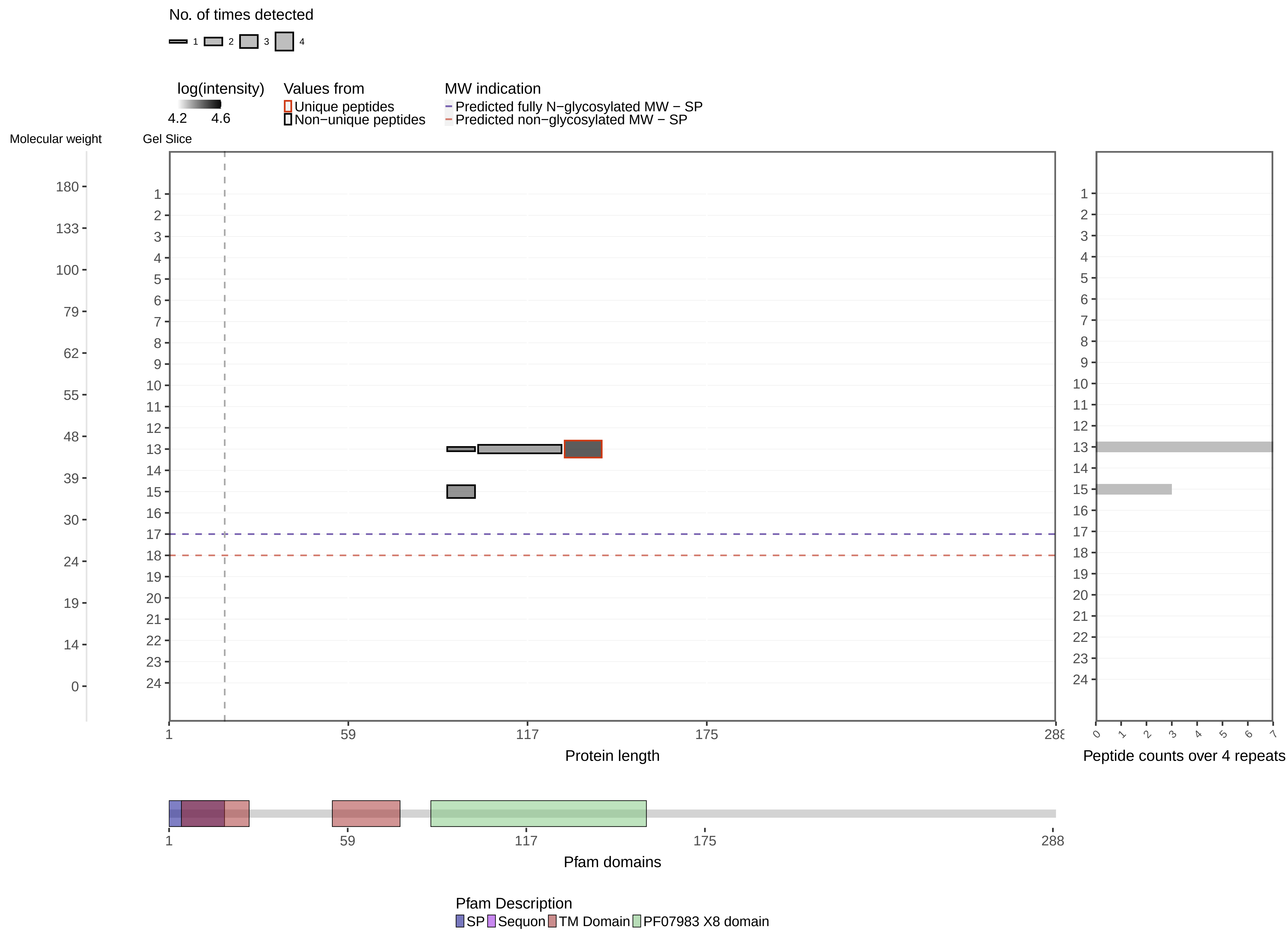


Protein sequence:

YAVNSYYQRKGAAGASCDFSGTATTSPNPPTTTGSGCVYQSTPGNTGGGSTTPTIAPPGSTTPTIVPPGSTTPTIAPPGTTIPTTAPPGTTTPGIGTGGTGTGIATTPGTNNPVFGPGPSGSGISNTDGNAAGHFKNSIFLTIALLAISLICLRA*

NbL04g05120.1 alpha−14−glucan−protein_synthase_[UDP−forming]_2−like [MW = 41.35 kDa]

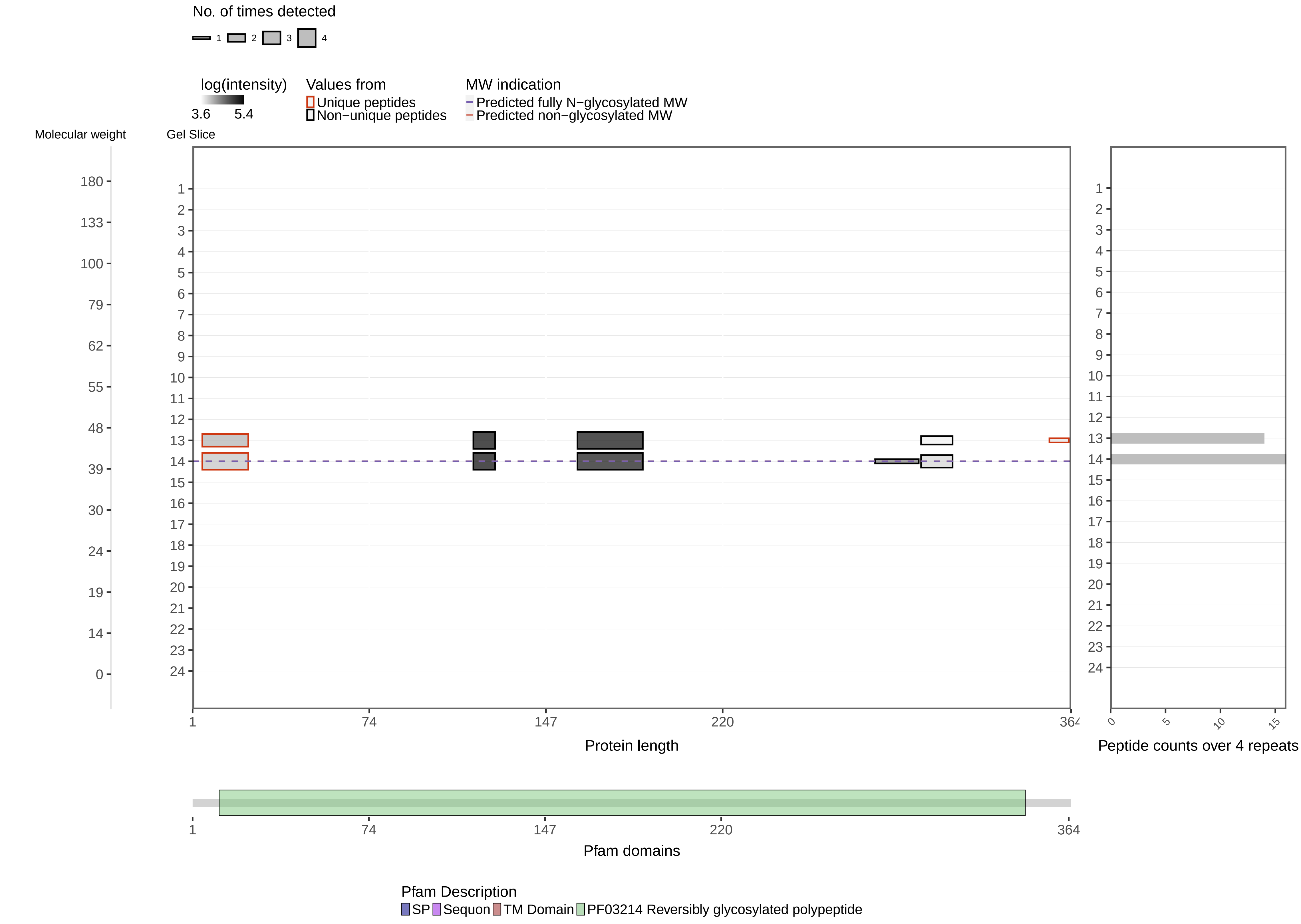


Protein sequence:

WLNIPDYDAPTQLVKPRERNTRYVDAVMTIPKGTLFPMCGMNLAFDRELIGPAMYFGLMGDGQPIGRYDDMWAGWCTKVICDHLGLGIKTGLPYIWHSKASNPFVNLKKEYKGIYWQEEIIPFFQSATLPKDCTSVQQCYIELSKQVKDKLSKVDPYFTKLADAMVTWIEAWDELNPTSEGLAKLSISNGPAK*

NbL04g05090.1 translin−associated_protein_X [MW = 33.65 kDa]

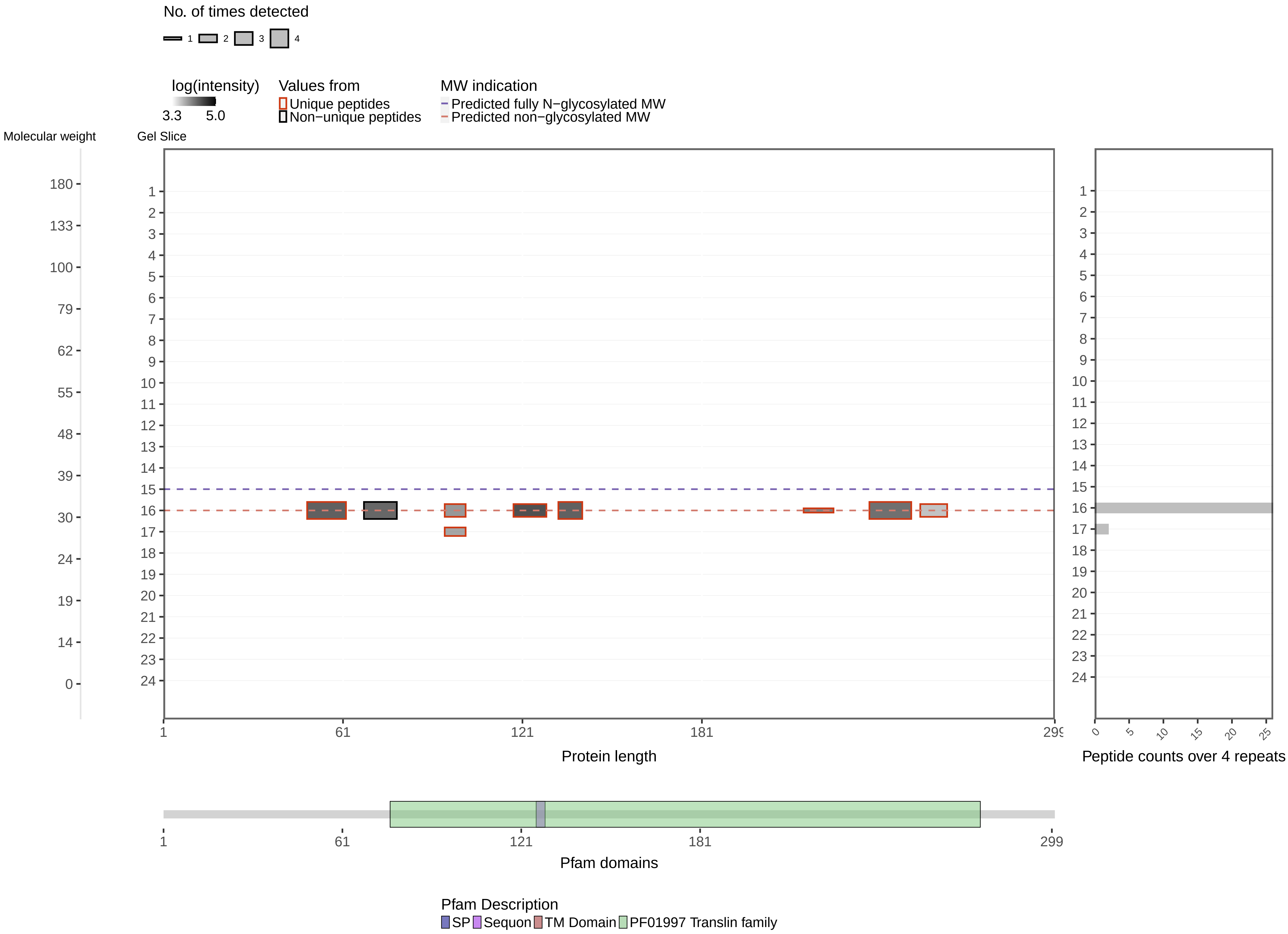


Protein sequence:
QGTDFWKLRRAYSPGVQEYVEAATLCNFCTTGTLLTLDEINAAFLPLSDPSVEPLQINILDYILGVADLTGELMRLAIGRISEGELDFAEKICSFVREIYRKLTLIAPEMDDPSDMKQKMETMLQSVMKIENACFSVHVRGSEYIPLLGSAEPSYPLLGMPDVE*

### NbL04g04980.1 protein_PHYLLO_chloroplastic [MW = 157.84 kDa]

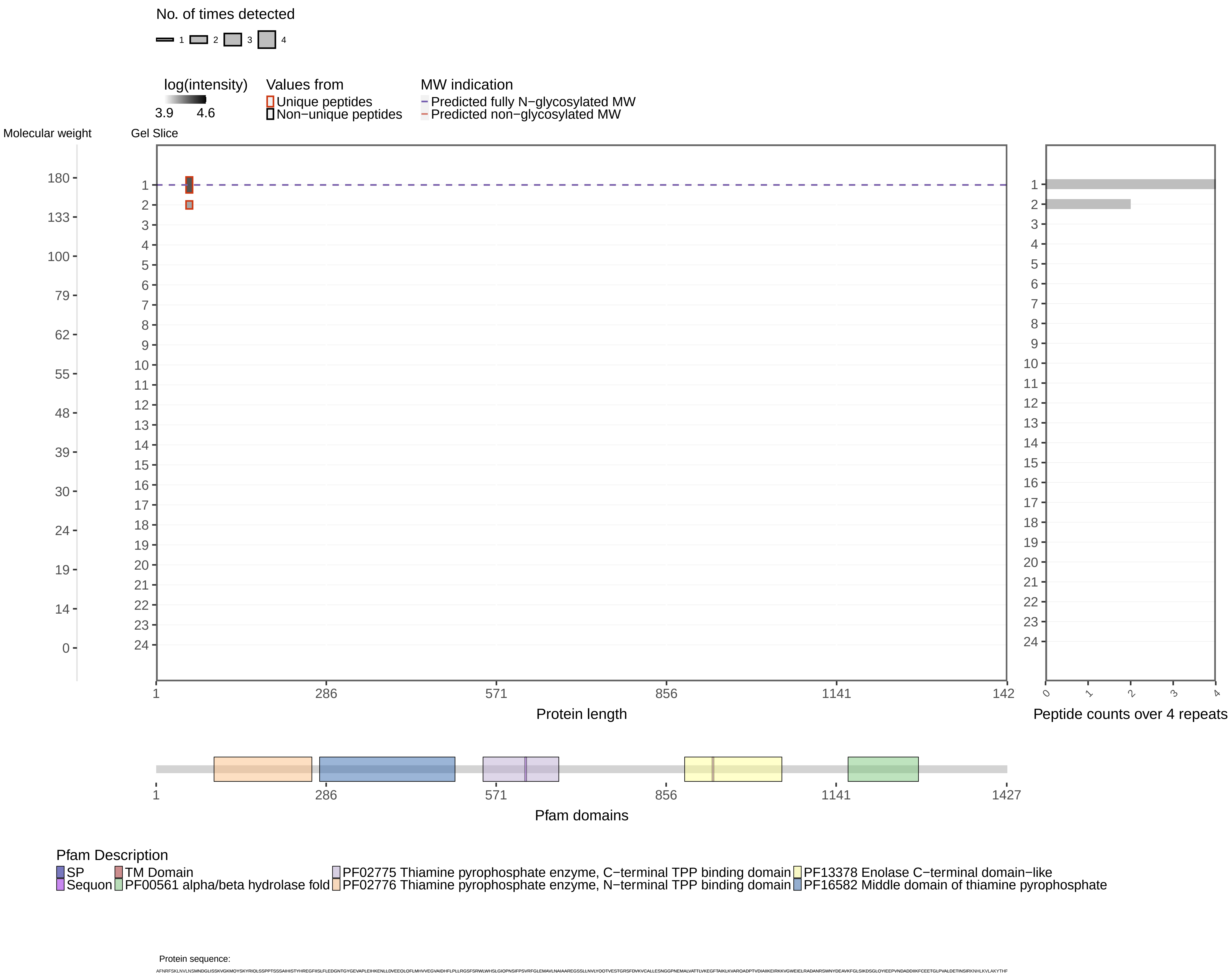

NbL04g04780.1 acyl−acyl_carrier_protein_thioesterase_ATL3_chloroplastic−like [MW = 24.1 kDa]

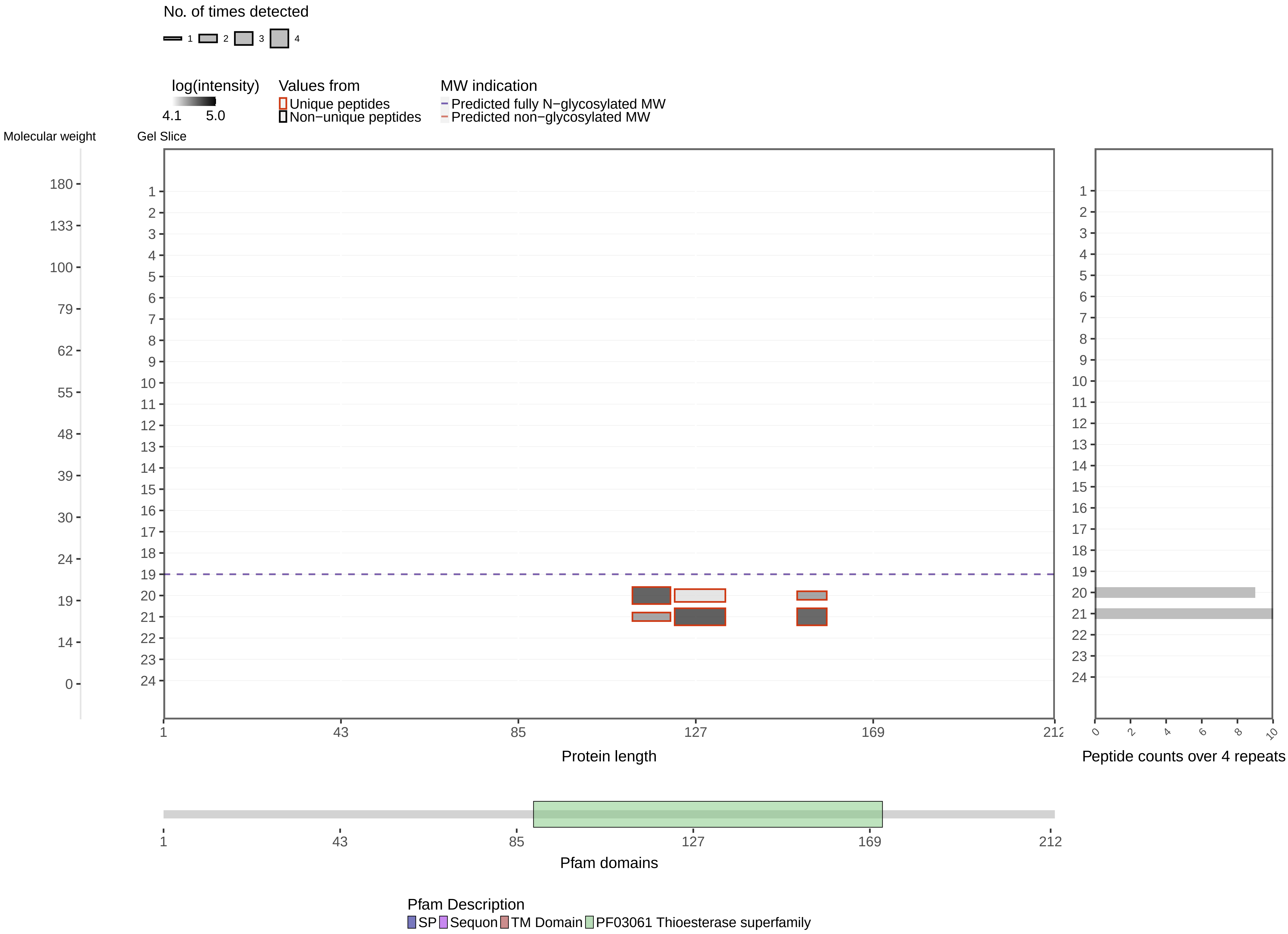


Protein sequence:

VNNAIYASYCQHCRHELLERLGVSADAVARNGDAIALTDLSLKYLAPLRSGDRFVVKVRISDSSAVRLFFEHFIFKLPDQQPILEATGTAVWLNKNYRPVRIPAEFRSKFVQFLRQEASK*

### NbL04g04700.1 acetylajmalan_esterase−like [MW − SP = 37.62 kDa]

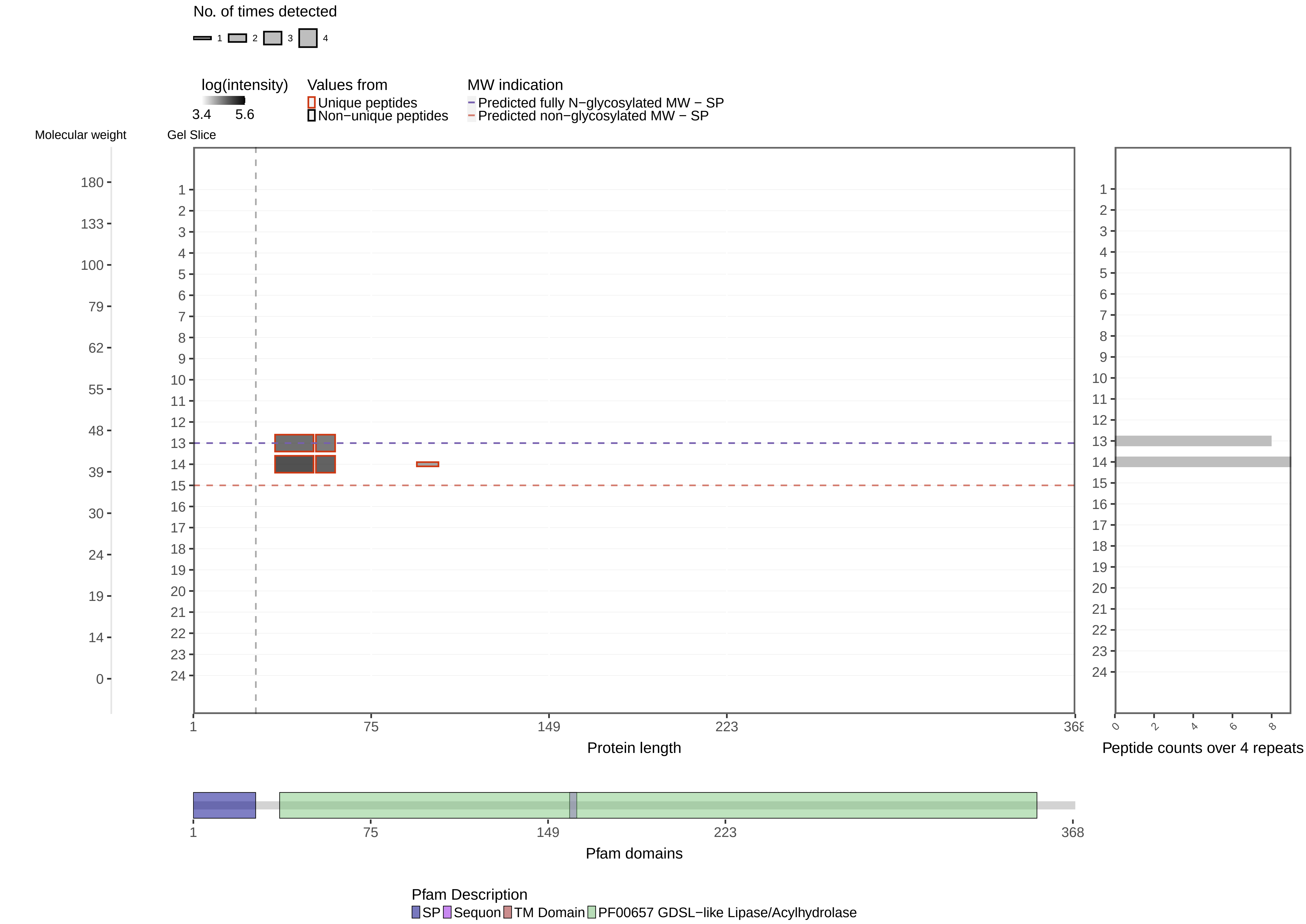


Protein sequence:

EWGGNDYYNGFFQRKQLPEVRTYVPFVVAGIMRSIKDVIQLGATRILVPGIYPLGCLPLYLTTFPDSNANGYDQLGCLRNYNDFASYLNRGLASLQRQFPNVRIVYGDYYGALLTLLRSPSSFGFNQNTLLTACCGTGGRYNFNFRTVCGSASVRACSNVAQYVHWDGIHLTDEAHRHMTDIVVKDLLSRFSCTV*

### NbL04g04560.1 ATP_synthase_subunit_beta_mitochondrial [MW = 59.81 kDa]

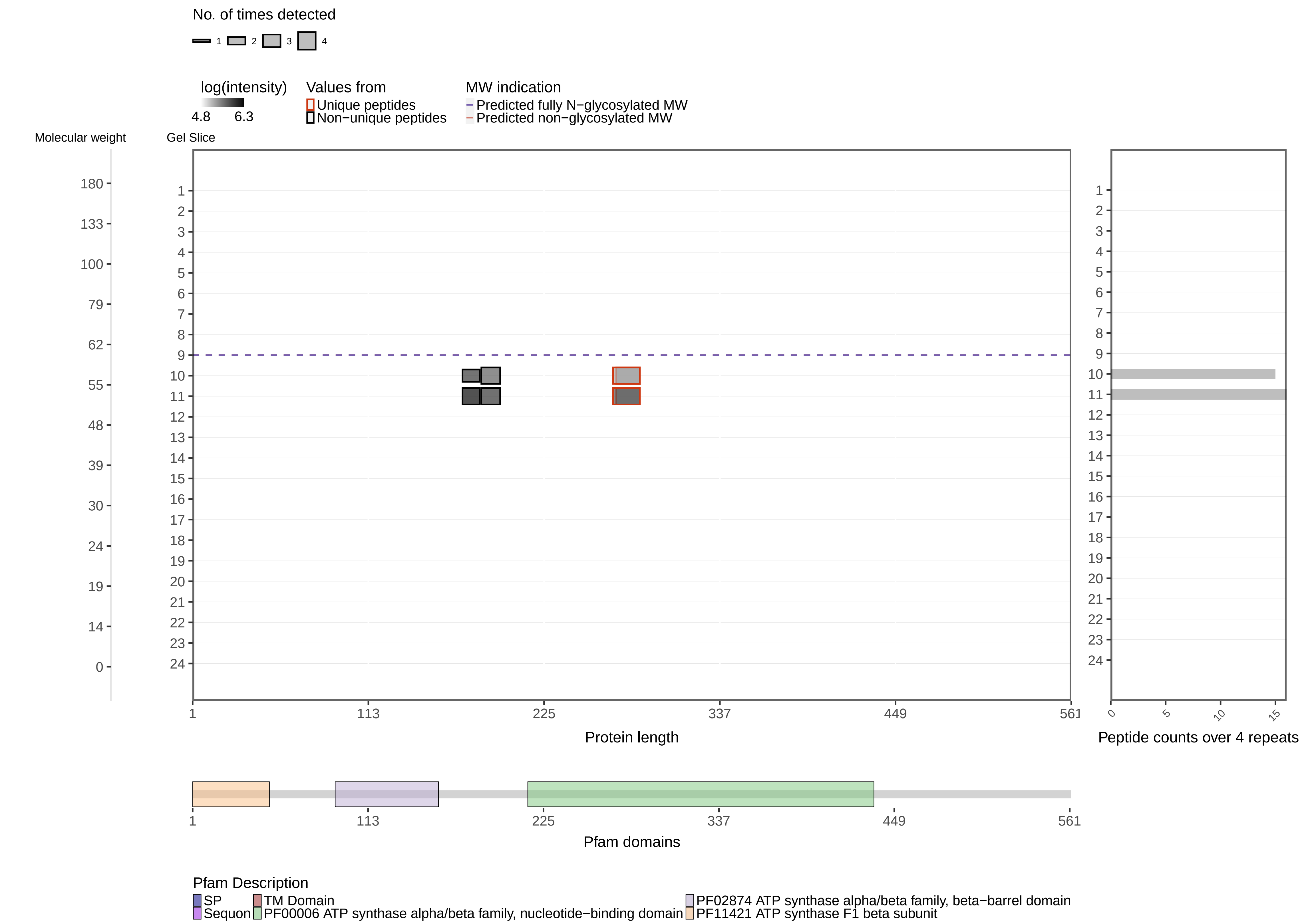


Protein sequence:

RTREGNDLYQEMIESGVIKLGEKQSESKCALVYGQMNEPPGARARVGLTGLTVAEHFRDAEGQDVLLFIDNIFRFTQANSEVSALLGRIPSAVGYQPTLATDLGGLQERITTTKKGSITSVQAIYVPADDLTDPAPATTFAHLDATTVLSRQISELGIYPAVDPLDSTSRMLSPHILGEDHYNTARGVQKVLQNYKNLQDIIAILGMDELSEDDKMTVARARKIQRFLSQPFHVAEVFTGAPGKYVDLKESINSFQGVLDGKYDDLSEQSFYMVGGIDEVIAKAEKIAKESAA*

### NbL04g04320.1 kirola−like [MW = 18.71 kDa]

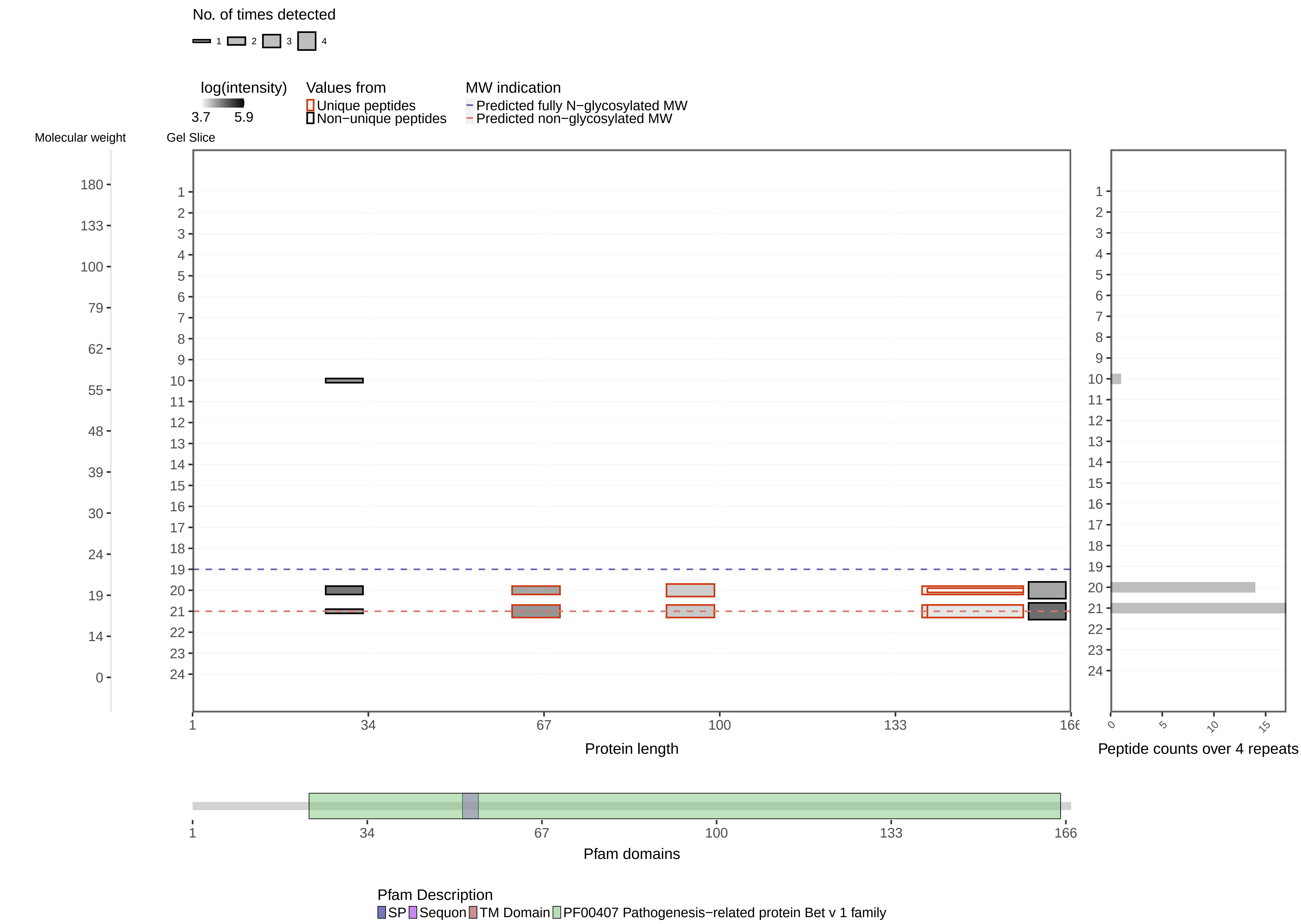


Protein sequence:

/GSVVSWKYNDDGKDKICKQVIEAVDHQKKSITWKVIGGDLLESYNSFTITTSHDHQWTTWTFVYEKKTEDTPEPLVPLGLVLHVTKDIEGHLLK*

### NbL04g04200.1 HMG−Y−related_protein_B−like [MW = 21.16 kDa]

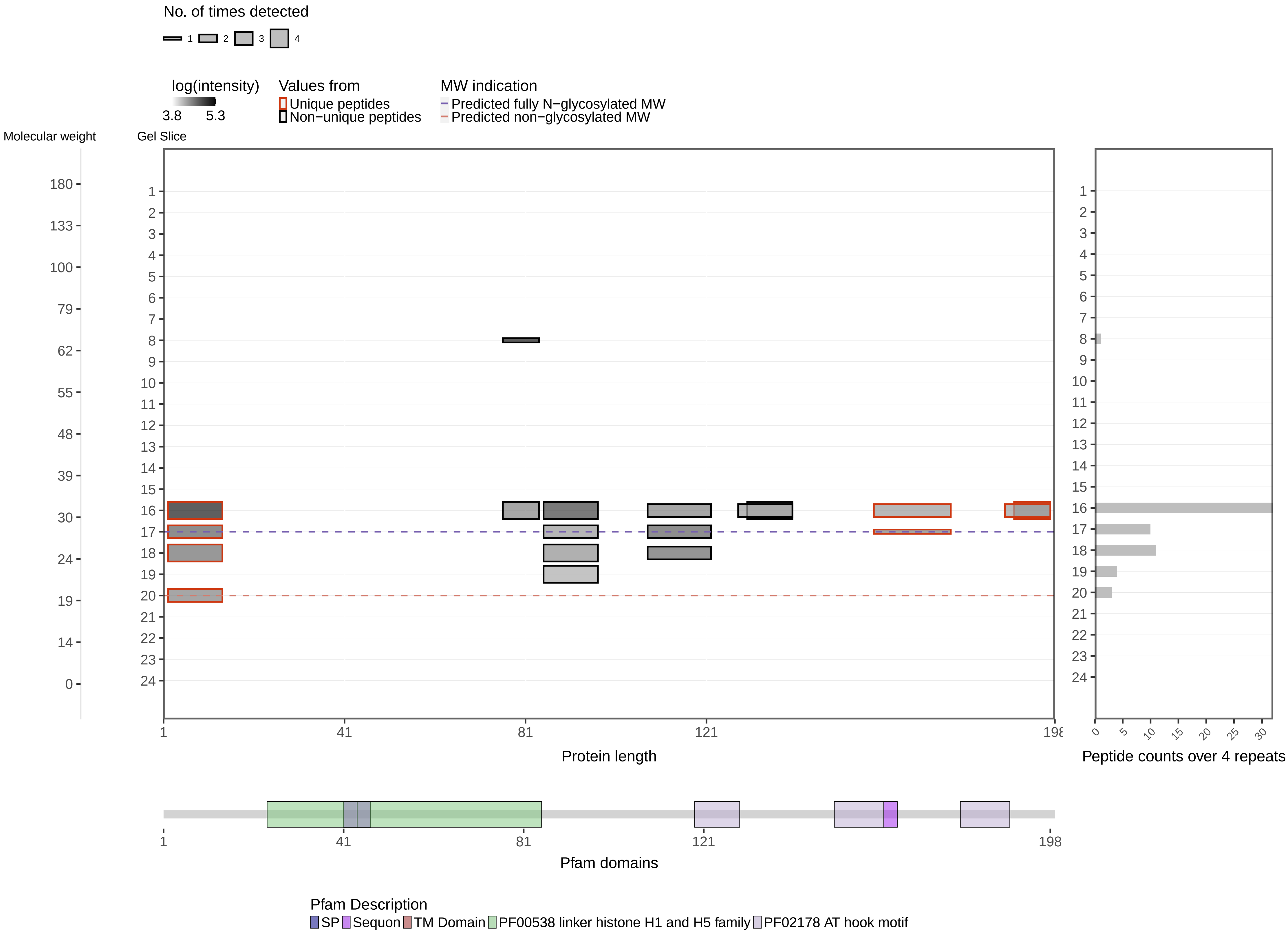


Protein sequence:

MKPDPNAPPRRGRGRPPKPKAPLPEGYVPPPPRPRGRPPKERDPHAPISVPMKVTSPSSGGTGKKRGRPRKYPNDTSSPSPAVTLPPRGTPRGRGRPPKVKTPVAAEVGA*

NbL04g04190.1 glucan_endo−13−beta−glucosidase_3−like [MW − SP = 52.07 kDa]

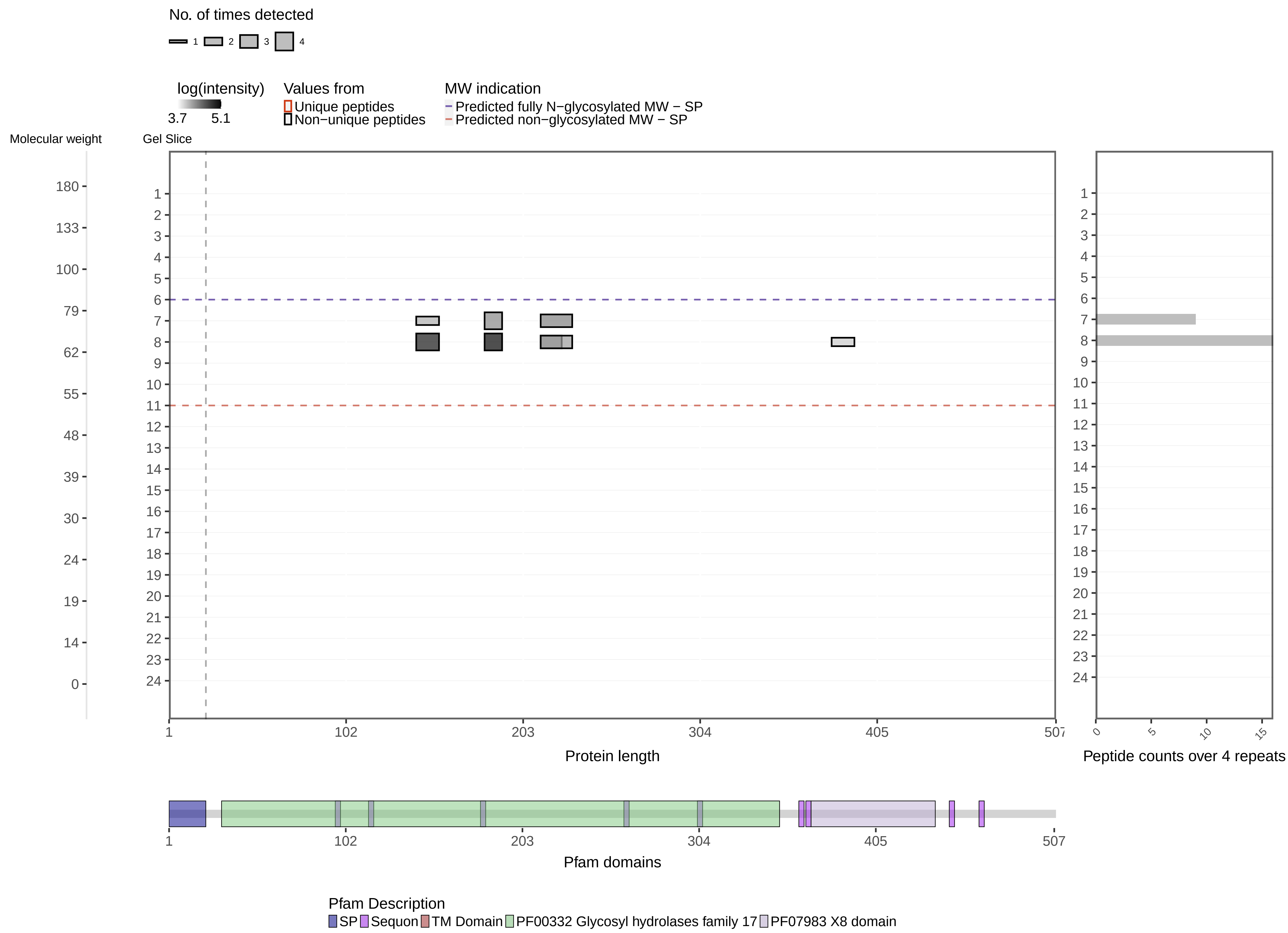


Protein sequence:

NVFDAVVDAAYFSMSYLNFTNIPIIVTESGWPSKGDSSEPEASLDNANTYNSNLIRHVLNNTGTPKHPGIAISTYIYELYNEDLRHDSISEKNWGLFDSSGMPVYTMHLTGSGTVLANDTTNQTYCVAKENAEKKMVQAALDWACGPGKVDCTPLLQGNPCYEPNTVIAHASYAFDAYYRKMGMADGTCNFNGVATVSTTDPSNGSCIYPGSGGRTGTLTNSSSHAPSSDSTASGCHSQYSHDVRTFLSSLIVGLVLCTAACL*

### NbL04g04180.1 cysteine−rich_repeat_secretory_protein_12−like [MW − SP = 27.31 kDa]

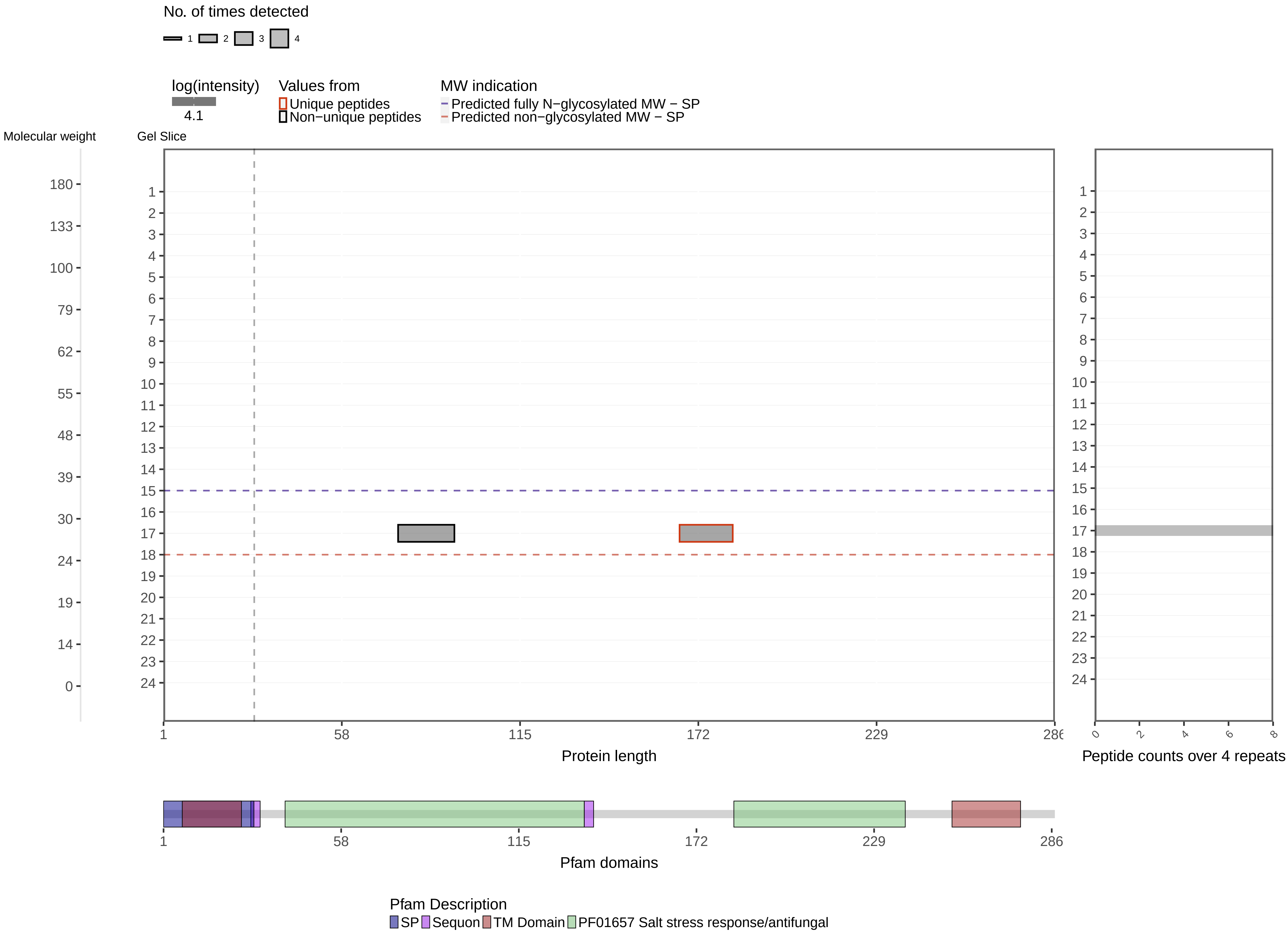


Protein sequence:

FVKYDNVSFLGVEDKGVVMHKCGPSIMDDNEMLTRNDAVLTYLGSVPQGQYFRVGGSGKIQAVAQCTQDLSVGECDDCLSEAIRQLKSECGSSPWGDMFMTKCYARFSERGFTSKSGDEDVEKTLAITIGLIAGVAVLIVFLSVLNKLCEKKGGK*

### NbL04g04140.1 1−aminocyclopropane−1−carboxylate_oxidase_homolog_4−like [MW = 49.16 kDa]

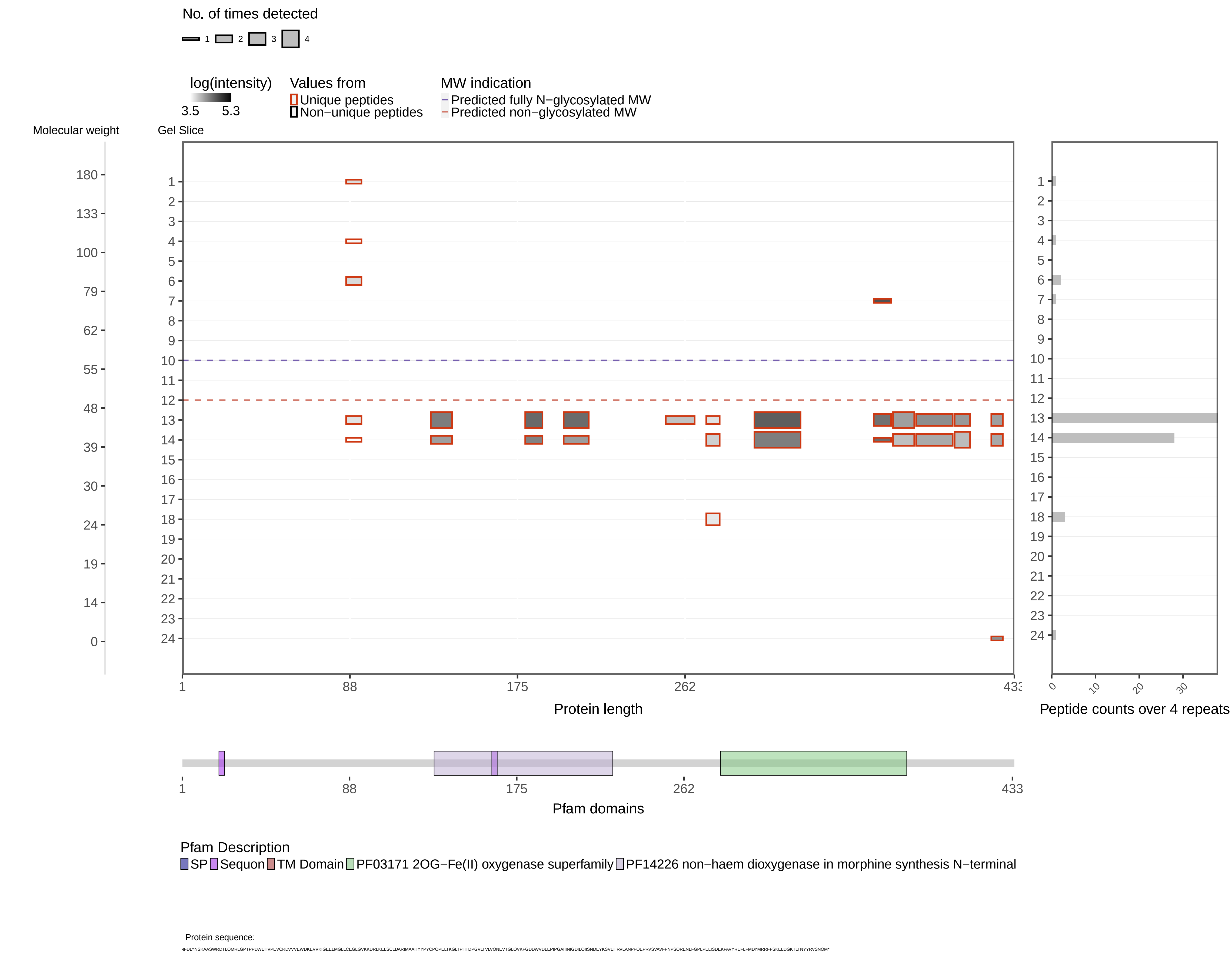

NbL04g03820.1 putative_methyltransferase_DDB_G0268948 [MW = 29.98 kDa]

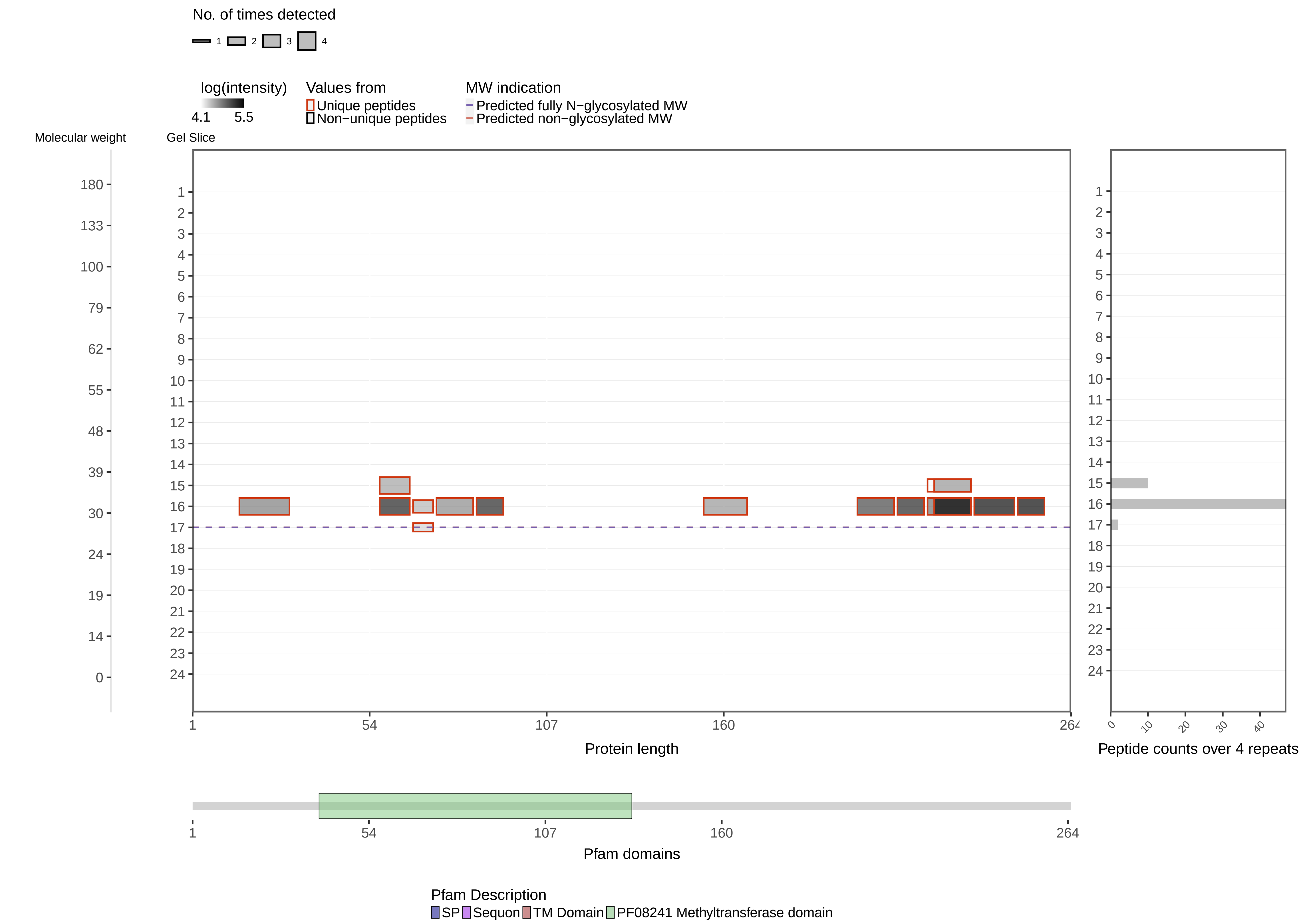


Protein sequence:

[illegible]QVKWLLKKPNGVIAAWCYTVPEVSNSVDPIFEKFYTVDAGPYWECPRKLVDERYKTIDFPFEPVSGYDHNGPFEFKIEKVMDLNSYFTYLKSWSAYQTAKEKGVELLTDDVVEKFTSVWNEDGESQKTVSFPIYLRIGKVGNL*

NbL04g03540.1 vegetative_cell_wall_protein_gp1−like [MW − SP = 8.7 kDa]

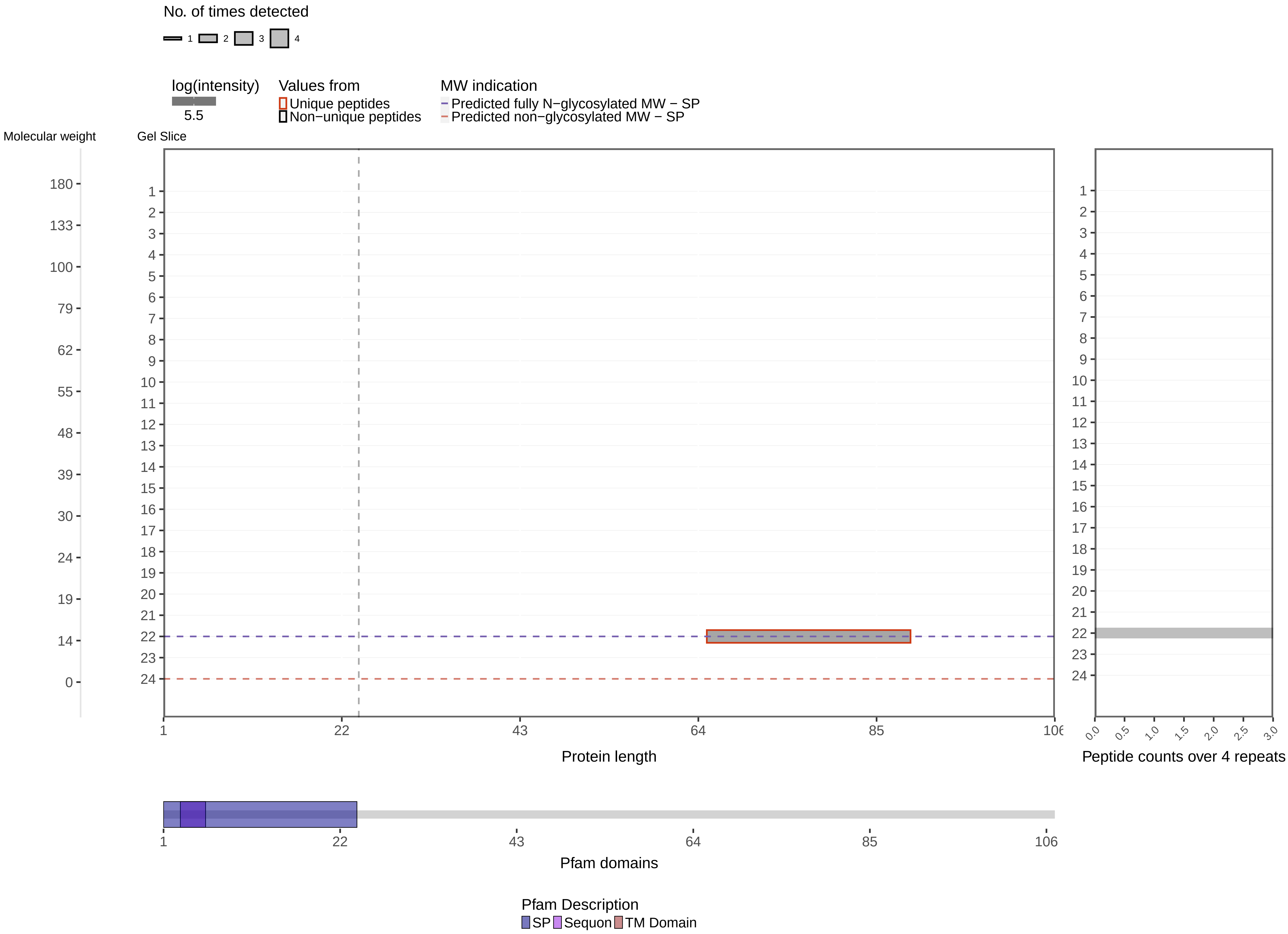


Protein sequence:

GLPKPEIPTLPQPGLPTLPTLPMPKLPLPQLPMPQLPTLPTPDLPVPIPKLPLPELPVPLPPLPSP*

### NbL04g03500.1 alpha−galactosidase_1−like [MW = 48.21 kDa]

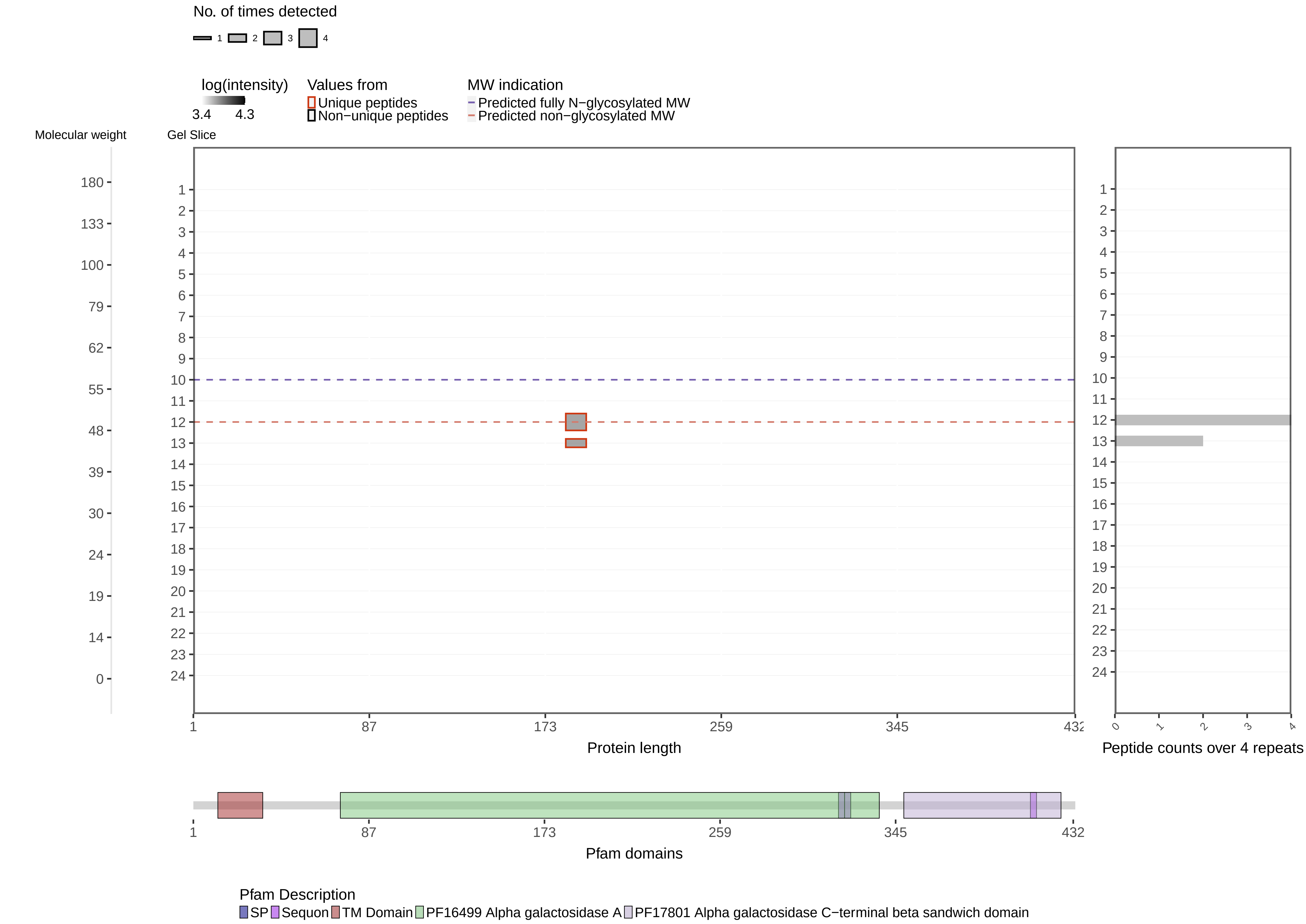


Protein sequence:

GTKPTARYPVMTRALMNAGRPIFFSLCEWGDLHPALWGGKVGNSWRNTIDIQDTWESMISRADQNEVYADLARPGGWNDPDMLEVGNGGMTKDEYIVHFSLWAISKAPLIIGCDVRNLSNDTKEILANKDVIAVNQDELGIQGKKVRMQGDLEVWAGPLSGYRVALVLLNRGSTKIEITALWEDIGIPPISVVVARDLWEHKTLETRFVGNLTATVMSHACKMYVLKPIAS*

### NbL04g03480.1 pyruvate_kinase_cytosolic_isozyme [MW = 55.06 kDa]

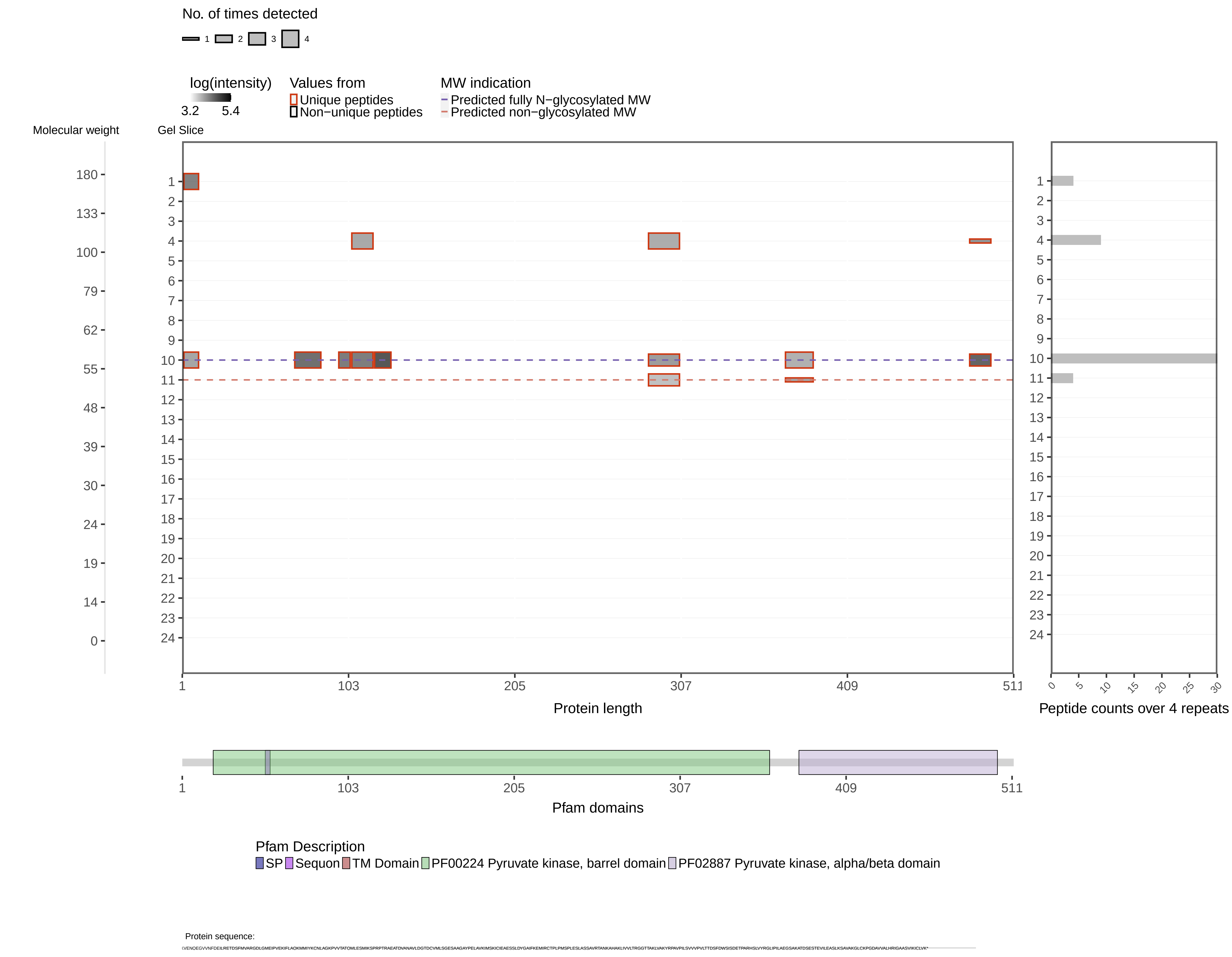

### NbL04g03430.1 probable_leucine−rich_repeat_receptor−like_protein_kinase_At1g35710 [MW − SP = 63.27 kDa]

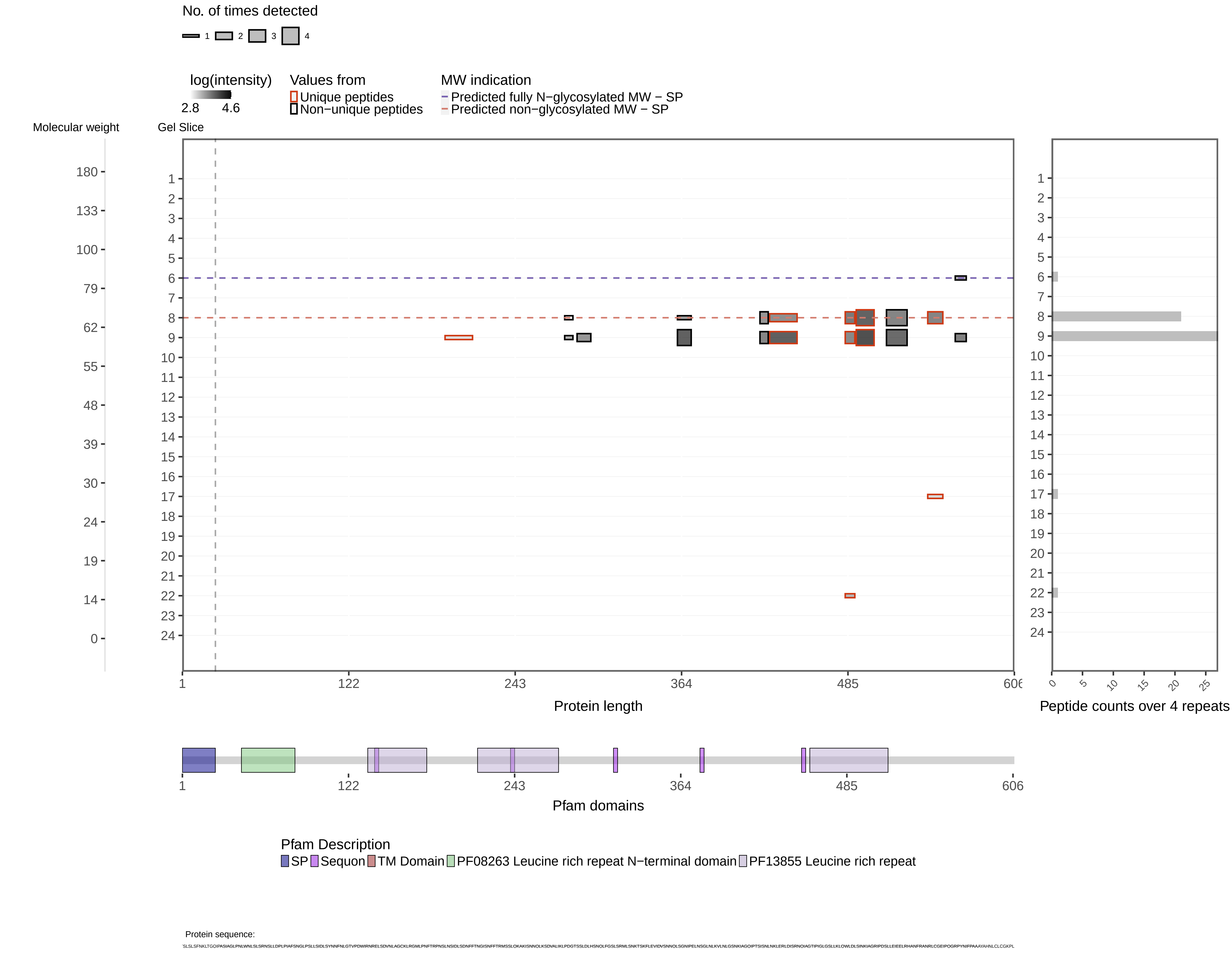

NbL04g03320.1 FAD_synthetase_2_chloroplastic−like [MW = 40.55 kDa]

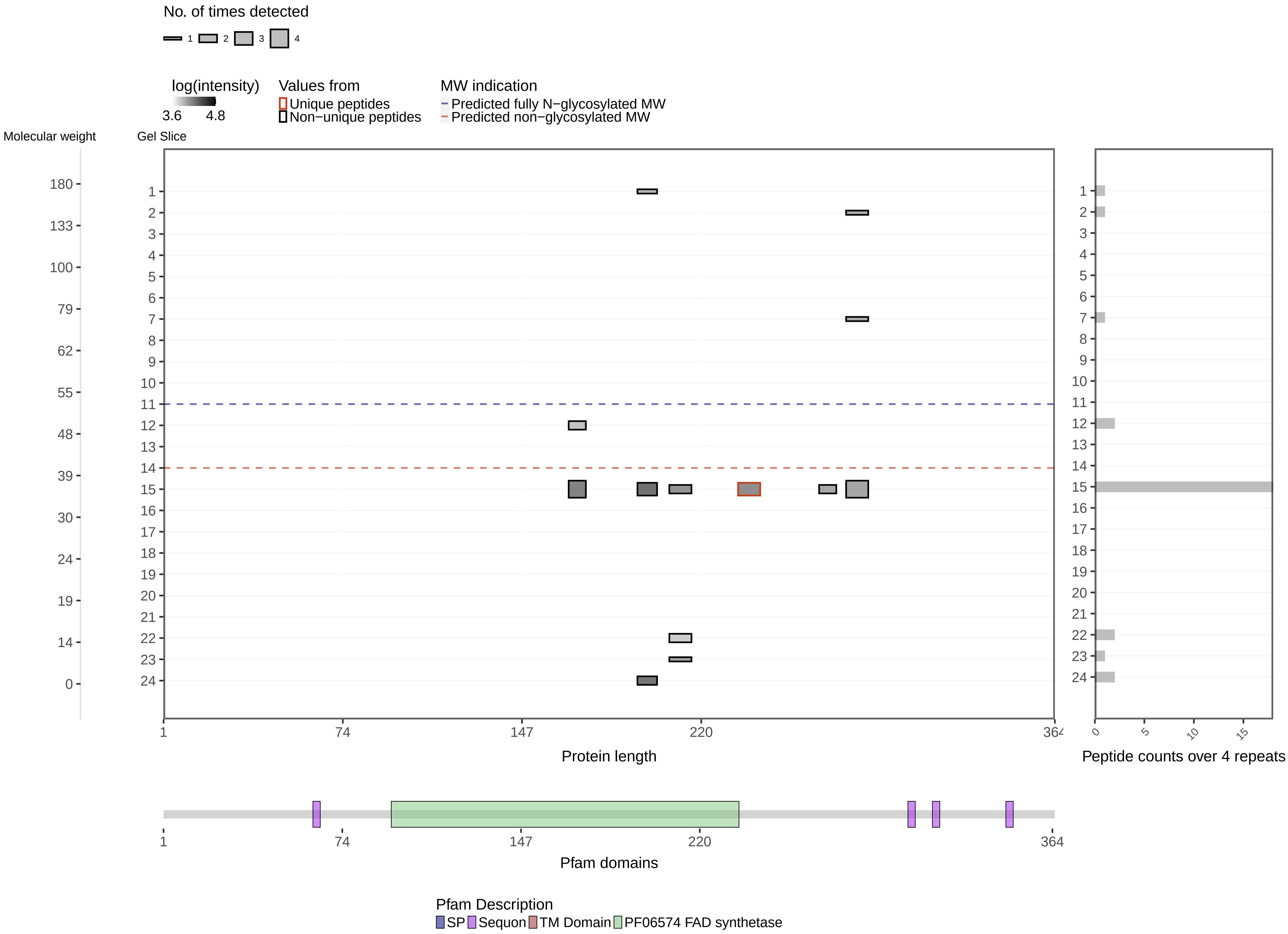


Protein sequence:

QIEFSKVRSLTPRQFVEKLSKELGVRGVVAGENYRFGYRASGDASDLVKLCEEYGLEAYIINSVMEKNQISEDLNSREKKERGQVSSTRVRYALDKGDMKYVSELLGRNHRLVLLMEDQERFTSEKSRLSAPKSCLLNLSPKEGLYENCSISIDENVVPCRVIVDTTDIHLECYEVANFTCITSRDIKILGIDFGS*

NbL04g03270.1 glyceraldehyde−3−phosphate_dehydrogenase_A_chloroplastic [MW − SP = 41.06 kDa]

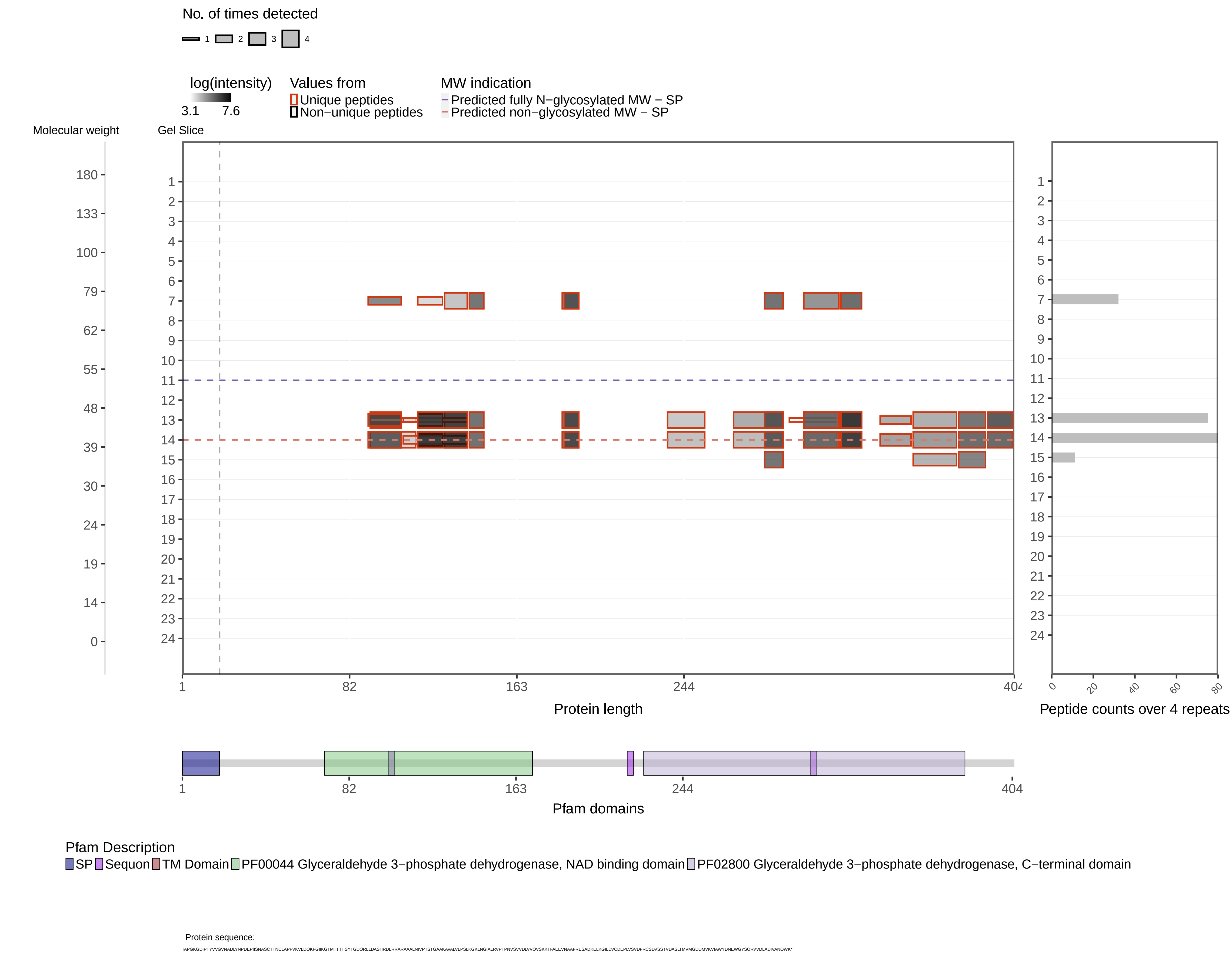

### NbL04g03230.1 glutamate−1−semialdehyde_21−aminomutase_chloroplastic [MW = 58.46 kDa]

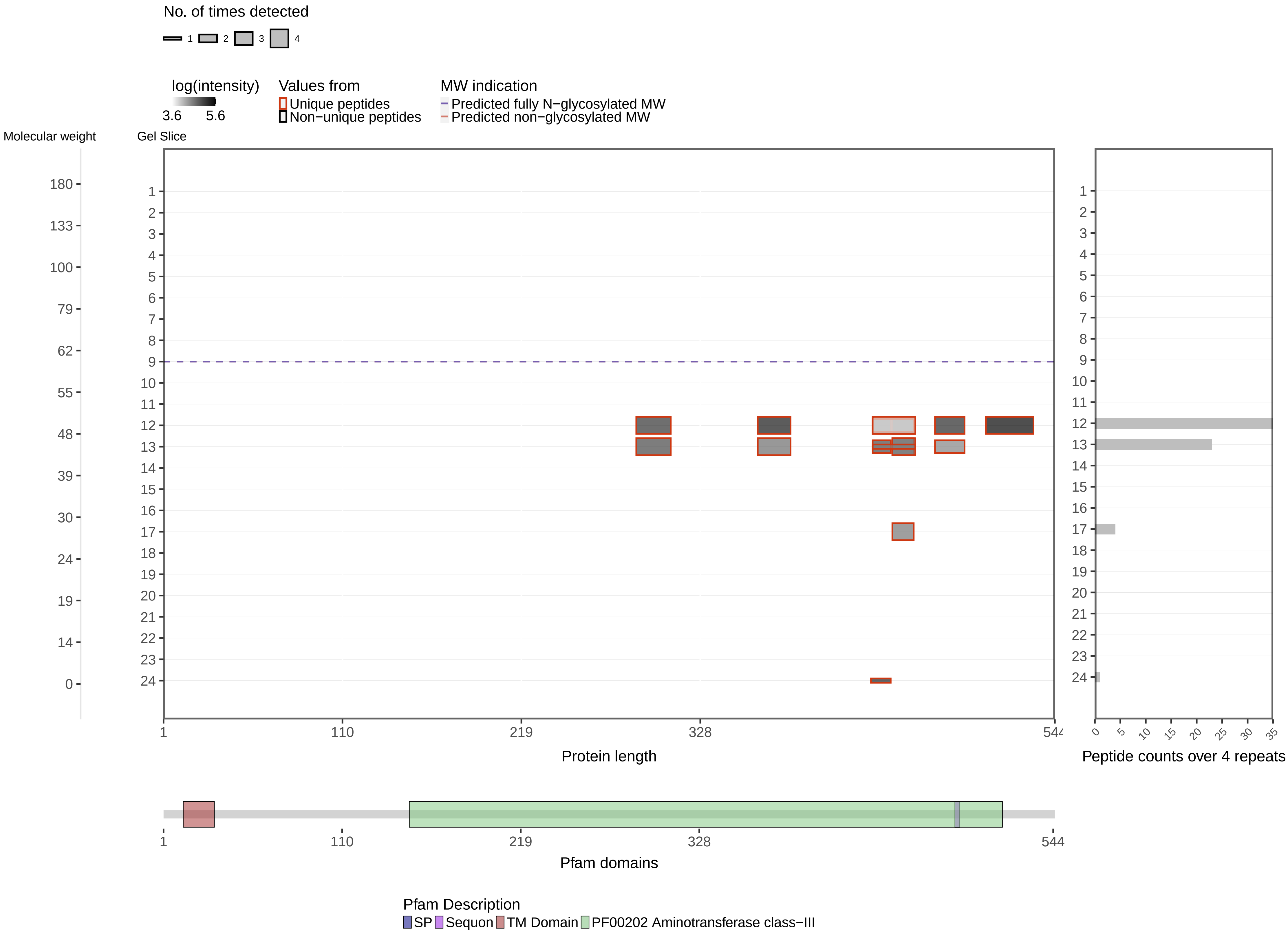


Protein sequence:

CYHGHADPFLVKAGSGVATLGLPDSPGVPKAATTDTLTAPYNDISAVASLFKEHKGEVAAIILEPVVGNAGFIQPNPDFLAAIRKITKENDALLIFDEVMTGFRLAFGGAQEYFGITPDLTTLGKIIGGGLPVGAYGGRRDIMEMVAPAGPMYQAGTLSGNPLAMTAGIHTLKRLQGPGTYEYLDKITGVLTQGILDAGKKTGHAMCGGYIRGMFGFFFAEGPIHNFSDAKKSDTEKFGRFYRGMLEEGVYFAPSQFEAGFTSLAHTSEDIQKTVAAAEKVLKQI*

NbL04g03110.1 lignin−forming_anionic_peroxidase−like [MW − SP = 31.82 kDa]

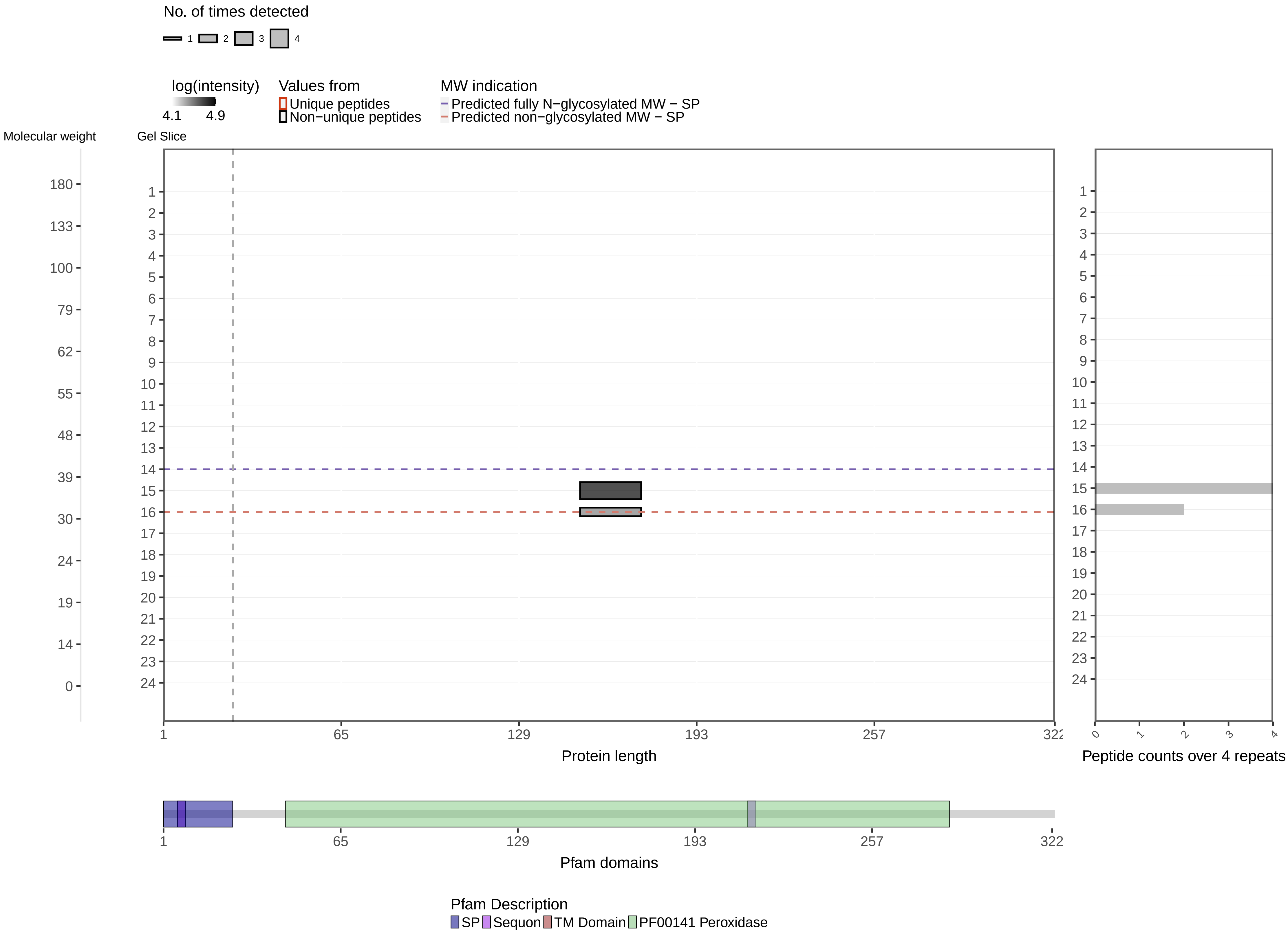


Protein sequence:

RRDSTTASHTLAETDLPGPFDPLNRLISGFASKGLSTRDMVALSGAHSIGQAQCFLFRDRIYGNGTDIDAGFASTRRRQCPKEDQNGNLAPLDLITPNQLDNNYFKNLRQRKGLLQSDQILLSGGSTDDIVSEYSNSPPAFASDFAAAMIRMGDISPLTGQNGIIRTVCGSIN*

NbL04g03100.1 lignin−forming_anionic_peroxidase−like [MW − SP = 31.79 kDa]

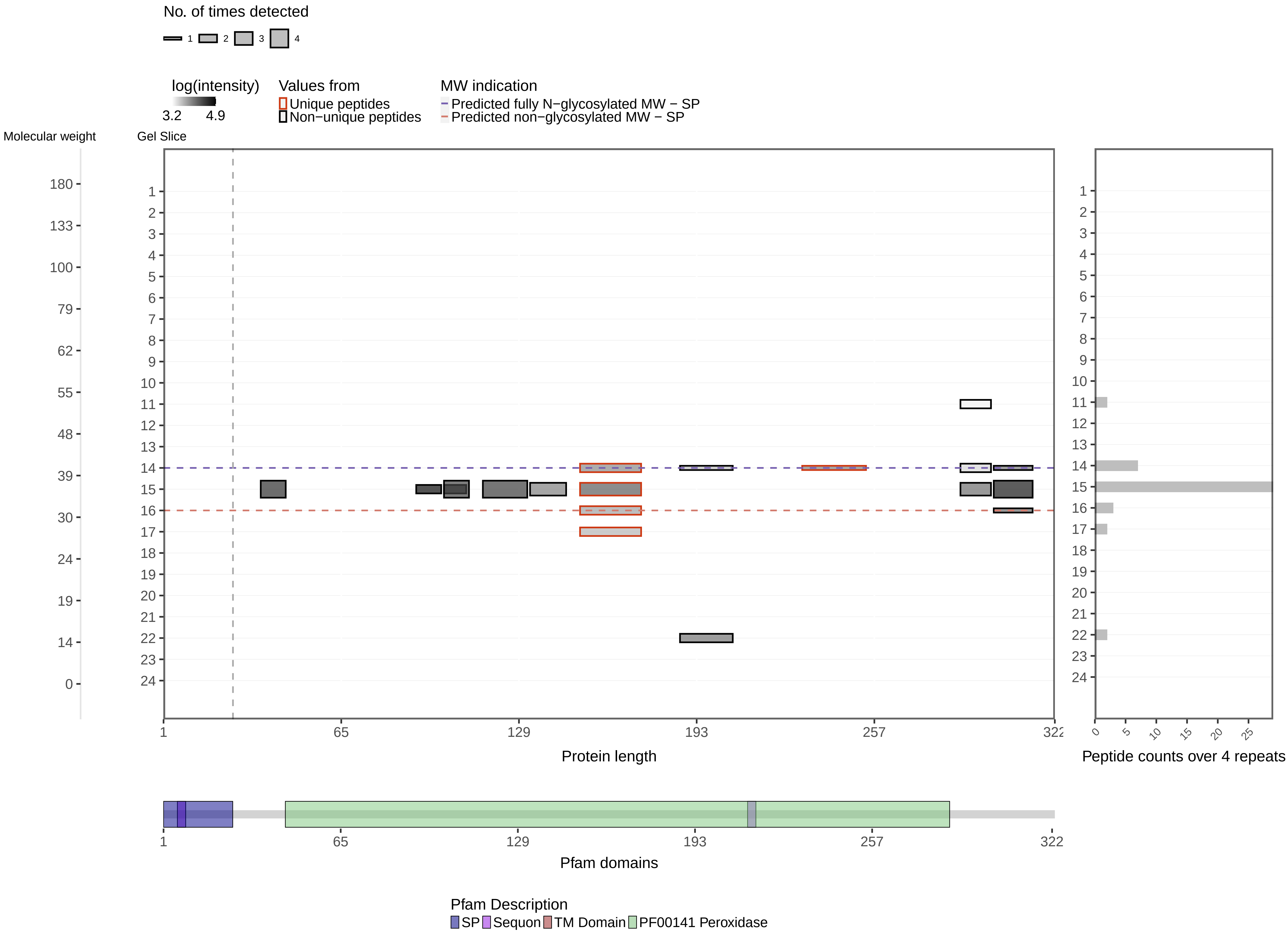


Protein sequence:

RDSTTASHTLAETGLPGPFDPLNRLISGFASKGLSTRDMVALSGAHSIGQAQCFLFRDRIYGNGTDIDAGFASTRRRQCPKEDQNGNLAPLDLVTPNQLDNNYFKNLRQRKGLLQSDQVLLSGGSTDDIVSEYSNSPRAFASDFAAAMIRMGDISPLTGQNGIIRTVCGSIN*

NbL04g03020.1 histidinol−phosphate_aminotransferase_chloroplastic [MW = 46.23 kDa]

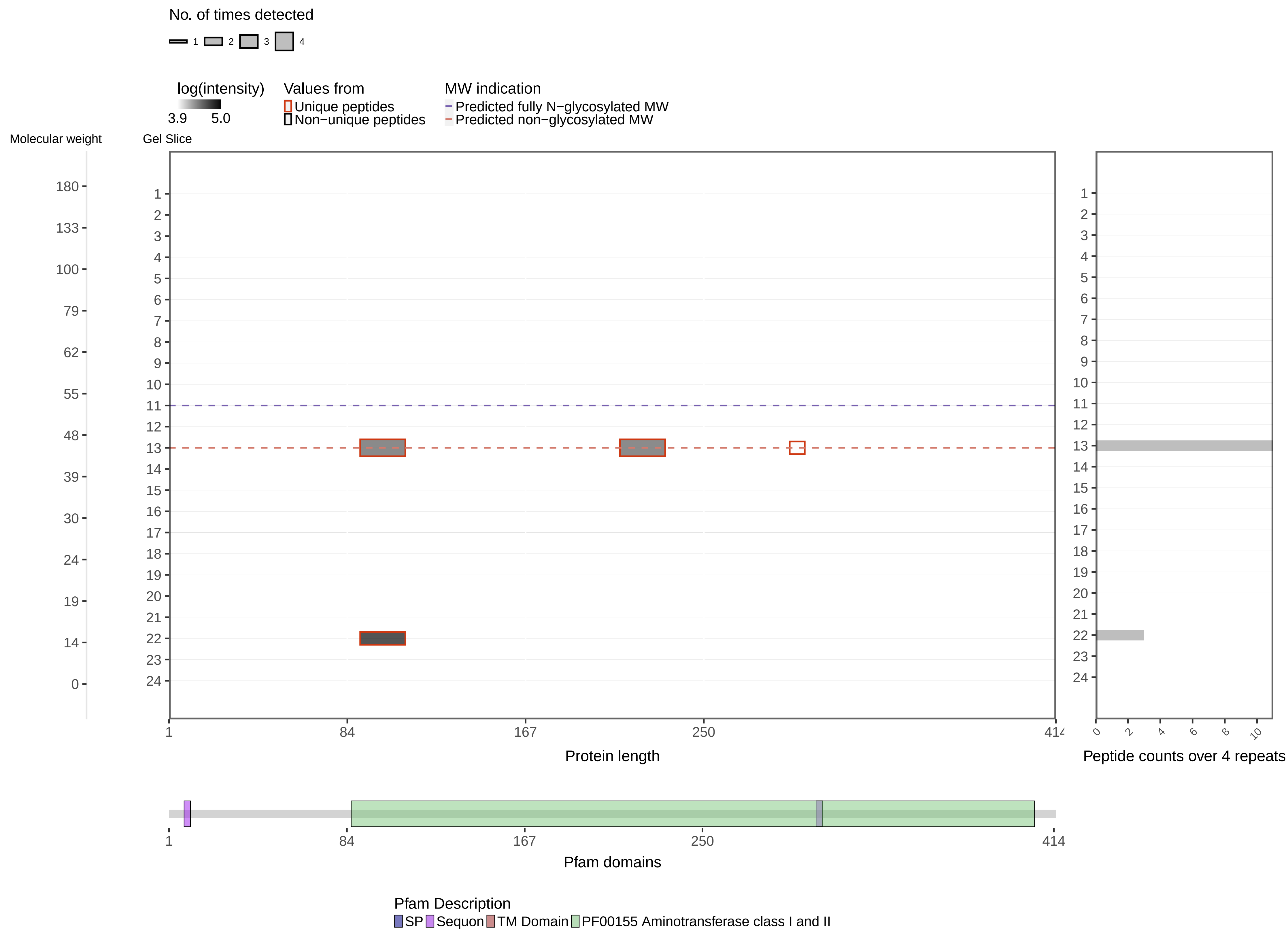


Protein sequence:

SLDVERIAEVVEHEKPKCIFLTSPNNPDGSIIDGETLLKILDLPILVILDEAYVEFSGMESKMKWVKKHENLIVLRTFSKRAGLAGLRVGYGAFPKSIIEFLWRAKQPYNVSVAAEVAACAALNNPAYLENVKVALVQERERLFNLLKEVPFLDPYPSYSNFILCKVTSGMDAKKLKEDLATMGVMIRHYNSKELKGYVRVSVGKPEHTEALMKCLKHFY*

NbL04g02980.1 DNA_repair_RAD52−like_protein_2_chloroplastic [MW = 23.13 kDa]

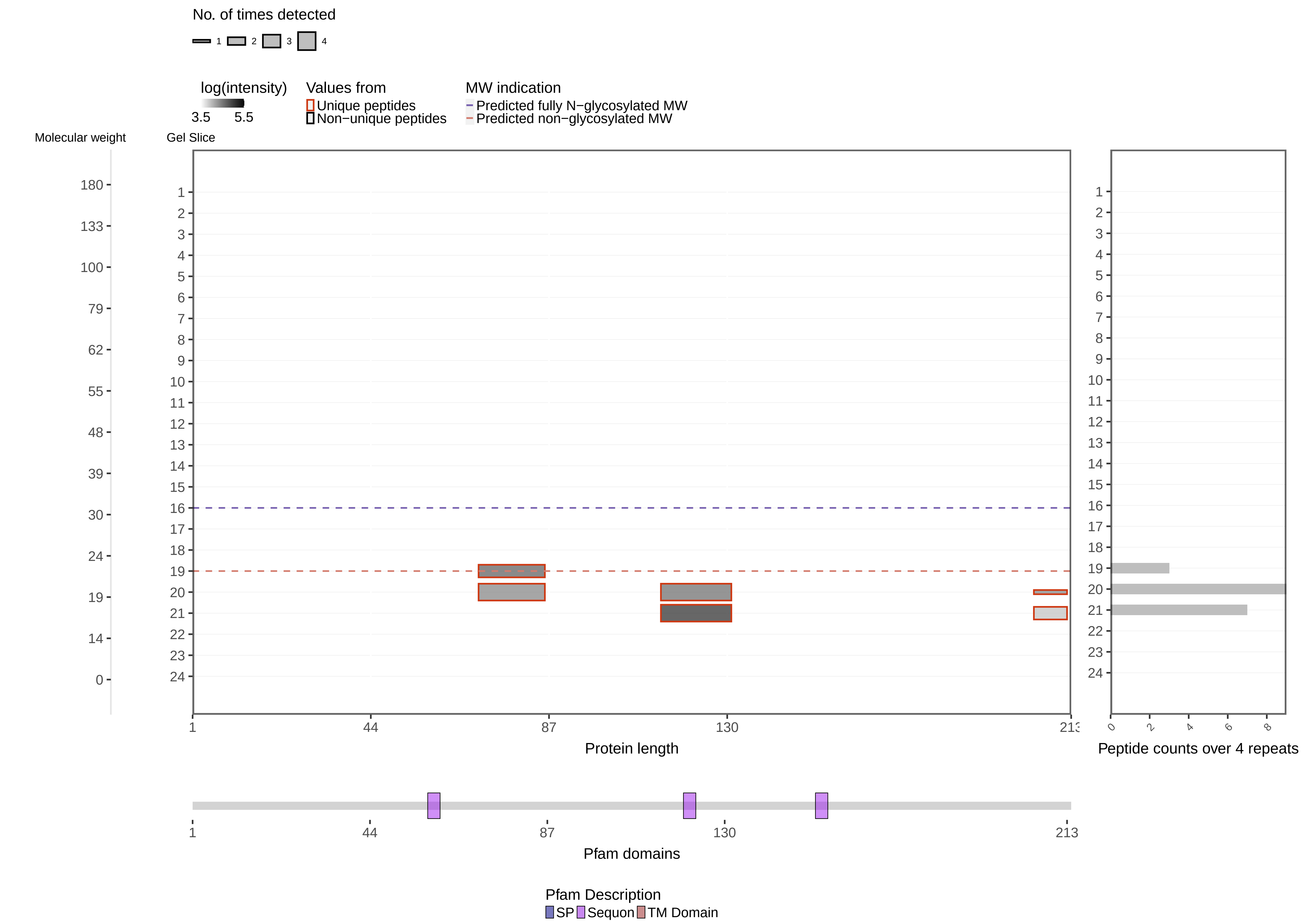


Protein sequence:

TRPLAEILRDLNKRIADNIIKANDGDDNSTFIPWYHANRMLSFYAPGWCGEVRDVTFSDNGSVTVVYRVTIRGSDGEAHRESTGTVSPGNGPSTDLVAAAEEIAFCRACARFGLGLYLYH*

### NbL04g02880.1 glycine−−tRNA_ligase_mitochondrial_1−like [MW = 81.33 kDa]

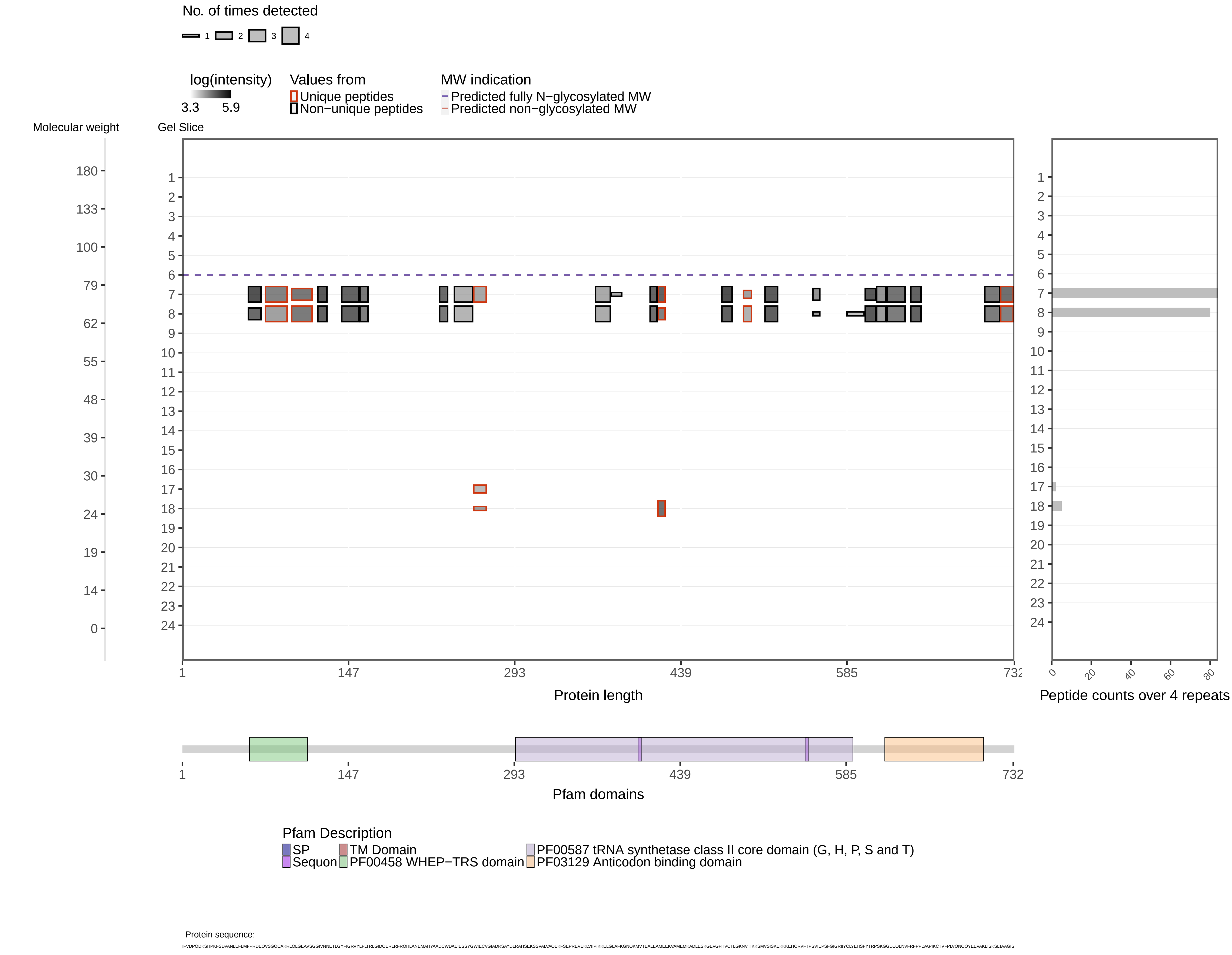

### NbL04g02860.1 proteasome_subunit_beta_type−3−A−like [MW = 22.69 kDa]

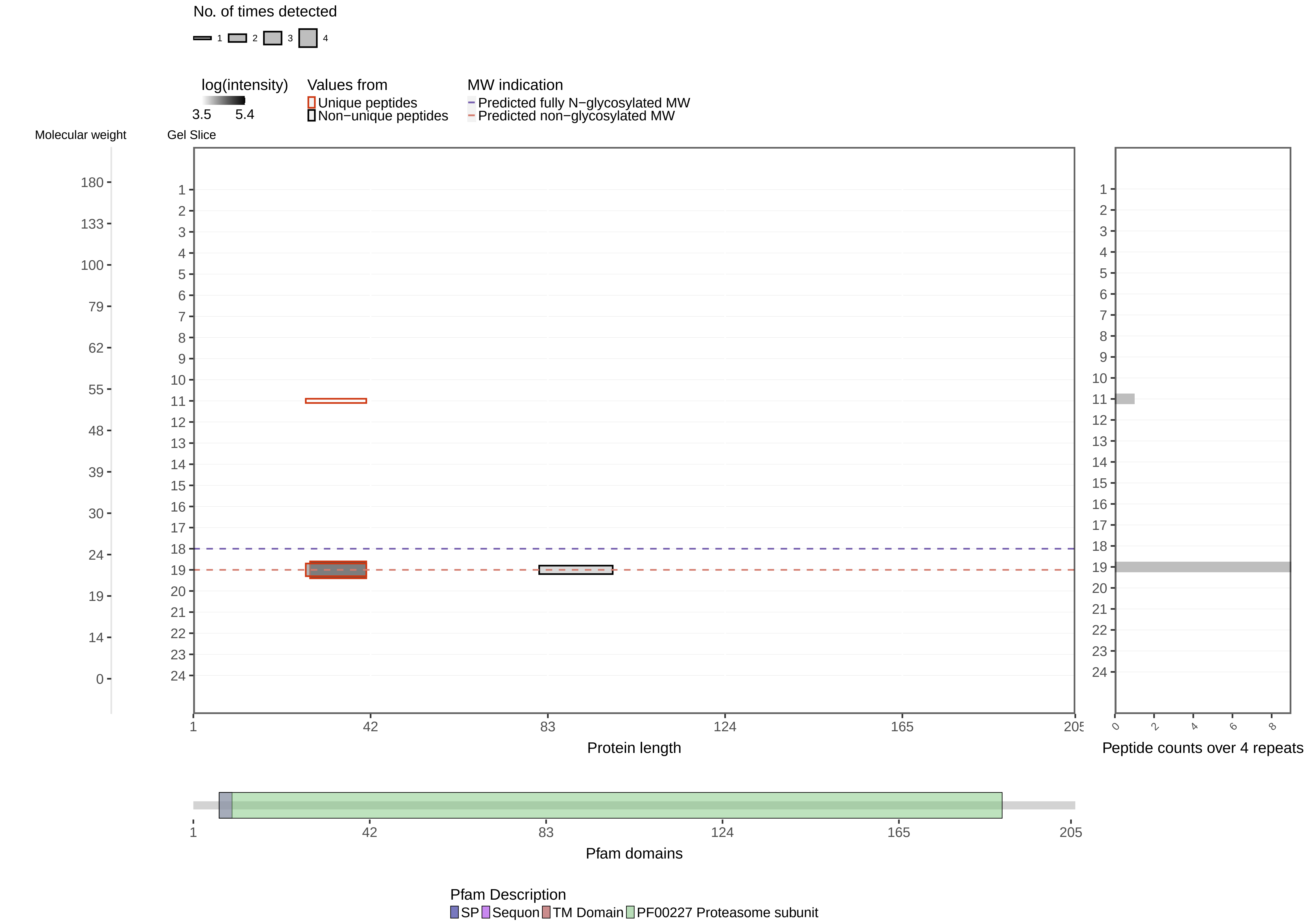


Protein sequence:

.VSAILYEKRFGPYFCQPVIAGLGDDDKPFICTMDAIGAKELAKDFVVAGTASESLYGACEAMFKPDMGPEELFETISQALLSSVDRDCLSGWGGHVYVVTPTEVTEKILKGRMD*

NbL04g02850.1 psbP_domain−containing_protein_2_chloroplastic−like [MW = 28.07 kDa]

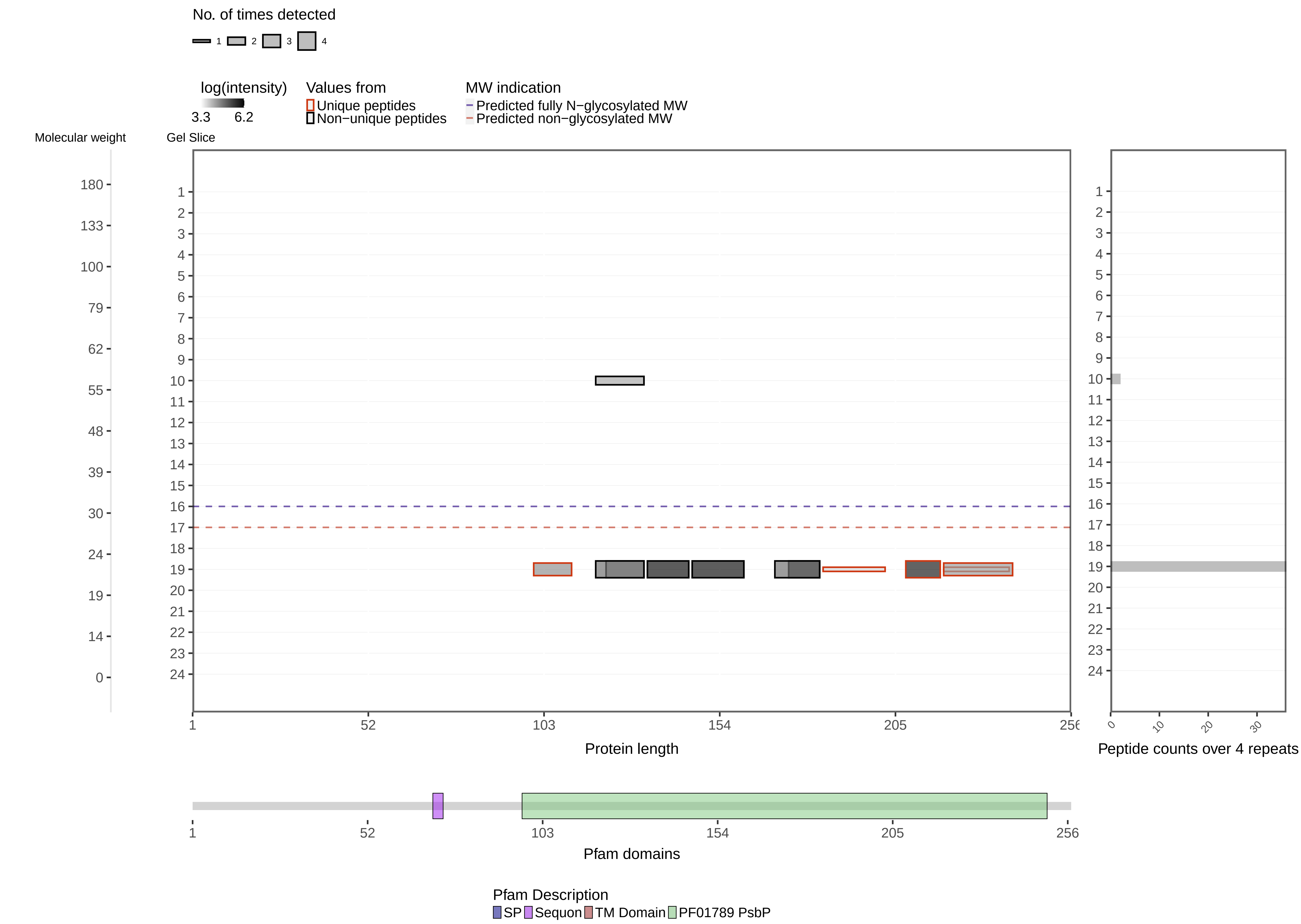


Protein sequence:

WIKVDKAGATVLFQDANKGSNNVGVVVIPVKITTLSQFGTPQFVADKLILAEKRKESTKEAEVVAVSERTGEGGLQVYEFEYTIDSTRGGKKRIFSAAFVASGKLYLLNIANSDGLESPIDPDRRNTLEQILHSFDTAPST*

### NbL04g02500.1 cell_division_cycle_5−like_protein [MW = 118.18 kDa]

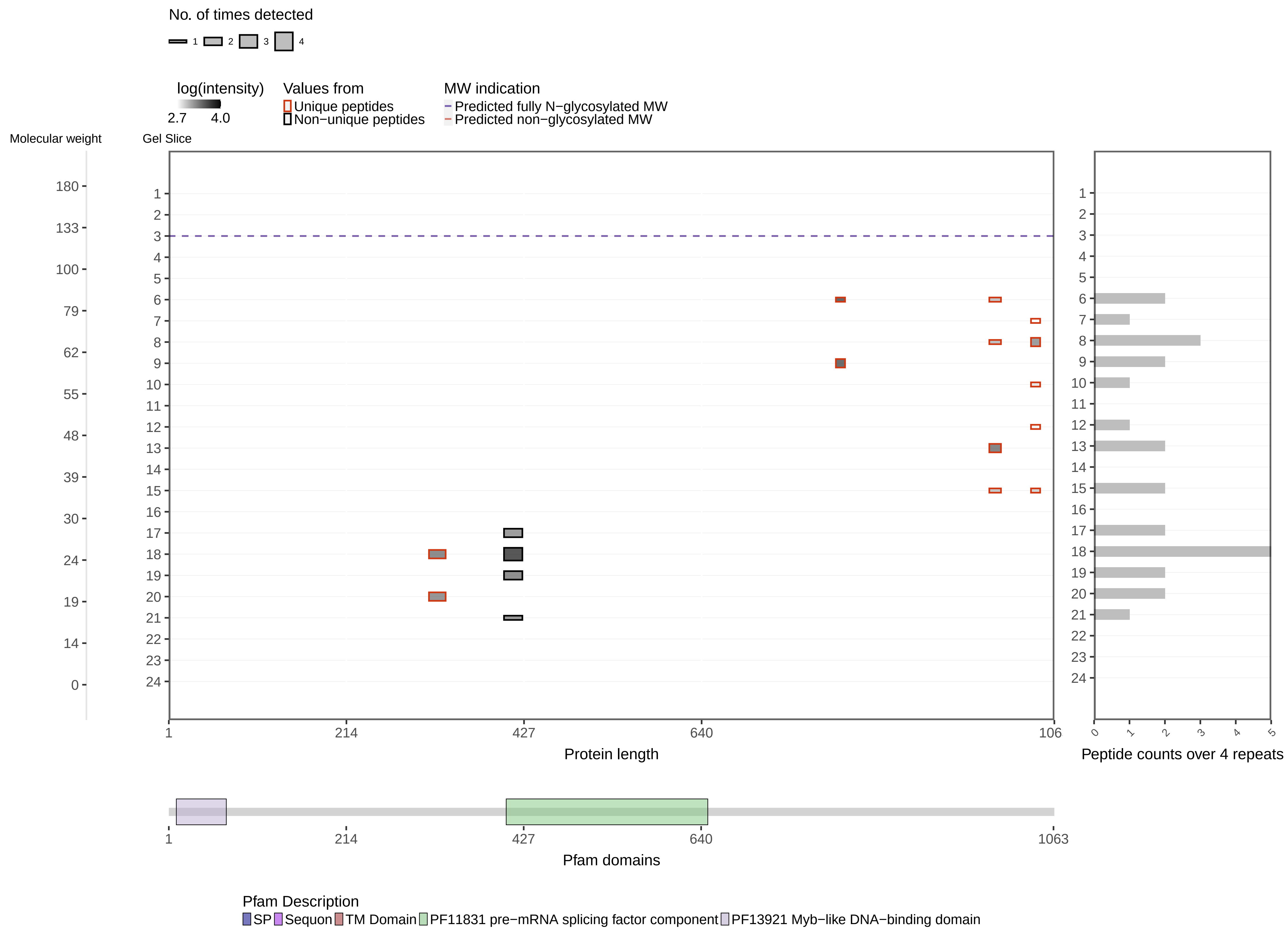


Protein sequence:

### NbL04g02300.1 splicing_factor_3B_subunit_3−like [MW = 134.62 kDa]

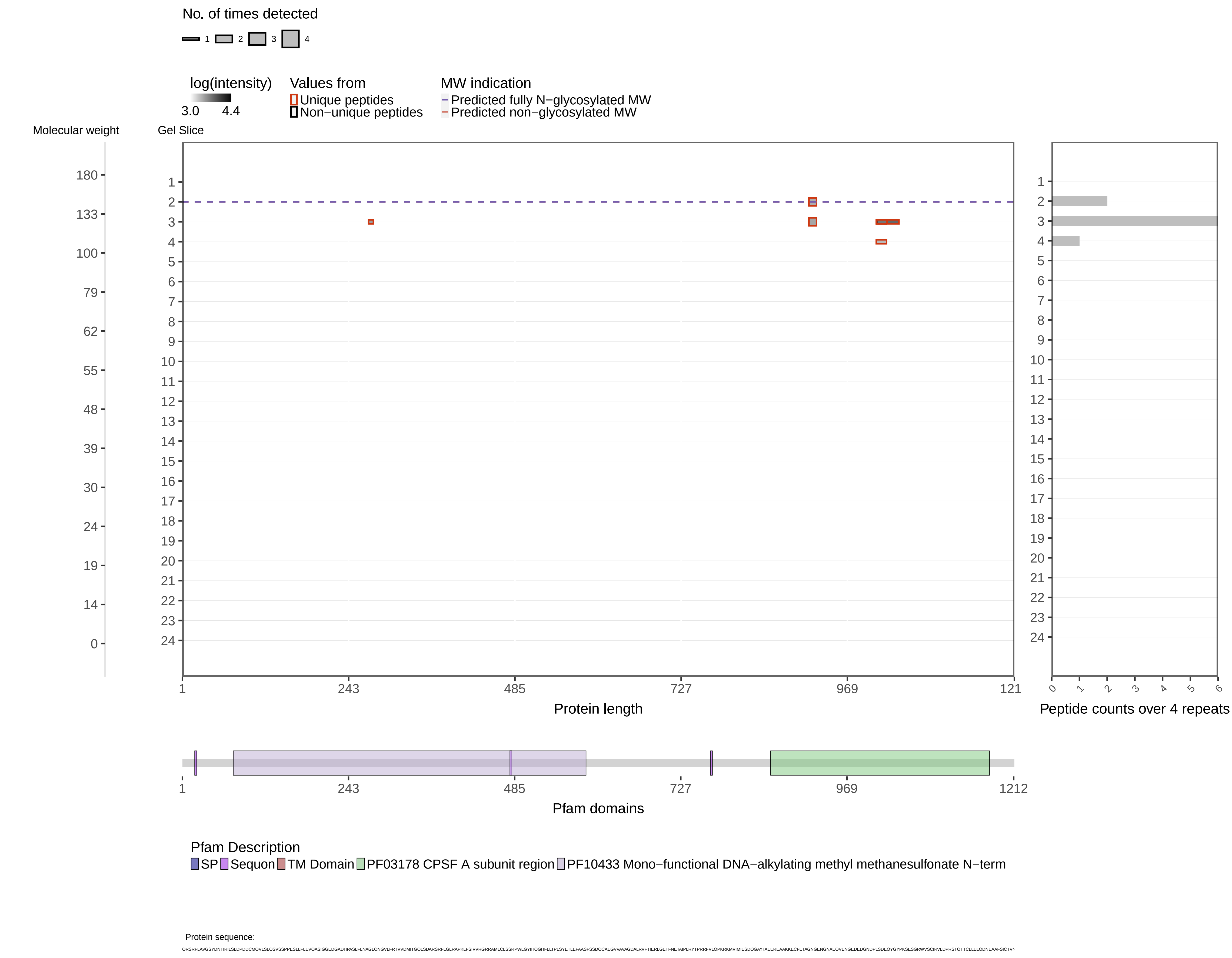

### NbL04g02270.1 histidine_biosynthesis_bifunctional_protein_hisIE_chloroplastic [MW = 31.45 kDa]

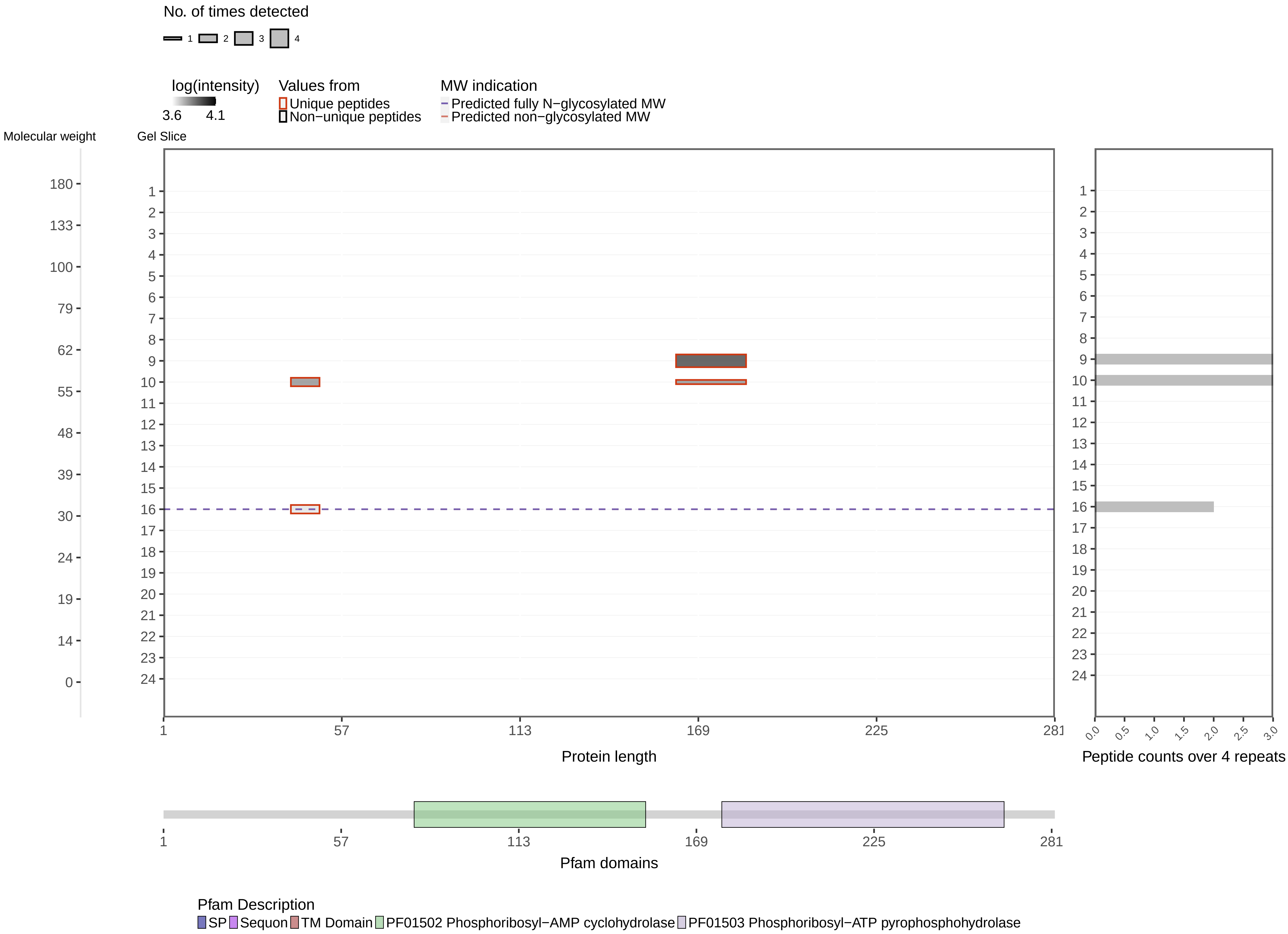


Protein sequence:

CDRDSIIYLGKPDGPTCHTGSETCYYTSVDGILKNTEVEKNELAMTTLYALESTINKRKAETSSSSSGKPSWTKRLLLDDKLLCSKIREEADELCRTLEENEDKGRTASEMADVLYHAMVLLSLRGVKIEEVLQVLRQRFSKSGIEEKNSRKS*

NbL04g02220.1 2−oxoglutarate_dehydrogenase_mitochondrial−like [MW = 115.56 kDa]

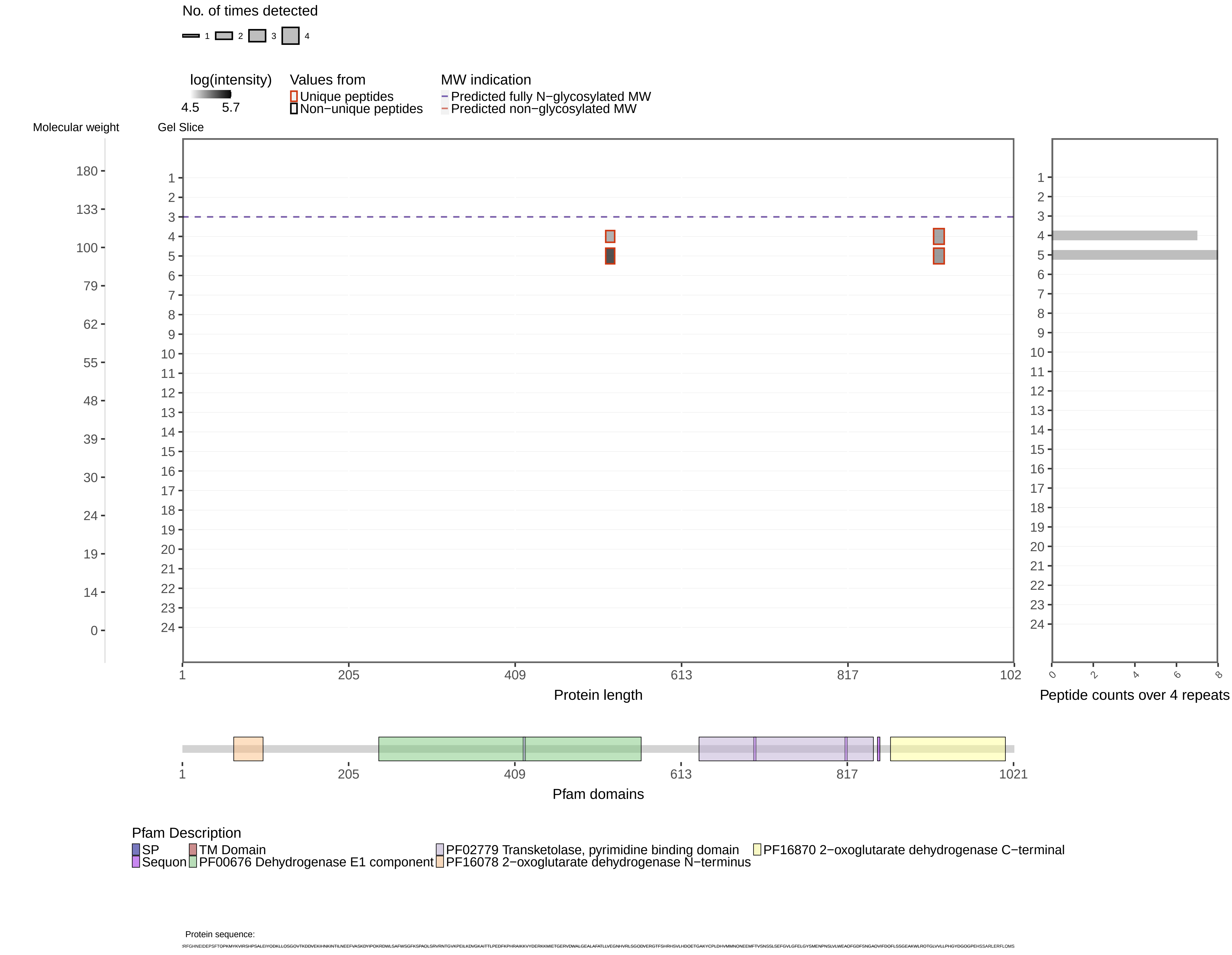

### NbL04g02200.1 actin−depolymerizing_factor_2 [MW = 23.93 kDa]

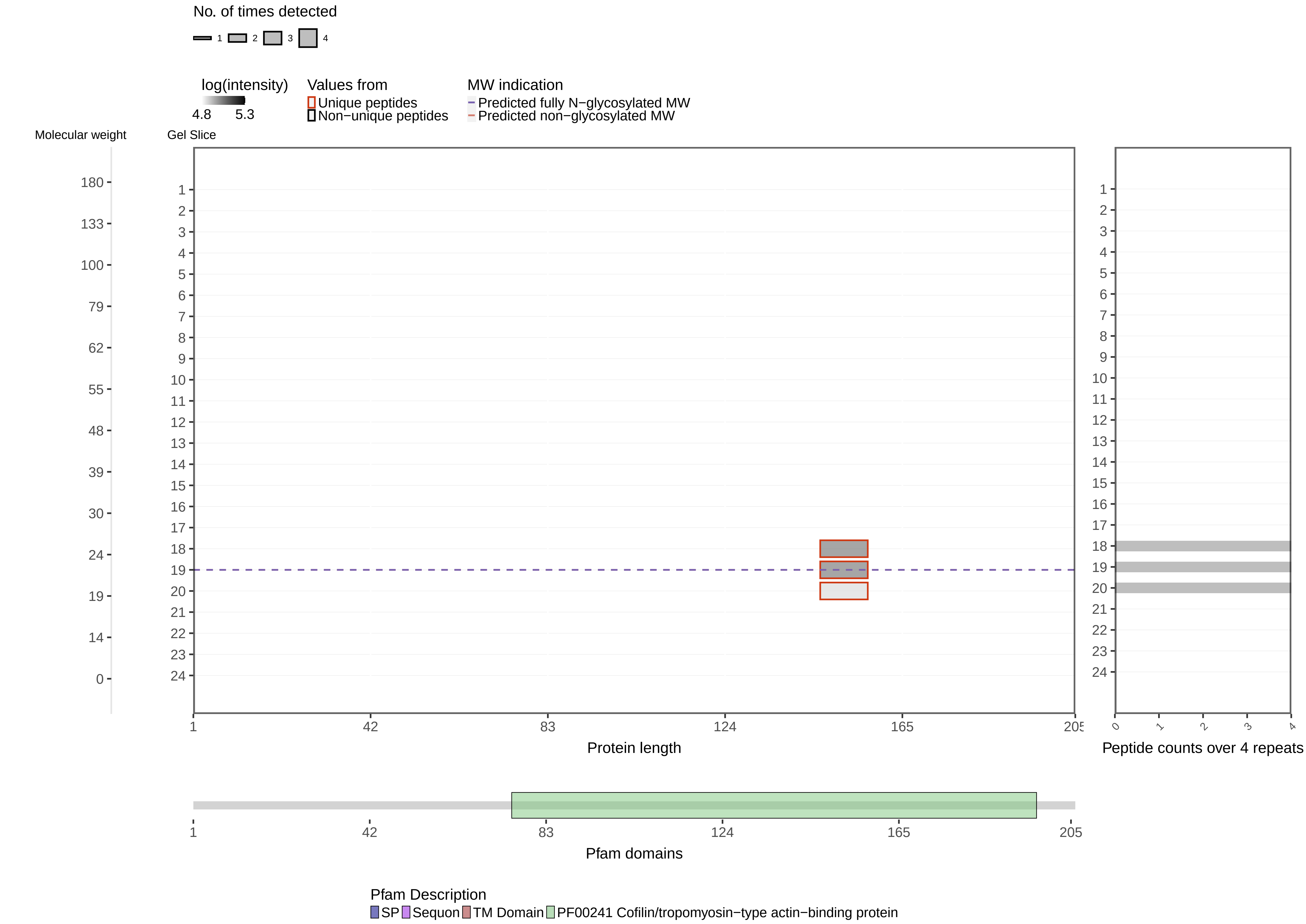


Protein sequence:

RFIIYKIEEKQKQVVVEKLGEPTESYEDFTAGLPADECRYAVYDFDFTTEENVQKSRIFFIAWSPDTARVRSKMIYASSKDRFKRELDGFQVELQATDPTEMGLDVFRSRVGGIEC*

NbL04g02180.1 heat_shock_cognate_70_kDa_protein_2−like [MW = 71.31 kDa]

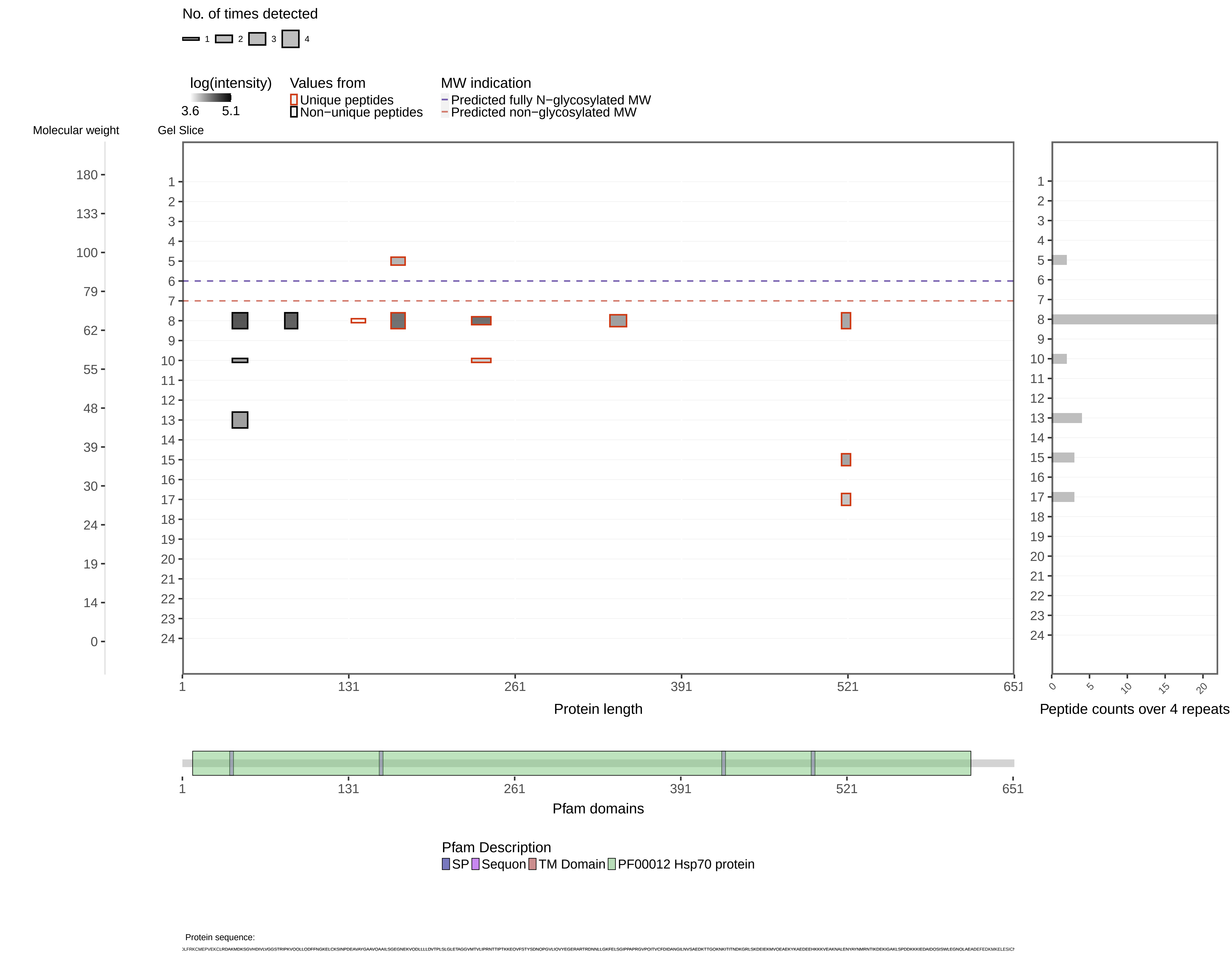

NbL04g02150.1 triosephosphate_isomerase_cytosolic [MW = 27.56 kDa]

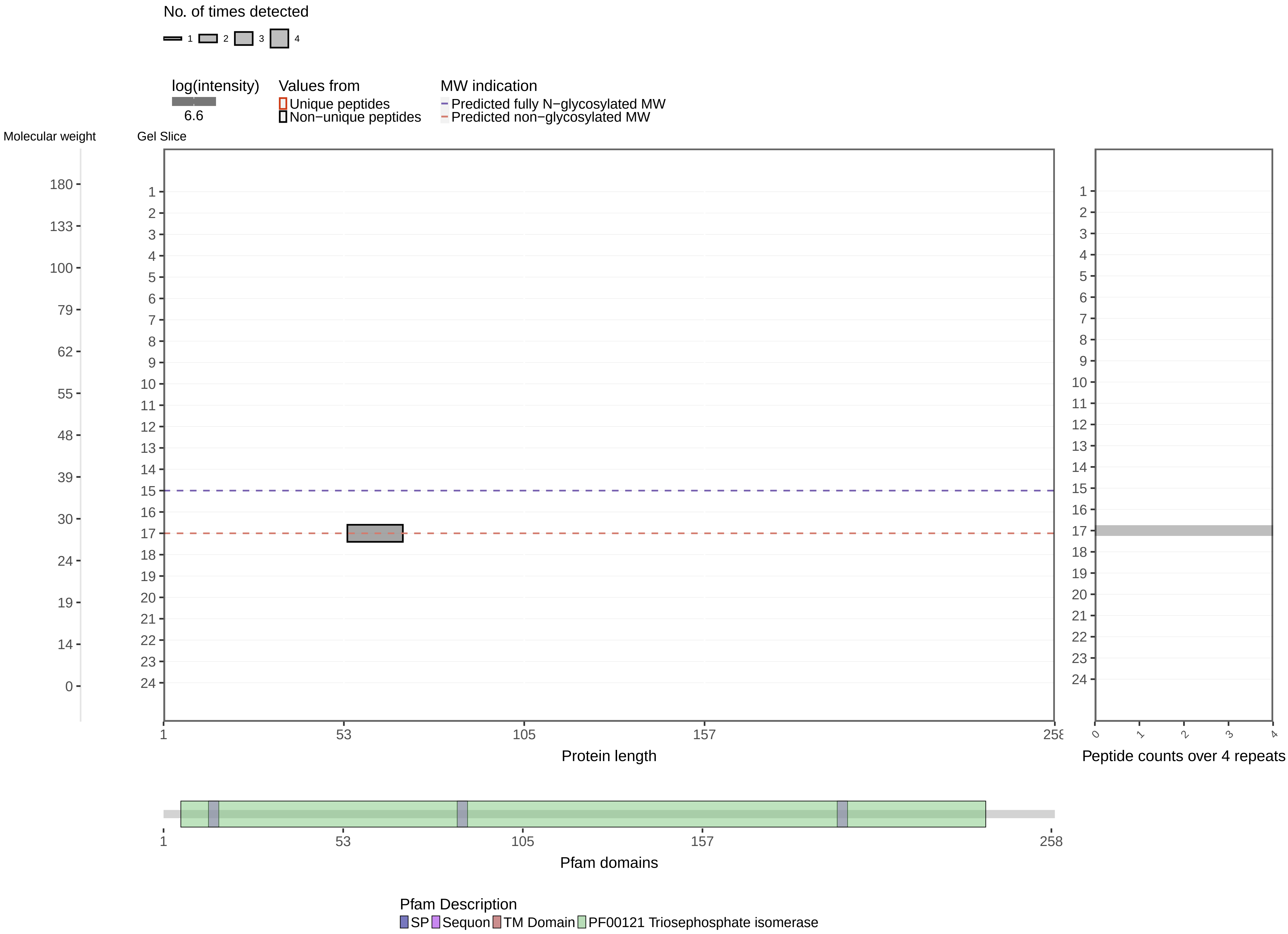


Protein sequence:

AYAISQGLKVIACVGETLEQRESGSTIDVVAAQTKAIADRVKDWTKVVIAYEPVWAIGTGKVASPAQAQEVHAELRKWLQVNVSAEVAASTRIIYGGSVNGANCKELAGQPDIDGFLVGGASLKVCENTLELLSSVYSWLNKQ*

### NbL04g02080.1 peptidyl−prolyl_cis−trans_isomerase_FKBP20−1 [MW = 20 kDa]

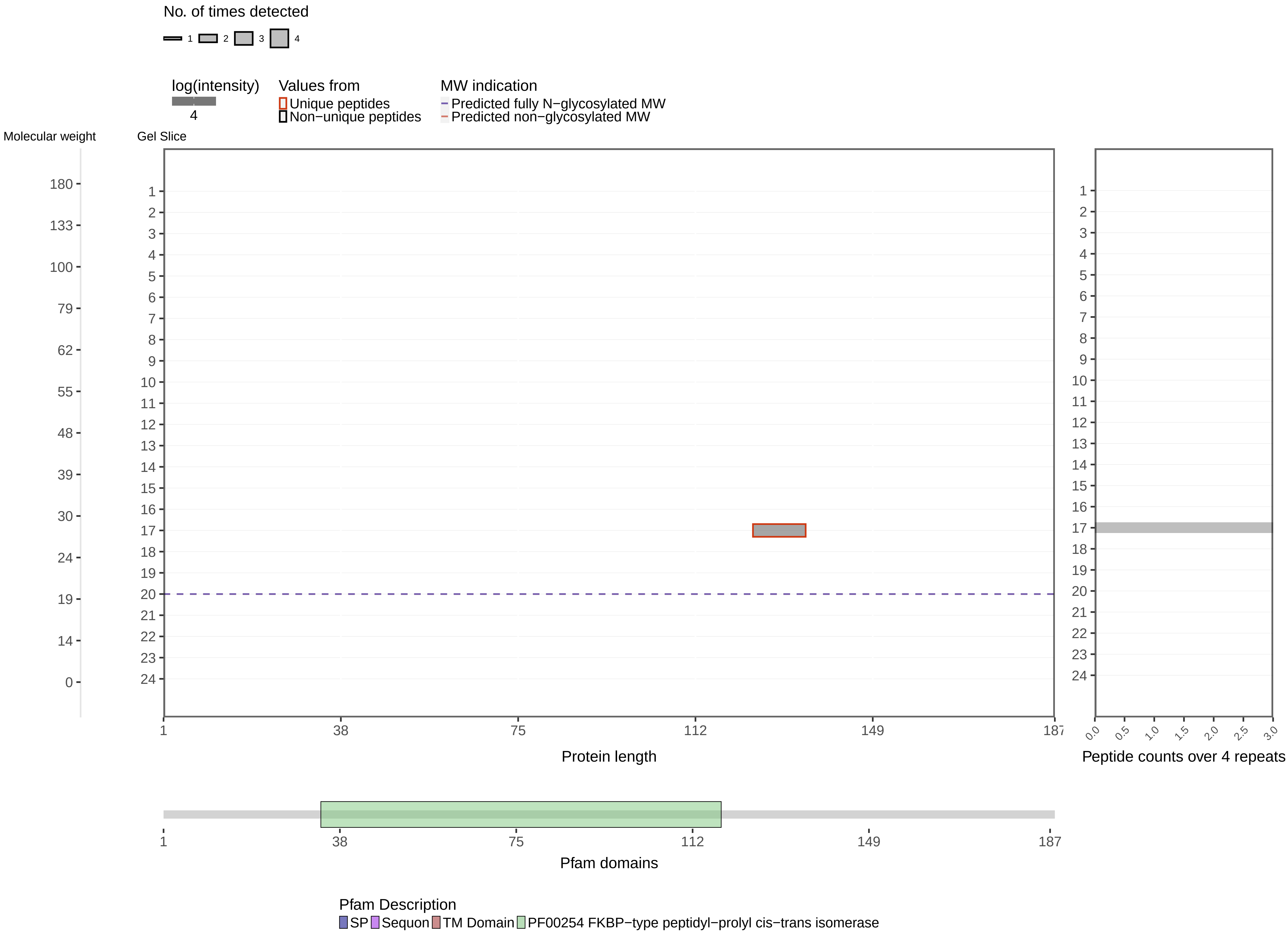


Protein sequence:

KVGEIAVITCKPEYAYGSAGSPPDIPPDATLVFEVELVACRPRKGCSISSVSDERARLEELKKQREMAAAAKEEEKKKREEAKAAAAARIQAKLEAKKGKGKGKAK*

NbL04g01920.1 uncharacterized_LOC107771755 [MW = 35.97 kDa]

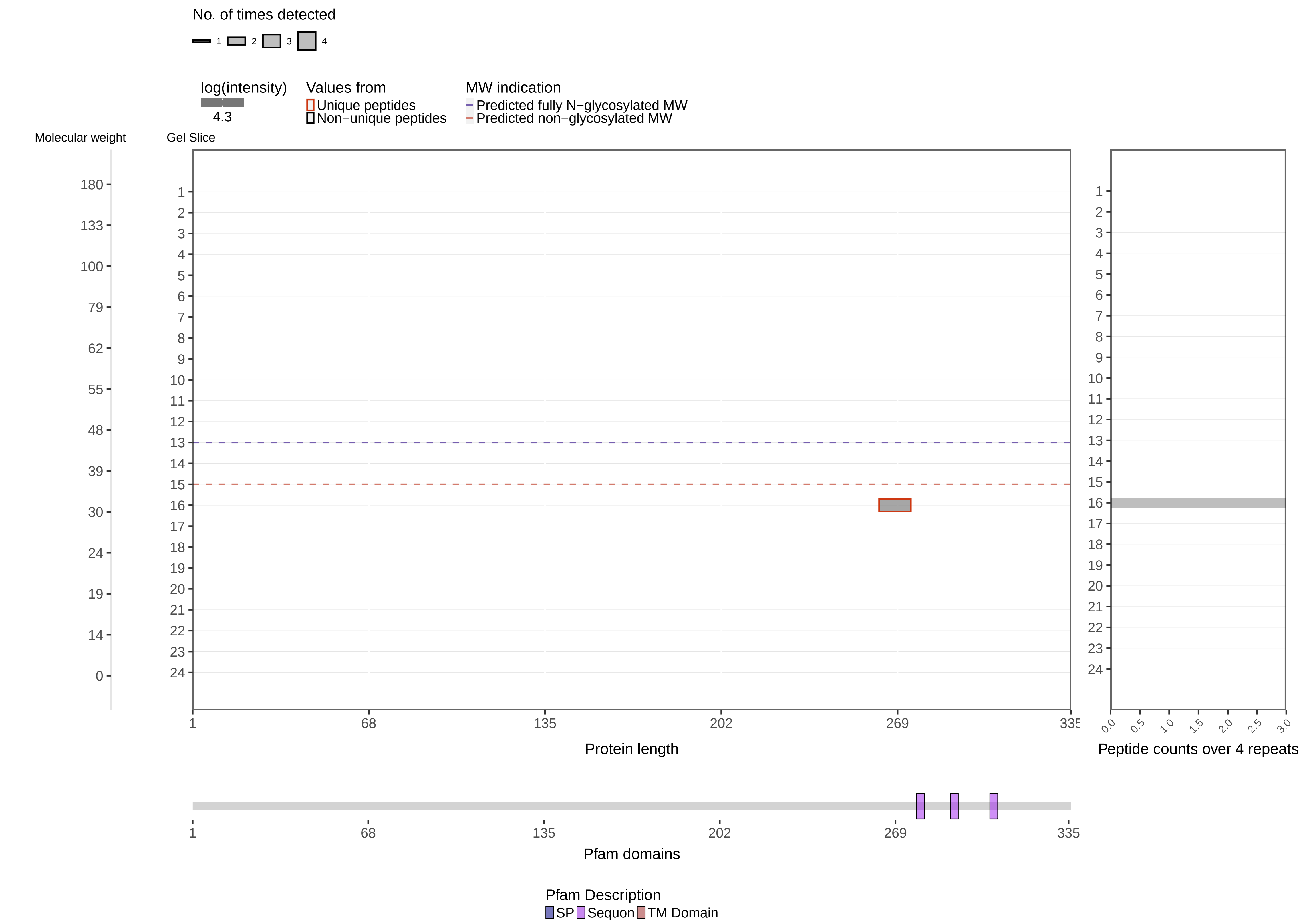


Protein sequence:

YLIGSISLVVLGIGSTSPGLLQAAIDGFSSFFPDYQERIARHEAAHFLVAYLIGLPILGYSLDIGKEHVNLVDERLEKLIYSGQLDSKELDRLAVVAMAGLAAEGLKYDKVIGQSADLFTLQRFLNRSKPSLSKDQQQNLTRWAVLFAGSLLKNNSKLHEALITAMTNKASVTECIEAIEKAA*

### NbL04g01860.1 acylsugar_acyltransferase_3−like [MW = 51.42 kDa]

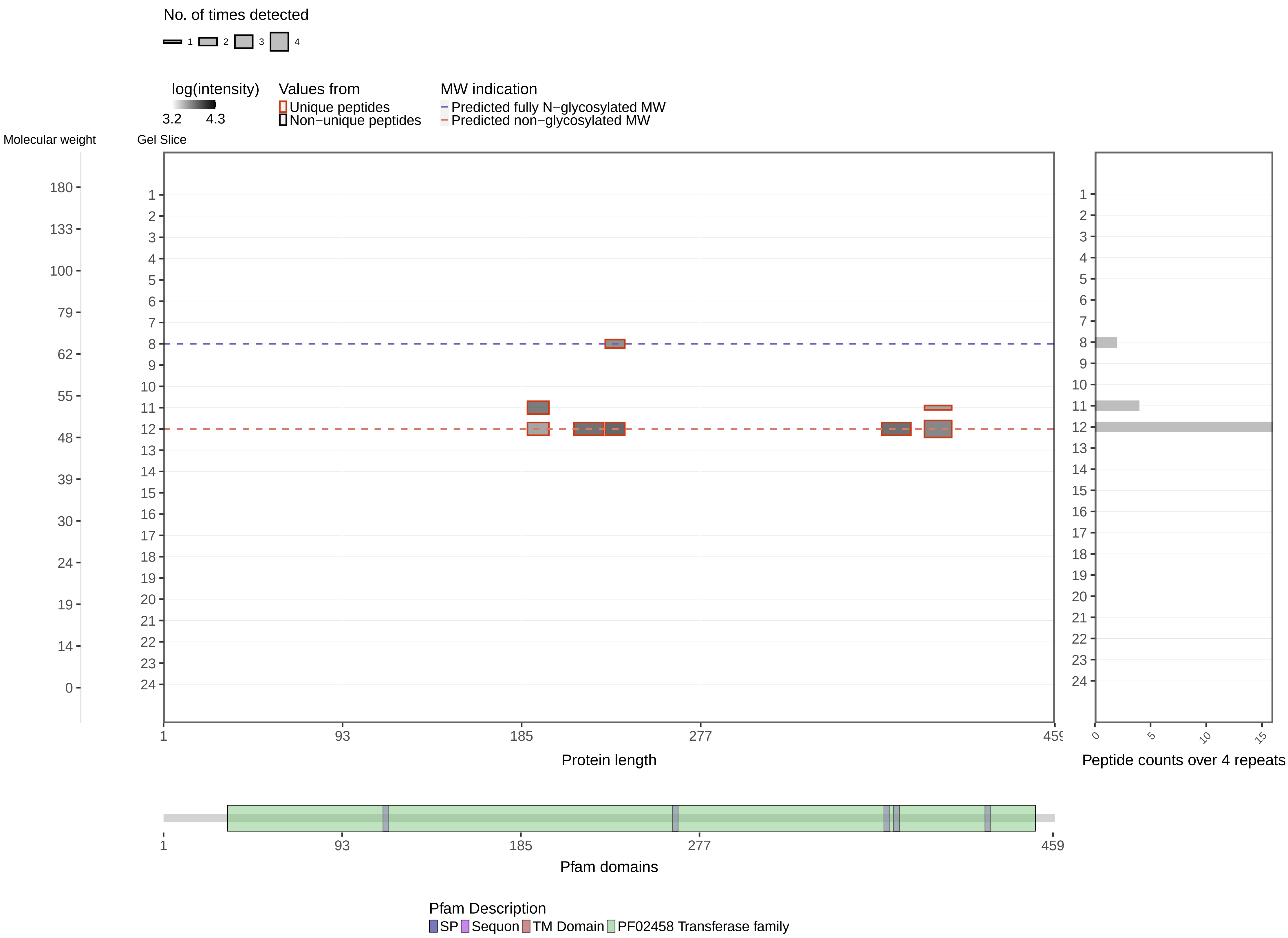


Protein sequence:

AASIYPPTKDMANLPEVVPKGEECSSKSFTFSSSKLAALRARVINNSNIQNPTTTEIVSAFIYQRAMATKKKSLGSICPSVLVQAMNLRPPLPKTLMGNIGSFFSVITTDEKEMDLPRLVGKLREAKEEVRRKYKHDKTDEFLPTTIELYKEANDSFFNNSCYDIYRFSSINKFPFYNVDFGWGKPEKVSSSSNGPVKNMLIVMDNKSGDGVEVVSYMKEQDMSALEKDEELLEFASPNM*

### NbL04g01850.1 acylsugar_acyltransferase_3−like [MW = 51.4 kDa]

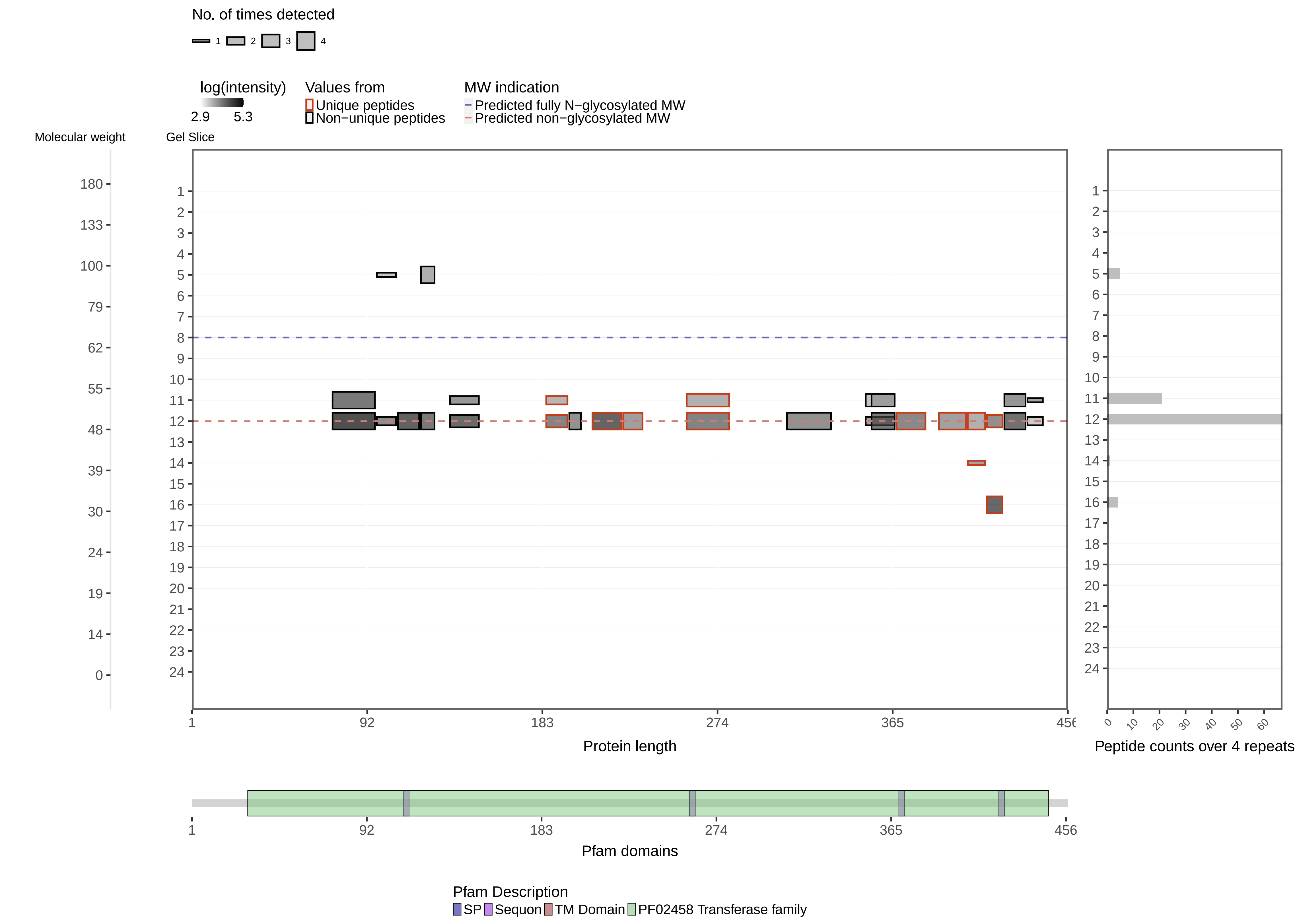


Protein sequence:

SIFPPTKDMINLLEVVPKGEECSSKSFTFSSSKLAALRARIINNSNIQNPTTTEIVSAFIYQRAMATKKKTSGSICPSVLVQAMNLRPPLPKTLMGNIGSFFSVITTDEKEMDLPRLVGKLREAKEEVRRKYKHDKTDEFLPTTIELYKEGNDSFYSNSCYDIYRFSSVHKFPFYNMDFGWGKPEKVSMSSNGPVKNTLMLMDNKSGDGVEVVSYMKEQDMSALEKDEELLEFASPNM*

NbL04g01810.1 reductase_large_subunit−like [MW = 91.34 kDa]

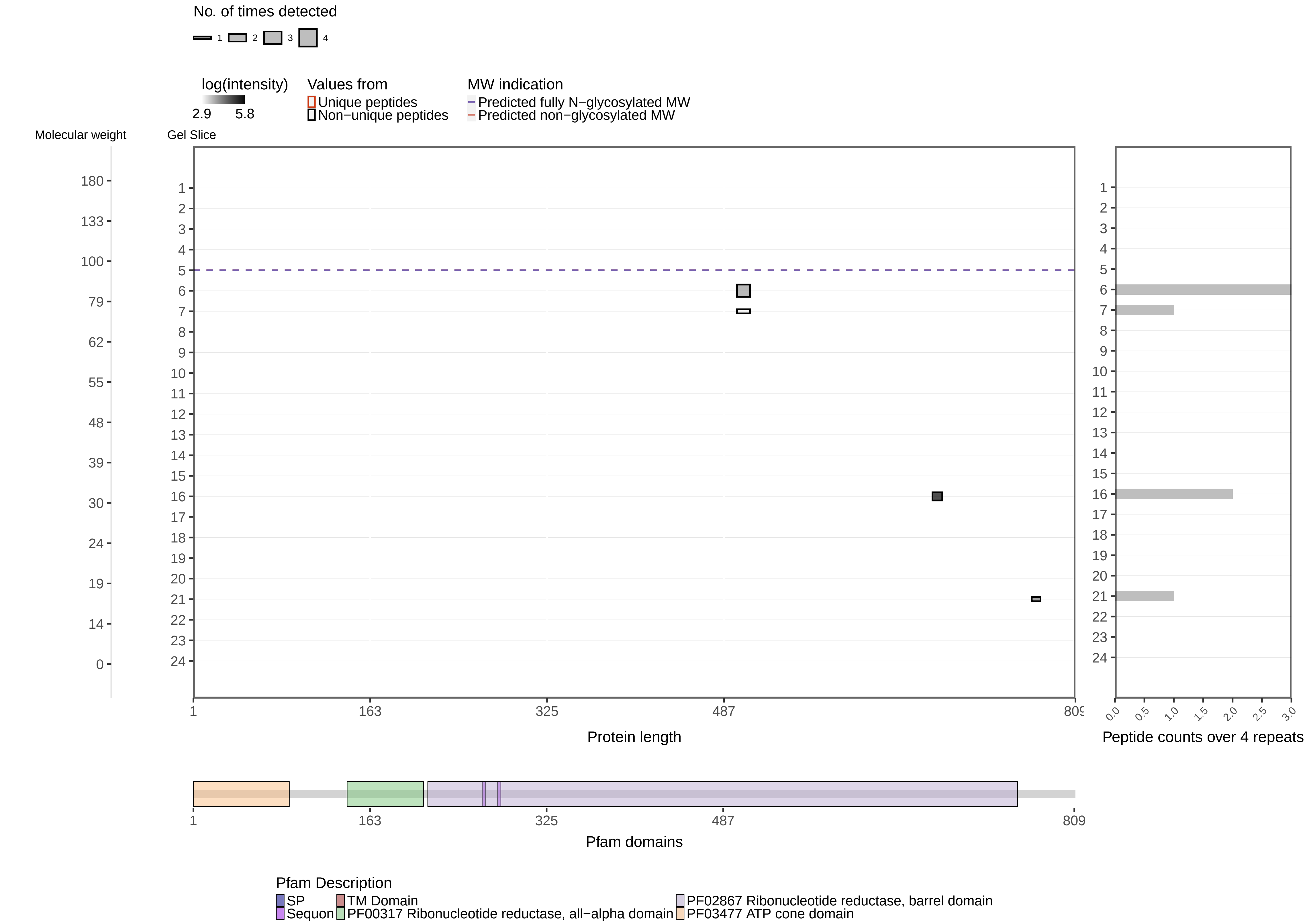


Protein sequence:

VFEILKSQIETGTPYMLYKDSCNRKSNQQNLGTIKSSNLCTEIIEYTSPTETAVCNLASIALPRYVREKEVPTESQPSKLVGSRGSKNRYFDFDKLAEVTALVSTNLNKIIDVNYYPVETAKRSNLRHRPIGLGVQGLADTFILLGMSFDSPEAQQLNKDIFETIYYHALKASSELAAKEGPYETYAGSPVSKGILQPDMWGVTPSNRWDWGVLREMIVKNGVRNSLLVAPMPTASTSQILGNNECFEPYTSNIYSRRVLSGEFVVVNKHLLHDLTEMGLWSPTLKNRIIYDDGSVQ

### NbL04g01790.1 ras−related_protein_Rab11D [MW = 24.59 kDa]

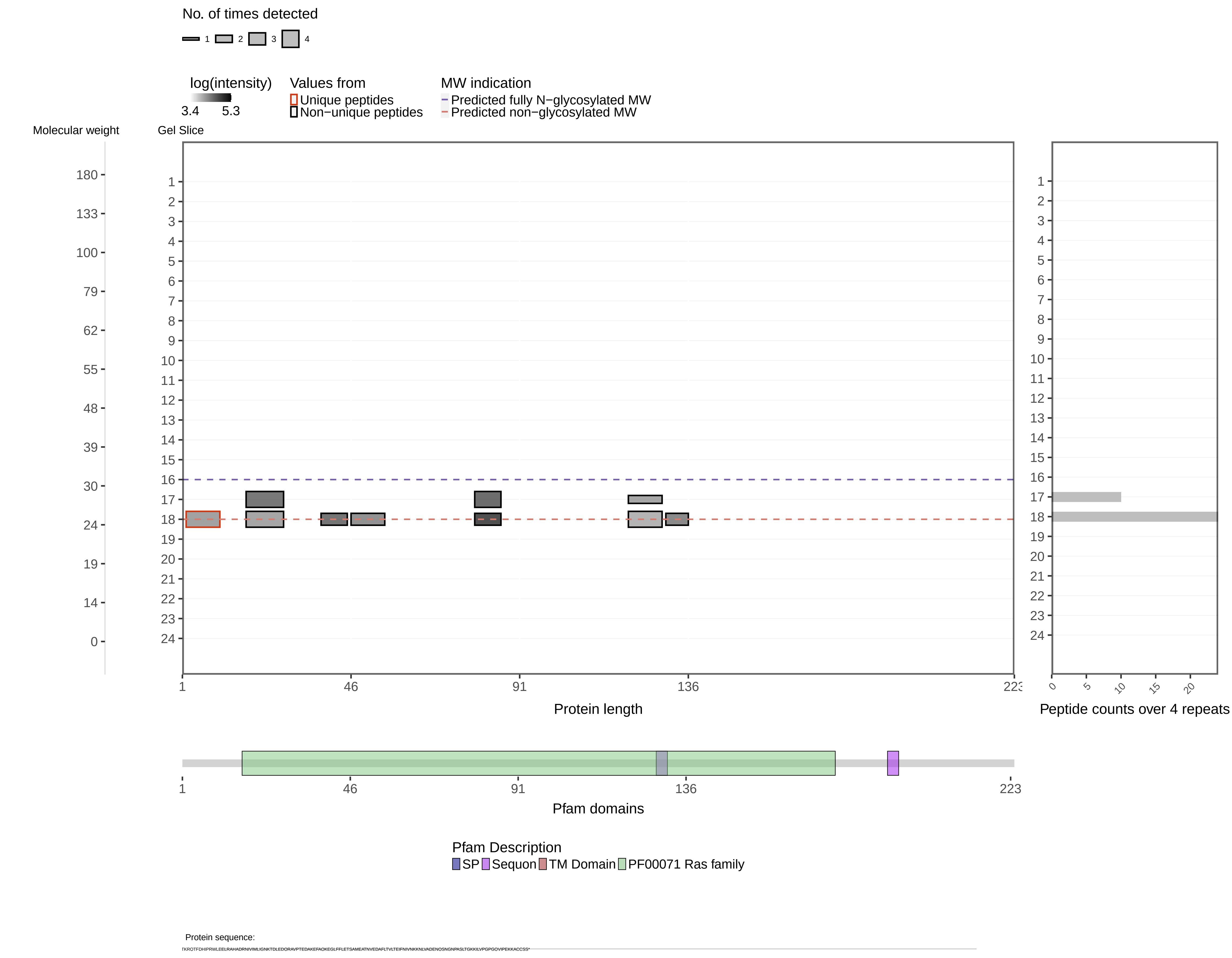

### NbL04g01770.1 14−3−3−like_protein_A [MW = 30.25 kDa]

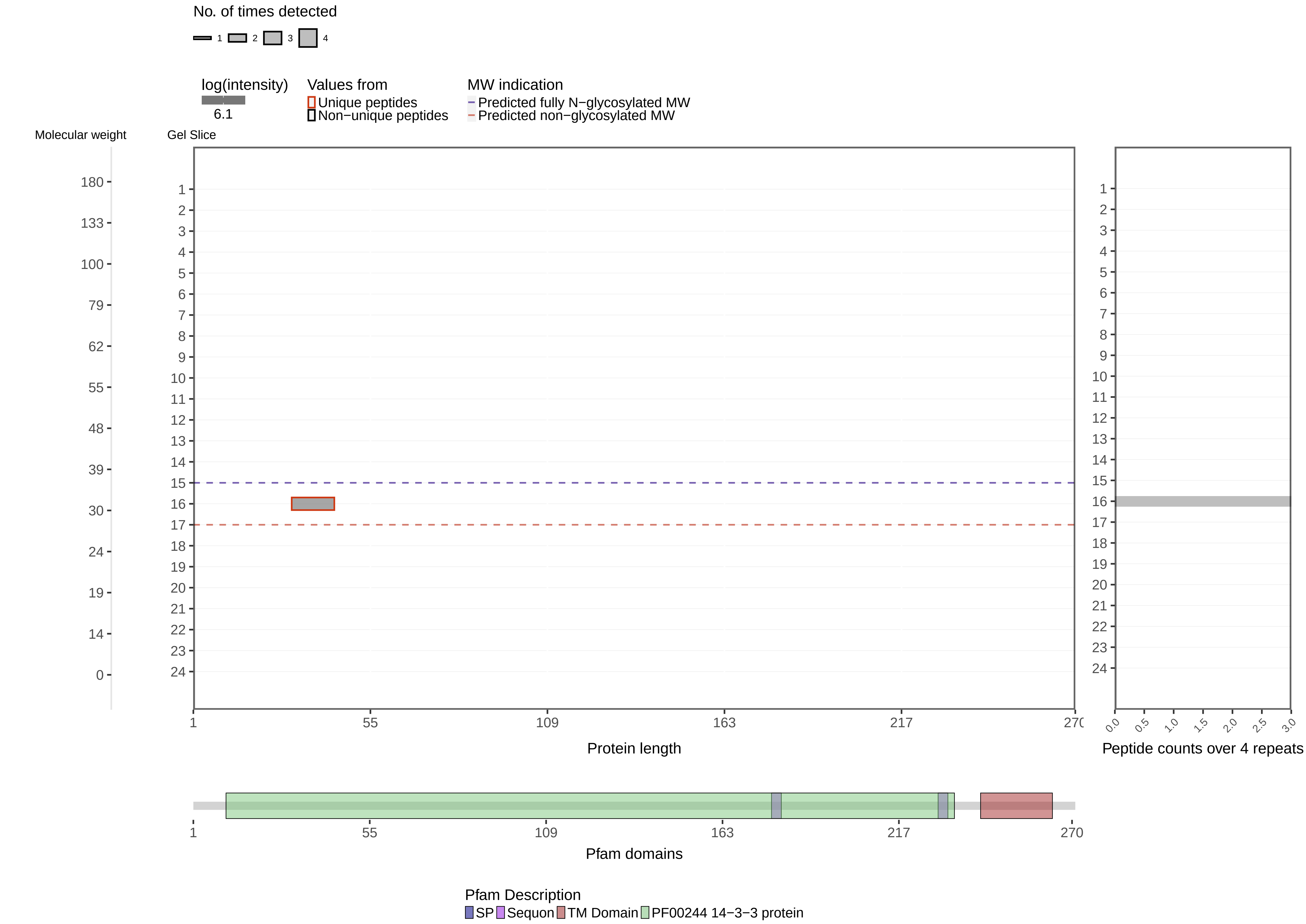


Protein sequence:

FYLKMKGDYYRYLAEFKTGAERKEAAENTLSAYKSAODIANVELAPTHPIRLGLALNFSVFYYEILNSPDRACNLAKOAFDEAIAELDTLGEESYKDSTLIMOLLRDNLTLWTSDMOVSSFVLLFIGLVYLFFDSLANPYLVGFSSSG*

### NbL04g01670.1 ATP−dependent_6−phosphofructokinase_5_chloroplastic [MW = 60.76 kDa]

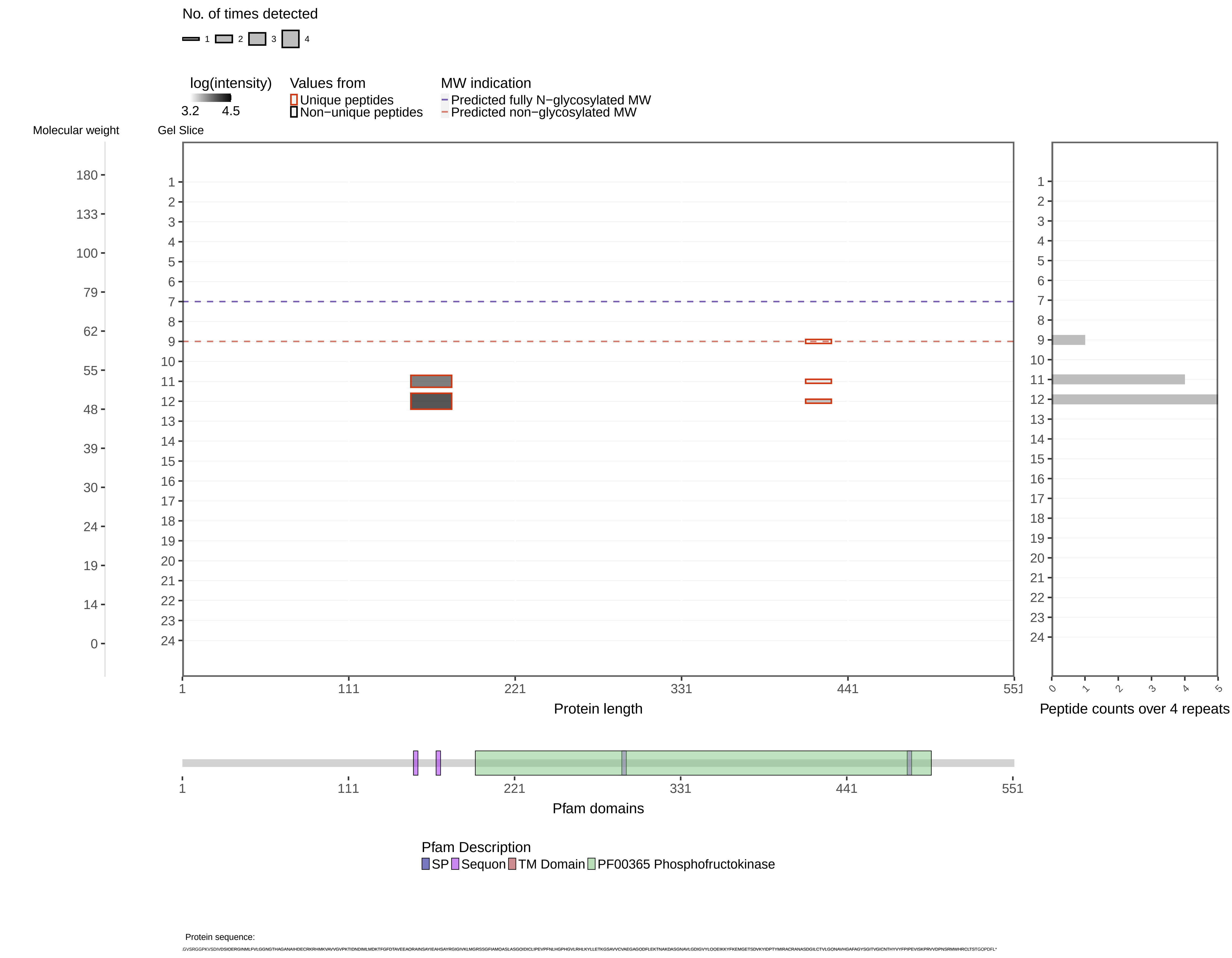

### NbL04g01590.1 54S_ribosomal_protein_L51_mitochondrial−like [MW = 14.53 kDa]

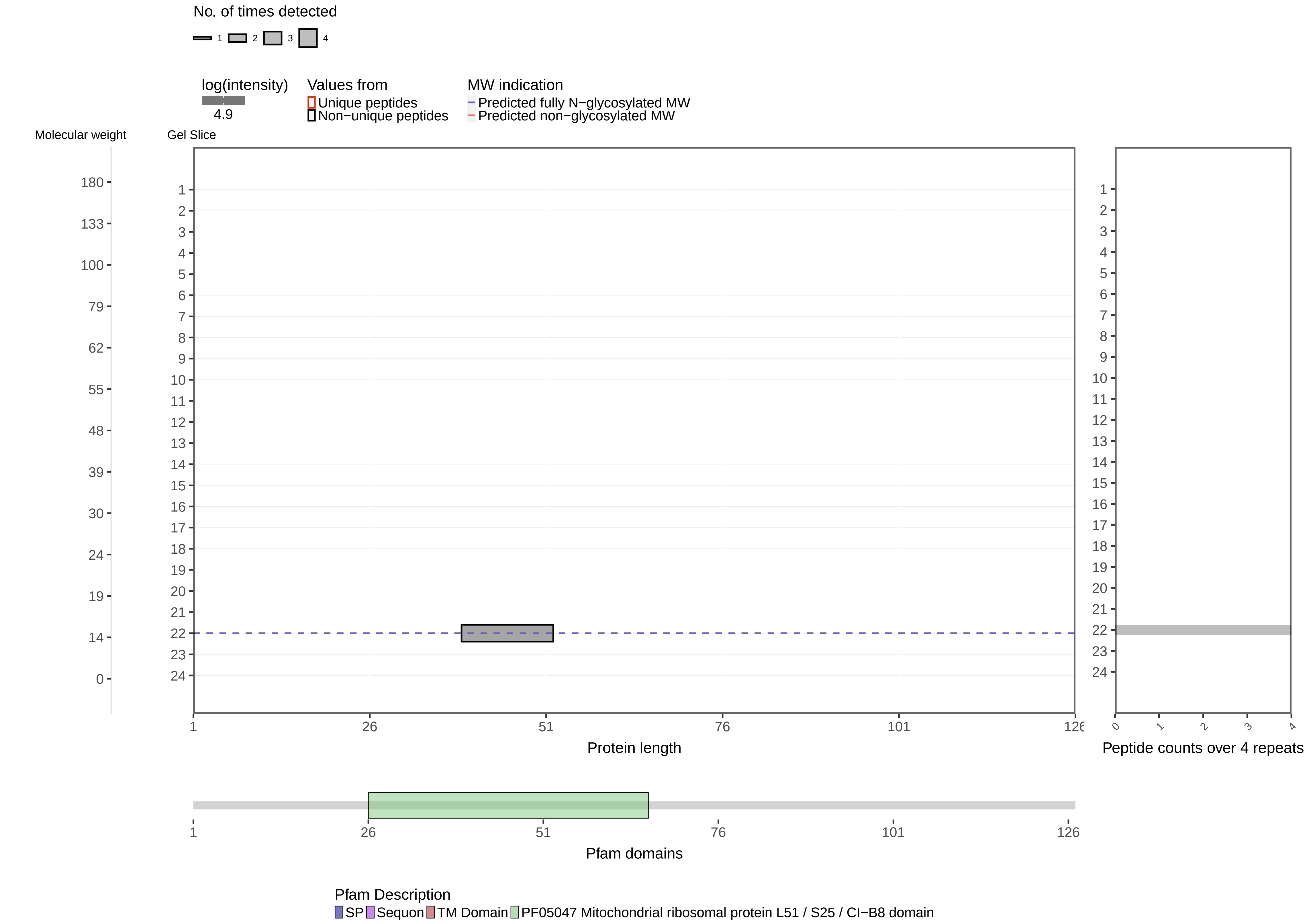


Protein sequence:

.NRGQHPFLKGLYKQERACCMCEKFESRRSTRSSNQAKEFAWQKGGEDEDSACYKTTKRSRYMVNSIEVISLQKFF*

NbL04g01440.1 glutamine_synthetase_cytosolic_isozyme_1−1−like [MW = 39.18 kDa]

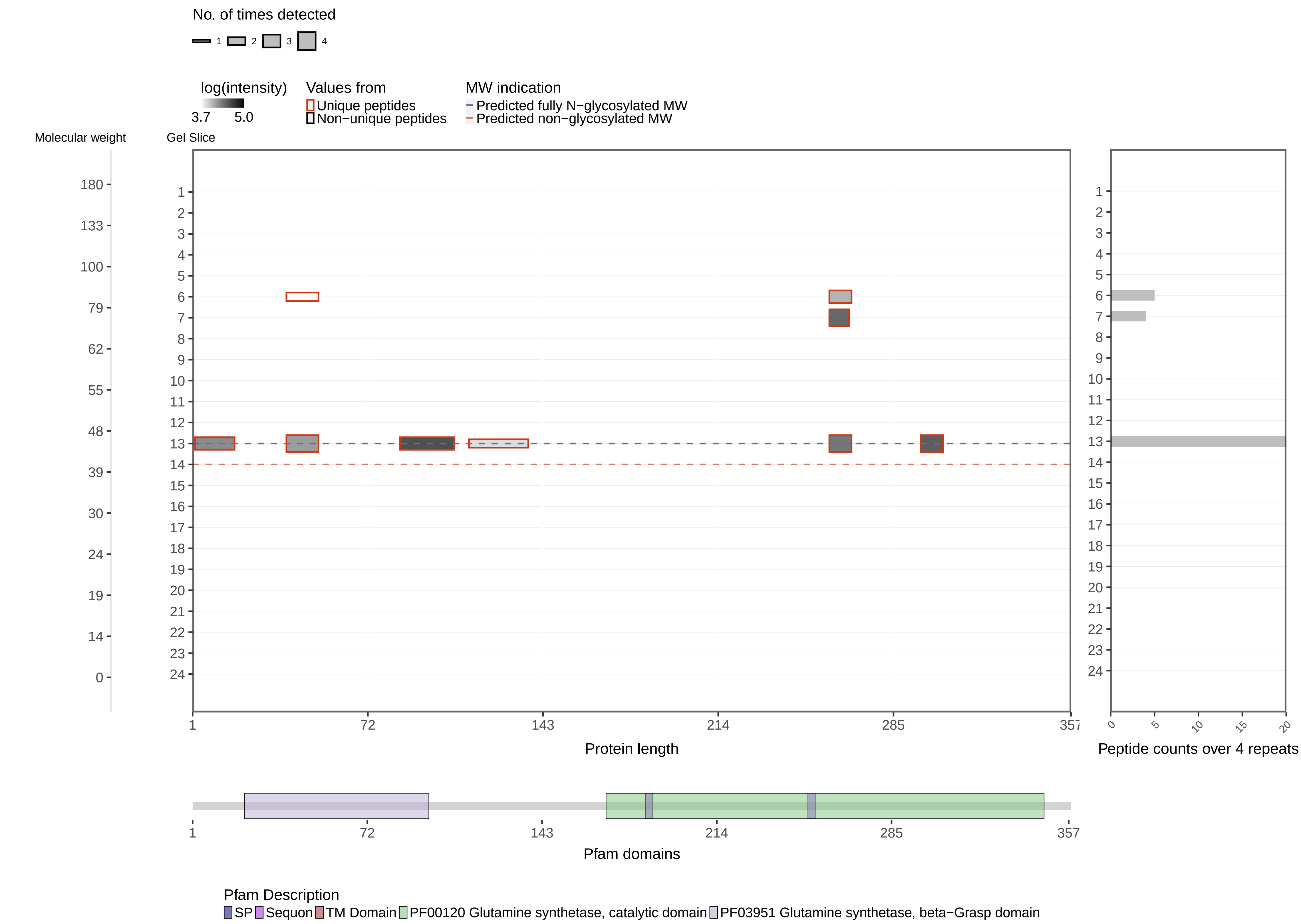


Protein sequence:

NAFGRDIVDSHYKACLYAGINISGINGEVMPGQWEFQVGPSVGISAADELWAARYILERITEIAGVVVSFDPKPIPGDWNGAGAHTNYSTKSMRNEGGYEVIKKAIQKLGLRHKEHIAAYGEGNERRLTGRHETADINTFKWGVANRGASIRVGRDTEKEGKGYFEDRRPASNMDPYVVTSMIAETTILYEL*

### NbL04g01370.1 universal_stress_protein_PHOS32 [MW = 17.82 kDa]

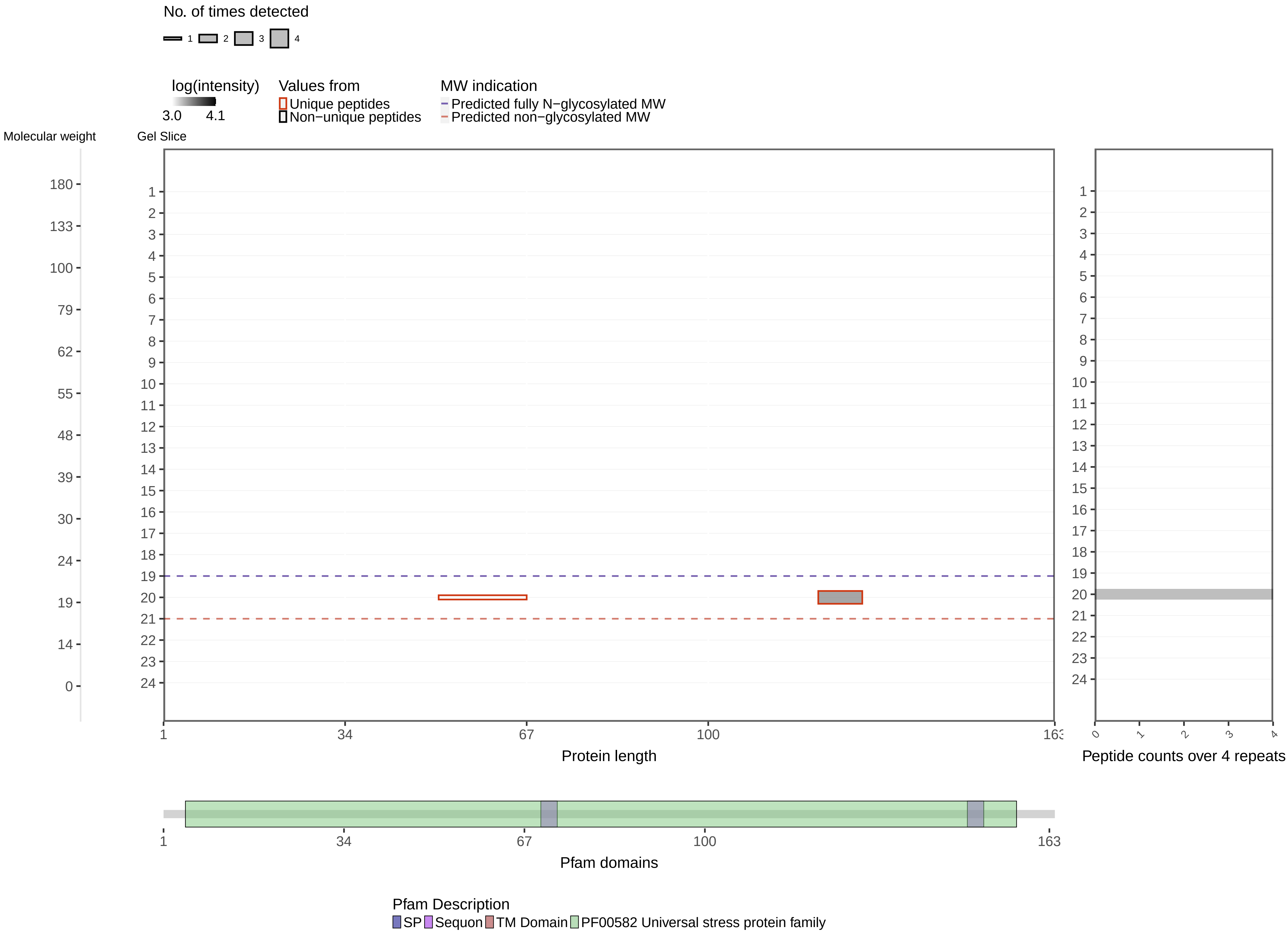


Protein sequence:

'SKYYGLDPDREVLDMLDTVSKLKKVIVVAKVYWGDAREKLCDAAEHLKLDSLVLGSRGLGVLKRMLLGSVSNHLVQNATCPVTVVKSSPPK*

NbL04g01330.1 tropinone_reductase−like_3 [MW = 26.95 kDa]

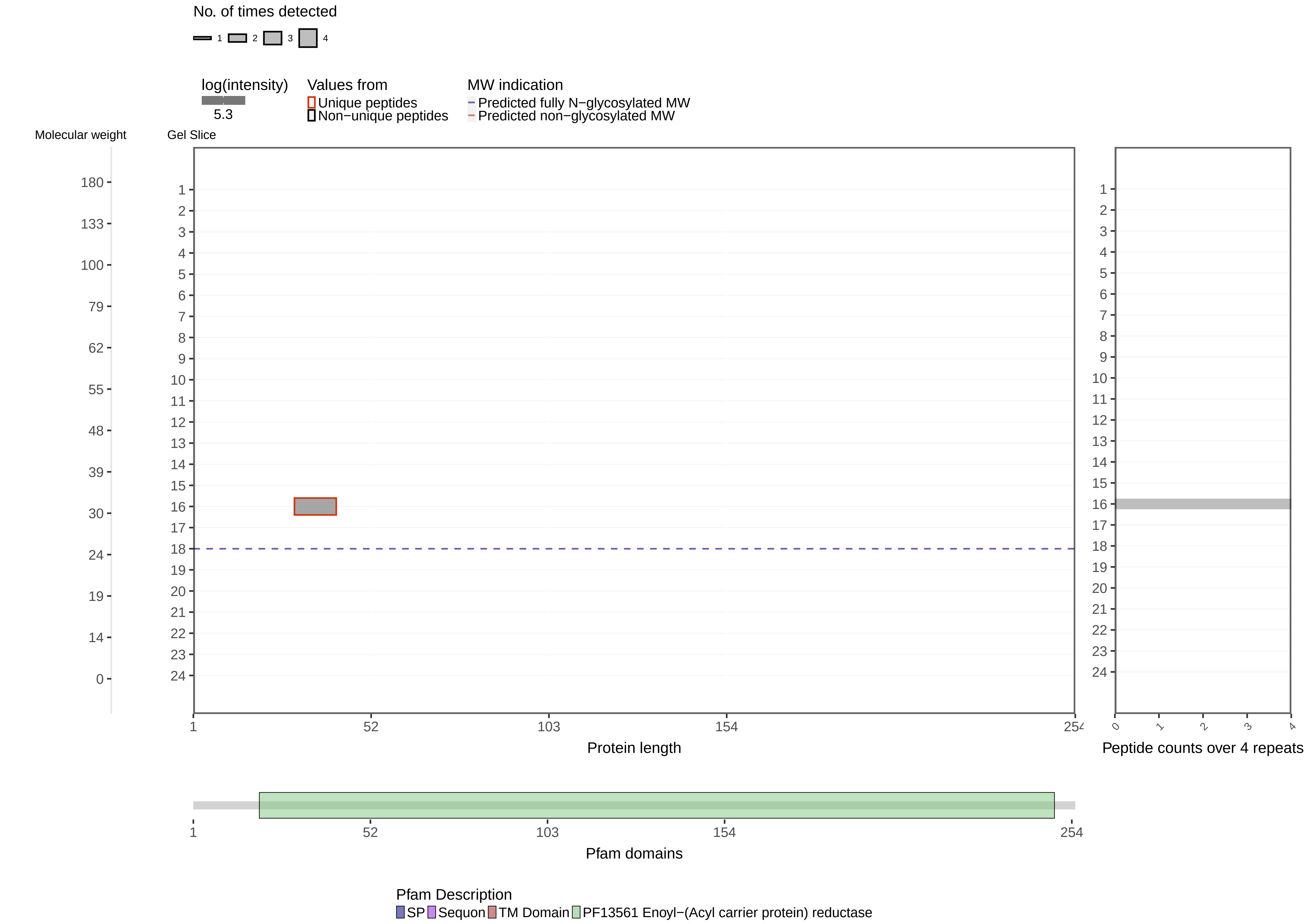


Protein sequence:

KLWDINVKASILLLQDAAPYLKKGSSVVLISSISAYSPPASMGMYGVTKTALLGLTKALASEMSPDTRVNCVAPGFVPTHIADLITQNEHVRREIEGKTLLNRLGTTQDMAAAVGYLASDDAAYVTGETLVVAGGMPSRL*

NbL04g01160.1 peptidyl−prolyl_cis−trans_isomerase_FKBP16−3_chloroplastic [MW = 27.14 kDa]

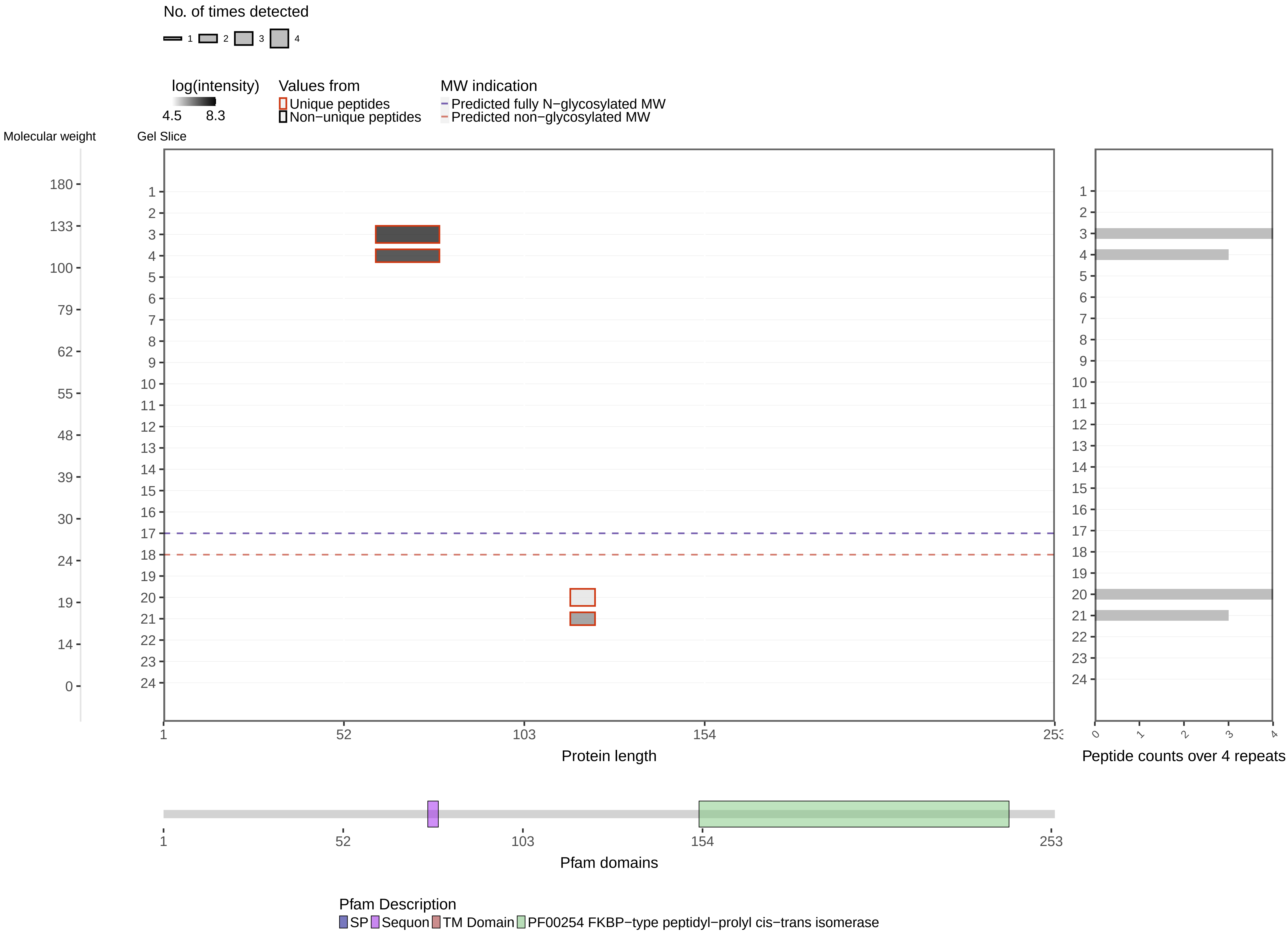


Protein sequence:

PKLCDATCAKELENVPMVTTESGLQYKDIKVGGGPSPPVGFQVAANCVAMVPSGQIFDSSLEKGQLYIFRVGAGQVIKGLDEGILSMKVGGKRRLYIPGSLAFPKGLTSAPGRPRVAPNSPVIFDVSLEYIPGLESDEE*

NbL04g01150.1 trafficking_protein_particle_complex_subunit_5 [MW = 24.39 kDa]

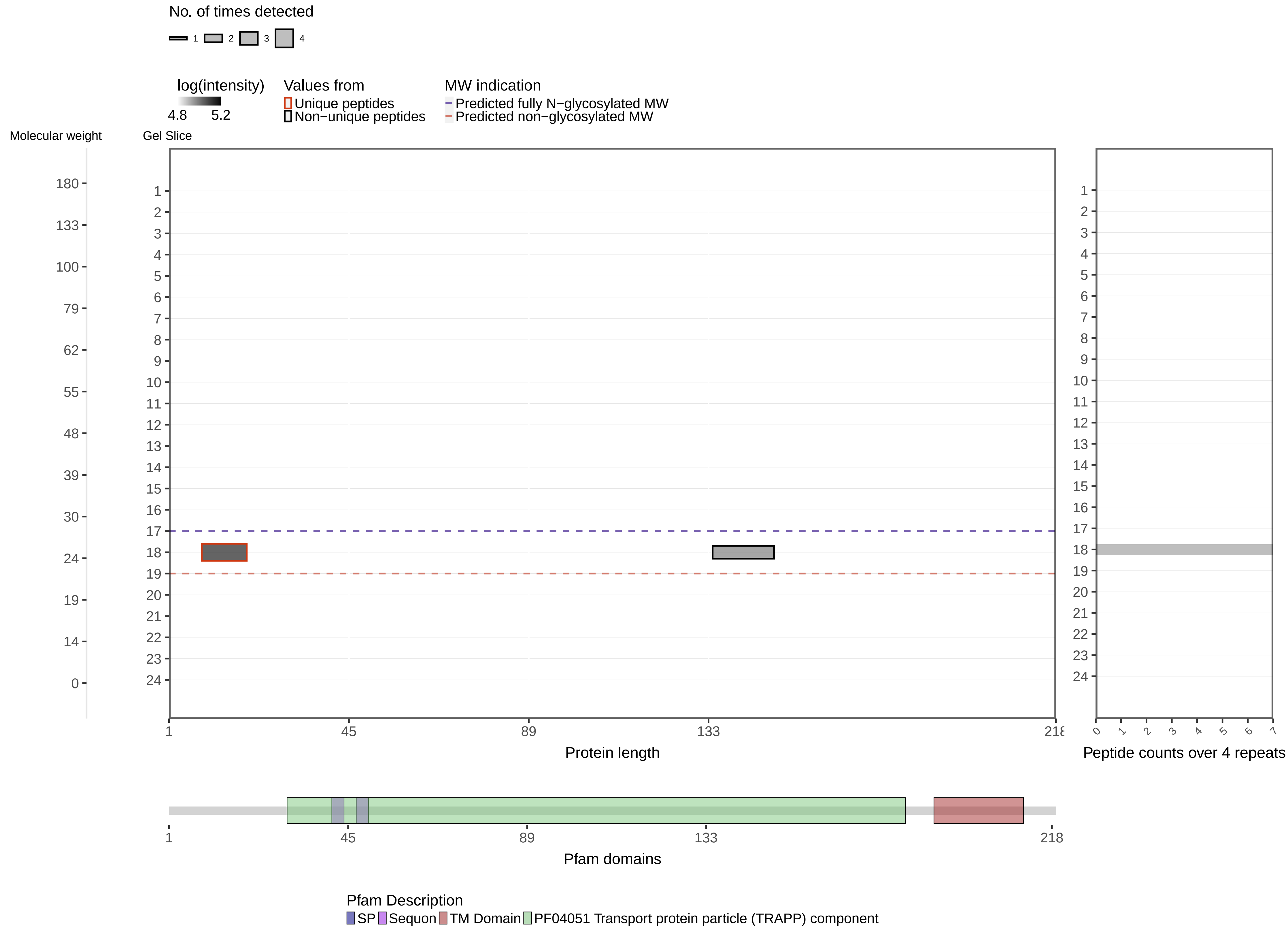


Protein sequence:

WKVLFGKVADSLEKGTEHEDEYMISEKELLVNRFISIPKDMGAFNCGAFVAGIVRGVLENAGFPAVVTAHFVPVEGOORPRTTILIKFAEEVNOFFLSFCSTLLVGCSTDFLPIAFTVTYFFT*

### NbL04g01110.1 acetyl−CoA_acetyltransferase_cytosolic_1−like [MW = 43.11 kDa]

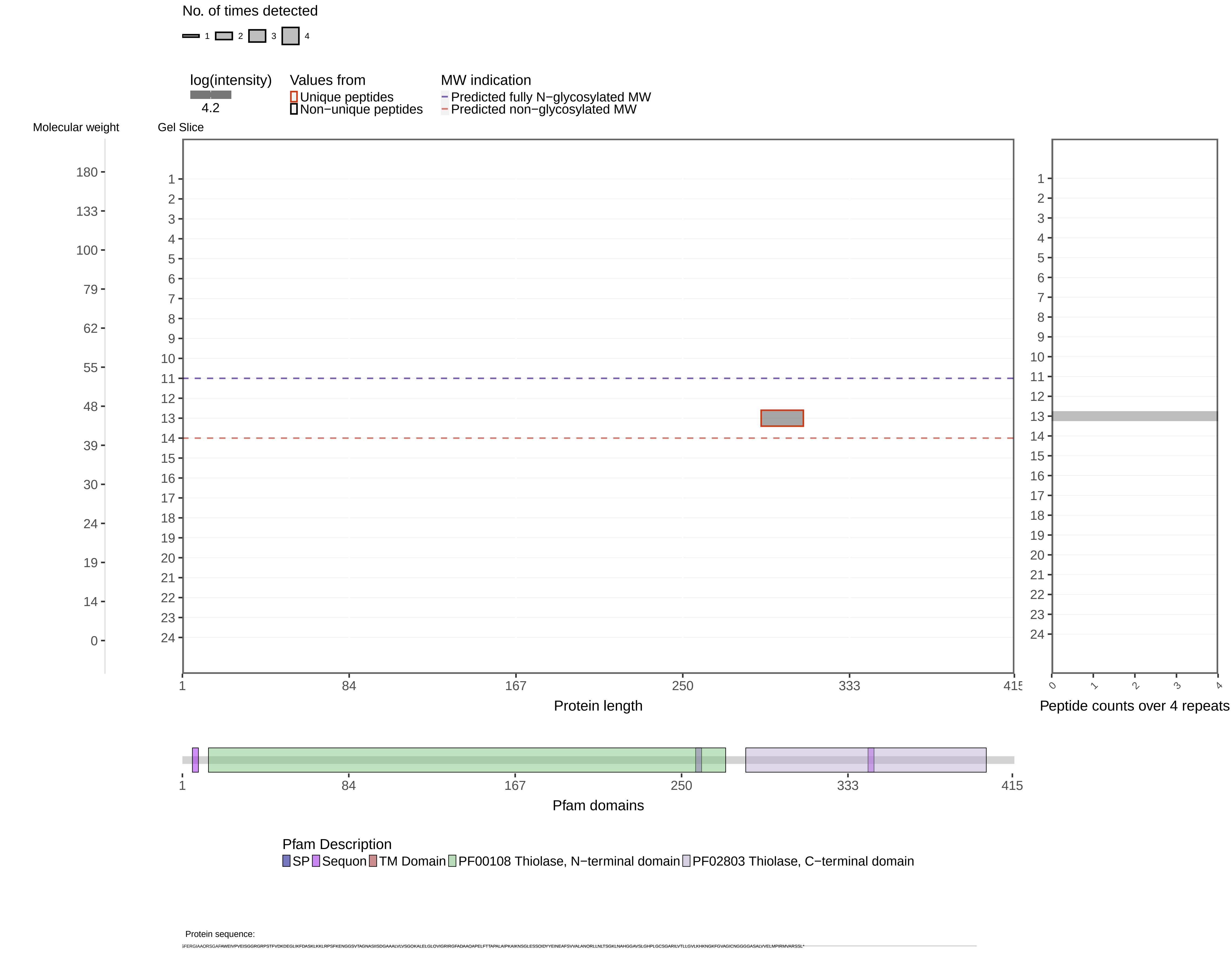

### NbL04g01060.1 phosphoglycolate_phosphatase_2−like [MW = 34.55 kDa]

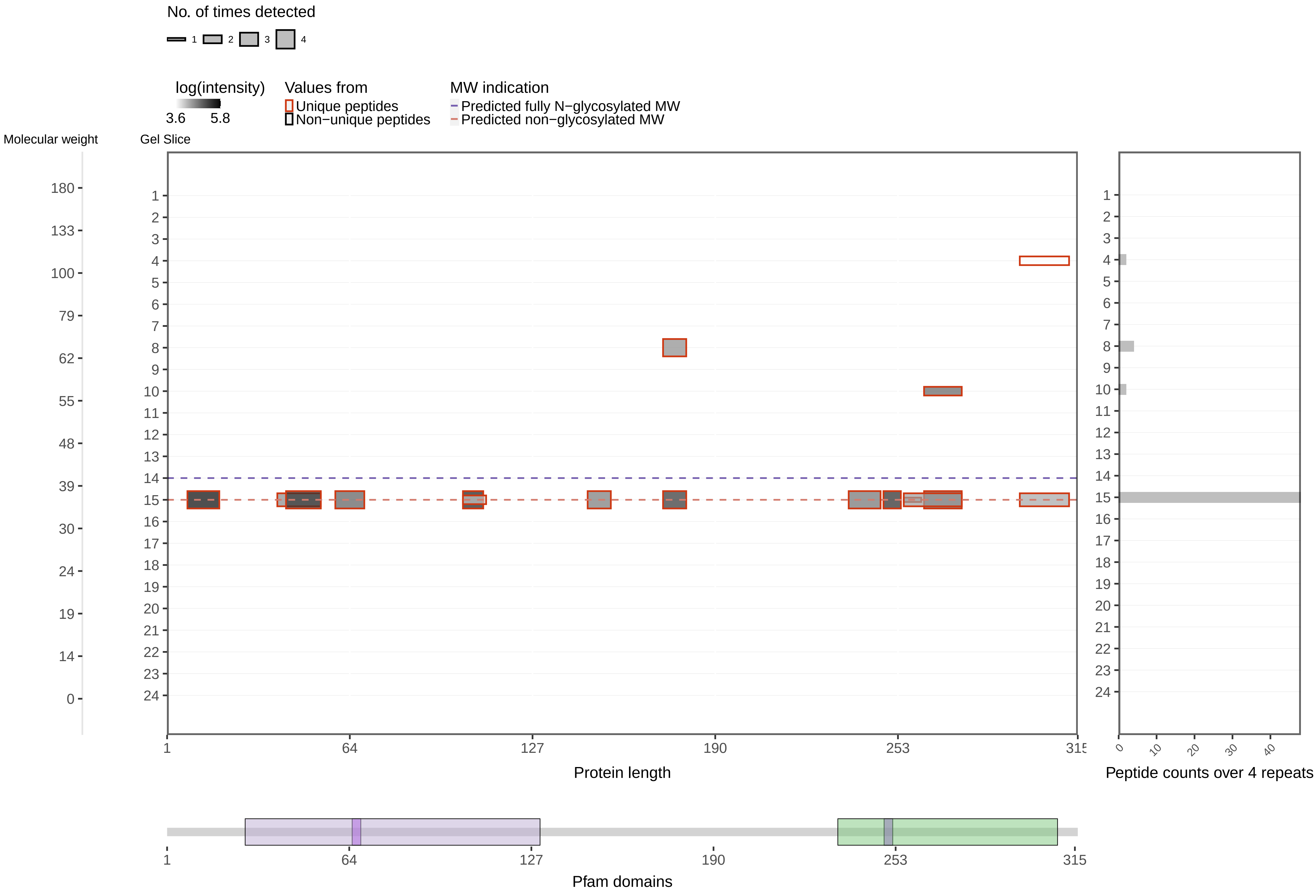


Protein sequence:

[illegible]SNCLFEHDKSVGAVVVGLDQYINFYKLQYGALCIRENPGCLFIATNRDASGHLTDLQEWPDTSFLVTVGAGCMVAAVCGATQKEPIVVGKPSTFLMDFLLQKYNITTSRMCMVGDRLDTDILFGQNAGCRTLLVFSGVVTESAFQDLSKELQVQPEYYTNSVSDILKYV*

### NbL04g00830.1 uncharacterized_LOC109239397 [MW = 35.51 kDa]

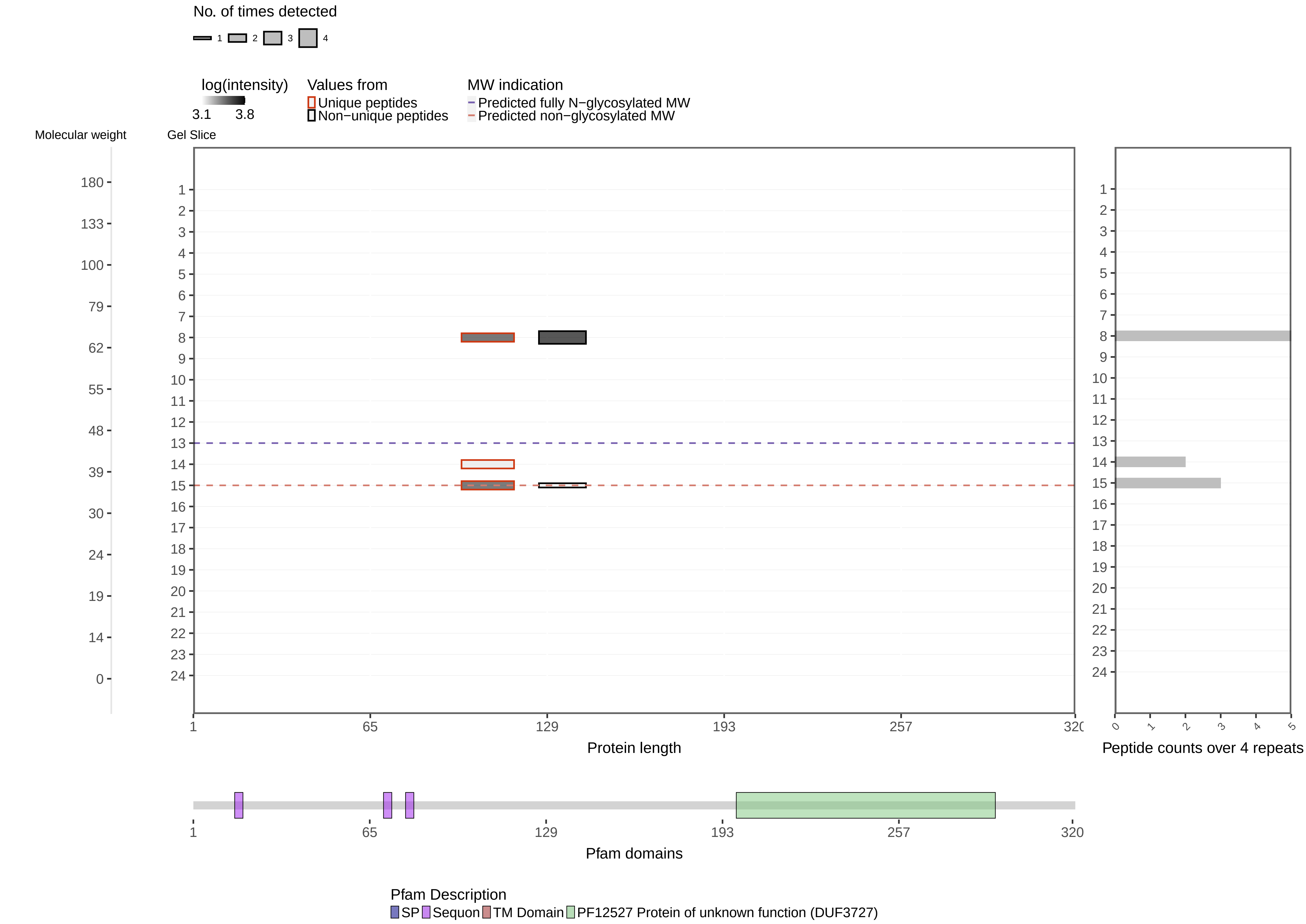


Protein sequence:

FRSLREDECLLLCPVDMPVQILKSTNVEGWSAVSDEEVEAILPTAAYALAKIHMHLVYSGFCYTARGGFCYTEDDIFEFRTDDGDDVDGLPSEGIEITCFHMDGSHYMIYTPSDPLLFVAVKGKDGLLQIADDELLEDPAIISAIDEETEFNALVEEEAALLESLLGERFKFYY*

### NbL04g00760.1 beta−glucosidase_BoGH3B−like [MW − SP = 66.09 kDa]

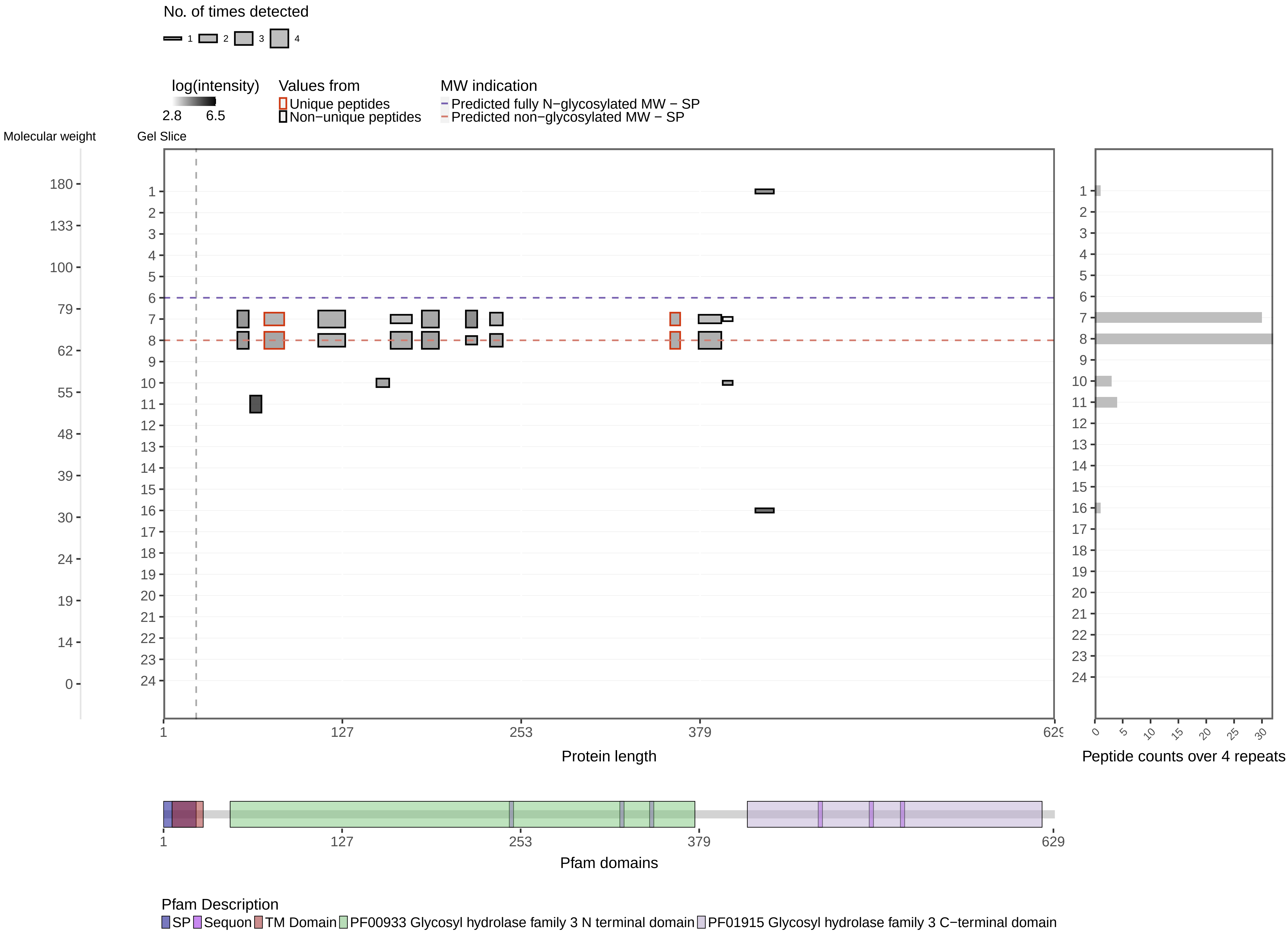


Protein sequence:

RFRGFVISDWAGIDKLTYPWHANYTYSILEGVNAGIDMVMLPYNTTEFIDGLTYLVKNNFVQMSRIDDAVKRILRVKFQMGLFESPLADYSMVKYLGSAEHRELAREAVRRSLVLLKNGANADEPVLPLPKKTSSILVAGIHANNMGYQCGGWTITWQGLSGNTTIGTTILSAIENTVDPETKVVFKENPDAEFIKSNNFSYAIVAVGETPYAEGSGDSLNLTIPAPGPDTMTTVCASVKCVVVLLTGRPVMIQPYLAQIDALVAAWLPGTEGQGIADVLFGDYGFTGKLARTWFKT

NbL04g00720.1 anthocyanidin_reductase_((2S)−flavan−3−ol−forming)−like [MW = 25.24 kDa]

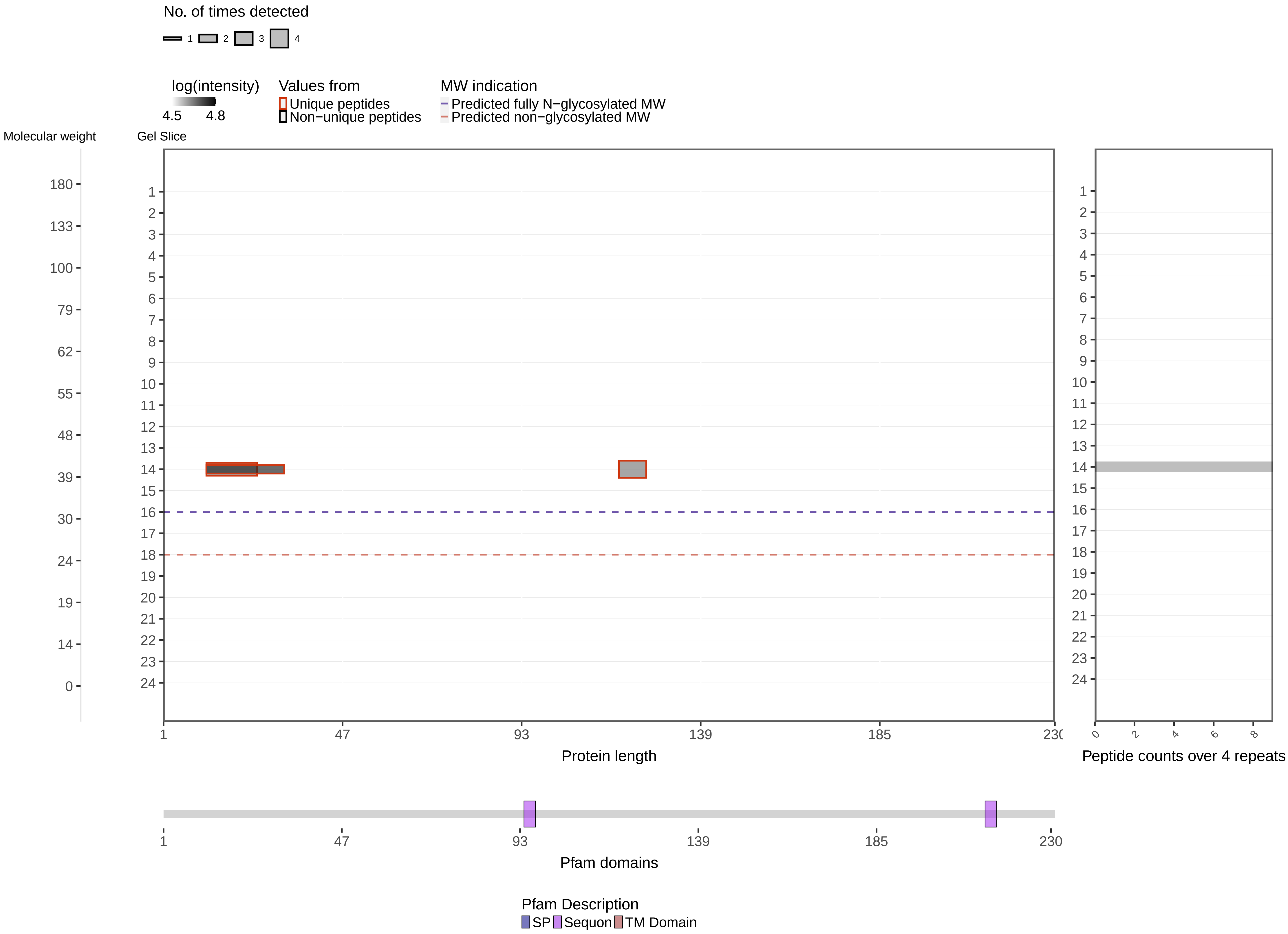


Protein sequence:

AILSVVTGDEISYQQLKFIEEVDGKAPIVHIEDVCEAHIFGMETASINGRFLCASAFVSSAEIASYYKEKYPEFHANQRYLDDPKREVKWGSKKLIDKGFVYKKDMKIILDDNISCAKEIGVLNLPPSA*

### NbL04g00660.1 insulin−degrading_enzyme−like_1_peroxisomal [MW = 112.06 kDa]

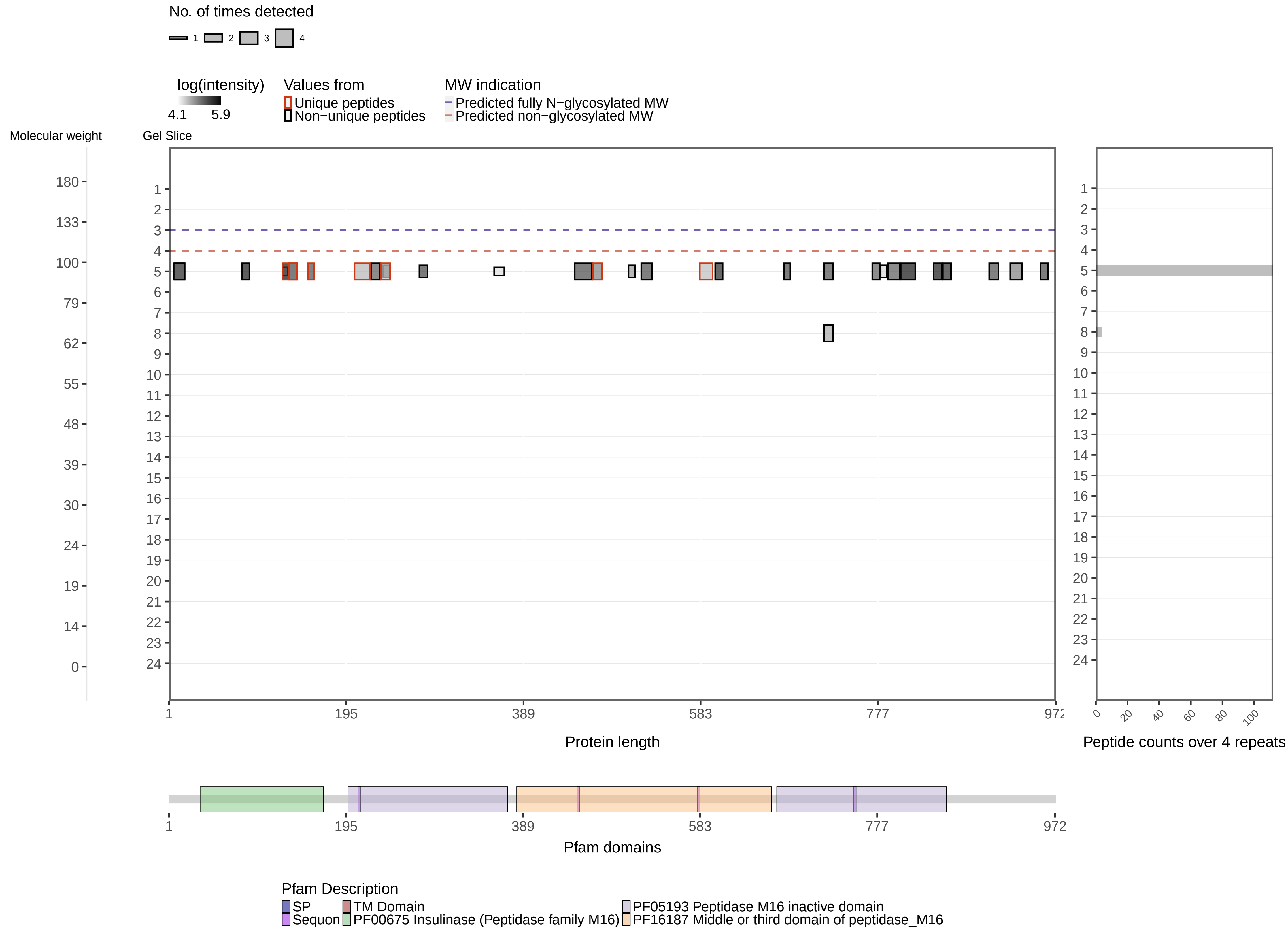


Protein sequence:

### NbL04g00620.1 magnesium−chelatase_subunit_ChlH_chloroplastic [MW = 153.89 kDa]

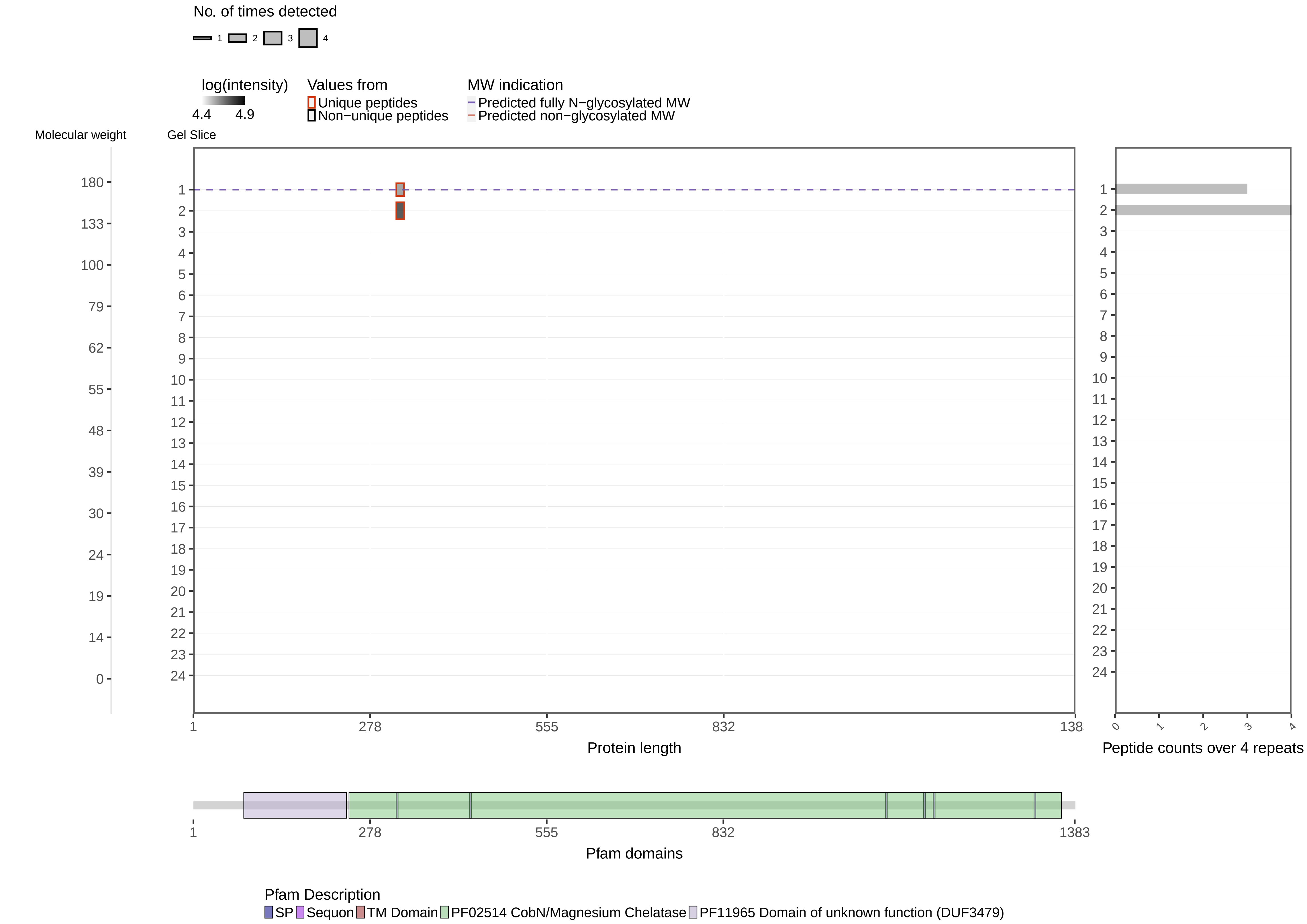


Protein sequence:

### NbL04g00610.1 villin−3−like [MW = 104.24 kDa]

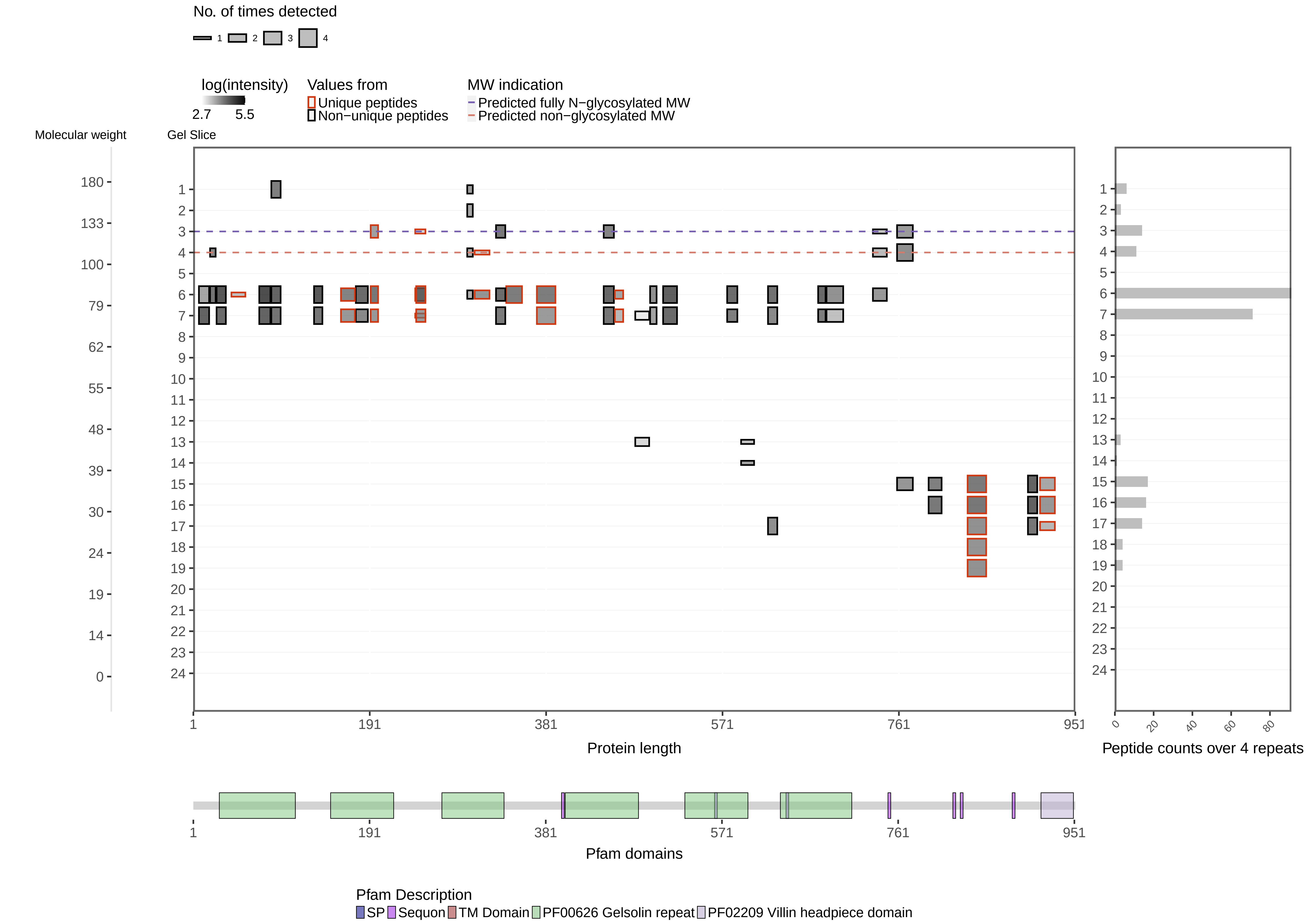


Protein sequence:

RPVLGRVFQGKEPPQFVAIFQPMLVLKGGLSSGYKSYIADKGLNDETYAADSVALIRLSGTSVHNNKAVQVDAVPASLNSNECFLLQSGSSLFSWHGNQSTYEQQQLAAKVAEFLKPGATVKHTKEGTESSAFWFALGGKQSYTSKKVAPEVSRDPHLFAYSFNKGKFEVEEIYNFSQDDLLTEDVLLLDTHAEVFVWIGQSADSKEKQSAFDIGQKYVEMAASLEGLSPNVPLYKVTEGNEPCFFTTFFSWDPAKASAHGNSFQKKVMLLFGVGHASENQQRSNGSGG

NbL04g00570.1 L−ascorbate_oxidase_homolog [MW = 19.51 kDa]

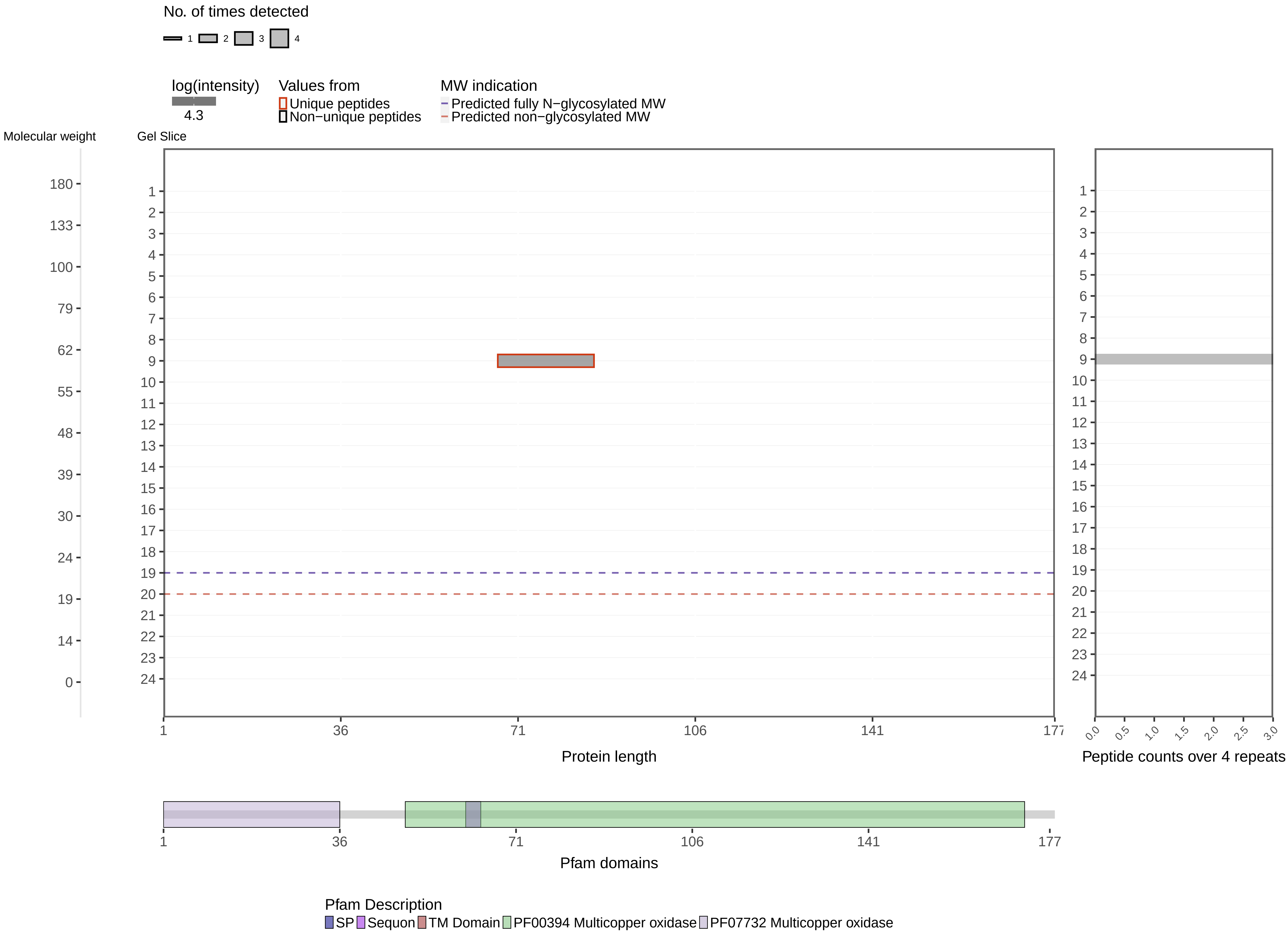


Protein sequence:

_PFPDGLLINGRGSNSLTFTVDQGRTYRFRISNVGLTTAINFRIQGHKMVLVEVEGTHTLQNTYESIDIHLGQSYSVLVTMDQPGQDYYIVASTRFTSQVLTA

NbL04g00540.1 dihydroneopterin_aldolase_1−like [MW = 14.25 kDa]

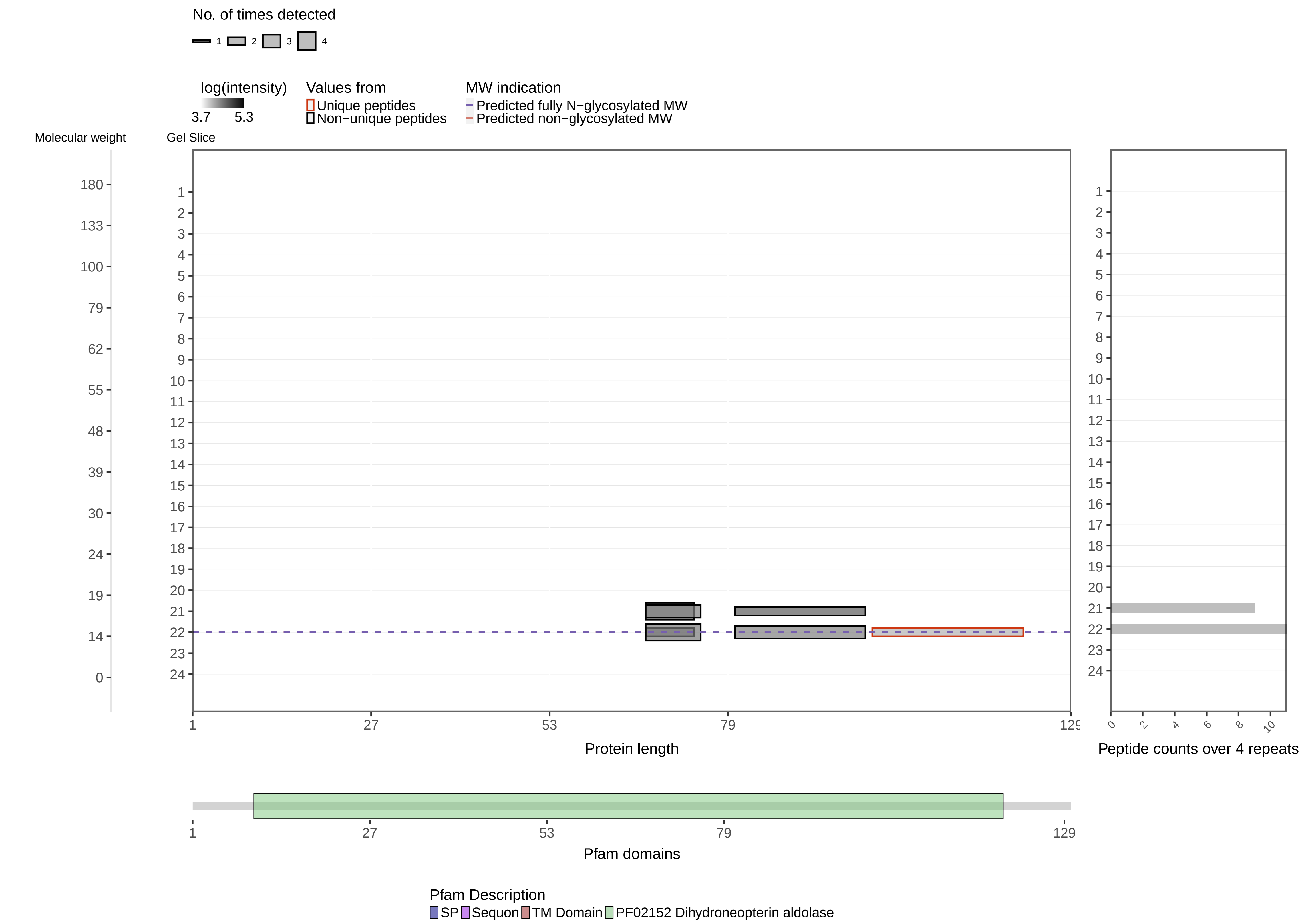


Protein sequence:

CLSDTLSYTDIYKIVKEIVEGPPRKLLEKVAELIASTTLDKYPQVSAVRVQVGKPHVAVQGSVDYLGVEIIRHRDLDG*

### NbL04g00380.1 DNA−(apurinic_or_apyrimidinic_site)_lyase_chloroplastic [MW = 59.58 kDa]

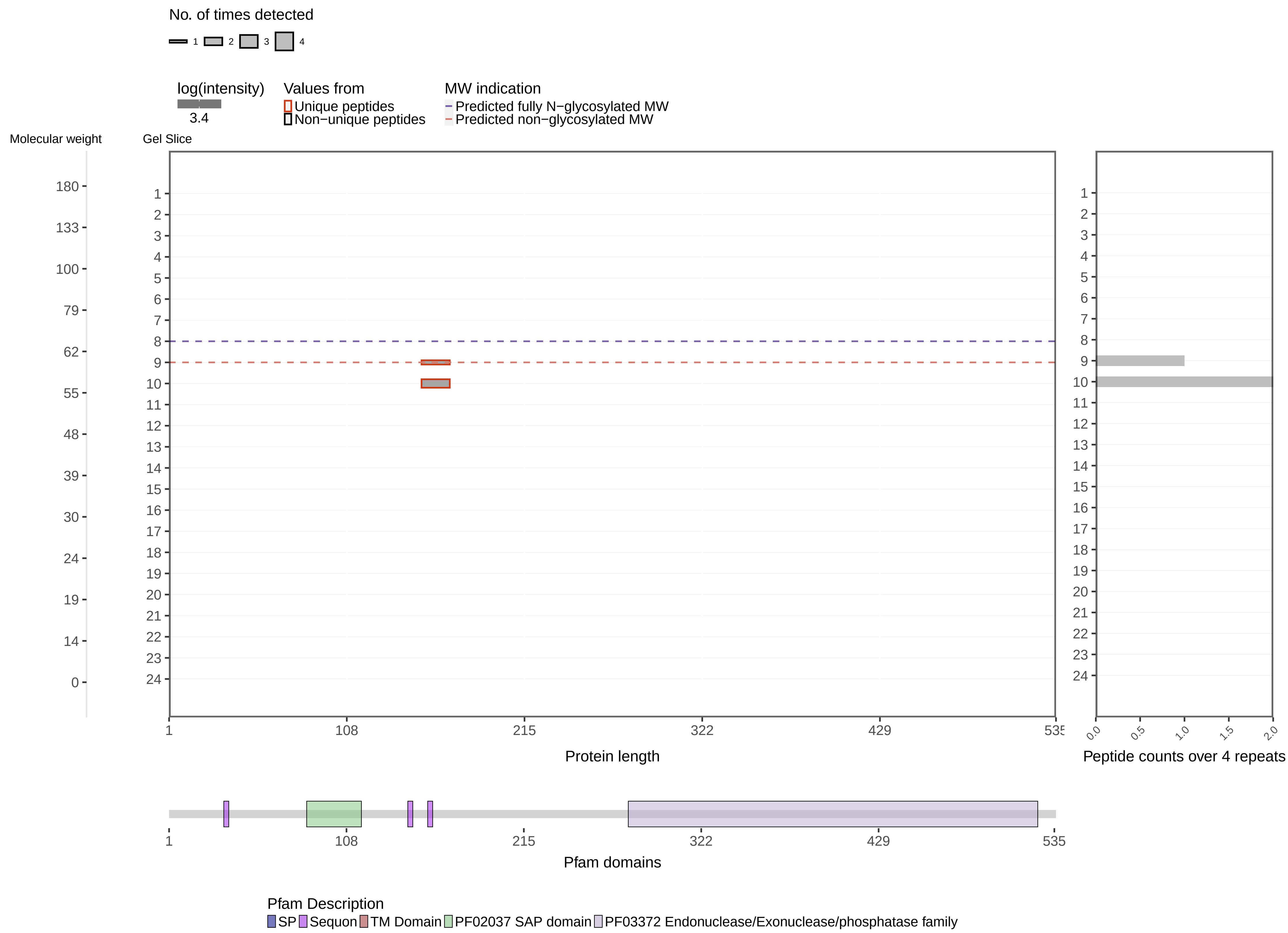


Protein sequence:

NbL04g00160.1 protochlorophyllide−dependent_translocon_component_52_chloroplastic−like [MW − SP = 64.46 kDa]

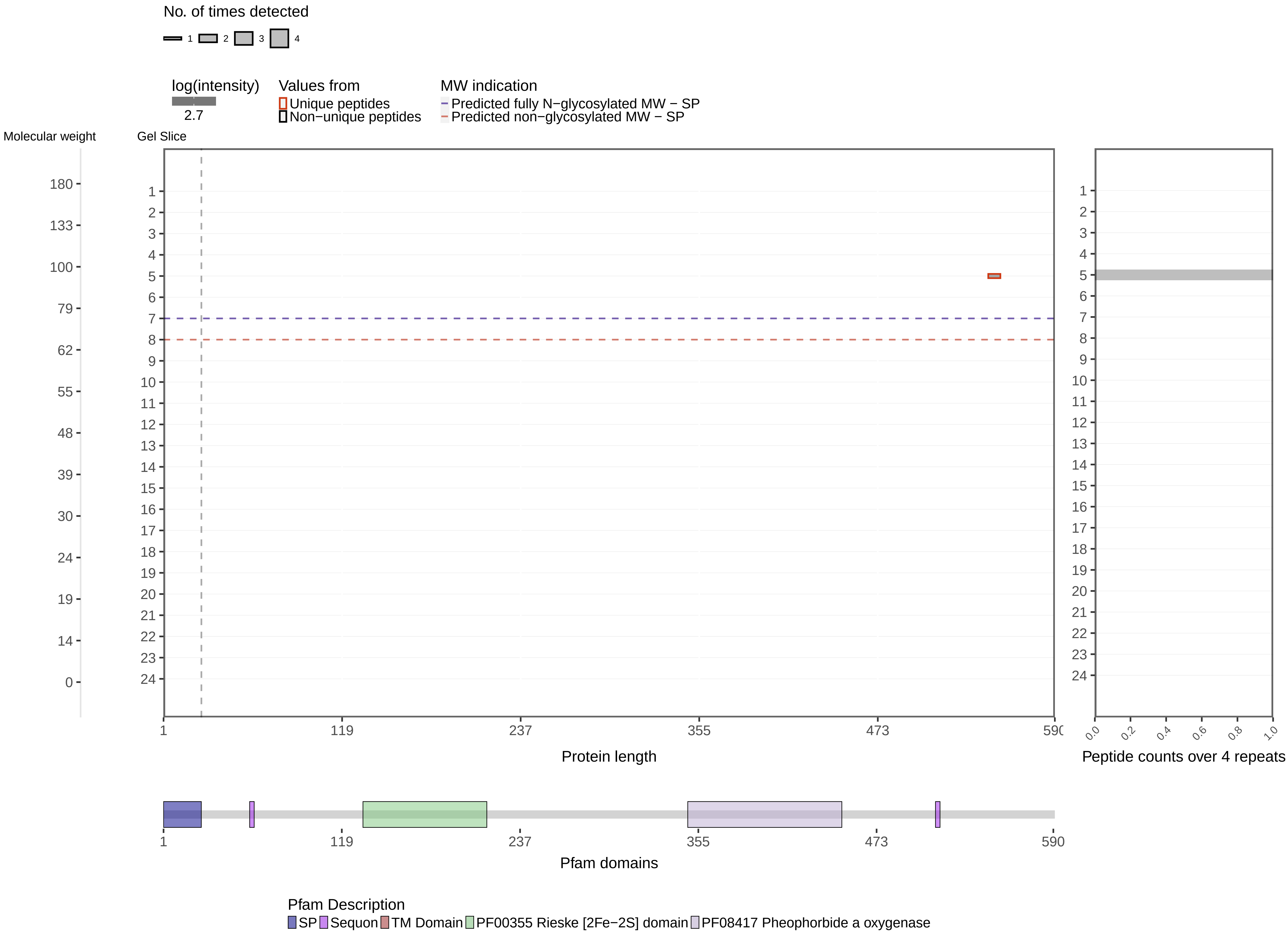


Protein sequence:

RDISYGYELLIENLMDPAHVQYSHYGIMNVPQPPKSVKADREGGRPLDITVTKLDVNSITANQGPGRNTFIAPCVYYSLFAFGGNQGKTSAVSSGTVQEKPSAEKQKKALLIFICIPVSPGHSRIIFASPRNFATWADRIIPRWIFHLGQNLILDSDLYLLHVEEHKLKEIGPYNWHNACYMPTKADAIVVAFRRWLNKYAGGQVDWRAKYNGDLPPTPPREQLLDRYWTHTVNCTSCNFAYKGLNGLEIILQIASIGMIGIVAAAKQGMLSMVMRYSLVSIALLCFVASRWLSHFIY

NbL03g24920.1 glutamyl−tRNA_reductase−binding_protein_chloroplastic [MW = 35.8 kDa]

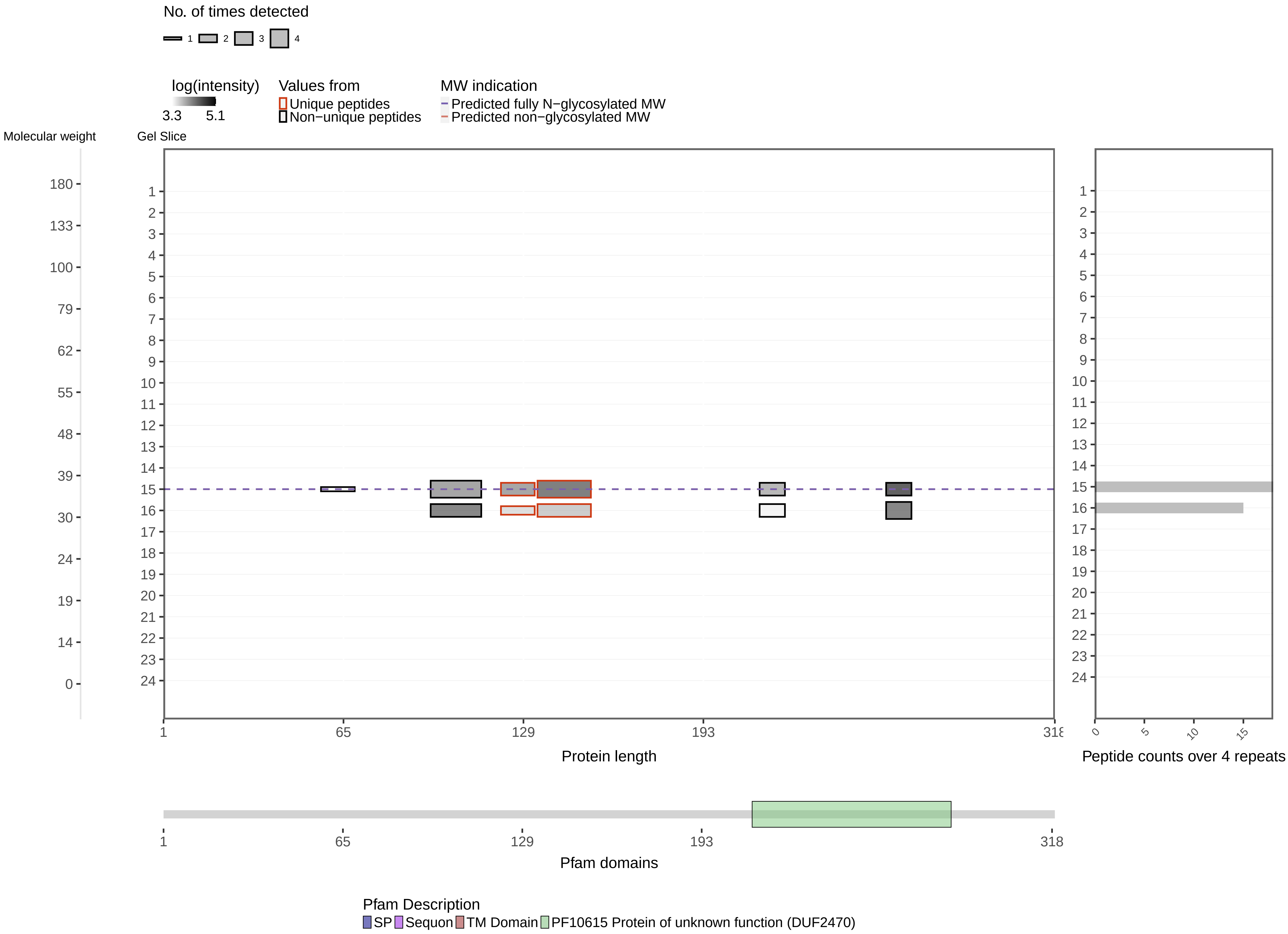


Protein sequence:

EDTTALKKLHSVWKKRFGHEIDEDHLFLLSVERVLQLEDFAEDGIWVTSSDYKLANPDPLRDFAERMIEEINTHNREDILRFCNIYIDLDFQVSDAKMLWVDRLGFDVRFRSPQNDVFEARIPFPREVTDEKGAKSSFNCMSQLAWEVEKNFHAAEFEKAKQLKKIEYTGL*

NbL03g24910.1 uncharacterized_LOC104100193 [MW = 28.76 kDa]

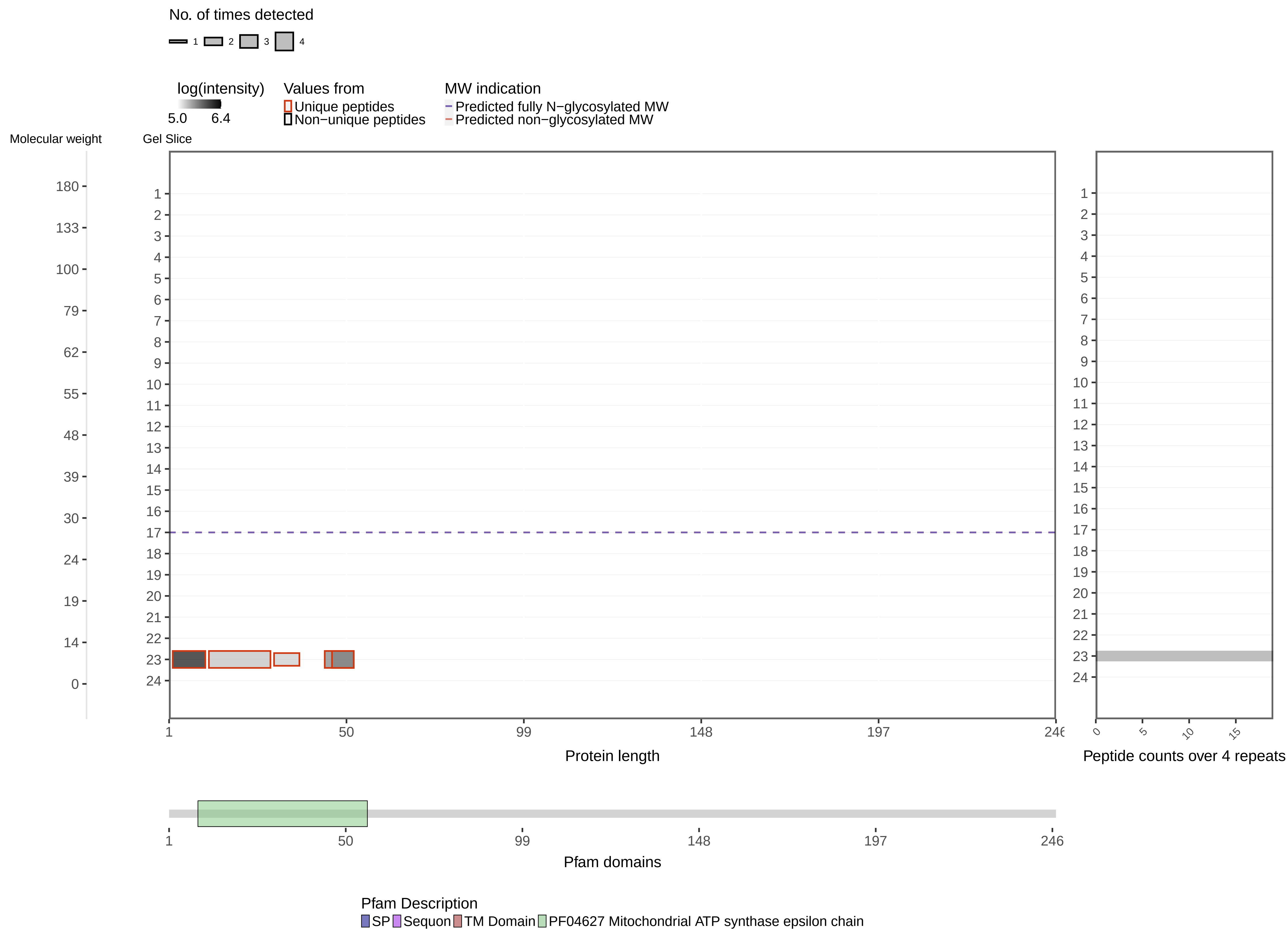


Protein sequence:
KFGAEPTEAVVEVRRYSHVIPKRGSCKYLRALVRPVMLYGTKCWPVKNSHIQKMKVAEMRMLRWMCGHTRMDKLKNEDIREKVGVAPMEDKMREARLRWFGHIQRRNTDAPVRRCERLAVVGTRKGRGRPKKY*

### NbL03g24880.1 cell_wall_integrity_protein_scw1 [MW = 35.03 kDa]

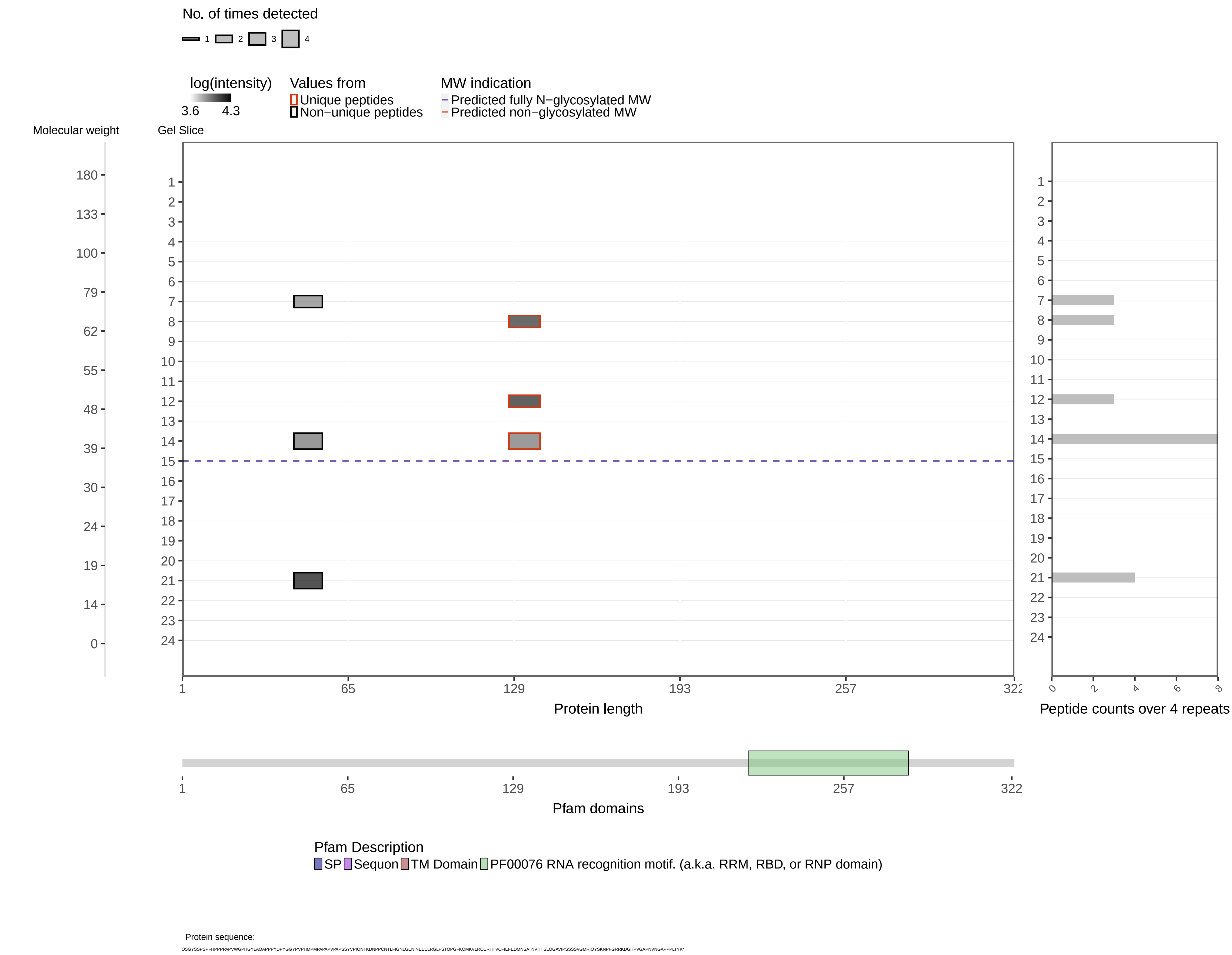

NbL03g24870.1 D−3−phosphoglycerate_dehydrogenase_2_chloroplastic−like [MW = 70.64 kDa]

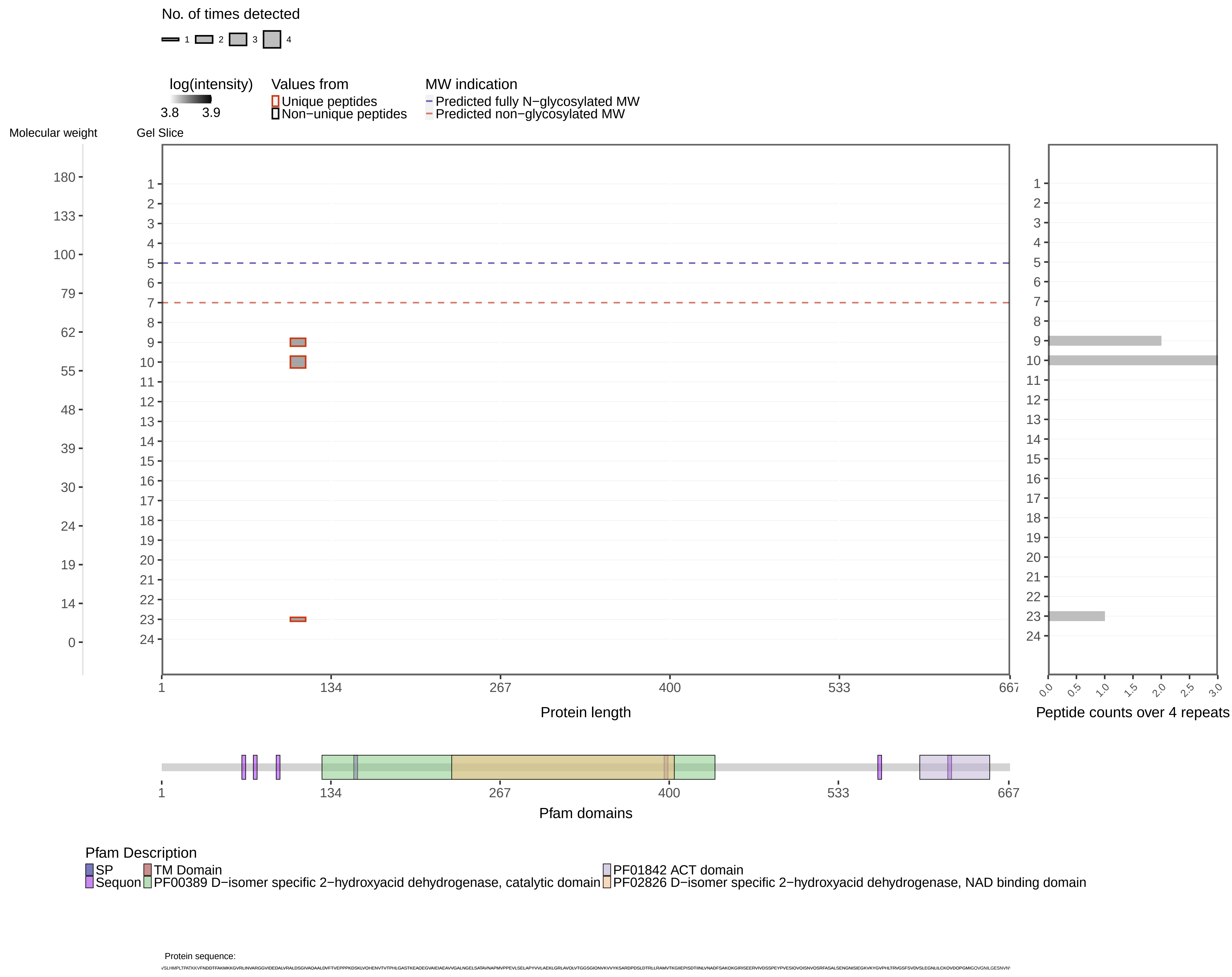

NbL03g24800.1 protein_STRUBBELIG−RECEPTOR_FAMILY_6 [MW − SP = 74.66 kDa]

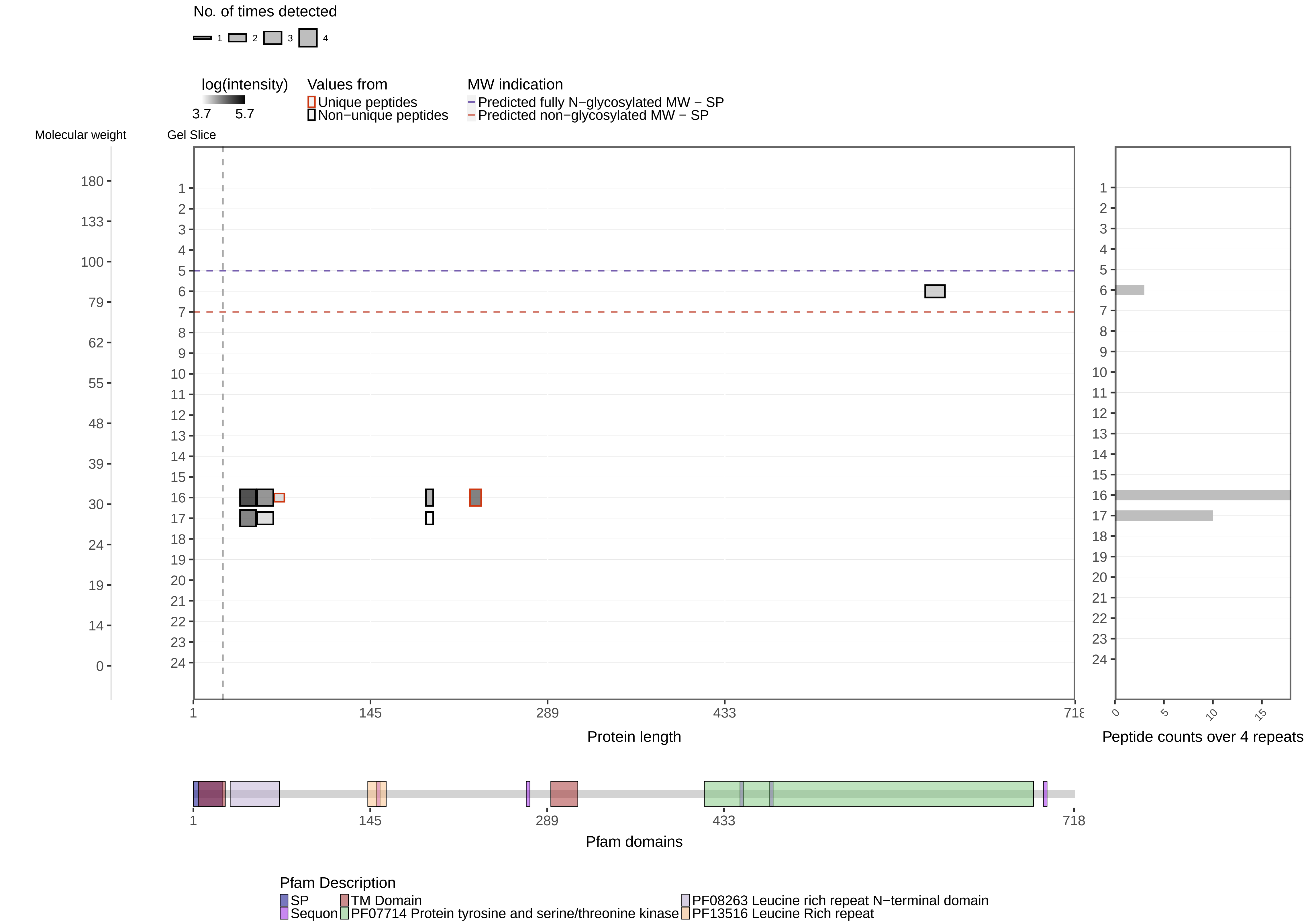


Protein sequence:

IKTNQASSAPSVKTFEAPAVVNLRPPPIERHKSFDEDEIAAKPIVPPKKVNTAKINARQYSIADLQMATDSFSVDNLIGEGSFGRVYRAQFDDGKVLAVKKINSSALHNPEDFLDIVSEISGLHHPNVTELVGYCSEHGQHLLVYEFHKNGALHDFLHLSDENSKPLTWNSRVKIALGTARALEYLHEVCSPSVVHKNIKSANILLDAELNPHLSDSGLASLIADVDQALNHNTGSGYGAPEVAMSGQFTIKSDVYSFGVVMLELLTGRKPFDGGRTRSEQSLVRWATPQLHDIDAL

### NbL03g24710.1 pectinesterase/pectinesterase_inhibitor_U1 [MW = 64.44 kDa]

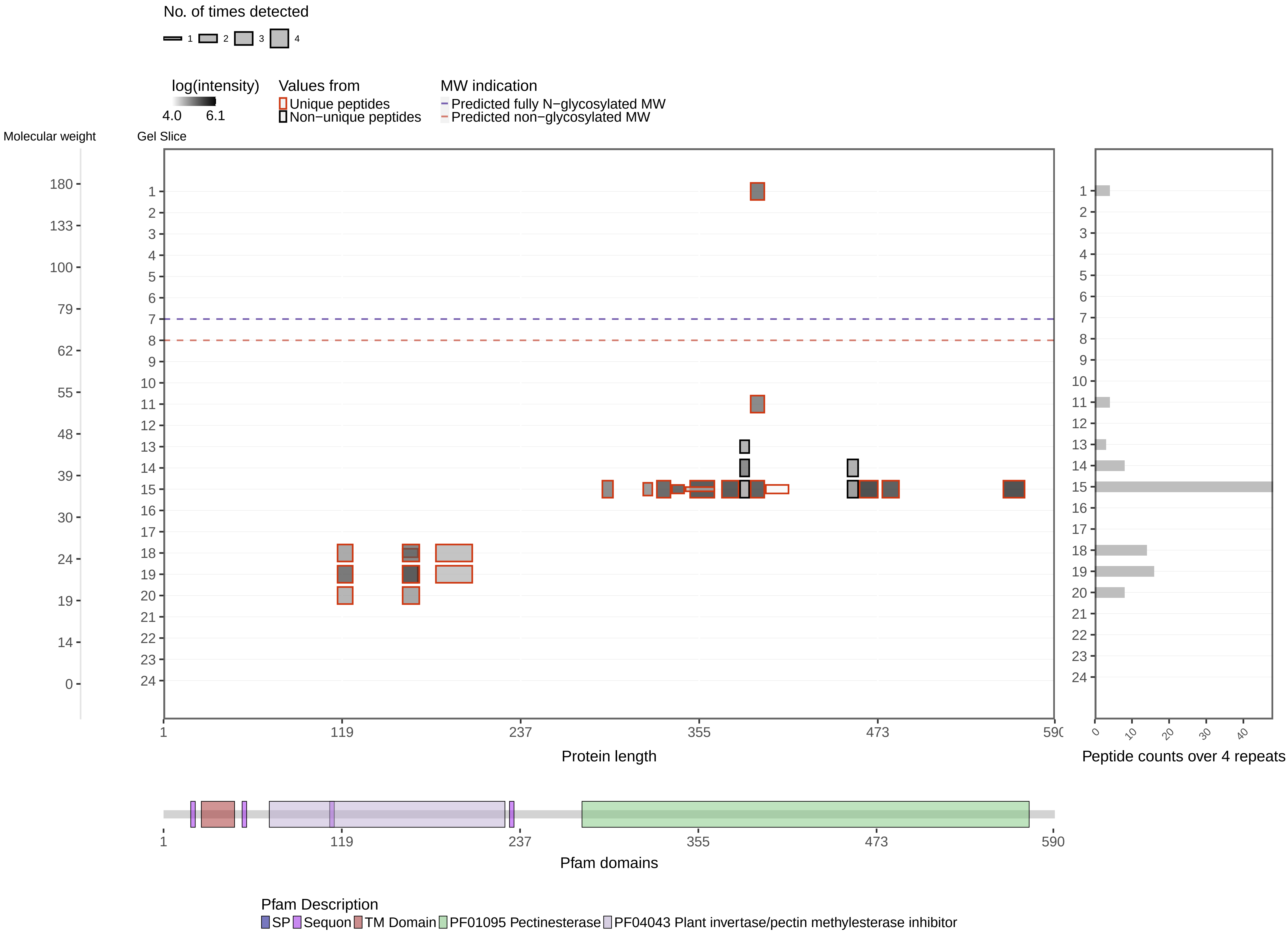


Protein sequence:

DGSGNFKTVSEAVARAPEKSSKRYVIRIKAGVYRENVDVPKKKTNIMFMGDGRSNTIITGSRNVKDGSTTFNSATVAAVGEKFLARDITFQNTAGAAKHQAVALRVGSDLSAFYRCDILAYQDSLYVHSNRQYFVQCLIAGTVDFIFGNAAAVLQDCDIHARRPGSGQKNMVTAQGRSDPNQNTGIVIQKCRIGATSDLRPVQKSFPTYLGRPWKEYSRTVIMQSSITDVINSAGWHEWNGNFALNTLFYGEYQNTGAGAGTSGRVKWRGFKVITSATEAQAYTPGRFIAGGS

### NbL03g24700.1 pectinesterase_3 [MW = 64.33 kDa]

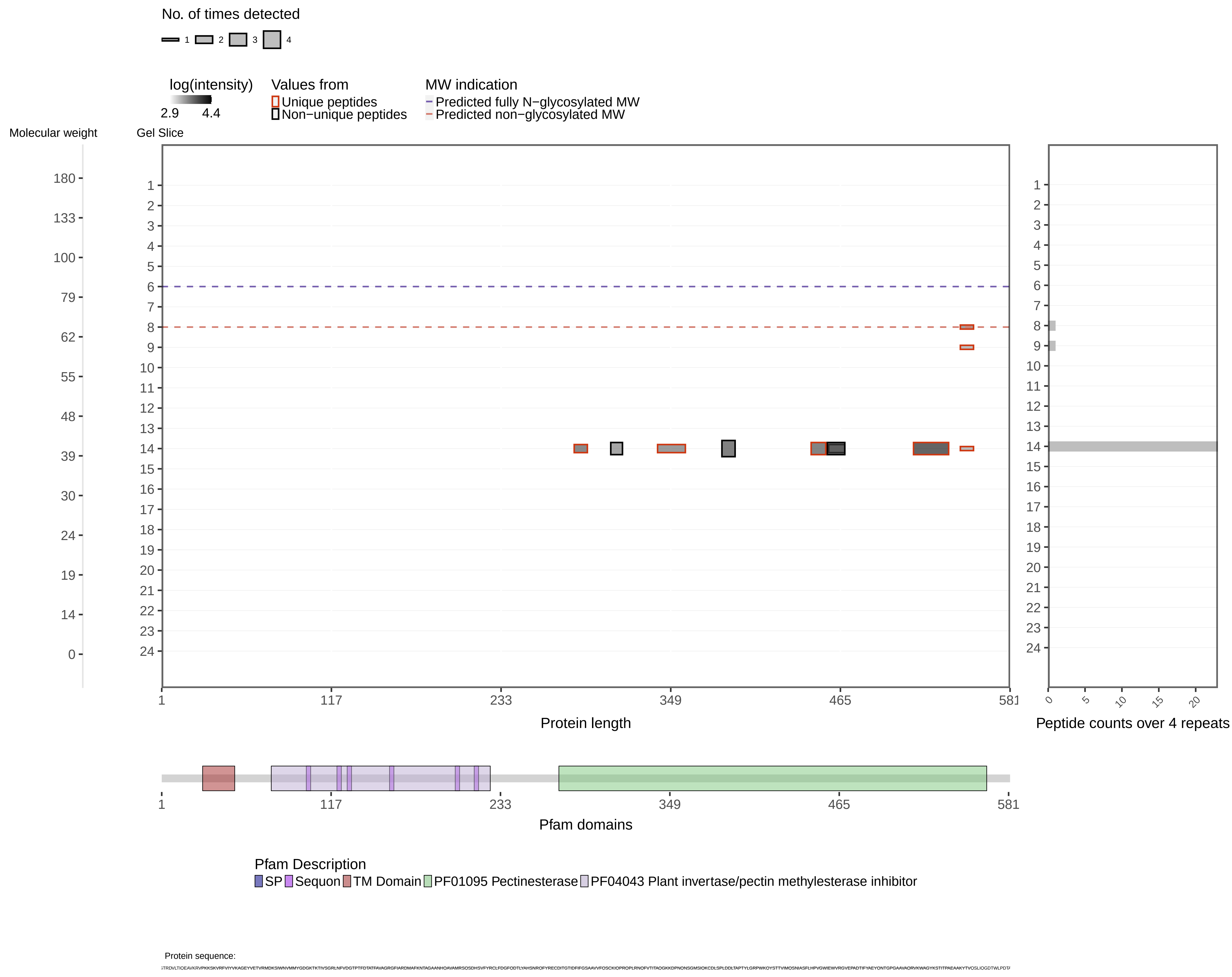

### NbL03g24680.1 alanine_aminotransferase_2−like [MW = 59.88 kDa]

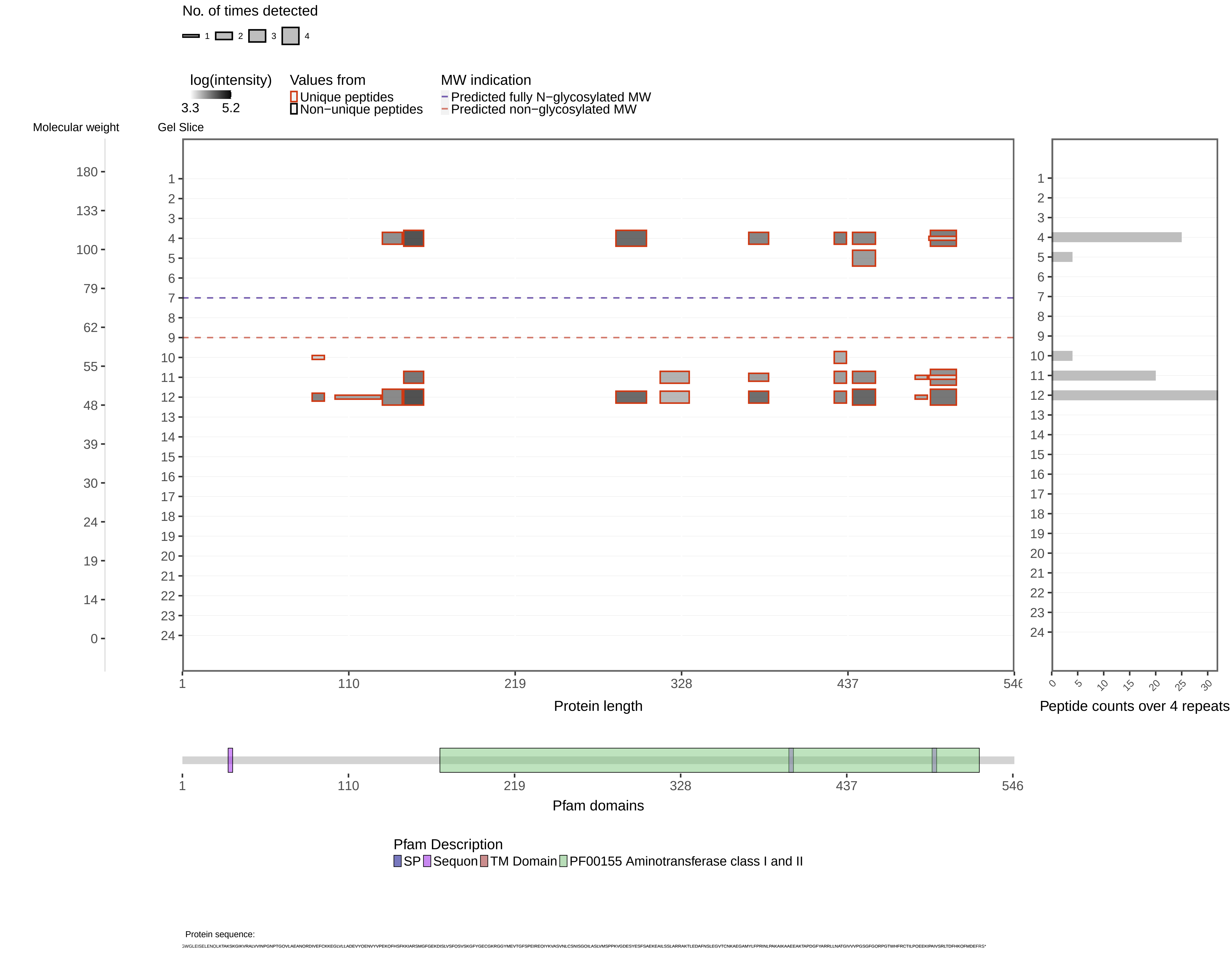

### NbL03g24600.1 serine/threonine−protein_kinase_Aurora−3−like [MW = 12.24 kDa]

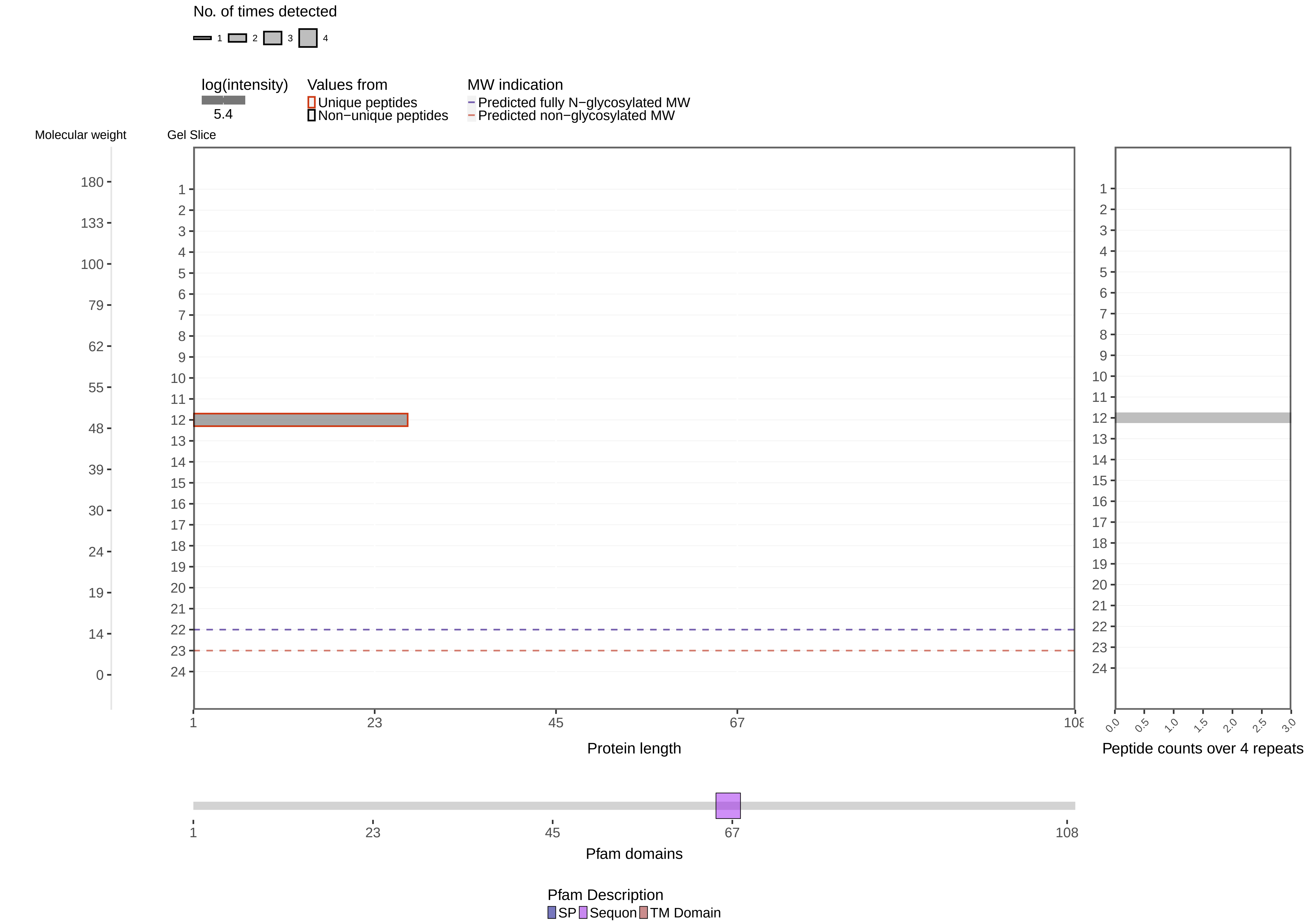


Protein sequence:

[illegible]DLEILEGPLLTGSFWWLFCGFRGVNLSFGSHFCKCLGKTASVIQFNMWREIGIVAALLRGIGYDPFVF

### NbL03g24550.1 subtilisin−like_protease_SBT1.5 [MW − SP = 80.57 kDa]

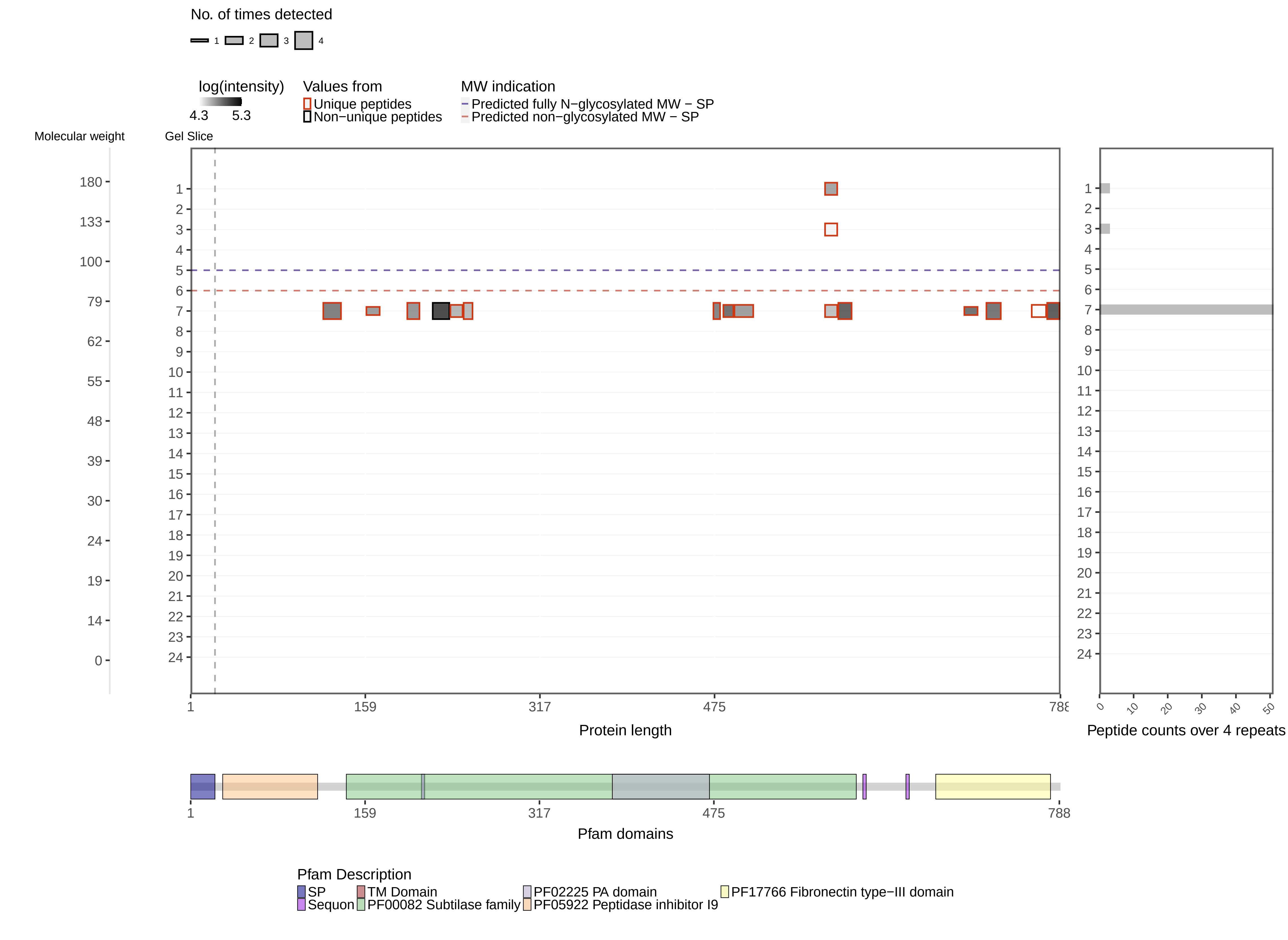


Protein sequence:

### NbL03g24390.1 linoleate_13S−lipoxygenase_3−1_chloroplastic [MW = 103.33 kDa]

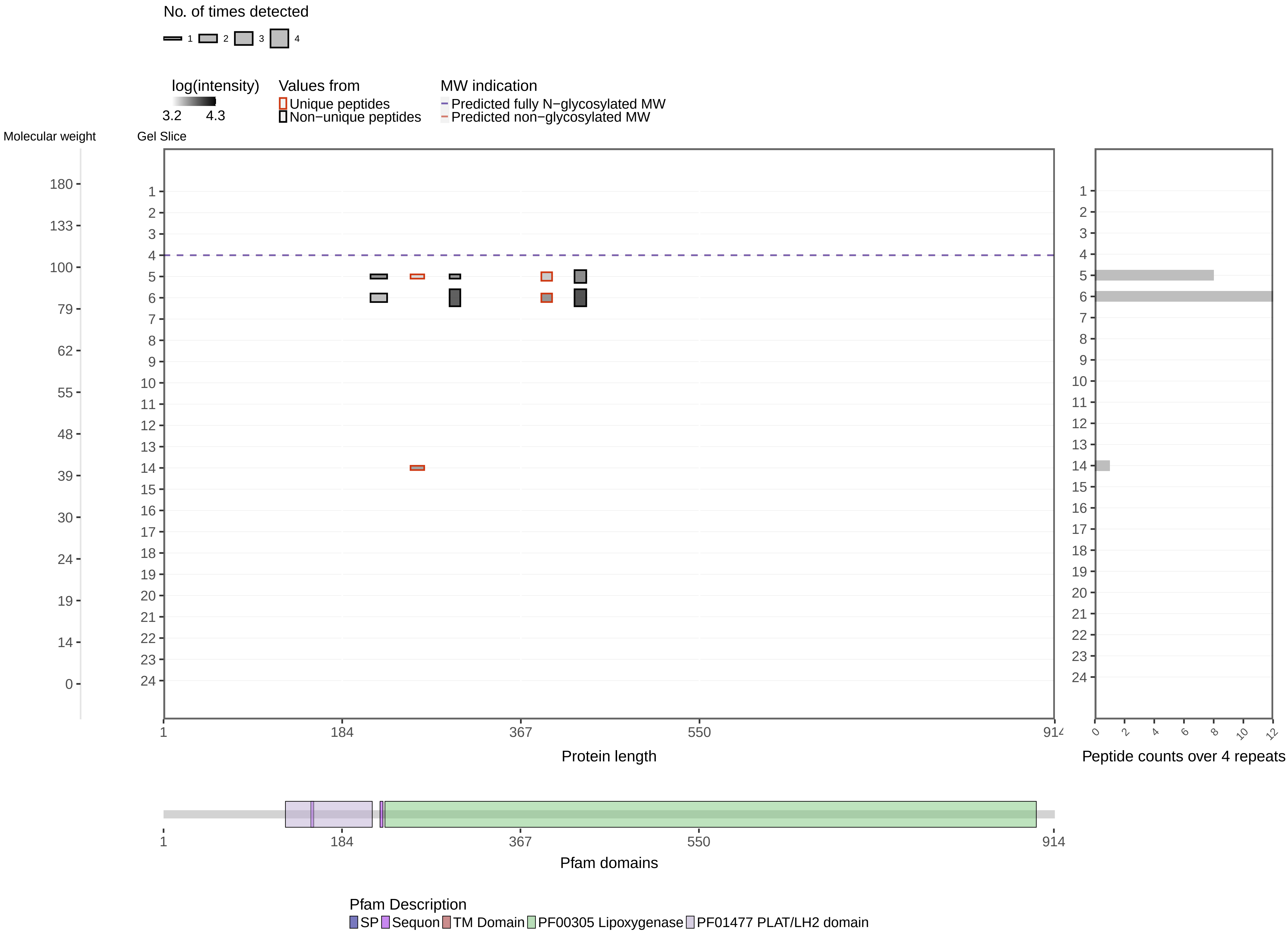


Protein sequence:

DPEIYGTQESAIKEEHILGHLNGMTVQEALDSNRLYIVDYHDVYLPFLDRINALDGRKAYATRTIFFLSDLGTLKPVAIELSLPQTGPSSRSKHVVTPPVDATGNWMWQLAKAHVCSNDAGVHQLVNHWLRTHACLEPFILAAHRQLSAMHPIYKLLDPHMRYTLEINALARQSLISADGVIEACFTPGRYCMEMSAAAYKNFWRFDLEGLPADLIRRGMAVPDPTQPHGLKLLIEDYPYAADGLMIWAAIEGWVRSYVNHYYPDSAQVCNDRELQAWYAESINVGHADLRNEE

NbL03g24370.1 aldehyde_dehydrogenase_family_7_member_B4 [MW = 54.63 kDa]

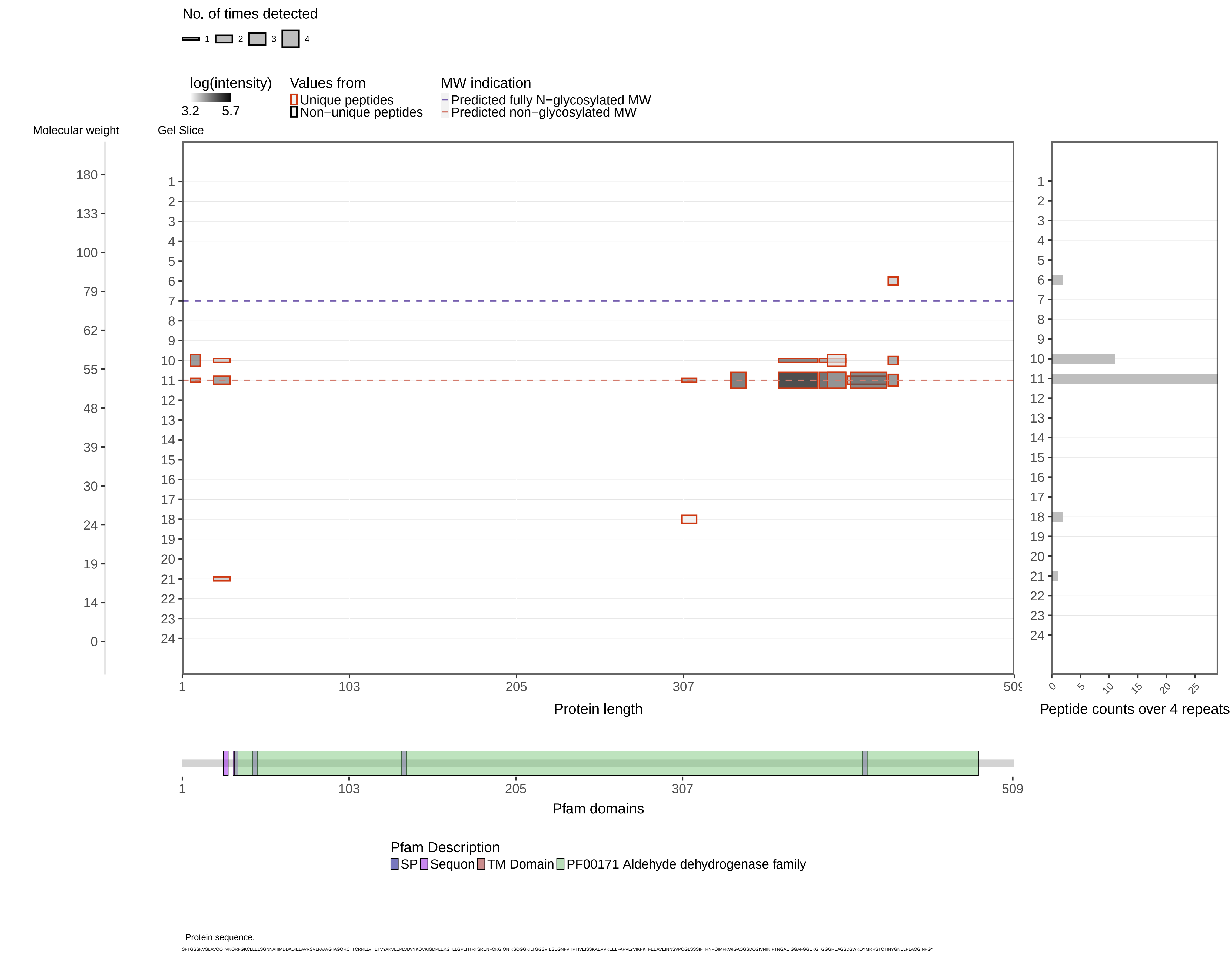

NbL03g24260.1 peroxisomal_(S)−2−hydroxy−acid_oxidase_GLO4 [MW = 39.73 kDa]

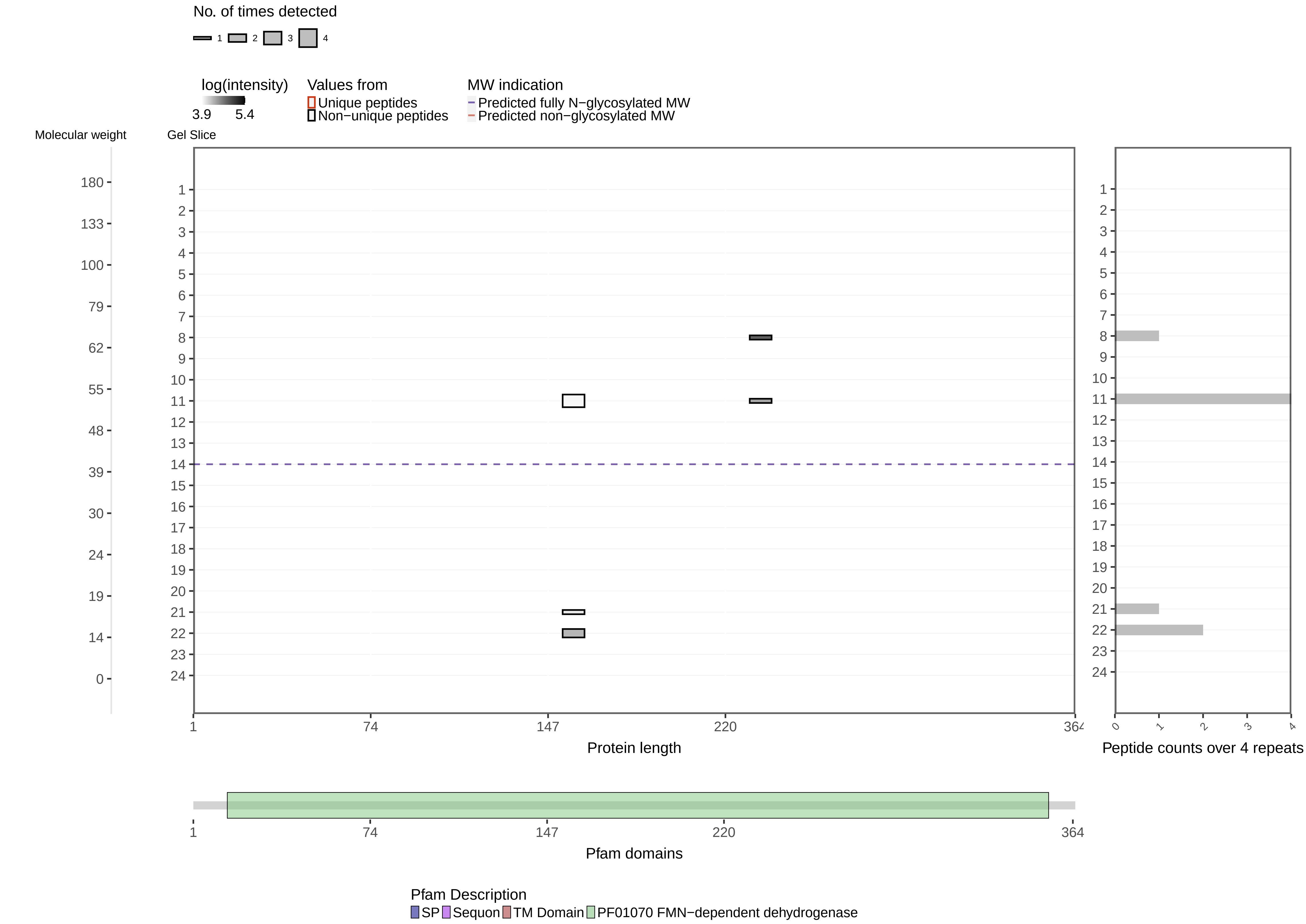


Protein sequence:

ADIKNKMVAPRLKNFEGLISTEYVNDKGSNIEALAAGTFDASLCWKDIPWLKSITKLPILIKGVLTSEDAIKAVEAGVEGIIVSNHGARQLNYAPATISVLEEVVHAVQDKVPVFFDGGISRGTDIFKALALGAKAVLIGRAVVYGLAAKGEHGVKQVIEMLKNELELTMALSGCCTINDITRSHVKTDKEIYCRM*

### NbL03g23940.1 glyoxylate/succinic_semialdehyde_reductase_2_chloroplastic−like [MW = 39.69 kDa]

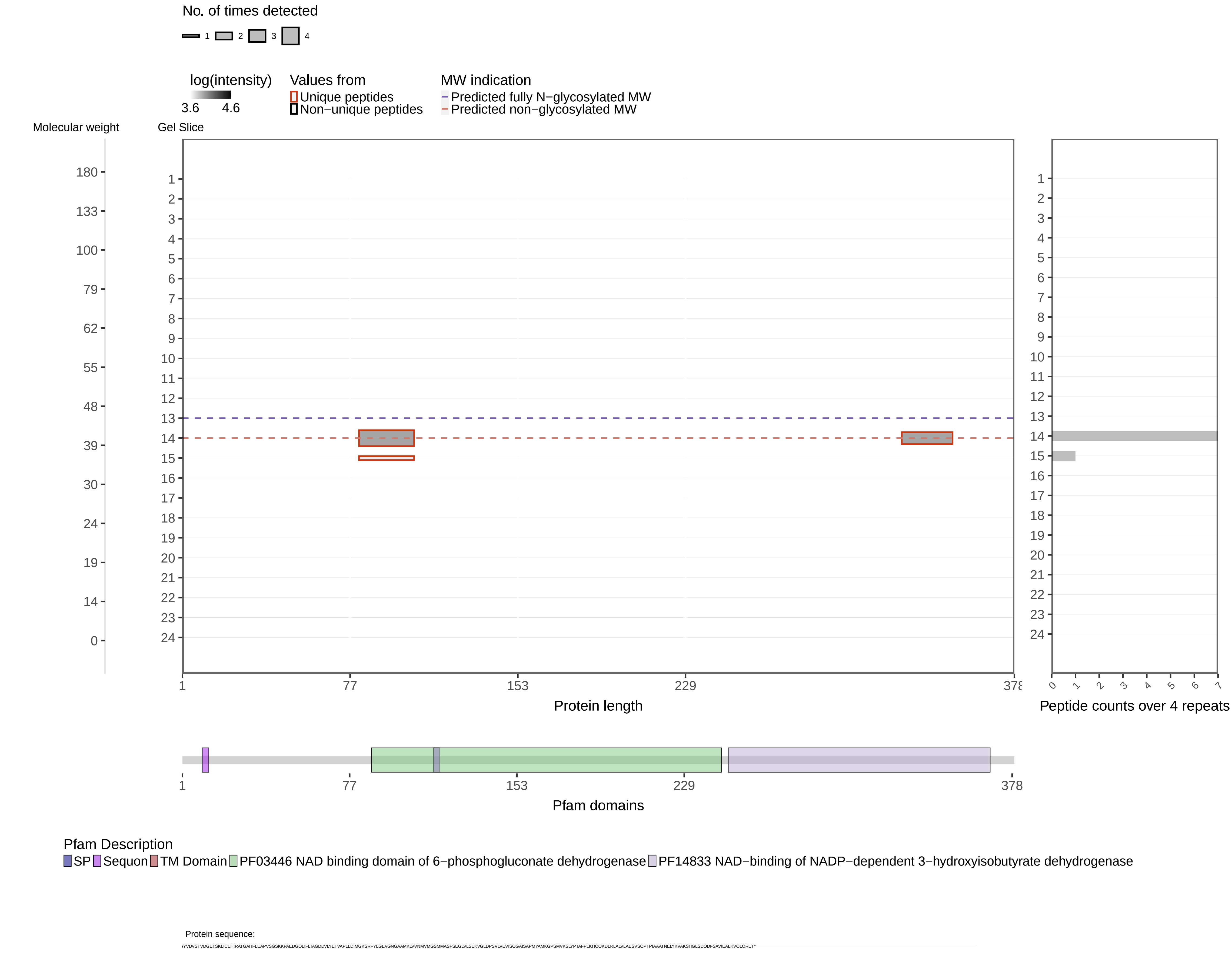

NbL03g23920.1 putative_hydrolase_C777.06c [MW = 39.39 kDa]

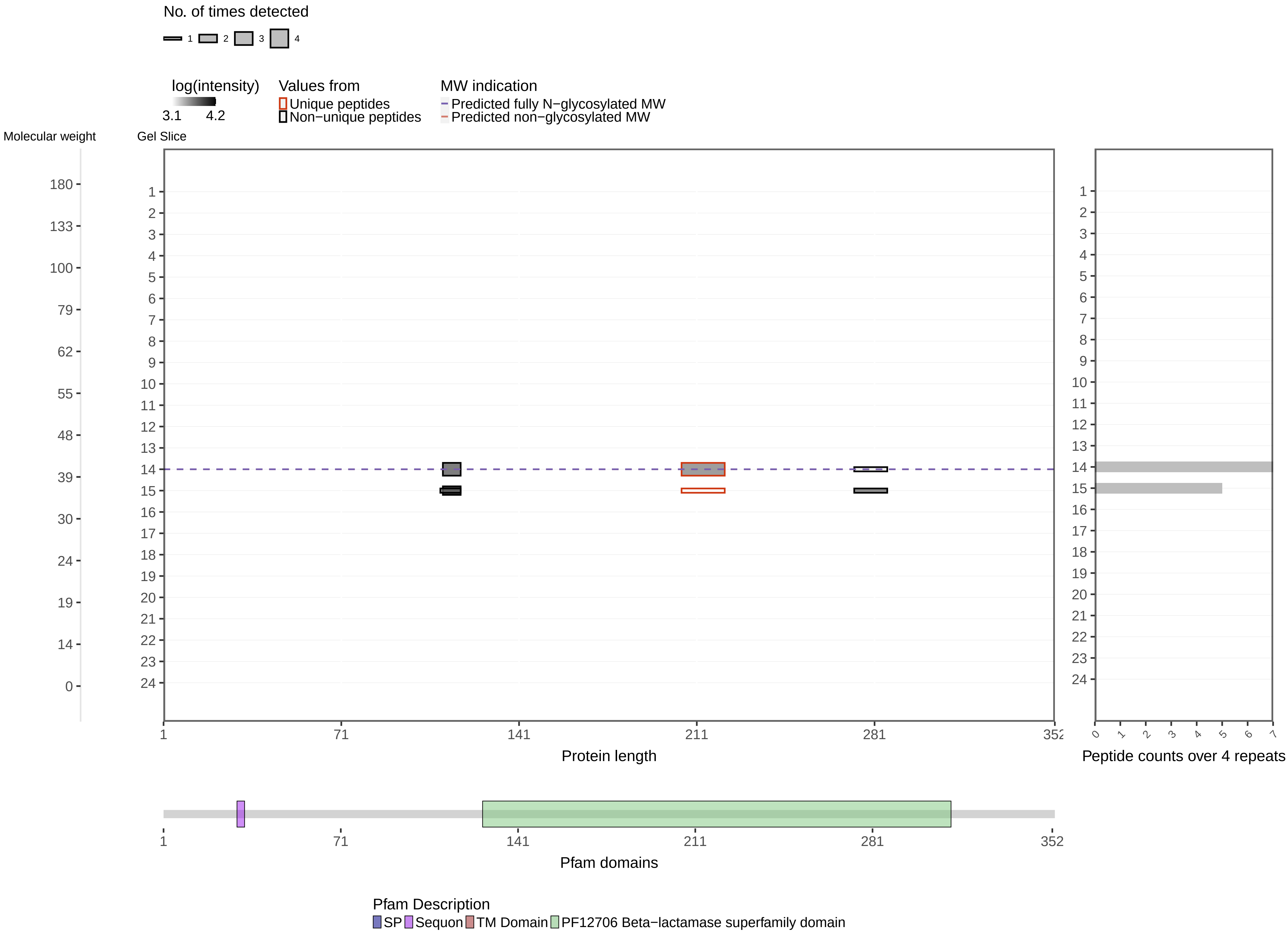


Protein sequence:

AIGGLDDLRDWTNNVQPSIPIYVASRDFEVMKKTHYYLVDRSGIIPGAAVSELQFNIIKEDPFIVHDLEIIPLPVWHGSGYRSLGFRFGDICYISDVSDIPEETYPLLKDCELLIMDALRPDRSSSTHFGLPRALEEVRKIKPKRTLFTGMMHLMDHEVVNERLLKLRETEGIDVQLSYDGLRVPVSL*

NbL03g23830.1 WD−40_repeat−containing_protein_MSI4−like [MW = 50.78 kDa]

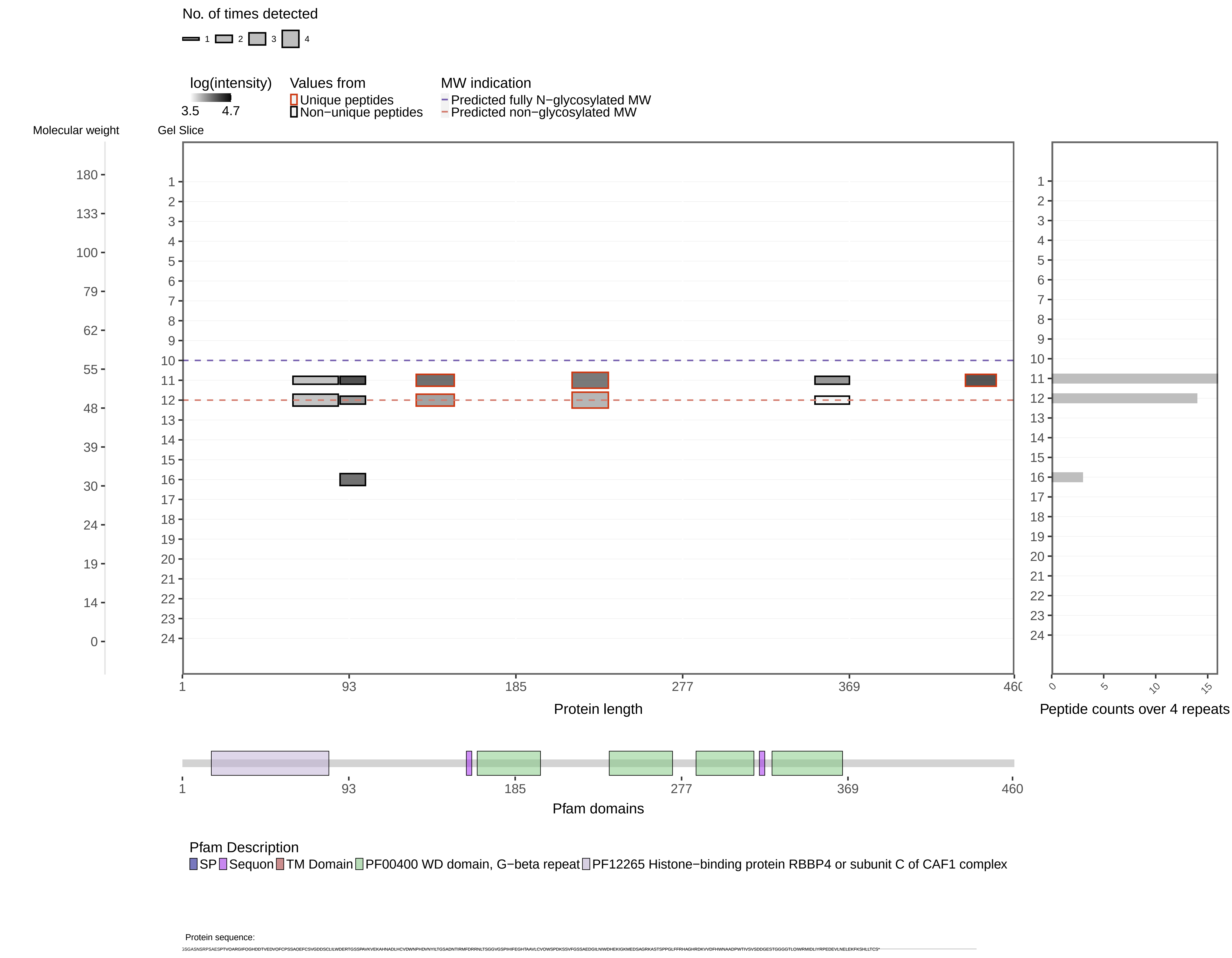

### NbL03g23770.1 eukaryotic_initiation_factor_4A−10−like [MW = 60.8 kDa]

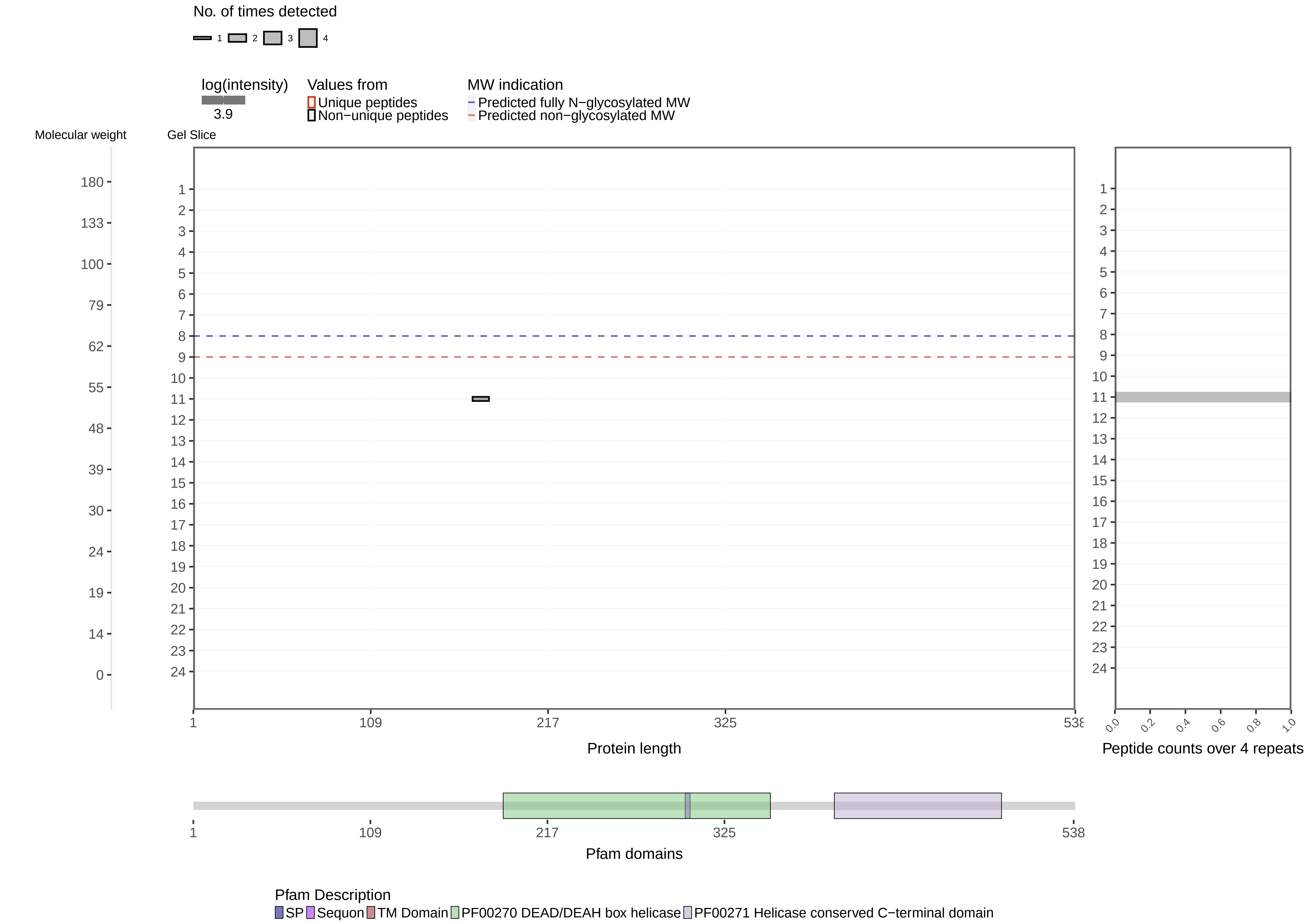


Protein sequence:
[illegible]

### NbL03g23740.1 PRA1_family_protein_F2−like [MW = 20.27 kDa]

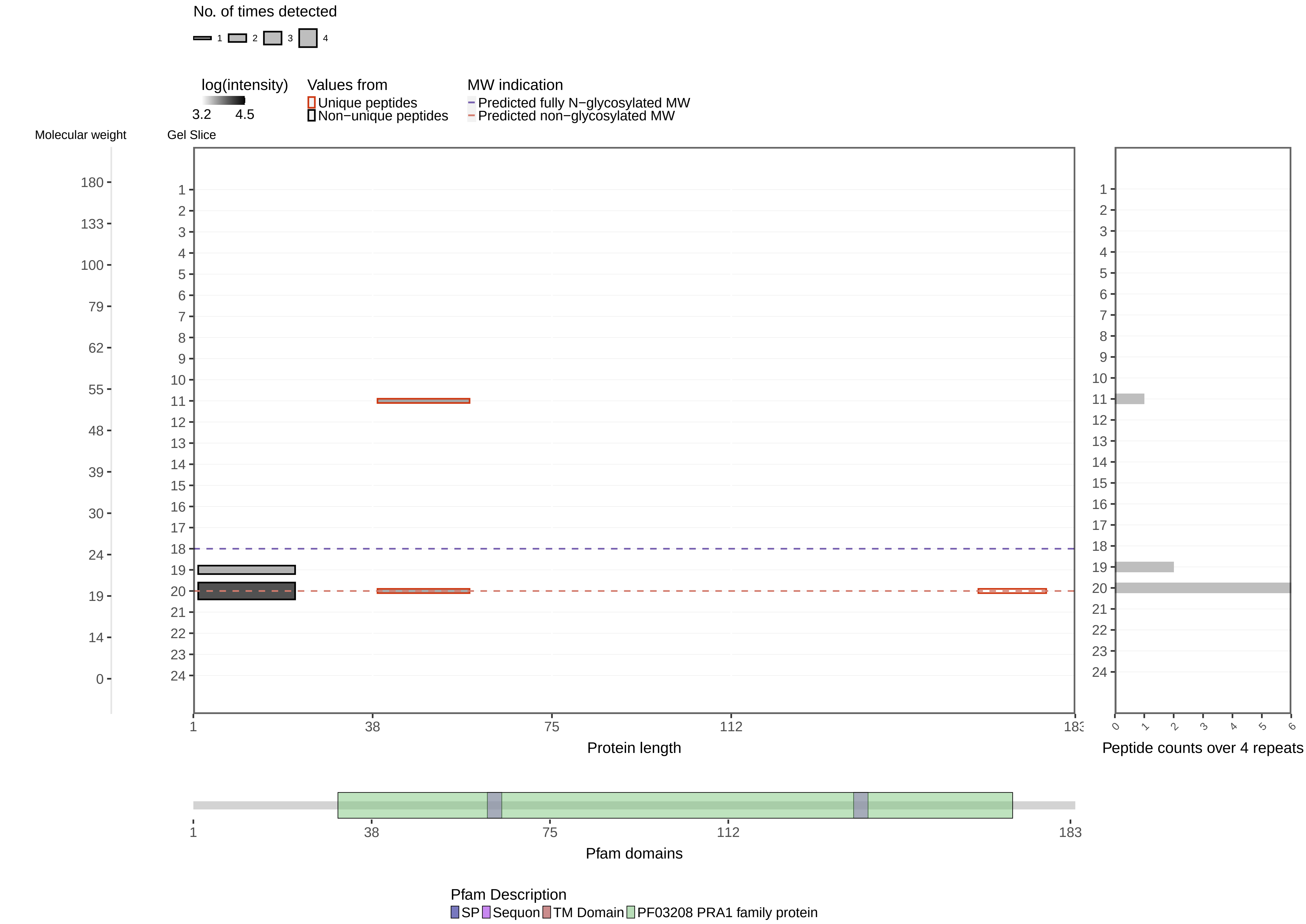


Protein sequence:

LGIVFLSLLWHPISLIVFLVLMAIWLFLYFLRDEPLVVFGRLISDHVVLIVLSVFTIGILLLTNATSNILISLAVAVFVVLIHASIRKTDDLFMDEEAAGFLRAPPSS*

### NbL03g23530.1 IAA−amino_acid_hydrolase_ILR1−like_4 [MW − SP = 45.63 kDa]

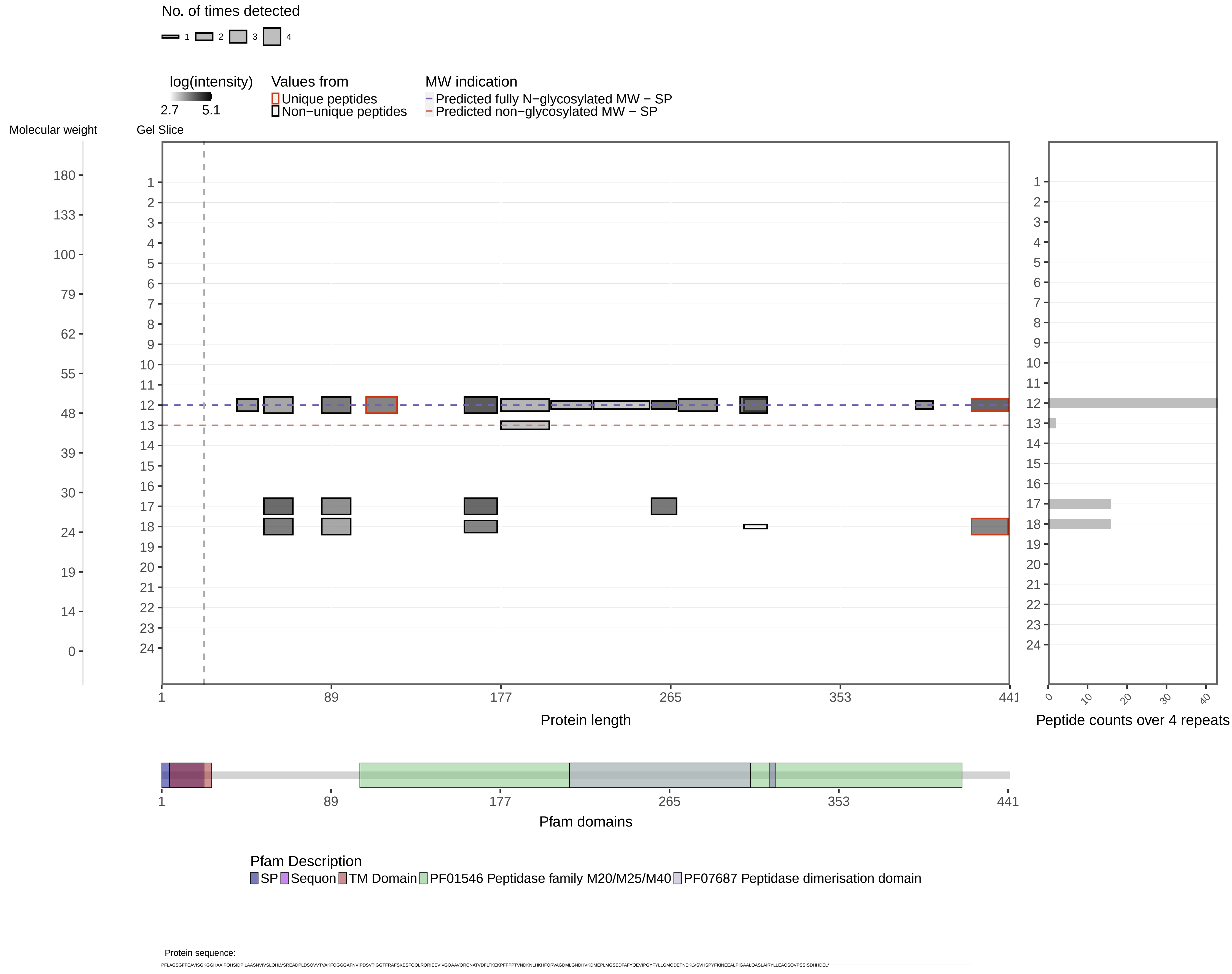

### NbL03g23450.1 diaminopimelate_epimerase_chloroplastic [MW = 39.69 kDa]

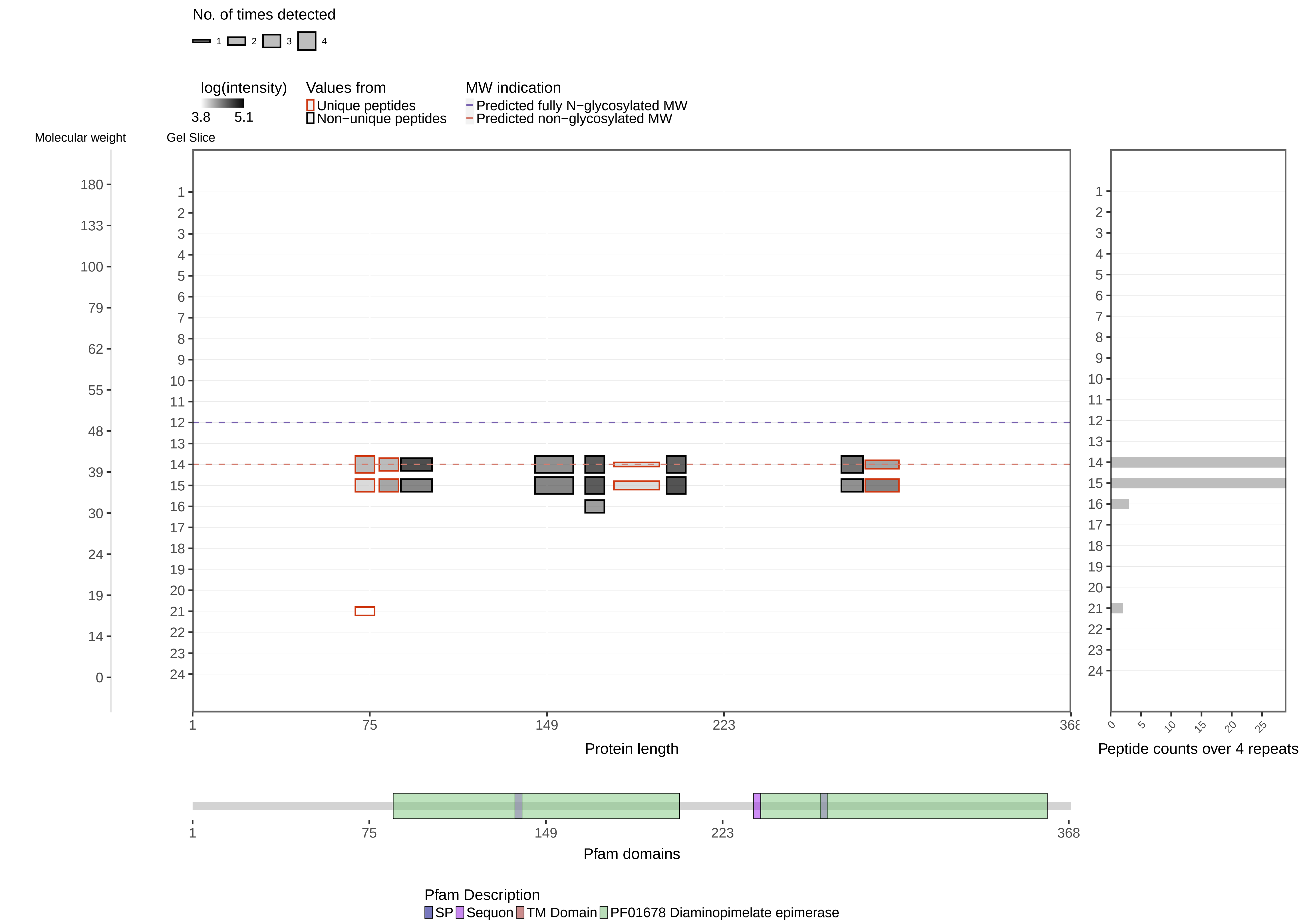


Protein sequence:

GKRSFTIHTGAGLIVPEIOEDGKVRVDMGEPILKASDVPTKLPPNKDOSVVKARLDVDGSAWNVTCVSMGNPHCVTFGTSOGODLOVDELNLSDIGPKFEHHVMFPARTNTEFVOVFSPTYLKMRVWERGAGATLACGTGACAVVVAAVLEGRAARWCTVDLPGGPLDIEWSEKDNHIYMTGPAEVVFYGSVPL*

NbL03g23220.1 protein_transport_protein_SEC13_homolog_B−like [MW = 19.23 kDa]

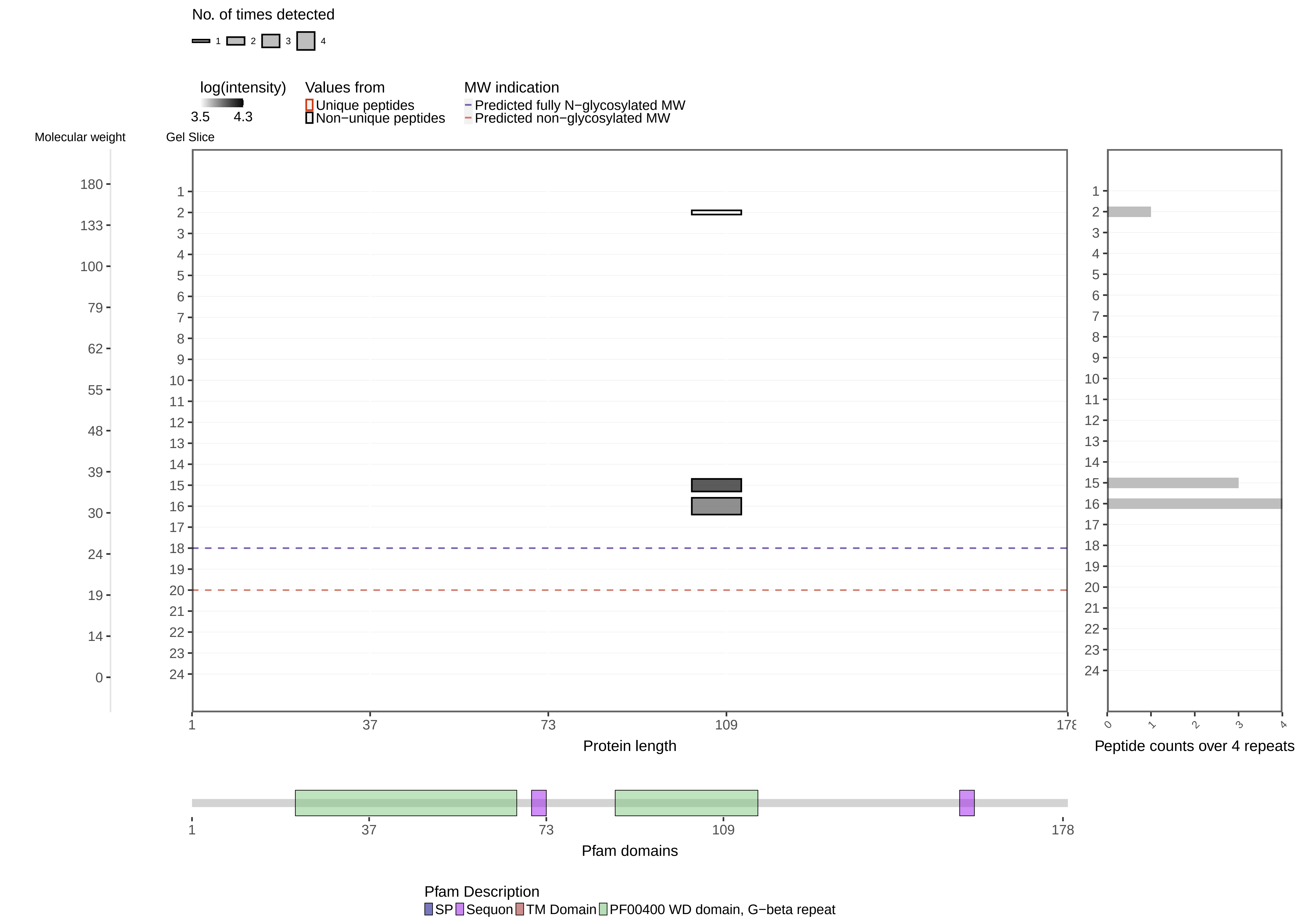


Protein sequence:

CFPALQMYTDWVRDVAWAPNLGLPKTTIASASEDGRVIIWTVAKEGDQWEGKVLKDFGSPVWRVSWSLTGNILAVADGNNNVTLWNEAVDGEWQPVTTVDA*

NbL03g23160.1 ruBisCO_large_subunit−binding_protein_subunit_beta_chloroplastic−like [MW = 64.44 kDa]

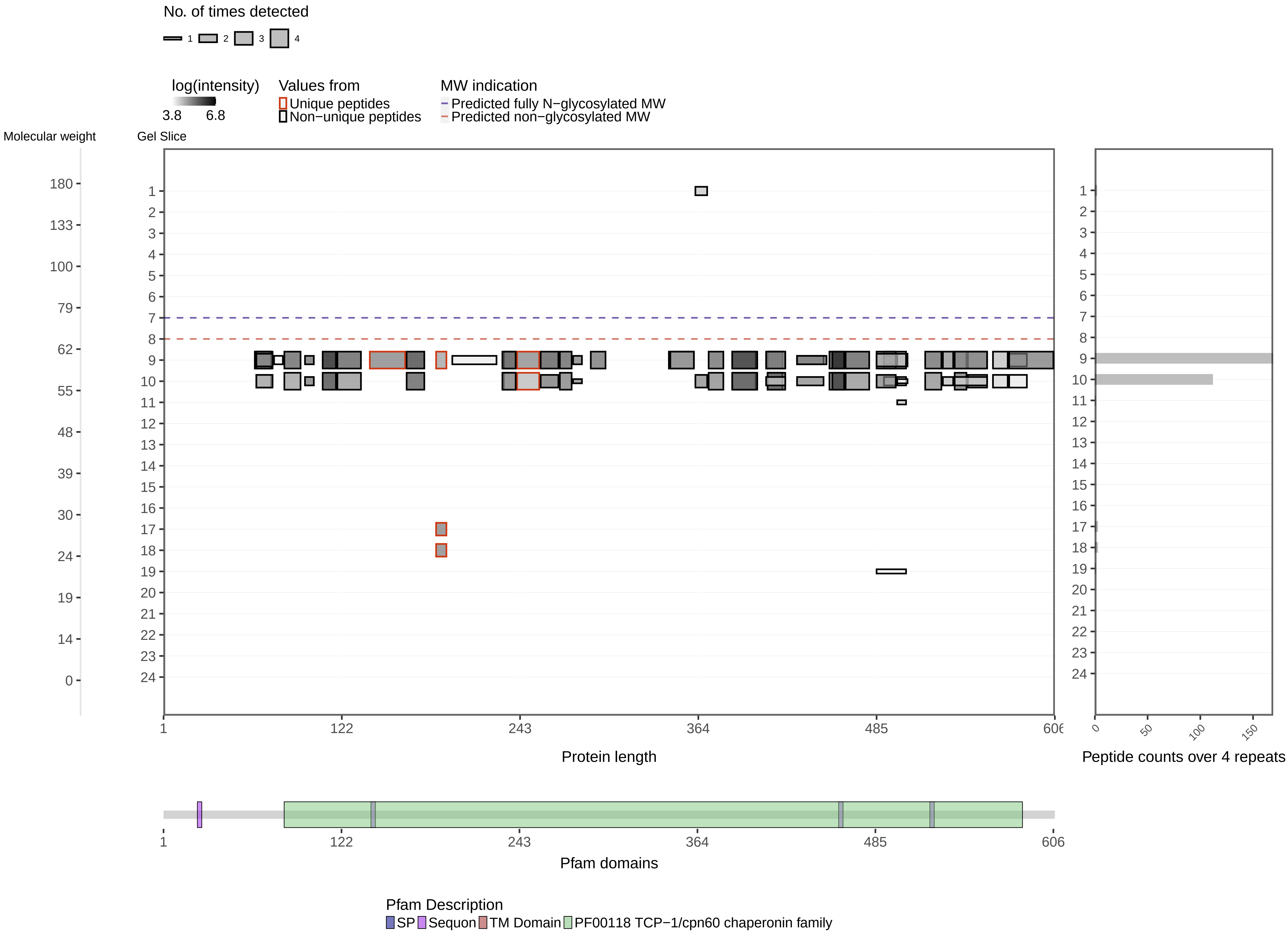


Protein sequence:

NARDLVNVLEEAIRGGYPILIIAEDIEQEALATLVVNKLRGALKIAALKAPGFGERKSQYLDDIAILTGGTVIREEVGLSLDRAGSEVLGHAGKVVLSKDSTTIVGDGSTQEAVNKRIAQIKRIVEATEQDYEKEKLNERIAKLSGGVAVIQVGAQTETELKEKKLRVEDALNATKAAVEEGIVVGGGCTLLRLAAKVEDIKGTLDDDEQKIGADIVKRALRYPMKLIAKNAGVNGSVVIEKVLSNDNPKYGYNAATGKYEDLMAAGIIDPTKVVRCCLEHAASVARTFLTSDAVVTENKE

### NbL03g23070.1 uncharacterized_LOC109224015 [MW = 17.81 kDa]

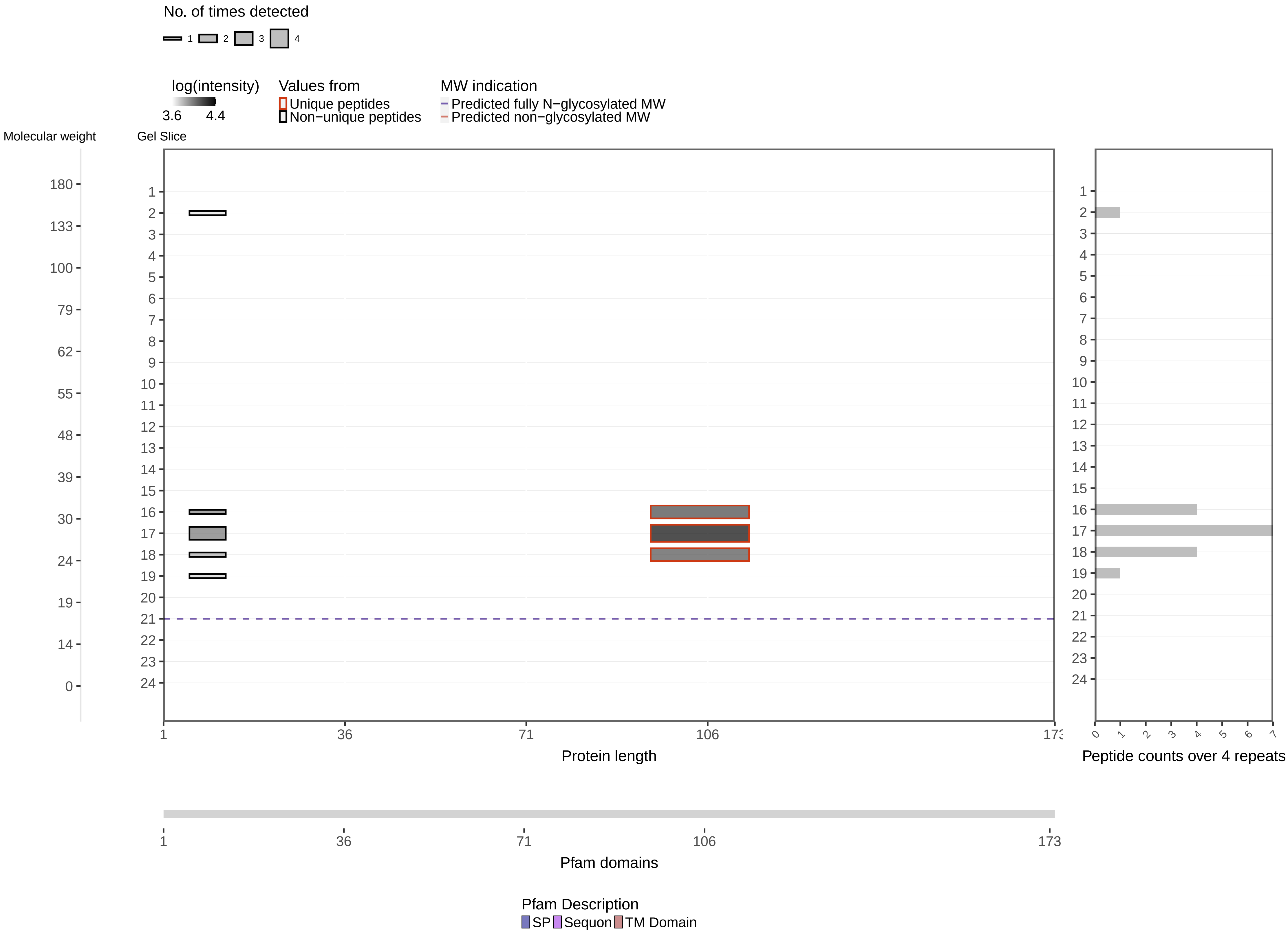


Protein sequence:

ALVGIAVAIEENLGKCITEKGSGGEVPEGTTDVVVVDGAKENSGDLVGEPSLPEGGESCVDGLQGLEIGGKGGKKKEVKEEEGEEGSGKPTLVEGYVV*

### NbL03g23030.1 hydroxyphenylpyruvate_reductase−like [MW = 33.78 kDa]

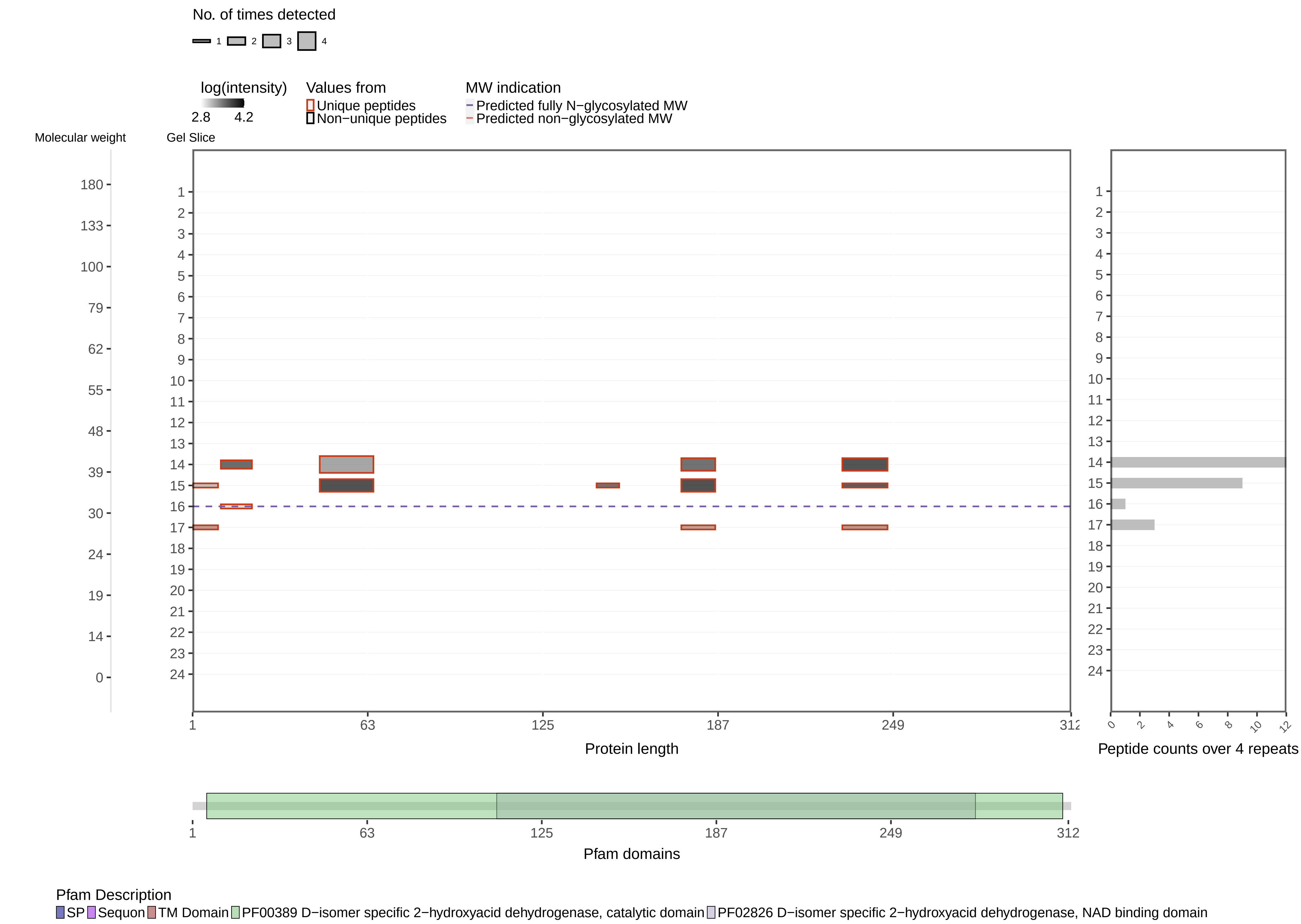


Protein sequence:

GKSVGIVGLGRIGSAIAKRAEAFGCSISYHSRSQKPESTYTYYPRVIDLASNCOILIVACALTDETHHIINREVIAALGPNGVVINIARGSHIDESELVSALAEGRLGGAGLDVLEYEPEAPVLLAGLDNVVLSPHVAASTLETRKDMADLVVANLEAHFSHKPLLTPVI*

NbL03g23020.1 hydroxyphenylpyruvate_reductase−like [MW = 34.16 kDa]

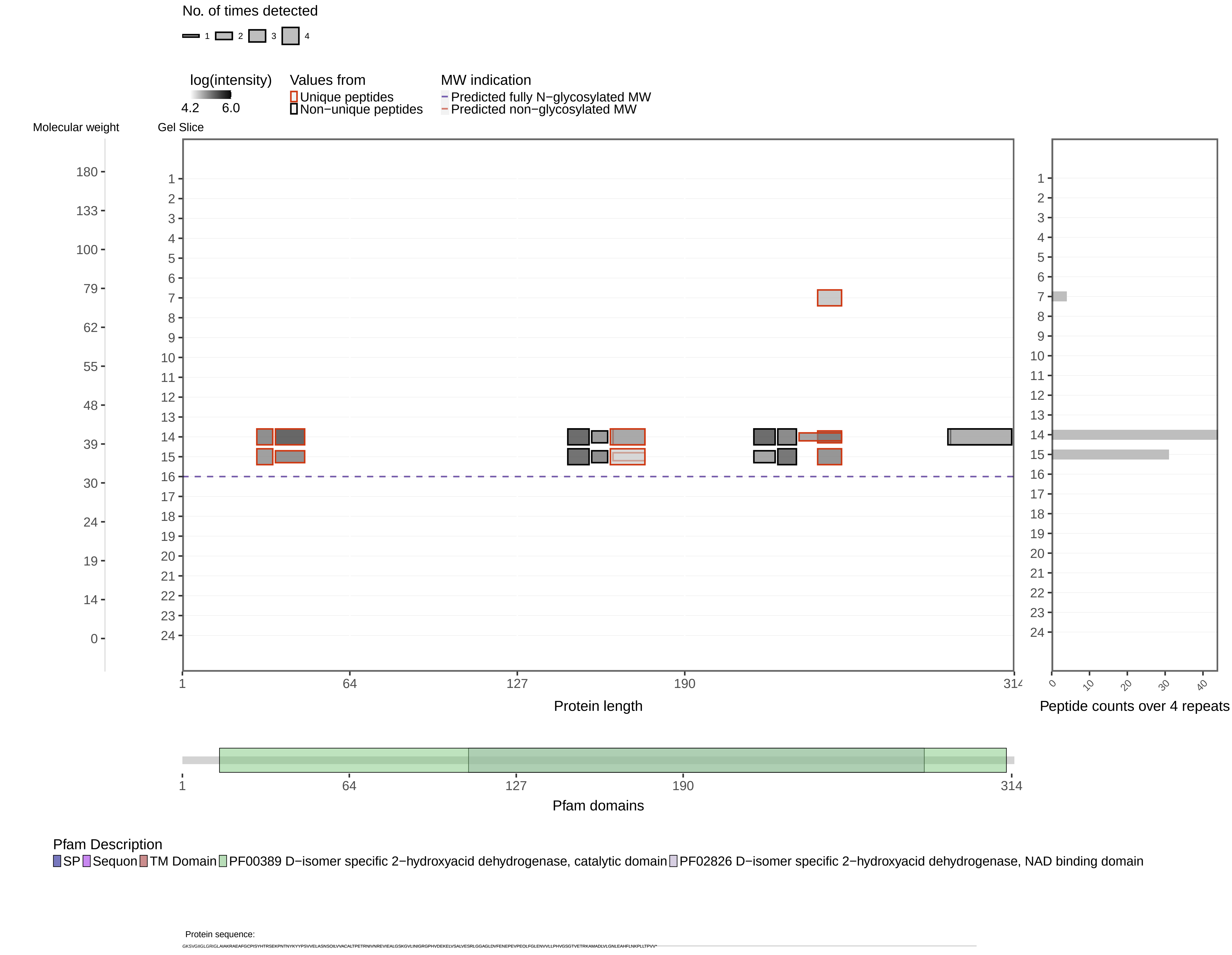

### NbL03g23000.1 30S_ribosomal_protein_S17_chloroplastic [MW = 17.59 kDa]

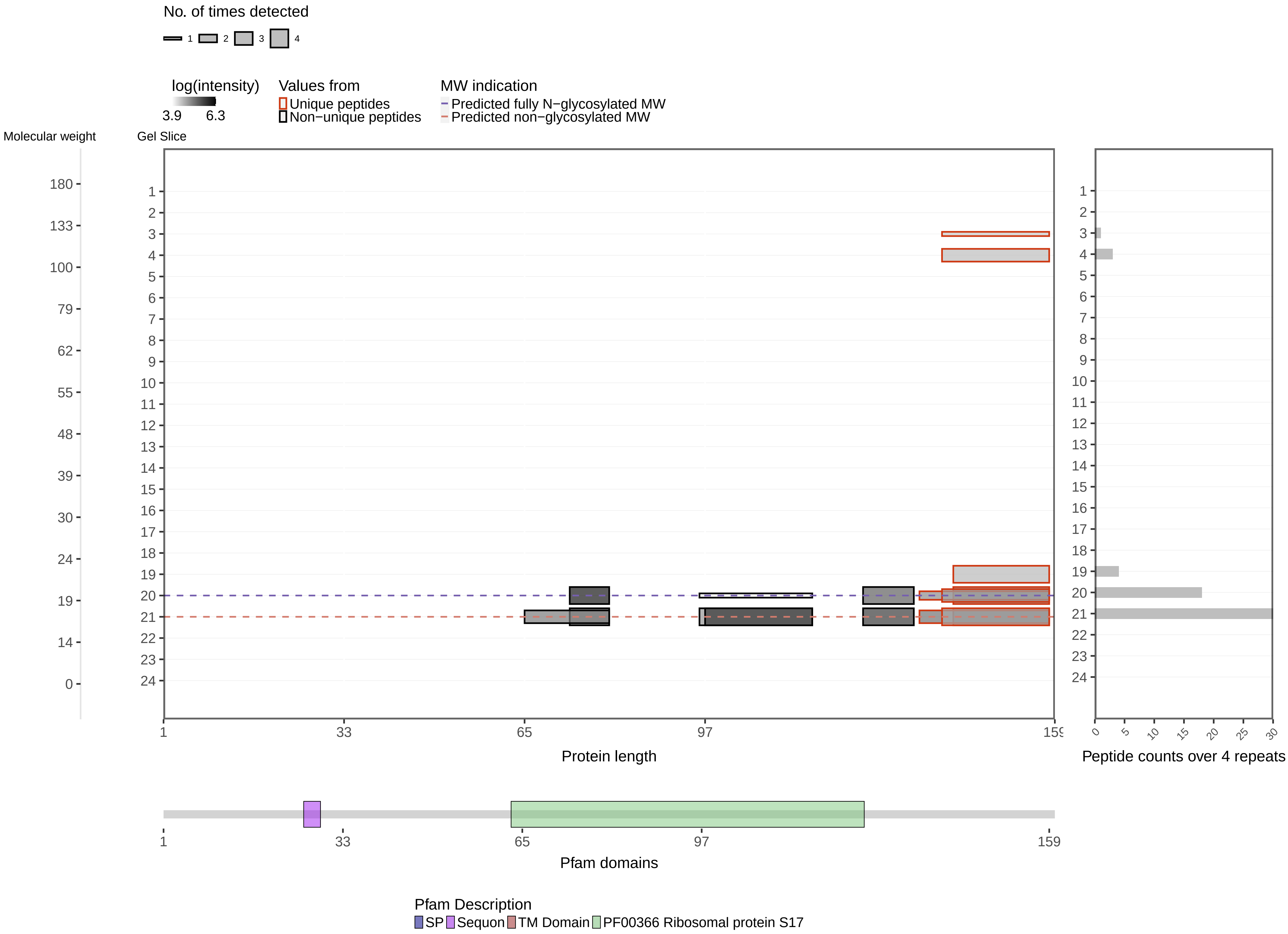


Protein sequence:

STNDKTVSVEVTRLAPHPKYKRRVRKKKKYQAHDPLNQFQVGDFVQLEKTAPISKTKTFLAVPVPPRNQPKPKEEENQELVLPLESSLQQS*

### NbL03g22950.1 malonate−−CoA_ligase [MW = 67.3 kDa]

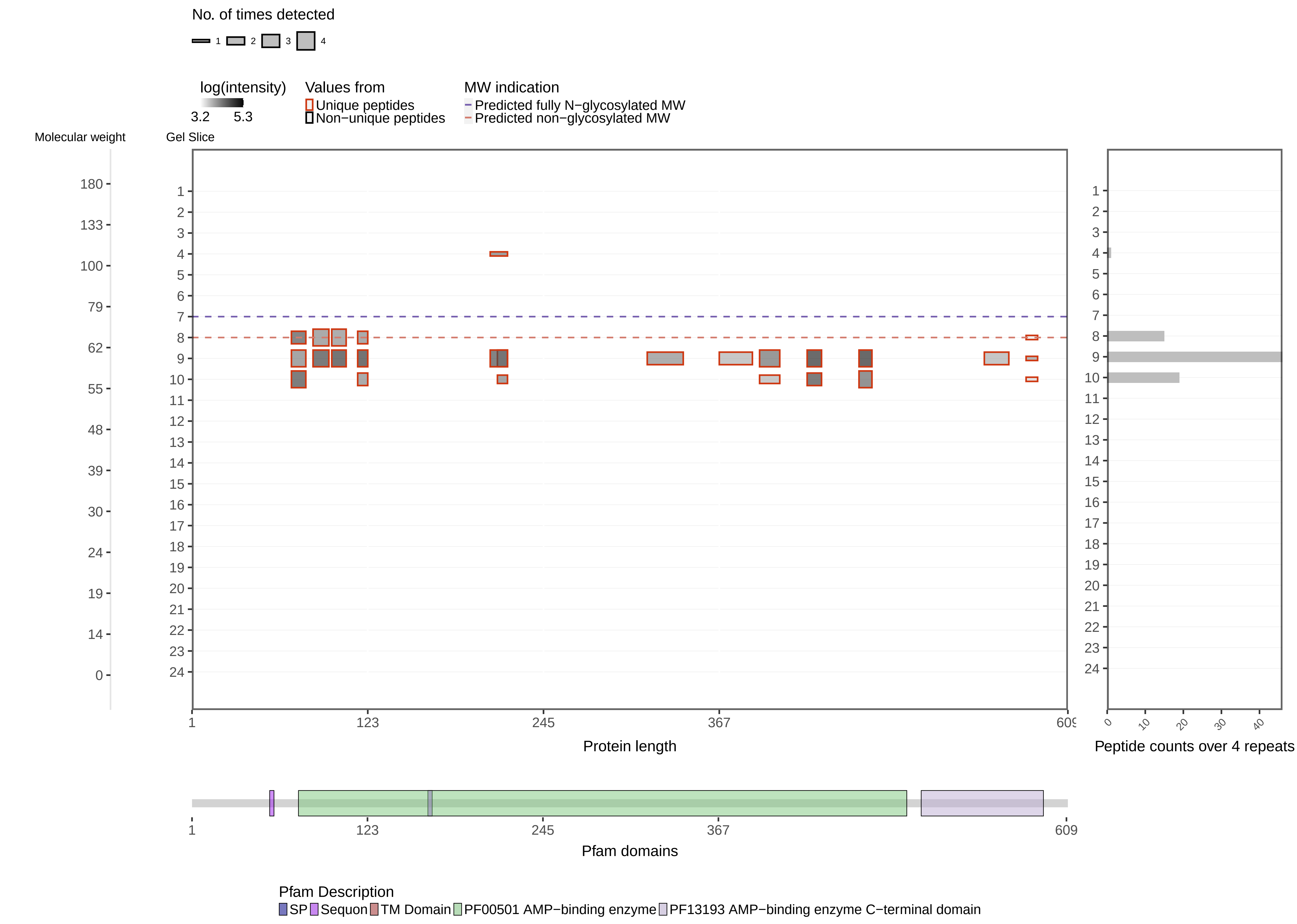


Protein sequence:

### NbL03g22850.1 aspartate_aminotransferase [MW = 43.39 kDa]

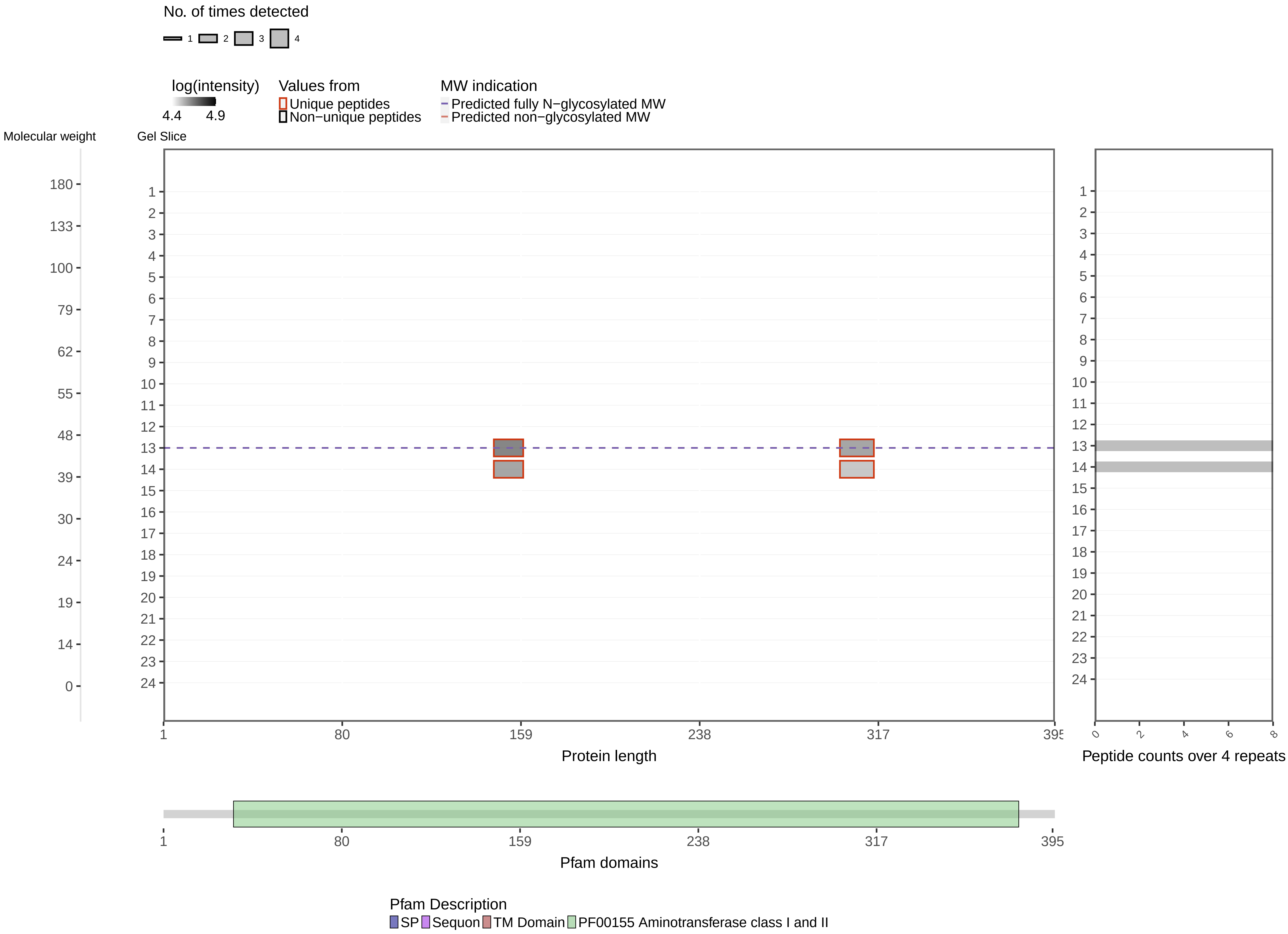


Protein sequence:

PESLLKRISDICKEAGCWLVIDNTYEYFMYDDRKHVCIEGNHIVNIFSFSKAYGMMGWRVGYIAYPSEVEGLAAOLLKVODNIPICASIISORLALYSMEMGPEWVTDOVKDLVKNRELLLEALSPLGEEAVKGGEGAIYLWAKLPDKYENDFEVVHWLAKRHGIVLIPGSSCGCPGHVRISFGGLIEKDCRVAAERLRKGLEELVNSGMIS*

NbL03g22830.1 D−glycerate_3−kinase_chloroplastic [MW = 51.1 kDa]

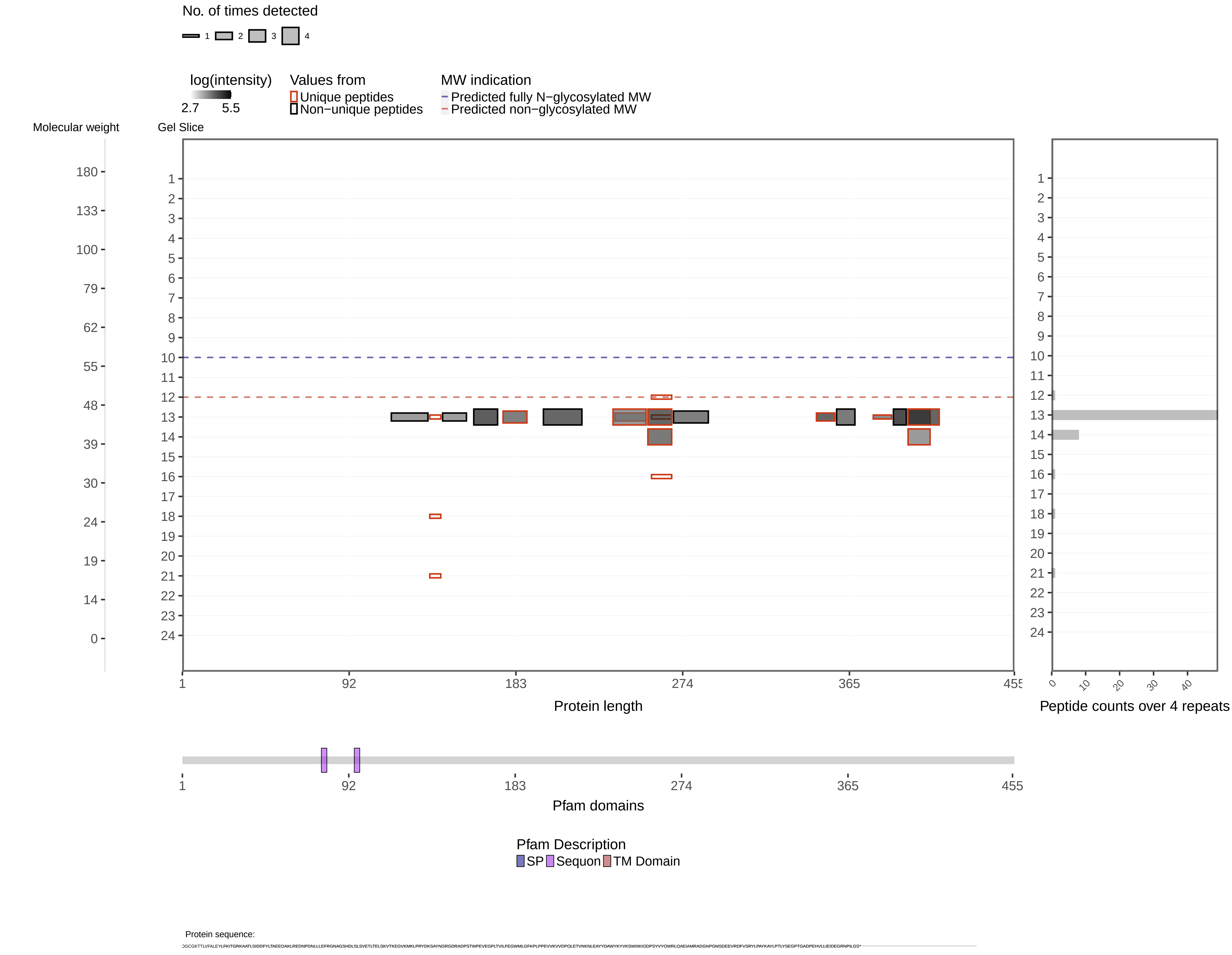

NbL03g22820.1 protein_CDI−like [MW = 31.98 kDa]

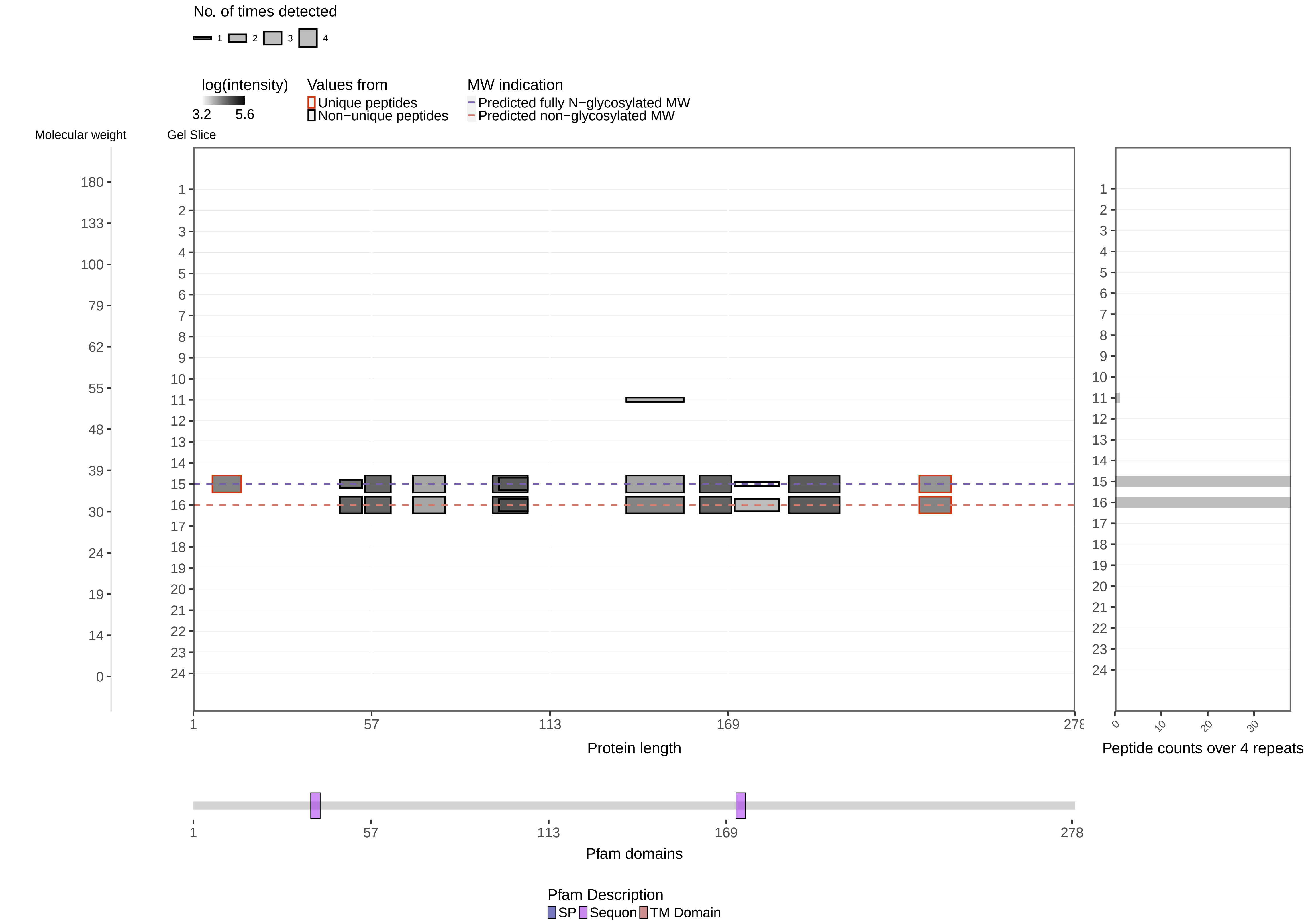


Protein sequence:

YLDDIKELRDLVDDKYALMCVQHDYTPKETTKMDGAVQTVYPRKNWSSMVLYNCGHPKNRALTPEVVNSESGAFLHRFMWLEDEEIGEVPFVWNFLVGHNKVVKDDPATFPKAIHYTLGGPWFEAWKDCEFGDLWLKELEECEKAKEKLG*

NbL03g22790.1 uncharacterized_LOC109210747 [MW = 83.38 kDa]

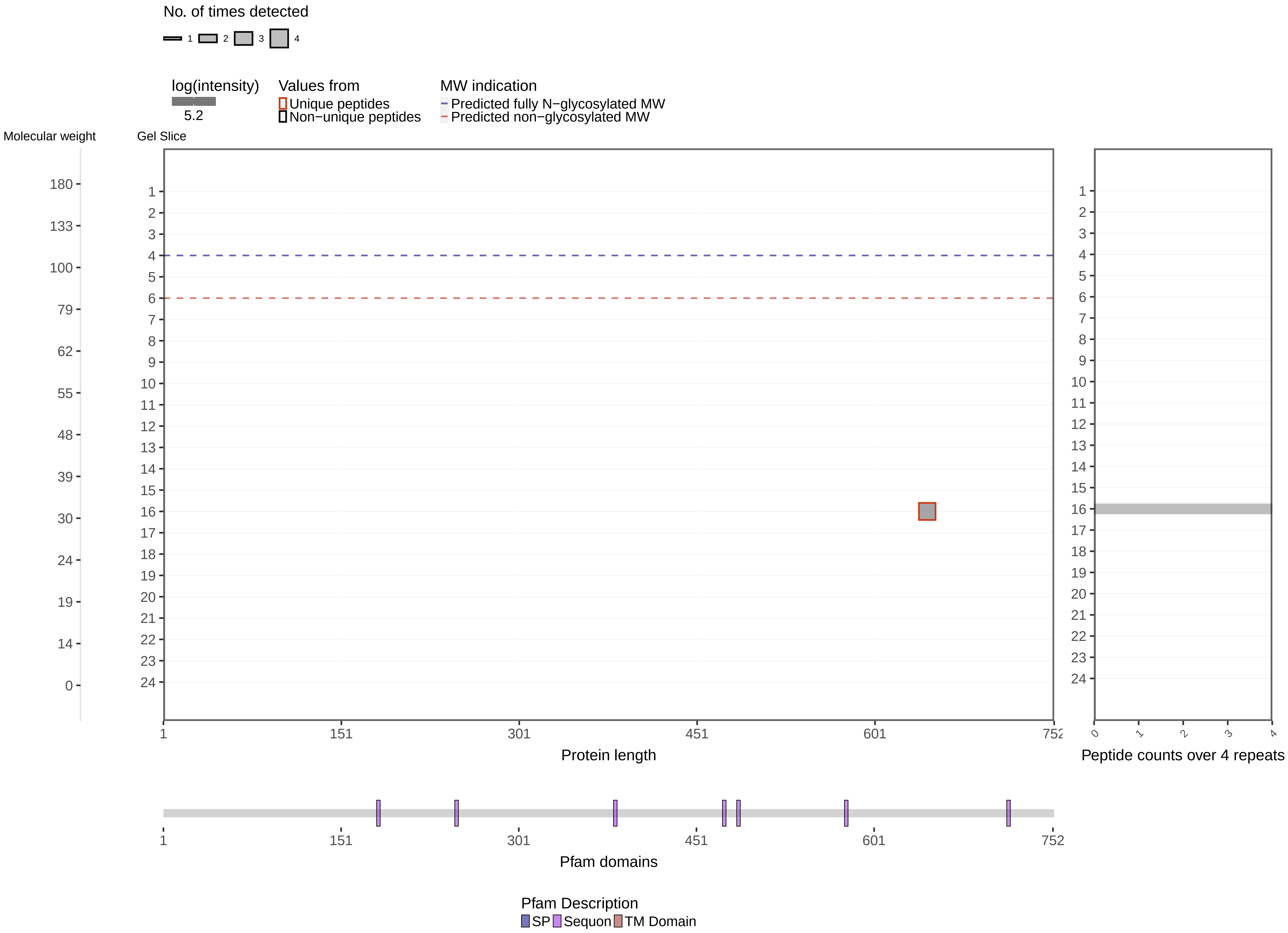


Protein sequence:

LRDPFSPLNIDLYNNDSQSDHPPDWSMKMNPQCISNNAGPTESHGETEYTAGSDNTNIMKMNPQCISNNAGPTESHGETENTAGSDNTNIIVPLSGSDFSHEQLMKNDSGKDNIKTGPNGSQSEKELENGYDDIDINNDINLNMRNMDSYYESGVLVKVNDVAVVNDVIKDQQGLQTESSISCQNMQDGEVLAETLSSPQAQRKADDTHNYSIETVAADSGSSEIDRQVDHMPLQREHSAEQDHYFEDSVKDVTSDQLRSVAVEVHSTEVRSVLSDMSPQHYAQAKDK

### NbL03g22780.1 N−alpha−acetyltransferase_15_NatA_auxiliary_subunit−like [MW = 52 kDa]

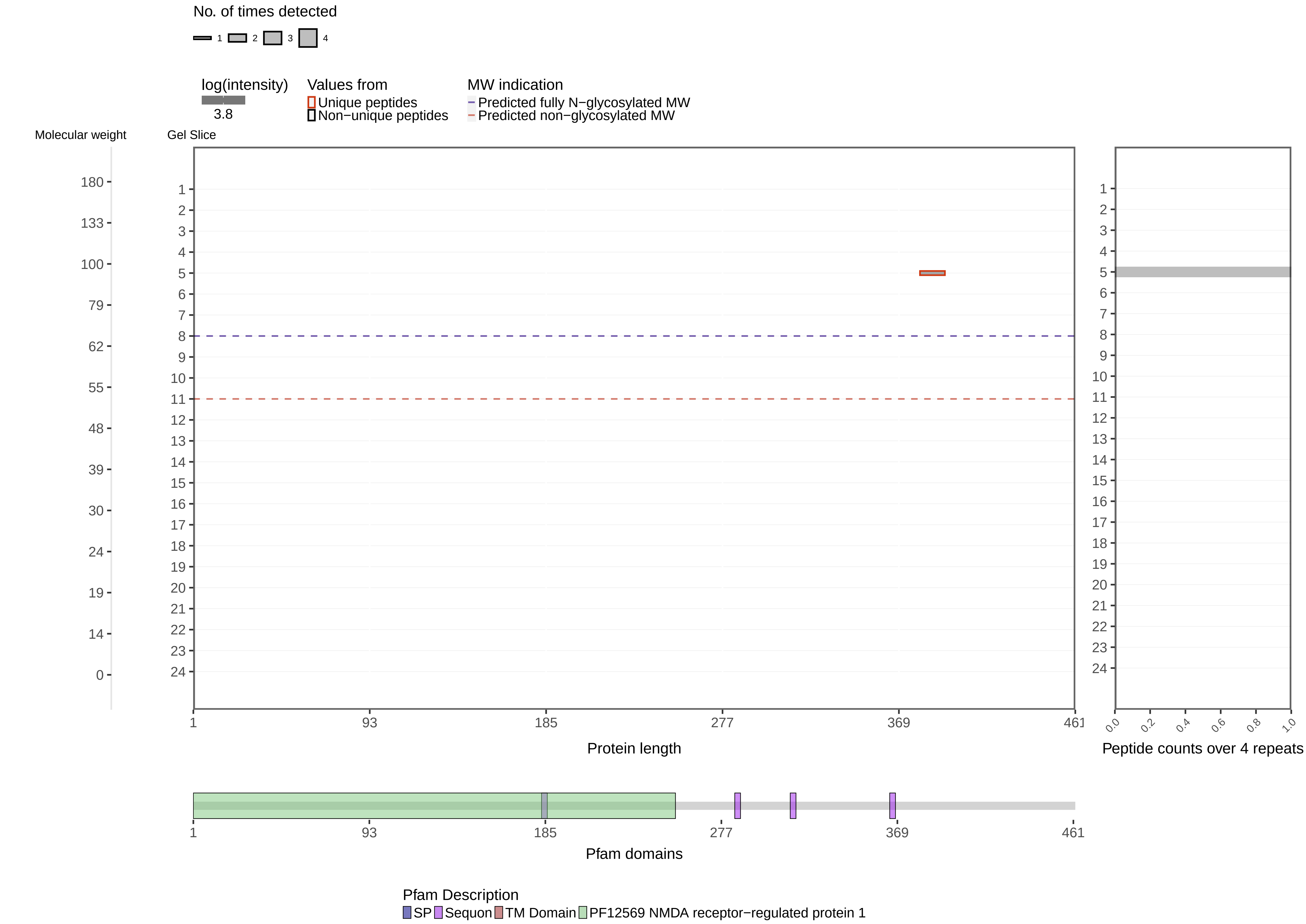


Protein sequence:

ASKYLKLLLKHSSDCLETHLLSFEVNMRKQKILLALQAIKYLMRLDADDPKSHLCLIKFFHKIDSLPTPVNDTEKLIWGVLEAERPTISQLHGKSLIDANNTFLEKHRESLMHRAAVAELLYVLEPNKKAEAVKLILDSTNDLVSIDGAQGNVSSWKLNDCITVHKLLETTLADSDAASRWKVRCAEYFSYSIHFGGFQSSAATSSANNQIQKSSANGALGLNADGDSSLSSNGRLDKLNALKDLHI*

### NbL03g22690.1 nuclear_pore_complex_protein_NUP50A−like [MW = 47.84 kDa]

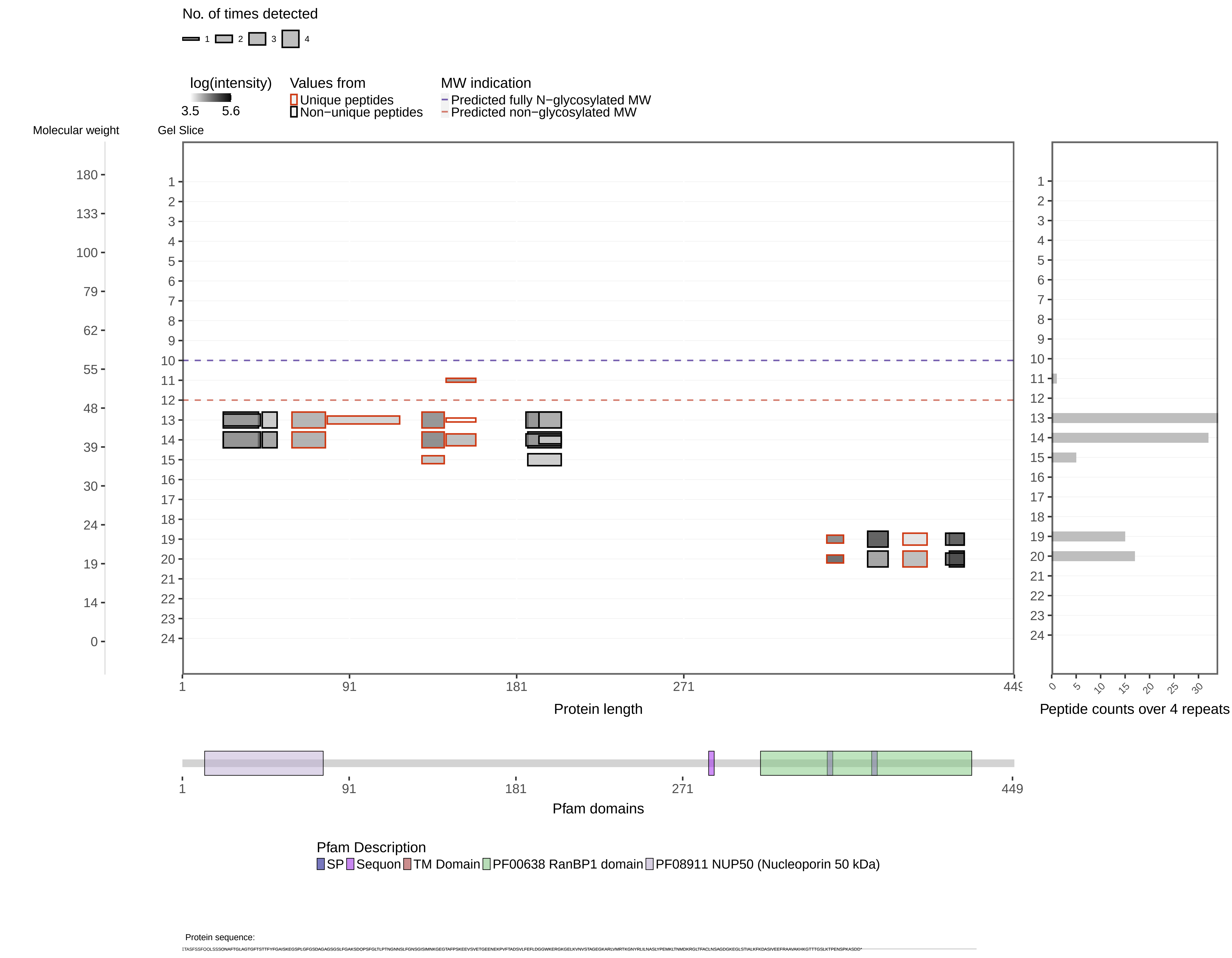

### NbL03g22650.1 MAR−binding_filament−like_protein_1−1 [MW = 82.35 kDa]

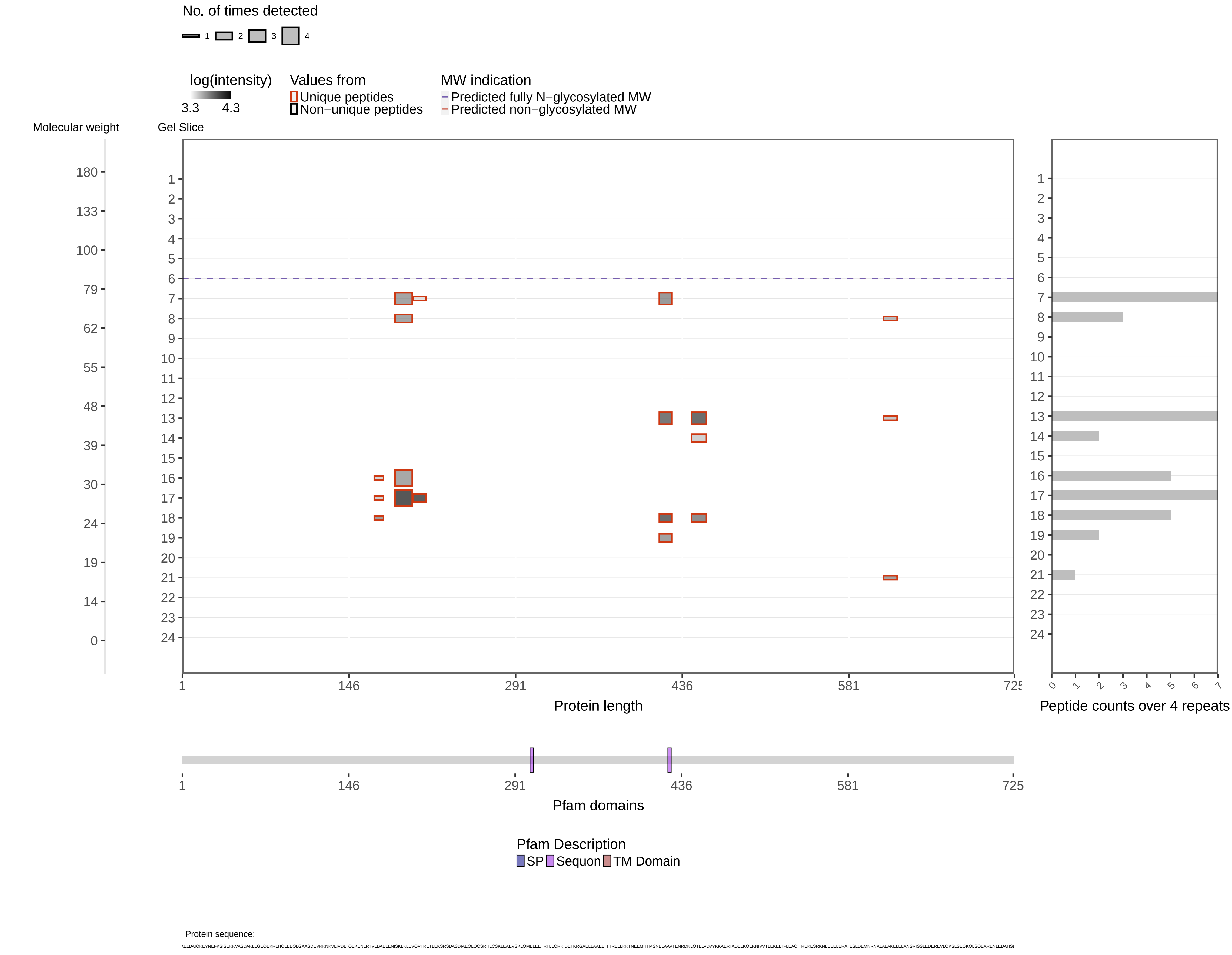

NbL03g22570.1 pyridoxal_5'−phosphate_synthase−like_subunit_PDX1.2 [MW = 32.88 kDa]

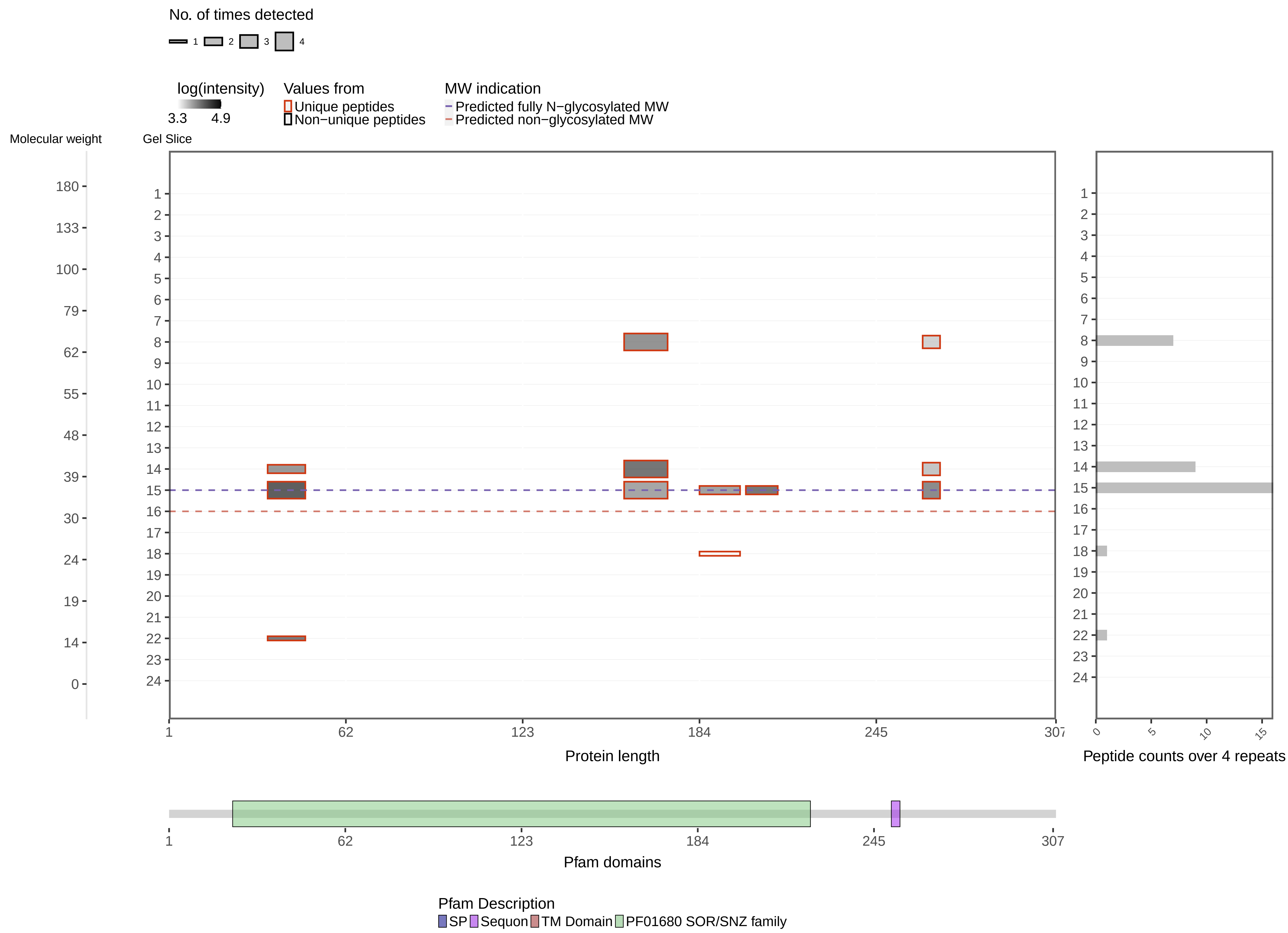


Protein sequence:
LGEALRRVREGAAMIRTQGDLLGTGNIVDTVRNVRKVMGDIRLLSNMDEDEVFTFSKKISAPYDIVAQTKQMGRLPVVHFAAGGIVTPADAALMMQLGCDGVFLGPDIFNCSNPYKKVRAIVQAVRNYNDPHILAAASSGLEEAMGGLNLNENRVERFVSAEETY*

### NbL03g22550.1 uncharacterized_LOC107817663 [MW − SP = 38.57 kDa]

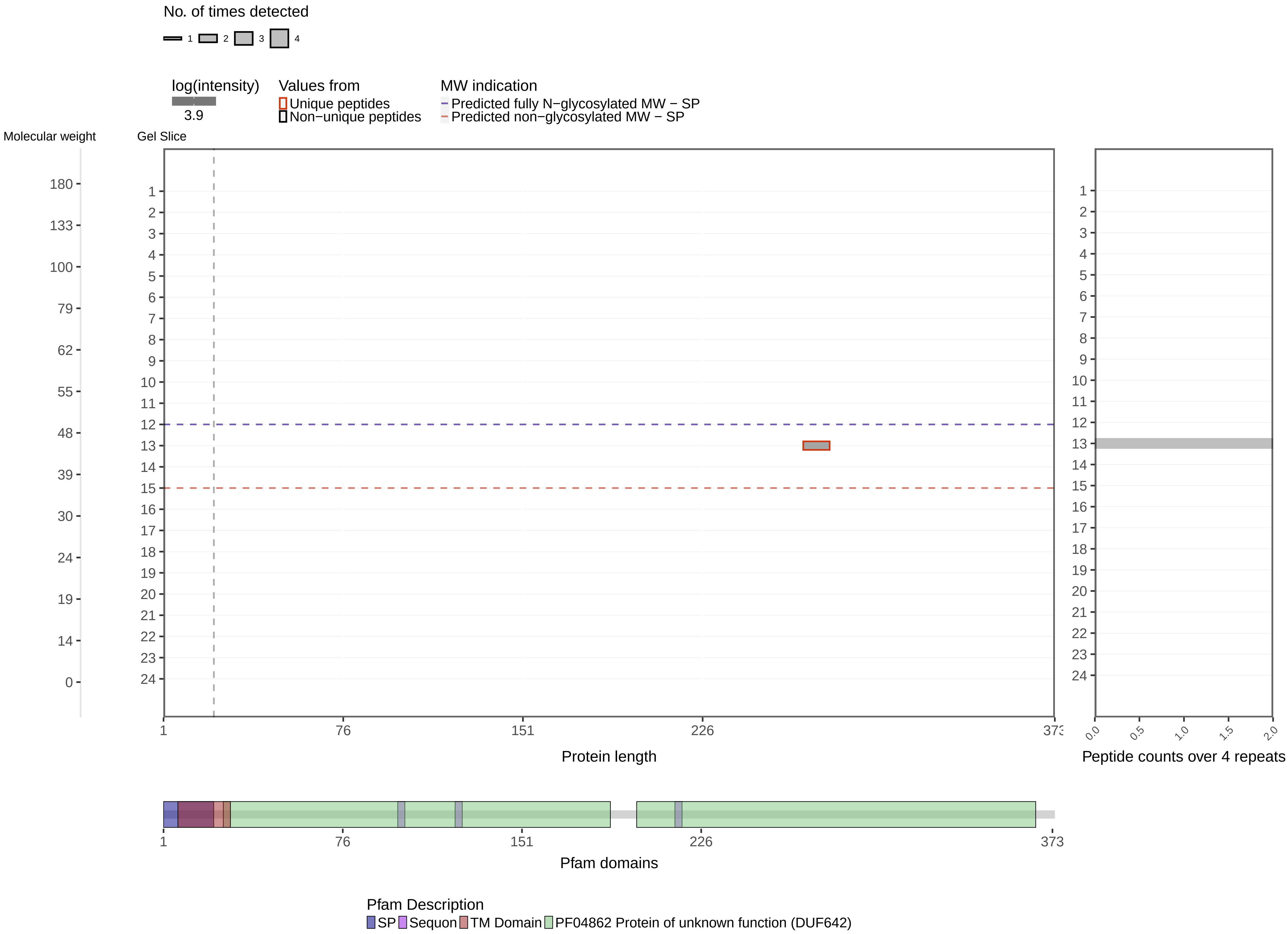


Protein sequence:

EKDPACGPLIDFIALKALKNPHRPKGNMLKNGNFEEGPYIFPNTSWGVLIPPNIEDDHSPLPGWMIESLKAVKYIDAAHFTVPEGKRAIELVAGRESTIAQQVITTKGKVYDLMFSVGDASNSCQGSMLIEAFAGQINLQVPYESTGNGGFKRAKLRFKAVSERTRVRFLSTYYHMKSDHSGSLCGPVVDDVRLVGVRHP*

### NbL03g22480.1 serine−threonine_kinase_receptor−associated_protein−like [MW = 38.43 kDa]

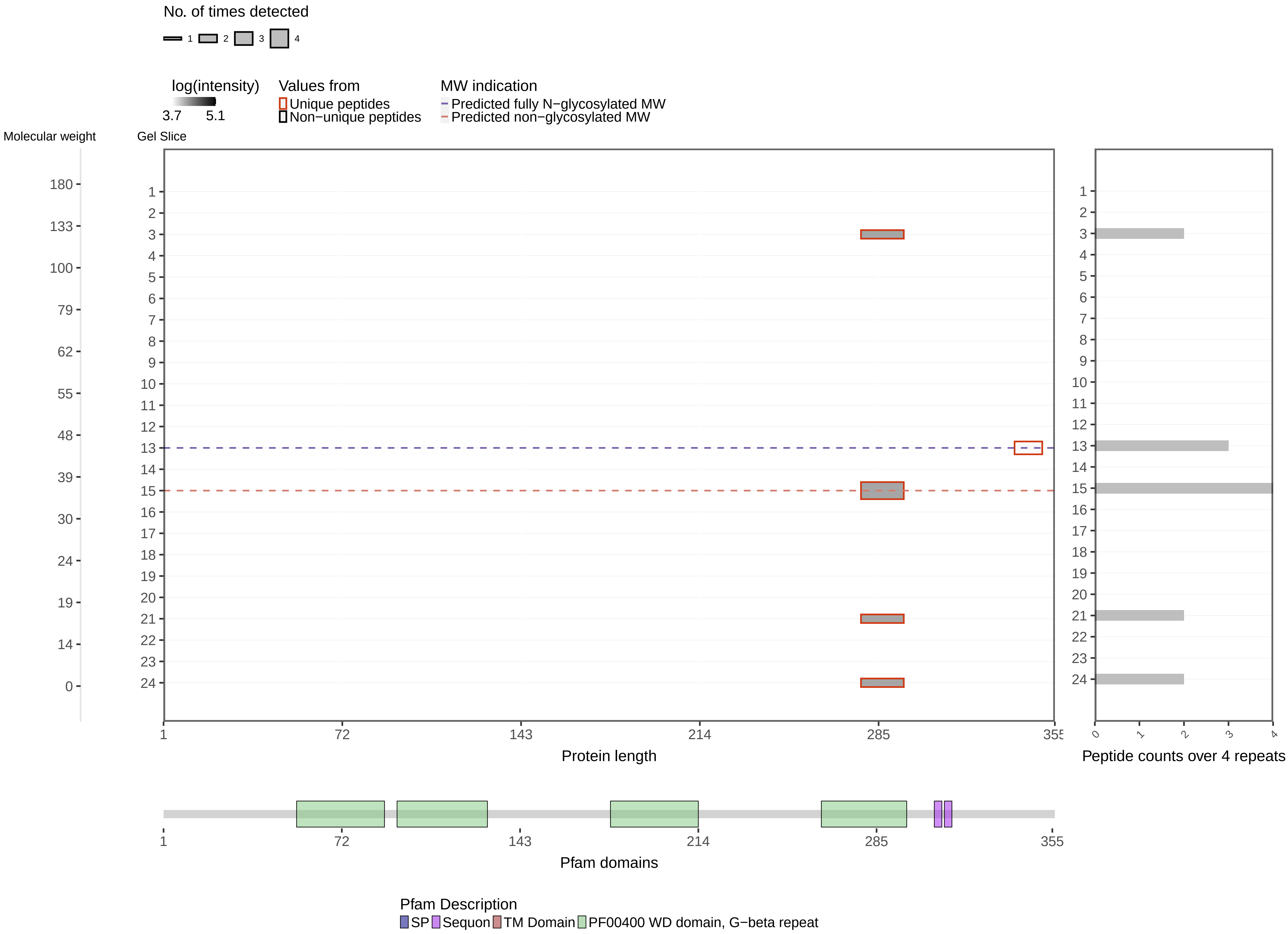


Protein sequence:

)AGGLRLWDVRSGKVVQTLETKFPVTSAEVSQDGRYITTADGSSVKFWDANHFGLVKSYDLPCKVESASLEPKFGNRFIAGGEDMWVHVFDFDTGEEIGCNKGHHGPVHCLRFSPGGESYASGSEDGTIRIWQLGPLGQNEDNSTANGSIANAKDDMGEVTQKIEELDVSEAKKTEETQIDGAVKVADA*

### NbL03g22400.1 beta−galactosidase_9 [MW = 104.99 kDa]

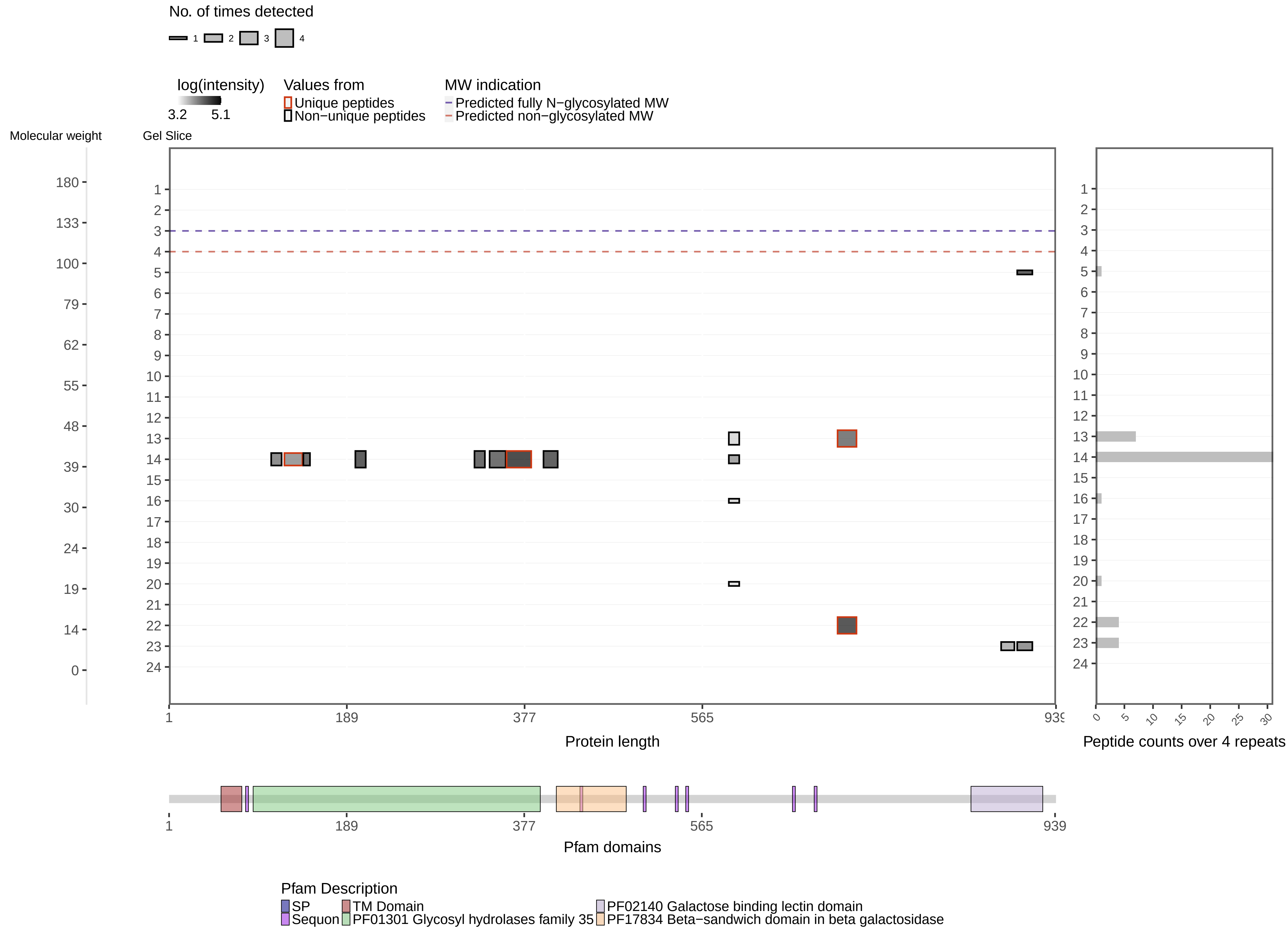


Protein sequence:

[illegible]

NbL03g22220.1 protein_ASPARTIC_PROTEASE_IN_GUARD_CELL_1−like [MW − SP = 49.16 kDa]

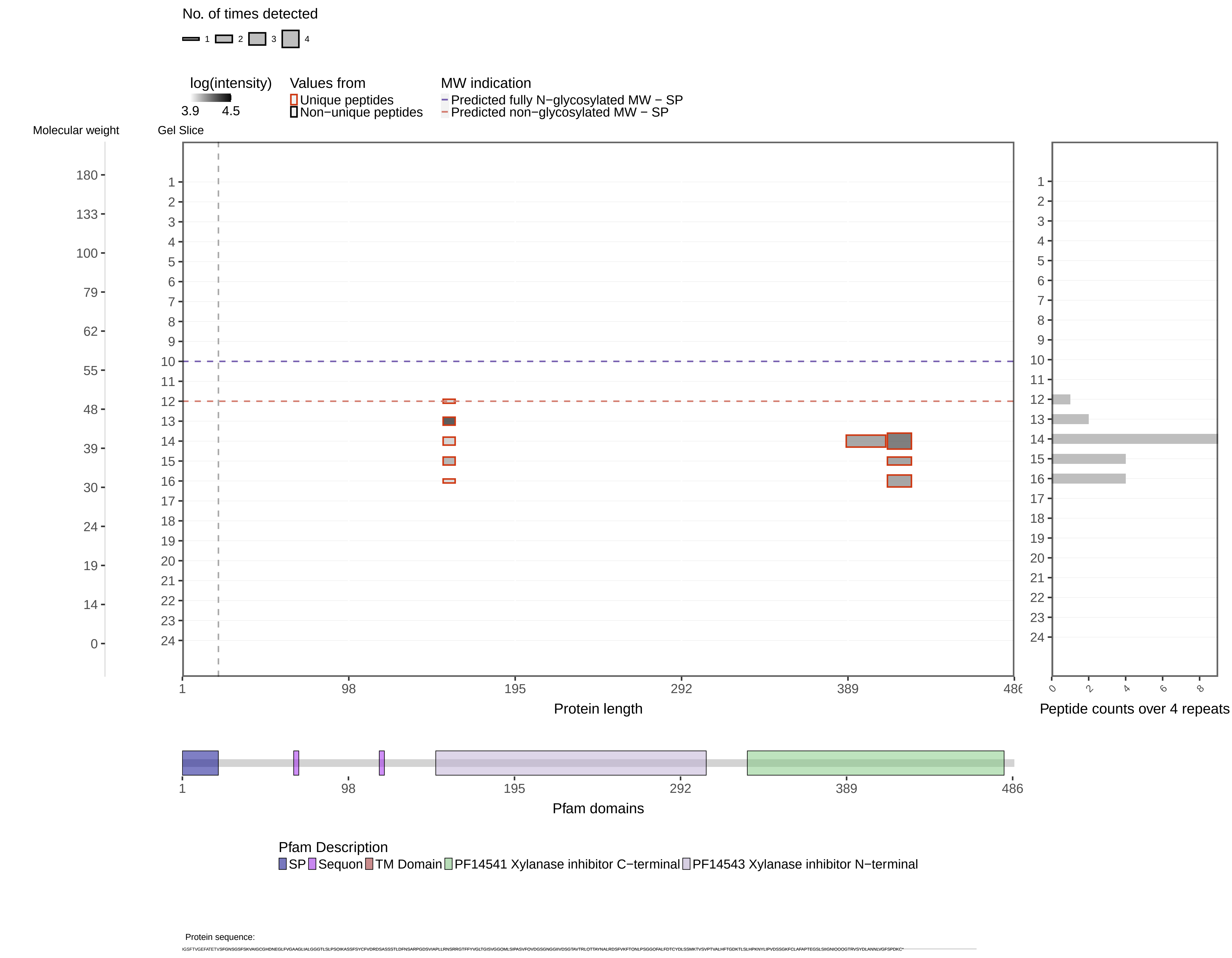

### NbL03g22110.1 phenylalanine−−tRNA_ligase_alpha_subunit_cytoplasmic−like [MW = 56.26 kDa]

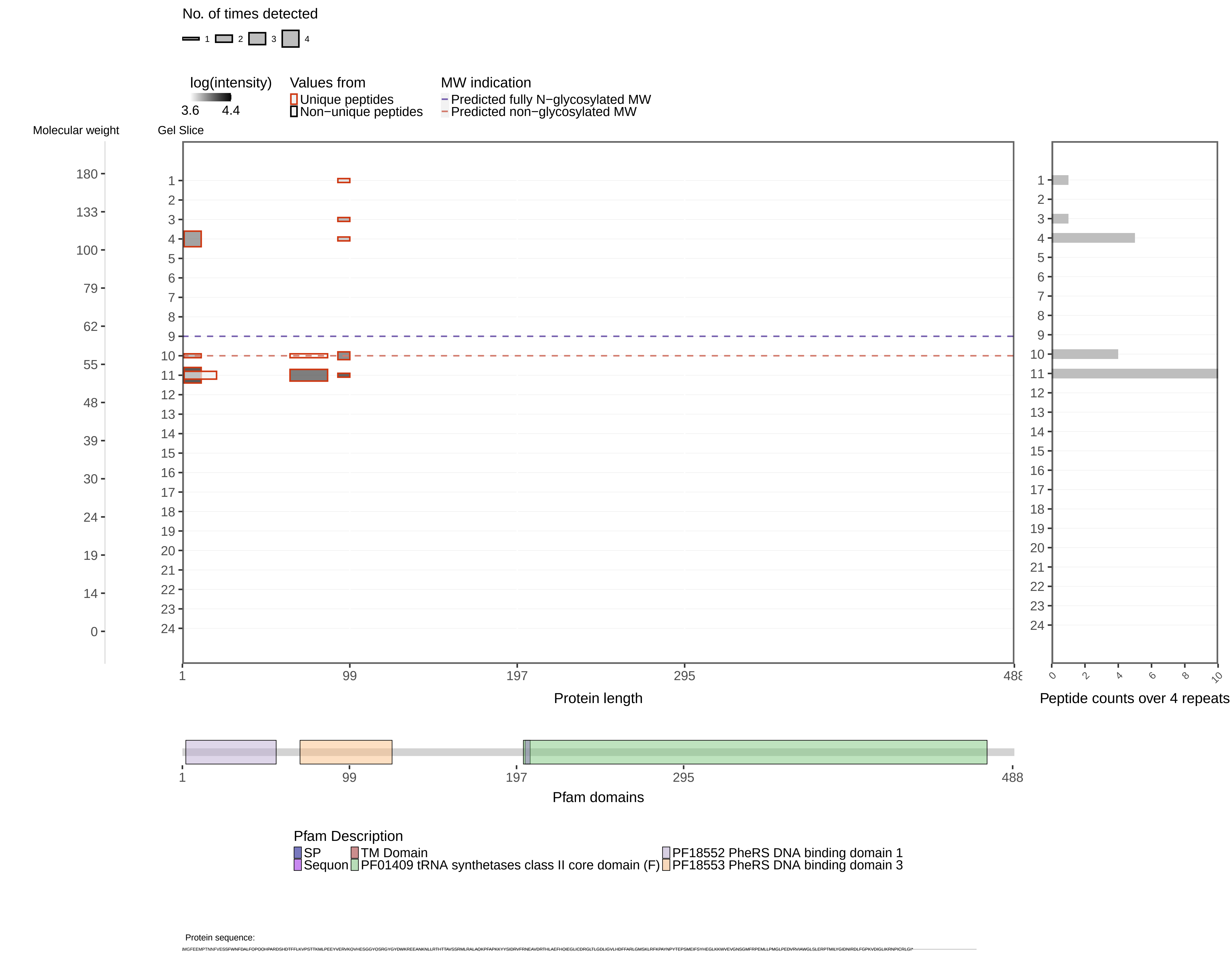

NbL03g21930.1 histidine−−tRNA_ligase_chloroplastic/mitochondrial [MW = 56.75 kDa]

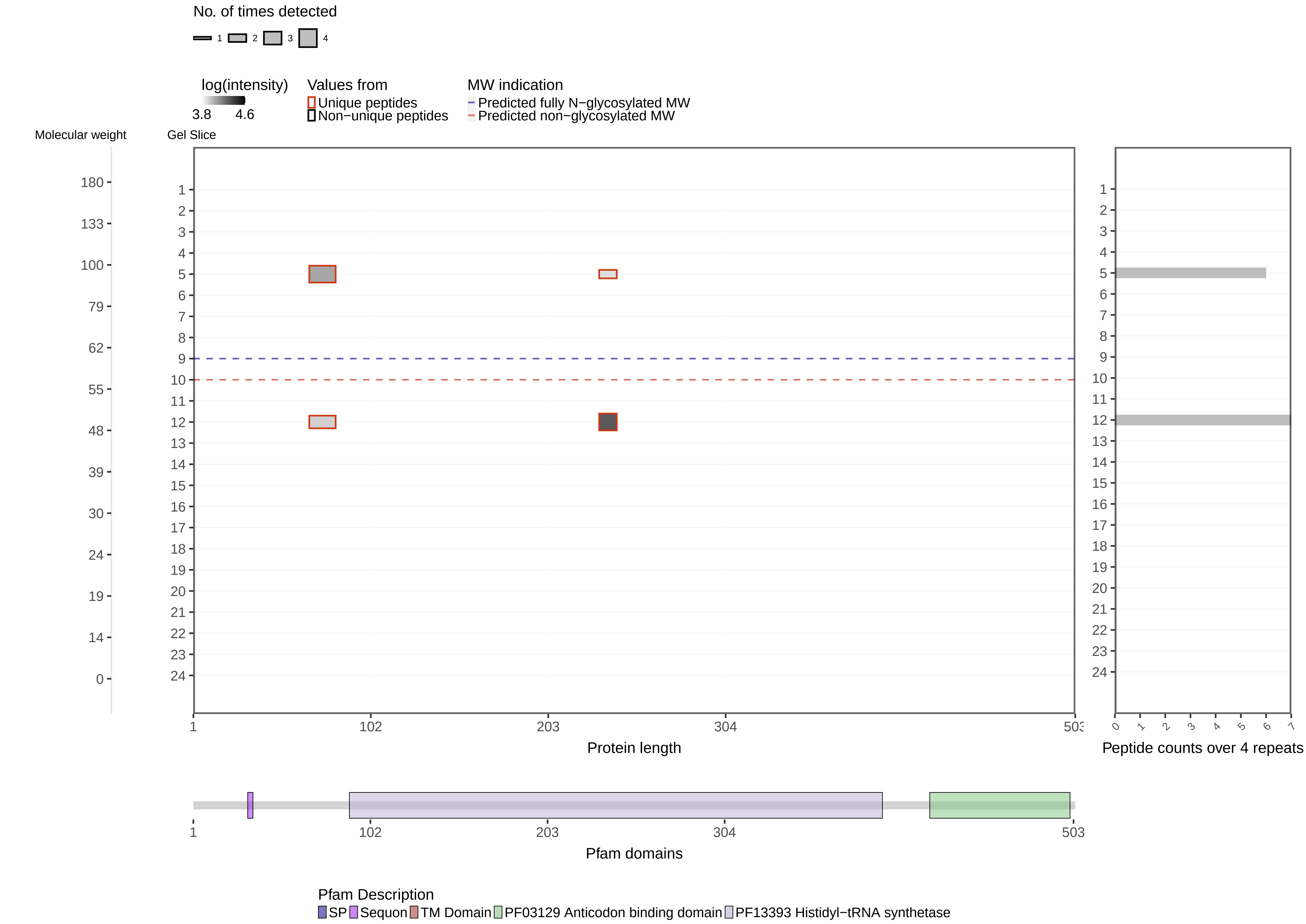


Protein sequence:

NbL03g21830.1 SH3_domain−containing_protein_C23A1.17−like [MW = 57.53 kDa]

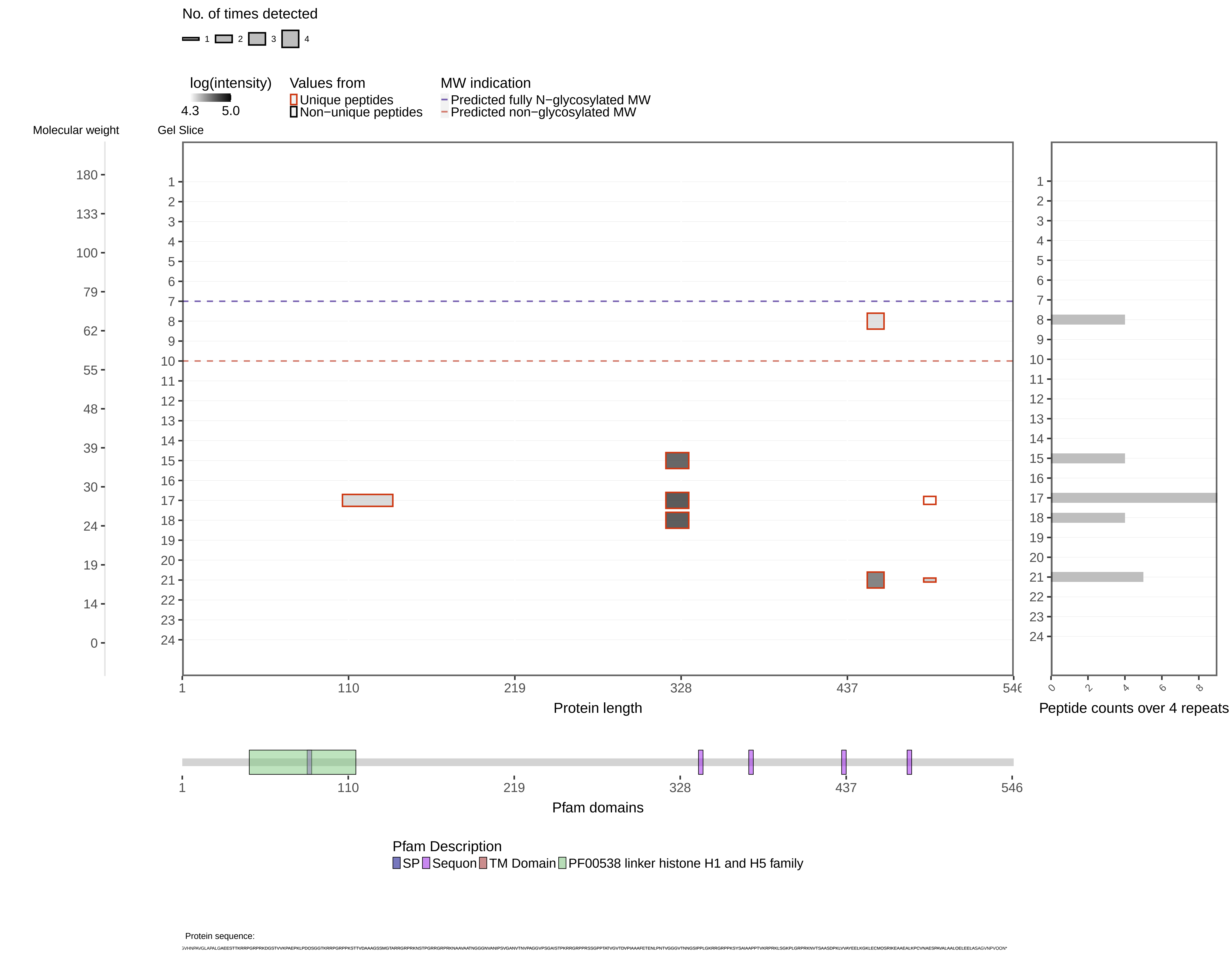

NbL03g21790.1 66_kDa_stress_protein−like [MW = 66.03 kDa]

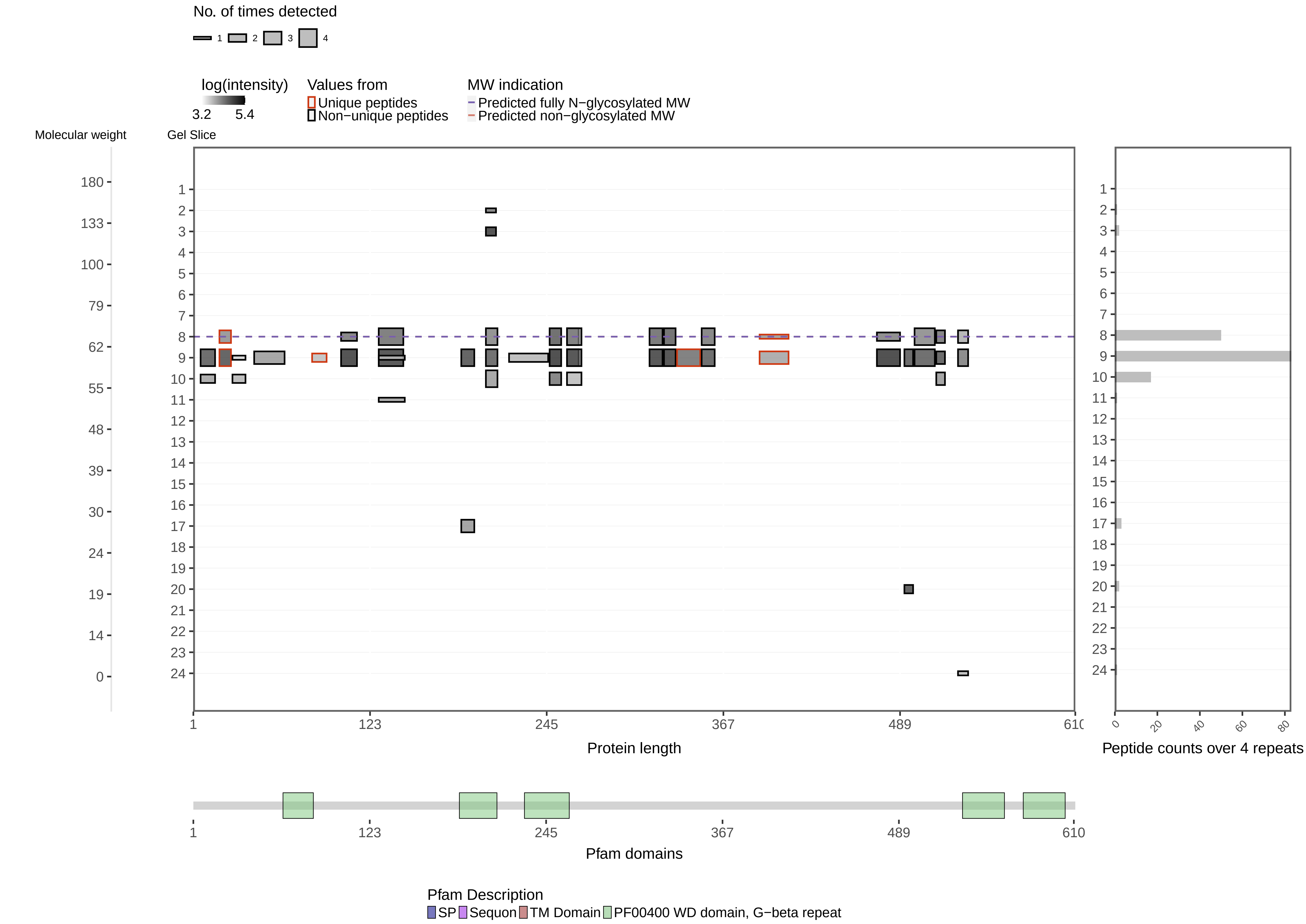


Protein sequence:

WONDHLVTVSLGGTISIFSASDLEKSPVSFSGHMKNVNSLAVLKSDPOIMLSSSYDGLIVKWIOGIGYSGKLERKVNSOIKCFEVVEGEIVSCGFDNKVWRVSLLGDOCGDANSIDVGNOPKDLSLALSSPEATLVSFDTGVILLRGTKVLSTIDLGFTVTASAISPDGSEAIVGGODGKLRLYSITGDTLNEEAVLEKHRGAITVIRYSPDVSMFASADVNREAVVWDRASREVKLKNMLYHTARINCLAWSPDNTMVATGSLDTCVIIYDISKPASSRITIKGAHLGGVYGLAFT

### NbL03g21780.1 beta−glucosidase_44−like [MW = 60.33 kDa]

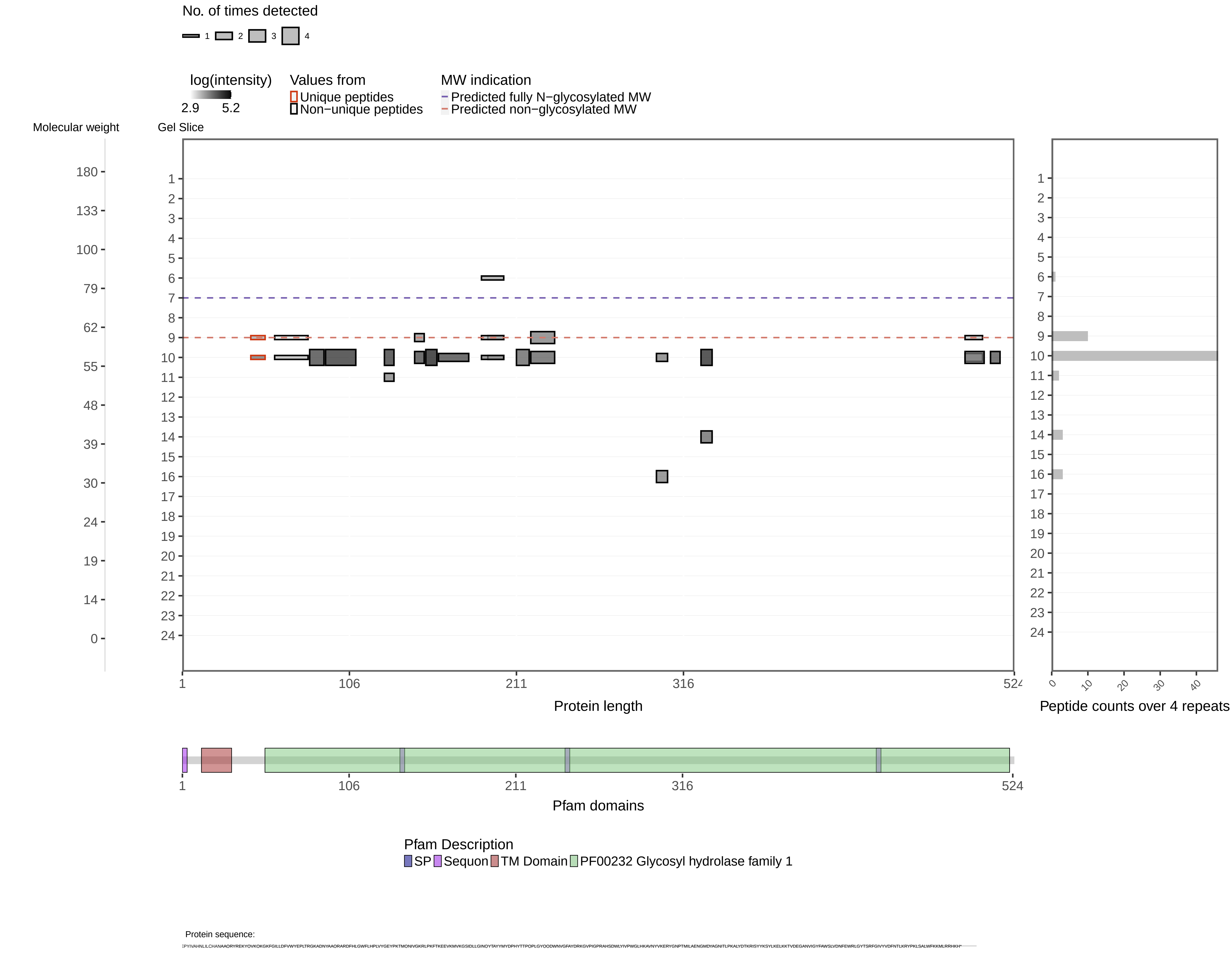

NbL03g21740.1 guanine_nucleotide−binding_protein_subunit_beta−like_protein [MW = 35.99 kDa]

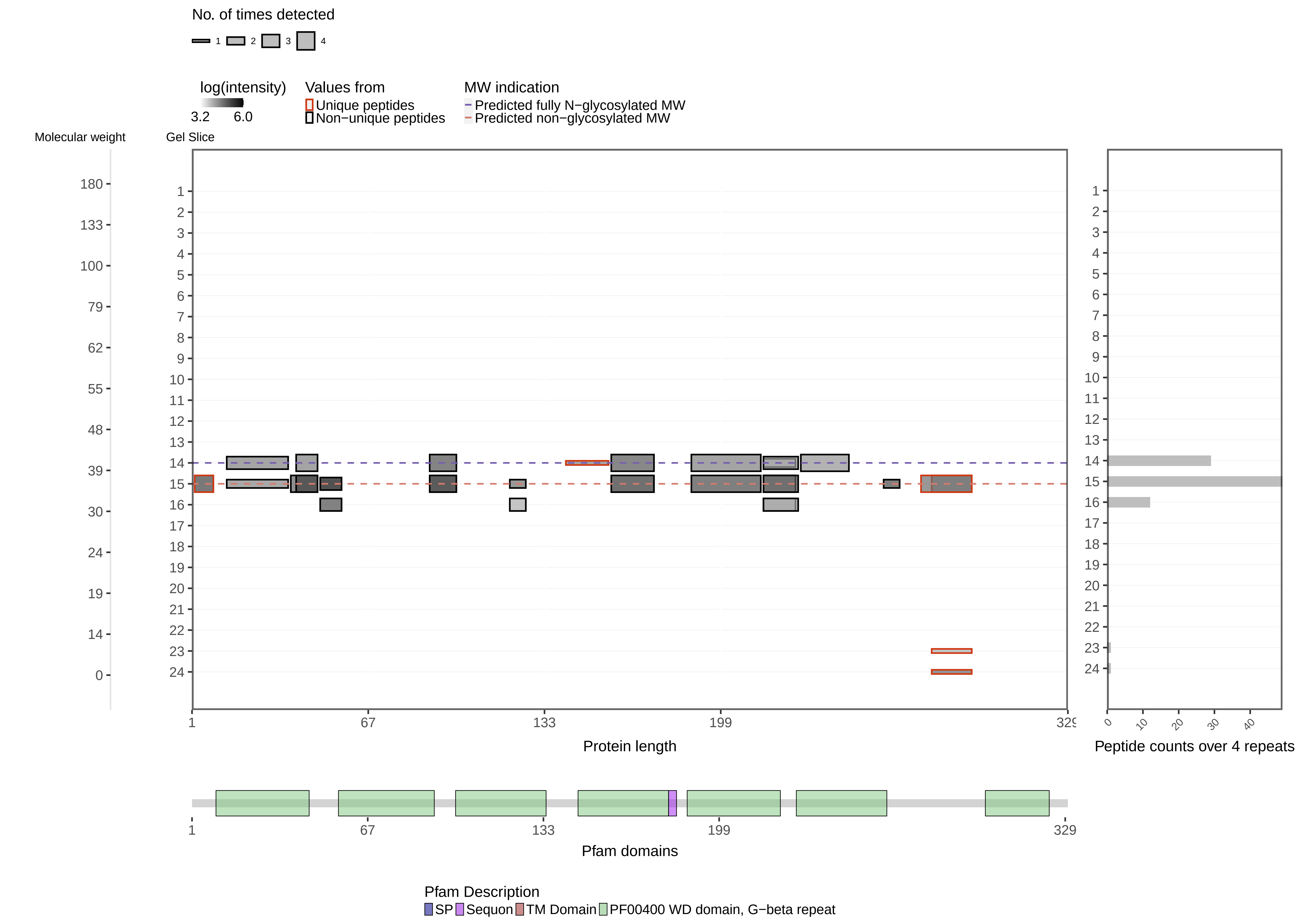


Protein sequence:

DWVSCVRFSPNTLQPTIVSGSWDRTVKIWNLTNCKLRSTLAGHAGYVNTVAVSPDGSLCASGGKDGTILLWDLAEGKKLYSLDAGSIIHALCFSPNRYWLCAATETSIKIWDLESKSIVVDLKVDLKQESEMATEGTTGSACKNKIMYCTCLSWSADGSTLFSGYTDGLIRVWGIGRY*

### NbL03g21700.1 eukaryotic_peptide_chain_release_factor_GTP−binding_subunit_ERF3A−like [MW = 57.42 kDa]

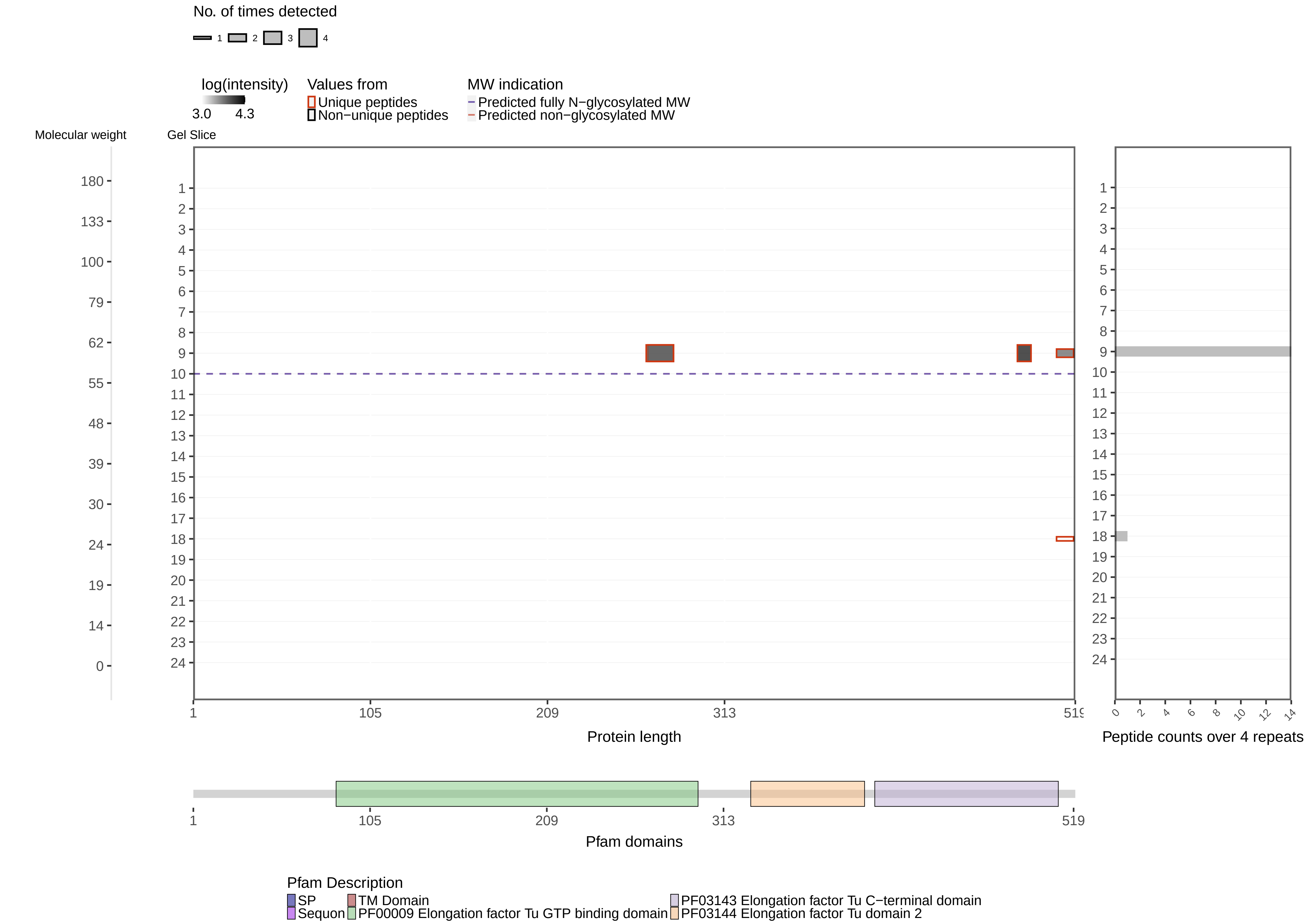


Protein sequence:

VAKERYDEIESKMIPFLKSSGYNVKKDVQFLPISGLLGSNMKNRVEKSVCSWWNGPCLFEALDAVEVPPRDPKGPLRIPIIDKFKDMGTVIMGKIESGSIREGDNLLVMPNKAAVKGLAIFCDEDRVRGAGPGENVRVRLSGIEEEDIMSGFVLCSVAQPIPAVTEFVAQLQILELLDNAIFTAGYKAVLHIHAVVEECEIVELIQQIDLKTKKPMKKKPLFVKNGAIILCRVQVNNLICIEKFSDFPQLGRFTLRTEGKTVAVGKVAALPSVADNA*

### NbL03g21620.1 probable_protein_phosphatase_2C_8 [MW = 24.86 kDa]

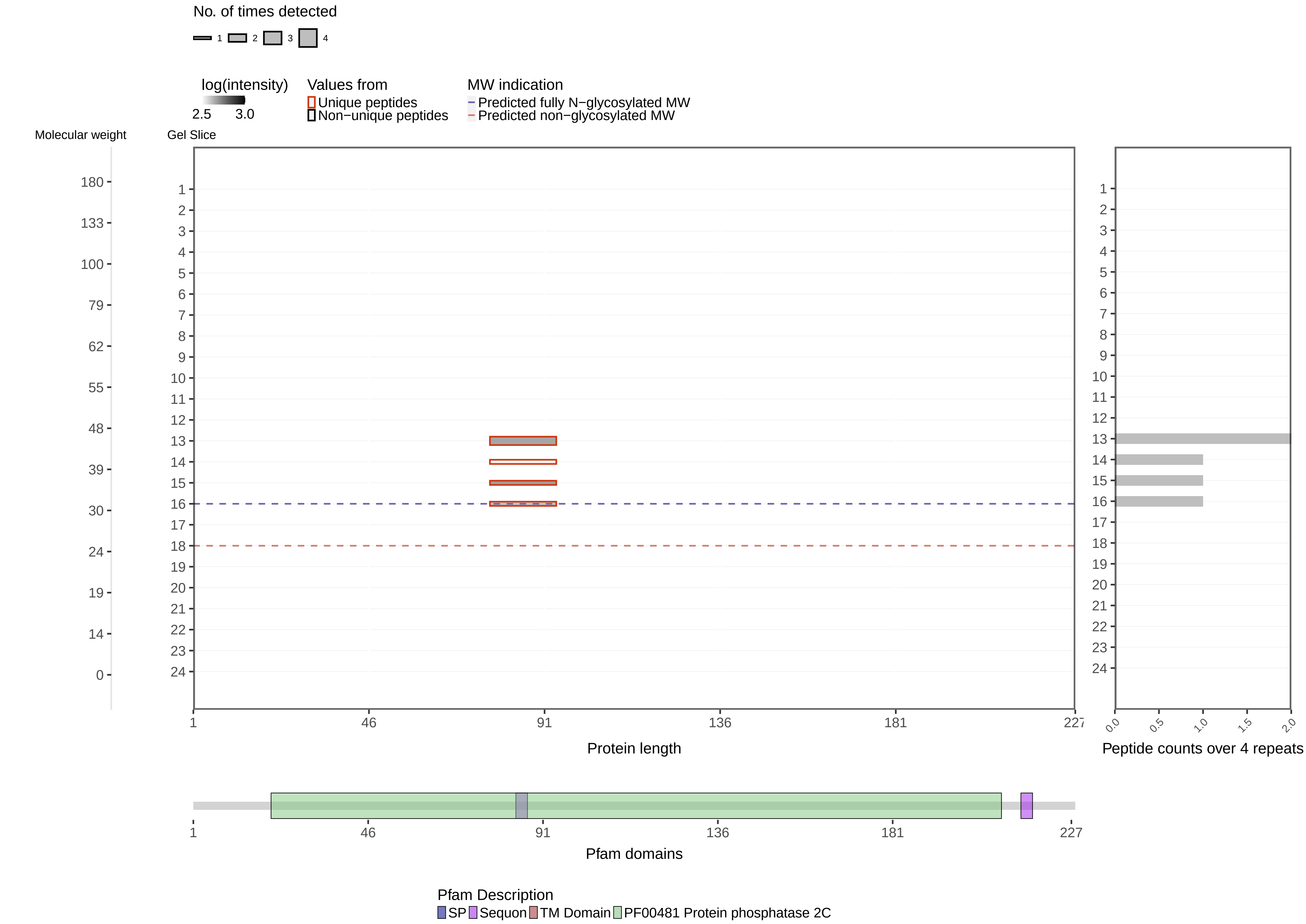


Protein sequence:

EHKAIYPQERARIQKAGGSVSSNGRLQARLEVSRAFGDRQFKKVGVIATPDVHSFDLTERDHFIILGCDGLWGVFGPSDAVDFVQKLLKEGLPVSVVSRRLVREAVRERQCKDNCSAVVIAFRKKQ*

### NbL03g21550.1 calcium−binding_allergen_Ole_e_8−like [MW = 18.74 kDa]

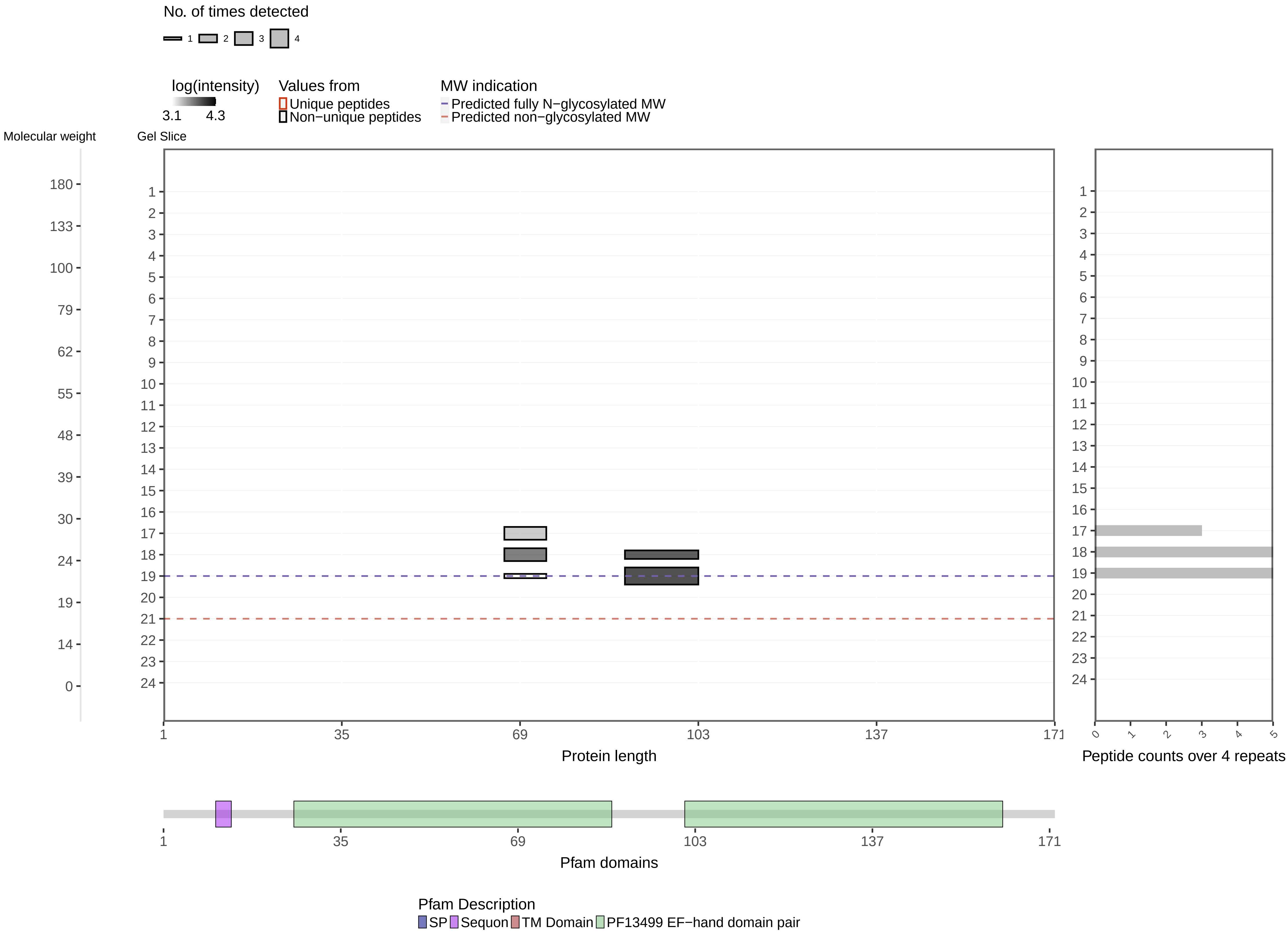


Protein sequence:

DRDGYINLDEFAEFCKGDVNGVGDGGVNELREAFDLYDQDNNGLISDAELHQILTRLGESCSVQDCTKMISSVDADGDGYVSFDEFKKMMTNNNFQKC*

NbL03g21410.1 uncharacterized_LOC109223953 [MW = 151.74 kDa]

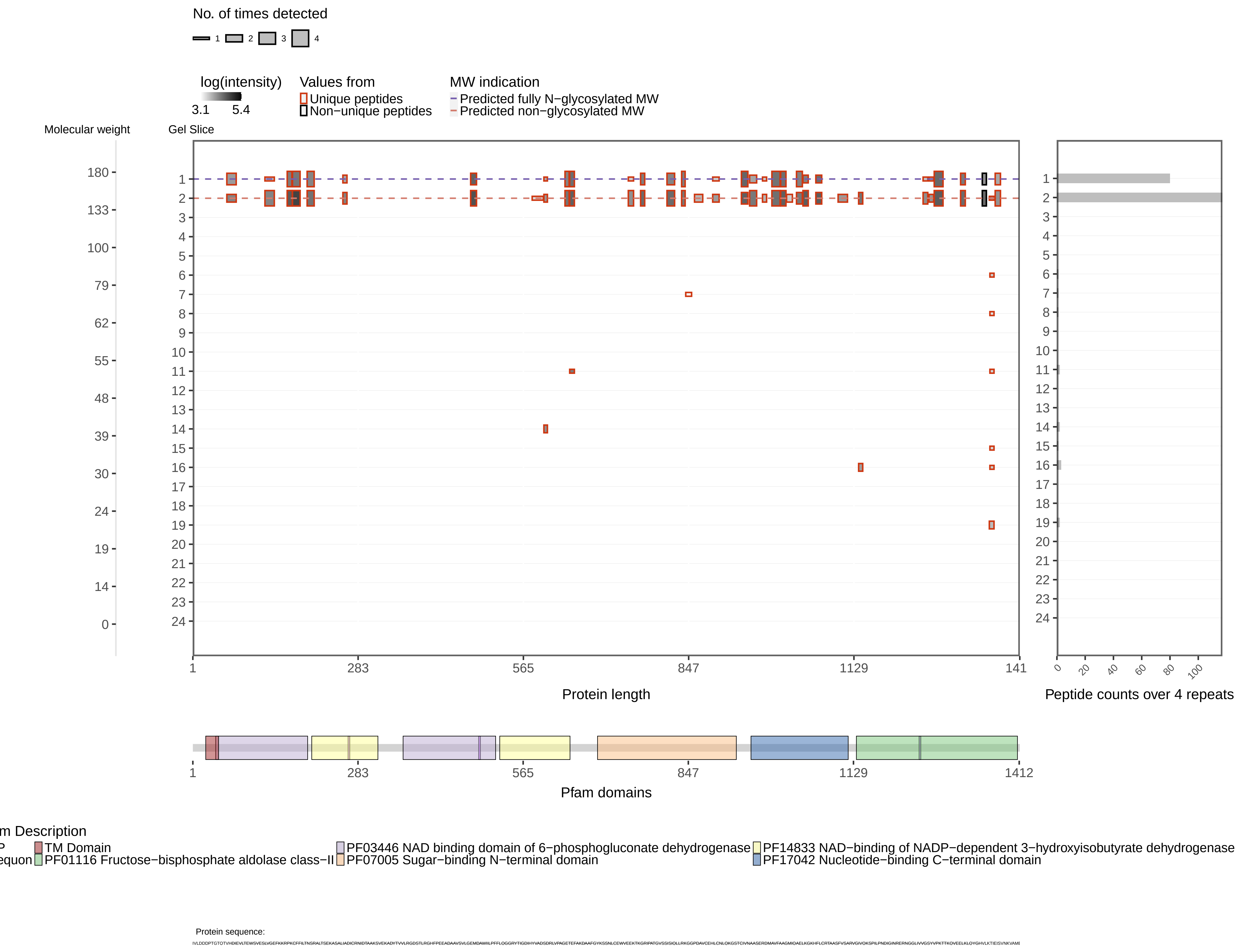

NbL03g21350.1 geranylgeranyl_transferase_type−2_subunit_beta_1−like [MW = 34.61 kDa]

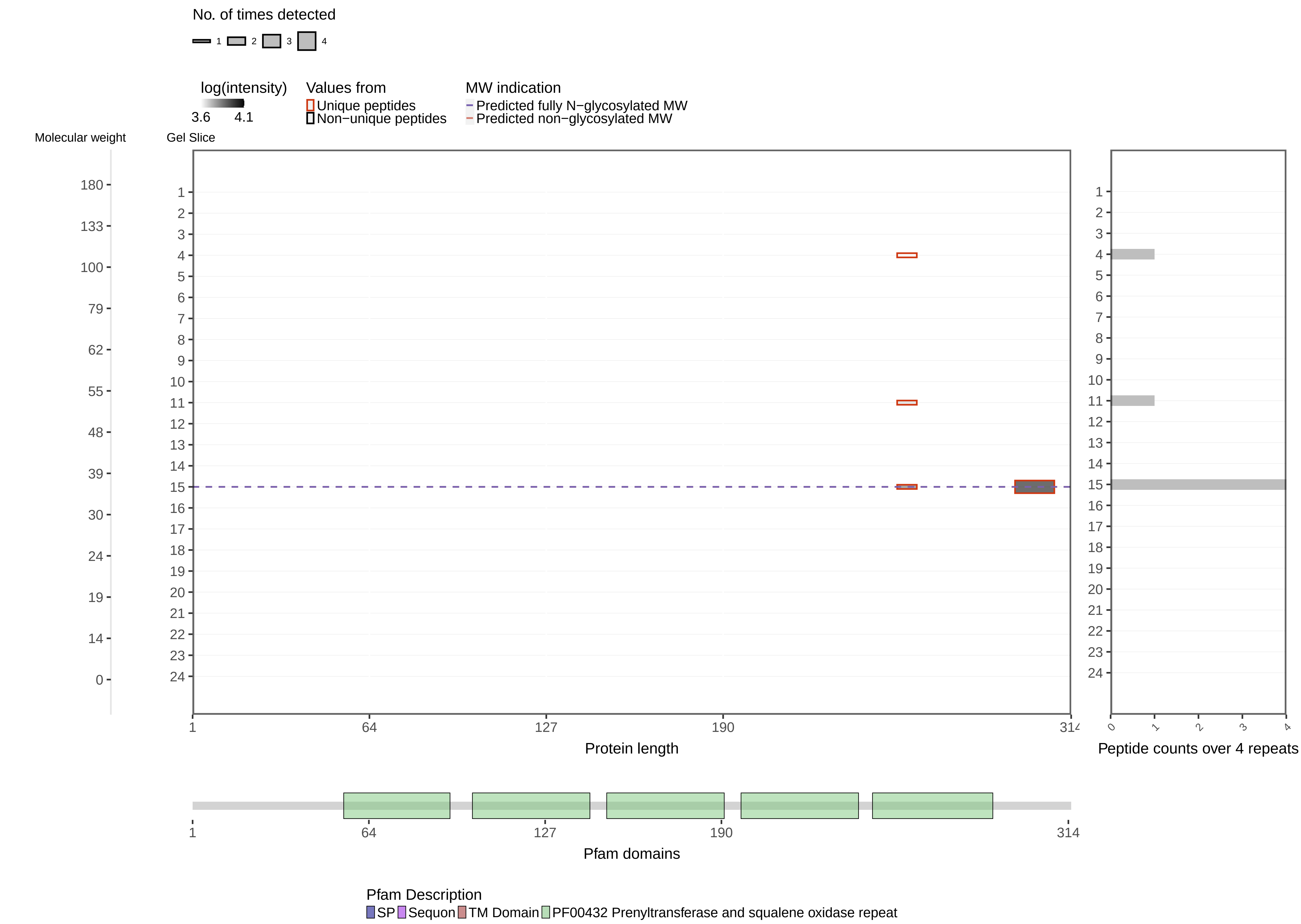


Protein sequence:

[illegible]LDKVDVGKAVKYILSCKNVDGGFGCTPGAESHAGQIFCCVAALAITGSLHHVDKDLLGWWLCERQVKSGGLNGRPEKLPDVCYSWWVLSSLIMIDRVHWIDKGKLVKFILDCQDKENGGISDRPDDAVDVFHTYFGVAGLSLLEYPGIKPIDPAYALPVDVVNRVILGR*

NbL03g21200.1 stromal_processing_peptidase_chloroplastic [MW = 116.49 kDa]

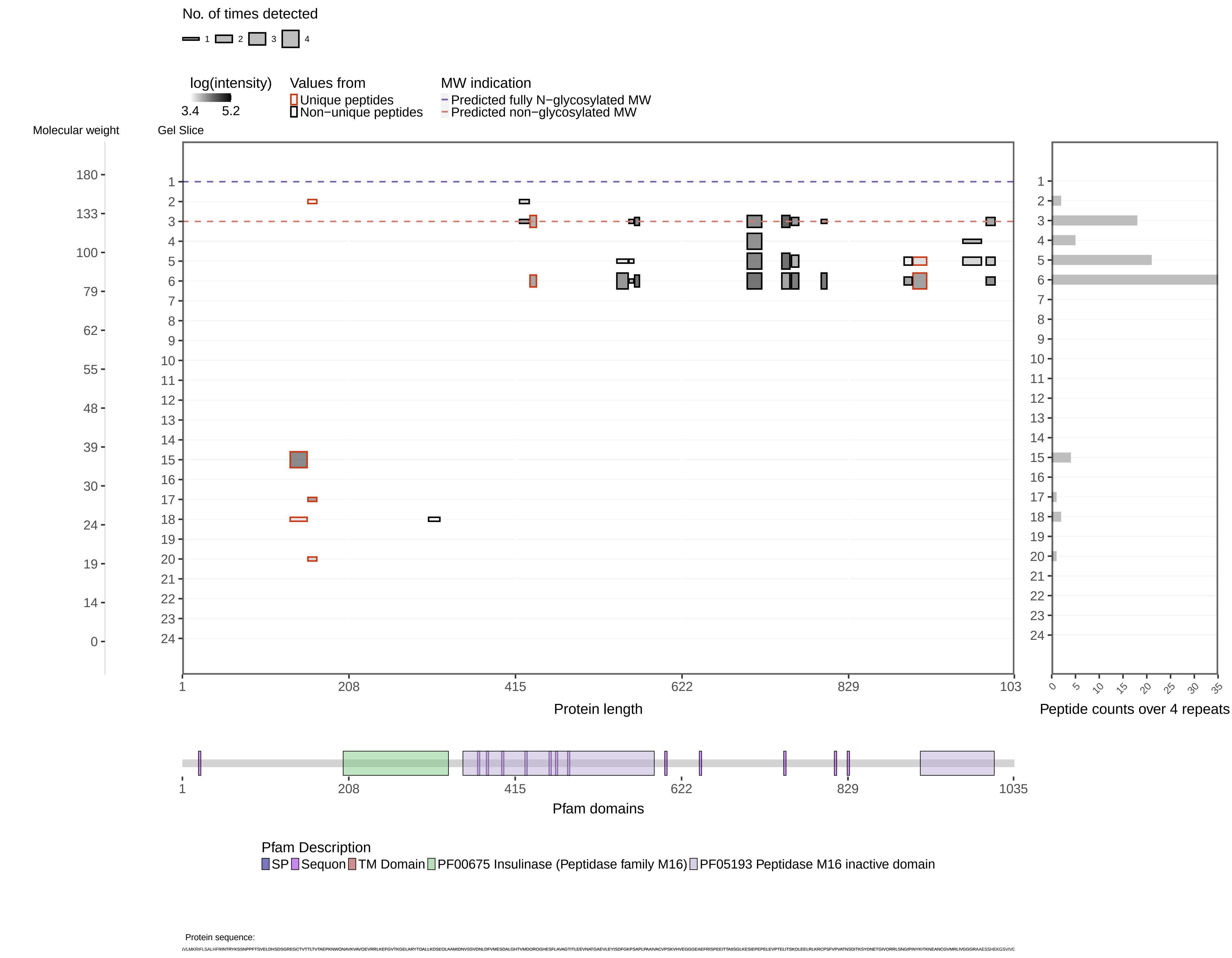

NbL03g21010.1 uncharacterized_protein_At4g14100−like [MW − SP = 22.88 kDa]

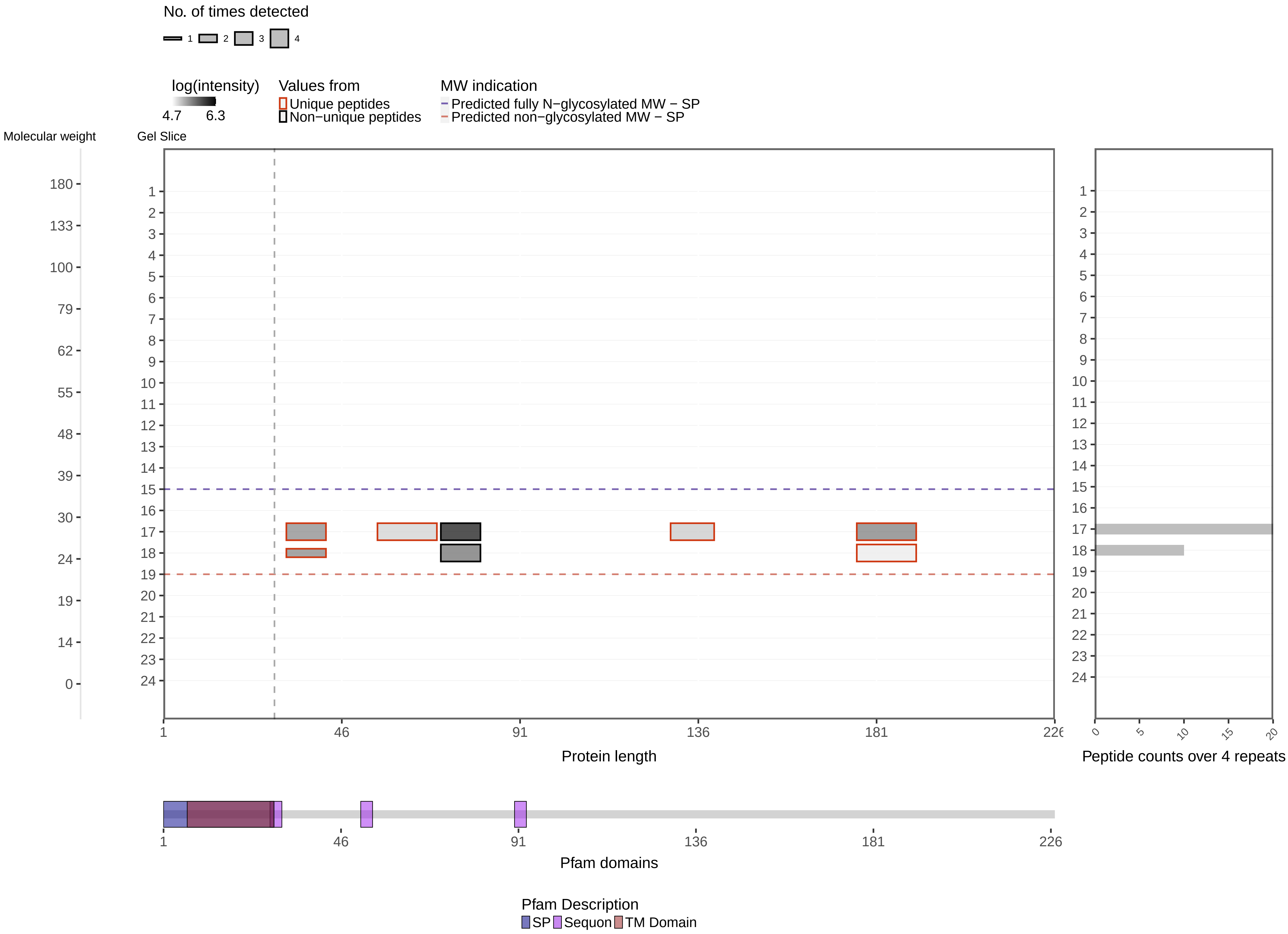


Protein sequence:

ECRVMHFPVGILRPNWLQGGKYLGQRYMDGFLCNVWEKVDFIWYYEDIVTKRPVYWAFFTGMIAHVMTFEVGKVLEDPNWQAPVYCFNKALEQEKSPVIENLGNLDASVSRLMGSVVDVSLLL*

NbL03g21000.1 succinate_dehydrogenase_assembly_factor_2_mitochondrial−like [MW = 21.03 kDa]

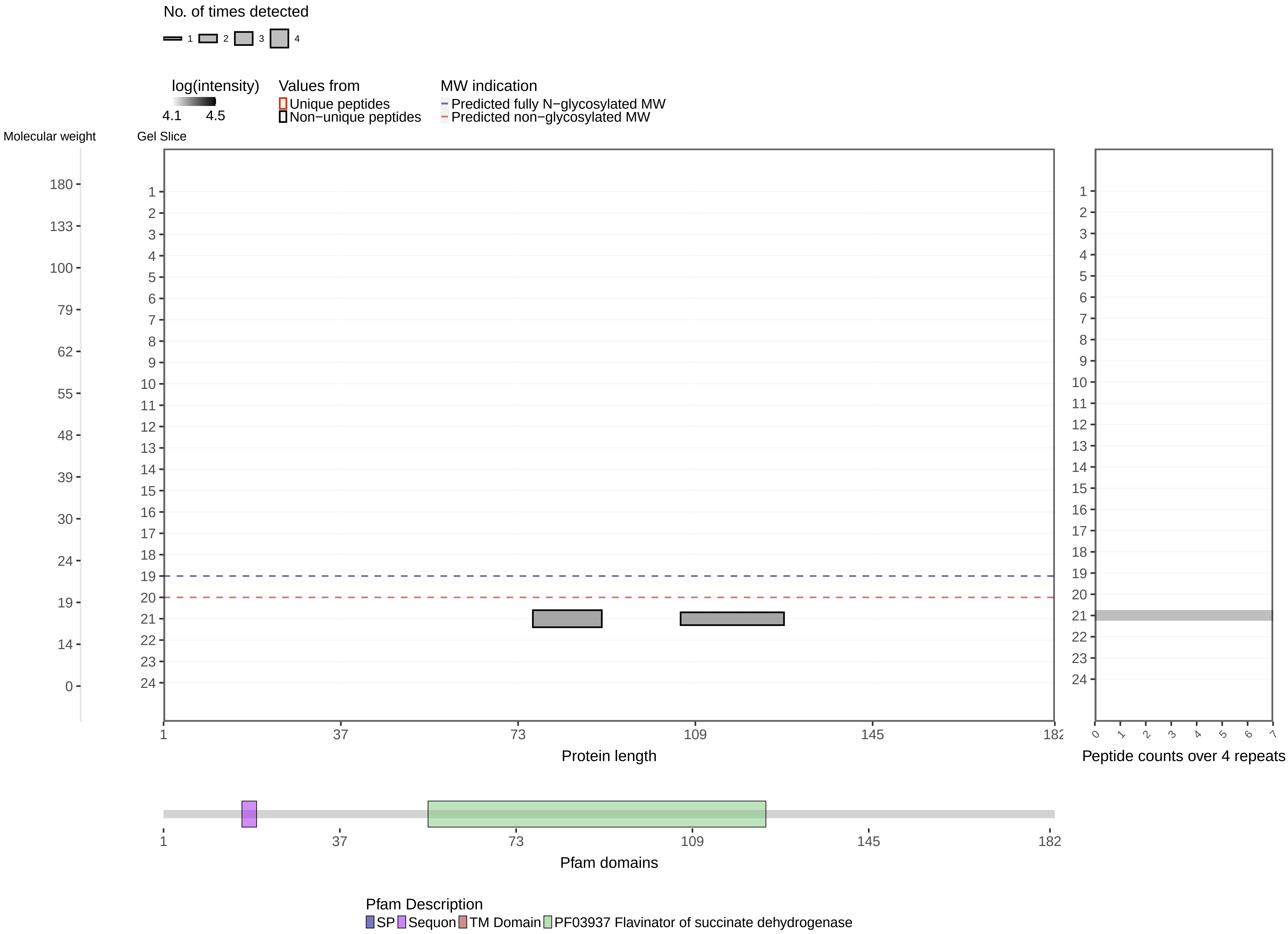


Protein sequence:

EHIQSMDENGIKFLVHVLDQENPDLWKWLTGQEQPPDAISTNPVFSAVREKVINNLDKHASPETRLFLENLGLEDGMISREVEMRVLLEISRSFYCWEASKNI*

### NbL03g20930.1 ribonuclease_TUDOR_1−like [MW = 107.67 kDa]

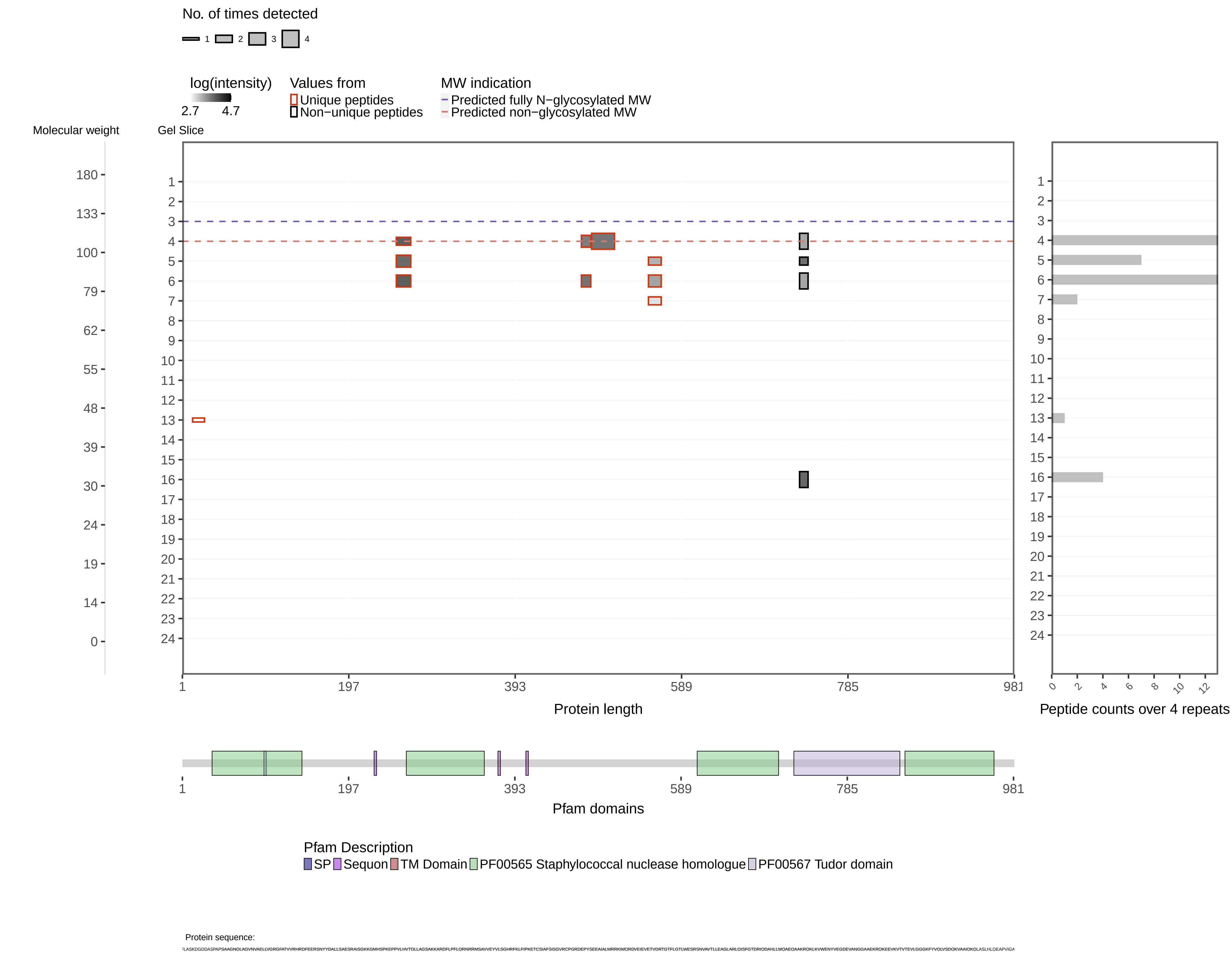

### NbL03g20890.1 respiratory_burst_oxidase_homolog_protein_C [MW = 105.92 kDa]

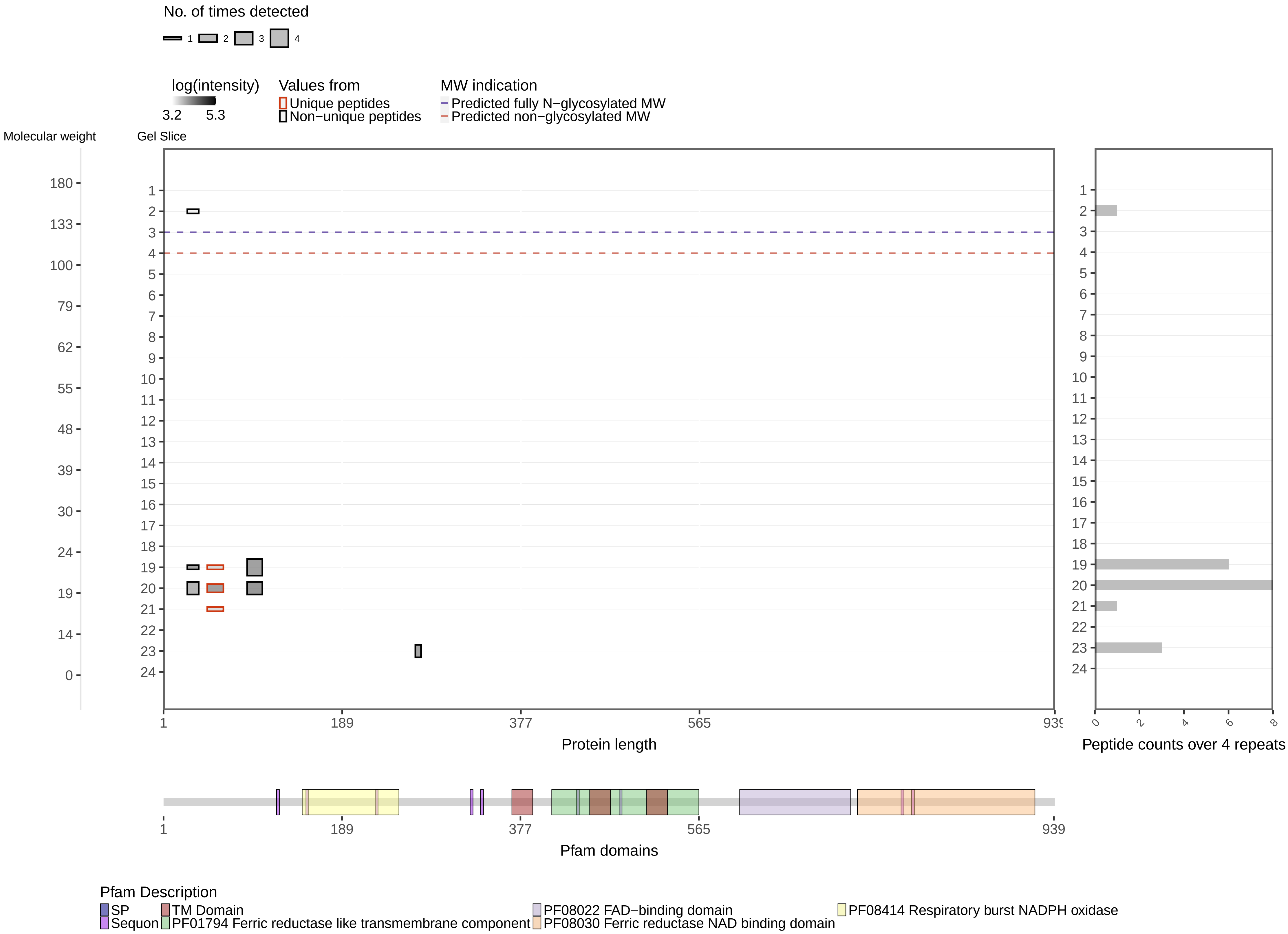


Protein sequence:

AVAIALGVGVHGLSHLTCDFPRLLNASEEEYEPMKYYFGDQPESYWWFIKGVEGVTGIIMVVLMAIAFTLATPWFRRNRVSLPKPFHKLTGFNAFWYSHHLFVIVYTLFIVHGEKLYITKDWYKRTTWMYLTIPIILYASERLIRAFRSSIKAVKILKVAVYPGNVLALHMSKPQGYKYKSGQYMFVNCAAVSPFEWHPFSITSAPGDDYLSVHIRTLGDWTRQLKTVFSEVCQPPPNGKSGLLRADYLQGENNPNFPRVLIDGPYGAPAQDYKKYEVVLLVGLGIGATPMISIVKDI

NbL03g20860.1 U2_small_nuclear_ribonucleoprotein_A'−like [MW = 32.63 kDa]

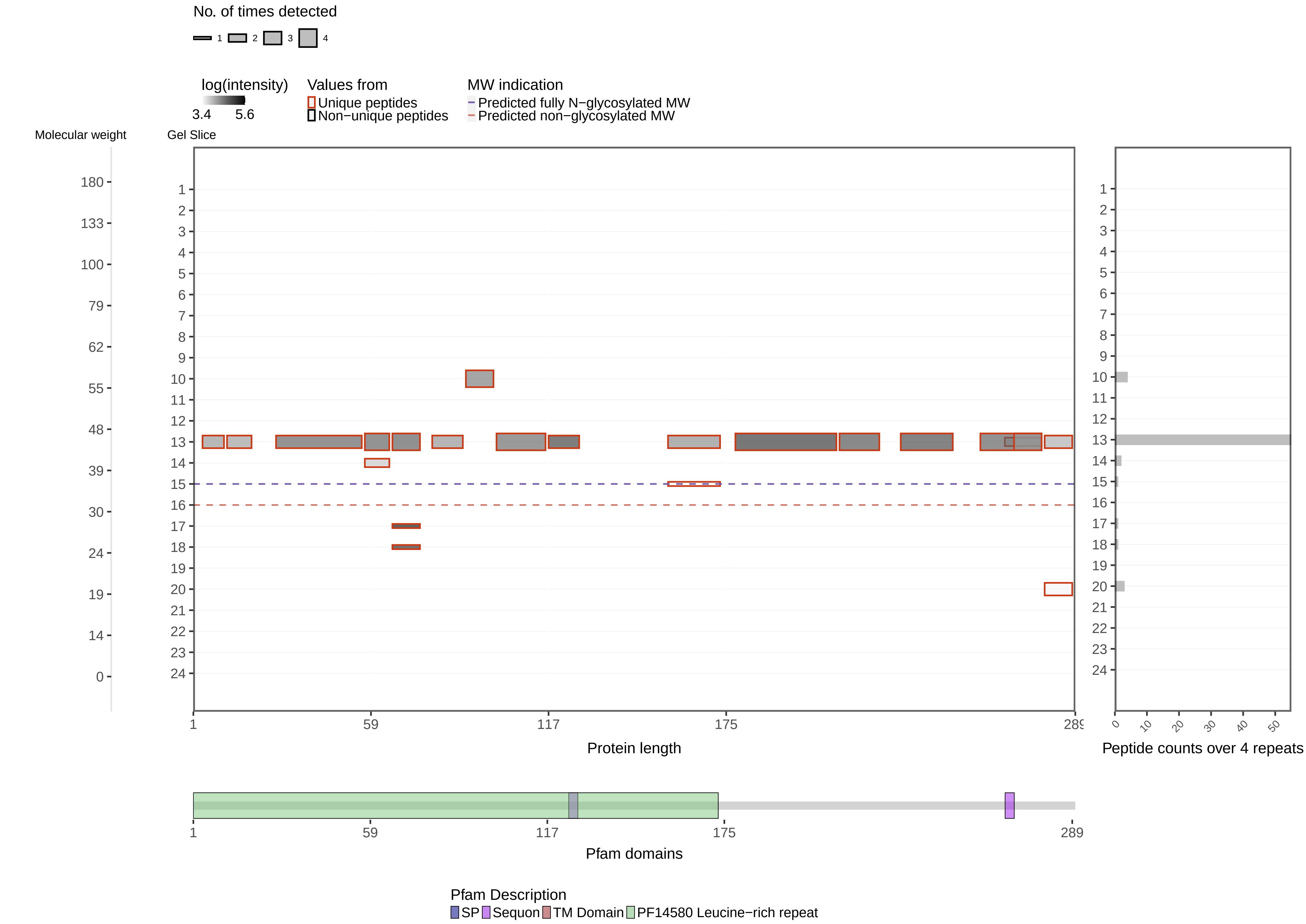


Protein sequence:

YVIHKLKSLRLLDFRKVKQKERLEASNLFASEEAEEEAKKESVKTFVPGEVPAAPEEPKEDEAPKPVAPTPEQIIAIKAAIVNSQTLEEVARLEQALRSGQLPADLNIGDYDIAAKKEDAKEDKMVTDGDDKANKTEEENIPEQKTDGPTDMEQE*

### NbL03g20830.1 ACT_domain−containing_protein_DS12_chloroplastic−like [MW = 31.04 kDa]

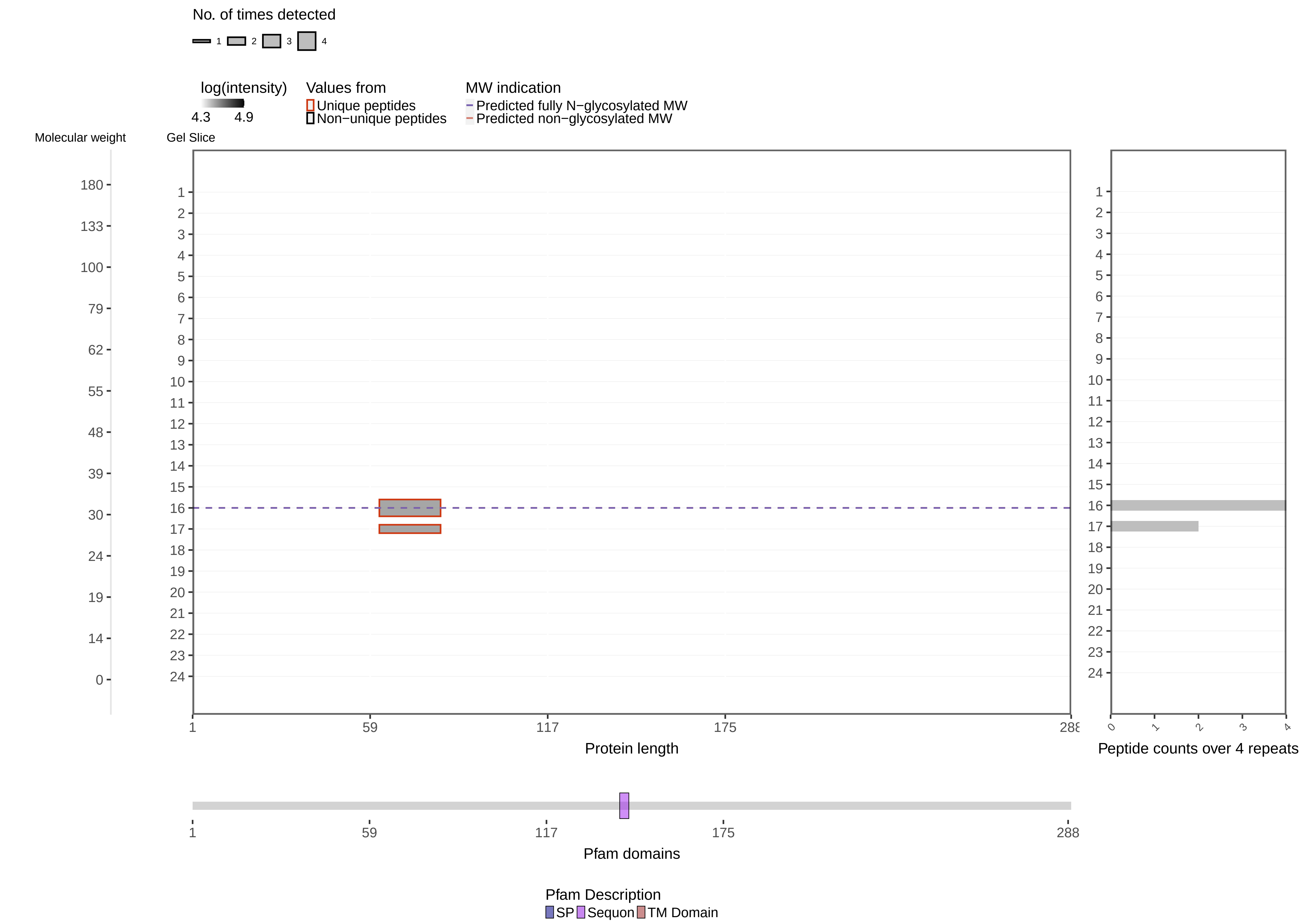


Protein sequence:

SGKHTTLCITNASTGRKVDDPEQLEAIRLTIINNMIEFHPESSAQLAMGEVFGAYQPNQKLDVDIATHIHVYDDGPDRSLLYVEAADRPGLIVDLVKIITEINIDVESGEFDTEGLLAKAKFHVSYKNKALIKPLQQVLANSLRYYLRRPTTEESSF*

NbL03g20820.1 4−coumarate−−CoA_ligase_1 [MW = 59.56 kDa]

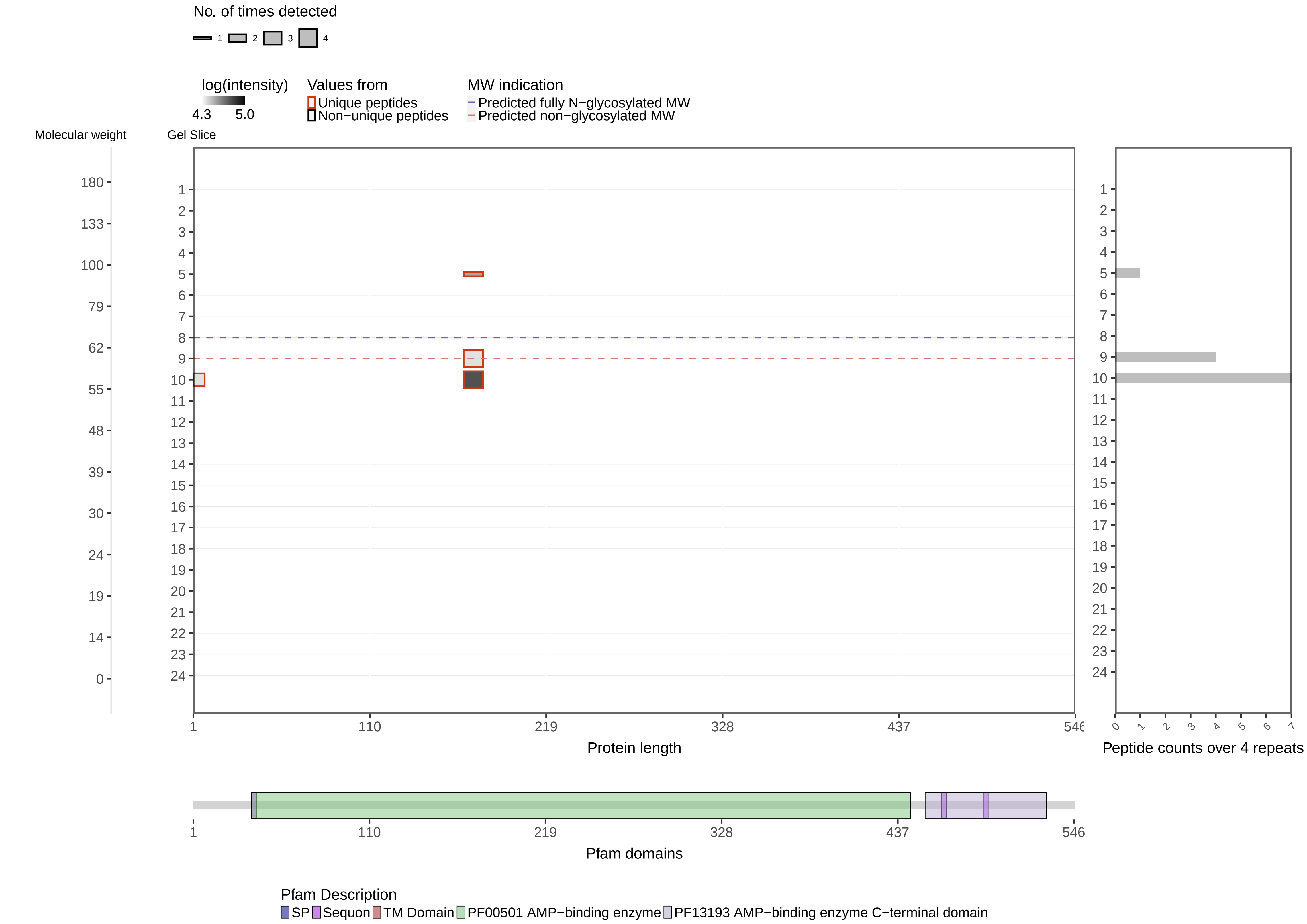


Protein sequence:

QKFDIAPFLELIQKYKVSIGPFVPPIVLAIAKSPIVDSYDLSSVRTVMSGAAPLGKELEDAVRTKFPKAKLGQGYGMTEAGPVLAMCLAFAKEPFDIKAGACGTVVRNAEMKIVDPDTGCSLPRNQPGEICIRGDQIMKGYLNDPEATTRTIDKEGWLHTGDIGFIDEDDELFIVDRLKELIKYKGFQVAPAEIEALLLNHPNISDAAVVPMKDELAGEVPVAFVVRSNGSAITEDEVKDFVSKQVIFYKRVKRVFFVETVPKSPSGKILRKDLRAKLAAGVPN*

NbL03g20730.1 ubiquitin_receptor_RAD23b−like [MW = 39.28 kDa]

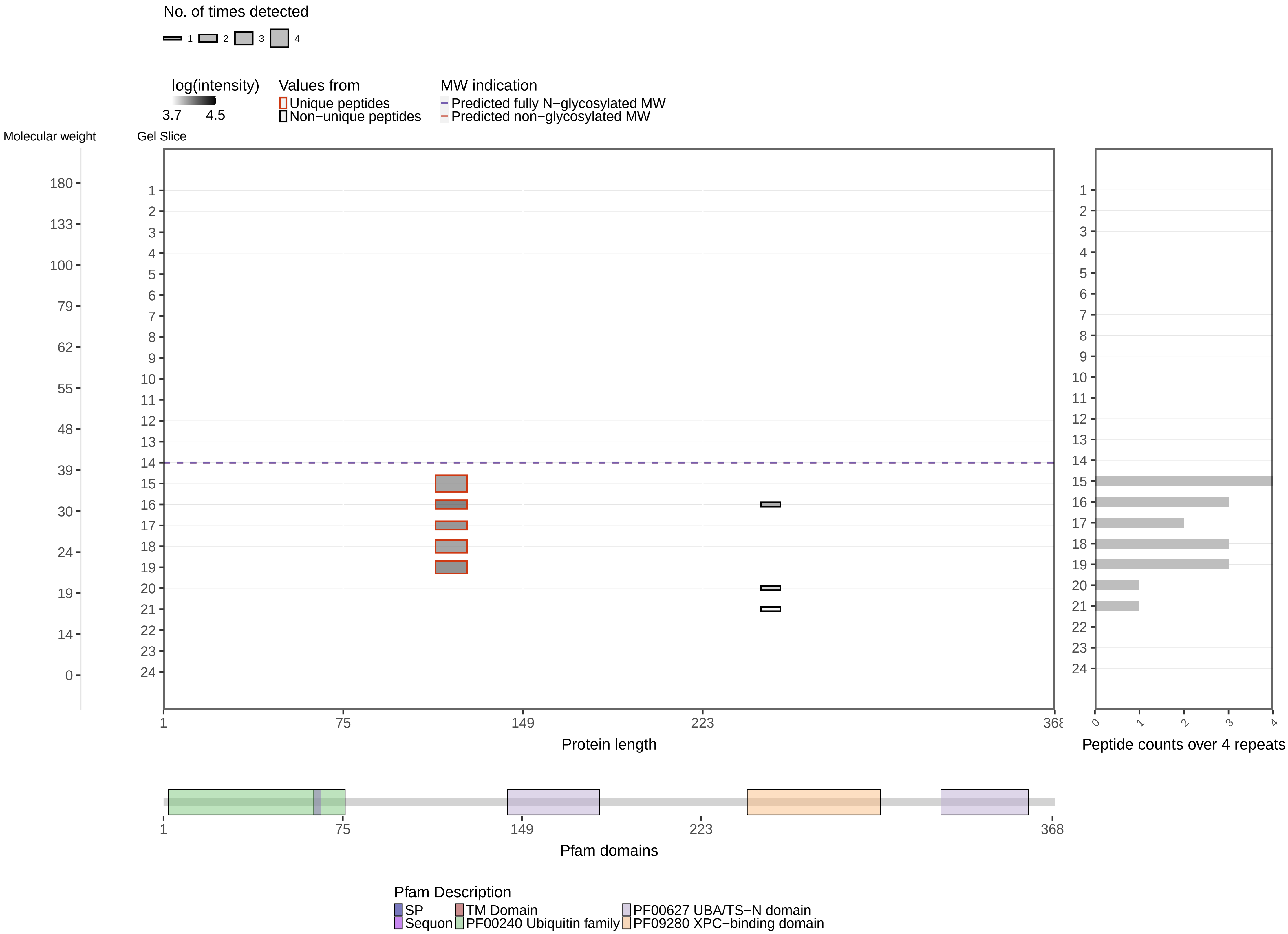


Protein sequence:

YNNPERAVDYLYSGIPETAEVAVPVAHGGVNSAAGTTAAPTAPSSGGPNSAPLNLFPQENVAGAGGAGLGSLDFLRNNQQFQALRSMVQANPQILQPMLQELGKQNPQLLRSIQEHHQEFLQLINEPVHGSEGDIFDHAEQEIPHTVSVTPEEQAVIERLEAMGFDRALVIEAFLACDRNEELAVNYLLEHAGDYED*

NbL03g20680.1 uncharacterized_LOC107804369 [MW = 33.66 kDa]

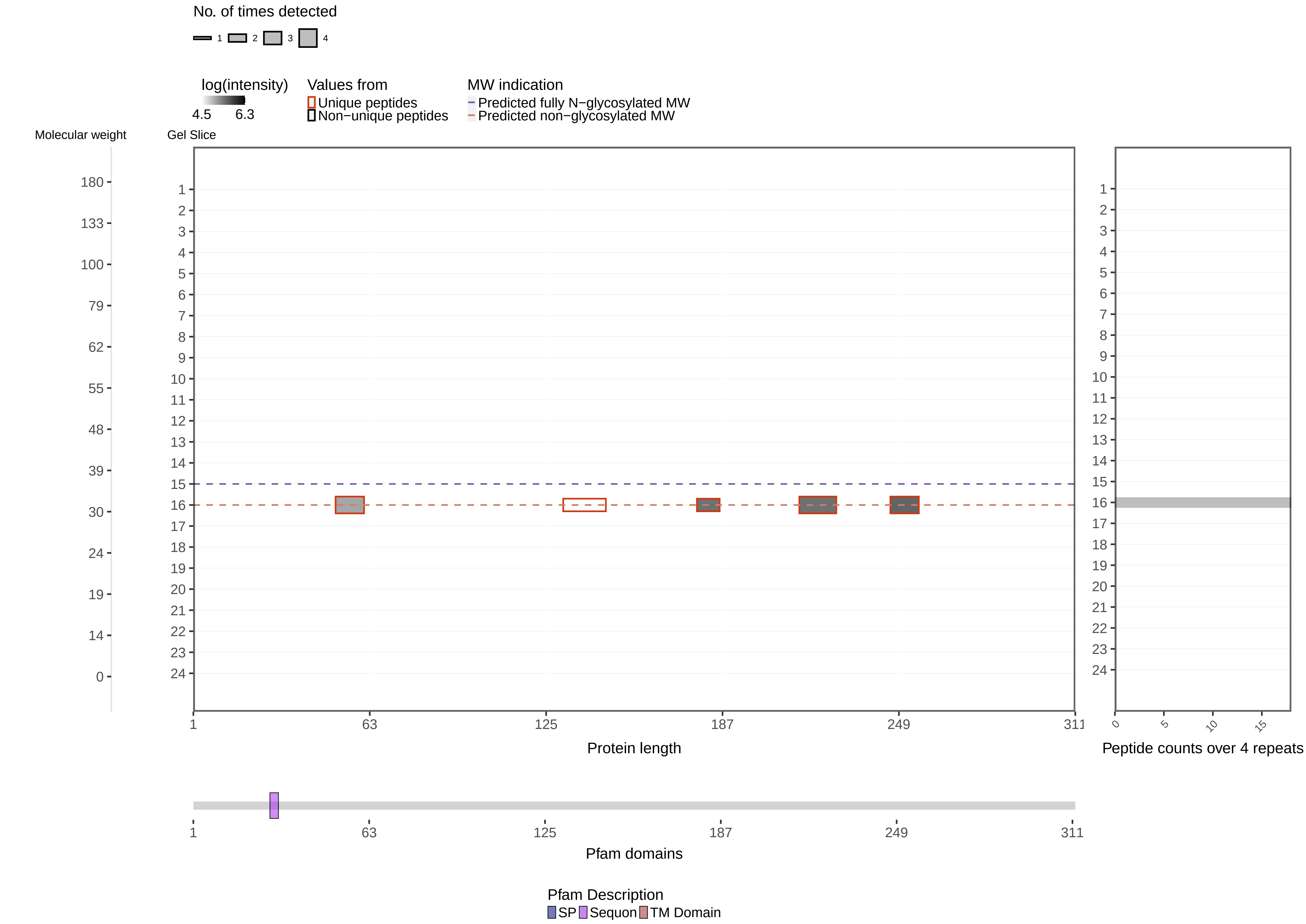


Protein sequence:

VSKLGEPPTDGKTDTNPEGLTAAYGKWASAVAARLQAGGLSCKVLEKEPFQKQMLEKLIWICAFMLVGARHPGATVGAVEEEYRSEVSQLIAELAAAAAAEKGLVFEEAMEDRLCAYSRAVAHFPTAVKEFKWRNGWFYSLSEKAIAEGKPDPCPLHTAWLKELKIV*

### NbL03g20580.1 heat_shock_70_kDa_protein [MW = 71.92 kDa]

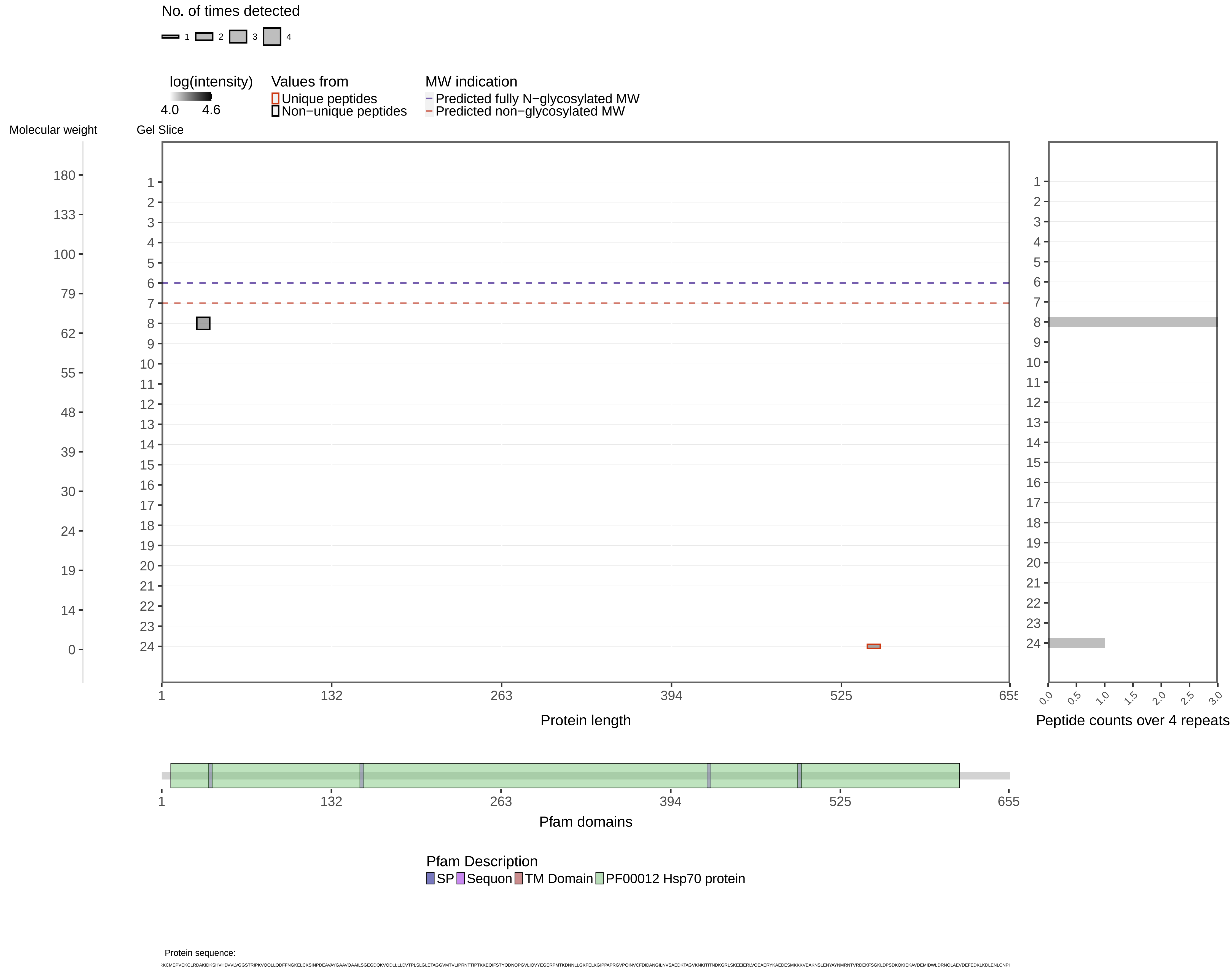

NbL03g20570.1 shikimate_O−hydroxycinnamoyltransferase [MW = 48.4 kDa]

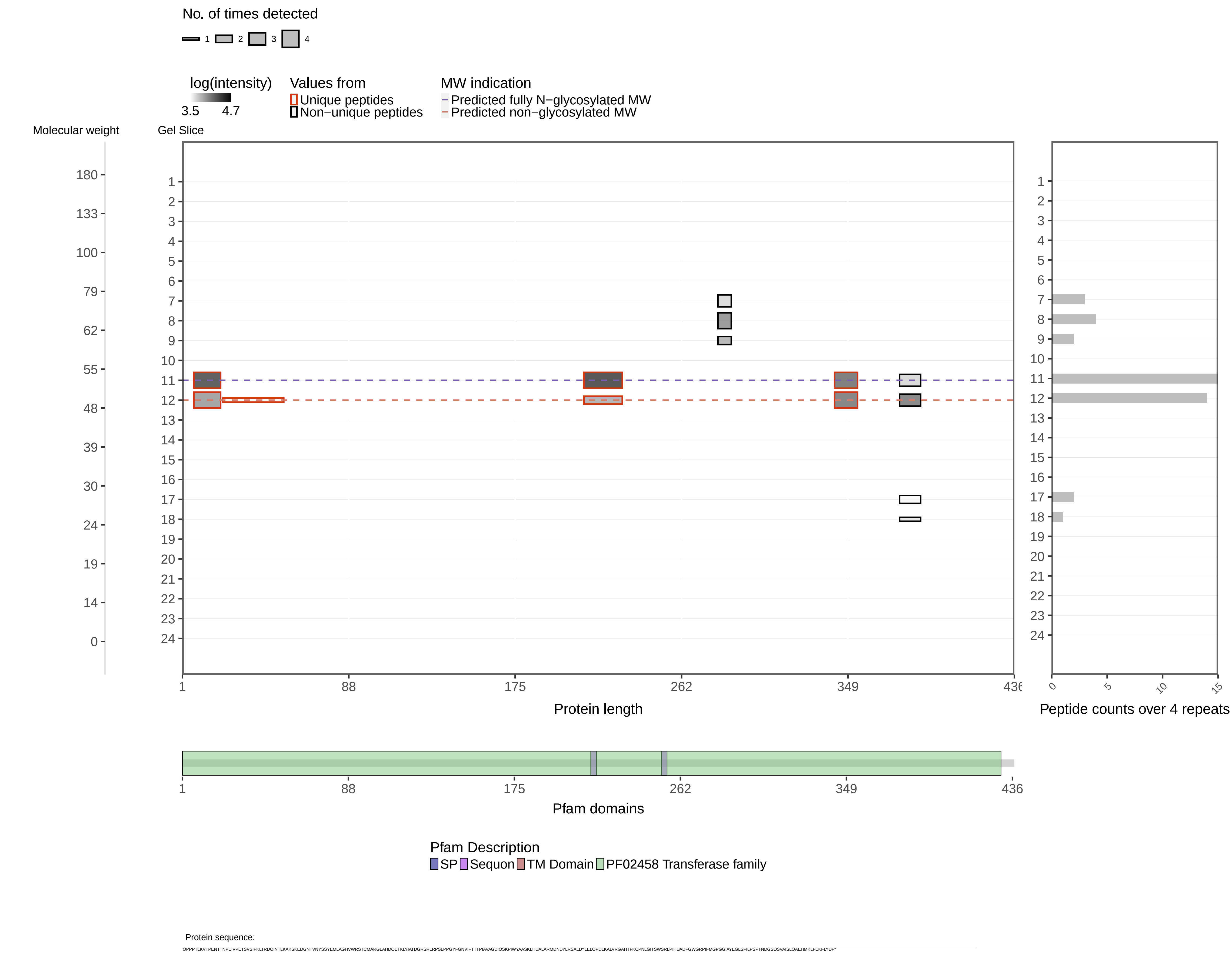

### NbL03g20470.1 glycerol_kinase [MW = 56.56 kDa]

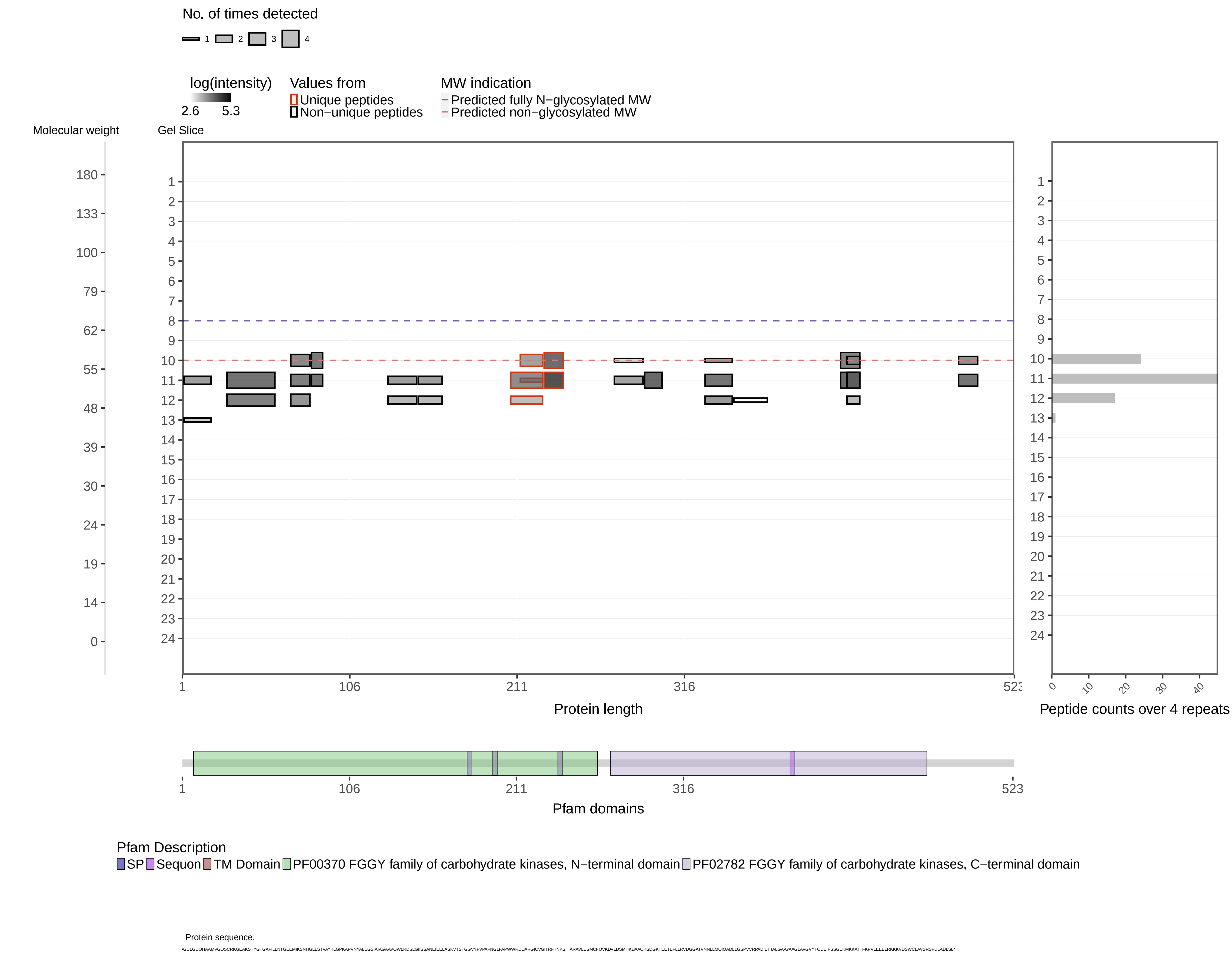

NbL03g20410.1 protein_TOPLESS−like [MW − SP = 126.47 kDa]

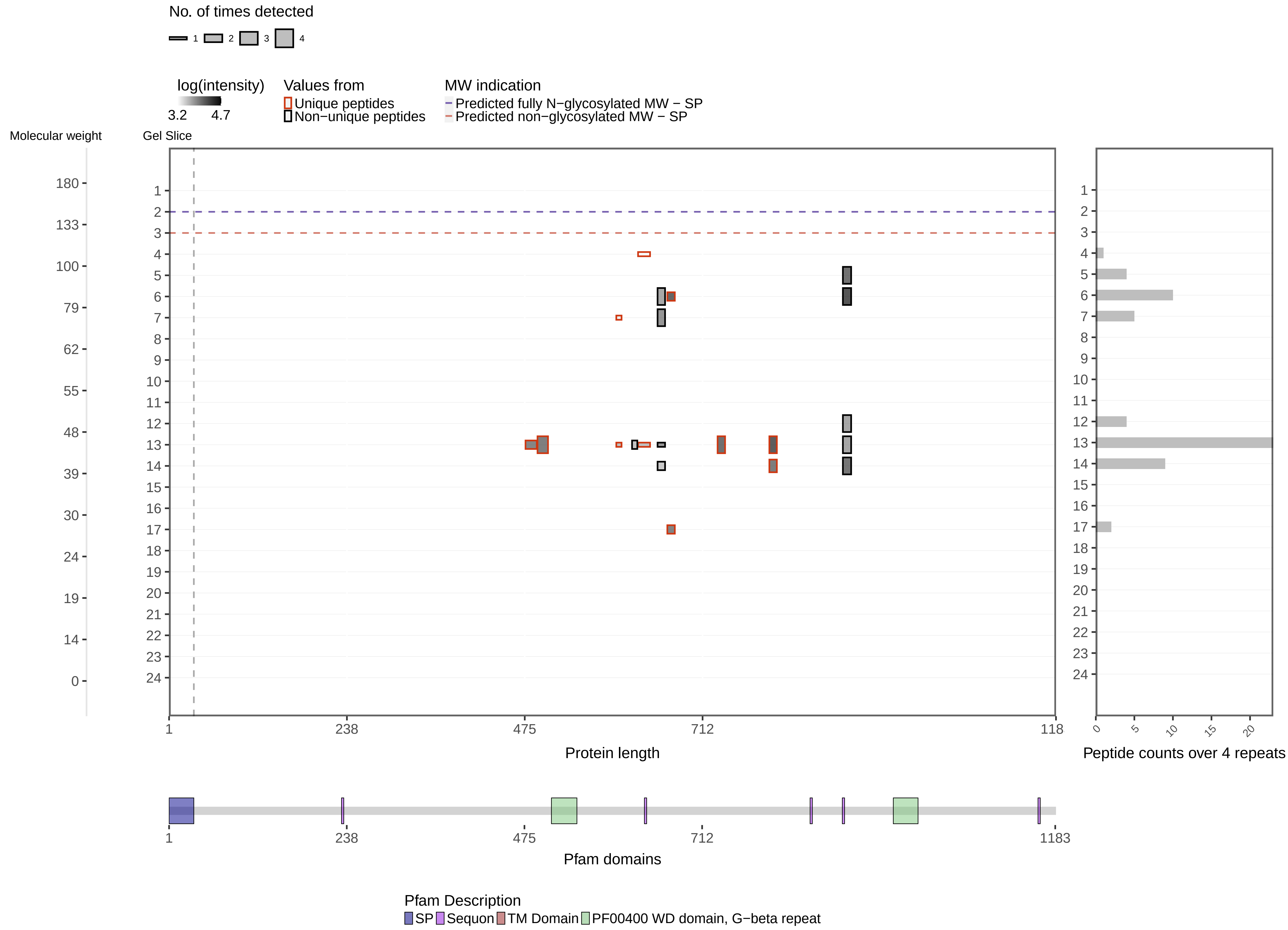


Protein sequence:

[illegible]

NbL03g20380.1 proline−−tRNA_ligase_cytoplasmic−like [MW = 58.76 kDa]

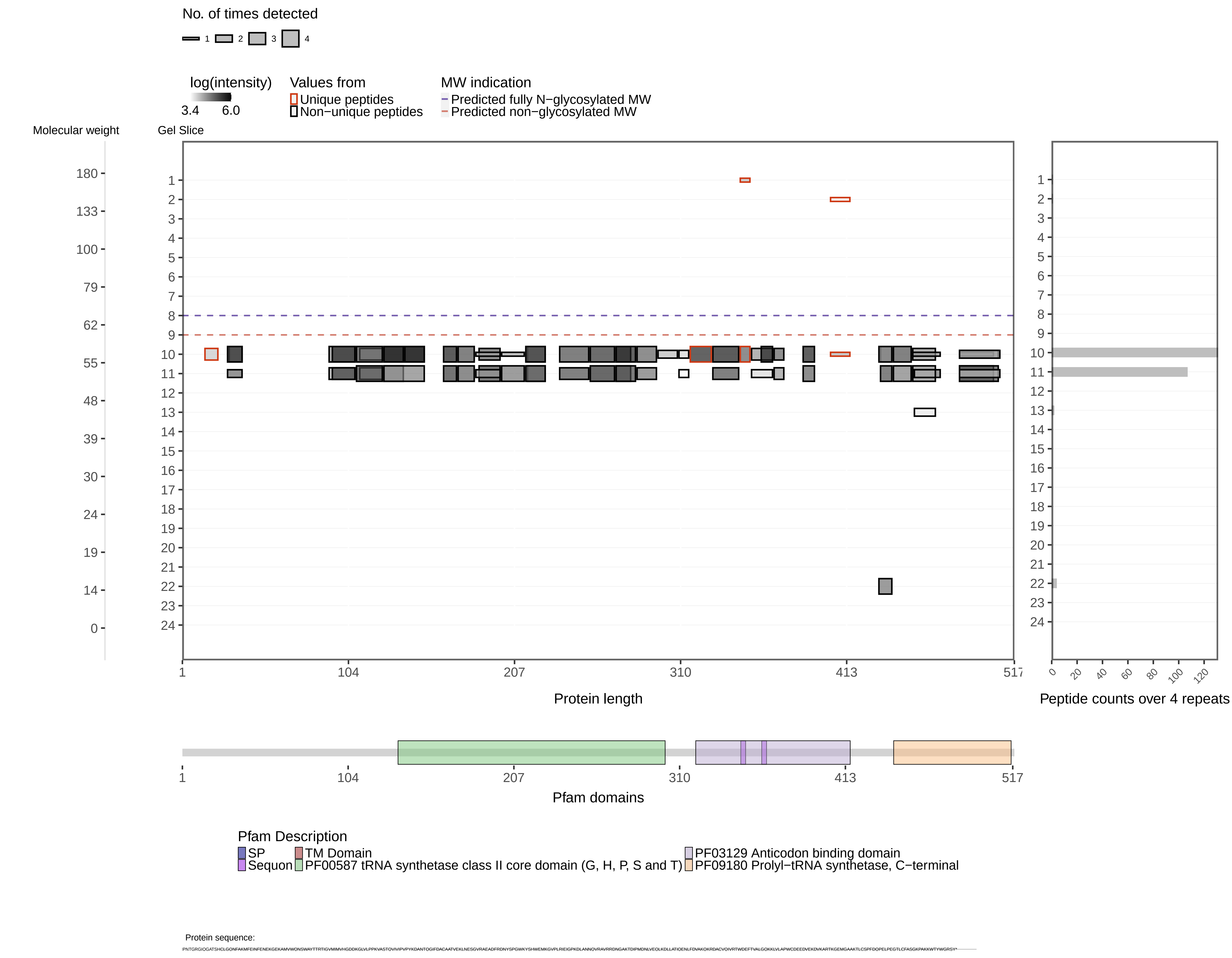

### NbL03g20320.1 uncharacterized_LOC109223891 [MW − SP = 55.22 kDa]

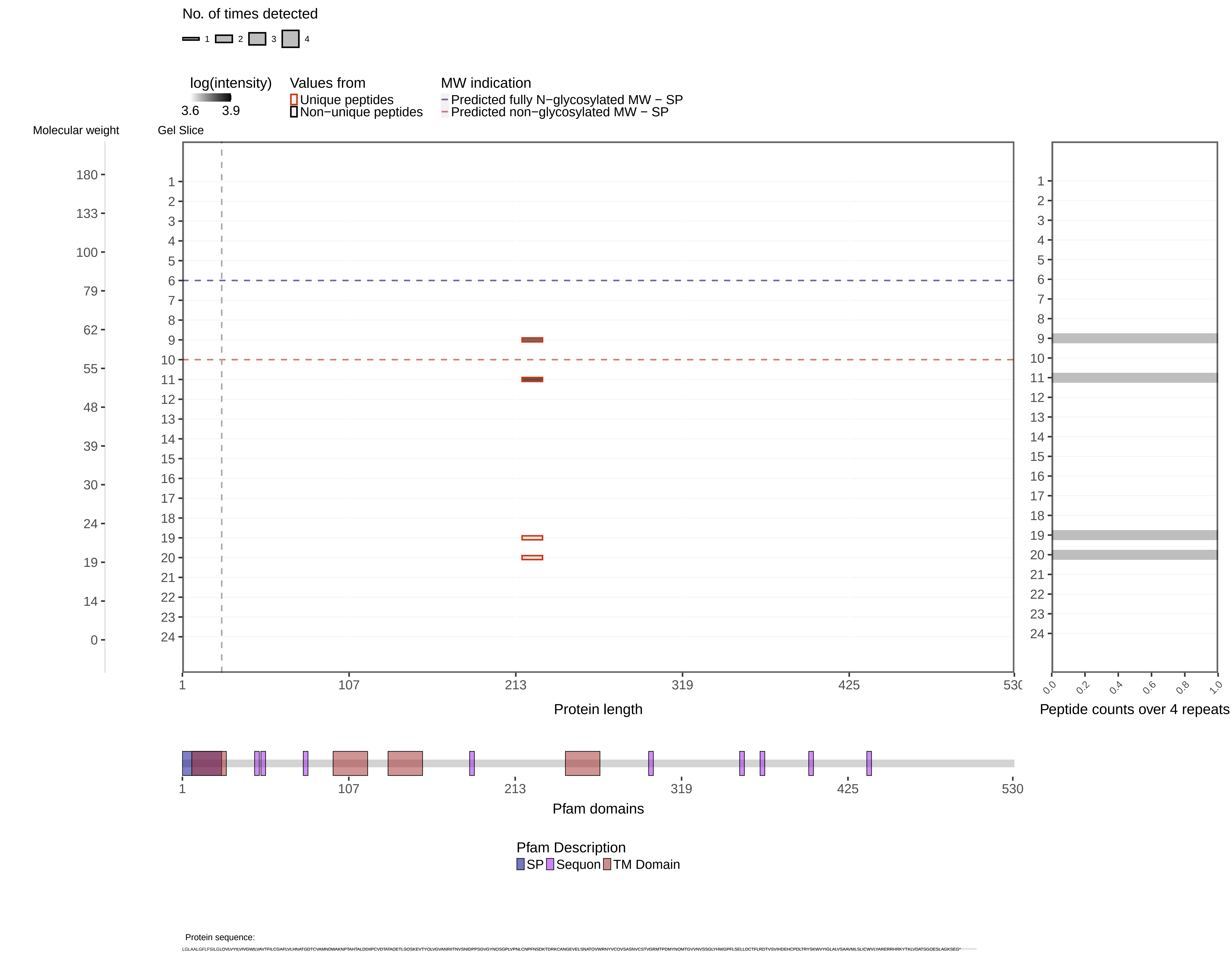

### NbL03g19970.1 protein_LURP−one−related_12−like [MW = 23.63 kDa]

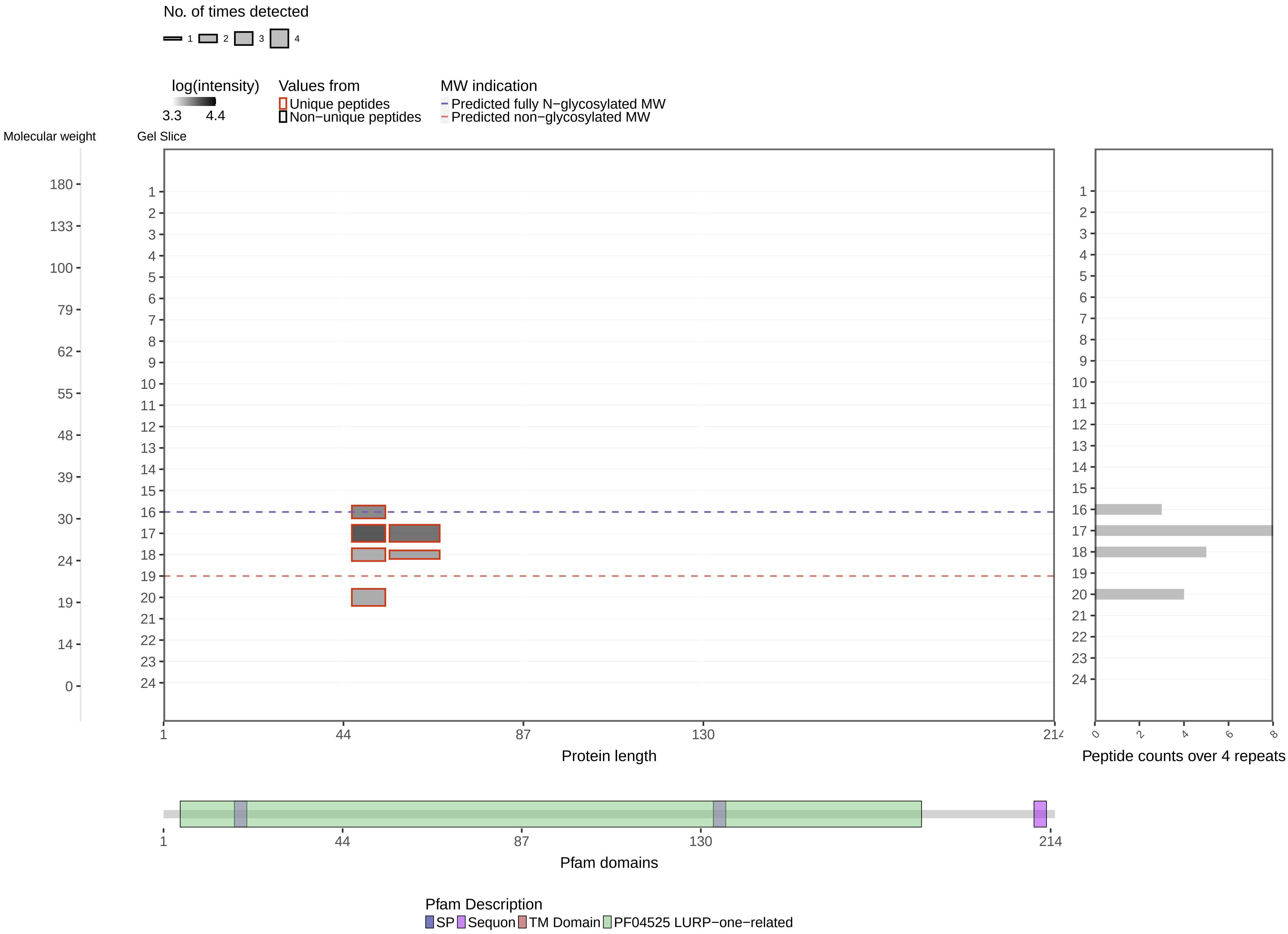


Protein sequence:

QKPIFSVRRSSIIGRSSVTAEVYSNPMEEYQIEGSFAQRNCTFFNADKVSVAEIRRKVDACANVVLGKDVFSLSVKPGFDGAFAMGLVLVLDQIQGDDYVNPNDNRVDQDPISDDPNLSS*

NbL03g19960.1 cucumber_peeling_cupredoxin−like [MW − SP = 16.34 kDa]

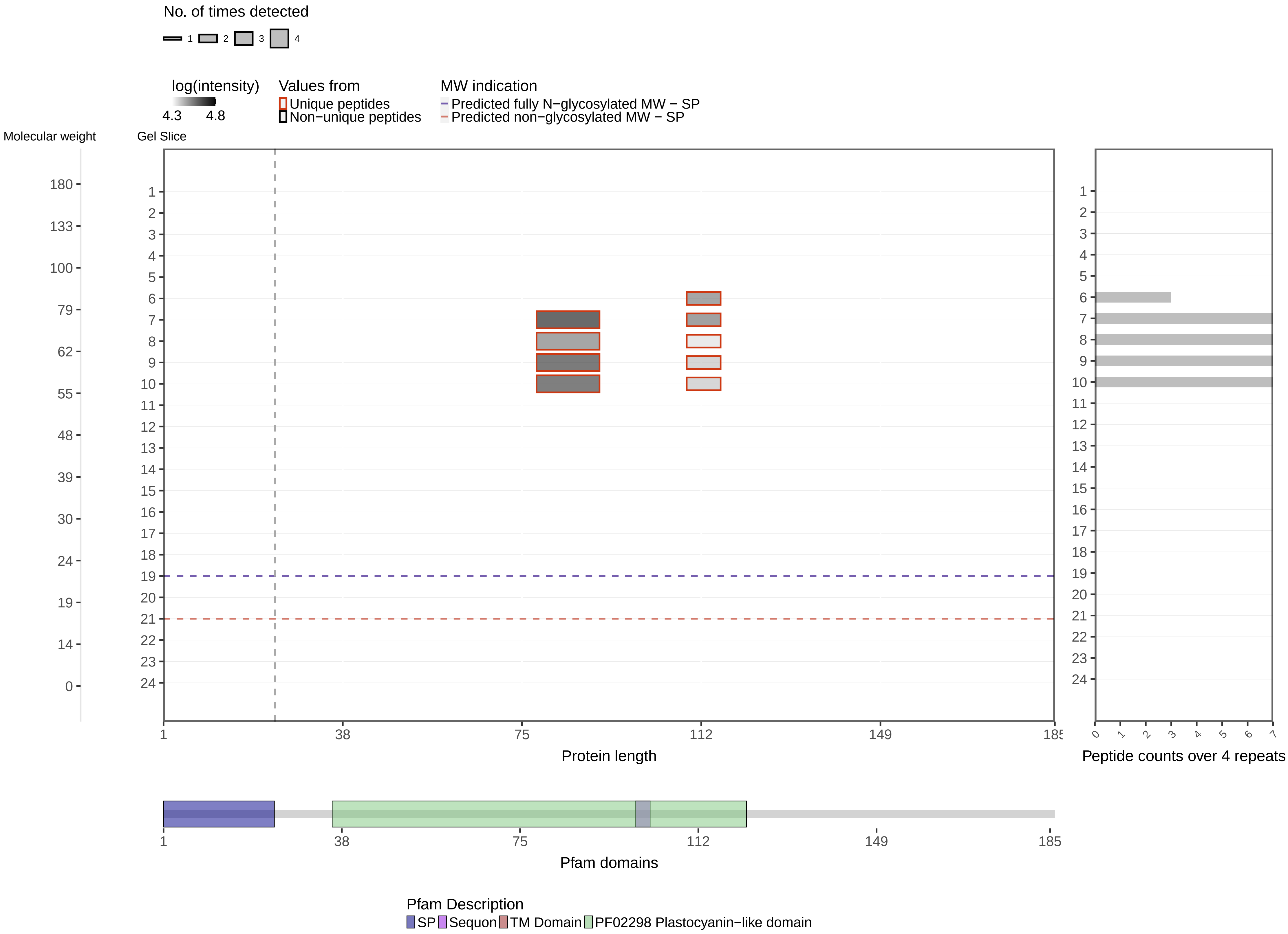


Protein sequence:

ASFDGCSSQNAIGRAIINGPANVTLDSAGERYYICTFGRHCQNGQKLAITVSSSTGTPGANPPTSFAAGPSGSVPGGIAPPPPPSSSTIVFASFLLSLSSIALAIFL*

### NbL03g19950.1 methyl−CpG−binding_domain−containing_protein_11−like [MW = 38.26 kDa]

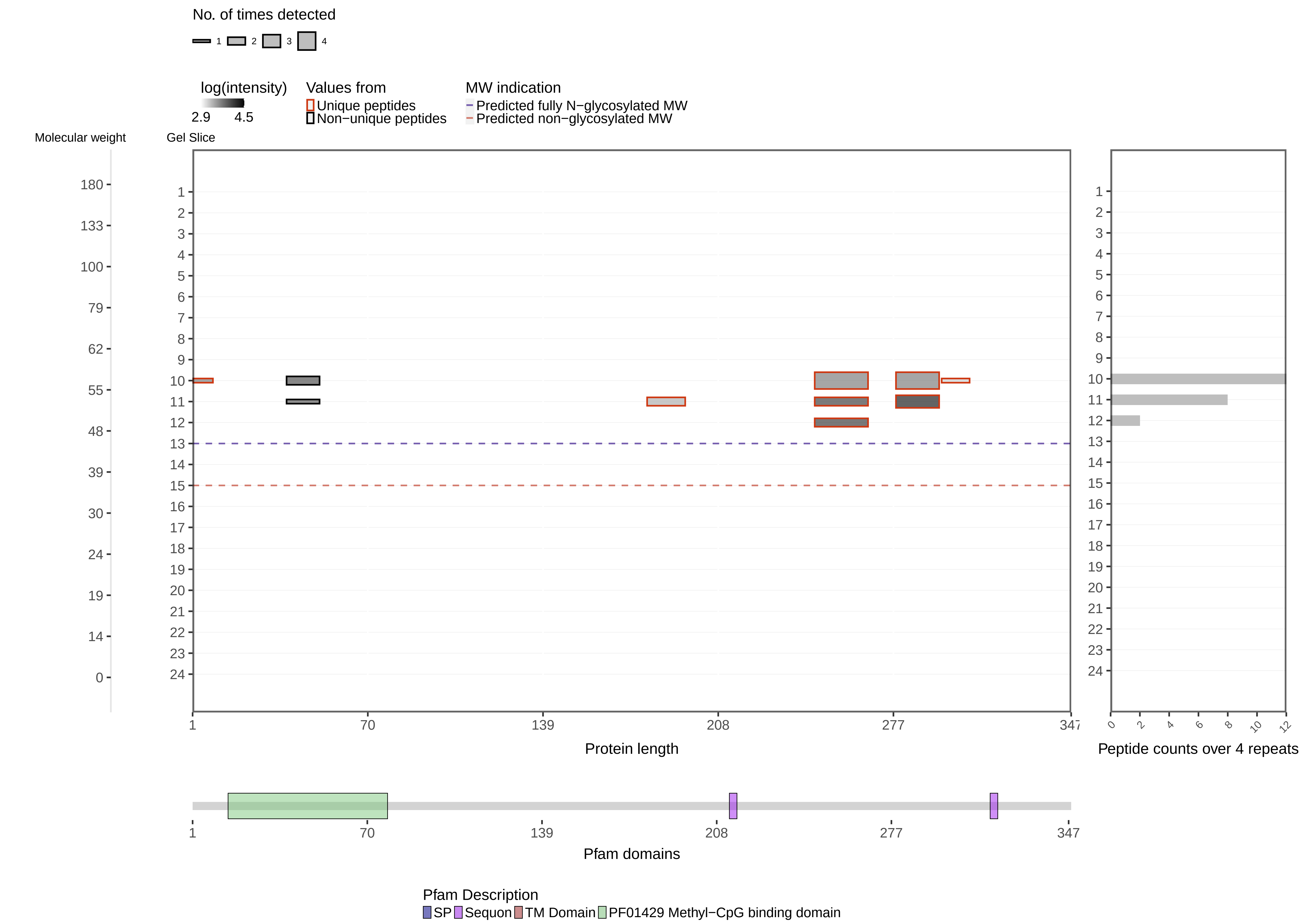


Protein sequence:

VTEHKKEEEMPDKDASEKKMEASDETEVDNGRDKMAGHGSEETIAIDVEVHNNSTEKDNFEGVVSDAAIGEANAAEAGLEVEEKHGSGEKPETQIDEDSGAGDVNRDVPDKVTPERSVAGDGTGNIQEPGLVGKATNSYANFGEYRSLEQEKEKNRTSLMMDNGEINQPGPAHTPQHQPAASISC*

### NbL03g19560.1 protein_RKD1 [MW = 16.14 kDa]

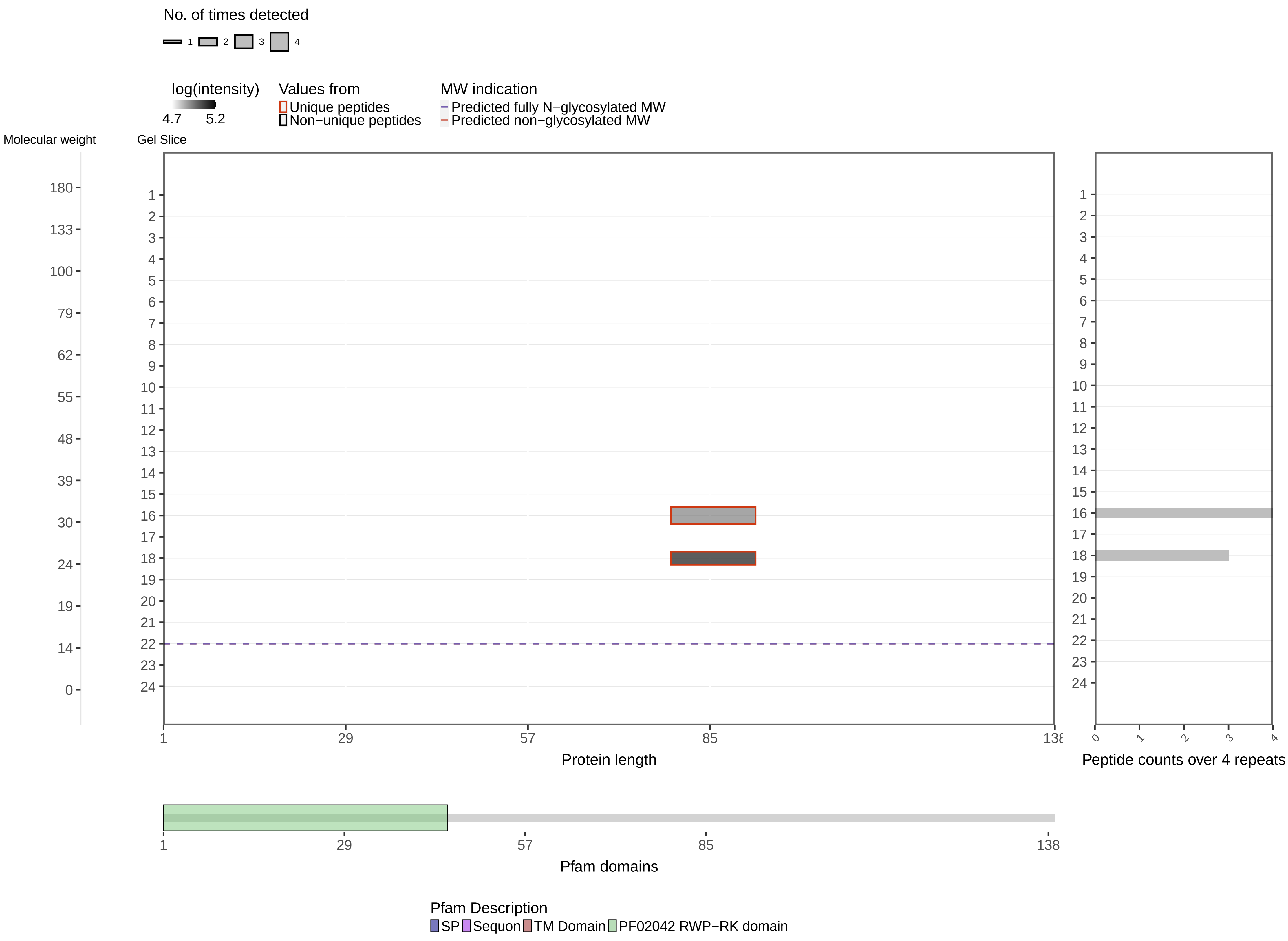


Protein sequence:

GGGSGMEQKLKDVIKLLEEEKKMLEEVPDMELEEKTKRLRQARFKANYKRRKLMSMTELQASFGSYCNSNPAVCIGYGHRE

### NbL03g19540.1 malate_dehydrogenase_chloroplastic [MW = 43.39 kDa]

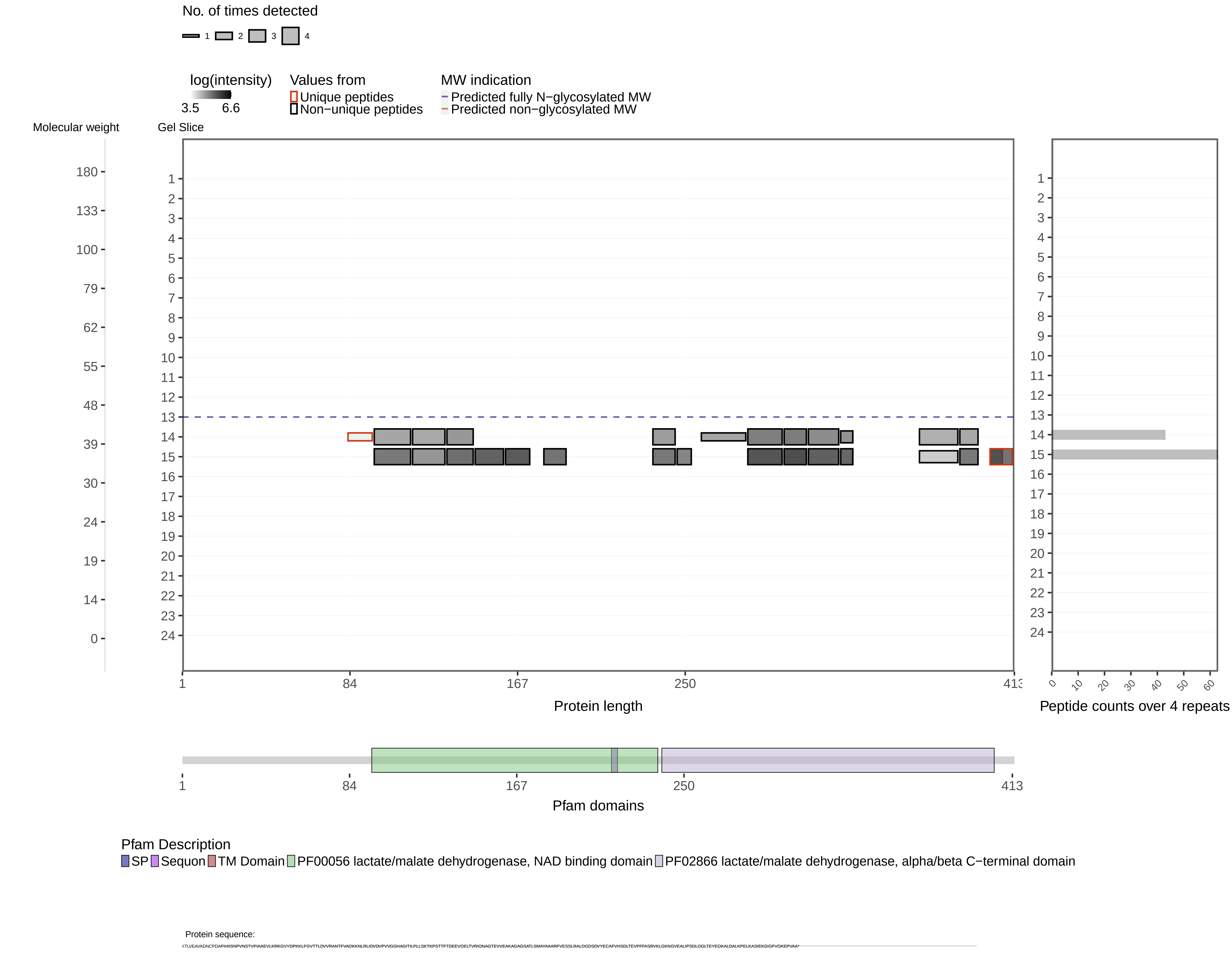

NbL03g19530.1 geranylgeranyl_diphosphate_reductase_chloroplastic [MW = 51.25 kDa]

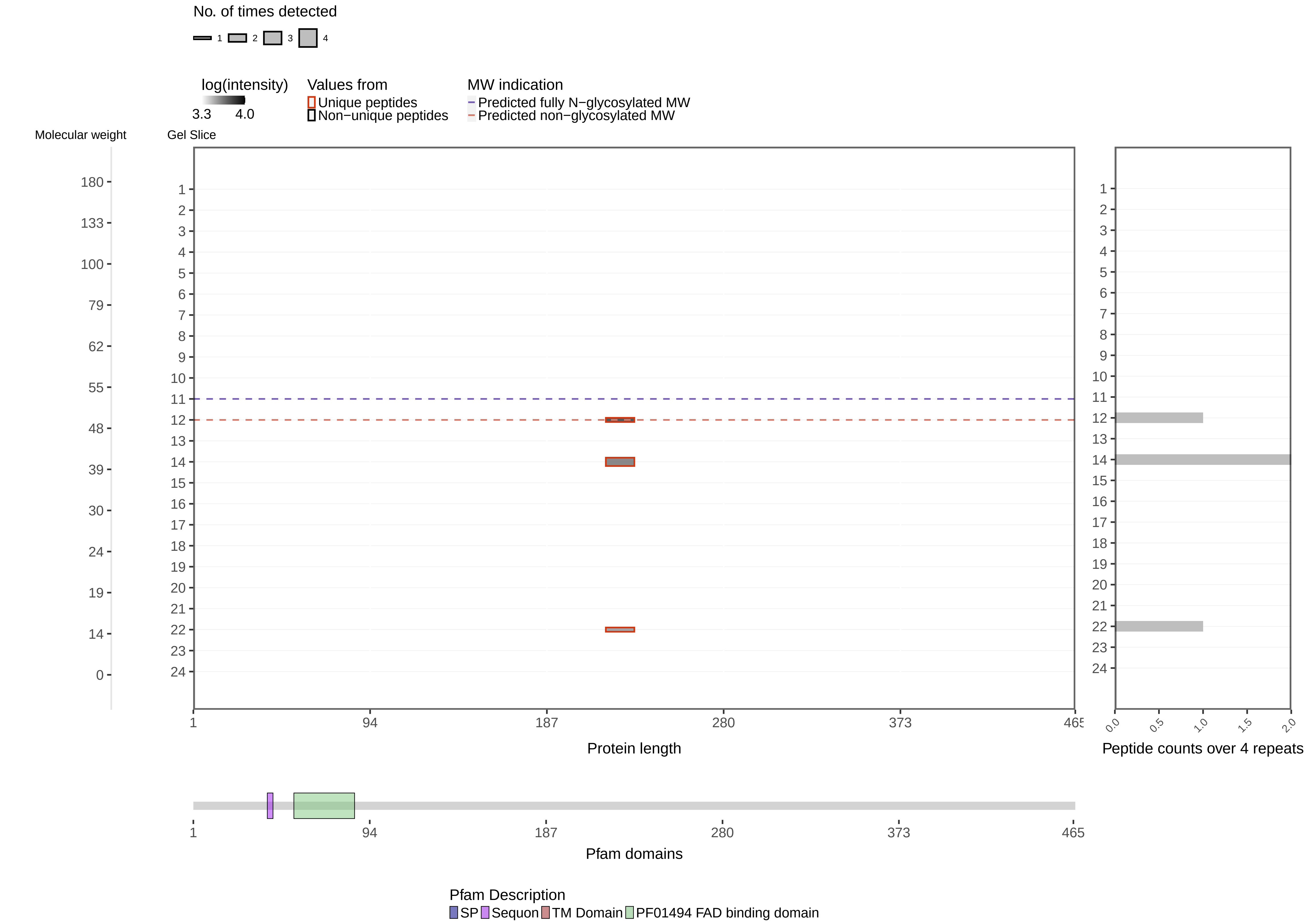


Protein sequence:

AGDYEYAIAFQERIRISDDKMKYYENLAEMYVGDDVSPDFYGWVFPKCDHVAVGTGTVTHKADIKKFQLATRLRADSKITGGKIIRVEAHPIPEHPRPRRLQDRVALVGDAAGYVTKCSGEGIYFAAKSGRMCAEAIVEGSEMGKRMVDESDLRKYLEKWDKTYWPTYKVLDILQKVFYRSNPAREAFVEMCADEYVQKMTFDSYLYKKVAPGNPIEDLKLAVNTIGSLVRANALRKEMDKLSV*

### NbL03g19430.1 ribulose−phosphate_3−epimerase_chloroplastic [MW = 30.6 kDa]

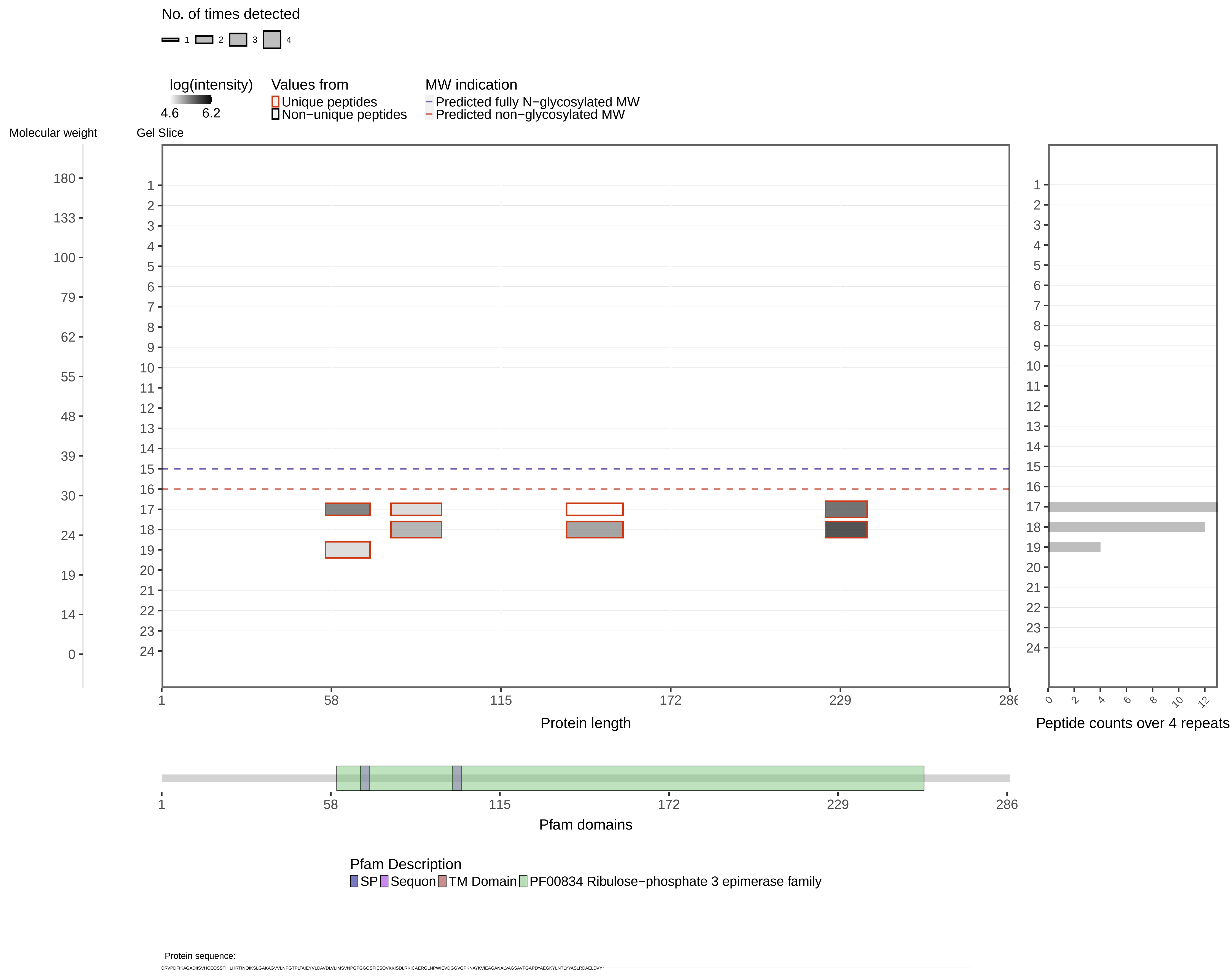

### NbL03g19370.1 vesicle_transport_v−SNARE_13−like [MW = 25.11 kDa]

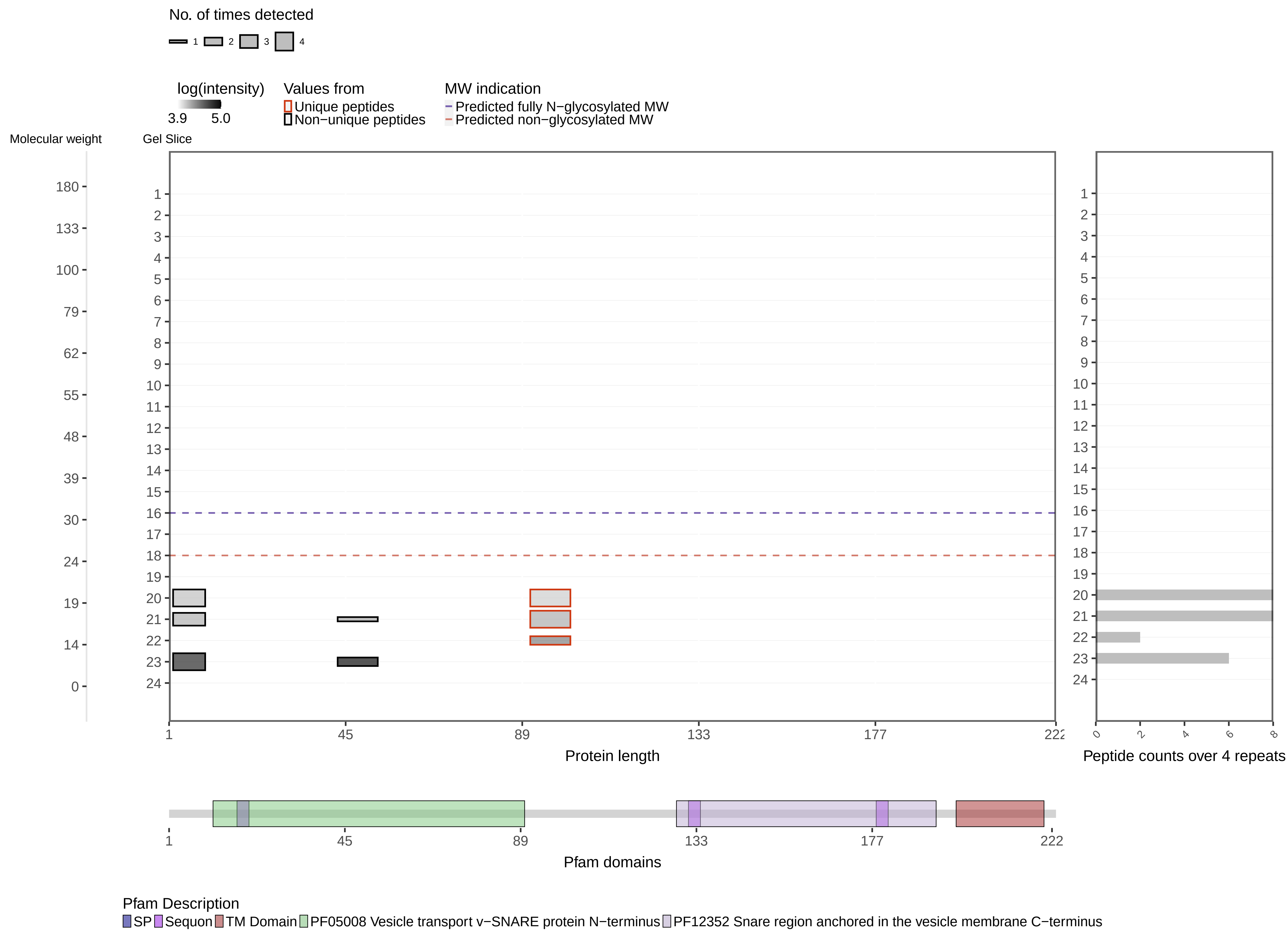


Protein sequence:

QAARDELLESGMVDAMMVSADQRQRLMMSTERLNHSSDRIRESRKTMLETEELGVSILQDLHQQRQSLLHAHDTLHGVDDNISRSKKILTTMSRRMSRNKWIIGSIIAVLVLAILLVLYFKLAH*

### NbL03g19290.1 eukaryotic_translation_initiation_factor_5A−1 [MW = 17.37 kDa]

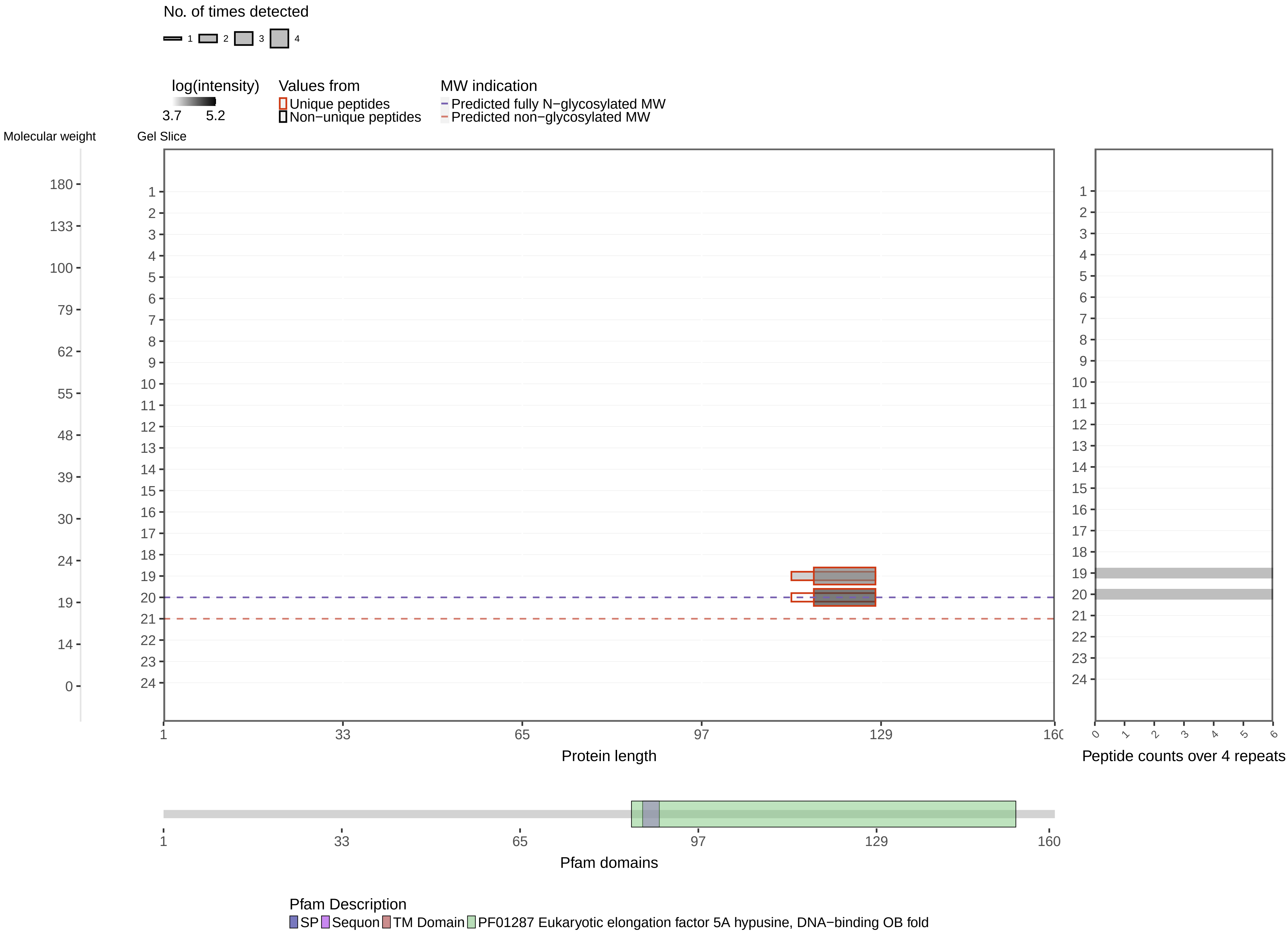


Protein sequence:

TGKKLEDIVPSSHNCDVPHVNRTDYQLIDISEDGFVSLLTENGNTKDDLRLPTDDNLLALIKDGFAEGKDLVLSVMSAMGEEQICGIKDVGPK*

### NbL03g19180.1 LYR_motif−containing_protein_4 [MW = 9.88 kDa]

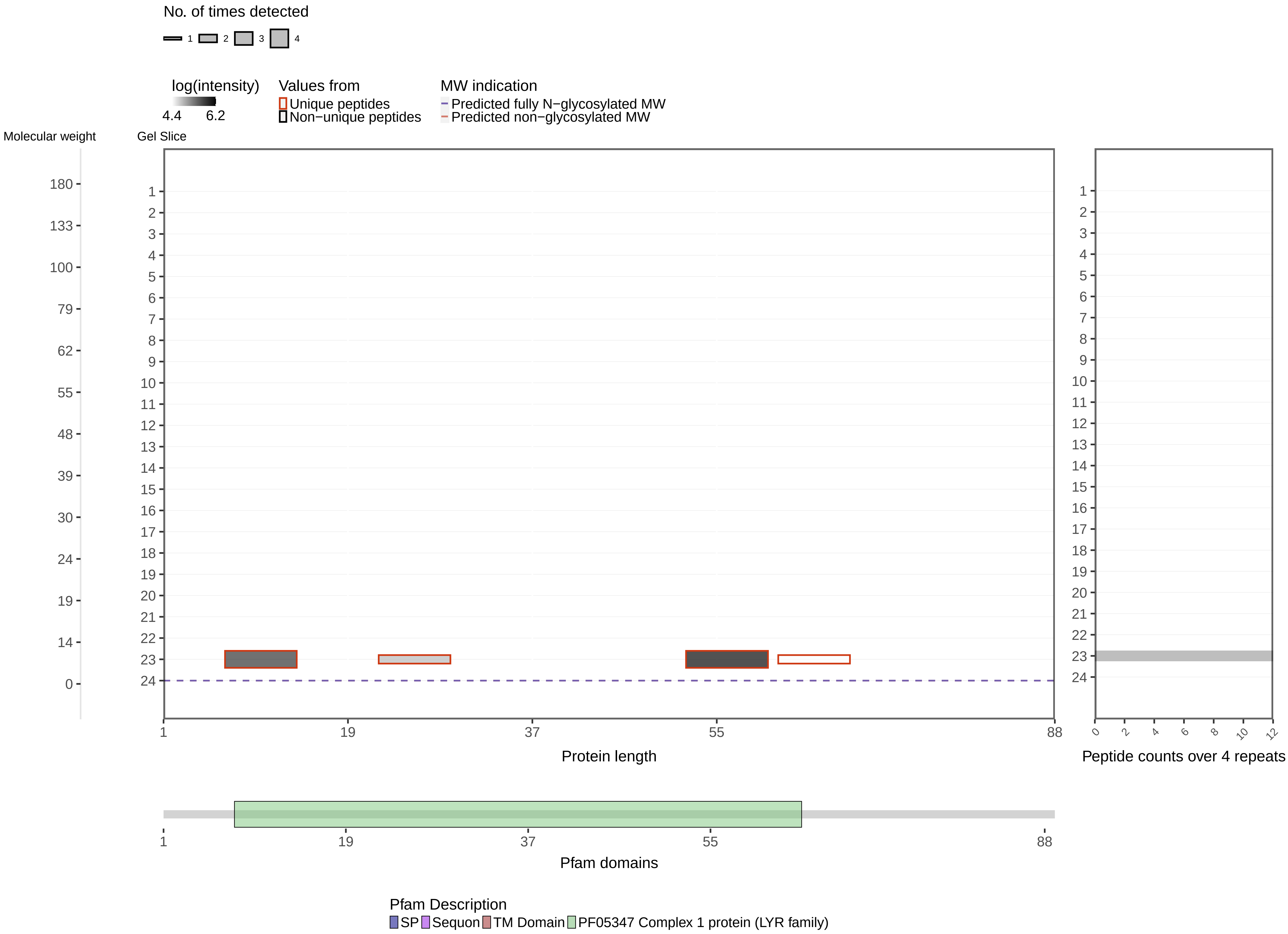


Protein sequence:

TKRRIIDGFRQNKGLSDPSKVAAAFSEGRSQLEVAKRQAVVYSLYAPKVKSVMELN*

### NbL03g19120.1 pre−mRNA−splicing_factor_ATP−dependent_RNA_helicase_DEAH1−like [MW = 119.68 kDa]

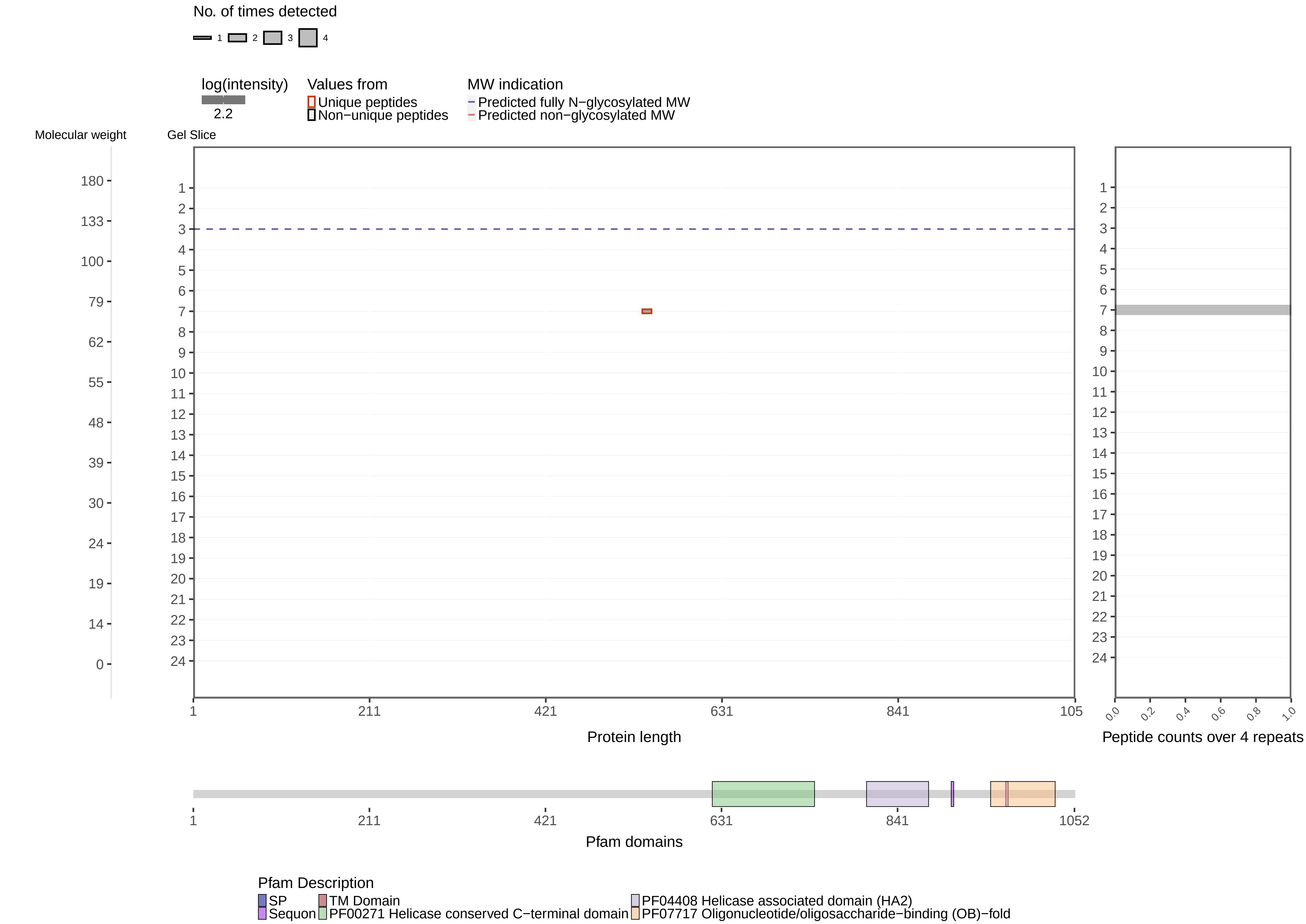


Protein sequence:

YMTDGMLLREFLGEPDLASYSVIMVDEAHERTLSTDILFGLVKDIARFRPDLKLLISSATLDAEKFSDYFDCAPIFKIPGRRFPVEIHYTKAPEADYLDAAVVTALQIHVTQPPGDGDILIFLTGQEEIETAEEIIKHRIKGLGTKIAELIICPIYANLPTELQAKIFEPTPEGARKVVLATNIAETSLTIDGIKYVIDPGFSKMKSYNPRTGMESLLVTPISKASANQRAGRSGRTGPGKCFRLYTAYNYMNDLEDNTVPEIQRTNLANVVLSLKSLGIHDLLNFDFMDPPPAEALLKALEMLF

### NbL03g19100.1 UDP−glucose_6−dehydrogenase_5−like [MW = 52.91 kDa]

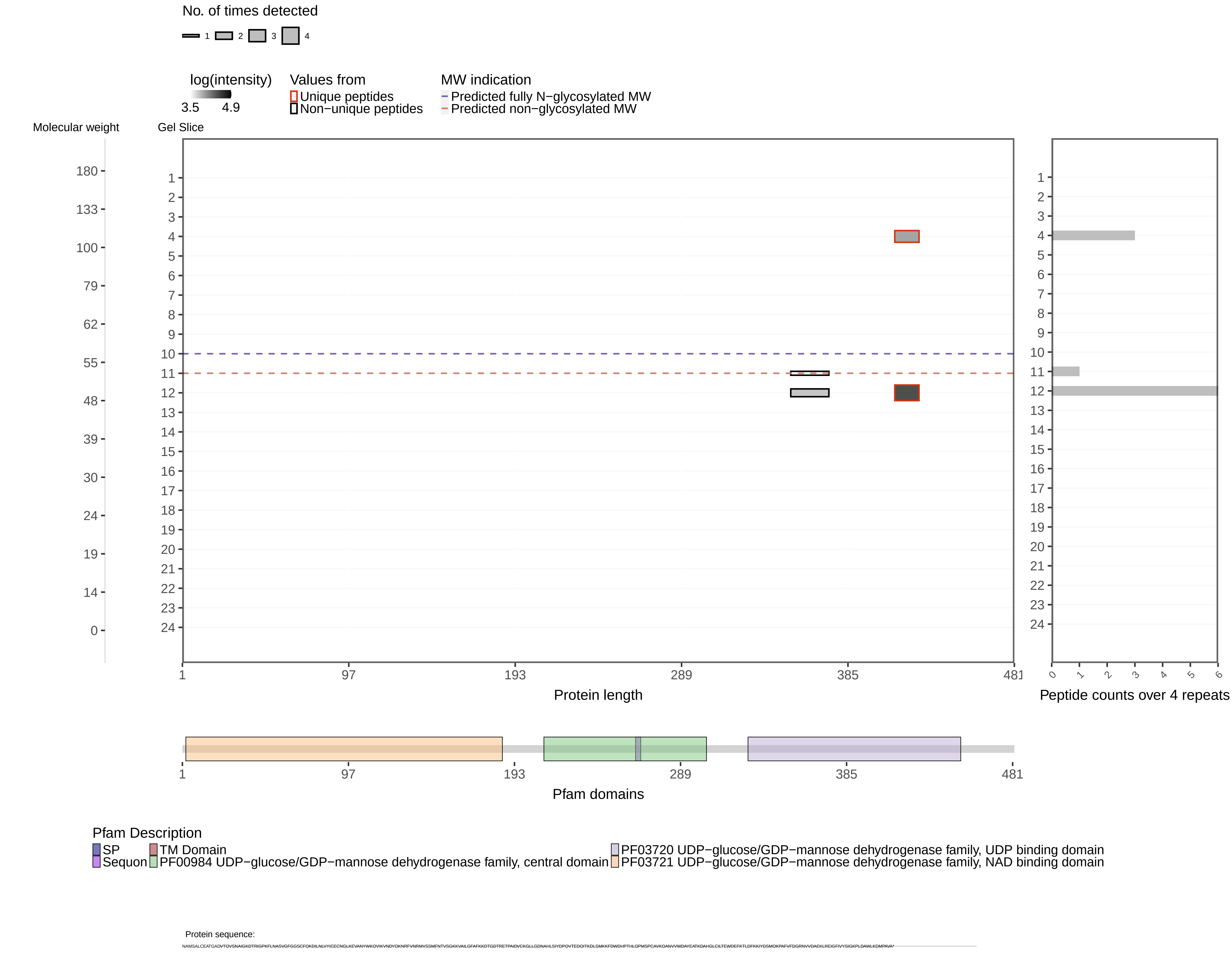

### NbL03g19080.1 40S_ribosomal_protein_S19−3−like [MW = 16.03 kDa]

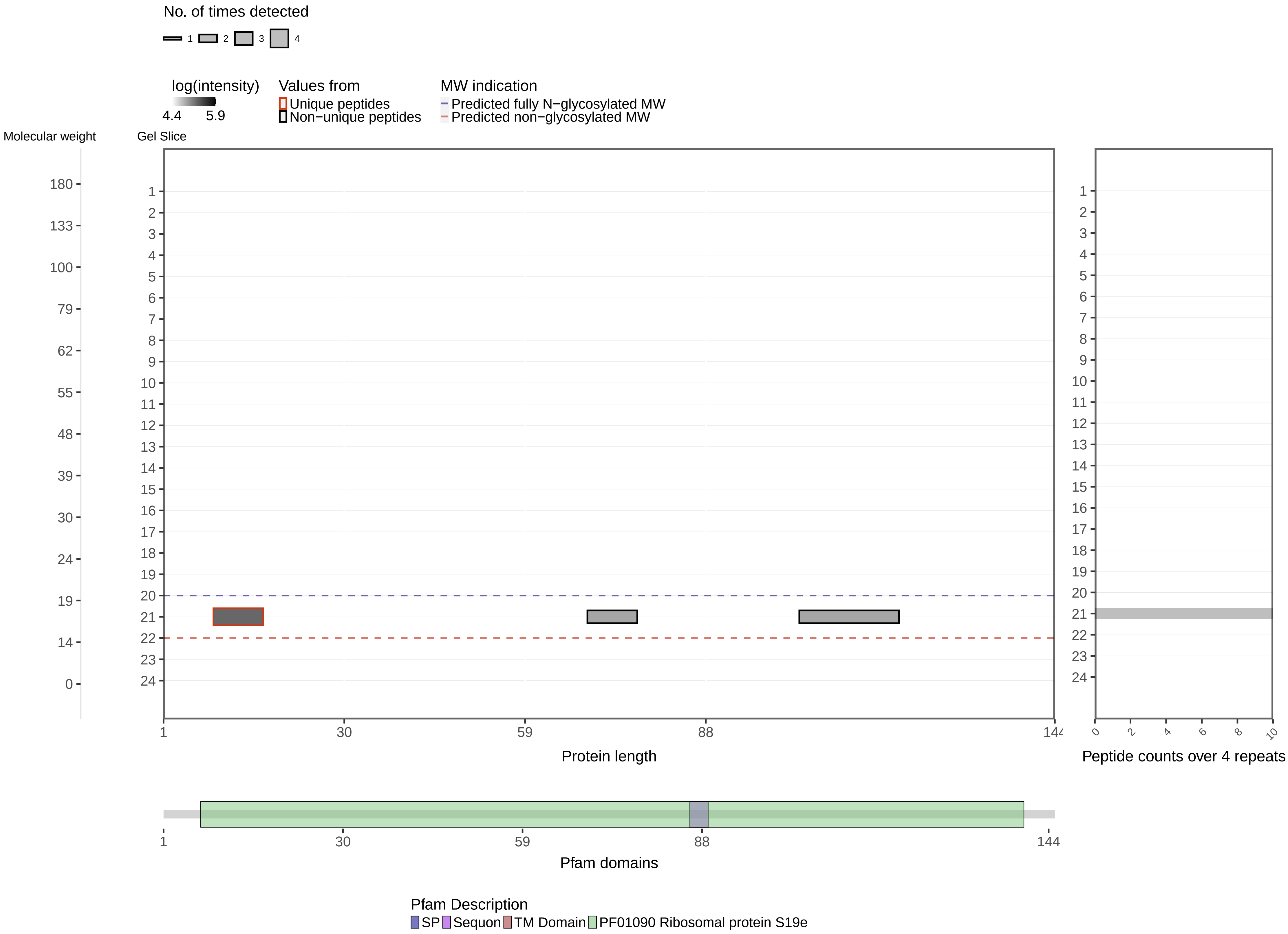


Protein sequence:

[illegible]ASMARKIYLRGGLGVGGFRRIYGGNQRNGSRPRHFCKSSGSVARNILQQLQNMNIIDFDPKGGRRITSNGQRDLDQVAGRIAVAI*

NbL03g18870.1 uncharacterized_LOC109223785 [MW = 18.77 kDa]

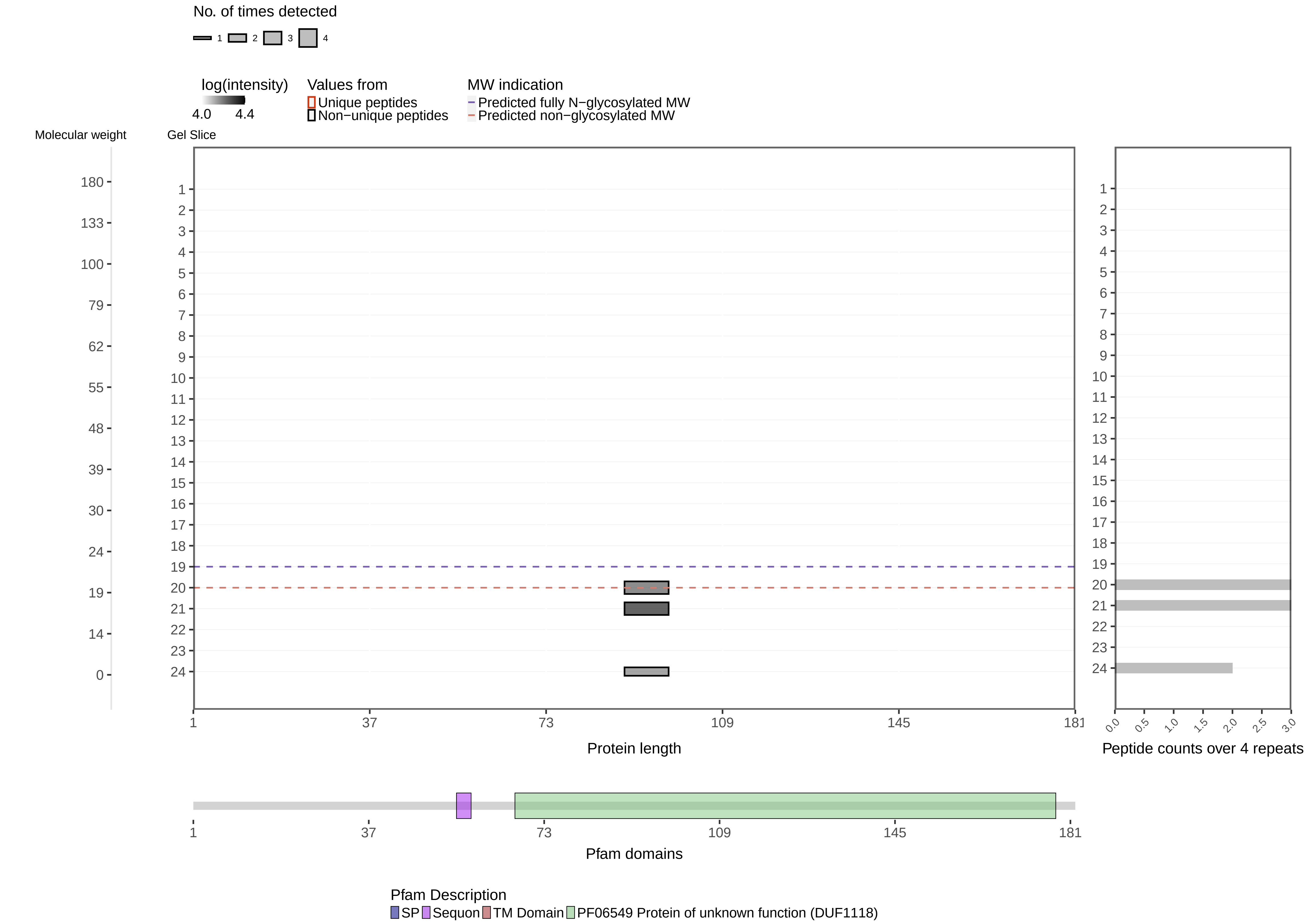


Protein sequence:

AEKAGLLSAAEKSGLSLSTIERLGLLSKAEEFGILSAATDPGTPSALFSLSLGLLVLGPACVYLIPEDYPWQIVVQVLVALLSVVGGSAAFGASNLVSNLQKLK*

### NbL03g18790.1 probable_UDP−arabinopyranose_mutase_2 [MW = 41.18 kDa]

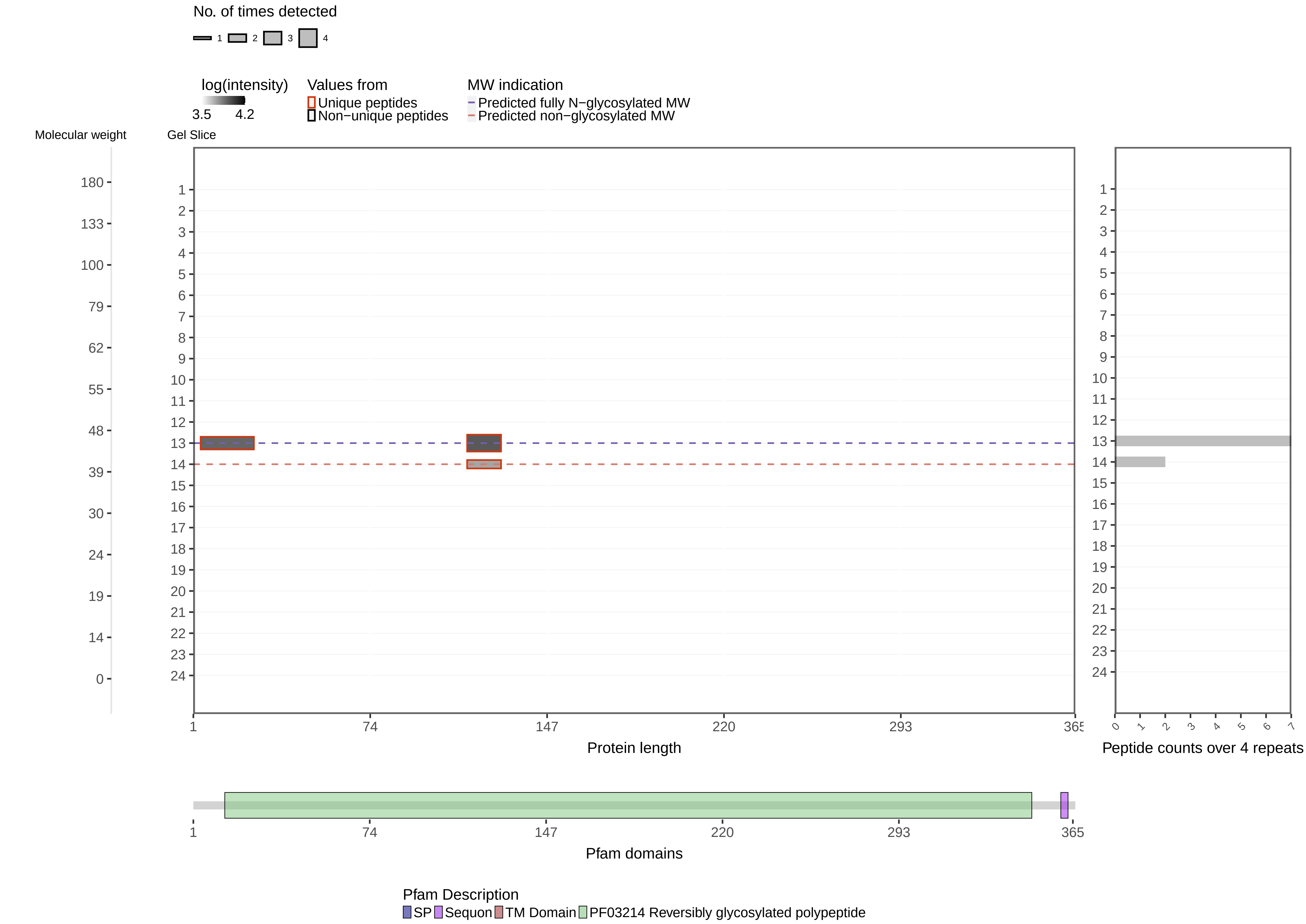


Protein sequence:

LWLNIPDYDAPTQLVKPLERNTRYVDAVMTIPKGTLFPMCGMNLAFDRELIGPAMYFGLMGDGQPIGRYDDMWAGWCTKVICDHLGLGVKTGLPYIWHSKASNPFVNLKKEYKGIYWQEELIPFFQSVTLPKDCTSVQKCYLELSKQVKAKLGKVDDYFNKLADAMITWIEAWDELNPSATATAVKIPNGSSK*

### NbL03g18730.1 flavonol_3−sulfotransferase−like [MW = 14.89 kDa]

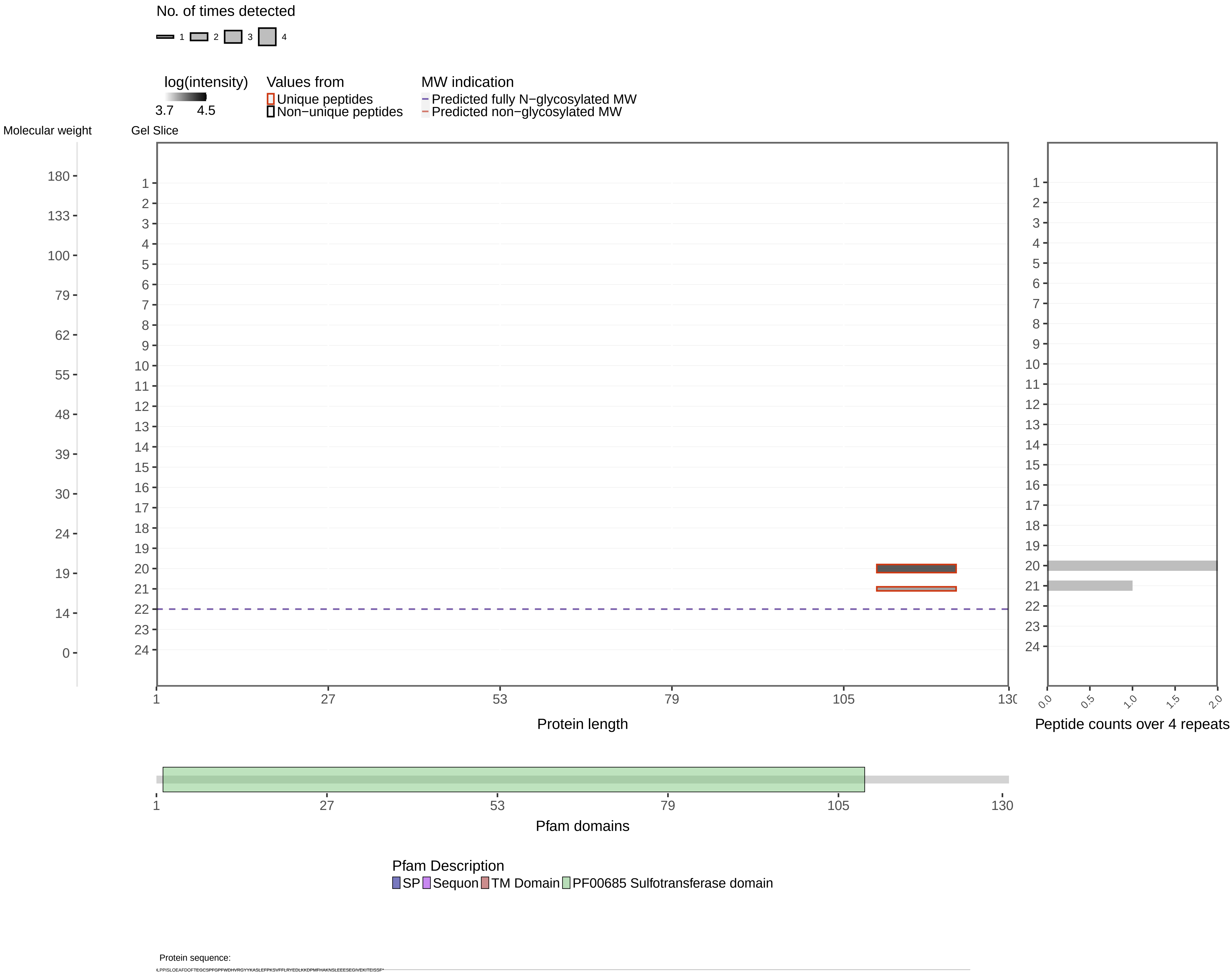

### NbL03g18610.1 UDP−glycosyltransferase_89B2−like [MW = 53.43 kDa]

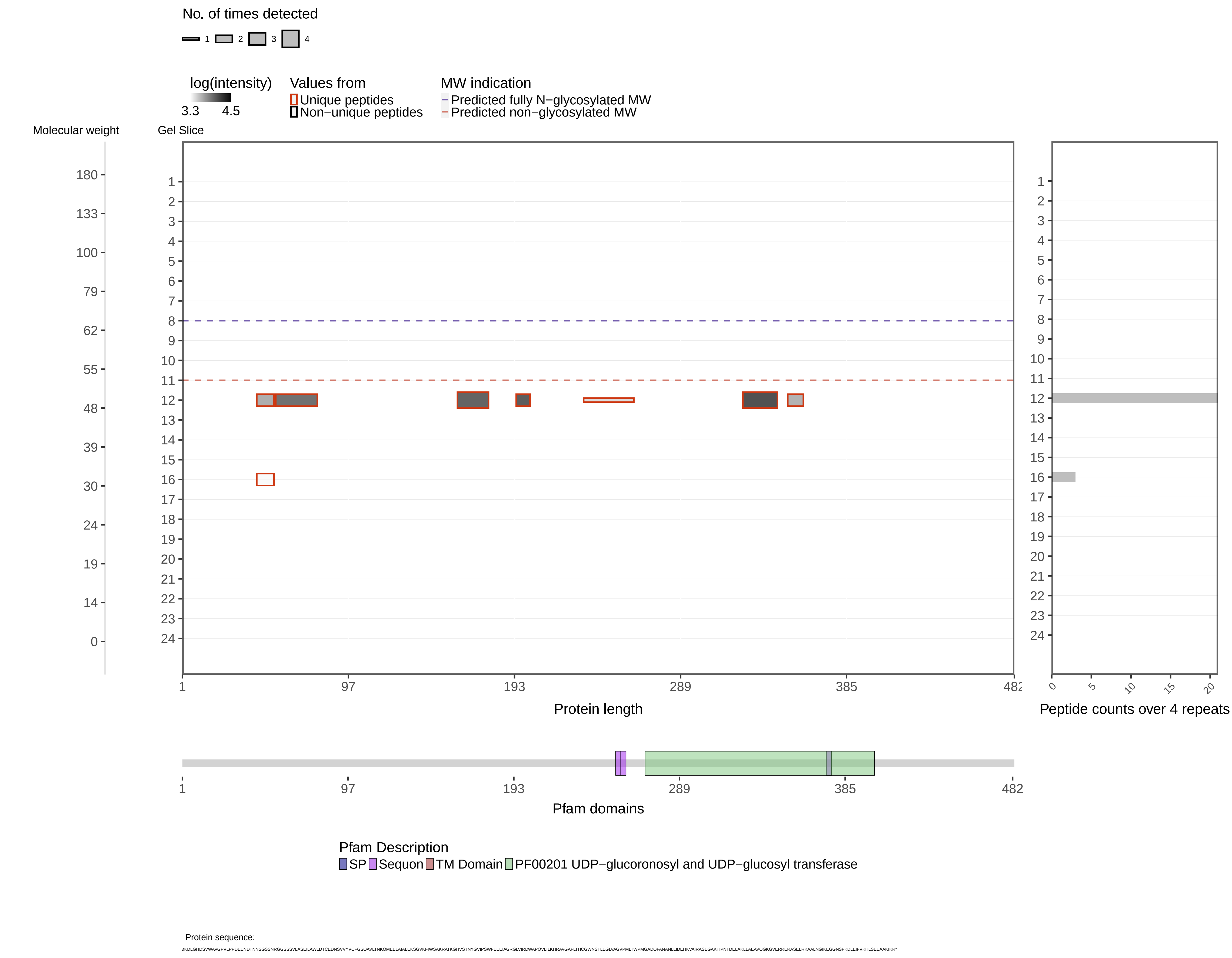

NbL03g18430.1 calcium_sensing_receptor_chloroplastic−like [MW = 41.87 kDa]

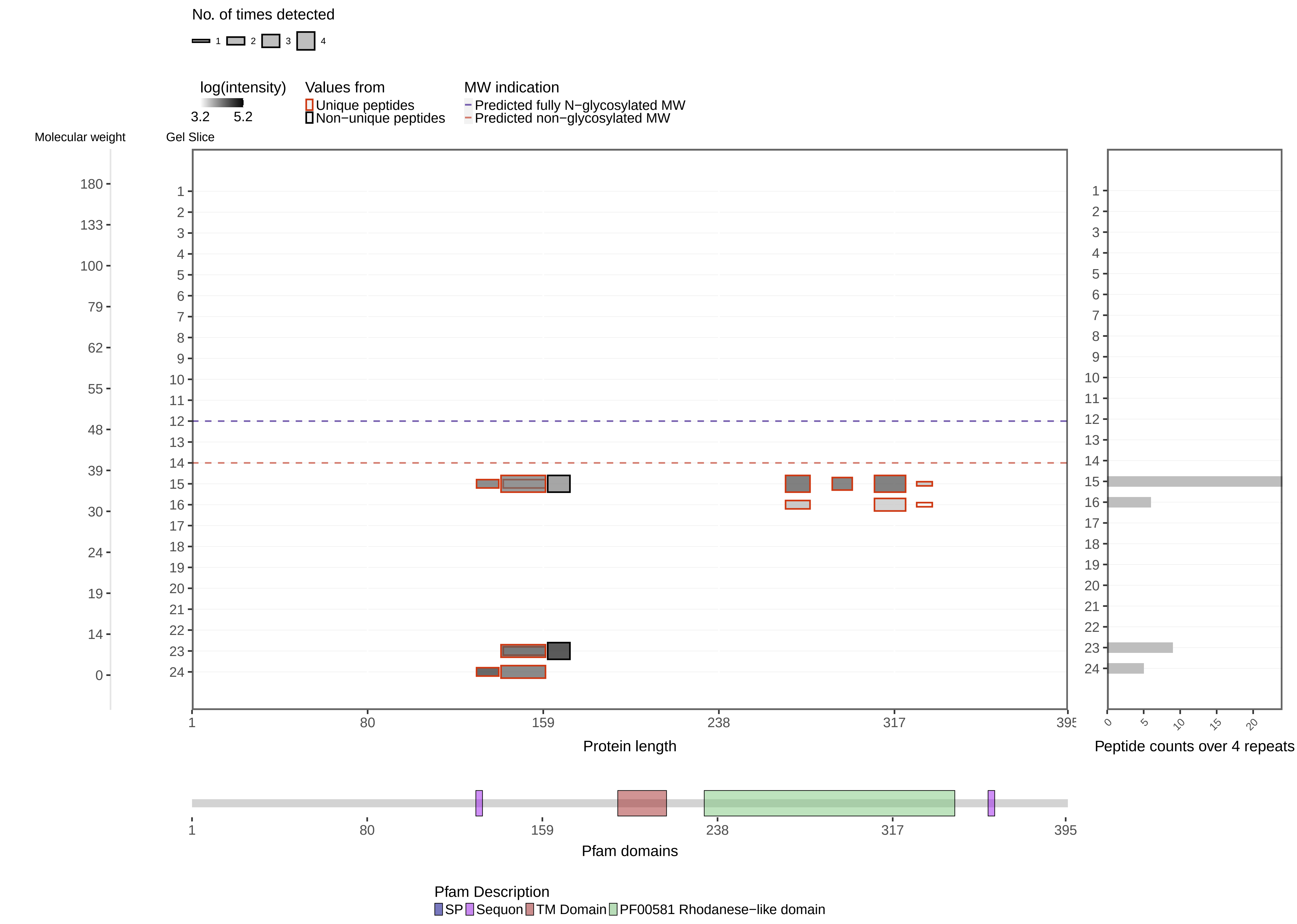


Protein sequence:

VETISSADPAIIAVAGGTLFLAYLLFPPVLSALSFSLRGYKGELTPAQTLDQMCSKNYVLIDIRTEKDKDKAGIPRLPSSAKNKMIQIPLEDLPSKLRSLVRNAKKVEAELVALKISYLKKINKGSNIVIMDSYSDSAKTVAKTLTSLGFKNCWIMTDGFSGGRGWLQSKLGTDSYNFSFAQVLSPSRVIPAAARRFGTTGTVKLLSGGSD*

NbL03g18370.1 1−deoxy−D−xylulose_5−phosphate_reductoisomerase_chloroplastic [MW = 51.29 kDa]

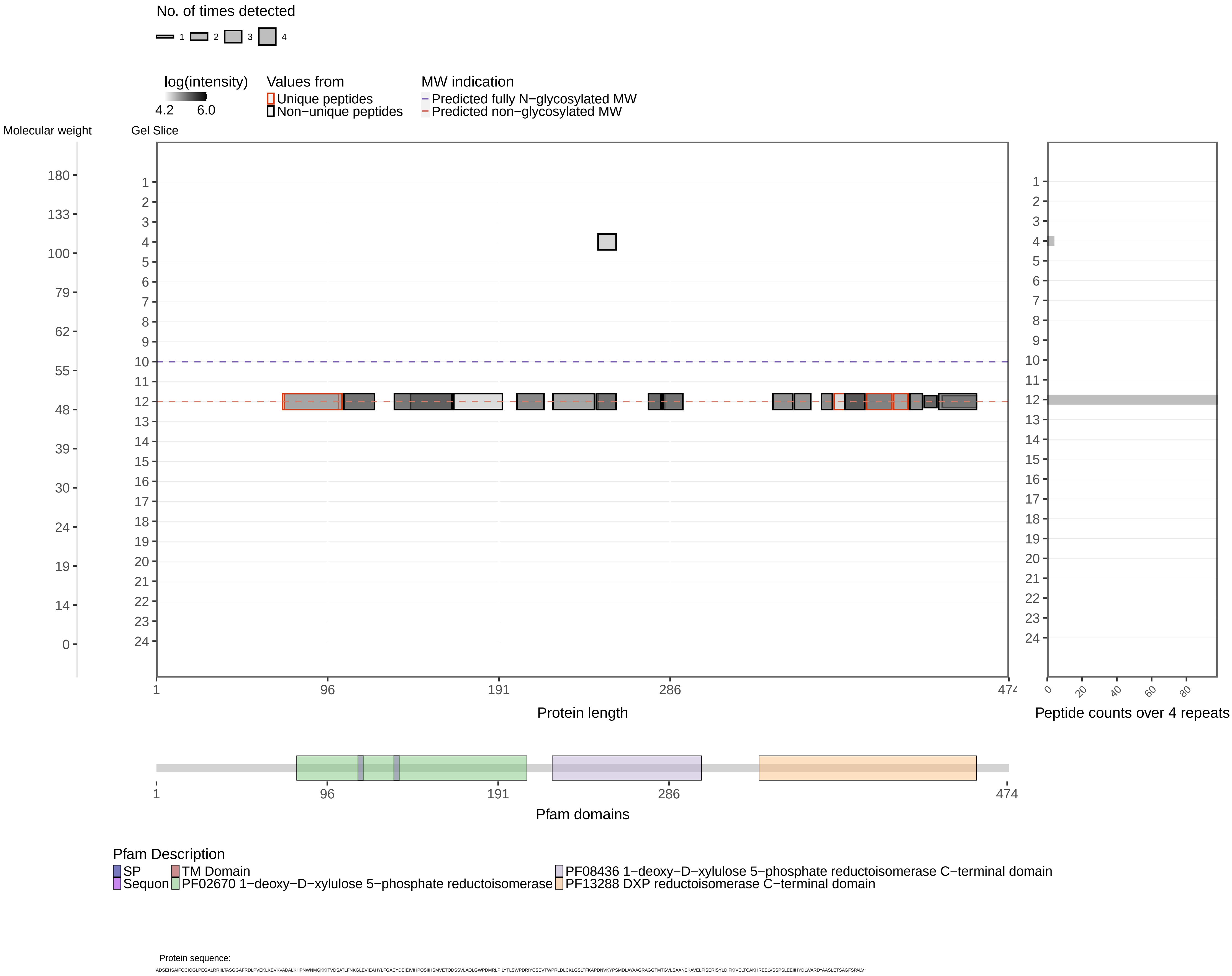

### NbL03g18270.1 uncharacterized_LOC109214664 [MW = 33.39 kDa]

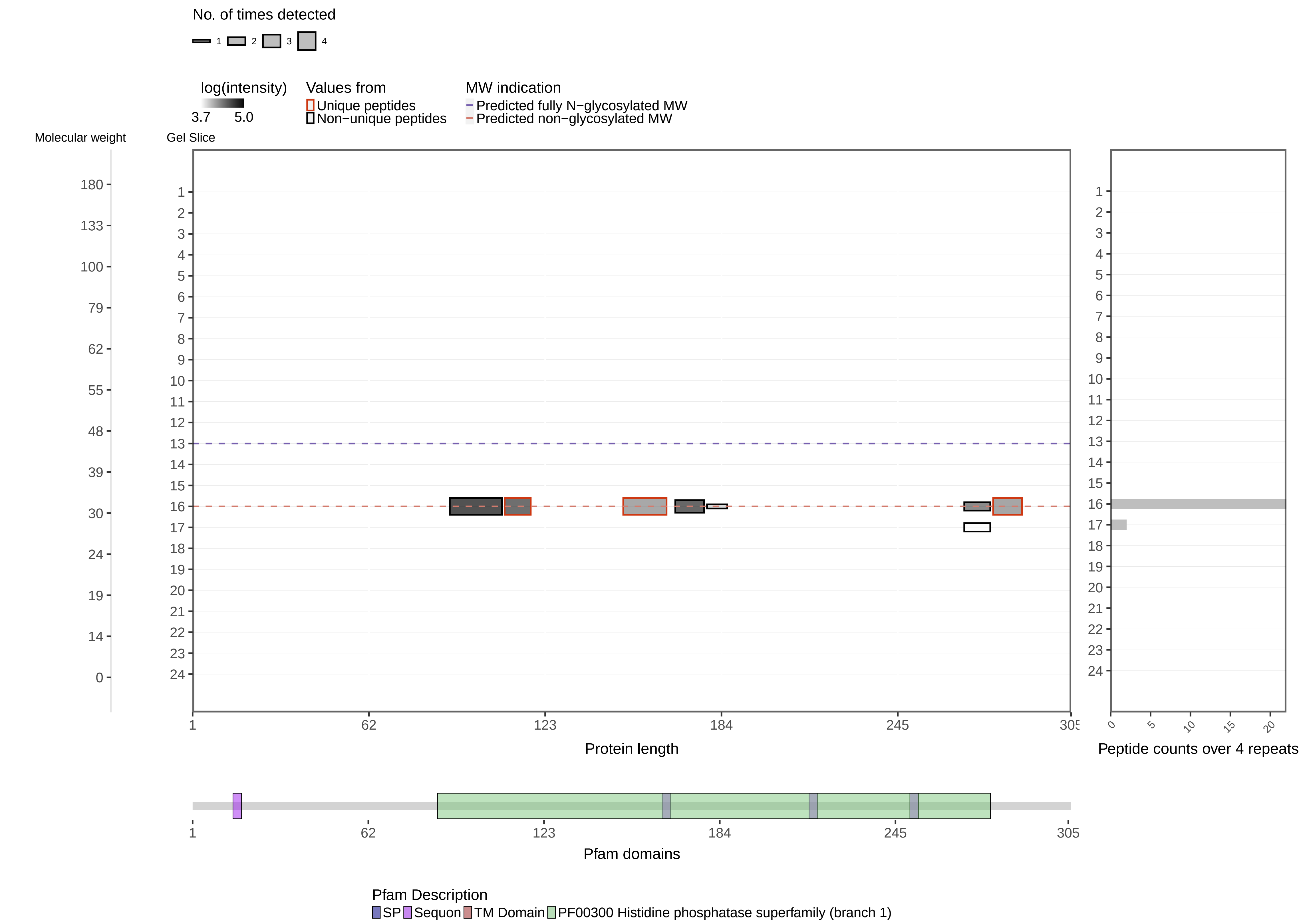


Protein sequence:

CWIWPSITQRAYQTAEIIAAVNGVNRSRIVPEYSFLDARGLGAYEGKSLDTLAEVYESDSLSPNIKPPPTDDGTPNESVSDVFVRVTQVMSVLETQYSEDTVIIVSPDSDNLTILQAGLTGLDLRRHRDLSFGPGEVRYVDGSSIPTYKQPASKVYKCLNPPSCT*

### NbL03g18210.1 UDP−glycosyltransferase_90A1−like [MW = 54.62 kDa]

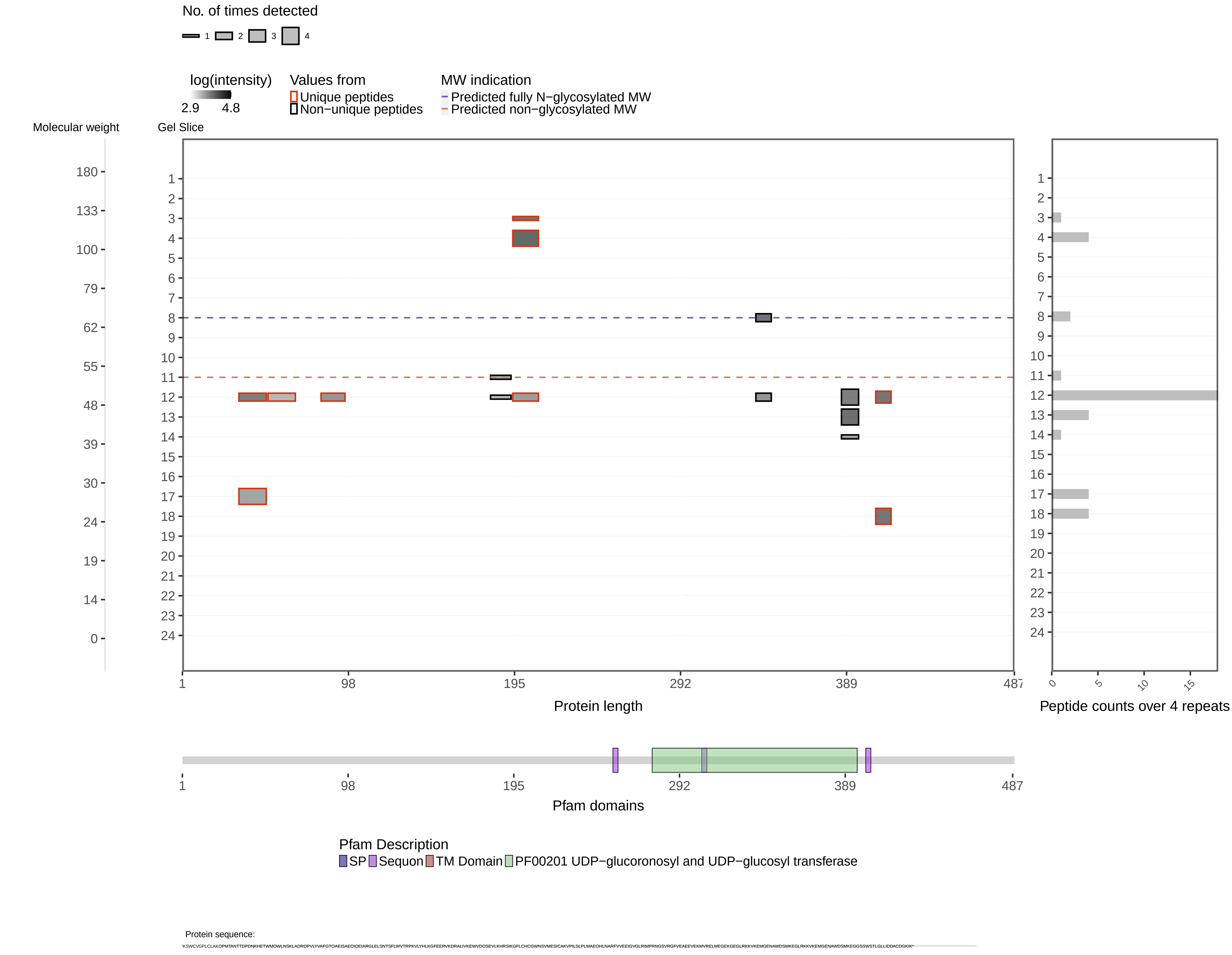

### NbL03g18170.1 aldehyde_dehydrogenase_family_2_member_B4_mitochondrial−like [MW = 58.53 kDa]

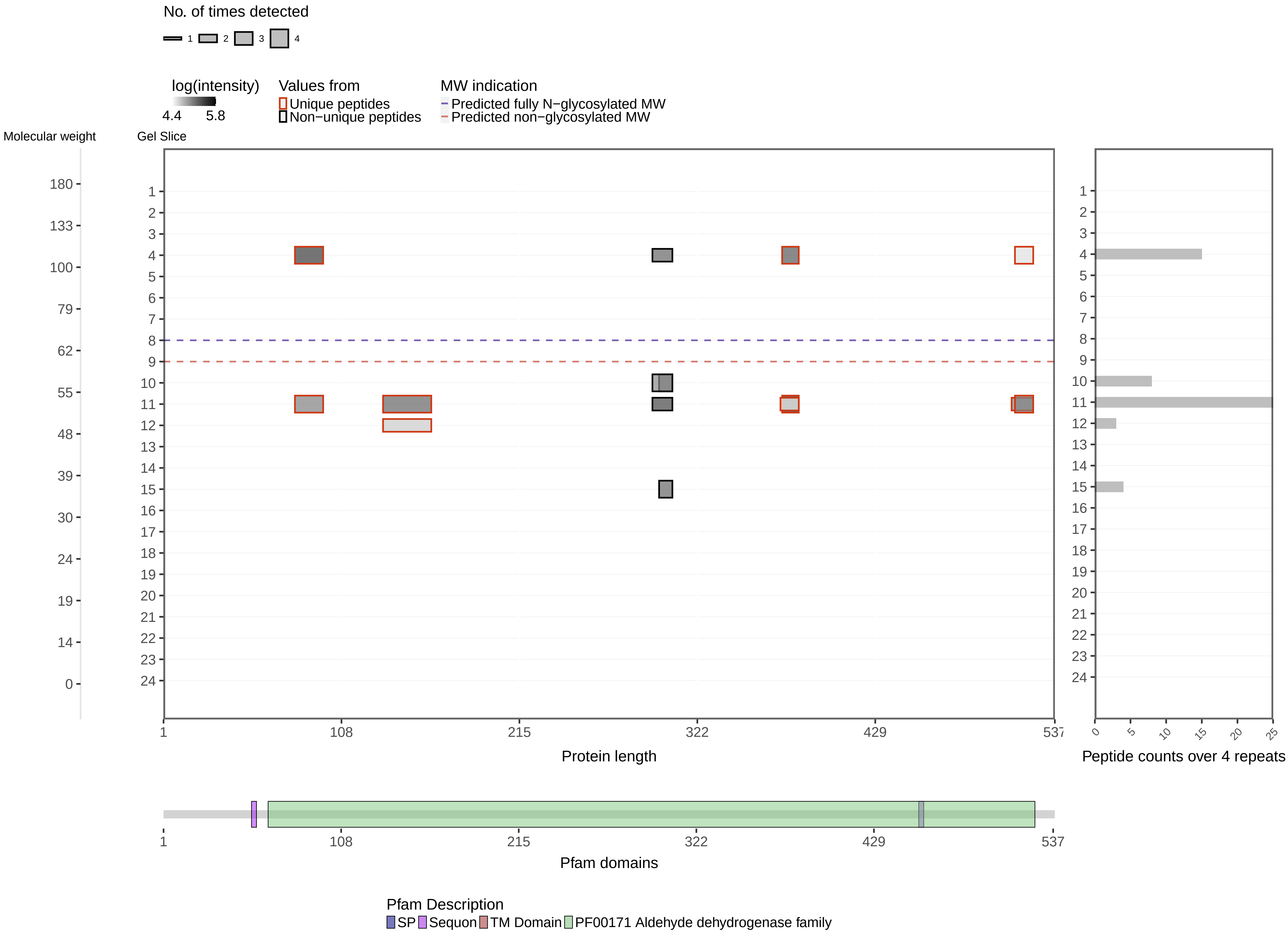


Protein sequence:

### NbL03g18140.1 protein_NUCLEAR_FUSION_DEFECTIVE_2 [MW − SP = 17.02 kDa]

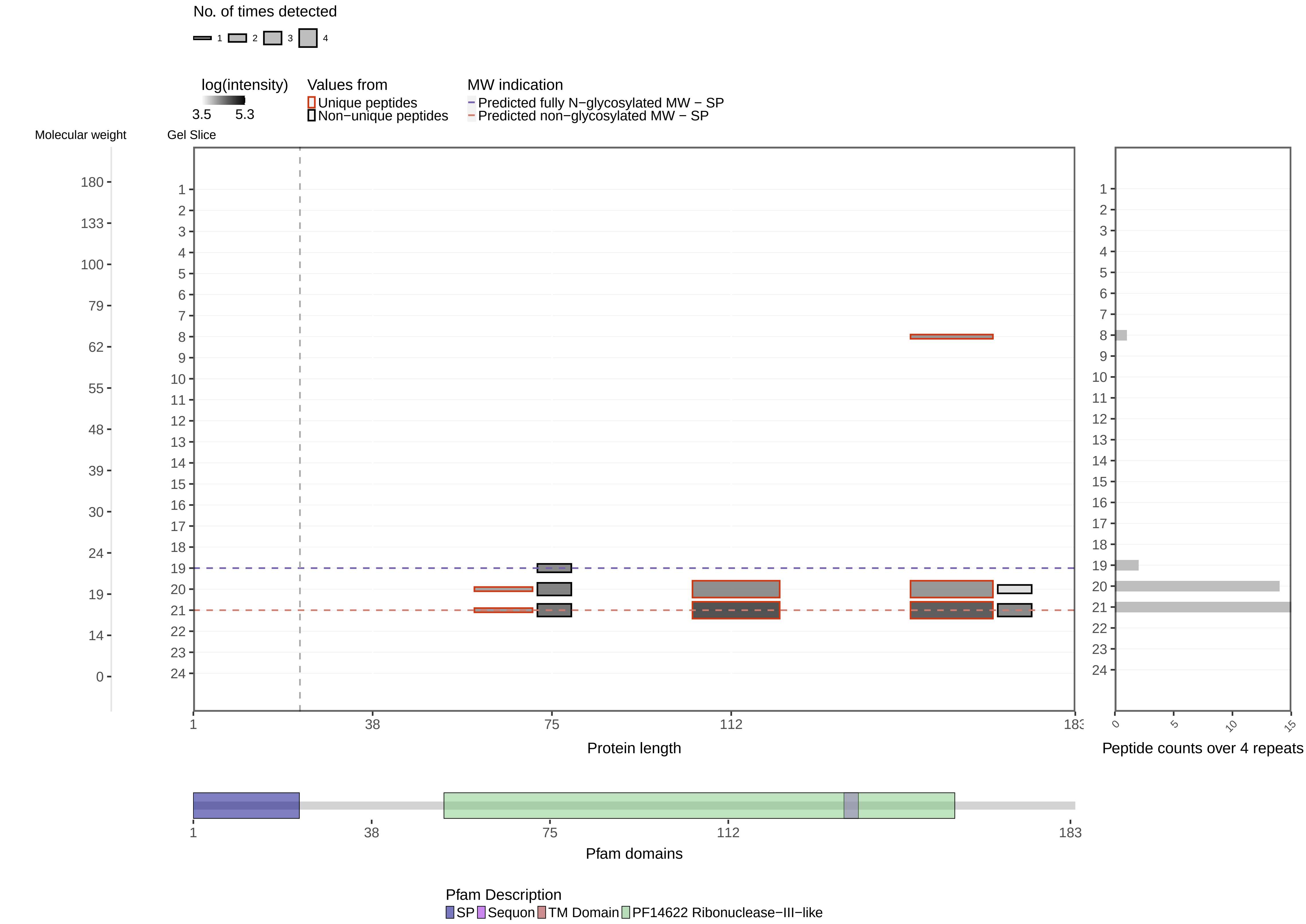


Protein sequence:
KAIEGSVSLQLLRKNIDMSPKDLNRAIAELSSNVVTSCAADGERLGLQKIVRVSHKTNSSAPAVVCGAFRAMFGAIAIDTSSLDNAGKVFFGIHGKGMEQAMAM*

NbL03g18090.1 39S_ribosomal_protein_L47_mitochondrial−like [MW = 18.97 kDa]

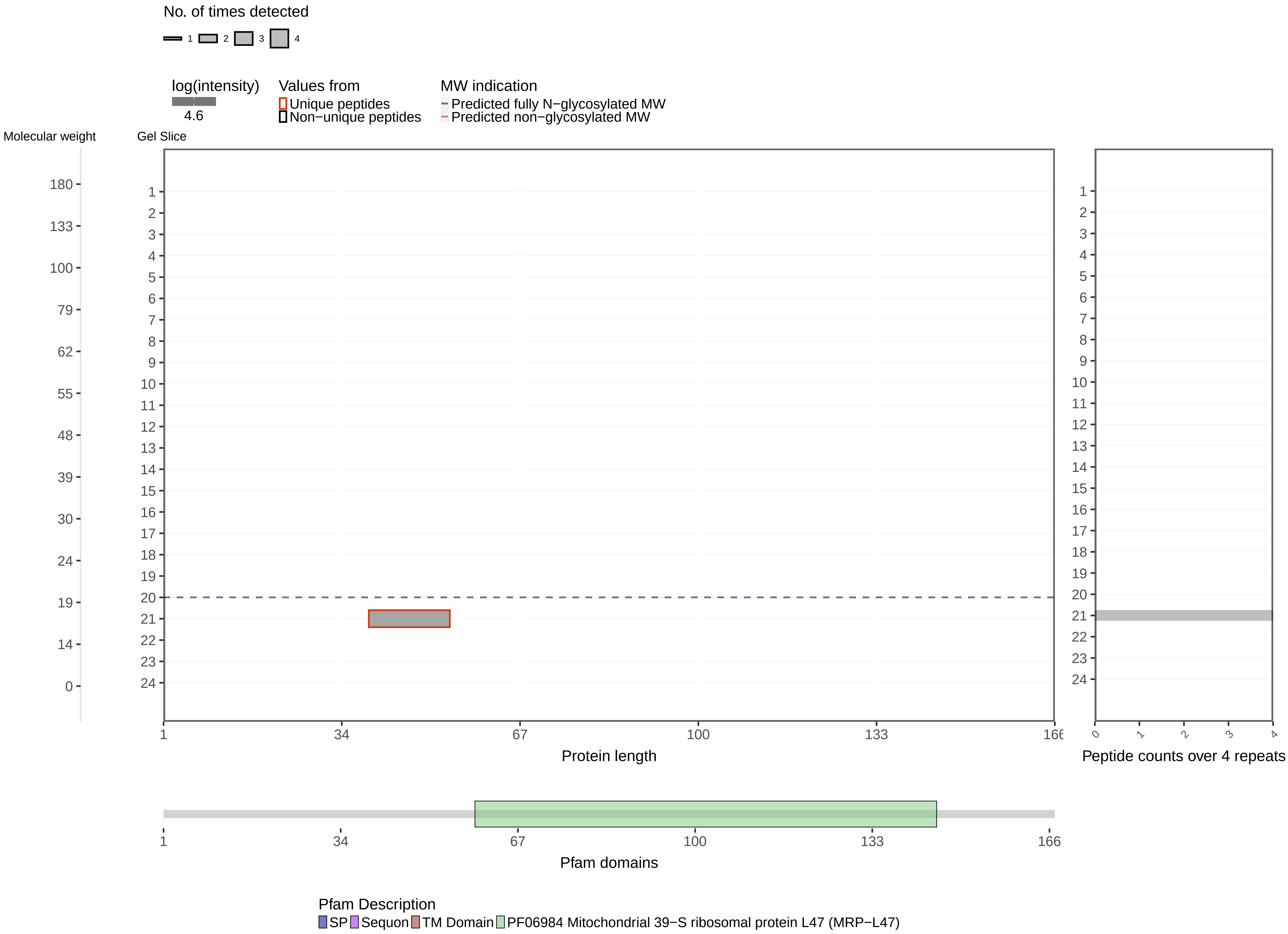


Protein sequence:

DKPVVYGRGWKASELRLKSWDDLHKLWYVLMKEKNMLMTQRQMLNAQNLRFPNPERISKVRKSMCRIKHVLTERAIDEADPRRSTEMKRMINAL*

NbL03g17910.1 betaine_aldehyde_dehydrogenase_chloroplastic−like [MW = 55.67 kDa]

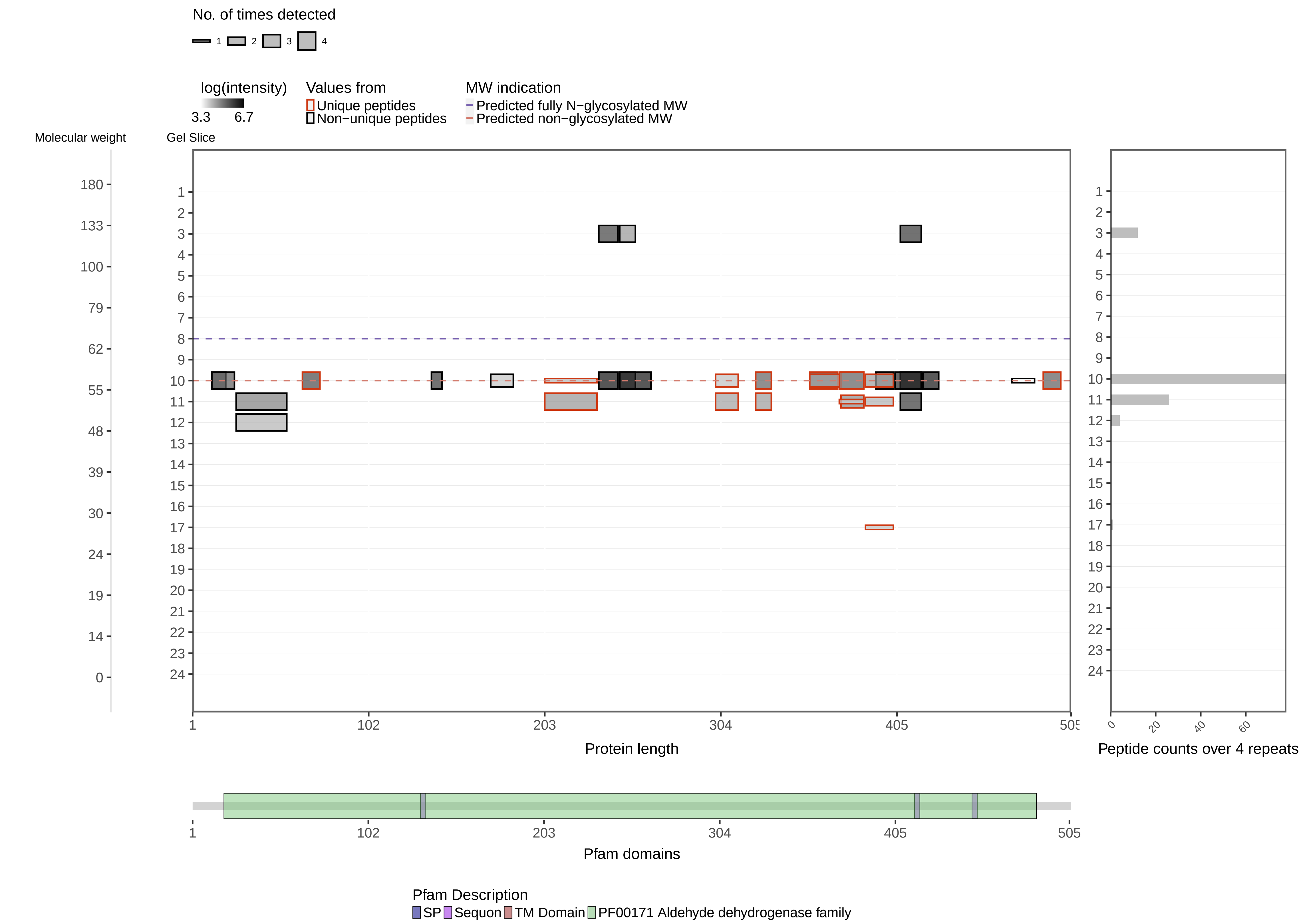


Protein sequence:

GPTGIKIMTAAAQLVKPVTLELGGKSPIVVFDDIPNIDITVEWTLFGCFWTNGQICSATSRLIIQDTIASQFLDRLLEWTKNIKISDPLGEDCKLGPVISRGQYEKVLKLISTAKNEGATILYGGDRPQHLKKGYYIQPTIITDVDRSMEIWKEEVFGPVLCVKTFKTEEEAIELANDTKYGLGAAILSKDLERCERFTKAFQSGIVWINCSQPCFWQPPWGGKKRSGFGRELGEWSLENYLNIKQVTRYVSDEPWAFYKSPSKL*

NbL03g17900.1 mannose−1−phosphate_guanyltransferase_alpha−B−like [MW = 45.58 kDa]

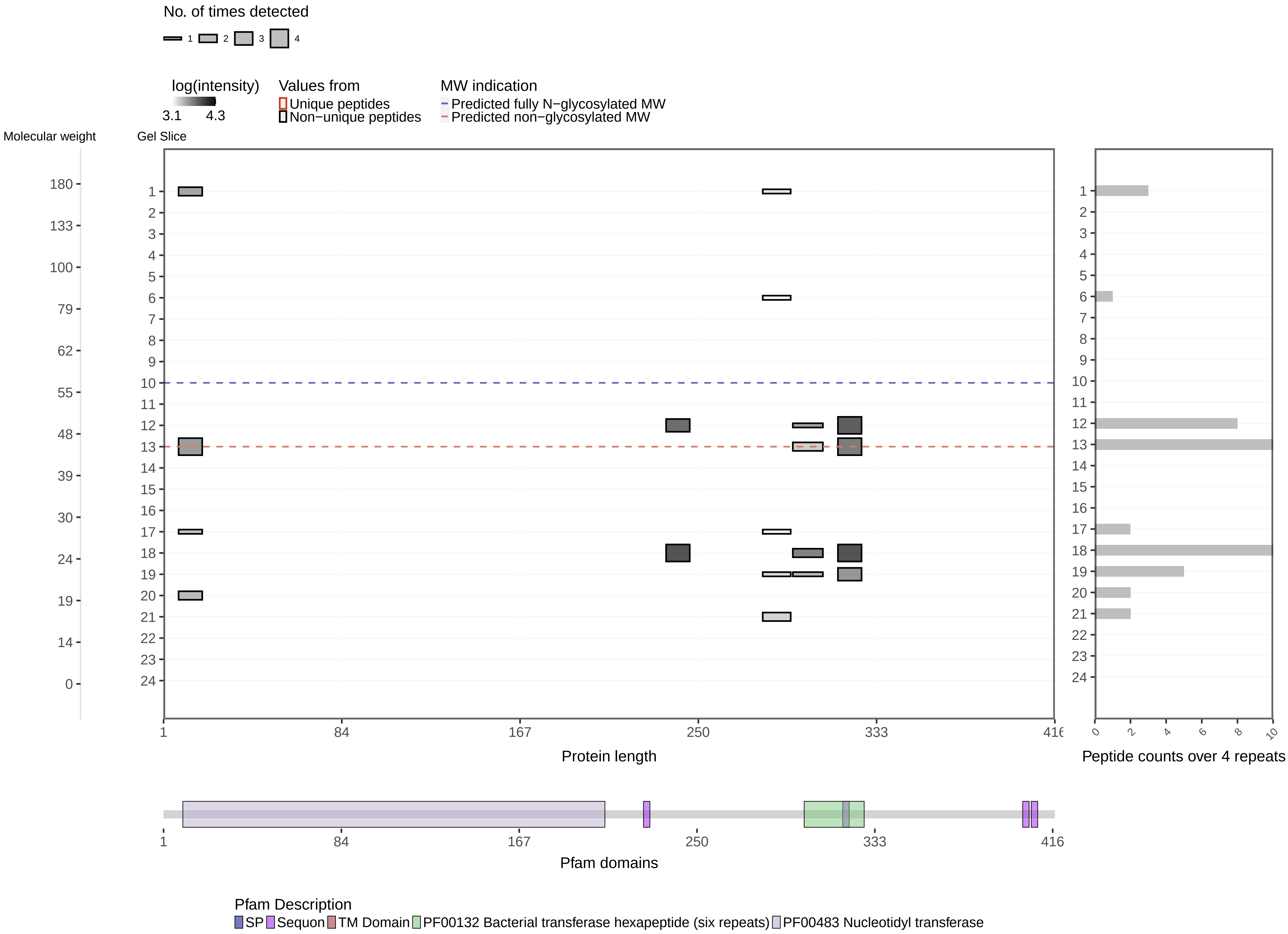


Protein sequence:

PDIFNAIQGVSSQRKDRANLRRVSSFEALQPVNRSLPTDFVRLDQDILTPLAGKKQLYTYETMDFWEQIKTPGMSLKCSGLYLALYRSTSPHLLASGDGSKTAAISGDVYIHPSAKVHPTAKIGPNVSISANARIGAGARLISCIILDDVEIKENGVVIHAIVGWKSSIGRWSRVQAAGDYNAKLGITILGESVSVEDEVVVINSIVLPNKTLNVSVQEEIIL*

NbL03g17890.1 pyridoxine/pyridoxamine_5'−phosphate_oxidase_1_chloroplastic−like [MW = 62.16 kDa]

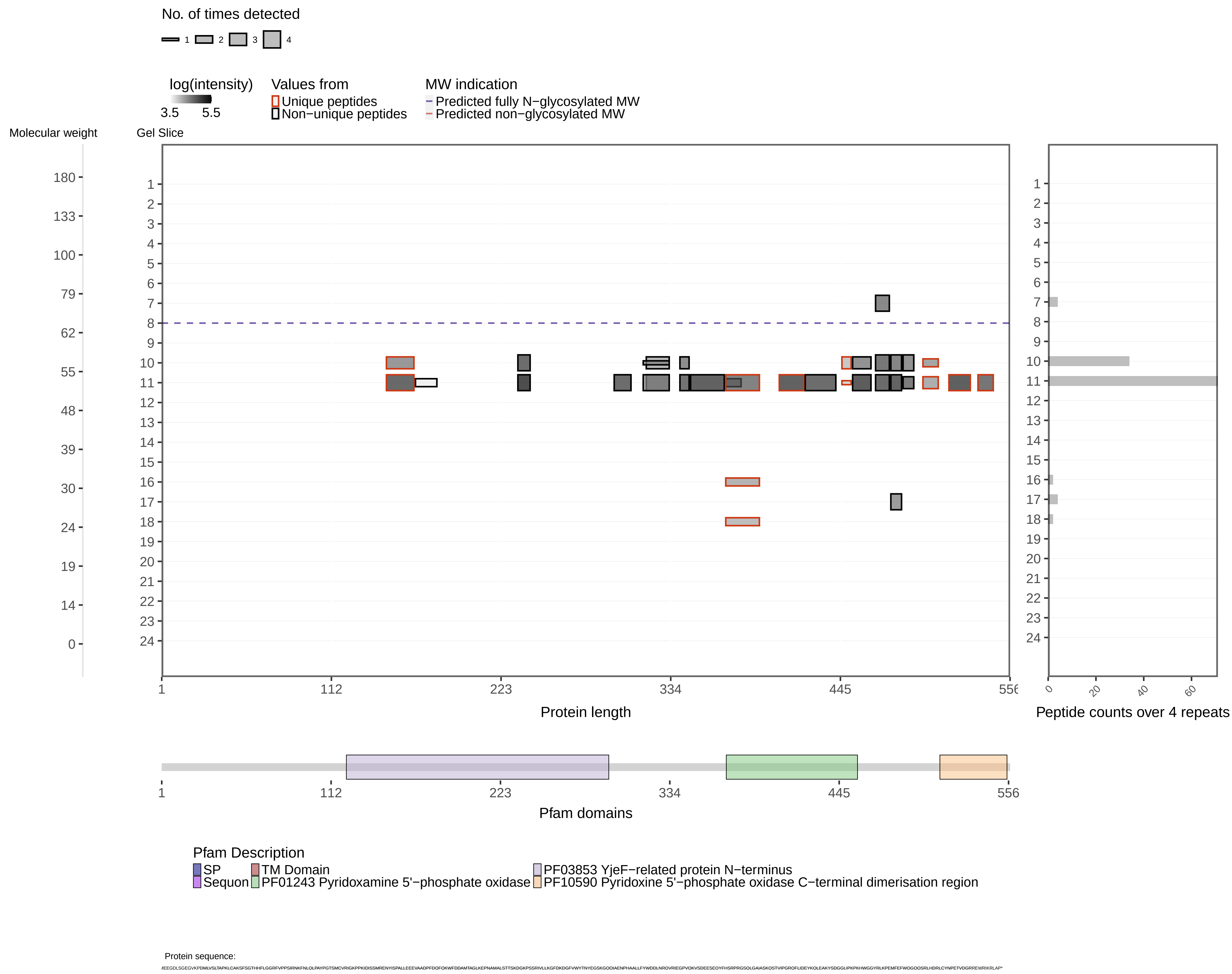

### NbL03g17770.1 5−formyltetrahydrofolate_cyclo−ligase_mitochondrial−like [MW = 31.68 kDa]

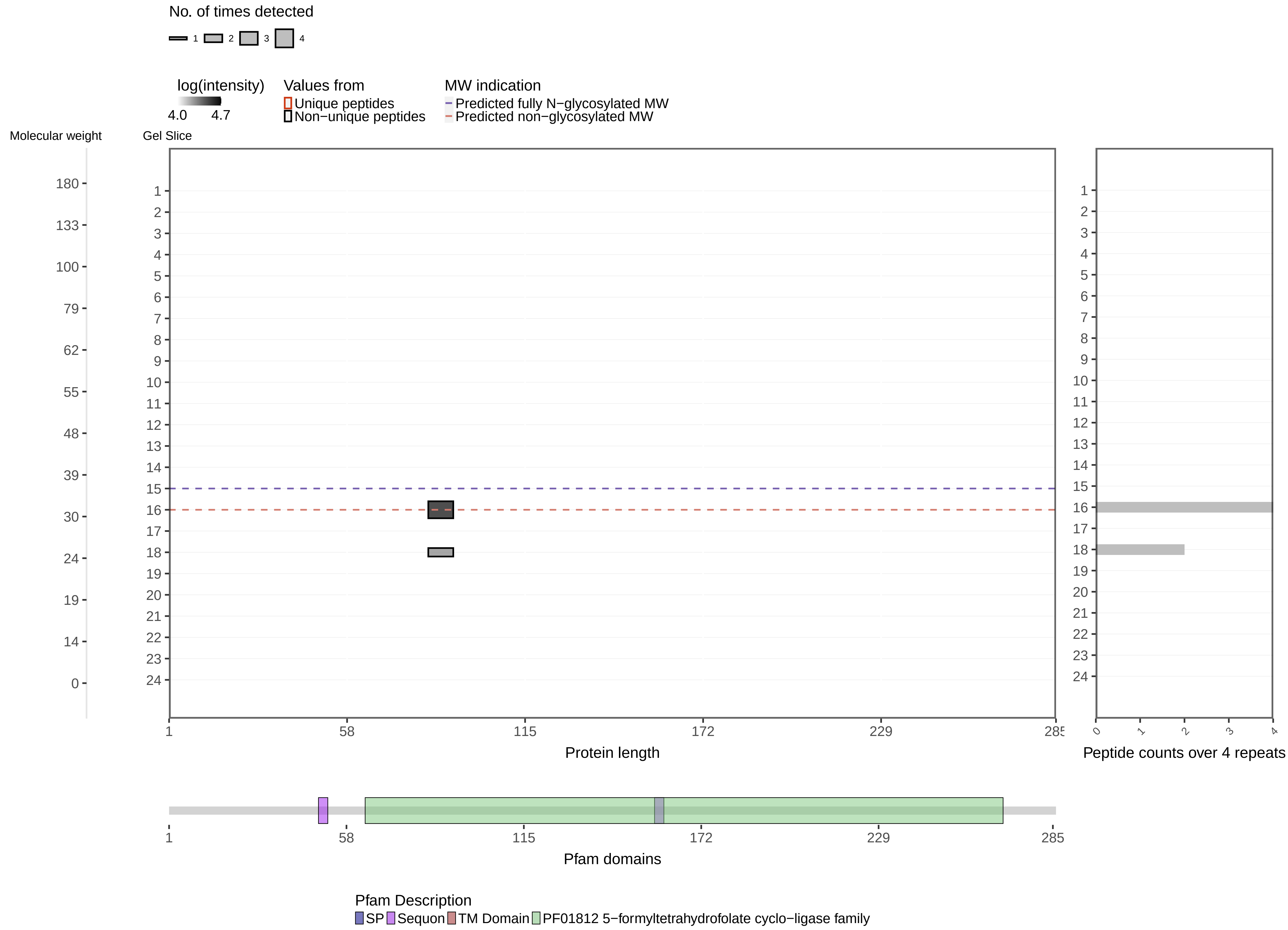


Protein sequence:

GSHPEDLQIQKKLFVPRVEDRNRNMRMLNISSTEDLIANSMDILEPAPVDADGNEREDVLFANEPVDLFLLPGLAFDKAGRRLGRGGGYYDTFLSRYQELAVDRNWKQPLKIALCYSMQIVDDGTIPLTPNDILVDALVSPSGVIPISPVALEICH*

NbL03g17740.1 HIPL1_protein−like [MW − SP = 72.57 kDa]

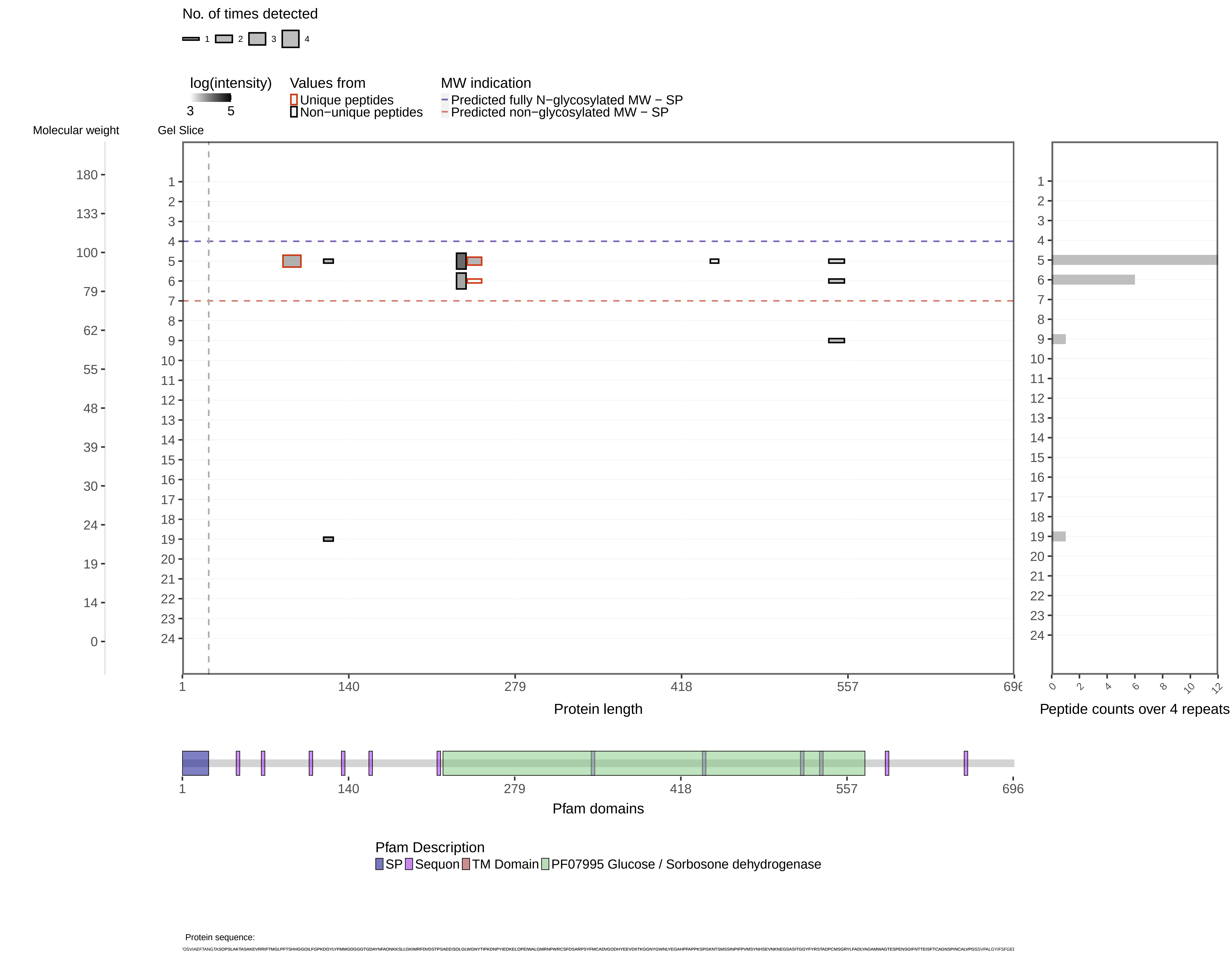

### NbL03g17690.1 heterogeneous_nuclear_ribonucleoprotein_1−like [MW = 34.84 kDa]

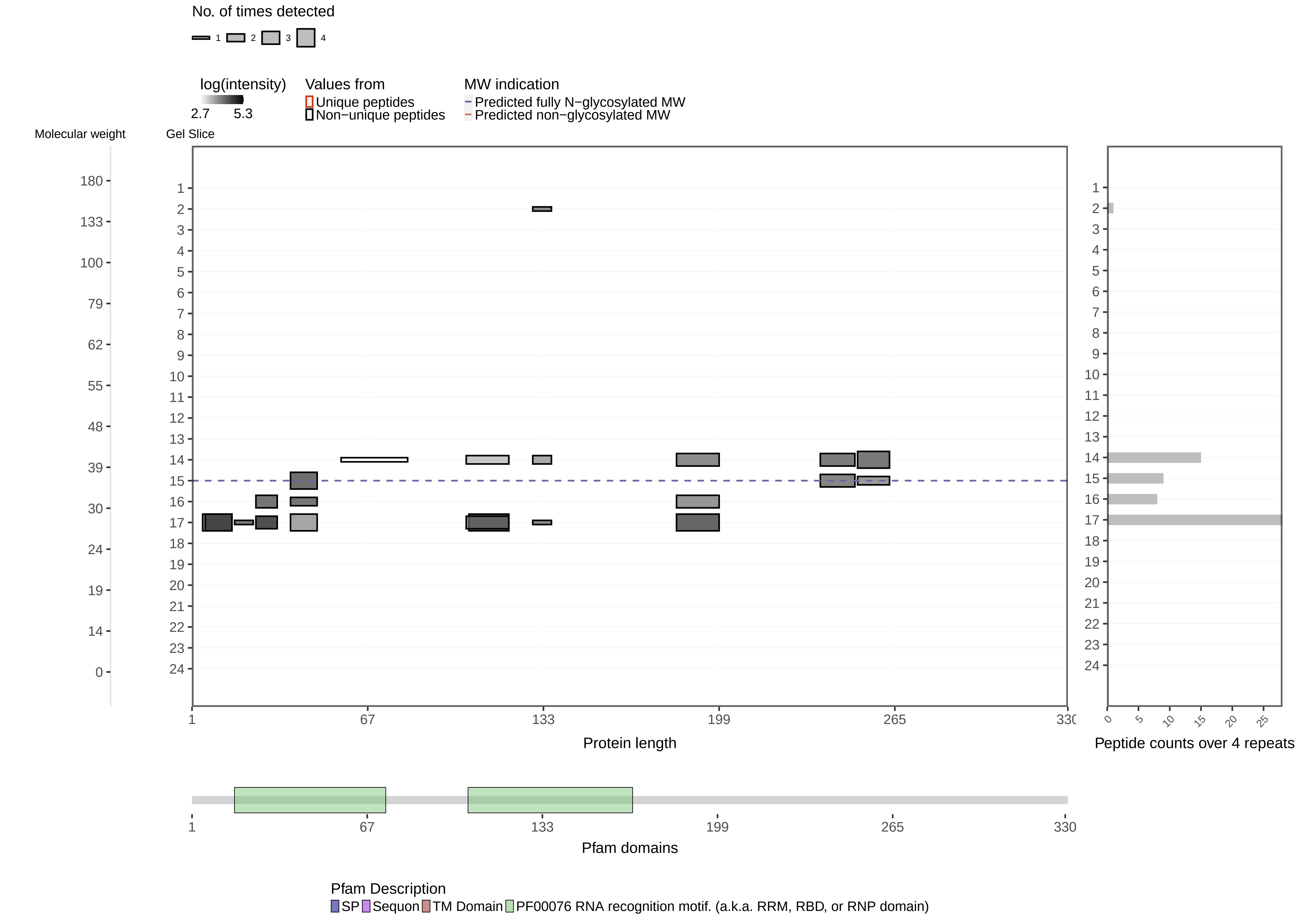


Protein sequence:

NVLSEGNMIDMLGTQVEIKKAEPKKPSNPASSGHGYGSESRGRGYSDSYGGFGNSYSSFGSGGFGPASYRSSGGMGGRFGDYGYGGGEYGGRYGDFGGGEFGGYRGDPSLGYSSRYGSYAGGFGGGAGGYGGSGLMAYGRGASYGGYGGAGSGAGYDSVLALVMVDQEIYM*

NbL03g17480.1 protein_QUIRKY [MW = 118.43 kDa]

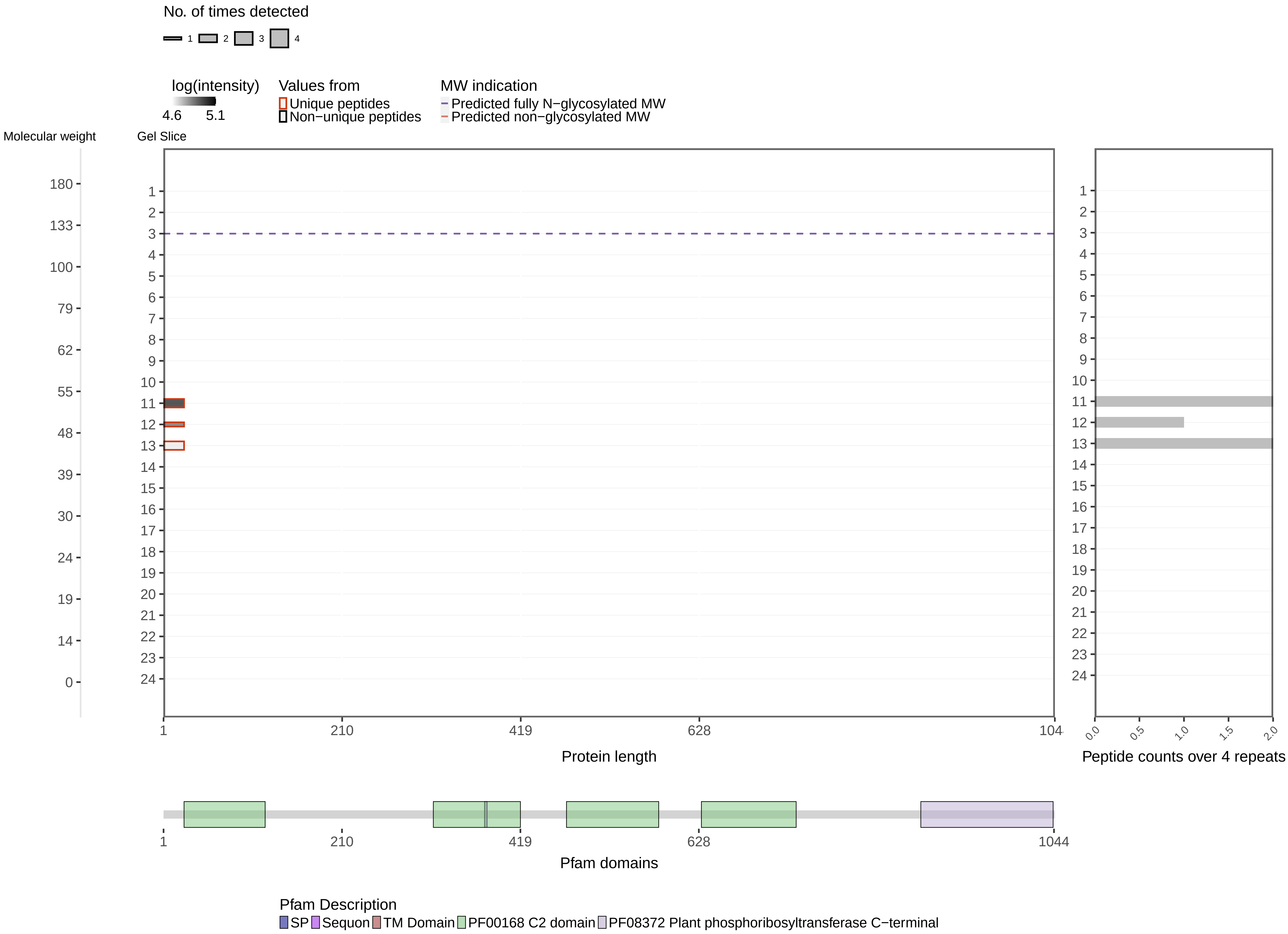


Protein sequence:

QLGFQSVRTRRGSMNHHSSAFHWNEDLIFVAGEPLEDGLILLVEDRTTKDPVLLGHIIIPVSSIEQQLDERLVSAKWFGLEGGPGGSYCGRVNLRMCLEGGYHVLDEAAHVCSDFRPTAKQLWKPPVGILELGILGARGLLPLKSKGPGKGSTDAYCVAKYGKKWVRTRTITDTFDPRWNEQYTWQVYDPCTVLTIGVFDNWHMFADAGDDKPDYPIGKVRIRVSTLENNKVYTNSYPLLVLLRSGLKKMGEIEVAIRFVCPSLLPETCAVYGQPVLPKMHYLRPLGVAQQE

### NbL03g17140.1 probable_3−hydroxyisobutyrate_dehydrogenase−like_1_mitochondrial [MW = 37.81 kDa]

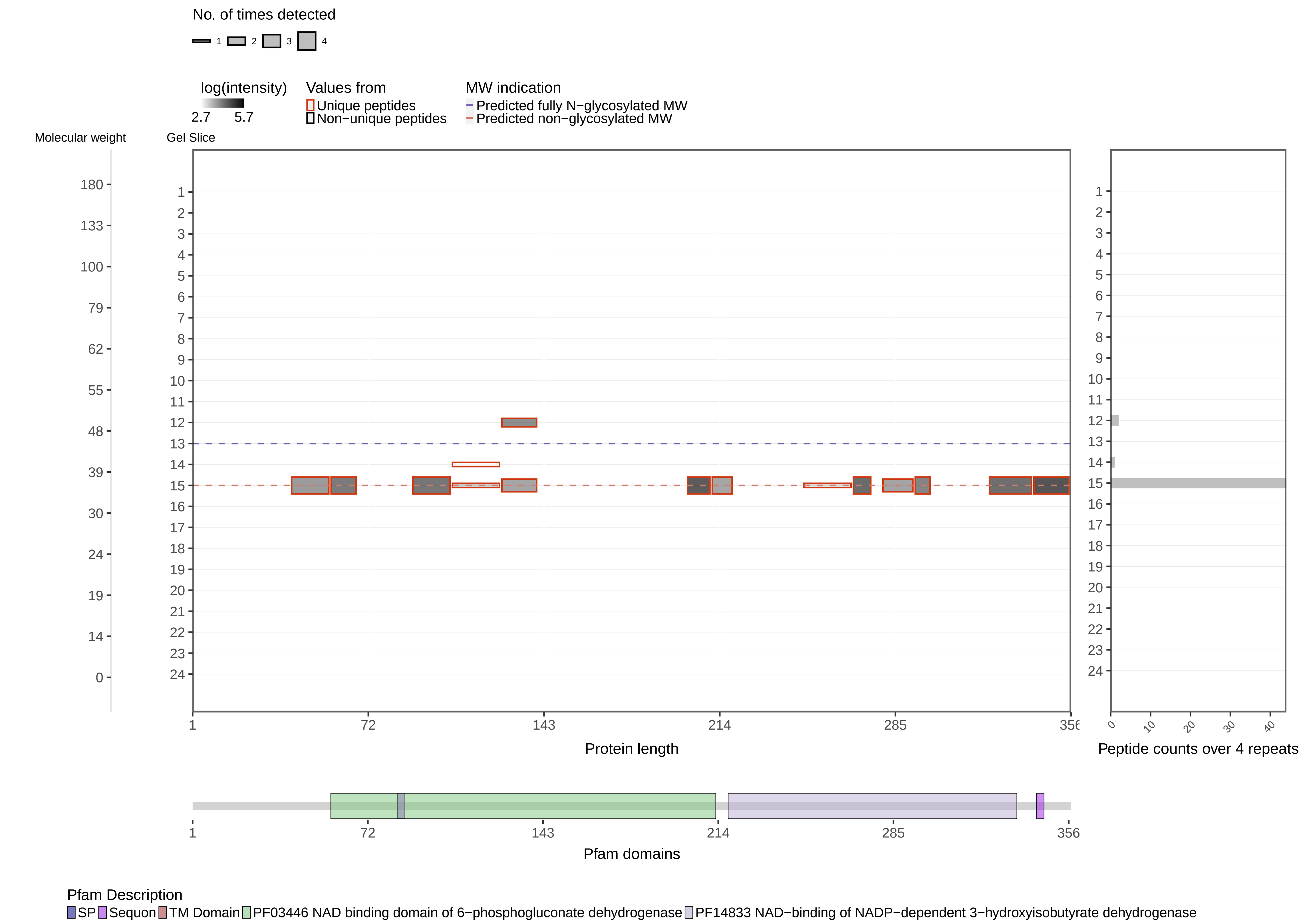


Protein sequence:

AAGCSAVDAPVSGGDLGARNAALSIFAGGEESIISRLNPLFNLLGRVYYMGAPGKGQFTKLGNQVVIASTMVGLCEGLIYAHKAGLDLNMFLNAISTGAAGSKSLDLYGSRIHCRDFEPGFYVNHFVKDLGICLRECQNMGLALPGLALAQQLYLSLKAHGEGNLGTQALILALERINNLSLASSSSVGKP*

### NbL03g16810.1 60S_ribosomal_protein_L27a−3 [MW = 16.46 kDa]

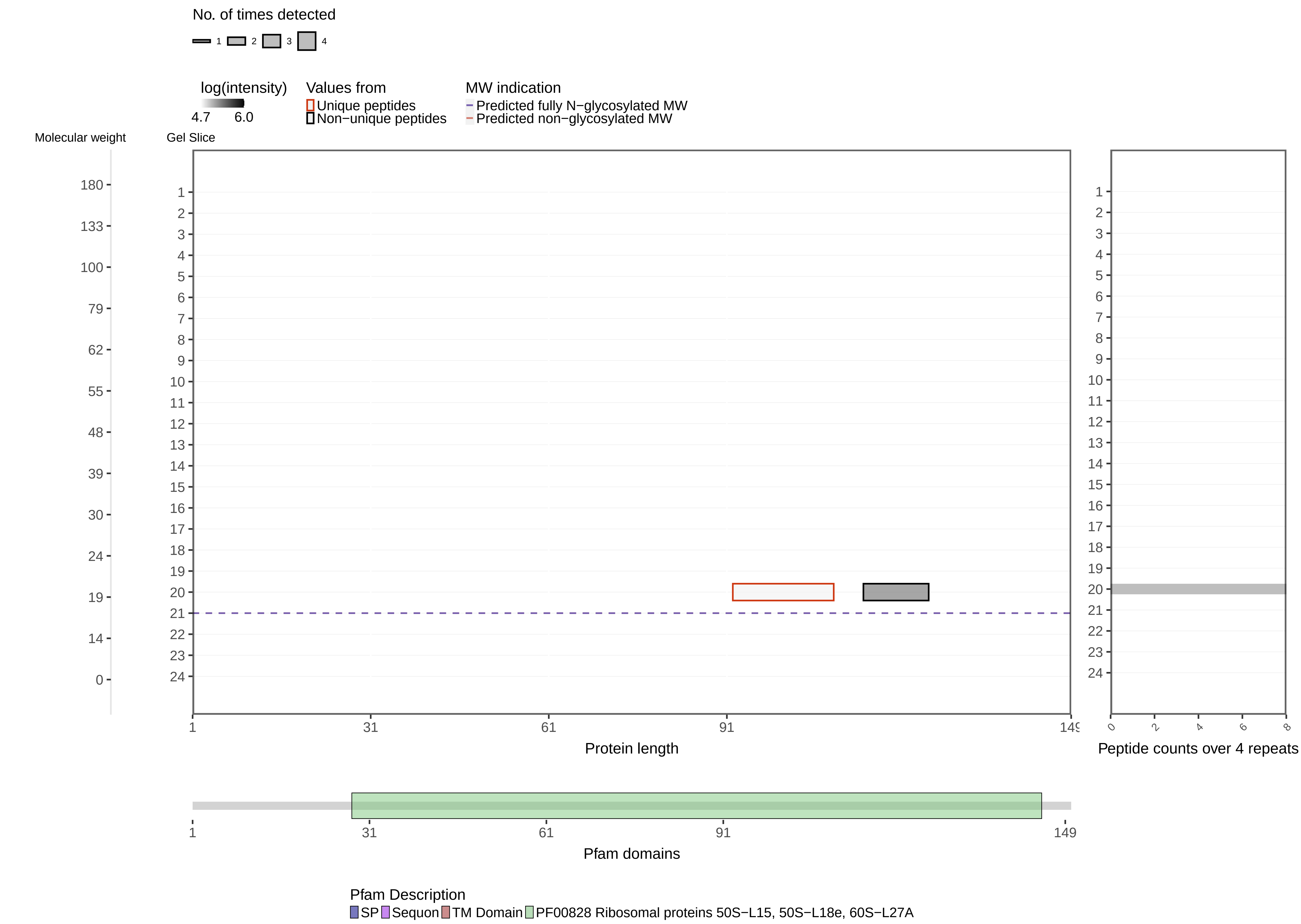


Protein sequence:

RYFHKLRNKFHCPTINIDNLWSLVPQEVKDKAKSSSGTAPLIDVTQYGYFKVLGKGVLPENQAVVVKAKLISKNAEKKIKEAGGAVVLTA*

NbL03g16740.1 histidinol_dehydrogenase_chloroplastic−like [MW = 53.14 kDa]

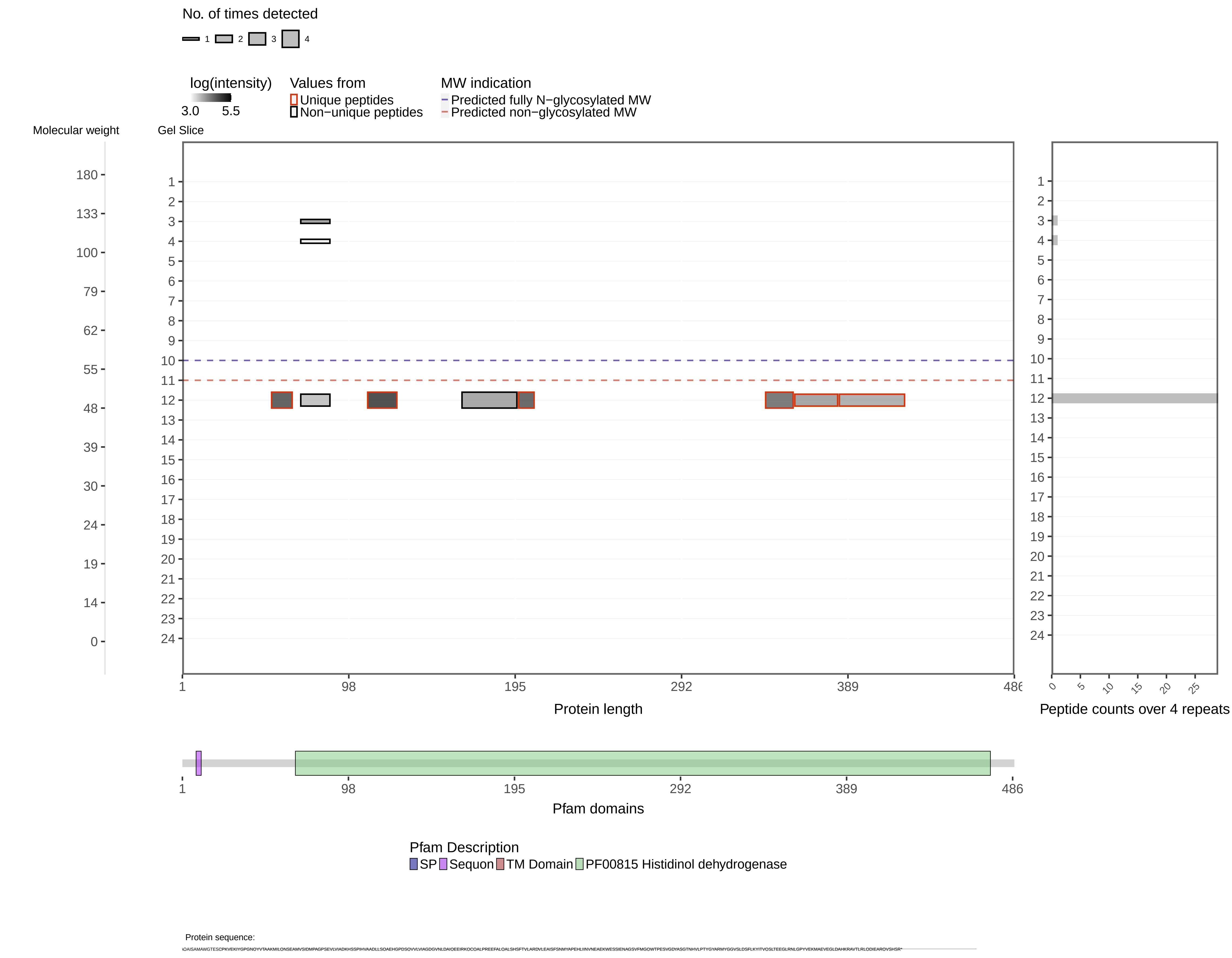

NbL03g16620.1 manganese−dependent_ADP−ribose/CDP−alcohol_diphosphatase−like [MW = 35.73 kDa]

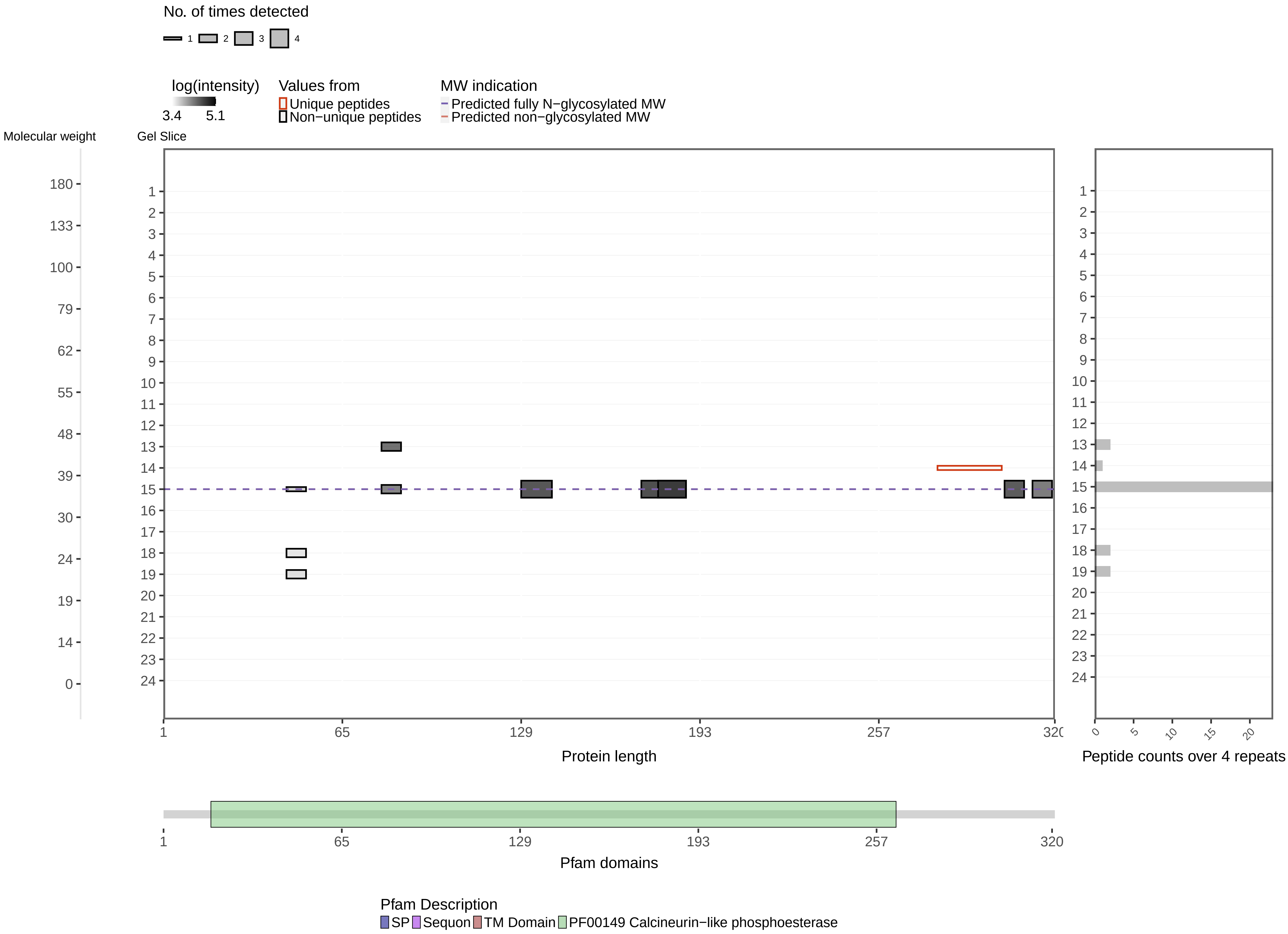


Protein sequence:

AIGWPEDHPNTLKALKVLQERNPNSDKNSPNGLVGLERRFLKFNGGVGKEQLEWLDHVLQEATELNQNVVVCCHLPLDPGASSFAALLWNYDEVMDVIHCYSCVKVCLGGHAHKGGQSVDSHGVHHRVLEAALECPPGTDAFGHIDAFSDRLSLFGNDRMKSTEMVFGH*

### NbL03g16530.1 indole−3−glycerol_phosphate_synthase_chloroplastic−like [MW = 45.03 kDa]

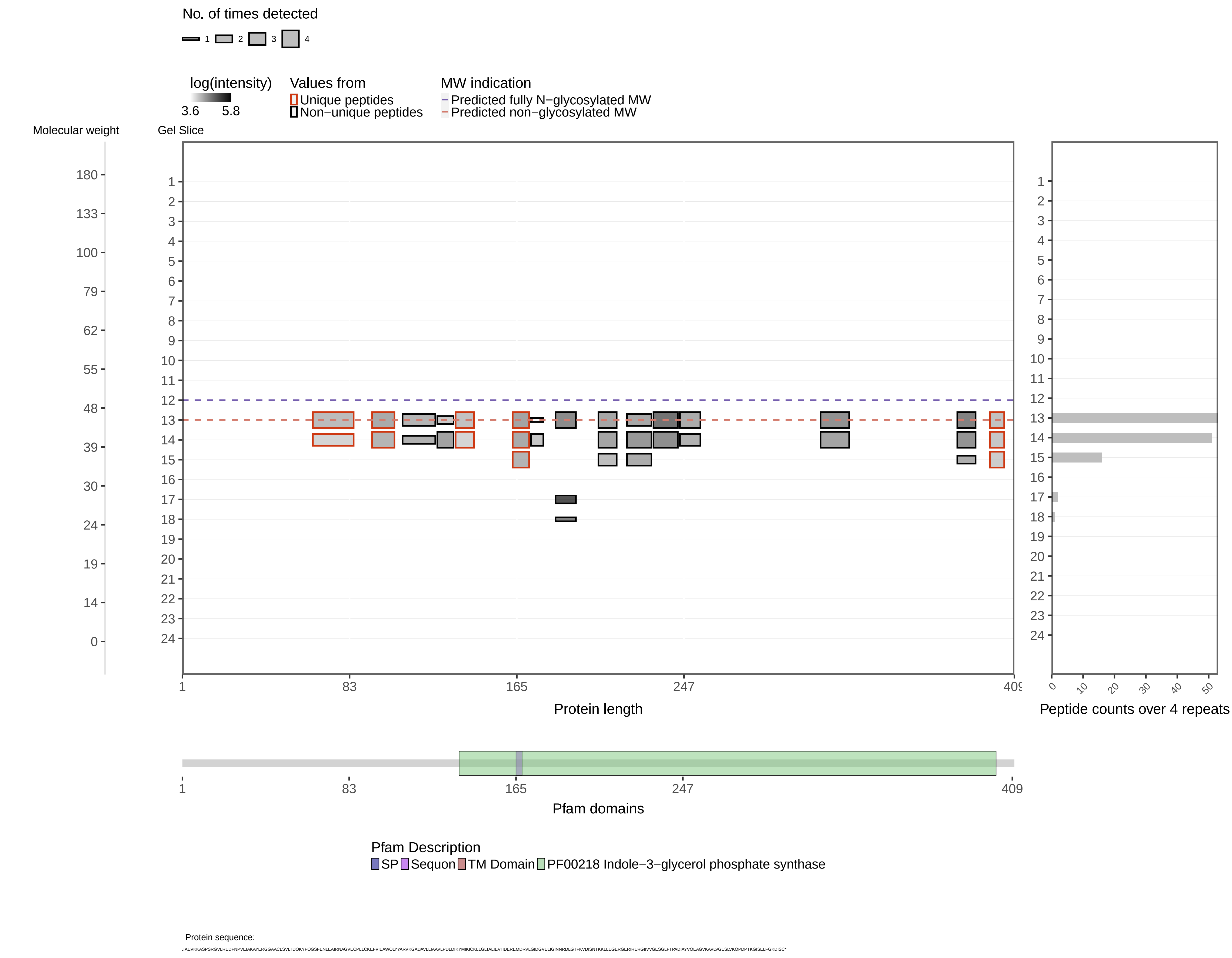

### NbL03g16510.1 28_kDa_ribonucleoprotein_chloroplastic [MW = 32.19 kDa]

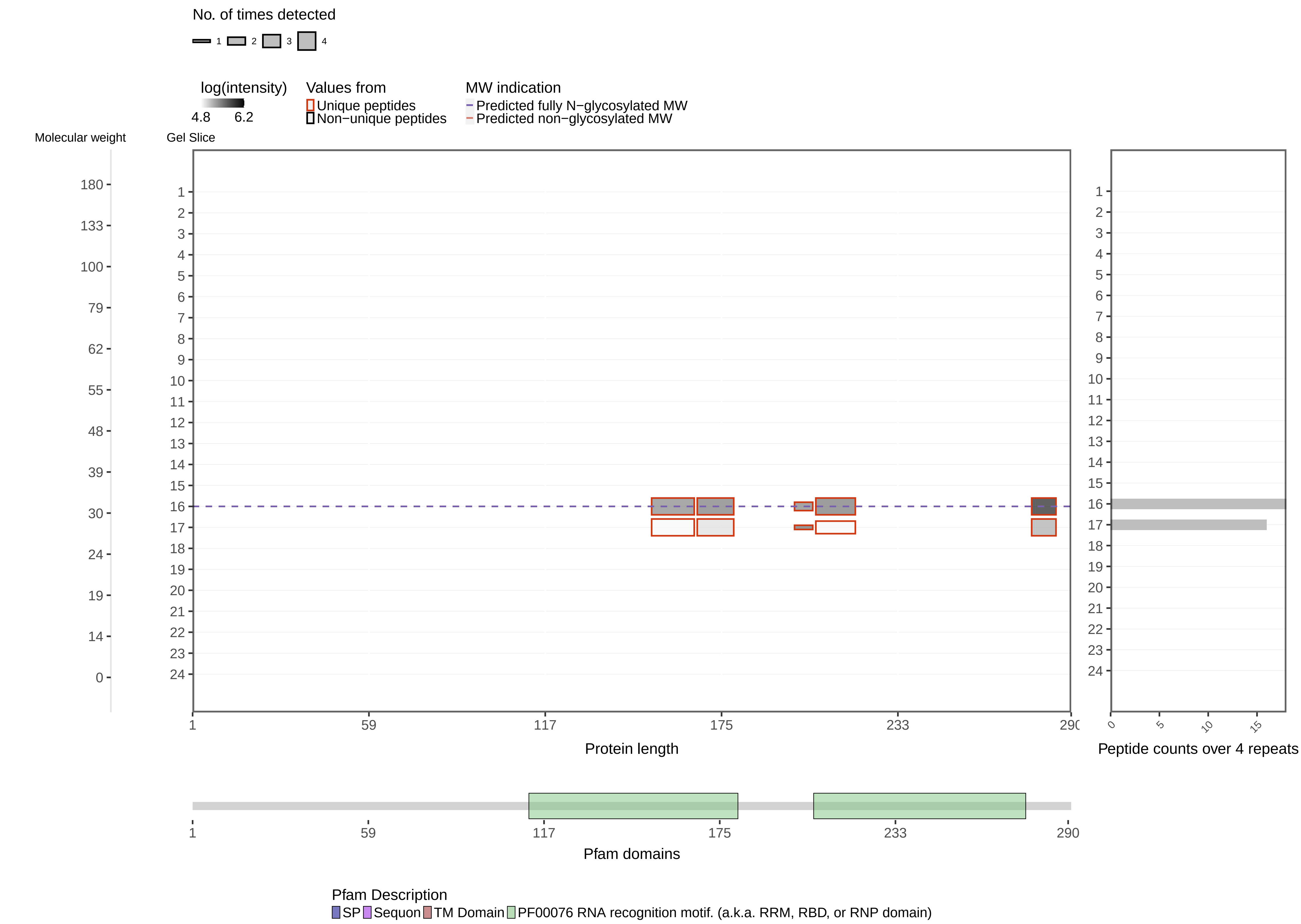


Protein sequence:

GVVEIAEVIYNRETDRSRGFGFVTMSTVEEADKAVELYSQYDLNGRLLTVNKAAPRGSRPERPPRTFQPTYRIYVGNIPWDIDDARLEQIFSEHGKVVSARVVFDRESGRSRGFGFVMMSSEAEMSEAIANLDGQTLDGRTIRVNAAEERPRRNTY*

### NbL03g16360.1 uncharacterized_LOC107819570 [MW = 116.33 kDa]

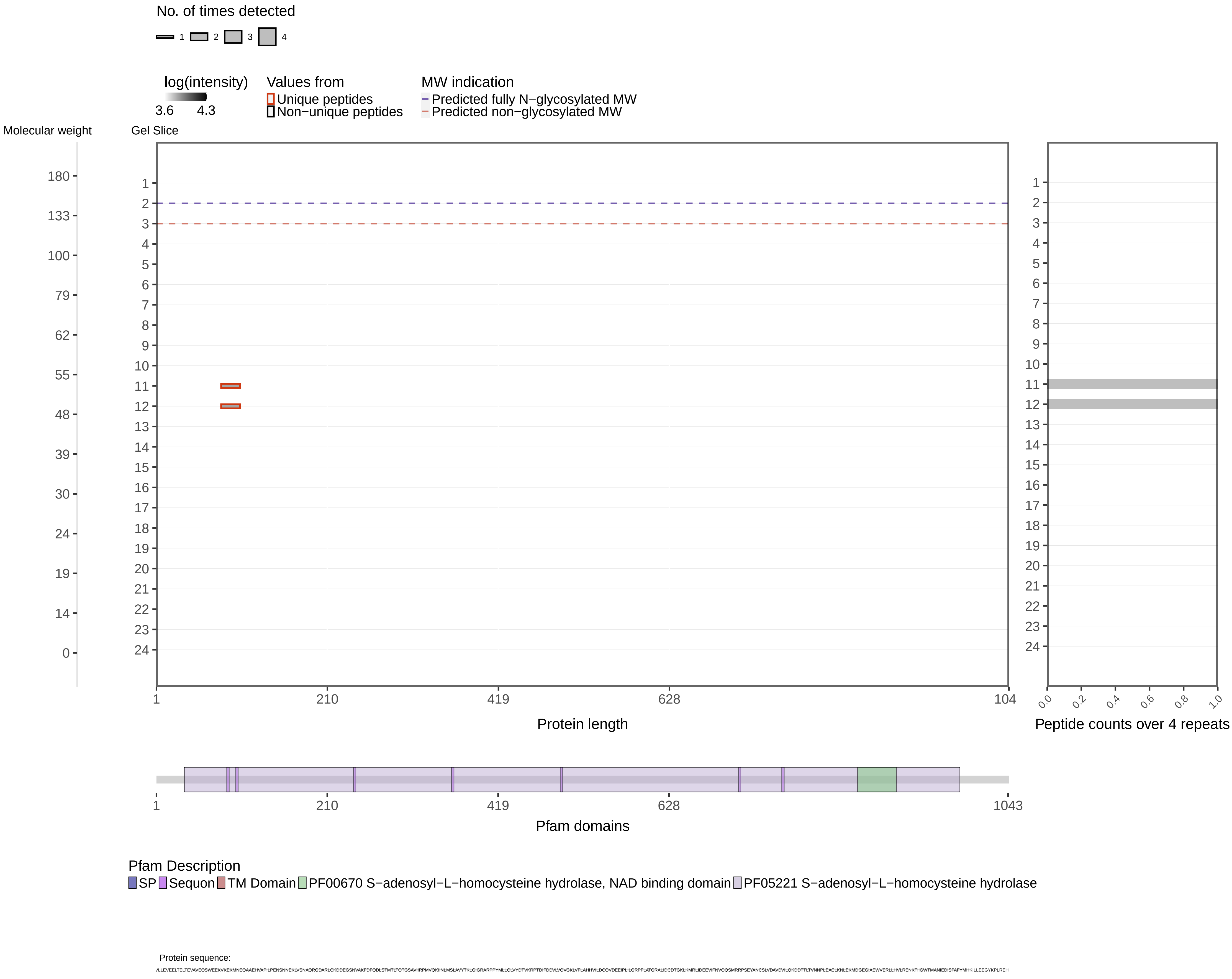

NbL03g16310.1 haloacid_dehalogenase−like_hydrolase_domain−containing_protein_At3g48420 [MW = 33.64 kDa]

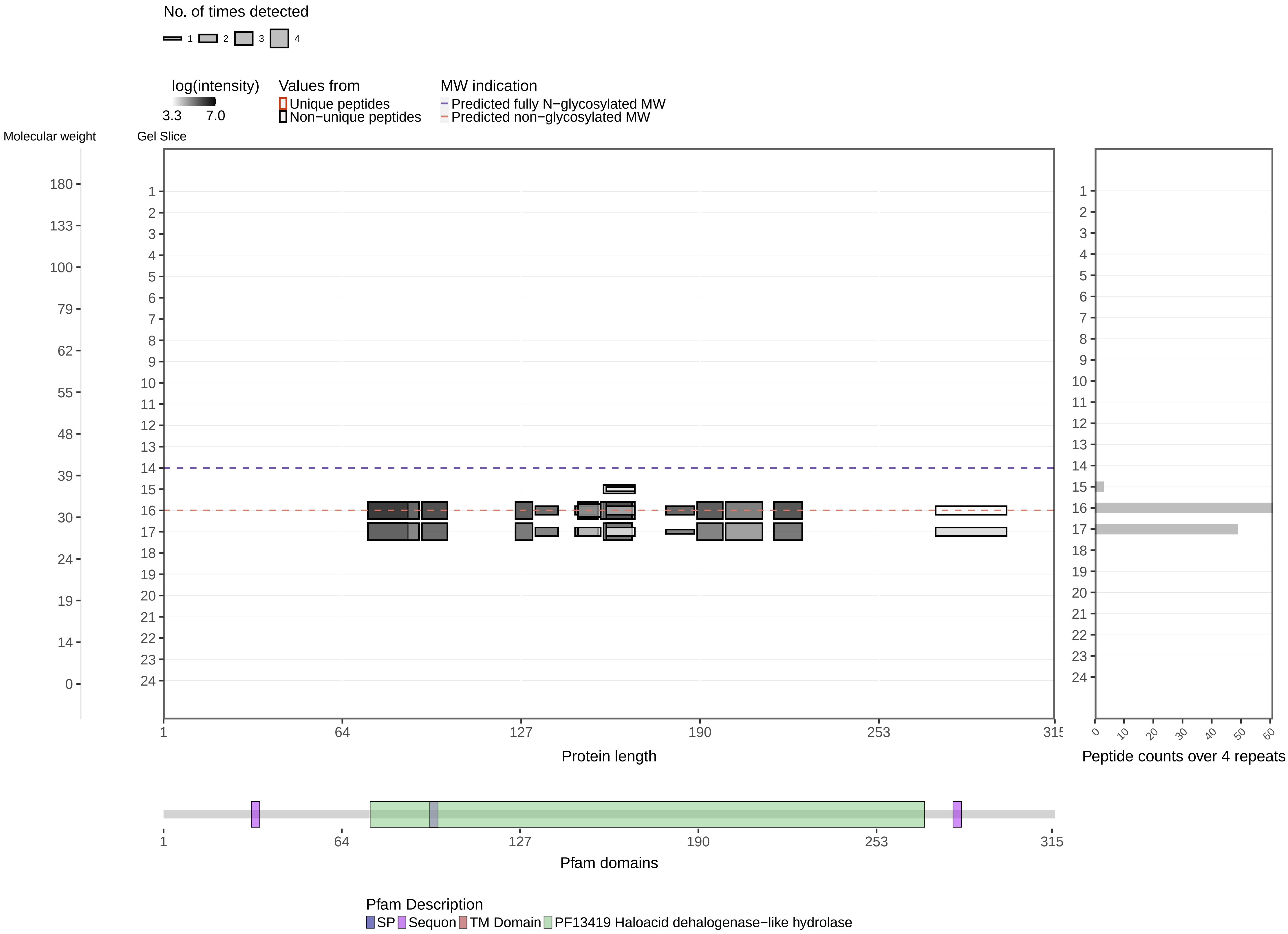


Protein sequence:
ERKQFIAGLHKRKTELFMALIEKKLLPLRPGVAKLVDQALGNGVKVAVCSTSNEKAVSAIVSCLLGPERAEKIQIYAGDVVPRKKPDPAIYLLAADTLGVDPSSCVVVEDSGIGLAAAKAAGMKCIVTKSGYTADENFSNADAVFDCIGDPPEERFDLAFCGSLLEKQYVS*

### NbL03g16220.1 ubiquitin−like_modifier−activating_enzyme_5 [MW = 46.66 kDa]

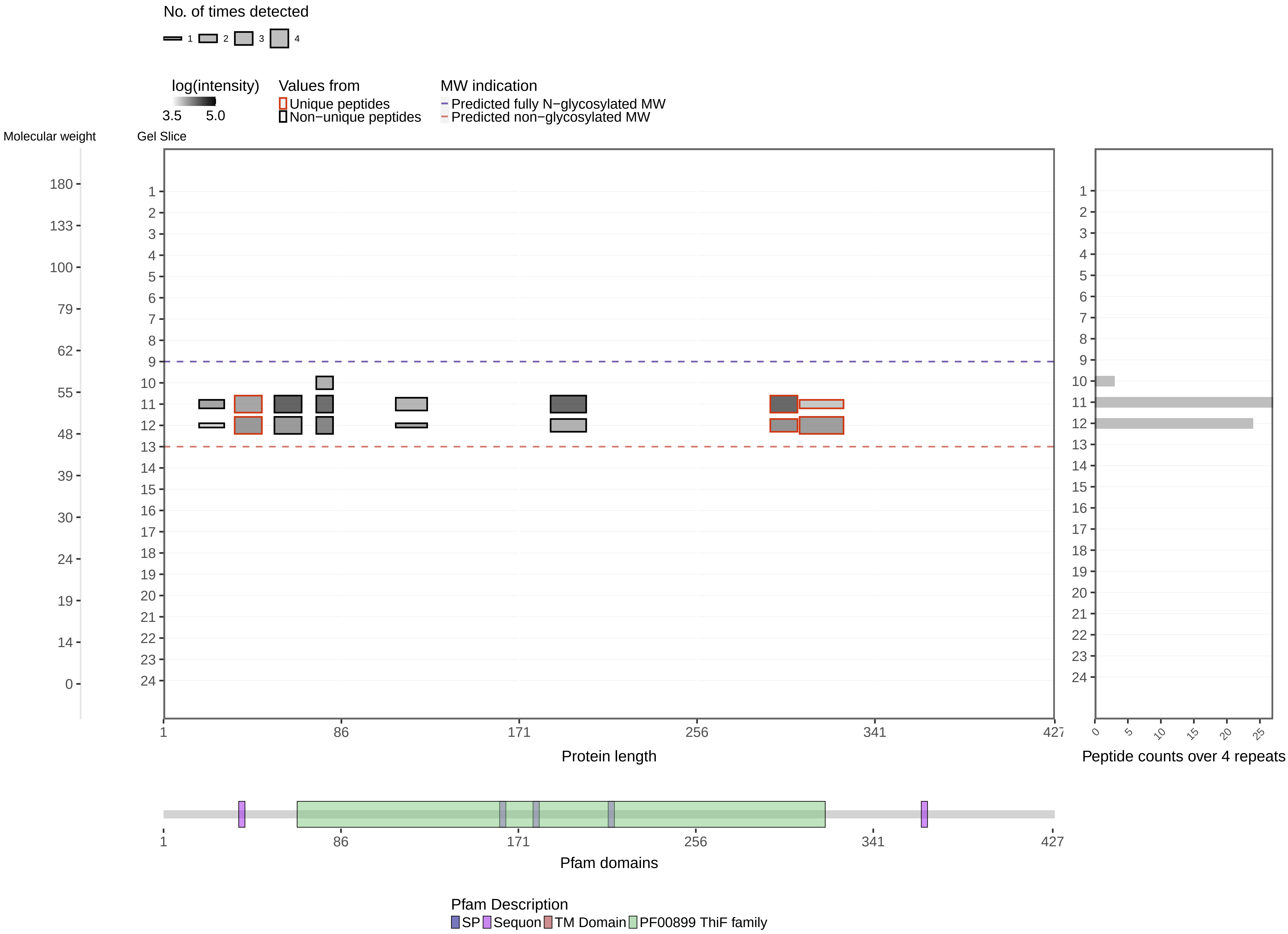


Protein sequence:

RMVVNQACNELNQTWMESGVSEDAVSGHIQLLIPGESACFACVPPLVVASGIDERTLKREGVCAASLPTTMGVVAGLLVQNTLKYLLKFGQVSPYLGYNALKDYFPTMEMKPNTQCSNAACLERQKEYILAKPTRDAAAKAKADKEALSAPENPVHADNEWNISVVDDNEVDGTVTGHSGVLPEGIVHELPVADEYPKPAITETVGPADDLEELRRQLEALNAE*

NbL03g16130.1 uncharacterized_LOC104224837 [MW = 93.85 kDa]

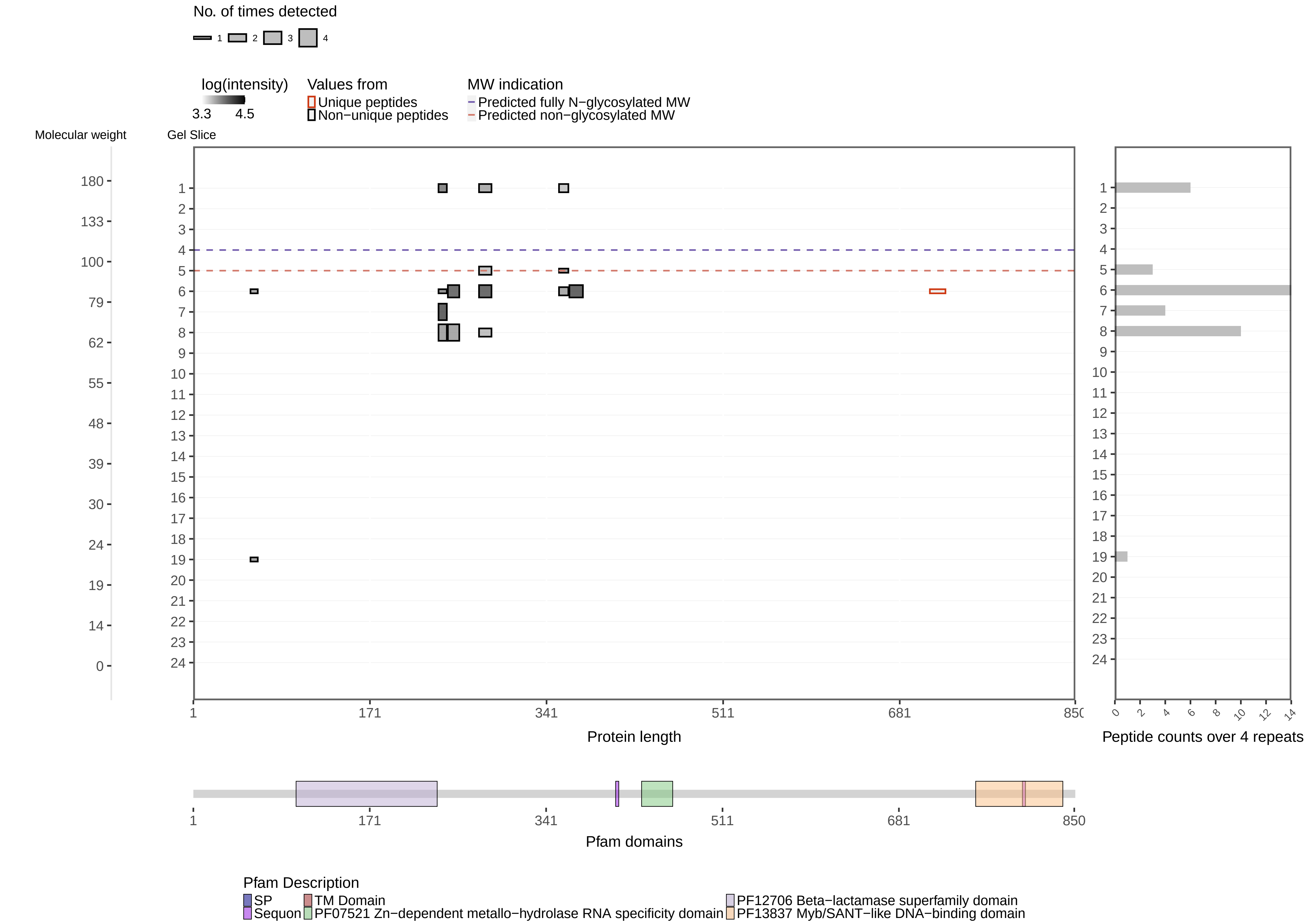


Protein sequence:

[illegible]

### NbL03g16090.1 probable_aspartyl_aminopeptidase [MW = 52.65 kDa]

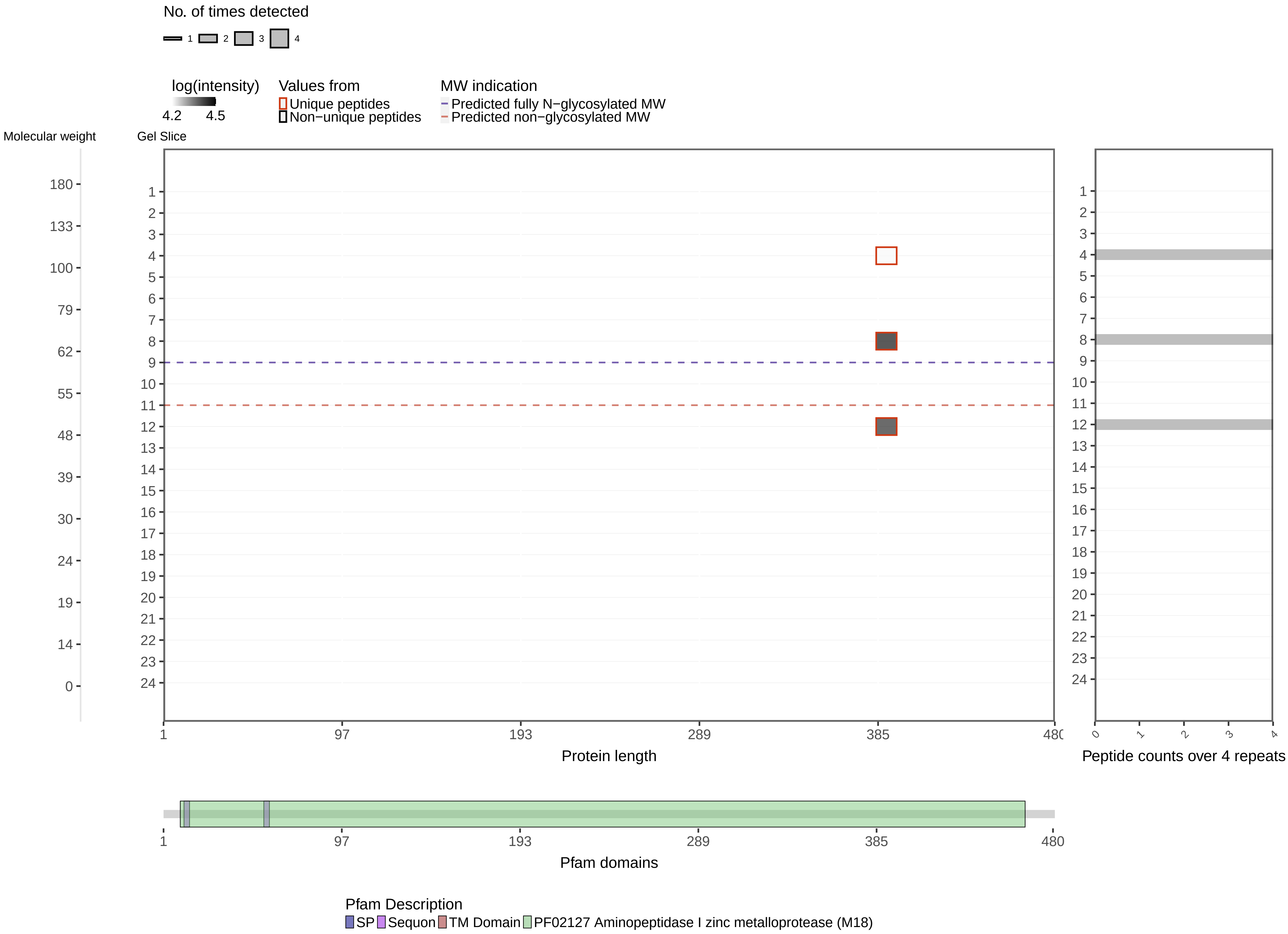


Protein sequence:

.AAQAGCQPSDICDFELQACDTQPSVIAGAMKEFVFSGRLDNLCSSFCSLKALIDATSQNSLEDEVGVRMVALFDHEEVGSDSAQGAGSPVMFDALSRITSTFSPDLKLIPKAIQRSFLVSADMAHALHPNYPDKHEENHQPKFHGGLVIKHNANQRYATNAVTSFLFREIAAKHNIPIQDFVIRNDMPCGSTIGPILASGIGIRTVDVGAPQWSMHSIREMCAVDDVKHAYHHFKAFFQDFSHLDAKIAVDV*

NbL03g16080.1 4−coumarate−−CoA_ligase−like_9 [MW = 59.66 kDa]

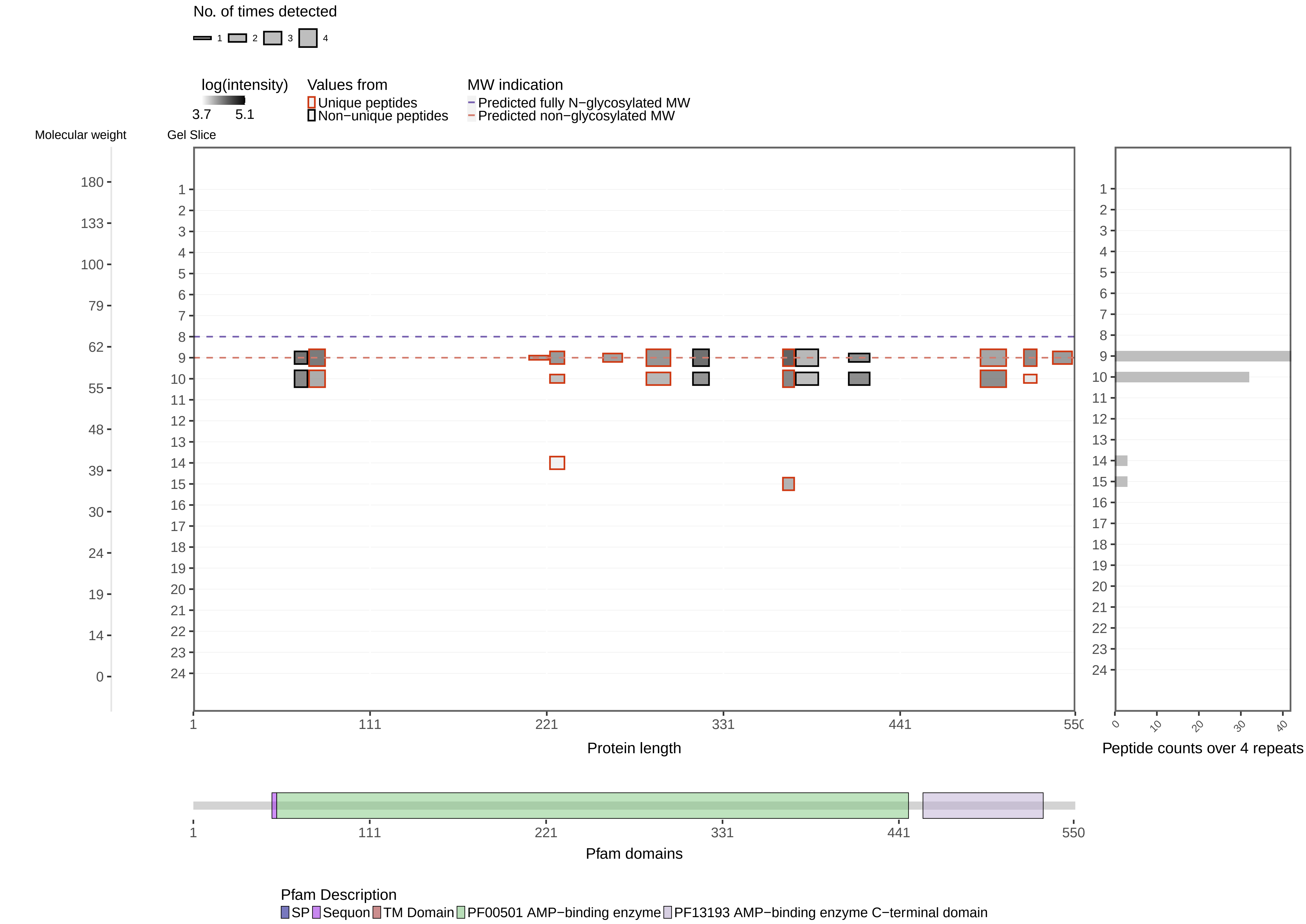


Protein sequence:

MIERFDFEKMLAAVEKYRVTYMPVSPPLVVAMAKSDLALKHDLSSLKLLACGGAPLGKEVAERFKRRFPNVEIIQGYGLTETTGGATGMSGPEESDRYGSAGRLAVNVEAKIVDPDNGETLPPGQRGELWIRGPTIMKGYVGDKQATSATLNSEGWLKTGDLCYFDADGFLYVVDRLKELIKYKAYQVPPAELEHLLQSIPDVADAAVIPYPDEEAGEIPMAYIVRRPGSTISEAQIIDAIAKQVAPYKKIRRVAFIDAIPKSPAGKILRRELVNHASSGASARL*

### NbL03g15940.1 nucleoside_diphosphate_kinase_2_chloroplastic [MW = 25.38 kDa]

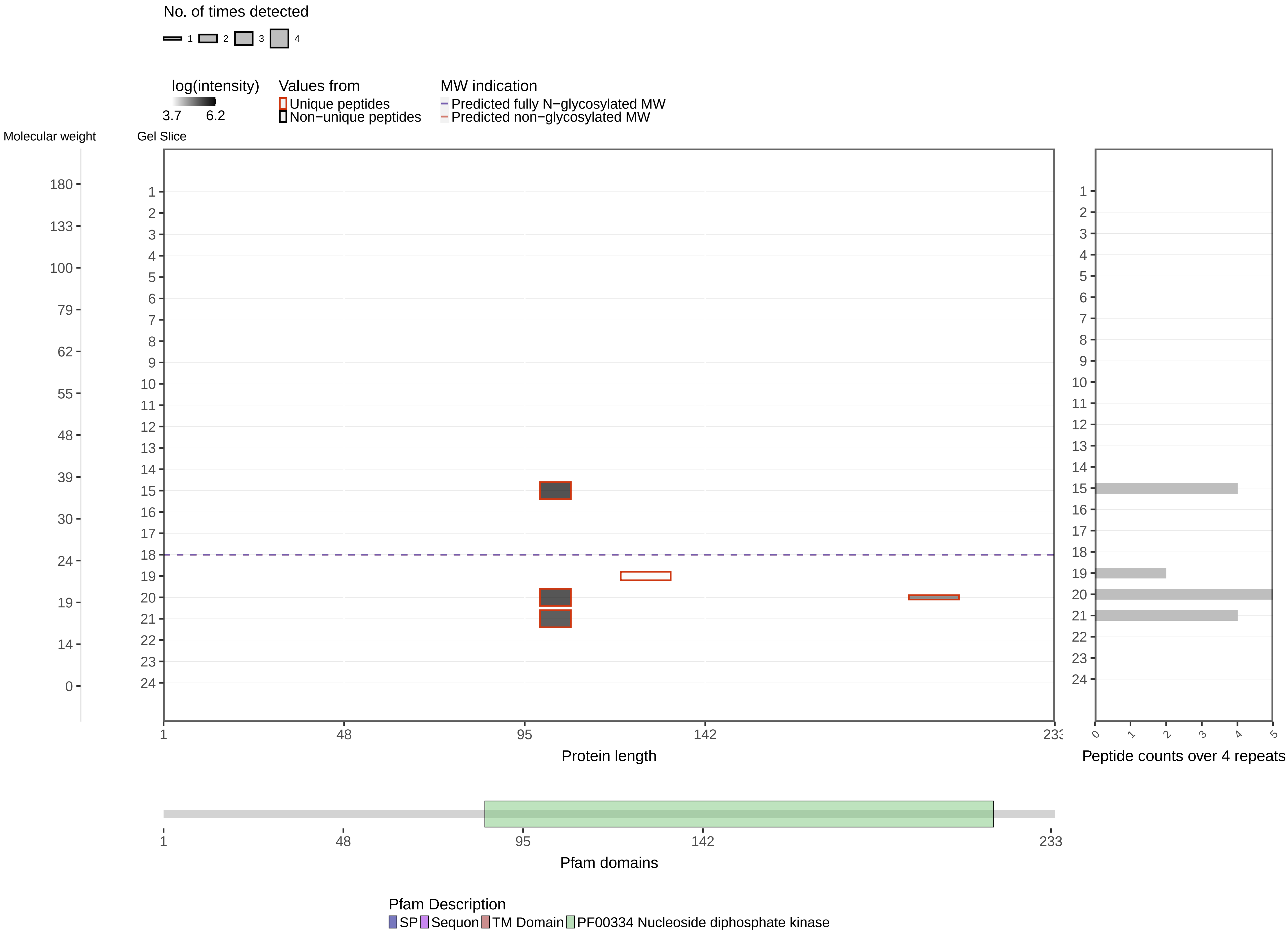


Protein sequence:

TSRFEKKGFKLTGLKLFECPKELAEEHYKDLQSKSFFPKLIDYITSGPVVCMAWEGVGVVASARKLIGATNPLNAEPGTIRGDLAVQTGRNVVHGSDSPDNGKREIALWFREGELCSWTAVQEPWLIE*

### NbL03g15780.1 gamma_carbonic_anhydrase−like_2_mitochondrial [MW = 33.12 kDa]

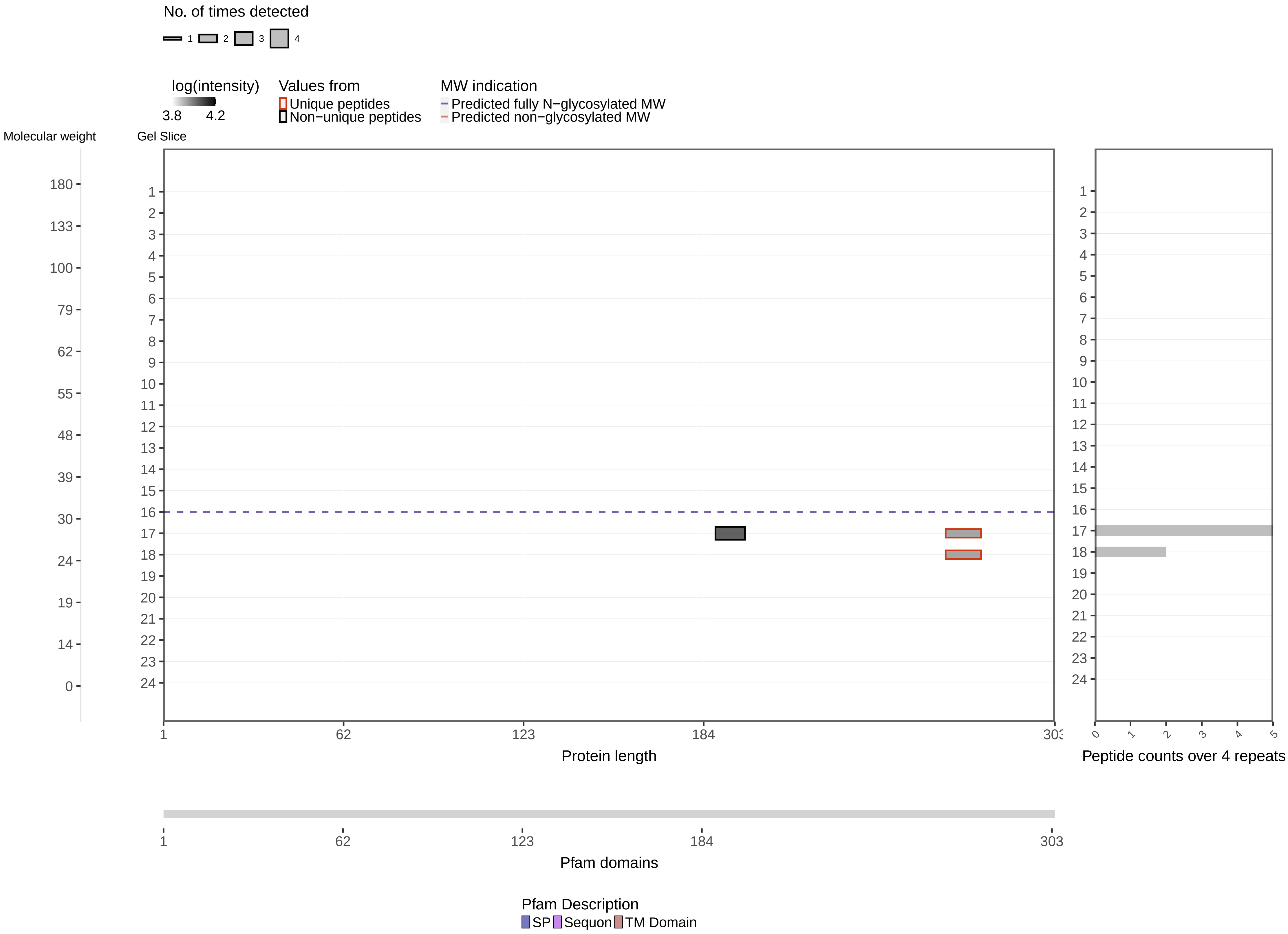


Protein sequence:

YDGASVWNGAVLRGDLNKITIGFCSSIQEKCVVHAAWSSPTGLPAETLVDRYVTVGAYSLLRSCTIEPECIIGQHSILMEGSLVETNSILEAGSVLPPGRRIPSGELWAGNPARFVRTLTHEEIKEIPRLSAAINDLANAHRSEFLPYSTVYLEVEKLKKSFGISI*

### NbL03g15750.1 alpha−14−glucan−protein_synthase_[UDP−forming]−like [MW = 35.11 kDa]

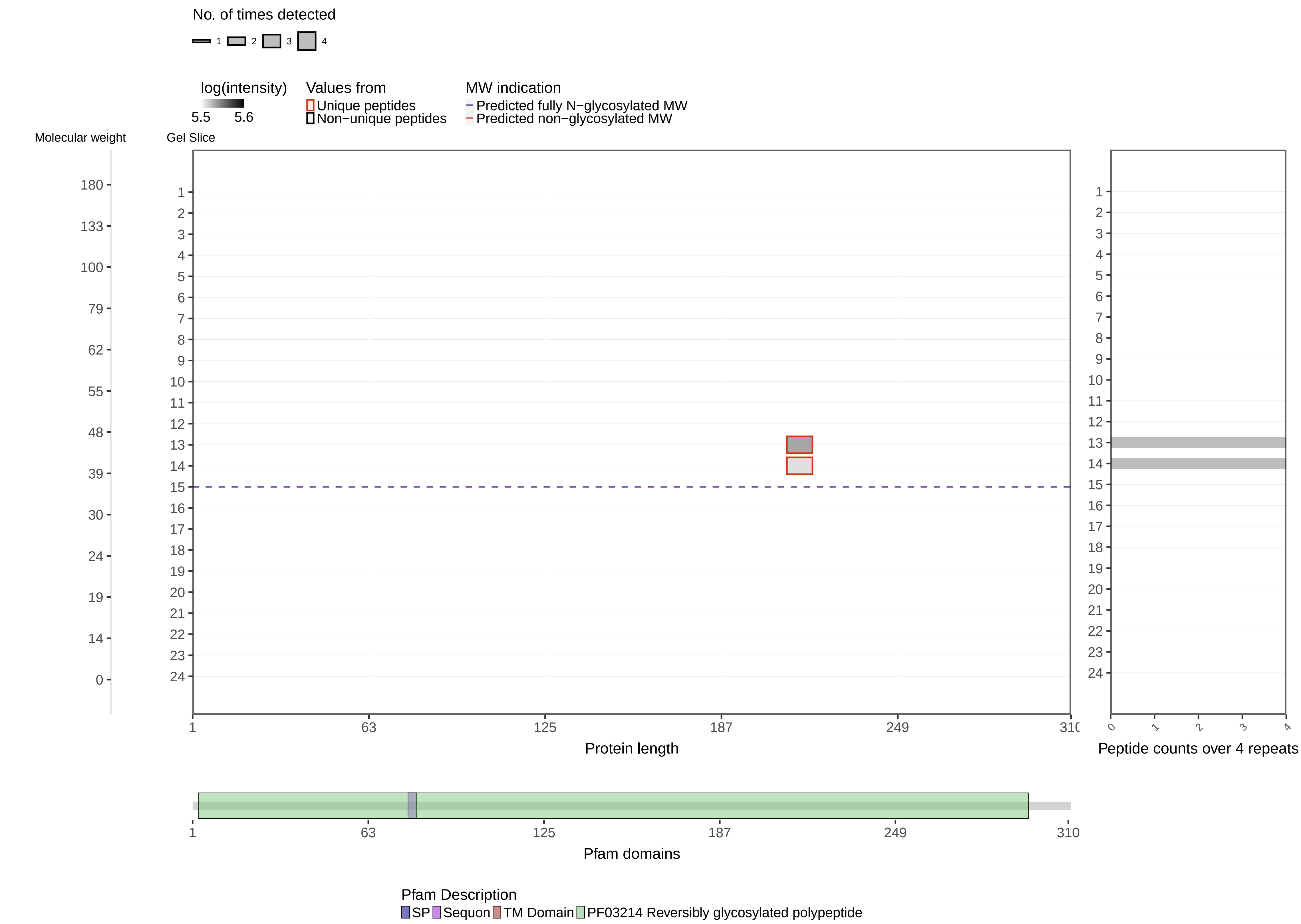


Protein sequence:

DAVMTIPKGTLYPMCAMNLAFNRELVGPAMYFGIMGDGQPLGRYDDMWAGWCSKVVCDHLGYGVKTGLPYLWHSKASNPFVNLKKEFKGLFWQEEMIPFFQSVVIPKECTNAQQCYLELAKMVKEKLGPIDPYFHNFAAAMVTWIEAWEEFNAPAKAKIDSVKSA*

### NbL03g15710.1 40S_ribosomal_protein_SA−like [MW = 32.49 kDa]

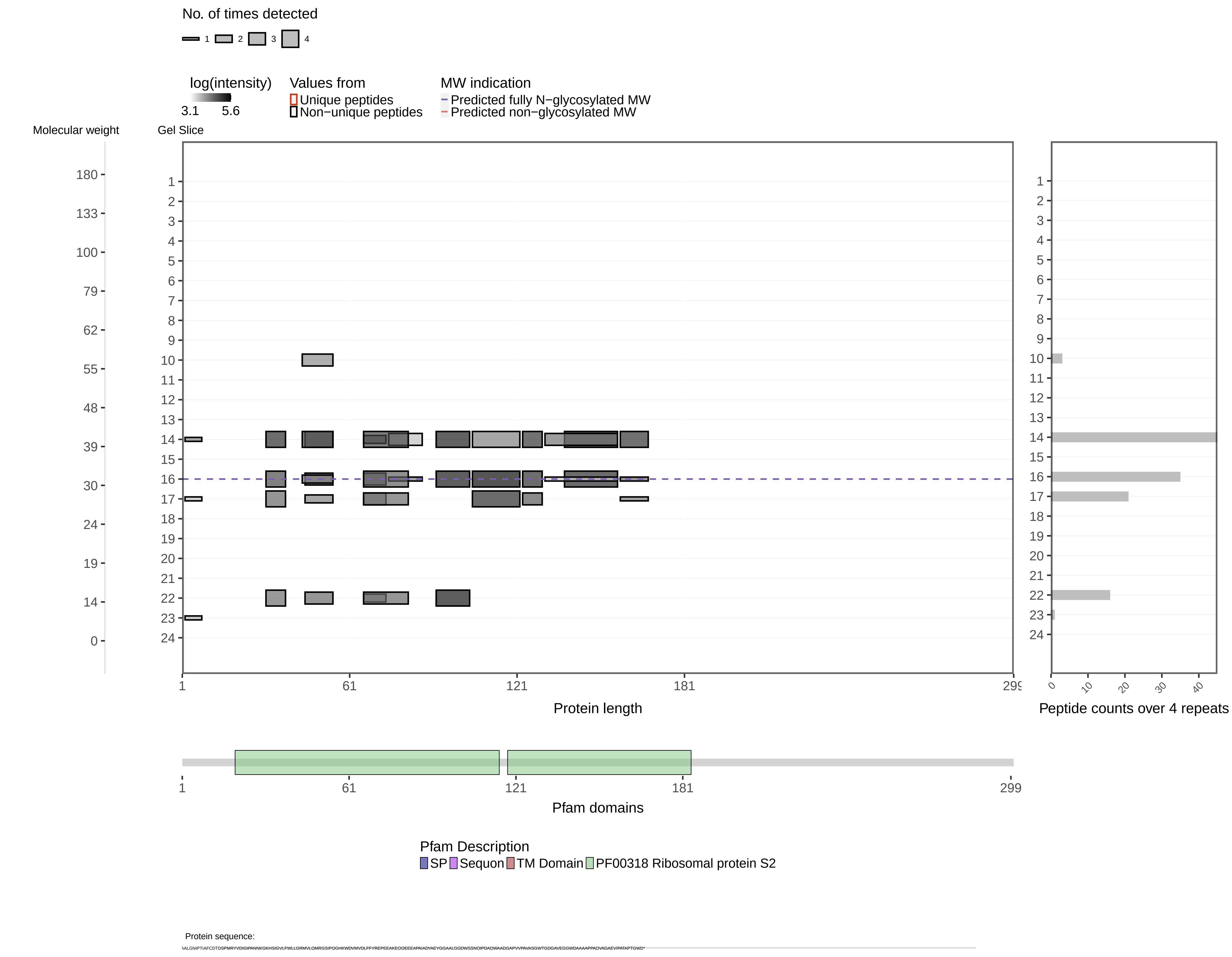

### NbL03g15700.1 alpha−galactosidase [MW − SP = 42.75 kDa]

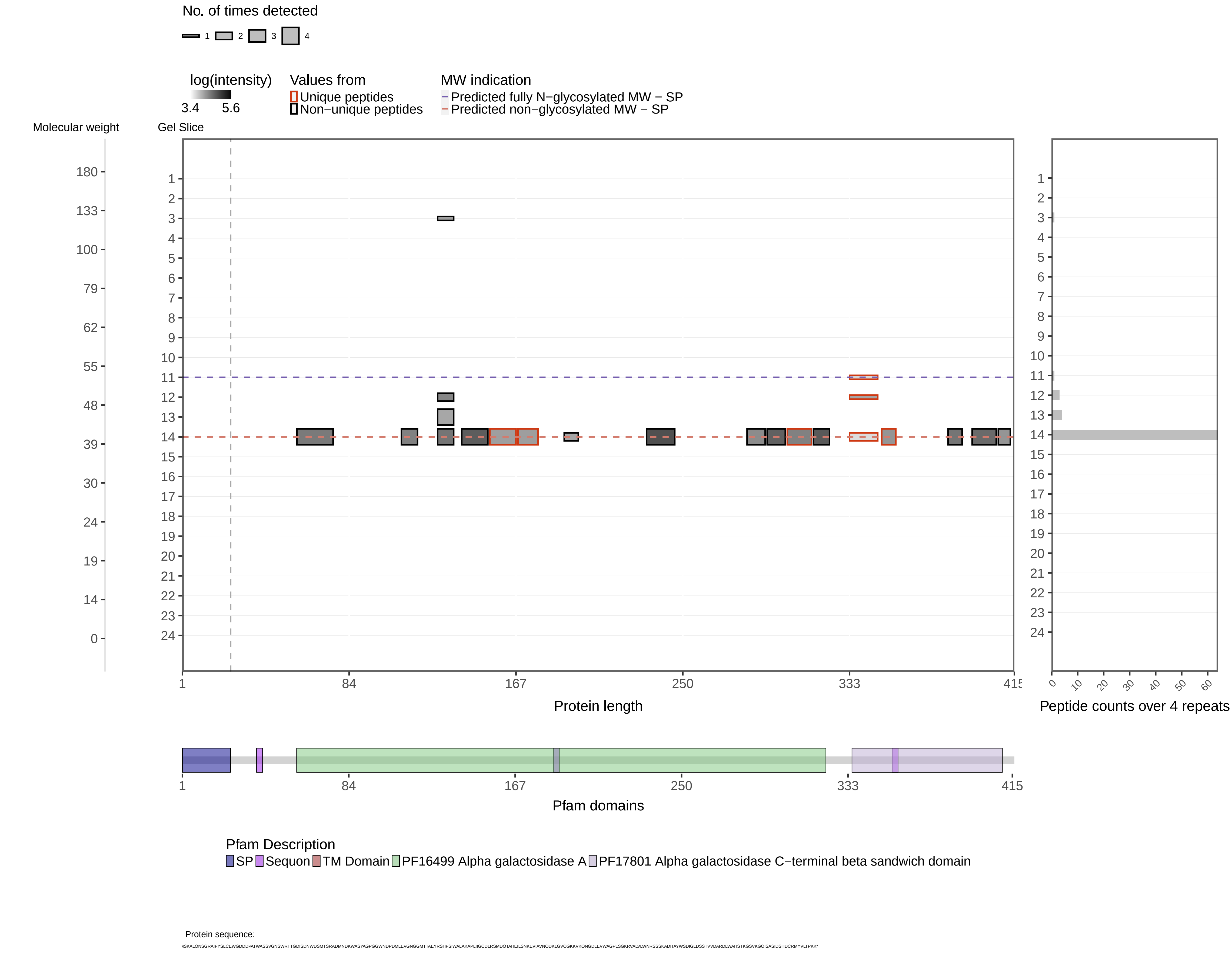

### NbL03g15610.1 beta−galactosidase_10 [MW − SP = 97.02 kDa]

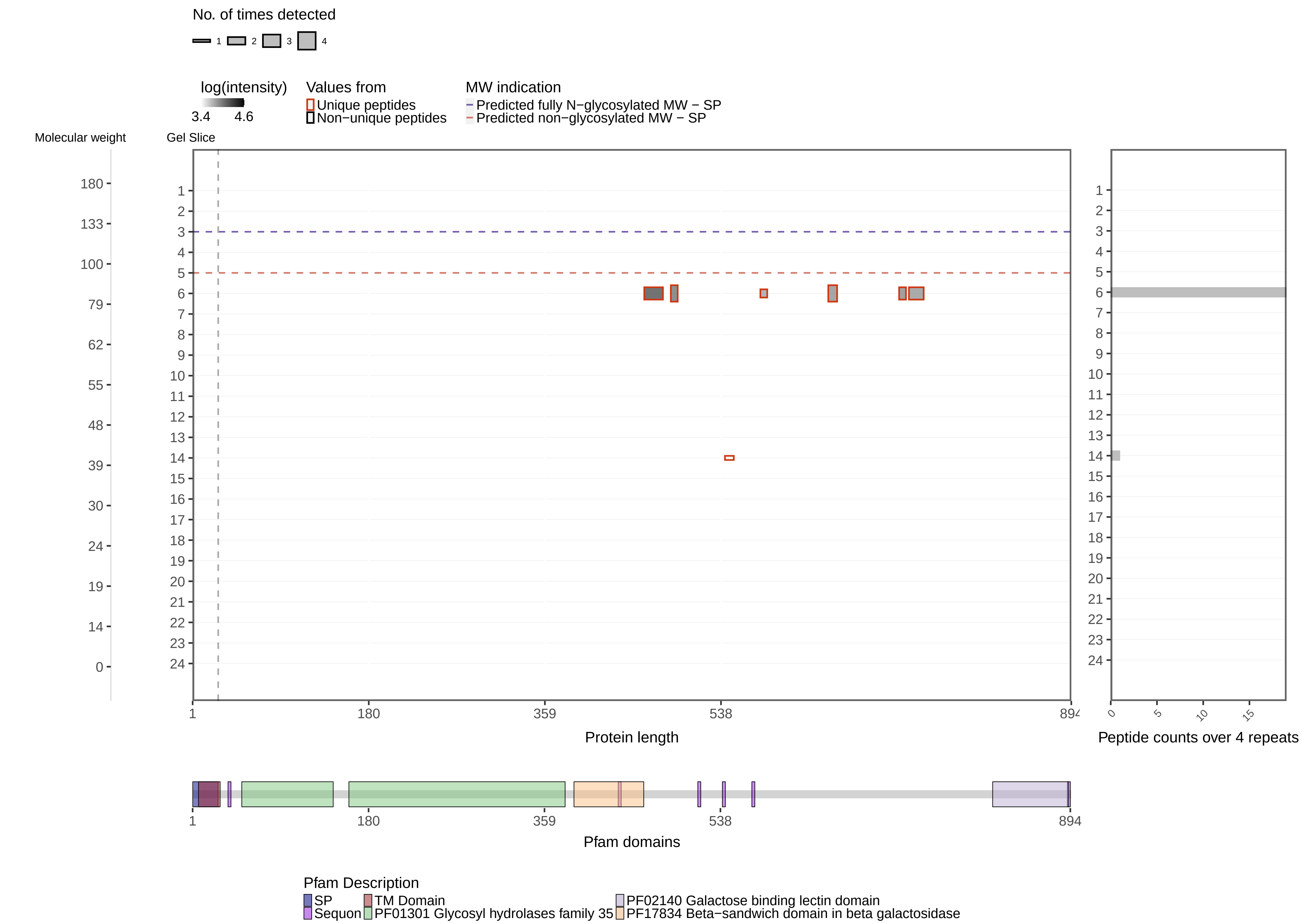


Protein sequence:

SYOLPAWSVSILPDCKNVVFNTAKVGCOTSIVNMEPVDLHPTVRSPKRDIKALOWEVFKETAGVWGVADFTKNGFVDHINTTKDTTDYLWYTTSIFVNEEEFLONKSSAMLFVESKGHAMHVFINHVLOASASGNGTVPHFKFGTPIVLKAGKNDIALLSMTVGLOTAGSFYEWIGAGPTSVKVEGFNSGTVDLSASTWTYKVGLHGEYLRIYESGGLNNKIWALTSEPPKOOPLTWYKAVVDVPPGDEPVALDMIHMGKGLAWLNGOEIGRYWPRKSSKLEKCVTOCDY

NbL03g15600.1 phytochrome−associated_serine/threonine−protein_phosphatase_3 [MW = 34.84 kDa]

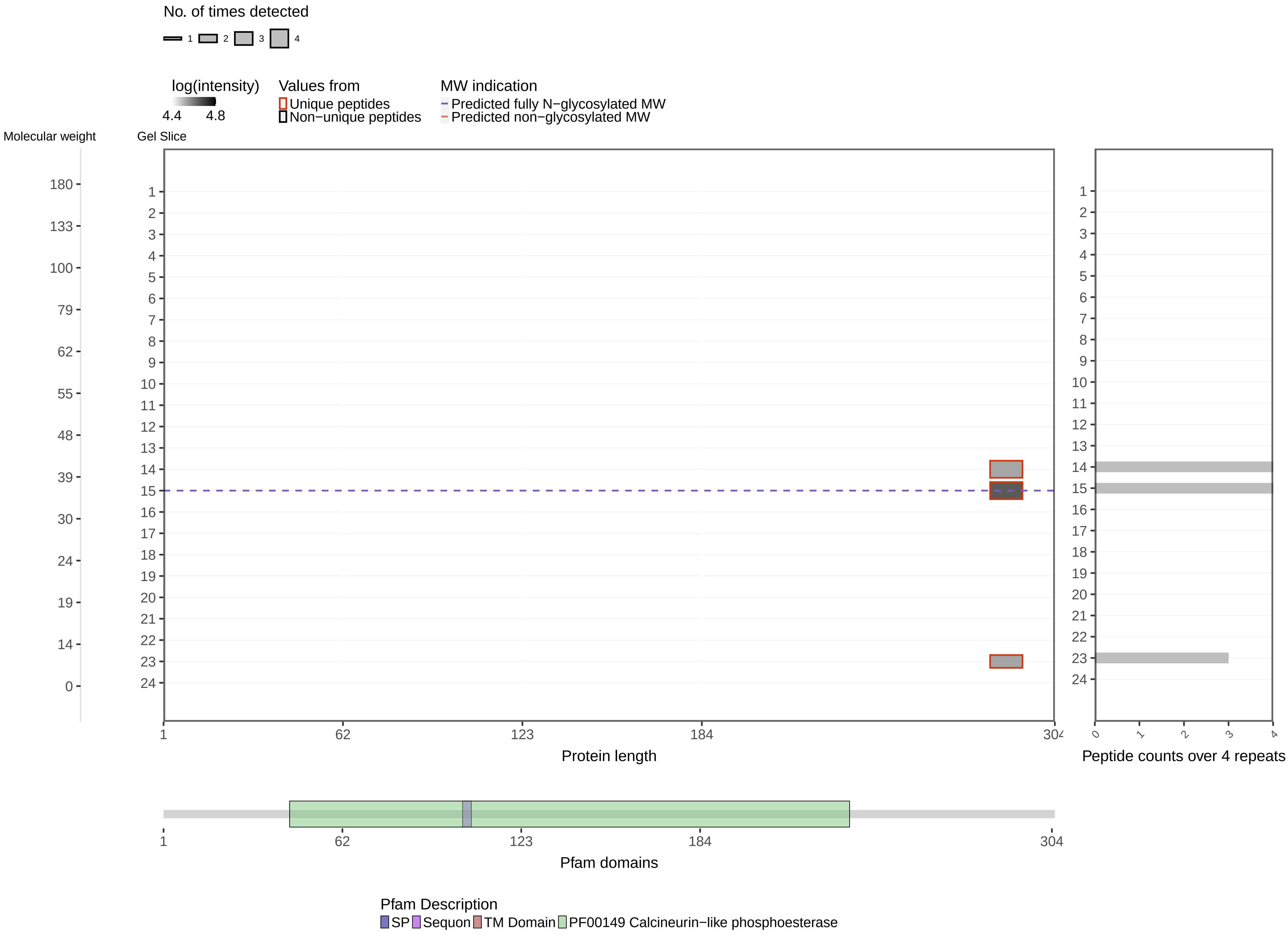


Protein sequence:
CTDVFDYLTLSAIIDGTVLCVHGGLSPDVRIIDQIRVIERNCEIPHEGPFCDLMWSDPEDIETWAVSPRGAGWLFGSRVTTEFNHINKLDLVCRAHQLVQEGLKYMFQDKGLVTVWSAPNYCYRCGNVASILSFNENMEREVKFFSETEENNQMRGPRTGVPYFL*

### NbL03g15570.1 asparagine−−tRNA_ligase_chloroplastic/mitochondrial−like [MW = 66.02 kDa]

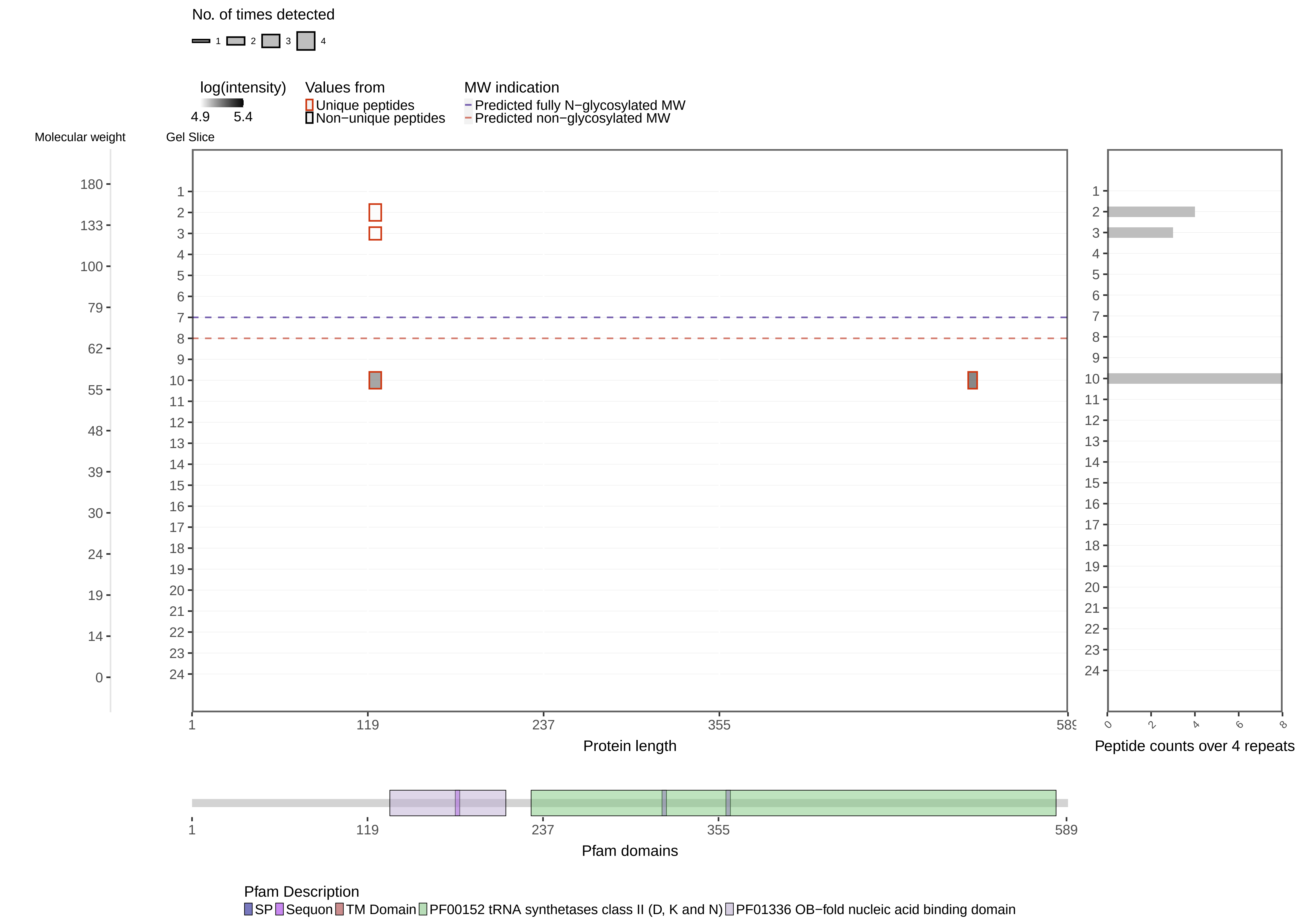


Protein sequence:

EGAGEQFRVTTLIPTTKEGGDSPVNAIPTTESGSVNWSQDFFGKQAFLTVSGQLNGETYATALSDIYTFGPTFRAENSNTSRHLAEFWMIEPELAFADLDDDMACATAYLQYVVQYVLENCKEDMDFFNTWIEKGIITRLSDVVEKNFVQLTYTDAVELLLKARKKFDFPVKWGCDLQSEHERYITEEAFGGCPVIIRDYPKDIKAFYMRQNDDGKTVAAMDMLVPRVGELIGGSQREERLGYLEERLDNLKLNKESFWWYLDLRRYGSVPHAGFGLGFERLVQFATGIDNIR

### NbL03g15530.1 serine_protease_inhibitor_1−like [MW − SP = 23.95 kDa]

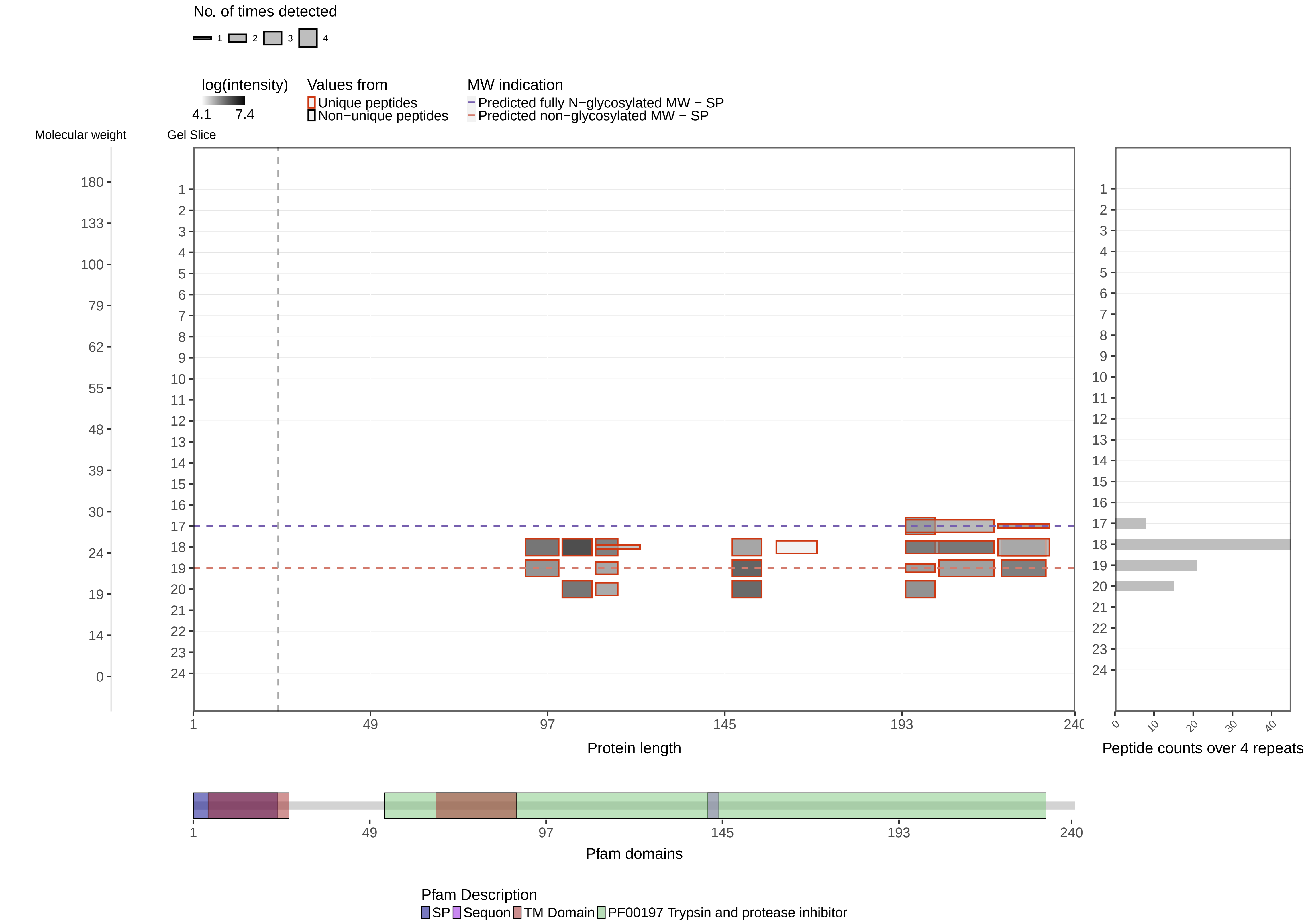


Protein sequence:

VKFFTTHPRPLNVVRENTDINIMFSVATTRLCVNETVWKVGDPEPIARLTRFVLTGGTLGNLRPENIDSWFKIERVTKQGPFYKLRYCPATIDSRPVVCSDVGVTHYFGVRRLALTNQPFVVVFRKFQKTDG*

NbL03g15280.1 protein_argonaute_1−like [MW = 76.89 kDa]

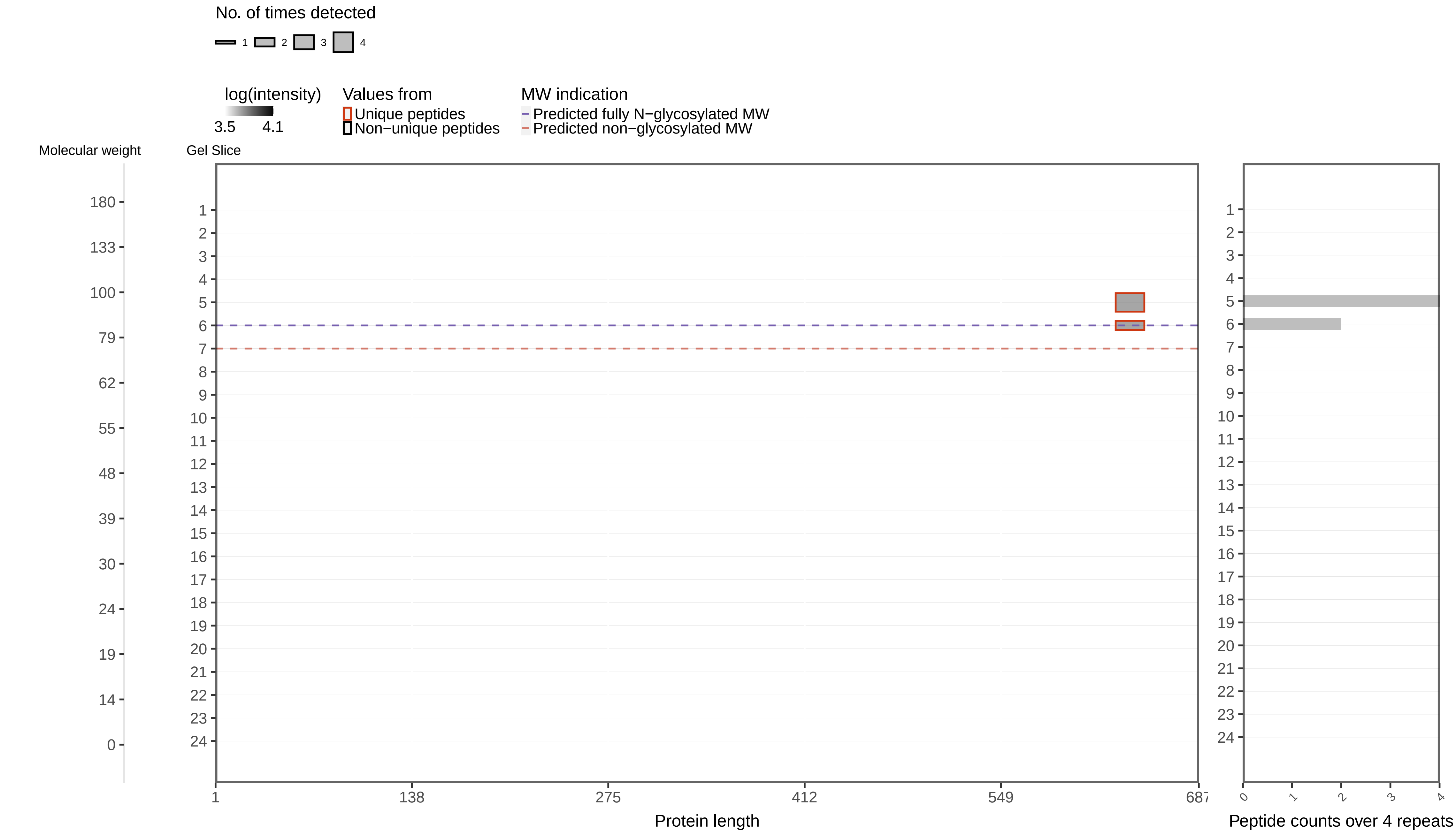


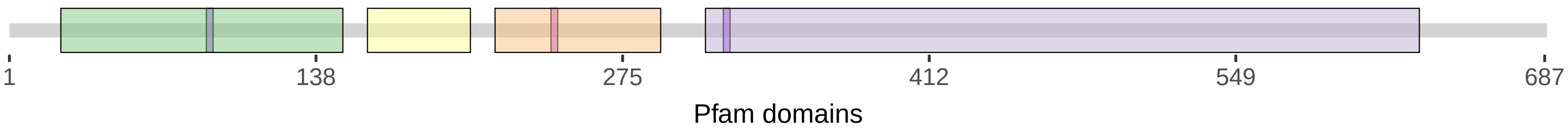


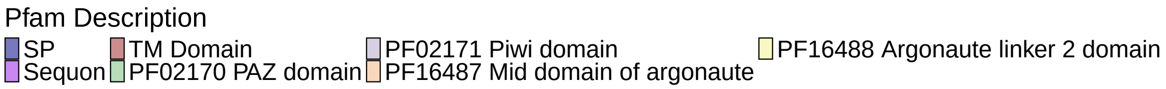


Protein sequence:

RICETELGVVSQCCLTKHVFKMSKQYLANVALKINVKVGGRNTVLVDAISRRIPLVSDRPTIIFGADVTHPHPGEDSSPSIAAVVASQDWPEITKYAGLVSAQAHRQELIQDLYTTRQDPVKGTVAGGMIKDLLISFRRATGQKPQRIIFYRDGVSEGQFYQVLLFELDAIRKACASLEPNYQPPVTFVVVQKRHHTRLFANNHRDRNAVDRSGNIIPGTVVDSKICHPTEFDFYLCSHAGIQGTSRPAHYHVLWDENKFTADALQSLTNNLCYTYARCTRSVSIVPPAYYAHLAAF

### NbL03g15240.1 glutamate_decarboxylase−like [MW = 58.69 kDa]

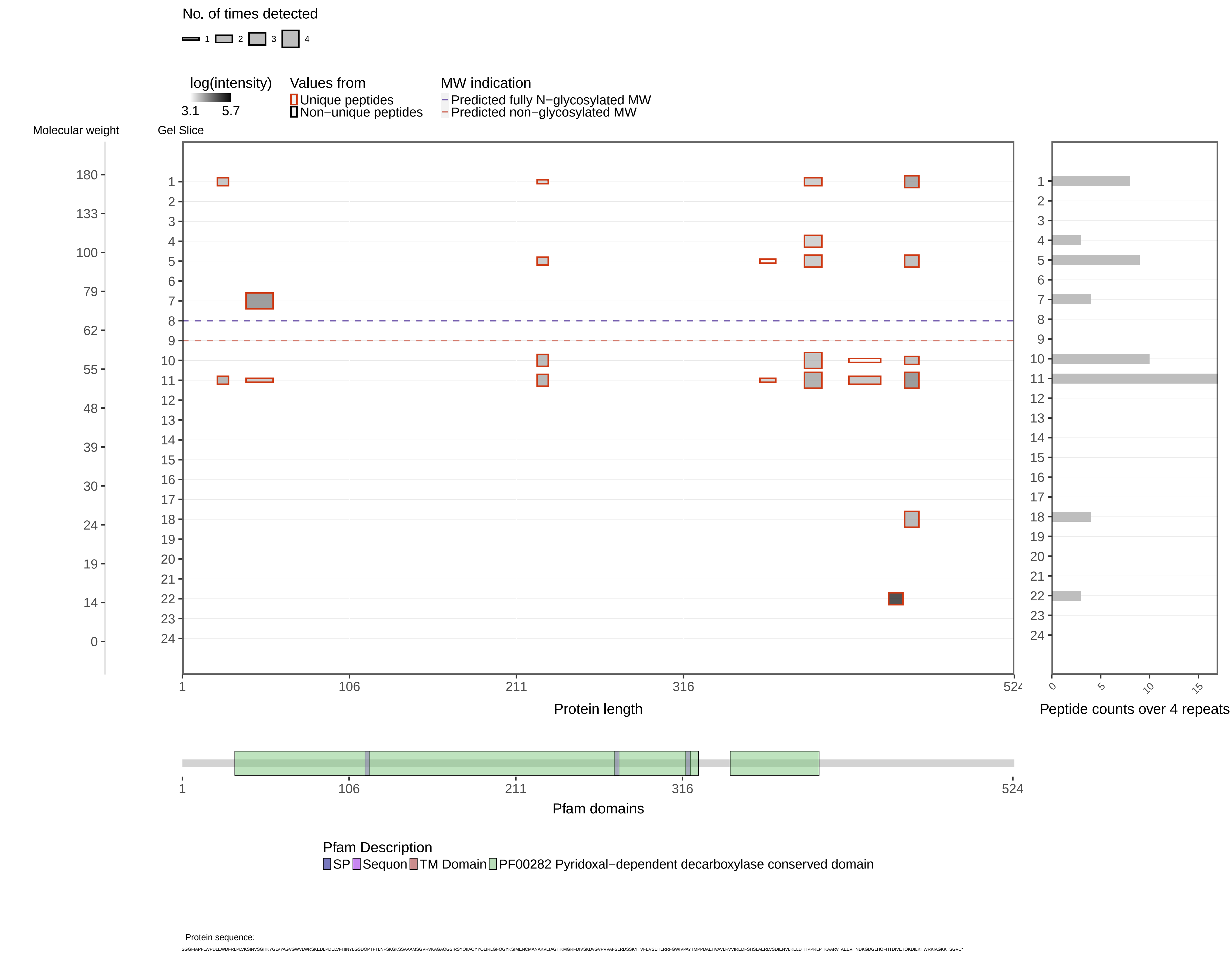

### NbL03g15140.1 uncharacterized_LOC109207500 [MW = 41.82 kDa]

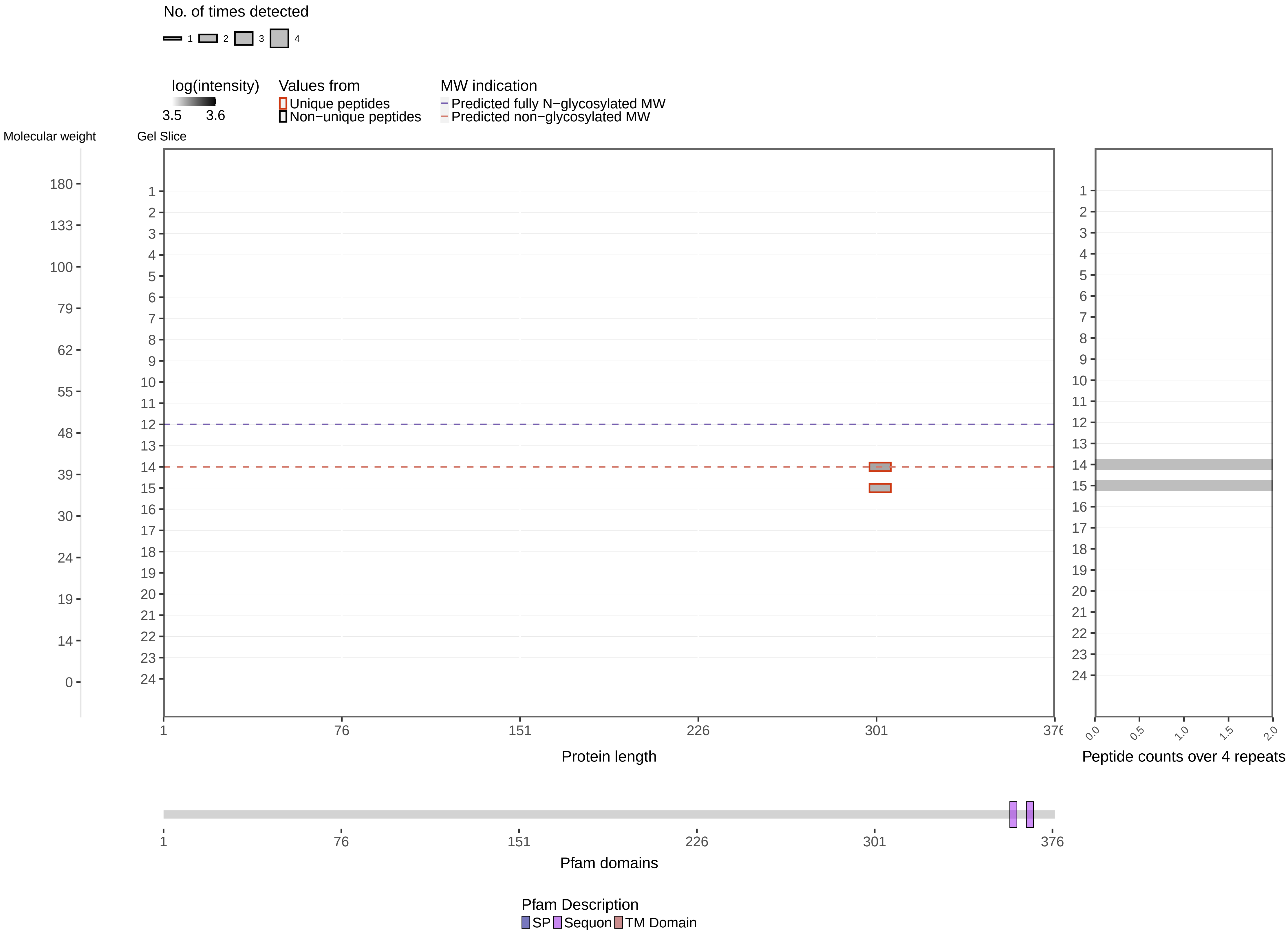


Protein sequence:

GDNNRAADVDALKIGEFDRVRRLFEYIEDRDLWRWKLPDSKAFSSGLDDCNIEYDINLNPSLFQQLLALDLKSLIEQGKISLSHKQKRIDEVLEKSFEIALGAGTFGCCLAVNADSLSELRSELGNQLAARSRNMKFRGIGAVVYKVPELENDQMLKVSLRSIGDEDTTSISQAFGGGGHRNASSFMLNSTEFENWKVK*

NbL03g14930.1 dihydropyrimidine_dehydrogenase_(NADP(+))_chloroplastic−like [MW = 47.15 kDa]

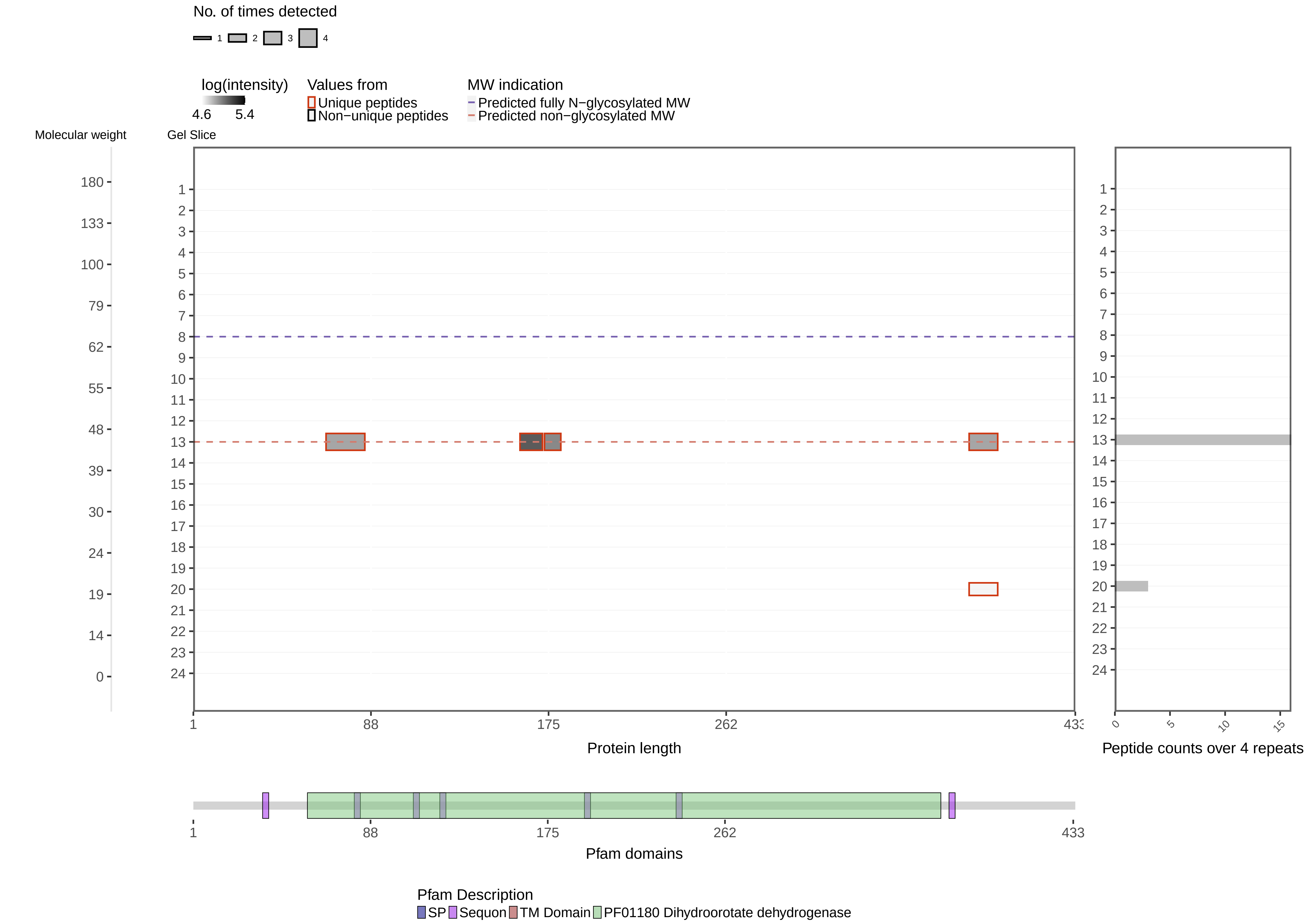


Protein sequence:

MGAAVGQDCDLLEEVCGWINAVATVPVWAKMTPNITDITKPARVSLNQGCEGVSAINTIMSVMGINLNTLRPEPCVEGYSTPGGYSAKAVHPIALAKVMNIARMMKSEFGDKDYSLSAIGGVETGGDAAEFILLGANTVQVCTGVMMHGYGLVKTLCSELKDFMRKHNFSSIEDFRGASLEYFTTHTDLVKRQQEAIRQRKAIKKGLQSDKDWTGDGFVQETESMVSN*

NbL03g14920.1 40S_ribosomal_protein_S3a−like [MW = 29.7 kDa]

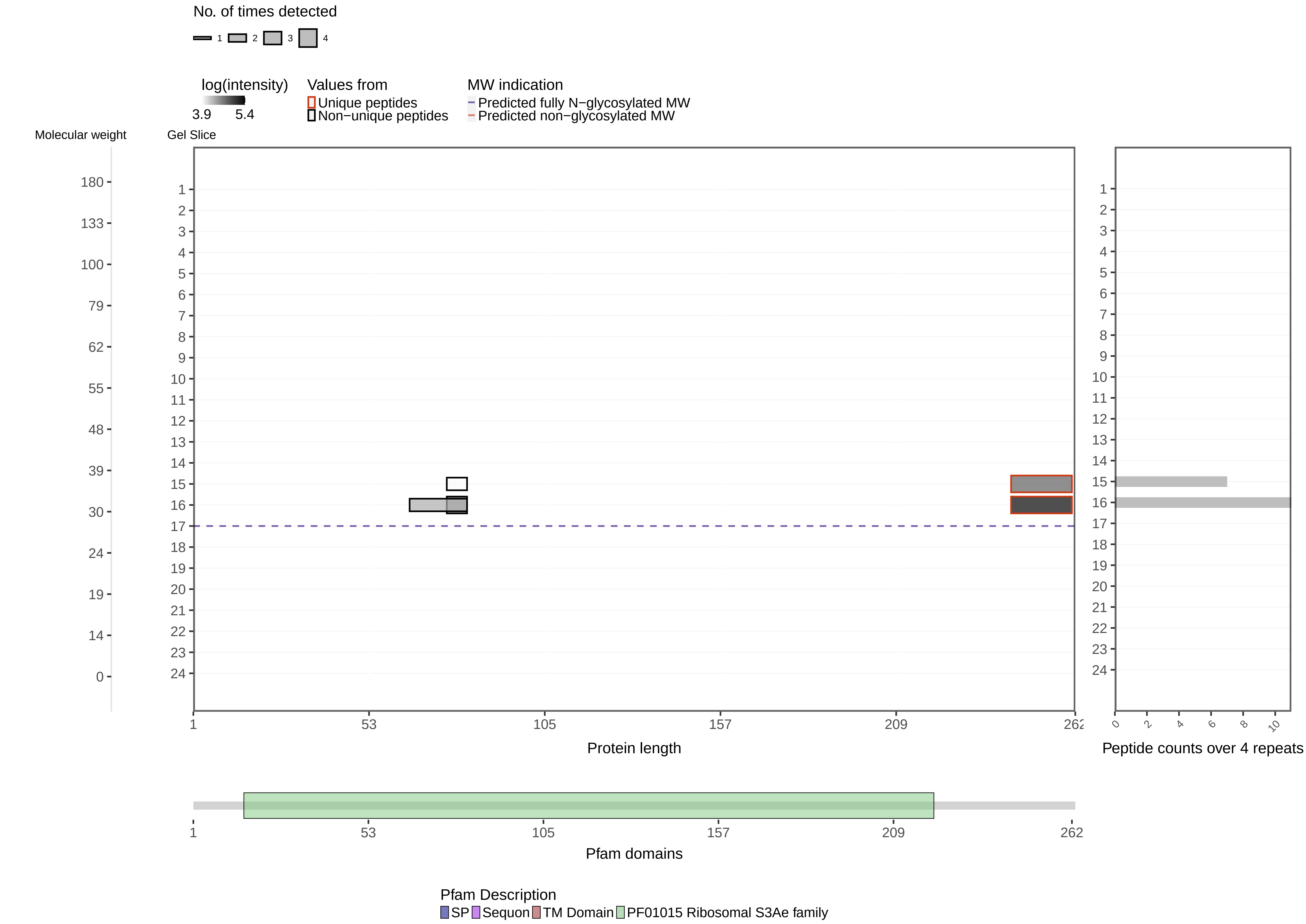


Protein sequence:

KWQTLIEAHVDVKTTDSYTLRMFCIAFTKKRPNQQKRTCYAQSSQIRQIRRKMVEIMRNQASSCDLKELVAKFIPESIGREIEKATSSIFPLQNVYIRKVKILKAPKFDIGKLMEVHGDYSEDVGVKLDRPADETAAEAEPEVPGA*

### NbL03g14870.1 V−type_proton_ATPase_subunit_C−like [MW = 42.92 kDa]

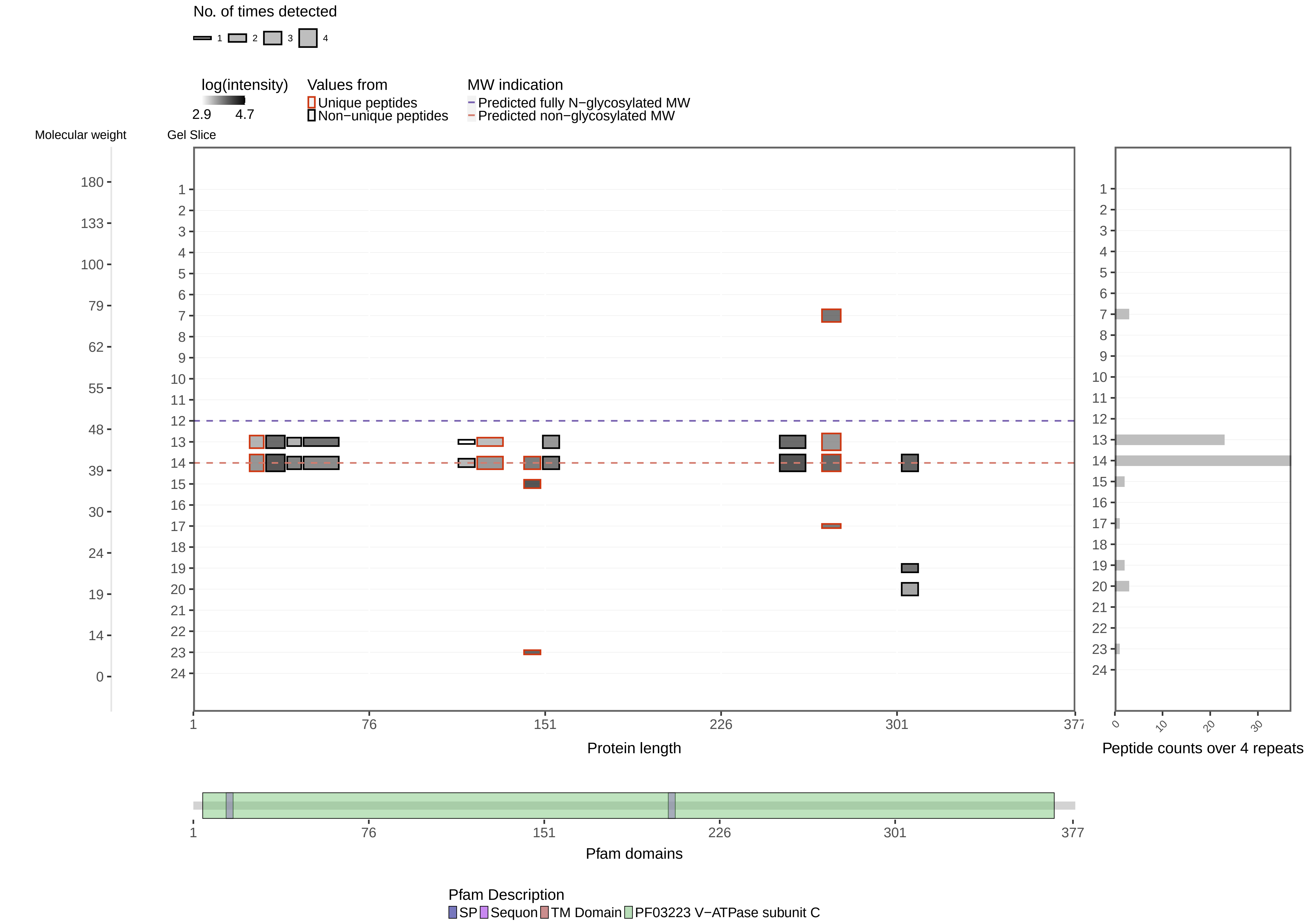


Protein sequence:

DIVTSEHLTTLLAVVSKYSOKDWLSSYENLTTYVVPRSSKMLYEDNEYALYTVTLFNRDADNFKIKARERGFOIRDFEHNPETOESRKOELEKLMODOETFRSSLLOWCYTSYGEVFSSWMHFCAVRIFAESILRYGLPPSFLSVVLAPSTKSEKKVRSILESLCDSSNSNFWKSEDEGGMAGFGGDTEAHPYVSFTINLV*

### NbL03g14600.1 3−oxoacyl−[acyl−carrier−protein]_synthase_3_A_chloroplastic−like [MW = 46.66 kDa]

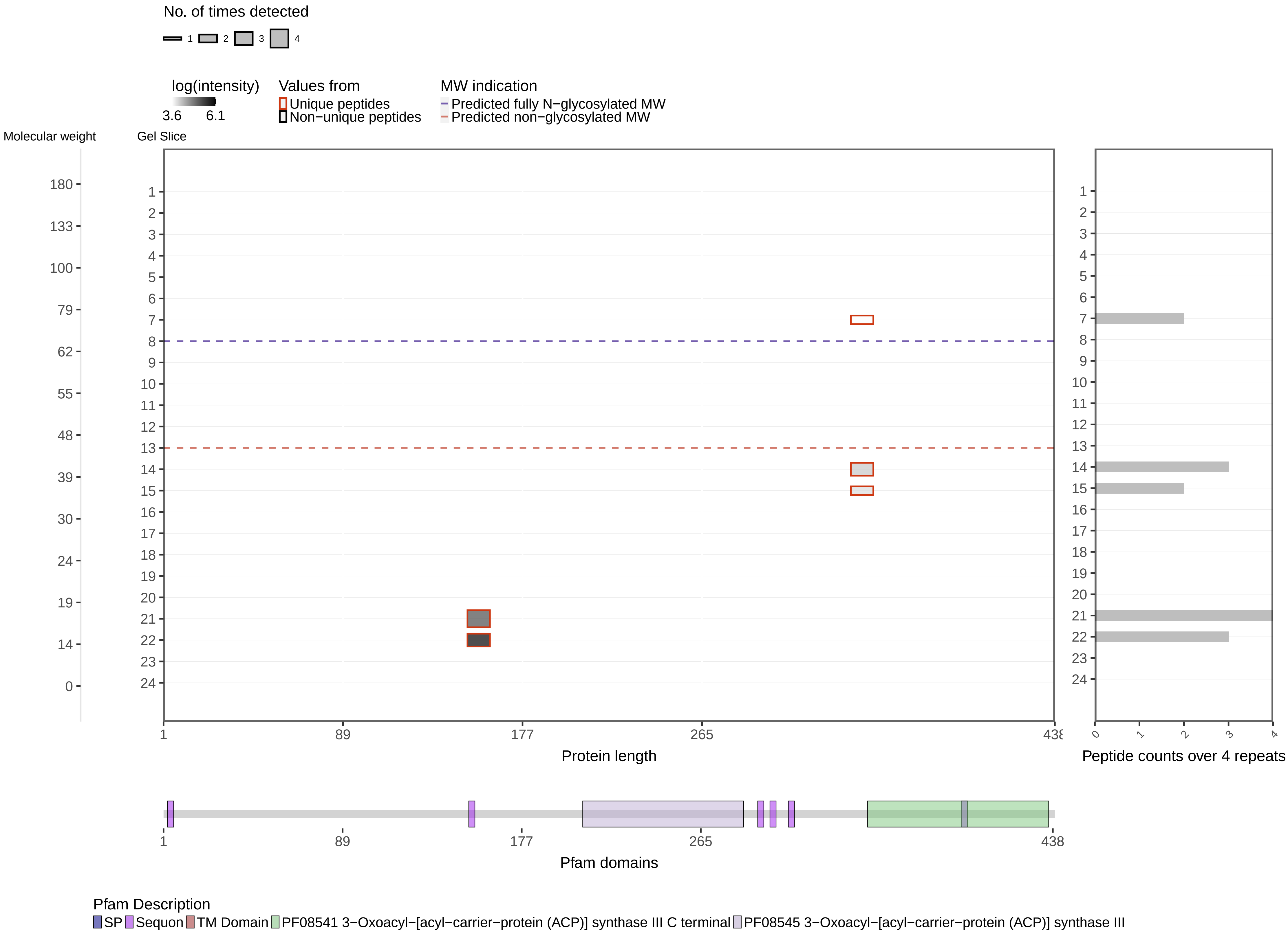


Protein sequence:

FDITAACSGFMLGLVSAACYIRGGVFKNVLVIGADALSRYVDWSDRGTCILFGDAAGAVVMQACDIGEDGLFGFDLHSDGEGQRHLNASIKENETDHAFGTNGSVIGFAPKGSSYSCIQMNGKEVFRFAVRVVPLSIEAALEKAGLPGSKIDWLLLHQANQRIIDGVATRLEVPSERVISNLANYGNTSAASIPLALDEAVRSGKVQAGHVIAAAGFGAGLTWGSAILRWG*

NbL03g14500.1 DNA−binding_protein_SMUBP−2 [MW = 108.1 kDa]

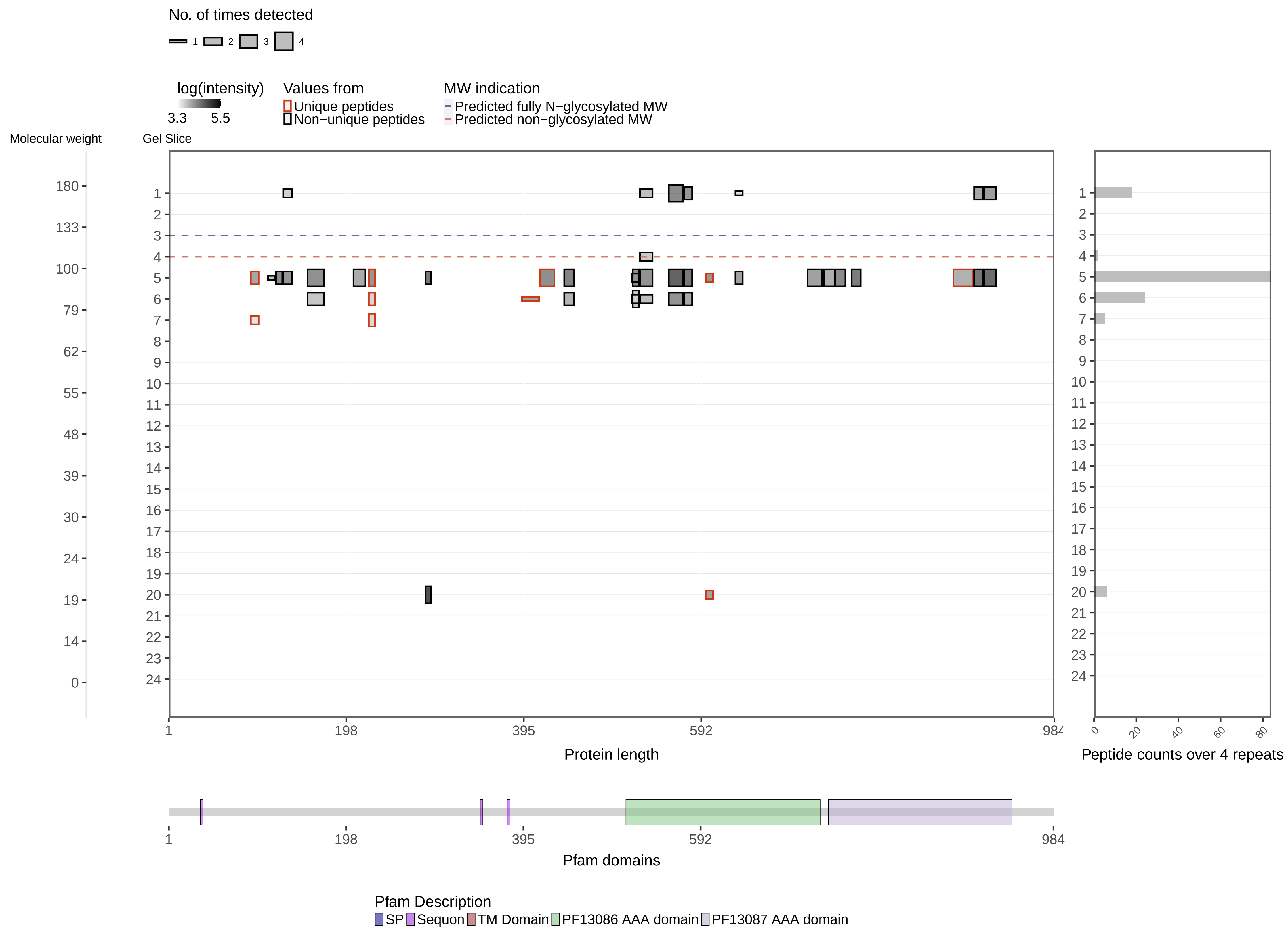


Protein sequence:

NbL03g14490.1 protein_BOLA4_chloroplastic/mitochondrial [MW = 17.23 kDa]

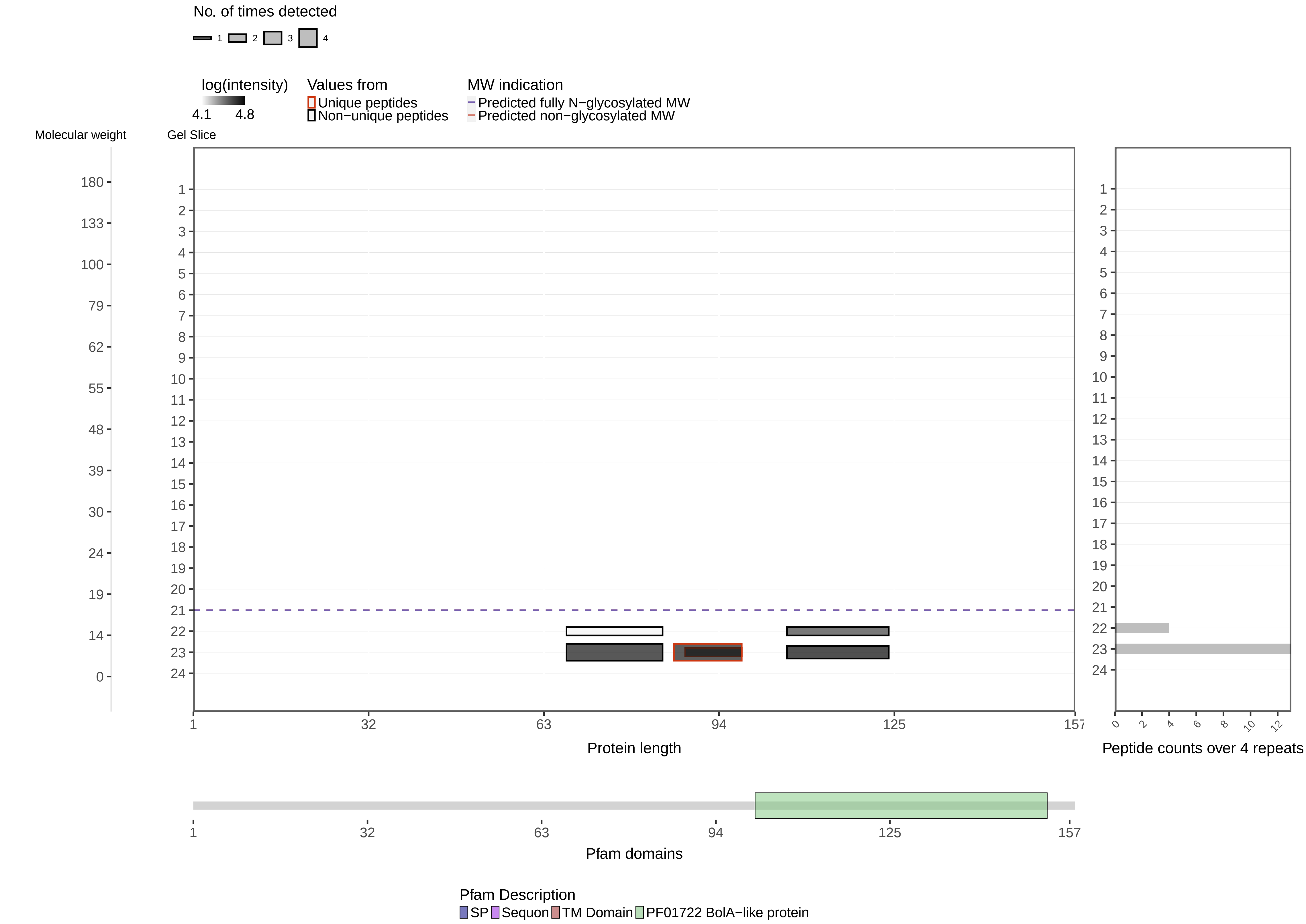


Protein sequence:

ATSDTGSIDSPLMESMQKKIKEQLNADSVIVKDAYGDGRHVSIDVVSSAFEGQSAVNRQRMVYKAIWEELQNTVHAVDQMTTRTPAEAGK*

NbL03g14430.1 cysteine_proteinase_inhibitor_6 [MW − SP = 29.48 kDa]

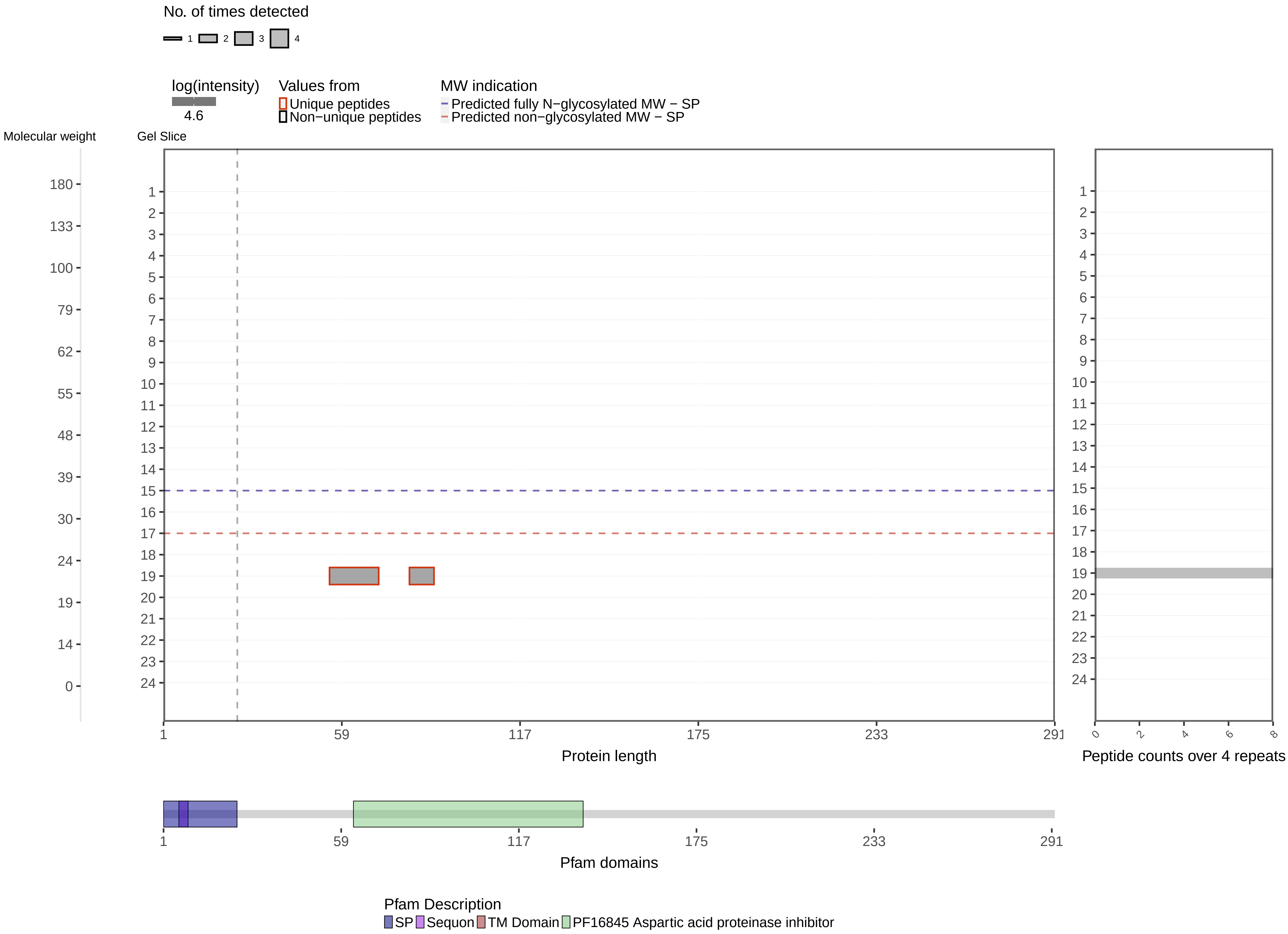


Protein sequence:

OEFTHVEDVPTLTSSDLGVKOGDMSLCOVCDYIOKKILDSISGVNRKKCSSSVLOIIVSAIYEEEGSGLKSVPVHDPVVOEAADHAIKTIOORSNSLLPYELOEIVHANAEVIGEDNMKLHMLIKTSRGGKOEKFKVOVHHNNEGAFOLNHMEPDHS*

### NbL03g14420.1 alanine−−tRNA_ligase−like [MW = 112.75 kDa]

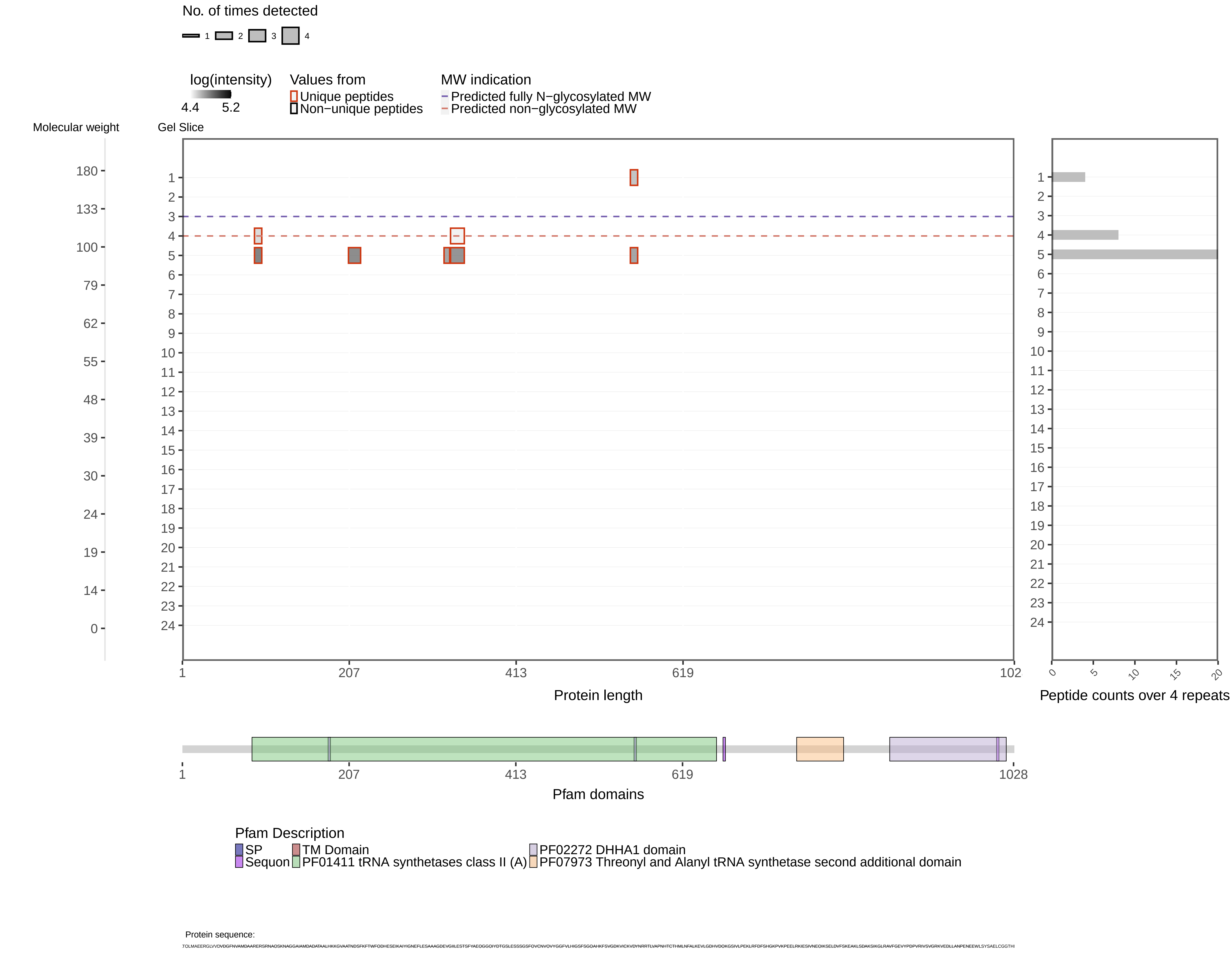

NbL03g14350.1 WEB_family_protein_At3g02930_chloroplastic−like [MW = 109.69 kDa]

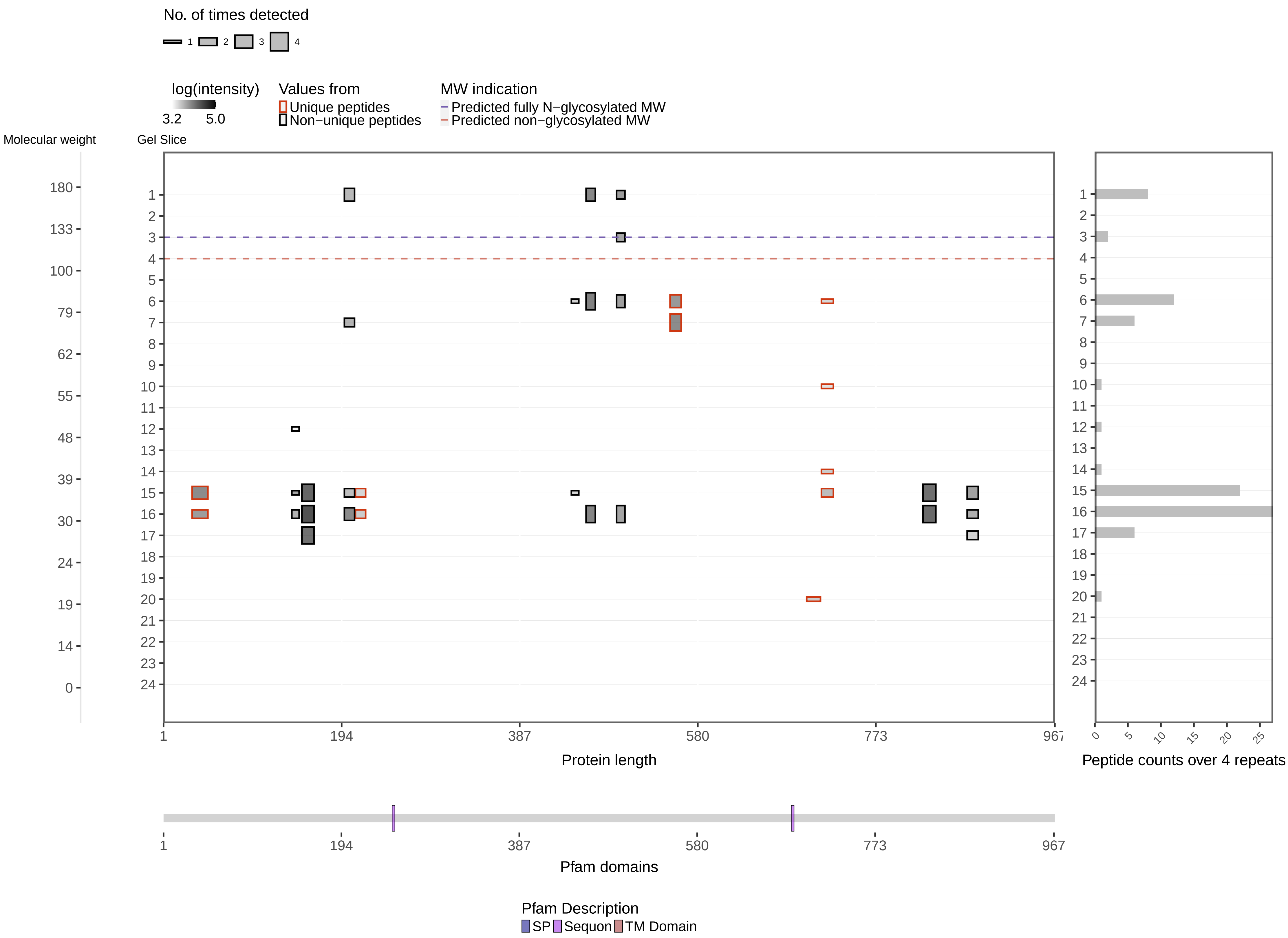


Protein sequence:

GSNDLLHDAESEIASLKEKVGLLEMSTTRQKGDLEESERRAQIAKEEASEMAKKVESLIAELETVKEEKTQAIEHEKLAAENVQSLLEEKNKLINELENSREEEEKSKKAMESLASALHEVSSEAREAKERWLSSQAEHEHYETQIEDLKLVLKATNEKYESLLDEAKEKIDDLTNSVEQSKNEHQISKAEWEDKELYLMNYVKKTEEENSSMEKEINRLVNLLKEAEQEASFEEEAVQLKNSLNEAESEVTYLKEVLGEAKGESMKLKESLLDKENEVQNILQENEELRSREA

NbL03g14250.1 binding_partner_of_ACD11_1−like [MW = 29.58 kDa]

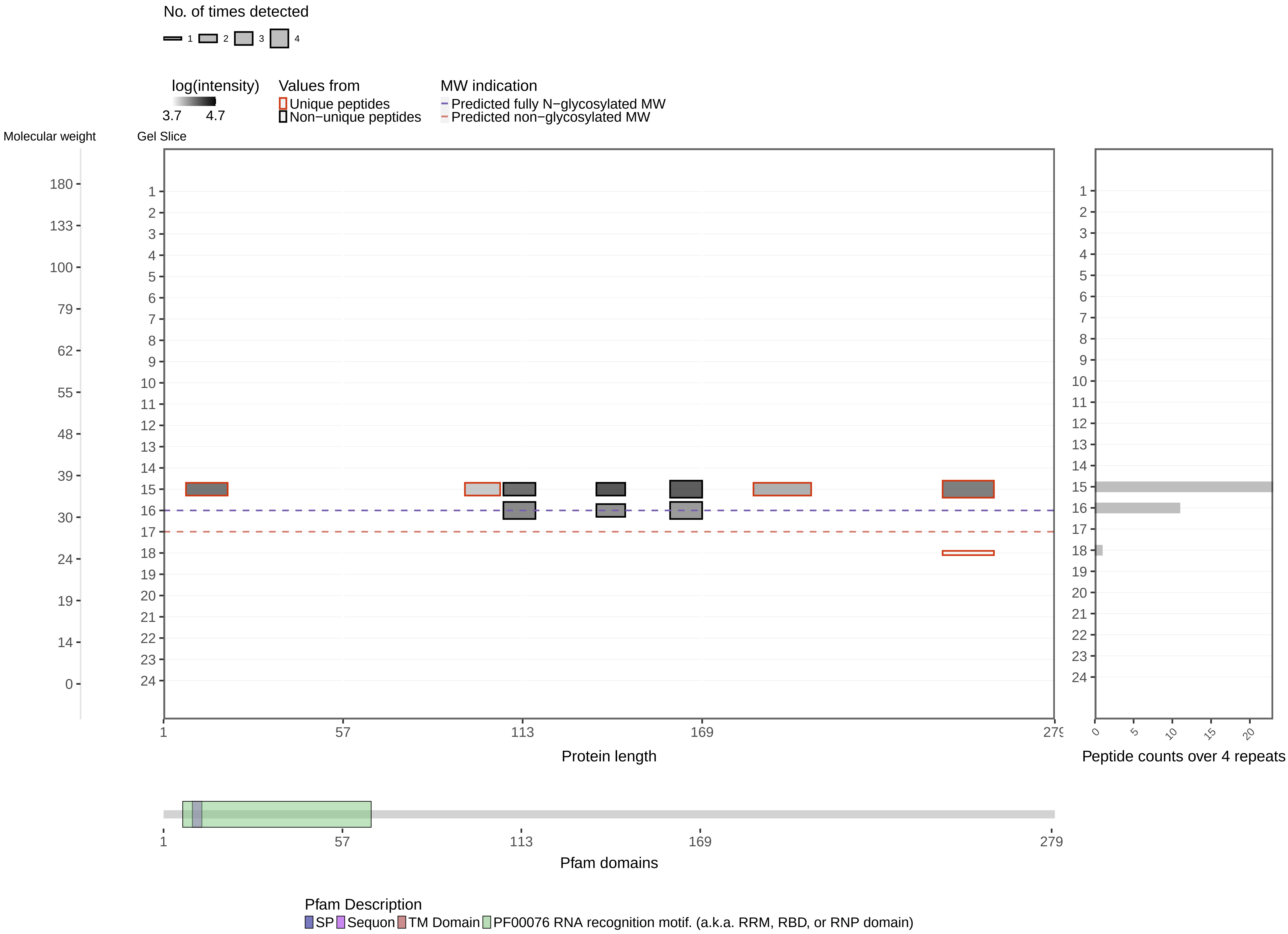


Protein sequence:

KAKTFDEKHQFISTASAKVASLDQKIGLSEKINMGTTIVNDKVKEMDQKFQVTEKTKSALAAAEQTVSTAGSAIMKNRYVLTGASWVTGAFNKVTKAAGEVGQKTKEKMAEEEQGRSSATGYVPIHAFSESPKASKTEEPAKPSSPKGLIL*

NbL03g14230.1 TOM1−like_protein_1 [MW = 43.75 kDa]

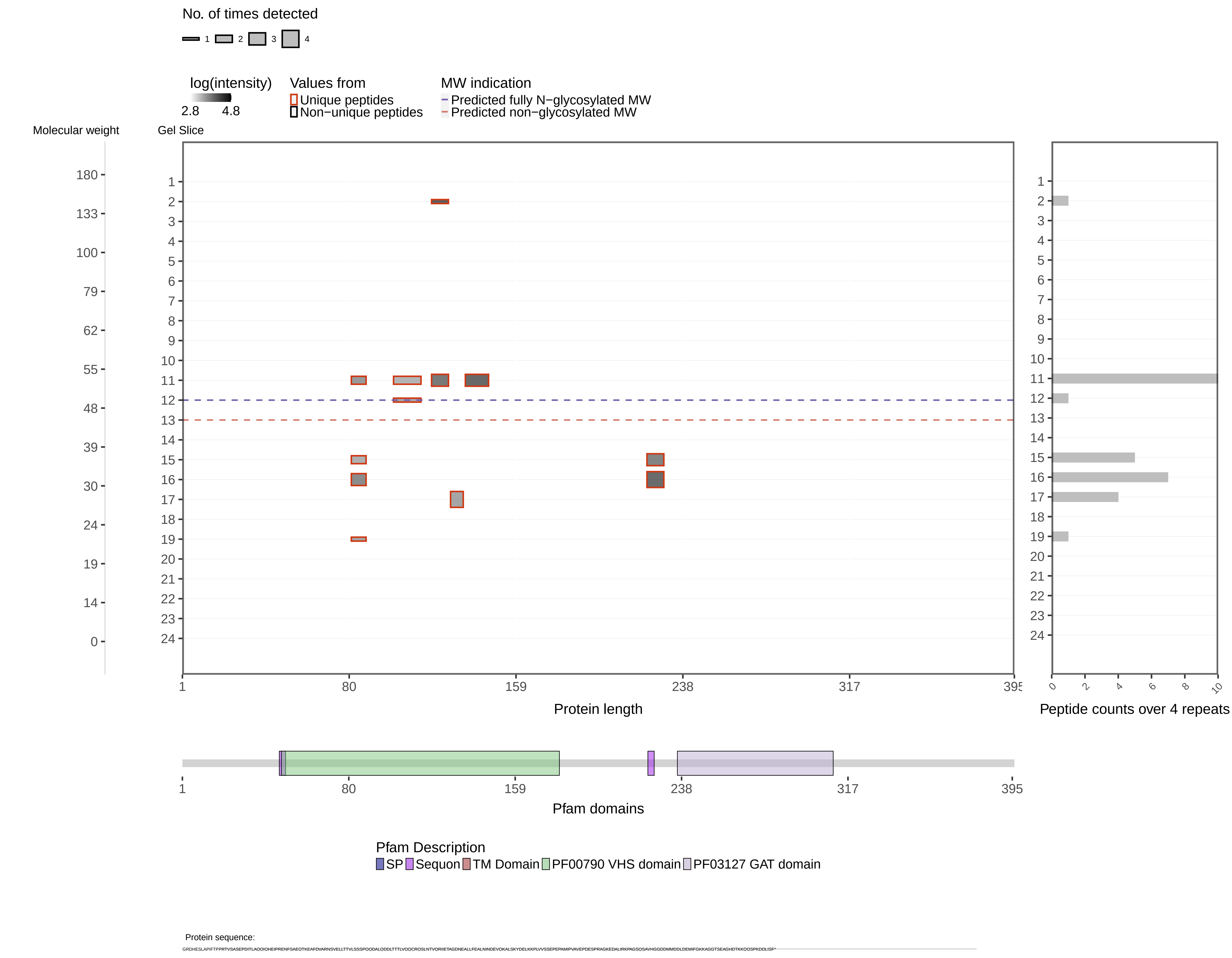

NbL03g14200.1 4−coumarate−−CoA_ligase_2 [MW = 67.08 kDa]

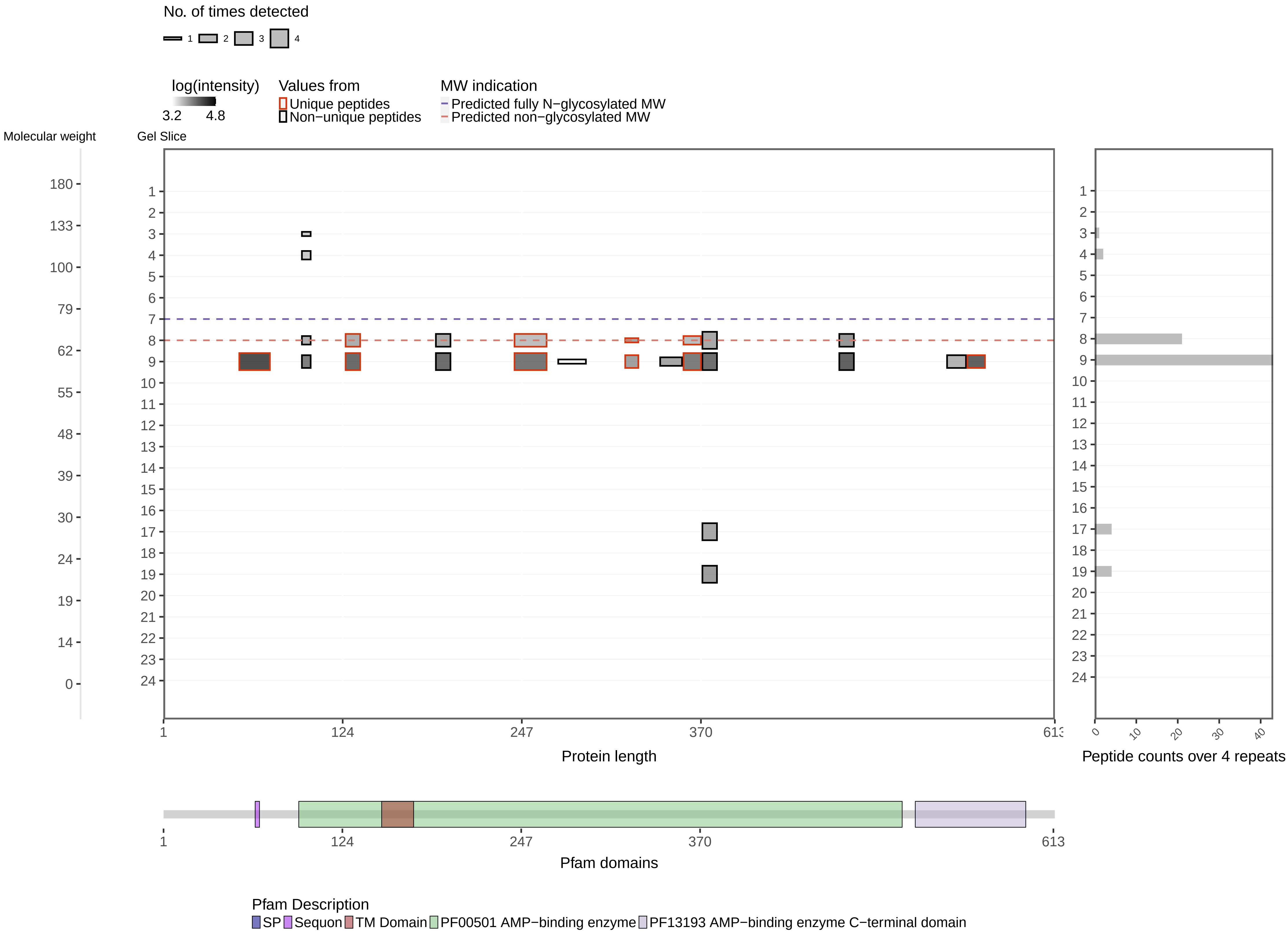


Protein sequence:

[illegible]

NbL03g14050.1 oleoyl−acyl_carrier_protein_thioesterase_chloroplastic−like [MW = 41.42 kDa]

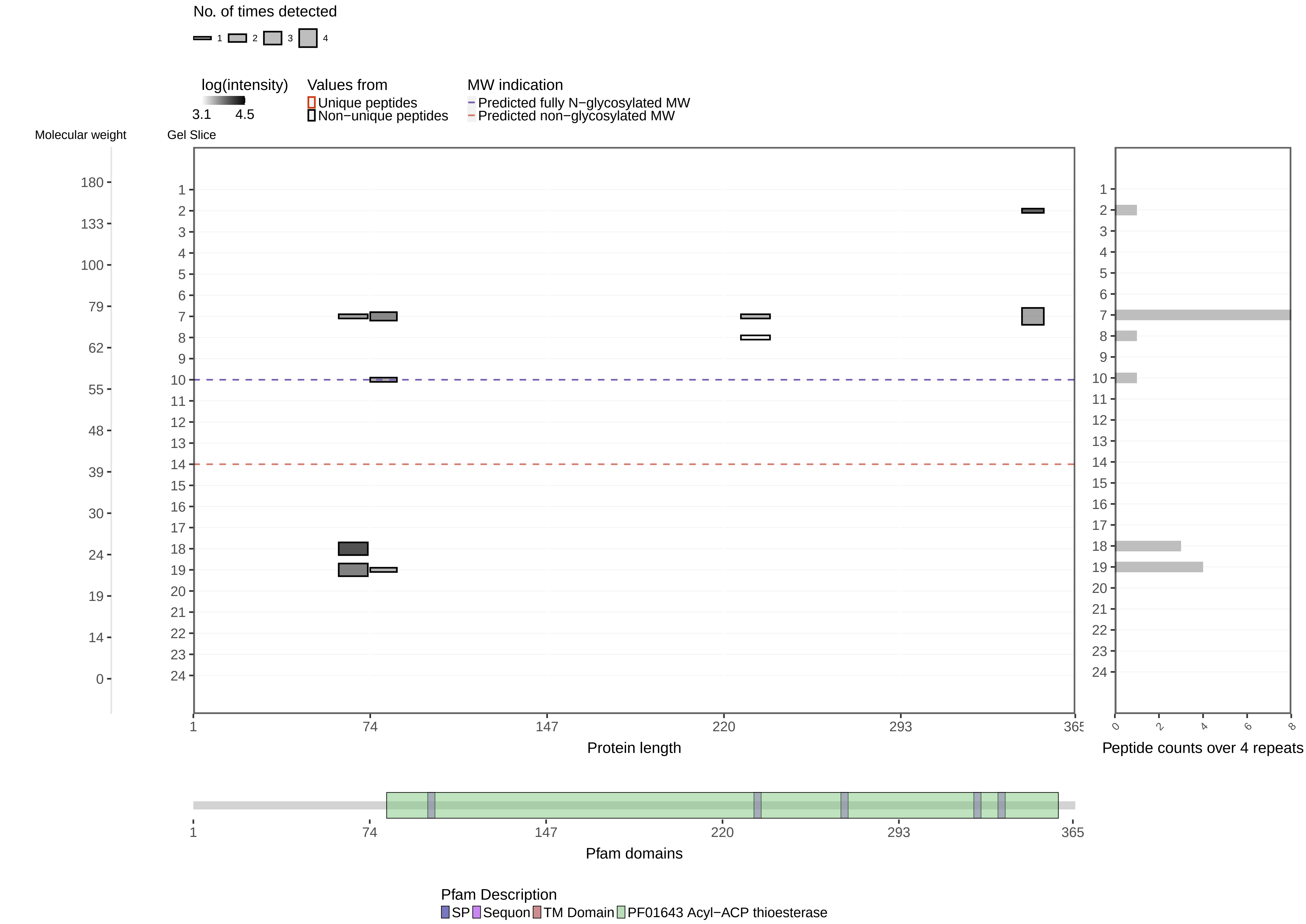


Protein sequence:

RIGTRRDWILKDYATGEFIGRATSKWVMMNQDTRRLQKVTDDVHEEYLVFCPKELRLAFPEENNSSLKKILKLEDPAHSSRLGLAPRRADLDMNQHVNNVTYIGWVLESMPQDILDTHELQTITLDYRRECQHDDVVDSLTSLESFEDSALGTNGSASASREINKSFLHLVRLSGDGLEINRGRTEWRKKSVKK*

### NbL03g14020.1 uncharacterized_LOC109209395 [MW = 29.13 kDa]

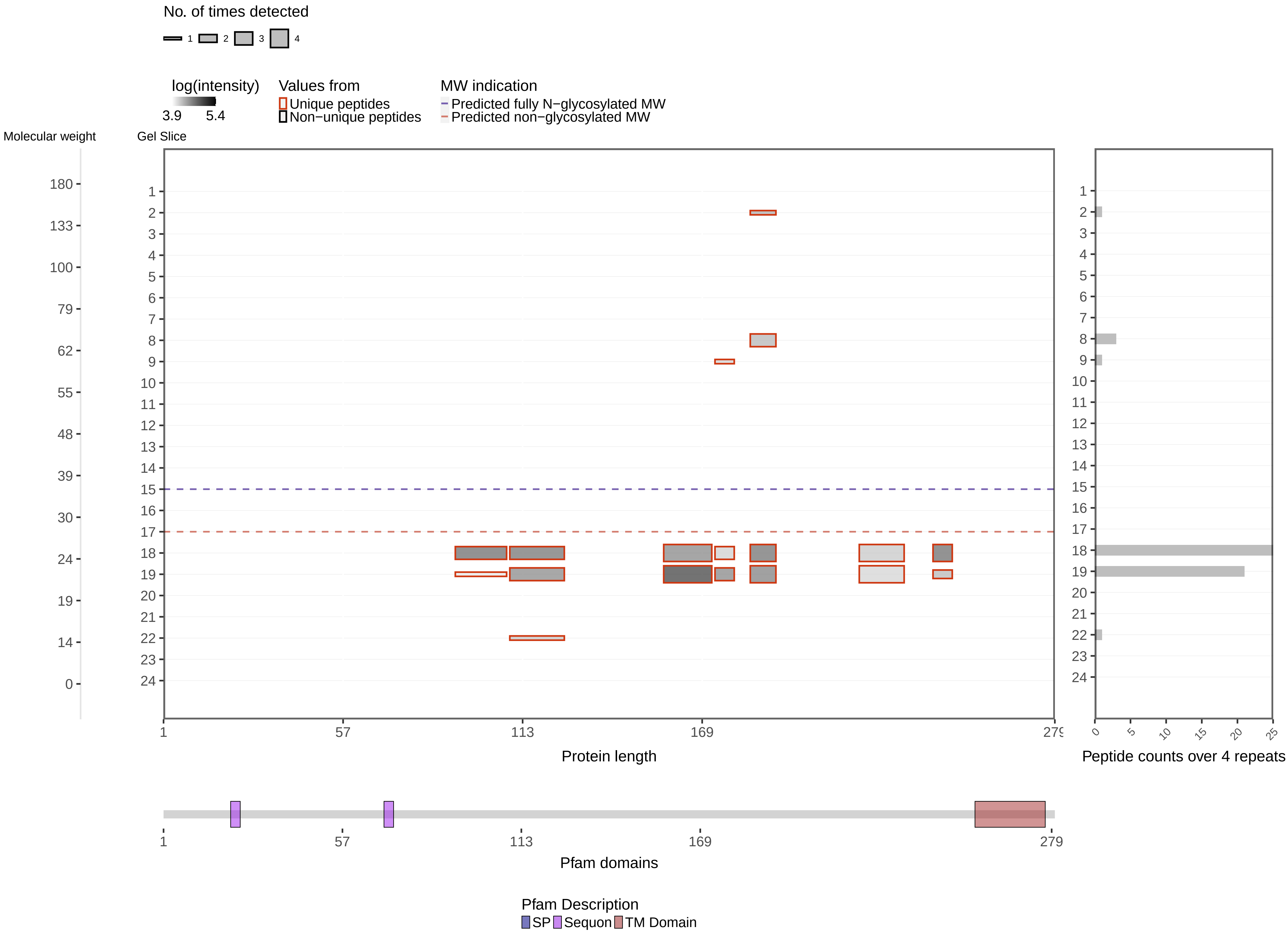


Protein sequence:

AGLKVGDQVVYTSSFFGDELWPADKLGFTKTAIQAKPDSVYFVVSRGVDVDVKRLPKRPAPPQFGRKLTDAQKARATHICLDCGYIYTLPKSFDDQPEDYACPQCRAPKKRFAKYDVNTGRAIGGGLPPIGVIIGLIAGIGGVGALLVYGLQ*

### NbL03g13970.1 membrane_protein_of_ER_body−like_protein [MW = 132.66 kDa]

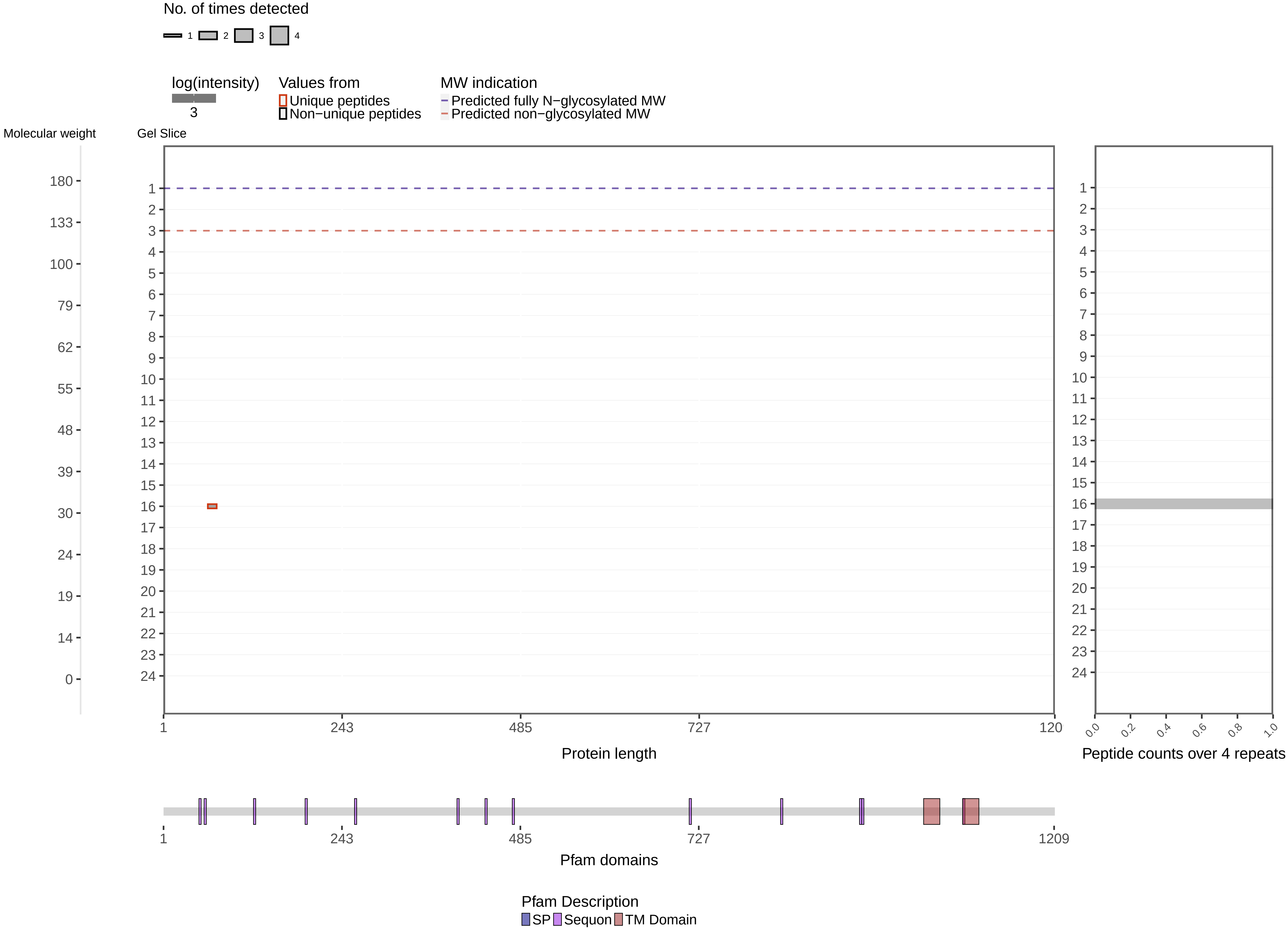


Protein sequence:

LSCFSFFIPTGNGFKLFRFFGEKGDKQNVKGEQIPTTSKNWFSSIFASDKGKASVEQGSGTGANAVKNDAGVLTPSDNFKDQSAKPLVIRESAPPAYSSSGTQEAGGKILGSENQDAKMKGKMAINTVGKSVNATKSQAGGSVFDIFEEGILNLPYVAASSEEVRGNDRNDELLRTAGRIQVSNGNLAQESLPASQQTGSKVLITSTKEESLTIEKSRTELKSDVMITSTKEESLTIEKSETDLKSDMTLLNKGPDNESLLSSATTSFALNGLDENGKPKFLTELPQEHEQNVKA

NbL03g13790.1 nascent_polypeptide−associated_complex_subunit_alpha−like_protein_2 [MW = 23.2 kDa]

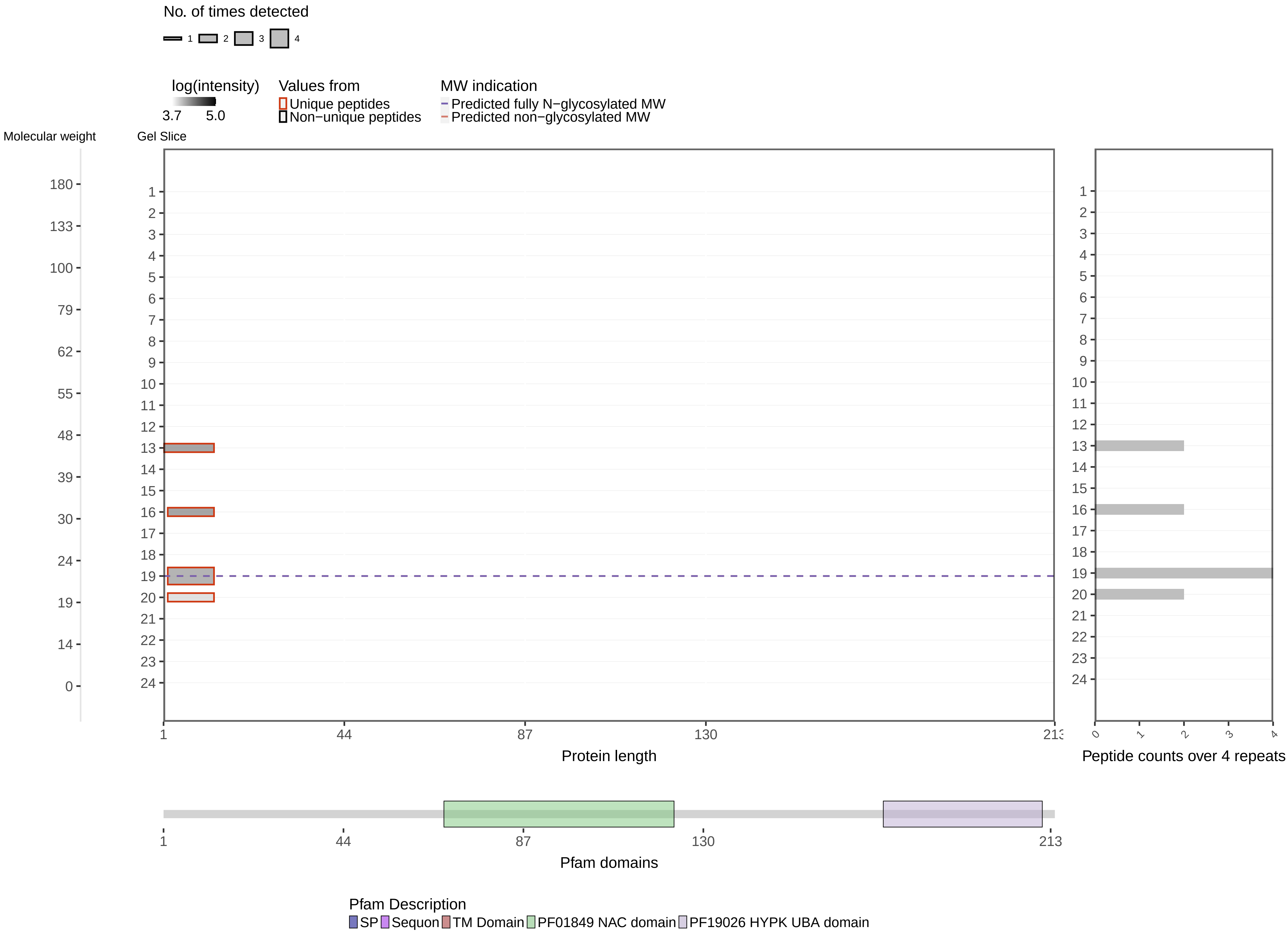


Protein sequence:

RTKNVLFFISKPDVFKSPNSETYVIFGEAKIEDLSSQLQTQAAQQFRMPDMGSVMAKSDVSPSTAAVEADEEEEEVDETGVEPRDIDLVMTQAGVSRSKAVKALKCHNGDIVSAIMELTT*

NbL03g13770.1 caffeic_acid_3−O−methyltransferase−like [MW = 39.6 kDa]

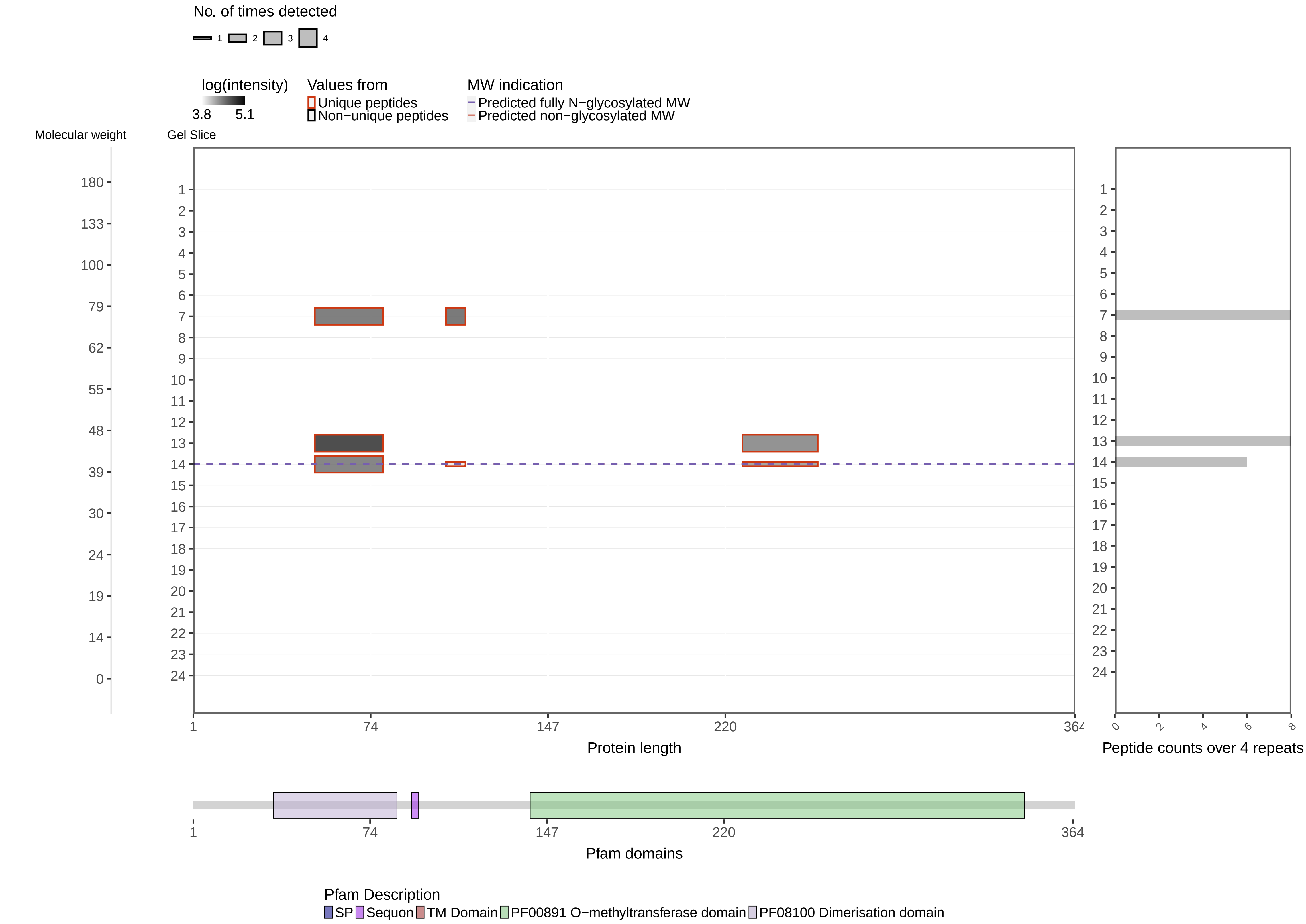


Protein sequence:

DPRFNKVFNRGMSDHSTMSMKKILEDYKGFEGLNSIVDVGGGTGATVNMIVSKYPSIKGINFDLPHVIGDAPAYPGVEHVGGDMFASVPKADAIFMKWICHDWSDEHCLKFLKNCYEALPANGKVIVAECILPEAPDTSLATKNTVHIDIVMLAHNPGGKERTEKEFOALAKGAGFTGFHKACCAYNTWVMEFNK*

### NbL03g13630.1 probable_6−phosphogluconolactonase_4_chloroplastic [MW = 37.26 kDa]

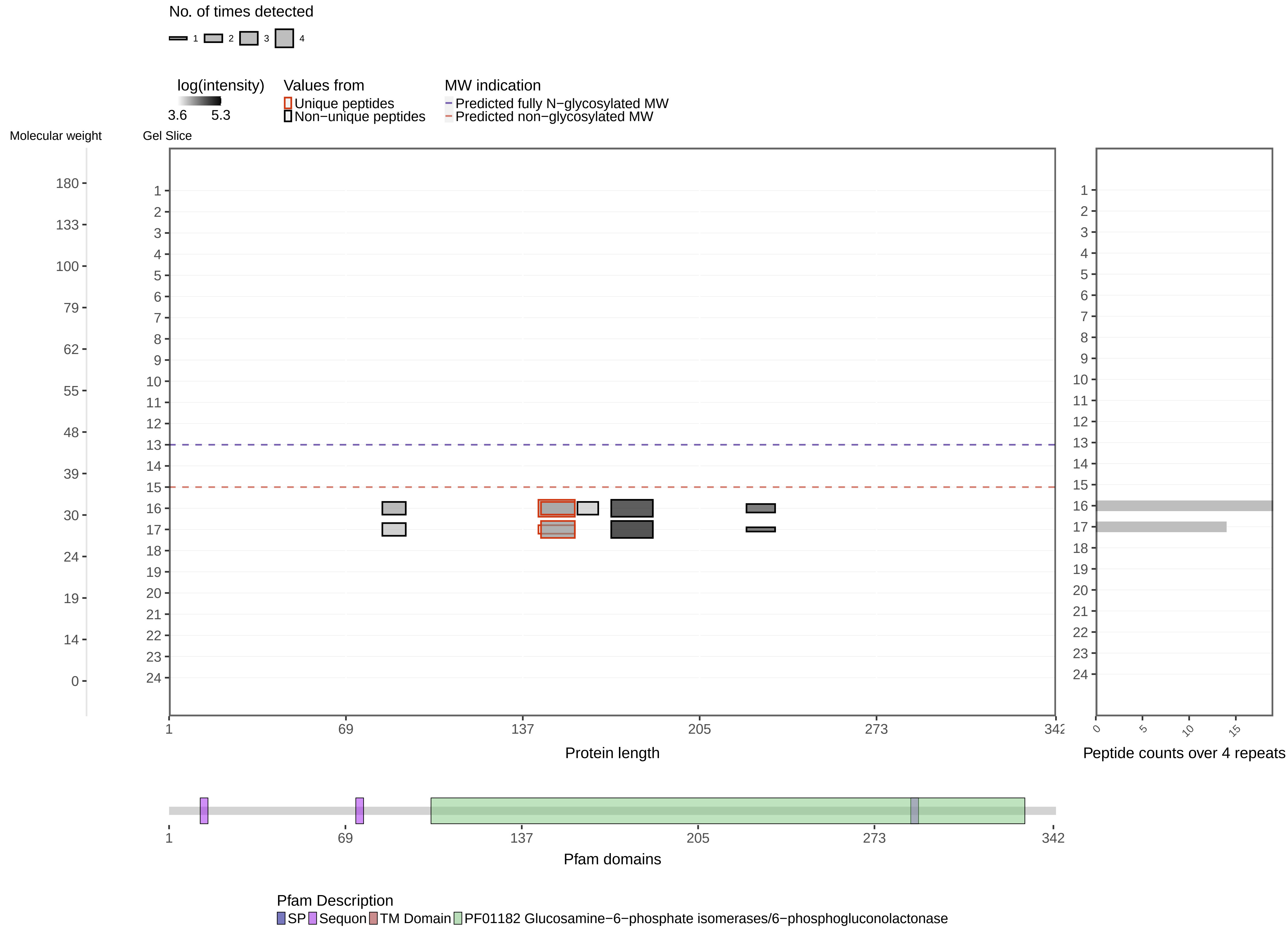


Protein sequence:

/HVFWVDERVVPKDHPDSNYLLAYDAFLSKIPIPSGNVYAINDALSAEGAADDYETCLRHLVKSKIVDISDATGFPKFDVMLLGMGPDGHVASLFPGHPLVHEKEKWVTFIKESPKPPPERITFTFPVINSSANIALVVAGAGKADVVHKSLGDSESSDLLPVOMVSPEGELIWFLDKAAASKL*

### NbL03g13390.1 protein_yippee−like_At4g27740 [MW = 12.01 kDa]

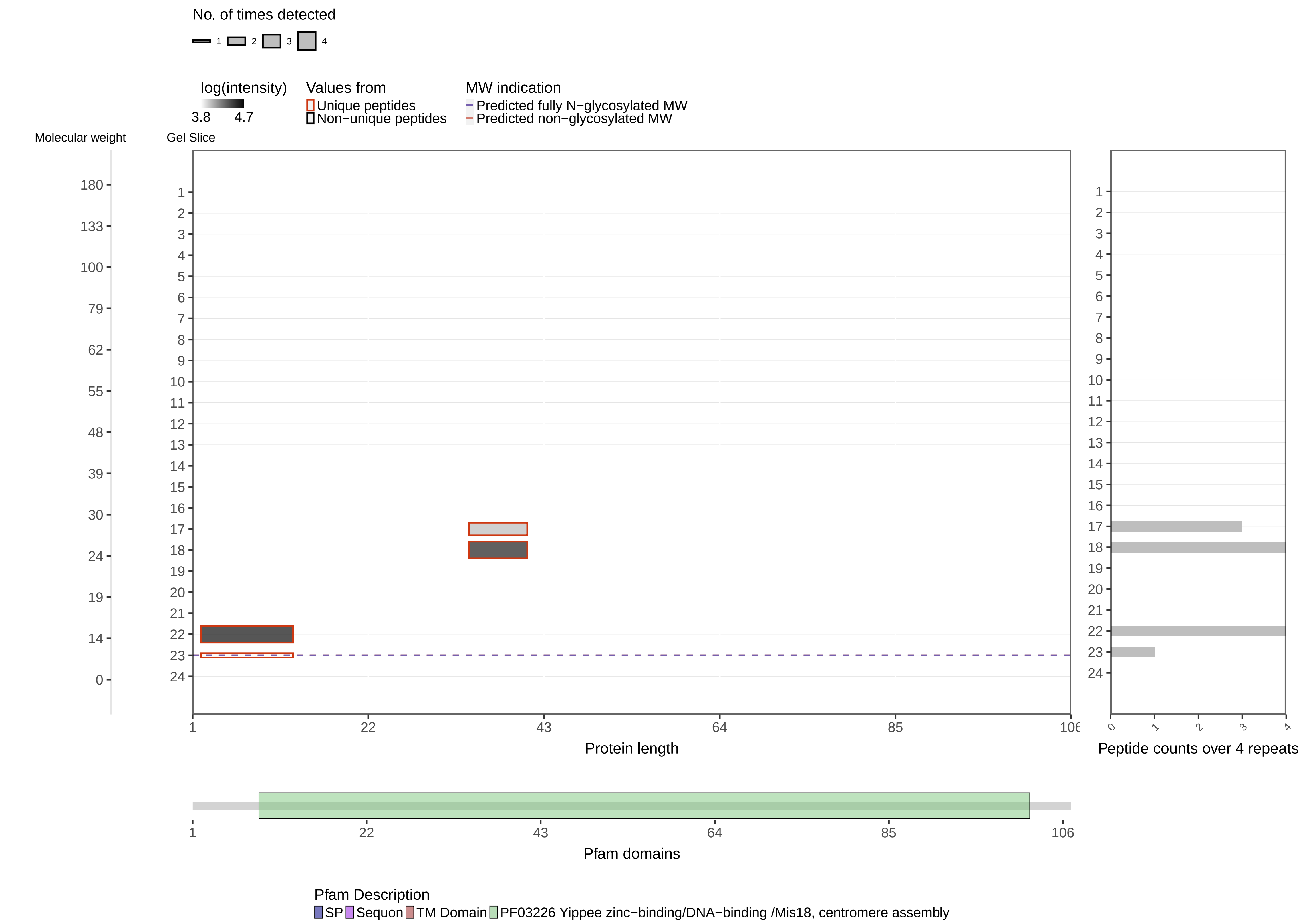


Protein sequence:

FKHAMNVVVGKKEDRQLLTGYFSVADIFCSKCGEELGWKYVKAYDPSQKYKEGRFIIETAKILKEY*

NbL03g13270.1 thylakoid_lumenal_17.4_kDa_protein_chloroplastic [MW = 27.68 kDa]

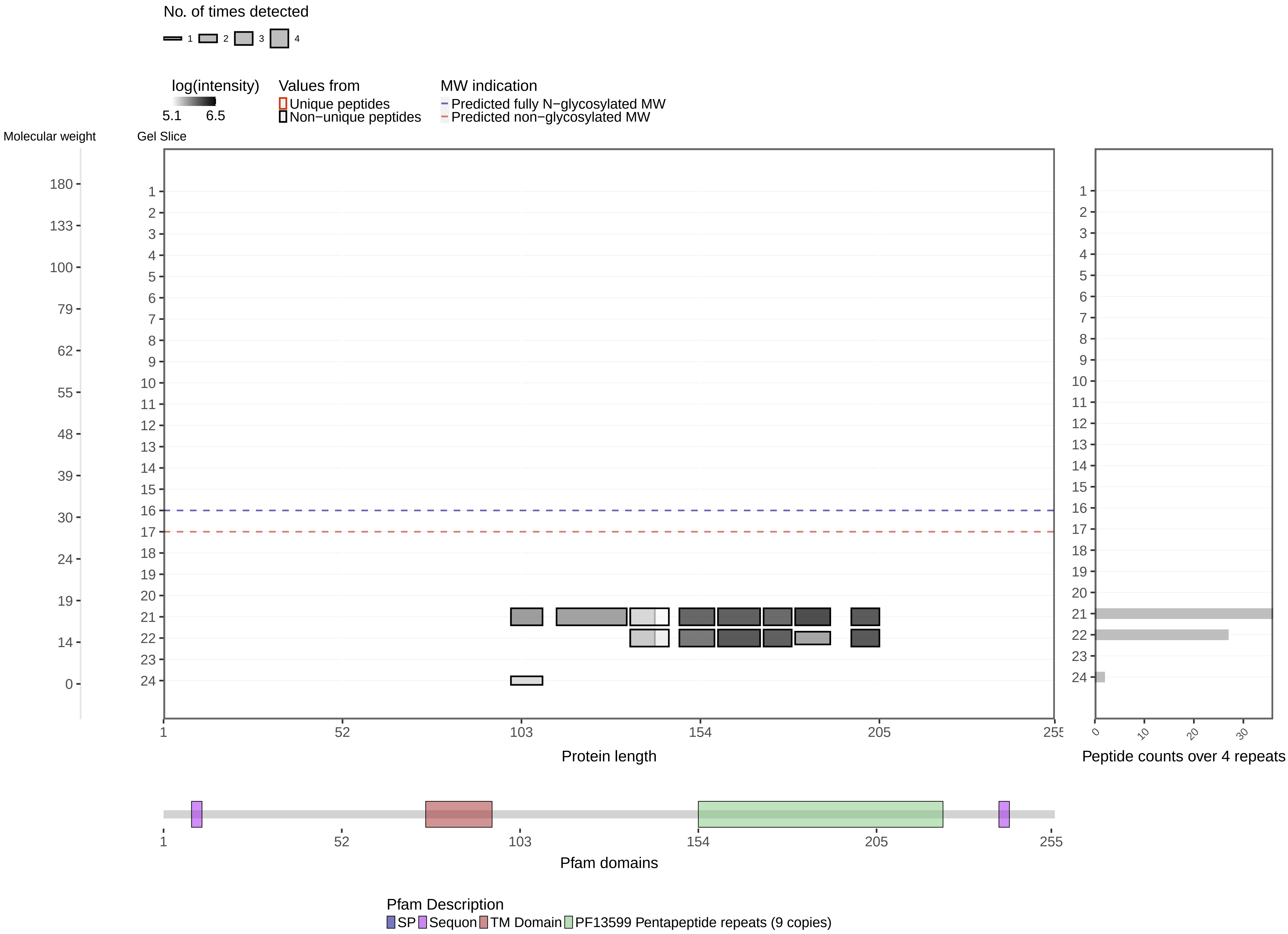


Protein sequence:

VGNTIGQANGVYDKPLDLRFCDYTNEKNNLKGKSLAAALMSGAKFDGADMTEVIMSKAYAVGASFKGTDFSNAVLDRVNFEKANLQGASFKNTVLSGSTFNDAQLEDADFEDTIIGYIDLQKICLNKTINEEGRVNLGCR*

### NbL03g13240.1 luminal−binding_protein_5 [MW − SP = 70.7 kDa]

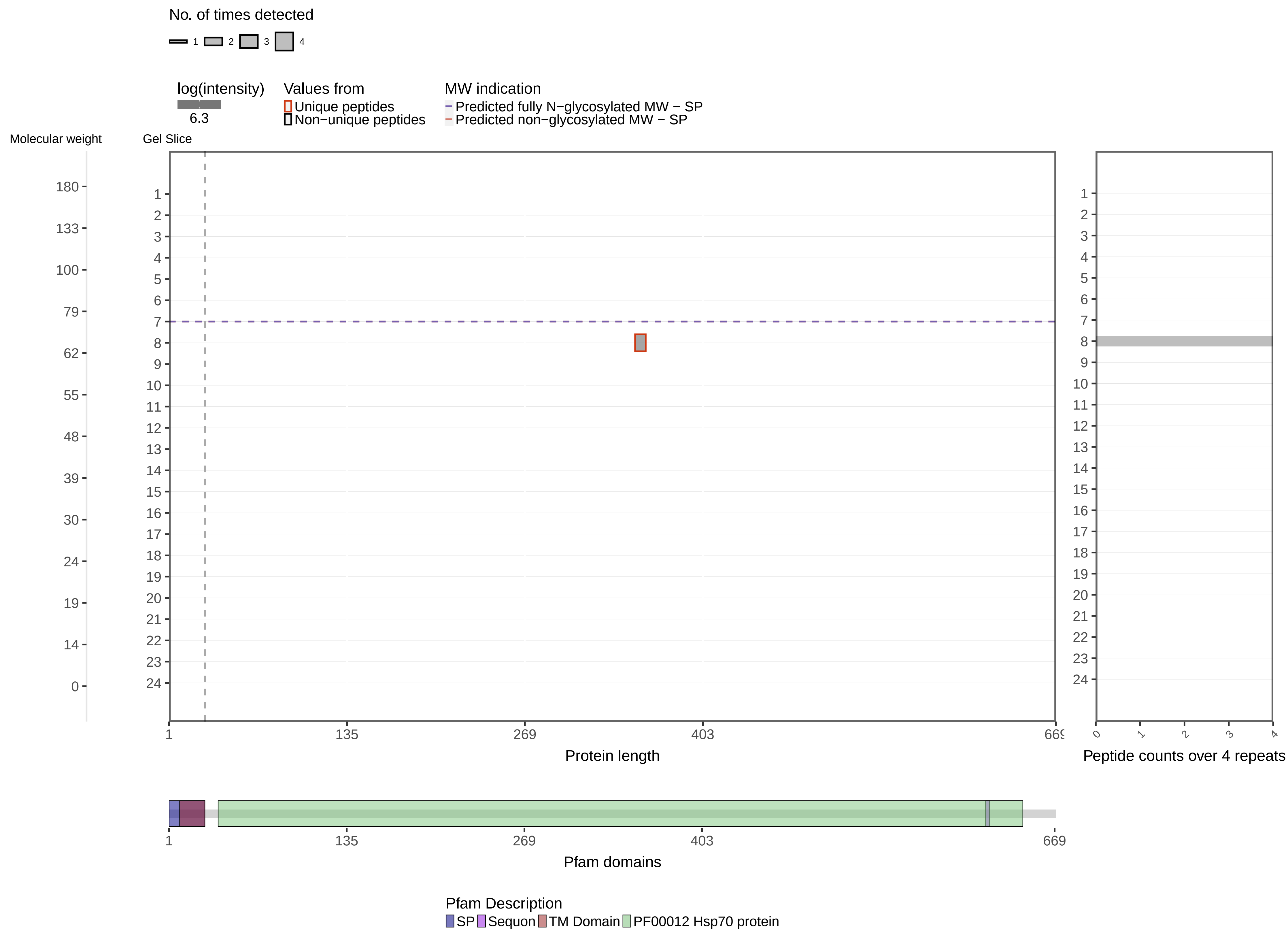


Protein sequence:

FSEPLTRARFEELNNDLFRKTMGPVKKAMEDAGLQKNQIDEIVLVGGSTRIPKVQQLLKDYFDGKEPNKGVNPDEAVAYGAAVQGGILSGEGGDETKDILLLDVAPLTLGIETVGGVMTKLIPRNTVIPTKKSQVFTTYQDQQTTVTISVFEGERSLTKDCRLLGKFDLTGIAPAPRGTPQIEVTFEVDANGILNVKAEDKASGKSEKITITNDKGRLSQEEIERMVKEAEEFAEEDKKVKERIDARNSLETYVYNMRNQINDKDKLADKLESDEKEKIETATKEALEWLDDNQSAE

NbL03g13200.1 KH_domain−containing_protein_At4g18375 [MW = 74.11 kDa]

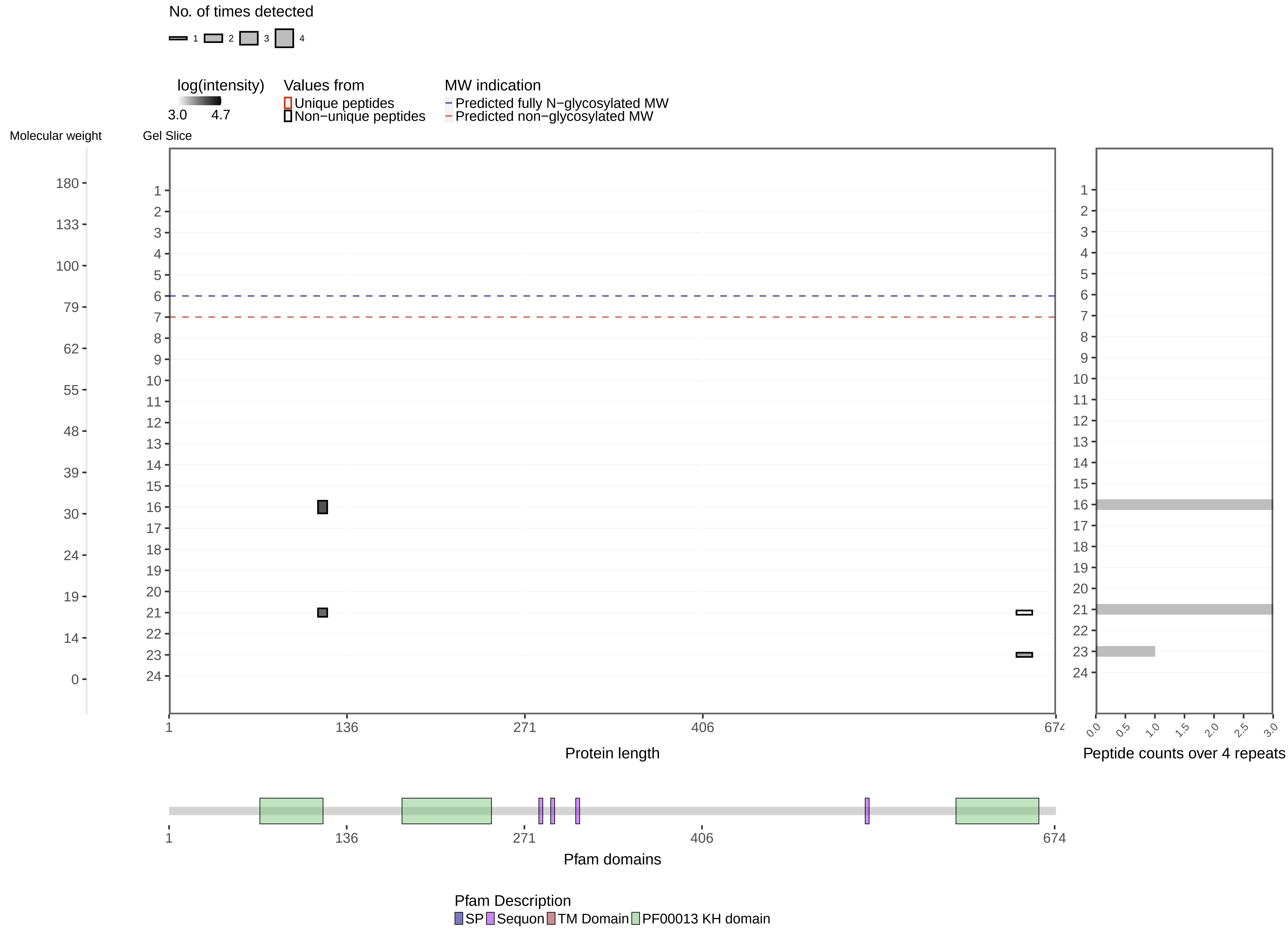


Protein sequence:

YAVESGKAPGNENGPVMYVEDLVFRILCPVDKVDTVAGESDGIIELLONEIGVDVKILNPVASSNEOVIIIMSDEGPDDELFPAOEALLHIOTRIVDLVPEKENVITTRLVVOTDEVECLGGRDGPLSDMOKITGATVOILPKEELPPCVSGTDEIIOIVGEIKAAREALVEVTSMLRSCTYREFFOKDIPSPAVPASSPARCNIGADKSSVNNTSPSOONYNVNDVOTSIYONAPAKLMTOPVKETGASASEIAKONENERCEDIPSGLNRMHVPLVTRSTLEVVIPPHAVPKLITKSF

### NbL03g12940.1 dipeptidyl_aminopeptidase_4 [MW = 87.25 kDa]

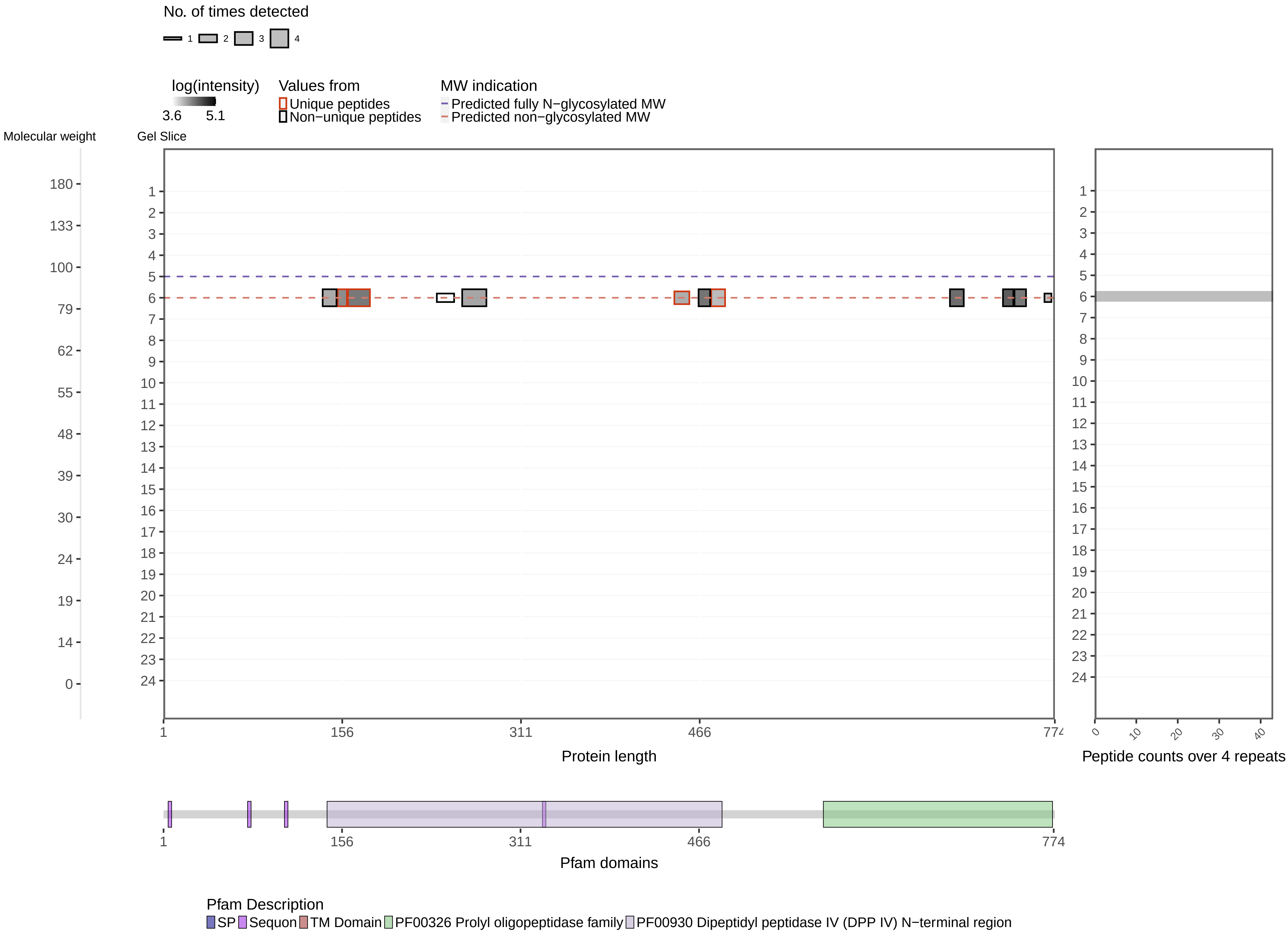


Protein sequence:

[illegible]

### NbL03g12880.1 protein_BOBBER_1−like [MW = 33.98 kDa]

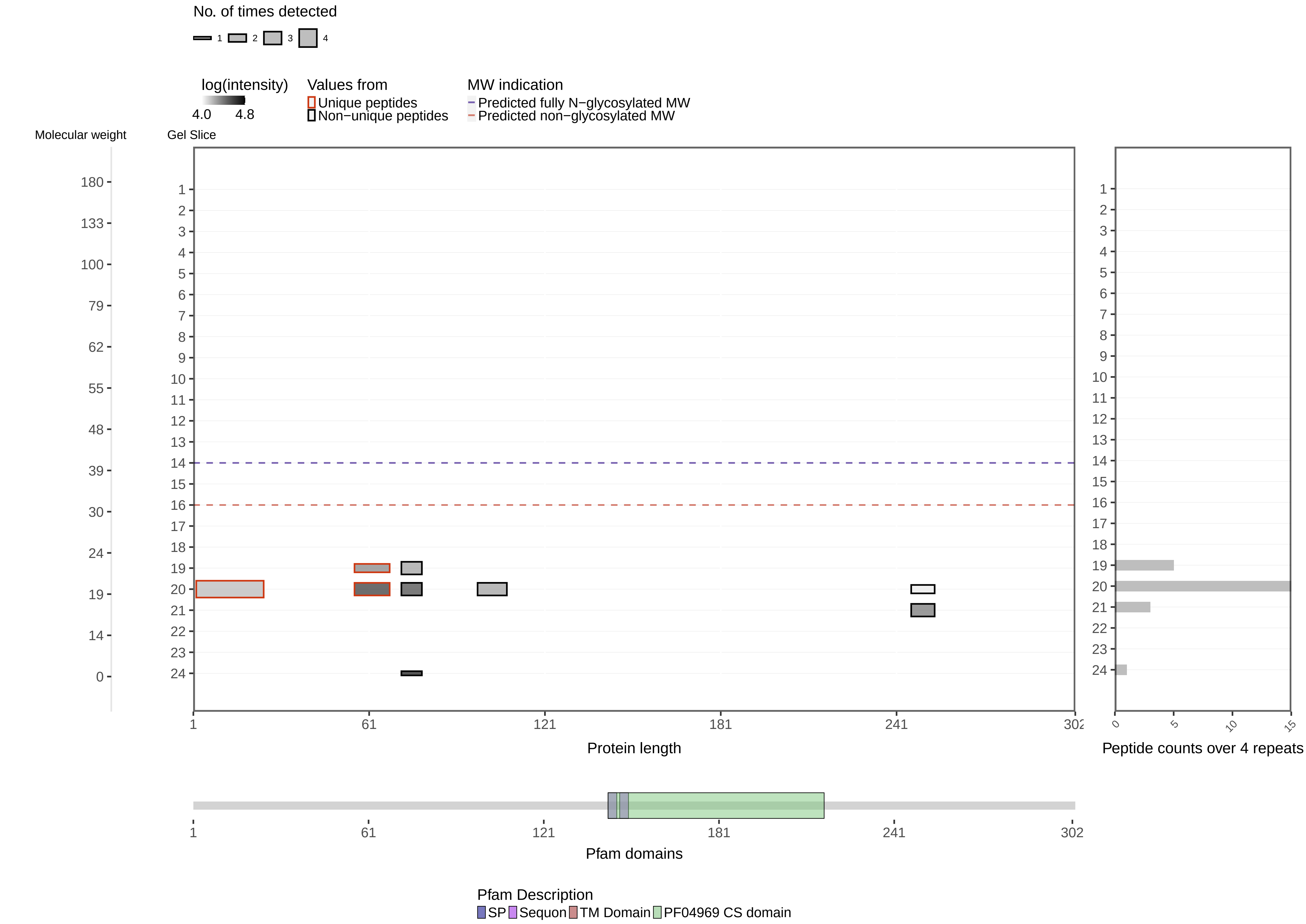


Protein sequence:

DLENYSWNQSLQEVNVNIPVPHGTKSRFIVCDISKNHLKVGLKGQPPIIDGELYRPVKVDDCFWSLEDQKSISVLLTKKDQMEWWKCCVKGEPEIDTQKAEPESSKLADLDPETRSTVEKMMFDQRQKSMGLPTSDETQKQEILKKFMAEHPEMDFSKAKIS*

### NbL03g12860.1 probable_plastid−lipid−associated_protein_14_chloroplastic [MW = 75.92 kDa]

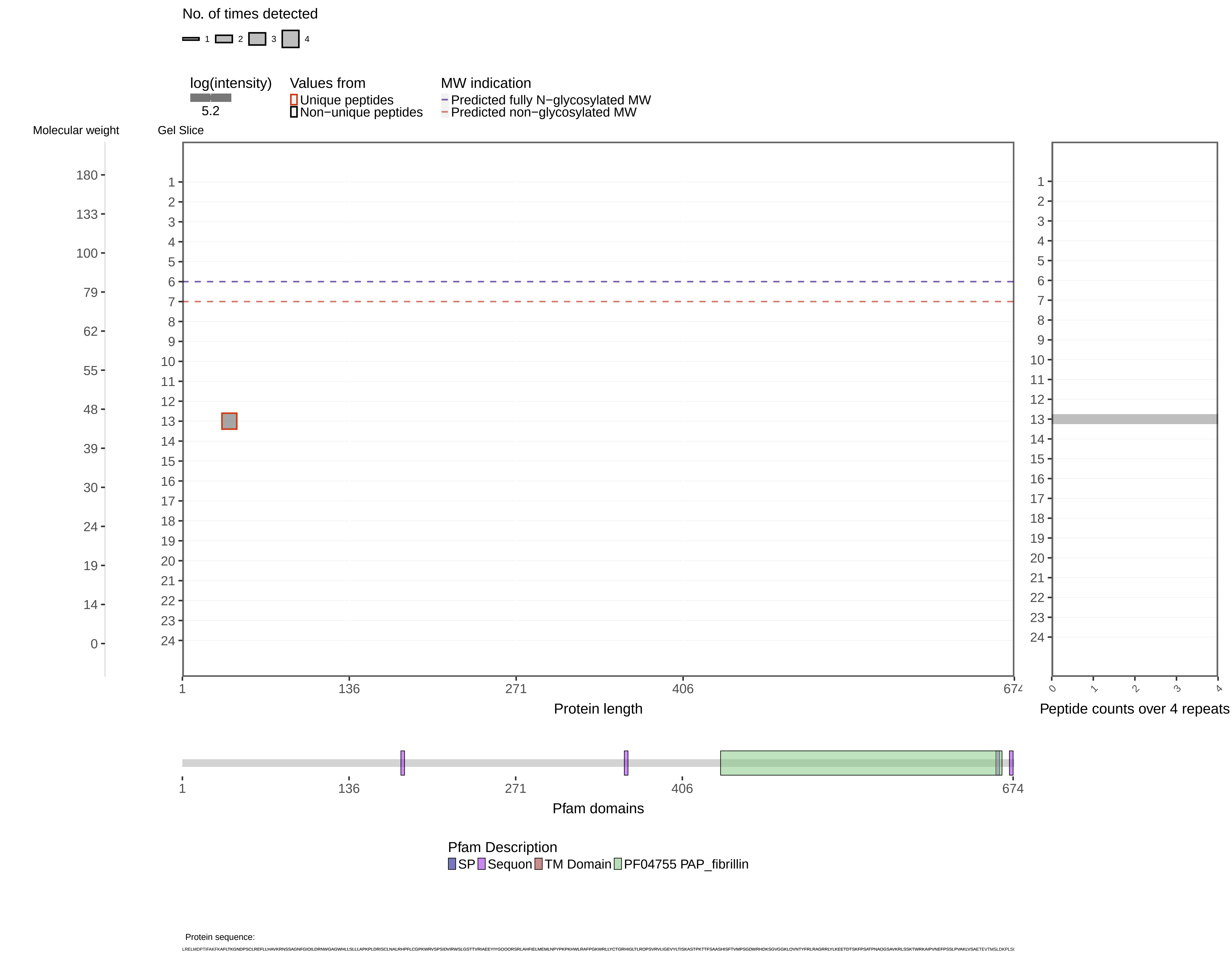

NbL03g12840.1 splicing_factor_3A_subunit_3−like [MW = 59.76 kDa]

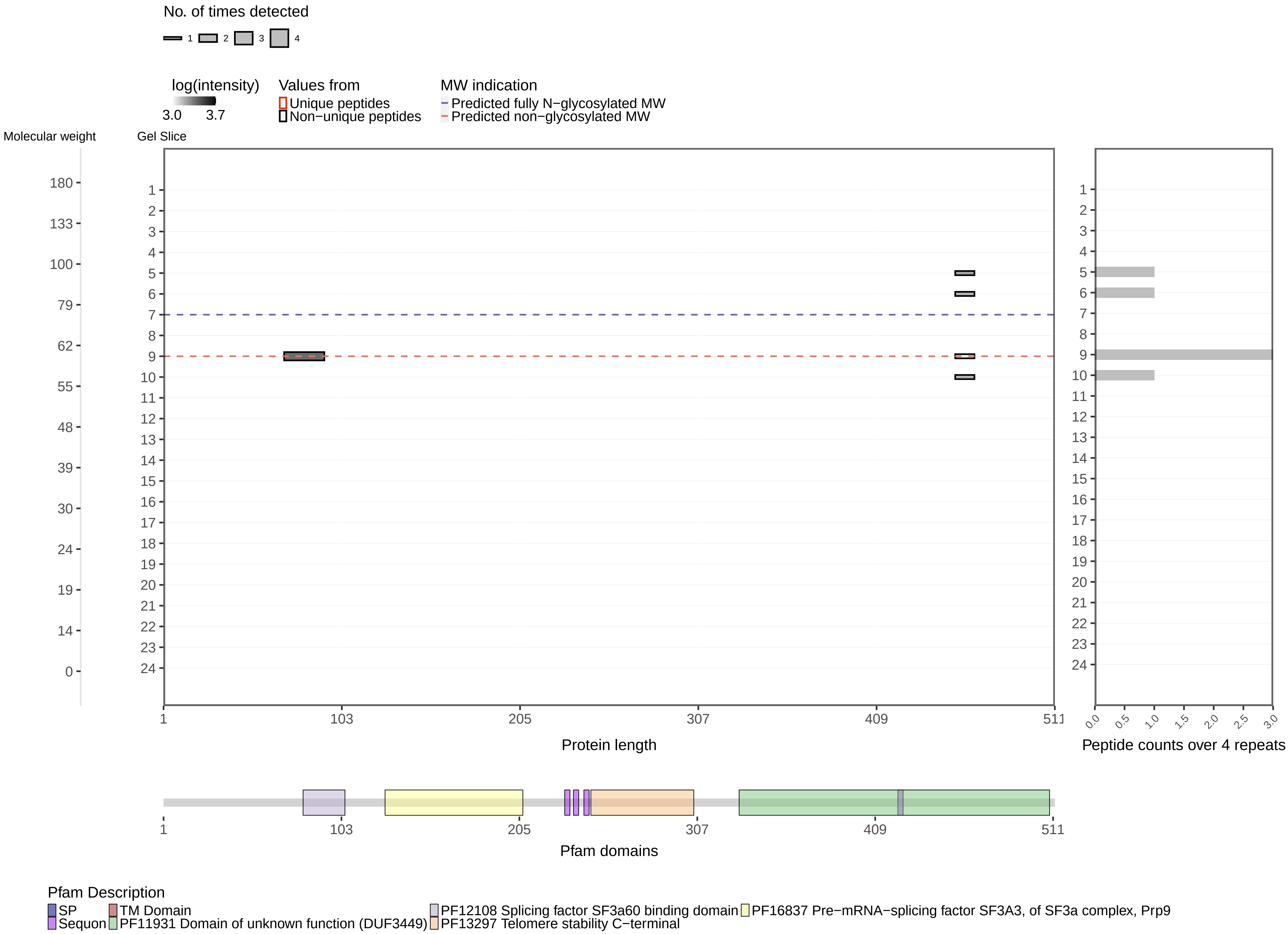


Protein sequence:

VIDLNSYSTVEELMIEGPDKLKEALGALGLKTGGTIQQRAERLFLTKHTPLEKLDRKHFAKGSRKPEQNGVMPAPQVSDDVKEIALMEAKIRKLCDLLDETIVRTKENVEKKQALTYEEMEAEREEDDVQAESESDEEDQQIYNPLKLPMGWDGKPIPYWLYKLHGLGQEFKCEICGNHSYWGRRAYERHFKEWRHQHGMRCLGIPNTKNFNEITSIEEAKQLWERIQEKQGLNKWRPDLEEEYEDQEGNIYNKKTYTDLQRQGLI*

NbL03g12820.1 glutamate_synthase_1_[NADH]_chloroplastic [MW = 243.02 kDa]

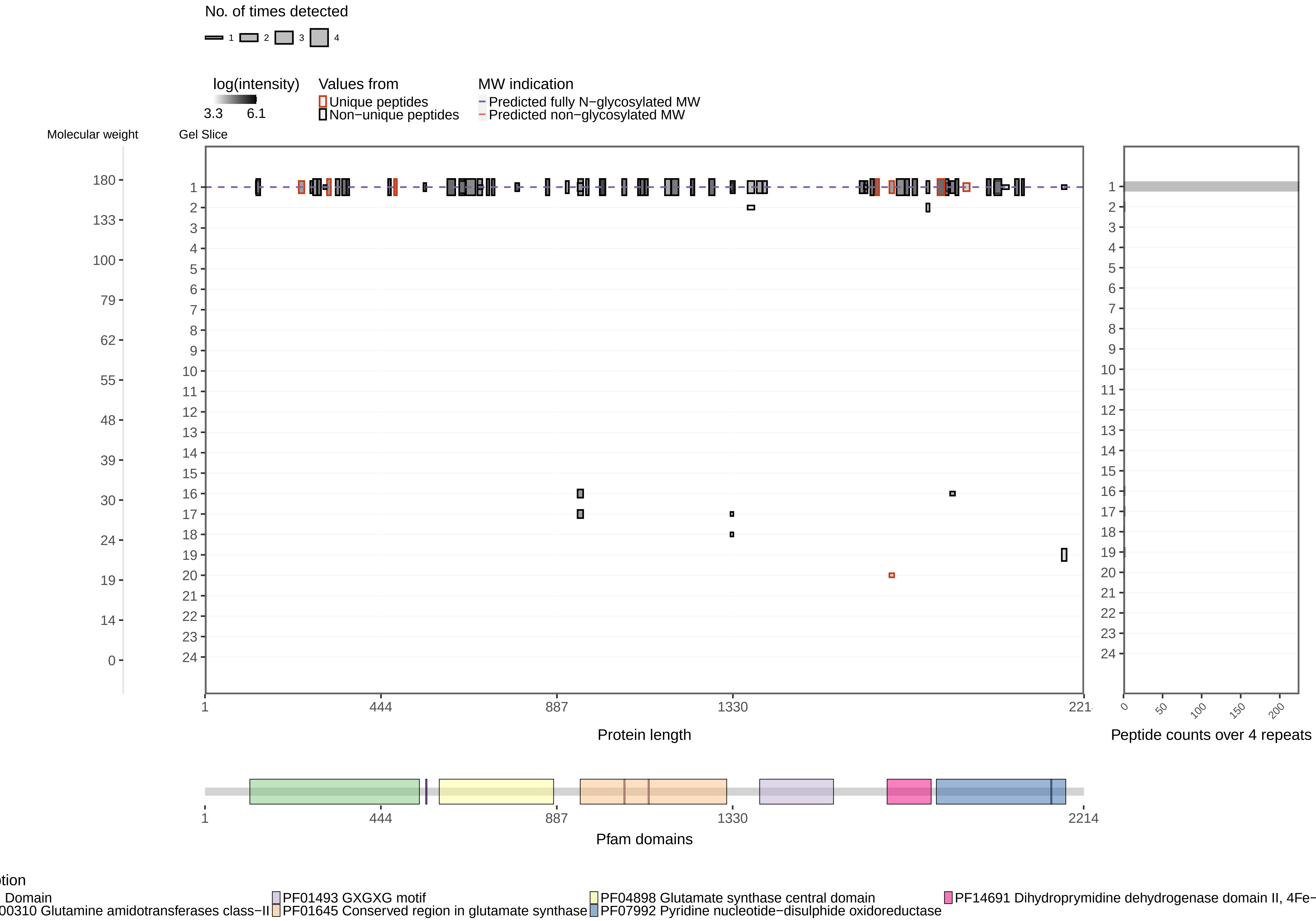


Protein sequence:

OGAKPGEGGELPGHKVIGDIAVTRNSTAGVGLISPPPHHDIYSIEDLAOLIHDLKNANPEARVSVKLVSEAGVGVIASGVVKGHADHVLISGHDGGTGASRWTGIKSAGLPWELGLAETHOTLVANDLRGRTTLOTDGOLKTGRDVAVAALLGAEEFGFSTAPLITLGCIMMRKCHKNTCPVGIATODPILREKFAGEPEHVINFFFMLAEEVREIMSHLGFRTLTEMVGRSDMLELDKDLTKNNDKLKNIDLSLLLRPAADIRPEAAOYCVOKODHGLDMALDNNLIALSKAALE

NbL03g12720.1 probable_protein_phosphatase_2C_76 [MW = 38.77 kDa]

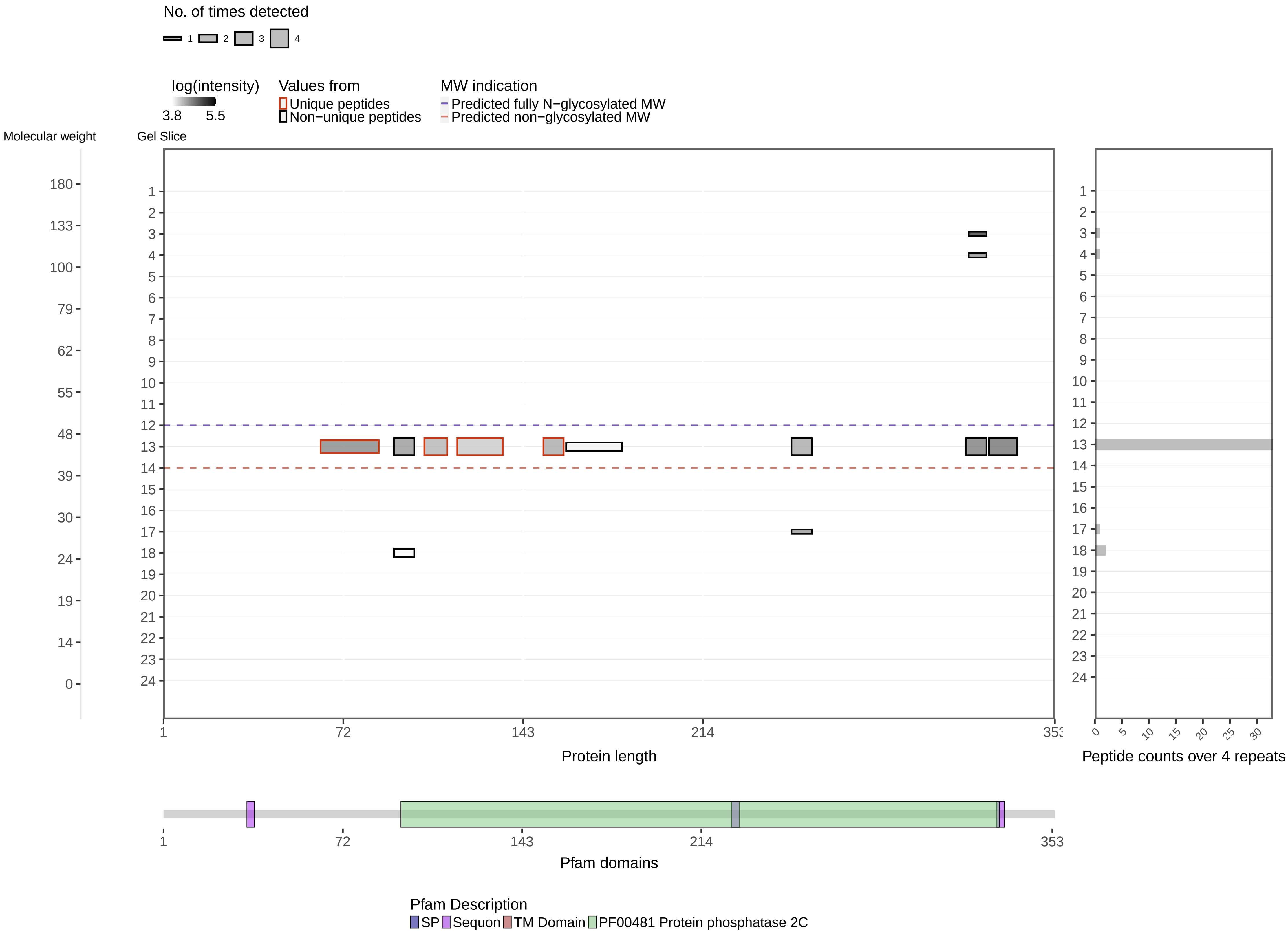


Protein sequence:

SETYOOTDMDFLDSEKDTFRDDGSTASTAVLVGNHLYVANVGDSRTIISKGGKAIALSEDHKPNRTDERKRIESAGGVVMWAGTWRVGGVLAMSRAFGNRMLKOFVVAEPEIODOEIDEEIELLVLASDGLWDVVPNEDAISLAEAEEEPEAAARKLTETAFTRGSADNITCIVVKFHHKKVEPEGSOOG*

### NbL03g12490.1 uncharacterized_LOC109206341 [MW = 19.05 kDa]

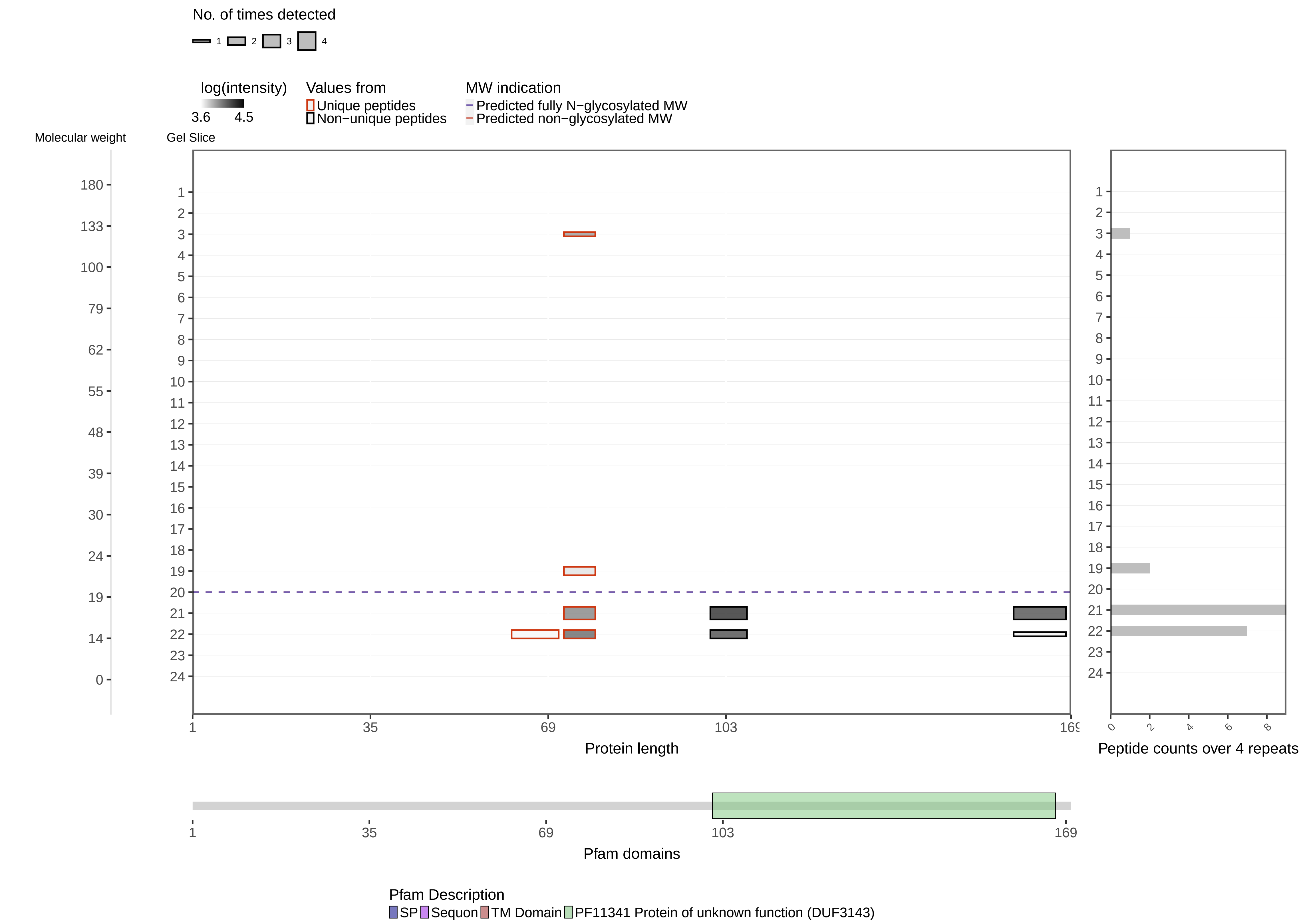


Protein sequence:

DEWLQKLPDKKKALYSHSLPCIEAWLRSLGFYQSRDDRALWFVEKPDWHAHLSLDITDIYIRYLKNGPGNLEKDIERRFSYALSREDIENAILGGP*

NbL03g12390.1 probable_NADH_dehydrogenase_[ubiquinone]_1_alpha_subcomplex_subunit_5_mitochondrial [MW = 25.77 kDa]

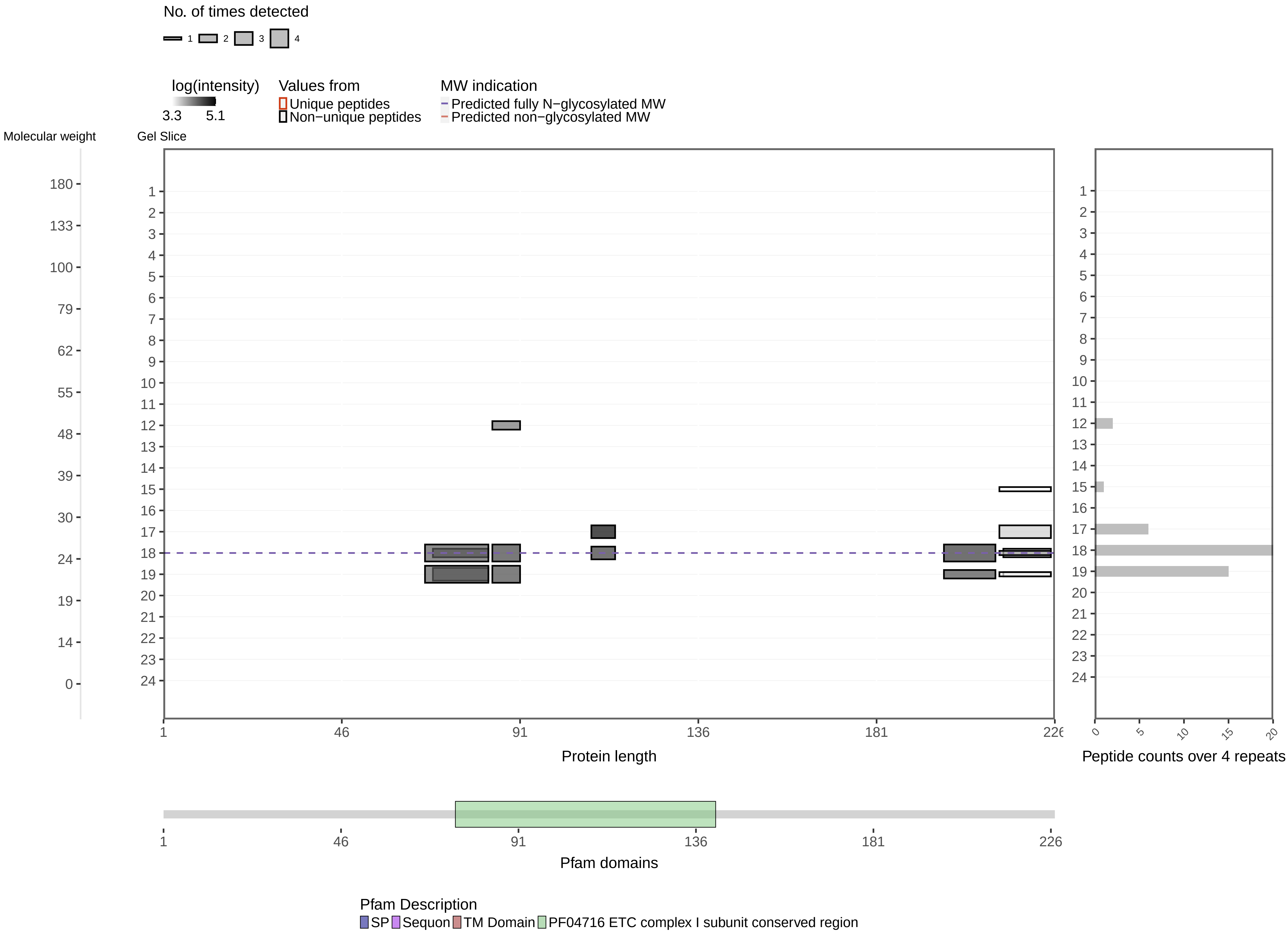


Protein sequence:

EDEGYRKAVESFTRHRLSVCQEEEDSEMIEKRLGCGQVEELIEEAQDELKLIGHMNEWKPWGVPDDYECEVIENDAPVPKHIPLHRPGPLPEEFYKTLEAVTSGTLETSKKDEPAIASGESQPK*

### NbL03g12380.1 protein_SGT1_homolog [MW = 47.56 kDa]

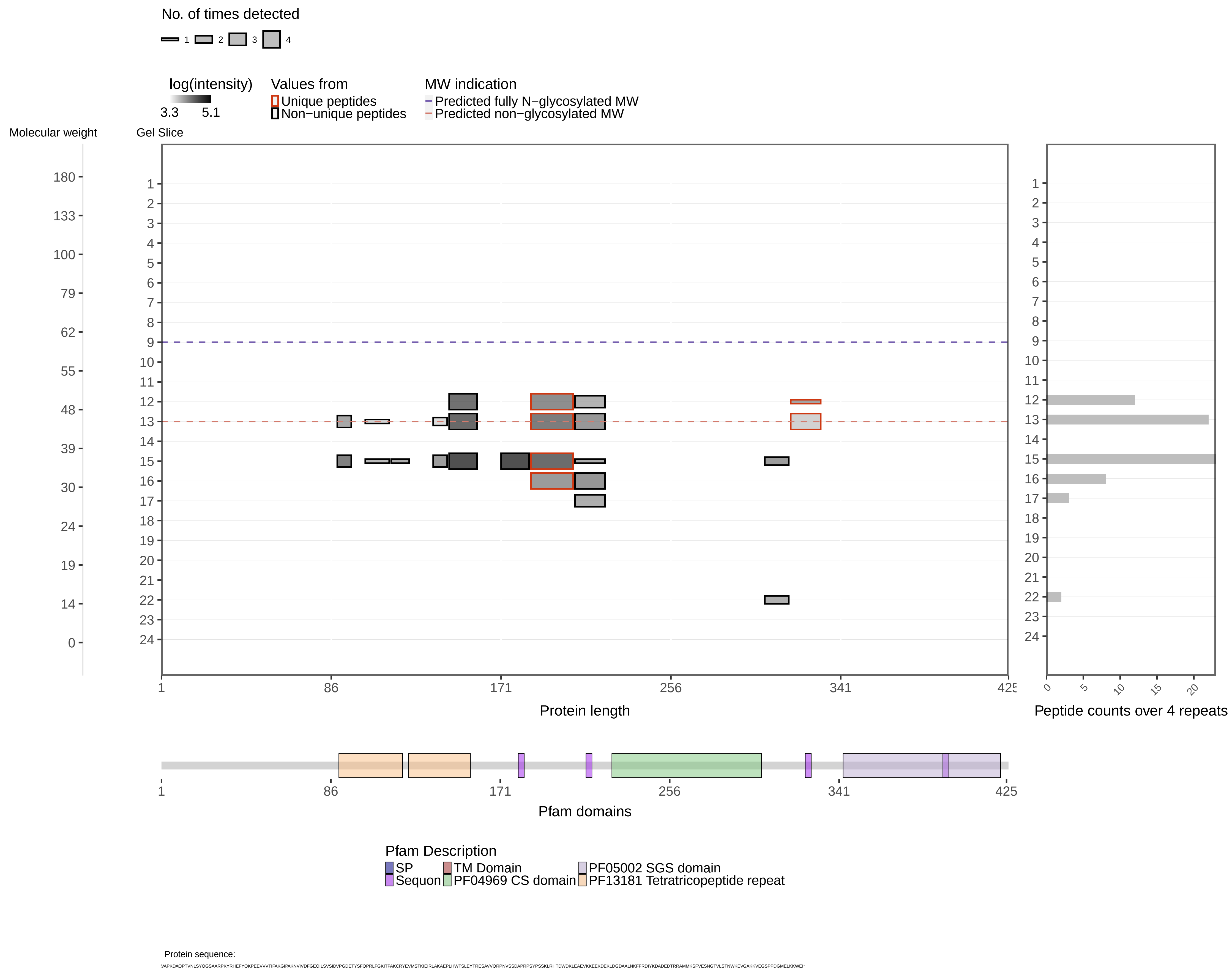

NbL03g12220.1 heat_shock_protein_82 [MW = 80.76 kDa]

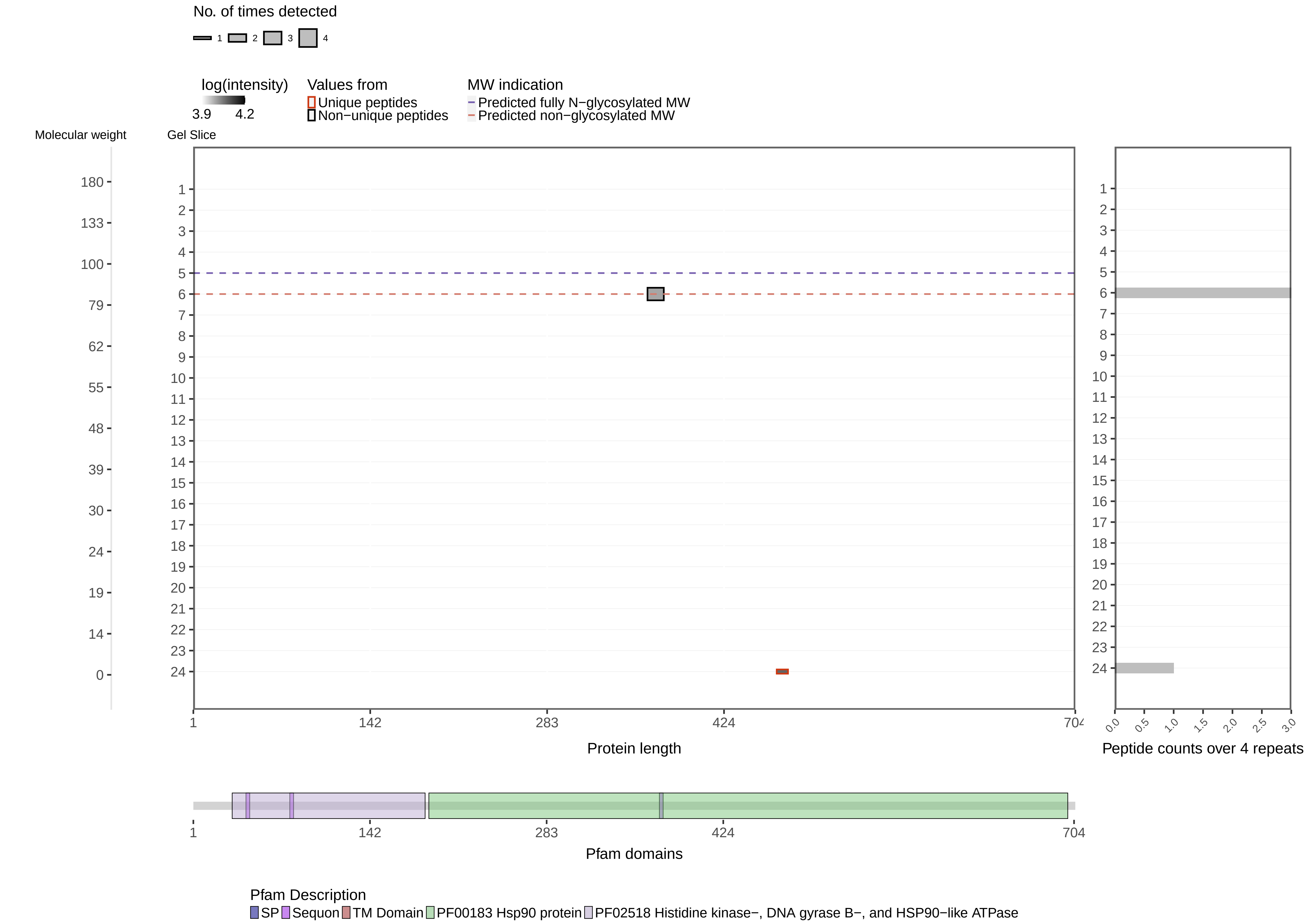


Protein sequence:

/RRVFIMDNCEELIPEYLGFVKGVVDSDDLPLNISREMLQQNKILKVIRKNLVKKCIEMFNEIAENKEDYNKFYEAFSKNLKLGIHEDSQNRAKLADLLRYHSTKSGDEMTSLKDYVTRMKEGQKDIYYITGESKKAVENSPFLERLKKKGYEVLYMVDAIDEYAVGQLKEYDGKKLVSATKEGLKLDDDSEEEKKKKEEKKKSFENLCKIIKDILGDKVEKVVVSDRIVDSPCCLVTGEYGWTANMERIMKAQALRDSSMSSYMSSKKTMEINPDNGIMEELRKRAEADKNDKS

### NbL03g12140.1 plastidial_pyruvate_kinase_2−like [MW = 66.32 kDa]

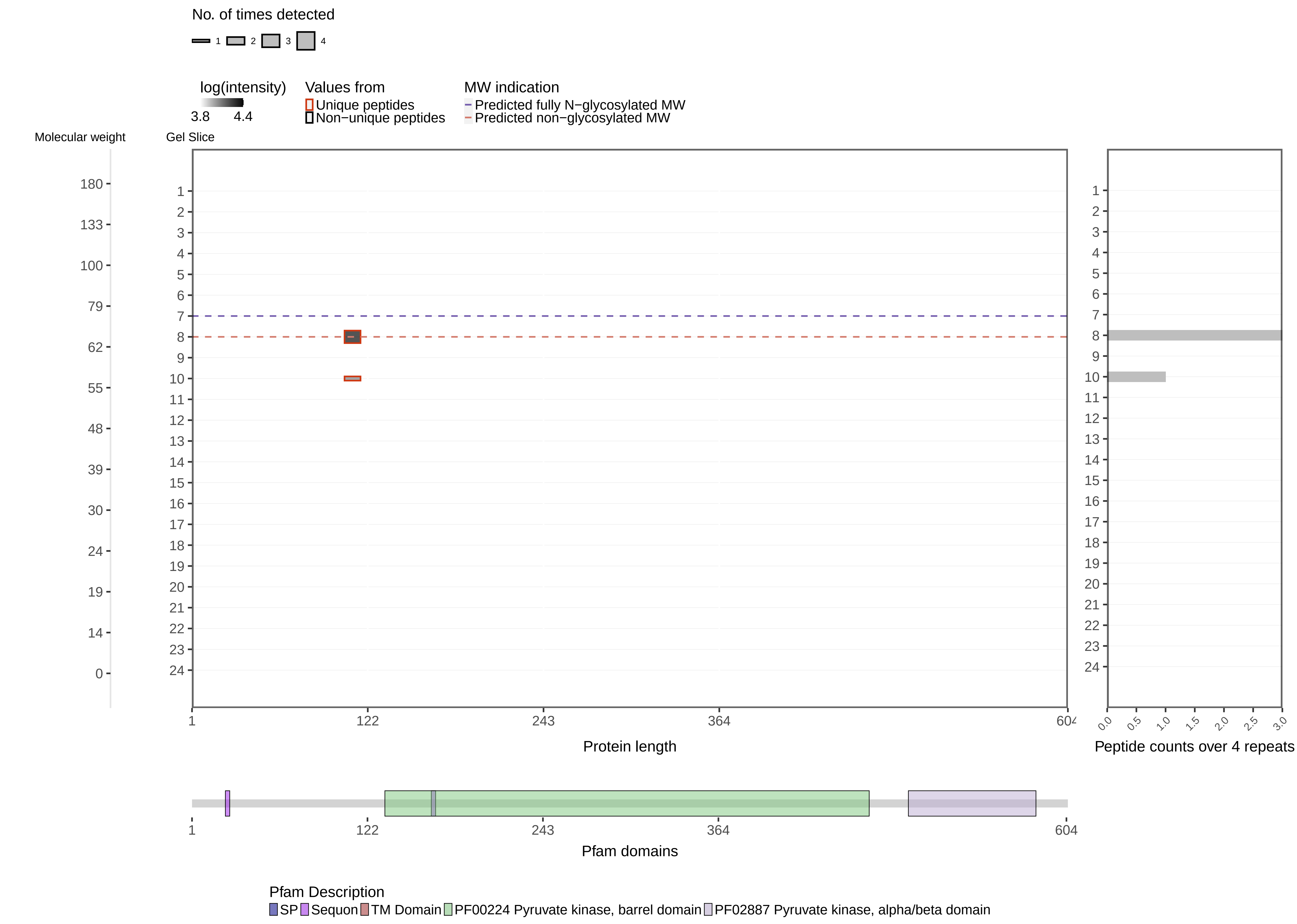


Protein sequence:

### NbL03g11990.1 sec−independent_protein_translocase_protein_TATB_chloroplastic [MW = 35.09 kDa]

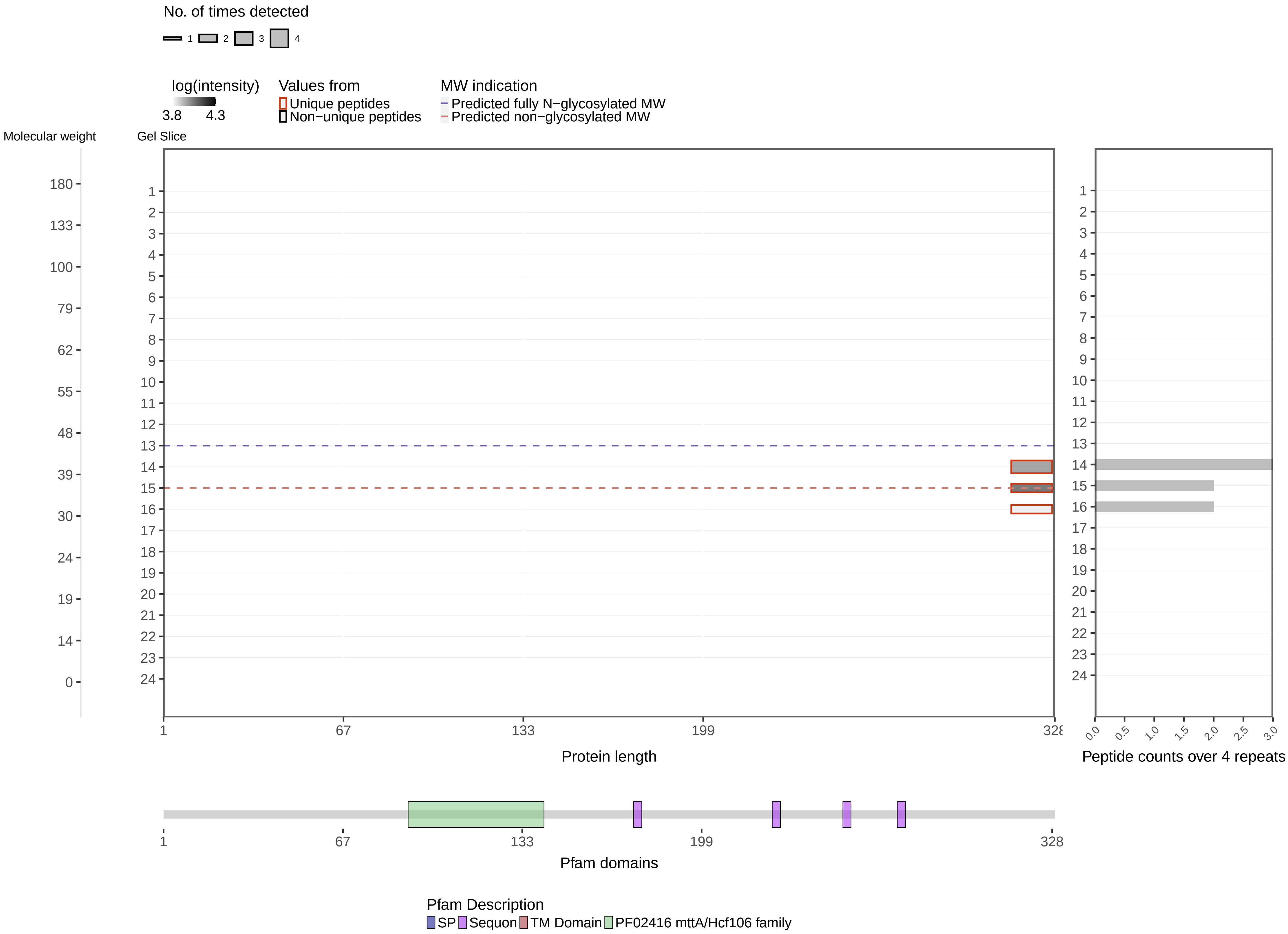


Protein sequence:

ↄIKGSGODTIKSSTTSROEVDPNGSPSPSIASSSEEDMERLLRIAEAEKOAEKDLASLLESOSESRTVSOVADNSSSAYSTEDYLKITEEOLKAAAOOONESSIPEOISSNAOSLSOADNSSSSDGAYSTDEVSKITEOOPSSPOOSLSNAOSOLEETLKEAASAIPSSOKPESET*

### NbL03g11960.1 protein_SRC2_homolog [MW = 34.9 kDa]

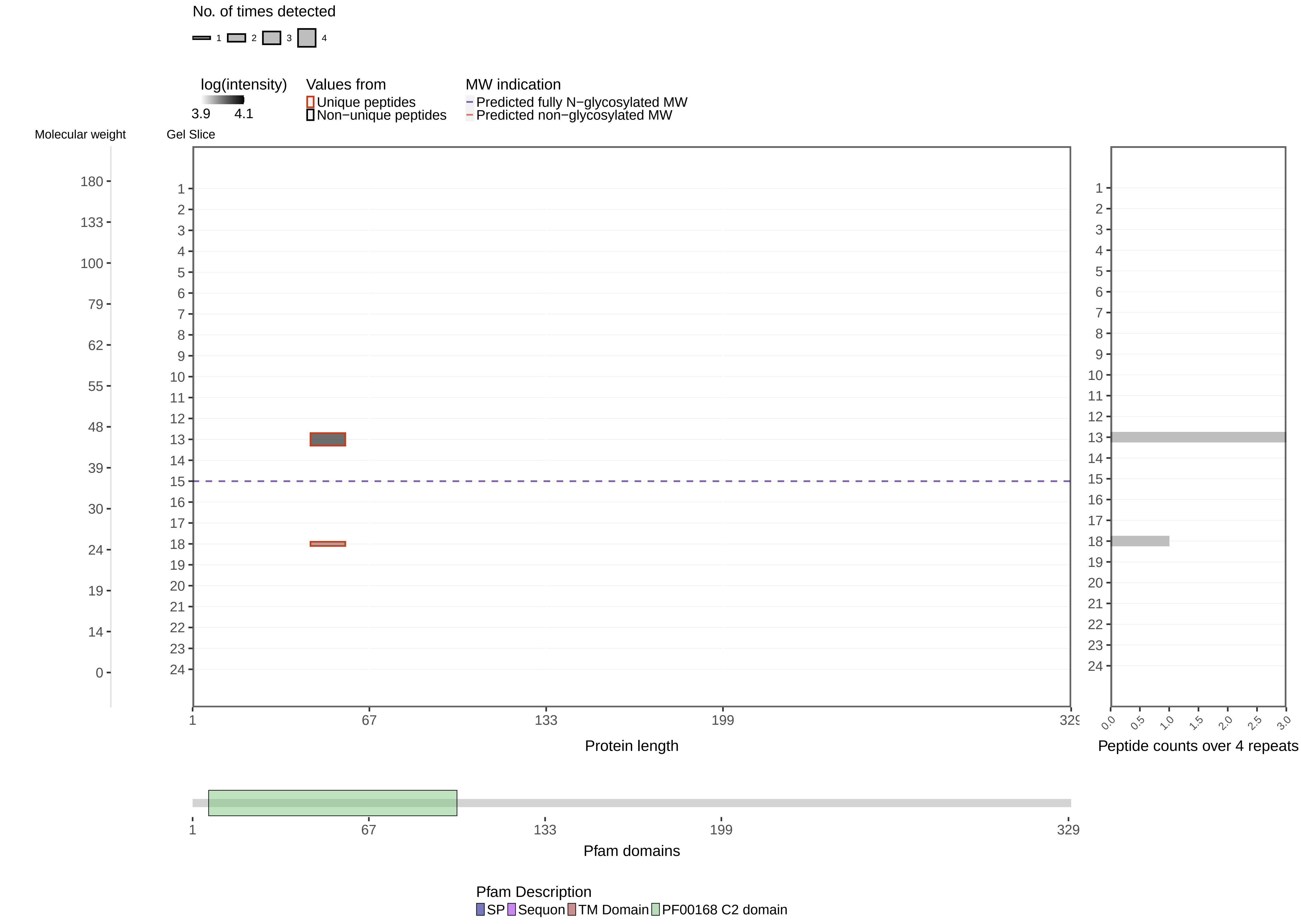


Protein sequence:

SRDYPAAPPQPYGGSYGAPSPAPSAAPPQPYGGSYGAPAPYAAPPSGYPYSAAPAPAPYGQQQPNYGAPPAPYSQGGSYGQPGYSMGTYGQQQGNYGQASYGQQSGYKCEEEKKKGKFGGMGLGAGLAMGAVAGALGGLAISEGIDHIEDNIADKAAEKVEDDLADDDYDGDDF*

NbL03g11930.1 short−chain_dehydrogenase_TIC_32_chloroplastic−like [MW = 39.32 kDa]

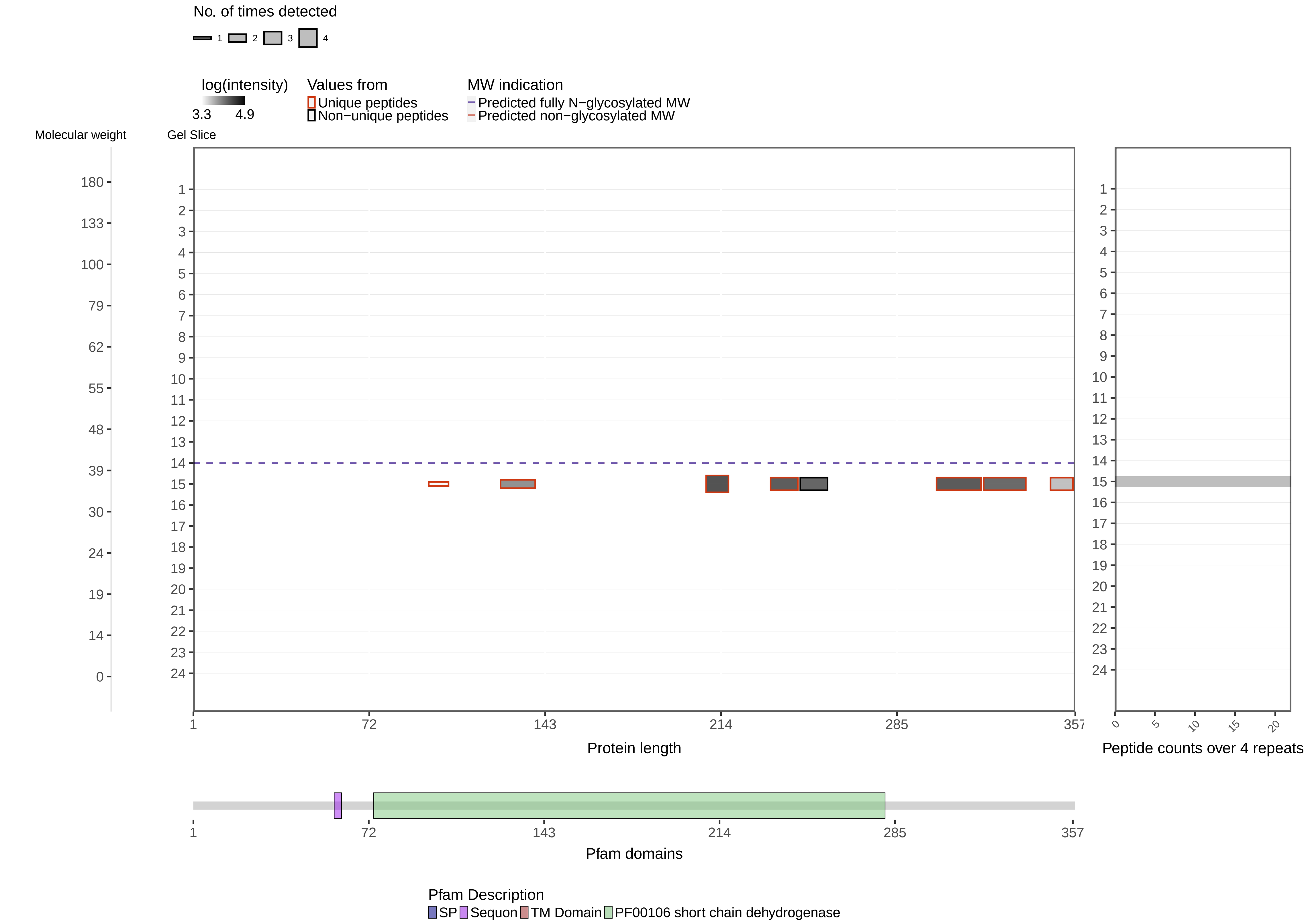


Protein sequence:

MLSKDNIELQFATNHLGHFLLTNLLLEKMKETTRKTKREGRIVNVASEAHRFAYREGVRFGKINDRKSYNSFGAYGQSKLANILHANELTRRLKEDGAEITANSLHPGMITTNLFPHMGVFDGLVSNLGKLVLKNVQQGASTTCYLALHPQVKGVSGEYFLDNNLATTTAKAKDMDLAKRLWDFSMDLVK*

### NbL03g11800.1 pectin_acetylesterase_9−like [MW − SP = 17.07 kDa]

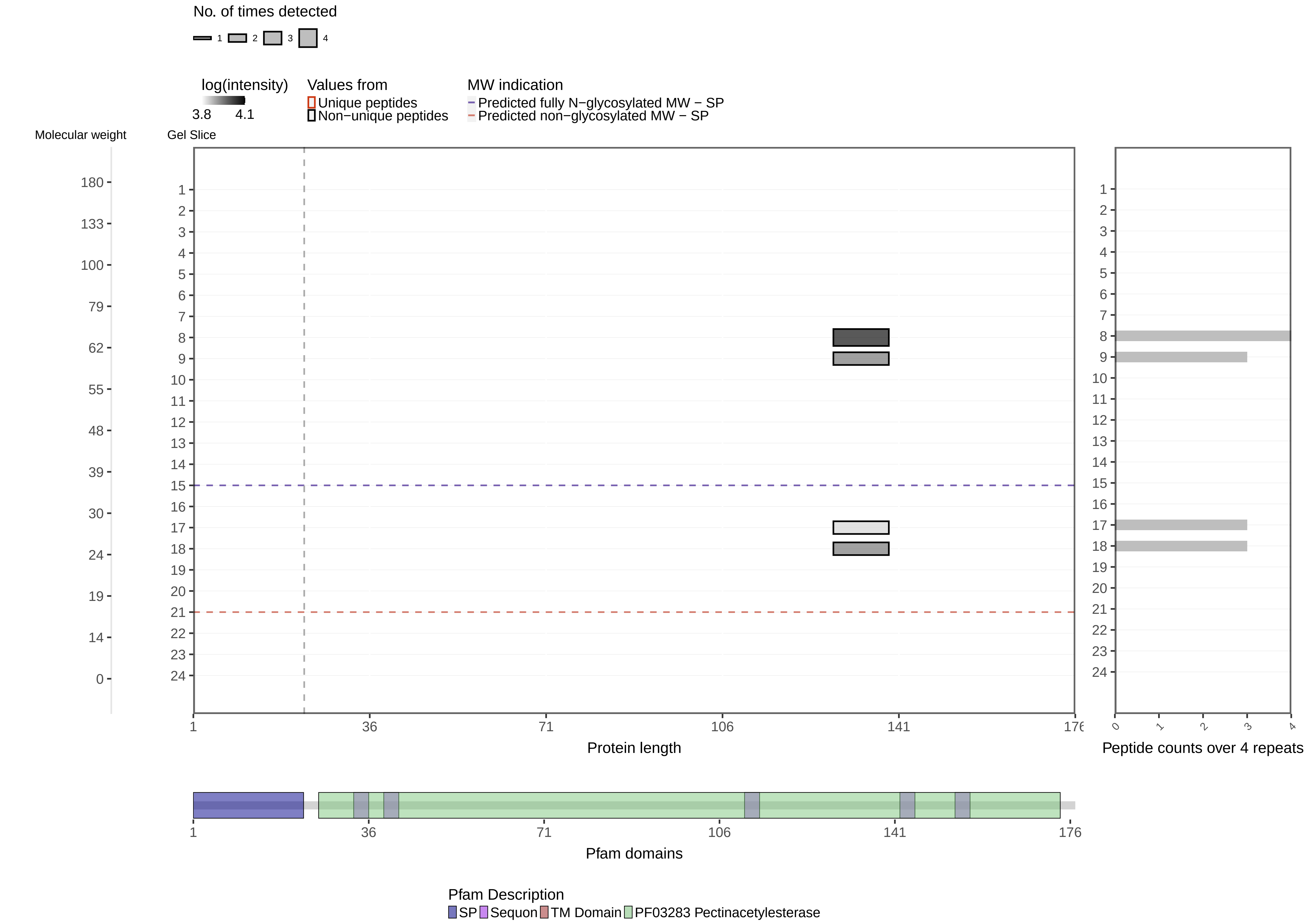


Protein sequence:

GWCNDIKSCLDRSTTHRGSTRYMNKWEVFSGILSNNASLNPDFYNWNRVKLRYCDGASFAGDSKFYNGTYSRSSSERNGSTLLYFRGQRIWKAIILDLLP*

### NbL03g11790.1 tripeptidyl−peptidase_2 [MW = 149.42 kDa]

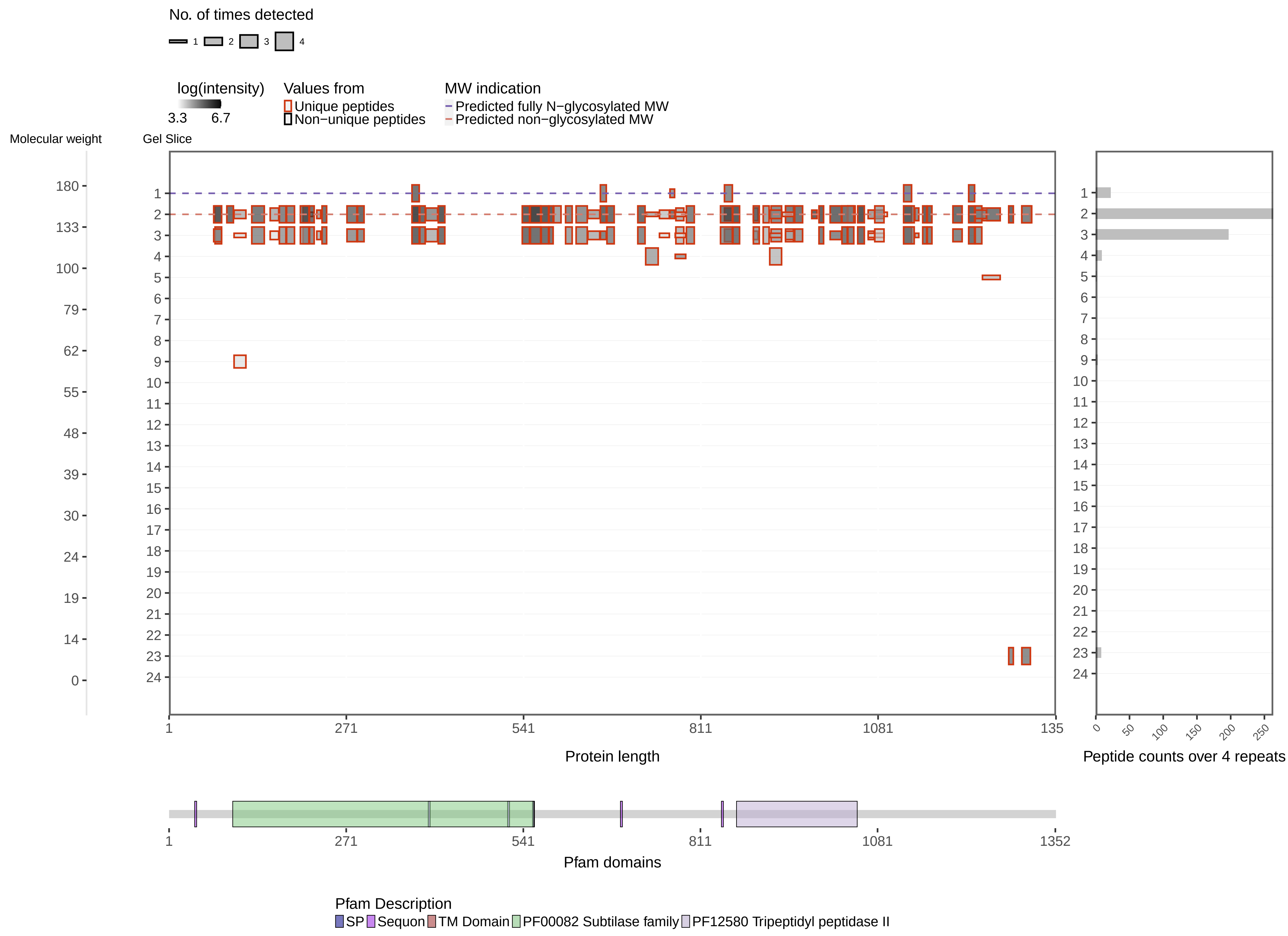


Protein sequence:

### NbL03g11750.1 oxalate−−CoA_ligase−like [MW = 56.21 kDa]

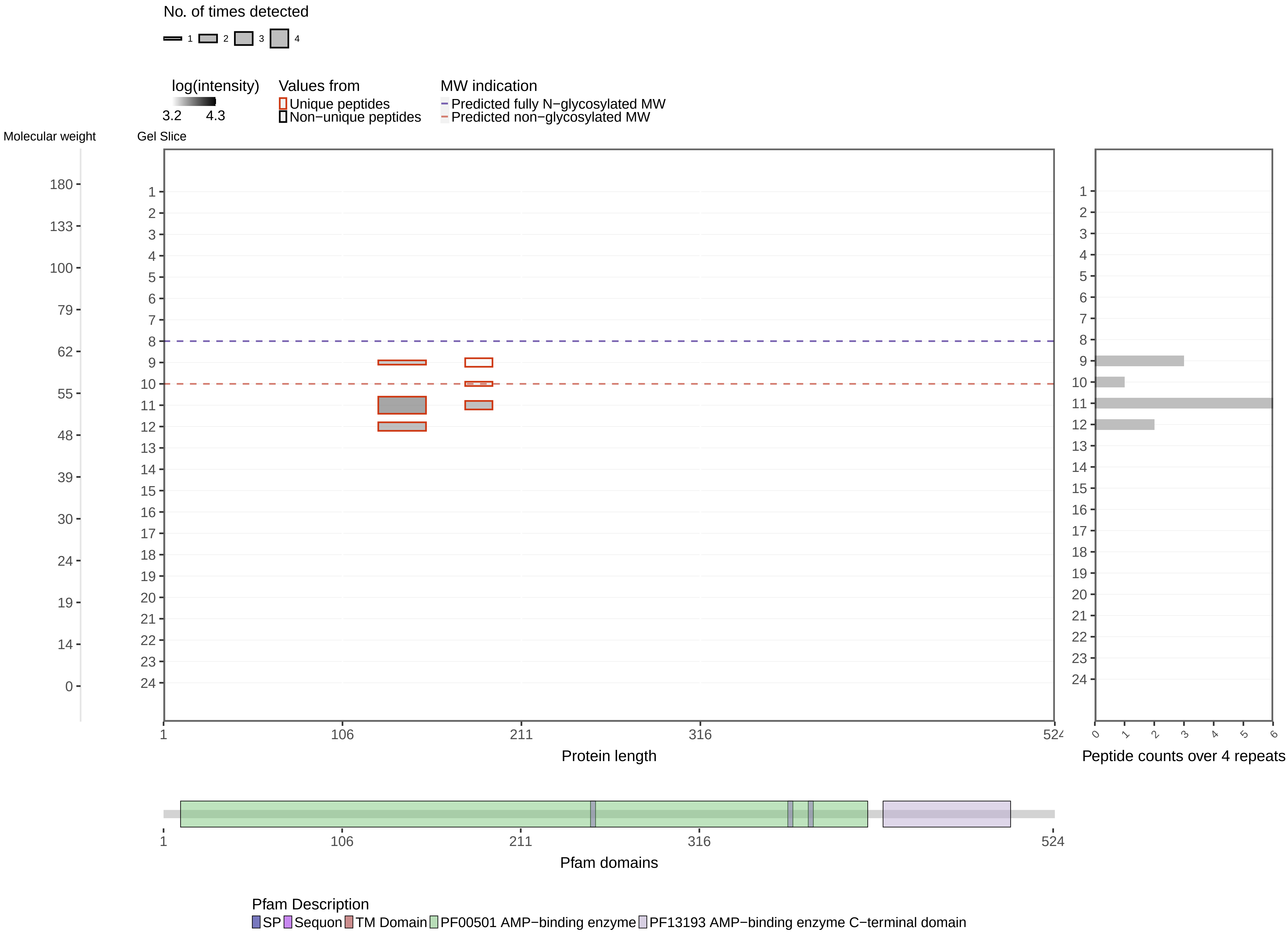


Protein sequence:

### NbL03g11650.1 remorin−like [MW = 23.17 kDa]

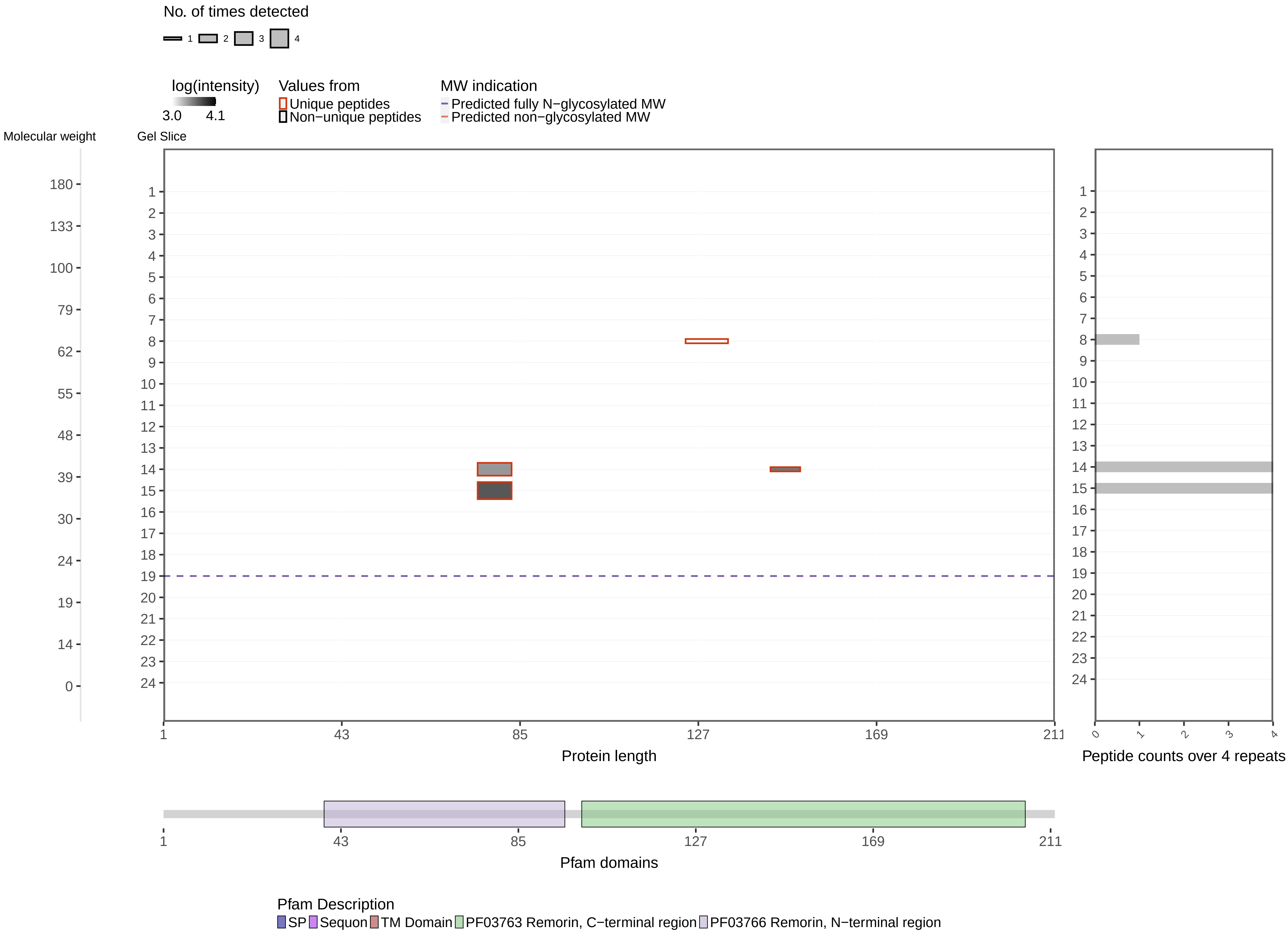


Protein sequence:

LARVATEKRLSLIKAWEESEKSKAENKAQKKVSAIAAWENSKKANLEAELKKMEEQLEKKKAEYIEKMRNKIALLHKEAEEKRAMIEAKRGEDLLKAEELAAKYRATGTAPKKLLGCF*

### NbL03g11550.1 3−hexulose−6−phosphate_isomerase [MW = 21.42 kDa]

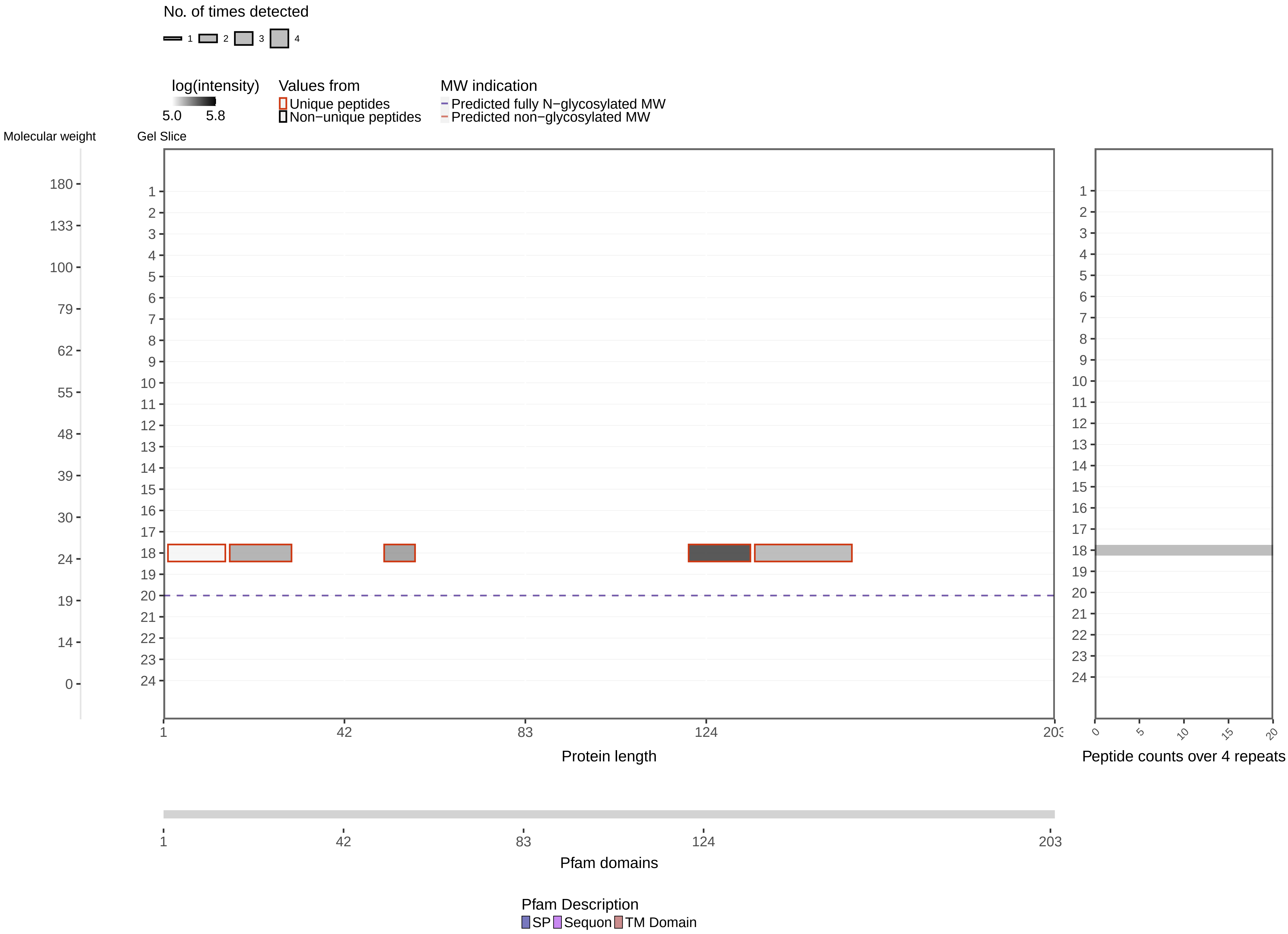


Protein sequence:

RPPDLLIASAGPGGFSTVDAICGVAKSNGARVLLLTAQPESGSSVKYASVVAYIPAQTMADDSAGDAERERPLLPMGSLYEGAMFVLFEMVVFKLADVLKQSPEAVRARHTNLE*

NbL03g11380.1 putative_4−hydroxy−tetrahydrodipicolinate_reductase_3_chloroplastic [MW = 26.96 kDa]

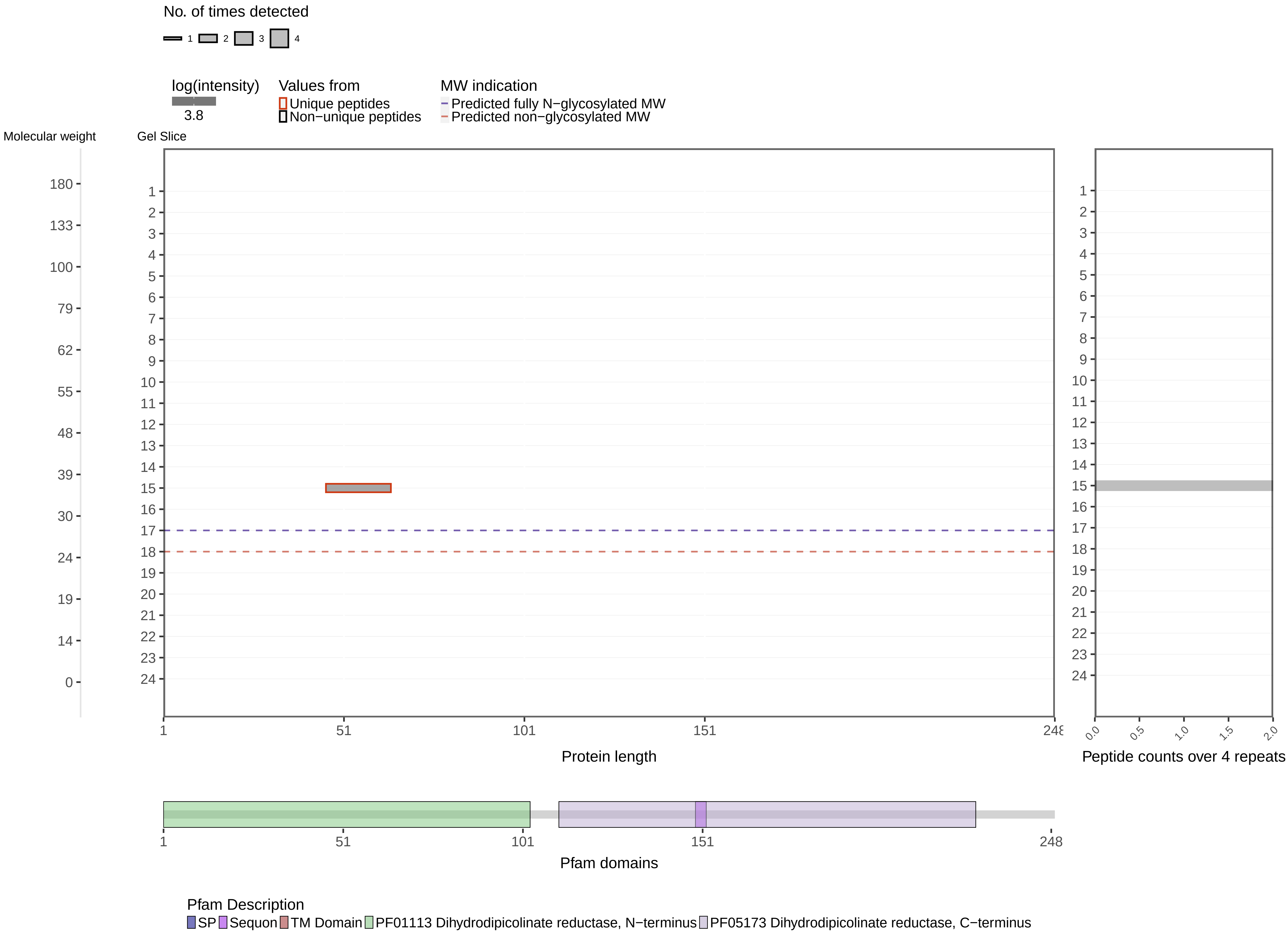


Protein sequence:

.LQQAAISASFHYNNVEIVESRANAVDFPTQDAVQIANNLSNLGQLYNREDISTDIPARGQILGEDGVRVHSLVLPGLPSSTSVYFSKPGEVYTLQHDITDVQCLMPGLILAIRKLFASRTWSMAWRSFYKKQTVRG*

NbL03g11080.1 ATP−dependent_Clp_protease_ATP−binding_subunit_CLPT1_chloroplastic−like [MW = 25.99 kDa]

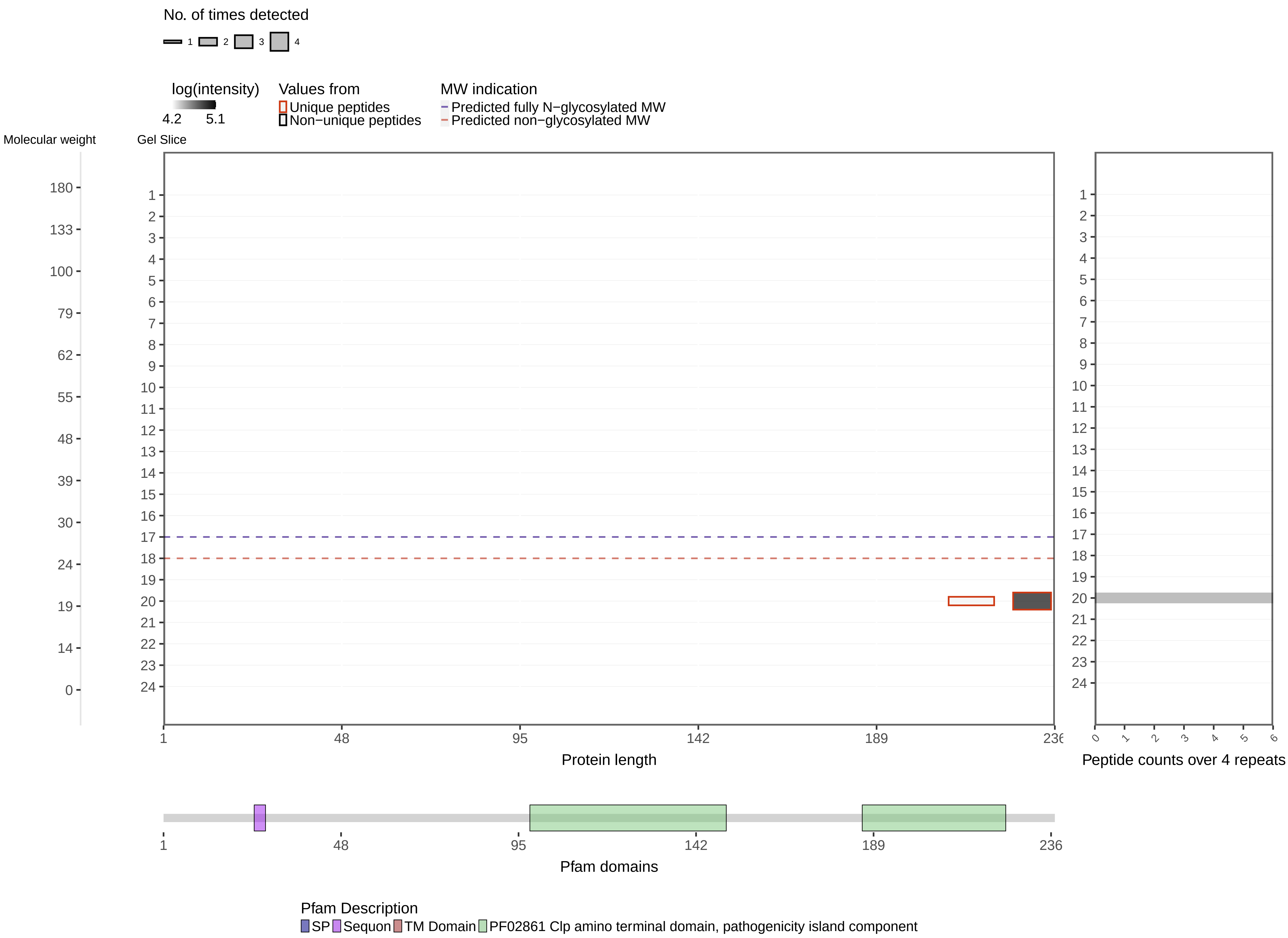


Protein sequence:

.KYPNTGTEALLMGILVEGTSLAAKFLRANGVTLLKVREETVKLLGKSDMYFFSPEHPPLTEPAQRALDWAVDEKLKSGESGEITIAHLALGIWSEKESAGHKIMATLGFDDEKARELAKSMDKDIEMSYT*

### NbL03g10900.1 UMP−CMP_kinase [MW = 26.78 kDa]

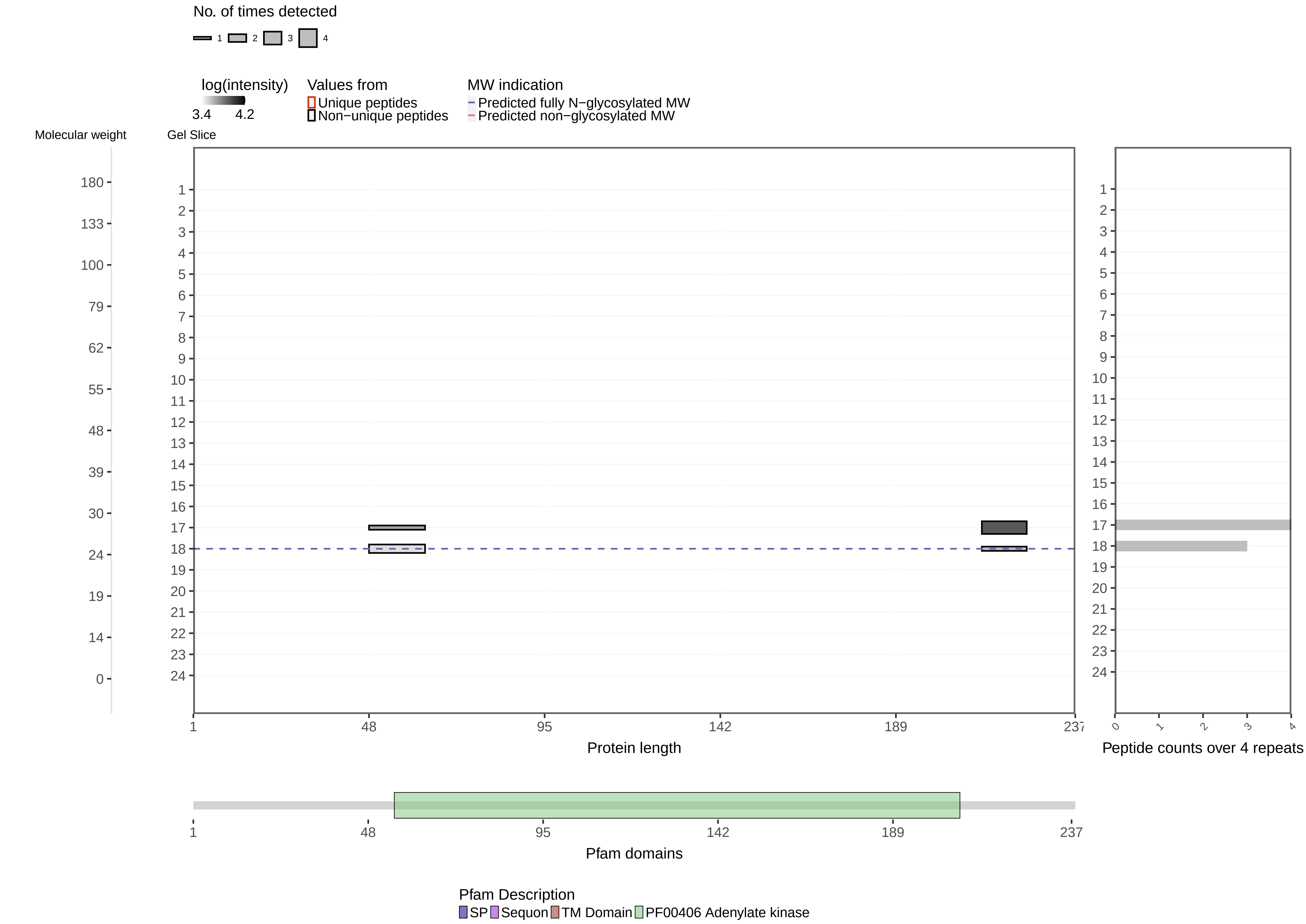


Protein sequence:

EGSIAPSEVTVKLIRKAIESAENHKFLIDGFPRSEENRMAYERIIGAEPNFVLFFDCPEEVMVKRVLNRNEGRVDDNEHTVKERLKVYKAVTLPVVNHYAKKGKLYKIDGTGTQEEIFERVRPIFSSLSSLV*

NbL03g10880.1 21_kDa_protein−like [MW − SP = 18.69 kDa]

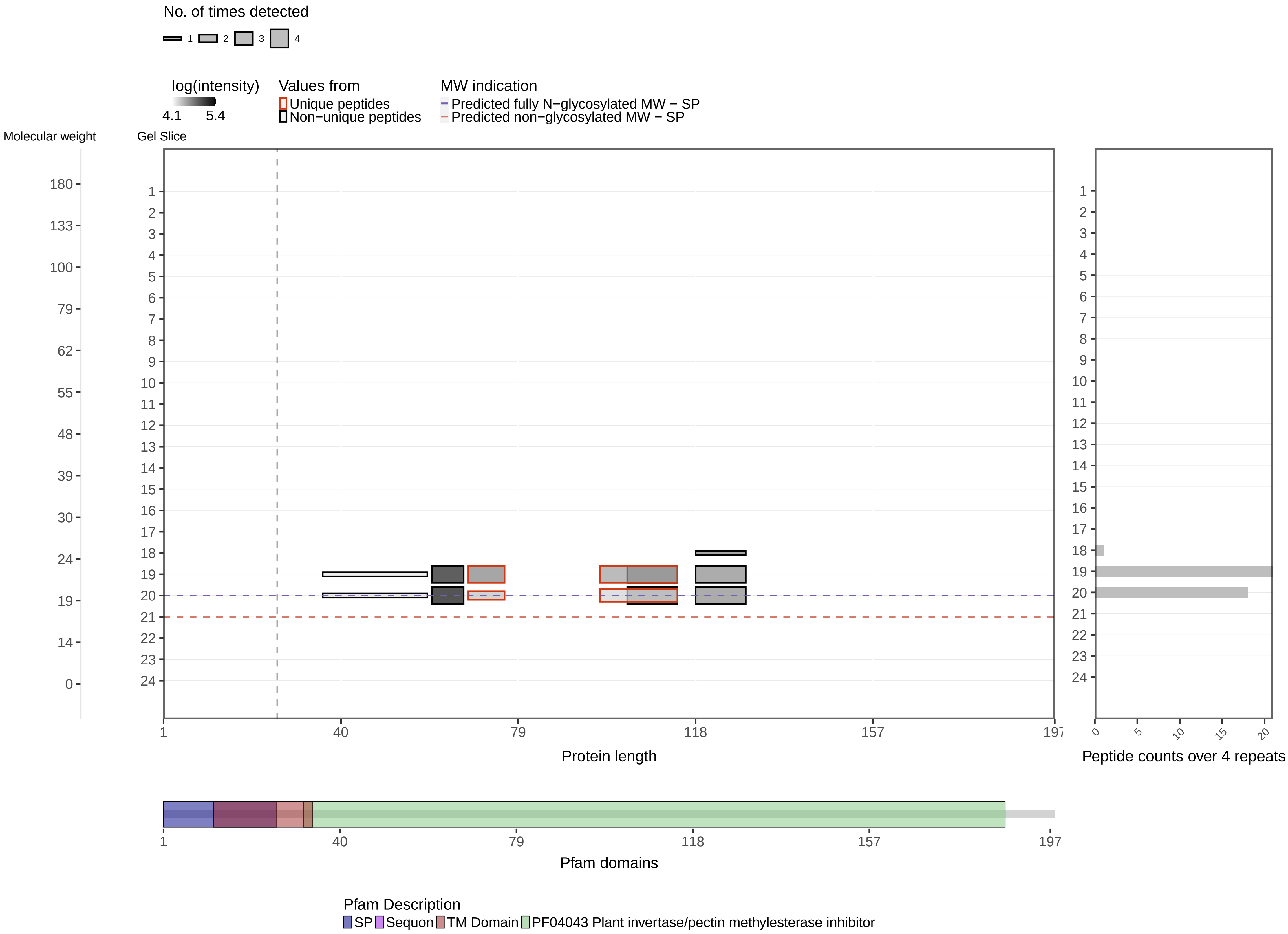


Protein sequence:

KLLKFKGLKPREYEAIKDCIEETSDSVDRLSKSVSELKEVDHSRGKDFLWHISNVETWVSAAITDENTCTDGFAGRALNGRIKASIRNRIRNLAEVTSNALGLINQYAAKH*

NbL03g10800.1 monocopper_oxidase−like_protein_SKS1 [MW − SP = 63.52 kDa]

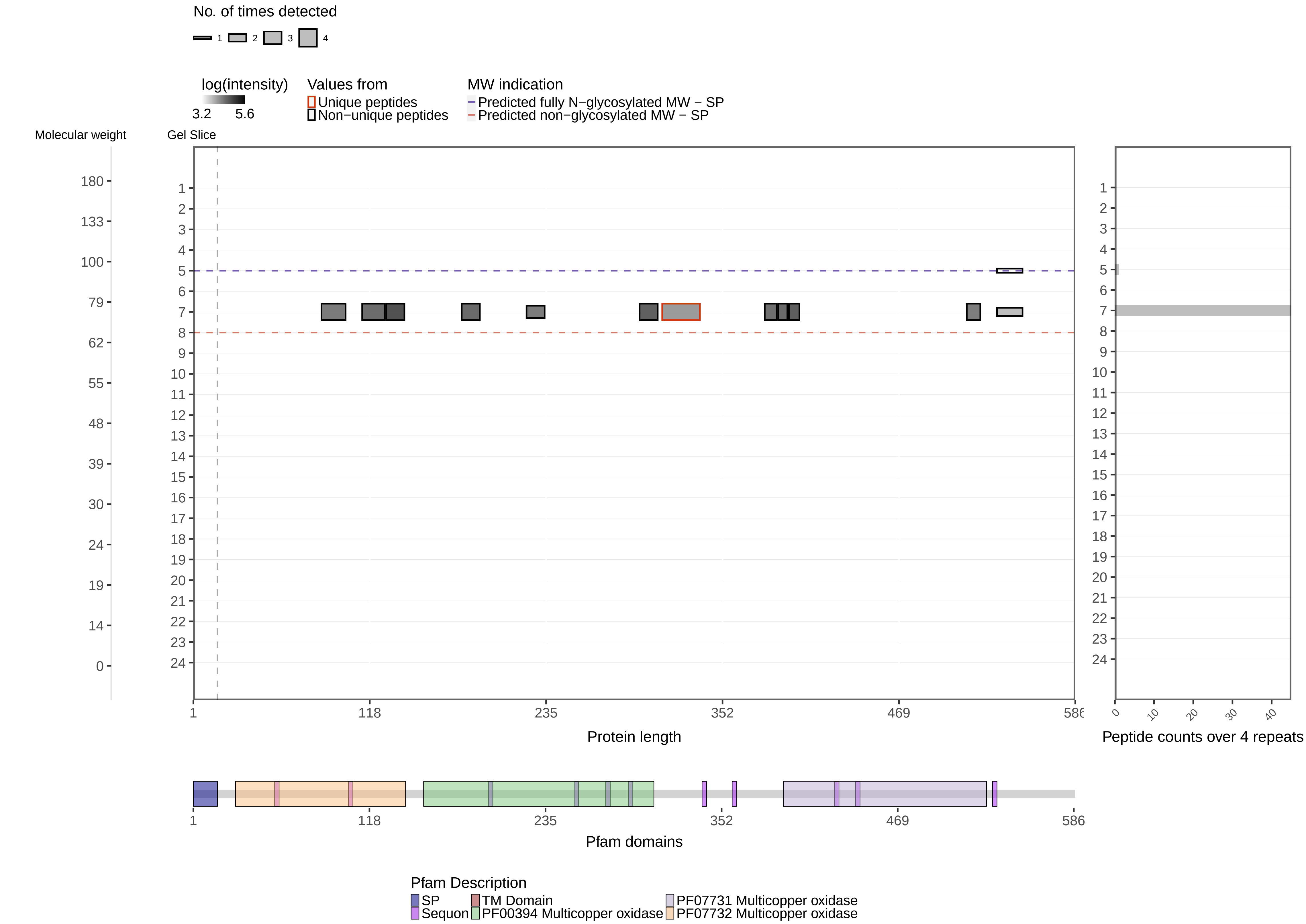


Protein sequence:

### NbL03g10790.1 acid_beta−fructofuranosidase_AIV−18−like [MW = 74.77 kDa]

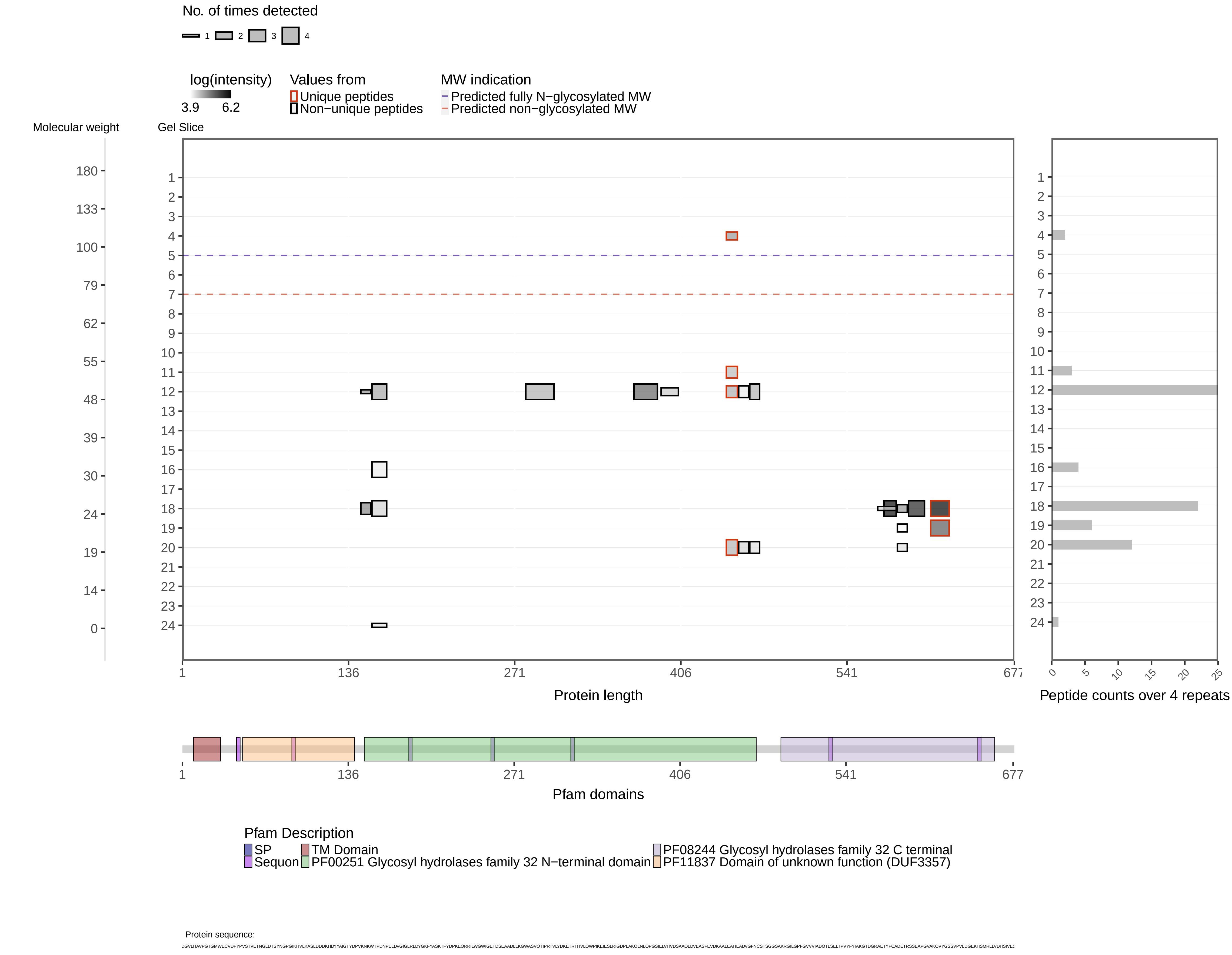

NbL03g10460.1 acid_phosphatase_1−like [MW − SP = 28.53 kDa]

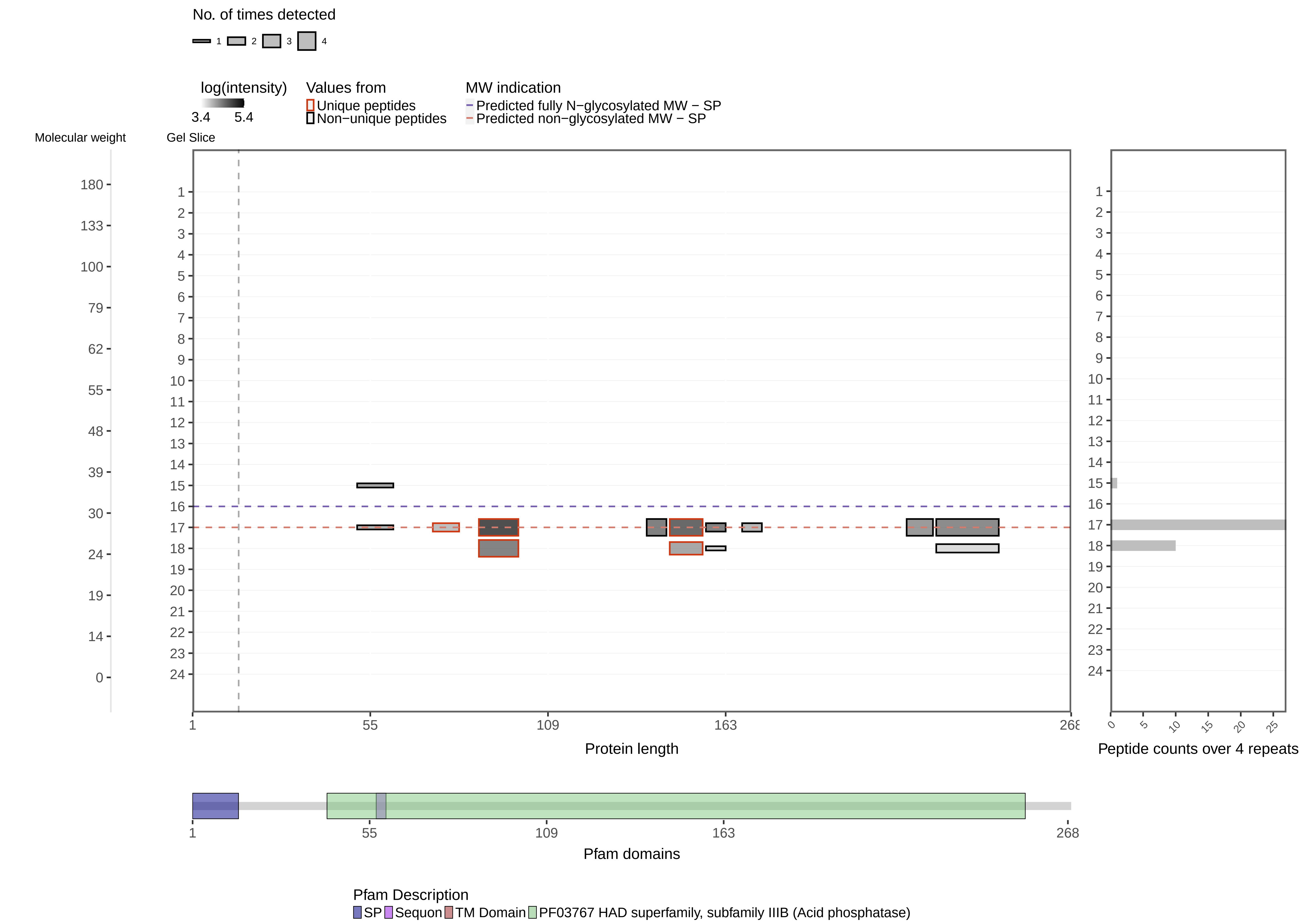


Protein sequence:

'YYSQHGYGLEVFDSVKFDEWVEKGVAPVIGSSLKLYQDVMRLGFKVFLLTGRSERHRIVTVENLMNAGFQDWDKLILRGSKDHGKSATIYKSEKRDEMVEEGFRIVGNSGDQWSDLLGSSASIRSFKLPNPIFSRTSSIGSGFFF*

### NbL03g10410.1 ATP−dependent_6−phosphofructokinase_4_chloroplastic [MW = 61.97 kDa]

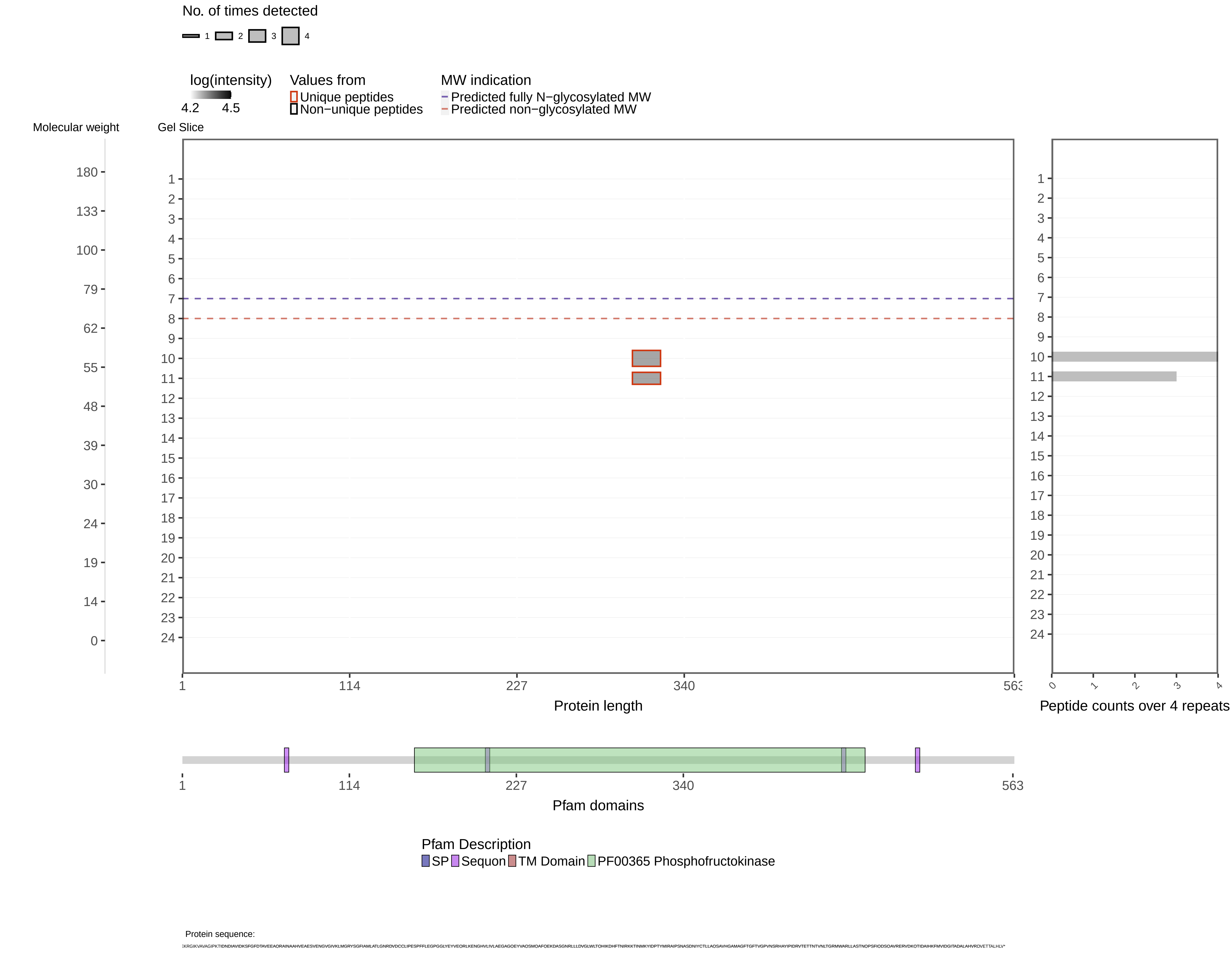

### NbL03g10290.1 protein_mago_nashi_homolog [MW = 17.83 kDa]

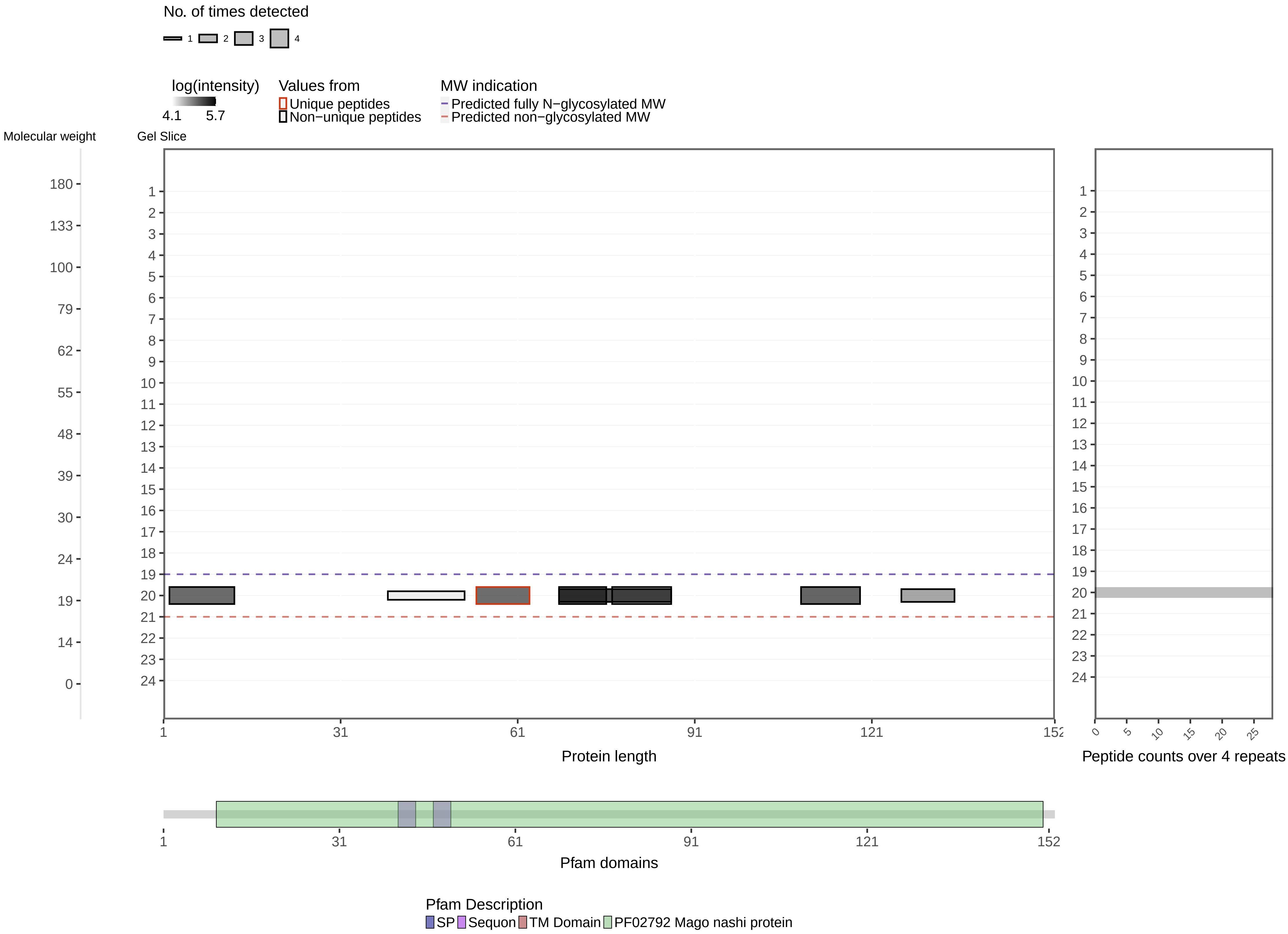


Protein sequence:

LRECRRIVADSEIMKEDDNNWPEPDRVGRQELEIVMGNEHISFTTSKIGSLMDVQTSKDPEGLRIFYYLVQDLKCFVFSLISLHFKIKPI*

### NbL03g10150.1 superoxide_dismutase_[Fe]_chloroplastic−like [MW = 34.8 kDa]

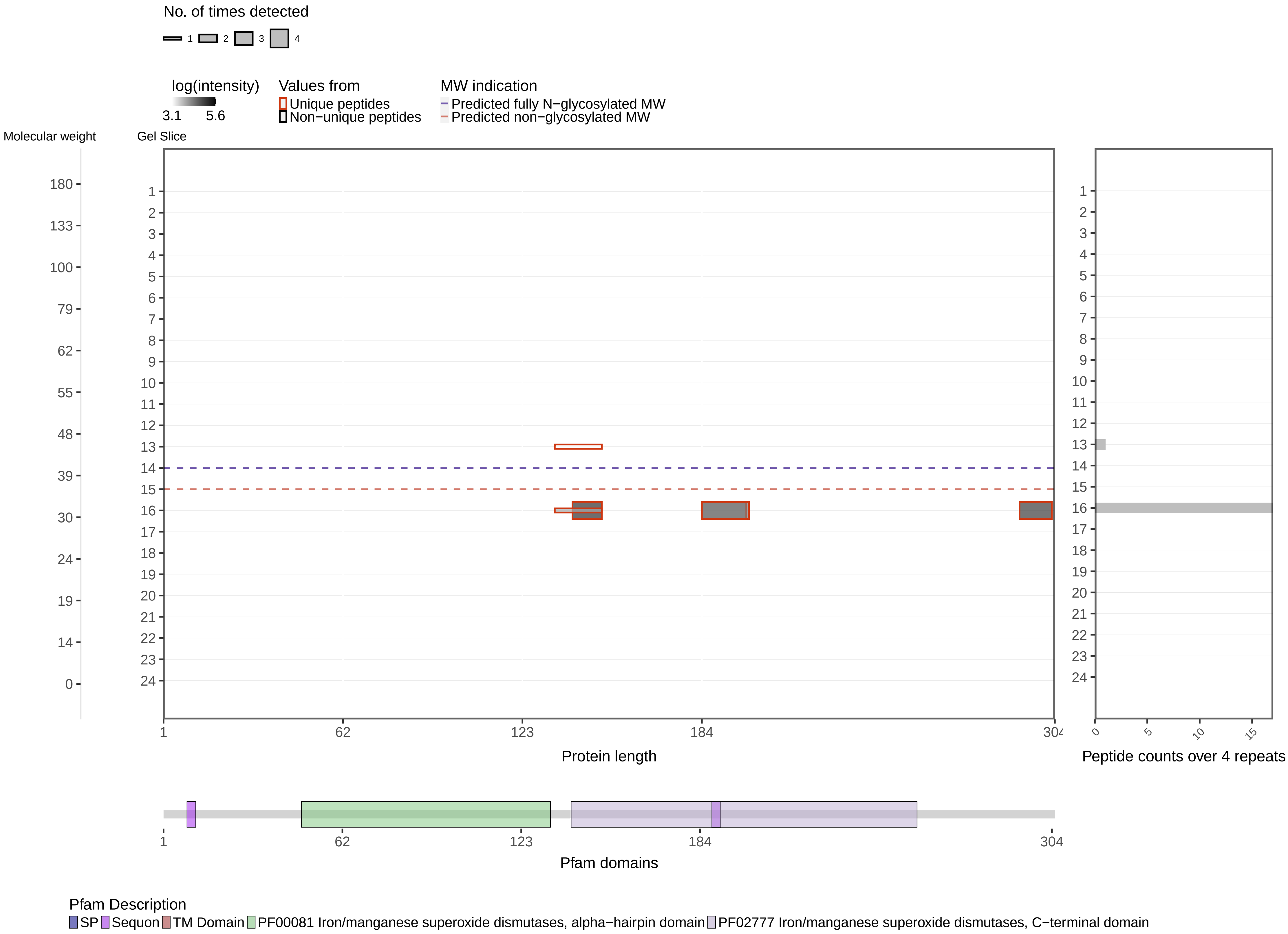


Protein sequence:

KPSGELLELINRDFGSFEAFVKEFKVAAATOFGCGWAWLAYKANRLNVGNASNPHPTDEDKKLVVVKTPNAVNPLVWDYSPLLTIDVWEHAYYLDFRNRRPDYISIFMEKLVSWEAVSSRLEVAKARVAEREEEEERKKREEEEEYKAGGEVREMYVESTHSDTE*

### NbL03g10050.1 quinone_oxidoreductase−like_protein_2_homolog [MW = 37.32 kDa]

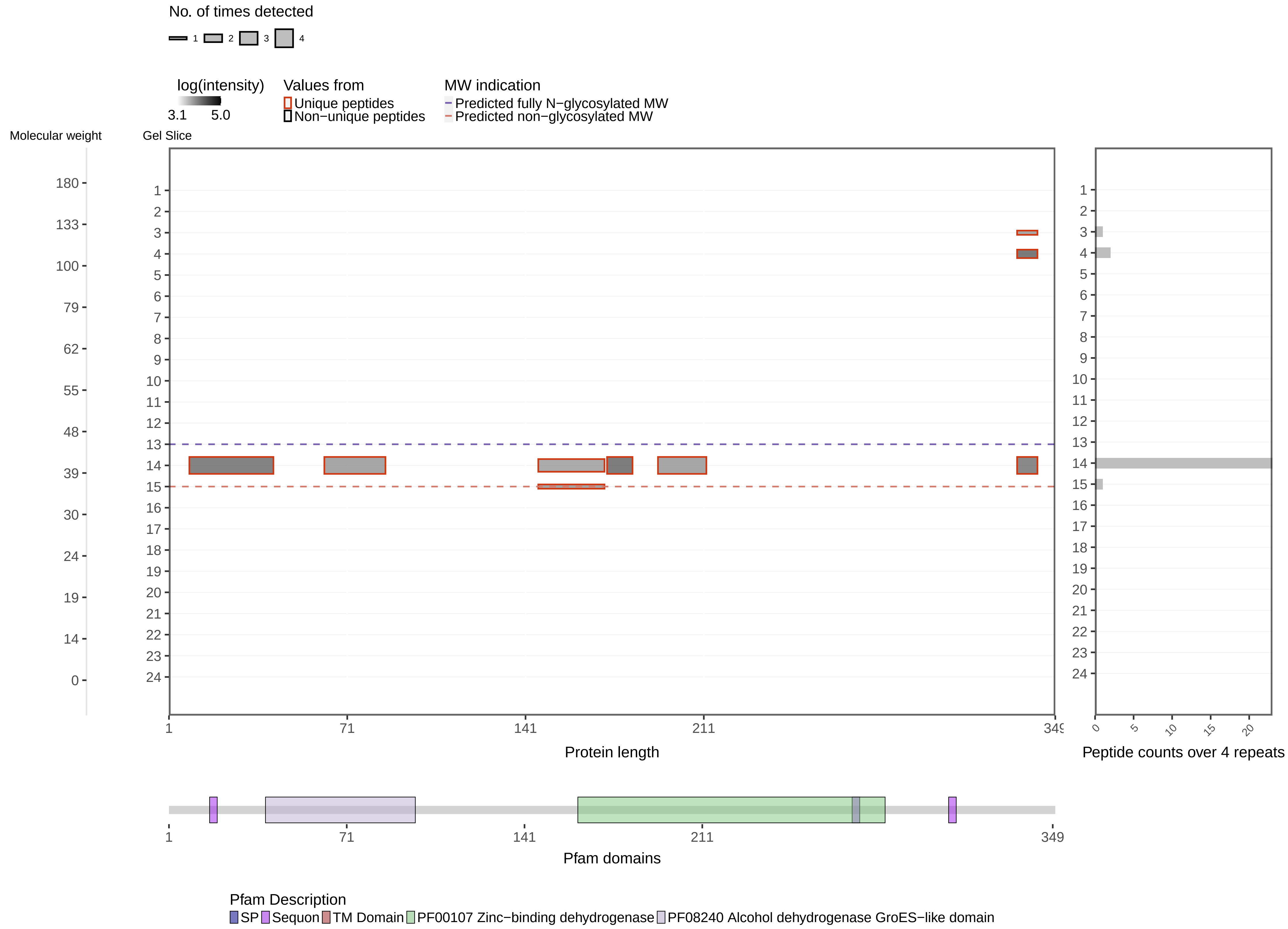


Protein sequence:

AGGVGLSAVQIGKICGATVIAVARGNEKVQFLKSLGVDHVVDLSNANVIESVKGFLKSRKLKGVDVLYDPVGGKLTKHSMKLLNWGAQILVIGFASGEVPVIPANIALVKNWTIHGLYWGSHKIYQPNVLGDSLNELLSWLSKGLITINISHTFSLAEAHLAFTALKDRRAIGKVMITFDEDRTLKSKL*

NbL03g09750.1 expansin−A4−like [MW − SP = 24.07 kDa]

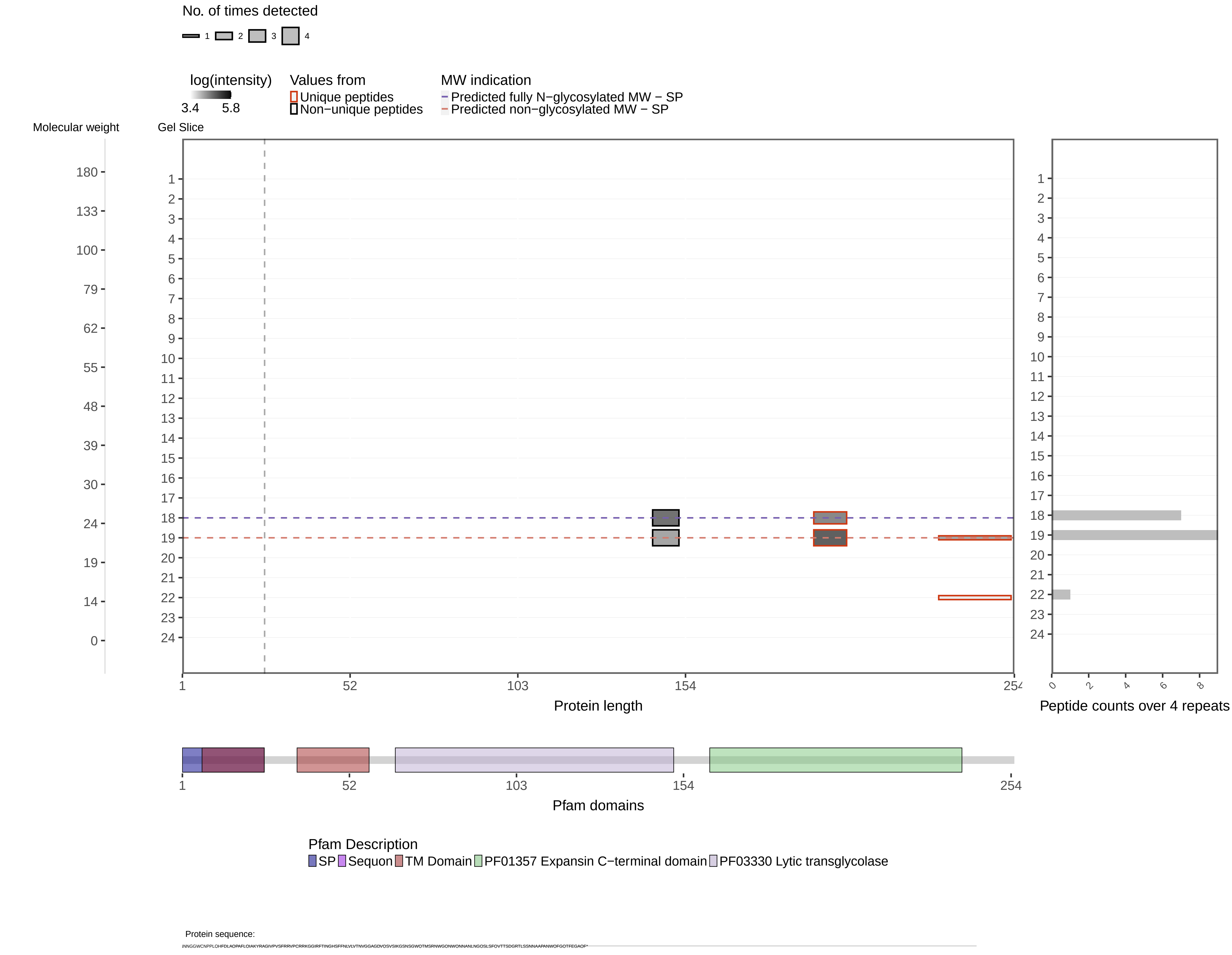

NbL03g09710.1 alpha−galactosidase−like [MW = 47.85 kDa]

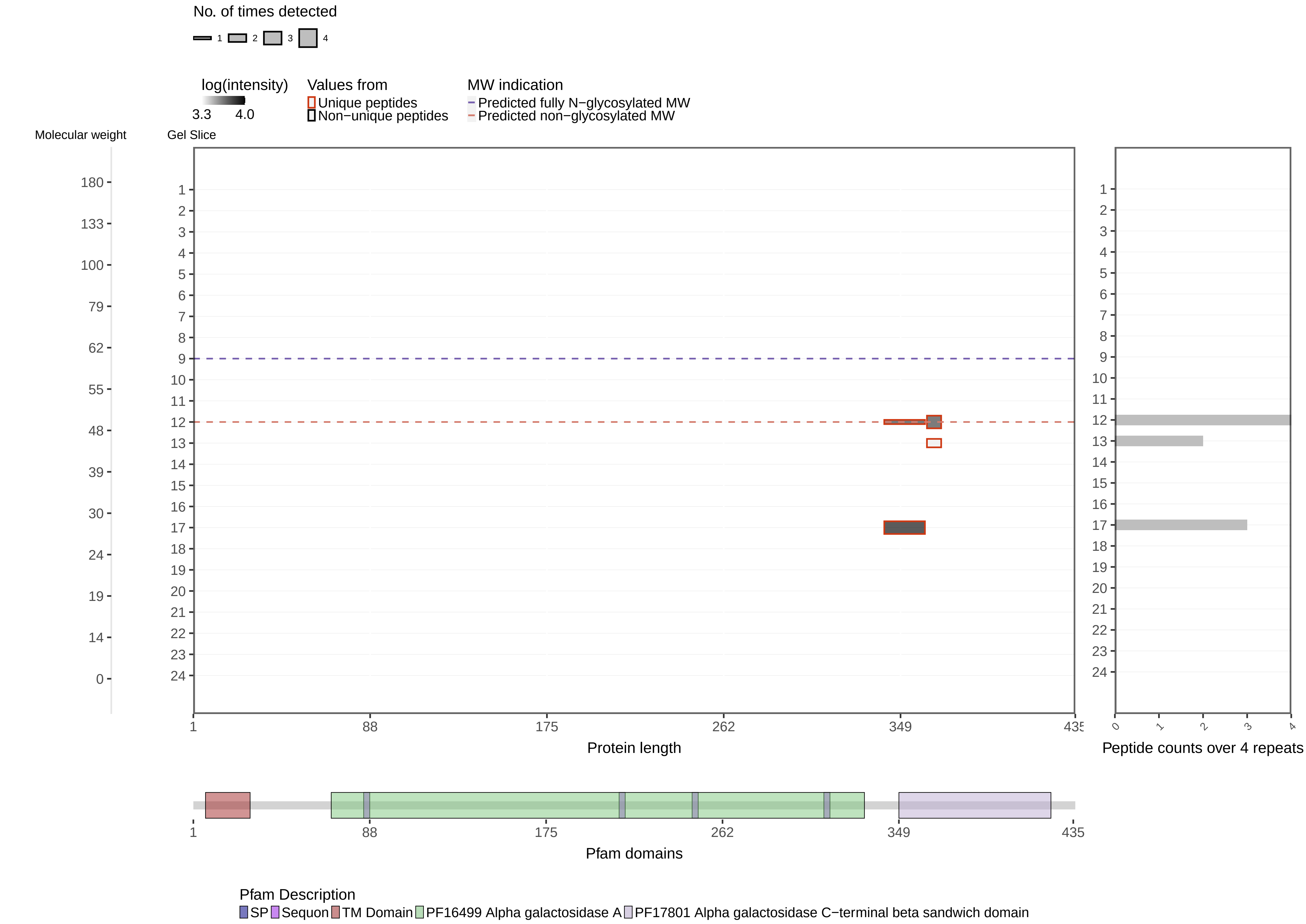


Protein sequence:

MRDALNATGRTIFYALCEWGVDDPAKWAGKIGNSWRTTDDINDTWASMTTIADINDKWASYAGPGGWNDPDMLEVGNGGMNYHEYKGHFSIWALMKAPLIIGCDVRNITAETLEIFSNKEVIAVDQDPLGVQGRRVYAAGPDGCEQVWAGPLSGNRLAIVLWNRCSKAASITAKWCAIGLQSSISVSVRDLWKHEVVSENTVGSFSARVDAHGCEMYILTPKRATRASI*

### NbL03g09700.1 uncharacterized_LOC107819606 [MW = 19.28 kDa]

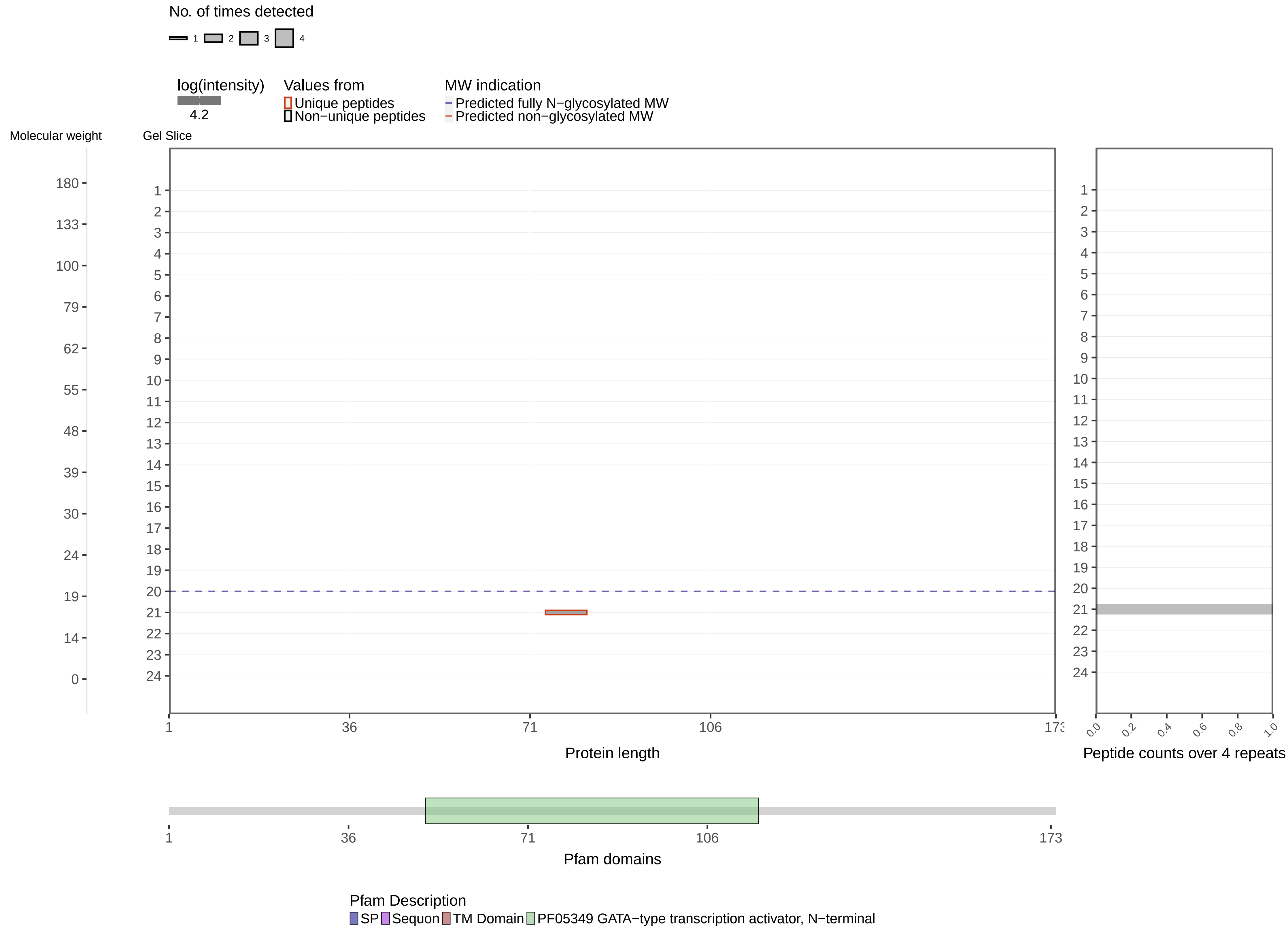


Protein sequence:

AELIDEAREATQSGVRMSREQYGALRRKIGGTYKDFFKSYVEVKGEYVEEGWVDKTCKVCKKDTRGEARQVDNFGRYAHVACLDKSKSGNFFTNLFSR*

NbL03g09650.1 uroporphyrinogen_decarboxylase_chloroplastic [MW = 43.33 kDa]

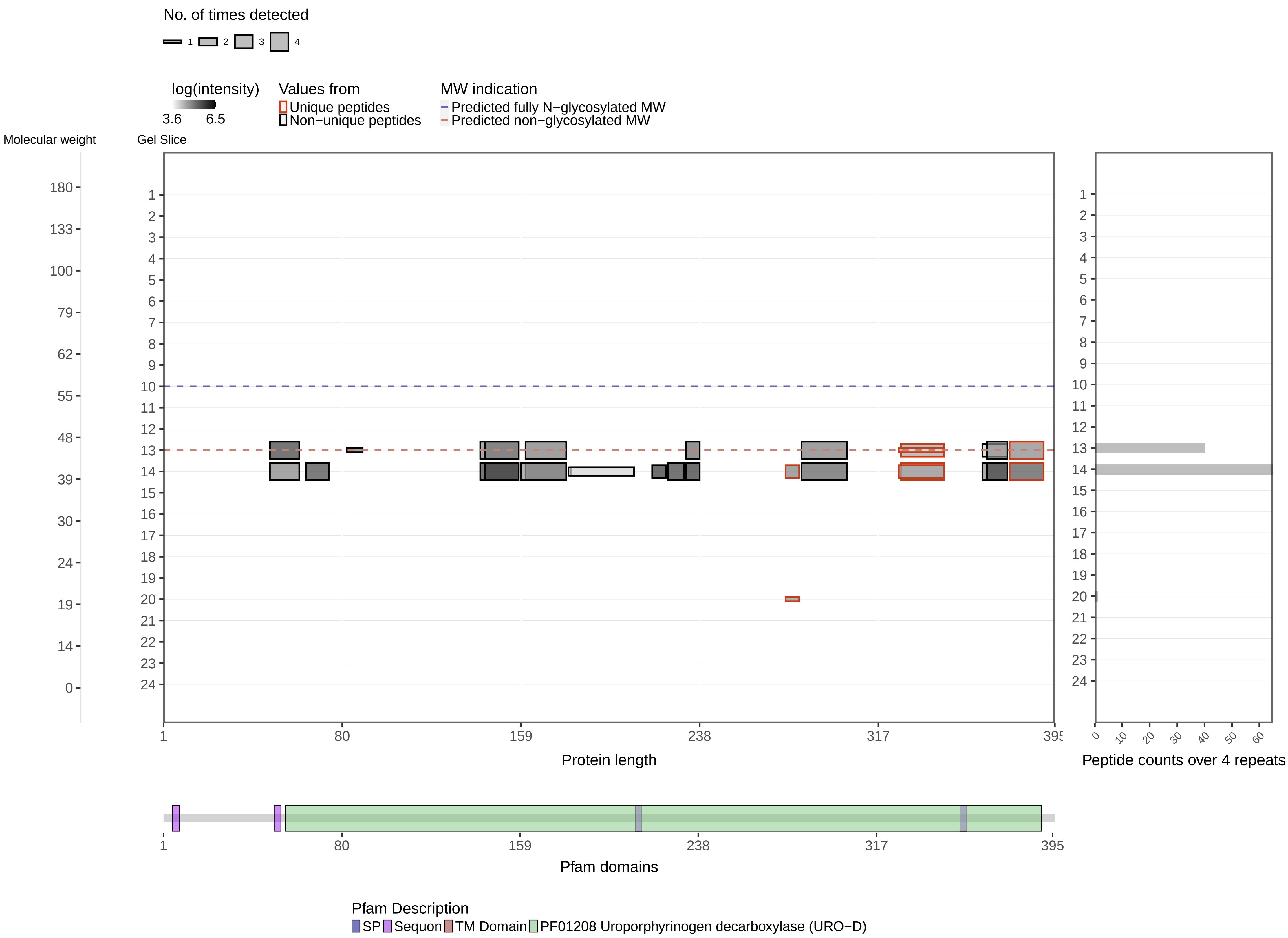


Protein sequence:
IQAAVLGFVGAPFTLASYVVEGGSSKNFTKIKRLAFAEPKVLHALLQKFATSMAKYIRYQADSGAQAVQIFDSWATELSPVDFEEFSLPYLKQIVDSVKLTHPNLPLILYASGSGGLLERLPLTGVDVVSLDWTVDMADGRRRLGPDVAIQGNVDPGVLFGSKEFITNRINDTVKKAGKGKHILNLGHGIKVGTPEENVAHFFEIAKGLKY*

NbL03g09540.1 photosynthetic_NDH_subunit_of_subcomplex_B_2_chloroplastic−like [MW = 38.79 kDa]

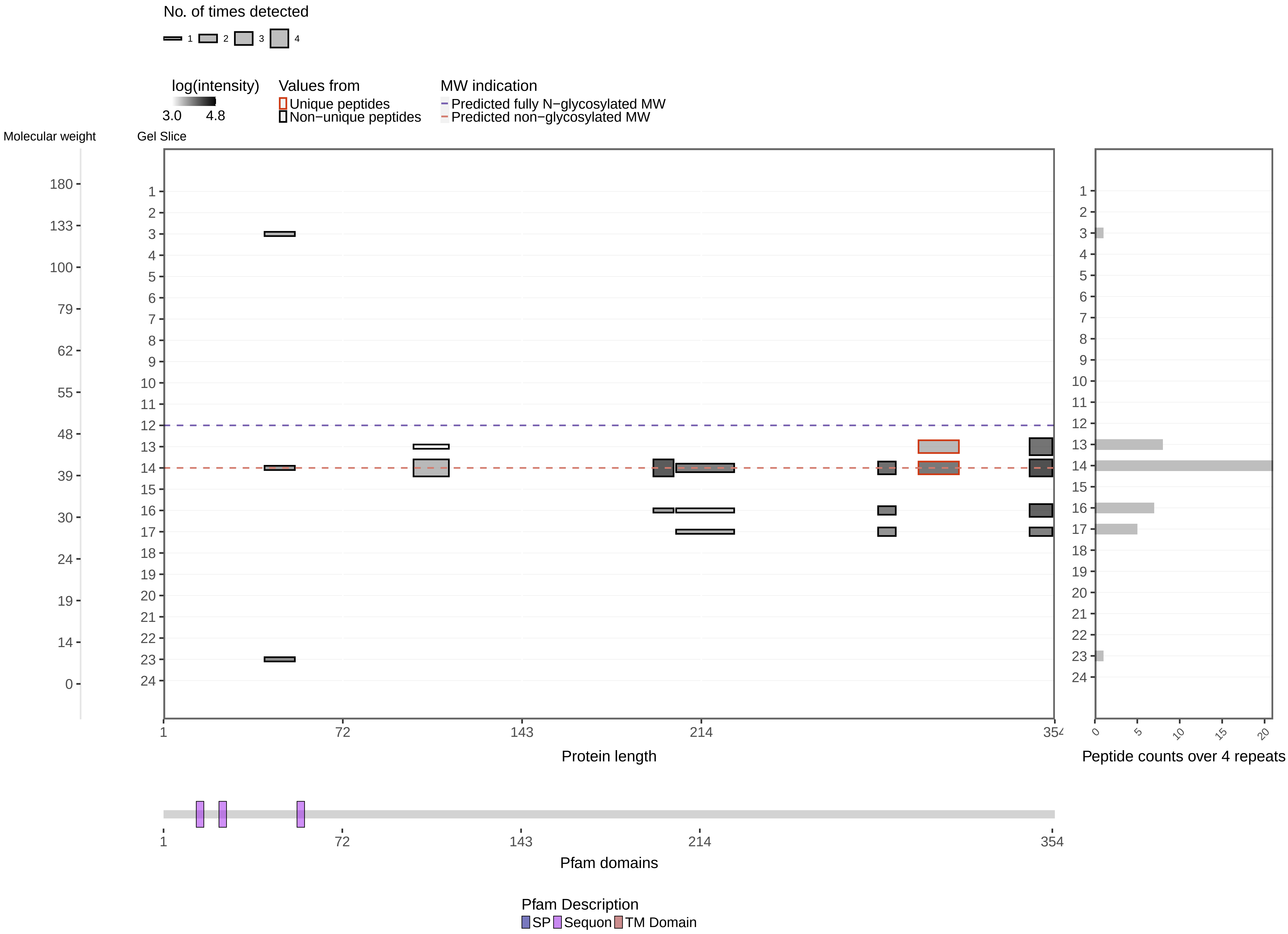


Protein sequence:

MATAVIVKNNGRKPVKLTSAILSHLKSKTRGGTGIQGLRSCTYCAHPPLSSPFEILSPGEAMKTEEPGLFSFGWEPELKPGIWSAQDVPITVLKHKLSRLYSVPPEERAKEFYNSIPSKYETIDQGRELFFRIIRMGFEDIYLSSPGSFSDKYGKDYFICTGPASMLIPLVVNPGEEWRGAQVIEHDNL*

### NbL03g09520.1 ADP−glucose_phosphorylase [MW = 37.84 kDa]

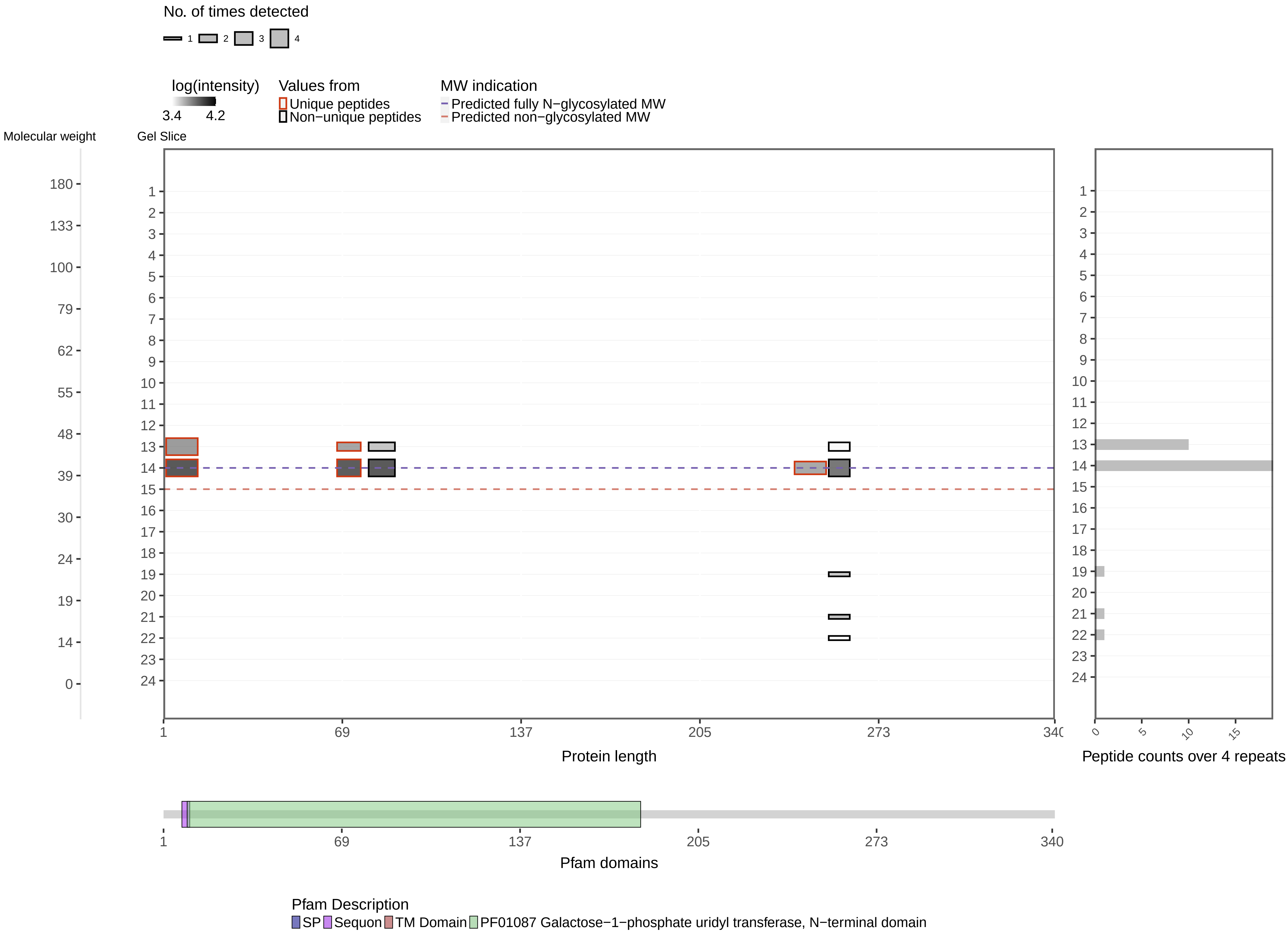


Protein sequence:

VQVFKNHGASAGASMSHSHSQMIALPIVPPTVSARLDSMREYYKQTGKCSLCDIQPNELLIDQSAHFISLVPFAATFAFEIWIIPRDHSSHFHQLDNEKANDLGGLLKLMLLKMSLQLNNPPFNLLIHTSPFQVDPSYASSTHWFLQIAPHLSVVGGFEIATGCHINPVFPEDAAKILRDVRILN*

### NbL03g09500.1 probable_phosphatase_SPAC5H10.03 [MW = 32.26 kDa]

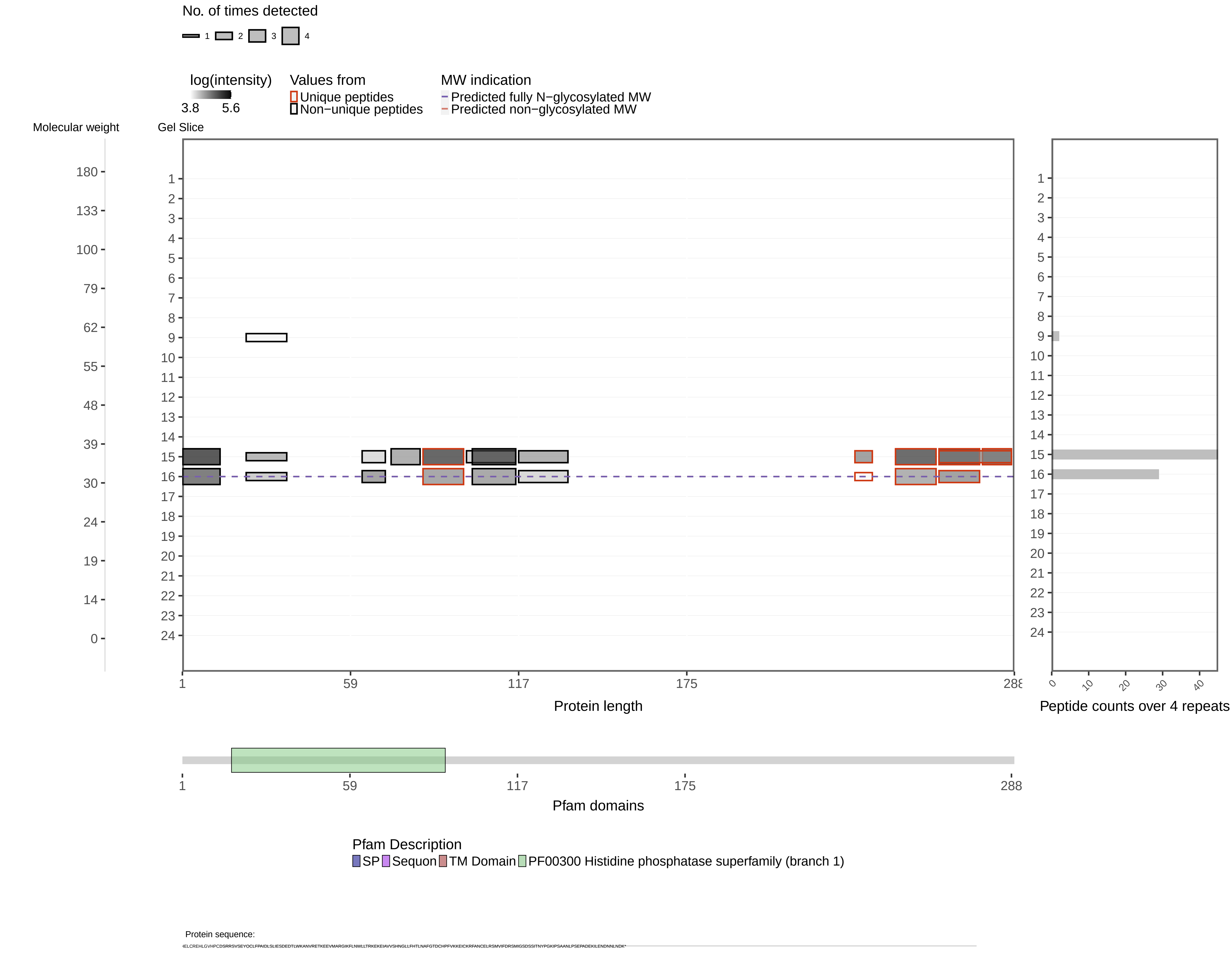

NbL03g09230.1 beta_carbonic_anhydrase_5_chloroplastic−like [MW = 33.78 kDa]

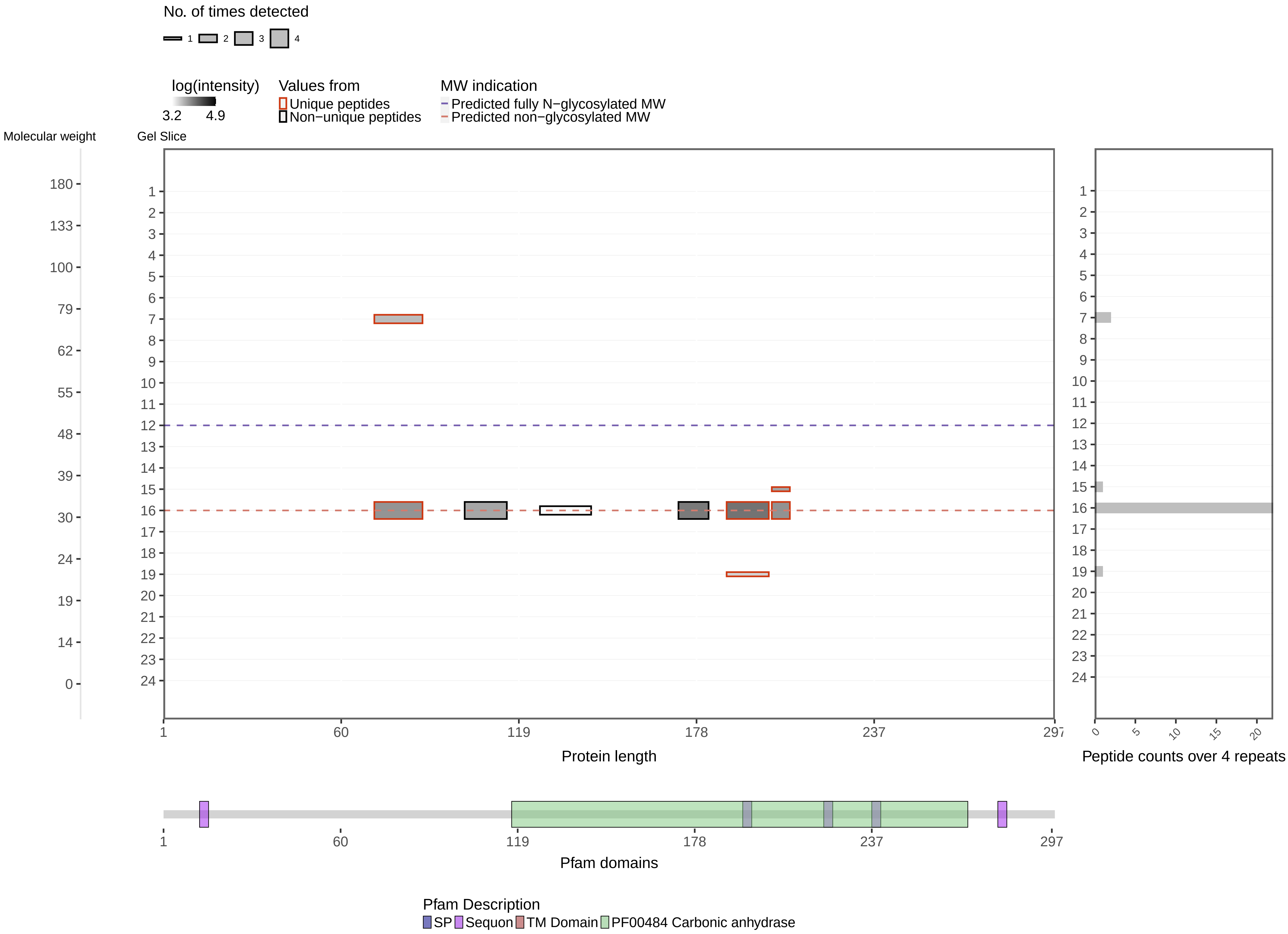


Protein sequence:

GEAFVVRNVANLVPPYENGPSETNAALEFAVNTLKVQNILVIGHSRCGGIRALMSMDNETPSSFIRSWVVLGKSARASTKAAASNLSFDQQCKHCEKESVNCSLMNLLTYPWIKDKVNKGELLIHGGYYDFVDCTFEKWTLENNSRVDNQVSIRNREFWY*

NbL03g09110.1 D−3−phosphoglycerate_dehydrogenase_1_chloroplastic−like [MW = 63.29 kDa]

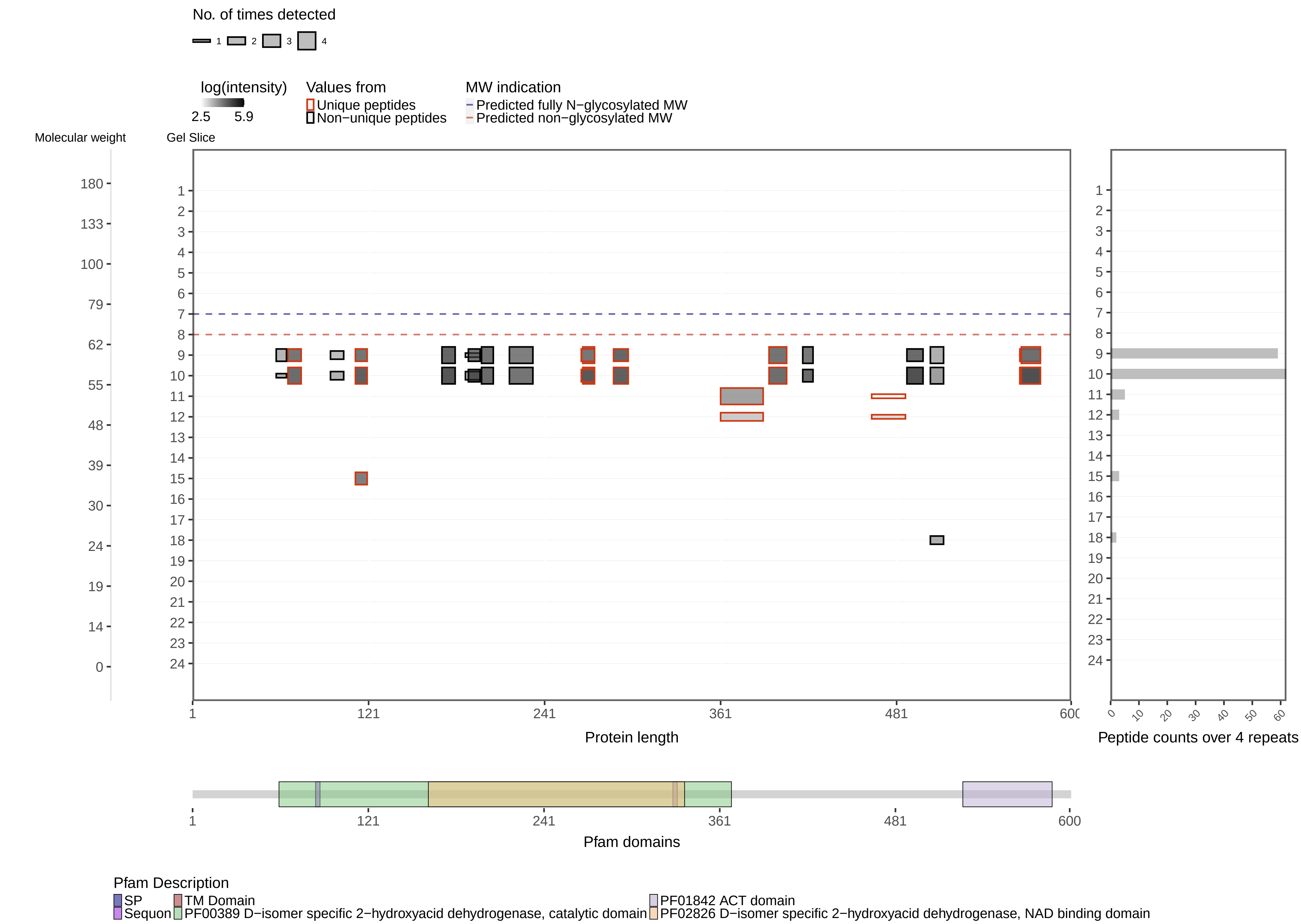


Protein sequence:

### NbL03g08920.1 glycine−rich_RNA−binding_protein−like [MW = 16.97 kDa]

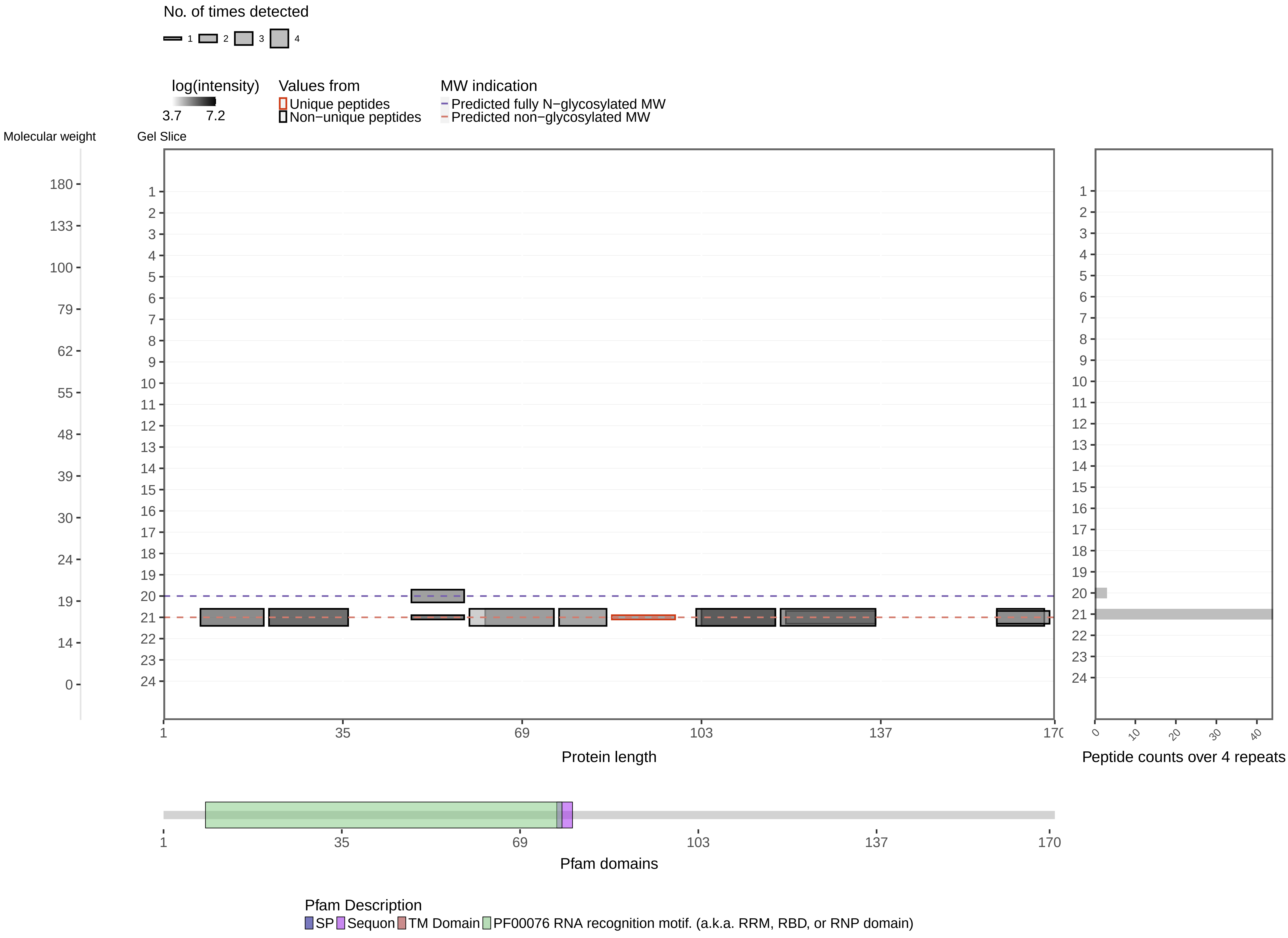


Protein sequence:

ITVNEAQSRGSGGGGGGGGGFRGGRREGGGGYGGGGYGGGRREGGGGGYGGGGGGYGGGRDRGYGGGGDRGYGGDGGSRYSRGGGDSDGNWRN*

NbL03g08910.1 probable_pinoresinol−lariciresinol_reductase_3 [MW = 34.6 kDa]

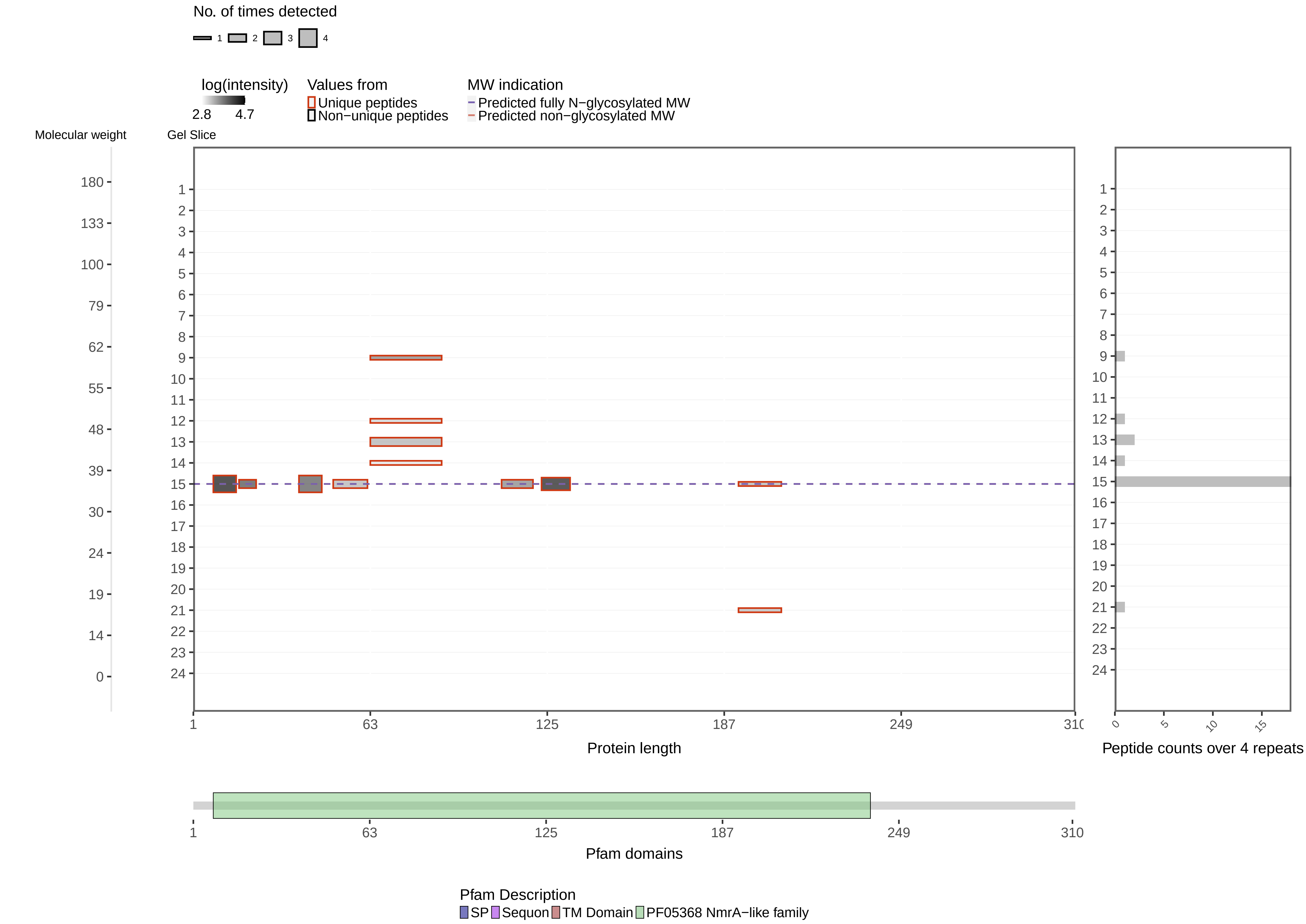


Protein sequence:

AEGIPYTYVCCNFFTSFFLPSLVQPGRKTPPRDEVSIFGDGTAKAVFVKENDAAAFAINTVDDLRTLNKVVHLRPPGNVYSMNELVDIWEGKIGKALKRNYFTEDELLKKIREMPYPENMELVFIYSAFVKGDQTYFSIDSSGGLEGTQIYPQITYTTISEFLDTLL*

NbL03g08870.1 bifunctional_dihydrofolate_reductase−thymidylate_synthase−like [MW = 52.67 kDa]

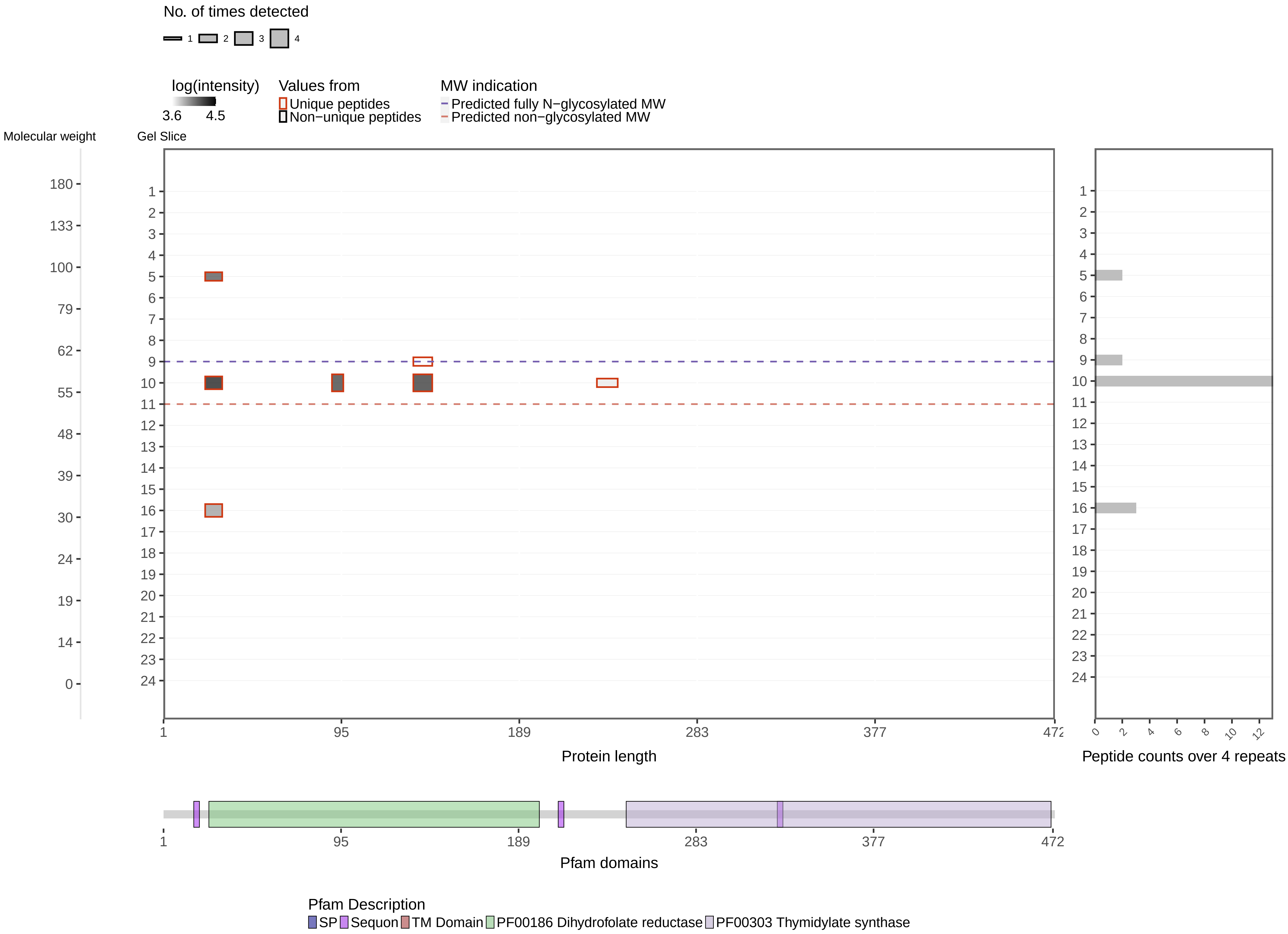


Protein sequence:

DEVKTFSFLPNMIFEKHEEYKYLRLVEEIISNGMPKDDRTGTGTLSKFGCQMRFNLRKSFPLLTTKKVFWRGVVEELLWFISGSTSSKVLQQKGIHIWDGNASRDYLDSIGLKDREEGDLGPVYGFQWRHFGARYIDMHTDYSGQGFDQLADVINKIKNNPDDRRIILSAWNPSDIKLMALPPCHMFAQFYVANGELSCQMYQRSADMGLGVPFNIASYALLTCMVAHVTDLVPGDFVHVIGDAHV*

NbL03g08820.1 purple_acid_phosphatase_2−like [MW − SP = 51.45 kDa]

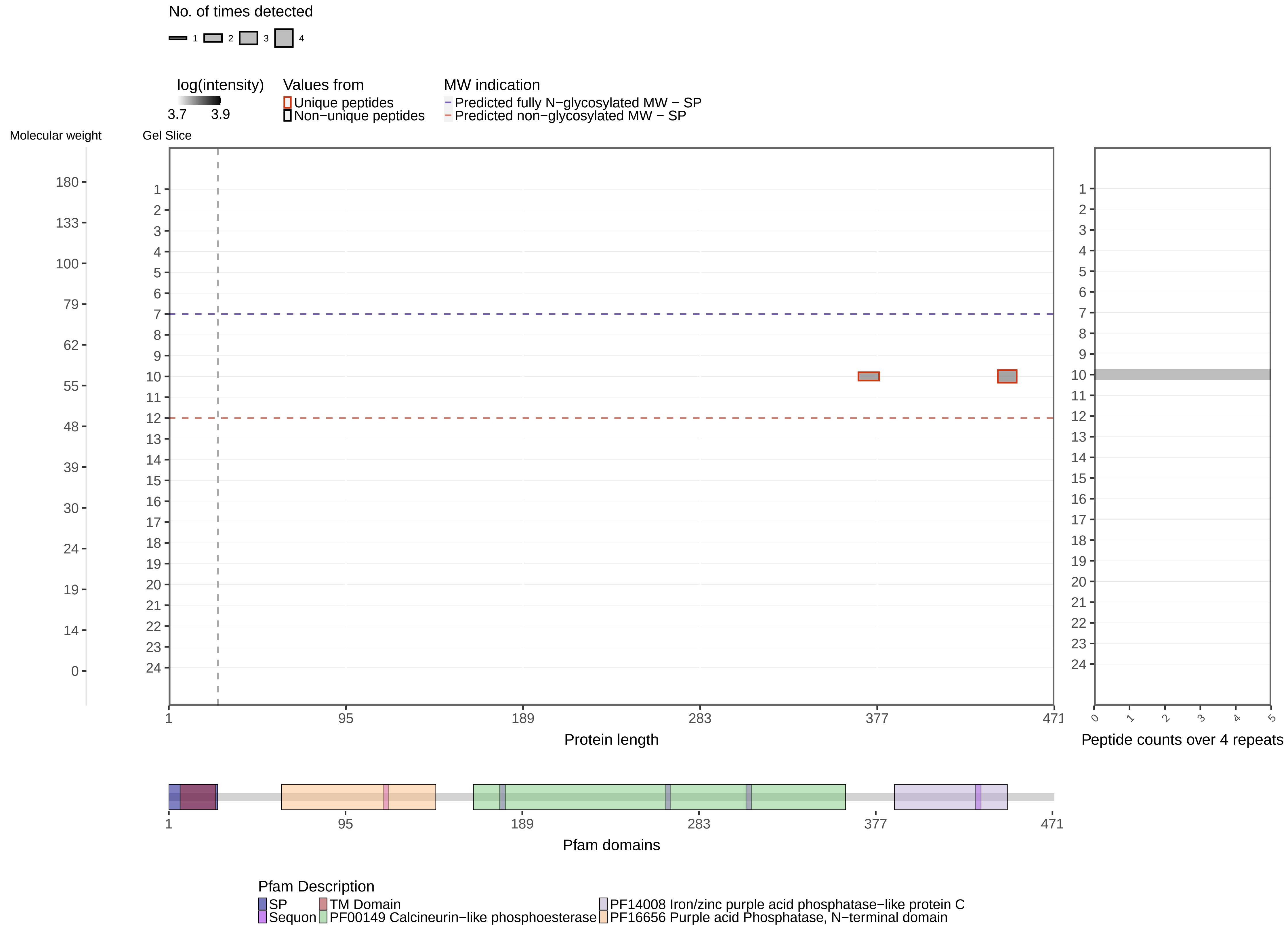


Protein sequence:

### NbL03g08800.1 xylulose_kinase−like [MW = 52.42 kDa]

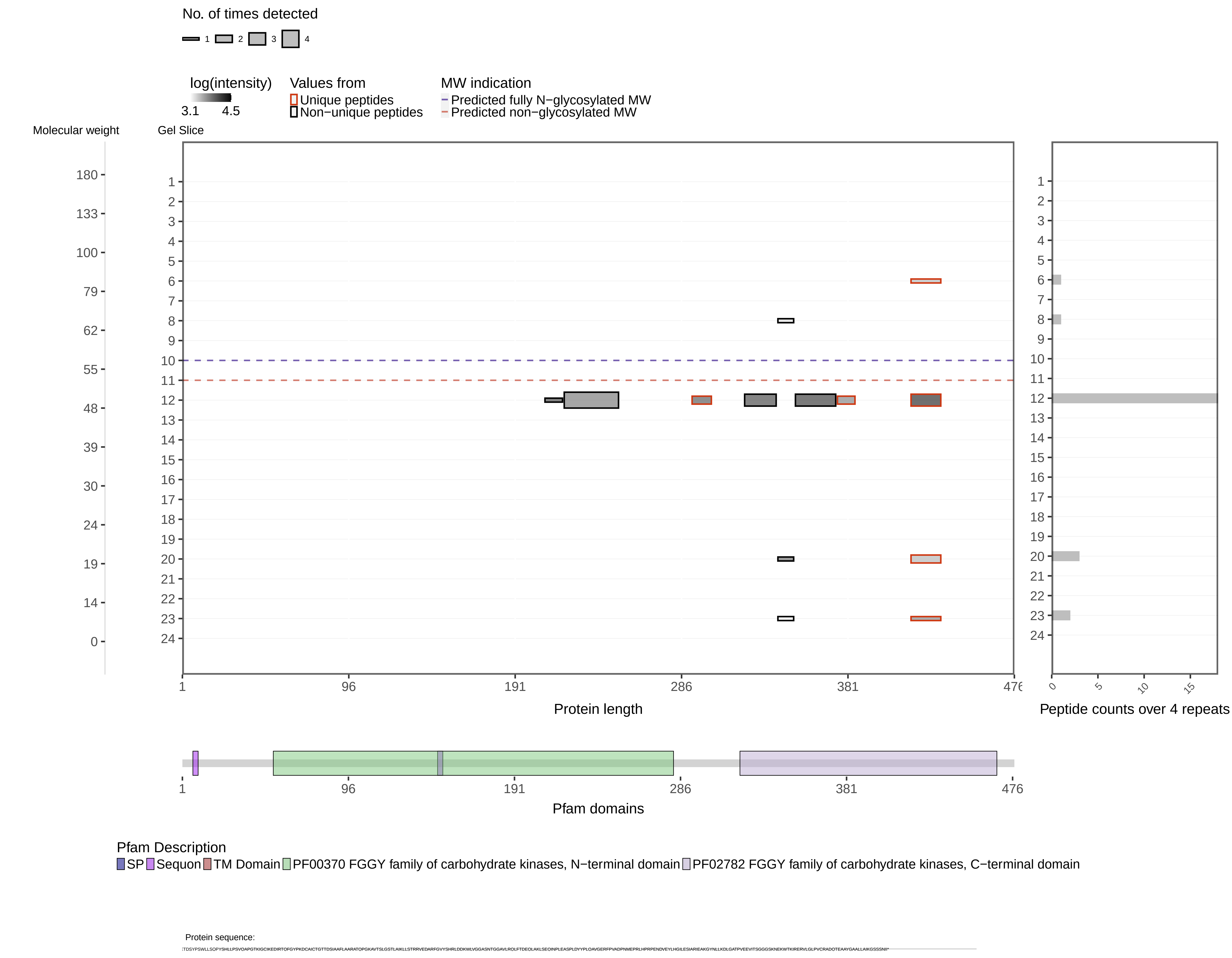

NbL03g08720.1 epimerase_family_protein_SDR39U1 [MW = 38.71 kDa]

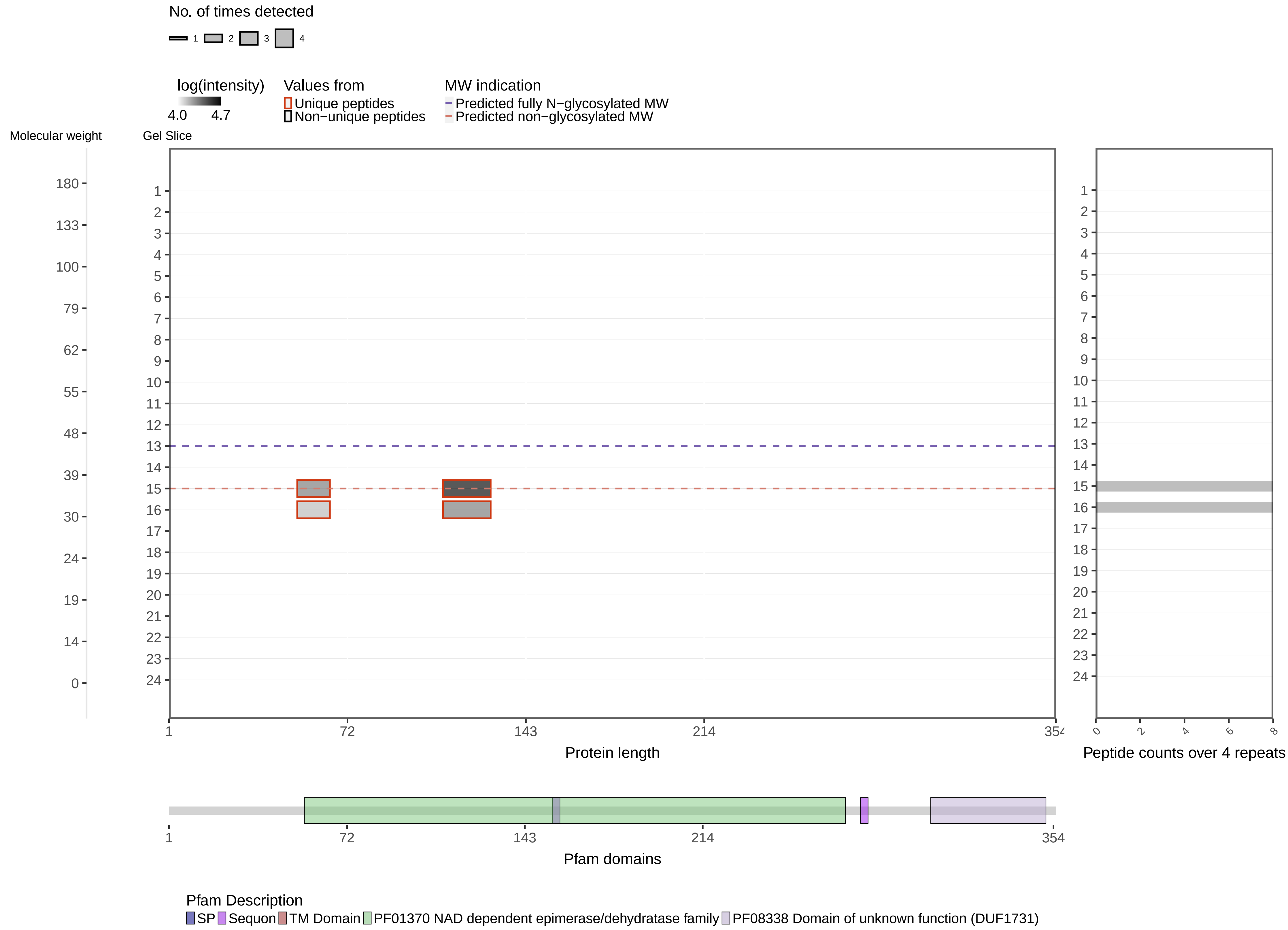


Protein sequence:

FISASAVGYYGISETRVFDEQSPSGNDYLAEVCREWEGAALKVNKDVRLALIRIGVVLGKDGGALAKMIPIFMMFAGGPLGSGNQWFSWIHLDDLVDLIYEALSNPSYKGVINGTAPNPVRLAEMCEQLGSVLGRPSWLPVPEIALKAVLGEGASVVLDGQKVVPTRAKELGFPFKYRYIKDALKAIMS*

### NbL03g08710.1 ubiquitin_fusion_degradation_protein_1_homolog [MW = 35.69 kDa]

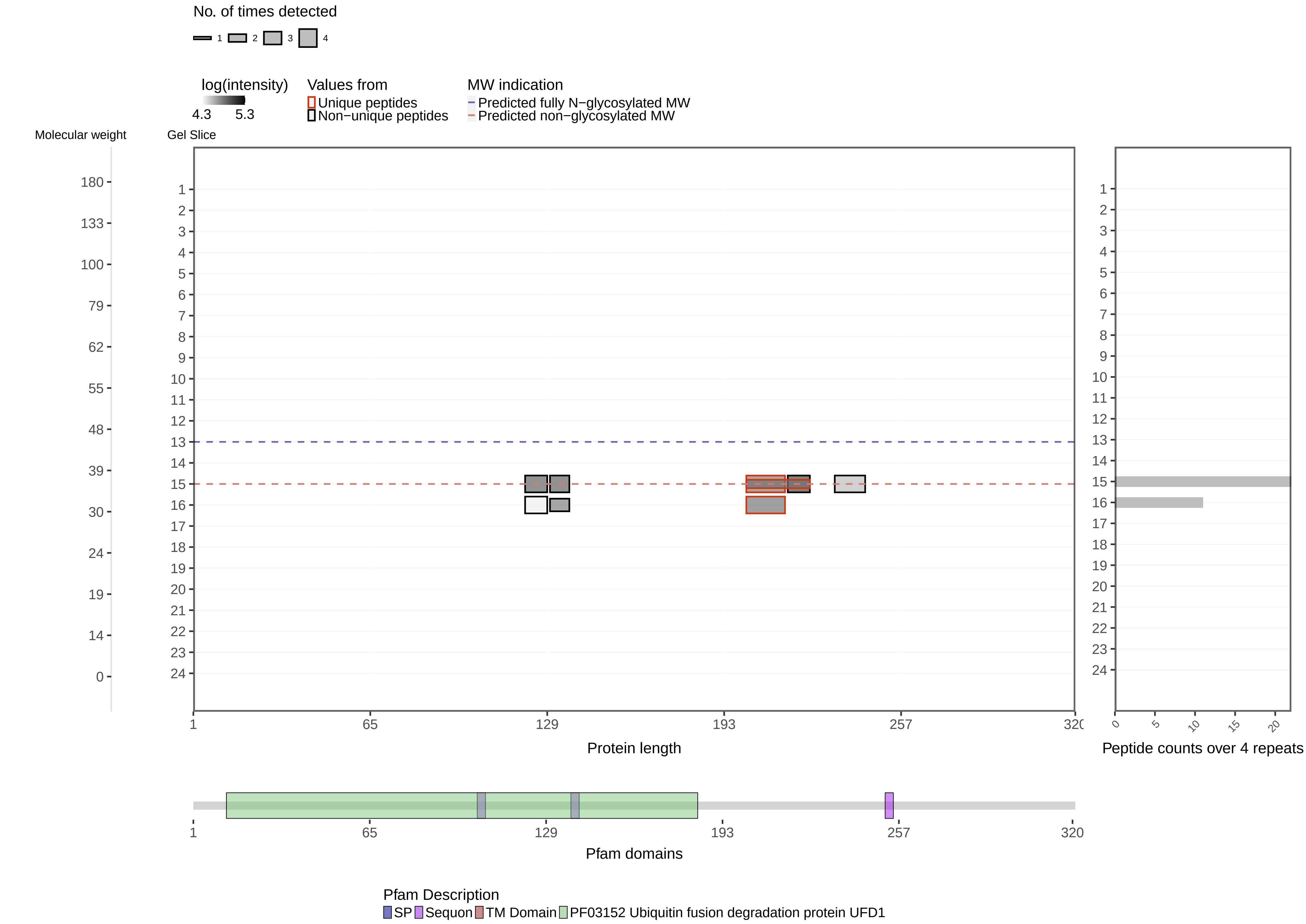


Protein sequence:

/VAYNNKKYYIDIVESKPSNAISIIETDCEVDFAPPLDYKEPERPVPSLPSKTPAEGEGAAAEEEPKFNPFTGSARRLDGKPLKQQPPPVSTSGSKDKRPDMANATGQSAAGSSSQSSNRQTQGKLVFGSNANRNLDKQKEPVKEPVKEEPQKREEPKFQAFTGKKYSLRG*

NbL03g08590.1 SKP1−like_protein_1A [MW = 17.51 kDa]

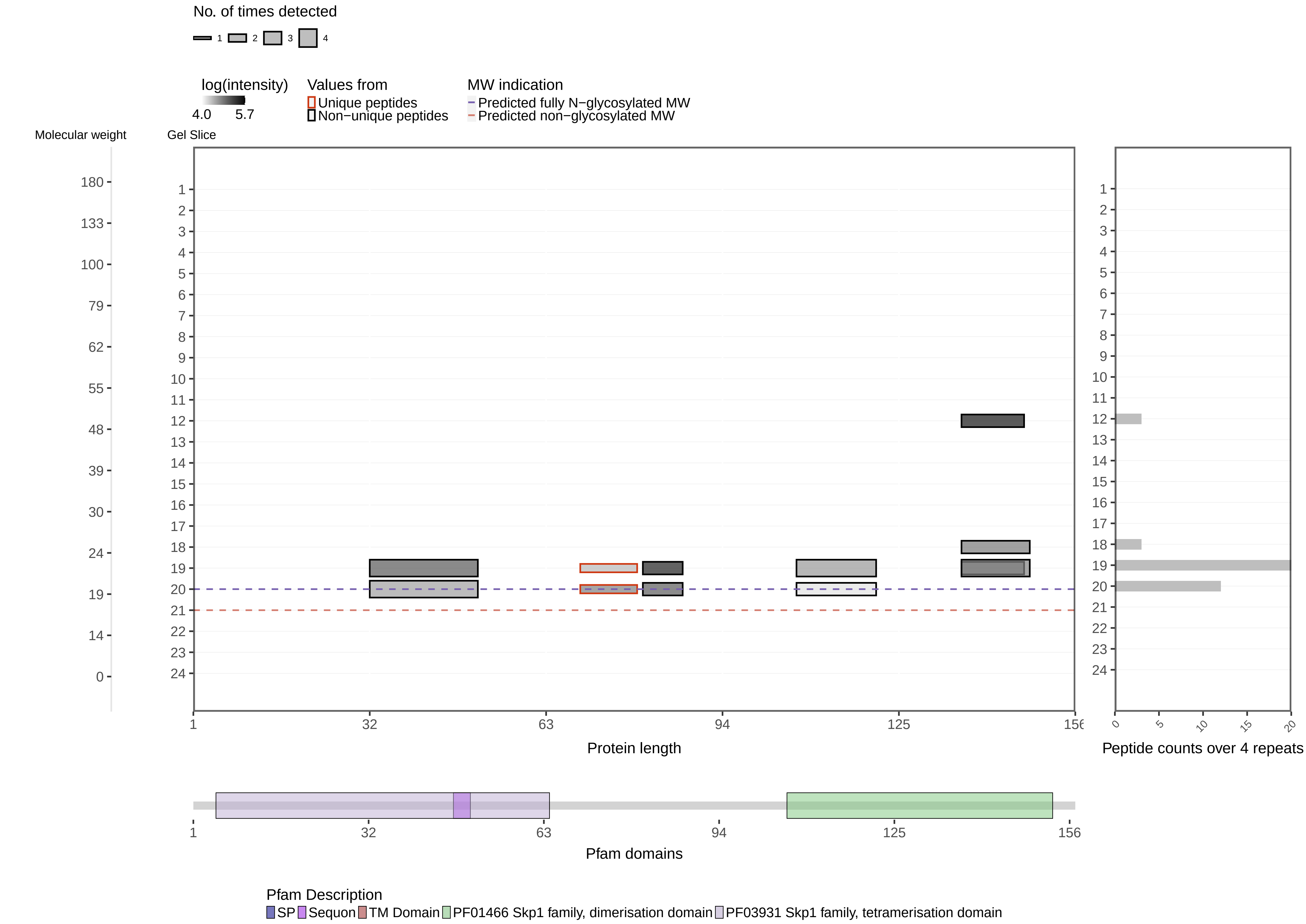


Protein sequence:

DAAKTEDKASEDELKAFDSDFVKVDQATLFDLILAANYLNIKSLLDLTCQTVADMIKGKTPEEIRKTFNIKNDFTPEEEEEVRRENAWAFE*

### NbL03g08470.1 expansin−like_A2 [MW − SP = 26.52 kDa]

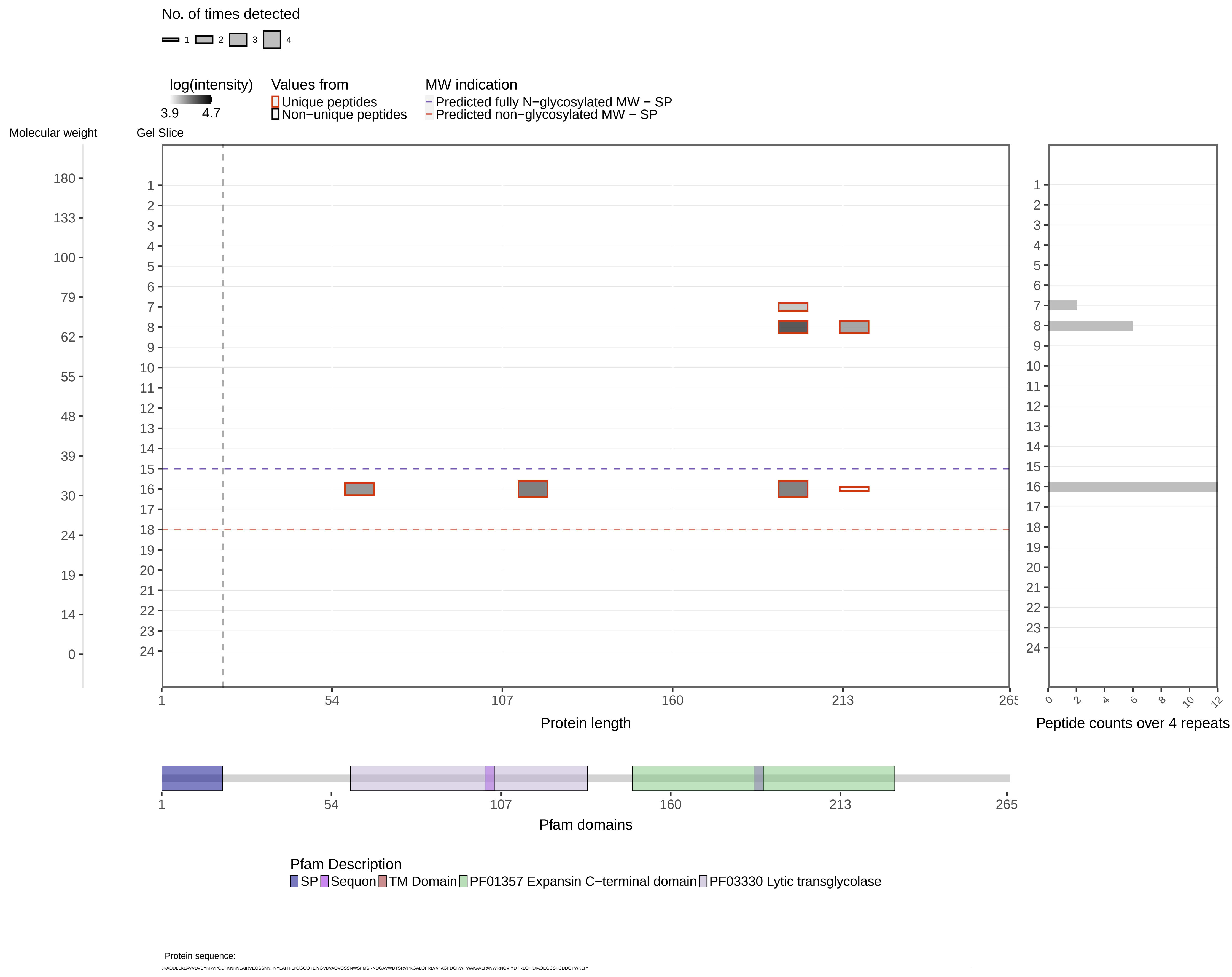

NbL03g08400.1 DNA_repair_RAD52−like_protein_1_mitochondrial [MW = 21.27 kDa]

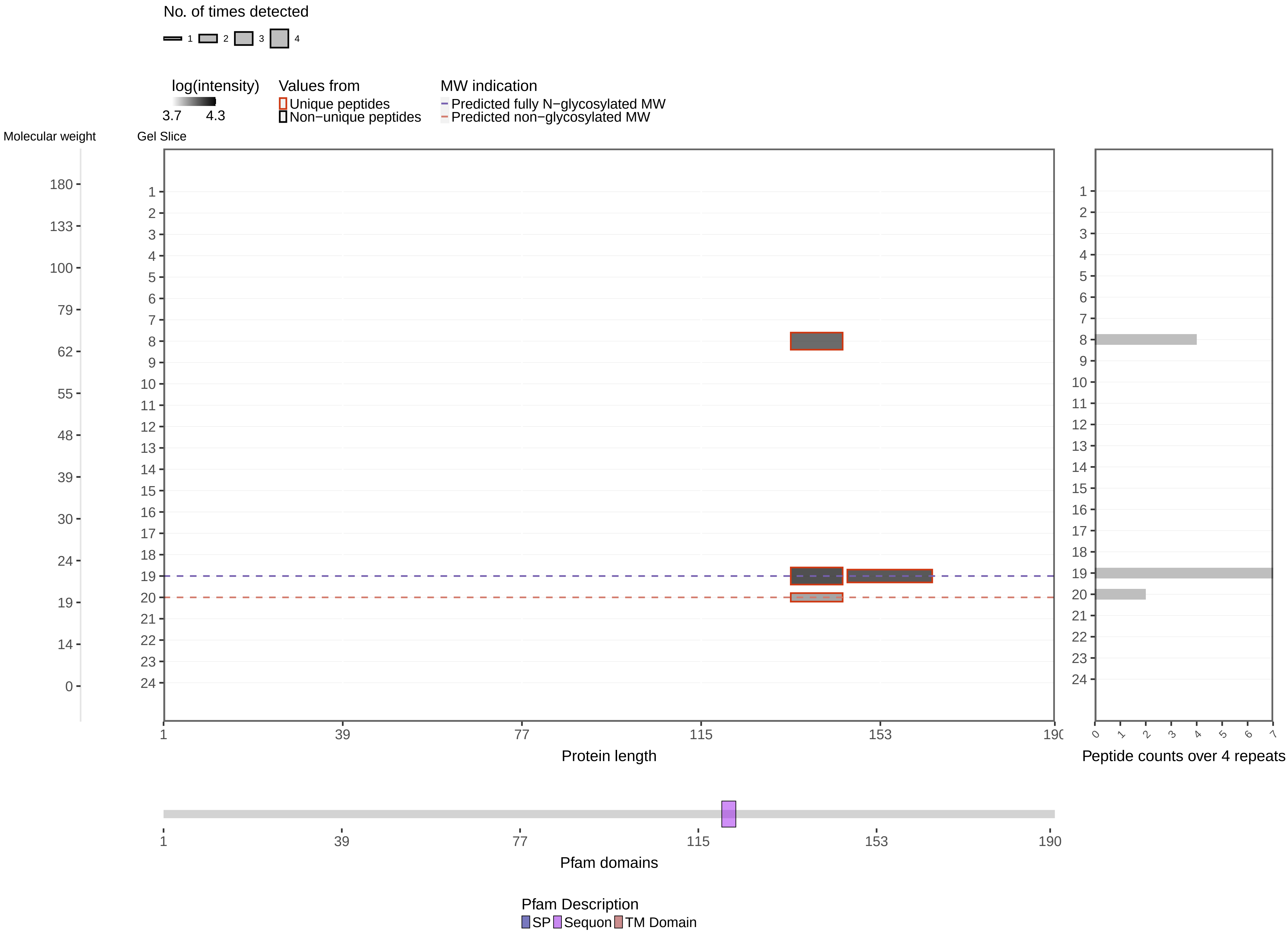


Protein sequence:

IKFRHEPNGISLKYIPWHIVNRILNLHAPEWSGEVRSINYSSDGKSVSVVYRVTLYGTDAEIYRESTGTASVDDPGYGDPVQKAEGMAFRRACARFGLGLHLYHEDMS*

### NbL03g08350.1 single−stranded_DNA−binding_protein_WHY2_mitochondrial [MW = 26.82 kDa]

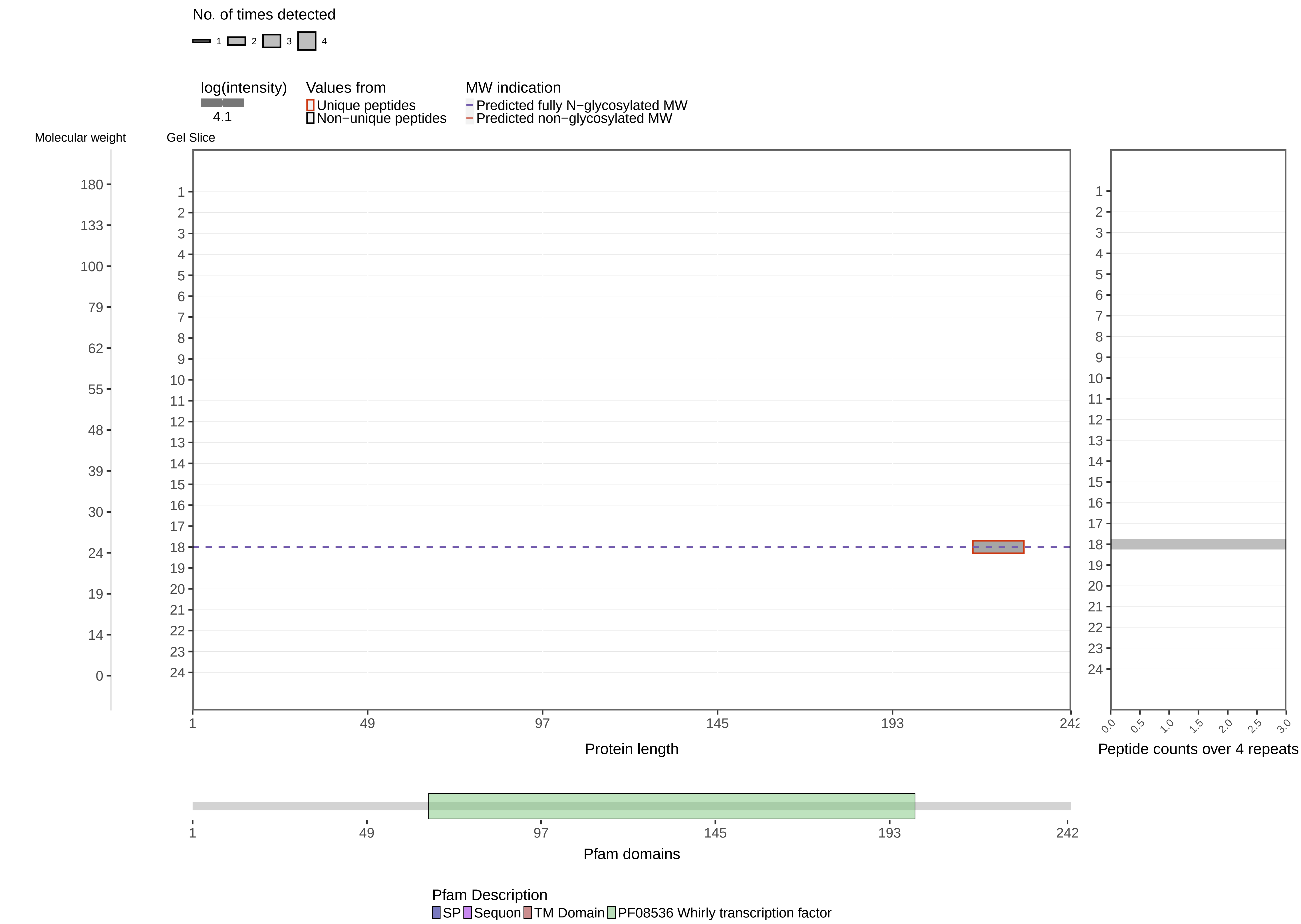


Protein sequence:

RKYDWEKRQLFALSATEVGSLISIGTRDSCEFFHDPSMLSSNAGQVRKSLSFKPNADGTGYFISLSVVNNNLKNNNRFTVPVTTAEFAVMRTAFSFALPHIMGWDRFTNQPSESISQTPSKVVPQLMETEWDR*

### NbL03g08280.1 60S_ribosomal_protein_L7a−2−like [MW = 29.35 kDa]

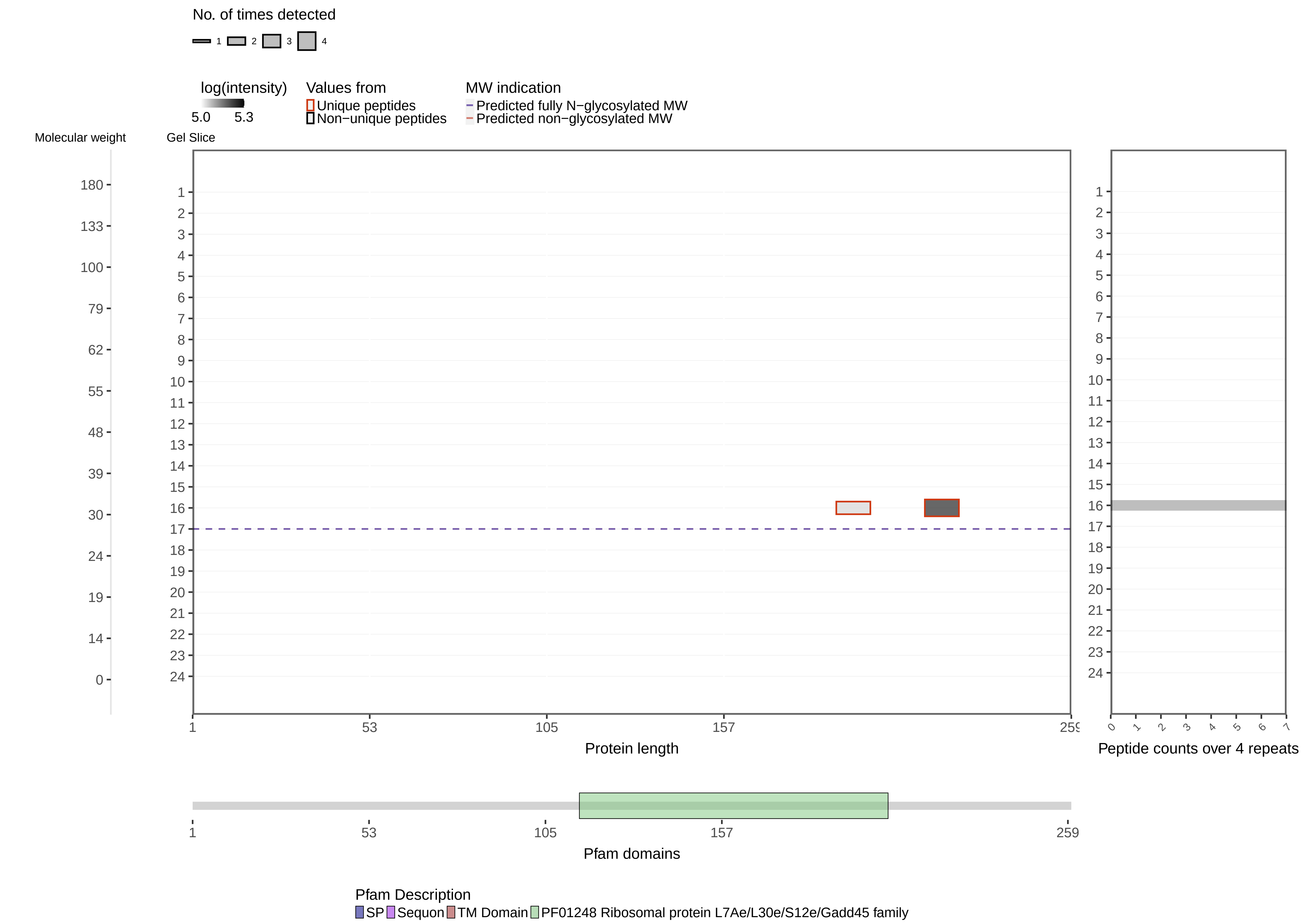


Protein sequence:

AEGKTPETKKPIIVKYGLKHITYLIEQNKAQLVVIAHDVDPIELVVWLPALCRKMEIPYCIVKGKARLGSIVHKKTASALCLTTLKNEDKMEFSRILEAIKANFNDKFEENRKKWGGGVMGSKSQARTKAKERVLAKEAAQRMN*

NbL03g08120.1 2−alkenal_reductase_(NADP(+)−dependent)−like [MW = 44.46 kDa]

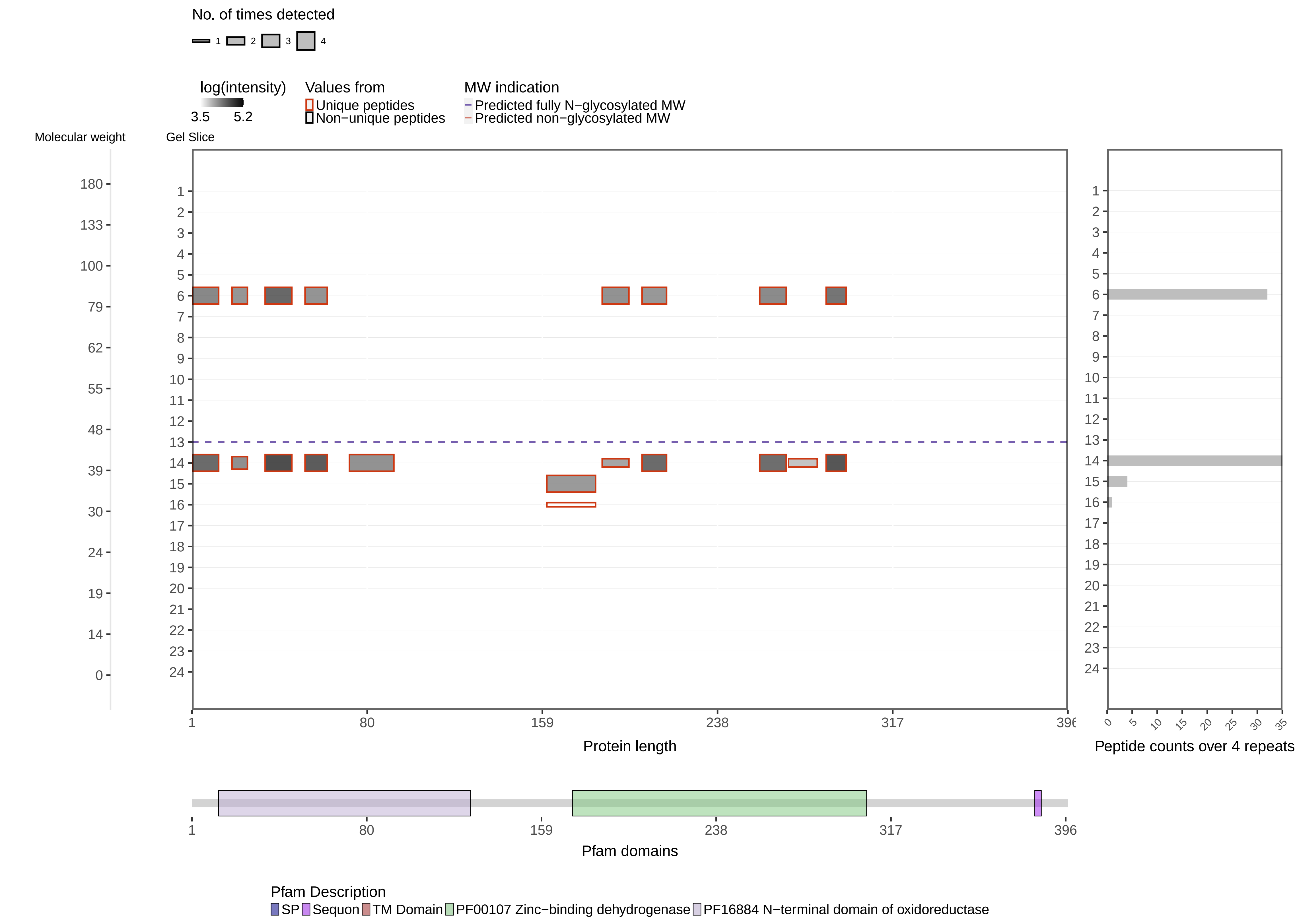


Protein sequence:

KGCYVVGSASTDEKVNLLKSELGFDDAFNYKEGDDFAGALKRHFPKGIDVYFENVGGNMLDEVLLHMNLYGRITVSGMISQYNLKKPDGIHNLFCLITKRLRMEGFSELDFRHKVPEYLEFVIQLIREKKLVFIEDIAEGLENAASAFVGIYHGRNVGKQIIQRLNRDKWSLRKNGFPAREFSWEIKVCLSSTRMYSNESSLFGVNIWLSR*

### NbL03g08110.1 UPF0301_protein_TC_0483 [MW = 53.28 kDa]

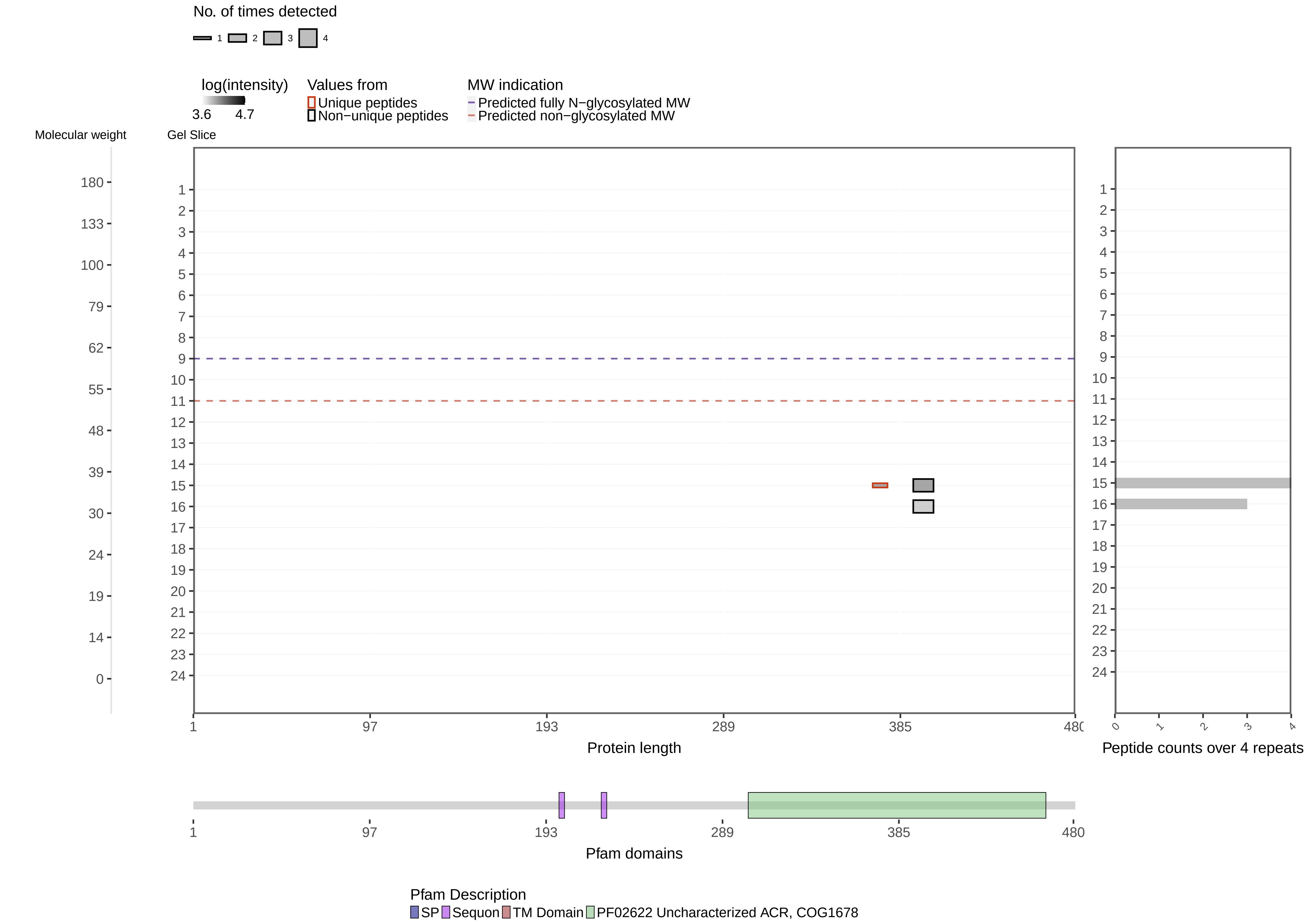


Protein sequence:

APKSHQKPLDWREFRASLYIQEQQAATAVSDTHKQEGTSFGPKALPLKWAHPISAPENGCVLVATEKLDGVRTFERTVVLLLRSGTRHPQEGPFGVVINRPLGKKIKHMKPTNLDLATTFADCSLHFGGPLEASMFLLRGSASSELPRFEEVIPGVCFGARNSLEEASVLVNKGVLKPQDFRFFVGYAGWQLDQLREEIESGYWYVASCSSNLIFAGSQTSSSESLWGEILQLMGGQYSEVSRKPKQDI*

NbL03g08020.1 uncharacterized_LOC104218509 [MW = 31.34 kDa]

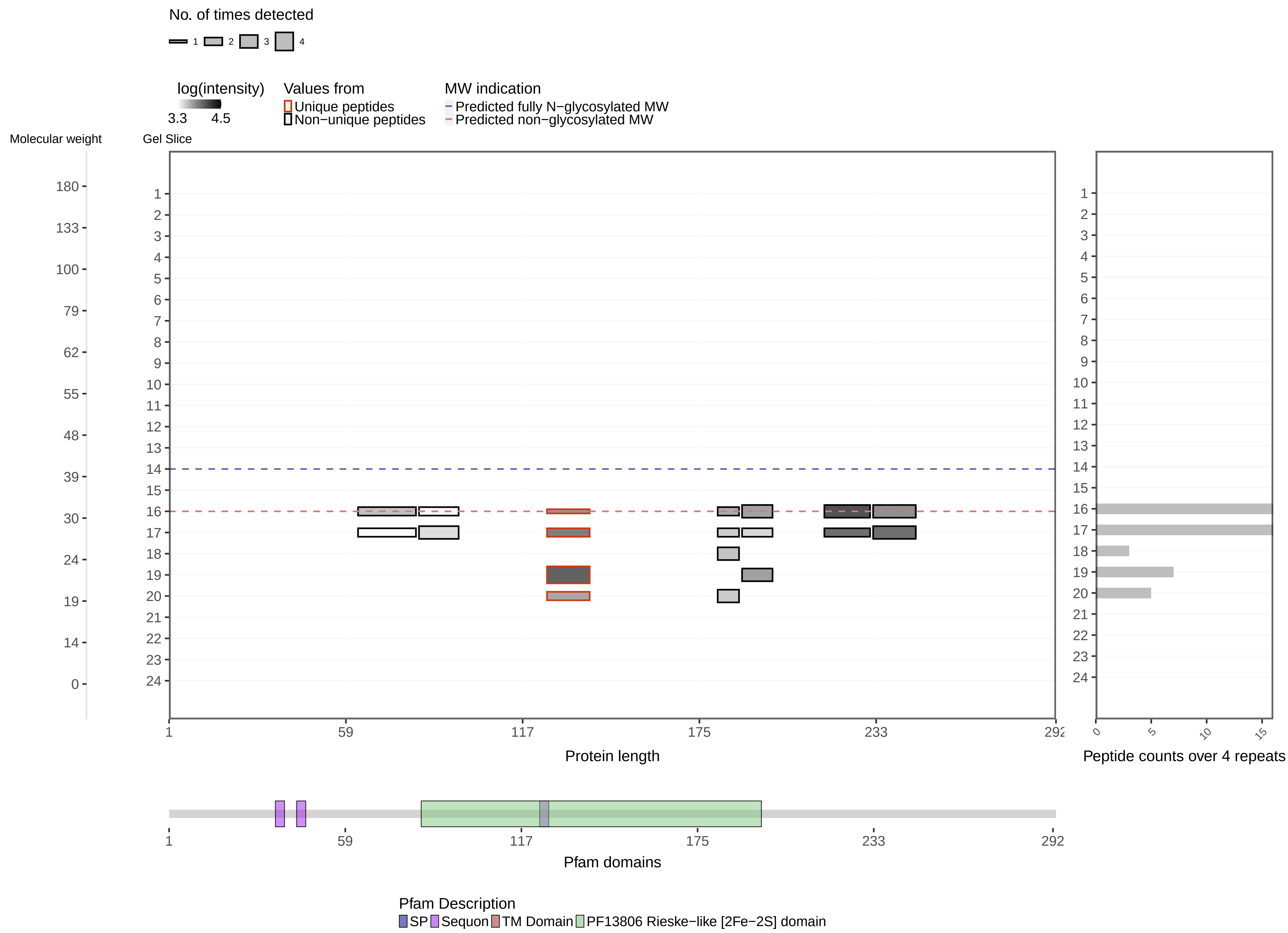


Protein sequence:

EGLLNAKLTQDGCIVCPTTDSTFFLQNGAIKEWYPNNPVLRVLTPALRNLFVYPVKTDDENIYISMRGGGKVDASAEIVFSGKAQPGVTASDVNVDEVRMVIDEGQEAFGFTGRNELINGKAAVIGFLLLLDFELLTGKGLLKGTGFLDFIYSVSDAFK*

### NbL03g07950.1 cytochrome_c−like [MW = 12.07 kDa]

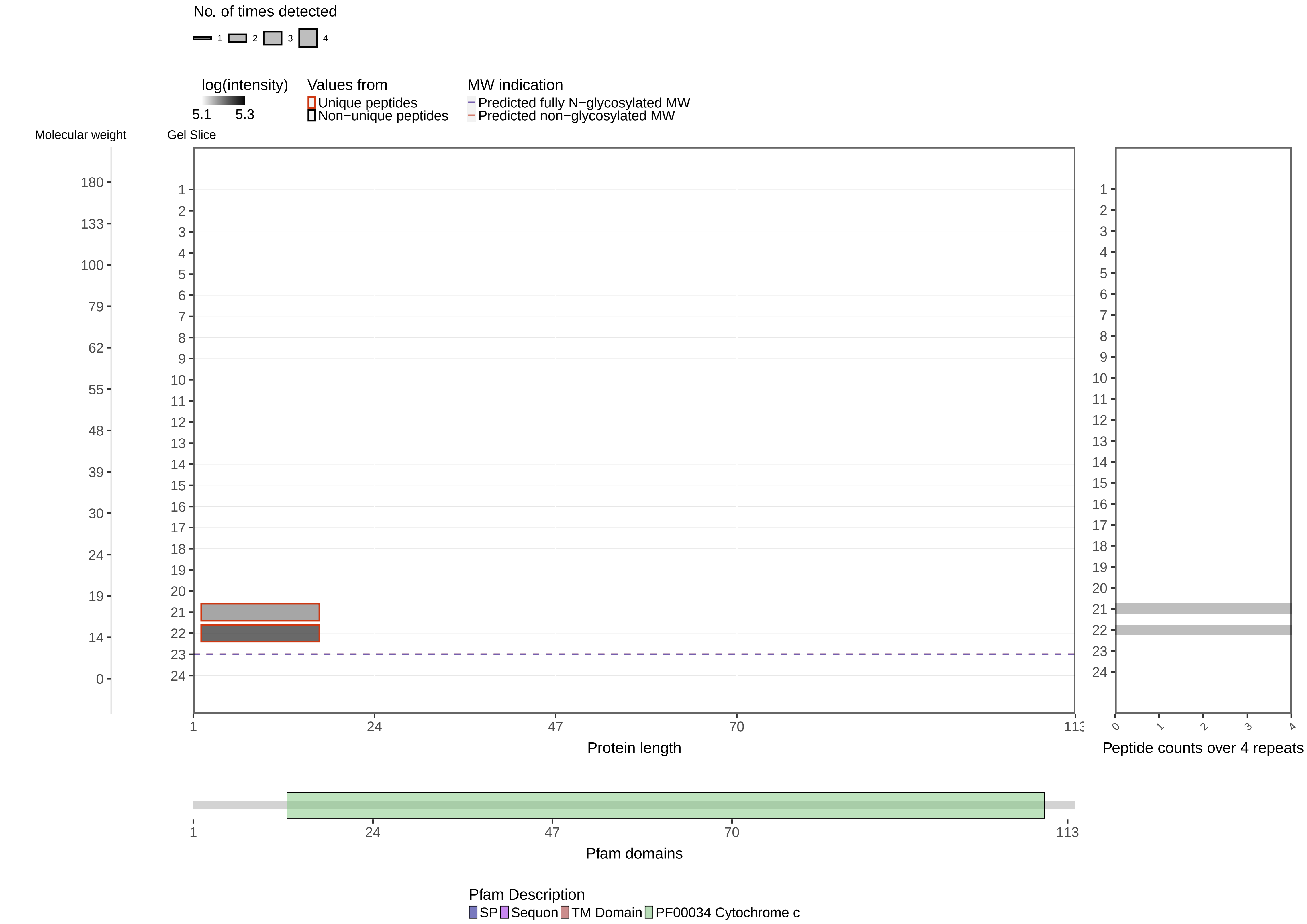


Protein sequence:

GLFGRQSGTTAGYSYSAANKNMAVMWEEKTLYDYLLNPKKYIPGTKMVFPGLKKPQDRADLIAYLKSATA*

### NbL03g07860.1 far_upstream_element−binding_protein_1−like [MW = 84.09 kDa]

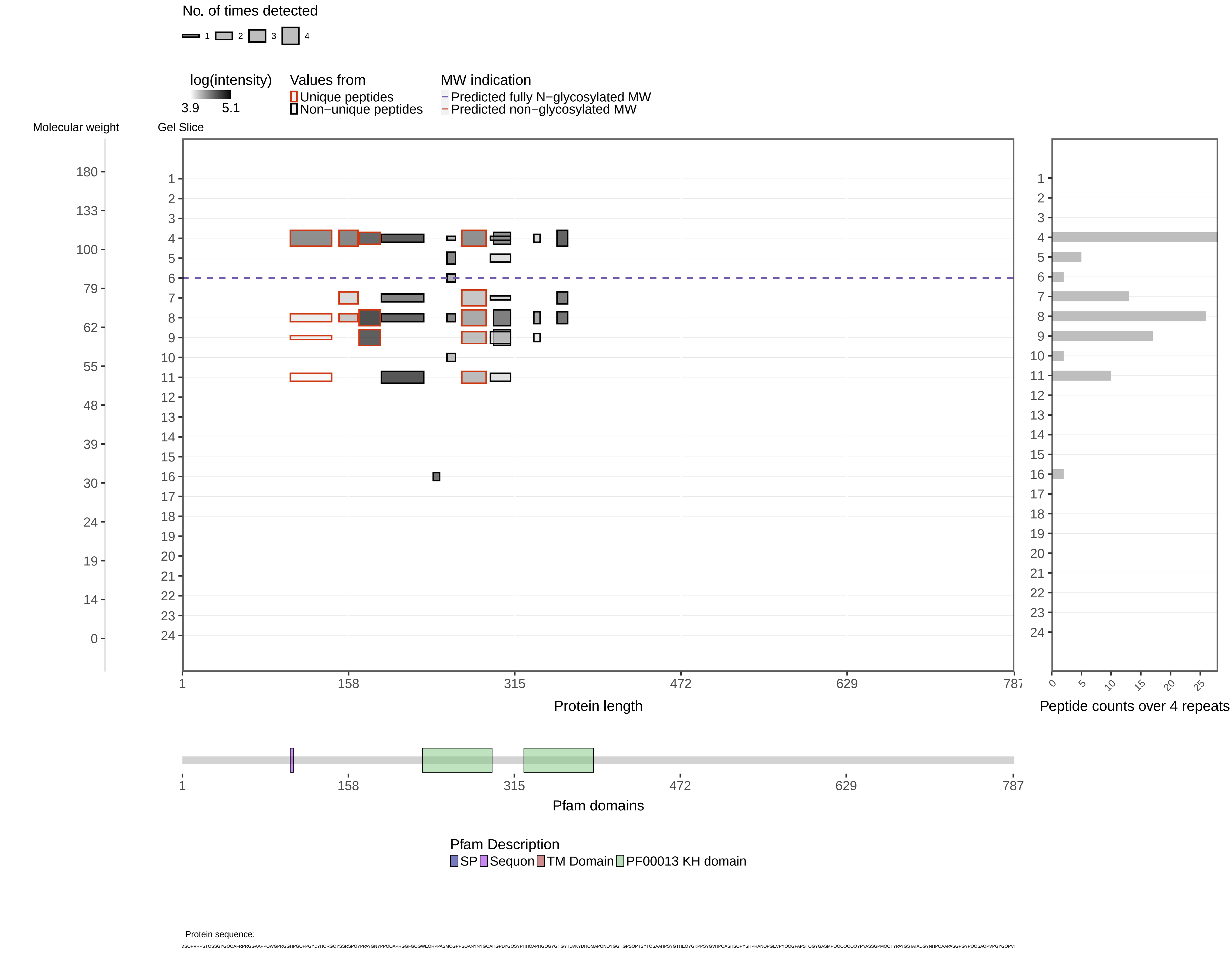

### NbL03g07800.1 bifunctional_aspartokinase/homoserine_dehydrogenase_chloroplastic−like [MW = 100.37 kDa]

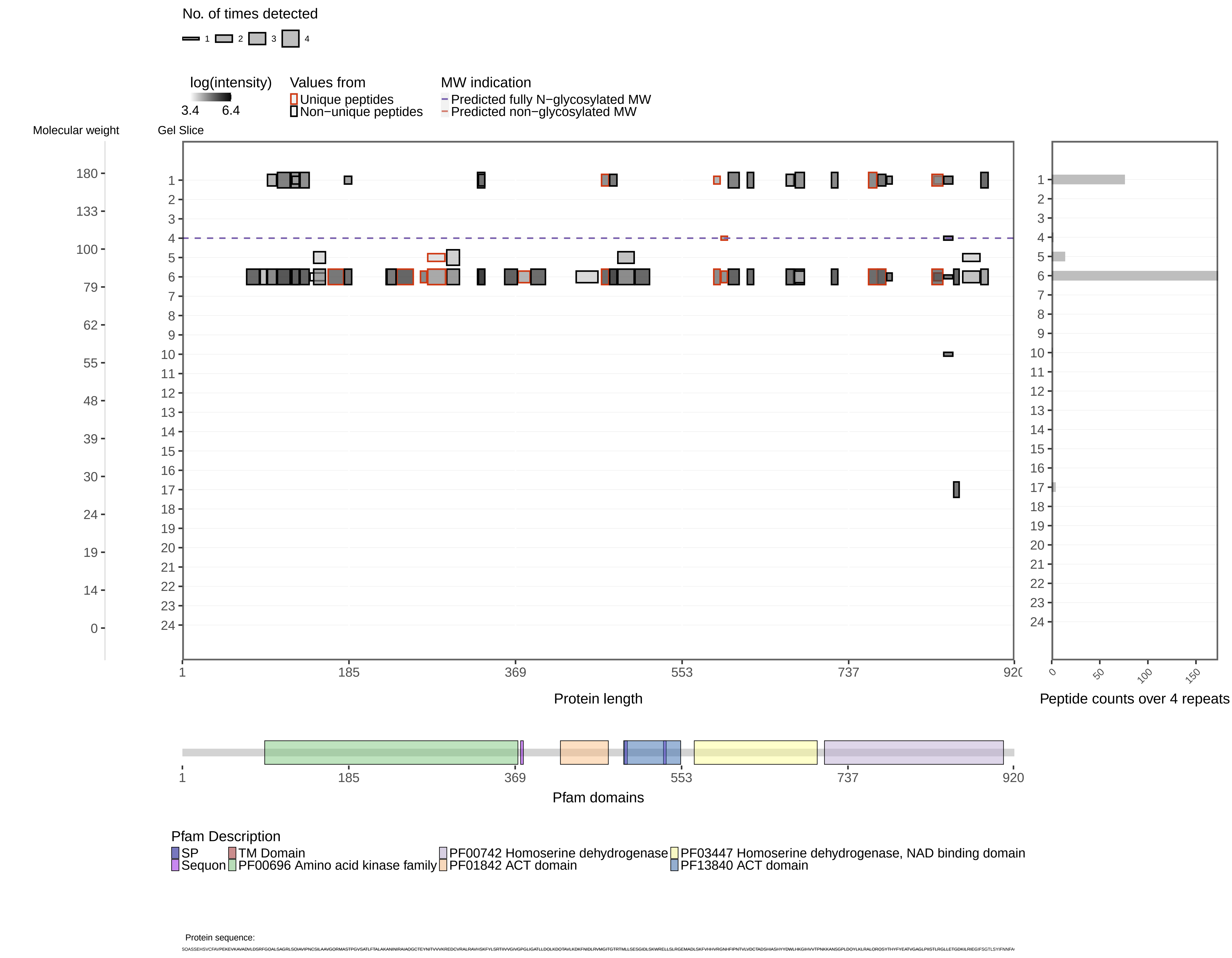

### NbL03g07790.1 histone_H2B−like [MW = 18.82 kDa]

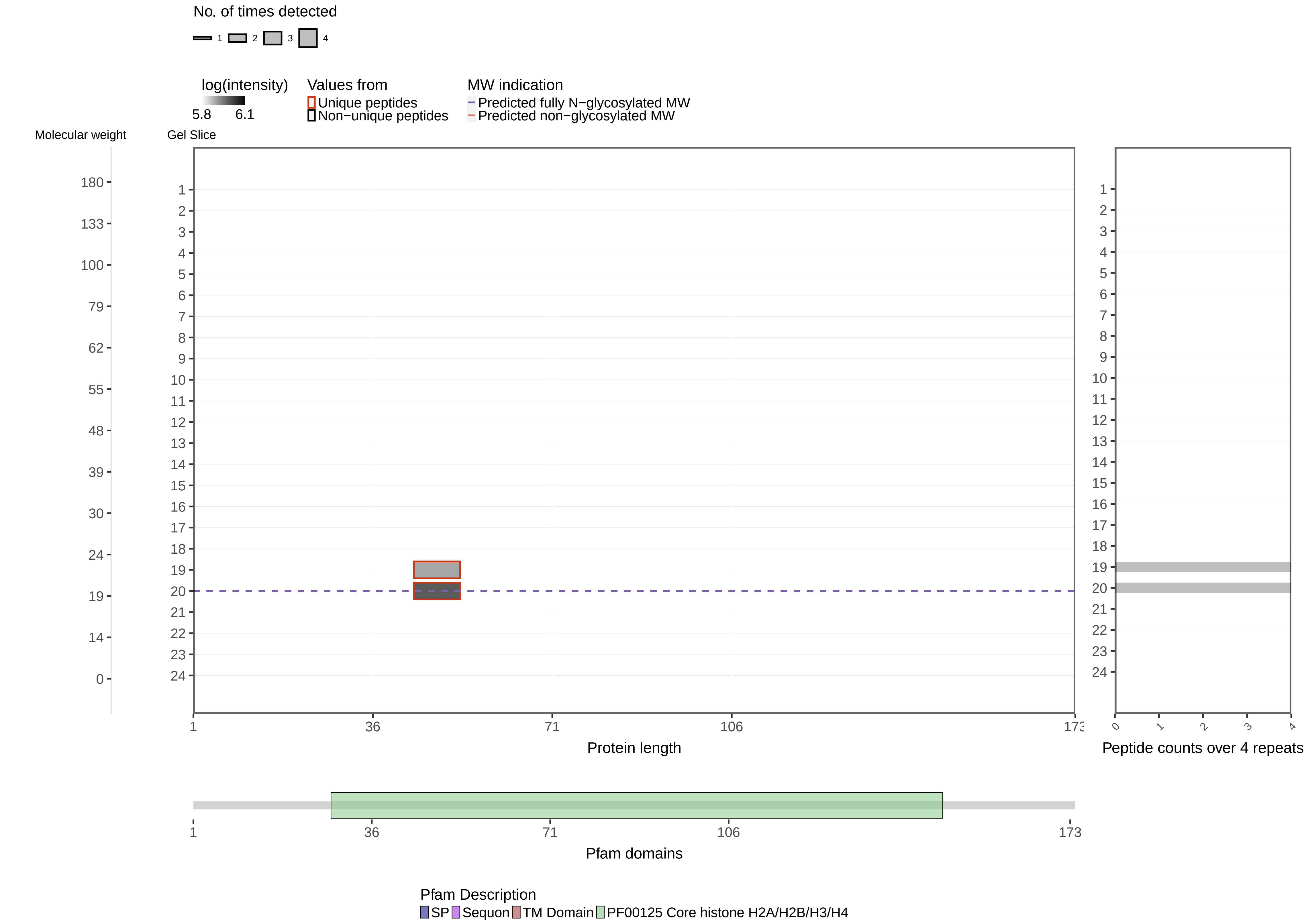


Protein sequence:

DKKKKRLKKSVETYKIYIFKVLKQVHPDIGISSKAMGIMNSFINDIFEKLAQESSRLARYNKKPTITSREIQTAVRLVLPGELAKHAVSEGTKAVTKFTSS*

### NbL03g07780.1 histone_H2B−like [MW = 19.72 kDa]

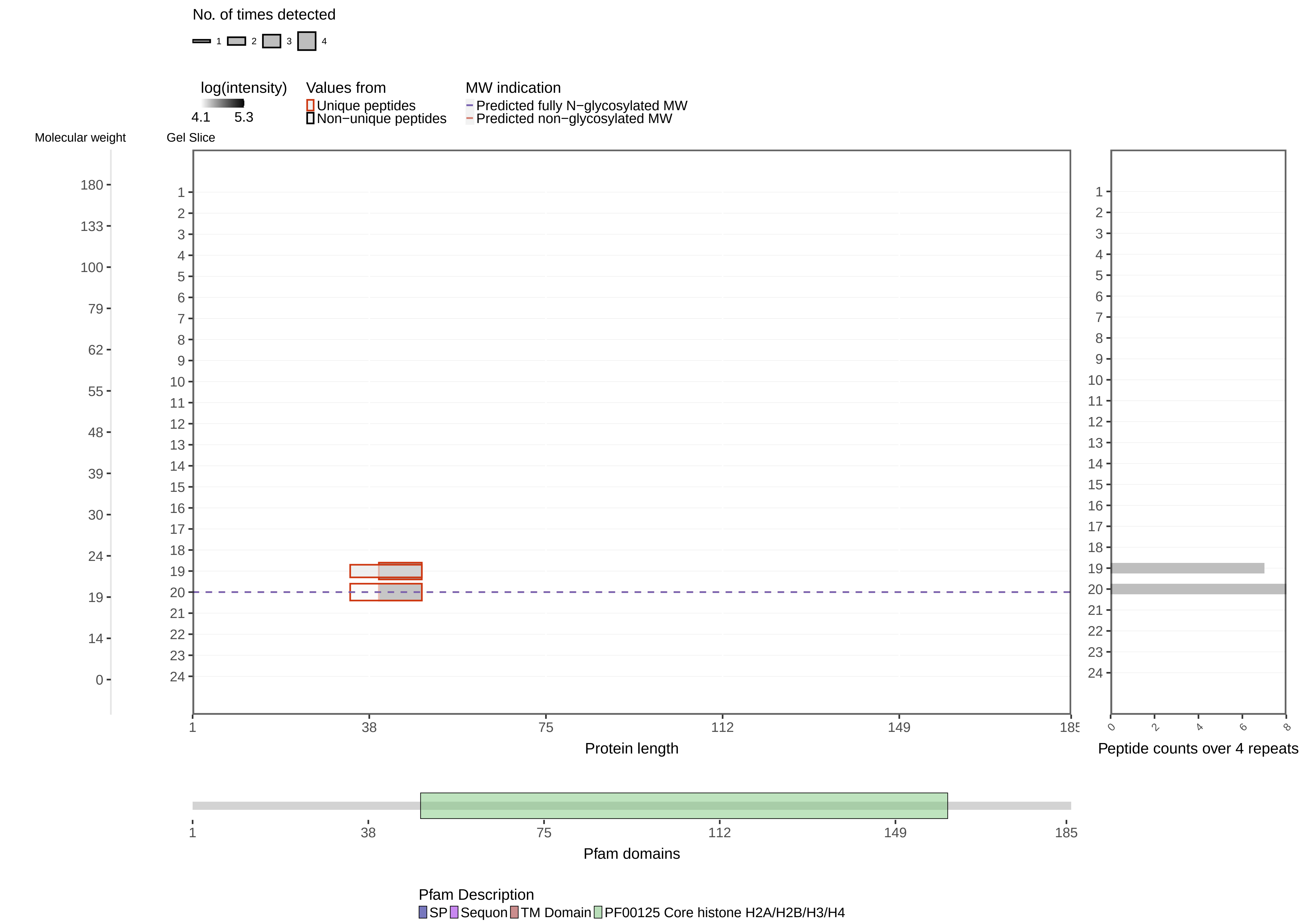


Protein sequence:

AGAAAGDKKKKRLKKSVETNKIYIFKVLKQVHPDIGISSKAMGIMNSFINDIFEKLAQESSRLARYNKKPTITSREIQTAVRLVLPGELAKHAVSEGTKAVTKFTSS*

### NbL03g07700.1 serine/threonine−protein_phosphatase_PP1_isozyme_3 [MW = 34.52 kDa]

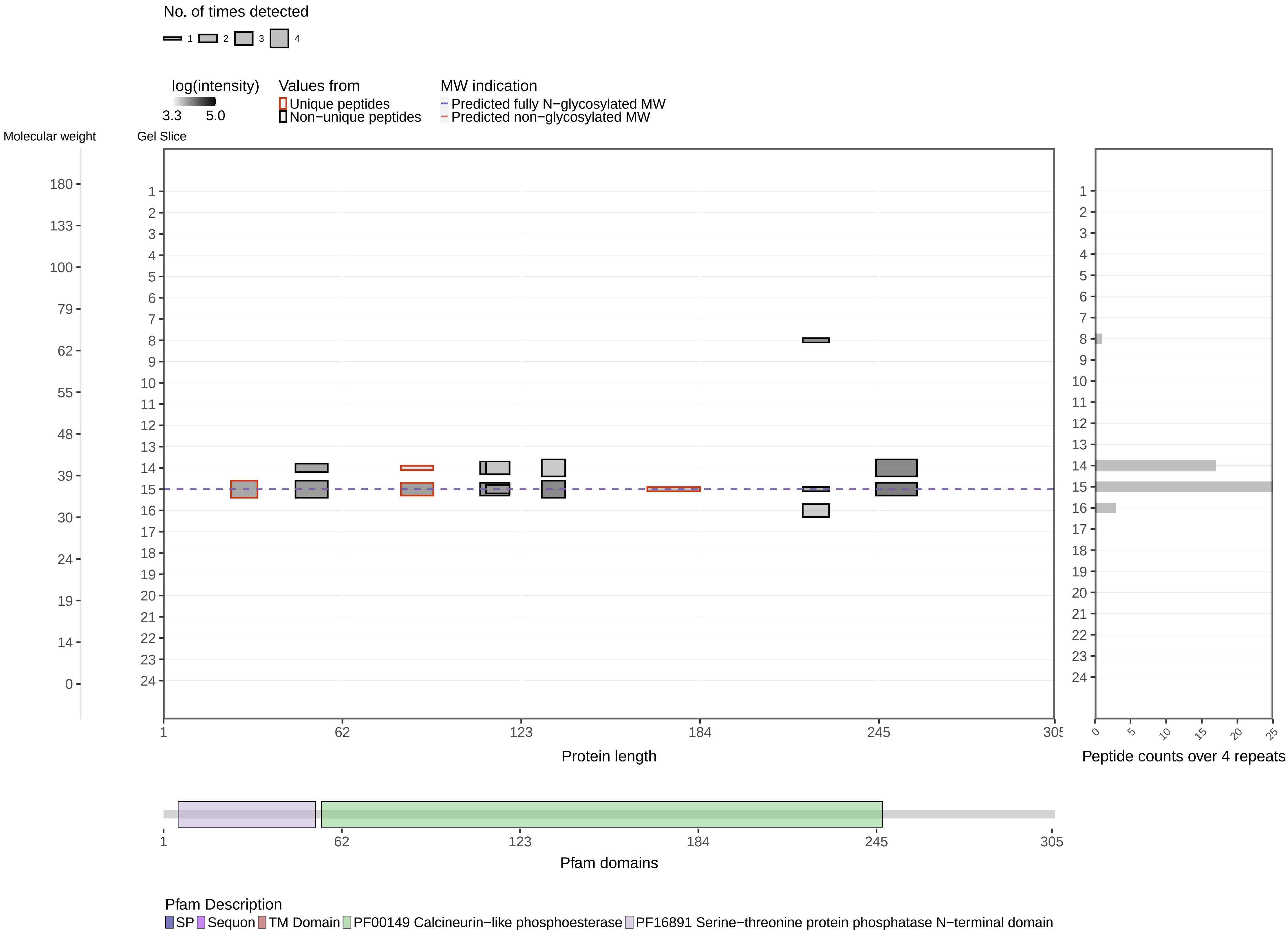


Protein sequence:

VKLWKSFTDCFNCLPVAALIDEKILCMHGGLSPDLSSLDQIRNLPRPTAIPDTGLLCDLLWSDPGKDVKGWGMNDRGVSYTFGPDKVSEFLSKHDLDLVCRAHQVVEDGYEFFAERQLVTIFSAPNYCGEFDNAGAMMSVDENLLCSFQILKPAEKKNKFMM*

### NbL03g07670.1 small_nuclear_ribonucleoprotein_E−like [MW = 10.31 kDa]

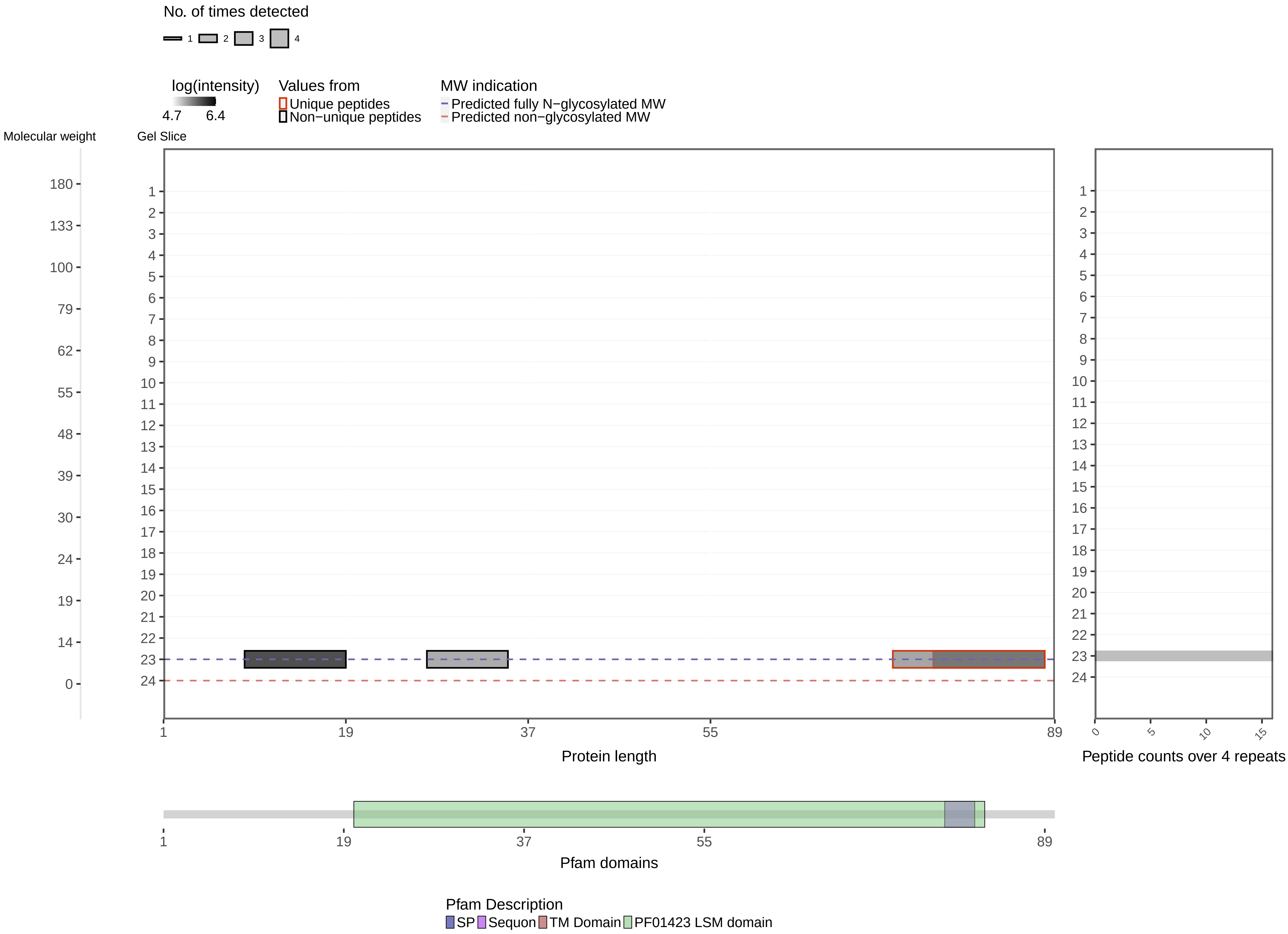


Protein sequence:

FEQKDLRIEGRIIGFDEYMNLVLDDAEEVSVKKKSRKQLGRILLKGDNITLMMNTGK*

### NbL03g07640.1 delta−1−pyrroline−5−carboxylate_synthase−like [MW = 77.87 kDa]

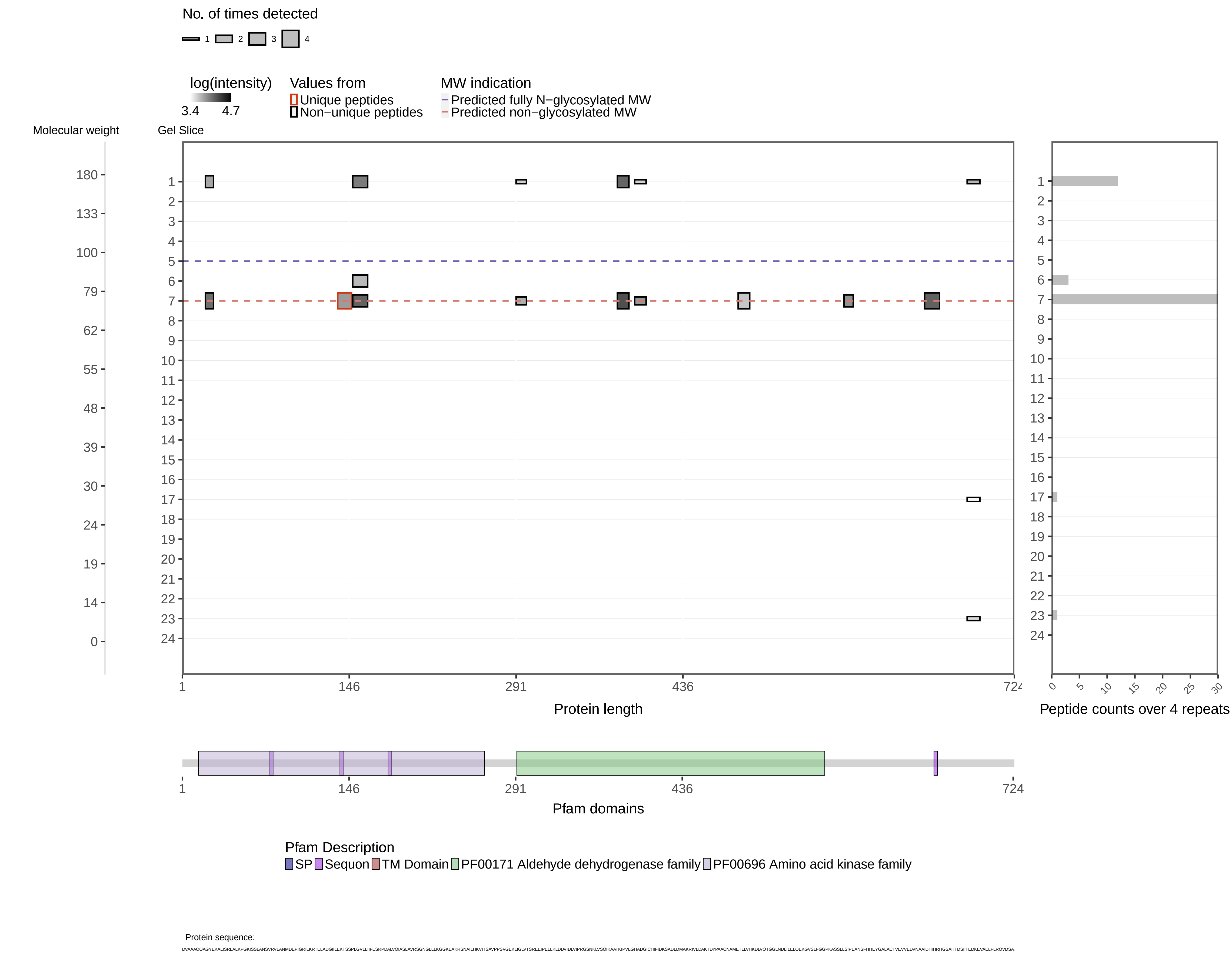

NbL03g07630.1 uncharacterized_protein_At2g39795_mitochondrial−like [MW = 28.9 kDa]

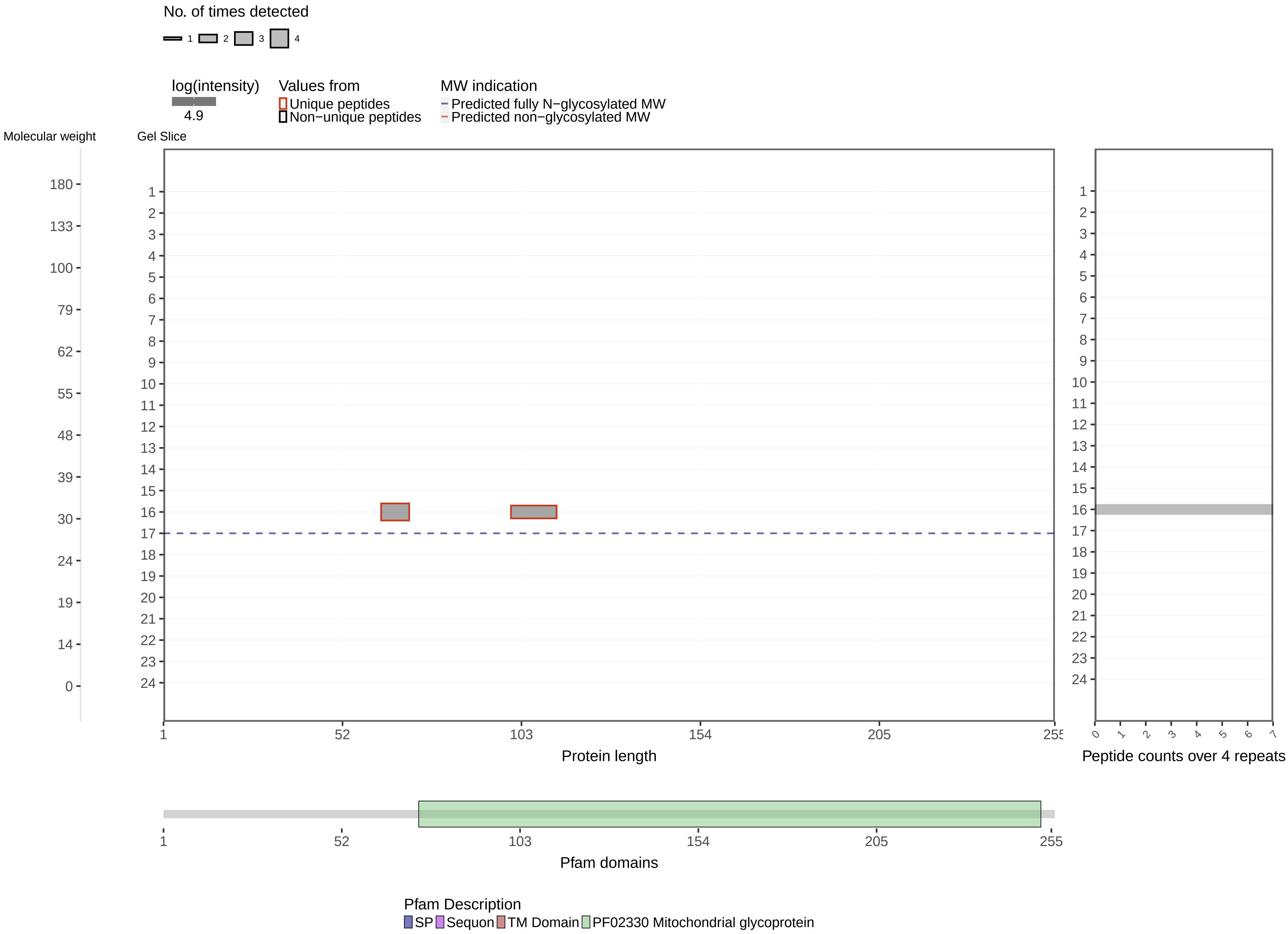


Protein sequence:
YQGETVNVEVHMPDLVTGDEDEDGNDNDDTERPAQSQLPLVVRVSKRNGPCLEFGCTAYPDEVVIDHLAIKDPDAADEQIAYEGPEFSDLDENLQKAFHKYLEIRGIKPSTTNFLHEYMINKDSREYLMWIKNVKKFIEA*

NbL03g07510.1 anthranilate_synthase_alpha_subunit_1_chloroplastic−like [MW = 60.3 kDa]

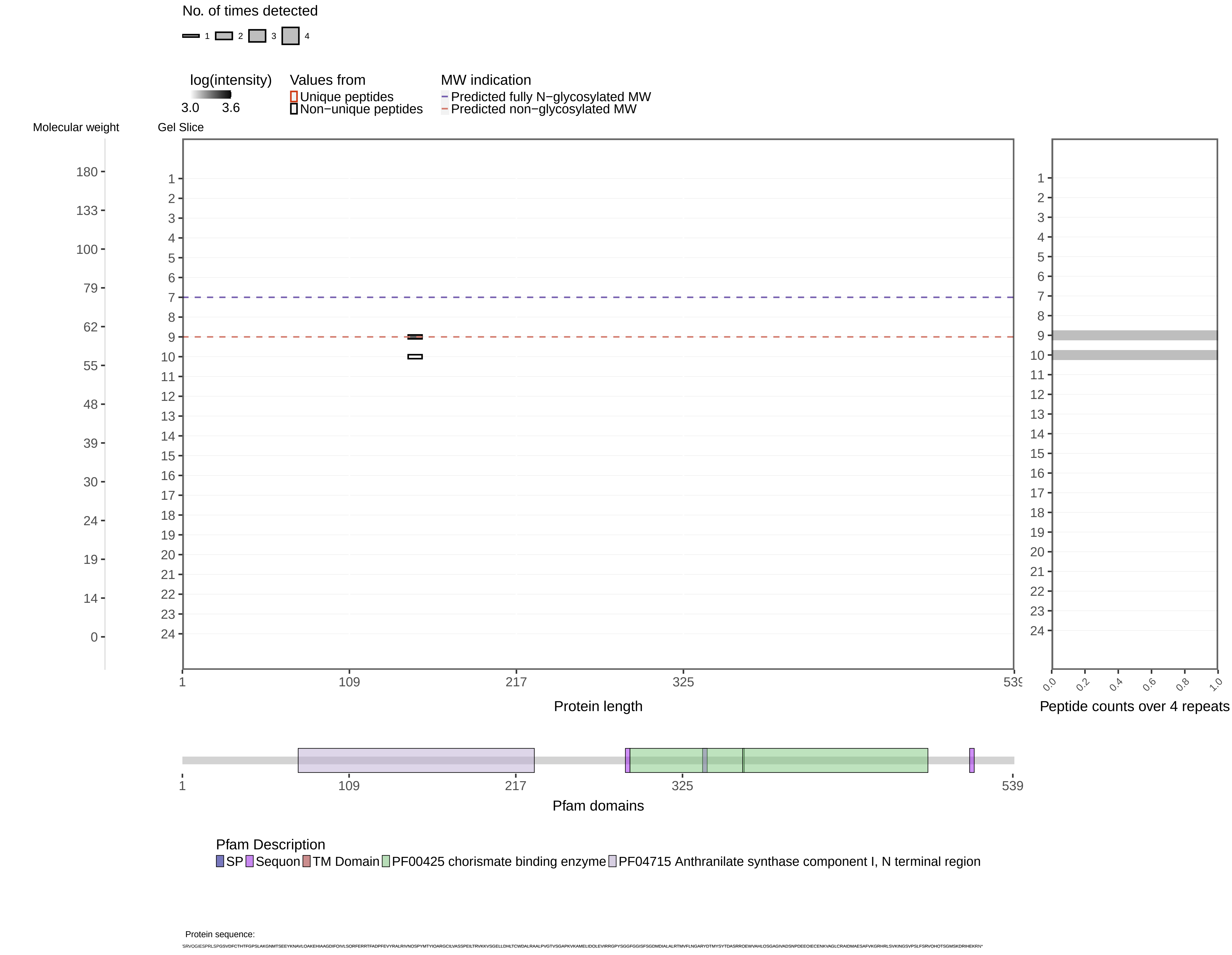

NbL03g07310.1 sedoheptulose−17−bisphosphatase_chloroplastic−like [MW = 42.43 kDa]

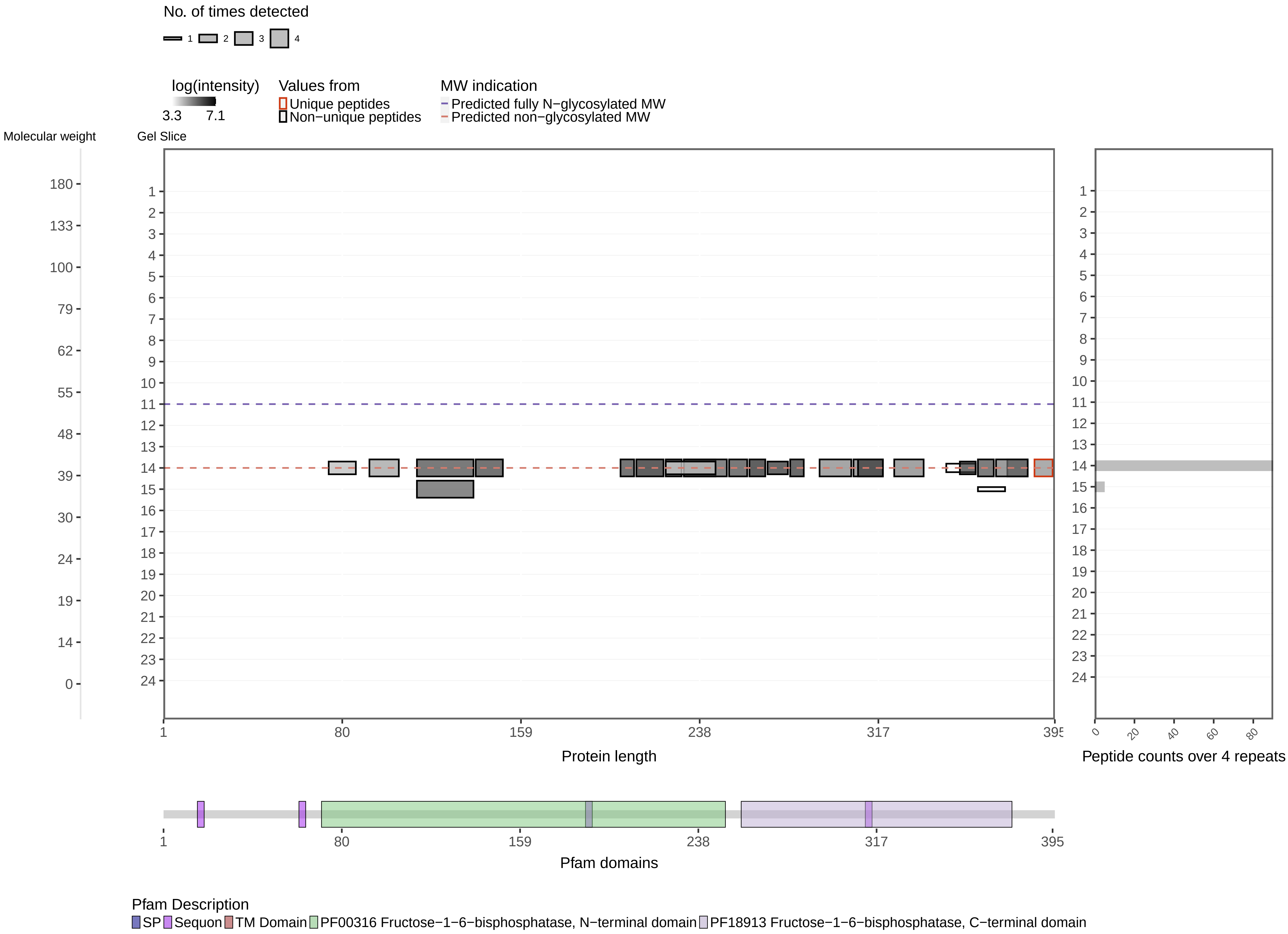


Protein sequence:

IVDTNFTVGTIFGVWPGDKLTGITGRDQVAAAMGIFGPRTTYVLALKDVPGTHEFLLLDEGKWQHVKDTTEIGEGKMFSPGNLRATFDNADYAKLIDYYVKEKYTLRYTGGMVPDVNQIIVKEKGIFTNVTSPTAKAKLRLLFEVAPLGFLIEKAGGYSSDGKQSVLDKVIGTLDERTQVAYGSKNEIIRFEETLYGSSRLKAAEPVGAAV*

NbL03g07090.1 GDSL_esterase/lipase_CPRD49−like [MW = 28.91 kDa]

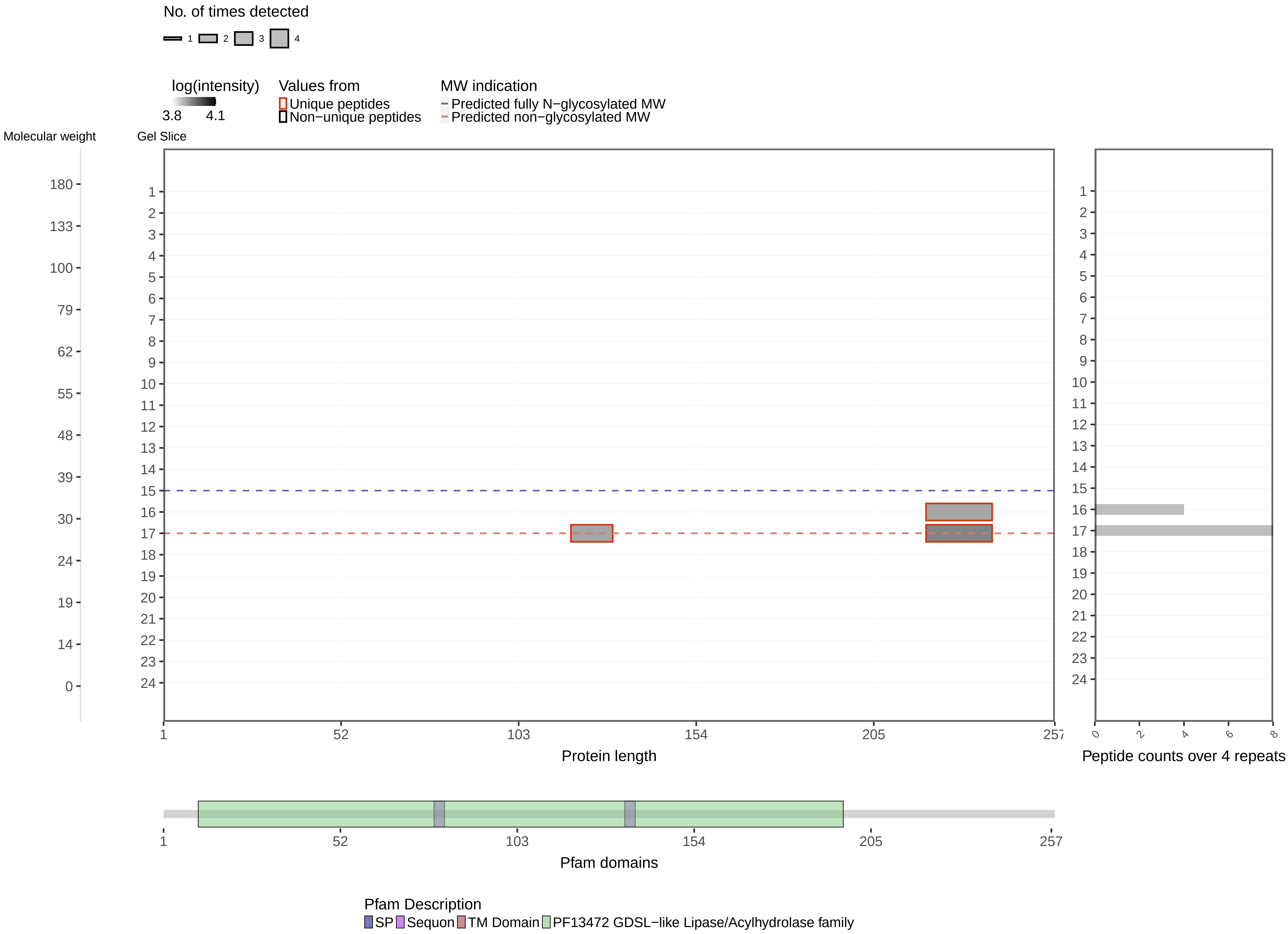


Protein sequence:

:NIRIIVLSCPPVDEARVRENTSSYFSELVRTNELCRQYSDACIELCNEMNLKVVDLWTALQKREDWLTACFTDGIHLAEAGSKIVVEEIIKVLKEAGWTPSLYWKSMPTEFAEDSPYDLVLADGKTTINPSDWTYHREIRWD*

### NbL03g06650.1 ferritin−1_chloroplastic [MW = 27.87 kDa]

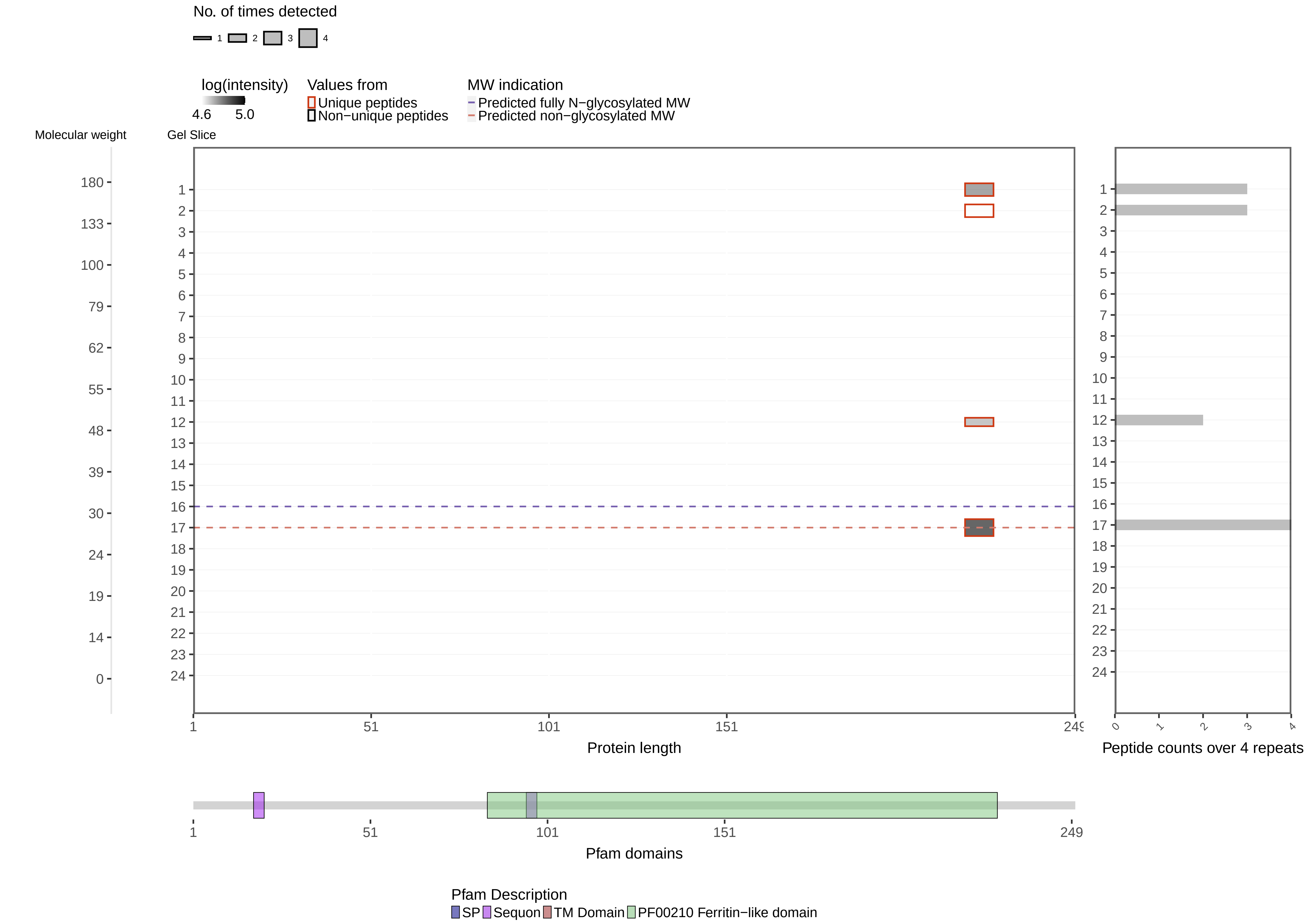


Protein sequence:

VALKGLAKFFKESSLEERDHAEKLMEFQNKRGGRVKLLSICAPPTEFDHCEKGDALYAMELALCLEKLTNQRLLNLHAVASRSNDVHLADFLESEFLVEQVDAIKKISEYVAQLRRVGQGHGVWQFDQMLLNEGAAA*

NbL03g06630.1 cytosolic_endo−beta−N−acetylglucosaminidase_1−like [MW = 81.99 kDa]

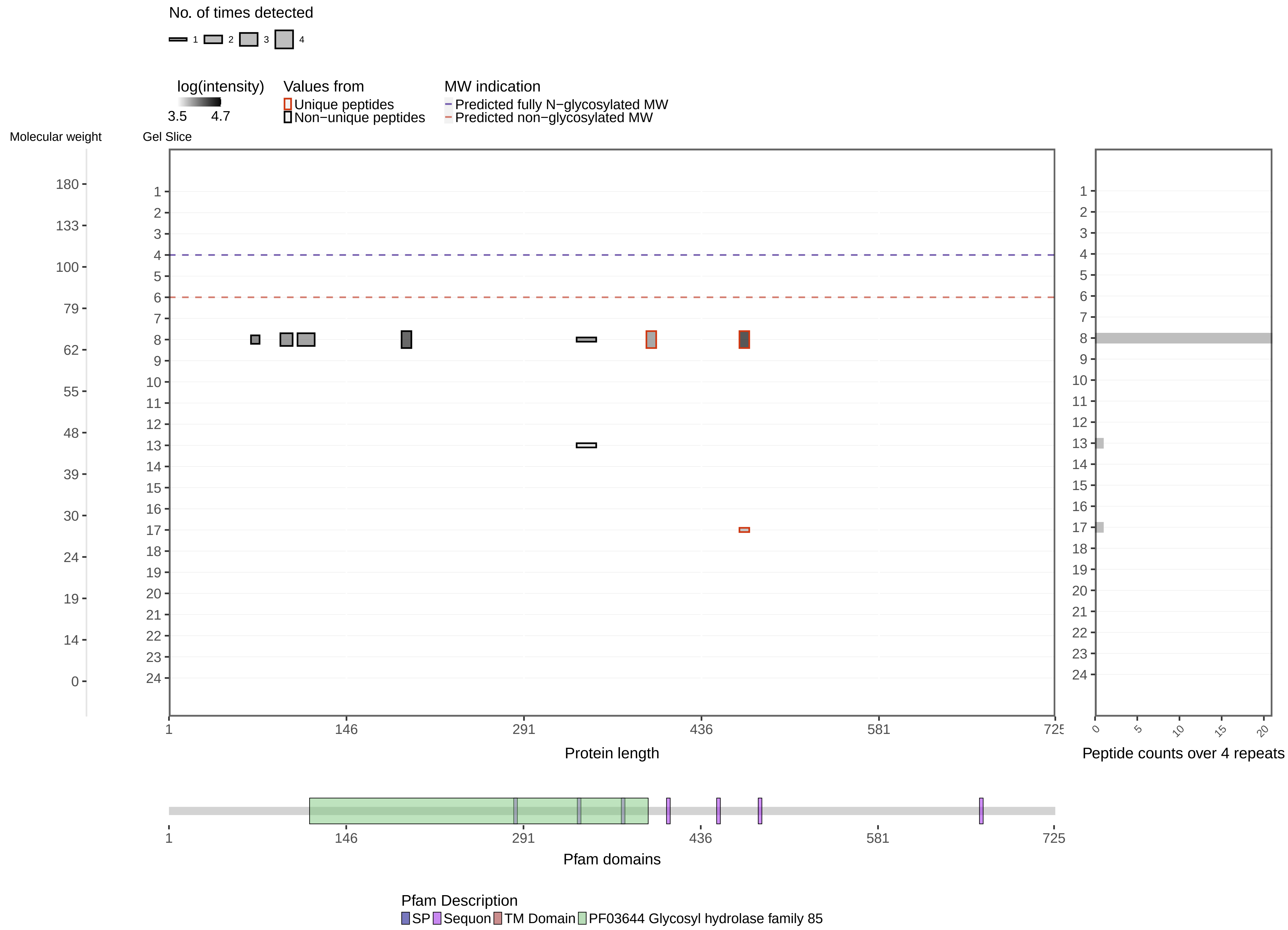


Protein sequence:

KQLPDFQTAQNRWWGLVERSWNISQNYPLTLPFYSNFDQGRGYHICVDGKQISQTPWNNISSQSFQPLLEFSGESTAGPLEVVVDLKEPSYNGGGNLAFNGTLDDDIHFSARLFEGKIQLADSPVRFTYSVKSNGSSLLGLSLEFNSAVRKQKSVLLASWGDSLLTMTRFVRQFDNVIMPRRVTKLETEPSWVIQESSVTMEGYMLTKIHAVCYKLRPEVHKSESQAKIMAFSPTEYHAVLGHLAIDSRTLNSDFPPSTSWLVDGNYIEWSFGSHGSRKLNVKLVWKLKD

NbL03g06530.1 biotin_carboxyl_carrier_protein_of_acetyl−CoA_carboxylase [MW = 30.03 kDa]

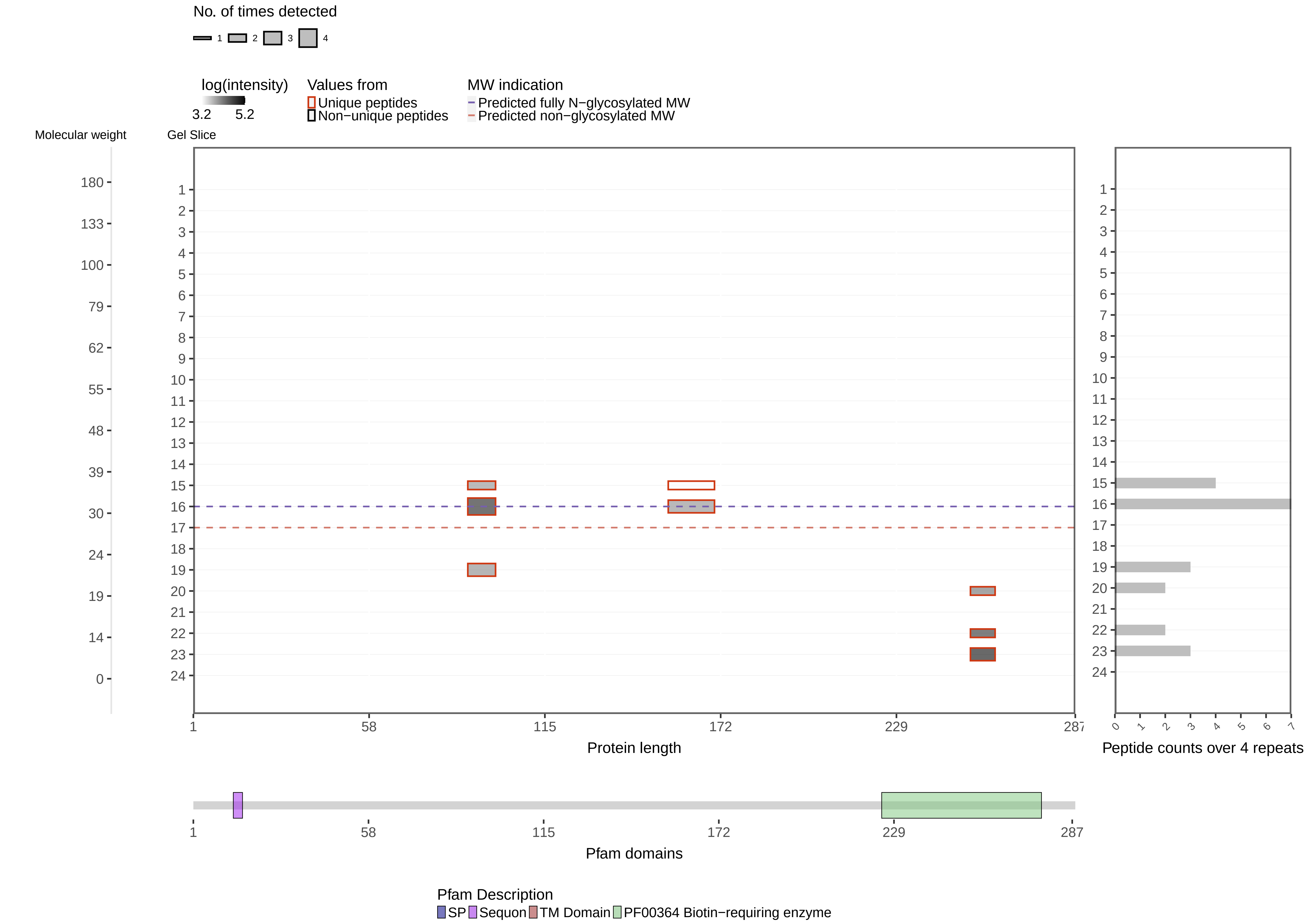


Protein sequence:

RDIEAPMVPAPVVSSPPPPPPSASKPSTASTAASPATSPGKSSSEKISPFTNVAAEKSAKLAALETSGASGYVLVSCPTVGSFRRARTLKGKKQPPACKEGDVIKEGQIIGFLDQFGTELPVRSDAAGELLKILFNDGEAVGYGDPLIAVLPSFRGI*

### NbL03g06460.1 alpha−soluble_NSF_attachment_protein [MW = 32.68 kDa]

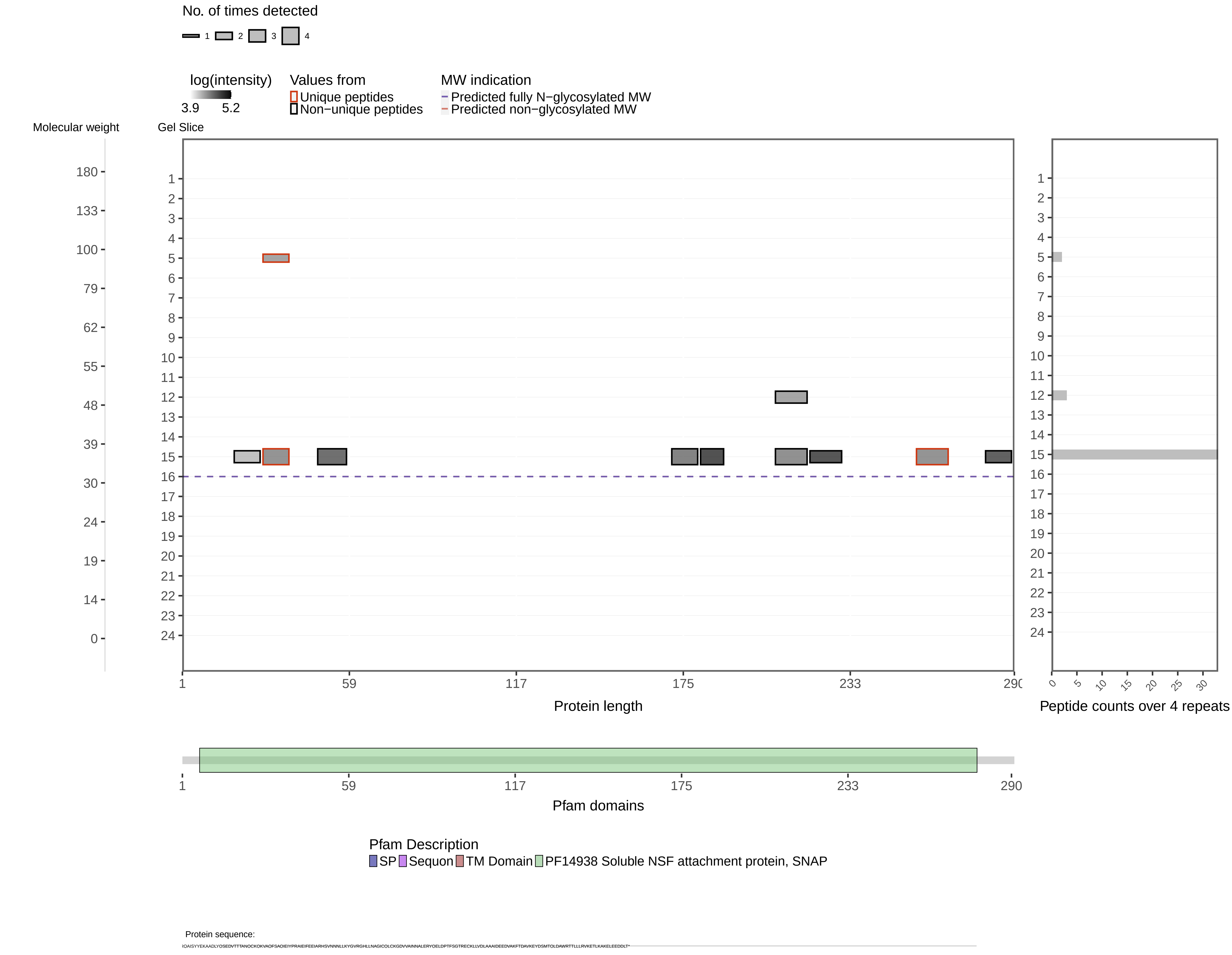

NbL03g06040.1 zinc_finger_CCCH_domain−containing_protein_14−like [MW = 30.88 kDa]

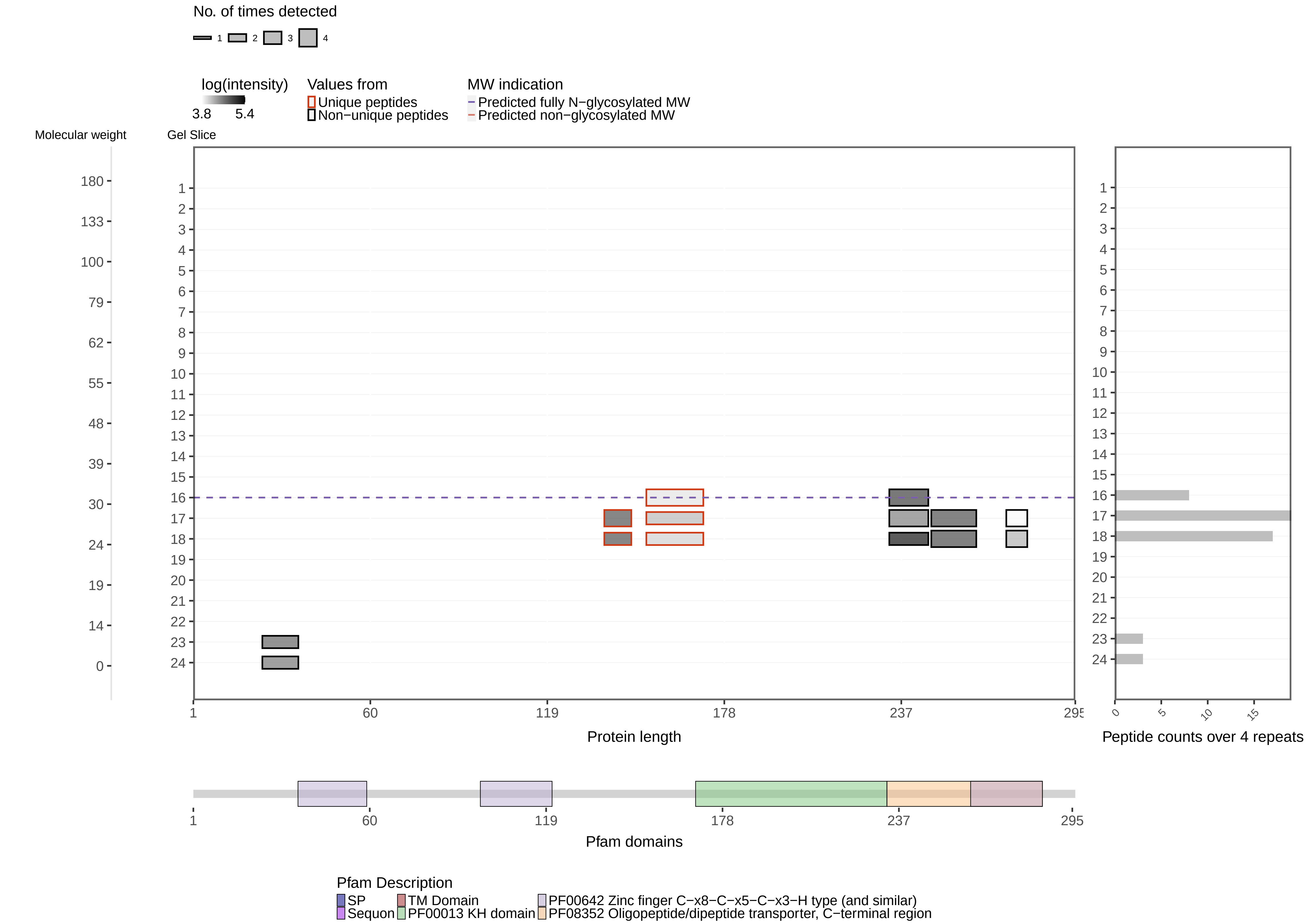


Protein sequence:

OEDPRAMGAGPIPGRFGGRMEPPPISGPAASFGTSATAKISVDASLAGAIIGKGGVNSKOICROTGAKLAIREHETDKNLRNIELEGTFEOISOASAMVRELISSLGSVGGPGRAPAVOGGPPPOMSNYKTKLCENFAKGSCTFGERCHFAHGAAELRKTGV*

### NbL03g06030.1 enoyl−[acyl−carrier−protein]_reductase_[NADH]_2_chloroplastic−like [MW = 41.35 kDa]

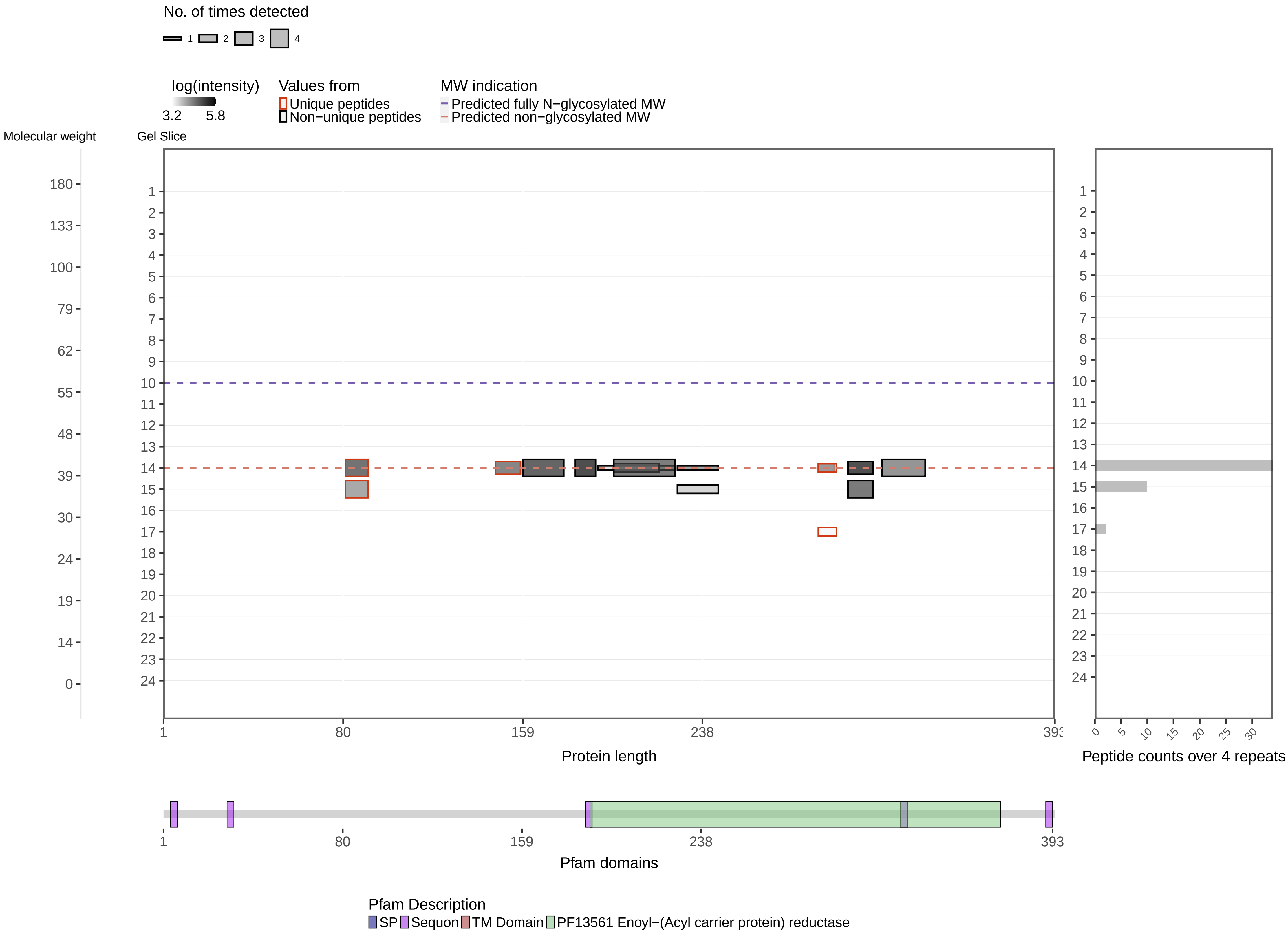


Protein sequence:

AGSSNWTVKEAVESVKQDFGTIDILVHSLANGPEVSKPLLETSRKGYLAAVSASSYSFVSLLRHFLPIINPGGASISLTYIASERIIPGYGGGMSSAKAALESDTKVLVFEAGRKNKVRVNTISAGPLGSRAAKAIGFIDMMINYSLENAPLQKELYAEEVGNAAAFLVSPLASAITGATIYVDNGLNAMGVGVDSPAFEGLDIPKDNKS*

### NbL03g06000.1 nudix_hydrolase_23_chloroplastic−like [MW = 31.85 kDa]

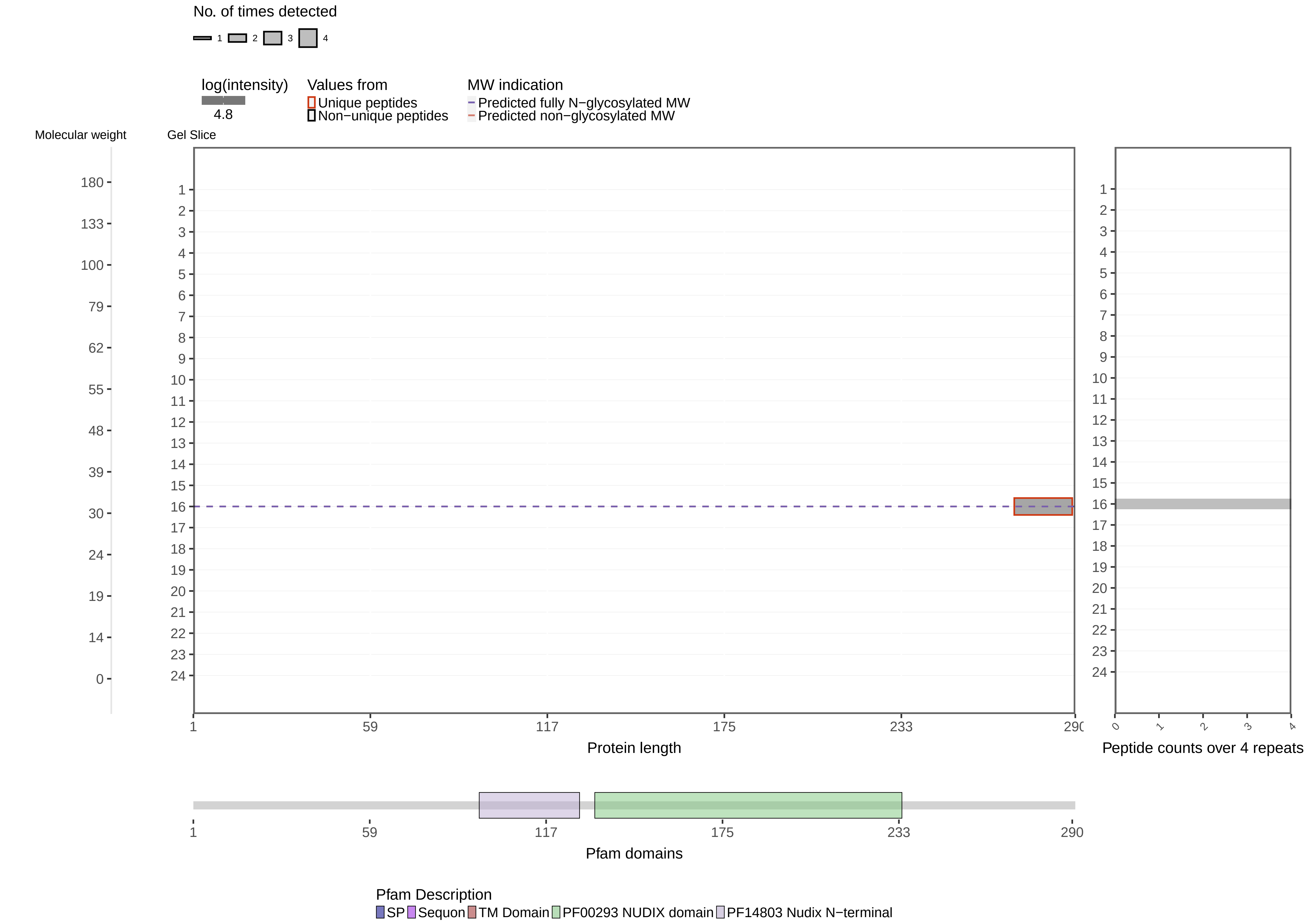


Protein sequence:

KMVVGCLIEHDKKILMCRRKIQPSSGLWTLPAGYMEIGESAAEGAIRETLEEANAEVEVESPFAQLDIPLIGQTYMIFLAKLKTPHFSPGPESSECRLYELDDIPFDSLAFSSMLVTLKLYIEDIKVGRPKFHYGIINKRPGTSPSEIHAYTLEYHMQS*

### NbL03g05910.1 probable_ribosomal_protein_S11_mitochondrial [MW = 26.13 kDa]

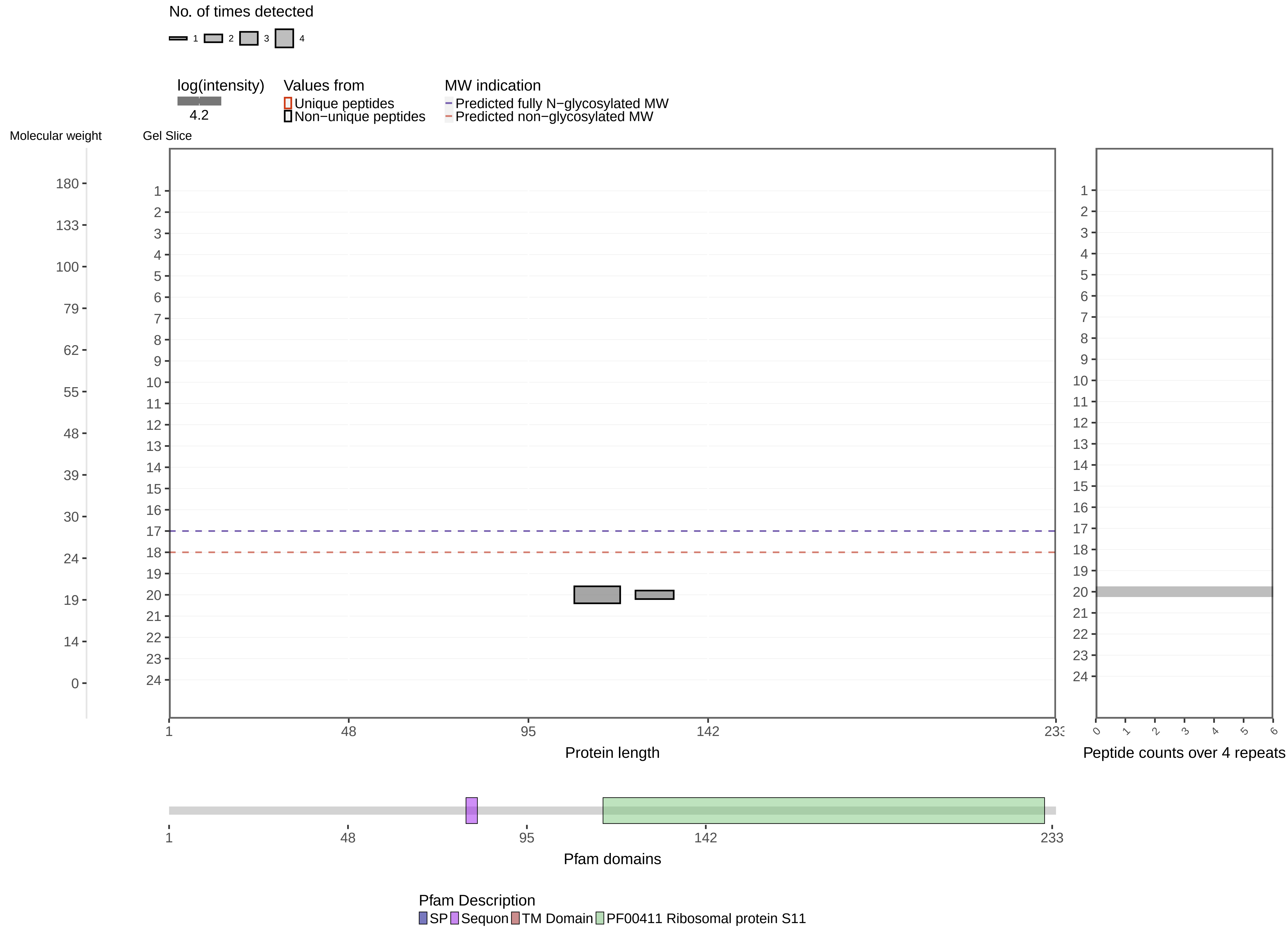


Protein sequence:

APLSRYHVEQDADIVHIKVLRNNAFVTVTDSKGNKKFGATAGKITGKGTKIARYAAESTAEHVGREARDRGLRSVVMKVNGFTFFKKKKQAILSFREGYTHSRGDKNPVVFIEDTTRRPHNGCRLPKKRRI*

NbL03g05820.1 uncharacterized_LOC107781405 [MW = 31.43 kDa]

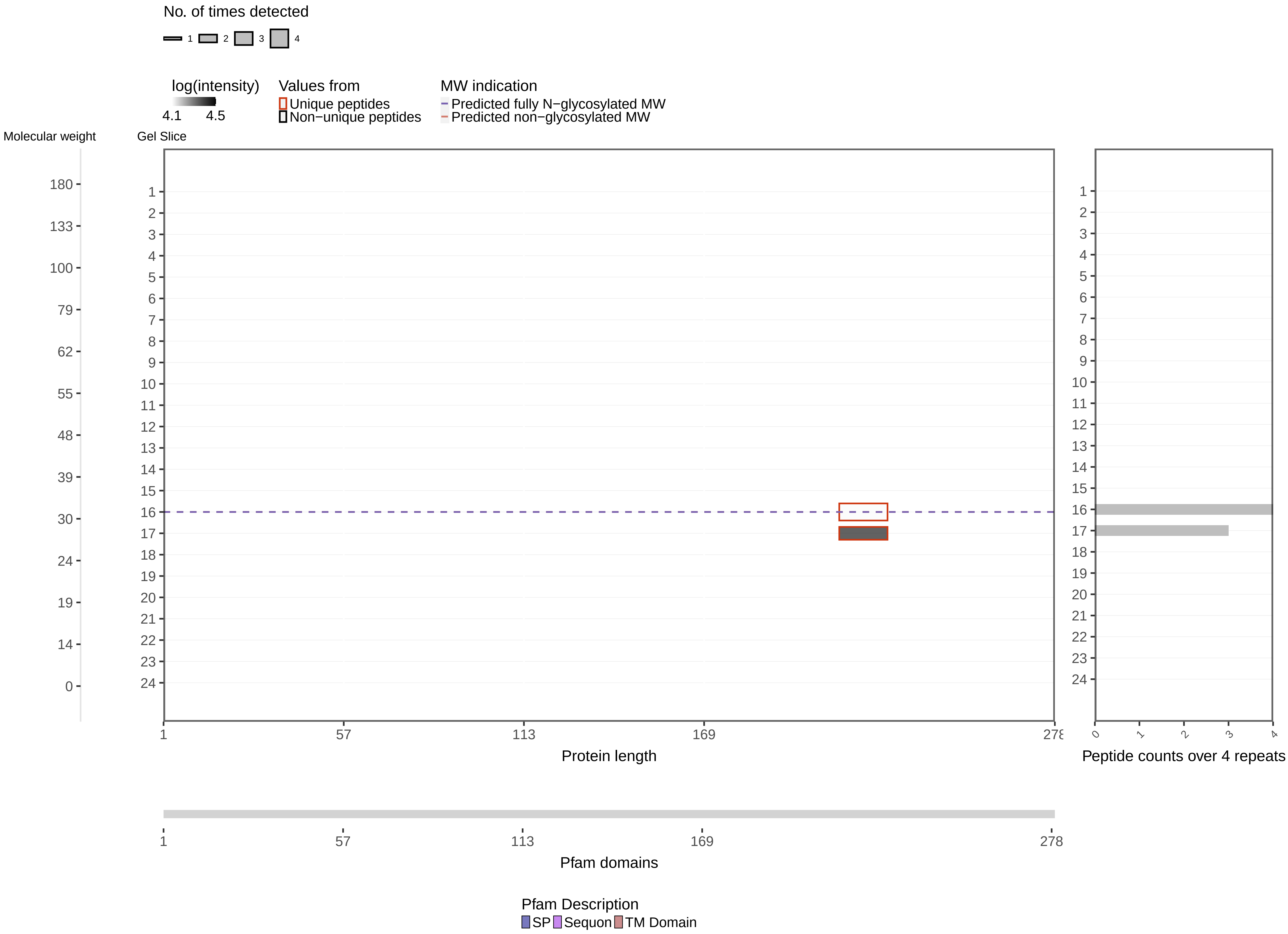


Protein sequence:

EEFVDRVVDSSNCROLLVVHDSLFMLACIKEKYDKVKCWOGELIYVPEKWTSFDVIFLYFLPALAFELGOIMEVLRKRCLPGARVVISHPOGSOVVEEORKLYPDVVVSNLPEKMILONVAAEHSFEVVKFTDEPGFYLAILKFEPENKND*

### NbL03g05600.1 uncharacterized_LOC107780098 [MW = 38.03 kDa]

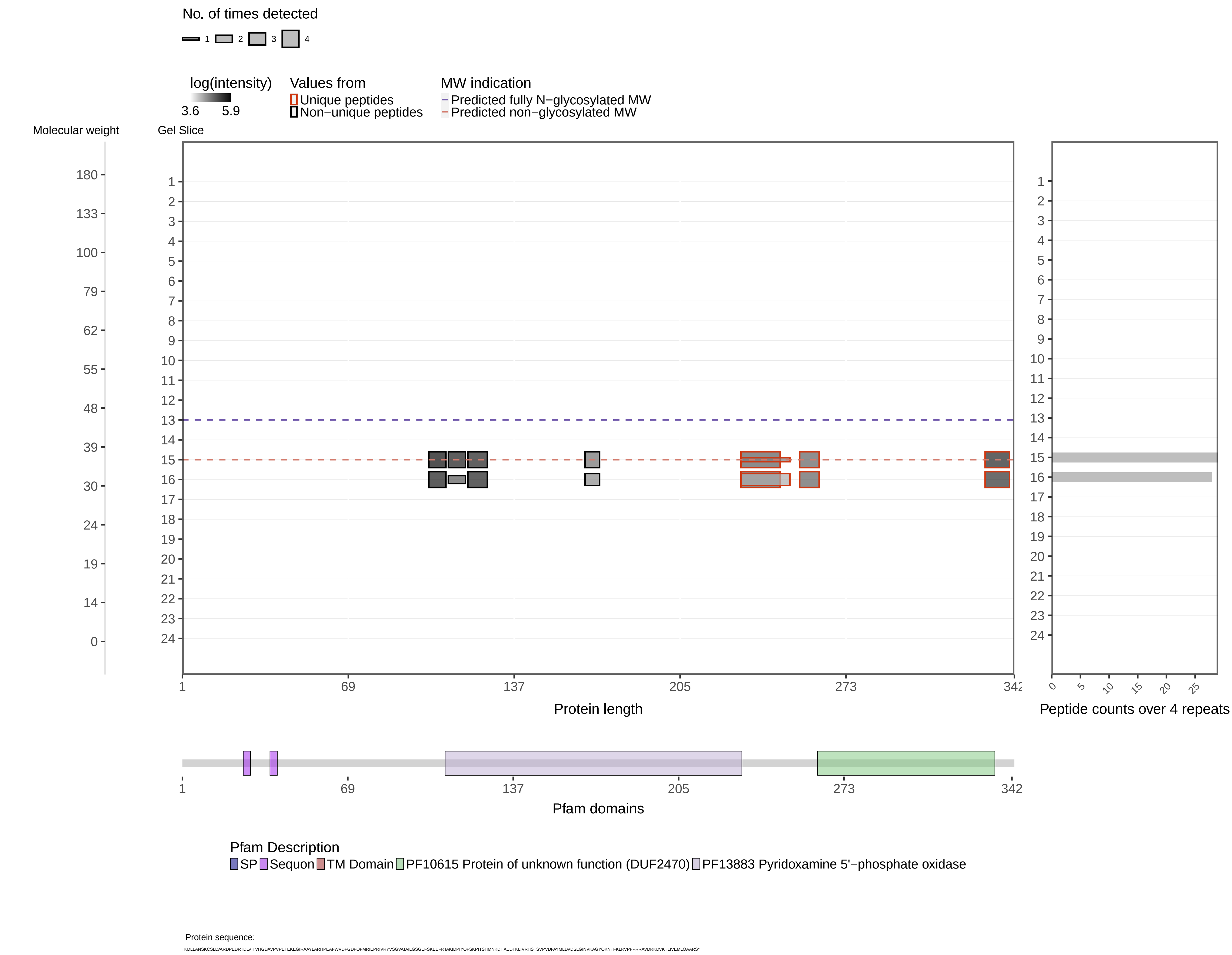

NbL03g05540.1 50S_ribosomal_protein_L18−like [MW = 26.06 kDa]

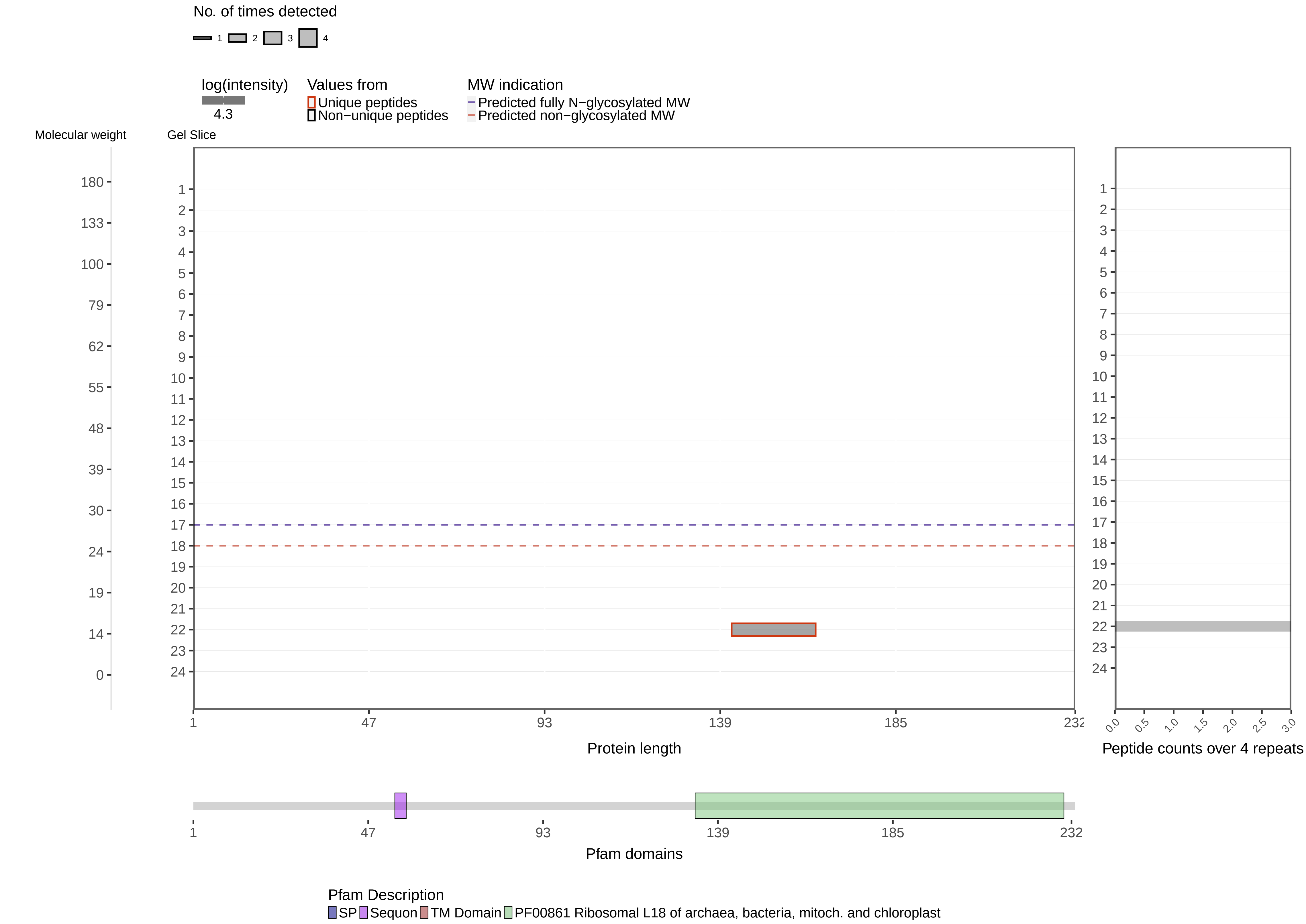


Protein sequence:

VKIASFNVEISRMVIPPPIRPDRVTKYLKPYVLRMHFTNKYVNAQVVHTPTATVAASASTQEKGLRLAMVEGKENTRDVAAAAKIGKLLGERLQAKGIPAISVFFKREQRYHGKVKAVIDSVKEVGIELV*

NbL03g05470.1 uncharacterized_LOC107828093 [MW = 72.43 kDa]

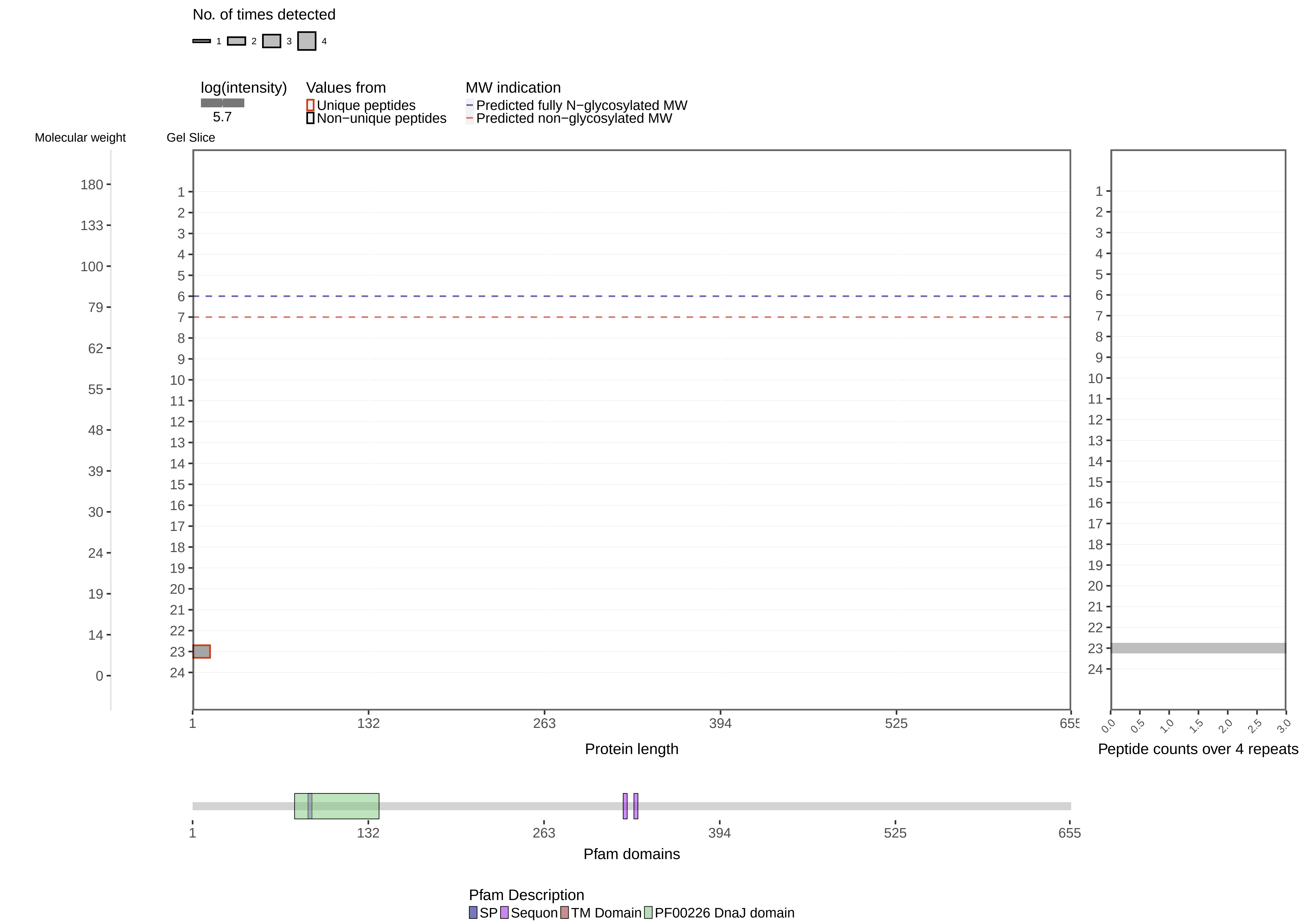


Protein sequence:

[illegible]

### NbL03g05330.1 adenylosuccinate_synthetase_chloroplastic−like [MW = 54.86 kDa]

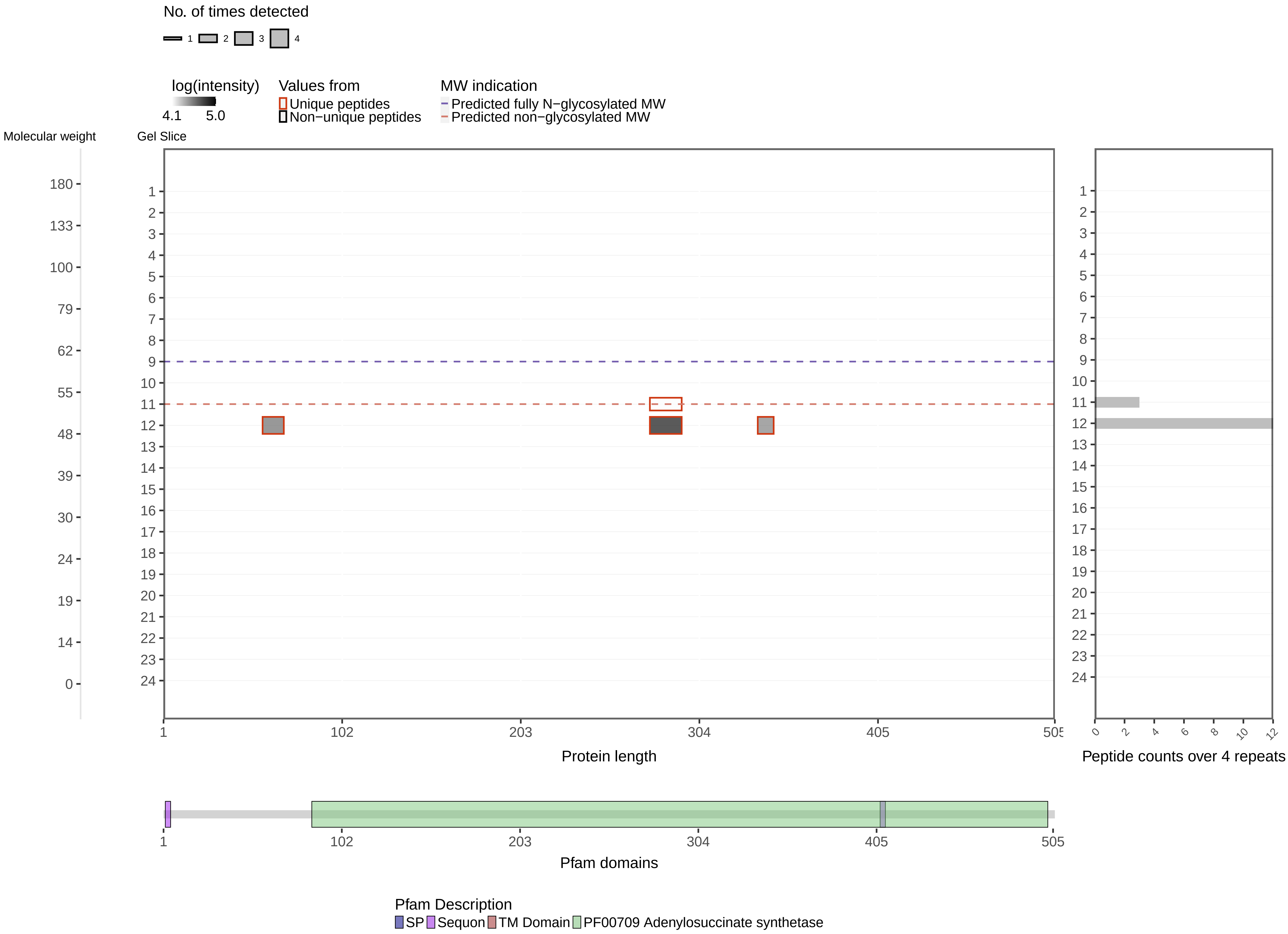


Protein sequence:

PQKLDLLLSDAASRFKGFNYGPDMLREEVERYKKFAERLEPFITNTVHFMNDAISQKKKILVEGGQATMLDIDFGTYPFVTSSSPSAGGICTGLGIAPRMVGDLIGVVKAYTTRVGSGPFPTEILGKGGDLLRFAGQEFGTTTGRPRRCGWLDIVALRFCCQINGFASLNLTKLDVLSDLSEIELGVSYRQPDGTPVESFPSDLRLLDQIKVEYEVLPGWQTDISSIRKYSELPKAARDYIERIEELVGVPIHYIGVGPGRDALIYK*

### NbL03g05170.1 aspartate_carbamoyltransferase_1_chloroplastic [MW = 45.34 kDa]

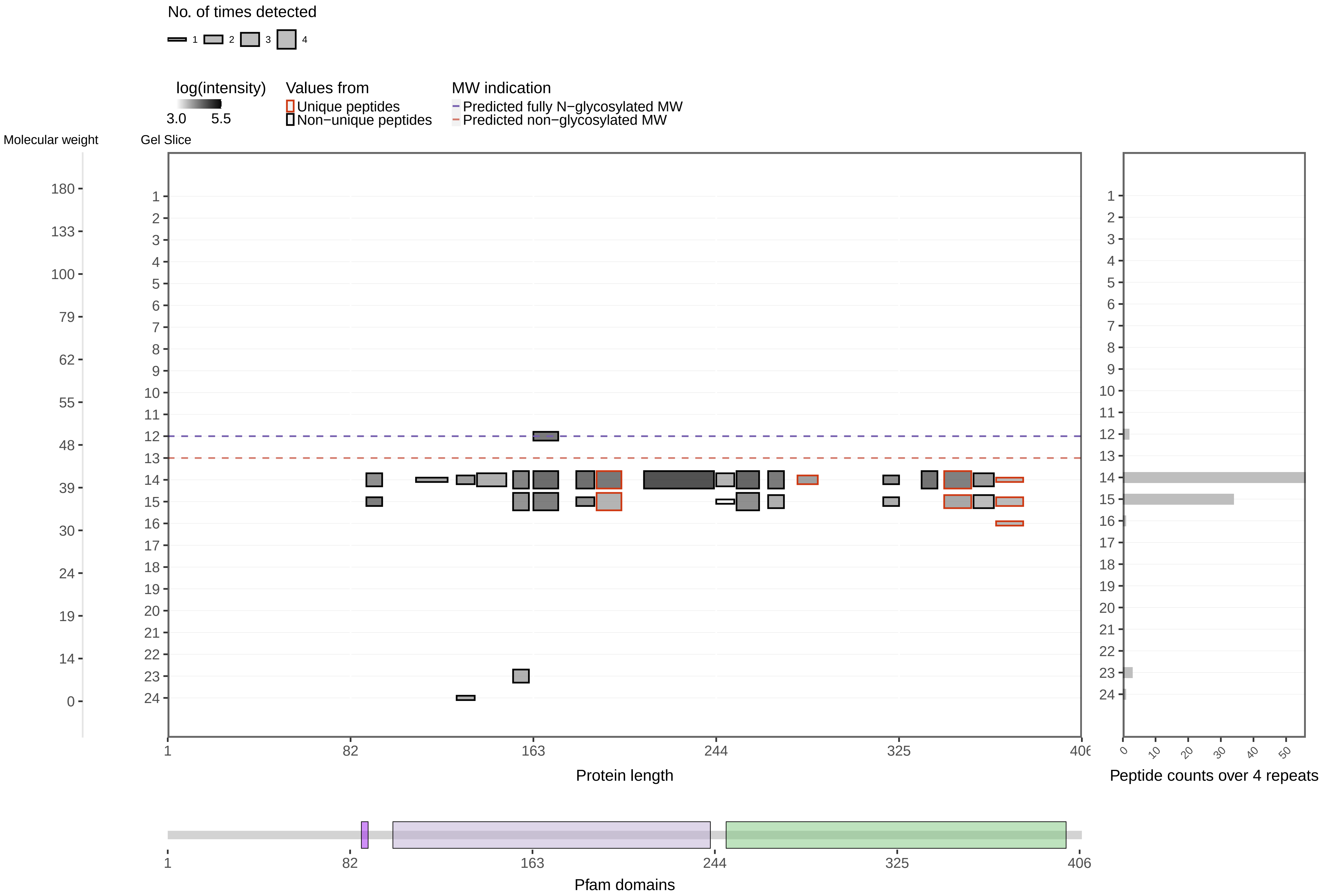


Pfam Description

SP Sequon TM Domain PF00185 Aspartate/ornithine carbamoyltransferase, Asp/Orn binding domain PF02729 Aspartate/ornithine carbamoyltransferase, carbamoyl−P binding domain

Protein sequence:

SIRTVEGYSDIIVMRHFESGAAKRAAATASIPIINAGDGPGQHPTQALLDVYTIEREIGKLDGIKVALVGDLAYGRTVRSLAYLLAKYQDVKIYFVSPDVVKMKDDIKDYLTSMGVQWEESADLVDVASKCDVVYQTRIQRERFGERVDLYEEARGKYIVDLGVLNAMQKHAVVMHPLPRLDEITIDVDEDPRAAYFRQAKNGLYIRMALLKLLLLGW*

NbL03g05150.1 eukaryotic_translation_initiation_factor_5B−like [MW = 152.47 kDa]

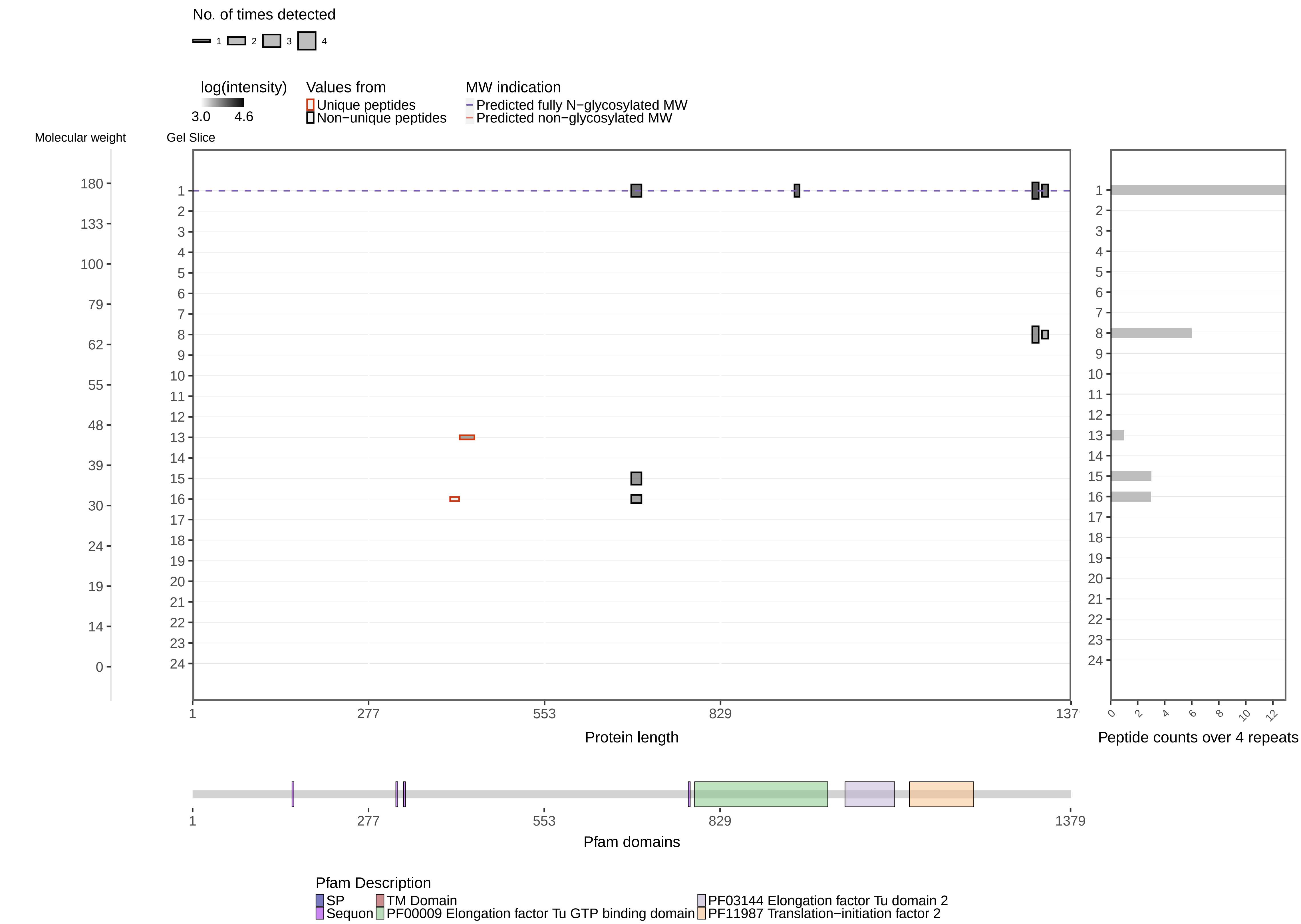


Protein sequence:

EEEDDDEEWDAKSWDDADLKLPGKSAFEDEEVDSEPQPITKKEIKTASLAAHDAATLPVAAKLVIPTQKAAATVAGLSKNDQSRKGEPEDRVADQNKQKGSPEDRVTVKHGVPPNQSEDNLRSPICCIMGHVDTGKTKLLDCIRGTNVQEGEAGGITQQIGATYFPAENIRERTKELKADATLKVPGLLVIDTPGHESFTNLRSRGSGLCDIAILVVDIMHGLEPQTIESLNLLKMRNTEFIVALNKVDRLYGWKVCKNAPIVKAMKQQSKDVQFEFNNRLTQVVTQFKEQGINT

### NbL03g05060.1 stearoyl−[acyl−carrier−protein]_9−desaturase_chloroplastic [MW = 44.88 kDa]

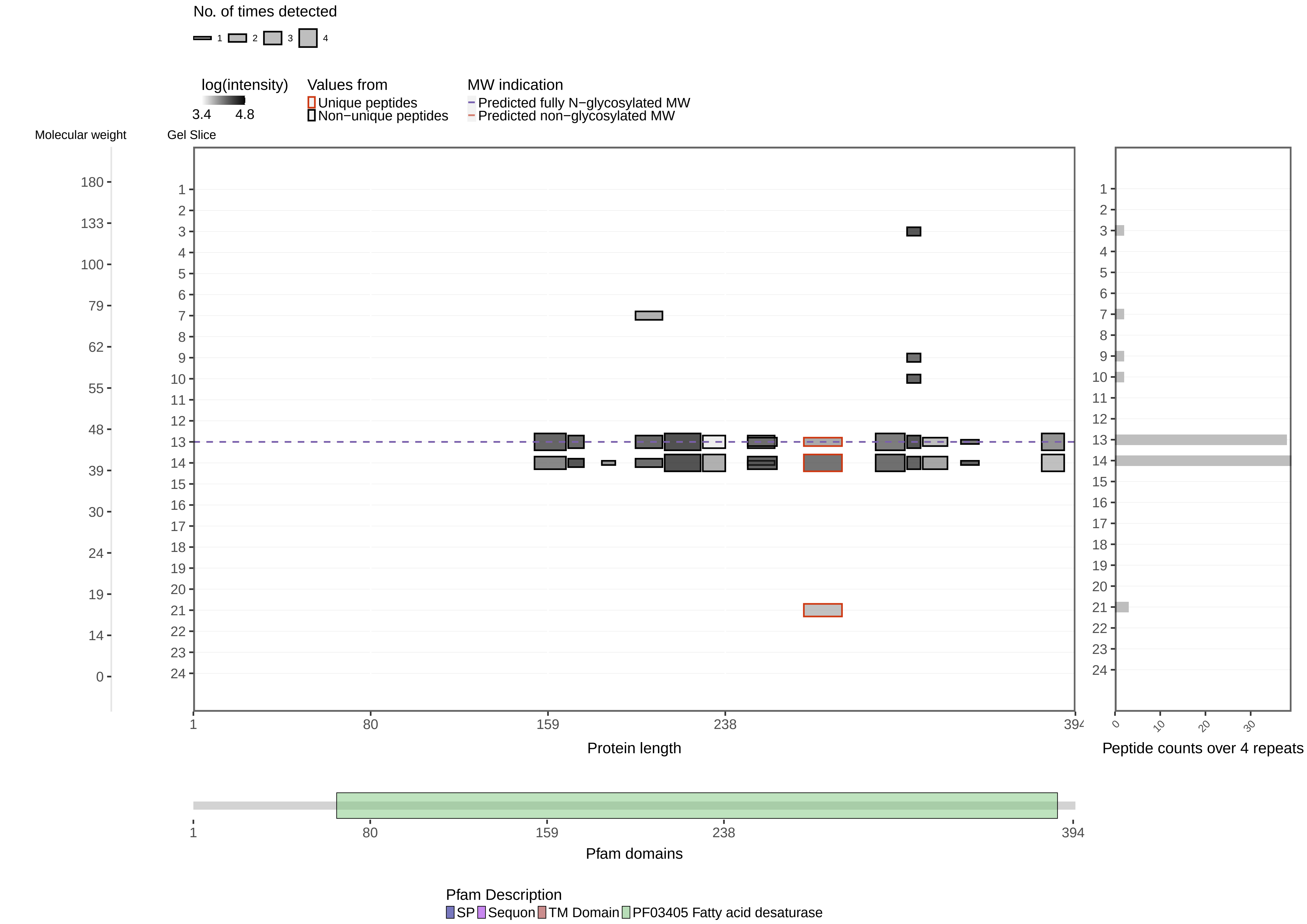


Protein sequence:

[illegible]YLYLSGRVDMRQIEKTIQYLIGSGMDPRTENNPYLGFIYTSFQERATFISHGNTARLAKEHGDLKLAQVCGIIAADEKRHETAYTKIVEKLFEVDPDGTVLAVADMMRKKISMPAHLMYDGRDDNLFEHFSAVAQRLGVYTAKDYADILEFLVGRWEMEKLTGLSGEGRKAQDYVCGLAPRIRKLEERAQARAKEKAPVPFSWVFGREIKL*

NbL03g04950.1 probable_LRR_receptor−like_serine/threonine−protein_kinase_At1g53430 [MW − SP = 109 kDa]

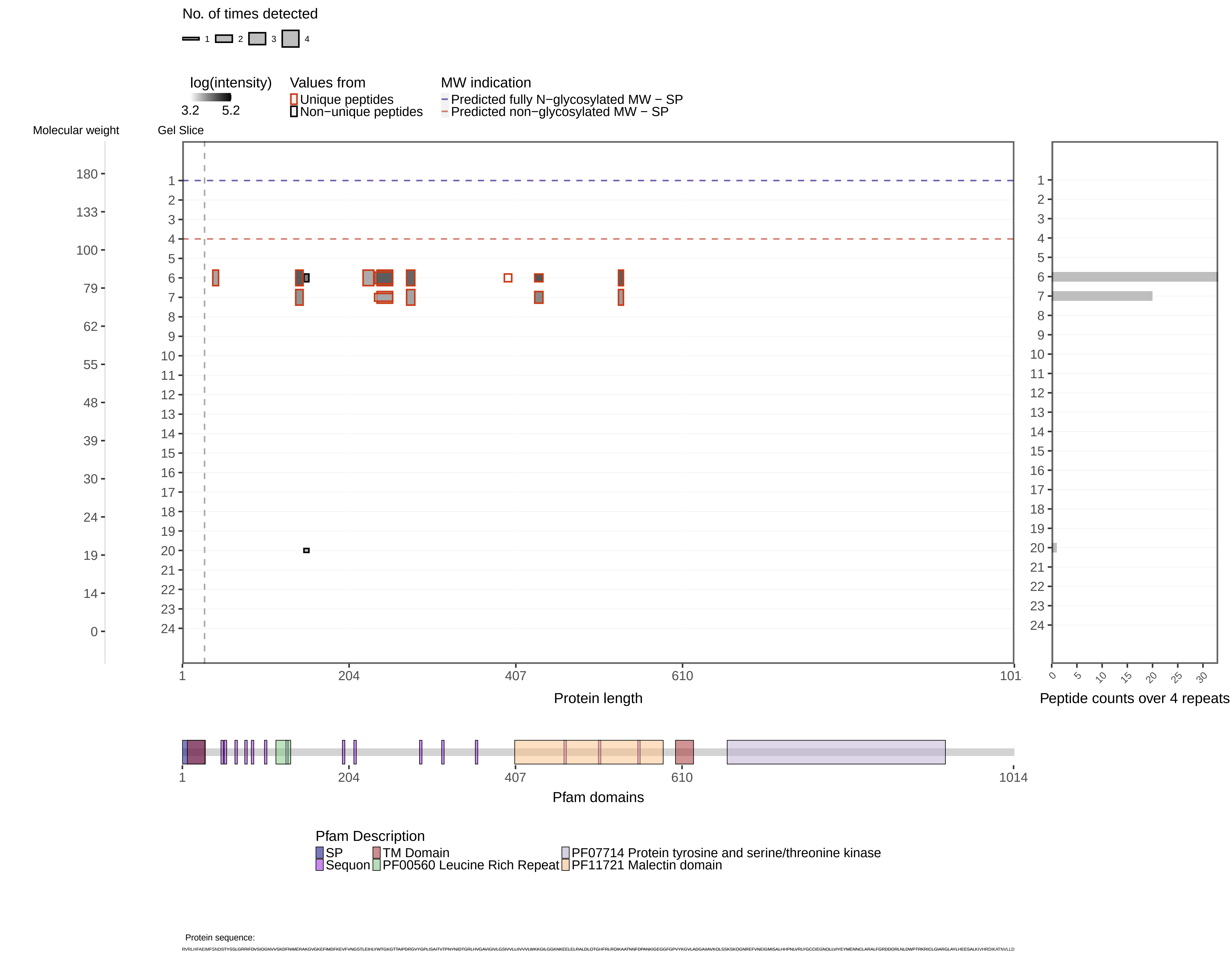

### NbL03g04820.1 peptidyl−prolyl_cis−trans_isomerase_CYP37_chloroplastic [MW = 50.71 kDa]

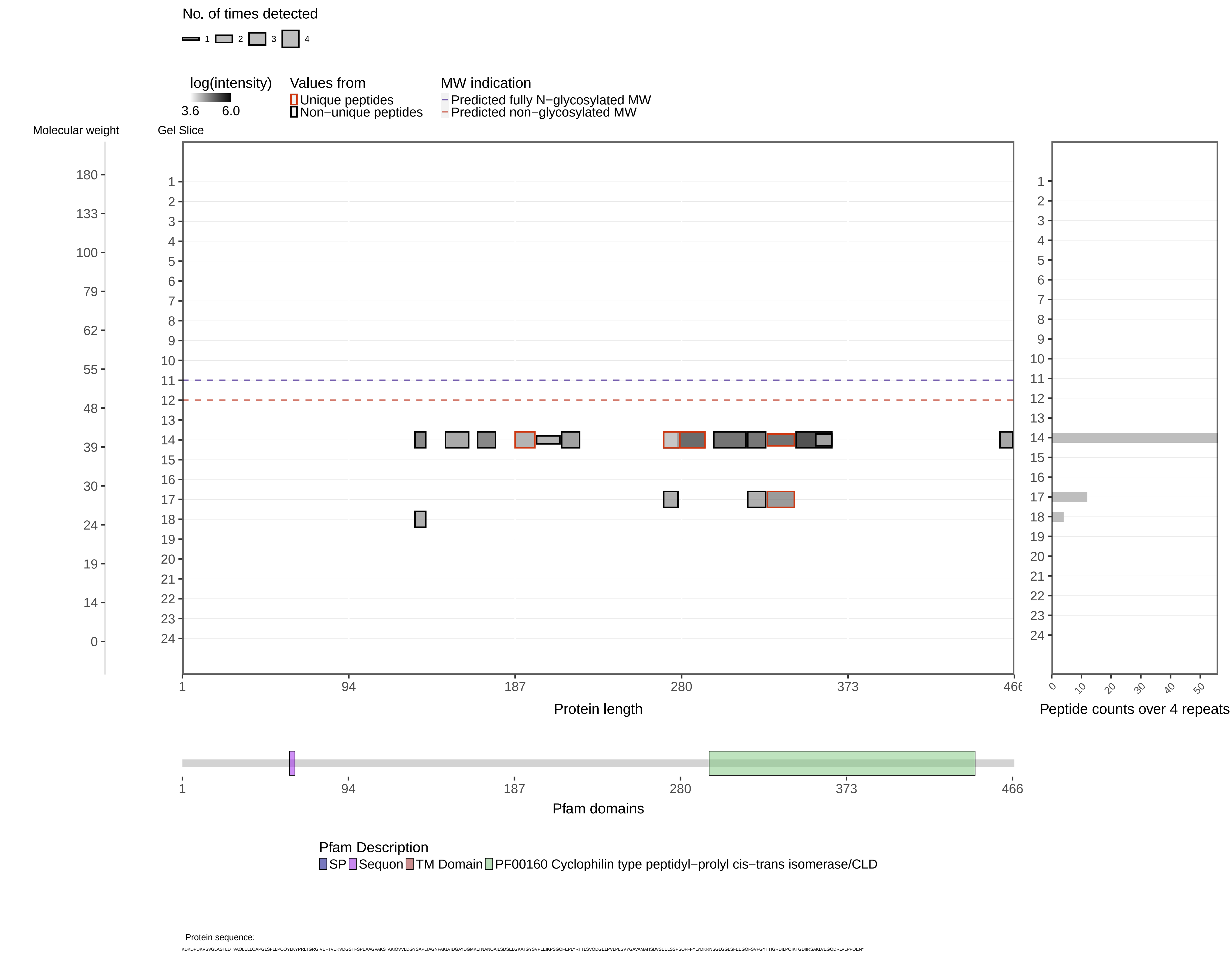

### NbL03g04700.1 thioredoxin−1−like [MW = 19.73 kDa]

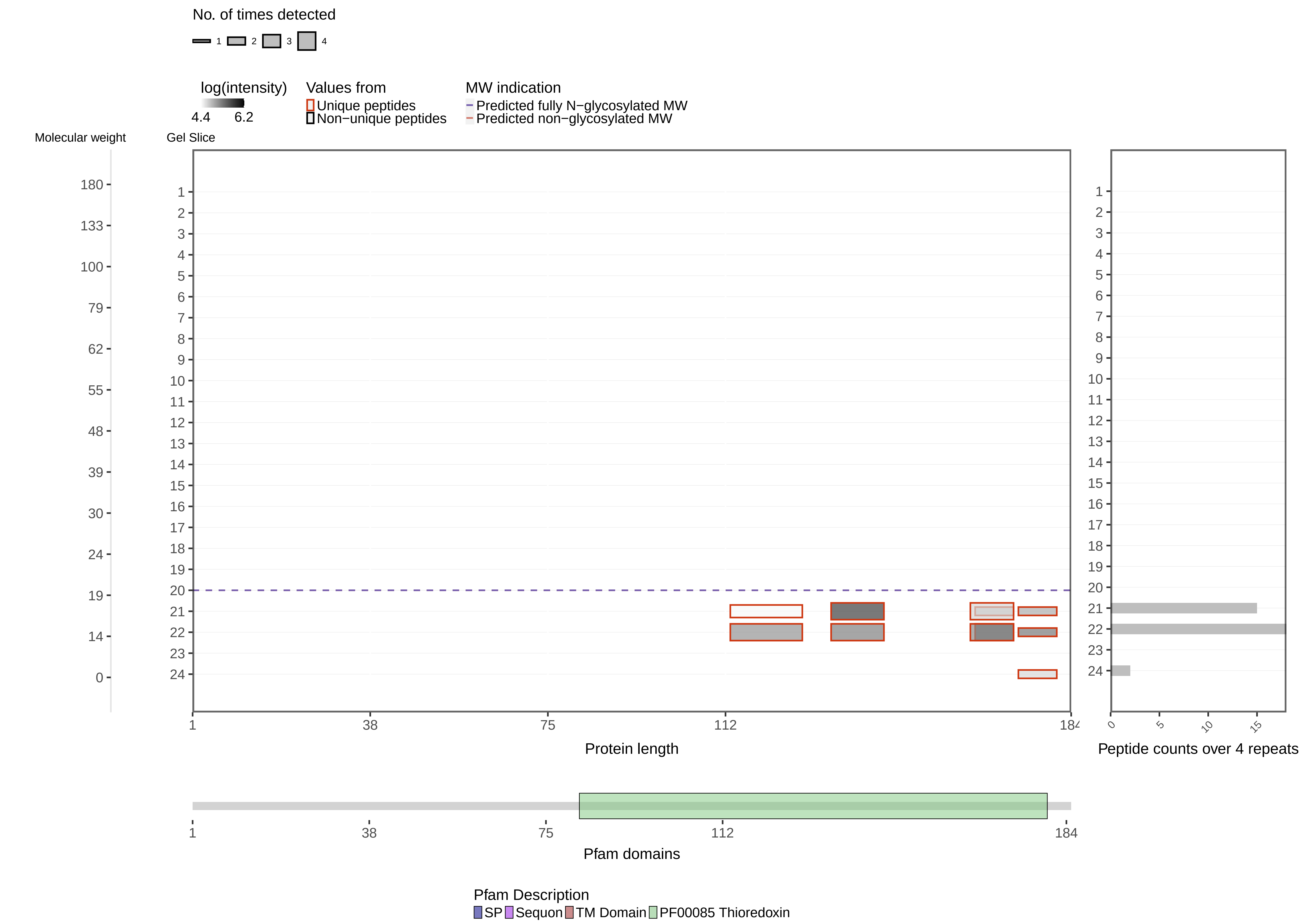


Protein sequence:

VLVPPVNDQTWESLVIGSDVPVLVEFWAPWCGPCRMIHPTVDELAQEFAGKFKFYKVNTDESPAIATRYGIRSIPTVMIFKDGEKKDTVIGAVPKTTLTASIEKFL*

### NbL03g04620.1 early_nodulin−like_protein_2 [MW − SP = 40.58 kDa]

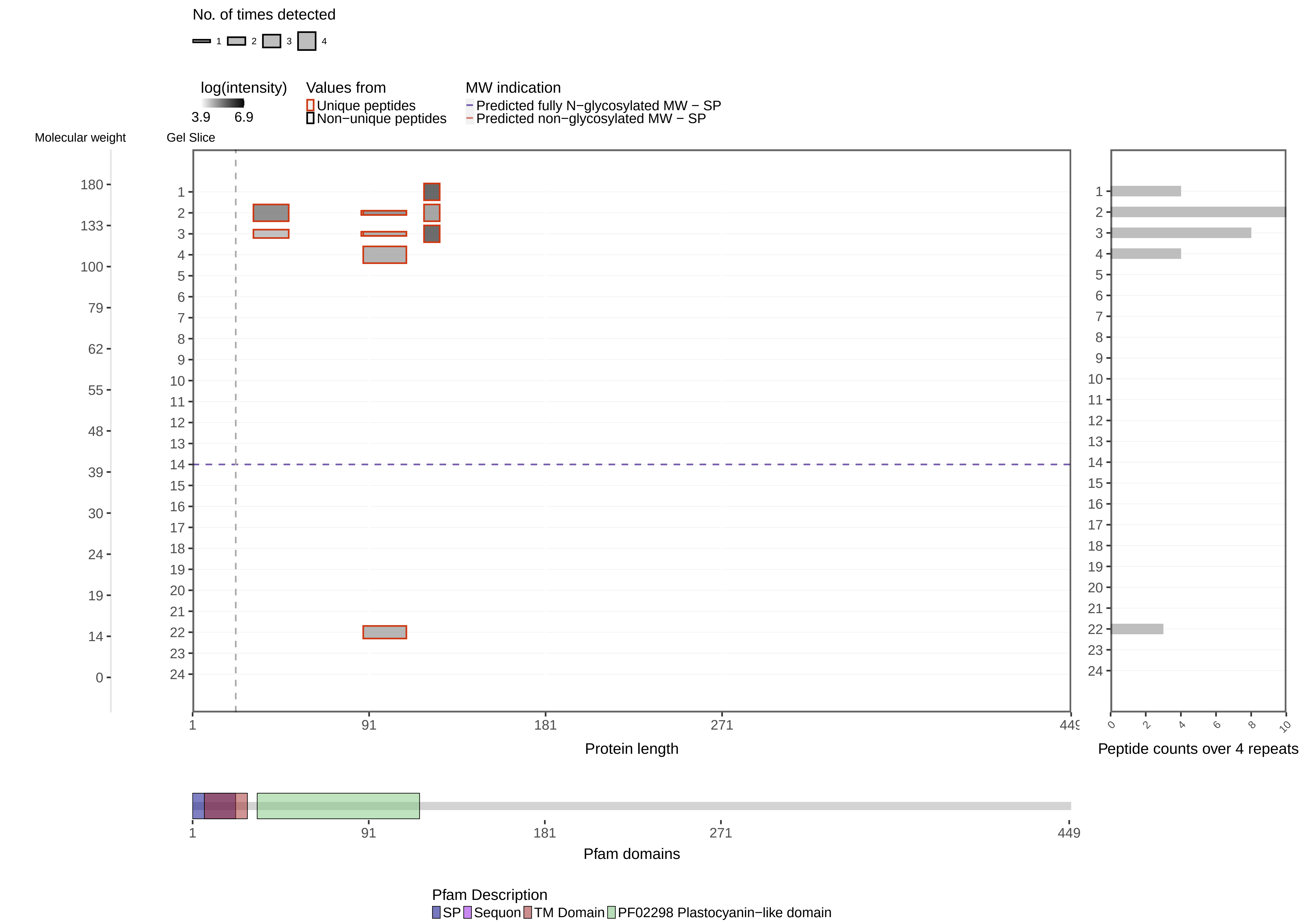


Protein sequence:

TPSGGSATPPTPSGGSATPPSPSGGSATPPNPSGGSTTPPAAGSPKGSSTPGTPSPAGVPSSPSAGSTTPPPTTSAPSATTPAGAPSKGSTTPDTSSPNGAPVVSPPAAGKSPTSSPTPAGSIVSPATSSMPPTGSPAPAGGASDLSPSPSSSSLAPGSSALSPGSIAPAGGPTSSMAPGGMAPGAGAGSTPADINSPAGAPSNPNSFAVKAFTPSVVLLSTVSLILTMAFGEFIISP*

### NbL03g04490.1 homogentisate_12−dioxygenase [MW = 51.69 kDa]

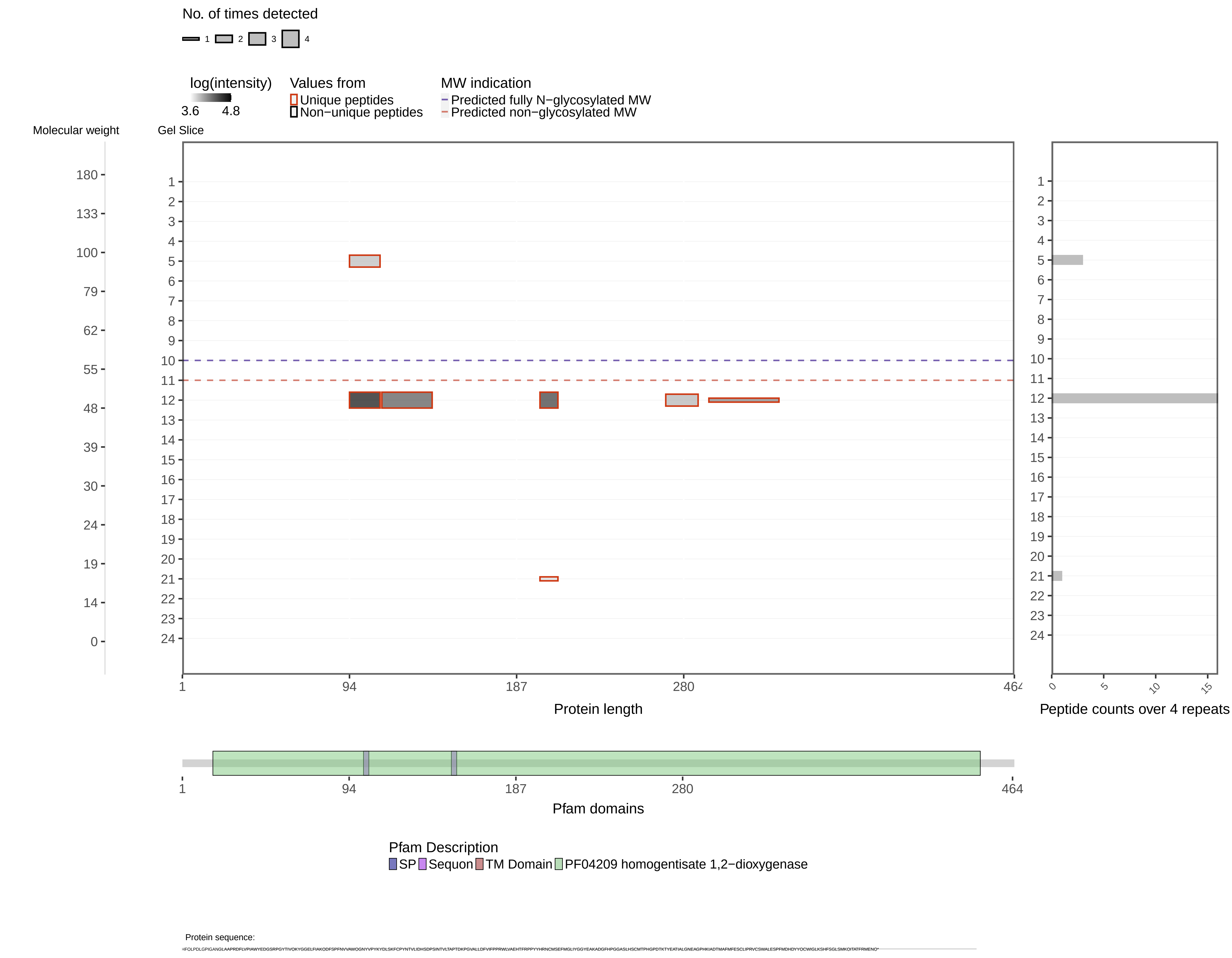

NbL03g04400.1 malate_dehydrogenase_2_mitochondrial [MW = 35.59 kDa]

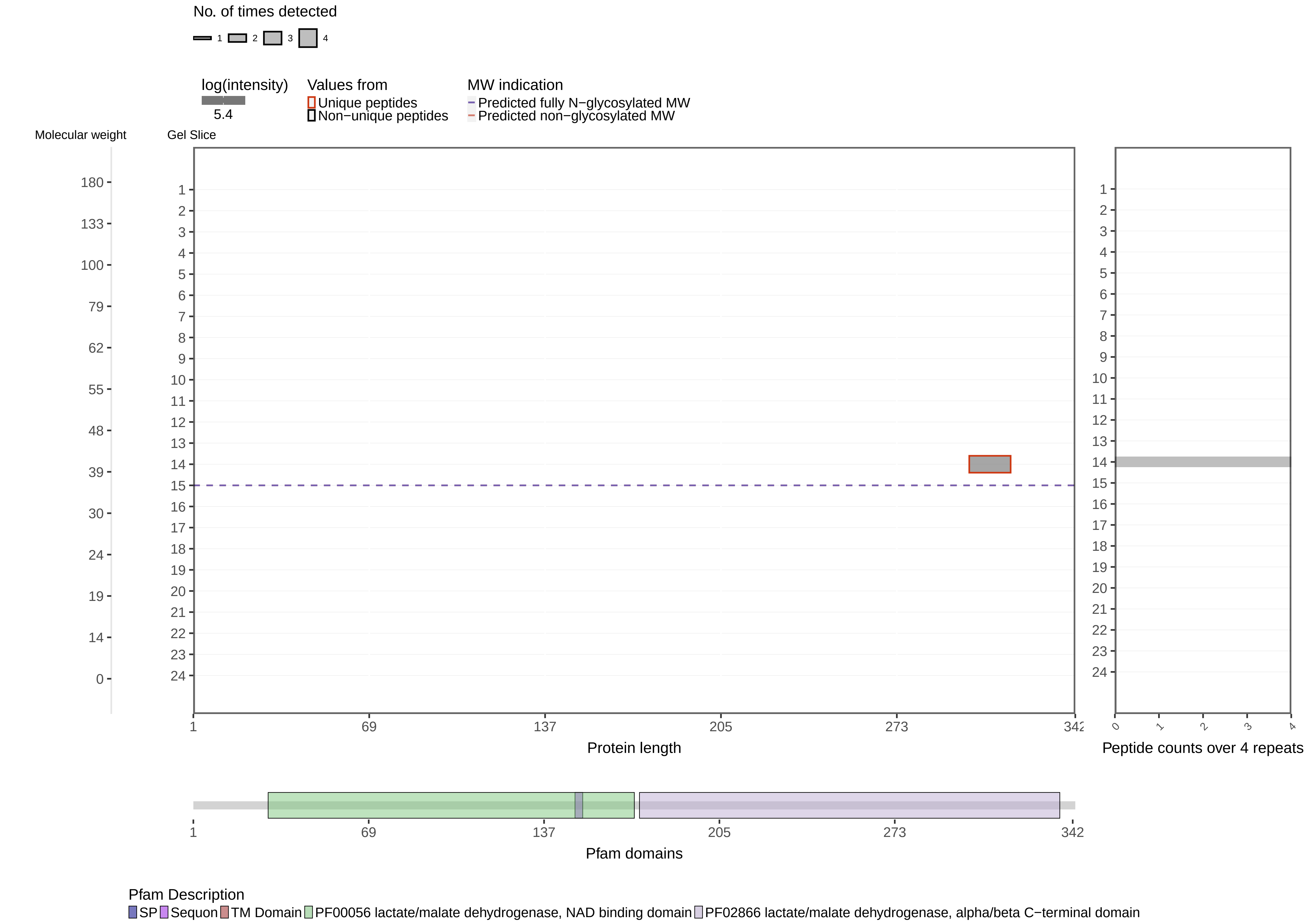


Protein sequence:

FKKAGTYDEKRLFGVTTLDVVRAKTFYAGKAKVNVADVIVPVVGGHAGITILPLFSQATPKANLGDEEIEALTKRTQDGGTEVVEAKAGKGSATLSMAYAGAIFADACLKGLNGVPDVVECSFVQSTVTDLPFFASKVRLGKNGVEEVLGLGPLSVFEKQGLEALKPELKSSIEKGIKFANDN*

### NbL03g04350.1 multiple_organellar_RNA_editing_factor_8_chloroplastic/mitochondrial [MW = 48.21 kDa]

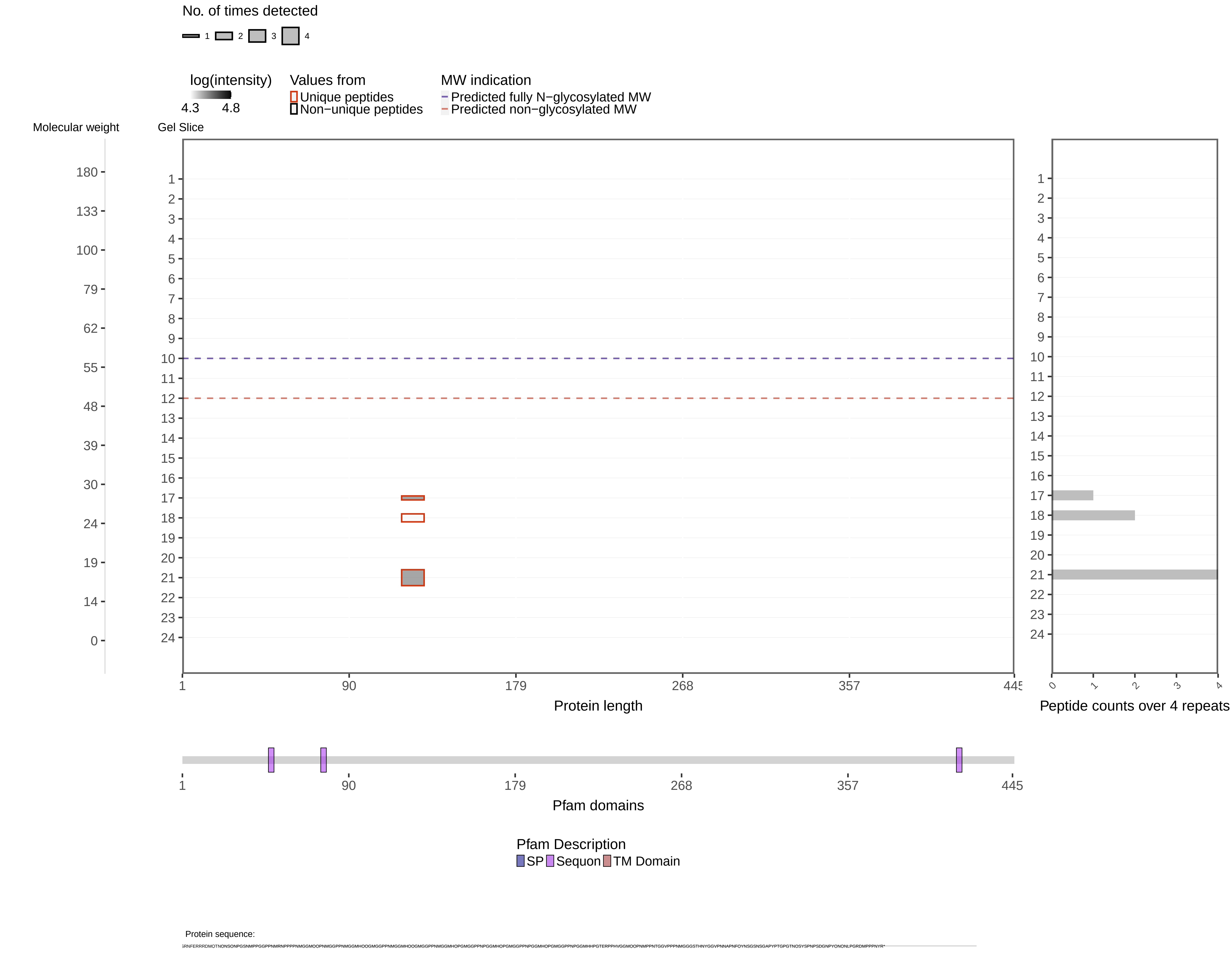

NbL03g04250.1 interactor_of_constitutive_active_ROPs_1−like [MW = 35.85 kDa]

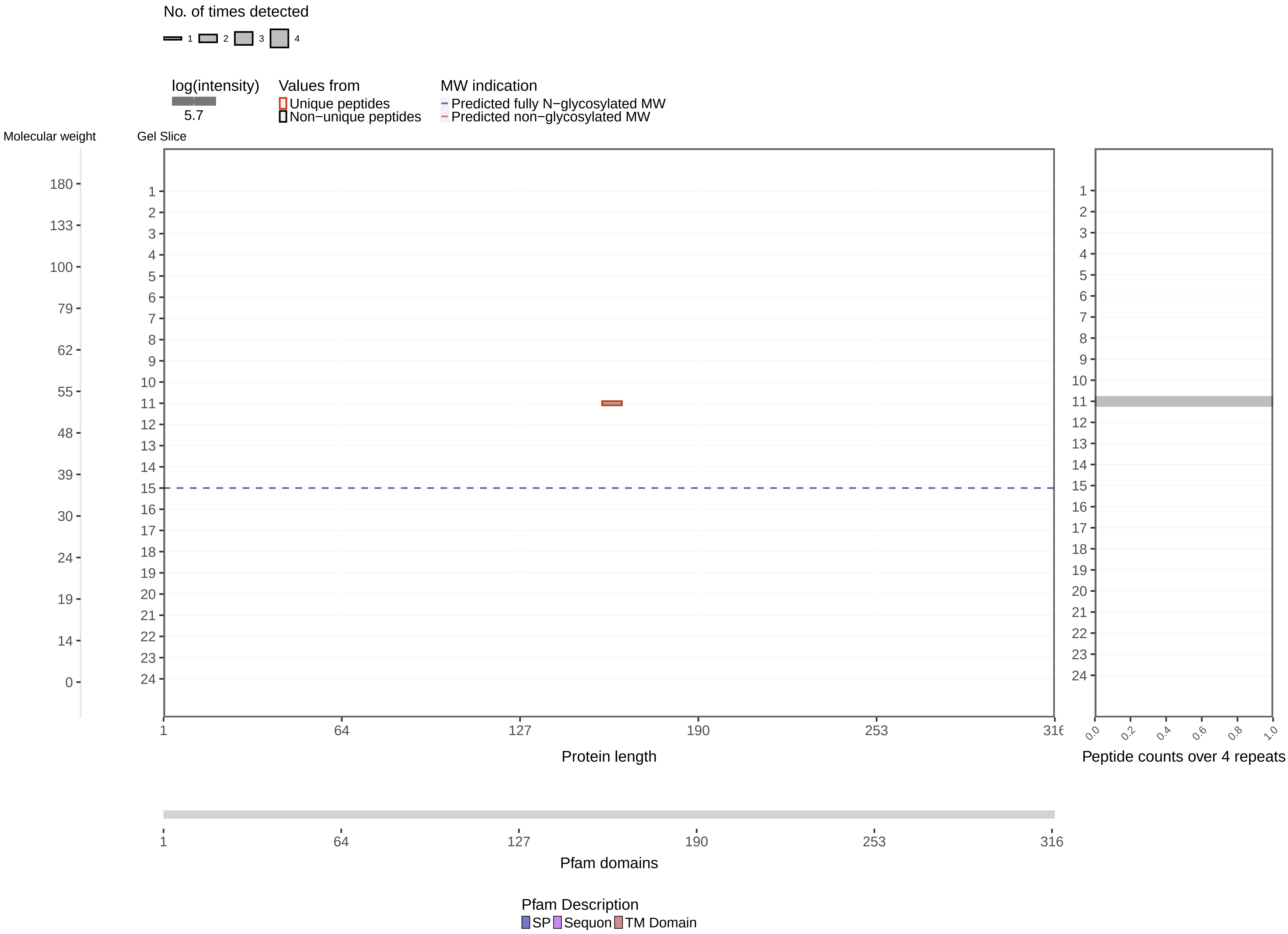


Protein sequence:

TALSYKELDLKNEEIIMLKAKLEEKEKEFQVFCQENENLKKELNDKTLEISSTKANTEEMNLKFNQVIQELETTKRNAFEINEKLEASEKAKDELESEMKKLRVQTEQWRKAADAAAAVLAGEVEVNGRRLSERCGSMDYKRYGNVFEPEVGGYGSYVGYPGLNDDSDDAFG

### NbL03g04160.1 elongation_factor_Ts_mitochondrial−like [MW = 43.07 kDa]

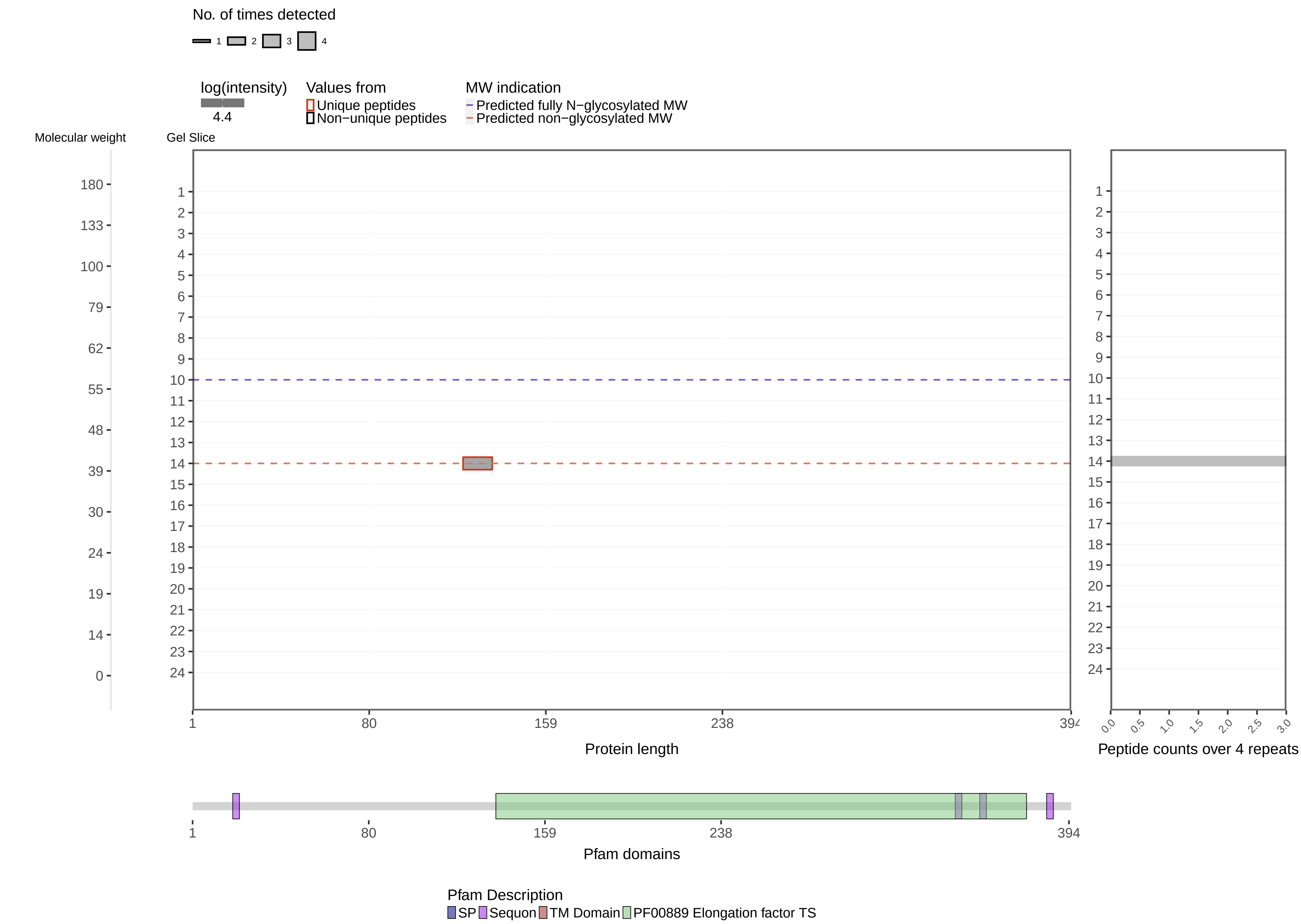


Protein sequence:

ILEDLKMNLDHPKLSGEKSVQNAITEVAAMMGENVKLRRGFAMSTPSIGVISTYLHTSPQPGVGRIAGILSLEVEDQNVPQDALQRVGSEIAMHVVAAKPLFLTKEDVSSDALSNERDILKSQAESSGRPQIAIEKMVEGRMRKYFEEVALMEQKFVVNDTINVKTLISNLSKEVGSPVKIGGFLRMEVGEGLQRLEASNESEPLANAA*

NbL03g04080.1 3−hydroxyisobutyryl−CoA_hydrolase−like_protein_5 [MW = 42.78 kDa]

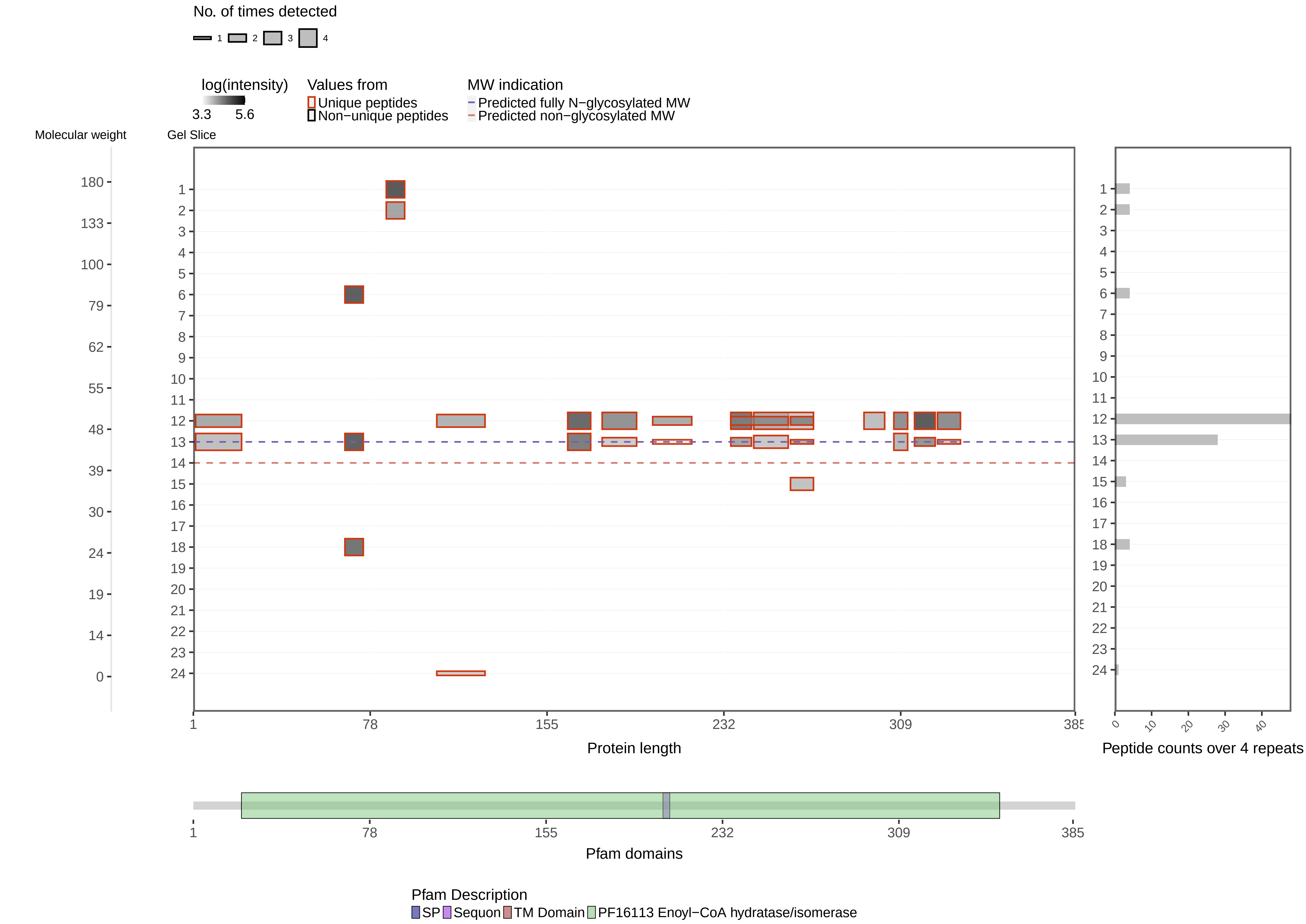


Protein sequence:

GKELVAAGLATHFVPLEKLPELEKHLLSLNTSDEAVVSSAIKEFSTDVQIDEESVLKKQAIIDECFSKDSVAEIISSFEAEAGKEGNSWIVPVLKGLKRSSPTGLKITLRSIREGRKQSLSECLKKEFRLTINILRTKISGDVYEGIRALTIDKDNSPKWDPTTLEKLDDERVNLVFGPFEEDLELKVSENEQCRWDGKYEDSAYCRQ*

### NbL03g03970.1 citrate_synthase_glyoxysomal−like [MW = 56.07 kDa]

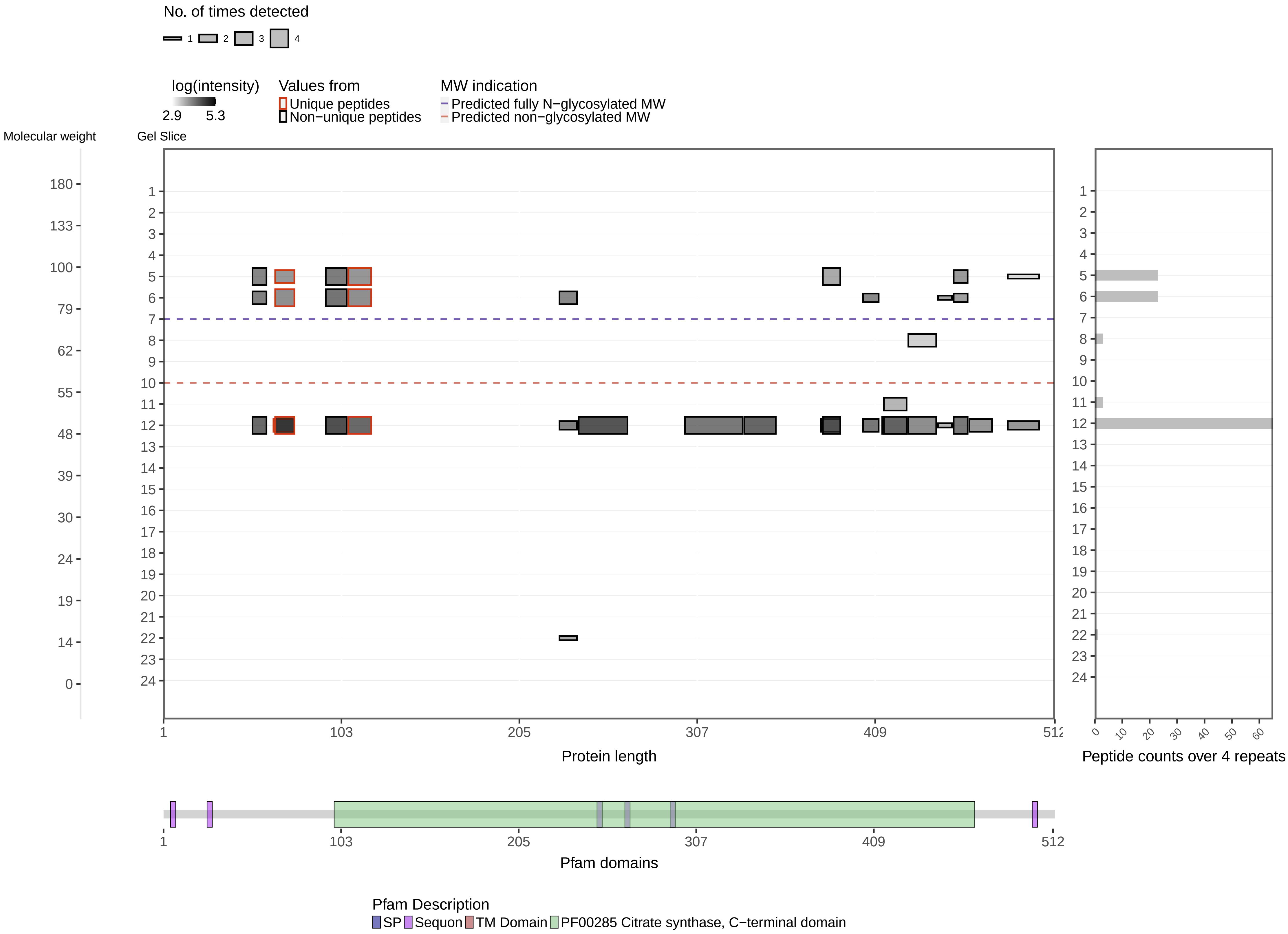


Protein sequence:

VLPSNNLSYAENFLYMLDSLGNRSYKPNPRLARVLDILFILHAEHEMNCSTAAARHLASSGVDVYTAIAGAVGALYGPLHGGANEAVLKMLSEIGSVENIPEFIEGVKNRKRKMSGFGHRVYKNYDPRAKVIKKLADEVFSIVGRDPLIEVAIALEKAALSDEYFVKRKLYPNVDFYSGLIYRAMGFPTEFFPVLFAVPRMAGYLSHWRESLDDPDTKIMRPAQVYTGVWMRHYMPLKERSPYTETDKLGQVSVSNATKRRLAGSGA*

### NbL03g03880.1 14−3−3_protein_9−like [MW = 28.55 kDa]

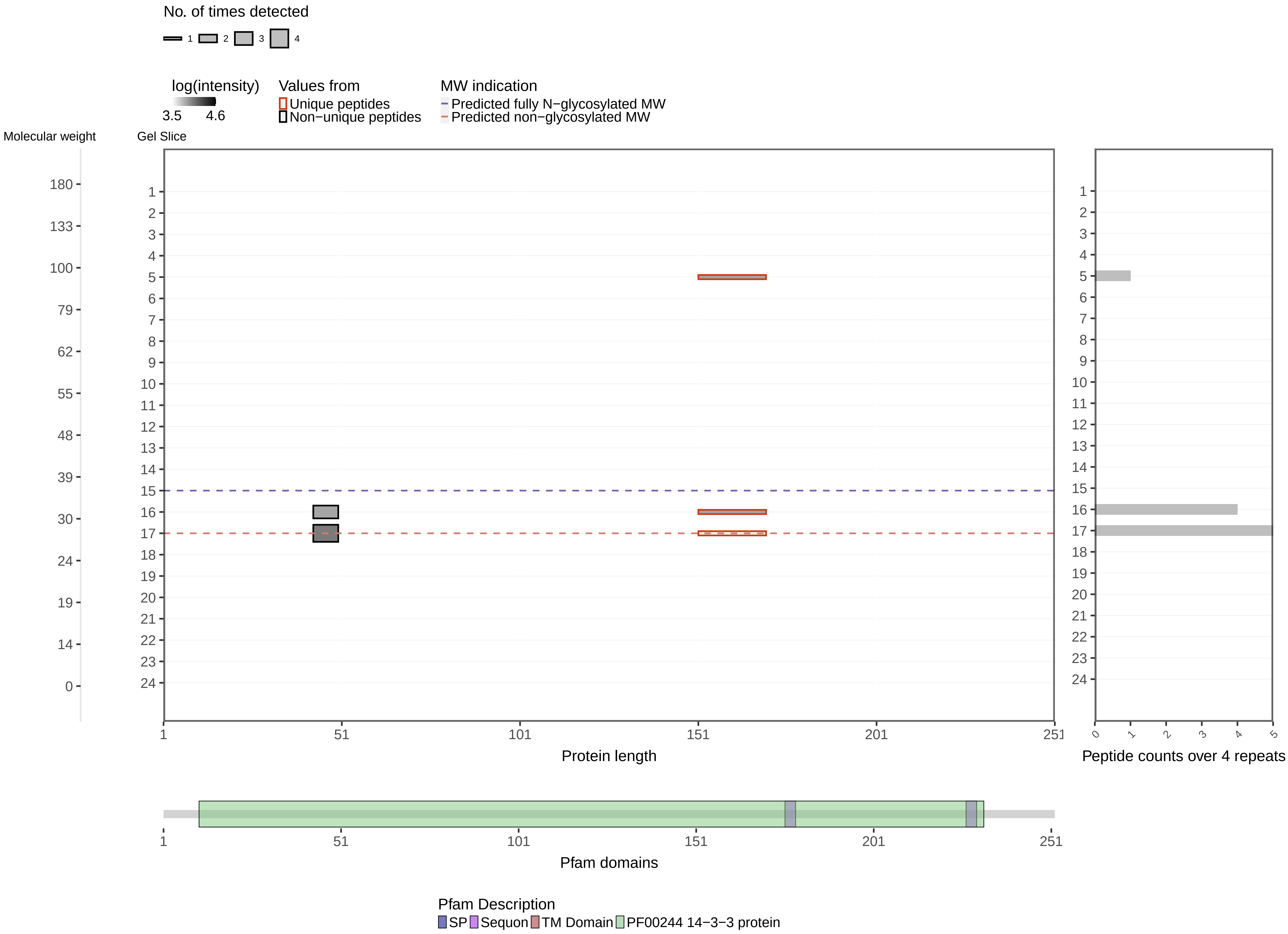


Protein sequence:

CTSGESSVFYYKMKGDYYRYLAELKTGNDKKEVSDLSLKAYQTATATAEAELSTTHPIRLGLALNFSVFYYEIMNSPERACHLAKQAFDEAISELDALNEDSYKDSTLIMQLLRDNLTLWTSDIPDDAGMCKAFSQEDA*

NbL03g03800.1 ethylene−responsive_transcription_factor_4−like [MW = 31.04 kDa]

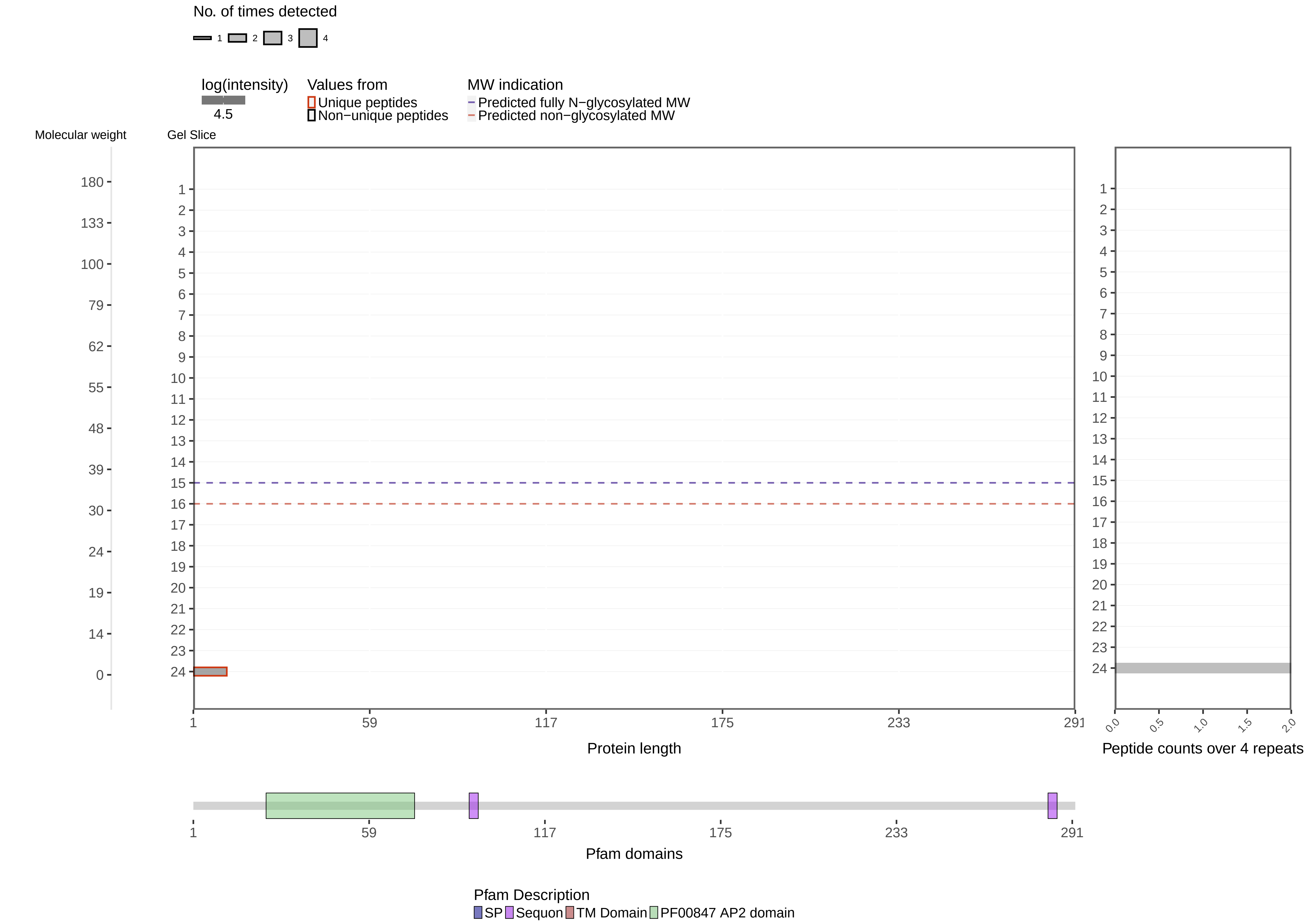


Protein sequence:

CKSEVGLVRNGFPIFHHQPVMGALTSGSSSPVESSSGETGVHAPLELDLTRRLGAVGEGGNHVRNGFPIFHQQPTVAVLPNGQPILLFDSNFVPTGVLNRPMPYRFEQVTVEFNGVSGRVVHSDSSSSTTVNENYYGGGGVSRRELNLDLNLSPPMEA*

### NbL03g03630.1 probable_cinnamyl_alcohol_dehydrogenase_6 [MW = 42.55 kDa]

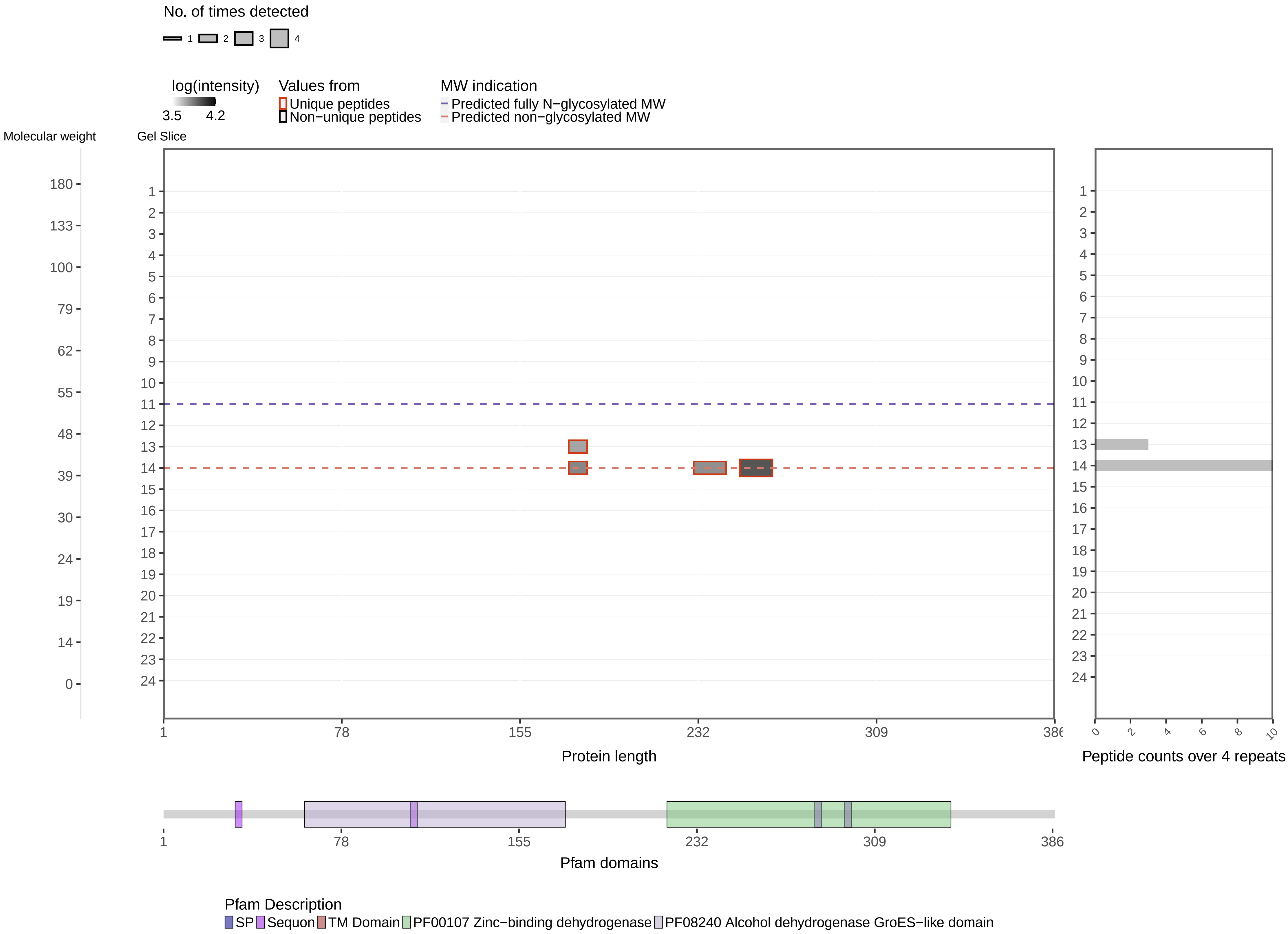


Protein sequence:

PDNLAMDRATPLLCAGITVYSPLKDNNLVETKGKRIGVIGLGGLGHVAVKFGKAFGHHVTVISTSPSKQQEAKHNLGADDFILSTNPKQIQARKRTLDFILDTVAANHSLGIYLELLKVNGTLVIVGAPEKPIDLPAFPMIFGKRNVKGSMIGSIEETQEMMDLCEKHNILSDIEIITTDQINEGLDRLAKNDVKYRFVIDIAGQSSNL*

NbL03g03540.1 cytochrome_b6−f_complex_iron−sulfur_subunit_2_chloroplastic [MW = 26.13 kDa]

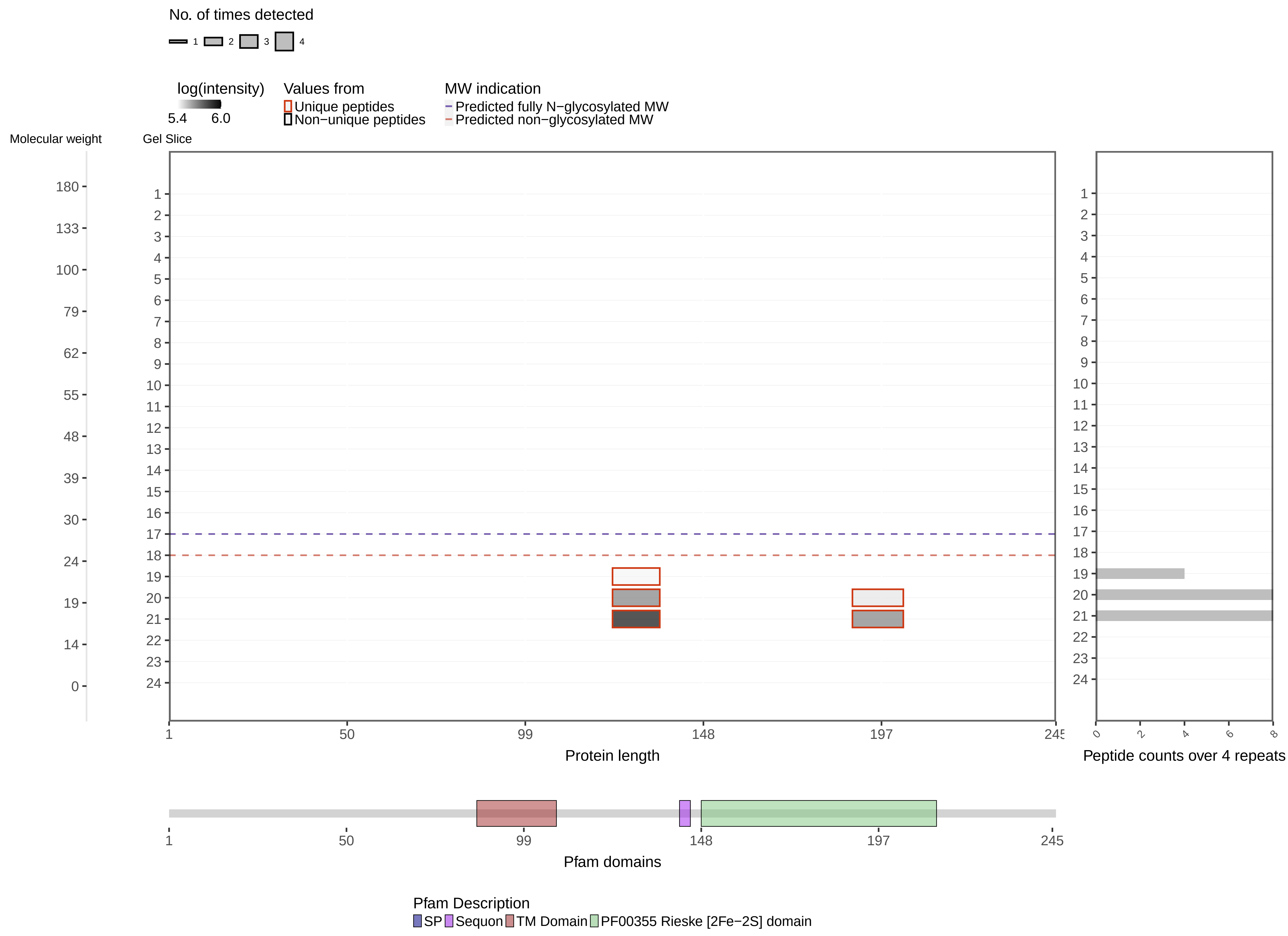


Protein sequence:

GGGGGGIPAKDALGNDVVASEWLKTHPPGNRTLTQGLKGDPTYLVVENDGTLATYGINAVCTHLGCVVPFNAAENKFICPCHGSQYNNQGRVVRGPAPLSLALAHADIDDGKVVFVPWVETDFRTGEAPWWA*

### NbL03g03390.1 uncharacterized_LOC107778869 [MW = 10.46 kDa]

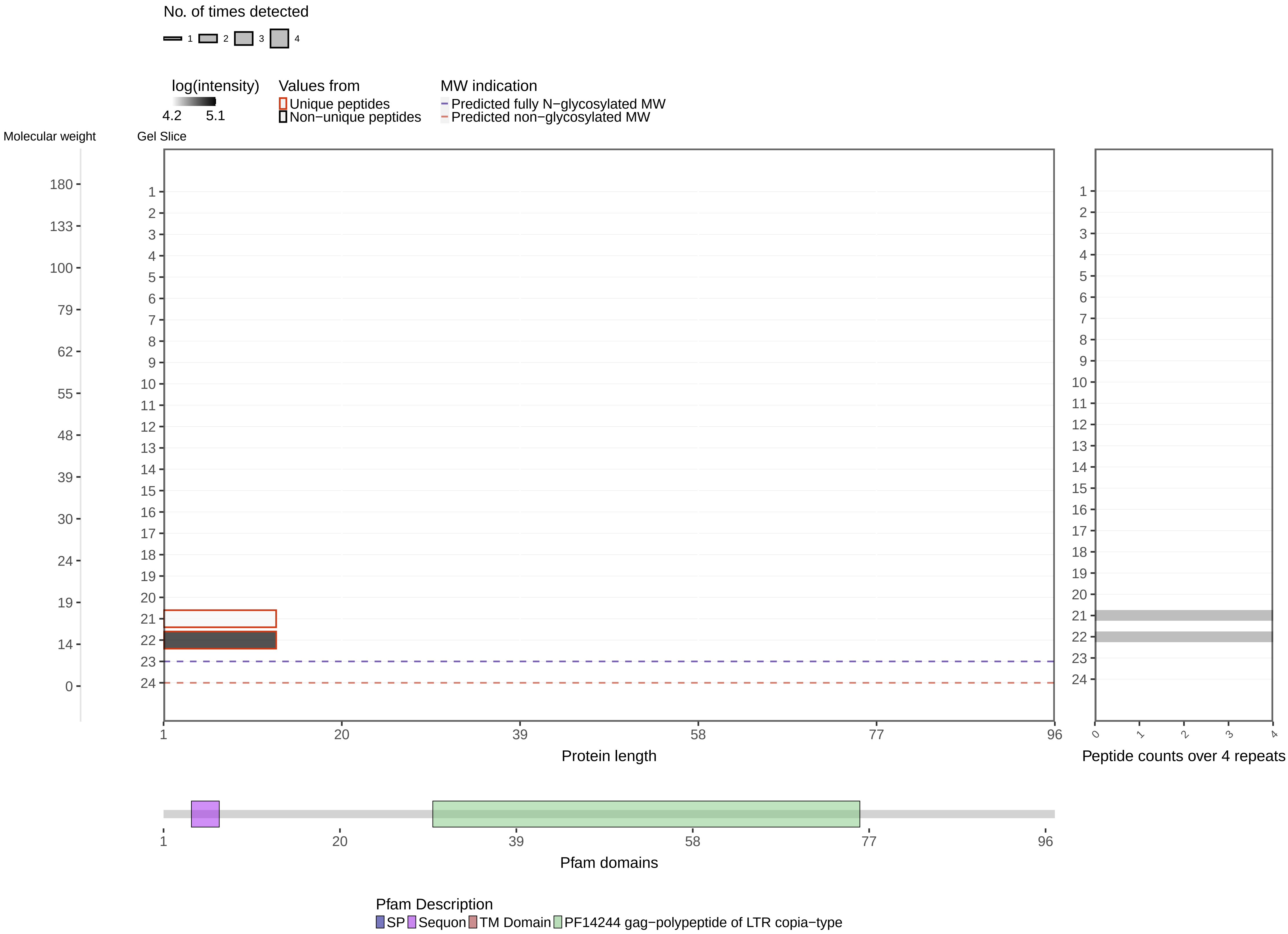


Protein sequence:
GMVLVNSVFDGKEYGGWNRAIVIALSSKNKLGFIDGTYFEPDASSTDFKQWNCCNDMVIS*

### NbL03g03170.1 ACT_domain−containing_protein_DS12_chloroplastic−like [MW = 30.88 kDa]

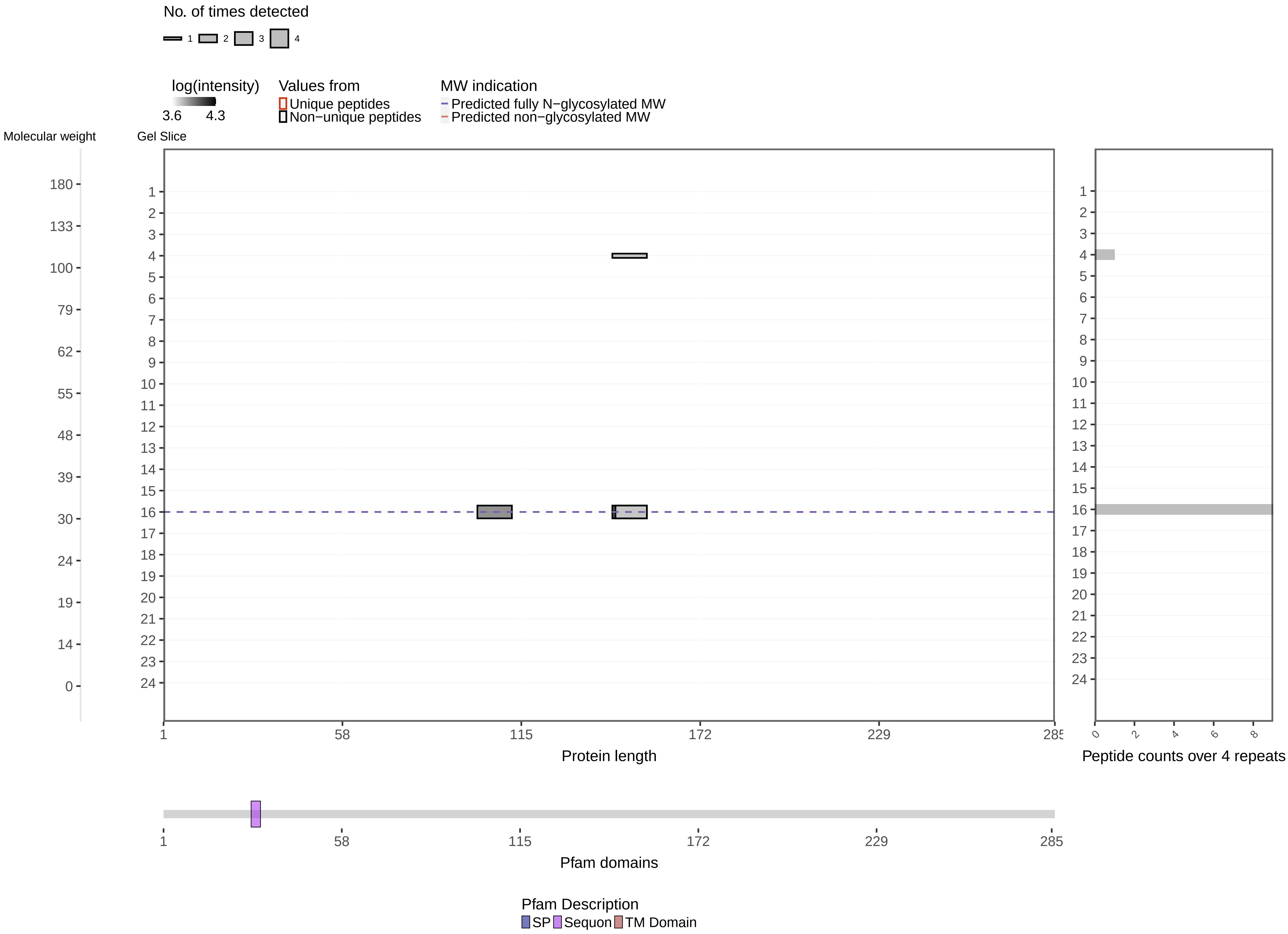


Protein sequence:

GKHNKFAITKASTGRKVDDPELLEAIRLTIINNMMEYHPESSSMLAMGEAFGVFQPYQKIDVDIATHIHIYDDGPERSLLCVETADRPGLIVDLVKIITDINVDVESGEFDTEGLLAKAKFHVSYKGKALIKPLQQVLANSLRYFLRRPTTEDASF*

### NbL03g03130.1 calnexin_homolog_1−like [MW = 65.27 kDa]

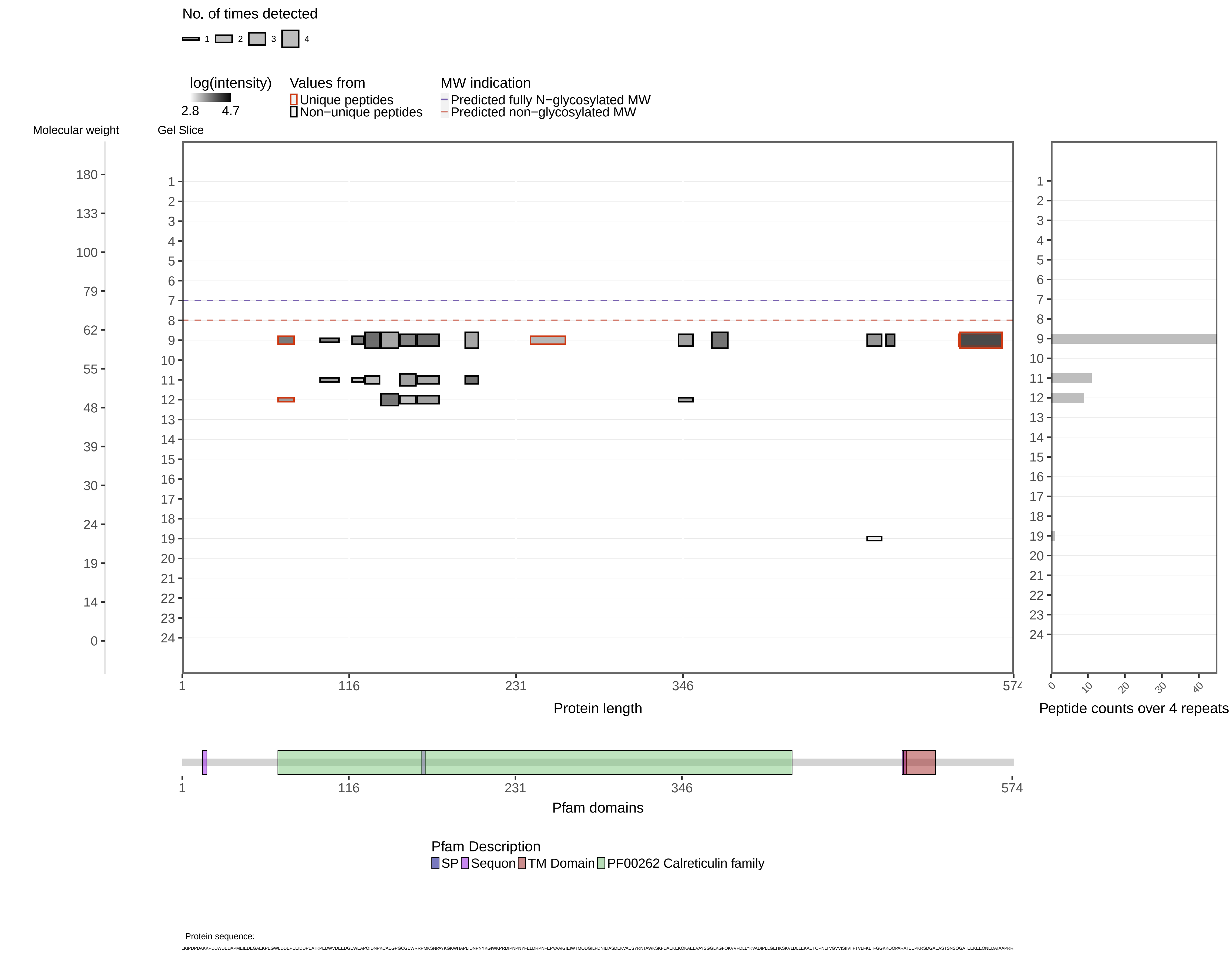

### NbL03g02970.1 lysosomal_alpha−mannosidase−like [MW − SP = 113.57 kDa]

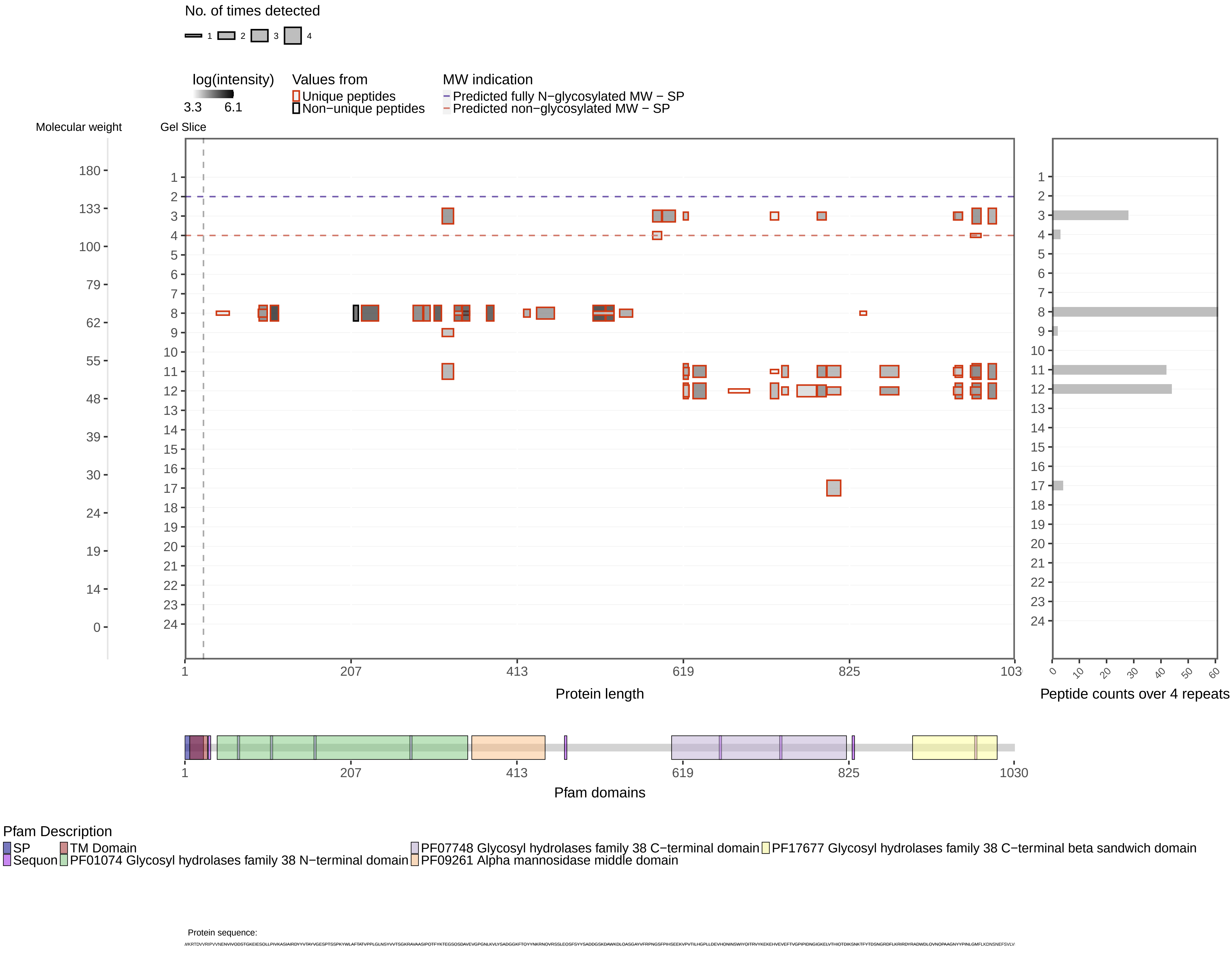

### NbL03g02850.1 ubiquitin−conjugating_enzyme_E2_variant_1C−like [MW = 16.77 kDa]

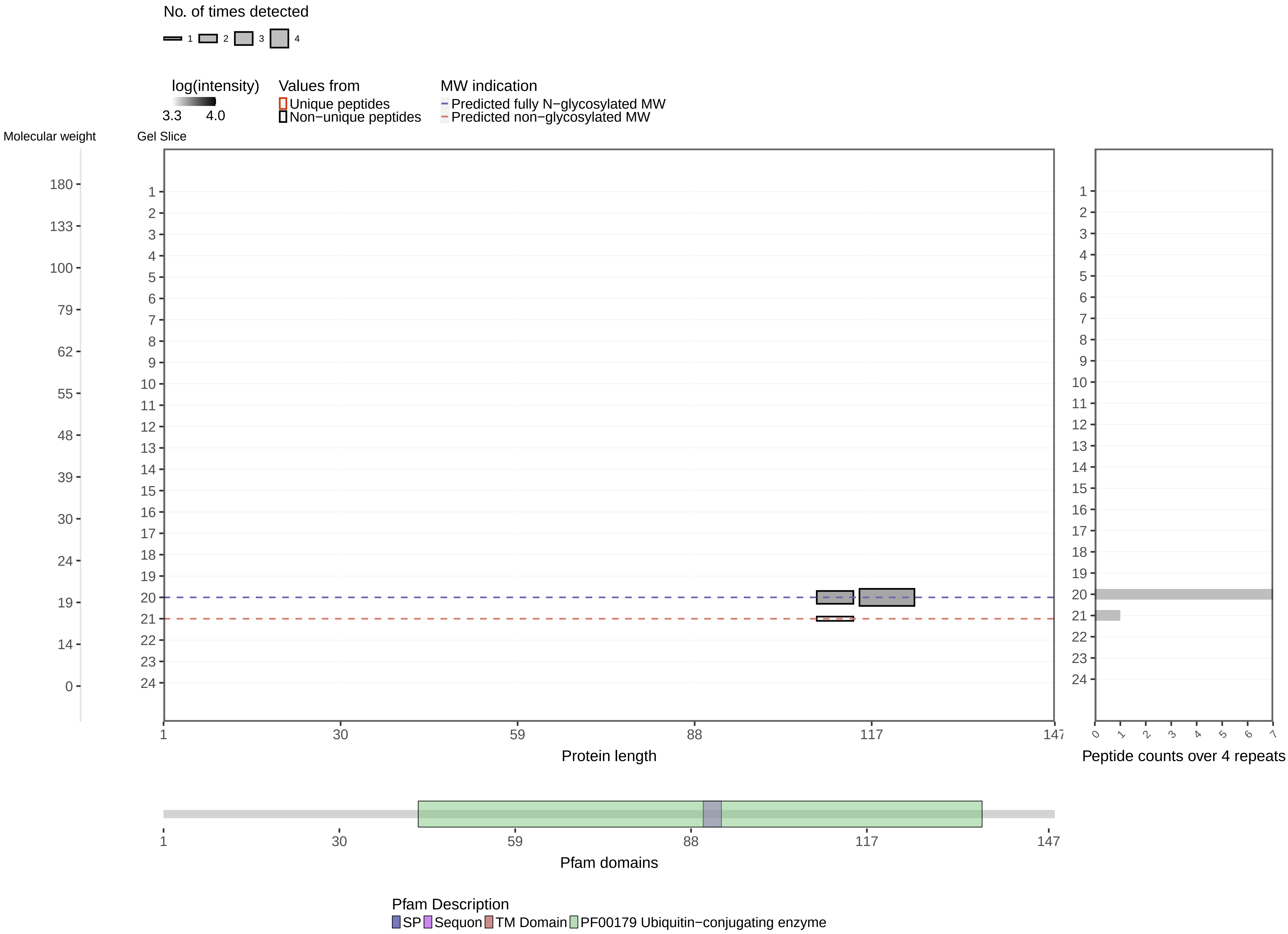


Protein sequence:

HQGRIYQLKLFCDKDYPEKPPSVRFHSRINMTCVNPESGVVDAKKFRMLENWQREYTMEDILTKLRKEMVAPQNRKLVQPREGTFF*

### NbL03g02720.1 serine/threonine−protein_phosphatase_BSL3 [MW = 108.97 kDa]

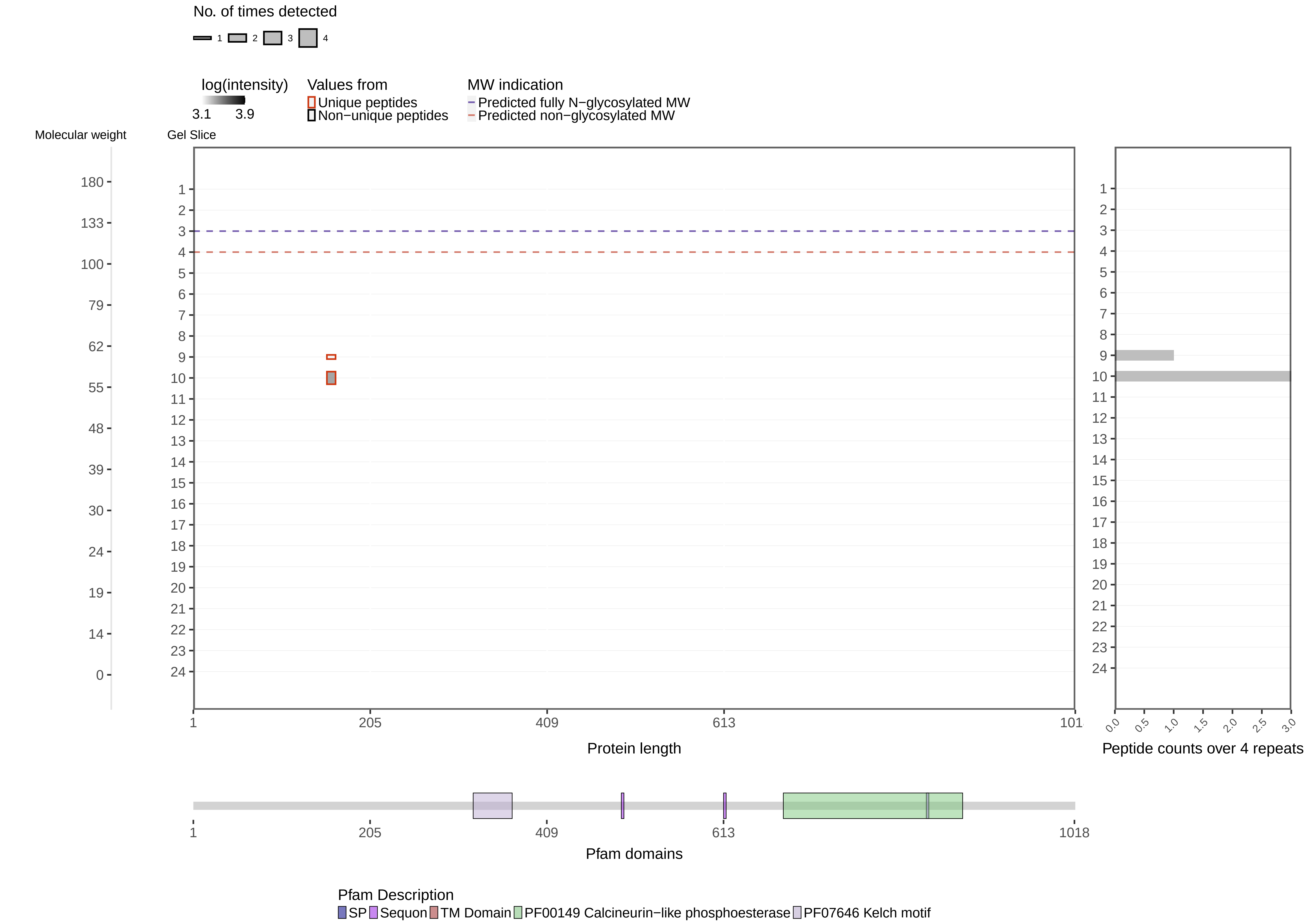


Protein sequence:

### NbL03g02670.1 succinate_dehydrogenase_subunit_6_mitochondrial−like [MW = 16.75 kDa]

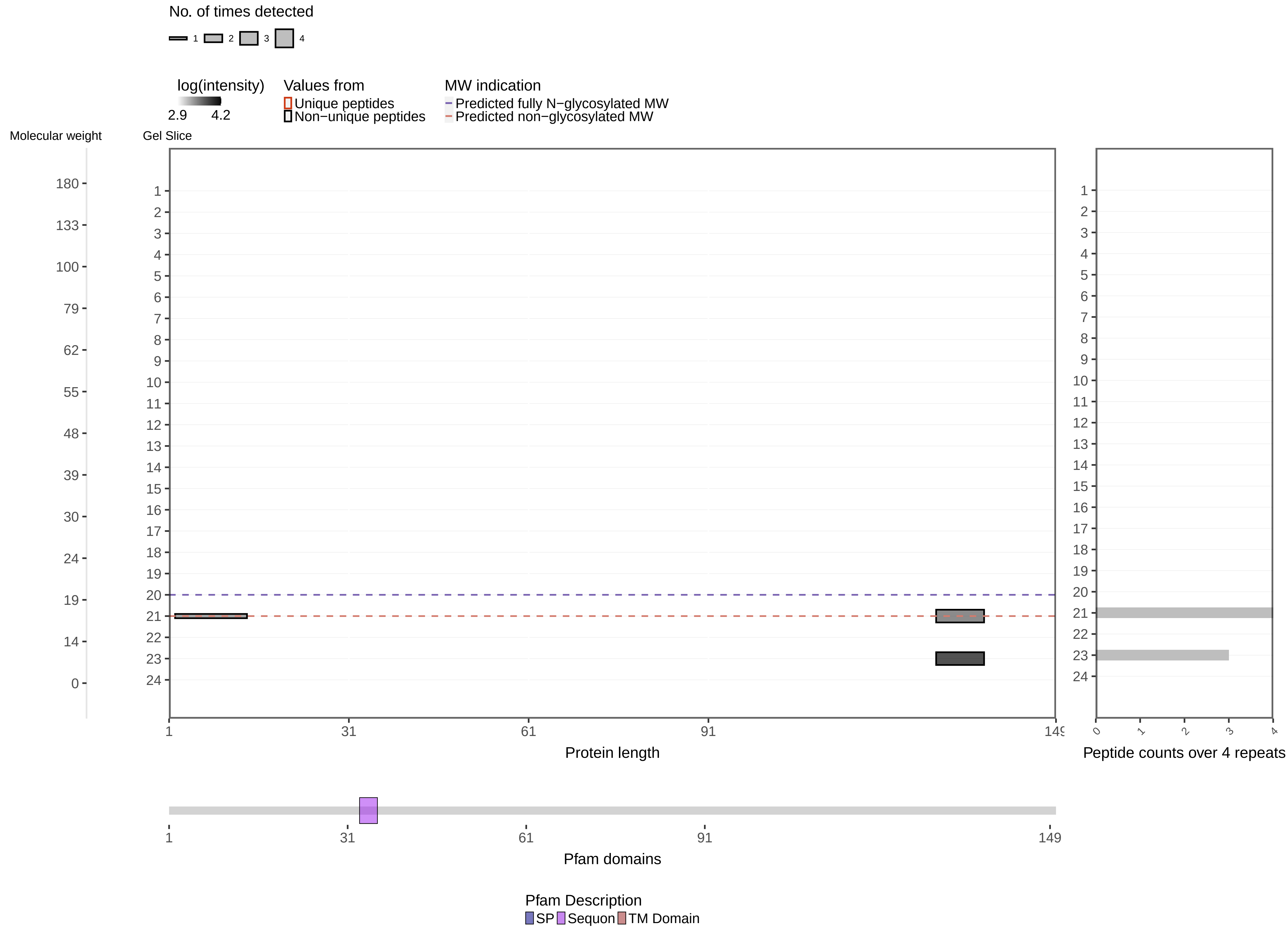


Protein sequence:

HGPTLRTAREAAKISAVGGIIGAVSTAGVAWKYSRSLHGTALSFGAGAVFGWTFGQEVANHWLQLYRLDTMAAQVKFMEWWQNKVEG*

### NbL03g02440.1 enoyl−CoA_delta_isomerase_2_peroxisomal−like [MW = 25.68 kDa]

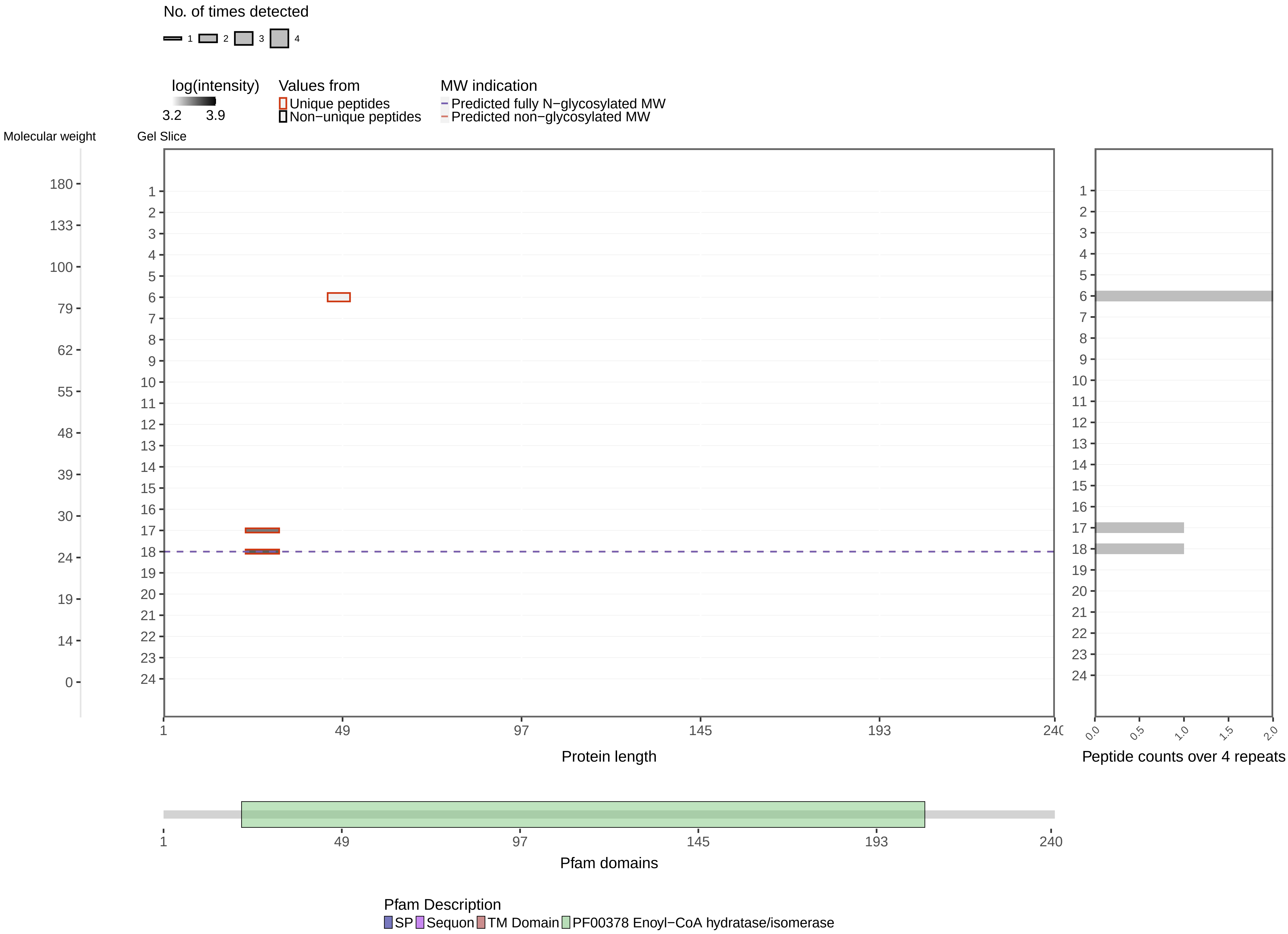


Protein sequence:

\AAAGLLLAICHDYVTMRSDKGVVYMSELDIGMTLPDYFTAMIRSKVGSAAARRNLVLKATKMKAEDAVAMGILDSAHPTAEEAVVAAVRLAEELGKKRWDGKTYAELRKSLVPEVCGVLGLKHTAVLPSRL*

### NbL03g02400.1 signal_recognition_particle_14_kDa_protein−like [MW = 14.22 kDa]

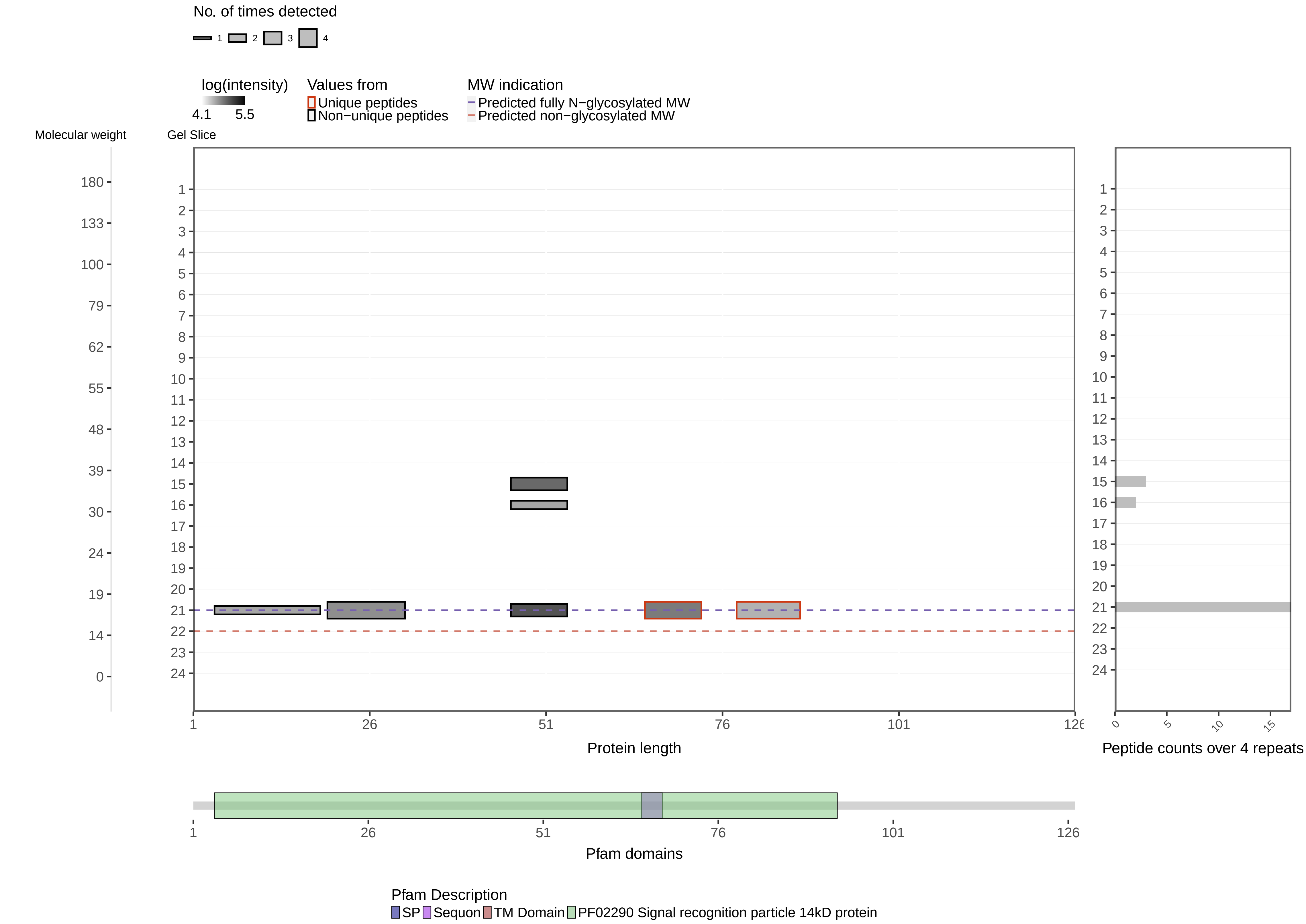


Protein sequence:

ENIEFKCLIRATDGKKNISTMVGAKDHQRFQASYAILLKARLTALKKRERKDKRKAADSDKKLEKSKKKSAAQKAST*

### NbL03g02340.1 2−isopropylmalate_synthase_B [MW = 65.79 kDa]

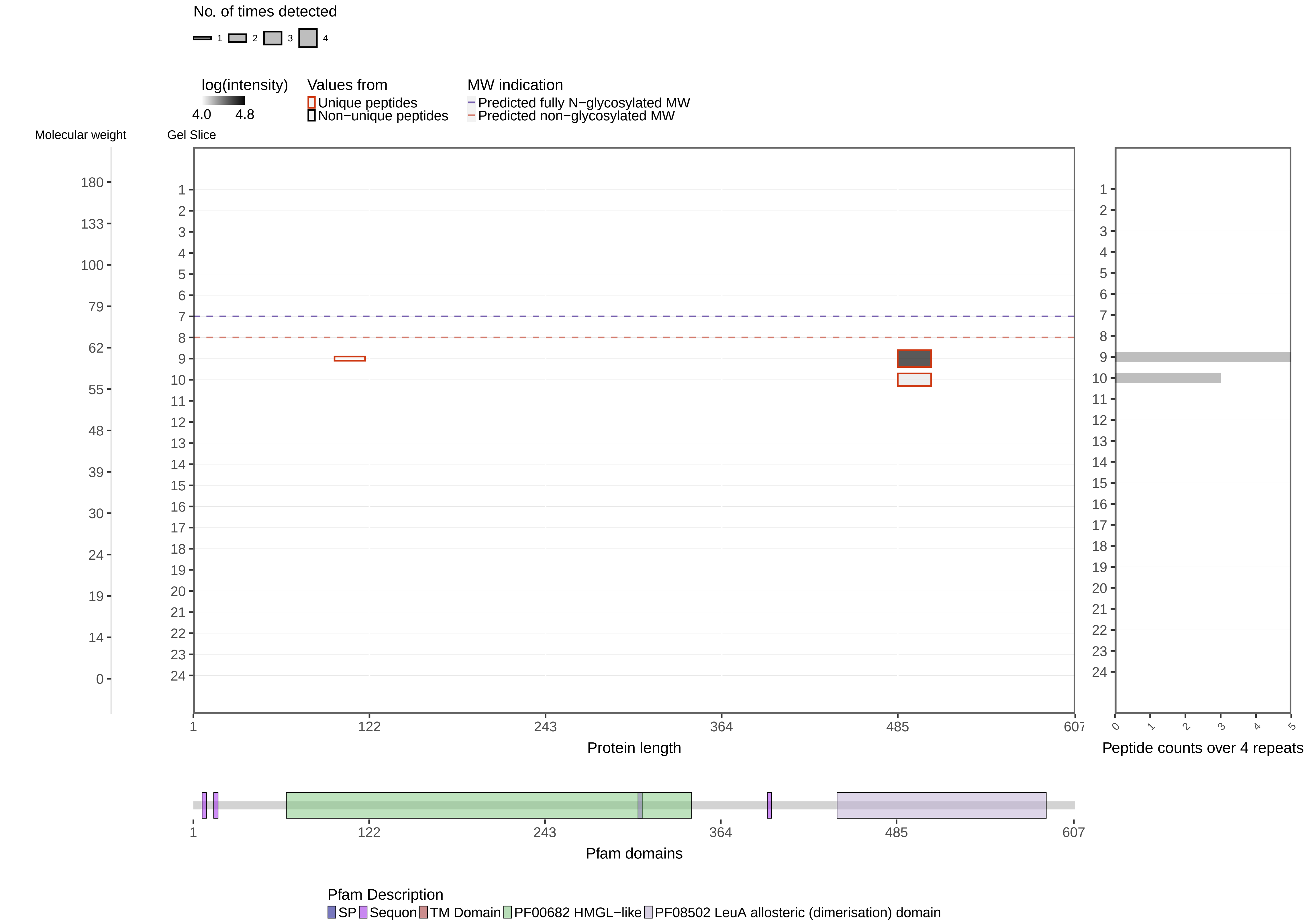


Protein sequence:

VEVTINGIGERAGNASLEEVVMALKCRGEQVLGGLYTGINTQHILMSSKMVEEYTGLHVQPHKAIVGANAFAHESGIHQDGMLKHKDTYEIISPEDIGLNRANESGIVLGKLSGRHALQAKMLELGYDIEGKELEDLFWRFKSVAEKKKKITDDDLIALMSDEVFQPQFVWQLENVQVTCGSLGLSTATVKLIDADGQEHVSCSVGTGPVDAAYKAVDLIVKVPVALLEYSLNAVTEGIDAIASTRVLIRGENGHTSTHALTGETVHRSFSGTGADMDIVISSVRAYIGALNKMLSI

### NbL03g02330.1 15_kDa_selenoprotein [MW − SP = 16.28 kDa]

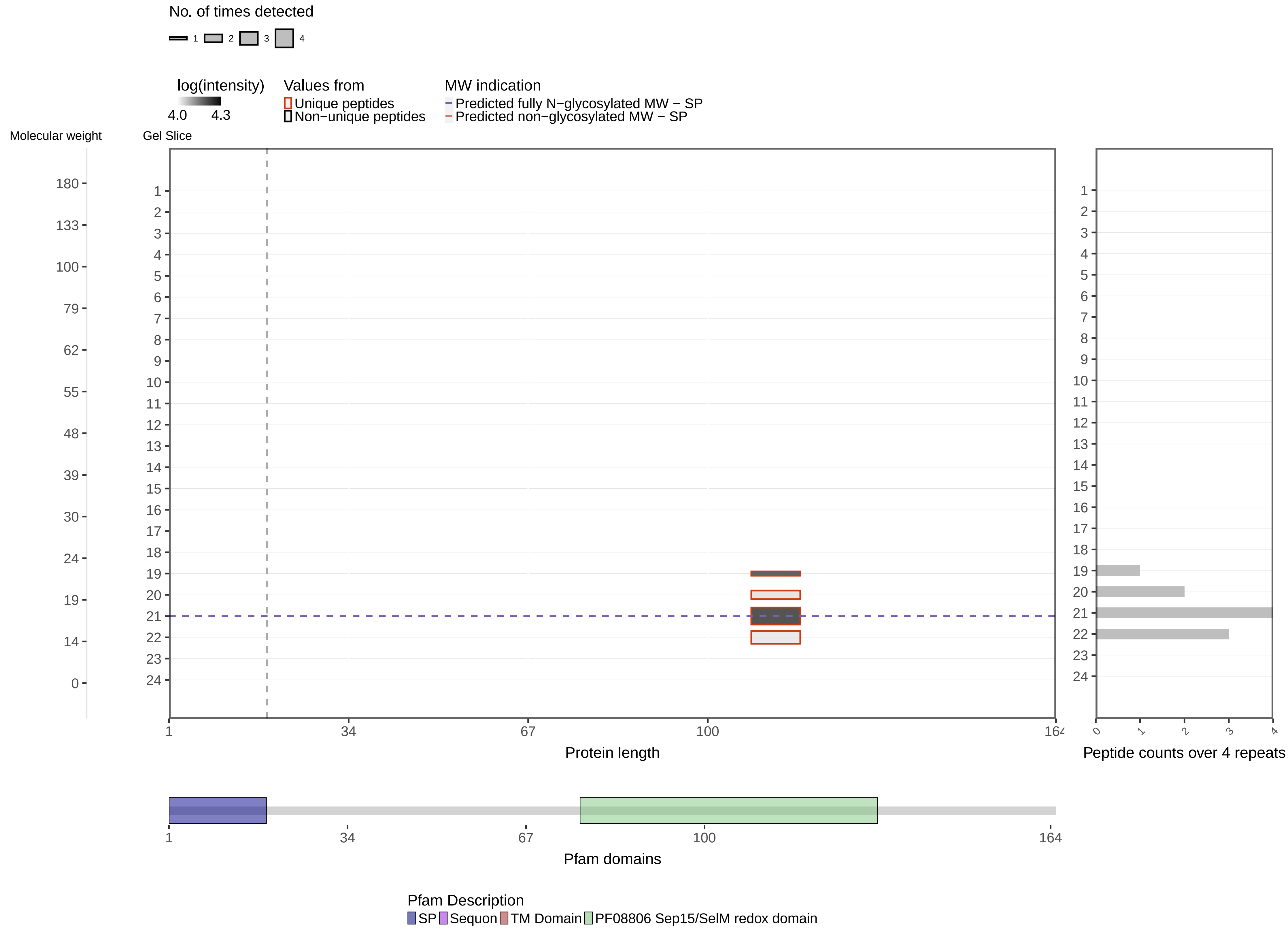


Protein sequence:

AMSKVVYSGAILEVCMRKLVFYPEVVGFIEEEKDKFPYVKVQYAYNSPPKLIMLDDHGEHKEAIRSSSCLLFPSFLCVMCISQPISYFYAVFIKVH*

NbL03g02270.1 probable_prefoldin_subunit_2 [MW = 16.19 kDa]

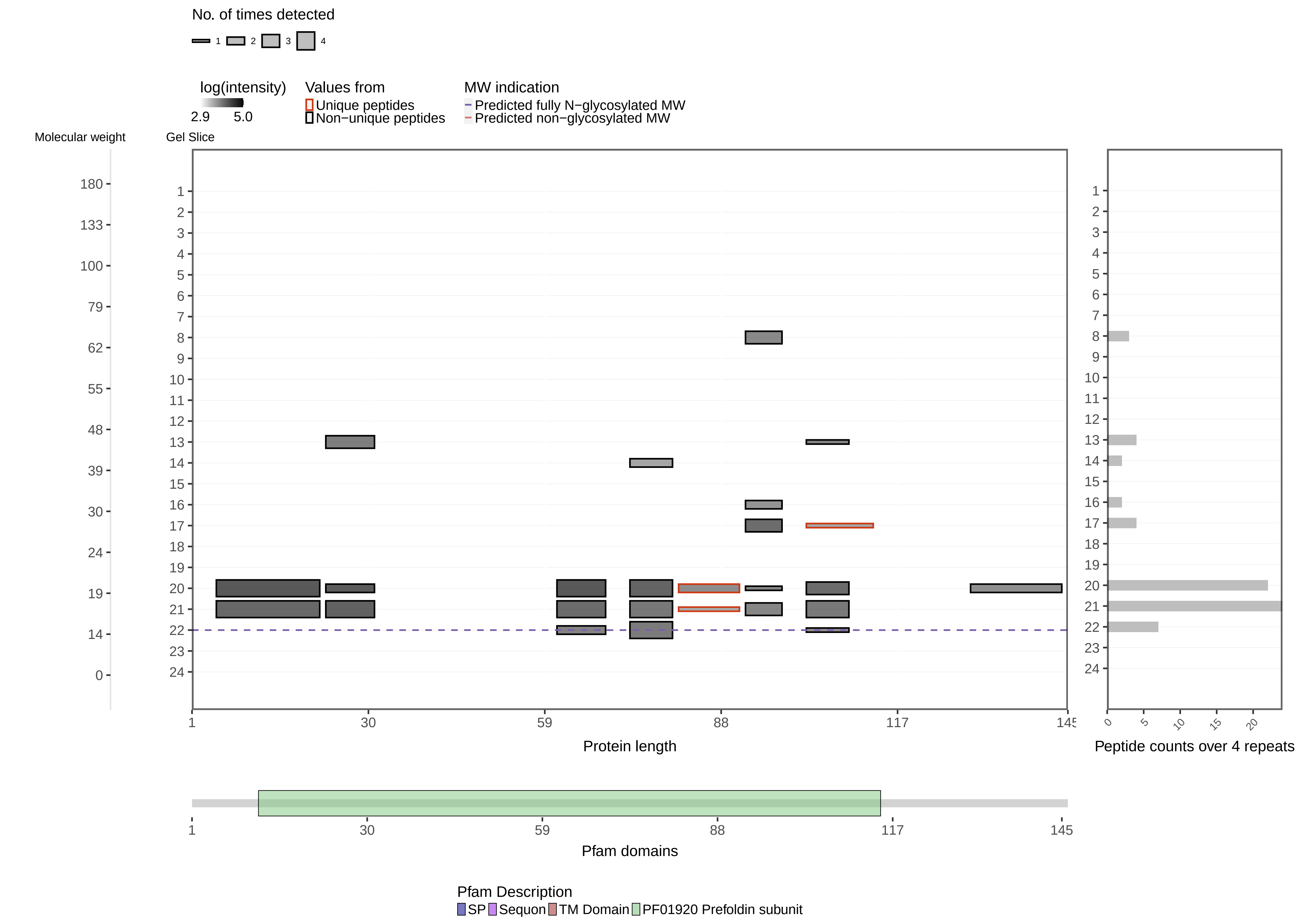


Protein sequence:

RMIGGVLVERTIKEVLPAVQRNKEGIEEVITRLNEALEKKKKEISDFETKYQIRMRKPDAEVKEESGRKEGSAQGVLVGPAGSNE*

NbL03g02250.1 stearoyl−[acyl−carrier−protein]_9−desaturase_chloroplastic [MW = 45.14 kDa]

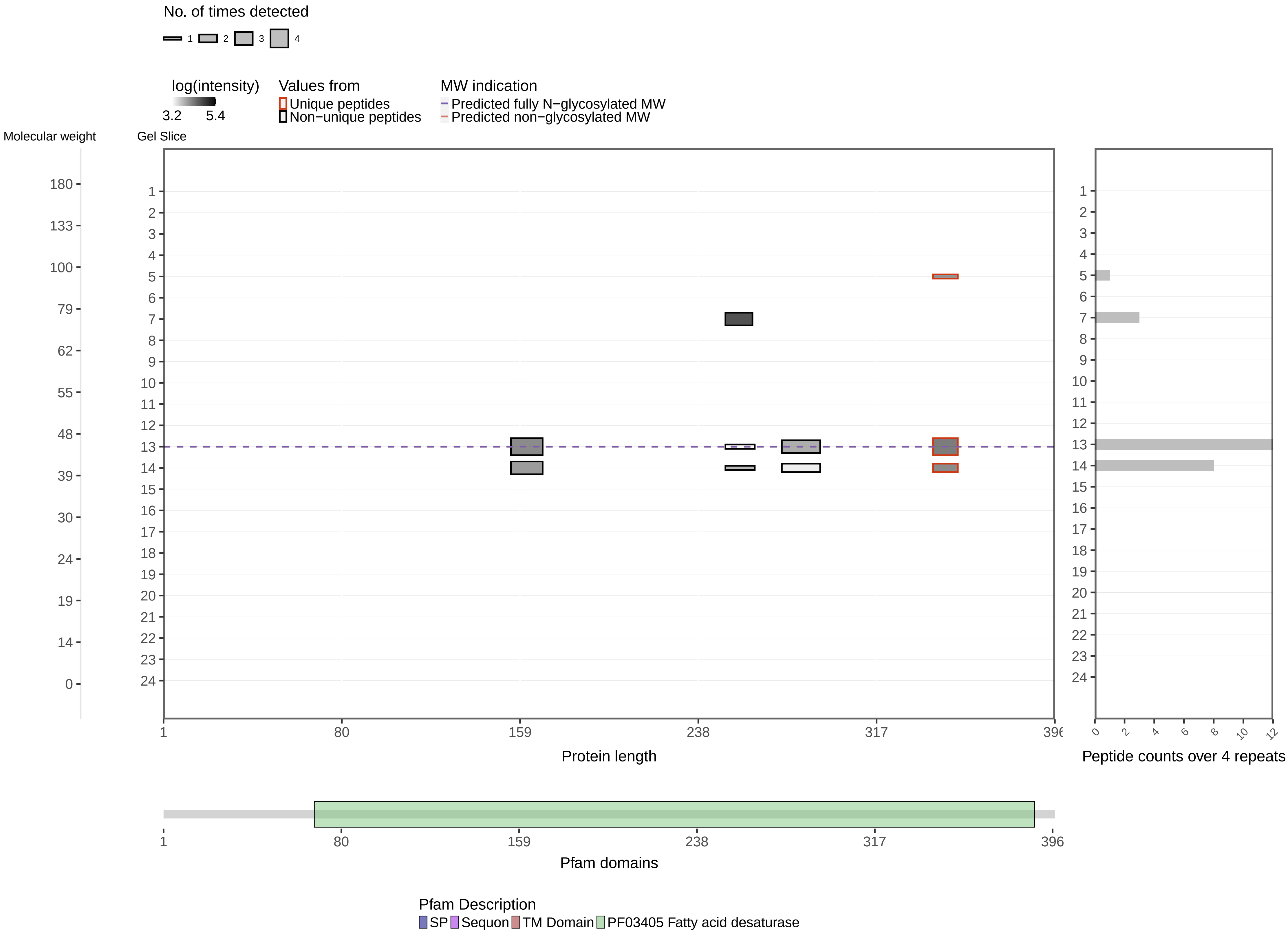


Protein sequence:

LNKYLYLSGRVDMRAIEKTIQYLIGSGMDPRTENNPYLGFIYTSFQERATFVSHGNTARLAKEHGDIKLAQICGIIAADEKRHEIAYTKIVEKLFEIDPDGTVLAFADMMRKKISMPAHLMYDGRDESLFDHFSAVAQRLGVYTAKDYADILEFLVGRWKVADLTGLSGEGRKAQEYVCGLPPRIRRLEERAQGRAKQGPIMPFSWIYDREVKL*

NbL03g02240.1 protein_FAM136A−like [MW = 17.09 kDa]

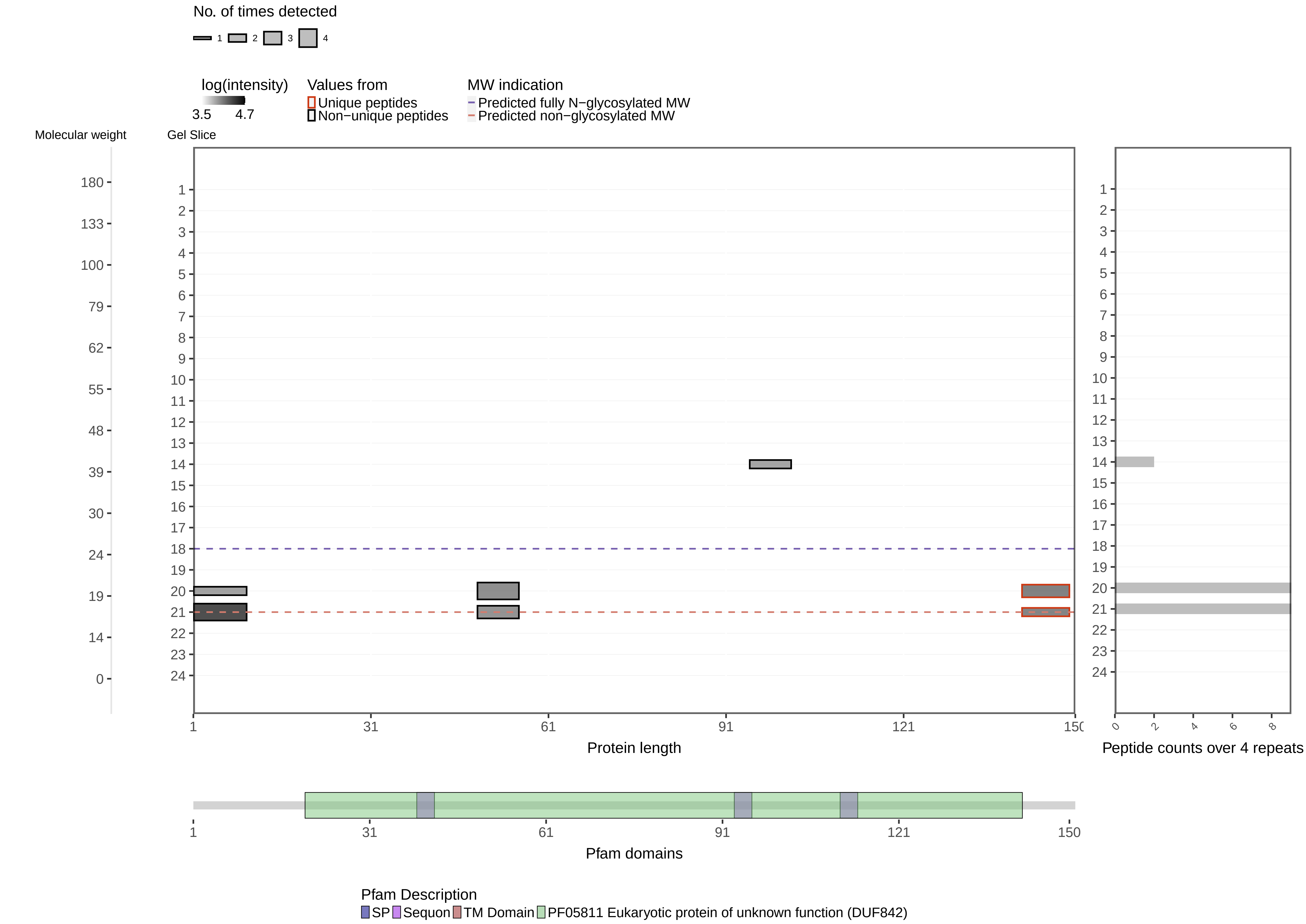


Protein sequence:

IGHCVEHCSVHVHNAQNLVQNEMAKFQERMNRSLMVCQDKYETAKLQKNRSDATKDLESCVDQSVHDSINTLPHLVARLKGSLGITN*

### NbL03g02160.1 nuclear−pore_anchor [MW = 229.78 kDa]

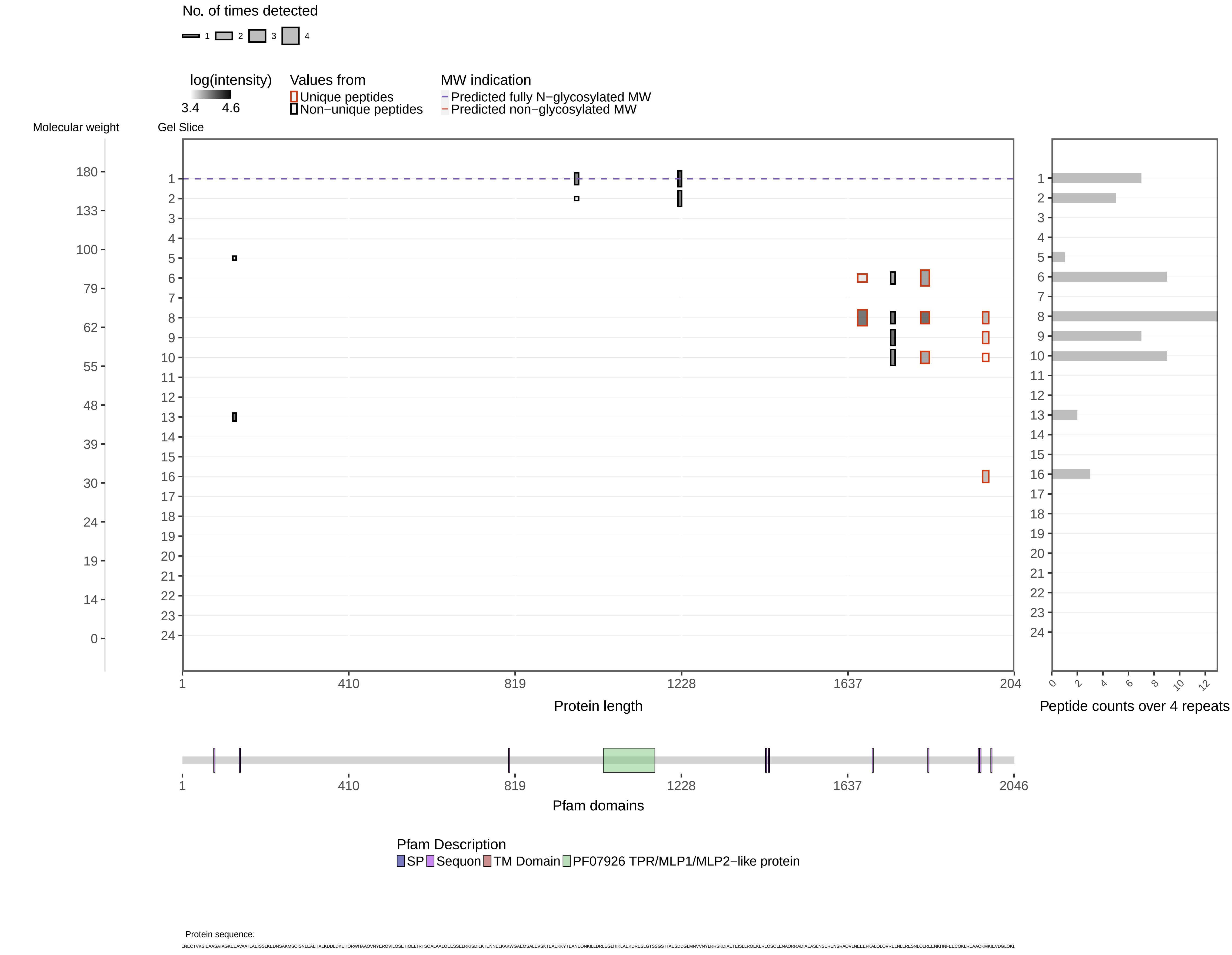

NbL03g02150.1 aldo−keto_reductase [MW = 47 kDa]

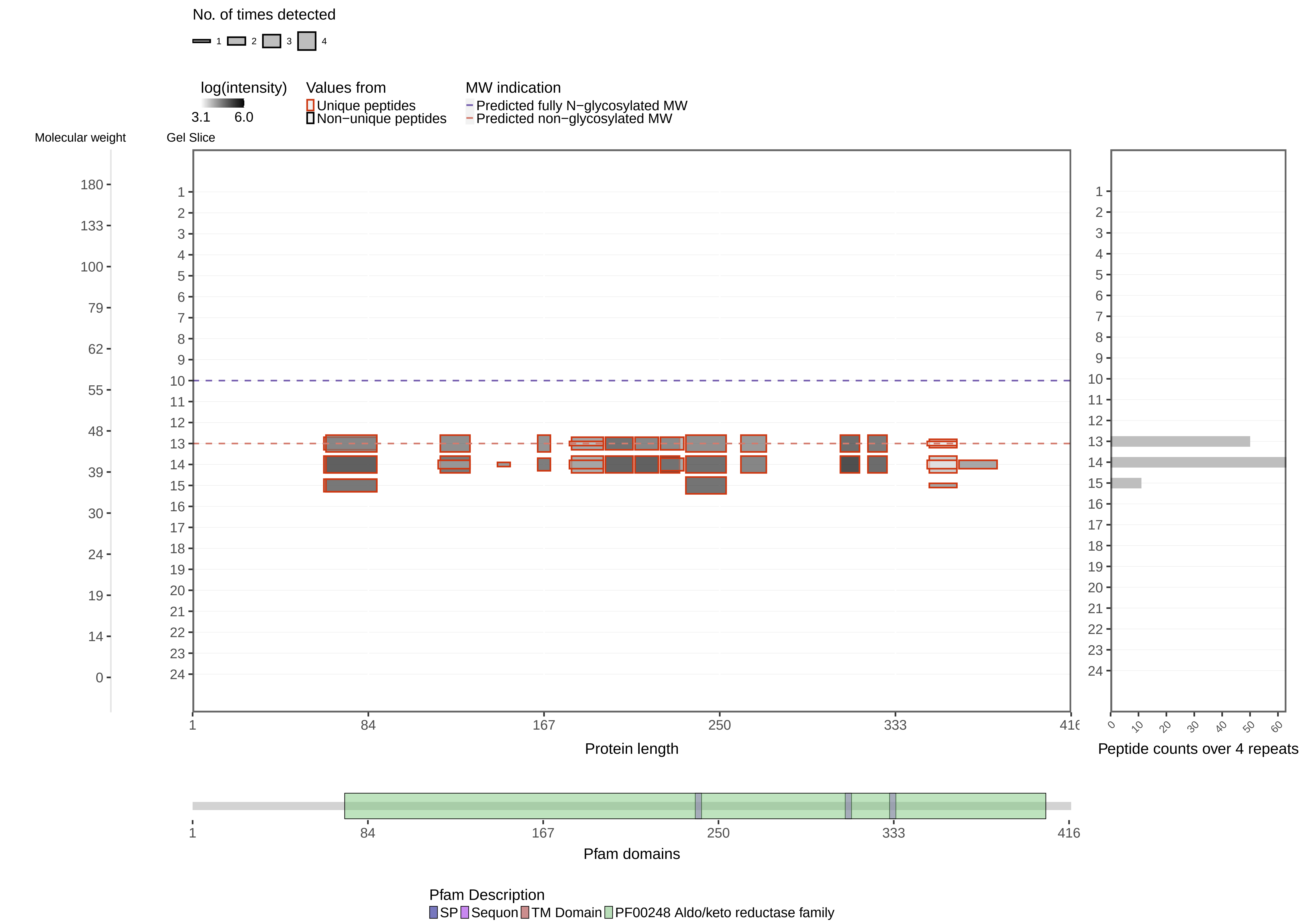


Protein sequence:

YVPLFGEFFYESSKWRPSVPFVEQLRAFQELIDEGKVRYIGVSNETSYGVMEFVHAAKVEGMPKIVSIQNSYSLLVRCHFEVDLVEVCHPNNCNIGLLSYSPLAGGTLSGKYLDNNSEAARKGRLNLFPGYMERYNKSLAKEATKKYEEVAKKHGLTLVELALGFARDRPFMTSSIIGATSVEQLKEDIDAFLTTERPLPPQVMDDIEDIFKRYRDPASR*

NbL03g02120.1 serine/threonine−protein_phosphatase_PP1_isozyme_2−like [MW = 36.7 kDa]

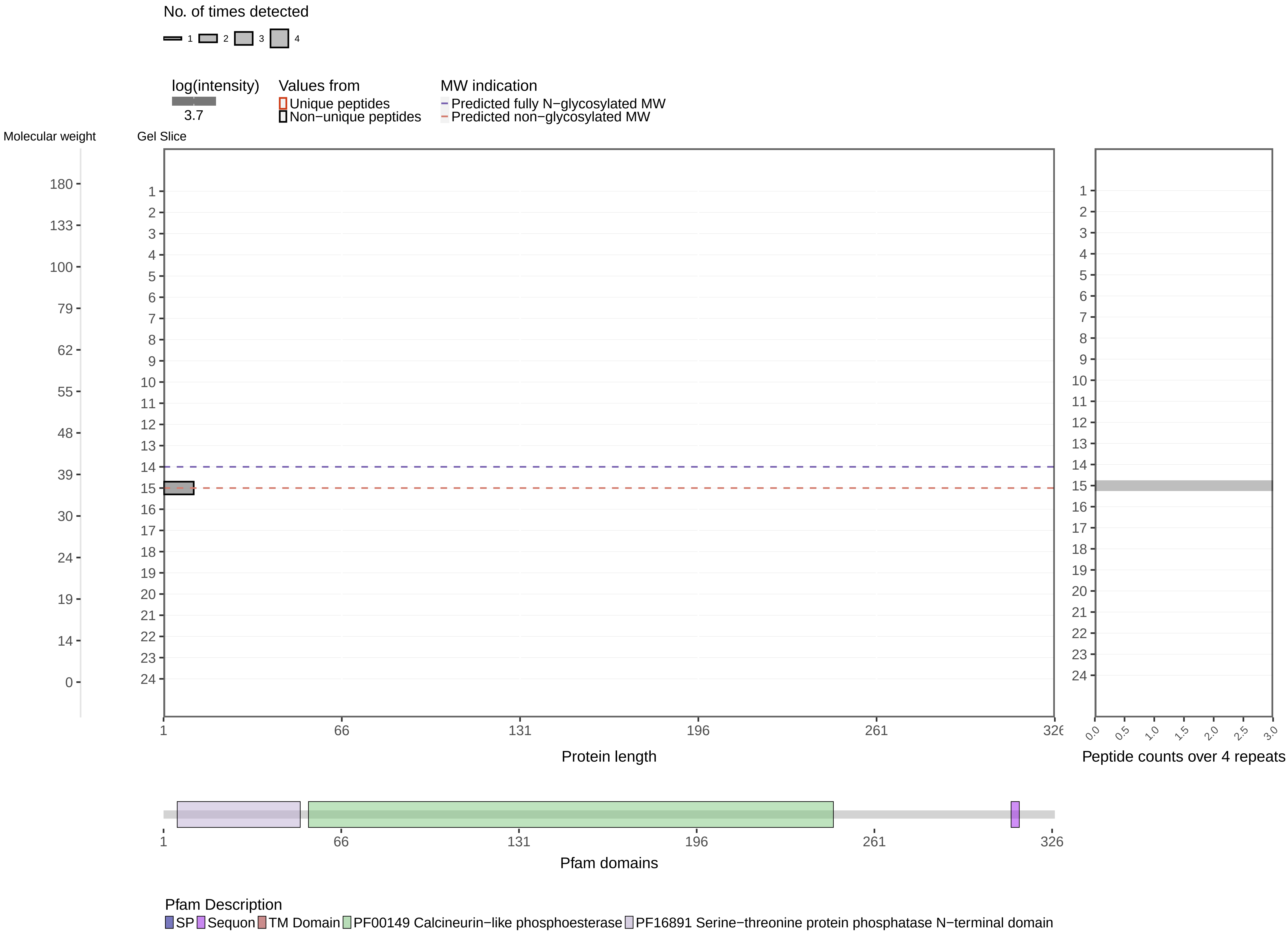


Protein sequence:

FNCLPVAALIDEKILCMHGGLSPDLNHLDOIRNLORPTDVPETGLLCDLLWSDPSKDVKGWGMNDRGVSSTFGPDKVTEFLOKLDLDLVCRAHOVVEDGYEFFADROLVTIFSAPNYCGEFDNAGAMMSVDETLMCSFOILKPAEKKPKFGFGSATSAKNATPTKSKSFLGKIG*

### NbL03g01910.1 probable_hydroxyacylglutathione_hydrolase_2_chloroplastic [MW = 36.22 kDa]

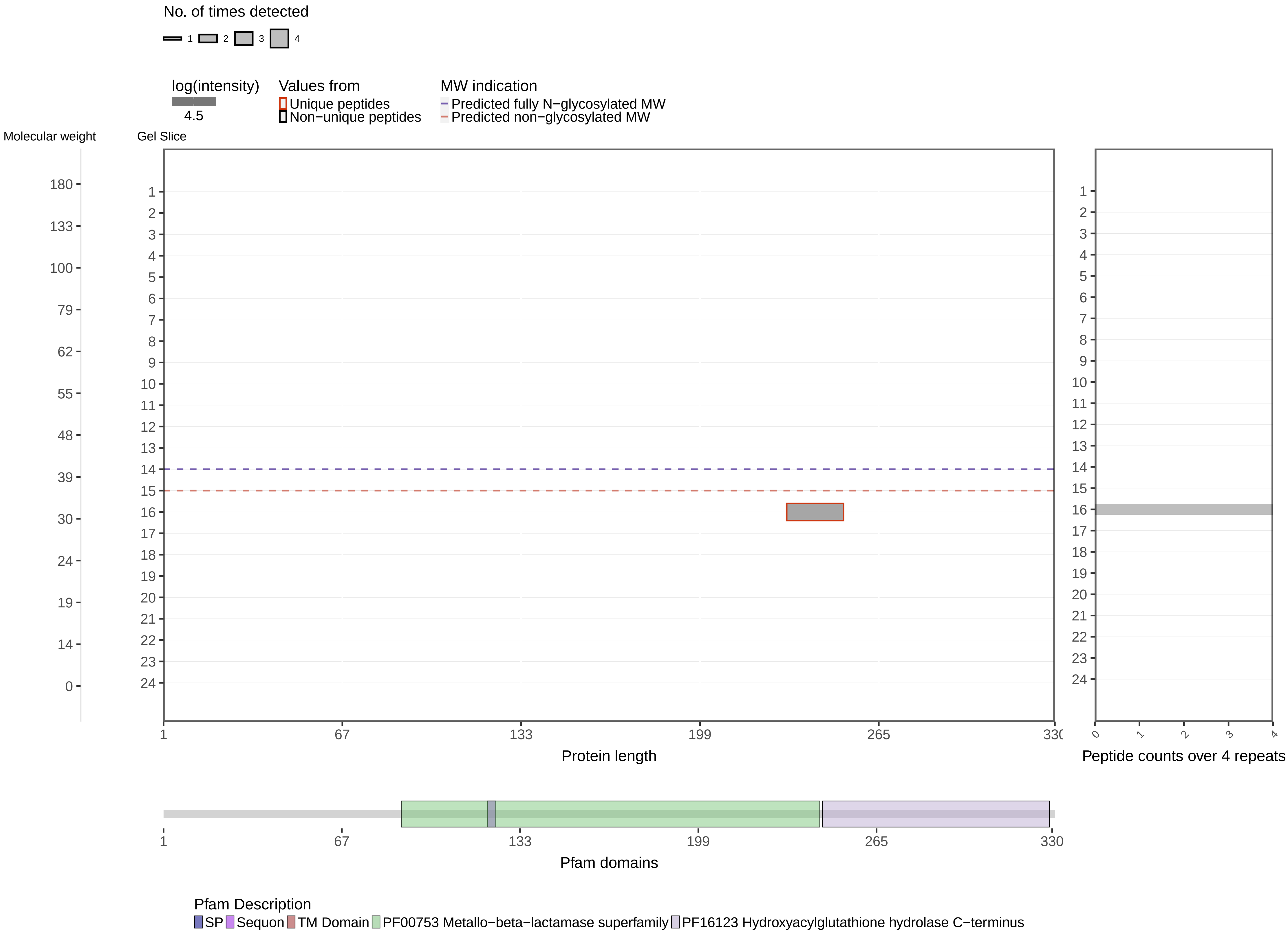


Protein sequence:

SGADSDRIPGIDIALNDGDQWMFAGHEVLVMETPGHTRGHISFYFPRSKAIFTGDTLFSLSCGKLFEGTPQQMLSSLRKIMSLPDDTNVYCGHEYTLTNSKFALSVEPGNEELQSYAAQVVNLCKKGLPTIPTTLKAEKLCNPFLRTSSTEIRKLLNIPPTADDAEALGAIRRAKDTF*

### NbL03g01870.1 30S_ribosomal_protein_S16−like [MW = 14.65 kDa]

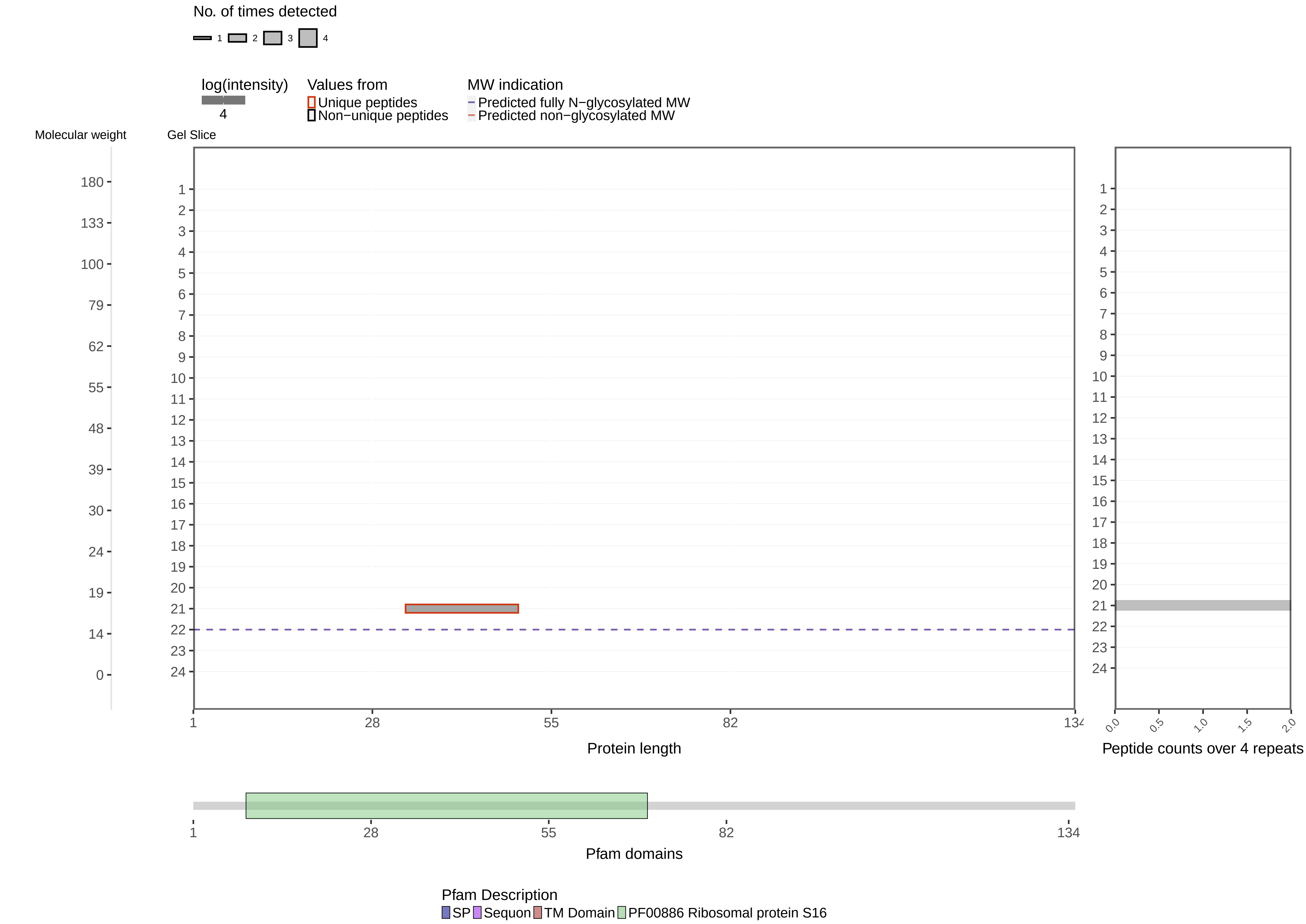


Protein sequence:

LNFDRVKYWLSVGAQPSEPVERLLFRAGVLPPPPMLAMGQKGGPRDTRPVHPMTGRVVAPESVKIADPKVSAKVDGDDKA*

NbL03g01820.1 methyl−CpG−binding_domain−containing_protein_11−like [MW = 37.48 kDa]

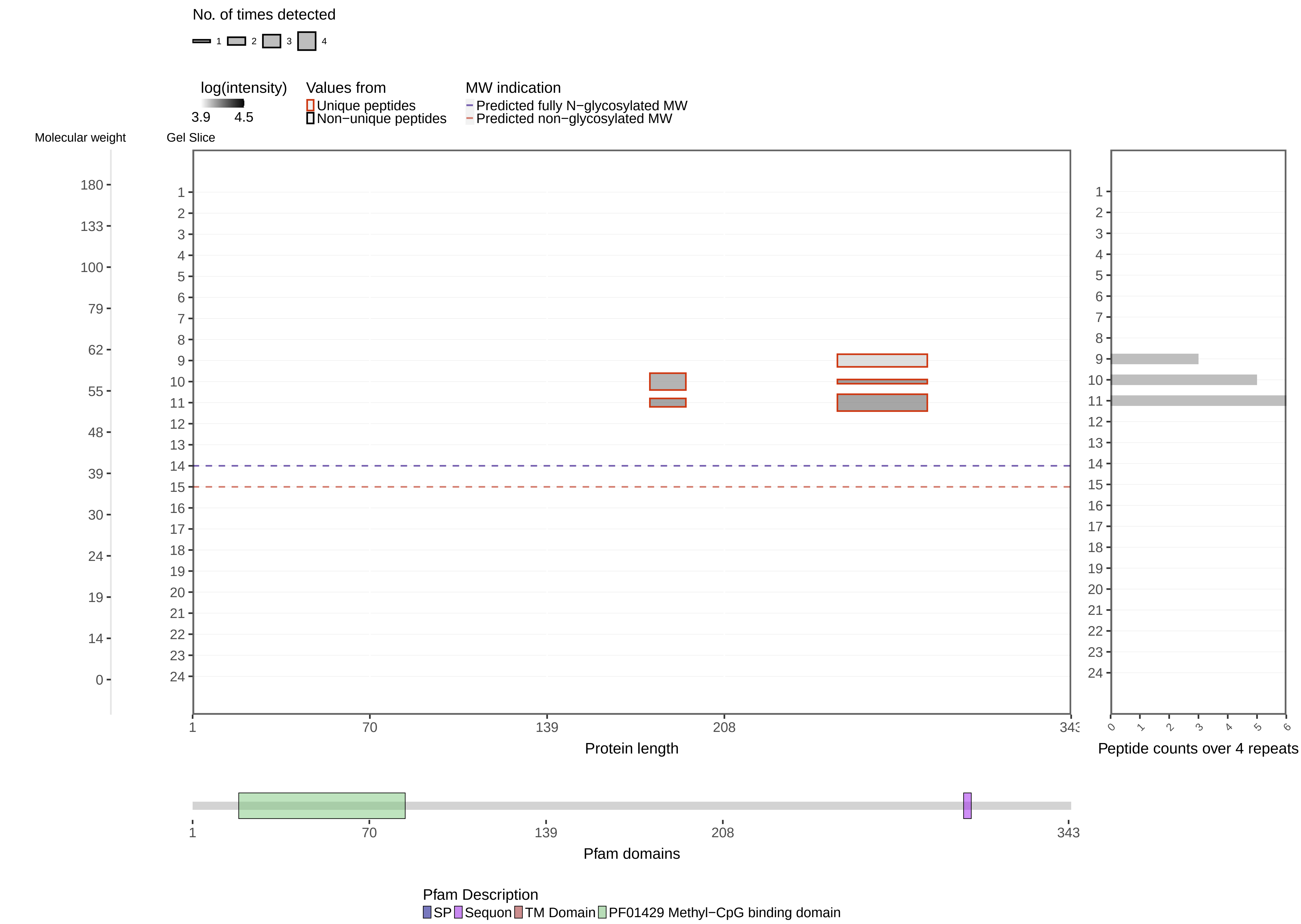


Protein sequence:

NKGKKEVEAADEEAEGMKGAELHPEEKPEMDDKGKKEVEAANEEAEGMKGAELHPEEKPESEPDKVDSADDGKQDKSDNAKIEDKLVSGEKLESLSDKESPAADVNENKPEVVTAEGINEAVGGSSKDVEHAGNGESNNGENRSQEEEKEMKGTDLVMGNNKINQPGPAHSQQHQSPAPISC*

### NbL03g01790.1 phospholipase_D_alpha_1−like [MW = 92.19 kDa]

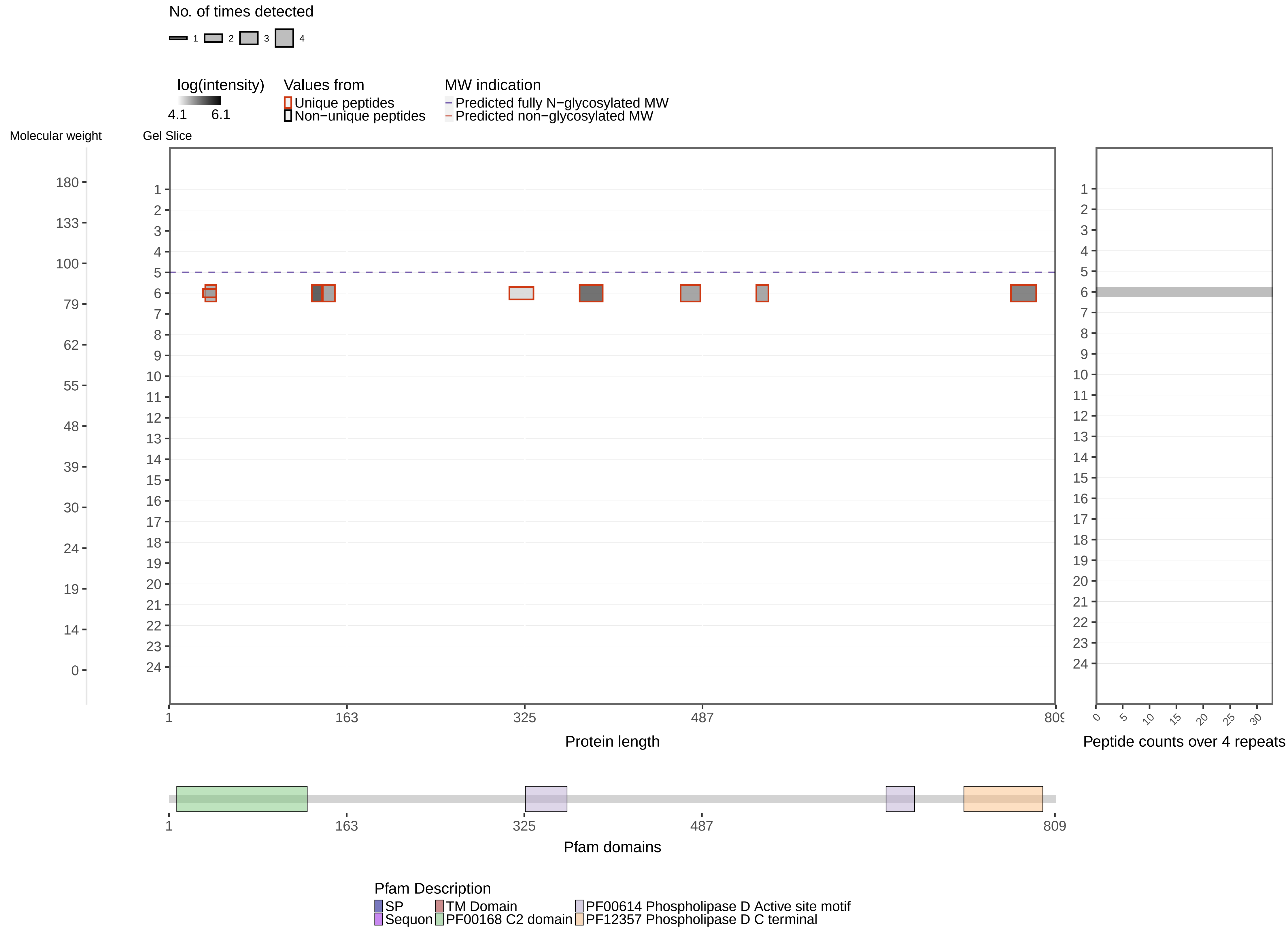


Protein sequence:

SAITKGGPREPWHDIHSRLEGPIAWDVLFNFEQRWRKQGGKDVLVNFRELDDIIIPPSPVMYLDDSETWNVQLFRSIDEGAAFGFPDTPEDAAKAGLVSGKDNIIDRSIQDAYIHAIRRAKNFIYIENQYFLGSSYDWQSDDIKVEEIGALHVIPKELALKIVSKIEAGERFTVYYVVVPMWPEGIPESASVQAILDWQRRTMEMMYKHIVQALNAKGIEEDPRNYLTFFCIGNREVKKSGEYEPSETPEPDSDYMRAQEARRFMIYVHSKMMIVDDEYIIVGSANINQRSMDGARD

NbL03g01770.1 probable_polygalacturonase_At1g80170 [MW = 49.3 kDa]

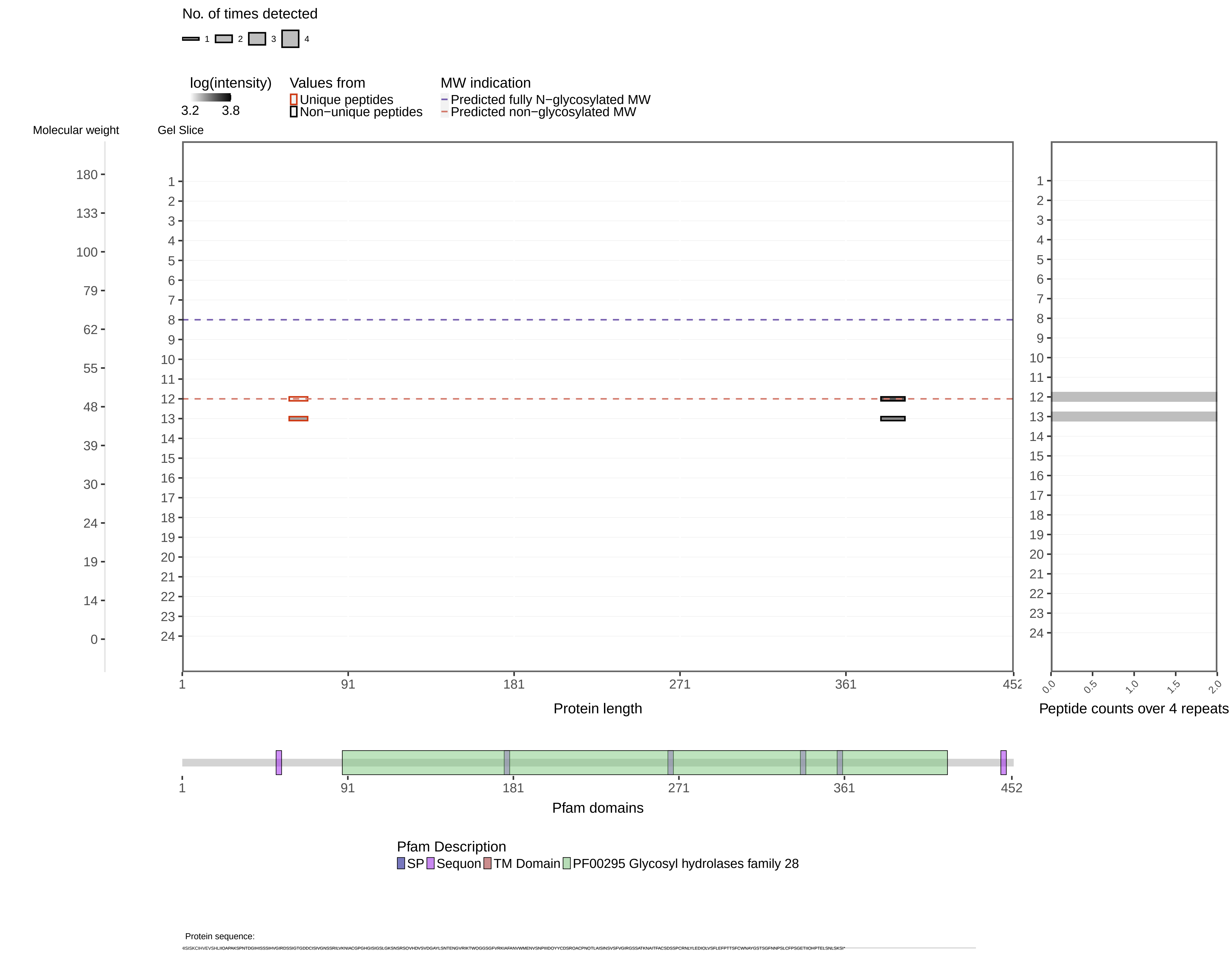

NbL03g01760.1 trafficking_protein_particle_complex_subunit_2−like [MW = 15.35 kDa]

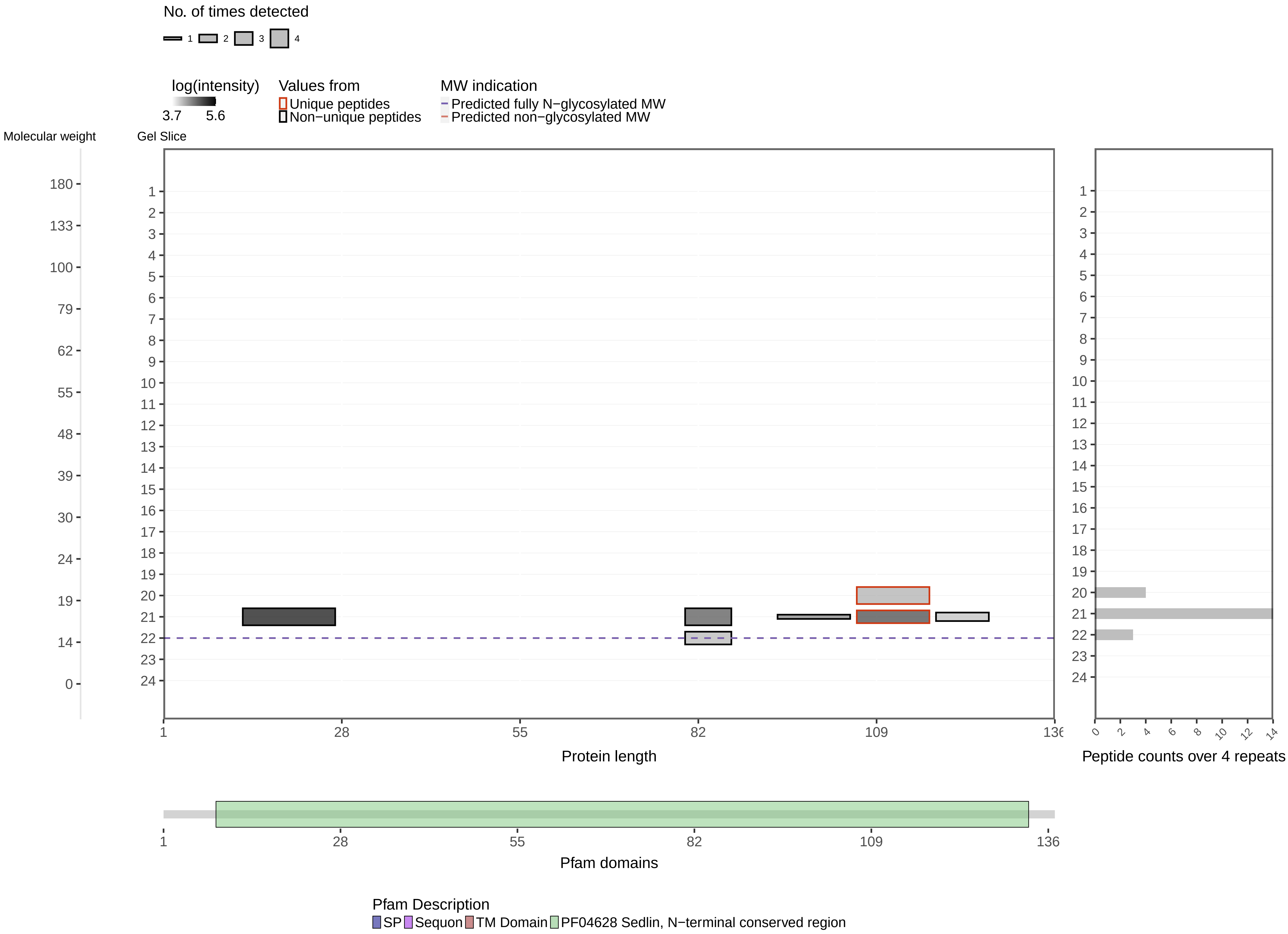


Protein sequence:

AMFLKSIDRFNDLVVSVYVTAGHTRLMLLHDSRNDDGIKSFFQEVHELYIKILLNPLYLPGSRITSSHFDTKVRALARKYL*

### NbL03g01750.1 biotin_carboxyl_carrier_protein_of_acetyl−CoA_carboxylase_1_chloroplastic−like [MW = 29.49 kDa]

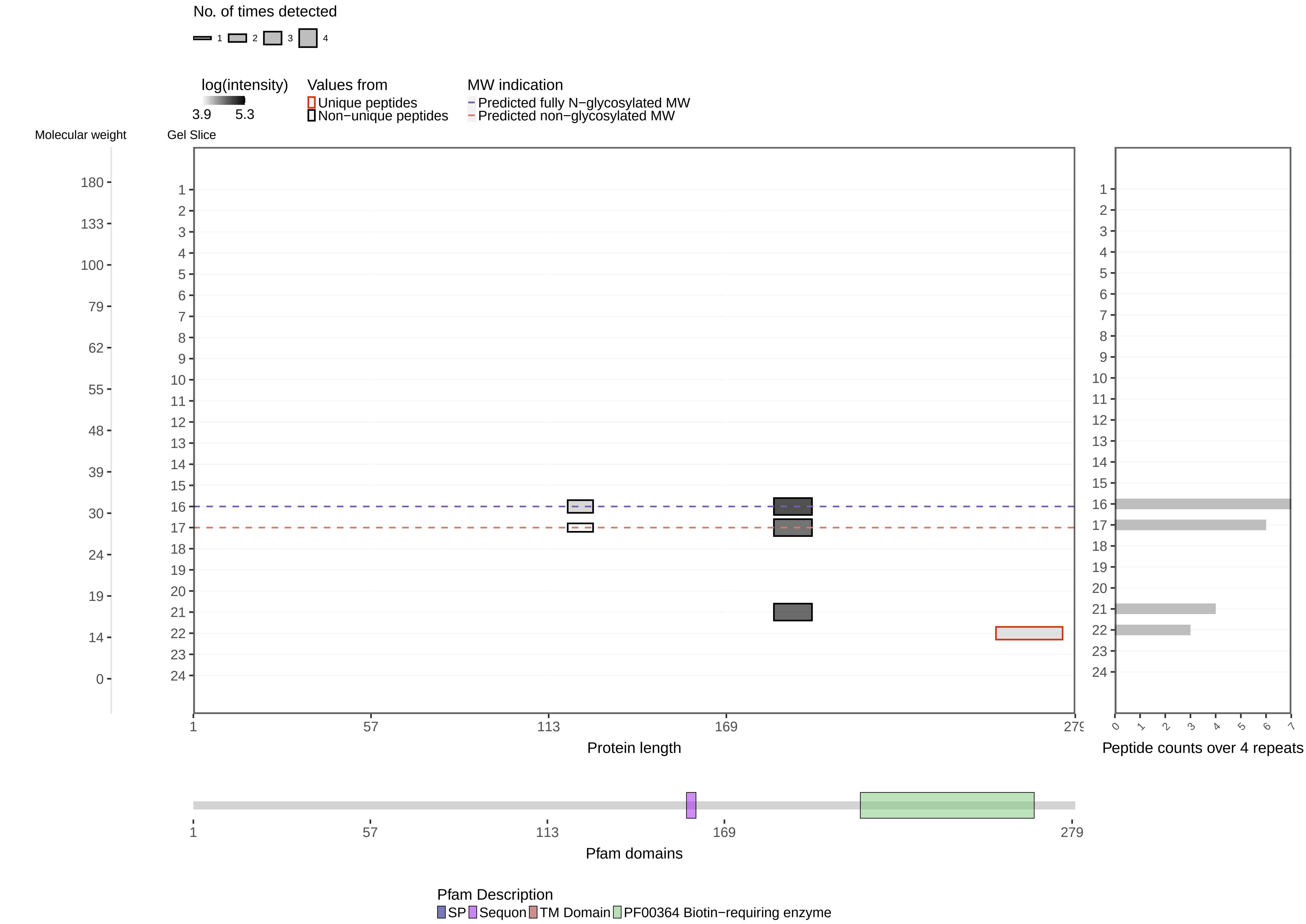


Protein sequence:

/KRDLTGPSTTSLPAISNPVNIHSSVEAADSNGSASSPSLAITKSSPPSDGIRTMIDKAADEGLVIIOSPRVGYFRRSRTIKGKRAPPSCKEKOOVKEGOVVCFIEOLGGELPVESDVSGEVIKILOKDGDPVGYGDPLISILPSFPGIKKLO*

NbL03g01710.1 monothiol_glutaredoxin−S15_mitochondrial−like [MW = 18.96 kDa]

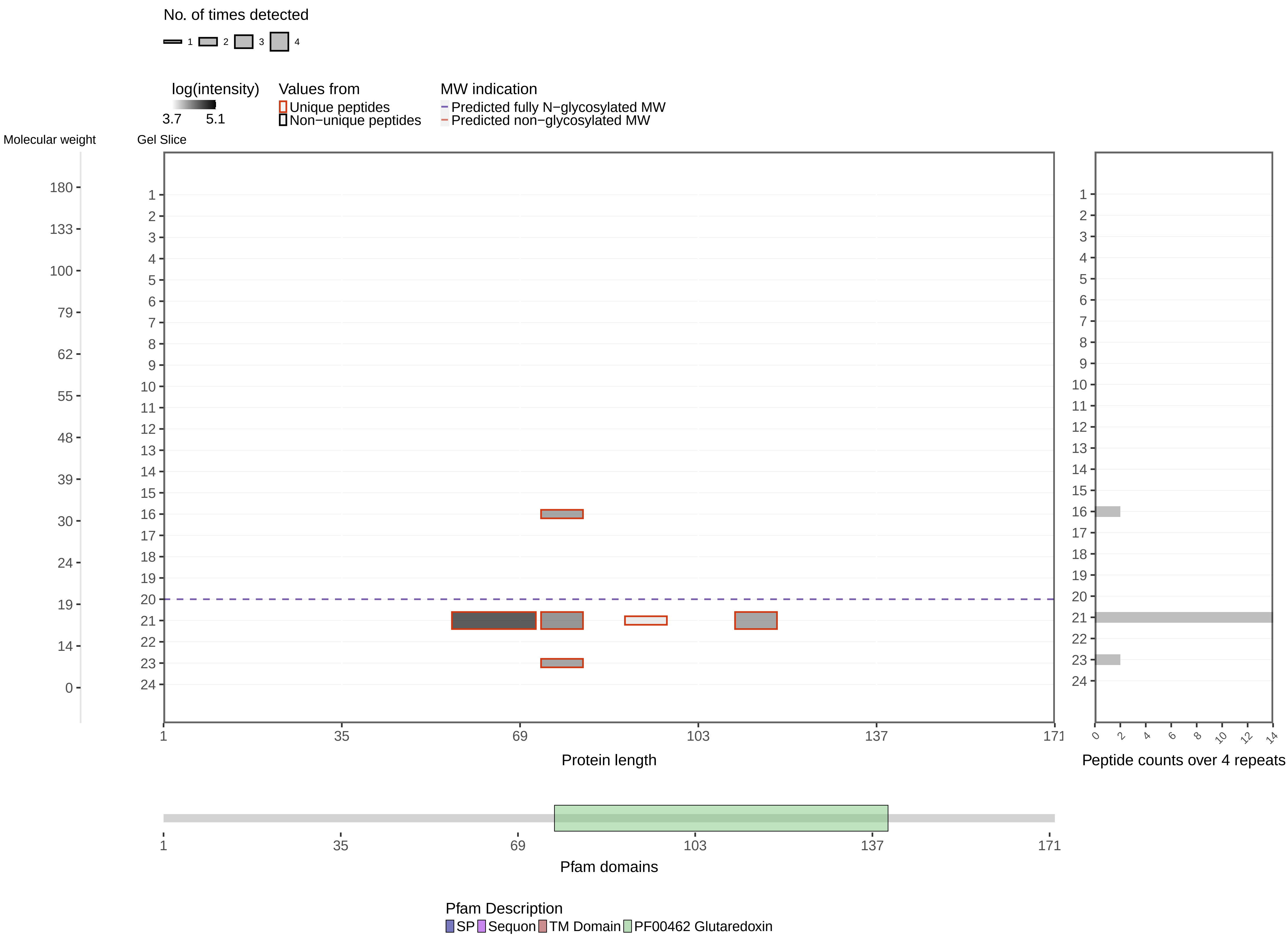


Protein sequence:
KENPVMIYMKGVPELPRCGFSSLAVRVLKEYSVPISARNILEDPELKNAVKAFSHWPTFPQIFINGEFIGGSDIILNMHQSGELKEKLKTIAANQGKAE*

### NbL03g01650.1 uncharacterized_LOC107814272 [MW − SP = 38.87 kDa]

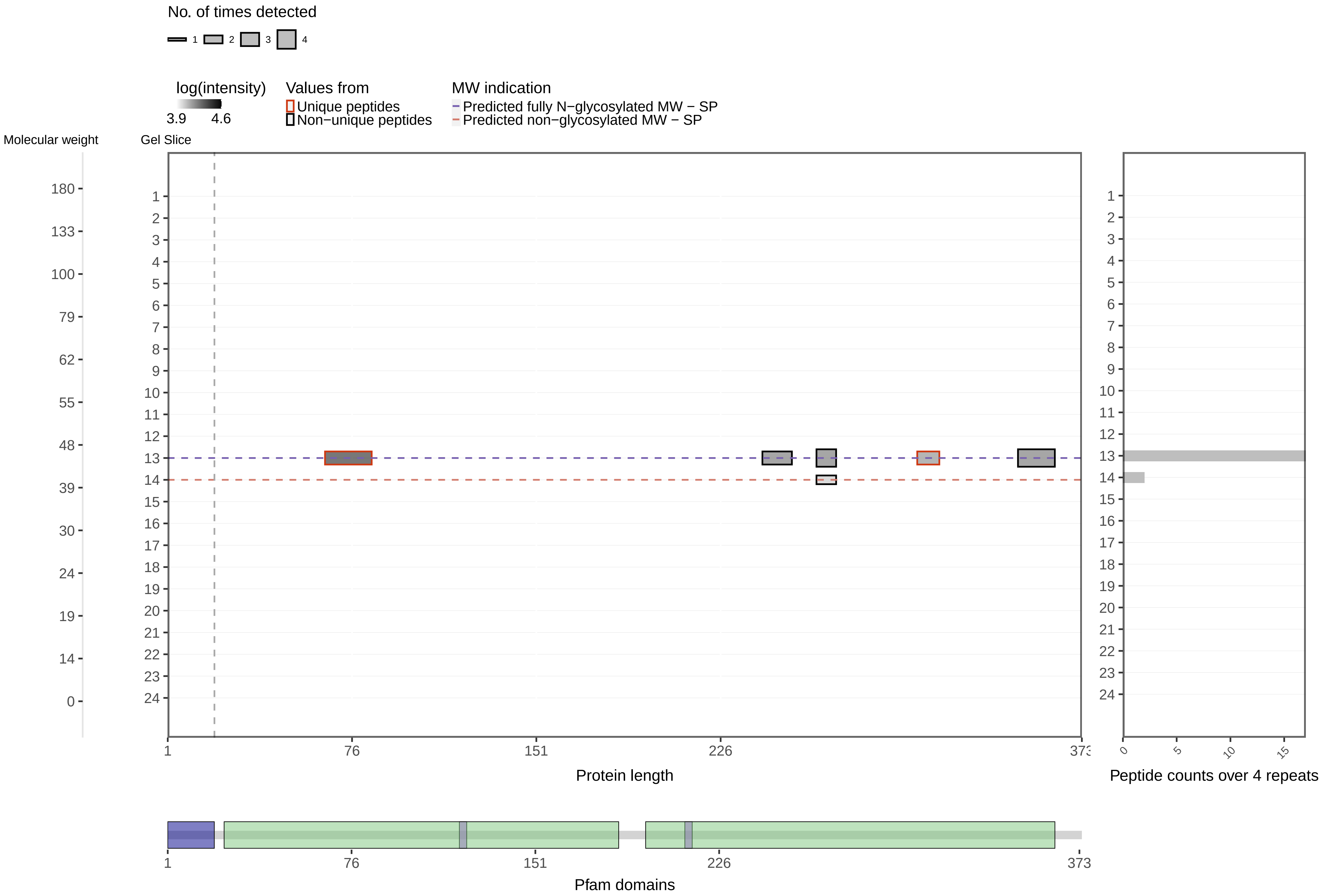


Protein sequence:

PACGPLIDFVALKALKTPKRPKGNMLKNGNFEEGPYIFPNTTWGVLIPPNIEDDHSPLLGWMIESLKAVKYIDAEHFTVPEGKRAVELVAGRESAIAQVVRTRPGKIYDLIFSIGDASNSCEGSMLVEAFAGKNTMQFPYESKGKGGFKRARLRFTAISPRTRVRFLSTYYHMKSDNSGSLCGPVVDDVRLVGVRNPRHP*

### NbL03g01570.1 heat_shock_70_kDa_protein_15−like [MW = 93.68 kDa]

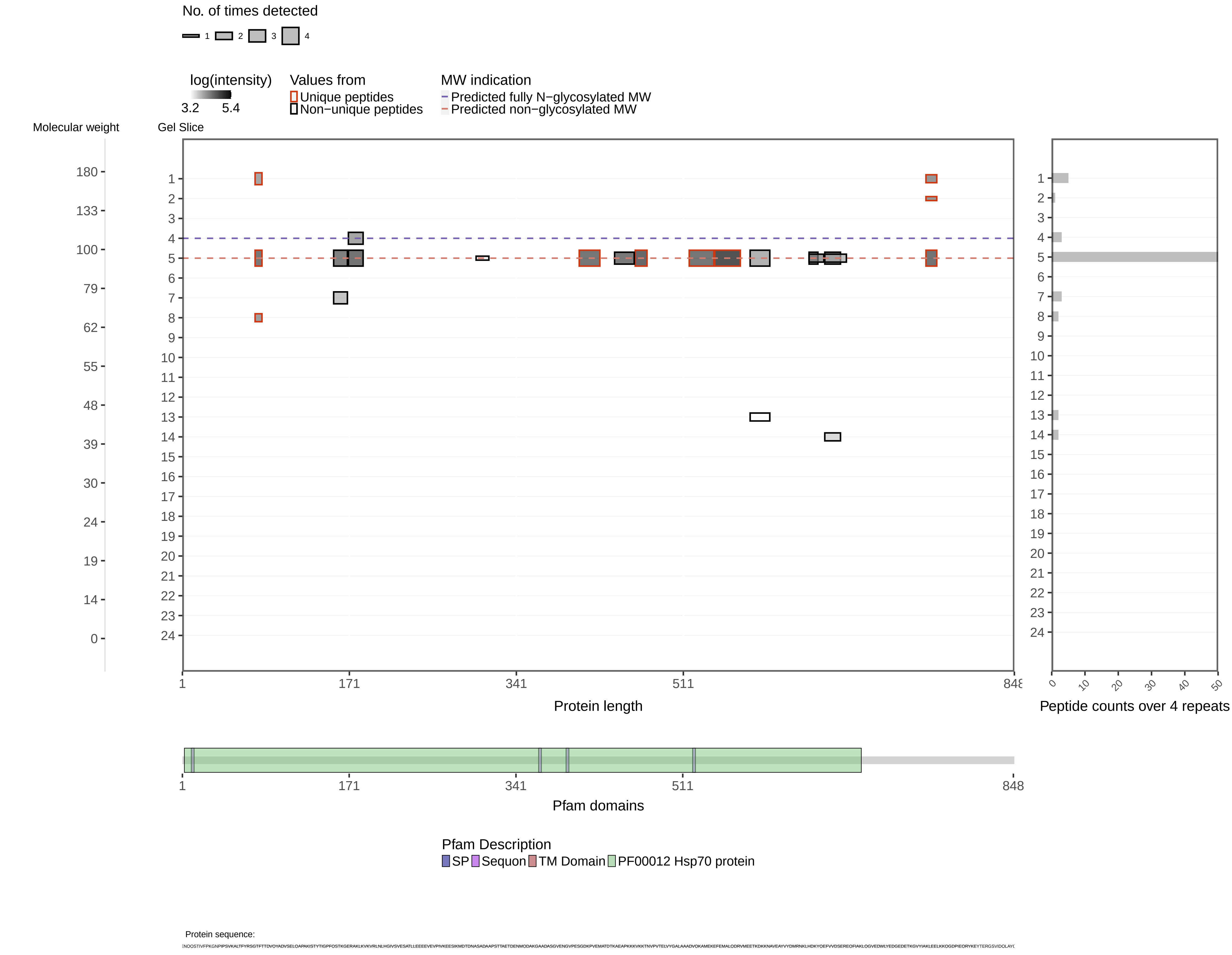

NbL03g01550.1 nuclear_pore_complex_protein_NUP50B−like [MW = 47.73 kDa]

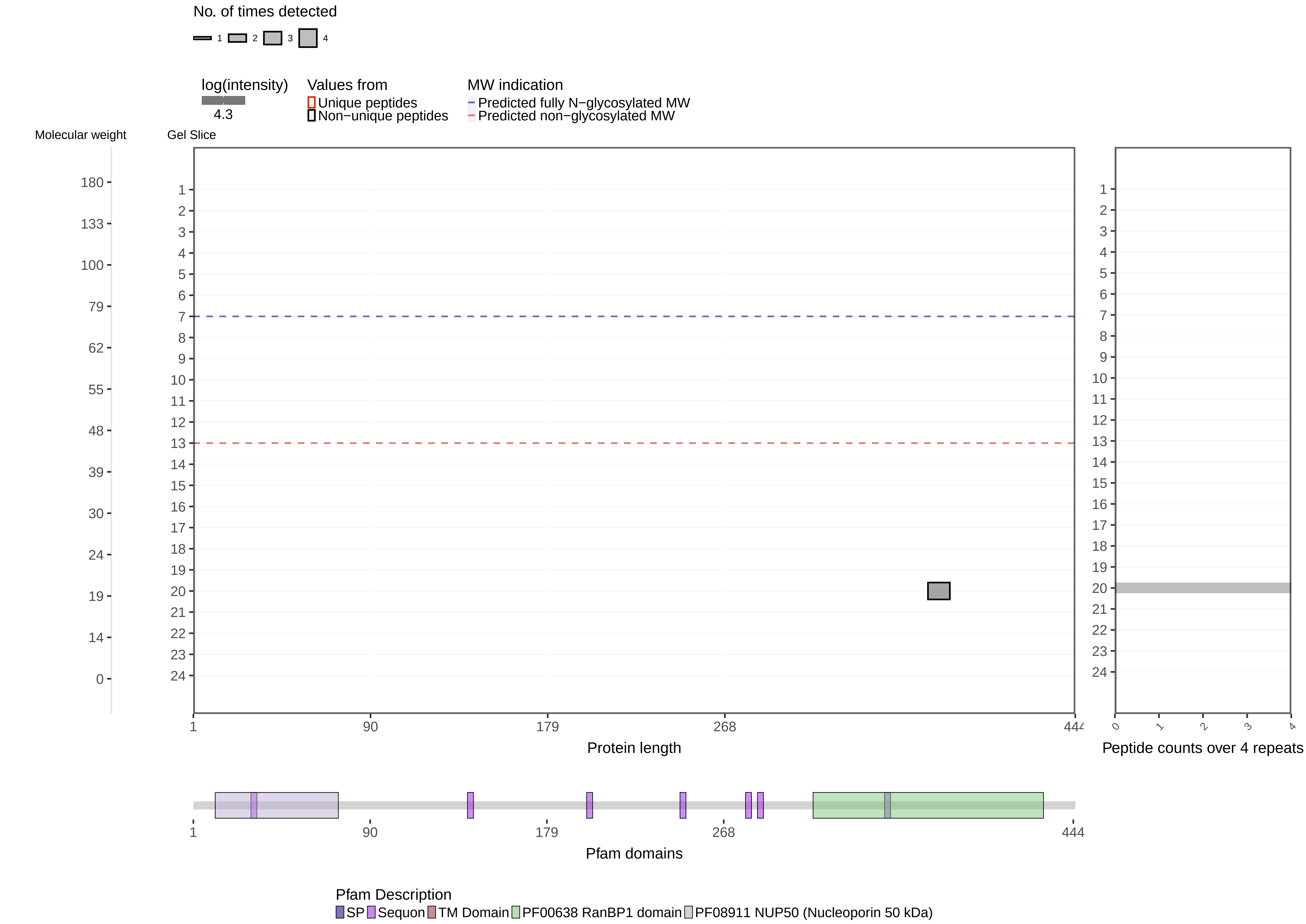


Protein sequence:

SDGENEKGAETASFSSFQQLSSSQNAFTGLAGTGFSNTTFSFGVSSKEGSPLGPGSLFGVKSDQSPFGLNLSTNGNASLFGNSGSSSVNKGEGIGIPSMQEVPVETGEENEKAIFTADSALFEFLNGGWKERGKGELKLNISSTETGKARLVMRTRGNYRLILNANLYPEMKLTNMDKKGVTFACMNSAGDGKDGLSTIALKFKDASIVEEFRAVVMEHKDKMPGSLKTPQNSP*

### NbL03g01480.1 N−alpha−acetyltransferase_16_NatA_auxiliary_subunit [MW = 102.98 kDa]

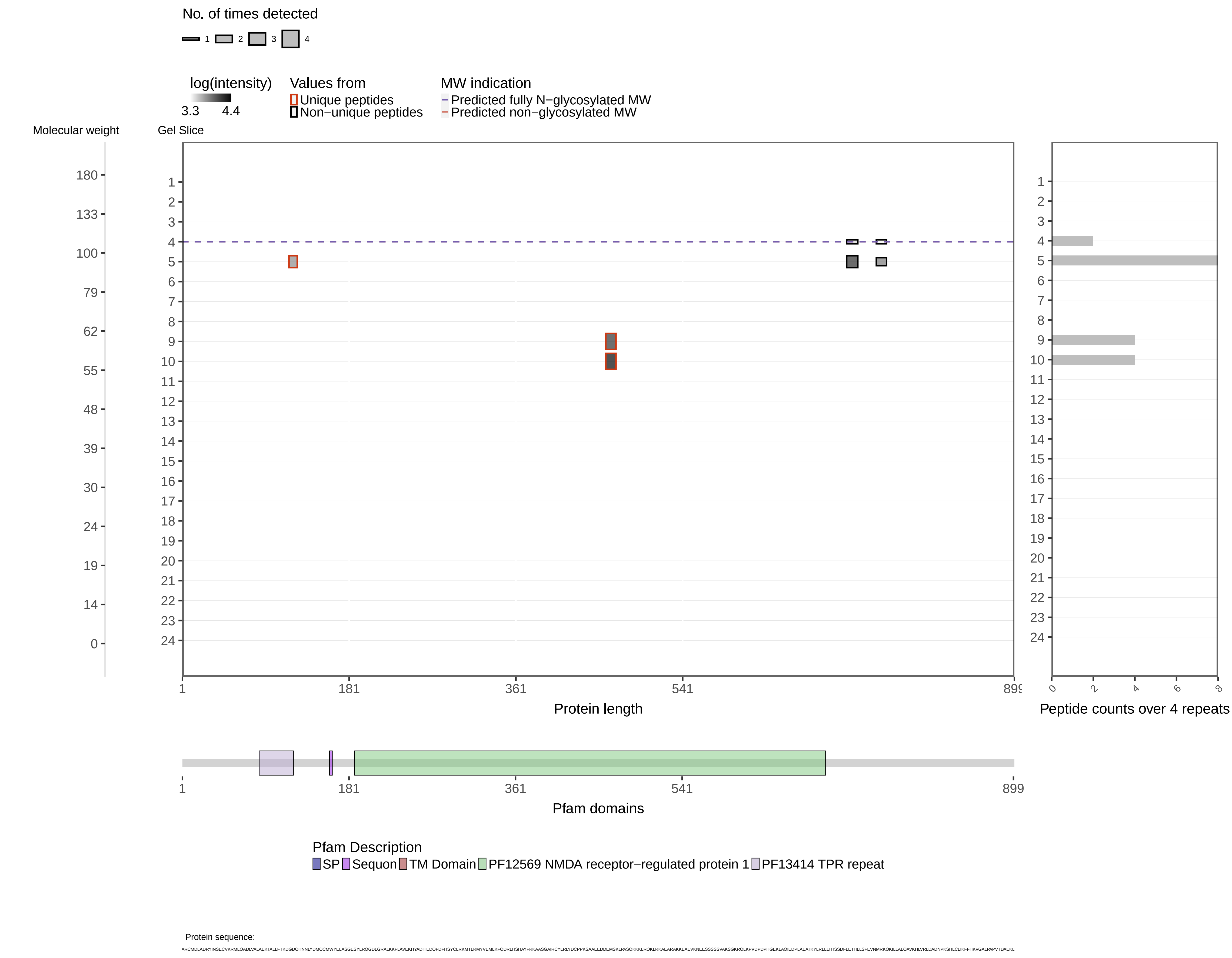

NbL03g01360.1 golgin_candidate_5 [MW = 114.88 kDa]

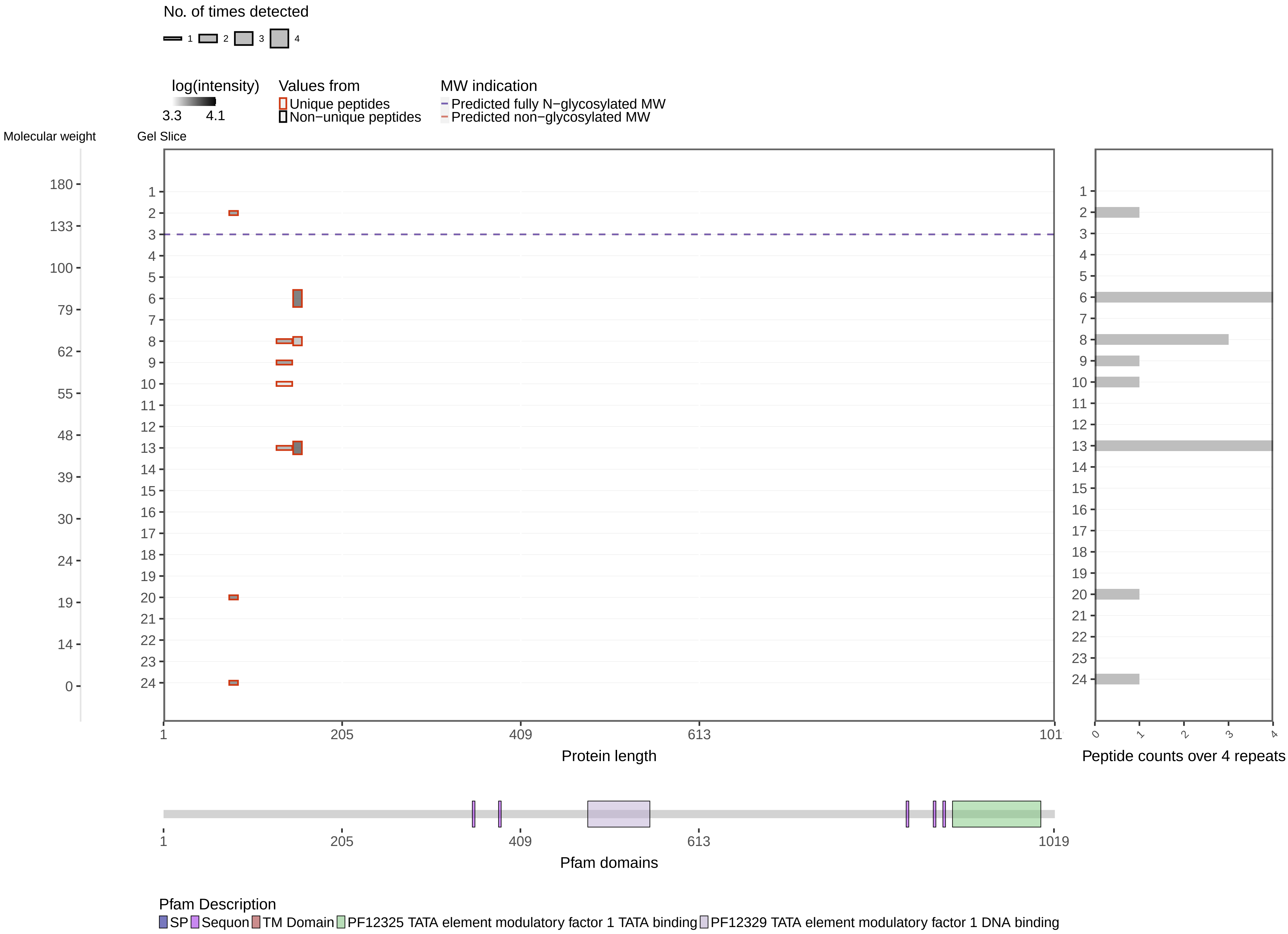


Protein sequence:

DEIITQVMAEGEQLSKKQAAQEAQMRKLRAQIRELEEEKKGLLTKLEVEENKVESIKRDKAATEKLLHETVEKHQAELATQKDYYTNALNAAREAEALAEARANNEARTQLEGRLREAEDREAMLVQALEELRQTLSRTEQQAVFREDMLRRDIEDLQKRYQASERRCEELITQVPESTRPLLRQIEAMQETNARKTEAWAAVERTLNSRLQEAEEKAATAEEKERSISERLSQTLSRINVLEAQISCLRAEQTQLTRSLDKERQRAGENRQEYLALKEEAESNEGRVNQLE

NbL03g01030.1 beta−galactosidase−like [MW = 93.86 kDa]

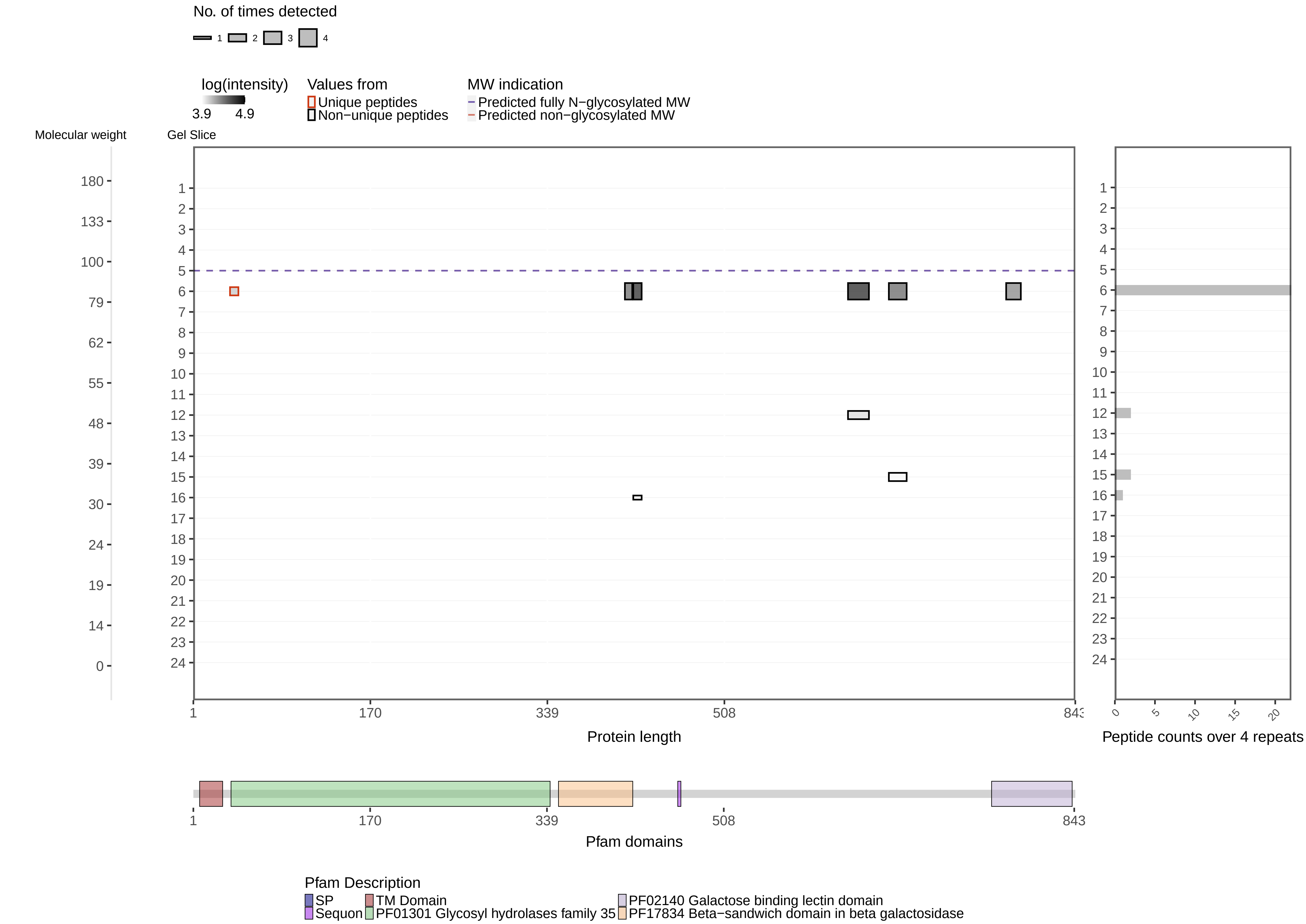


Protein sequence:

[illegible]

### NbL03g01000.1 protein_HOTHEAD−like [MW − SP = 63.77 kDa]

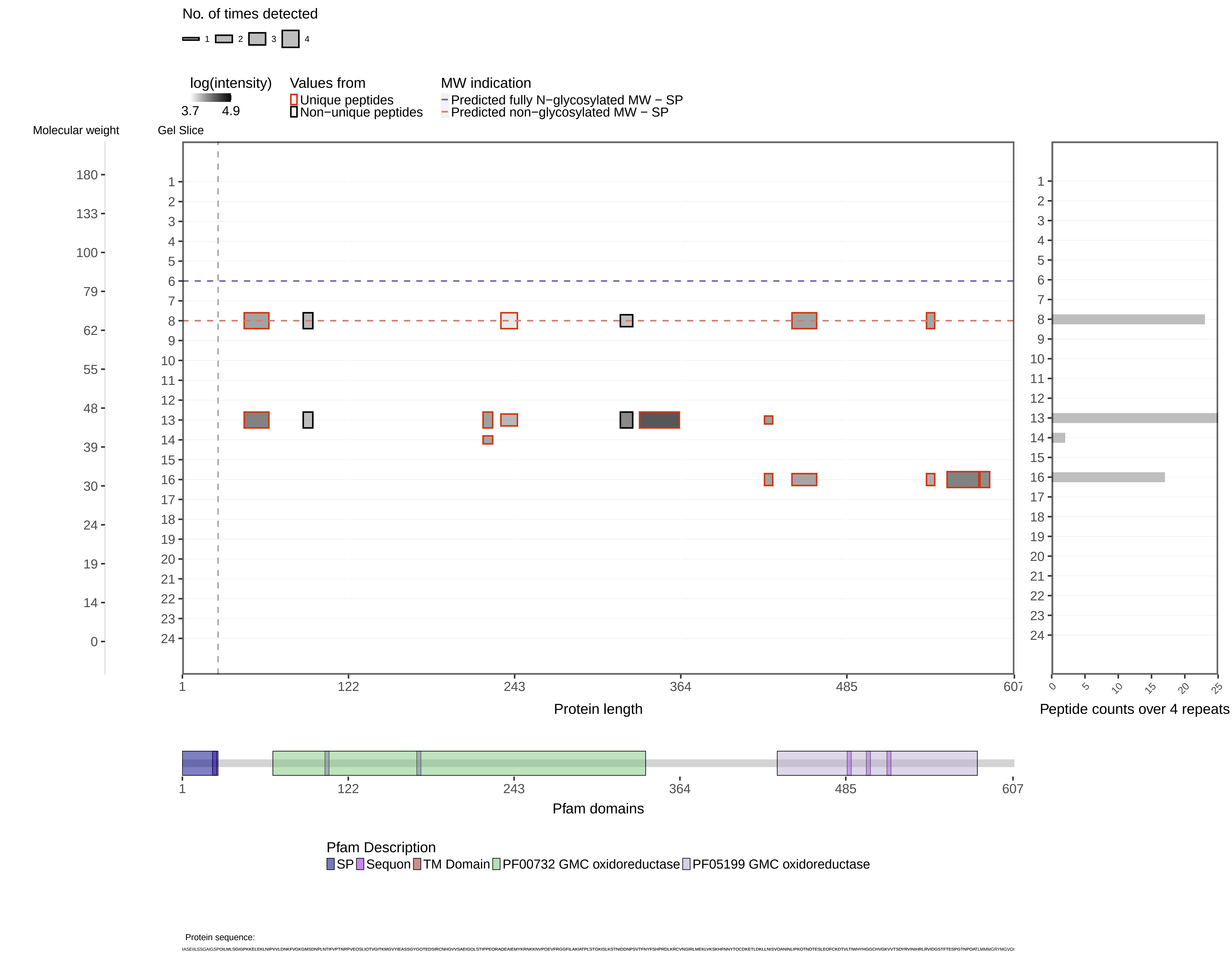

### NbL03g00970.1 beta−galactosidase_17 [MW = 80.87 kDa]

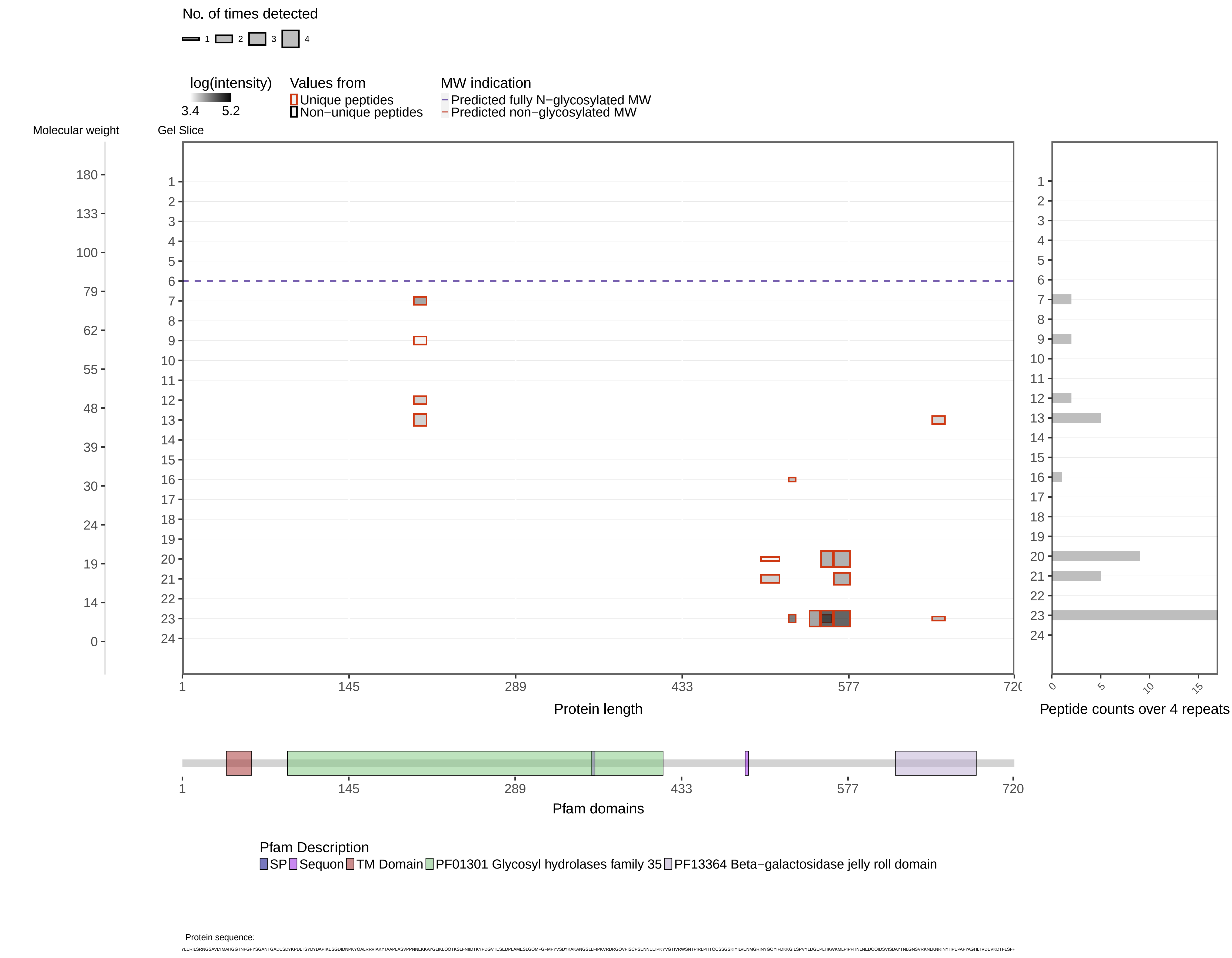

NbL03g00820.1 threonine_synthase_chloroplastic−like [MW = 61.9 kDa]

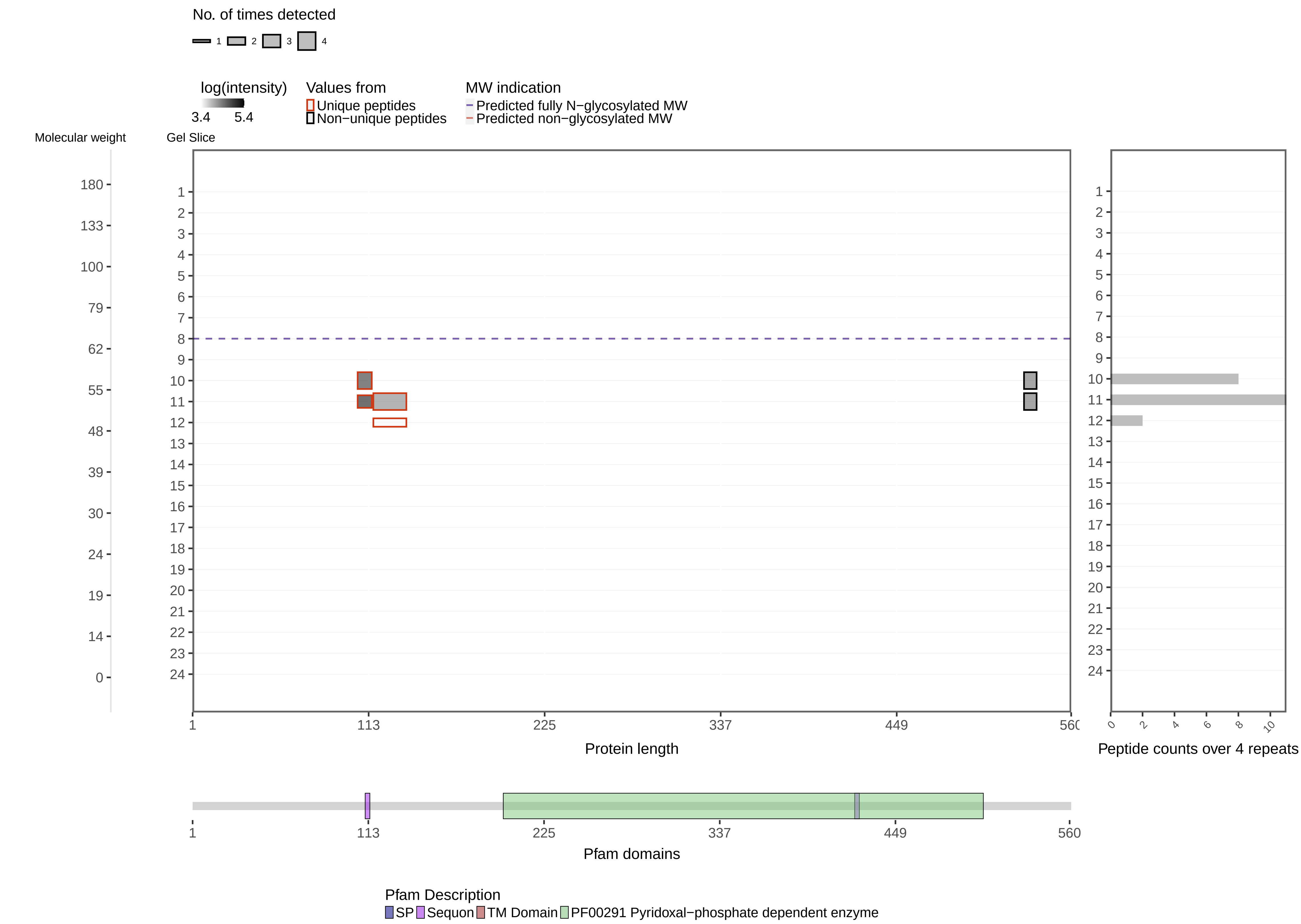


Protein sequence:

STGDTSAALSAYCASAGIPSIVFLPANKISMAQLVQPIANGAFVLSLDTDFDGCMQLIREVTAELPIYLANSLNSLRLEGQKTAAIEILQQFDWEVPDWVIIPGGNLGNIYAFYKGFQMCKELGLVDRIPRLVCAQAANANPLYLHYKSGWKEFKSVKANTTFASAIQIGDPVSIDRAVYALQNSNGIVEEATEEELMDAMAQADSTGMFICPHTGVALTALSKLRKTGVIRPTDRTVVVSTAHGLKFTQSKVDYHSKEIKNMECRFANPPVQVKADFGSVMDVLKKYLLSKNSKF*

### NbL03g00730.1 subtilisin−like_protease_SBT1.4 [MW − SP = 79.85 kDa]

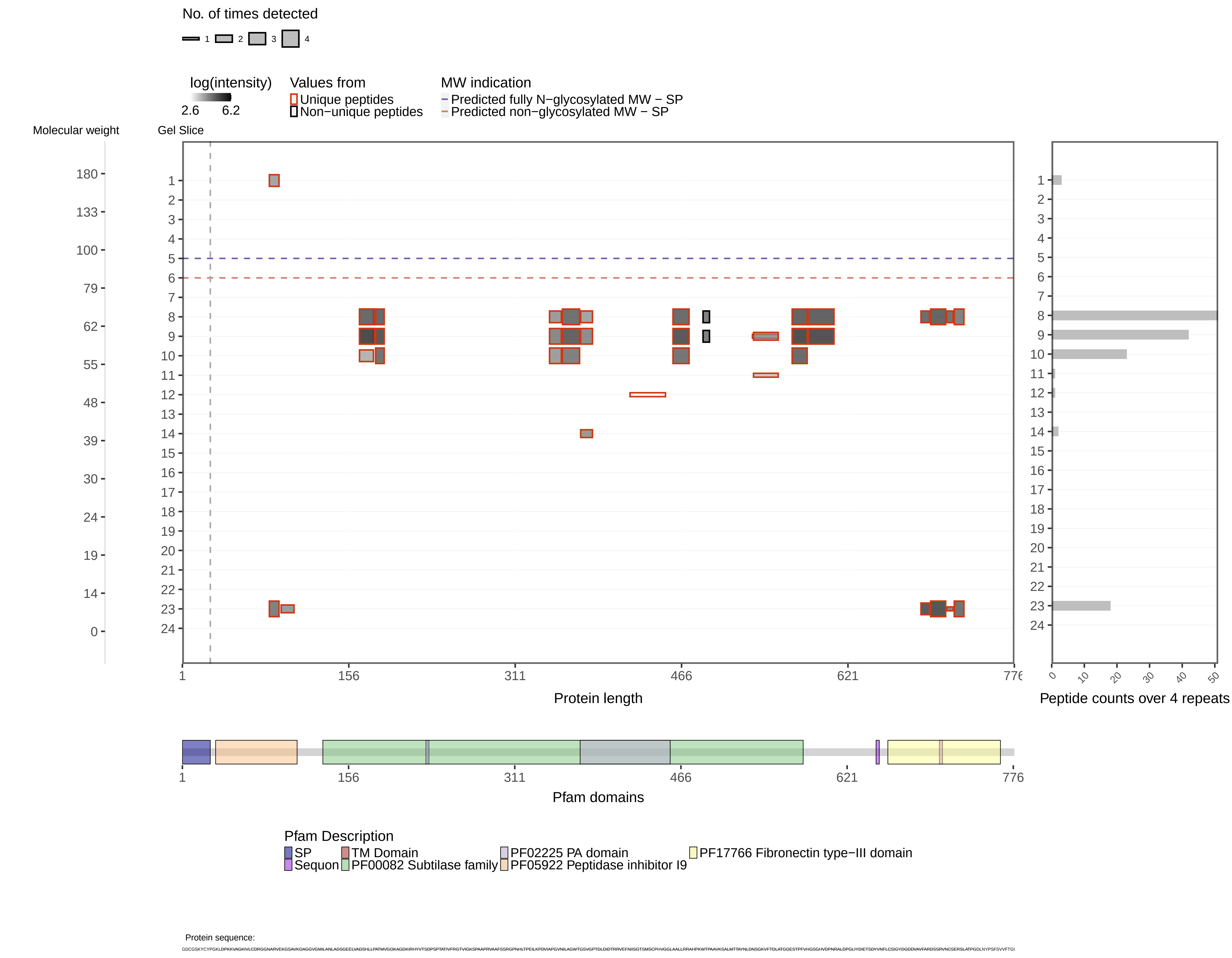

NbL03g00600.1 alanine_aminotransferase_2−like [MW = 58.35 kDa]

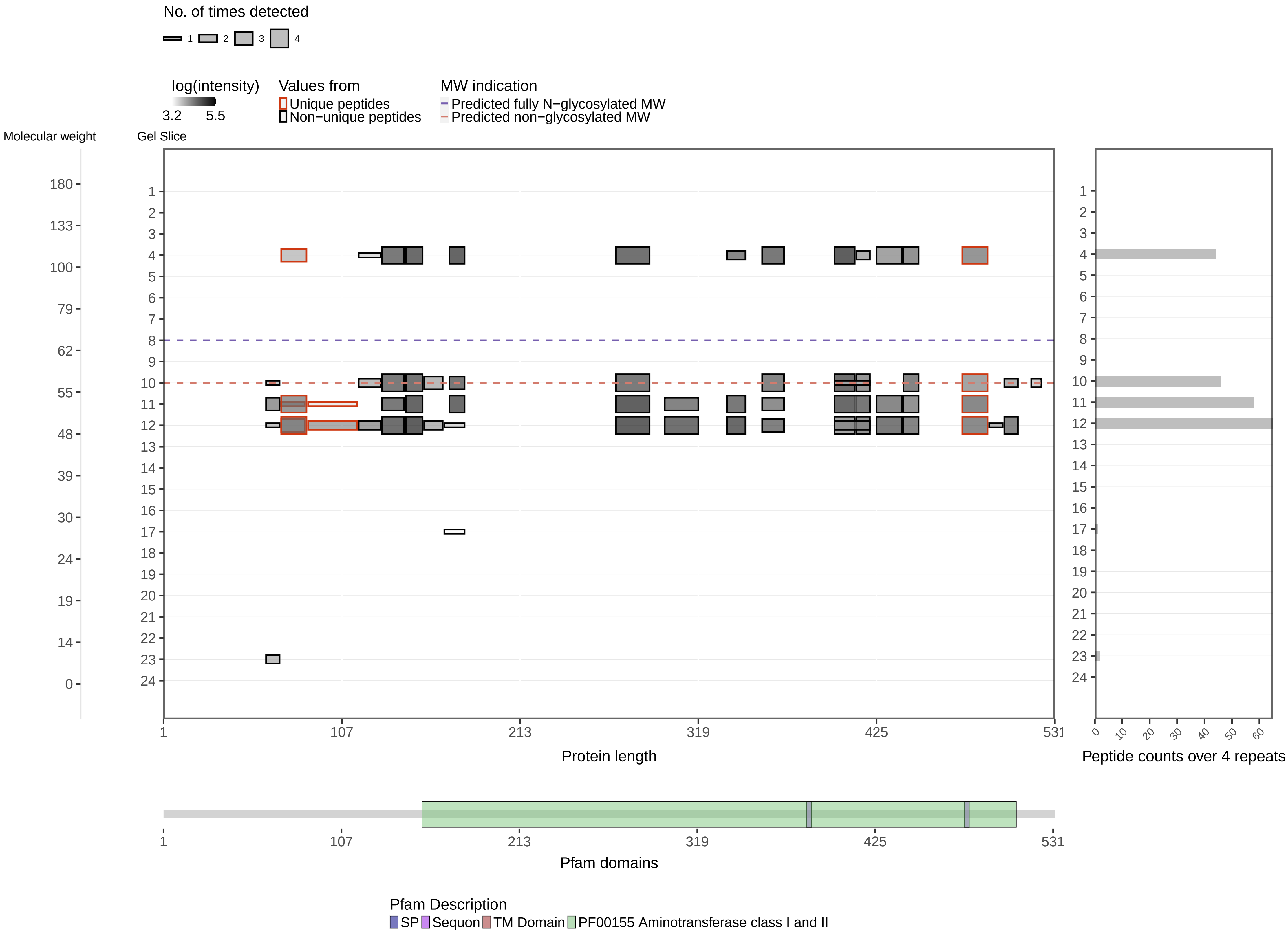


Protein sequence:

EQQLKTAKSKGIDVRALVVINPGNPTGQVLGEANQREIVEFCKREGLVLLADEVYQENVYVPDKKFHSFKKVSRSMGYGEKDISLVSFQSVSKGFYGECGKRGGYMEVTGFSPEVREQIYKVASVNLCSNISGQILASLIMSPPKVGDESYESFSAEKEGILSSLARRAKTLEDALNSLEGVTCNRAEGAMYLFPRINLPDKAIKAAEAAKTAPDAFYAKRLLNATGIVVVPGSGFRQVPGTWHFRCTILPQEEKIPAIVSRLTEFHEKFMNEFRG*

NbL03g00450.1 uncharacterized_LOC107810895 [MW = 44.37 kDa]

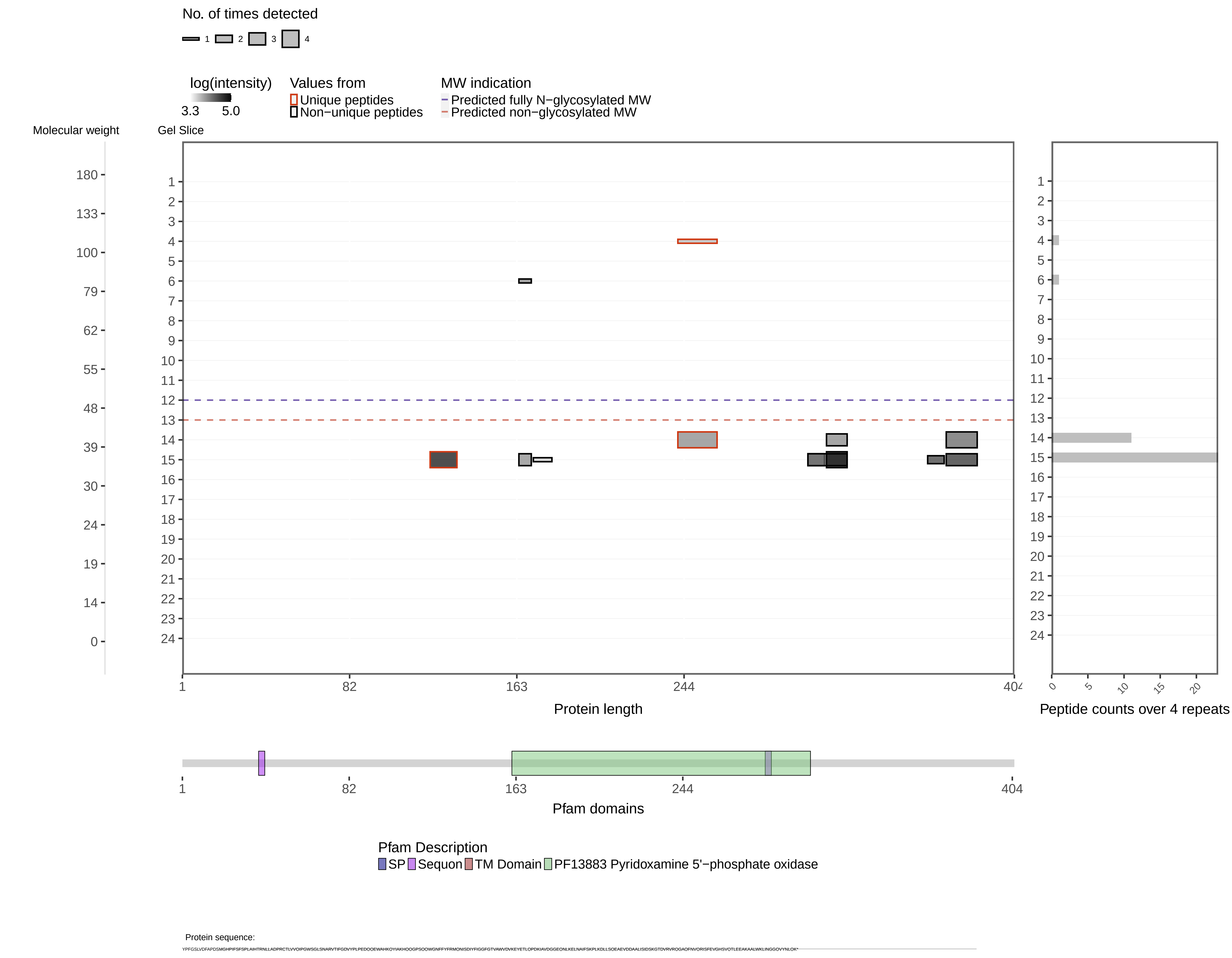

### NbL03g00400.1 V−type_proton_ATPase_catalytic_subunit_A [MW = 49.81 kDa]

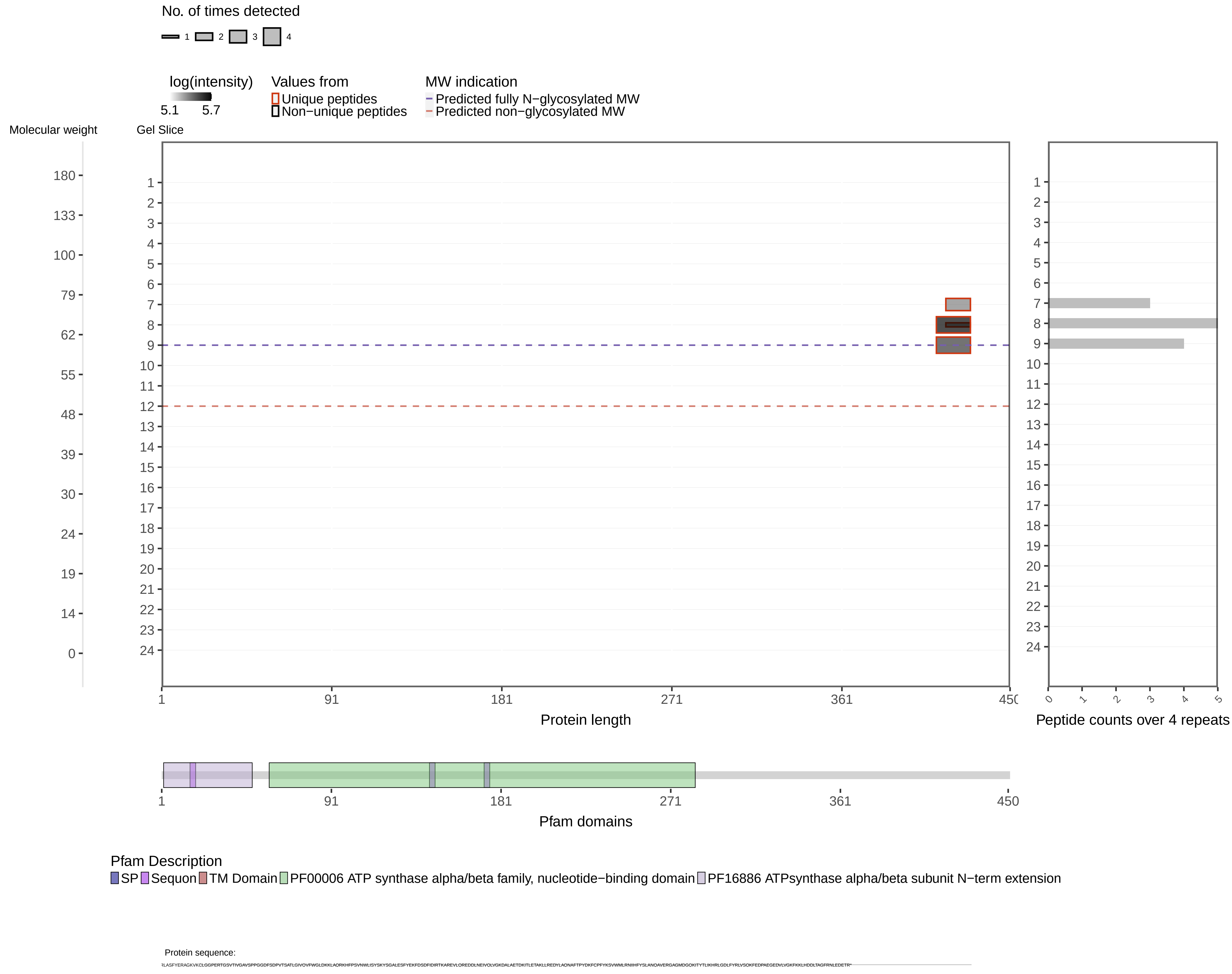

NbL03g00220.1 60S_ribosomal_protein_L15−like [MW = 24.19 kDa]

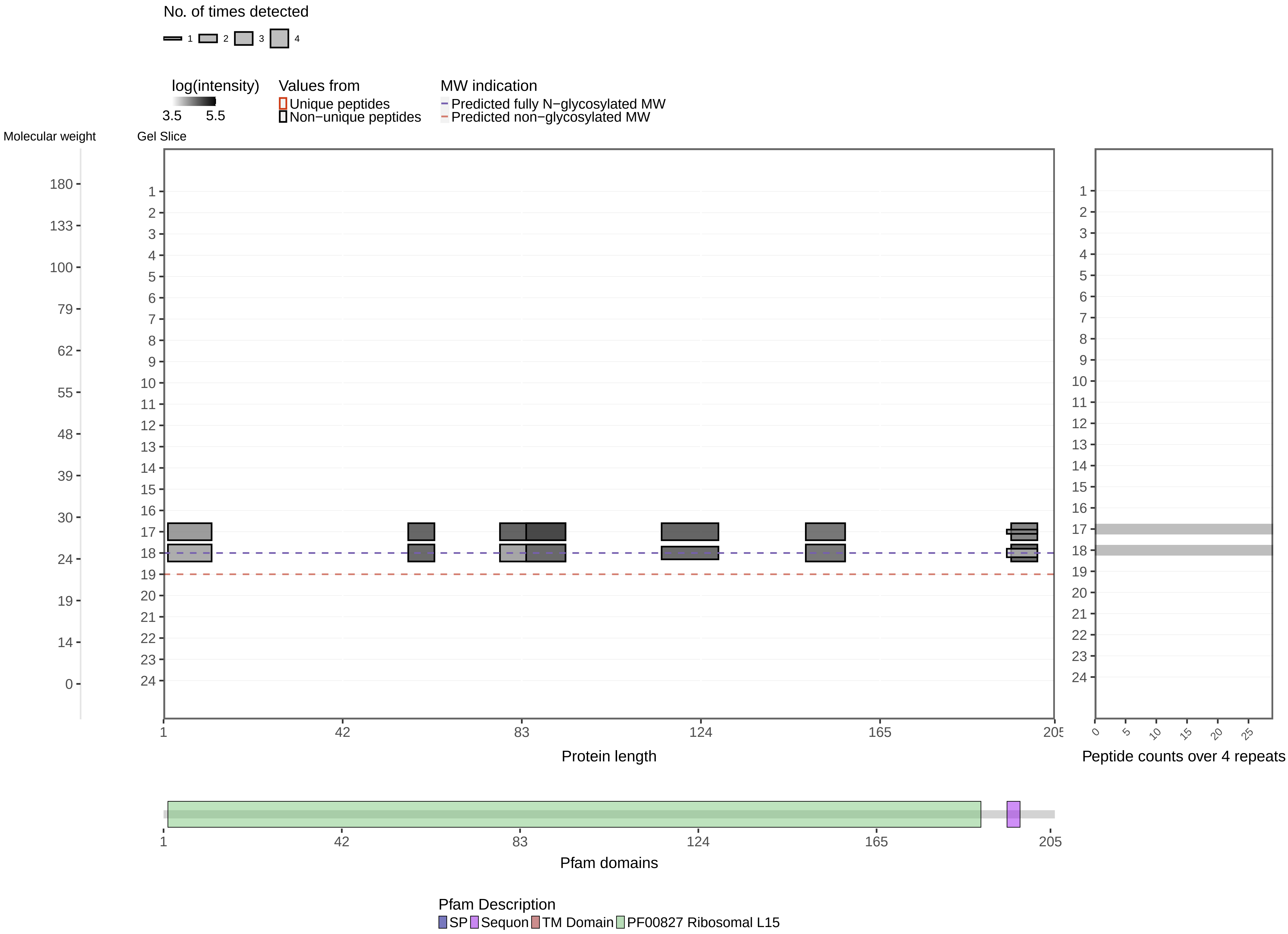


Protein sequence:

VTQLKFQRSKRSVAEERAGRKLSGLKVLNSYWINEDSTYKYFEVILVDAAHAAIRNDPRINWICNPVHKHRELRGLTSAGKKYRGLRGKGHLYHKNRPSRRGTWKRNQTLSLRRYR*

### NbL03g00210.1 protein_ABHD11 [MW = 33.15 kDa]

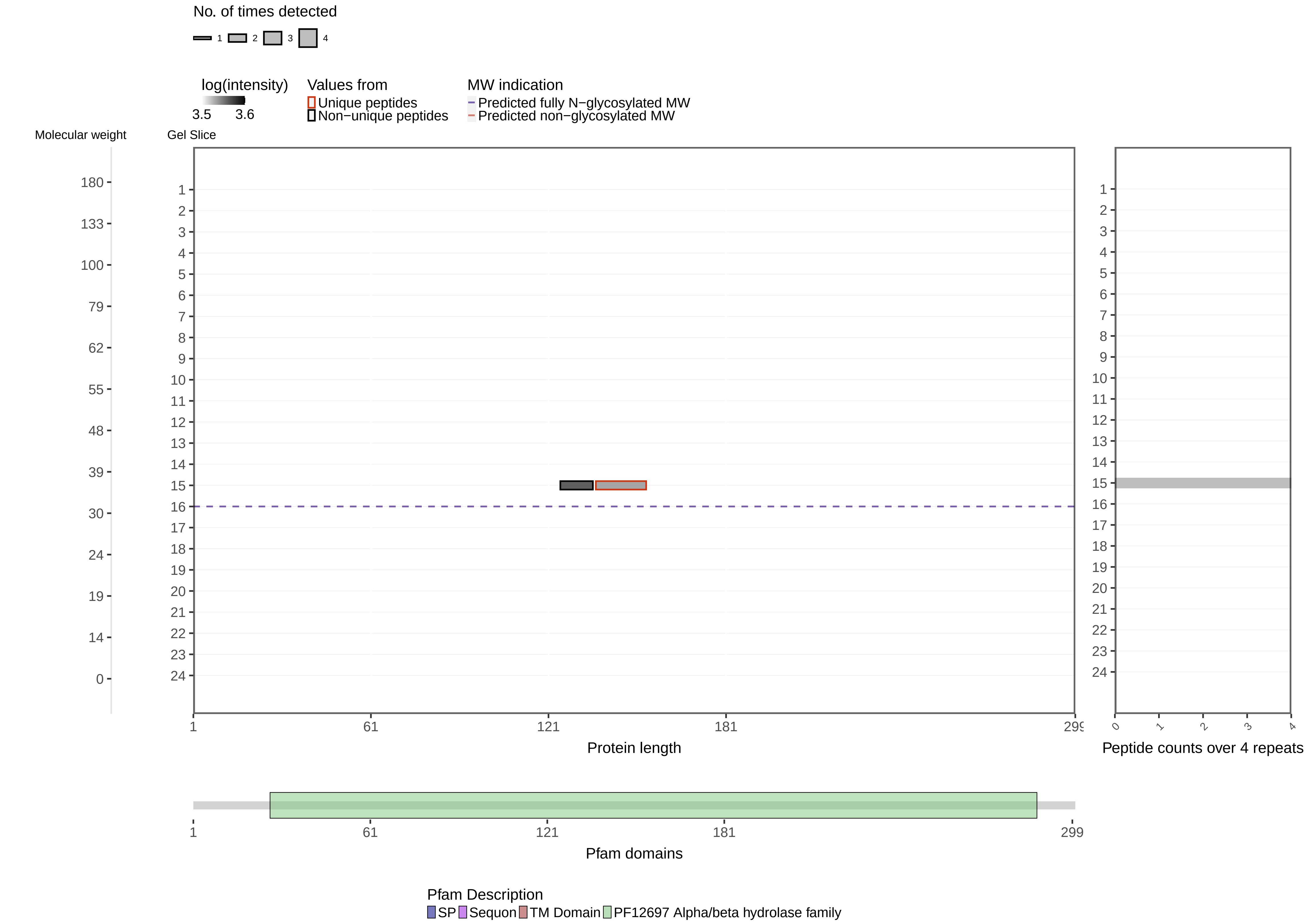


Protein sequence:

[illegible]AGADGDDHPAELISFLSKLPEEVSSKRDIVEALILEGFSRDVAQWVVTNLRQTNPAGSSPSHLSWVFDLKGIAEMYQSYEDTNLWKIVEDVPRGVHVNFLKAERSLHRWALEDIRRIHVAEEQAVEEGGGVEMHVLEDAGHWVHADNPDGLFKILSFSFQGF*

### NbL03g00170.1 uncharacterized_LOC107816116 [MW = 15.55 kDa]

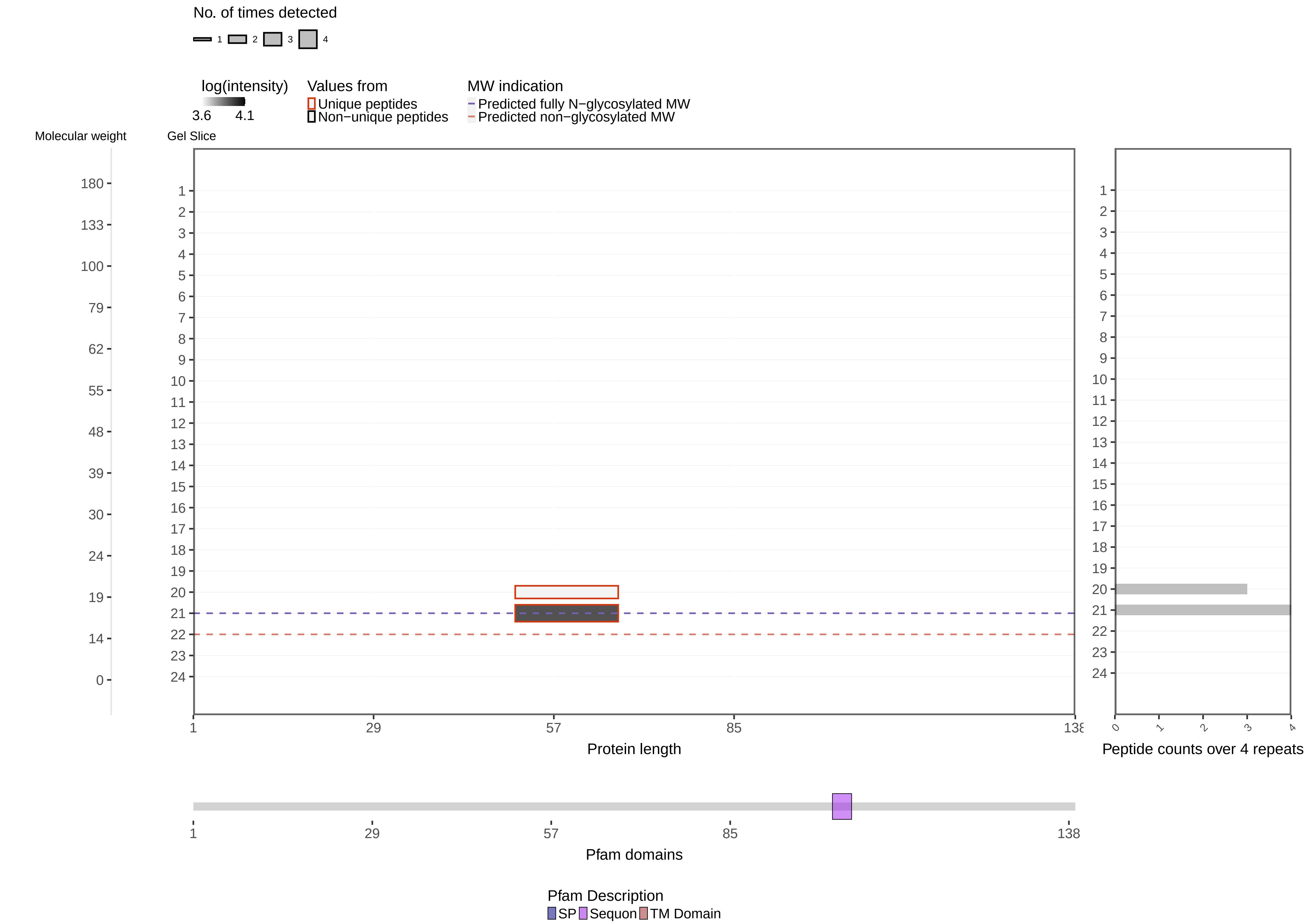


Protein sequence:

IEADDSLEIDPELRYSFQRNFQFLQRVFSIDTVVKPLPPAMAYNVARNLTFFTSIFTQFFDPEGIANAQKSLGLGQEEKVRKVR*

NbL03g00060.1 patellin−3−like [MW = 64.31 kDa]

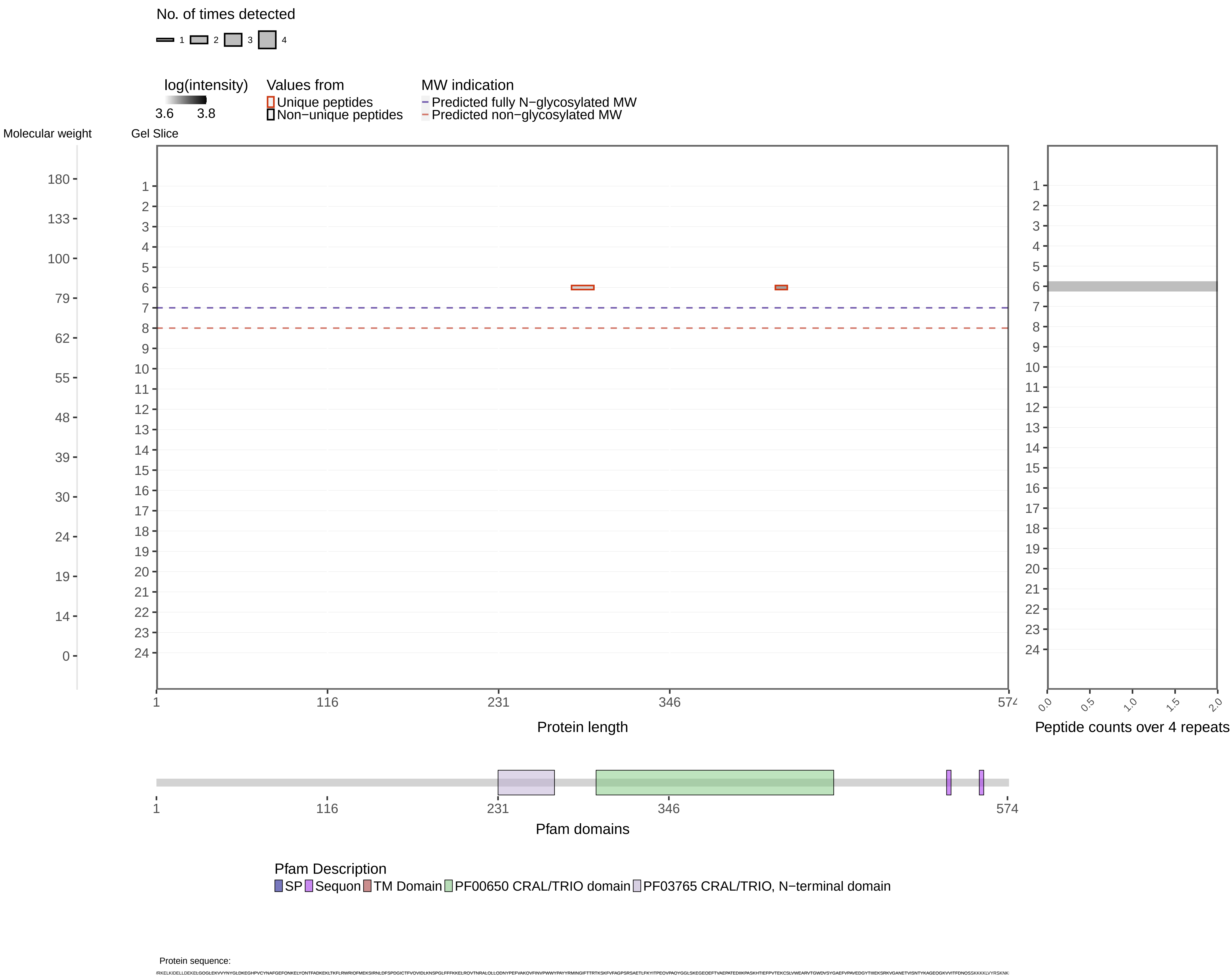

### NbL02g26240.1 nuclear_transport_factor_2−like [MW − SP = 17.75 kDa]

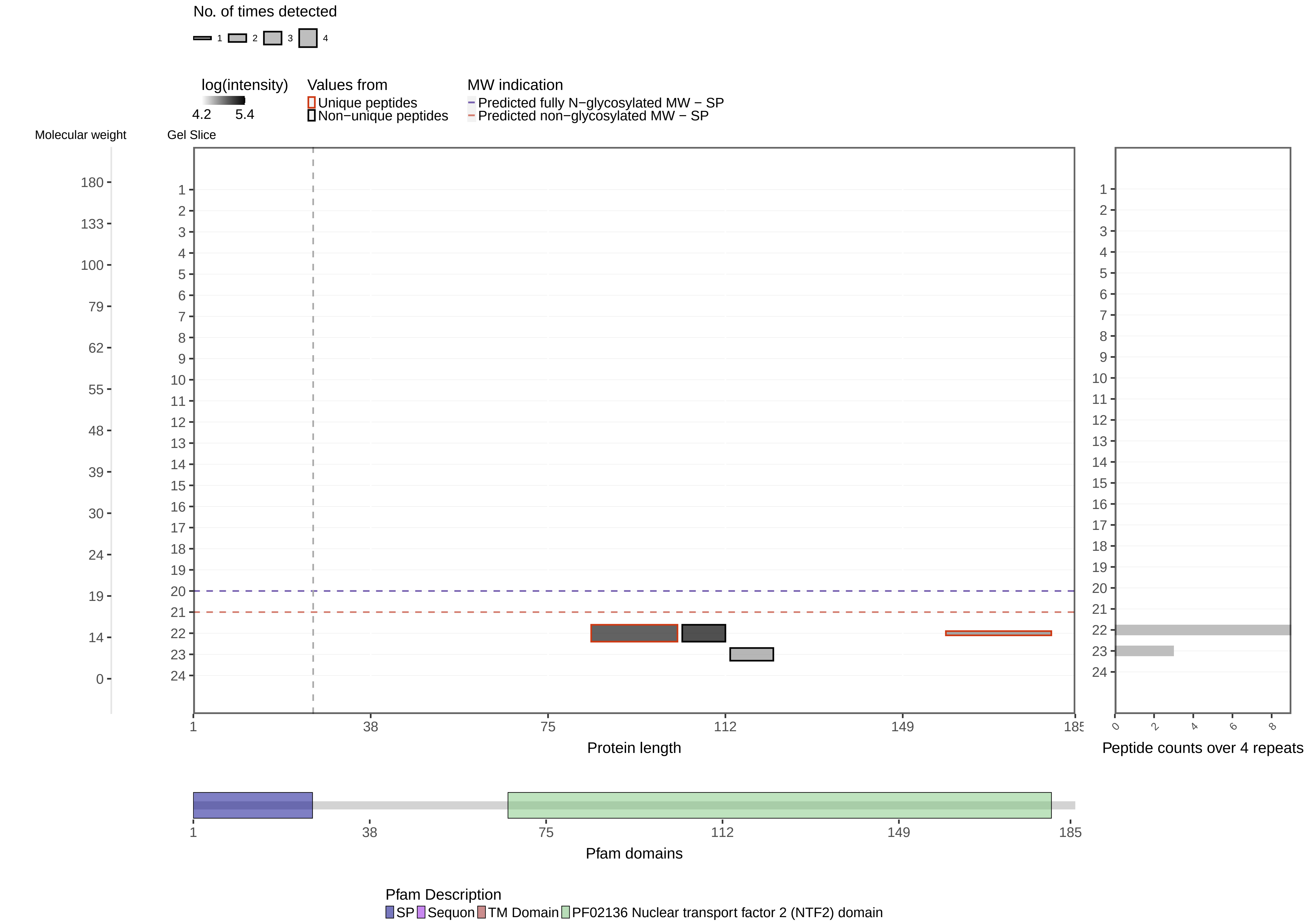


Protein sequence:

NRAGLANLYQENSMLSFEGVRIQGAQHIVAKLTSLPFQQCKHQINTVDCQPSGPIGGMLVFVSGNLQLPGEEHVLKFSQMFHLMPTQQGSFYVMNDIFRLNYA*

NbL02g26180.1 sulfite_oxidase [MW = 43.51 kDa]

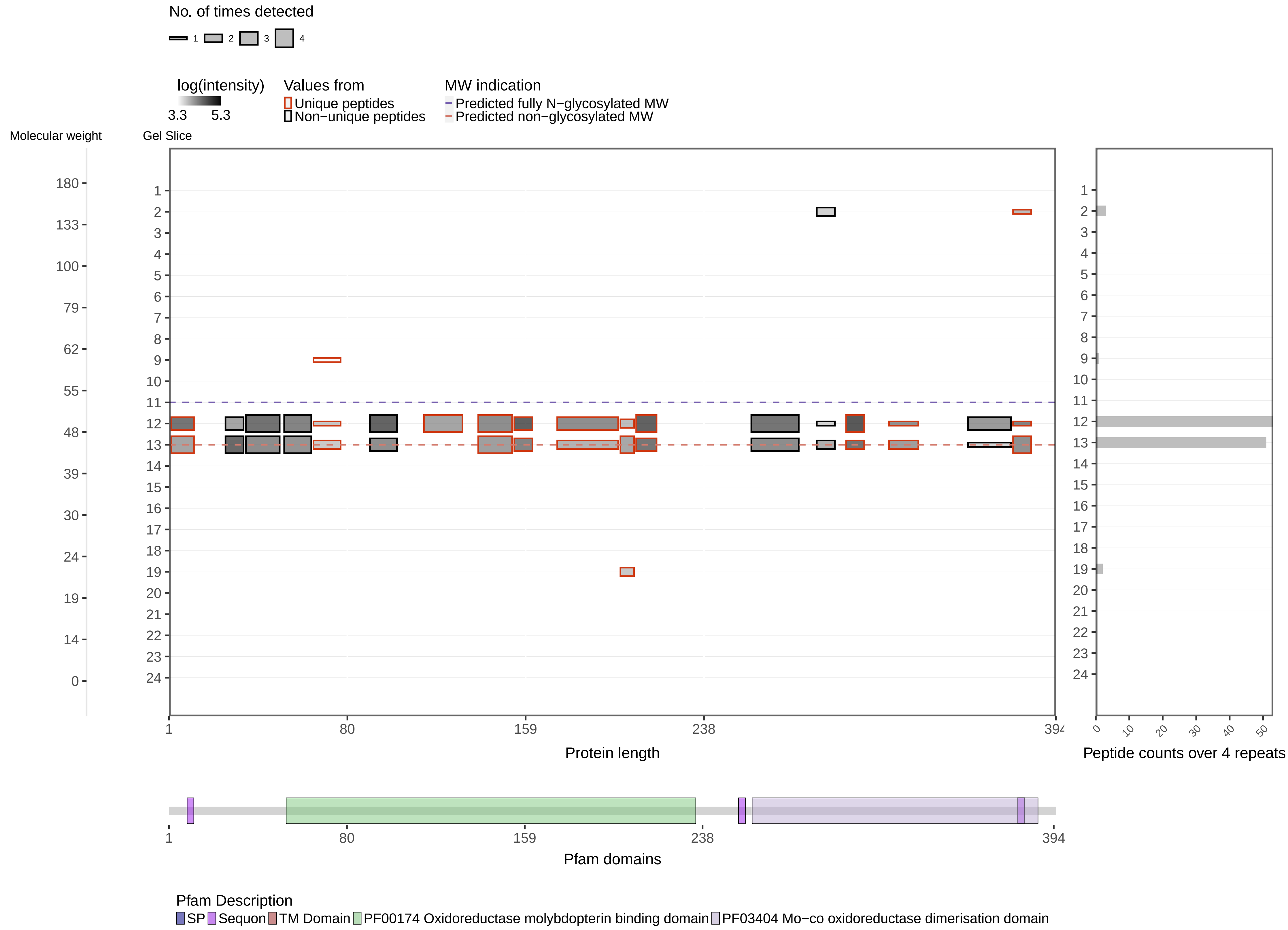


Protein sequence:

[illegible]

### NbL02g25910.1 glucose−6−phosphate_1−dehydrogenase_cytoplasmic_isoform [MW = 58.7 kDa]

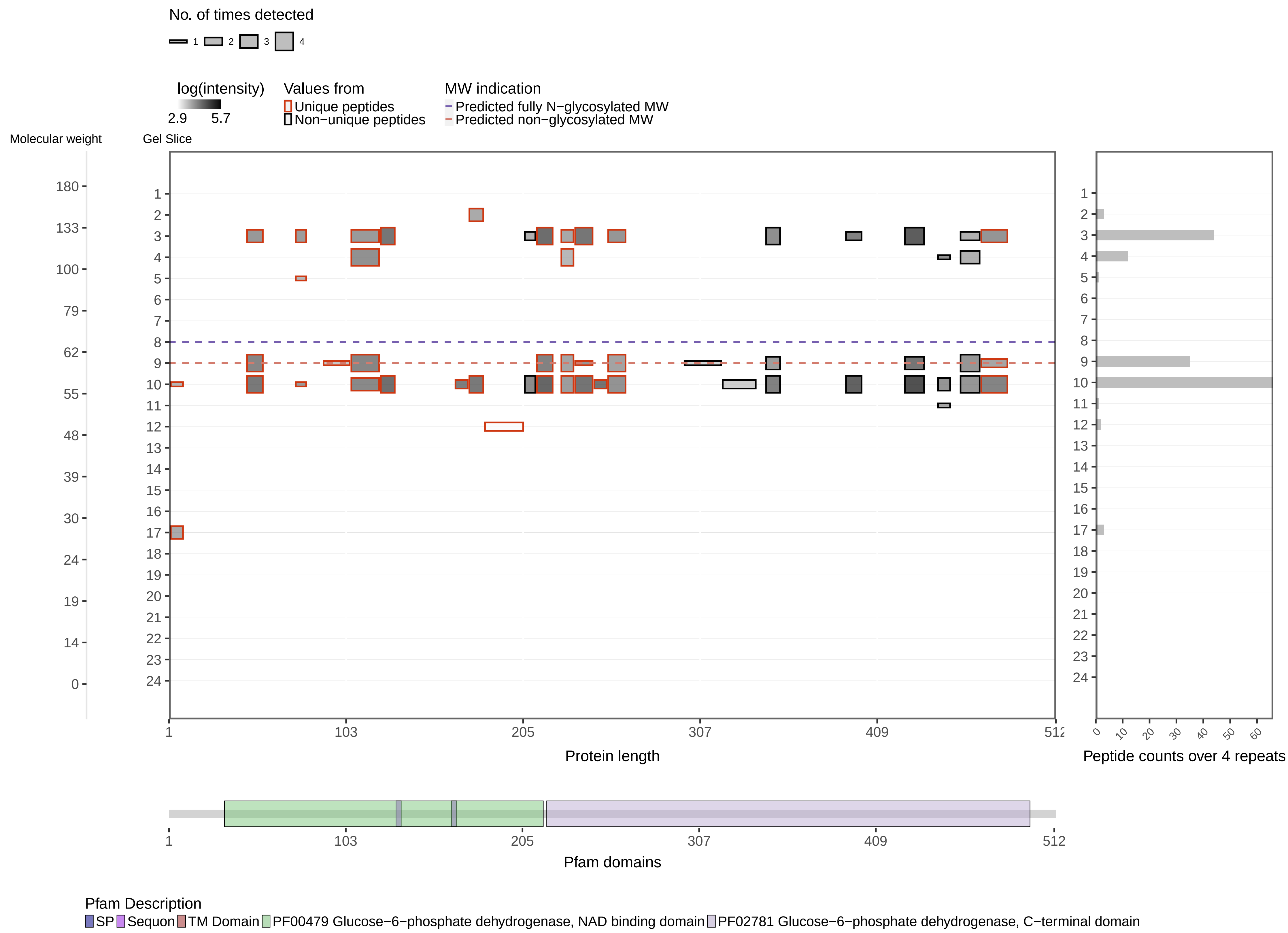


Protein sequence:

REDFGTEGRGGYFDEYGIIRDIIQNHLLQVLCLVAMEKPVSQKPEHIRDEKVKVLQSMLPIKDEEVVLGQYEGYKDDPTVPDNSNTPTFATMVLRIHNERWEGVPFIMKAGKALNSRKAEIRVQFKDVPGDIFRCKKQGRNEFVIRLQPSEAMYMKLTVKQPGLEMSTVQSELDLSYRQRYQGVVIPEAYERLILDTIRGDQQHFVRRDELKAAWEIFTPLLHRIDDGEIKPIPYKPGSRGPAEADELLQNVGYVQTHGYIWIPPTL*

### NbL02g25500.1 glucuronokinase_1−like [MW = 39.95 kDa]

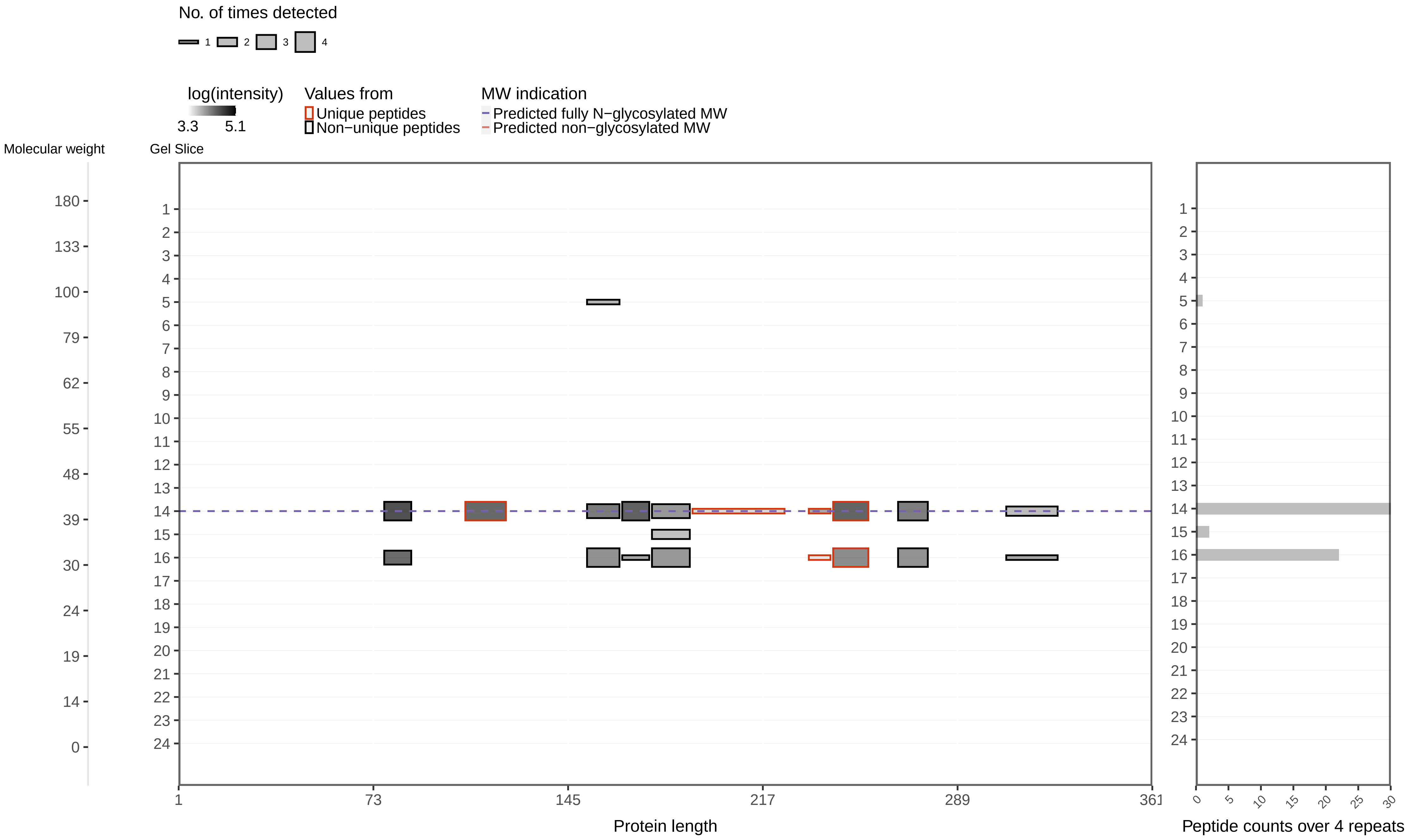


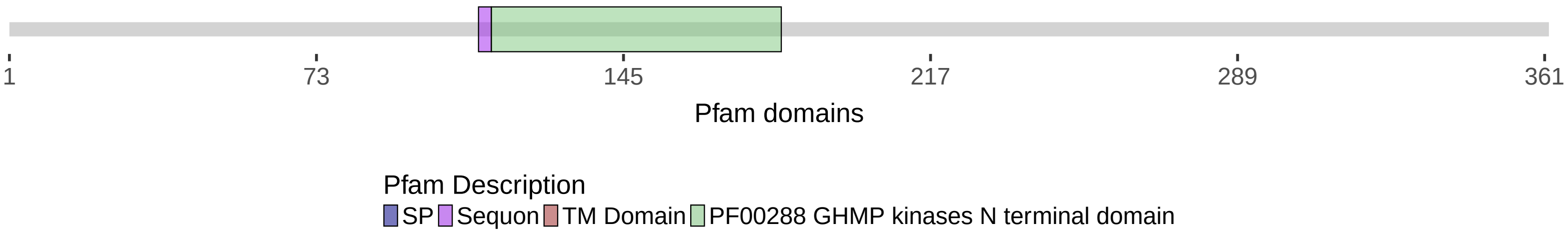


Protein sequence:

AGLQDRVAQVYGGLVYMDFGKNHMDELGHGIYTPMDVDLLPPLYLIYAENPSDSGKVHSTVRQRWLEGDEFIRSTMEEVANIAVEGRKTLLEKDYIKLAALMNHNFDLRRRMFGDDALGAMNIEMVEIGRRIGAASKFTGSGGAVVVFCPDGPSQVKQLEDACHTAGFTFQPIKVMPSFLSEVDLQTLGSN*

### NbL02g25330.1 titin [MW = 427.49 kDa]

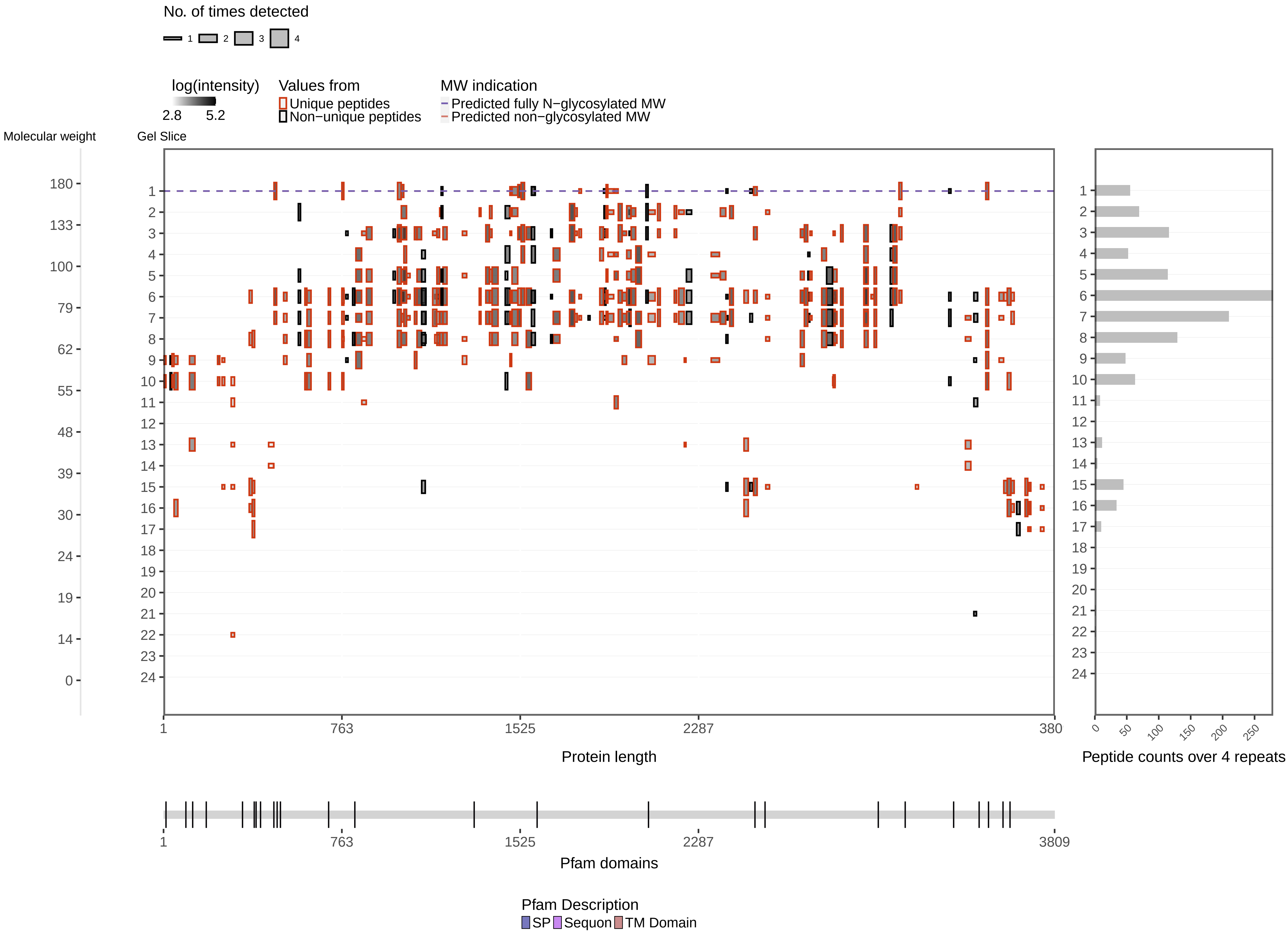


Protein sequence:

EPAAEKKPEEPVATEKPEEPVATENPEEPAKEKADEPAVLEKPEEPAVEKKPEEPVATEKPEEPAIKEKDDEPAVQEKPEEPTVEKKPEEPAVEEKFDEPAVGEKSEEPVVEQTPAGPAWEETPKELAFEEKPEEPAVEETPEVLAVEKNPEKPAVGEKHEKVTLLEKNIPIIELNEKNDTATPCNTNNDVADYLAVEEKPEEPAIEKKPEEPMAKEKLKEPNVEDKPEKSVVEEKSEKLATEETPEEQAEEETPEALAVEENPDESAVEETVEELAIEQKPKTTATEKEKPEAVTLLI

NbL02g25190.1 isocitrate_dehydrogenase_[NADP] [MW = 53.9 kDa]

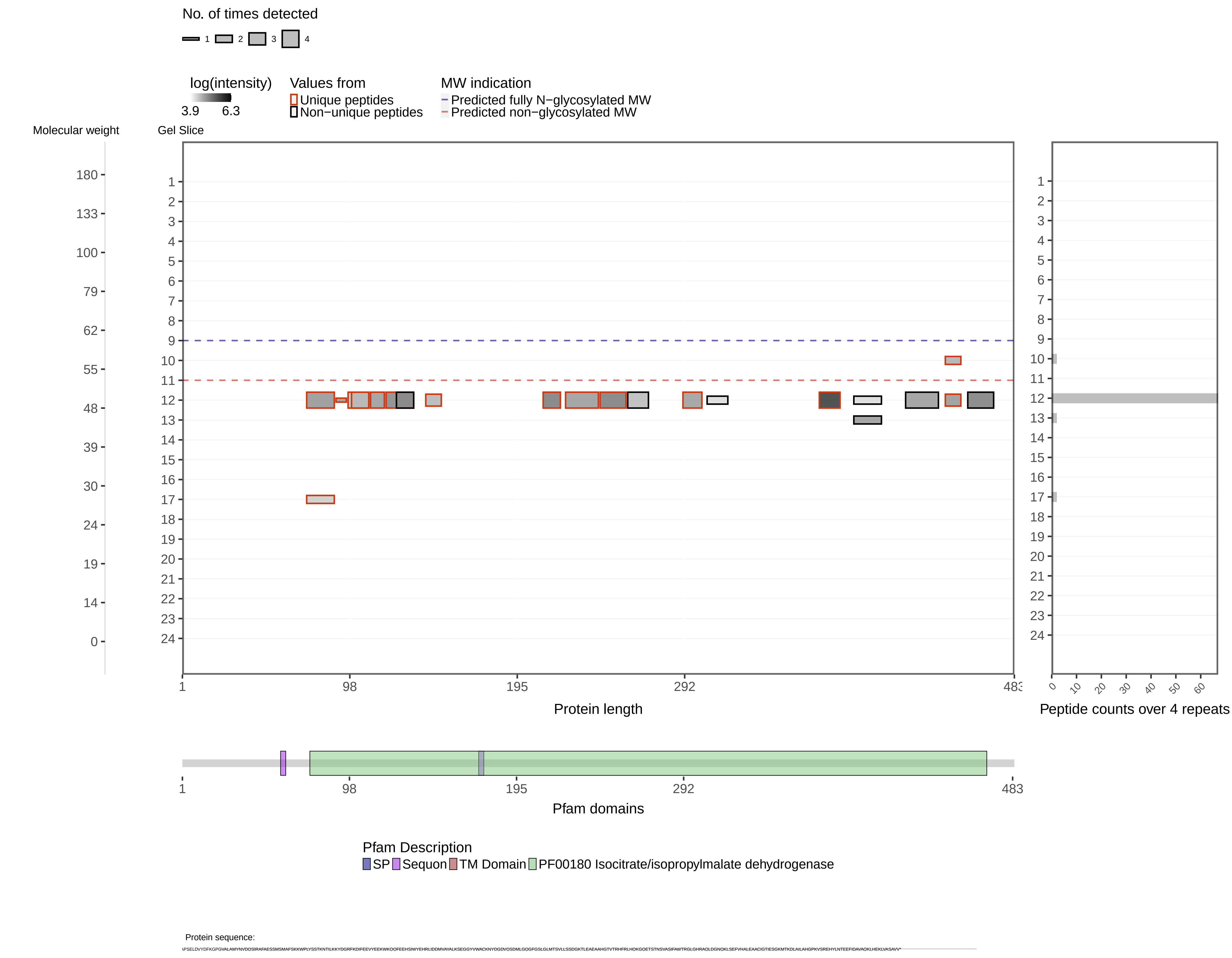

NbL02g25170.1 tRNA_(adenine(58)−N(1))−methyltransferase_catalytic_subunit_TRMT61A [MW = 37.99 kDa]

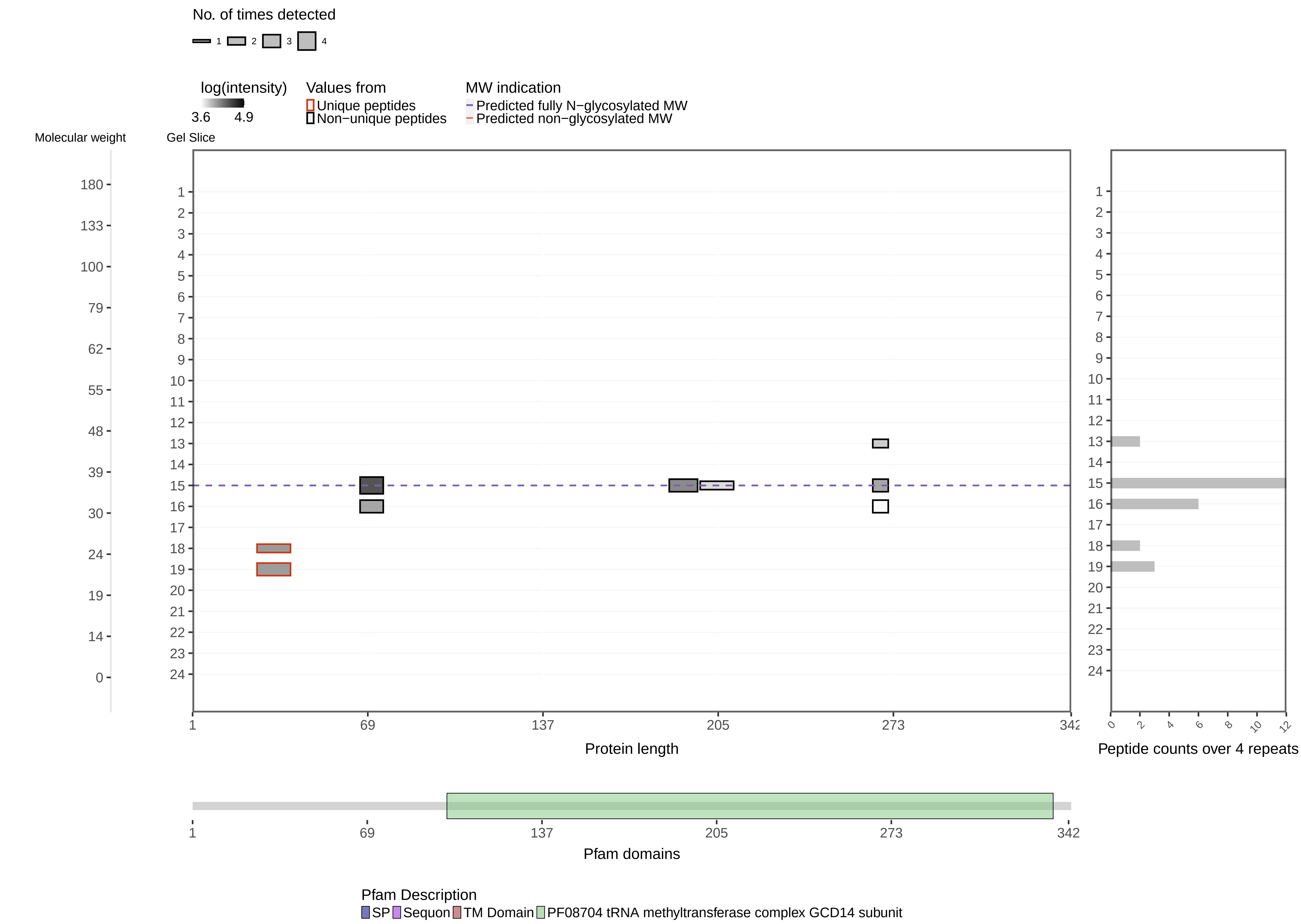


Protein sequence:

PTGHVYTFDFHEQRAASAREDFERTGLSNLVTVGVRDIQGEGFPDEFCGRADSVFLDLPQPWLVIPCAGKMLKPDGVLCSFSPCIEQVQRSCEALKSHFTDIRTFEVLLRTYDVRQRIVESCPVNEGGSNERPPRKRRQGCAEGNNGIECTSSPTVMIRPSGEARGHTGYLTFARLKCFA*

### NbL02g25140.1 peptide_deformylase_1B_chloroplastic [MW = 30.9 kDa]

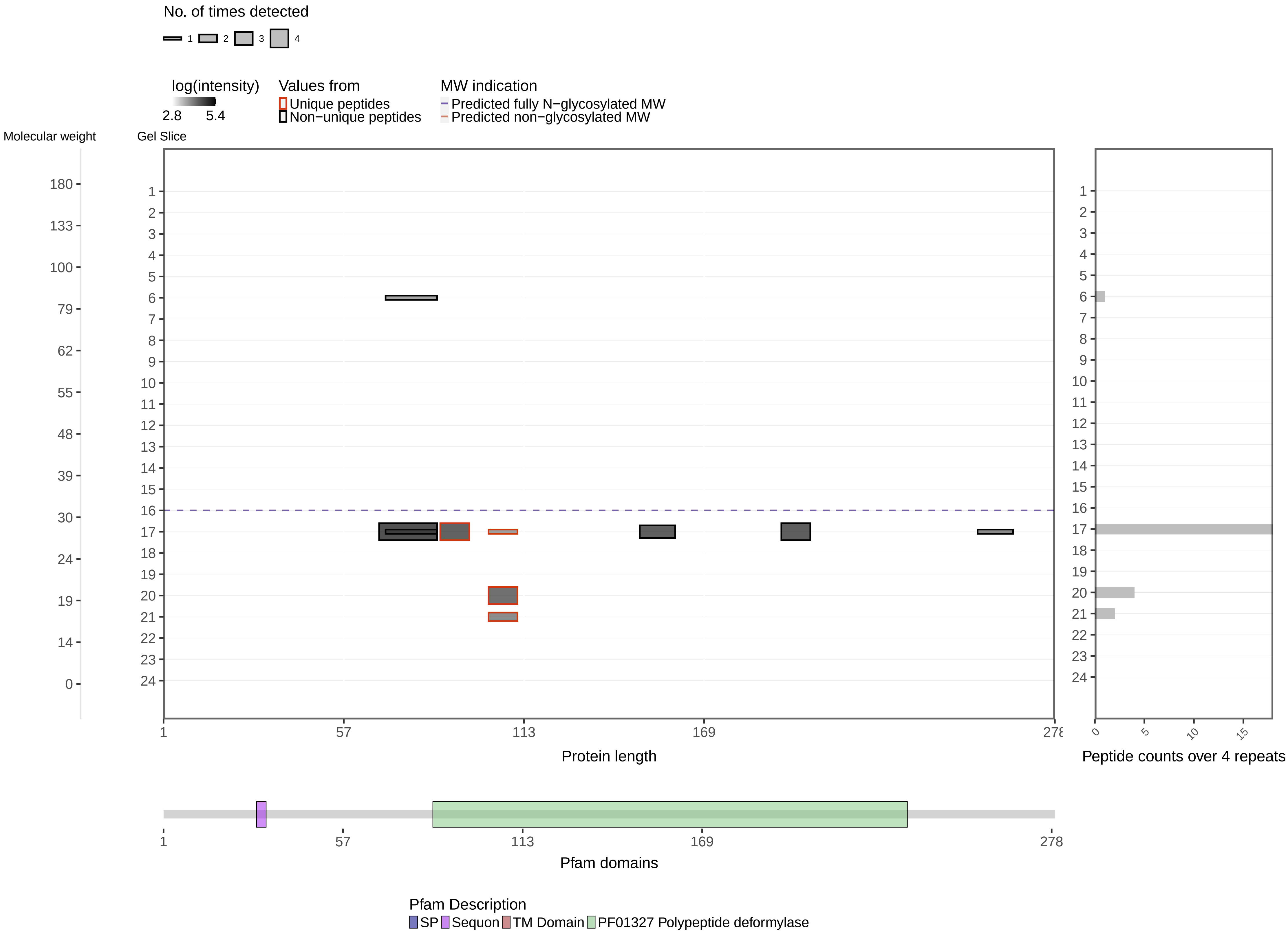


Protein sequence:

IGLSAPQVGMNVQLMVFNAAGERGEGEEIVLVNPRVSRYSRRIIPYDEGCLSFPEIYGDVERPDSVKVDAQDINGARFELTLTALPARVFQHEFDHLQGVLFFEKMTDEVLDTIREDLVALEKKFEEKTGLPTPESINTRKIKKAAVGFGKS*

NbL02g25100.1 50S_ribosomal_protein_L12_chloroplastic [MW − SP = 20.26 kDa]

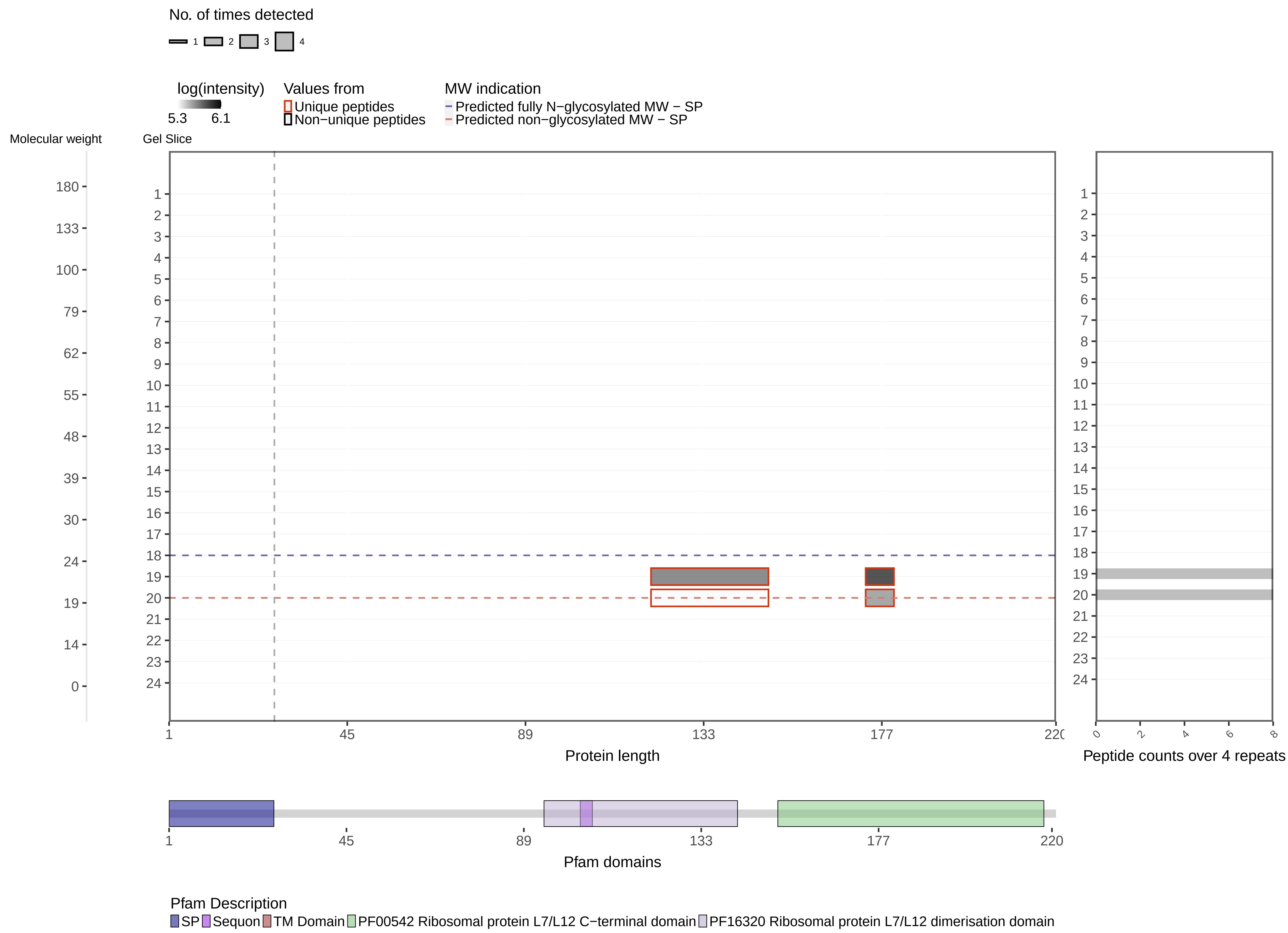


Protein sequence:

LGDEISNLTLADAQKLVEYLQDKLGVTAASFAPAAAVAAPGAAAEAPAVVEEKTEFDVVIDEVPSNARIATIKAVRALTSLALKEAKELIEGLPKKFKEGVSKDEAEDAKKQLEEAGAKVSIA*

### NbL02g25020.1 protease_Do−like_1_chloroplastic [MW = 45.29 kDa]

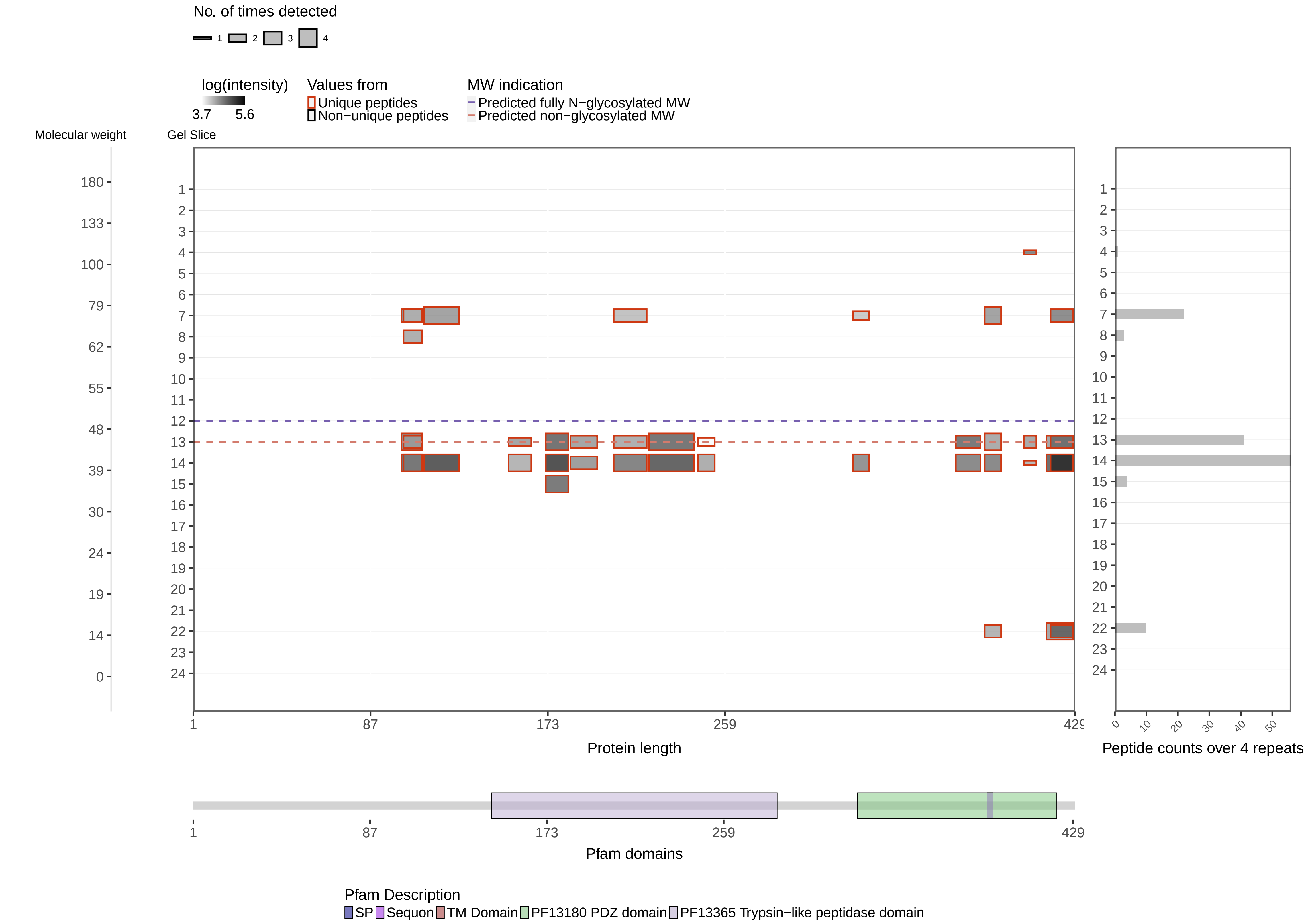


Protein sequence:

DAPKDKMRPIPIGVSADLLVGQKVFAIGNPFGLDHTLTTGVISGLRREINSAATGRPIQDVIQTDAAINPGNSGGPLLDSSGNLIGINTAIYSPSGASSGVGFSIPVDTVSGIVDQLVKFGKVTRPILGIKFAPDQSVEQLGVSGVLVLDAPPNGPAGKAGLLPTKRDSYGRLILGDIITSINGKKVSNGTDLYRILDQCKVGEKVIVEVLRGDQKEKIPVLLEPKPEES*

NbL02g24980.1 formate_dehydrogenase_mitochondrial [MW = 42.47 kDa]

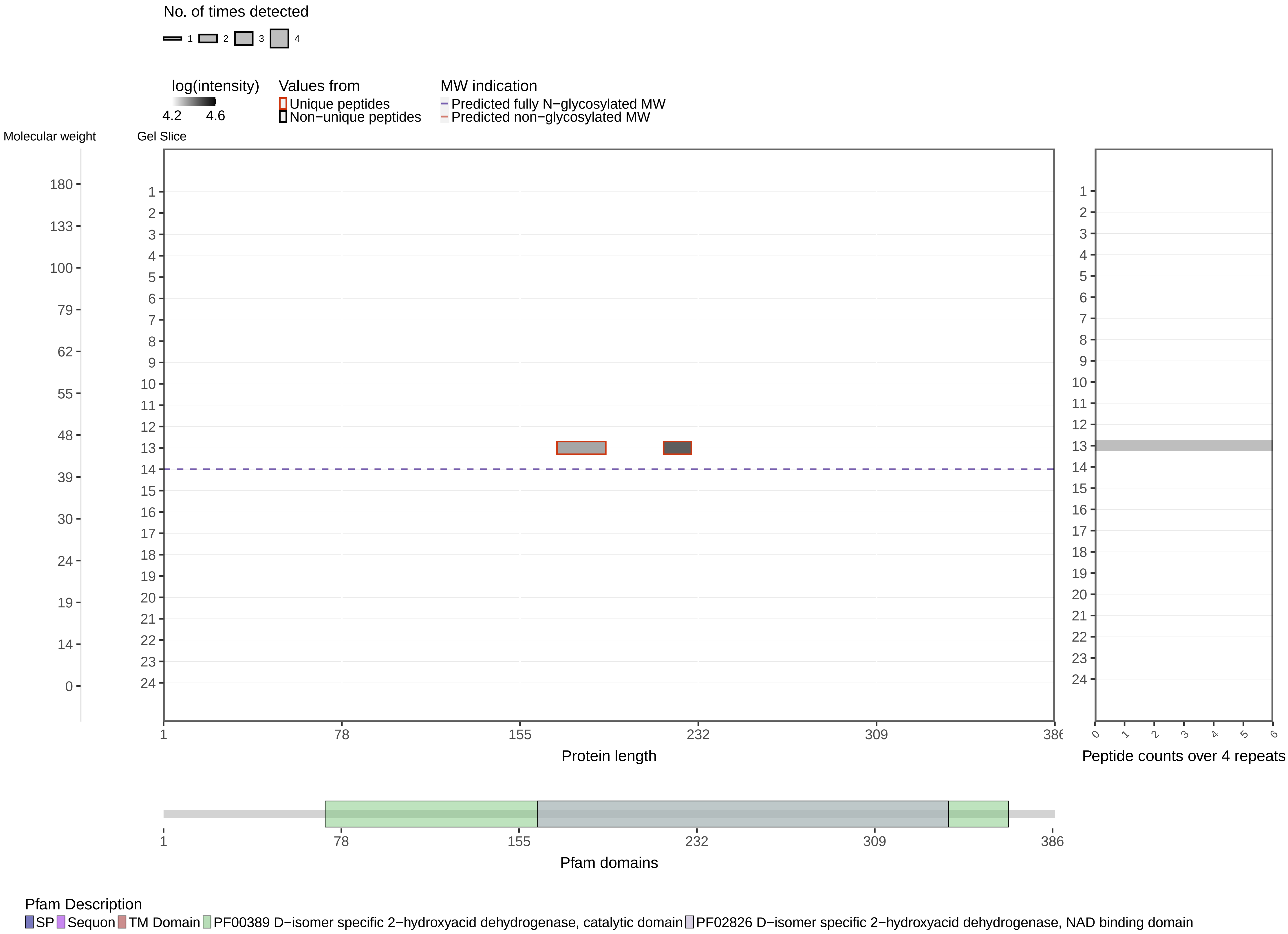


Protein sequence:

EWNVAGIAYRAYDLEGKTVGTVGAGRIGRLLLQRLKPFNCNLLYHDRVKMDSELENQIGAKFEEDLDKMLSKCDIVVVNTPLTEKTKGMFDKERIAKMKKGVLIVNNARGAIMDTQAVVDACNSGHIAGYSGDVWYPQPAPKDHPWRYMPNQAMTPHISGTTIDAQLRYAAGTKDMLDRYFKGEDFPAQNYIVKDGELAPQYR*

NbL02g24960.1 peptidyl−prolyl_cis−trans_isomerase_CYP38_chloroplastic [MW = 49.39 kDa]

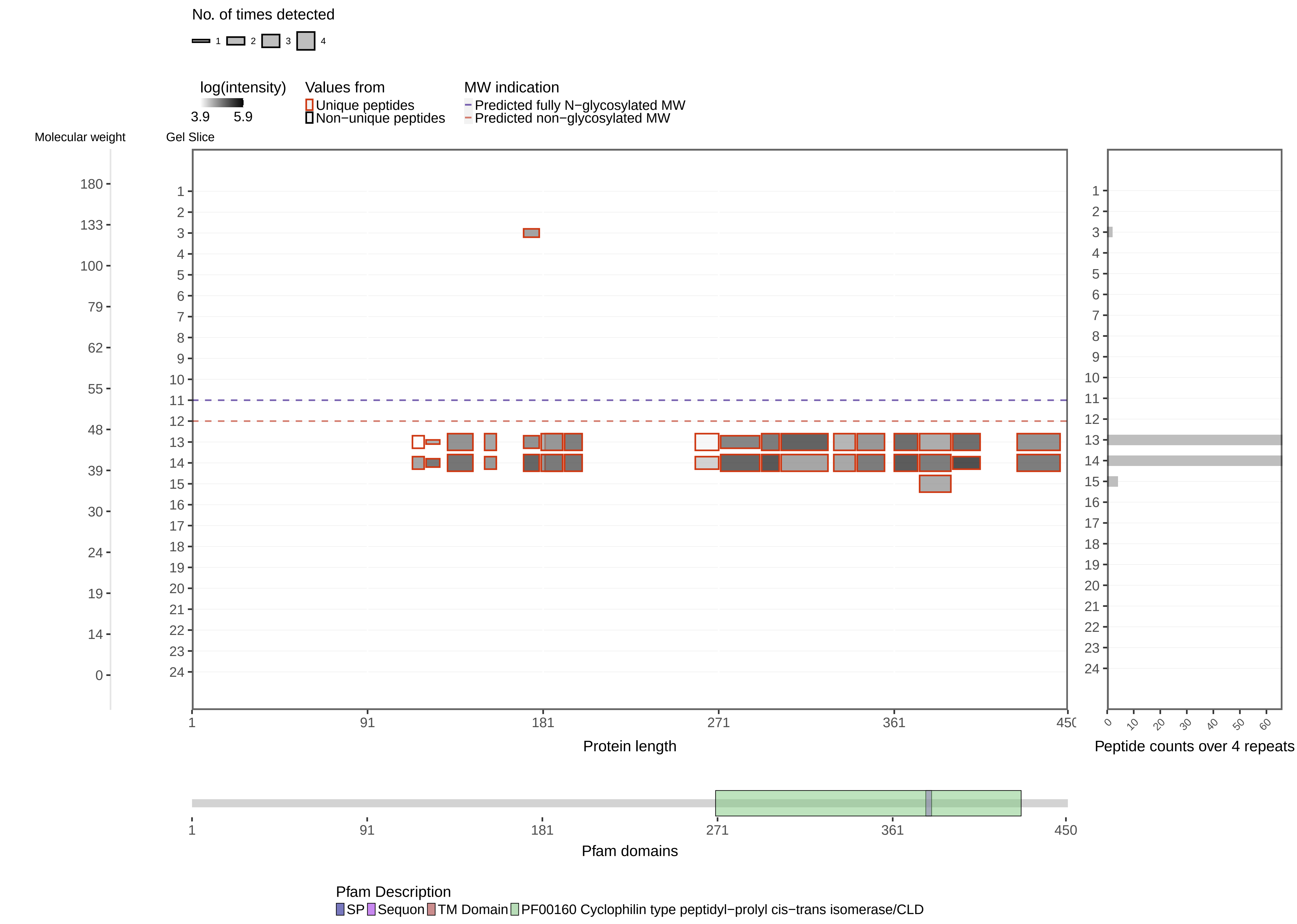


Protein sequence:

QKELLNYVGGVEEDMVDGFPYEVPEEYQNMPLLKGRATLDMKVKVKDNPNVEECVFRIVLDGYNAPVTAGNFIDLVERHFYDGMEIQRADGFVVQTGDPEGPAEGFIDPSTEKIRTVPLEIMVDGEKVPFYGETLEDLGLYKAQTRLPFNAFGTMAMAREEFENNSASSQVFWLLKESELTPSNANILDGRYAVFGYVTENEDFLADLKVGDVIDSIQVVSGLDNLVNPSYKIAG*

NbL02g24790.1 uncharacterized_LOC107798474 [MW = 24.91 kDa]

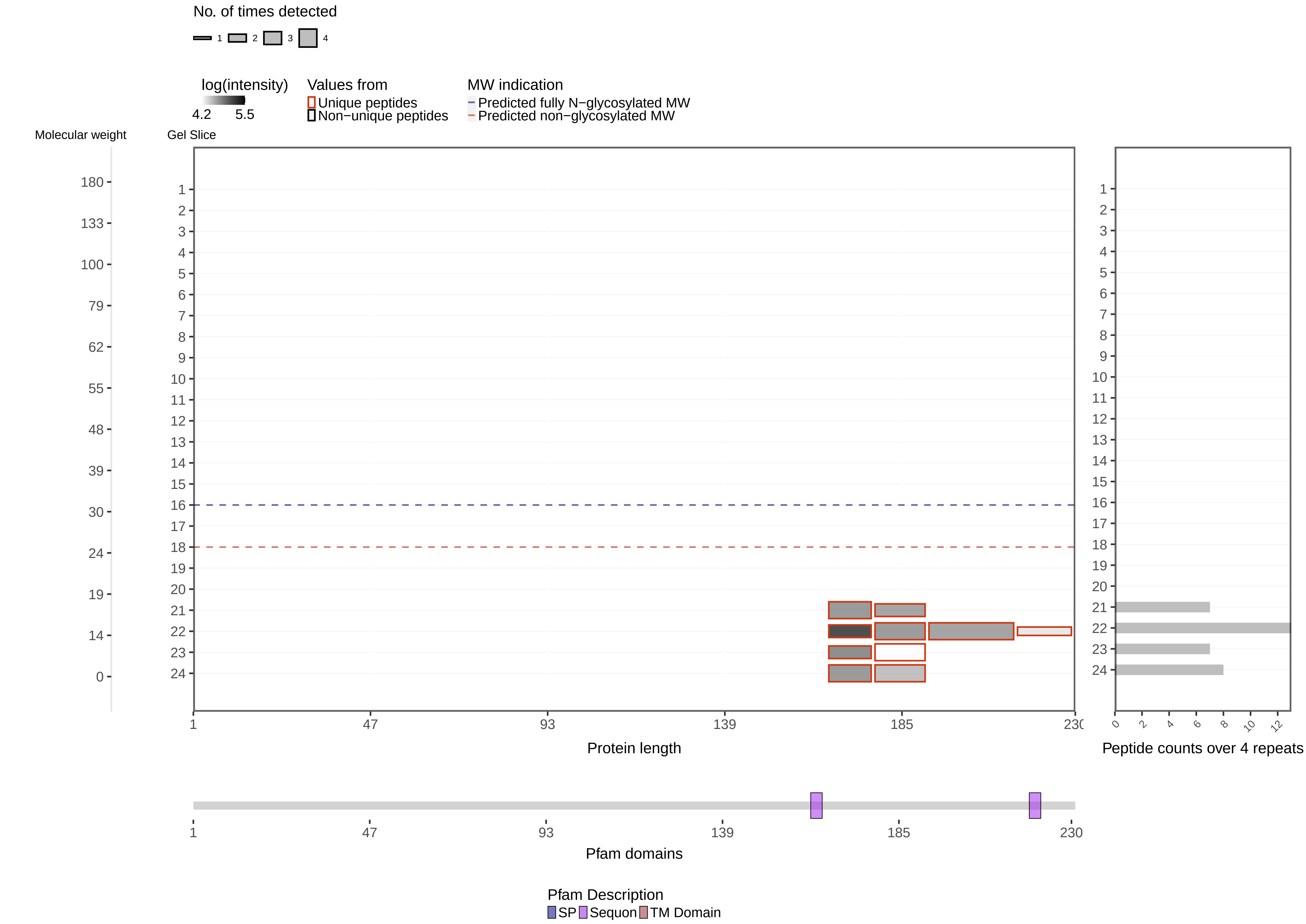


Protein sequence:

NTRIWSATEEEGATAVVDEASAALTTTVDOTVSVSVSPSDVLTMFFOSOAEGTMNEAAVPNVTKALEEIEGITDLKVOVVEGIASVELTKOTTIOATGVSSSLVETIOGSGFKLOTLNLSFODEEDTS*

### NbL02g24770.1 translation_initiation_factor_IF−1_chloroplastic [MW = 16.98 kDa]

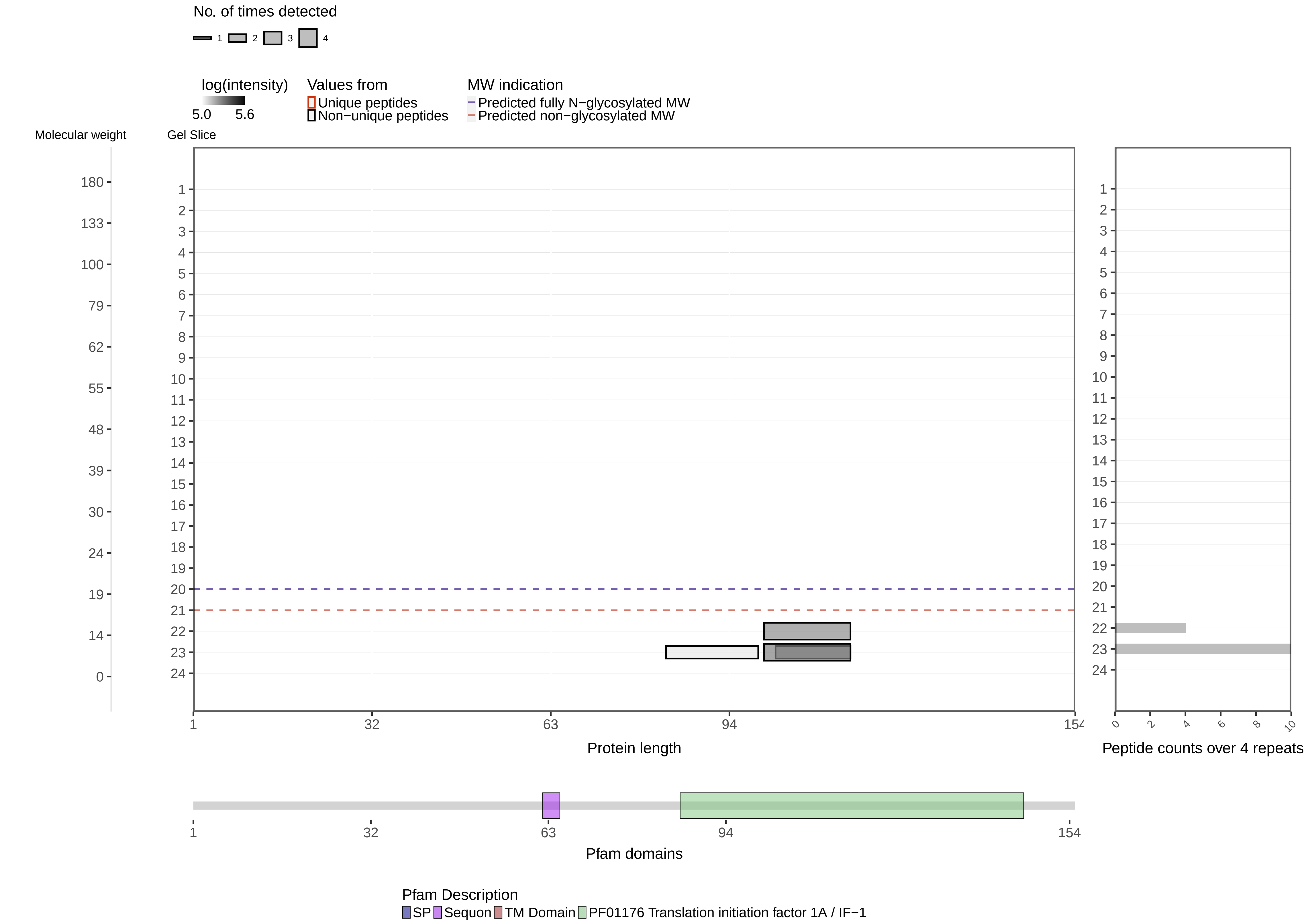


Protein sequence:

HSFTNSRSGTSIQCLSQEQKWTHEGSITESLPNGMFRVKLDNADVVLGYISGKIRKNFIRLLPGDRVKIEVSRYDSTKGRIIYRLRGGREG*

NbL02g24750.1 peroxidase_63−like [MW − SP = 34.25 kDa]

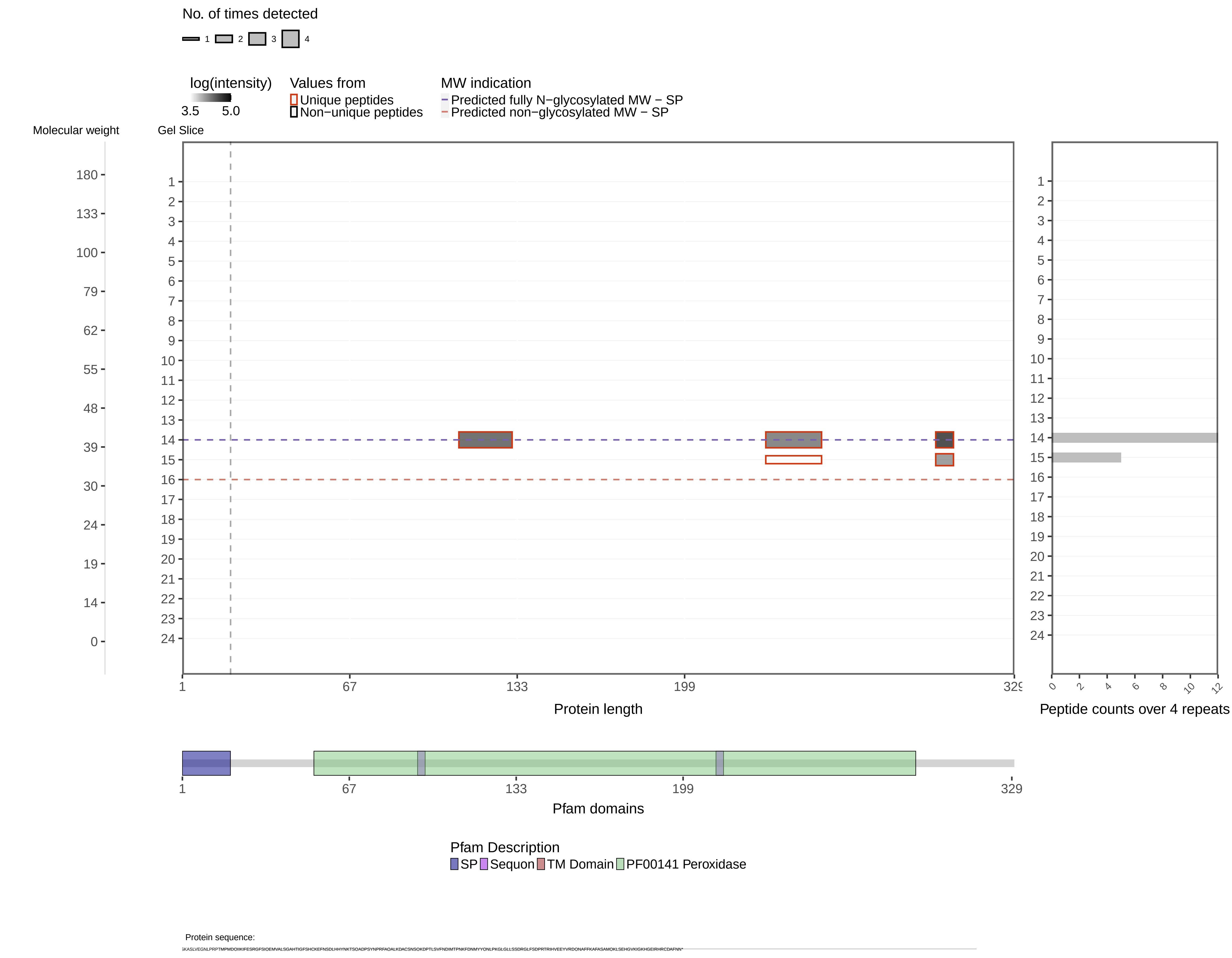

### NbL02g24730.1 ferredoxin−thioredoxin_reductase_catalytic_chain_chloroplastic [MW = 17.1 kDa]

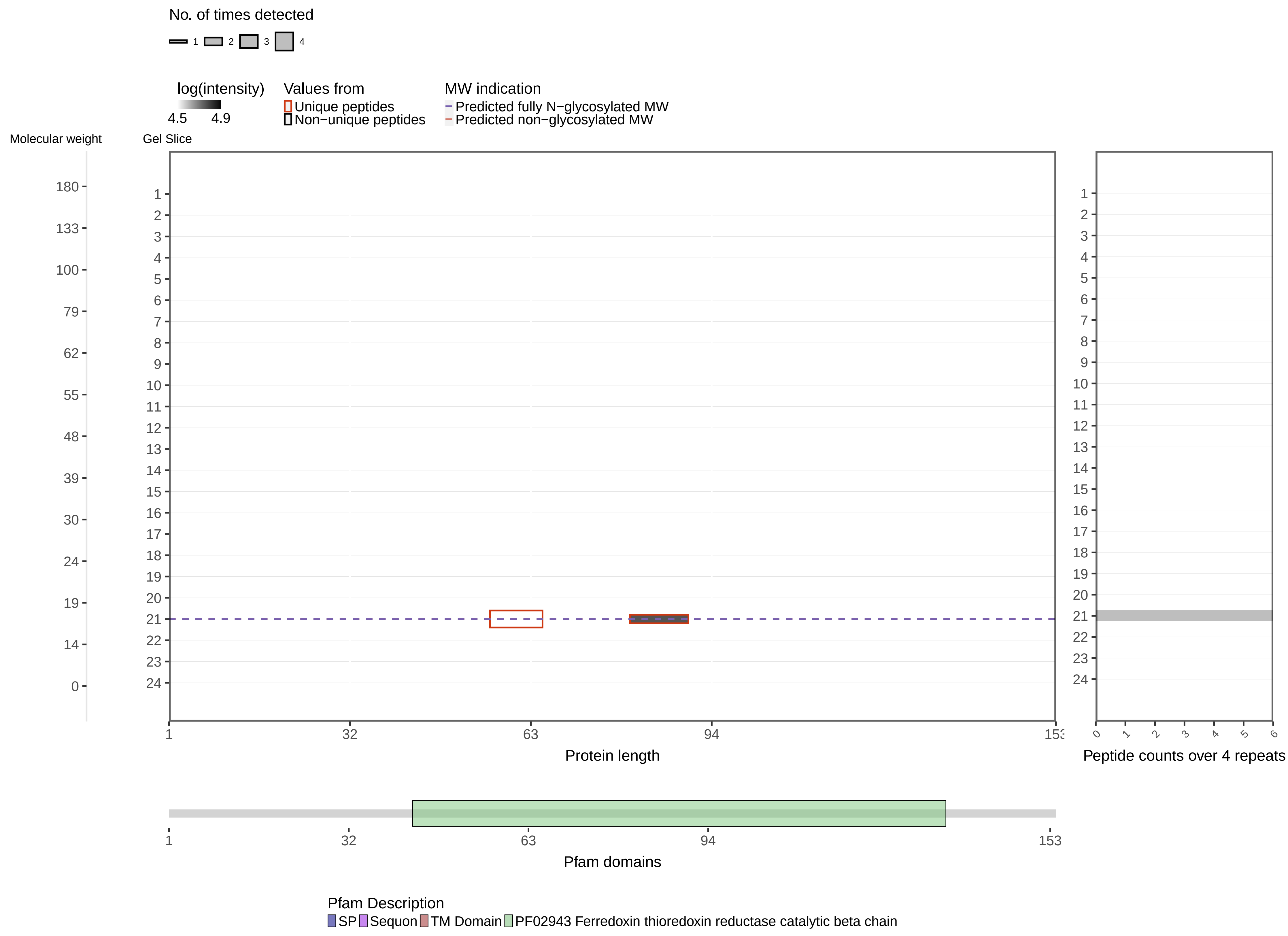


Protein sequence:

)KGVTSVVIKGLAEHKDILGAPLCPCRHYDDKAAEAQQGFWNCPCVPMRERKECHCMLFLTPDNDFAGEEQVYANMRSLVSNYLLRLGI*

### NbL02g24690.1 protein_CREG1 [MW − SP = 20.31 kDa]

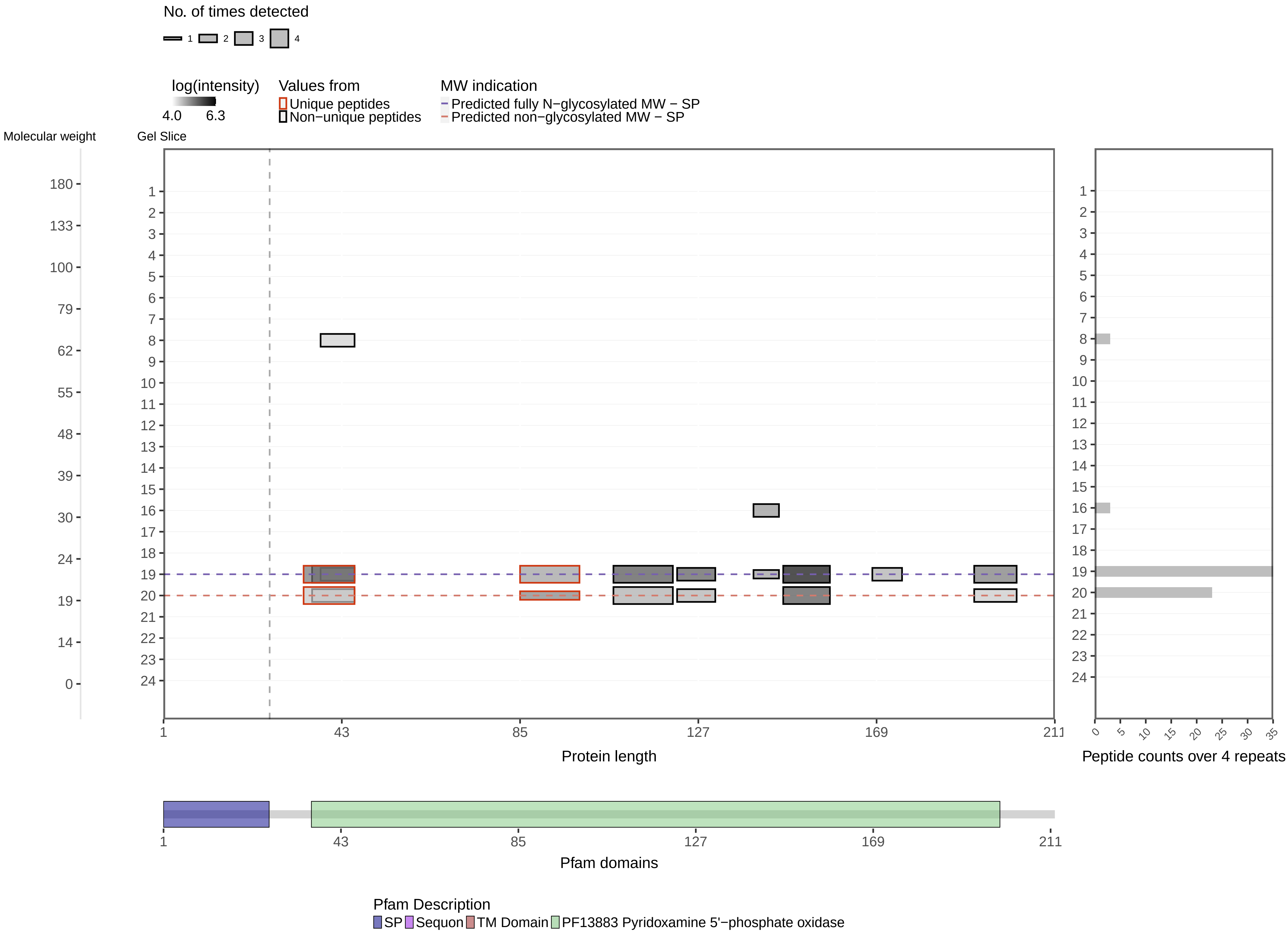


Protein sequence:

.TTLDPTARNALKDQRSSLTISEYAIGTCGKTDPENPSCAKITLIGKLKLLNGDPKETSFAQTALFTKHPEMKGWPKGHNFQIFKLEIEEIFMINWFGGPKPLTVDQYLQAKMDSRTAIA*

NbL02g24670.1 alpha−mannosidase_2 [MW = 132.39 kDa]

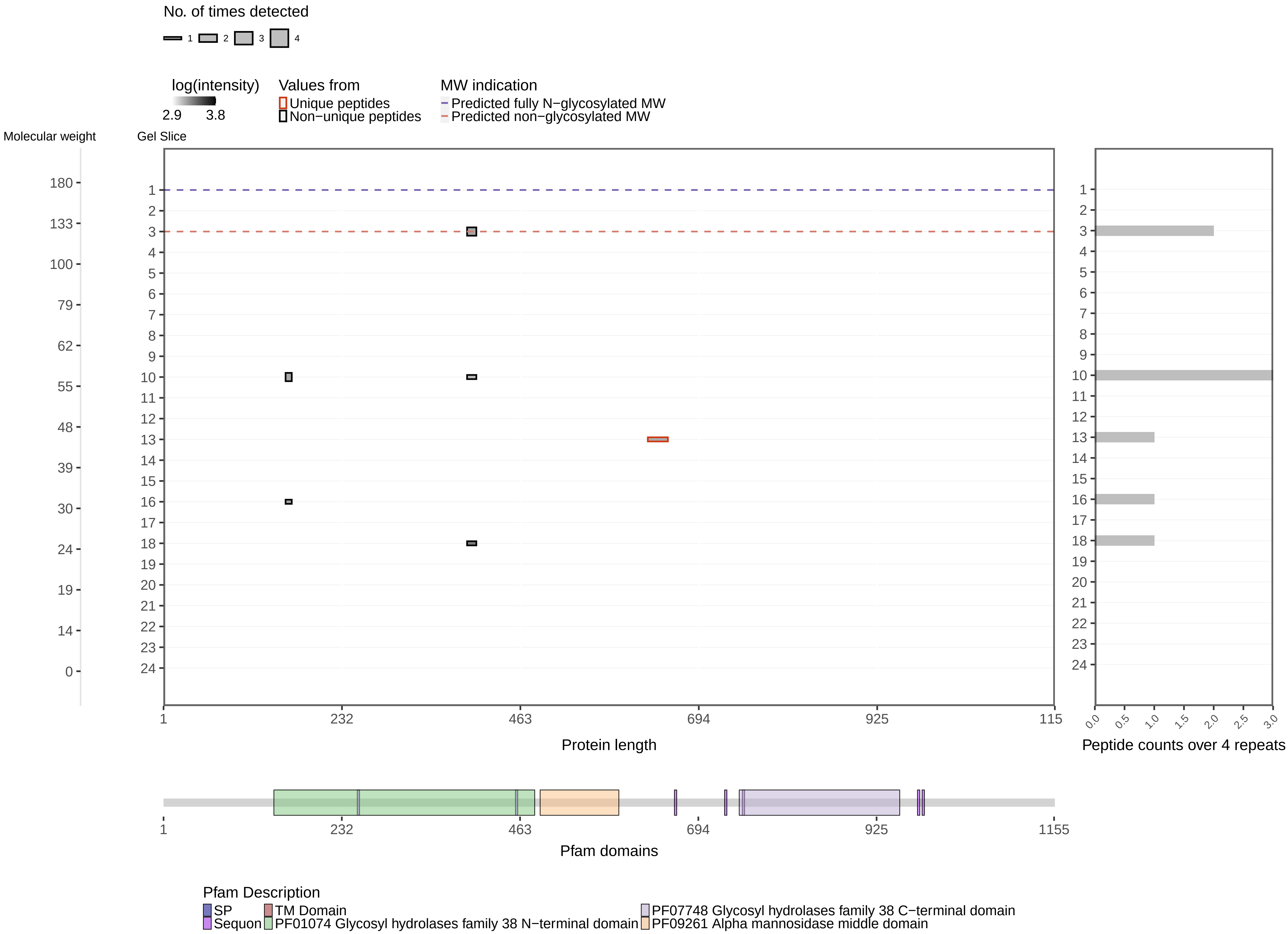


Protein sequence:

DHVVQDYGTRMHMALQDLQIFMSKAIEVLLGIRHEKNDQHPSQFEPAQVRSKYDAQPVVKVISAREGTVQTVVLFNPSEQTINEVVMVTVERPDVTILDSNWTCIKSQISPELSHDKENTFTRRHRVYWKASVPAMGLQTYYVANGFAGCEKAIPAQLRILVPSGNFSCPAPYTCSKLDSNEAAIQNENLTVTFSTKFGLLQKVSHSDGRQNVMDEEIDMYSSSGGAYLFKPEGEAEPIIQAGGIMVVSEGHLVQEVYSYPMTAWDKSPISHSTRIYNGHNTIQEHIIEKEYHV

### NbL02g24660.1 protein_transport_protein_SEC13_homolog_B [MW = 32.63 kDa]

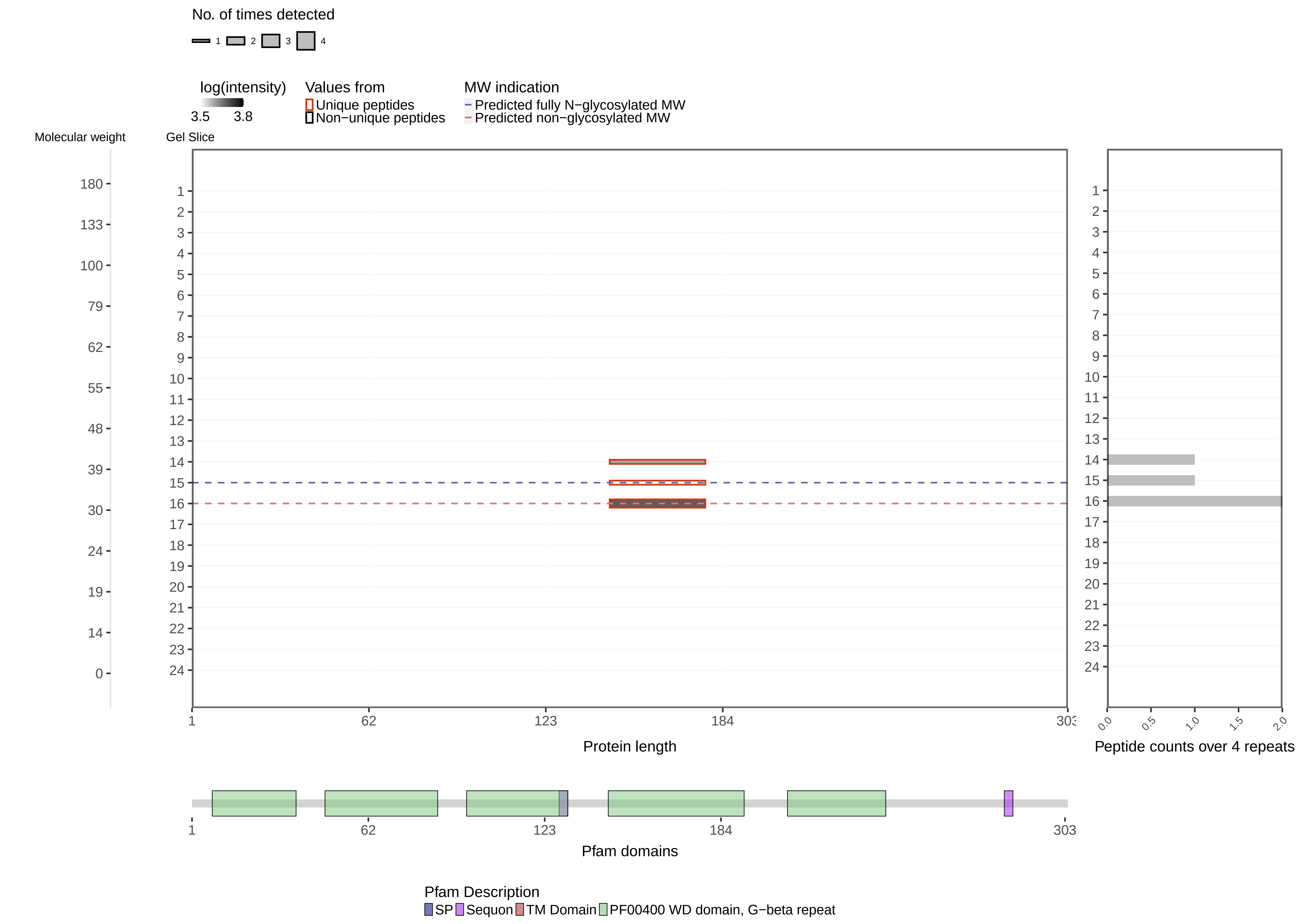


Protein sequence:

/DTTRIDOAHPVGVTSVSWAPSMAPGALVGSGVLEPVOKLASGGCDNTVKVWKLYNGIWKMDCFPALOMHTNWVRDVAWAPNLGLPKSTIASASEDGTVVIWTVAKEGDOWEGKVLKDFKTPVWRVSWSLTGNLLAVAAGDNNVTLWKEAVDGEWOOASTVDO*

### NbL02g24600.1 signal_recognition_particle_43_kDa_protein_chloroplastic [MW = 45.36 kDa]

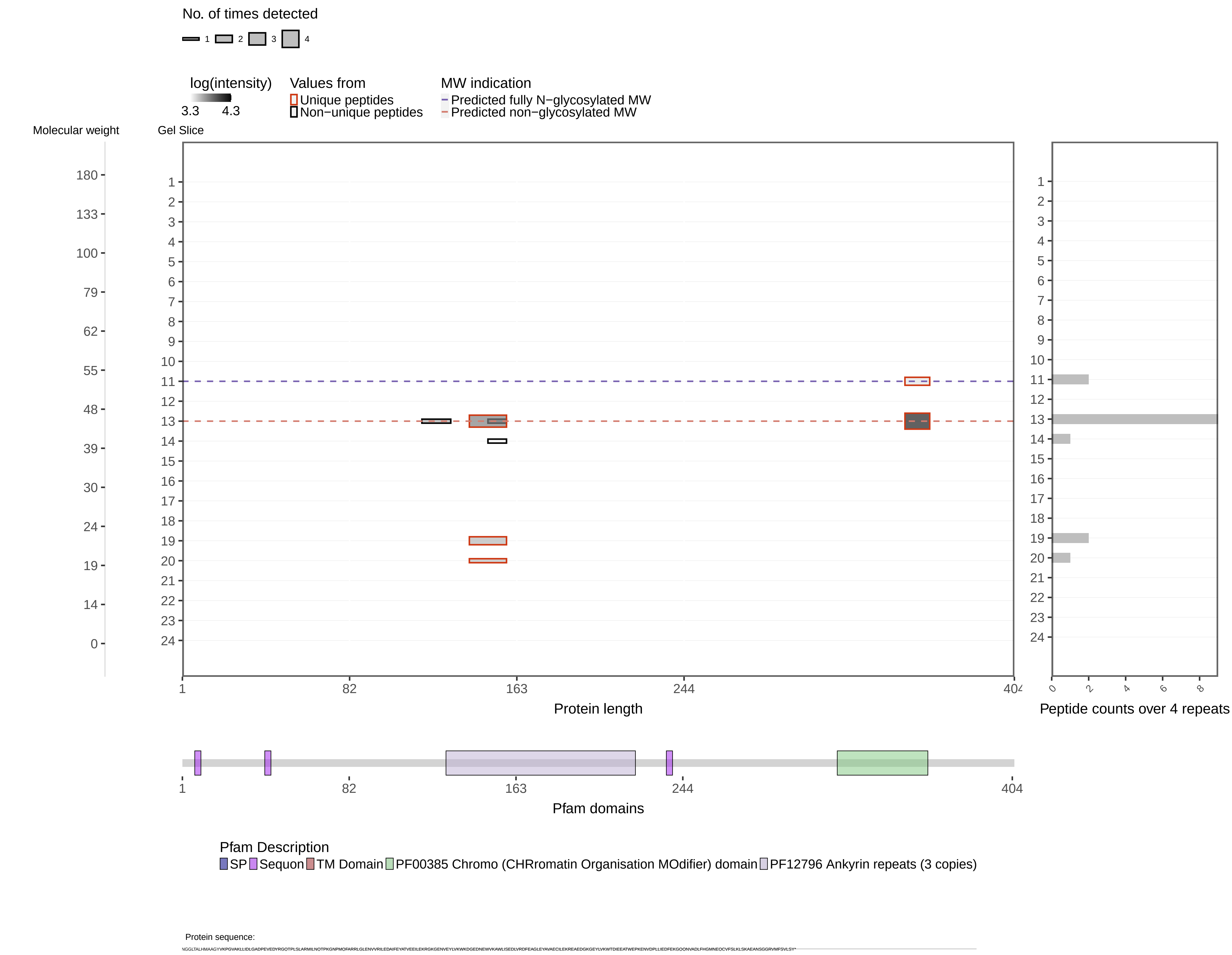

NbL02g24520.1 inactive_LRR_receptor−like_serine/threonine−protein_kinase_BIR2 [MW − SP = 63.66 kDa]

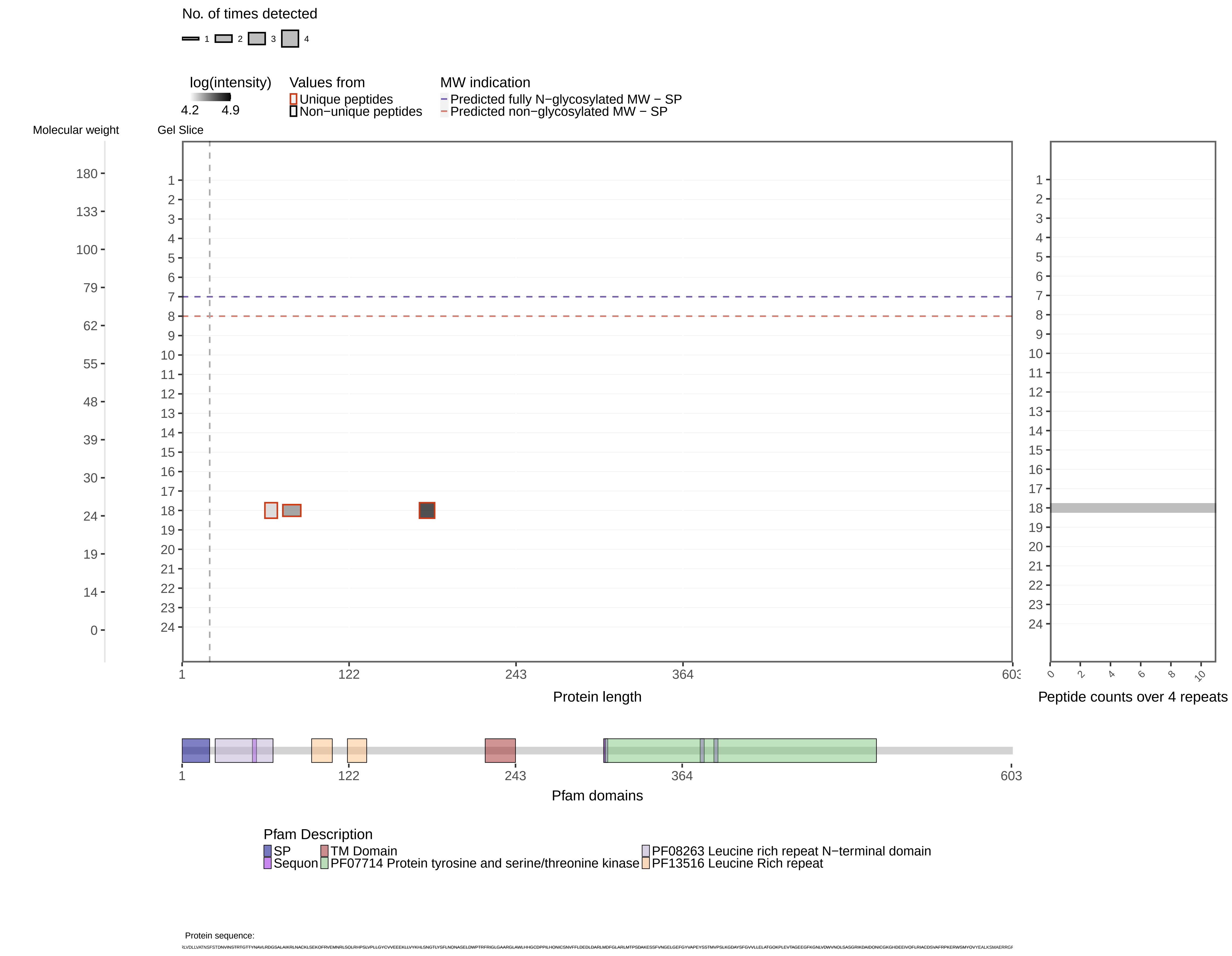

### NbL02g24440.1 inositol_hexakisphosphate_and_diphosphoinositol−pentakisphosphate_kinase_2−like [MW = 95.13 kDa]

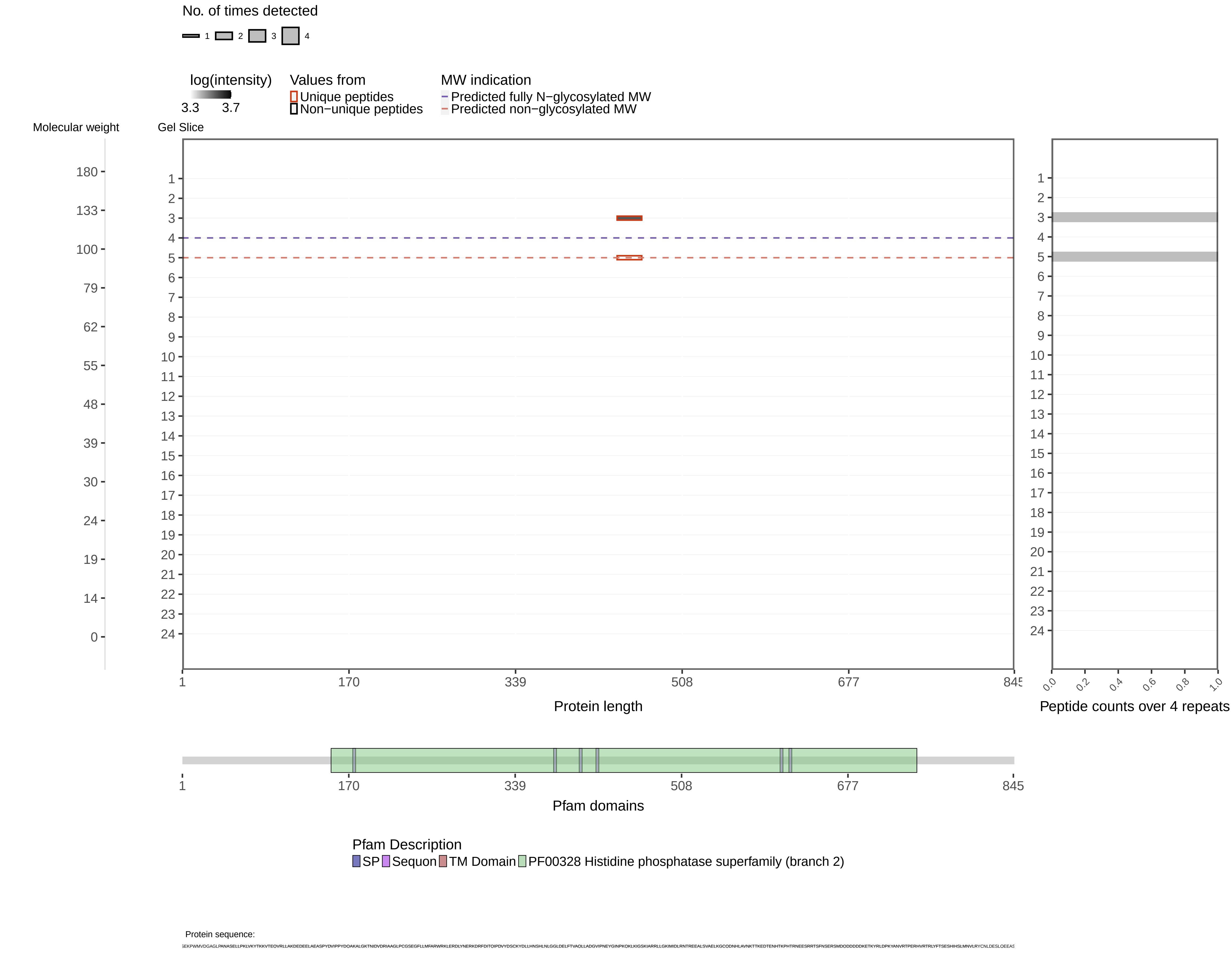

### NbL02g24410.1 FACT_complex_subunit_SSRP1 [MW = 70.83 kDa]

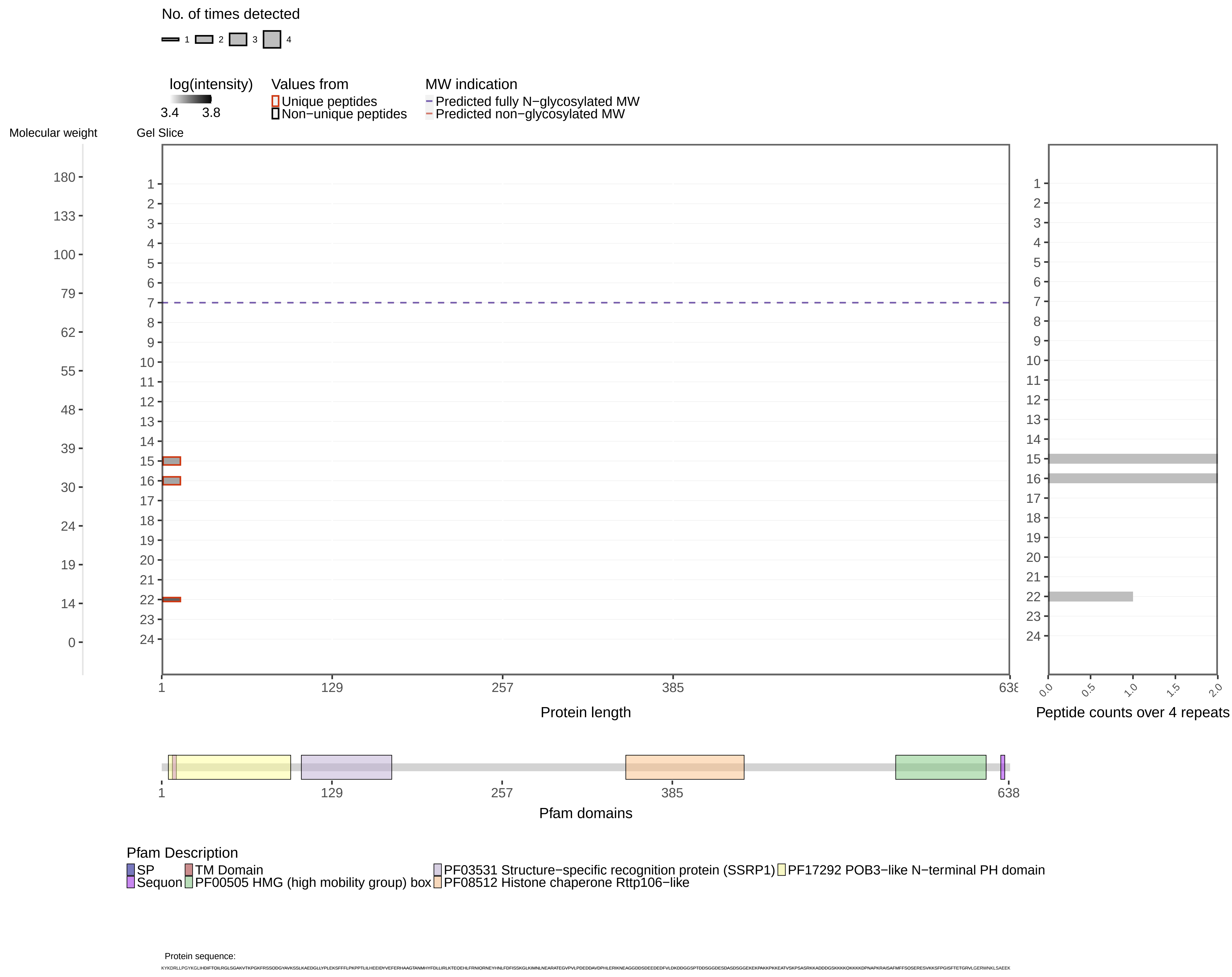

### NbL02g24350.1 ubiquitin−conjugating_enzyme_E2_2 [MW = 17.37 kDa]

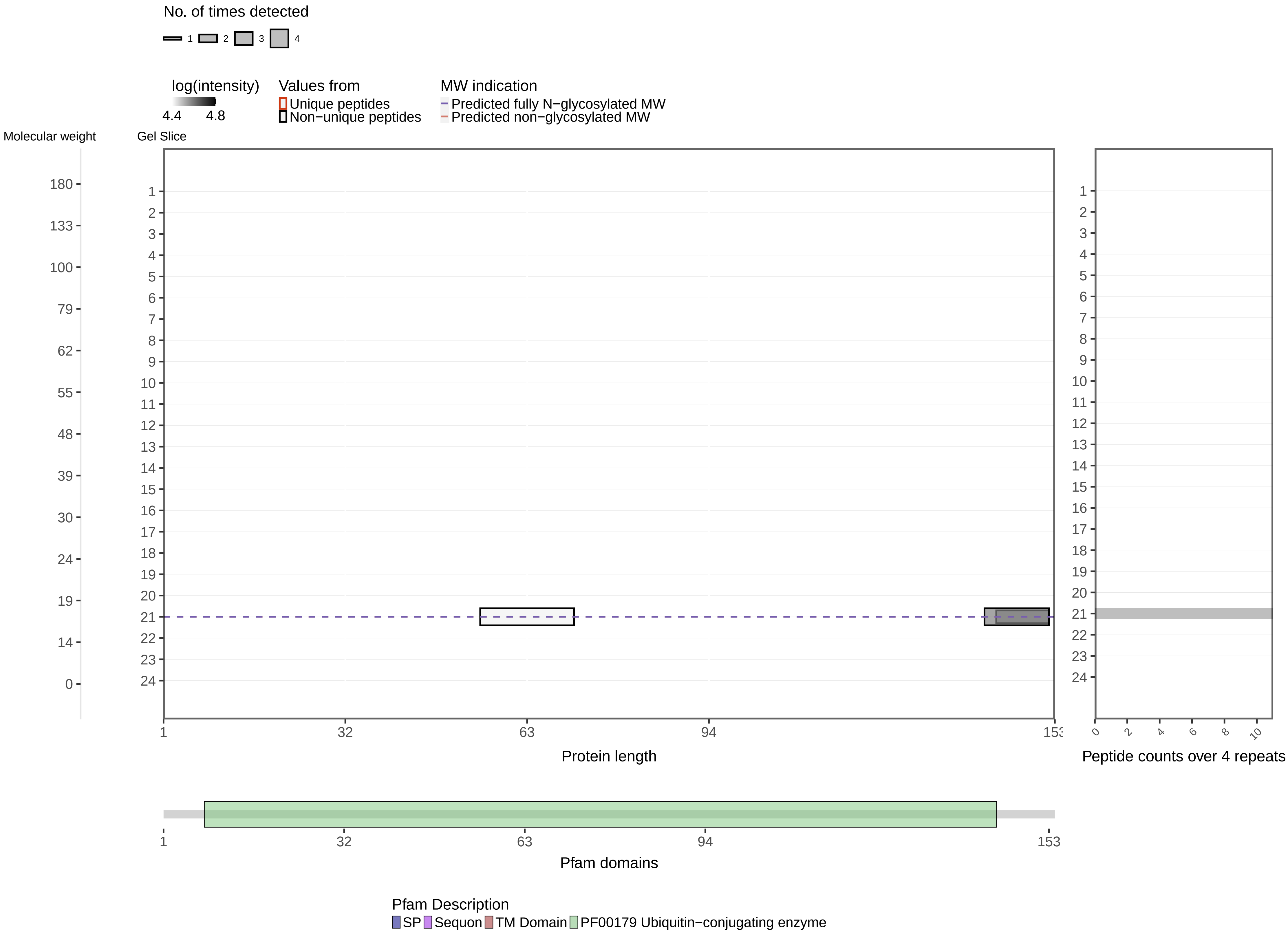


Protein sequence:

)YPNKPPTVRFISRMFHPNIYADGSICLDILQNQWSPIYDVAAILTSIQSLLCDPNPNSPANSEAARMFSENKREYNRKVREIVEQSWTAD*

NbL02g24320.1 aldose_1−epimerase−like [MW − SP = 37.17 kDa]

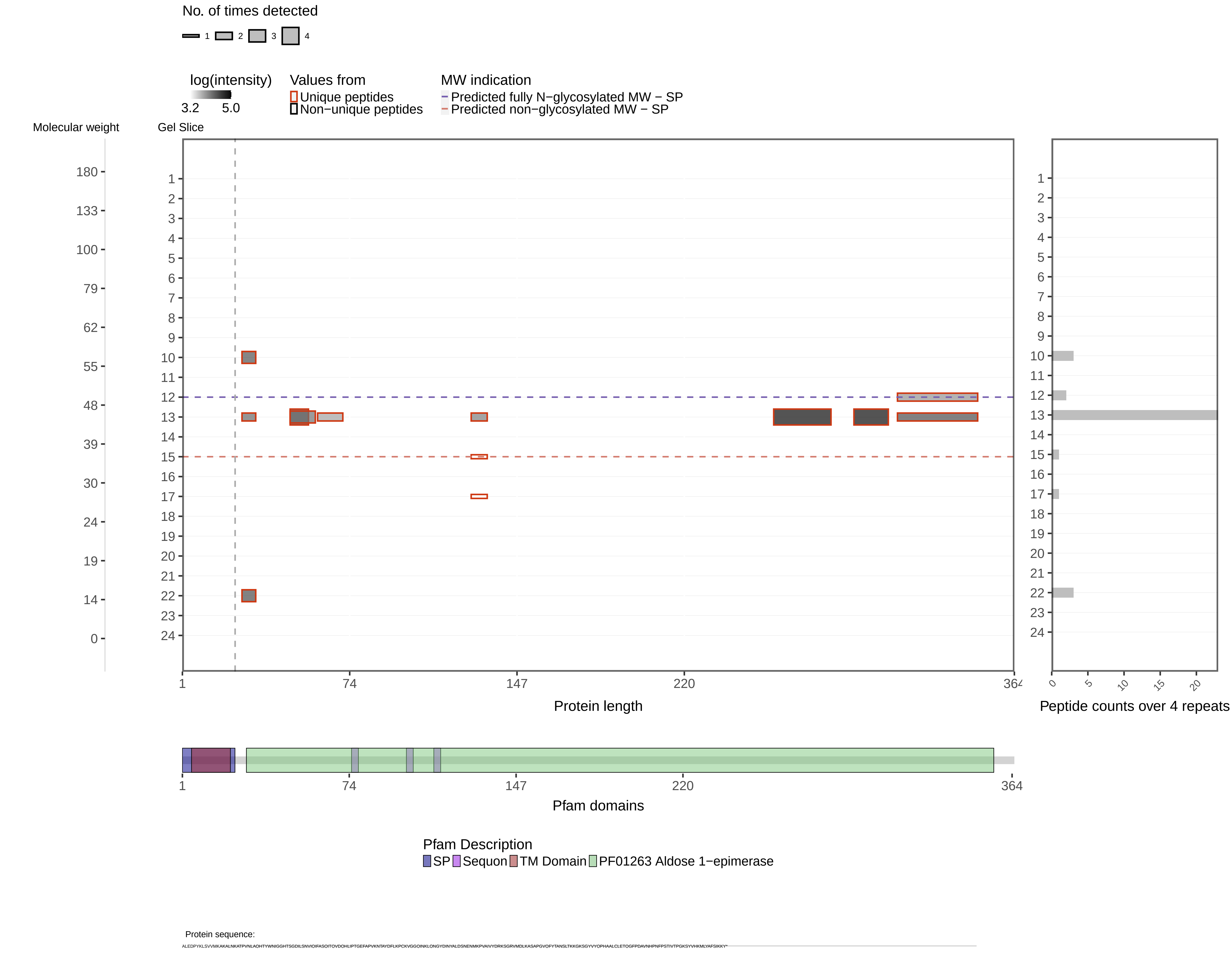

NbL02g24300.1 aldose_1−epimerase−like [MW = 40.23 kDa]

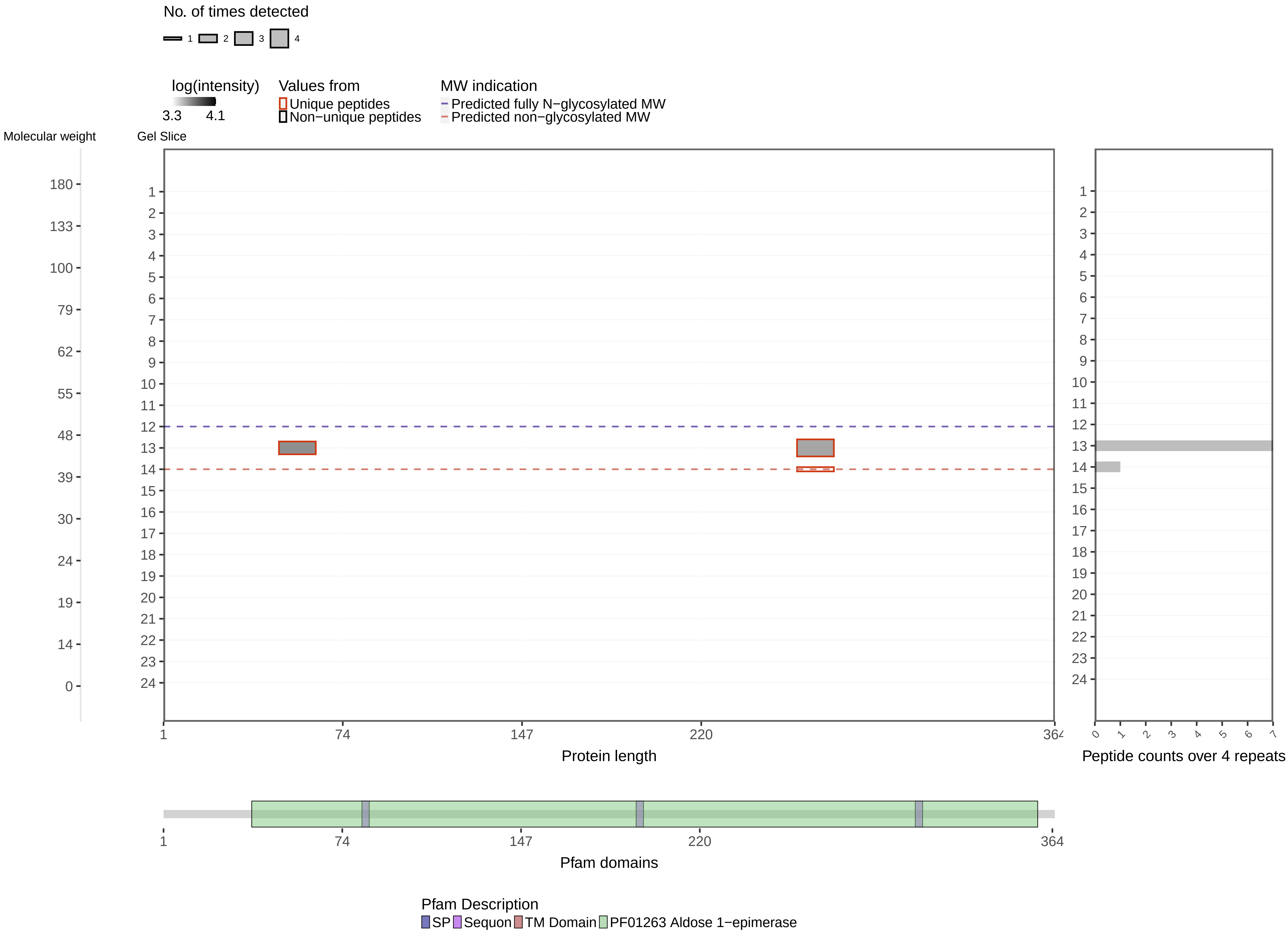


Protein sequence:

YTLKDPYKLSVVFRAKALNKATPINLSHHPYWNIGGHNSGDVLSQVLQIFGSHITLVDKELIPTGEIAPVKNTPYDFLKPRKVGSRINKLQNGYDINYVLDSNKKMKPVGLVYDKKSGRVMDVQASAPGVQFYTANFVNNTKGKGGFVYQPHSALSLETLVFPDAVNHPNFPSTIVNPGDKYVHSVLYTFSIKK*

### NbL02g24260.1 tubulin_alpha_chain−like [MW = 49.63 kDa]

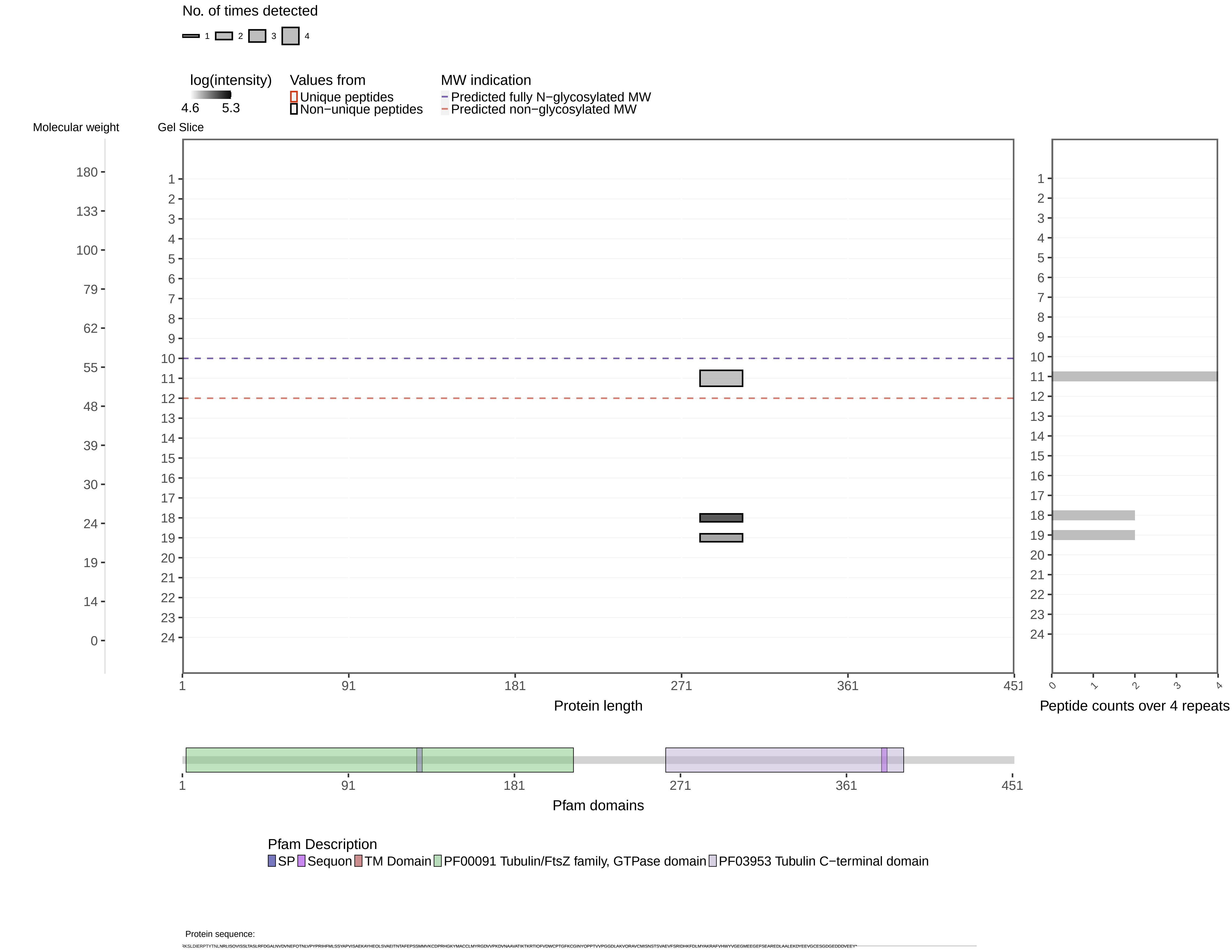

### NbL02g23980.1 cystathionine_gamma−synthase_1_chloroplastic−like [MW = 57.98 kDa]

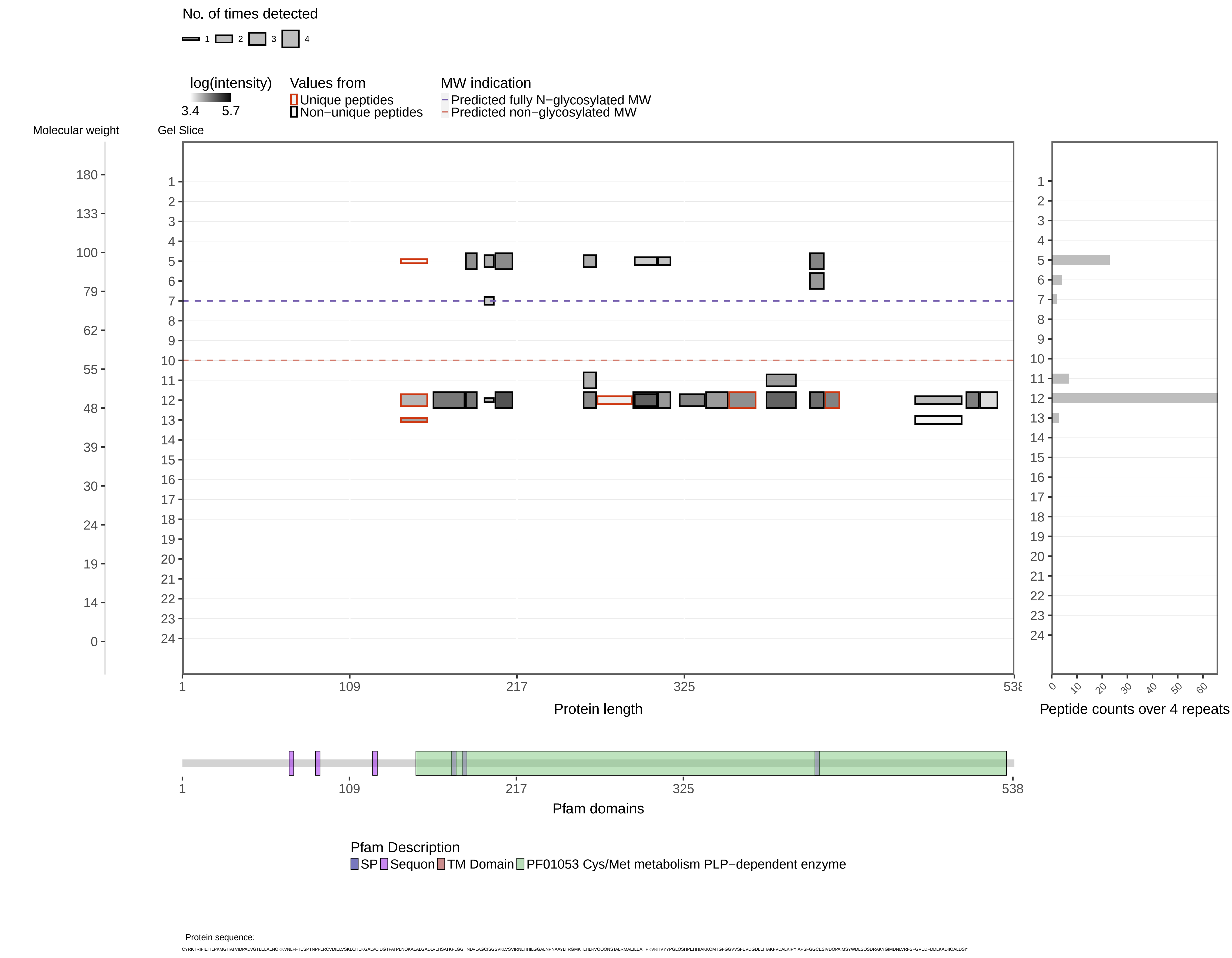

### NbL02g23860.1 uncharacterized_LOC109219811 [MW = 46.51 kDa]

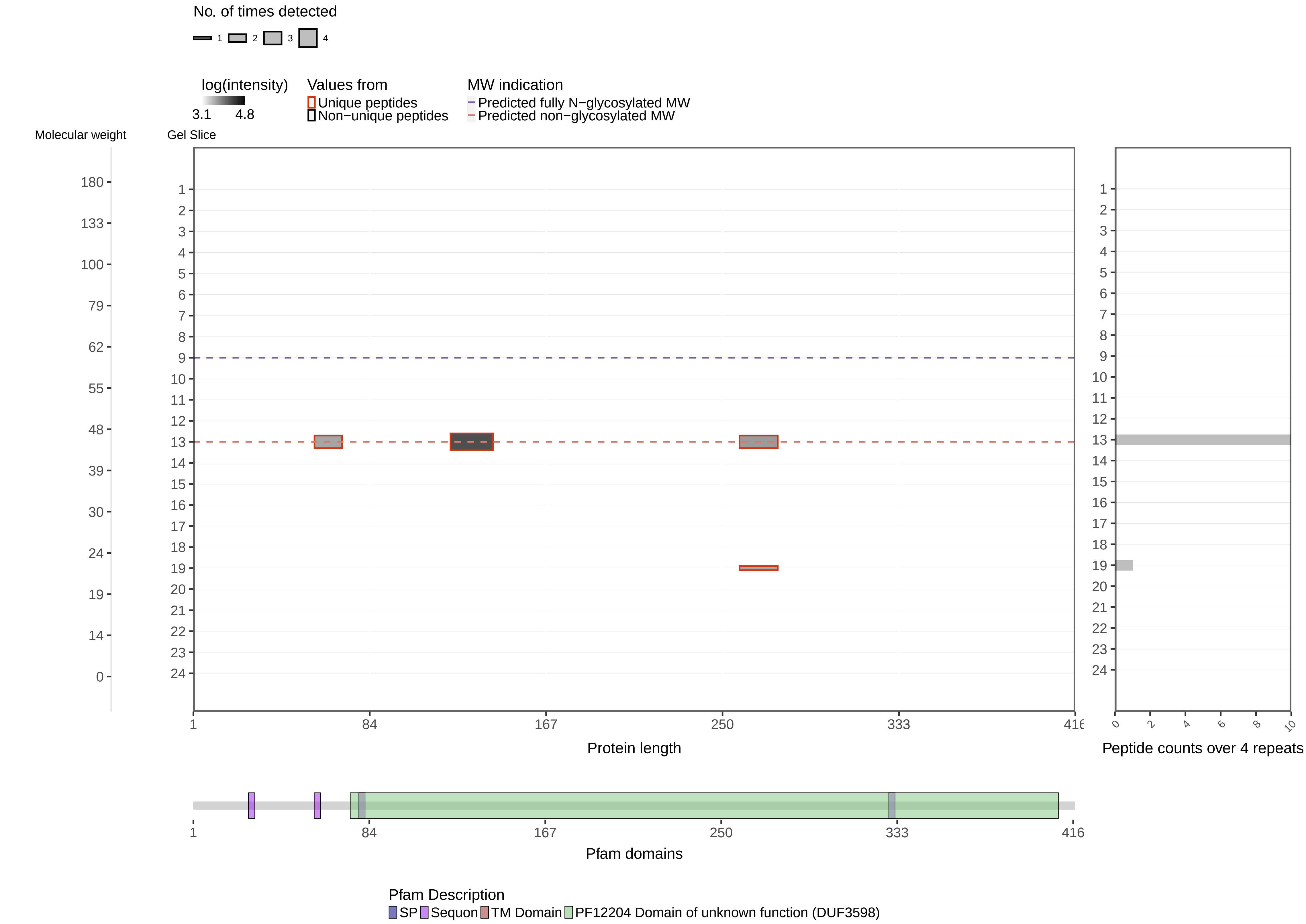


Protein sequence:

[illegible]GVLGEDDIGEESPRNLKLPSKKPSIVCENCLYSLEKDKRVRAFHIMDPKGILEMLLVFLEERGNGDVIPPSLDNFKEDSERIMPHLGTWKGHSLTKRTGVYGATITEAETTAVLEMNKDGOLVODITSTSGASNVTTNVHWTGTMSGNLVIFDGGFOLTLLPGGIYMGYPSDVANSVOESKSFHLELCWLEAPGKRORLIRTYDVEGLAVSSTYFIESKI*

### NbL02g23840.1 chorismate_mutase_1_chloroplastic−like [MW = 36.15 kDa]

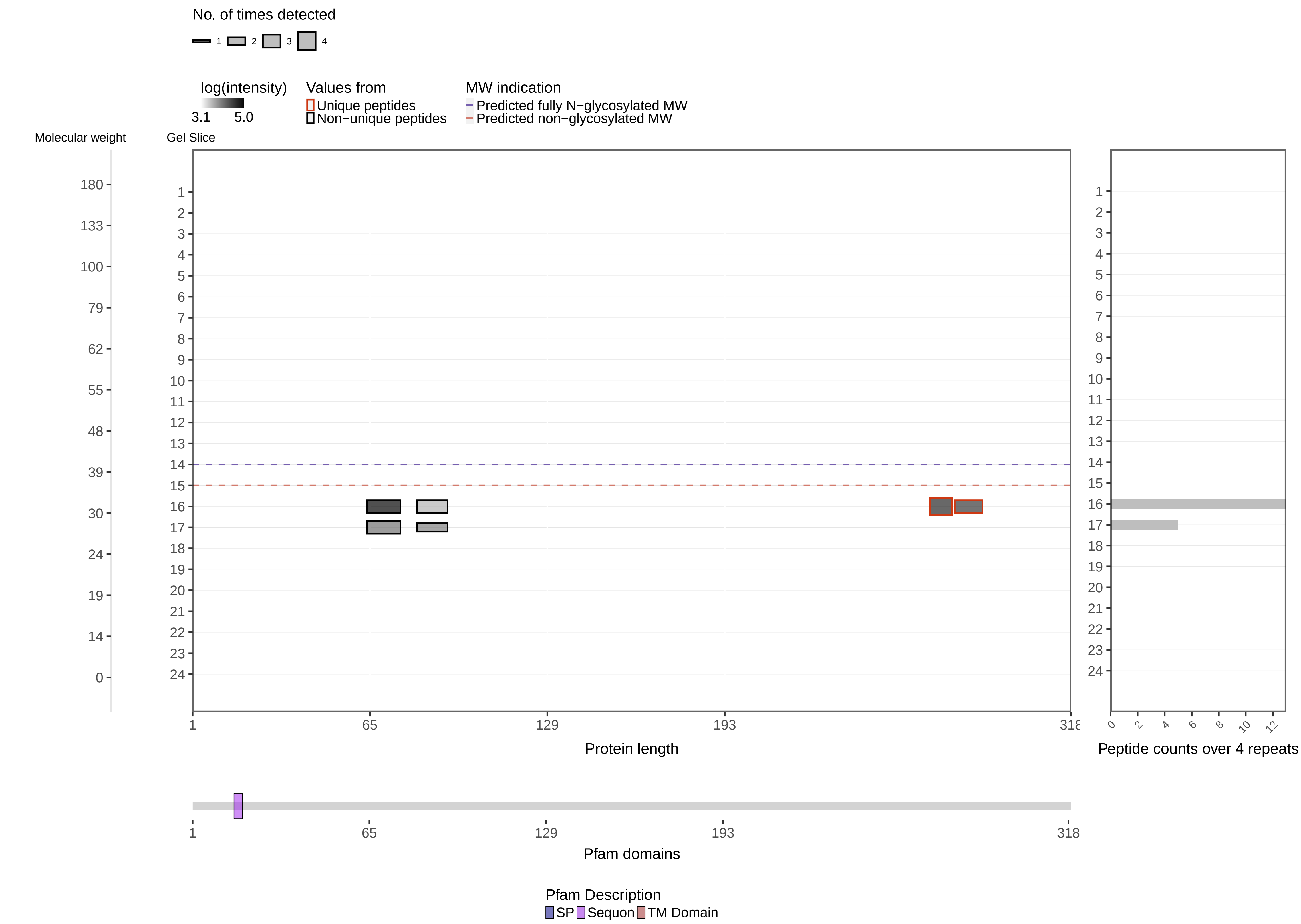


Protein sequence:

ELPEPMLPPMQFPKVLHSVADSININVKIWEMYFKNLLPRLVKEGDDGNFGSTAVCDTICLQVLSKRIHYGKFVAEAKFRASPDVYKAAIRAQDRNGLMDLLTYPTVEEAITRRVEMKTRTYGQEFNIKGPENGGDPVYKIKPSLVAELYGDWIMPLTKEVQVEYLLRRLD*

### NbL02g23780.1 probable_ubiquitin_conjugation_factor_E4 [MW = 117.79 kDa]

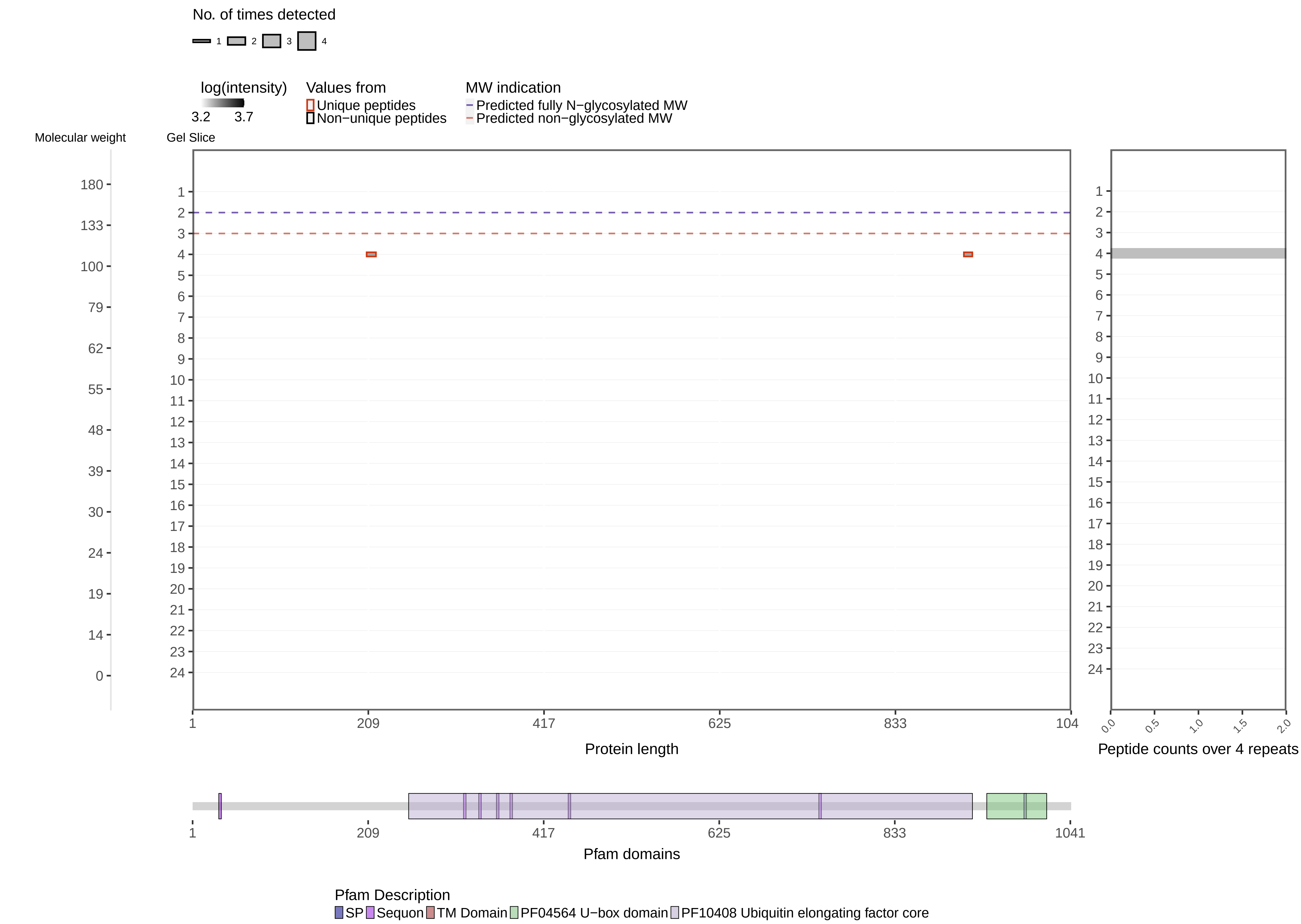


Protein sequence:

EDHLSTMKTMLEOAPSPOLOOEISRLEKELELYSOEKLCYEAOILRDGGLLORALSFYRLMVIWLVGLVGGFKMPLPSPCPMOFASMPEHLVEDAMELLIFASRIPRALDGVLLDDFMNFIIMFMASPEYIRNPYLRAKMVEVLNCWMPRRSGSTATSTLFEGHOLSLEYLVRNLLKVYYVDIEFTGSHTOFYDKFNIRHNIAELLEYLWOVPSHRNAWROIAKEEEKGVYLNFLNFLINDSIYLLDESLNKILELKELEAEMSNTVEWERRPAOEROERTRLFHSOENIIRIDMKL

### NbL02g23710.1 chaperone_protein_ClpB3_chloroplastic [MW = 109.64 kDa]

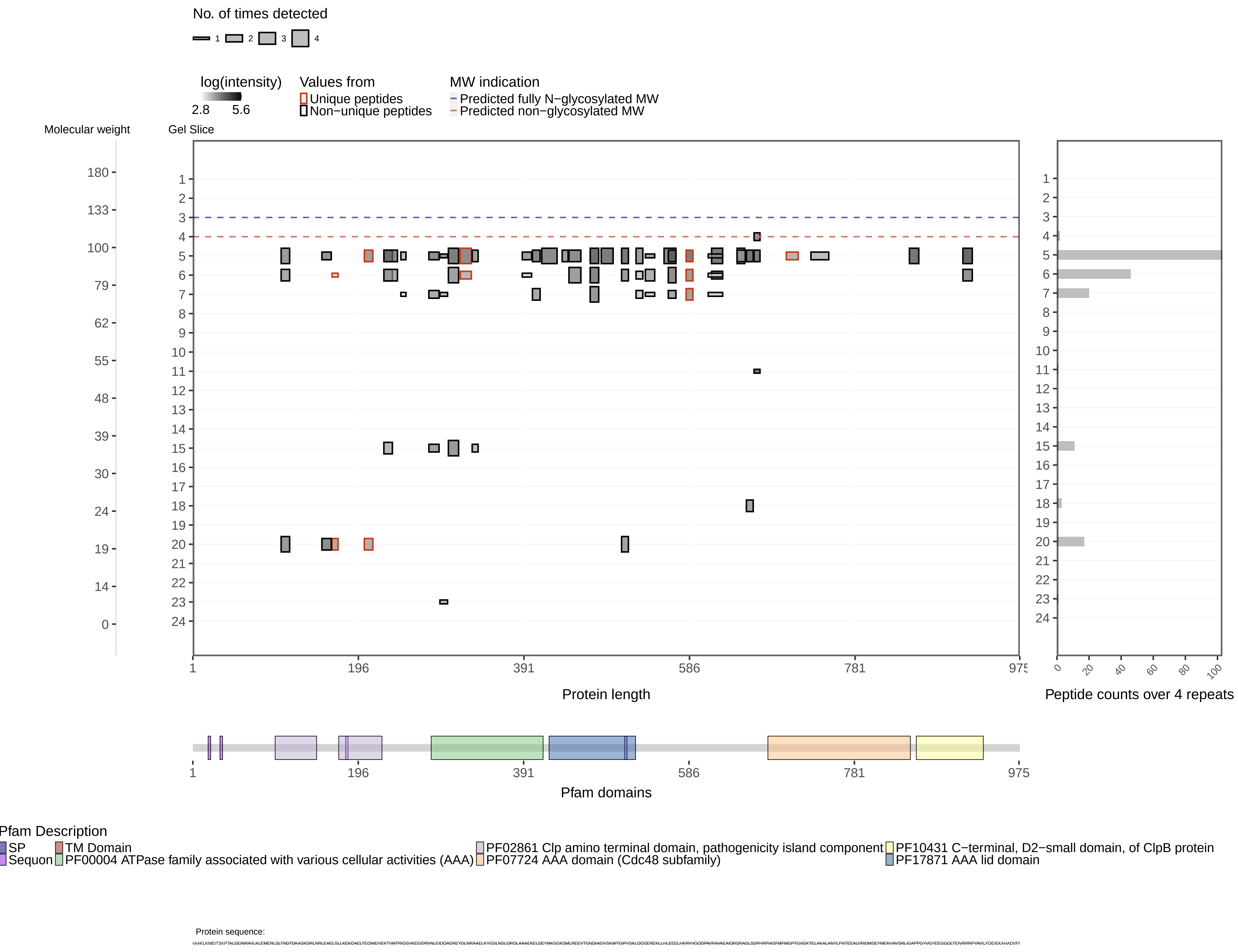

### NbL02g23660.1 UDP−glucose_6−dehydrogenase_3−like [MW = 53.07 kDa]

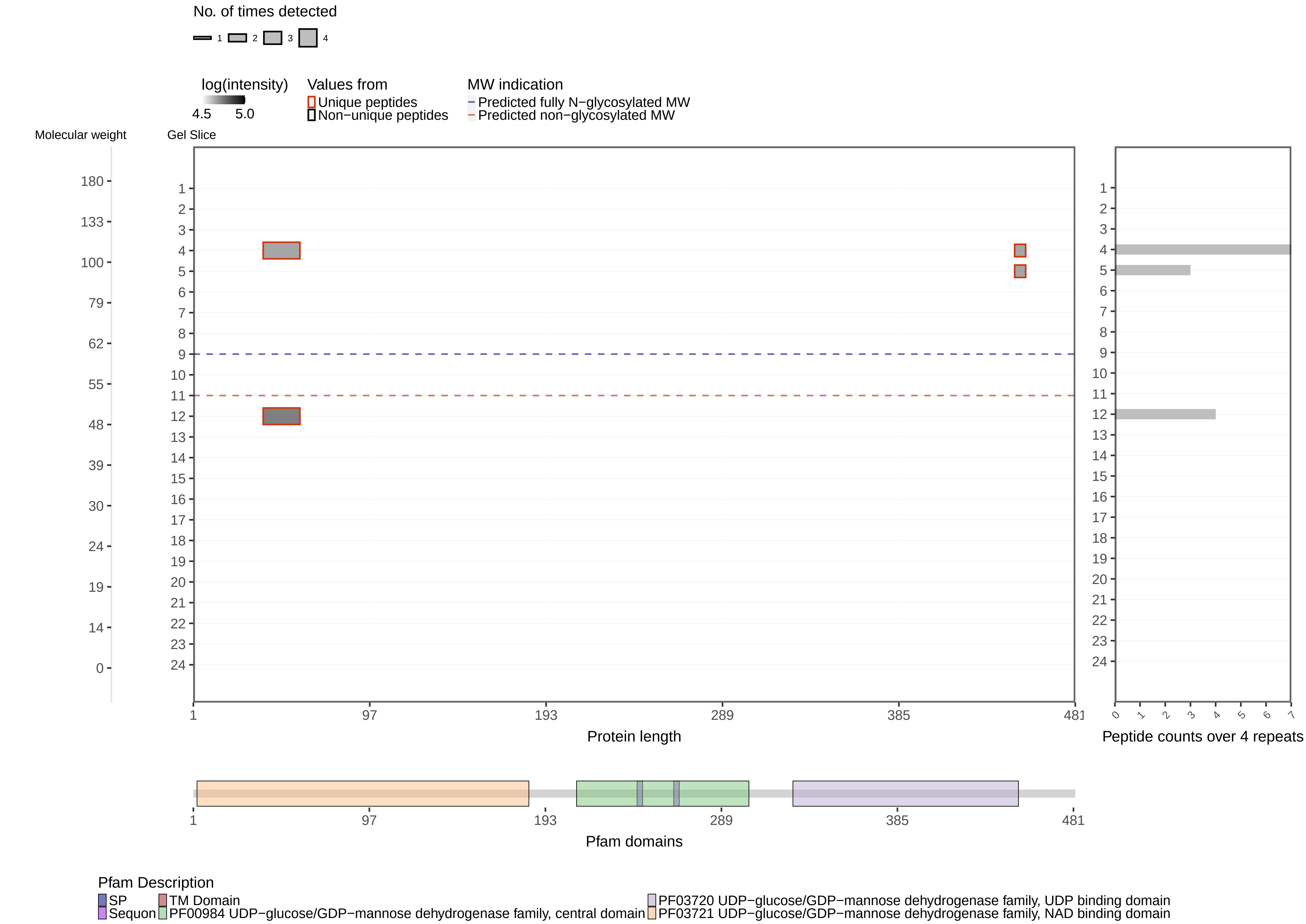


Protein sequence:

VNAMSALCEATGANVSQVSYAVGKDSRIGPKFLNASVGFGGSCFQKDILNLVYICECNGLPEVAEYWKQVIKINDYQKSRFVNRVVASMFNTVSNKKVAILGFAFKKDTGDTRETPAIDVCKGLLGDKARLSIYDPQVQEDQIQRDLSMNKFDWDHPLHLQPMSPTTKKLVSVTWDAYEATKDAHAVCILTEWDEFKTLDYQRIYENMQKPAFIFDGRNIVNVDKLREIGFIVYSIGKPLDAWLKDMPAVA*

### NbL02g23640.1 probable_mitochondrial−processing_peptidase_subunit_beta_mitochondrial [MW = 59.07 kDa]

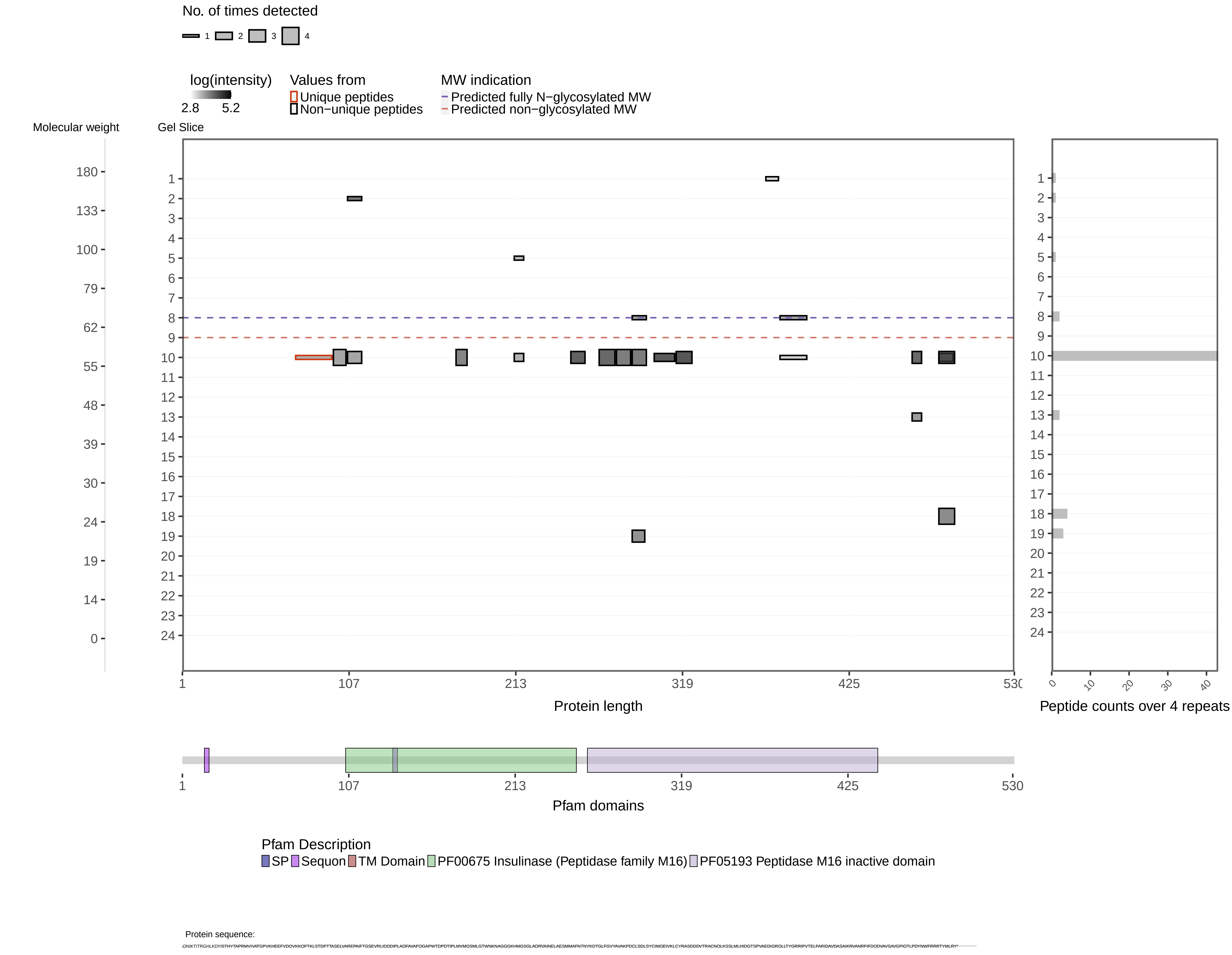

### NbL02g23580.1 glycine−rich_RNA−binding_protein_3_mitochondrial−like [MW = 31.21 kDa]

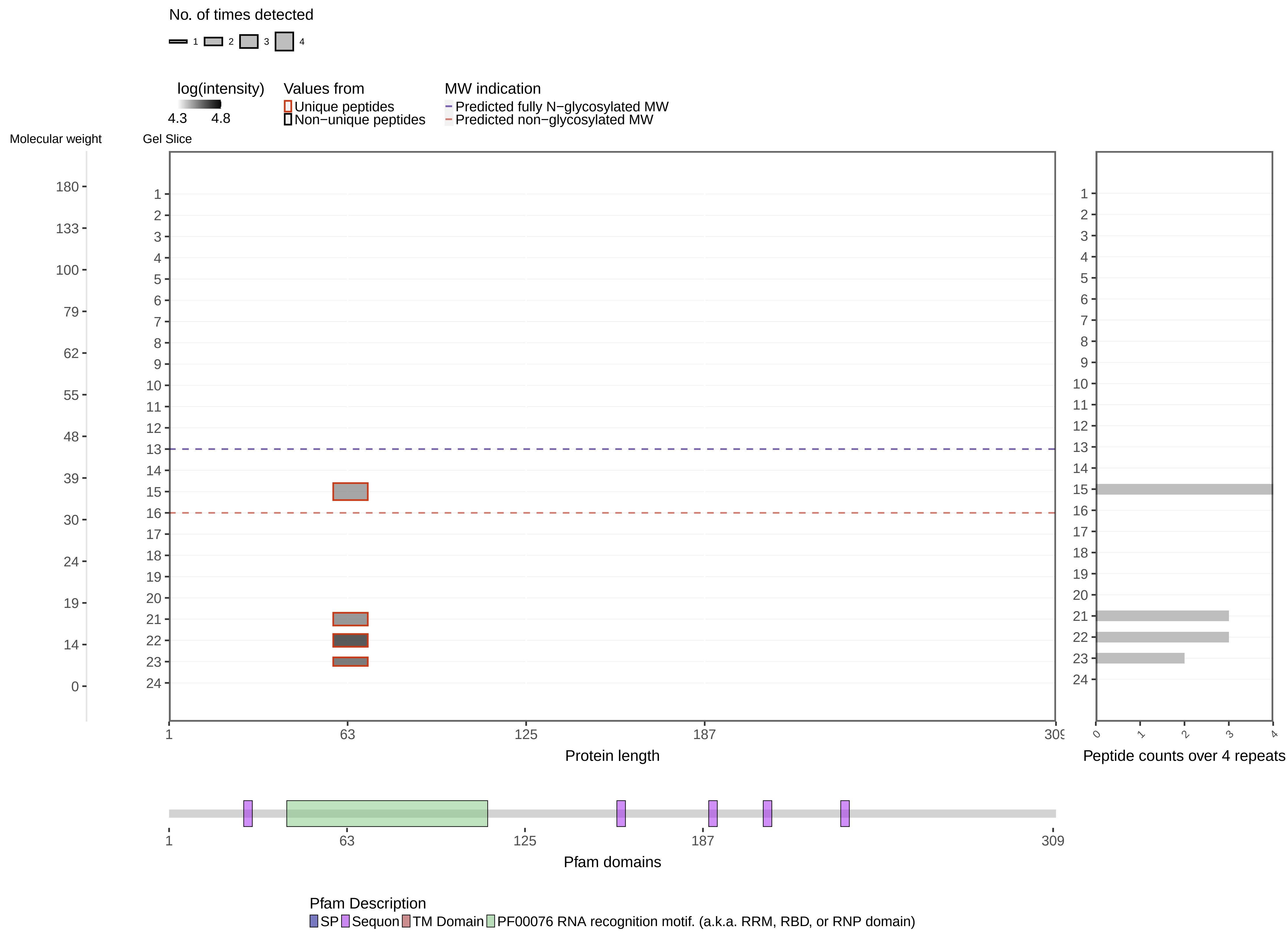


Protein sequence:

NYGGGGGGYGNSSGFNYGGEGGNSAGAGGYPTSNYGGGFSGGNSSTGGYGSYGGGSDYGNDNSSPVEGGNFGGGVGYGSGNYGNTPNSNYSGQGNSFSGGYDSGSSGNGFGESPGASFNGGQEQVSVDQGSQSLNEDIGQETSEGNYRDDDDEPKDFANTRG*

### NbL02g23560.1 serine_carboxypeptidase−like_25 [MW − SP = 50.53 kDa]

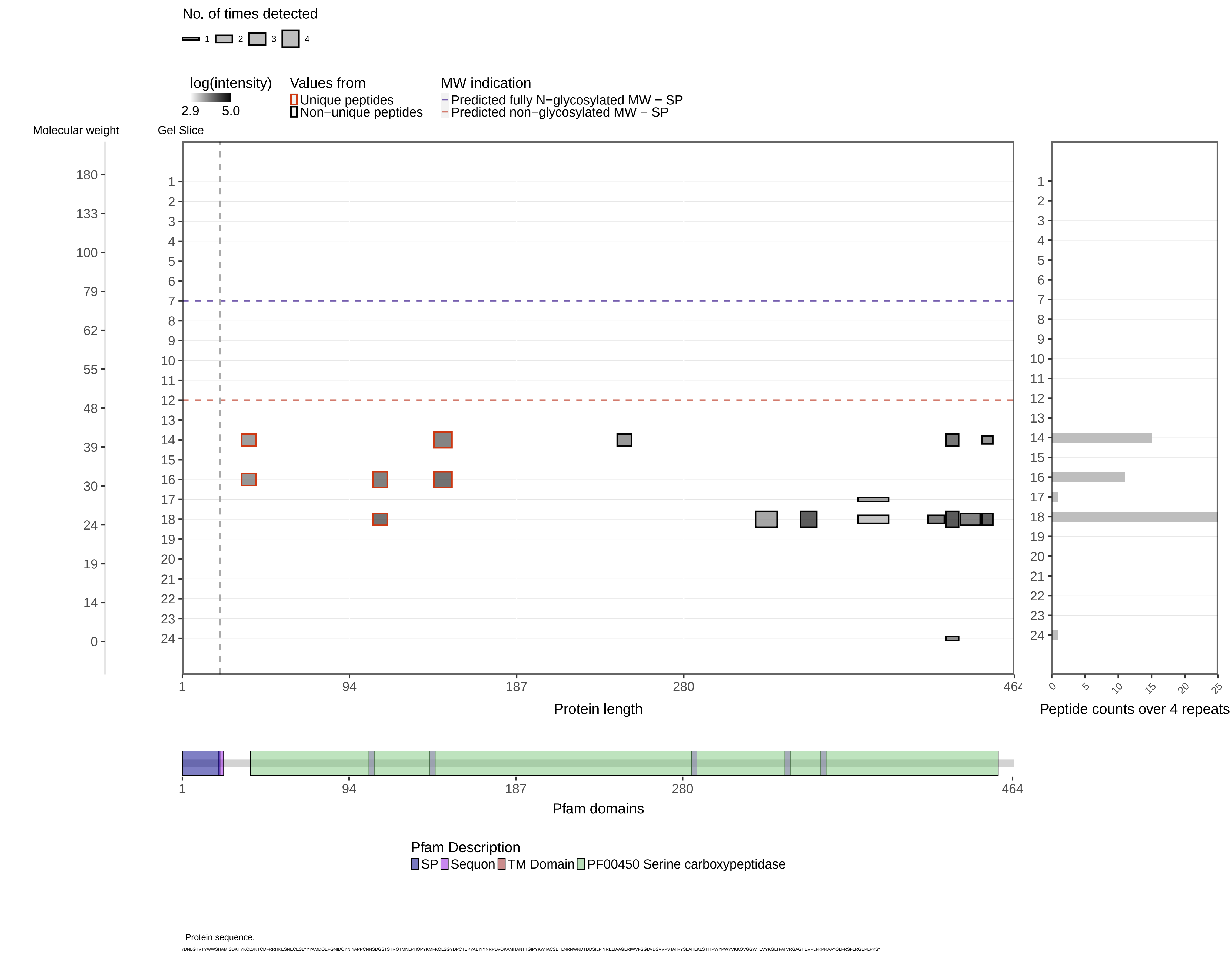

### NbL02g23450.1 uncharacterized_LOC109206144 [MW = 20.51 kDa]

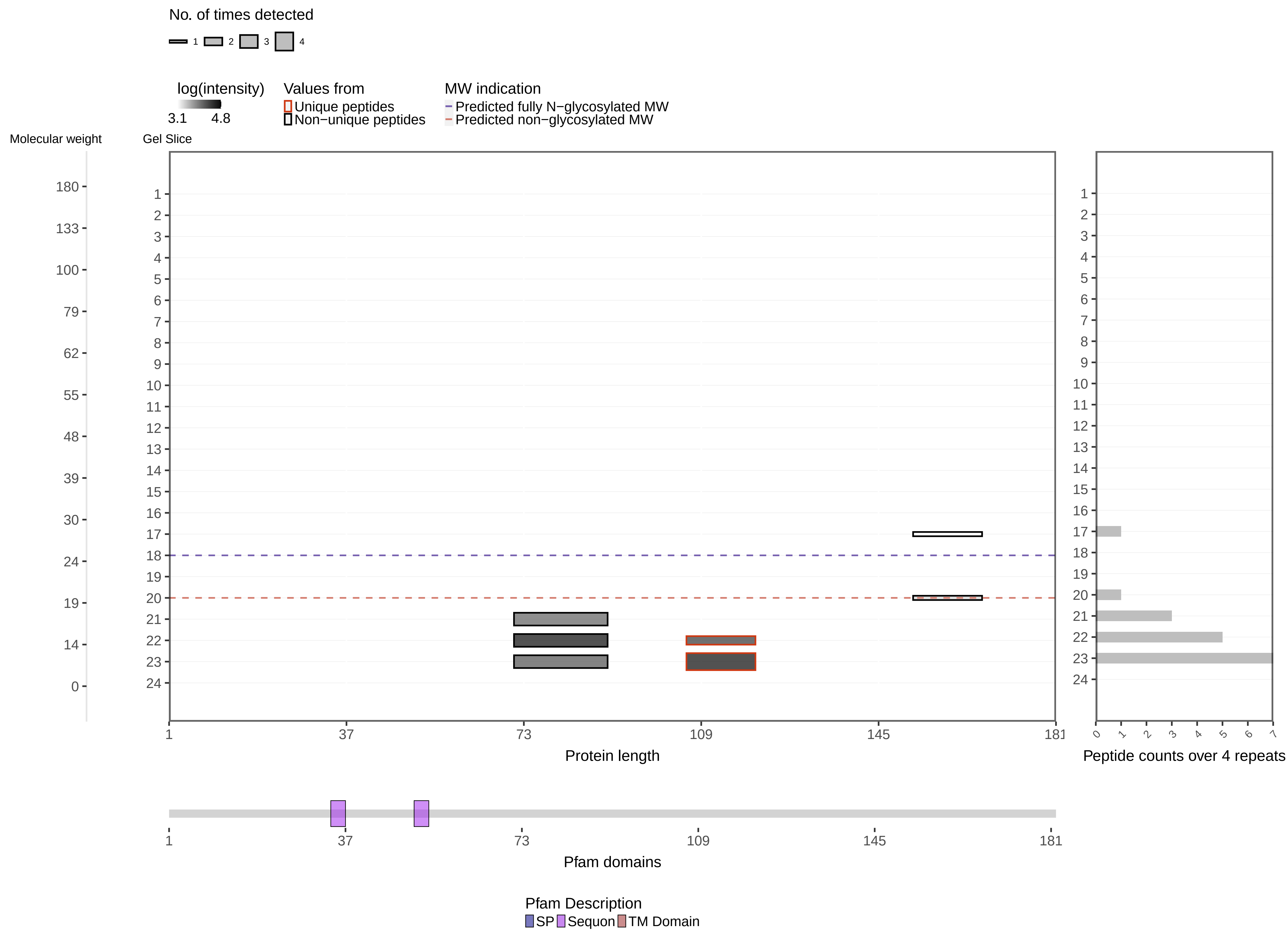


Protein sequence:

;POTSPEYLSAVPKATDYRLFCPATAOAKAIVPTSDPETVFDIKYFSRDORRNRPPIRRTFLNKADVEKMMMEKKTFDVNDFPOPYLTAKIEEDMDAIGGGYOK*

NbL02g23340.1 protein_COBRA−like [MW − SP = 46.99 kDa]

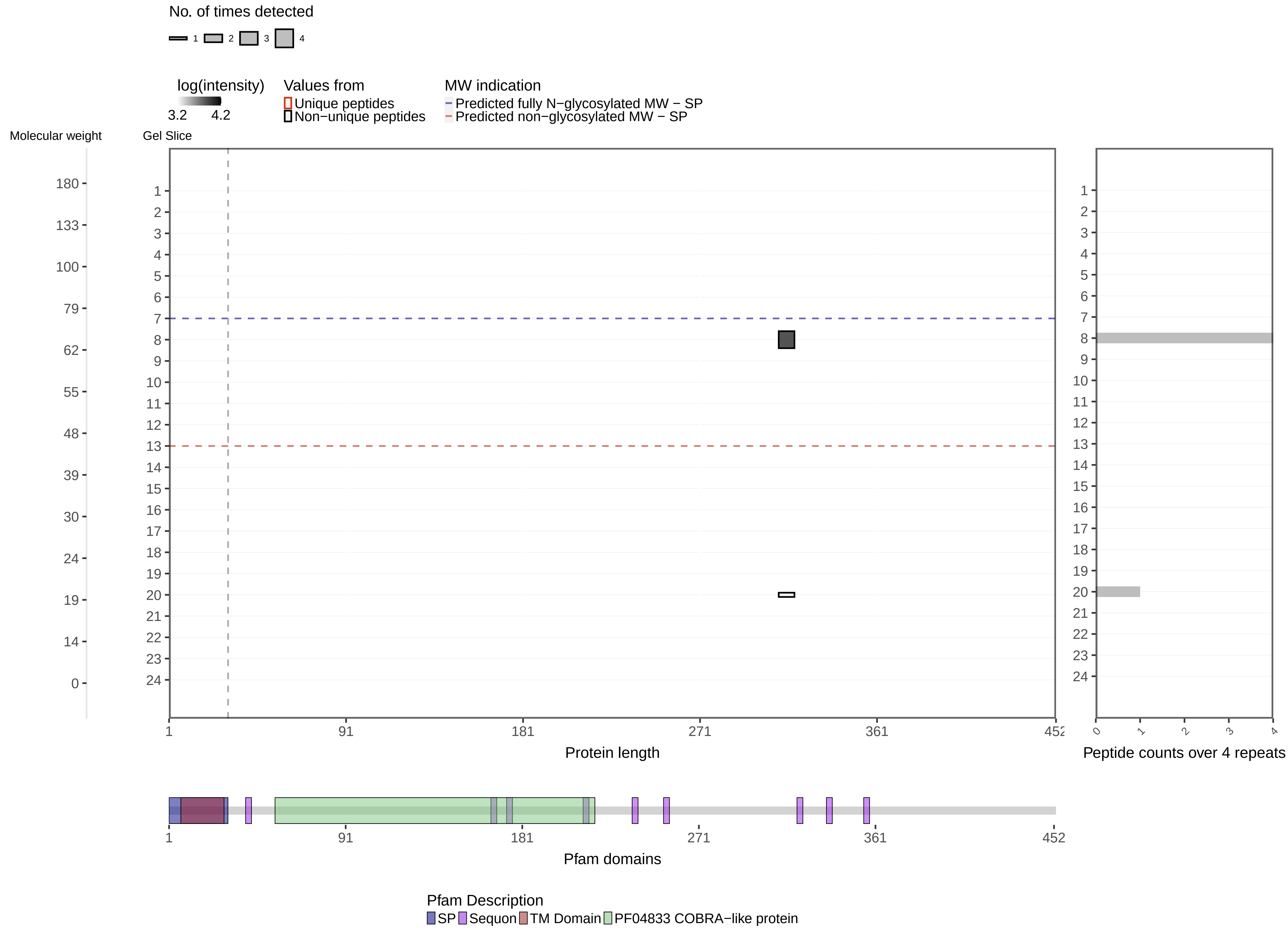


Protein sequence:

'TCTYSQFLAQKTPTCCVSLSSFYNDTIVPCPTCTCGCQTNGTQRGNCVDSKTPHLASVVSDHGKTNYVPLVQCTNHMCPIRIHWHVKVNYKEYWRVKITITNFNYRMNYTQWNLVVQHPNFDNLTTLFSFNYKSLTPYGAINDTAMLWGVKFYNDLLMQAGPLGNVQSELLFRKDASTFTFEKGWAFPRRIYFNGDNCVMPPPDAYPYLPNDGSRWGVSLLKLLVTLISSTAIVLAYV*

### NbL02g22940.1 putative_endo−13(4)−beta−glucanase_2 [MW = 75.97 kDa]

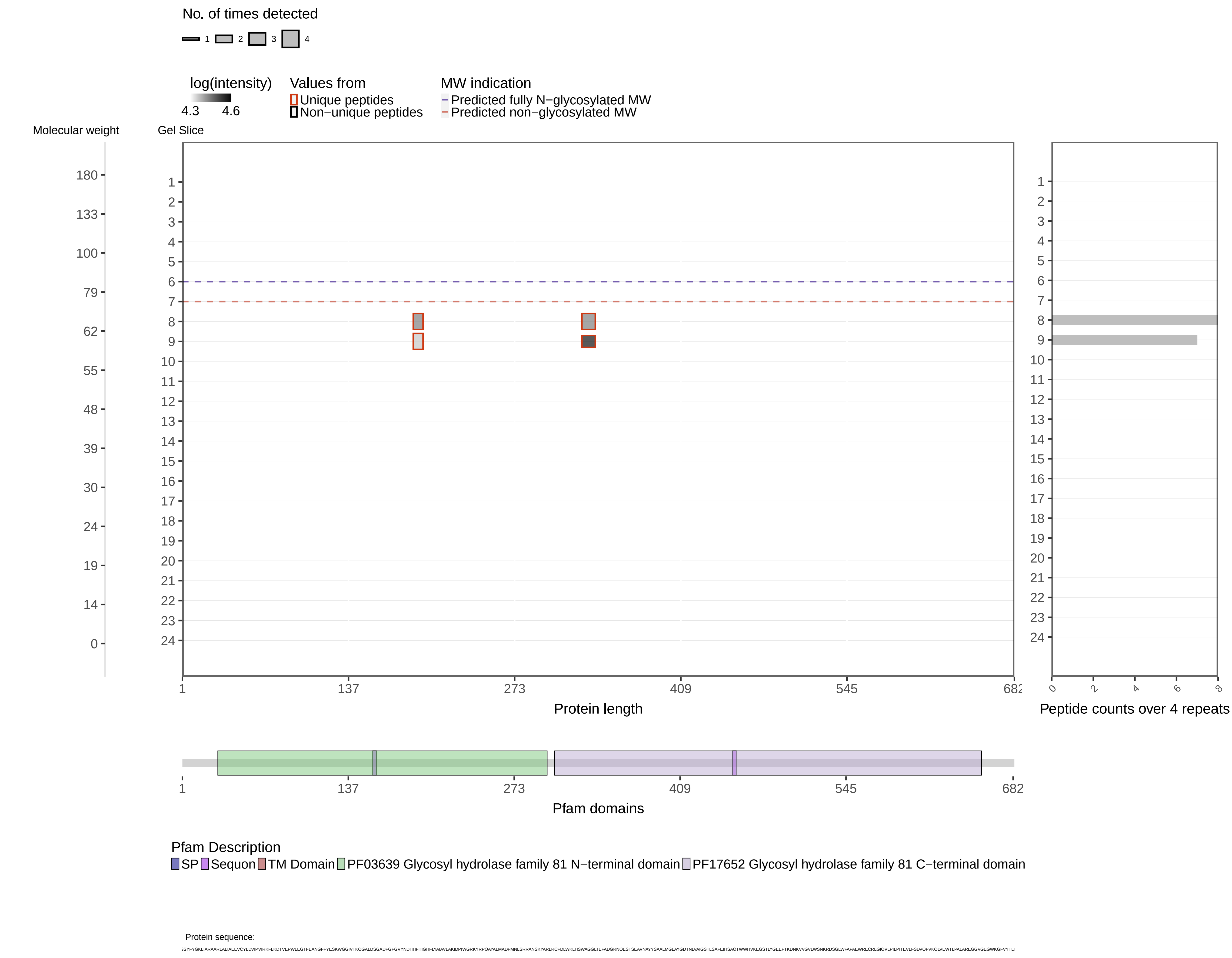

### NbL02g22710.1 protein_DJ−1_homolog_D [MW = 41.79 kDa]

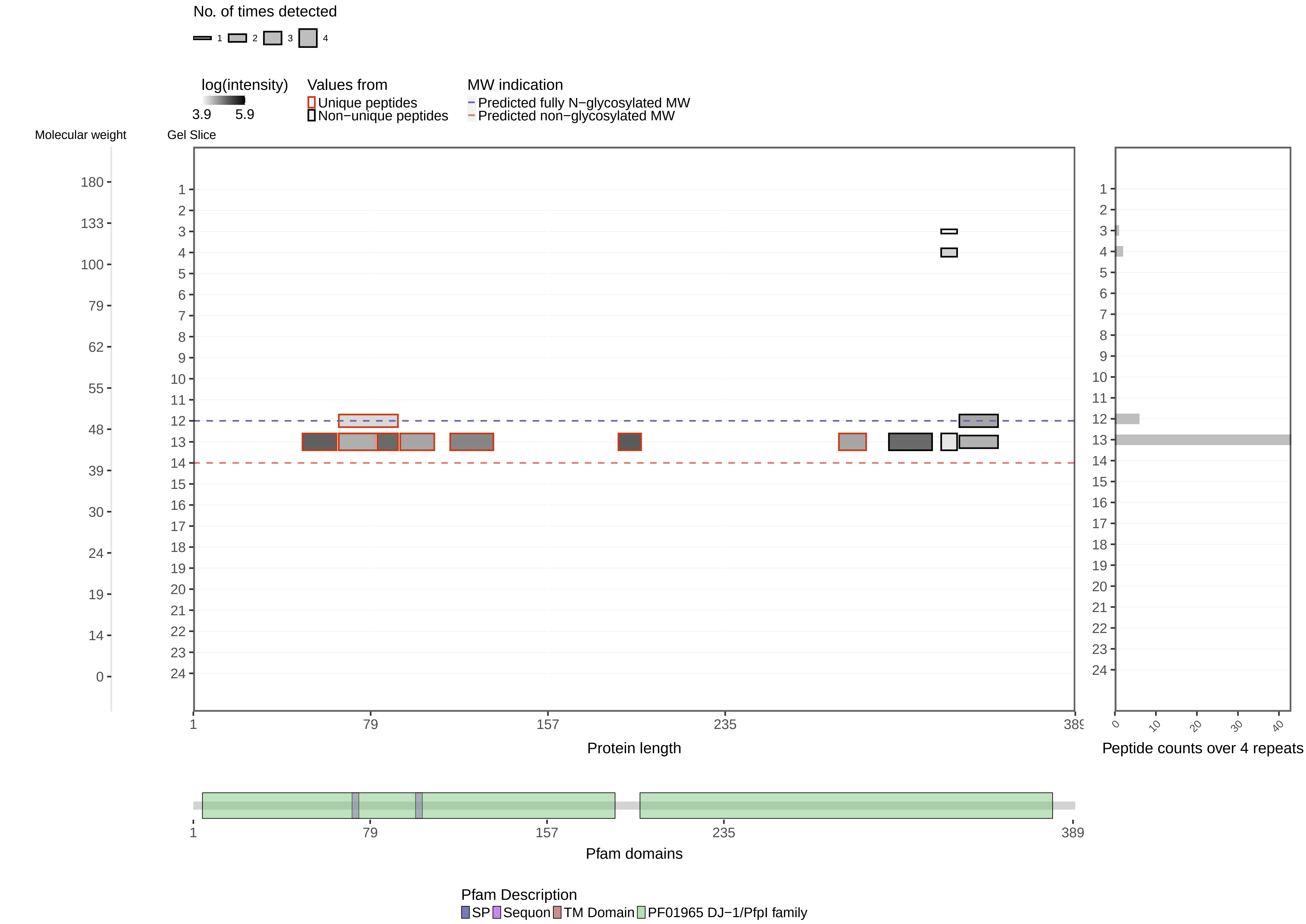


Protein sequence:

RLFVKALGGSIVGSGKRILFLCGDFMEDYEVMVPFQSLQALECHVDAVCPRKKAGDKCPTAVHDFEGDQTYSEKPGHDFTLNADFESVDSSSYDGLVIPGGRAPEYLALDDAVIKLVKEFMQSKKPVASICHGQQILSAAGVLKGKKCTAYPAVKLNVVLGGGTWLEPEPIDRCFTDGHLVTGAAWPGHPEFISQFMALIGVHVKF*

### NbL02g22690.1 sucrose_synthase_7−like [MW = 98.82 kDa]

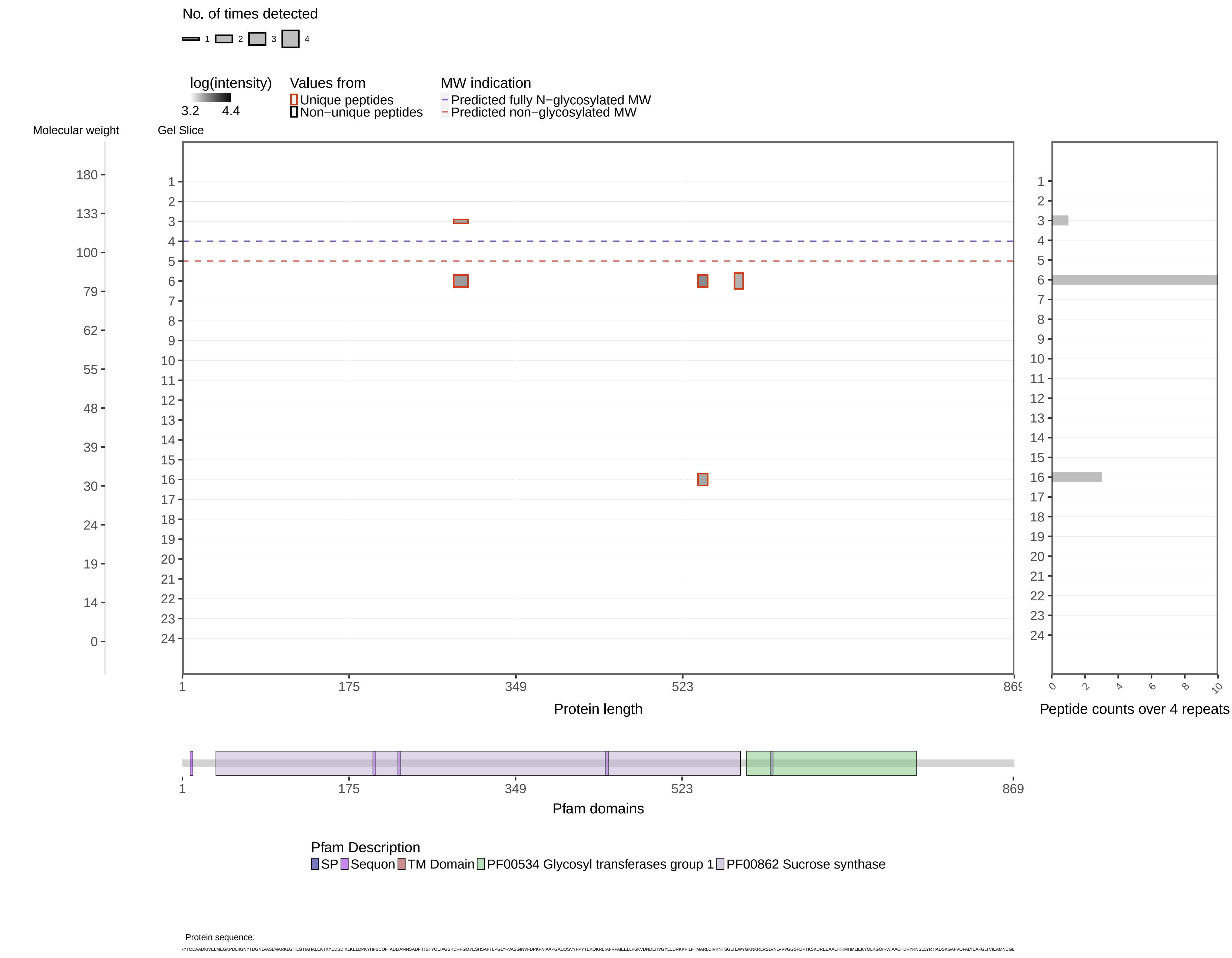

NbL02g22630.1 HSP−interacting_protein−like [MW = 87.91 kDa]

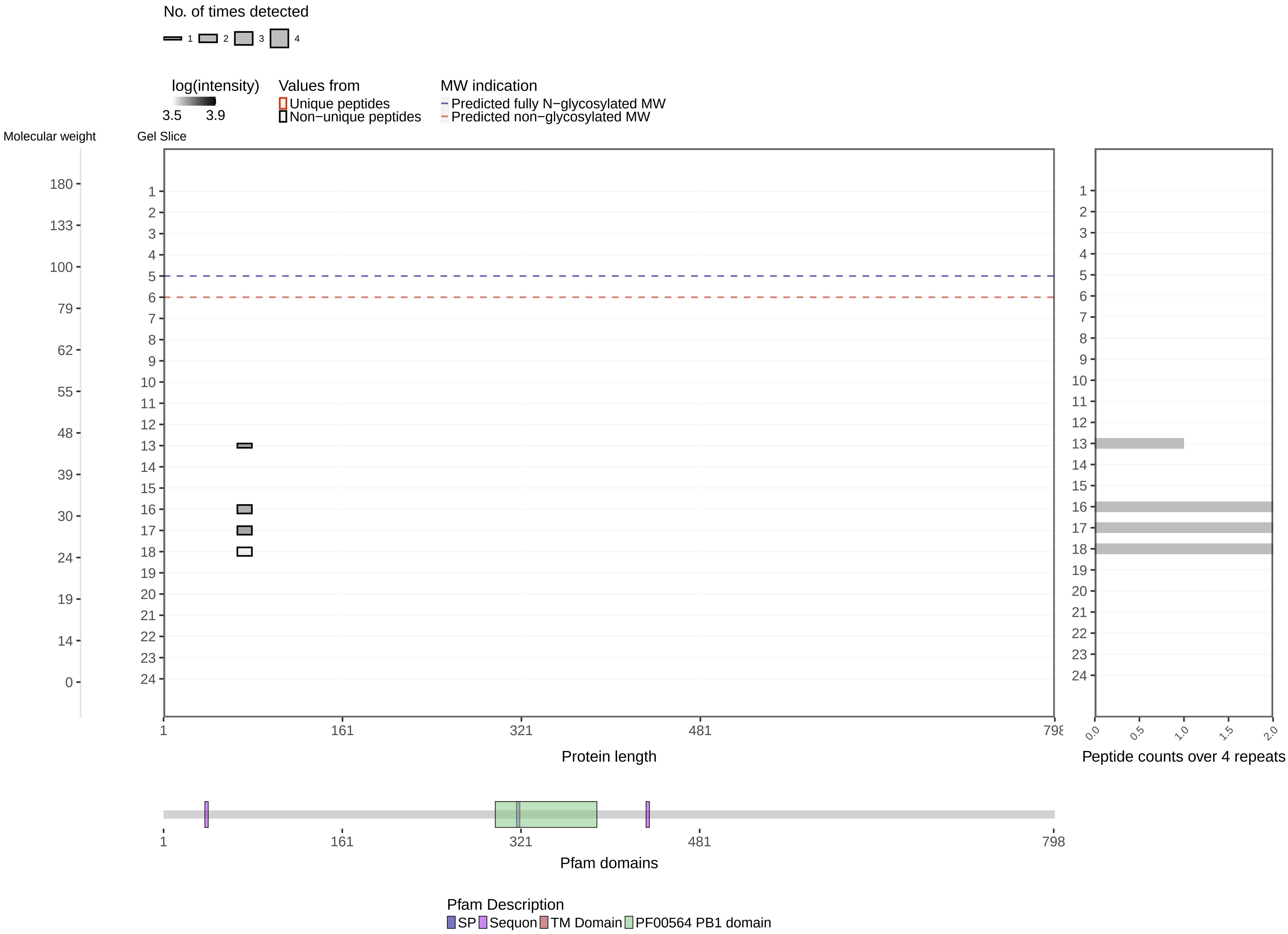


Protein sequence:

LHVVEVSPEQEPALLEEEEEKLVESEPSKGDDSGSHSSISDSVVETVDNESNKAEKDATKEKVATTENPDCKEVEMDDWLFEFAQLFRTHVGIDPDAHIDLHELGMELSSEALEETVTSEEAQALFDKAALKFQEVAALAFFNWGNVHMCAARKRIPIDDSASKEMMATQLQAAYDWVKEKYSLAKEKYEEALLIKPDFYEGLLALGQQQFEMAKLYWSFILAKKEDLLNWDPAETLALFDSAEEKMKAATEMWEKLEEQRAKELKDPSASKKDELLRRRKKQGSGPEGE[illegible]

### NbL02g22620.1 1−aminocyclopropane−1−carboxylate_oxidase−like [MW = 36.19 kDa]

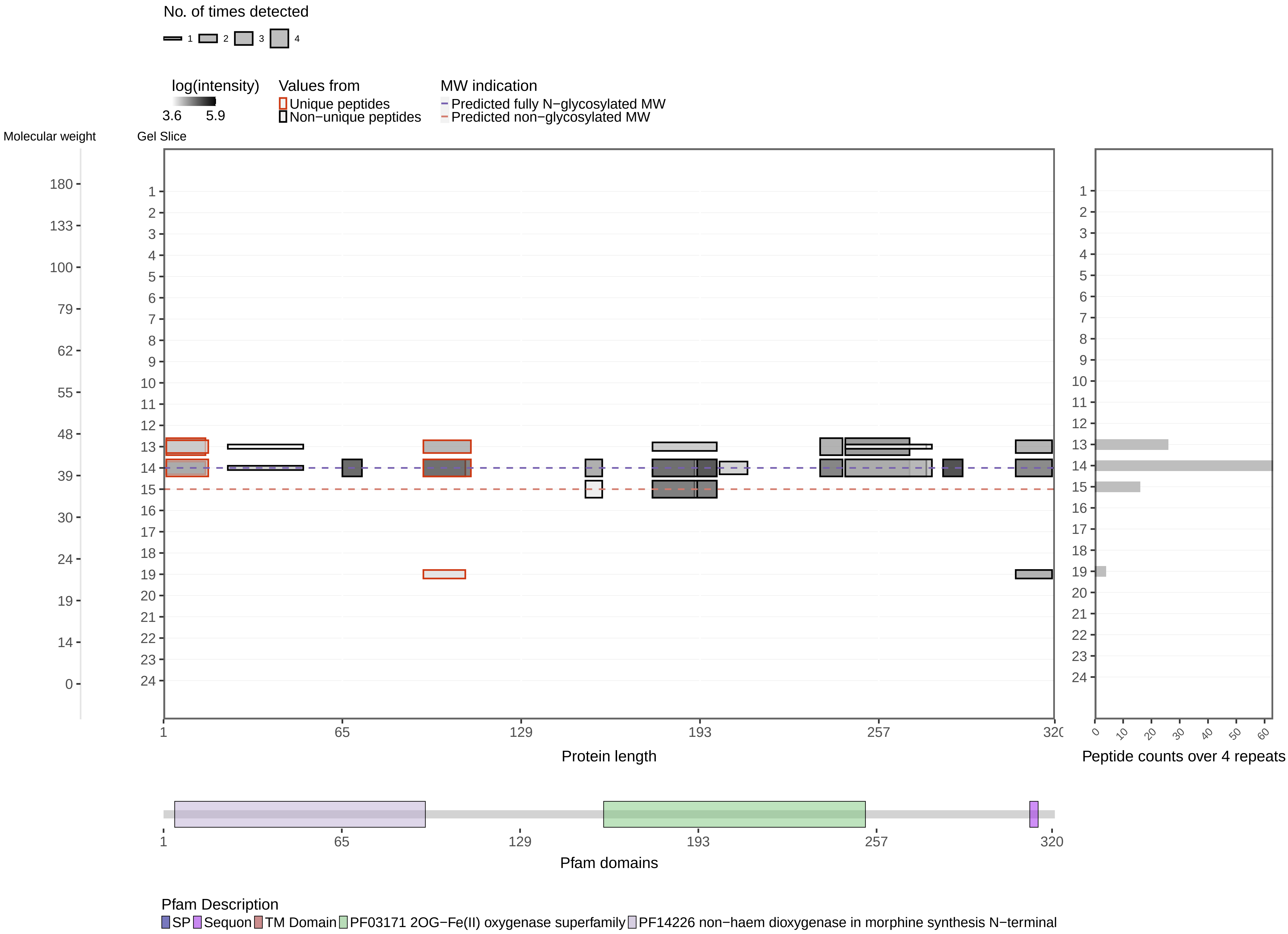


Protein sequence:

### NbL02g22600.1 light−induced_protein_chloroplastic [MW = 34.95 kDa]

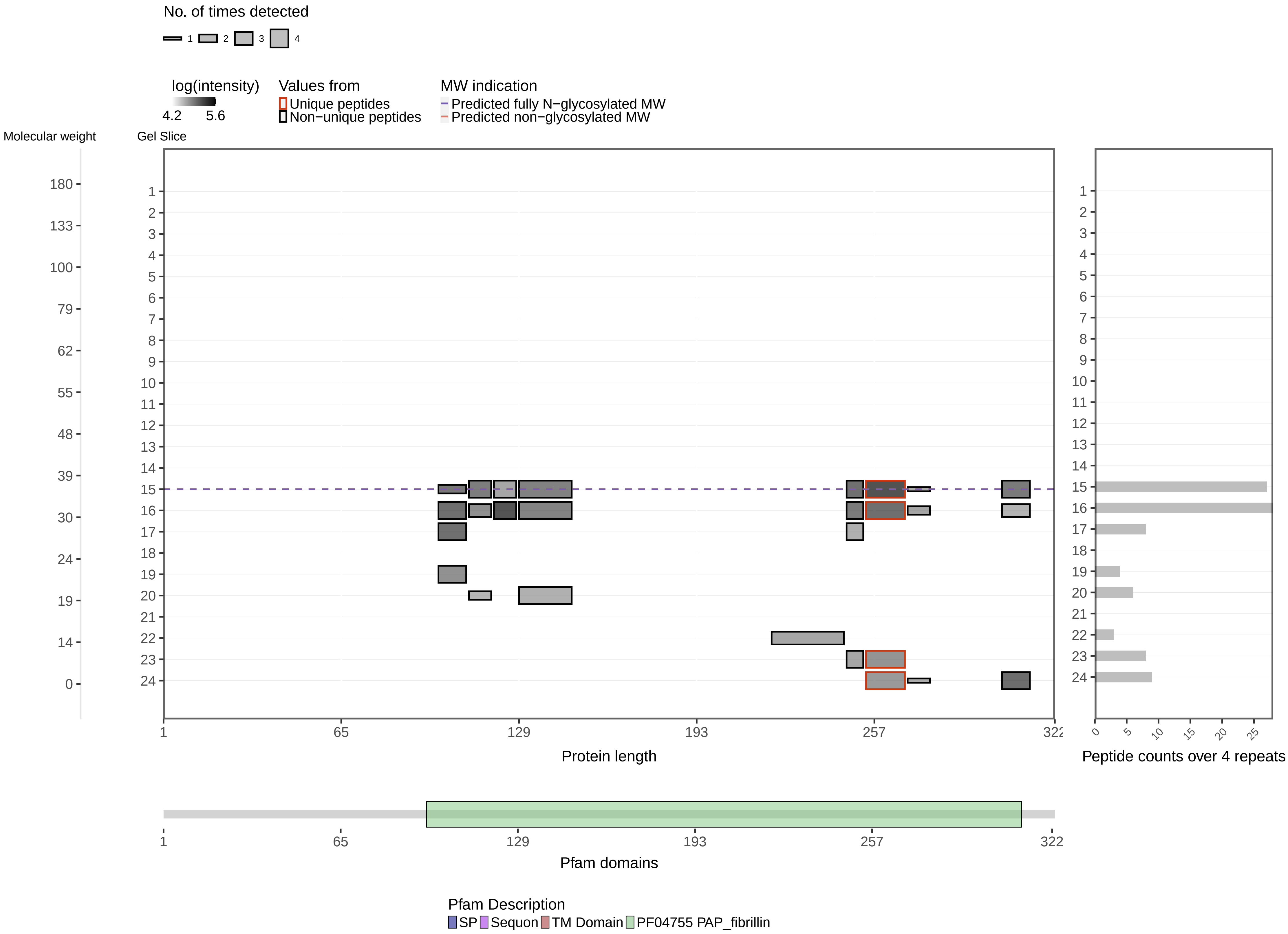


Protein sequence:

GKWILAYTSFSGLFPLLSRGTLPLVRVEEISOTIDSEAFAVONSVVFAGPLATTSITTNAKFEVRSPKRVOIKFDEGIIGTPOLTDSIELPENVEFLGOKIDLSPFKGLVNSVODTASSVAKSISSOPPIKFPISNSNAOSWLLTTYLDDELRISRGDGGSVFVLIKEGSPLLKP*

NbL02g22590.1 pyrophosphate−−fructose_6−phosphate_1−phosphotransferase_subunit_beta [MW = 61.78 kDa]

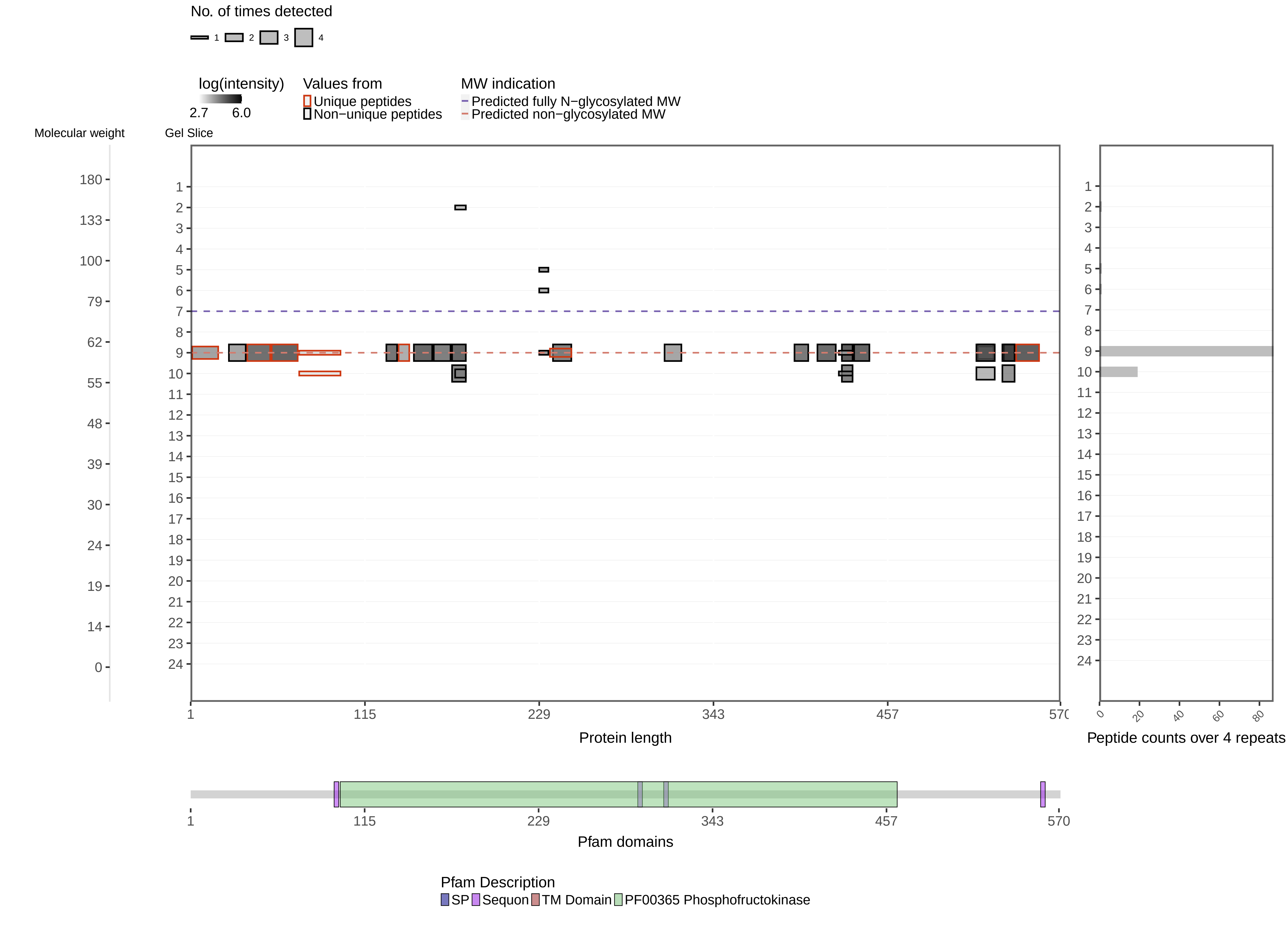


Protein sequence:

### NbL02g22570.1 UBA_and_UBX_domain−containing_protein_At4g15410−like [MW = 32.69 kDa]

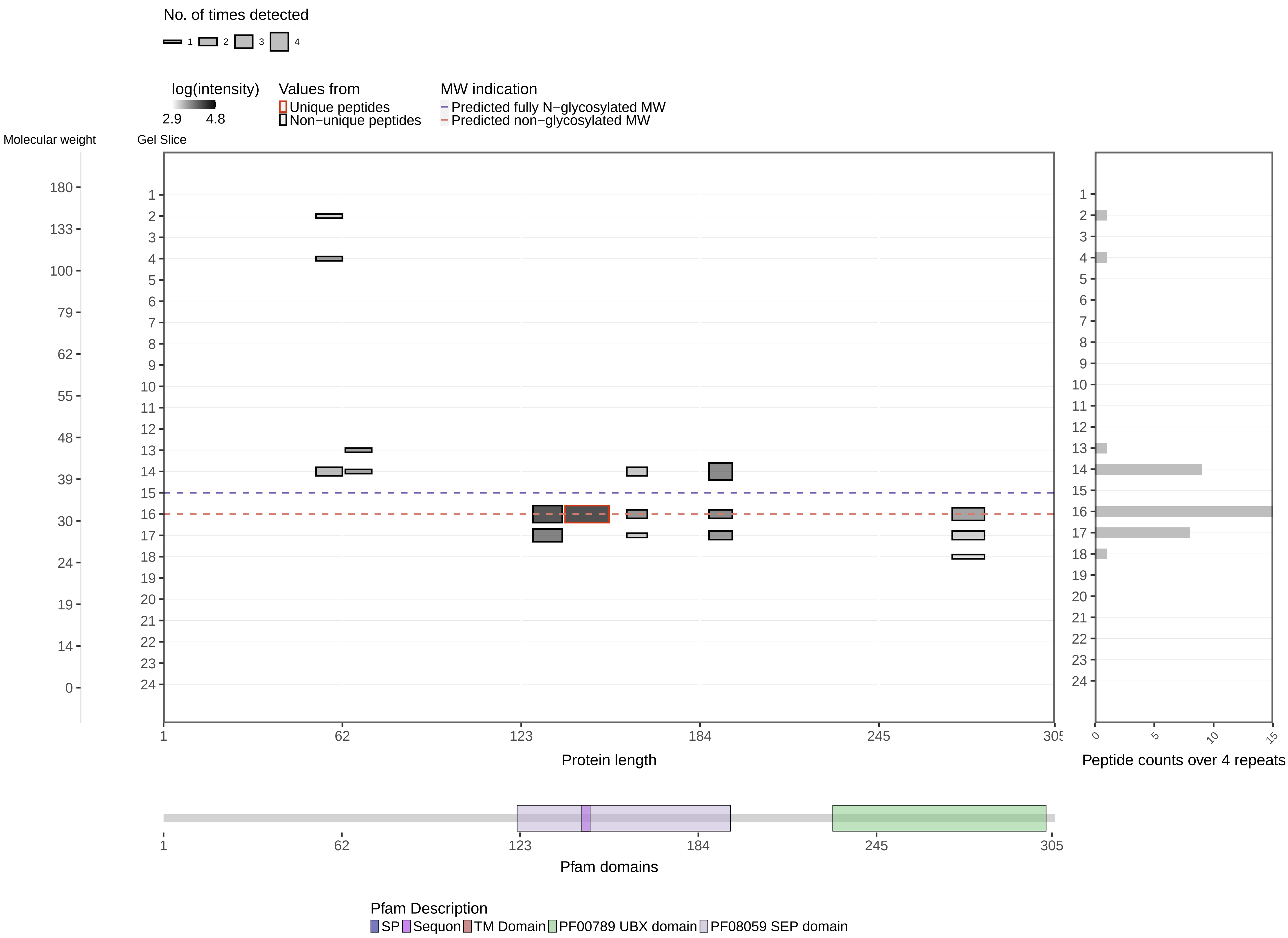


Protein sequence:

.RSLDDPENASFLESITKSECPRELAPADTRSQVHCNLIRRDENCPEPEKRQVPFQGVGRTLGSSSIPATSEPSASVPISTASAPTSILDDSLPSTSLQIRLADGTRVVAHFNFQQTVGDIRAFIDASRPSDTRAYLLQTVGFPPRILCDTTQTIEQAGLANSVVIQKF*

NbL02g22460.1 leucine−−tRNA_ligase_chloroplastic/mitochondrial [MW = 110.28 kDa]

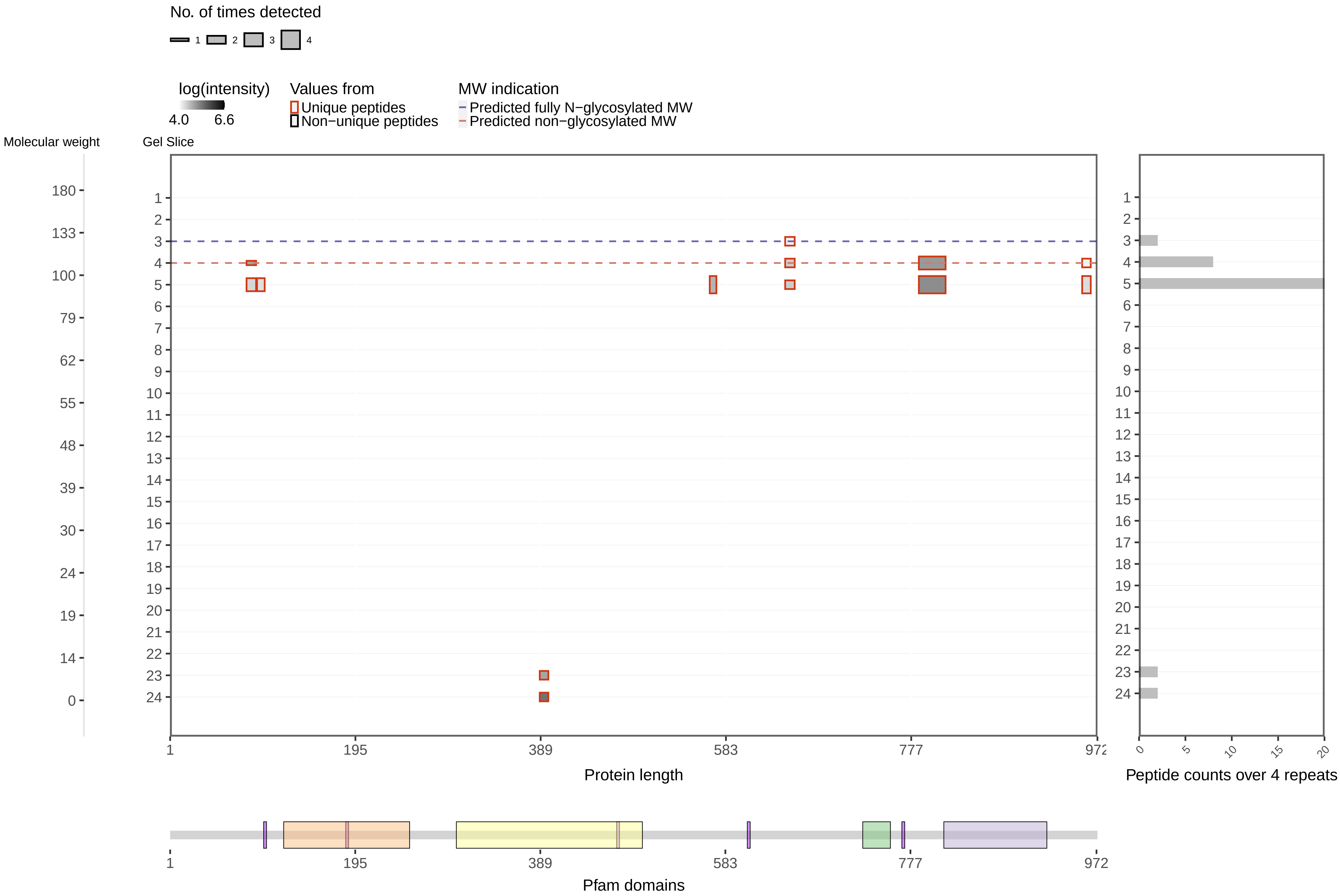


Pfam Description

SP | TM Domain | PF08264 Anticodon−binding domain of tRNA ligase | PF13603 Leucyl−tRNA synthetase, Domain 2
Sequon | PF00133 tRNA synthetases class I (I, L, M and V) | PF09334 tRNA synthetases class I (M)

Protein sequence:

NbL02g22340.1 aminomethyltransferase_mitochondrial [MW = 44.24 kDa]

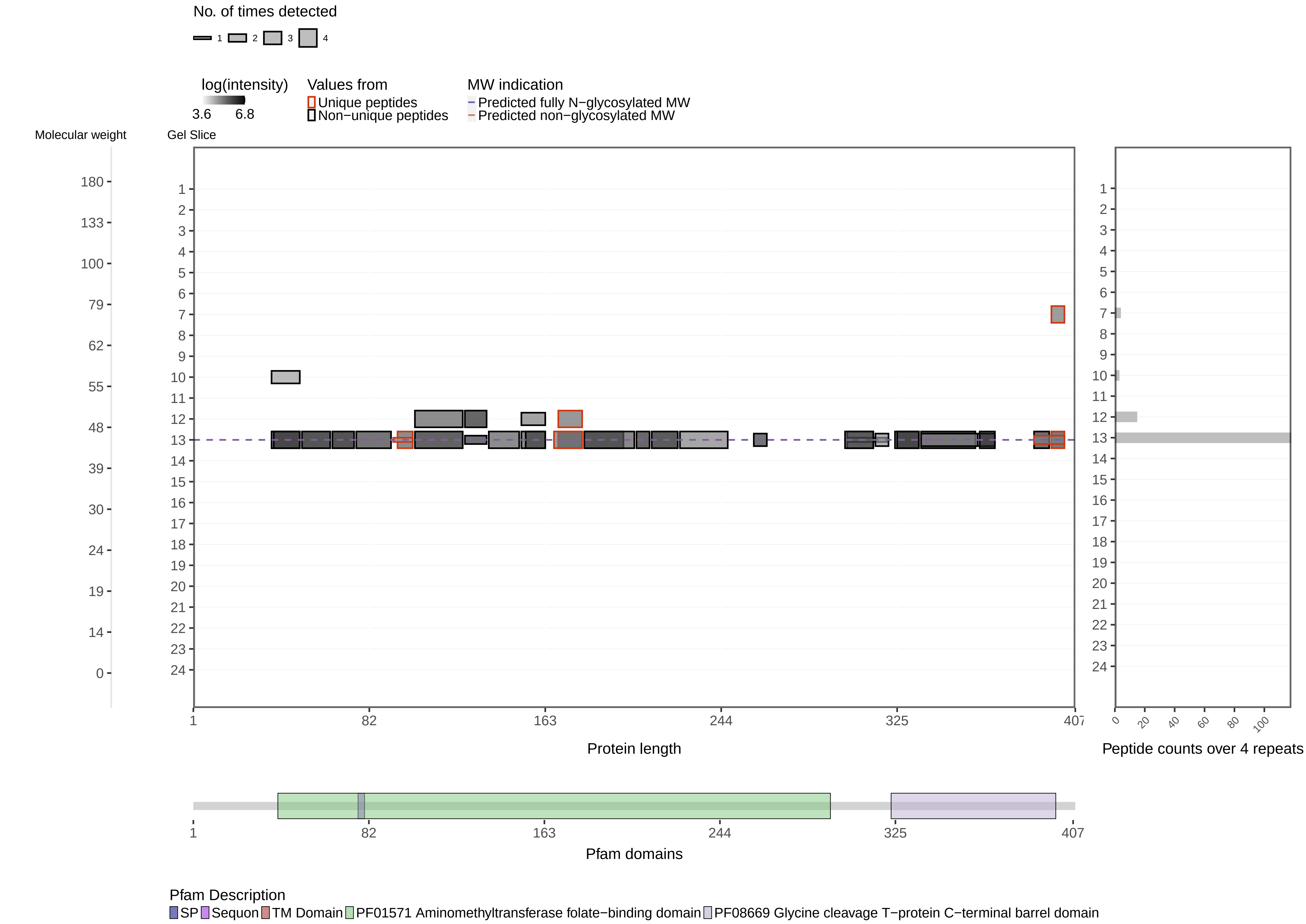


Protein sequence:

AAPVLQHLTKDDLSKMYFGEFRVLDINGASCFLTRTGYTGEDGFEISVPSENAVDLAKAILEKSEGKIRLTGLGARDSLRLEAGLCLYGNDMEQHITPVEAGLTWAIGKRRRAEGGFLGAEVILKQIEEGPKIRRVGFFSSGPPPRSHSEIQDSNGQNIGEITSGGFSPCLKKNIAMGYVKTGNHKAGTNVKIVIRGKSYDGAVTKMPFVPTKYYKP*

### NbL02g22310.1 glucan_endo−13−beta−glucosidase_1 [MW − SP = 54.35 kDa]

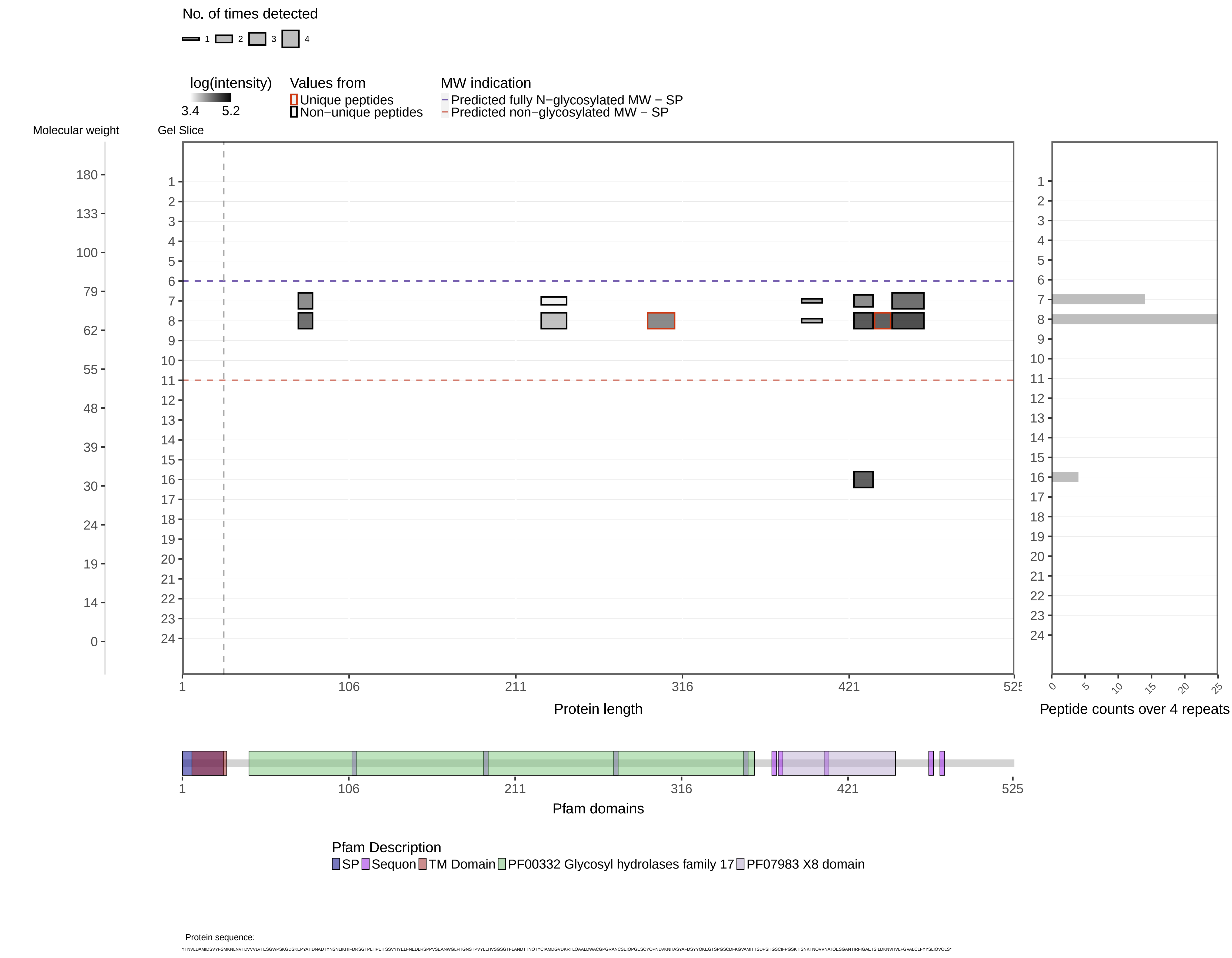

NbL02g22290.1 5'−adenylylsulfate_reductase_1_chloroplastic−like [MW = 51.18 kDa]

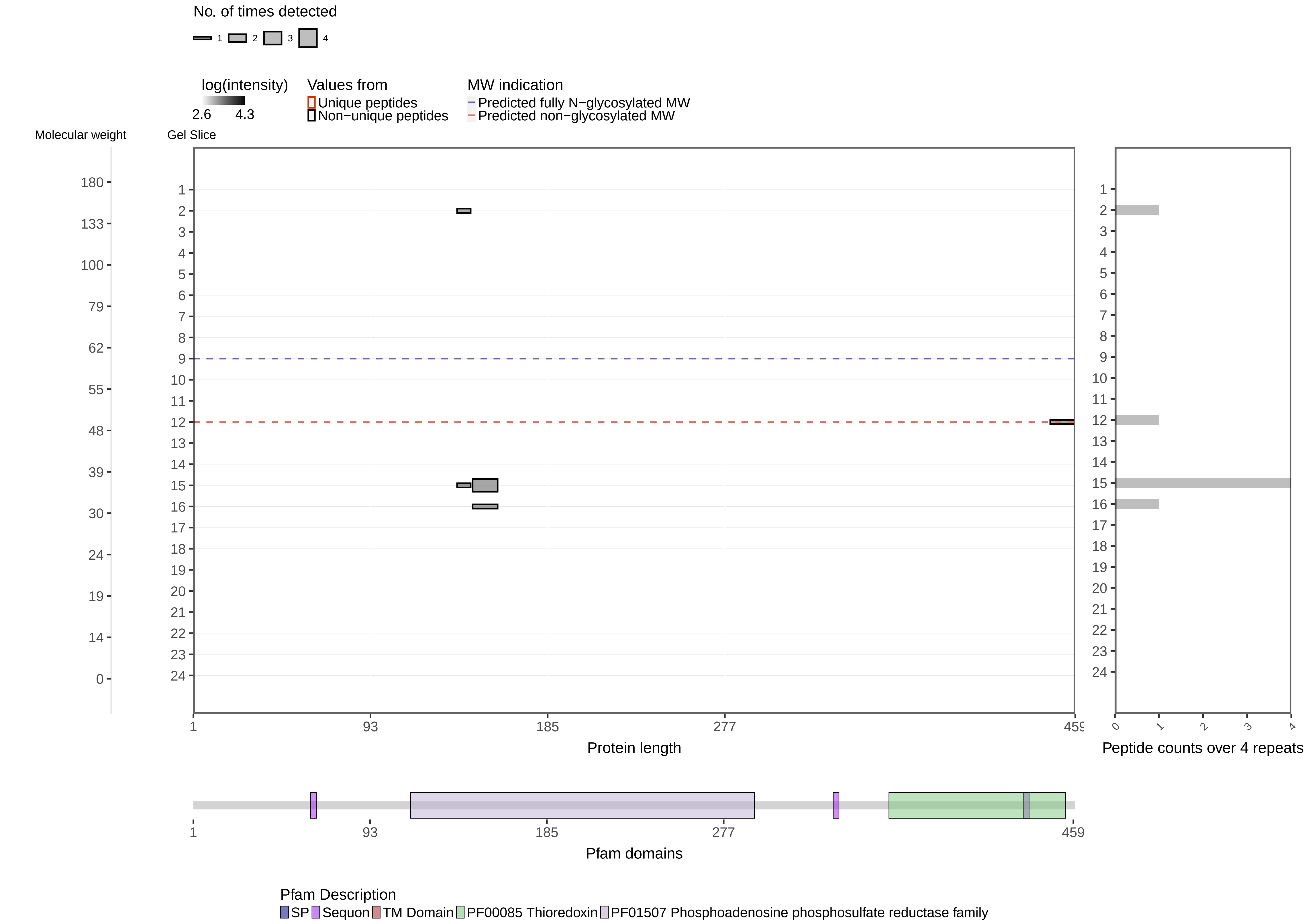


Protein sequence:

RKDQSPGTRSEIPVVQVDPSFEGLDGGAGSLVKWNPVANVDGKDIWNFLRAMNVPVNSLHSQGYVSIGCEPCTRPVLPGQHEREGRWWWEDAKAKECGLHKGNIKDETVNGAVQTNGTATIADIFDTKDIVTLSRPGVENLLKLDDRREPWLVVLYAPWCQFCQAMEGSYVELAEKLAGWGVKVGKFRADGEQKAFSQQELQLGSFPTILFFPKNSSKAIKYPTEKRDVDSLLAFVNALR*

### NbL02g22280.1 lactoylglutathione_lyase_GLX1 [MW = 32.86 kDa]

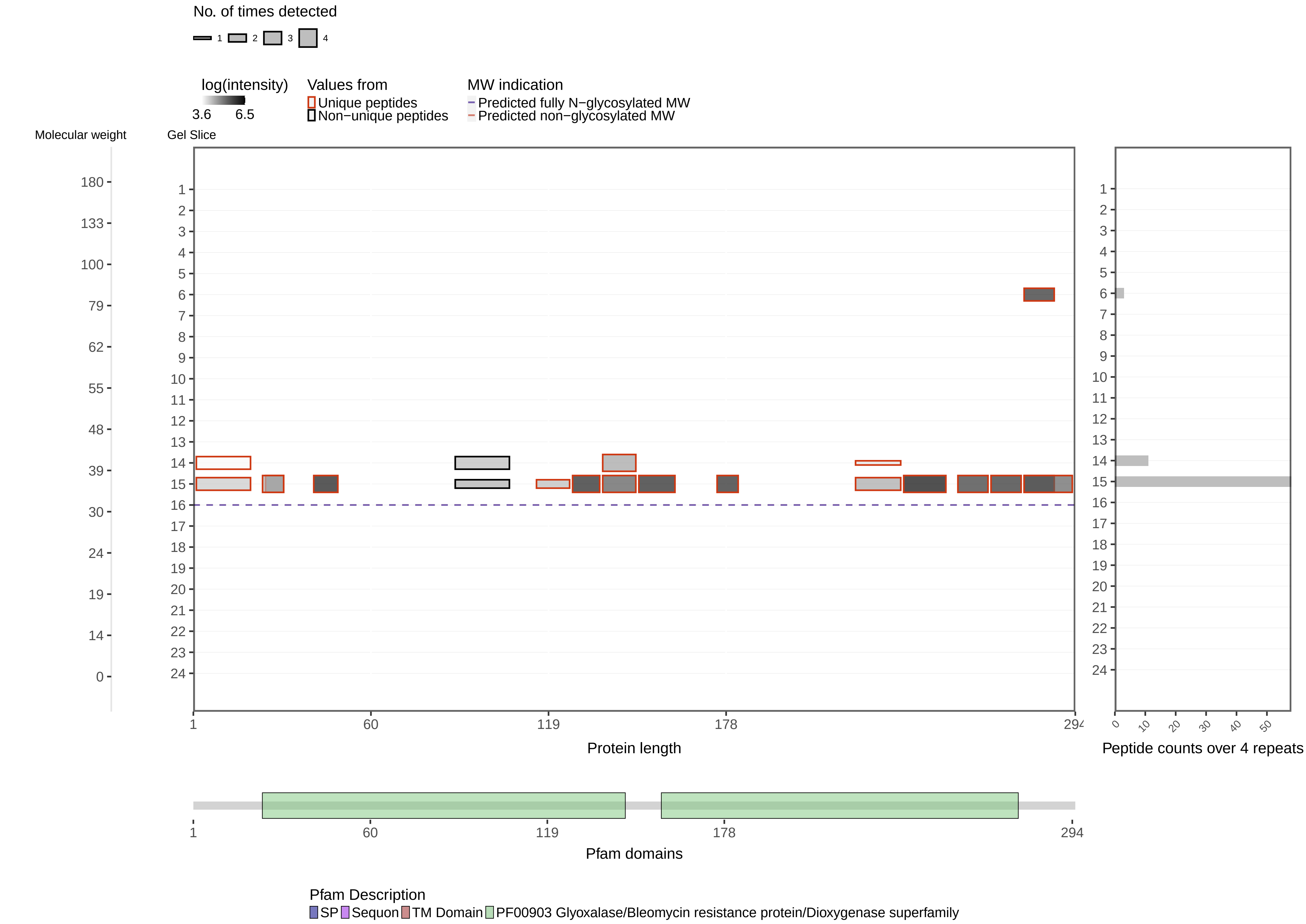


Protein sequence:

VKDPDGYLFEILQRASTPEPLCQVMLRVGDLERSIKFYEKALGMQLVKKVDRPEQKYSIAMMGYAPEHETVVLELTYNYGVTEYTKGNAYAQVAISTDDVYKSAEVVNLVTQELGGKITRQPGPIPGLNTKITSFLDPDGWKTVLVDNQDFLKELESSK*

NbL02g22190.1 soluble_starch_synthase_3_chloroplastic/amyloplastic [MW = 141.42 kDa]

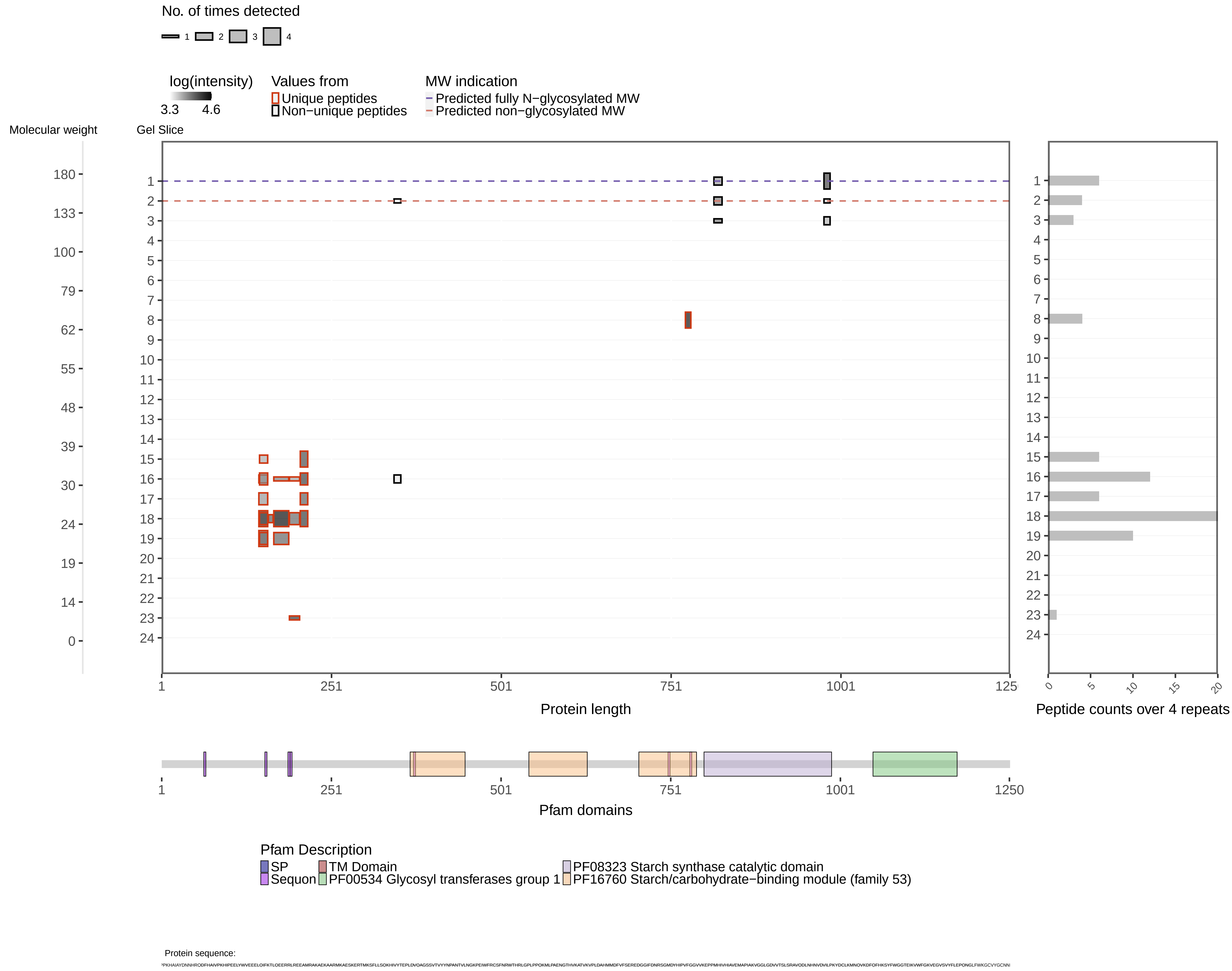

### NbL02g22160.1 ATP_synthase_gamma_chain_chloroplastic [MW = 41.43 kDa]

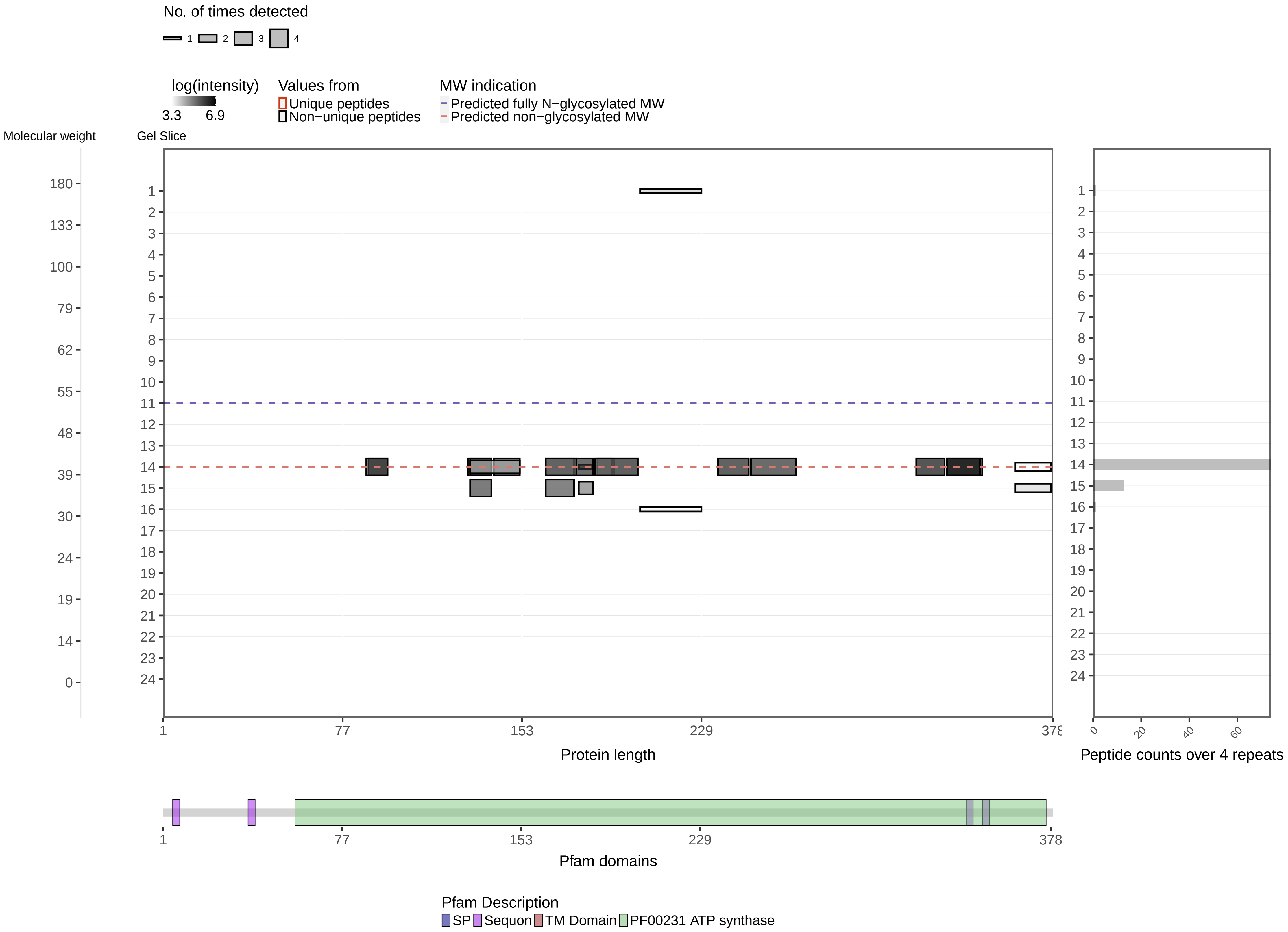


Protein sequence:

KKGNSYFLRRPYIPVDKFLEGSNLPTAKDAQAIADDVFSLFVSEEVDKVELLYTKFVSLVKSEPVIHTLLPLSPKGEICDINGNCVDAAEDEFFRLTTKEGKLTVERDIMRTKTTDFSPILQFEQDPVQILDALLPLYLNSQILRALQESLASELAARMSAMSAATDNASELKKNLSRAYNRQRQAKITGEILEIVAGADALV*

NbL02g22140.1 nucleolar_protein_56−like [MW = 53.86 kDa]

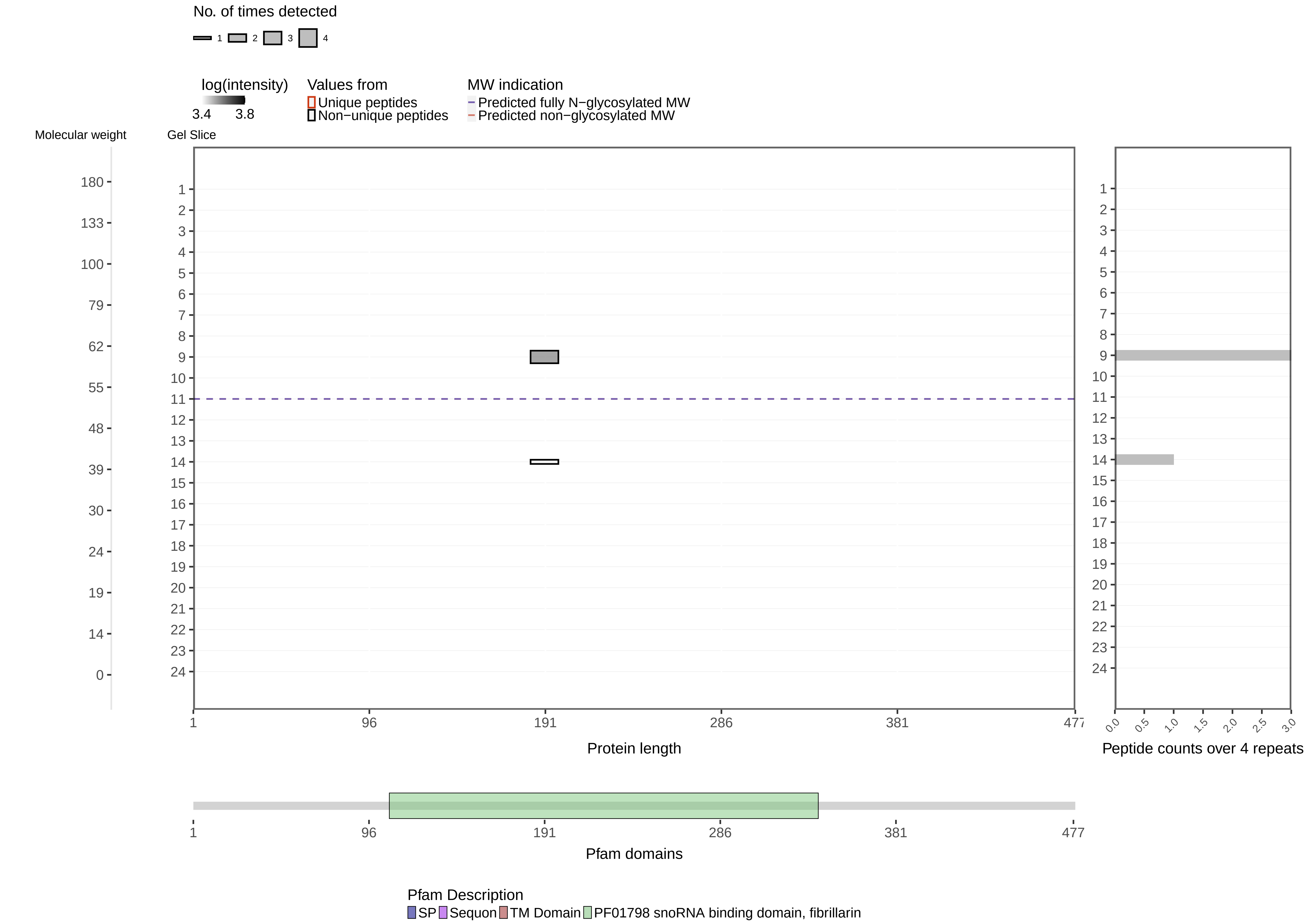


Protein sequence:

NLAALIGEVVGARLISHAGSLTNLAKCPSSTLQILGAEKALFRALKTKGNTPKYGLIFHSSFIGRASARNKGRMARYLANKCSIASRLDCFLDTSTTTFGEKLREQVEERLDFYDKGVAPRKNIDVMKSAIETVEGKDAEMDVDEPSAMKSKKKKSKVEVVEYAQPMIEDQPLVTNVDASEEPKSEKKKKKKKRKTEQEREHEEDQTNGAPEDGTAKKKKKSKEENGEELQAASEVKKKKKKSKVQDDE*

NbL02g22060.1 zinc_finger_MYND_domain−containing_protein_15 [MW = 70.51 kDa]

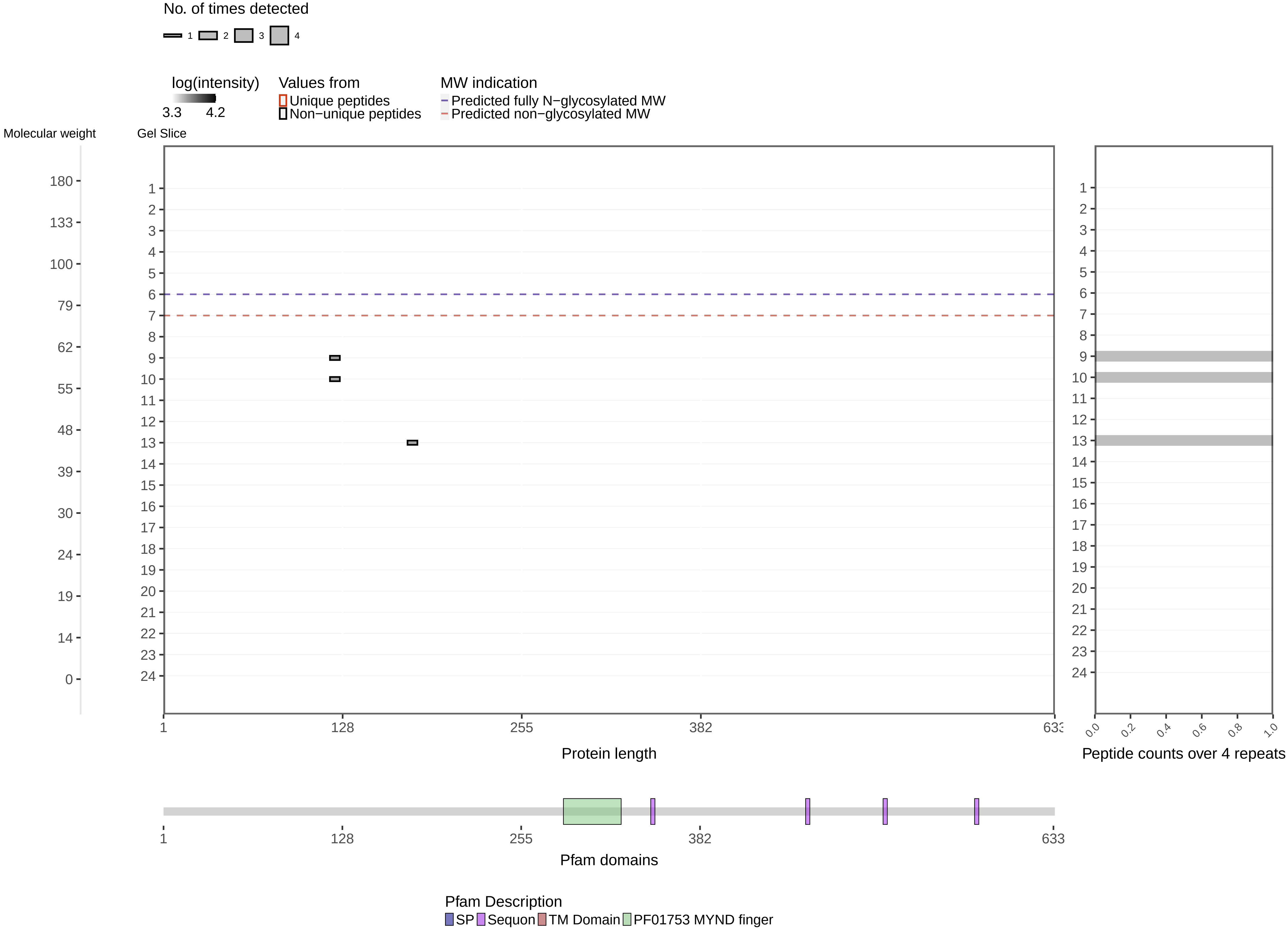


Protein sequence:

CSAVVYCSSVCQKQHWKEAHKDVCGLYRAMMEREEELAMKIFIFNCSAEQPCKWLESLGIHQKGMWRRKCNCYSHCPFGLLPVPGGLWNSWGGLDENEYPHDSPFHNHFRDGISSSVFLSGWLEYYSLRSLPLTSPVADILSHPLTIYYMLTALNISSKNLLLKNKEVILHYIGPESELDWMPAFAEIGHLLHGMGNIHIIMVGPEVPTNLSGTTSGIGSRVRVNLVRGLYQDEASYLPTPHVIMSLNCGLGSYSCWDGALDLIKSMNVPAYFTNESELSCSSSKQVLRAAG

### NbL02g21830.1 pectinesterase−like [MW = 58.73 kDa]

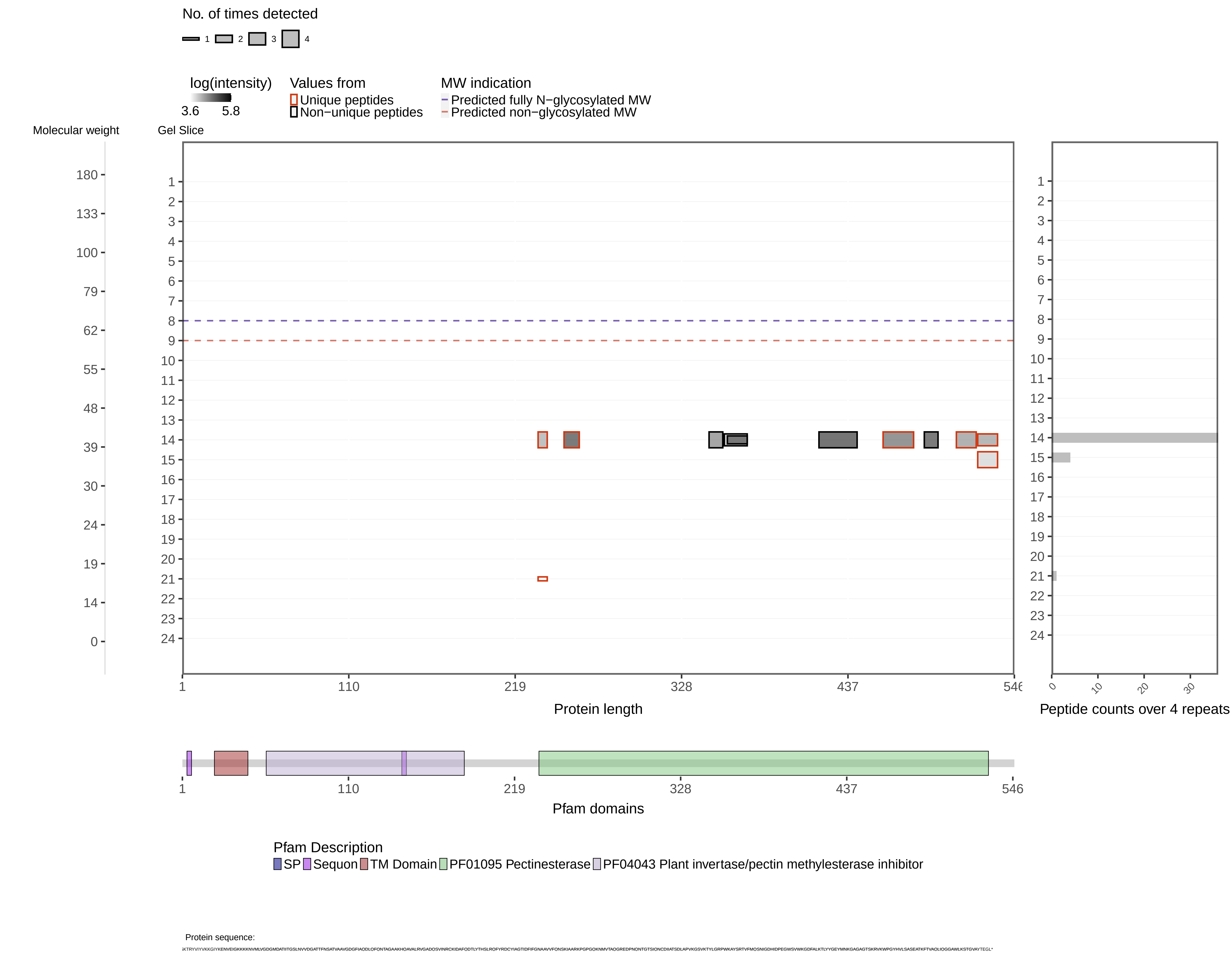

NbL02g21770.1 putative_receptor−like_protein_kinase_At4g00960 [MW − SP = 72.45 kDa]

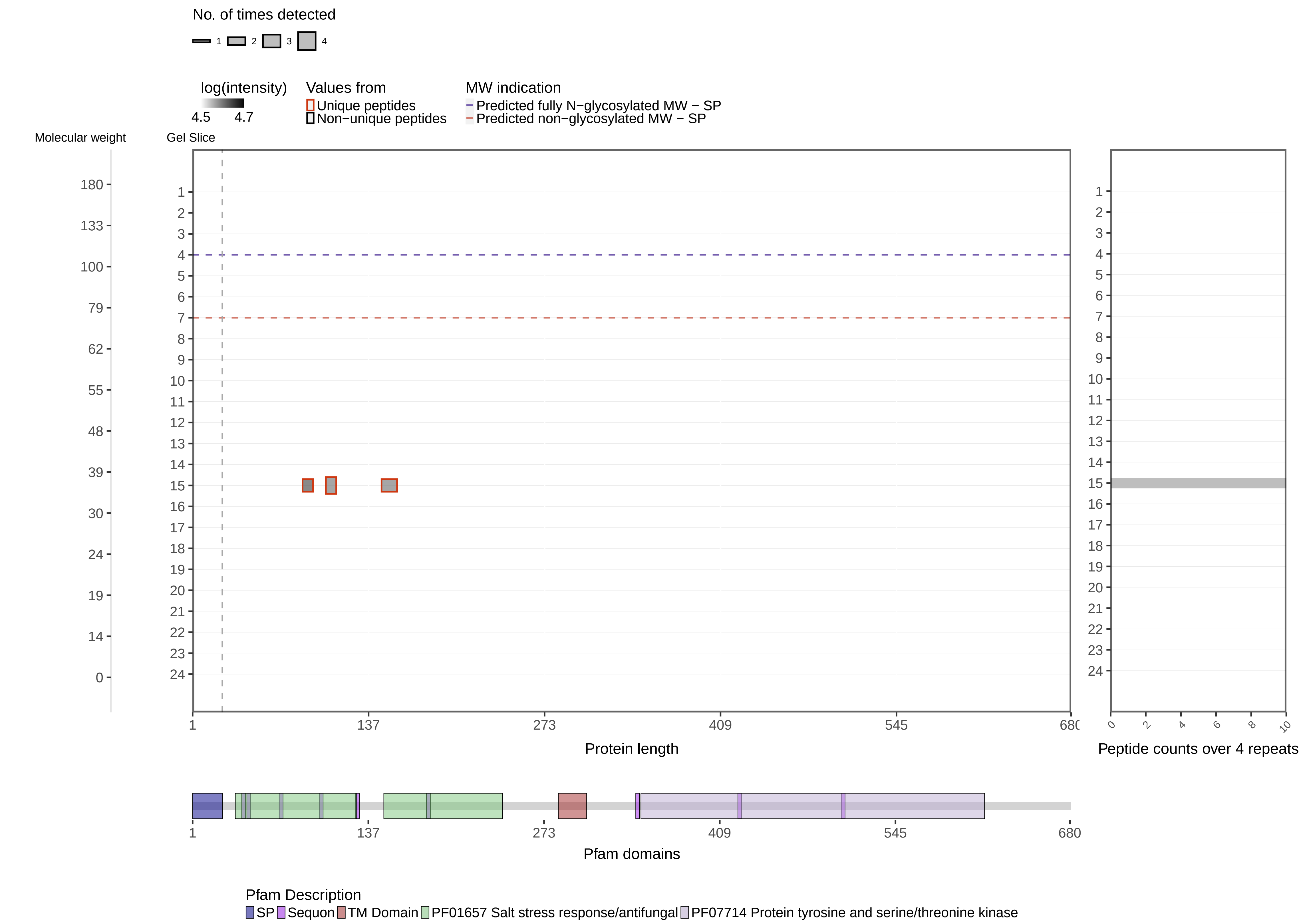


Protein sequence:

TAESLQYDFSTIRAATDNFSNANKLGQGGFGPVYKGKLSNGQEVAVKRLSADSGQGDLEFKNEVVLVSRLQHRNLVRLLGFCLDGAERLLVYEFVPNASLDQFLFDPVKRRQLDWERRSRIIGGVARGILYLHEDSRLRIIHRDLKASNVLLDAGMNPKISDFGMARLFTLDETQGNTSRIAGTYGYMAPEYAMHGQFSVKSDVFSFGVLVLEILSGQKNTCFRNGESVEDLLSYAWSNWRDGTATNLIDPMLRGSTVLVRDILRSIHIALLCVQENVADRPTMAAVVLMLSSL

NbL02g21630.1 probable_NAD(P)H_dehydrogenase_(quinone)_FQR1−like_1 [MW = 21.6 kDa]

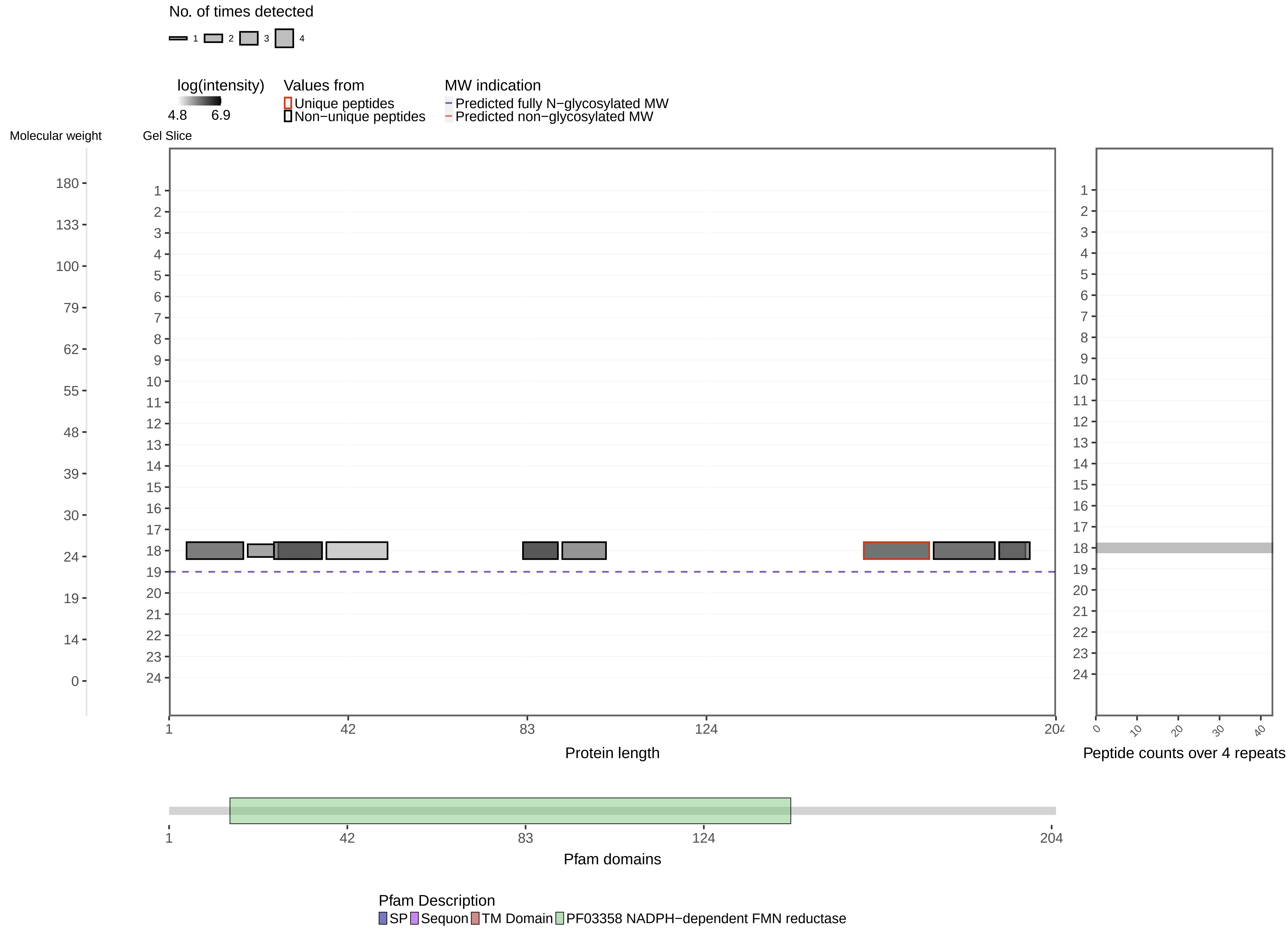


Protein sequence:

FKAFLDATGGLWRTQQLAGKPAGLFYSTGSQGGGQETTALTAITQLVHHGMIFVPIGYTFGGGMFEMENIKGGSPYGAGTYAGDGSRQPTELEMEQAFHQGKYIATITKKLKGSA*

### NbL02g21580.1 G−type_lectin_S−receptor−like_serine/threonine−protein_kinase_At1g11410 [MW − SP = 90.65 kDa]

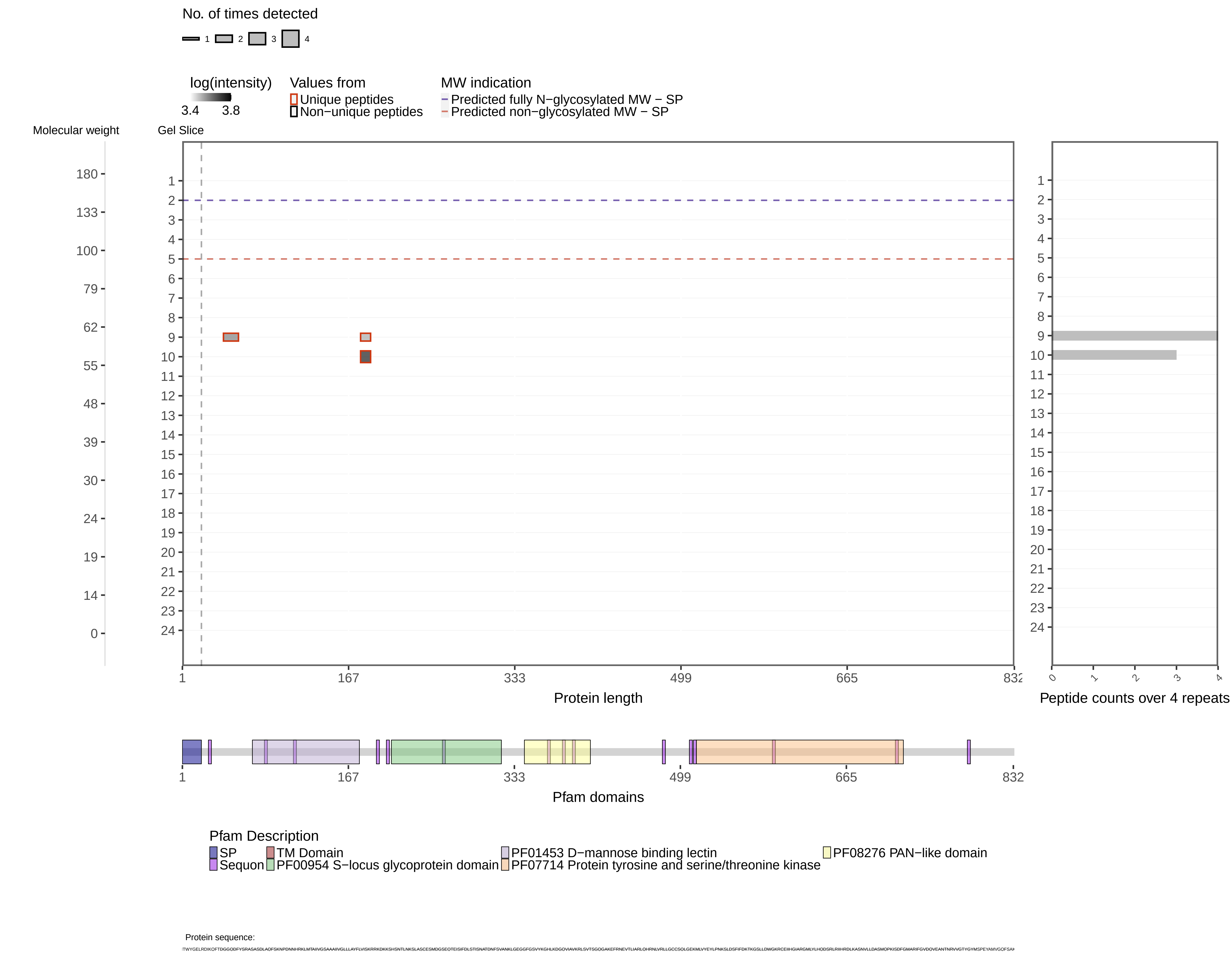

### NbL02g21450.1 universal_stress_protein_PHOS32−like [MW = 26.82 kDa]

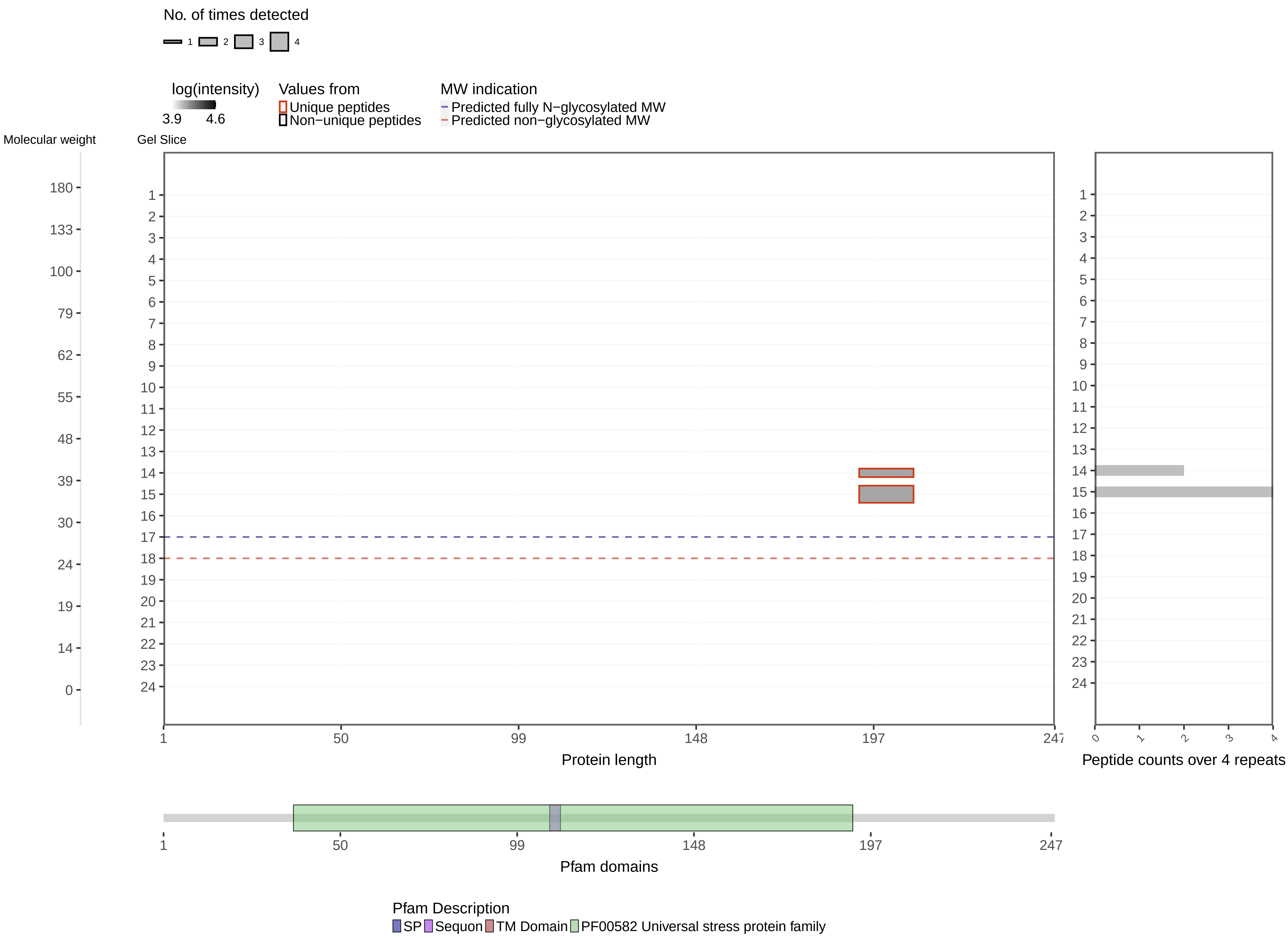


Protein sequence:

TTKANDLAQPLVEGNIPFKIHIVKDHDMKERLCLEVERLGLSAVIMGSRGFGASRRSIKGGRLGSVSDYCVHHCVCPVVVVRYPDDKDGVVADDANERSVKKETDGVLHPVPEDEPIYHDAEDKAAVTGQRKGLPI*

NbL02g21370.1 putative_PAP−specific_phosphatase_mitochondrial [MW = 21.92 kDa]

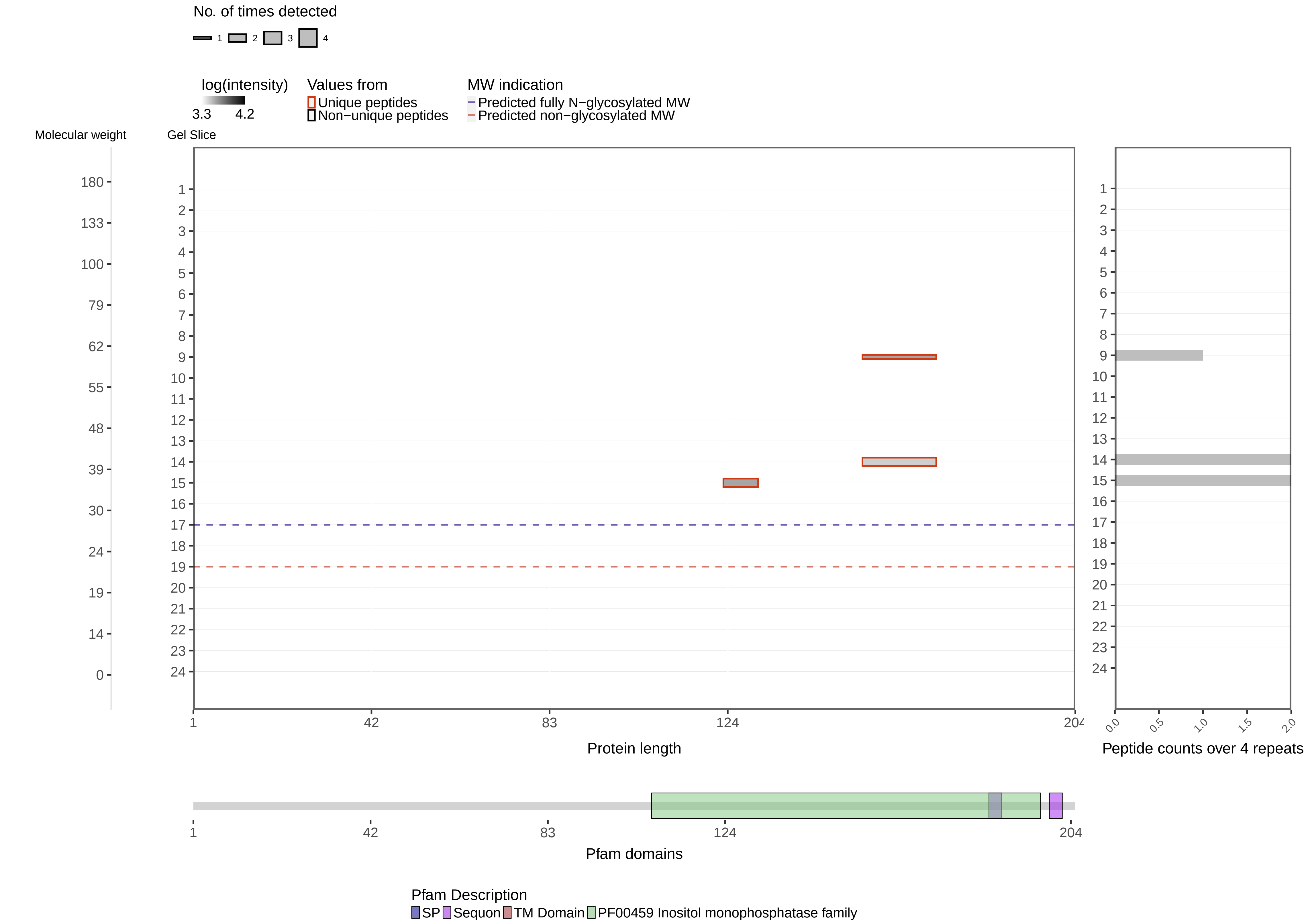


Protein sequence:

SFDAERDSENVGEGNILLLSACCGSLCKYLLVASGRASVYIQGPKATSTIKAWDHAVGIICVHEAGGKVTDWEGSSLDFAADQSERRLIFPSGGVLVTNGSLHSKIIGMISSNTSVL*

### NbL02g21190.1 eukaryotic_translation_initiation_factor_3_subunit_J−A−like [MW = 25.2 kDa]

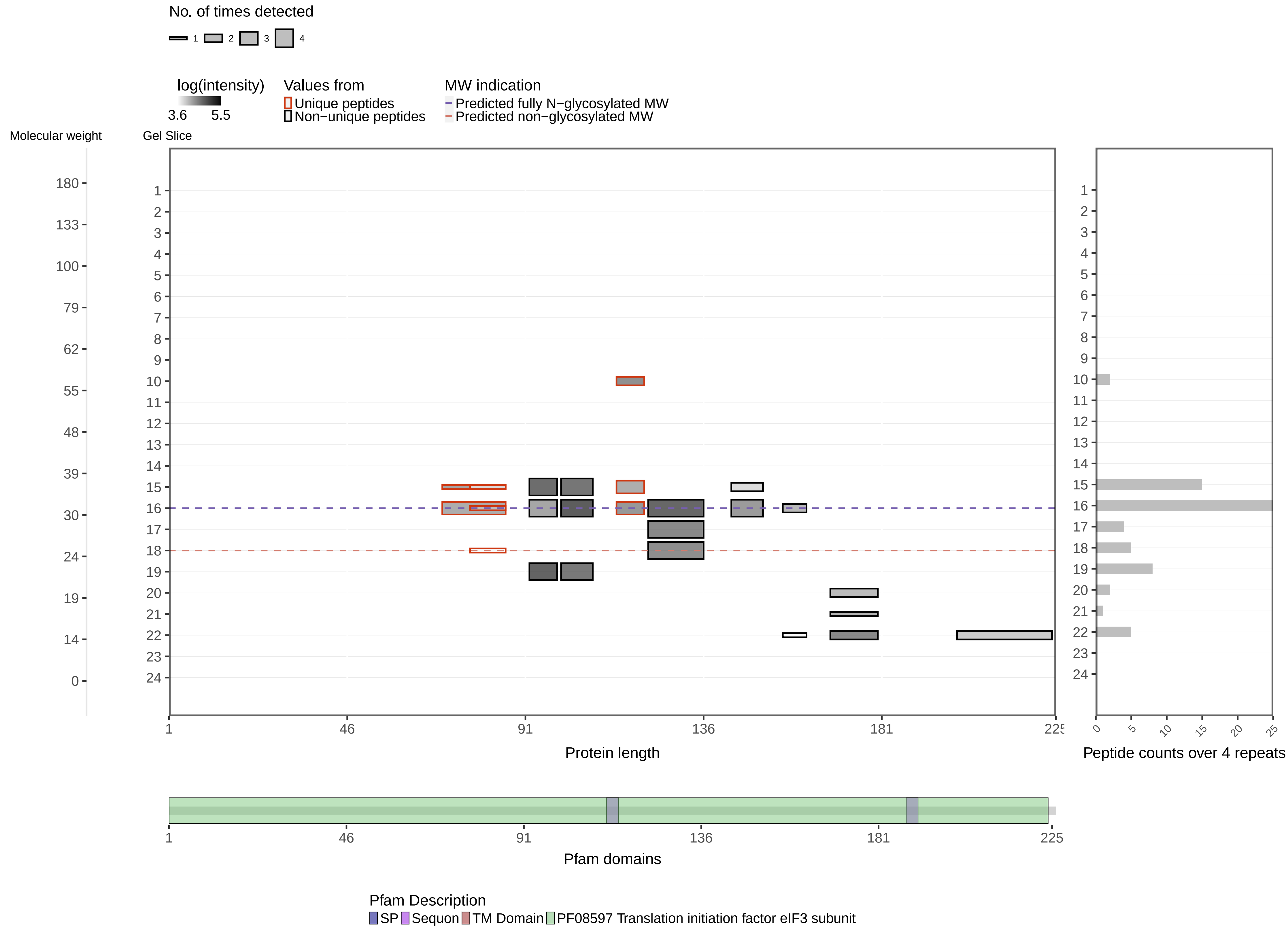


Protein sequence:

DYKSTAELFATKGGDNKTVDNFIPKSENDFLEYAELISHKLRPYEKSYHYIGLLKAVMRLSMTALKGADAKEIASSVTAIANEKIKAEKEANASKKKAGAKKKQIHVDKADDEAVVNAYDGYDDYDFM*

NbL02g21100.1 beta−galactosidase_8−like [MW = 19.95 kDa]

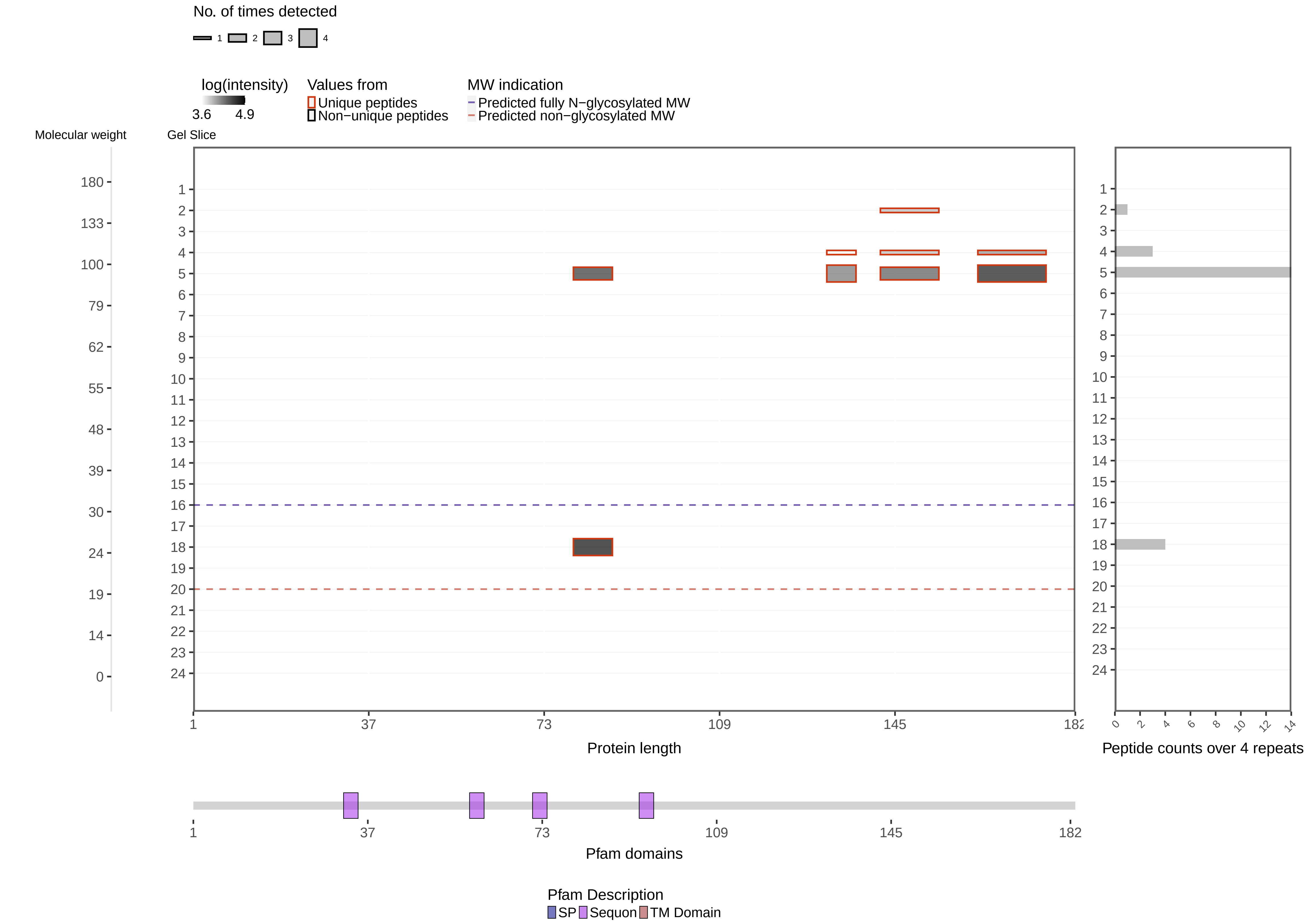


Protein sequence:

SFSPPEGKGPVSLNLSSMGKGQAWVNGQIVGRYWSSYRSPSTGCSNNCDYRGAYNSWKCLKKCGQPAQVLYHVPRSWLTPGKNLLVLHEELGGDPSKISFSS*

NbL02g21070.1 quinone_oxidoreductase_PIG3 [MW = 34.57 kDa]

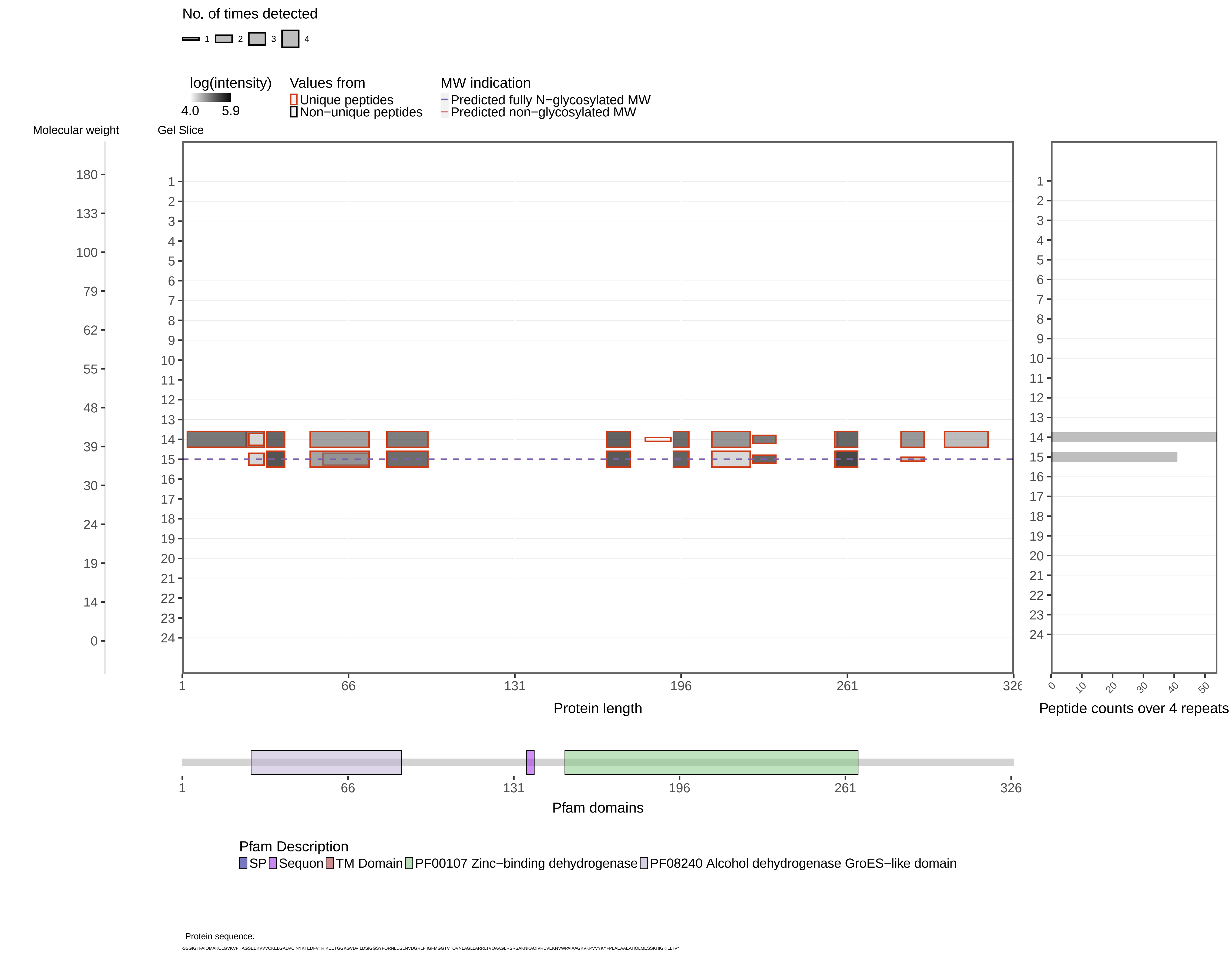

NbL02g20940.1 glucose−induced_degradation_protein_8_homolog [MW = 27.71 kDa]

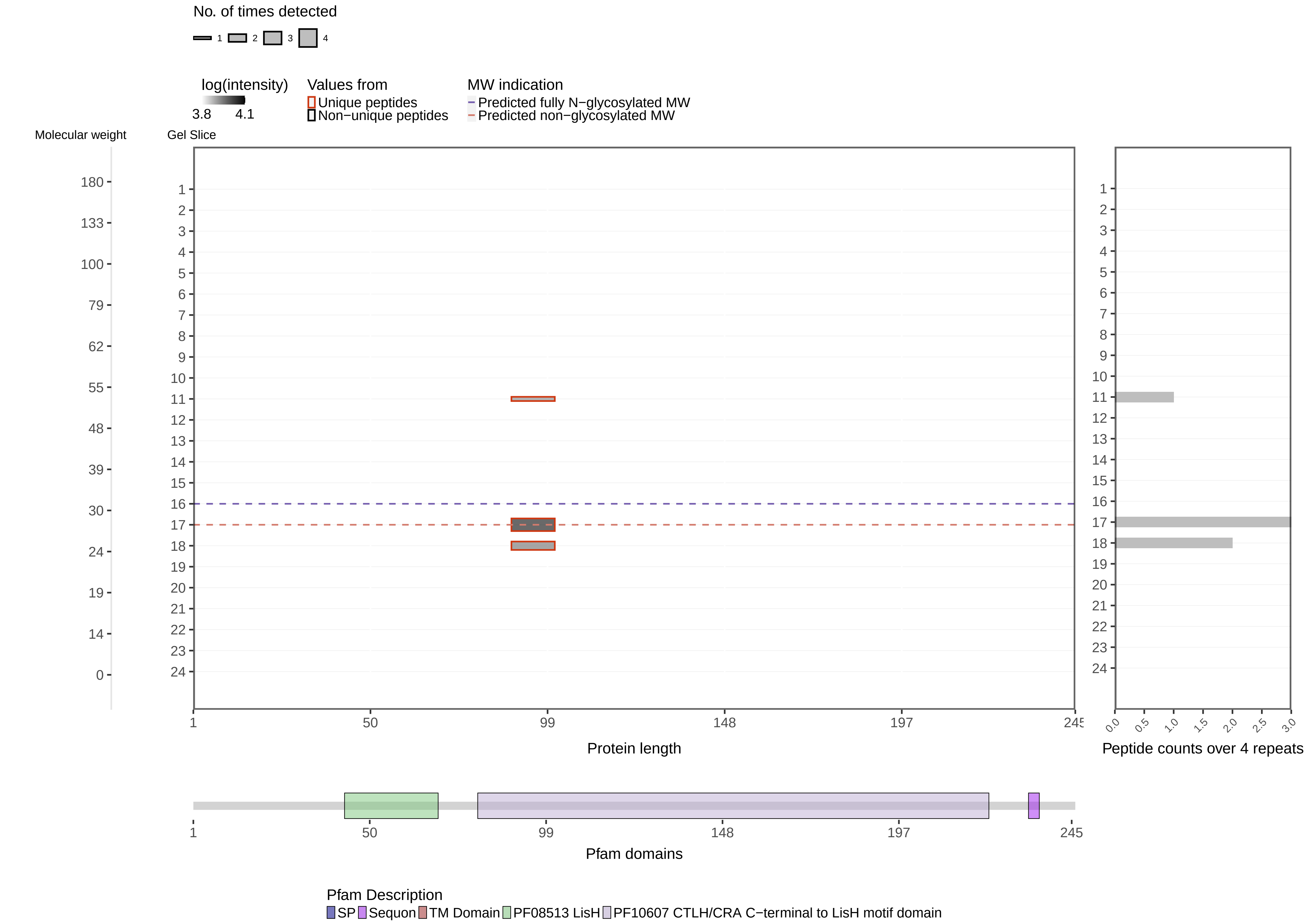


Protein sequence:

ILDTNPQLFFHLQQQRLIELIRNGKVEEALEFAQEELAPRGEENQAFLEELEKTVALLAFEDVSNCPVGALLDVSQRLKTASEVNAAILTSQSHEKDPKLPSLLKMLIWAQNQLDEKAVYPRITNLSTASLEDPAV*

NbL02g20810.1 bifunctional_epoxide_hydrolase_2−like [MW = 25.24 kDa]

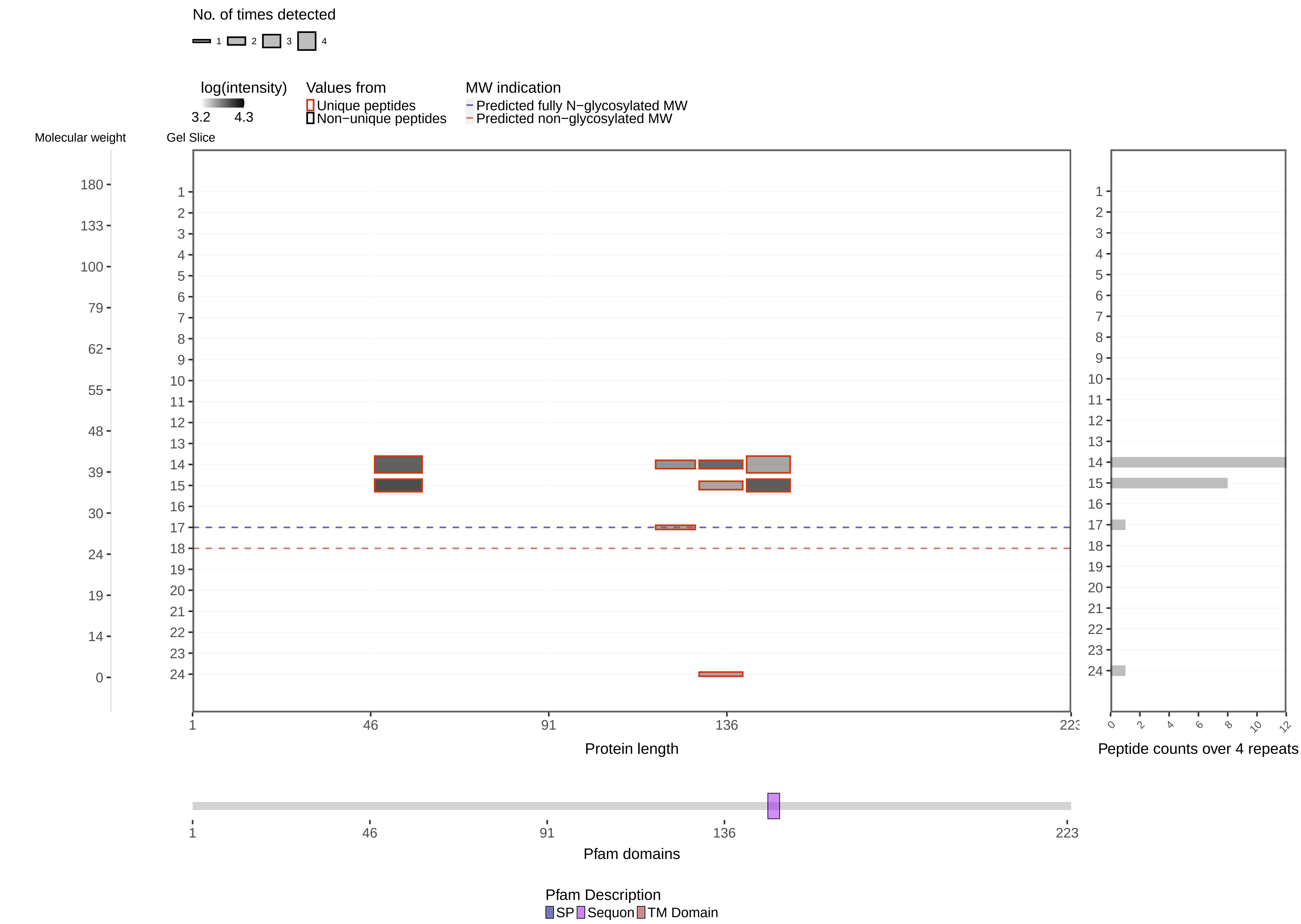


Protein sequence:

EIMDIVQPSTPLPPWFSEKDLETYGALYEKSGFQTALQVPYRSLLEQANITDPRVHVPALFIVGEKDYFLKFPGVRDYISGGLKSLVPNLQMVNLLEGTHFVQEQLPDEVNQLVLDFLTRNSQS*

### NbL02g20780.1 clathrin_light_chain_1−like [MW = 35.48 kDa]

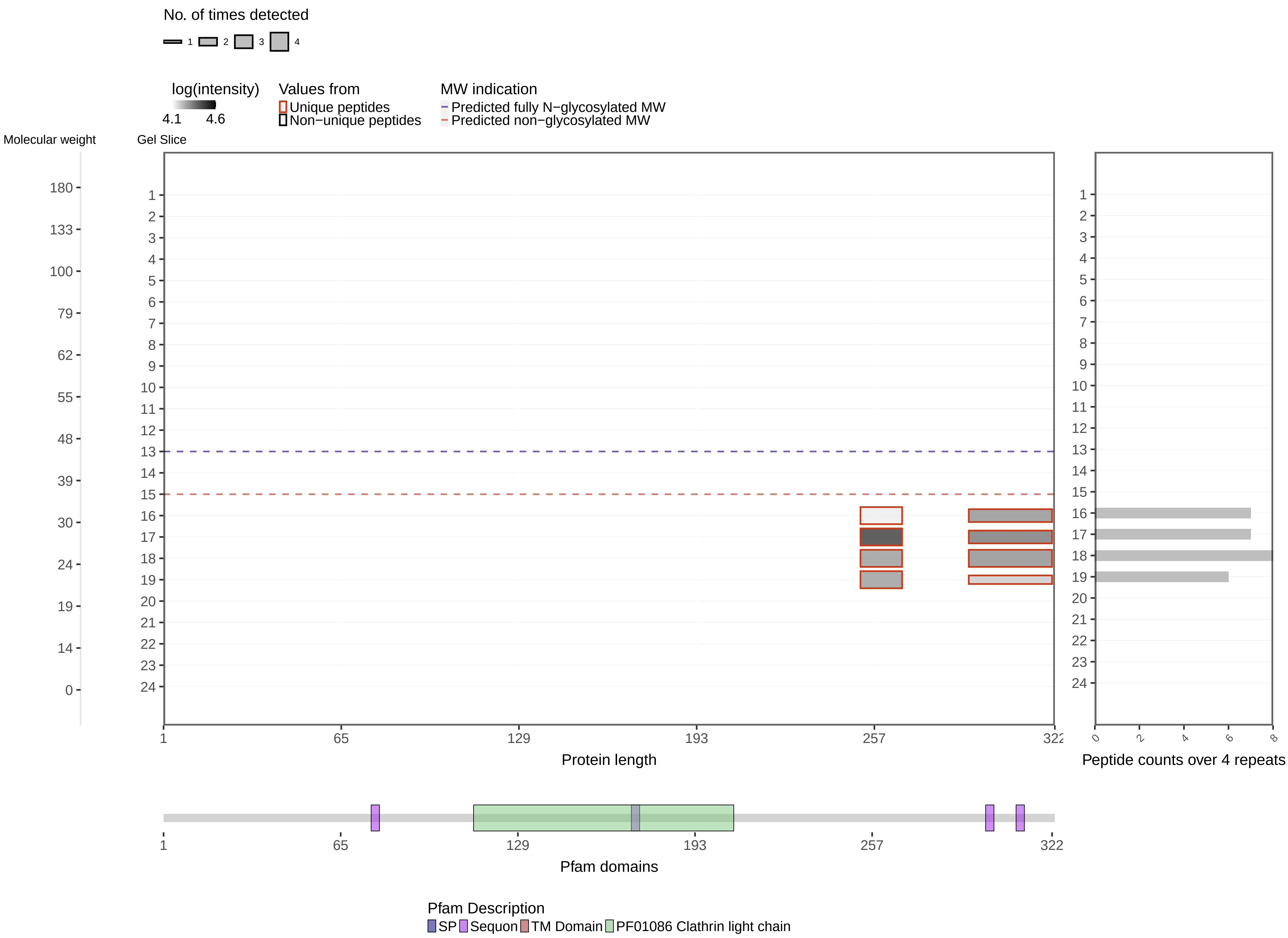


Protein sequence:

QIIEEGEEYKKAFYEKRKLTVETNQSTNREKEKLYLSNQEKFHKEADKQYWKAIAELIPKEVATIEKKGRKKDQEKKPSVTVVQGPKPGKPTELSRMRHILLKLKHSPPPHMIPPPPPPAKDAKAGKDGKDTKDGKDTKGKDAKNAKDEAPNGTSDAPPSDANVTQTSEEPAAAE*

### NbL02g20660.1 14−dihydroxy−2−naphthoyl−CoA_thioesterase_1−like [MW = 18.96 kDa]

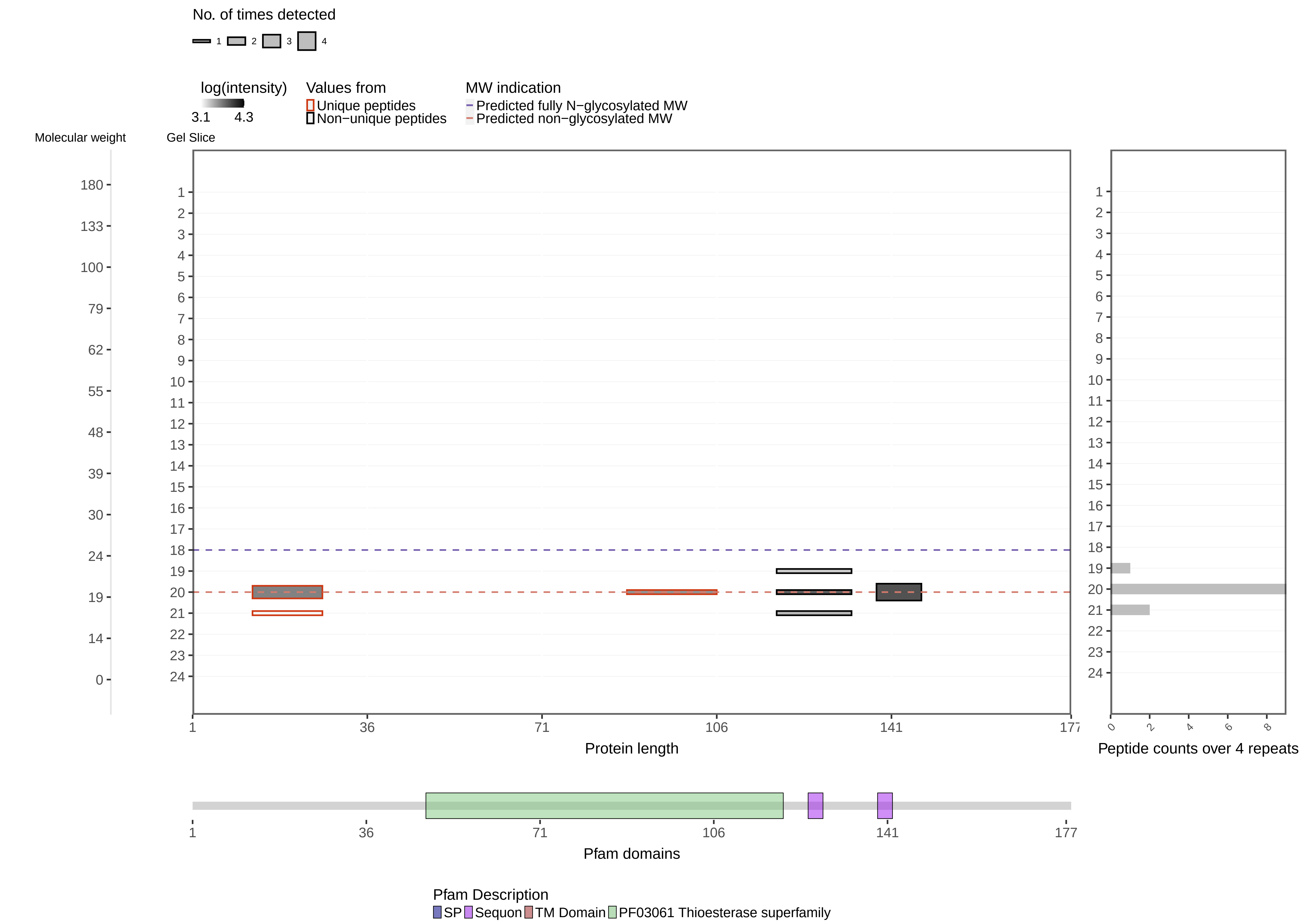


Protein sequence:

RVAGVHLSIHHLKSAHIGDVVFAEAAPVNIGKTIHVWEVCIWKIDPNNEENKTLIASSRVTLKANMSVPENAKDAAVNLKKYAKKVLPAVQGVARWSFSYDI*

NbL02g20550.1 DNA−directed_RNA_polymerase_II_subunit_RPB2 [MW = 135.26 kDa]

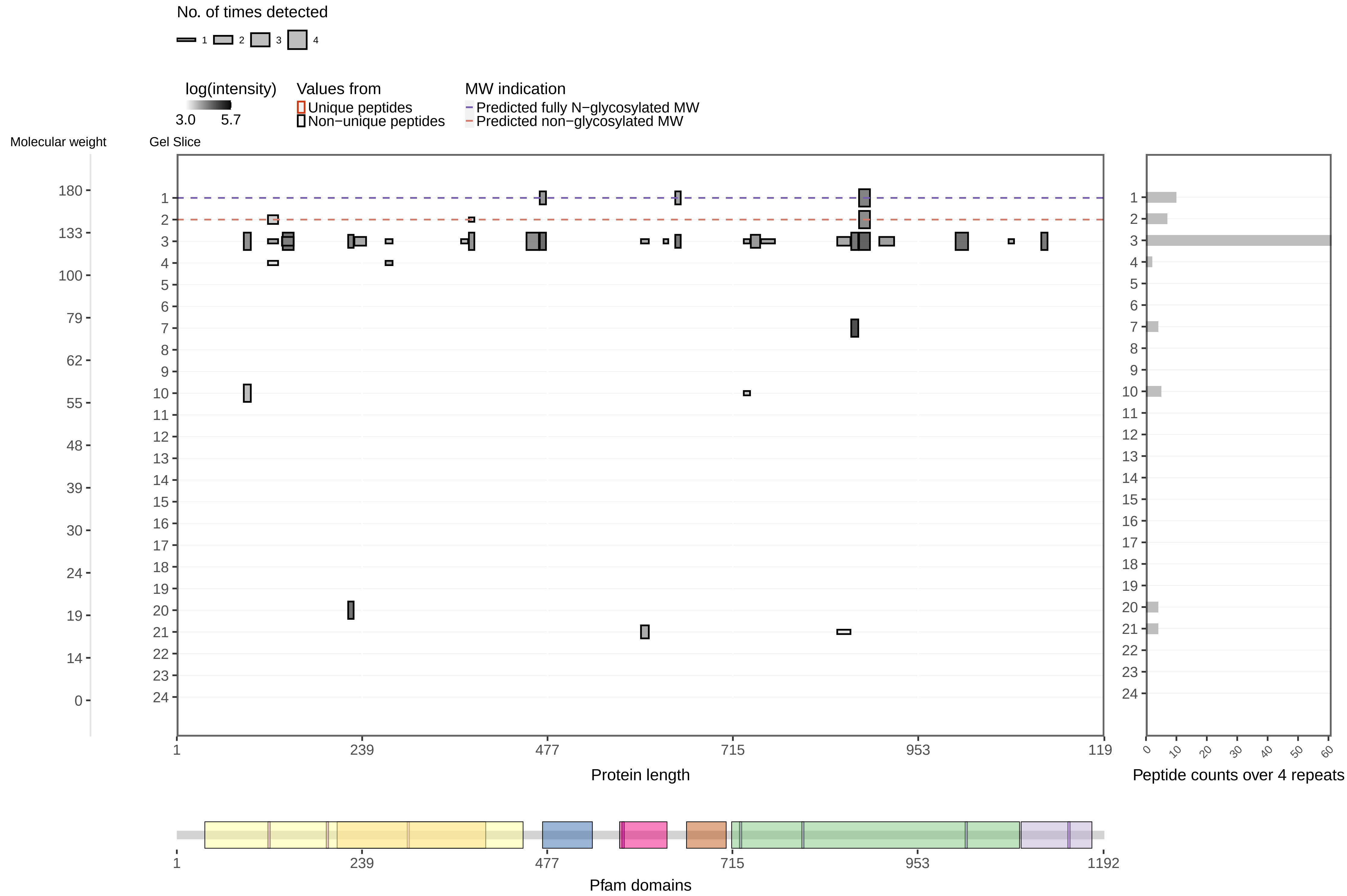


Pfam Description

- SP
- Sequon
- TM Domain
- PF00562 RNA polymerase Rpb2, domain 6
- PF04560 RNA polymerase Rpb2, domain 7
- PF04561 RNA polymerase Rpb2, domain 2
- PF04563 RNA polymerase beta subunit
- PF04565 RNA polymerase Rpb2, domain 3
- PF04566 RNA polymerase Rpb2, domain 4
- PF04567 RNA polymerase Rpb2, domain 5

Protein sequence:

NbL02g20540.1 metal−independent_phosphoserine_phosphatase−like [MW = 32.97 kDa]

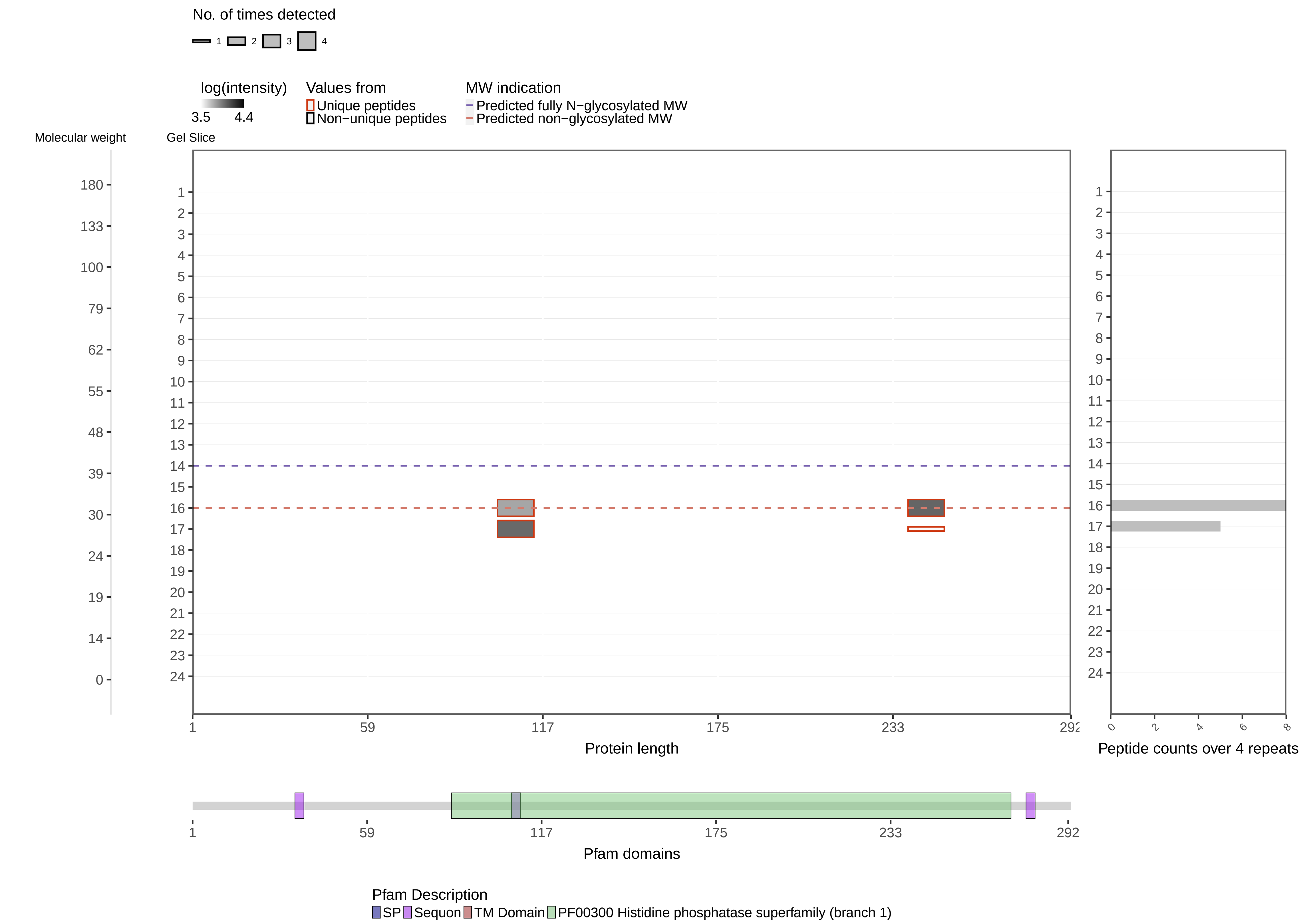


Protein sequence:

CFSSPISRAKTTAEIIWQGKEEPLVFLDSLKEAHLYFLEGLRNEDAKRIYPKEYTTWREDPSNFCVDGVYPLQKLWGRAYEAWEEILLTPGENFLVVTHKSILRALICTPLGLGPERFRSVDINNGGLCVFKFNKQGEAMLHSLNMTAHMYTDHIYHY*

NbL02g20340.1 30S_ribosomal_protein_S5_chloroplastic [MW = 35.44 kDa]

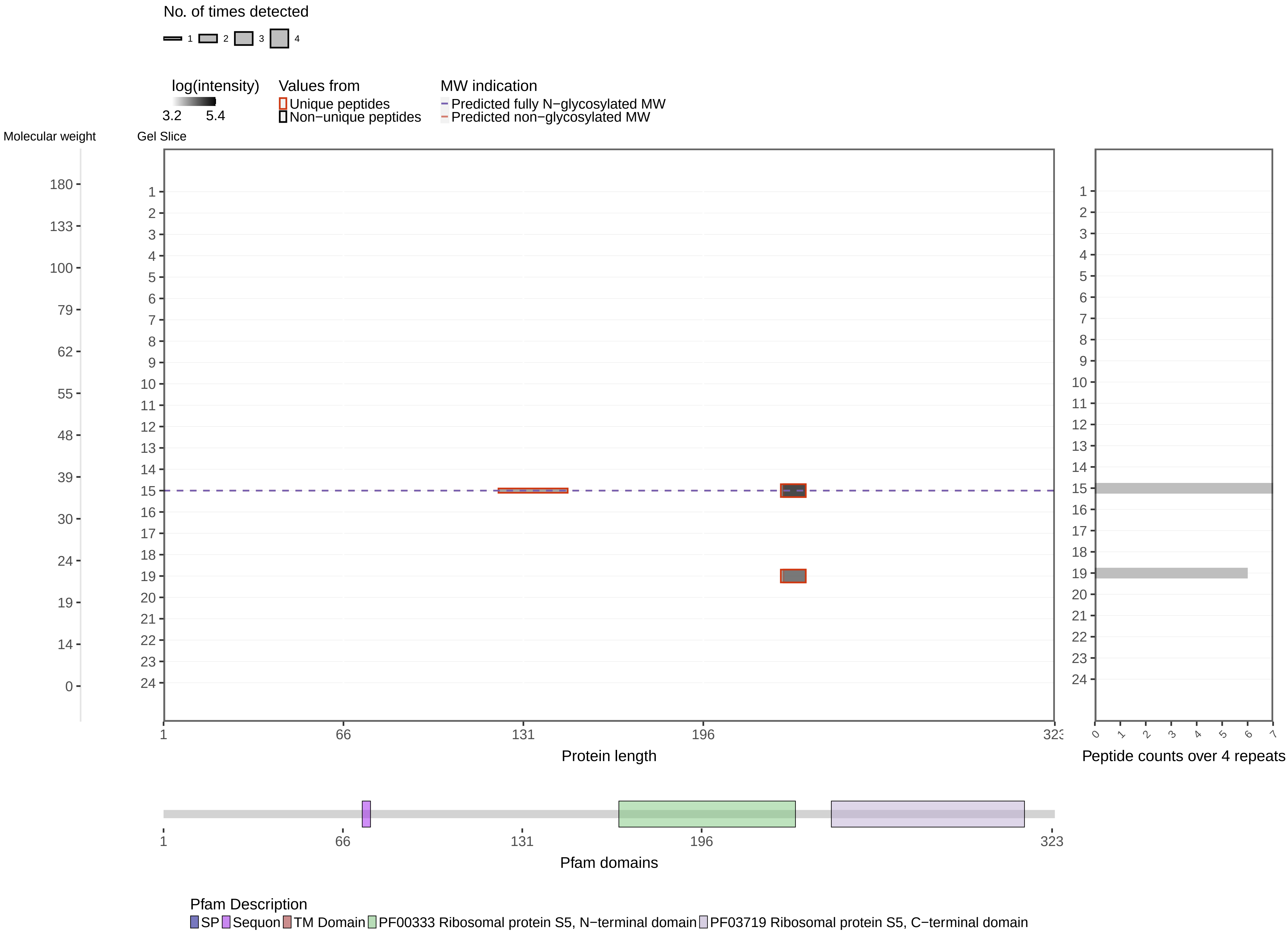


Protein sequence:

AFGKTKKEKAKDGFDERVVOVRRVTKVVKGGKOLHFRAIVVVGDKKGOVGVGVGKAKEVIAAVOKSAVNARRNVITVPMTKYLTFPHRSEGDFGAARVMLRPAAPGTGVIAGGAVRIVLEMAGVENALGKOLGSNNALNNARATVVAVOOMROFSEVAOERGIPMEELWK*

### NbL02g20130.1 homeobox−DDT_domain_protein_RLT1 [MW = 188.29 kDa]

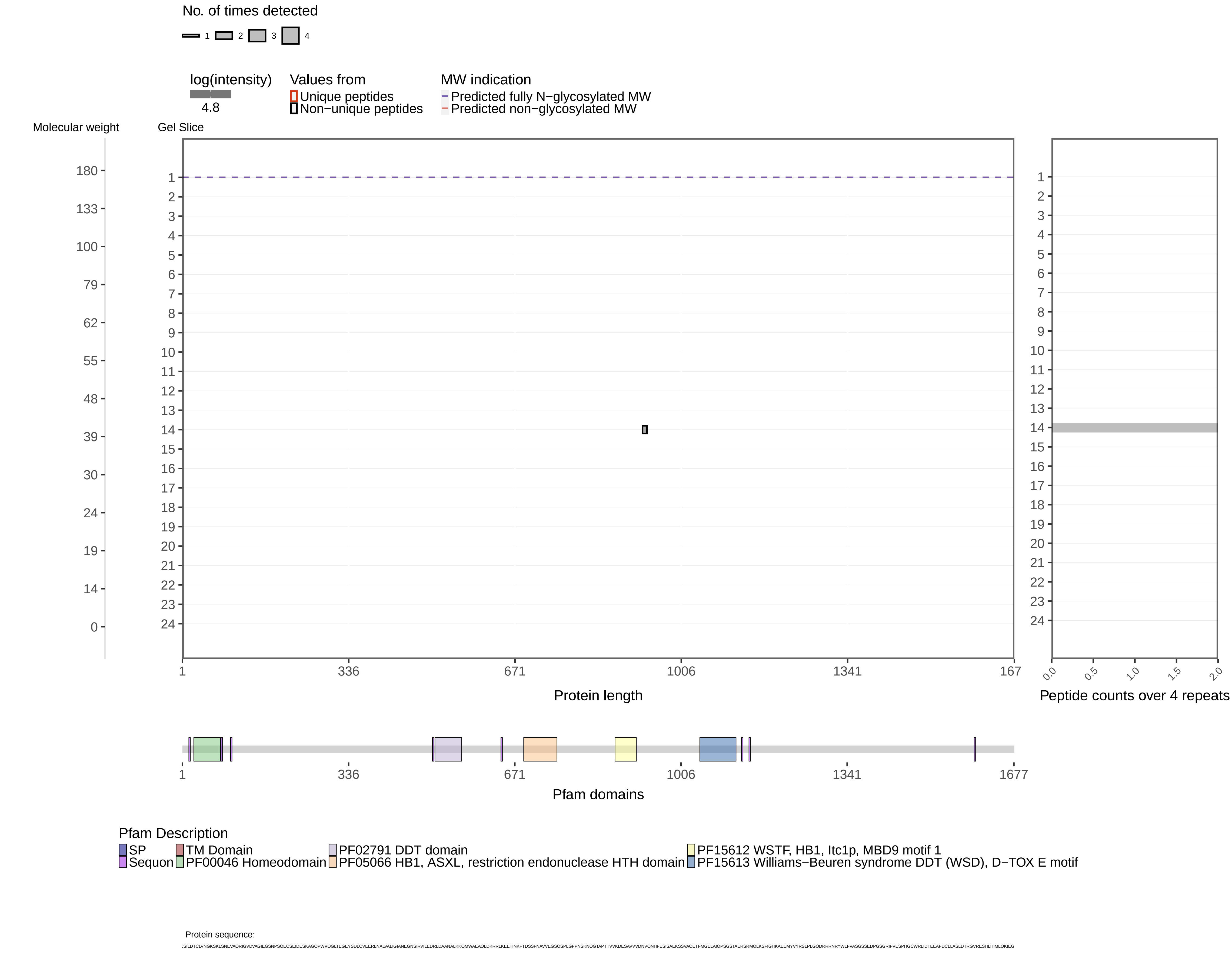

### NbL02g19760.1 GDSL_esterase/lipase_At5g45920 [MW = 25.97 kDa]

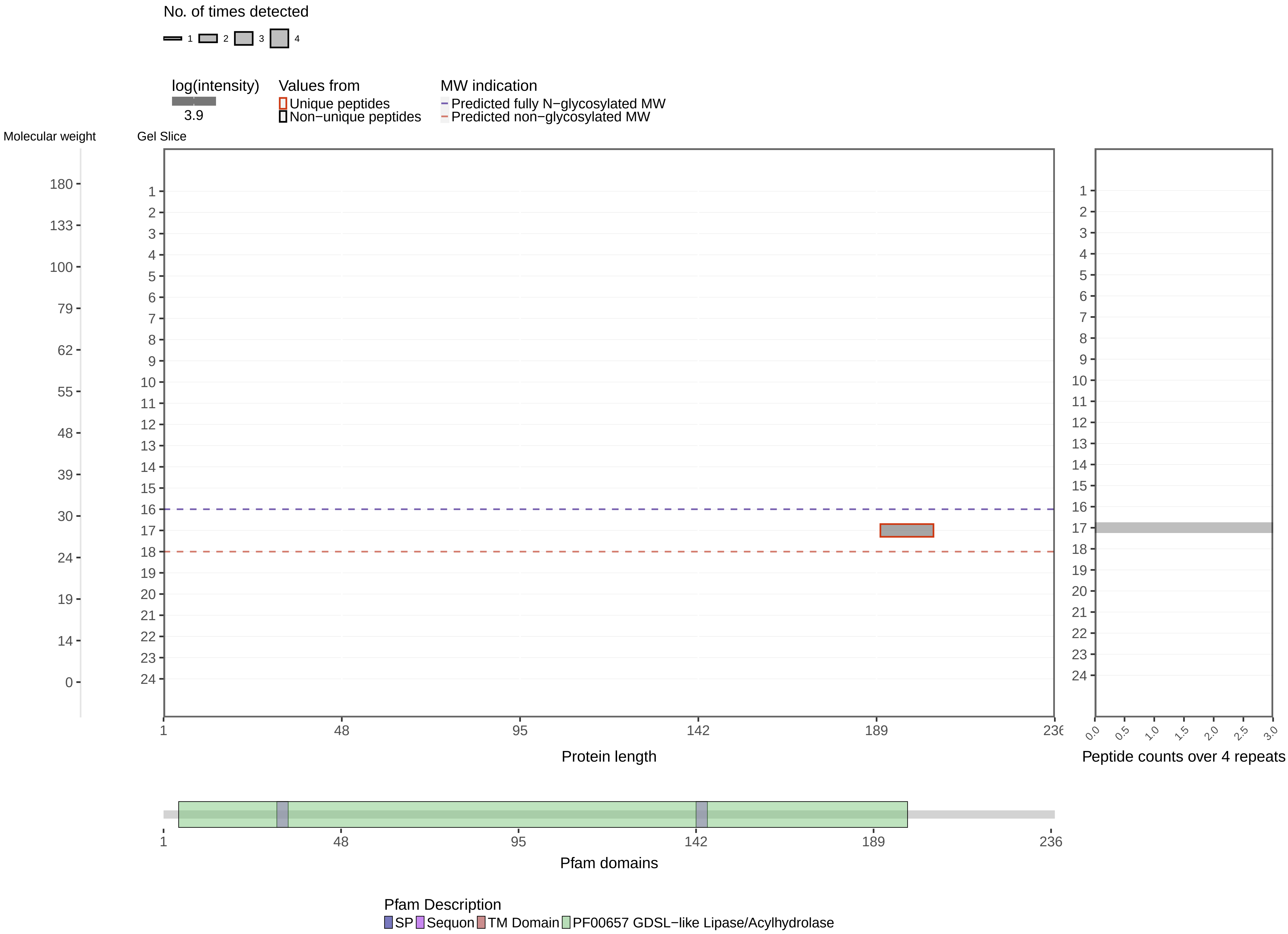


Protein sequence:

GRWPTTQIVLISPPPIDEPTRLLYPFMENKLGLSERTNETAGNYAKASLAVAAECGVLAVDLWTRMQQIPGWQTACLSDGLHLSKTGNEIVFEEVVAALKKKGLSVEALPVDLPVINEIDPNDPLKSFEGI*

### NbL02g19500.1 low−temperature−induced_cysteine_proteinase−like [MW − SP = 53.42 kDa]

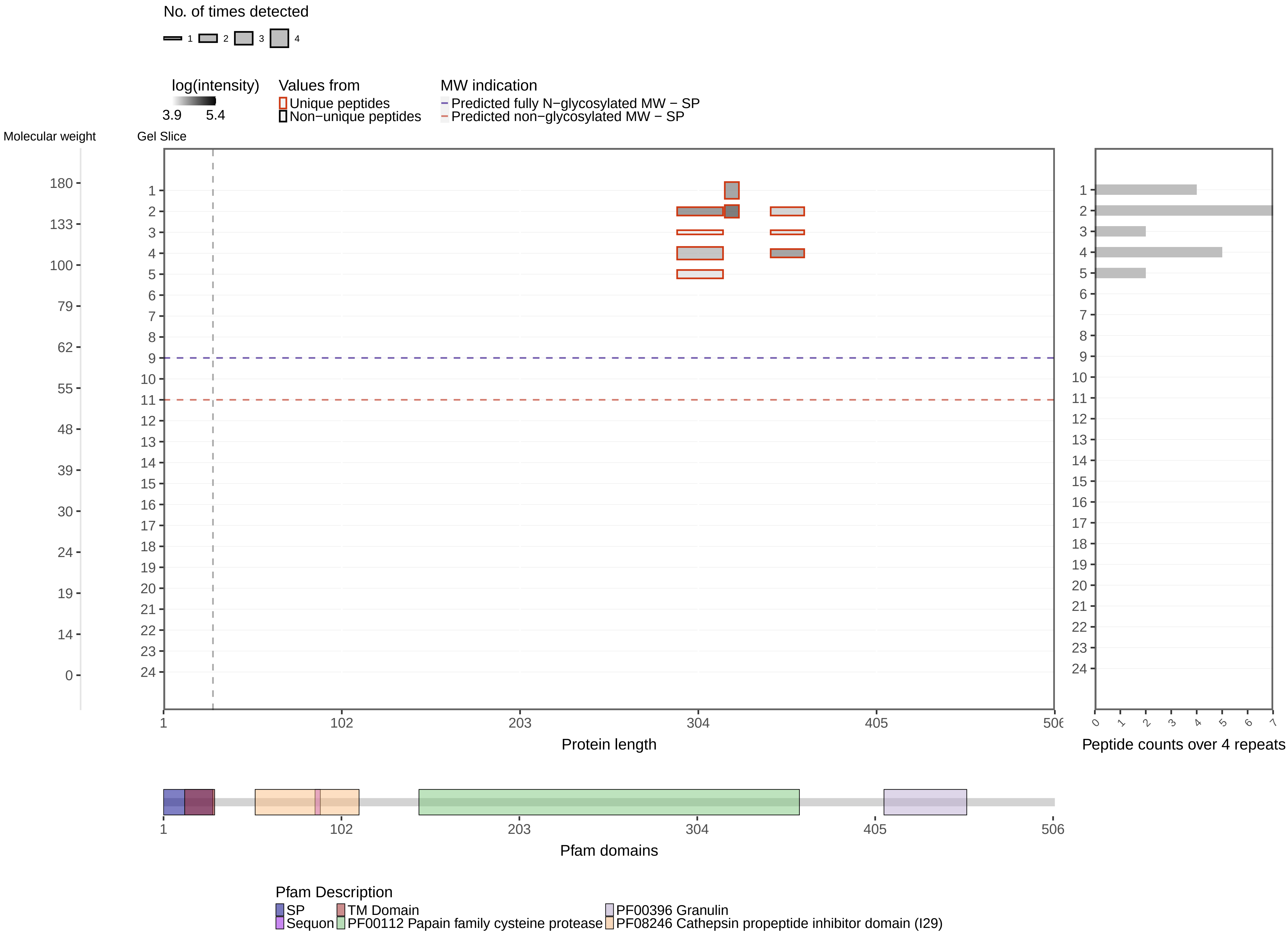


Protein sequence:

CNYDKVNHKVVTVDGYQDVPEDENALLCAVAQQPVSVAIDGSSLDFQLYRGGIYDGECSSNPDDLSHAVLIVGYGSKGDDDYWIIKNSWGTTWGMEGYAYIRRNTYLPYGVCAINSLASYPTKEASTAPSPYPSPAVPPPPPPPSPPPPPPPSPKPSECGDFYYCPEDQTCCCELYLFGICFIQGCCPYENGVCCHGSEFCCPADYPICDVYEGLCLKDYRDNIGVSARKRRMAKHKLPWRANTEPTEERDQTLKWKRNHVAAMR*

### NbL02g19230.1 protein_HIGH_CHLOROPHYLL_FLUORESCENCE_PHENOTYPE_173_chloroplastic [MW = 66.16 kDa]

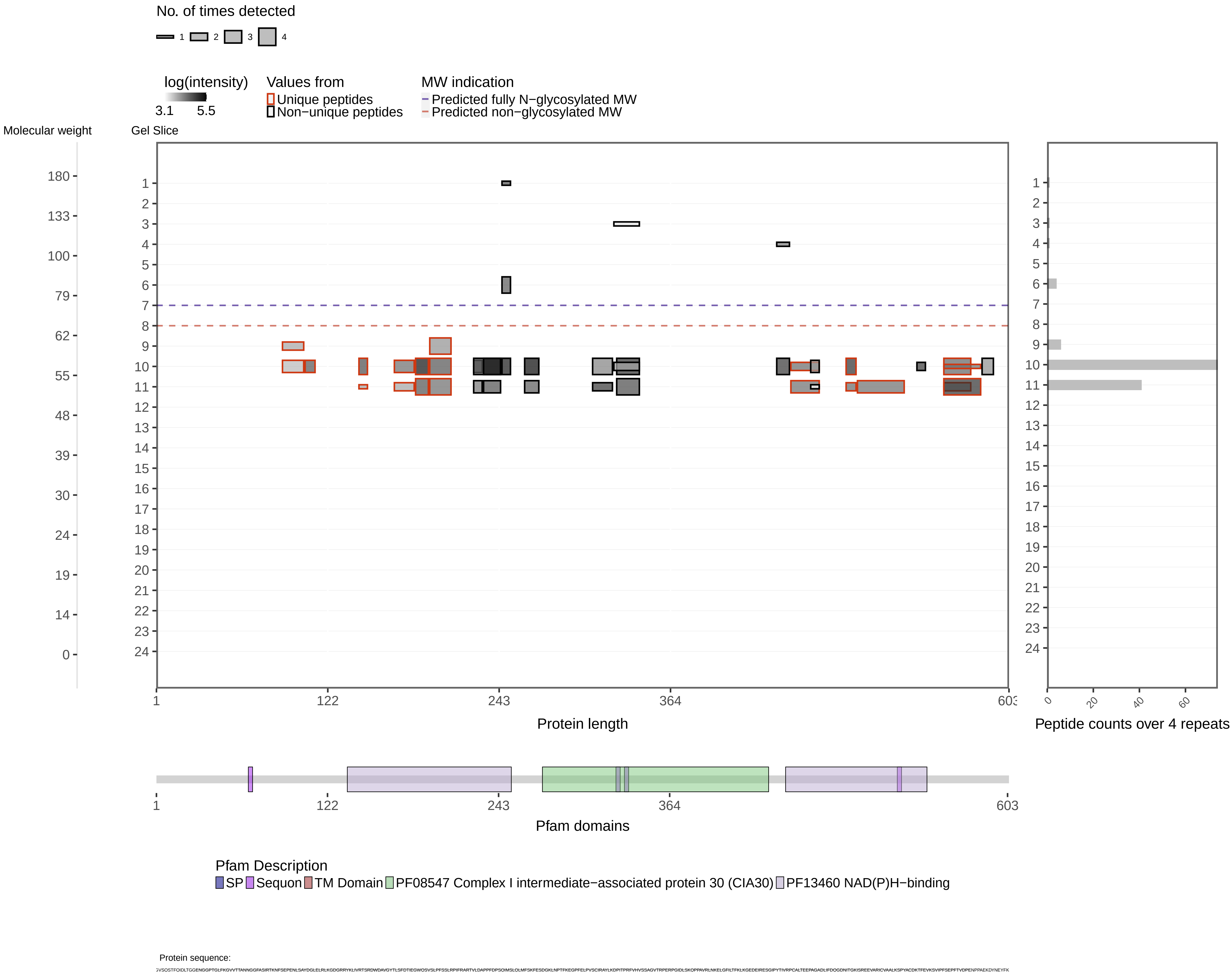

NbL02g19210.1 probable_alphaalpha−trehalose−phosphate_synthase_[UDP−forming]_7 [MW = 96.77 kDa]

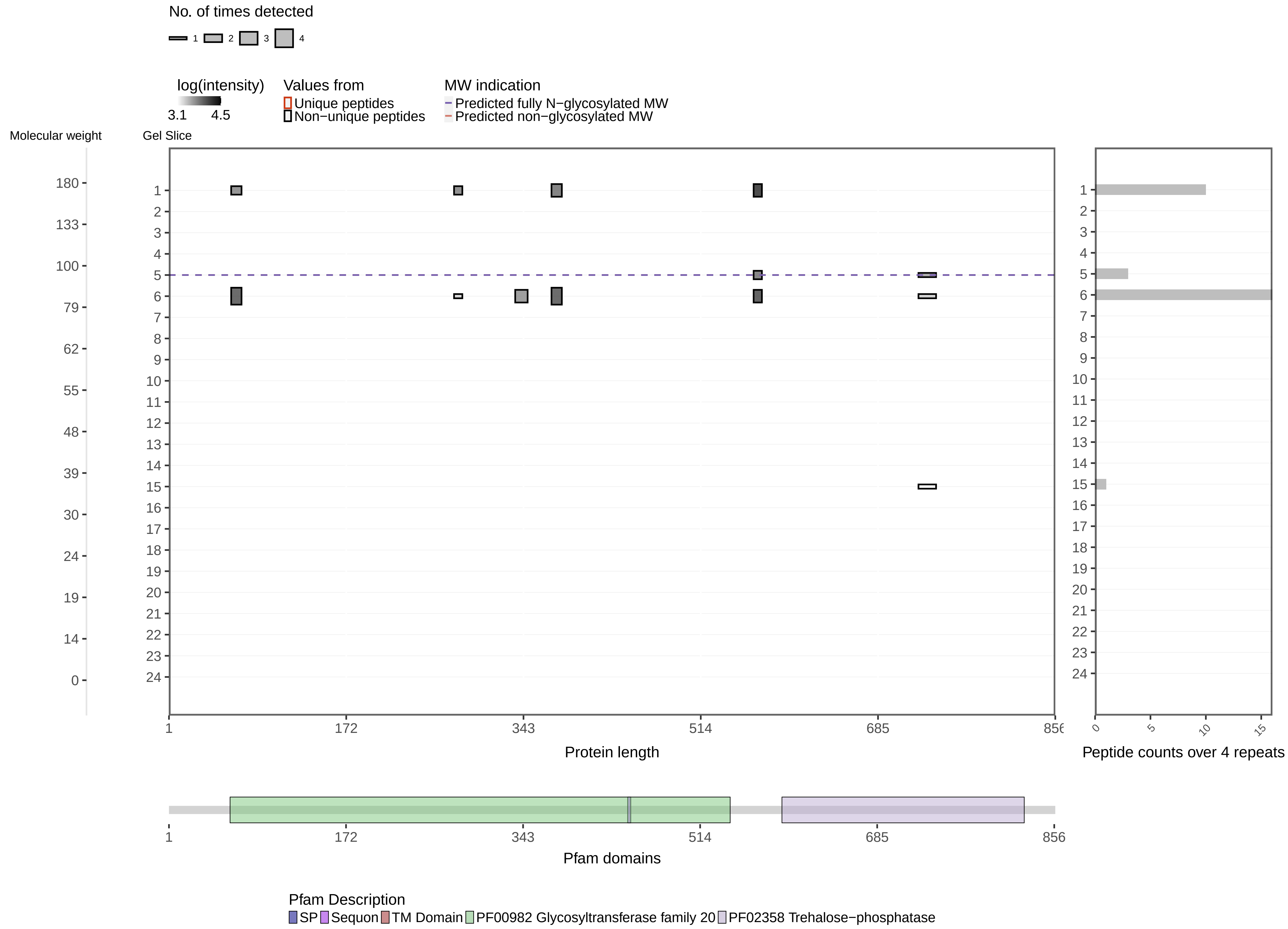


Protein sequence:

PVSLSERTAYYTVAECVVVTAVRDGMNLTPYEYIVSRQGIPGSDCTAESNGQKKSMLVVSEFIGCSPSLSGAIRVNPWNVEATAEALNEAISMADAEKVLRHEKHYKYLSTHDVAYWSRSFFQDLERTCKDHFRRRCWGIGLSFGFRVVALDPNFRKLSMDNIVSSYSKAKNRAILLDYDGTLMPQTSINKVPSPEVISIINTLCGDERNTVFLVSGRGRDSLGRWFSPCEKLGIAAEHGYFLRWSAHKEWEVCGQNSDFQWMQIAEPVMKQYTEATDGSYIETKESALVWH

NbL02g19160.1 dihydroorotase_mitochondrial [MW = 43.11 kDa]

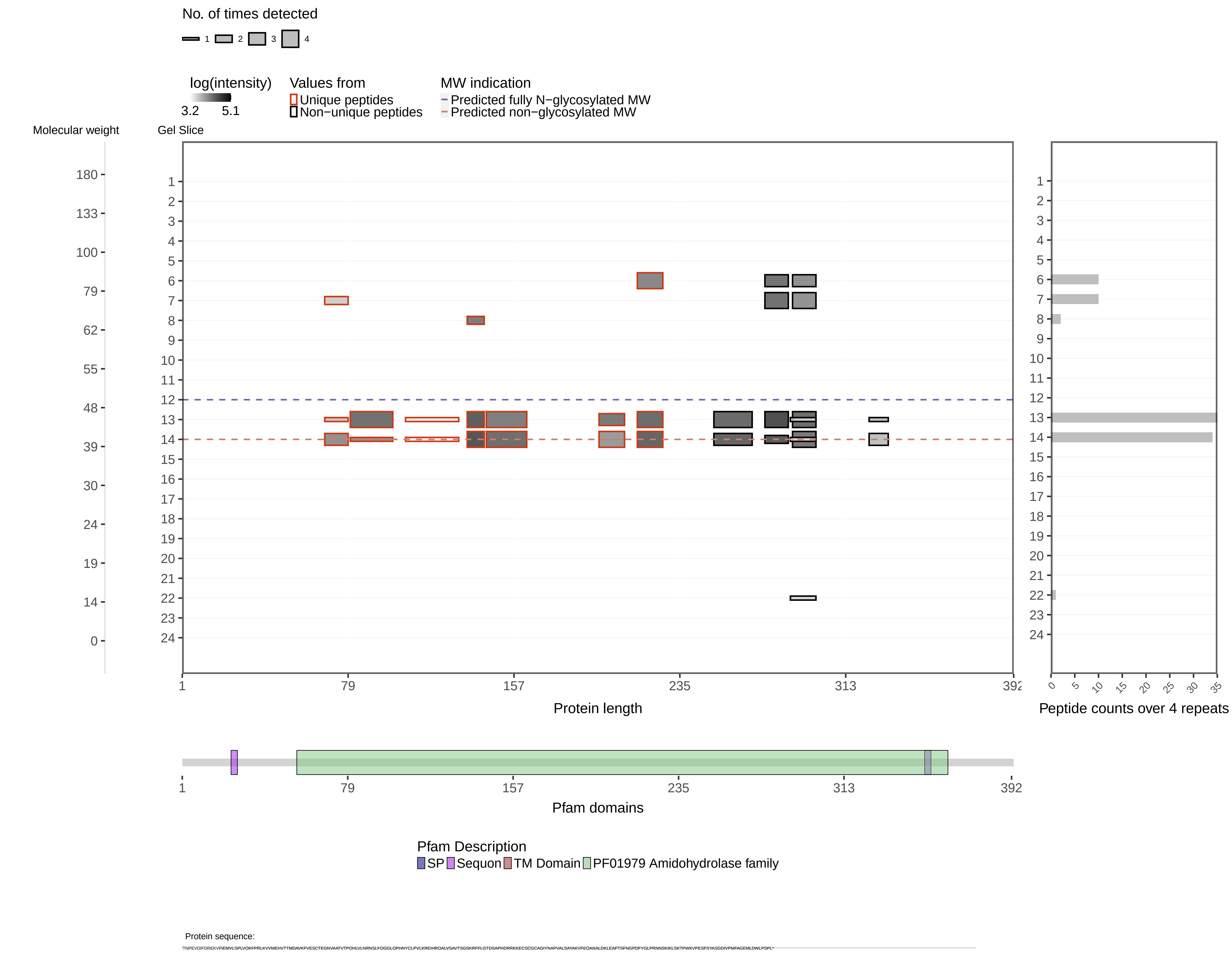

### NbL02g18960.1 histidinol_dehydrogenase_chloroplastic [MW = 52.75 kDa]

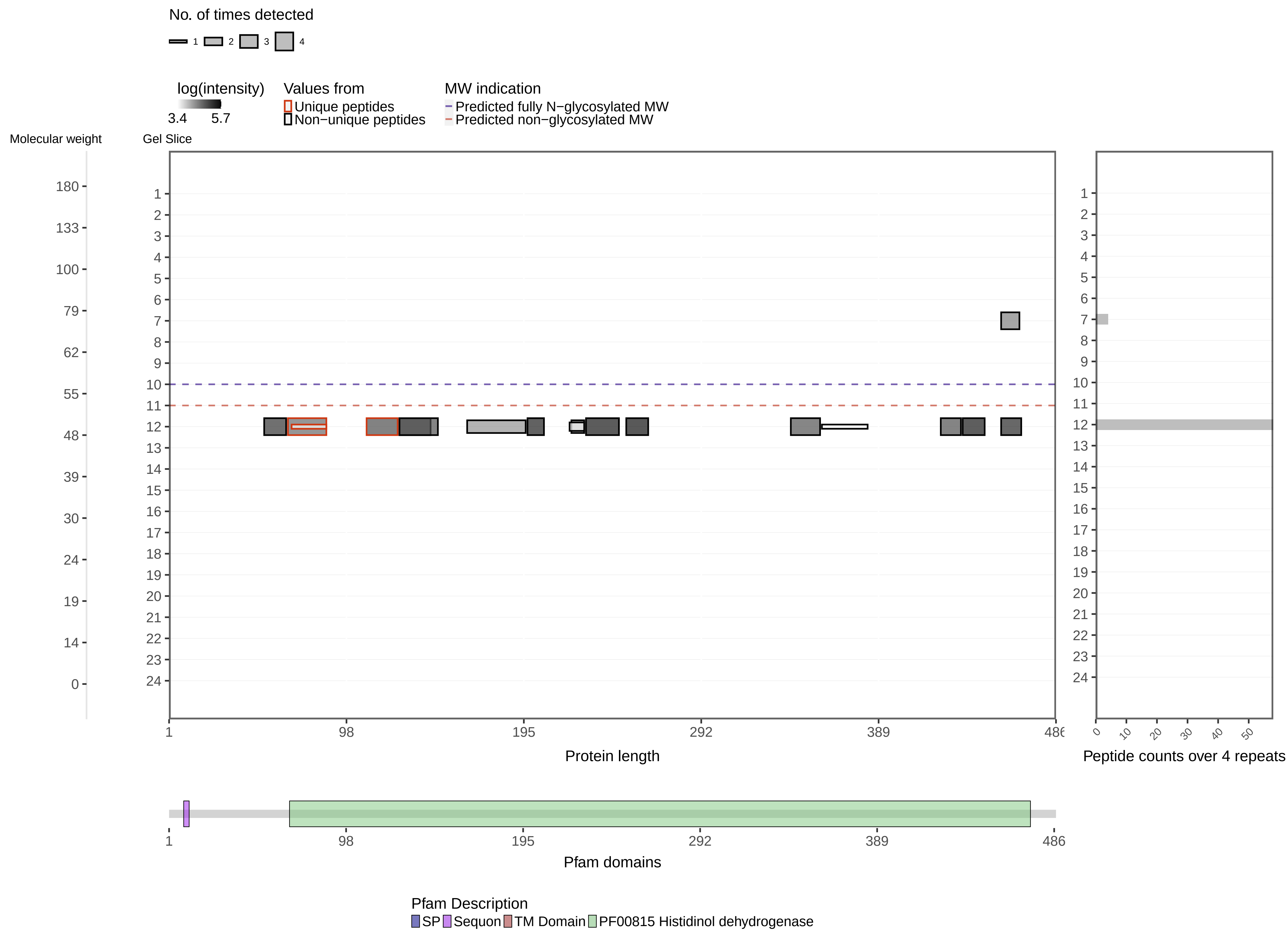


Protein sequence:

AQAISAMAWGTESCPKVEKIYGPGNQYVTAAKMILQNSEAMVSIDMPAGPSEVLLIADKHSSPVHVAADLLSQAEHGPDSQVVLVIAGDDVDLNAIQEEIRKQCKALPRGEFALQALSHSFTVLARDMLEAISFSNMYAPEHLIINVNEAEKWESLIENAGSVFMGQWTPESVGDYASGTNHVLPTYGYARMYGGVSLDSFLKYITVQSLTEEGLRNLGPYVEKMAEVEGLDAHKRAVTLRLQDIEARQISNSR*

NbL02g18950.1 probable_LRR_receptor−like_serine/threonine−protein_kinase_RFK1 [MW − SP = 109.82 kDa]

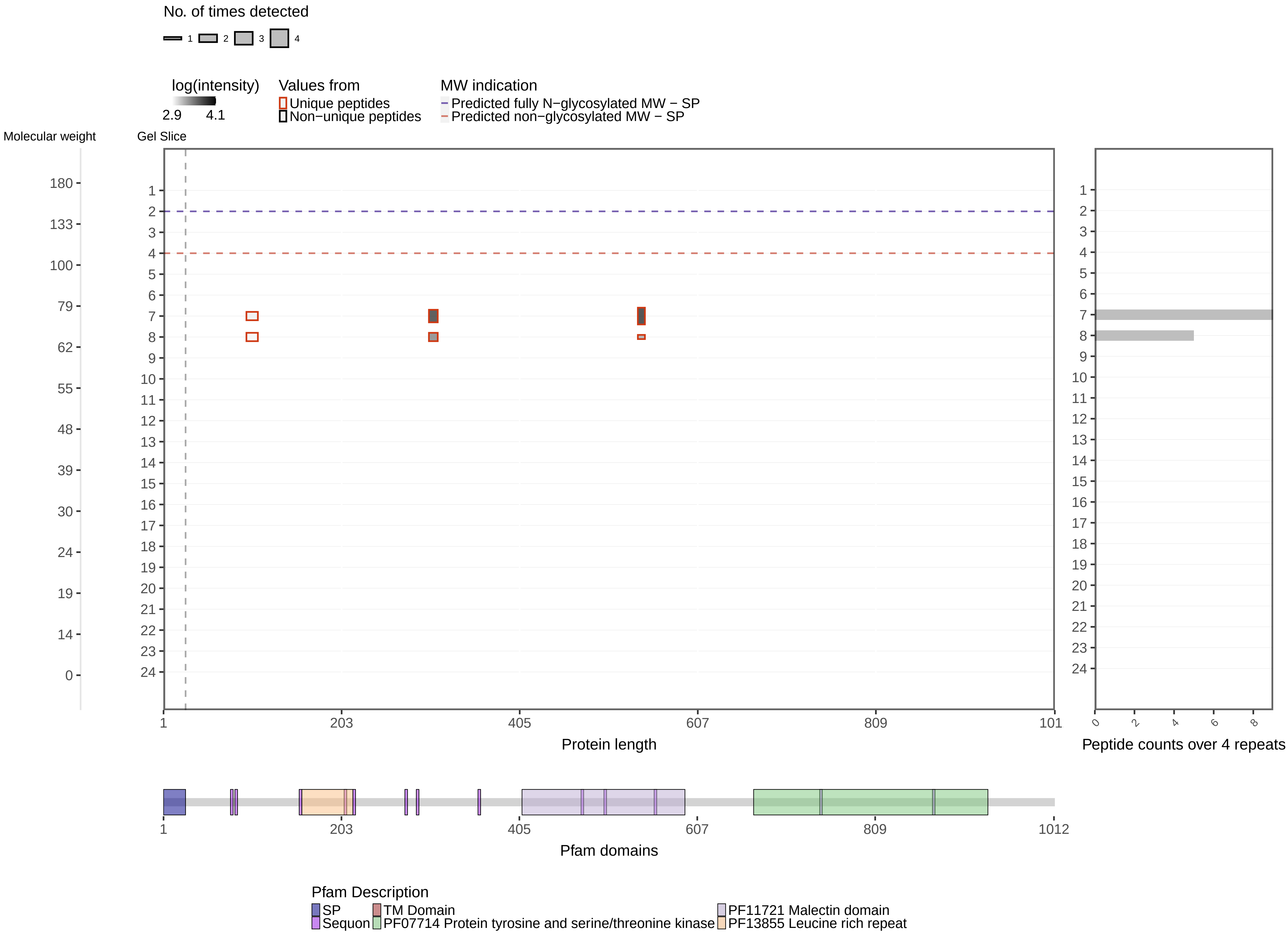


Protein sequence:

YCLENGNYSVSLHFAEIMFKEDHTYYNLGRRVFDIYIQEKLVWKDFNIEEEARGVKRPVIRRFNATVTDSVLEIRLYWGGKGTTRIPLRGHYGALISAISVDSNFKFCSNEDRKTVIIYVTVGVLAACITFFVLSILWWKGYLCRERKRRDLNGVELQMVCFTLKQIKAATKNFDASNKIGEGGFGPVYKGQLPDGTLVAVKQLSSQSKQGNREFLNEISTISCLQHPNLVKLHGCCIEADQLLLVYEYLDNNSLAGALFQDSRLTLDWATRFRICLGIAKGLAFLHEESSLKIVHRD

NbL02g18840.1 GDSL_esterase/lipase_At1g29670−like [MW = 40.83 kDa]

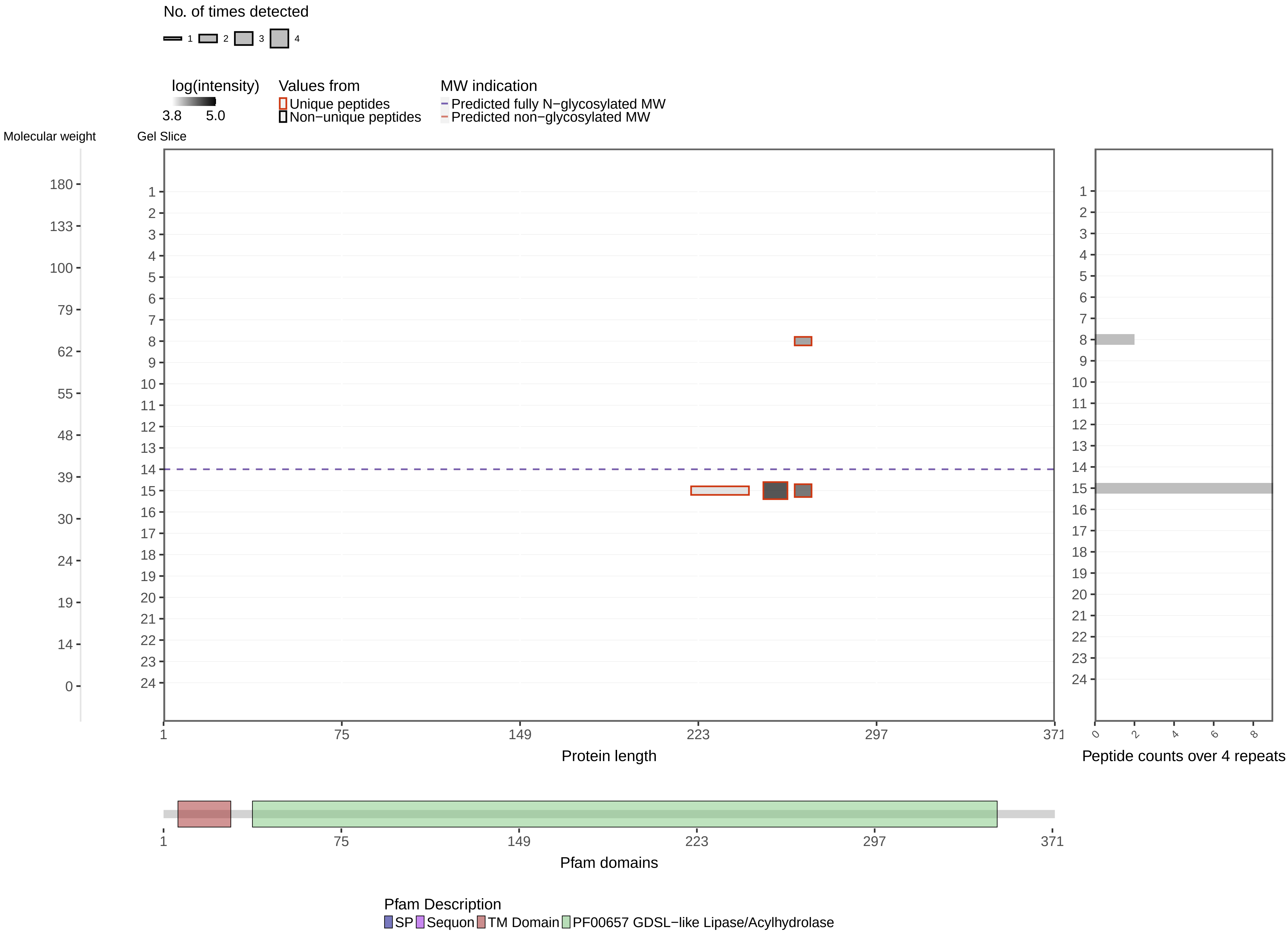


Protein sequence:

GSNDYLNNYFMPLYYPTSRQFTPQQYANVLIQQYTQQLRILYNNGARKFVLIGVGQIGCSPNALAQNSPDGRTCVQRINVANQLFNNKLKALVDNLNRNTPDAKFIYIDAYGIFQDLIDNPSAFGLSVTNAGCCGVGRNNGQITCLPFQRPCRNRNQYLFWDAFHPTEAANIVVGRRSYRAQKSSDAYPFDISRLAQL*

### NbL02g18830.1 GDSL_esterase/lipase_At5g45670−like [MW = 46.52 kDa]

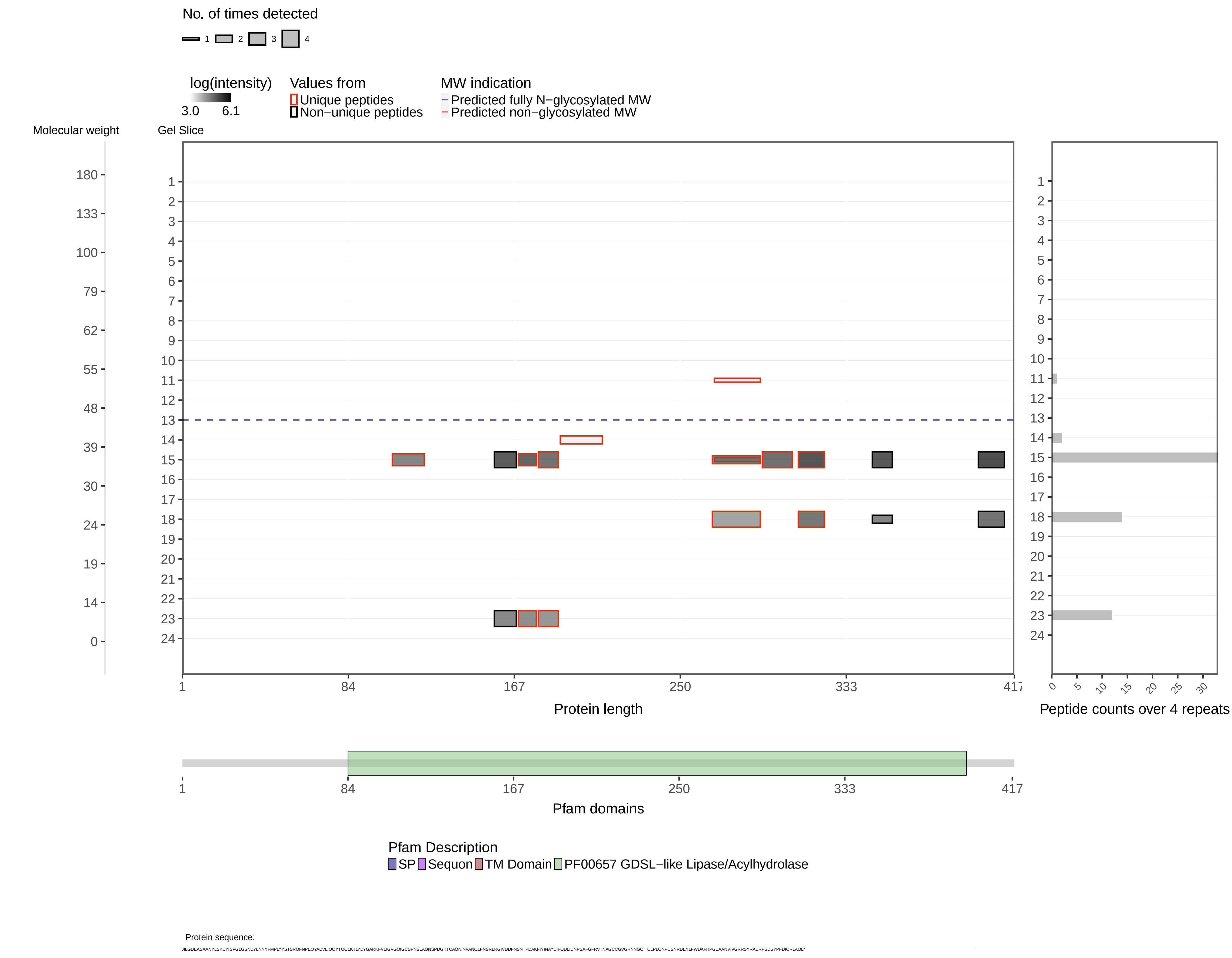

NbL02g18800.1 subtilisin−like_protease_SBT5.3 [MW = 86.41 kDa]

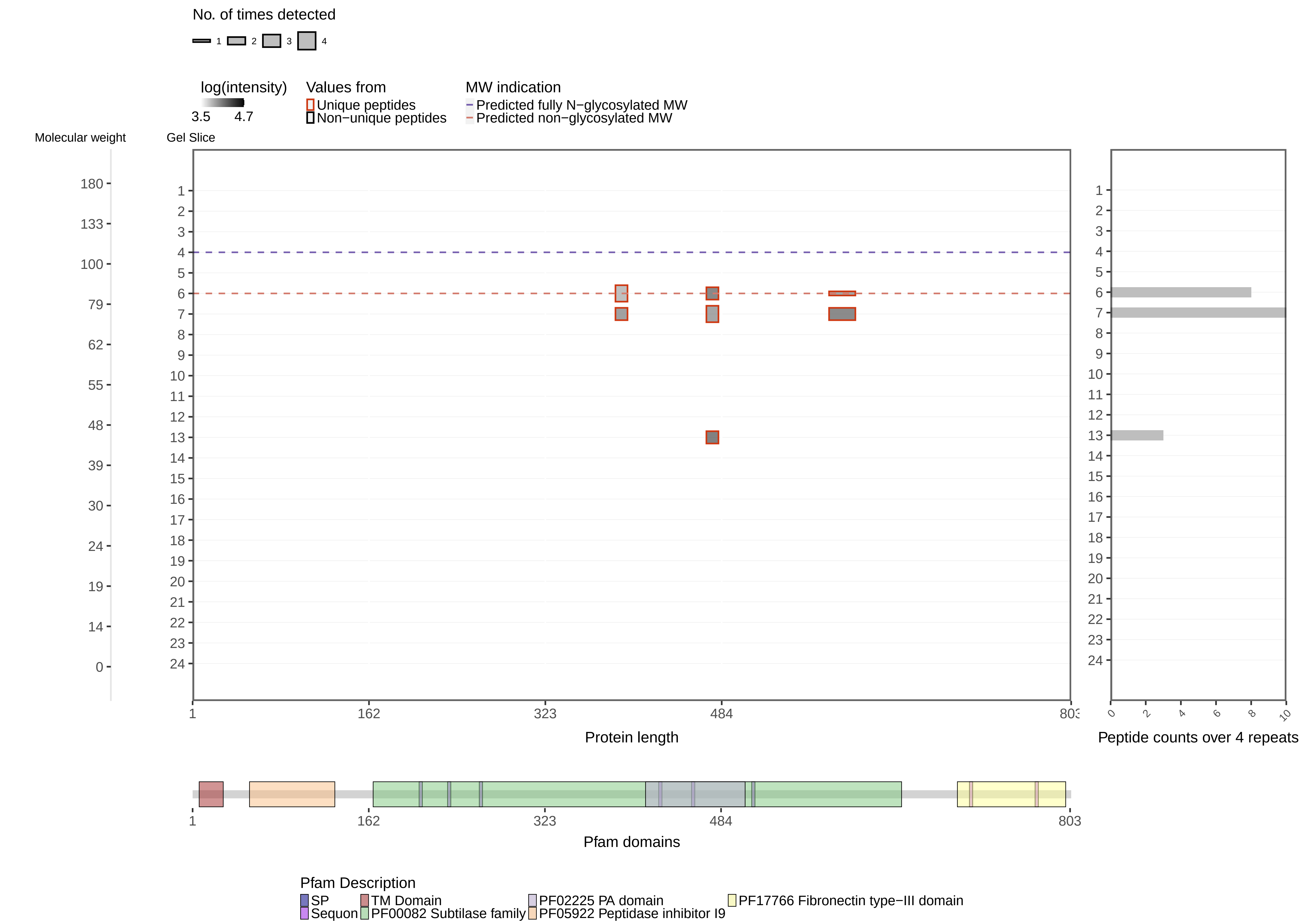


Protein sequence:

[illegible]

### NbL02g18780.1 subtilisin−like_protease_SBT5.3 [MW = 70.09 kDa]

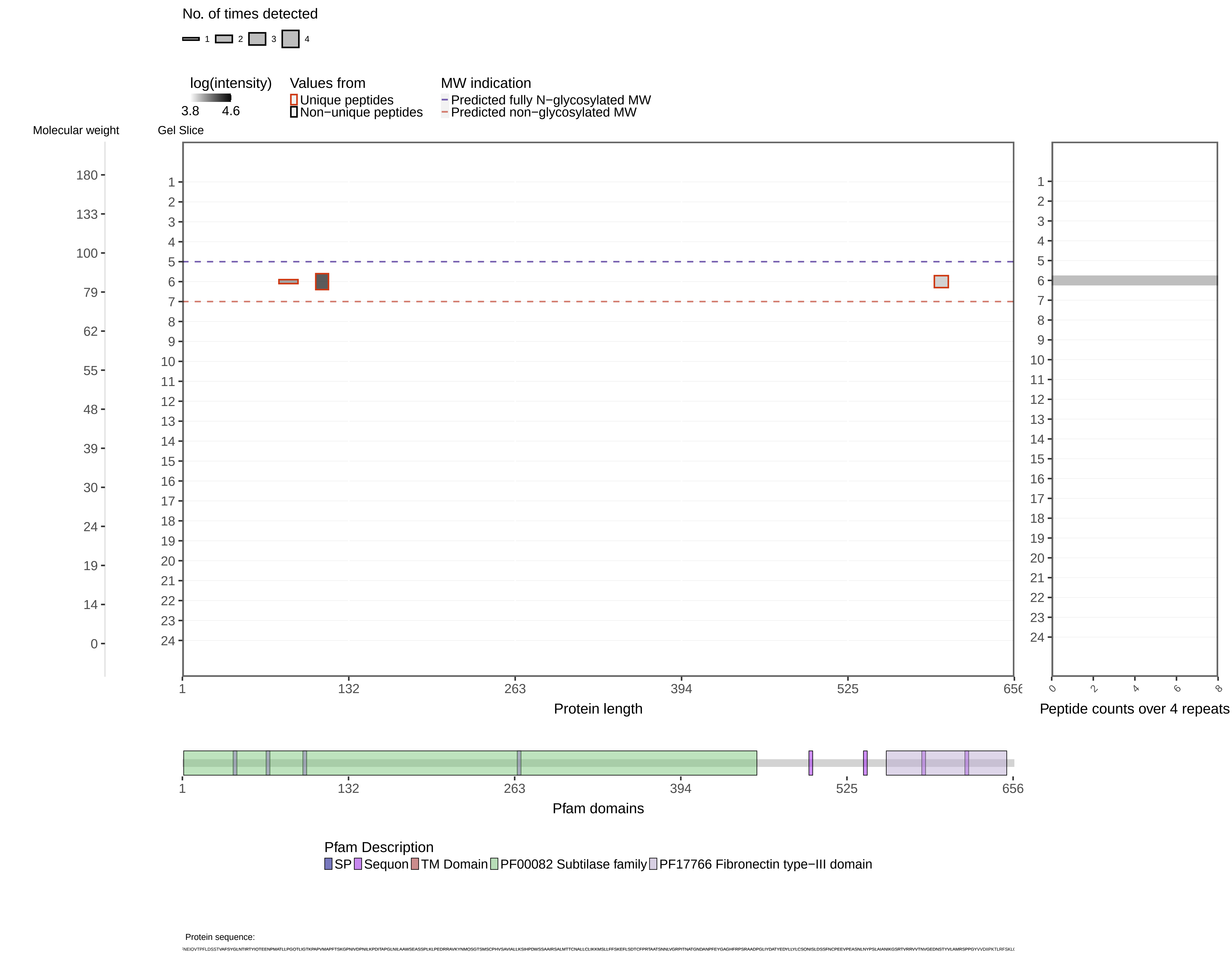

### NbL02g18680.1 protein_SRG1−like [MW = 41.9 kDa]

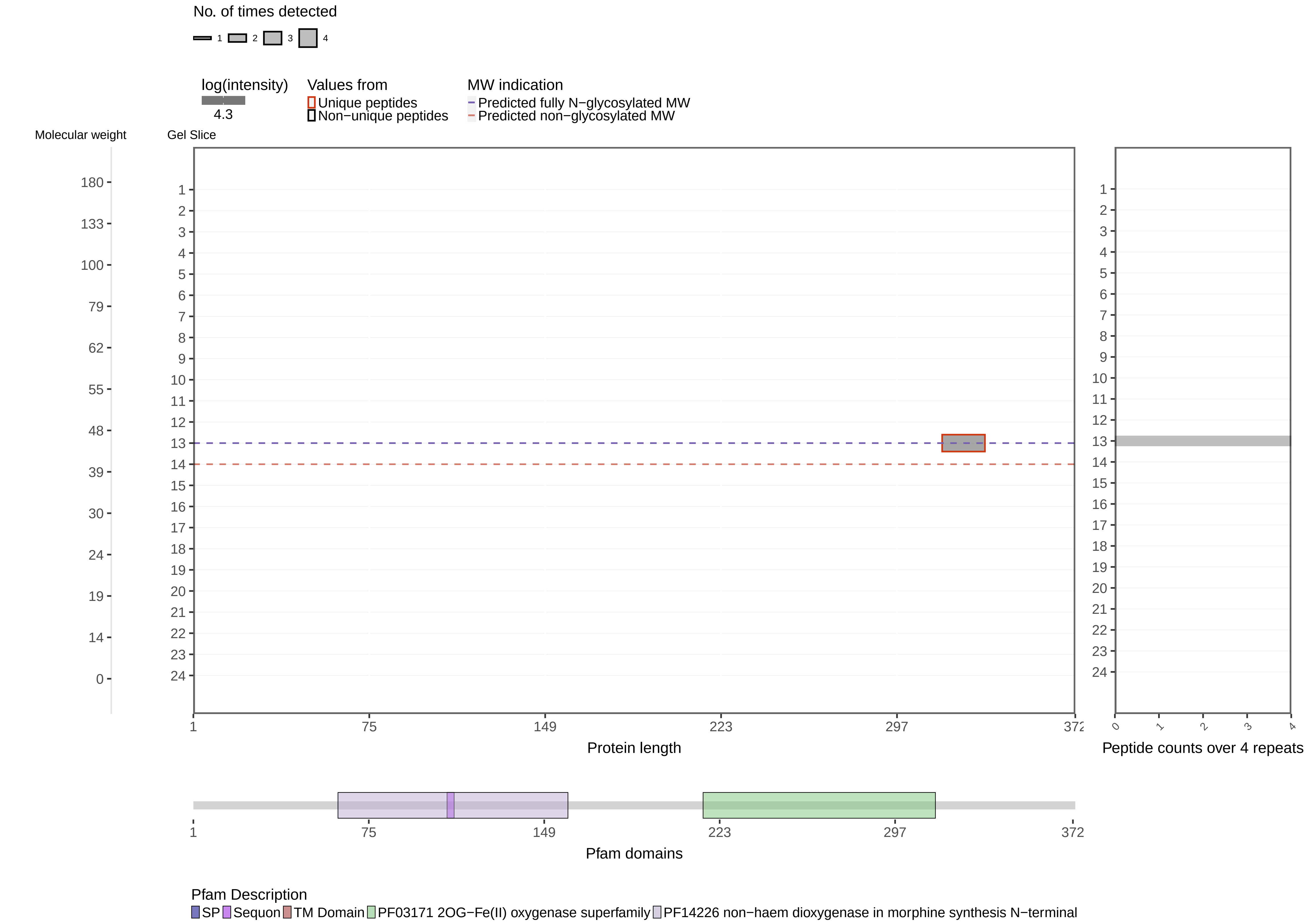


Protein sequence:

LPLPFRETVELYAAELKTLALOIINFIAKALNIDEEYMSELFGEGMOAMRMNYYPPCPEPEKVIGLSPHSDGVGLTILLOLNOMEGLOIKKDGLWIPISPLPNAFIINIGDILEIVTNGAYRSIEHRATVNNKKERLSIATFYNPKLDGOIGPAPSIISSENTAKFKNIGAADYFRGLFARKLDKKSYLDVLRIENYDNPAS*

### NbL02g18630.1 60S_ribosomal_protein_L32−1−like [MW = 19.71 kDa]

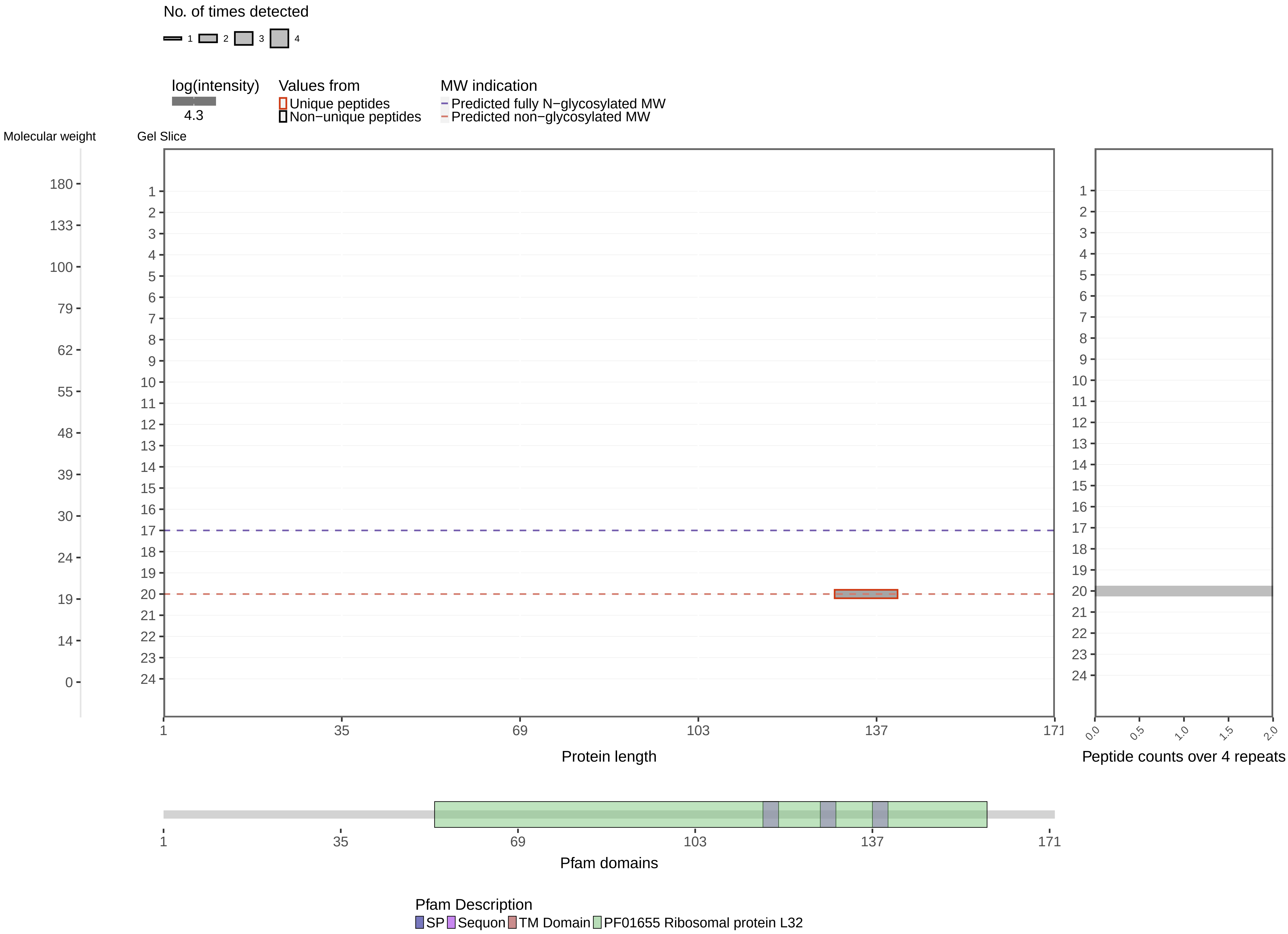


Protein sequence:

KGIDSRVRRKFKGCVLMPNIGYGSDKKTRHYLPNGFKKFVVHNASELDILLMHNRTCCAEIAHNVSTRKRKEIVERAAQLDVVVTNKLARLRGQEDE*

NbL02g18620.1 uncharacterized_LOC109210634 [MW = 80.67 kDa]

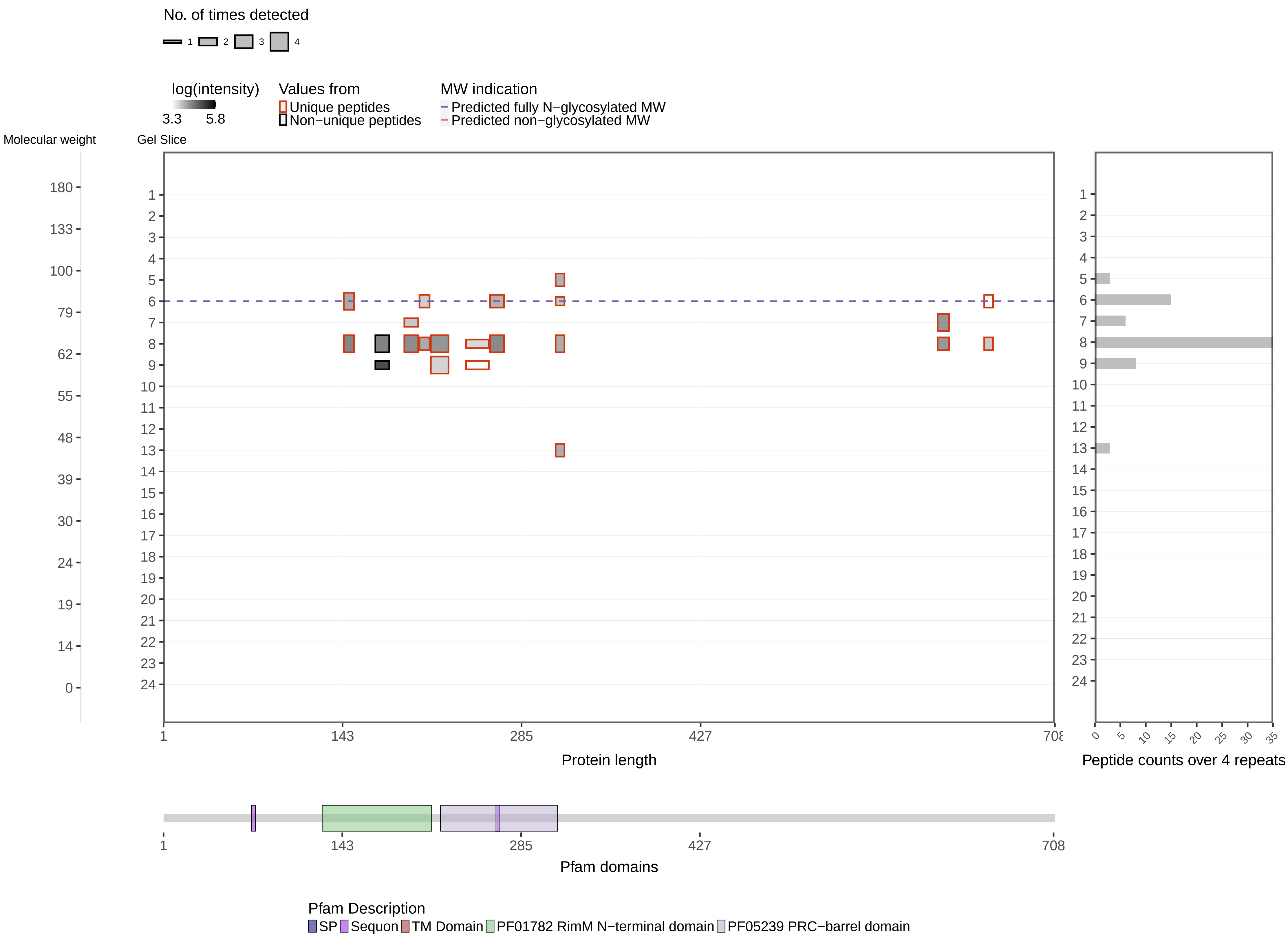


Protein sequence:

[illegible]

NbL02g18490.1 DNA−binding_protein_DDB_G0278111−like [MW = 14.8 kDa]

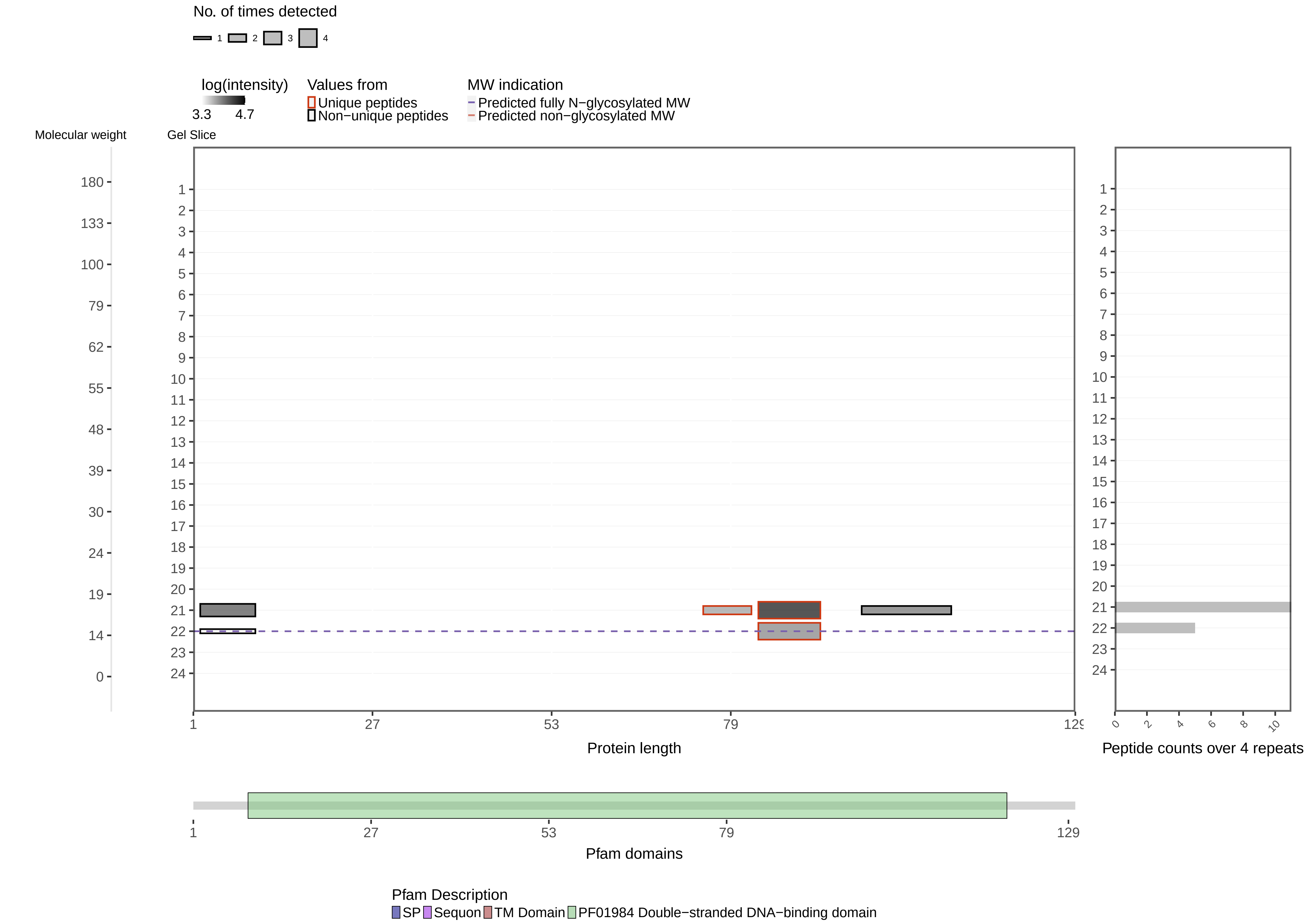


Protein sequence:
MLTQILSSEARARVARIALVKPDKARGVEDVILRAAQYGQITEKVSEEKLIQLLEQINTQTTKQTKVTIQRRRNVLEDDD*

### NbL02g18320.1 uncharacterized_protein_ycf23−like [MW = 33.91 kDa]

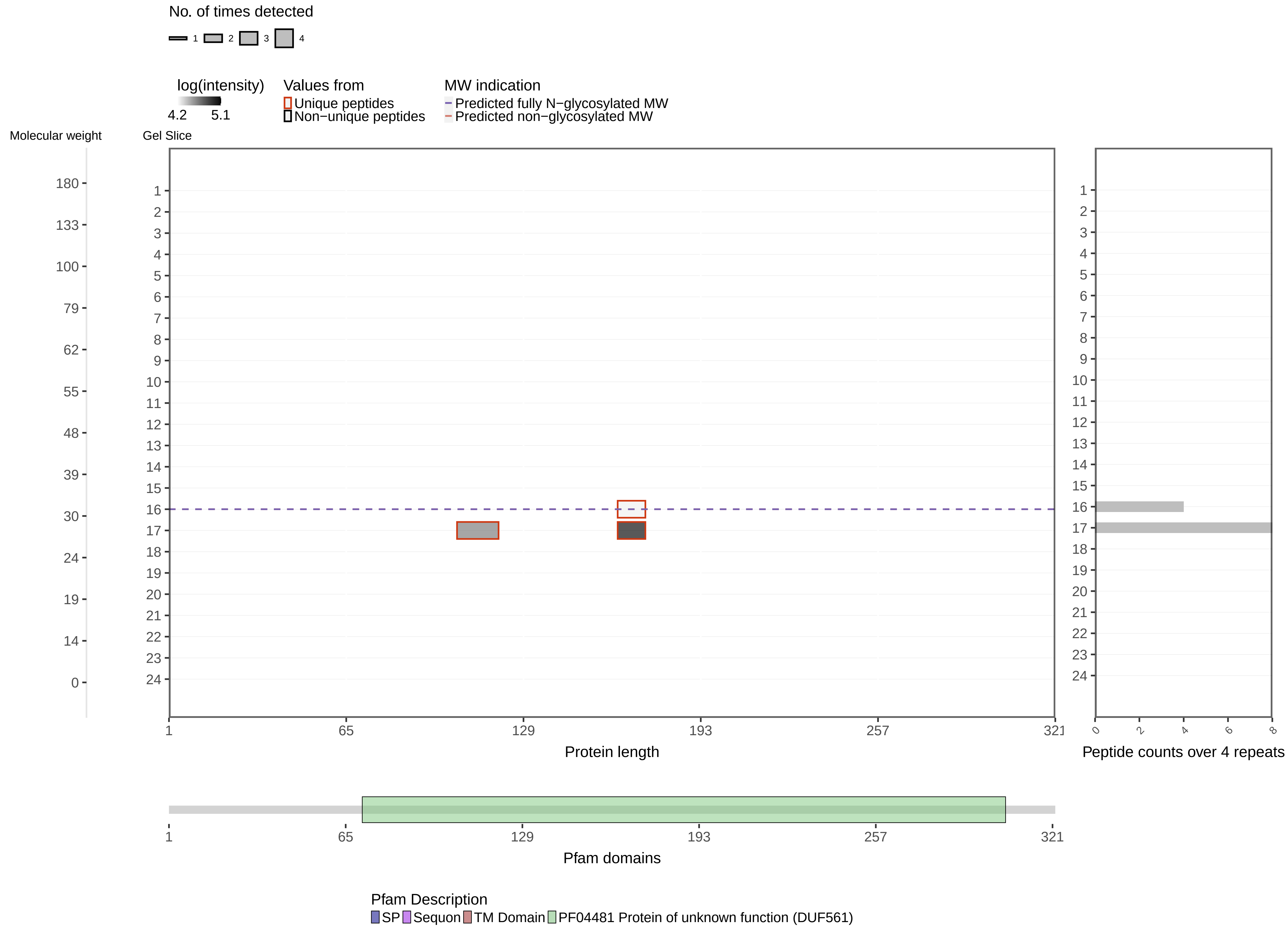


Protein sequence:

ALMVEIGNYDSFYEKGLVFSPEQILKLTKETKRILPLVTLSVTVPHTLSLPDQVKLAELLEQEGVDIIQTEGGKFSSPSKAGVLGLIEKATPTLAAAYSISRAVNIPVMCSSGLSAVTAPMAIAAGAAGVGVGSAINKLNDQVAMIAEVKSIAHSLGLSTKTQNLFEGSSLRL*

NbL02g18300.1 3−oxoacyl−[acyl−carrier−protein]_synthase_I_chloroplastic [MW = 49.98 kDa]

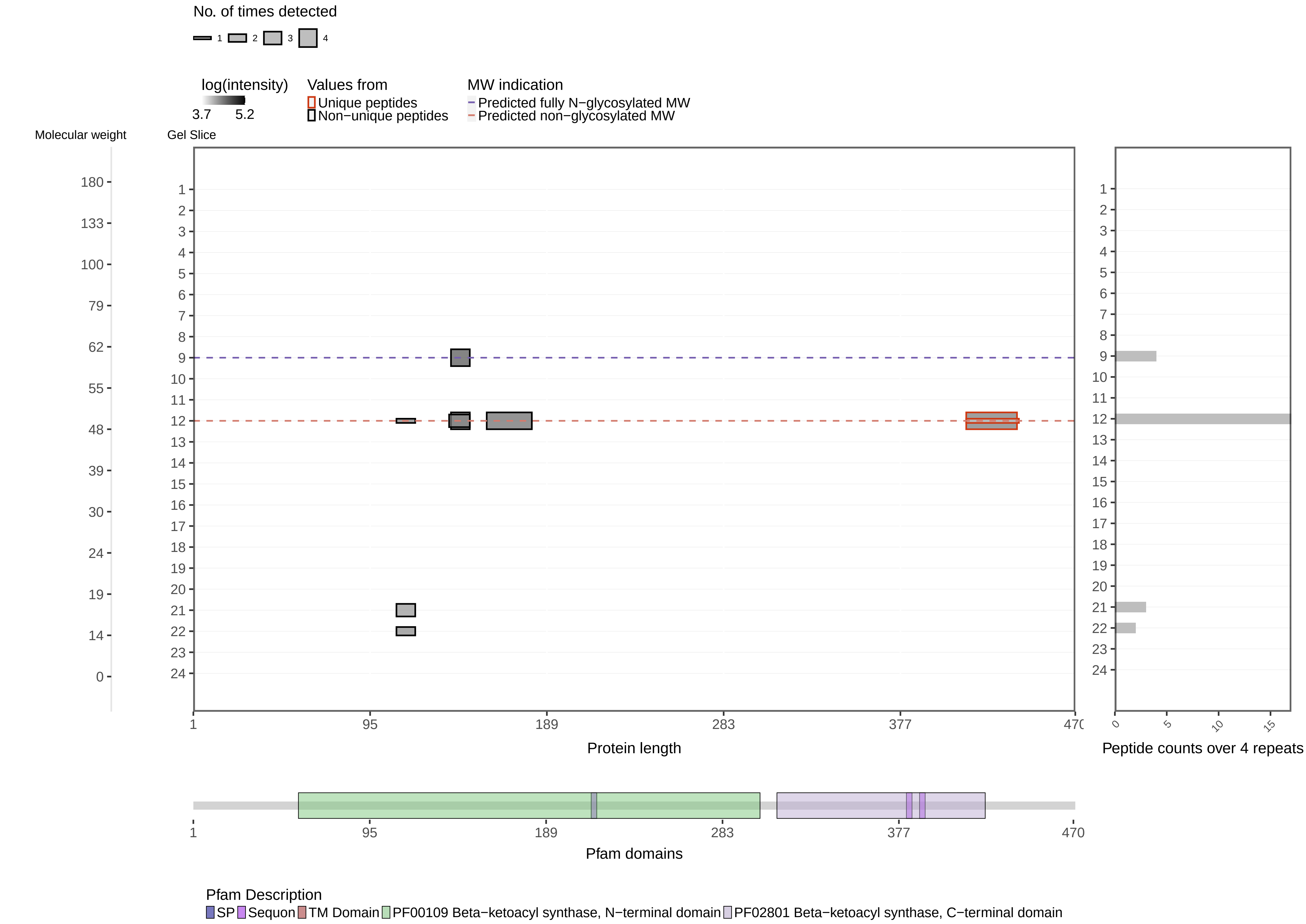


Protein sequence:

NYCFYAAANHIRRGEADLMLAGGTEAAIIPIGLGGFVACRALSQRNDDPQTASRPWDKERDGFVMGEGAGVLVMESLEHAMKRGAPIIAEYLGGAVNCDAHHMTDPRADGLGVSSCIQSSLEDAGVSPEEVNYINAHATSTIVGDIAEVNAIKKVFKNTSGIKMNATKSMIGHCLGAAGGMEAIATVKAITTGWVHPSINQFNPEPSVEFDTVPNKKQQHEVNVGISNSFGFGGHNSVVAFSAFKP*

NbL02g18280.1 uncharacterized_protein_At2g34460_chloroplastic−like [MW = 27.29 kDa]

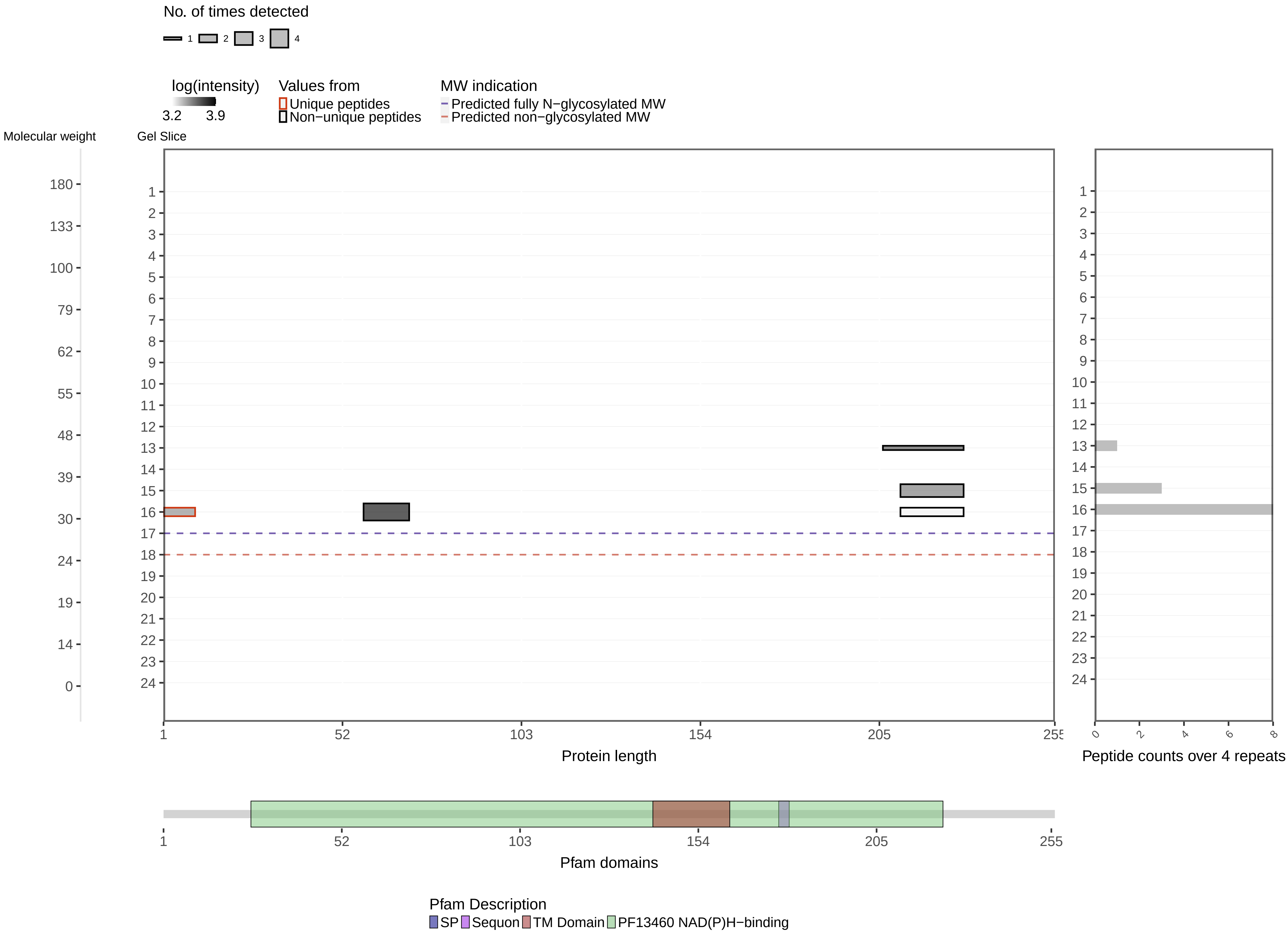


Protein sequence:

FGTVNLVDACQKLGIDRFILISSILVNGAAVGQLFNPVYIILNVMGLVLVAKLQAERYIRKSGINYTIIRPGGLKNDPPQGNIVMQPEDTLFRGSISRDQVAEVAVEALLHPESHYKVVEIVAGTDIPKRPFEELFGLIKQQ*

### NbL02g18210.1 60S_ribosomal_protein_L18a−like [MW = 21.41 kDa]

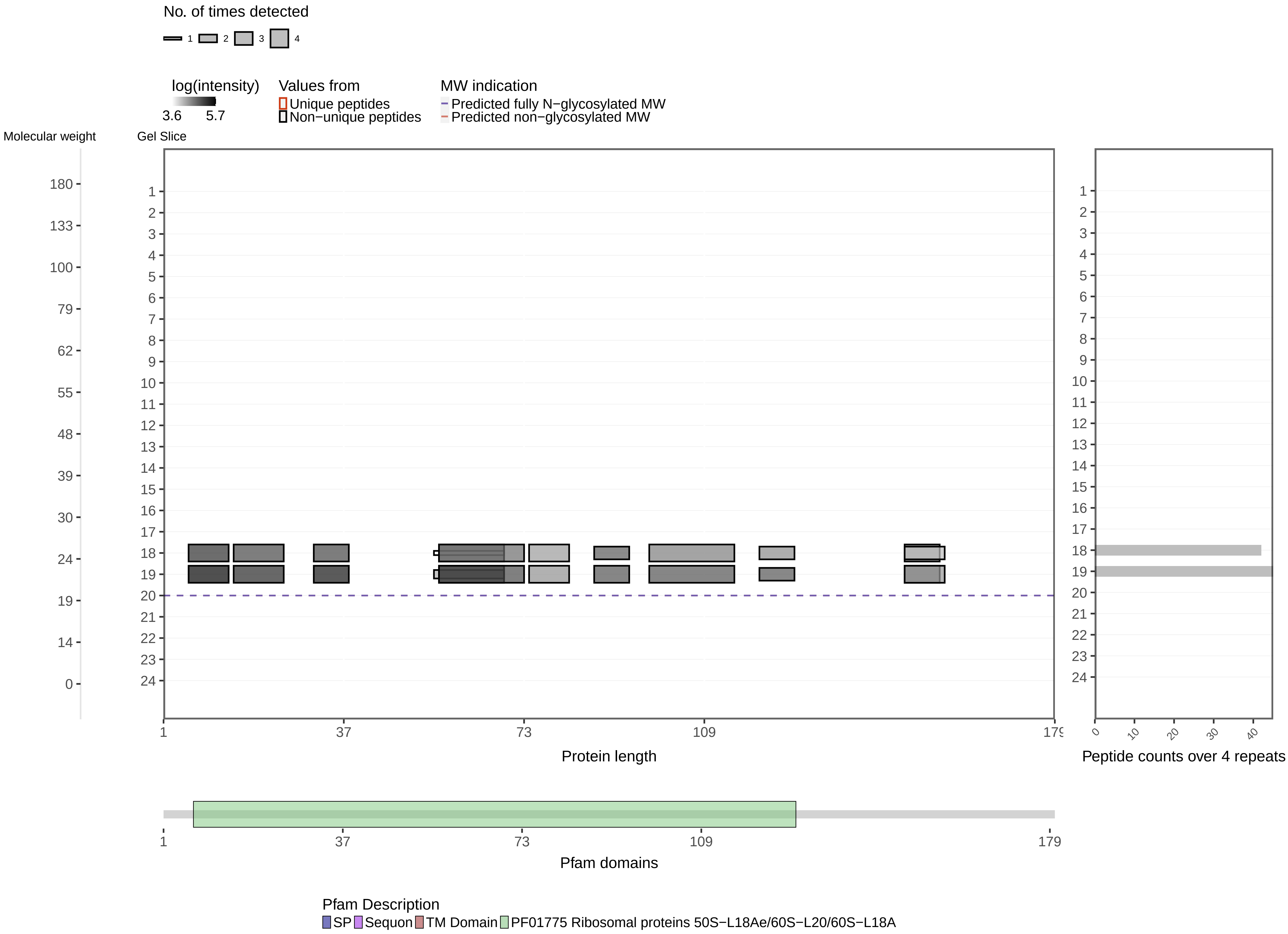


Protein sequence:

NYGIWLRYQSRTGYHNMYKEYRDTTLNGGVEQMYTEMASRHRVRHHCIQIIKTATIPAKLCKRESTKQFHDSKIKFPLVFKKVRPPSRKLKTTYKATRPNLFM*

### NbL02g18080.1 proteasome_subunit_alpha_type−5−like [MW = 26.07 kDa]

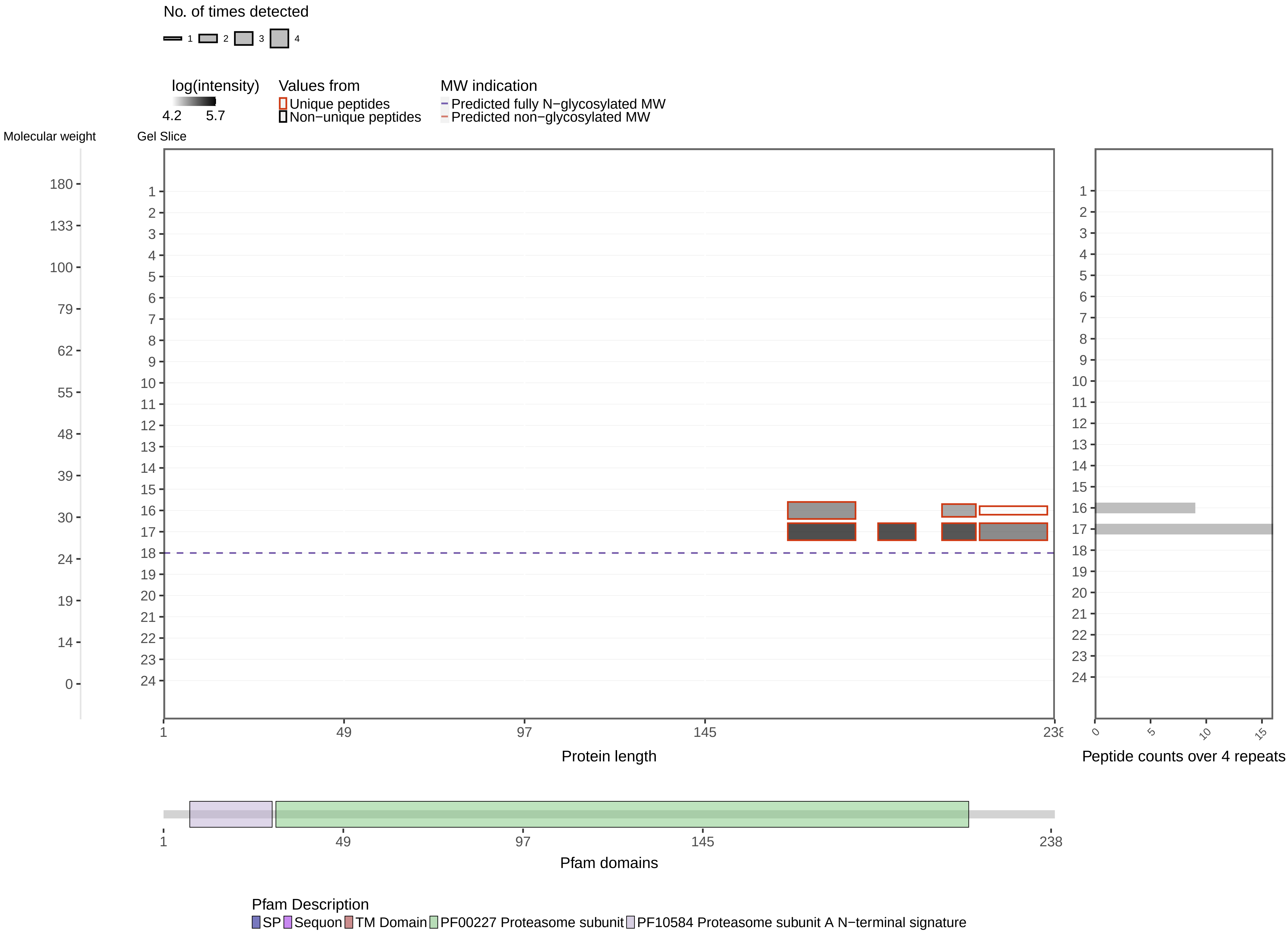


Protein sequence:

MTVESTTQALCDLALRFGEGDEESMSRPFGVSLLIAGHDENGPSLYYTDPSGTFWQCNAKAIGSGSEGADSSLQEQHNKDLTLKEAETIALSILKQVMEEKVTPDNVDIARVSPTYHLYSPSEVEEVISRL*

NbL02g18070.1 uncharacterized_LOC109223617 [MW = 16.75 kDa]

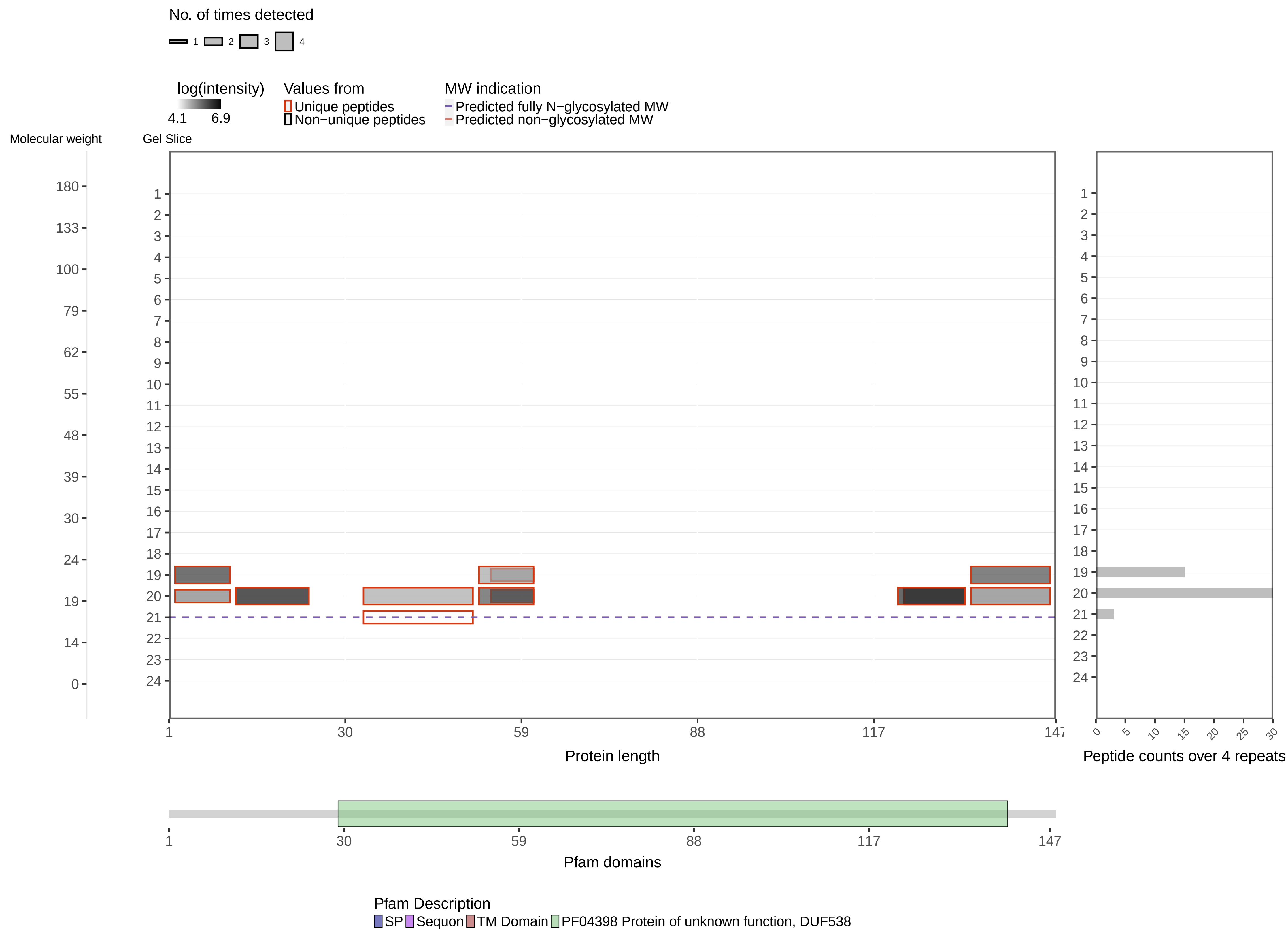


Protein sequence:

LKQKKAKEHKFKKIGKLVWYDTEVTAIVEDRRMKKLTGVKSKELLLWVTLSDISIQDPESKKITFATPAGLSRAYPVSAFEEEEVEK*

### NbL02g18000.1 40S_ribosomal_protein_S4 [MW = 34.99 kDa]

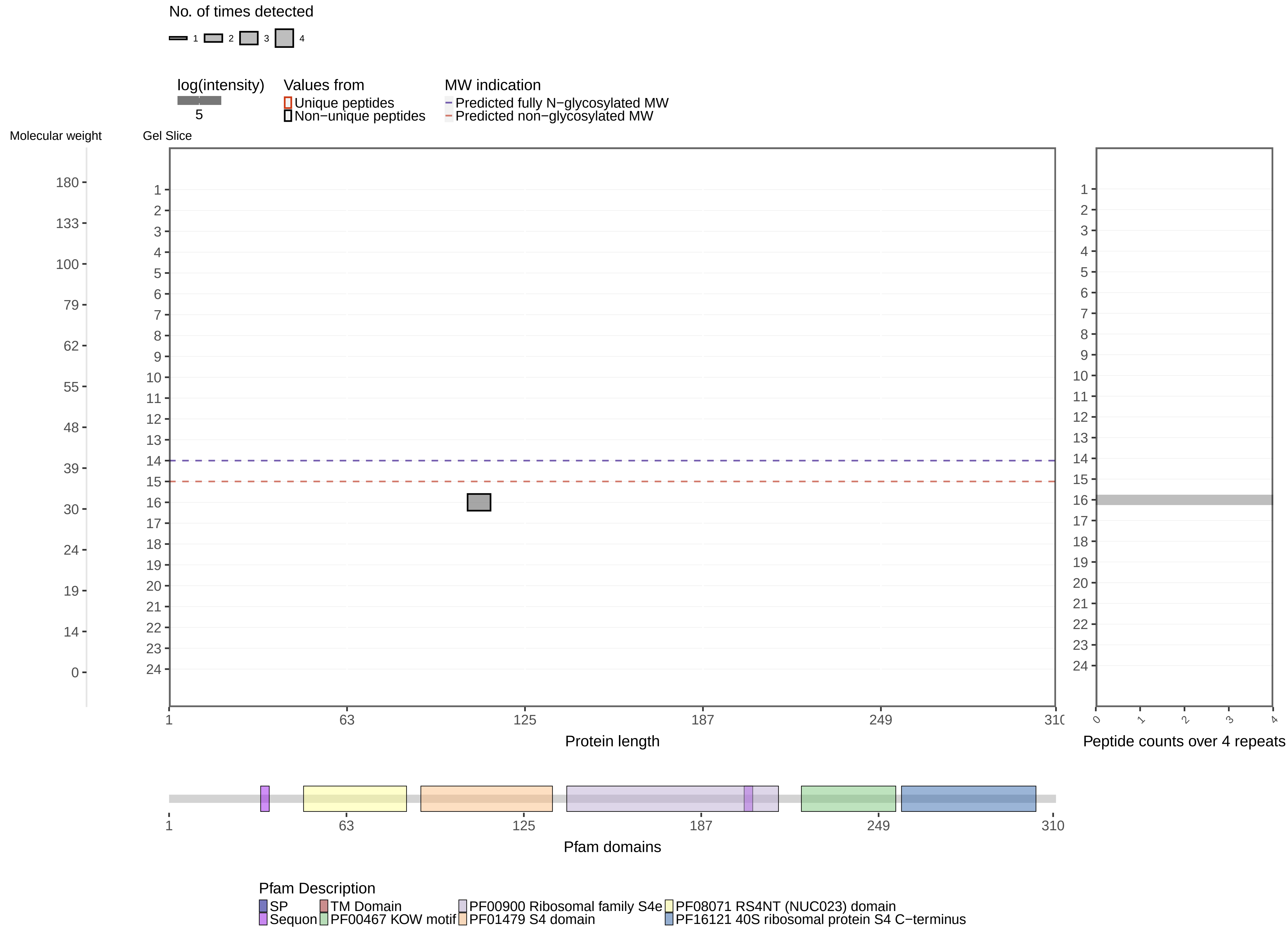


Protein sequence:

NENFRLLYDTKGRFRLHSLRDEEAKFKLCKVRSVQFGQKGIPYLNTYDGRTIRYPDPLIKANDTIKLDLESNKIVDFIKFDVGNVVMVTGGRNRGRVGVIKNREKHKGSFETVHIQDALGHEFATRLGNVFTLGKGTKPWVSLPKGKGIKLSIIEEARKRLAAQSATAA*

### NbL02g17860.1 berberine_bridge_enzyme−like_13 [MW − SP = 58.95 kDa]

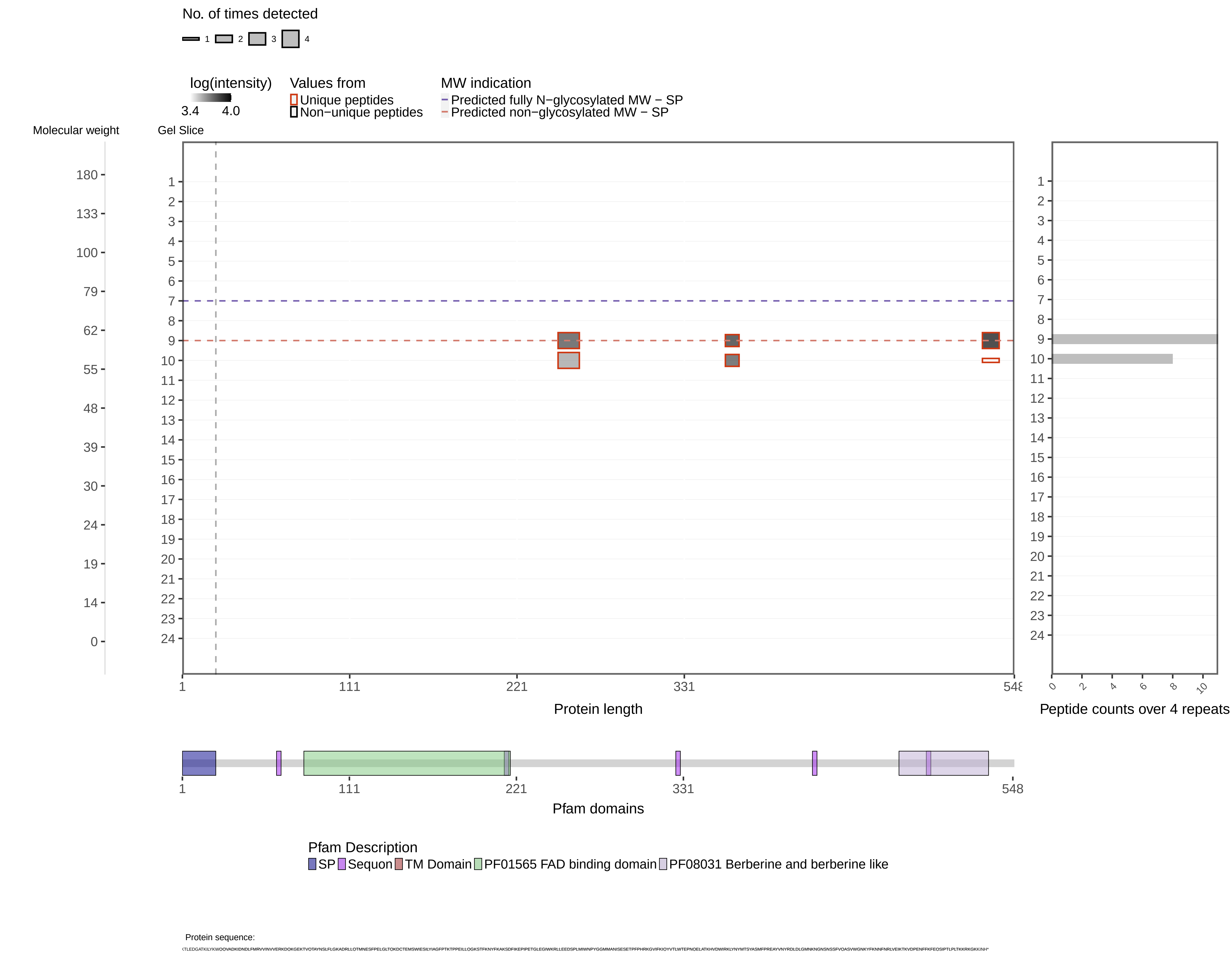

### NbL02g17790.1 probable_LRR_receptor−like_serine/threonine−protein_kinase_At4g20940 [MW − SP = 113.27 kDa]

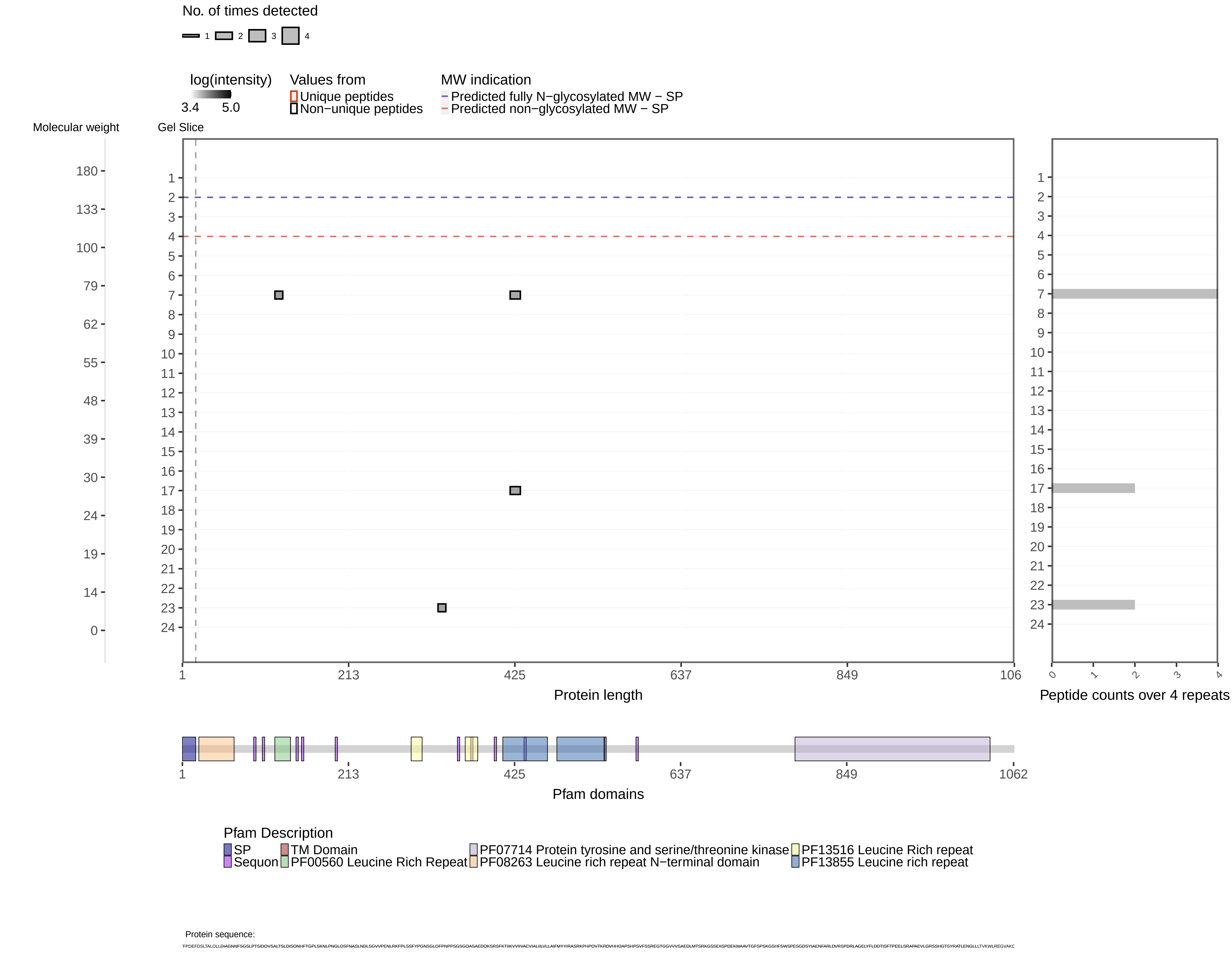

NbL02g17710.1 probable_carboxylesterase_17 [MW = 37.61 kDa]

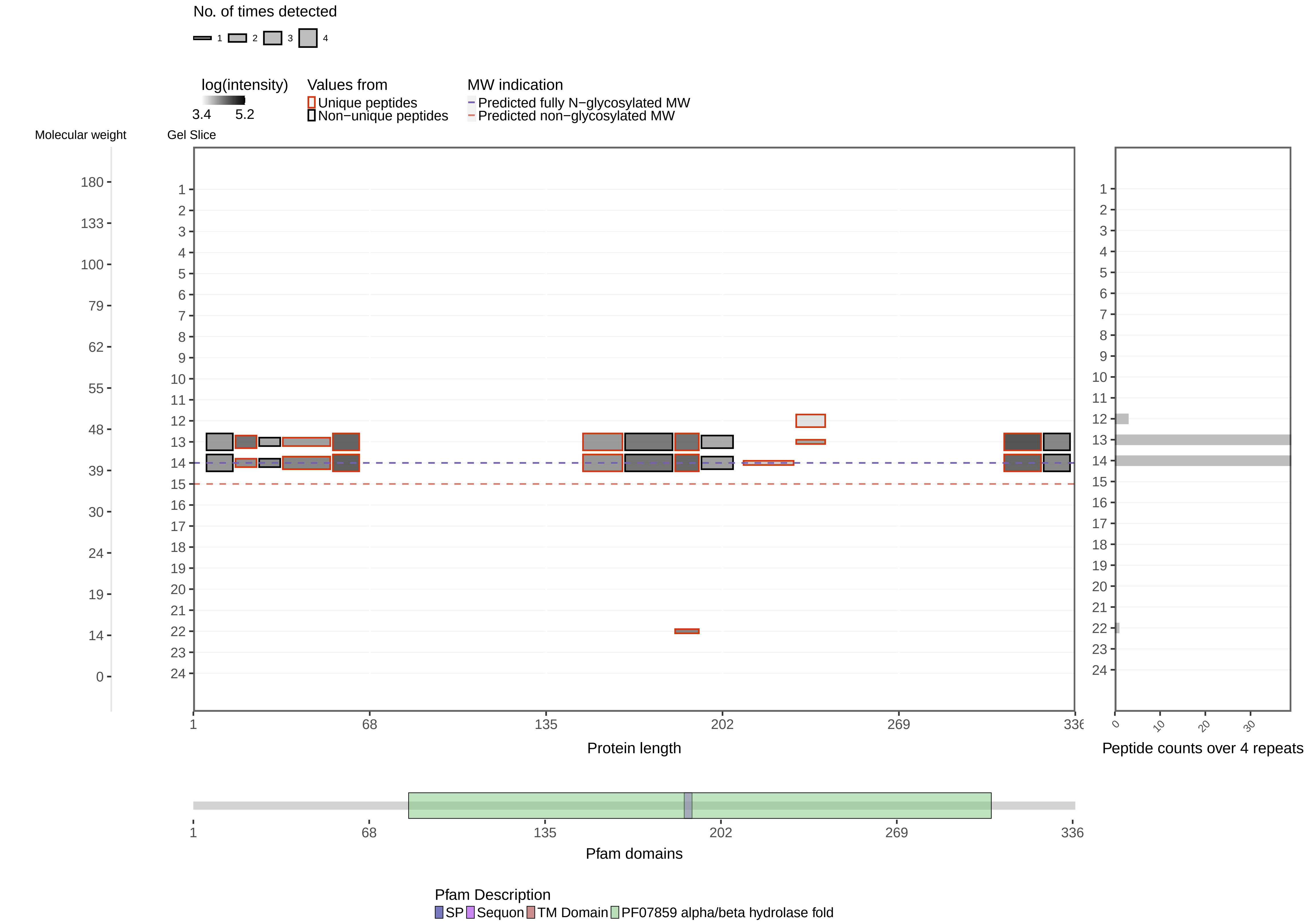


Protein sequence:

PWLNDYADFNRVFLIGDSSGGNIVHQVAVRAGEENLSPMRLAGAIPIHPGFVRSYRSKSELEQEQTPFLTLDMVDKFLGLALPEGSNKDHPITCPMGEPAQAVEELKLPPYLYCMAEKDLIKDTEMEFYEAMKNGKKDVELFINNGVGHSFYLNKIAVEMDPVTASETEKLYEAVAEFISKH*

NbL02g17670.1 probable_choline_kinase_2 [MW = 40.55 kDa]

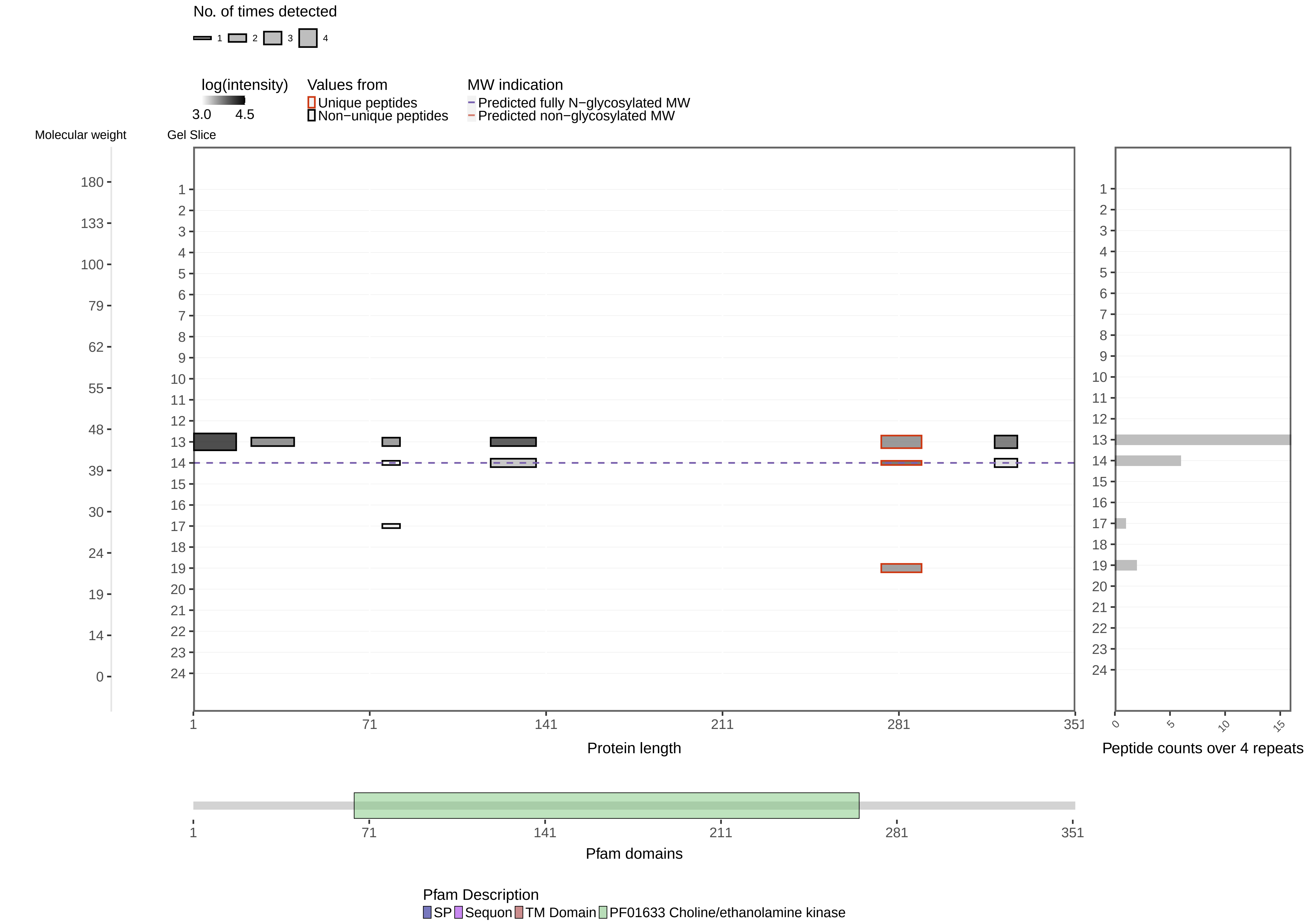


Protein sequence:

NVAKRLASAEEAKDFOLNHLEVEISLLEKSLAGNHLSMGFSHNDLOYGNIMMDEEIRSITFIDYEYAGYNHVAFDIANHFCEMVADYHTETPHIMDFSKYPGLEERKRFLRAYLSSSGGSPSEOELGKLAOEVEKYTLASHLFWGLWGIISHHVNKIDFNYLEYARORFOPYWSKKPELLGSSGPKKG*

NbL02g17620.1 flavin−dependent_oxidoreductase_FOX2−like [MW = 62.15 kDa]

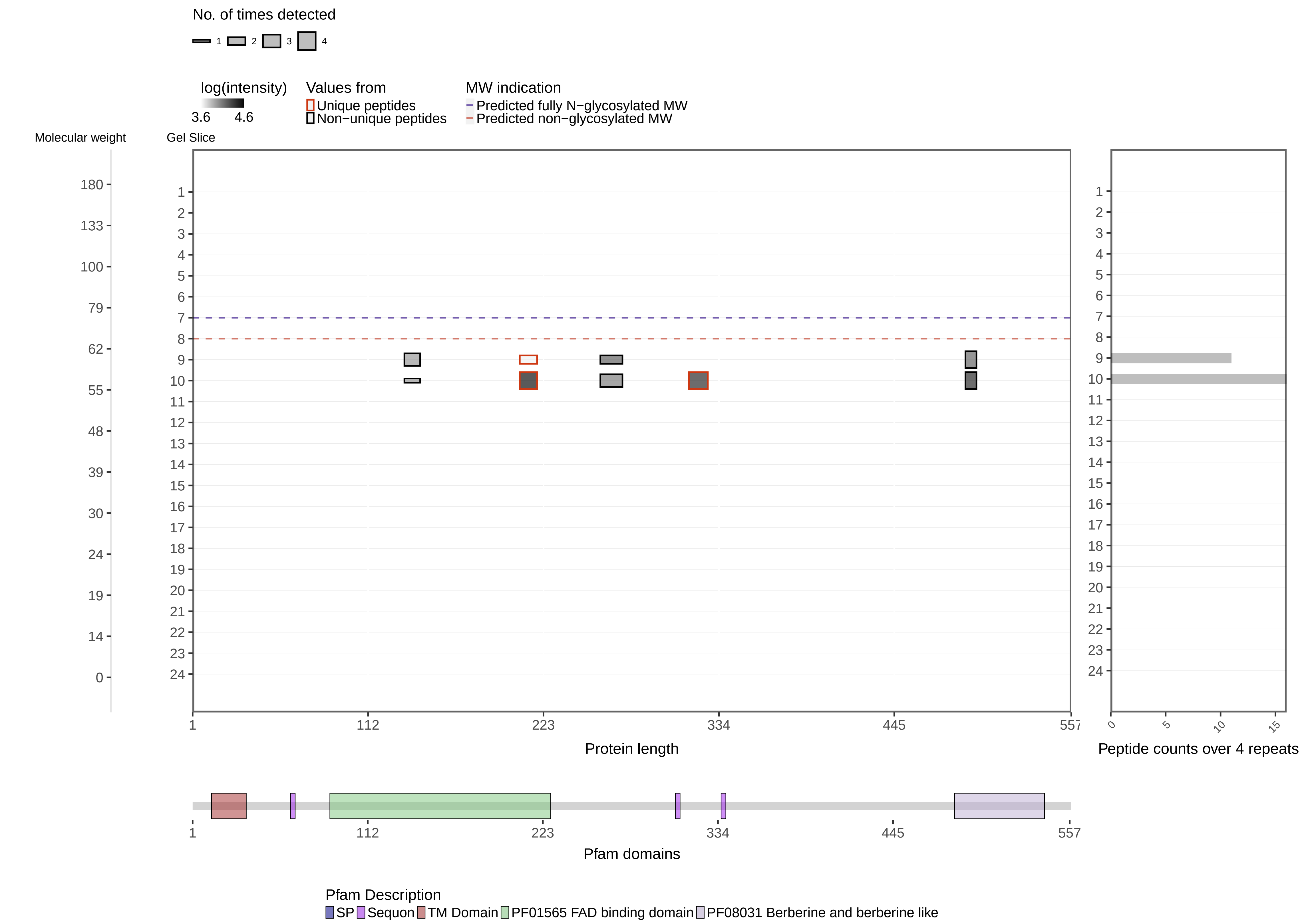


Protein sequence:

TVPKTLETGATKVLNKWQHVAHKIDEDLFVRVLITTANSTKTGKKTVQTAYQALFLGKTDRLLHVMNHSFPELGLTGKDCVEMSWIESVIYIAGYPSSIKPEFLLQGKSLFPPVNFKAKSDFLRVPVPETGLDGMWKKFLEDDSPMMIWNPYGGMMGKISESSIPFPHRKGVICKIQYLTSWTEGNKKSADKHINWIRGLYEYMGTFVSKFPREAYVNYRDLDLGMNRNANSNFLEASVWGKKYFKNNYDRLVLVKTKVDPDNFFWHEQSLPILPLKVGKDGKSLIH*

### NbL02g17590.1 probable_ribose−5−phosphate_isomerase_4_chloroplastic [MW = 31.49 kDa]

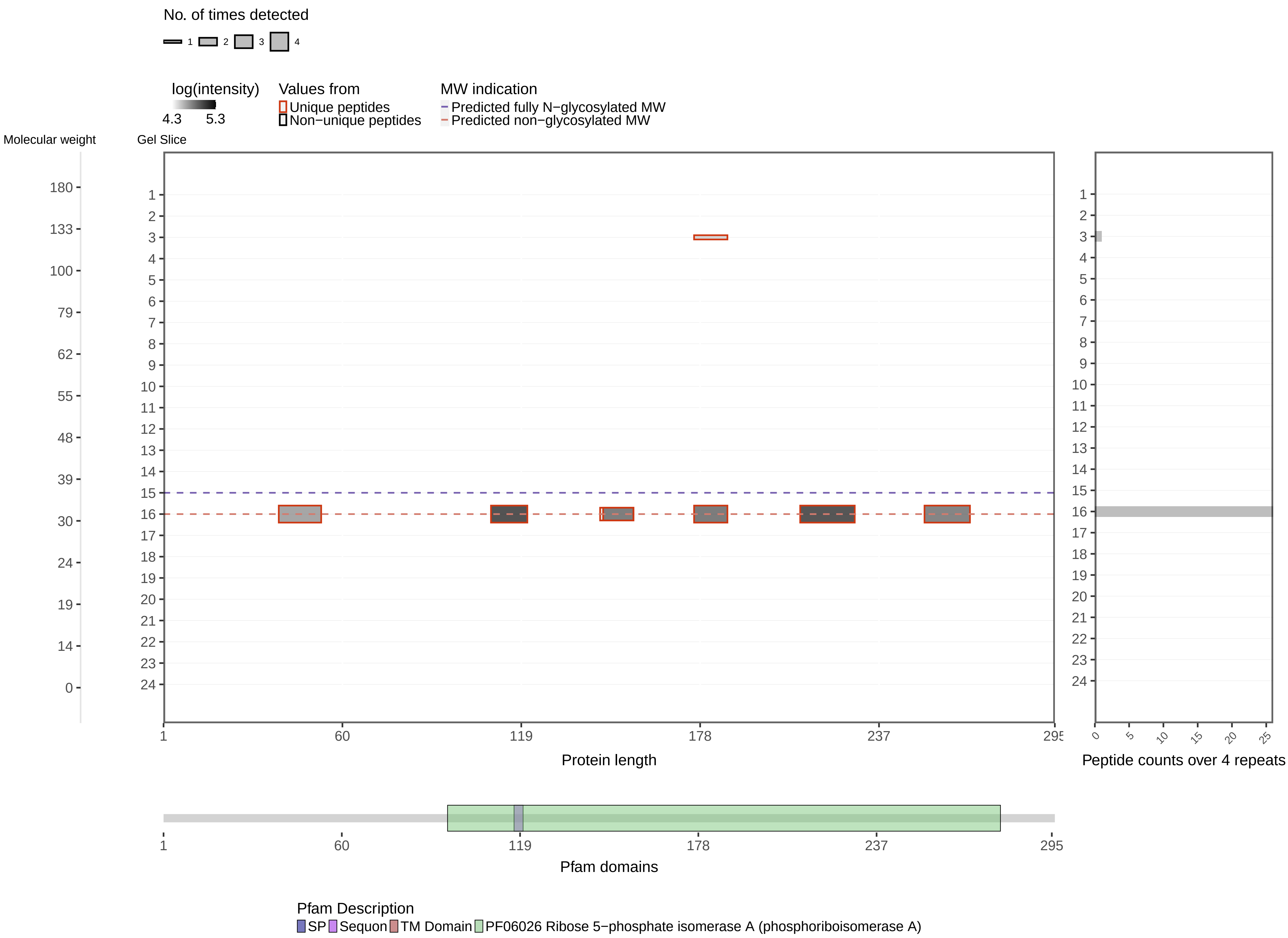


Protein sequence:

TMEEETLAAIIGRRKLQGDDSIIQEKIILEAADKLILIIKEKQFKGSVDGSIPVLIKSVNWLETAEEIDDLFLGDAEVWRRPAIGYAGPLGGDFPLVTKEGHNILDVIFTSPMESLVEVAKTLDQIDGVVEHGVIFRNPCTAIIVSENGLQIADNTIEMEASNL*

### NbL02g17580.1 coatomer_subunit_epsilon−1−like [MW = 31.94 kDa]

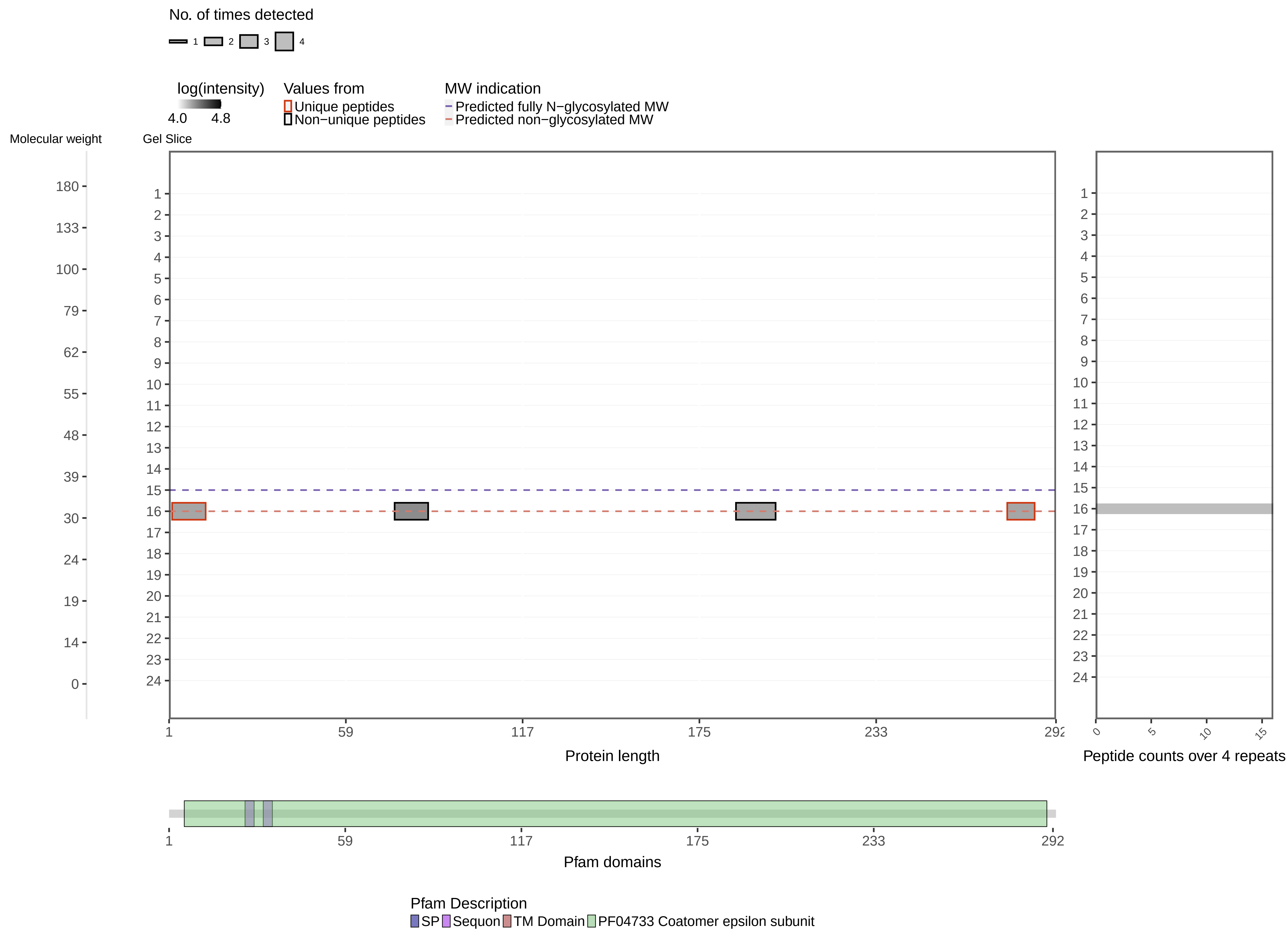


Protein sequence:

MELHALNVQILIKMHRSDYAEKQLRIMQQVDEDHTLTQLANAWLNLAVGGSKIQEAYLIFQDFSEKYQMTSLILNGKAVCCMHMGNFDEAETLLLEALNKDAKDPETLANLIVCSLHLGKPSARYLNQLKLSHPDHALVKRAASAEESFDRAVQSVA*

### NbL02g17470.1 photosystem_I_reaction_center_subunit_III_chloroplastic−like [MW = 24.98 kDa]

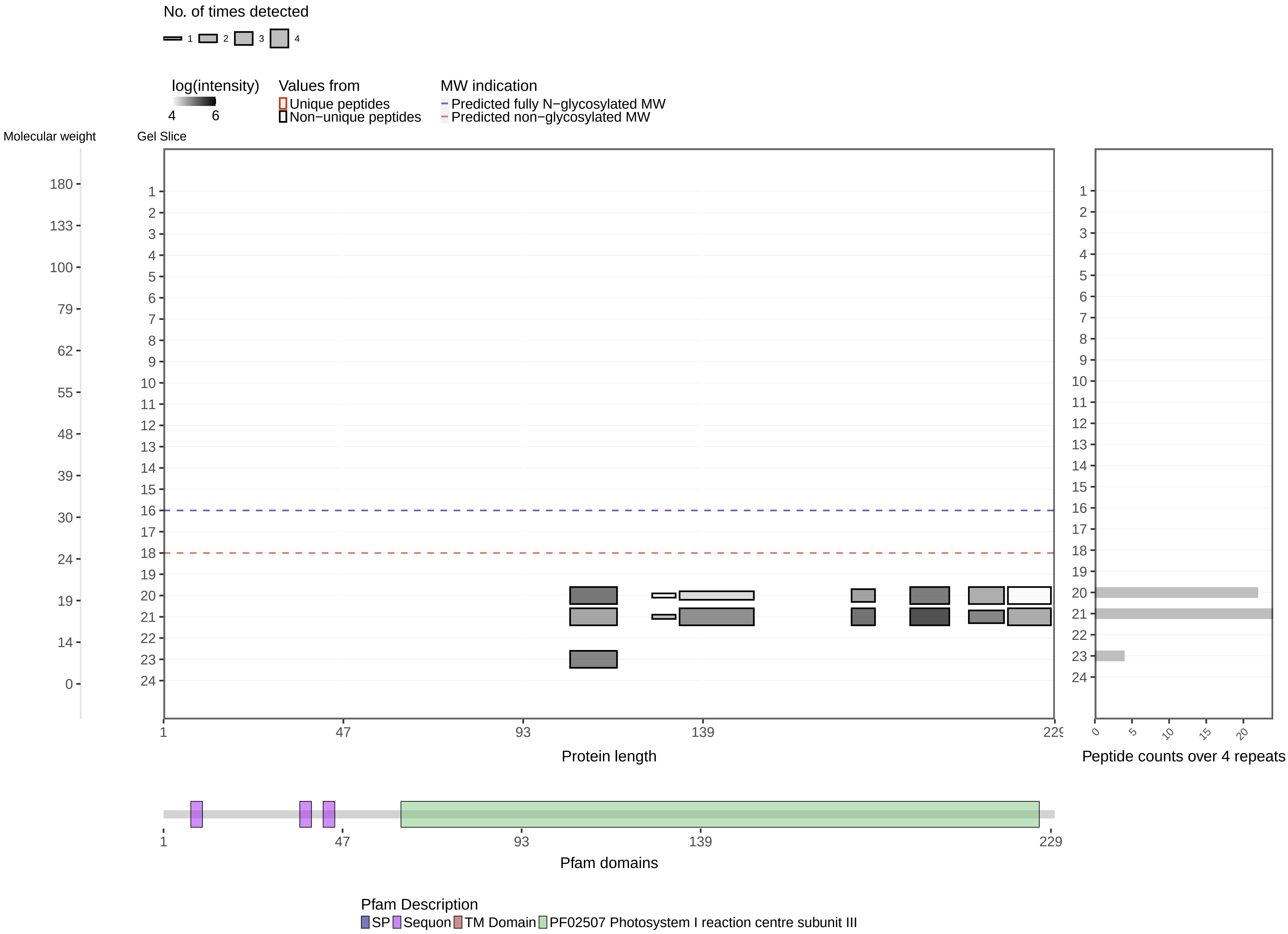


Protein sequence:

SLKLYAPDSAPALAIKATVEKTKRRFDNYGKQGLLCGSDGLPHLIVSGDQRHWGEFITPGILFLYIAGWIGWVGRSYLIAVRDDKKPTMKEIIIDVPLANKLIWRGFSWPVAAYREFLNGELTDPNF*

NbL02g17430.1 haloacid_dehalogenase−like_hydrolase_domain−containing_protein_At3g48420 [MW = 41.48 kDa]

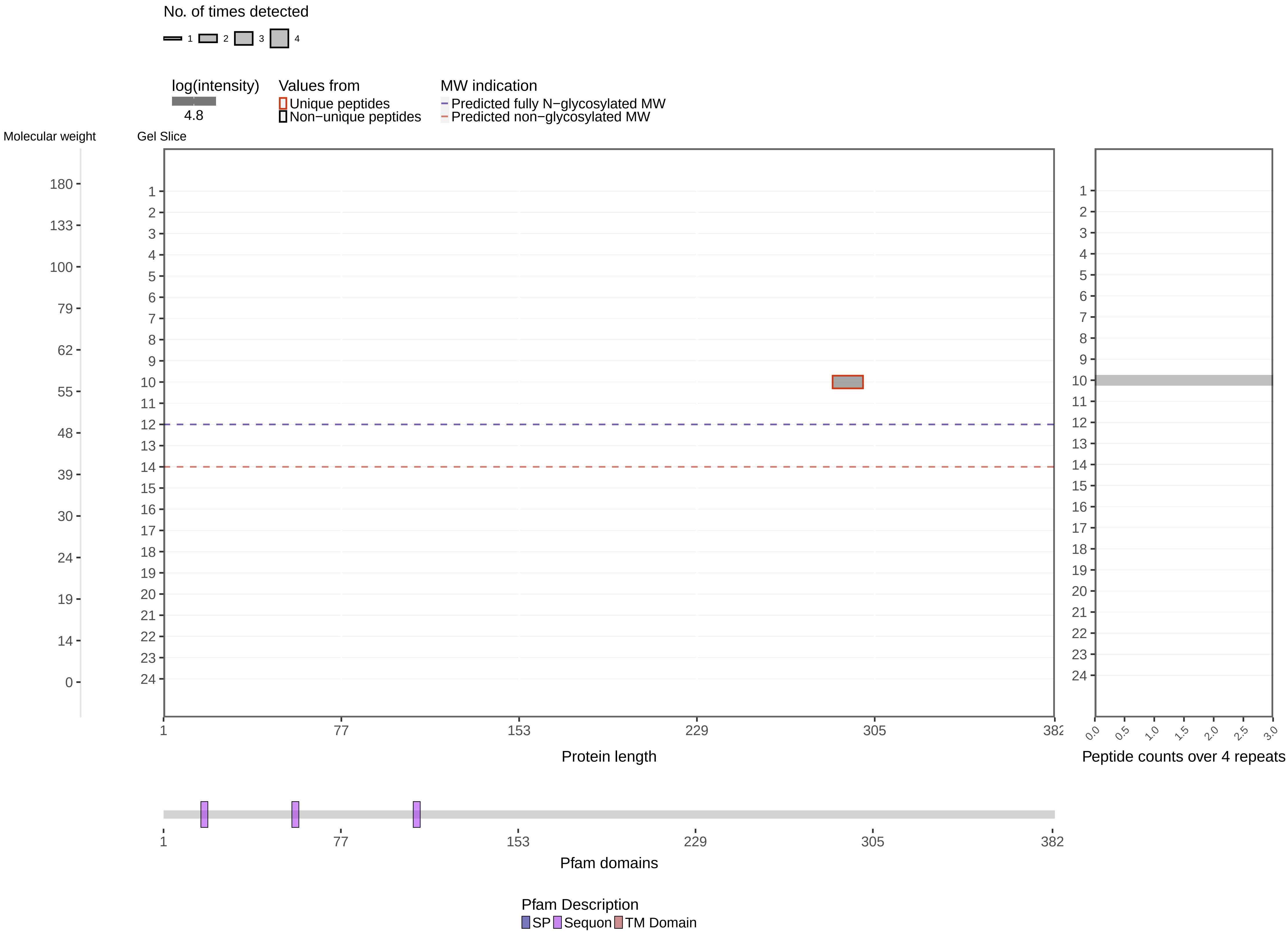


Protein sequence:

MSKSLPLRPGVEEFIDEAYEEGVPVVMLTSYCKCGEKPARSIIEKLGNDKMARIKILGMEEVKQSFYGQLVFGGGVASDLGEQLAKEARKAVSAEEQRIAKEVASMLKLSVDIHTTSSEEMKNVVAALRAGAEYANVPVDNCVLVAGSQSGVAGAERIGMSCVVVRSSSTARAEFPGAKAIMDGFGGGDLSISRLRQIQGLDL*

NbL02g17230.1 cathepsin_B−like [MW − SP = 36.66 kDa]

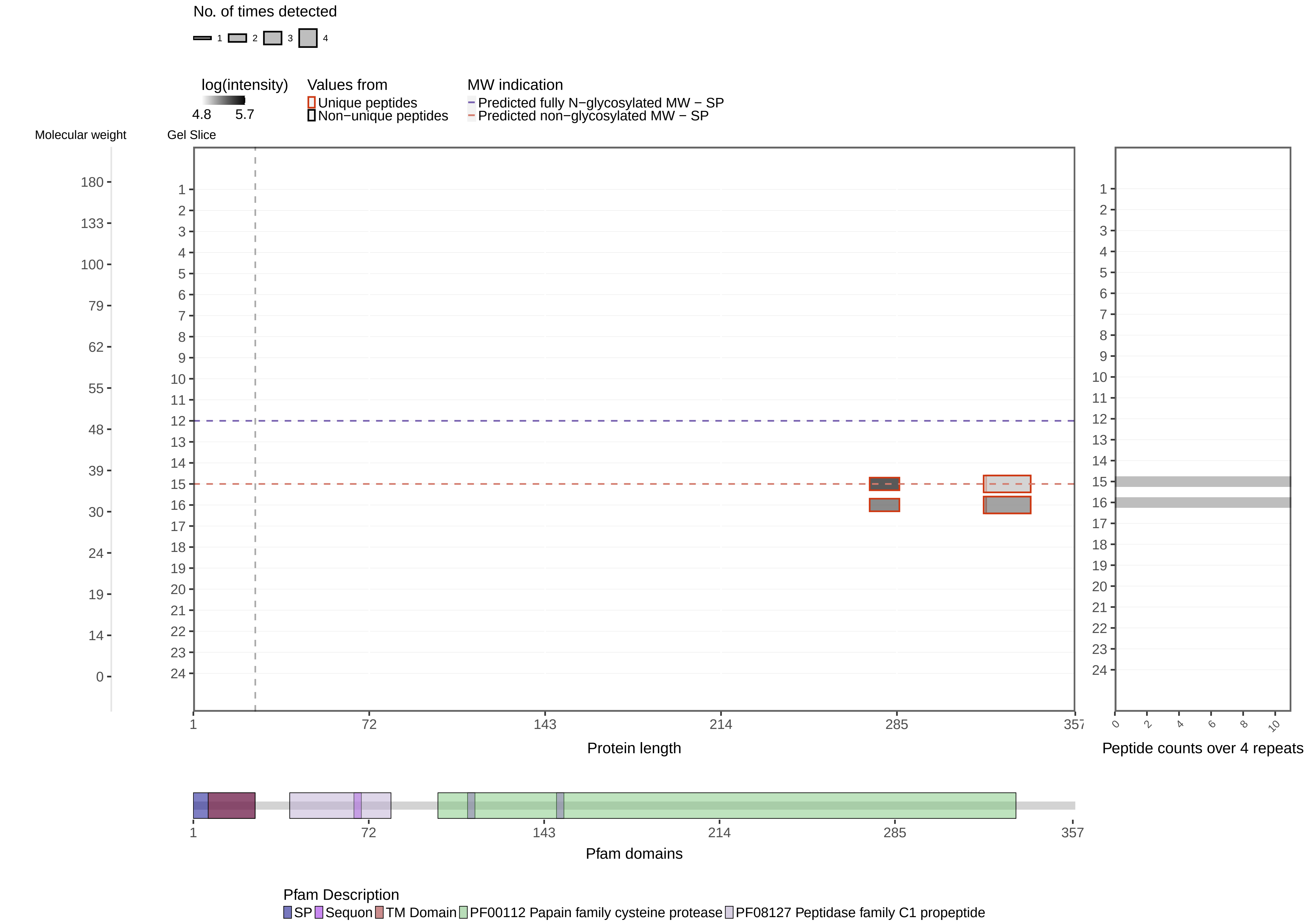


Protein sequence:

)GGYPLQAWKYFVRKGVVTDECDPYFDNEGCSHPGCEPAYPTPKCHRKCVKQNLLWSKSKHFGVNAYMISSDPHSIMTELYKNGPVEVSFTVYEDFAHYKSGVYKHVTGDVMGGHAVKLIGWGTSEDGEDYWLLANQWNRGWGDDGYFKIRRGTDECEIEDEVVAGLPSARNLNMELDVSDAFLDAAM*

NbL02g17190.1 phosphatase_IMPL1_chloroplastic−like [MW = 39.43 kDa]

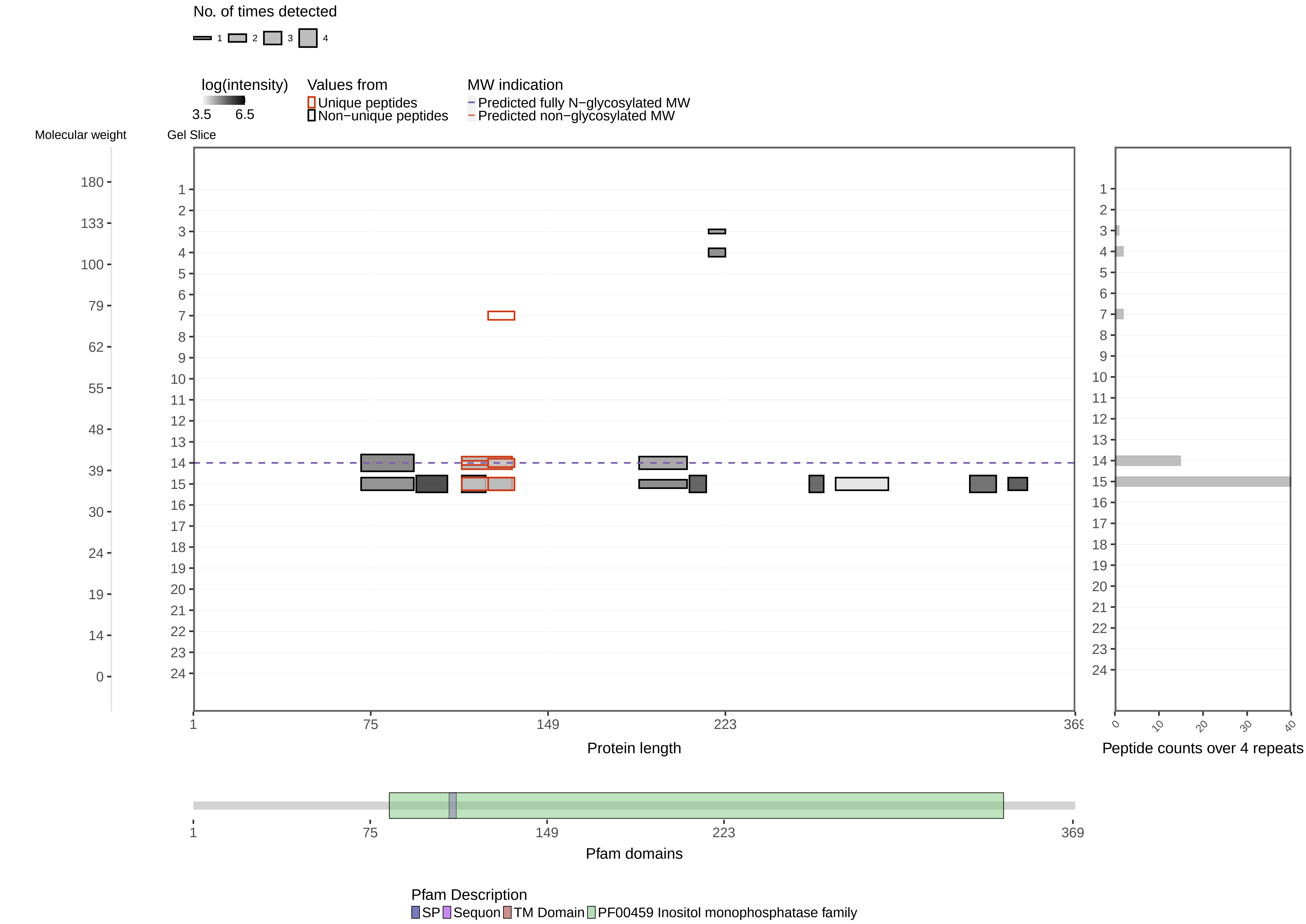


Protein sequence:

YPSFAVSVGVLFRGKPAAATVVEFVGGPMCWNTRIFSAAAGKGAFANGEKIHVSGTDKVEQSLLVTGFGYDHDDPWATNMELFKDFTDVSRGVRRLGAAAVDMCHVALGIAEAYWEYRLKPWDMAAGVLIVEEAGGTVSCMDGGKFSVFDRSVLVSNGLLHAKLLEKIGPATENLKAKGIDFSVWFKPENYHTEV*

NbL02g17160.1 cytochrome_c6_chloroplastic [MW = 25.78 kDa]

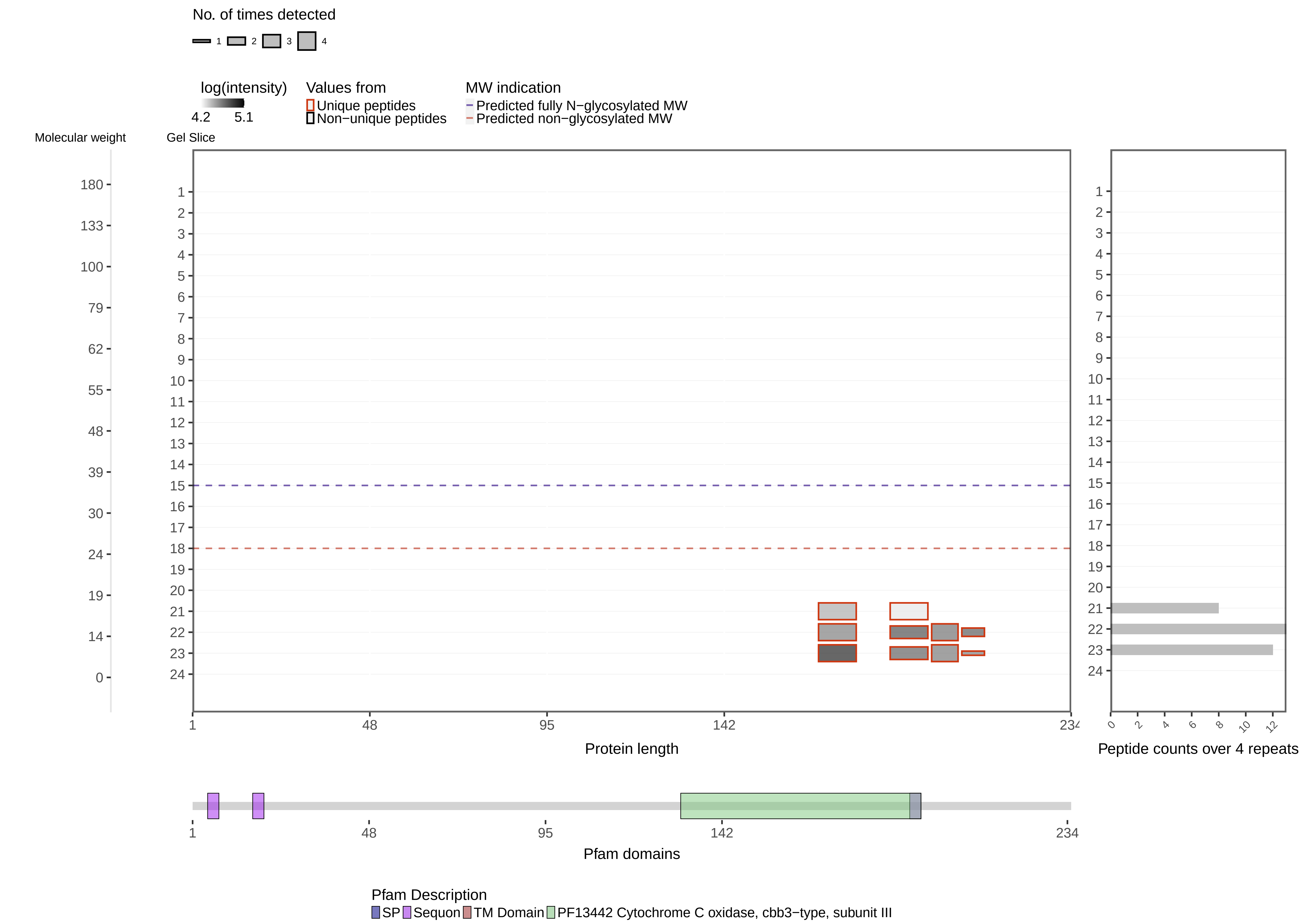


Protein sequence:

.APPLMTALIAFSPIVNPPVSVGOTIEVOKGAALFSKACIGCHYAGGNIIOPGATLFLKDLERNGADTEEEIYRITYYGKGRMPGFGONCTPRGOCTFGPRLODDEIKLLVEFVKSOADOGWPKIENSGD*

### NbL02g17130.1 glutathionyl−hydroquinone_reductase_YqjG−like [MW = 41.25 kDa]

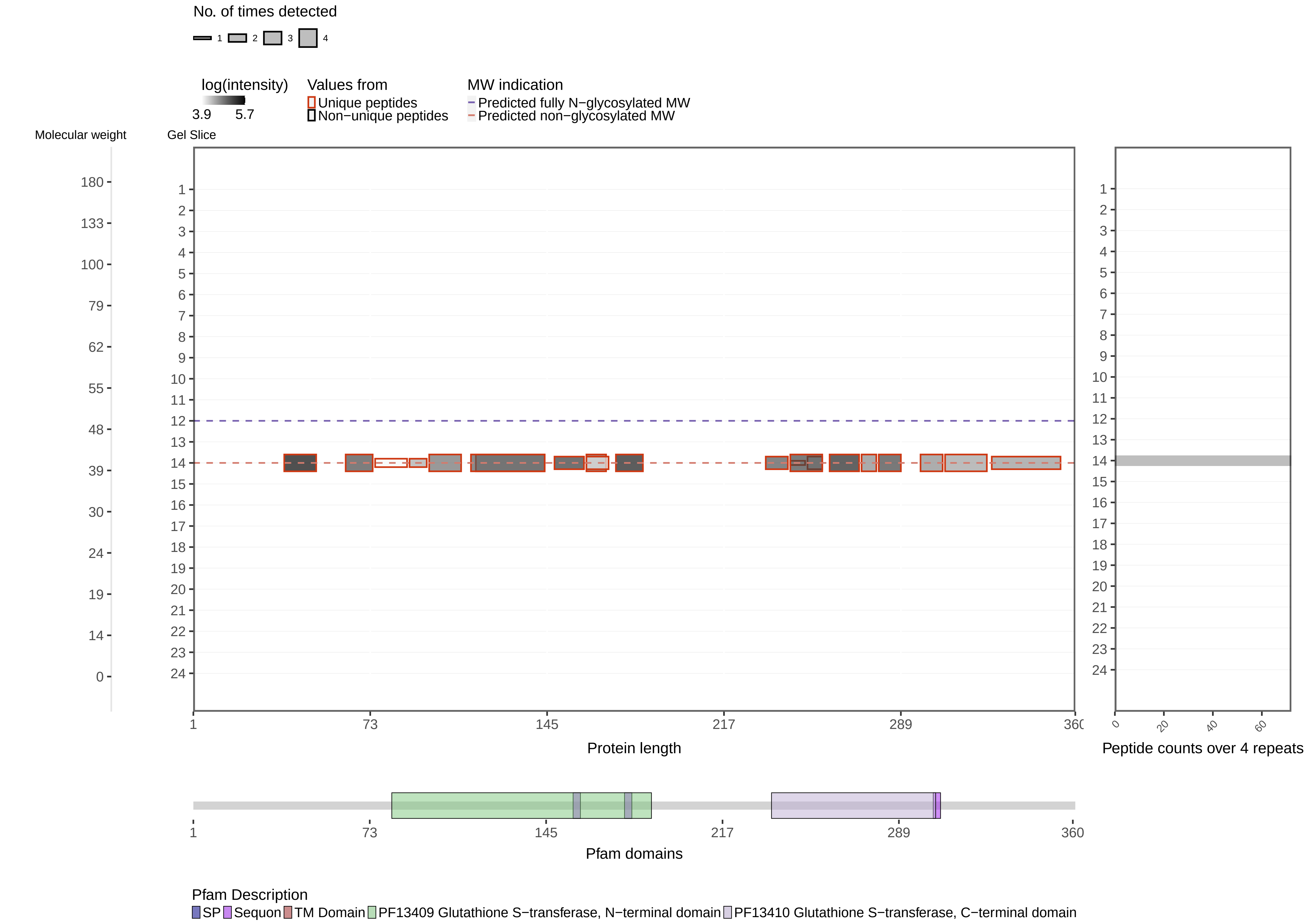


Protein sequence:

/LWDKKLKTIVNNESSDIIRMFNSEFNDISGNAALDLYPPHLQSQINEINEWIYDGINNGVYRCGFATKQEPYDEAVQKLYETLDKCEMILSKQRFLCGDQVTEADIRLFVTLIRFDEVYAVHFKCNKKLLREFPHLFNYTKDIFQIPGMSSTVNMEHIKKHYYGSHPTINPYGIIPQGPNIDYSSPHDREKFSK*

### NbL02g17080.1 50S_ribosomal_protein_L30−like [MW = 12.39 kDa]

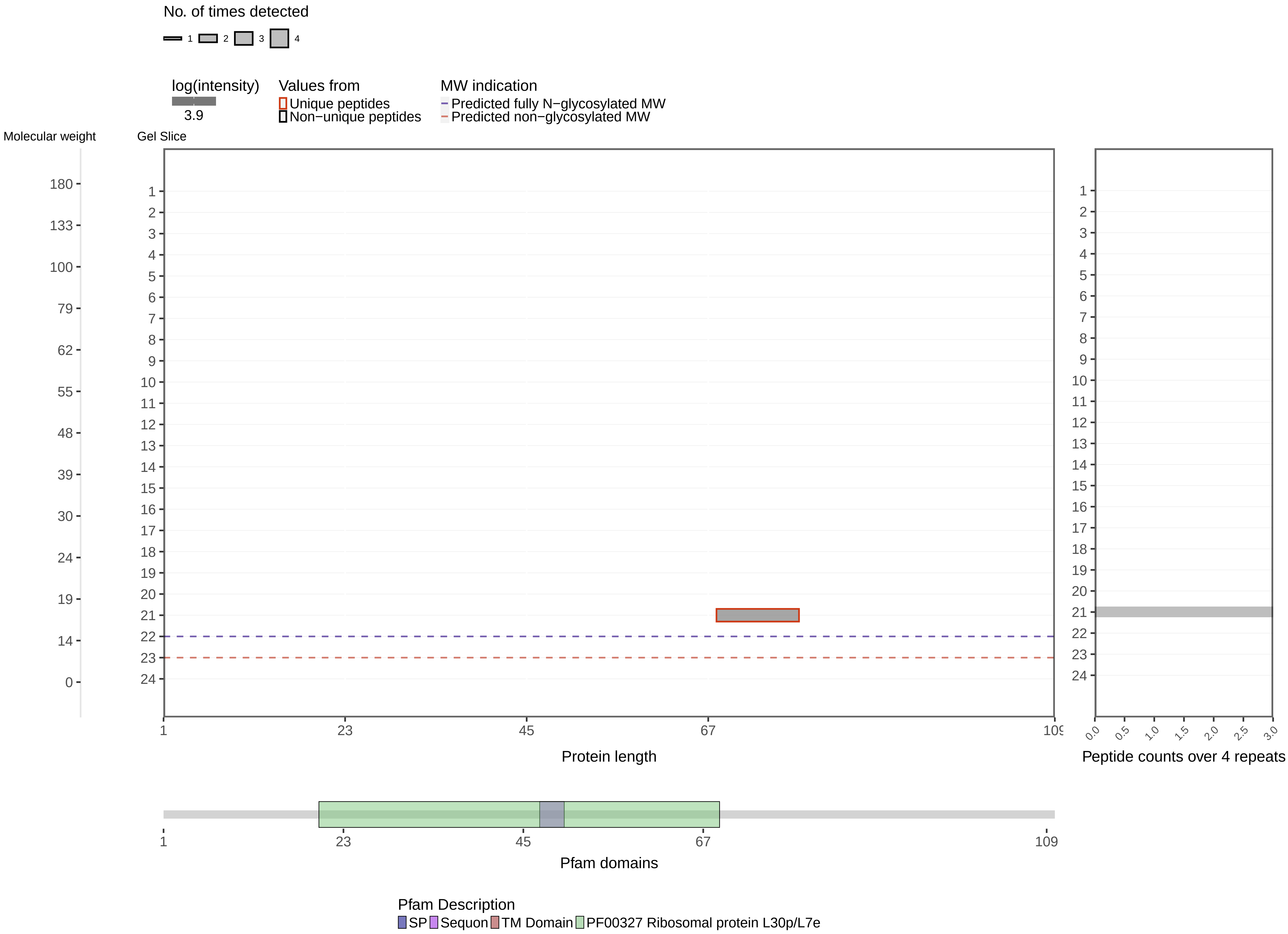


Protein sequence:

LRKCNRTVMRWNTPTVRGMLQQVKRLVVVETEEMYKARKEKLADHKAIRPPLVVNHHAAPAADSVL*

NbL02g17040.1 14_kDa_zinc−binding_protein [MW = 19.69 kDa]

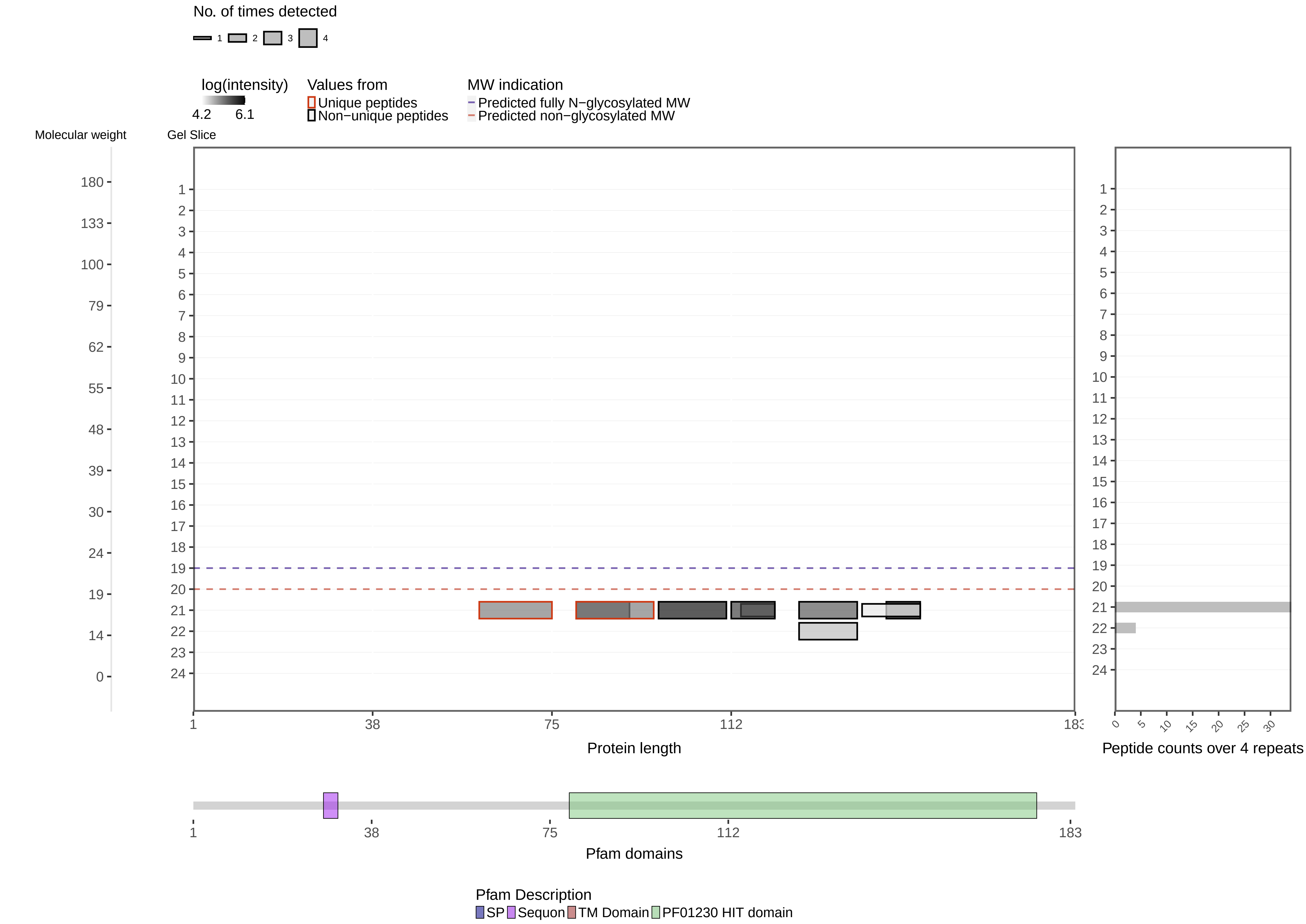


Protein sequence:

KEIPASVVYEDEKVLAFRDINPQAPVHVLVIPKLRDGLTELGKAEQRHEDILGNLLYAAKIVAEKEGIVDGFRVVINSGASACQSVYHLHLHVLGGRQLKWPPG*

NbL02g17020.1 tRNA_(guanine−N(7)−)−methyltransferase−like [MW = 32.8 kDa]

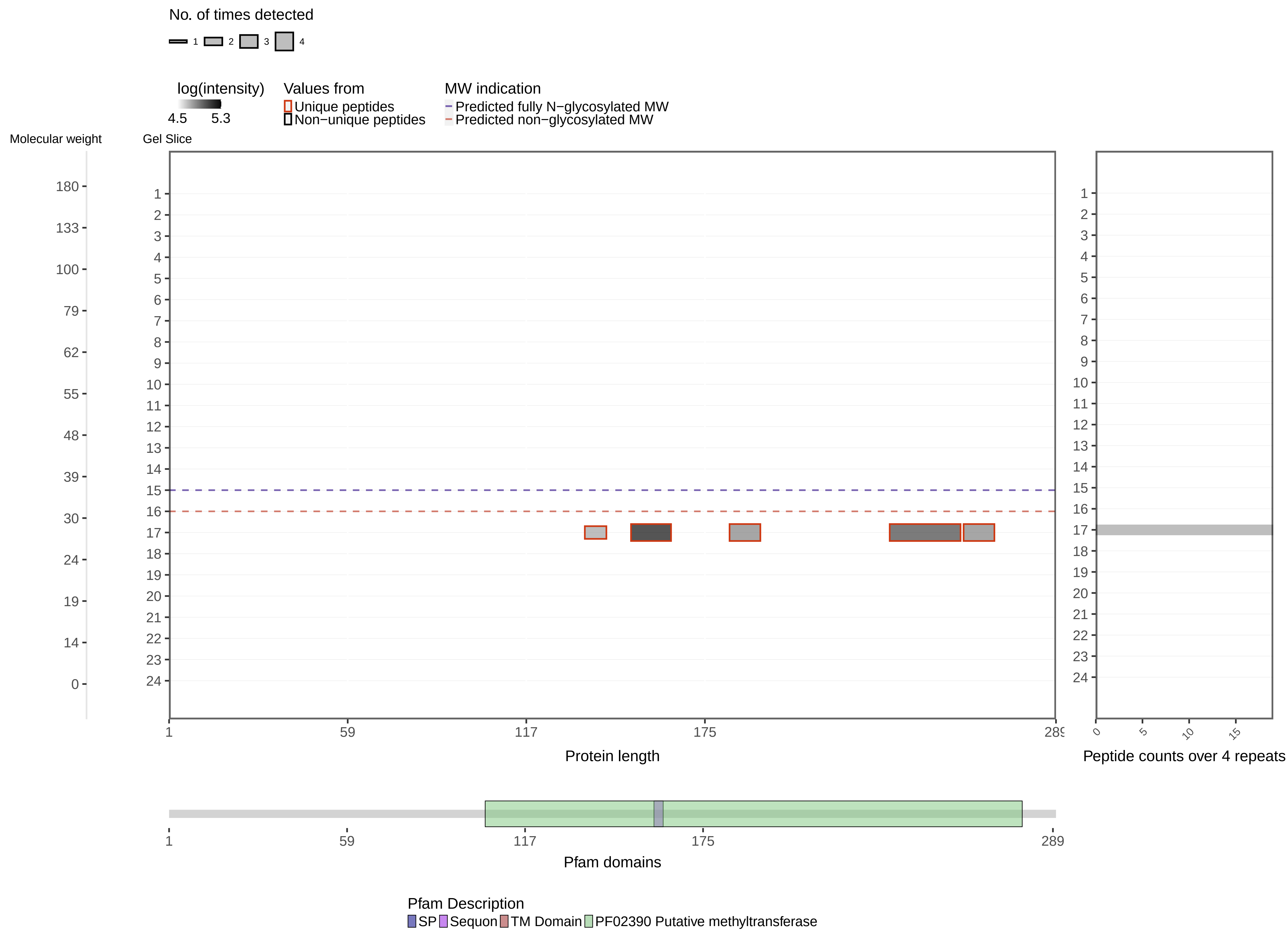


Protein sequence:

MELRDKVTEYVKERILGLRTTNPGQYQNVSVVRTNSMKYIPNYFGKGQLKKMFFLFPDPHFKEKNHRRRVISPFLLDEYAYVLAVGGIIYTITDVEELGEWMKSCLEEHPMFEPLTTEELEADQVVKLLSTATEEGQKVARNDGQTFRAVYRRIATH*

NbL02g17010.1 pyrroline−5−carboxylate_reductase [MW = 28.93 kDa]

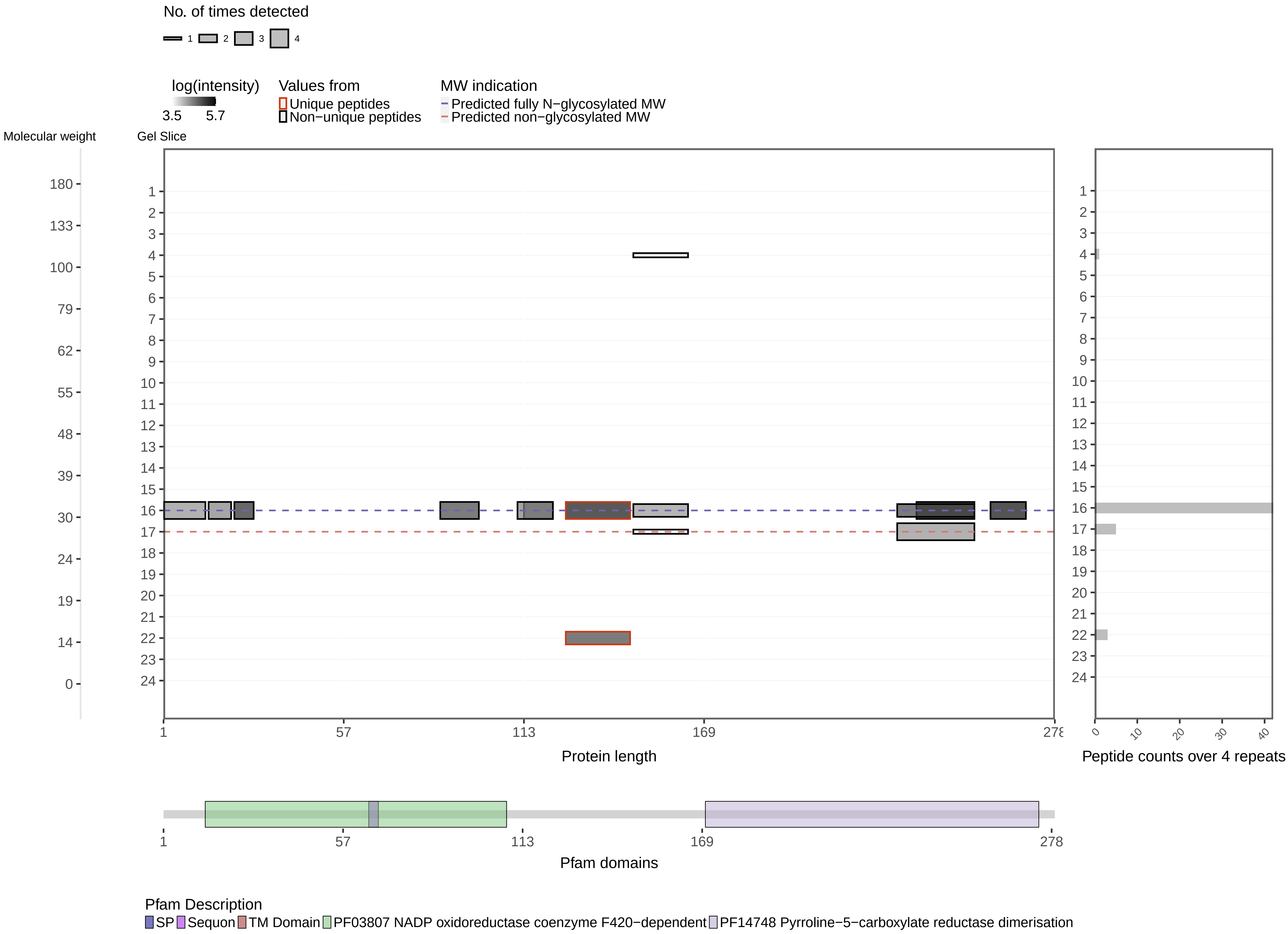


Protein sequence:

MPNTPSAVGDAATVFTLGEKATTEDGELISQLFGAIGKVWRADEKLFDAVTGLSGSGPAYIFLAIEALADGGVAAGLPRELALGLASQTVLGAASMVKGMAKHPGQLKDDVASPGGTTIAGIHELEKAGFRGILMNAVVAAAKRSRELSQN*

### NbL02g16830.1 bifunctional_purine_biosynthesis_protein_PurH [MW = 66.14 kDa]

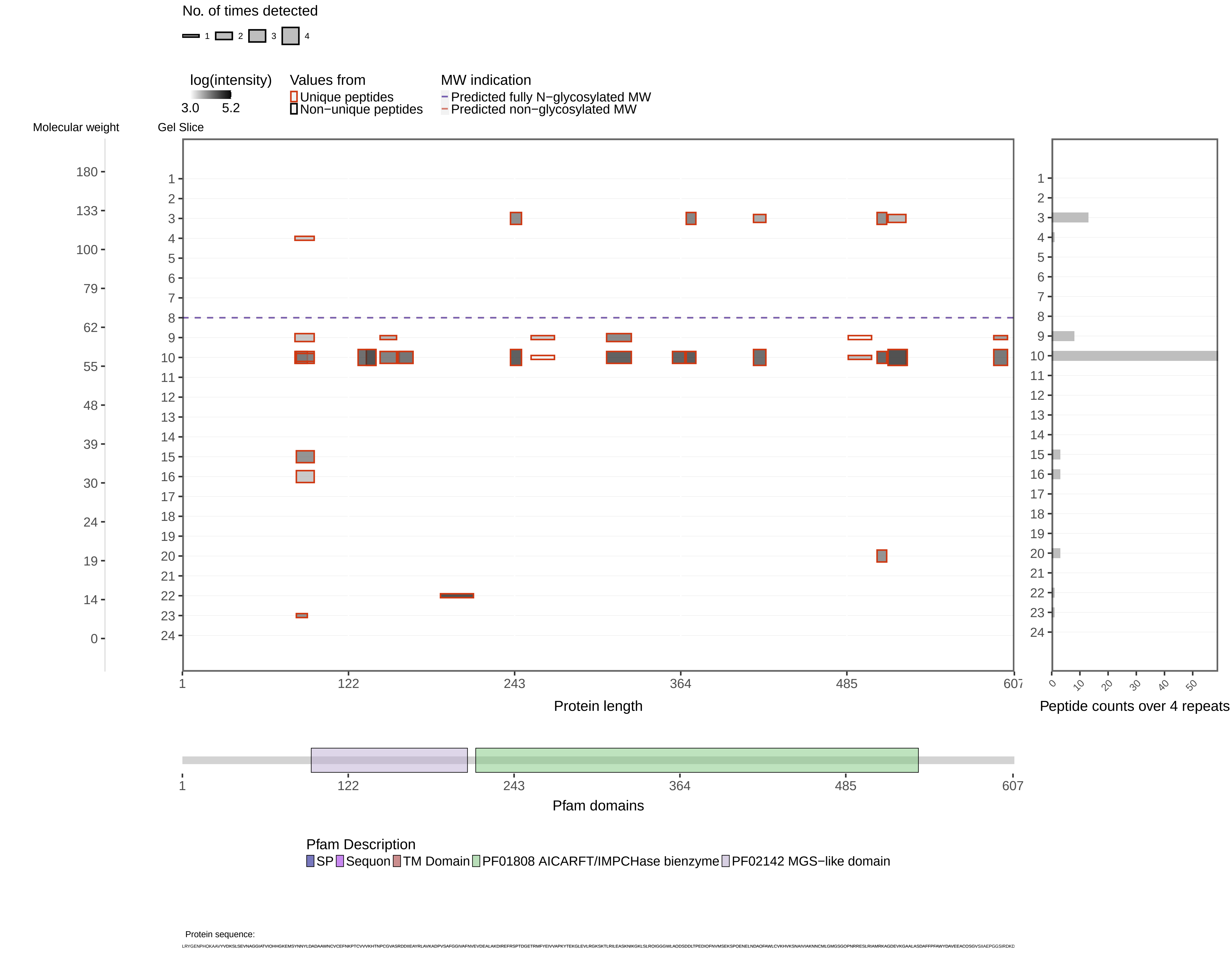

NbL02g16820.1 UDP−N−acetylglucosamine_diphosphorylase_1 [MW = 54.33 kDa]

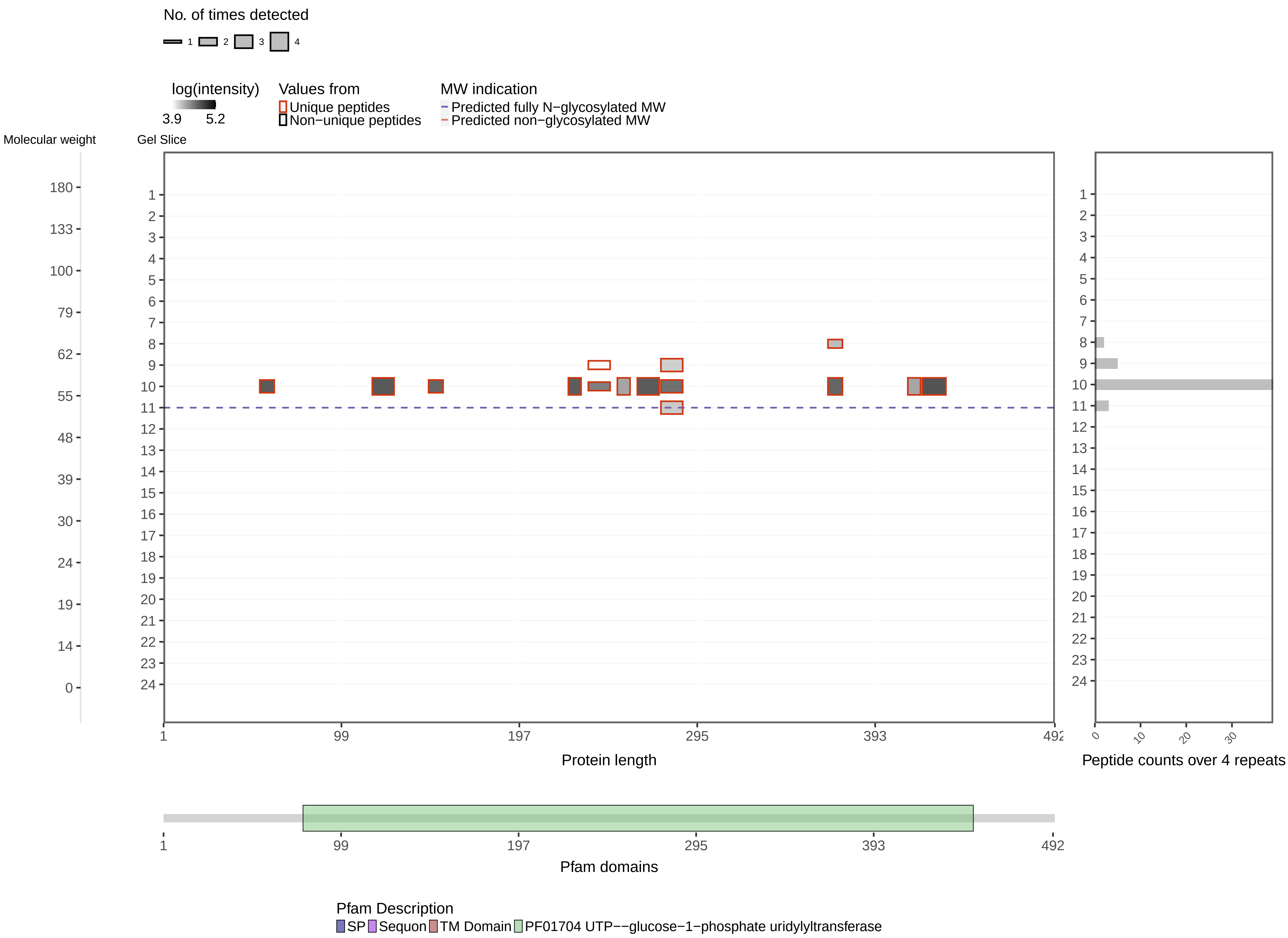


Protein sequence:

KSPDGNGGVYTALKYSRLLEDMNSRGIKYVDCYGVDNALVRVADPTFLGYFIDKGVSAAAKVVRKAYPQEKVGVFVRRGKGGPLAVVEYSELDPSLCSAVNQETGRLRFCWSNVCLHMFSLDFLNQVANGLEKDSIYHLAEKKIPSIHGHTMGYKLEQFIFDAFPYTPSTALFEVVREEEFAPVKNANGANFDTPDSARLLVLRLHTRWVVAAGGFLTHSVPLYATGVEVSPLCSYAGENLEAICRGRTFHAPCEISF*

### NbL02g16800.1 thioredoxin_O2_mitochondrial−like [MW = 21.88 kDa]

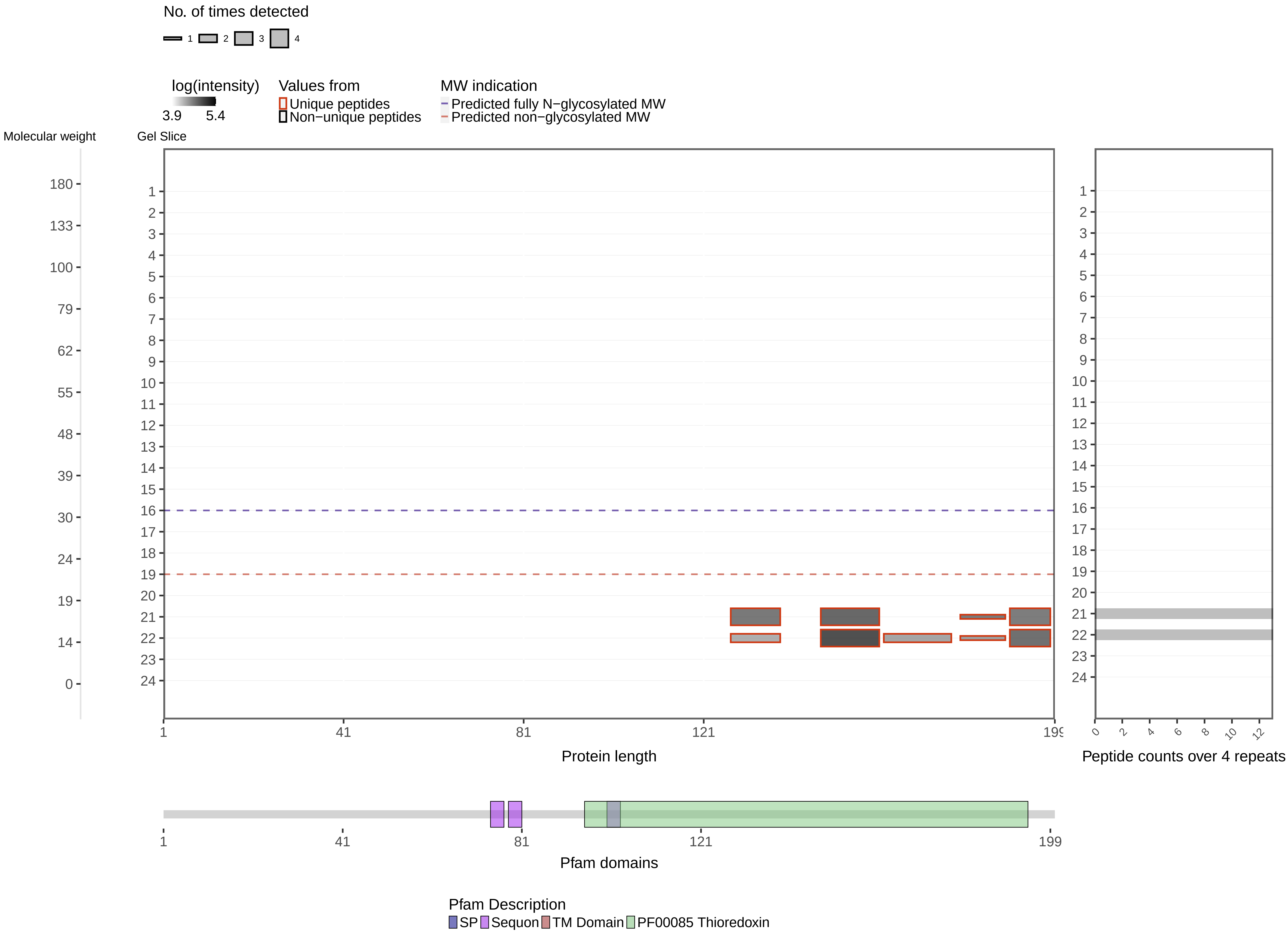


Protein sequence:

PSNIVSIESEEQFNTSLRKVQDESLPAIFYFTAVWCGPCRLLSPVIGQLSEKYPHVTTYKVDIDKEGLGNALSKLNIHSVPTLHFFQNGKKTSEVIGADVQRLKDTMEELYK*

### NbL02g16530.1 acidic_endochitinase_Q [MW − SP = 25.06 kDa]

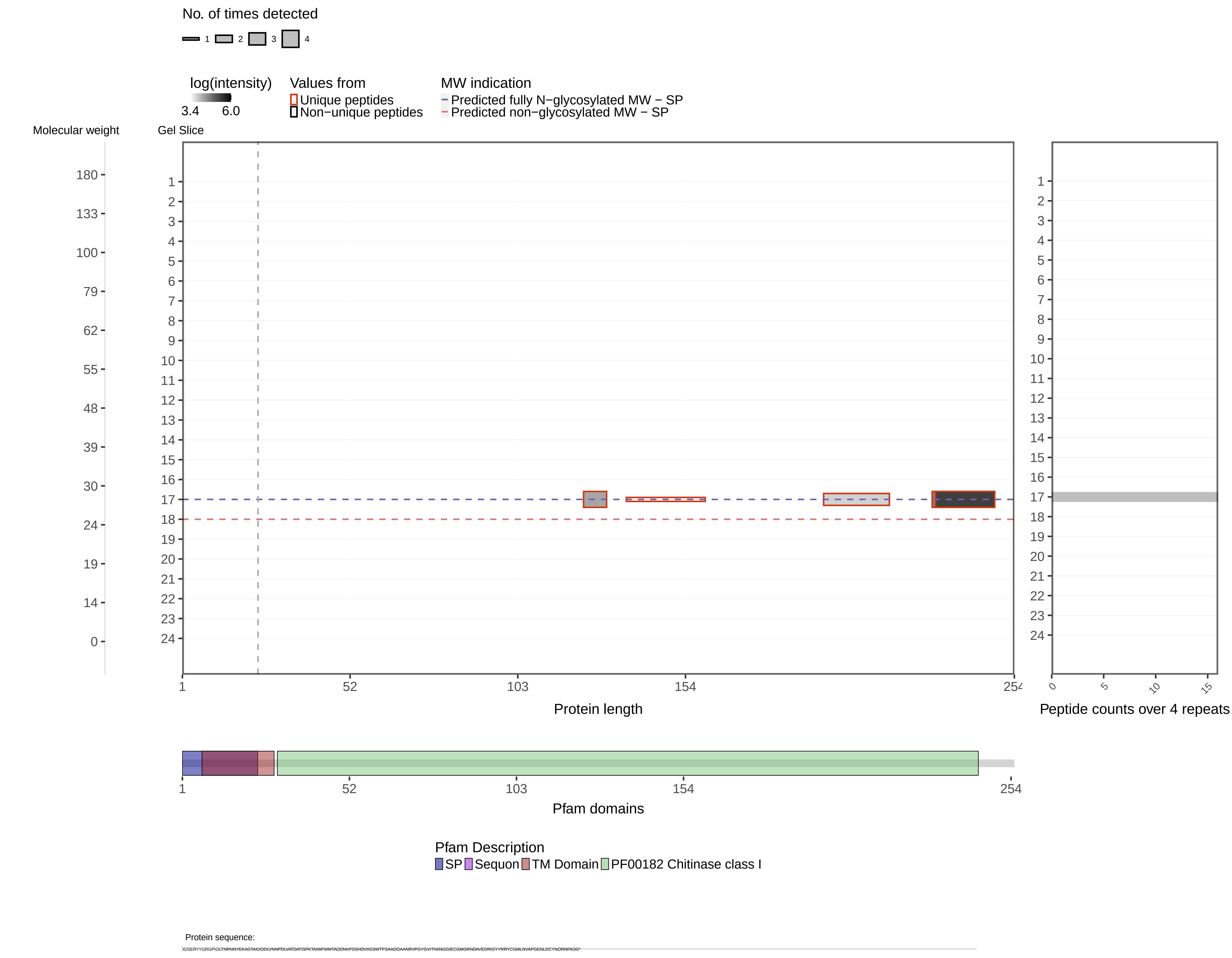

### NbL02g16460.1 phosphoserine_aminotransferase_2_chloroplastic−like [MW = 50.78 kDa]

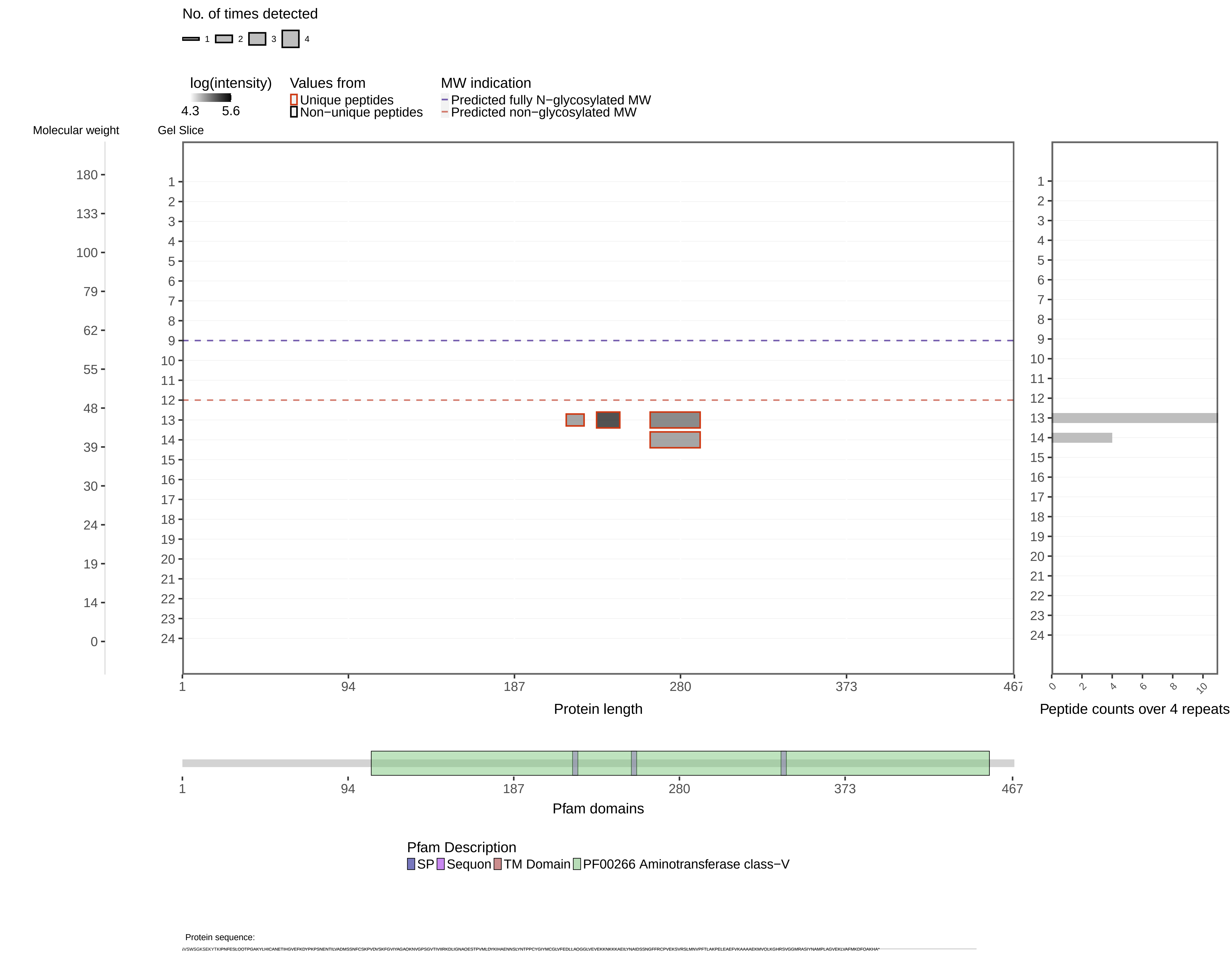

NbL02g16430.1 ubiquitin_domain−containing_protein_DSK2b−like [MW = 60.17 kDa]

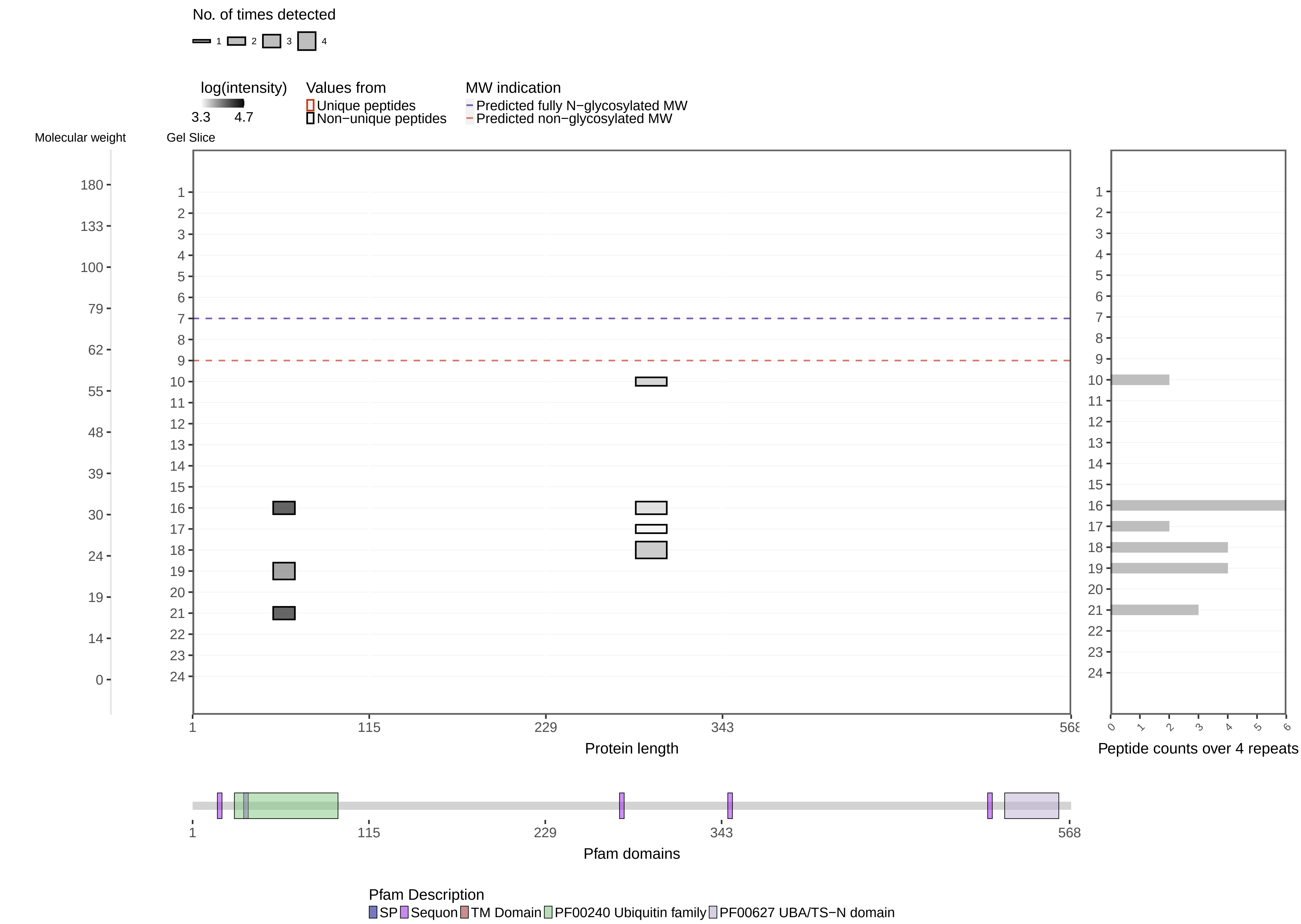


Protein sequence:

EPFLNATTMAGDARNDVGSNPFAALLGAQGAGQGRVQSTNPPITGSETTANTPVPNTNPLPNPWASAGAGAGQTNTTARSNAAGETRAPSPAGLGGLALPDLEGMLGGMQGGMPDAASLNQMMQNPAISQMMQSLLSNHQYMNQYLGVQILGLNPQLRNMLDSNSHLREMMQNPEFIRQLTSPETMQQLMTFQQGLLSQLGREQTNQPGQNAGGAALENNIGMEMLMNMFGGLGSAGLGGVTNRSNVPPEELYATQLVQLQDMGFFDTQENIRALVATAGNVH

NbL02g16410.1 catalase_isozyme_1 [MW = 56.91 kDa]

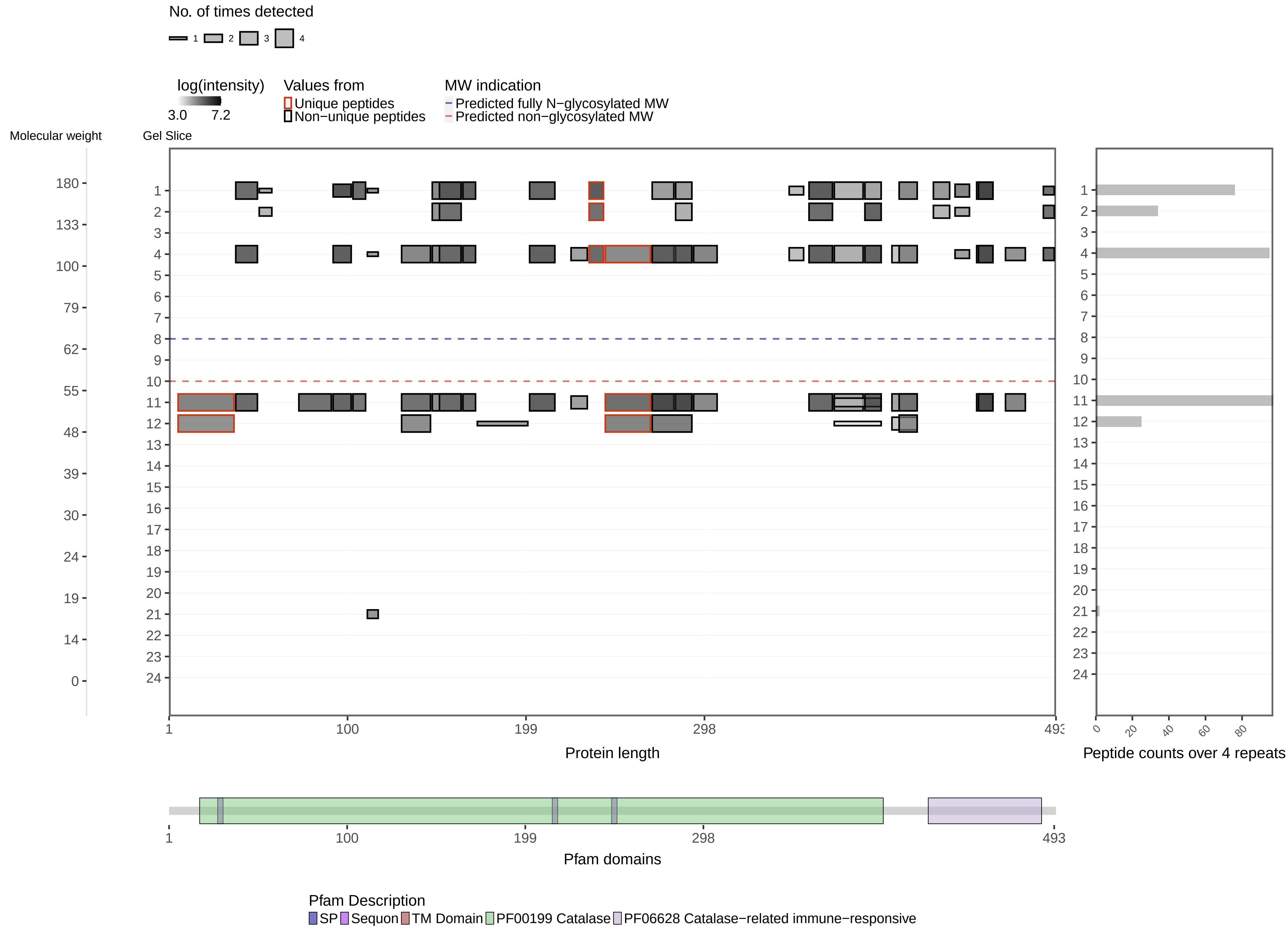


Protein sequence:

SLLEEEAARIGGANHSHATODLYDSIAAGNYPEWKLFIOTMDPDHEDRFDFDPLDVTKTWPEDILPLOPVGRLVLNKNIDNFFNENEOLAFCPSIVVPGVYYSDDKMLOTRIFSYSDTORYRLGPNYLOLPANAPKCAHHNNHYDGSMNFMHRDEEIDYFPSRYDPVRHAEKYPIPSTMCTGKREKCVIOKENNFKOPGERYRSFTPDROERFIRRWVEALSDPRITYEIRSIWISYWSOADKSLGOKLASRLNVRPSI*

NbL02g16360.1 uncharacterized_LOC109214258 [MW = 115.58 kDa]

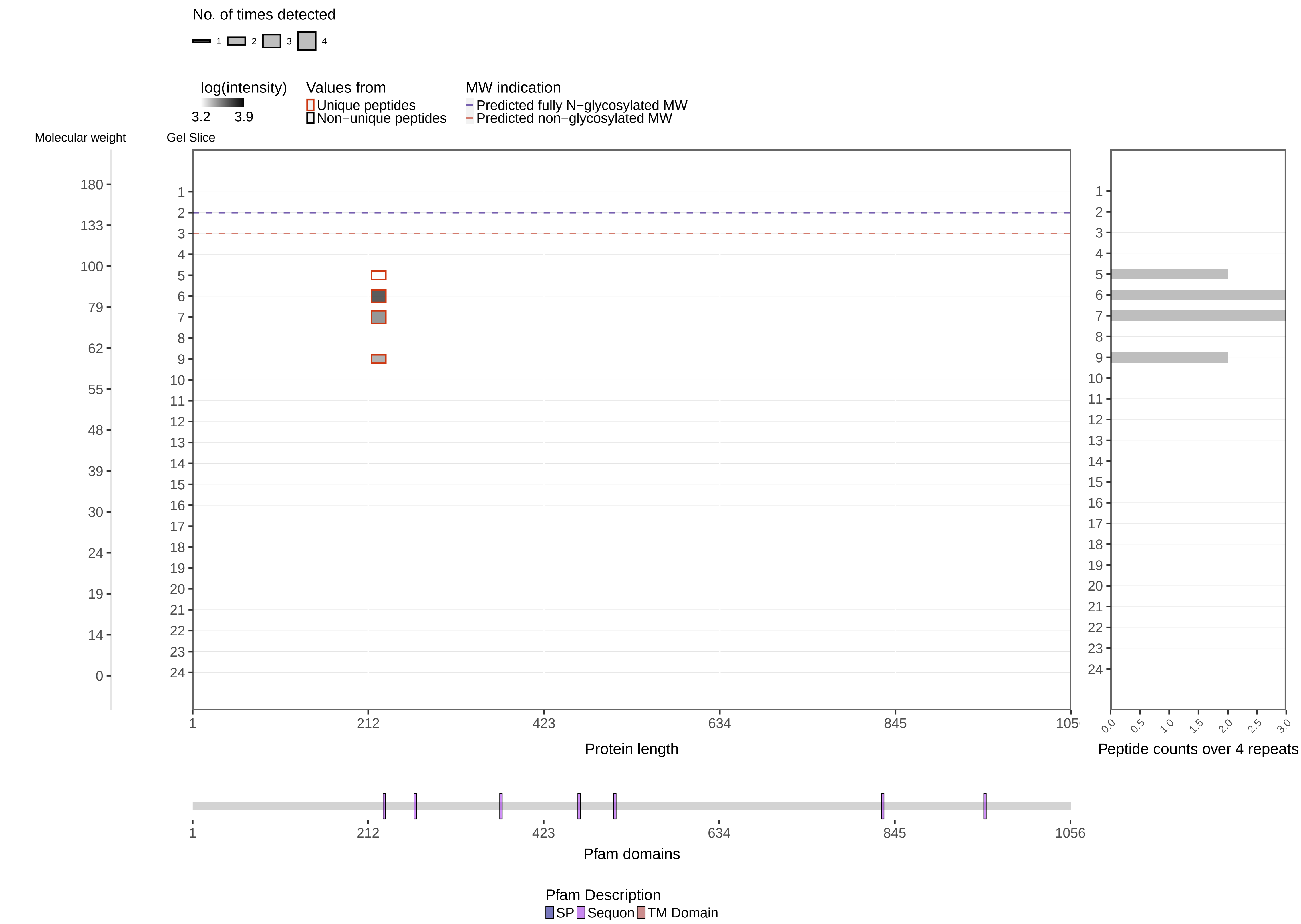


Protein sequence:

LCLTSDLHILISGDQSGTVRIYKFKTEFFAPDTSFLSFQVGSKKGSNPIQSVKLVKVNGAVLSISTSQESKYFAVGSDQGYVVLIDSDSKTILYQTHIASEICAGVISMQFNMCSVHGFDKNILVVATKDSSVLALETETGNILSPSMVHPKKPSRALFMQILDGQELSGRGLSISDGINMIKGNSDNSASKQPVVLLCSEKALYVVYSLLHIVQGIKKVYYKKKFHSSLCCWASTSDSPEAGLMLLFSNGKIEIRSLPELSLLKETSVRGLMLSPPKANSIADHSICASKNGELIVVDRD

NbL02g16260.1 epidermal_growth_factor_receptor_substrate_15−like_1 [MW = 130.55 kDa]

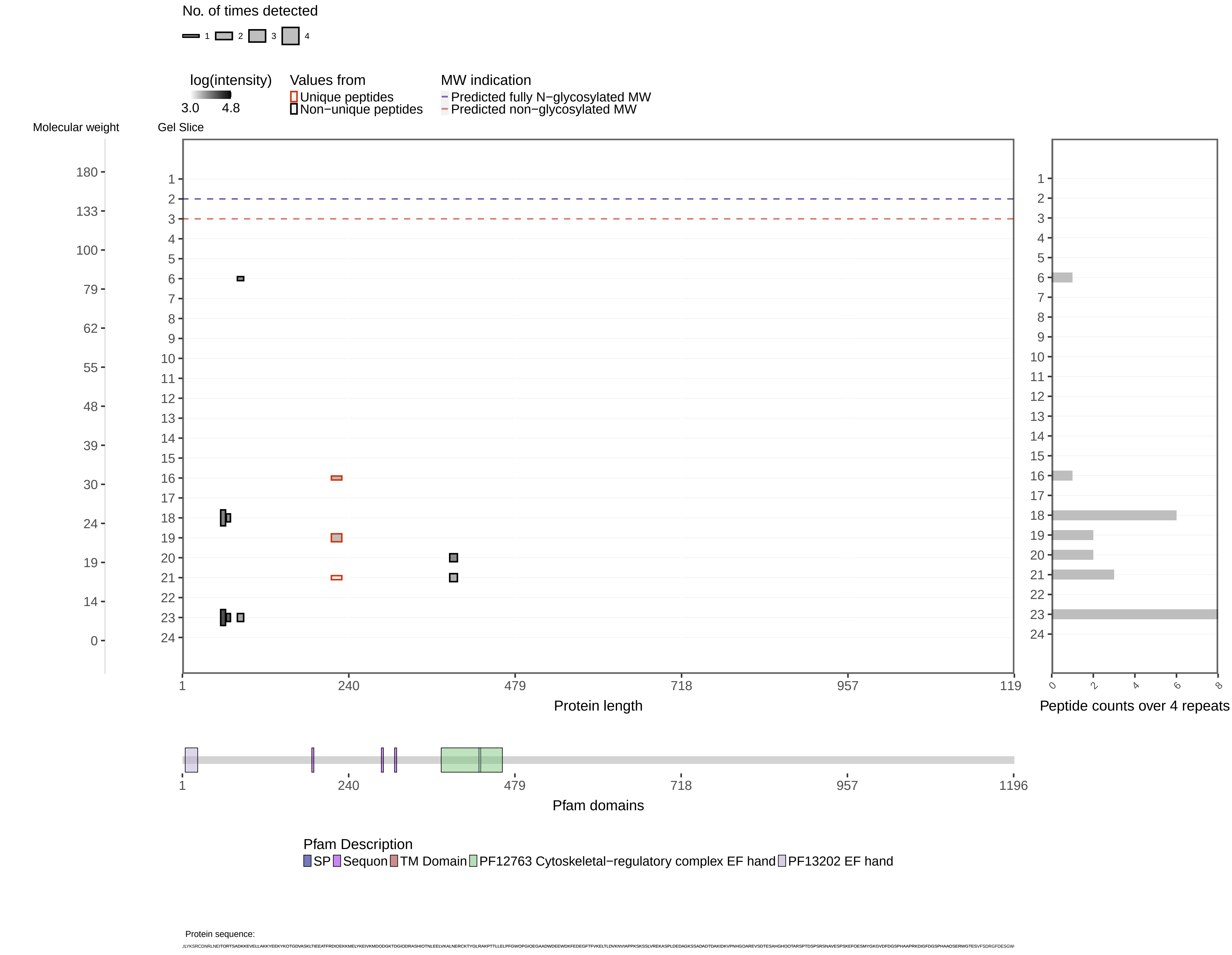

### NbL02g16160.1 thioredoxin_reductase_NTRB−like [MW = 40.23 kDa]

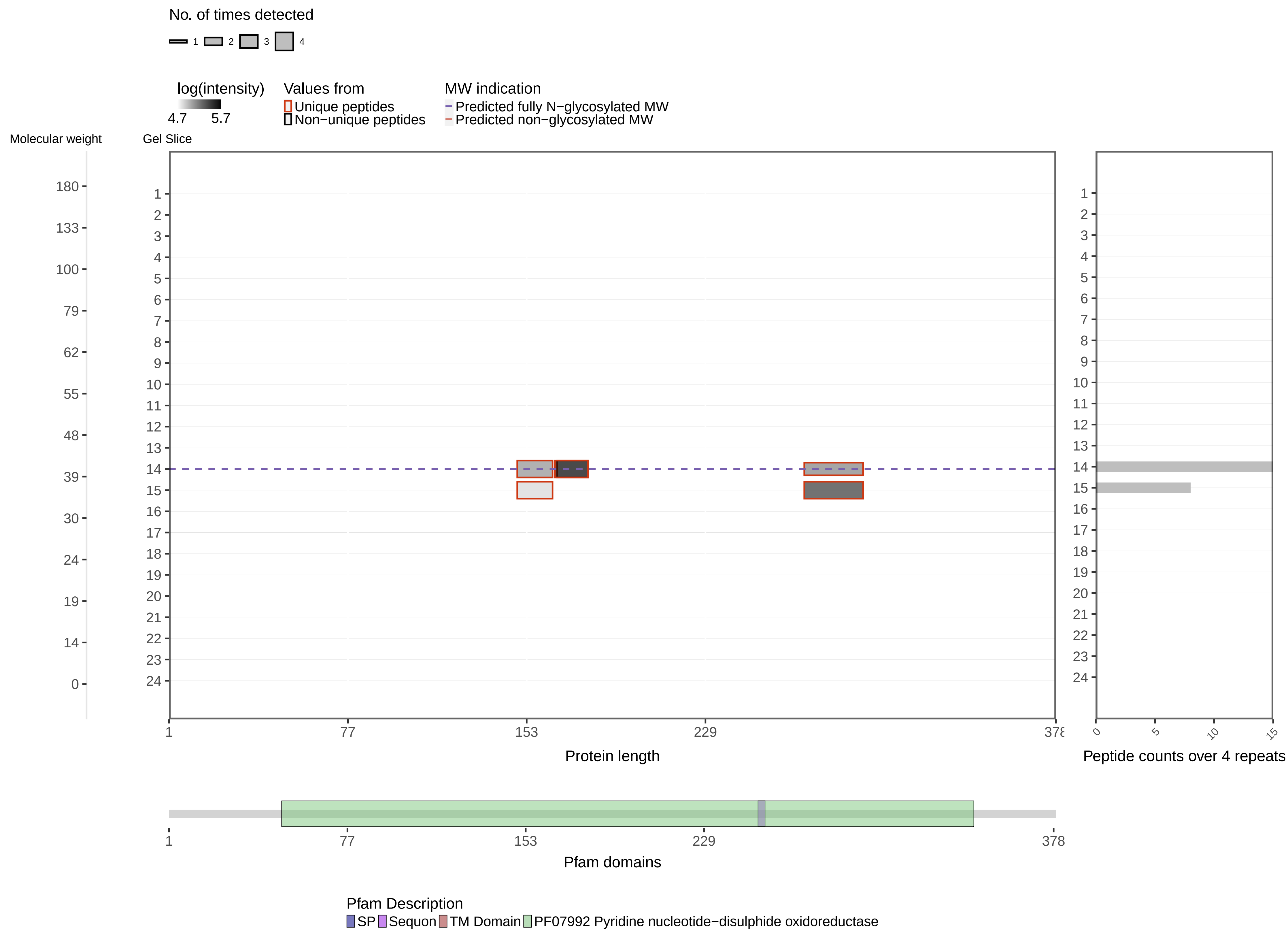


Protein sequence:

/NRGISACAVCDGAAPIFRNKPLAVIGGGDSAMEEANFLTKYGSKVYIIHRRDEFRASKIMQSRALSNPKIEVIWNSSVVEAYGEKLLGGLKVKNVVTGQVSDLQVSGLFFAIGHEPATKFLDGQLELDSDGYVVTNPGTTLTSVRGVFAAGDVQDKKYRQAITAAGSGNSLPSTCAPRGLVGALRDSDTRCKSKEKNYVF*

### NbL02g16000.1 uncharacterized_LOC107769872 [MW = 19.96 kDa]

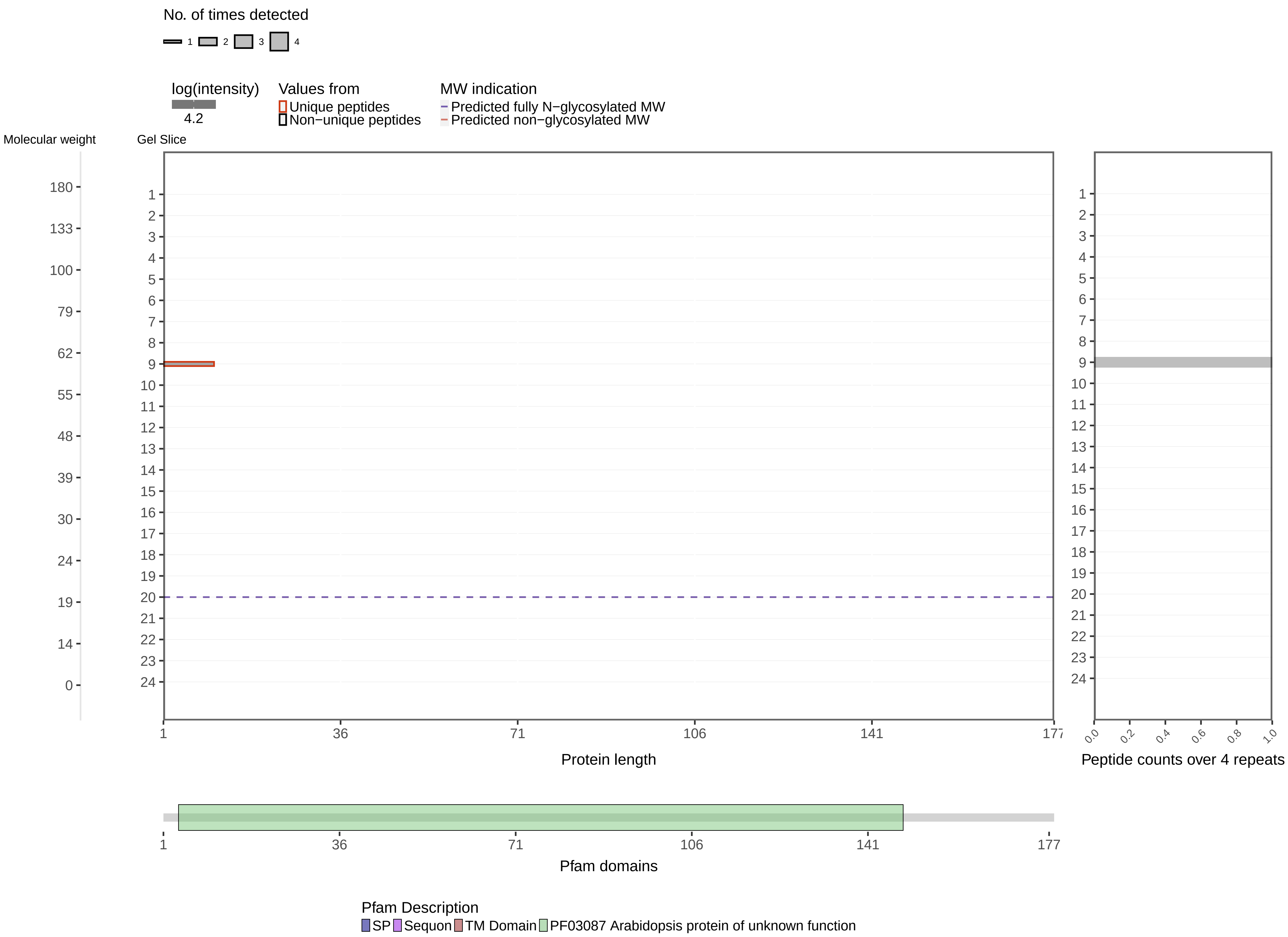


Protein sequence:

LTTDDSVTRFTSFSKKIKRDAKRLALNLKQMDQAAAVPVLLDADQDTIVAIRALREANAVCISTFQMLLSFLCVPLDSFENALEATFRCLIRSRSLLLNVLSC*

NbL02g15860.1 DNA−directed_RNA_polymerase_II_subunit_1 [MW = 204.78 kDa]

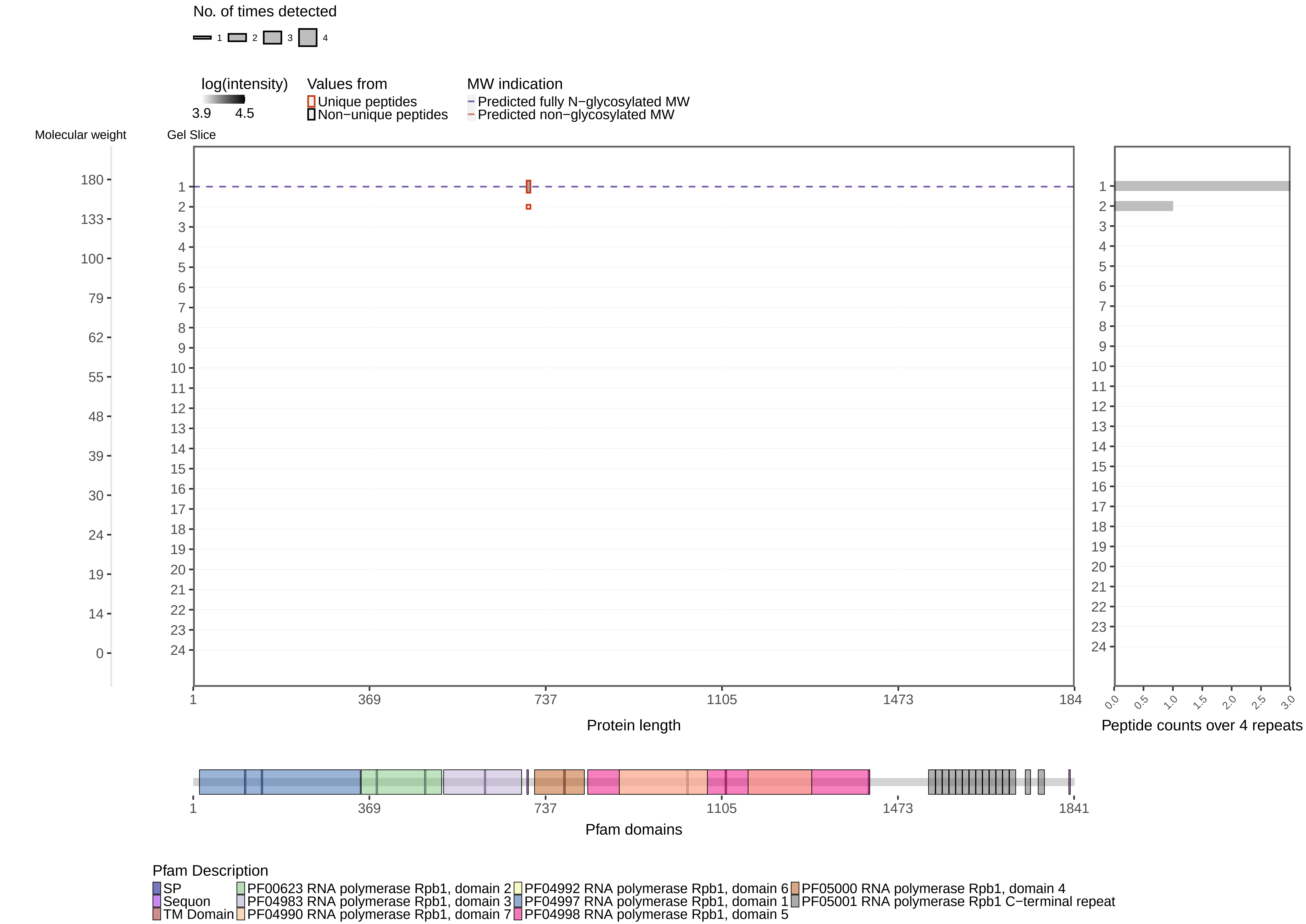


Protein sequence:

KAKKSTFDDLYAYEIDDPNWNPSYMLPEAIEDLKTIREIRSVFDAEVQKLQADRKQLGTEIAVTGDNSWPLPVNIQRLVLNAQKTFKIDFRRRSDMHPMEIVEAVDKLQERLKVVPGDDYLSMEAQKNATLFFNILLRSALASKRVLKEYRLSREAFDWVIGEIESRFLQSLVAPGEMIGCVAAQSIGEPATQMTLNTFHYAGVSAKNVTLGVPRLREIINVAKKIKTPSLSVYLKPEVGKTKERAKTVQCALEYTTLRSVTQATEVWYDPDPMSTLIEEDVEFVKSYYEMPDEEIDP

### NbL02g15790.1 ribulose_bisphosphate_carboxylase_large_chain [MW = 19.8 kDa]

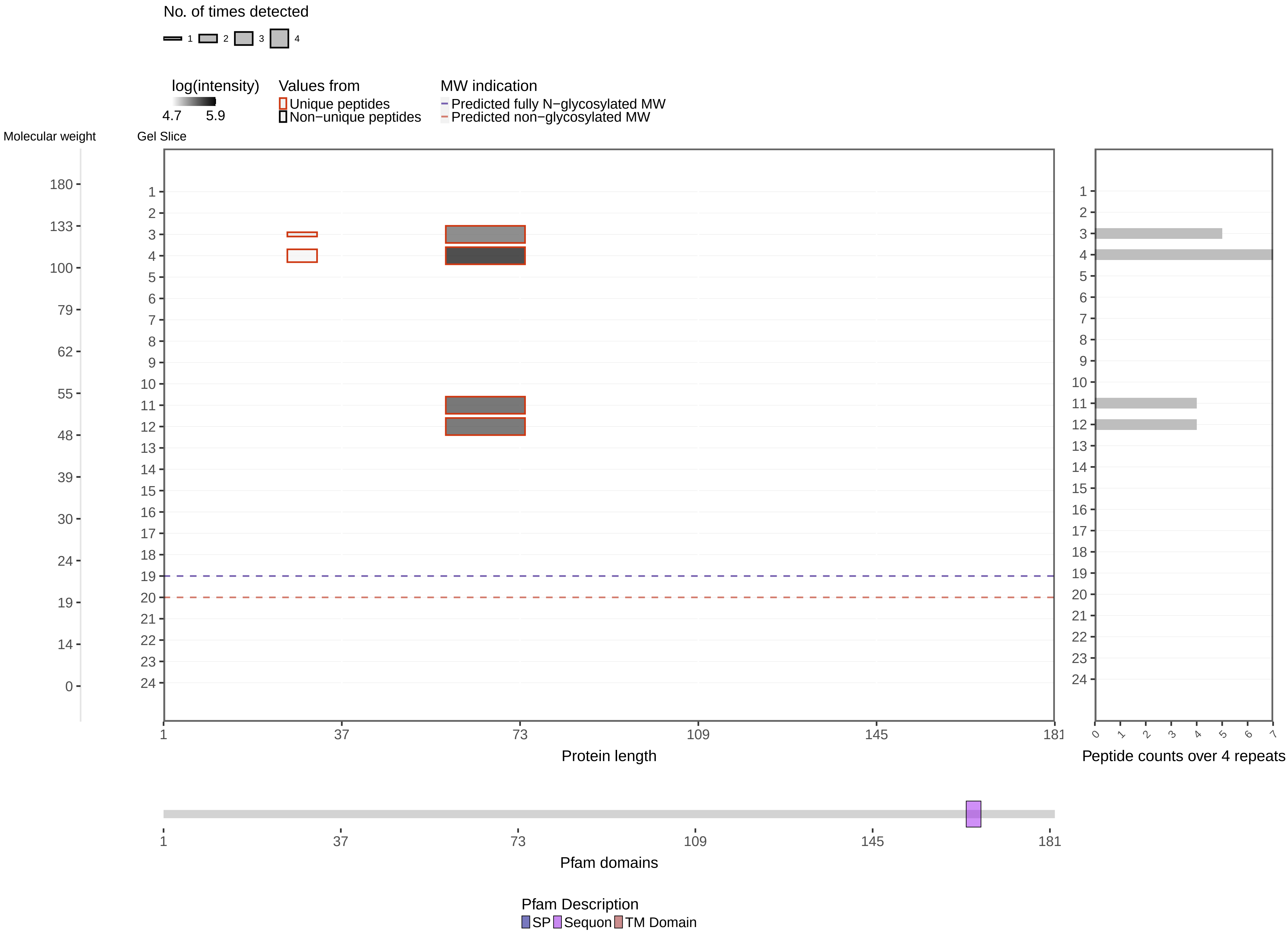


Protein sequence:

FNFAAVDVLDKSSILYRGRLESIPNQYSYPSNRREGGQKKVRIWQKHLLDRMGFCDKKCFSNGDSNPFPAPKMTGAKRSAEAVGCKNTSVGEHSALERSLRAR*

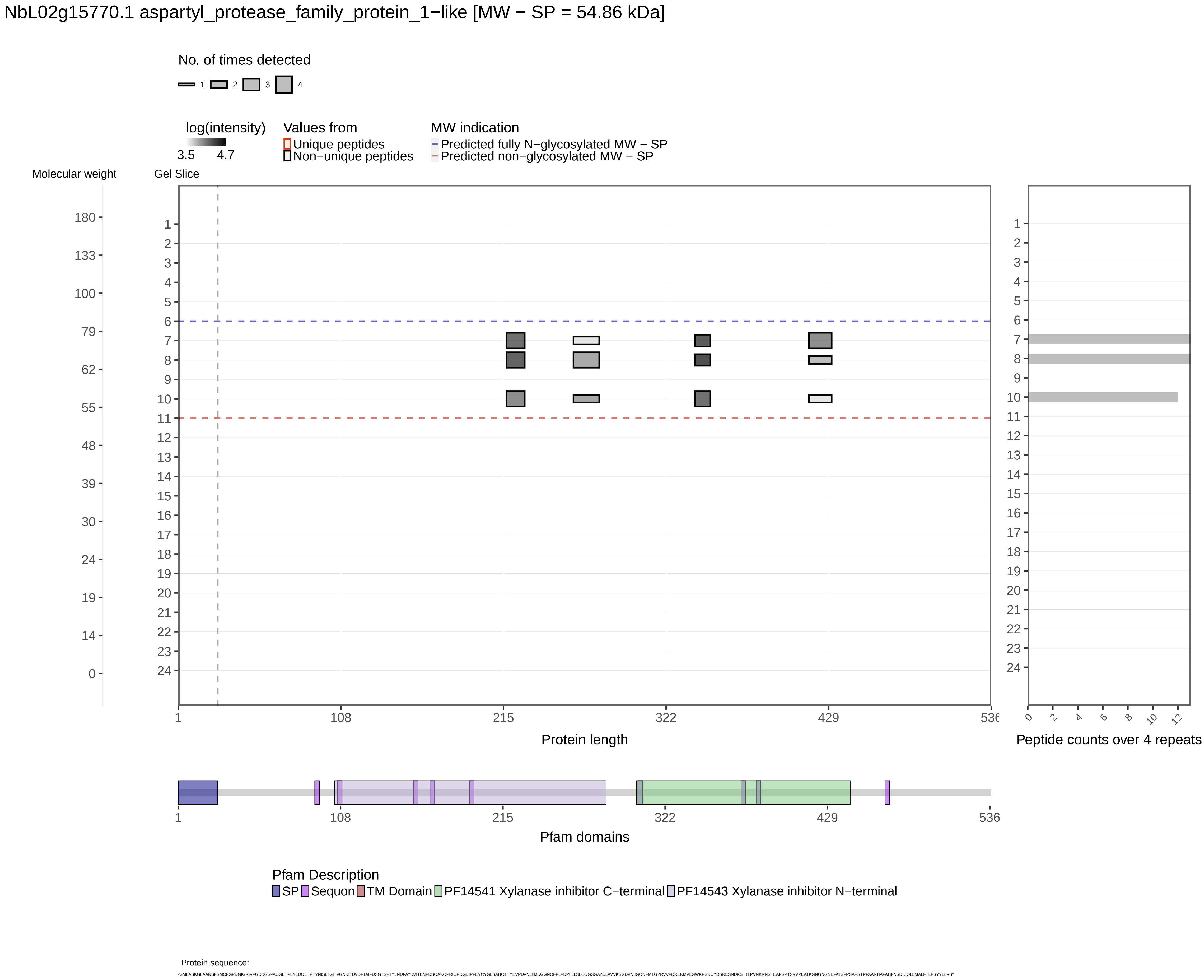
NbL02g15770.1 aspartyl_protease_family_protein_1−like [MW − SP = 54.86 kDa]
No. of times detected
1 2 3 4
log(intensity)
3.5 4.7
Values from
Unique peptides
Non−unique peptides
MW indication
Predicted fully N−glycosylated MW − SP
Predicted non−glycosylated MW − SP
Molecular weight
Gel Slice
180 133 100 79 62 55 48 39 30 24 19 14 0
1 2 3 4 5 6 7 8 9 10 11 12 13 14 15 16 17 18 19 20 21 22 23 24
1 108 215 322 429 536
Protein length
0 2 4 6 8 10 12
Peptide counts over 4 repeats
Pfam domains
Pfam Description
SP
Sequon
TM Domain
PF14541 Xylanase inhibitor C−terminal
PF14543 Xylanase inhibitor N−terminal
Protein sequence:

### NbL02g15750.1 uncharacterized_LOC109237051 [MW = 23.26 kDa]

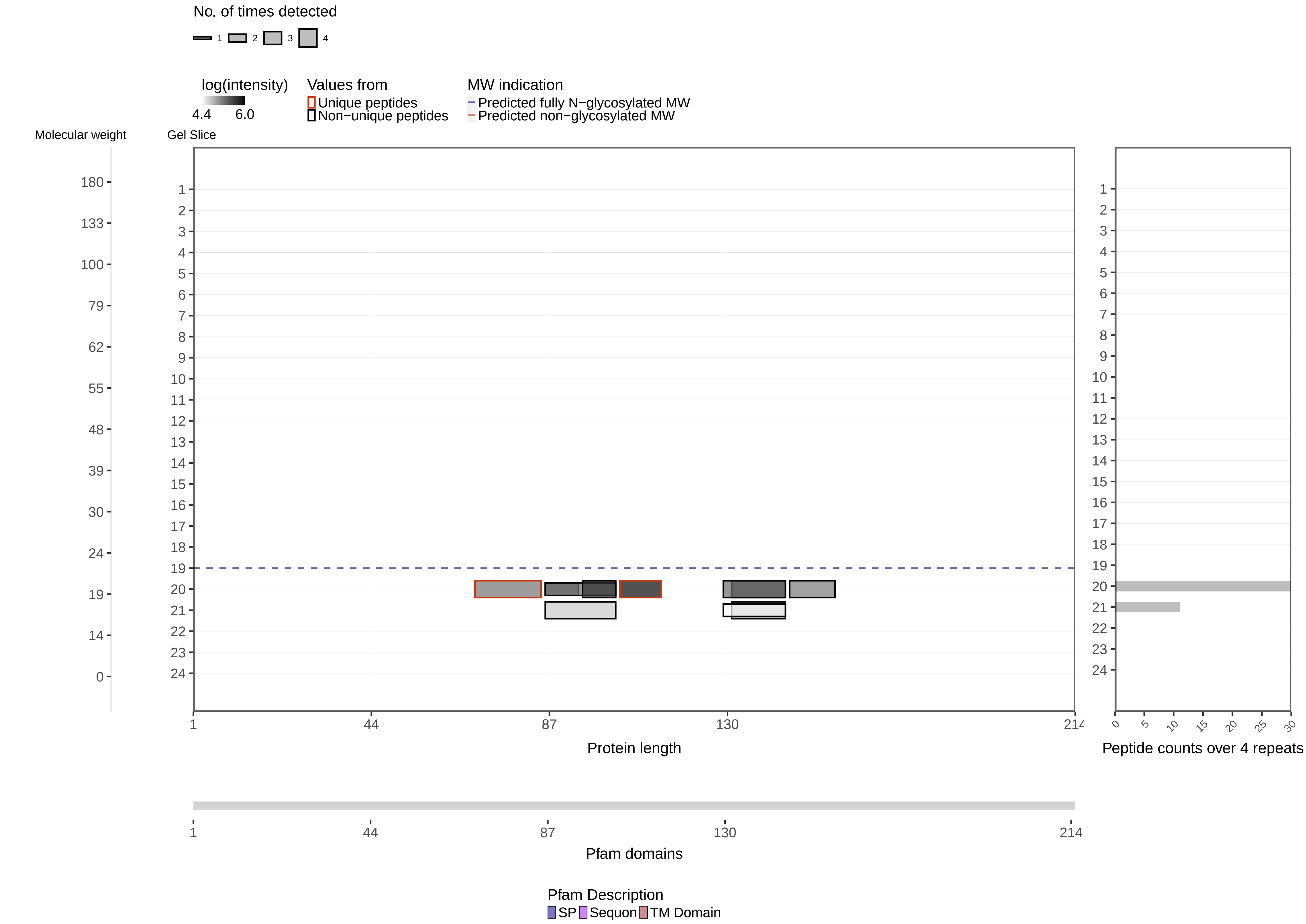


Protein sequence:

RNPYLSEGSRQAAATATALAKKYGADITVVVIDEKEKEAVPEHETQLASIRWHLSEGGYQEFKLLERLGEGSKPTAIIGEIADEMNLDVVIMSMEAIHSKHVDANLLAEFIPCPVLLLPL*

### NbL02g15660.1 methionyl−tRNA_formyltransferase−like [MW = 39.08 kDa]

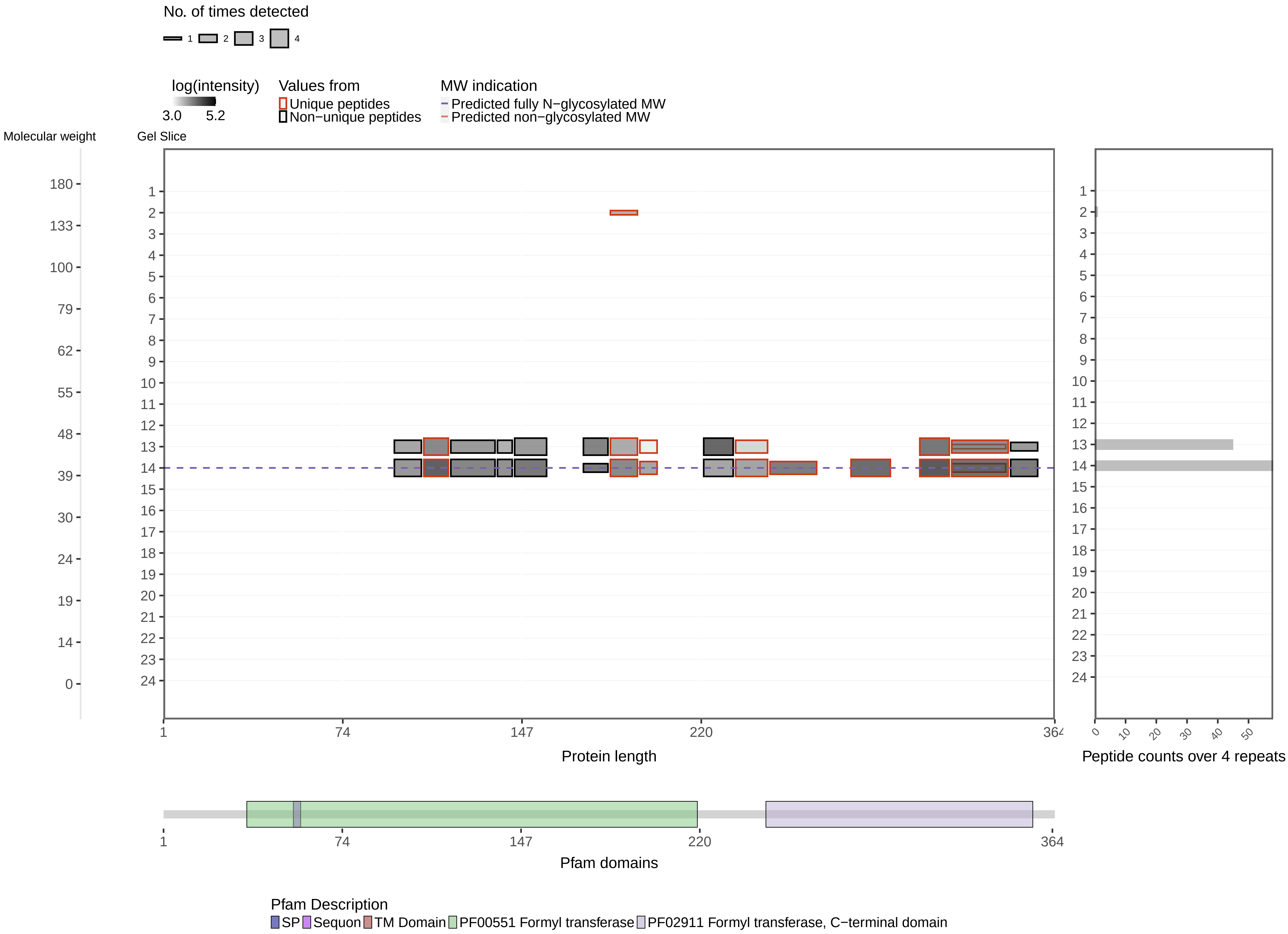


Protein sequence:

DGVKETGVSLAYTVRKLDAGPVIALERIKVDDQIKAPDLLDLLFALGSKLLIRELPSIFDGSASGRAEDQDDSKATLAPKITPEESWLSFDEDAQVLHNKVRAFAGWPGTRARLIVIDPSSGESSSIELKIITTRIYTGIRSQDIEANDVLFTKGSLVIPCGGDTALEVLEVQLPGKRAVNAASFWNGLRGQILKK*

### NbL02g15650.1 ATP−dependent_Clp_protease_proteolytic_subunit_3_chloroplastic−like [MW = 36.41 kDa]

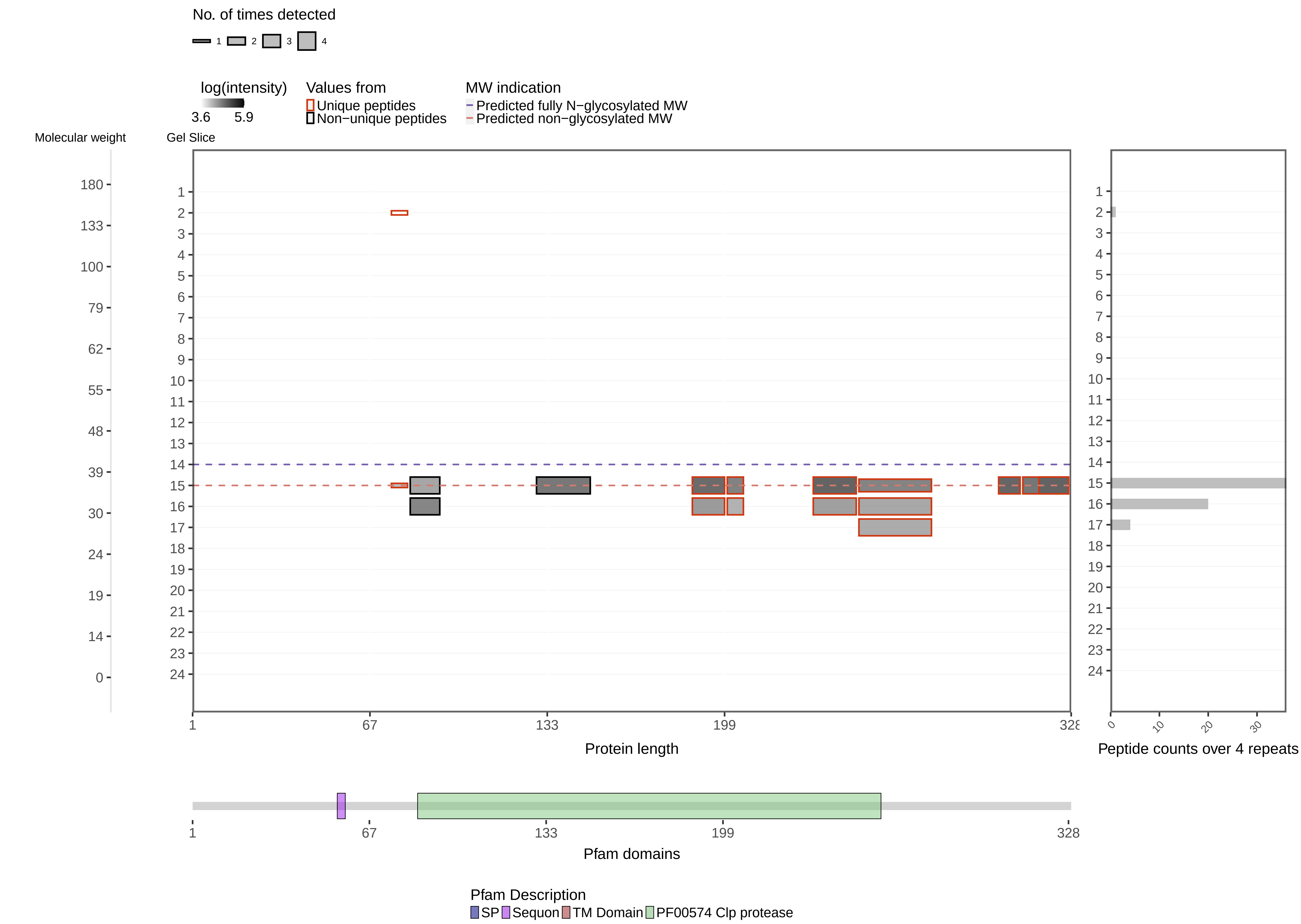


Protein sequence:

ADVSTICMGLAASMGAFLLASGSKGKRYCMPNARVMIHQPLGTSGGKATEMSIRIREMAYHKIKLNKILSRITGKPEEKIEEDTDRDNFMNAWEAKEYGLVDAVIDDGKPGLVAPTADATAPPKTRVWDMWKIEGSKKAKKNLPSEERMLQTGYVSRQDDEQKSNEQKDEAPSPL*

### NbL02g15540.1 2−hydroxyacyl−CoA_lyase [MW = 61.45 kDa]

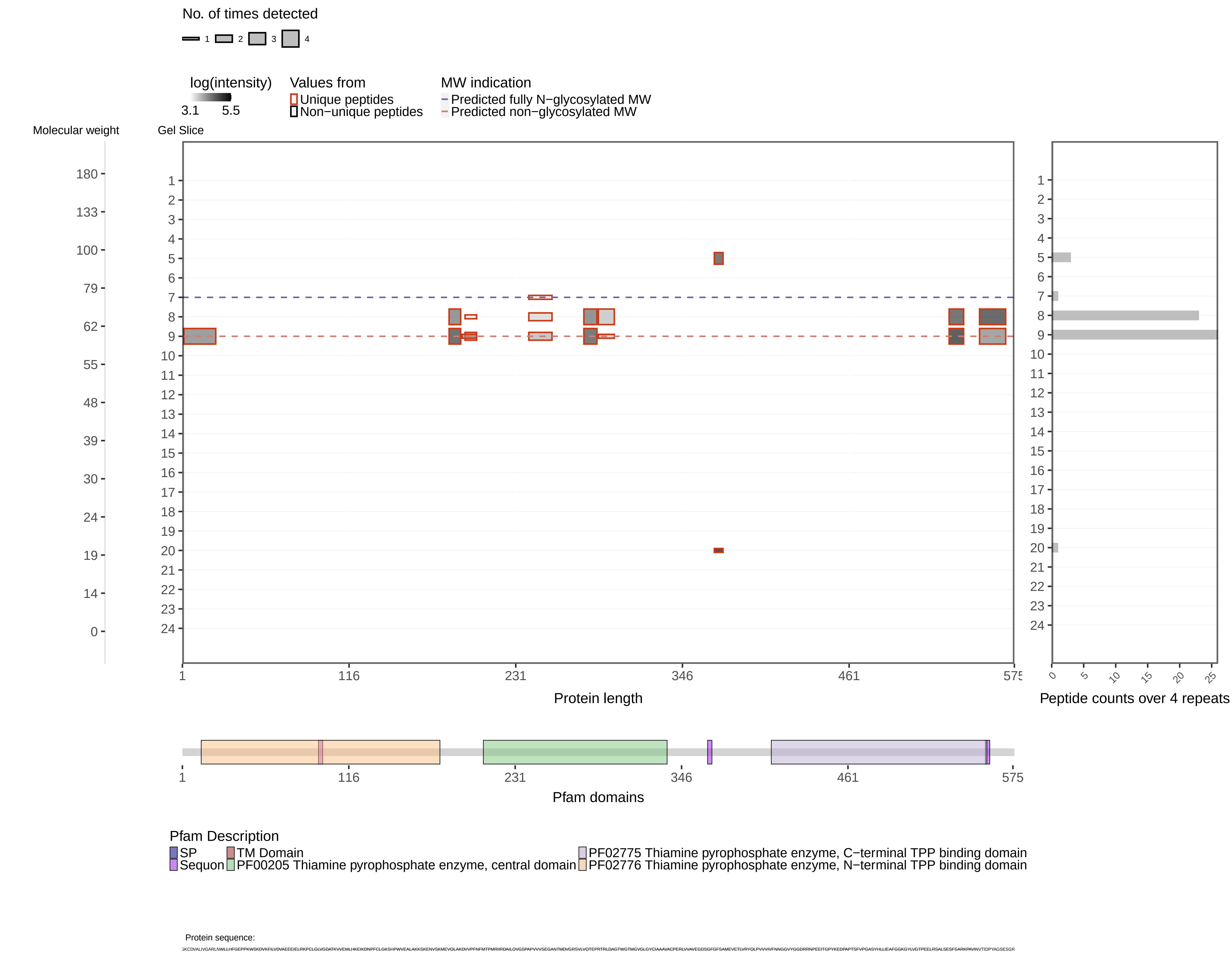

### NbL02g15180.1 ribulose_bisphosphate_carboxylase_small_chain_S41_chloroplastic−like [MW = 20.18 kDa]

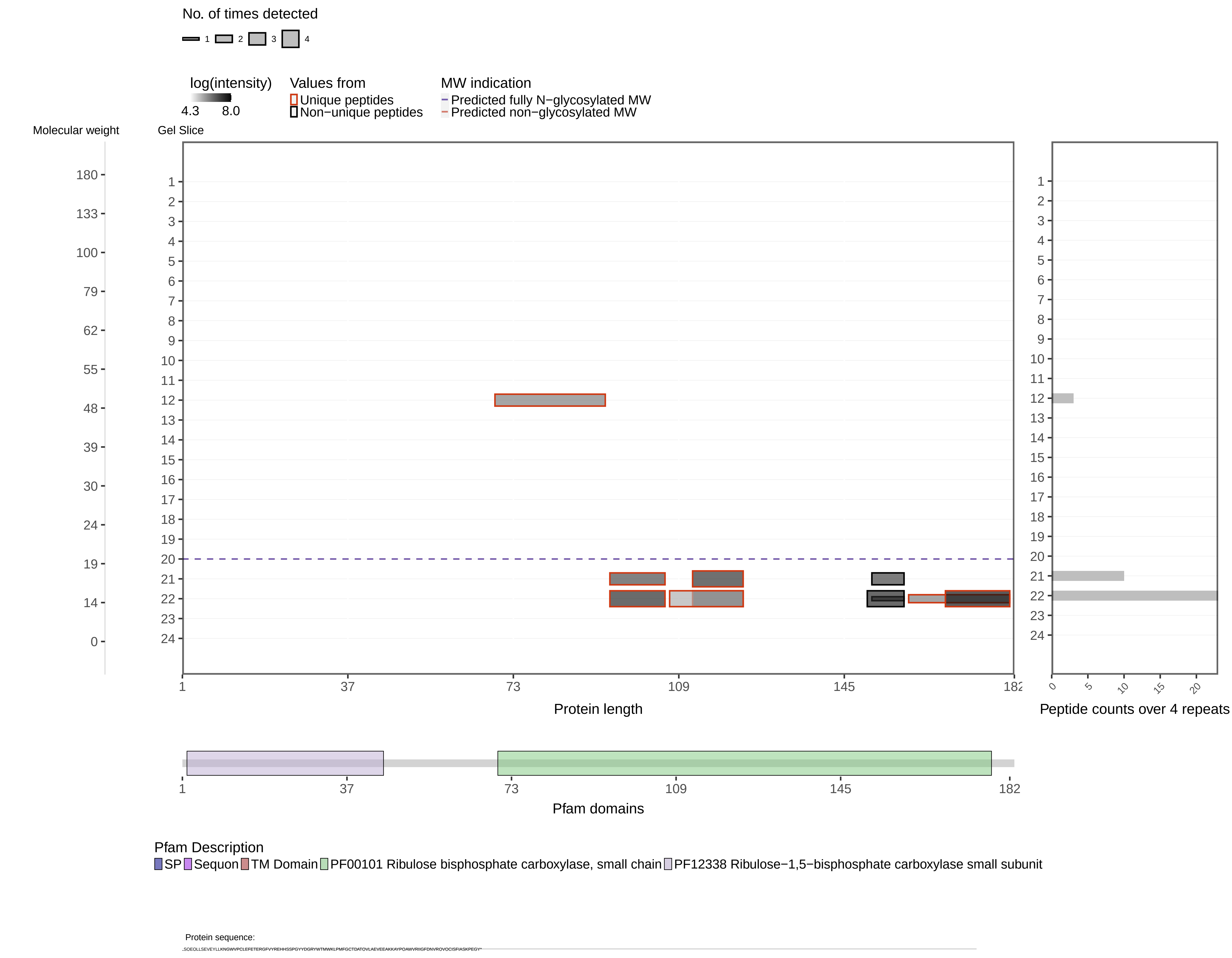

### NbL02g15130.1 ubiquitin_receptor_RAD23d−like [MW = 41.21 kDa]

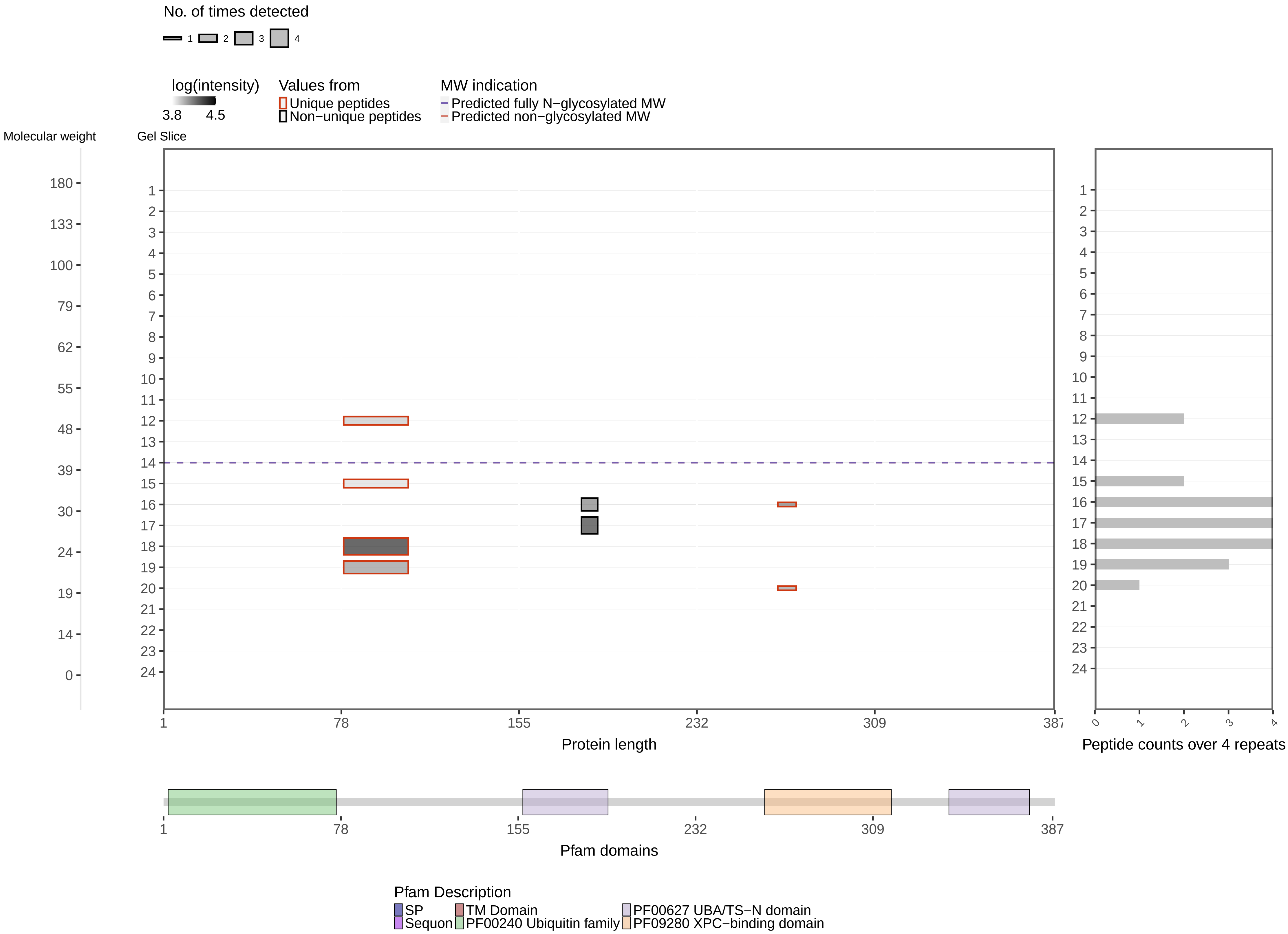


Protein sequence:

RAAYNNPERAVDYLYSGIPEQAEIPPFAPTAAATASGQAVNPSAQEASQPAVPSSGPNANPLDLFPQVGLPNVGSNAGSGNLDFLRNSPQFQALRTMVQANPQILQPMLQELGKQNPHLMRLIQEHQPDFLRLINEPVEGEGSALGQMAGAIPQAVTVTPEEREAIERLEAMGFDRALVLEVYFACNKNEELAANYLLDHMHEFDE*

NbL02g15100.1 HEAT_repeat−containing_protein_5B [MW = 251.17 kDa]

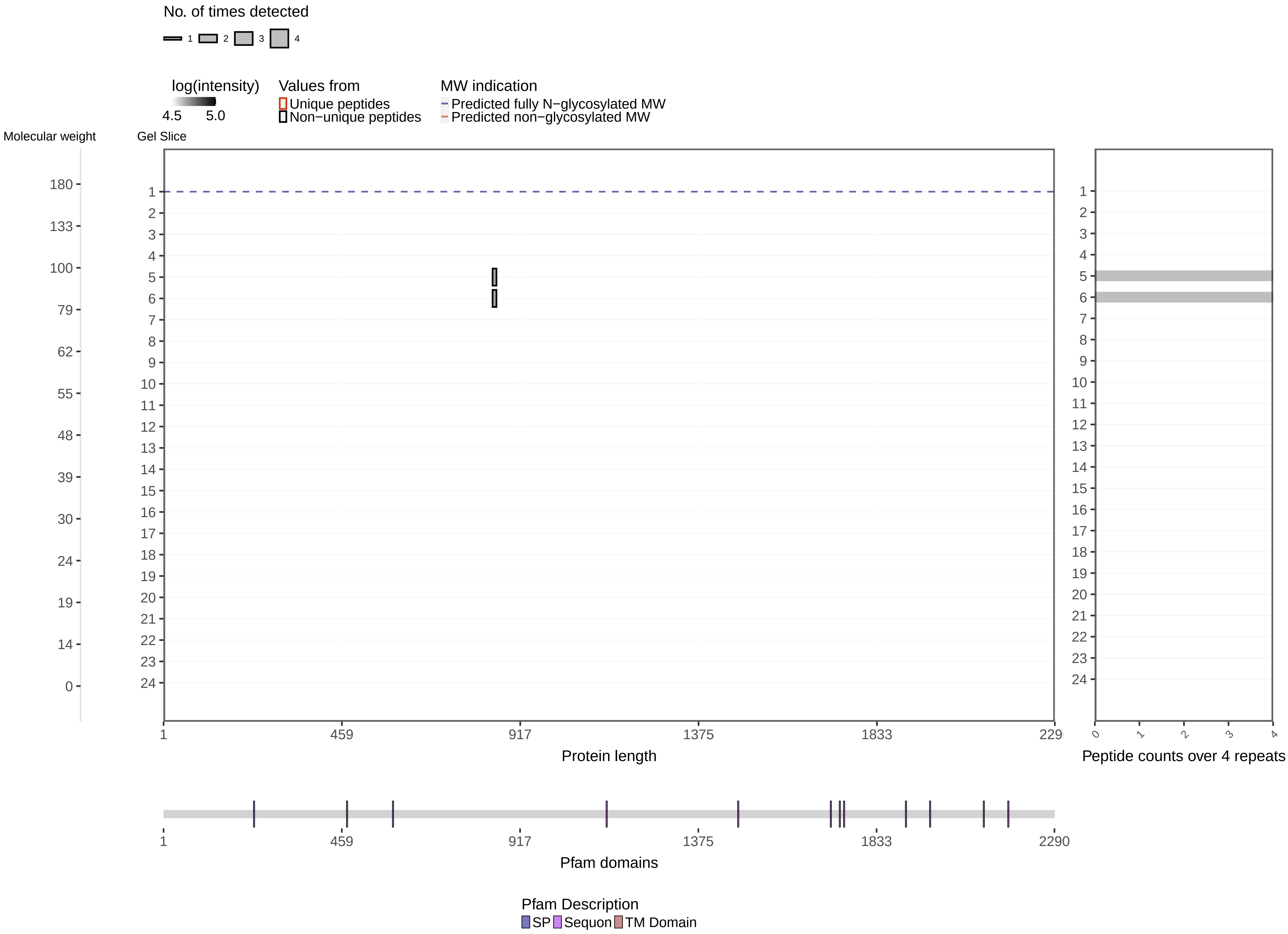


Protein sequence:

SQNDSSSGLDGEAGLNYGDDDEDMVSSSQNPTFQGYRSSHPIAYLPRNKHLRYRTRVFAAECLSHLPAAVGKNPAHFDVALARKQPGSGPSSGDWLVLQLQELVSLAYQISTIQFENMRPVGVNLLSTIIDKFGTLDPEFPGHLLLEQYQAQLVSAVRTALDSSSGPALLEAGLQLATKILTCKIVSRDQLAVKRIFSLISRPLNEFNDLYYPSFAEWVSCKIKVRLLTAHASLKCYTFAFLKNQQKEVSDEYLALLPLFSVSSKILGIYWLCLLKDYSYIRTQSFPKENWKPFLDGI

### NbL02g15050.1 ultraviolet−B_receptor_UVR8−like [MW = 41.71 kDa]

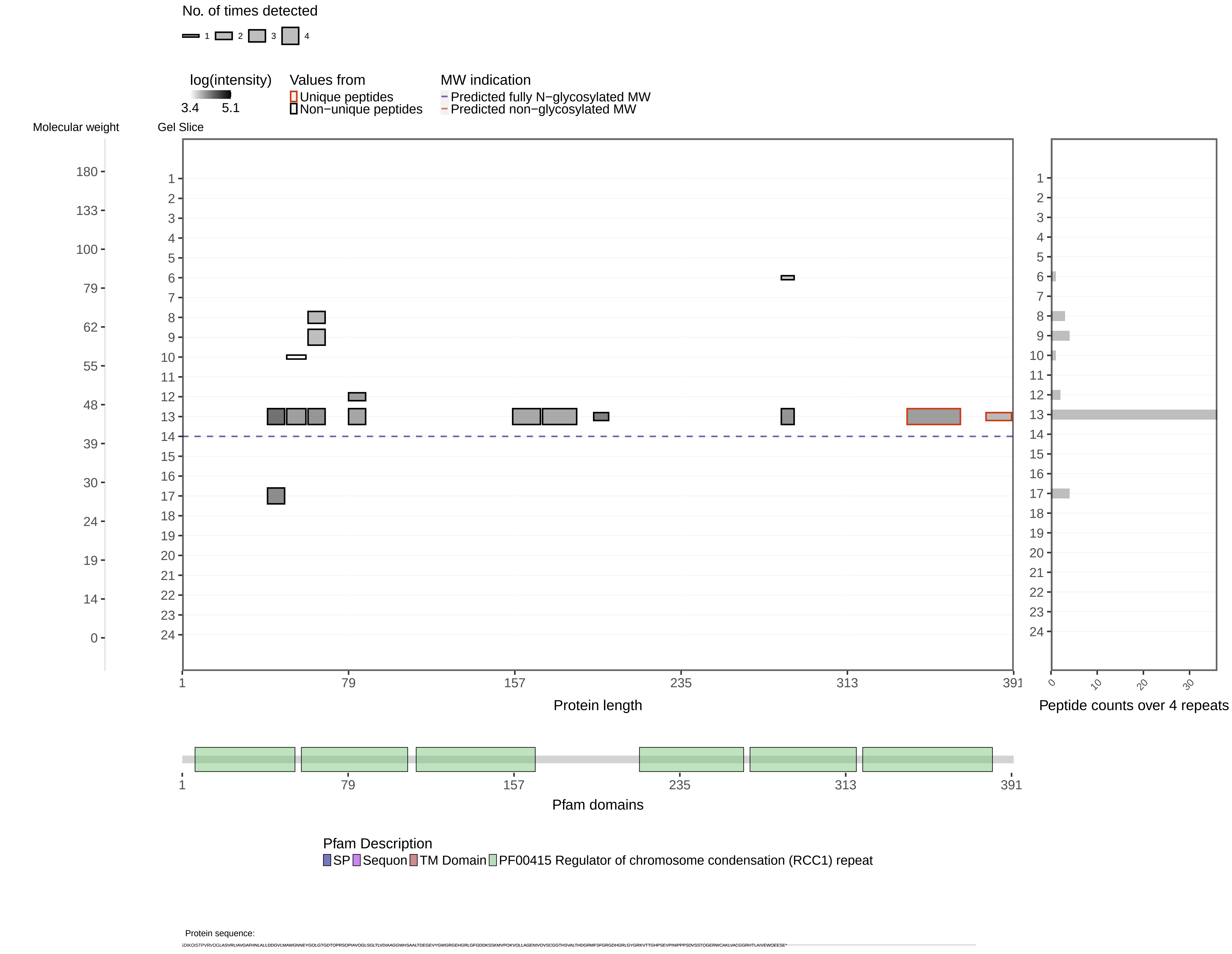

NbL02g15000.1 allene_oxide_cyclase_chloroplastic−like [MW = 26.18 kDa]

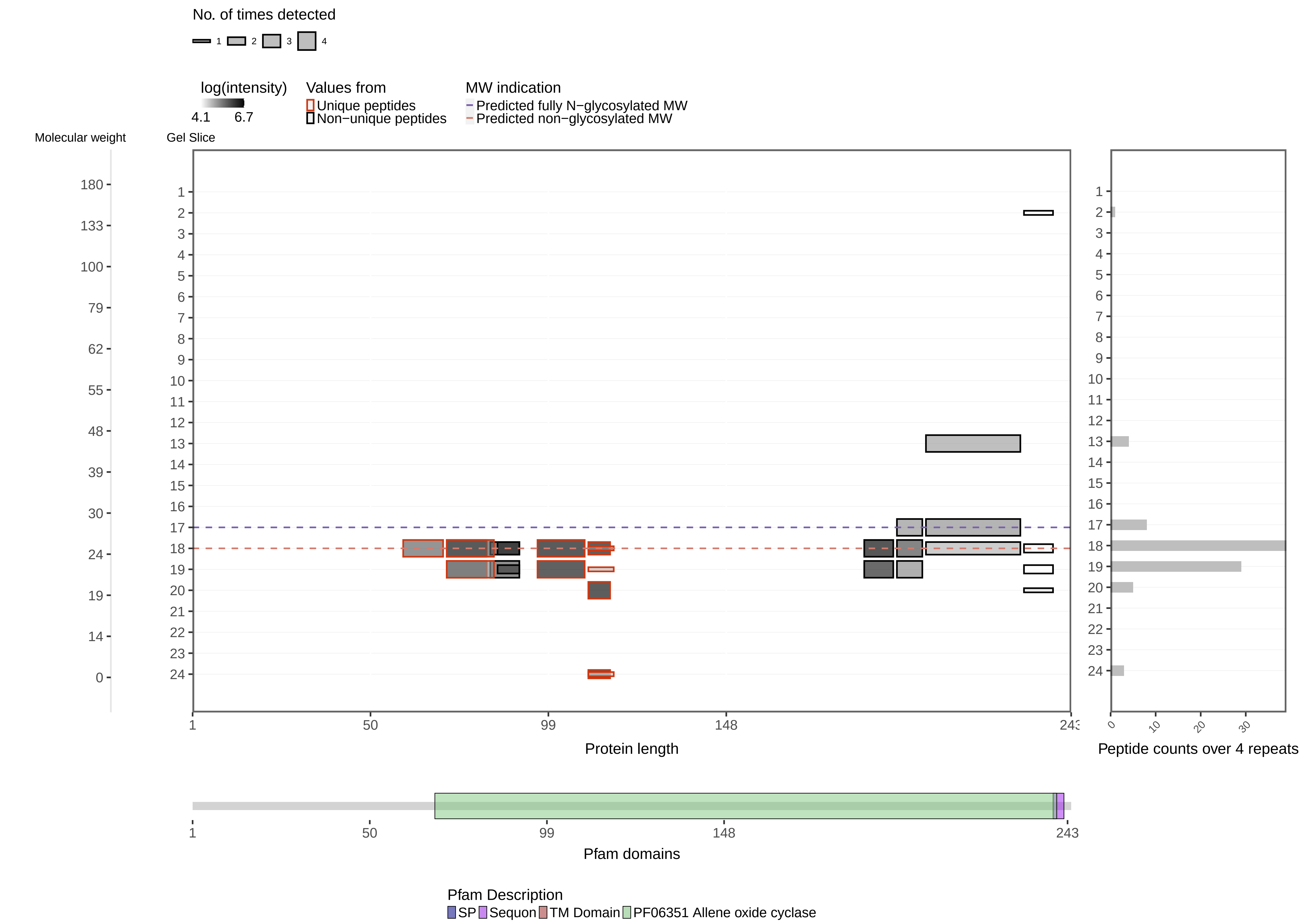


Protein sequence:

SNKLYTGDLKKRIGITAGLCILIKHEEEKKGDRYEAIYSFYFGNYGHIAVQGSYLTYEDSYLAVTGGSGIFAGVSGQVKLQQIIFPFKLFYTFYLKGIPDLPSELLVTAVPPSPTVEPAPEAKACEAGATVKNFTN*

### NbL02g14970.1 geranylgeranyl_pyrophosphate_synthase_chloroplastic−like [MW = 39.56 kDa]

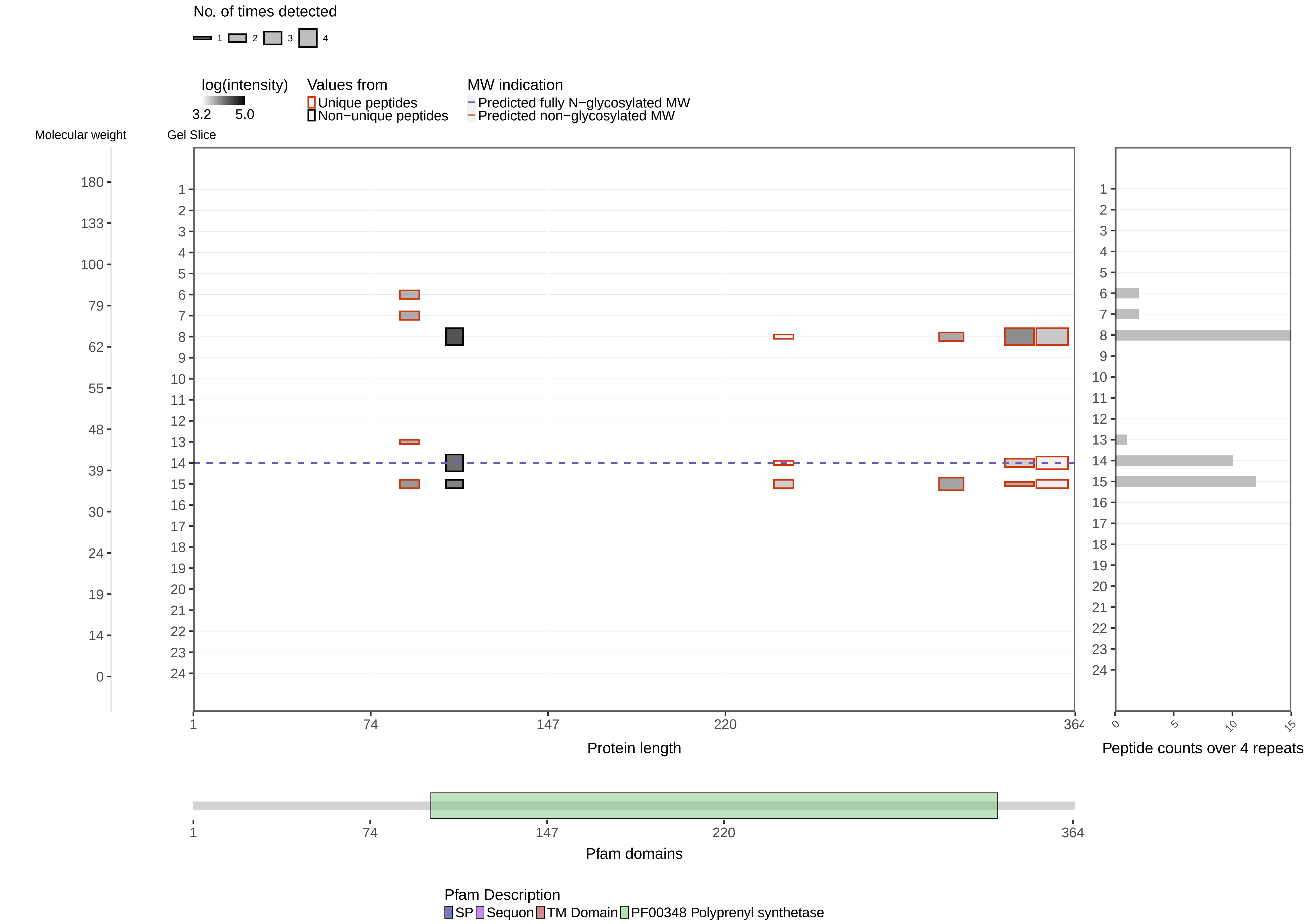


Protein sequence:

TNHKIYGEDVAVLAGDALLALAFEHIATQTKGVPSHRIVRVVGELAKCIGAEGLVAGQVVDIISEGVSDVDLKHLEFIHLHKTAALLEGSVVLGAILGWADDEEVEKLRKFARCIGLLFQVVDDILDVTKSSQQLGKTAGKDLVADKVTYPKLIGIEKSREFAEELNKEAQAQLAGFDQGKAAPLIALANYIAYREN*

### NbL02g14740.1 F−box_protein_At5g49610−like [MW = 40.18 kDa]

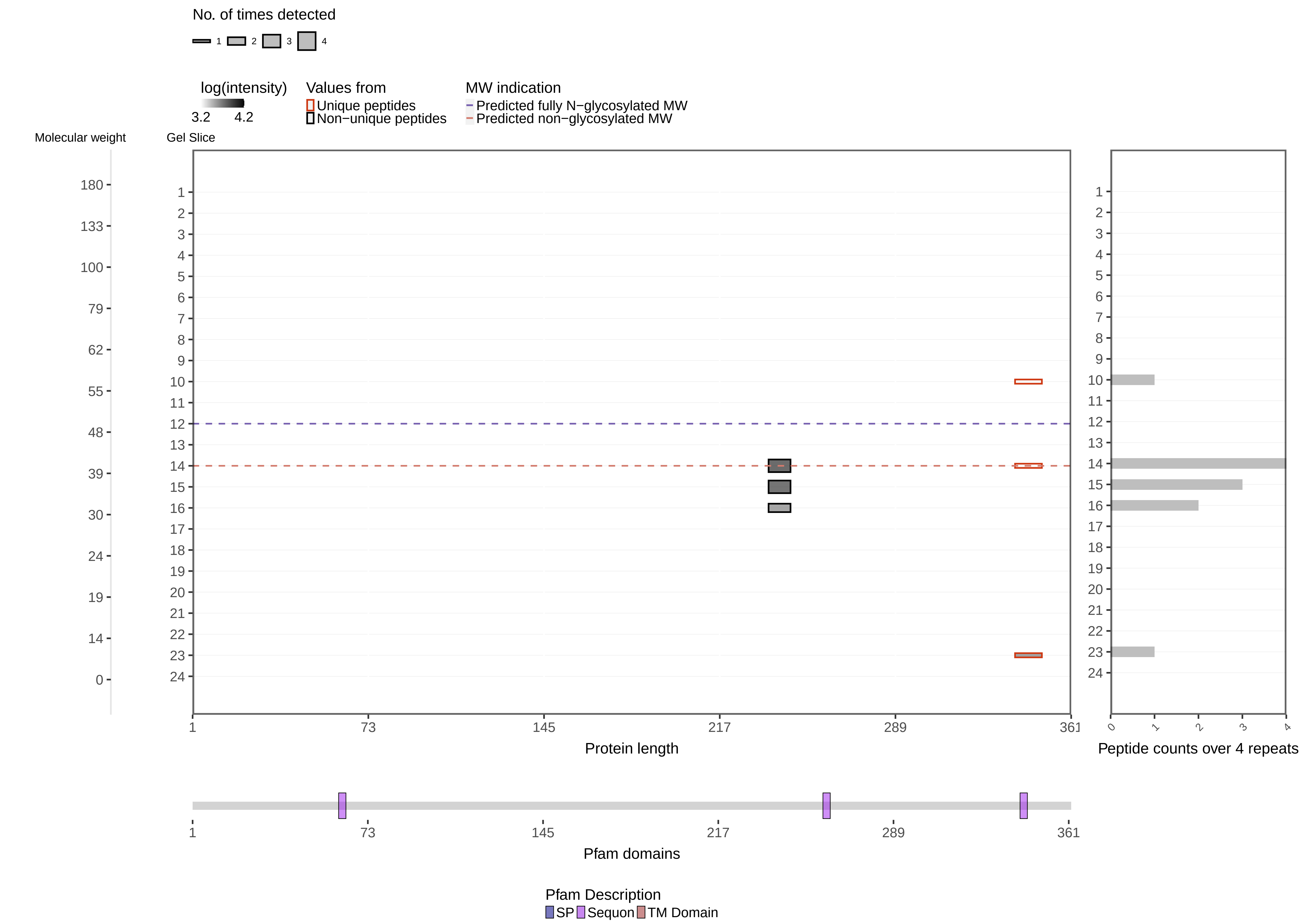


Protein sequence:

YKIICAFPSTDFDEATEFEIYSSKEGTWEIAGEICFGAWRVIPKSGVHVNGVVYWMTTGSILAFDLTKERSQLLQDYVARGILGVFSGKLCKANVSGNSIIIYVLTNAHSNTMQMGSNTRMWSEKQQIVLDSKIVGDVARDYSVLHVDGDIMVVHSRERMYSYDFKSRATKAMSNVSDIYNMRCLPYVNNLVSL*

NbL02g14680.1 succinate_dehydrogenase_[ubiquinone]_flavoprotein_subunit_1_mitochondrial [MW = 69.19 kDa]

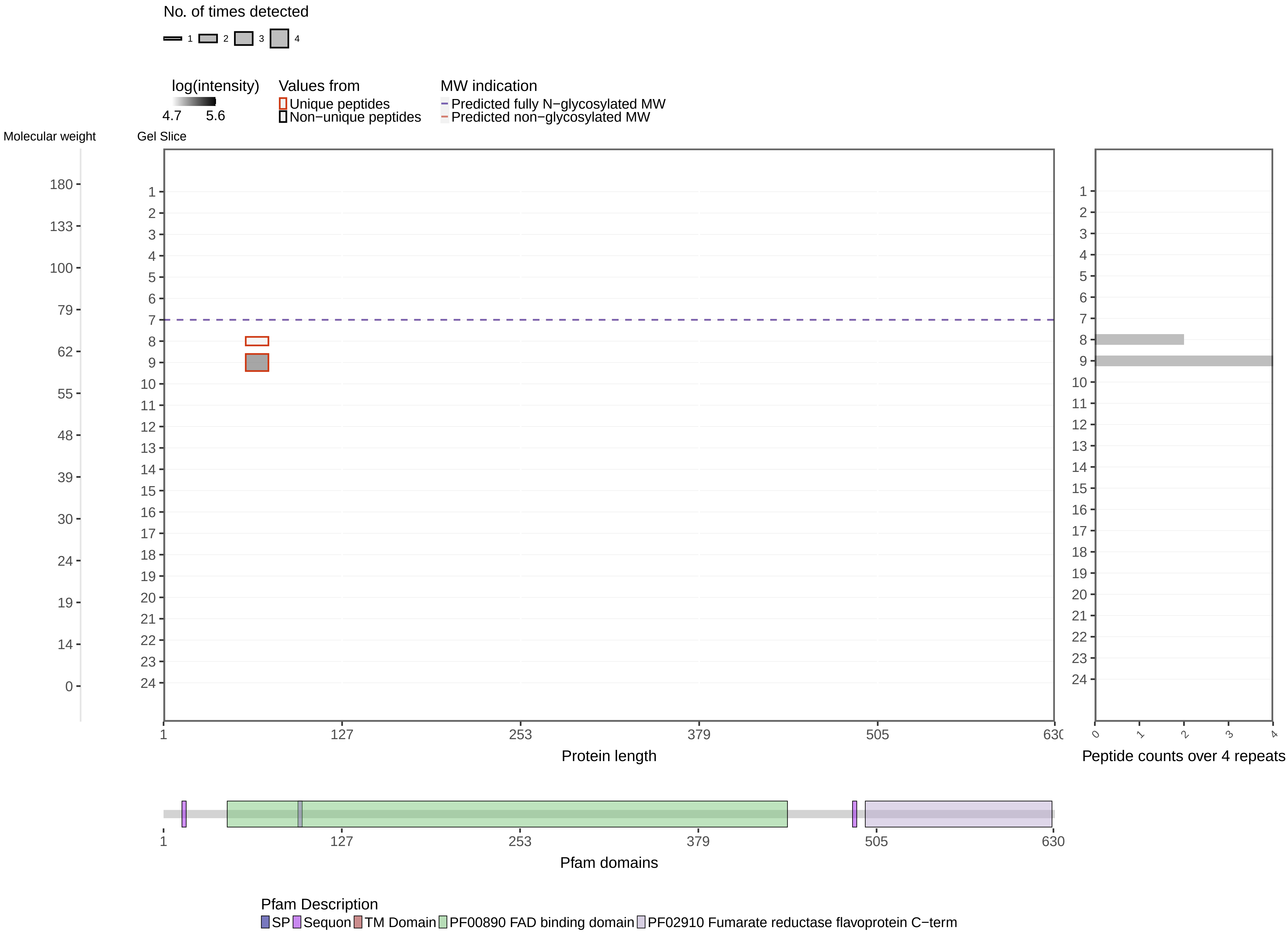


Protein sequence:

GILRNSEGERFMERYAPTAKDLASRDVVSRSMTMEIREGRGVGPLKDHIYLHLNHLPPEVLKERLPGISETAAIFAGVDVTKEPIPVLPTVHYNMGGIPTNYHGEVVTIKGDNPDAVVPGLMAAGEAACASVHGANRLGANSLLDIVVFGRACANRVAEIQRPGENQKPLEKDAGERTIAWLDKIRNSNGSLPTSKIRLNMQRVMQNNAAVFRTQETLEEGCQLIDKTWESFHDVQLKDRSLIWNSDLIETIELENLLINACITMHSAEARKESRGAHAREDFTKRDDEKWMK

NbL02g14600.1 pyridine_nucleotide−disulfide_oxidoreductase_domain−containing_protein_2 [MW = 61.22 kDa]

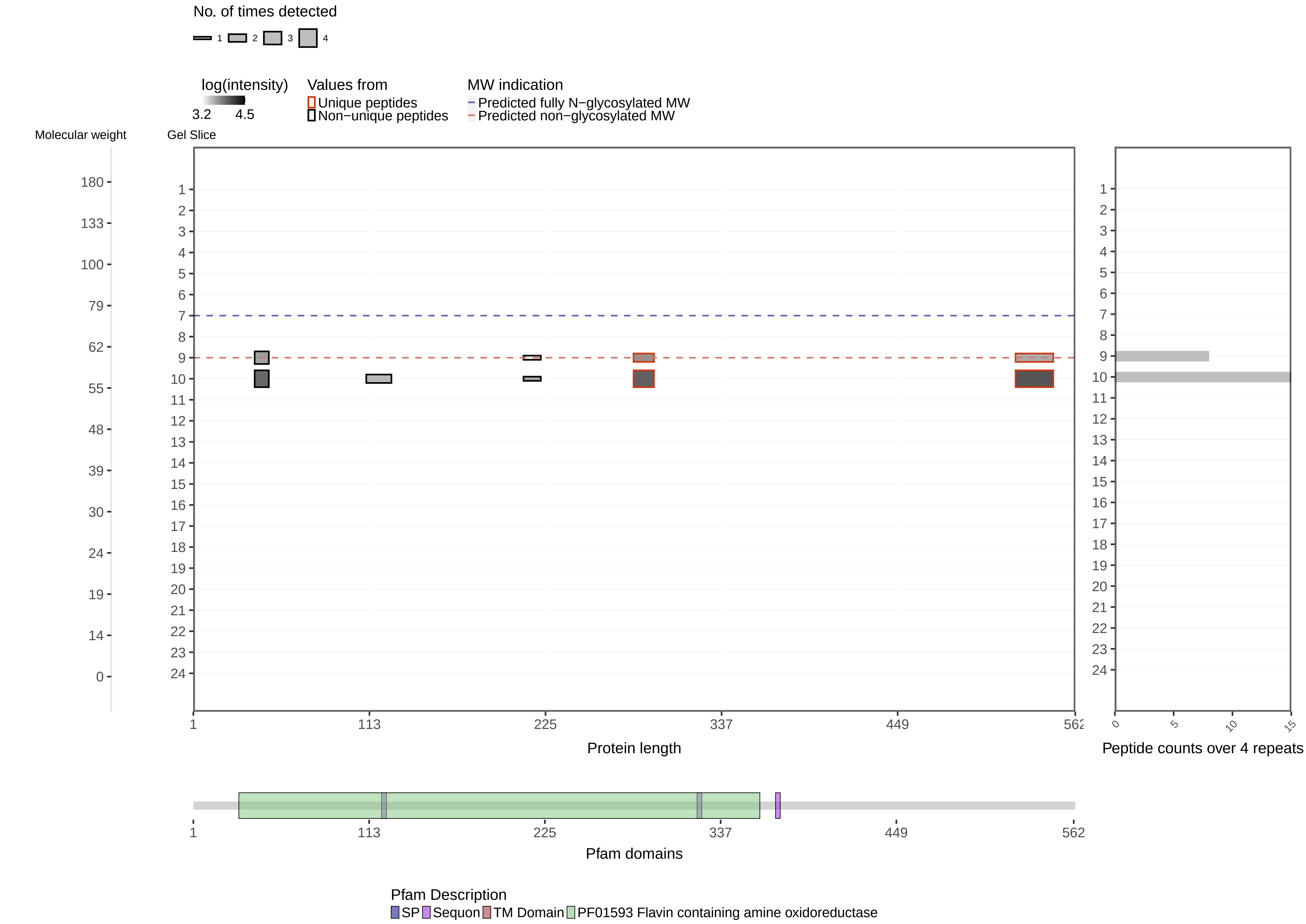


Protein sequence:

MGSVSMAIAAAAKEAGATILTNAEVSKLMIEDTGRVHGVMLADGTILHSSVVLSNATPFKTFMDLVPDDVLPHDFHNAIKSSDYRSATTKINLAVDGLPQFQCCNISHPDAGPQHSGTIHIGPESMEEMHLAAQDAENGLPSRRPIIEMTIPTVFDKTISPSGKHVIGLFIQYTPYKPSDGSWENPAYRESFAQRCFSMIDEYAPGFSSSVIGYDMLTPPDLEREIGLTGGNIFHGSMGLDSLFLMRPVKGWSNYRTPVEGLYLCGSGAHPGGGVMGAPGRNAAHVVIEDLKKS

NbL02g14440.1 uncharacterized_LOC109242336 [MW = 84.18 kDa]

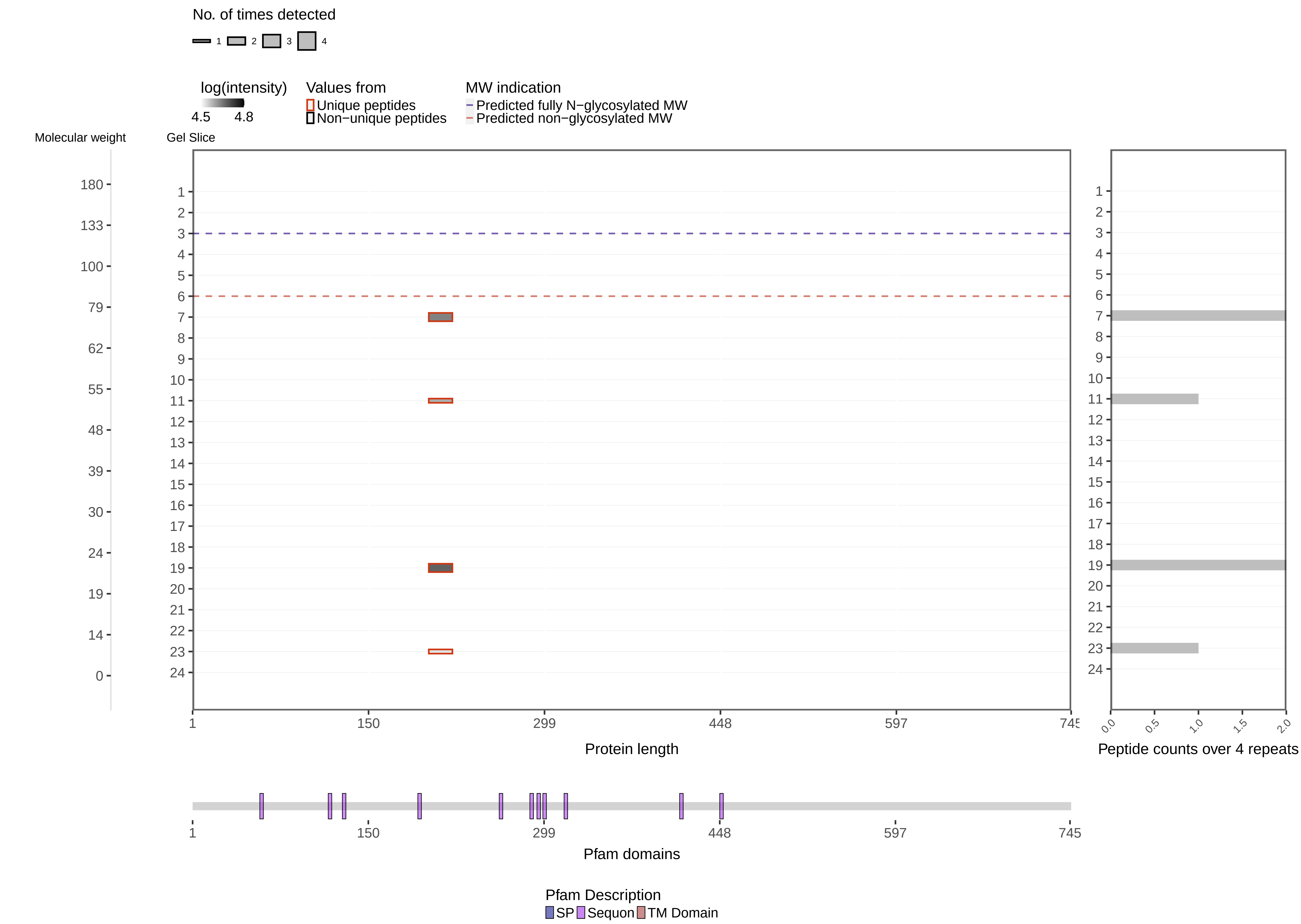


Protein sequence:

GSCRTSRRESPERGRPIDEALVIAKASIVESLRPFWADSYQPASLEEFTCHKKEALLLKDLVSSNETFPQVLFMGPSGSGKRTLAIALLREIYGDAICNISHDLKYFQIKETRPMQVVVPVSSSPHHIELNVQLEPNARYAIMAFVKQISSEYALTPEISRVNMKADYKVIVLYNVDKAAENIQHLIKWIMDCYSDACKLILCCEDDVSTLDSVKSRTKVVELAAPVTHEIMKVLIEIARKEDFELPMEFAAKIAAKSKQNLRRAIMALEACKAHNYPFAEDQPISIGWEDIVIELAAEILAD

NbL02g14360.1 beta−galactosidase_3 [MW − SP = 91.02 kDa]

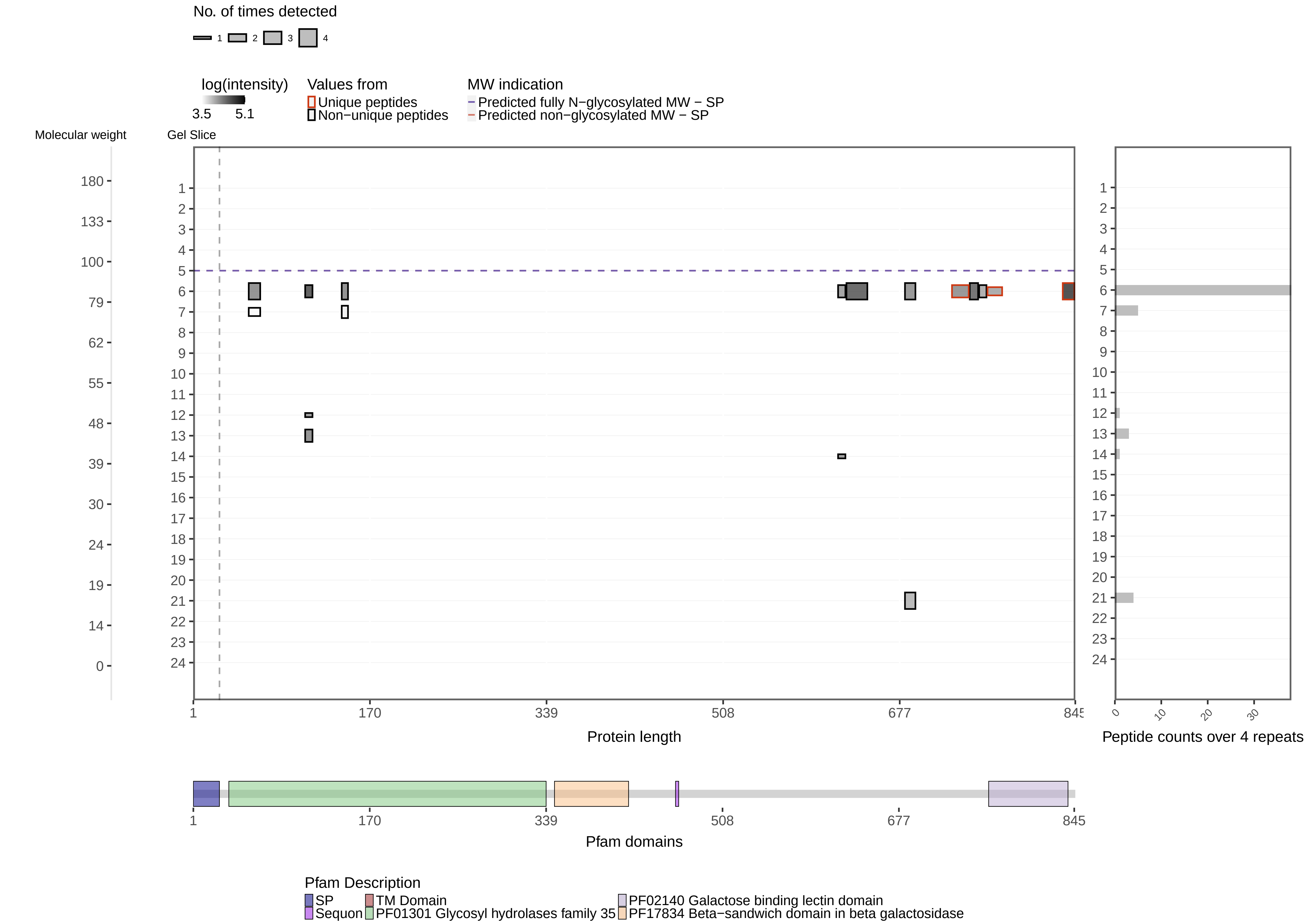


Protein sequence:

[illegible]

### NbL02g14320.1 probable_ubiquitin−conjugating_enzyme_E2_18 [MW = 22.16 kDa]

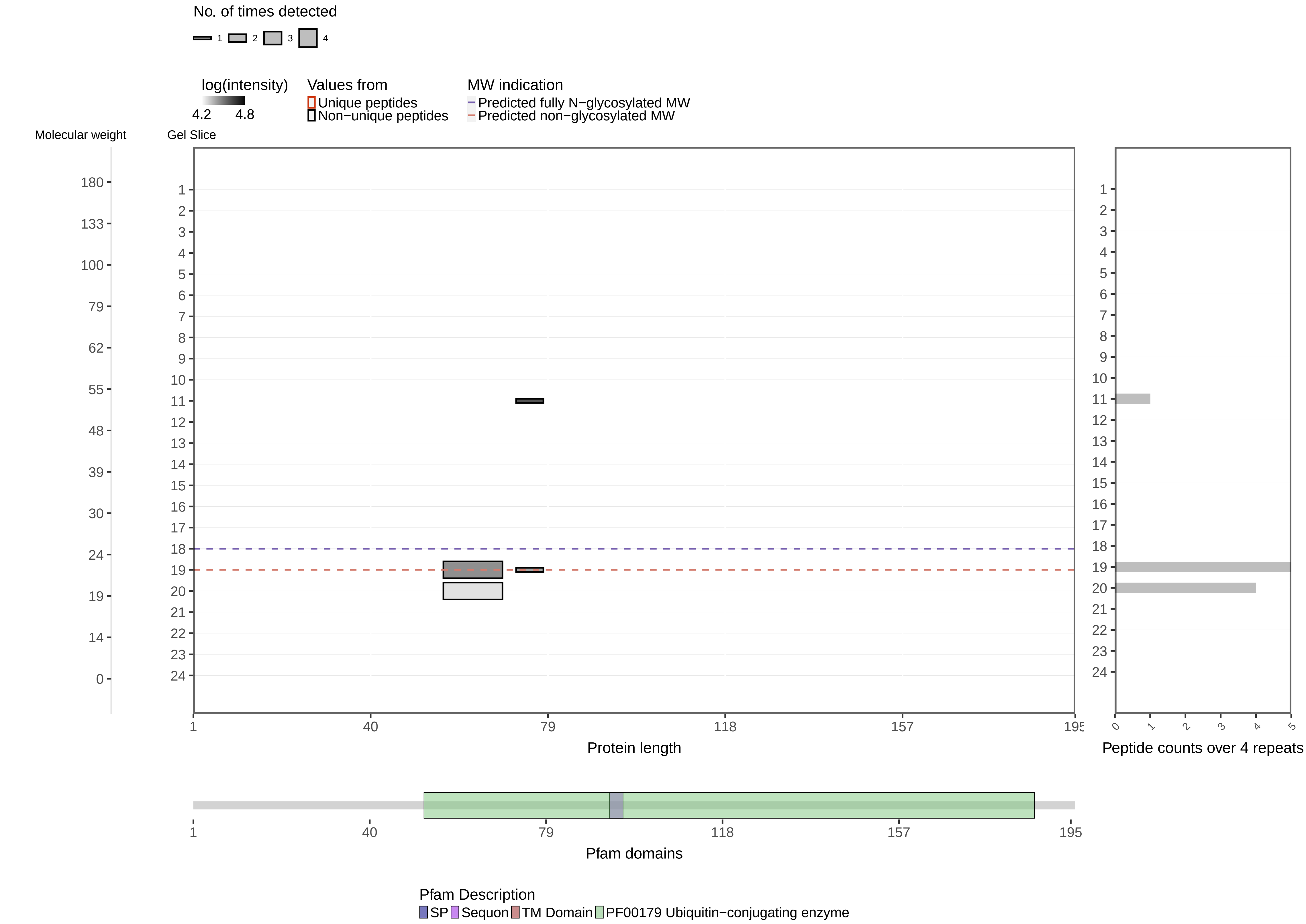


Protein sequence:

APGTLYANETYQLQVDFPEHYPMEAPQVIFVPPAPLHPHIYSNGHICLDILYDSWSPAMTVSSICISILSMLSSSTVKQRPEDNDRYVKNCRNGRSPKETRWWFHDDKV*

NbL02g14240.1 abscisic_acid_and_environmental_stress−inducible_protein_TAS14−like [MW = 22.14 kDa]

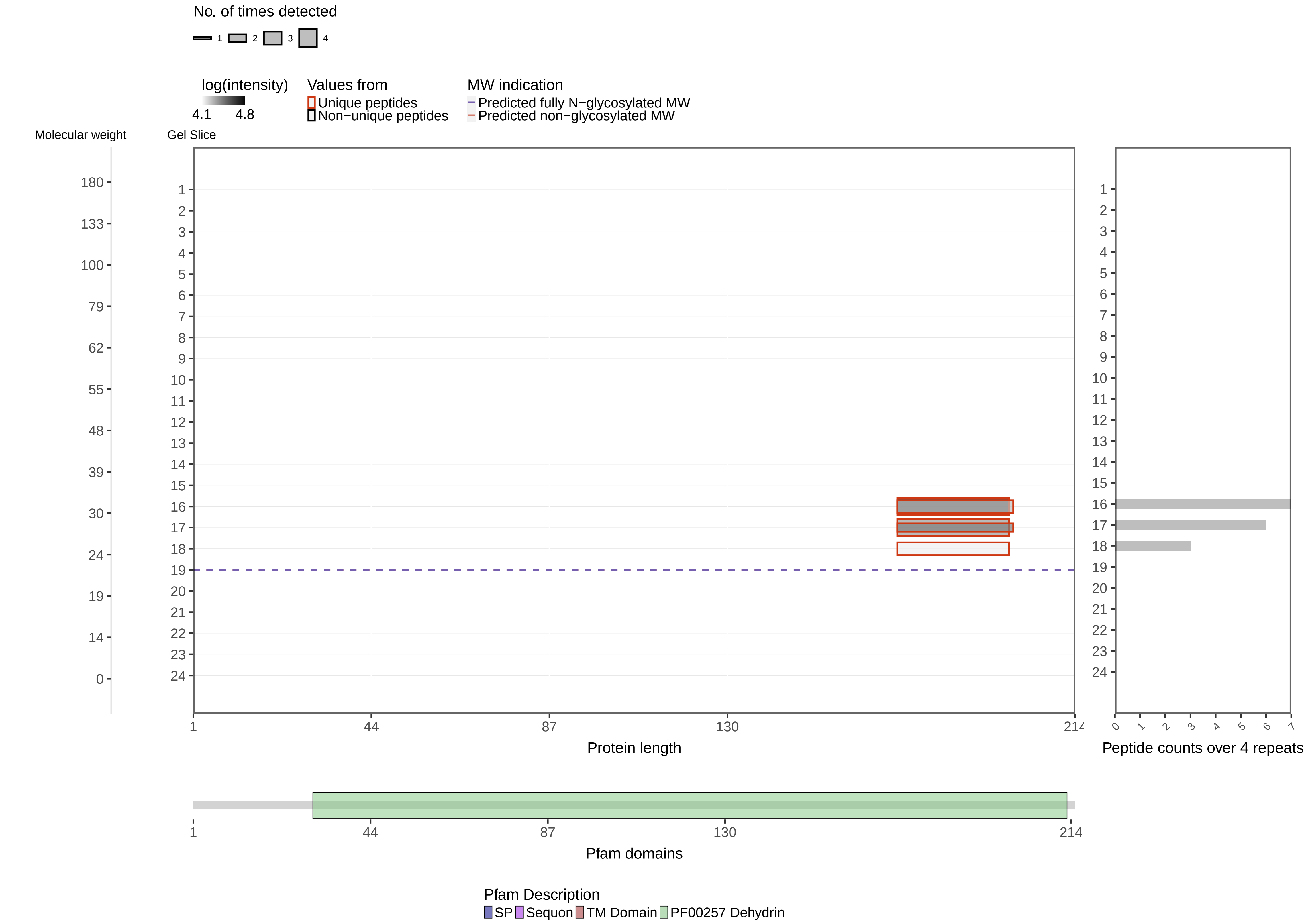


Protein sequence:

MGMGGEAYGTQGGTGMGAMGGEYATQGTGMGMGTGGMHTQHHEGQQQLRRSDSSSSSEDDGEGGRRKKGIKEKIMEKMPGGHAQQEGDQYNEQAQTTYSTEGGEKKGMMDKIKDKIPGMH*

### NbL02g13980.1 fructose−bisphosphate_aldolase_1_chloroplastic−like [MW = 42.75 kDa]

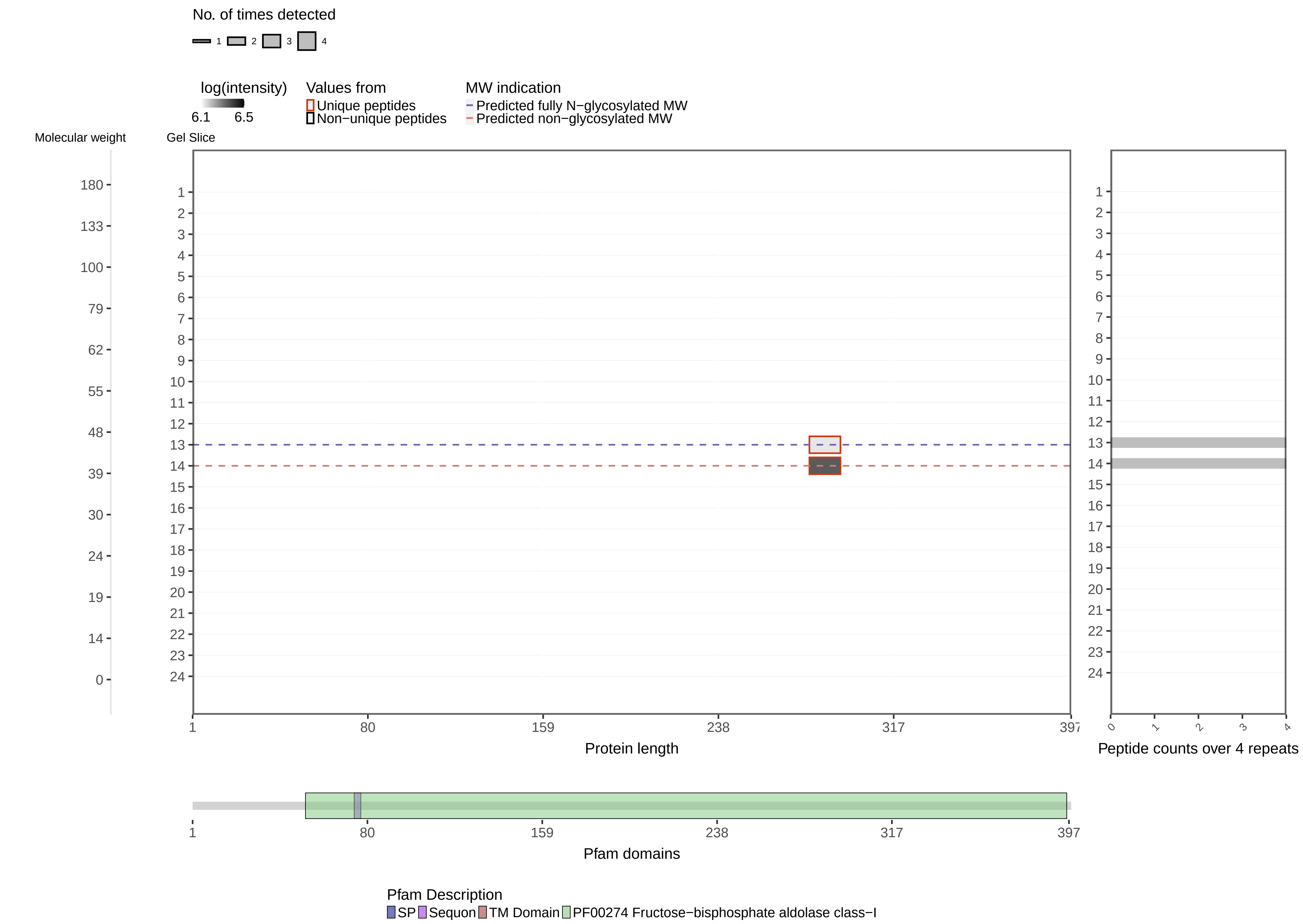


Protein sequence:

/VSIPNGPSALAVKEAAWGLARYAAISQDNGLVPIVEPEILLDGEHNIDRTFEVAKKVWAEVFFYLAENNVMFEGILLKPSMVTPGAECKERATPQQVADYTLNLLRQRIPPAVPGIMFLSGGQSEVEATLNLNAMNQGPNPWHVSFSYARALQNTCLKTWGGRPENVQAAQEALLVRAKANSLAQLGKYTGEGESEEAKKGMFVKGYVY*

NbL02g13710.1 naringenin2−oxoglutarate_3−dioxygenase [MW = 46.82 kDa]

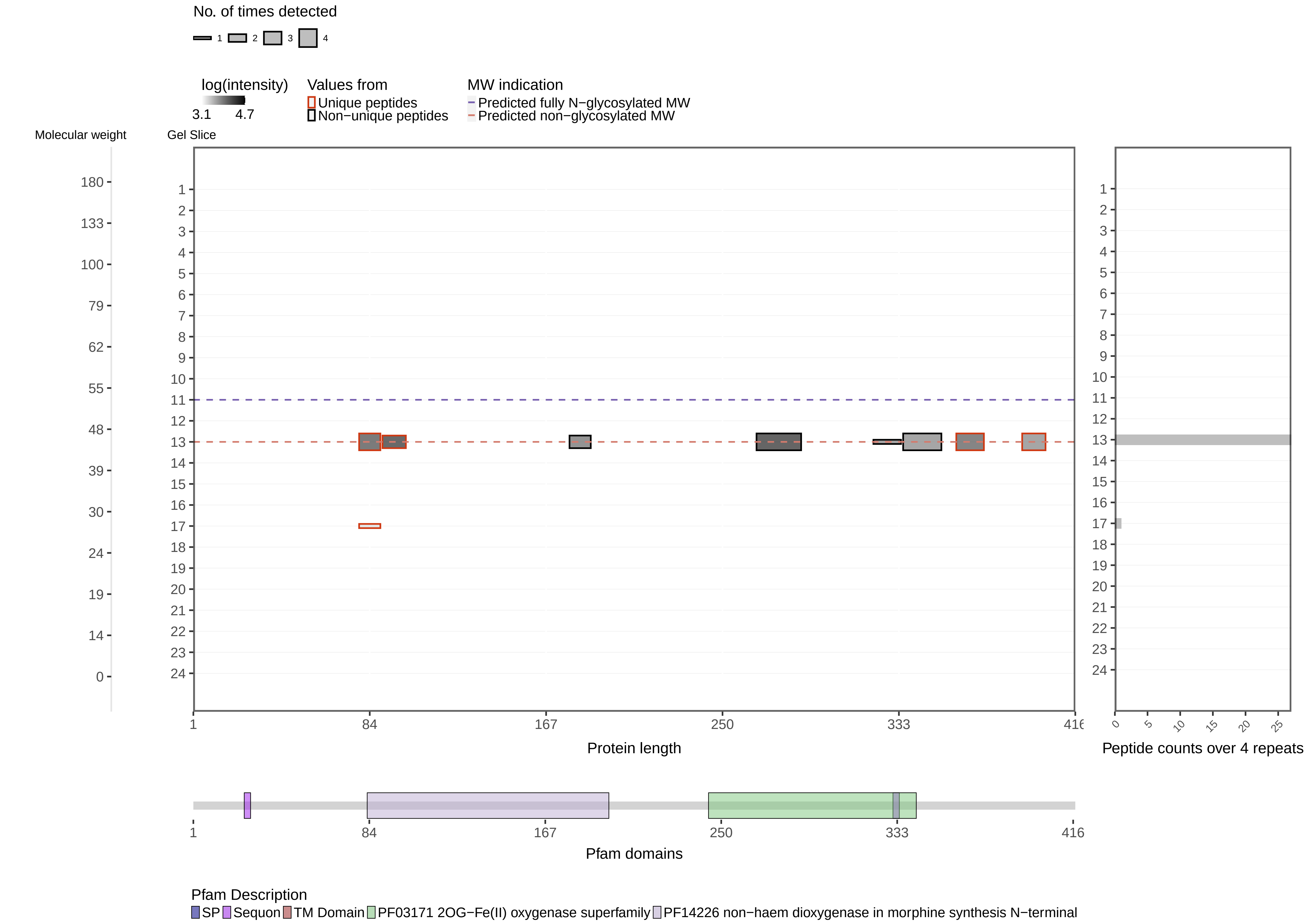


Protein sequence:

KPEGWIAVTQKYSEKLMELACKLLEVLSEAMGLEKEALTKACVDMDQKVVINFYPKCPQPDLTLGLKRHTDPGTITLLLQDQVGGLQATKDNGKTWITVQPVEGAFVVNLGDHGHFLSNGRFKNADHQAVVNSNSSRLSIATFQNPAPEAIVYPLKIREGEKAVMDEPITFAEMYRRKMSKDLELARLKKIAKEQQIQAEEAAEKAKLETKPIEEILA*

NbL02g13640.1 ferredoxin−−NADP_reductase_leaf−type_isozyme_chloroplastic [MW = 29.03 kDa]

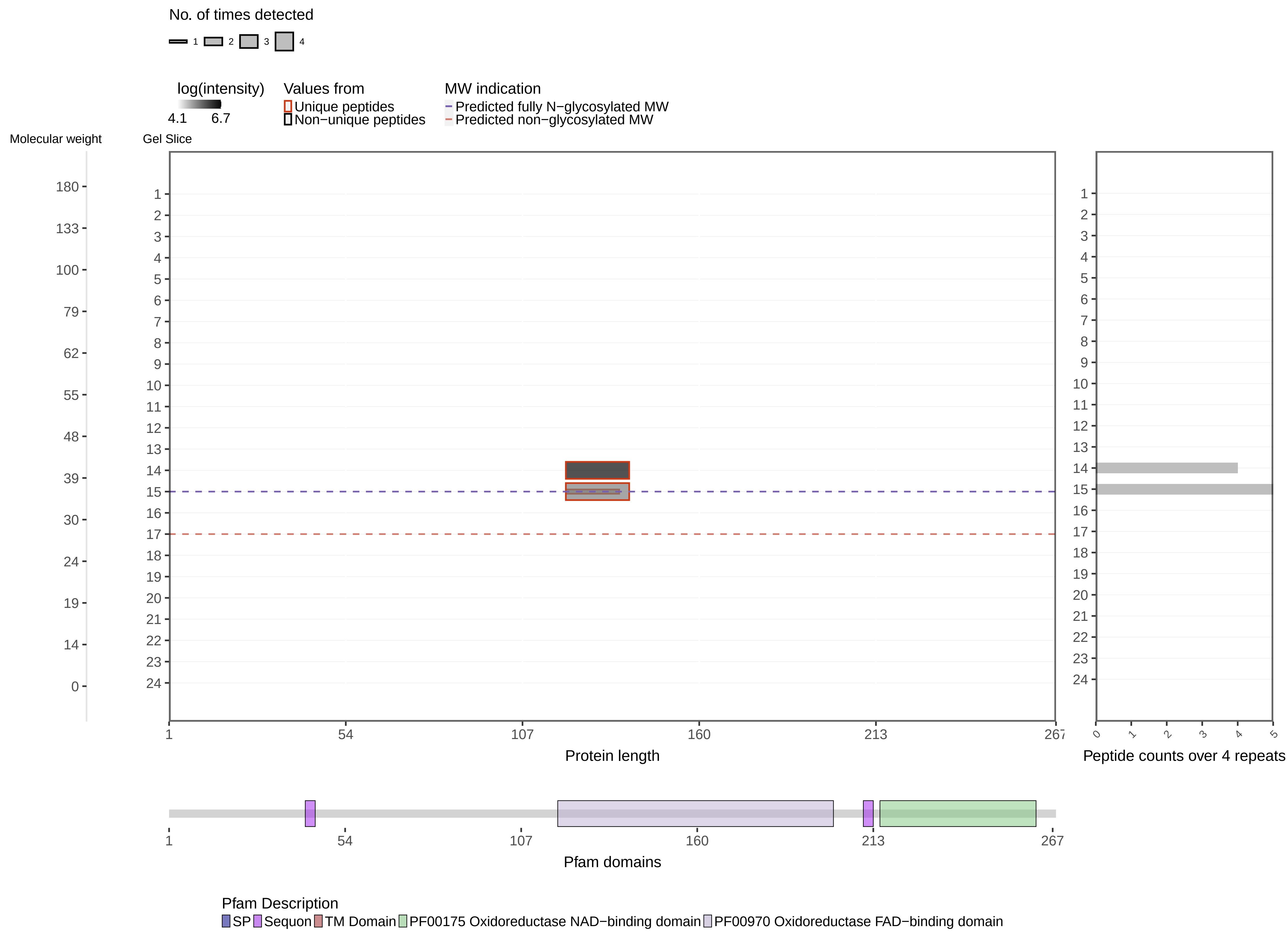


Protein sequence:

GQSIGVIADGVDANEKPHKLRLYSIASSALGDFGDSKTVSLCVKRLIYTNDKGEEVKGVCSNFLCDLKPGAEVKITGPVGKEMLMPKDPNATIVMLATGTGIAPFRSFLWKMFFEKHEDYKFNGLAWLFLGVPTSSSLLYKEVALL*

### NbL02g13580.1 26S_proteasome_non−ATPase_regulatory_subunit_4_homolog [MW = 42.39 kDa]

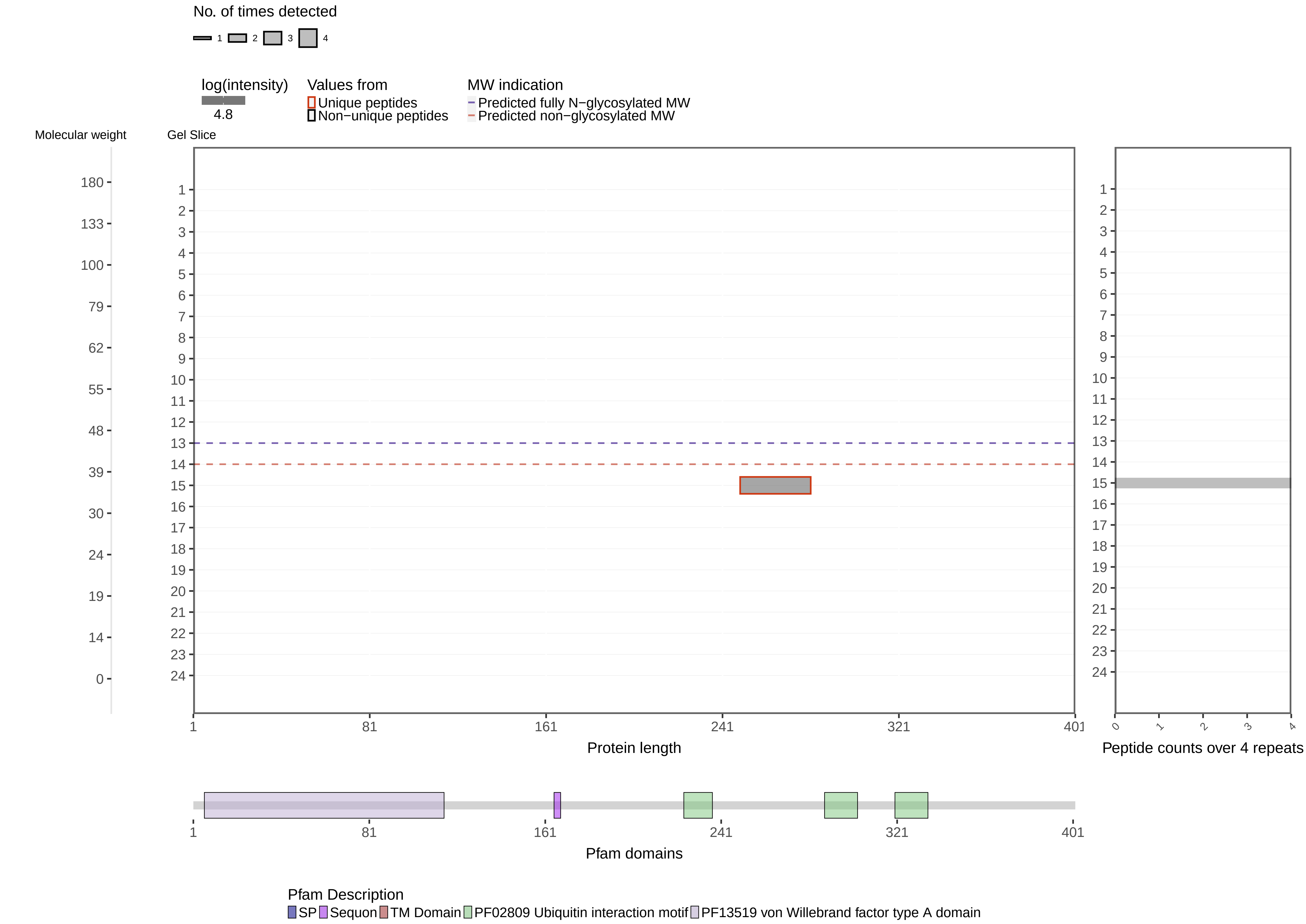


Protein sequence:

GDGEGGSGFAAAAAAAAAGGVSGFDFGVDPNLDPELALALRVSMEEERAROEAAAKKAAEETATOEKGETSODVTMAENVNAGTPEPENKATDLMEDENALLOOALAMSMDDASSNVATRDTDMSEAASEDODLALALOLSVODSTNDOSSOTDMSKLLADOSFVSSILASLPGVDPNDPSVKDLLASMOGOSEKKDEDTDKEOKEEKK*

### NbL02g13570.1 alpha−mannosidase [MW − SP = 13.24 kDa]

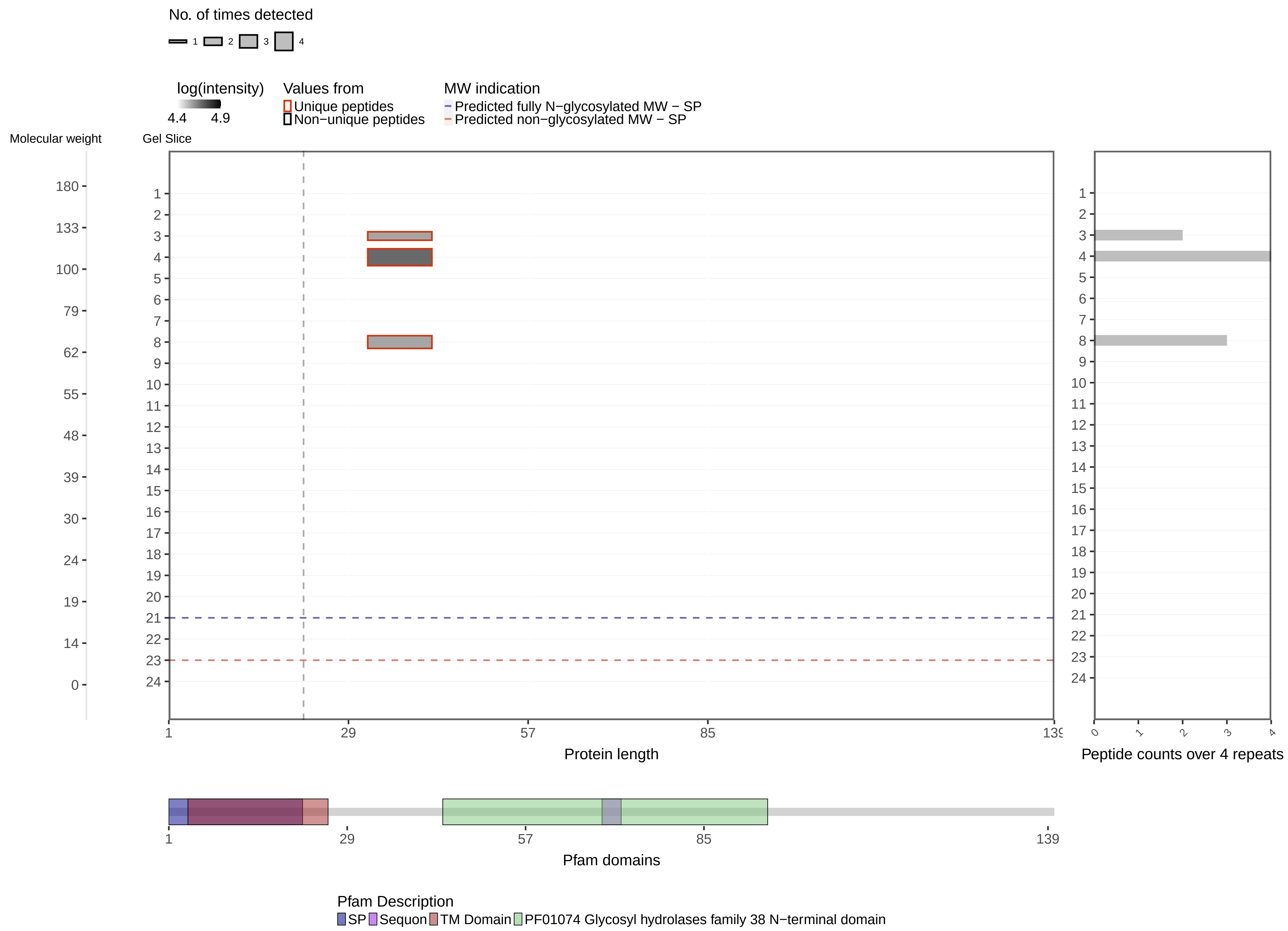


Protein sequence:

/LKTIDQYYVGSNNTIQGACVENVLDSVVMSLRRDPNRKLSLLKWHFSIVVDTAKPRNSRRSEKPCSFWSAGIHKWWLVYAR*

### NbL02g13470.1 3−dehydroquinate_synthase_chloroplastic [MW = 55.36 kDa]

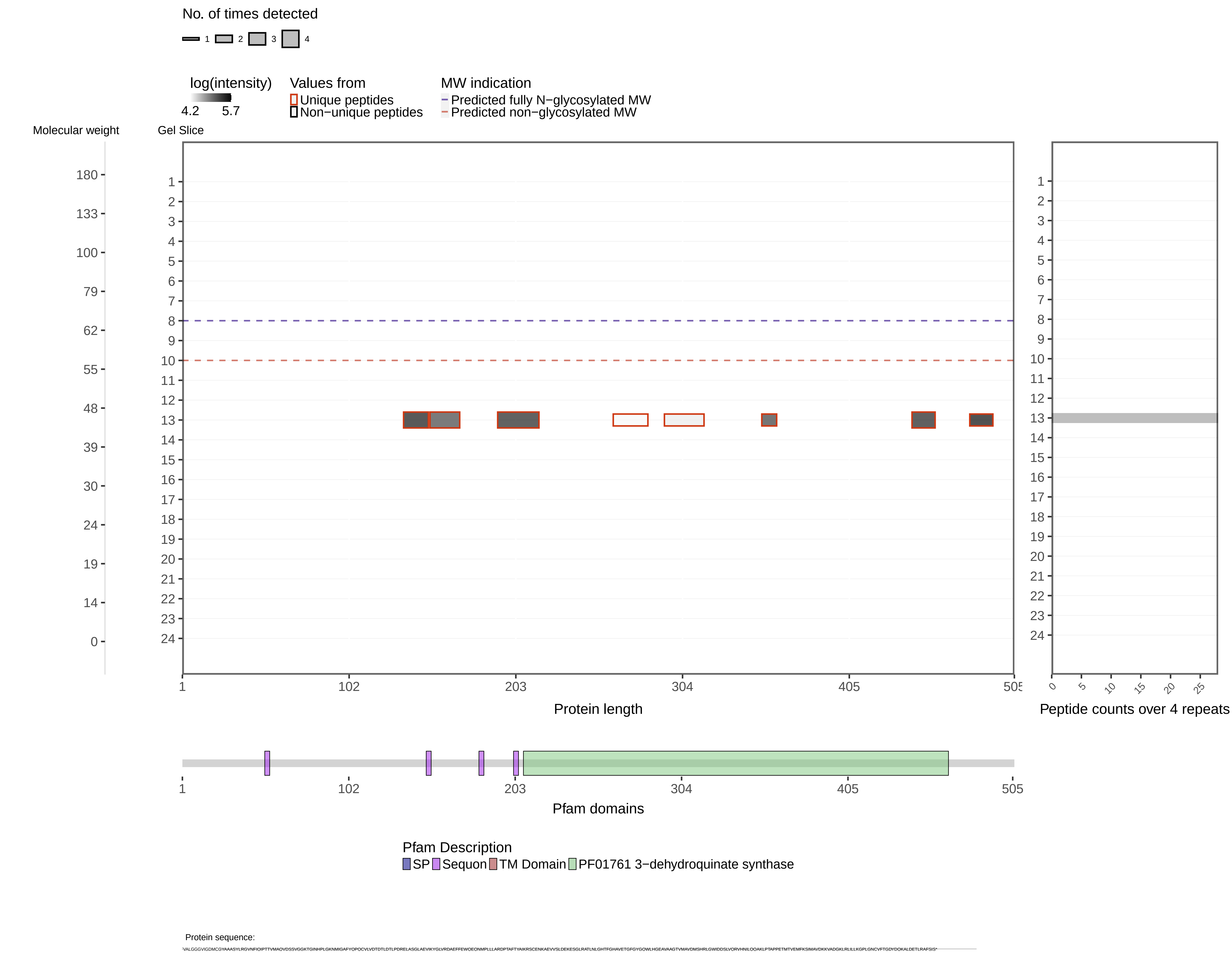

### NbL02g13210.1 serine_hydroxymethyltransferase_mitochondrial−like [MW = 57.36 kDa]

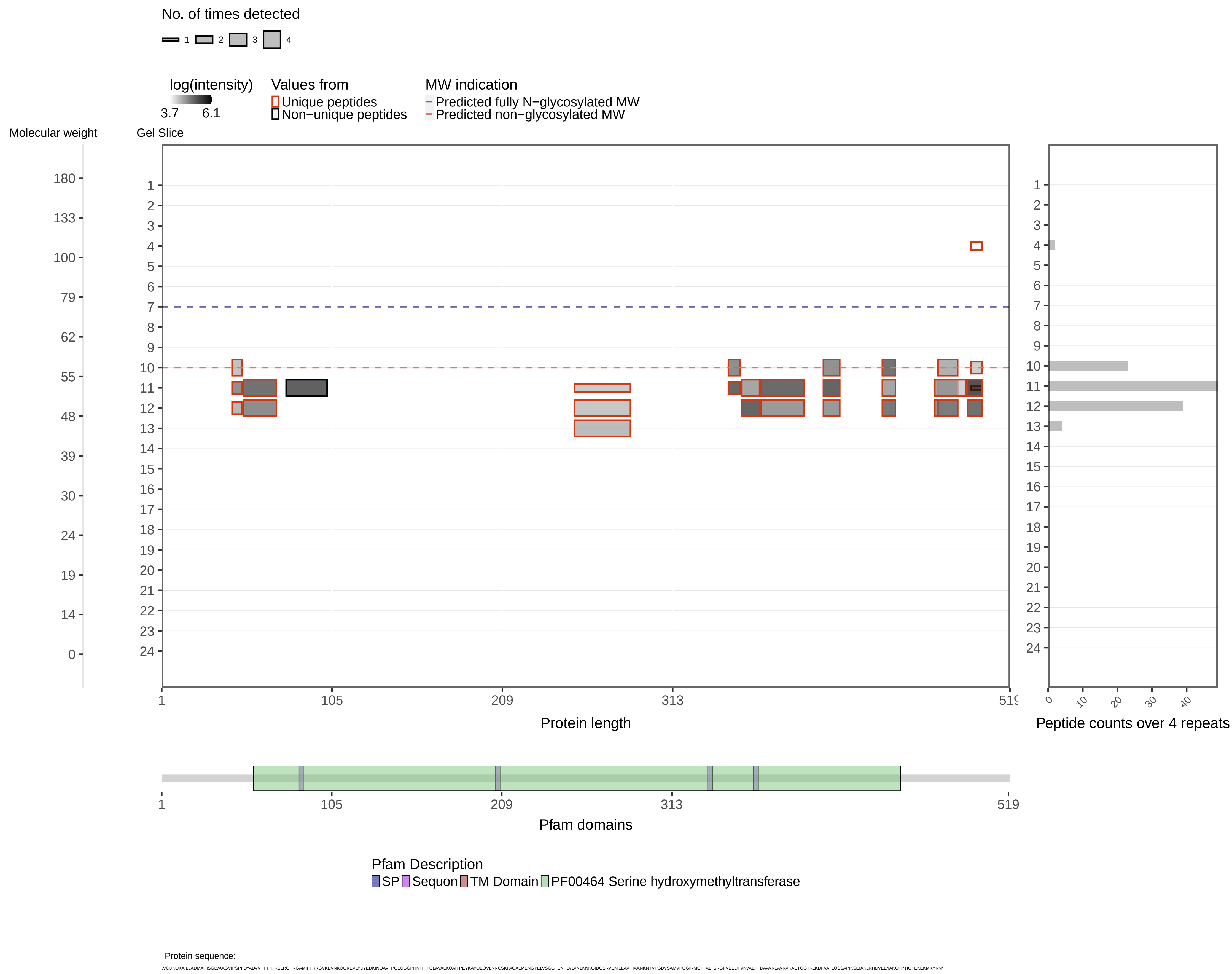

NbL02g13200.1 probable_cytosolic_oligopeptidase_A [MW = 94.36 kDa]

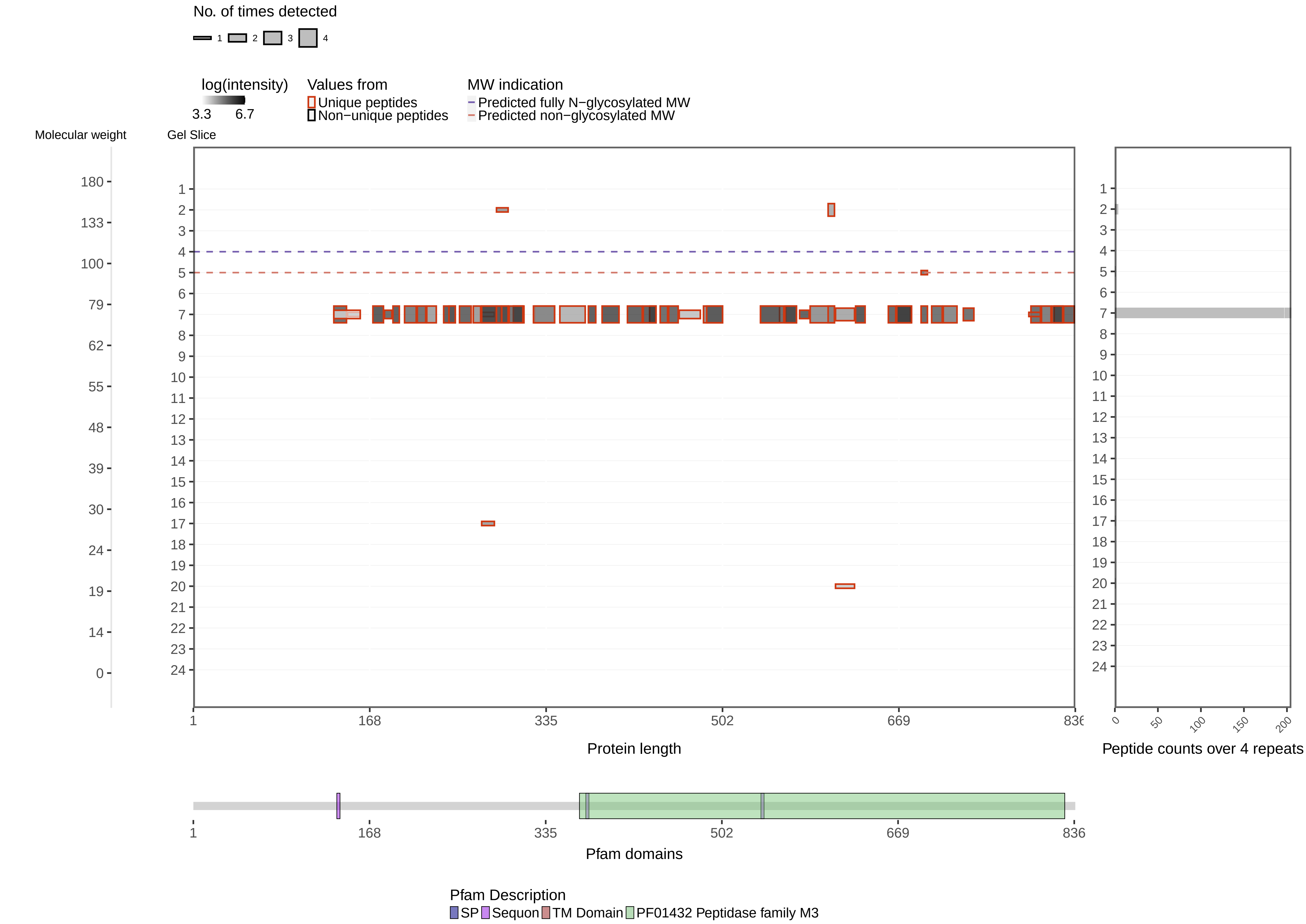


Protein sequence:

LLGYNNYAEVSMAMKMATVDKAEELLEKLRRSSWDPAVKDLEDLKEFCKSQGAPEADDLNHWDITFWSERLRESKYELNEEELRPYFSLPRVMDGLFSLVNLLFGINVEPADGLAPVWNNDVRFYRVNDSSGSPIAYFYFDPYSRPSEKRGGAWMDEVVSRSRVLSLDGASARLPVAHMVCNQMPPVGEKPSLMTFREVETVFHEFGHALQHMLTKQDEGLVAGIRGVEWDAVELPSQFMENWCYHRDTLMGIAKHHETGESLPEDIYKKLLTARTFRAGTLSLRQLF

### NbL02g13020.1 glutathione_S−transferase [MW = 23.8 kDa]

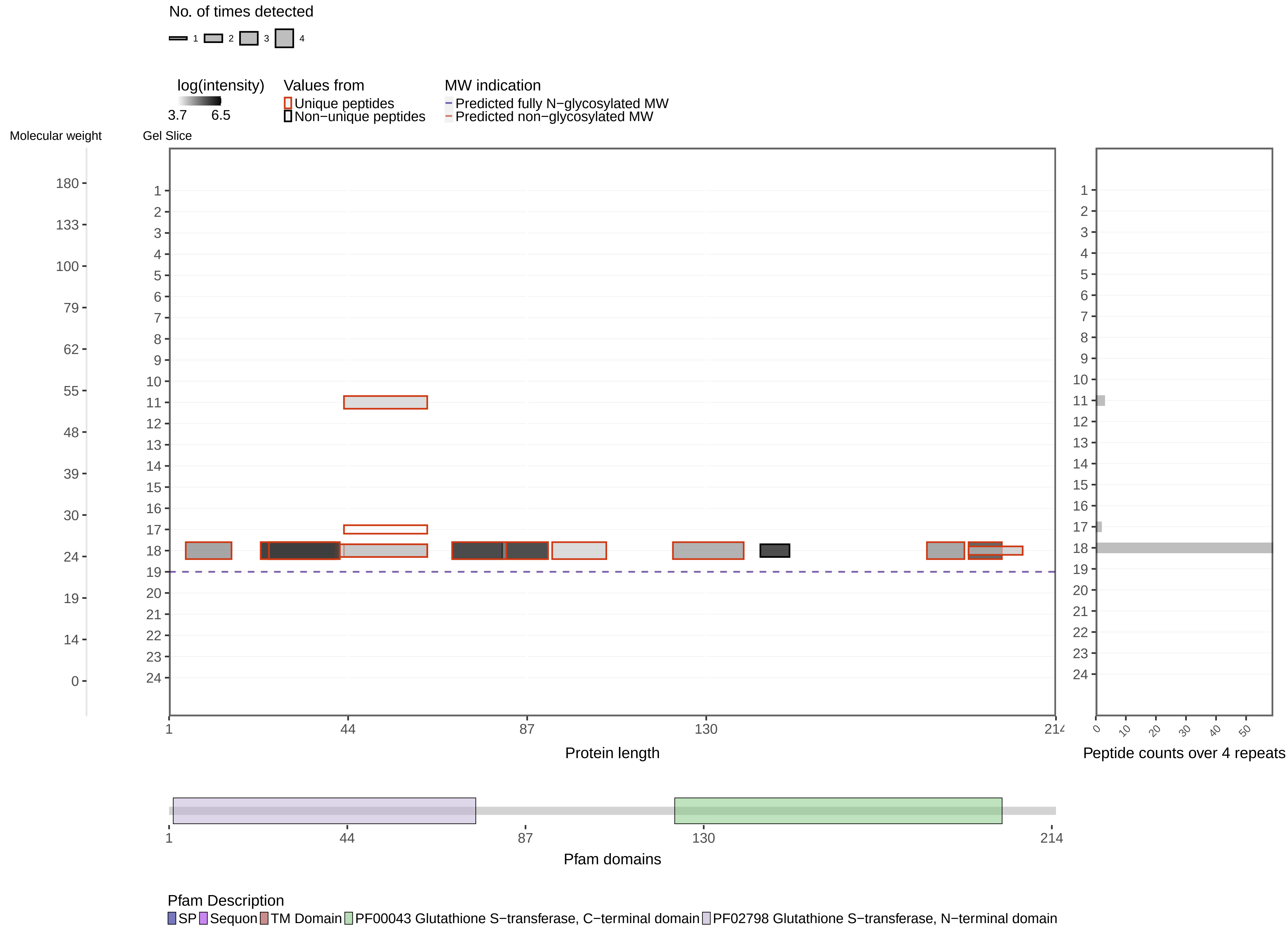


Protein sequence:

IMSVWMEVESTKFDPIGSKLCFELAVKPMLGMVTDDAAVAENEEKLGKILDVYESRLKDSKYLGGESFTLADLHHAPVVHYLMGTKVKSLFDARPHVKAWCADIVGRPAWSKALELHKQ*

### NbL02g13010.1 uncharacterized_protein_OsI_027940 [MW = 21.58 kDa]

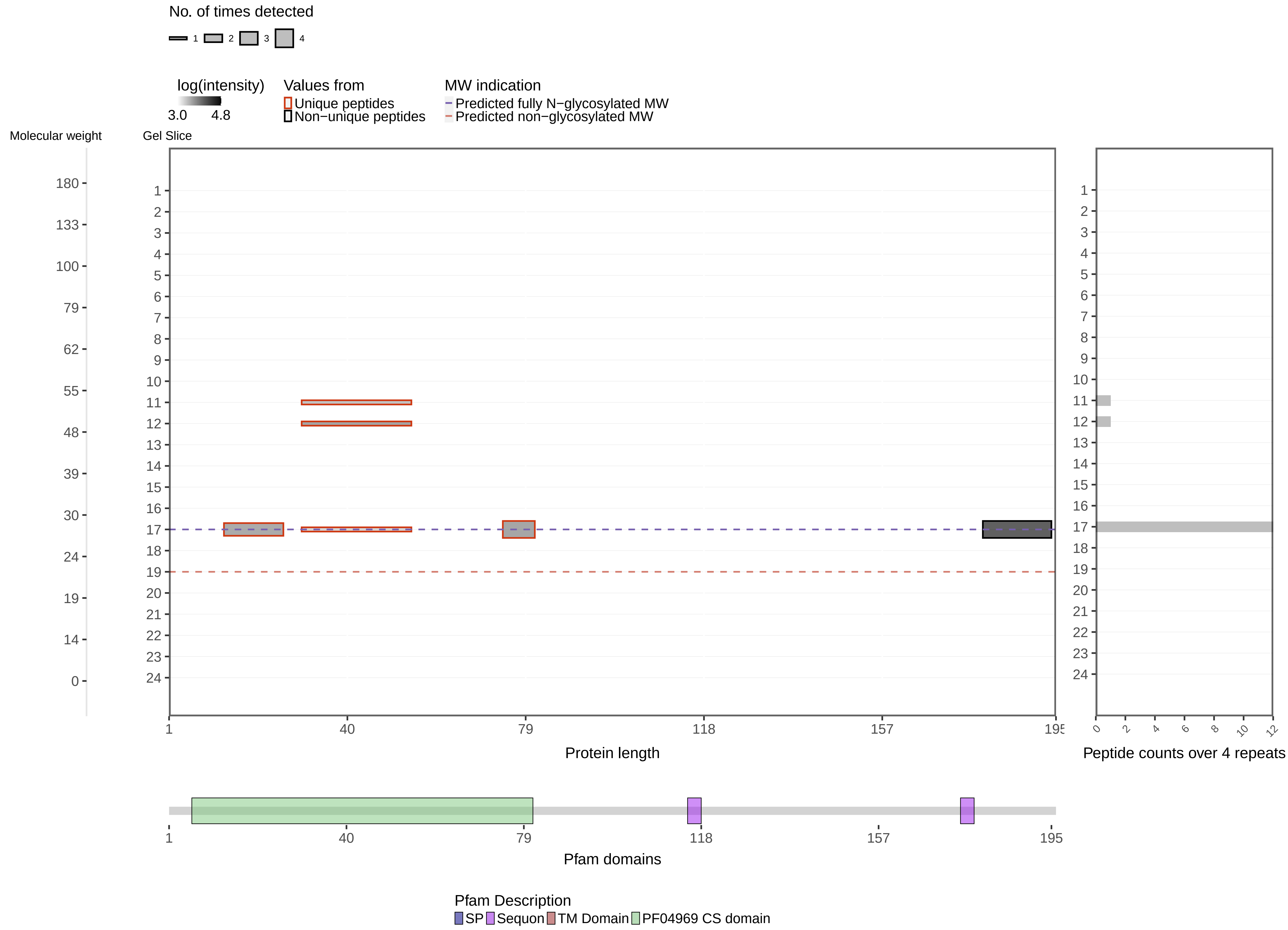


Protein sequence:

NKLLHGDEKAPHYVKVDWDKWVDEDDDNDTAAPGDFDMNSMDFSKFGDMGGMGLGDDAMGDDLDDSDDEEEEVTKPSETVEGEPKEQNNSKVGGEAAEGKAEAAPSS*

### NbL02g12890.1 oxygen−evolving_enhancer_protein_1_chloroplastic−like [MW = 35.34 kDa]

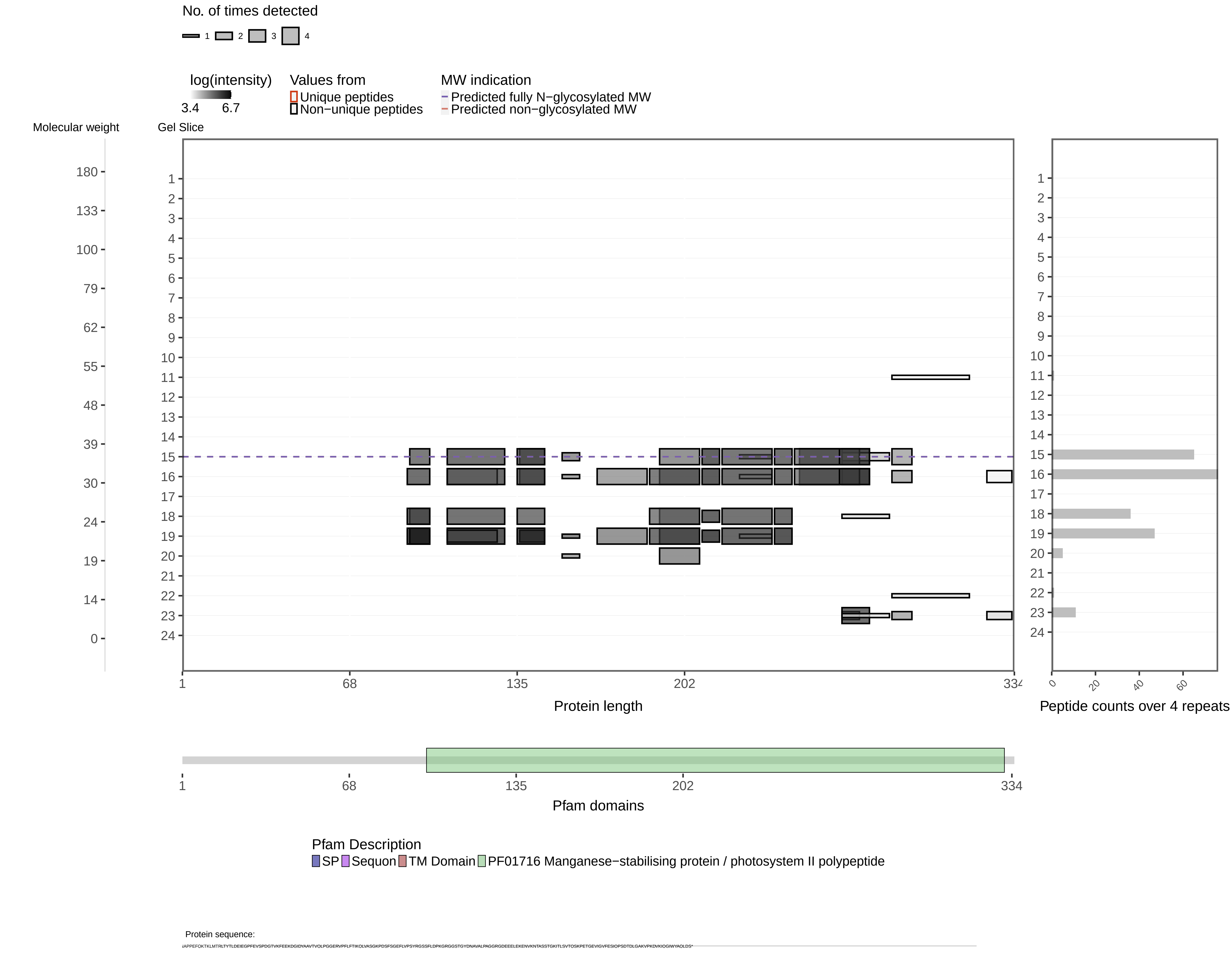

### NbL02g12820.1 5−oxoprolinase−like [MW = 137.99 kDa]

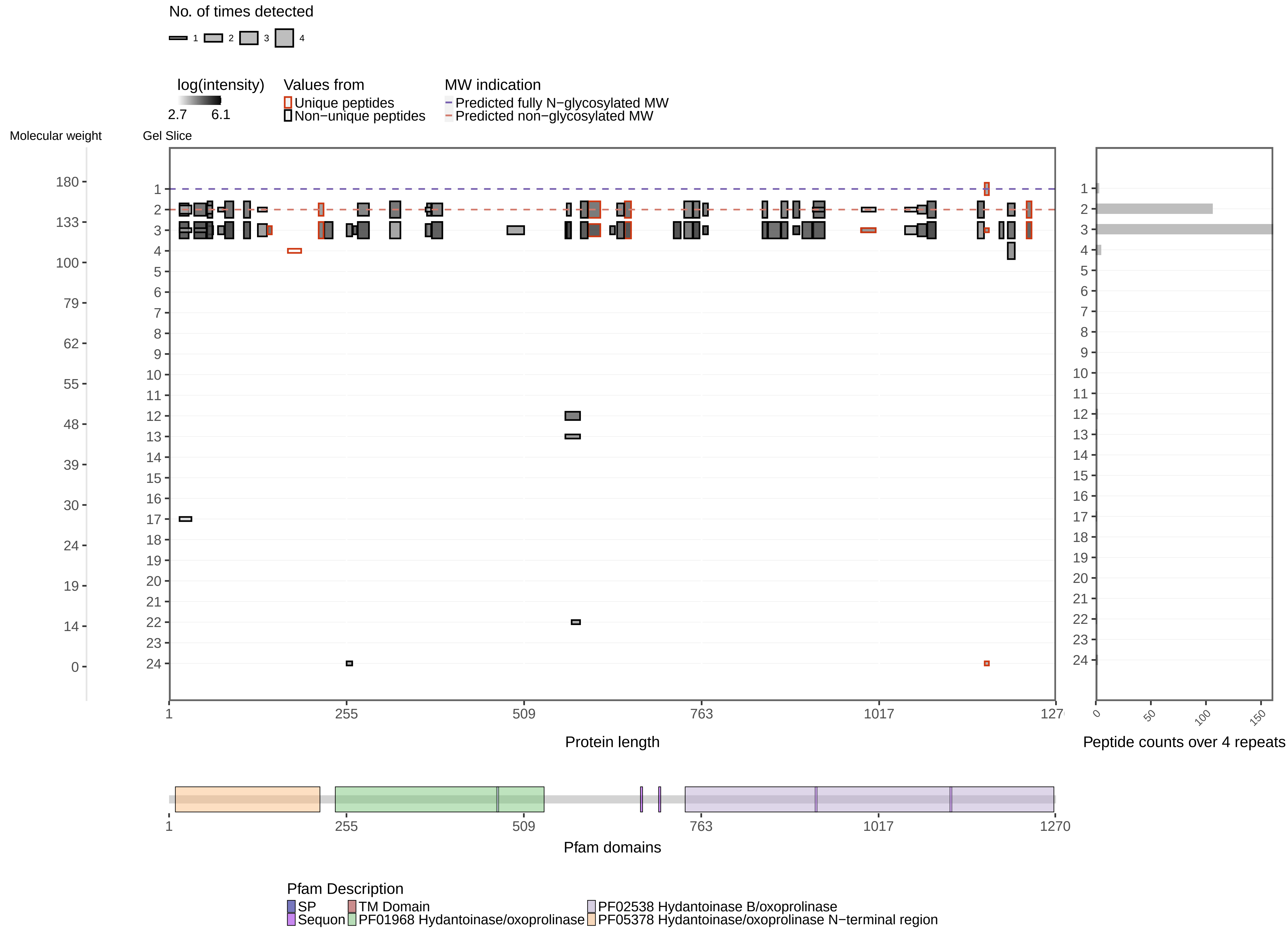


Protein sequence:

### NbL02g12380.1 uncharacterized_monothiol_glutaredoxin_ycf64−like [MW = 19.08 kDa]

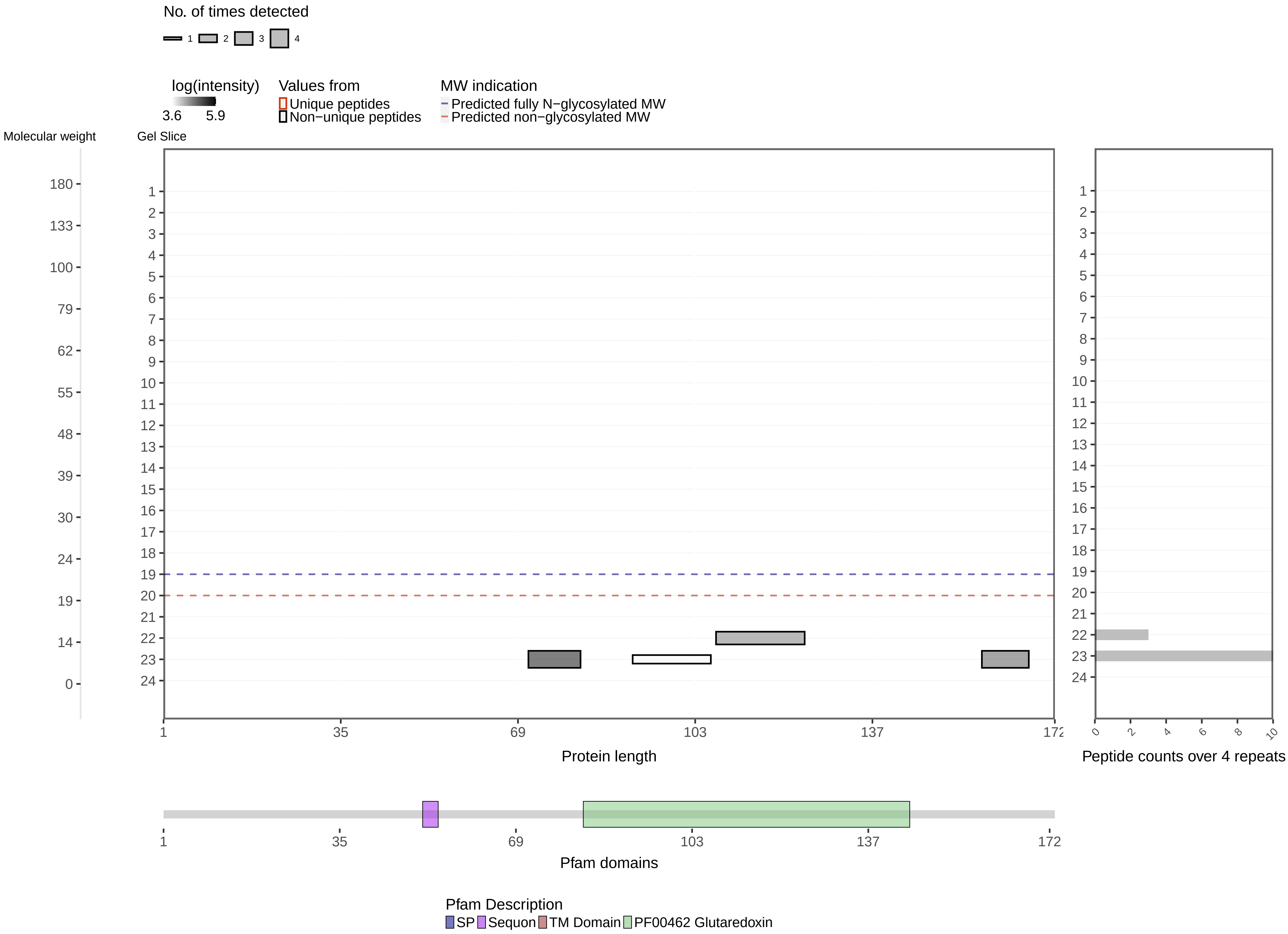


Protein sequence:

DKVVTSQKIVLFMKGTKDFPQCGFSNTVVQILKSLNAPFETINILENEALRQGLKEYSSWPTFPQLYIDGEFFGGCDITVEAYKSGELQELLERTLCS*

### NbL02g12270.1 uncharacterized_LOC109215327 [MW = 22.24 kDa]

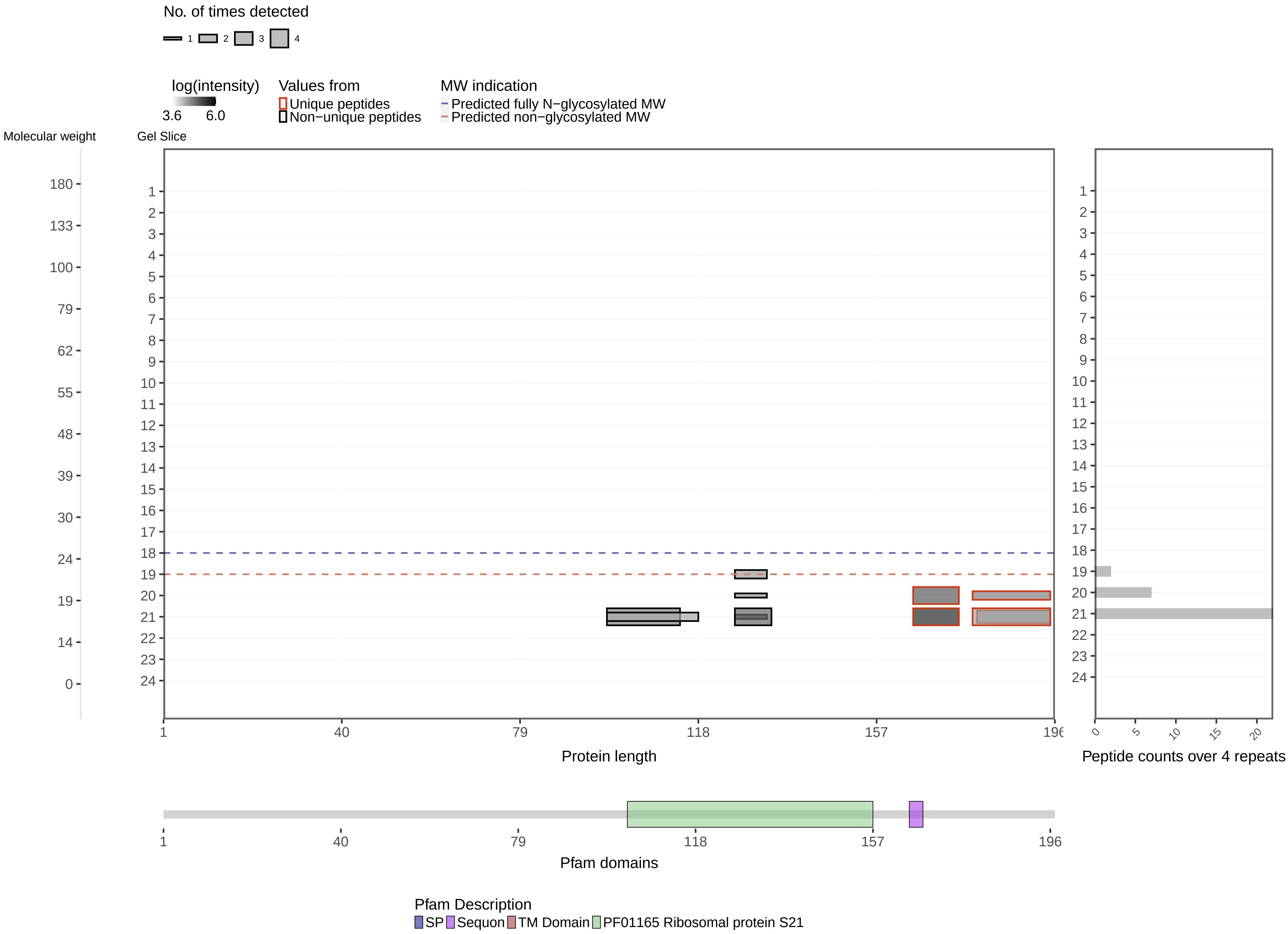


Protein sequence:

.ANANTLYFRSGYNVQVIVDDNEHEEKLVGRFRREVMRAGVIQECKRRRYFENKHDKLKRKSREAAKRNRKRRGPPRNFSDDKQEASKNKRDDEGEDNWELPDGGLPY*

### NbL02g12220.1 actin−histidine_N−methyltransferase [MW = 57.45 kDa]

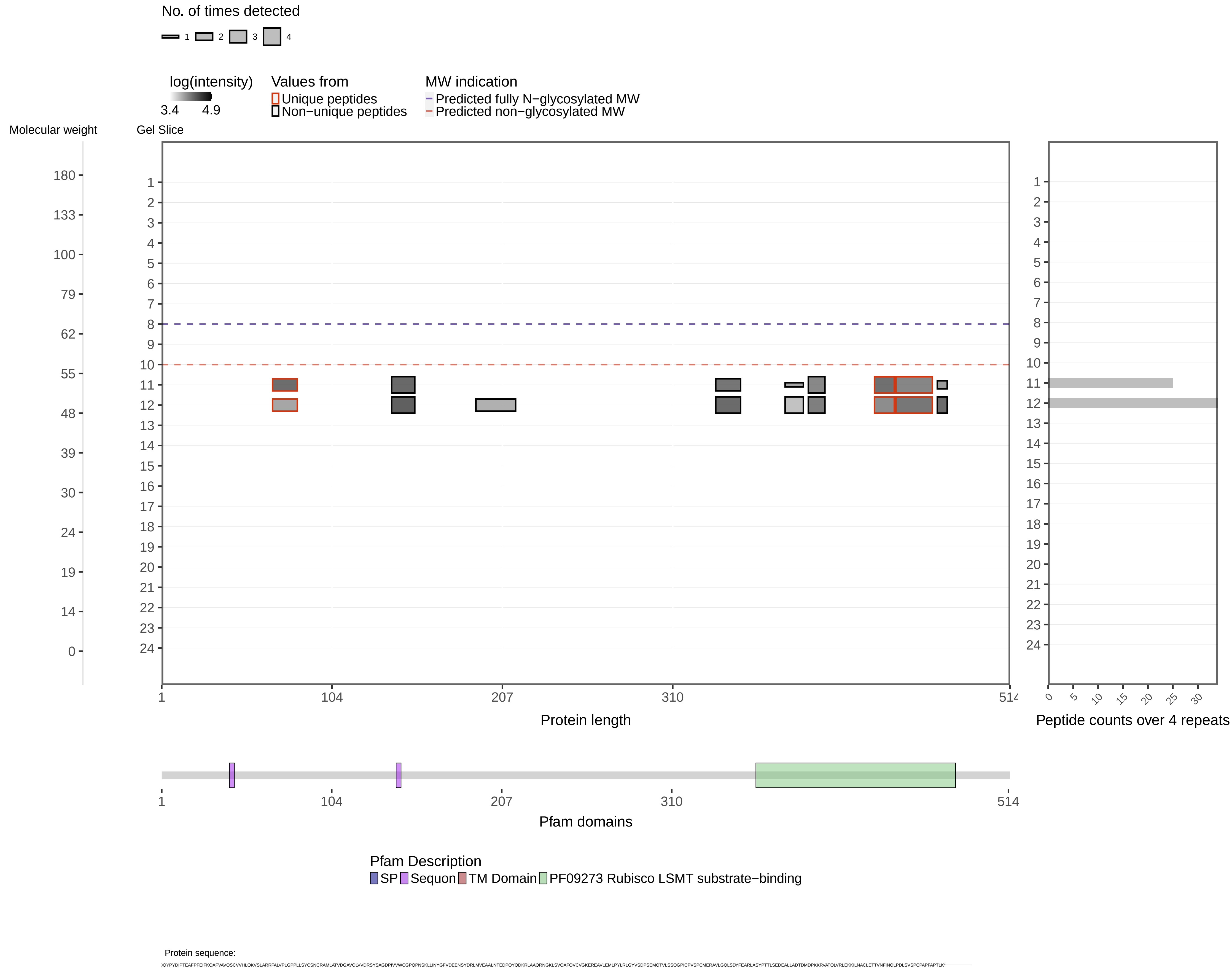

### NbL02g12080.1 carbonic_anhydrase_chloroplastic−like [MW = 28.63 kDa]

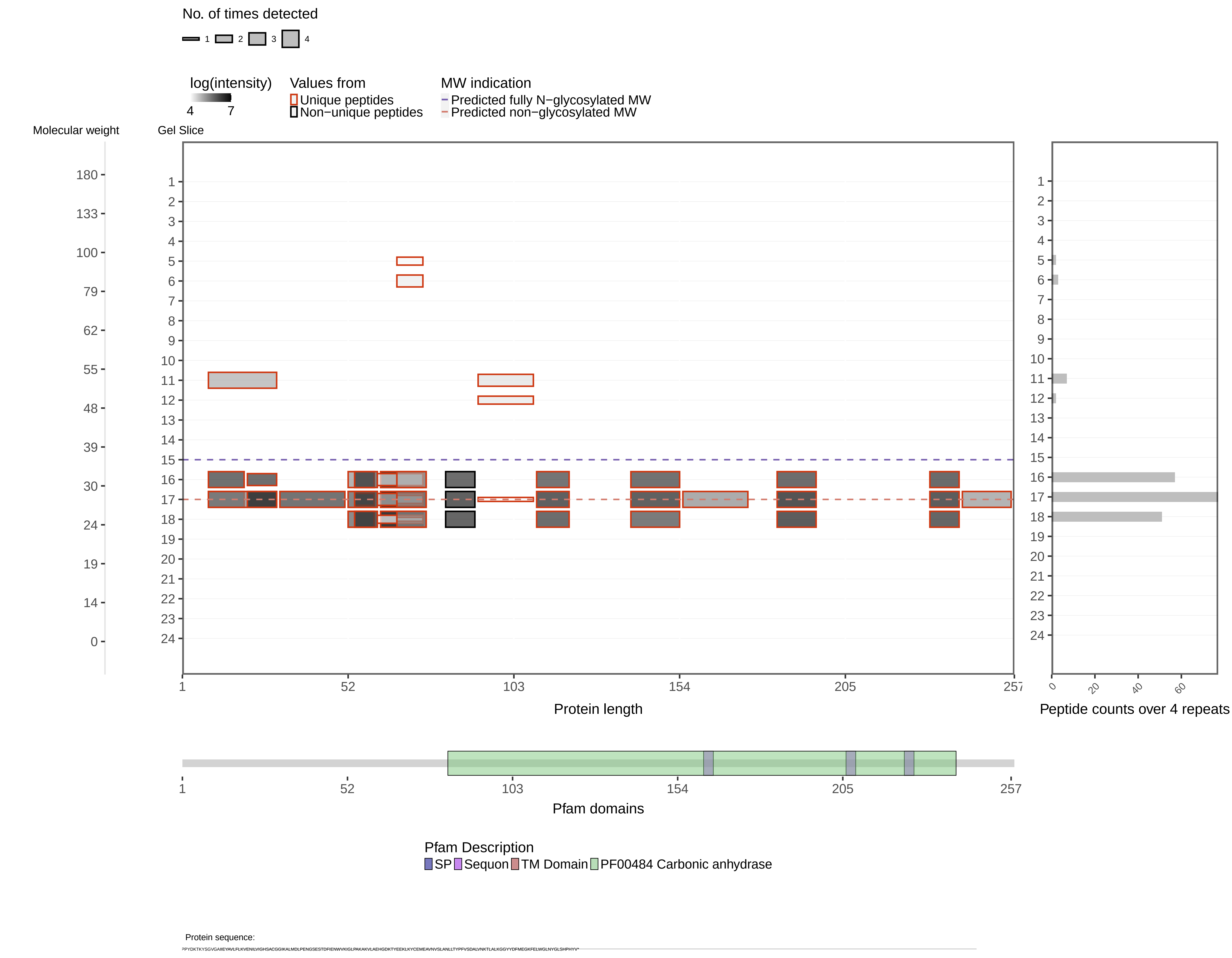

### NbL02g12060.1 putative_glucose−6−phosphate_1−epimerase [MW = 34.78 kDa]

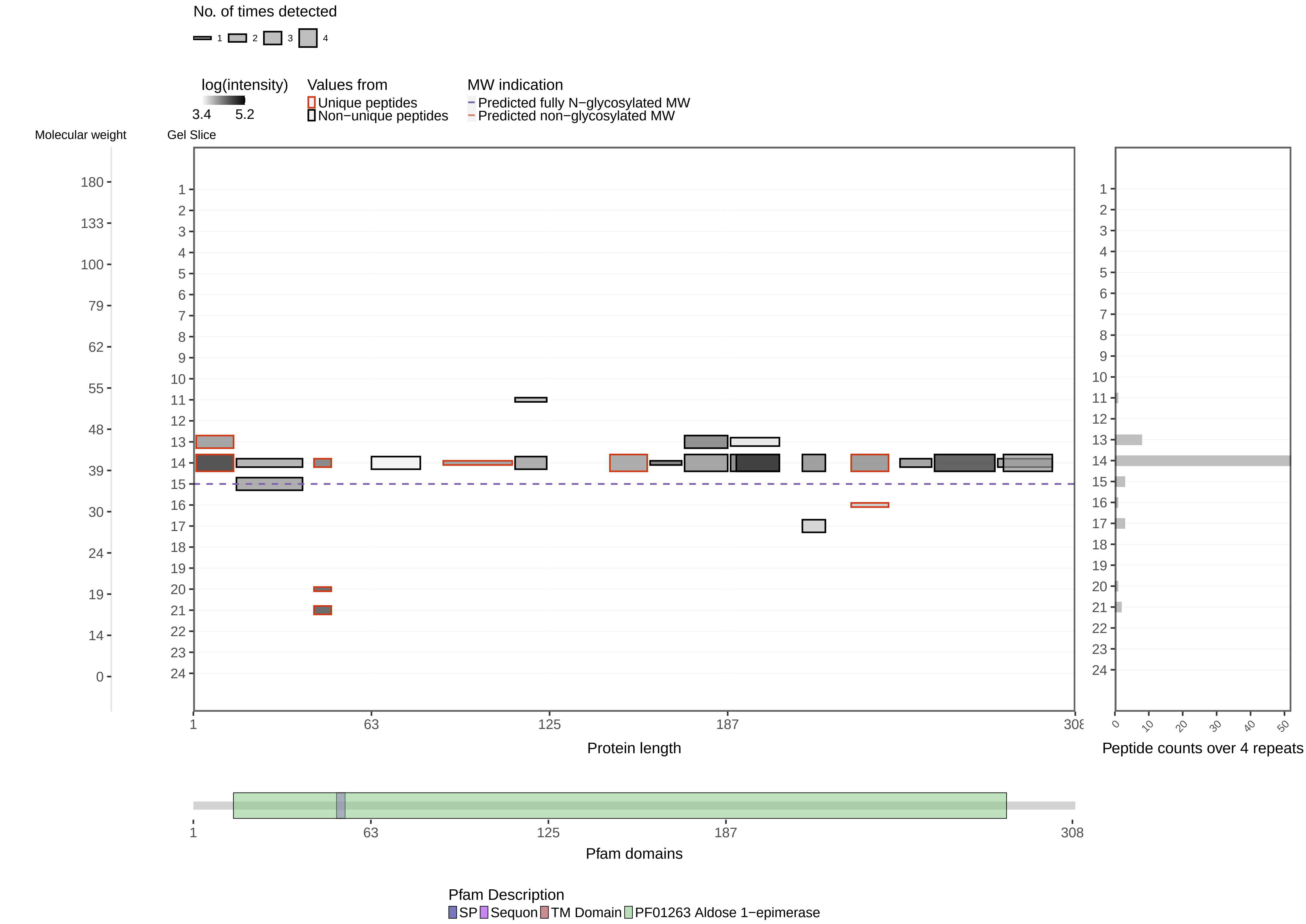


Protein sequence:

IPRVRNIDNKPFSFTISLRNYLSVSDISEVRVEGLETLDYFDNLQQRERYTEQADAITFDSEIDRVYLSTPKKIAIIDHEKKRTYVIQKEGMLDAVVWNPWNKKAKSLPDLGDEDYKKMLCVDSAAVETPIVLKPSEEWKGRQELSTVSSSYCSGQLDPRKVLYGLA*

### NbL02g11580.1 uncharacterized_LOC104228188 [MW = 20.53 kDa]

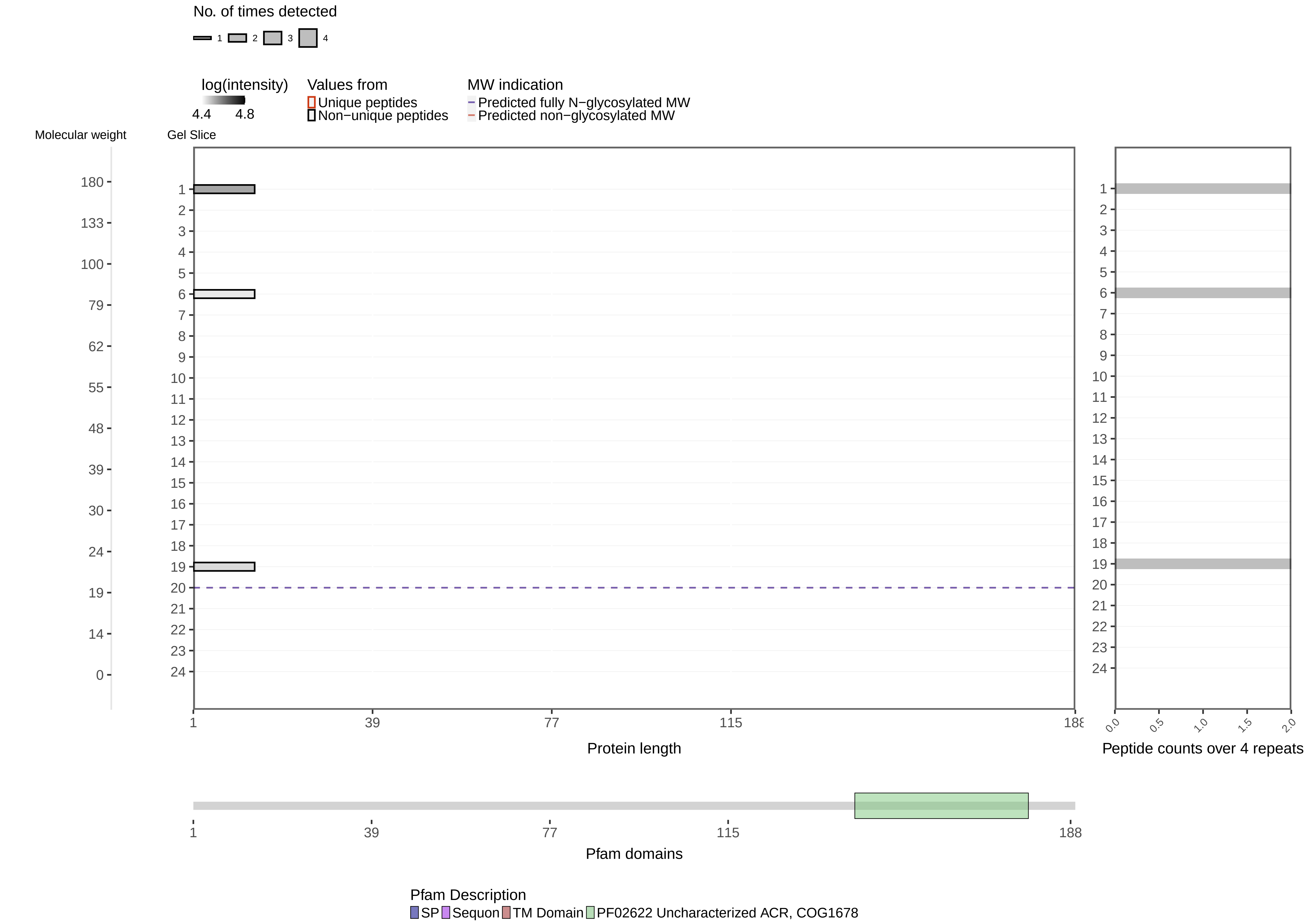


Protein sequence:

[illegible]ARLVAGEKALRSEEPSSVVNPDTVDDLPPPPAVTIGSKWAHTIHEPEKGCLLIATEKLDGVHIFERTVVLLLSMGPIGPTGVILNRPSLVSIKEMRSSVLGRVRDLCE*

NbL02g11570.1 UDP−glucose_6−dehydrogenase_3 [MW = 58.41 kDa]

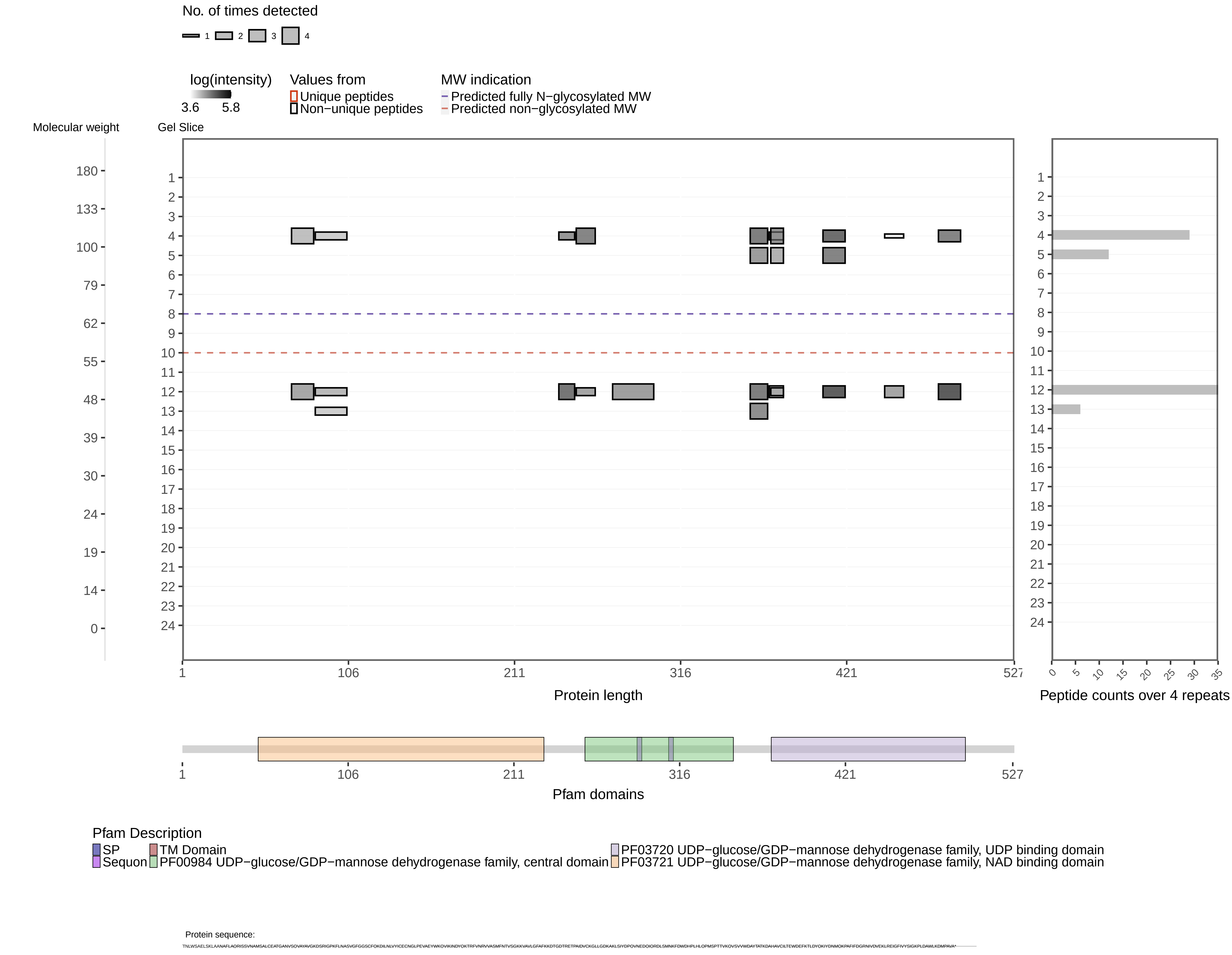

NbL02g11490.1 ras_GTPase−activating_protein−binding_protein_2−like [MW = 50.32 kDa]

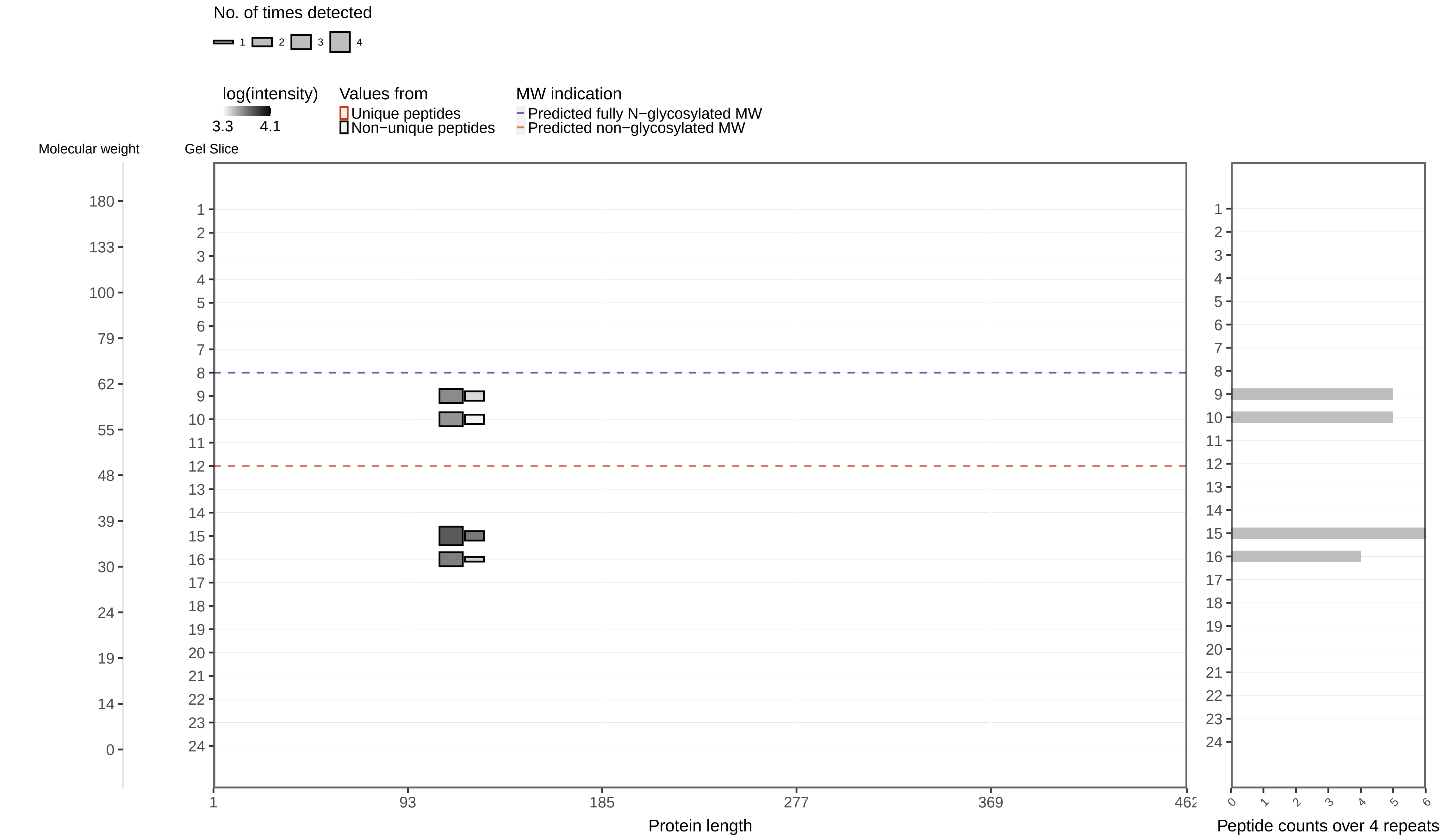


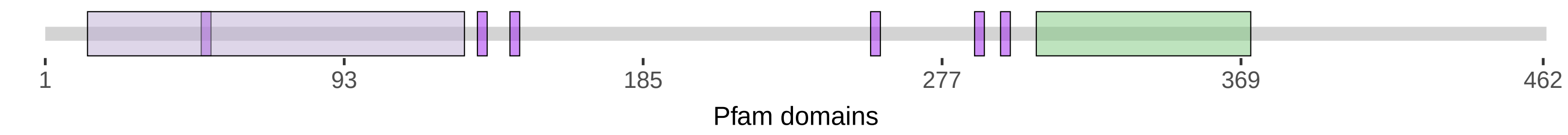


Pfam Description

SP Sequon TM Domain PF00076 RNA recognition motif. (a.k.a. RRM, RBD, or RNP domain) PF02136 Nuclear transport factor 2 (NTF2) domain

Protein sequence:

ENSVAQEDAPKKSYASIVSSQTKKGPTKIYVPTNTSRMAPAKAVKQPVAAVAQTPAPEASHPTVPNGTNVPETNDTEDEAEGHSIYVRNLPLDVTVAQLETEFKKFGPIKQGGVQVRSNRQQGFCFGFVEFEDVSSMHSAIQASPITIGGRQAVVEMKRTTTRVGSARGRFPPGRGGFRNDNFRGRGNFGGGRSYGRNEFGGRDFSGRGRGQGGRAGEGYQQVRGRGGRRGGPGQSSAPA*

### NbL02g11370.1 protein_FLX−like_2 [MW = 51.25 kDa]

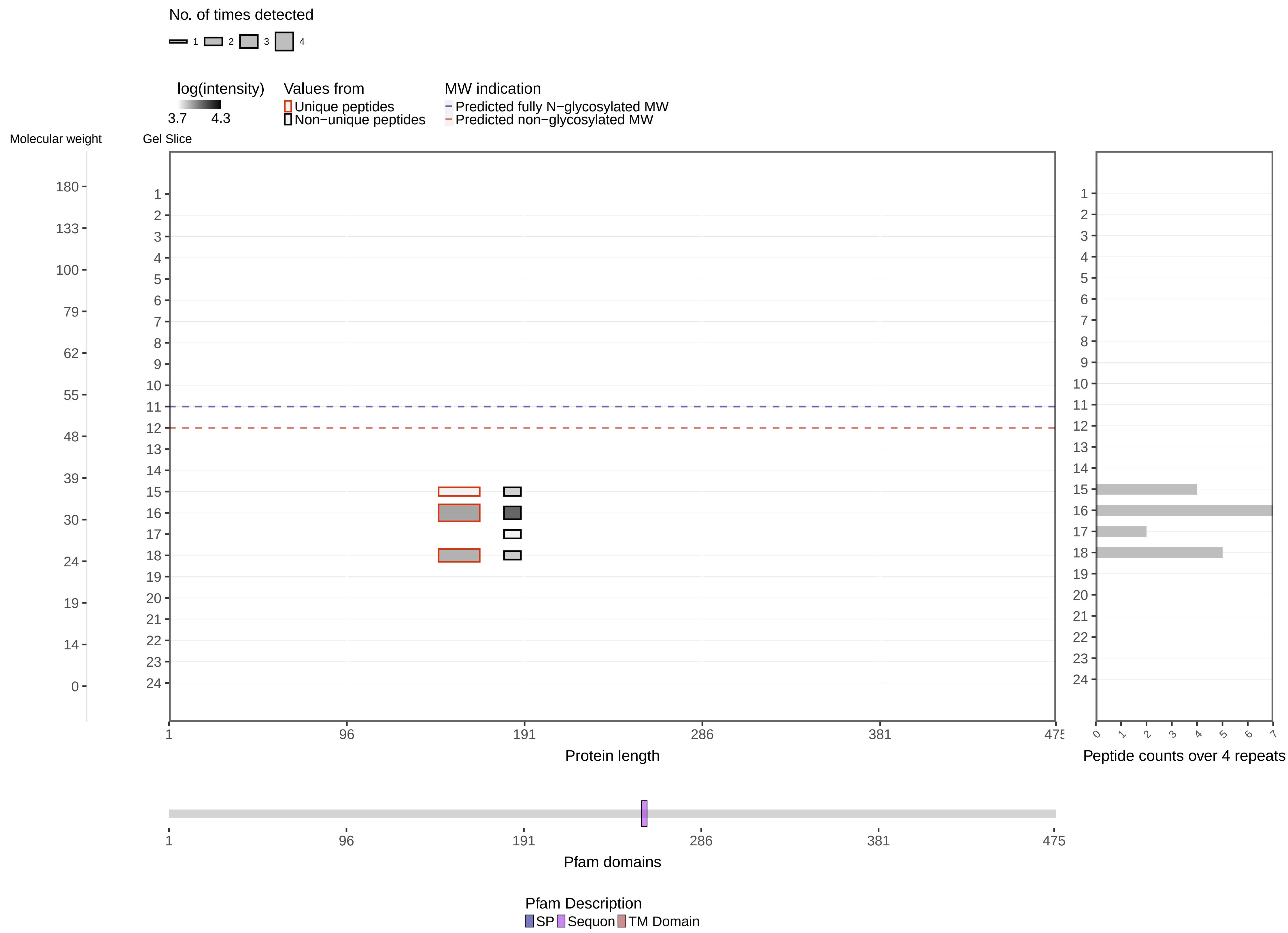


Protein sequence:

SLQVMEKNYMTMSREVEKLRAELNTSNSERRTGGPYGGATGYNENDAANNYAIGQNIYGDGYGIYQGRGPLPTATNVGGVAVVTSPRVGAQSVPPSNRPPYDSSNMPGYDAQRGMVPTGPGYDAQKGSALAAYEAQRGHGYDTMKGPGYDAQRAAGYETYGGPGYDGYGAPVYNAQRGSGYDVQRGGNYDPAKASNYDASNKGGVATQGQVAPMSNNPHGSSTPPGHAGPGYDAAAQGGNPARR*

### NbL02g11210.1 oxygen−evolving_enhancer_protein_1_chloroplastic [MW = 35.2 kDa]

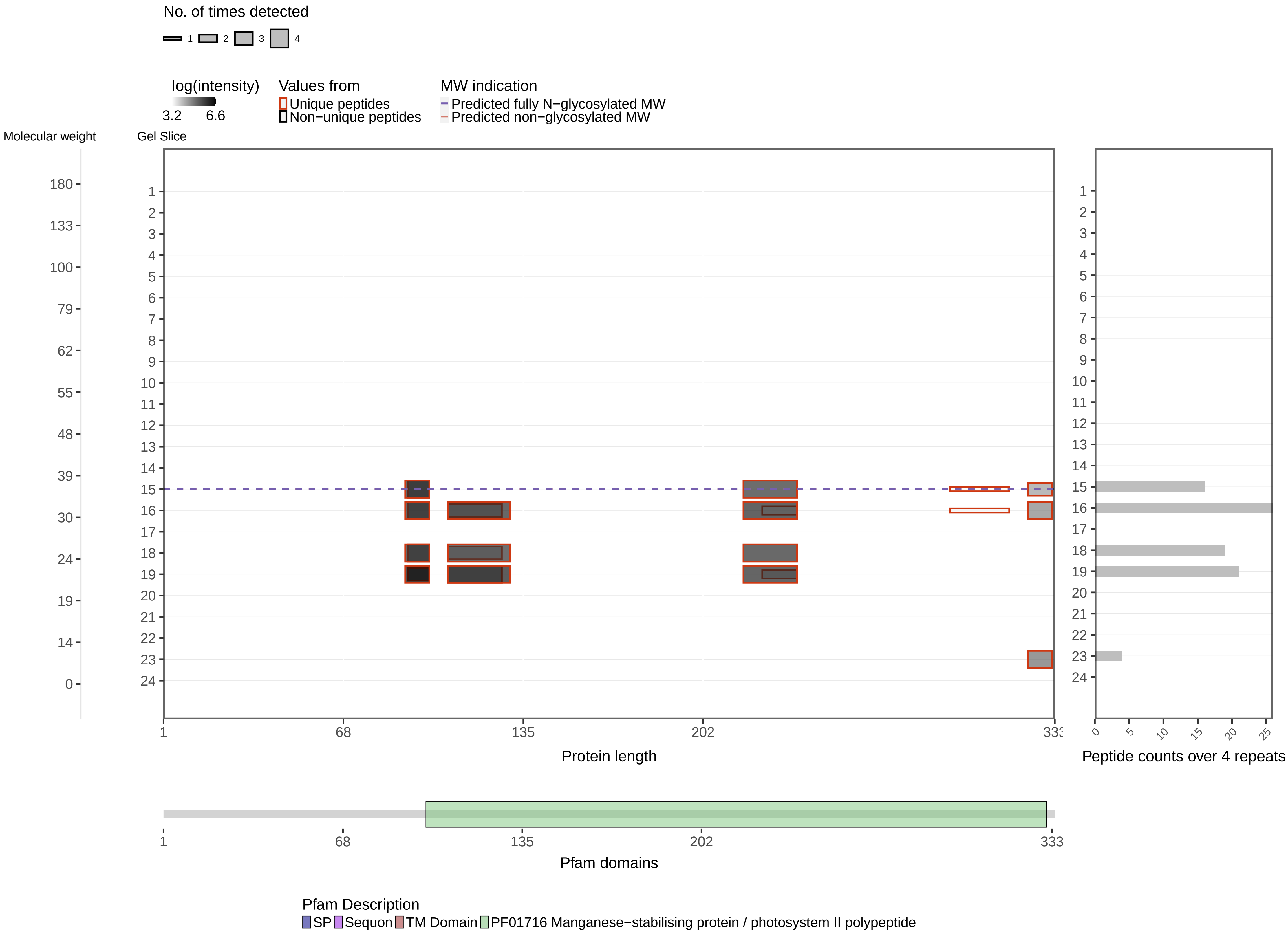


Protein sequence:

NAPPDFQKTKLMTRLTYTLDEIEGPFEVSSDGTVKFEEKDGIDYAAVTVQLPGGERVPFLFTIKQLVASGKPESFSGEFLVPSYRGSSFLDPKGRGGSTGYDNAVALPAGGRGDEEELEKENVKNTASSTGKITLSVTQSKPQTGEVIGVFESIQPSDTDLGAKVPKDVKIQGIWYAQLE*

NbL02g11120.1 patatin−like_protein_2 [MW = 54.01 kDa]

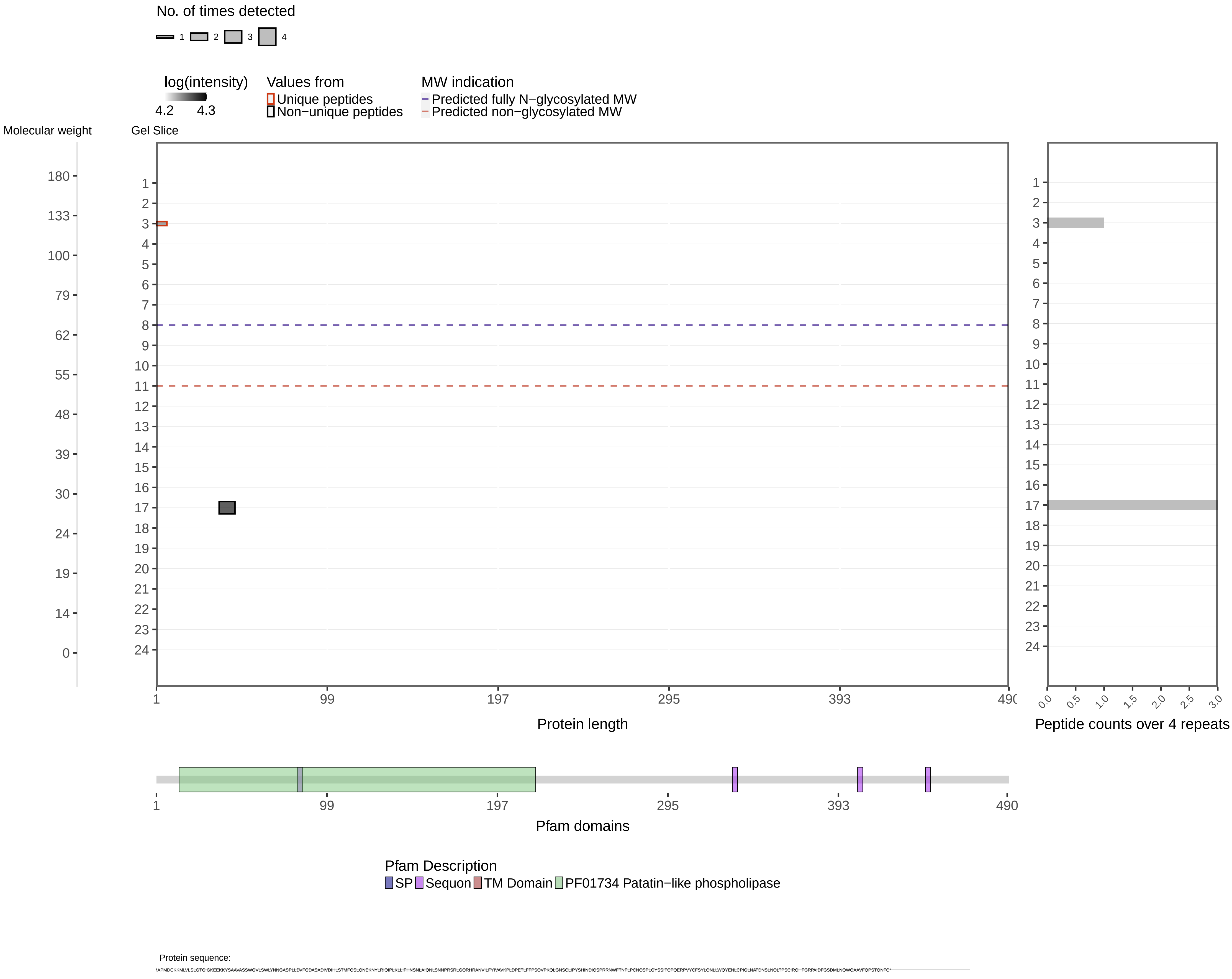

### NbL02g11050.1 L−ascorbate_oxidase_homolog [MW − SP = 57.39 kDa]

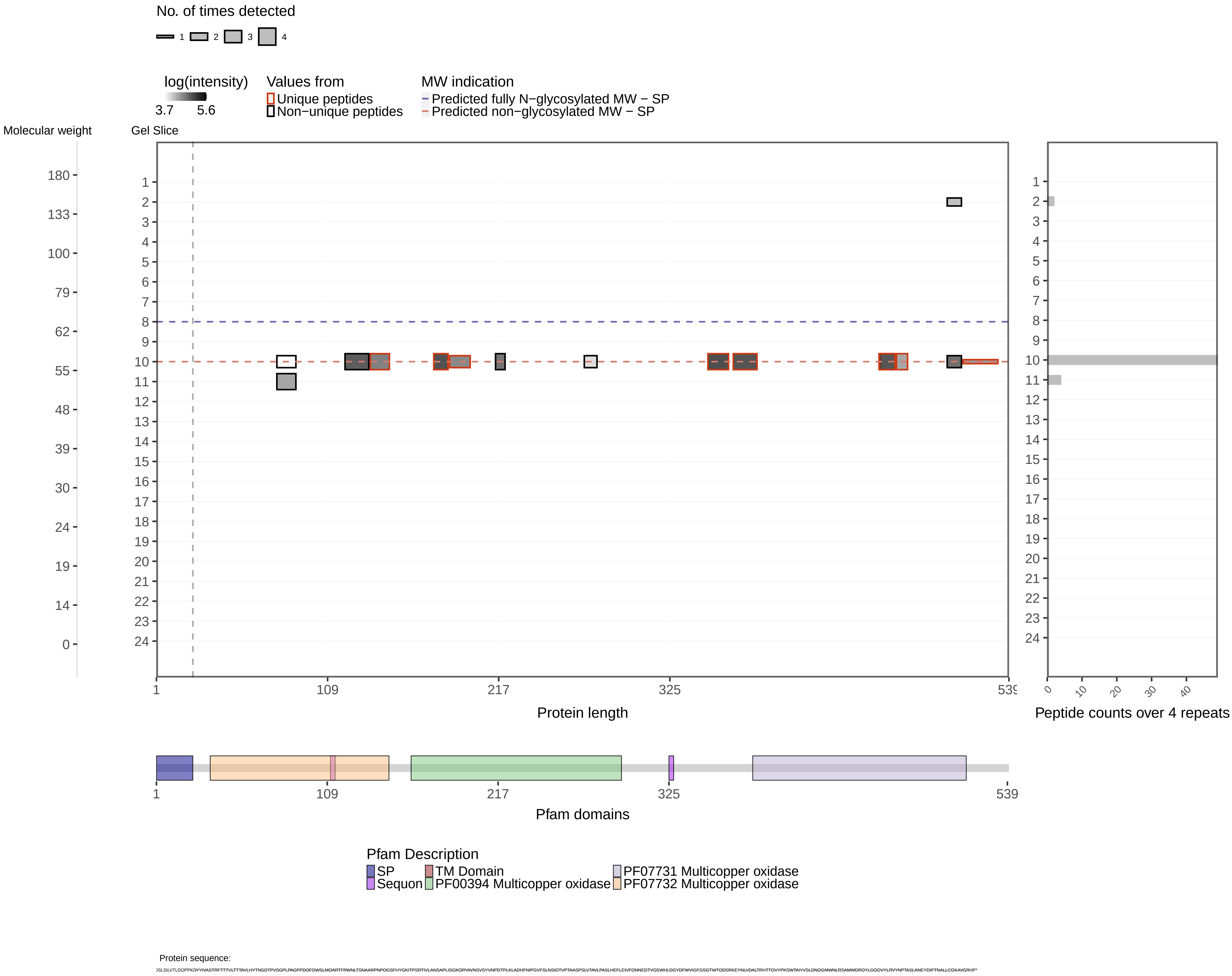

NbL02g10900.1 probable_esterase_KAI2 [MW = 30.33 kDa]

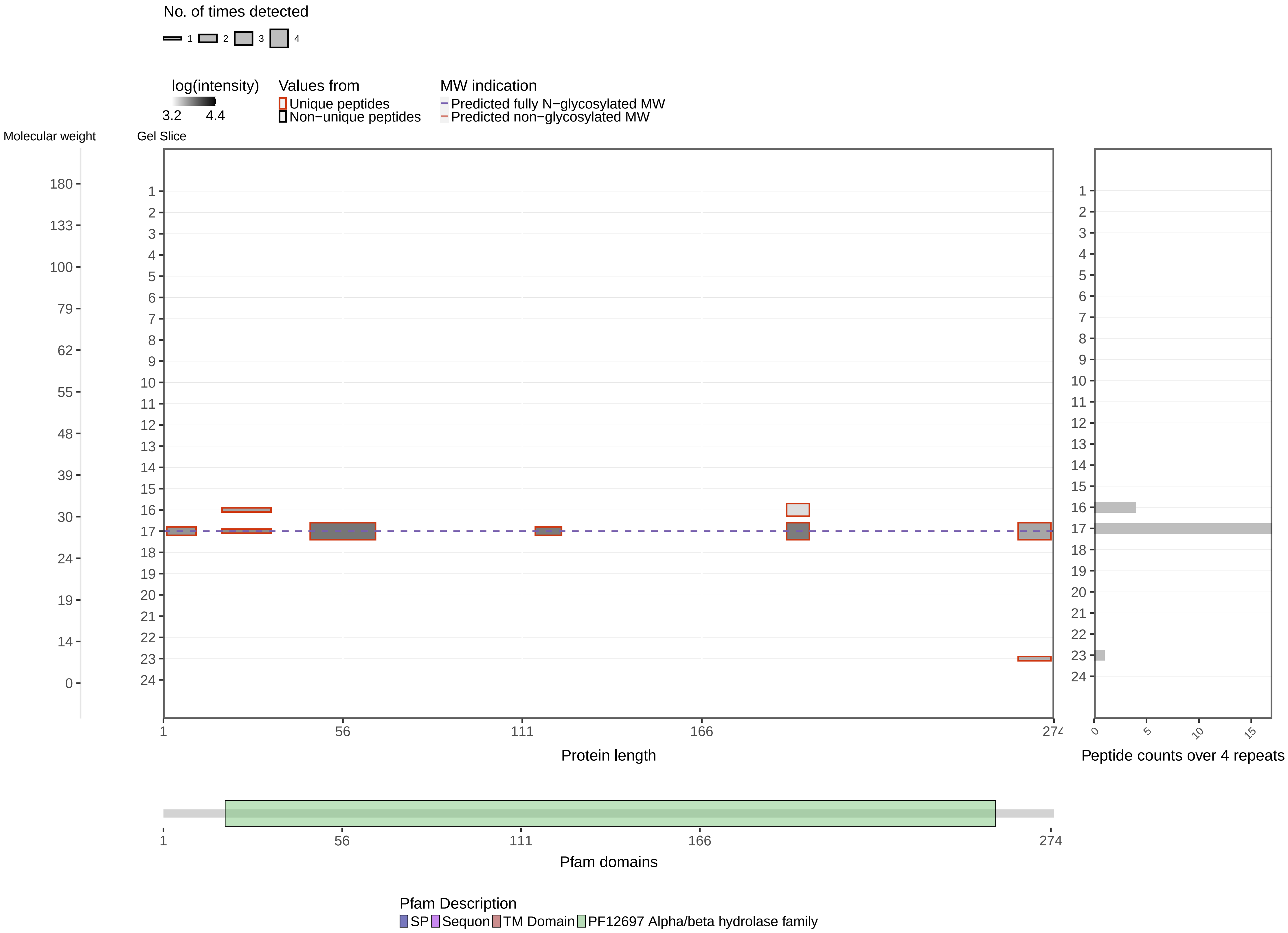


Protein sequence:

RFINTDDYYGGFEKEDIDQLSQAMESNYKSWIDGFAPLVIGGDMDSVAVQEFSRTLFNMRPDIALSVFRTIFTFDLRHFLSRVTVPCHIIQSSKDLAVPVAVSEYIHQNLGGKSIVEVISTEGHLPQLSAPEVTVPVLLRHISHDISSDTV*

### NbL02g10850.1 casein_kinase_II_subunit_alpha−2−like [MW = 49.57 kDa]

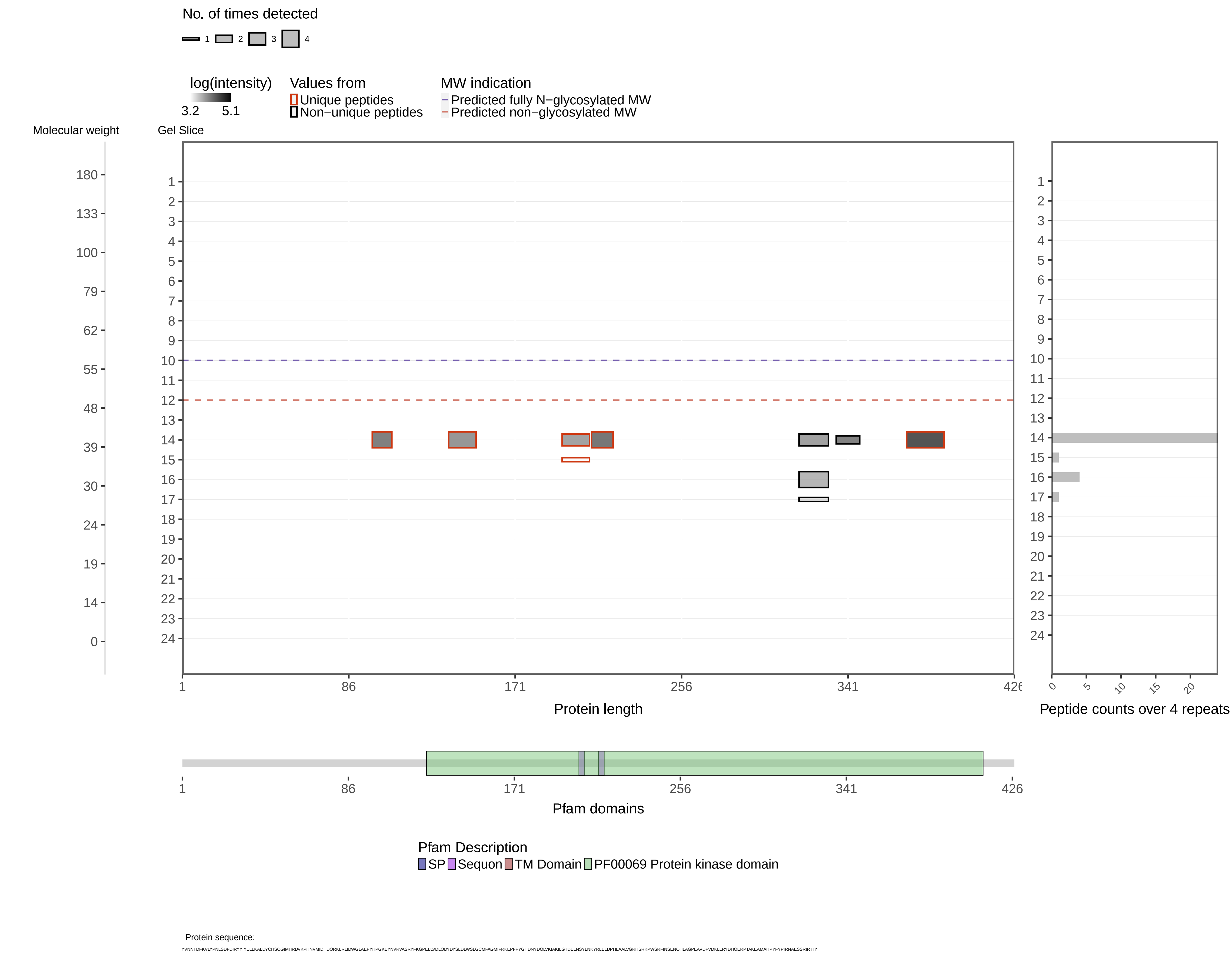

NbL02g10670.1 malate_dehydrogenase_glyoxysomal−like [MW = 37.95 kDa]

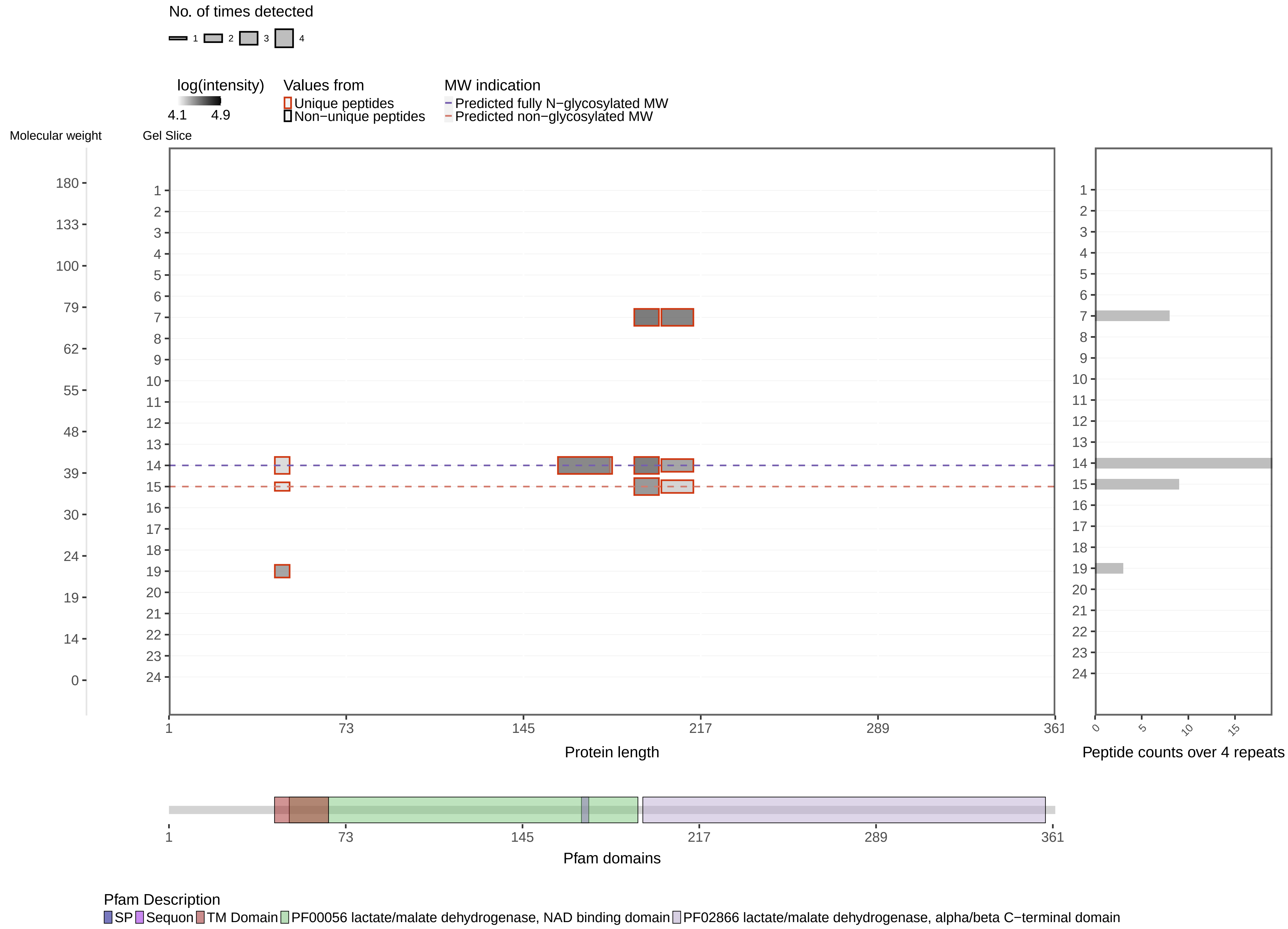


Protein sequence:

VNSTVPIAAEVFKKAGTFDPRRLLGVTMLDIVRASTFVAEVLGLDPREVDVPVVGGHAGVTILPLLSQVKPPCSFTPEETEYLTSRIQNGGTEVVEAKAGAGSATLSMAYAAVKFADACLHGLRGDAGIVECAFVSSQVTELPFFASKVRLGRNGVEEIYPLGPLNEYERSGLEKAKKELATSIQKGVNFVKK*

### NbL02g10610.1 serine_hydroxymethyltransferase_mitochondrial [MW = 57.17 kDa]

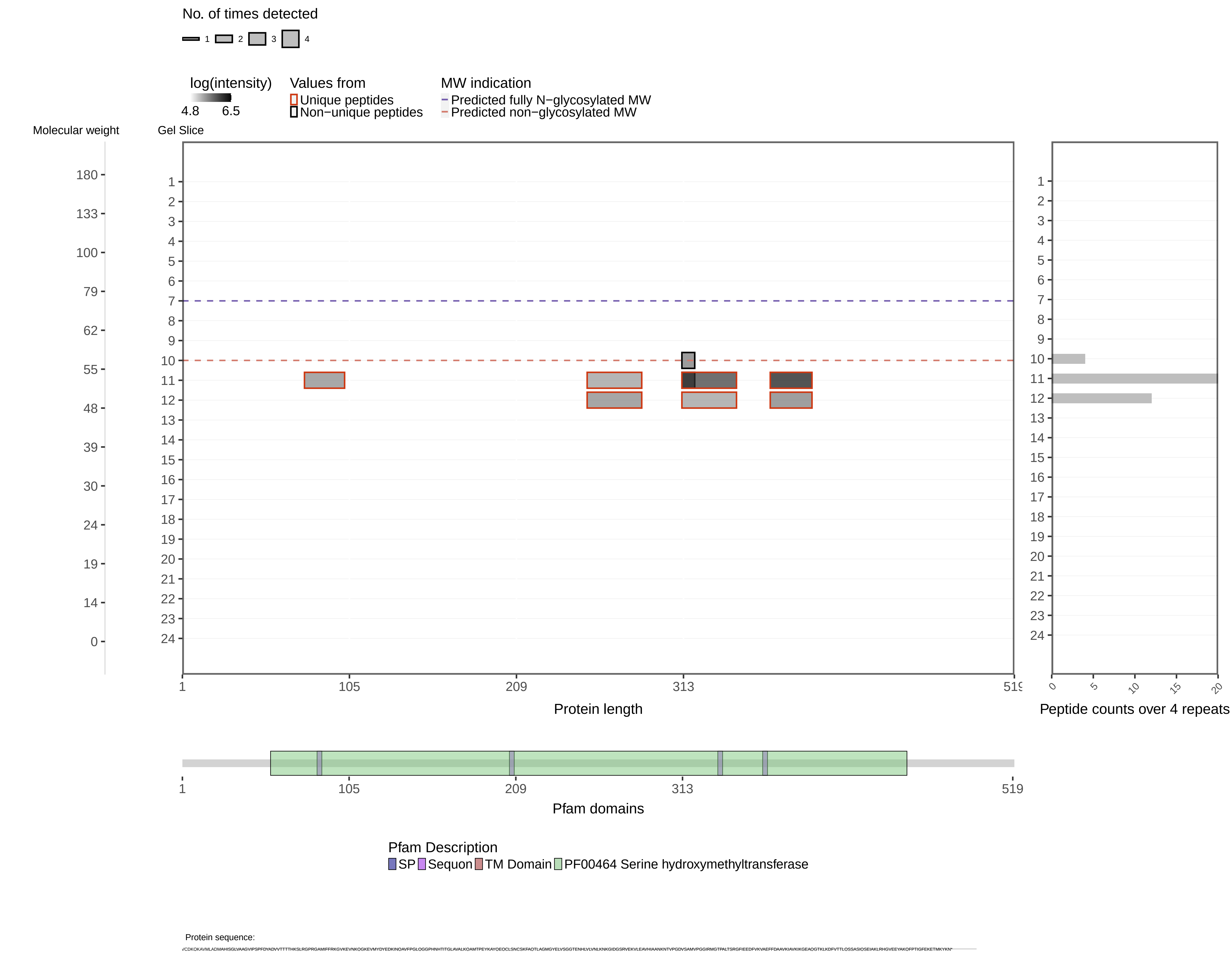

### NbL02g10390.1 ubiquitin_receptor_RAD23d−like [MW = 45.51 kDa]

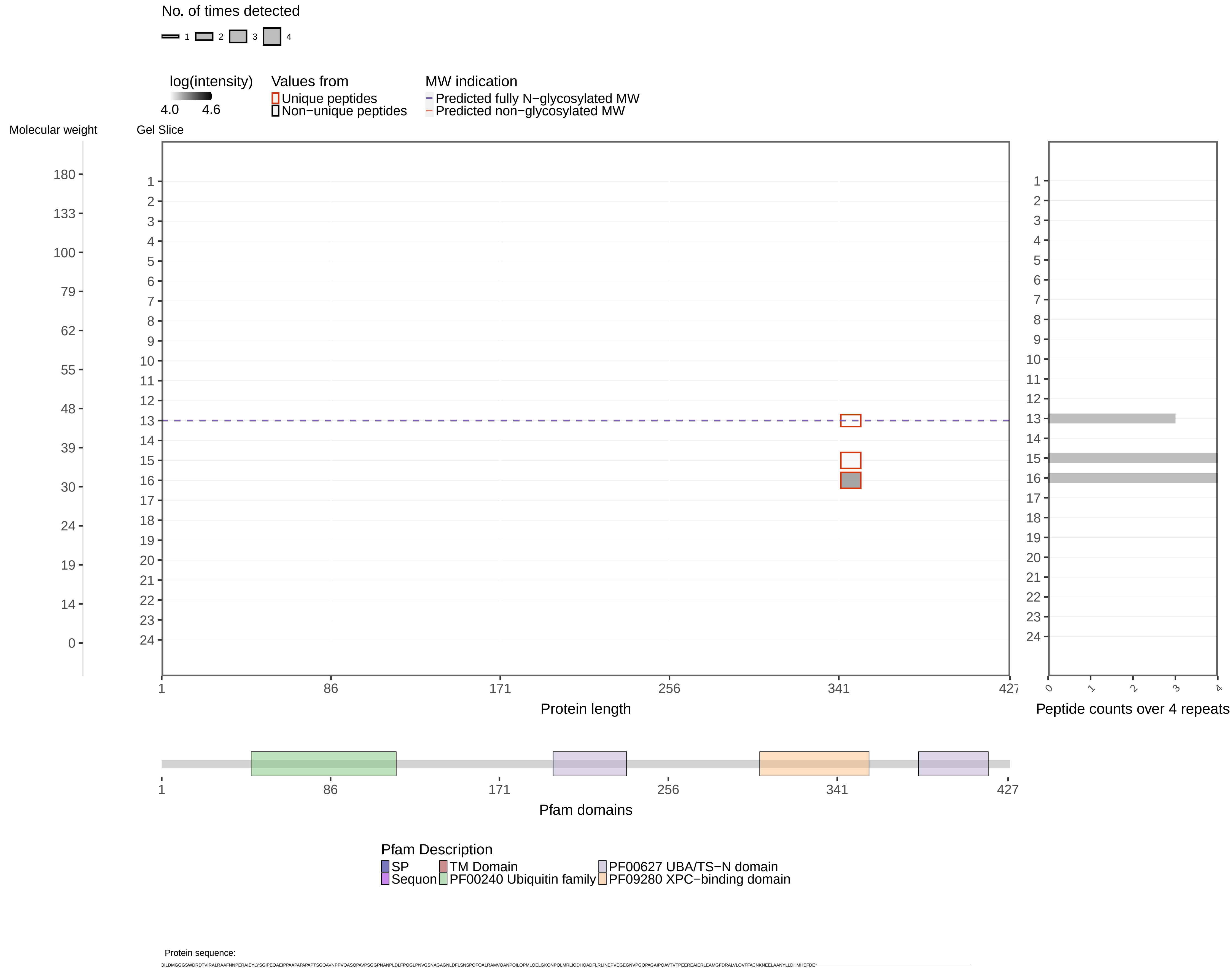

NbL02g10380.1 T−complex_protein_1_subunit_zeta_1−like [MW = 58.84 kDa]

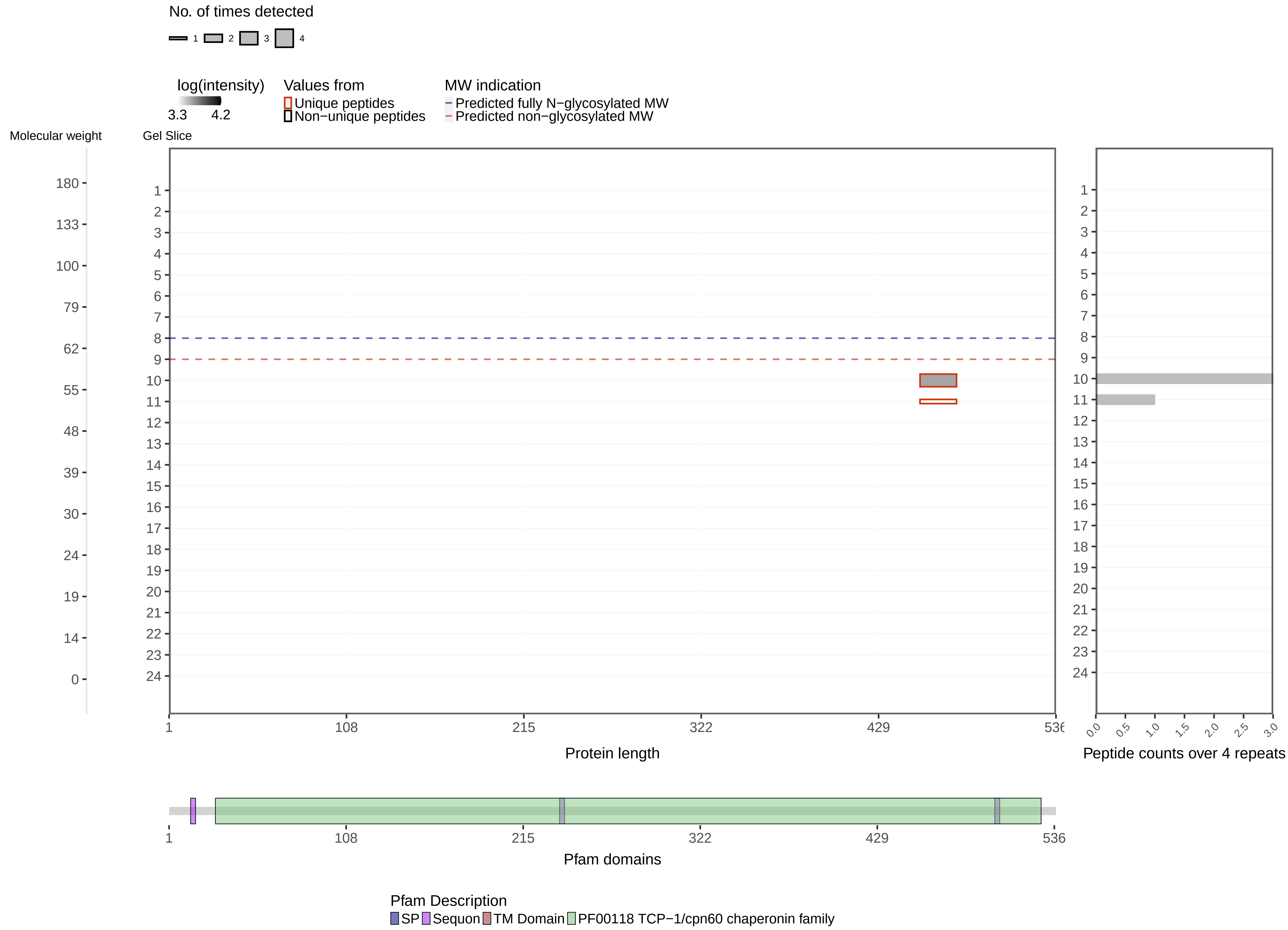


Protein sequence:

NAEQREAMVAAERRSVDERVQKIIDLKNKVCAGNDNNFVIINQKGIDPPSLDILARAGIIALRRAKRRNMERLVLACGGEAVNSVDDLTPDCLGWAGLVYEHVLGEEKYTFVENVKNPHSCTILIKGPNDHTIAQIKDAVRDGLRAVKNTIEDEAVVLGAGTFEVAARQHLINEVKKTVKGRAQLGVEAFANALLVIPKTLAENSGLDTQDVILALTGEHDKGNVVGLNHHTGEPIAPQMEGIFDNYSVKRQIVNSGPVIASQLLLVDEVIRAGRNMRKPN*

NbL02g10360.1 14−3−3_protein_4 [MW = 29.43 kDa]

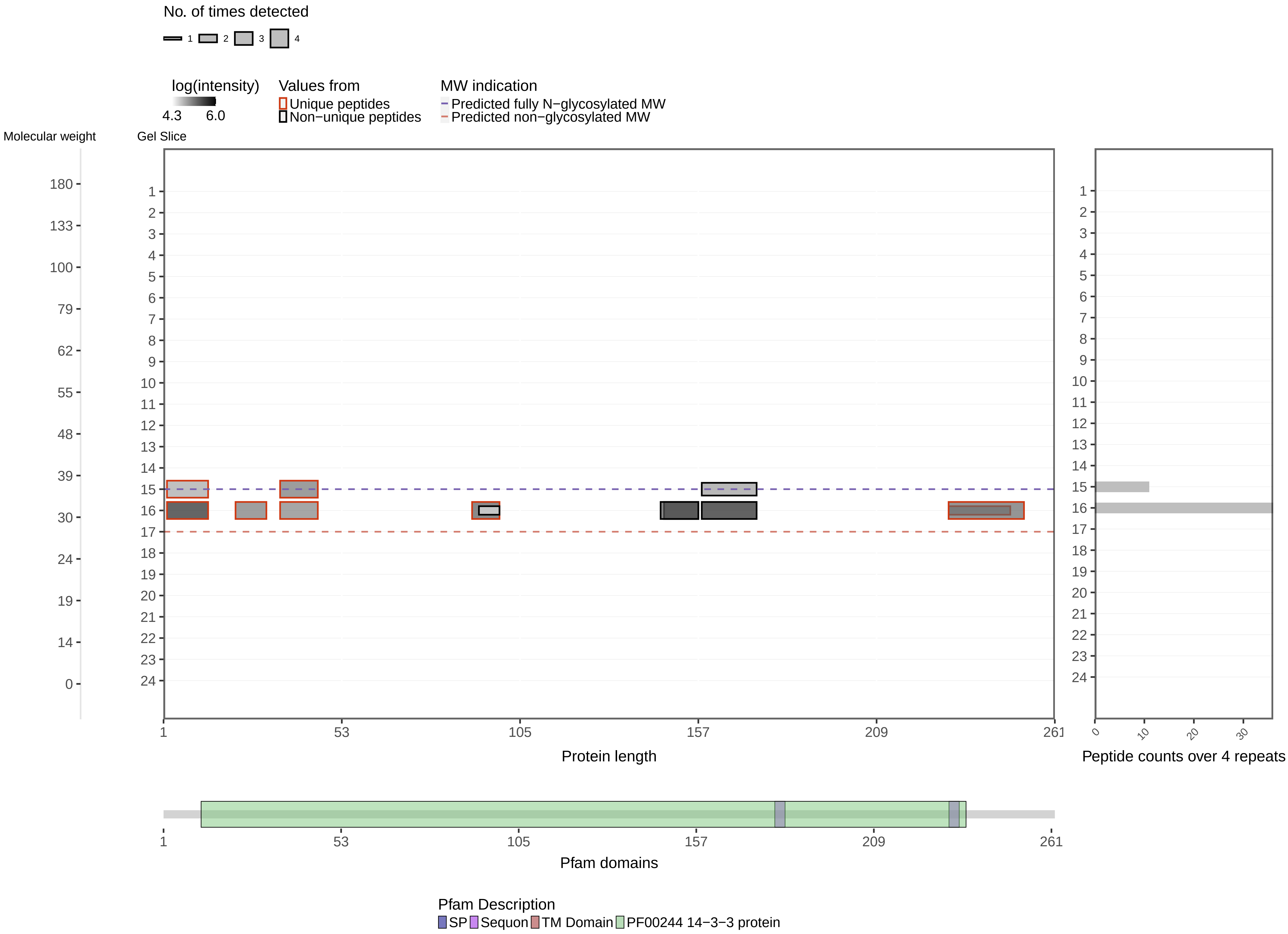


Protein sequence:

TAESKVFYLKMKGDYHRYLAEFKTGSERKEAAENTLLAYKSAQDIALTELAPTHPIRLGLALNFSVFYYEILNSPDRACNLAKQAFDEAISELDTLGEESYKDSTLIMQLLRDNLTLWTSDNAEDAGDDIKEAPKHESGEGQQ*

### NbL02g10210.1 glycine_cleavage_system_H_protein_2_mitochondrial−like [MW = 16.91 kDa]

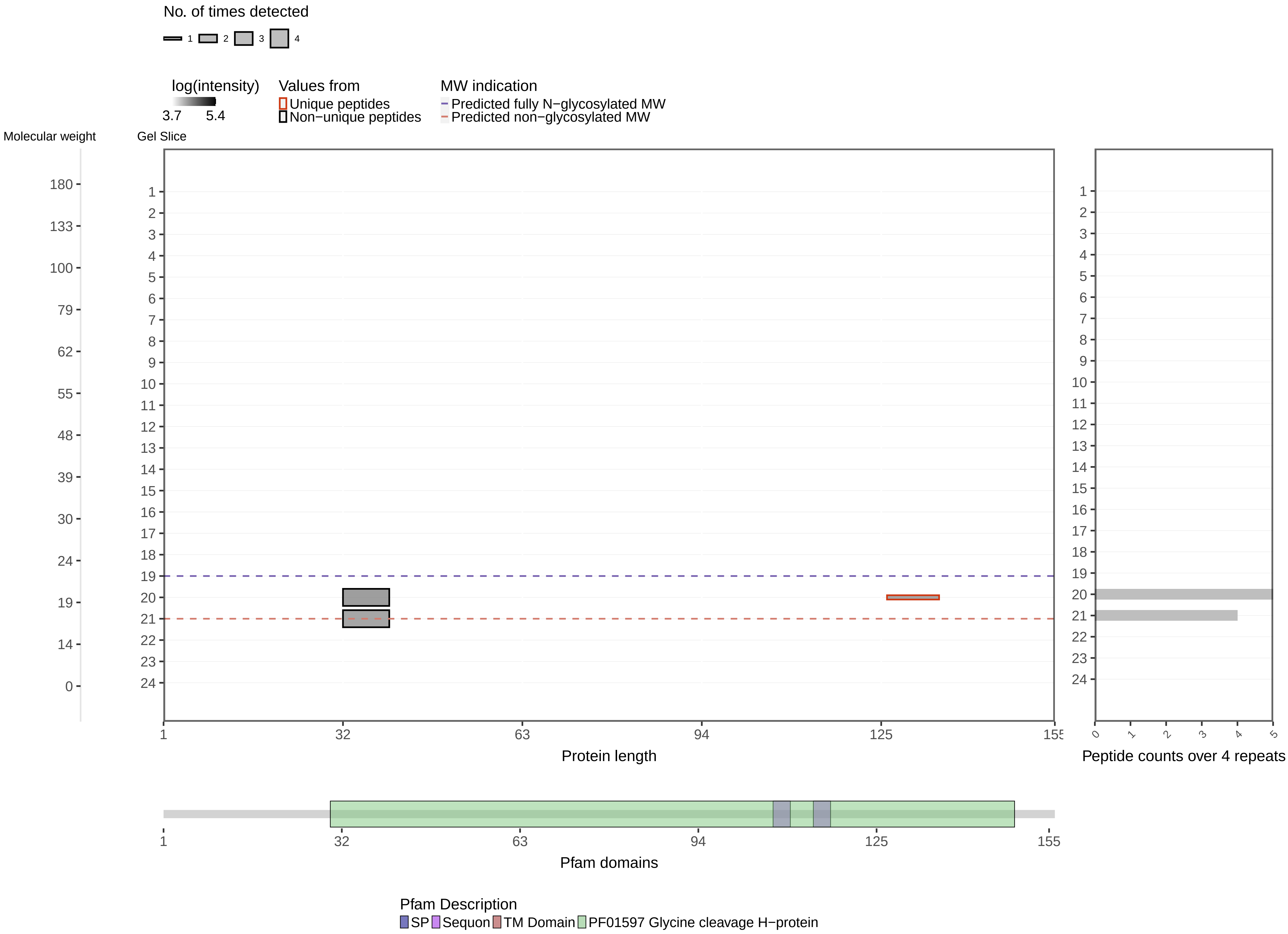


Protein sequence:

ELPEVGASVTQSGSFGAVESVKATSDINSPVSGKVVEVNEKLNDTPGLVNESPYQDGWIIKVEINNNDELKSLMDHDQYSKFCEEEDAKH*

### NbL02g09970.1 aspartic_proteinase−like [MW = 59.72 kDa]

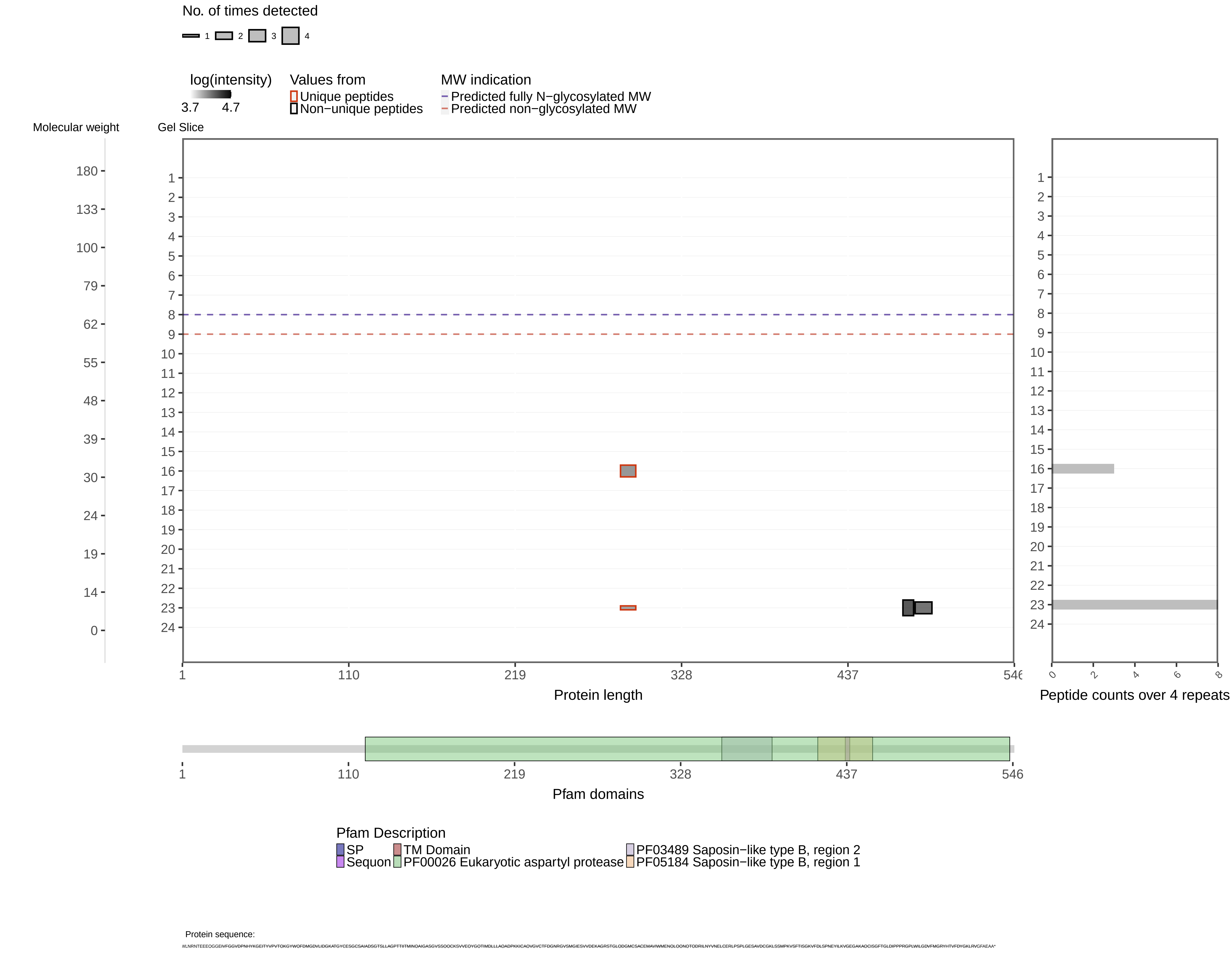

NbL02g09920.1 1−aminocyclopropane−1−carboxylate_oxidase [MW = 25.4 kDa]

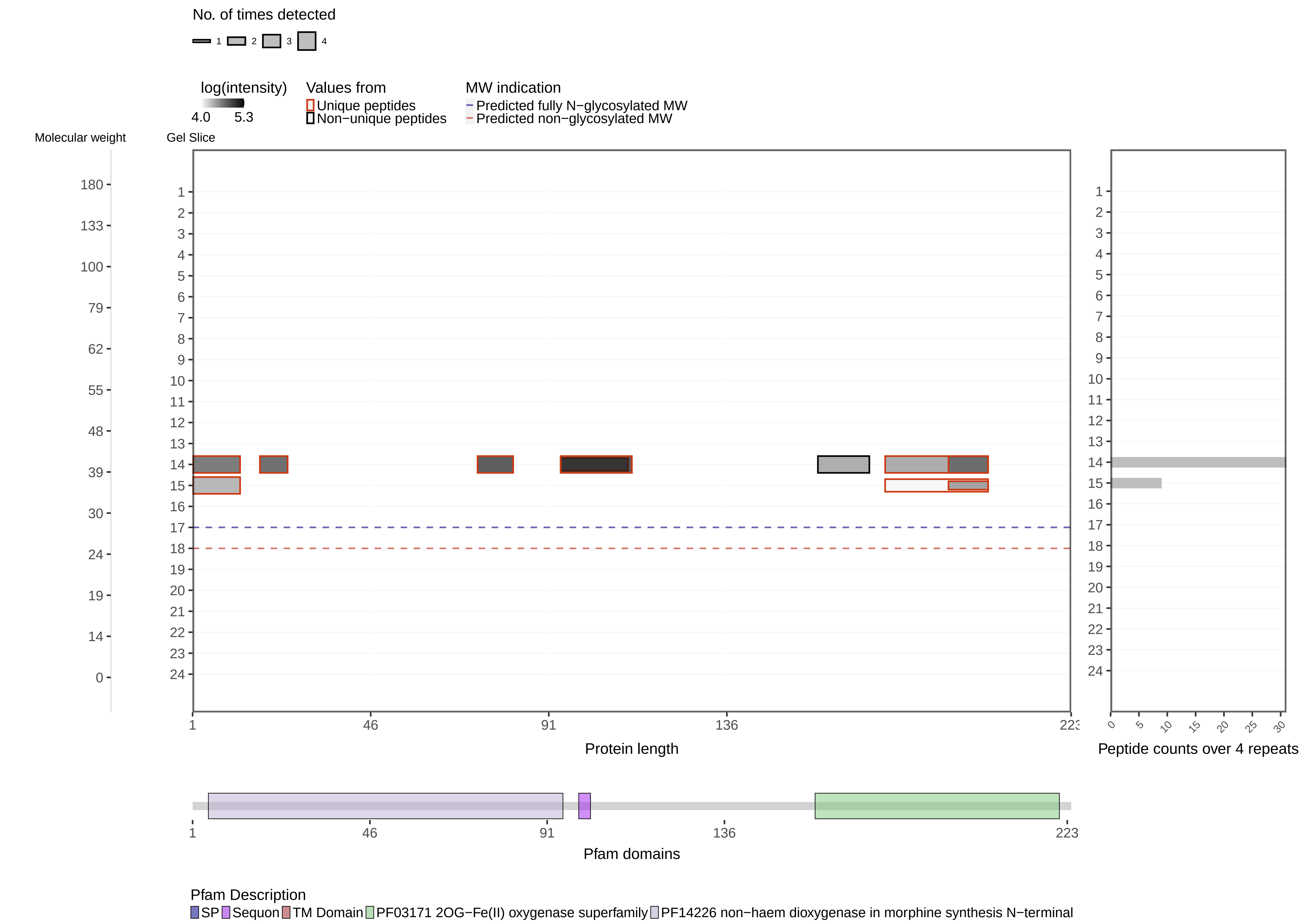


Protein sequence:

/SNISEVPDLEHEYRKIMKEFAEKLEKLAEQLLDLLCENLGLEQGYLKKAFYGSKGPTFGTKVSNYPPCPKPDLIKGLRAHTDAGGIILLFQDDKVSGLQLLKDDKWIDVPPMRHSIVINLGDQLR*

NbL02g09570.1 RNA_polymerase_II_transcriptional_coactivator_KIWI−like [MW = 11.67 kDa]

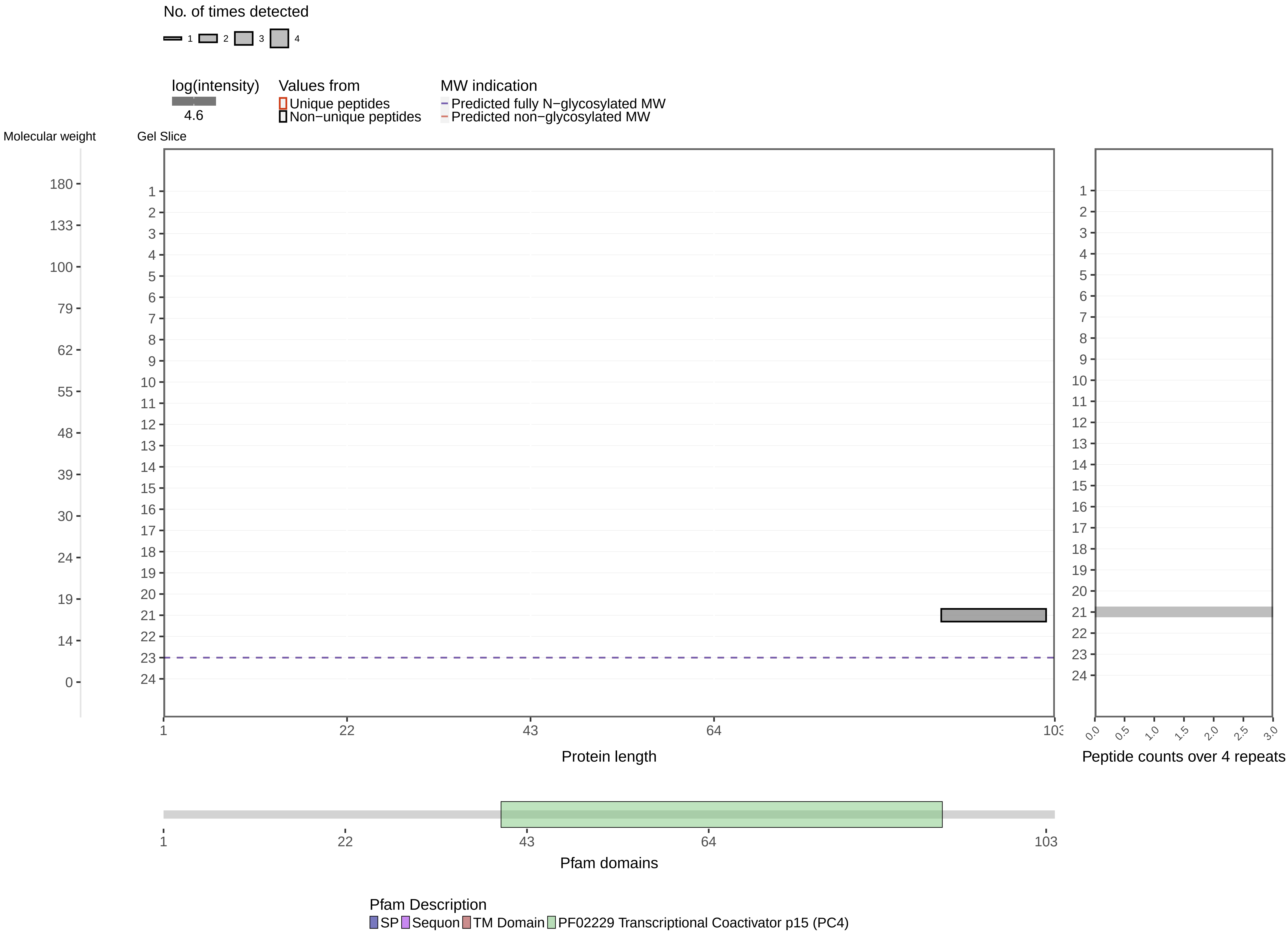


Protein sequence:

VCEISKNRRVSVRSFQGKIMVDIREFYVKDGKQMPGRKGISLSMDQWNVLRDHADEIDELVANS*

### NbL02g09300.1 phosphopantetheine_adenylyltransferase [MW = 19.52 kDa]

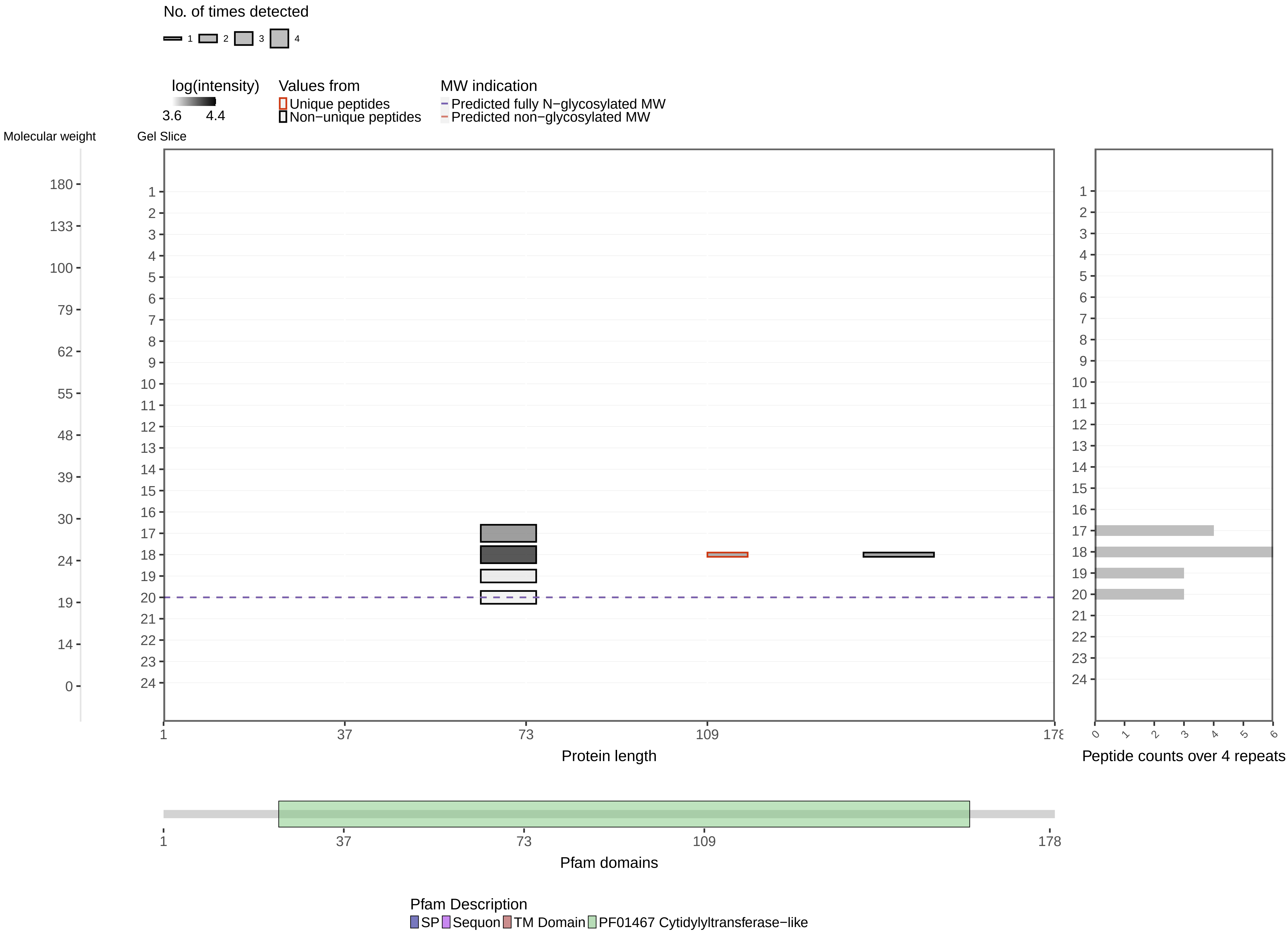


Protein sequence:

MQNVENYIKSLKPELVVEAVPIVDPYGPSIIDKNLEAIVVSKETLPGGLSVNRKRTERGLSQLKIEVVDLVSEESSGDKLSSTALRKLEAEKLRDHHPESQS*

### NbL02g09220.1 proteasome_subunit_beta_type−1 [MW = 24.69 kDa]

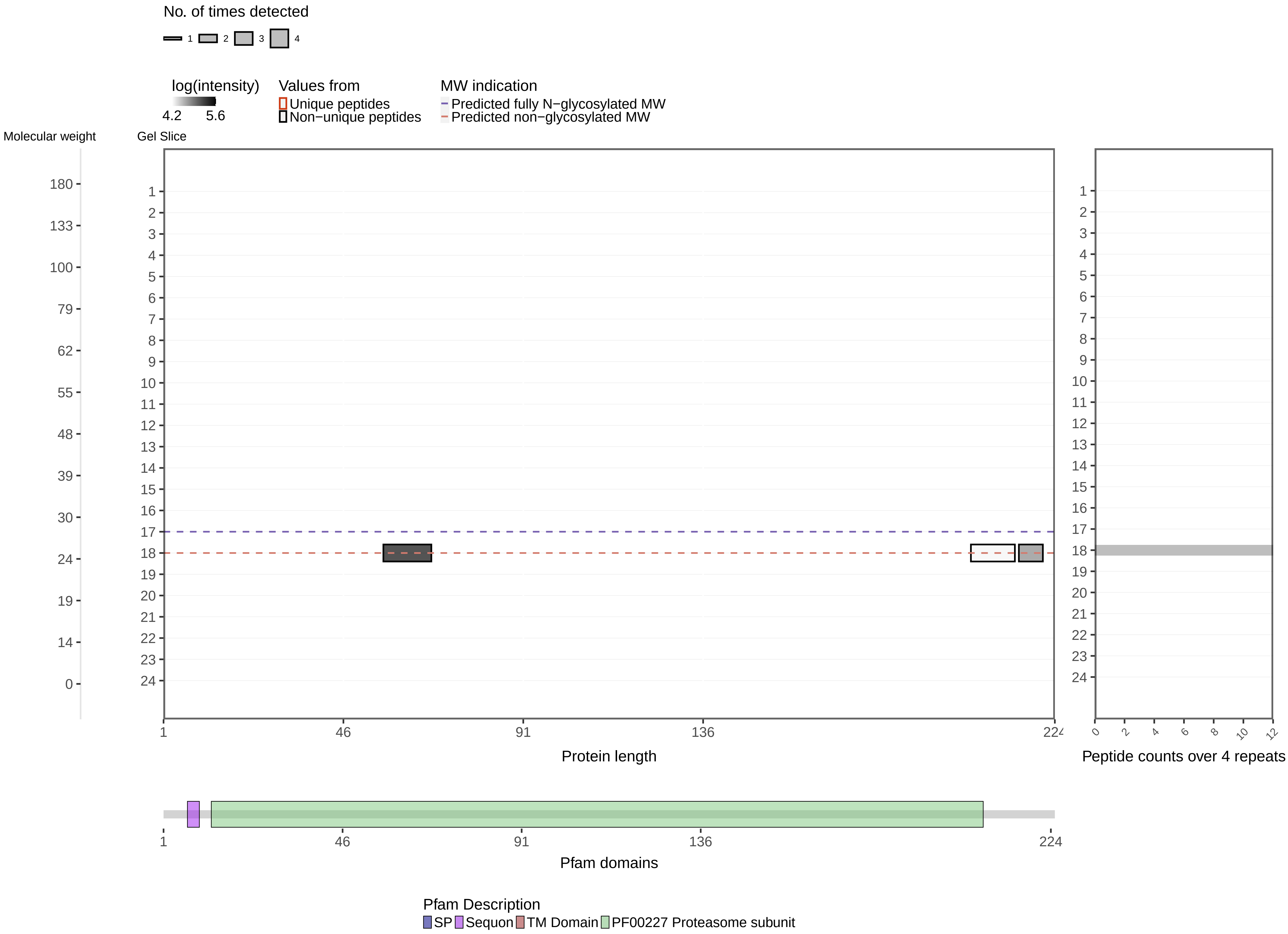


Protein sequence:

LLSNTLYYKRFFPYYSFNVLGGLDSEGKGCVFTYDAVGSYERVGYSSQGSGSTLIMPFLDNQLKSPSPLLLPAKDAVTPLSESEAIDLVKTCFASATERDIYTGDRLEIVILNANGIRREEMELRKD*

### NbL02g09170.1 fructose−bisphosphate_aldolase_1_chloroplastic [MW = 42.55 kDa]

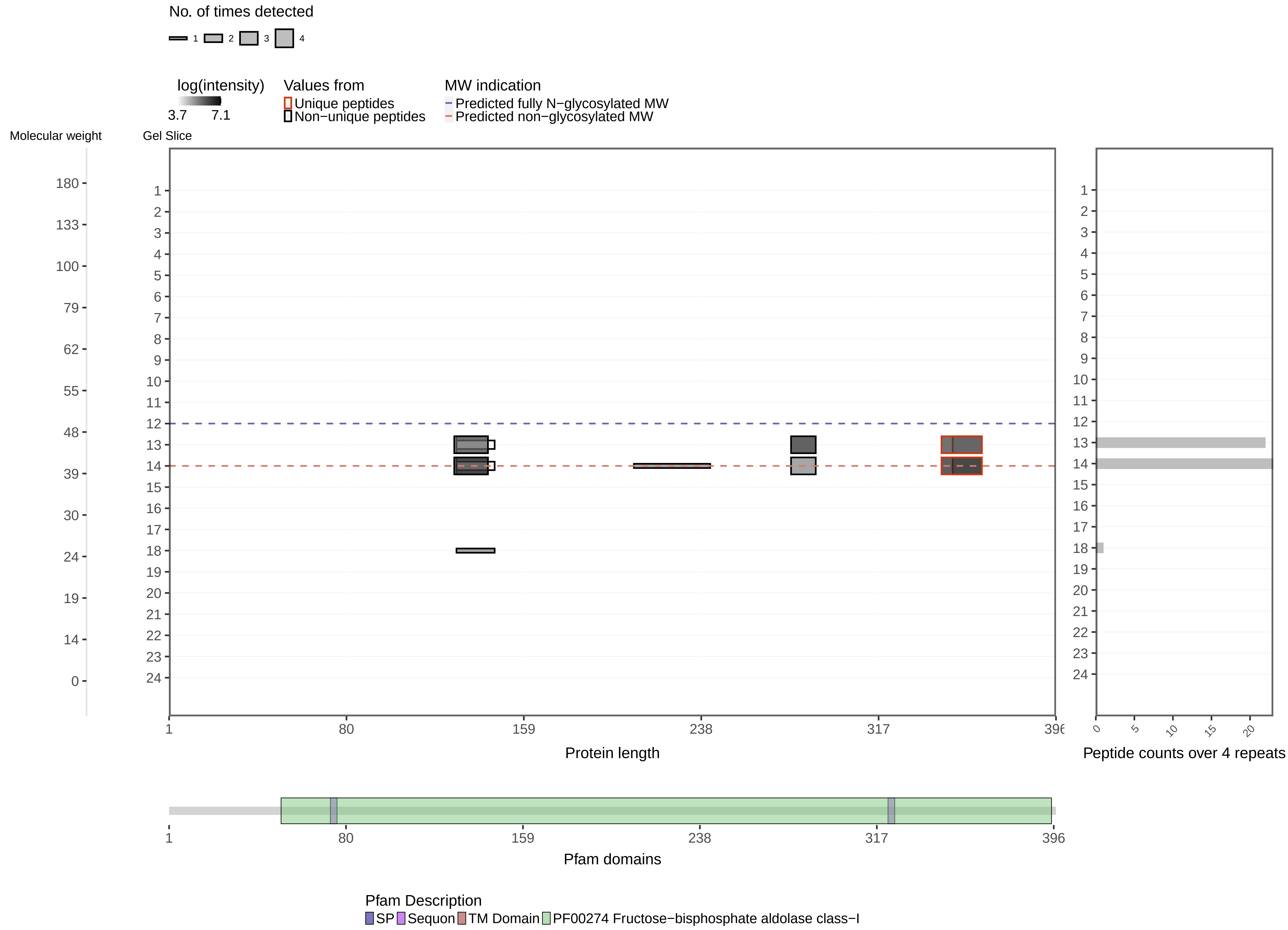


Protein sequence:

/VSIPNGPSALAVKEAAWGLARYAAISQDNGLVPIVEPEILLDGEHNIDRTFEVAQKVWAEVFFYLAENNVMFEGILLKPSMVTPGAECKERATPDQVADYTLKLLQRRIPPAVPGIMFLSGGQSEVEATLNLNAMNQSPNPWHVSFSYARALQNTCLKTWGGRPENVQAAQEALLIRAKANSLAQLGKYTGEGESEEAKKGMFVKGYVY*

### NbL02g09100.1 nucleolin−like [MW = 135.26 kDa]

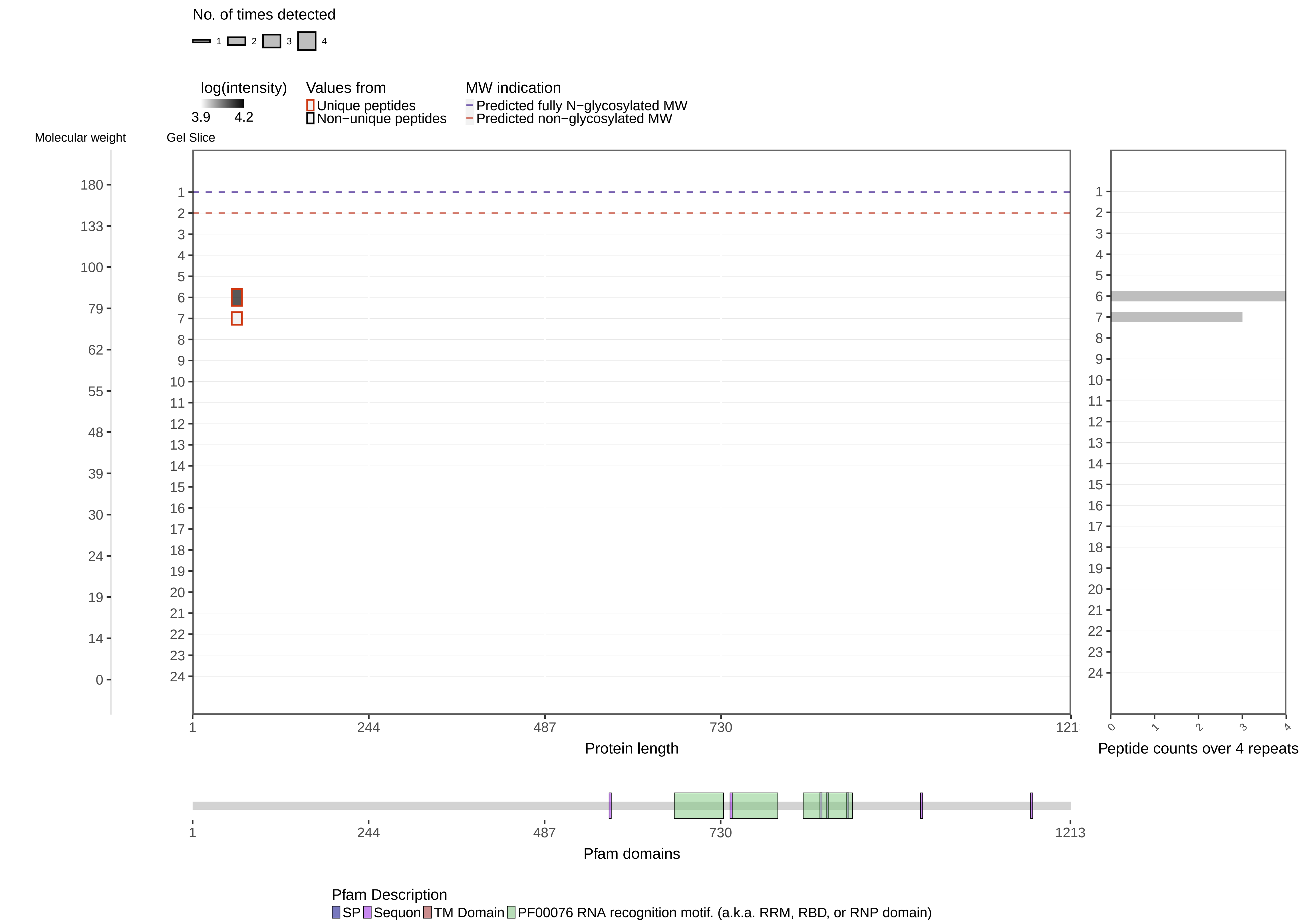


Protein sequence:

### NbL02g09080.1 peptidyl−prolyl_cis−trans_isomerase_Pin1−like [MW = 13.61 kDa]

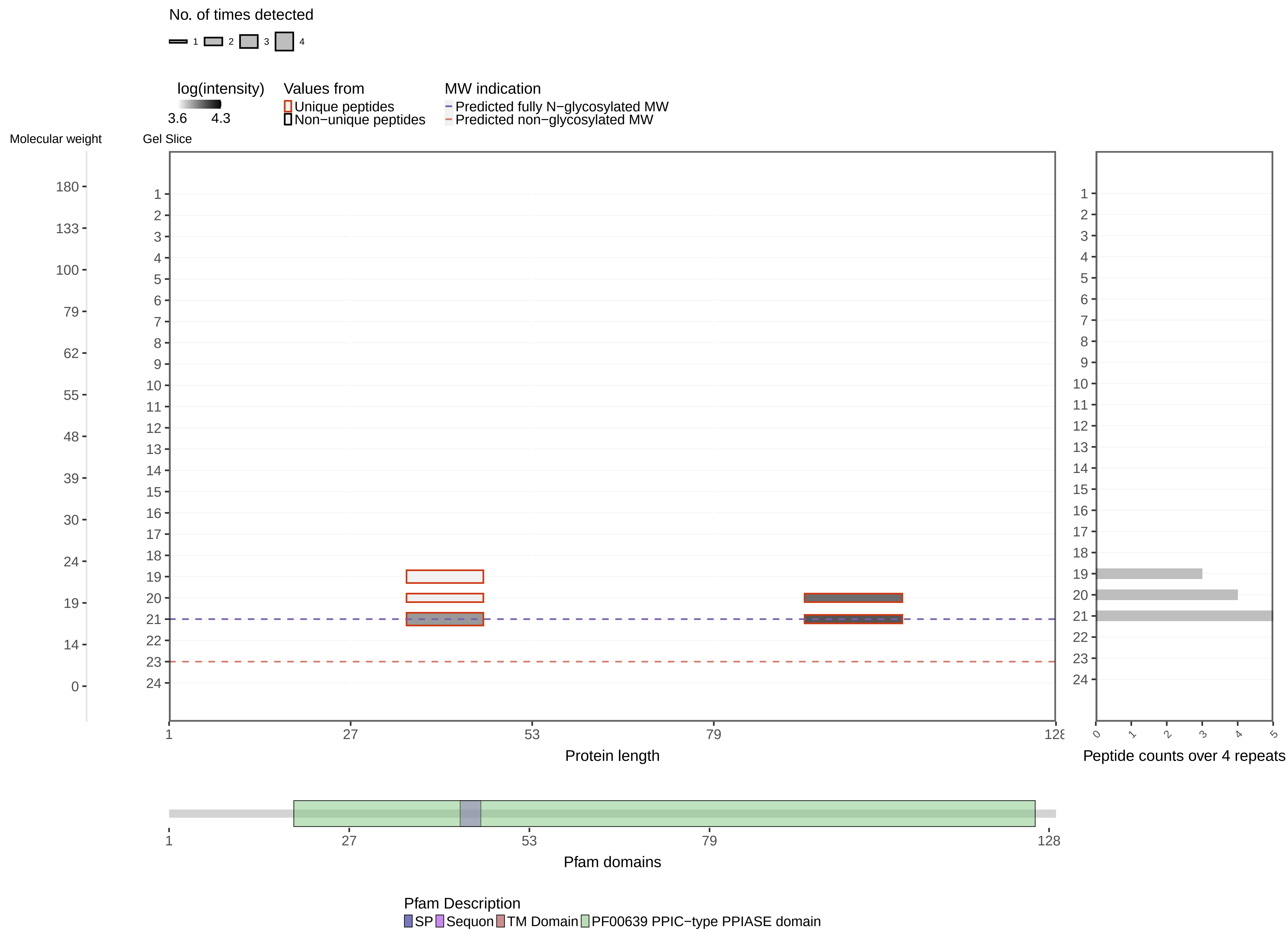


Protein sequence:

/SQLKSLREDIVSGKAKFDDVAATHSDCSSAKRGGDLGPFGKGQMQKPFEEATYALKVGEISDIVETDSGVHIIKRTA*

NbL02g08840.1 anthocyanidin_3−O−glucosyltransferase_5−like [MW = 55.33 kDa]

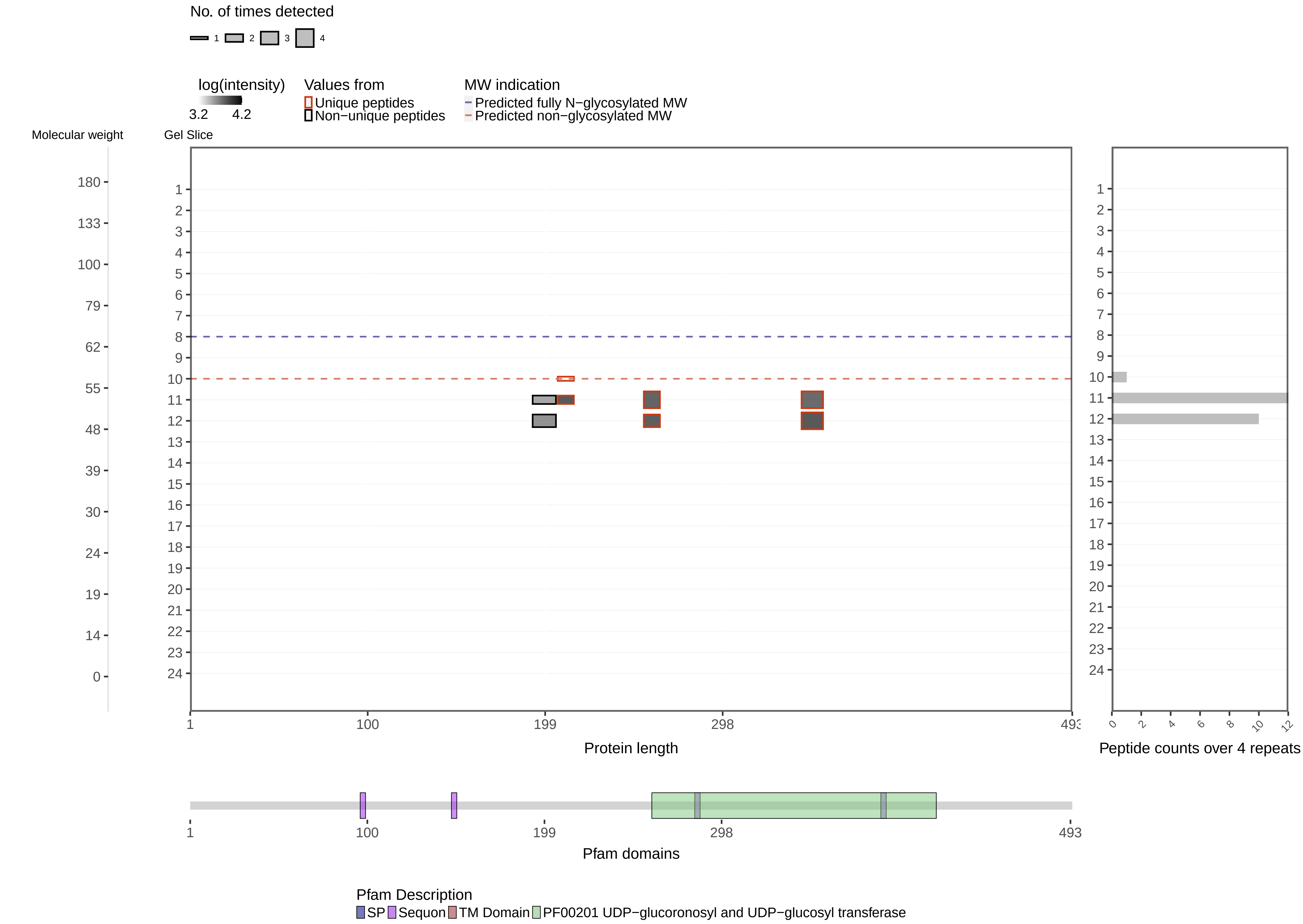


Protein sequence:

ALRYNEKLRLLLKVPVFPIGPLRRKVETTLNDEVIQWLDKQNNESVLFVSFGSGGTLSTKQMTELAWGLELSQQKFVWVVRPPSDGDADSAYLNSAGKDTRDMSEYLPEGFLTRTKDMGLVVPMWANQVEILSHSSVGGFLTHCGWNSTVESLTNGVPMIAWPLHAEQKMNAAMLTEELGVAIRPAVLPTKKLVKREEIQGMVRILMQTKEGKRIKEKAKKLKKSAENALSDGGSSYNSICELVKDIRSREL*

NbL02g08390.1 protein_plastid_transcriptionally_active_16_chloroplastic−like [MW = 54.4 kDa]

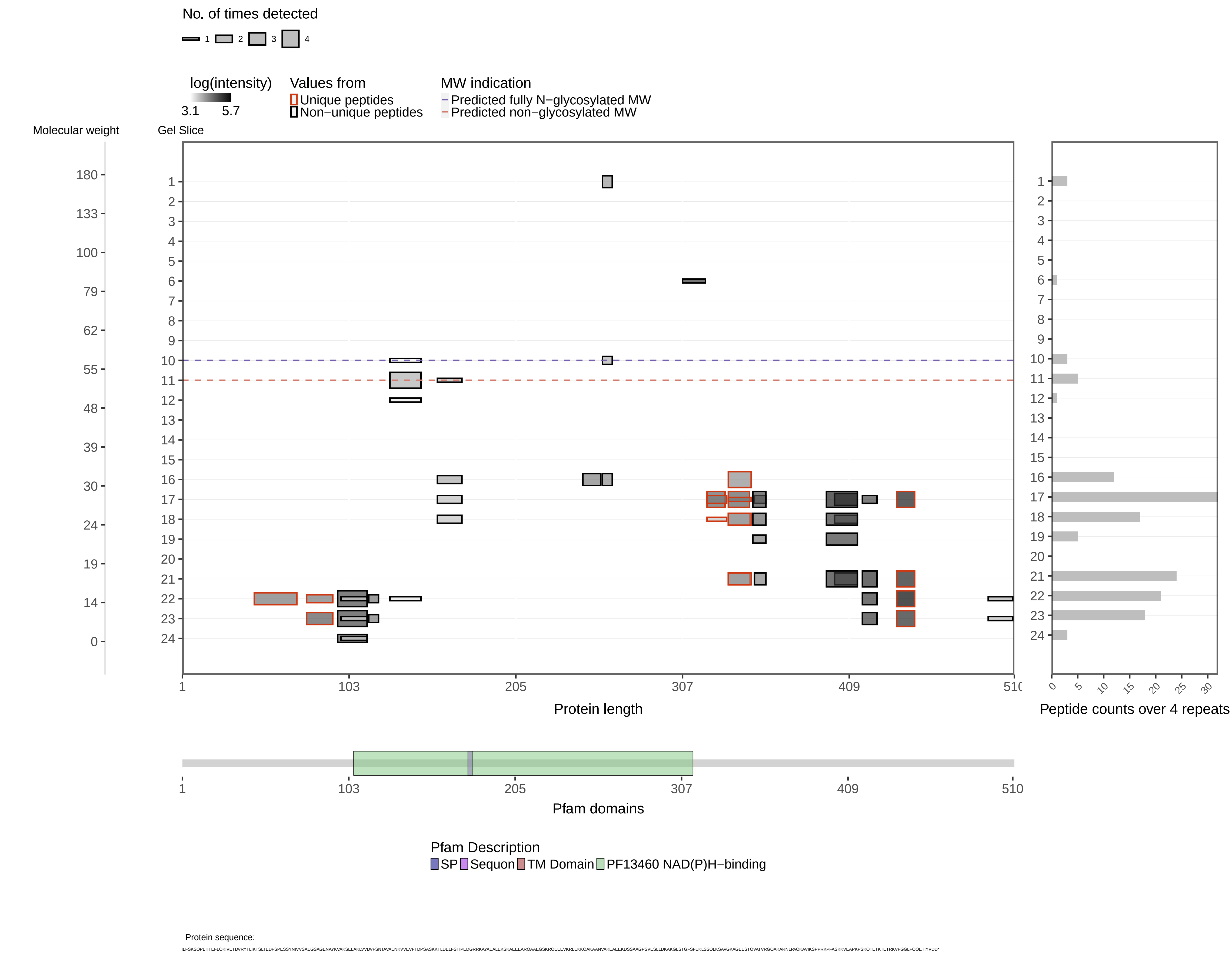

NbL02g08250.1 40S_ribosomal_protein_S4−like [MW = 29.91 kDa]

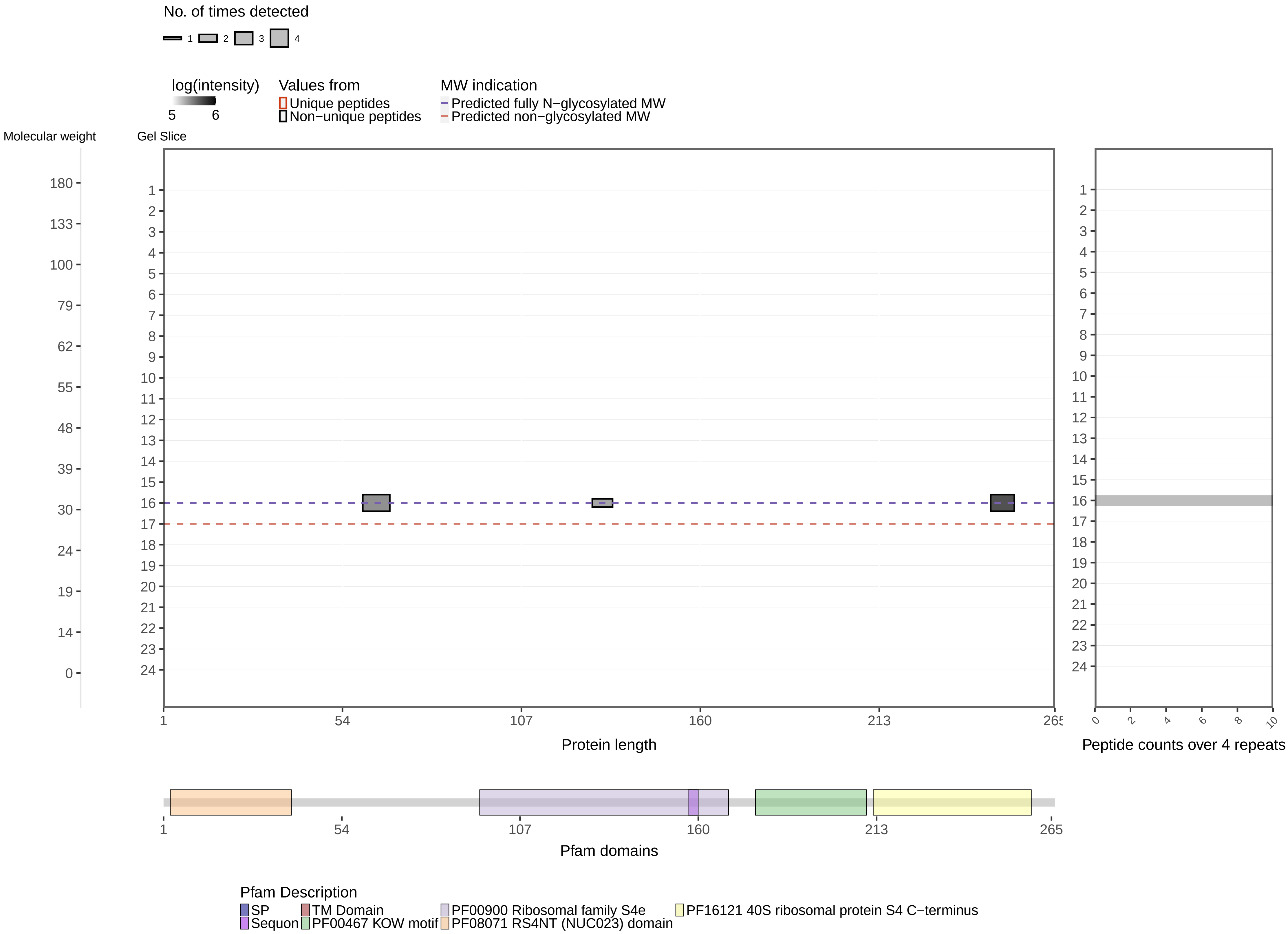


Protein sequence:

AKFKLCKVRSVQFGQKGIPYLNTYDGRTIRYPDPLIKANDTIKLDLESNKIVDFIKFDVGNVVMVTGGRNRGRVGVIKNREKHKGSFETVHIQDALGHEFATRLGNVFTLGKGTKPWVSLPKGKGIKLSIIEEARKRLAAQSATTA*

NbL02g08130.1 ERBB−3_BINDING_PROTEIN_1−like [MW = 39.64 kDa]

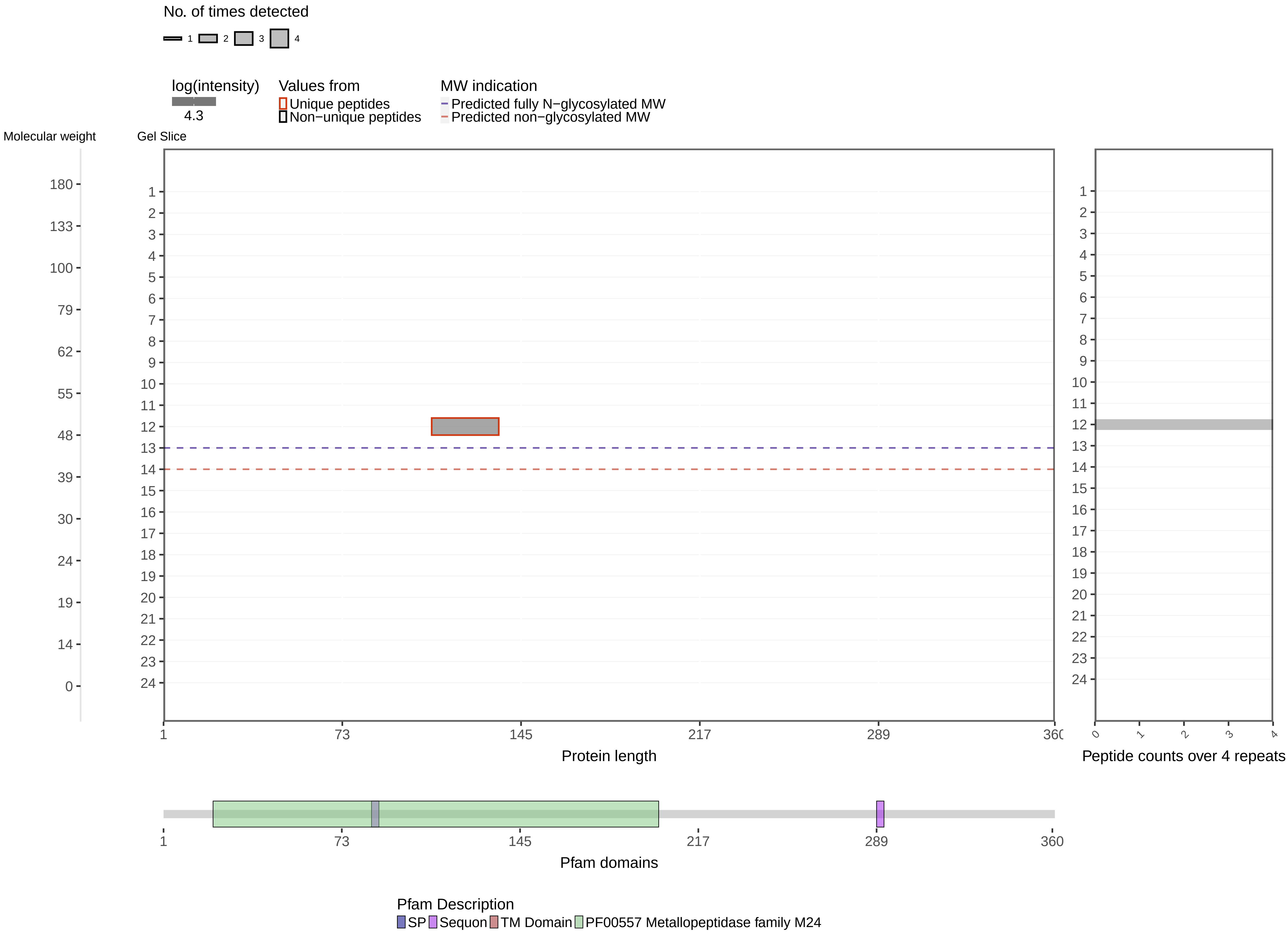


Protein sequence:
AIQKVAAAYDCKIVEGVLSHQMKQFVIDGNKPKLLDEKQTTIYKRAVDKSYNLKMKASRFIFSEISQKFPVMPFTARDLEEKRARLGLVECVNHELLQPYPVLHEKPGDLVAHIKFTVLLMPNGSDRITTHTLQELKPAKTIDDEPEIKTWLALPVKSKKKGGGKKKKAKKSEKTEDSSQAEPMEGESNGAES*

NbL02g08110.1 ran−binding_protein_1_homolog_c−like [MW = 24.04 kDa]

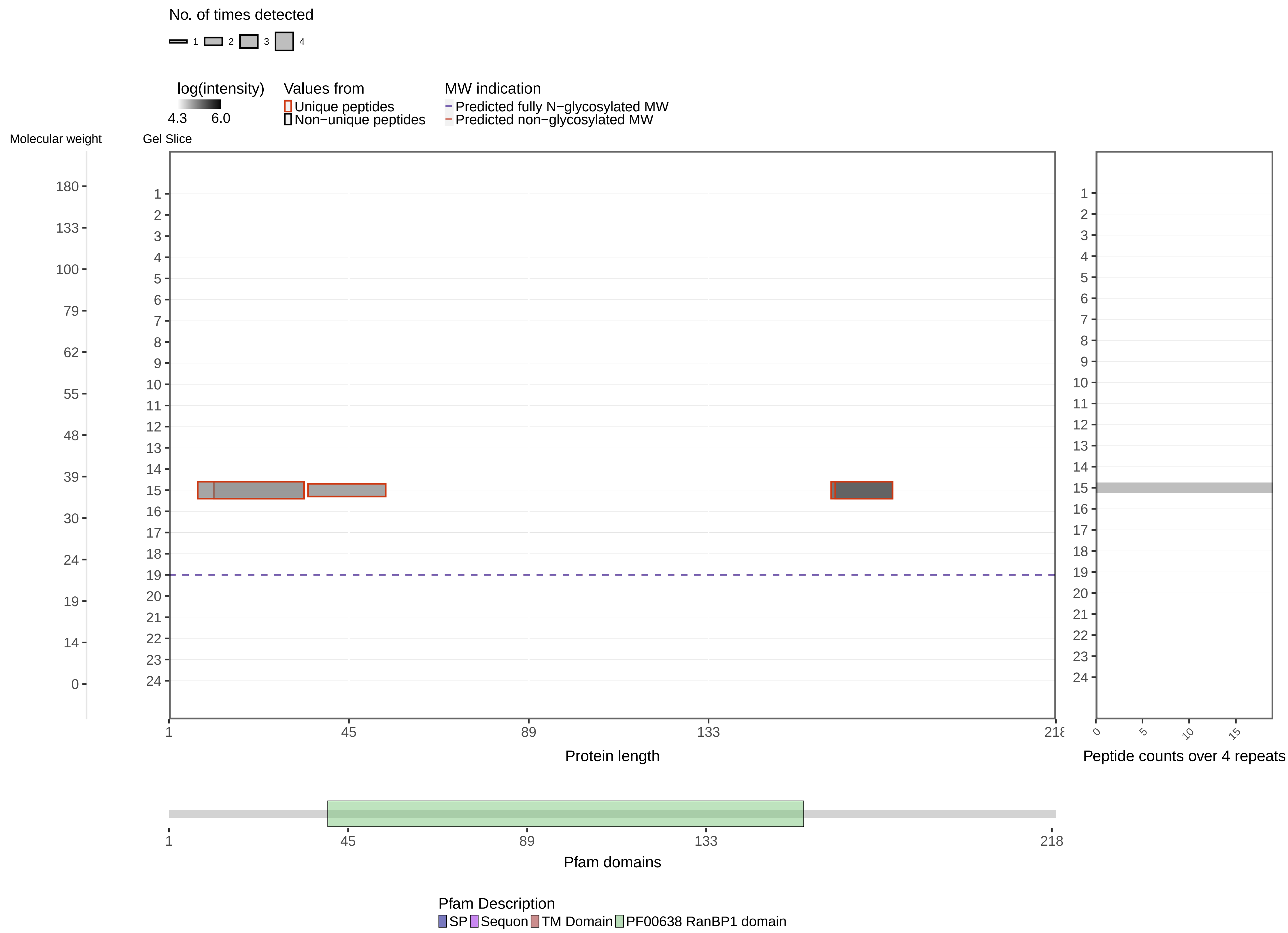


Protein sequence:

KICANHLVLPTMTLQEHAGNEKSCLWHAPDFADGELKDELFCLRFASIENCKSFMEMFQEVAESQKKVENEDGSTAAGLLEKLSVEDKKSEEKSEDKTKEGDGKGESPSKDTEKKDESASA*

### NbL02g08000.1 40S_ribosomal_protein_S26−3−like [MW = 14.51 kDa]

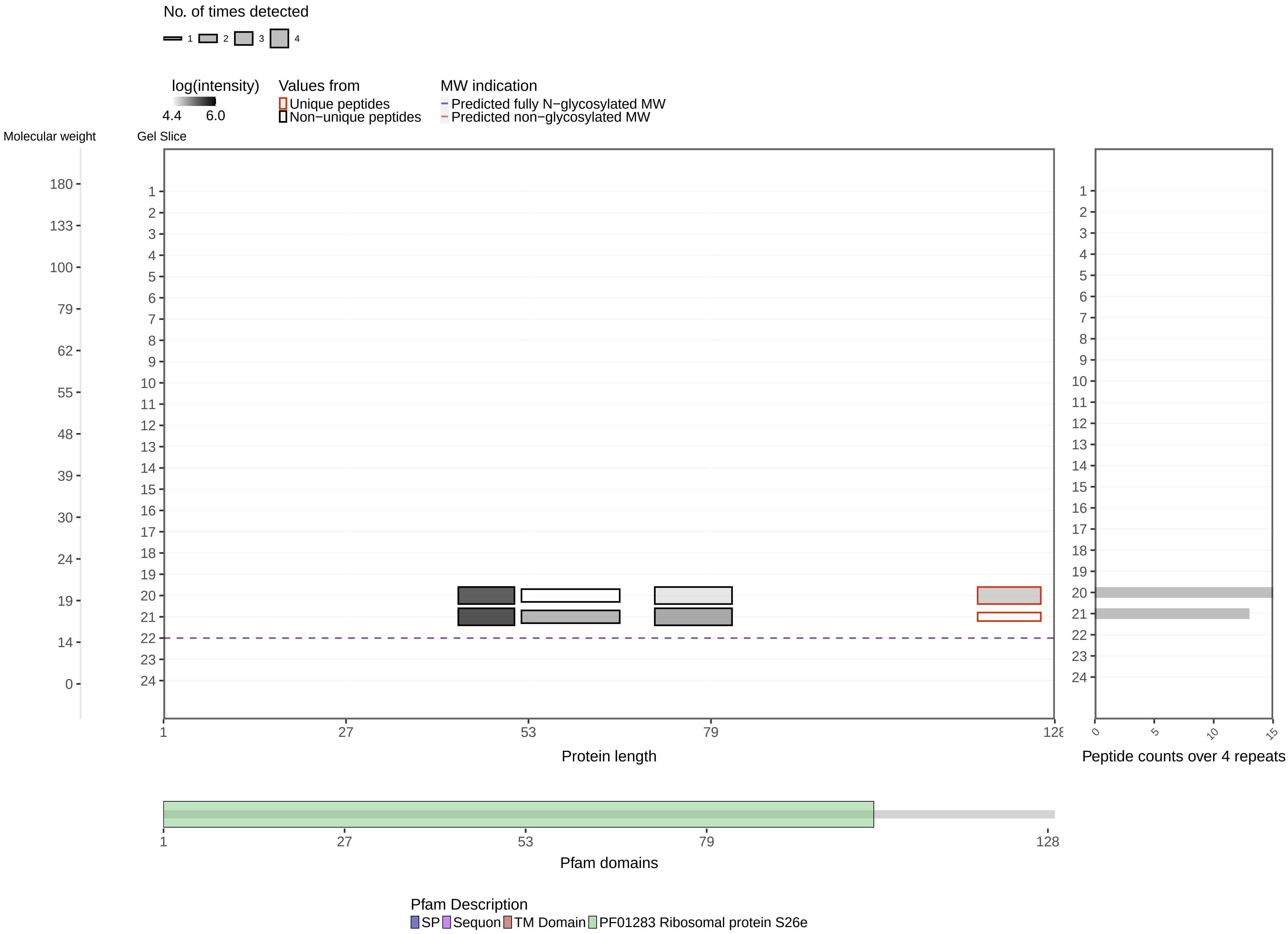


Protein sequence:

[illegible]DVQEACAFETYTLPKLYLKMQYCVSCAIHSKVVRVRSVTDRRVREPPQRFRRPRDDLPKPGQAPRPTGGAPAAPRT*

### NbL02g07960.1 60S_ribosomal_protein_L10−like [MW = 24.78 kDa]

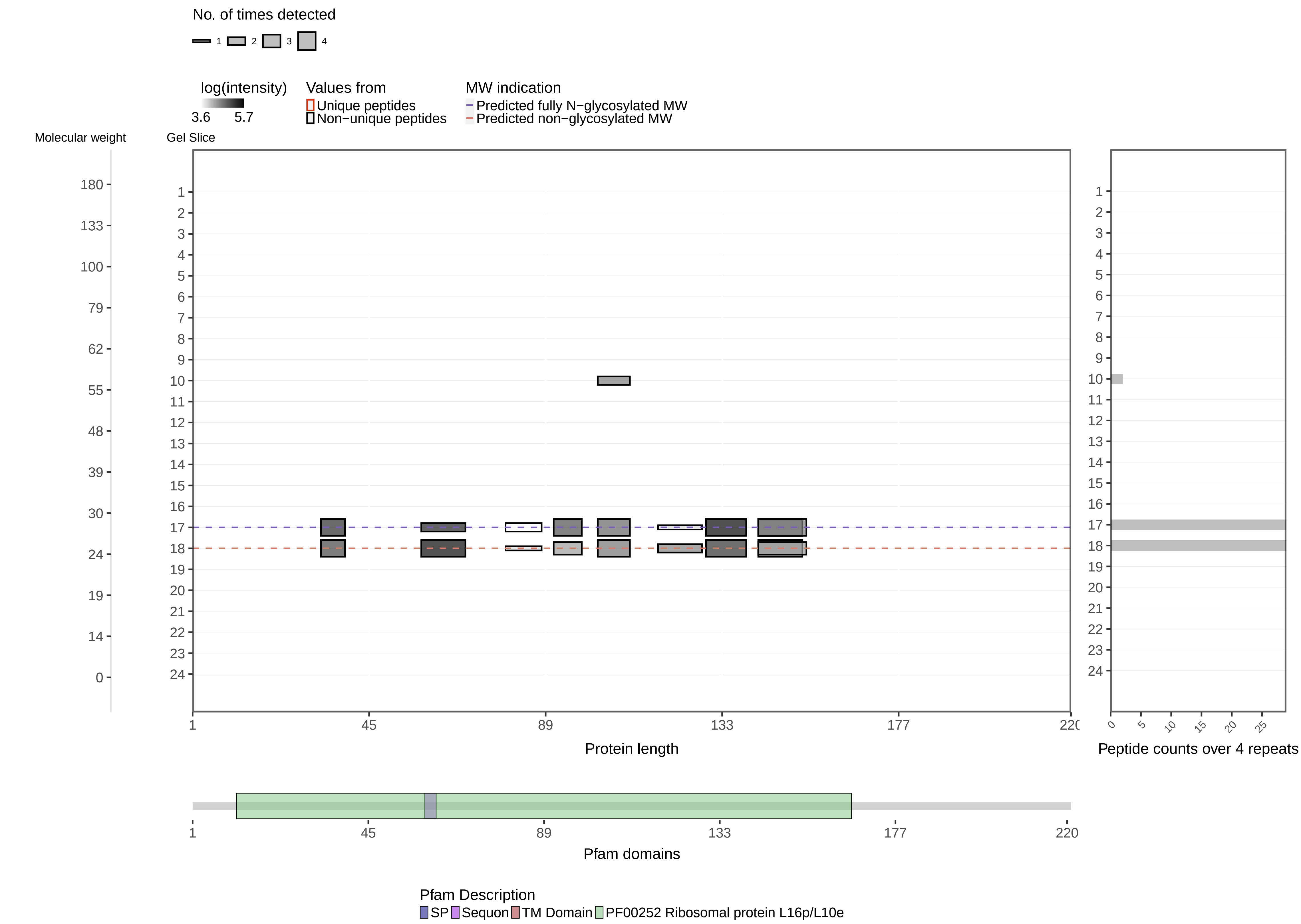


Protein sequence:

RINKMLSCAGADRLQTGMRGAFGKPQGVCARVAIGQVLLSVRCKDGNANHAQEALRRAKFKFPGRQKIIVSRKWGFTKFSRTDYLKYKSENRIVPDGVNAKLLGCHGRLAAROPGRAFLEAV*

NbL02g07830.1 60S_ribosomal_protein_L5−like [MW = 34.45 kDa]

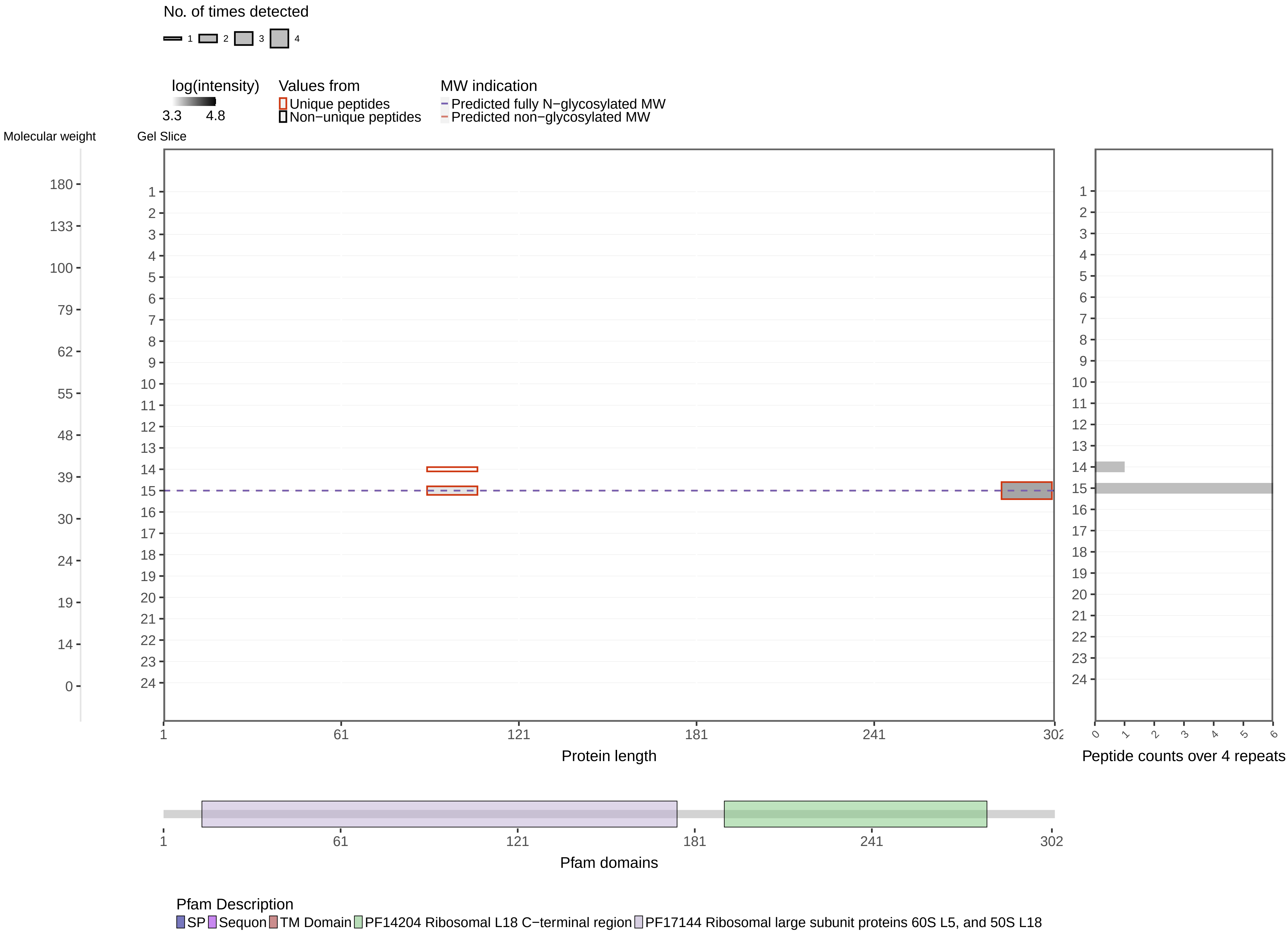


Protein sequence:

[illegible]PFRALLDVGLVRTTTGNRVFGALKGALDGGIDIPHSEKRFAGFGKDSKQLDAGVHRKYIYGGHVSAYMKTLMEDEPEKYQSHFSEYIKTGLEADDLEEMYKKVHAAIRADPSSKKSDKQPPKQHKRYNLKKLTYDERKARLIERLNALNAAAGNDDDEEDDD*

NbL02g07730.1 cytochrome_b5−like [MW = 14.99 kDa]

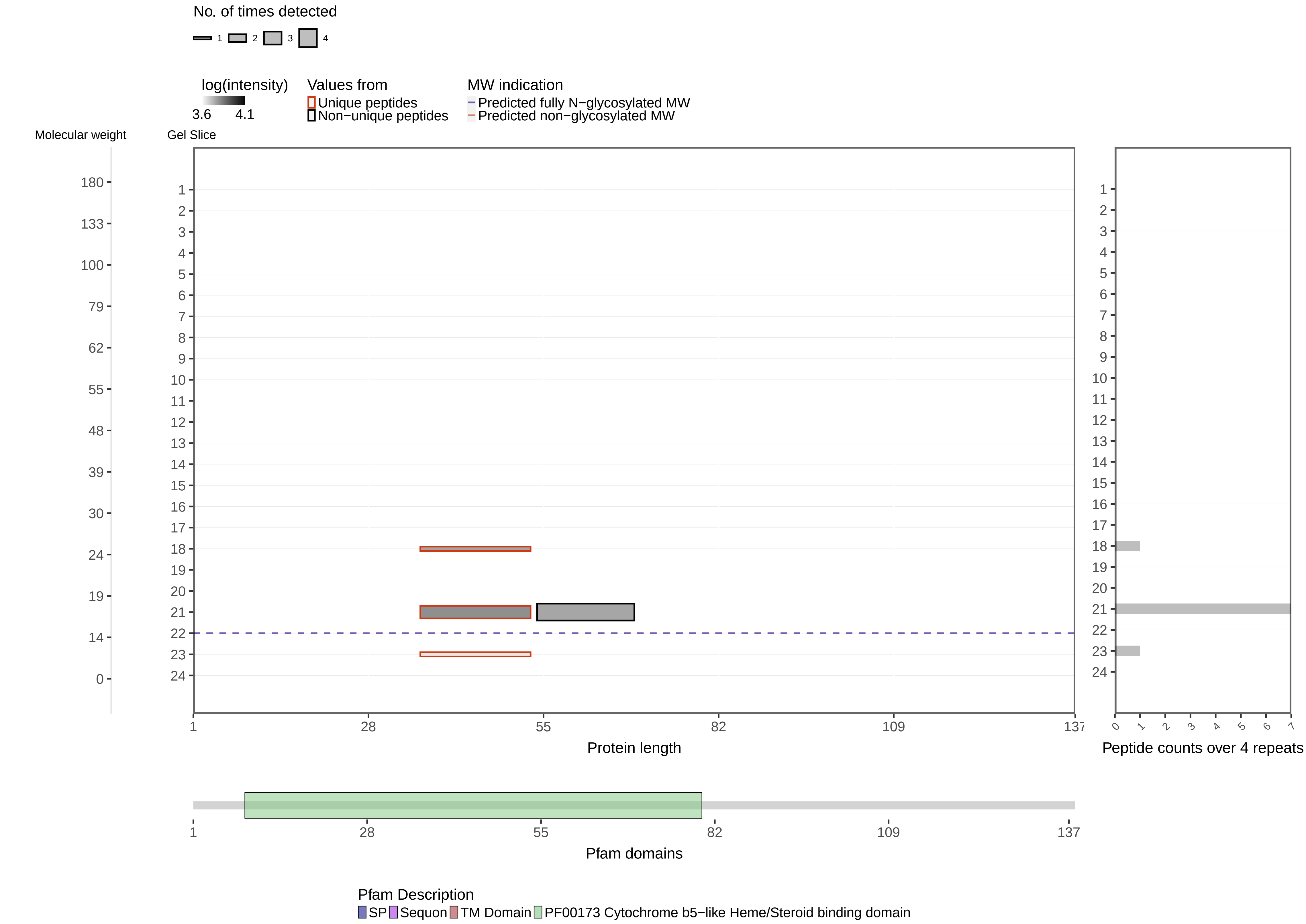


Protein sequence:

DATDDFEDVGHSTSARAMLDEYYVGNIDSATIPTKTKYTPPNQPHYNQDKTSEFVIKLLQFLVPLIILGVAVGIRFYTKQQSA*

### NbL02g07590.1 uncharacterized_LOC104215819 [MW = 16.63 kDa]

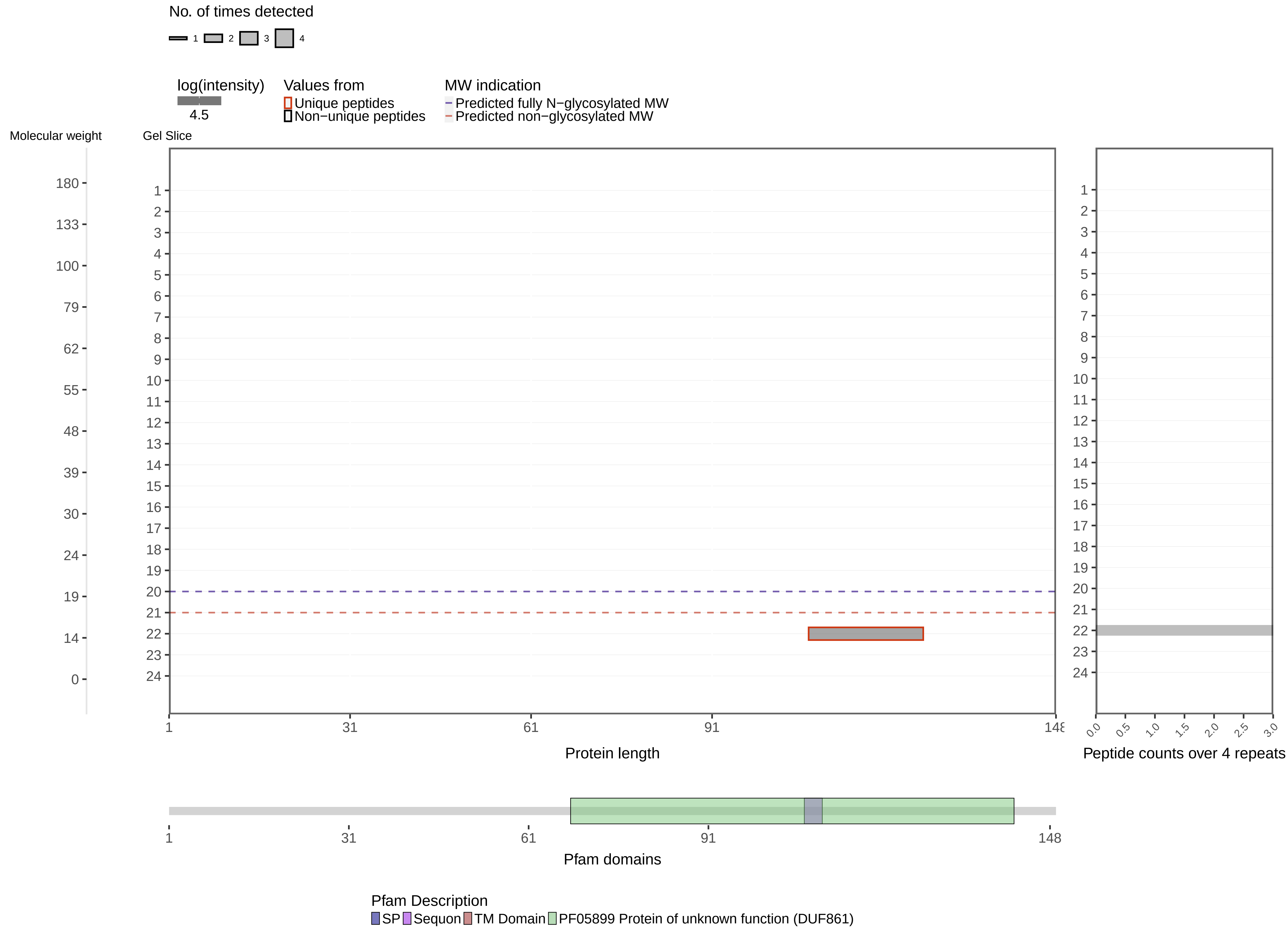


Protein sequence:

ESQLNDLGIKSWPKWGCSPGKYQLKFDAEETCYLLRGKVKVYPKNNSSDETVEFGAGDLVIIPKGLSCTWDVSVAVDKHYKFDSS*

### NbL02g07570.1 mitochondrial_import_receptor_subunit_TOM9−2 [MW = 11.13 kDa]

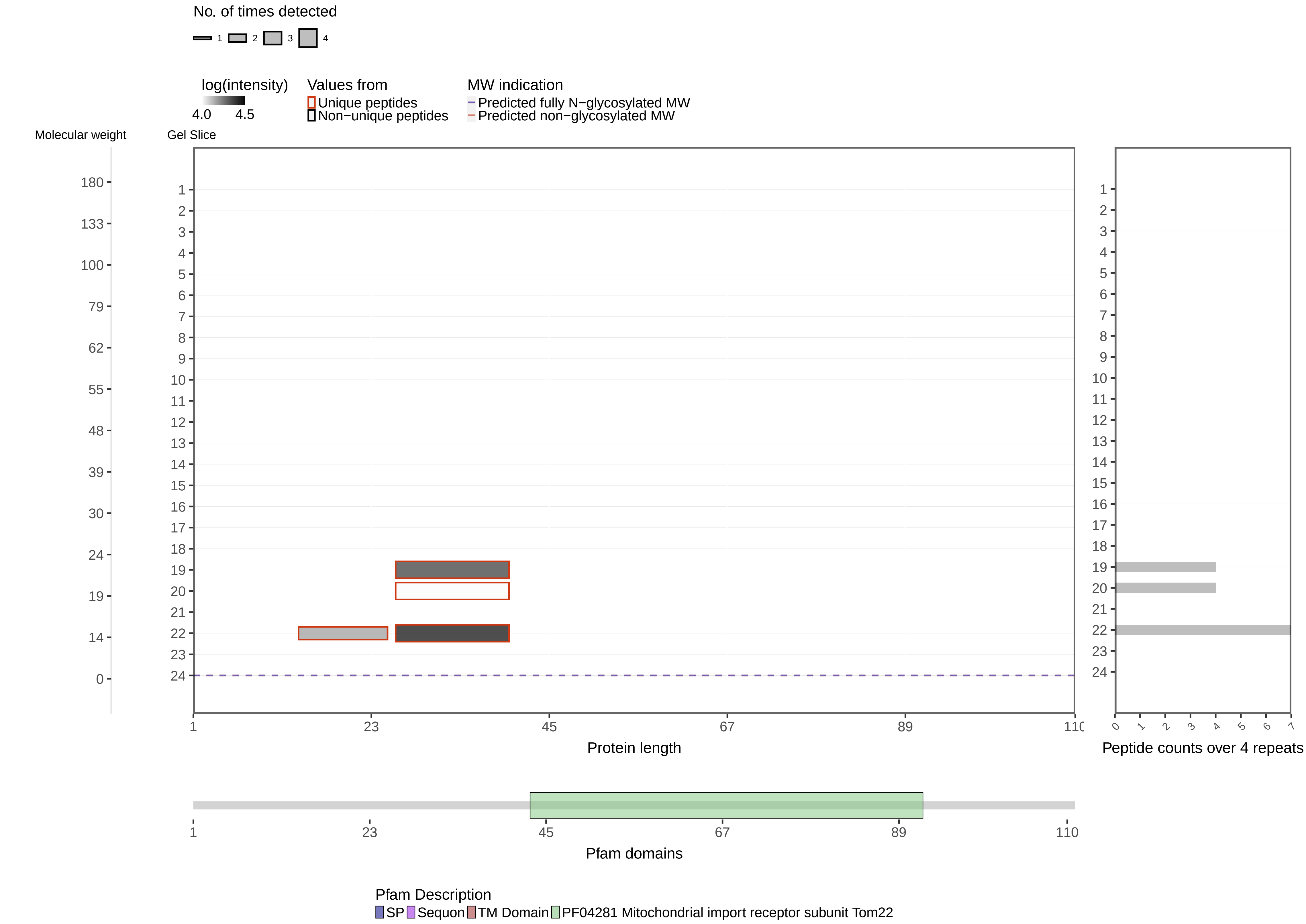


Protein sequence:

RAASDAALIAKKLLKSTGKAAWIAGTTFLILVVPLIIEMDRDAQLTELELQQASLLGTPPPPSAAGGPK*

NbL02g07080.1 leucine−rich_repeat_receptor−like_serine/threonine−protein_kinase_BAM1 [MW − SP = 108.87 kDa]

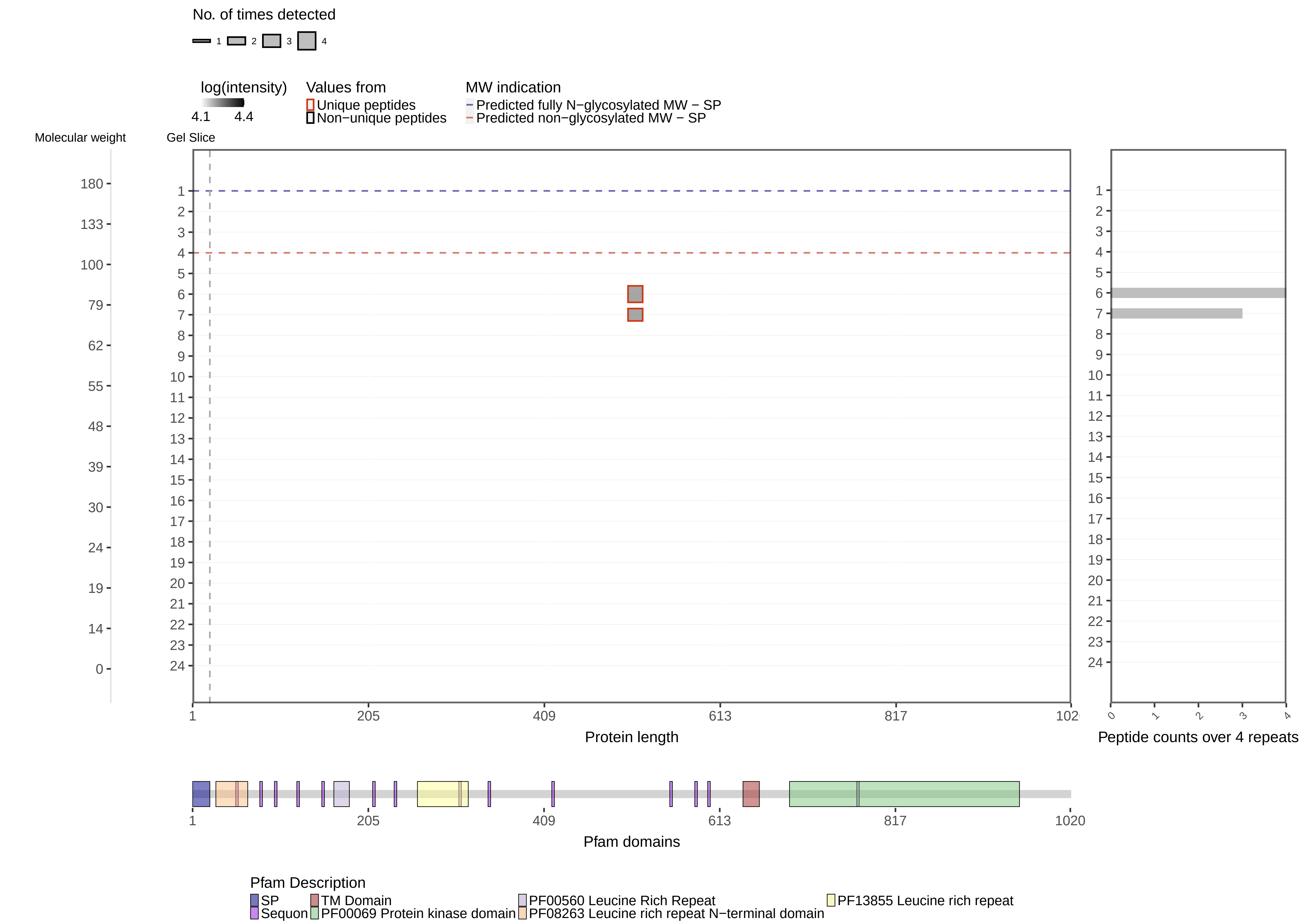


Protein sequence:

OLSKIDFSHNDFSGPIVPEISKCKLLTYVDLSRNOLSGEIPTEITGMRILNYLNLSRNHLVGSIPAPISSMOSLTSVDFSYNNFSGLVPGTGOFSYFNYTSFLGNPDLCGPYLGPCKEGVVDGVSOPHORGALSPSMKLLLVIGLLVCSIVFAVAAIIKARSLKKASEARAWKLTAFORLDFTCDDVLDSLKEDNIIGKGGAGIVYKGVMPSGEHVAVKRLPAMSRGSSHDHGFNAEIOTLGRIRHRHIVRLLGFCSNHETNLLVYEYMPNGSLGEMLHGKKGGHLHWDTRYKIAV

NbL02g07010.1 probable_xyloglucan_endotransglucosylase/hydrolase_protein_6 [MW = 34.85 kDa]

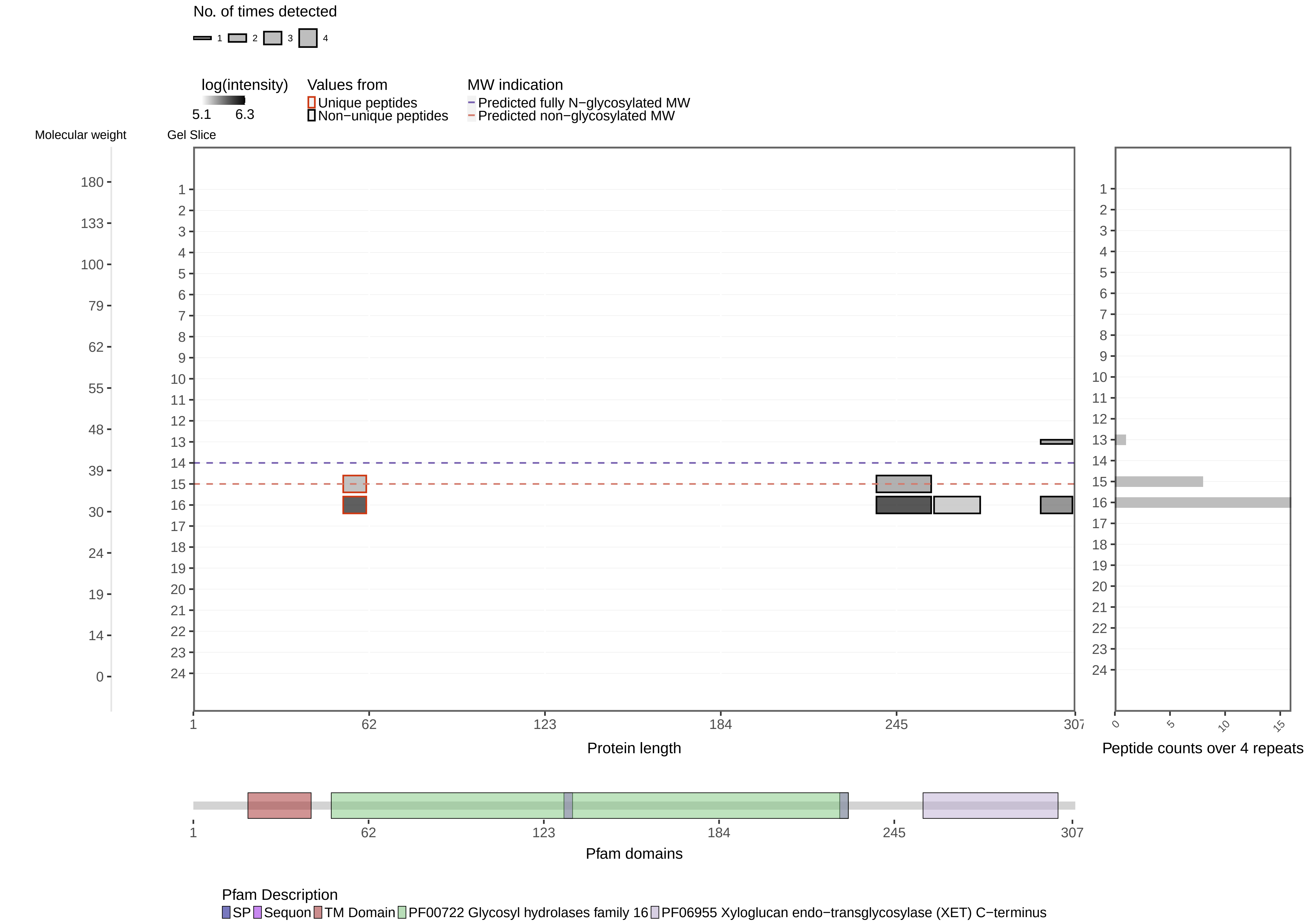


Protein sequence:

### NbL02g07000.1 probable_xyloglucan_endotransglucosylase/hydrolase_protein_7 [MW − SP = 30.21 kDa]

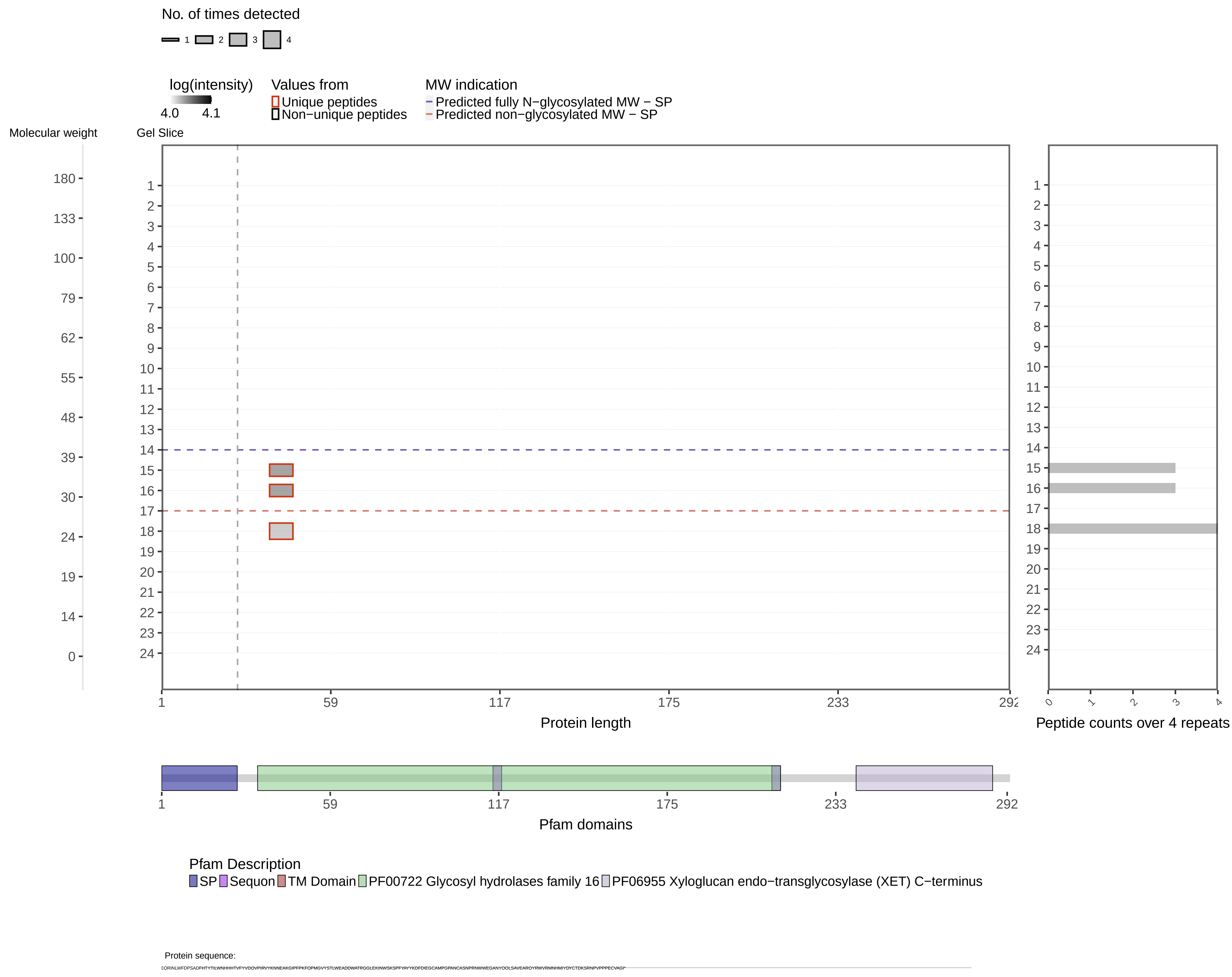

NbL02g06680.1 ADP−ribosylation_factor−like_protein_8a [MW = 20.52 kDa]

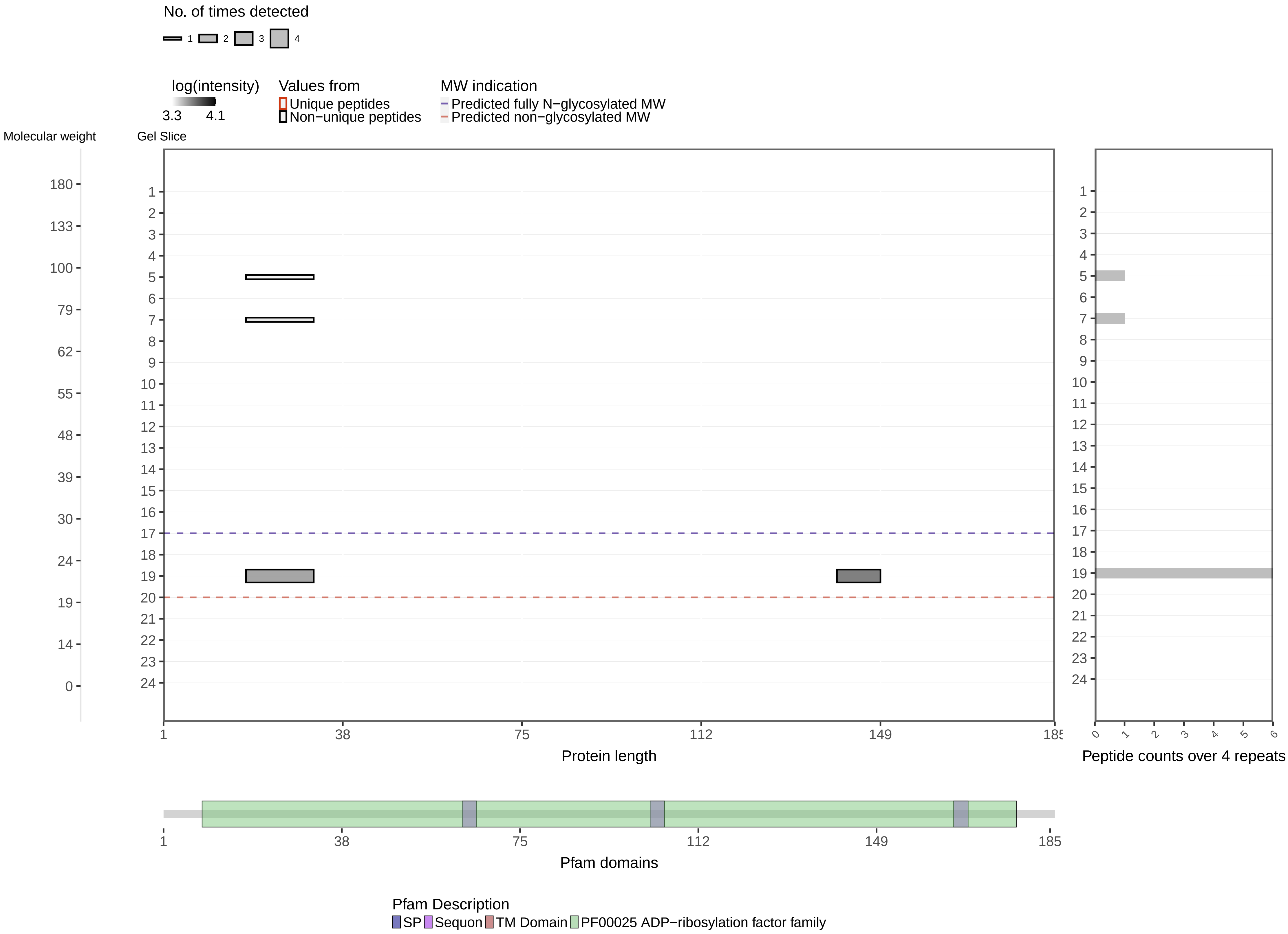


Protein sequence:

SMWERYCRAVSAIVYVVDAADPDNLSISSRELHDLLSKPSLSGIPLLVLGNKIDKPGVQSKQALTDQMGLKSITDREVCCYMISCKNSTNIDSVIDWLVKHSKSKS*

NbL02g06470.1 peroxidase_51−like [MW − SP = 33.23 kDa]

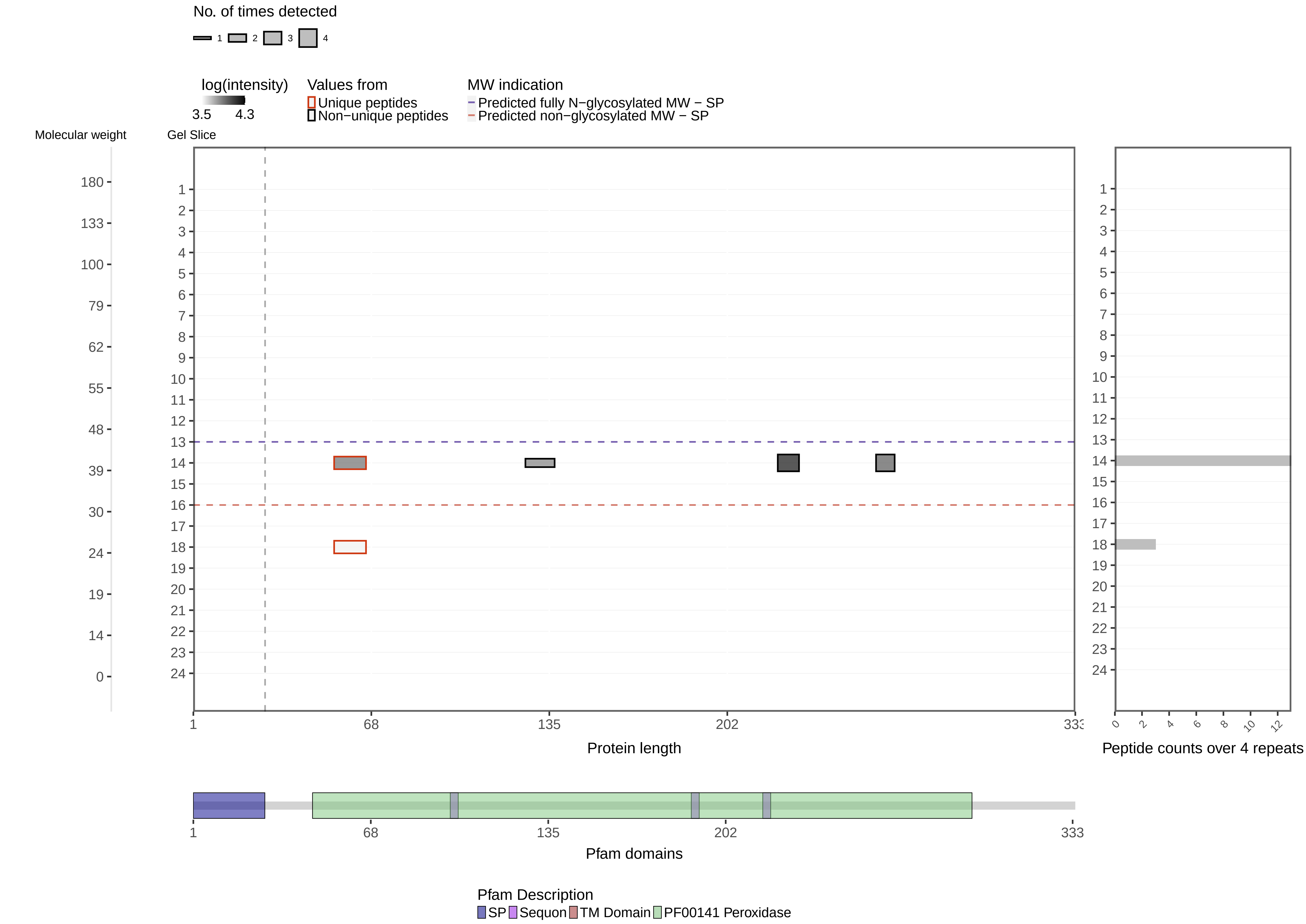


Protein sequence:

.DGFTSKASNVEGKLPKPTFNLNQLNSMFASHGLNQSDMIALSAAHSVGFSHCSKFANRIYNFSPKNPIDPTLNKQYAAQLQGMCPRNVDPRIAINMDPKTPRTFDNAYFKNLQQGMGLFTSDQVLYTDGRSKGTVDIWASNSKAFQNAFVSAMTKLGRVGVKTGSNGNIRFDCGRFN*

NbL02g06410.1 subtilisin−like_protease_SBT1.7 [MW = 81.95 kDa]

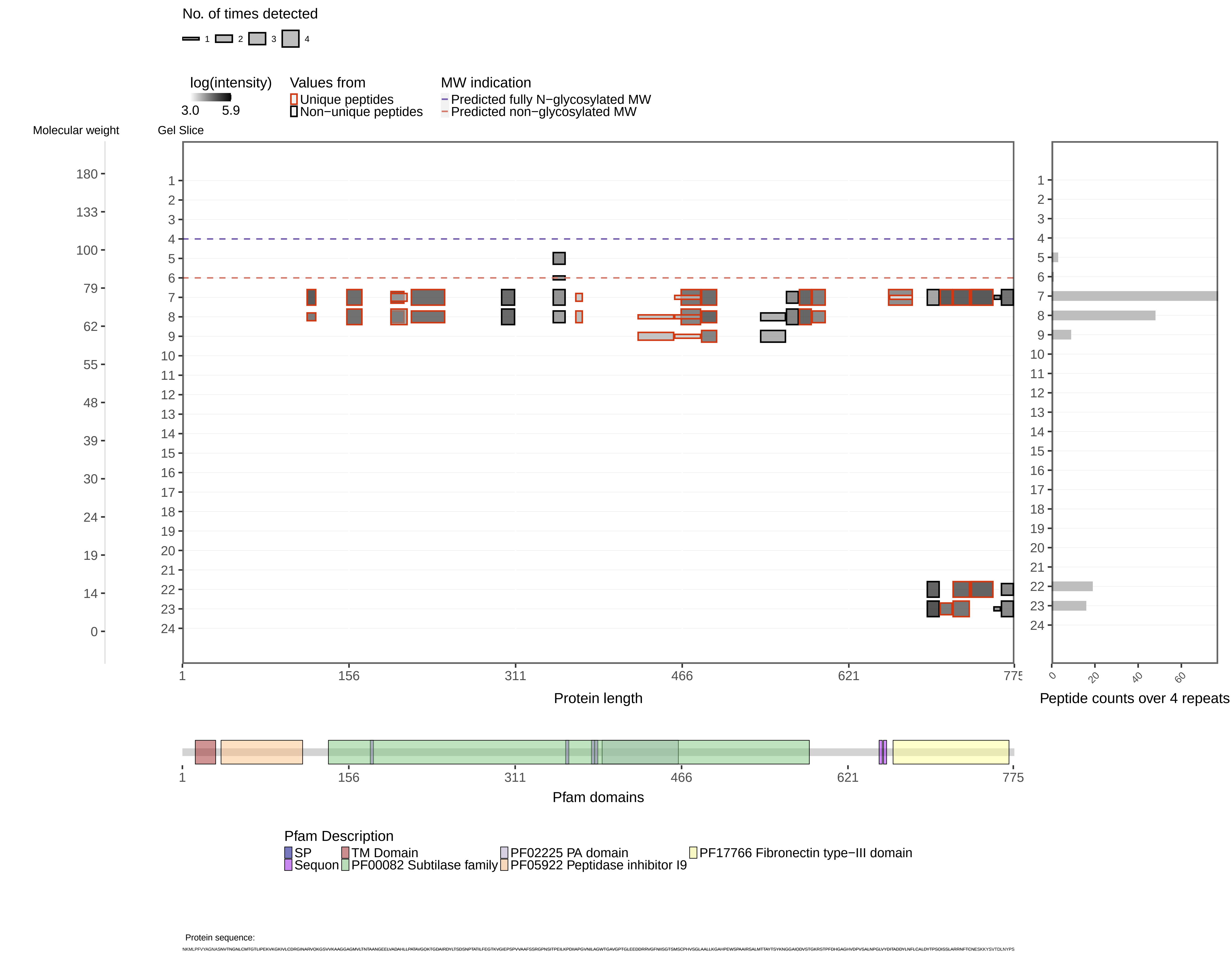

### NbL02g06340.1 probable_esterase_KAI2 [MW = 29.81 kDa]

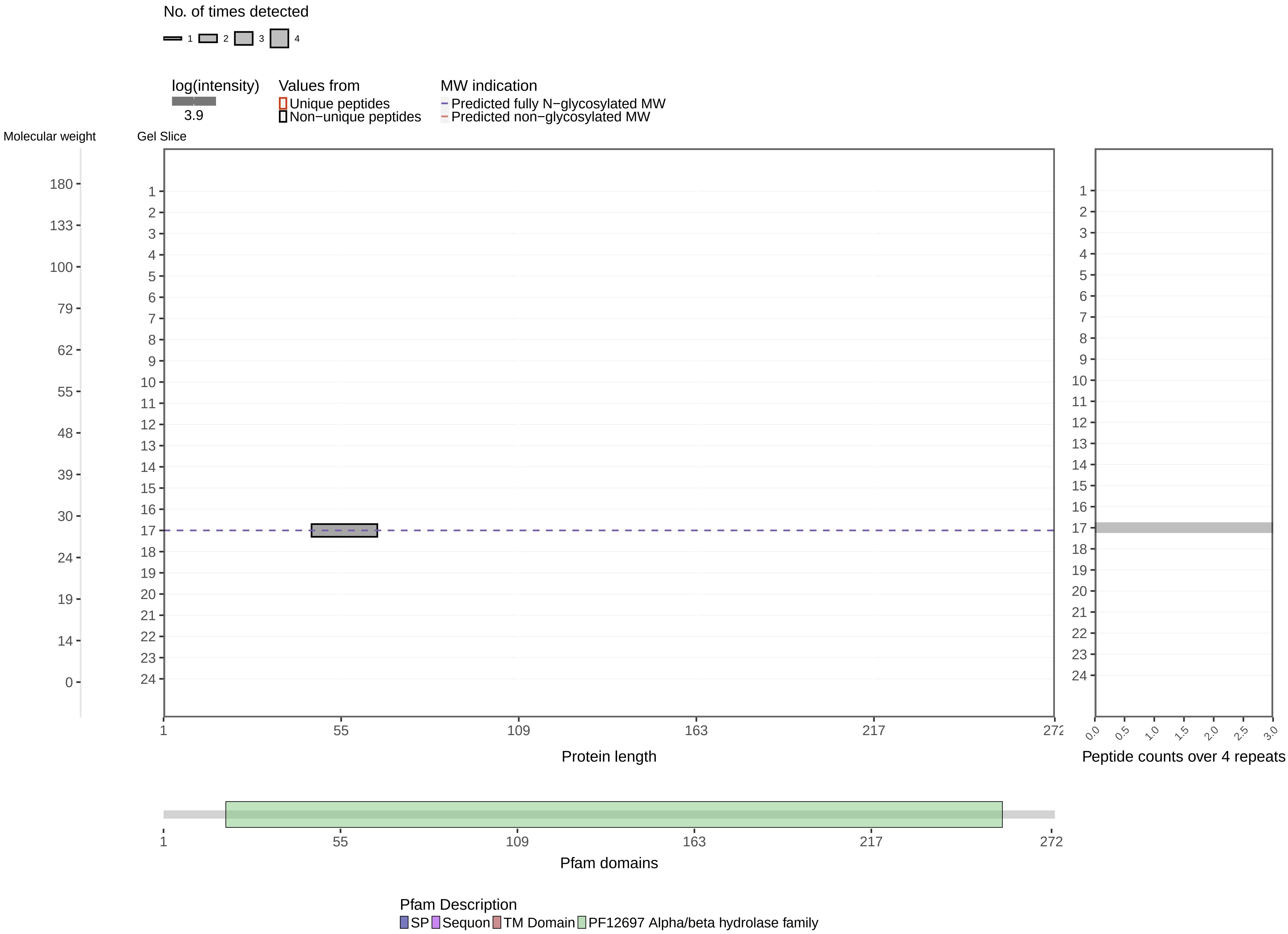


Protein sequence:

YLNDVDYYGGFEQEDLDQLFEAMTSNYKAWCSGFAPLAVGGDMDSVAVQEFSRTLFNMRPDIALSVLQIIFQSDLRHLLPHVTVPCHIIQSMKDLAVPVVVSEYLHQNLGGGSIVEVMSTDGHLPQLSSPDVVIPVLLRHIRYDIAVD*

NbL02g06000.1 caffeoyl−CoA_O−methyltransferase_4 [MW = 27.31 kDa]

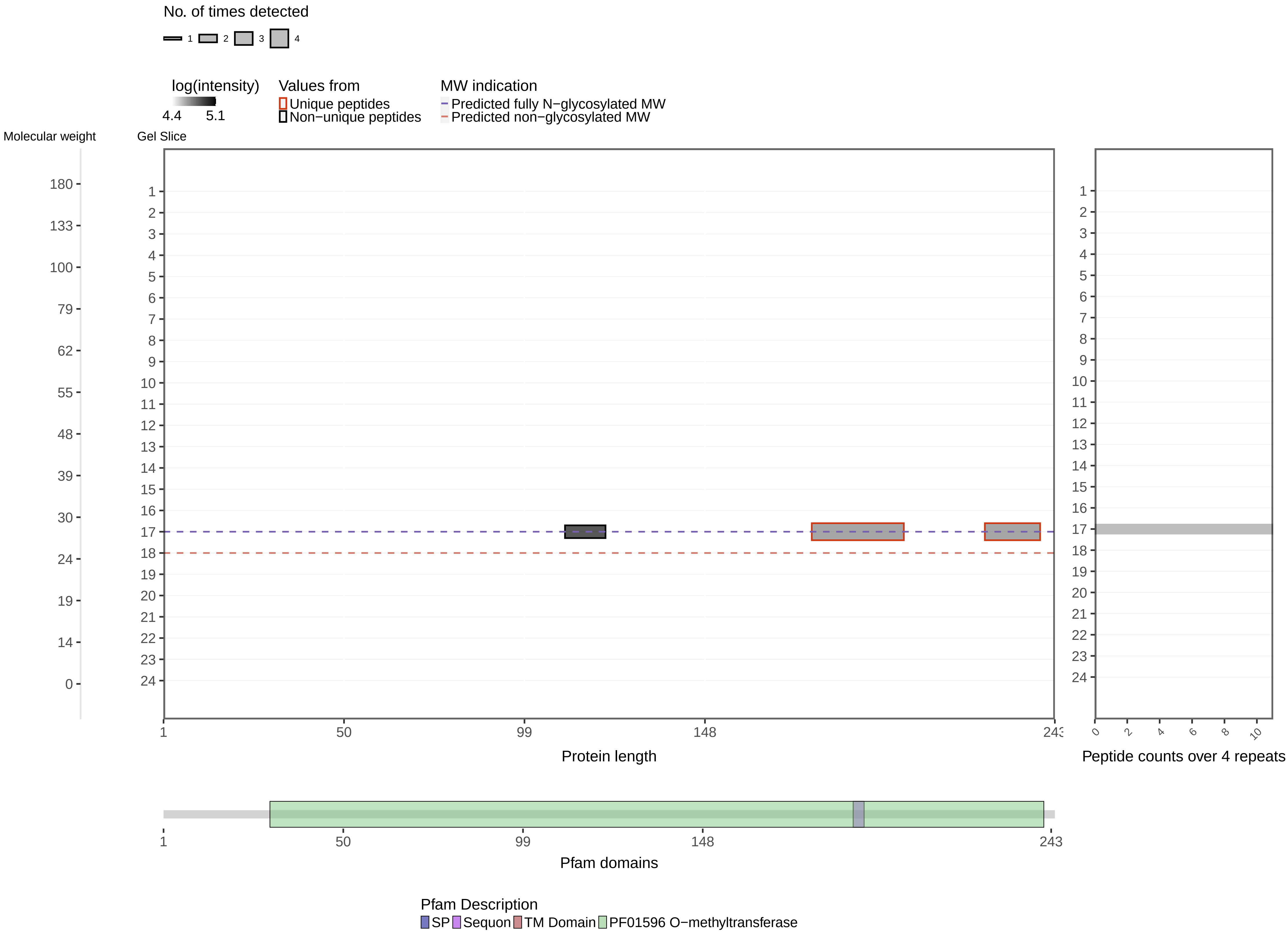


Protein sequence:

INRENYEIGLPVIEKAGLAHKIEFKEGPALPVLDRMIEDGKYHGSYDFIFVDADKDNYLNYHKRLIDLVKIGGLIGYDNTLWNGSVVAPPDAPLRKYVRYYRDFVLELNKALAADSRIEICQLPVGDGITLCRRIT*

### NbL02g05960.1 nicotinate_phosphoribosyltransferase_2−like [MW = 62.91 kDa]

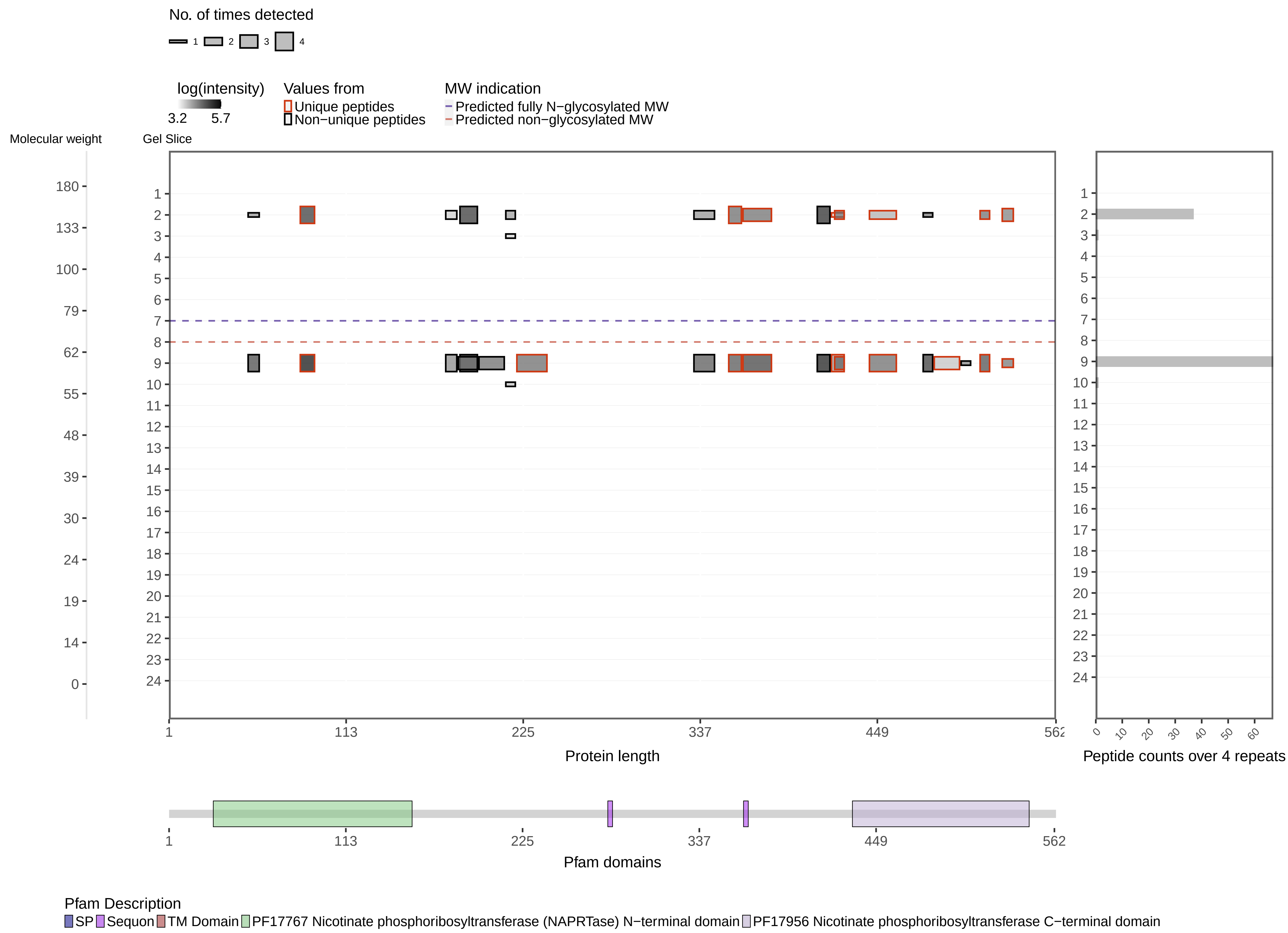


Protein sequence:

ILLGGIFSETNOSELAAFTSYALAFPGSFLALVDTYDVMRSGIPNFCAVALALNDLGYKAVGIRLDSGDLAYOSCEARKFFRAIEKEFGLPGFGKMNITASNDLNEETLDALNKOGHEVDAFGIGTHLVTCYAOAALGVVFKLVEINNOPRIKLSEDVSKVSIPCKKRCYRLYGKEGYALVDIMTGENEPOPKVGERILCRHPFSESKRAYVVPORVEELLECFWPGNSDOTREDLPPLKEIRERCIEOLDKMRPDHMRRLNPTPYKOVSVSAKLYDFIHFLWLNEAPVGELO*

NbL02g05940.1 PI−PLC_X_domain−containing_protein_At5g67130−like [MW − SP = 42.09 kDa]

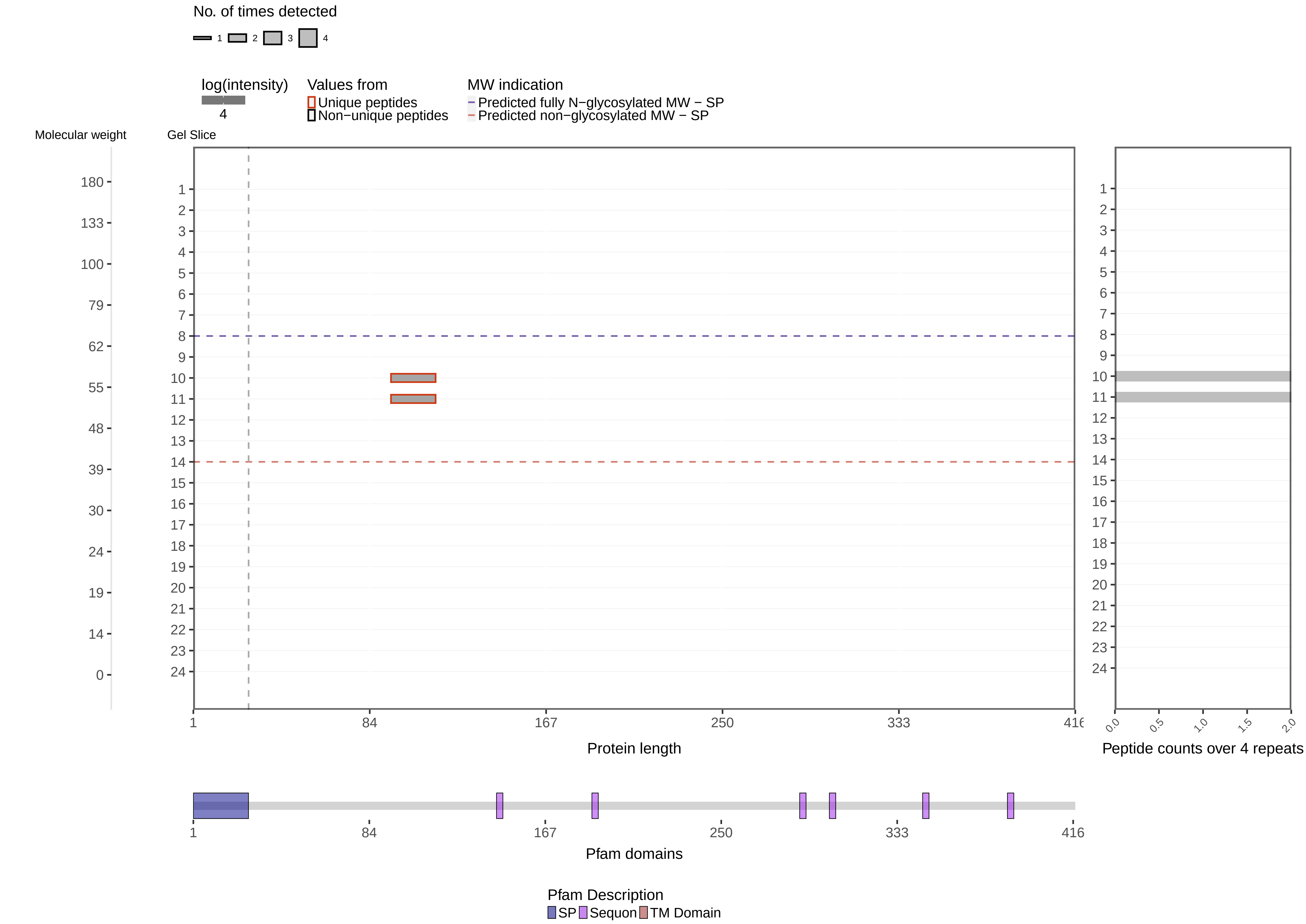


Protein sequence:

PVSRMPKKREDWPTVDDMVKQNQRLVVFTSKSSKEASEGIAYEWRYVVENQYGNGGMKVGSCPSRSESSPMSTSSISLVLQNYFPDNPNITAACVDNSATLINMTKTCFEAAGKRWPNFIAVDFYQKSDGGGAPEAVDEANGQLACNCSSIAYCKASGTSHKCEAPMLSPPPPAQATSIGTSQDGNNESNDNSHSSRSKPLHWLIGTIFVAALLLHA*

### NbL02g05860.1 zeaxanthin_epoxidase_chloroplastic [MW = 72.59 kDa]

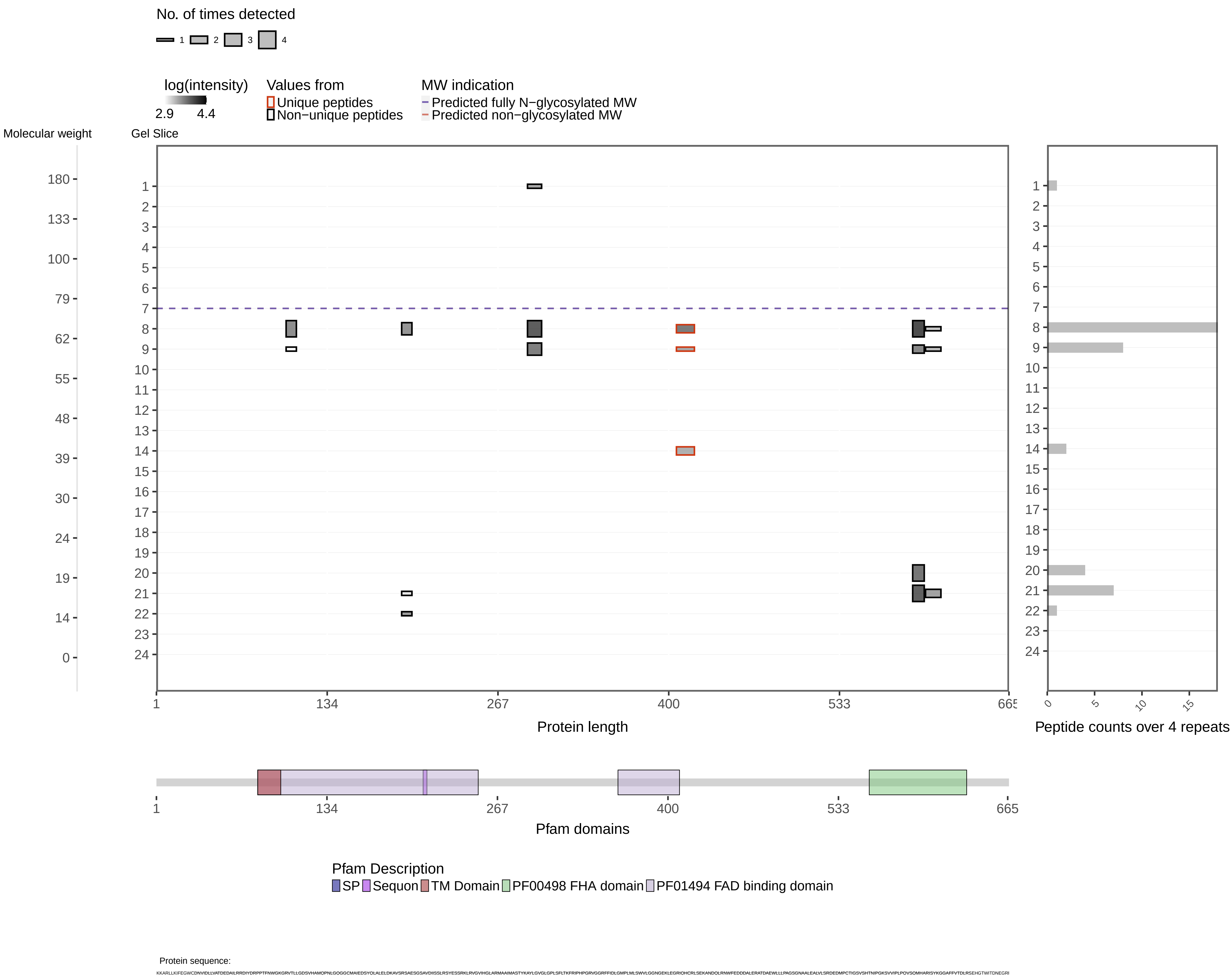

### NbL02g05810.1 high_mobility_group_B_protein_14 [MW = 18.56 kDa]

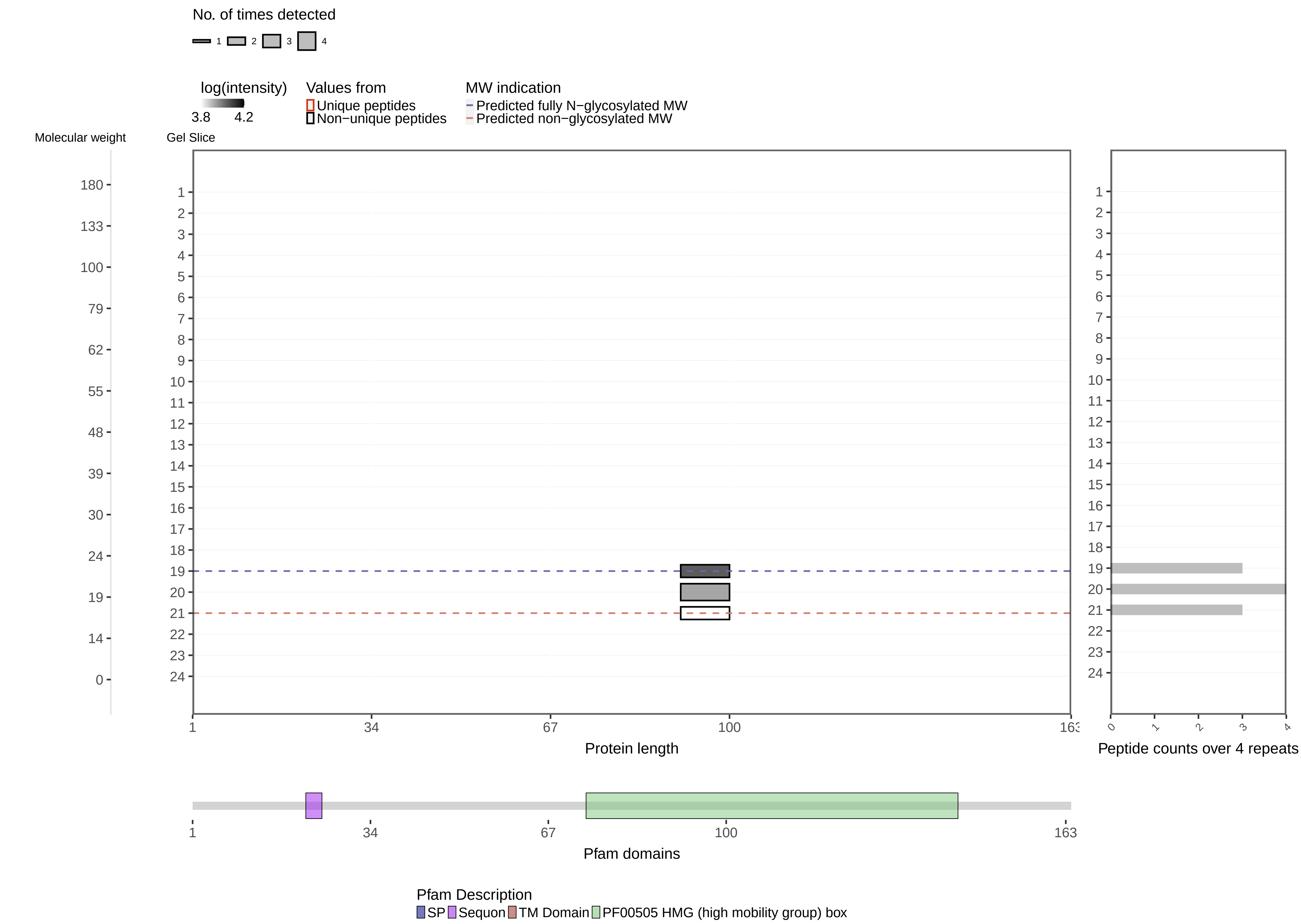


Protein sequence:

DAMKPKKPPTAFFYFLEDFRKEFQEQNPDVKSMRDVGKACGEKWKTMTYEEKVHYYDIATEKRREFDRAMADFNKRKESGEFQEFEEDSDFDE*

### NbL02g05550.1 DNA_damage−binding_protein_1 [MW = 122.19 kDa]

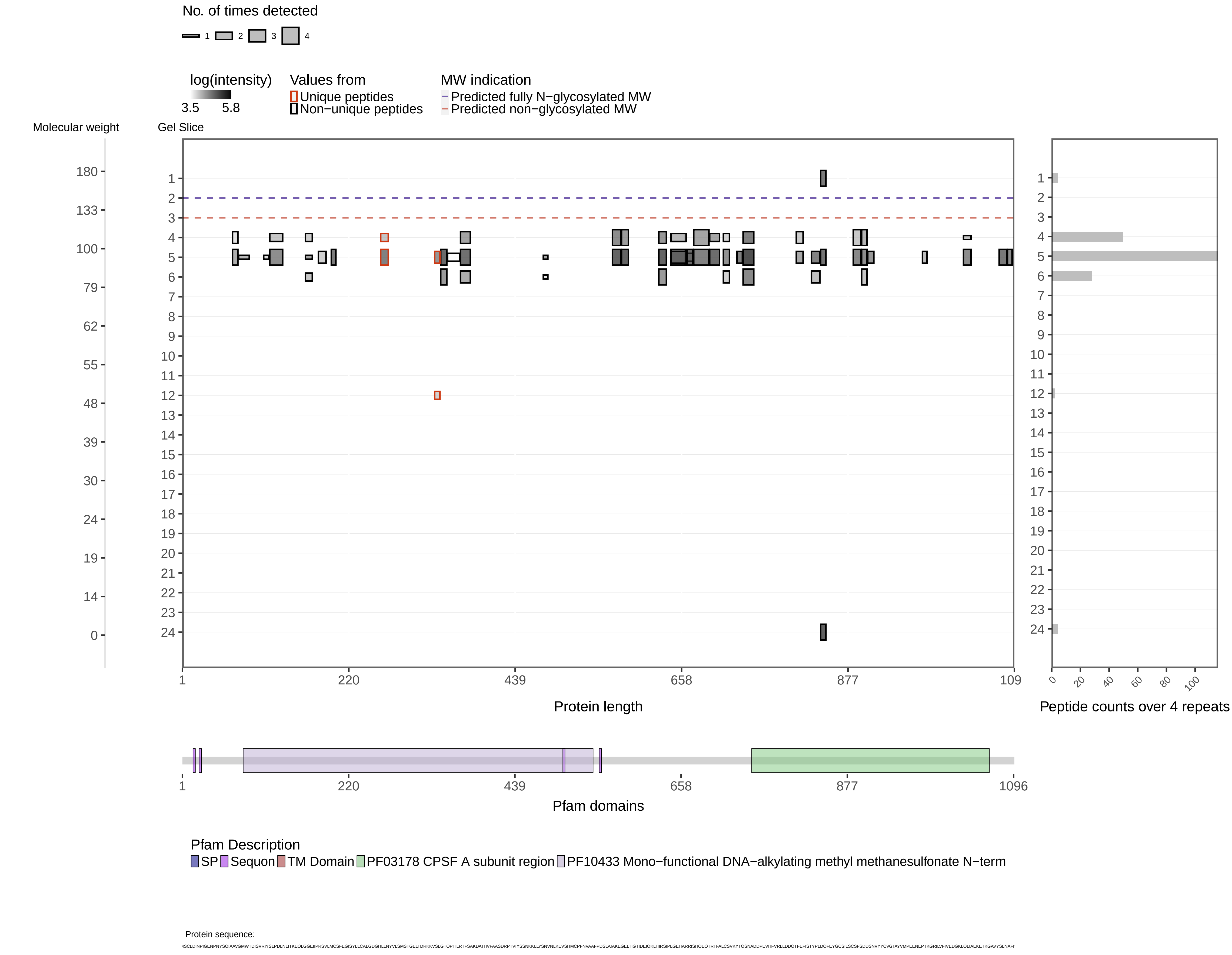

NbL02g05500.1 40S_ribosomal_protein_S4−like [MW = 29.88 kDa]

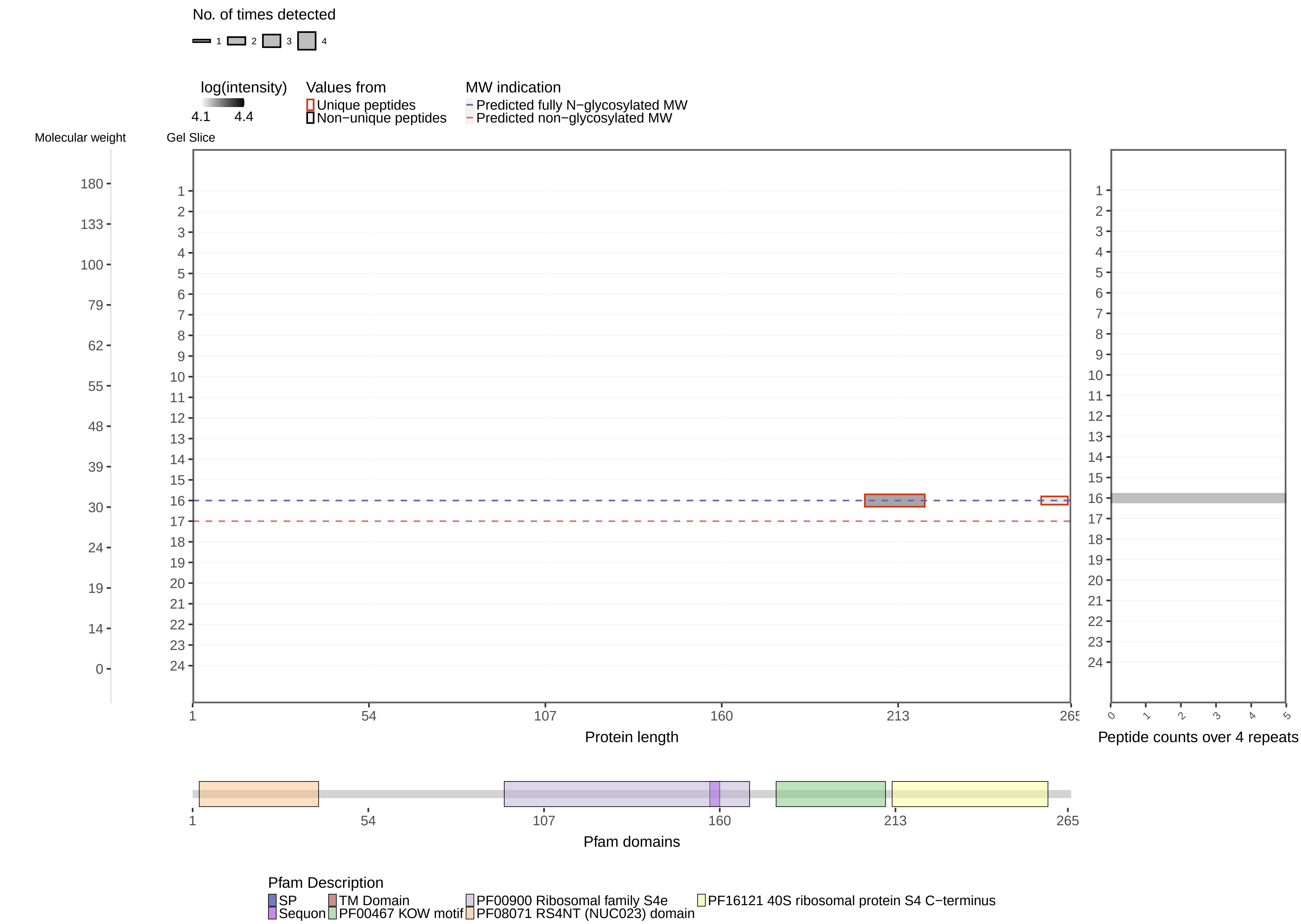


Protein sequence:

:AKFKLCKVRSVQFGQKGIPYLNTYDGRTIRYPDPLIKANDTIKLDLENNKIVDFIKFDVGNVVMVTGGRNRGRVGVIKNREKHKGSFETVHIQDAQGHEFATRLGNVFTLGKGTKPWVSLPKGKGIKLSIIEEARKRLAAQSATTT*

### NbL02g05280.1 tubulin_beta_chain [MW = 57.69 kDa]

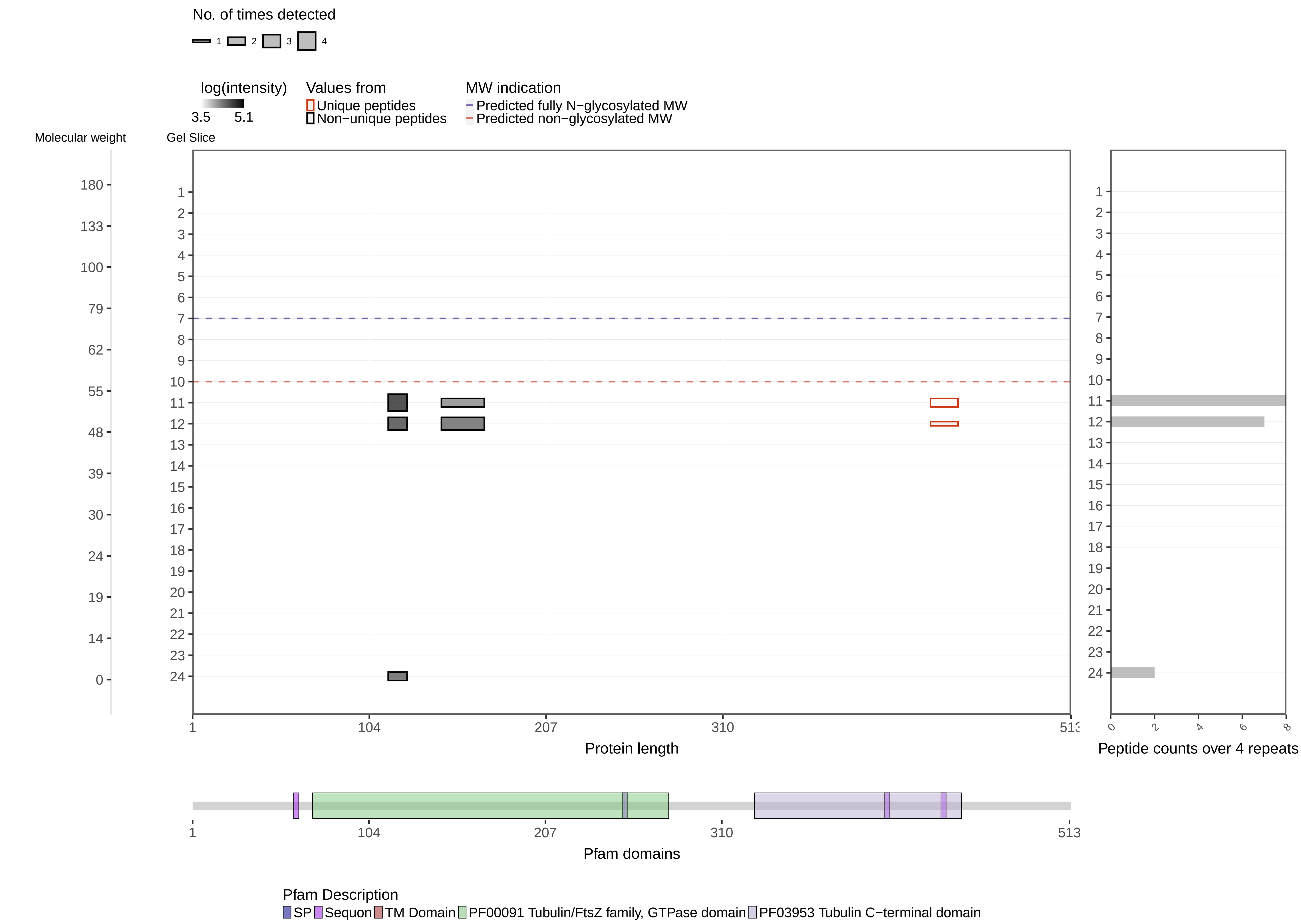


Protein sequence:

DTVVEPYNATLSVHQLVENADECMVLDNEALYDICFRTLKLSTPKFGDLNHLISATMSGVTCCLRFPGQLNSDLRKLAVNLIPFPRLHFFMVGFAPLTSRGSQLYRALTVPELTQQMWDAKNMMCAADPRHGRYLTASAMFRGKMSTKEVDEQMLNVQNKNSSYFVEWIPNNVKSSVCDIPPKGLKMASTFIGNSTAIQEMFRRVSEQFTAMFRRKAFLHWYTGEGMDEMEFSEAESNMNDLVAEYQQYQDATADDEEYEEEEEDVAA*

NbL02g05270.1 probable_3−hydroxyisobutyrate_dehydrogenase_mitochondrial [MW = 43.88 kDa]

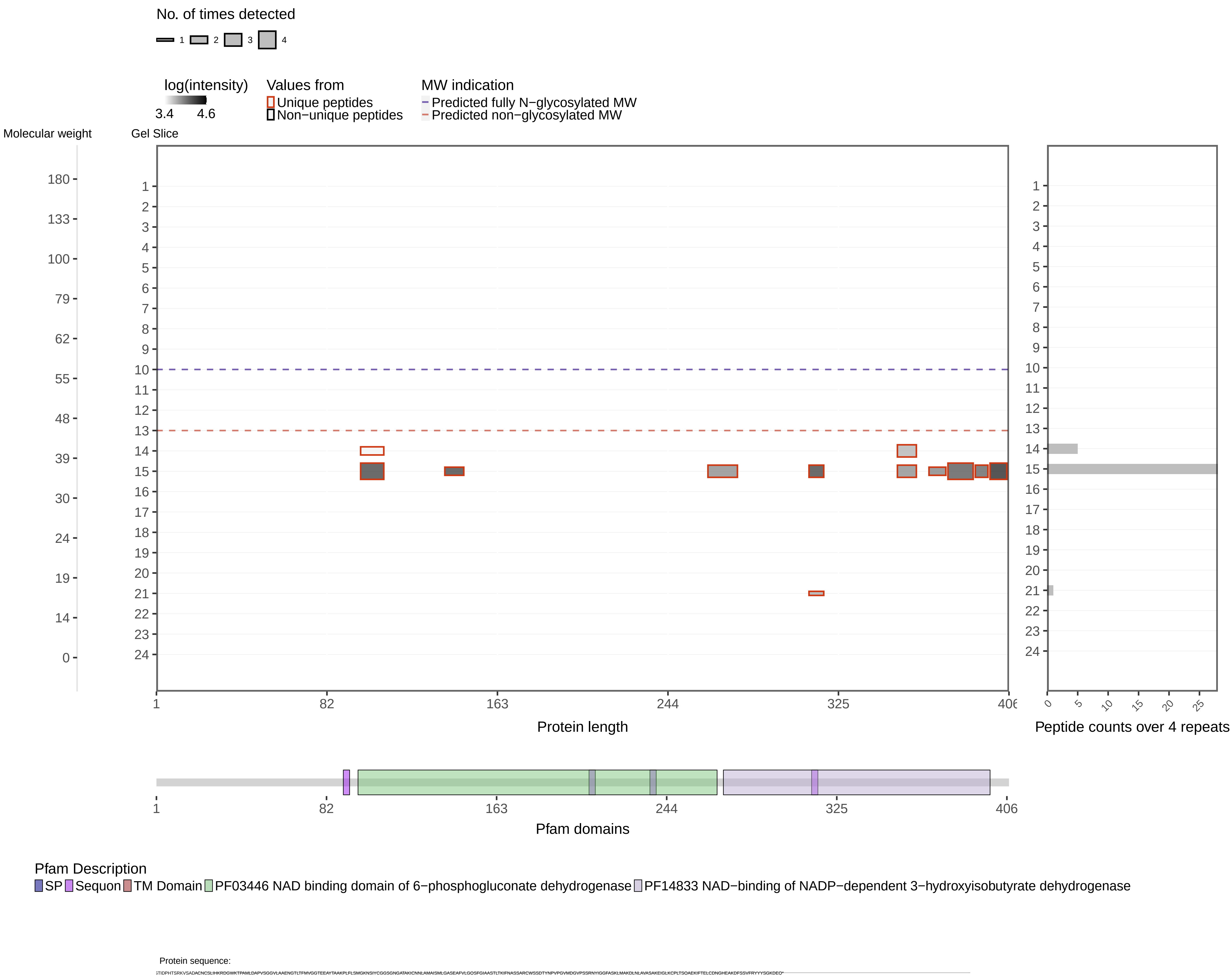

NbL02g05260.1 60S_ribosomal_protein_L35 [MW = 14.32 kDa]

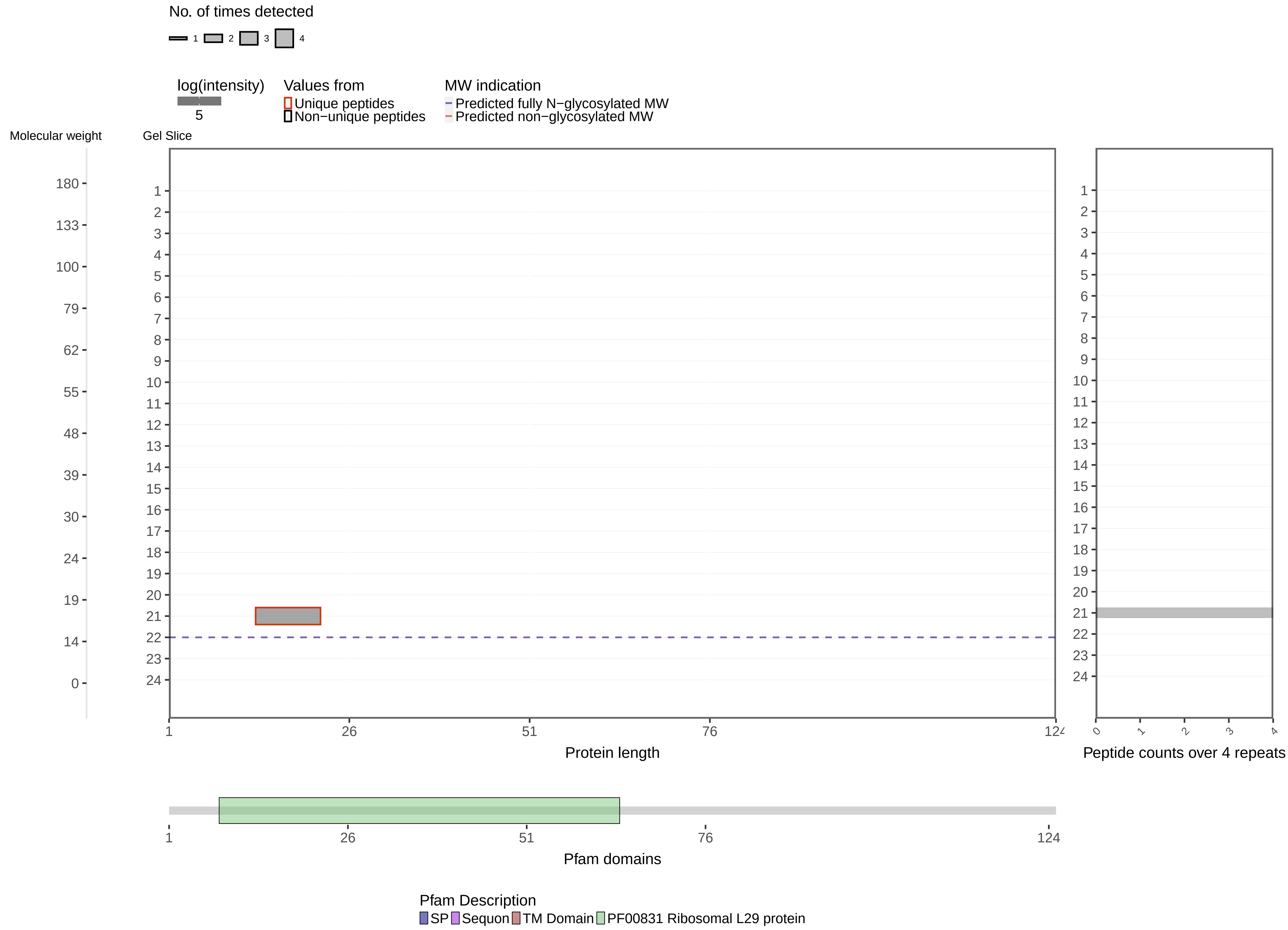


Protein sequence:

VVRLSIAQVLTVISQKQKTALREAYKNKKYLPLDLRPKKTRAIRRRLTKHQASLKTEREKKKEMYFPLRKYAIKV*

### NbL02g05220.1 actin−depolymerizing_factor_2−like [MW = 16.11 kDa]

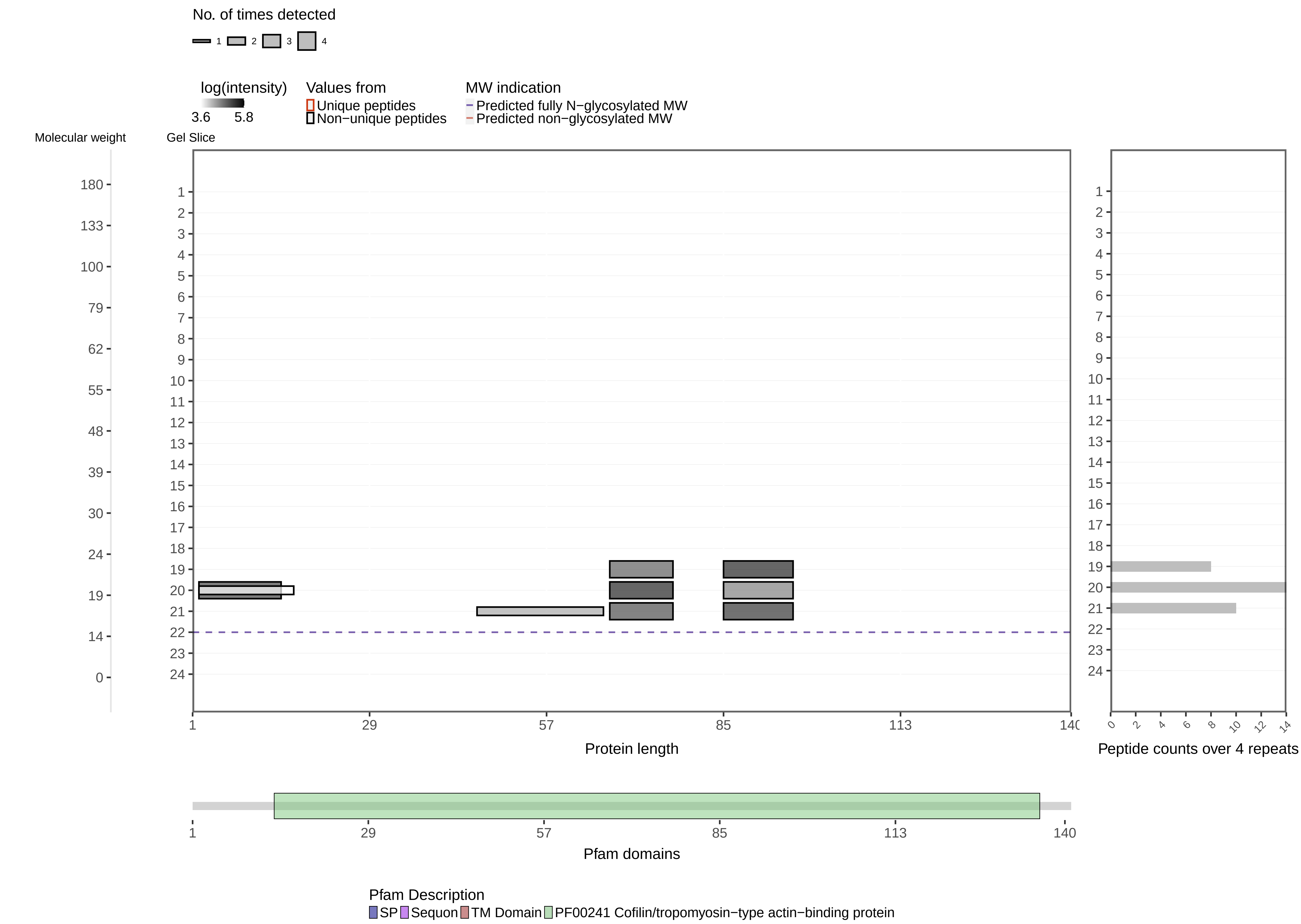


Protein sequence:

CTHLPADECRYAVYDFDFLTKESVPKSRIFFIAWSPDTSKVRSKMIYASSKDRFKRELDGIQIELQATDPTEMGLDVFKSRAN*

NbL02g05130.1 ferredoxin−−NADP_reductase_root−type_isozyme_chloroplastic−like [MW = 41.91 kDa]

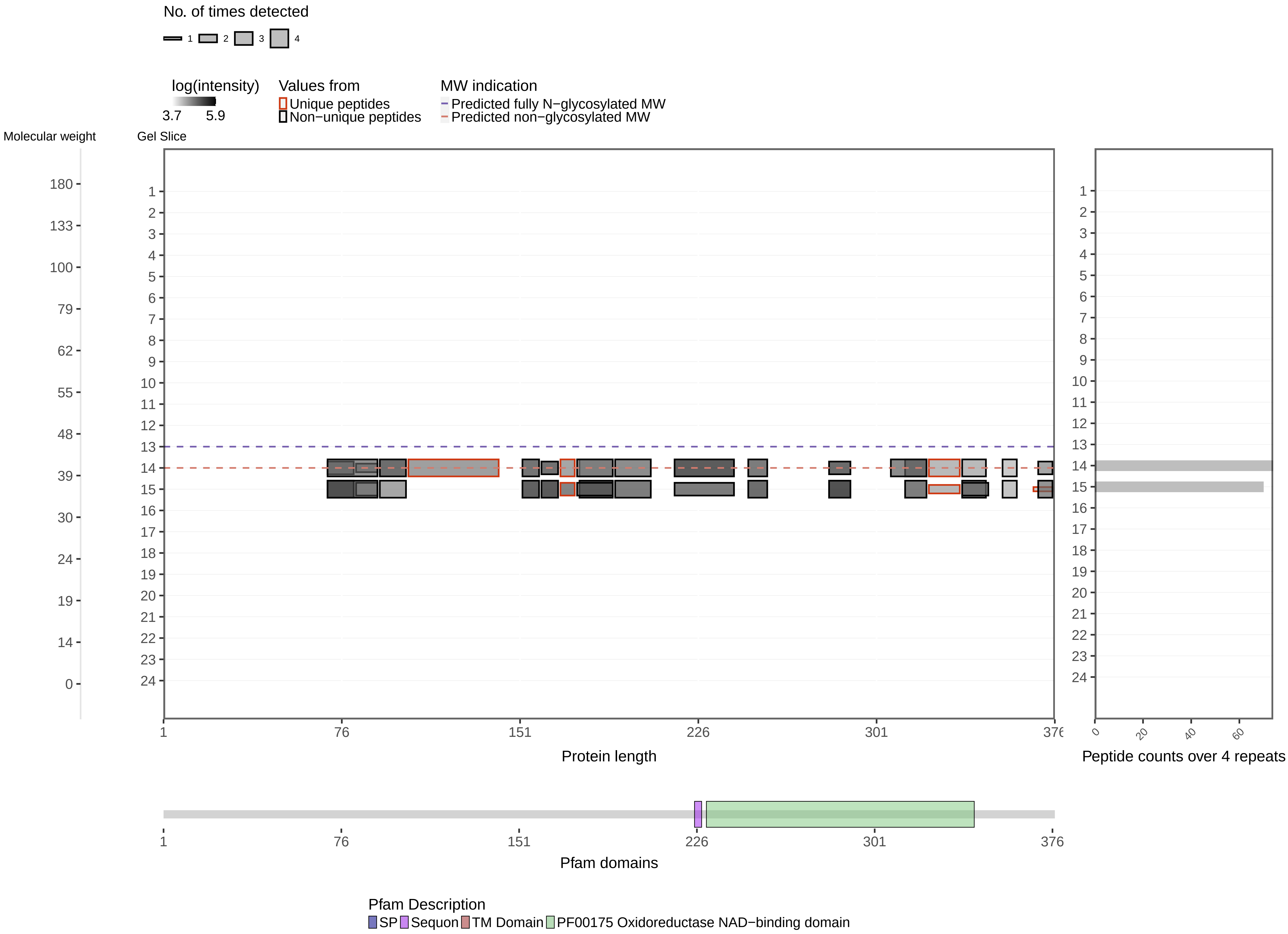


Protein sequence:

YYDPETGKEDPSKNGVCSNFLCDSKPGDKVKITGPSGKIMLLPEDNPNATHIMIGTGTGVAPFRGYLRRMFMESVPTKFNGLAWLFLGVANTDSLLYDDEFTKYLNDYPGNFRYDRALSREOKNNKGGKMYVODKIEEYSDEIFKLLDEGAHMYFCGLKGMMPGIODTLKRVAERRGESWEOKLSOLKKNKOWHVEVY*

### NbL02g05080.1 C−factor [MW = 32.48 kDa]

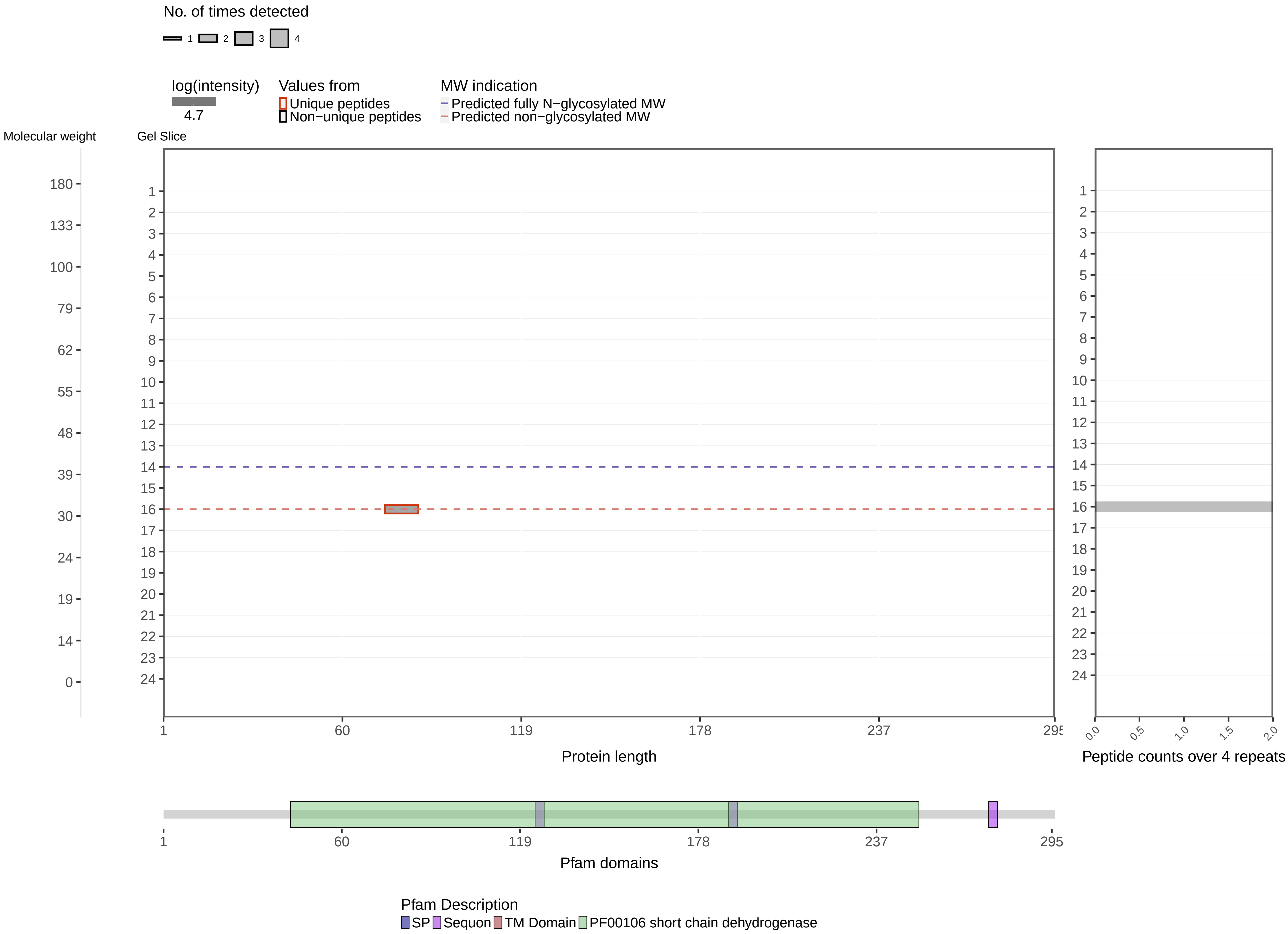


Protein sequence:

VMQPETTLSKVQRSSLLLAYDINAVGPILVIKHMWPLLKAGGGSGTDKDFAVVANLSARVGSIGDNILGGWHSYRASKTALNQLTKNVAVEFARKKDPIICILLHPGTVDTDLSKPFQRNVPKEKLFTKEYSVQTLLSIINNTKRSDNGKFVAWDAQQIPW*

NbL02g05060.1 obg−like_ATPase_1 [MW = 31.11 kDa]

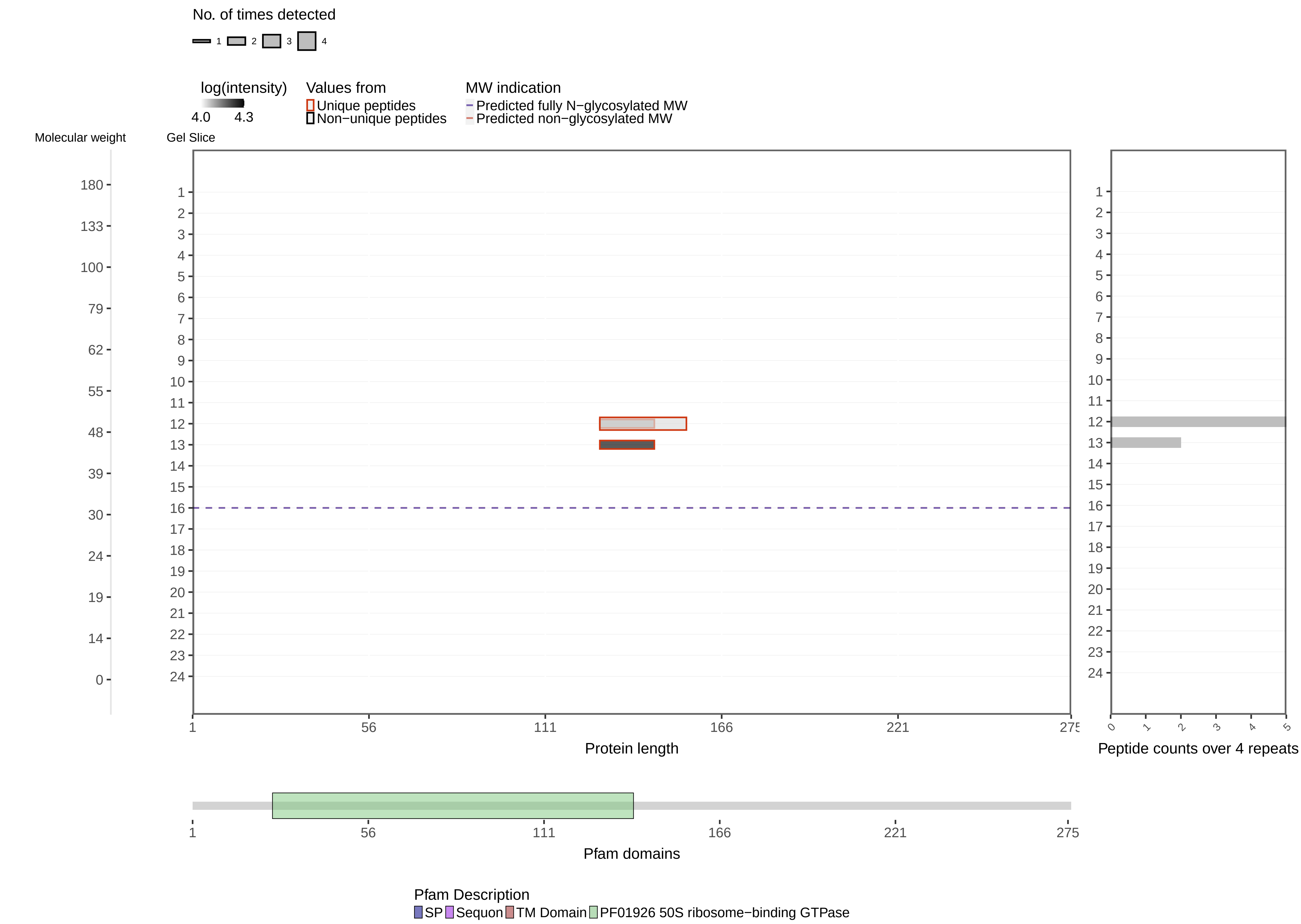


Protein sequence:

HVLRAFEDPDIIHVDDNVDPVRDLEIISEELRLKDIEFMERRIEDLEKSMKRSNDKQLKIEHELSLRVMASLKEGKDIRLGDWKAADVEFLNTFQLLTAKPVVYLVNMNEKDYARKKNKFLPKIHAWVQEHGGEIIIPFSAAVRETLLICH*

NbL02g05050.1 LRR_receptor−like_serine/threonine−protein_kinase_GHR1 [MW − SP = 112.79 kDa]

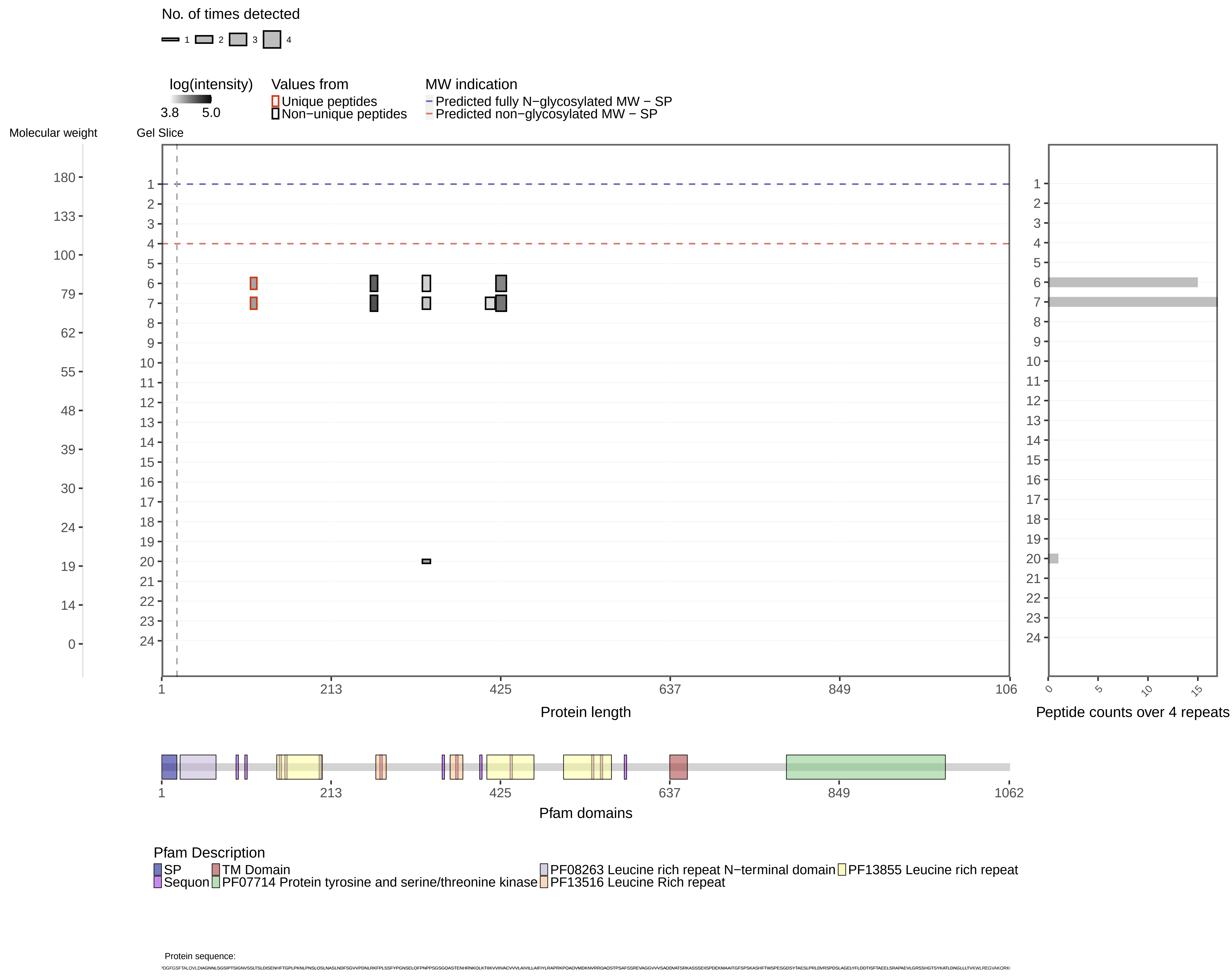

NbL02g04970.1 L−ascorbate_peroxidase_2_cytosolic [MW = 27.41 kDa]

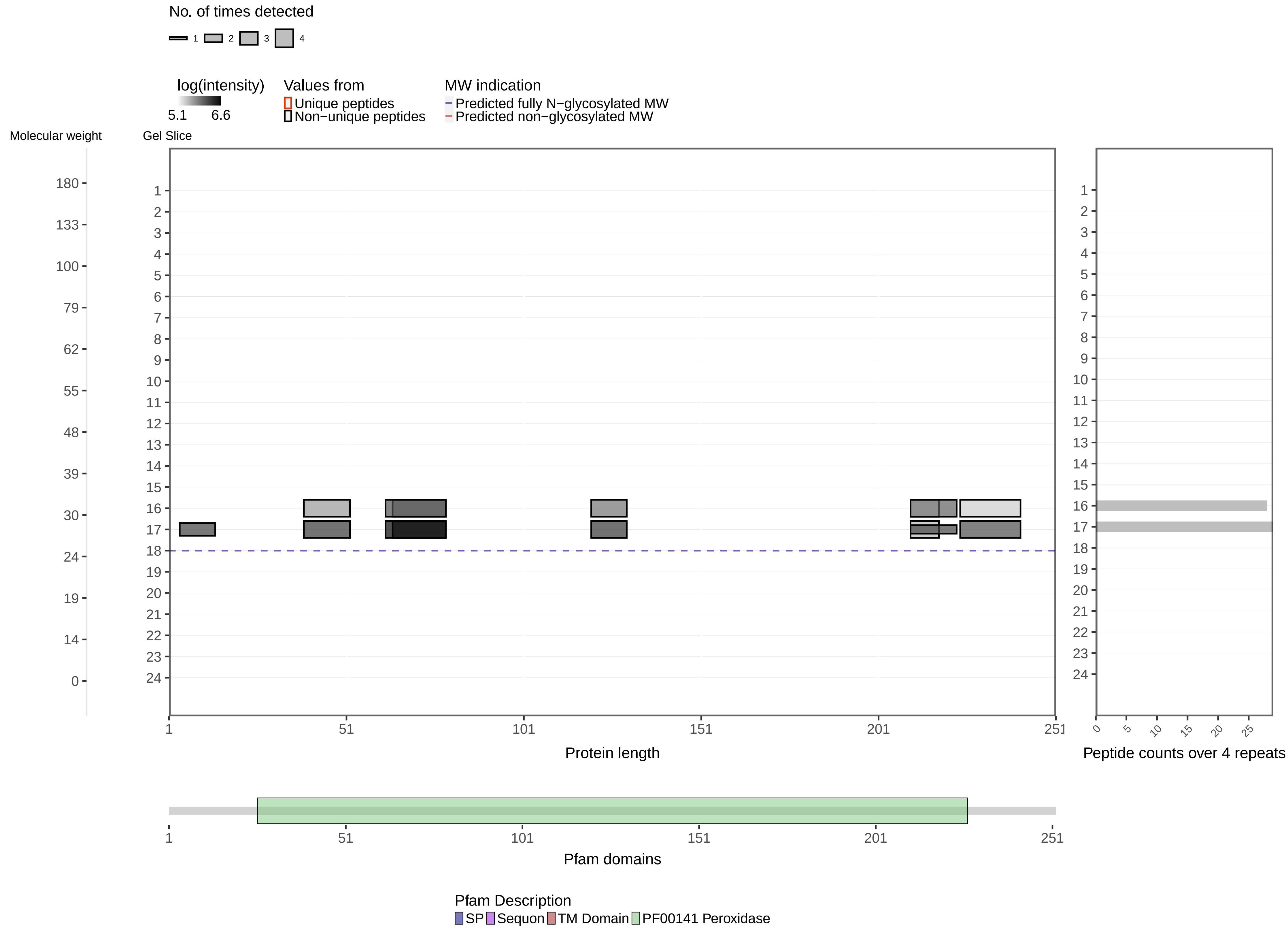


Protein sequence:

VPFHPGREDKTEPPVEGRLPDATKGSDHLRDVFVKQMGLSDKDIVALSGGHTLGRCHKERSGFEGPWTTNPLIFDNSYFTELLSGEKEGLLQLPSDKALLSDPAFRPLVEKYAADEDAFFADYAEAHLKLSELGFAEA*

NbL02g04860.1 elongation_factor_1−alpha−like [MW = 49.3 kDa]

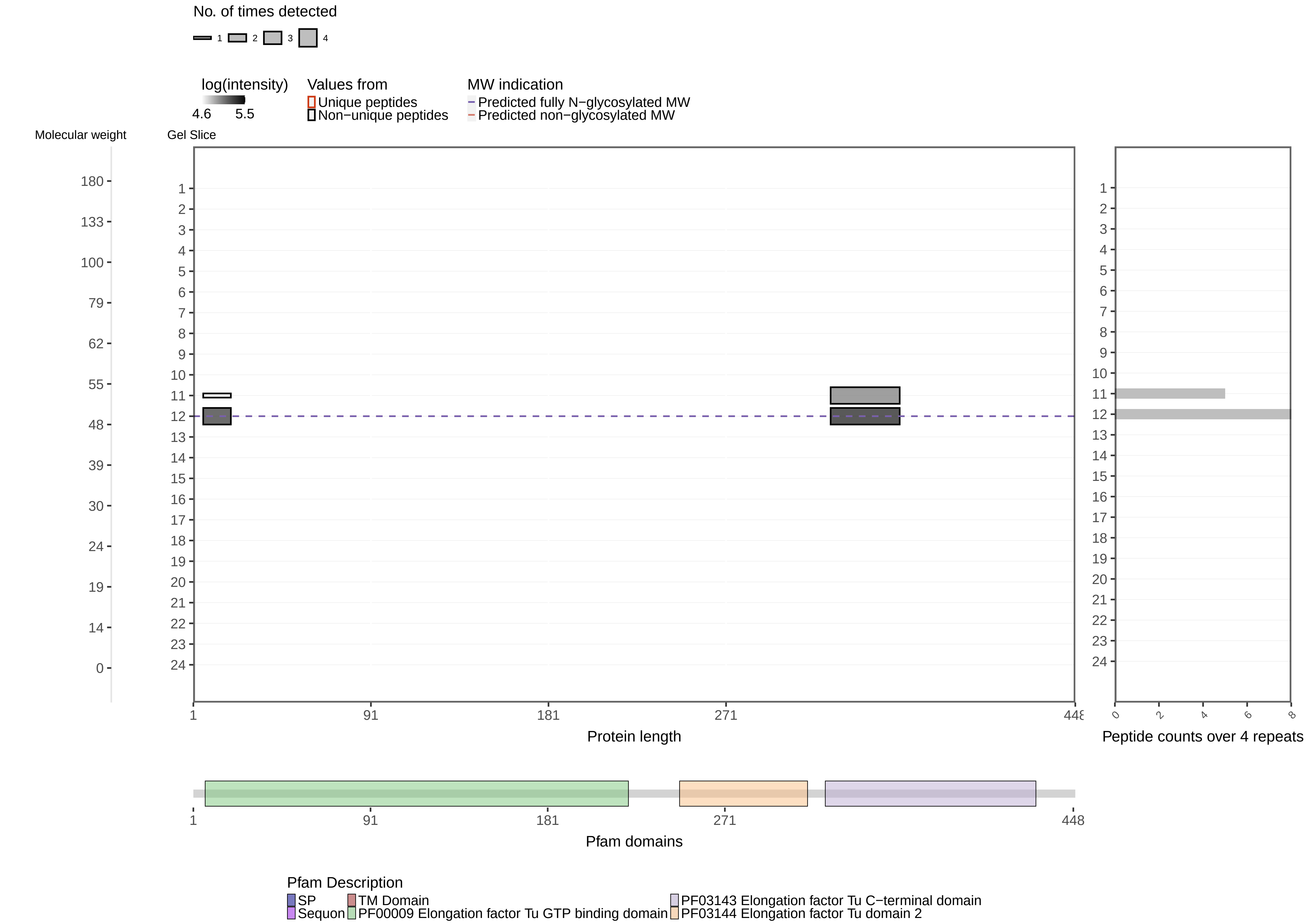


Protein sequence:

GPTLLEALDQINEPKRPTDKPLRLPLQDVYKIGGIGTVPVGRVETGVLKPGMVVTFGPTGLTTEVKSVEMHHEALQEALPGDNVGFNVKNVAVKDLKRGFVASNSKDDPAKGASSFTSQVIIMNHPGQIGNGYAPVLDCHTSHIAVKFAEILTKIDRRSGKELEKEPKFLKNGDAGMVKMIPTKPMVVETFSEYPPLGRFAVRDMRQTVAVGVIKNVDKKDPTGAKVTKAAQKKK*

### NbL02g04820.1 proteasome_subunit_beta_type−4 [MW = 27.88 kDa]

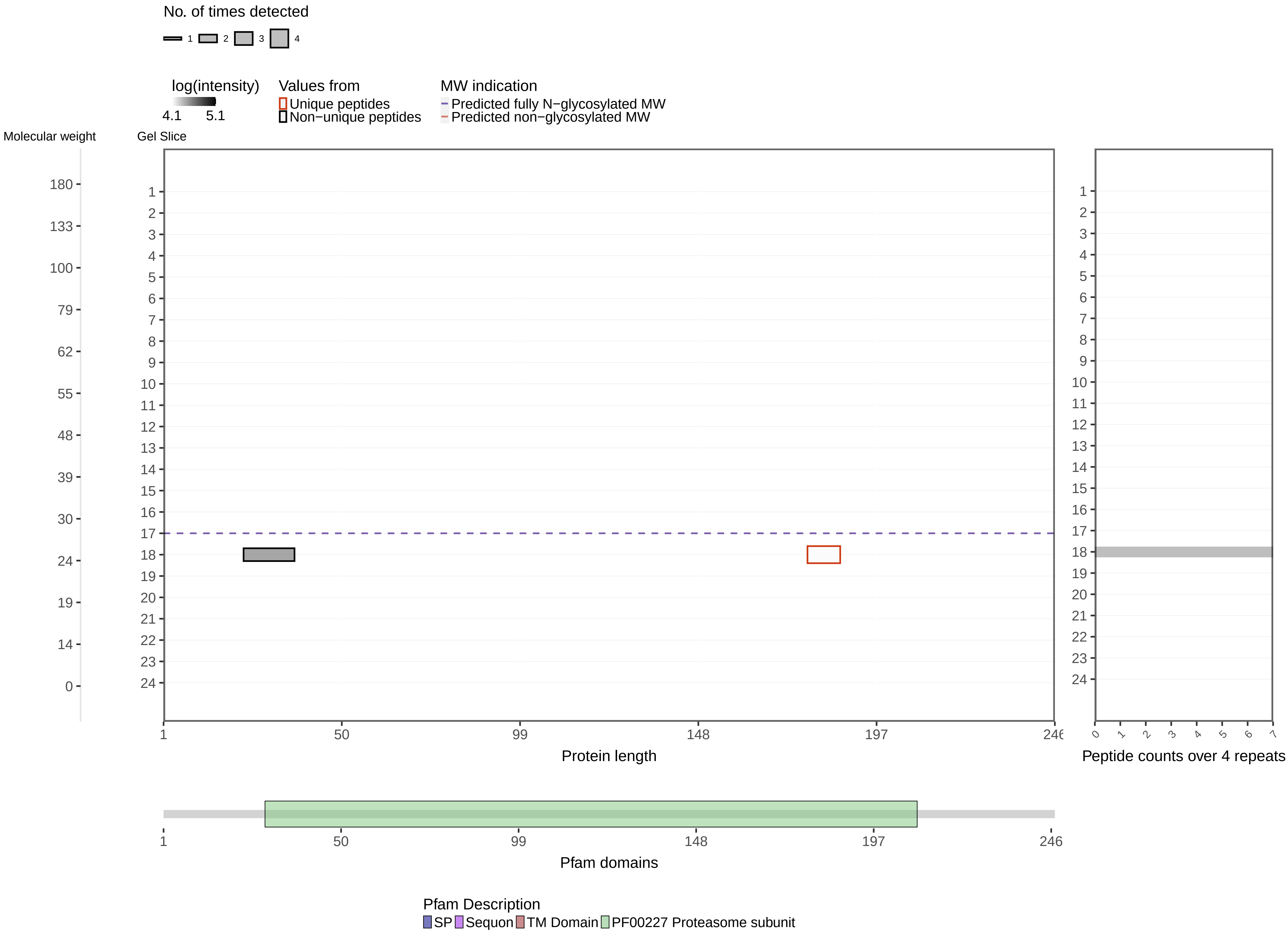


Protein sequence:

INYLTRMMYNRRNKFNPLWNSLVLGGVKNGQKYLGTVSMIGVHYEDNHVATGFGNHLAKPIIHDEWREDMSYEEGVKLLEKCMRVLLYRDRSAVNKLQMLDMLQLLLAETCQTASEFDETSFYESQIMIWTPTMW*

### NbL02g04680.1 PIP2−4−like [MW = 30.73 kDa]

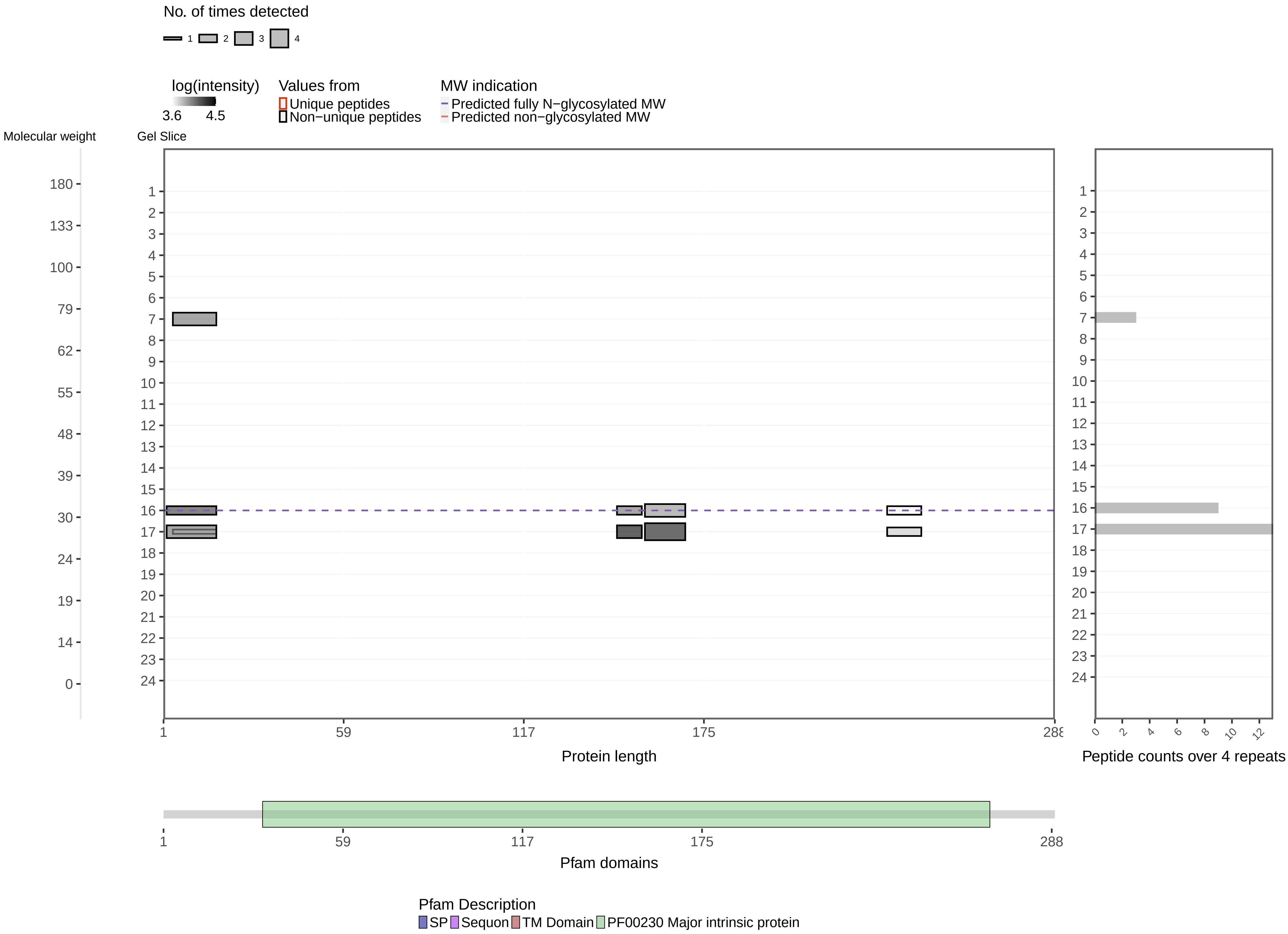


Protein sequence:

AQCLGAICGVGFVKAFQSAYYVRYGGGANVMAPGHTKGVGLAAEIIGTFVLVYTVFSATDPKRNARDSHVPVLAPLPIGFAVFMVHLATIPITGTGINPARSFGAAVIYNHDKAWDEHWIFWVGPFIGAFAAAVYHQFILRAGAIKALGSFRSNA*

### NbL02g04530.1 polyadenylate−binding_protein_8−like [MW = 70.46 kDa]

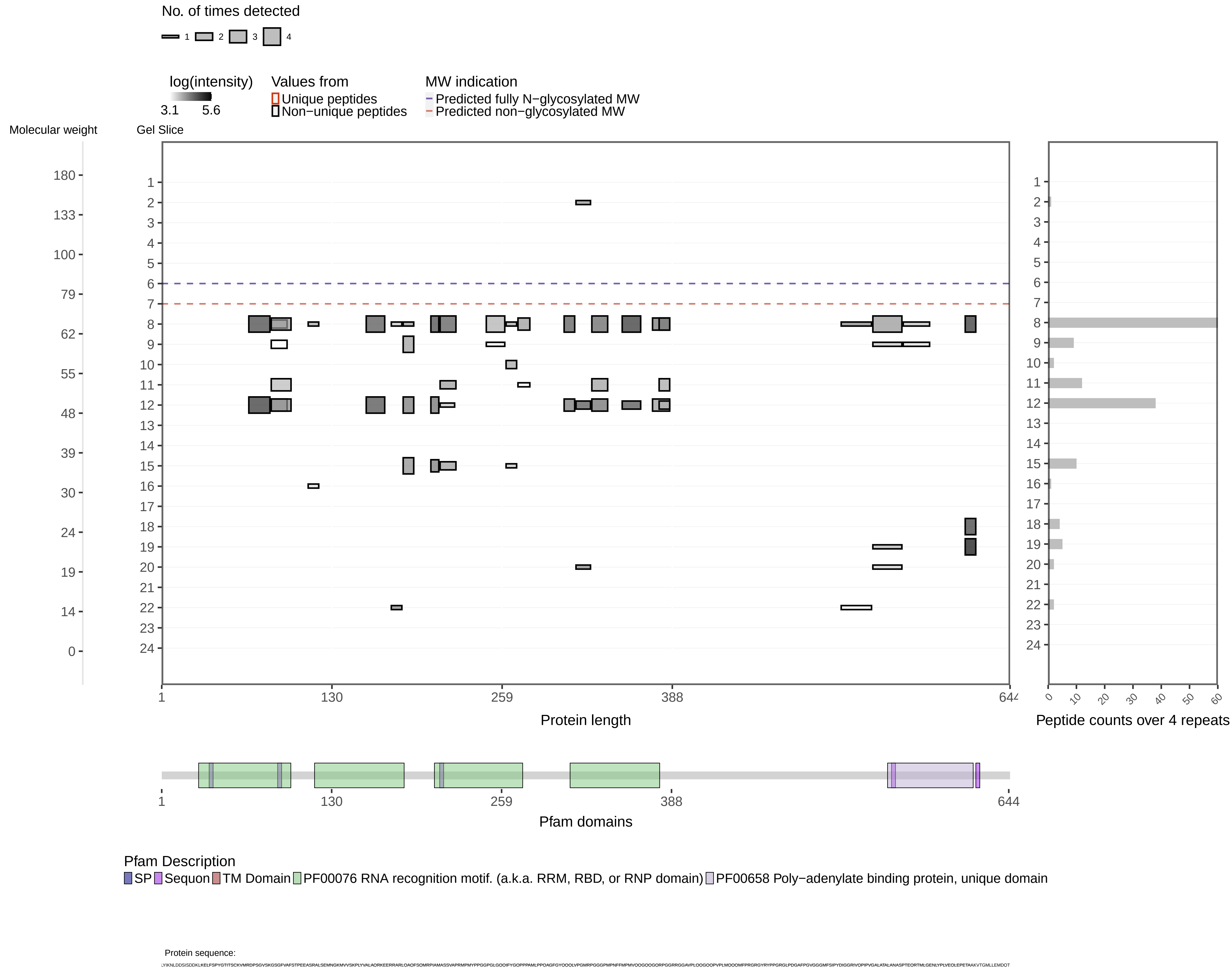

NbL02g04500.1 serine_carboxypeptidase−like [MW − SP = 53.97 kDa]

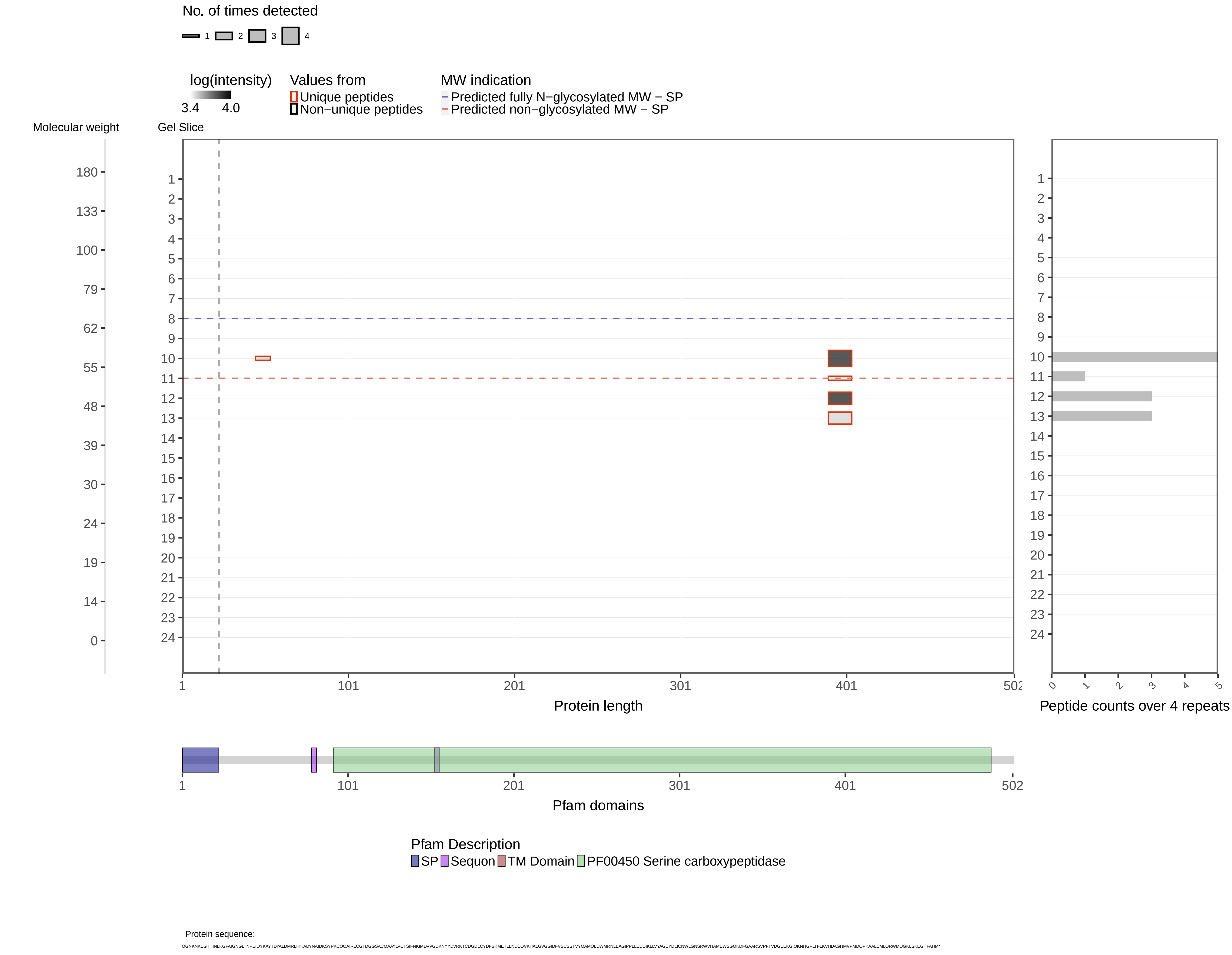

NbL02g04360.1 aminopeptidase_M1 [MW = 99.26 kDa]

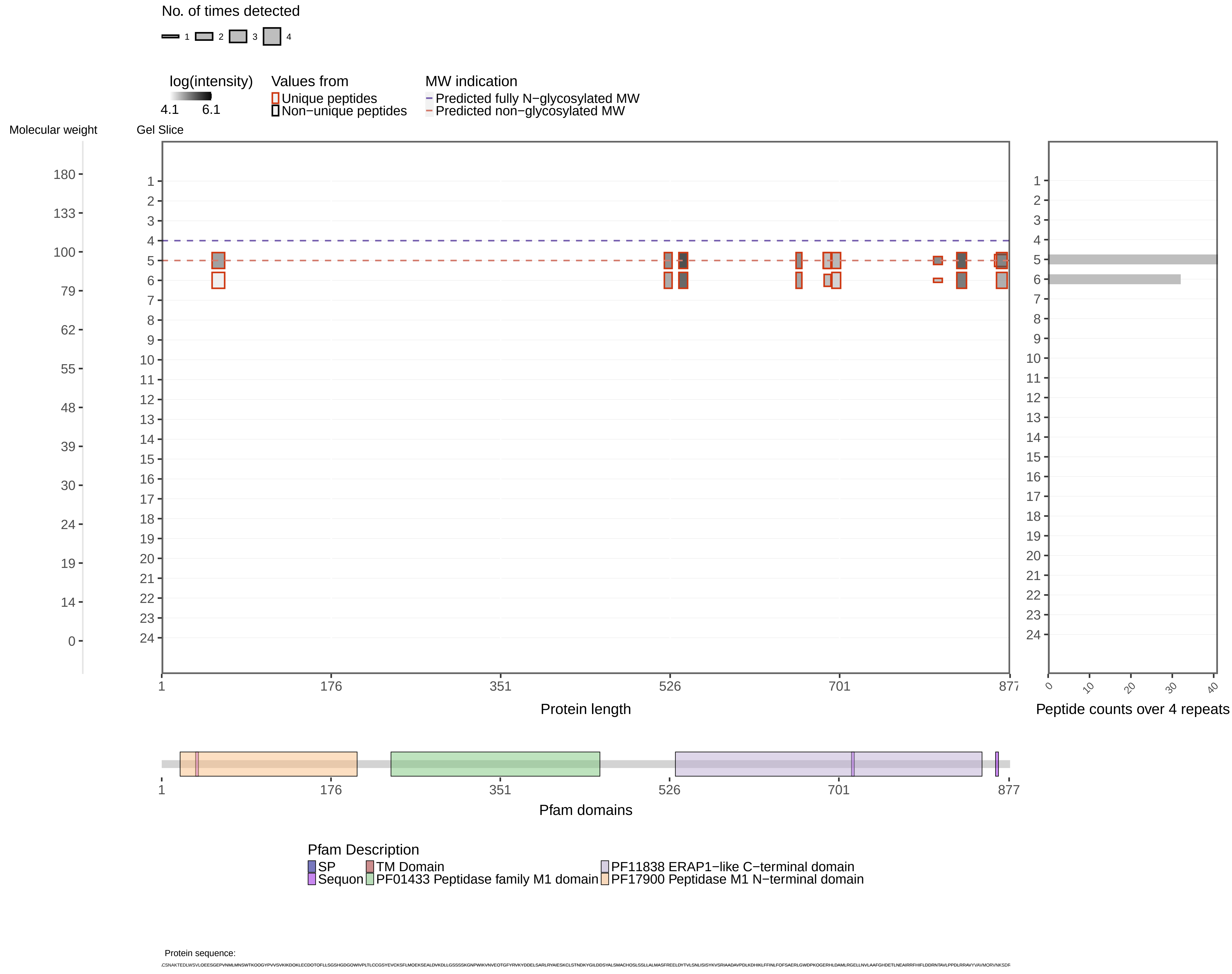

NbL02g03980.1 21_kDa_protein−like [MW − SP = 19.1 kDa]

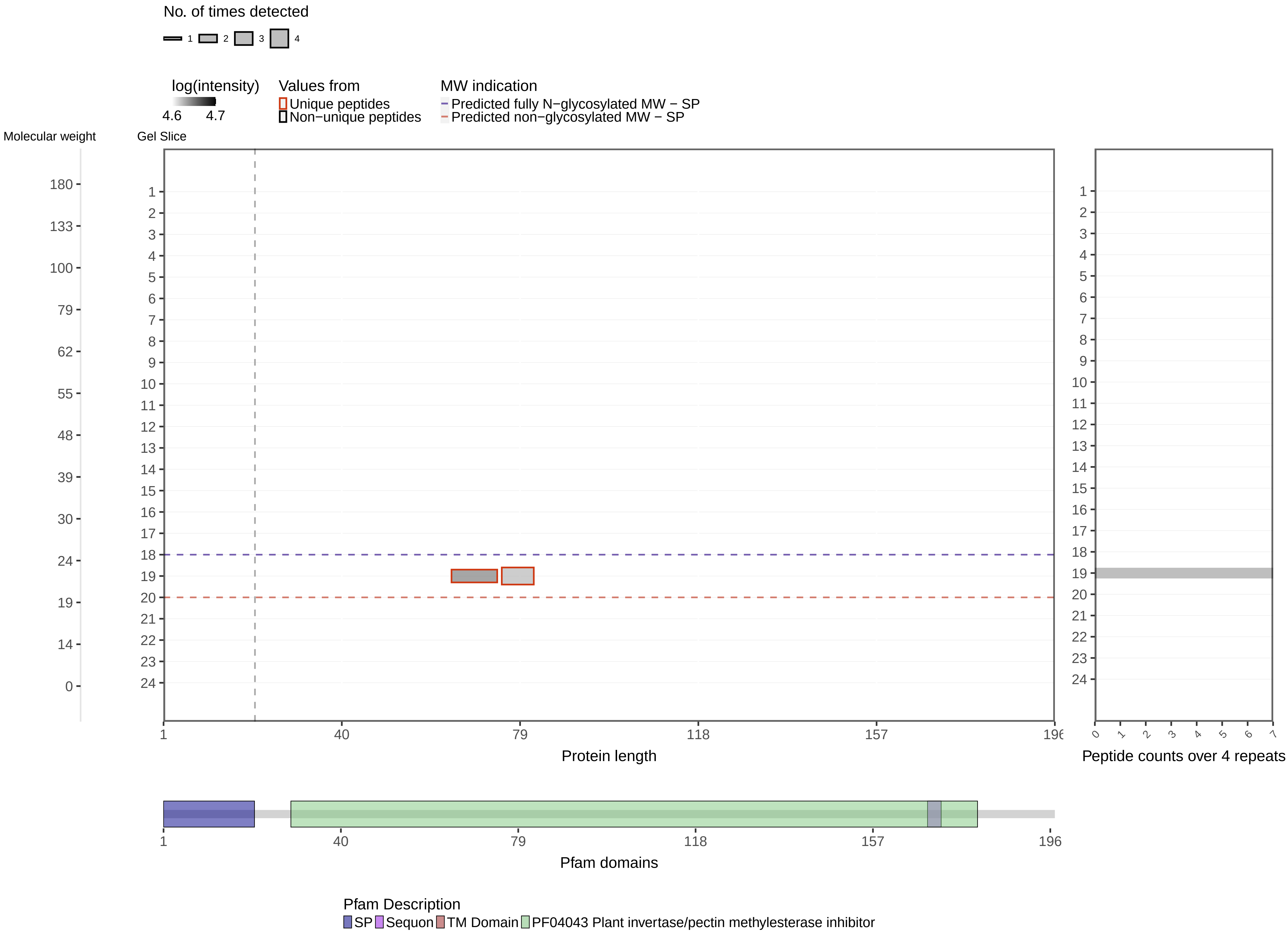


Protein sequence:

INSLKRRGRVALLDCVECFQDALDNLHNSLGVLRKLSSQTFNDQMGDVTTWLSAALTDEDTCLGGFEDQRRRKQVKLLLNRVTNVSYMTSNALALVNKLATTGPECLQNS*

NbL02g03910.1 protein_MET1_chloroplastic [MW = 36.05 kDa]

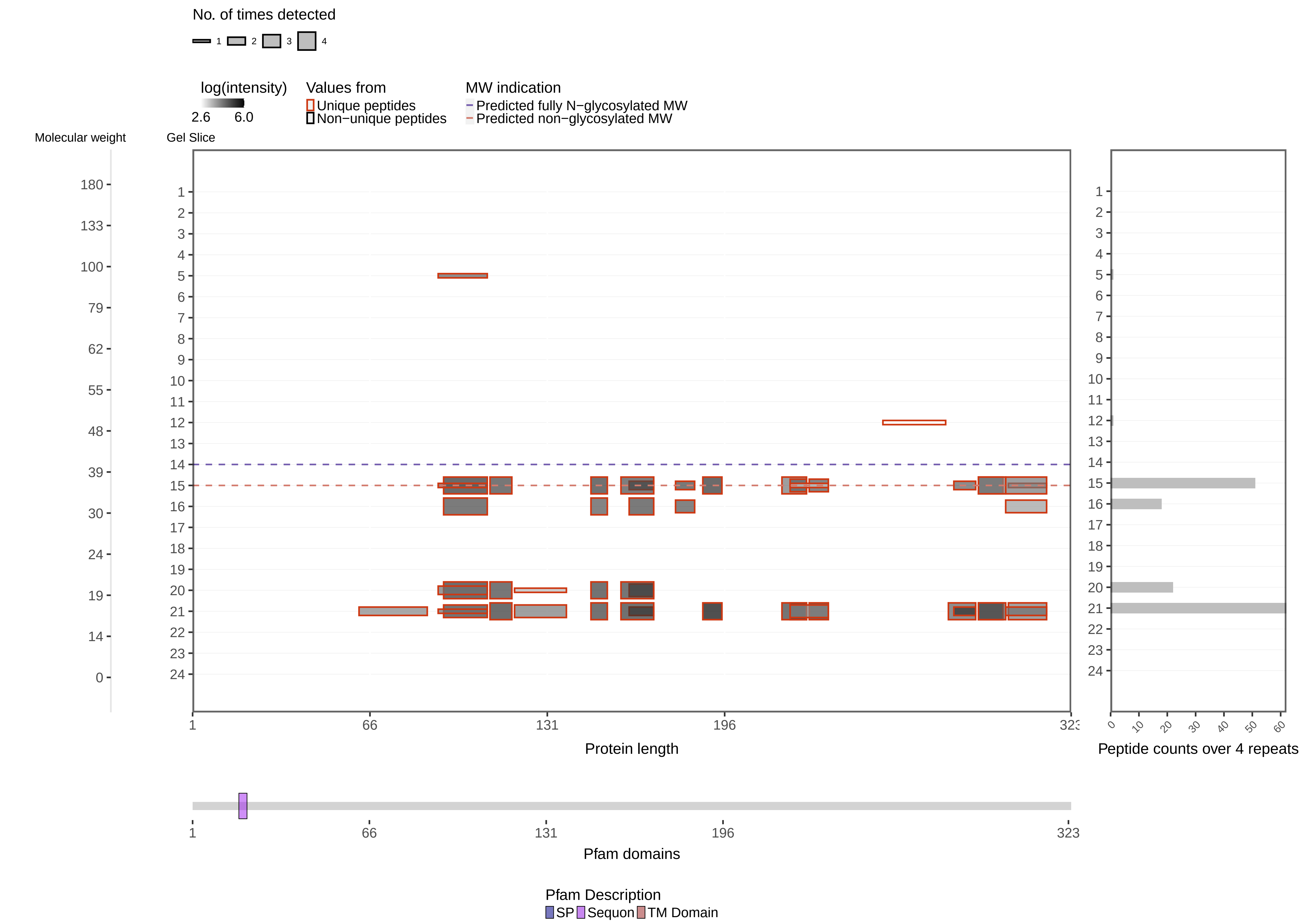


Protein sequence:
SPLLMKMQKRYGKVEEAGELTEKEIIRAERNAGFISDRVREIQLTNARRKKEQKEKREKDLREGLQLSKNGKYEEALEKFESVLGSRPTGNEAAVASYNVACCYSKLNQLQAGLSALEDAMDAGFDDFKRIRTDPDLANLRTSADYEPLMKKFDESFINENAINAIKSLFGFNKK*

### NbL02g03890.1 lysine−−tRNA_ligase_chloroplastic/mitochondrial [MW = 67.63 kDa]

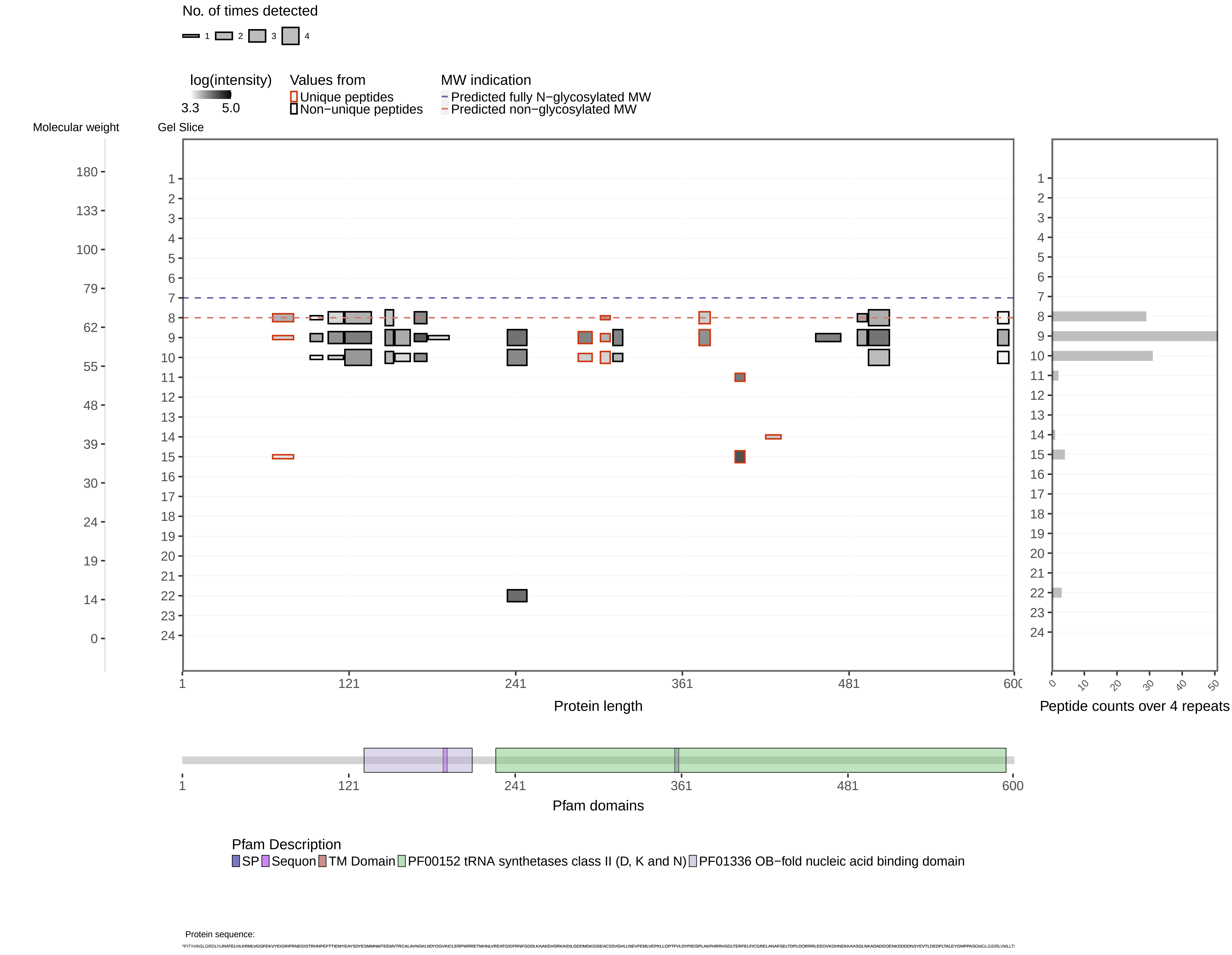

### NbL02g03870.1 isocitrate_dehydrogenase_[NADP] [MW = 46.78 kDa]

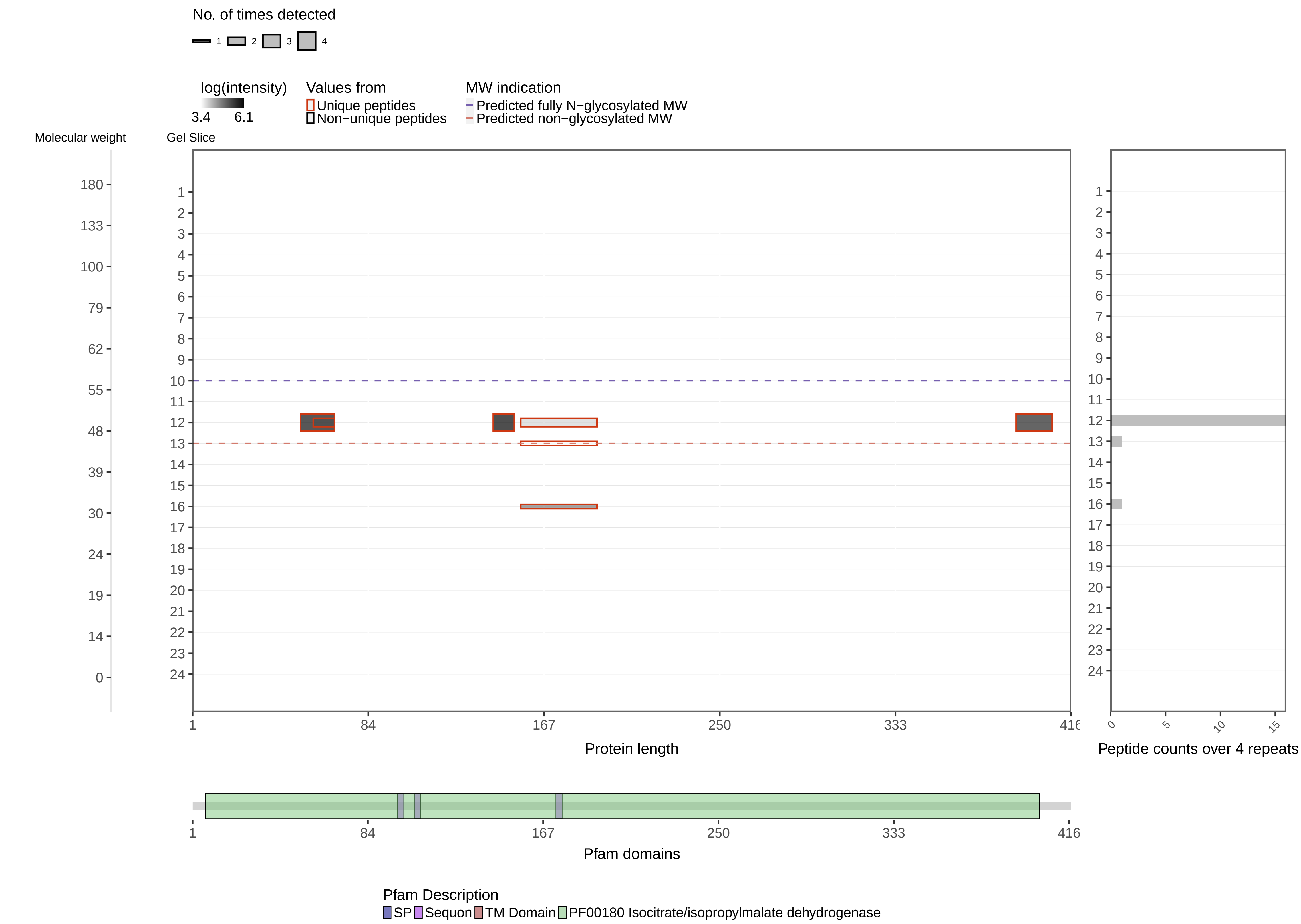


Protein sequence:

ASMNMAYQKKWPLYLSTKNTILKKYDGRFKDIFQEVYEANWKSKYEEAGIWYEHRLIDDMVAYALKSEGGYVWACKNYDGDVQSDFLAQGFGSLGLMTSVLVCPDGKTIEAEAAHGTVTRHYRVHQKGGETSTNSIASIFAWTRGLAHRATLDKNERLLDFTEKLEAACIGAVESGKMTKDLALIIHGSKLSREHYLNTEEFIDAVADELKARLLKAKA*

NbL02g03800.1 UPF0061_protein_azo1574−like [MW = 43.09 kDa]

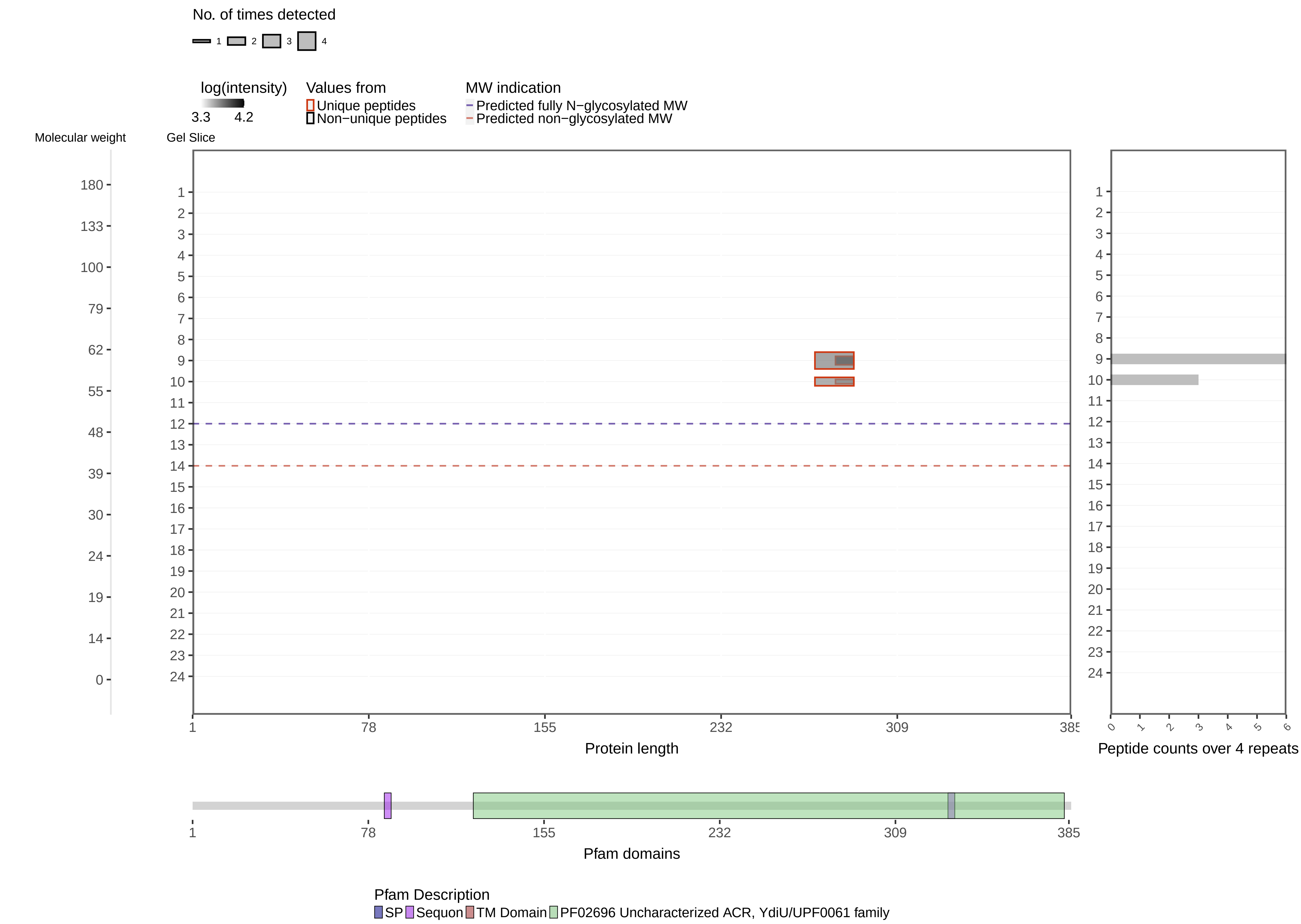


Protein sequence:

'AONYGGHOFGTWAGOLGDGRAITLGEVLNSKSERWELOLKGAGKTPYSRFADGLAVLRSSIREFLCSEAMHSLGIPTTRALCLVTTGKDVTRDMFYDGNPKDDPGAIVCRVAOSFLRFGSYOLHASRGKKDLEIVRTLADYAIRHHFPHLENMSKSESISFSTGEEDDSAVDLTSNKYAAWAVEVAERTASMIARWOGVGFTHE*

### NbL02g03750.1 enoyl−[acyl−carrier−protein]_reductase_[NADH]_chloroplastic−like [MW = 41.9 kDa]

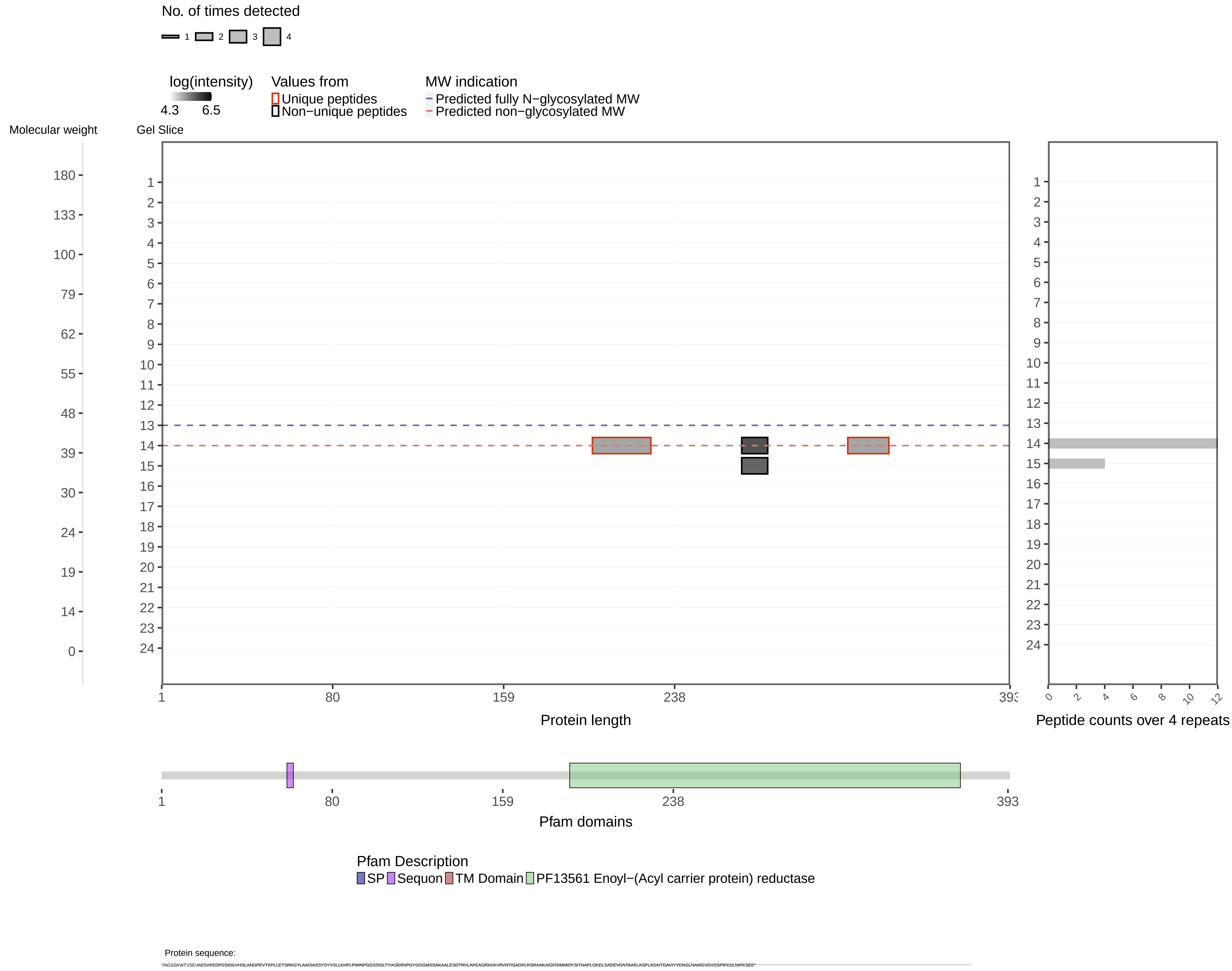

### NbL02g03640.1 vacuolar_protein_sorting−associated_protein_26A [MW = 35.61 kDa]

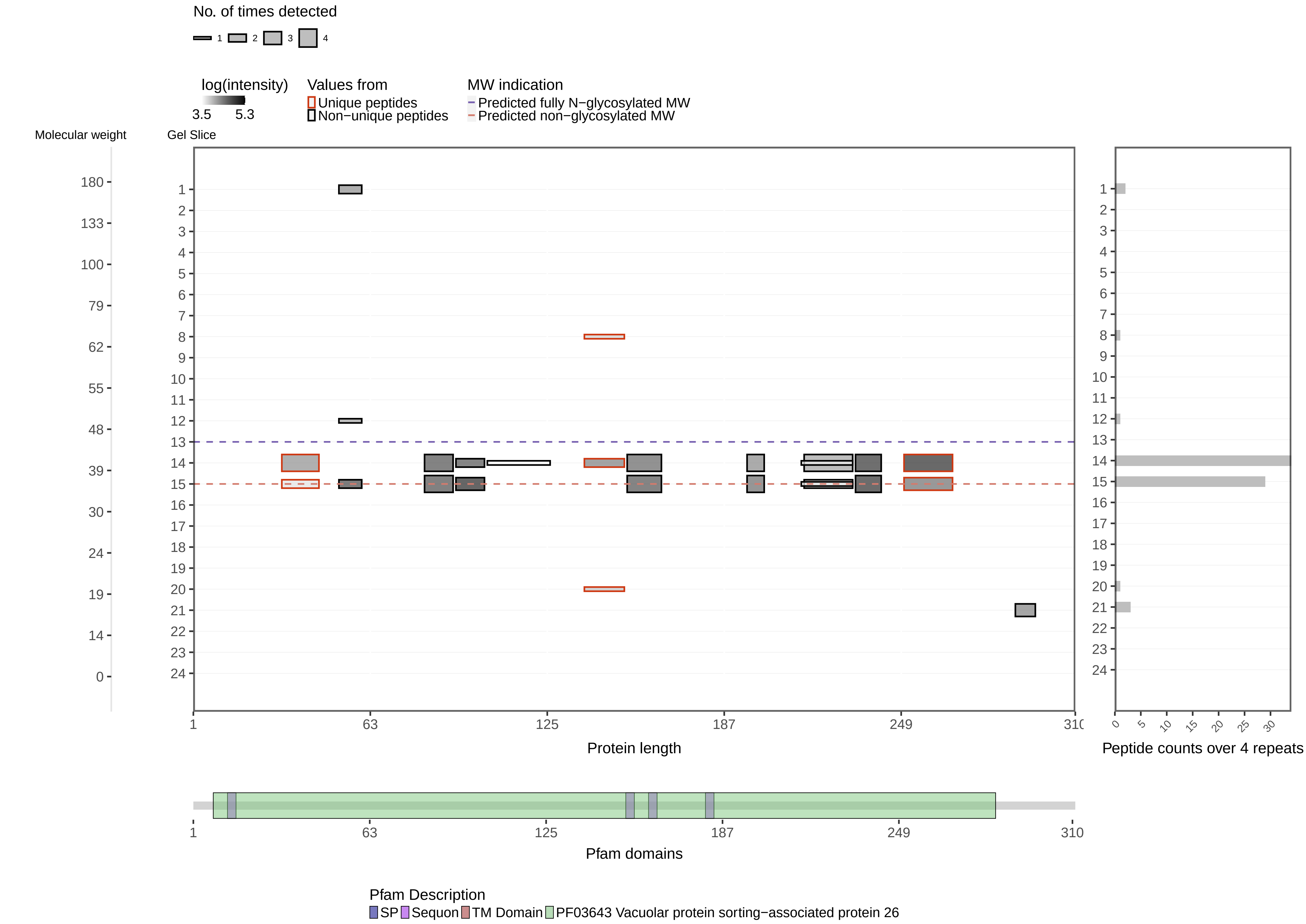


Protein sequence:

GSIVEYQDFVVRNYSPPPSINNSIKMEVGIEDCLHIEFEYNKSKYHLKDVIIGKIYFLLVRIKIKNMDLEIRRRESTGSGANTHVETETLAKFELMDGAPVRGESIPIRLFLSPYELTPTYHNINNKFSVKYYLNLVLVDEEDRRYFKQQEITMFRLAETSVINQAVGA*

NbL02g03290.1 ATP−dependent_6−phosphofructokinase_6−like [MW = 28.98 kDa]

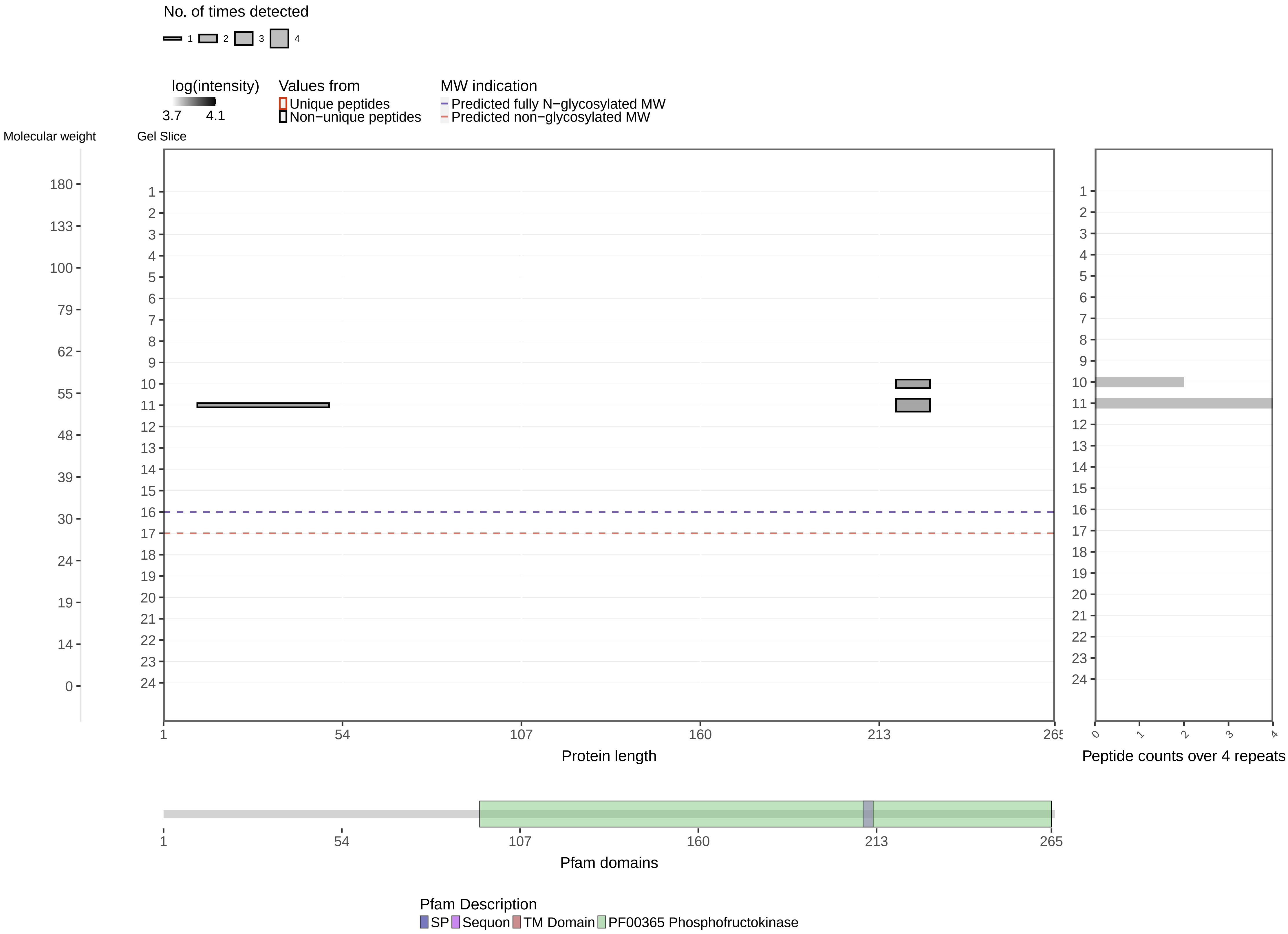


Protein sequence:
HHMYGVKRVMGIDGGYRGFYSKNIIPLTPKVVNDIHKRGGTILGTSRGGHVTKKIVDSIQDRGINQVYIIGGDGTQRGAAVIFEEIRRRGLNVSVAGIPKTIDNDILVIDRSFGFDSAVEEAQRAISAAHVEATSFENGIGLVKLMGR

### NbL02g03270.1 acid_phosphatase_1_pseudogene [MW − SP = 27.87 kDa]

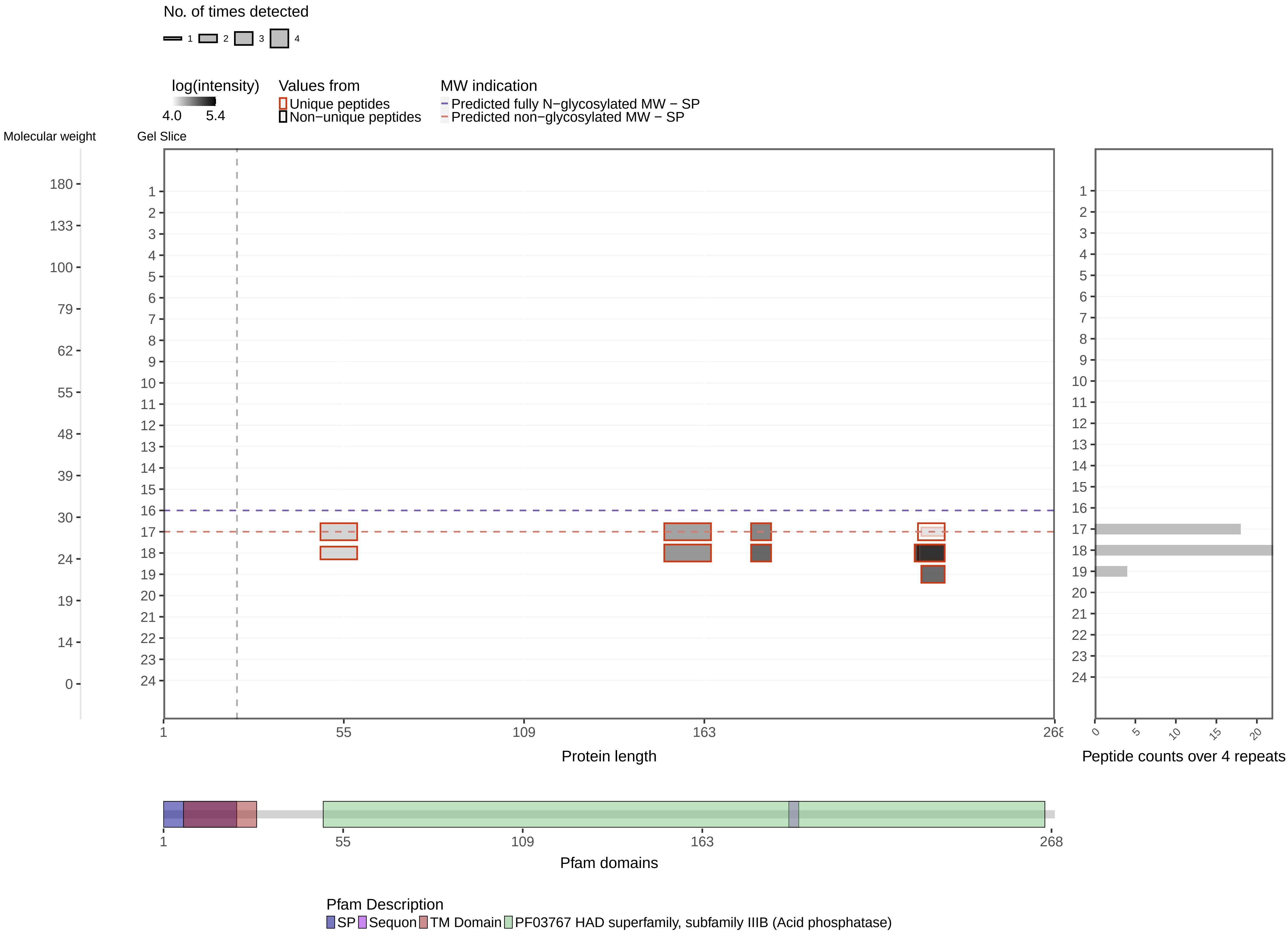


Protein sequence:

DIDETLLSNLPYYVTHGFGSEIFDEIAFDKWVNEAEAPAIPASLKLYKELQQRGFKIFLLTGRSELQRNYTEKNMVYAGYSNWERLILRGPSDKGKLATQYKSEKRKELEDEGYRIHGSSGDQWSYLNGFAVATRSFKLPNPMYYIA*

### NbL02g03250.1 phospholipase_D_alpha_1−like [MW = 92.3 kDa]

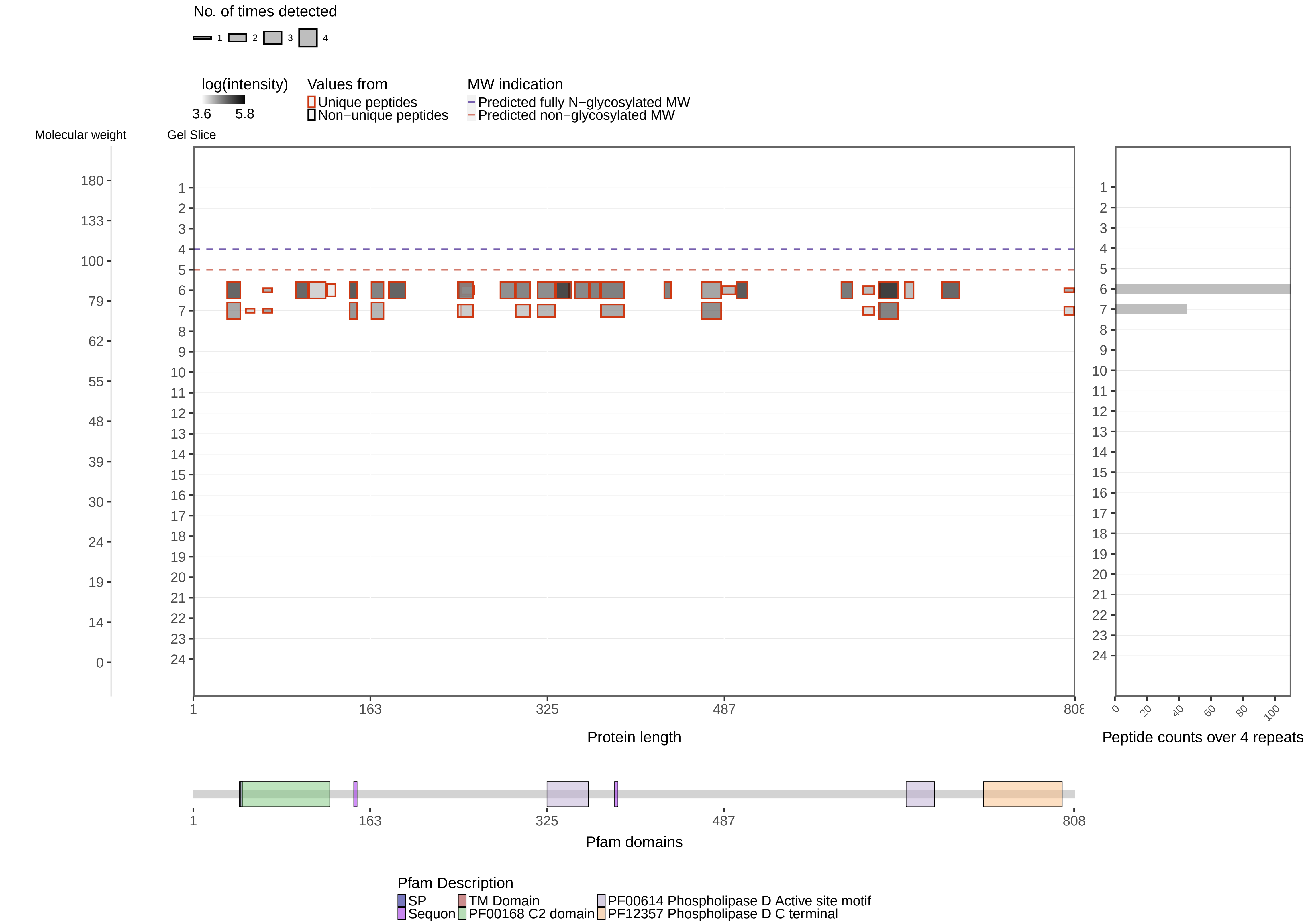


Protein sequence:

[illegible]

### NbL02g03210.1 arginine−−tRNA_ligase_chloroplastic/mitochondrial−like [MW = 73.54 kDa]

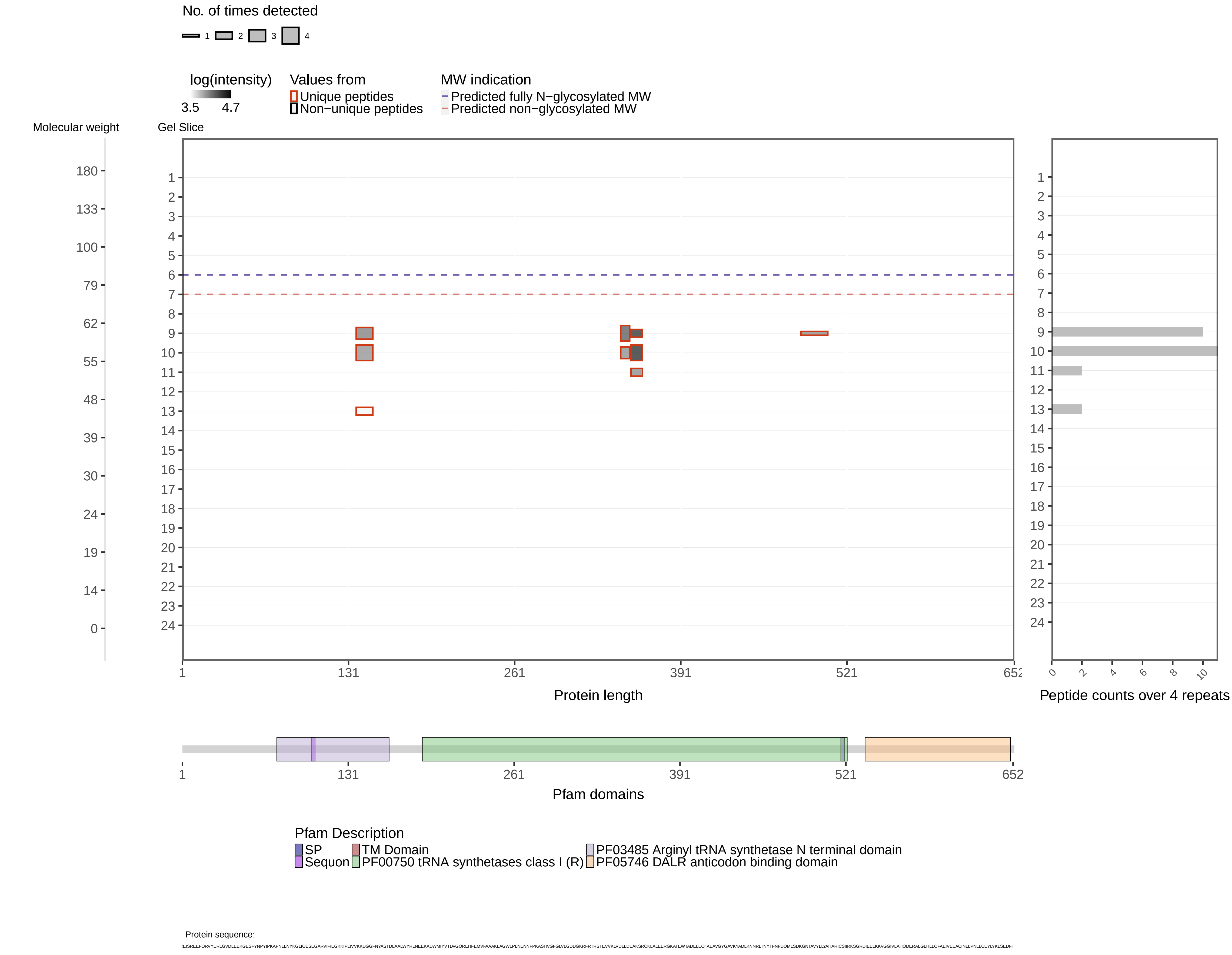

### NbL02g03060.1 uncharacterized_LOC104114303 [MW = 302.25 kDa]

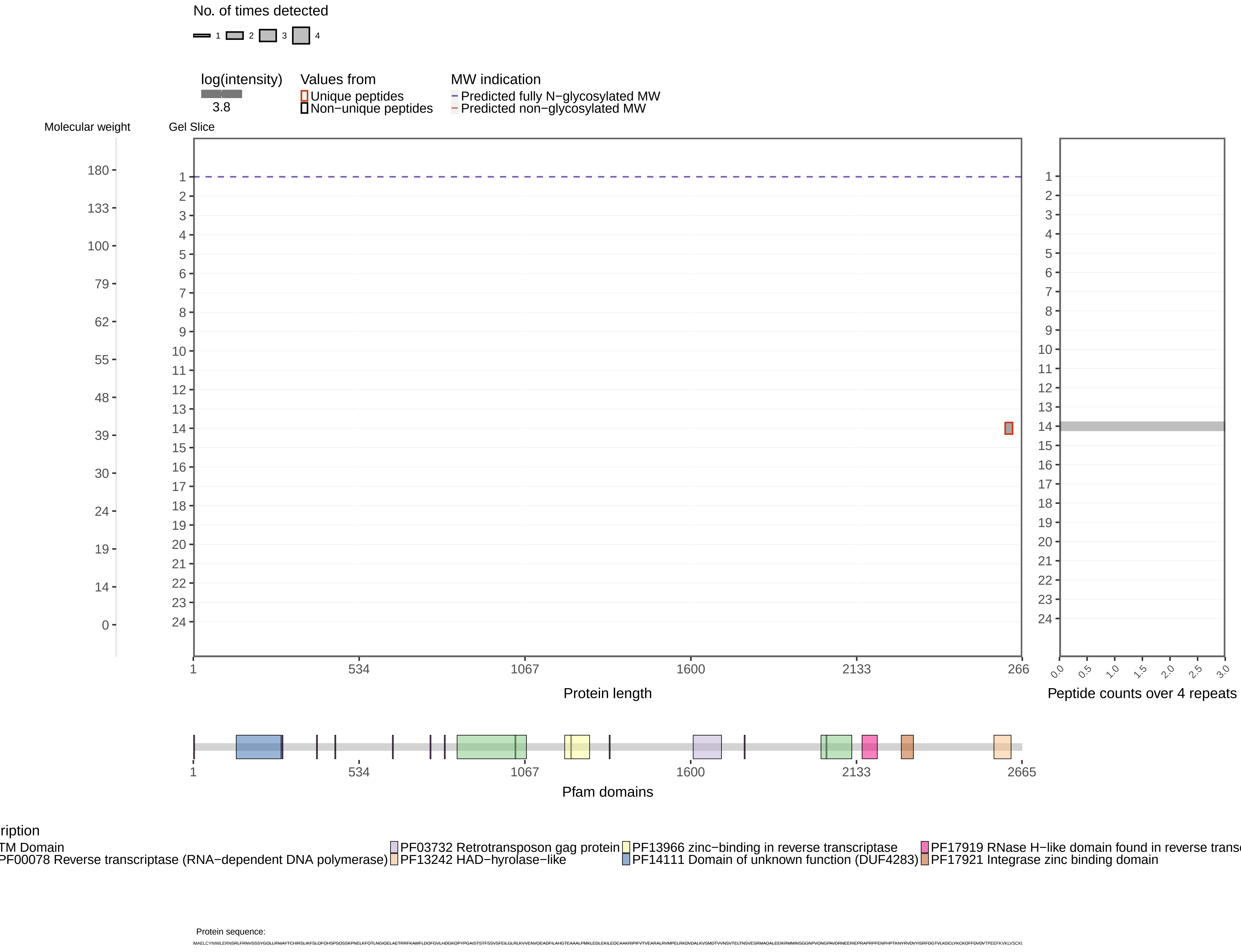

### NbL02g03030.1 plastocyanin [MW = 17.01 kDa]

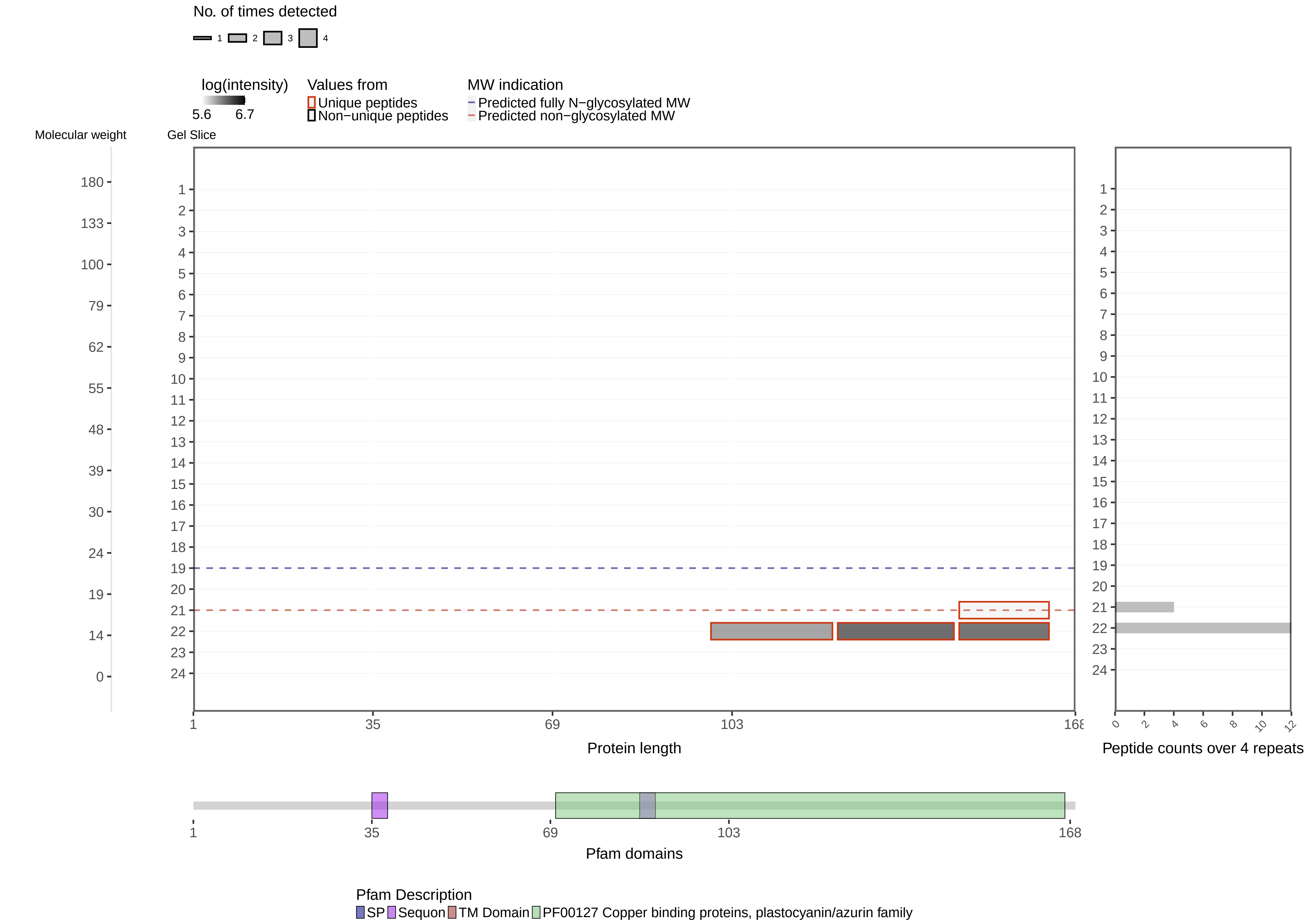


Protein sequence:

LGGDDGSLAFVPGNFSVSAGEKITFKNNAGFPHNVVFDEDEIPSGVDASKISMSEEDLLNAPGETYSVTLSEKGTYSFYCSPHQGAGMVGKVTVN*

NbL02g03010.1 uncharacterized_LOC109239937 [MW = 29.04 kDa]

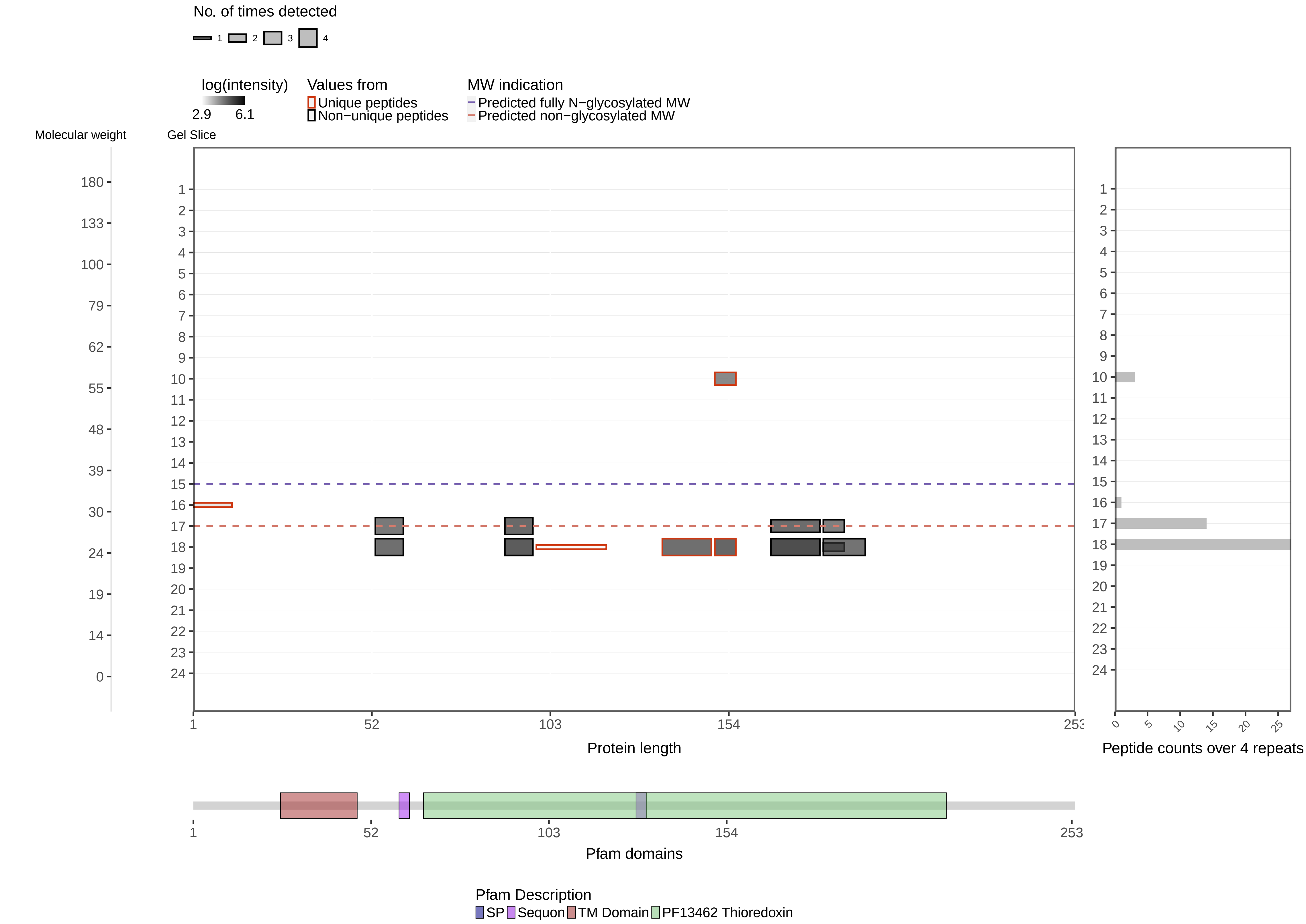


Protein sequence:

FLTSRALHVVNKLNASATYKLLESFFDHQEDFYNKATFDLSRASVVEKVARFSTNAIGNSNYDVIKAGFTDYKTDQSTRLSFKYGCVKGVYGTPFFFVNGFPLPDAGSALDYKGWRRVLDPLVHQQDTGRNYLLHFFL*

### NbL02g02920.1 uncharacterized_LOC109211680 [MW − SP = 16.95 kDa]

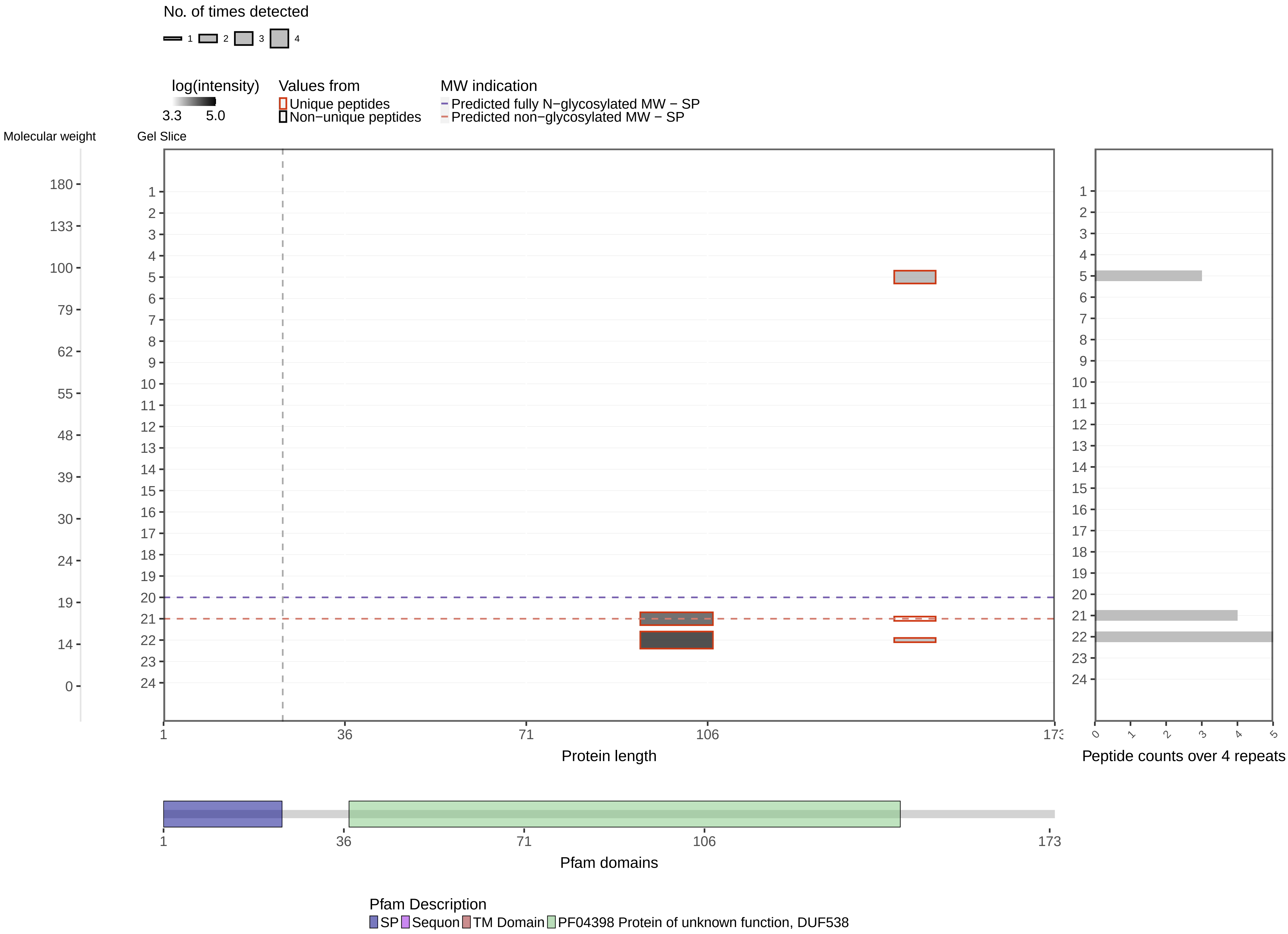


Protein sequence:

CYVQFQDQLVYYQKEIKGKMSYGVVSDVSGIQAKKLFVWVPVTGINVDEESHMIEFHVGFLSEKLPAKDFETIPTCKNKGAKILSFLQYDEDEWSSEVL*

### NbL02g02830.1 endoplasmin_homolog [MW − SP = 89.72 kDa]

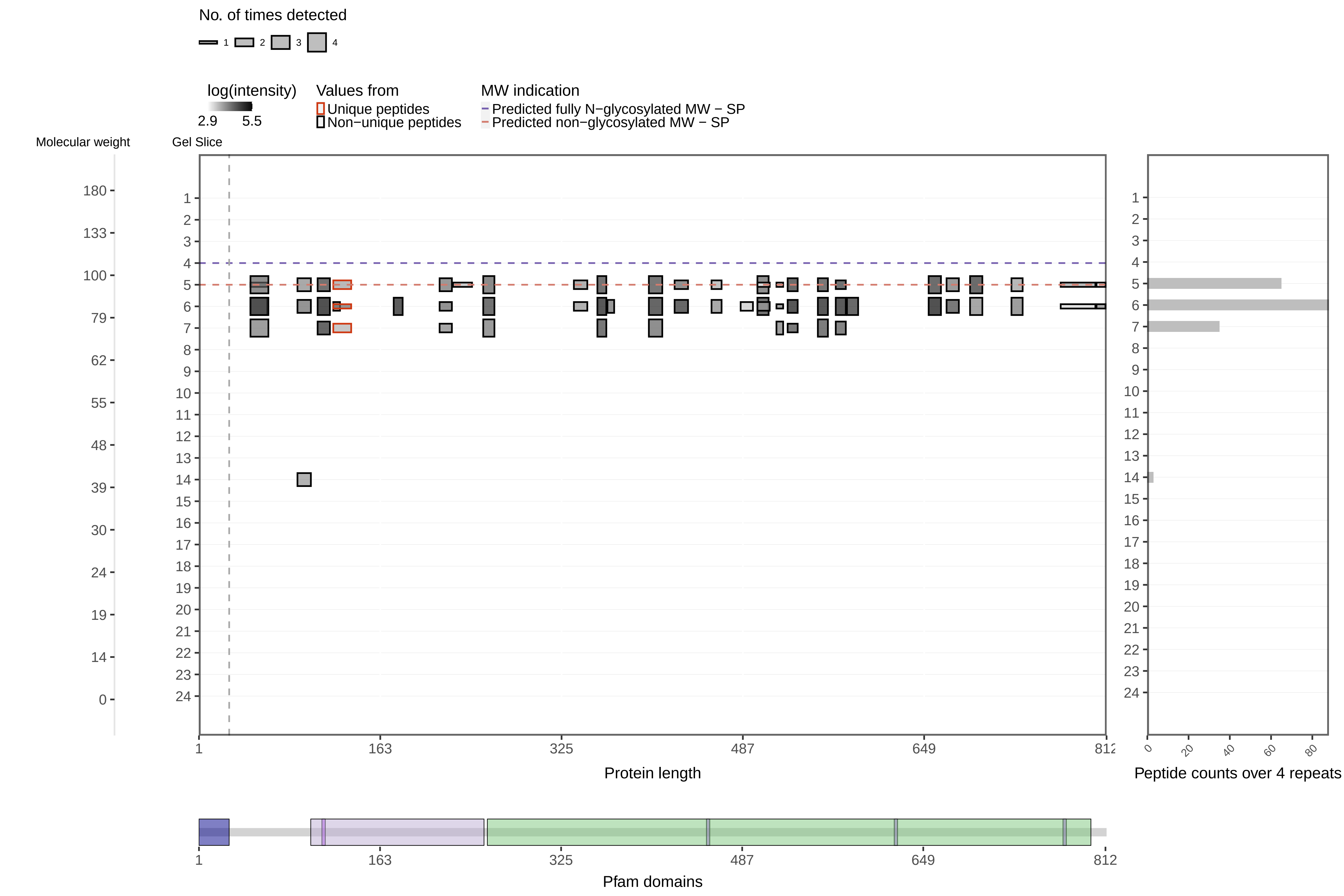


Pfam Description
SP Sequon TM Domain PF00183 Hsp90 protein PF02518 Histidine kinase−, DNA gyrase B−, and HSP90−like ATPase

Protein sequence:

AVLFVPPKAPHDLYESYYNSKKSNLKLYVRRVFISDEFDELLPKYLSFLMGLVDSDTLPLNVSREMLQQHSSLKTIKKKLIRKALDMIRKIADEDPDESNDKDKKDIEESSADNEKKGQYAKFWNEFGKSVKLGIIEDATNRNRLAKLLRFETSKSDGKLTSLDQYISRMKAGQKDIFYITGASKEQLEQSPFLERLTKKNYEVIFFTDPVDEYLMQYLMDYEDHKFQNVSKEGLKLGKDSKAKELKESFKELTKWWKGALASDNVDDVKISNRLADTPCVVVTSKYGWSANME

### NbL02g02760.1 prolyl_endopeptidase−like [MW = 23.48 kDa]

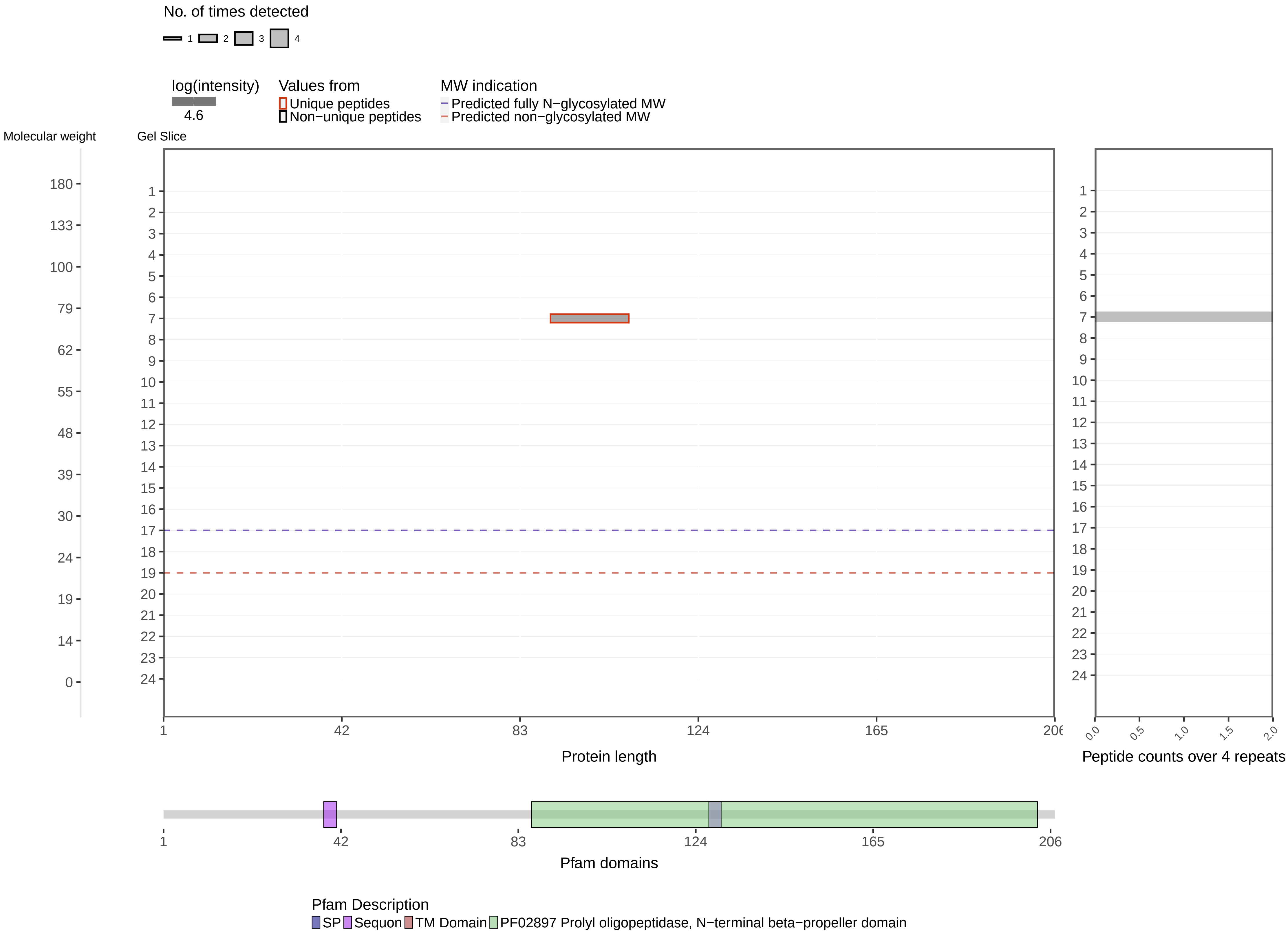


Protein sequence:

ESVVDNYHGVSVPDPYRWLEDPDSEETKEFVEKQVNLTDSVLKTCETREKLREKITKLYDFPKYEVPFRAGDKYFYFHNSGLQPQKVLYVQDSLDGEPEVLLDPNTLSETGQLH*

NbL02g02750.1 ATP−dependent_zinc_metalloprotease_FTSH_chloroplastic [MW = 76.59 kDa]

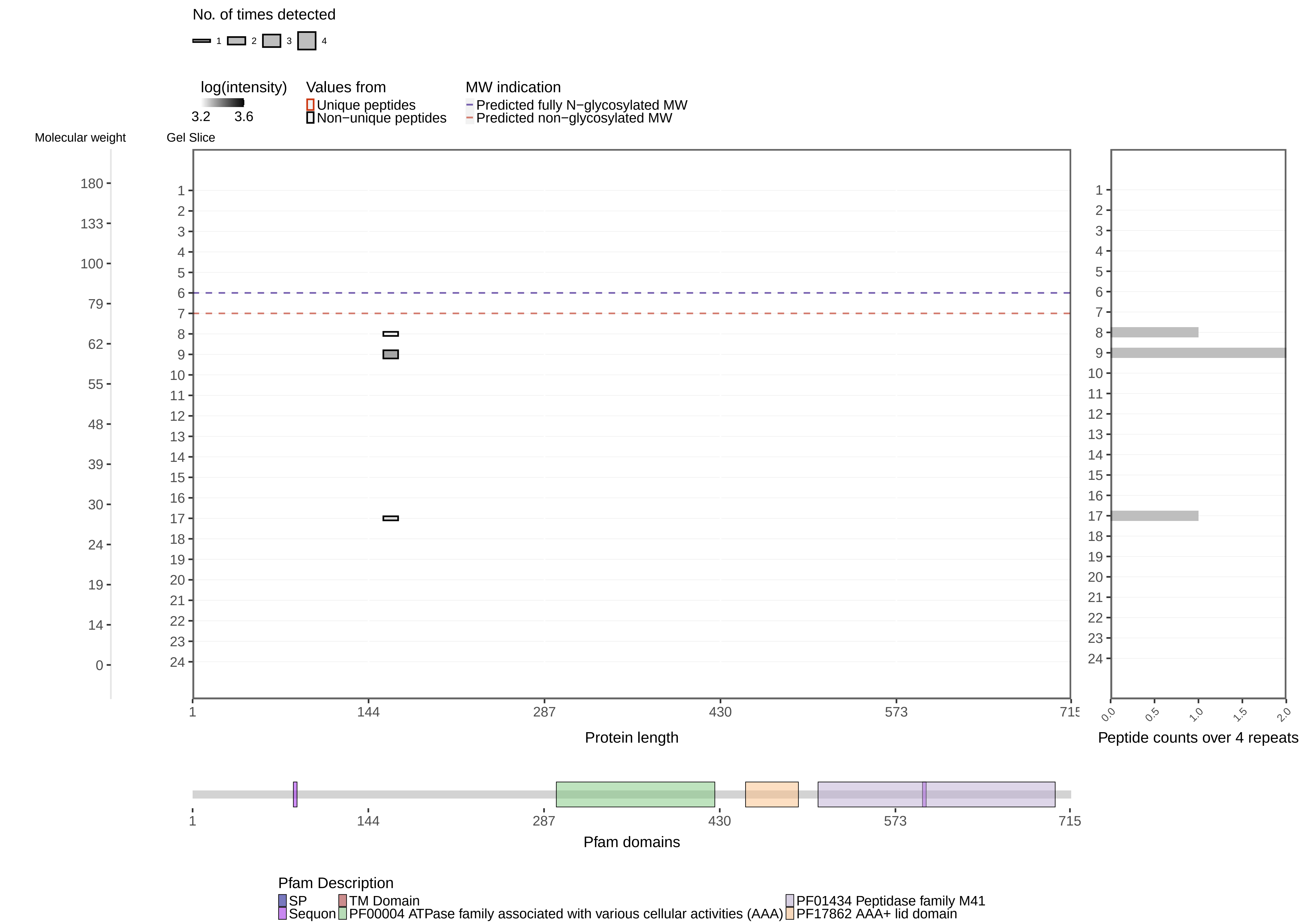


Protein sequence:

DLFEKAKSKAPCIVFIDEIDAVGRQRGAGLGGGNDEREQTINQLLTEMDGFSGNSGVIVLAATNRPDVLDSALLRPGRFDRQVTADRPDVAGRVKILQVHSRGKALAKDVDFDKIARRTPGFTGADLQNLMNEAAILAARRDLKEISKDEISDALERIIAGPEKKNAVVSDEKKKLVAYHEAGHALVGALMPEYDPVAKISIIPRGQAGGLTFFAPSEERLESGLYSRSYLENQMAVALGGRVAEEVIFGQDNVTTGASNDFMQVSRVARQMVERLGFSKKIGQVAIGGGGGNPF

### NbL02g02620.1 protein_TIC110_chloroplastic [MW = 112.44 kDa]

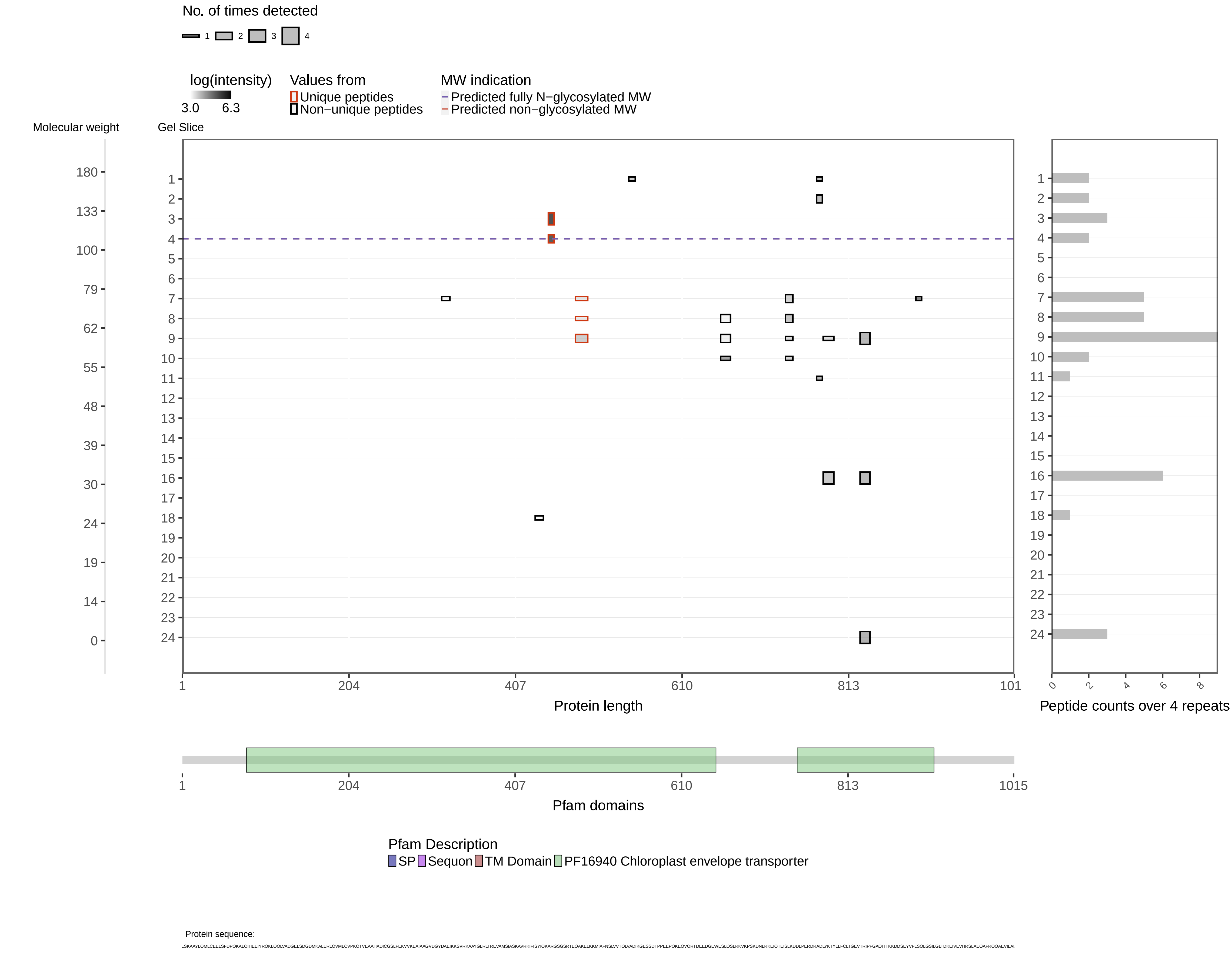

### NbL02g02550.1 CBS_domain−containing_protein_CBSX6 [MW = 15.2 kDa]

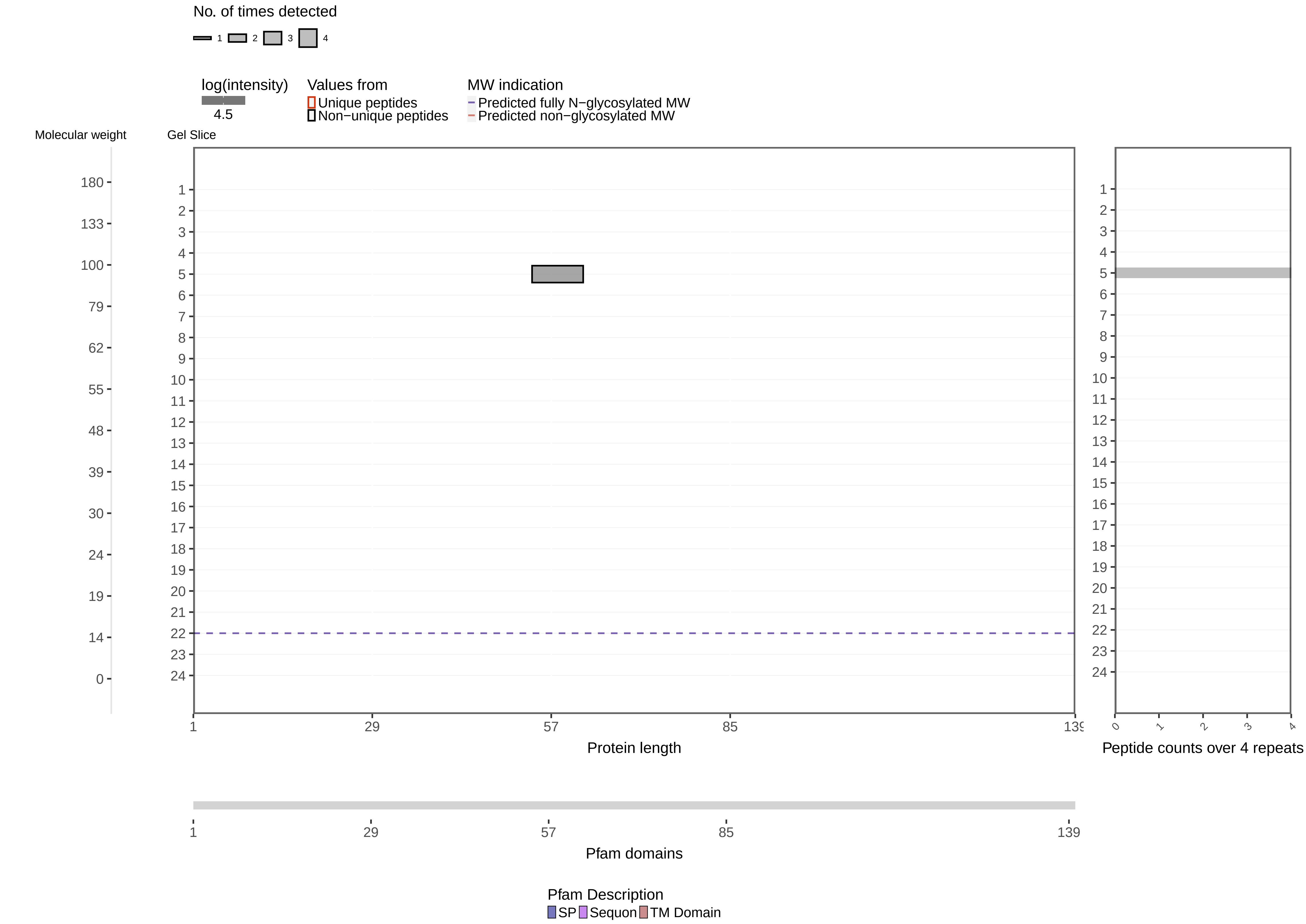


Protein sequence:

ENSEMRAKRFVGILTSLDIVAFLAREECLVDQEKAMKTPVSEVVVPDPSLLKELDPATRYINCARSSSSLLCQSFMFDQGVT*

### NbL02g02390.1 violaxanthin_de−epoxidase_chloroplastic−like [MW = 54.59 kDa]

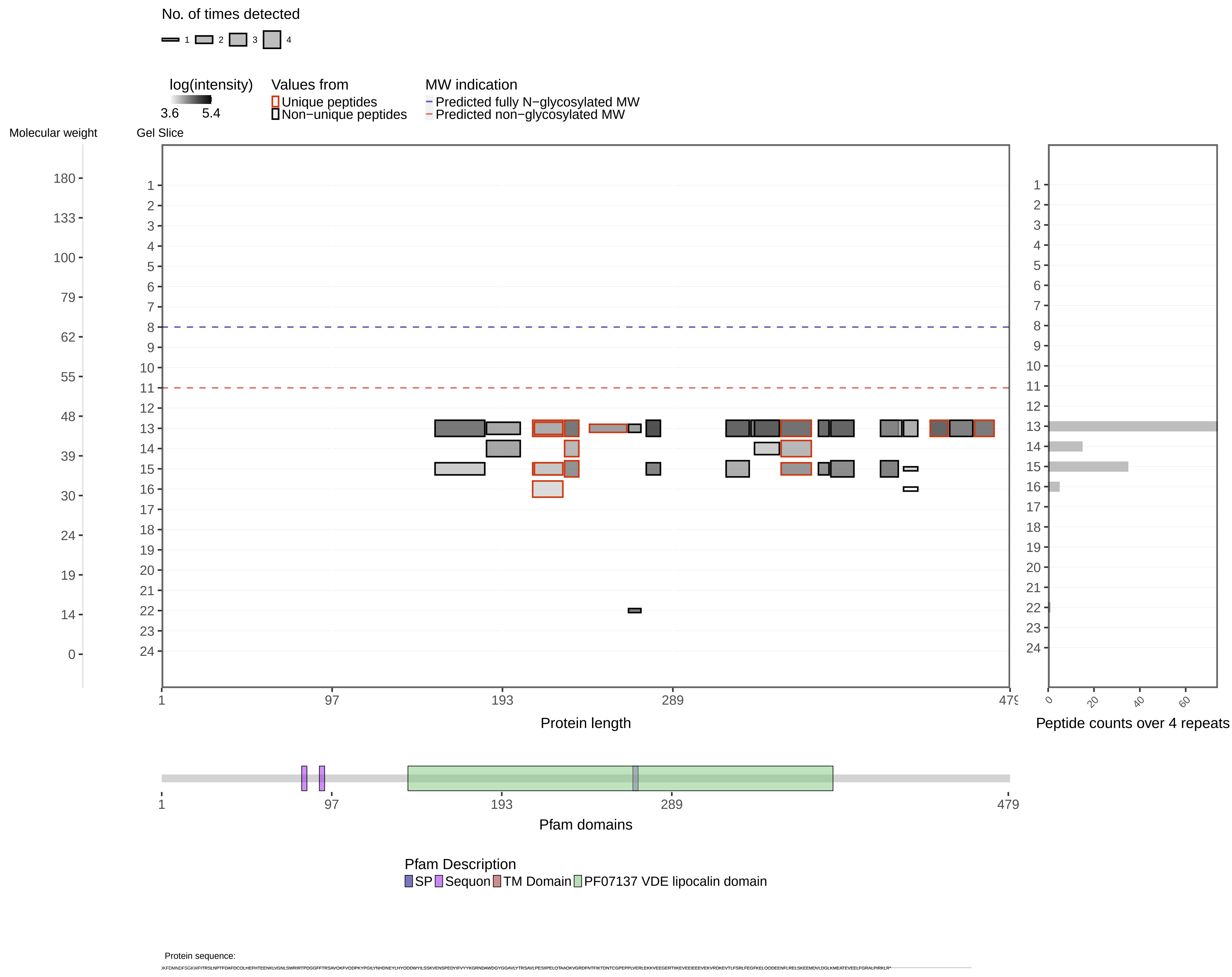

NbL02g02200.1 leukotriene_A−4_hydrolase_homolog [MW = 68.66 kDa]

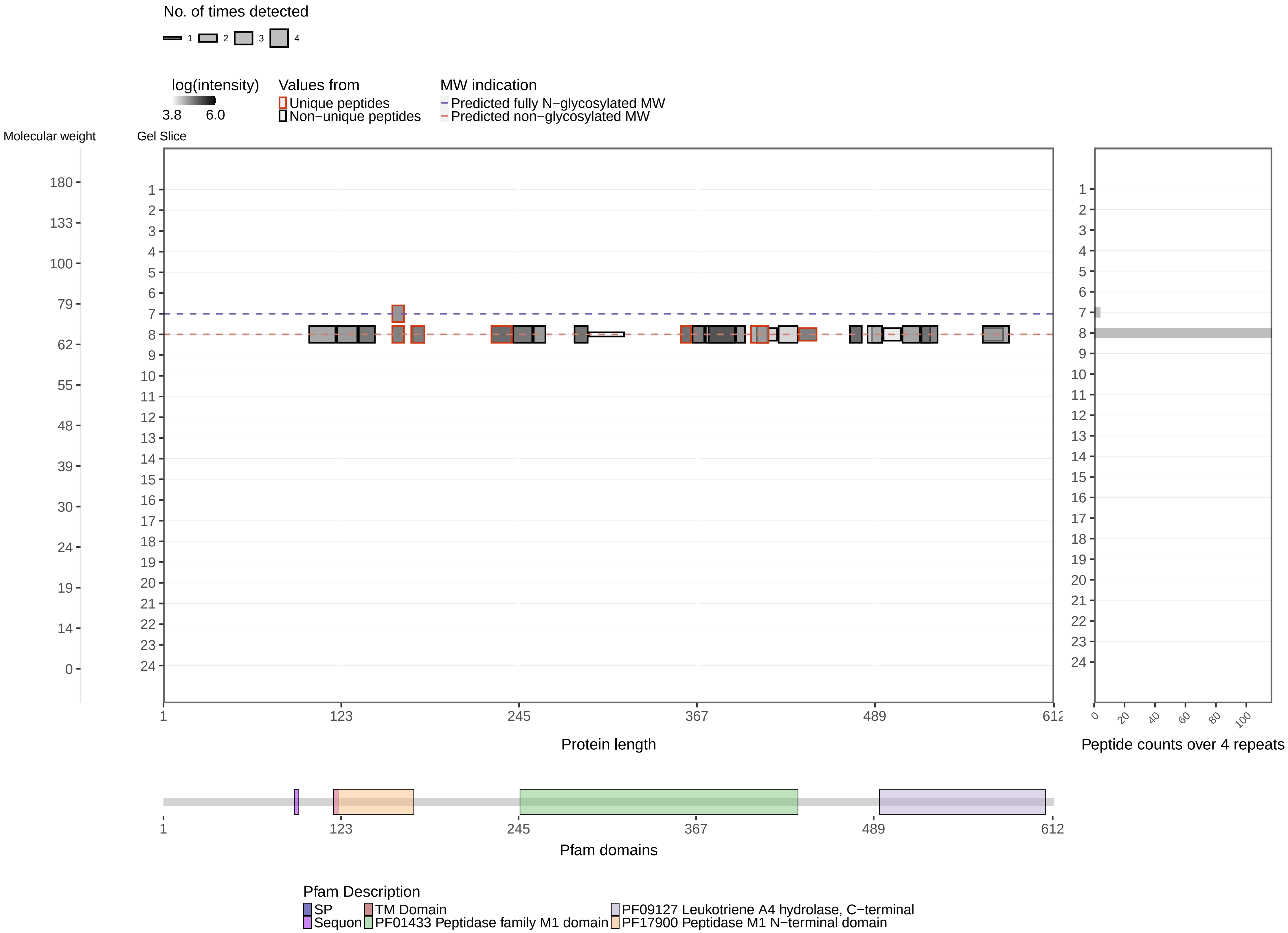


Protein sequence:

ASGAQVVAHELAHSWTGNLITNKNNDHFWLNEGFTTYAERRIVEVVQGEDIAALNIGIGWRGLIEEMERFKDNMEFTKLKTNQAGVDPDDVYSQVPYEKGFQFLWRIERQIGRPAFDDFLKKYIATFKFQSIDTDMFLNFLKADVPGIENEIDLKLWTEGTGIPPDAMEPVSNIYSKIVSLANEFKLGRMPREDEVADWKGQEWELYLENLPKSVEASQVTALDARYHLSESKDYEVRVAFLQLAISARCRDYYGEVEKTLKEVGRMKYLRPLYTALVQGAGKEEEKAFARRV

NbL02g02090.1 uncharacterized_LOC107785953 [MW = 29.08 kDa]

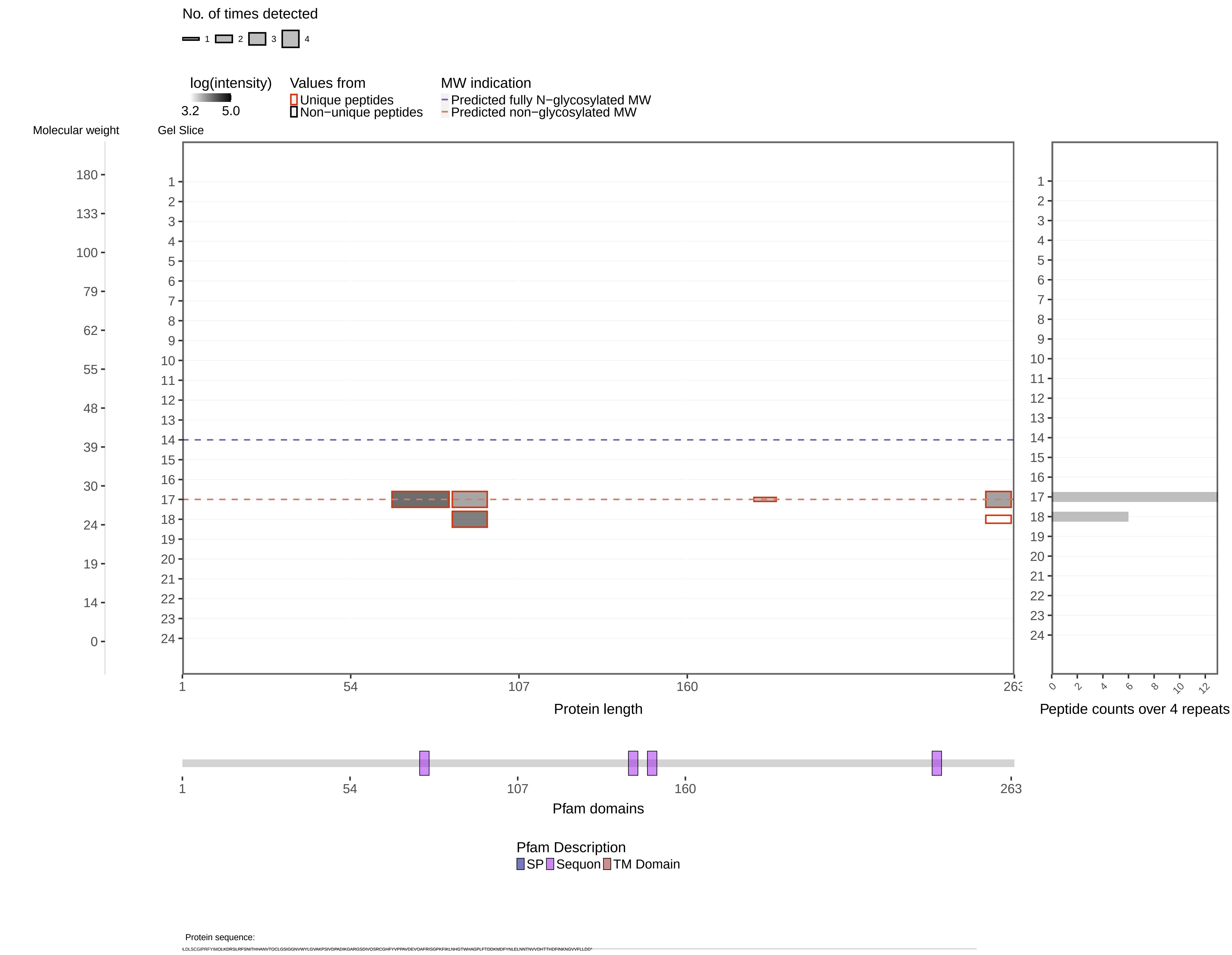

### NbL02g02070.1 uncharacterized_LOC104220881 [MW = 184.43 kDa]

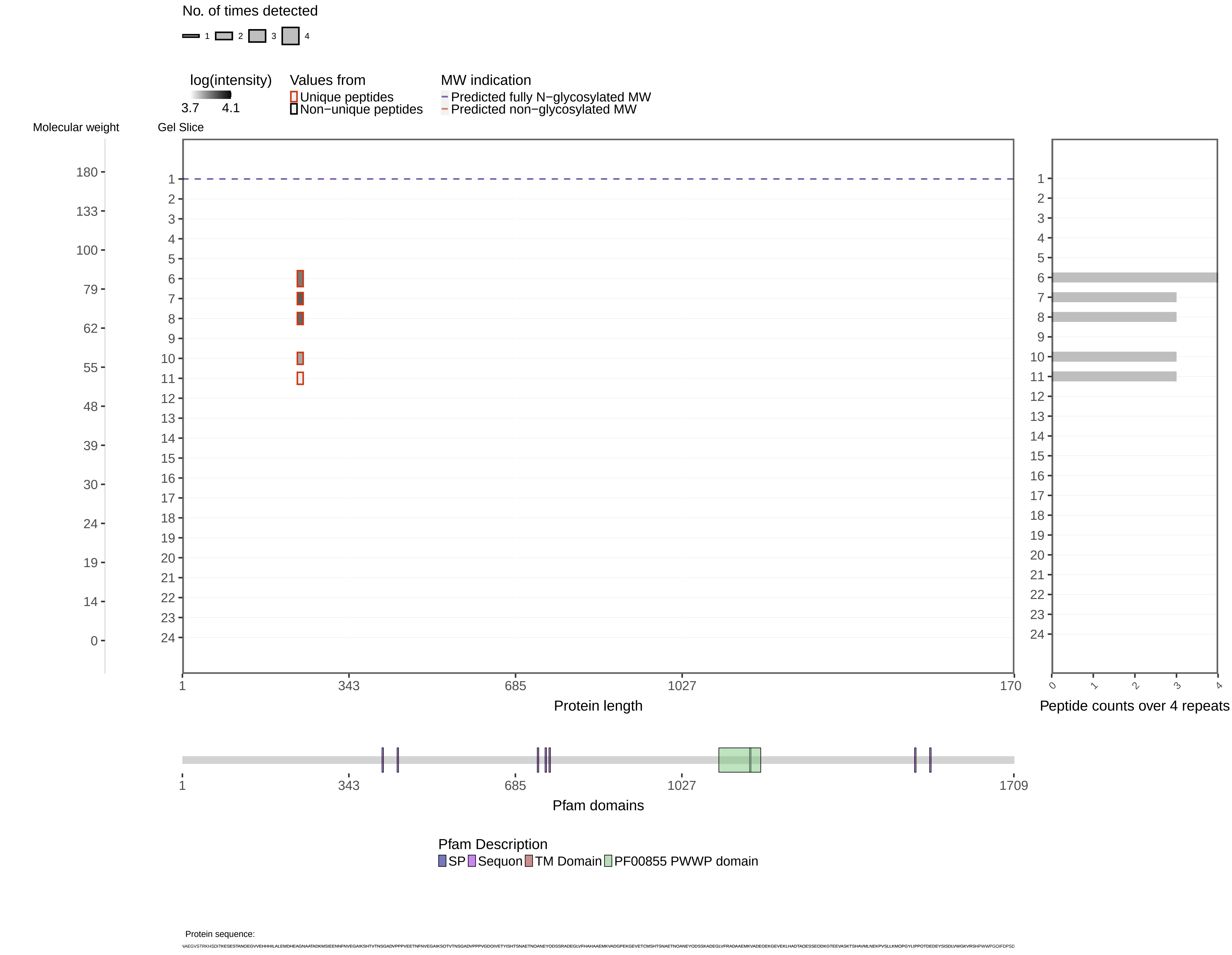

### NbL02g01760.1 isopentenyl−diphosphate_Delta−isomerase_I−like [MW = 33.19 kDa]

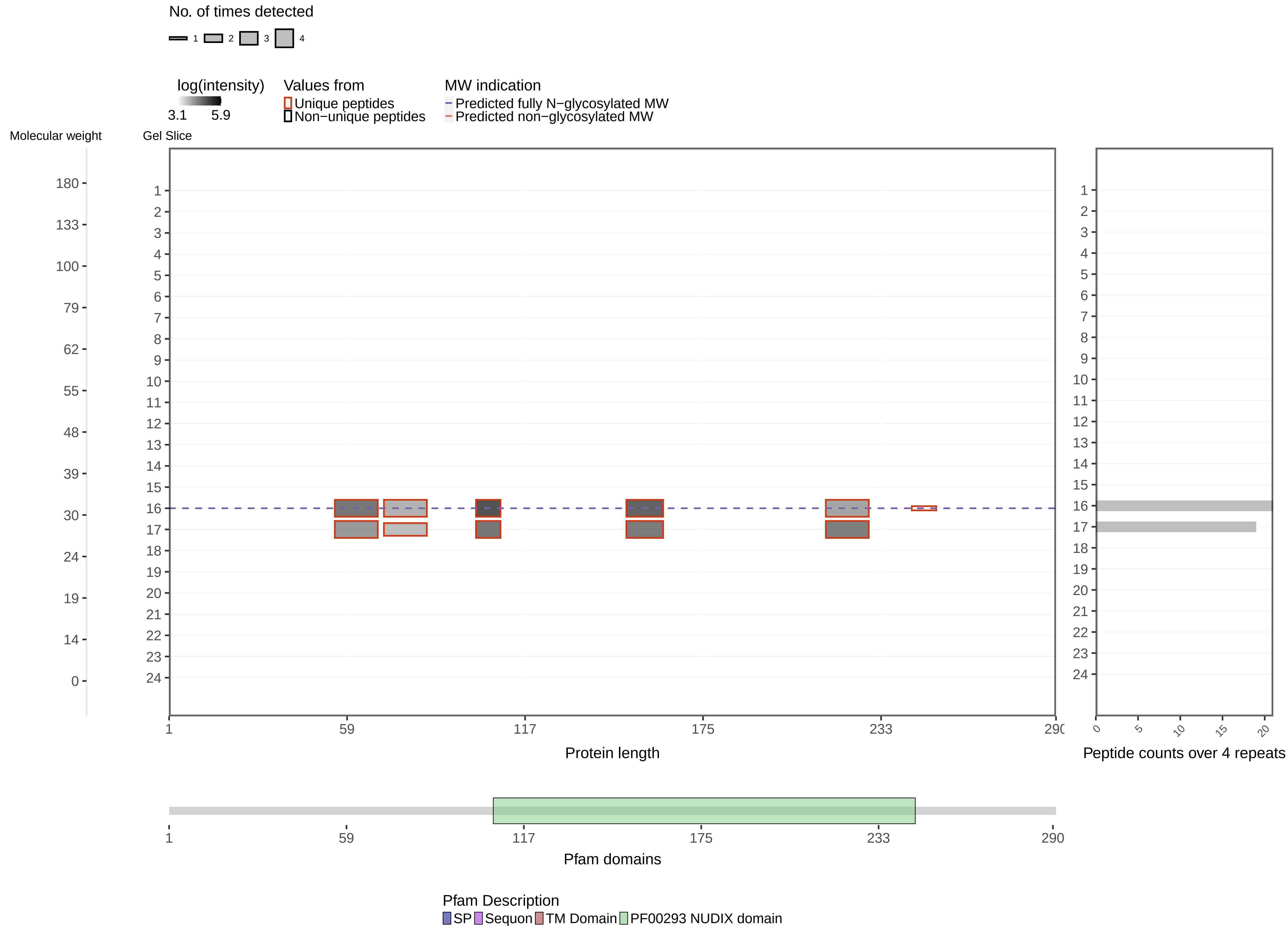


Protein sequence:
TFPLVWTNTCCSHPLYRESELIEENVLGVRNAAQRKLLDELGIPAEDVPVDQFTPLGRILYKAPSDGKWGEHELDYLLFTVREVNMKPNPDEVADVKYVNREQLKELLRKADAGEGGLKLSPWFRLVVDNFLFKWWDHLEKGTLKEVIDMKTIHKLT*

NbL02g01570.1 aspartate_aminotransferase_chloroplastic [MW = 50.72 kDa]

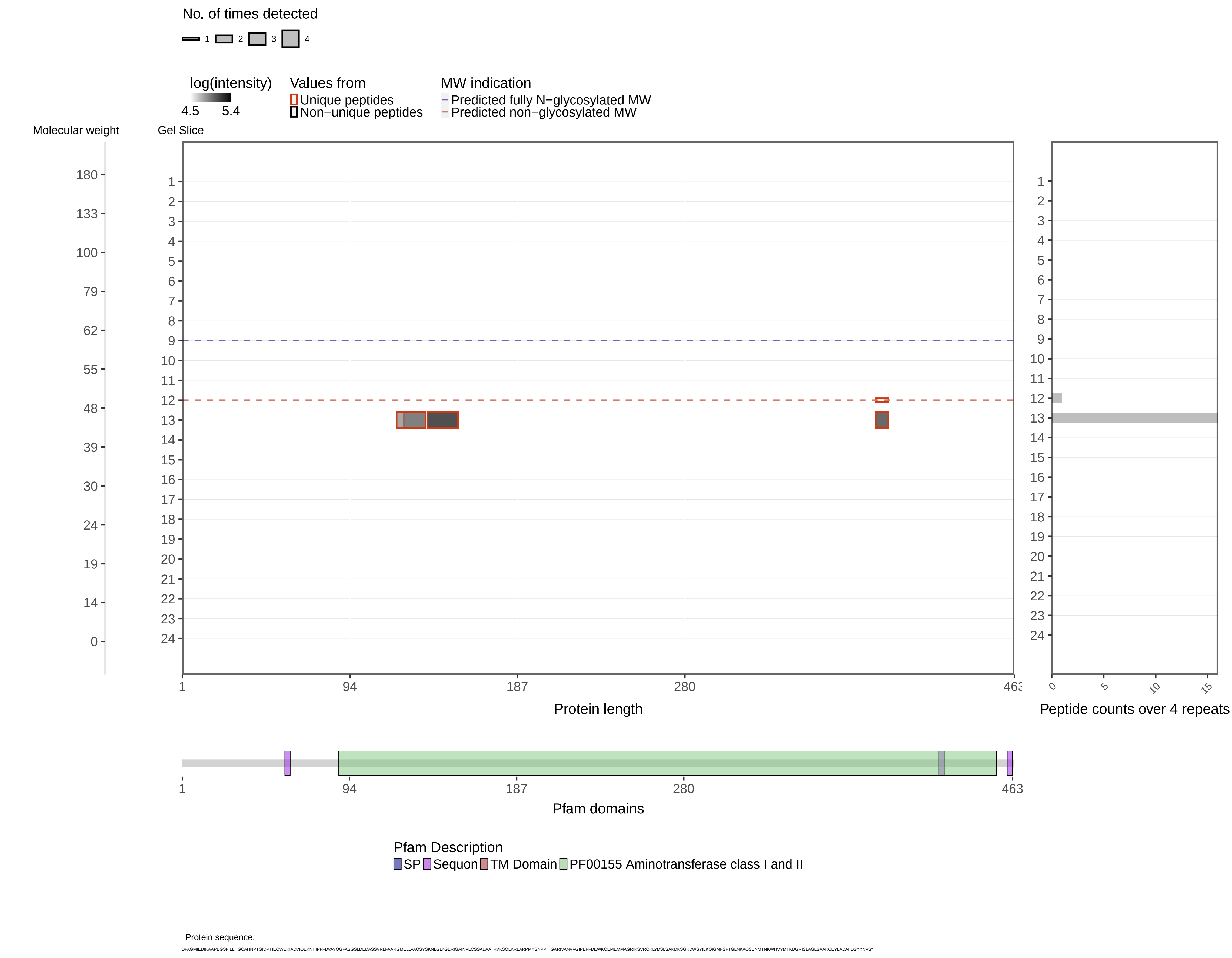

NbL02g01550.1 alanine−−glyoxylate_aminotransferase_2_homolog_1_mitochondrial [MW = 26.13 kDa]

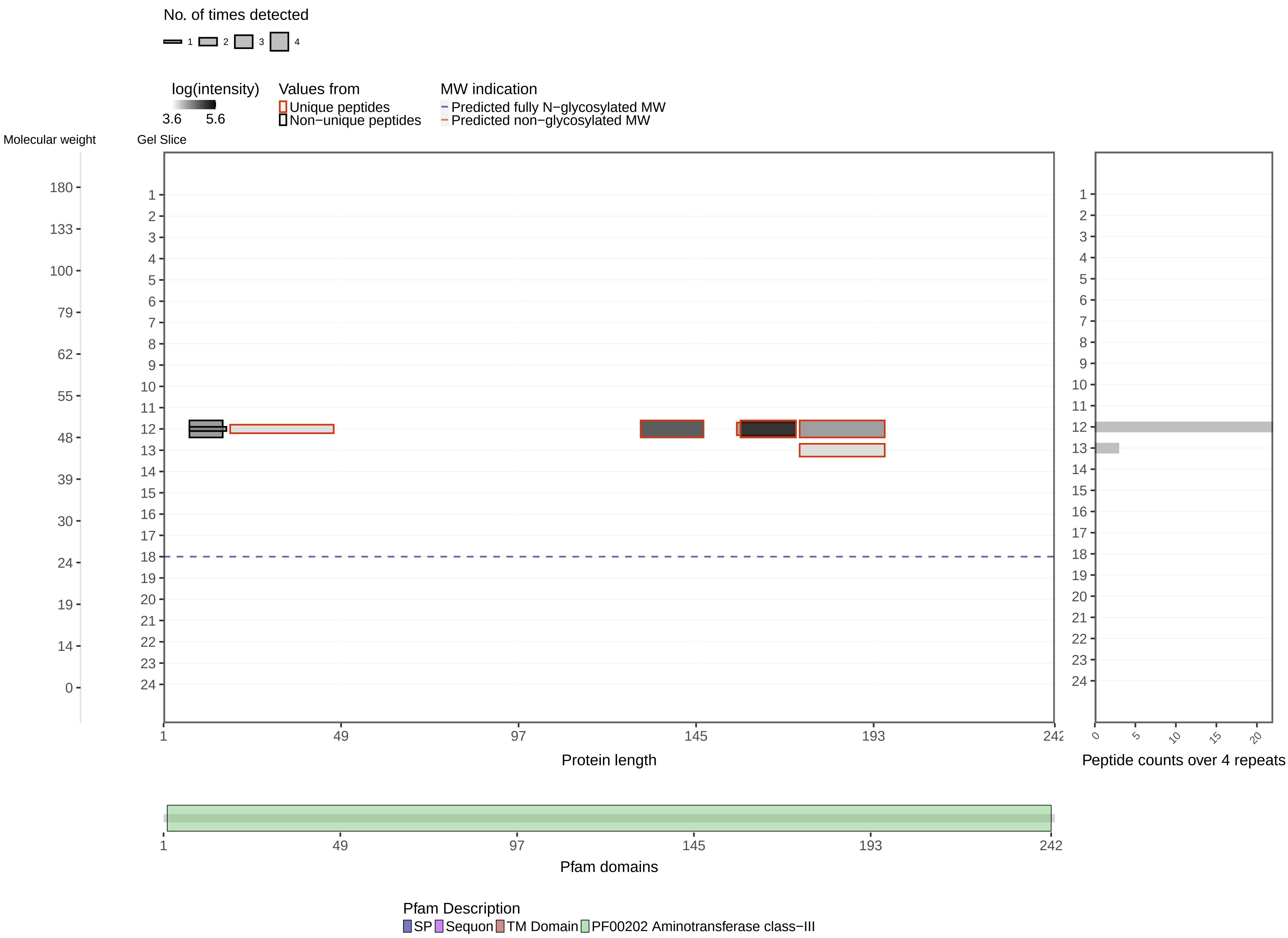


Protein sequence:

RNAYHGGSANTIGLTALNTWKYPIPQGEIHHVVNPNPYRGVFGSDAKRYAEDVQDHIDHGTSGKVAGFIAETIQGAGGAVELAPGYLKLVYDIARKAGGVCIADEVQTGFGRTGSDYWGFQTQGVIPDIVTMAK

NbL02g01540.1 alanine−−glyoxylate_aminotransferase_2_homolog_1_mitochondrial [MW = 23.32 kDa]

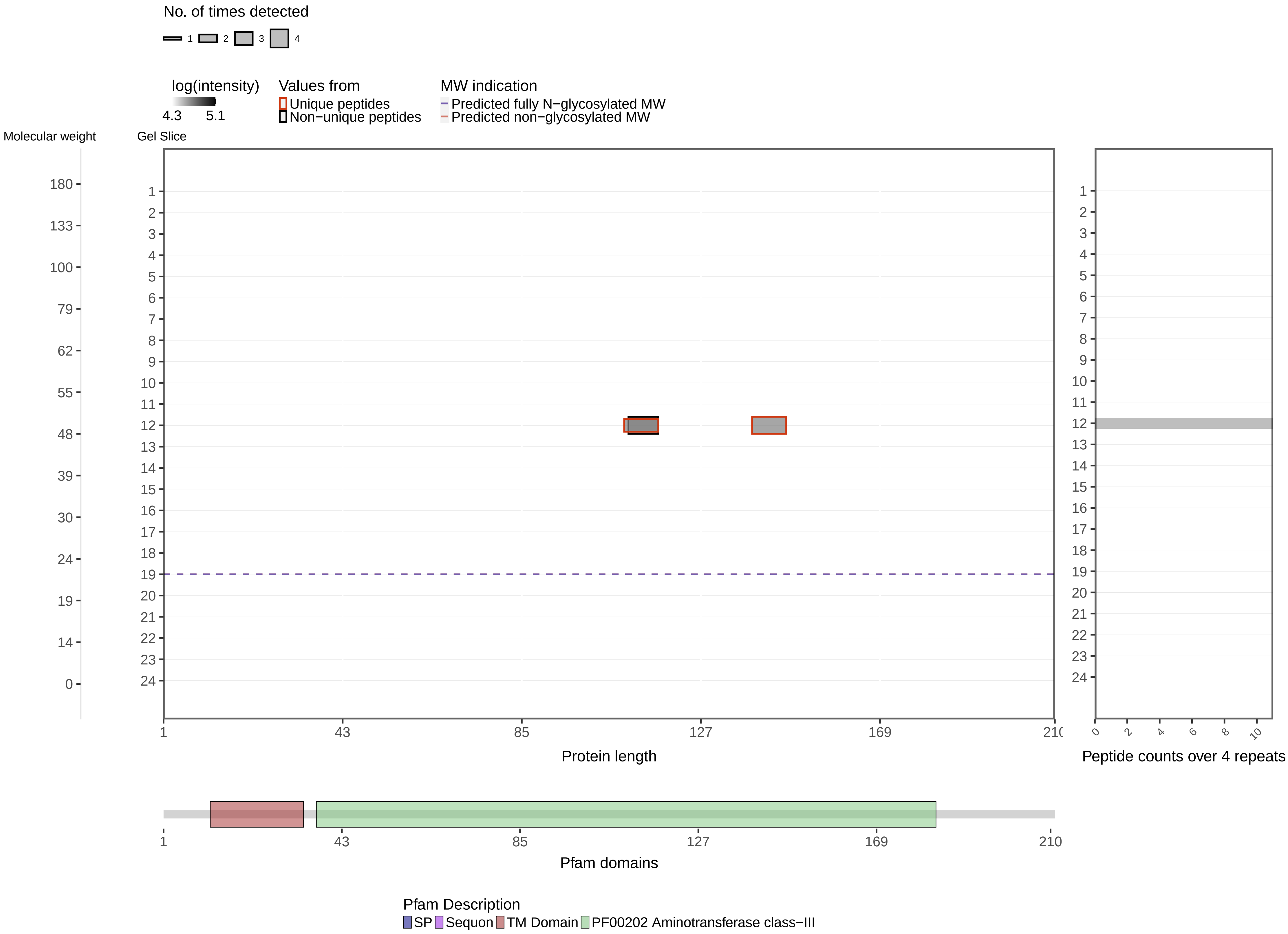


Protein sequence:

ICAEVGSHLIGRLRDLQKRHEIIGDVRGRGLMVGIELVTDRKEKTPAKAETGVLLEKLREFGVLVGKGGLHGNVFRIKPPMCFGKDDAGNLFTHWFNILRIFSWIIPSDLISIVENHFY*

### NbL02g01370.1 L−ascorbate_oxidase [MW − SP = 61.46 kDa]

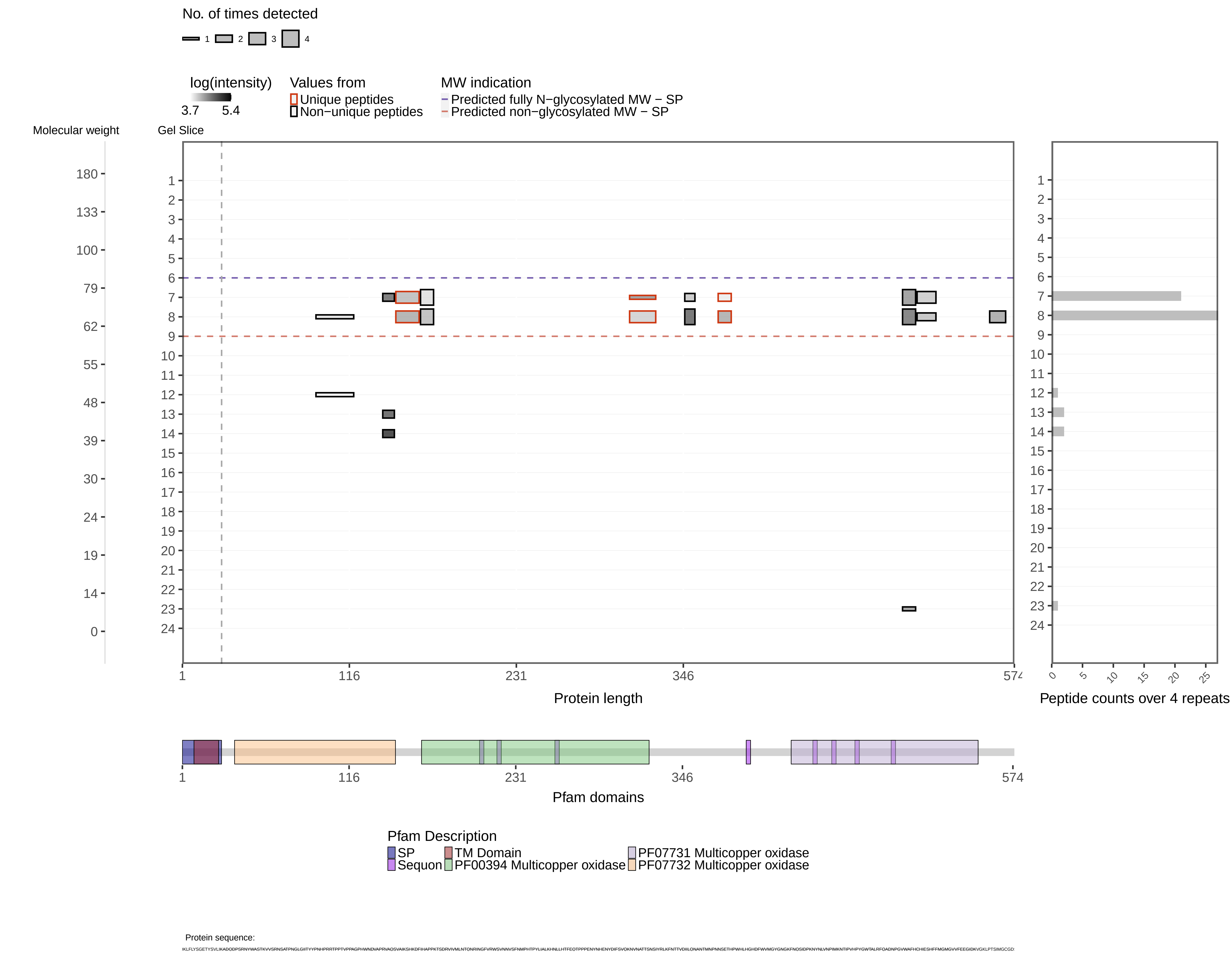

NbL02g01350.1 bifunctional_aspartate_aminotransferase_and_glutamate/aspartate−prephenate_aminotransferase−like [MW = 51.59 kDa]

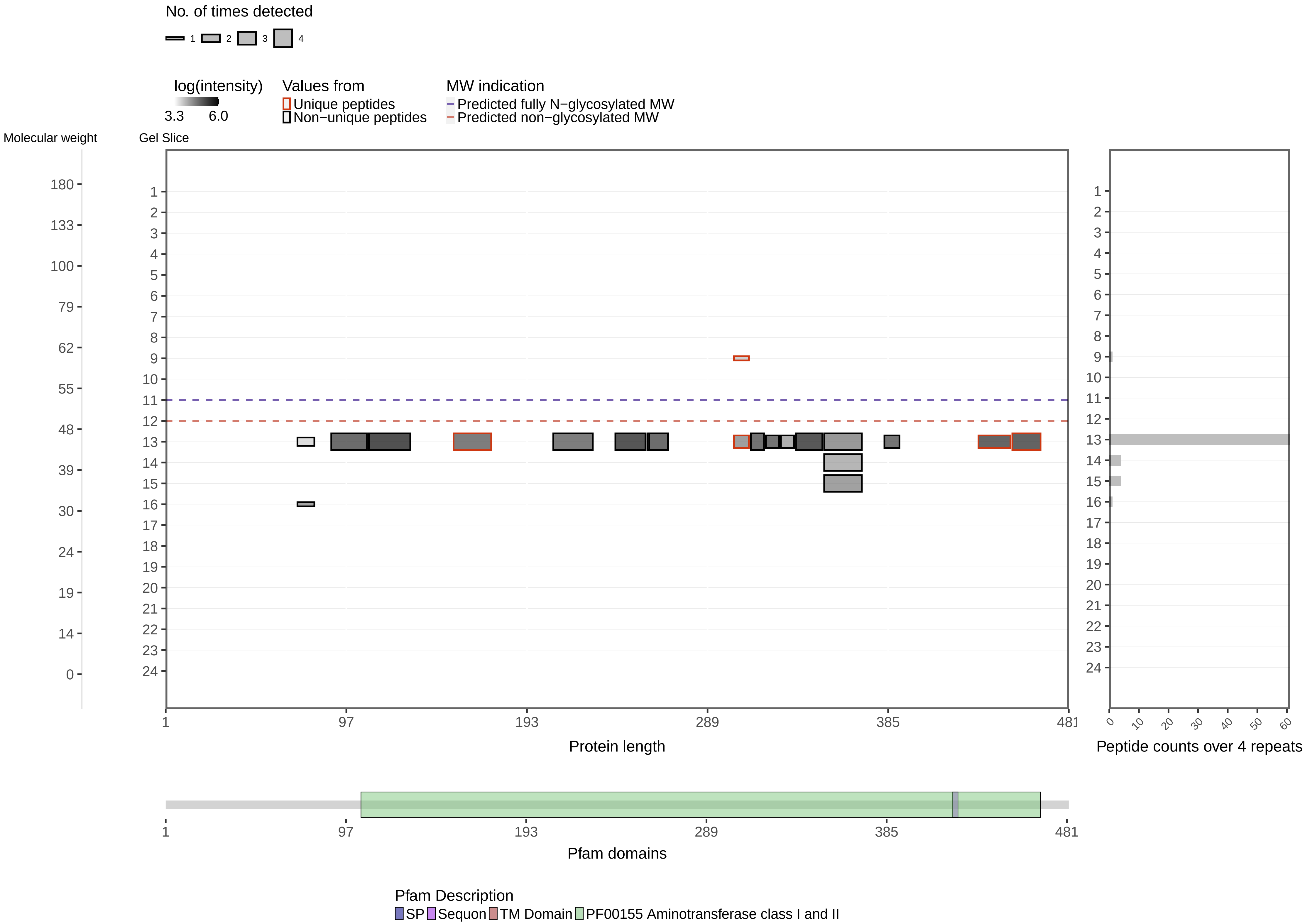


Protein sequence:

KLLESKLSEKSRLLILCSPSNPTGSVYPRKLLEEIAEIVARHPRLLVISDEIYEHIIYAPATHTSFASLPGMWDRTLTVNGFSKAFAMTGWRLGYIAGPKHFVSACNKIQSQFTSGASSISQKAAVAALGLGYAGGEAVATMVKAFRERRDYLVKSFGEIDGVKISEPRGAFYLFIDLSSYYGVEVDGFGSINDSESLCRYLLDKAQVALVPGDAFGDDTCIRISYAASLSTLQAAVERIKKALVTLRSHVPV*

### NbL02g01320.1 inositol−3−phosphate_synthase [MW = 56.39 kDa]

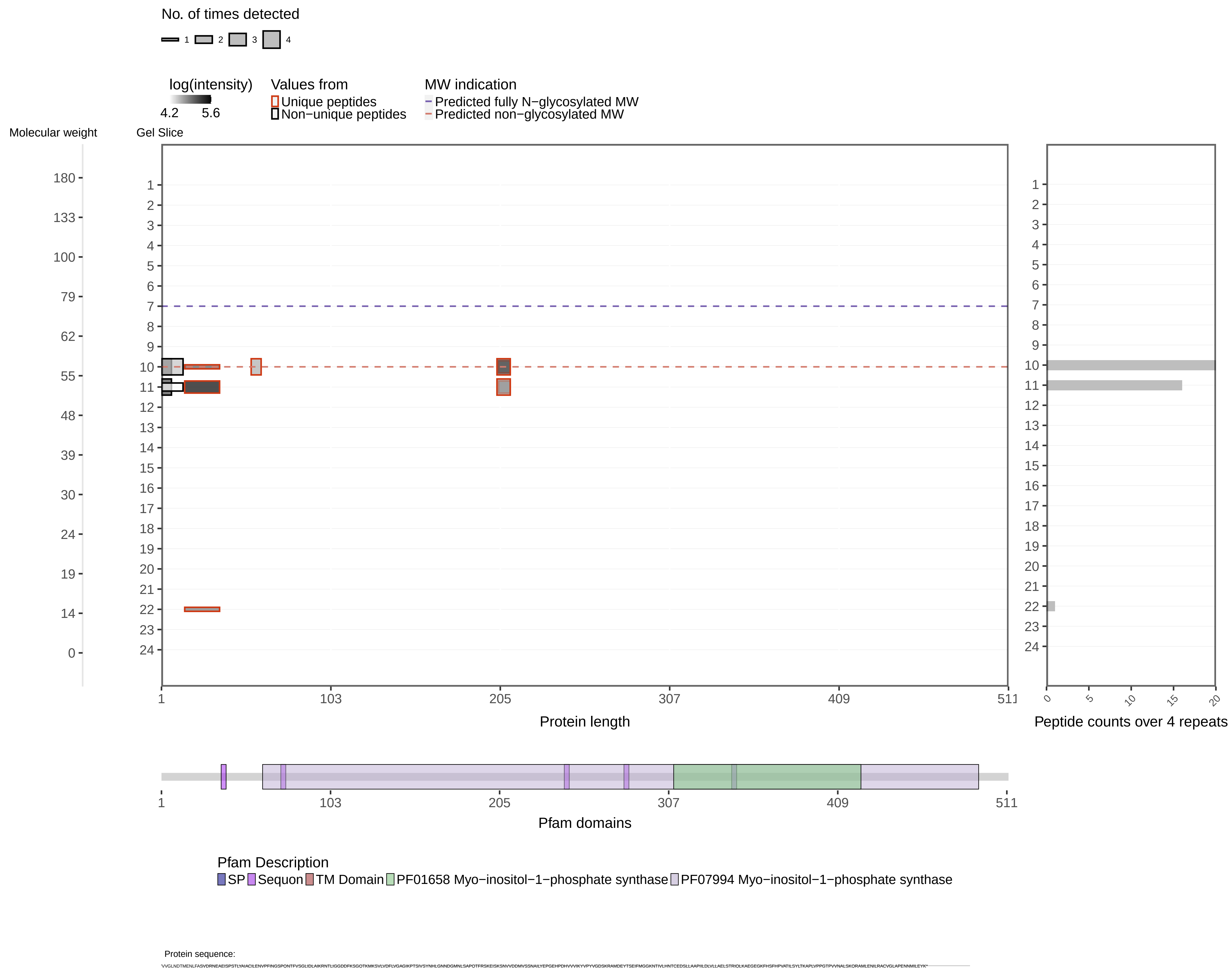

### NbL02g01210.1 asparagine_synthetase_[glutamine−hydrolyzing]_2 [MW = 70.87 kDa]

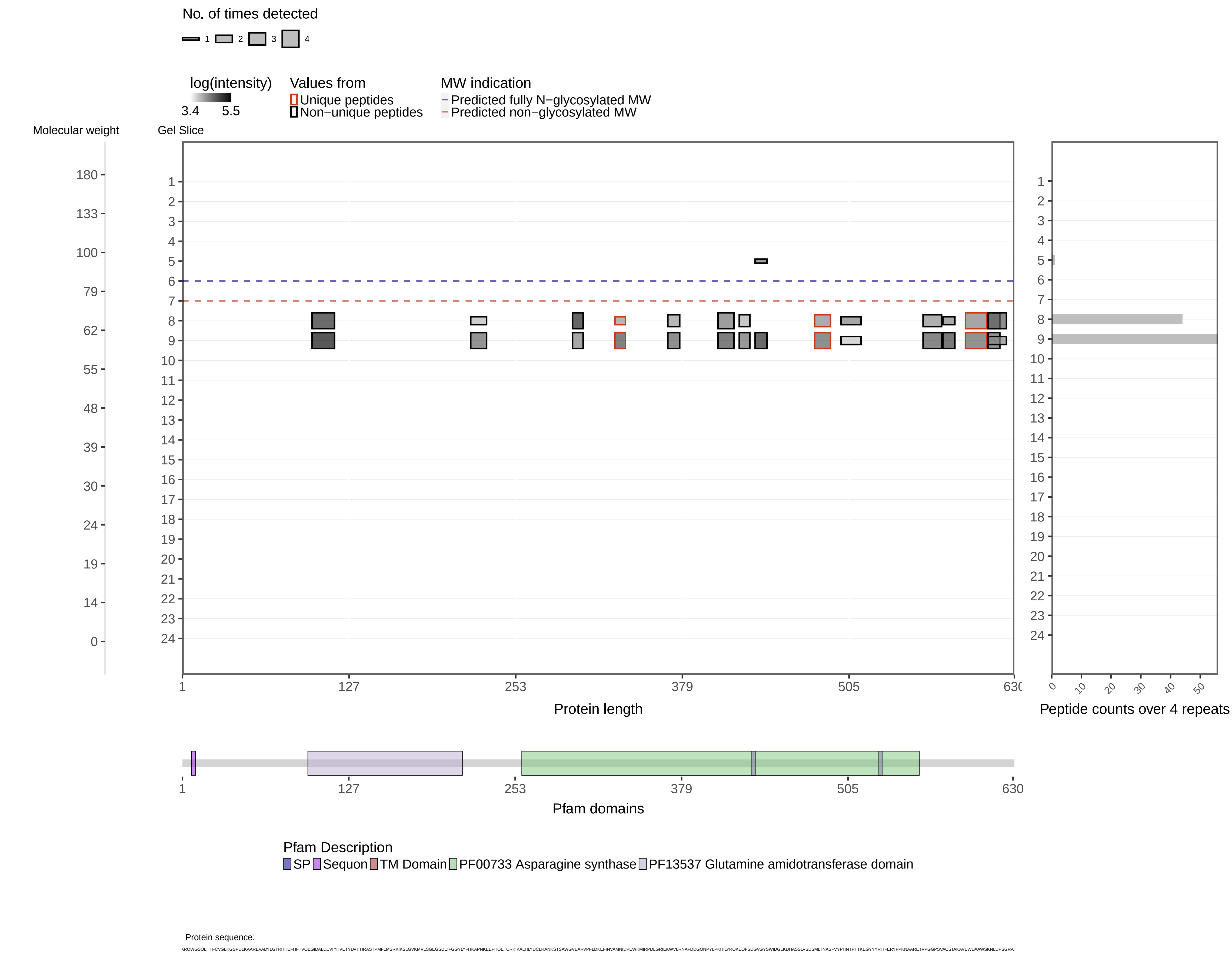

### NbL02g01180.1 annexin_D2−like [MW = 35.91 kDa]

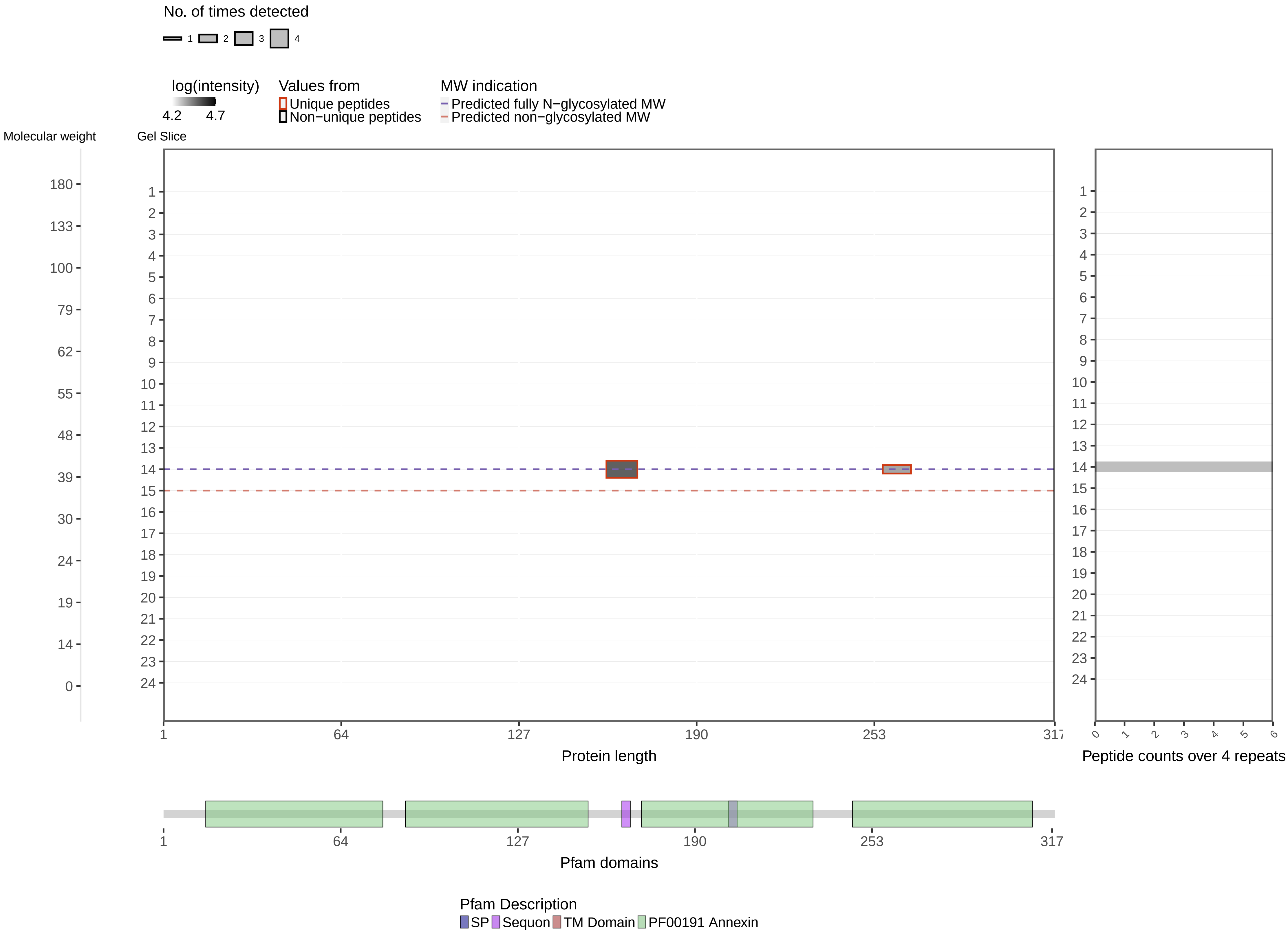


Protein sequence:

FRKVLVPLVTAFRYEGEEVNMTLARKEANILHEKISDKAYNDEELIRIISTRSKAQLNATFNHYLDQHGNEINKDLETDSDDEYLKILSAAIECLKTPEKYFEKVLRLGIKGAGTDEWDLTRVVTTRAEVDMERIKEEYQKRNSVPLDRAIAGDTSGDYERMLLALIGHGDA*

NbL02g00700.1 transketolase_chloroplastic−like [MW = 79.86 kDa]

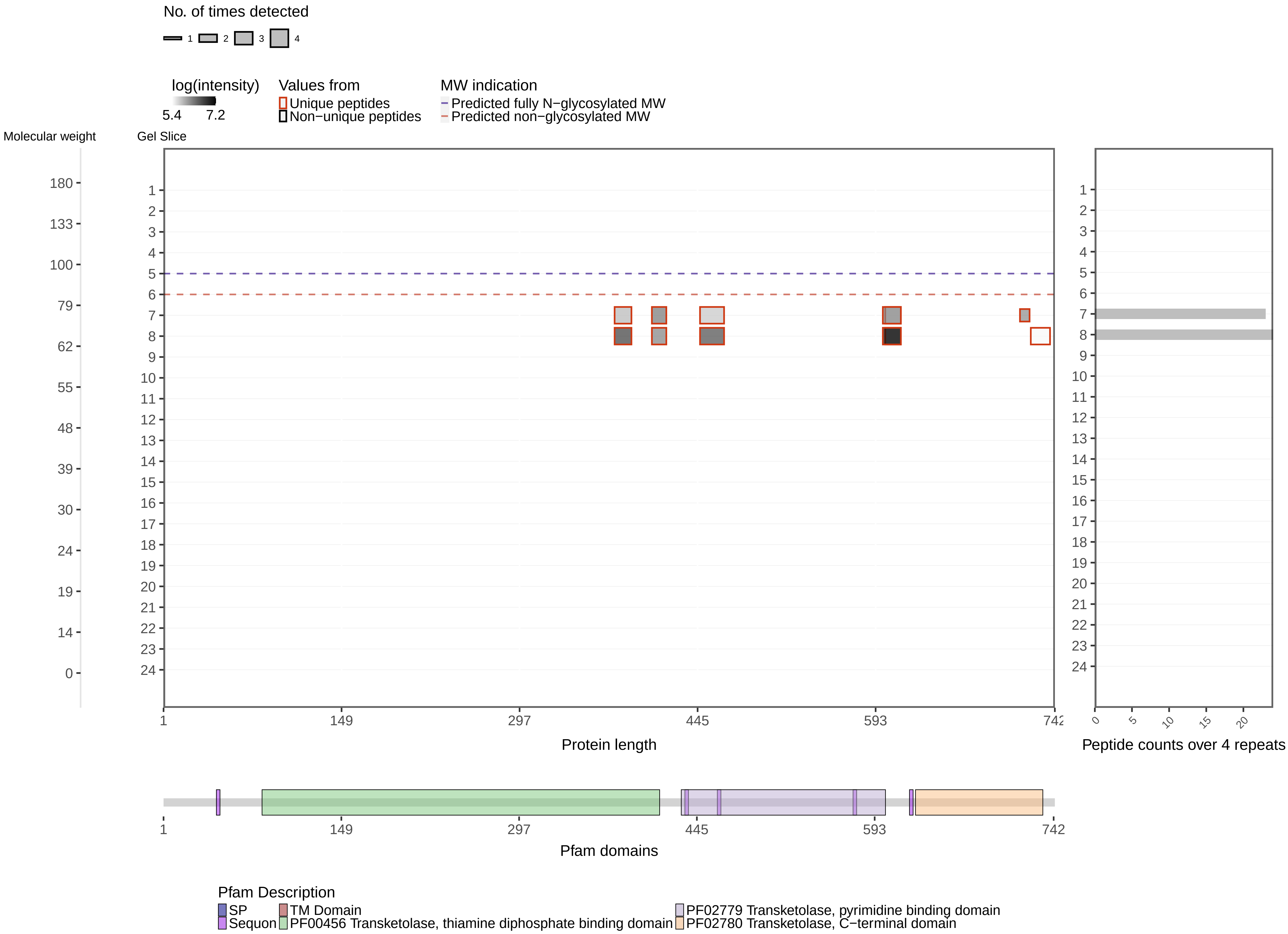


Protein sequence:

### NbL02g00300.1 uncharacterized_LOC109205733 [MW = 16.69 kDa]

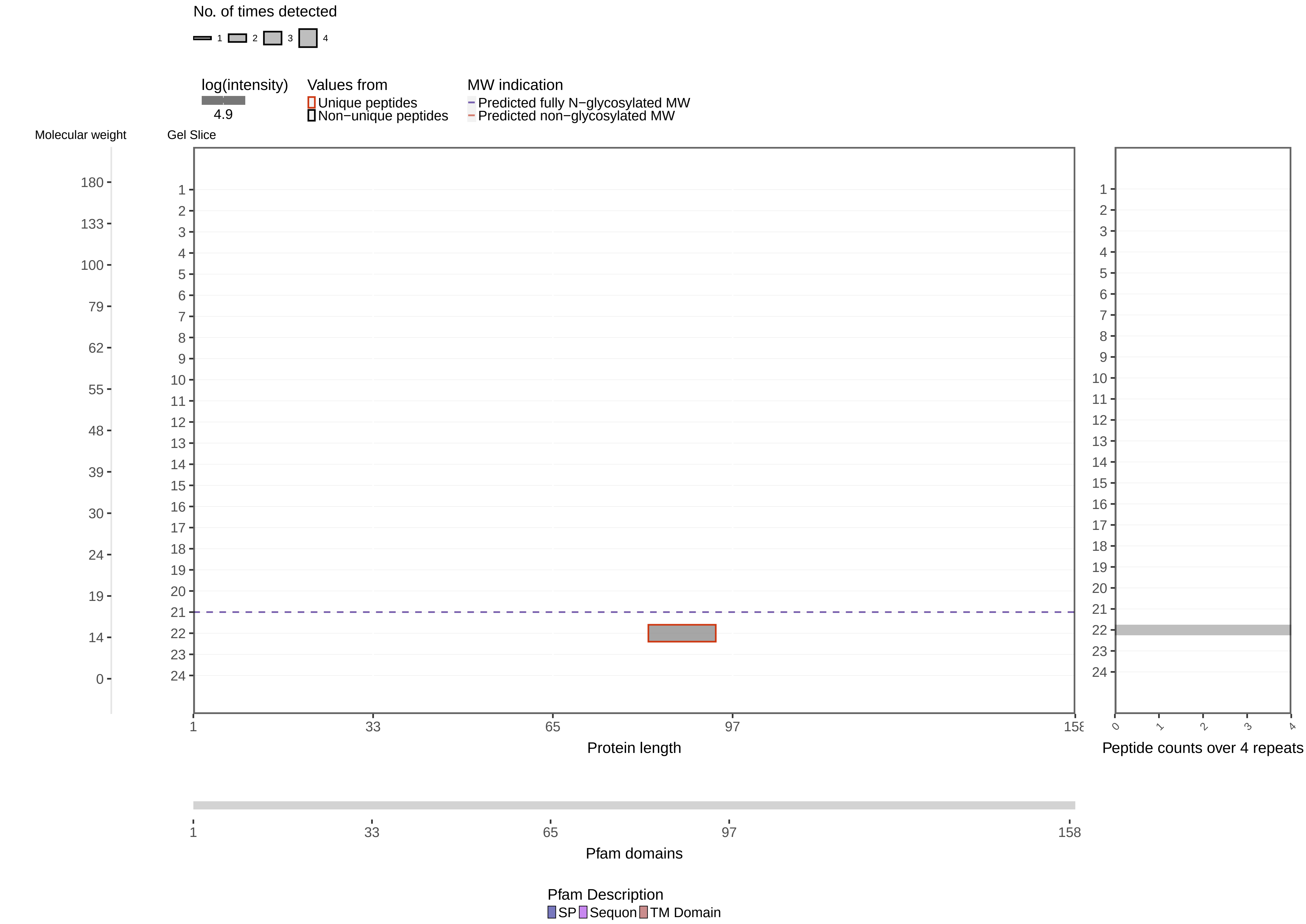


Protein sequence:

RCAEGQTQTVTREQSTITVAPVQGKEKSPALDDGGSGFPPRDDDGGGGGGGGGGGNWKGGFFFFGFLAFLGFLKDQEDEGPYRDQRRR*

NbL02g00260.1 4−alpha−glucanotransferase_chloroplastic/amyloplastic [MW = 65.29 kDa]

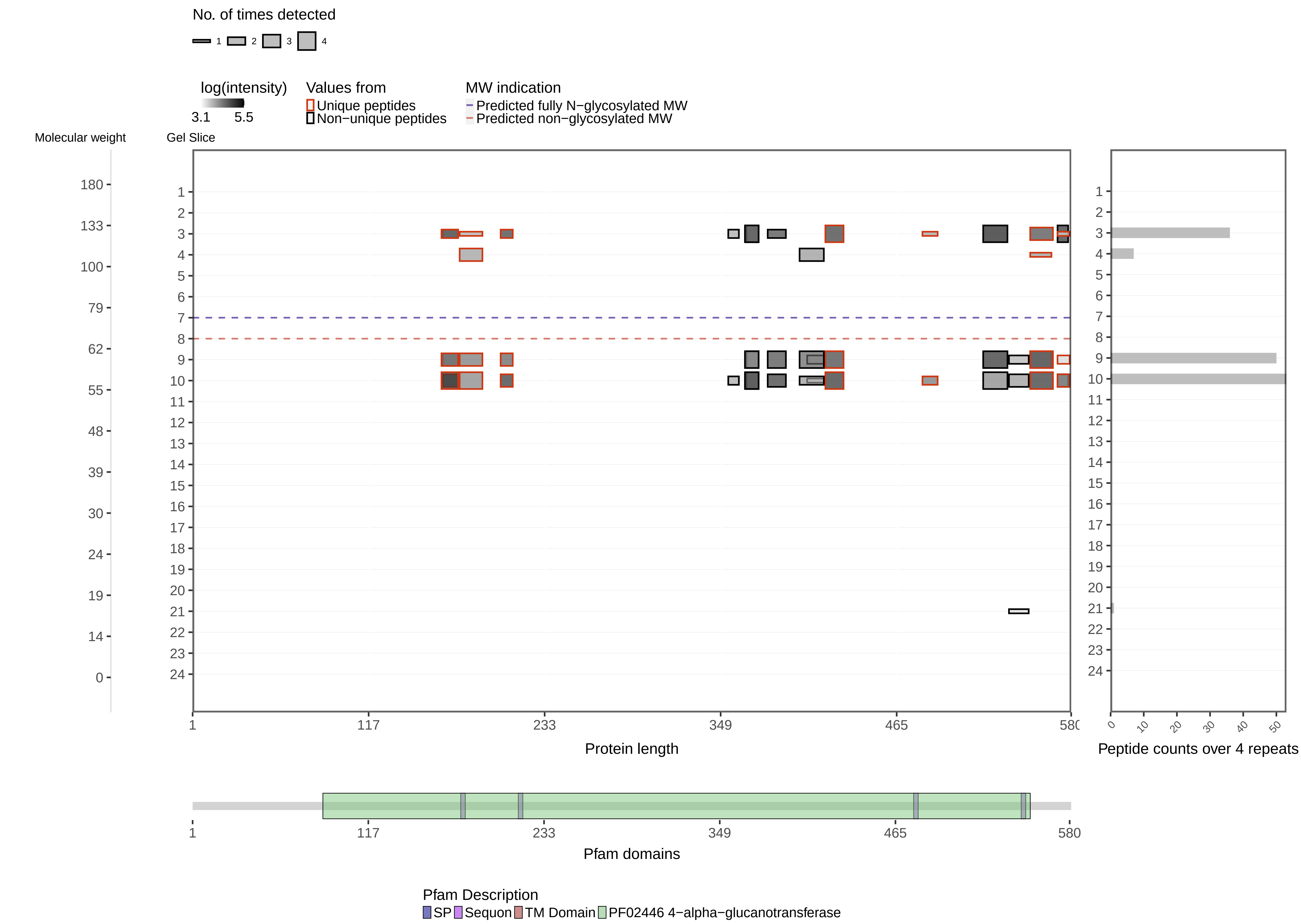


Protein sequence:

WQKVRDYARSKGISIMGDMPIYVGYHSADVWANKKQFLLNRKGFPLIVSGVPPDAFSETGQLWGSPLYDWKAMEKDGFSWWIRRIRRATDLFDEFRIDHFRGFAGFWAVPSEAKVAMLGRWKVGPGKPLFDAIFQAVGKINIIAEDLGVITKDVVQLRKSIEAPGMAVLQFGFGSDAENPHLPHNHEQNQVVYTGTHDNDTTRGWWNILPQEEKSNVLMYLSNIEEEEISWGLIEAAVSSVARIAIVPMQDVLGLGSESRMNIPATQFGNWSWRIPSSTSFDSLDSEAKKL[illegible]

### NbL02g00220.1 2−hydroxyisoflavanone_dehydratase−like [MW = 35.4 kDa]

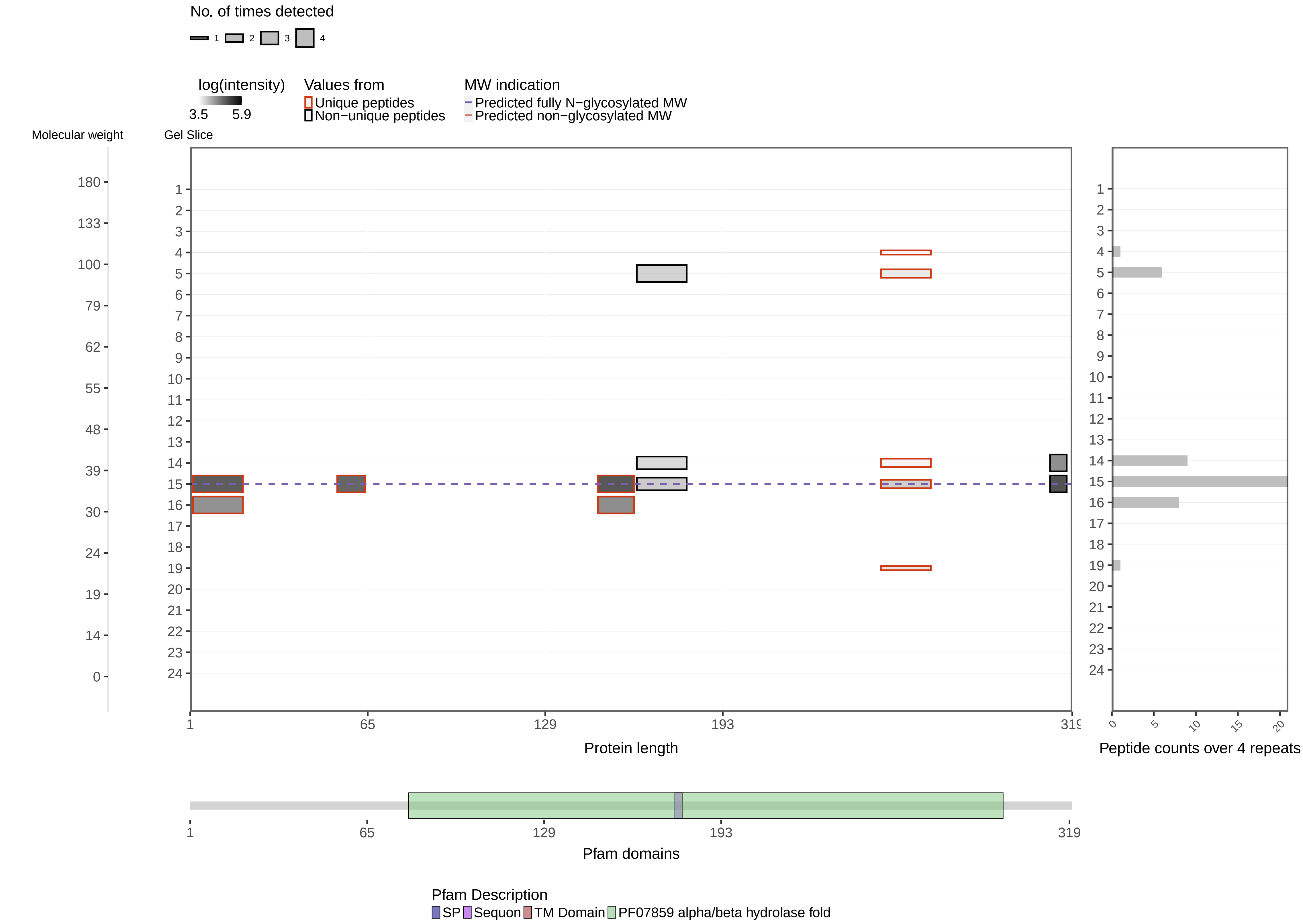


Protein sequence:

KDAEPWIANHGDFDRLFIAGDSAGGNIVFNMTMRVGRESLTGGVKLVGAIFSFPFFLIPSVENIEGYKLWNTICPPSEQGINSPMINPVSENGPSLSELGCSRLFLCAGEKDELIPSEVVSRFAEAVKNSGWKGELEYIEVEGEGHCFQAANPETEKSKHLIKRMASFIQRN*

NbL02g00180.1 uncharacterized_LOC109214761 [MW = 44.02 kDa]

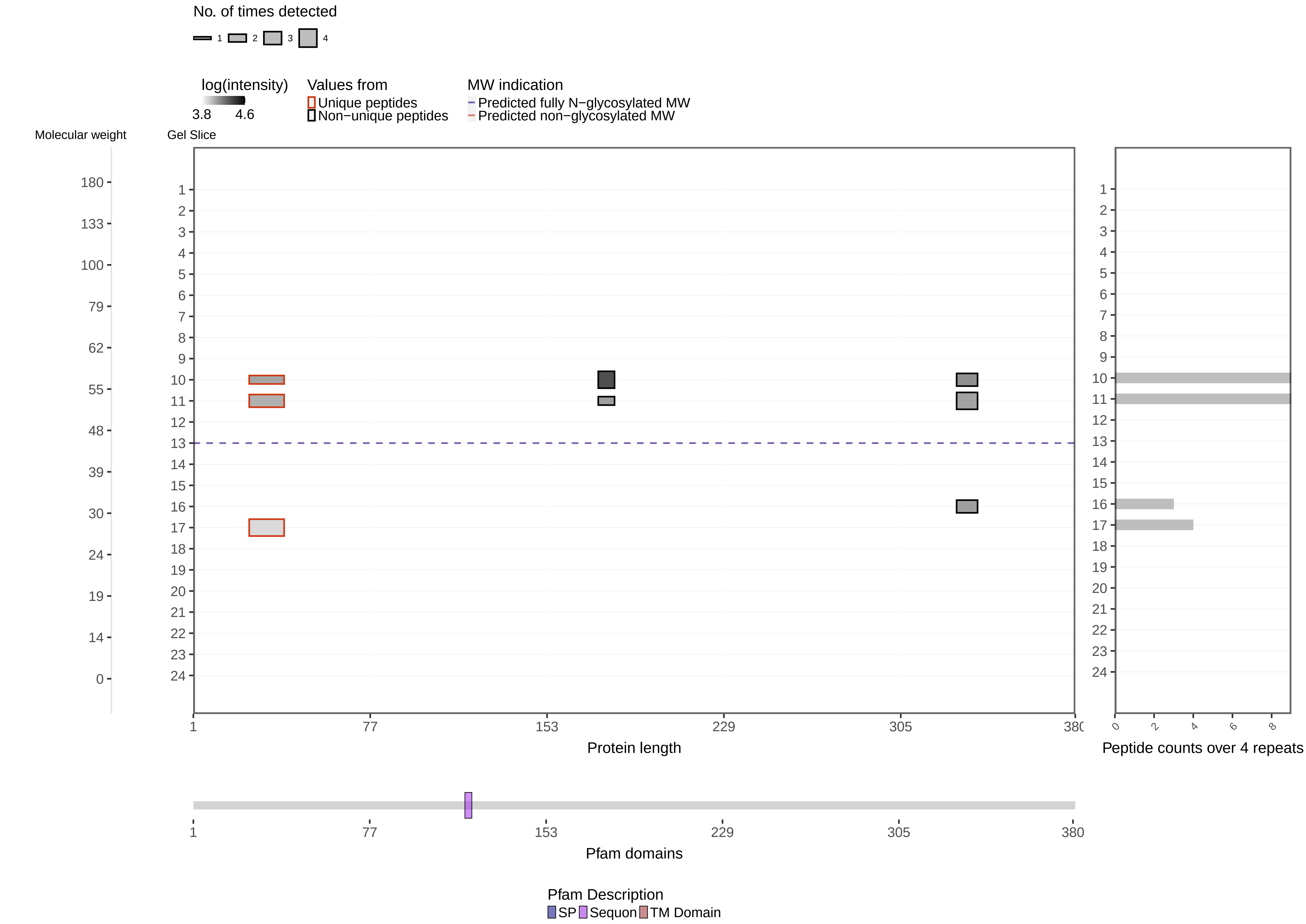


Protein sequence:

IGINSVQREAEEILPMKQMKMDWVPYIPLENRESQVERLKSQIFILRCTQRRAGLKHLKLERVKKFEYCLPYFYQPFAEDELEQSTIVQILFPTEPPVFCEFDWEFDEPEEMADKLIEAEELVADKKDEFMEFVKEKVREGKKNNREAREARKKAIQEMSEEAKAAFQSMKFYKFYPMPSPDAPDVSQVKSPFINRYYGKAHTVF*

NbL02g00150.1 probable_inactive_receptor_kinase_At5g10020 [MW − SP = 112.58 kDa]

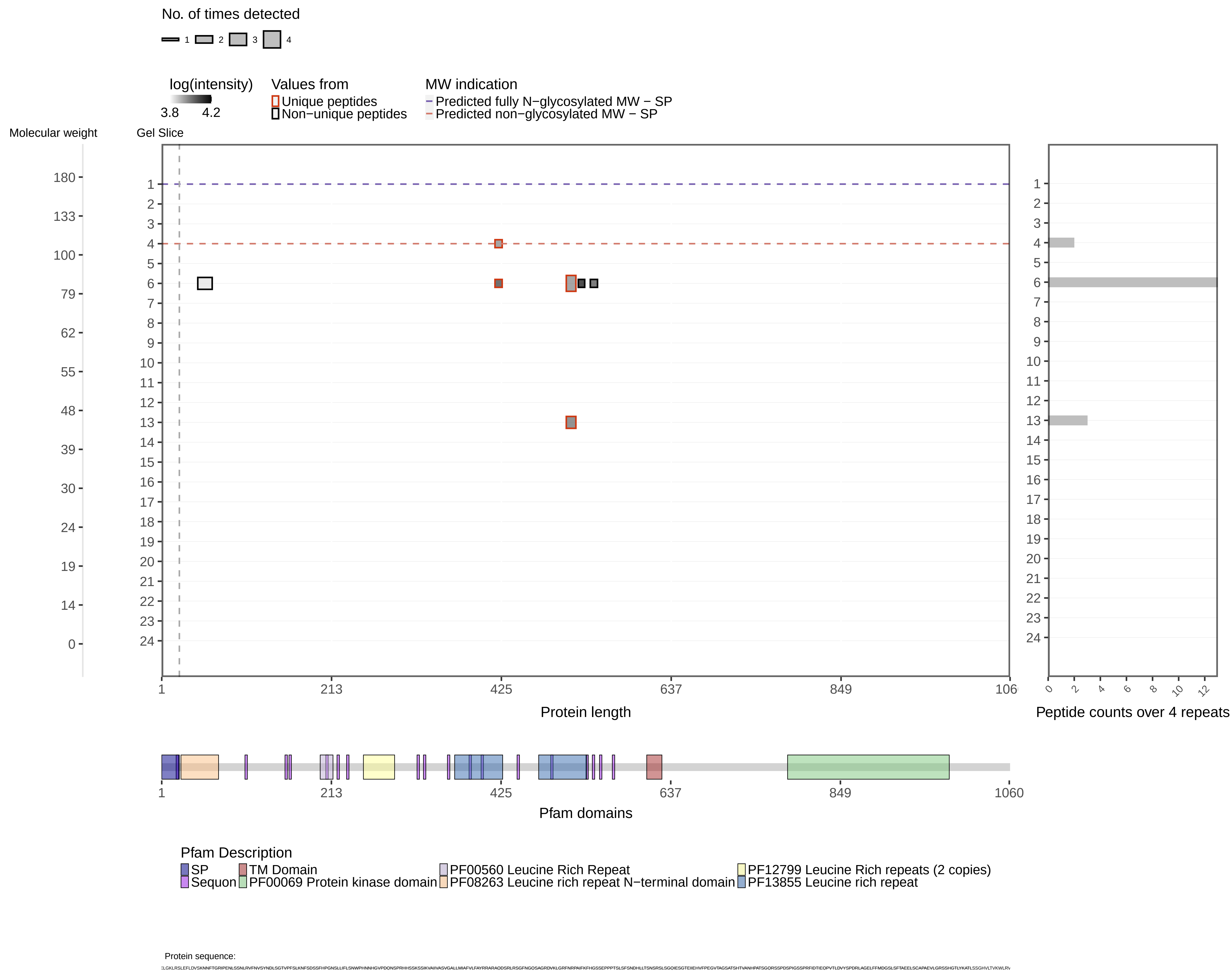

### NbL01g25400.1 chitinase−like_protein_1 [MW − SP = 33.32 kDa]

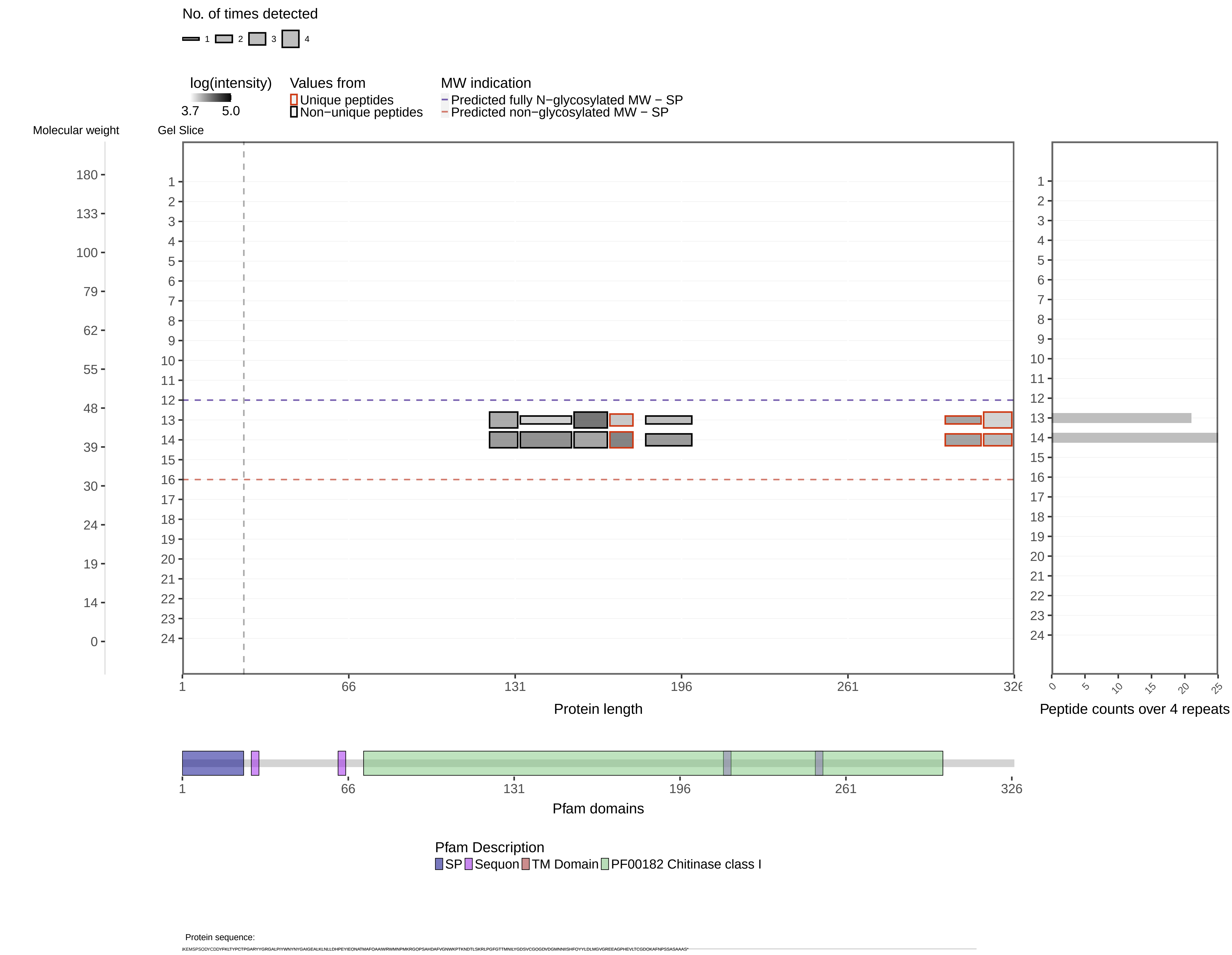

### NbL01g25140.1 pirin−like_protein [MW = 36.71 kDa]

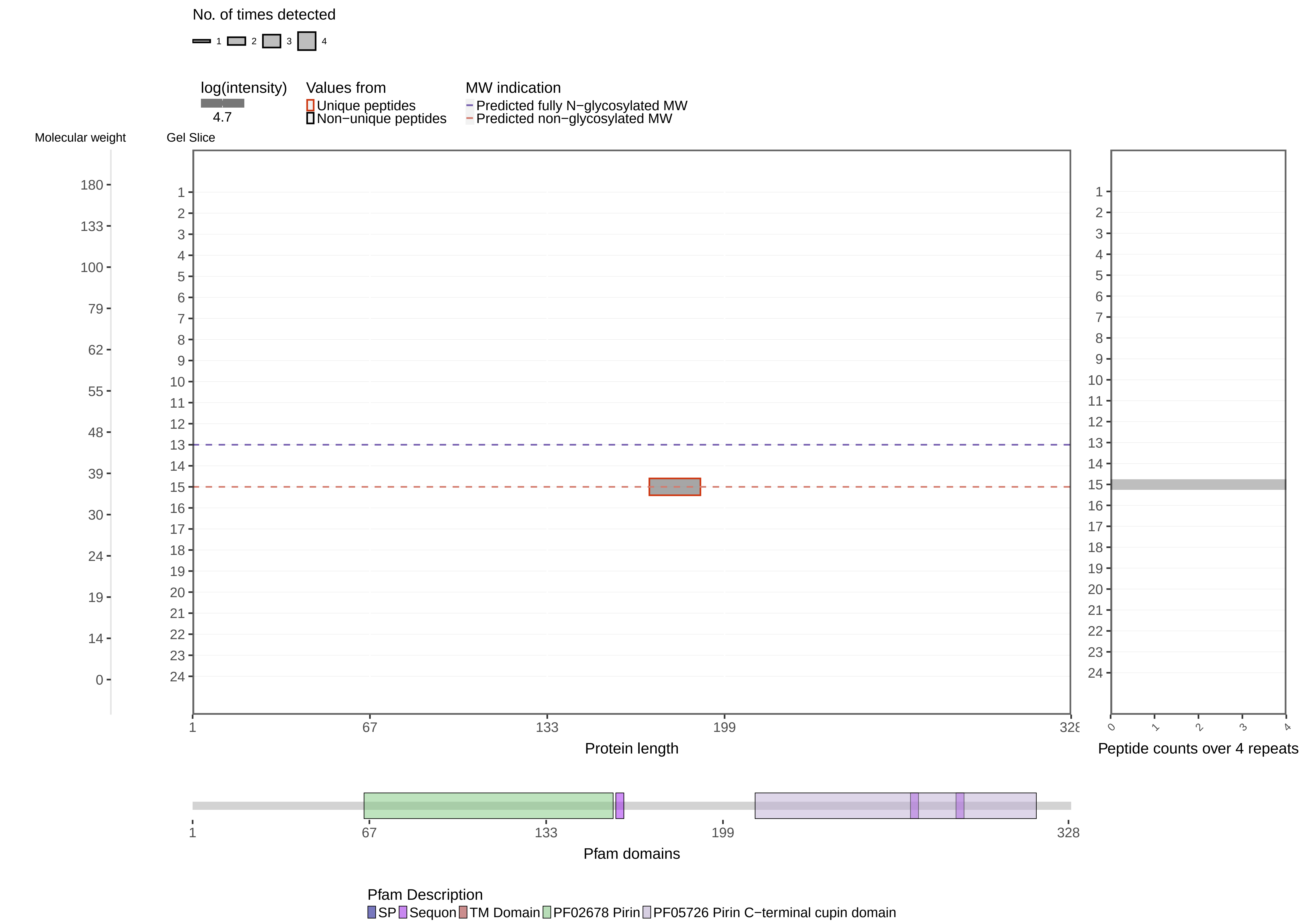


Protein sequence:

GLQLWINLSSRDKMIEPRYQELLNEDIPTAENEGVEVKIIAGEAMGVQSPVYTRTPTMYLDFTLQPTSYYHQPTPESWNAFVYIVEGEGVFGIPNSGPVSAHHCLVLGPGEGLSVWNKSSKPLRFVLLGGQPINESVVQHGPFVMNSQAEIDQTFEDYQYCKNGFENARYWRSGH*

NbL01g25120.1 pirin−like_protein [MW = 32.34 kDa]

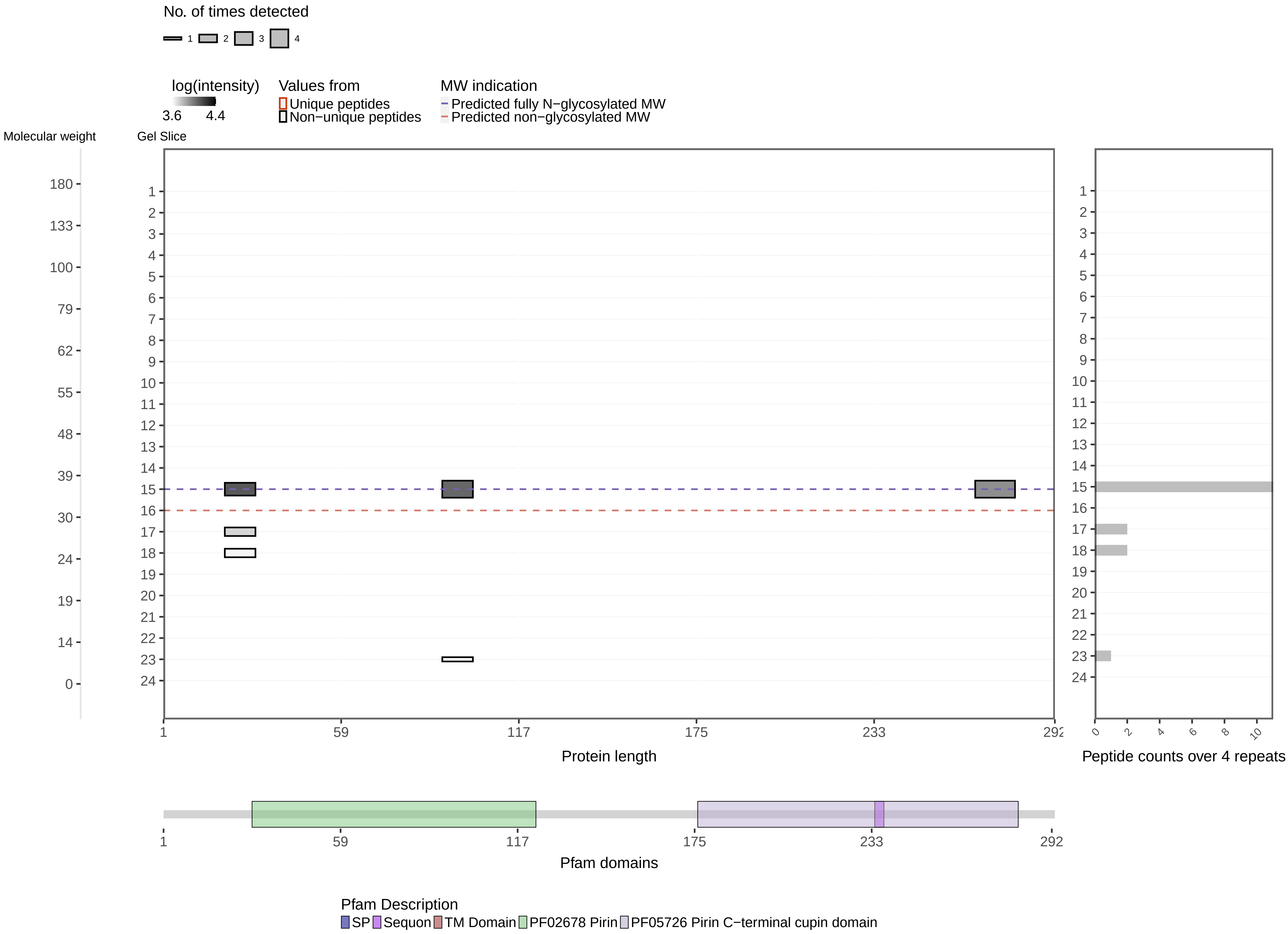


Protein sequence:

EPRYQELLHENIPKAEKYGVSVTIIAGEAMGVKSQVFTRTPTMYLDFTLKPASEHHQPIPESWNSFVYIIEGEGVFGSSDSTPTPAHHCLVLGPGEGLSVWNKSSKPLRFVLIGGQPINEPVAQYGPFVMNTKSEIMQAYQDYQLGKNGFERARHWNSK*

### NbL01g25010.1 auxin−induced_protein_PCNT115 [MW = 38.51 kDa]

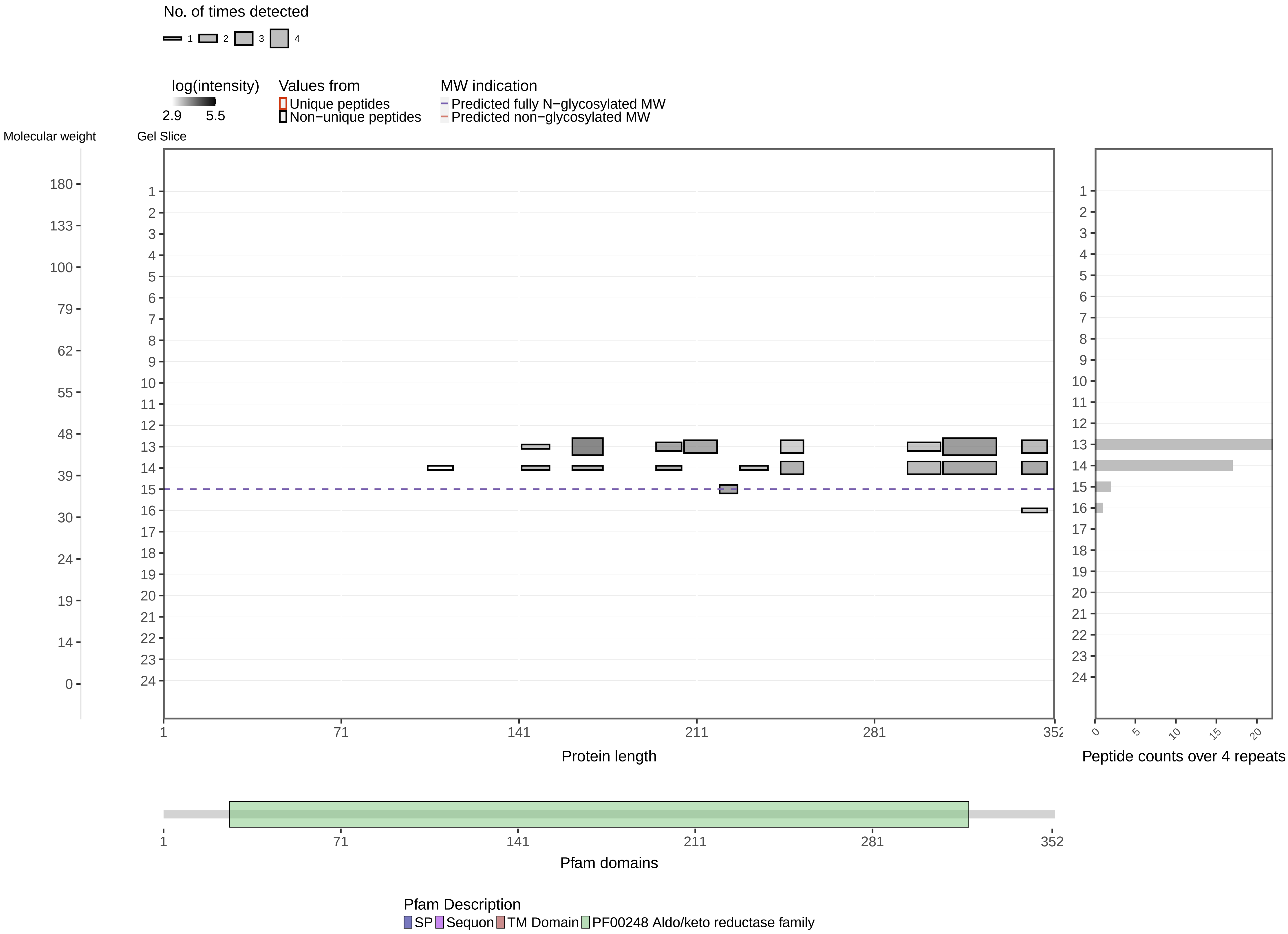


Protein sequence:

IGLSEASASTIRRAHAVHPITAVOLEWSLWSRDVEEEIIPTCRELGIGIVAYSPLGRGFLSSGPKLLEDMSNEDYRKHLPRFOAENLEHNKNLYERICOMAARKGCTPSOLALAWVHHOGNDVCPIPGTTKIENLNLNIGALSIKLTSEDMAELESIASDNAVOGDRYIPGMSTYKNSETPPLSAWKTT*

### NbL01g25000.1 auxin−induced_protein_PCNT115−like [MW = 41.41 kDa]

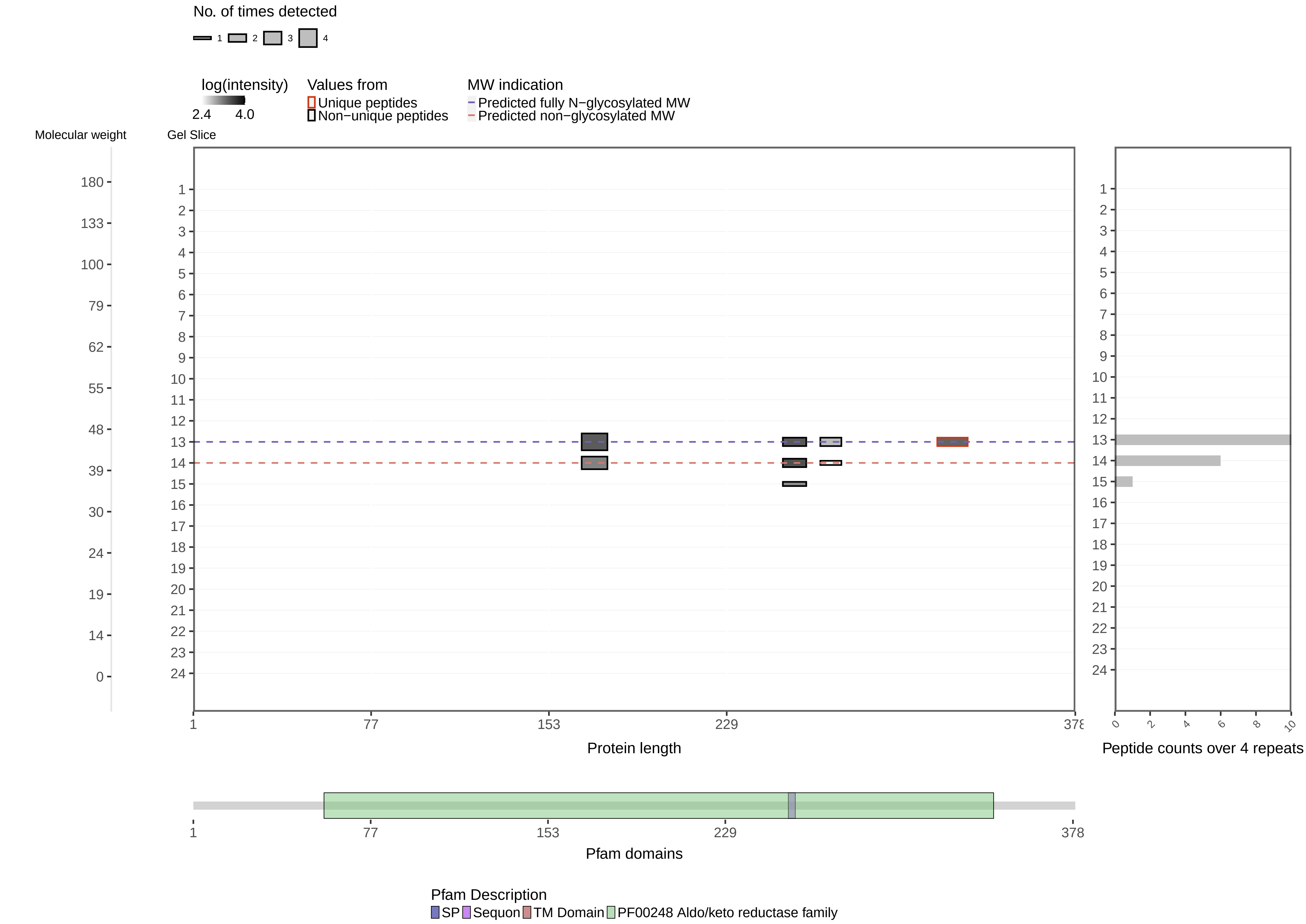


Protein sequence:

.KKLVEEGKIKYIGLSEASASTIRRAHAVHPITAVQLEWSLWSRDVEEEIIPTCRELGIGIVAYSPLGRGFLSSGPKLLENLSNQDFRKHLPRFQSDNLEHNKILYDRICQIAAKKGCTPSQLALAWVHHQGNDVCPIPGTTKIENLNQNIEALSIKLTSEDMVELESIASANAVQGDRYSSGASTYKDSDTPPLSAWKATE*

NbL01g24750.1 30S_ribosomal_protein_S1−like [MW = 42.62 kDa]

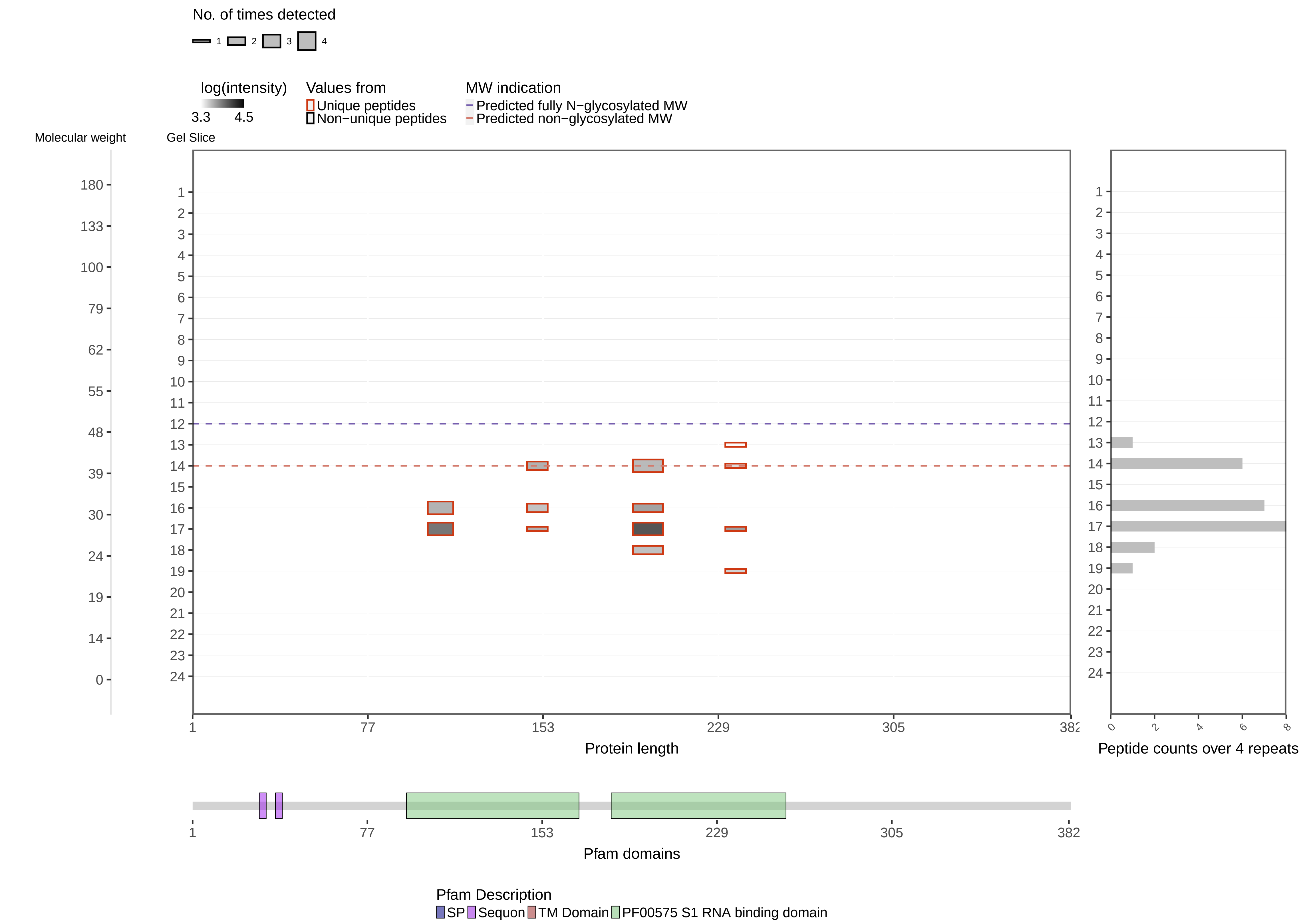


Protein sequence:

SKFSYQINVGDIFQARVGSVEDYGAFVHLRFPDGSYHLTGLVHVSEVSWDLVHDVRDILTEGDDVRVKIINIDREKSRITLSIKQLEEDPLLETLDKVIHKEASVDTNSLDSDGNFVIEPLSGLETIIEELMQEDGIYDVTITRQGFEKRVVSQDLQLWLSNAPASGDQFTLLARAGRQVQEIQLTTSLDQEGIKRALQRVLERVP*

NbL01g24560.1 uncharacterized_LOC104096711 [MW = 101 kDa]

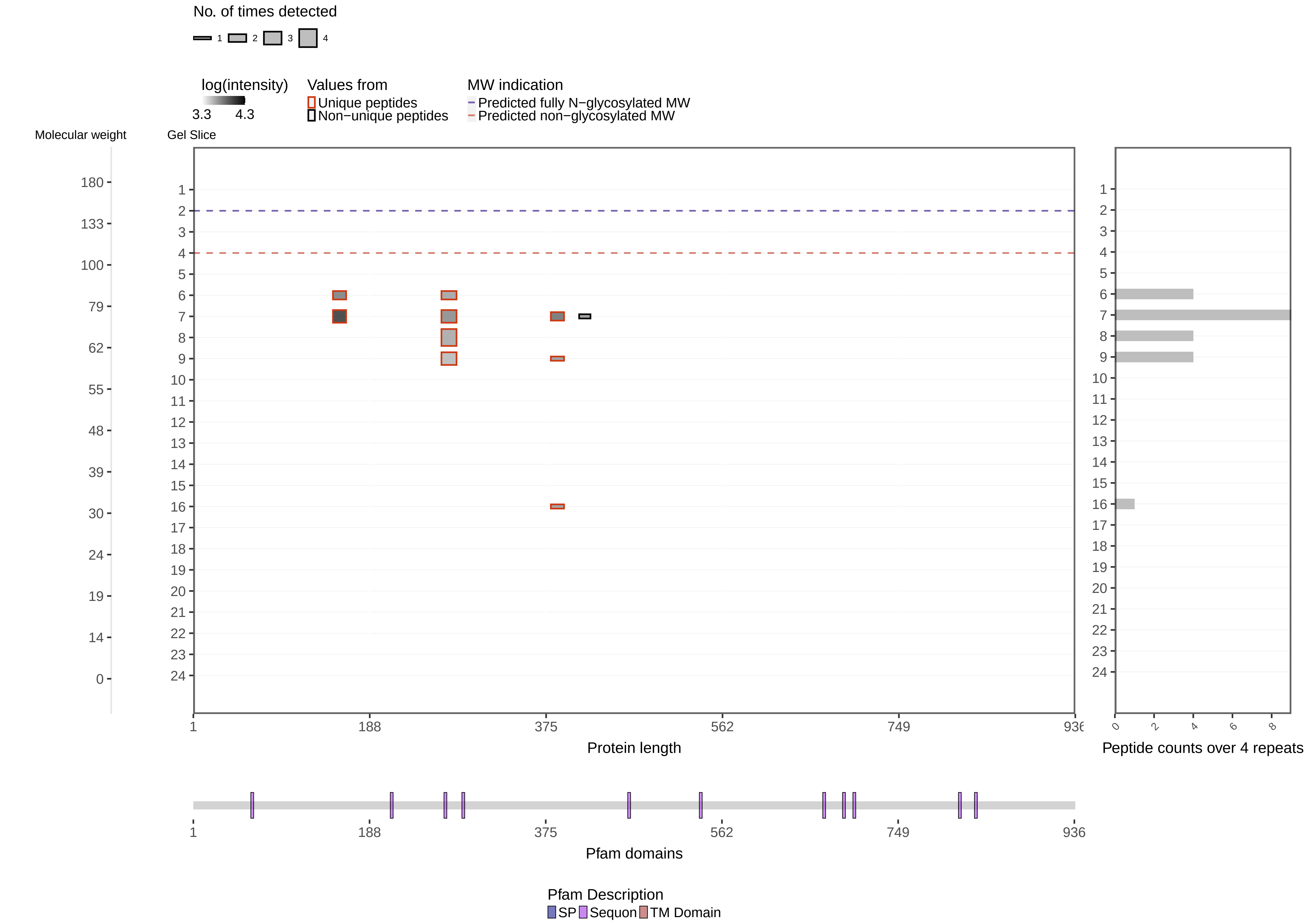


Protein sequence:

SIDGDSLNNSLPSGSPTDHLDMNTITSSVKEEAKQTTKVDDPVIERTETIFSMEEENKDGHPENELLANKETTVVDTSCLSEANQTTVTLGGSDHGEHEKARTEQPIVVGGETAKERSEEKRGVFEGDKTICPVGGHSDRAIVTDTTVHESRDEIFHEASEDSTVSNNAMPTSATGLDSKIISDCDVKEPAGLVVDTDLKGRVVDNSHAGDGIKNATGVISSASLNEGNDKLIKQNNTVSAVDVSTNSSSRADSLDANWGSVSVLSTQSESTAVPDAEATDTREKLEPDLQKPT

### NbL01g24550.1 protein_TIC_22−like_chloroplastic [MW = 36.45 kDa]

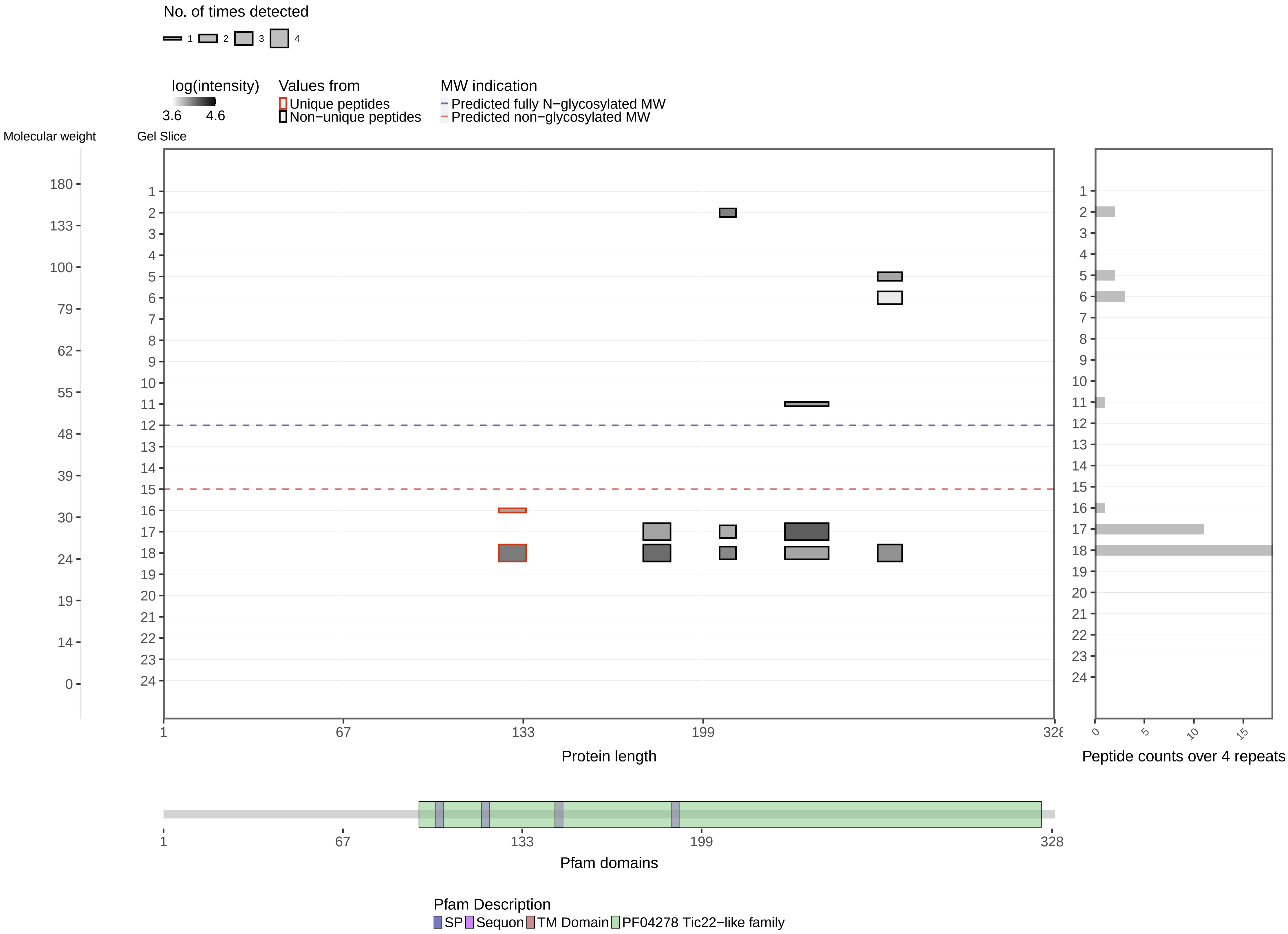


Protein sequence:

SGVSTGKNLGLFCFSEADAEALROOMESMDPTMRNGSRVVPVALNKVFOLKVDGVAFRLIPEASOVKNAMKERERTGTSDESFYGVPVFOSRSLILRSONKRYRPVFFRKEDLENSLVRASOOOGRLNPALKGDIOVAVLEDIIOGMKDTSTSKWDDVVFVPPGFDVSTDHSOK*

### NbL01g24500.1 crocetin_glucosyltransferase_chloroplastic−like [MW = 52.95 kDa]

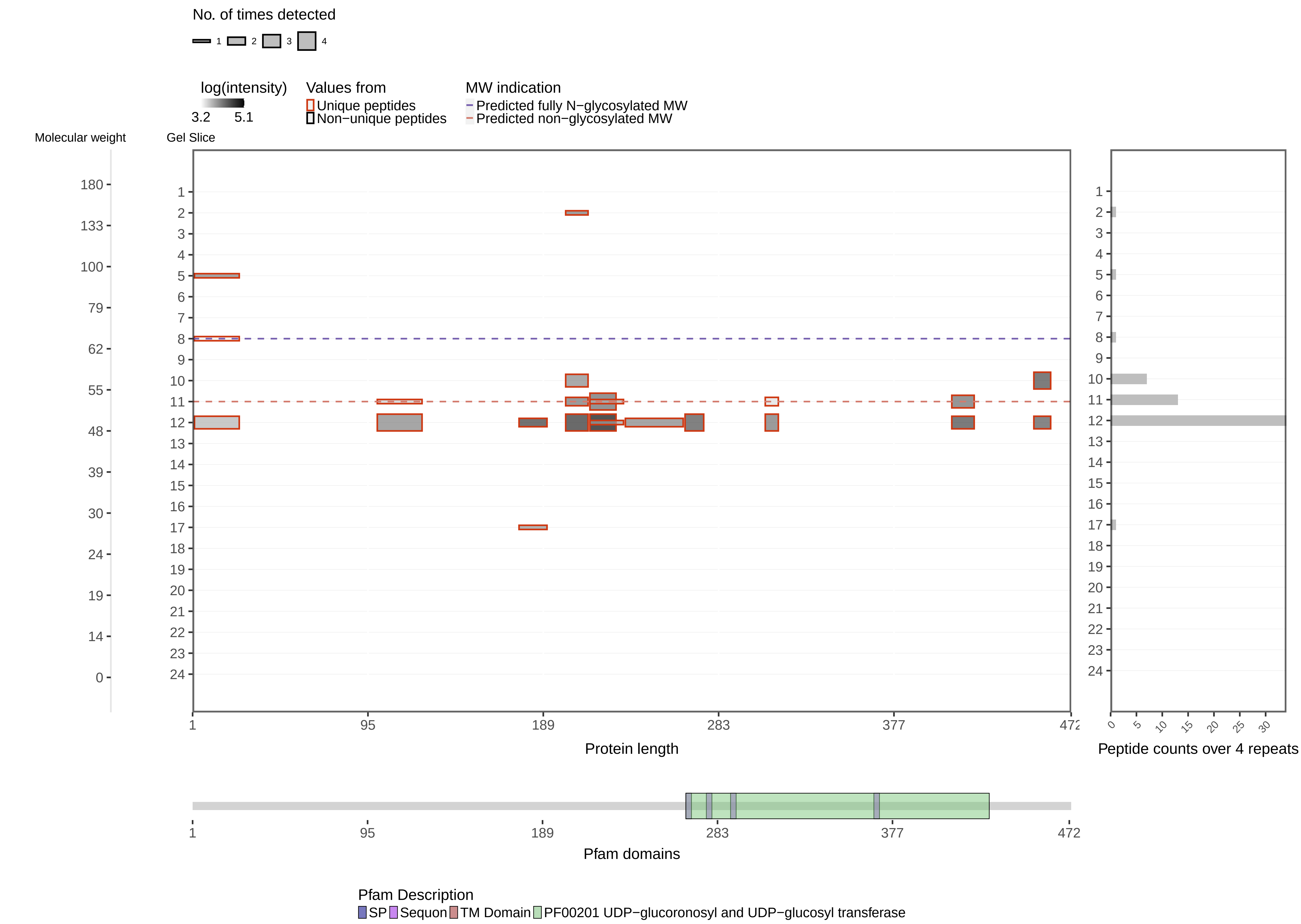


Protein sequence:

ELKAIEKYNLIGIGPLIPSTFLDGKDPLDSSFAGDLFQKSNDSIEWLNSKANSSVVYISFGSLLNLSKNQKEEIAKGLIEIKRPFLWVIRDQENGKGDEKEEEKLSCMMELEKQGKIVPWCSQLEVLTHPSLGCFVSHCGWNSTLESLSTGVPVVAFPHWTDQGTNAKLIEDVWKIGVRLKKNEDGVVESEEIKRCIEMVMNGGEKGEEMRRNAQKWKELAREAVKEGGSSYMNLKAFVQEVGKGC*

### NbL01g24450.1 receptor−like_kinase_TMK4 [MW − SP = 98.53 kDa]

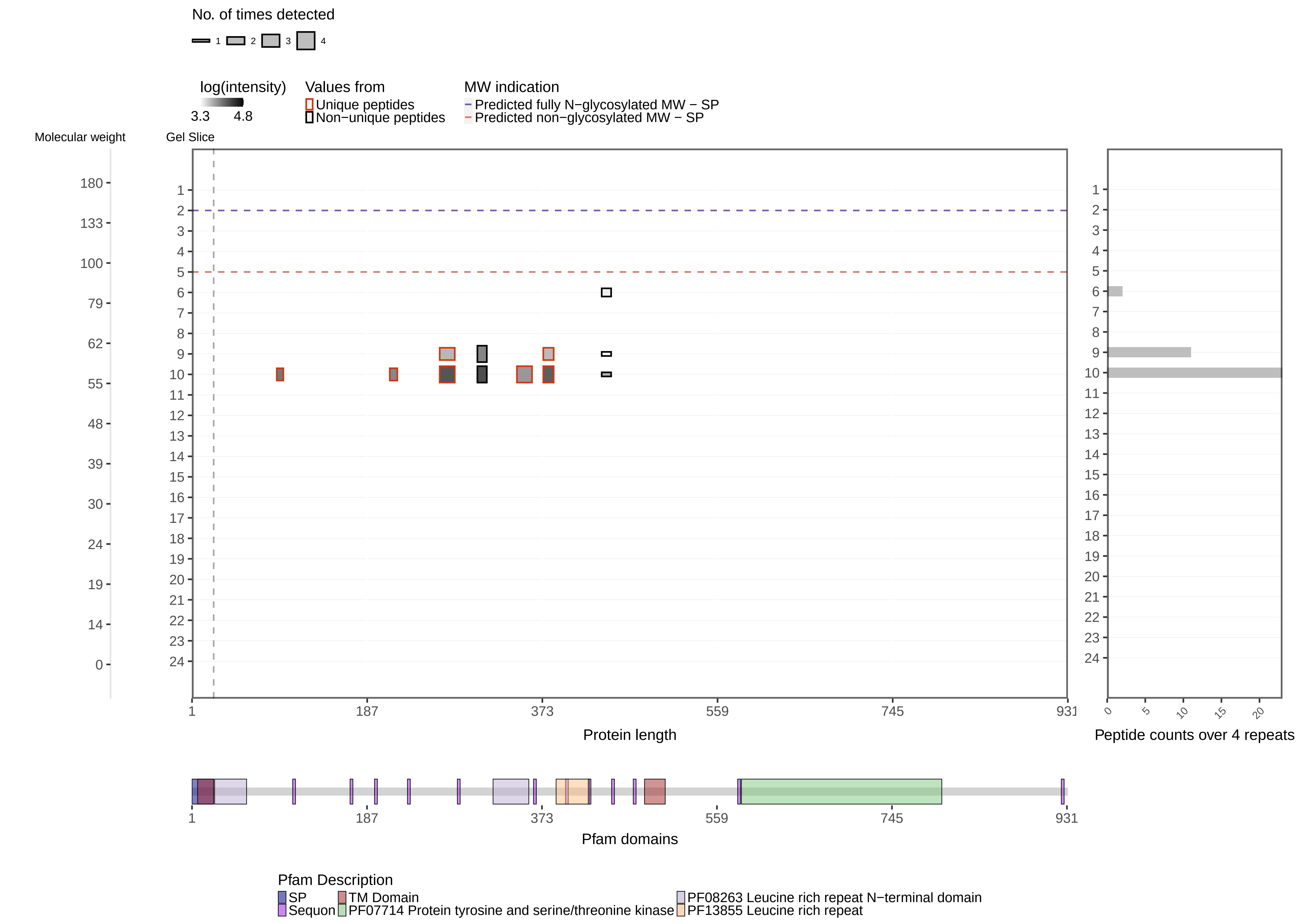


Protein sequence:

### NbL01g24440.1 probable_acyl−activating_enzyme_16_chloroplastic [MW = 80.49 kDa]

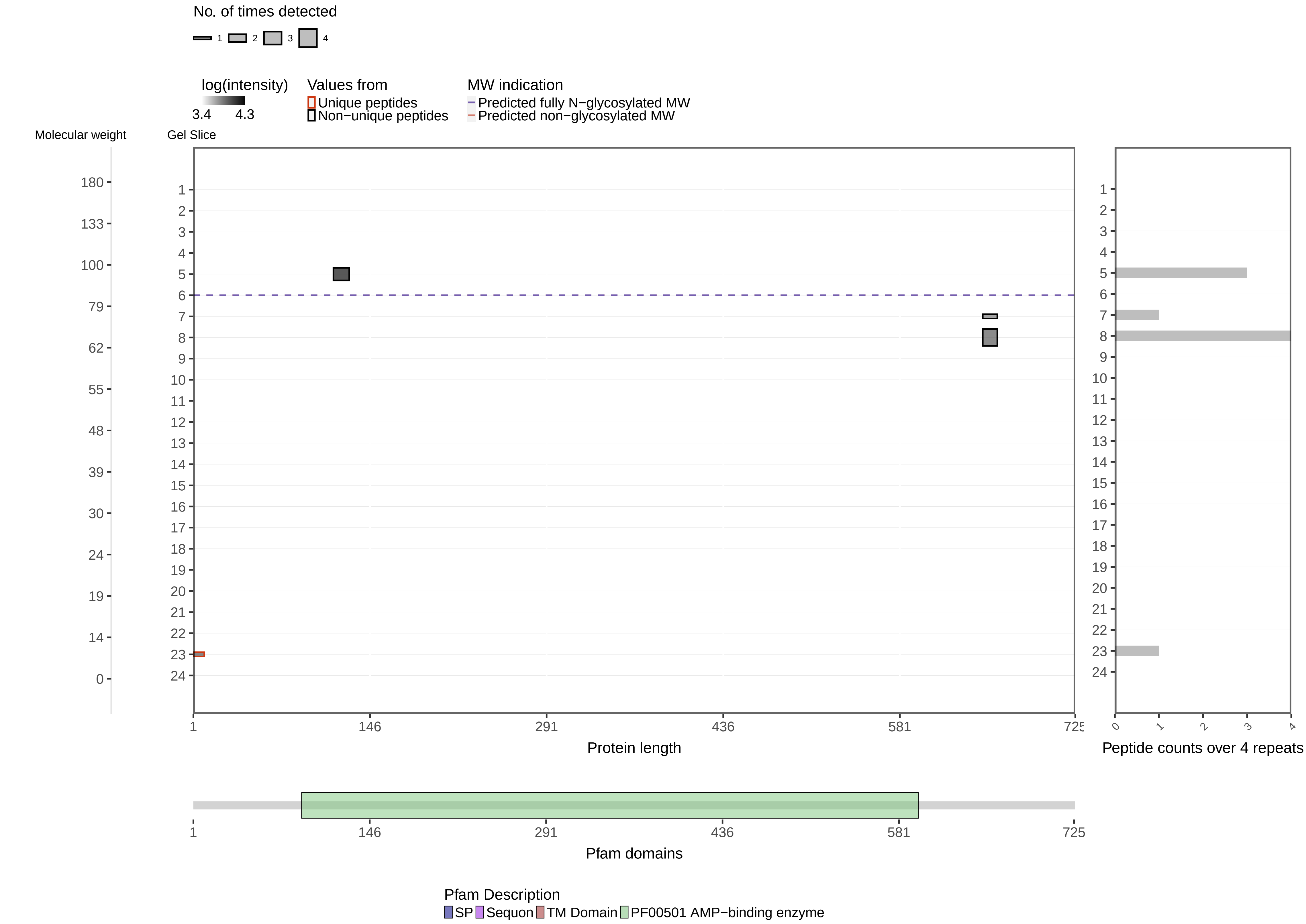


Protein sequence:

EDLRRYQPNYLISVPLVYETLYSGILKQINSNSAARKLIALLFLRISMAYMEAKRIYEGKCLMKDTKQPSYVVSLLDWLWARTIAAILWPLHMLAKKIVYSKIHASIGISKAGVSGGGSLSSRVDKFFEAIGIKIQNGYGLTESSPVISARHPACNVLGSVGHPIRHVEVKIVNAETDEVLPPGSRGIVKARGPLVMKGYYKNPLATKQAIDENGWLNTGDLGWIAPDHSVGRSRKSGGVIVLEGRAKDTIVLSTGENVEPSEIEEAATGSSLIQQIVVIGQDQRRLGAIIVPNKEELLLA

### NbL01g24400.1 selenium−binding_protein_1−like [MW = 53.62 kDa]

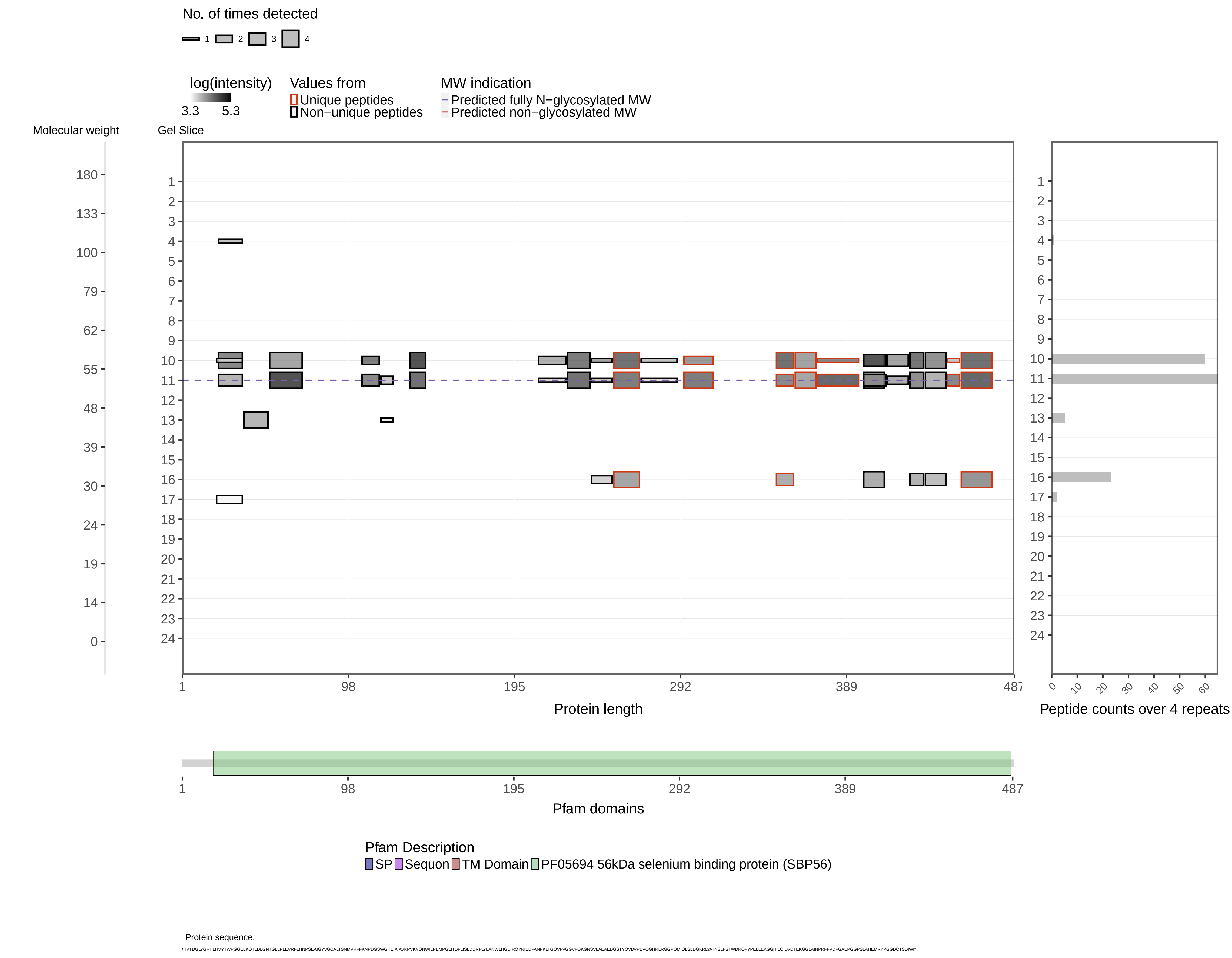

### NbL01g24330.1 NA [MW = 53.12 kDa]

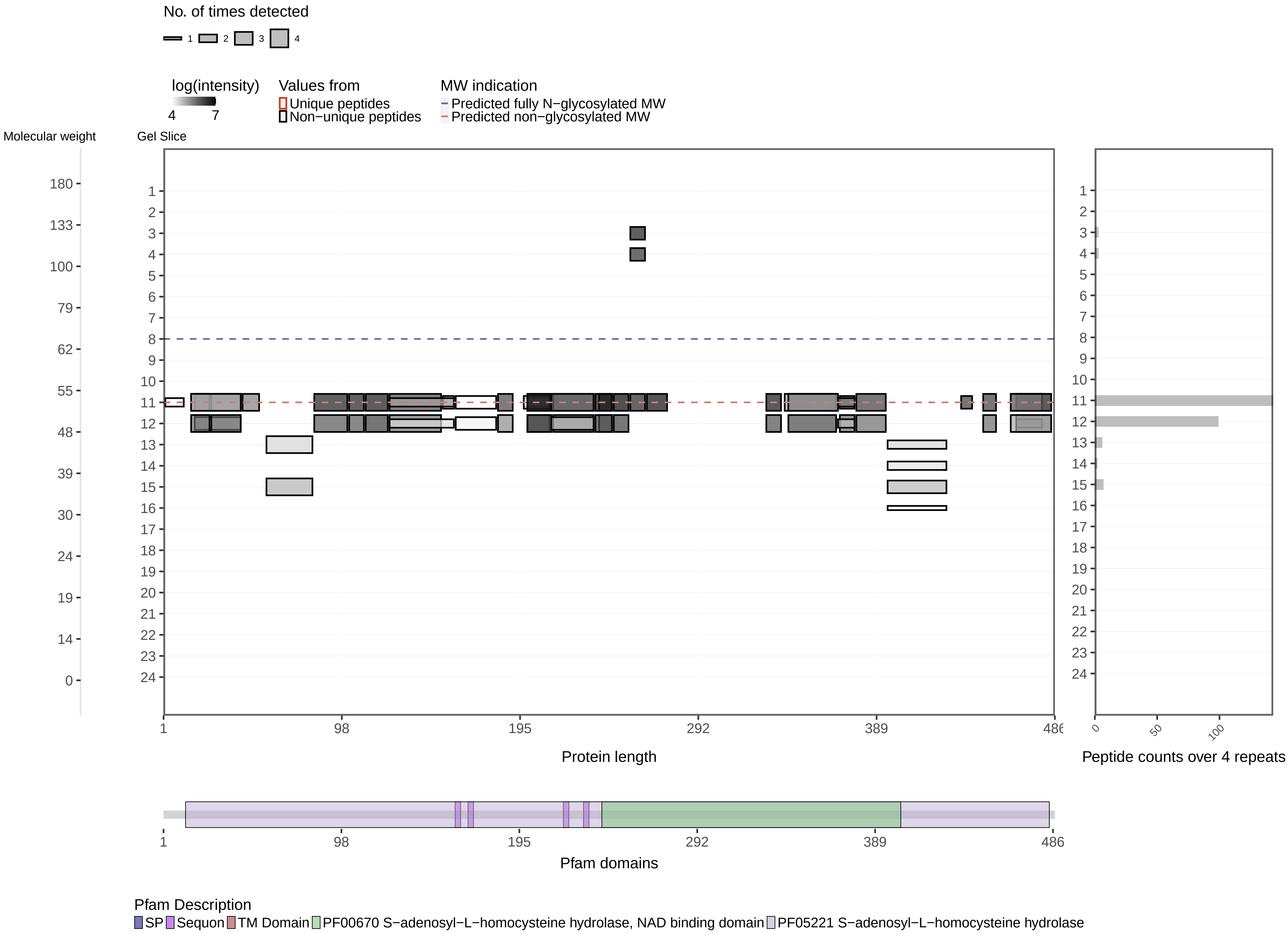


Protein sequence:

NDSVTKSKFDNLYGCRHSLPDGLMRATDVMIAGKVALVAGYGDVGKGCAAALKQAGARVIVTEIDPICALQATMEGLQVLTLEDVVSDVDIFVTTTGNKDIIMVDHMRKMKNNAIVCNIGHFDNEIDMLGLETYPGVKRITIKPQTDRWVFPDTNSGIIVLAEGRLMNLGCATGHPSFVMSCSFTNQVIAQLELWNEKSSGKYEKKVYVLPKHLDEKVAALHLGKLGAKLTKLSKDQADYISVPVEGPYKPAHYRY*

NbL01g24260.1 glycine−rich_RNA−binding_protein_4_mitochondrial−like [MW = 15.14 kDa]

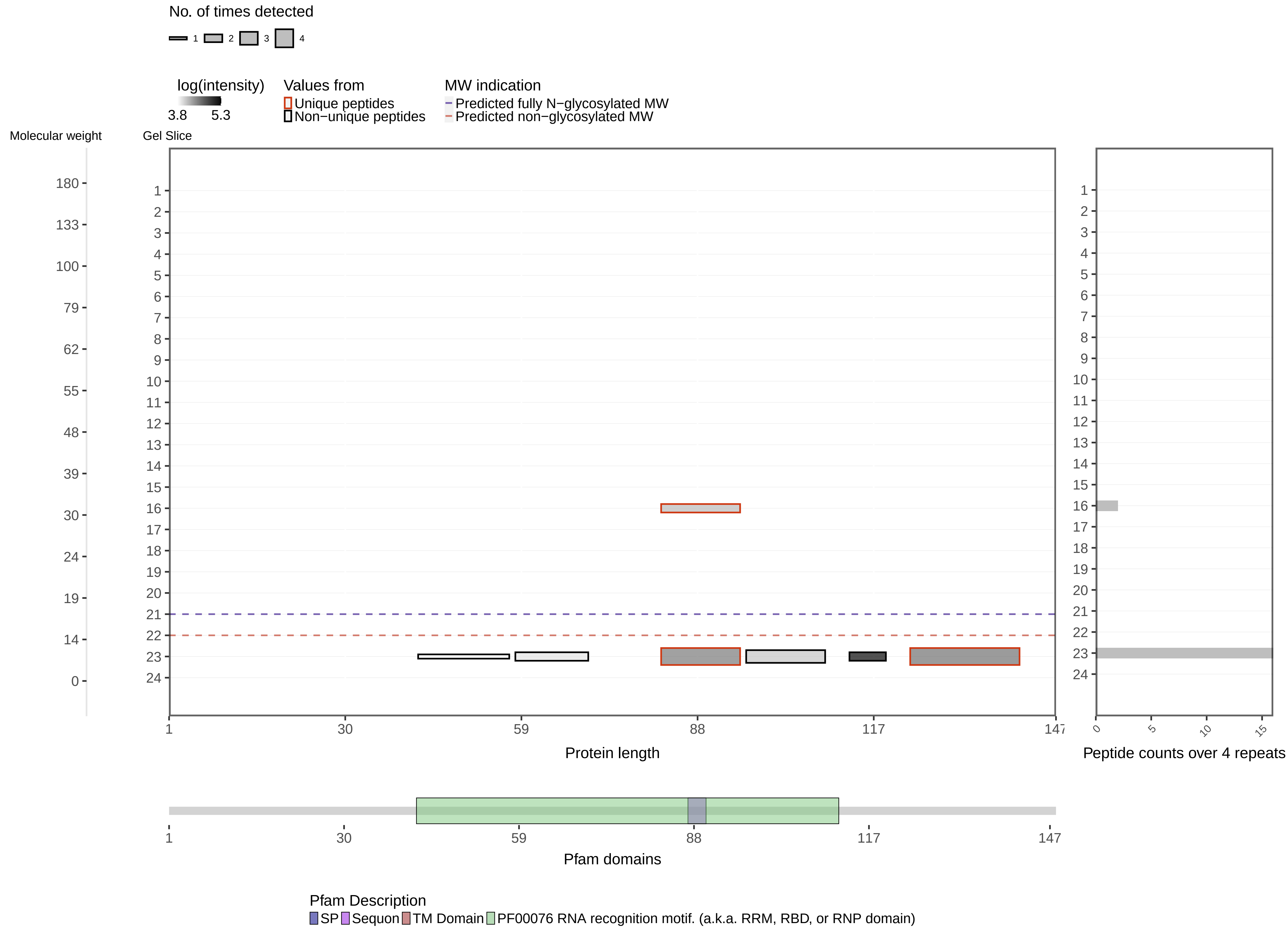


Protein sequence:

FGDVVDSKVITDRDTGRSRGFGFVNFTDGESAKEAMTAMDGQELNGRNIRVSLAQERAPRSGGFGSGGGGYGGGYGGSRGNDGF*

NbL01g24250.1 protein_ECERIFERUM_26−like [MW = 47.9 kDa]

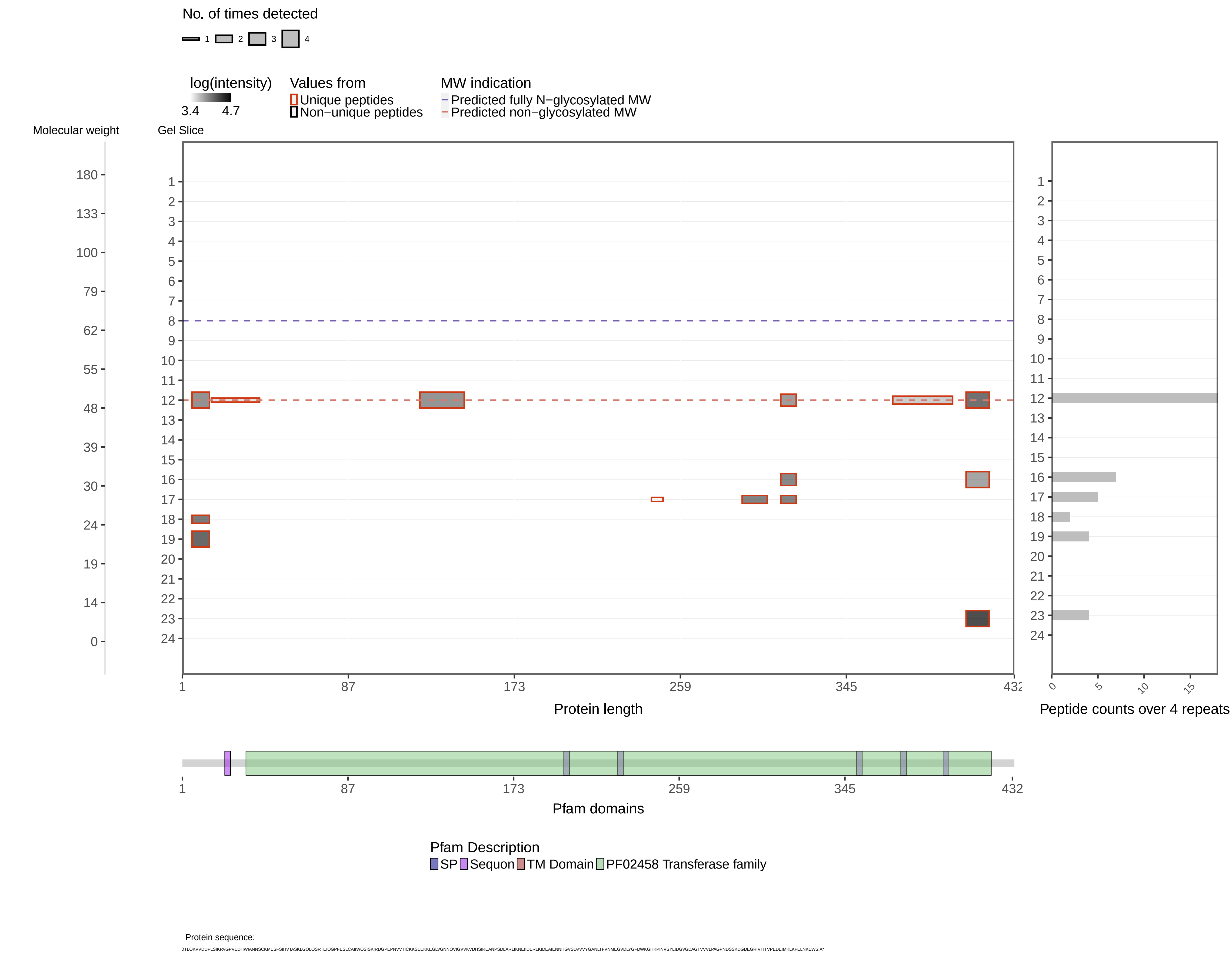

### NbL01g24060.1 probable_sucrose−phosphate_synthase_2 [MW = 119.69 kDa]

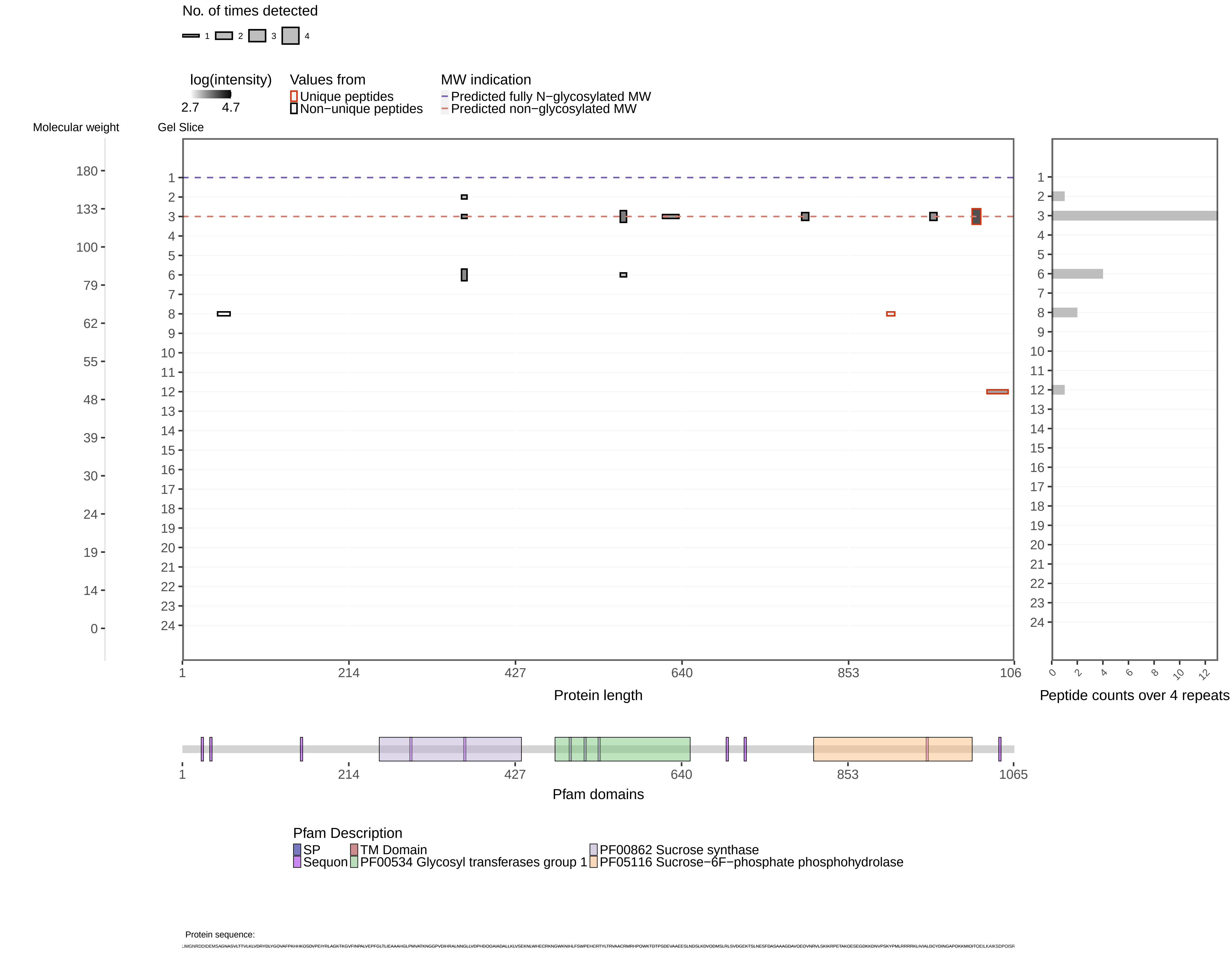

### NbL01g23940.1 purple_acid_phosphatase_15 [MW − SP = 59.96 kDa]

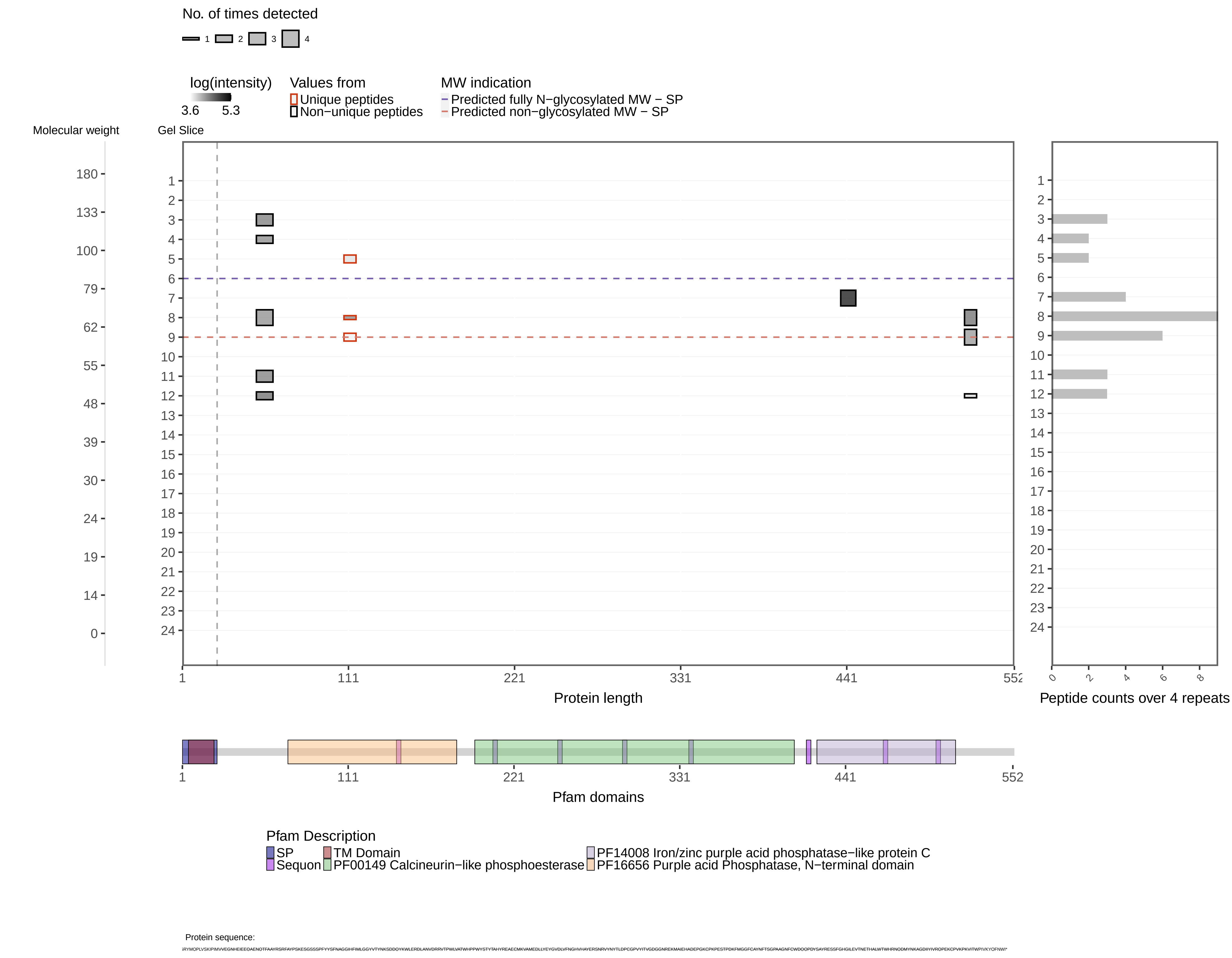

NbL01g23810.1 glutathione_reductase_chloroplastic [MW = 58.52 kDa]

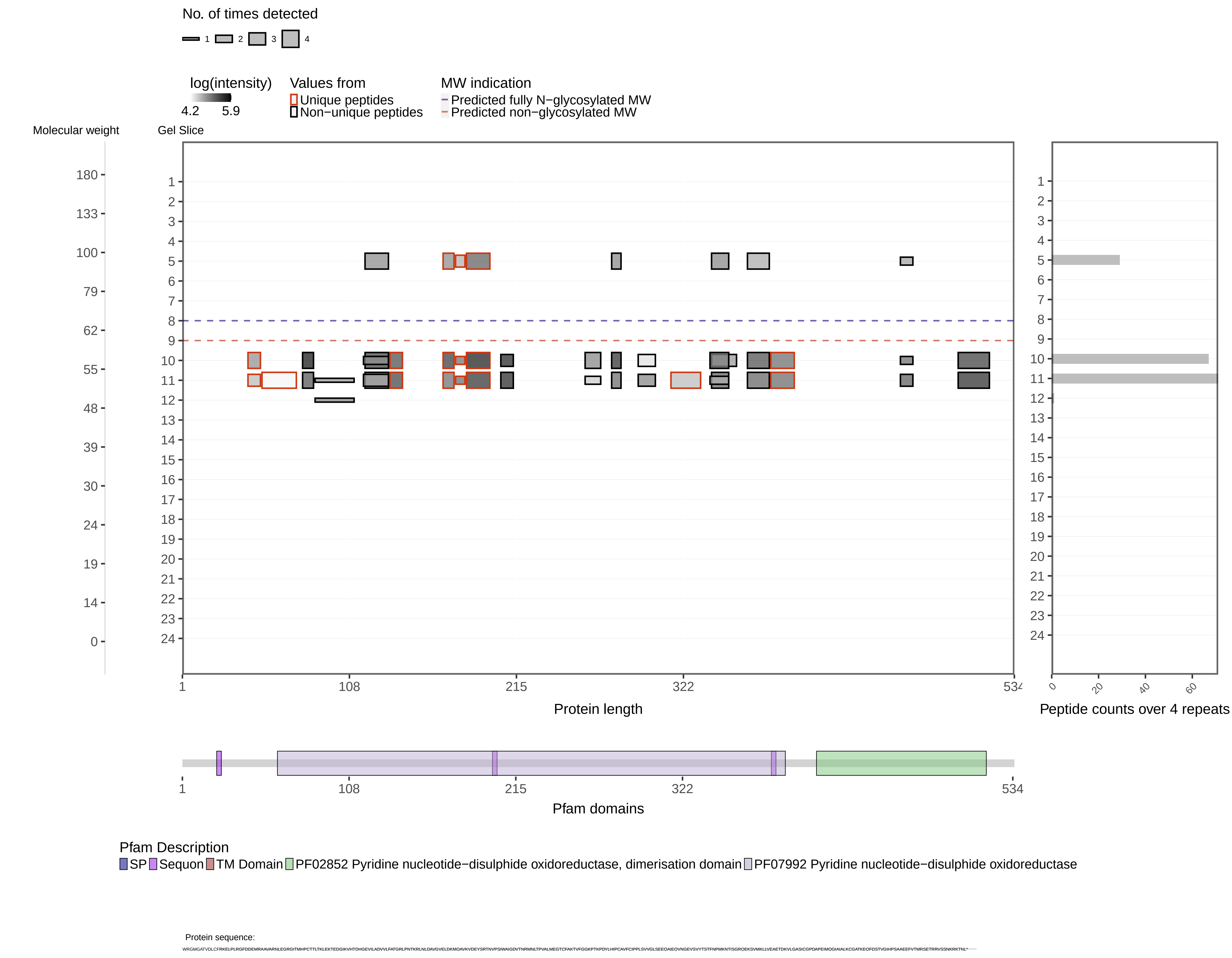

NbL01g23710.1 26S_proteasome_non−ATPase_regulatory_subunit_1_homolog_A−like [MW = 108.01 kDa]

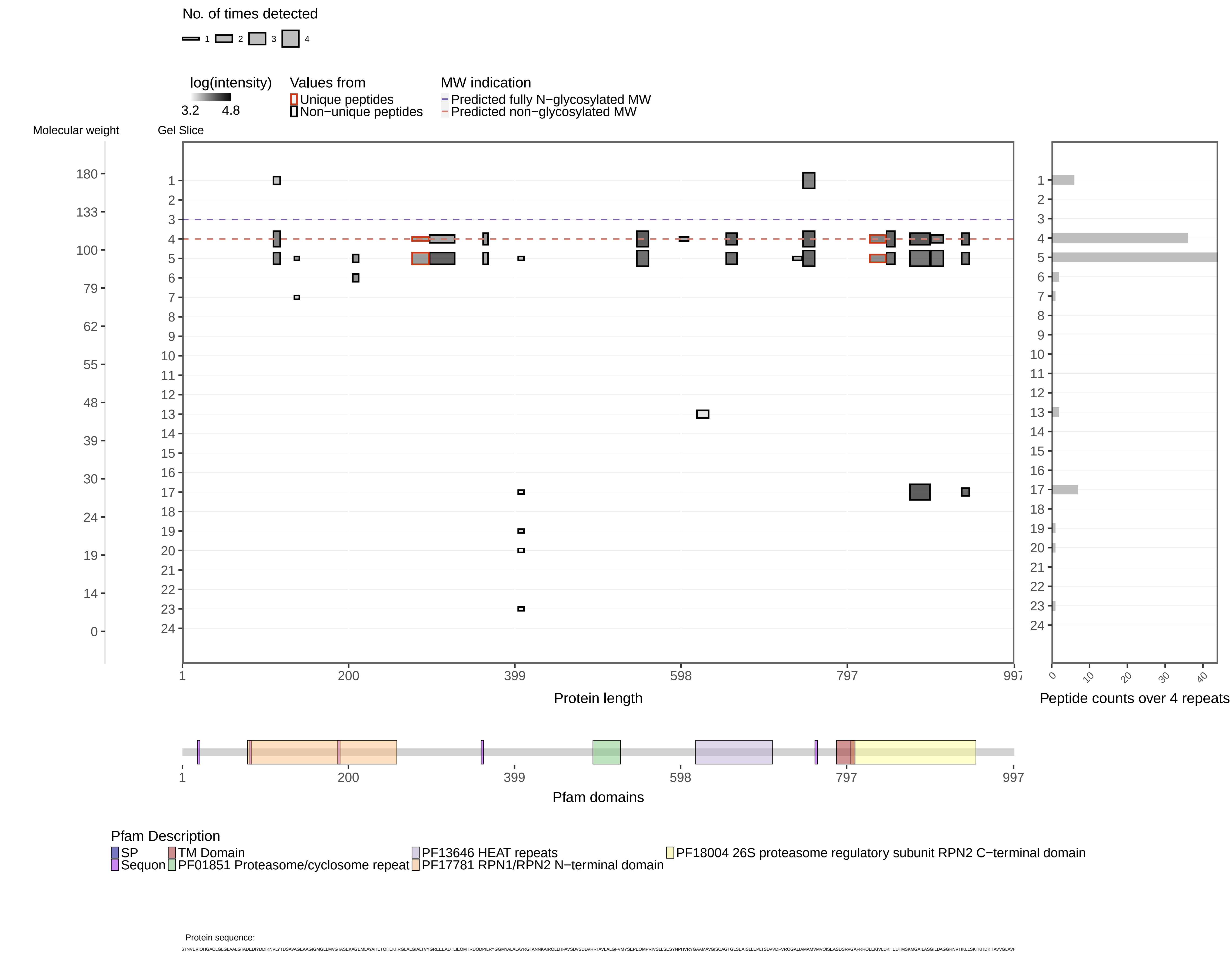

### NbL01g23610.1 pyruvate_kinase_isozyme_A_chloroplastic−like [MW = 65.22 kDa]

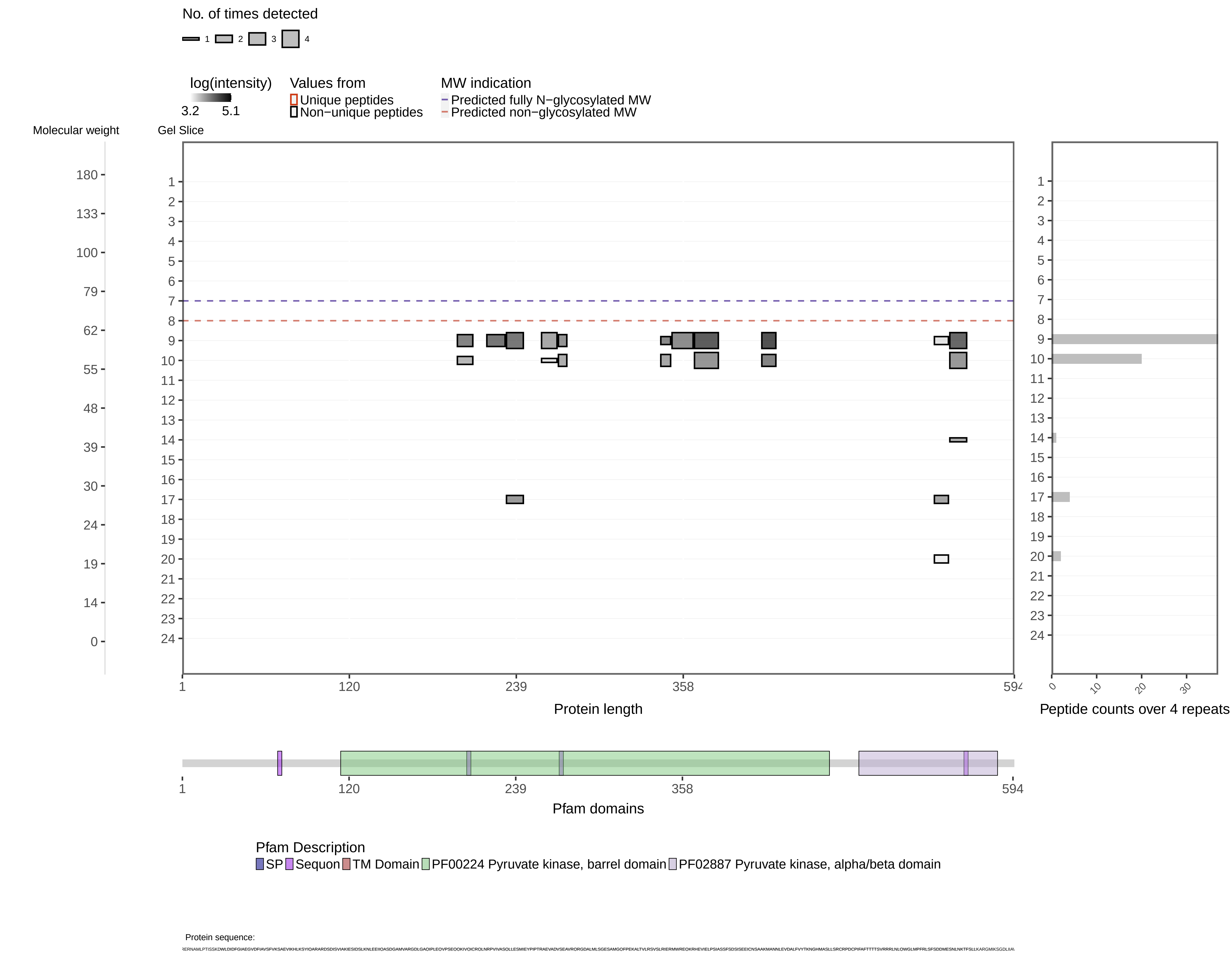

NbL01g23580.1 GDP−mannose_35−epimerase_1 [MW = 42.92 kDa]

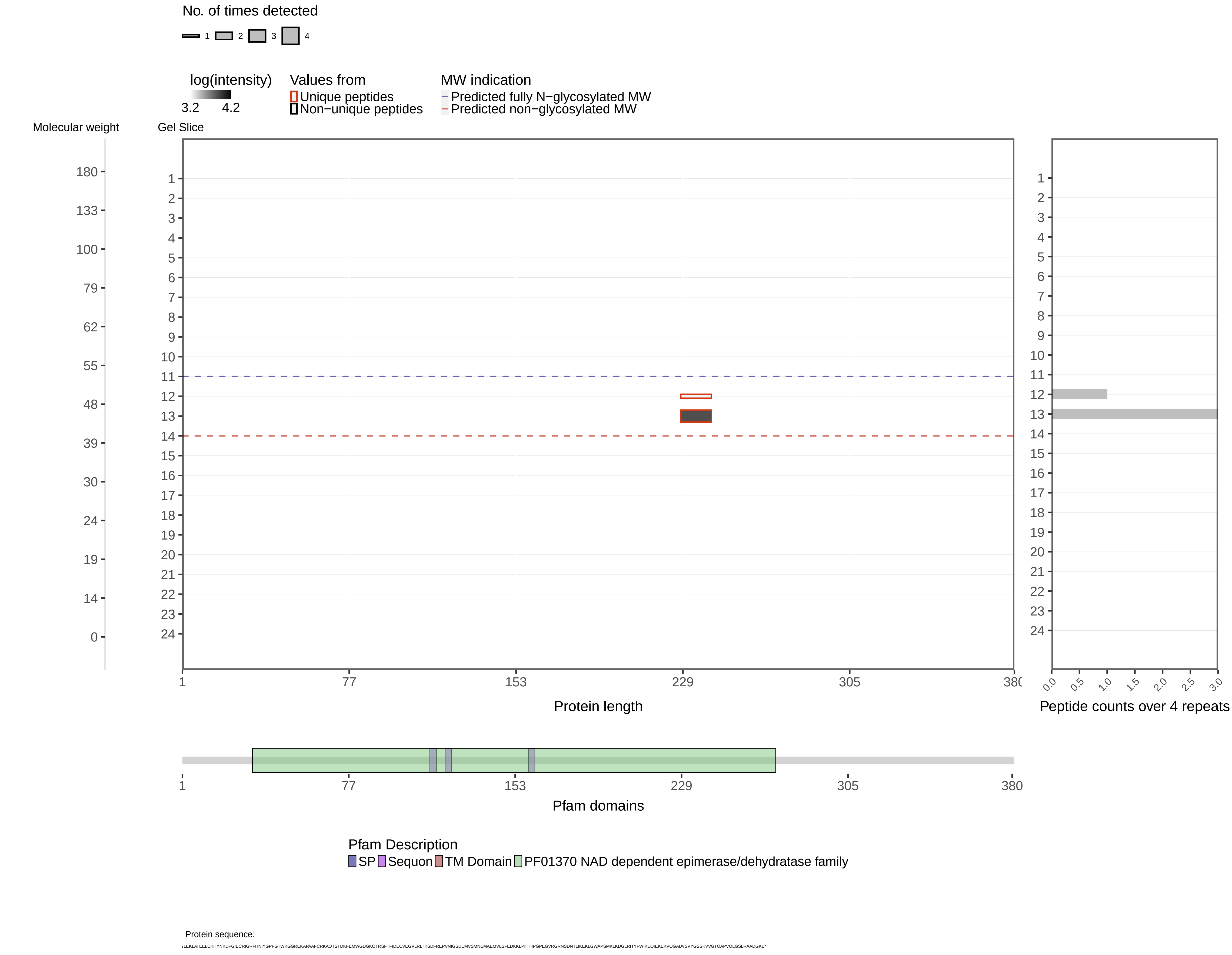

### NbL01g23460.1 NAD(P)H−quinone_oxidoreductase_subunit_N_chloroplastic [MW = 28.17 kDa]

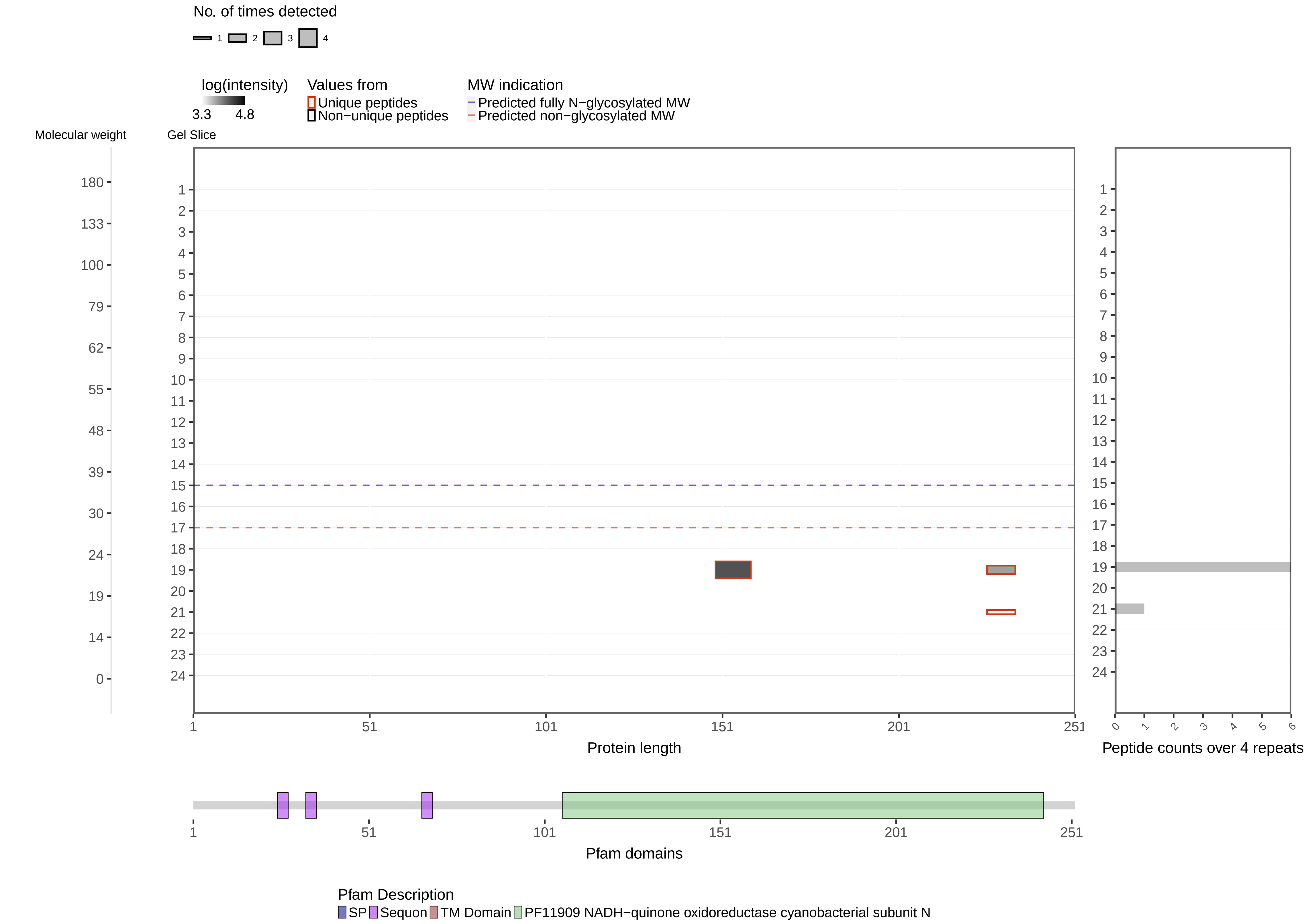


Protein sequence:

EHKALAIYSPHEGGYEGRYLTRLKYQGYHFLDLSARGLGDPETTLTKFHPVCPAHVGKQPIARWYFPPEVDYRLSLLPPNAKGLVVWIIEAKVLSKAELQFLALLPSLRPKVKVIAECGNWRKFMWKPLKEIAGLSQS*

### NbL01g23340.1 electron_transfer_flavoprotein_subunit_beta_mitochondrial−like [MW = 28.04 kDa]

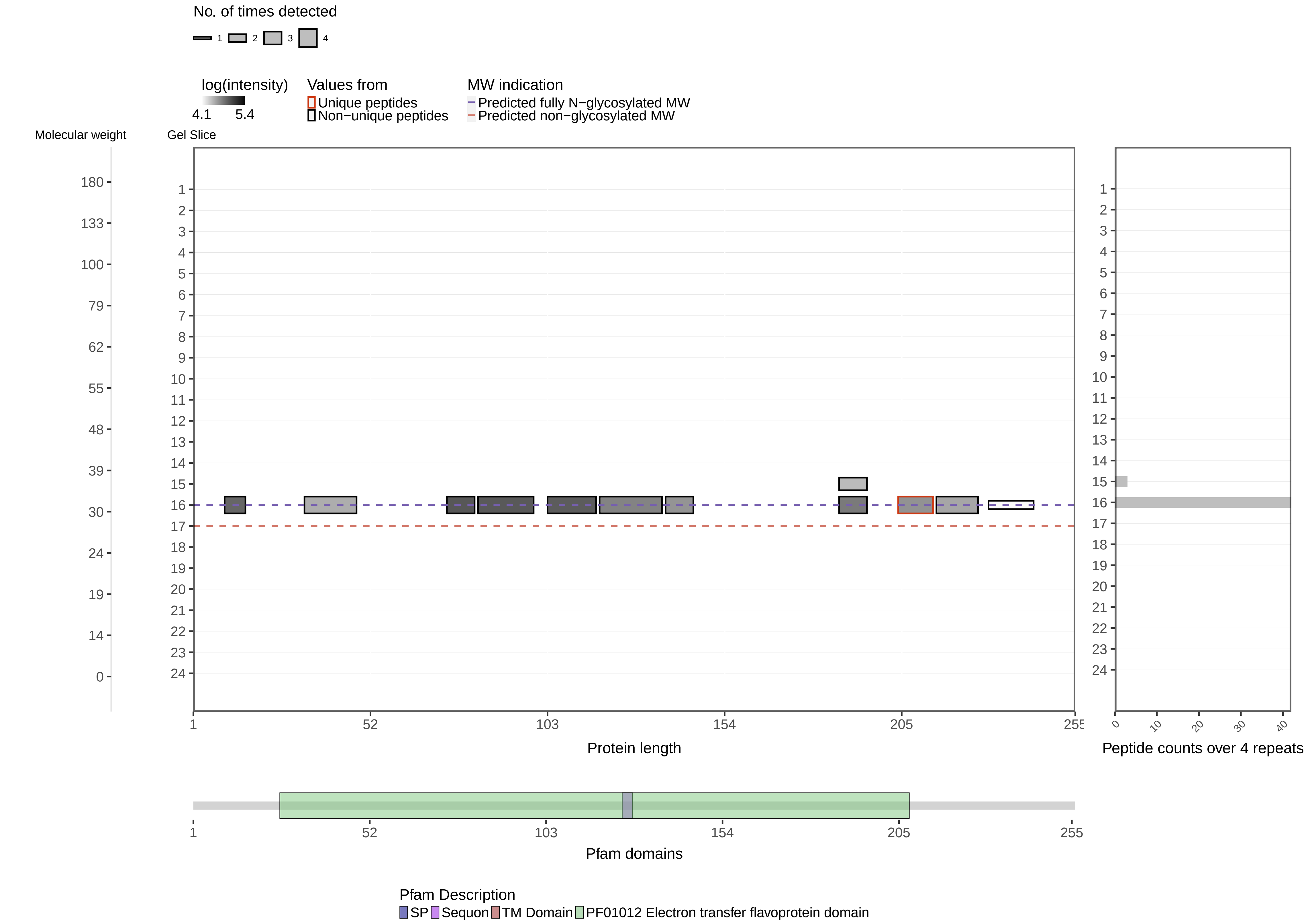


Protein sequence:

KQAIDNDQNQTGQMVAGLLKWPQGTFASKVVLDEEKQVATVDREVDGGIETLSLDLPAVITTDLRLNQPRYATLPNIMKAKSKPIKKFTPQDLNVEIKSDFEVVEVTEPPKRKAGLILSSVDELIDKLKNEARVIWVY*

NbL01g23220.1 DAG_protein_chloroplastic−like [MW = 28.61 kDa]

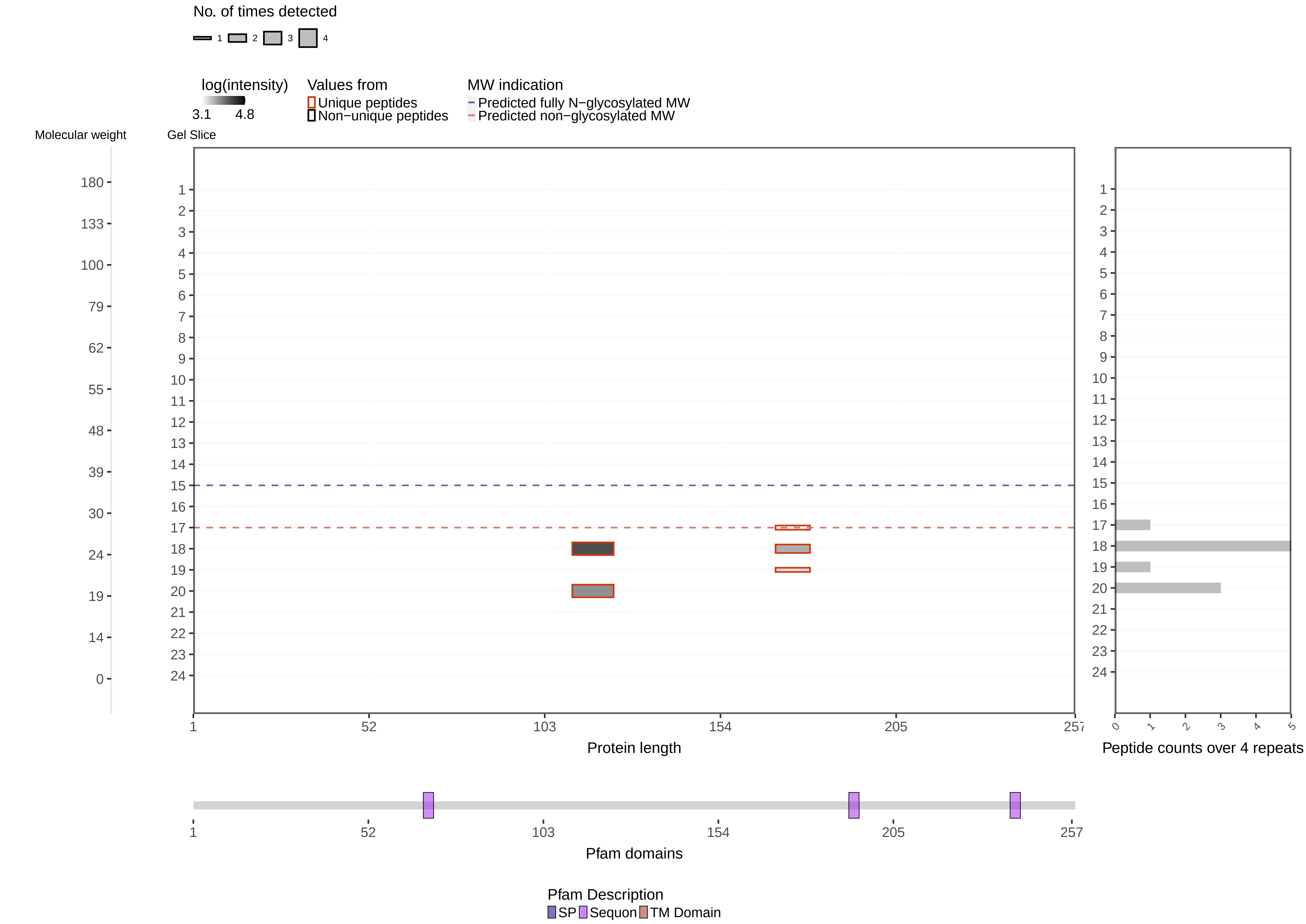


Protein sequence:

EEEAKKKIYSVSTTTYTGFGALISEELSYKVKGLPGVLWVLPDSYLDVPNKDYGGDLFVDGKVIHRPQYRFNQNQSSRPRPRPRYDRRREKMQVERREPIQRGAWTPNQQPPAQSVSQGQNFSHGTGGGPAGSQGNSG*

### NbL01g23210.1 uncharacterized_transmembrane_protein_DDB_G0289901−like [MW = 52.7 kDa]

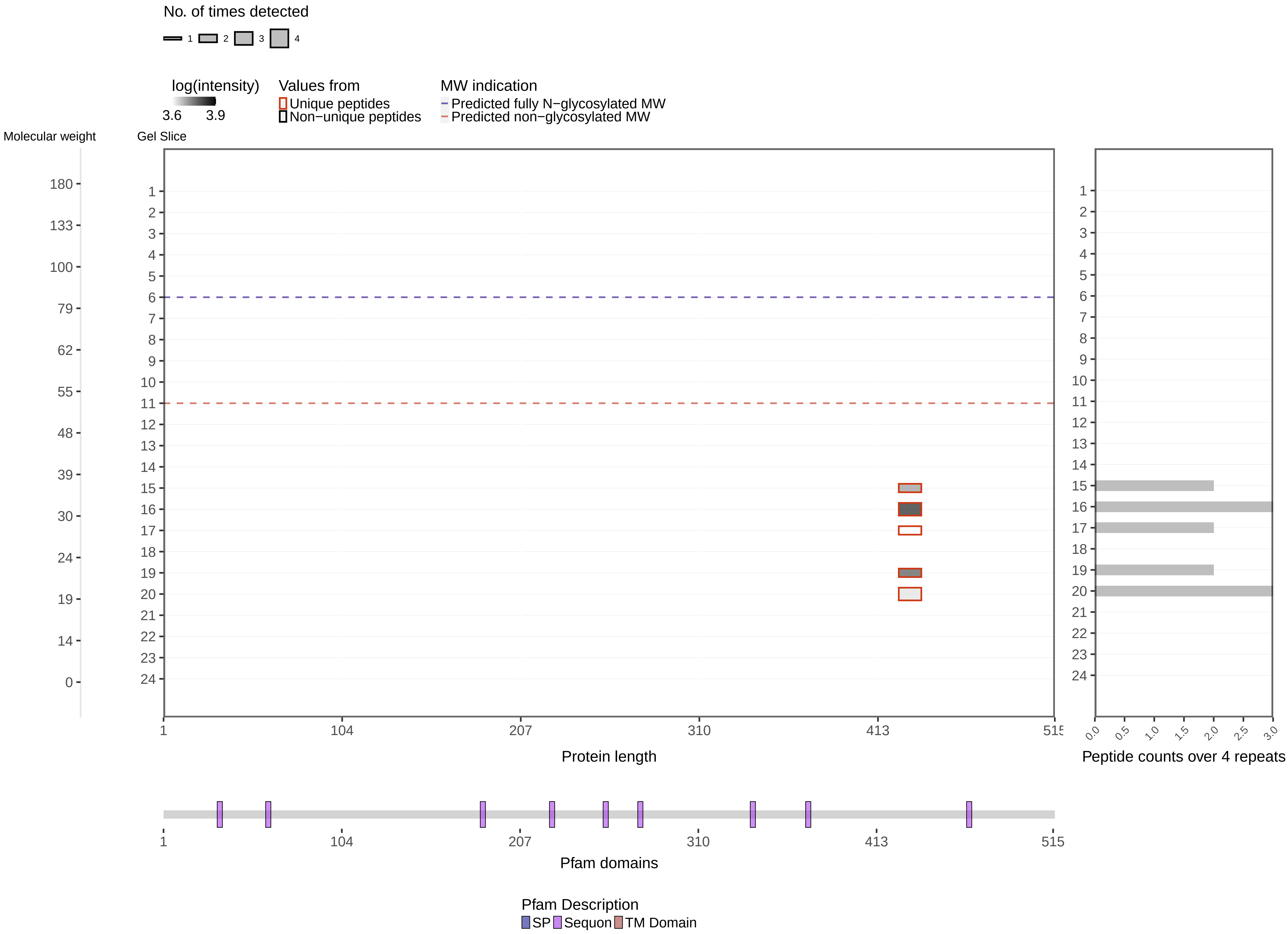


Protein sequence:

SSGDNVFGDFQNATKSGGSGFQSDPFPTSNVNASTGSNSGGAYSKVDDFGFTNTQSQASATQSSGVGDFGFTNTQSQPSATQSSGVDDFDSLFSSTNASFGGAAGGGSENQQFTGGDDWGFESEFVGANDSSGTTELEGLPPPPAGVSASAAKNKGMDNHKQGQYADAIKWLSWAVILLEKAGDGAGAMEVLSSRASCYKEVGEYKKAVADCTKVLEQDGKNVSVLVQRALLYESMEKYKLGAEDLRTVMNIDPGNRVARSTIHRLNKMAG*

NbL01g22830.1 ureidoglycolate_hydrolase [MW = 53.21 kDa]

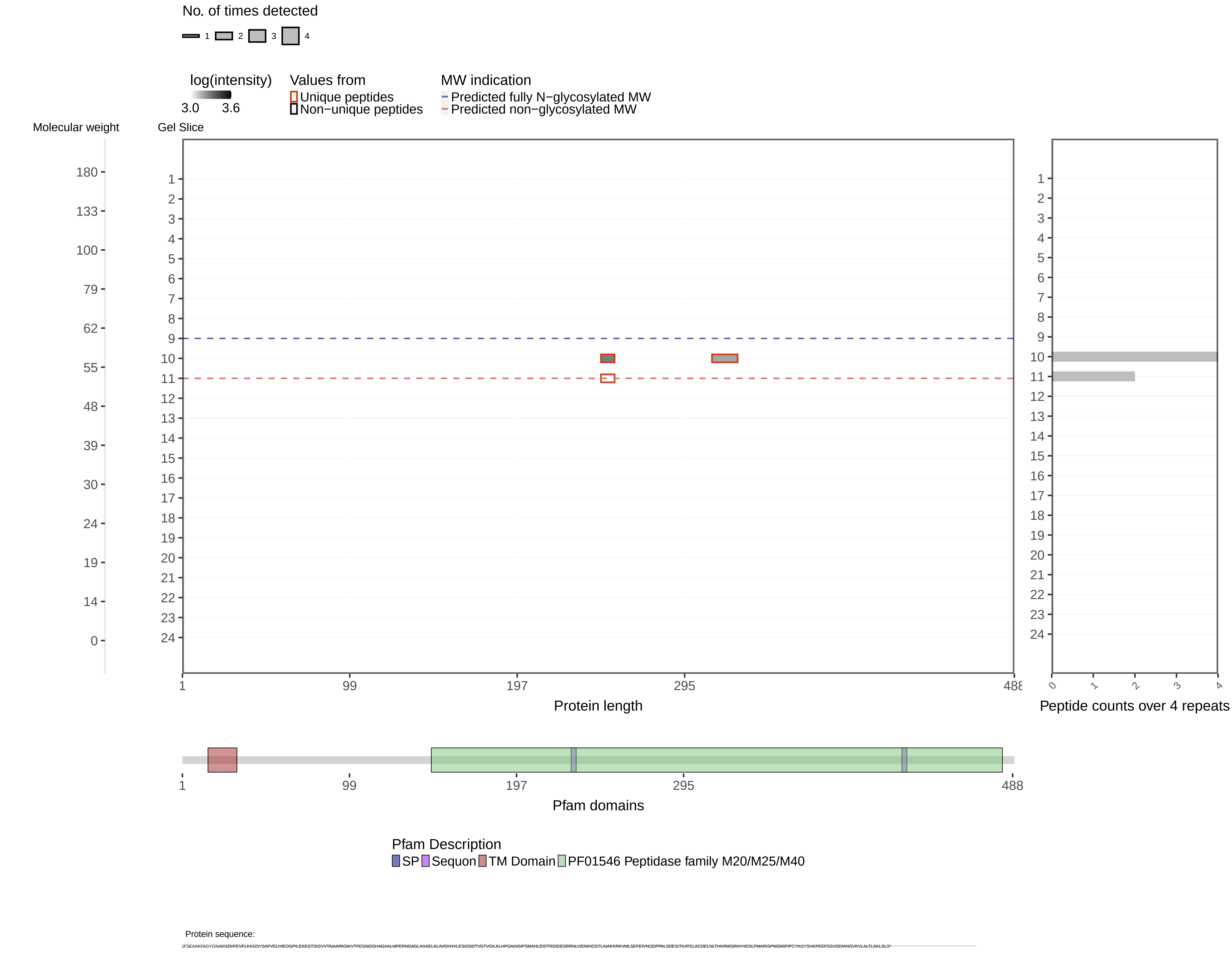

NbL01g22740.1 pleiotropic_drug_resistance_protein_1 [MW = 161.54 kDa]

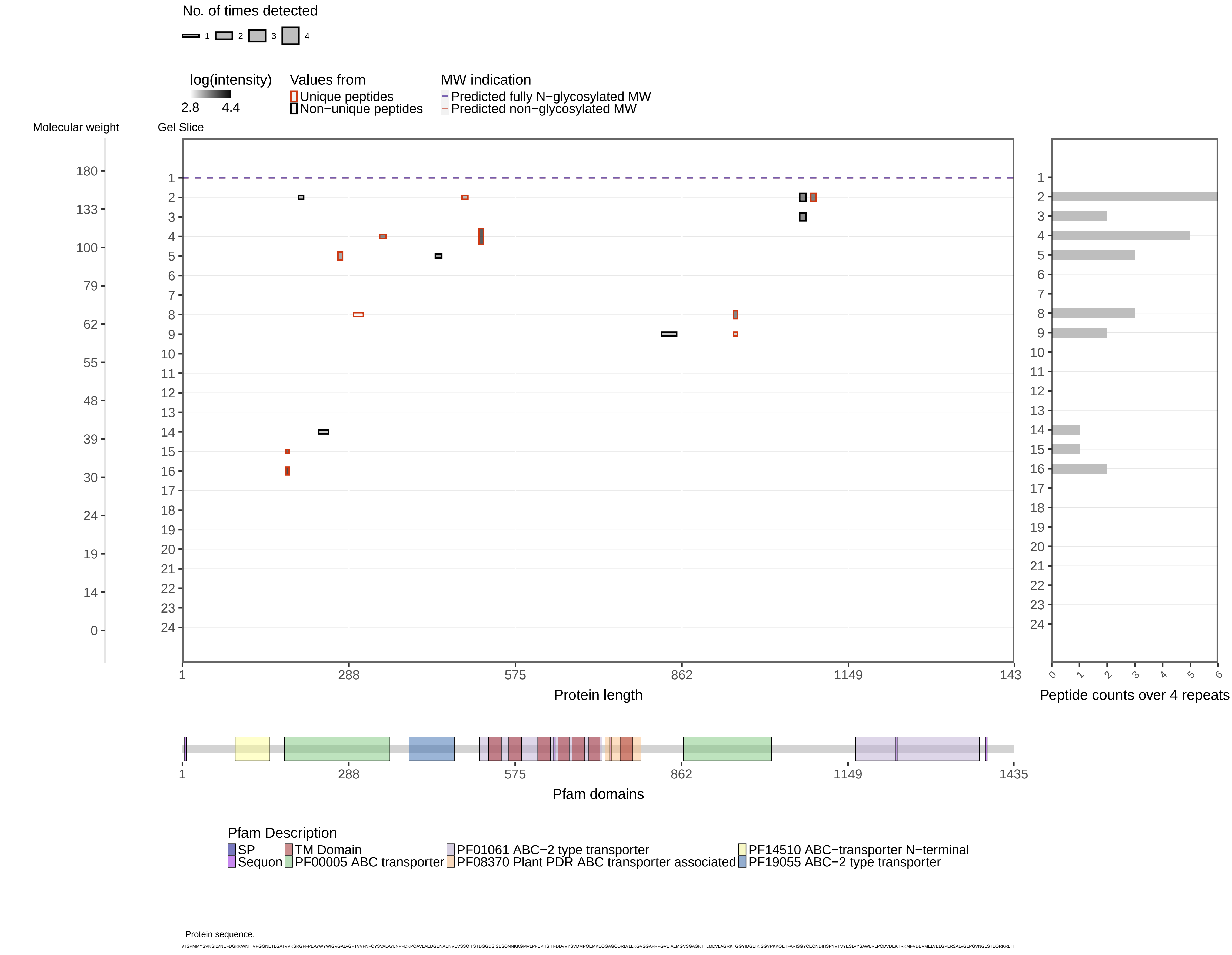

### NbL01g22720.1 pleiotropic_drug_resistance_protein_1 [MW = 84.52 kDa]

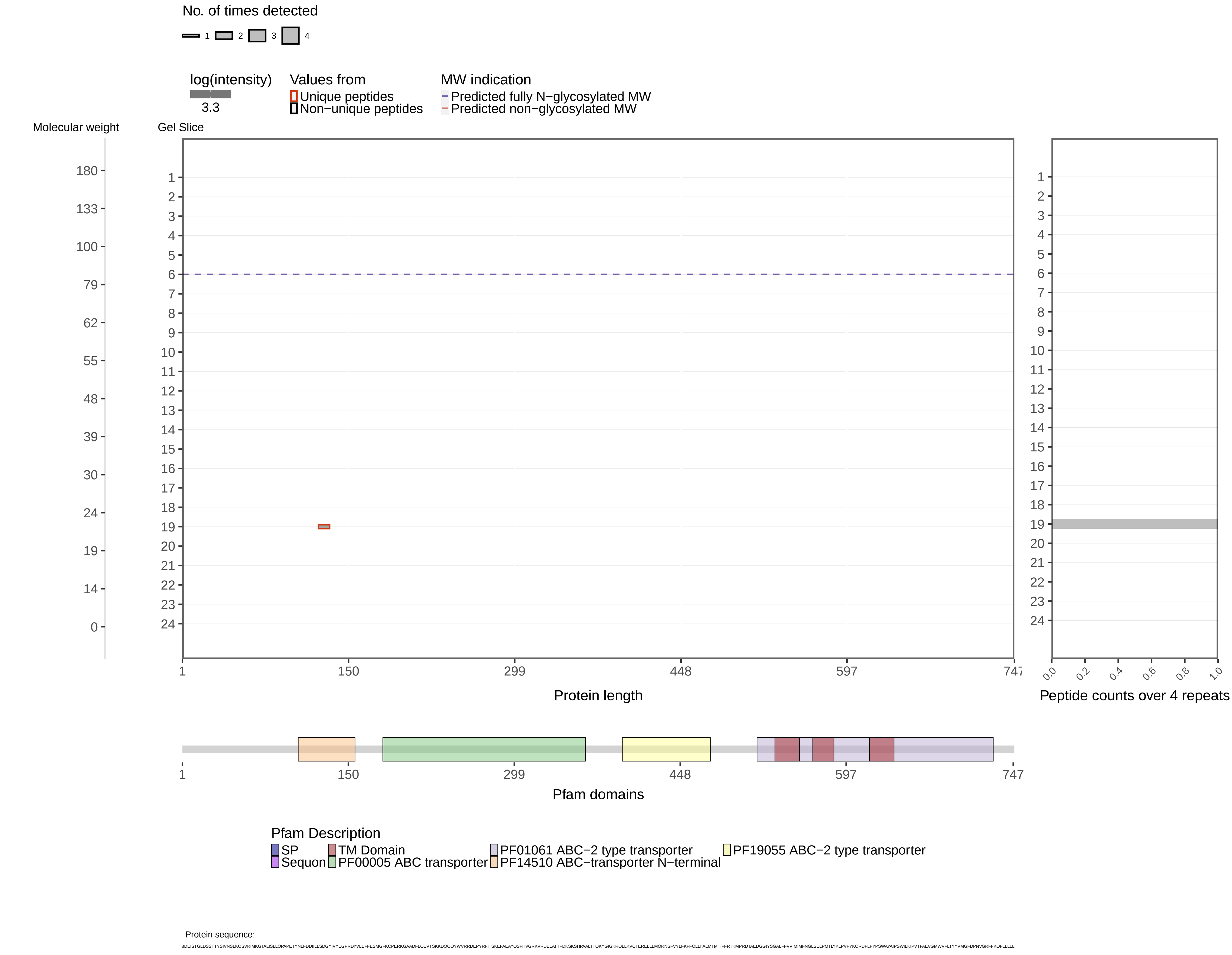

### NbL01g22630.1 sm−like_protein_LSM5 [MW = 5.94 kDa]

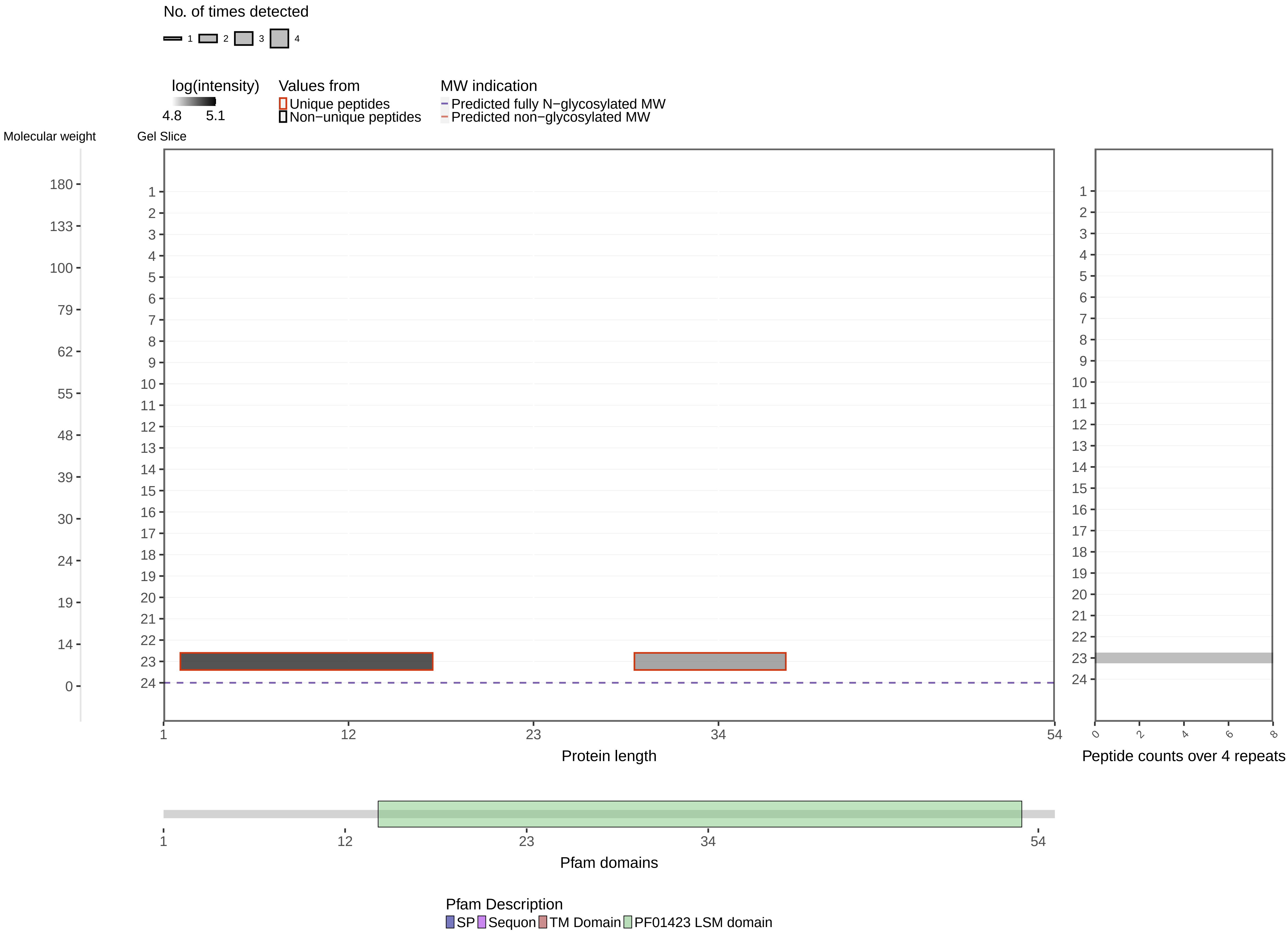


Protein sequence:

LIDRCIGSKIWVIMKGDKELVGTLRGFDVYVNMVLEDVTE*

NbL01g22610.1 3−ketoacyl−CoA_thiolase_2_peroxisomal [MW = 48.84 kDa]

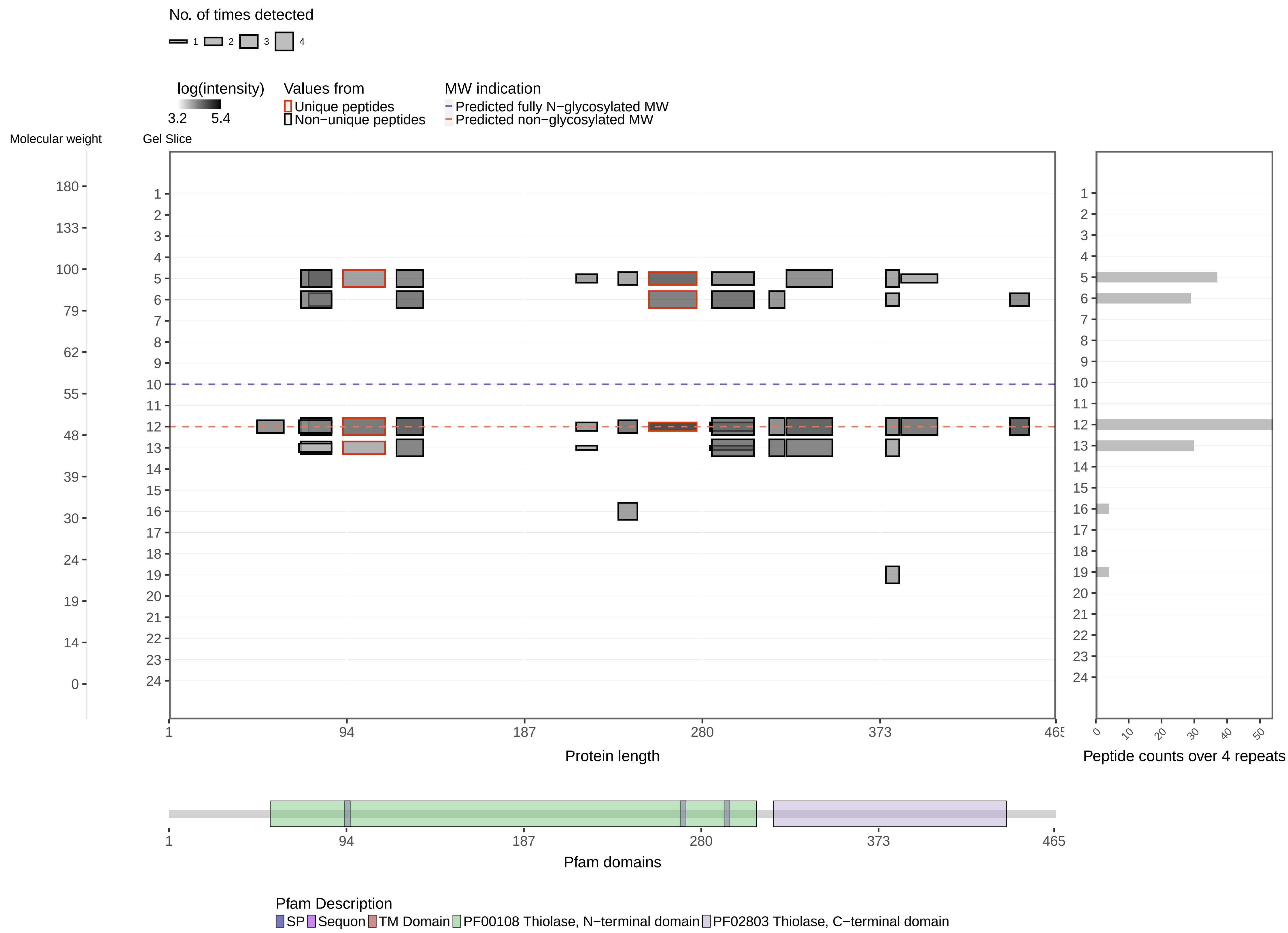


Protein sequence:

QAAVDSHRKAAAASASGKFKDEIIPVPTKIVDPKTGDETPVTISVDDGIRPNASVSDLAKLKPVFKKSGTTTAGNSSQVTDGAGAVLLMKRSLAMQKGLPILGVFRTFAAVGVDPAIMGIGPAVAIPAAVKSAGLELEDIDLFEINEAFASQYVYCRKKLELDPEKINVNGGAMALGHPLGATGARCVATLLHEMKRRGKDCRFGVVSMCIGTGMGAAAVFERGDSCDELCNARKIGSHNLLSKDAL*

### NbL01g22550.1 probable_pectate_lyase_8 [MW − SP = 46.54 kDa]

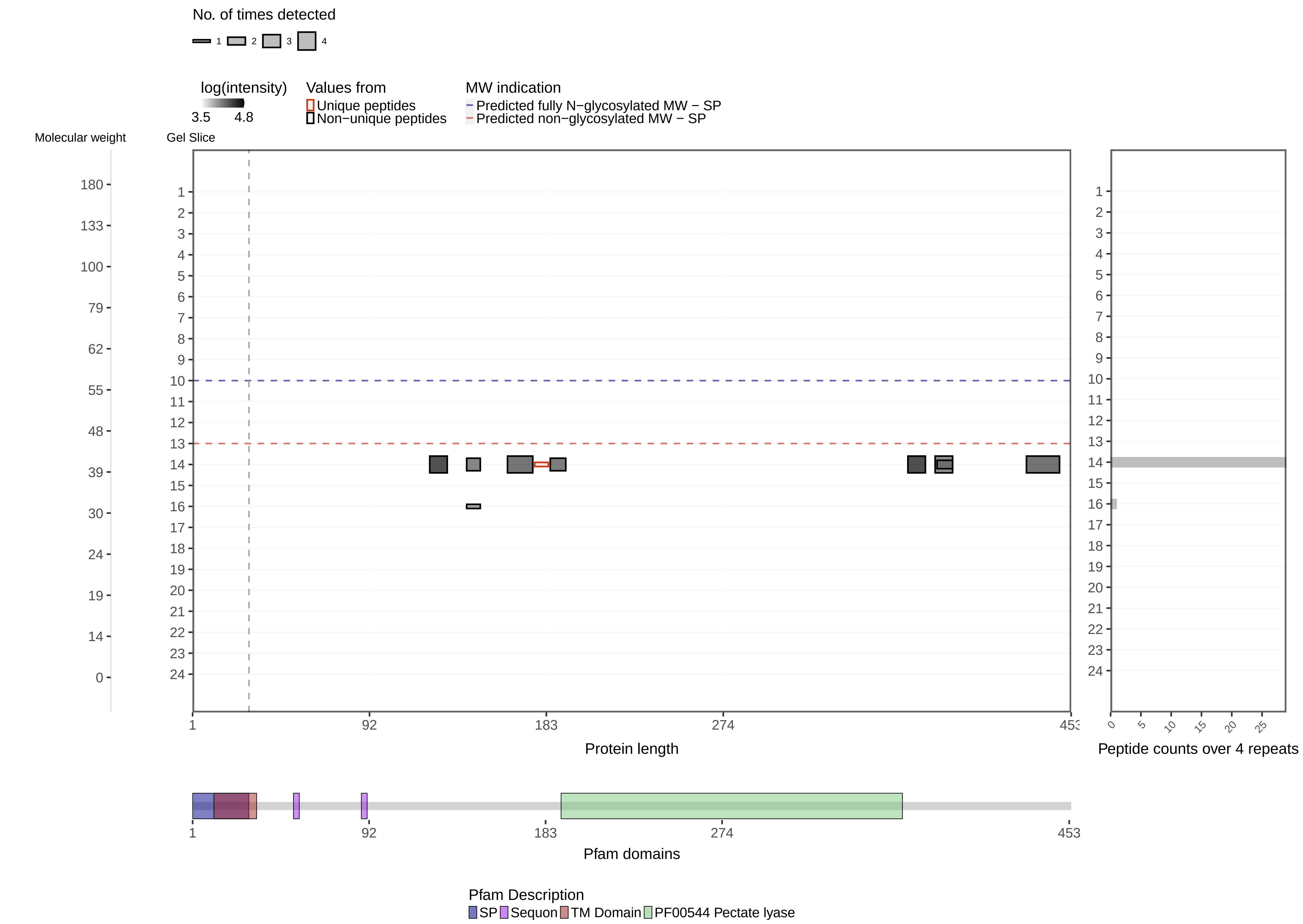


Protein sequence:

'TNIIIHGLHIHDCKPTGNAMVRSSPTHFGWRTMADGDAVSIFGSSHIWIDHNSLSHCADGLVDAVMGSTAITISNNHFAHHNEVMLLGHSDSYTRDKOMOVTIAYNHFGEGLIORMPRCRHGYFHVVNNDYTHWEMYAIGGSANPTINSOGNRYLAPTNPFAKEVTKRVDTAAGOWKGWNWRSEGDLMLNGAYFTPSGAGASASYARASSLGAKSSSMVGAITSGAGPLACRRSRMC*

### NbL01g22310.1 chaperonin_CPN60−2_mitochondrial [MW = 61.29 kDa]

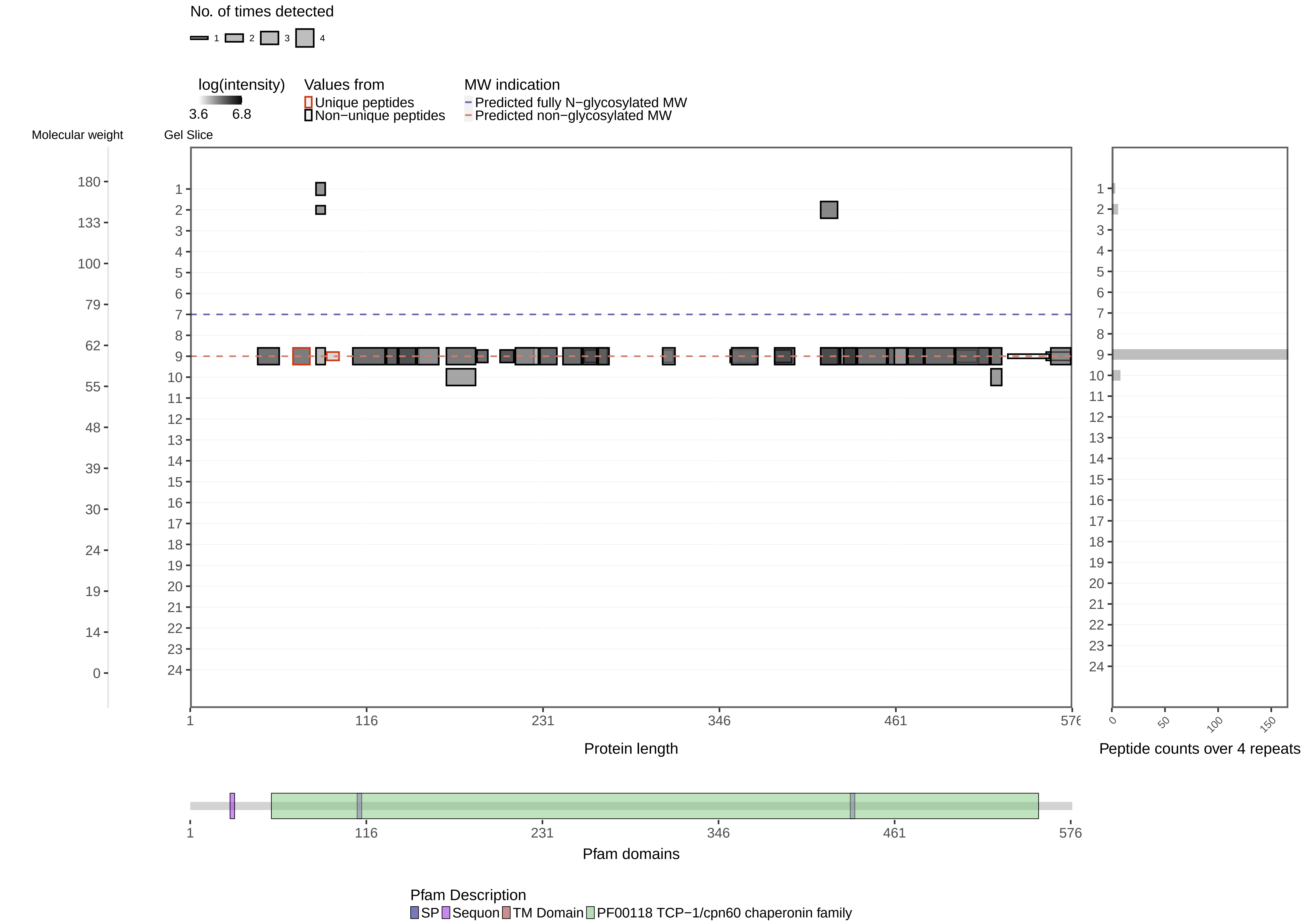


Protein sequence:

KRQRPLLIVAEDVDSEALATLILNKLRAGIKVCAIKAPGFGENRKANLQDLAILTGGQVVTEELGLNIENVELEMLGSCKKVAISKDDTVILDGAGEKKSIEERCEQIRSTIELSTSDYDKEKLQERLAKISGGVAVLKIGGASEAEVGEKKDRVTDALNATKAAVEEGIVPGGGVALLYAAKELDKLETANFDQKIGVQIIQNALKTPVHTIASNAGVEGAVVVGKLLEQDDTDLGYDAAKGEYVDMVKAGIIDPLKVIRTALVDAASVSSLLTTTEAVVVELPKDEKETPAMGGGMGG[illegible]

NbL01g22100.1 1_chloroplastic−like [MW = 64.38 kDa]

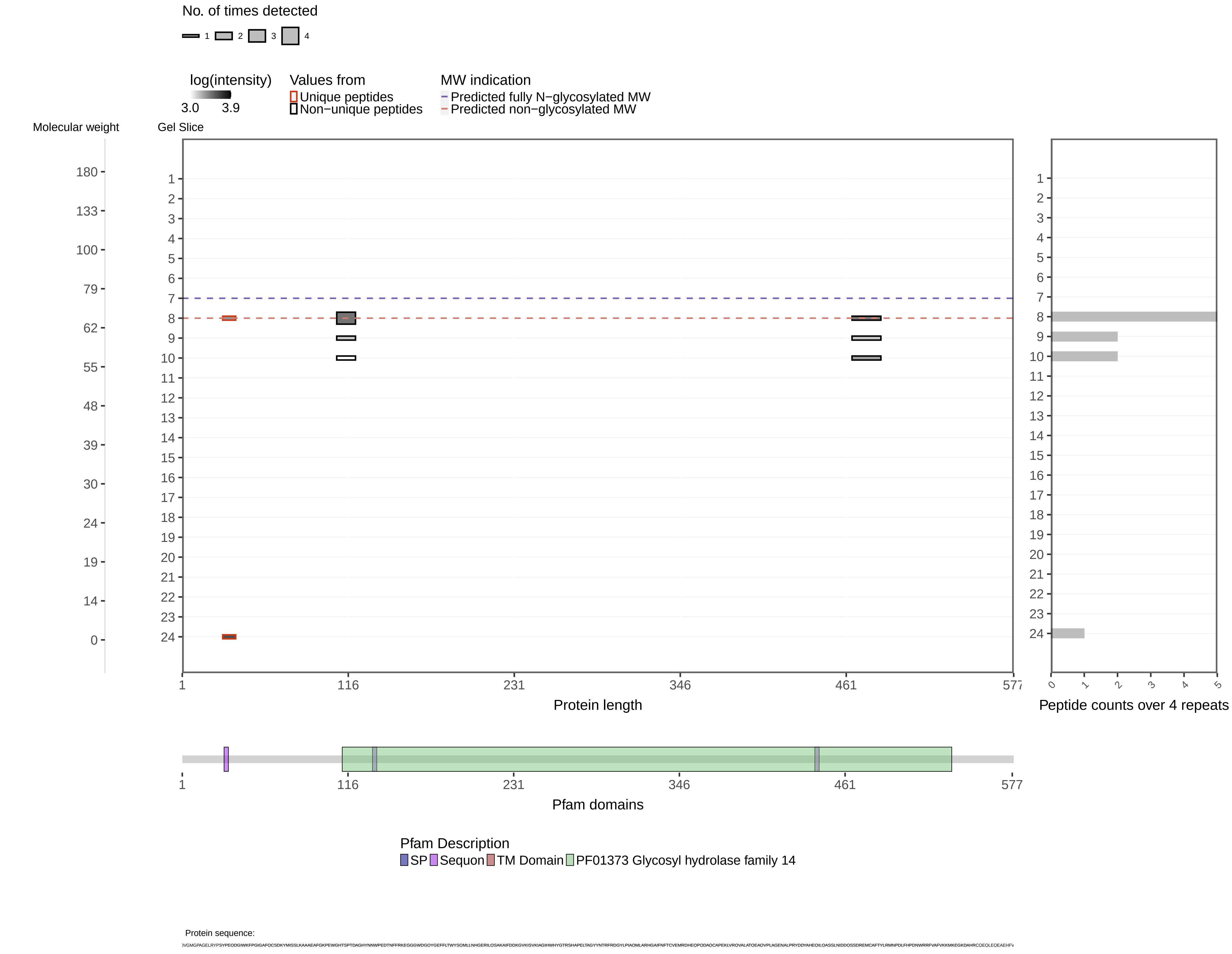

### NbL01g21920.1 glutamyl−tRNA(Gln)_amidotransferase_subunit_B_chloroplastic/mitochondrial [MW = 60.7 kDa]

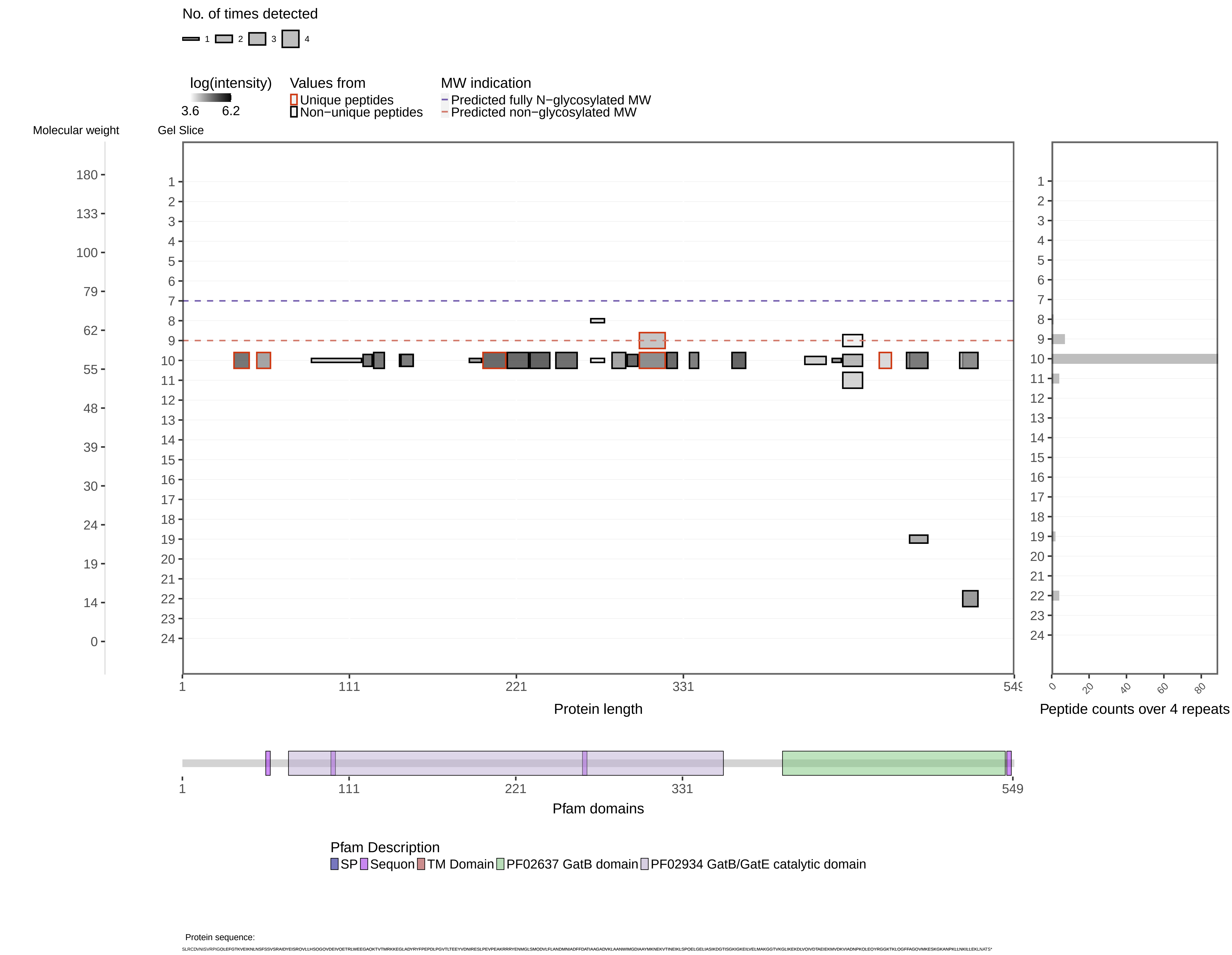

### NbL01g21890.1 3−isopropylmalate_dehydratase_large_subunit_chloroplastic−like [MW = 55.26 kDa]

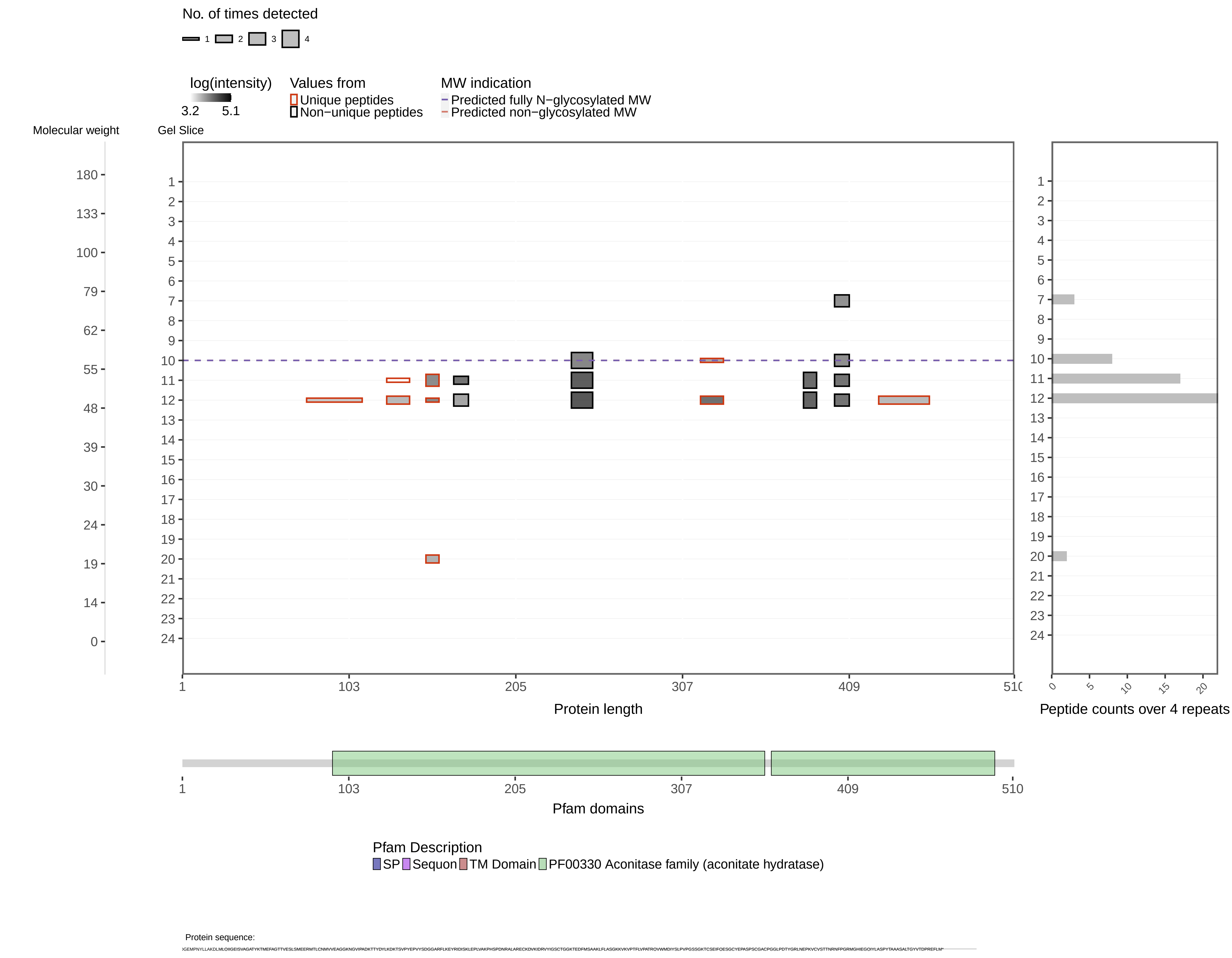

### NbL01g21880.1 3−isopropylmalate_dehydratase_large_subunit−like [MW = 57.34 kDa]

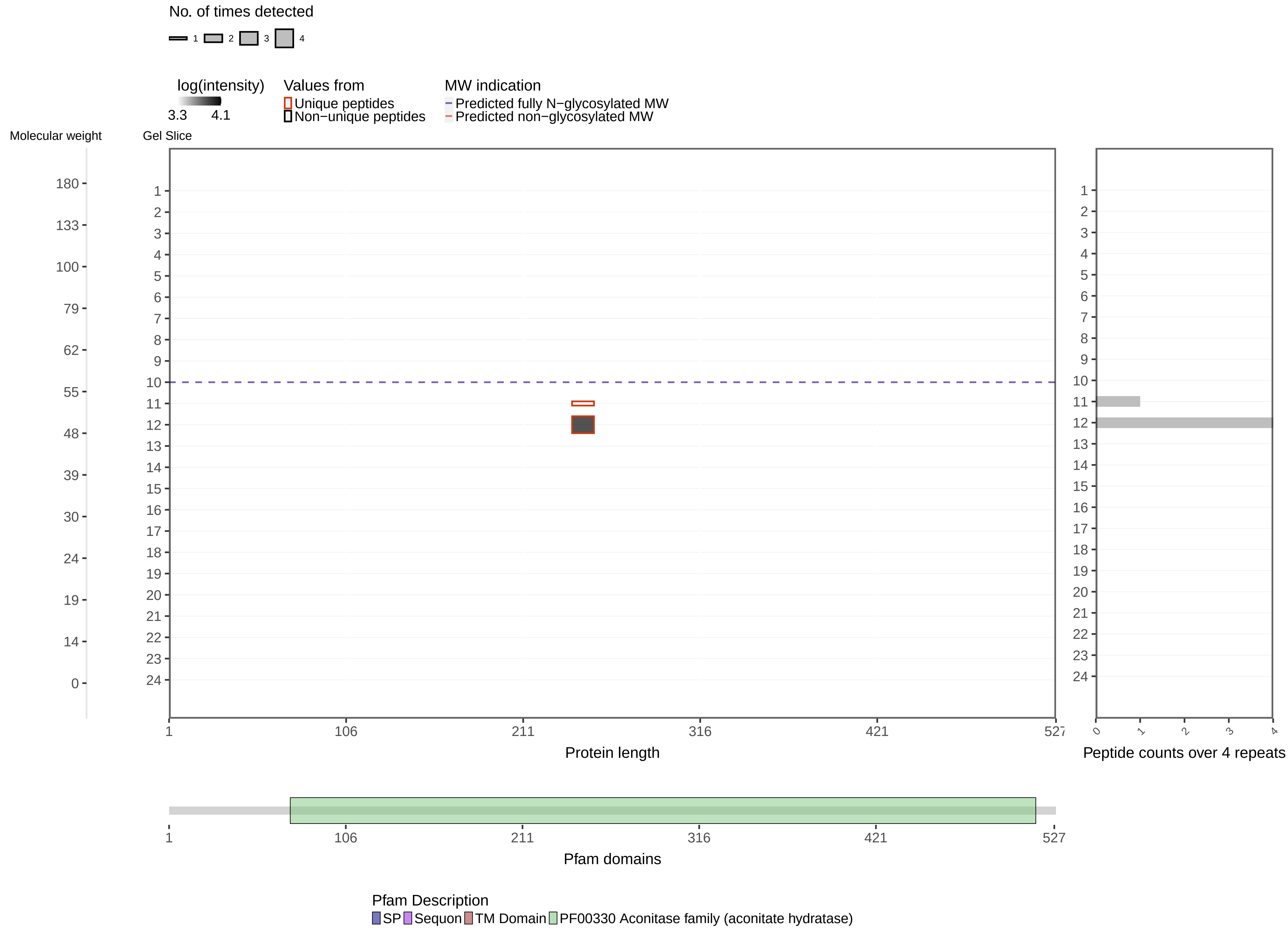


Protein sequence:

LLAKDLMLQIIGEIFVSGATYKTLEFVGTTVESLSMEERMTLCNMVVEAGGKNGVIPTDKTTYDYLKDKTSVPYEPVYSDEGARFLKEYRIDISKLEPLVAKPHSPDNRALARECKDVKIDRVYIGSCTGGKTEDFMAAAKVFLASGKKIKVPTFLVPATQKVWMDVYTLPVPGSGGKTCSQIFEEVGCETPASPSCAACLGGPQDTYARLNEPKEITSSSRGNSLLQKCRVCVSTTNRNFPGRMGHVEGQIYLASPYTAAASALTGKVTDPREFLM*

### NbL01g21830.1 RNA−binding_protein_CP33_chloroplastic [MW = 31.63 kDa]

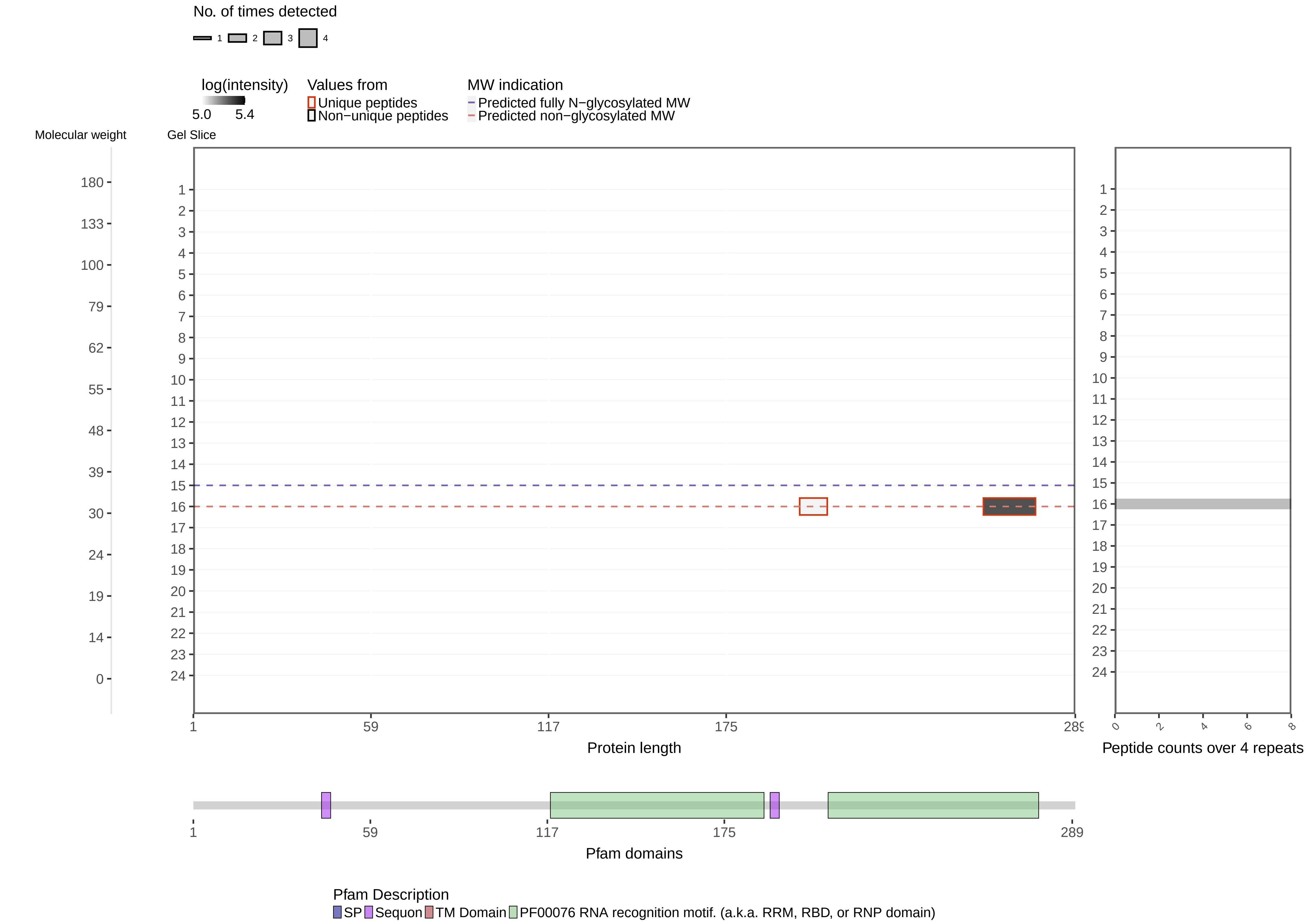


Protein sequence:

AGIVQDYATPELVEVLYDRETGKSRGFAFVTMSSLEDCKTVIKNLDRREYGGRTLRVNFSDKPKPKEPLYPETEHKLFVGNLAWSVTSESLAQAFQEYGTVVGARVLYDGETGRSRGYGFVSYENREQLENALQNLNGVELDGRAVRISLAQGKKQ*

### NbL01g21770.1 succinate−semialdehyde_dehydrogenase_mitochondrial−like [MW = 55.47 kDa]

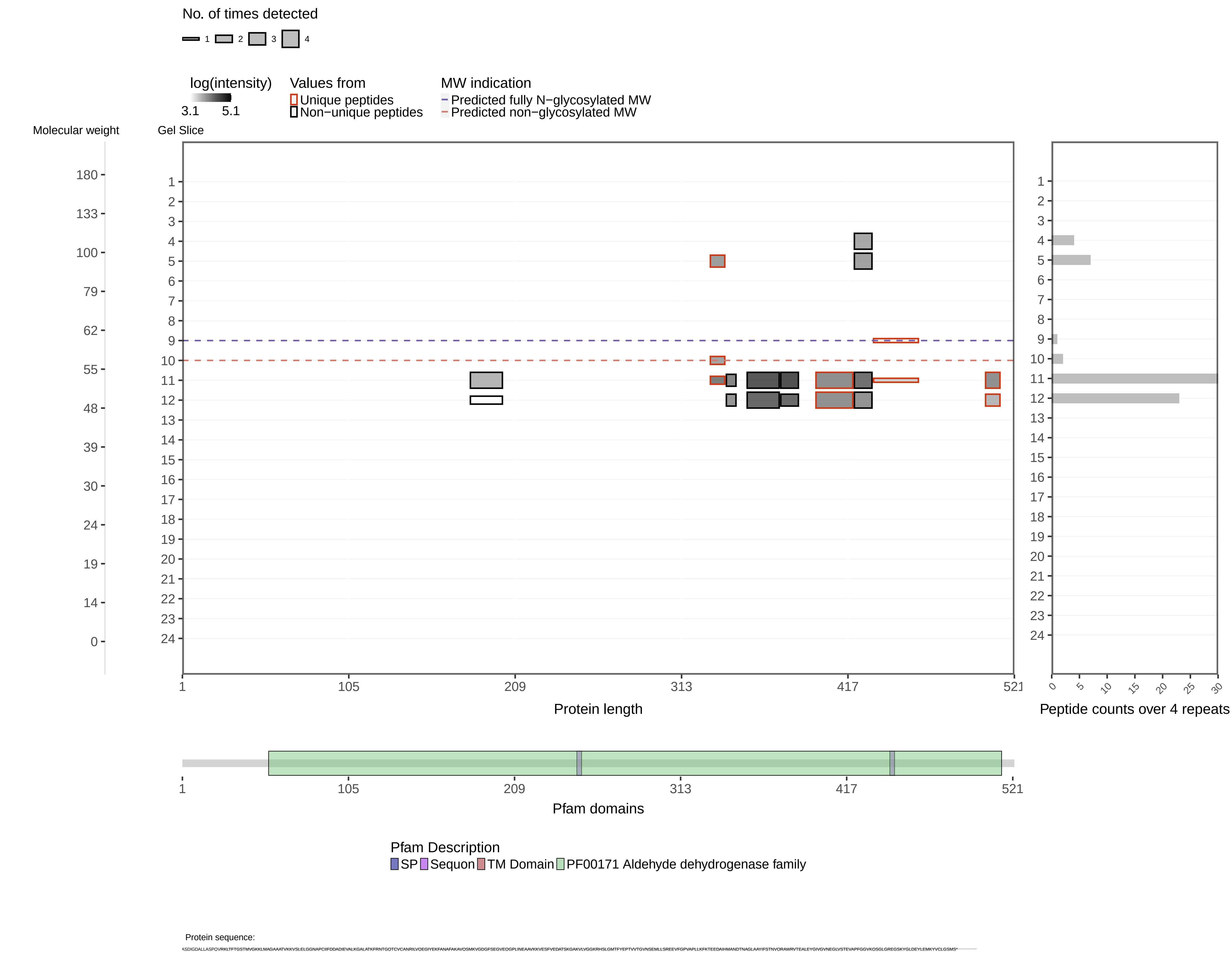

### NbL01g21750.1 glucose−6−phosphate_isomerase_cytosolic [MW = 63.67 kDa]

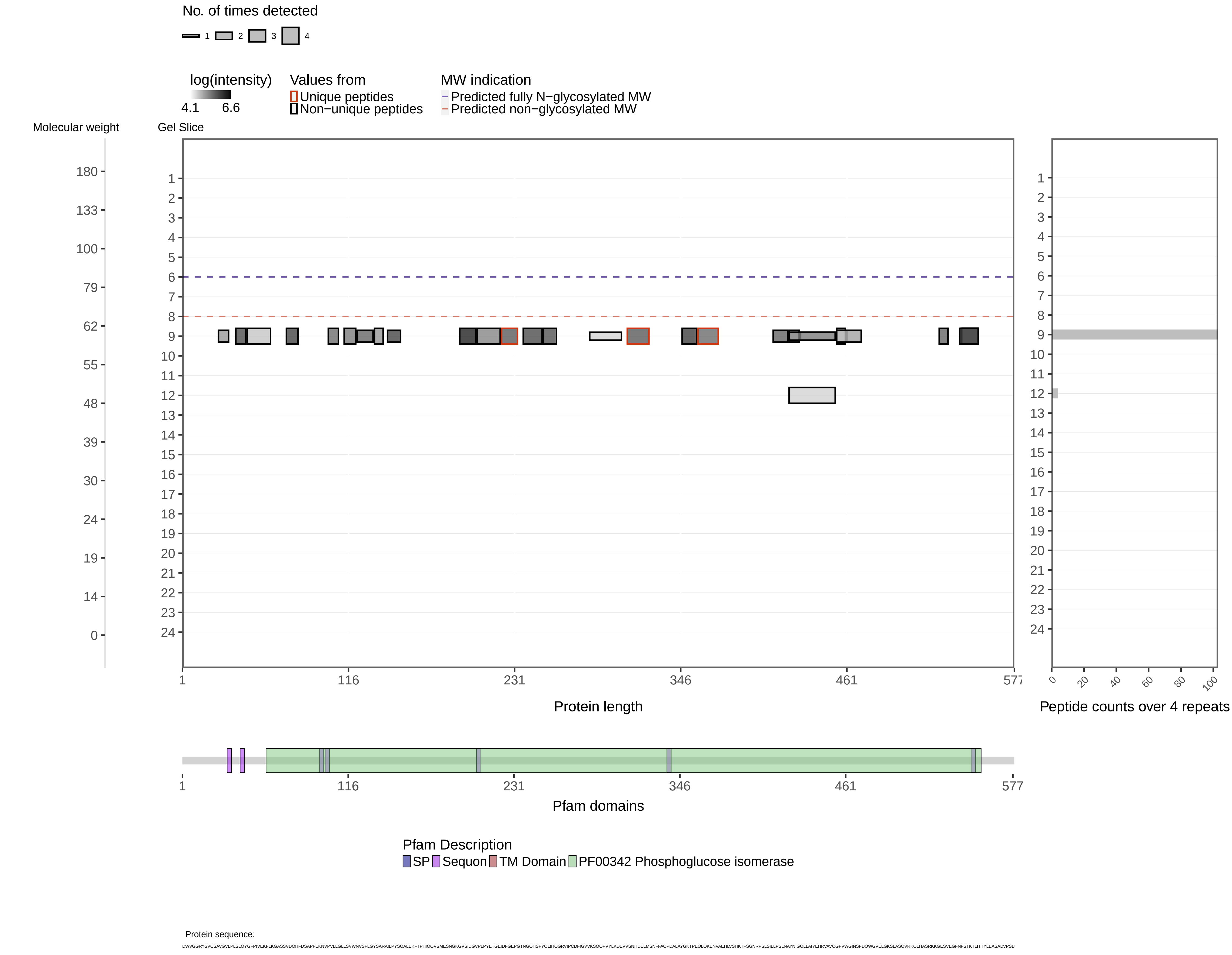

NbL01g21720.1 pre−mRNA−processing_factor_19_homolog_1−like [MW = 56.56 kDa]

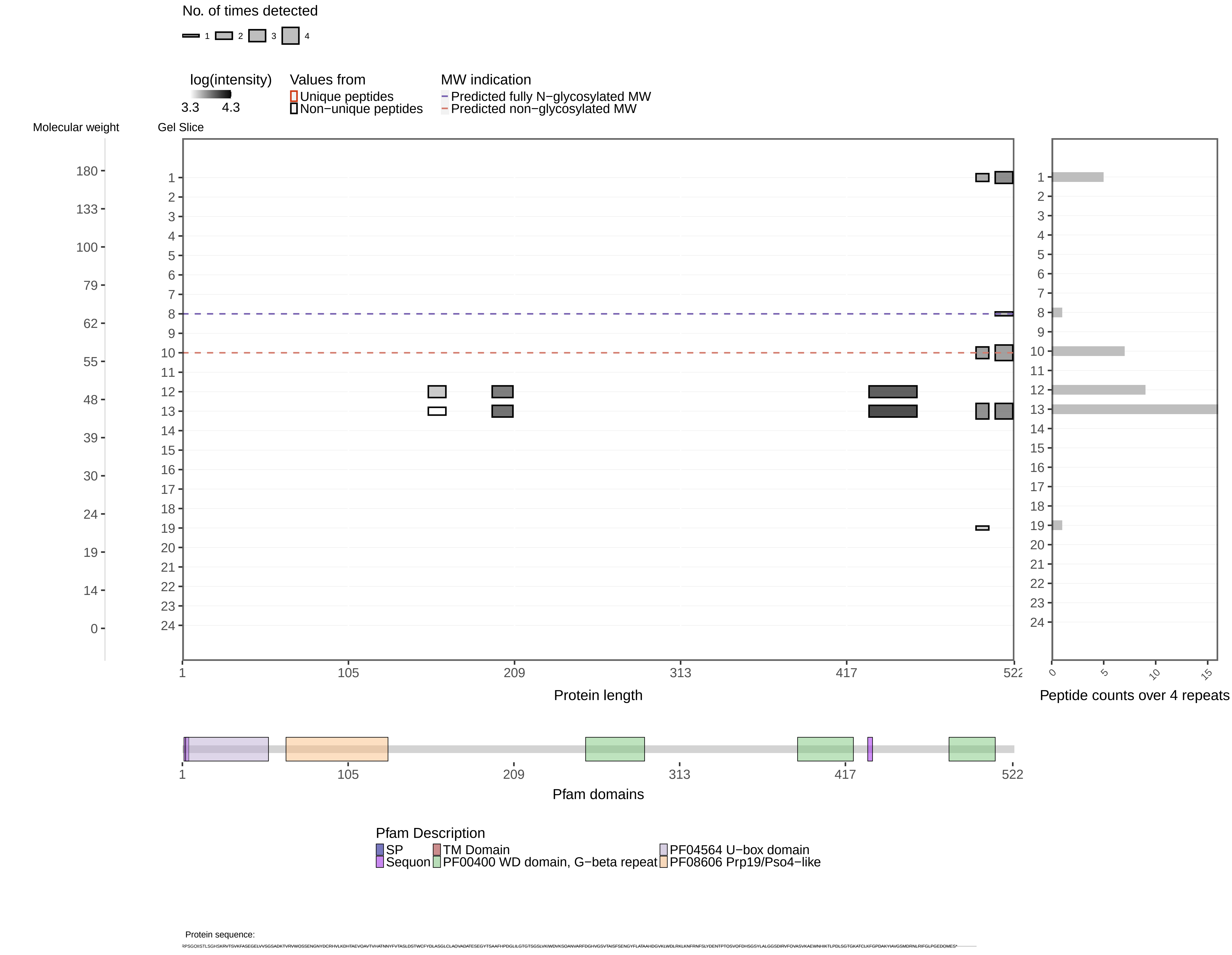

NbL01g21660.1 heterogeneous_nuclear_ribonucleoprotein_1−like [MW = 47.46 kDa]

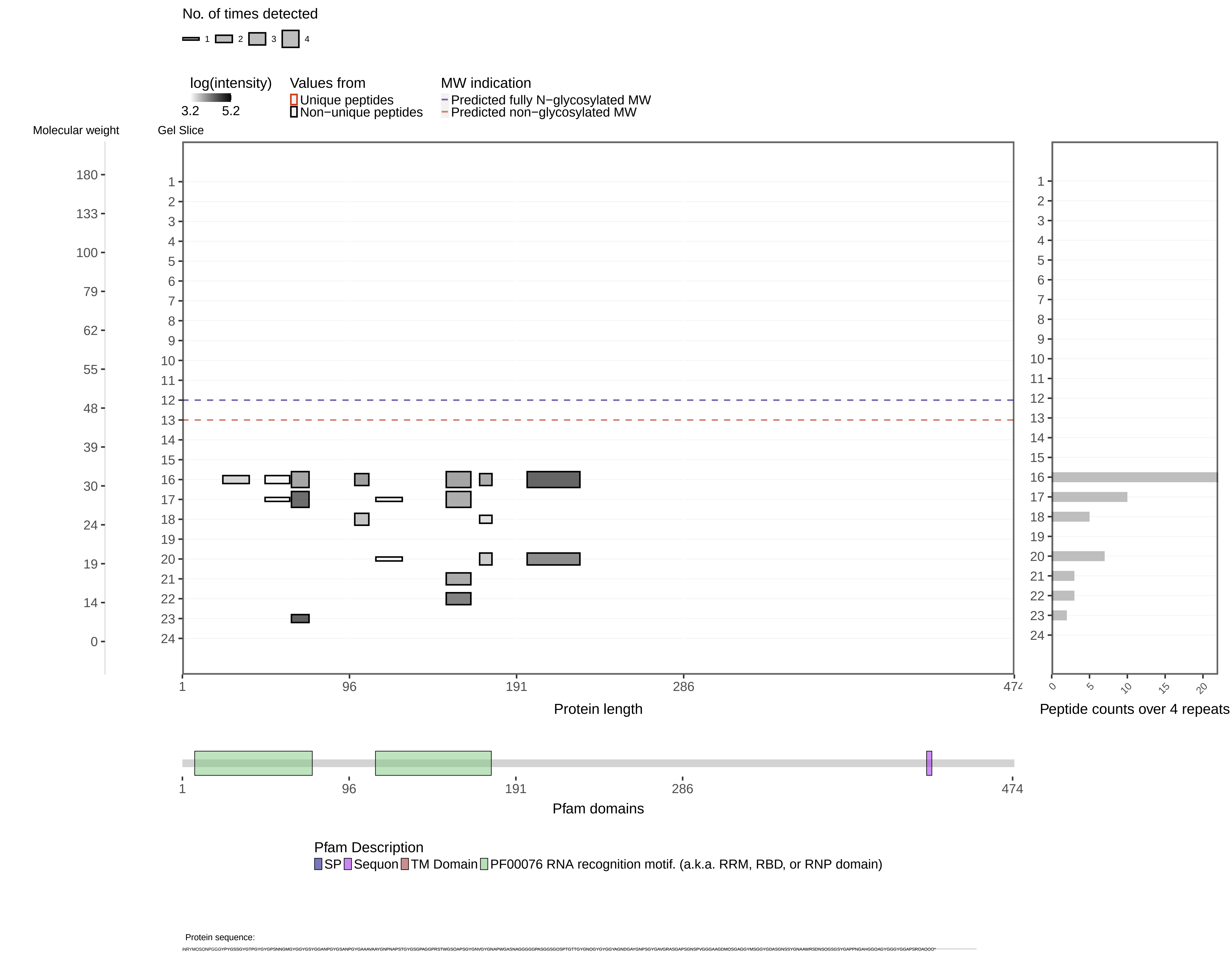

NbL01g21610.1 protein_PROTON_GRADIENT_REGULATION_5_chloroplastic−like [MW = 12.91 kDa]

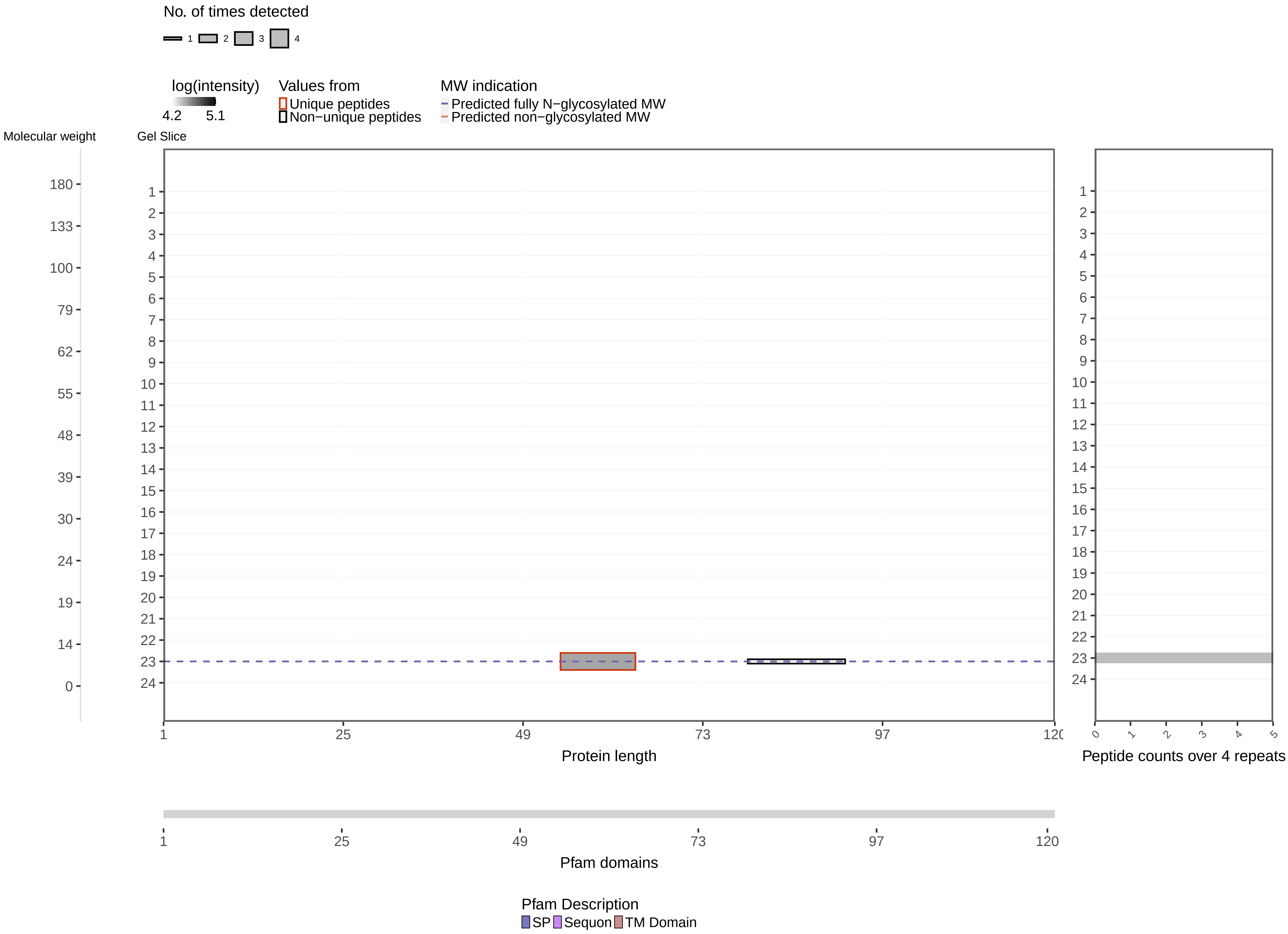


Protein sequence:

IGNINDGKGLFAPIVVVTRNIVGKKRFNQLRGKAIALHSQVITEFCKSIGADQKQRQGLIRLAKKNGERLGFLA*

### NbL01g21570.1 acid_phosphatase_1−like [MW − SP = 27.01 kDa]

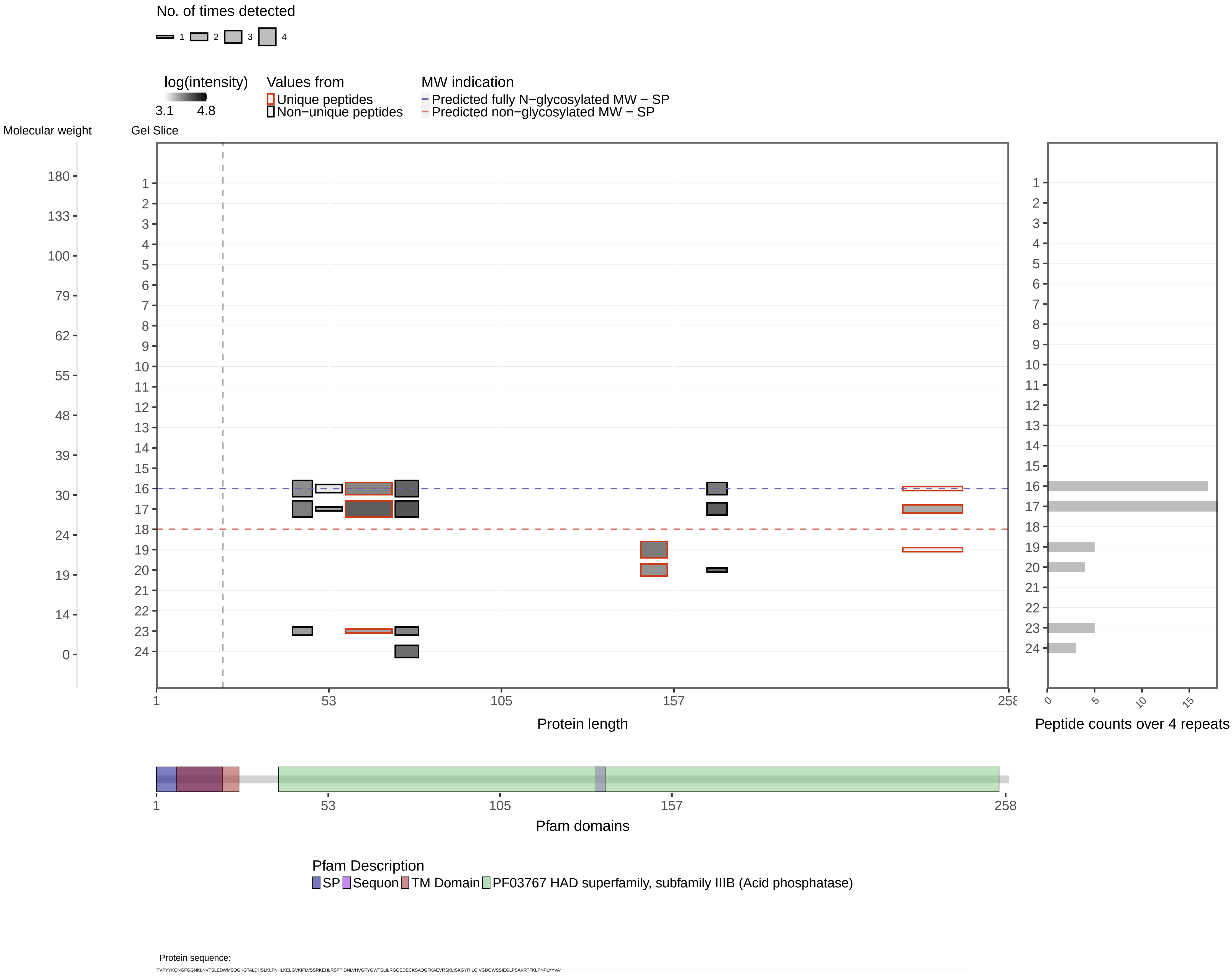

### NbL01g21430.1 glycine−rich_cell_wall_structural_protein [MW = 42.54 kDa]

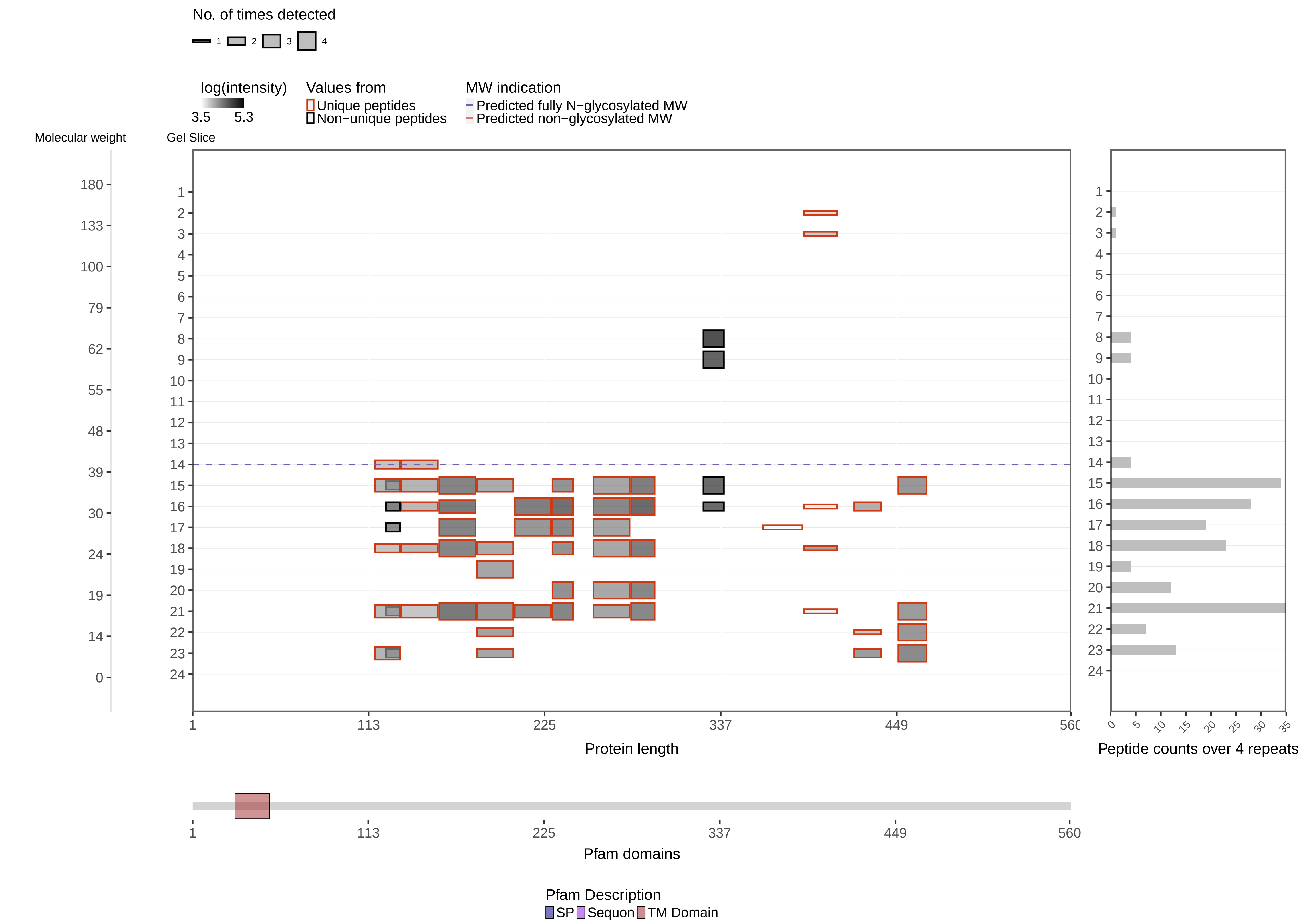


Protein sequence:
[illegible]

### NbL01g21050.1 putative_germin−like_protein_2−1 [MW = 25.82 kDa]

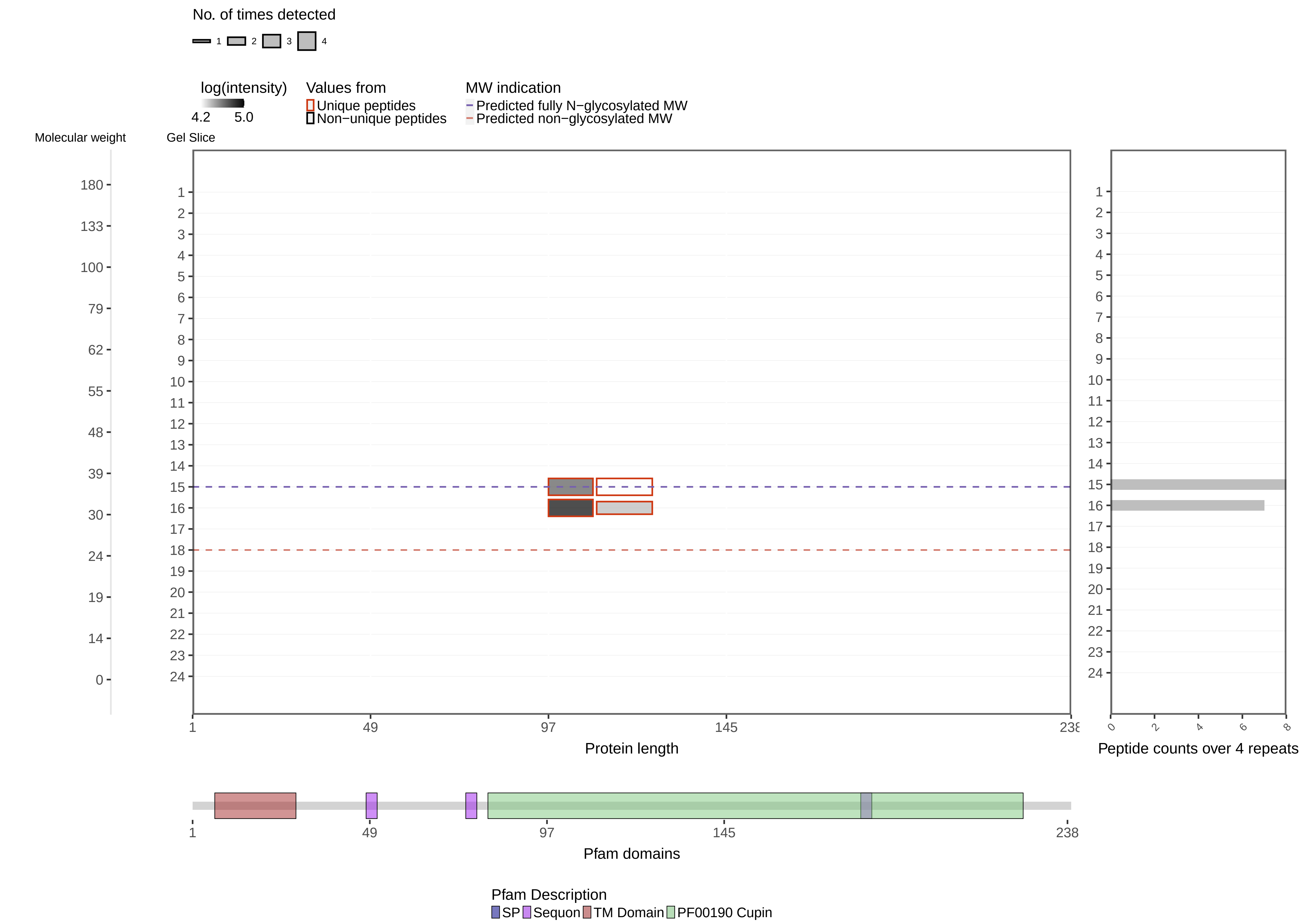


Protein sequence:

IVRADVEPFGLISPHIHPRGNELILVLEGILYVAFTIPDPGNPFKSRLFEKILNPGDVFVIPQSLVHVQYNLGTTNTTALSCFNSQNPGVHMIPFQLFGSNPPILDDVLVKGFRLDKKVVEHLRAQFSSKS*

### NbL01g20750.1 RNA−binding_protein_BRN1 [MW = 46.84 kDa]

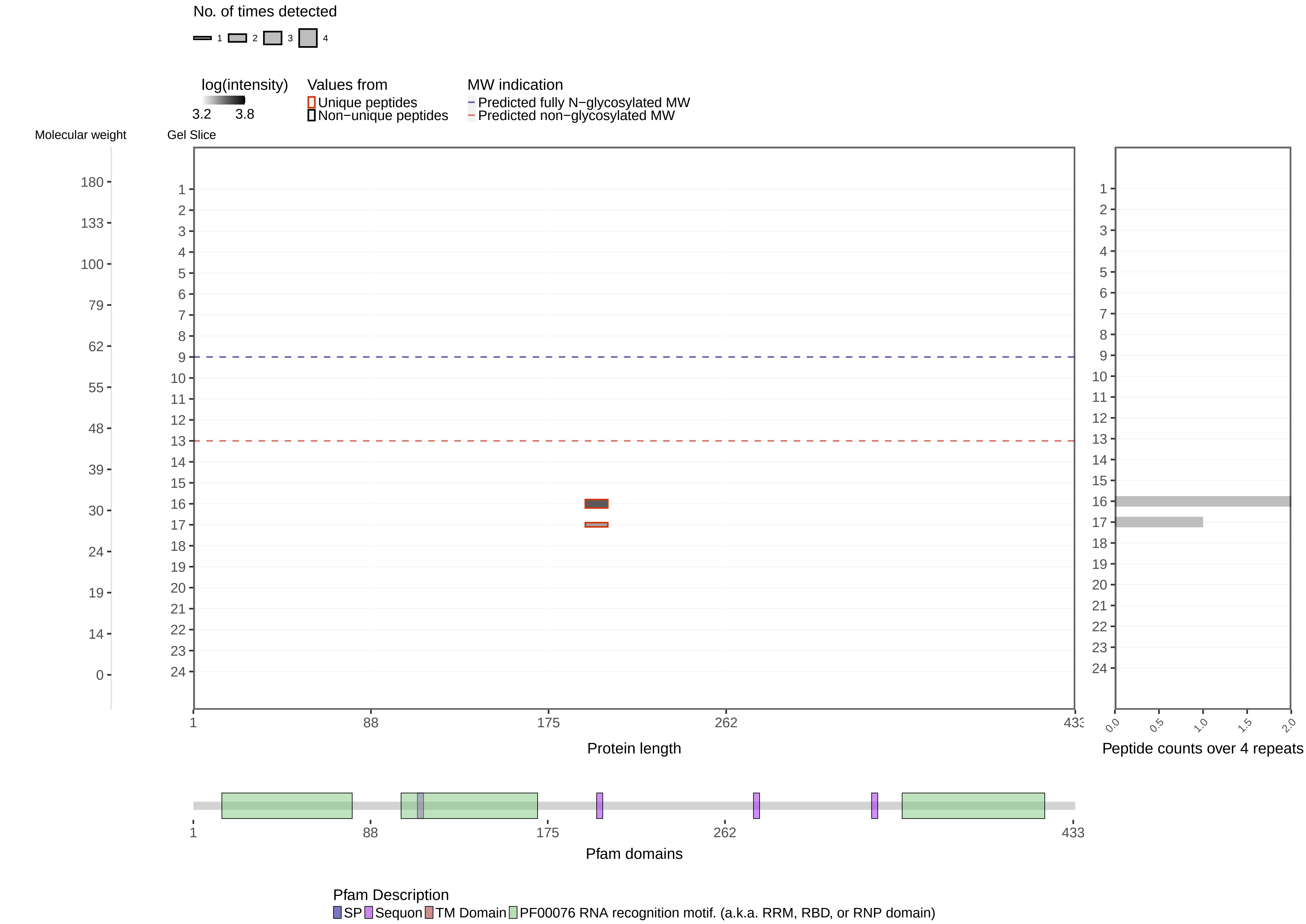


Protein sequence:

HPSLFGALPMGYMPPYNGYGYQTPGAYGLMQYRLPSVQNQPAFQNIVPPINQASALRGGAPDLSPGISPRNYSMSPGSYVGSAYPAVPRLQYPMPYAGGVMNARPPSGSPGSMPPSTANSHSAASSSVNSSTGGQVEGPPGANLFIYHIPQEFGDQDLANSFQPFGRVLSAKVFVDKATGVSKCFGFVSYDSPAAAQTAINMMNGCQLGSKKLKVQLKRDNKQNKHY*

NbL01g20720.1 protease_Do−like_2_chloroplastic [MW = 68.01 kDa]

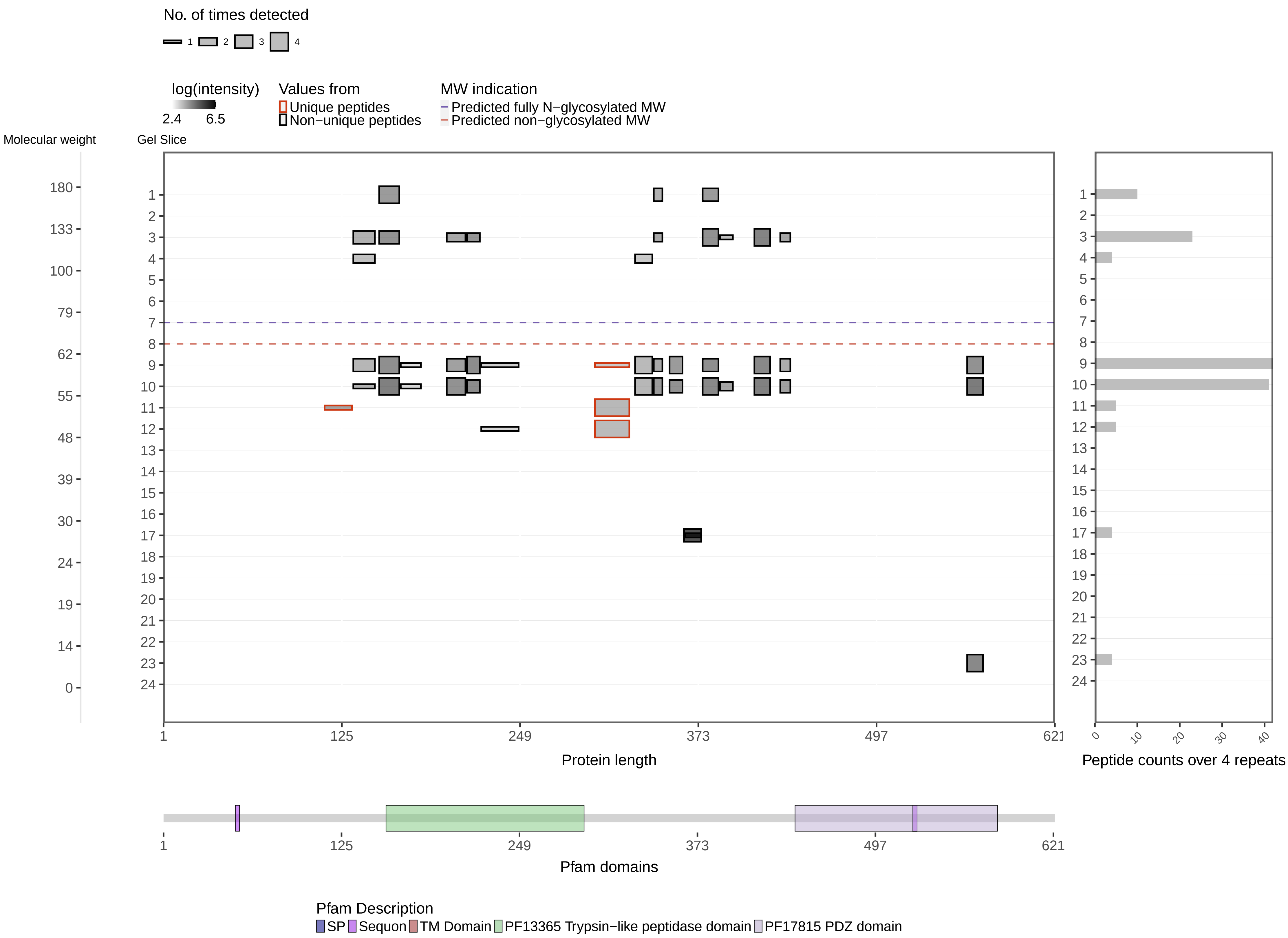


Protein sequence:

IGVAFQVYRSDETENIGYVIPTTVVSHFLEDYEKNGKYSGFPCLGVLLQKLESPALRACLKVPSNEGVLVRKVEPTSDVSNVLKEGDVIVSFDGVRIGCEGTVPFRSSARIAFRYLISQKFTGDVAELGIIRAGEFMKVQAVLKPRVHLVPYHIECGQPSYLIVAGLVFTPLSEPLIGEECEDSIGLKLLTKARYSLAKFEGEQIVVLSQVLANEVNIGYEDISNEQILKLNGTRIKNIHHLAHLVDSCKDKYLVLEFEDNFLVVLEREAAVSASASILKDYGIPAERSSDLLEPYIDPIRPD

NbL01g20600.1 NA [MW = 25.95 kDa]

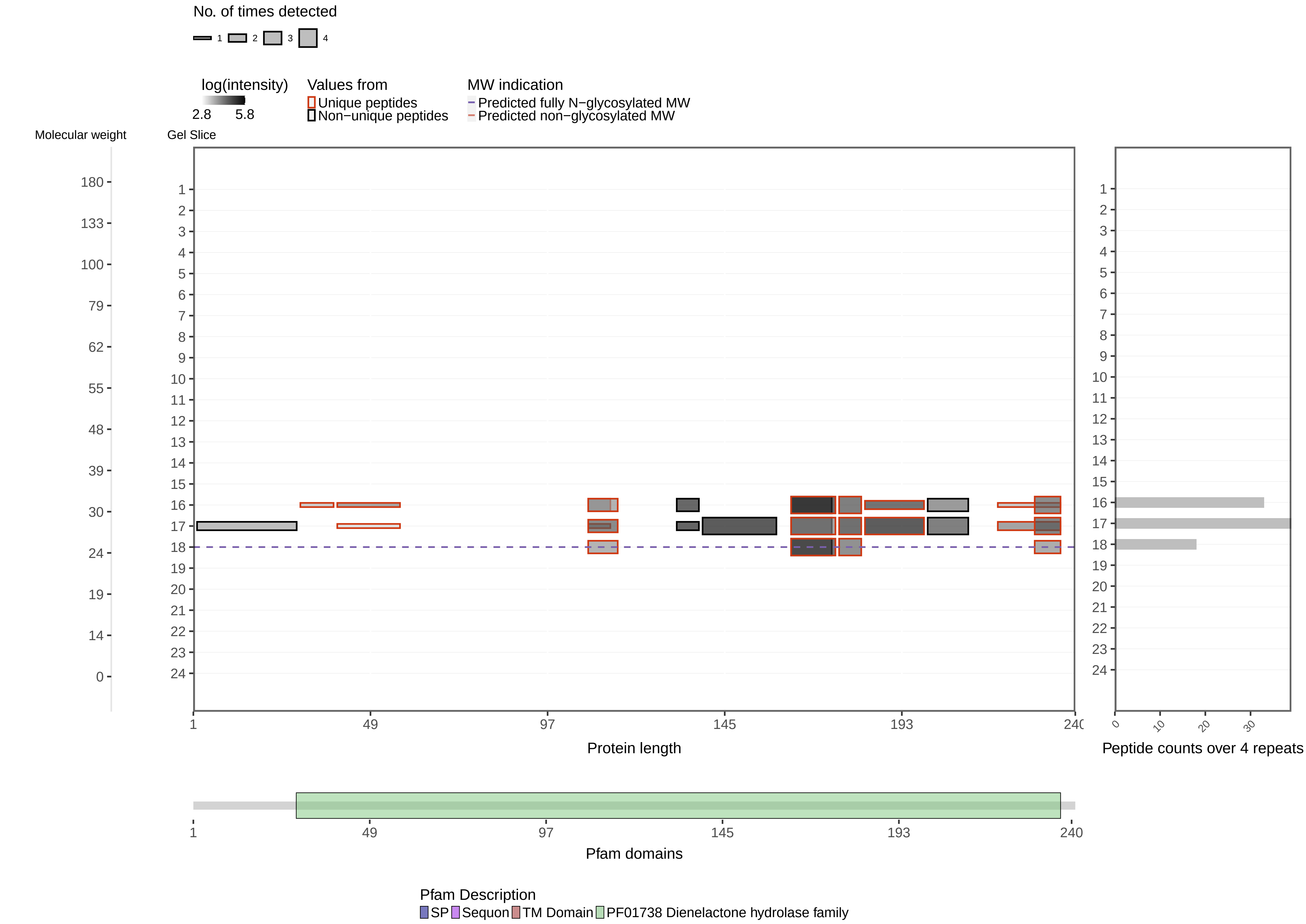


Protein sequence:

AKQVVSALKDKGISAVGAAGFCWGAKVVVQLAKSDDIQAAVLLHPSFVTVDDIKEVKAPIAILGAEIDKRSPPELIKQFEEILSSKPEVDNFVKIYPGVAHGWTVRYDVENKKAVESAEEAHRDMLEWFTKHVK*

### NbL01g20580.1 sm−like_protein_LSM2 [MW = 14.5 kDa]

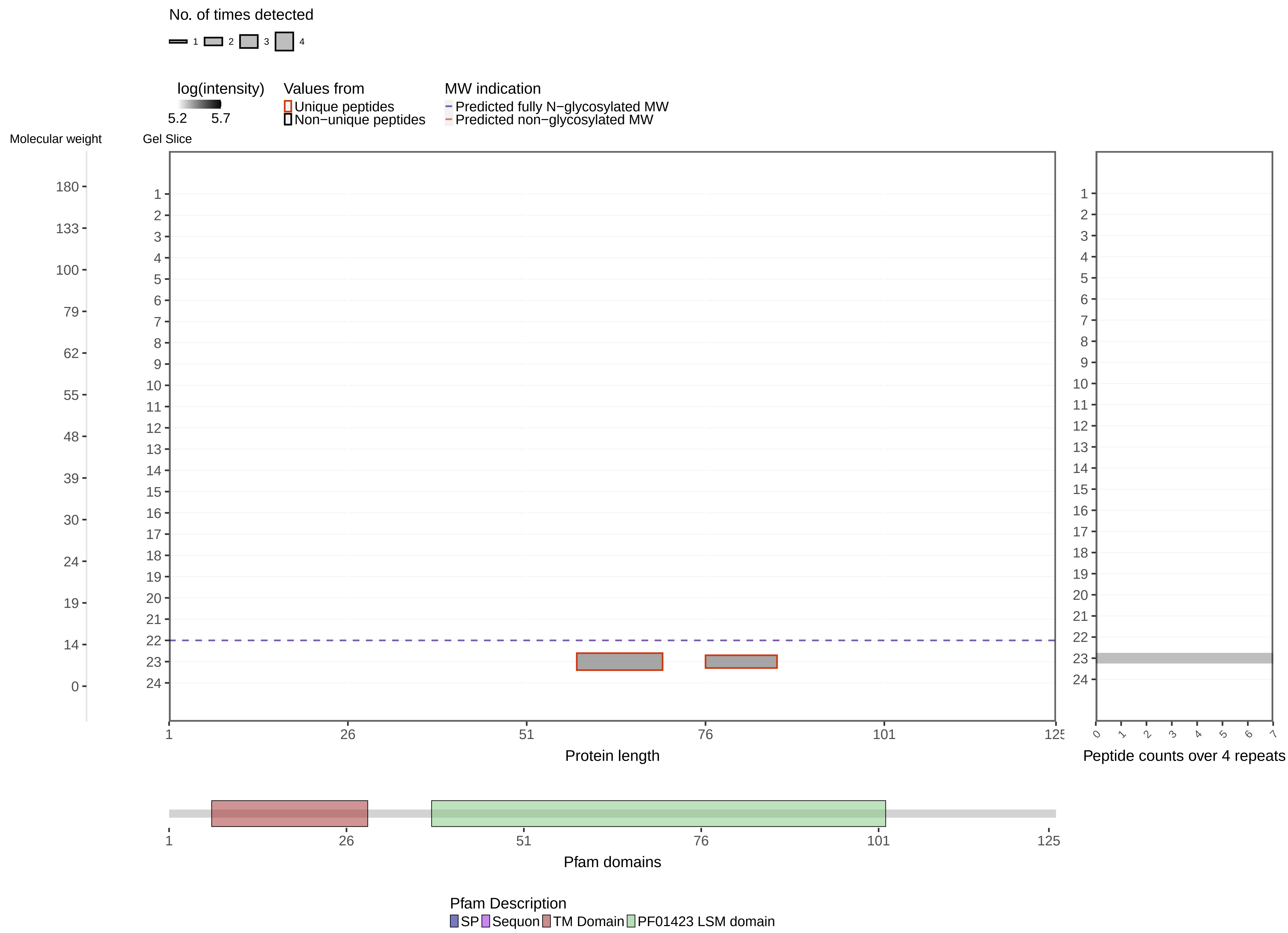


Protein sequence:

KNDLAIRGTLHSVDQYLNIKLENTRVVDEDKYPHMKSVRNCFIRGSVVRYVQLPPDGVDIELLHDATRREARGG*

### NbL01g20550.1 elongation_factor_Tu_mitochondrial [MW = 49.23 kDa]

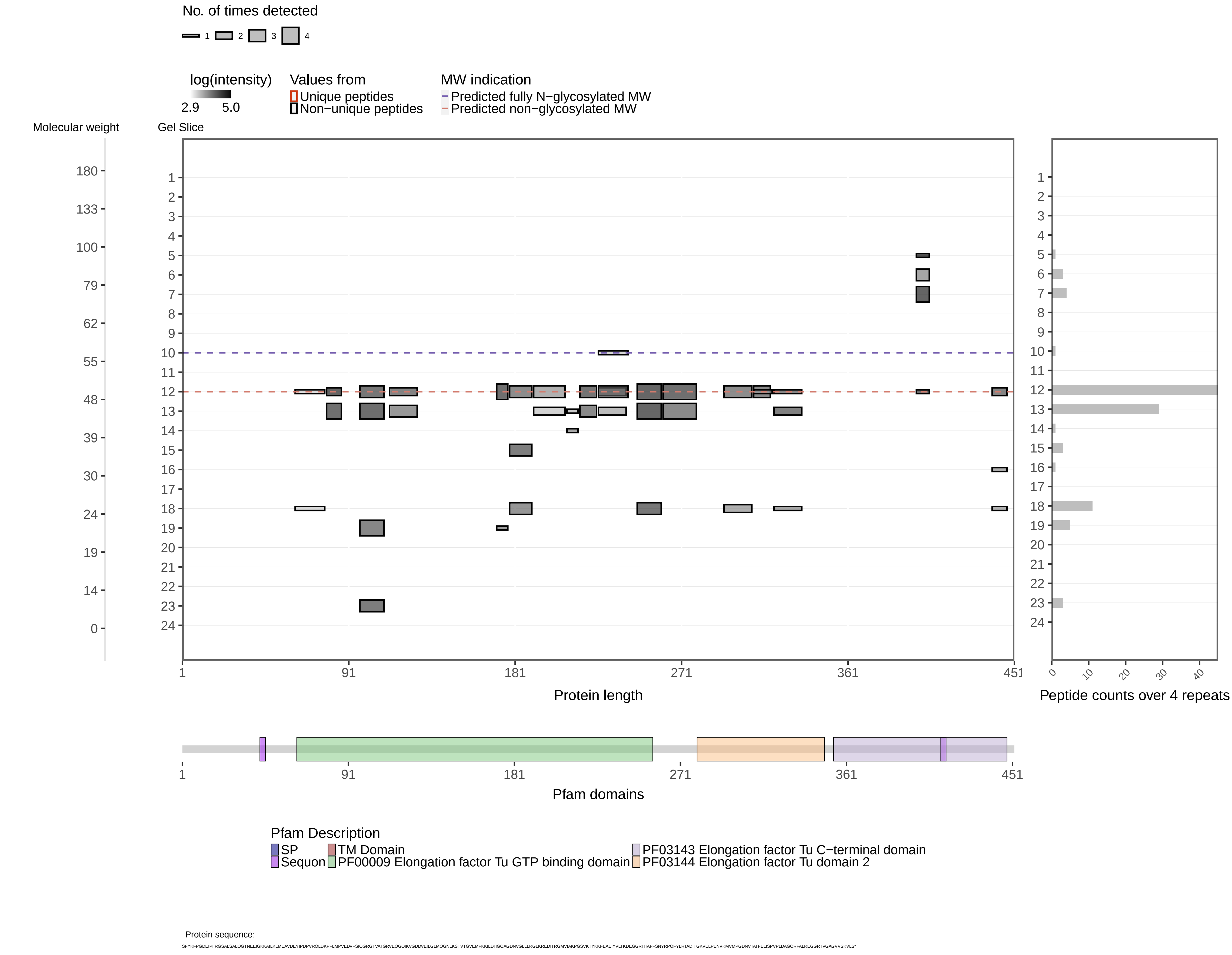

### NbL01g20300.1 probable_transmembrane_GTPase_FZO−like_chloroplastic [MW = 102.28 kDa]

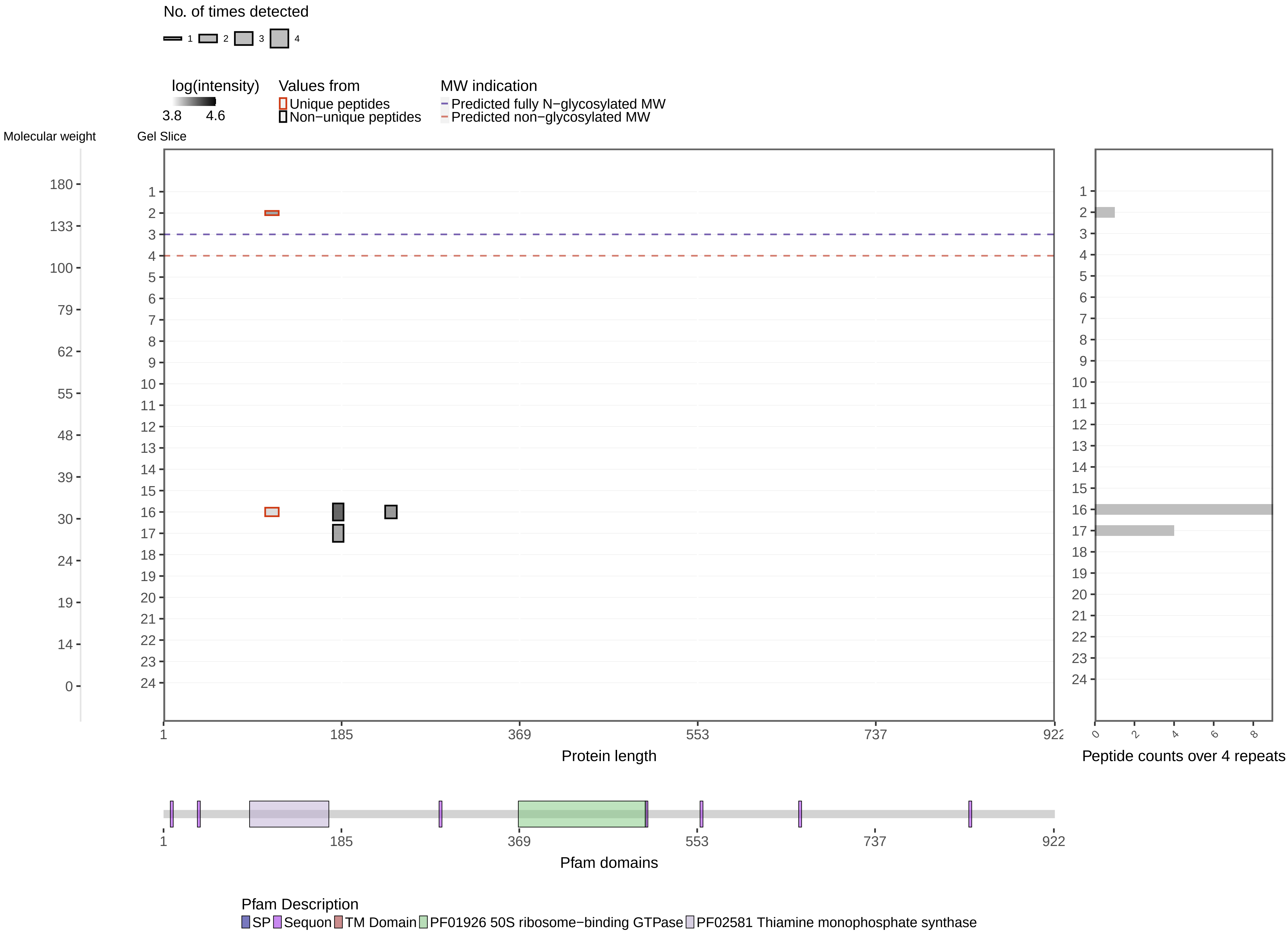


Protein sequence:

QRQQKLTEEFVPRADLLLFVMSADRPLTESEVNFLRYTQQWSKKVVFVLNKSDIYQNKGELEEAIGFIRENTQKLLNTDCVTLYPVSARLALEAKLSTFDGALSQSNRSSNDNFHWKTKGFYELEKYLYSFLDASTSTGIERMKLKLGTPIAIAEQLVSACQGLVRQECQQAKQDLLFVEDLVNSVKECTKKLEIDSILWKRQVLSMINSTQARVVRLVESTLQLSNVDLVTSYVFRGEKSTQMPATTSIQNDILGPAVLEGQNLLGEYTKWLQSKRDQEGQFYKQSFEKKWTS

### NbL01g20290.1 chloroplast_stem−loop_binding_protein_of_41_kDa_a_chloroplastic [MW = 44.17 kDa]

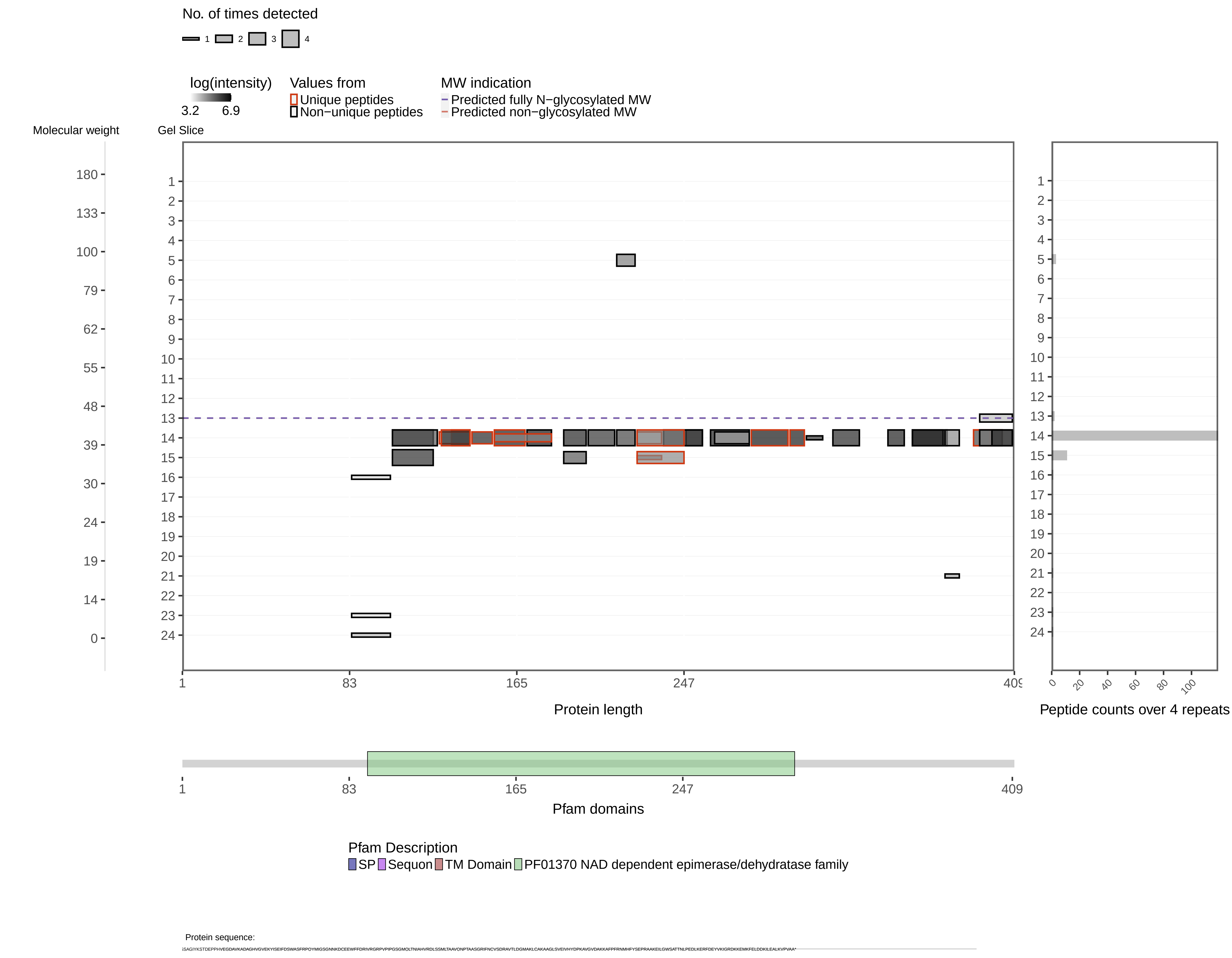

### NbL01g20280.1 RAN_GTPase−activating_protein_1−like [MW = 58.63 kDa]

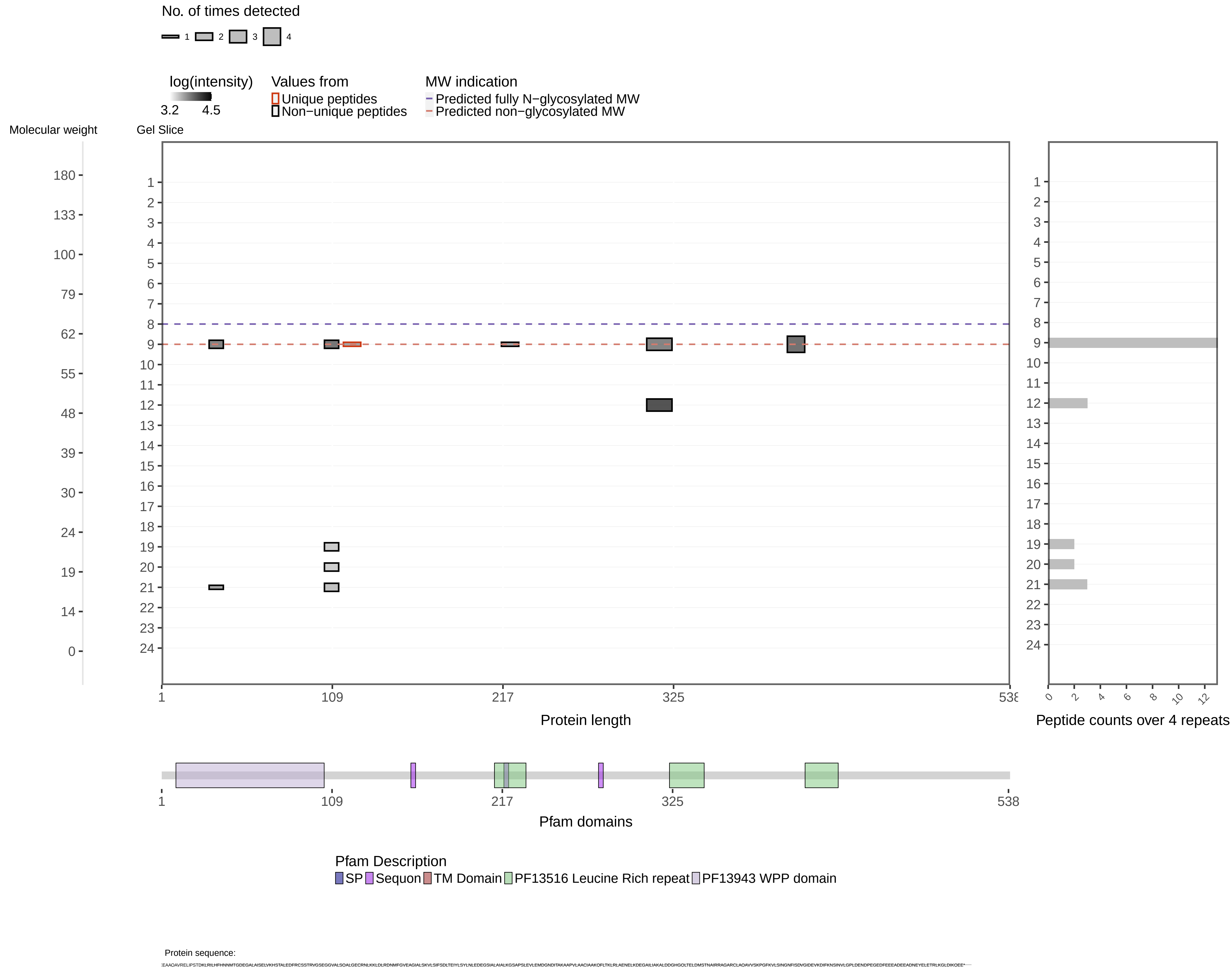

NbL01g20230.1 ribosome−recycling_factor_chloroplastic−like [MW = 35.05 kDa]

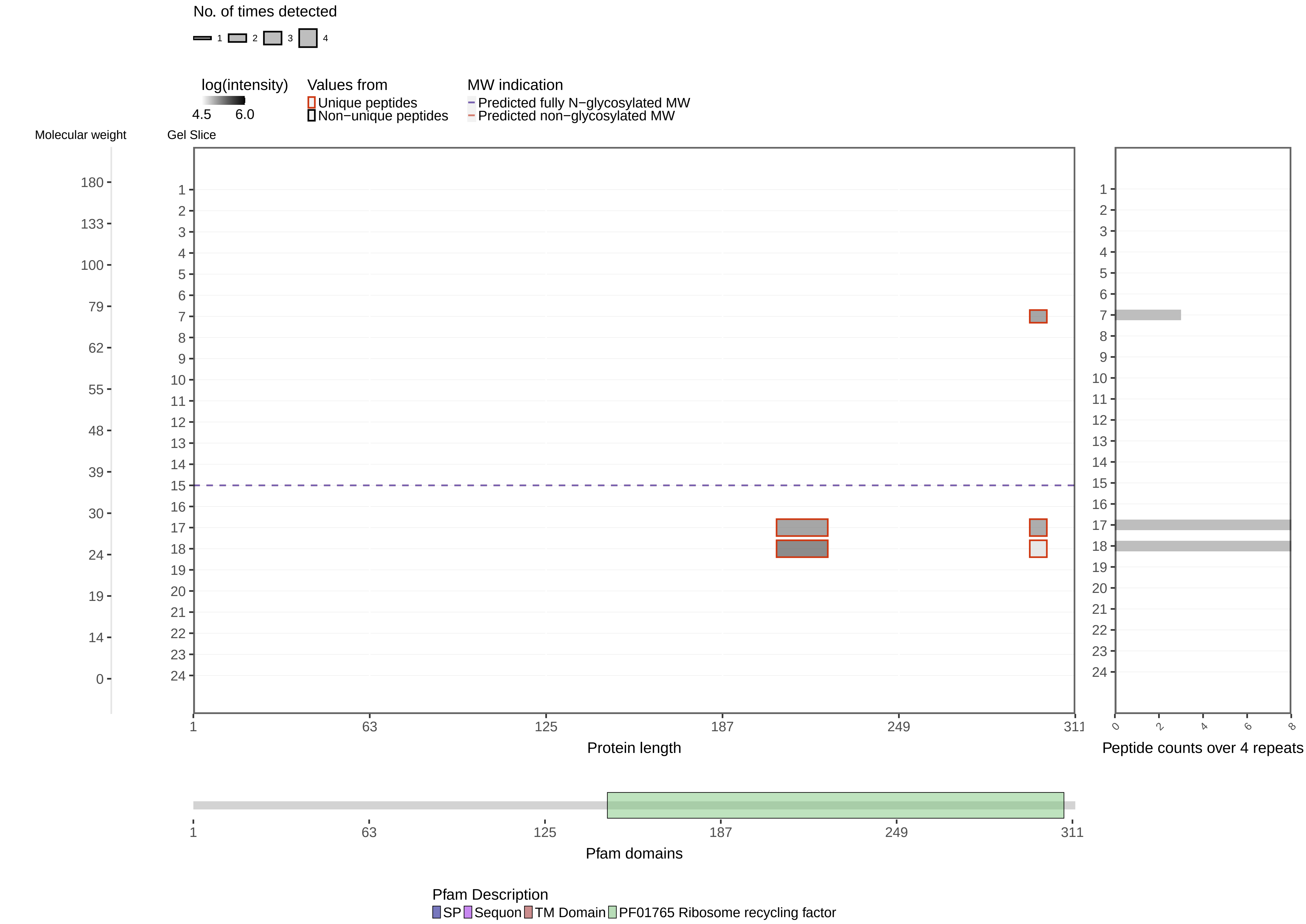


Protein sequence:

ENVRSNFNSIRTGRASPAMLDKIEVDYYGTPVSLKSIAQISTPDSSSILVQPYDKSSLKAIEKAIVGSDLGMMPNNDGEVIRMSVPQLTSDRRKELSKIVSKQAEEGKVALRNIRRDALKAYEKLEKEKKLSEDNVKDLSGDLQKVTDEYMKKVDTVFKQKEKELLTV*

### NbL01g20180.1 40S_ribosomal_protein_S24−1 [MW = 15.38 kDa]

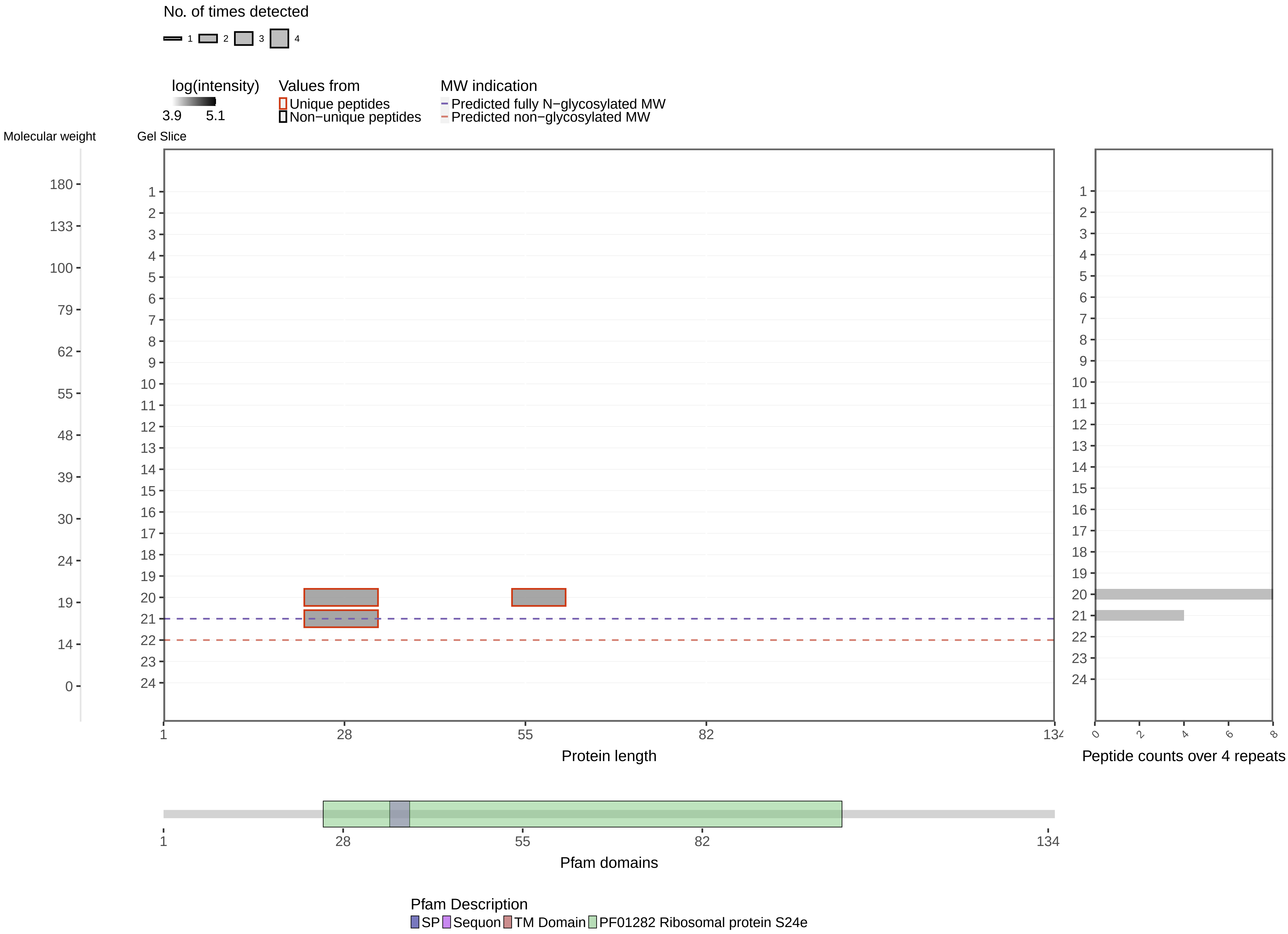


Protein sequence:

NAIFVFKFRTHFGGGKSTGFGLIYDSVENAKKYEPKYRLIRNGLDTKVEKSRKQMKERKNRAKKVRGVKKTKAGDAKKK*

### NbL01g19940.1 translocase_of_chloroplast_159_chloroplastic [MW = 158.11 kDa]

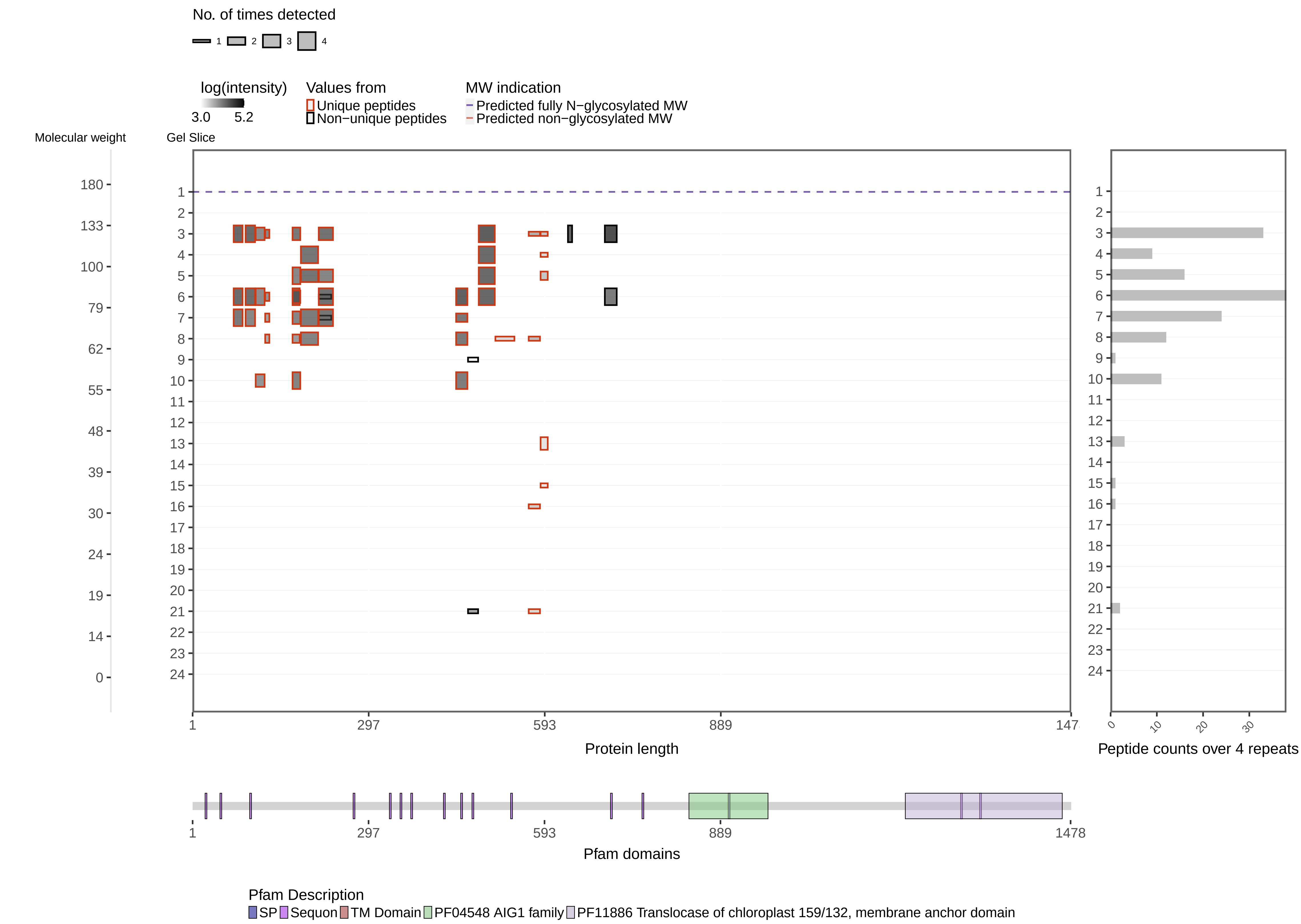


Protein sequence:

GLGSSLRSLRPAPQPNRPNLFTPSSLQNSGESENNLTEEEKKKLEKLQQIRVKFLRLIHRLGLSSDESIAAQVLYRLALIARRQNSPLFSLEAAKMRALQLEAEGEDDLDFSVNIQVIGKSGVGKSATINSIFGEEKTPINAFGPATTSVKEISGVVEGVRIRVFDTPGLKSSVMEQGFNRSVLSSAKKFTKKNPPDIFLYVDRLDAQTRDLNDLPMLKTITSCLGPSIWRSAIVTLTHGASAPPDGPSGSPLSYEVFVTQRSHVVQQSIGQAVGDLRMMSPSLMNPVSLVENHPS

### NbL01g19930.1 uncharacterized_protein_OsI_027940 [MW = 24.09 kDa]

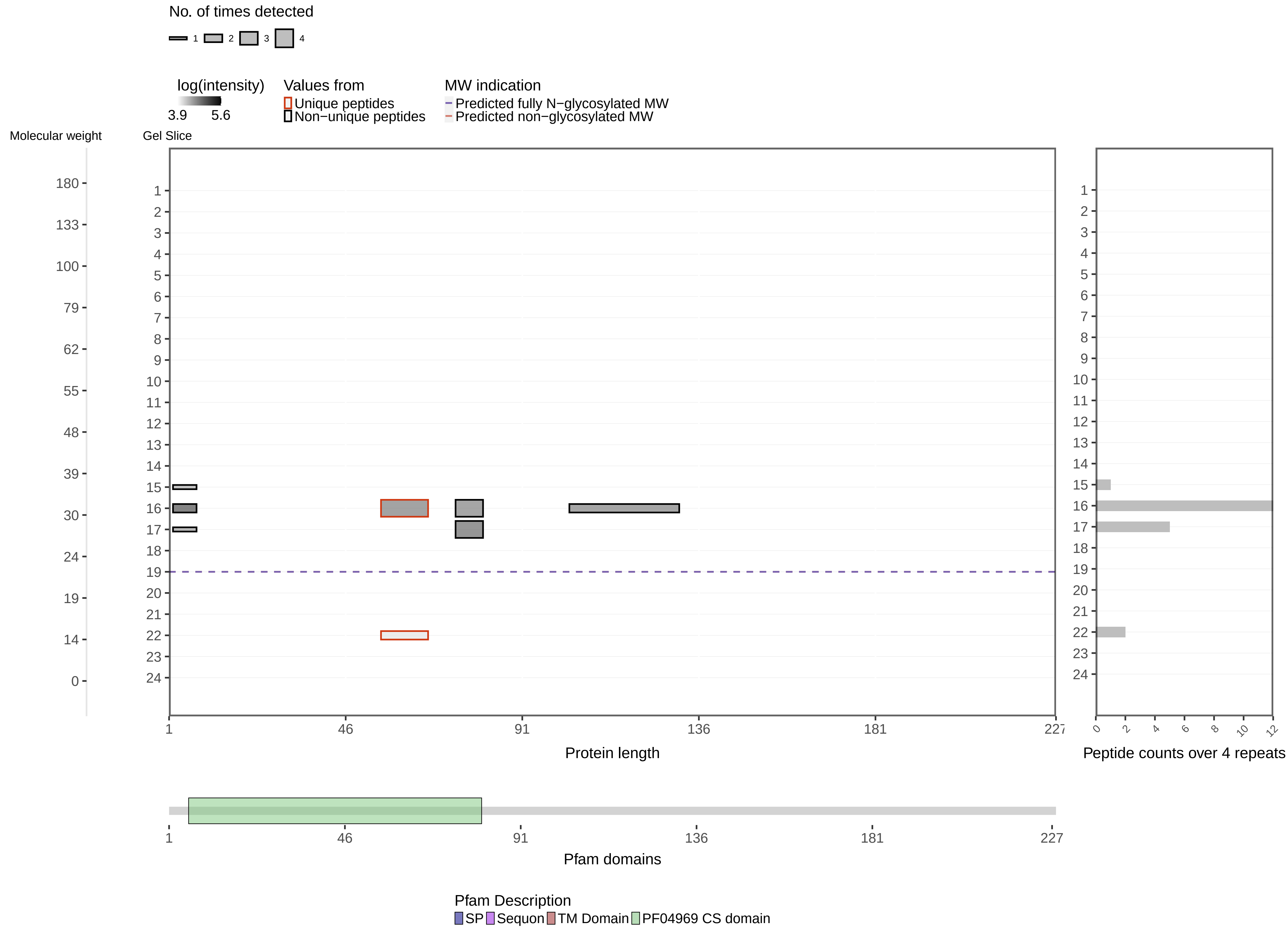


Protein sequence:

:WVDEDDEADAGPGDMDMNSMDFSKFGDMGGMMGGMGGDMAGMMGGMGGMGGMMGGMGGMMGGMGGDMAGMMGGMGGDMGDDLDDSDDEEQEVTKPSAKADEKVDGGPKEEGSKVADEAK*

### NbL01g19750.1 UDP−glucuronic_acid_decarboxylase_2−like [MW = 49.23 kDa]

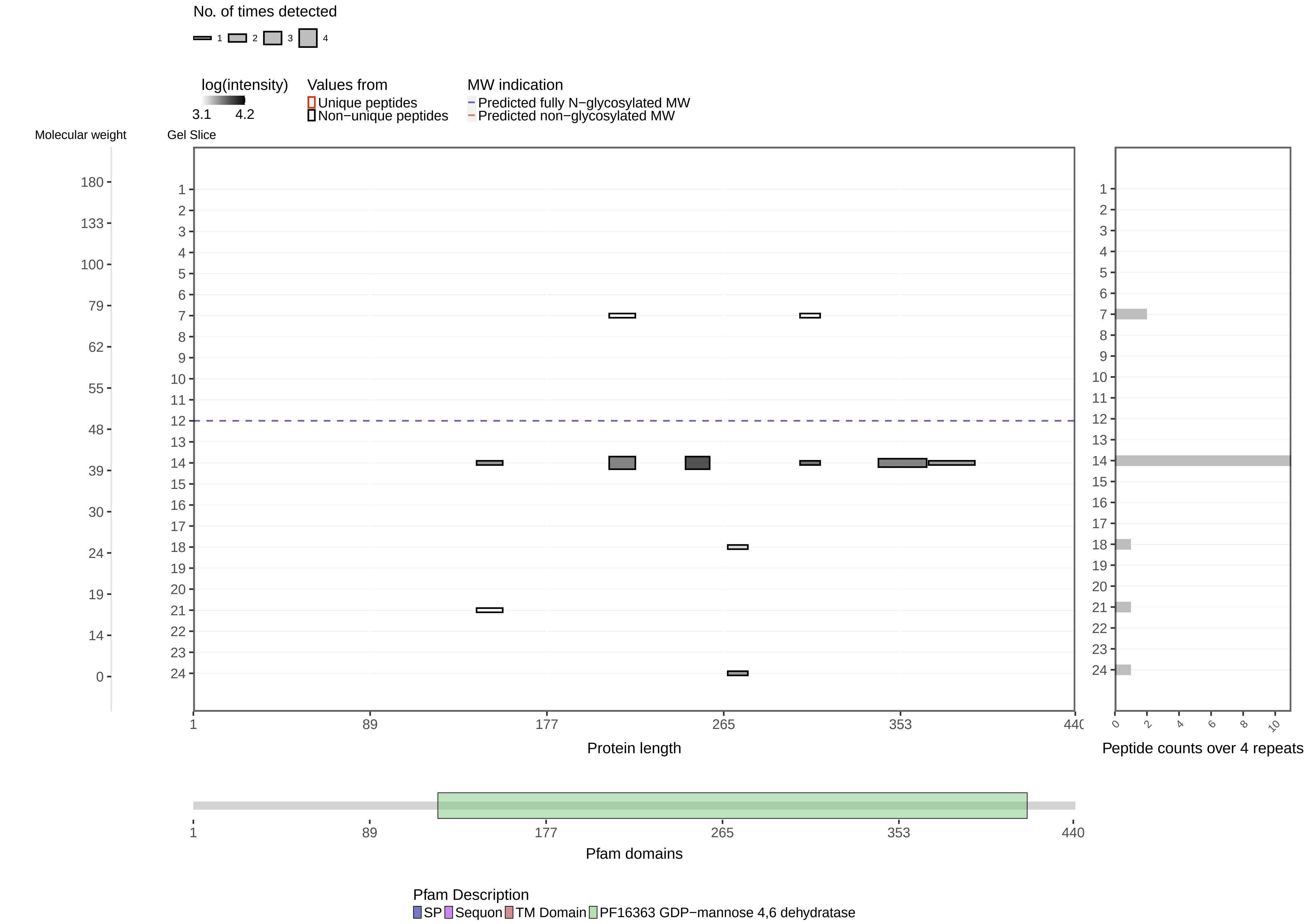


Protein sequence:

/VGTLNMLGLAKRVGARFLLTSTSEVYGDPLEHPQKETYWGNVNPIGVRSCYDEGKRTAETLTMDYHRGAGVEVRIARIFNTYGPRMCIDDGRVVSNFVAQALRKMPLTVYGDGKQTRSFQFVSDLVEGLMRLMEGEHVGPFNLGNPGEFTMLELAKVVQETIDPNAQIEFRPNTEDDPHKRKPDISRAKELLGWEPKVALRKGLPMMVQDFRQRIFADHKEDTSSATIA*

### NbL01g19610.1 pectinesterase−like [MW = 64.5 kDa]

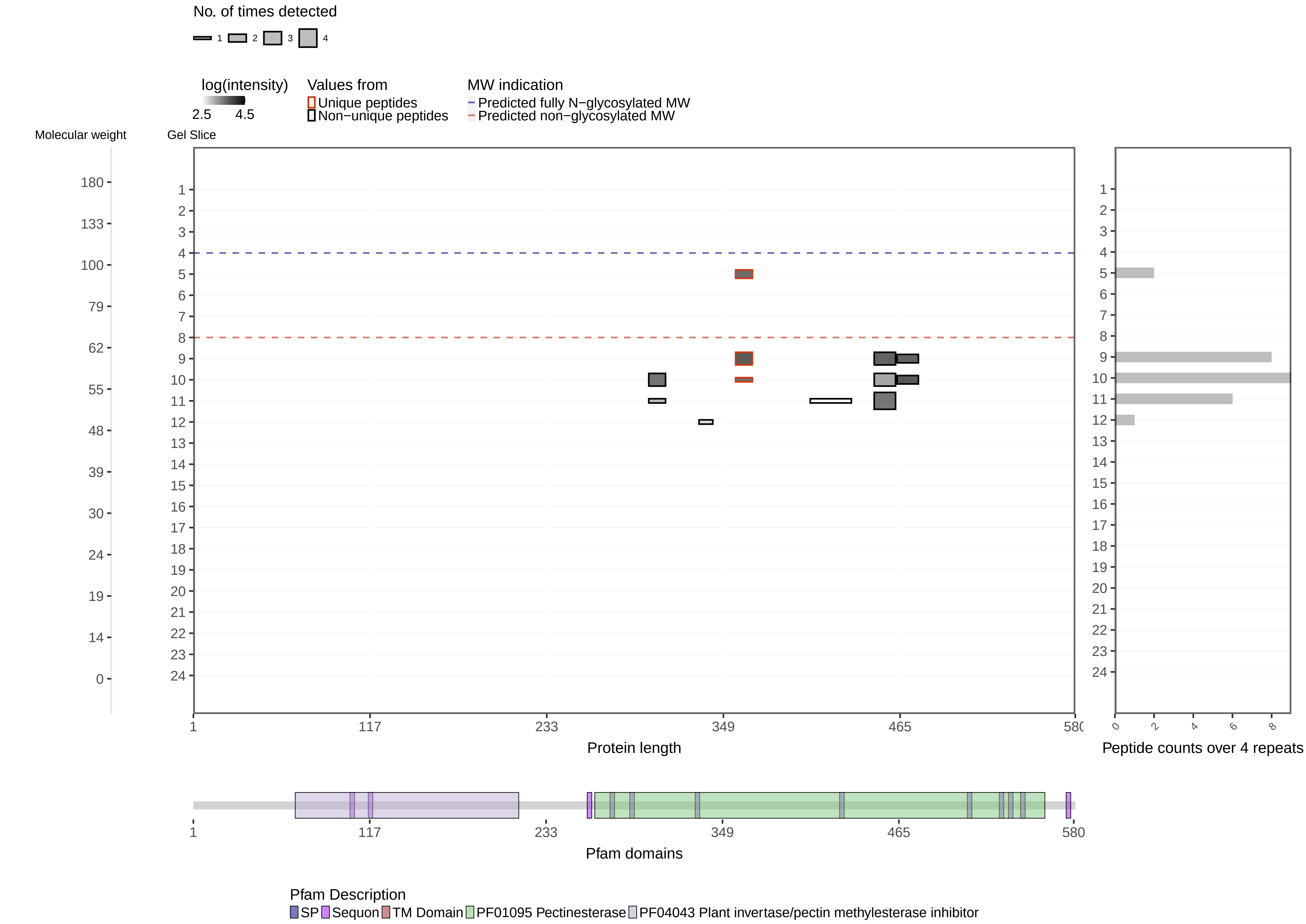


Protein sequence:

NEALSAAPNASRTRFVIHIKAGAYYEYIEVVRKKPMIMFLGDGIGKTLIKGNRSVVDGWTTFRSATVAVVGNGFIAKGITFENFAGASKHQAVAVRSGSDLSAFYQCSFVAYQDTLYVHSLRQFYRDCDVYGTVDFIFGNAAVVFQNCSLYARKPNENQKNIFTAQGREDPNQNTGISILNSKIAAAADLIPVQSSFKNYIGRPWKEYSRTVIMLTHIGSLIEPAGYLEWNGTFALDTLYYGEYKNRGPGSNTSARVNWSGYRVINNSAEANQFTVGNFIEGKEWLPDTGFPFYLN

### NbL01g19560.1 60S_ribosomal_protein_L19−1 [MW = 24.91 kDa]

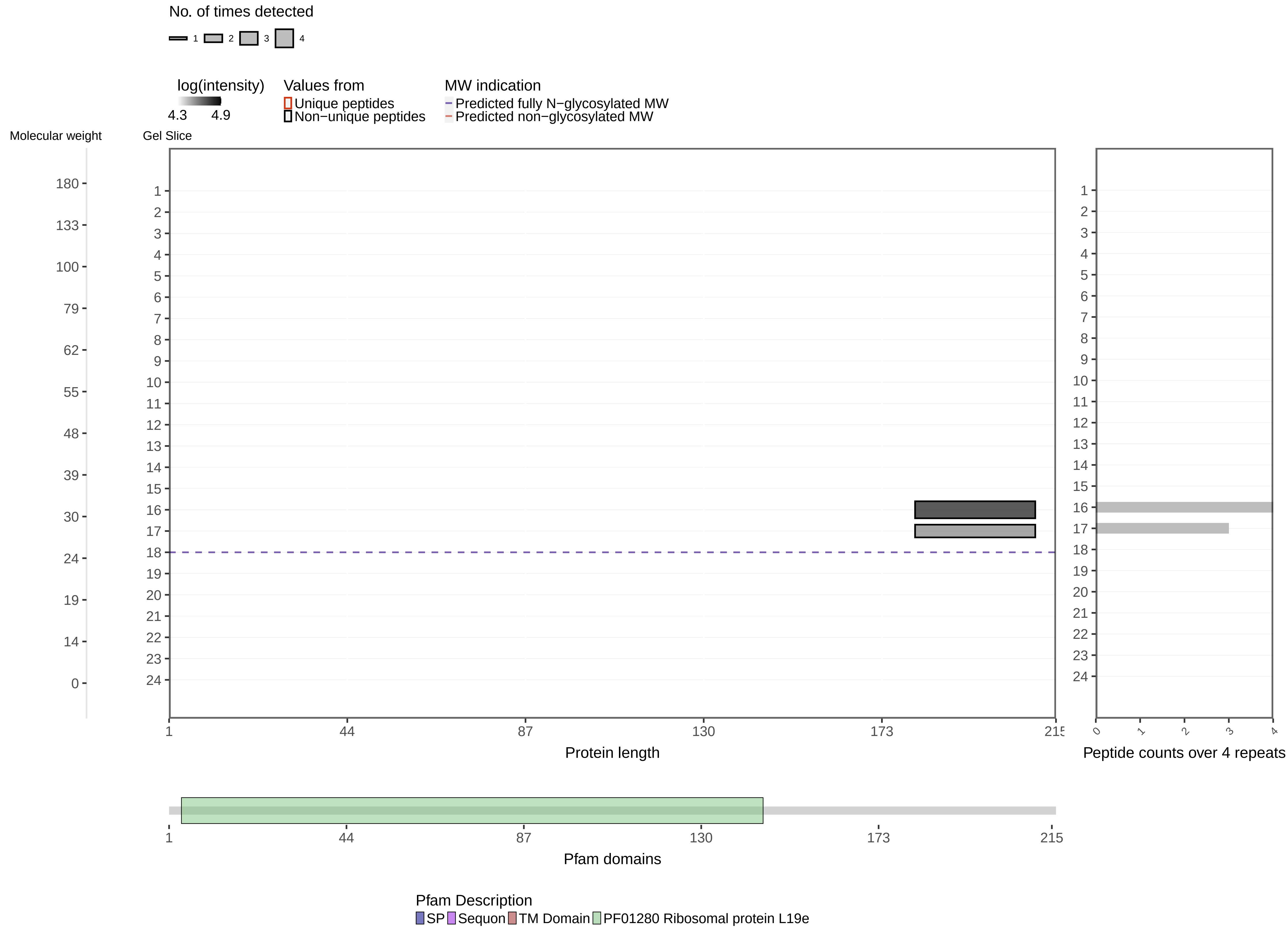


Protein sequence:

/MRRMRVLRRLLRKYRESKKIDRHMYHDMYMKVKGNVFKNKRVLMENIHKTKAEKAREKTLSDQFEARRAKNKASRERKFARREERLAQGPGERPVQPAAPAASATPAQTAQGGSKKSKK*

NbL01g19540.1 probable_polygalacturonase [MW − SP = 47.54 kDa]

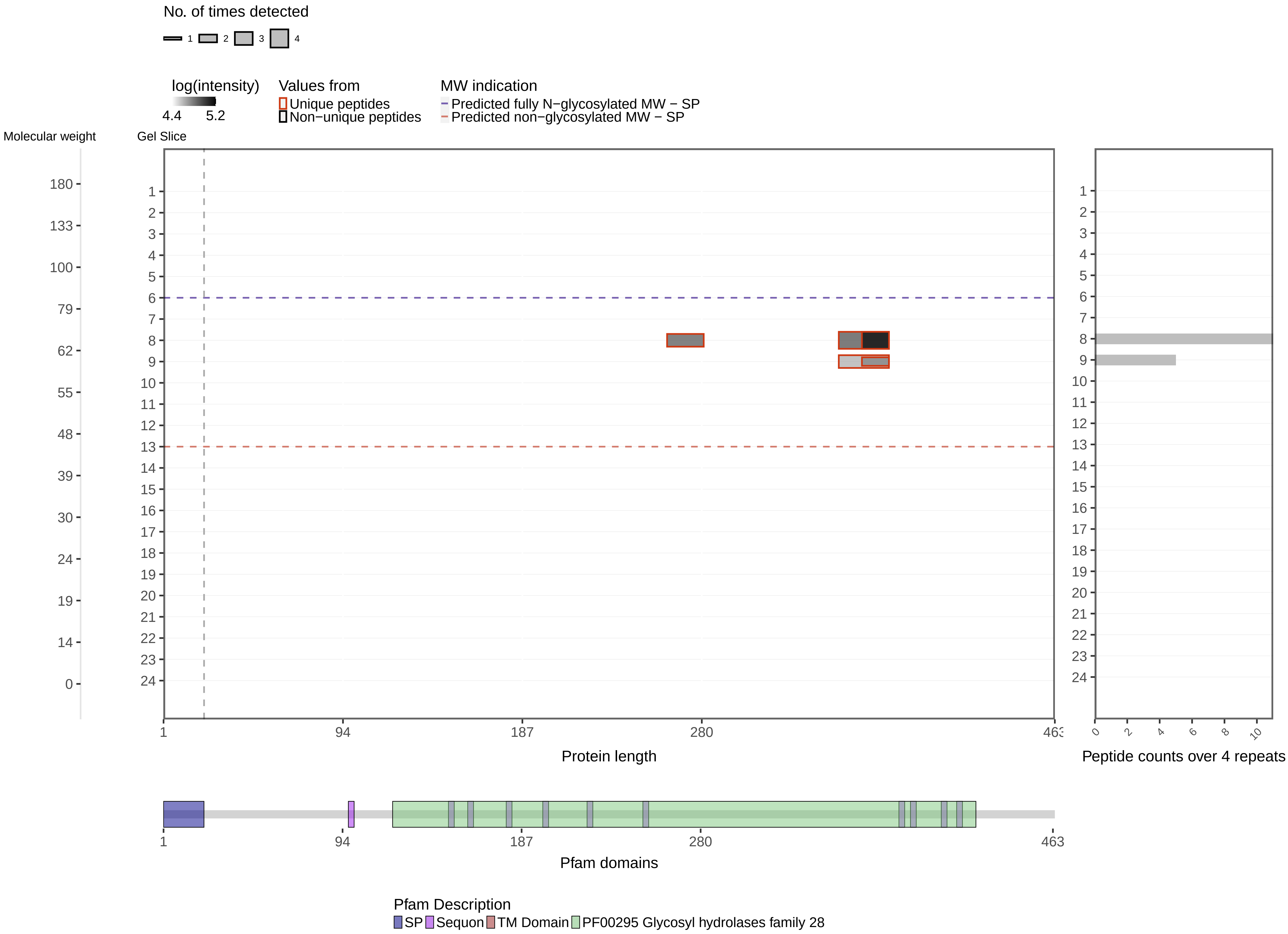


Protein sequence:

VIQNLTVHSPQDSPYTNGIVPDSSEHVCIENSNITIGYDAVVLKSGWDEYGISYGKPTSNVHIRGVRLQSSAGSAVAFGSEMSGGISNVLVEHISLHDSLFGIDLKTARGRGGYIKDILIMNVVMENVQVGIKATGYYDSHPDEKYDPSSLPVVSGITFKDIVGTNISIAGNFTGLSESPFTSICLSNISFSISSNLSTPWLCSDVSGSSENVFPEPCPELQSSFSSSSSACFALPDPYTEVAVL*

NbL01g19170.1 tubulin−folding_cofactor_A [MW = 13.75 kDa]

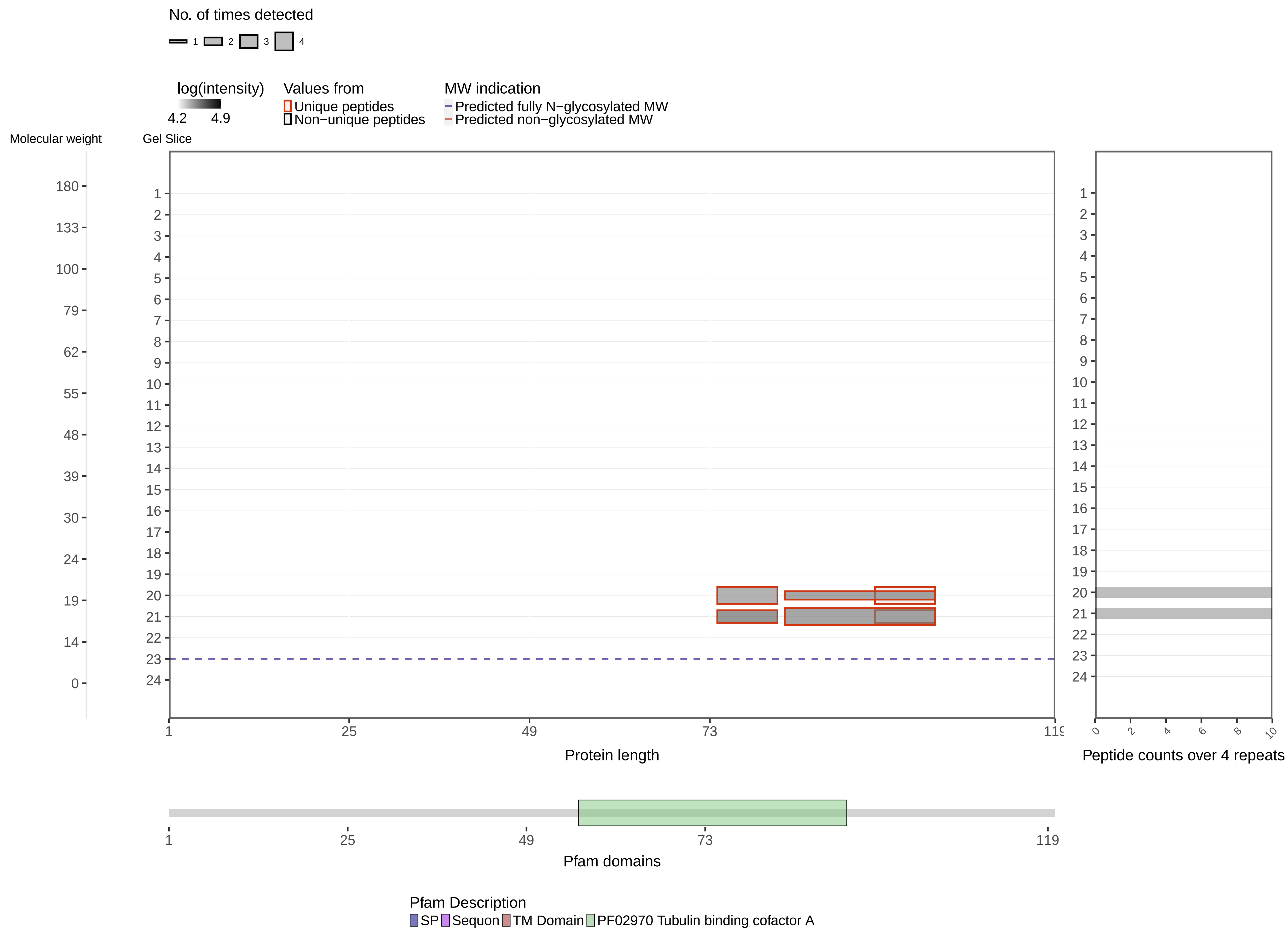


Protein sequence:

LFSSCFWVRQENVLAESRMMVPDCRKRLEAALDDLKGALVELEETDQKDGPEFEEARTIIADVEKLFESSEV*

NbL01g18990.1 40S_ribosomal_protein_S16−like [MW = 20.78 kDa]

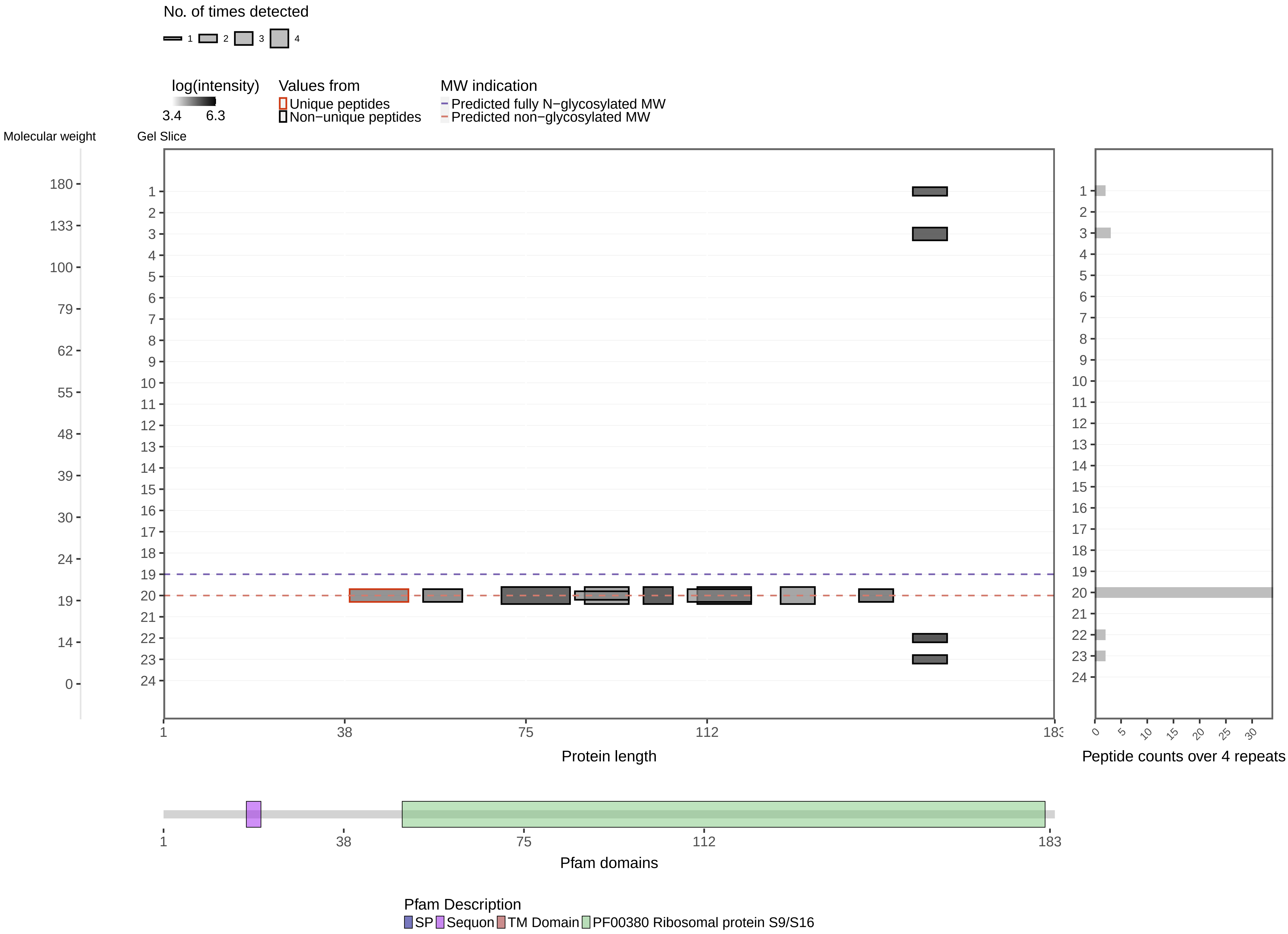


Protein sequence:

/QPEILRYKAFEPILLLGRHRFAGVDMRIRVKGGGHTSQIYAIRQSIAKALVAFYQKYVDEQSKKEIKDILVRYDRTLLVADPRRCEPKKFGGRGARARFQKSYR*

NbL01g18940.1 uncharacterized_LOC107810622 [MW = 32.17 kDa]

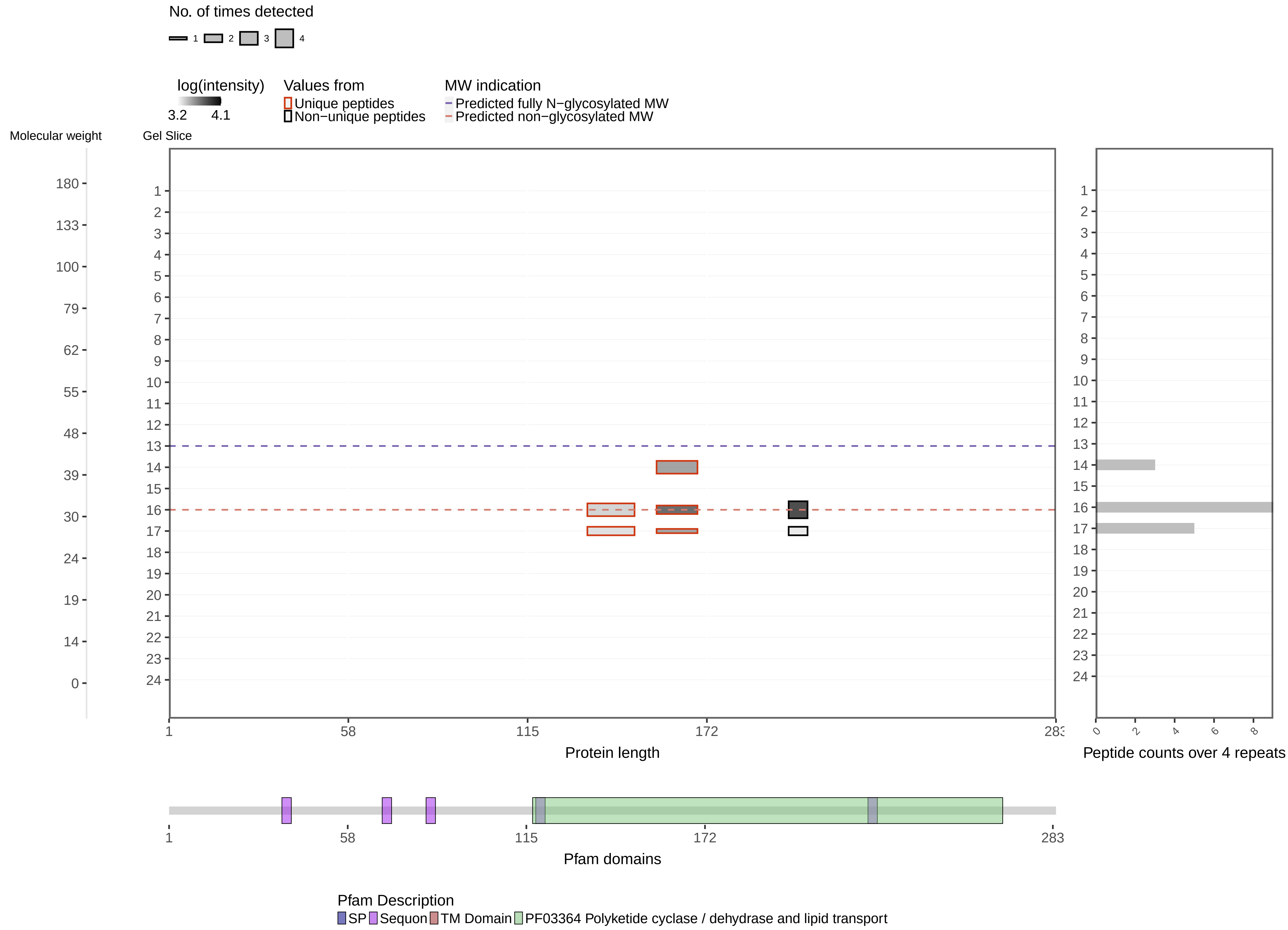


Protein sequence:

.TDYERLADFIPGLAVSQLLQKEPNFARLLQIGEQDLAFGLKFNAKGIVDCYEKDLEDLPIGQRRDIEFKMVEGDFKLFEGKWSIEQCNPGSVKEHNSSAGQPFHTTVFYIVDVEPKLWLPVRVIEGRLCKEIKTNLRSVREEAQKVYLTSLSSY*

### NbL01g18930.1 protein_transport_protein_SEC23−like [MW = 95.02 kDa]

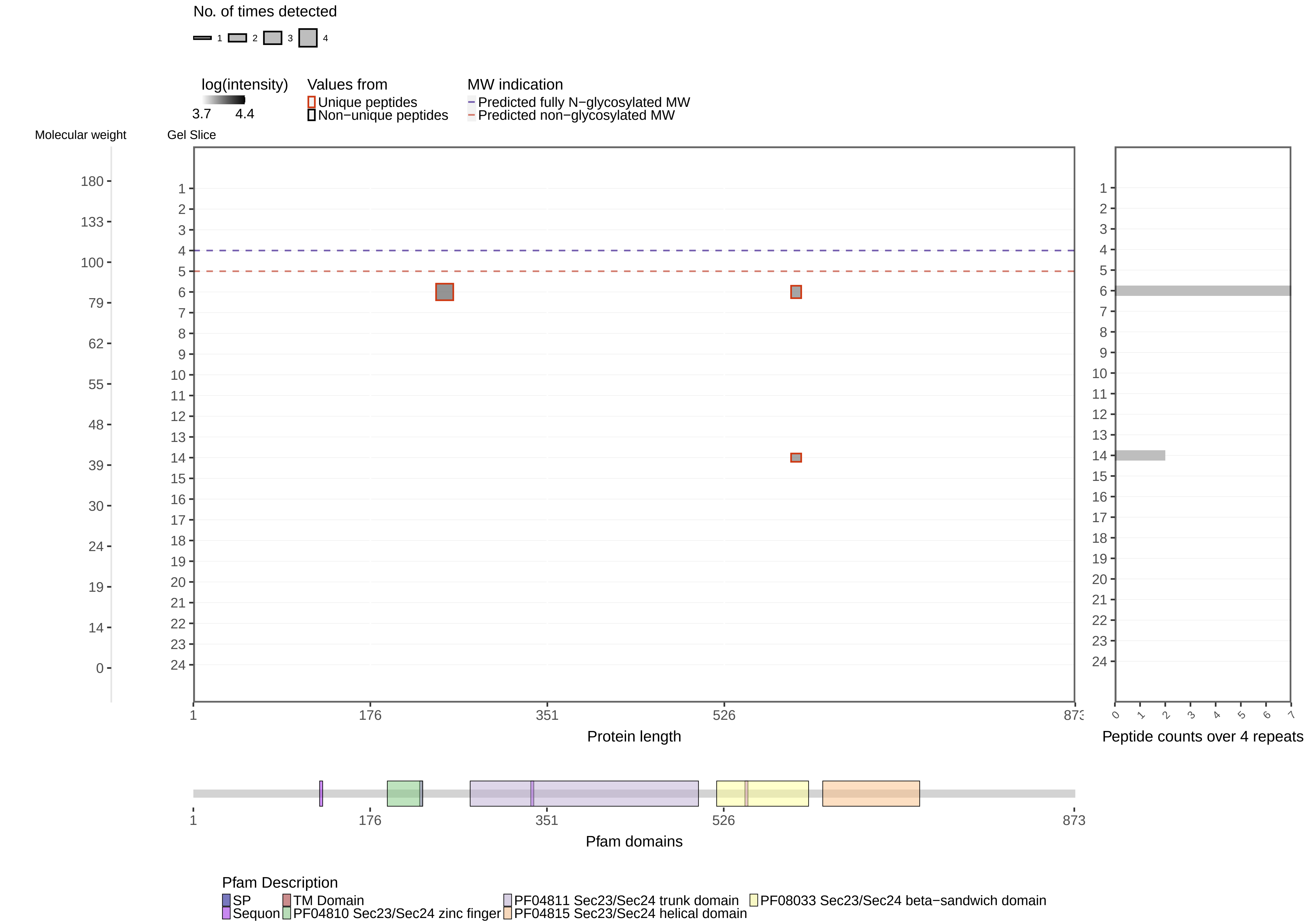


Protein sequence:

GPNTCGPGSVPYSFSHPNYAHMEKISLKWMETLGREAFRKNTVIDILCAGTCPVRVPVLQPLAKASGGVLILHDDFGEAFGVNLQRASGRAAGSHGLLEVRCSEDIFVSQVIGPGEEAHVDSNETFKNDDALVIQMLSVEETQSFALSMETKRDIKRDFVYFQFAFKFSDVYQSDITRVITVRLPTVDSVSSYLESVQDEVAAVLIAKRTLLRAKYANDALDMRVTVDERIKDITSKFGSQMPKSKLYRFPRELSVLPELLFHLRRGPLLGNILGHEDERSVLRNLFLNAGFDL

NbL01g18920.1 protein_translocase_subunit_SECA1_chloroplastic [MW = 115.22 kDa]

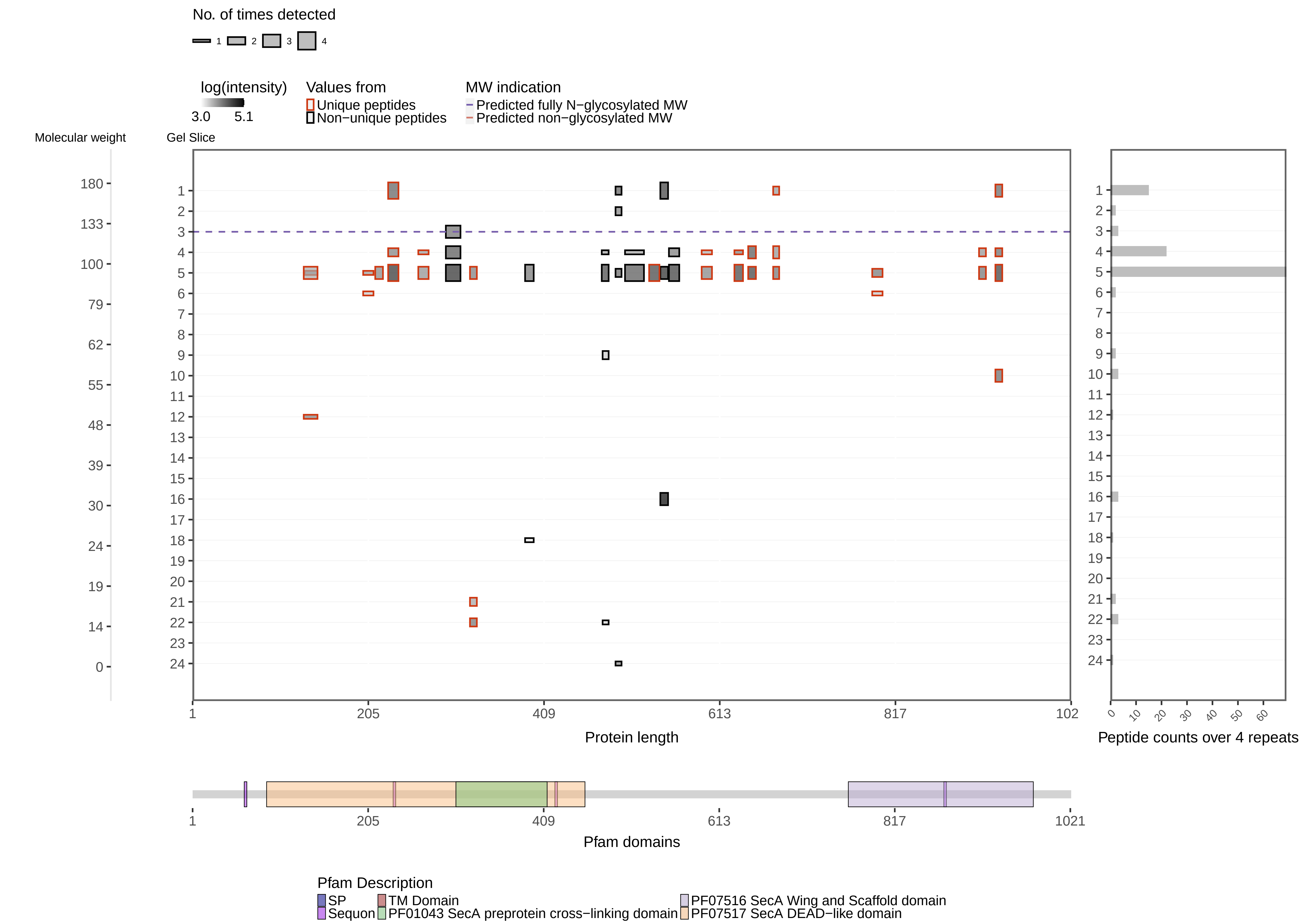


Protein sequence:

### NbL01g18890.1 protoporphyrinogen_oxidase_chloroplastic [MW = 59.65 kDa]

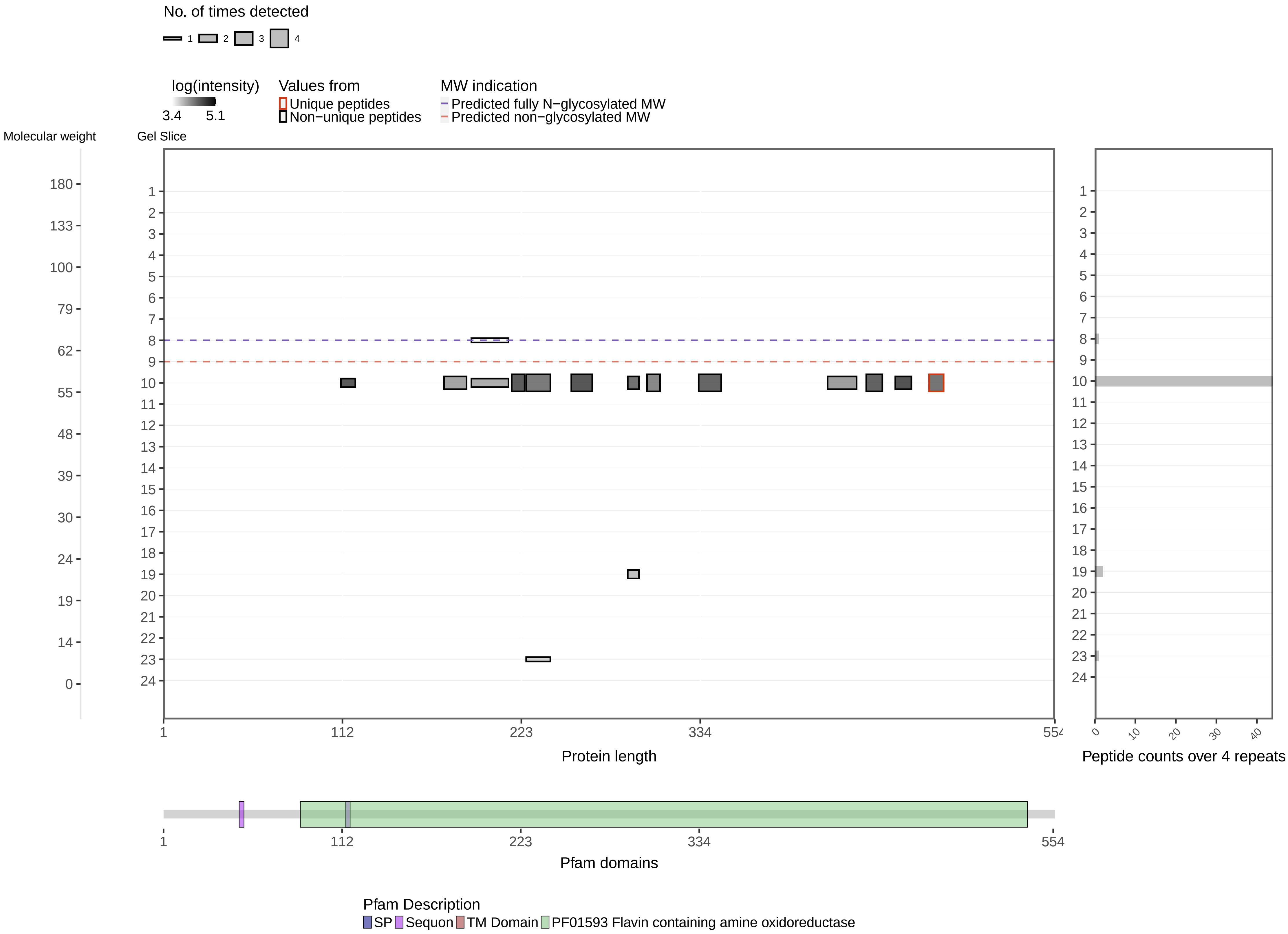


Protein sequence:

IGGTFKAIKERSSTPKAPRDPRLPKPKGQTVGSFRKGLRMLPDAISARLGSKLKLSWKLSSITKSEKGGYRLTYETPEGVVSLQSRSIVMTVPSYVASNILRPLSVAAADALSNFYYPPVGAVTISYPQEAIRDERLVDGELKGFGQLHPRTQGVETLGTIYSSSLFPNRAPKGRVLLLNYIGGAKNPEILSKTESQLVEVVDRDLRKMLIKPKAQDPLVVGVRVWPQAIPQFLVGHLDTLSTAKAAMSDNGLEGLFLGGNYVSGVALGRCVEGAYEVASEVTGFLSRYAYK*

### NbL01g18700.1 probable_pre−mRNA−splicing_factor_ATP−dependent_RNA_helicase_DEAH2 [MW = 82.08 kDa]

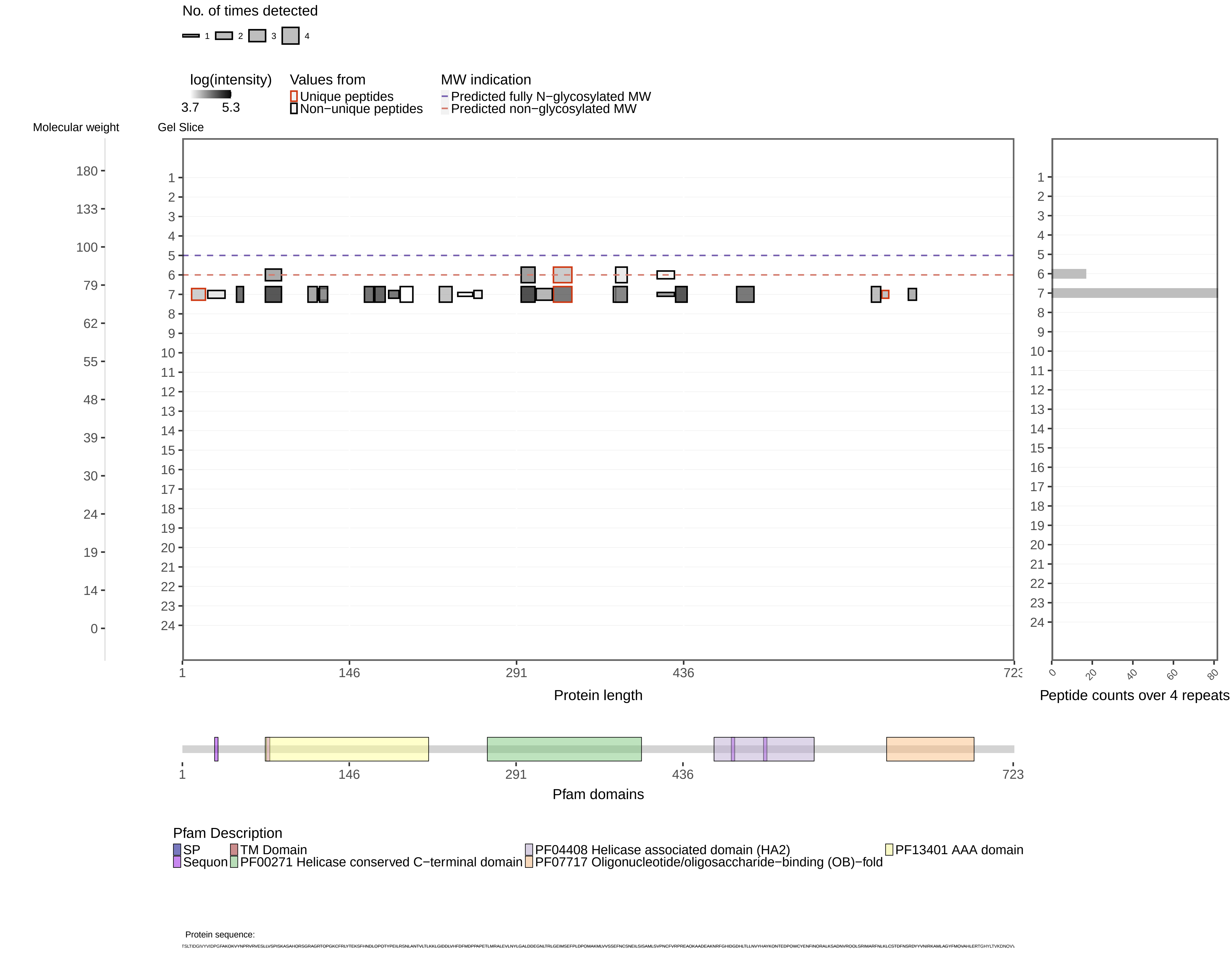

### NbL01g18690.1 long_chain_acyl−CoA_synthetase_1 [MW = 75.15 kDa]

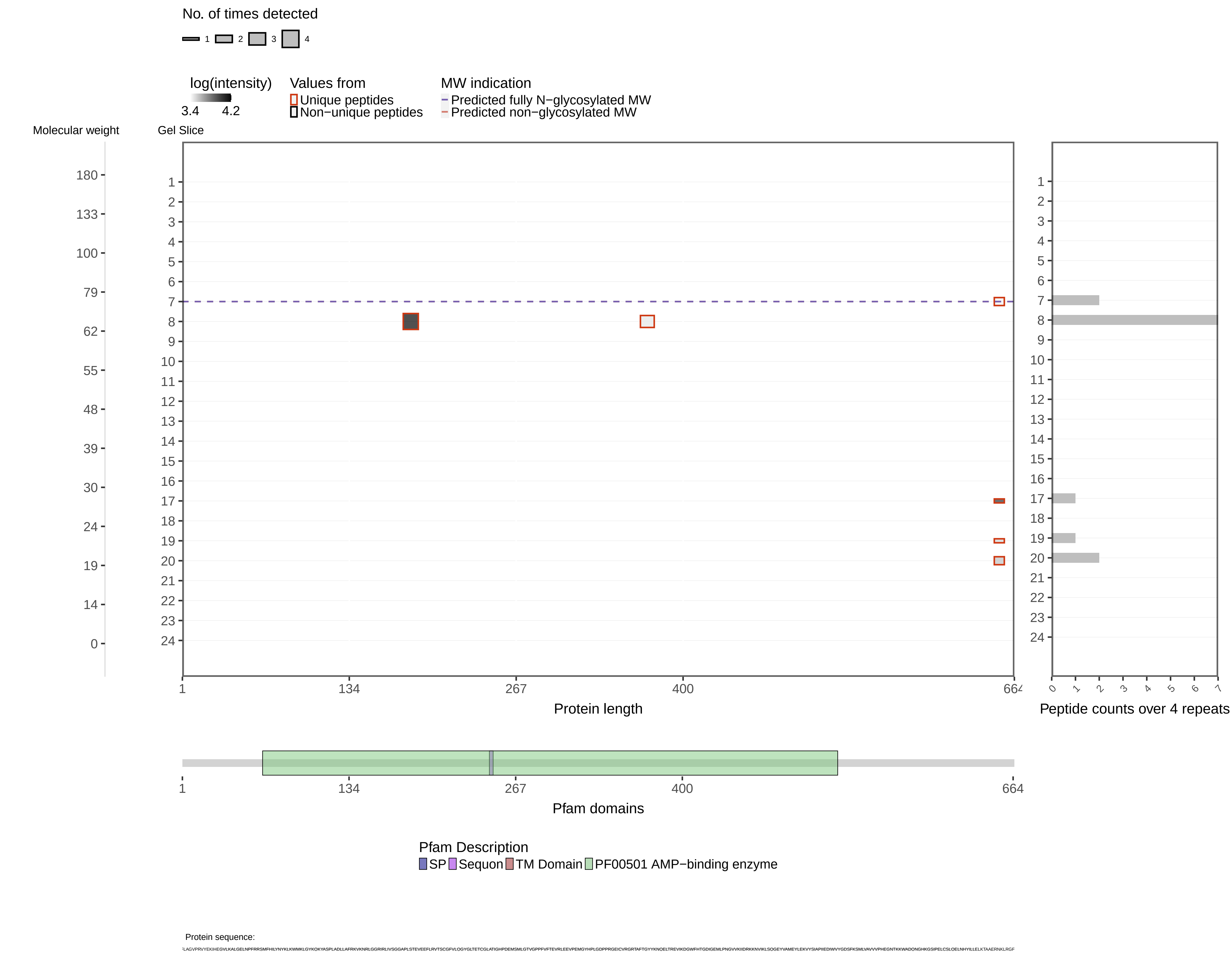

### NbL01g18620.1 ubiquitin−40S_ribosomal_protein_S27a [MW = 17.7 kDa]

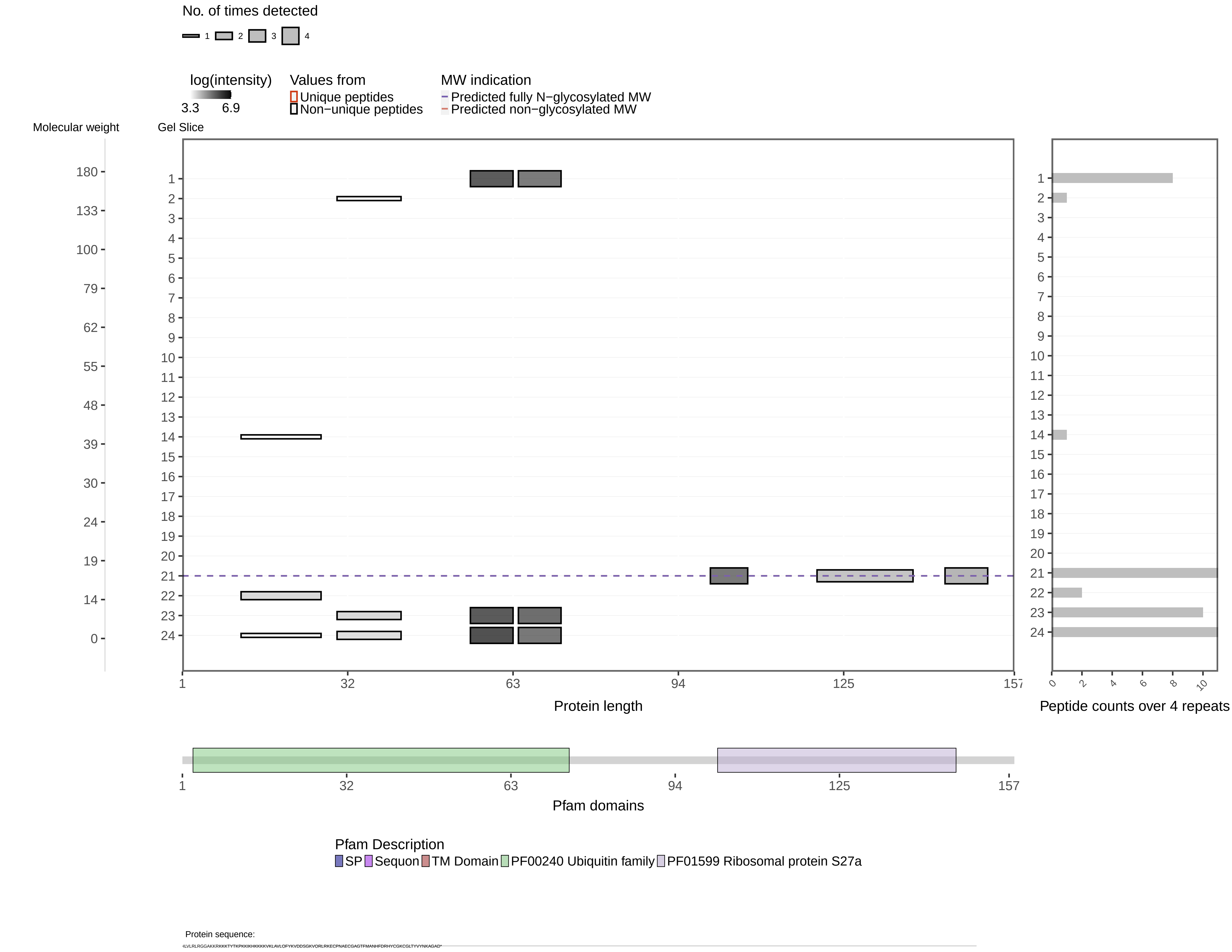

### NbL01g18510.1 probable_glutamyl_endopeptidase_chloroplastic [MW = 108.47 kDa]

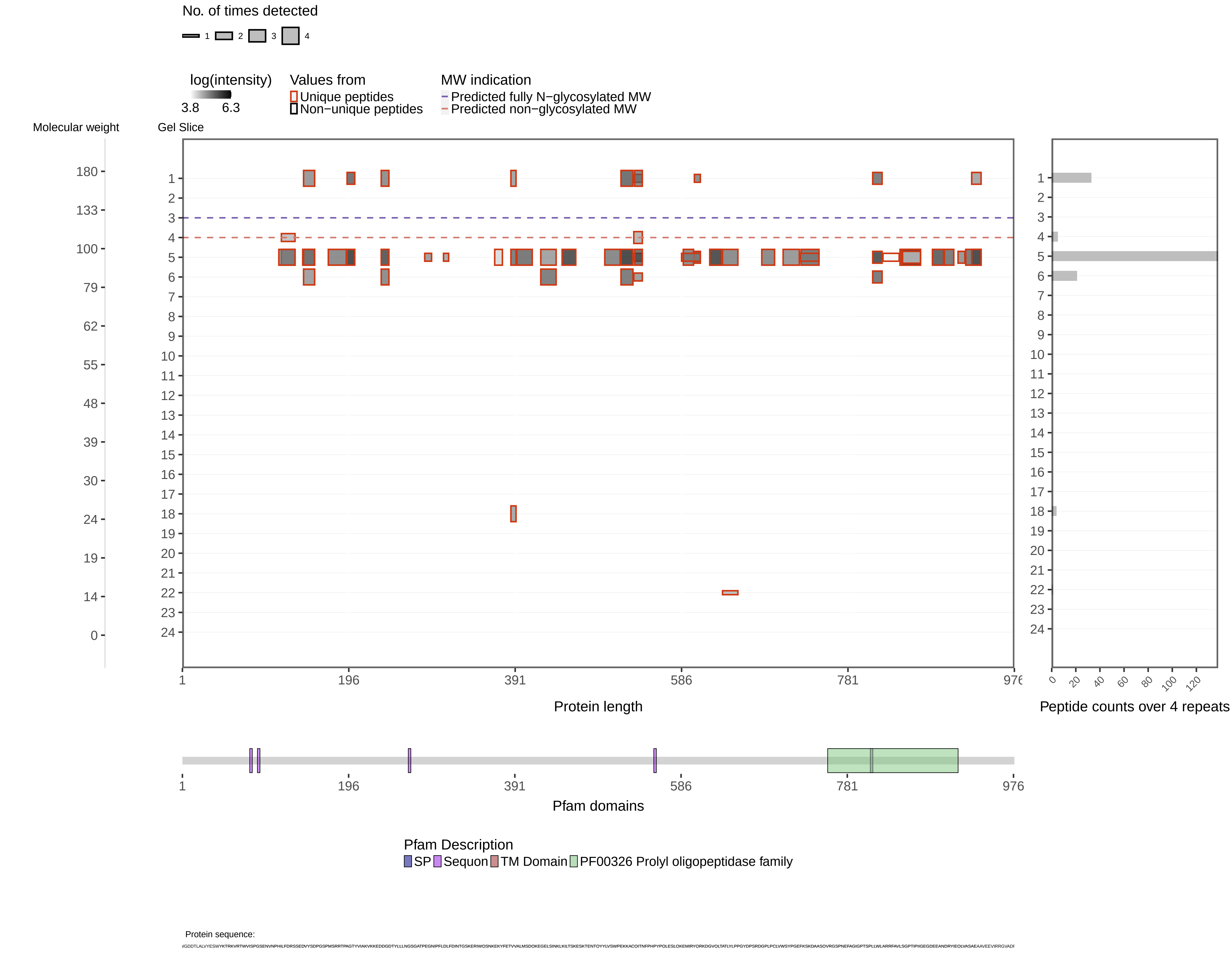

### NbL01g18490.1 calvin_cycle_protein_CP12−2_chloroplastic−like [MW = 14.06 kDa]

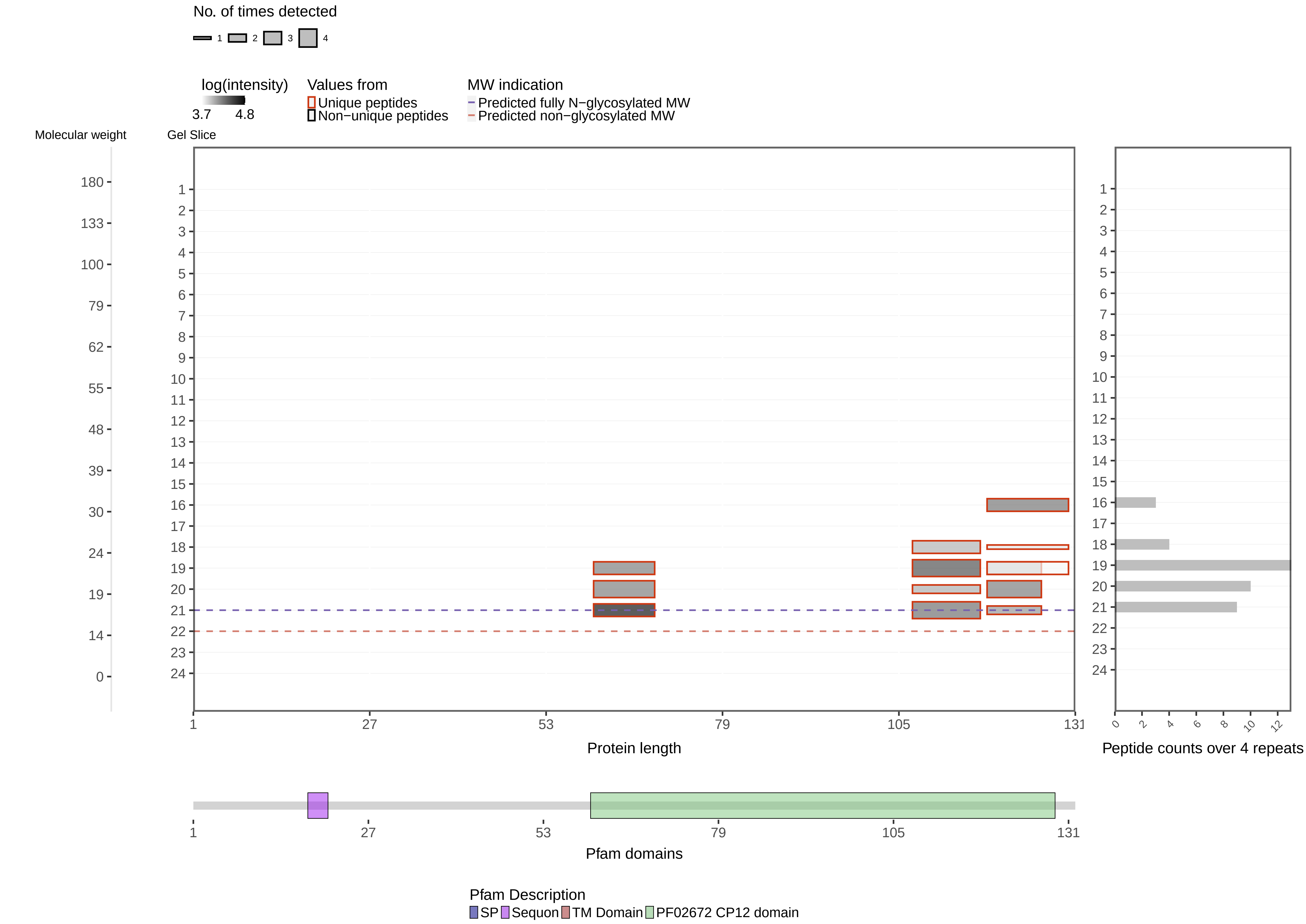


Protein sequence:

SAAPDKLSDLVADSVKGAEEACSENPASGECAAAWDVVEETSAAASHARDKQKQNSDPLEGYCKENPETDECRVYDN*

### NbL01g18390.1 uncharacterized_LOC107787269 [MW = 18.71 kDa]

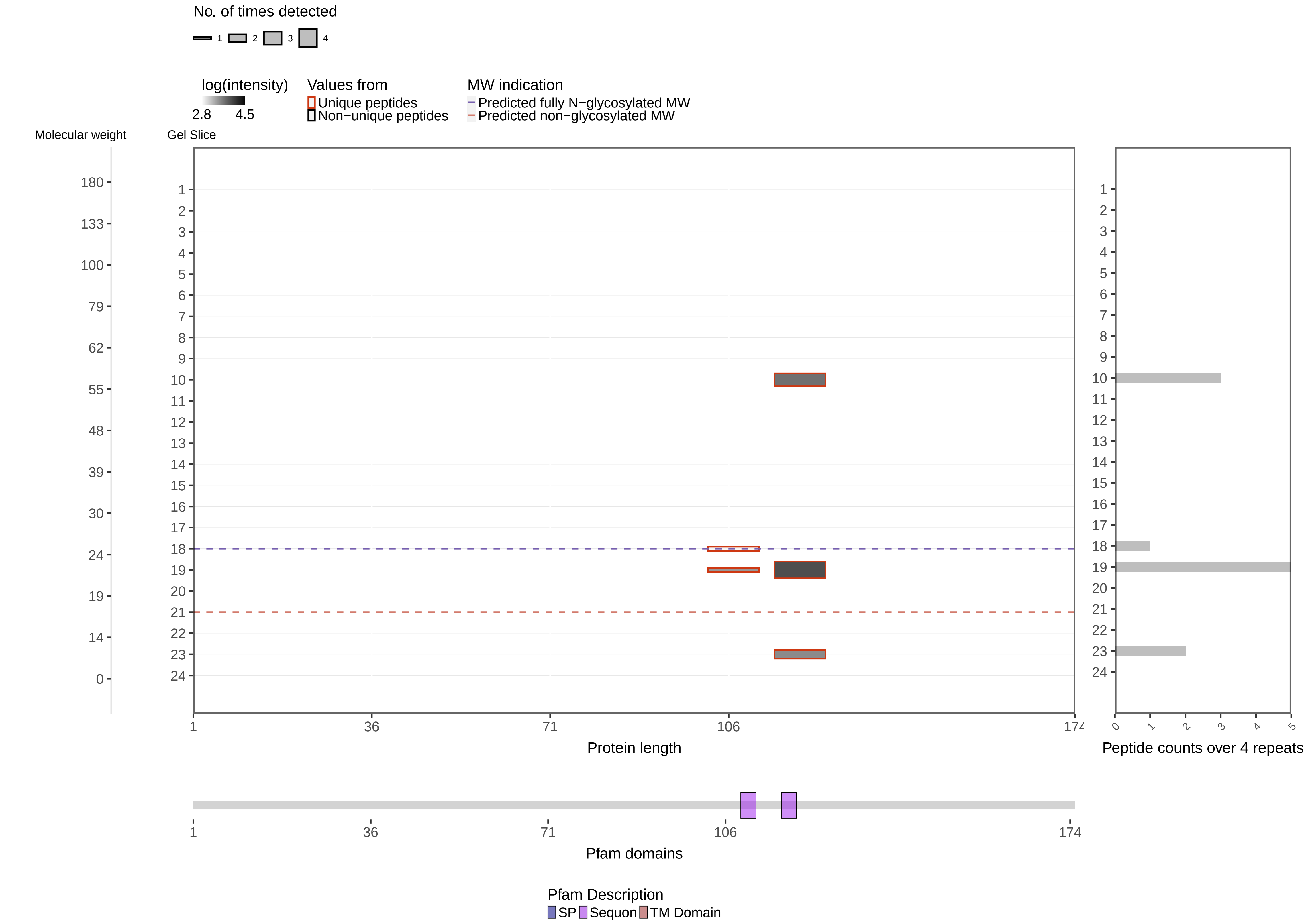


Protein sequence:

SELKLPDASVLLNSPSVAVGYMTDHSSRVAAAMAENASRKRDINVSASTYPRSKIPKGTLPHMKNVPETGSGLLVPPQLAGRSNVVTEDISKLFVKKTPS*

### NbL01g18300.1 protein_transport_protein_Sec31A−like [MW = 121.97 kDa]

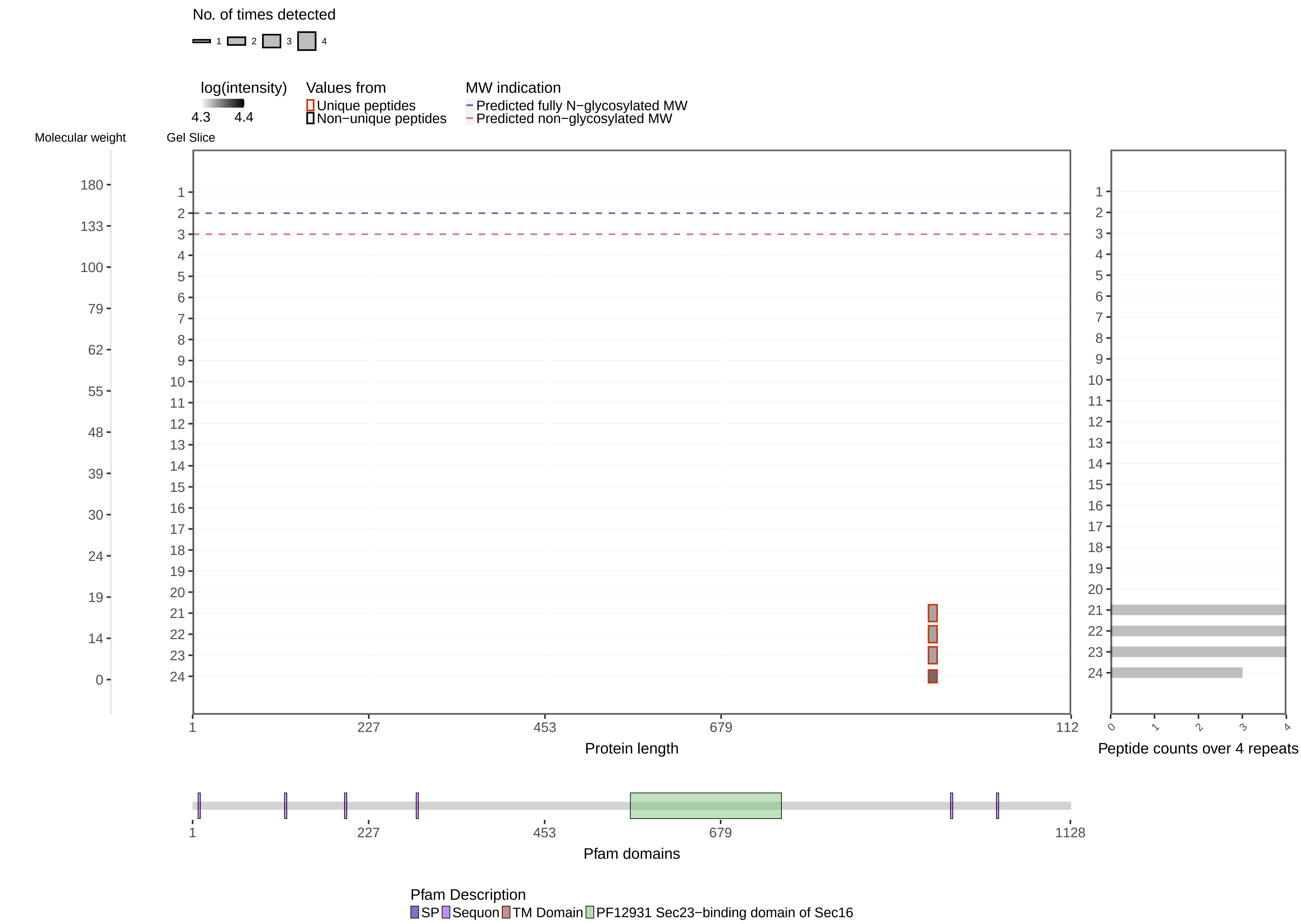


Protein sequence:

[illegible]

### NbL01g18230.1 serine_carboxypeptidase−like_40 [MW = 33.34 kDa]

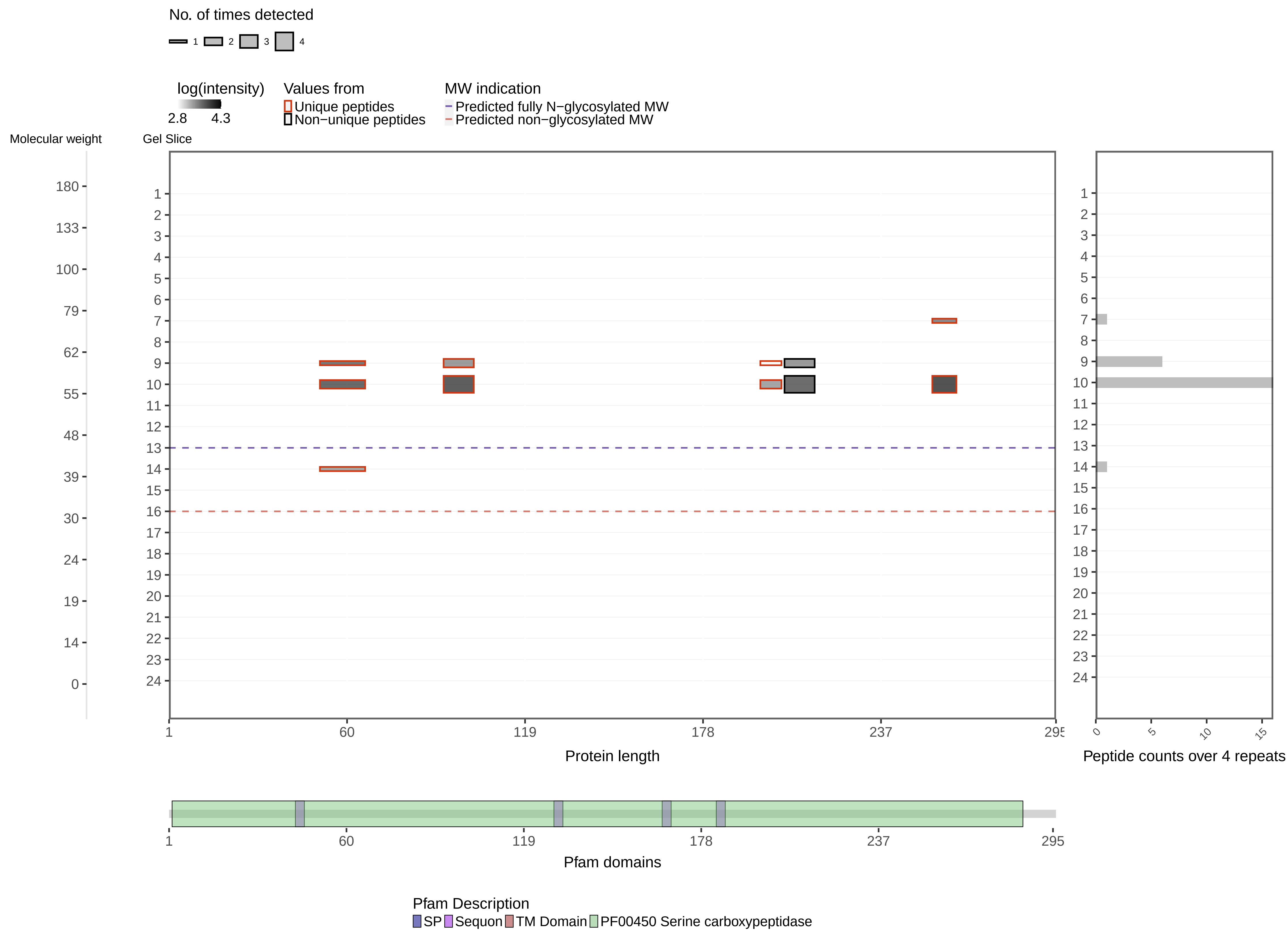


Protein sequence:

PSLVIDPCSDYYTIAYMNRPDVQNALHANVTNIKYSNWQPCSDVLTNWTDSSSTVIPLLREFMANGLRVWIFSGDTDGRVPVTSTKNSIKIMKLPIKTAWHPWYLSAEVGGYTQVYKGDMTFATVRGTGHQVPSYEAARSLSLVMHFLAGTNLPDKHT*

### NbL01g18210.1 N−alpha−acetyltransferase_11−like [MW = 21.2 kDa]

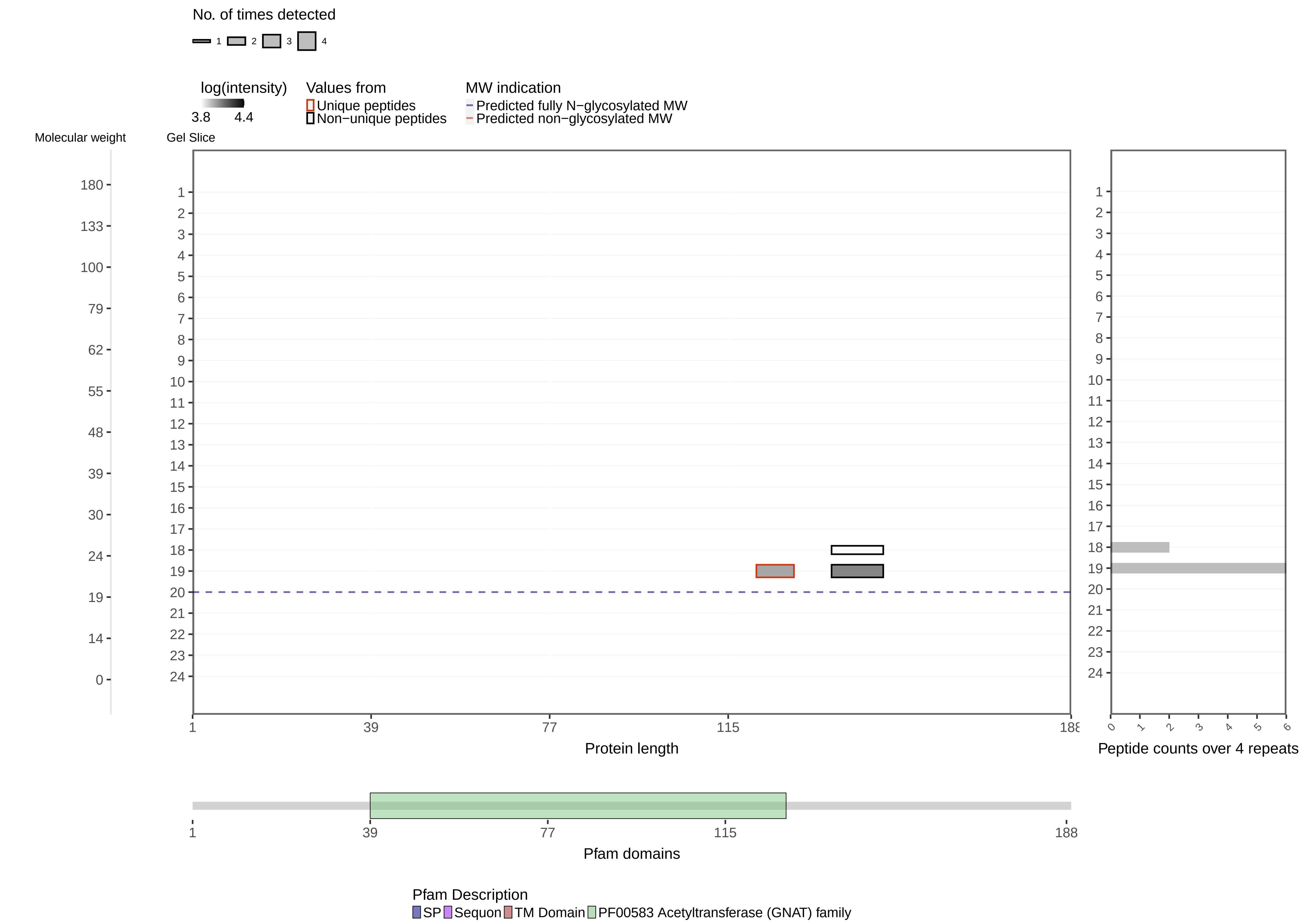


Protein sequence:

KLGIATKLMTAAQNAMEQVFGAEYVSLHVRKSNRAAFKLYTETLGYKIHDMEAKYYADGEDAYDMRKELKGKKHHQHHHHHHGGGCCSGEAKIGAKEGAAEAKAA*

### NbL01g18140.1 subtilisin−like_protease [MW − SP = 79.9 kDa]

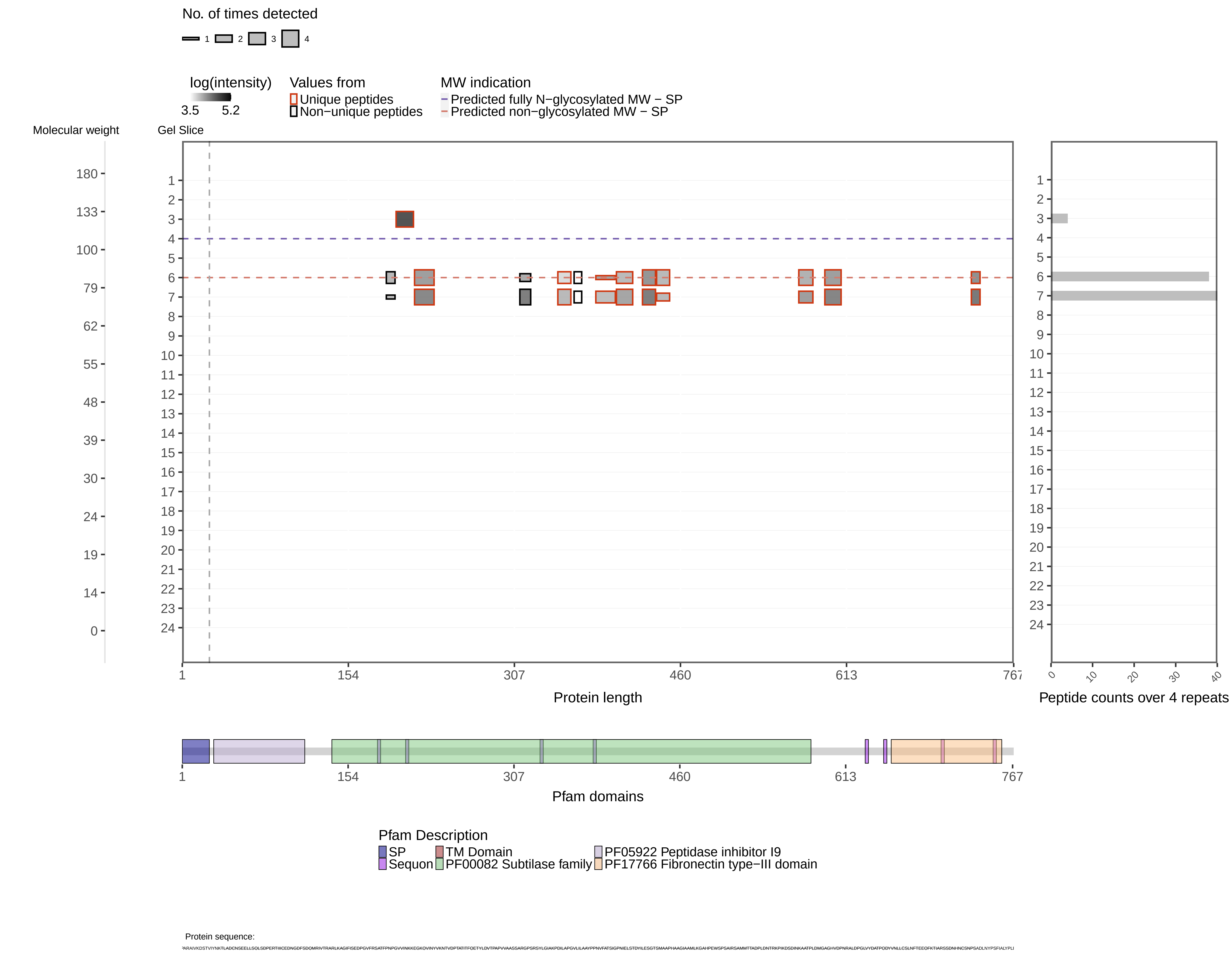

### NbL01g18130.1 protein_RETICULATA−RELATED_4_chloroplastic−like [MW = 41.04 kDa]

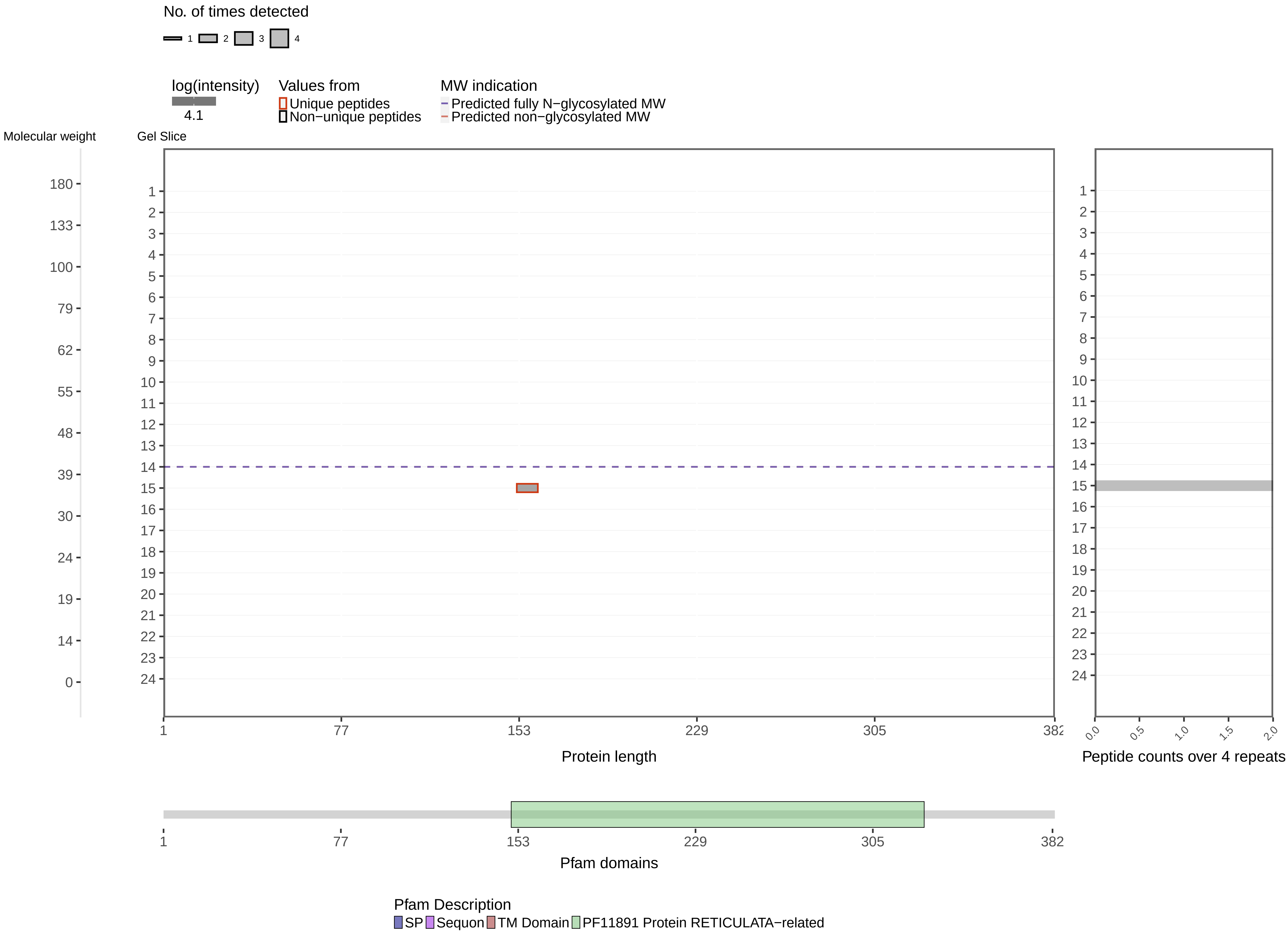


Protein sequence:

ELEKRRENFKNELDFVCADVVMAIVADFMLVWLPAPTVILRPSLAIGAGPLSRIFYNCPDNAFOVVLAGTSYTFLORIGAIVRNGAKLFAVGSGASLIGTGVTNGLINARKAIDKTFAGEAEDLPILPTSAAYGVYMAVSSNLRYOILAGVIEORILEPLLHKHKIILGALCFVFRTGNTFLGSLMWVDFARWIGIORSREOKET*

### NbL01g18110.1 50S_ribosomal_protein_L1_chloroplastic [MW = 37.1 kDa]

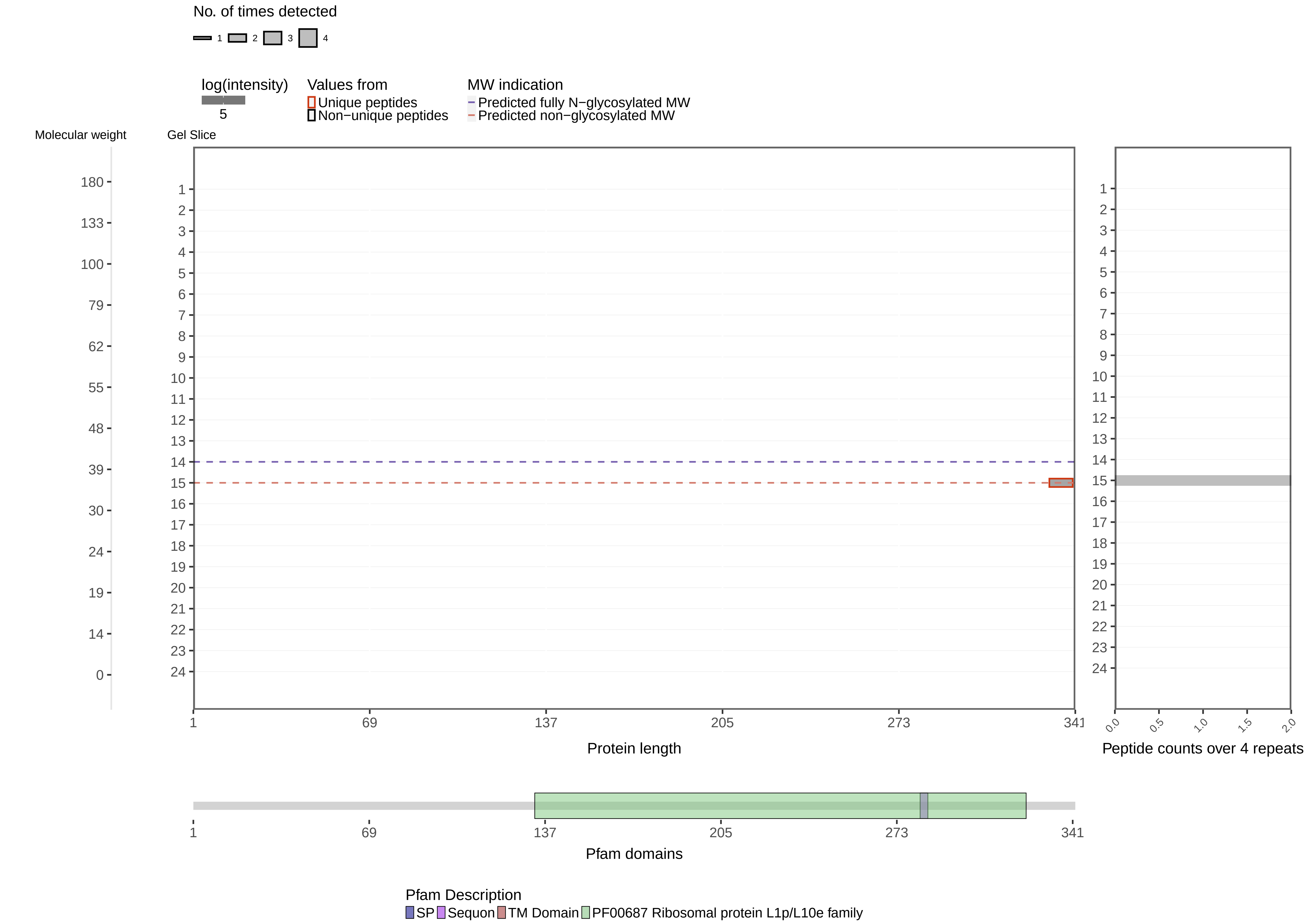


Protein sequence:

QQLRATVNLPKGTGQTVKVAVLTQGEKFDEAKNAGADLVGGEDLIEQIKGGFMDFDKLIATPDMMPKVASLGRILGPRGLMPNPKAGTVTTNIPQAIEEFKKGKVEYRADKTGIVHLPFGKANFSEEDLIINFIAAVKSIEANKPSGAKGVYWKSAHVCSSMGPSIRLNVREMLEYKLPNT*

### NbL01g17920.1 uncharacterized_LOC104211372 [MW − SP = 37.65 kDa]

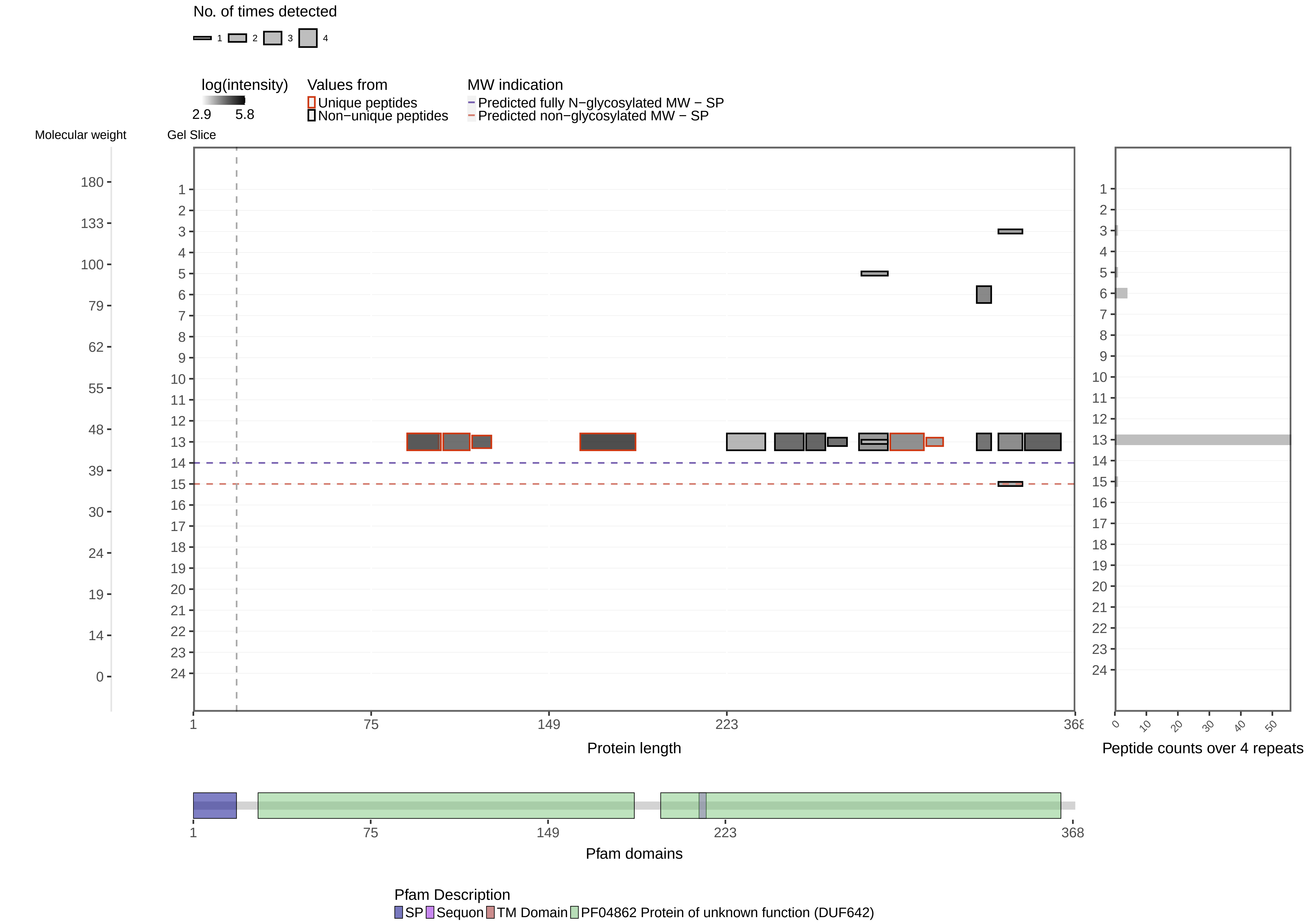


Protein sequence:

QEDPACGPLLDAVAIKELFPPQPTRVNLVKNPGFEEGPHLLINSSHGVLLPPKQEDLTSPLPGWIIESLKAVKFLDSAHFNVPFGRAAVELLAGRESAIAQIIRTVPKKVYAFAFTVGDAKNGCHGDMMIEAFAAKETFKVPFKSQGKGSFKTVSFKFTAIADRTRITFYSSYYHTKINDYGSLCGPVVDEVKVTPVA*

### NbL01g17560.1 thylakoid_lumenal_19_kDa_protein_chloroplastic−like [MW = 26.68 kDa]

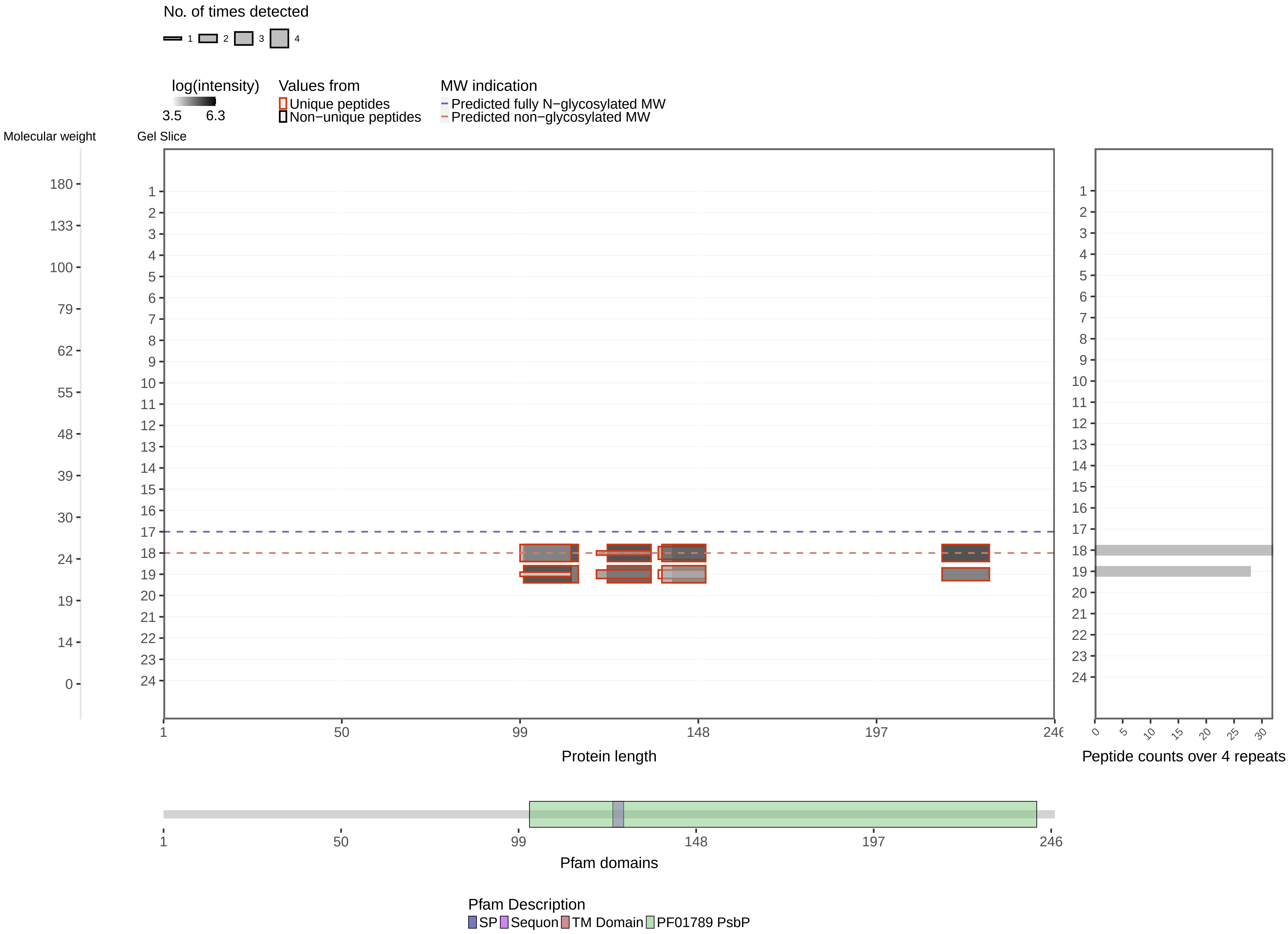


Protein sequence:

VKERLVSKVEKGTNGTDSEFYNPKKKTEKEYLTYLAGFRQLAPKDAVLNNLALSDVNLQDIIANADSVTSEERKDENGQLYYDYEIDGVAAHSLITVTCAKNKLYAHFVNAPAPEWKKDQDTLRHIHQSFKTVG*

NbL01g17470.1 uncharacterized_LOC109241047 [MW = 76.43 kDa]

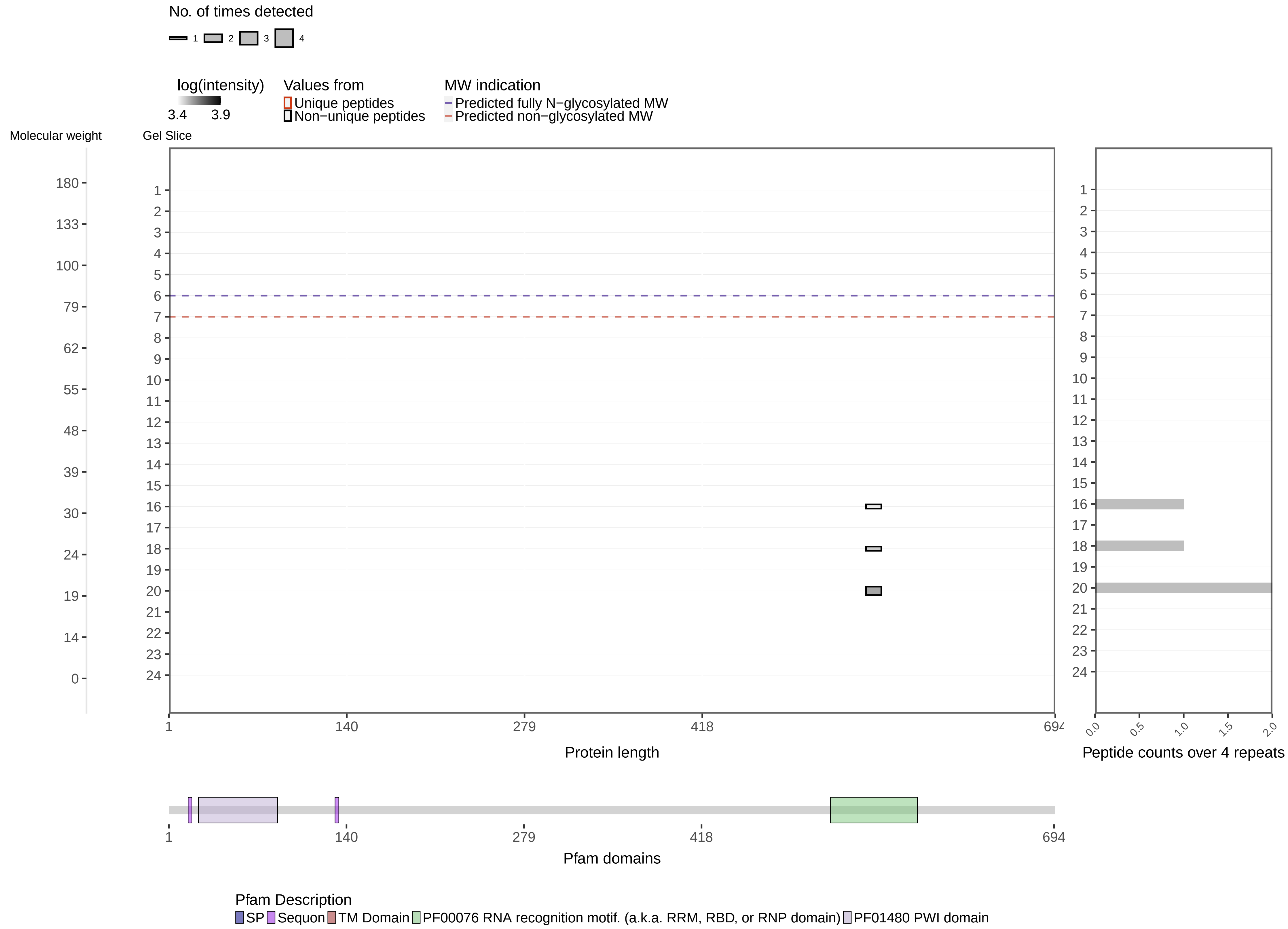


Protein sequence:

SNYRGKRIGKSTLNDDTAVLYDSALDEEADHGINIVNKGAIDASRTGACFKNIGETPLVVEYGVTVNANDILHKSQKDQDRPTFAPGAPHQPMNVPLNVNTWKSPQYQEARQALEREKQRSTLSSEGLANEPDMLLTKESTNPIAVTNGNVKPYADTQKESQKTASINQGLYSTGPPTEDADSRTIFVNNVHFAATKDSLSRHFNKFGEVLKVVILTDAATGQPKGSAYVEFMRKESAVNALSLDGTSFMSRILKVVRKSSATPEAASVAVWPRVTRAGPFAVSRFGRVPFPR

### NbL01g17430.1 DNA−damage−repair/toleration_protein_DRT100−like [MW − SP = 36.63 kDa]

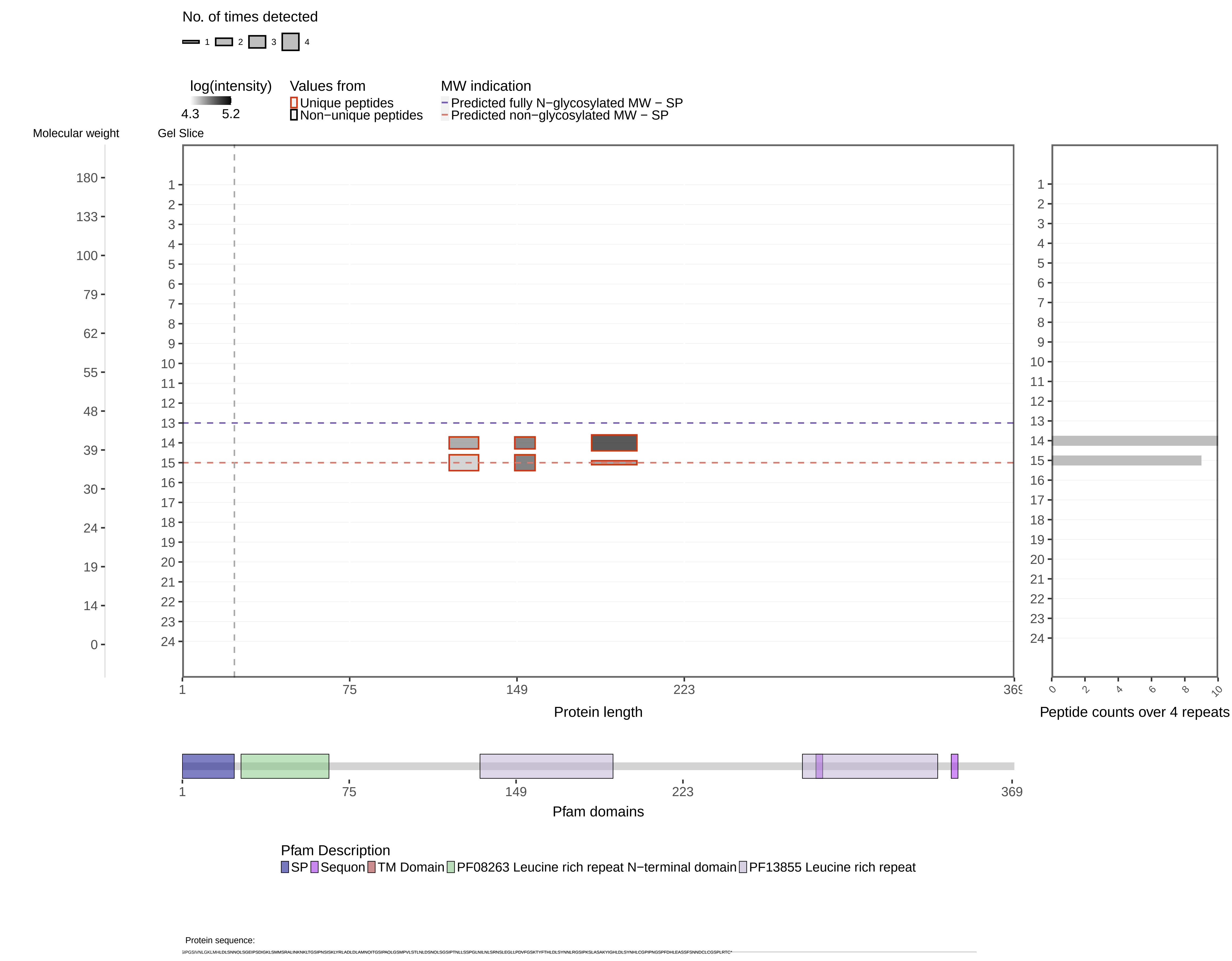

### NbL01g17240.1 siroheme_synthase [MW = 40.79 kDa]

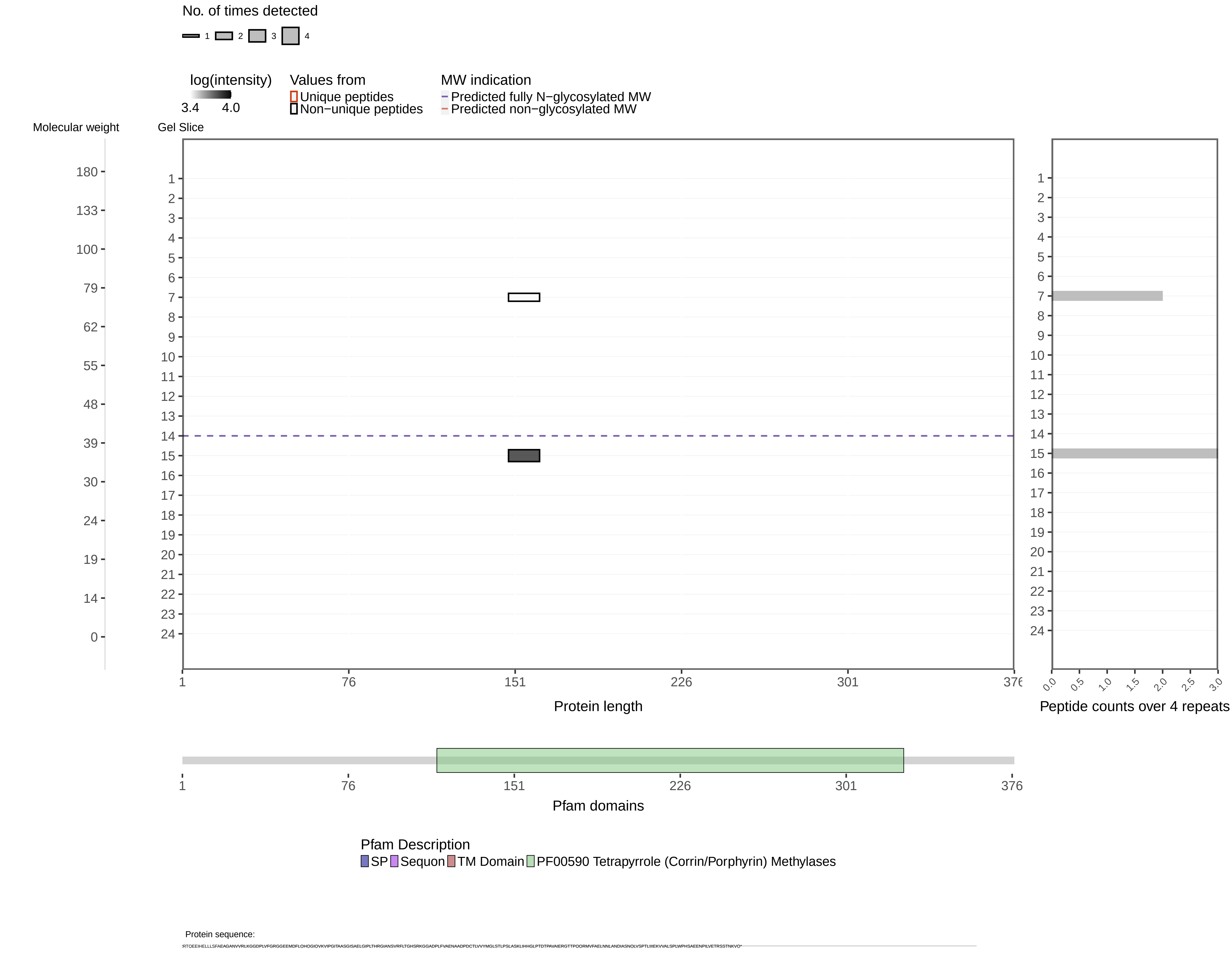

NbL01g17060.1 citrate_synthase_mitochondrial [MW = 29.12 kDa]

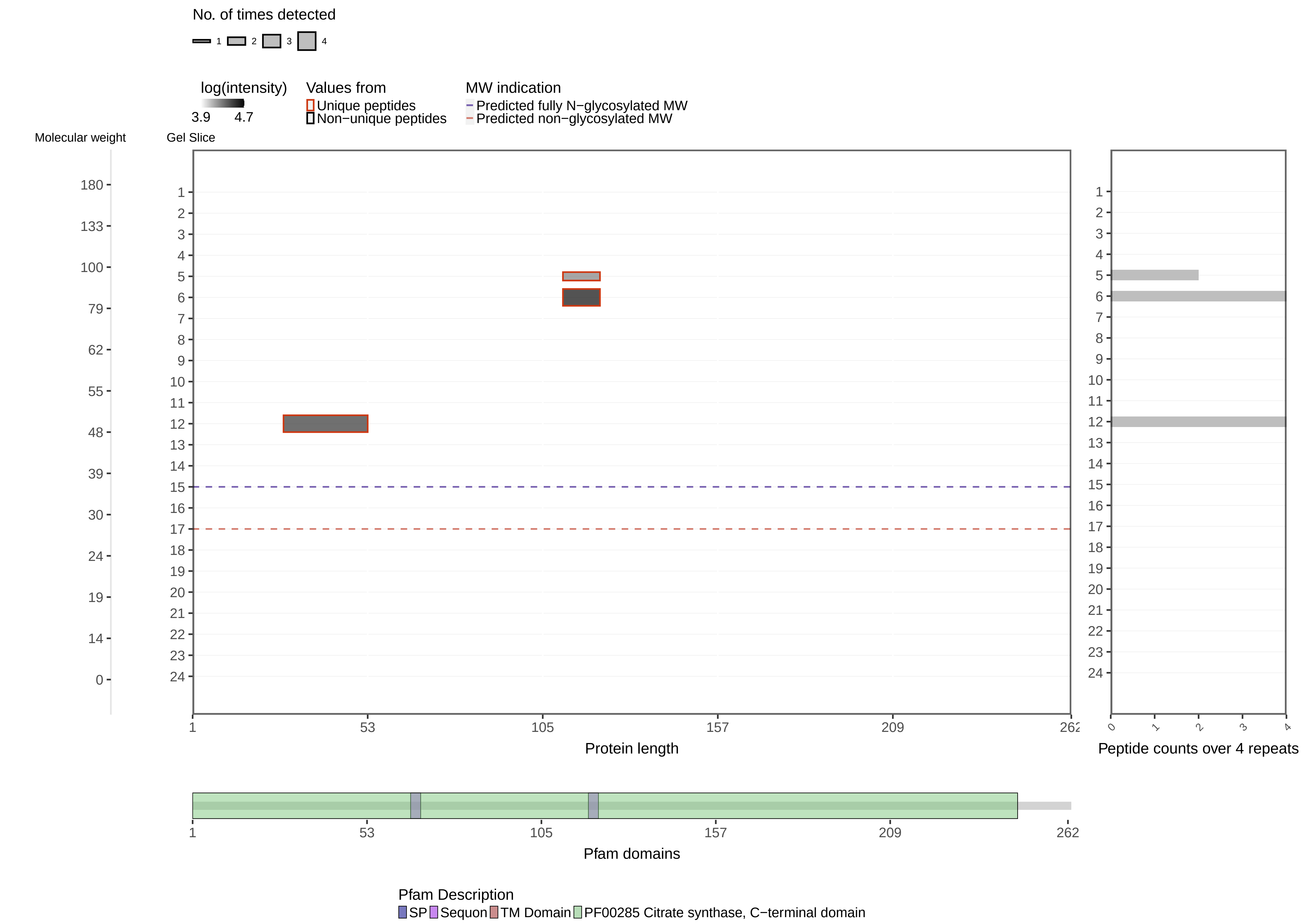


Protein sequence:

ENISKEQLKDYVWKTLKSGKVVPGFGHGVLRKTDPRYTCQREFALKHLPEDPLFQLVAKLYEVVPPILTELGKVKNPWPNVDAHSGVLLNYYGLTEARYYTVLFGVSRALGICSQLIWDRALGLPLERPKSVTMEWLENHCKKA*

NbL01g17020.1 uncharacterized_LOC104121022 [MW = 15.28 kDa]

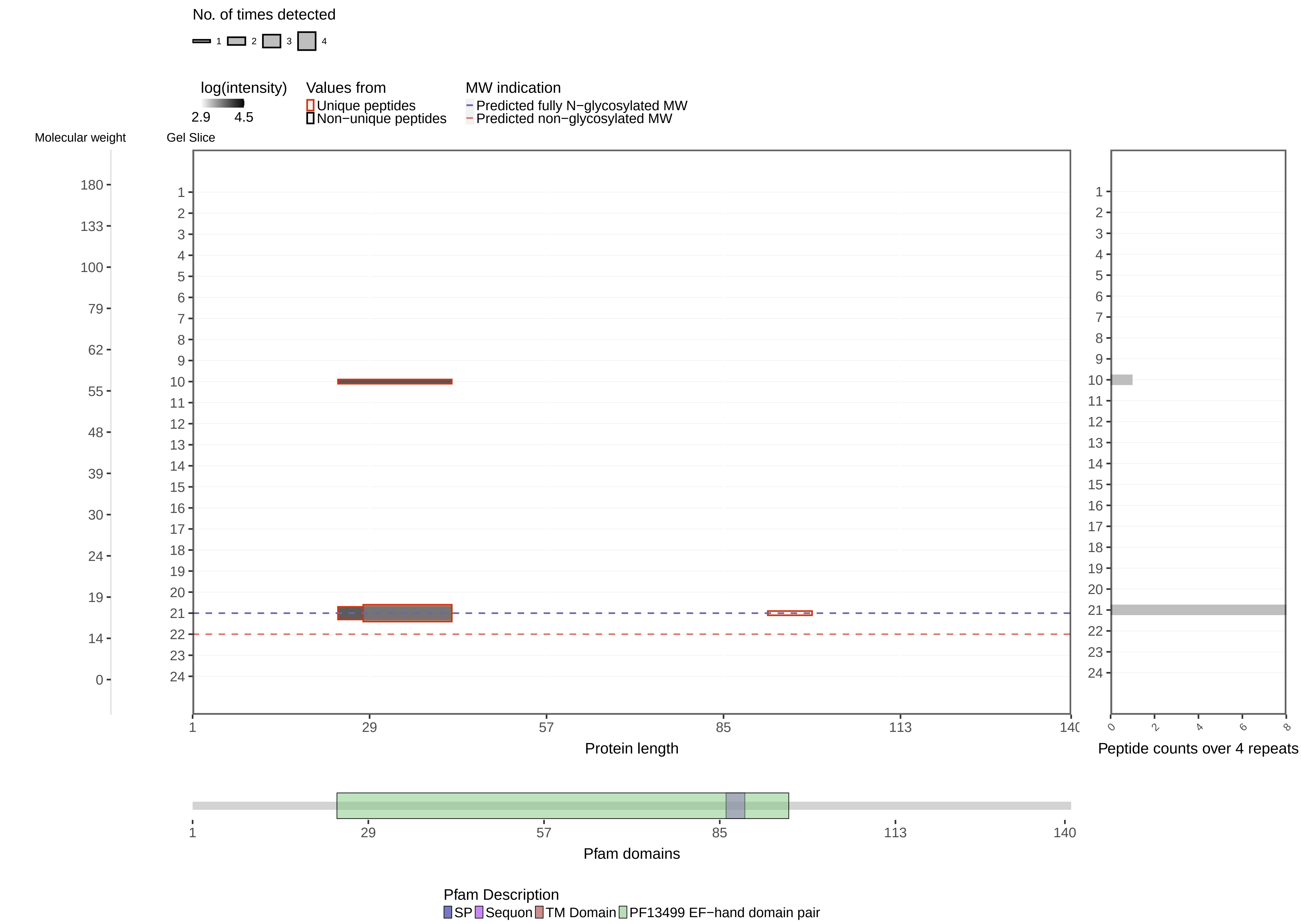


Protein sequence:

HFGVDVAASPEELTRLYDSIFEKFDCDHNGTVDRQEFGNEMRKIMLAIADGLGSSPIQMALDDSDQNLIKQAADLEASKLPA*

### NbL01g16910.1 beta−hexosaminidase_2 [MW = 67.11 kDa]

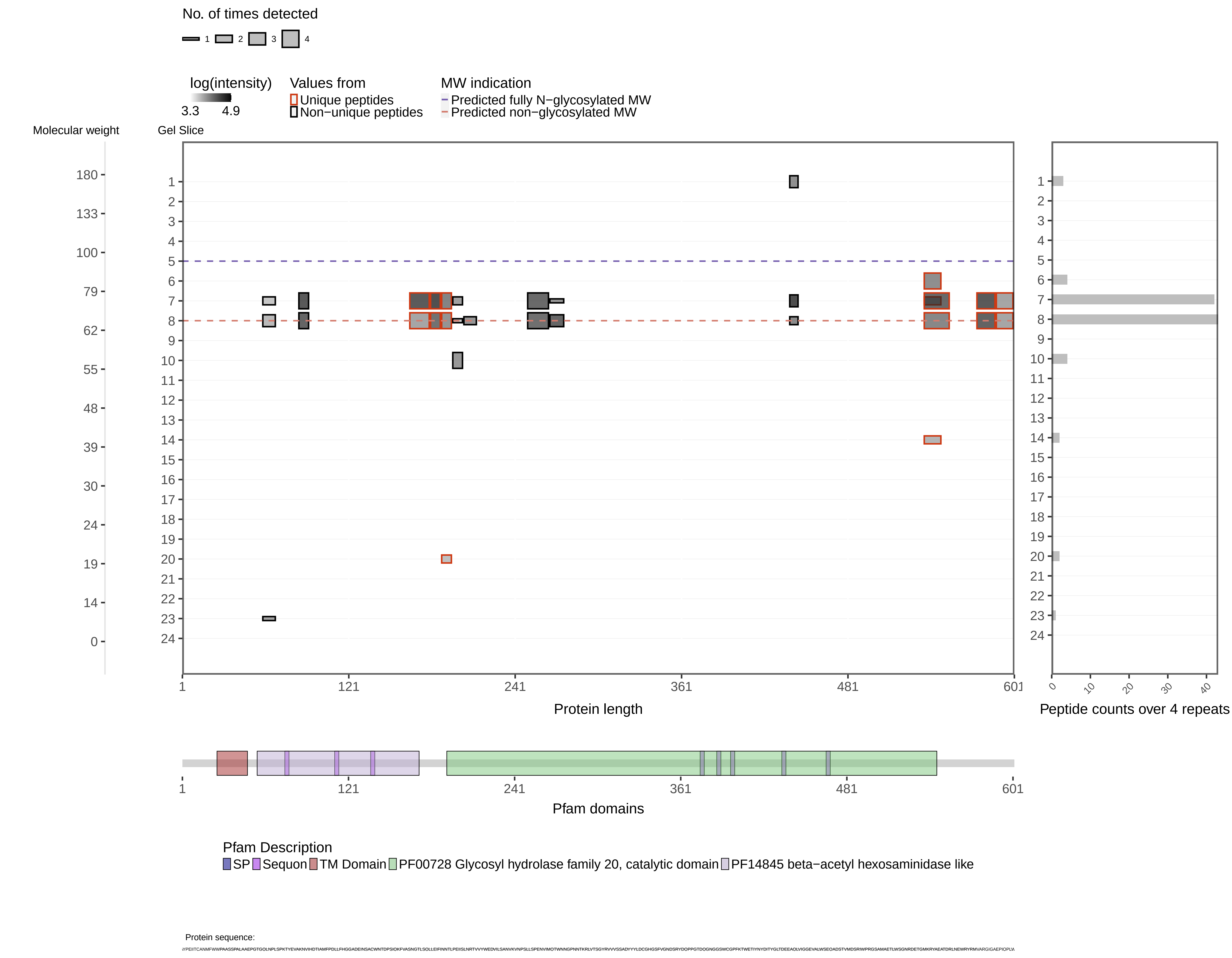

### NbL01g16900.1 non−specific_lipid−transfer_protein−like [MW − SP = 12.67 kDa]

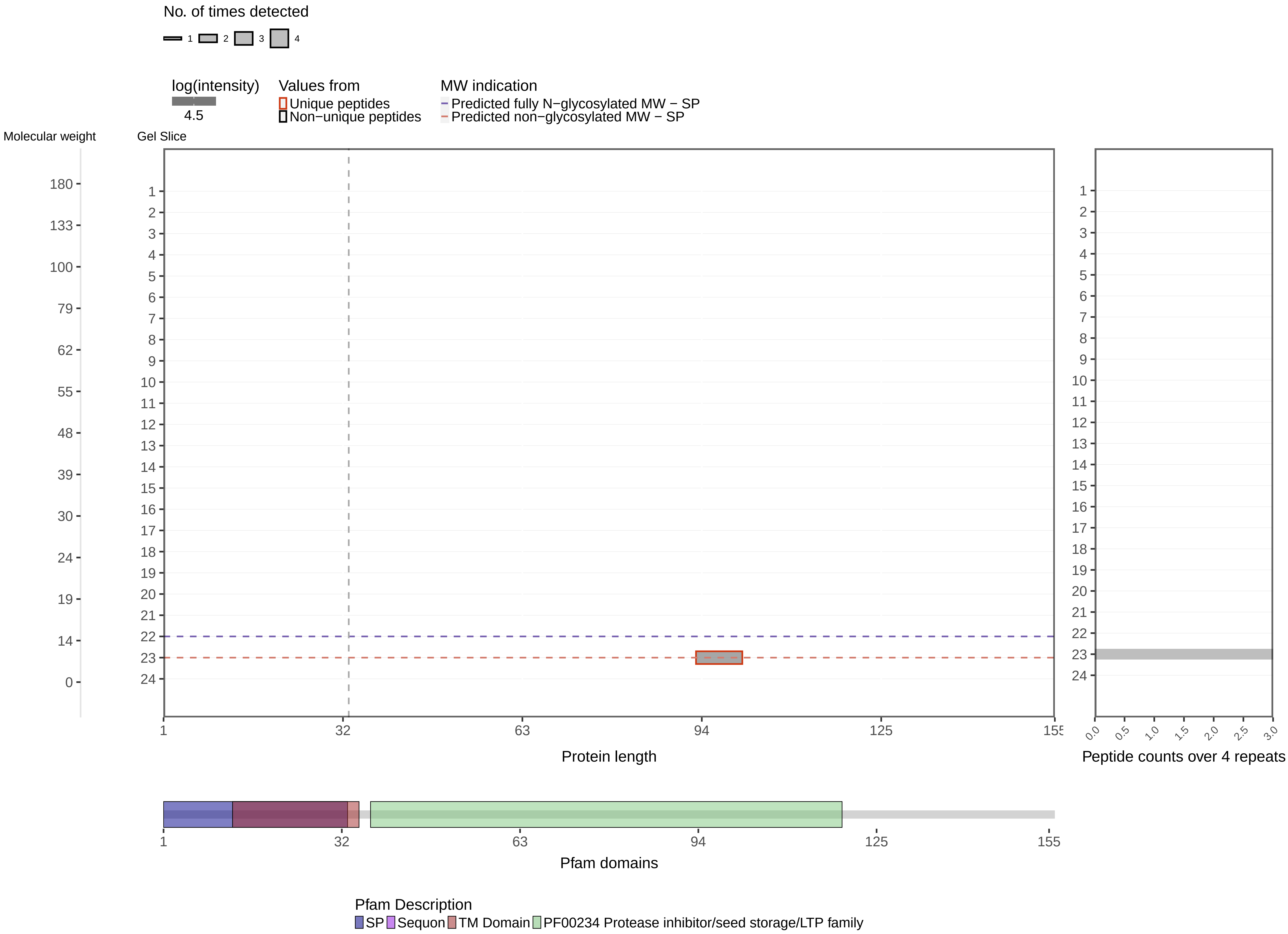


Protein sequence:

CQSLQSATSTTQDRQAACTCLKNAAQKVNINAQLAKGLPQSCGISLSFPISTSVDCTKYYSSPPPPNPQKITPLSSSYLALWQKNFGLS*

### NbL01g16880.1 uncharacterized_LOC109216616 [MW − SP = 44.3 kDa]

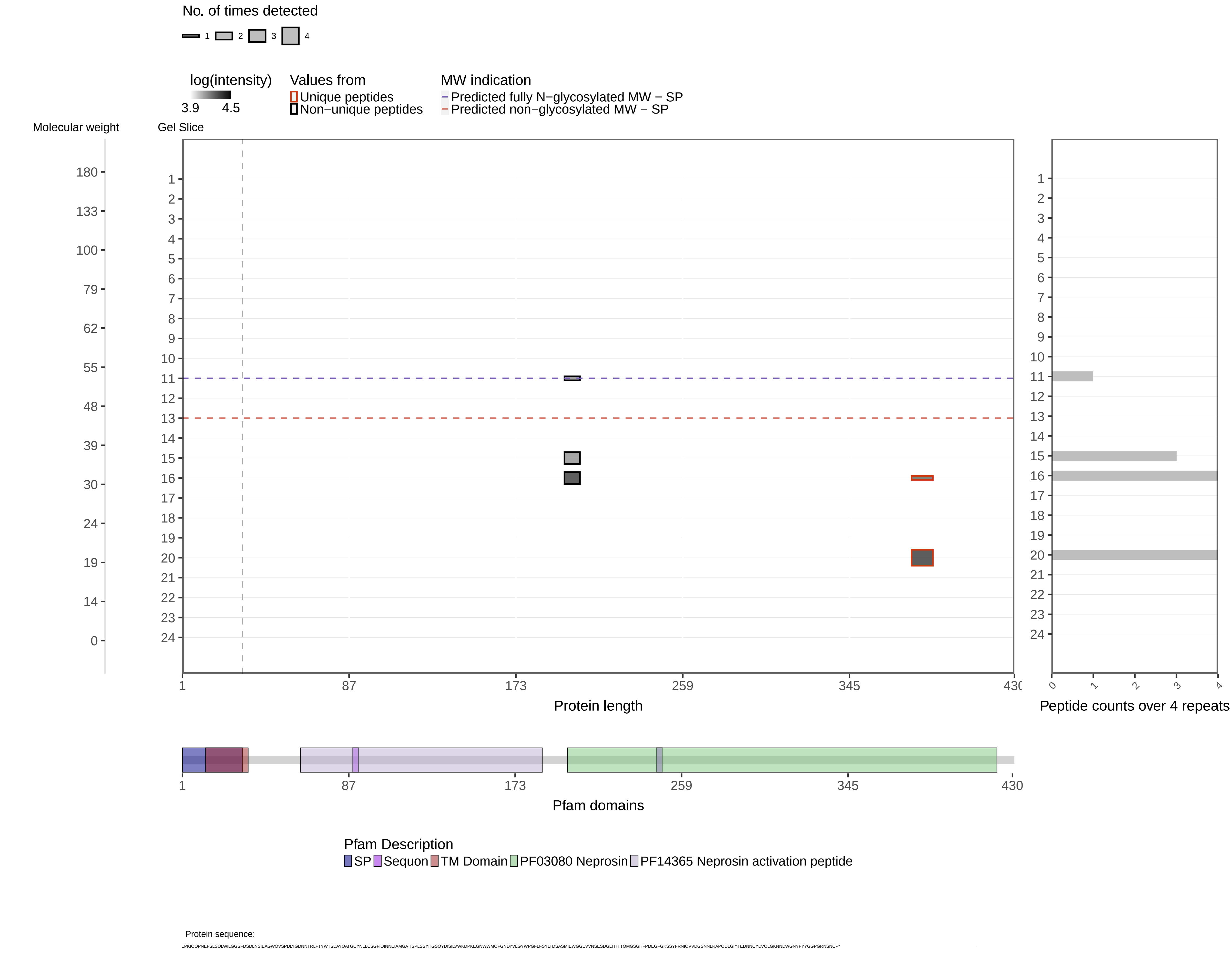

### NbL01g16810.1 probable_3−hydroxyisobutyrate_dehydrogenase−like_2_mitochondrial [MW = 31.65 kDa]

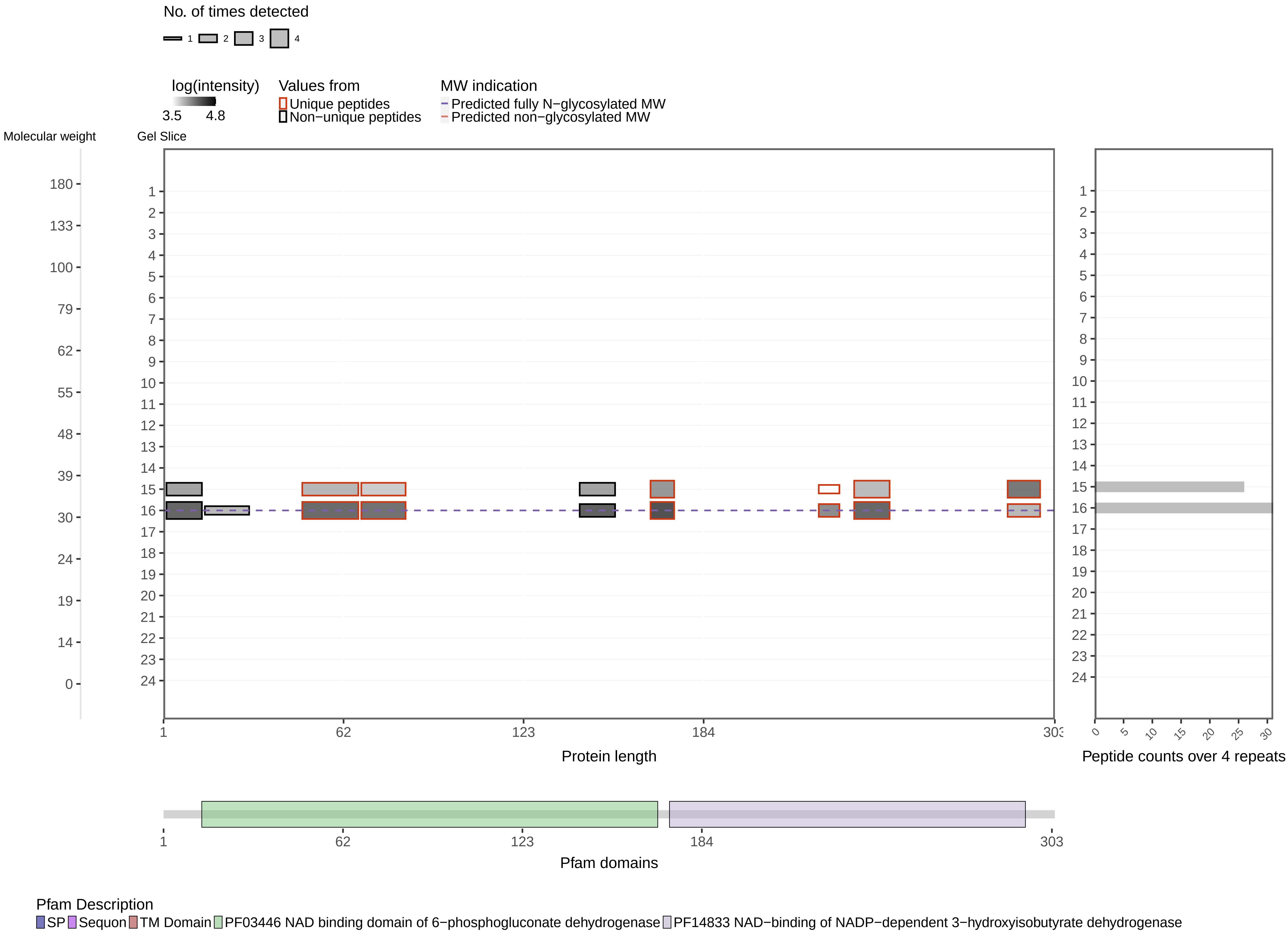


Protein sequence:

GKLAIFAGGNEDVVKWLKPLFDVLGKVTFVGGPGKGQSCKIANQIVVGGNLLGLSEGLVFADKAGLDKMKFVEAIKGGAAGSKVMELFGERMIGRDFRPGGFTDYMVKDLGMGLDVGAEQGDDVVVLPGAALSKQLFSAMVANGDGKLGTQGLITVIERINGK*

### NbL01g16640.1 glutathione_transferase_GST_23−like [MW = 25.99 kDa]

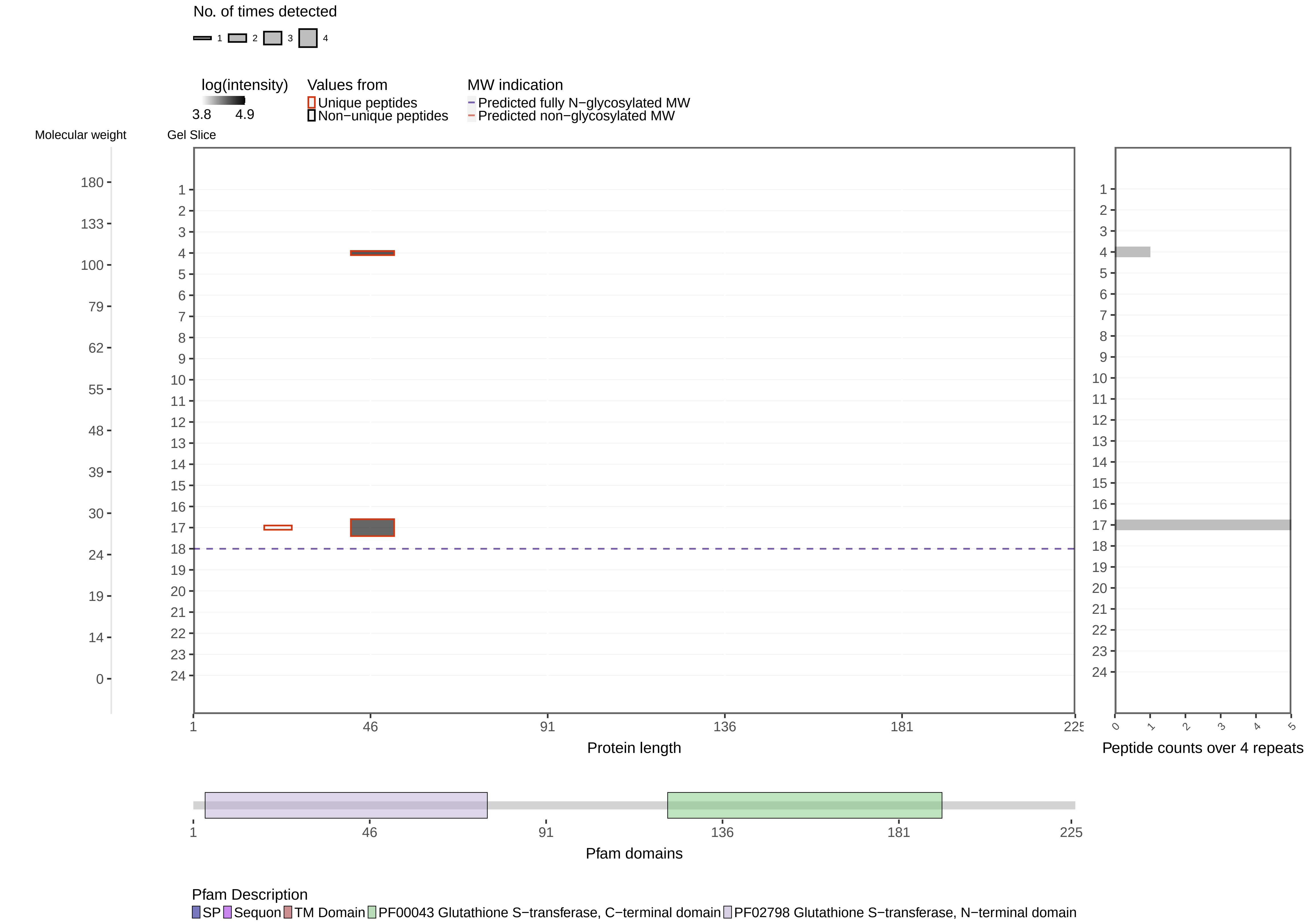


Protein sequence:

[illegible]GEEKLLPSMWSVFTGQGNEKNKALGPAVQNLELLEEQLKEKKFFGGEEIGYLDLSLGWLAYLLDVFEEVIDLKLFDAAKLPLLSAWMNNFYDAPIIKEHWPPRDKLVTKFQLLHENYQKAFTAK*

### NbL01g16540.1 DNA−directed_RNA_polymerases_II_IV_and_V_subunit_8B−like [MW = 16.84 kDa]

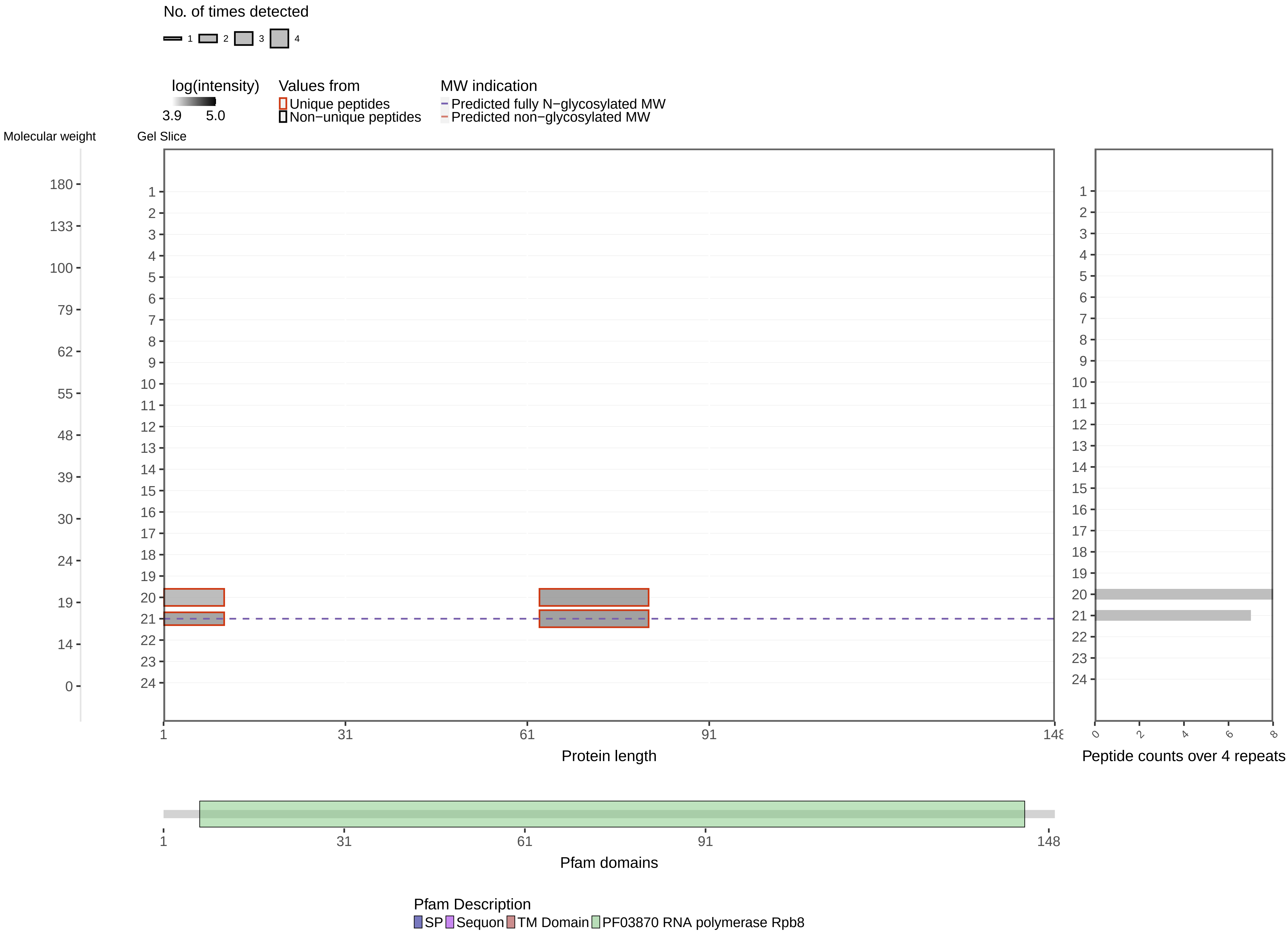


Protein sequence:

[illegible]KTLNMDGSAVTSHPPEGEQKSLADKFEYIVHGLVYKVSMDTSGPDKKVVVYVSFGGLQLMLKSDPLKVQKFKLDEKLFLLLRKMTK*

NbL01g16510.1 uncharacterized_LOC109245038 [MW = 10.57 kDa]

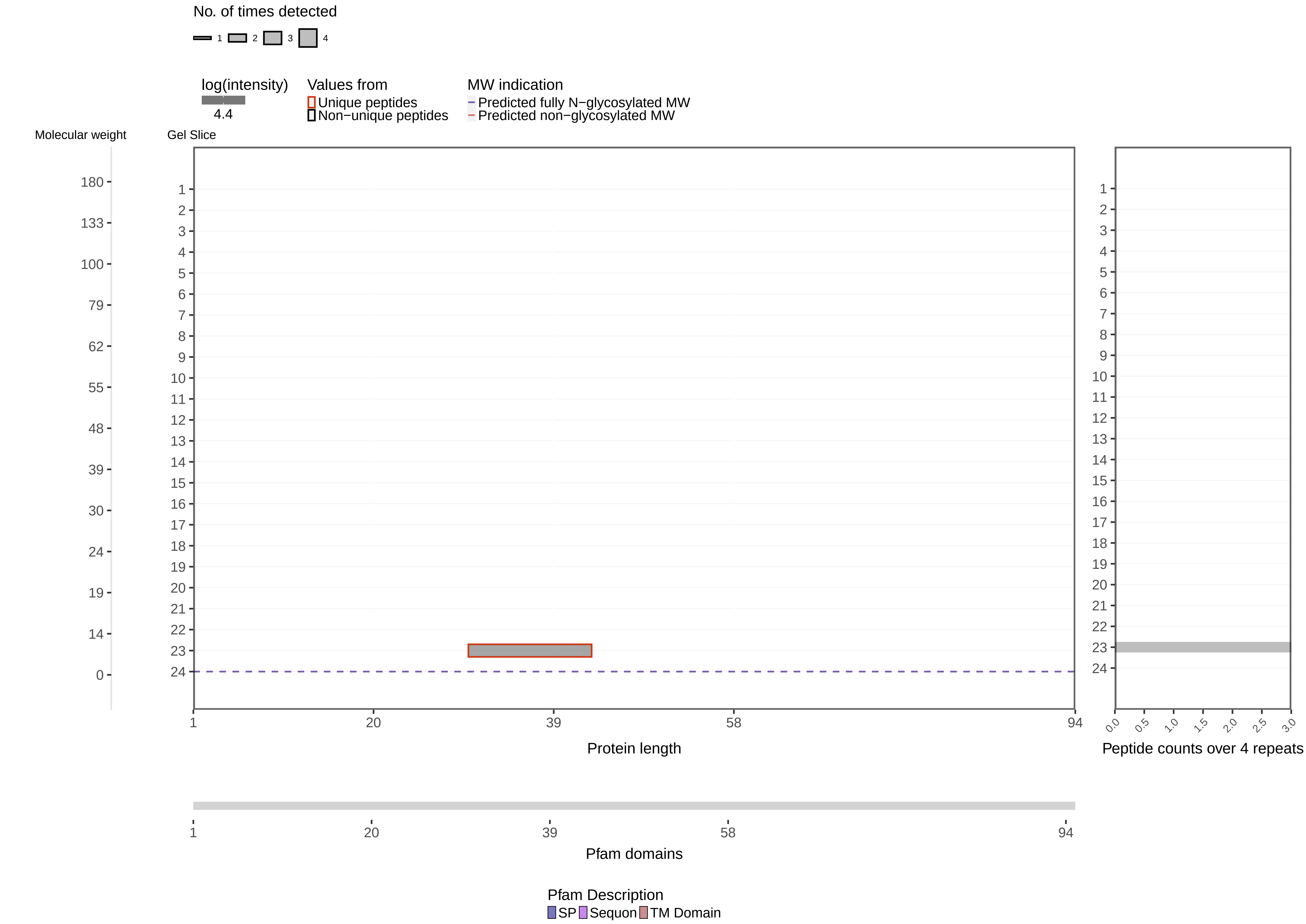


Protein sequence:

PALPLNLAARLQKKAEVRAMFMLGSEERSLHNIQDLIDAAEYSLSLLRKGEIPKHIQVYQ*

### NbL01g16400.1 protein_SCAI [MW = 69.82 kDa]

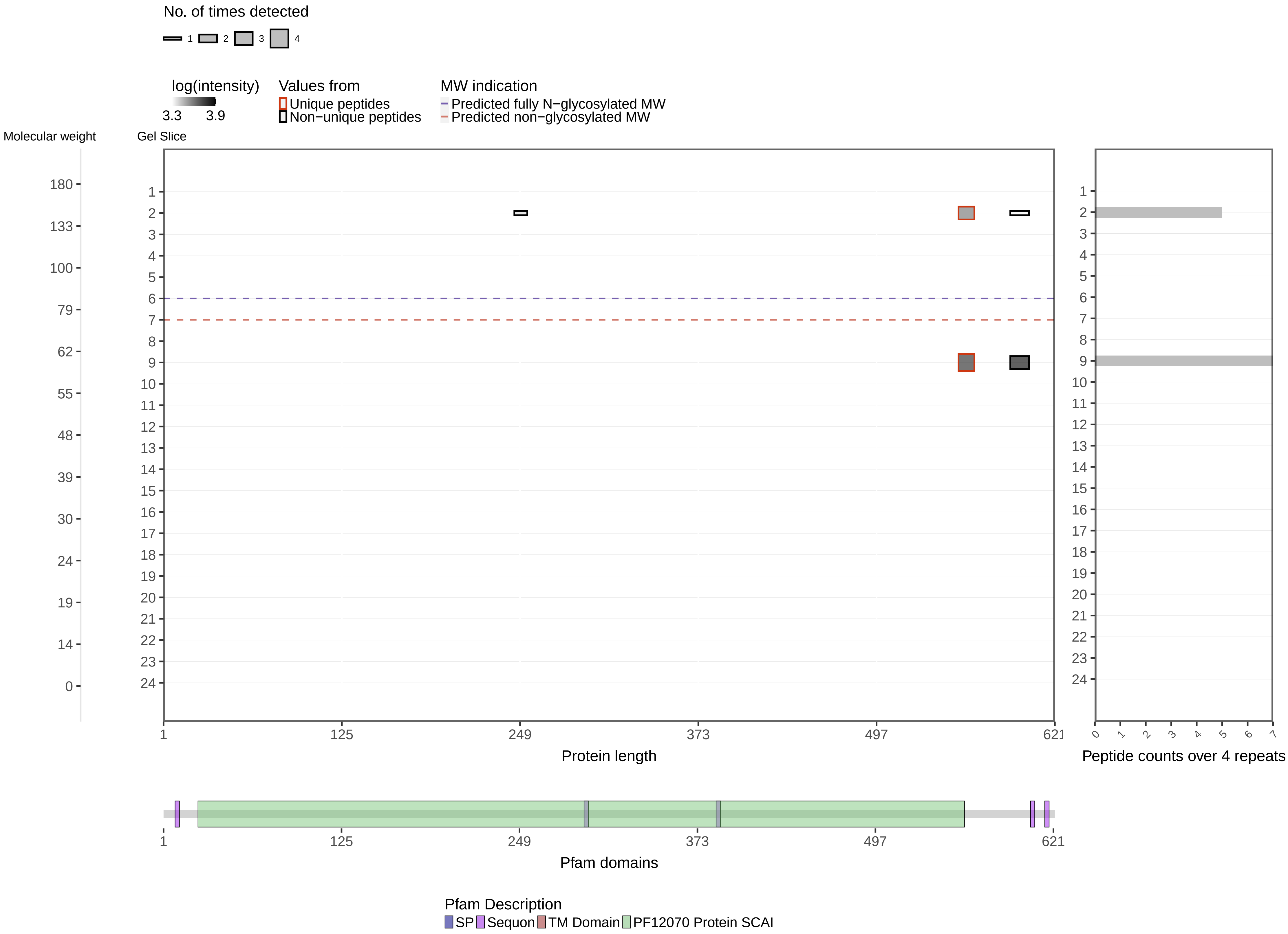


Protein sequence:

RVNYSQDISDPTLPPNPRKTILYRPSVTNFLAVLATICEELSHDGVLLLYLSAAGSSQNFFSSPSHGAGRSFMSRAVDLEATPTSSLISDTEIANTTSTREDCLTEQSSCIYIGSRGTGGVNGIYPCDLLPFTRRPLFLVIDSDGSKAFTALSGSEKGEPVAMLLSPTISLPISAMDSSRQPSGSLFTSFLTVPLQAFILMLGFTVSDIDMDMYGKAEKLLSSSLNEWGLLLAASDNLNPVWSQTLGDPFLRRLILRFIFCRTALALYAPTFNKIEFLPECIPCLPDAVLPSSAACQRTVL

### NbL01g16380.1 nucleolar_GTP−binding_protein_1−like [MW = 77.8 kDa]

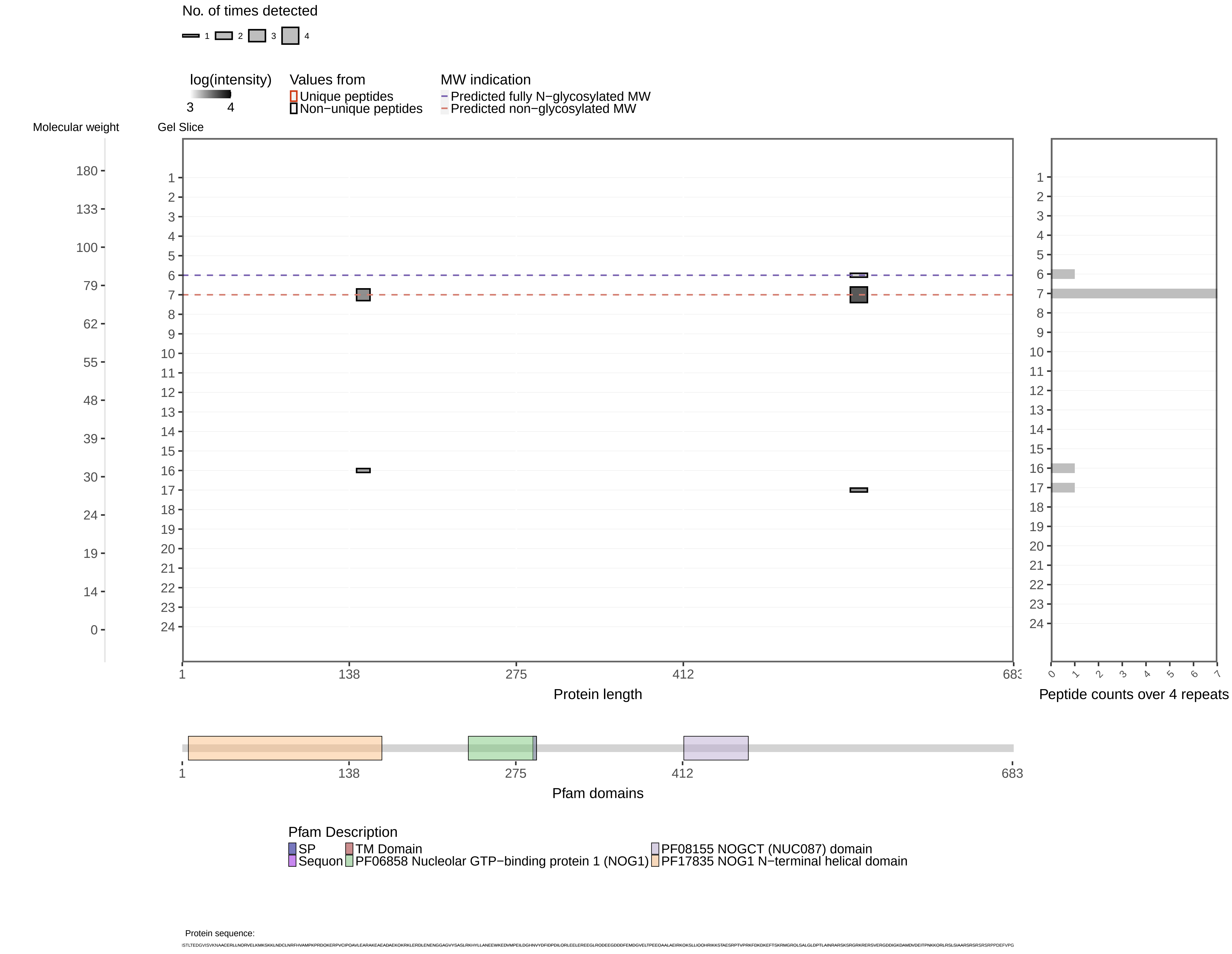

NbL01g16360.1 uric_acid_degradation_bifunctional_protein_TTL [MW = 37.3 kDa]

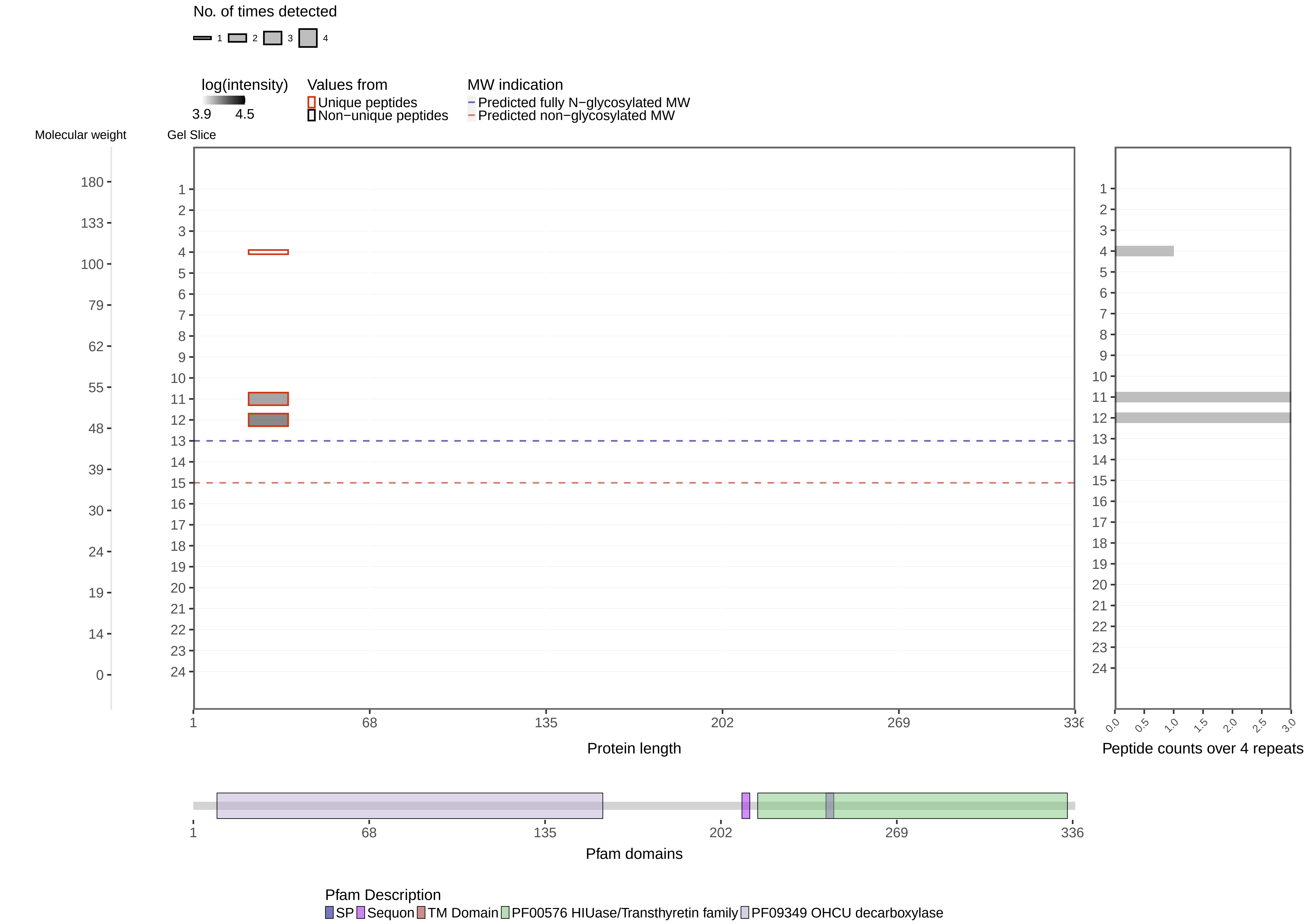


Protein sequence:

RLAKLFADKADTAQTIKSLARTNIARKAEEDRVNIIGAHLTAATGDAGAKPAHIPNRTRPPITTHVLDTSRGCPANGIEVQLEMWKDNQTRPQFGKANIGGWKVLGSSATDKDGRSGQLMNMVEALAPGIYRISFNTGEYIPDGFFPYVSIVFEIRESQKWDHFHVPLLLSPFSFSTYRGS*

NbL01g16350.1 3−hydroxyisobutyryl−CoA_hydrolase_1−like [MW = 42.61 kDa]

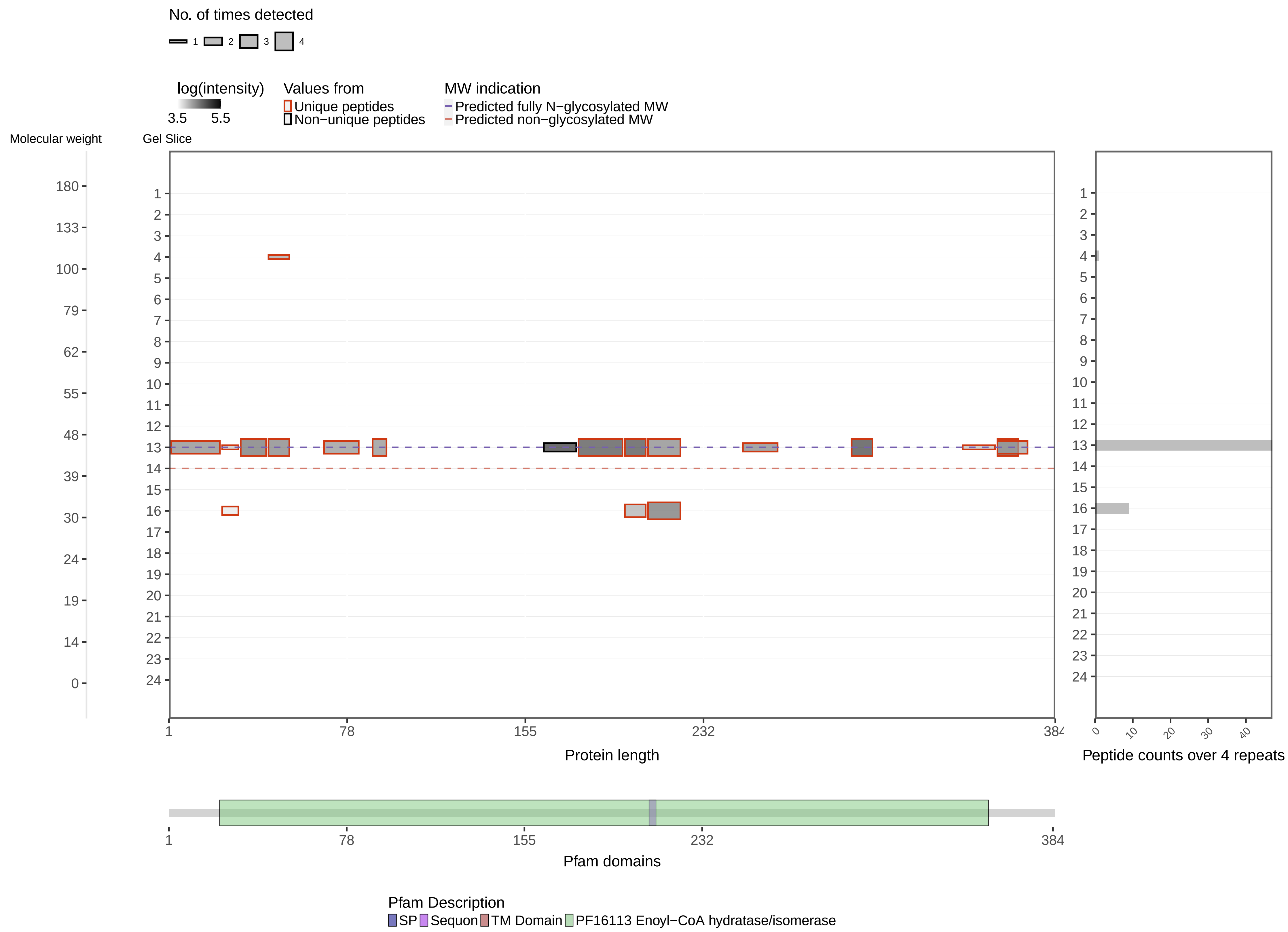


Protein sequence:

RLDGAEMLACGLATHFVPSERLSLLEQALVKVNTSDPDVISTIISRFSHIPKLKEGSSYHKMKIIDRCFSRRTVEEIISALESEALNKNDDWVSSTIQSLKKASPTSLKISLRSIREGRLQGVGSCLTREYRMVCHVLMGEFSKDFFEGCRALLIDKDRNPKWEPSRLDLIRDDDVDCYFSKIDDEDWEDLKLSPRSNLPPYAIAKL*

### NbL01g16140.1 UMP−CMP_kinase_3 [MW = 22.98 kDa]

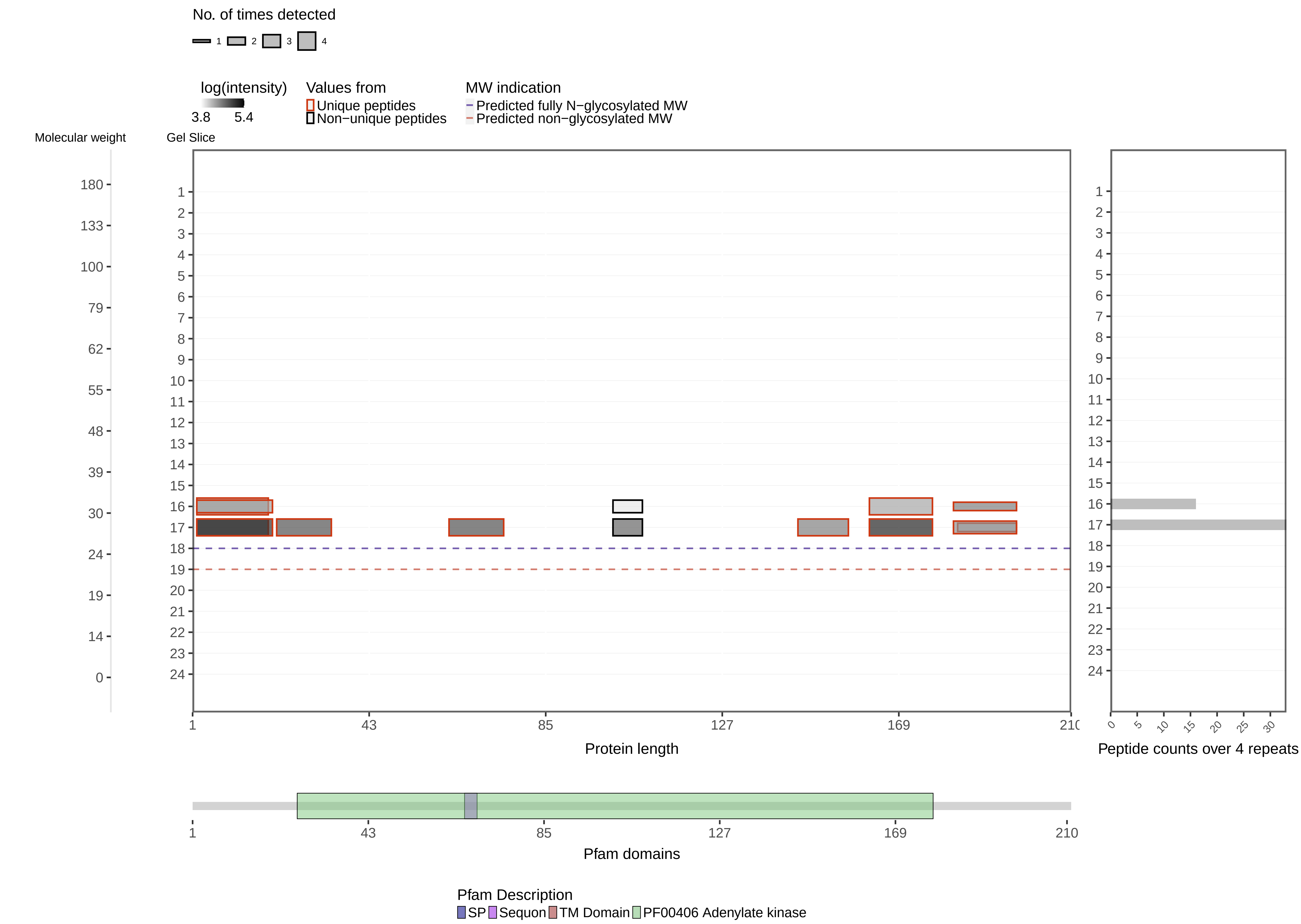


Protein sequence:

AIQENGNDKFLIDGFPRNEENRAAFESVTGIEPEFVLFFDCPEEEMERRLLGRNQGREDDNIDTIRKRFKVFLESSLPVIEYYNSKGKVRKIDAAKPVGEVFEAVKAVFAPAVEEVKY*

### NbL01g16040.1 acyl_carrier_protein_1_mitochondrial [MW = 13.75 kDa]

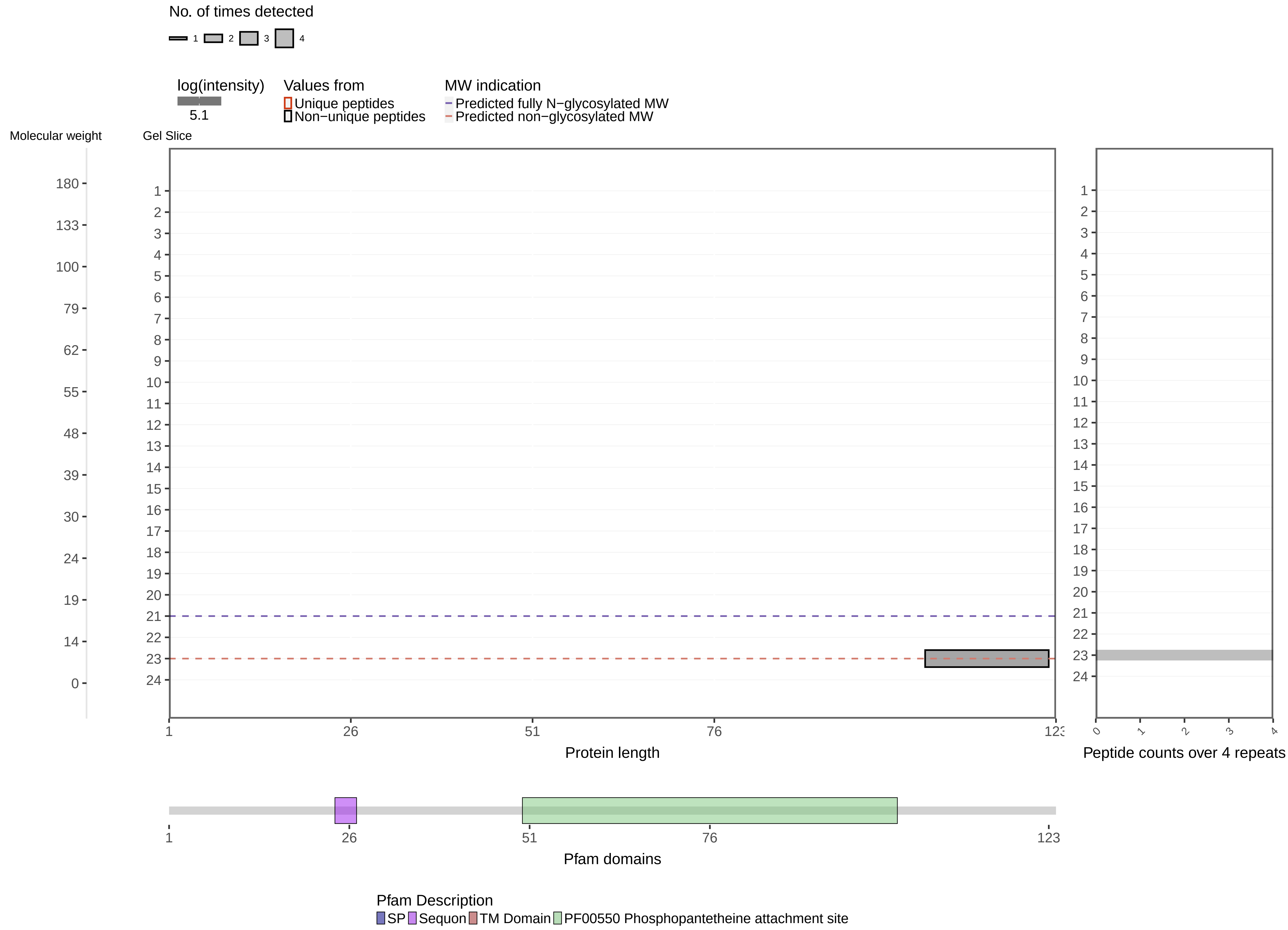


Protein sequence:

IRVLDVVKCFPKVDPSKVSPEVHFEKDLGLDSLDTVEIVMALEEEFKLEIPDKEAVRIESCEHAIEYIYNHPMSS*

NbL01g15560.1 ras−related_protein_RABD2a−like [MW = 23.24 kDa]

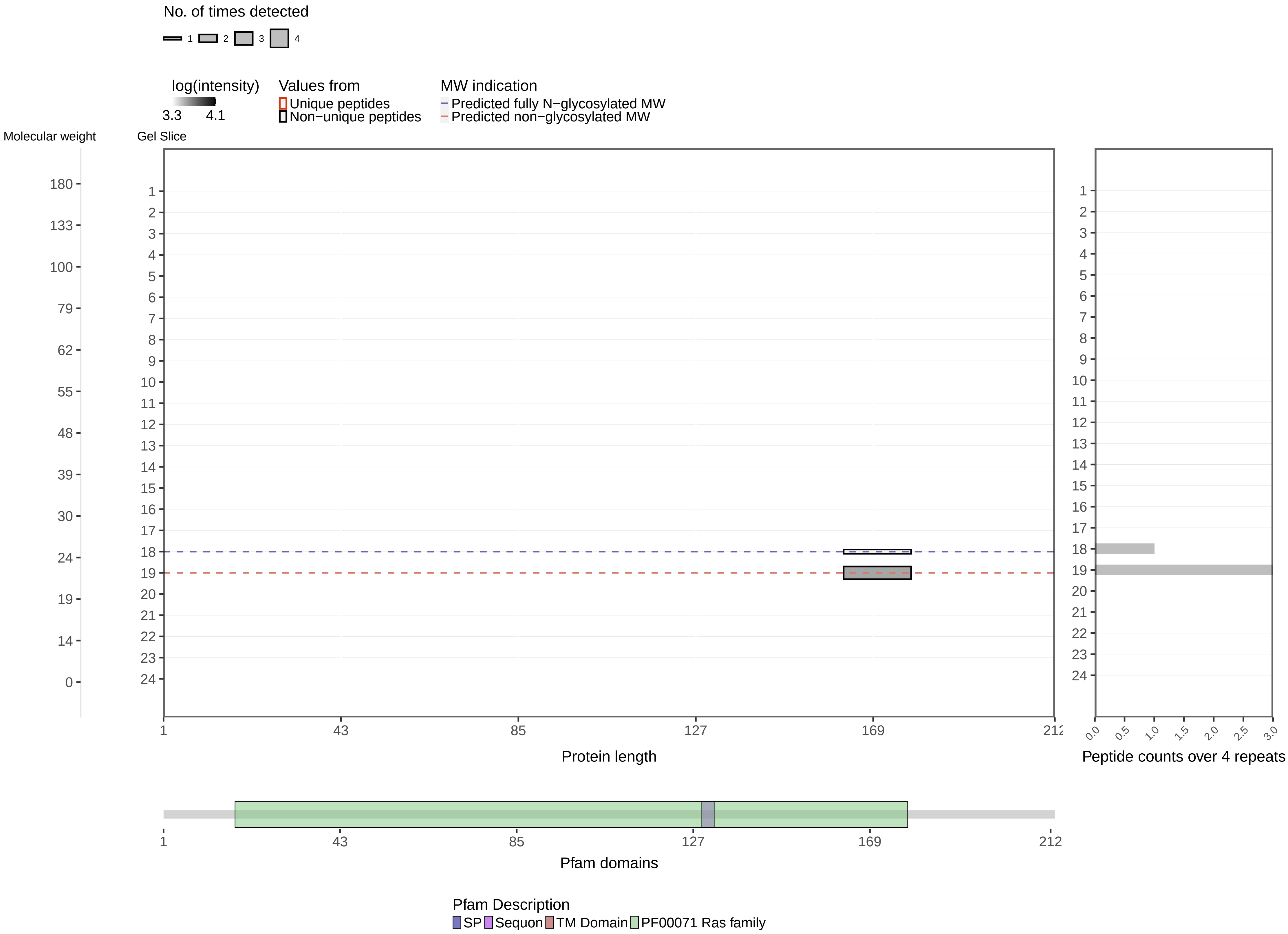


Protein sequence:

YDVTDQESFNNVKQWLNEIDRYATPNVNKILVGNKSDLTANRVVSYETAKAFADEIGIPFLETSAKDASNVEQAFMAMTSAIKNSMASQPATNTAKPPTVNIRGQPVAQNSGCCSS*

NbL01g15550.1 plasminogen_activator_inhibitor_1_RNA−binding_protein−like [MW = 38.73 kDa]

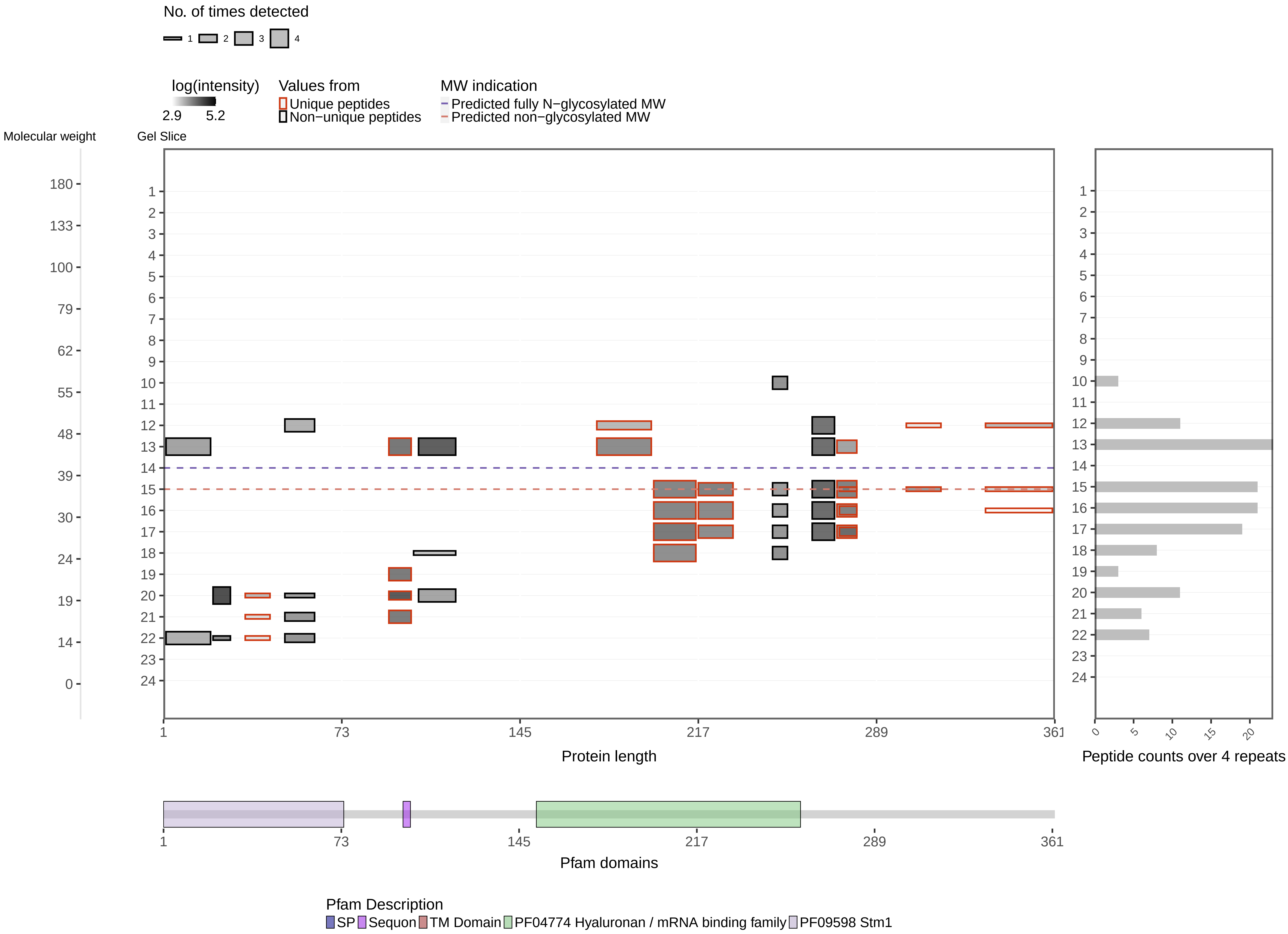


Protein sequence:

EIKREGAGRGNWGTQADEVTQISDEVANEGEKNLNAEKPSADGEAGDDKKENPAAEAEDKEPENKEMTLEEYEKVLEEKRQALQALKTEERKVDTKEFESMQQLSKKSSDDIFIKLGPKDKRKESAEKEEKAKKAVSINEFLKPAEGERYYAPGGRGRGGRGRGSRGYGGADTTSNVEAPPIEDPGHFPTLGGK*

### NbL01g15540.1 glycine−−tRNA_ligase_chloroplastic/mitochondrial_2 [MW = 119.77 kDa]

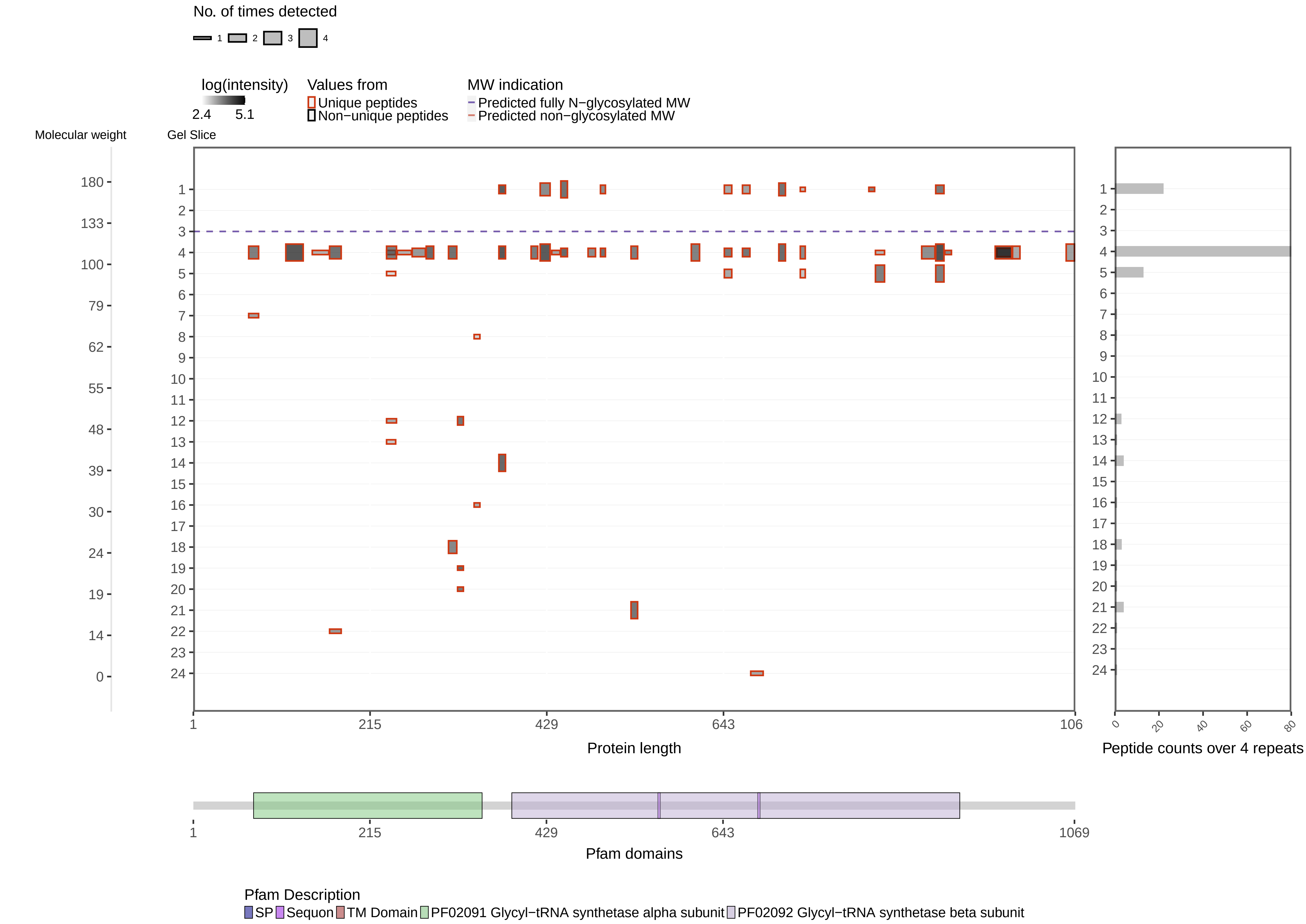


Protein sequence:

### NbL01g15490.1 tRNA_wybutosine−synthesizing_protein_4 [MW = 38.9 kDa]

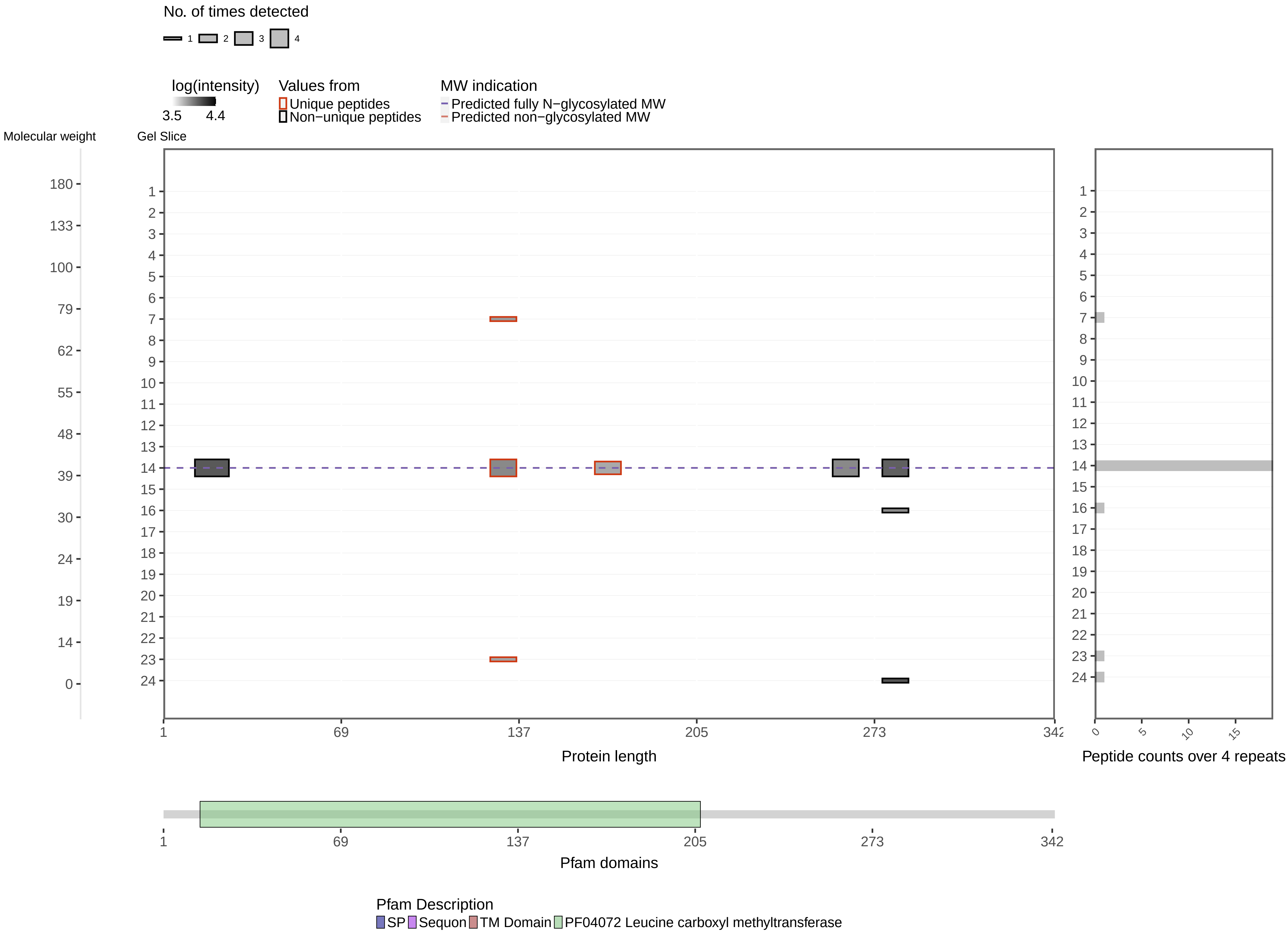


Protein sequence:

HYKLLPIDLRNIDDLNDIIARANLDPSLPTFIIAECVLIYLDPDSSRTIVGWASRTFTTAIFFLYEOIHPDDAFGOOMIRNLESRGCGLLGIYATPTLOEKENLFLDOGWOKAVAWDMLRVYSDFIEAOERRRIERLEIFDEFEEWYMMOEHYCVTYAINDAMGLFKEFGFPNTOPMTDTSIATST*

NbL01g15190.1 nuclear_pore_complex_protein_NUP62−like [MW = 73.5 kDa]

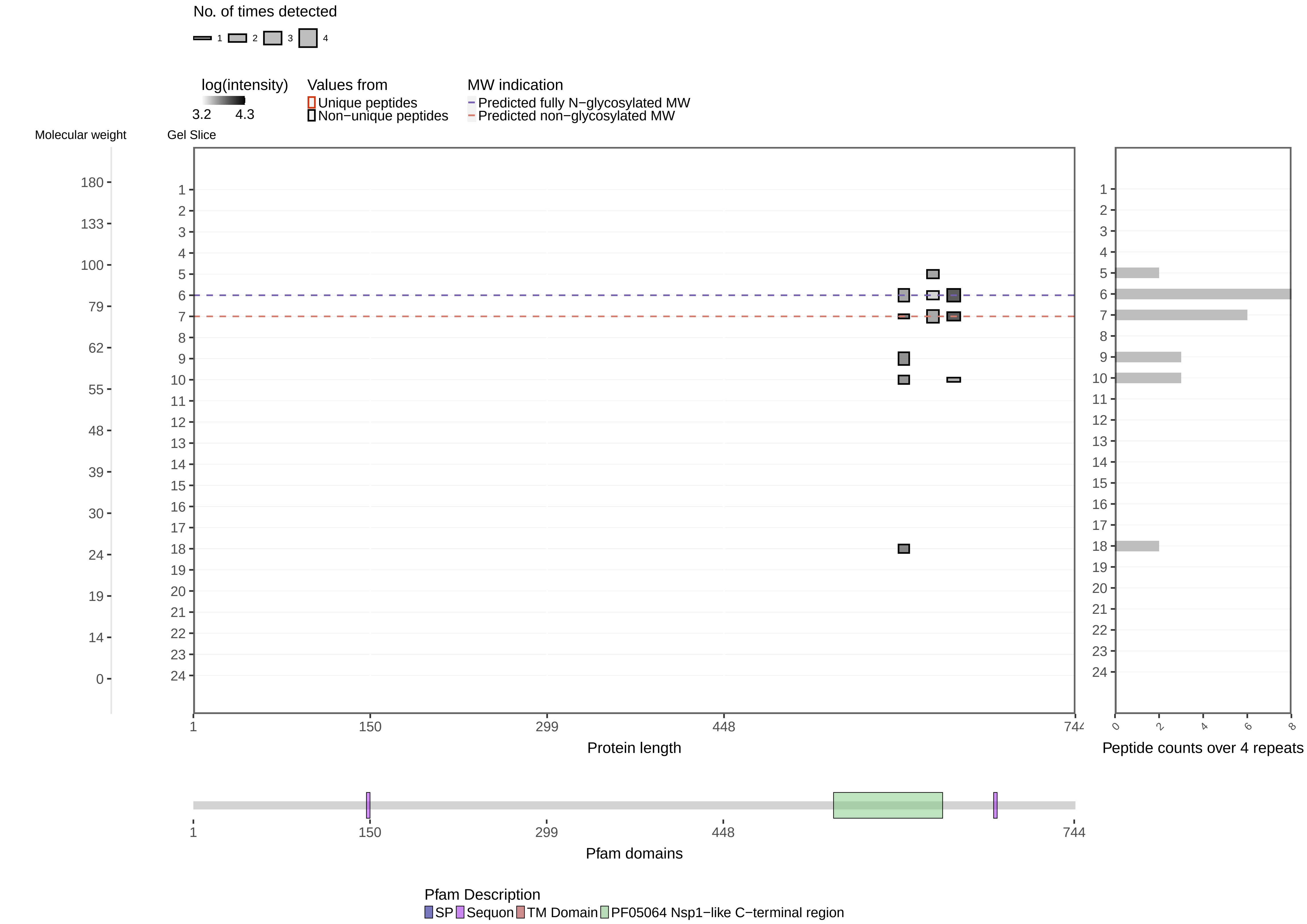


Protein sequence:

STSLSFSVTKGTGSSAAASSAPSLTTAFPFSTSSSAASSSATPVSKPASTGFSIASSPLFSTVTTTTTTGISTTATATGTSSAVSTTSGTGTGSSLTFPAFNAAPSSSASTTSAAAPTTSFSLSSKTSASTAALPSFSVPSTTSGAATSSAAPTSSALTVTASSGTTSTSTVAATSTLKLPSESTGKSVEEIIKEWNAELQERSAKFRKQANAIADWDKRILQNRDILLRLESEVGKVVETQTSLERQLELIETHQEEVDKALQSMEEEADRIYKDERGVILDDEAASTRDAMYEQAEF

NbL01g15180.1 elongation_factor_G−2_mitochondrial−like [MW = 87.4 kDa]

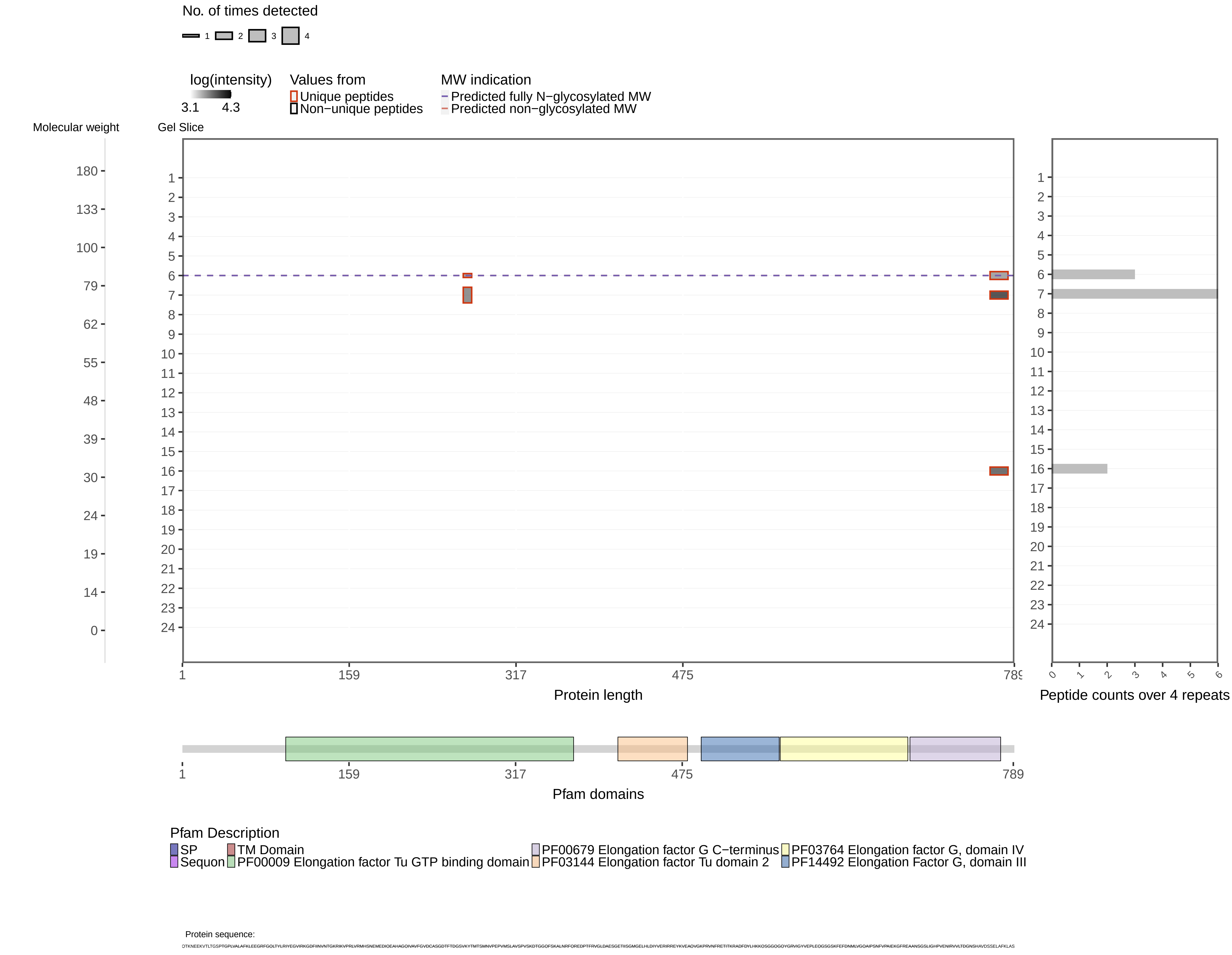

NbL01g15170.1 3−hydroxyisobutyryl−CoA_hydrolase−like_protein_1_mitochondrial [MW = 45.06 kDa]

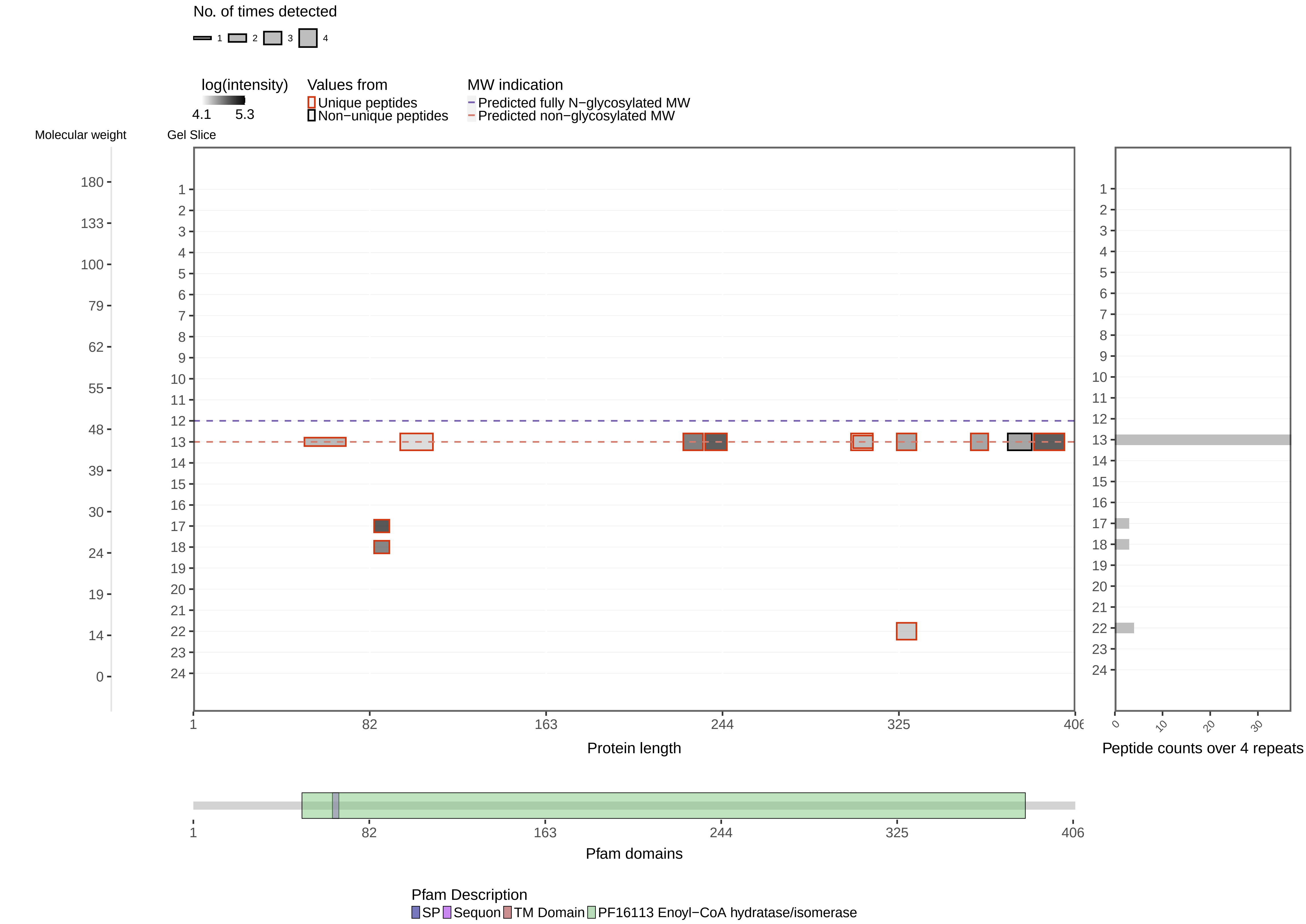


Protein sequence:

LPGYLGEYLALTGDKINGAEMISCGLATHYLHSAKLPLIEEQLGKLMTDDPSVIERSLENCGEIVHPDPTSVLHRIETLNKCFSHDTVEEIIDALESEAAKKQDAWCVSTLRKLQETAPLSLKVSLRSIREGRHQTLDQCLIREYRMSVQAFSGQITNDFCEGVRARLVDRDFAPKWDPPSLDKVTDDMVDQYFSRLTAFEPELELPTQQREAFT*

NbL01g15160.1 malate_dehydrogenase_cytoplasmic−like [MW = 35.84 kDa]

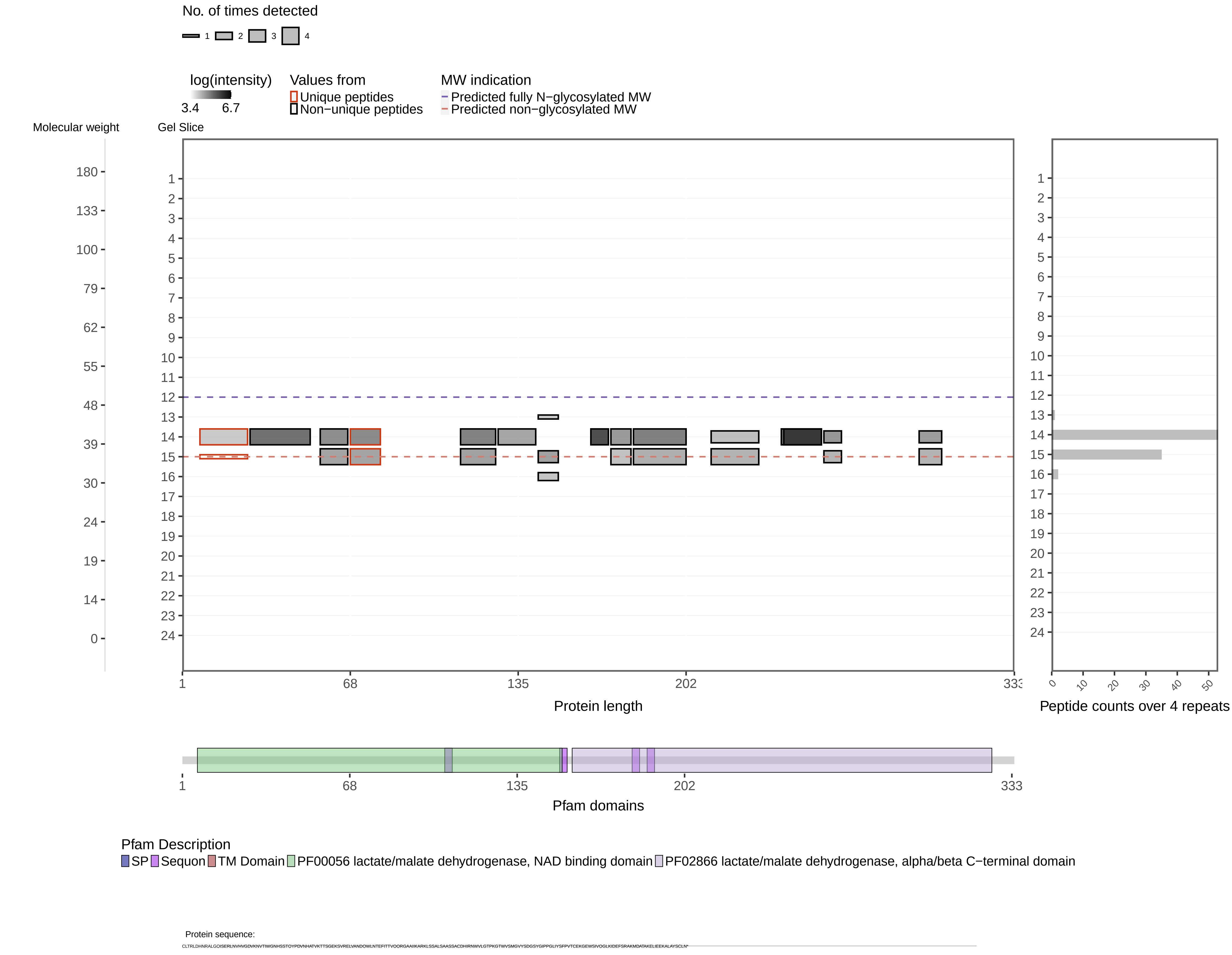

### NbL01g15120.1 T−complex_protein_1_subunit_alpha [MW = 59.14 kDa]

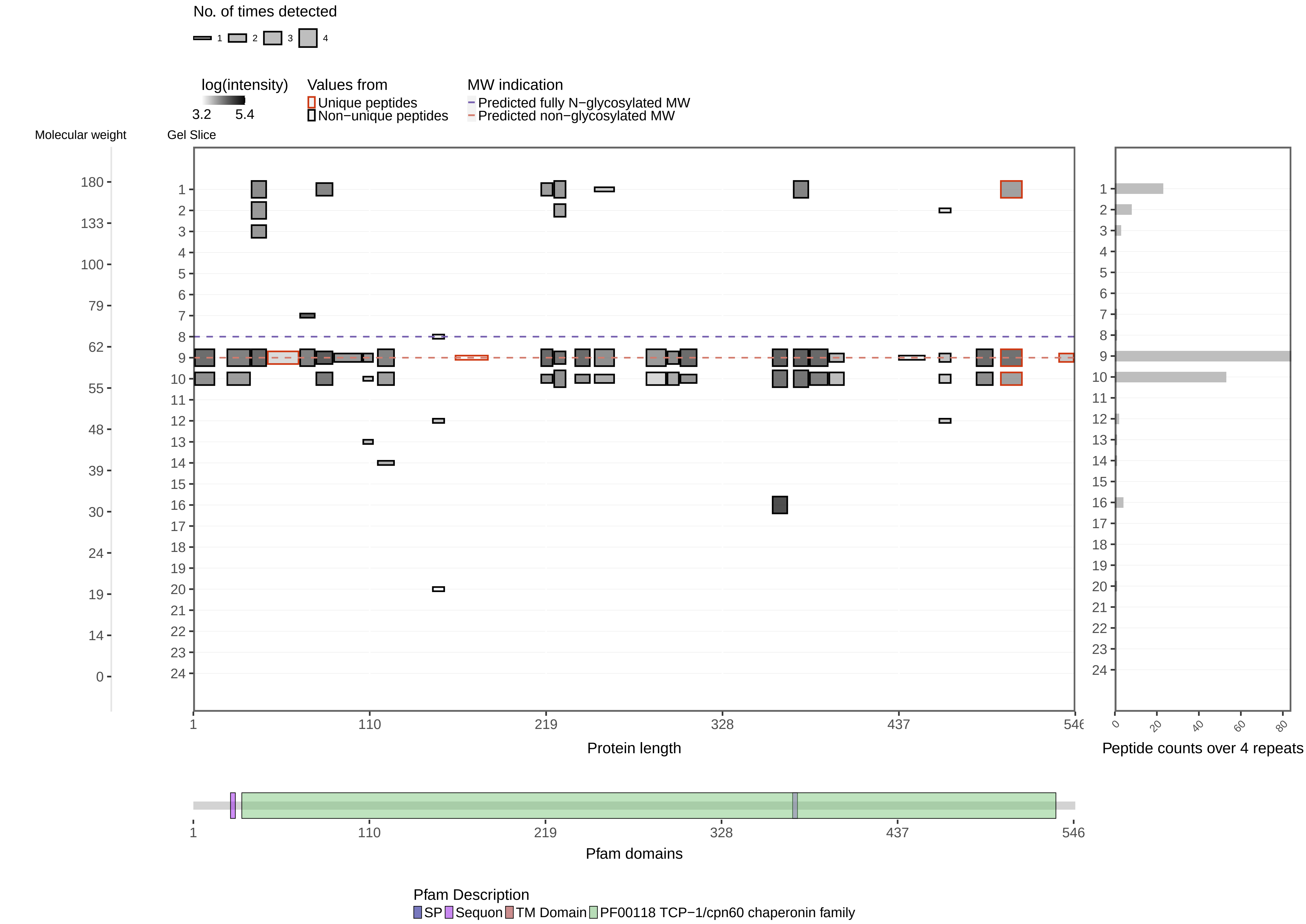


Protein sequence:

DPRELERIRQREADMTKERIQKLLSAGANVVLTSKGIDDMALKYFVEAGAIAVRRVRKEDLRHVAKATGATVVSTFADMEGEETFEPSLLGHAEEVVEERIADDDVIMVKGTKNSSSVSLILRGANDFMLDEMERALHDALCIVKRTLESTTVVAGGGAVEAALCVYLEYLATTLGSREQLAIAQFAESLLVIPKVLANNAAKDSSDLVSKLRSCHYLAQTKADKKHLSSMGLDLAKGTVRNNLEAGVIEPAMSKVKIIQFATEAAITILRIDDMIRLVKDENQNDD*

### NbL01g14870.1 ATP_synthase_subunit_O_mitochondrial−like [MW = 26.42 kDa]

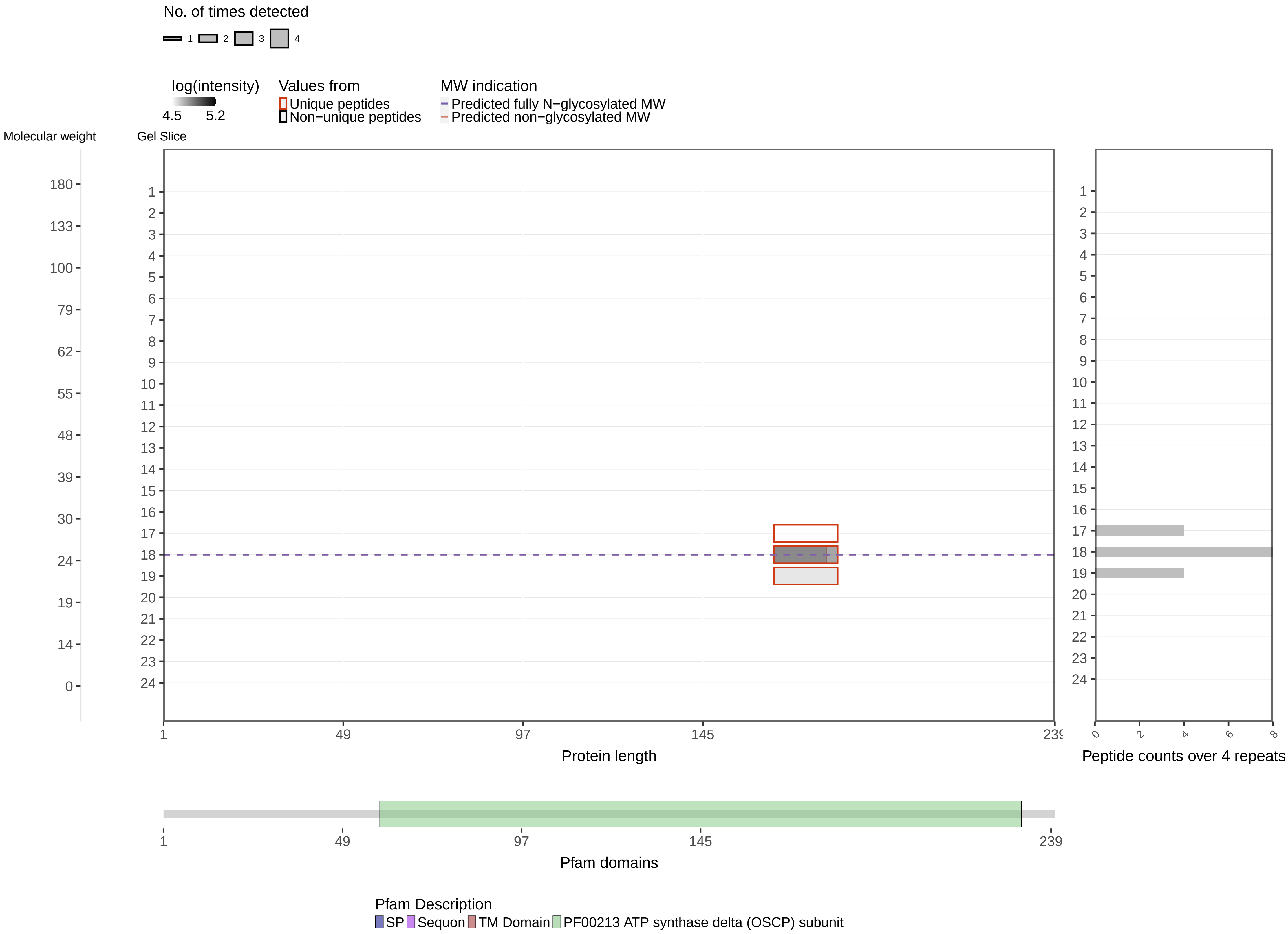


Protein sequence:

VPTDTRVKAISDICAQAKFGDLTRNFLVVLAENGRLKHIDAIAKRFAELTMAHKGELKATVTTVIPLPPEEEKELKETLQDVMGKGKKVNIEQKIDPSILGGLVIEFGQKVFDMSIRTRARQMERLLKEPLNF*

NbL01g14860.1 polyadenylate−binding_protein_2−like [MW = 22.23 kDa]

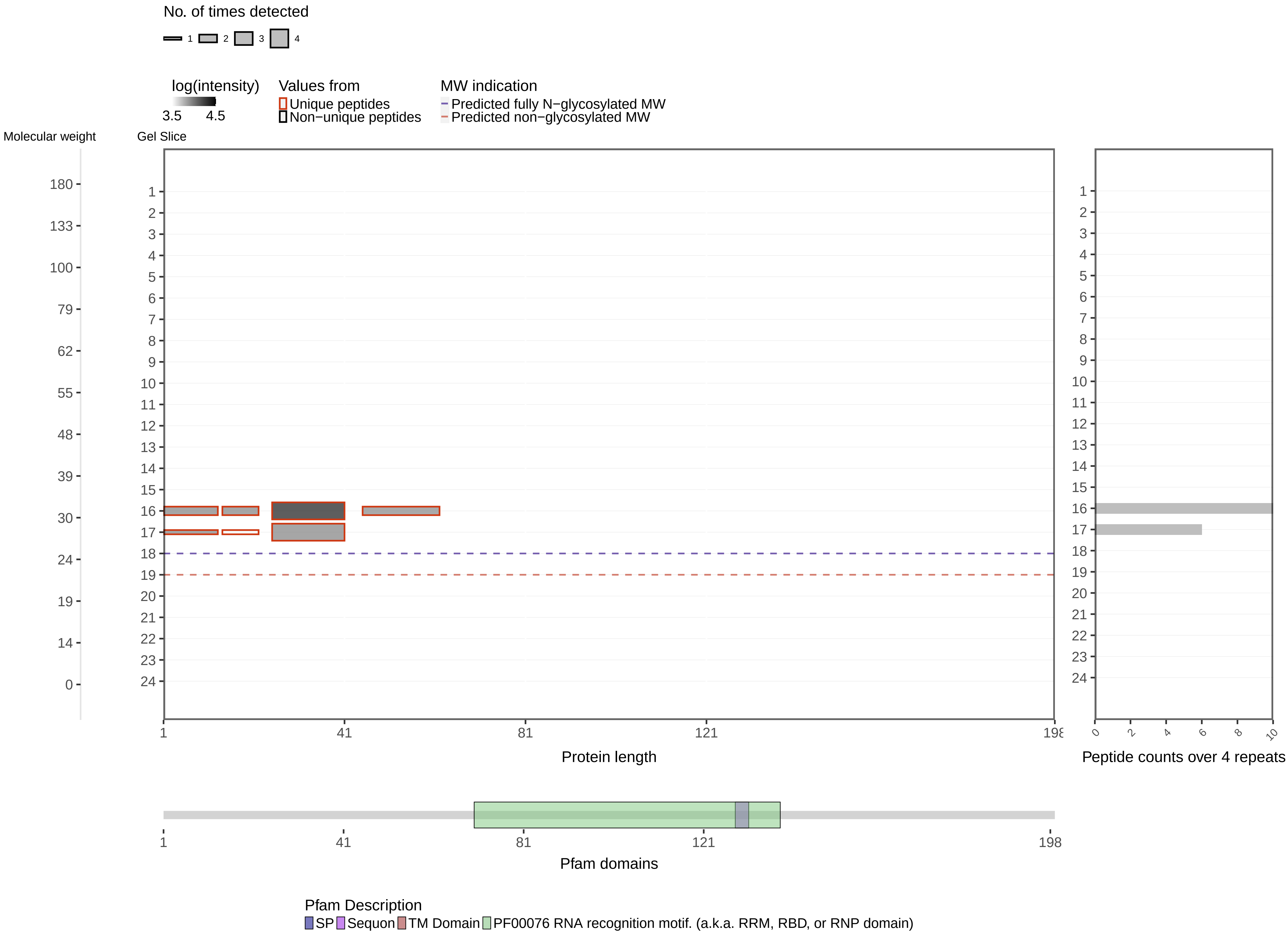


Protein sequence:

QHFQACGTVNRVTIRTNKFGQPKGYAYVEFLEAEAVQEALLLNESELHGRQIKVSAKRTNVPGMKQFRARRPNPYMGFRGQTPFIPAPYFYSPYGYGKVPRARAPMRYSPYF*

### NbL01g14840.1 protein_EXORDIUM−like_2 [MW − SP = 29.18 kDa]

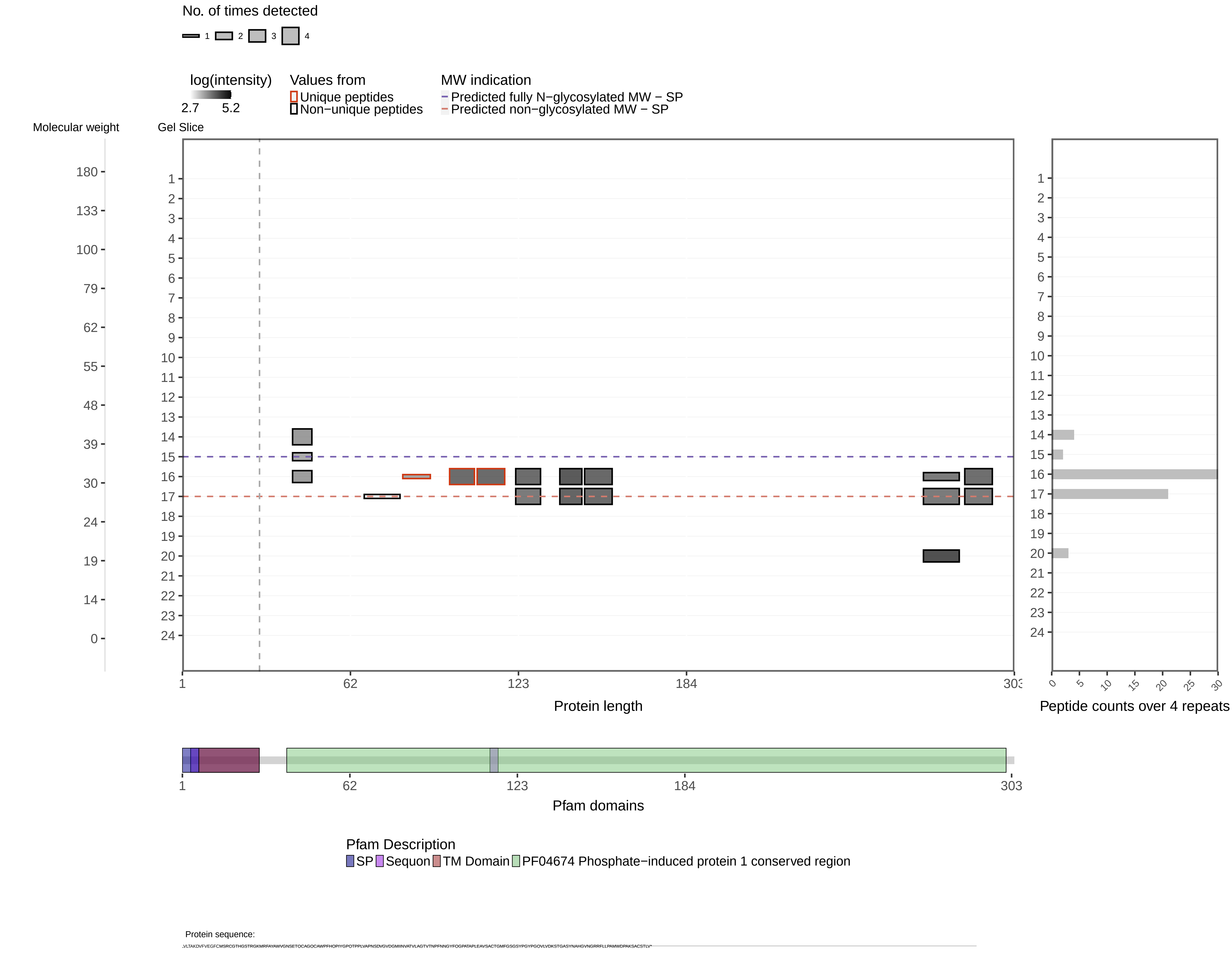

### NbL01g14600.1 peroxidase_N1 [MW = 38.4 kDa]

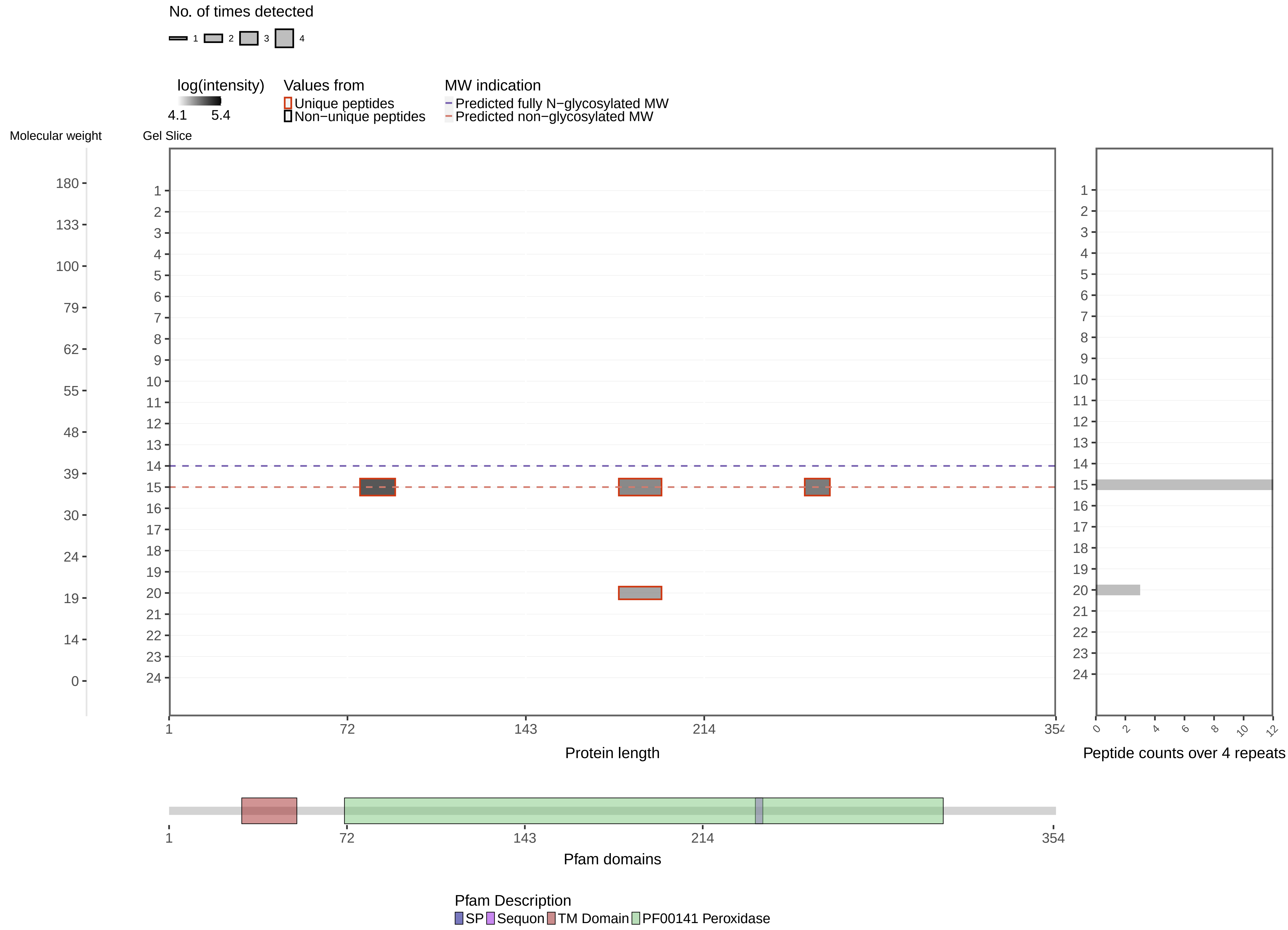


Protein sequence:

SVPTGRRDGRVSRAADAANLPAFFDSVDIQKQKFLEKGLNTQDLVALTGAHTIGTAGCAVIRDRLFNFNSTGGPDPSIDATFLPQLRALCPQNGDASRRVGLDTGSVNNFDTSYFSNLRNGRGVLESDQKLWTDASTRVFVQRFLGVRGLLGLTFGVEFGRSMVKMSNIEVKTGTNGEIRKVCSAIN*

### NbL01g14580.1 uncharacterized_LOC104221519 [MW = 18.92 kDa]

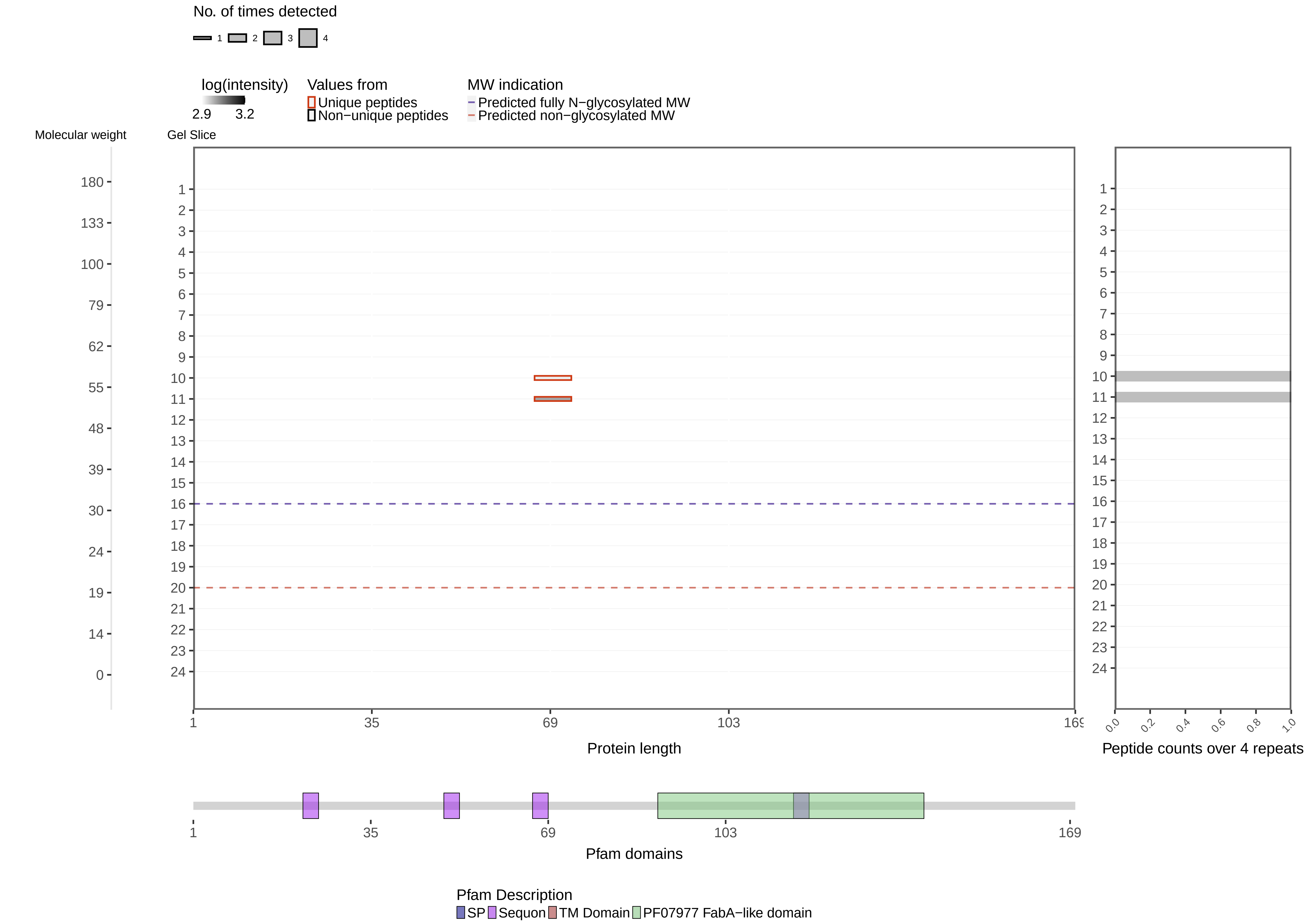


Protein sequence:

IELRYPAFPTVMDINQIREILPHRFPFLLVDRVIEYNPGVSAVAIKNVTINDNFFPGHFPERPIMPGVLMVEIGEVLTLQELLLLSLGFQYLGRVLLRY*

### NbL01g14560.1 probable_protein_phosphatase_2C_62 [MW = 86.59 kDa]

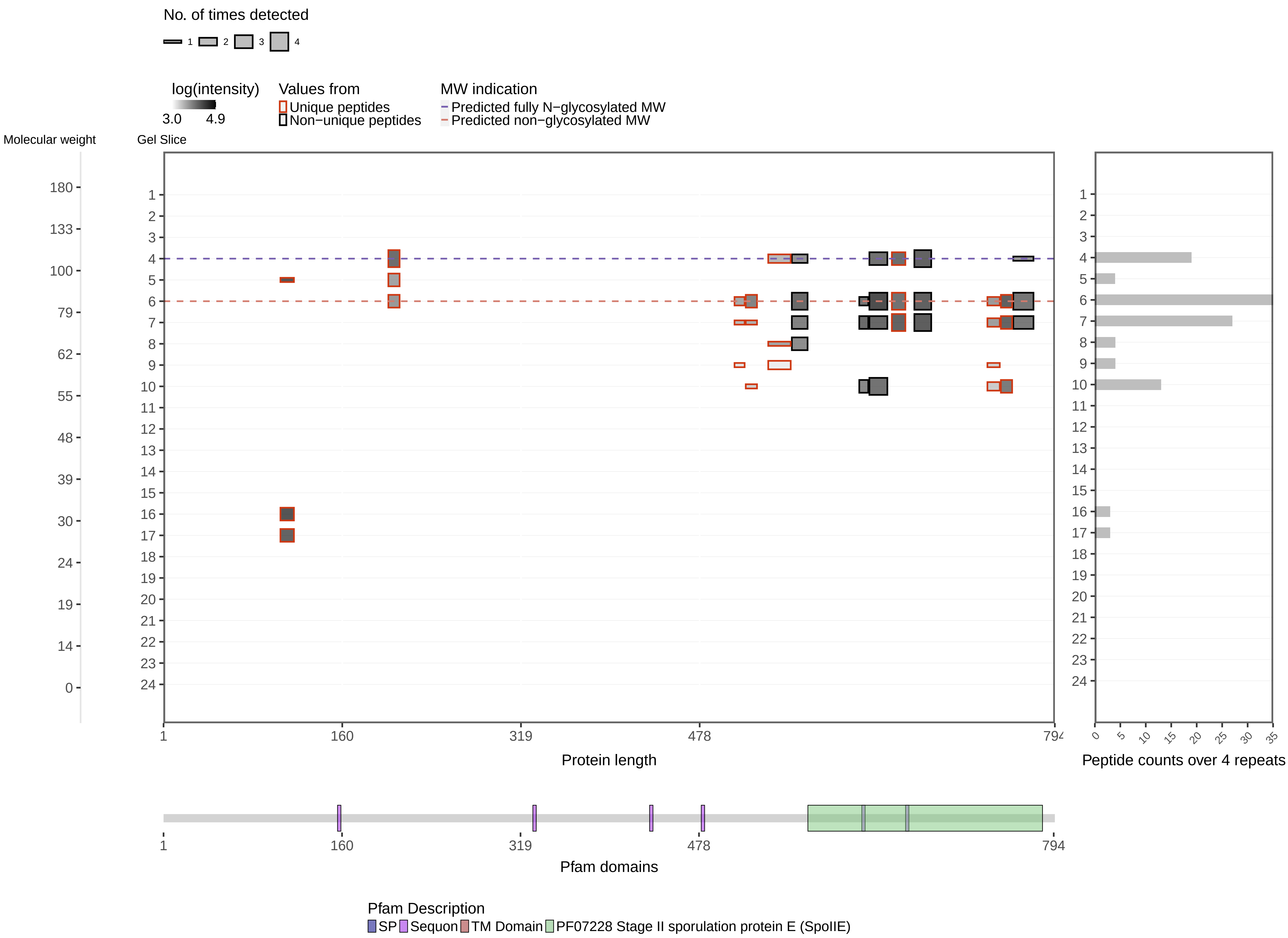


Protein sequence:

VPEEINLENIETVVDEISPENADIRHLSIDAEDLSTQVLSLENDDQTELTNFSEDTEDOASQKGTFKDSDEANVIEVMPLSPMLAAEPILDEEVDHNLTSESNEEGESPLSPEENLPSLSVEAEKEANAENSEVREISGHEVVEDRSSEAIGSSKEAMPAELILSSGATLLEYPSKALTGGHEAYFVACGKWLGVADGVGSWSLEGSDKGVYAKELMONSESIVSQCDNDILNDPKQVLNLSVSKTDSPGSSTALIAHFDGKDLRVANIGDSGFIIVRNGSVYRKSSPMLHEFNLI

NbL01g14500.1 cytochrome_f [MW = 24.69 kDa]

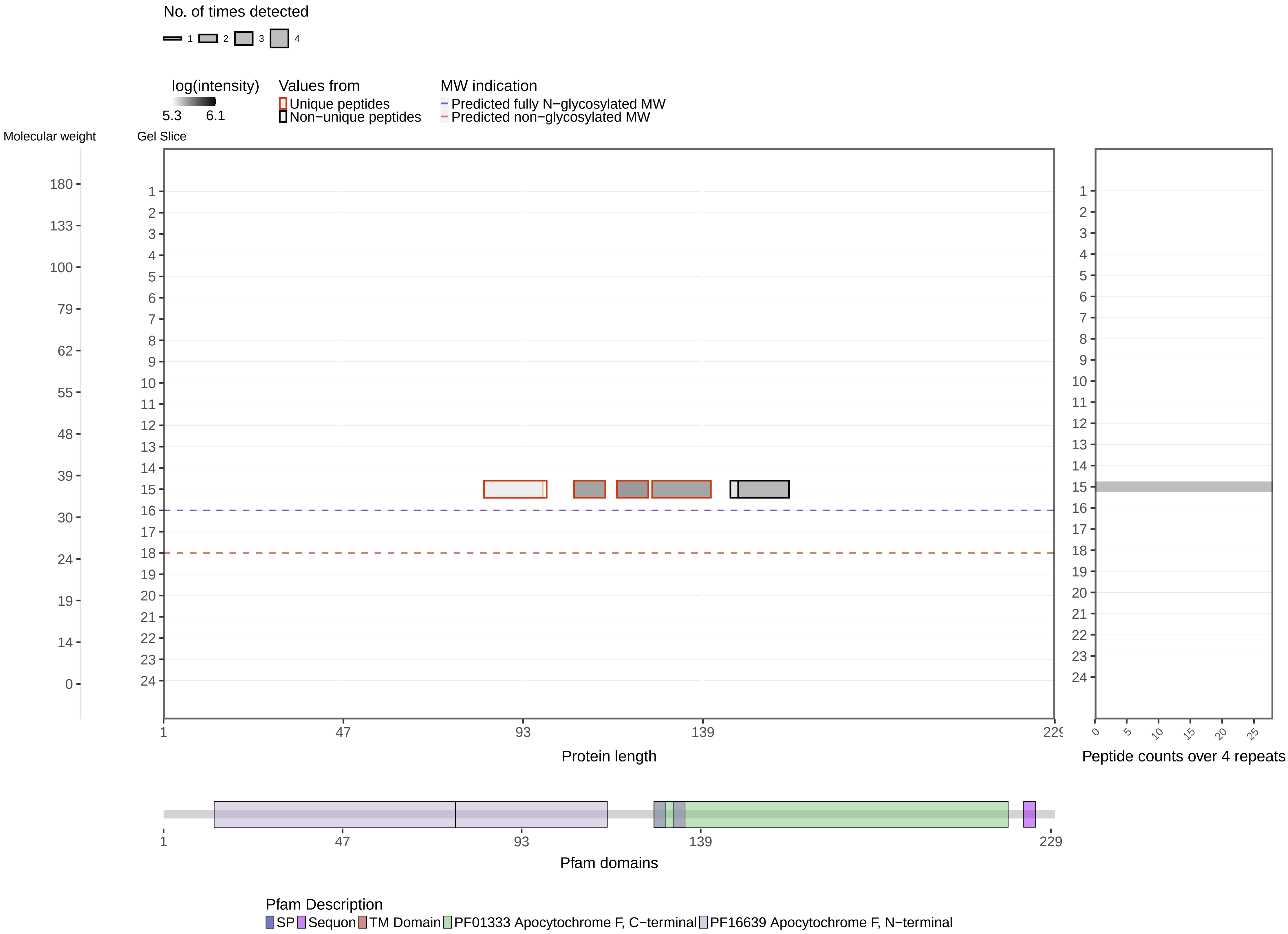


Protein sequence:
/HFLKYPIYVGGNRGRGQIYPDGSKSNNTVYNATAAGIVSKIIRKEKGGYEITITDASDGRQVVDIIPPGPELLVSEGESIKFDQPLTSNPNVGGFGQGDAEIVLQDPLRVQGLLFFTGPHNYSSDYF*

NbL01g14480.1 beta−xylosidase/alpha−L−arabinofuranosidase_2−like [MW − SP = 80.98 kDa]

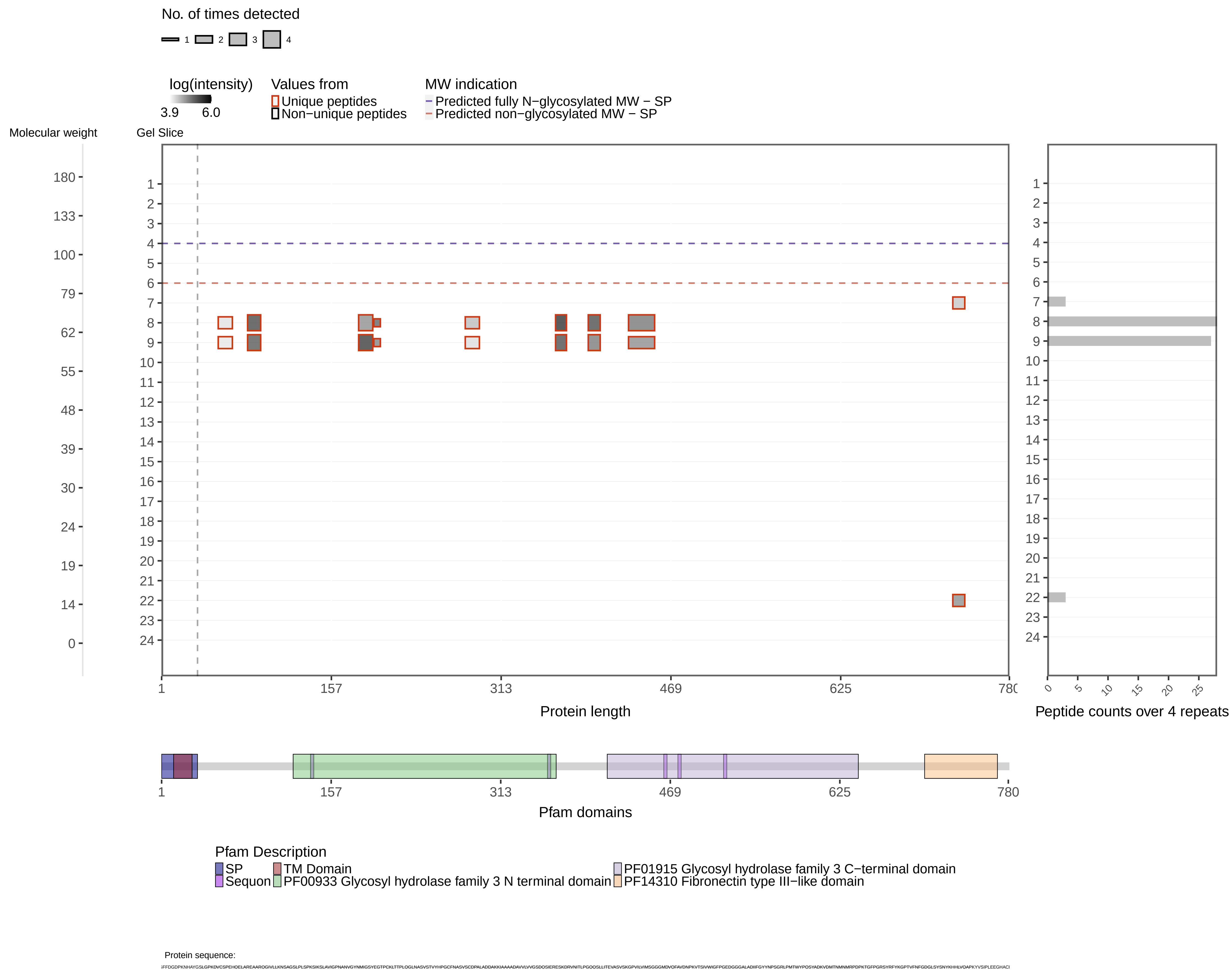

NbL01g14440.1 phosphatidylserine_decarboxylase_proenzyme_2−like [MW = 71.83 kDa]

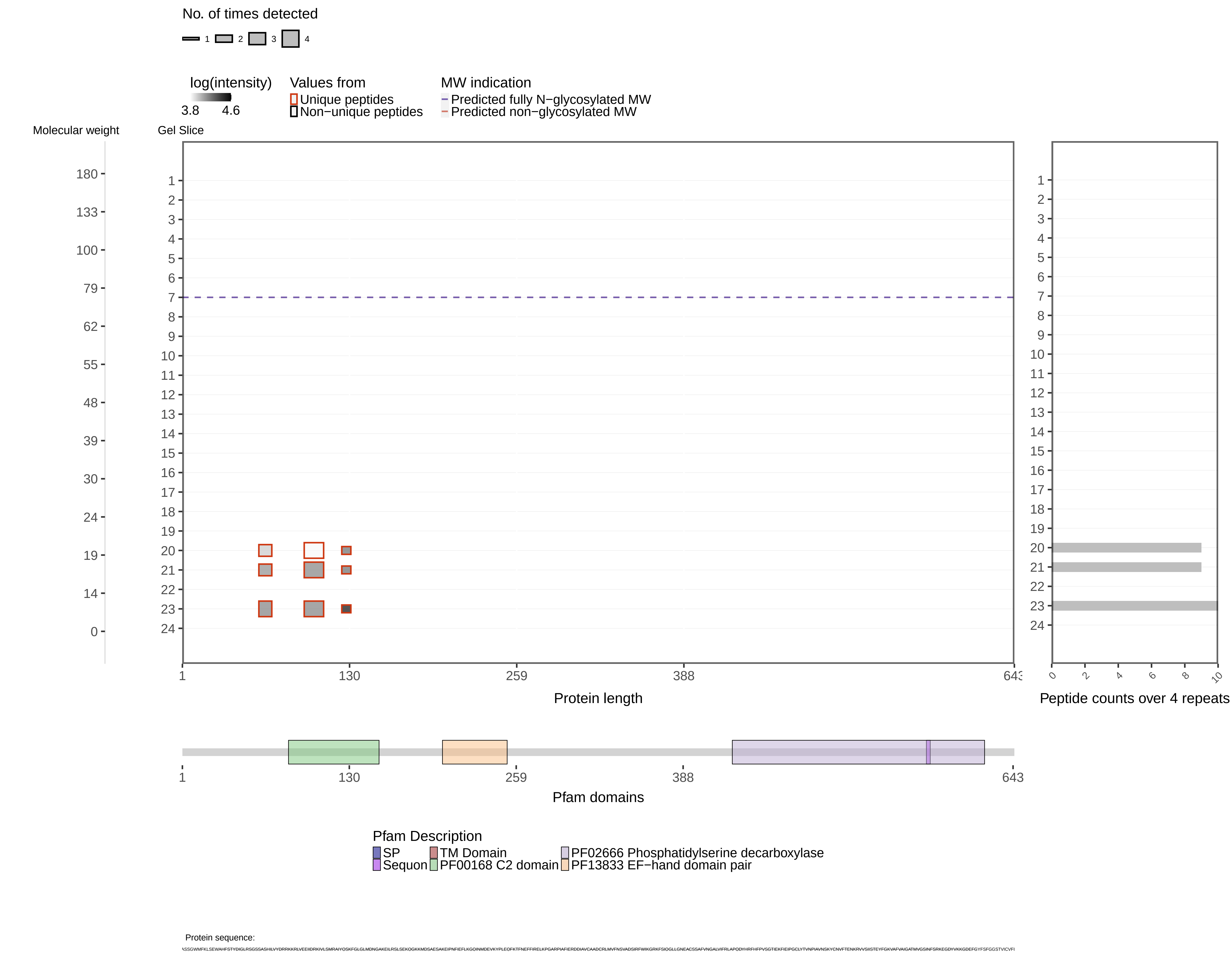

### NbL01g14370.1 glycine−rich_RNA−binding_protein_RZ1A−like [MW = 22.3 kDa]

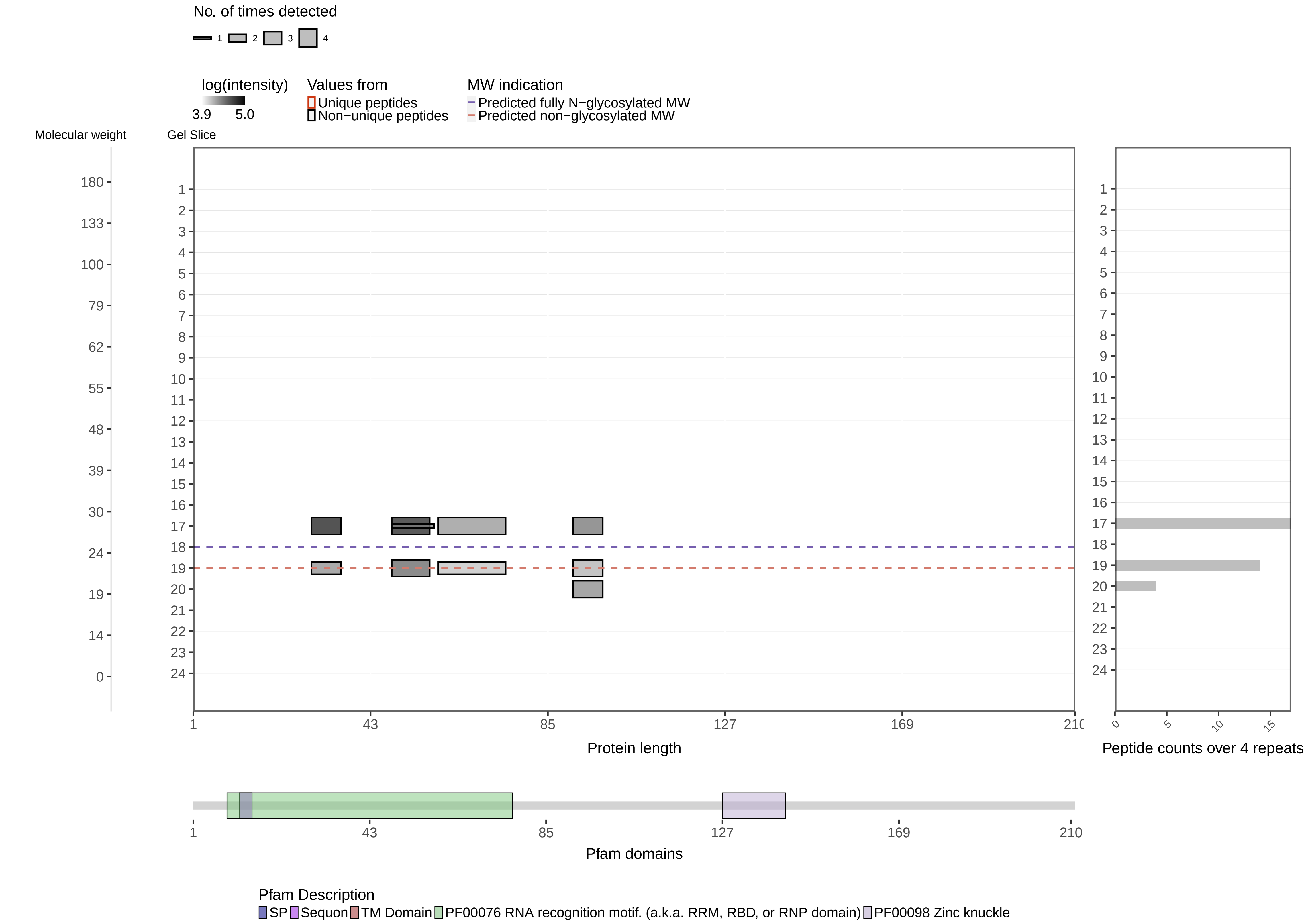


Protein sequence:

RPRDRDRDRGRDRDRGSRDYGGGRGSGGGGDCFNCGKPGHFARECPSEGGRGGRYGGGGGGGGSRSSGYGPDRNGDRYGSRSGRDGGGRGGGERFSRDRSGPYERRSSGGSRAG*

### NbL01g14330.1 actin−101 [MW = 41.66 kDa]

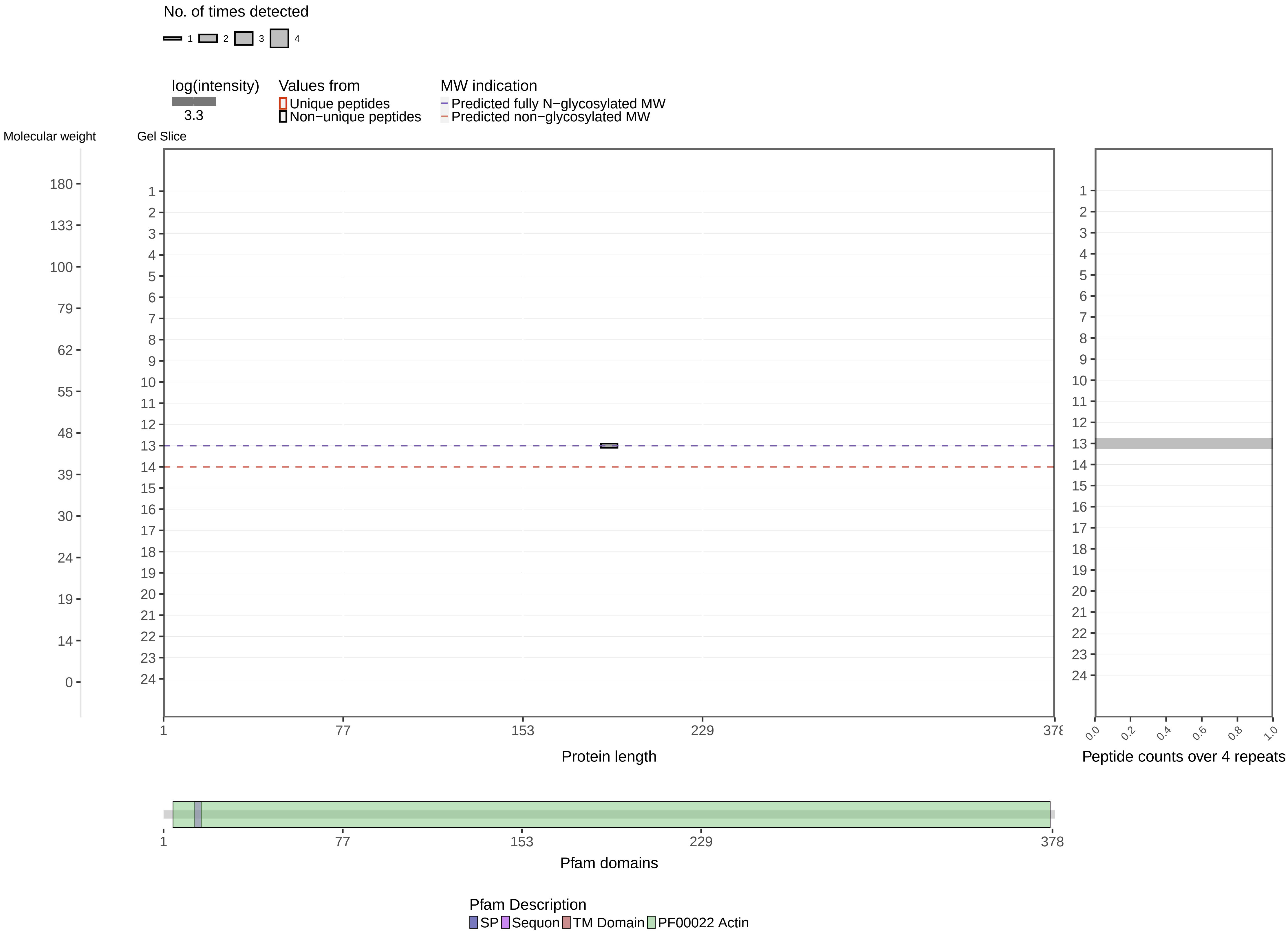


Protein sequence:

PHAILRLDLAGRDLTDCLMKILTERGYSFTTSAEREIVRDVKEKLAYVALDYEQELETAKSSSSVEKSYELPDGQVITIGSERFRCPEVLFQPSLVGMEAAGIHETTYNSIMKCDVDIRKDLYGNIVLSGGTTMFPGIADRMSKEITALAPSSMKIKVVAPPERKYSVWIGGSILASLSTFQQMWITKGEYDESGPSIVHRKCF*

### NbL01g14180.1 60S_ribosomal_protein_L3 [MW = 44.45 kDa]

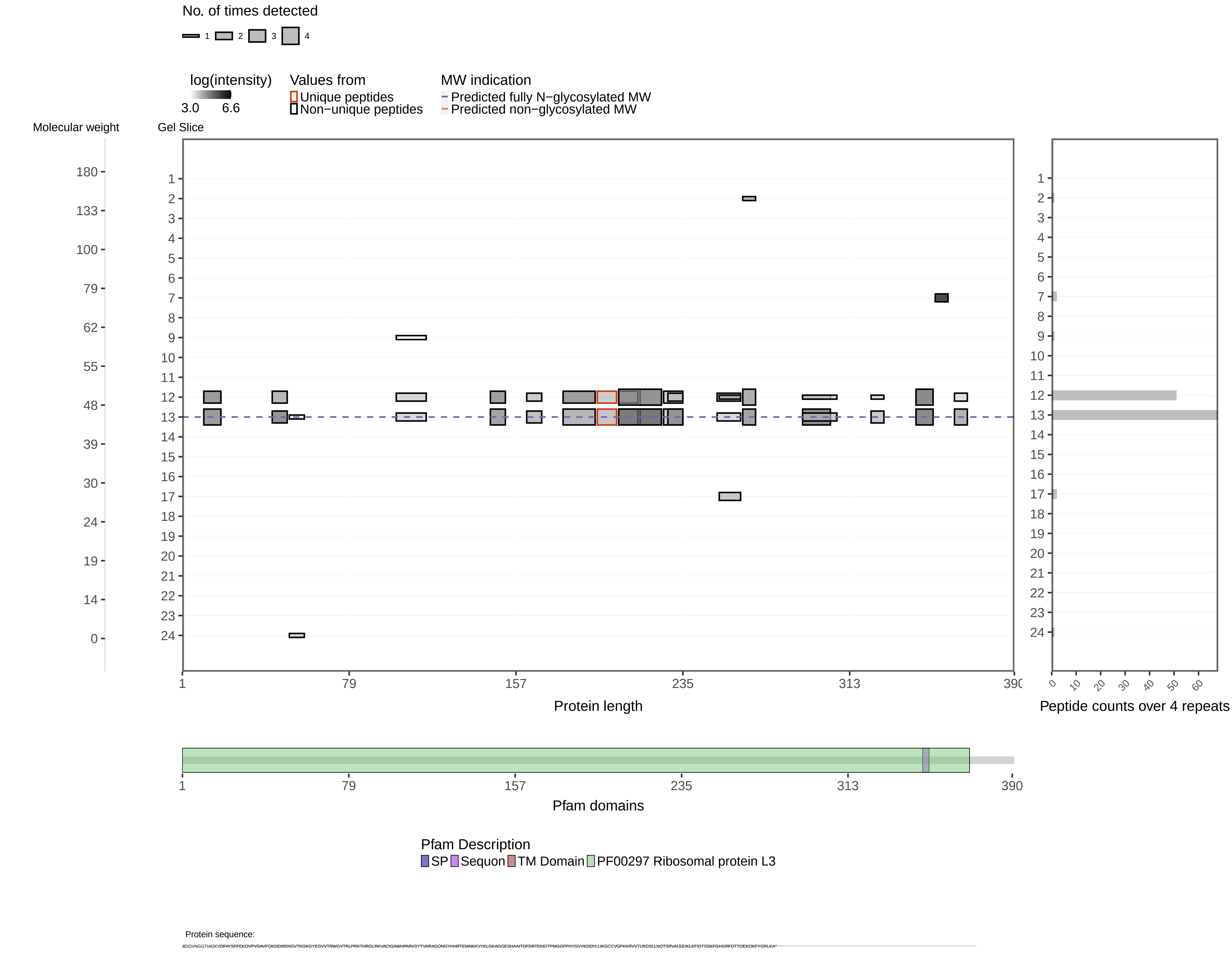

### NbL01g14110.1 WD−40_repeat−containing_protein_MSI1 [MW = 48.37 kDa]

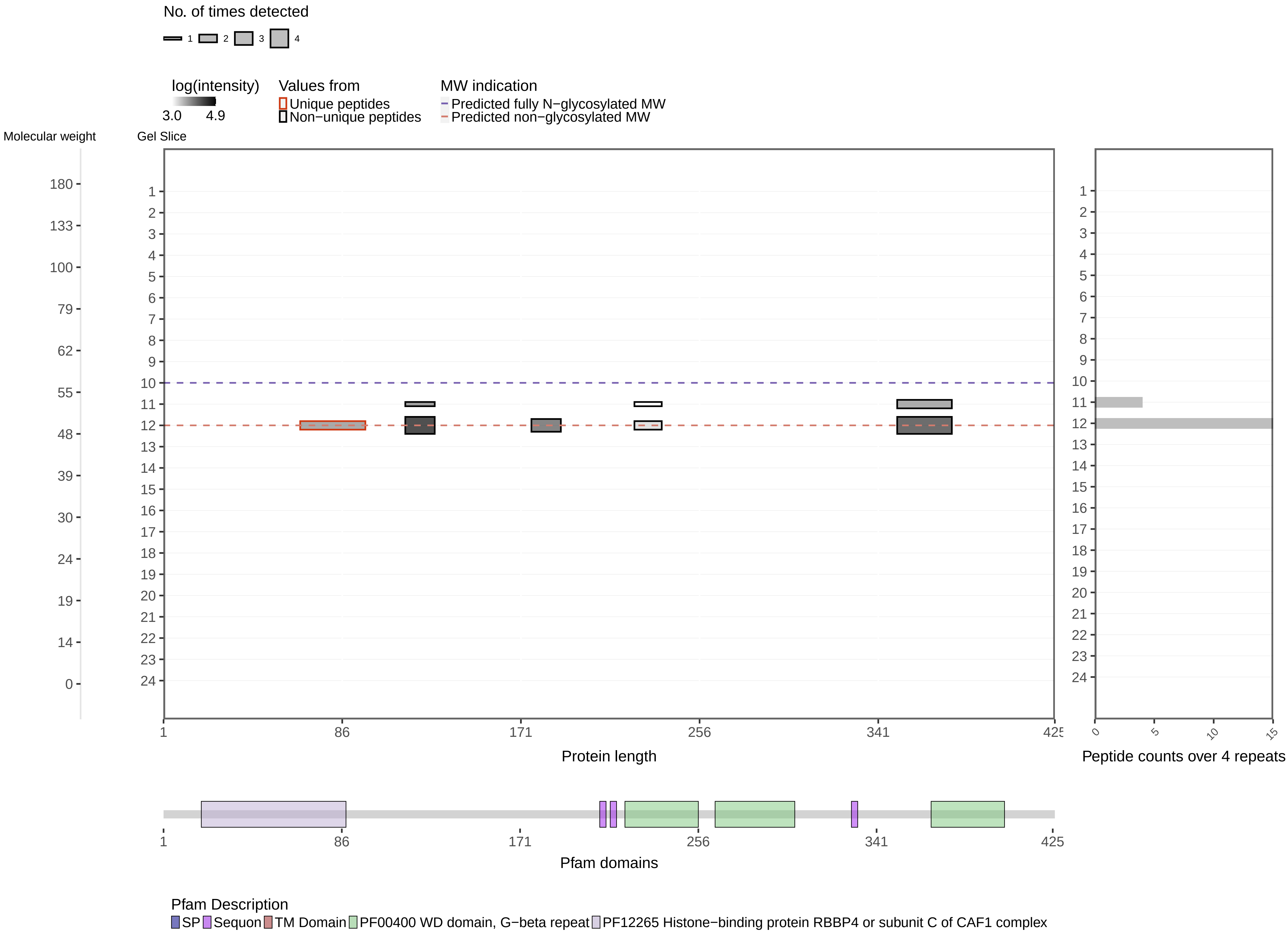


Protein sequence:

DDAQICLWDINATPKNNSLDAMQIFKIHEGVVEDVAWHLRHEYLFGSVGDDQYLHIWDLRTPSVTKPIQSVVAHQSEVNCLAFNPFNEWVVATGSTDKTVKLFDLRKISTALHTLDCHKEEVFQVGWNPKNETILASCCLGRRLMVWDLSRIDEEQTPEDAEDGPPELLFIHGGHTSKISDFSWNPCEDWVVASVAEDNILQIWQMAENIYHDEDDLPGDDATKAP*

### NbL01g14050.1 basic_blue_protein [MW − SP = 10.4 kDa]

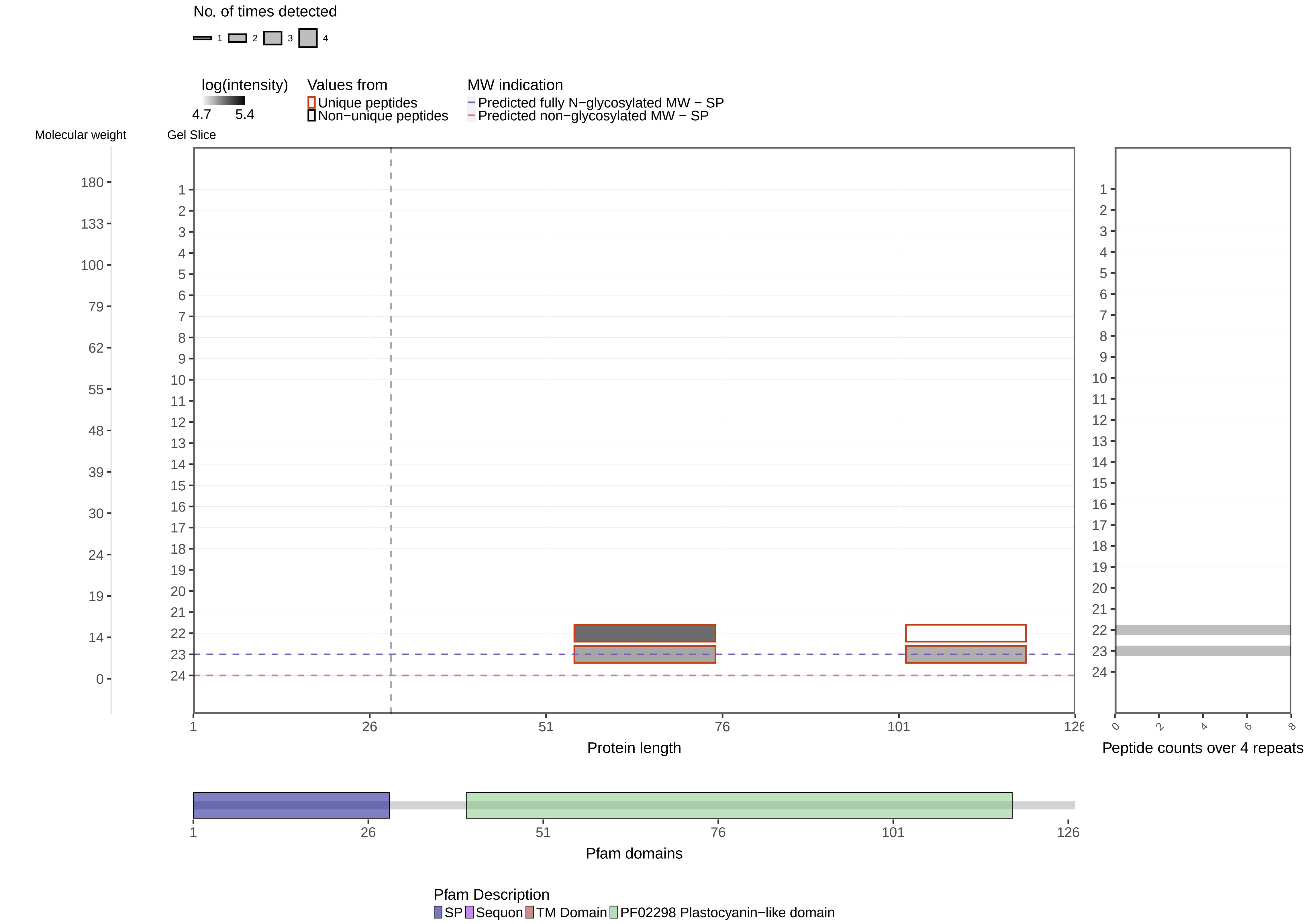


Protein sequence:

GKRFRAGDILAFNYPSGAHNVVVVNKAGYNSCRTPRGSKVYNKGGDKIKLVKGQNYFICNFAGHCESGMKISVFAS*

### NbL01g13880.1 ankyrin_repeat_domain−containing_protein_2B−like [MW = 37.19 kDa]

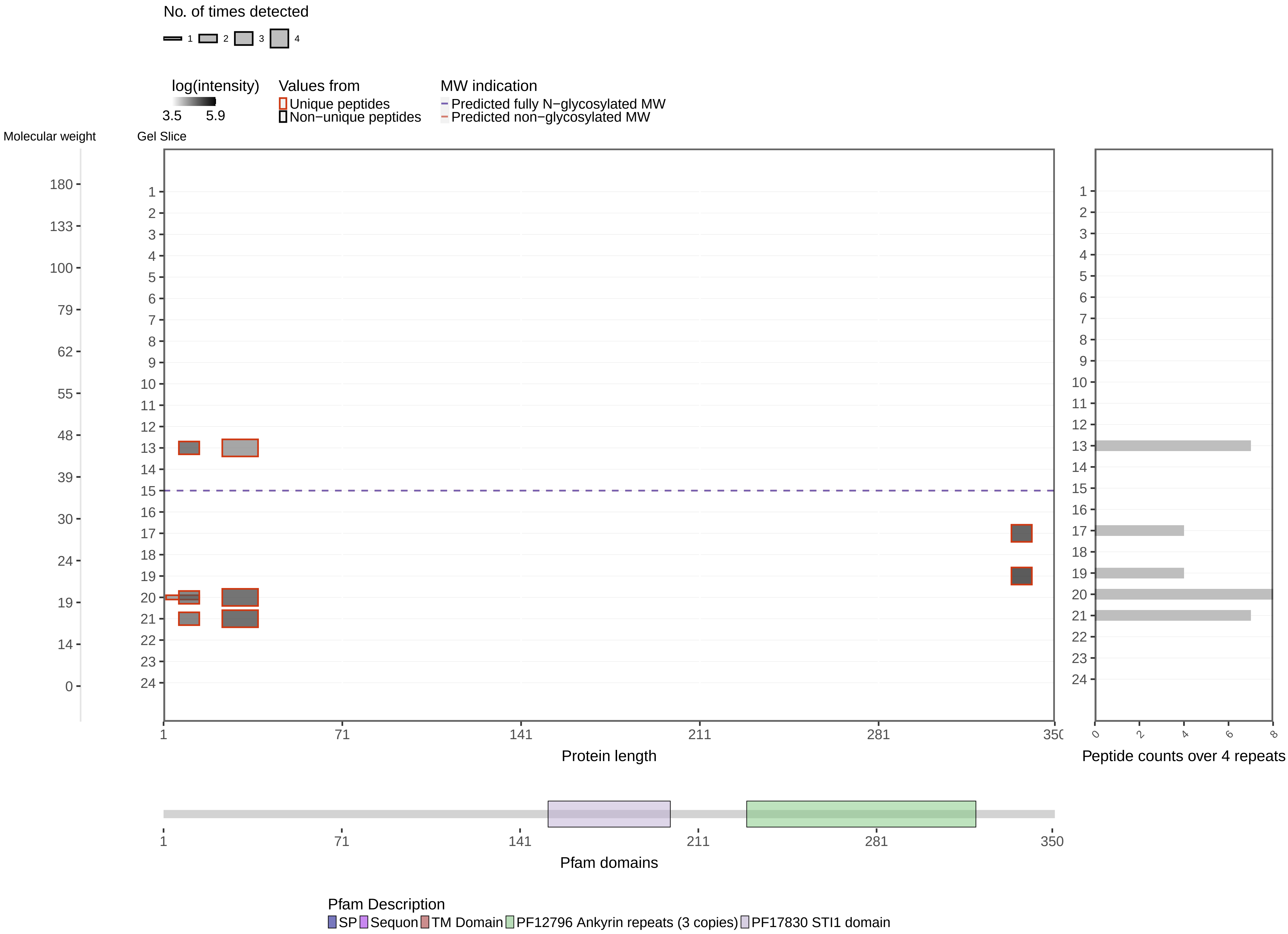


Protein sequence:

RIKEDPSLKPILEEIESGGPAAMMRYWNDQETLKKIGEAMGFAAGGEGATSSAIPGTDETEEANEDESVVHQCASVGDAEGLKAALTAGADKDEEDSEGRTALHFACGYGEVKCAQILLEAGAKVDALDKNKNTALHYAAGYGRKECVALLLENGAAVTLQNLDGKTPIDVAKLNNQEEVLKLLEKDVFL*

NbL01g13780.1 UDP−glucose:glycoprotein_glucosyltransferase [MW − SP = 182.79 kDa]

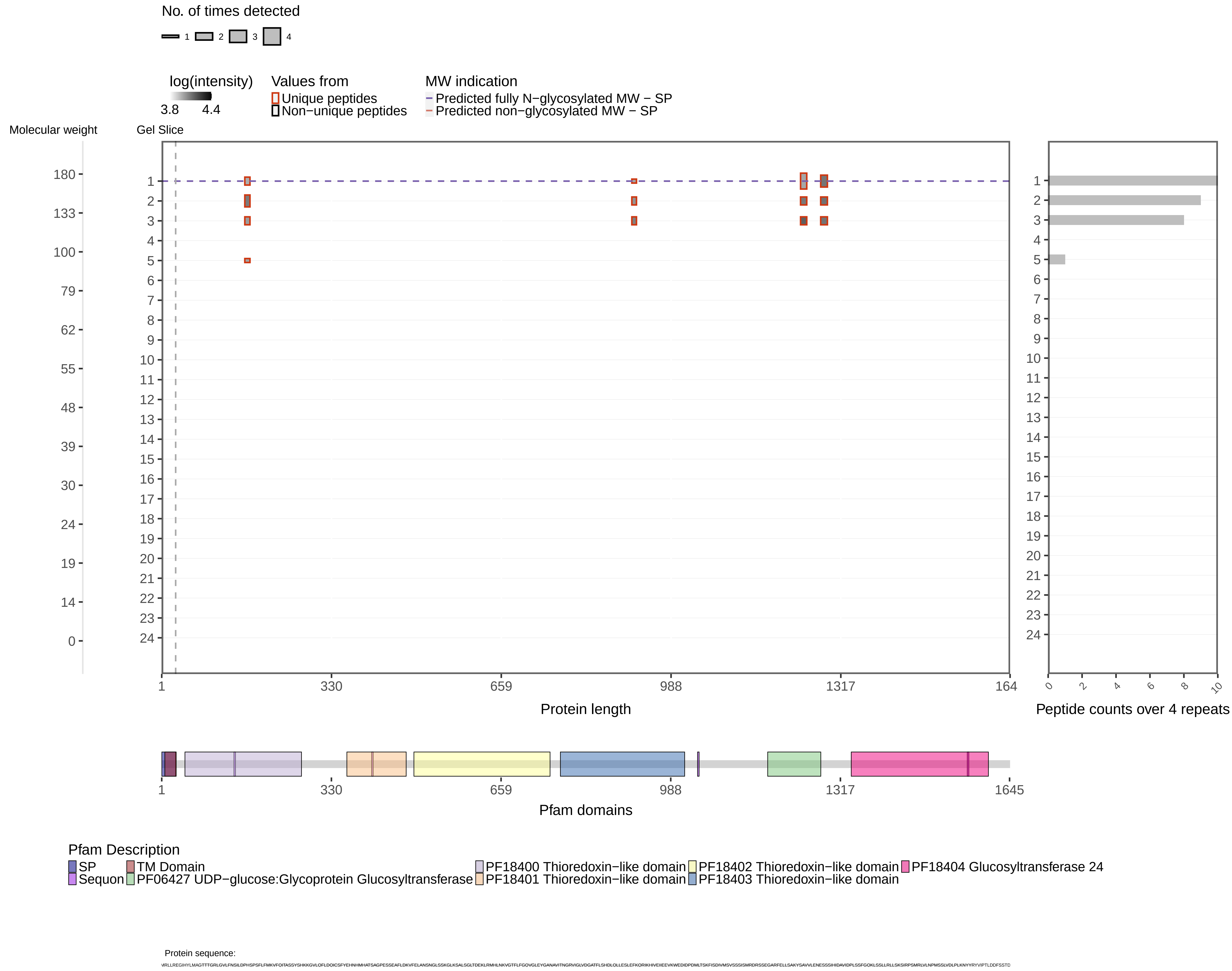

NbL01g13750.1 serine_hydroxymethyltransferase_7−like [MW = 65.53 kDa]

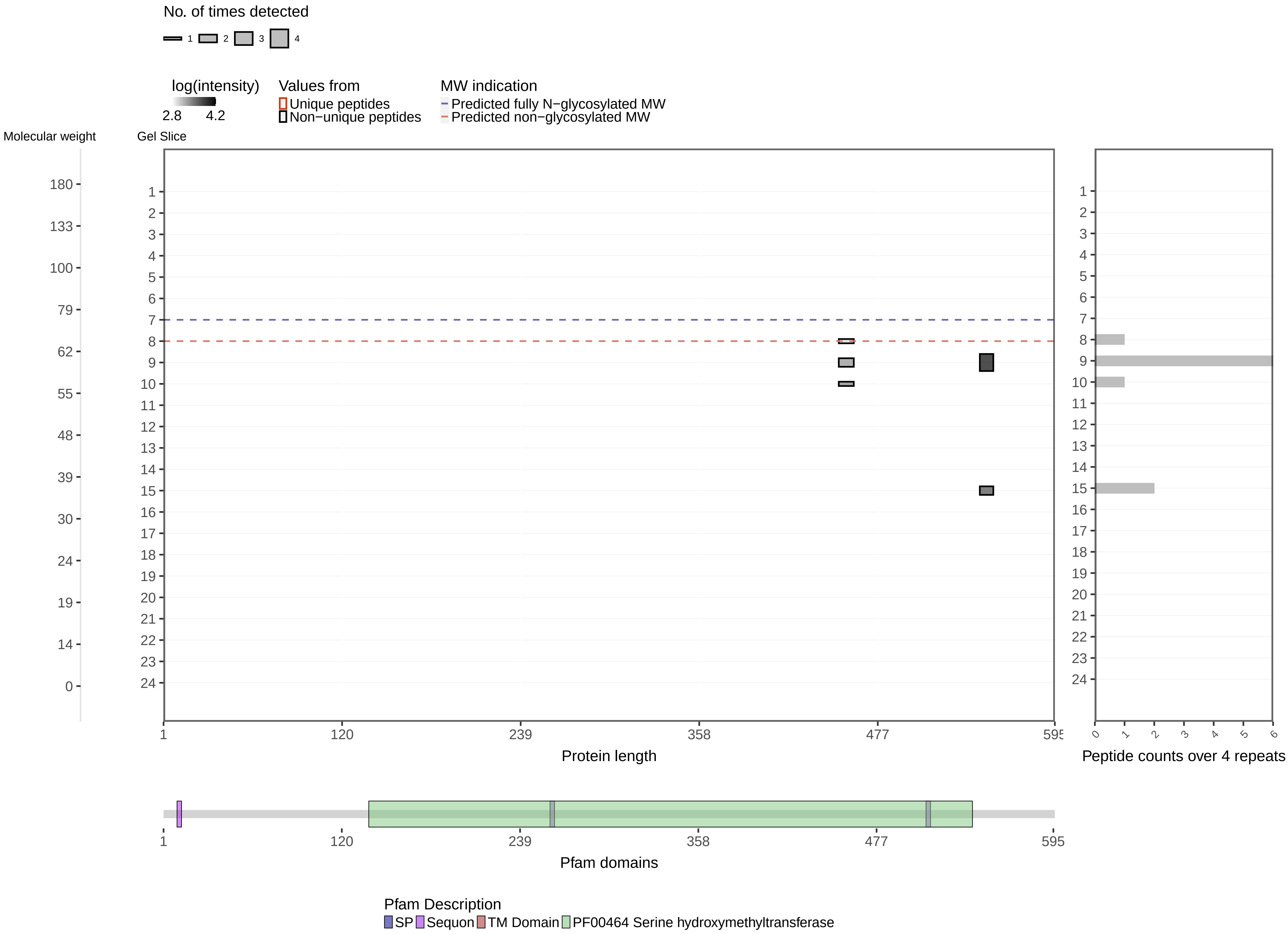


Protein sequence:

SYKVDPQTGYVDFDKLEERALDFRPKILICGGSSYPREWDYAKFRQIADKCGAVLLCDMAQISGLIAAKECASPFDYCDIVTSTTHKSLRGPRGGIIFYRKGSKPRKRGLLLNQGDGGDKYDFEEKINFAVFPALQGGPHNNHIAALAIALKQVATPEYKTYMQQVKRNAQALAAALLRRNCRLVTGGTDNHMILWDLRNLGLTGKNFEKVCELCHITLNKVMVFDDNGSITPGGVRIGTPAMTSRGCIENDFETITDFLLKAAQITNSVQREHGKLVKAFLKGLENNKDVIELRT

### NbL01g13700.1 thiamine_biosynthetic_bifunctional_enzyme_TH1_chloroplastic−like [MW = 60.72 kDa]

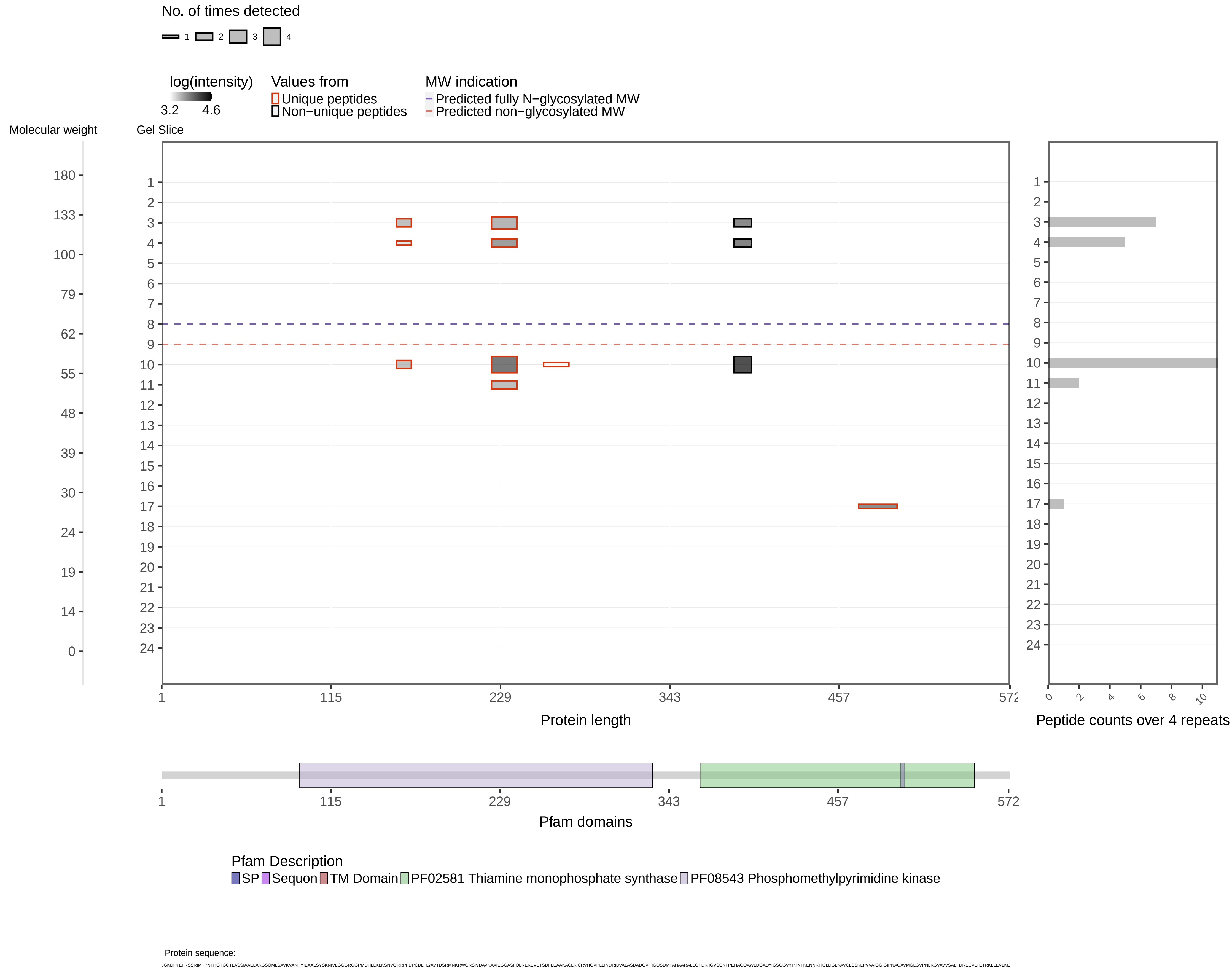

NbL01g13690.1 kinase_SRK2A [MW = 40.88 kDa]

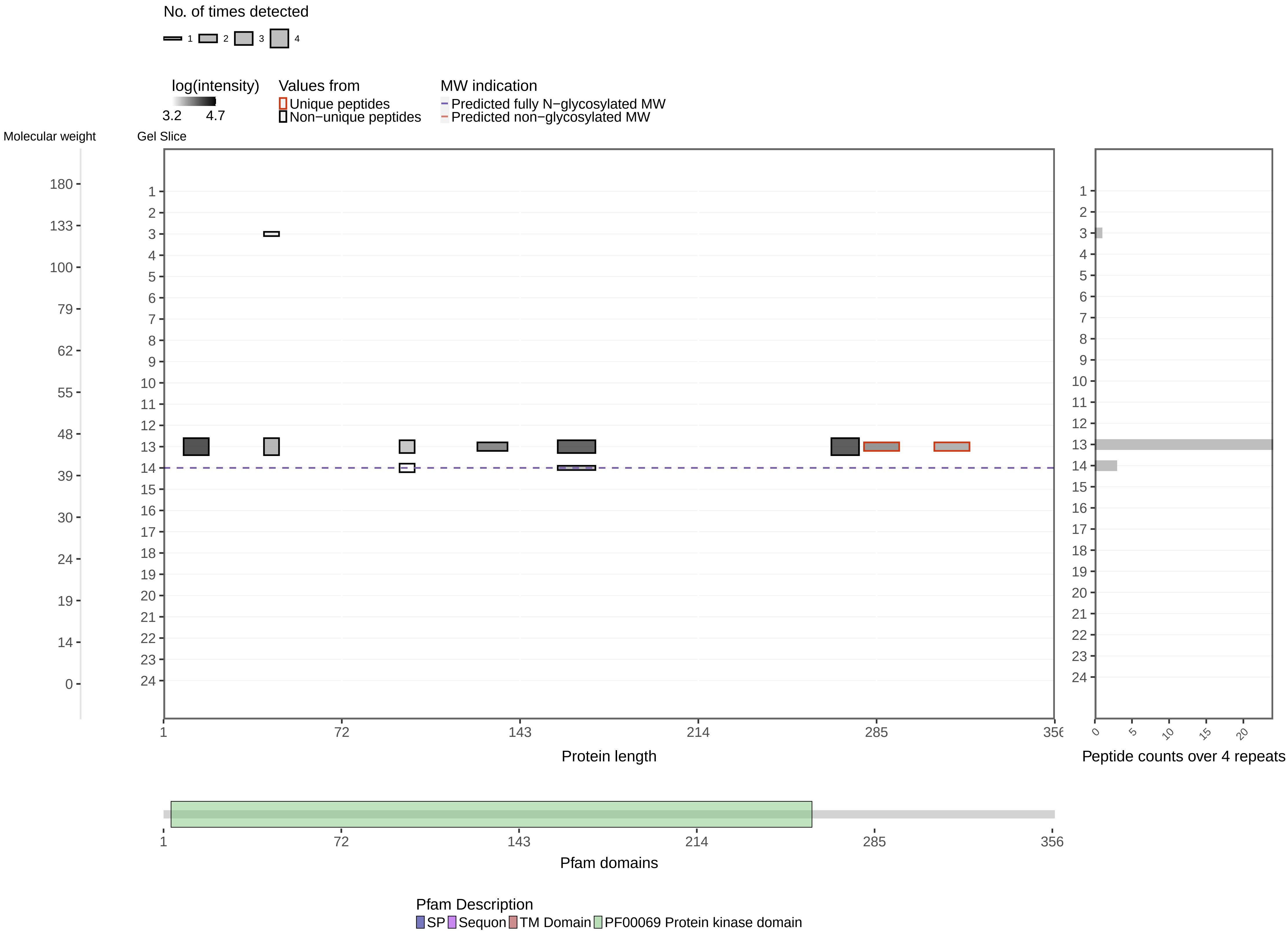


Protein sequence:

APEVLSRREYDGKLADVWSCGVTLYVMLVGAYPFEDQEDPKNFRKTIQRIMAVQYKIPDYVHISQDCRHLLSRIFVANPARRITIKEIKSHPWFLKNLPRELTEAAQAAYYRRENPTFSLQSVEEIMKIVEEAKTPAPASRSVSGFGWGGEEEEEKEGDVEEEGEDEEEEDEYEKQVKQAHESGEVRLT*

NbL01g13610.1 ubiquitin−like_domain−containing_CTD_phosphatase [MW = 38.74 kDa]

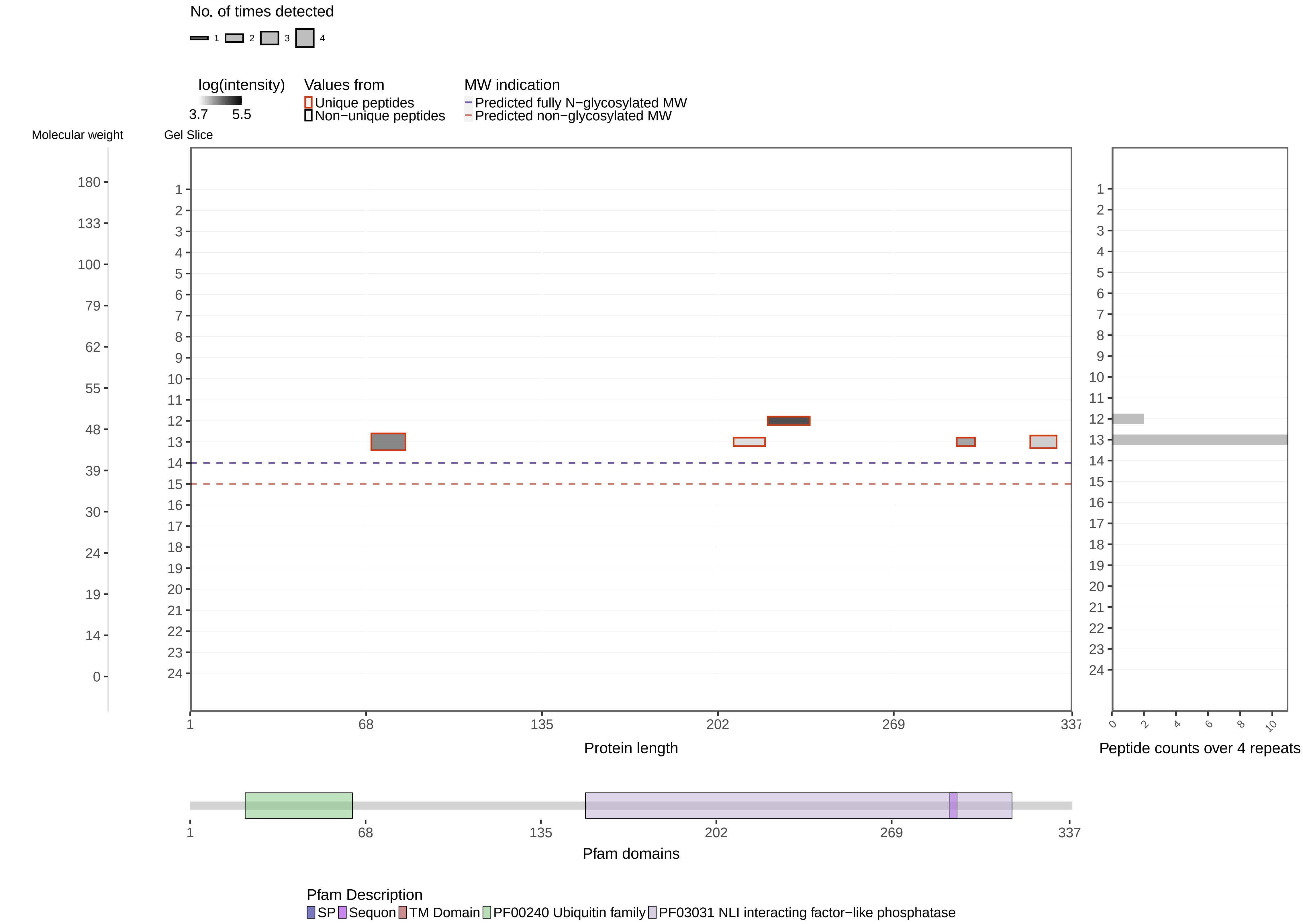


Protein sequence:

IDYTLFDHRSTAENPLQLMRPYLHEFLSAAYAEYDIMIWSATSMKWVELKMGQLGVLDNPHYKITALLDHMAMITVQSDRYGVFDCKPLGLIWAHFPEFYSPKNTIMFDDLRRNFVMNPQNGLVIKPFRKAHANRSTDQELMKLTQYLLAIADLDDVSVLDHKNWESFNEDNFKRRRHA*

### NbL01g13550.1 ATP_phosphoribosyltransferase_2_chloroplastic−like [MW = 43.01 kDa]

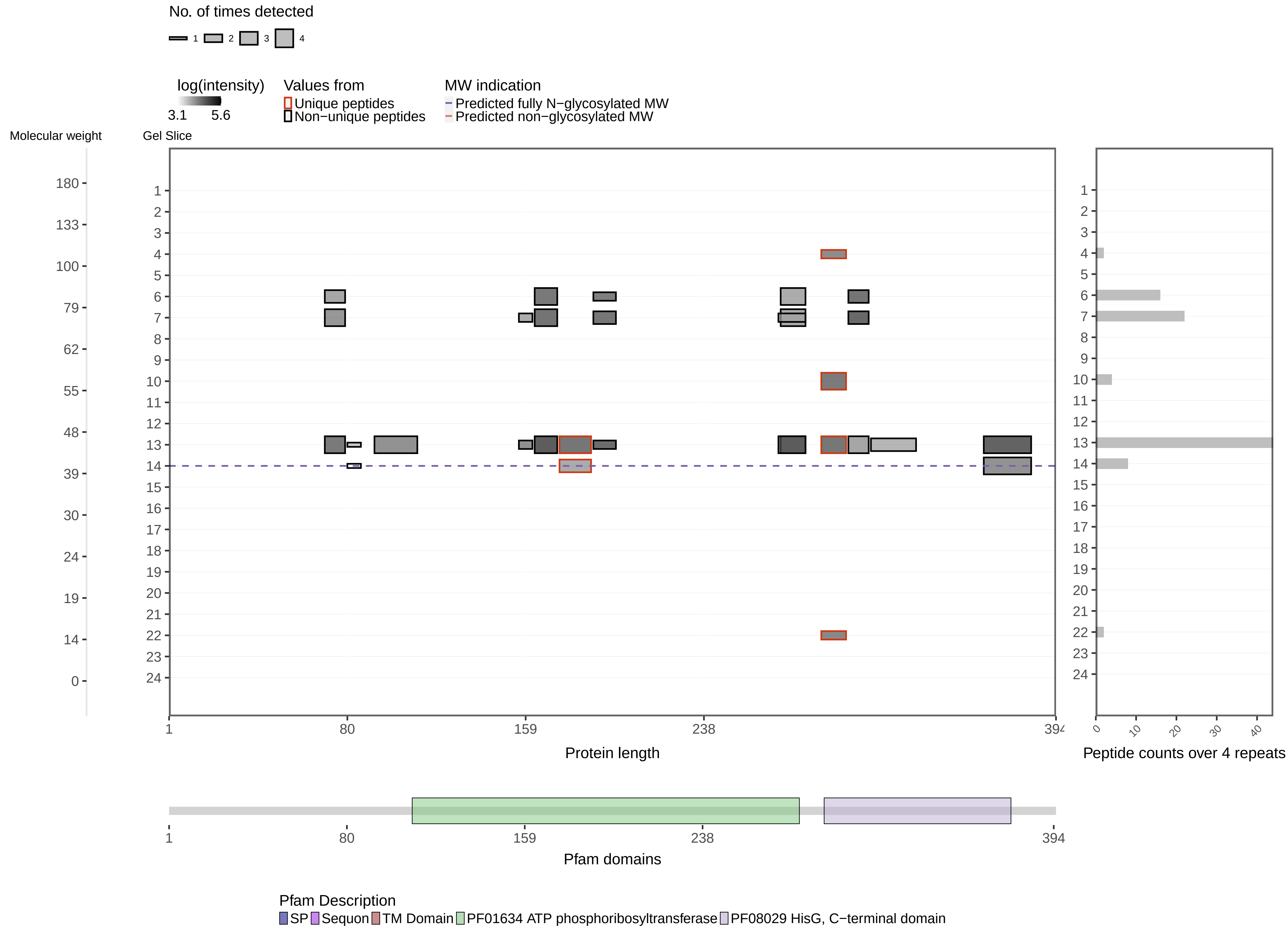


Protein sequence:

TPERPLRVATGFTYLGPKFMKDNGLKHVTFSTADGALEAAPAMGIADAIVDLVSSGTTLRENNLKEIEGGIILESQAVLVAGRKSLMQRKGVLDITHEMLERLEAHLRAIGQFTVTANMRGSSAEEVAERILSQTSLSGLQGPTVSPVFCKRDGRVTADYFAIVICVPKKALYKSVQQLRAIGGSGVLISPLTYIFDEETPRWRQLLSMLGL*

NbL01g13530.1 nifU−like_protein_2_chloroplastic [MW = 26.35 kDa]

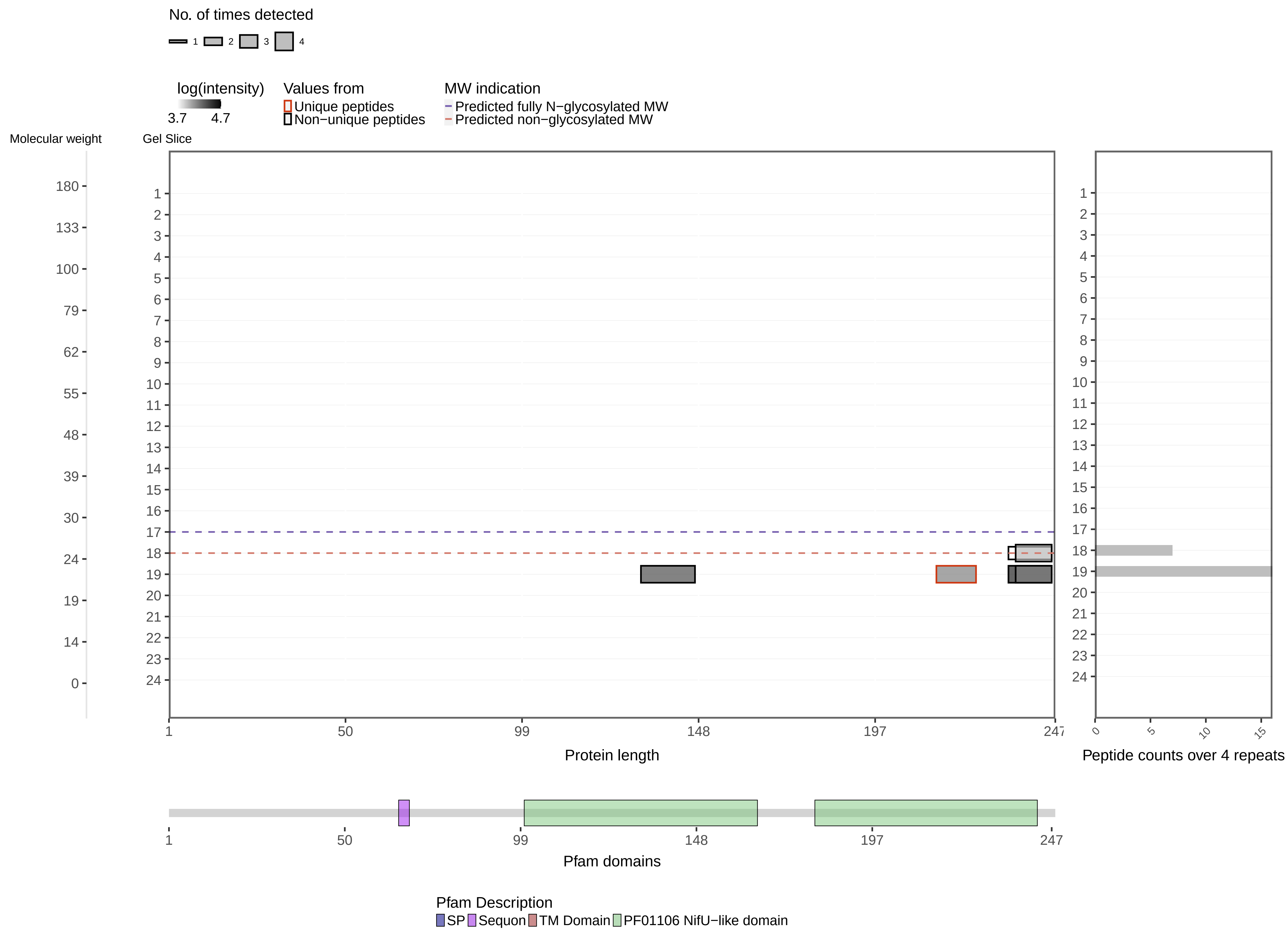


Protein sequence:

PYLIADGGNVALHKIDGNVVKLKLQGACGSCPSSVTTMKLGIERRLMEKIPEIVAVEAVPDEETGLELNEENIEKVLEEIRPYLVGAAGGSLELVAIEEPIVKVRITGPAAGVMTVRVAVTQKLREKIPAIAAVQLLQ*

### NbL01g13380.1 60S_ribosomal_protein_L3−2 [MW = 44.4 kDa]

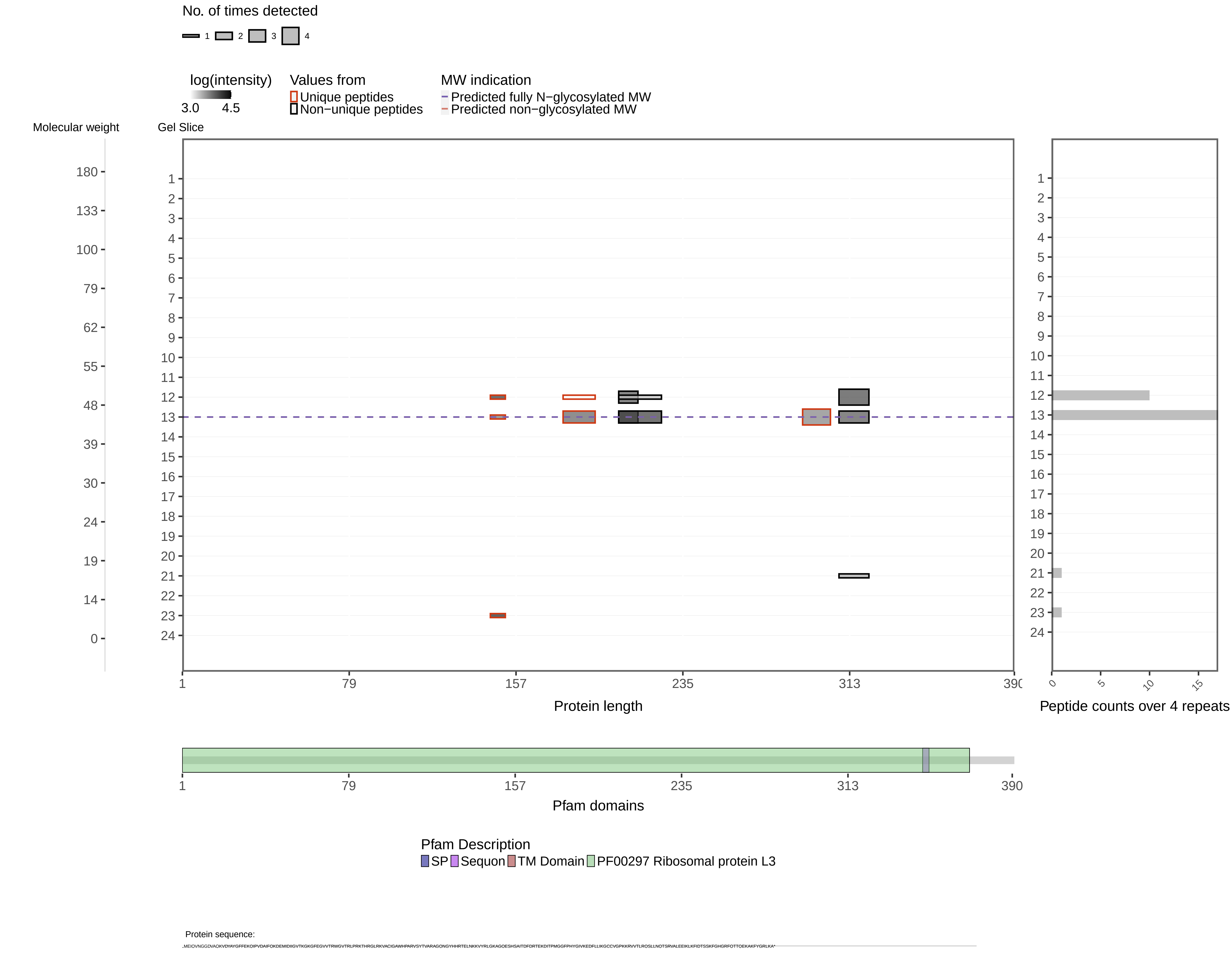

NbL01g13370.1 50S_ribosomal_protein_L4_chloroplastic [MW = 34.37 kDa]

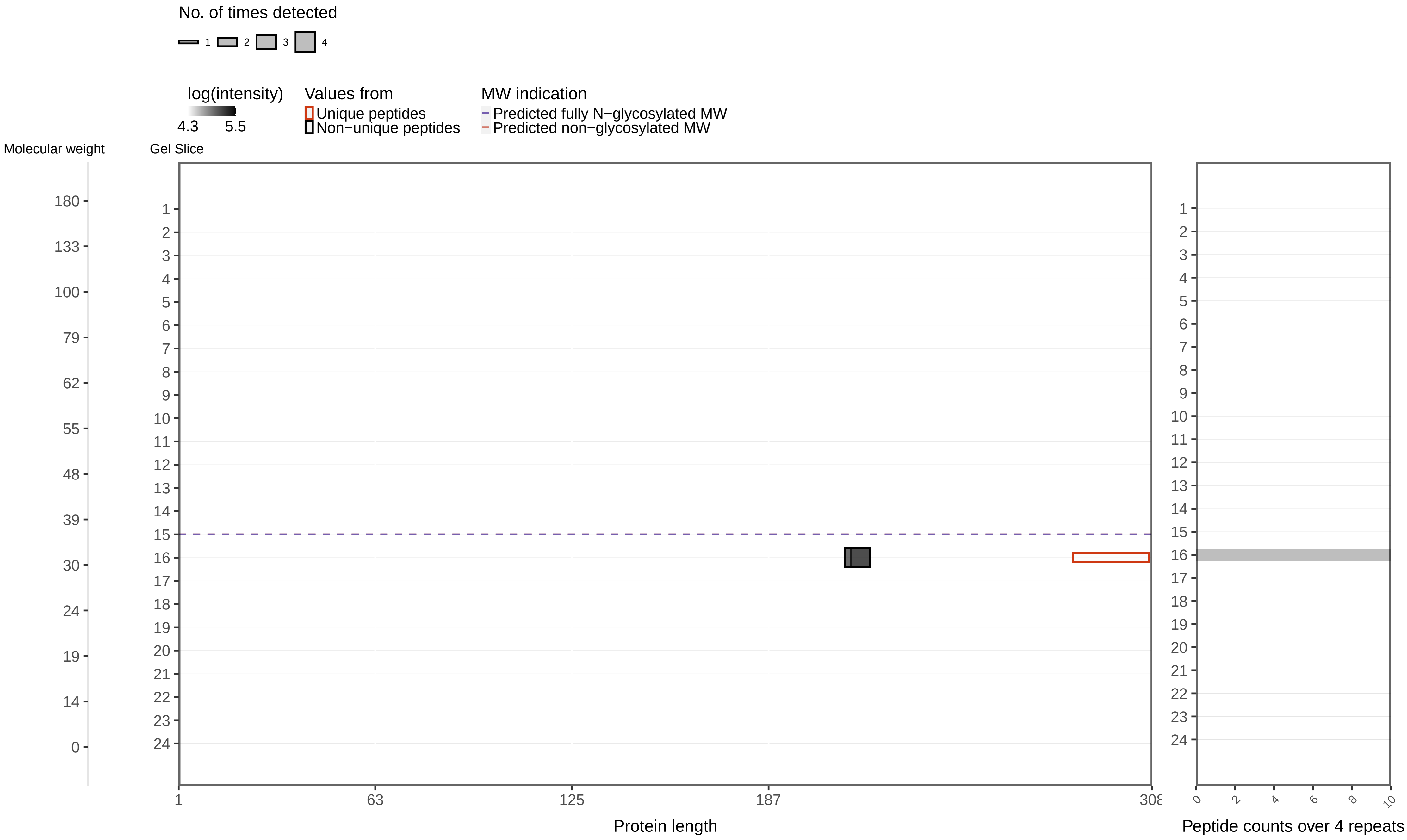


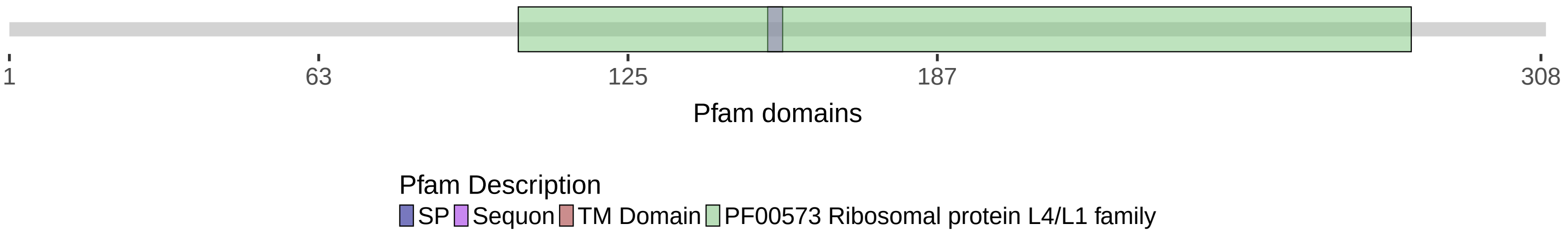


Protein sequence:

:KTGRARRGSNRTPLRPGGGVVFGPKPKDWSVKINKKEKRLAISTALSSASENTIVVEEFNDKFEKPKTKEFIELMRRWGLDPKEKSLFLMTEVSDNVILSSRNIGTLKMLTPRTLNLFDILDSEKLVLTKSAVDFLNERYGDENEWEDEGQEDNDGGEAEESSE*

### NbL01g13340.1 coatomer_subunit_delta−like [MW = 58.02 kDa]

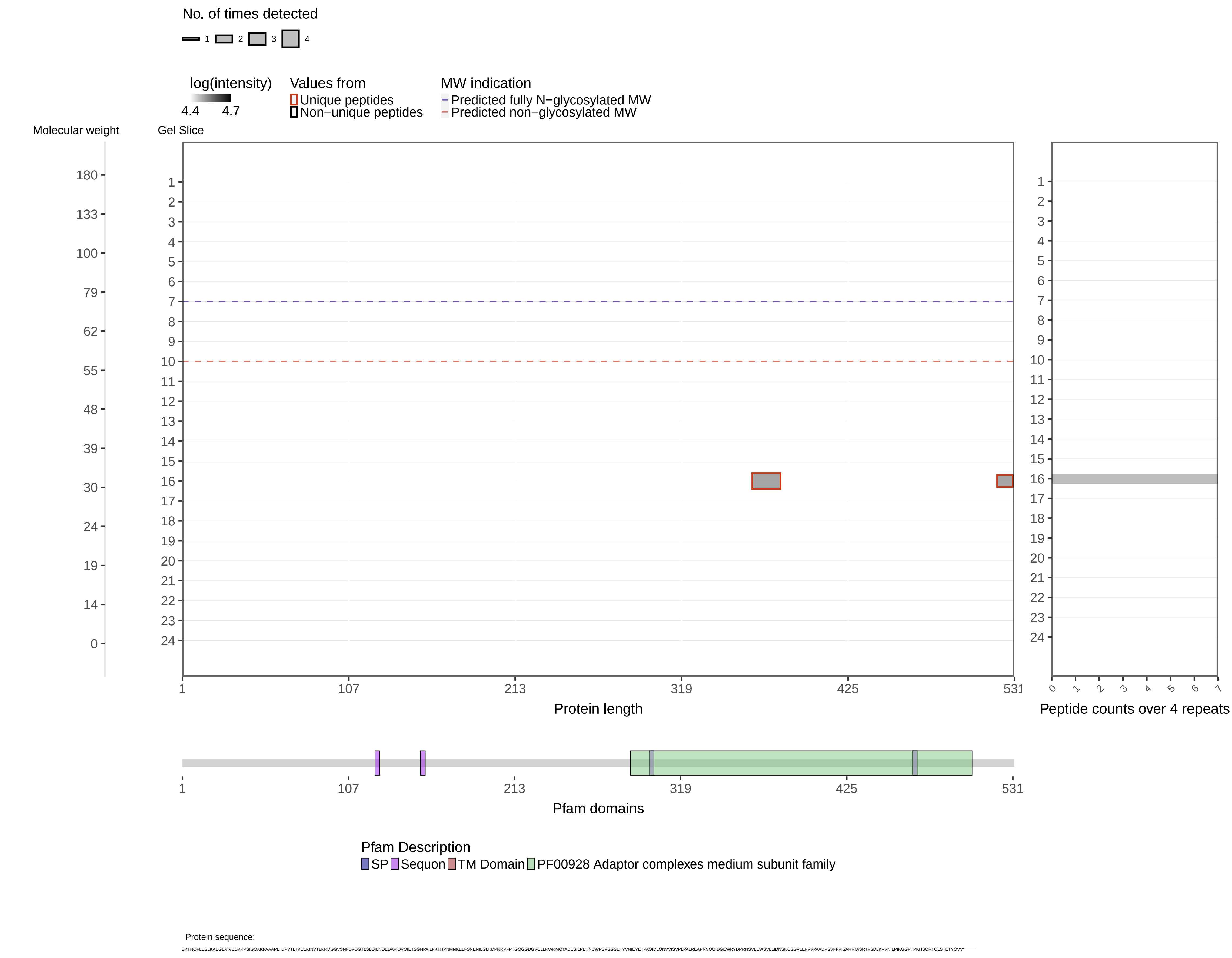

NbL01g13320.1 stromal_70_kDa_heat_shock−related_protein_chloroplastic [MW = 75.32 kDa]

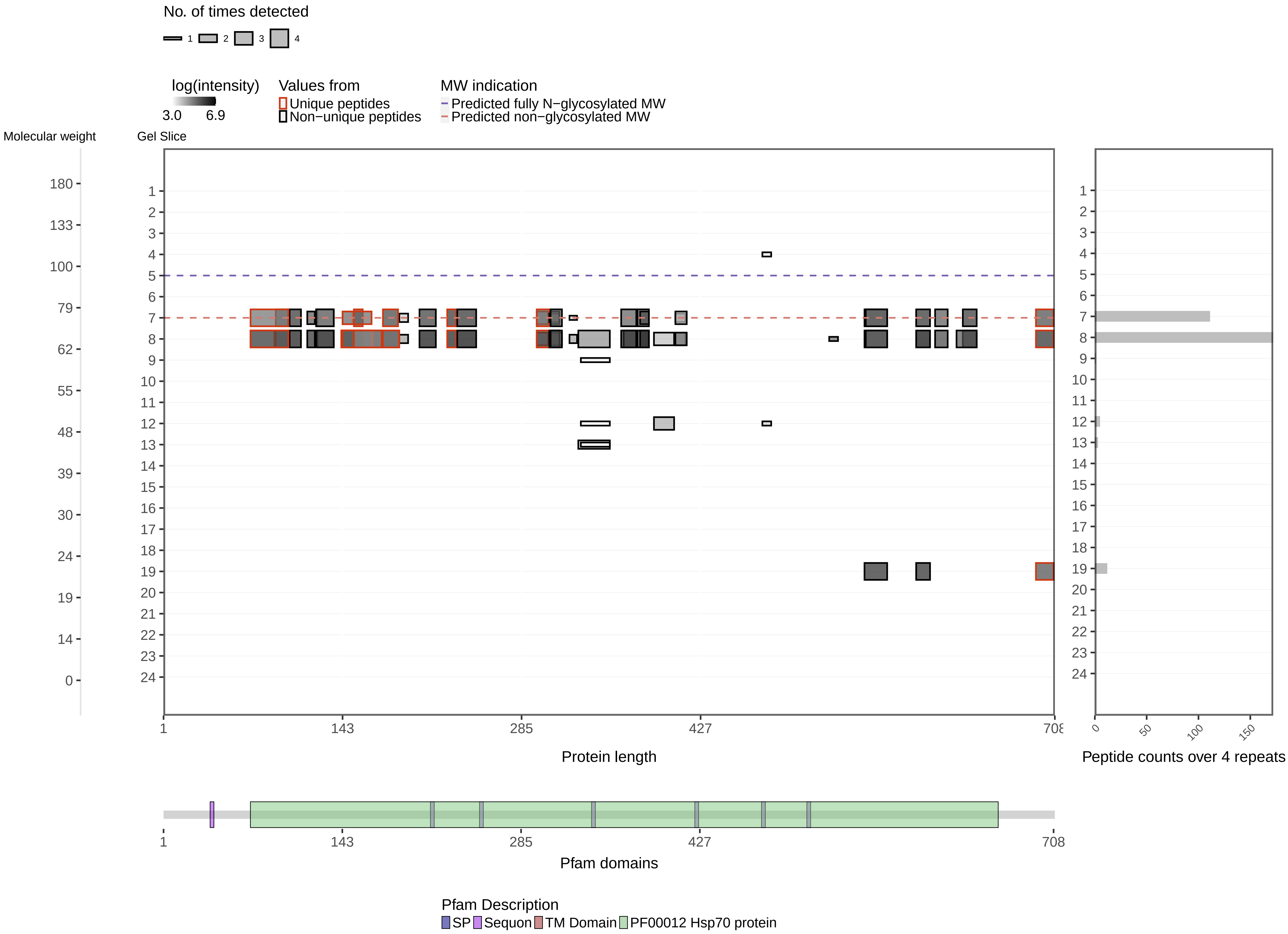


Protein sequence:

TNISLPFITATADGPKHIETTITRGKFEELCSDLLDRLKTPVQNSLRDAKLSFSDIDEVILVGGSTRIPAVQELVKKLTGKDPNVTVNPDEVVALGAAVQAGVLAGDVSDIVLLDVTPLSIGLETLGGVMTKIIPRNTTLPTSKSEVFSTAADGQTSVEINVLQGEREFVRDNKSLGSFRLDGIPPAPRGVPQIEVKFDIDANGILSVTAIDKGTGKKQDITITGASTLPGDEVERMVKEAERFAQEDKEKRDAIDTKNQADSVVYQTEKQLKELGDKVPGPVKEKVEAKLGELKEAISGG

### NbL01g13260.1 acetolactate_synthase_1_chloroplastic [MW = 59.9 kDa]

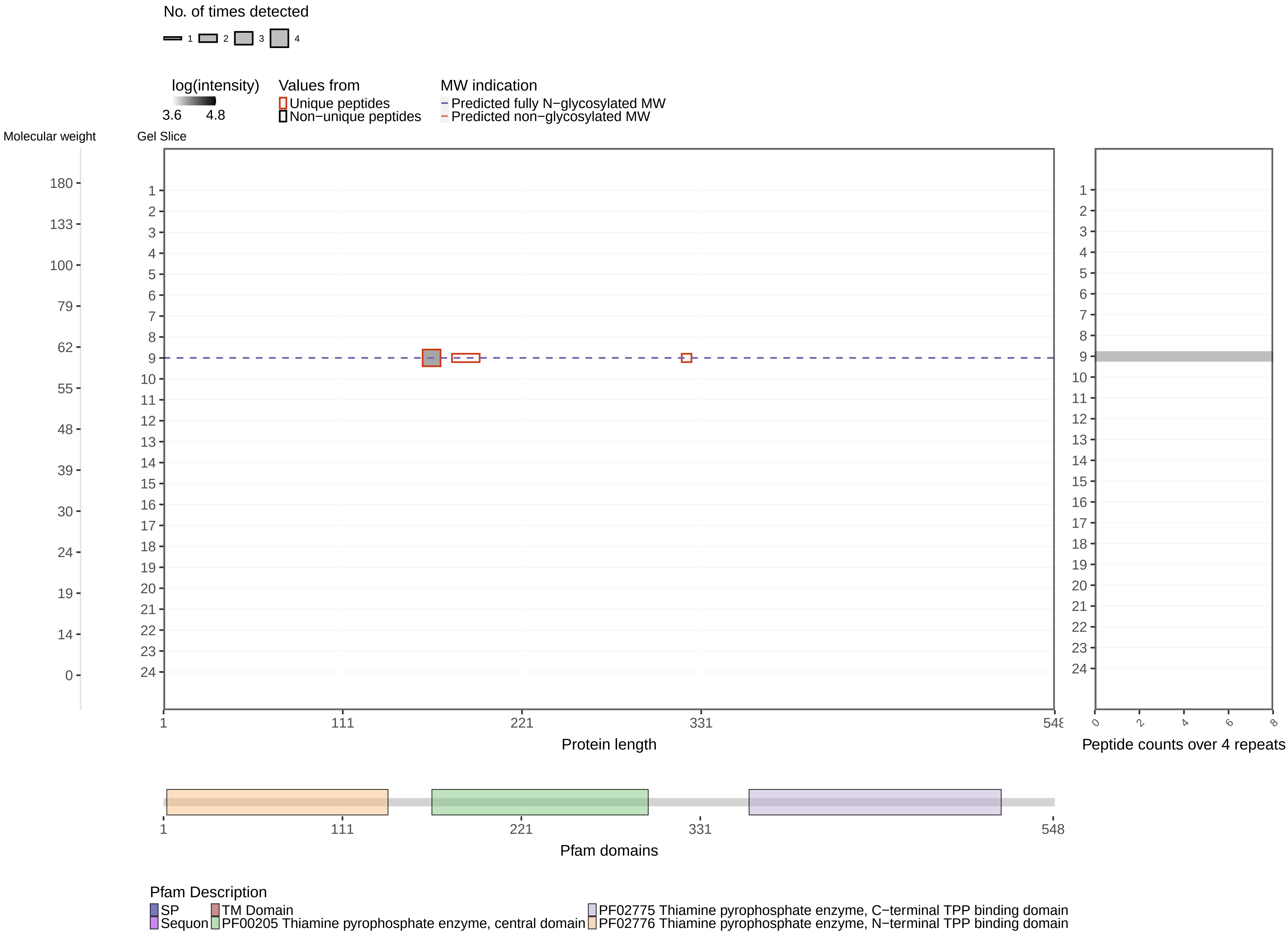


Protein sequence:

[illegible]

### NbL01g13060.1 uncharacterized_LOC107796198 [MW = 52.54 kDa]

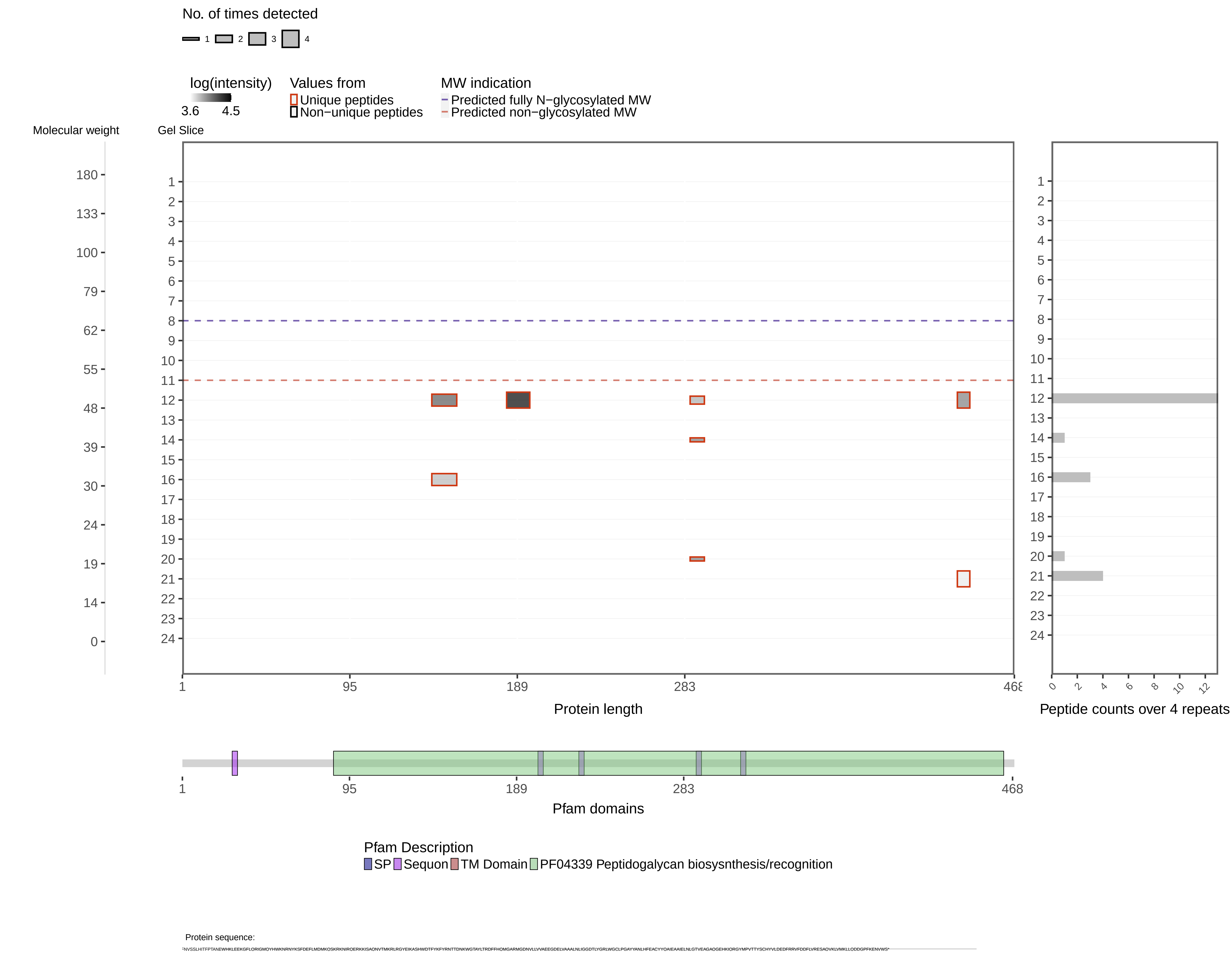

NbL01g13000.1 survival_of_motor_neuron−related−splicing_factor_30 [MW = 33.32 kDa]

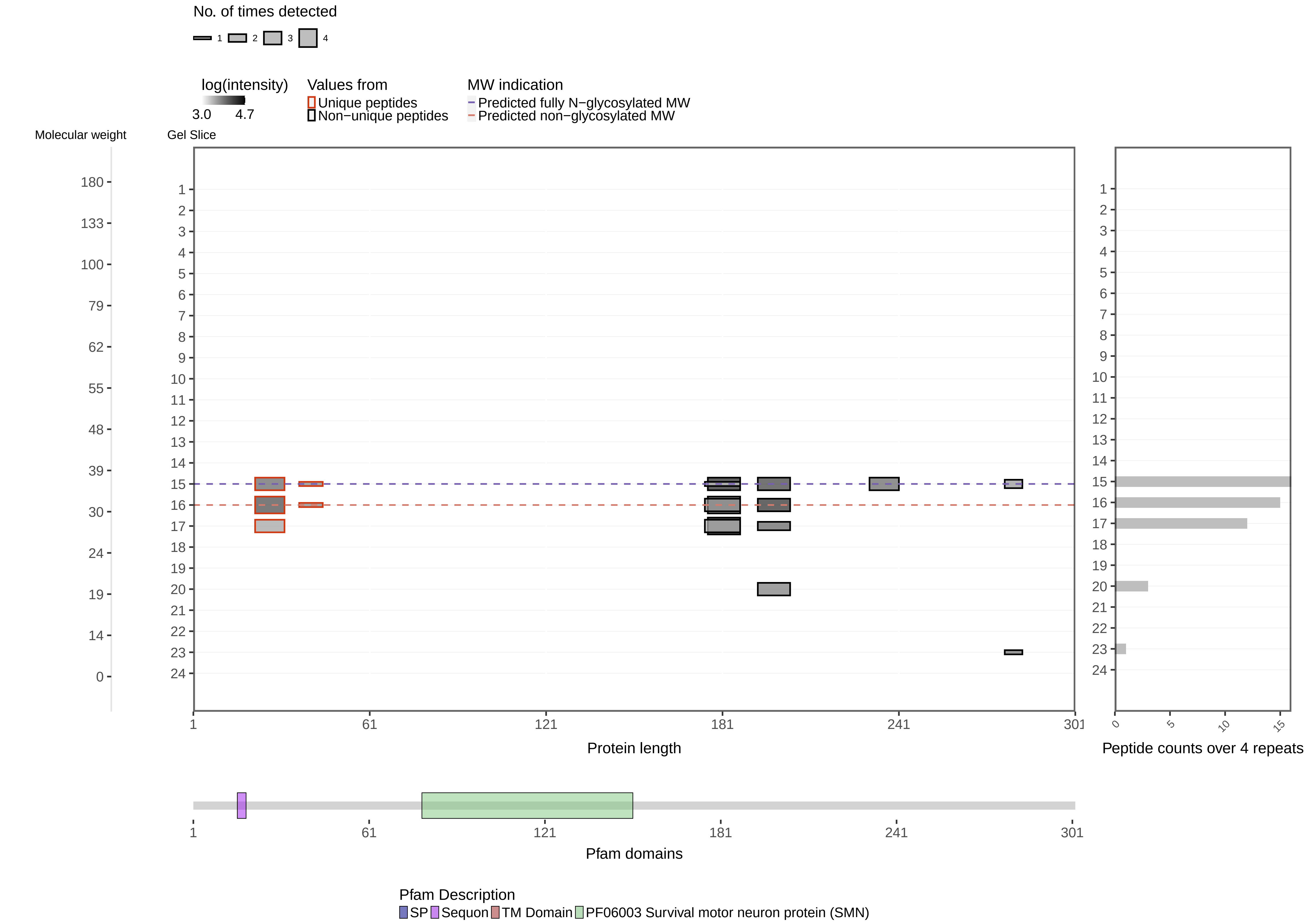


Protein sequence:
/GNKEEVDHANIRPLOEGPVNPLVEAEKIAEATKOALKRKIAOAAASDVOSRSLPAKLLIDPNDPEDVKAAKRKKIHAFKSKVRIEOLEVAONKRONAWOOFOSTKGOTKKVGFFSGRKRESIFKSPEDPHGKVGVTGSGKGLTDFORREKHLHLKGANAEASDE*

NbL01g12920.1 methionine_S−methyltransferase−like [MW = 121.29 kDa]

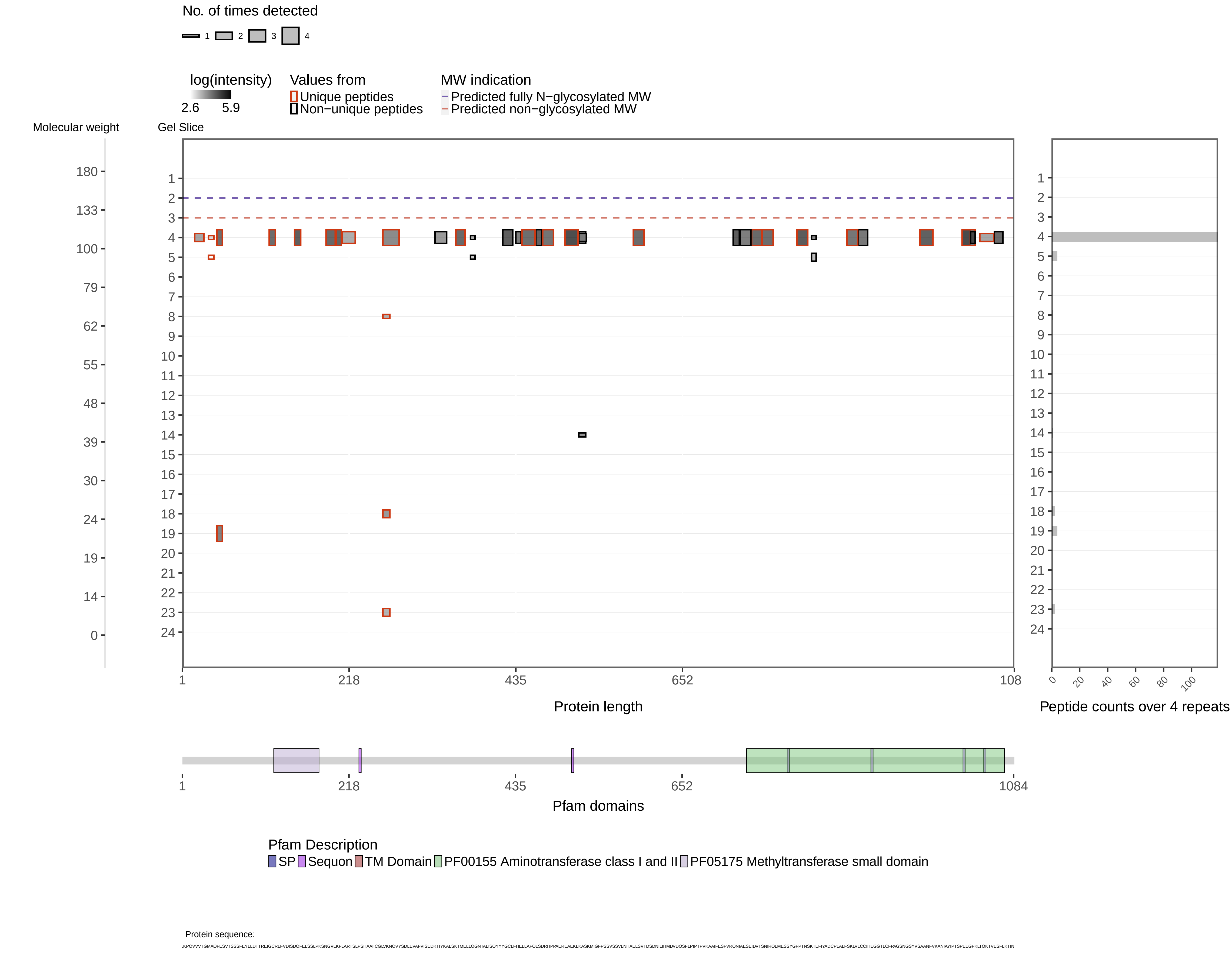

NbL01g12820.1 2−C−methyl−D−erythritol_4−phosphate_cytidylyltransferase_chloroplastic [MW = 34.25 kDa]

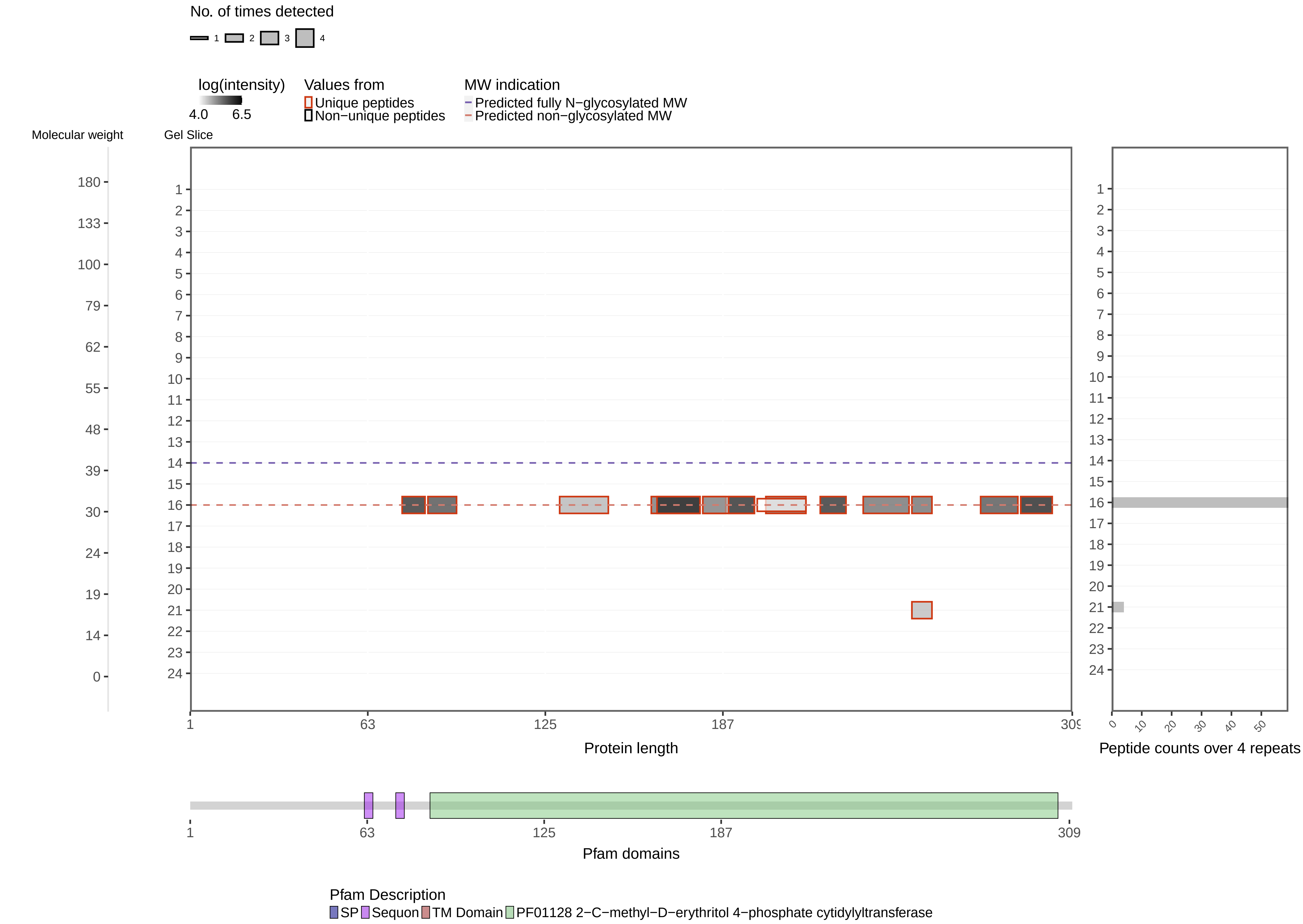


Protein sequence:

DIFEDAKENIQVKLKFALPGKERQDSVYSGLQEVDSNSKLVCIHDSARPLVWTEDVEKVLKDGWLVGAAVLGVPAKATIKEANSESFVVKTLDRKTLWEMQTPQVIKPELLKKGFELVNREGLEVTDDVSIVEHLKHPVYITEGSYTNIKVTTPDDLLLAERILNTEE*

### NbL01g12760.1 cinnamoyl−CoA_reductase_1 [MW = 35.66 kDa]

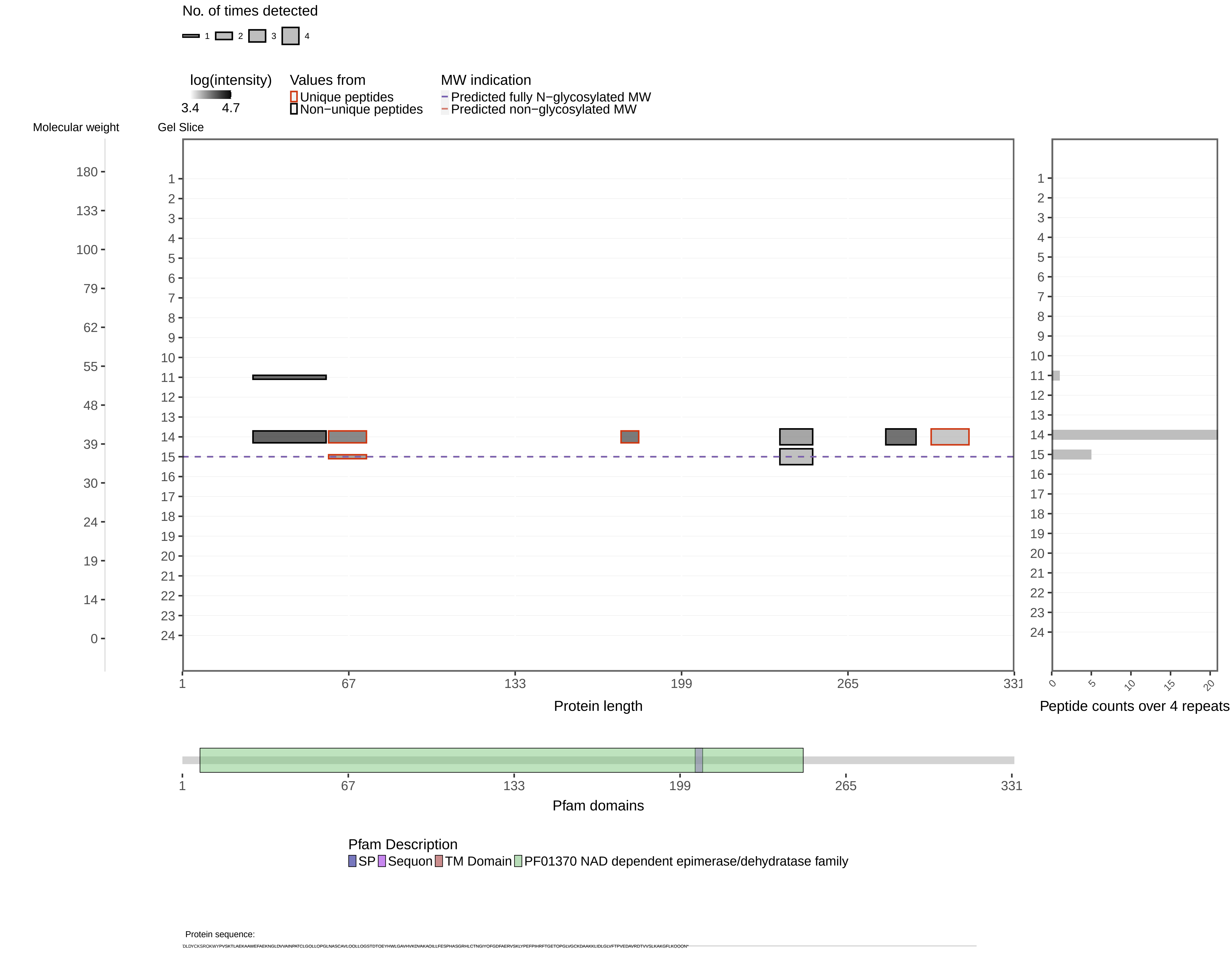

### NbL01g12560.1 NADPH:adrenodoxin_oxidoreductase_mitochondrial [MW = 57.74 kDa]

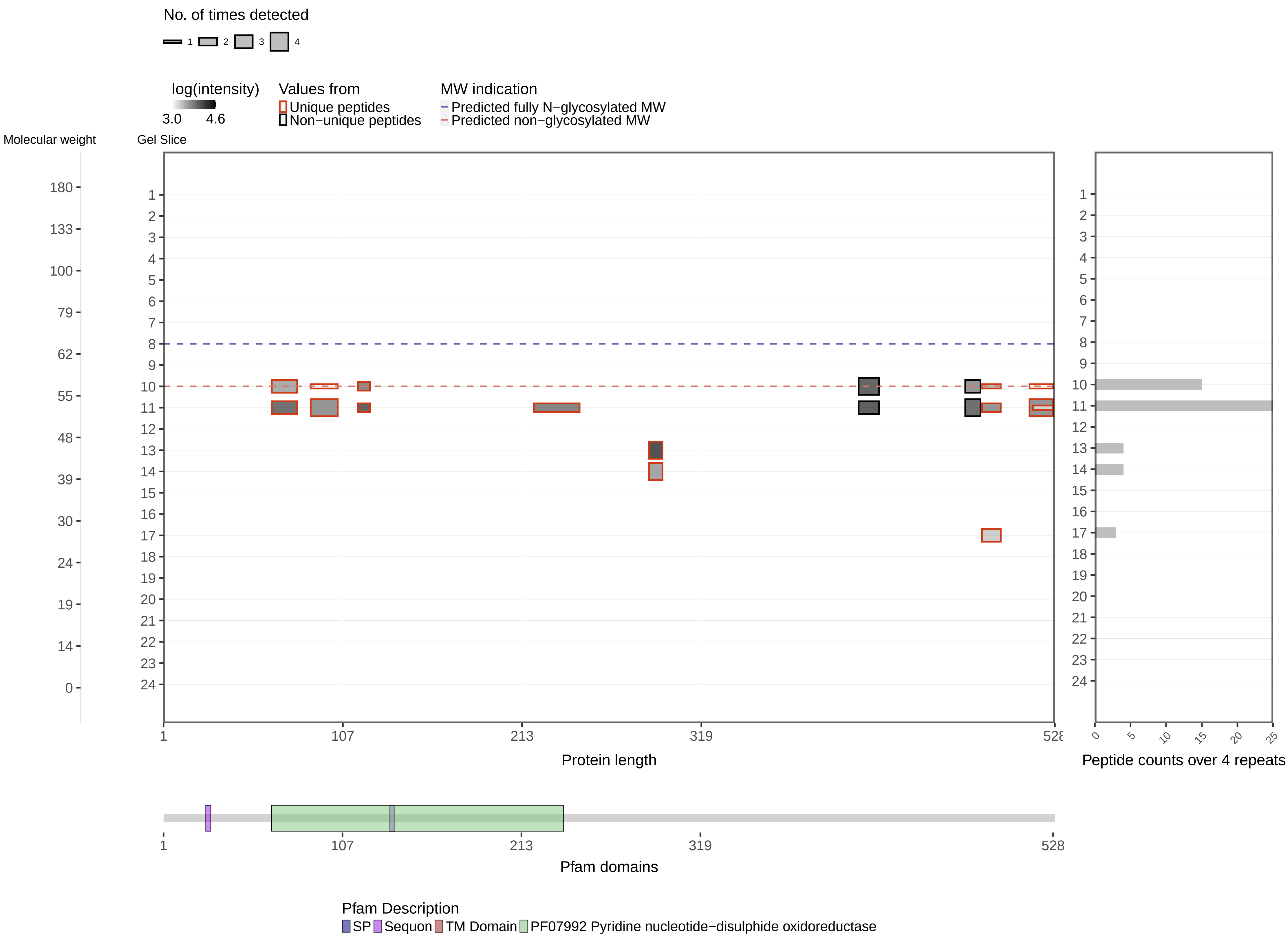


Protein sequence:

GRRGAAQAAFTAKELREVLGIKDLFIEIQQADLSKTPADAEELKSNRIKRRIHELLSKAATSAASVCSPGQKELHFVFFRKPDRFLESDCRSGHVAGLRLERTILKEDAGSGRQIAVGTELFDEMECGLVLKSVGYKSRPVDGLPFDHHKGIVPNVQGRVLSEASGDALVEKGLYVCGWLKRGPTGIIATNLYCAEETVASISEDVQGGLVTSSSGLSKPGREGLLQLLDSRNVNIVPFGGWEKIDSEEKRRGSLKNKPREKLTAWQELLEVATK*

NbL01g12380.1 cyprosin−like [MW − SP = 52.45 kDa]

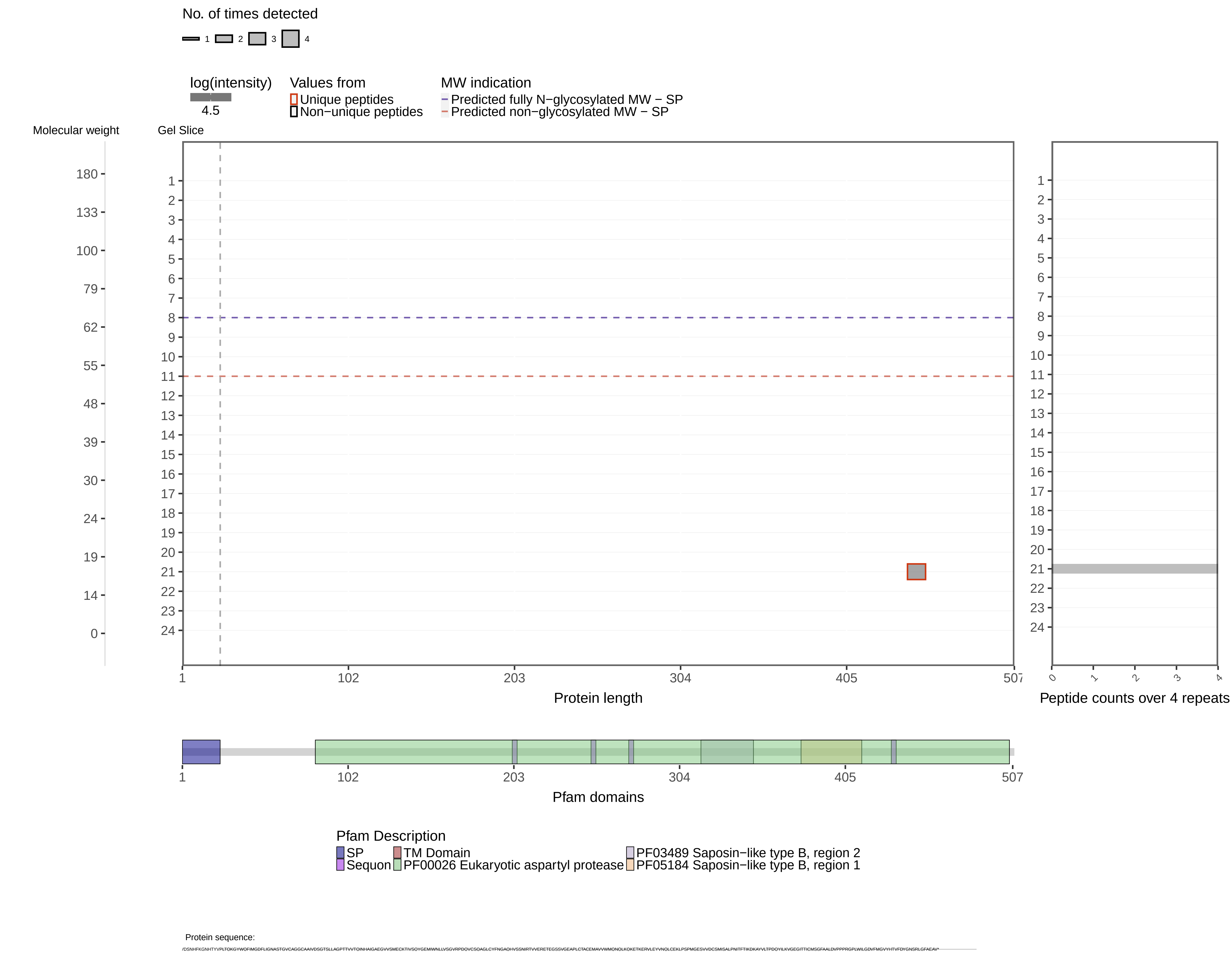

NbL01g12350.1 pentatricopeptide_repeat−containing_protein_At3g53700_chloroplastic−like [MW = 82.13 kDa]

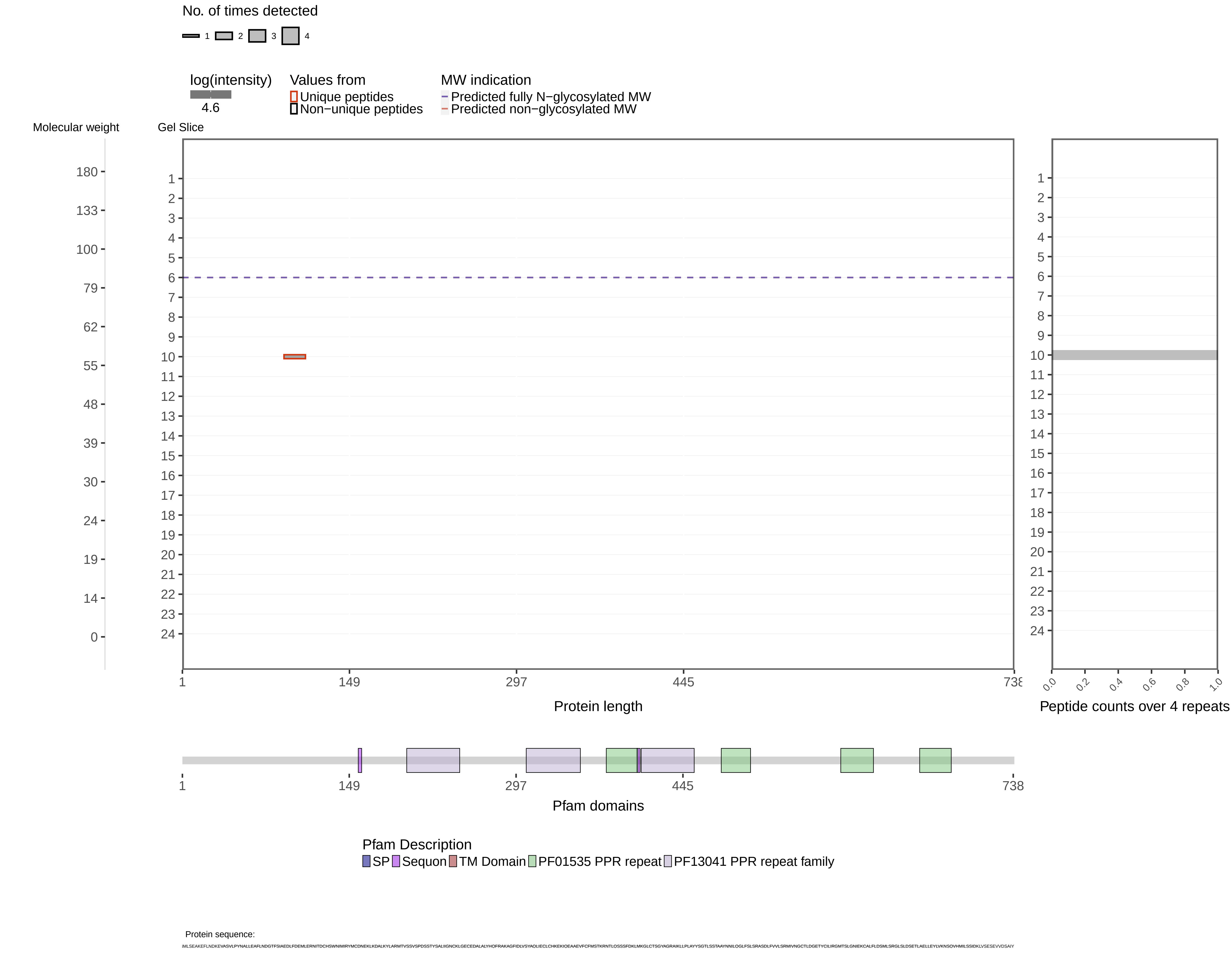

### NbL01g12230.1 ATP−citrate_synthase_alpha_chain_protein_2 [MW = 46.85 kDa]

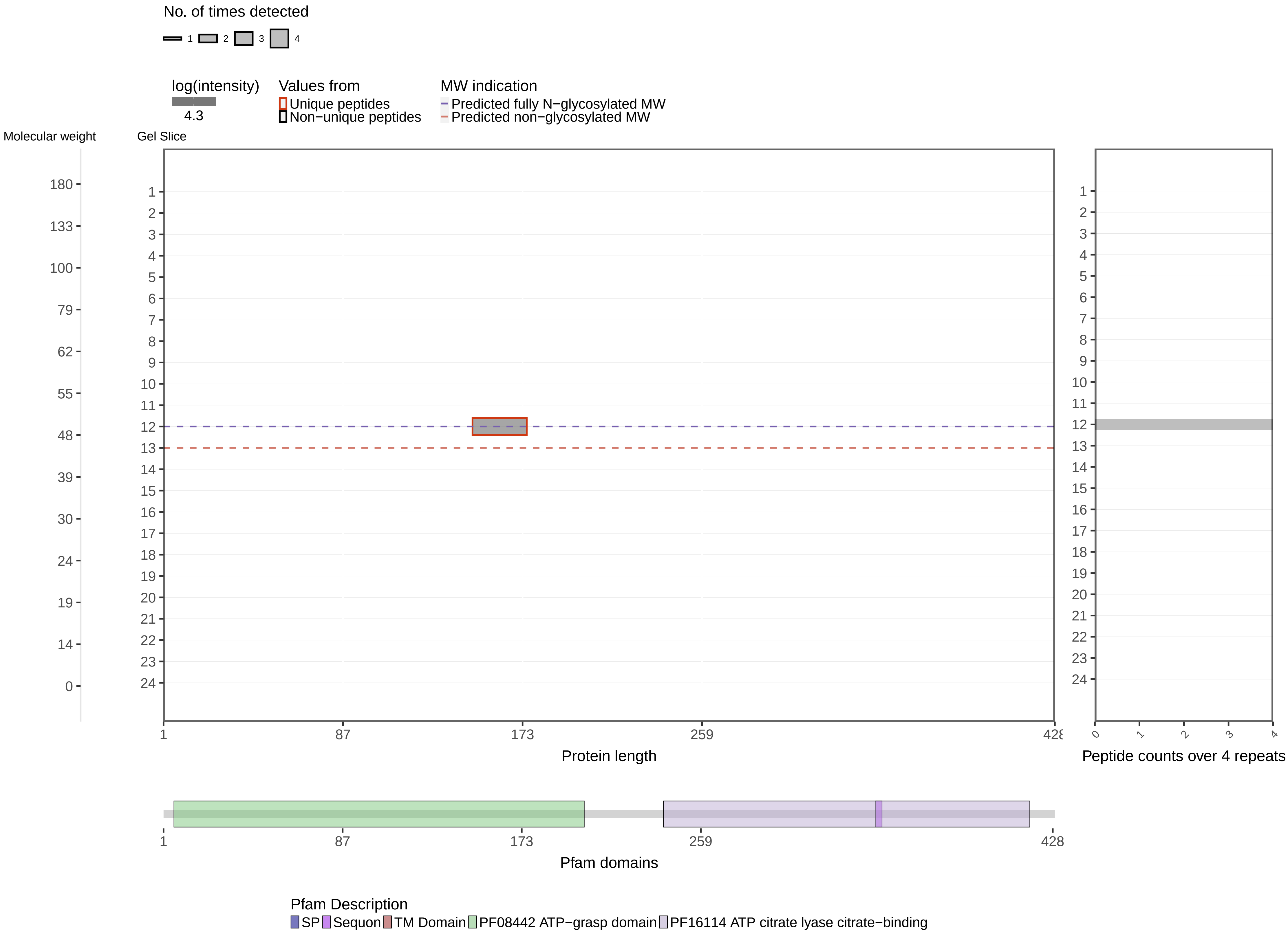


Protein sequence:

TLVNGDPYPLDMRGELDDTAAFKNFKKWGSIEFPLPFGRVLSPTESFIHSLDEKTSASLKFTVLNPKGRIWTMVAGGGASVIYADTVGDLGYASELGNYAEYSGAPNEEEVLOYARVVLDCATANPDGRKRALIVGGGIANFTDVAATFNGIIRALREKEAKLKAAGMHIYVRRGGPNYOTGLAKMRALGEELGVPLEVYGPEATMTGICKRAIDCIMSEASVCF*

### NbL01g12150.1 pyrrolidone−carboxylate_peptidase−like [MW = 23.92 kDa]

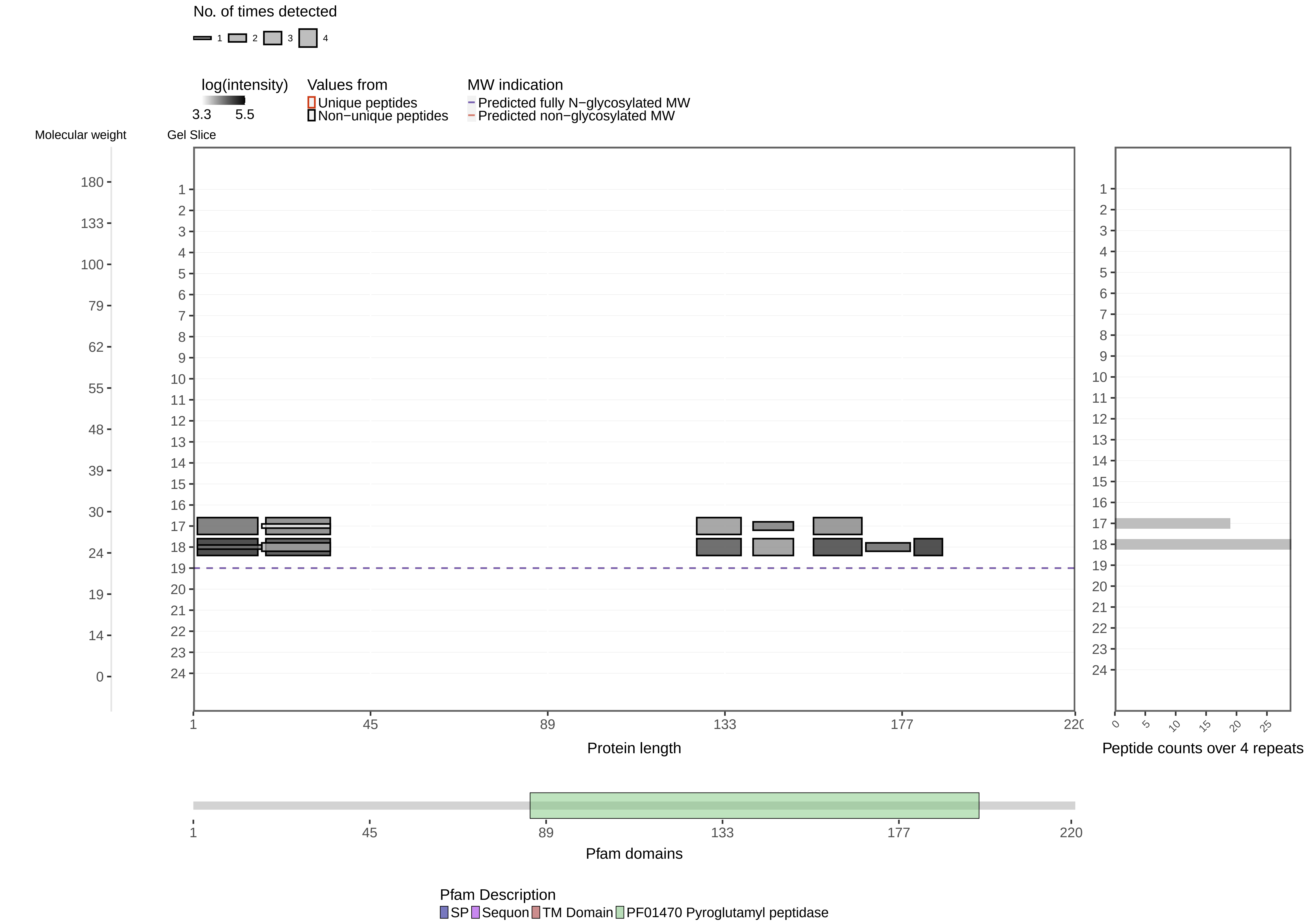


Protein sequence:
SGATMFAIENQAVNEATFRCPDEMGWKPQKVPIVHADGAISRKRETALPVEEITKVLAKMGYEVMTSDDAGRFVCNYVYYHSLRFAEQNGIRSLFVHVPLFLTIDEETQMQFAASLLEVLASLY*

NbL01g12120.1 GDSL_esterase/lipase_At1g09390 [MW − SP = 37.13 kDa]

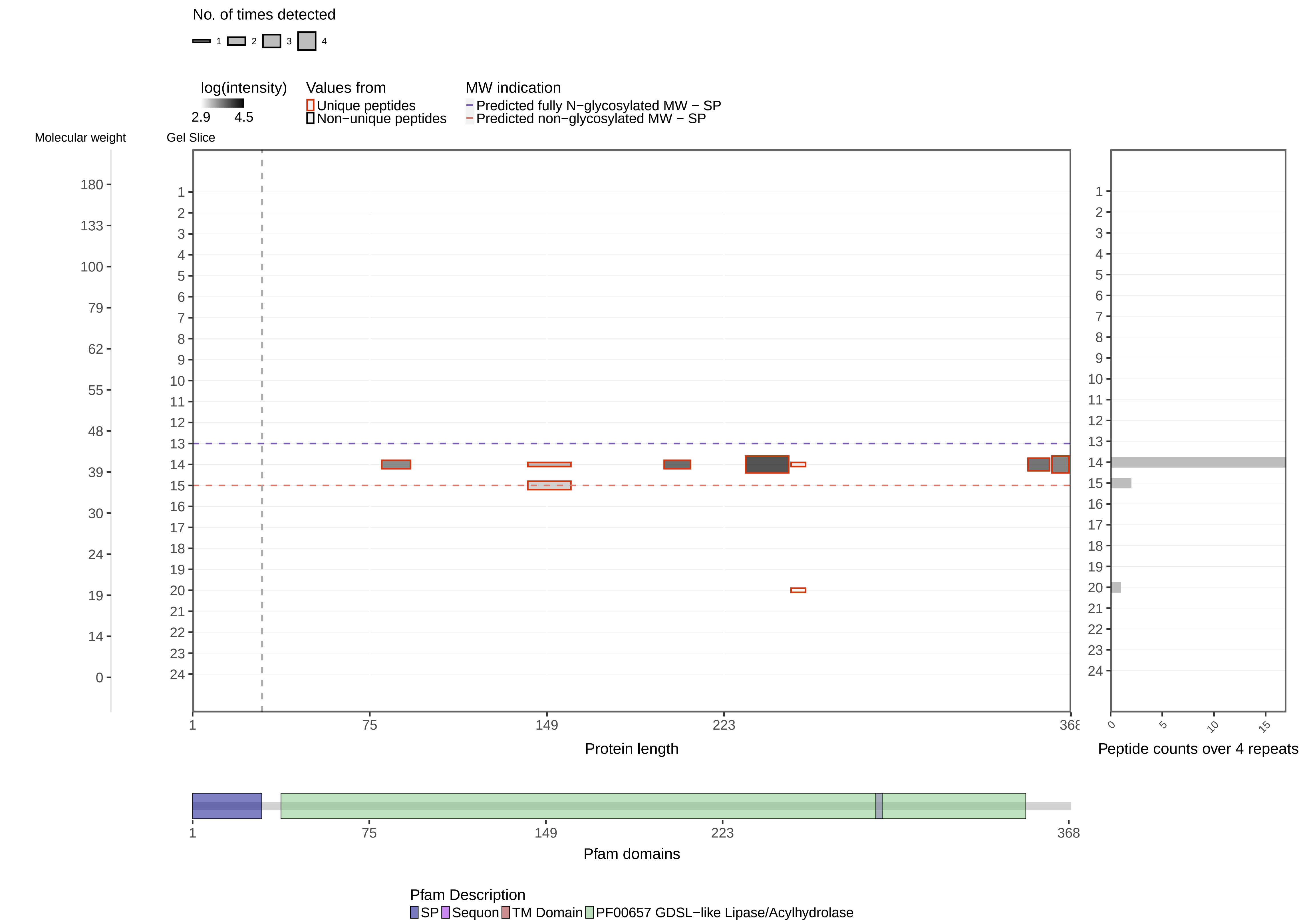


Protein sequence:

IADSFAKNMPYIEVVKMIPSFITEIKNAIQVIYSQGGRKFWVHNTGPLGCLPQKLTLVQKVSSDLDPHGCLASYNSAAKLFNEGLRHVCQELGSEMKDATIVYVDIYAIKYDLIANSSNYGFSNPLMACCGYGGPPYNYRKGVTCGHPGYQVCSEVSEFISWDGVHYTETANRVIASKILSTEYSTPRIGFDFFCH*

NbL01g11980.1 NHL_repeat−containing_protein_2 [MW = 117.18 kDa]

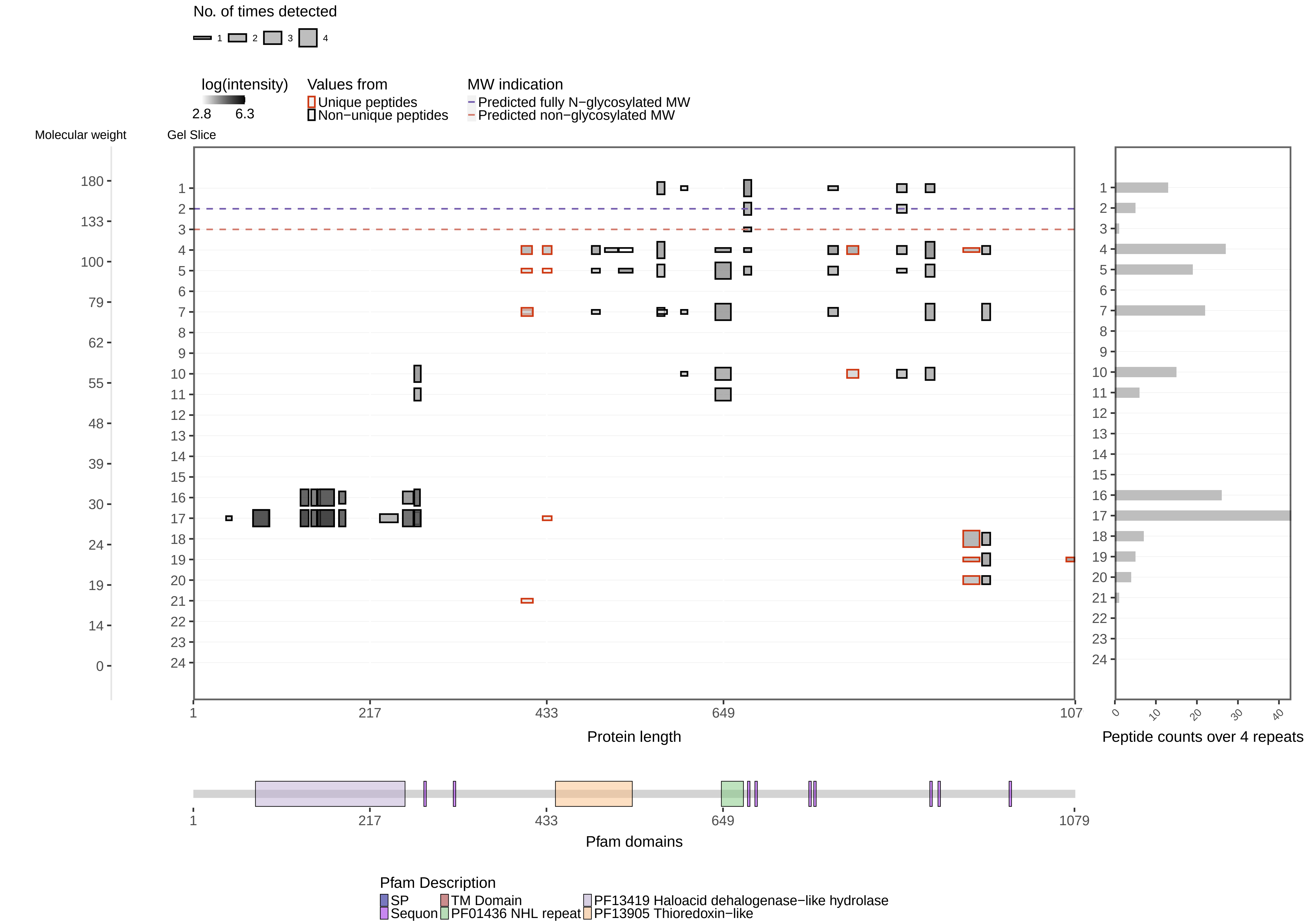


Protein sequence:

WPTFALVGPNGKLLAQIAGEGHRKDLDYLVEAALLFYGRKKLLDSTPIPLRLEKNNDPRLLTSPLKFPGKLAVDILNNRLFISDSNHNRIVVTDLEGNFLVQVGSTGAEGLRDGNFDDATFNRPQGLAYNAKKNLLYVADTENHALRVIDFVNETVRTLAGNGTKGSDYEGGGTGTAQLLNSPWDVCFEPENGIIYIAMAGQHQIWVHETSGGVTRAFSGNGFERNQNGSSSTNTSFAQPSGISLSRDLKEAYIADSESSSIRAVDLRTGGSRLLAGGDPIMAENLFKFGDHDG

NbL01g11880.1 uncharacterized_LOC109210159 [MW = 20.11 kDa]

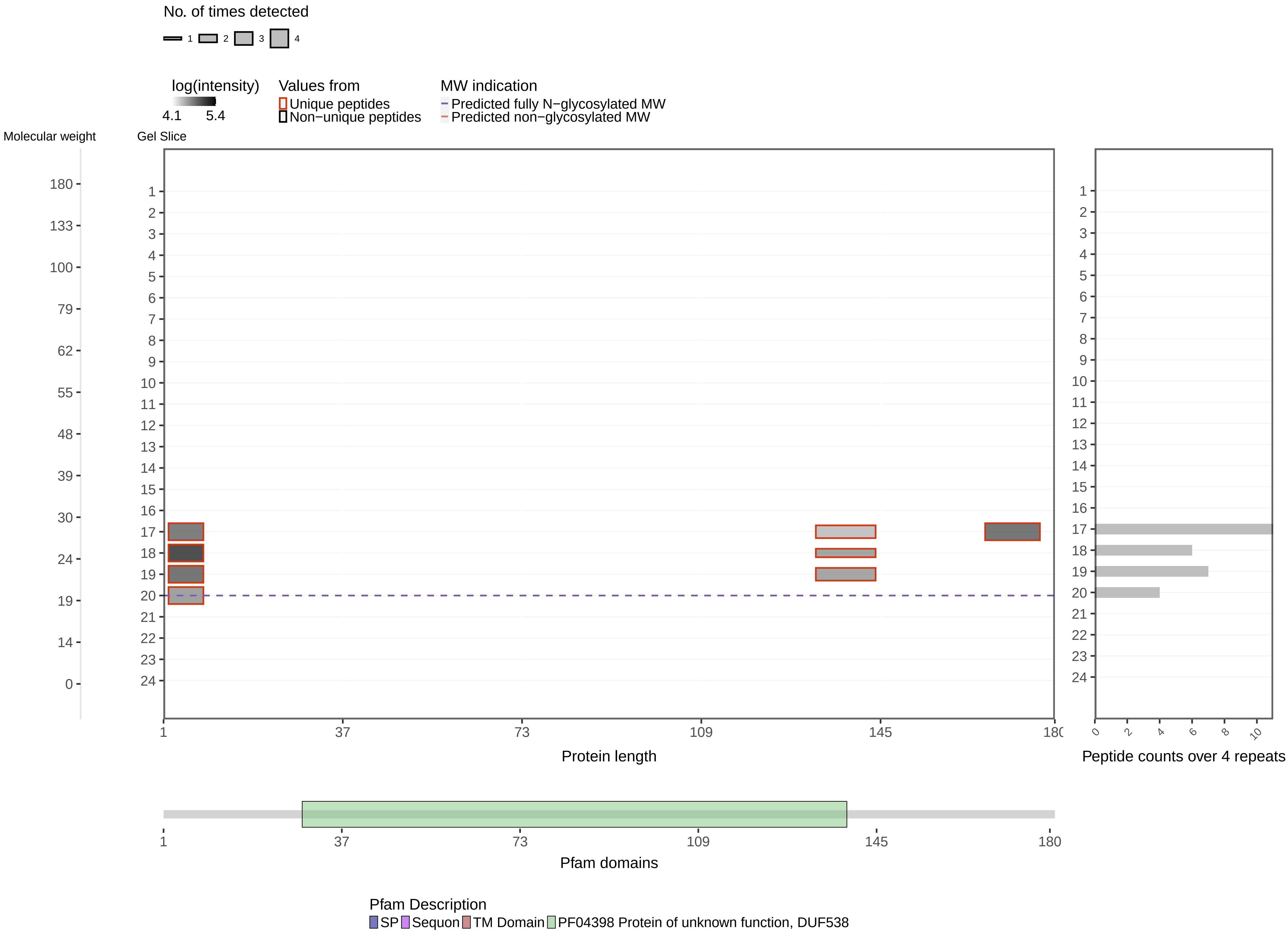


Protein sequence:

VQYGTEVTAIVEQNKIKKLTGVKAKELLIWLTLTEICVDDPPTGKIHFKATSGLSRTFPVSAFEIDVTKKEEVKEEVKEEVNEVVNKENKEVIPAAAAVEVKEV*

### NbL01g11870.1 uncharacterized_LOC107830650 [MW = 37.8 kDa]

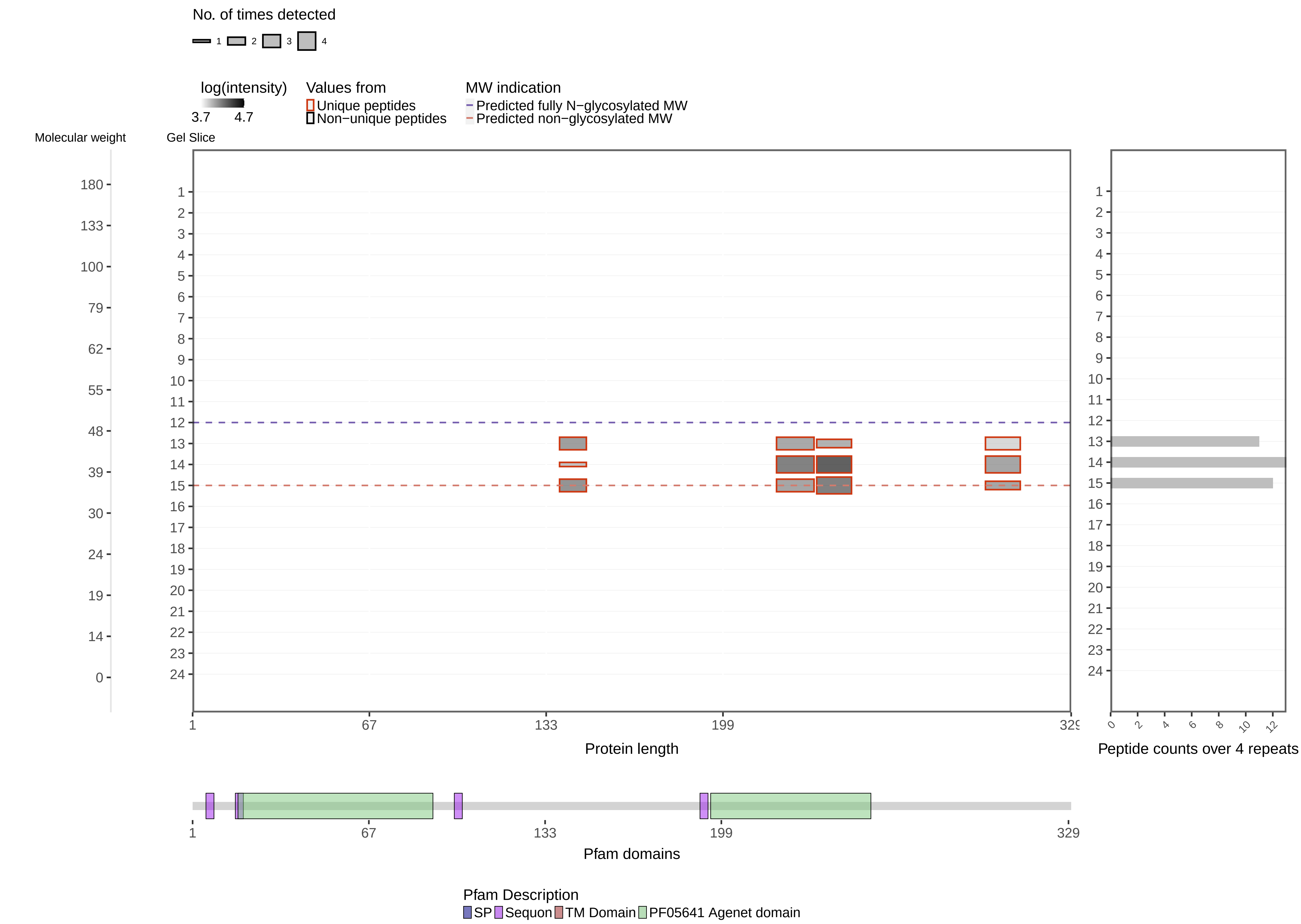


Protein sequence:

EWANGKWTPPLEQGGEGDEDEVEETLASTDIKPSNDAAENFSEGSLVEVSSDEEGFEGAWFAATVVKLLDNGNYLIEYQNLRNDDDTAFLQEEADRLYVRPRPPDIGPFKSFTVLEEVDALYNDGWWVGVVSKVLKGQRYRVQFKGNNEELEFNHADLRLHLDWINGKWVTPSQALK*

### NbL01g11820.1 trafficking_protein_particle_complex_subunit_4 [MW = 16.02 kDa]

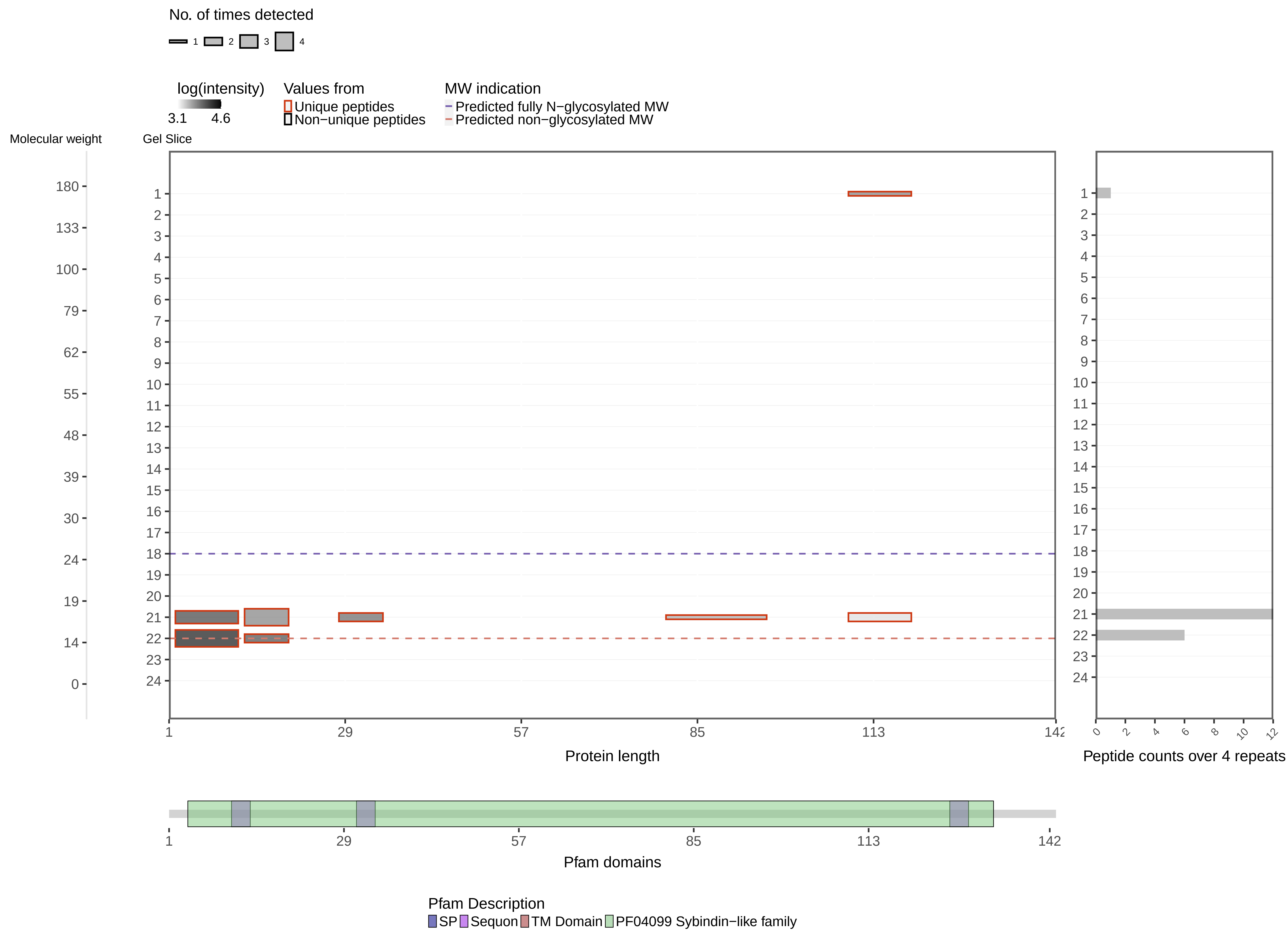


Protein sequence:

GIELLQADNFDLHCFQSLTGTKFFVVCEPGSLHMESLLKYVYELYTDYVLKNPFYEMEMPIRCELFDINLSQAVQKDRVALLGR*

### NbL01g11810.1 ABC_transporter_I_family_member_20−like [MW = 36.67 kDa]

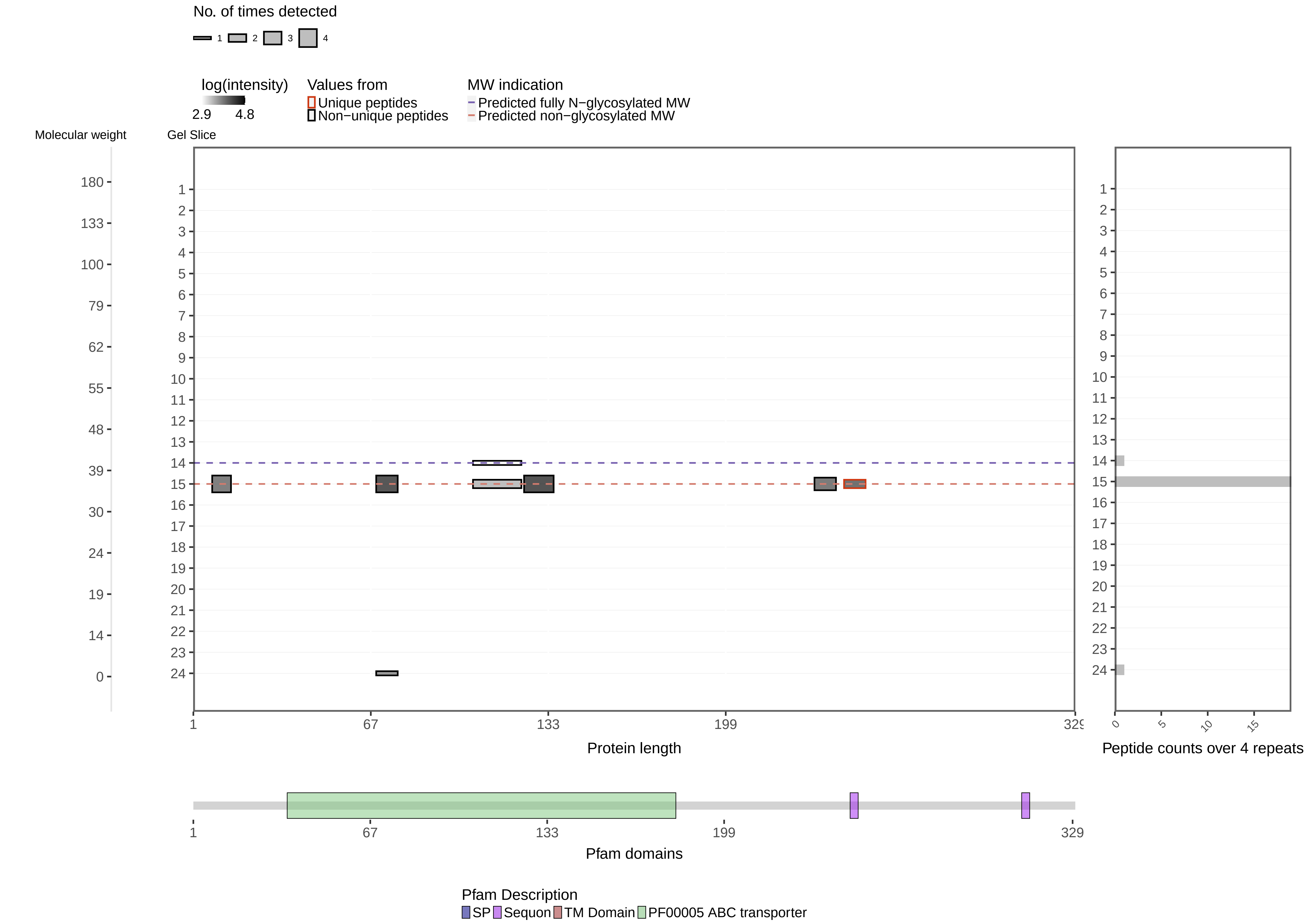


Protein sequence:

KASDGQRRRVQICMGLLKPFKVLLLDEITVDLDVLARADLLKFLSKECEERGATIIYATHIFDGLENWPSHMLYVAHGKLQLAMPMDKVKEISNMSLMRTVESWLRKERDEERKRRKERKAKGLPEYGKQVEGSRVVGDPARAAARPLNNGWAGGRLNSTIAGEENFVLSSNRVLR*

### NbL01g11760.1 60S_ribosomal_protein_L36−2−like [MW = 12.36 kDa]

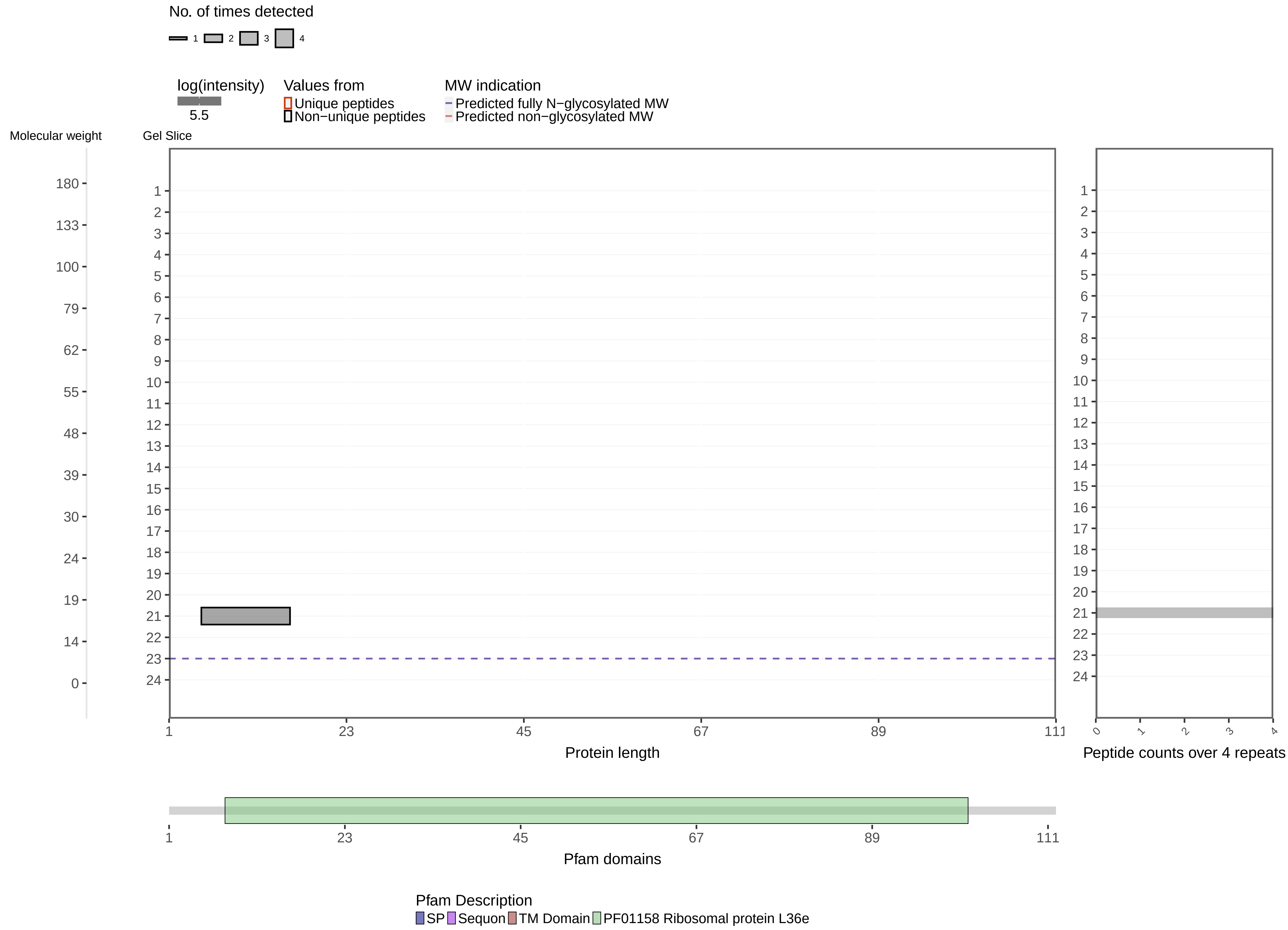


Protein sequence:

FVRSLIREVAGFAPYEKRITELLKVGKDKRALKVAKRKLGTHKRAKKKREEMSSVLRKMRATGGGEKKK*

### NbL01g11620.1 iron−sulfur_assembly_protein_IscA−like_2_mitochondrial [MW = 16.95 kDa]

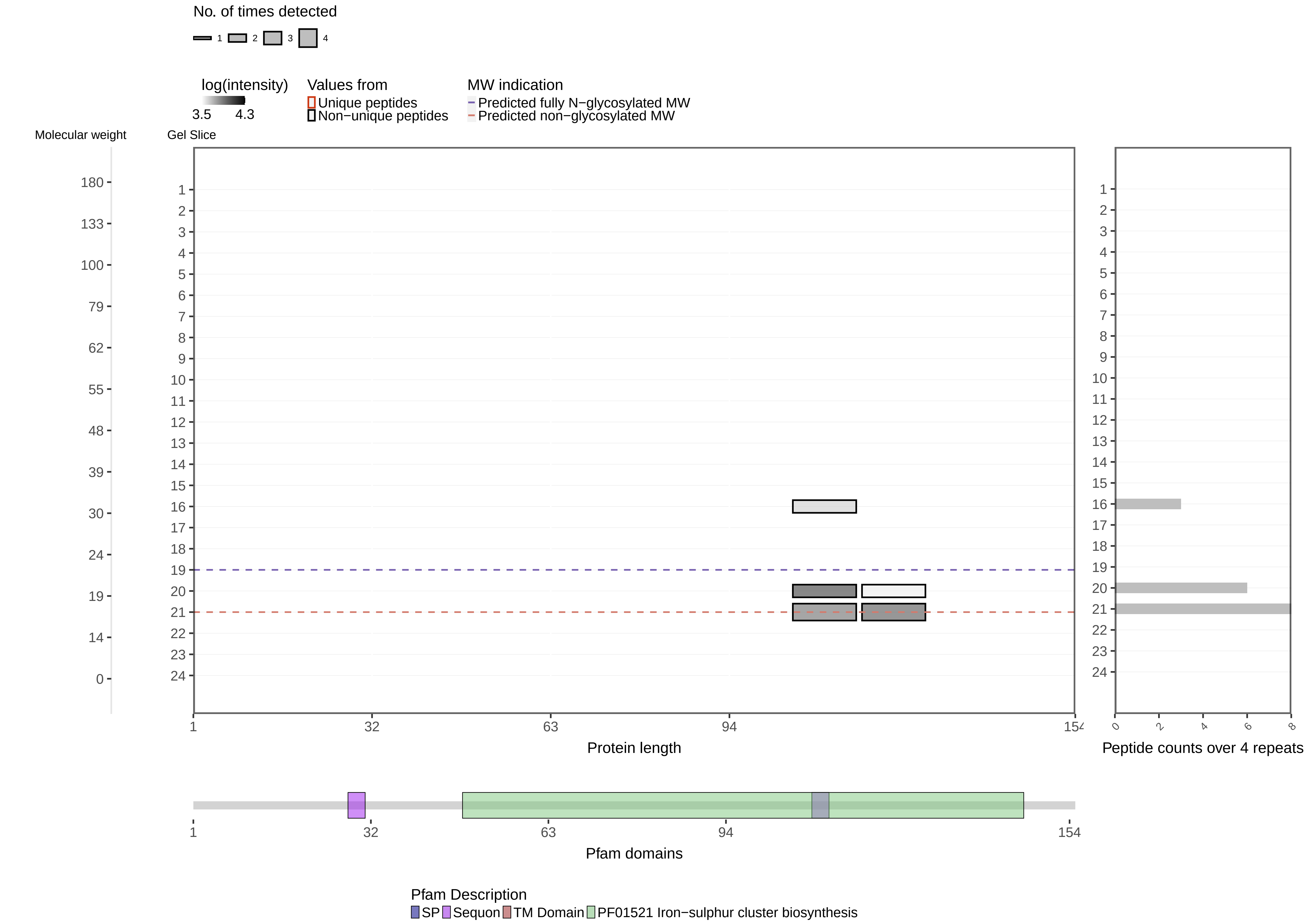


Protein sequence:

KEKMLRLSVEAGGCSGFQYEFSLDDQTNSDDRIFERAGVKLVVDNVSYDFVKGATVDYVEELIRSAFQVSTNPSAVGGCSCKSSFMVKQ*

### NbL01g11590.1 calreticulin [MW − SP = 44.24 kDa]

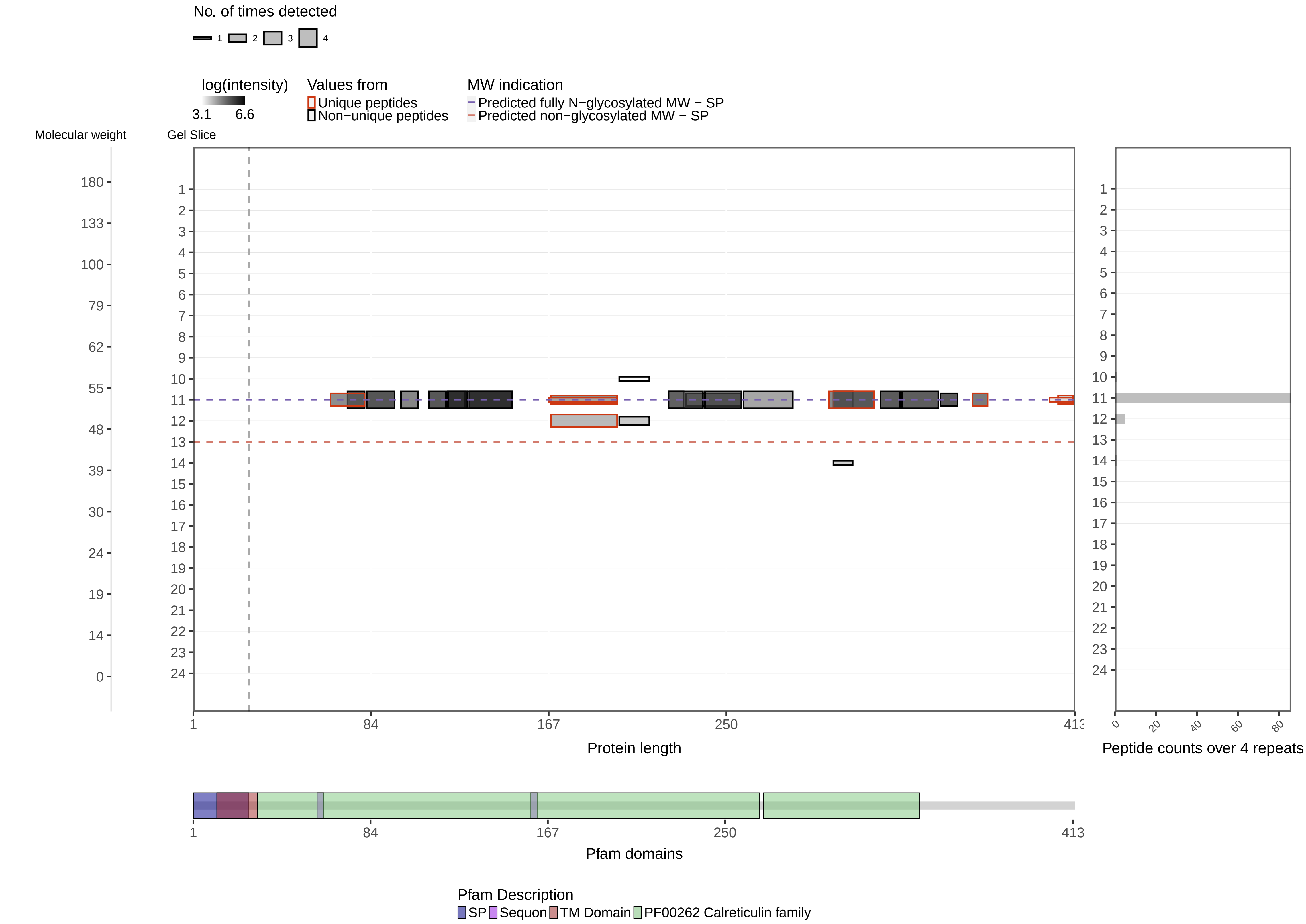


Protein sequence:

DNVEKQSGSLYSDWDLLPPKTIKDPSAKKPEDWDEKEFIDDPEDKKPEGYDDIPEEITDPDAKKPEDWDDEEDGEWTAPTIPNPEYKGPWKPKKIKNPNYKGKWKAPLIDNPDFKDDPDLYVFPKLKYVGVELWQVKSGTLFDNIVICDDPEYAKAIAEETWGKQKDAEKAAFEEAEKKREDEESKDAPADSDAEEDDDSDDADDKSESKDDEAHDEL*

NbL01g11550.1 dihydrolipoyl_dehydrogenase_2_chloroplastic−like [MW = 60.59 kDa]

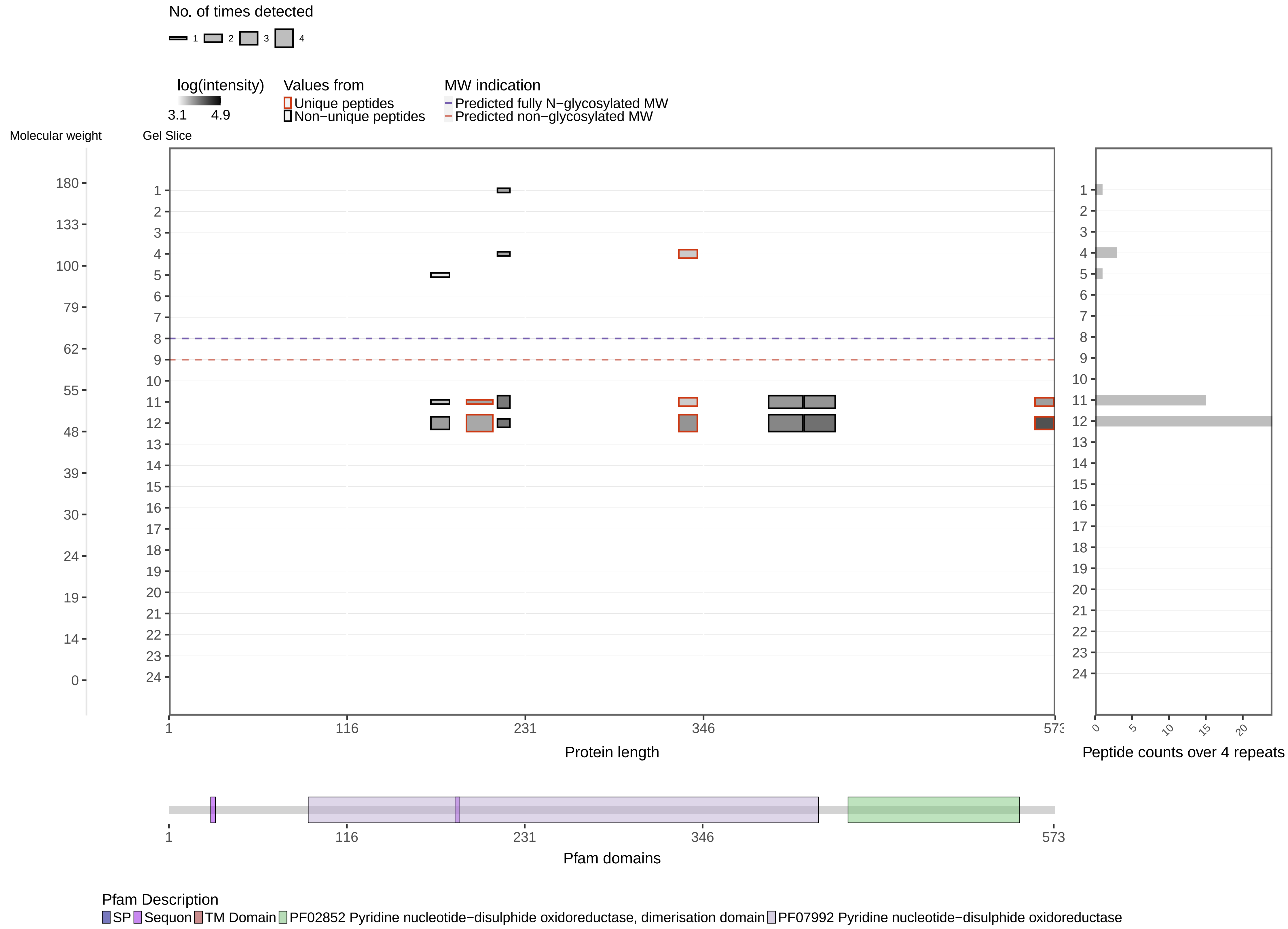


Protein sequence:

EFSDVYTALGSEVTFVEALDQLMPGFDPEIGKLAQRVLINPRKIDYHTGVFASKITPAKDGKPVTIELIDAKTKELKDTLEVDAALIATGRAPFTQGLGLENINVQTQRGFIPVDERMCVIDANGELVPHLYCIGDANGKMMLAHAASAQGISVVEQVTGKDHILNHLSIPAACFTHPEISMVGLTEPQAREKAEKEGFEIGIAKTSFKANTKALAENEGEGIAKMIYRPDNGEILGVHIFGMHAADLIHEASNAIAMGTRIQDIKFAVHAHPTLSEVIDELFKAAKVKAHTSSPVTEPVA

NbL01g11510.1 probable_protein_disulfide−isomerase_A6 [MW − SP = 37.02 kDa]

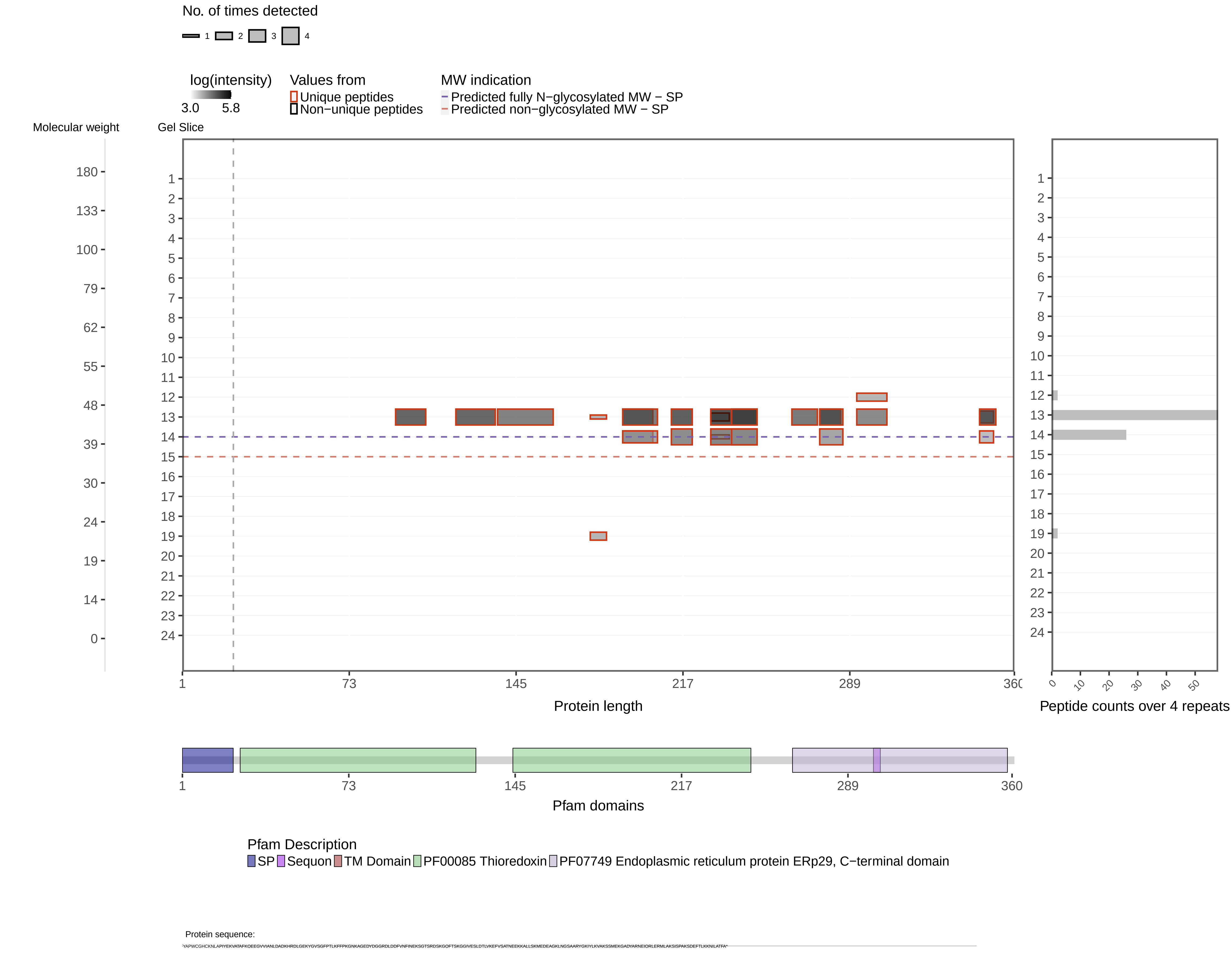

NbL01g11210.1 topless−related_protein_3−like [MW = 124.46 kDa]

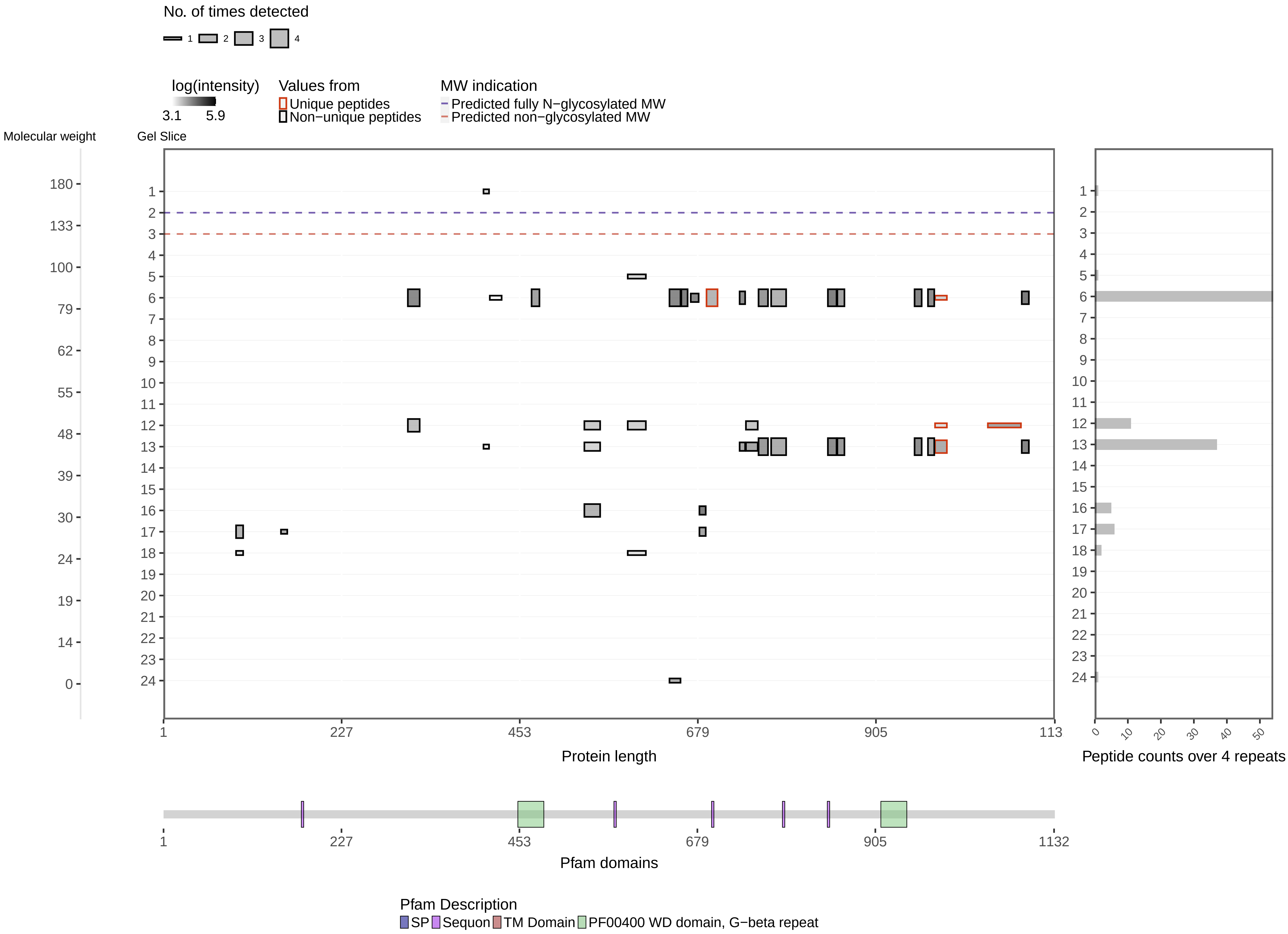


Protein sequence:

NbL01g11190.1 deoxyuridine_5'−triphosphate_nucleotidohydrolase [MW = 24.67 kDa]

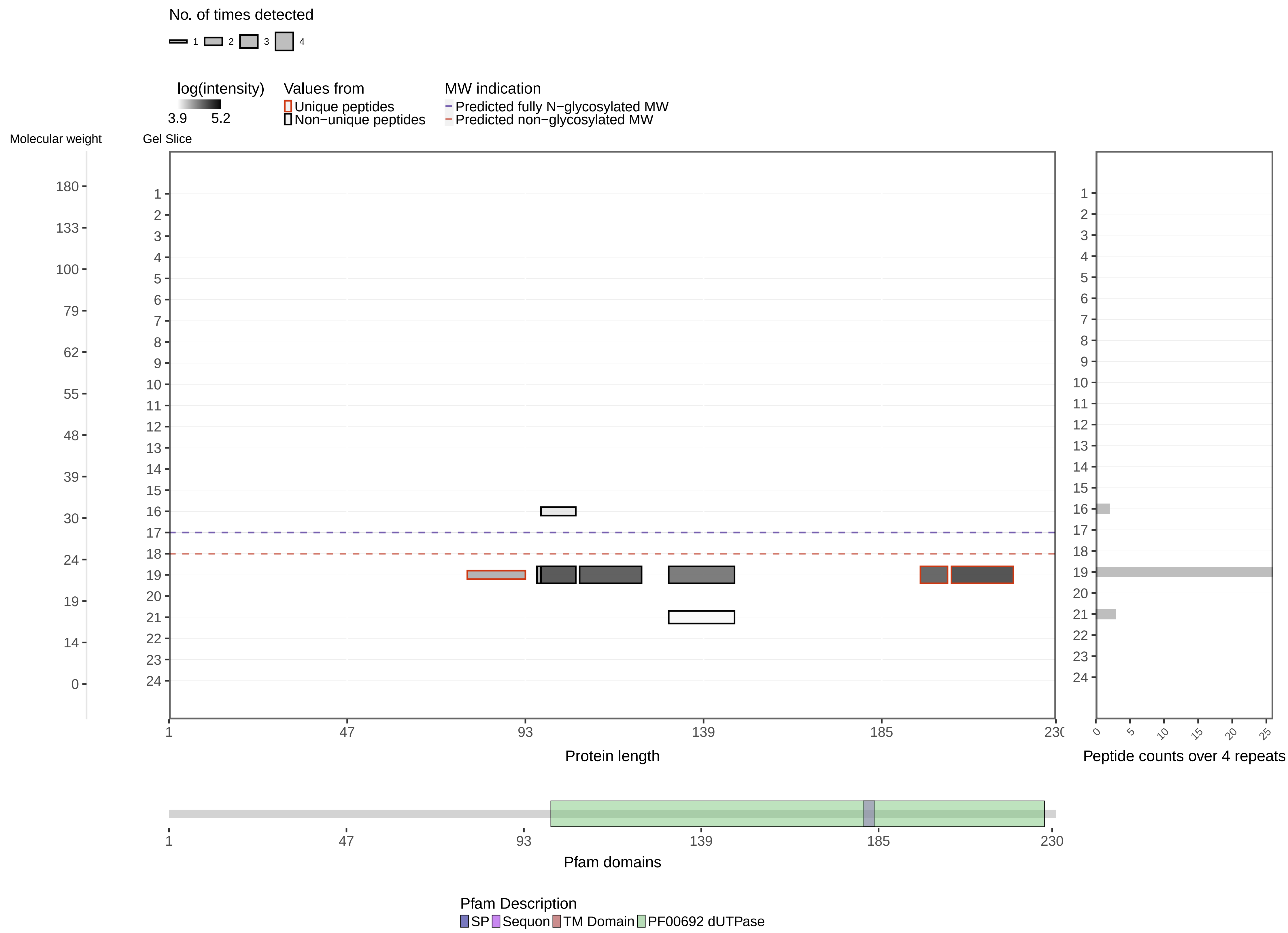


Protein sequence:

VLPSRASPLSAGYDLSSAAEAKVPARGKALIPTDLSIAVPQGAYARIAPRSGLAWKHSIDVGAGVIDADYRGPVGVVLFNHSDVDFEVKAGDRIAQLIIQKIVTPDVEEVDDLDSTVRGSGGFGSTGV*

NbL01g11080.1 bifunctional_epoxide_hydrolase_2−like [MW = 35.58 kDa]

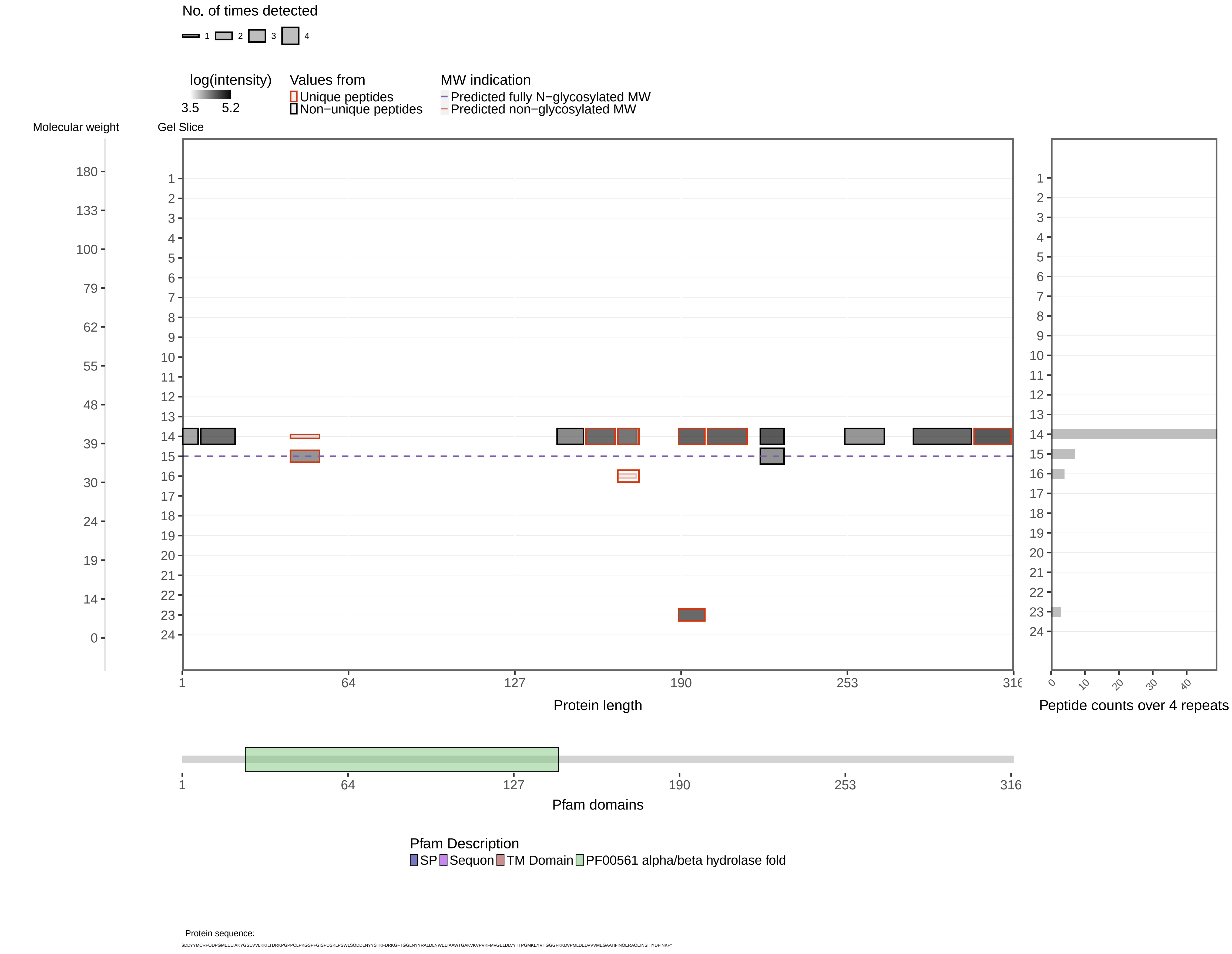

NbL01g11070.1 60S_ribosomal_protein_L18−2−like [MW = 26 kDa]

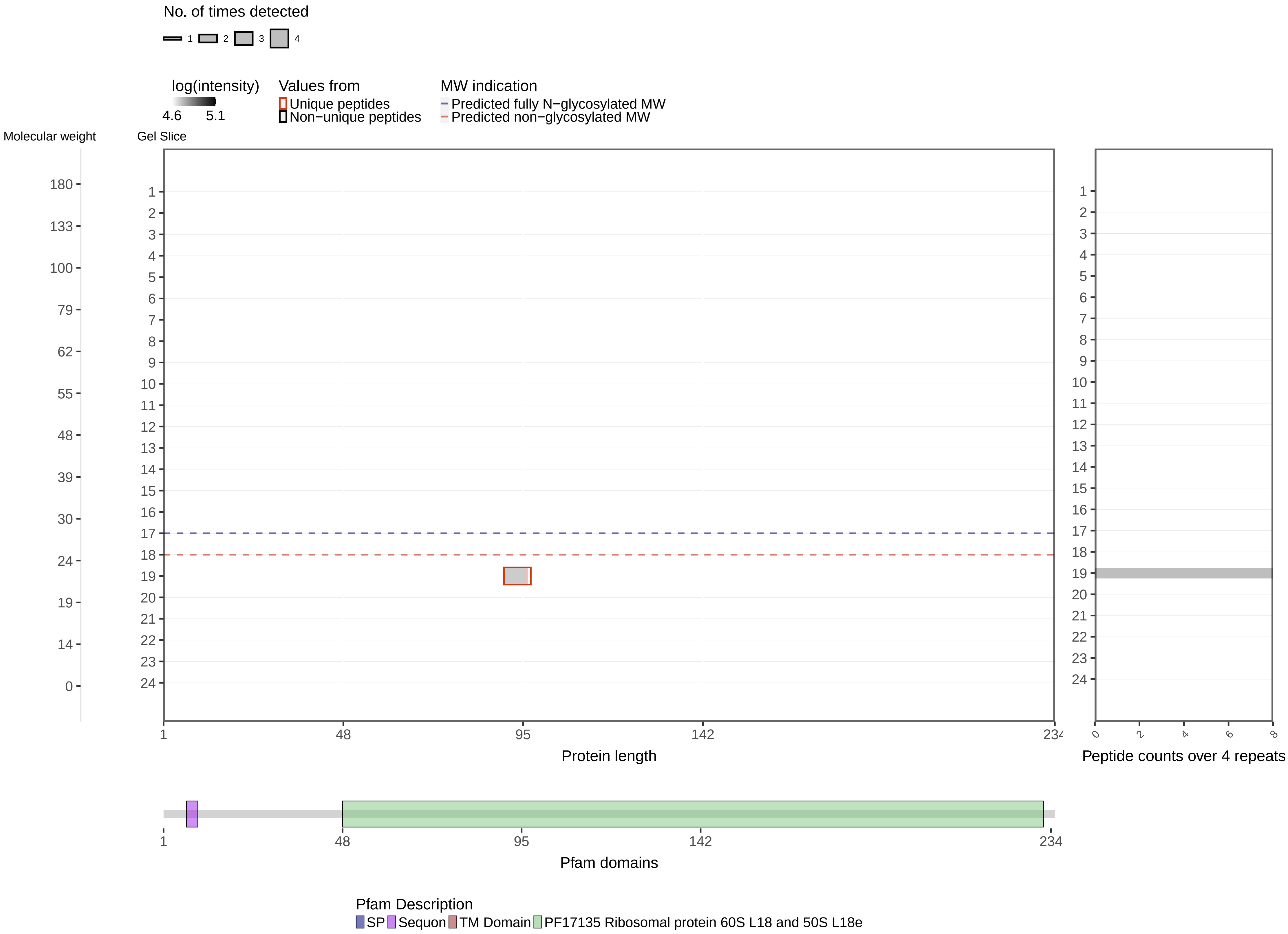


Protein sequence:

KPPLSLSRLISYAKGKEDKVVVIVGTITDDVRAYEVPALKVCALRFTKTARARIEKAGGECLTFDQLALRAPLGQNTLLLRGPKNAREAVKHFGPAPGVPHSHTKPFVRSKGRKFERARGRRNSRGYKA*

NbL01g11060.1 malignant_T−cell−amplified_sequence_1_homolog [MW = 21.49 kDa]

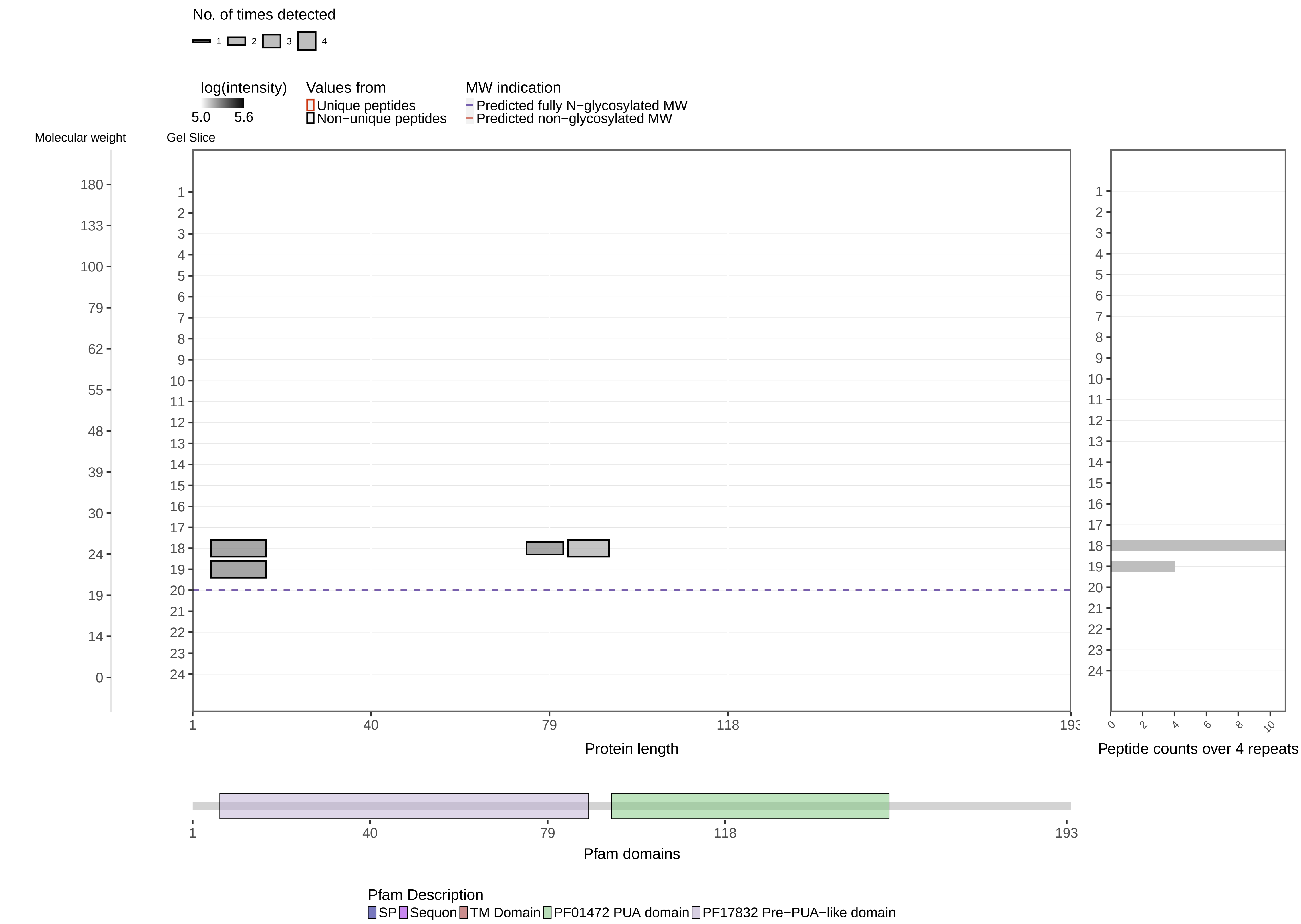


Protein sequence:

.HQYPNIMKKLQVDRGAIKFVLSGANIMCPGLTSPGGALDDEVGAETPVAIMAEGKQHALAIGFTKMSAKEGQSTRELVWITCIILMMASGRWSDWIELEWKTMFTFCK*

### NbL01g11000.1 auxin−repressed_12.5_kDa_protein−like [MW = 13.71 kDa]

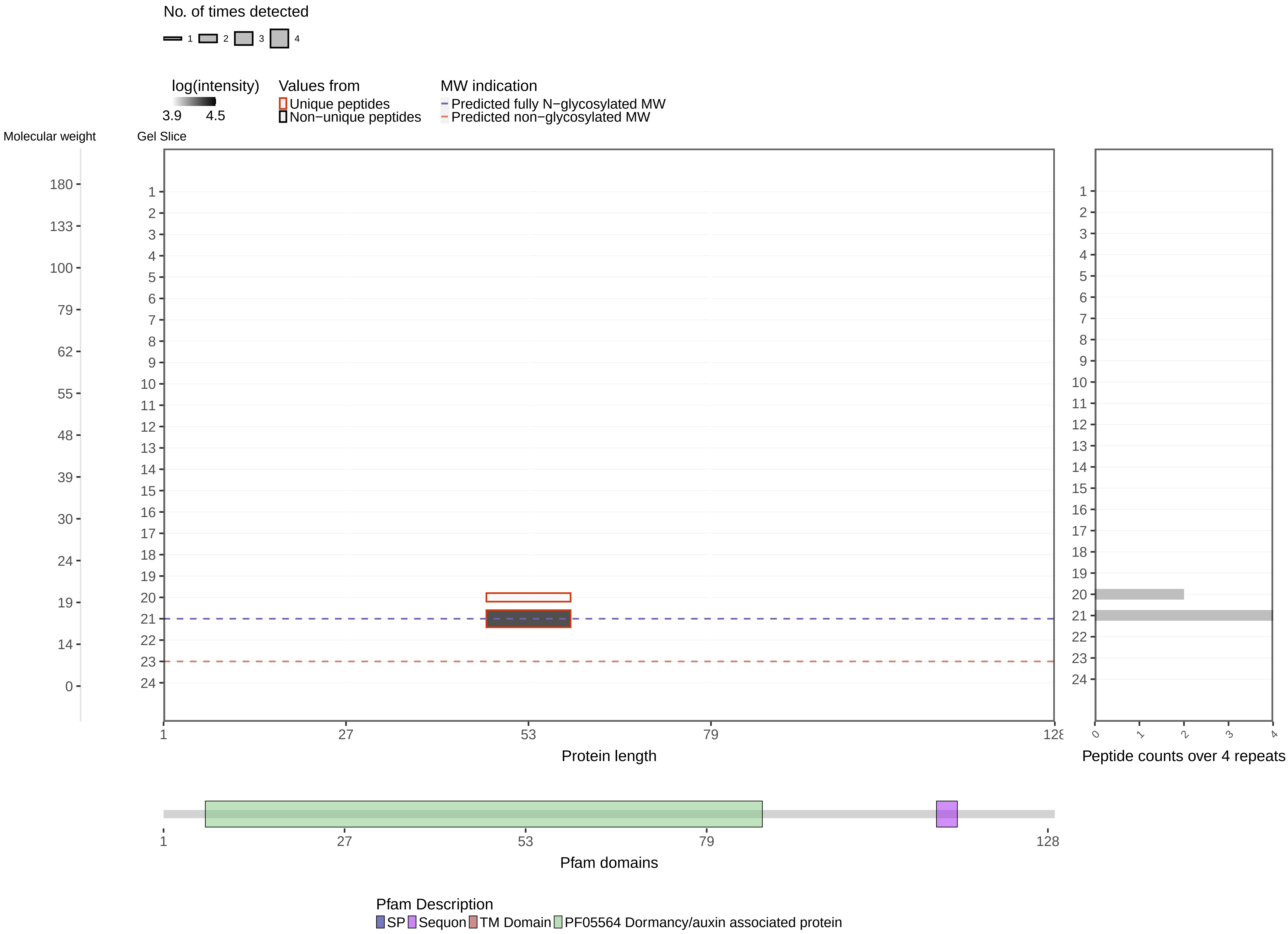


Protein sequence:

EASSDDAVRVTRSIMIVKPPGSQNRDSPPVSPAGTTPPVSPFAGAGGREAYWSRRRSFAYENASGVGPRSPRPPYDL*

NbL01g10880.1 ATP−dependent_Clp_protease_proteolytic_subunit−related_protein_3_chloroplastic [MW = 35.77 kDa]

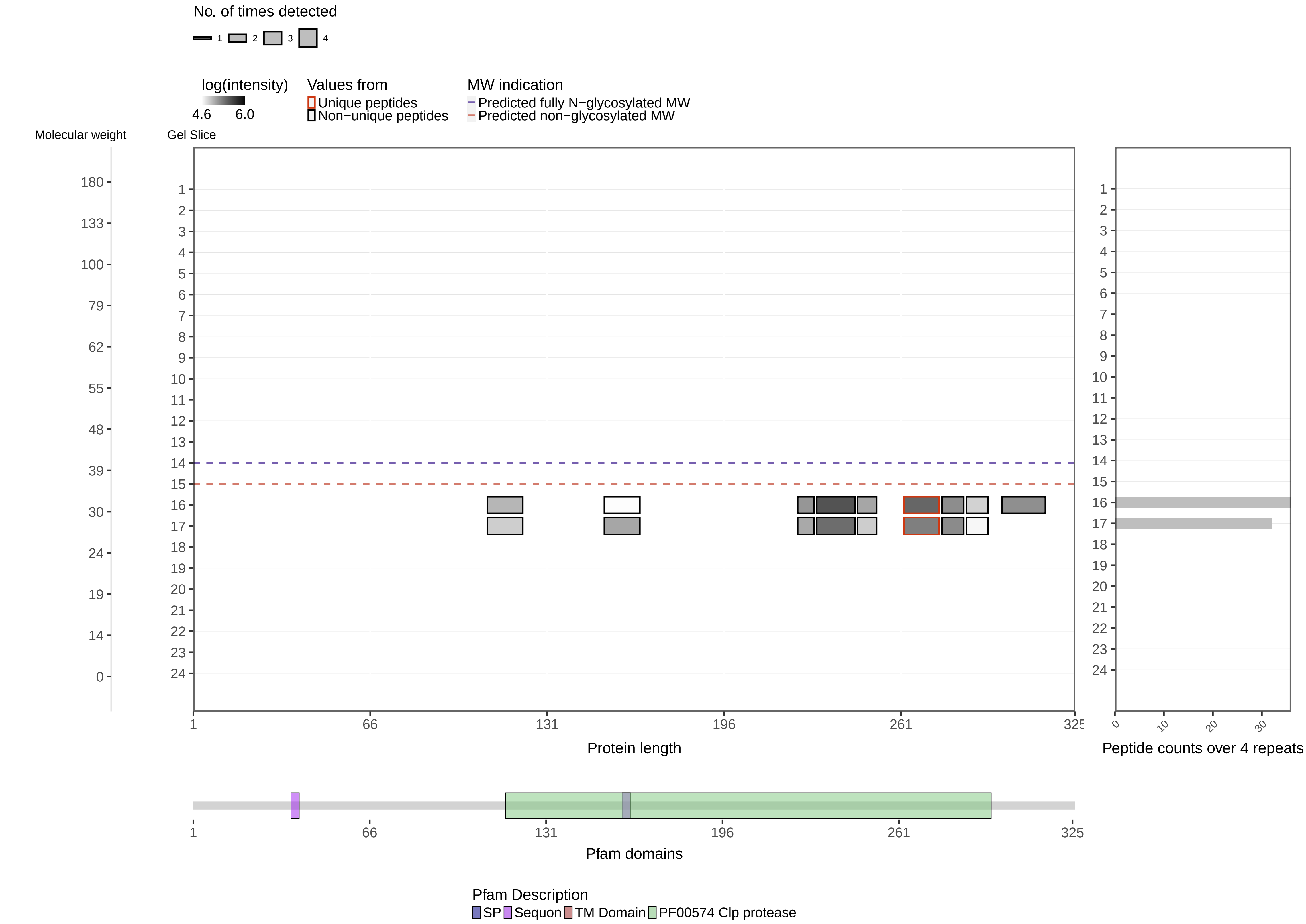


Protein sequence:

DPKEPIYLYINSTGTTRDDGETVGMETEGFAIYDAMMQLKNEIHTVAVGAAIGQACLLLAAGSKGKRFMMPHAKAMIQQPRVPSSGLMPASDVFIRAKEVIINRDILVKLLAKHTENSEEAVANVMRRPFYMDATRAREFGVIDKILWRGQEQIMADVSAPEQWDKNAGIKVADAI*

### NbL01g10850.1 40S_ribosomal_protein_S10−1−like [MW = 20.06 kDa]

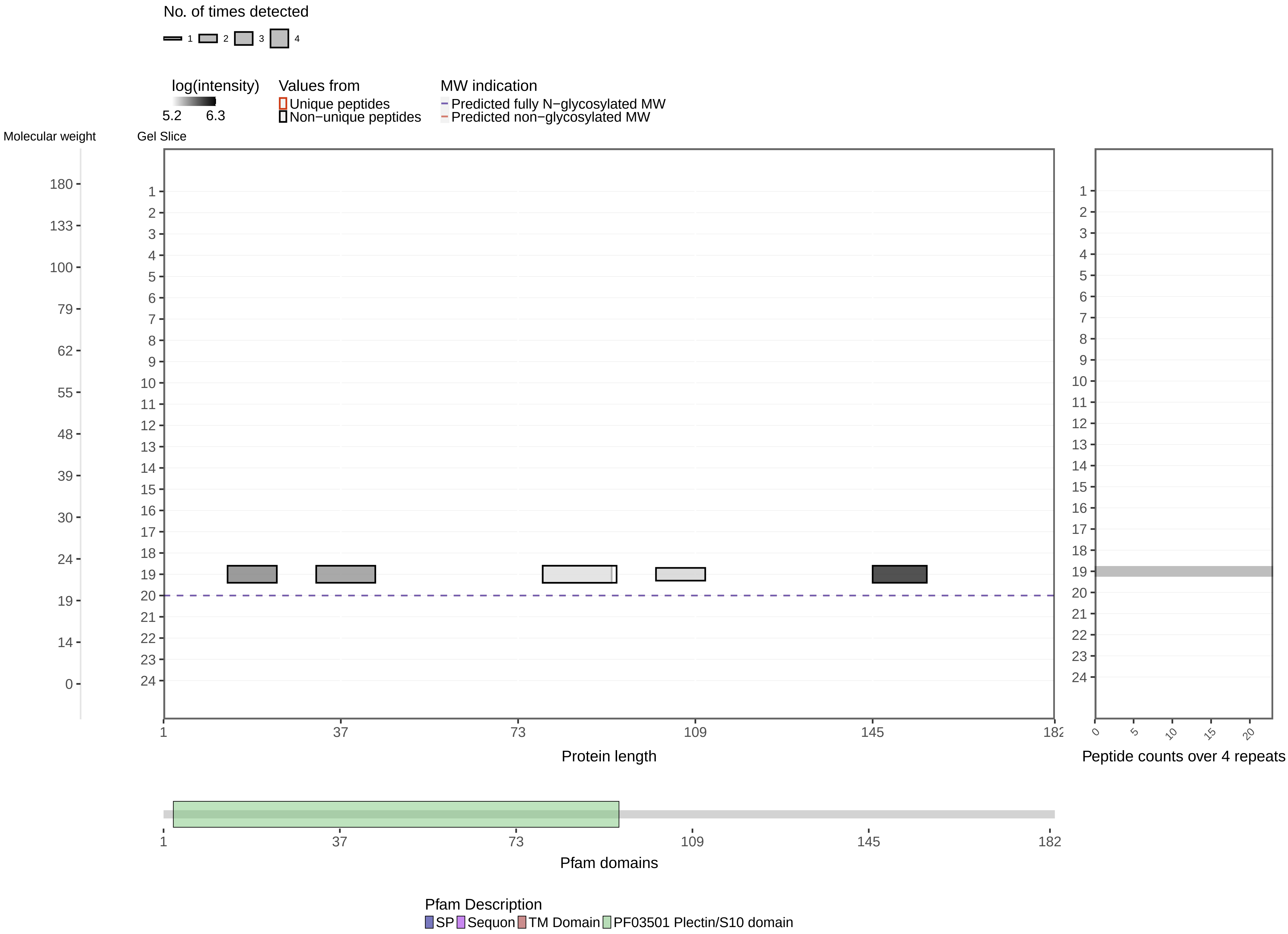


Protein sequence:

LPSEIVPATLKKSAKPLGRPMGGPPGDRPRGPPRFEGDRPRFGDREGYRAGPRGPPGEFGGEKGGAPADYQPAFRGSGGRPGFGRGSGGFGGAPPSSSFS*

NbL01g10820.1 xyloglucan_endotransglucosylase/hydrolase_protein [MW − SP = 31.68 kDa]

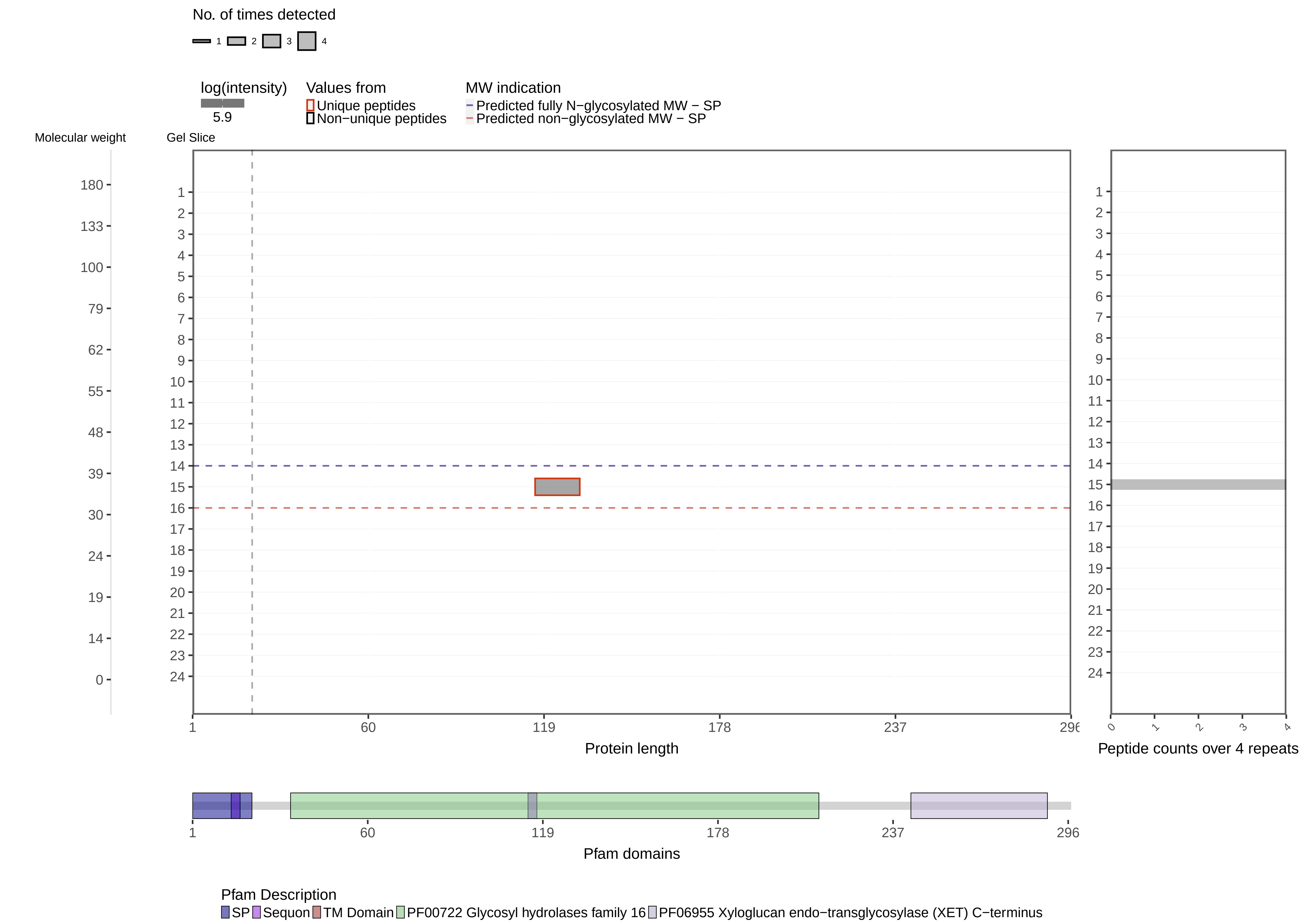


Protein sequence:

RIYLWFDPTKGYHSYSVLWNTFQIVIFVDDVPIRAFKNSKDLGVKFPFNQPMKIYSSLWDADDWATRGGLEKTDWSNAPFTASYTSFHVDGCEAATPQEVQVCNTKGMRWWDQKAFQDLDALQYRRLRWVRQKYTIYNYCTDRKRYPTLPPECTKDRDI*

### NbL01g10780.1 glutathione_S−transferase_U7−like [MW = 25.67 kDa]

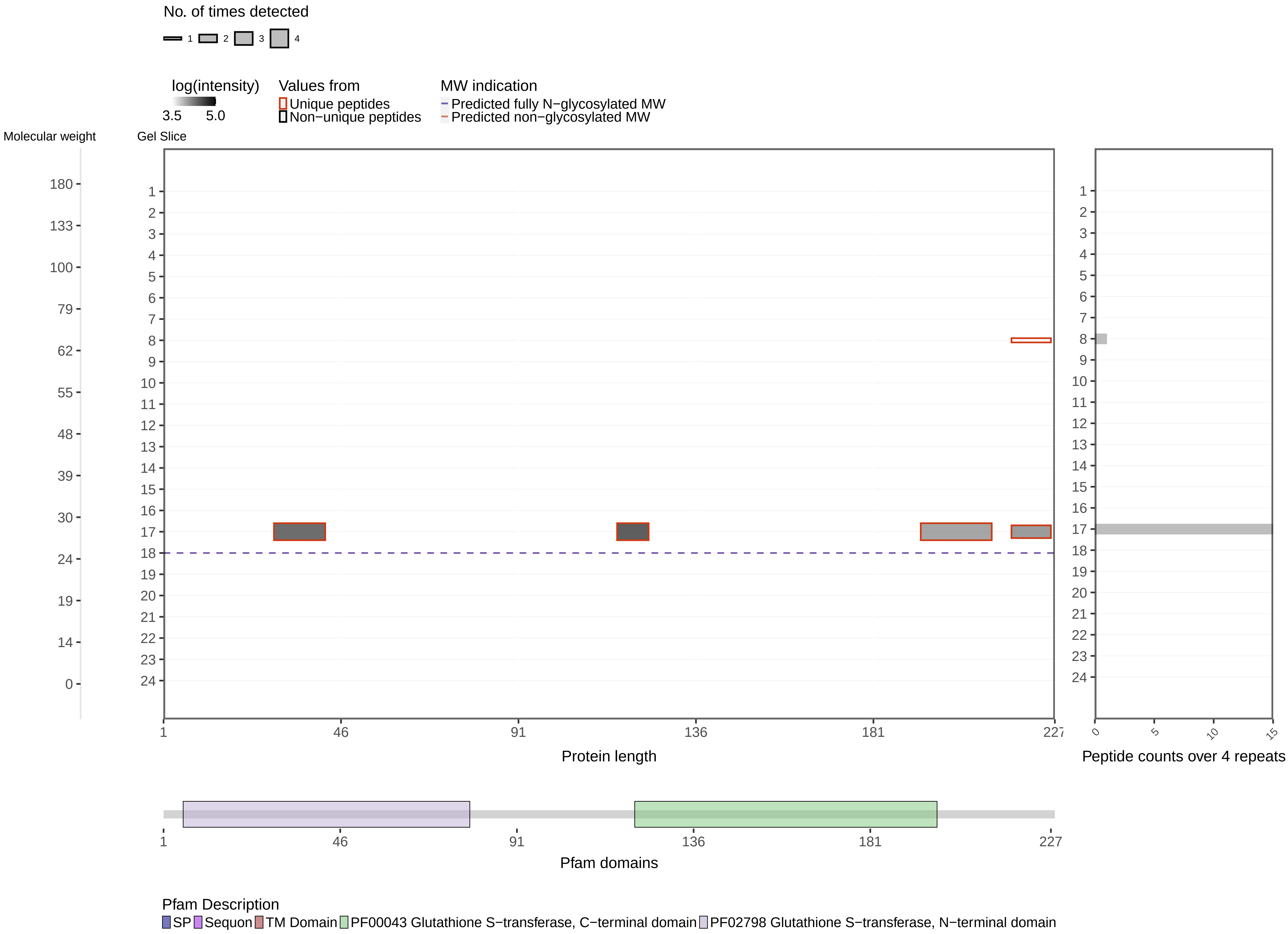


Protein sequence:

RFIDGKFYEAAKKAFFSSGESKAEGVDSVAEGLQLLEGQITGKKFFGGEKIGYVDIVVGWIAYWFQYVEEVGEFKAMDSKEYPCLNAWIKNFIQVPLIQQSLPKPDDVKAVFKGFKDAALADTN*

### NbL01g10730.1 histone_H2A.1 [MW = 15.53 kDa]

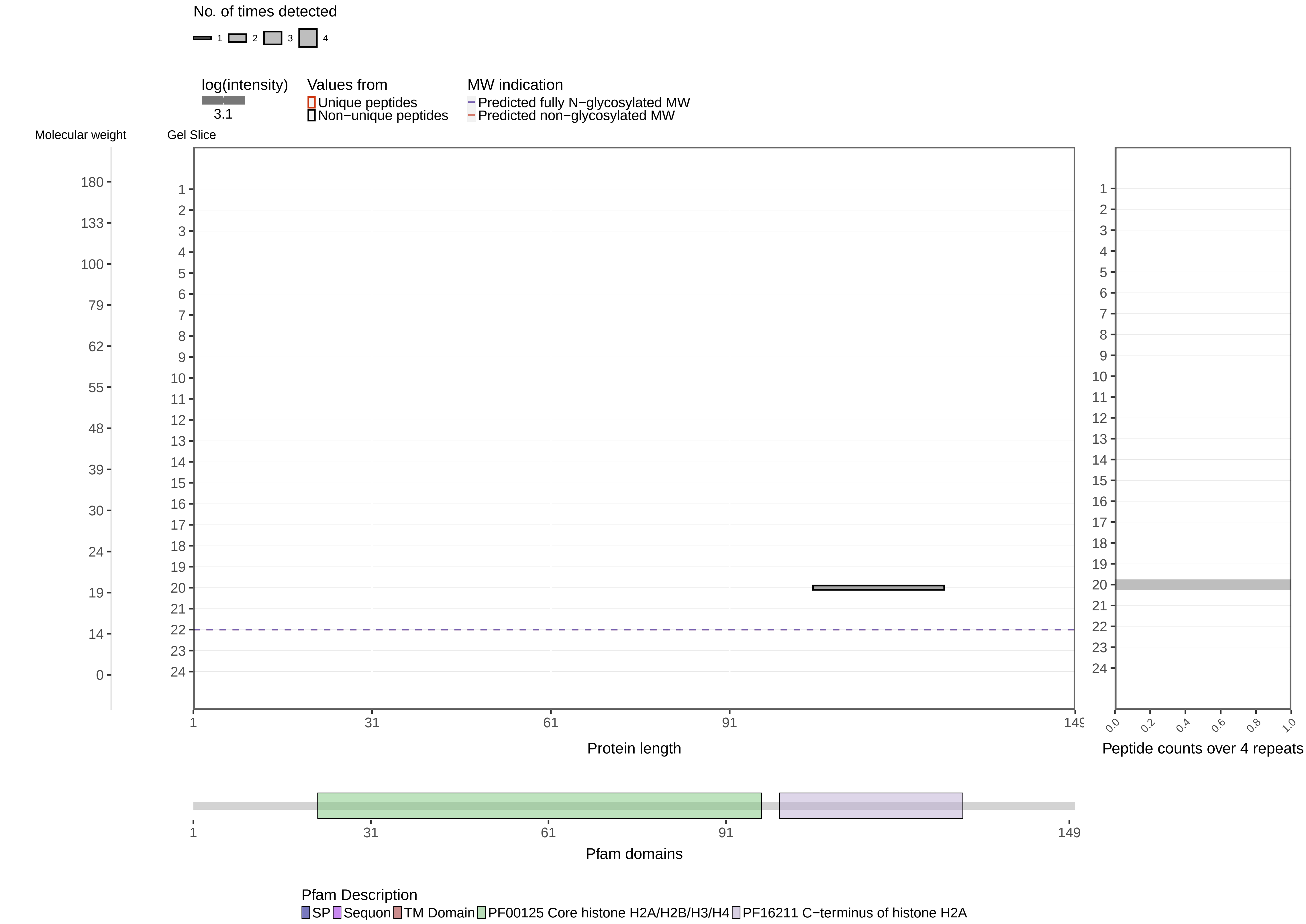


Protein sequence:

.AAVLEYLAAEVLELAGNAARDNKKSRIIPRHVLLAVRNDEELGKLLSGVTIASGGVLPNINPVLLPKKTAAVEEKASKPKATKSPKKA*

### NbL01g10720.1 autophagy−related_protein_18a−like [MW = 48.98 kDa]

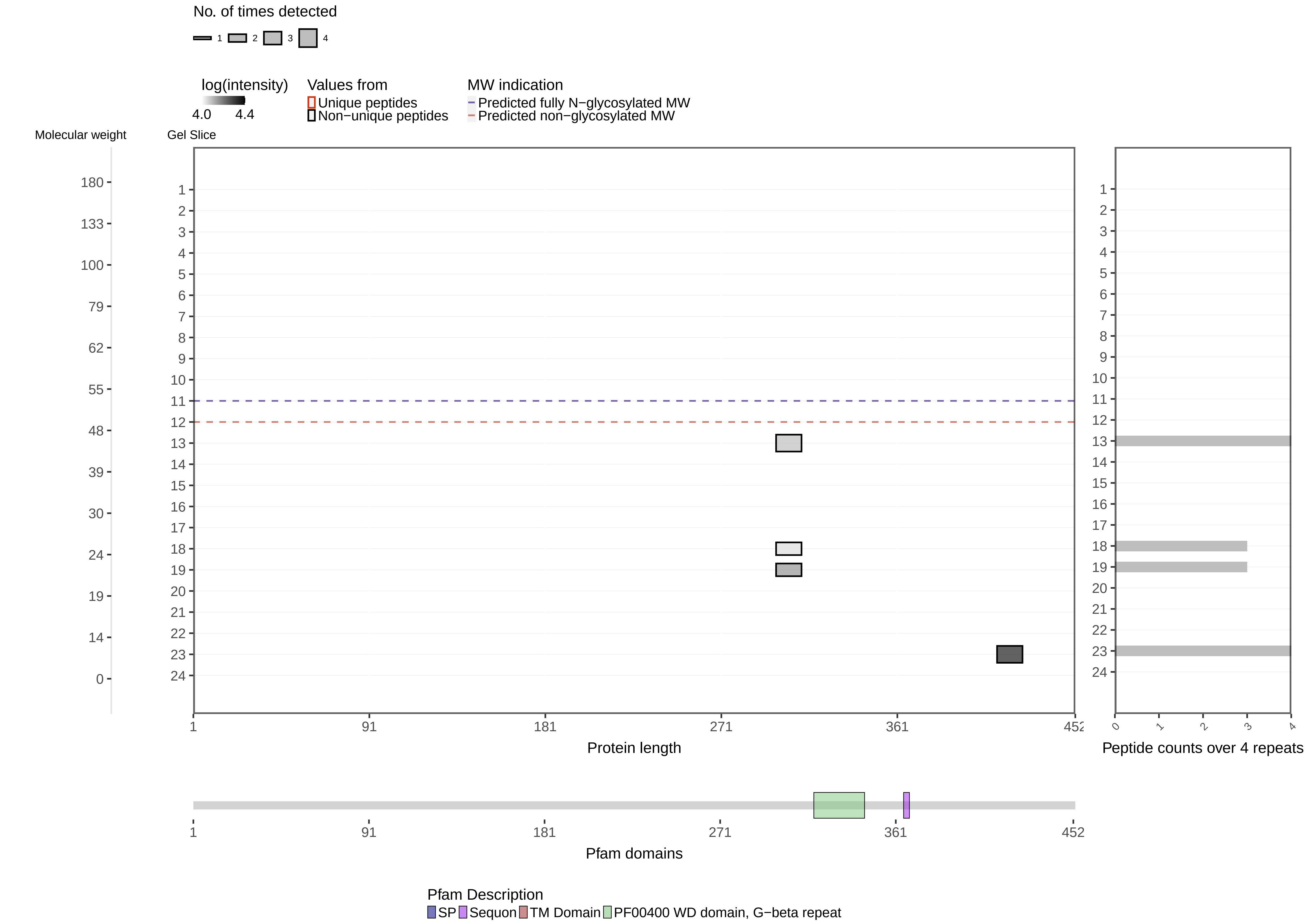


Protein sequence:

FADLKLLHQIETVANPKGLCEVSQVSGSMVLVCPGLQKGQVRVEHYASKRTKFIMAHDSRIVCFALTNGGRLLATASSKGTLVRVFNTLDGSLVQEVRRGADRAEIYSLAFSSTAQWLAVSSDKGTVHVFNLKVDSAAVGVDRPRGASESNATSPSAVSHLSFIRGVLPKYFSSEWSVAQFRLPEGLQHIVAFGHQKNTVVILGMDGSFYRCQFDAAAGGEMTLLEHHSFLKPEENF*

NbL01g10540.1 probable_linoleate_9S−lipoxygenase_5 [MW = 97.51 kDa]

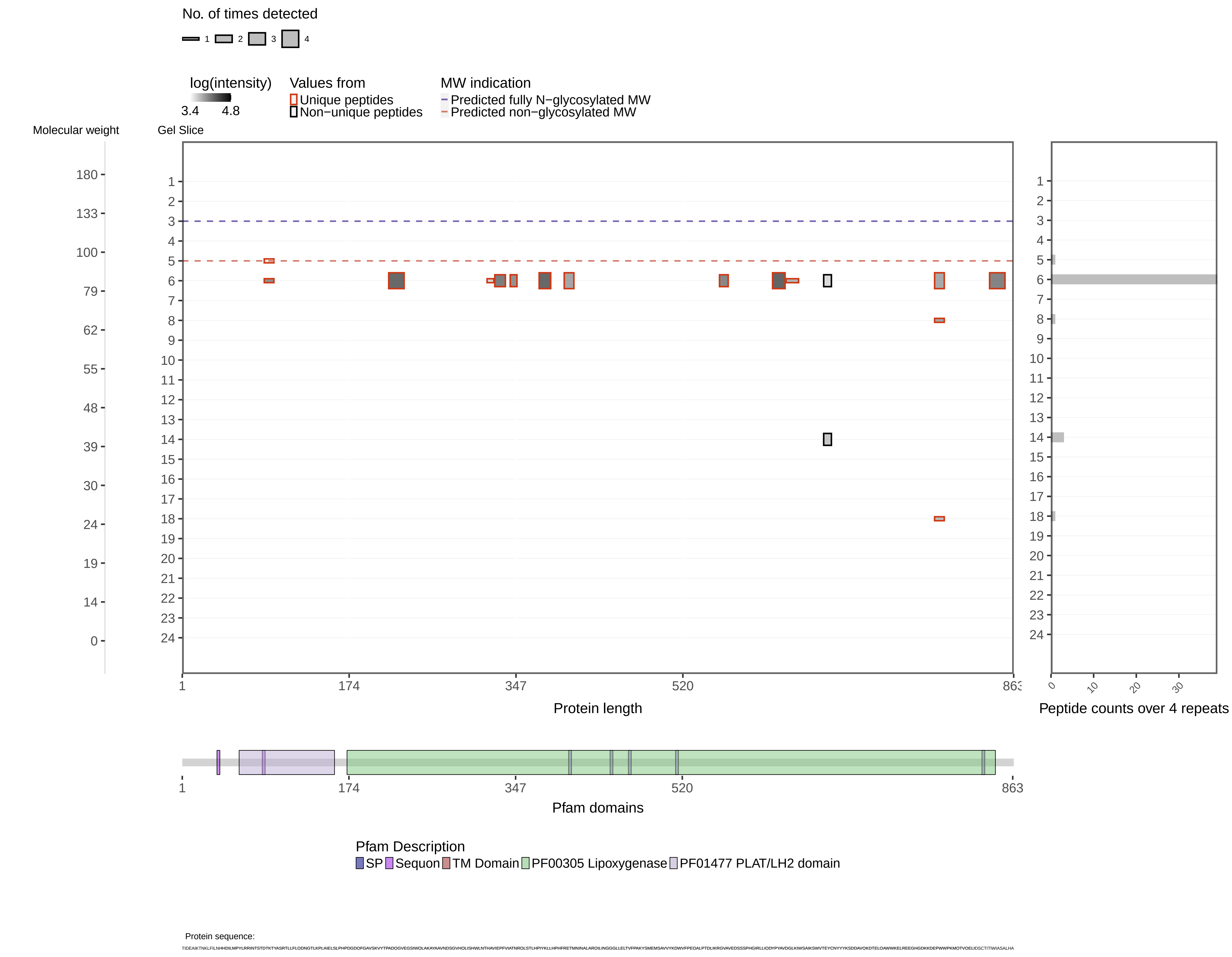

NbL01g10510.1 probable_Xaa−Pro_aminopeptidase_P [MW = 80.29 kDa]

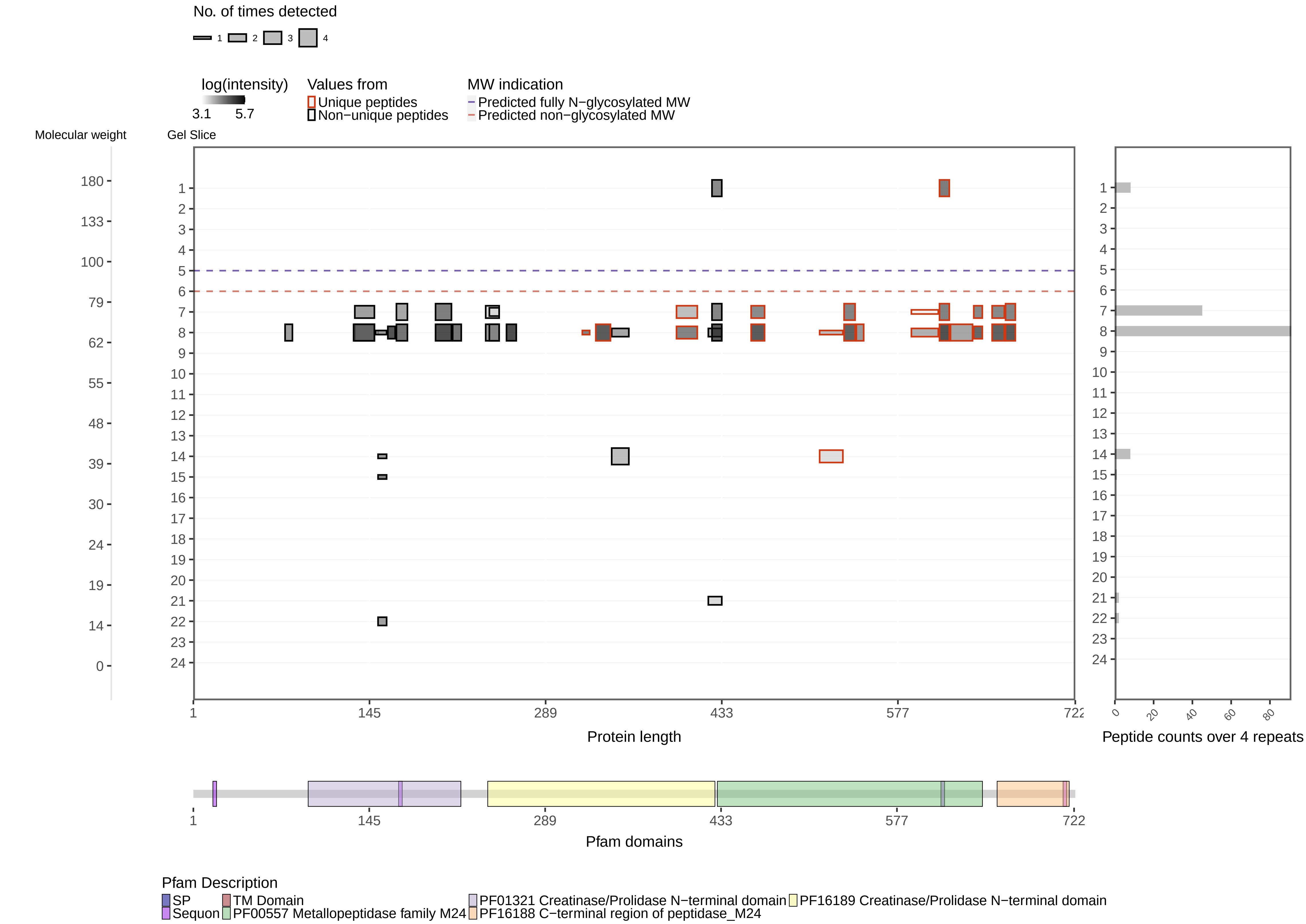


Protein sequence:

[illegible]

### NbL01g10500.1 long_chain_acyl−CoA_synthetase_6_peroxisomal [MW = 77.1 kDa]

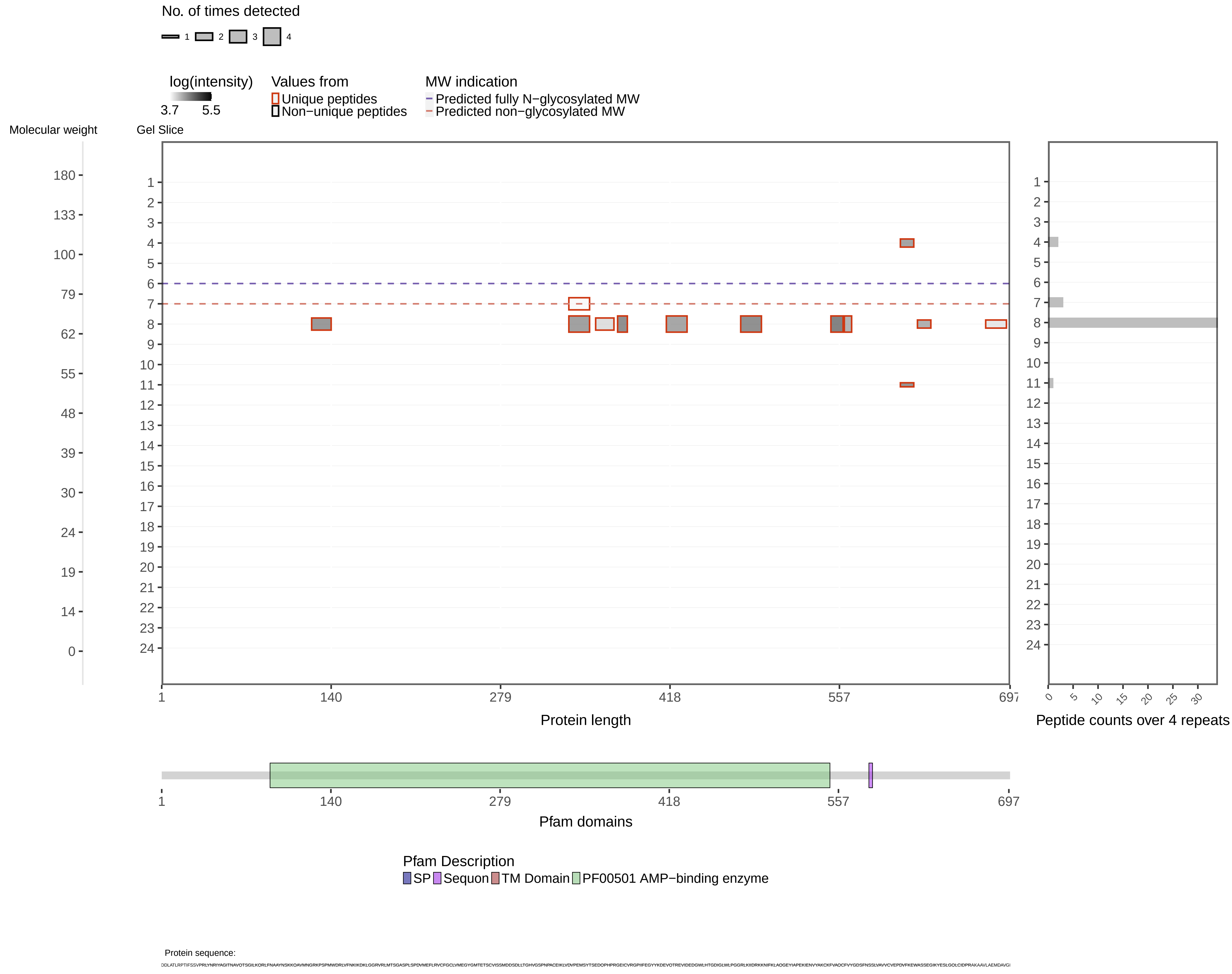

NbL01g10480.1 mannosylglycoprotein_endo−beta−mannosidase [MW = 110.56 kDa]

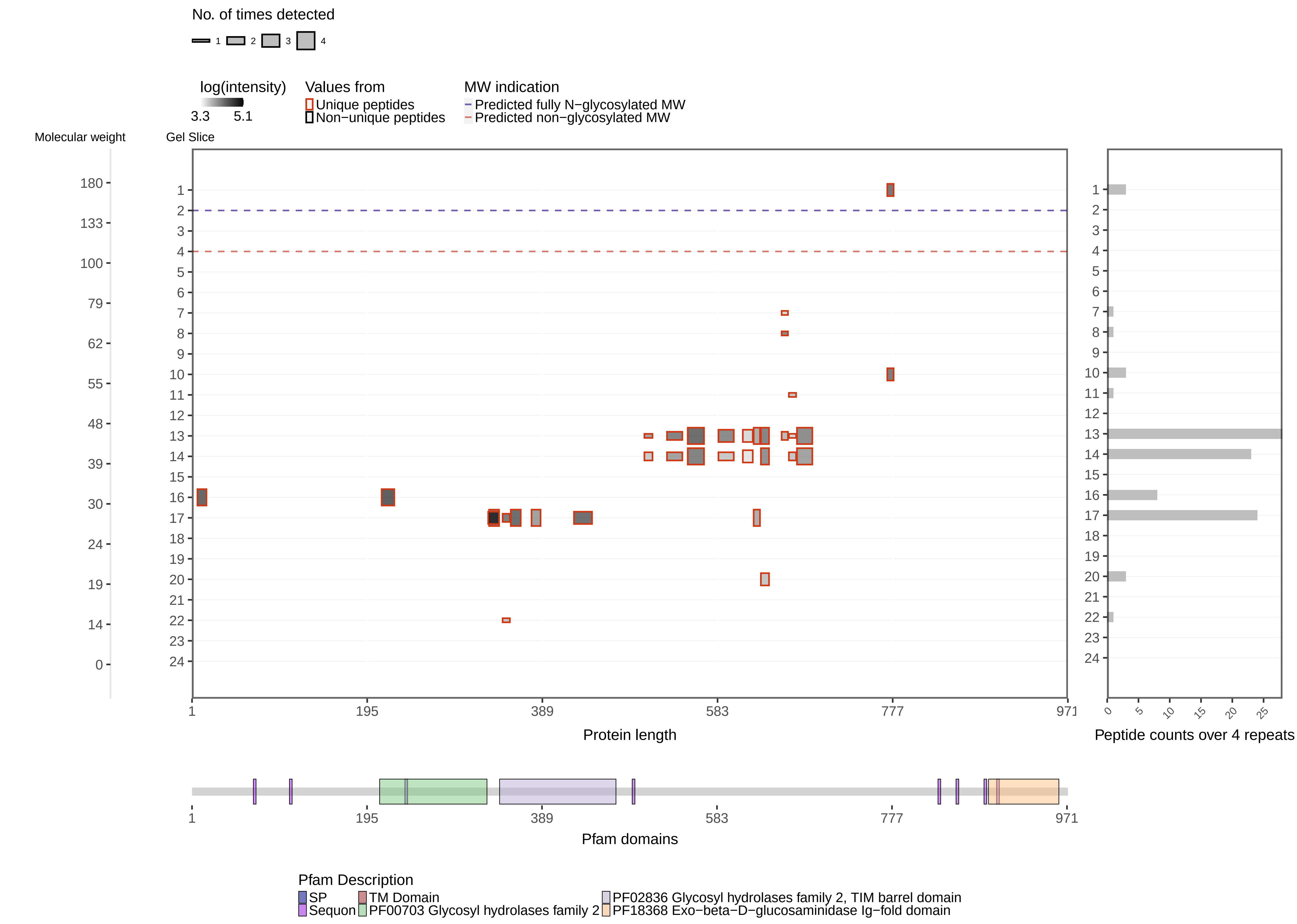


Protein sequence:

NAALKNDLQLHPYFMKSNDSGISAITPVLKDPSQYLDGTRVYIQGSMWDGFADGKGDFTDGPYEIQNPEDFFKHDYYKYGFNPEVGNVGMPVAATIRATMPPEGWQIPLFKKLSDGYIEEVPNPIWNYHKYIPYSKPGKVHDQILSYGTPKDLDDFCLKAQLVNYVQYRALLEGWTSQMWSKYTGVLIWKTQNPWTGLRGQFYDHLHDQTAGFYGCRSAAEPIHVQLNLATYSIEVVNTMSEELPNVAIEASIWDLEGECPYYKTSEKLTVPPKKTISTFEMKYPKSKNPK

NbL01g10330.1 serine−−tRNA_ligase−like [MW = 51.07 kDa]

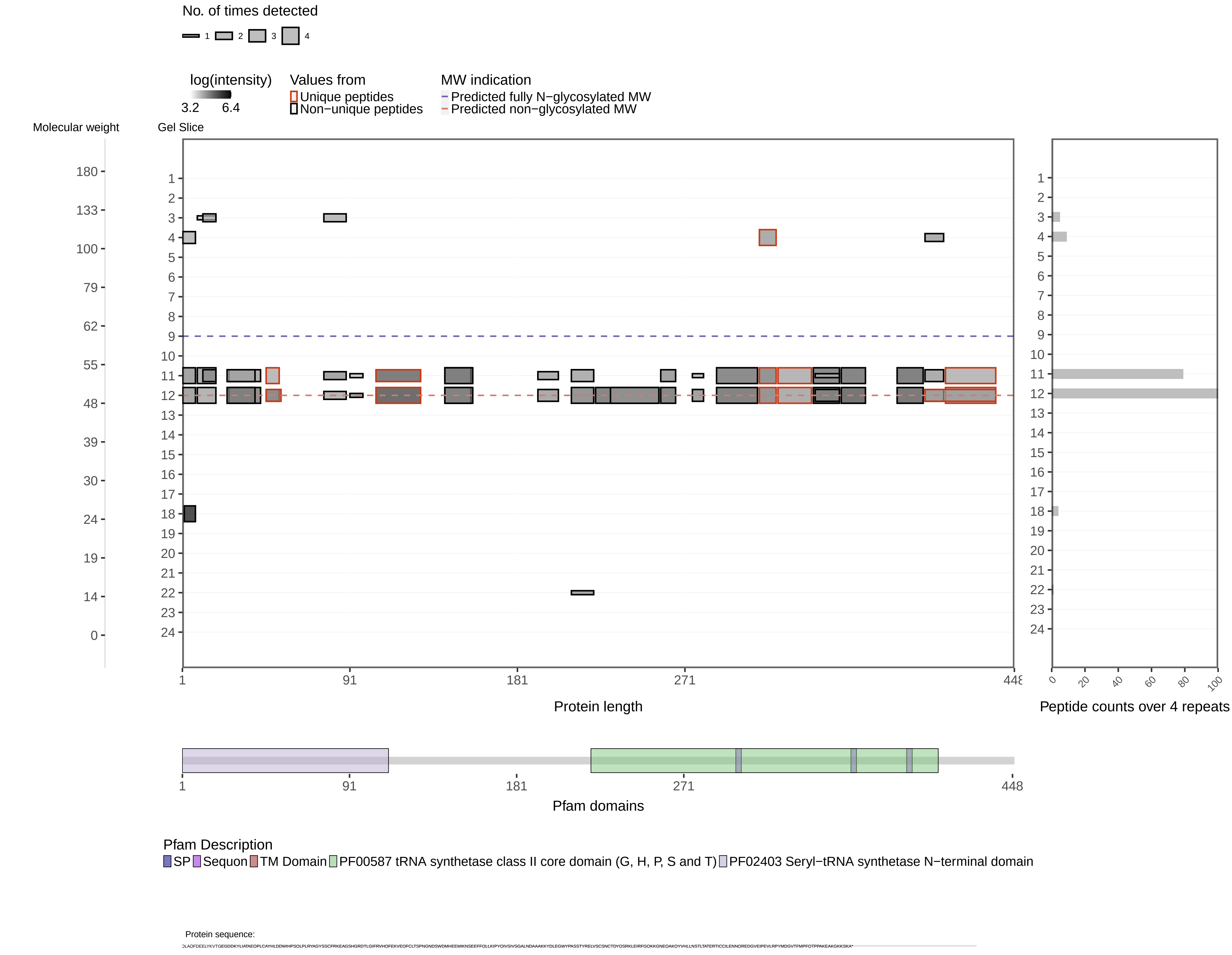

NbL01g10290.1 glucose_and_ribitol_dehydrogenase_homolog_1 [MW = 31.73 kDa]

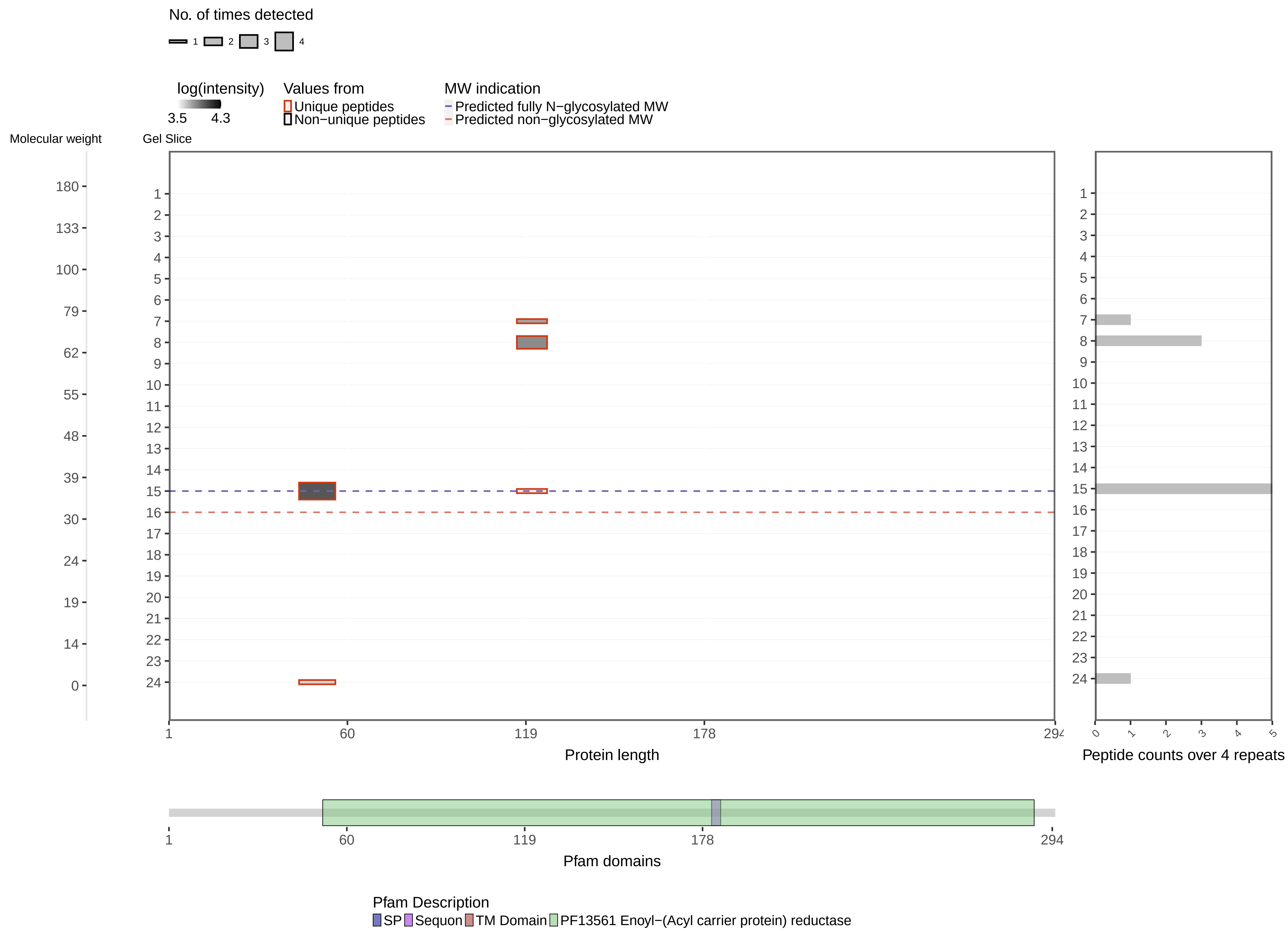


Protein sequence:

VNNAAEQYEATSVEEINEERLERVFRTNIFSYFFVTRHALKHMKEGSSIINTTSVTAYKGNAKLLDYTATKGAIVAFTRGLALQLVERGIRVNGVAPGPIWTPLIPASFTEEECANFGKQVPMKRAGQPIEVAPSYVFLASSPESSYISGQVIHPNGGTIVNA*

### NbL01g10140.1 branched−chain−amino−acid_aminotransferase−like_protein_1 [MW = 66.4 kDa]

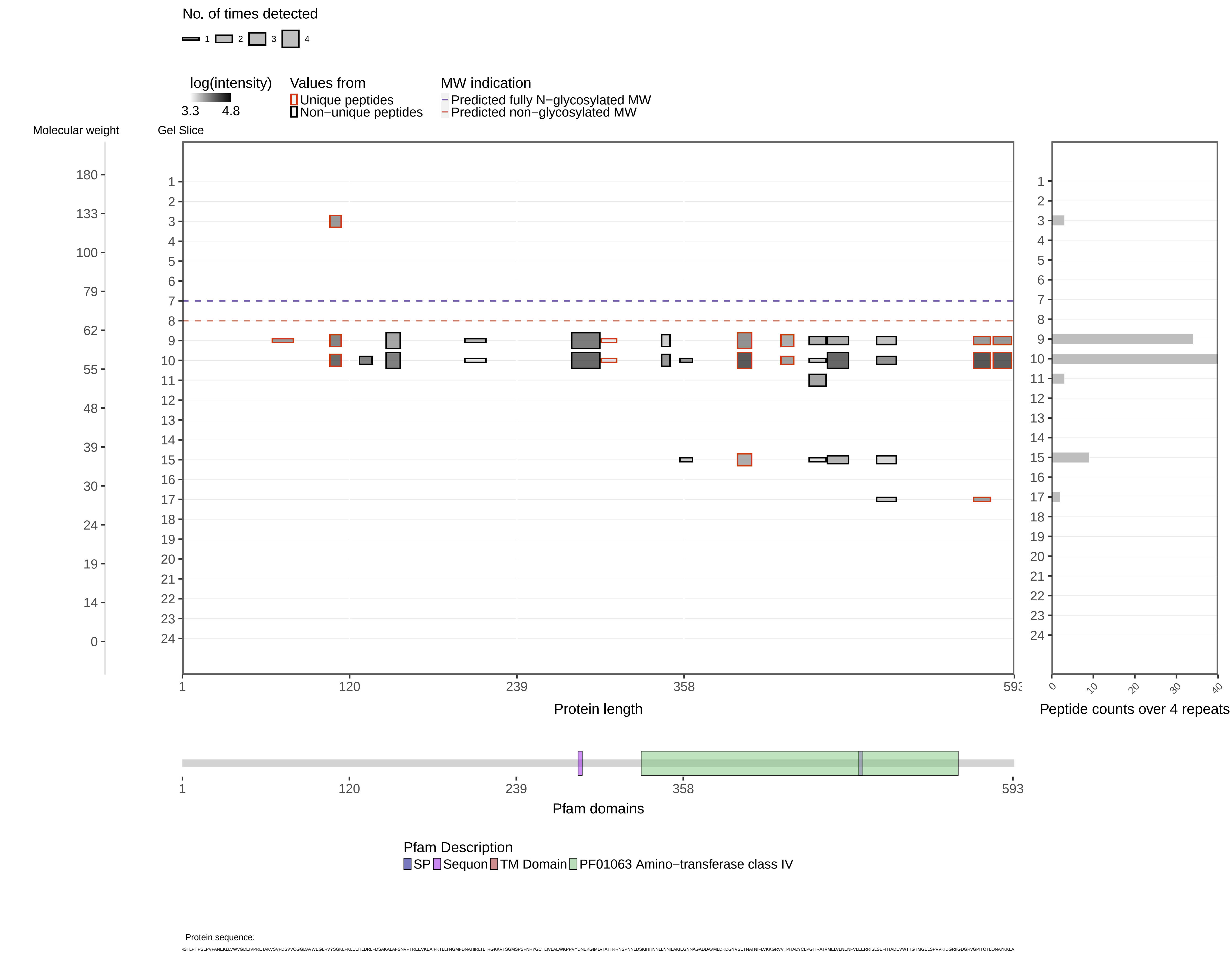

### NbL01g10120.1 GDSL_esterase/lipase_At1g54790 [MW − SP = 42.82 kDa]

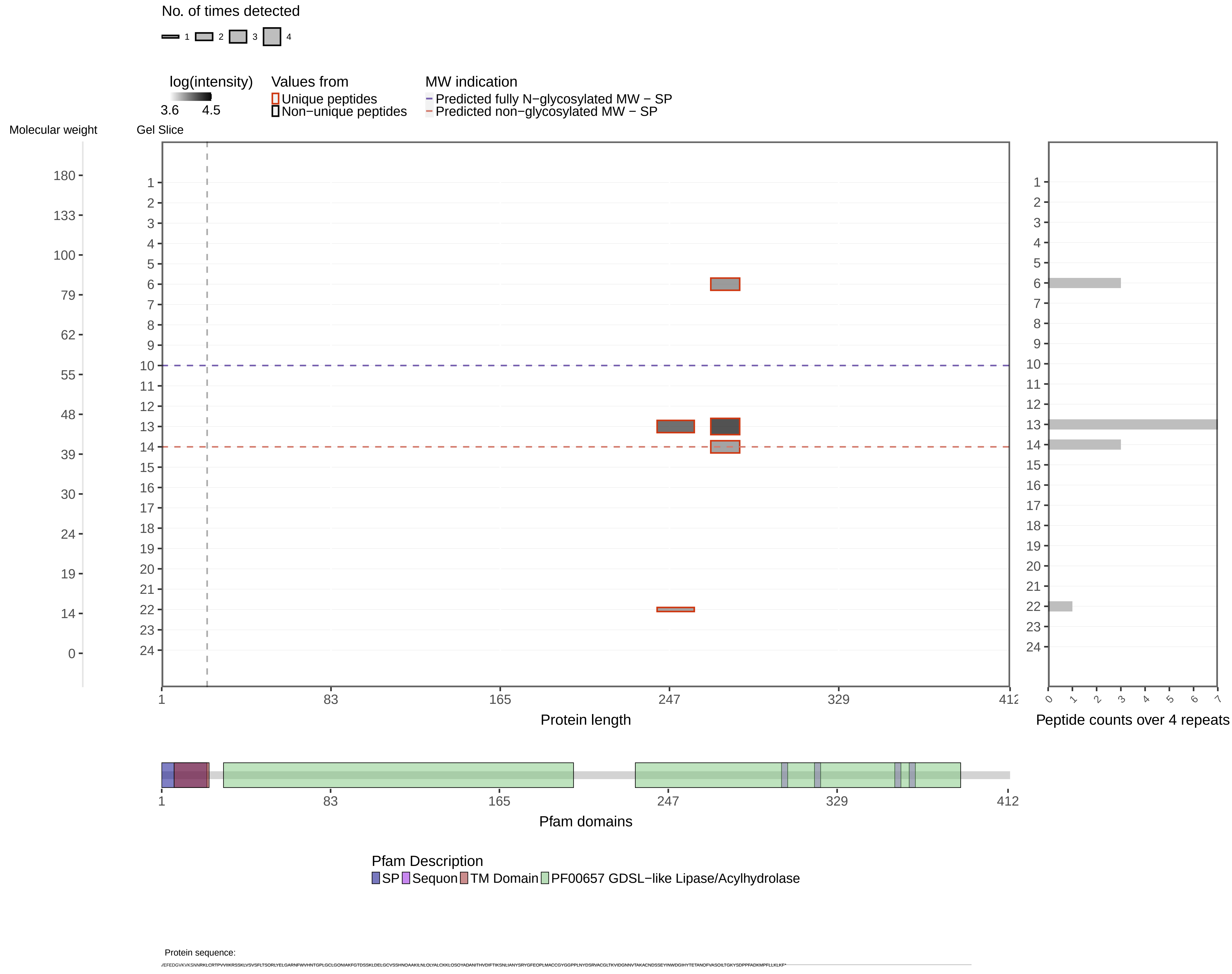

NbL01g10080.1 glutathione_synthetase_chloroplastic [MW = 64.25 kDa]

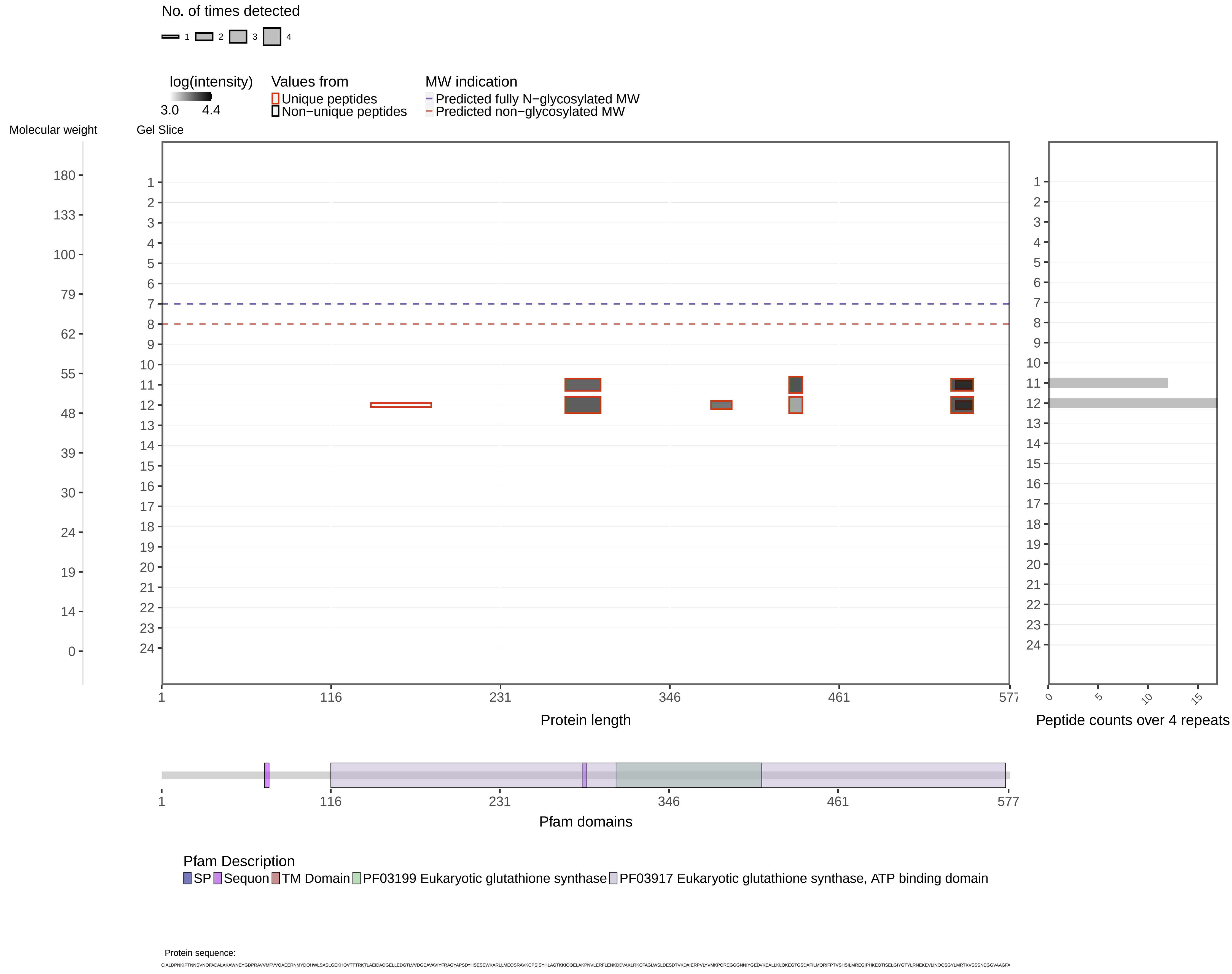

### NbL01g10030.1 broad−range_acid_phosphatase_DET1−like [MW = 34.49 kDa]

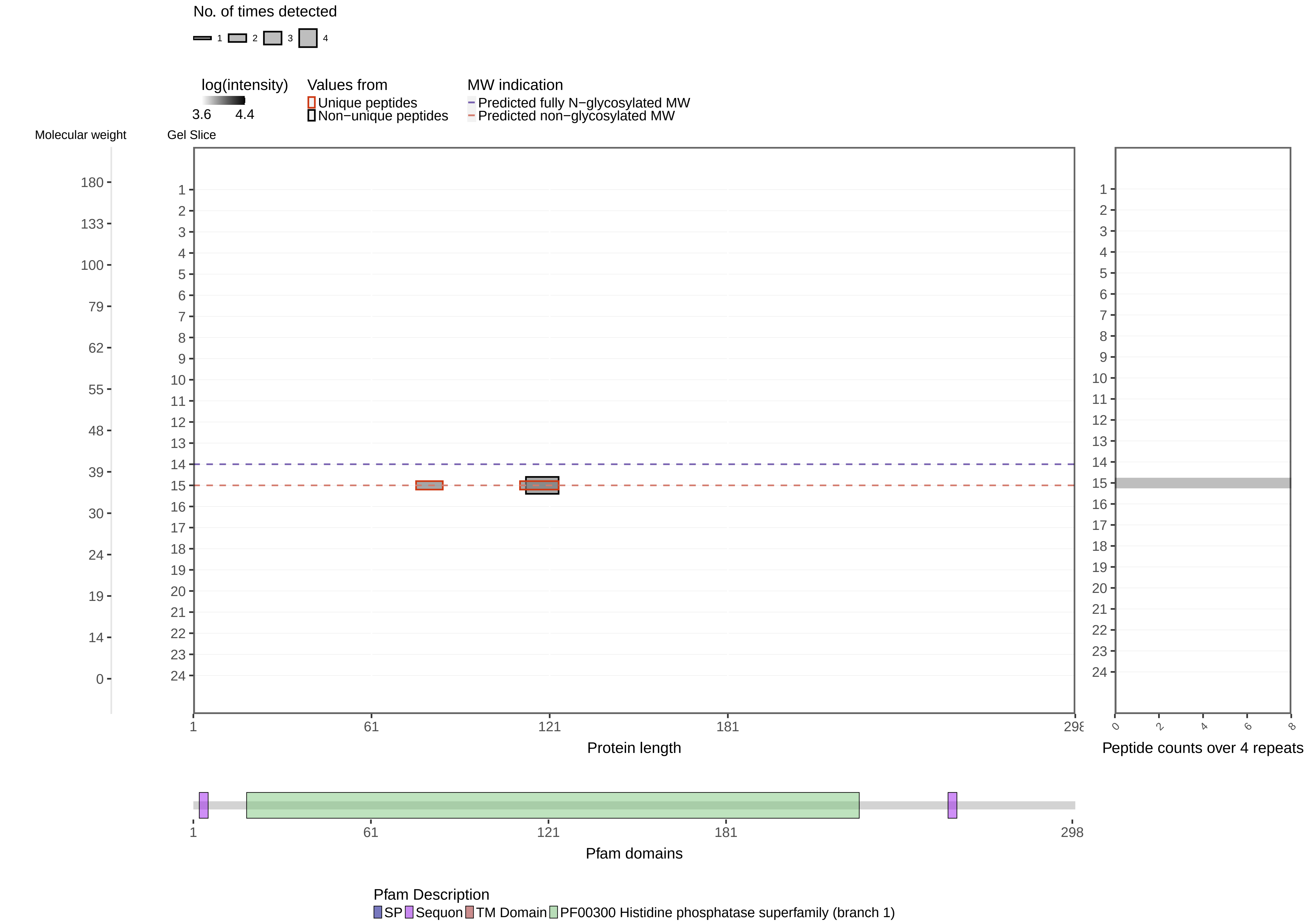


Protein sequence:

FYRFPEGESAADVYDRVSHFLESLWRDVDMNRHHHDPNDELNLVIISHGLASRVFLMKWFKWTVEQFEYLNNLGNCEFRVMELGAGGEYSLAVHHSDEEMLEWGLSPEMIADQKWRANASRGSLNDHCPTYLDAFFDHLTDSNDEDGEDVKSNNNLLCP*

NbL01g09900.1 4−hydroxy−tetrahydrodipicolinate_reductase_1_chloroplastic−like [MW = 36.26 kDa]

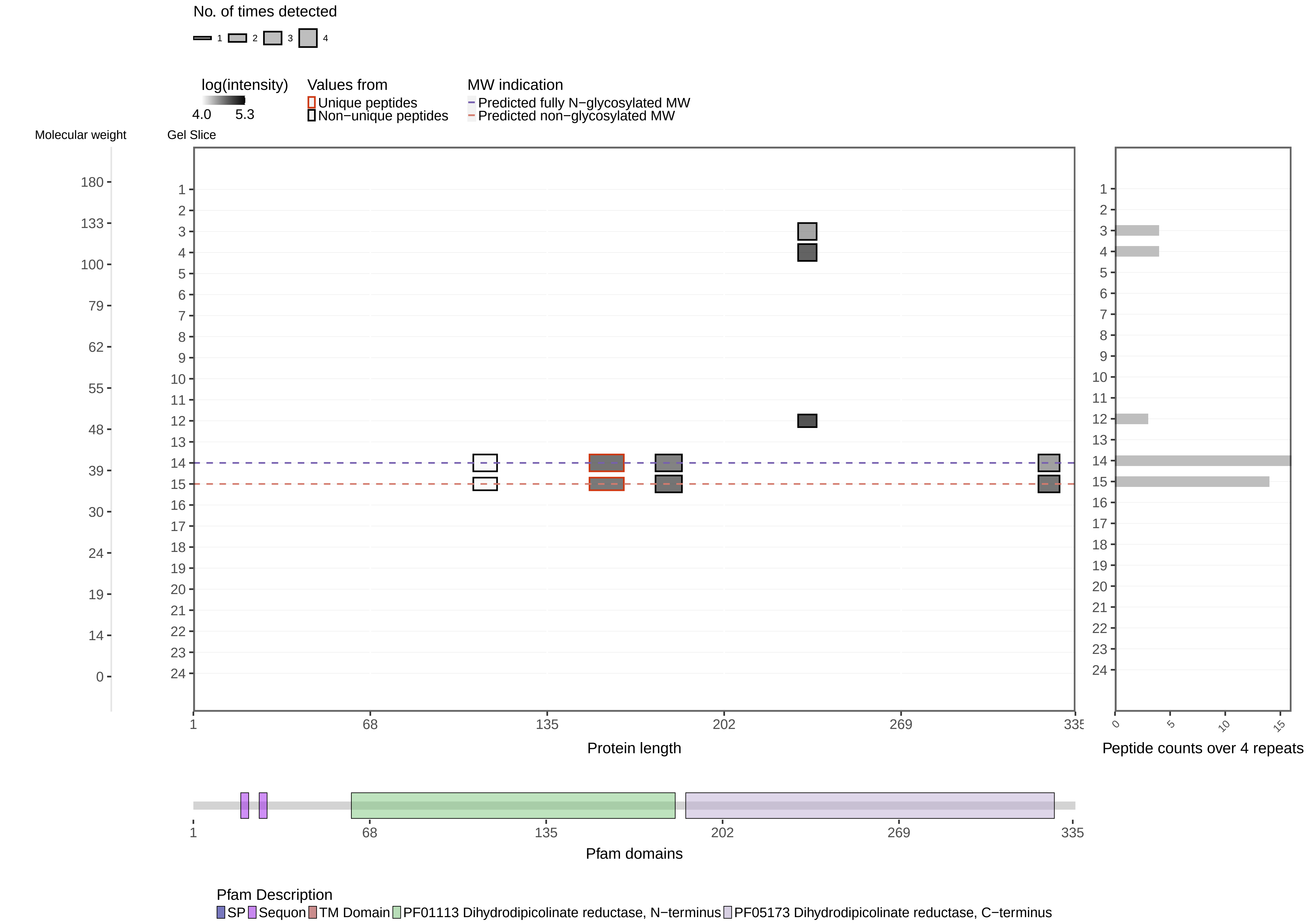


Protein sequence:

VMGTTGGDREKLYKTVADSKVYAVISPQMGKQVVAFLAAMEIMAEQFPGAFSGYTLQVMESHQASKLDTSGTAKAVISCFQKLGVSFDLDEVQLIRDPKQQVEMVGVPEEHLLGHAFHMYHLTSPDGSVSFEFQHNVCGRSIYAEGTVDAILFLAKKVKLKEEKRIYDMIDVLREGNMR*

NbL01g09760.1 translation_initiation_factor_eIF−2B_subunit_epsilon [MW = 81.99 kDa]

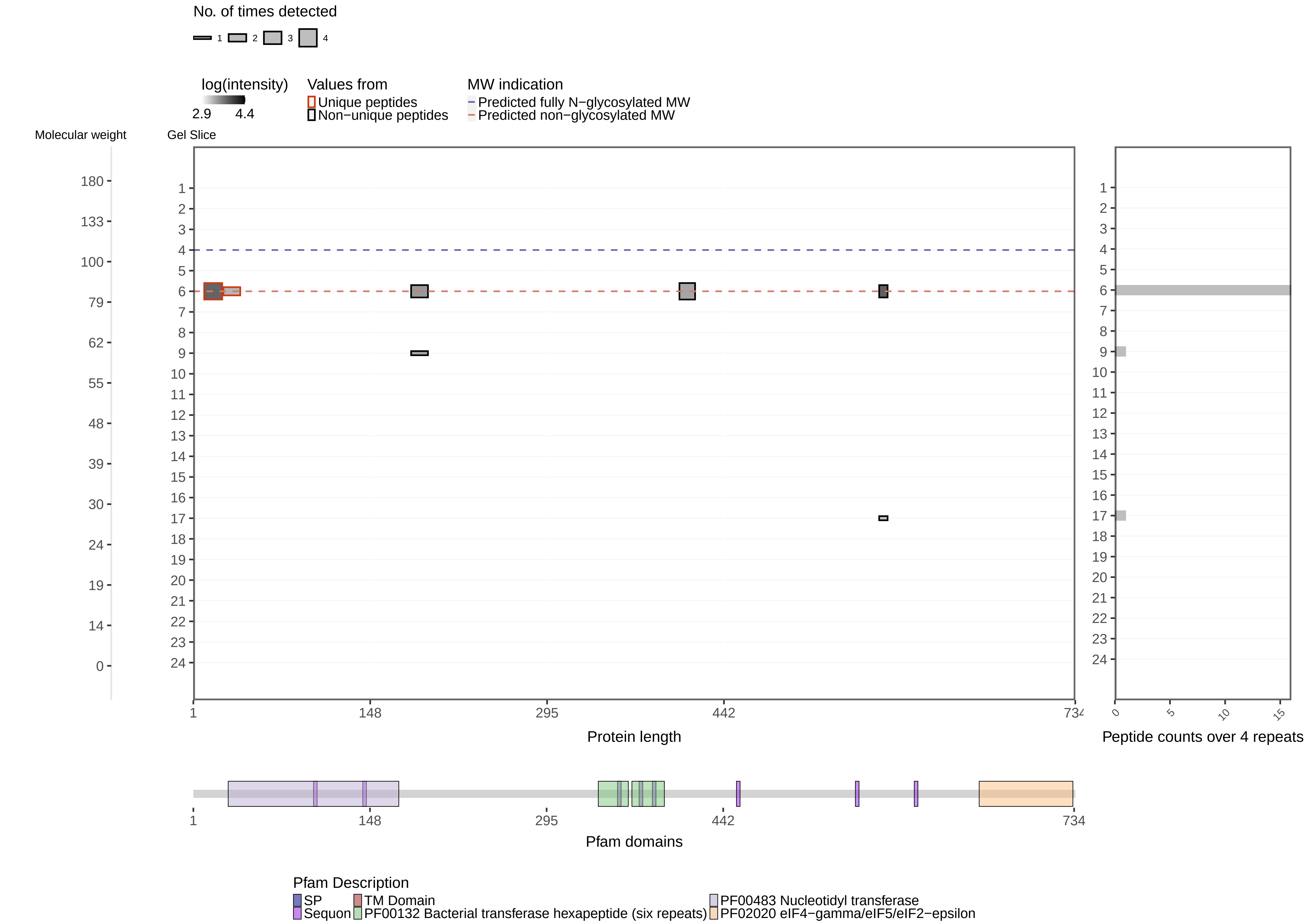


Protein sequence:

NbL01g09670.1 translation_initiation_factor_IF−2−like [MW = 62.24 kDa]

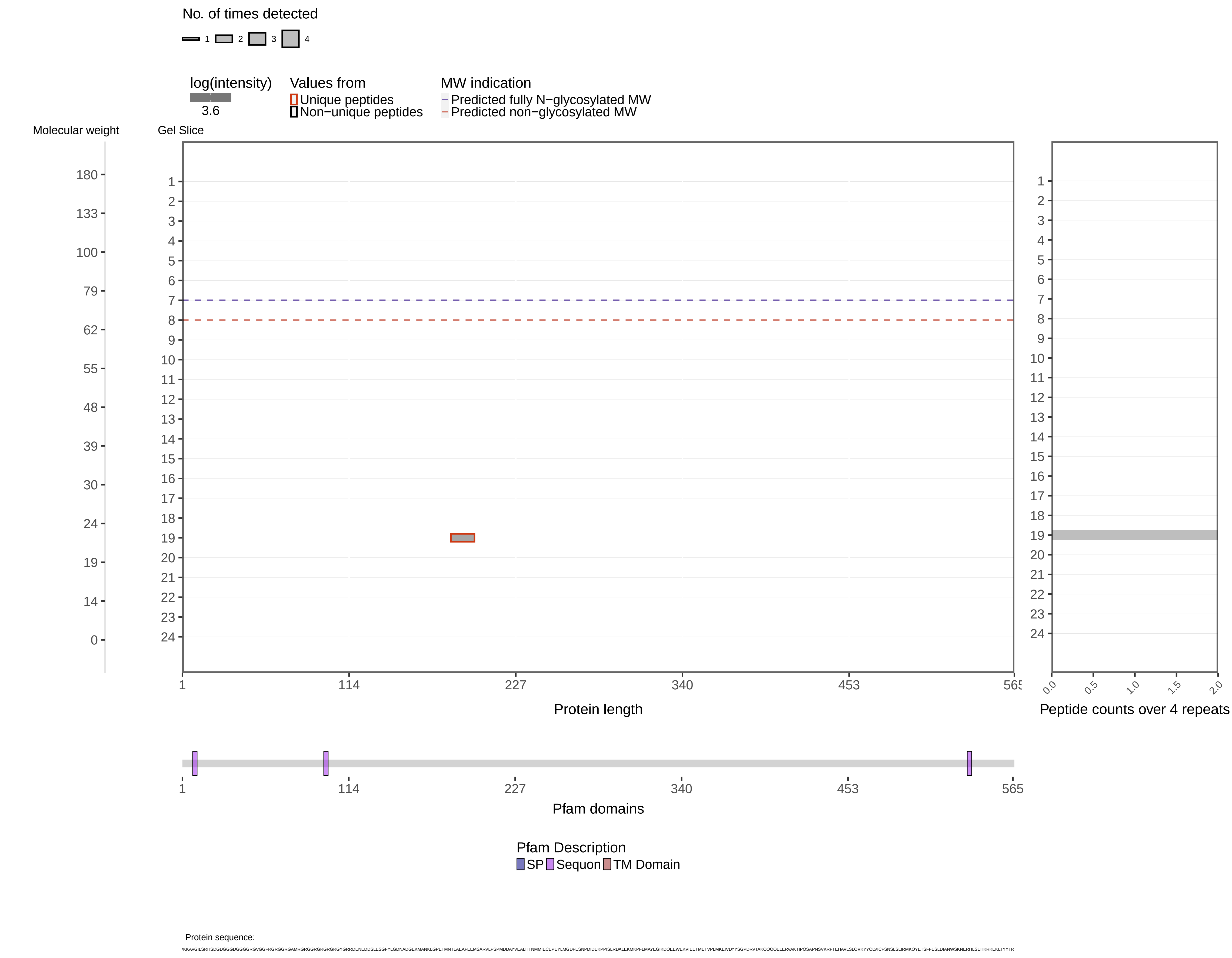

NbL01g09660.1 uncharacterized_N−acetyltransferase_p20−like [MW = 19.37 kDa]

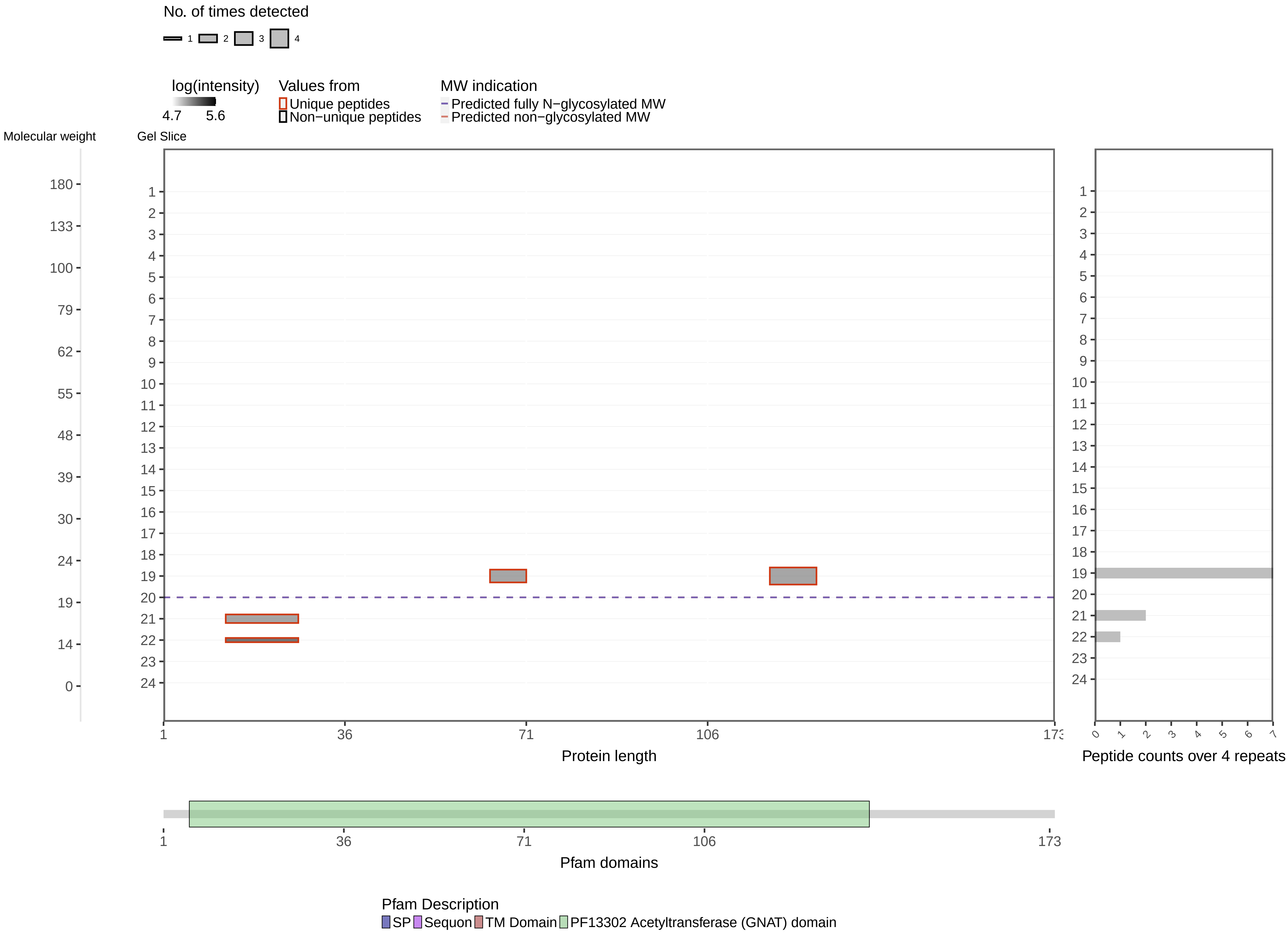


Protein sequence:

SDDDKCRAEIGYALAAEYWGQGITPKAVKMAIPLIFSDFPEIVRLQALSDANNRASHRVLEKAGFVKEGMFRKYFYFKGEIKDVVLYGLLSTDPIPLHP*

NbL01g09640.1 deoxyuridine_5'−triphosphate_nucleotidohydrolase−like [MW = 18.36 kDa]

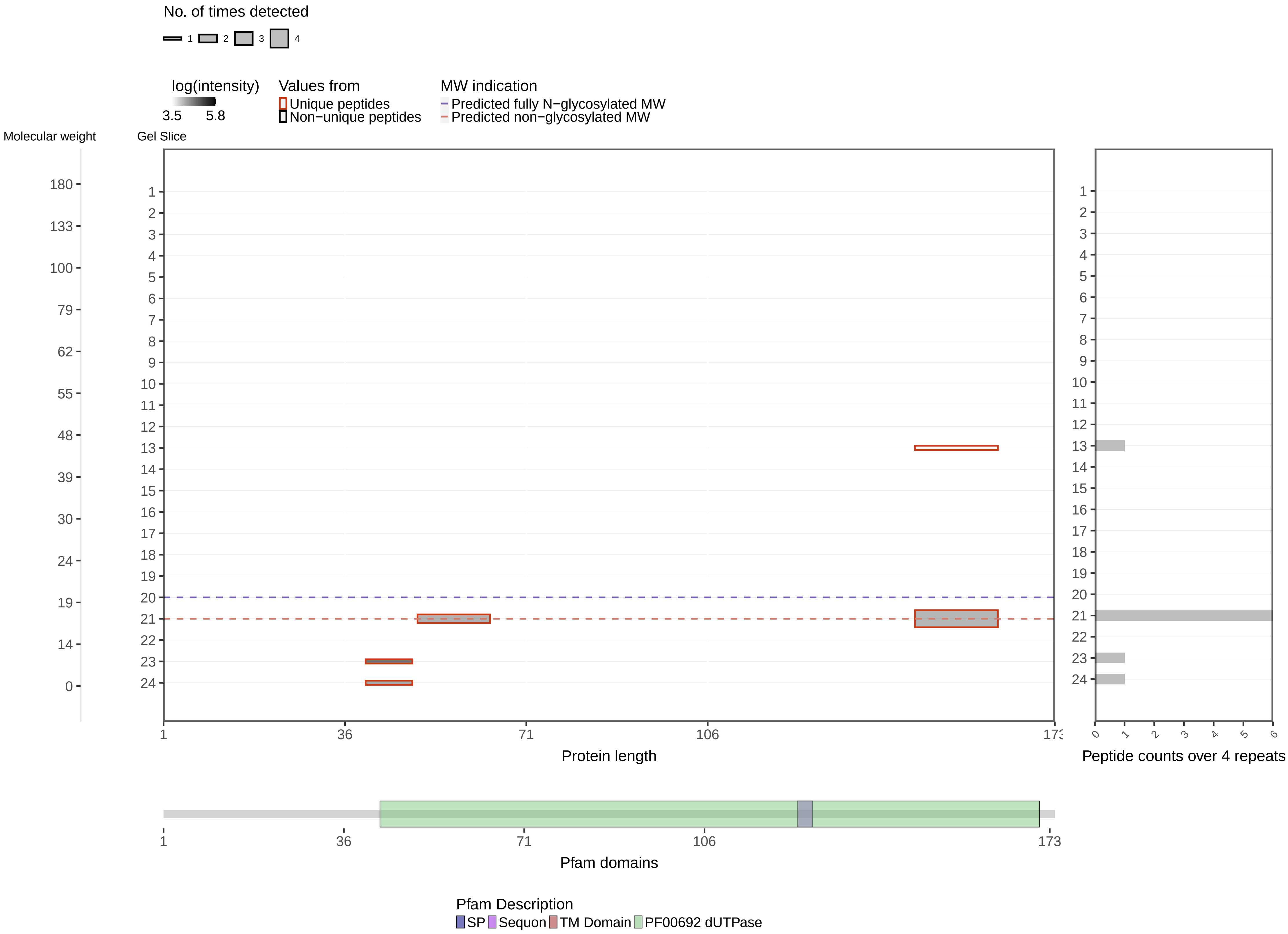


Protein sequence:

KALVPTDLSIAVPEGTYARIAPRSGLTWKHSIDVGAGVIDADYRGPVGVILFNHSDVDFEVKIGDRIAQLIIEKIIVPDVEEVDDLDSTVRGSGGFGSTGV*

NbL01g09630.1 cysteine_synthase−like [MW = 34.57 kDa]

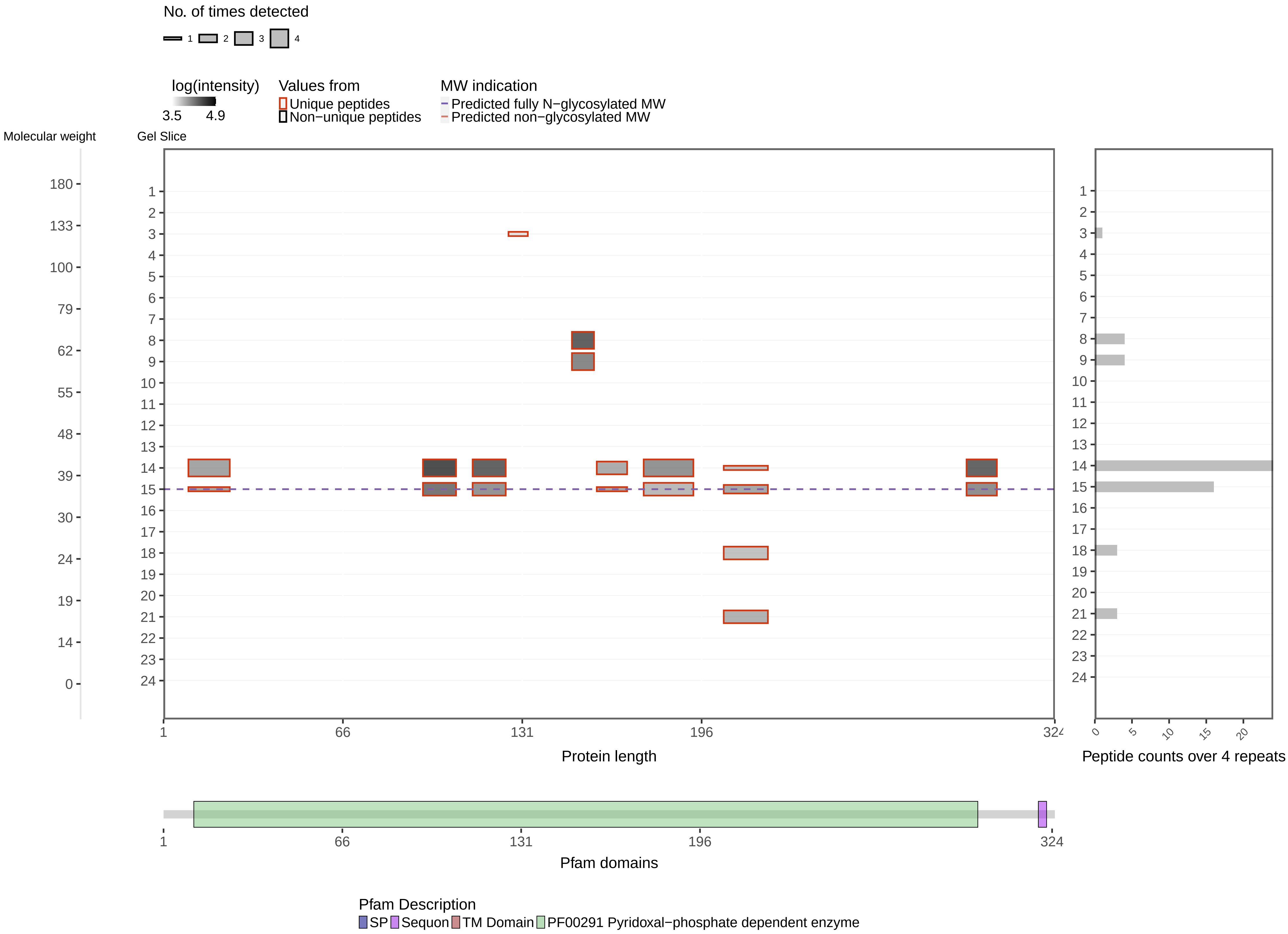


Protein sequence:

ƆFENPANPKIHYETTGPEIWKSTGGKVDIFIAGIGTGGTVTGAGRFLKEKNPEMKVYGIEPTENALLNGGQKGPHKIQGIGAGFVPAVLDLSILDEVLQVSSDEAIEMTKLLAVKEGLLVGISSGAAAAAAVKVAKKPENAGKLIAVIFPSFGERYLSTVLFDSIREEVENMTFE*

### NbL01g09600.1 guanine_deaminase−like [MW = 20.33 kDa]

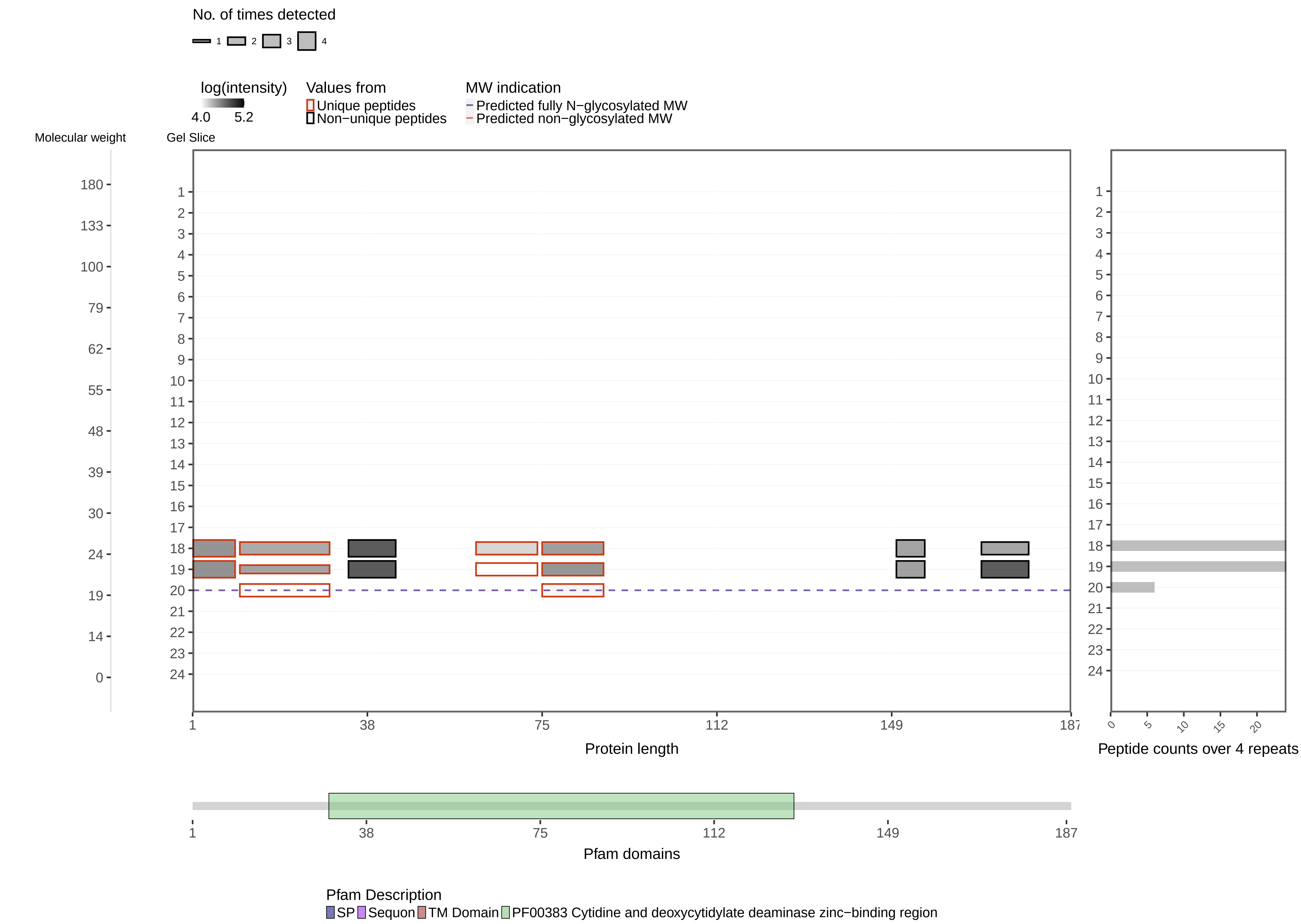


Protein sequence:

PTAHAEVTAVREACKKLNRFELSDCEIYASCEPCPMCFGAIHLSRIKRLVYGAKAEAAIAIGFDDFIADALRGTGFYQKAQLEIKKADGKGALIAEQVFEKTKEKFTIY*

NbL01g09550.1 nudix_hydrolase_20_chloroplastic−like [MW = 40.69 kDa]

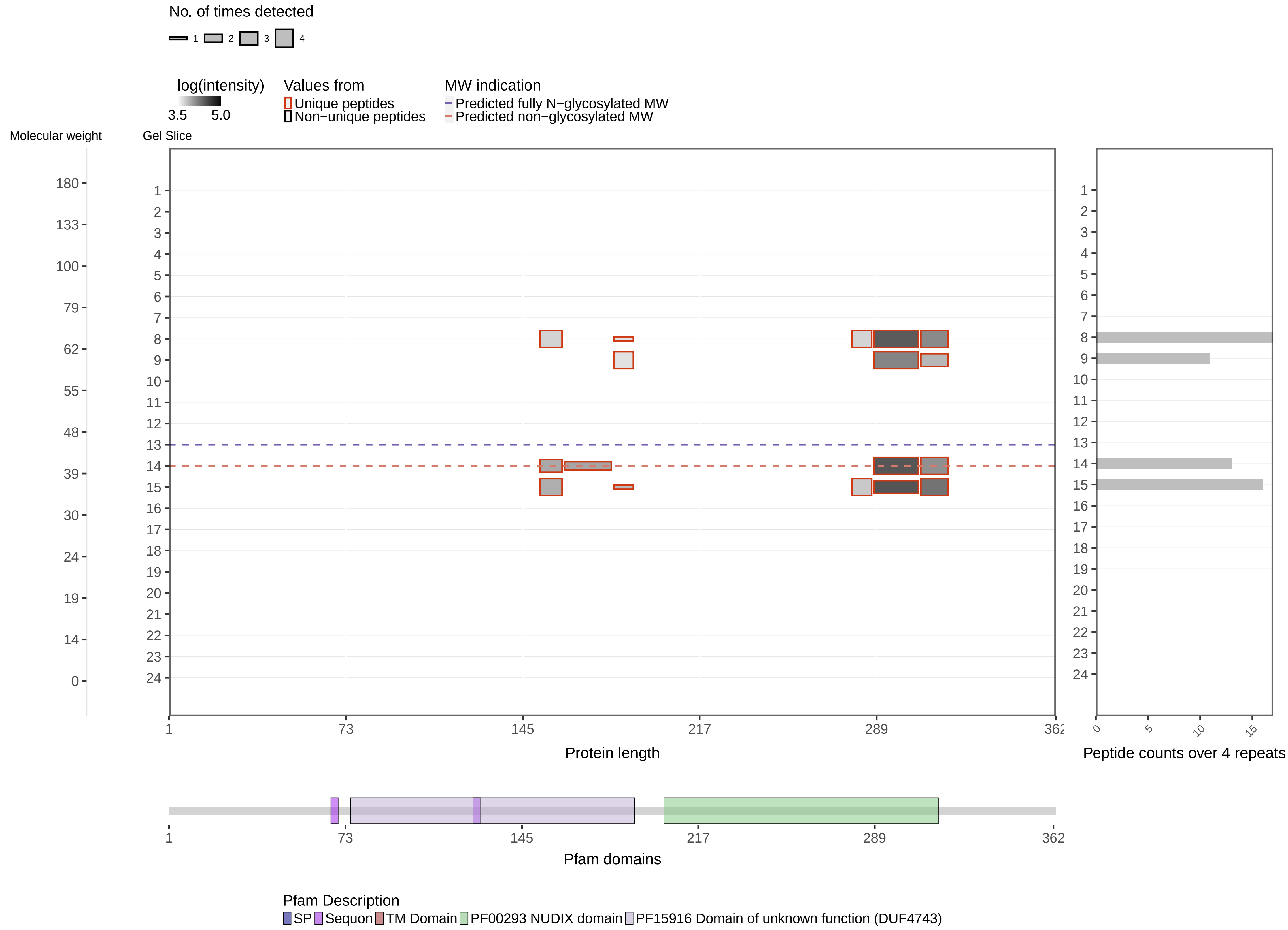


Protein sequence:

SAFGEPVFFSLERAAAPYFGIKAYGIHMSGYVEKNDOKFLWLAKRSATKSTYPGMLDHLVAGGLPHGISCGENVVKECKEEAGIPRSISROAVPVGAVSYIDIEGYRMKRDVLFCYDLKLPAEGFVPHNEDGEVESFRLVPVTOVANVIRSASFFKANCNLVIIHFLFRHGYIRPEEFGYLRLLOSLKSGCCS*

### NbL01g09540.1 zeta−carotene_desaturase_chloroplastic/chromoplastic [MW = 59.06 kDa]

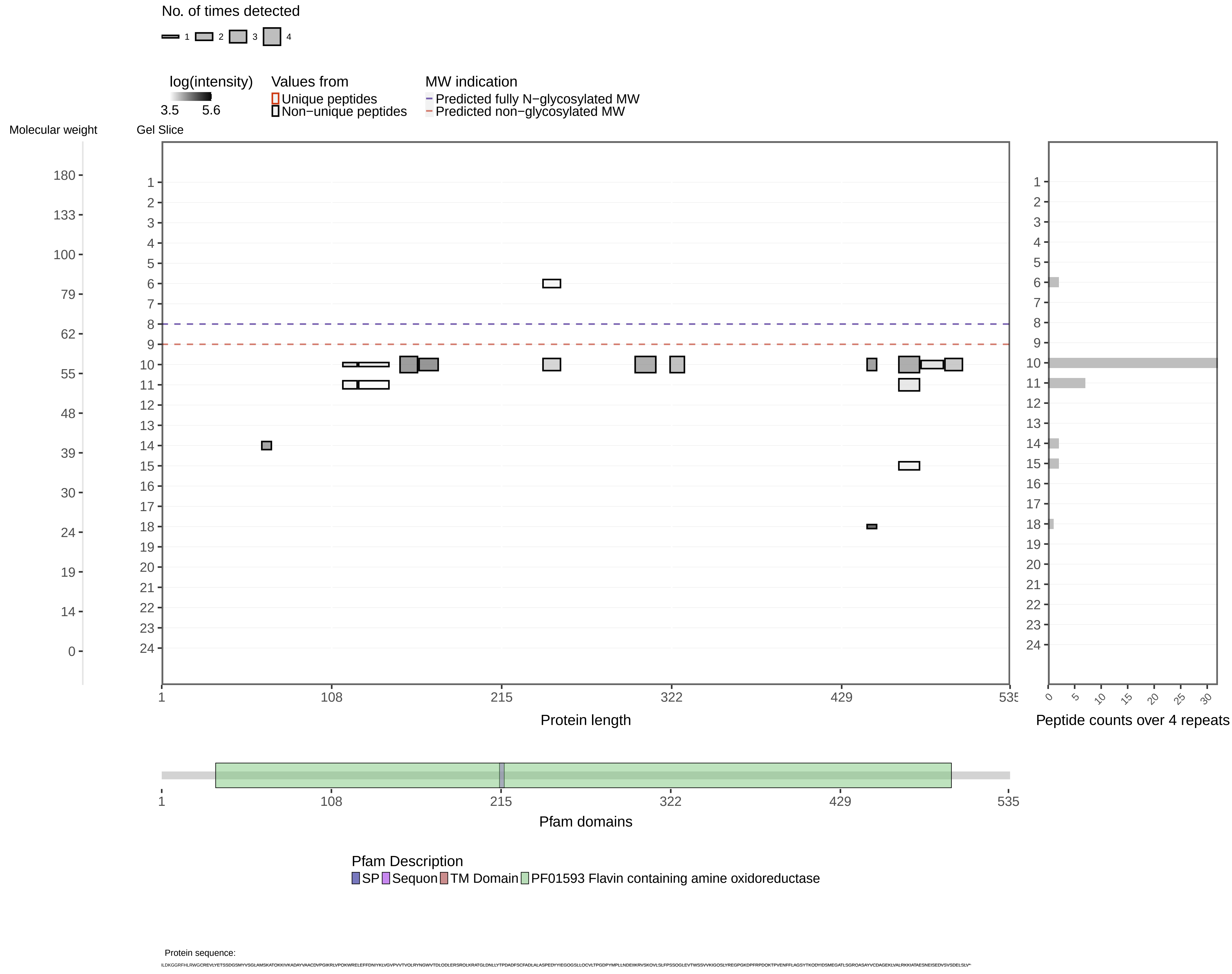

### NbL01g09530.1 phototropin−2−like [MW = 107.8 kDa]

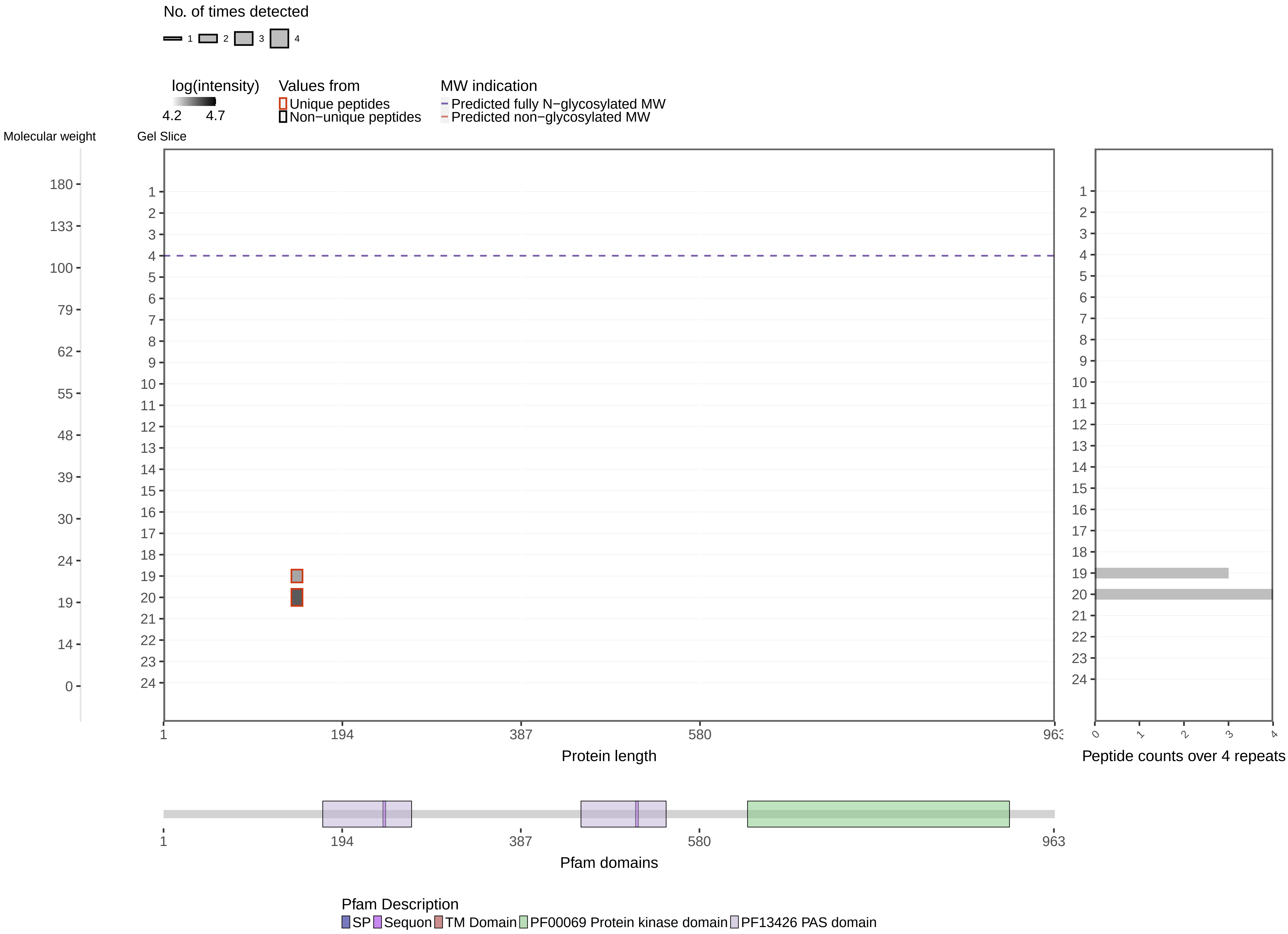


Protein sequence:

[illegible]

NbL01g09450.1 multisubstrate_pseudouridine_synthase_7 [MW = 83.47 kDa]

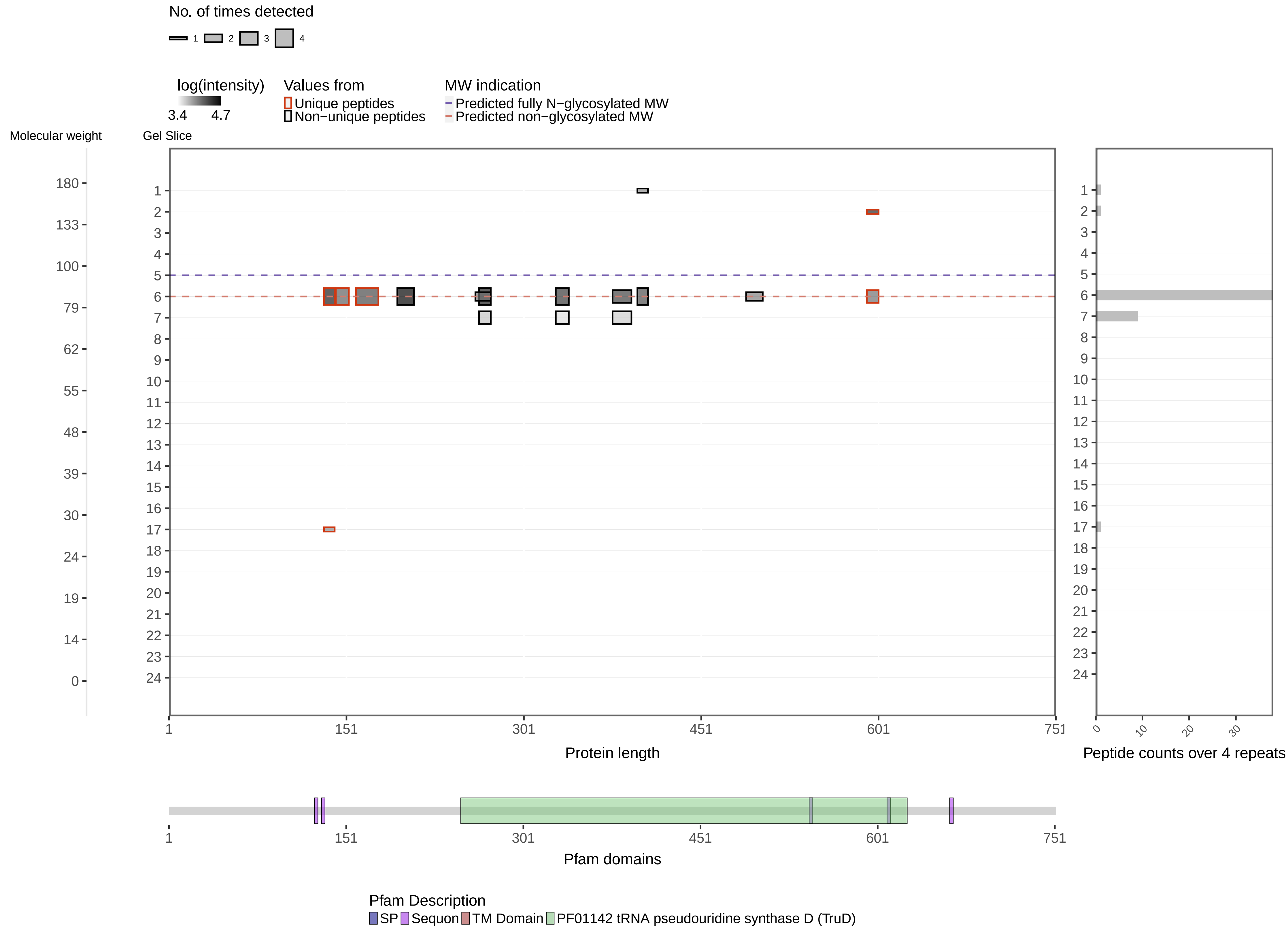


Protein sequence:

EHGFINYFGLQRFGSSSVPTHLIGAALLRGEWKAAVSLILDPREGKKNAISTVREYYKESGDTDGTLRQLPRHLVAERAILQCLKKSPGNCLQALKGIPRTLRMMYVHSYQSYLWNHAASMRVQKHGFDQVVLGDLVYSKEQPKEKETFVAACEDVNGNEMDDYSNLDETSETDPPEEKNISVKAINEEDLISGTYTIEDVLLPLPGSRVIYPSNDIGEVYCDLAQKDGISLTETAHKIKEFSITNMTGAYRRVFQKPKDFEWELLSYTDGTIPLAETDFDAIAKSKKKSKVREE

### NbL01g09390.1 probable_ribose−5−phosphate_isomerase_3_chloroplastic [MW = 30.36 kDa]

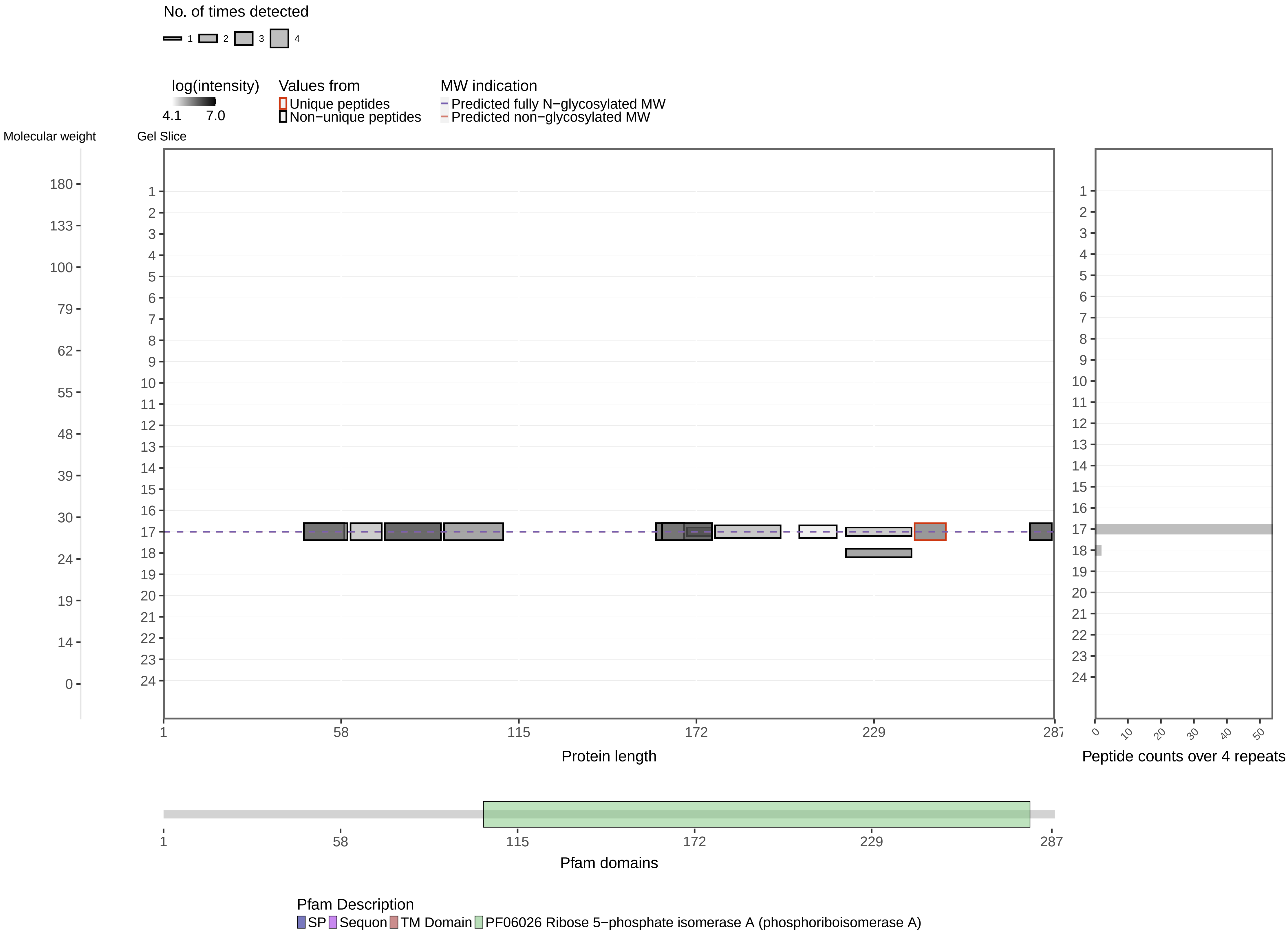


Protein sequence:

DLAIDGADEVDPNLDLVKGRGGALLREKMVEAASDKFVVVVDDSKLVSGLGGSGLAMPVEVVQFCWKYNLVRLQELFKAEGVDAKLRVDGNGKPYVTDNSNYIVDLYFKTPIRDSAAAGKEIAALEGVVEHGLFLDMTTAVIIAGKEGVSVQSK*

NbL01g09380.1 PITH_domain−containing_protein_At3g04780 [MW = 19.65 kDa]

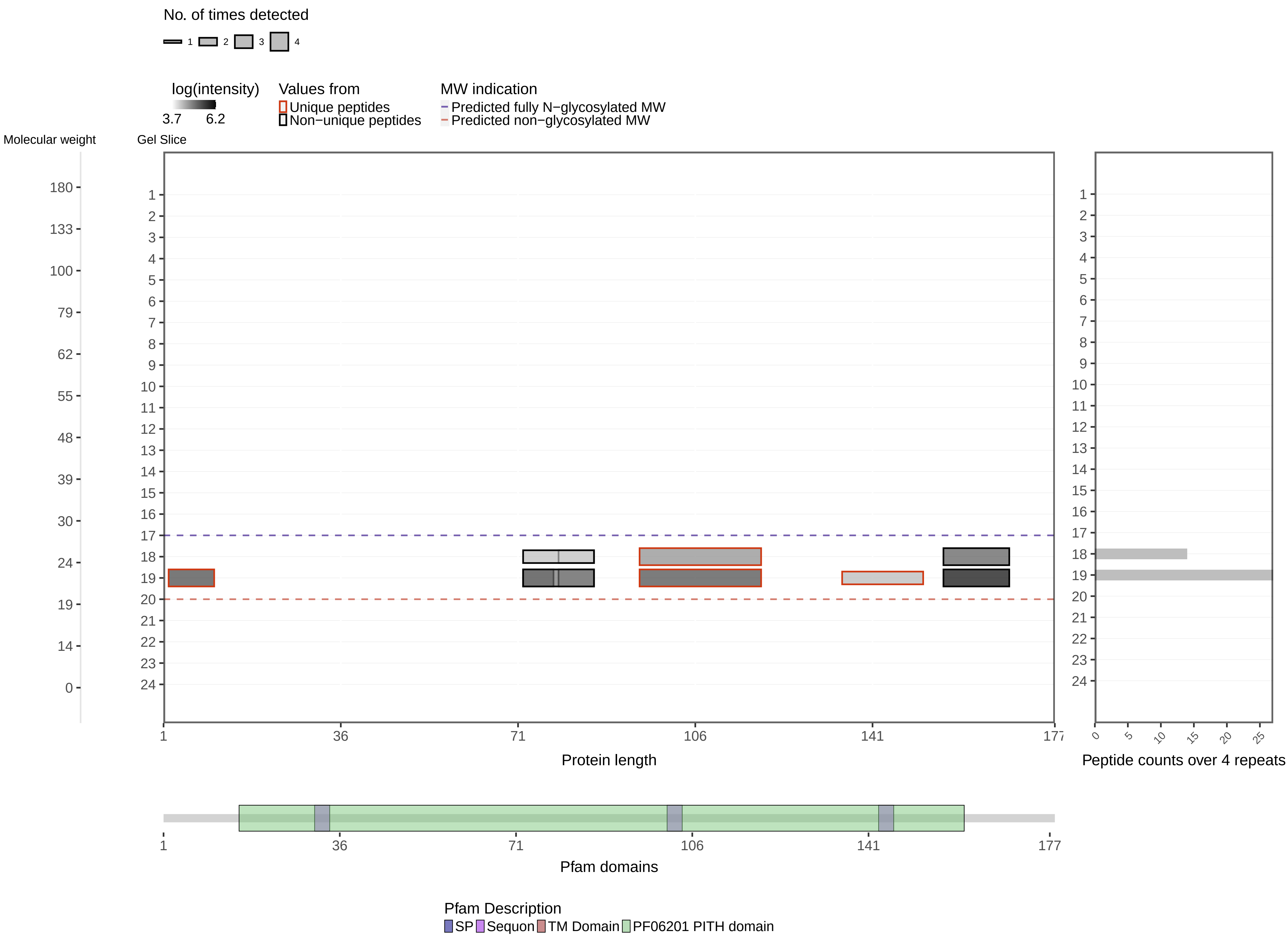


Protein sequence:

FVIKGPEEEGPKTVKLFSNKEHMGFSNVSDYPPSDTAVLSEDNLKGKPVILKYVKFQNVRSVTIFIEDNQSDSELTKVQKIVLYGTTVETTDMKGLKKIEDH*

### NbL01g09320.1 uncharacterized_LOC104217073 [MW = 36.25 kDa]

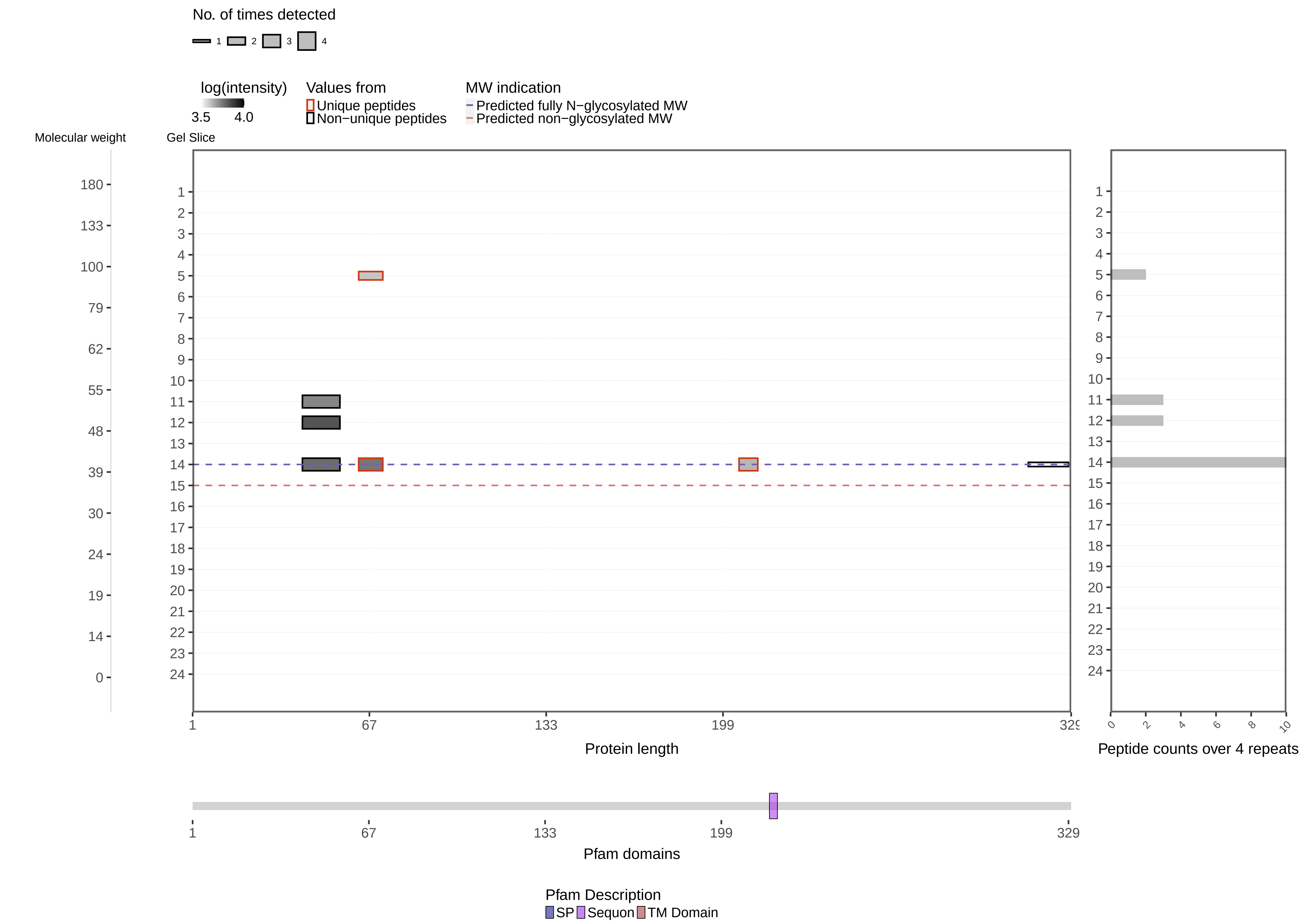


Protein sequence:

MGVEMGIPPLSEFPLLSDILKSHGAEGEDELGQAQFAHLLQPVLQELADALAKNPVVVVQKIKINNGSKLRKVLADEKQLRETVEKIMQEKQDEKDSPSNKDVIRCYLEKNGASLGLPPLKNDEVVILLYDIVLGDMENGKTDAASDKDEILVFLKDILEKFAAQLEVNPTFHDLDN*

### NbL01g09310.1 GDP−mannose_35−epimerase_1−like [MW = 42.53 kDa]

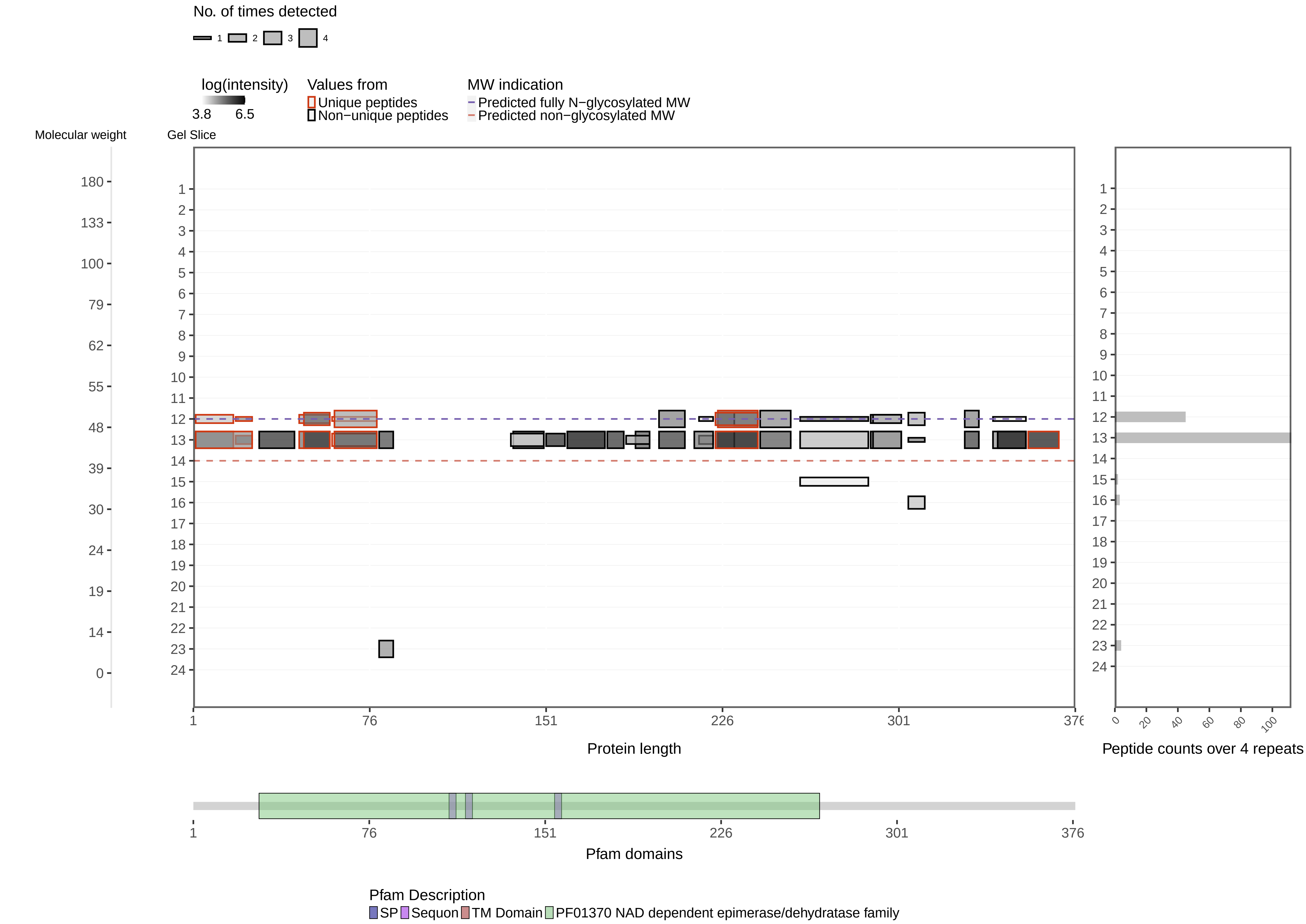


Protein sequence:

KLATEELCKHYNKDFGIECRIGRFHNIYGPFGTWKGGREKAPAAFCRKAQTAVDKFEMWGDGLQTRSFTFIDECVEGVLRLTKSDFREPVNIGSDEMVSMNEMAEMVLSFEDKKLPIHHIPGPEGVRGRNSDNTLIKEKLGWAPTMRLKDGLRITYFWIKEQIEKEKSQGVNIANYGSSKVVGTQAPVELGSLRAADGKE*

NbL01g09280.1 sec−independent_protein_translocase_protein_TATA_chloroplastic−like [MW = 16.27 kDa]

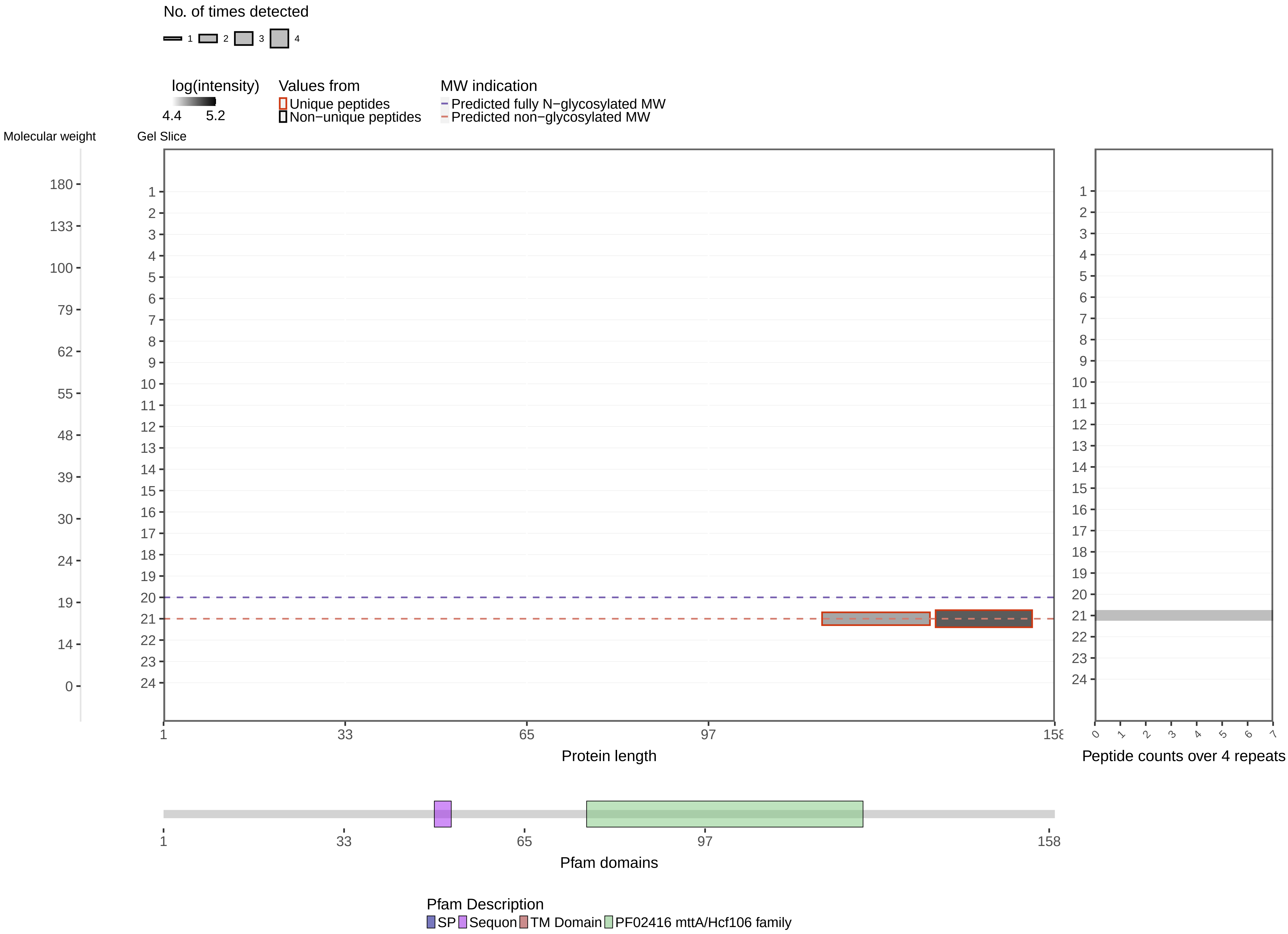


Protein sequence:

LSCNCLFGLGVPELVVIAGVAALVFGPKQLPEVGRTIGKTVKSFQQAAKEFETELKKEPEASAEPTAEKAMEASQEEKQDATVSSTKESS*

### NbL01g09270.1 probable_prefoldin_subunit_4 [MW = 14.83 kDa]

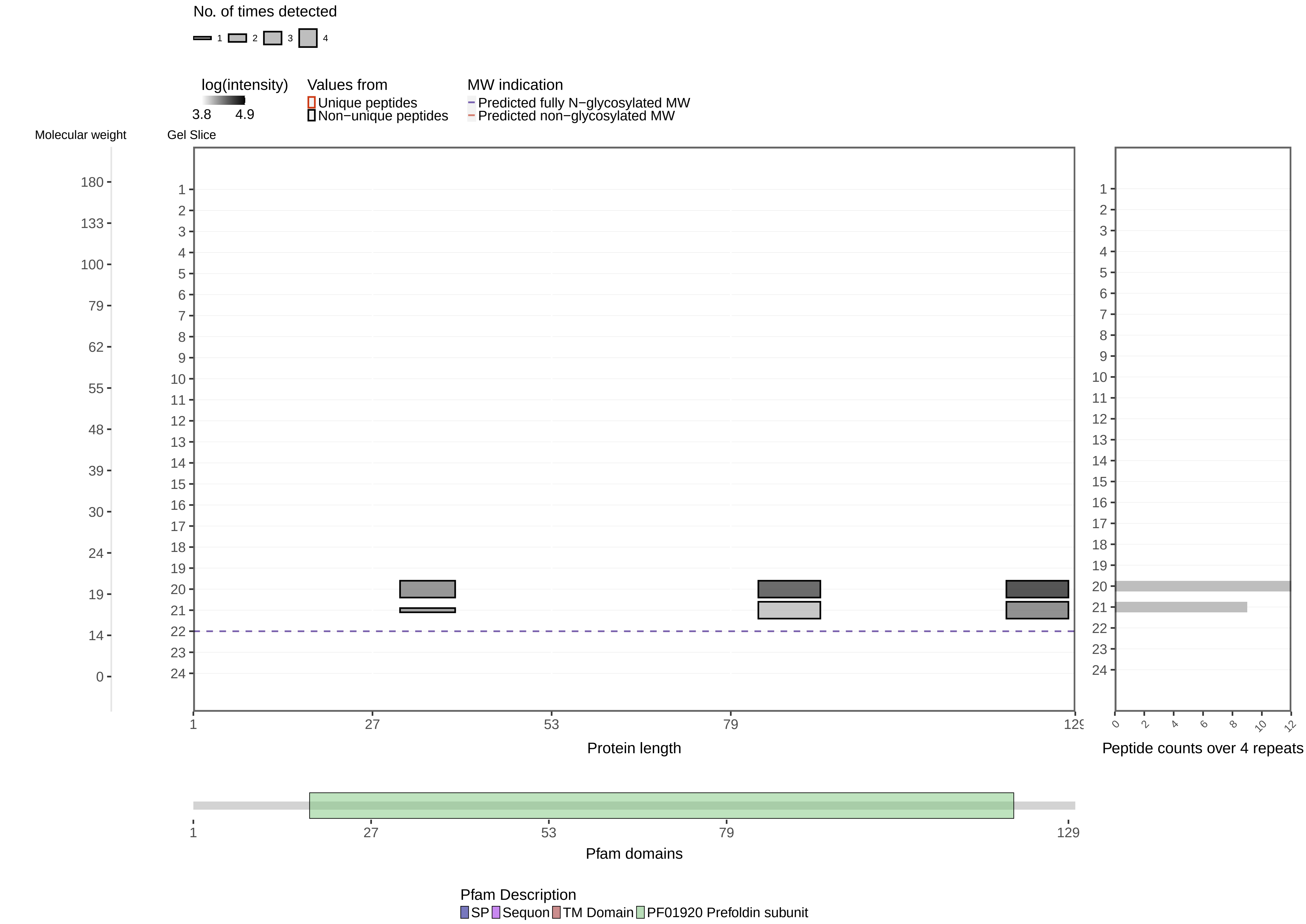


Protein sequence:

DASNELILTDEEIVRFQIGEVFAHVPKEEVESRIENMQEVTAKNLEKLEEEKETVVSQMAELKKILYGKFKDSINLEED*

### NbL01g09230.1 wound−induced_protein_WIN1 [MW − SP = 19.69 kDa]

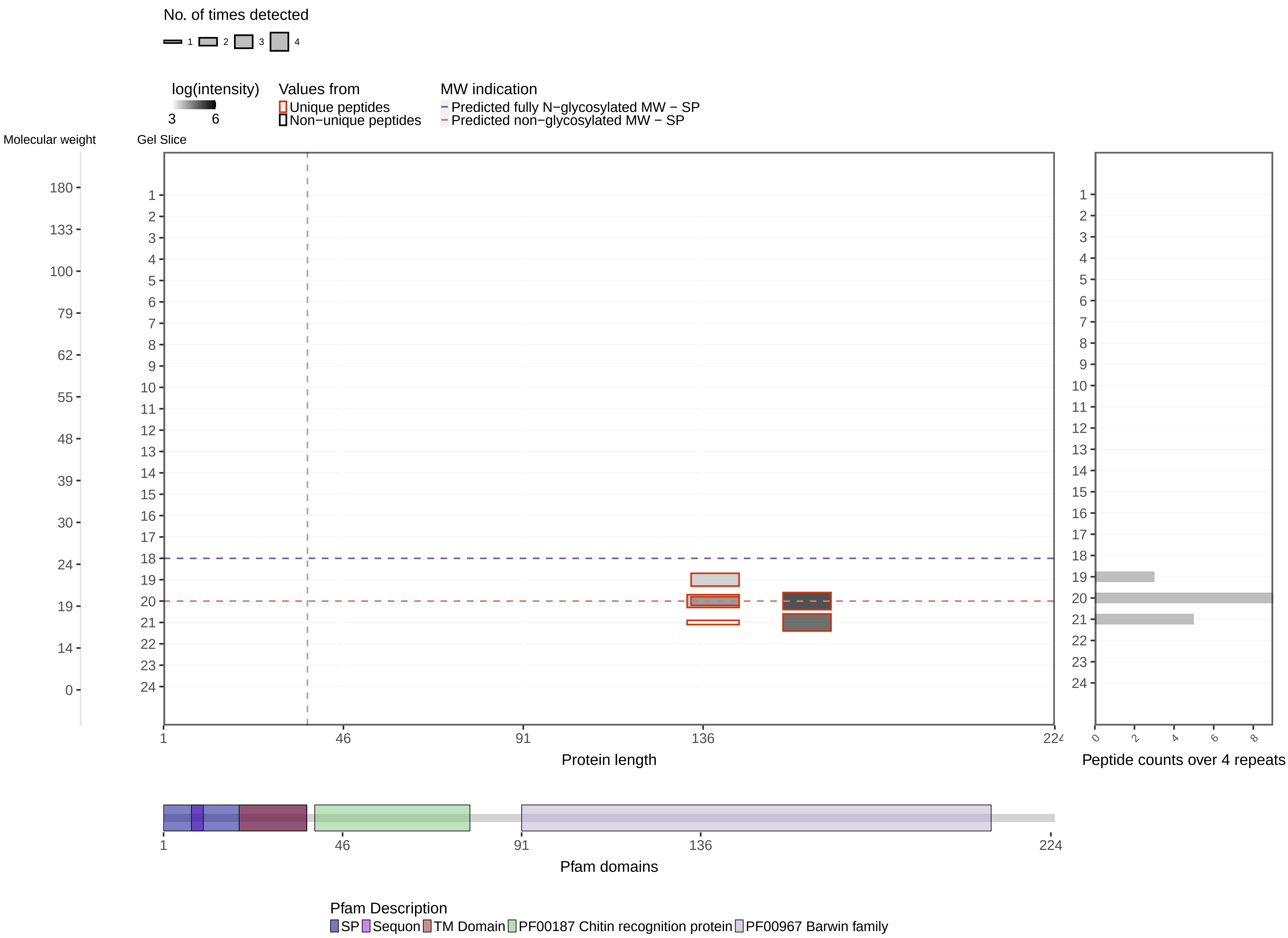


Protein sequence:

YHIYNPQNVGWDLYAVSAYCSTWDGNKPLAWRRKYGWTAFCGPVGPRGRDSCGKCLRVTNTGTGAQTTVRIVDQCSNGGLDLDVNVFRQLDTDGRGNQRGHLIVNYEFVNCGDNMNVLVSPVDKE*

NbL01g09210.1 pathogenesis−related_protein_PR−4A [MW − SP = 13.63 kDa]

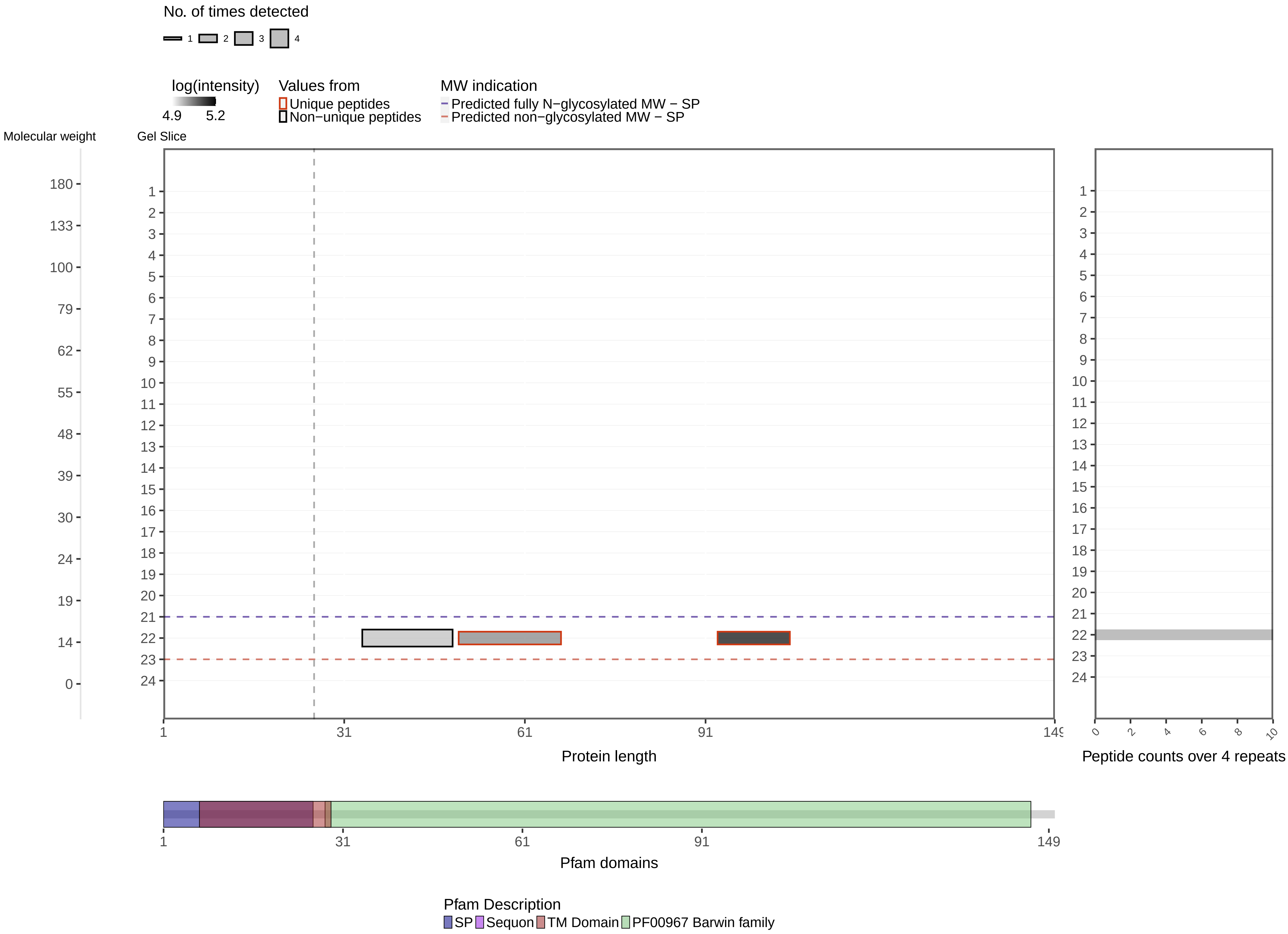


Protein sequence:
PLTWRQKYGWTAFCDAAGPQGQDSCGRCLRVTNTGTGTQTTVRIVDQCRNGGLDLDVNVFNQLDTNGVGYQQGHLIVNYEFINCDD*

NbL01g09160.1 uncharacterized_LOC104210129 [MW = 53.57 kDa]

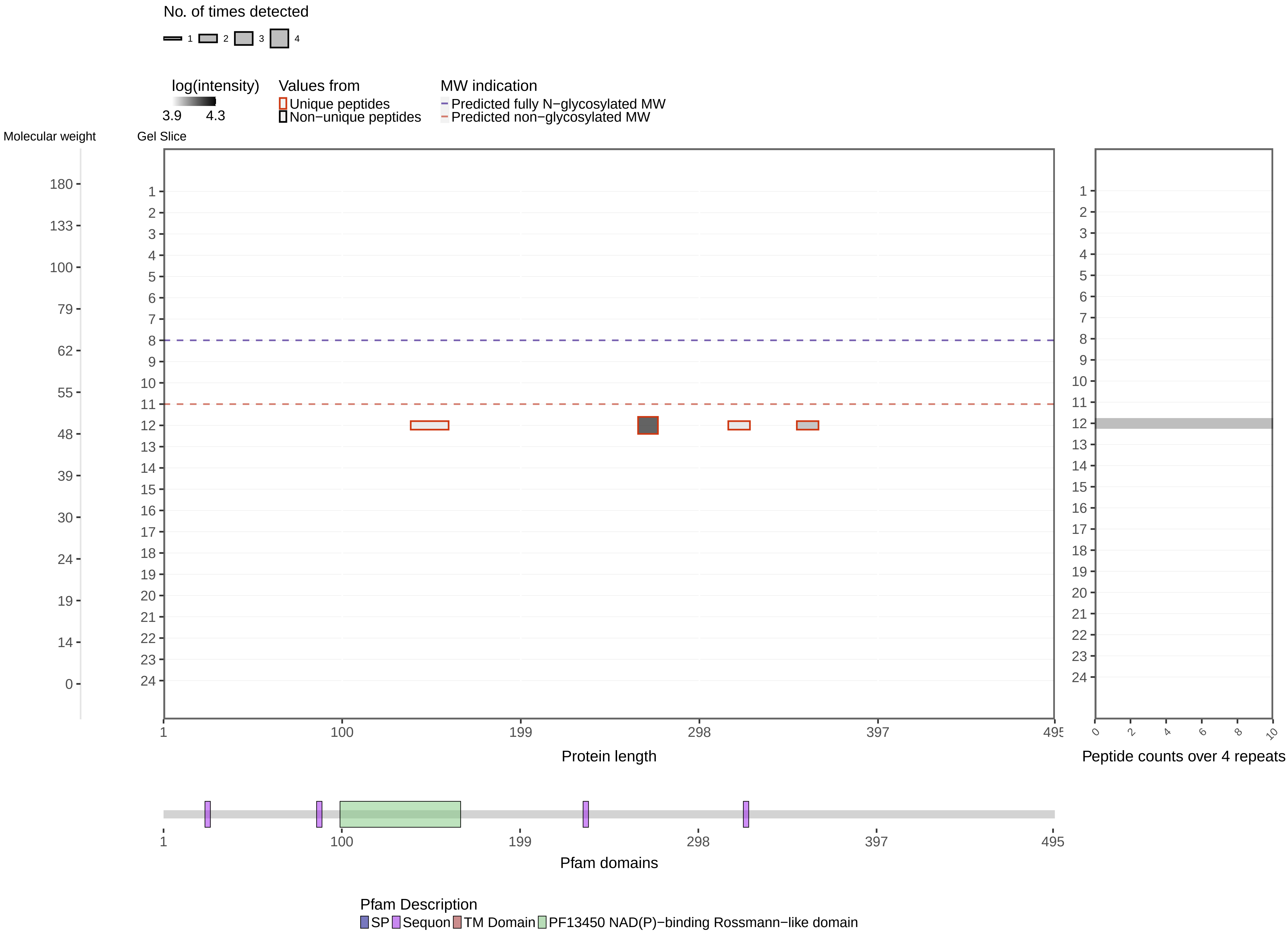


Protein sequence:

GTWHLSEKEKPCGQFDAIVIAHNGKCANRLLASSGLPLIARQMKRLELSPIWALLAAFEDPLPTAVNAESLPFEGAFVKGVESVSWMANNTKKLLGRDSGQQCWTFFSTAAFGKQNKVPQESIPVATAERVKEAMLEGVEKALGLSKNSLQRPFYTRLQLWGAALPTNTPGVPCIFDPRGRAGICGDWLLGSSVEAAAISGIALADHIADYFEQGGPCSDEFAVGLHDDYKPLGGHDIGQFPGLEYVDQTTKPPALTLAT*

NbL01g09120.1 Fanconi_anemia_group_D2_protein_homolog [MW = 166.81 kDa]

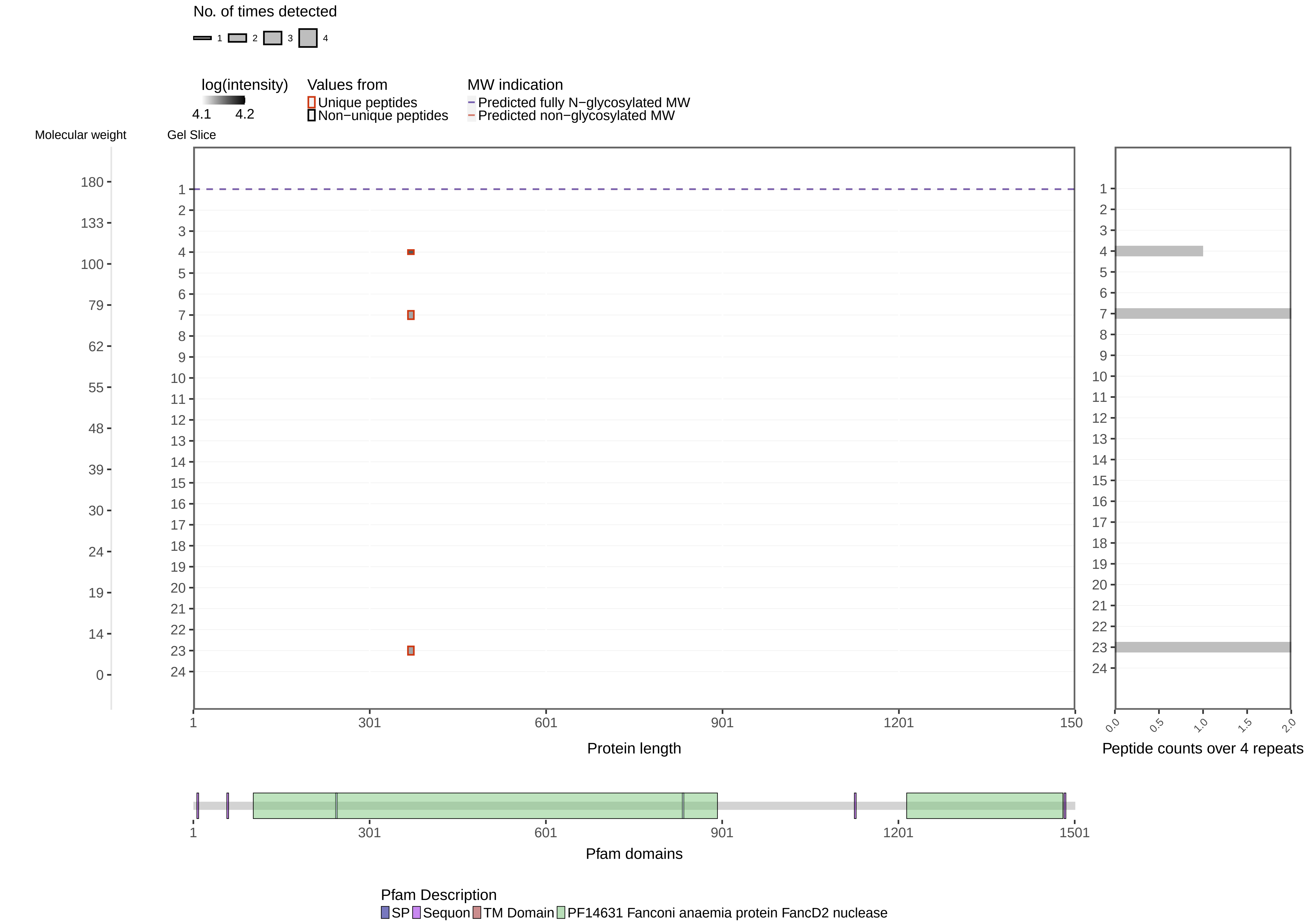


Protein sequence:

SLAGIDALLGCPIHLPSSKFFSEPLWGSLSGKQKQIIILSLYYAANWLRELLNAFCTQAVNECDGFSQATREEITLKLFKRLRNLLFLESLLNSSLRQCSLSLPELQPHLESLSLNQLDHNRDQEIKNEHGNGNGGLSQKKKEKKTSQASLTDGKLRQPTIMDVLRKAGAVPSQEPSVSPSGTCSKGSAPEPSGNQLRNSKIPANVDVSTAVEHVEAQRYKFRPLLLDCFAILAFQKTQDSCCADPASELPLCLYILRDLNRKLDYFSPRRHILVRRMSVPPAFGEMKVIEFISKIQ

NbL01g09040.1 flowering_locus_K_homology_domain−like [MW = 53.82 kDa]

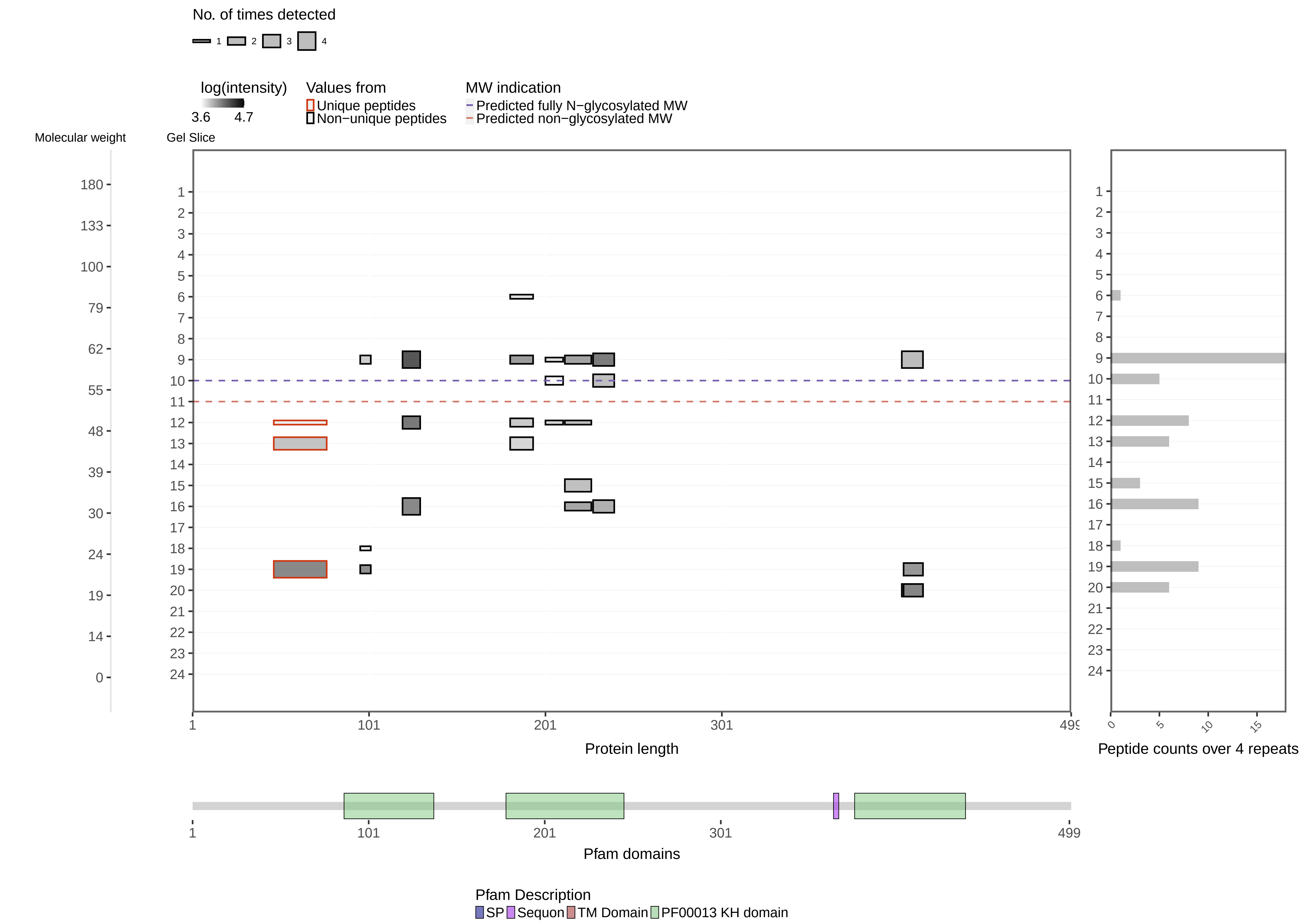


Protein sequence:

### NbL01g09000.1 retinoblastoma−binding_protein_5_homolog [MW = 58.07 kDa]

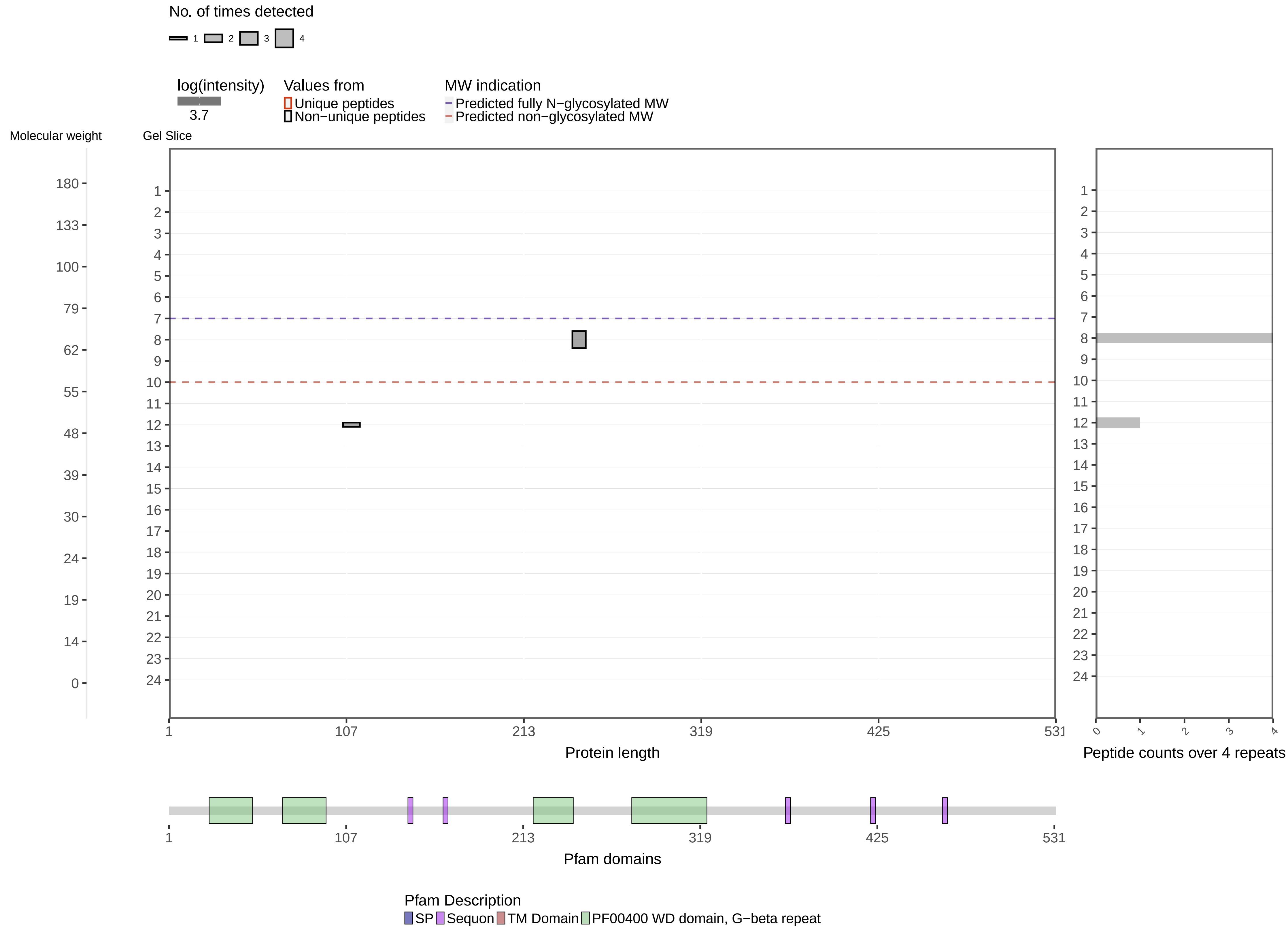


Protein sequence:

DEATNDLNELEGVEKLKAVGAQCLALFREFQDSVTRVHWKAPCFSGDGEWVVGGSANKGEHKIYIWDRAGHLVKILEGPKEAIIDLAWHPVHPIVVSVSLAGVVYIWAKDYTENWSAFAPDFKELEENEEYVEREDEFDLNPEIEKVKESDANEDEDIDIVGVENDSTFSDSDMSQGEICFLPADPIPDAPEQQDKCVGSTSKLGESNQSGSPISEEAGQNGQVNHESSPFEGEDTGTTCVKRKRKPSEKVLELQAEKVKKPSQKTKPSGKHQN*

### NbL01g08920.1 proline−−tRNA_ligase [MW = 60.88 kDa]

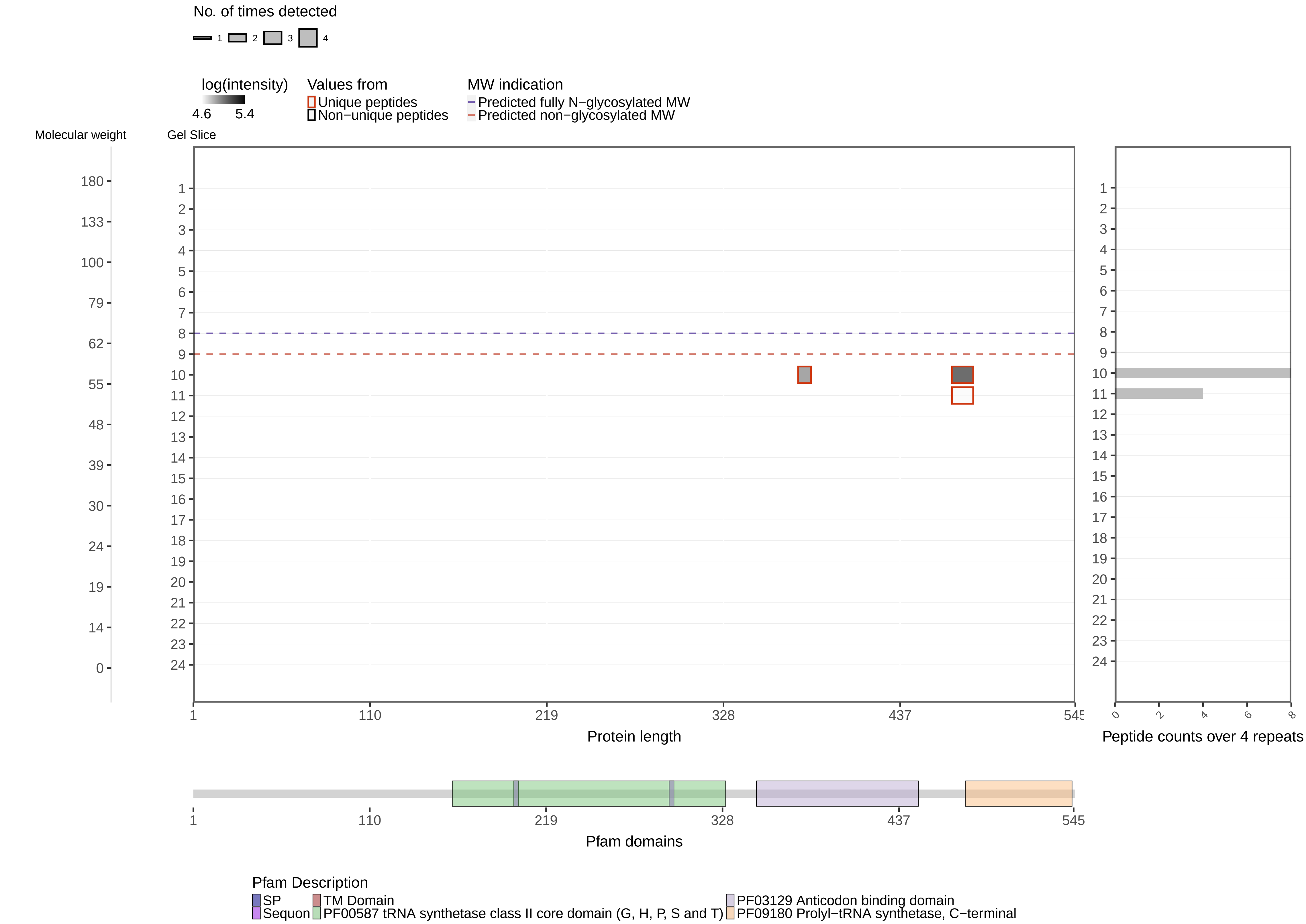


Protein sequence:

[illegible]

### NbL01g08750.1 protein_MAINTENANCE_OF_PSII_UNDER_HIGH_LIGHT_1 [MW = 25.79 kDa]

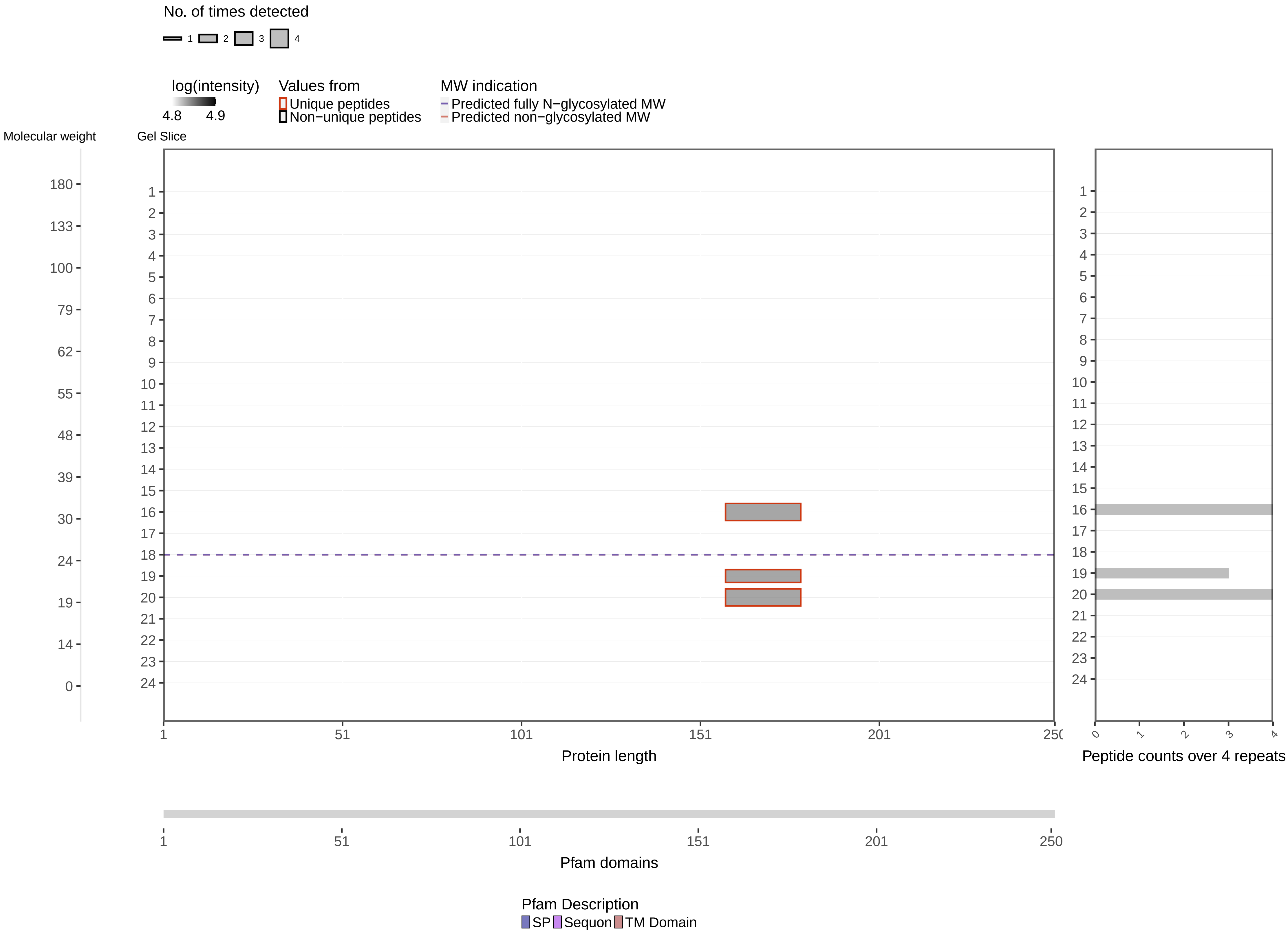


Protein sequence:

AVYVAESAISSGSAGSTSDGAENTAAIAAGIALISLAAASSILLQVGKNSPPPVQTMEYSGPSLSYYINKFKPVEIVQASVAPETEAPTAPEVAQASVAAETEAPTAPEVESSALEAPAPQVEVQSEAPQDTSSSSSDVS*

### NbL01g08710.1 40S_ribosomal_protein_S20−2−like [MW = 13.73 kDa]

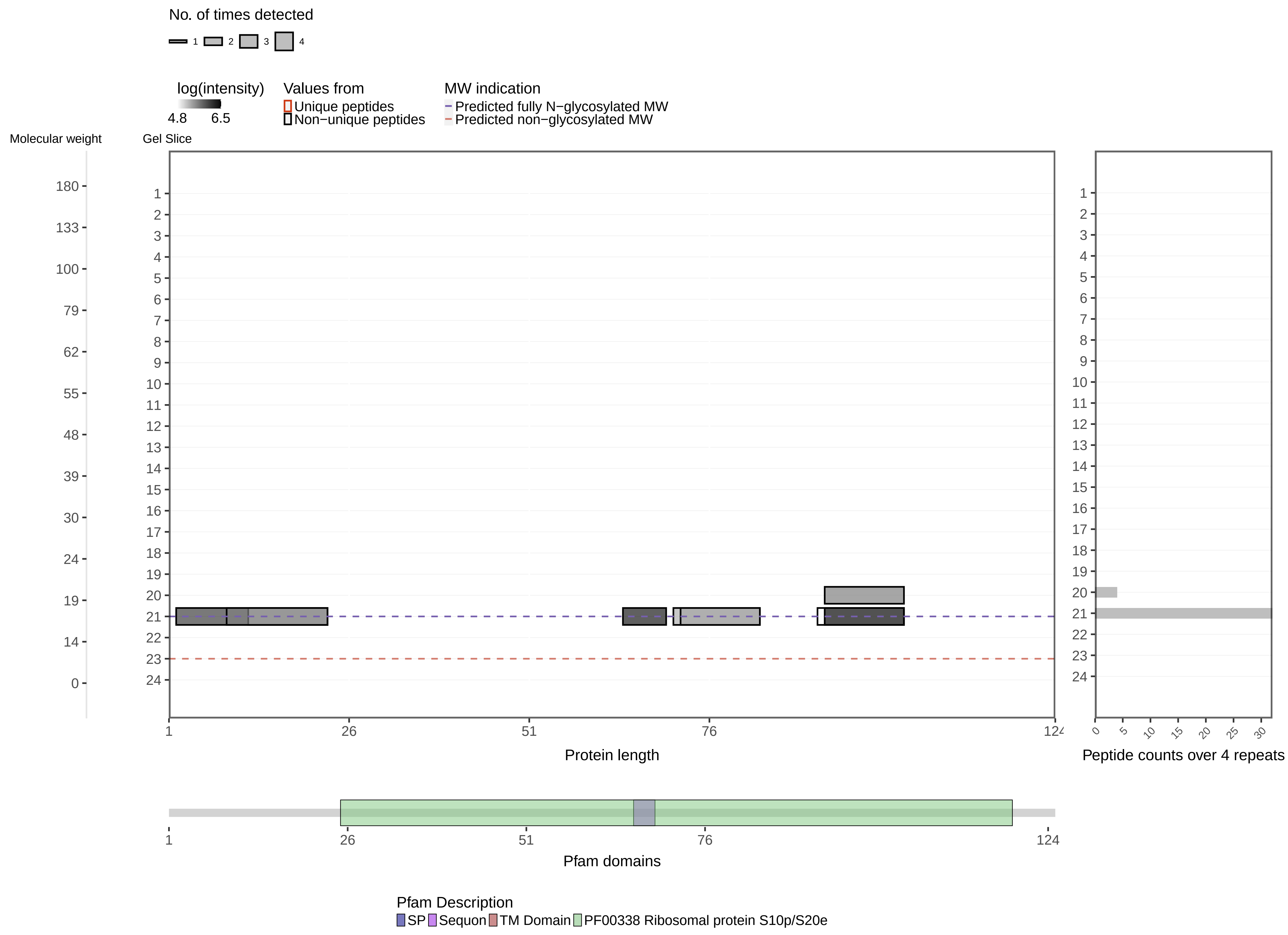


Protein sequence:

DKRLRVKGPVRMPTKVLNITTRKSPCGEGTNTWDRFELRVHKRVIDLFSSADVVKQITSITIEPGVEVEVTIADS*

### NbL01g08660.1 peptidyl−prolyl_cis−trans_isomerase_CYP22−like [MW = 20.59 kDa]

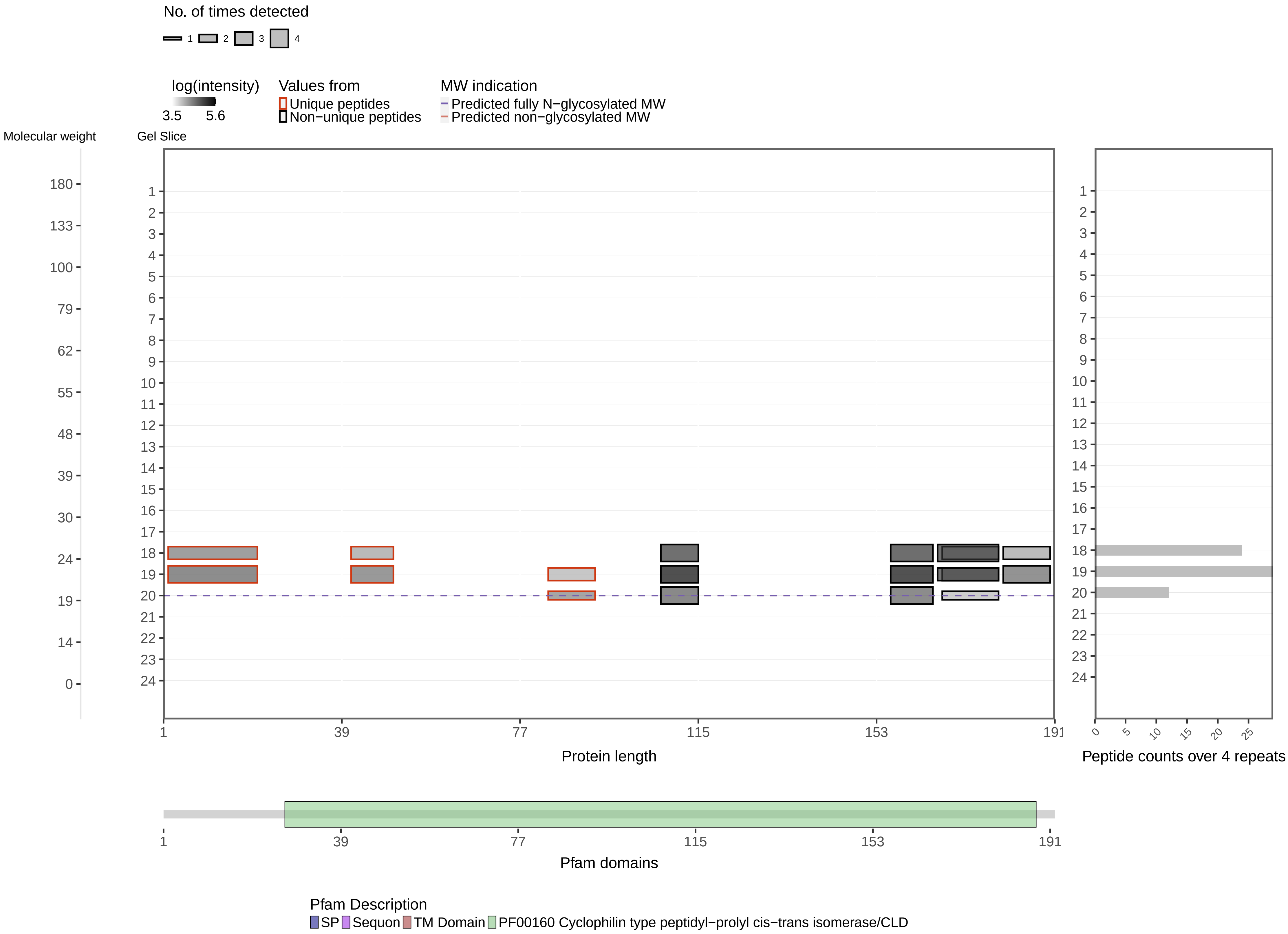


Protein sequence:

FMIQGGDFLKGDGSGCVSIYGSKFEDENFIAKHTGPGLLSMANSGPNTNGCQFFVTCAKCDWLDNKHVVFGRVLGDGLLVVRKIENVAVGANNKPKLACMIAECGEM*

### NbL01g08600.1 aspartyl_protease_AED3−like [MW = 49.6 kDa]

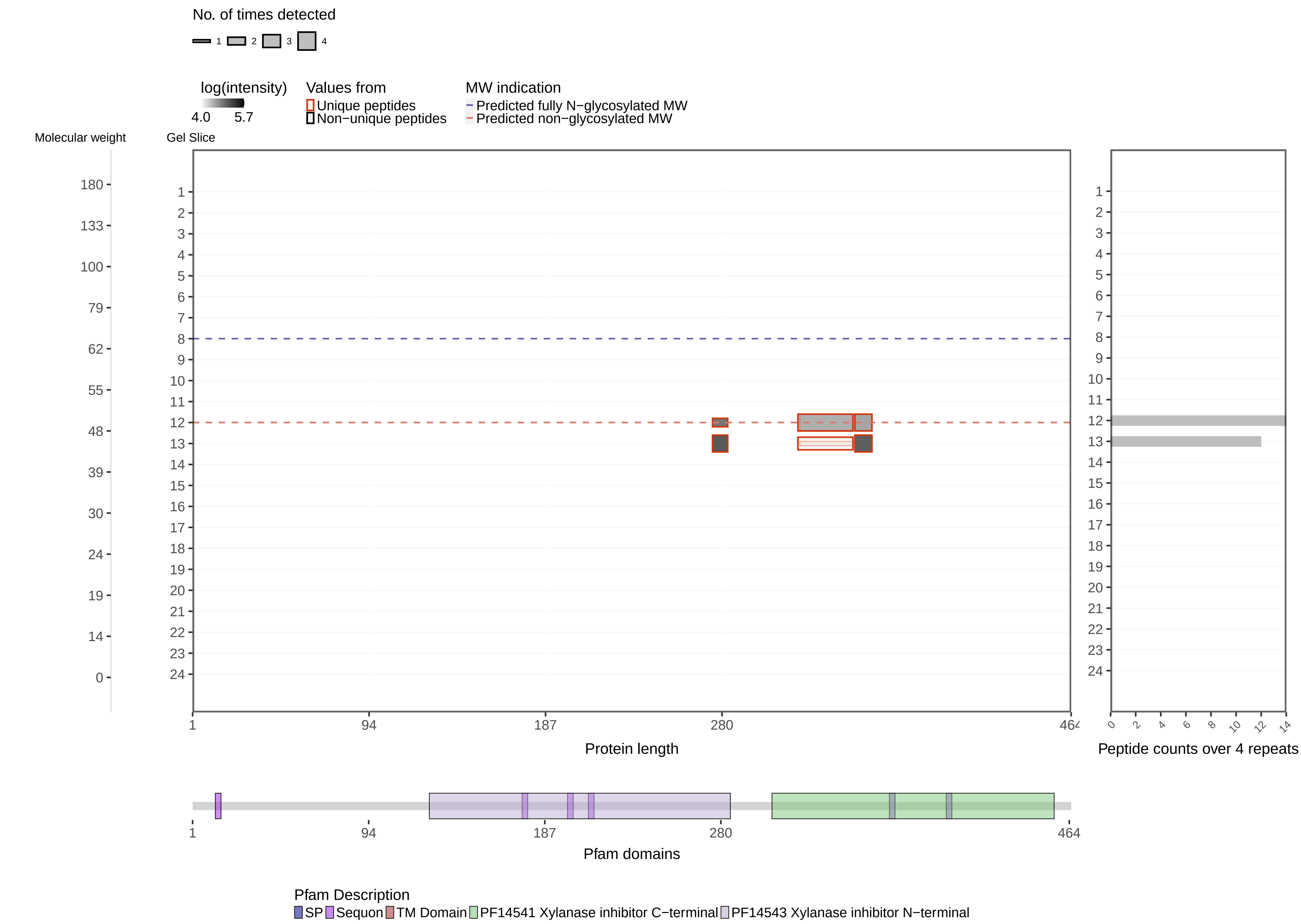


Protein sequence:

STDAVPSYTFGCVQKATGSSAPPQGLLGLGRGPLSFLSQTQSHYQSTFSYCLPSYKSPNFAGTLRLGPNGQPKRIKTTQLLRNPRRSSFYYVNLVGVKVGRRIVDIPPSALAFNPTTGAGTIIDSGTVFTRLVESAYTAVKNEFRRRMGRNTTVTSLGGFDTCYTVPITIPTITLMFAGMNVTLPQDNFLIRSSSSSTTCLAMAASPADPVNSVLNVIANWQQQNHRFLFDVPNSKLGVARETCS*

### NbL01g08580.1 NAD(P)H:quinone_oxidoreductase−like [MW = 21.68 kDa]

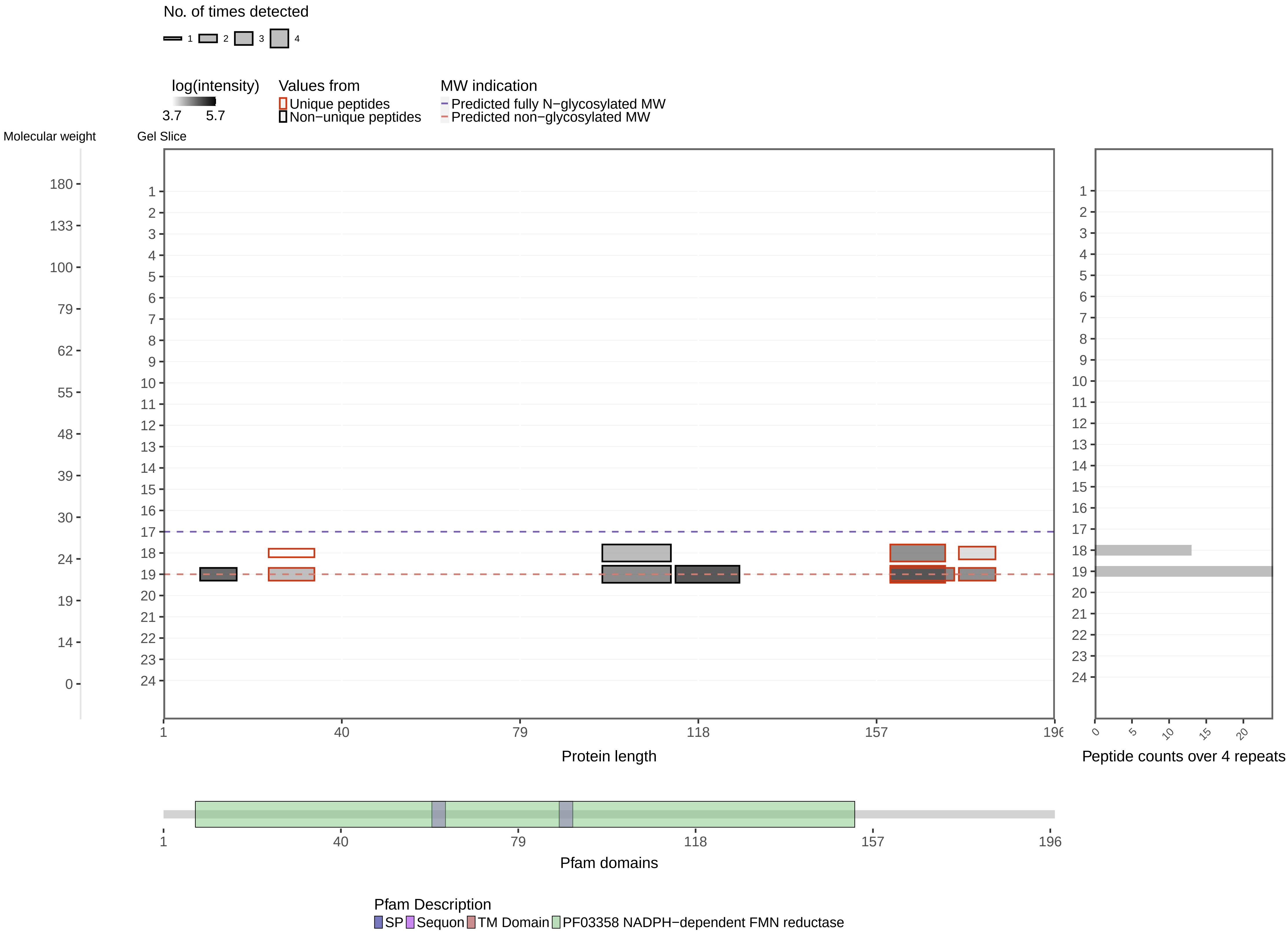


Protein sequence:

EYNYSVTGPLKNAIDWASRPPNVWGDKAAAIVSAGGNFGGGRSHYHLRQIGIYIDLHFINKPEFFLNAFELPPKFDDQGVLTDEESKNRLKEVLLVLKAFSLRLKGKLSE*

NbL01g08560.1 NAD(P)H:quinone_oxidoreductase [MW = 21.94 kDa]

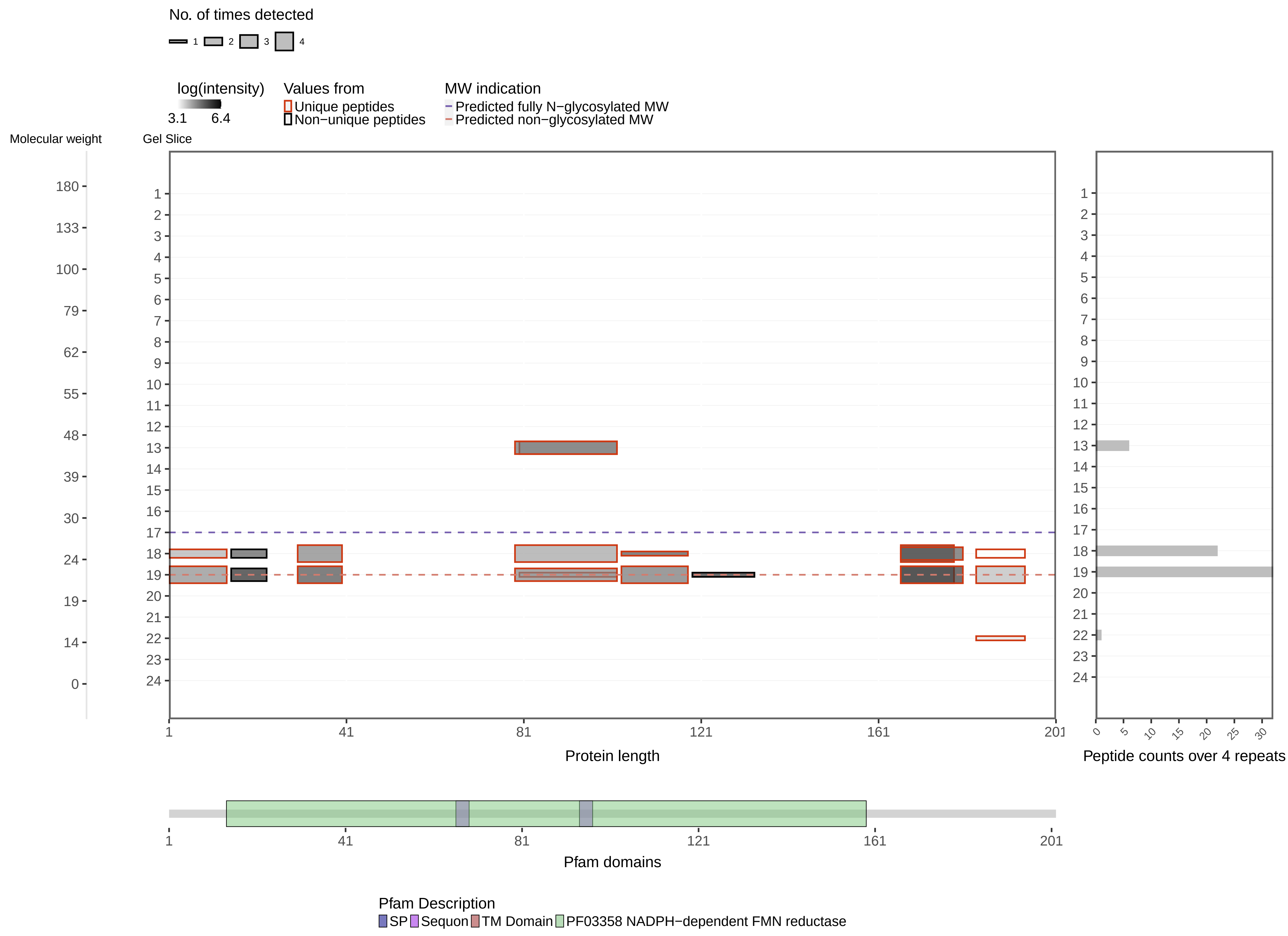


Protein sequence:

ASPEYNYSVTGPLKNAIDWASRPPNVWADKAAAIVSAGGGFGGGRAQYHLRQIGVYLDLHFINKPEFFLNAFQPPSKFDSDGVLTDEETKQKLKAILLSLQAFTLRLKGKCE*

### NbL01g08040.1 clustered_mitochondria_protein_homolog [MW = 209.8 kDa]

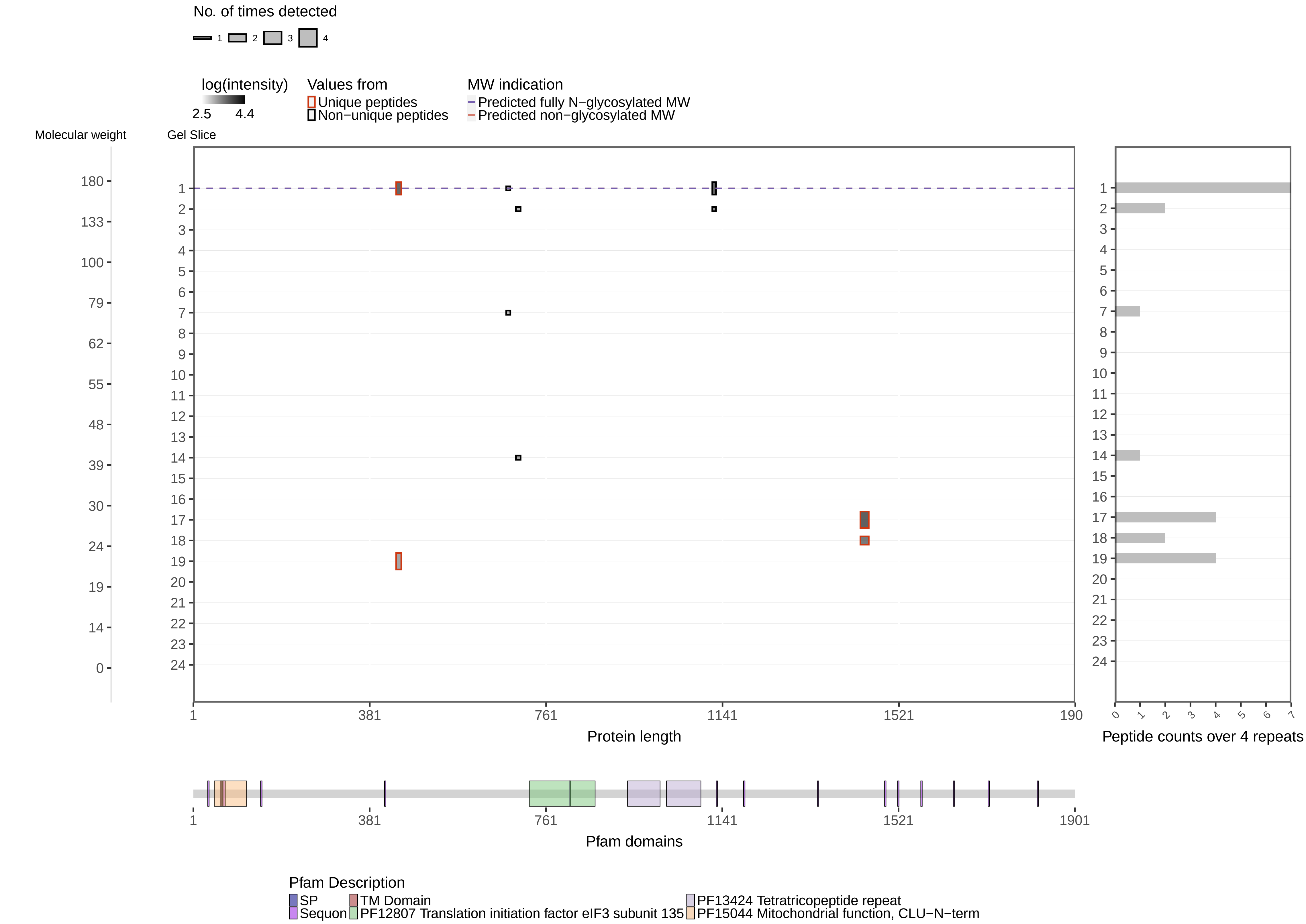


Protein sequence:

[illegible]

NbL01g07800.1 jasmonic_acid−amido_synthetase_JAR1−like [MW = 67.33 kDa]

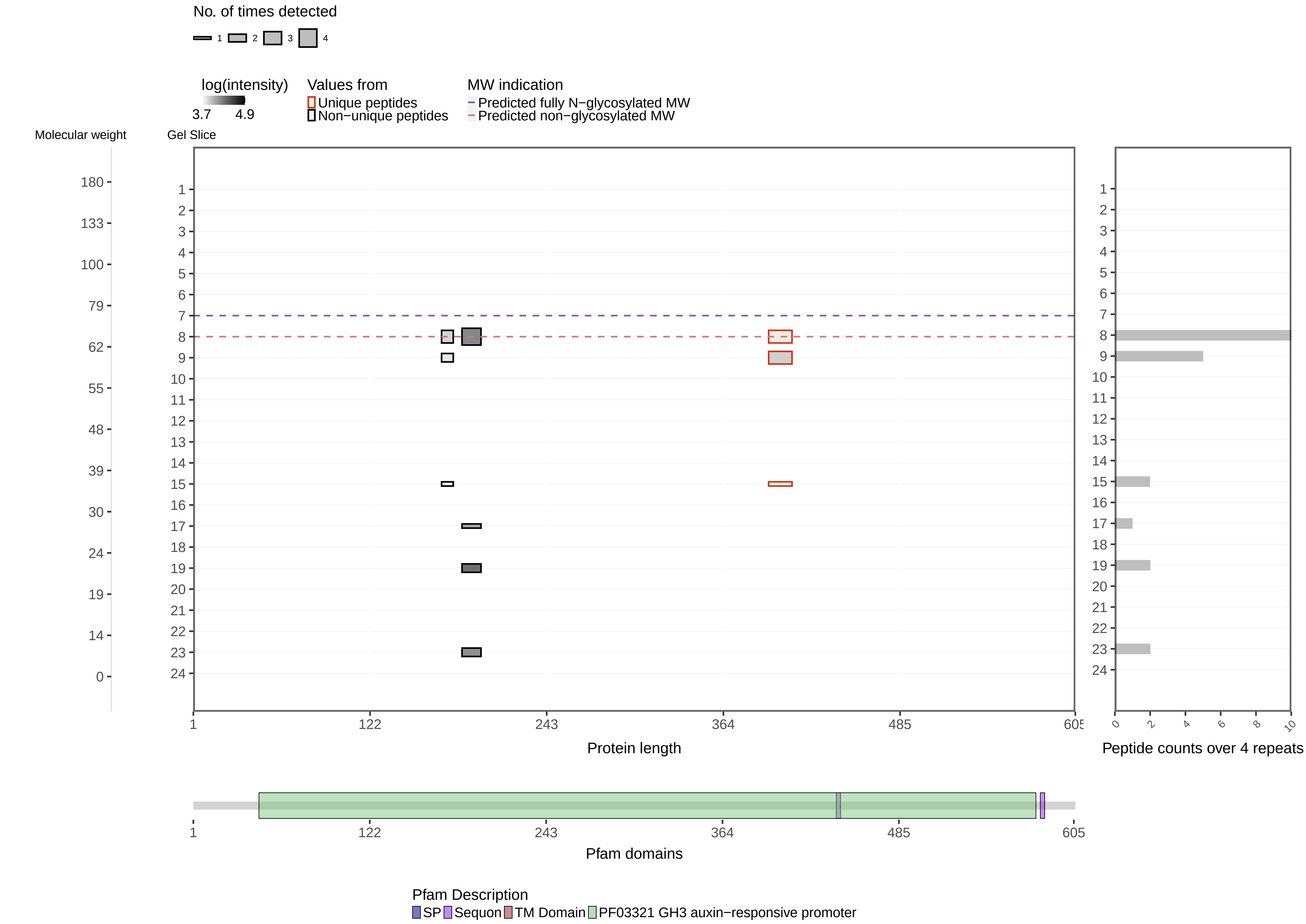


Protein sequence:

SKLLKPDPEQADTIFRKCSGLSNWYGLIPVLFPNAKYIYGIMTGSMEPYLKKLRHYAGELPLQSADYGSSEGWIGANVNPKLPPELVTYAVLPNIGYFEFIPLRGDLDGVEPTPVGLAEVKLGEEYEIVVTNFAGLYRYRLGDVVKIKGFHNATPELQFICRRNLLLSINIDKNTEKDLQLAVEAAAKLLVNEKSEVVDFTSHVNVAADPGHYVVFWELSGEASDETLQECCNCLDKSFVDAGYLGSRKANAIGALELRVVKRGTFHKILDHFVGLGGAVSQFKTPRCVGPNNSTI

NbL01g07710.1 exosome_complex_component_RRP41_homolog [MW = 26.62 kDa]

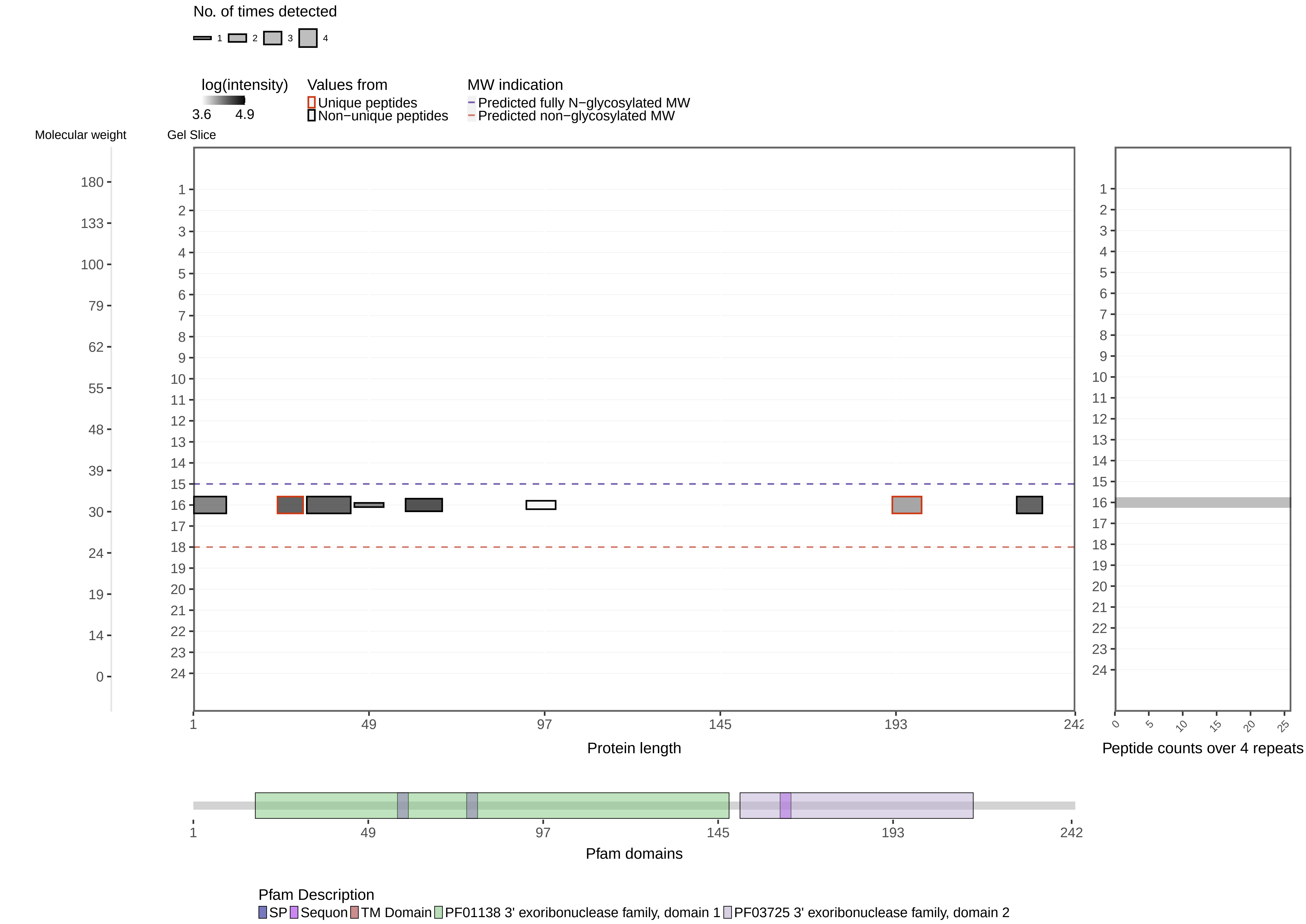


Protein sequence:

ACILTHLMPRSQIDIFVQVLQADGGTRSACINAATLALADAGIPMRDLVTSCSAGYLNSTPLLDLNYLEDSAGGPDVTVGILPKLDKVTLLQMDAKLPIDTFETVMQLAVEGCKAVESYIREILLENTKKLEIRRGV*

NbL01g07670.1 uncharacterized_LOC107820362 [MW = 27.55 kDa]

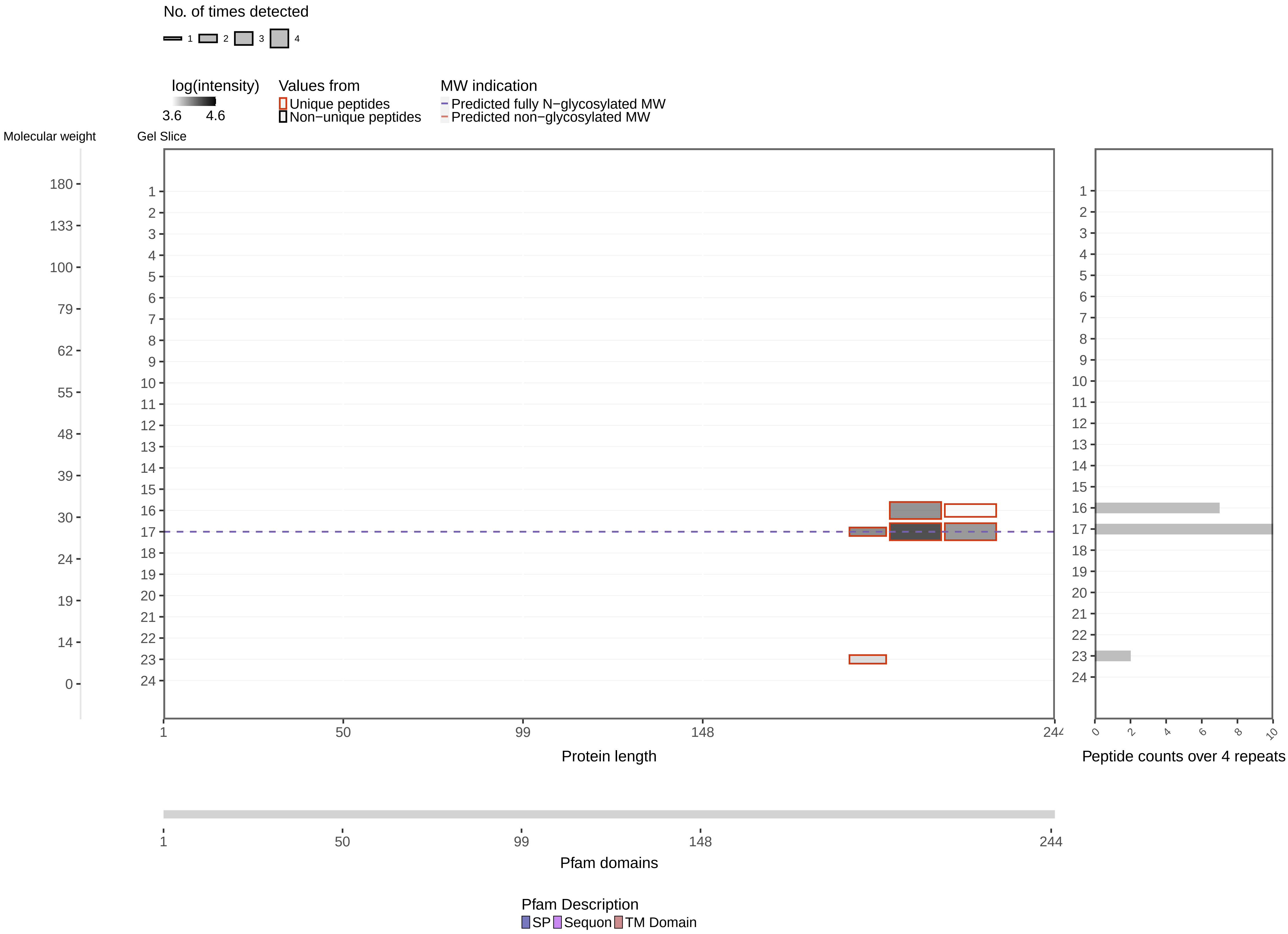


Protein sequence:

RDEELKKMKKMKKEFANPWPIGPPGLDLISLGLVDVEKLPKYEMTVEDGRRLAKEYSRVLMRKHRQRQAAESTLLRLKKEAIEALPENLKAAALIPDLTPFPVNRFMATLTPPIEGYIEKVNEAAKKSSAKEKLR*

### NbL01g07660.1 eukaryotic_translation_initiation_factor_1A−like [MW = 16.32 kDa]

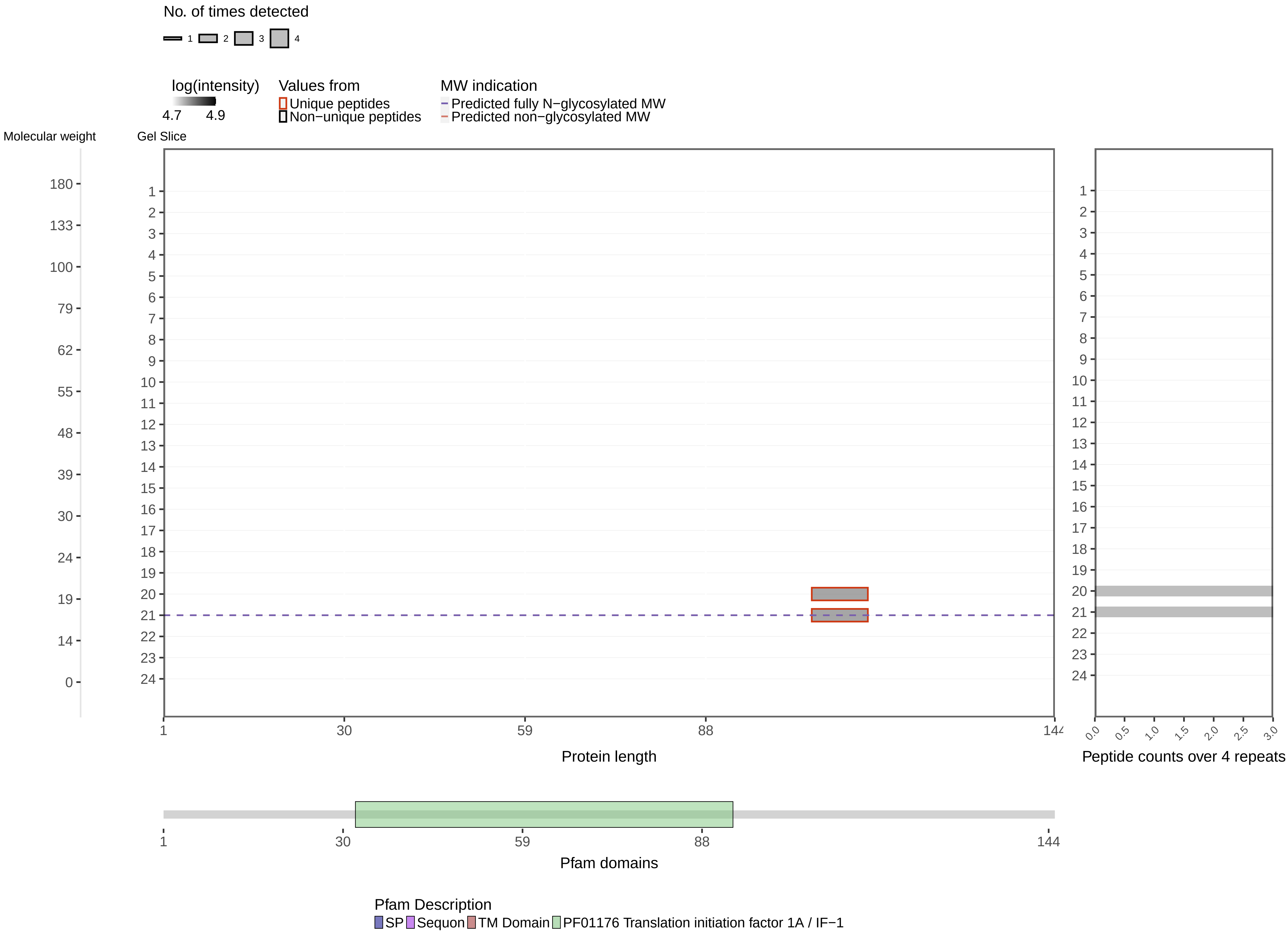


Protein sequence:

RLCHIRGKMHKKVWIAAGDIILVGLRDYQDDKADVILKFMPDEARLLKAYGELPESTRLNEVNGLDEEDDGDGDDYIEFEDEDIDKI*

### NbL01g07470.1 desiccation_protectant_protein_Lea14_homolog [MW = 18.52 kDa]

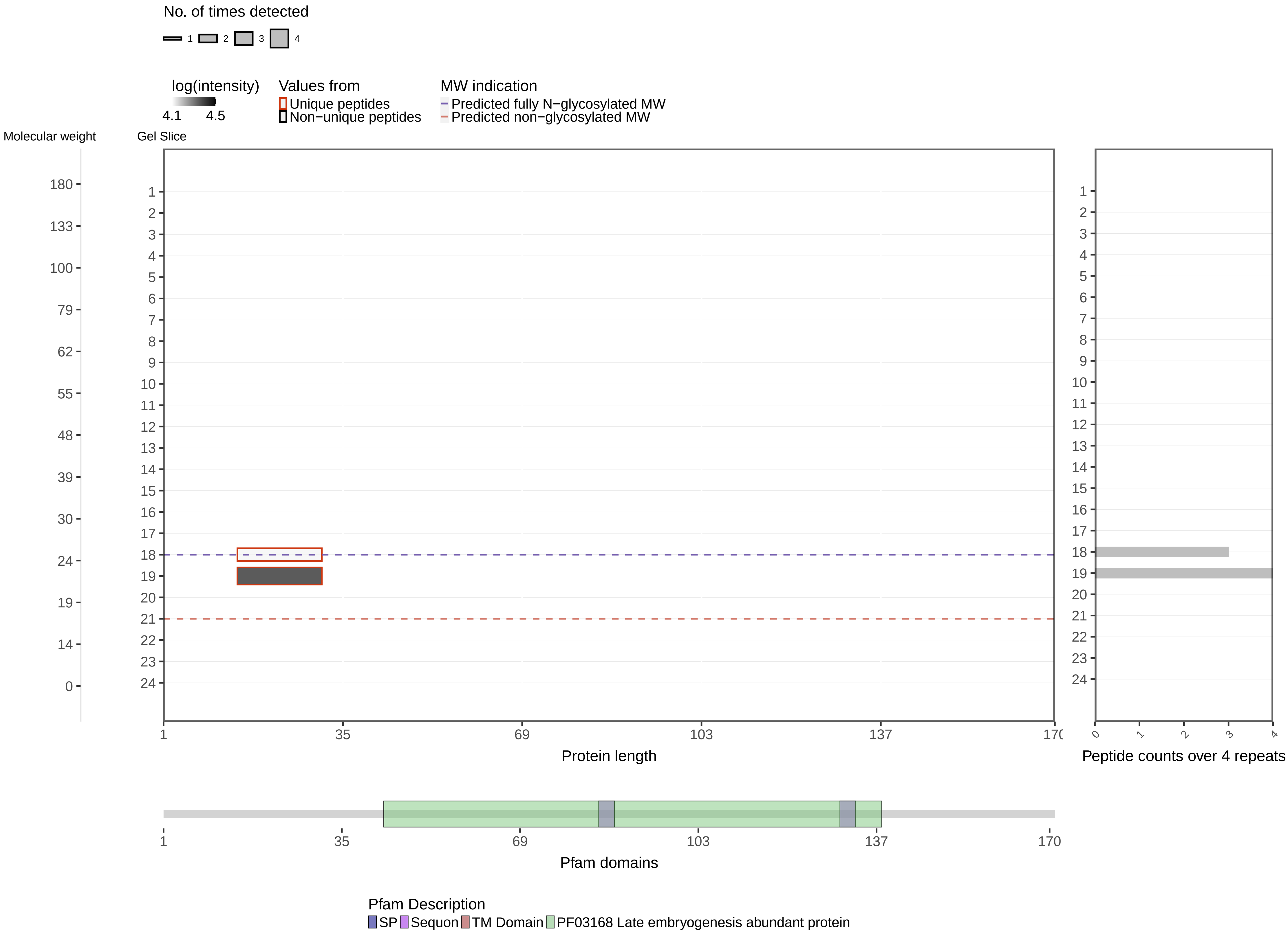


Protein sequence:

GRIPDPGNIKANDSTMLDVPVQVPHSVLLSLVKDLGGDWDIDYTLELGLIIDLPVIGNFTIPLSYSGEYKLPTLSDLWKGTGKEEDEKEDPTKVTEI*

NbL01g07460.1 proline_iminopeptidase−like [MW = 57.49 kDa]

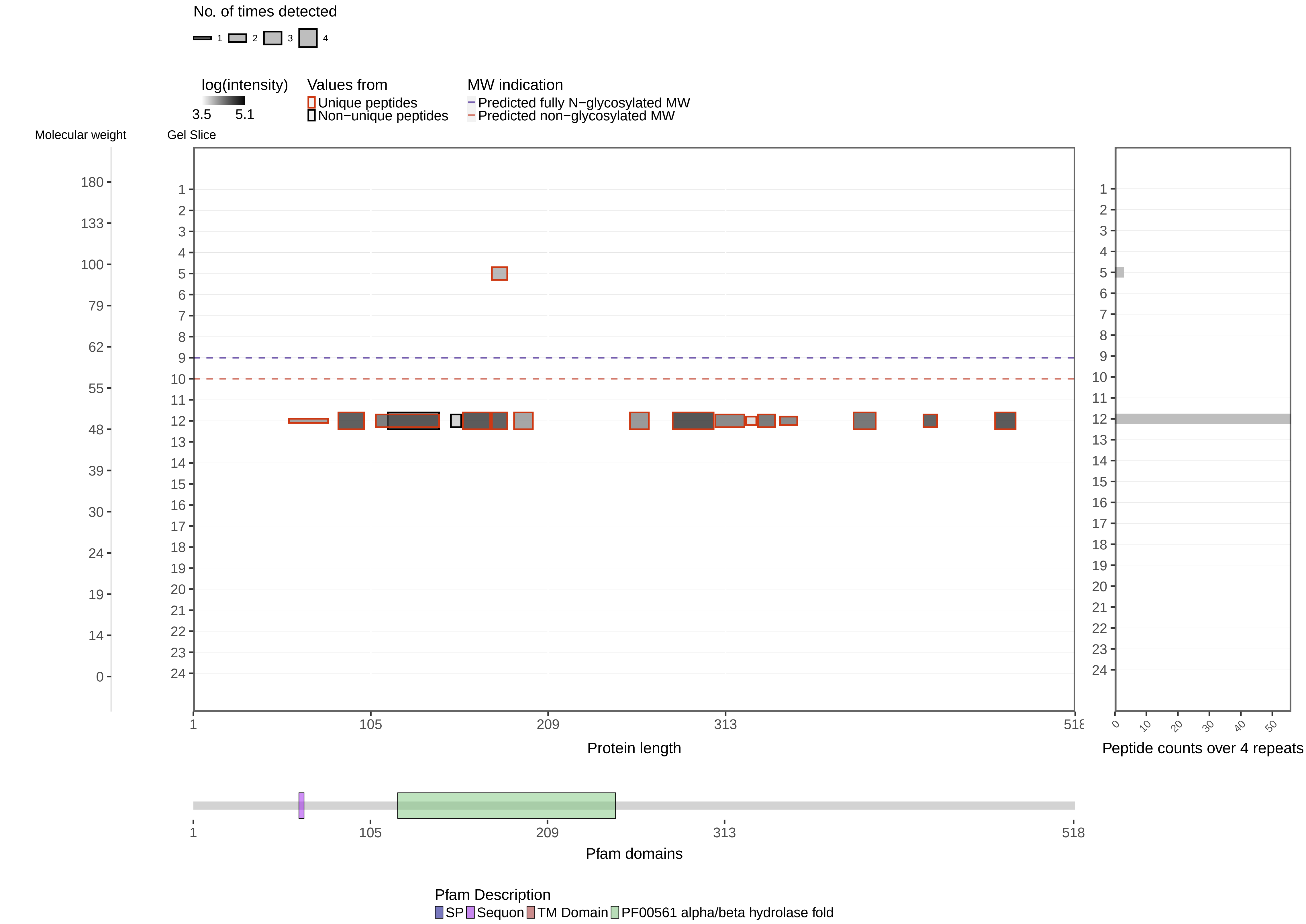


Protein sequence:

KGCTADAVYRAGFEQLVHQNEKYYKRFPQDIEIVRDVVNYLAESEGGGLPLPSGGILTPRGLQLLGLSGLGSSNGFERLHYLFERVWDSVIVQGAKRRISDYFLNAYERWIAFDANPLYALLHESIYCQGASSLWSAHRIRAENENKFDAIEAAKGSSPVLFTGEMIFPWLFDEVHALRPFKDAAQLLAEKKDWPSLYDIAALKDNKVPVAAAVYYEDMYVNFKLSMETASQIAGIRLWITNEYMHSGLRDAGSKVLDHLLAMLNGKKPLF*

NbL01g07280.1 succinate_dehydrogenase_[ubiquinone]_iron−sulfur_subunit_2_mitochondrial−like [MW = 31.15 kDa]

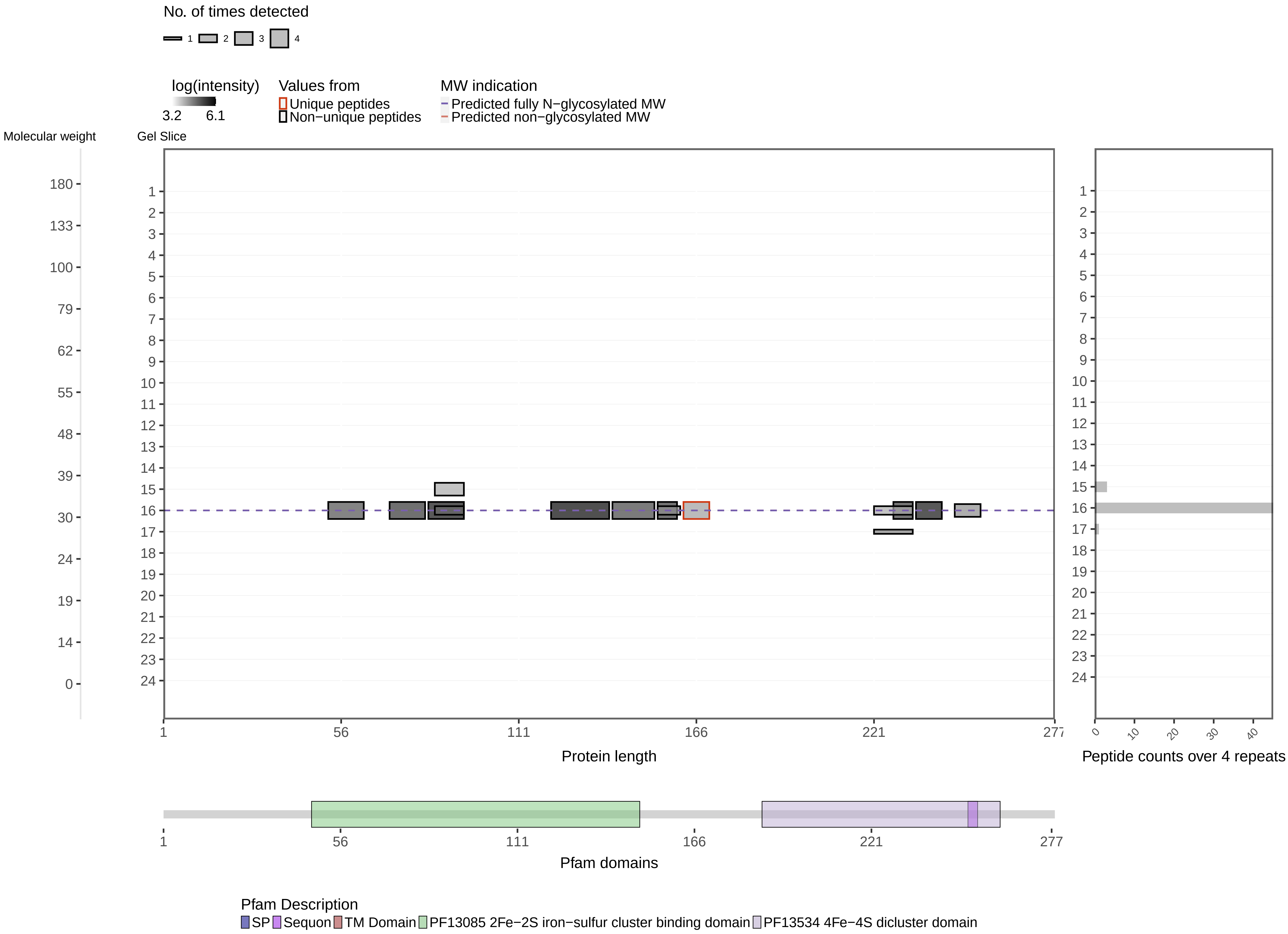


Protein sequence:

STITPLPHMFVIKDLVVDMTNFYNQYKSIEPWLKRKTEPPTPGKEIPQSKSDRAKLDGMYECILCACCSTSCPSYWWNPESYLGPAALLHANRRWIMDSRDEYTQERLDAVNDEFKLYRCHTILNCSRACPKGLNPGKHIQNIKRLELAS*

NbL01g07120.1 bifunctional_L−3−cyanoalanine_synthase/cysteine_synthase_2_mitochondrial [MW = 39.41 kDa]

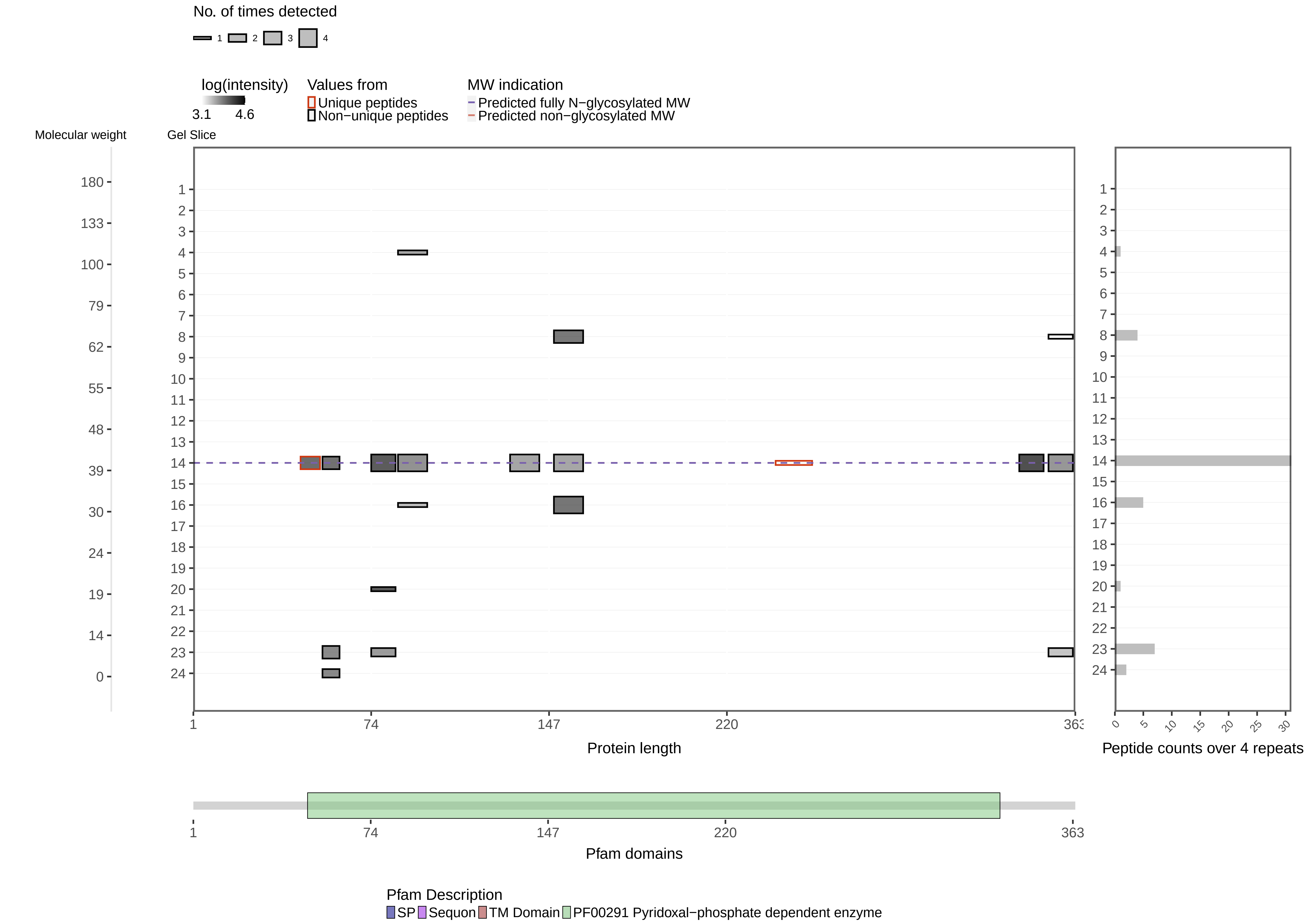


Protein sequence:

:AYDLLESTPNAFMLQQFSNPANTQIHFETTGPEIWEETQGNVDIFVMGIGSGGTVSGVGQYLKSKNPNVKIYGVEPTESNVLNGGKPGPHHITGNGVGFKPDILDMDIMEEVLMVSSEDAVNMARELAVKEGLMVGISSGANTVAALQLAQKPENKGKLIVTIHASFGERYLSSVLYQDLRKEAENMQPVSVD*

NbL01g06840.1 small_nuclear_ribonucleoprotein−associated_protein_B'−like [MW = 28.91 kDa]

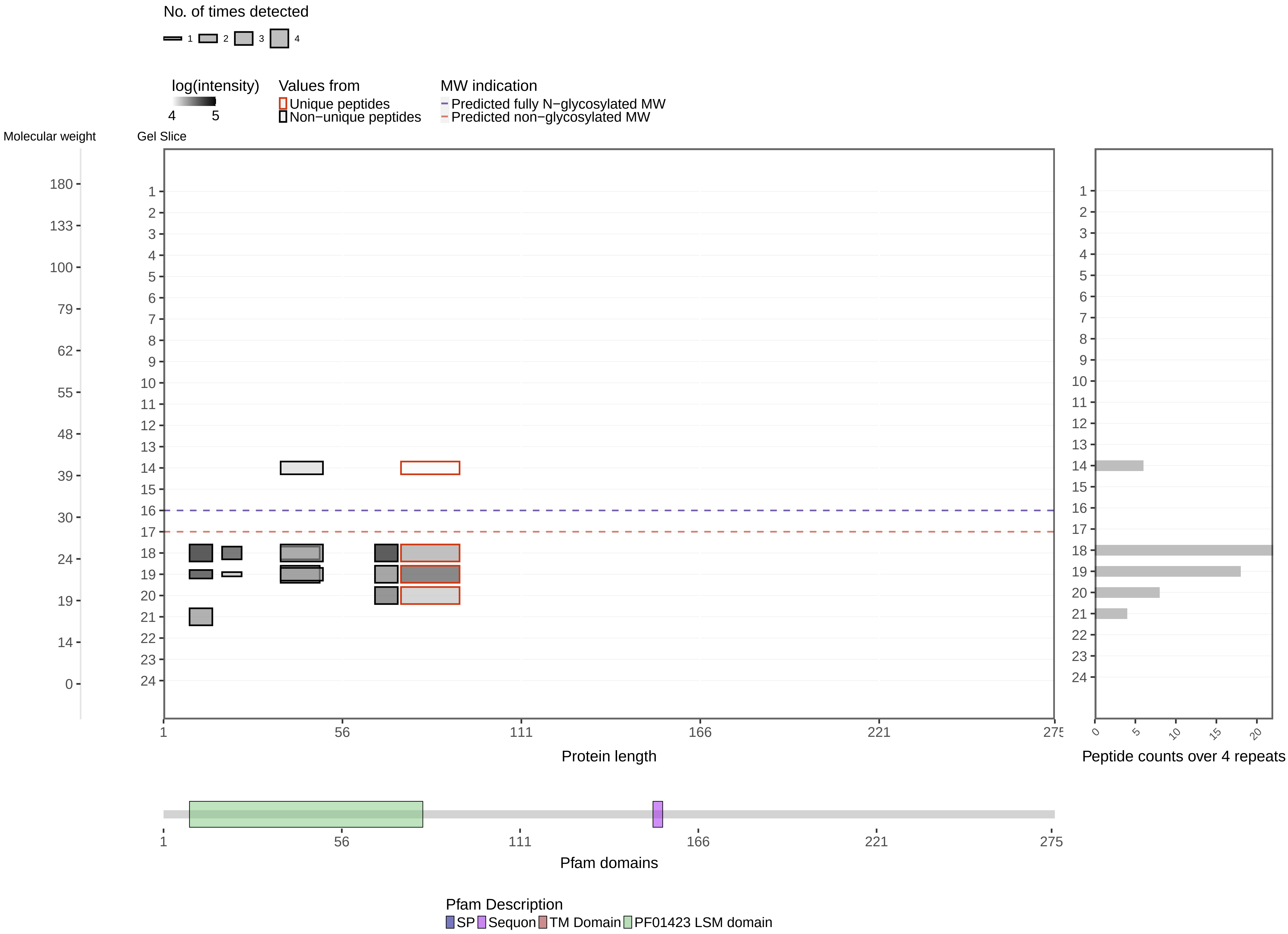


Protein sequence:

VRGVGGPAPGMMQPQISRPPMPNVSAPPMNYPPPVMRPPGGGPPMQMQMQRPGGPPPPMAPPPQFRPGGGPPQFPPQFGQRPMVPPPPMMRGPPPPGAPPRPGMPGGPPGQPPRPGMPPPPPGGQVPVYGPPRPGMPPPPNQQNQ*

### NbL01g06830.1 acyl−CoA_thioesterase_2−like [MW = 49.12 kDa]

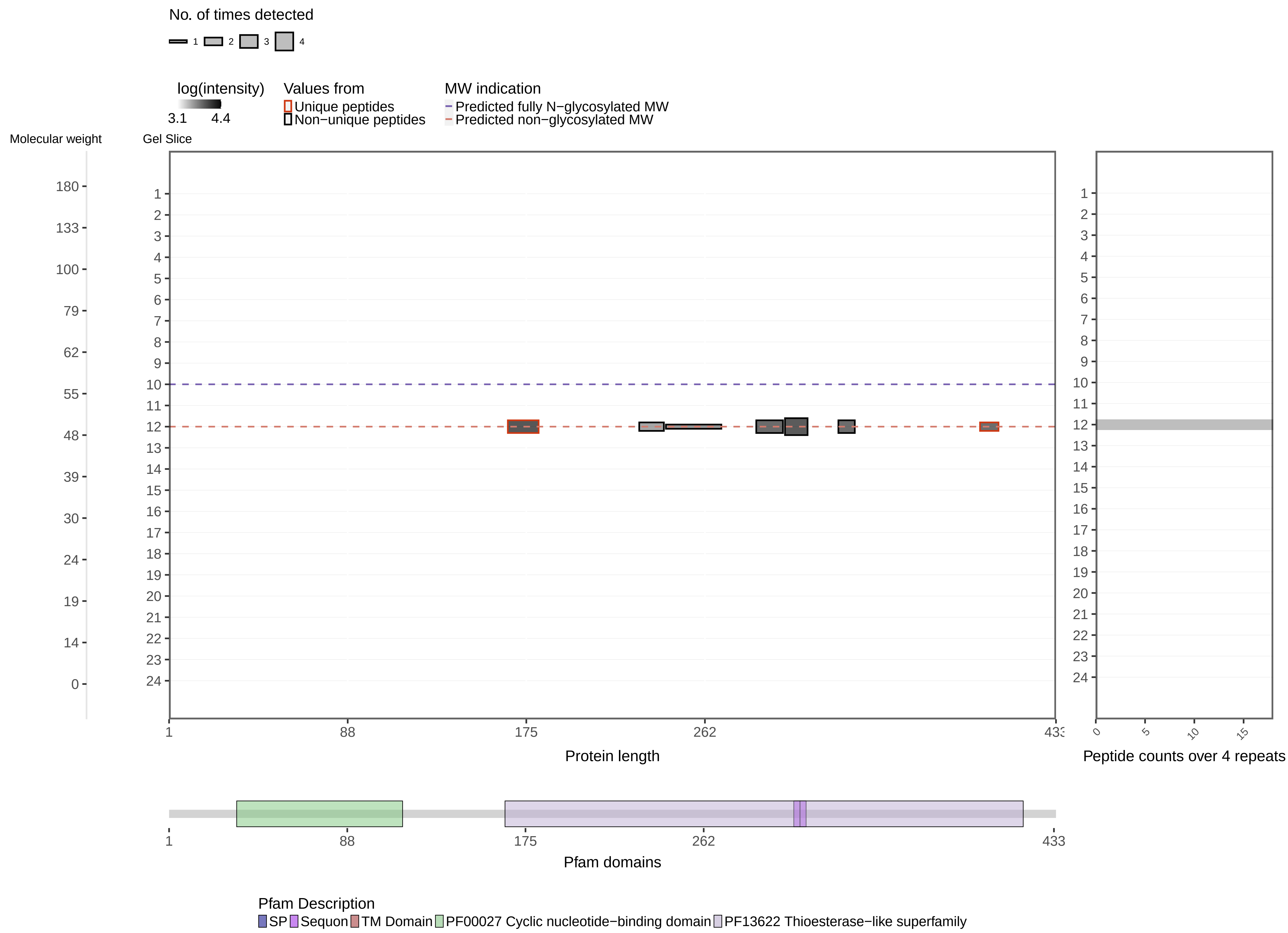


Protein sequence:

PIIYOVYRVRDGKSFATRRVDAIOKGNVIFTLLASFOKDEDGFDHOOAOMPDVPDPETLLSMEDLREMRKTDPRLPRTYRNKVATANFVPWPIEIRFCEPNNATNOTKSPPSLRFWFRAKGKLSDDOALHRCVAAFASDLIFLTVSLNPHRKKDFKTSSVSLDHSMWFHRPFRADDWILFVITSPAAYNARGFVSGEMFNRKGELVLSLTOEGLLRPFRKPPSVRPKL*

NbL01g06800.1 riboflavin_biosynthesis_protein_PYRD_chloroplastic [MW = 47.85 kDa]

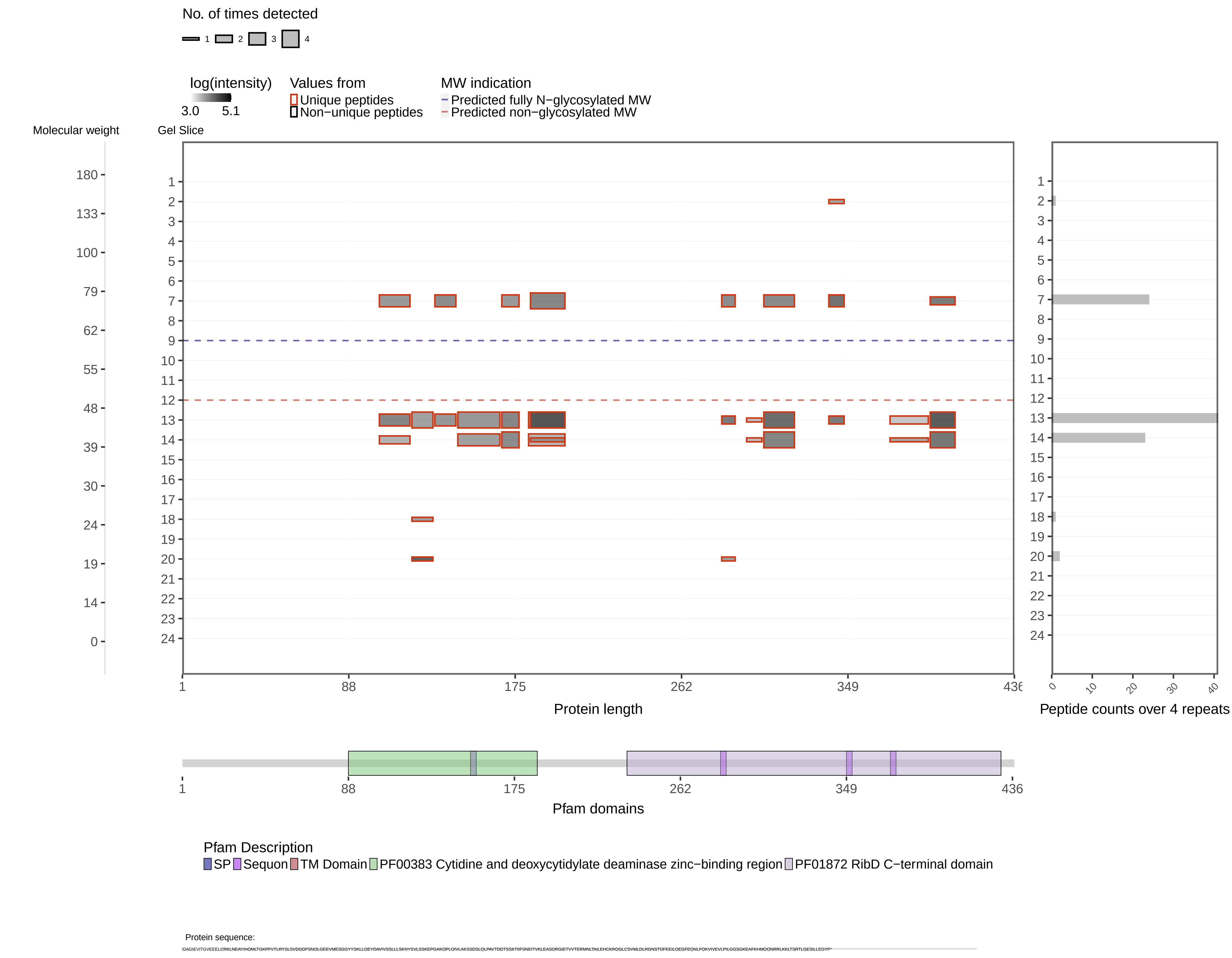

NbL01g06780.1 uncharacterized_LOC107784826 [MW = 74.58 kDa]

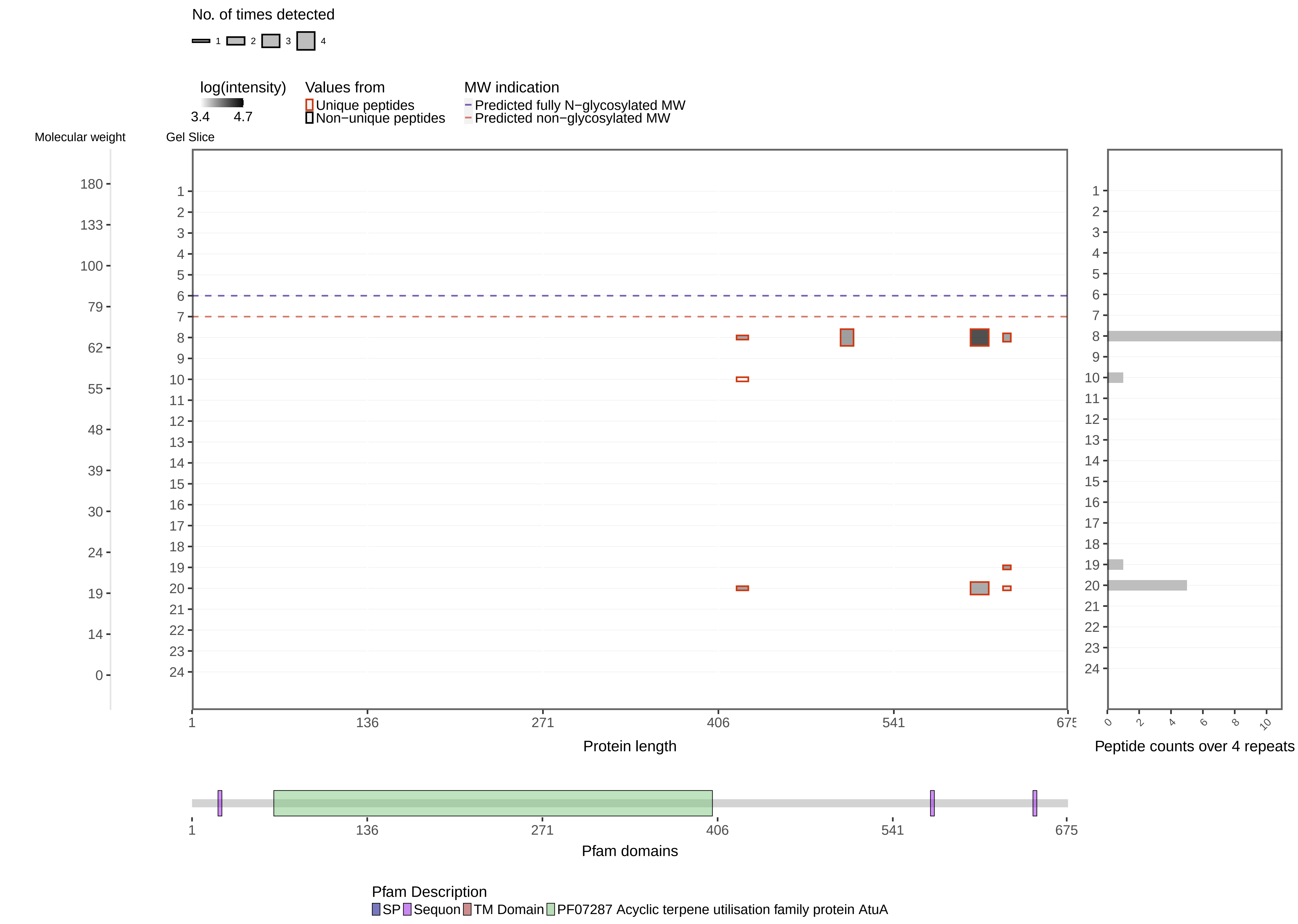


Protein sequence:

NbL01g06650.1 protein_pleiotropic_regulatory_locus_1−like [MW = 53.4 kDa]

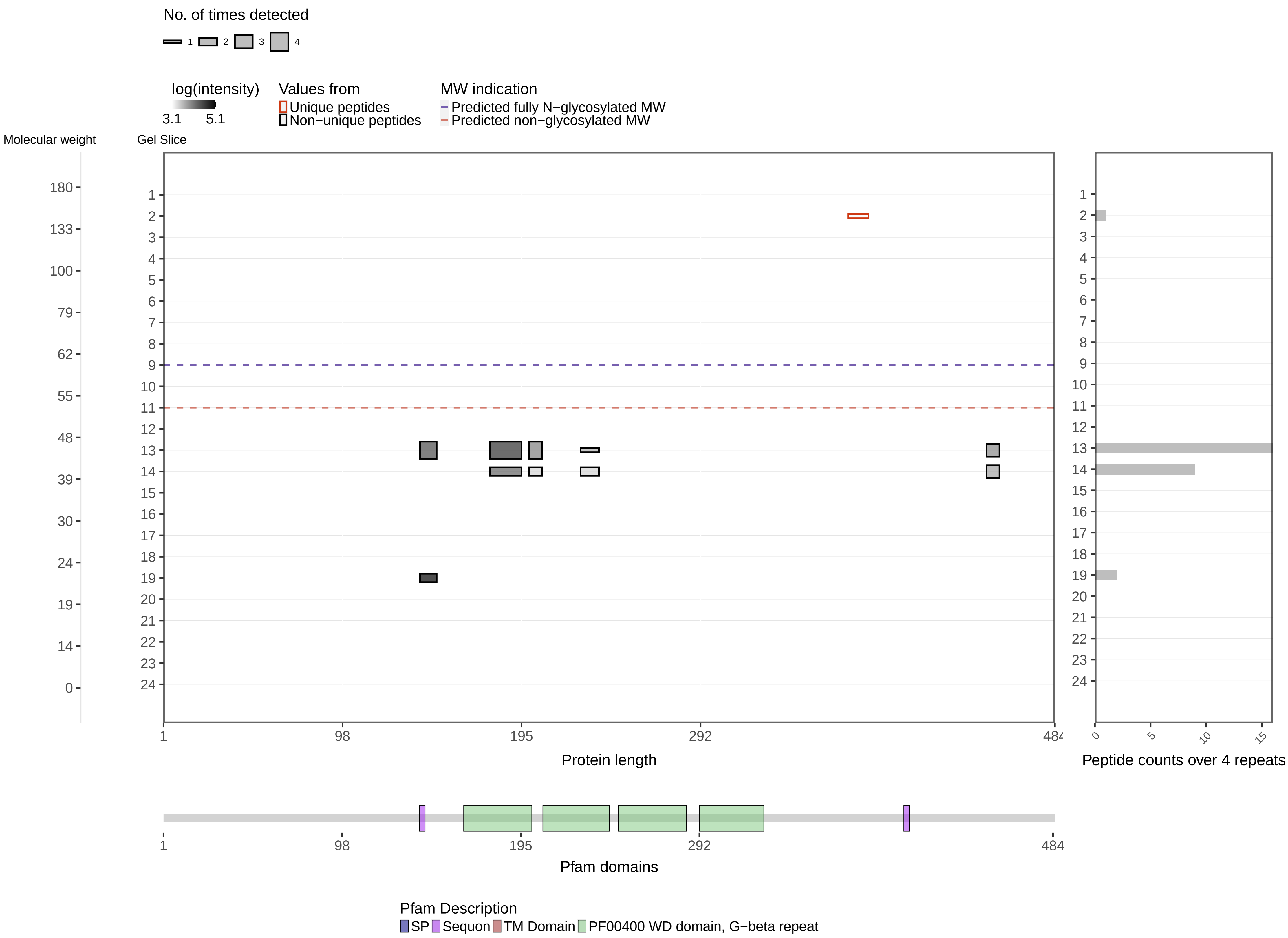


Protein sequence:

MFSAGDDKLVKCWDLEQNKVIRSYHGHLSGVYCLALHPTIDILFTGGRDSVCRVWDIRSKMQIHALSGHDNTVCSVFTRPMDPQVITGSHDSTIKFWDLRYGKTMATLTHHKKSVRAMAQHPTEDCFASASADNIKKFNLPKGEFLHNMLSQQKTIVNAMAVNDDGVMATGGDNGSMWFWDWISGHNFQQSQTIVQPGSLDSEAGIYALTYDMTGSRLISCEADKTIKMWKEDETATPETHPLHFKPPKDIRRF*

NbL01g06600.1 K(+)_efflux_antiporter_2_chloroplastic−like [MW = 130.34 kDa]

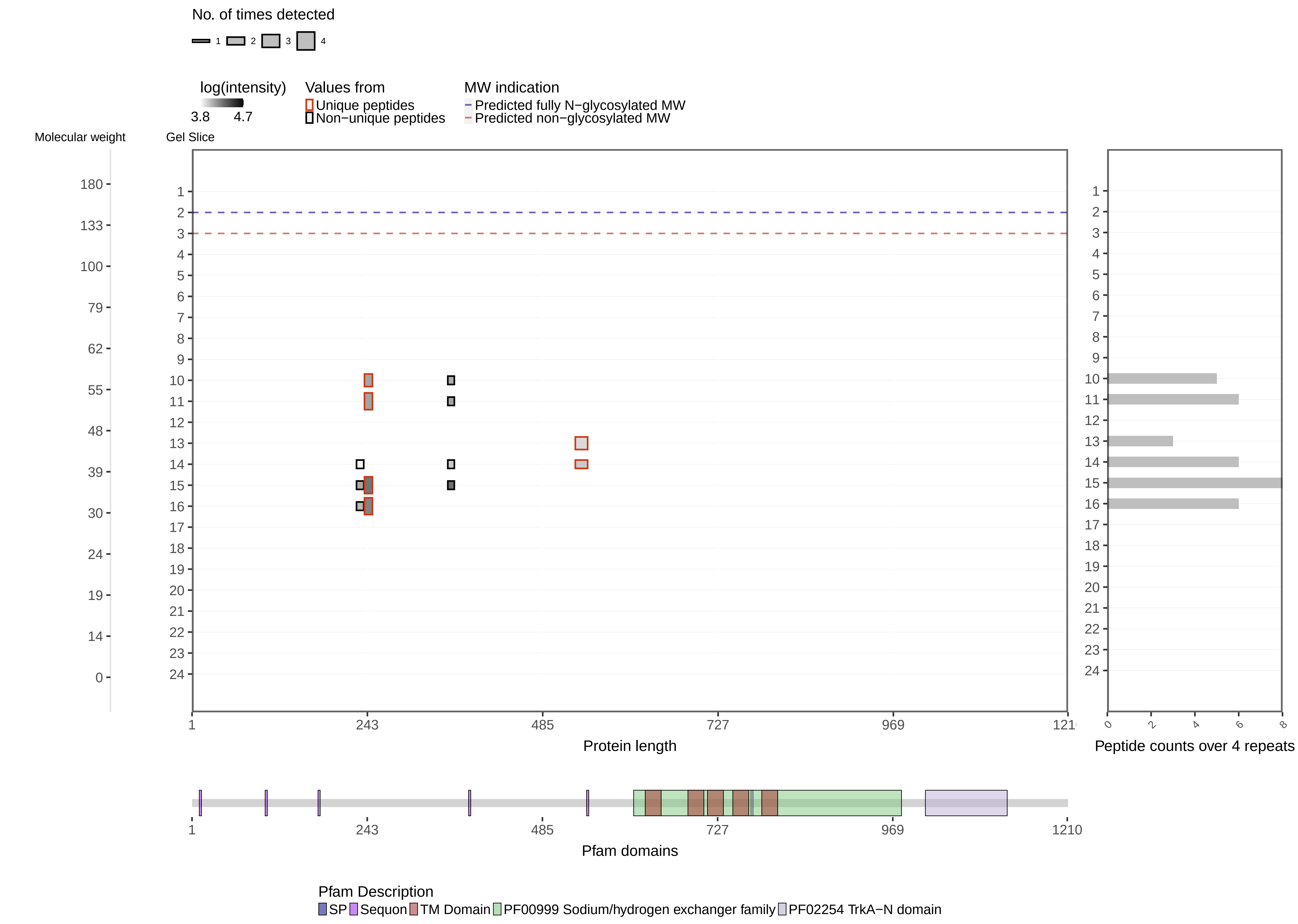


Protein sequence:

KLPEKLKTLMEILPHQEINEEEASLFDMLWLLLASVIFVPIFQKIPGGSPVLGYLAAGILIGPYGLSIIRHVHGTKAIAEFGVVFLLFNIGLELSVERLSSMKKYVFGLGTAQVLVTAVVVGLVAHFVAGQAGPAAIVIGNGLALSSTAVVLQVLQERGESTSRHGRATFSVLLFQDLAVVVLLILIPLISPNSSKGGVGFGAIAEALGLAAVKAIVAITAIIAGGRLLLRPIYKQIAENQNAEIFSANTLLVILGTSLLTARAGLSMALGAFLAGLLLAETEFSLQVESDIAPYRGLLLGLFFMTV

### NbL01g06540.1 (+)−neomenthol_dehydrogenase−like [MW = 33.37 kDa]

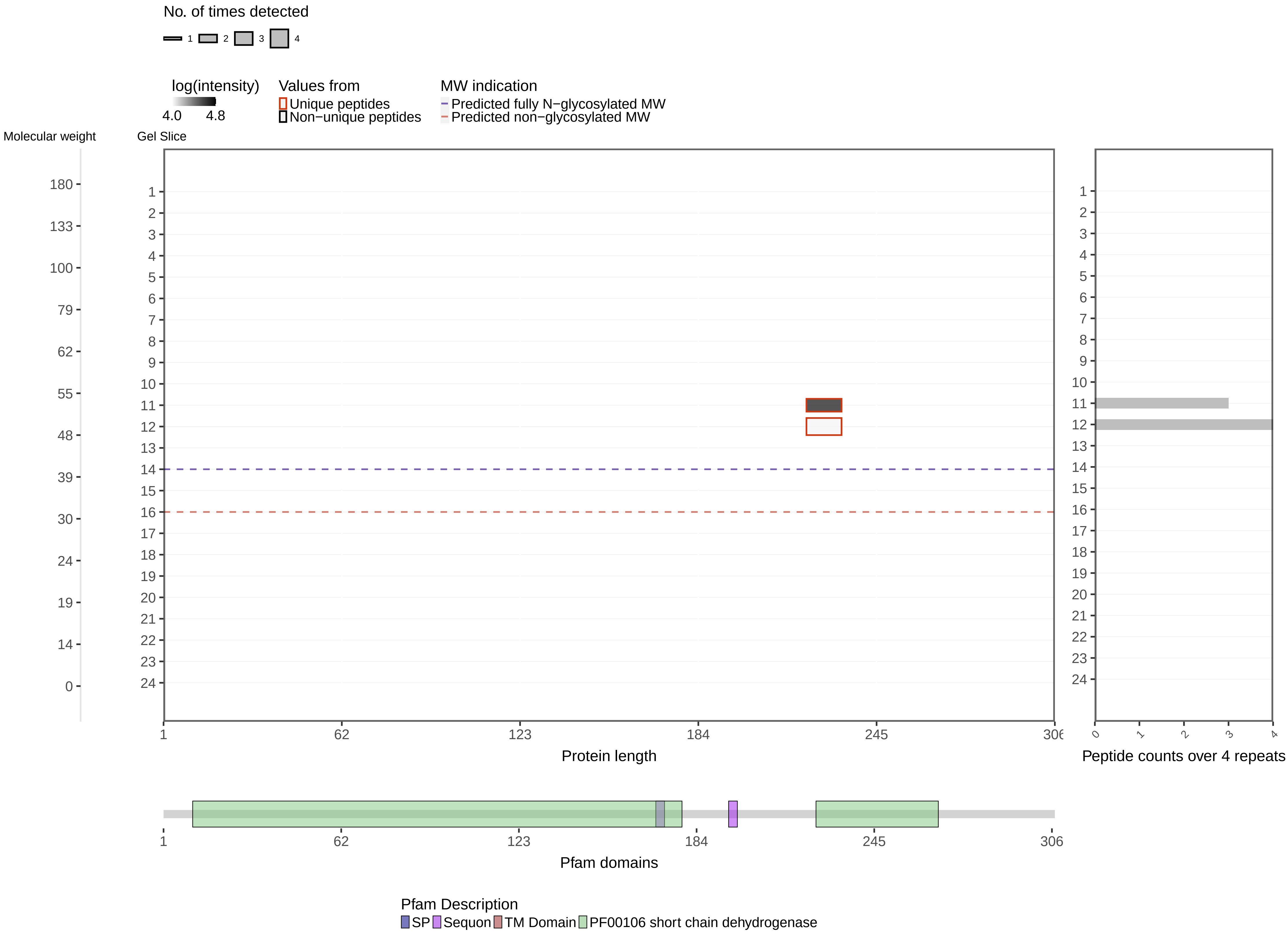


Protein sequence:

CLQTNYYGVKRMTEAFIPLLKLSKSPRIVNVSSSMGRLKNLKHEWAKGVLSDSENLTEEKIQEVIEQYLRDYKENALQAKDWPSLMSAYILSKAAMNAYSRVMAKKHMSIQINCICPGFVKTDMTYNSGILSIEQGAESPVMLALQPHGGPSGFFFVRKKVSSSL*

### NbL01g06520.1 (+)−neomenthol_dehydrogenase−like [MW = 34.8 kDa]

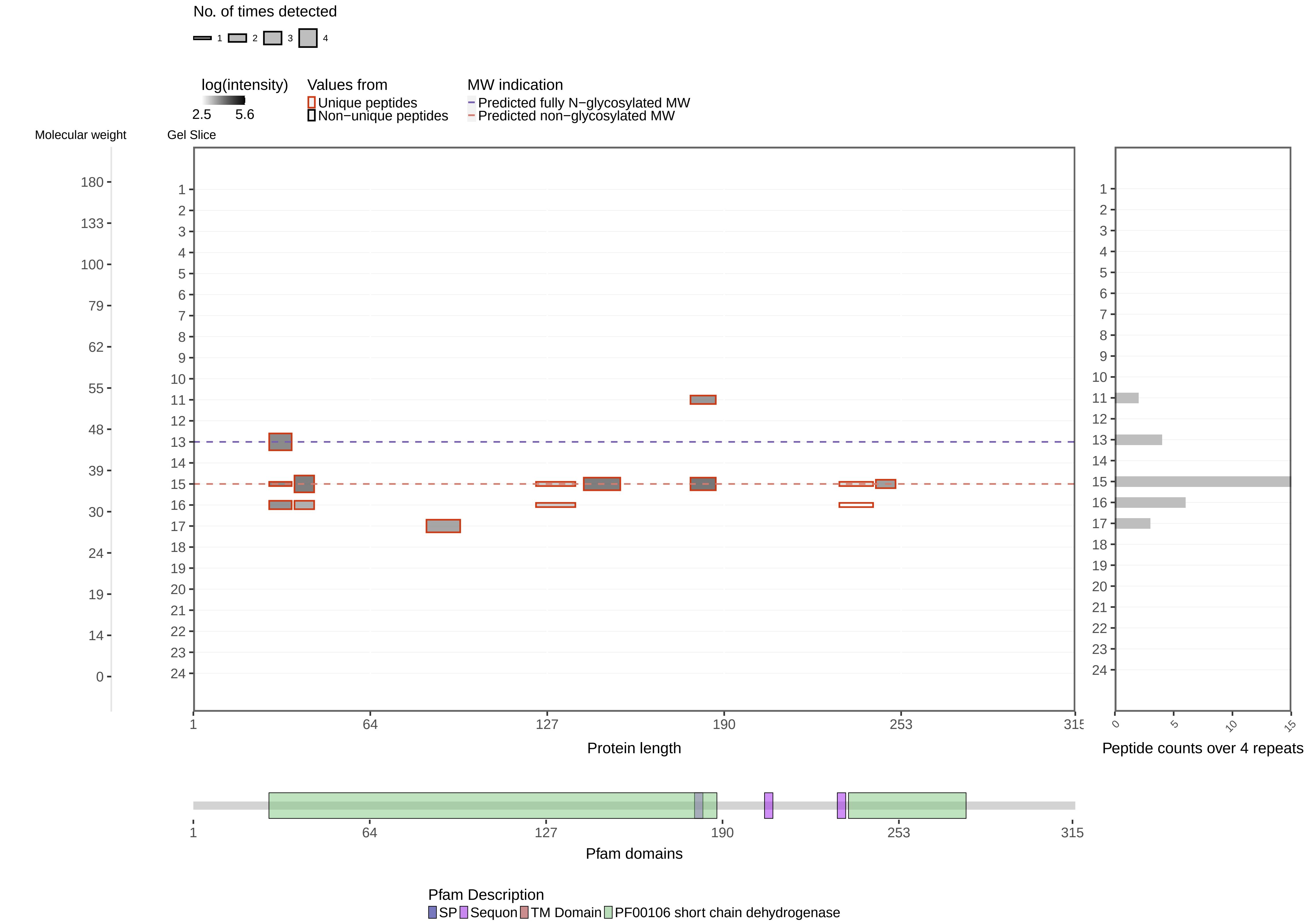


Protein sequence:

AOECLKTNYYGAKOMIOELLPLLOFSDSPRIVNVSSIVGKLEHVRNEWAVGVLSDADNLTEERVDEVLNAFLODLKDGLMETKNWSLIFPAYTLSKAAMNAYTRILAKKYPSLLINCVCPGYVKTDMTINCGKLSVEEGAESPVWLALLPKGGPSGKYFNRKEVSPF*

### NbL01g06490.1 NAD−dependent_malic_enzyme_59_kDa_isoform_mitochondrial [MW = 66.31 kDa]

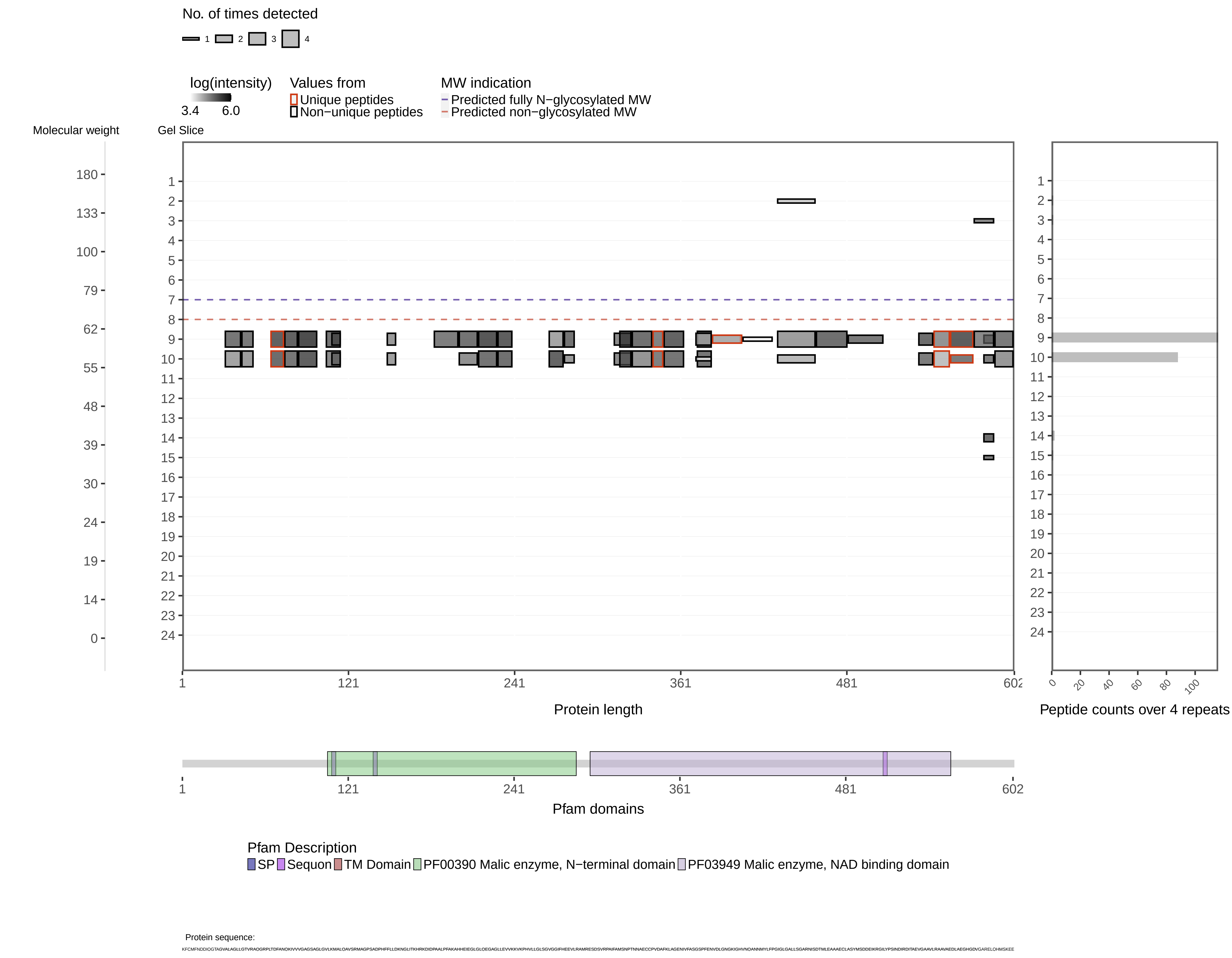

### NbL01g06380.1 synaptotagmin−4 [MW = 55.92 kDa]

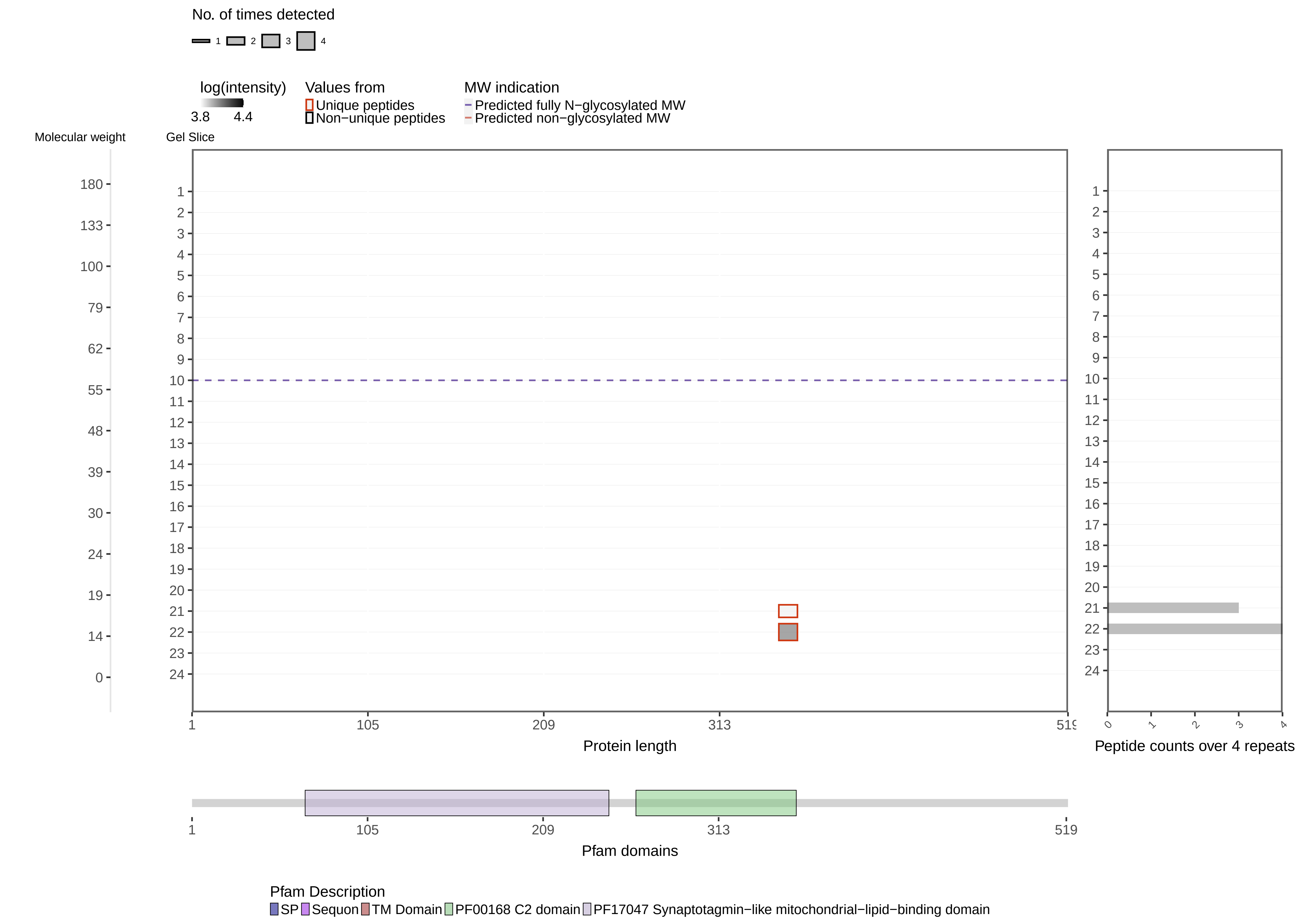


Protein sequence:

GVPVDTSDLELKPQGKLIVTVVKANGLKNHEMIGKSDPYAVVYIRPLFKVKTKTIDNNLNPVWDQTFELIAEDKETQSLIVEVFDKDVGQDQRMGVAKLPLNELVAEAAKEIELRLLPKLDMLKVKDKKDRGTITIKVLYHEFNKEEQLAALEAEKAILEERKKLKSEGVIGSTMDALDGAASLVGSGVGLVGTGLGAGVGLVGTGVGAGVGIVGSGFGAVGSGLSKAGKFMGRTFTSSSKKSGSSTPVNSIQENGGAKPLKTVLANTD*

### NbL01g06310.1 histone_deacetylase_complex_subunit_SAP18−like [MW = 17.76 kDa]

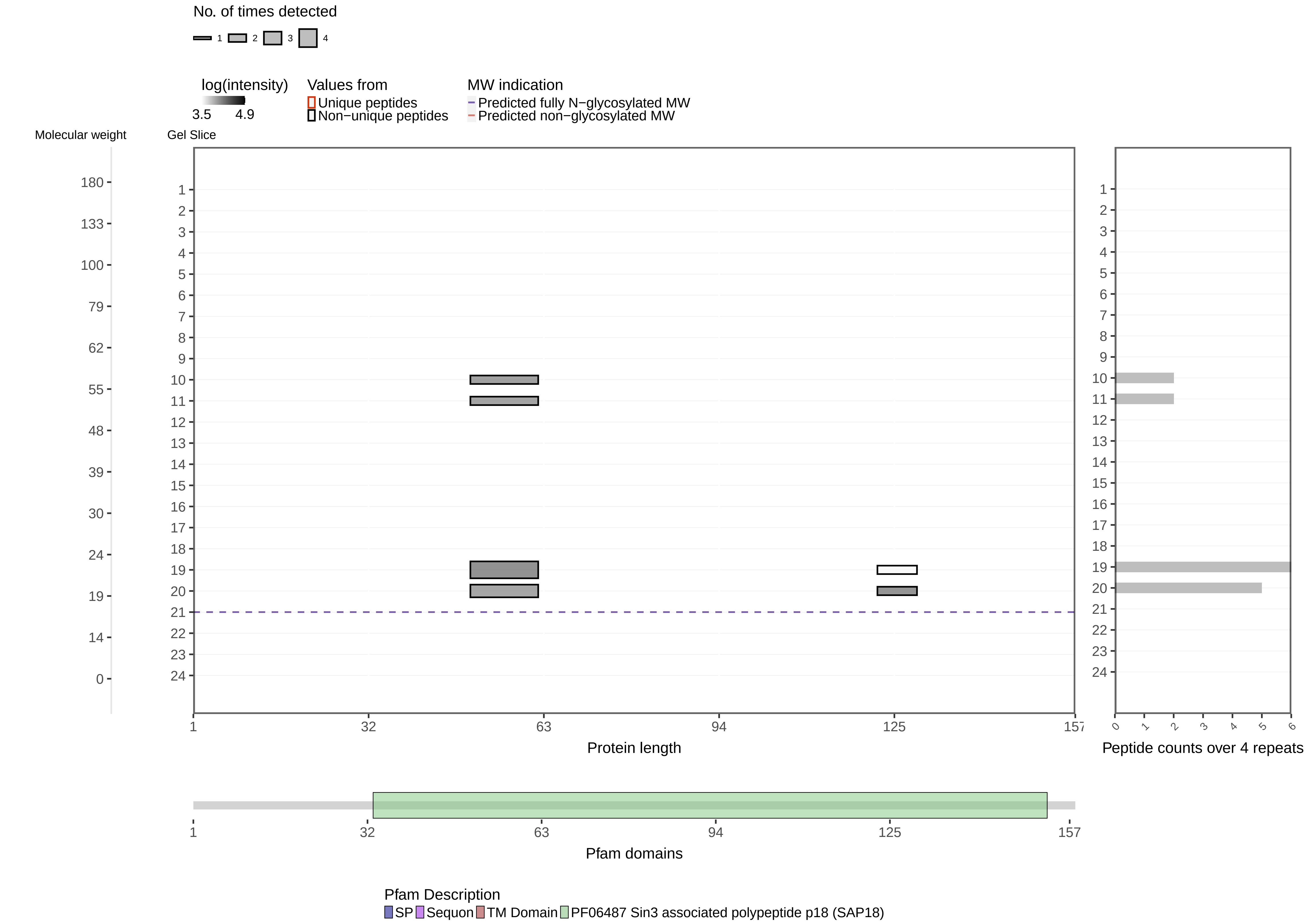


Protein sequence:

EPKDEVQIYTWMDATLRELTDLVKEVAPEARRRDASLSFAFVYPDKRGRFVVREVGRTYSYPNGRRPDSGSKSLCELKFQIGDYLDVAITFQ*

### NbL01g05960.1 COP9_signalosome_complex_subunit_1 [MW = 50.28 kDa]

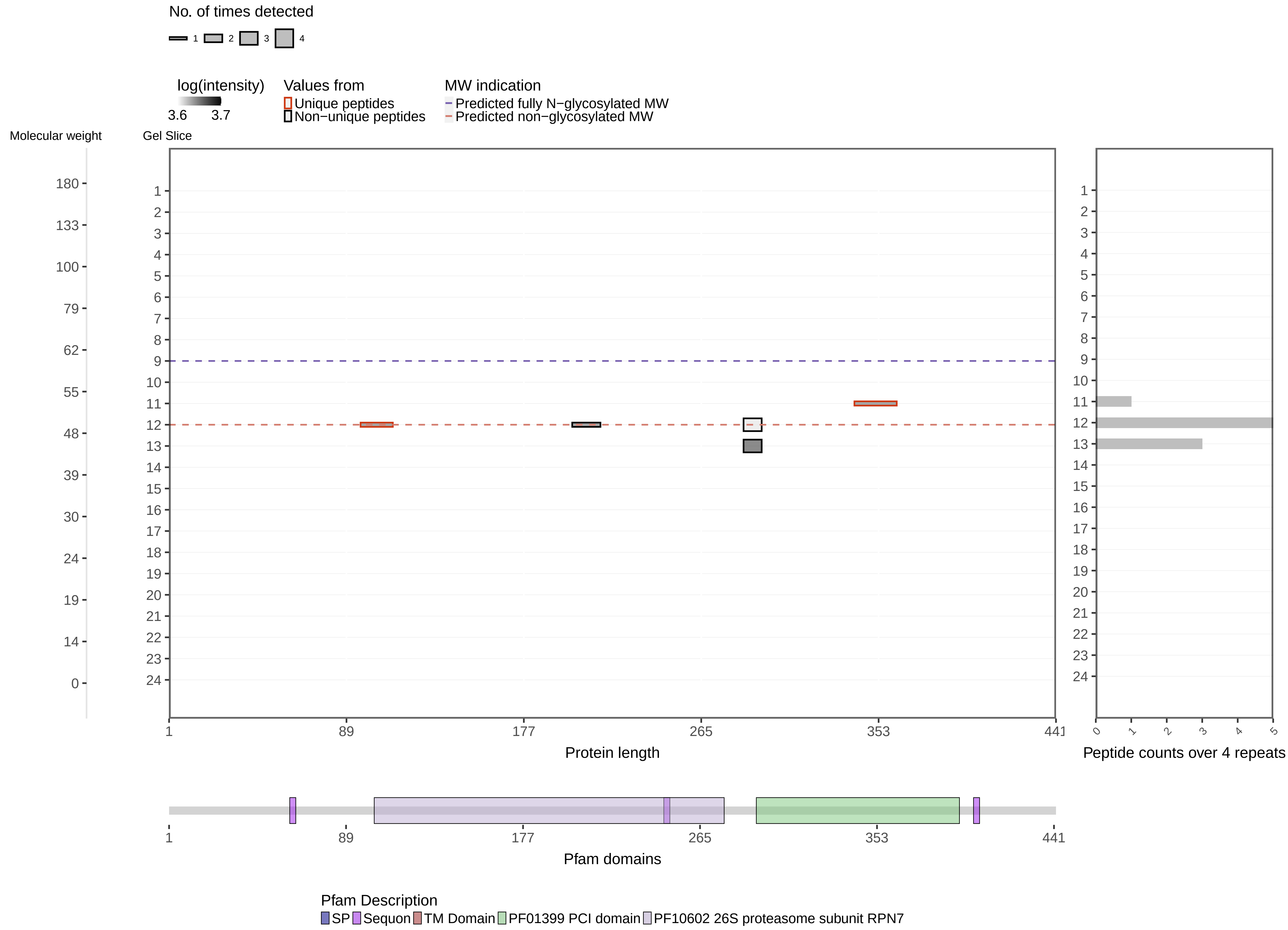


Protein sequence:

NbL01g05840.1 uncharacterized_LOC107827105 [MW = 19.53 kDa]

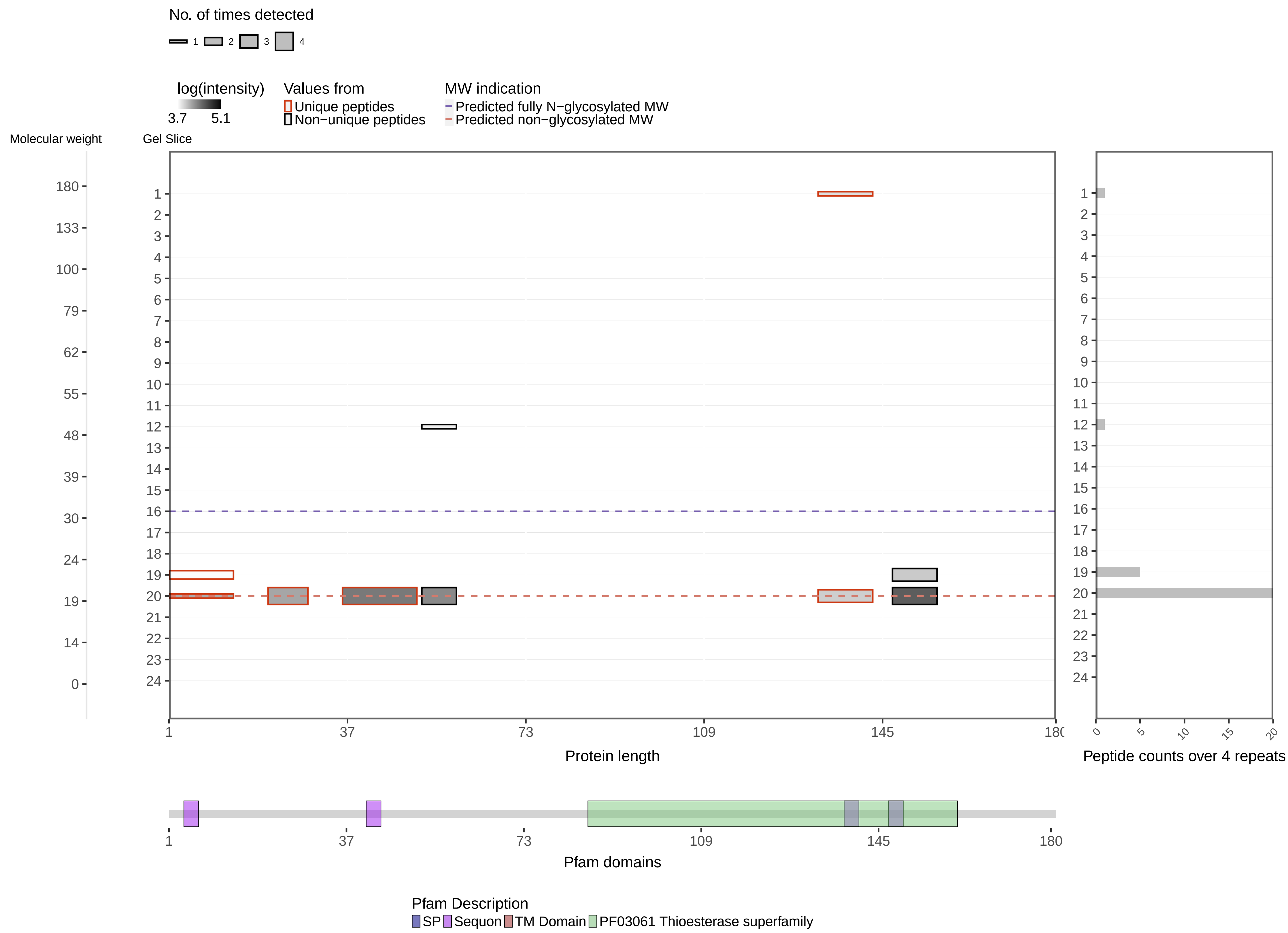


Protein sequence:

.SVKPPIMNIYGSLHGGAVGDVTVRIATACARTIVGKDKELFLGELSISYLSGASRNAEVIVNATVIRSGRNLTVVAVDFRVKDSEKLSYISHATFYHMPVASL*

### NbL01g05810.1 fasciclin−like_arabinogalactan_protein_8 [MW − SP = 41.49 kDa]

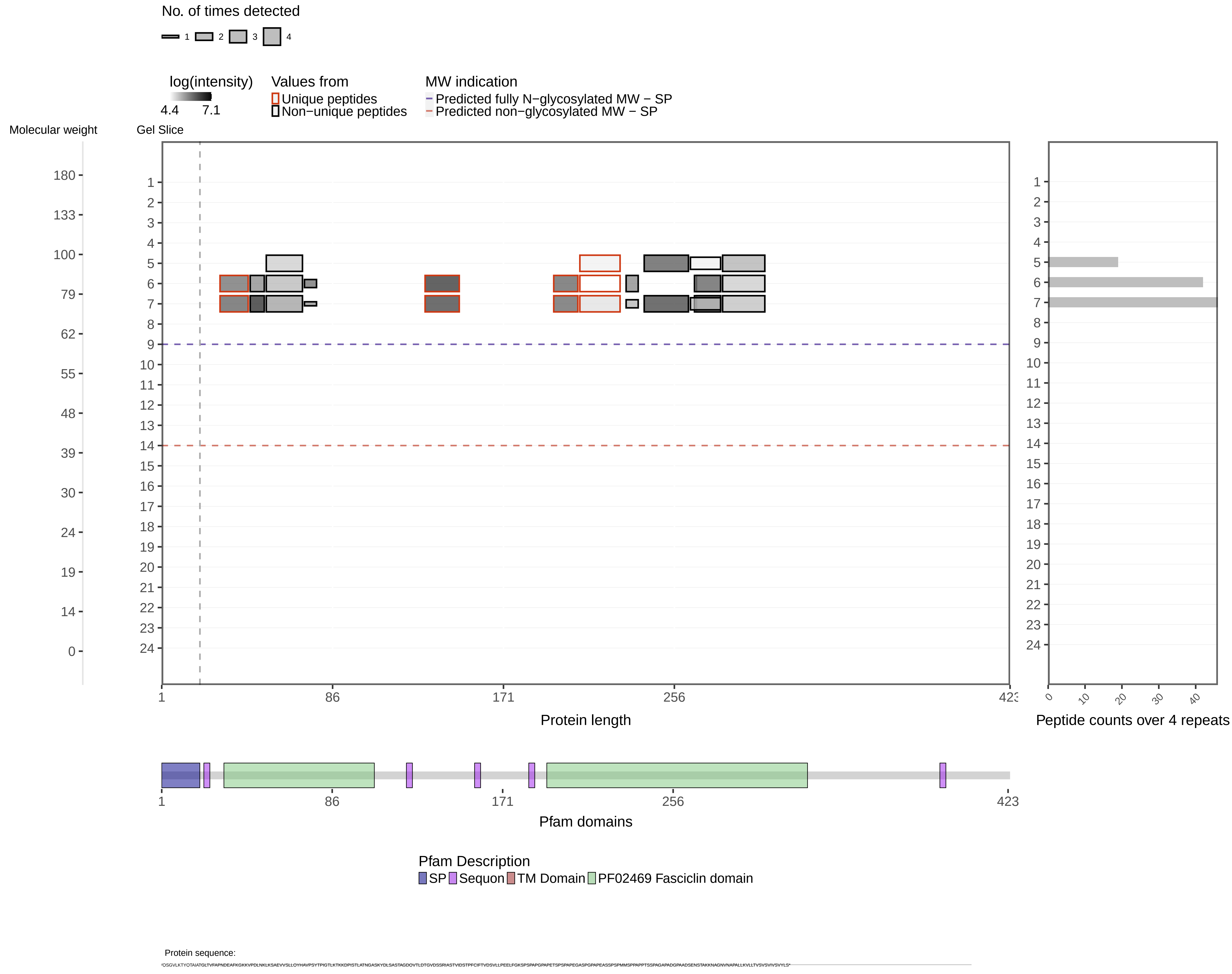

### NbL01g05620.1 uncharacterized_LOC107773275 [MW = 25.32 kDa]

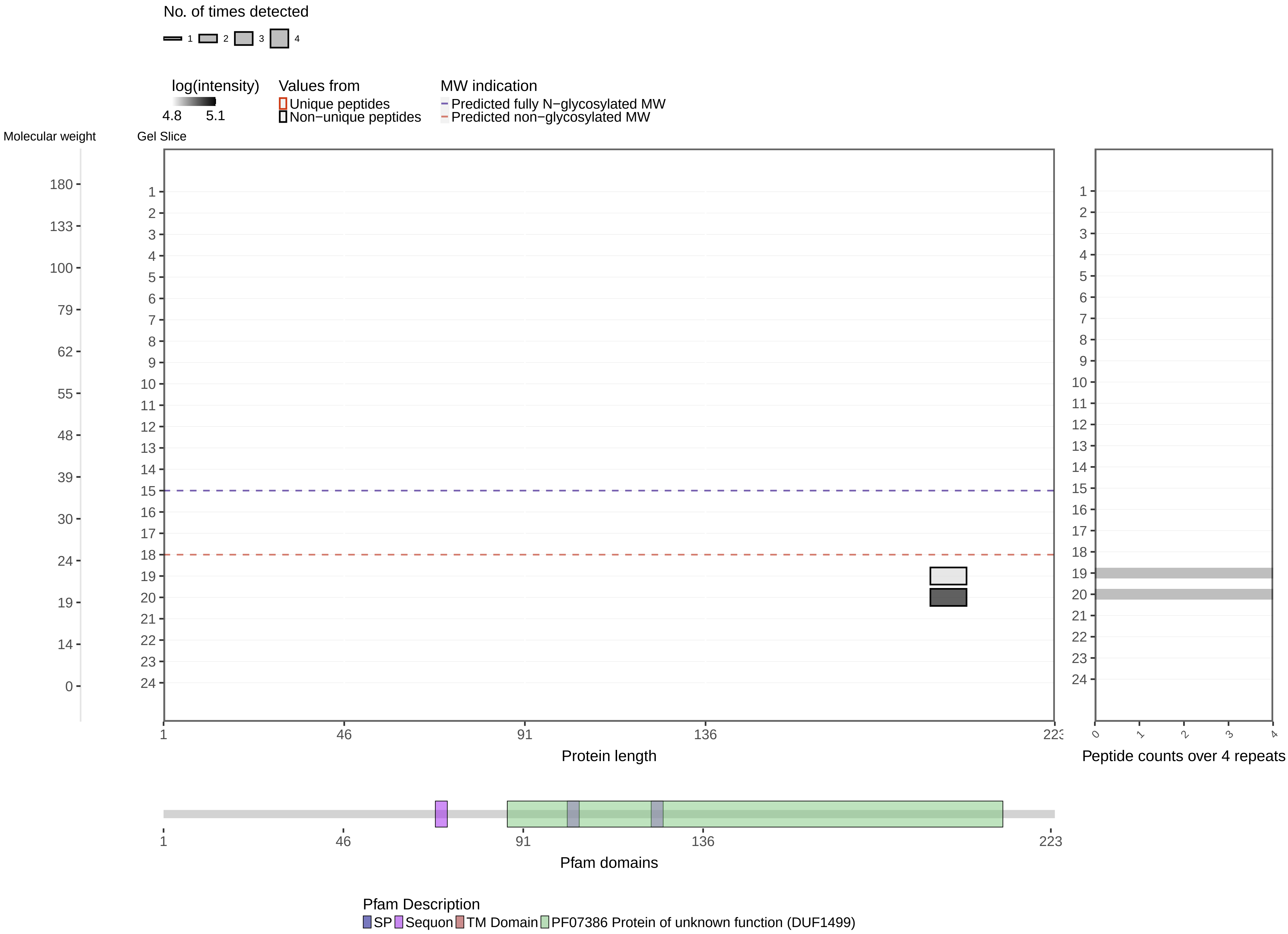


Protein sequence:

TSENISDSAHYAPPWNYNPEGKRGNVSREKAMEELLQVLKSTKPDKFSPKIAEMKDDYVRVEYESPILGFVDDMEFWFPPGKKSIVQYRSASRLGNYDFDANRKRIKALRLQLEKKGWASEDTV*

### NbL01g05470.1 tRNA_2'−phosphotransferase_1−like [MW = 33.61 kDa]

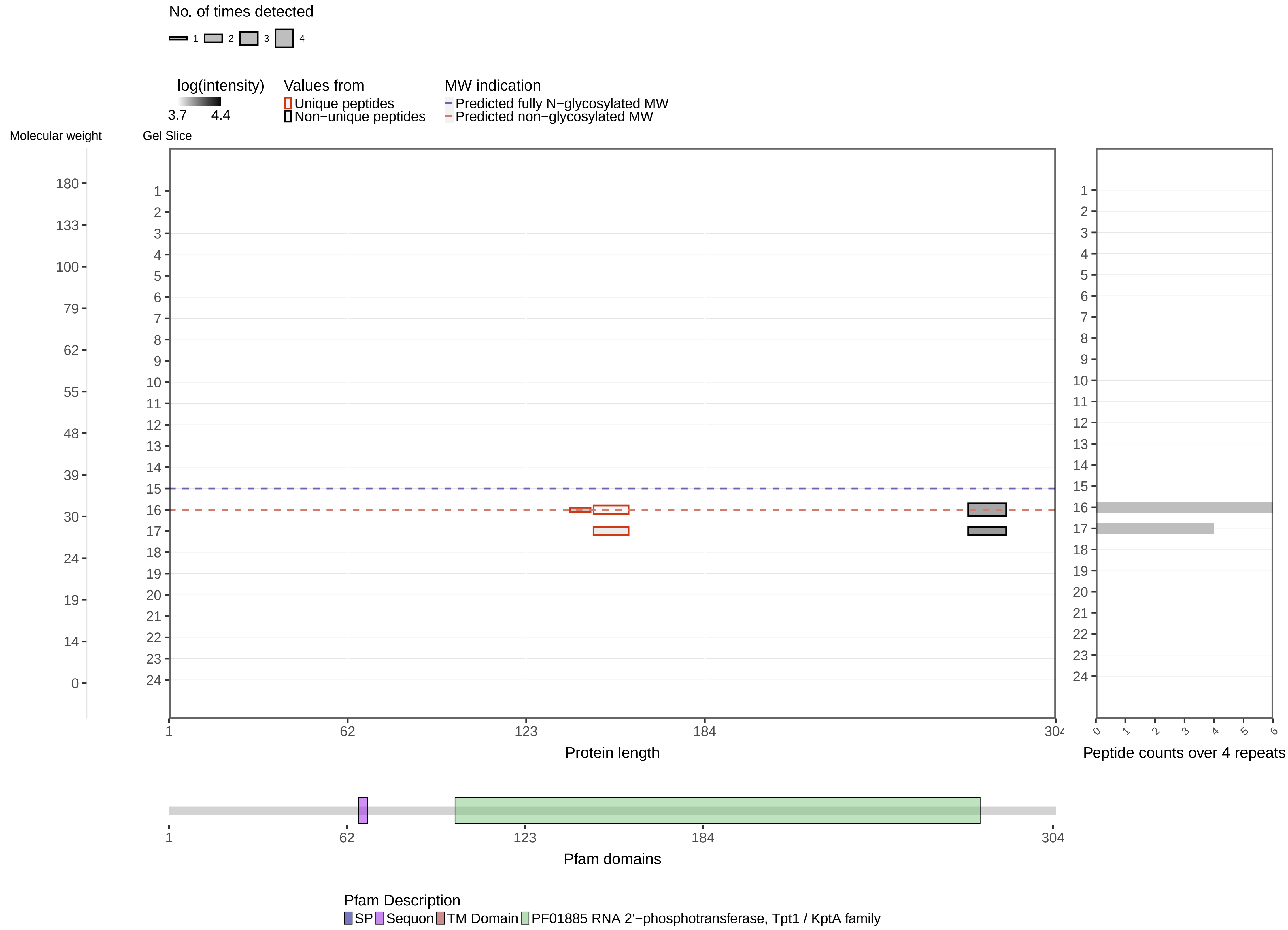


Protein sequence:

TFANVPLRSHTVDDVKEAVQKDNKQRFGLLEENGELLIRANQGHTVKIVETESLLKPILSADEVPVCVHGTYKKNLESILEHGLKRMKRLHVHFSCGLPTDGEVISGMRRDVNVLIFLDVRRALEDGMKLYISDNRVILTEGFDGAVPVKYFEKVESWPDRKPLPL*

### NbL01g05450.1 3−phosphoshikimate_1−carboxyvinyltransferase_1_chloroplastic [MW = 55.79 kDa]

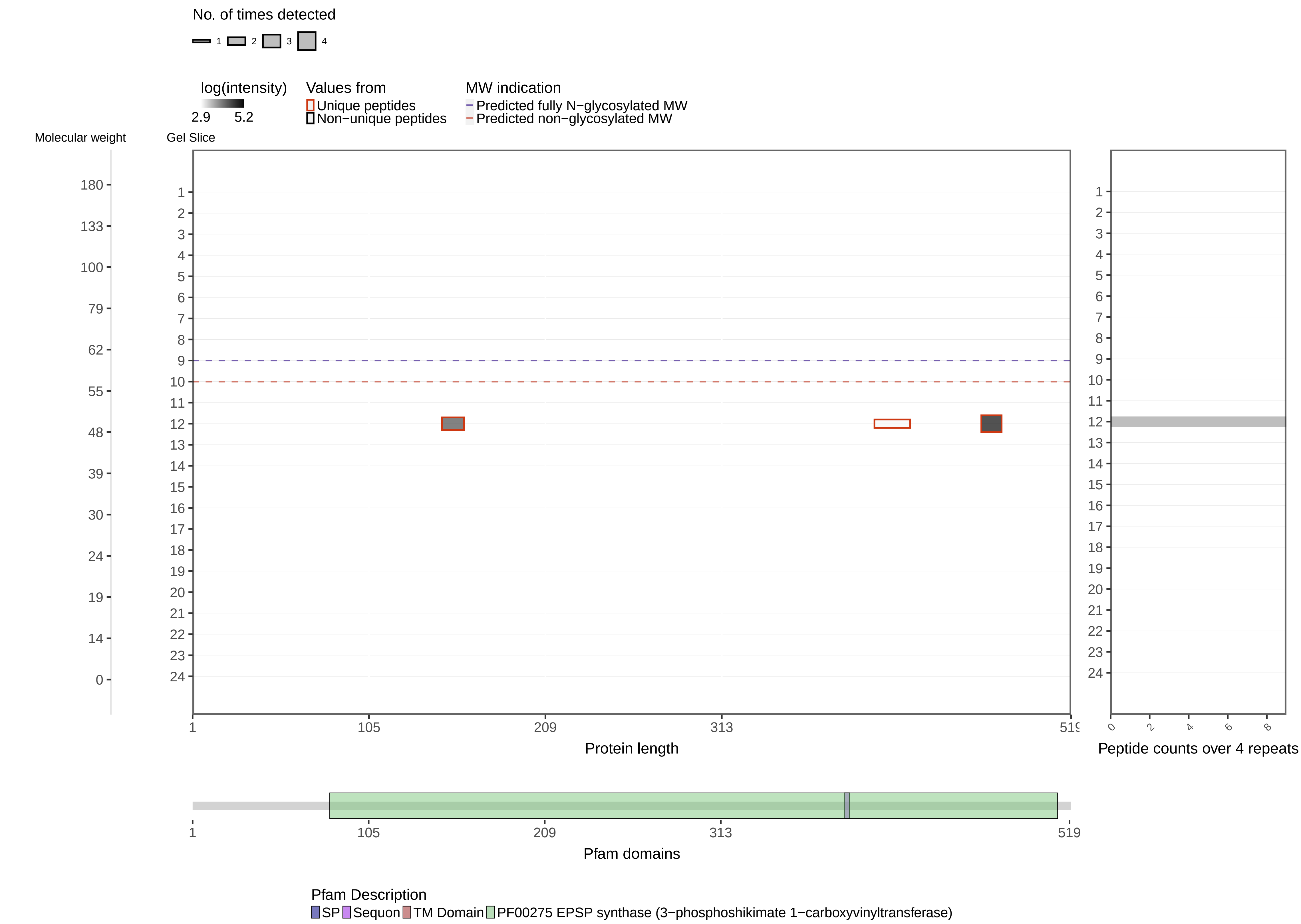


Protein sequence:

SGSISSQYLTALLMAAPLALGDVEIEIIDKLISVPYVEMTLKLMERFGVSVEHTSSWDKFLVRGGQKYKSPGKAYVEGDASSASYFLAGAAVTGGTVTVEGCGTSSLQGDVKFAEVLKKMGAEVTWTENSVTVKGPPRNSSGMKHLRAVDVNMNKMPDVAMTLAVVALFADGPTAIRDVASWRVKETERMIAICTELRKLGATVVEGSDYCIITPPEKLSVTEIDTYDDHRMAMAFSLAACADVPVTIKDPGCTRKTFPNYFDVLQQYSKH*

### NbL01g05350.1 methionine_aminopeptidase_1A−like [MW = 43.49 kDa]

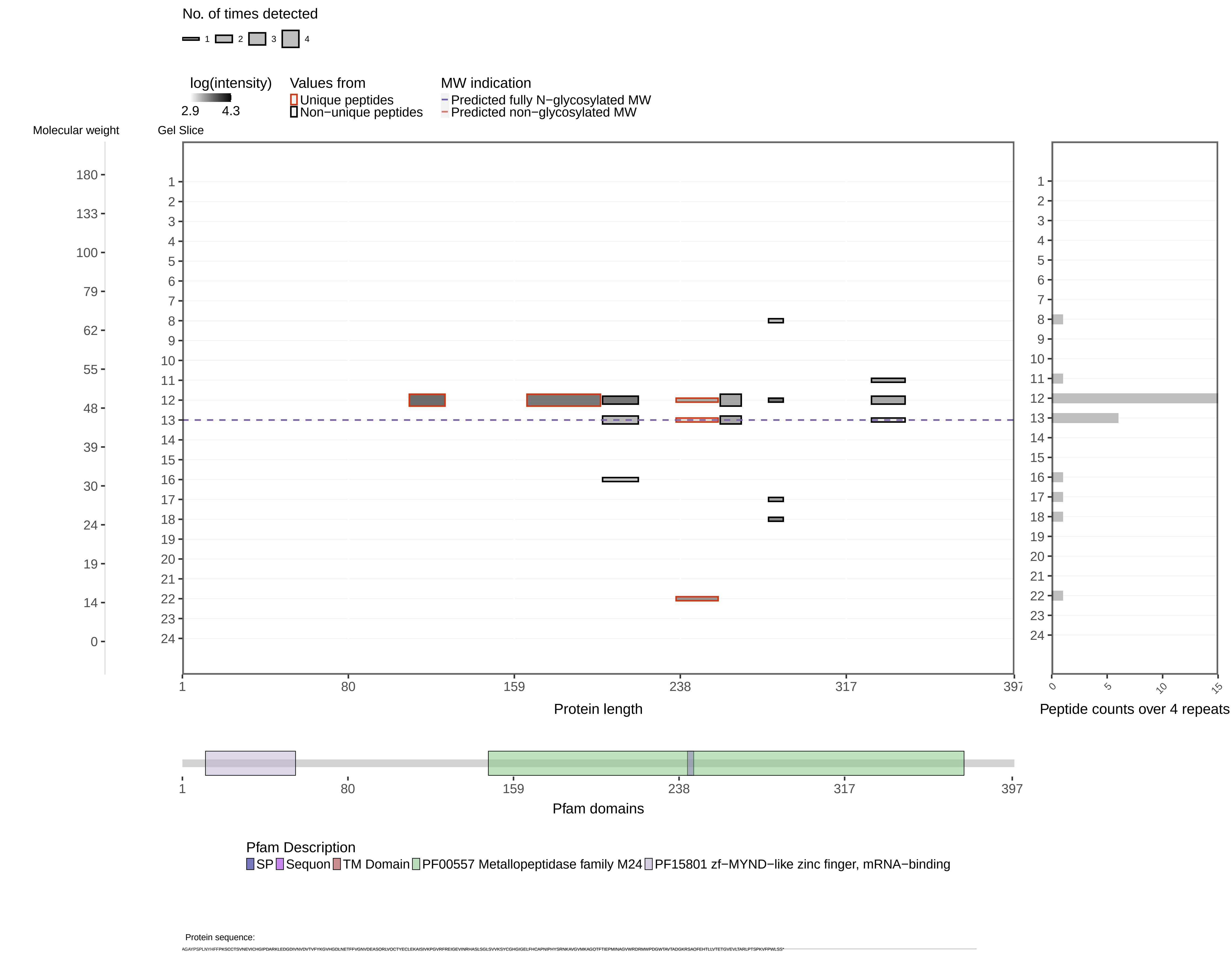

NbL01g05070.1 uncharacterized_LOC104217986 [MW = 27.44 kDa]

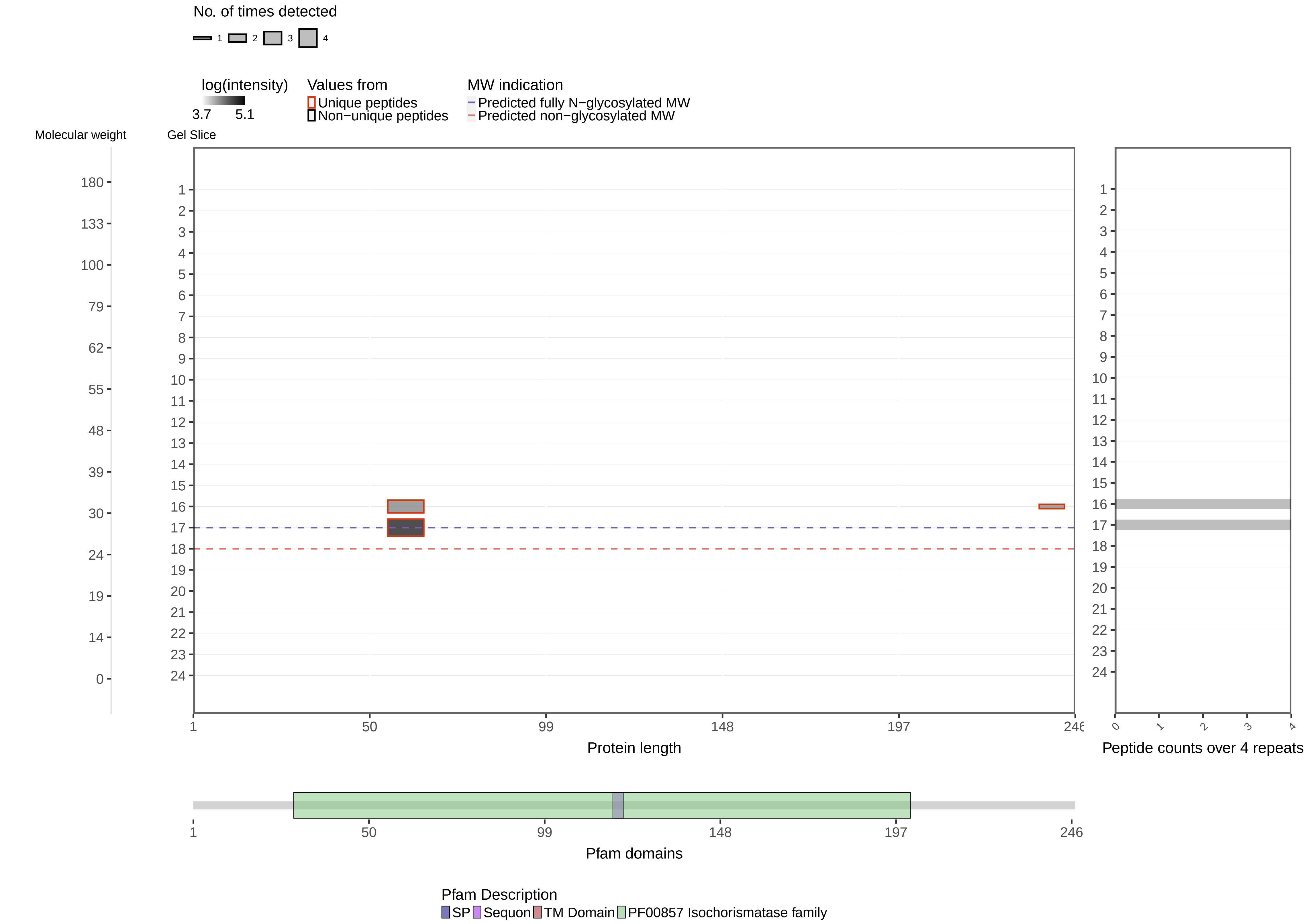


Protein sequence:

/LEKEQNVTIRRKDCYDGYIGSFQEDGSNVFVDWVKNNKIQTLLVVGACTDICVLDFVCSTLSAKNRGFLGPLEEVVVYSQGCATFDFPASVARNTKDISPHPQELMHHVGLYMAKERGAKIAREVSFNSLKKP*

### NbL01g05060.1 fructose−16−bisphosphatase_class_1−like [MW = 44.81 kDa]

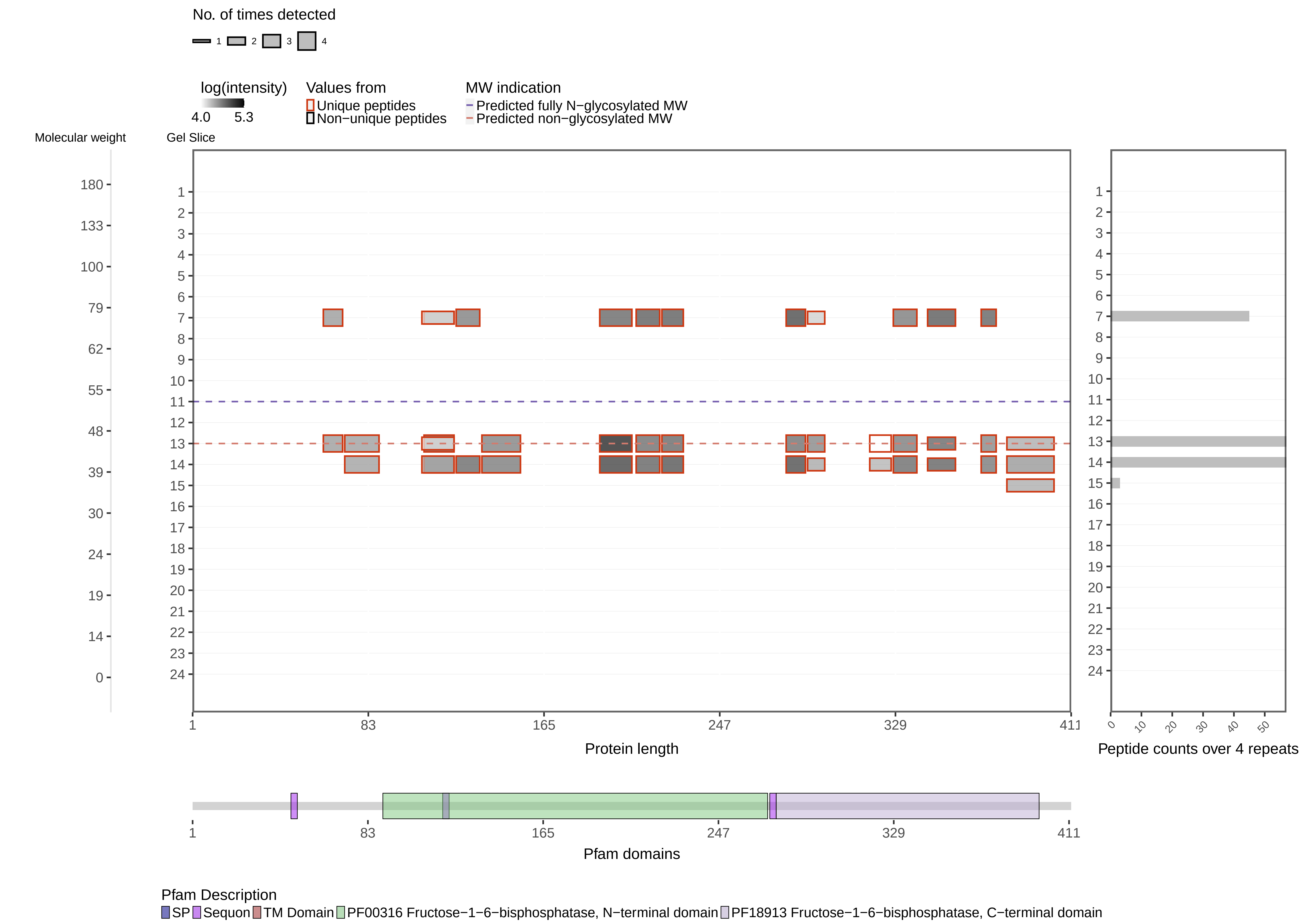


Protein sequence:

NIDASIPTGTIFGIYKRLVEIDHLPVEEKASLNCLOSGAKLIAAAYVLYSSATILCASFGSGTHAFTLDHTTGDFLLTHPNISIPPRGOIYSVNDARYFDWPEGLROYIDTIROGKGKHPKKYSARYICSLVADFHRTLLYGGVAMNPRDHLRLVYEANPLSFLAEKAGGRGSDGKNRILSIOPVKLHORLPLFLGSSDDIEELOSYGDVOOKVNPGYDV*

### NbL01g04990.1 methylmalonate−semialdehyde_dehydrogenase_[acylating]_mitochondrial [MW = 56.83 kDa]

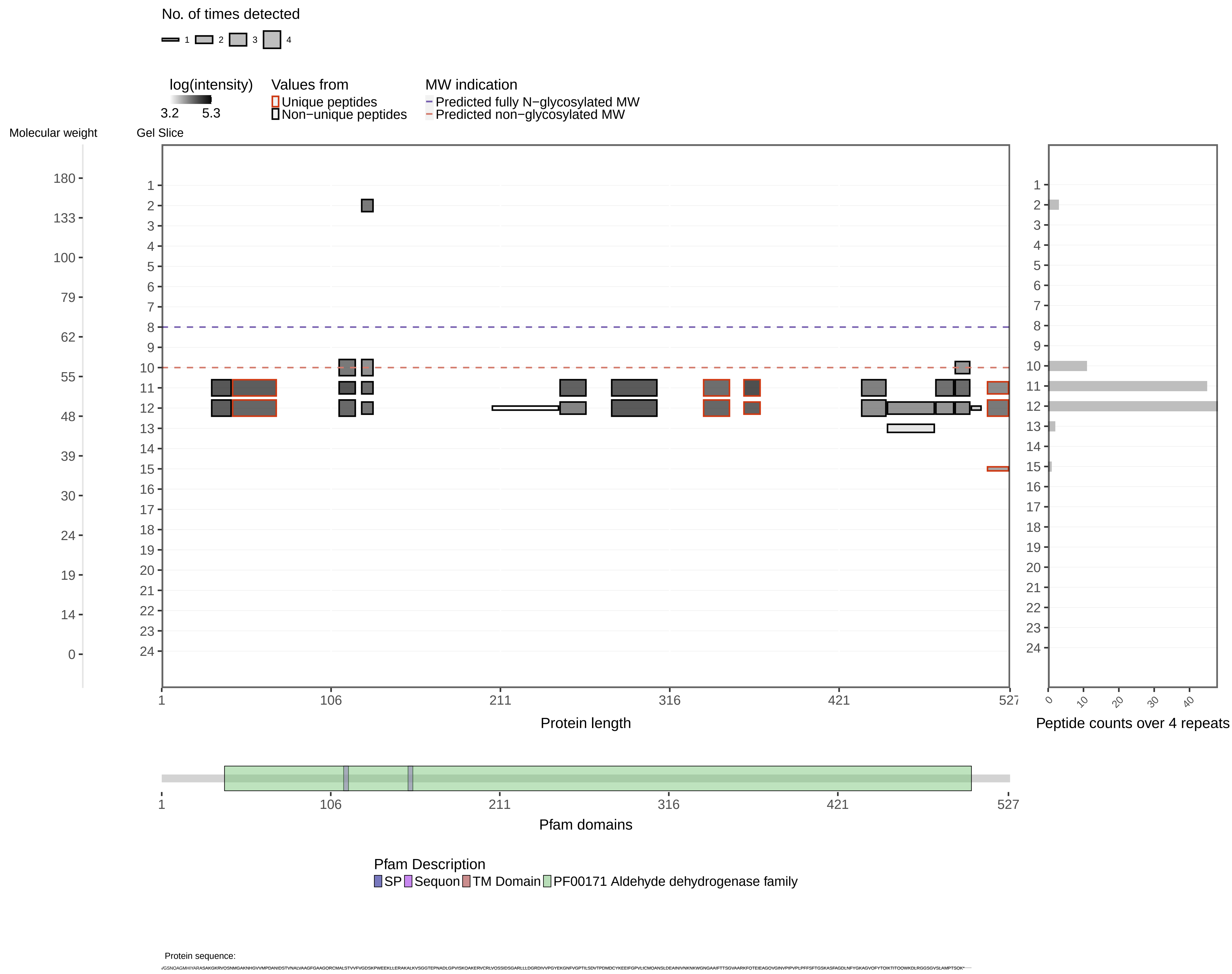

NbL01g04970.1 psbP_domain−containing_protein_1_chloroplastic−like [MW = 38.28 kDa]

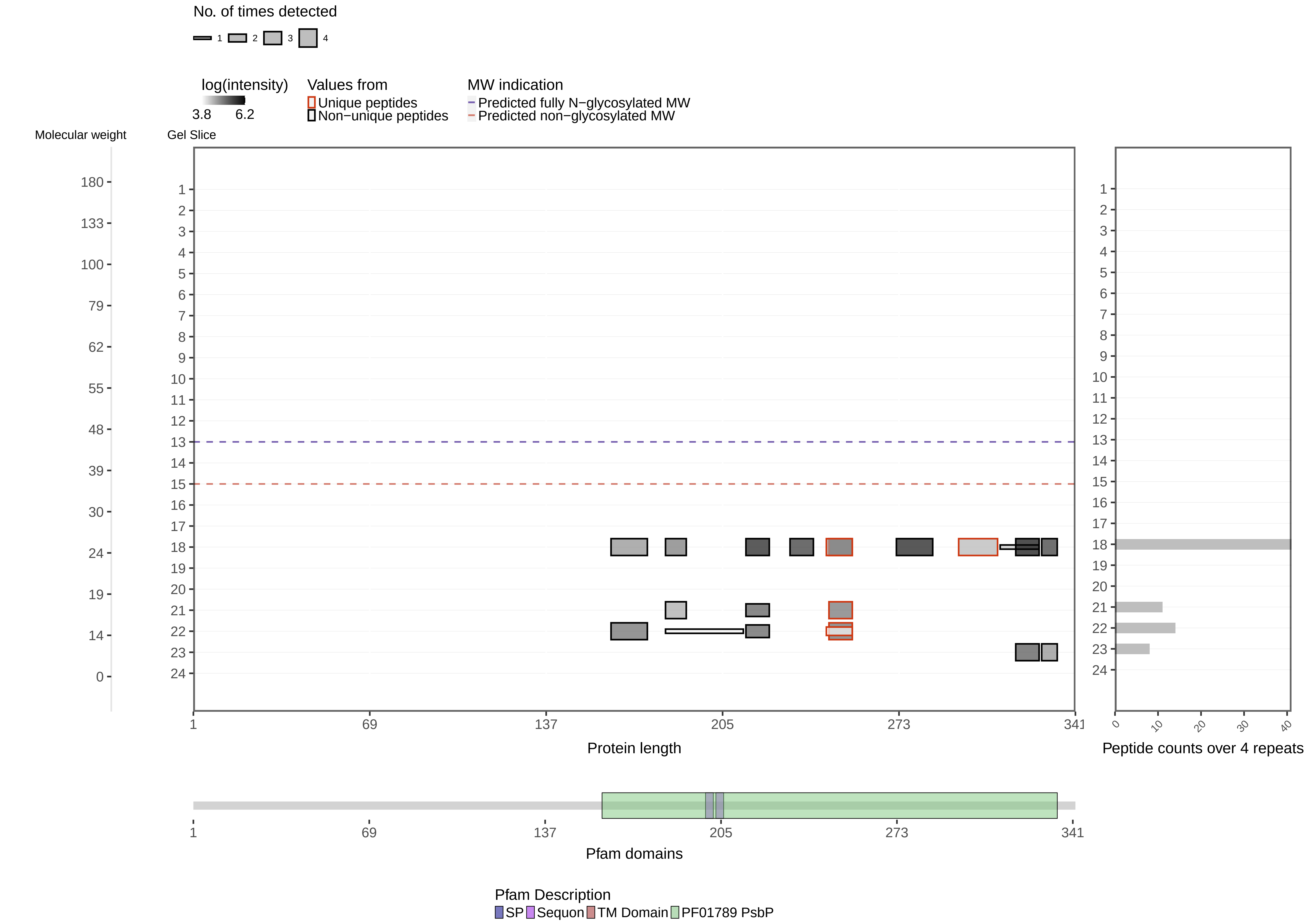


Protein sequence:

EFREYIDTFDGYSFMYPRNWIQVRGAGADIFFRDPLVLDENLSMNISSPSSSAYKSVEDLGPPEKAGKGVLRQYLTEFMSTRLGVRRESSVLSASSRVADDGKLYYEVEVNIKSYANNNELAVMPQDRVARLEWDRRYLSVLGVENNCLYELRIQTPEKTFADEESDIRAIMDSFRVNKIEV*

### NbL01g04860.1 glutamate−−tRNA_ligase_chloroplastic/mitochondrial [MW = 63.37 kDa]

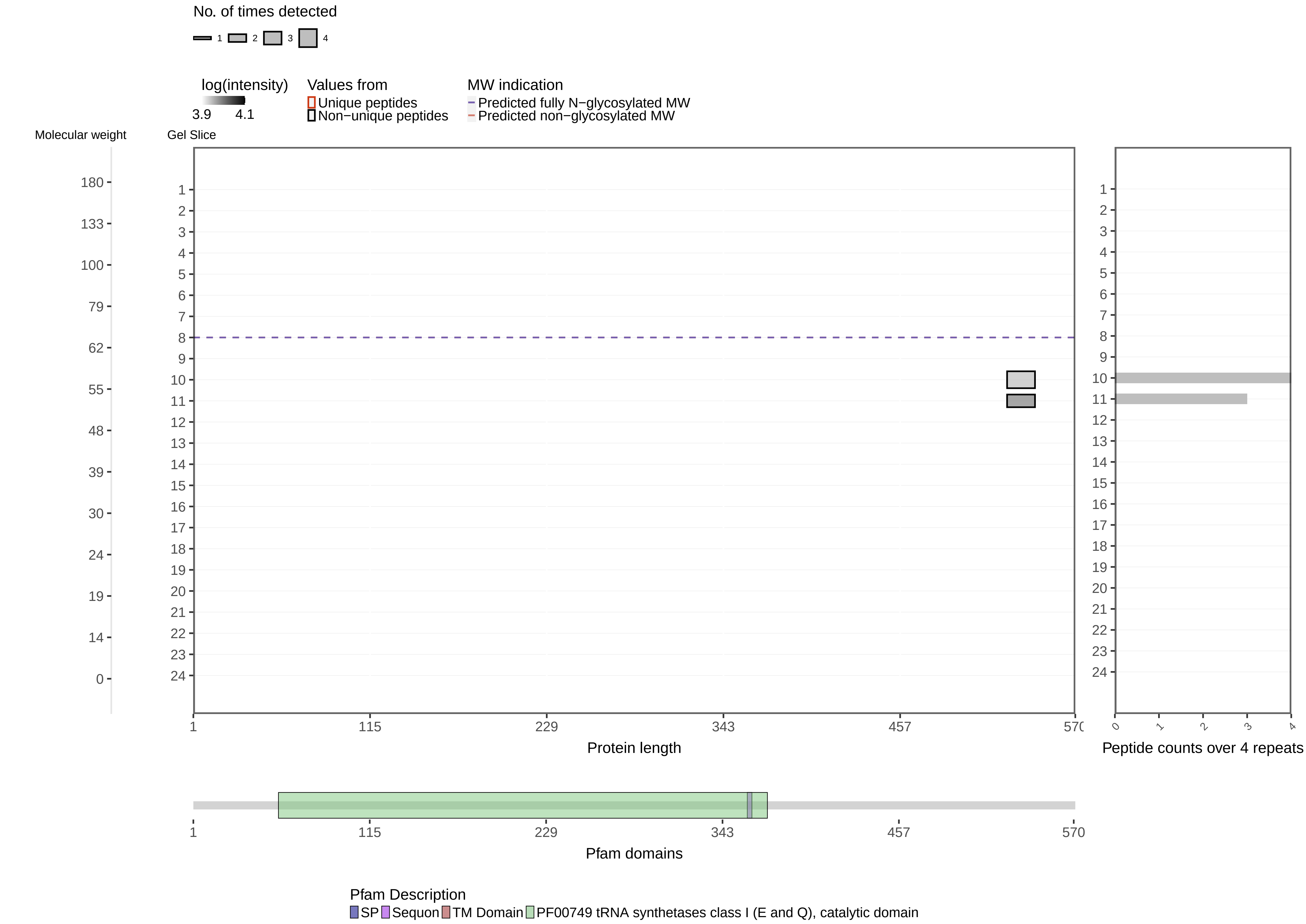


Protein sequence:

.PNTLRQALIYKALGFPMPHFAHVSLILAPDRSKLSKRHGATSVGQFRDMGYLPQAMVNYLALLGWGDGTENEFFTLEQLVEKFTIERVNKSGAIFDSTKLRWMNGQHLRSLPSEELNRIIGERWKGAGITTESQGIFIQDAVLLLKDGIDLITDSEKALSSLLSYPLYETLASAEGKPILEDGVSEVAKSLLAAYDSGELSGALAEGQPGWQKWVKNFGKLLKRKGKSLFMPLRVLLTGKLHGPDIGATTVLLYKAGTSGSVVPQAGFVTFDERLKILREVQWESFSTDVPLSA

### NbL01g04850.1 aminoacylase−1 [MW = 52.27 kDa]

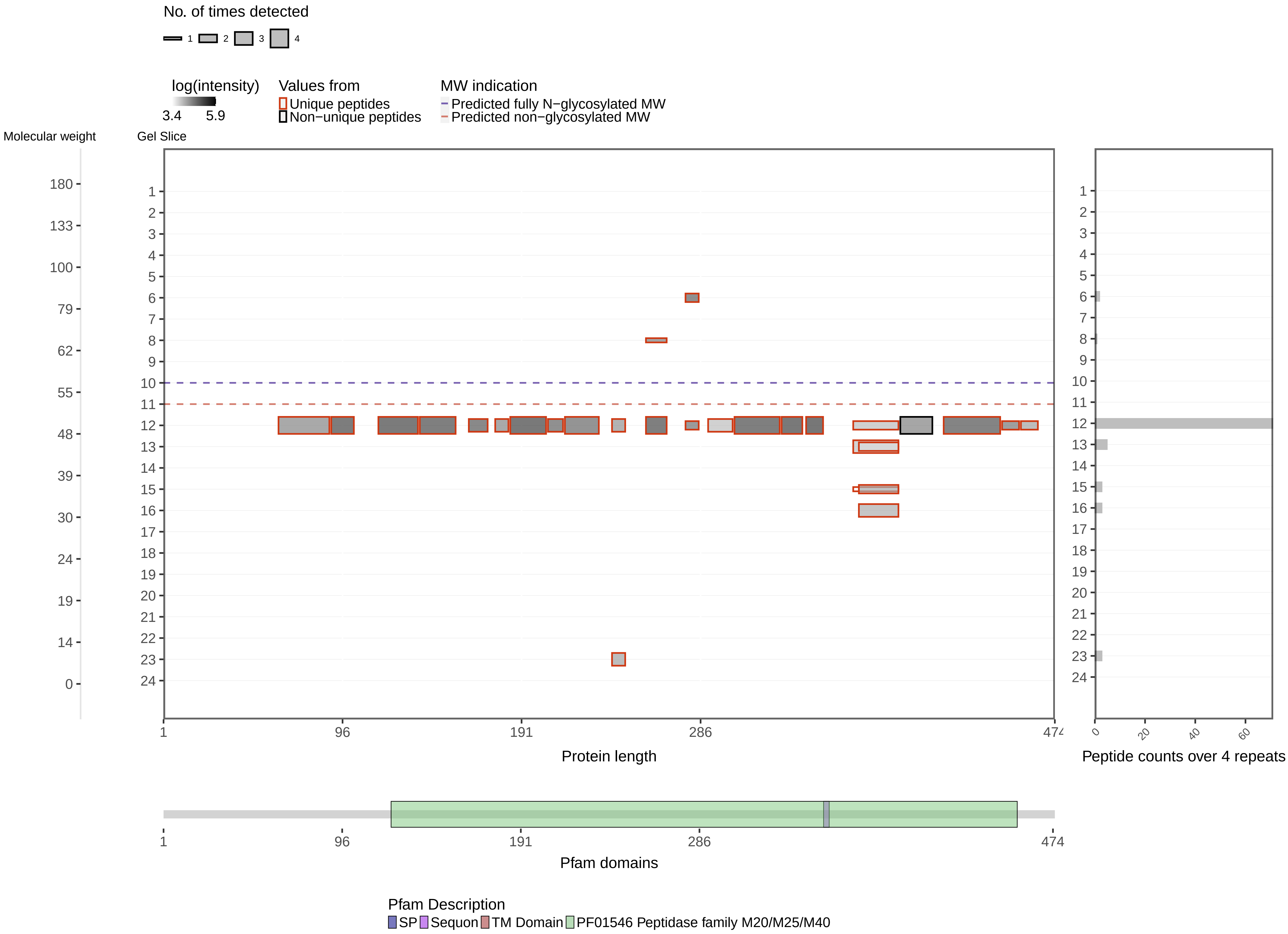


Protein sequence:

PSPTENYRAFYGERSPWWLVVKAVGAPGHGAKLYDNSAMENLLKSVETIRRFRAAQFDLVKAGLKAEGEVISVNMVFLKAGTPSPSGFVMNLQPSEAQAGFDIRVPPTADQASLEKRIAEEWAPASRNMTFEFKQKVSVNDKFGRPAITAVDSSNIWWALLEESIVKANAGVGKPEIFPASTDARYFRERGLPAIGFSPMANTPILLHDHNEFLNKDEYLKGIDVYGSIIKTYASYIEHPRDEASREEL*

NbL01g04840.1 uncharacterized_protein_At2g17340−like [MW = 40.69 kDa]

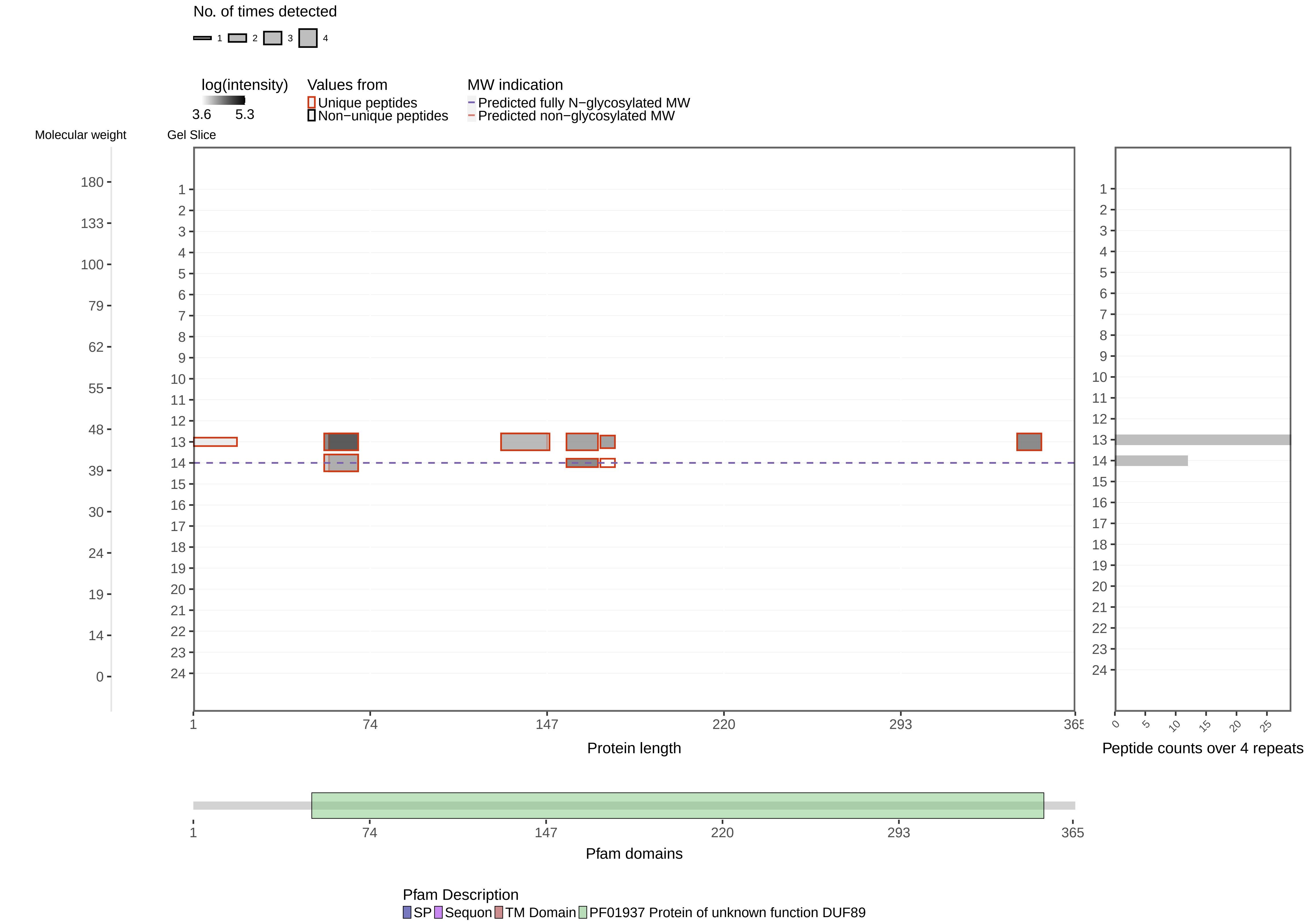


Protein sequence:

AKLAELFSRDGISFLASCENLVPRPWVIDDMDTFITKLSRKSWKKAVIFVDNSGADIILGILPFARELLRHGTQIILAANDLPSINDVTYHELVEIISKLKDEHGKLVGVDTSNLLVANSGNDLPVIDLMGISPELAYLASDADLVILEGMGRGIETNLYAQFKCDSLKIGMVKHPEVAEFLGGRLYDCVFKYNEIME*

### NbL01g04800.1 homoserine_kinase−like [MW = 38.42 kDa]

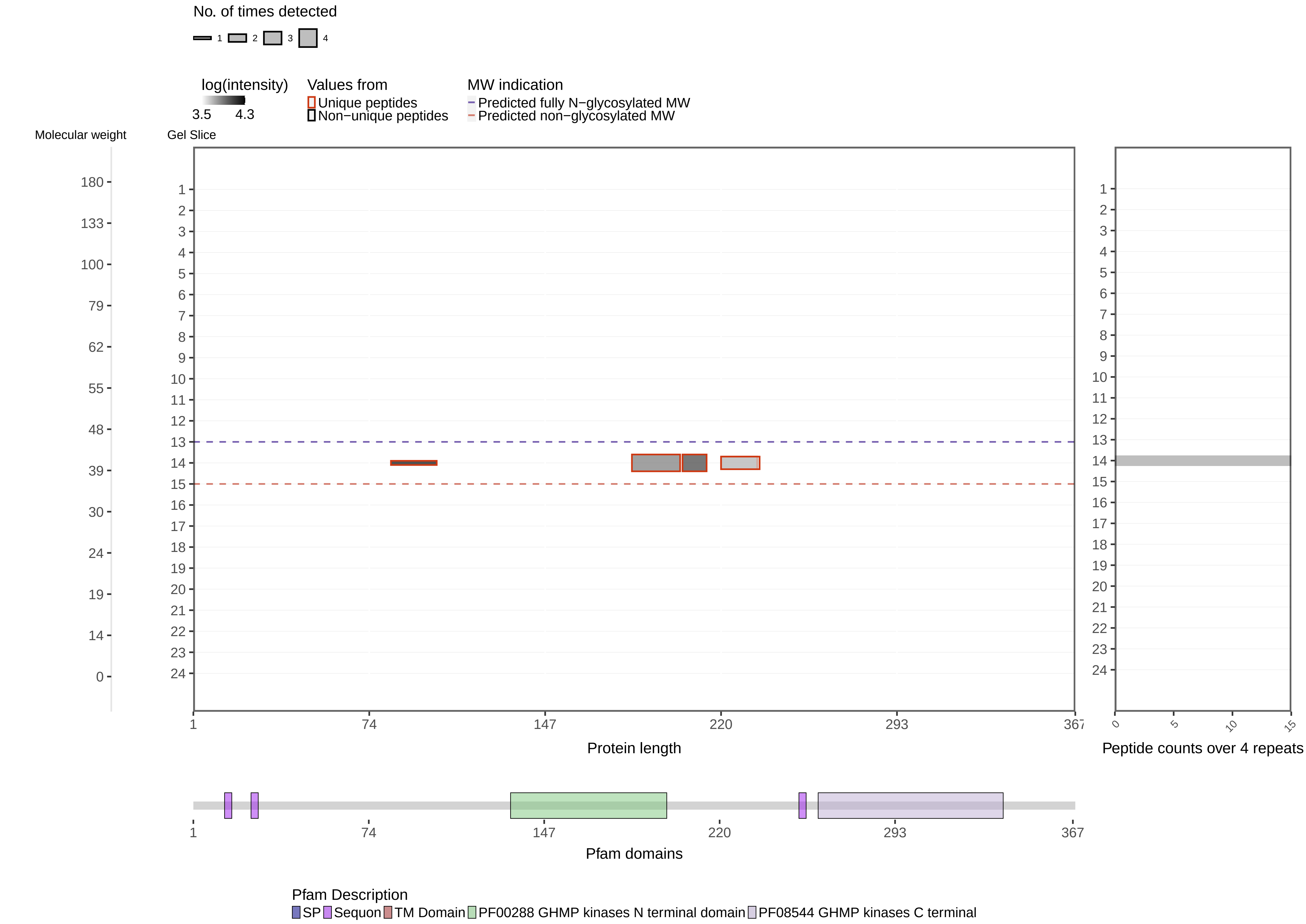


Protein sequence:

ILVLAGLESESKVSGYHADNVAPAIMGGFVLIRSYDPLELIQLKFPHEKELFFVLVNPEFEAPTKKMRAALPQQITMSHHVWNCSQAAALVAAVLQGDLSVLGKALSSDKIVEPRRAPLIPGMEGVKKVAMEAGAFGCTISGAGPTAVAVTDNEEKGREIGQRMVEAFLKEGNLKSSAMVKKLDRVGARLVSSDPR*

NbL01g04700.1 metal−independent_phosphoserine_phosphatase [MW = 25.64 kDa]

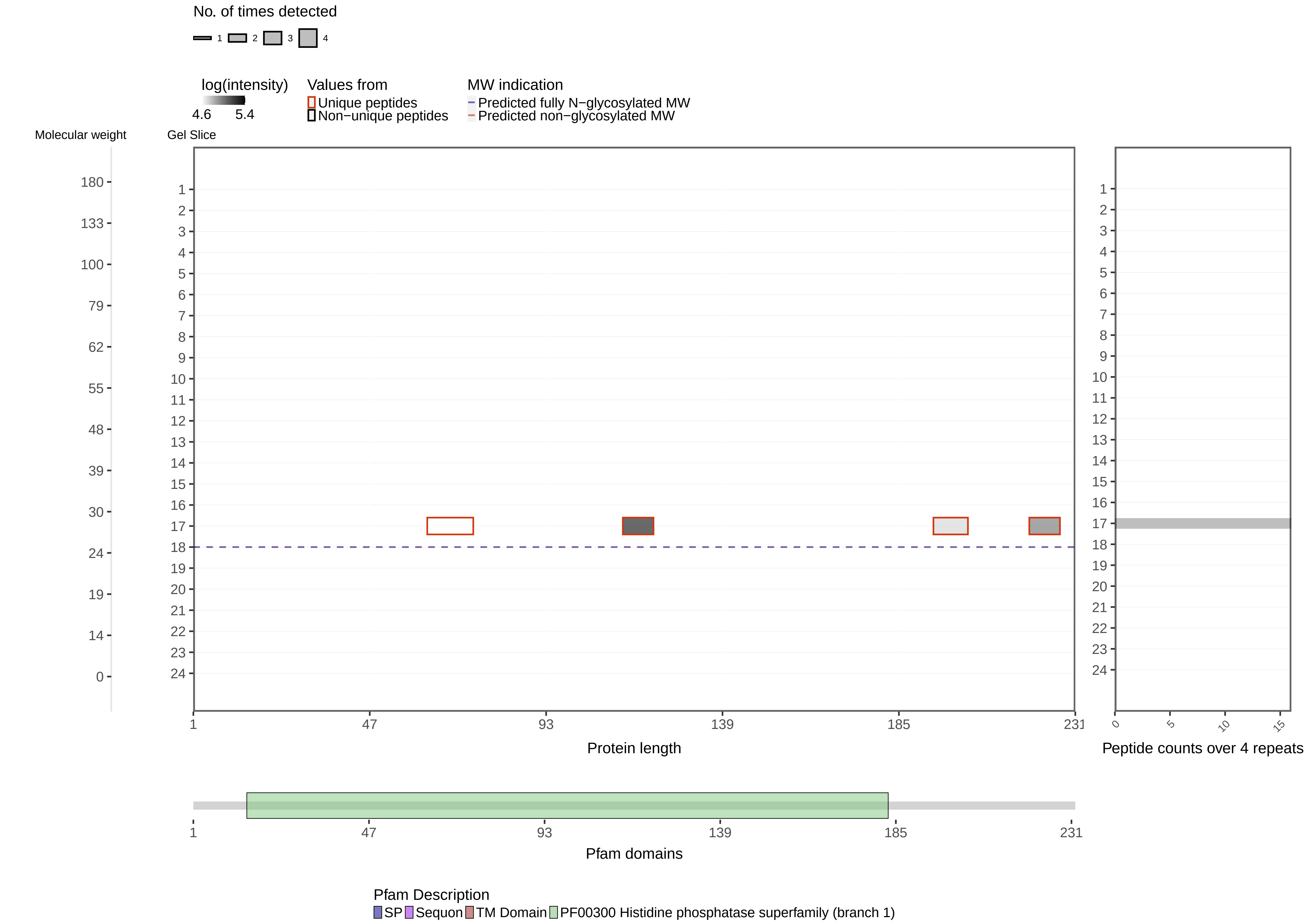


Protein sequence:

CMVLEDLRERFFGPSYELKSHDKYPDIWALDEKDPFMQPDGGESVADVVSRLTSALINMESQFQGCTVLVVSHGDPLQIFQTVLNAAKDYKGCCENDLTSRVQAVKIPCVLSQHRRFALQTGELRSVV*

### NbL01g04690.1 SEC1_family_transport_protein_SLY1−like [MW = 69.54 kDa]

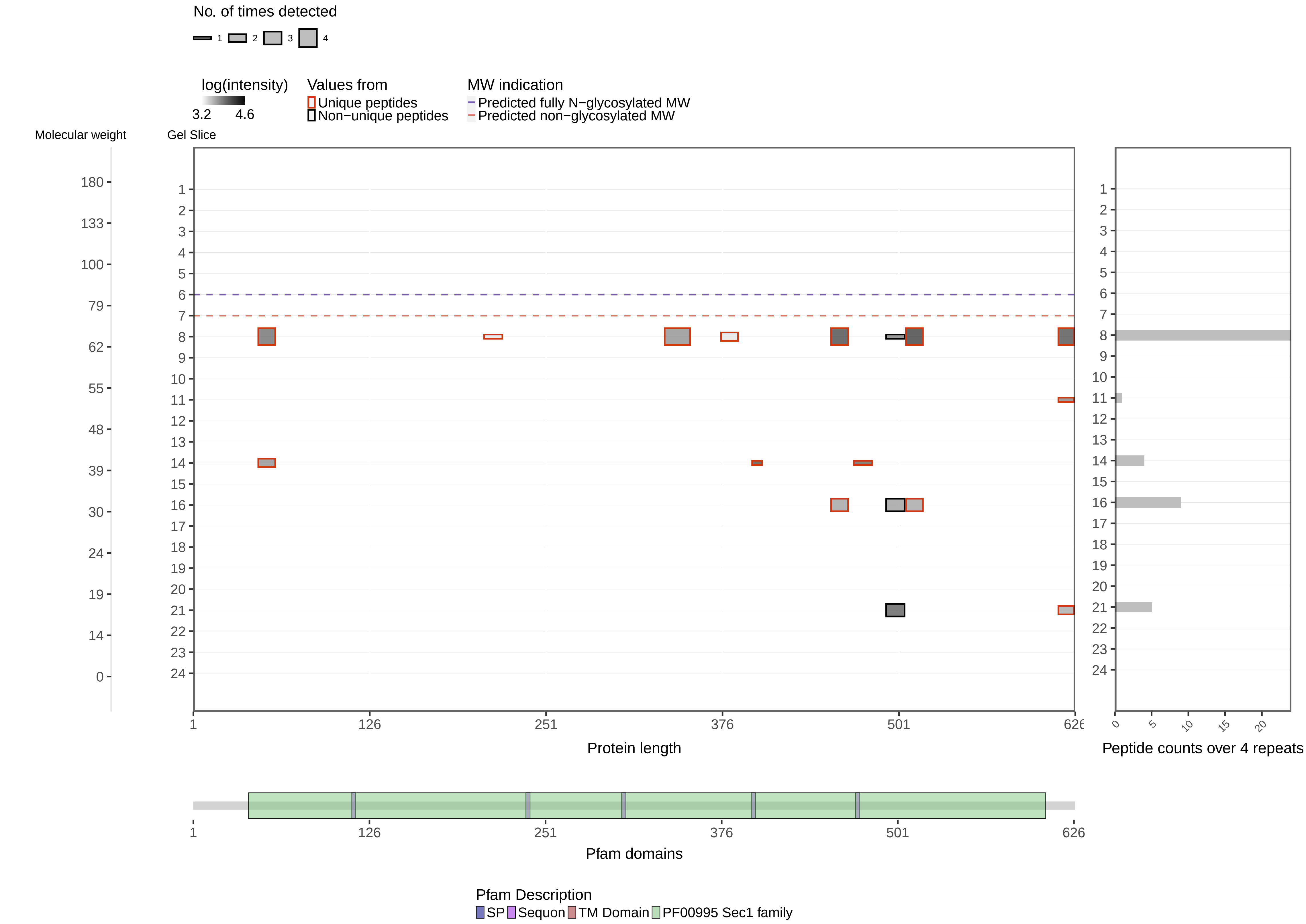


Protein sequence:

[illegible]

NbL01g04650.1 lysM_domain−containing_GPI−anchored_protein_2 [MW − SP = 35.32 kDa]

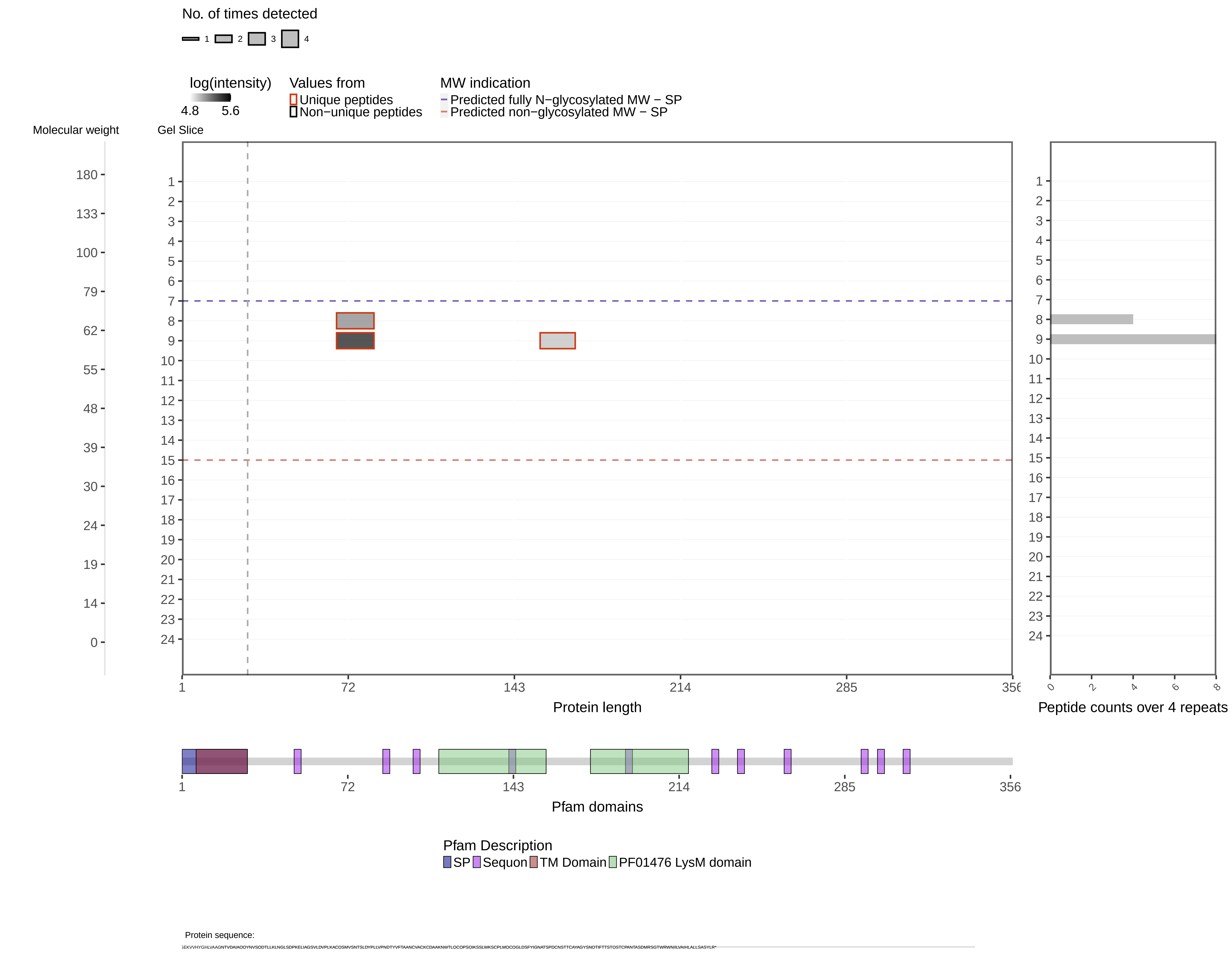

NbL01g04600.1 uncharacterized_protein_ycf39−like [MW = 43.89 kDa]

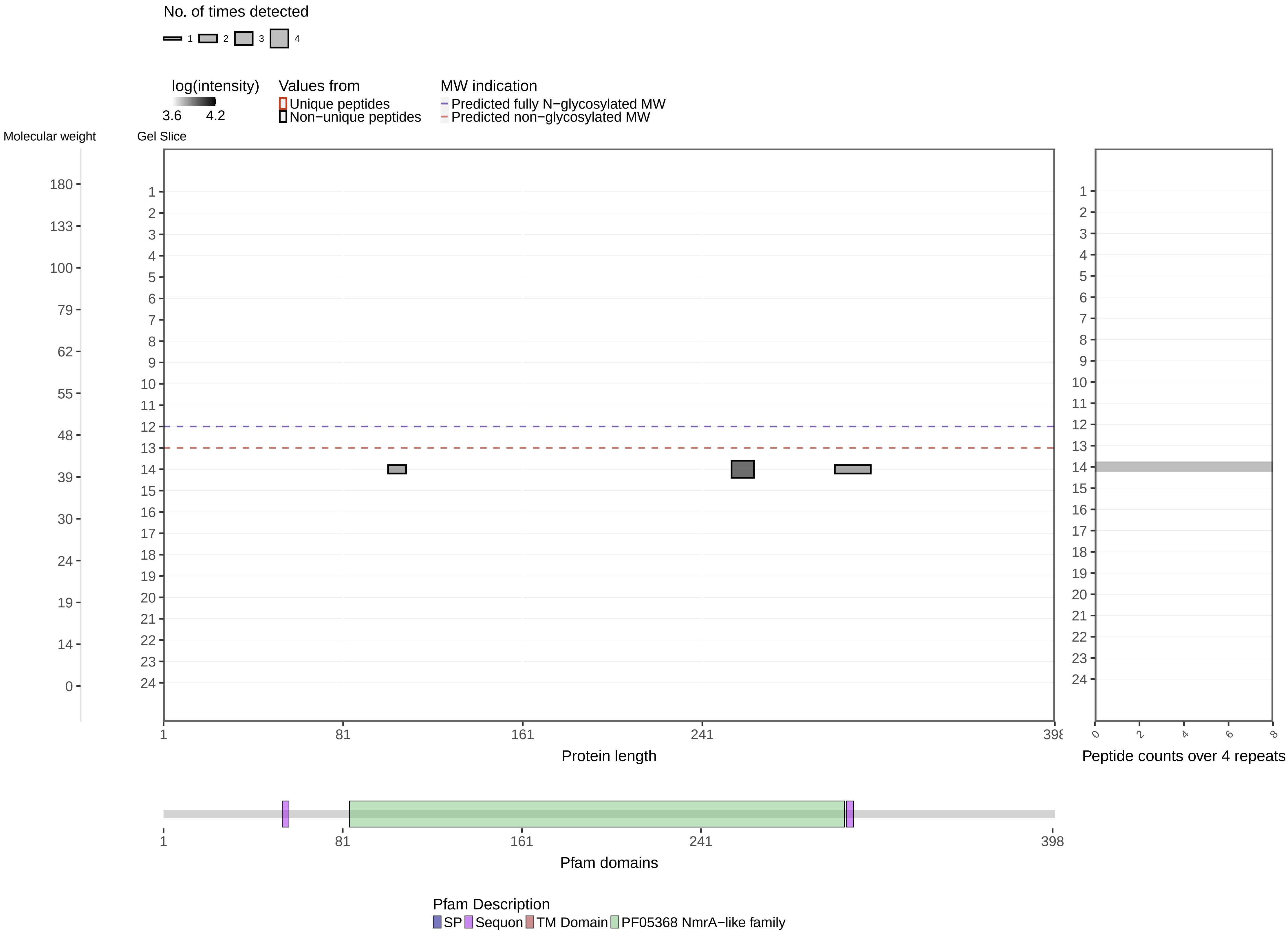


Protein sequence:

FIFFSIHNCDKHPEVPLMEIKYCTEKFLRDSGLNHVVIRLCGFMOGLIGOYAVPILEEKSVWGTDAPTRIAYMDTODIARLTFIALRNENINGKLLTFAGPRAWTTOEVITLCERLAGODANVTTVPVSVLRLTROLTRLFEWTNDVADRLAFSEVLTSDTVFSVPMTETYSLLGVDAKDVSSLEKYLODYFTNILKKLKDLKAOSKOTDIFF*

NbL01g04530.1 alanine−−tRNA_ligase_chloroplastic/mitochondrial [MW = 109.18 kDa]

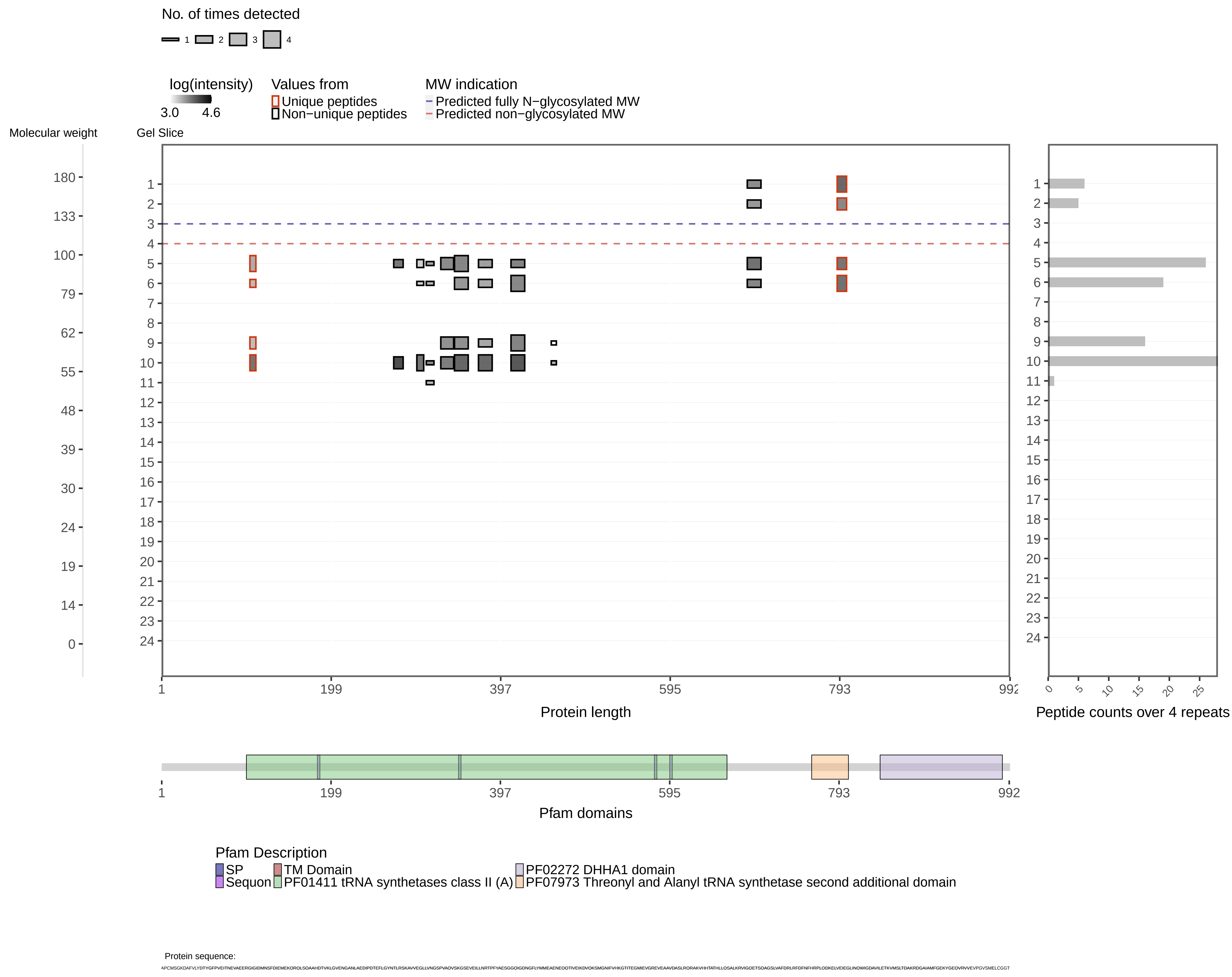

NbL01g04470.1 coiled−coil_domain−containing_protein_124−like [MW = 27.1 kDa]

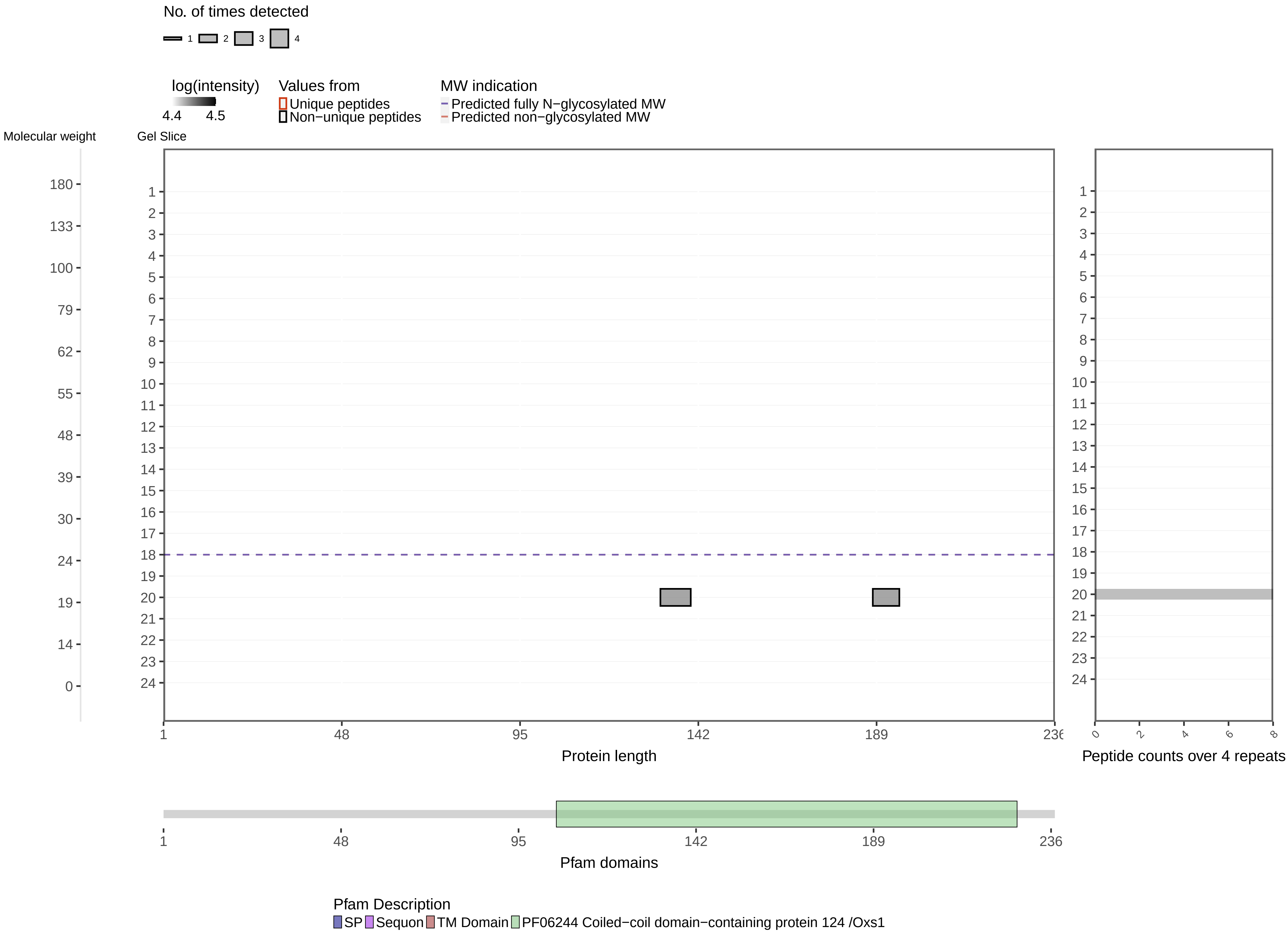


Protein sequence:

TEAELRRRKEEEQVALAKKAEEEKRRQSRMAAEEEYEKTVLVTNRNRDDSIIEASTVDDAIASLAISDSLPPDRHPEKRLRAAFKAFEEAELRVLKEEKPGLTHTQYKDLIWKLWKKSPDNPLNQINLRCTLC*

### NbL01g04130.1 SKP1−like_protein_1A [MW = 17.6 kDa]

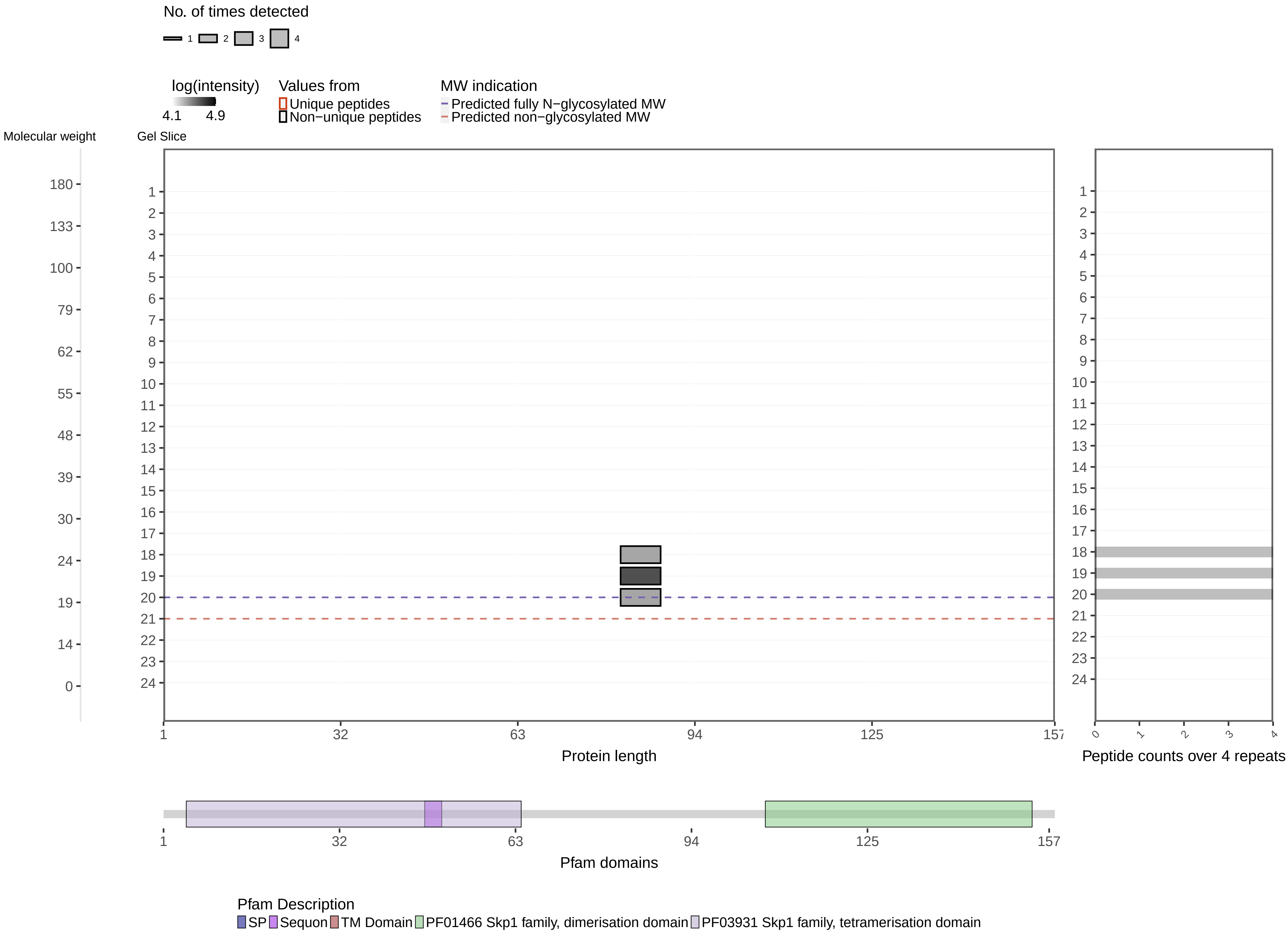


Protein sequence:

AVAKTDDKASEDDLKSFDADFVKVDQATLFDLILAANYLNIKSLLDLTCQTVADMIKGKTPEEIRKTFNIKNDFTPEEEEEVRRENAWAFE*

### NbL01g04120.1 UDP−glycosyltransferase_73C2−like [MW = 54.83 kDa]

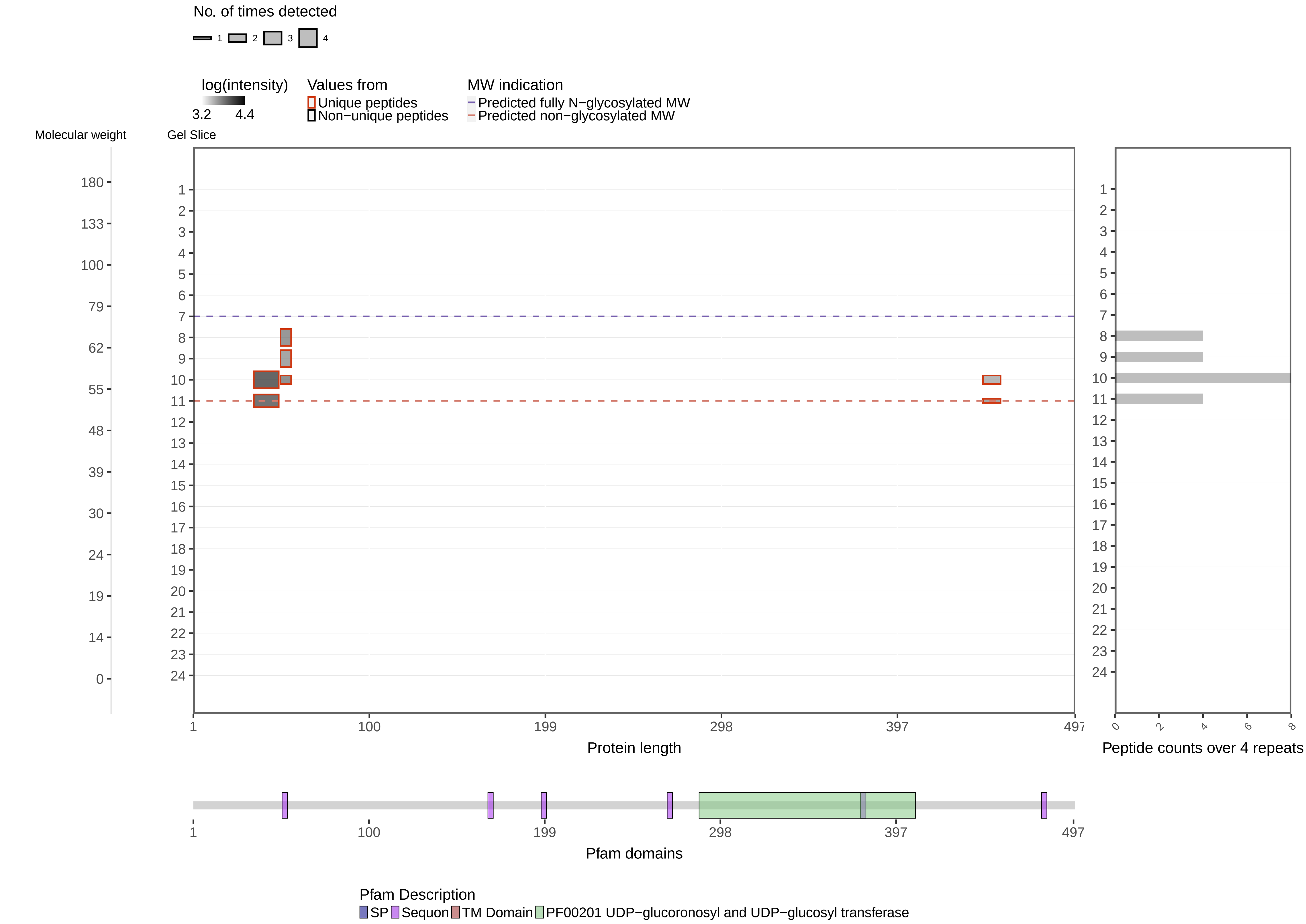


Protein sequence:

KARGKKVWCVGPVSLCNKENEDLVTRGNKTAISNQDCLKWLDGFETESVVYVSLGSLSRLTVLQMVELGLGLEESNRPFVWVLGGSDKLNDLEKWILENGFEERIKERGVLIRGWAPQVLILSHPAIGGVLTHCGWNSTLEGISAGLPMVTWPLFAEQFCNEKLVVQVLKIGVSLGVKVPVKWGDEENVGVLVKKDDVKKALDKLMDEGEEGQVRRTKAKELGELAKKAFGEGGSSYVNLTSLIEDIIEQQNHKDK*

### NbL01g04110.1 glycerate_dehydrogenase [MW = 42.3 kDa]

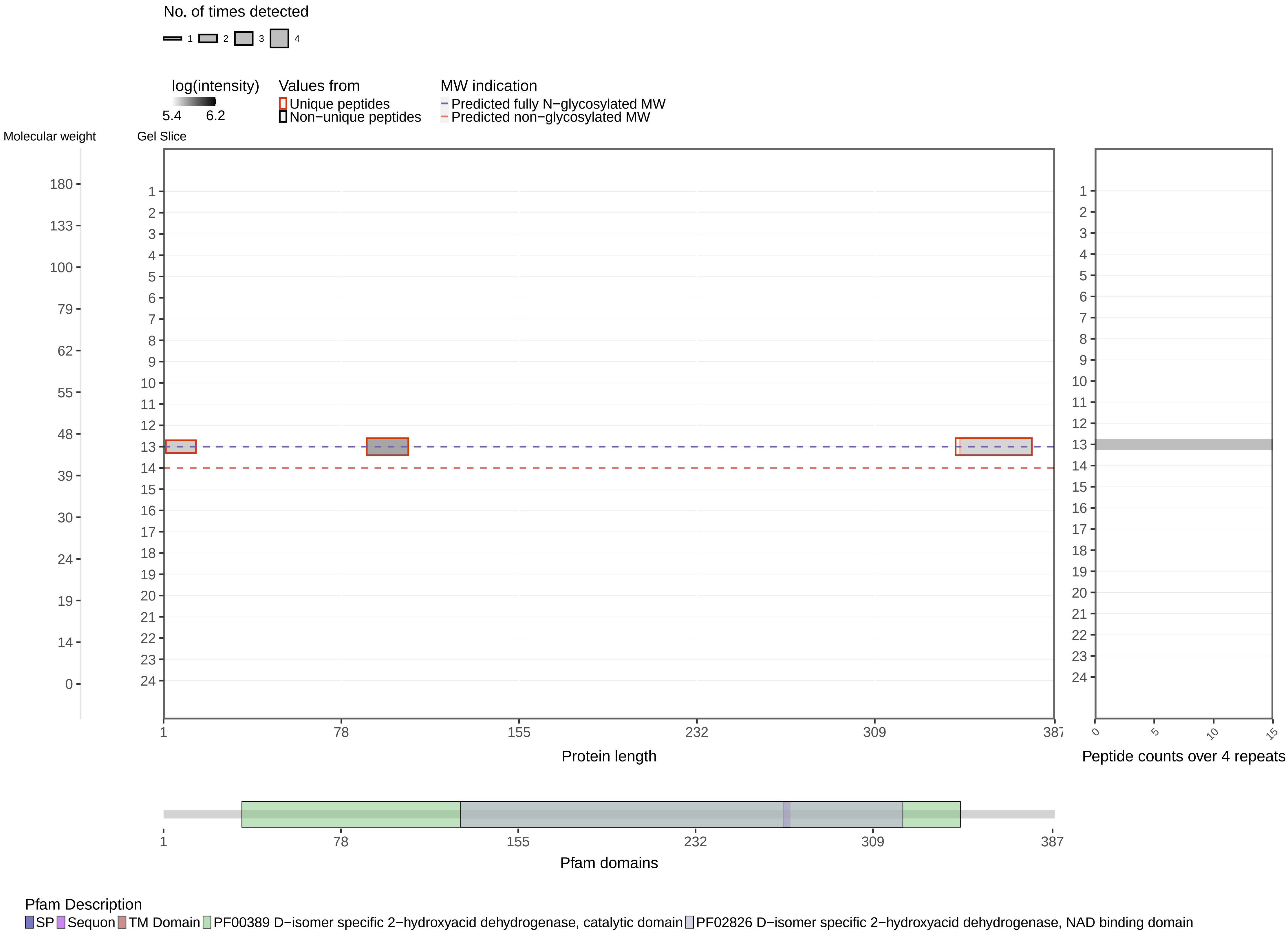


Protein sequence:

ARMMIEGFKMNLIYYDLYQATRLEKFVTAYGQFLKASGEQPVTWKRAASMEEVLREADVISLHPILDKTTYHLVNKERLAMMKKEAILVNCSRGPVIDEVALVEHLRENPMFRVGLDVFEDEPYMKPGLADMKNAVVVPHIASASKWTREGMATLAALNVLGKIKGYSVWGNPNSVDAFLNENSQPPAACPSIVNSKTLGLPVSKL*

### NbL01g04040.1 WD−40_repeat−containing_protein_MSI2−like [MW = 45.1 kDa]

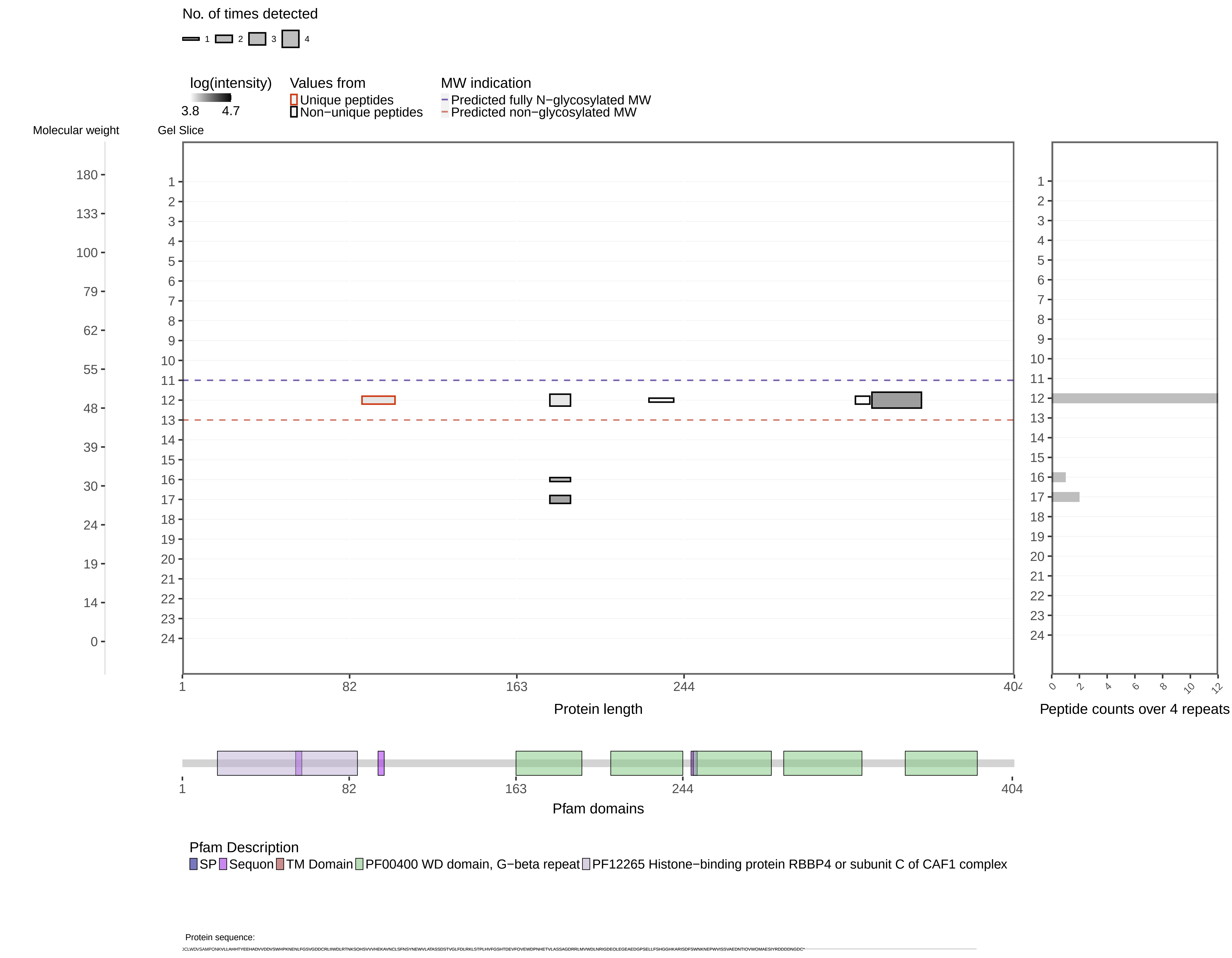

### NbL01g03750.1 vacuolar−sorting_receptor_3−like [MW − SP = 66.85 kDa]

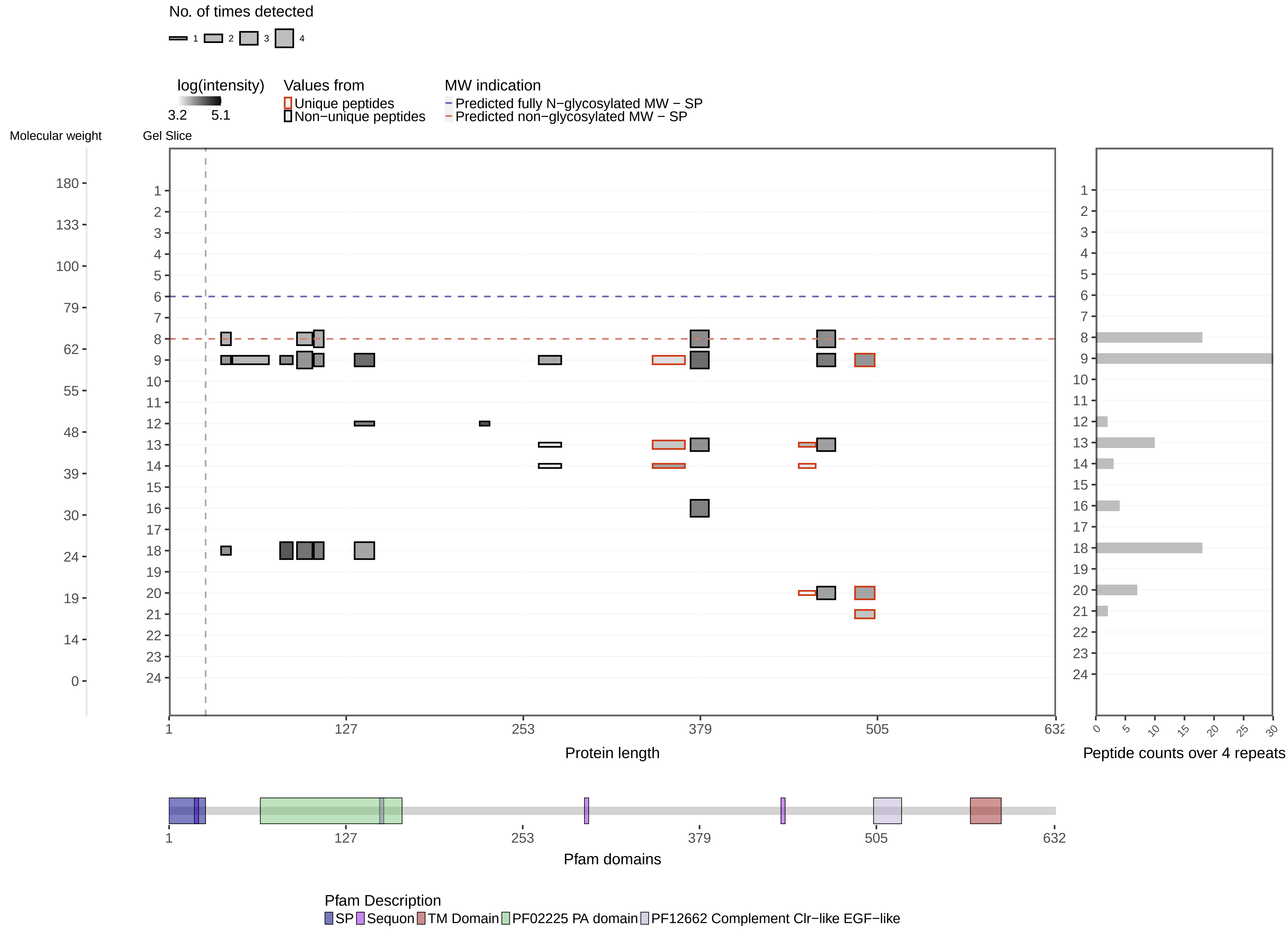


Protein sequence:

### NbL01g03710.1 mitotic_checkpoint_protein_BUB3.1−like [MW = 37.8 kDa]

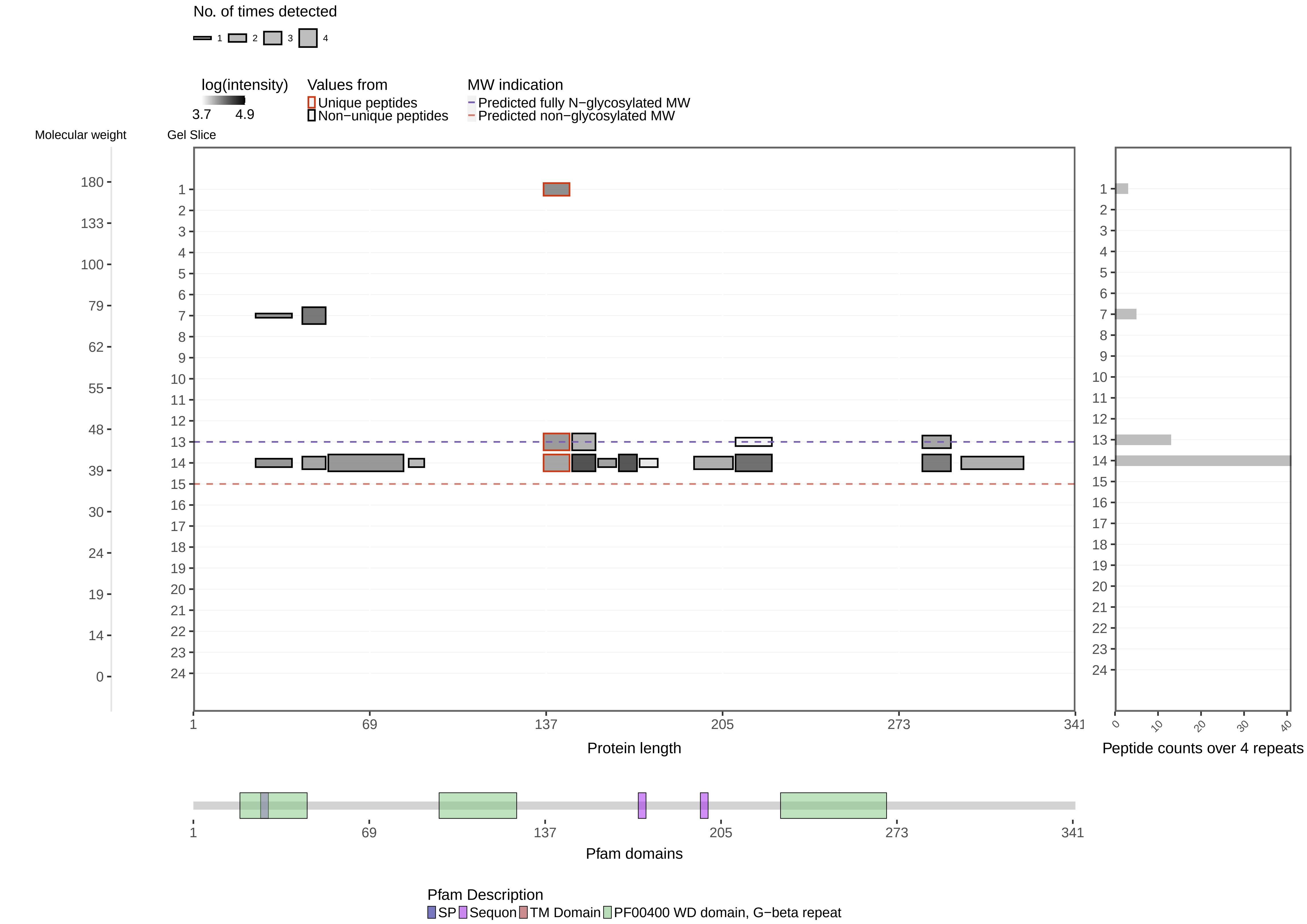


Protein sequence:

LVVATAGRHVNVYDLRNMSQPEQRRESSLKYQTRCVRCYPNGTGYALSSVEGRVAMEFFDLSEAGQSKKYAFKCHRKTEAGRDIVYPVNAIAFHPIYGTFATGGCDGYVNVWDGNNKKRLYQYPKYPSSIAALSFSRDGRLLAVASSYTFEEGEKPHEPDAIIVRSVNEVEVKPKPKVLPNPTT*

NbL01g03620.1 somatic_embryogenesis_receptor_kinase_2−like [MW = 31.7 kDa]

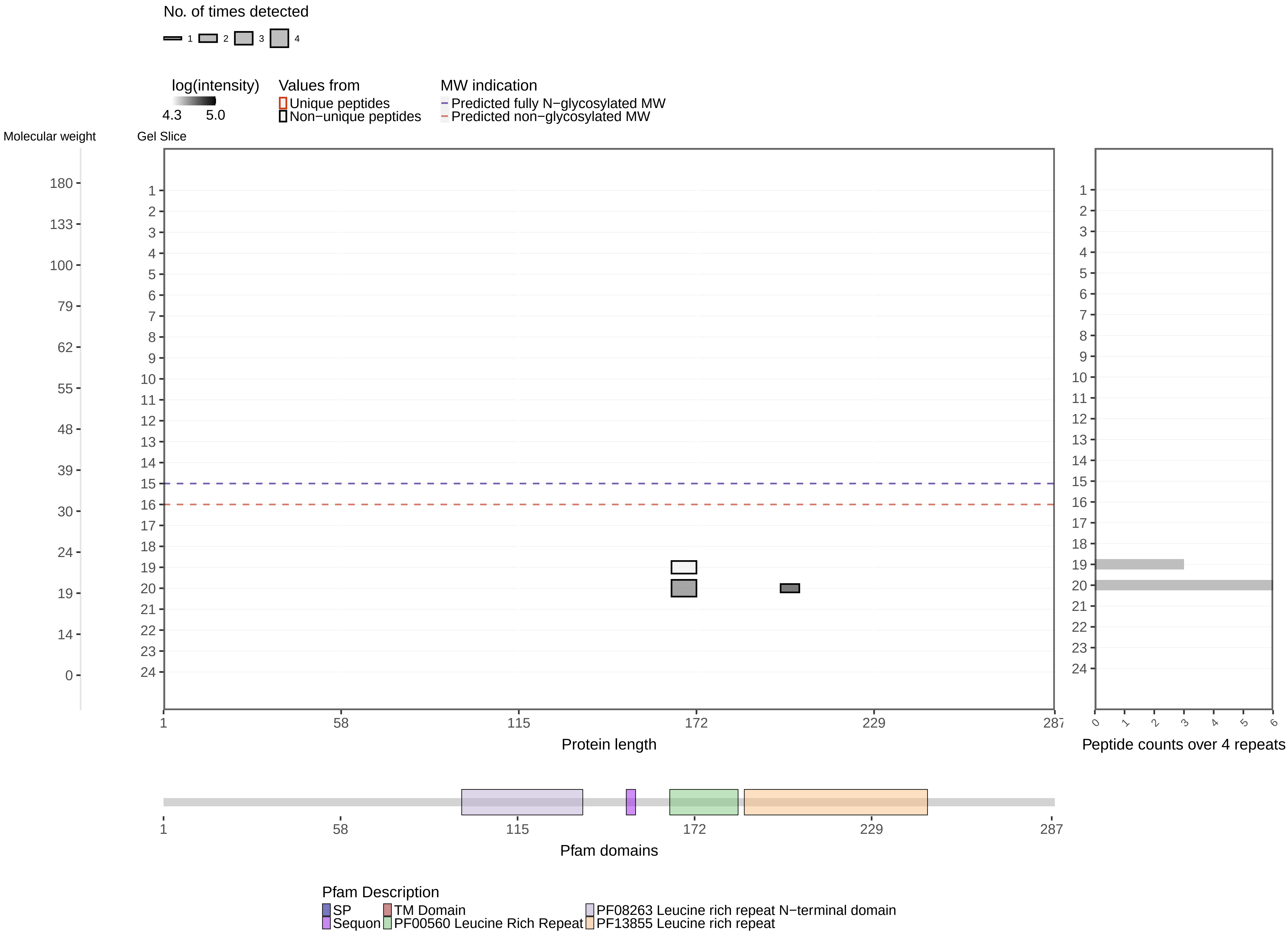


Protein sequence:

HITCDSNNHVIRLDLGNTNISGTLGPELGELKNLQYLELYRNNIGGKIPKELGNLENLISMDLYGNRFEGNIPKSFAKLKSLRFLRLNDNKLTGSIPRELTTLPNLKVFDVSNNDLCGTIPVDGPFGSFPMEGFAHNRLNGPELKGLAPYDFGCK*

NbL01g03580.1 protein_TRI1−like [MW = 21.96 kDa]

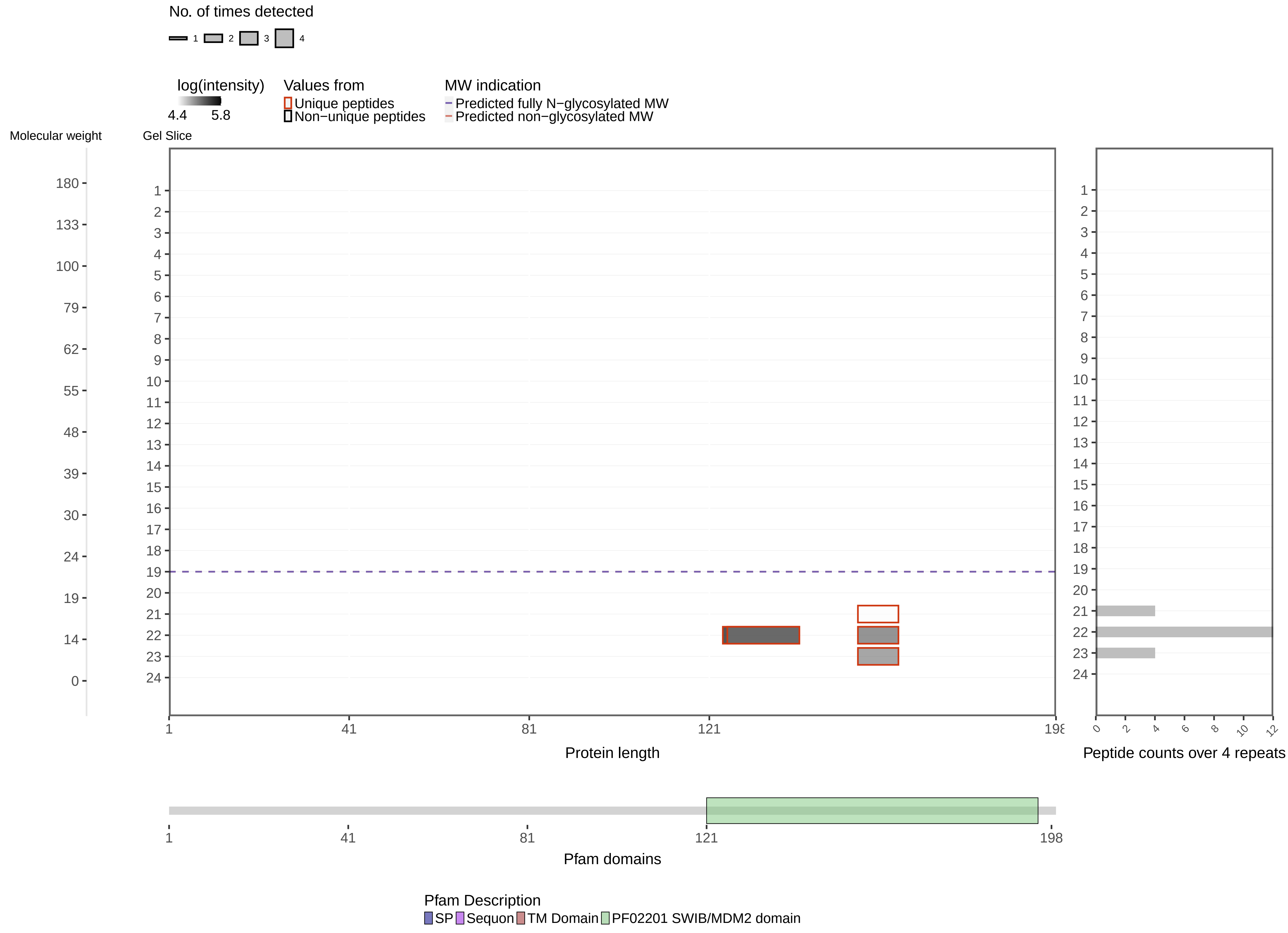


Protein sequence:

.AAHHANLRTVRMVTFATTSKPVATEPKKREPRGIMKPRRVSPEMQAFLGGIAEIPRTQALKLIWAHIKENNLQDPQNKKVIICDEKLKNIFPGKDRVGFLEIAGLISPHFLK*

### NbL01g03470.1 TIP41−like_protein [MW = 34.62 kDa]

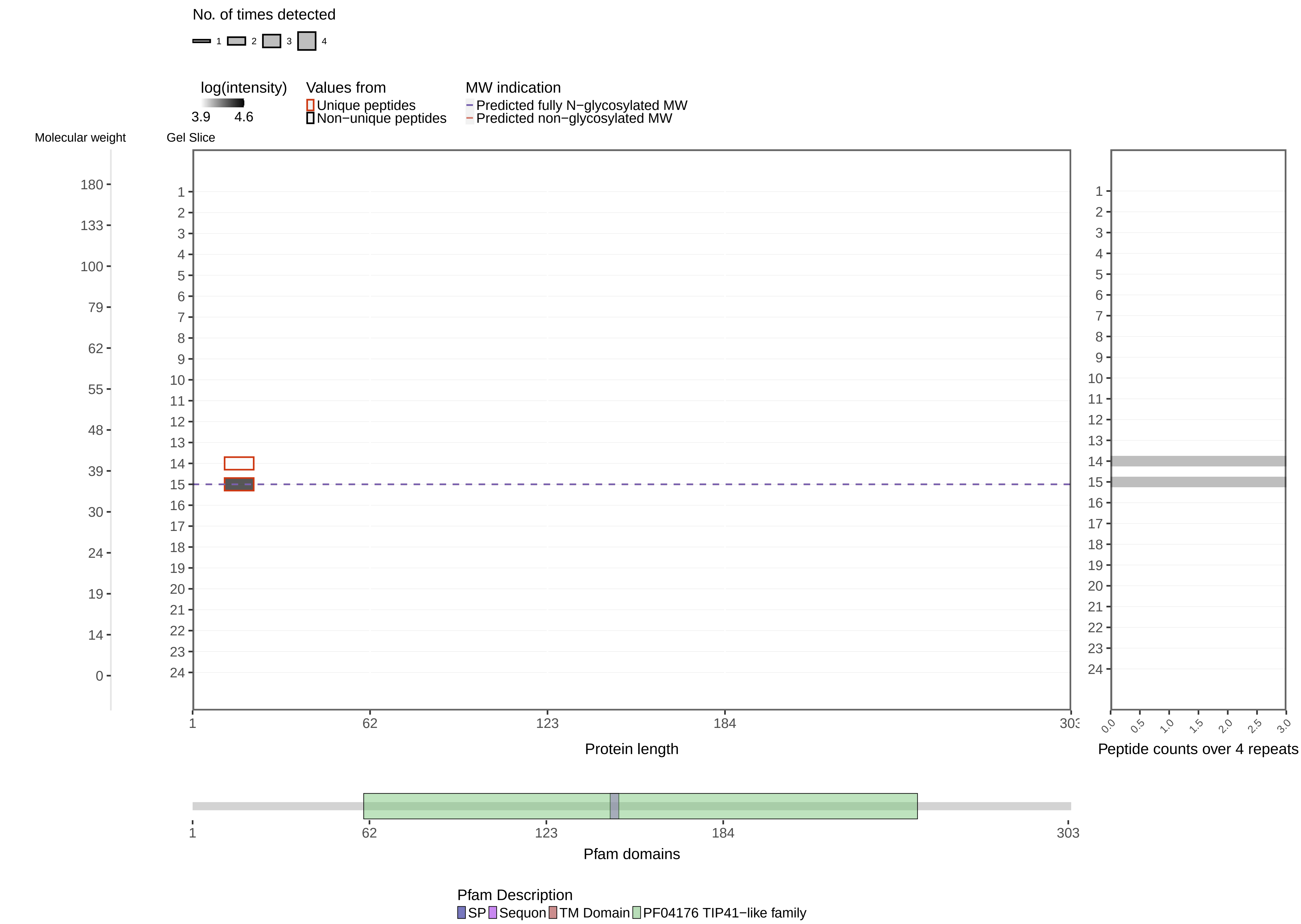


Protein sequence:

AERVRGLSNESSCSLQWEDCTEKIDLVALASKEPILFYDEIILYEDELADNGISLLTVKVRVMPSGWFLLLRFWLRVDGVLMRLRDTRLHCLFGDHMKSVILRESCWREATFQALSSKGYPSDCAAYSDPSTISERLPIIMQKTQKLSIEIPCVVGLYGDTDQKDS*

NbL01g03400.1 methylthioribose_kinase [MW = 47.81 kDa]

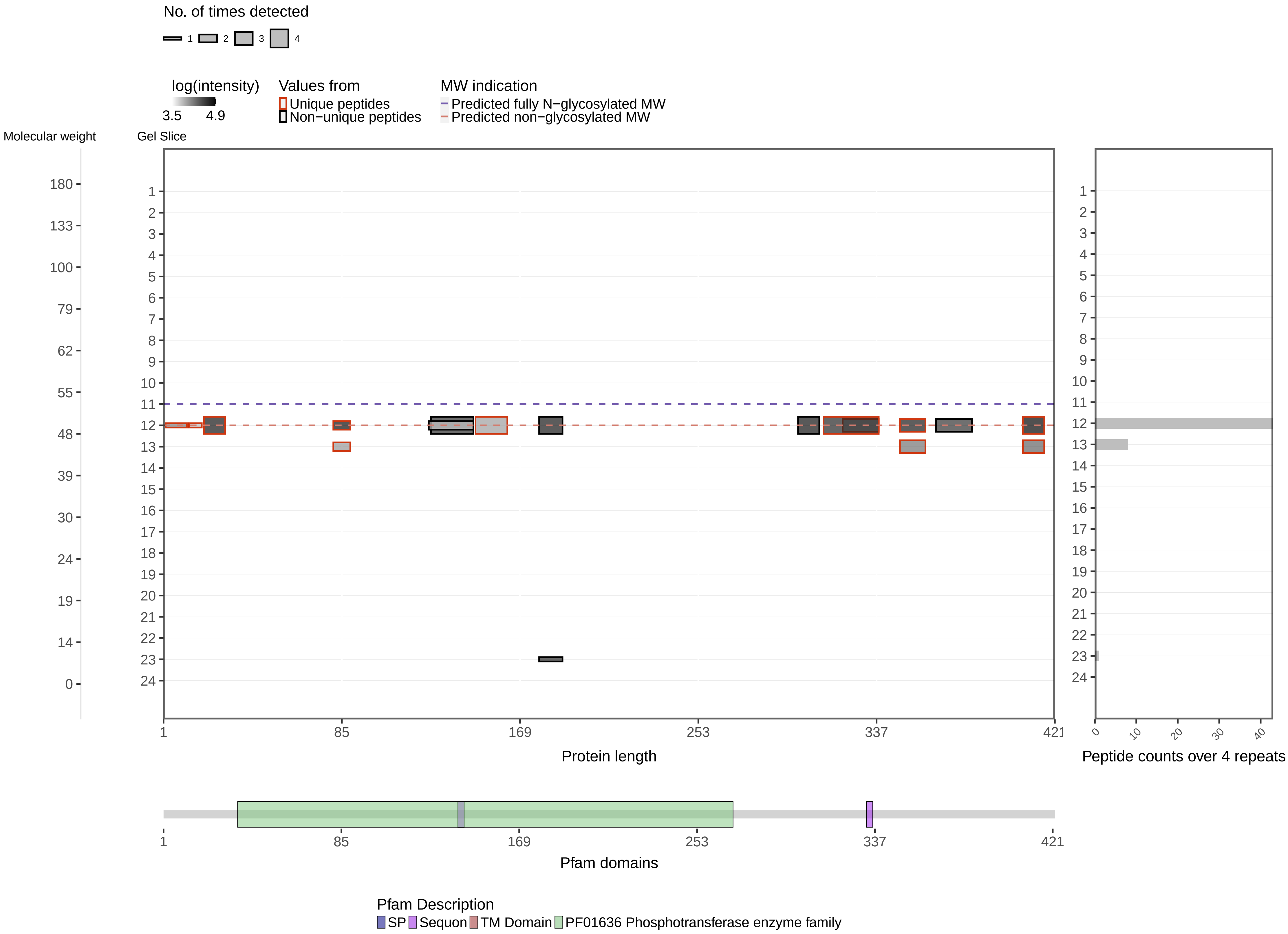


Protein sequence:

PYLDADAEAVRKDNGLKIEVAELKSKFCERAQALIHGDLHTGSLMVTQDSTQVIDPEFAFYGPMGFDIGAFIGNLILAYFSQDGHADQANDRKSYKVWILKTITETWNLFHKKFLALWDEHKDGPGEAYLPEIYNNTELRELVQQKYMKELFEDTLGFGGAKMIRRIVGVAHVEDFESIKDAAKRADCERRALNCAKKLLKERRNVKSIEEVVSTIEKVHLQ*

### NbL01g03340.1 probable_cinnamyl_alcohol_dehydrogenase_2 [MW = 38.77 kDa]

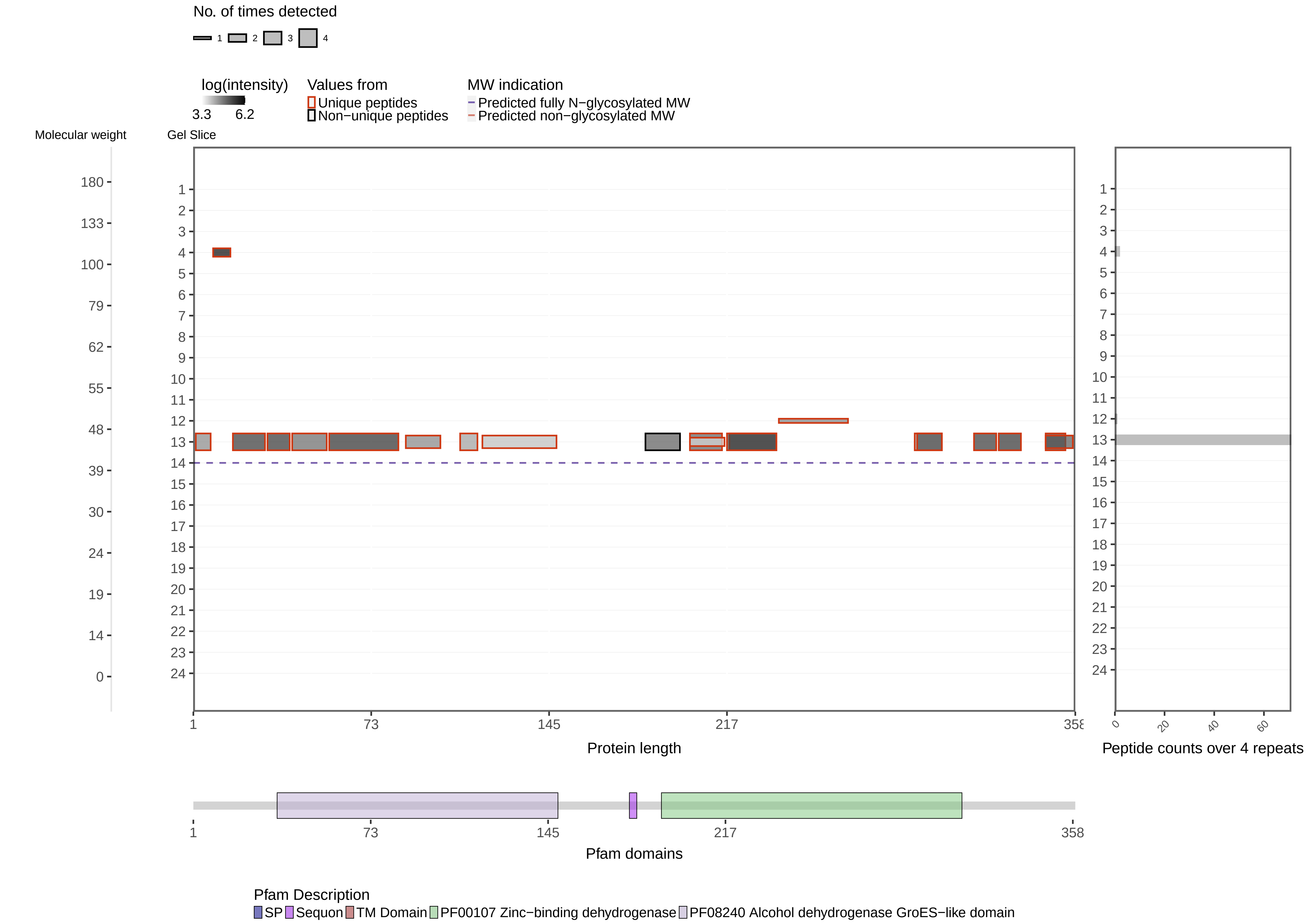


Protein sequence:

CAGITVYSPLNHFGFNOSGLRGGILGLGGVGHMGVKIAKAMGHHVTVISSSNKKRQEALEHLGADDYLVSSDTDKMOEAADSLDYIIDTVPVGHPLEPYLSLLKIDGKLILMGVINTPLOFISPMVMLGRKSITGSFIGSMKETEKMLEFCKEKGVTSOIEIVKMDYINTAMERLEKNDVRYRFVVDVAGSKLDO*

NbL01g03260.1 cyclase−like_protein_2 [MW − SP = 28.11 kDa]

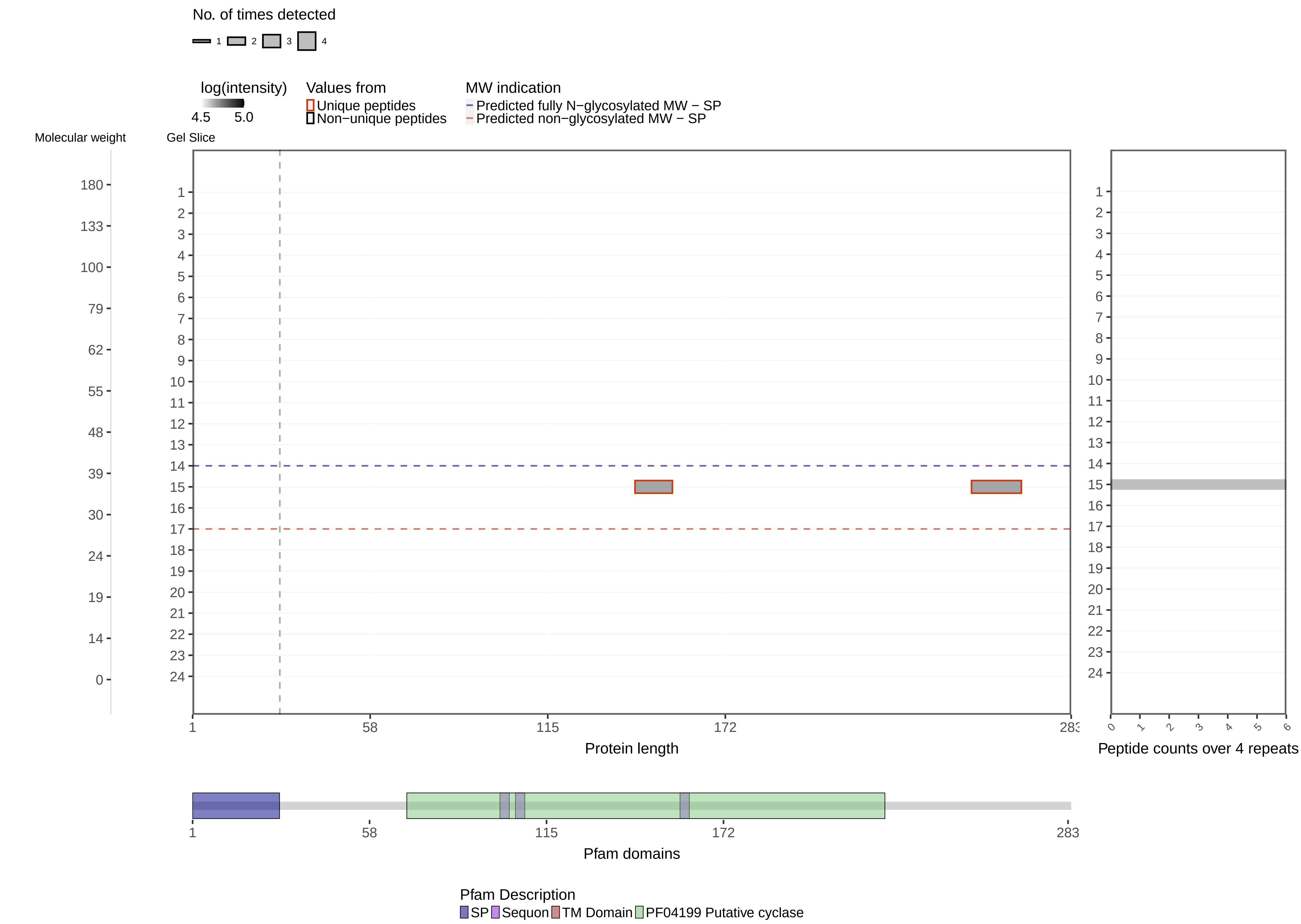


Protein sequence:

IYAEYYDAGFDVDSLDLRVLNGPALVVDVPRDKNLTADVMQSLHIPKGVKRVLFRTLNTDRRLMWKKAFDTSYVGFMKDGAQWLVDNTDIKLVGIDYLSVAAFDDLIPSHLVFLKSREIILVEGLKLDDIETGIYHVHCLPLRLLGAEGSPIRCILIK*

### NbL01g03160.1 glucosyltransferase [MW = 53.56 kDa]

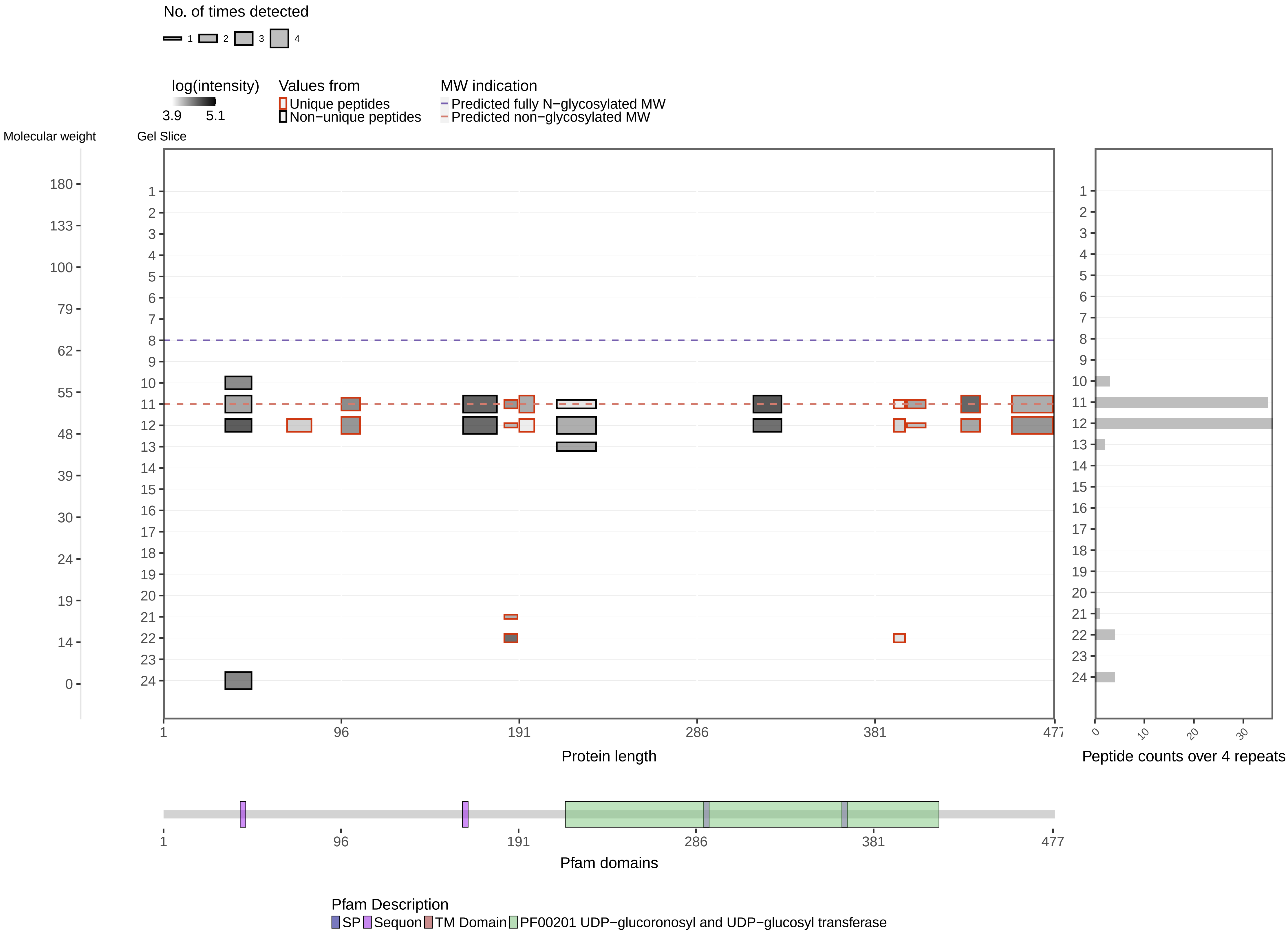


Protein sequence:

HYTKVLGRRAWAIGPLSMCNRDIVDKAERGKKSSIDKHECLKWLDSKKPSSVVYICFGSVANFTASQLHELAMGIEASGQEFIWVVRTELDNEDWLPEGFEERTKEKGLIIRGWAPQVLILDHESVGAFVTHCGWNSTLEGVSGGVPMVTWPVFAEQFFNEKLVTVVLKTGAGVGSIQWKRSASEGVKREAIAKAVKRVMVSEEADGFRNRAKAYKEMATKAIEKGGSSYTGLTTLLEDISTYSSTDH*

### NbL01g03130.1 polyadenylate−binding_protein_8−like [MW = 71.81 kDa]

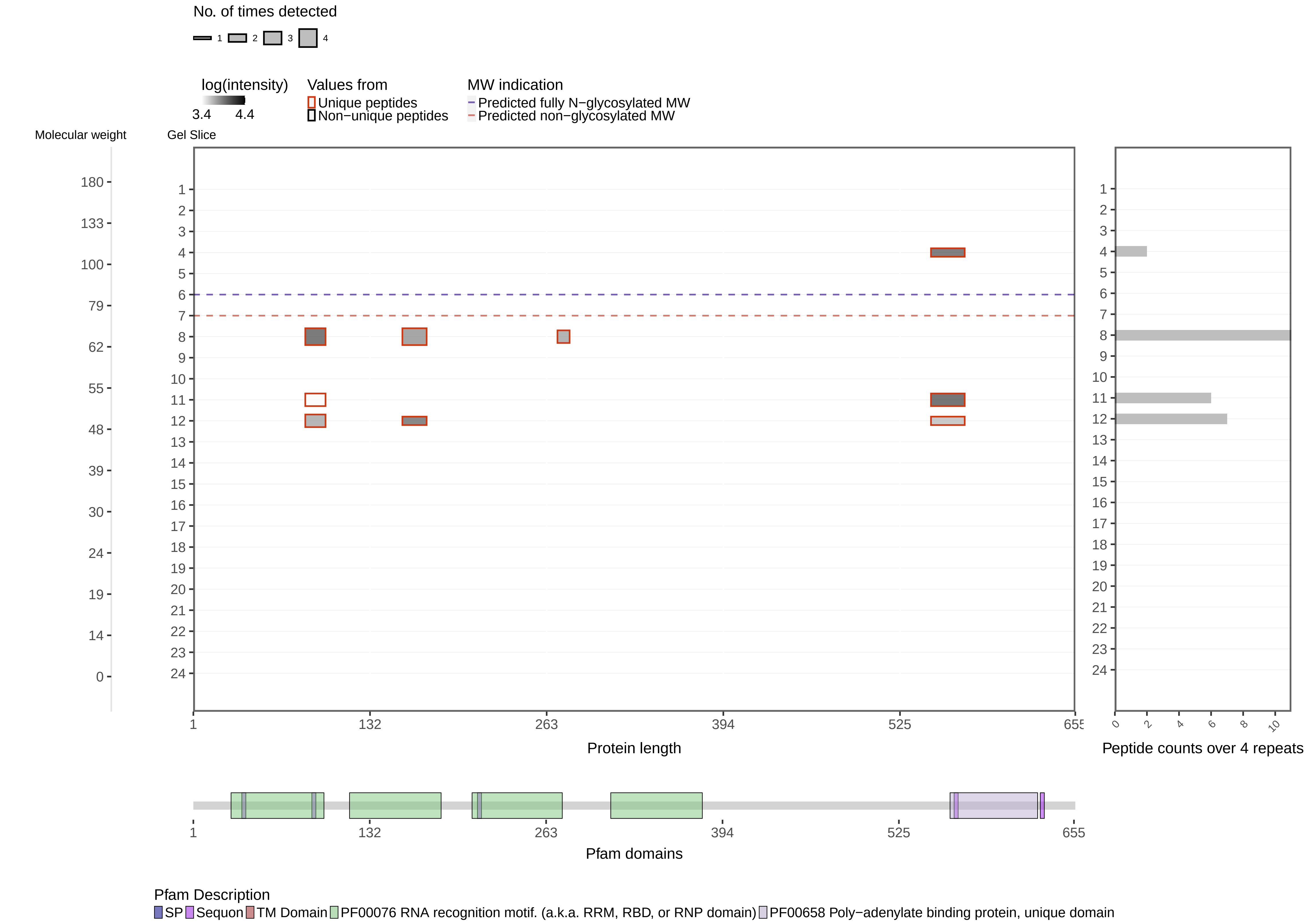


Protein sequence:

DSISDEKLKELFSPYGTITSCKVMRDPSGVSKGSGFVAFSTPEEASRALSEMNGKMVVSKPLYVALAQRKEERRARLQAQFSQMRPIAMASSVAPRMPMYPPGGPGLGQQIFYGQPPPAMLPPQVINRRAGFGYQQQLVPGMRPGGGPMPNFFMPMVQQGQQGQRPGGRRGGAVPLQQGQQPVSLMQQQMFPRGRGYRYPPGRGLPDGGFPGVGGGMFSIPYDMGGMPVRDAGIVQPIPVGALATALANASPTEQRTMLGENLYPLVEQLEPETAAKVTGMLL

### NbL01g03090.1 filament−like_plant_protein_7 [MW = 96.15 kDa]

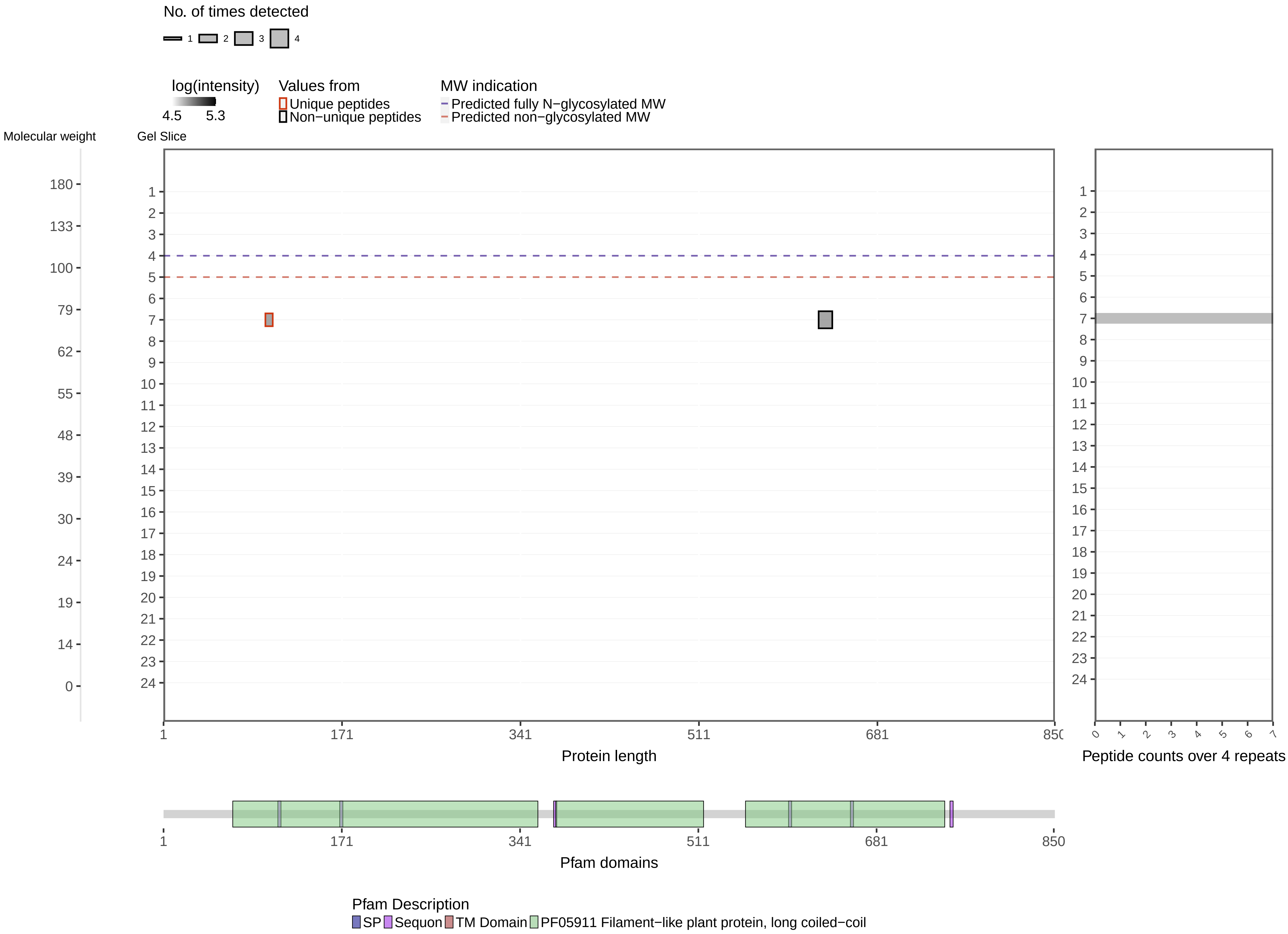


Protein sequence:

LAIVAVDSTIGSSYPPSDTSLDLSDSSRIETHGHQIDSTGKELVKVGQYDLYDLGMEVQGQDLLSKRSPNWLHEILKTIMEQSHVSKKGIDELIEDIRVALHSVTPPCTGPSELLPISGFITWKSPVTSPKMQSLTTEPGSMQSVISELEKIHLILQEENERLKAEINSMKSSNKDVQVKLQVVKNKSENLANELKQSQQSIVSLRAELEEVKESKRMMEDLMENQKSINEDLDTQLTVTKVKLNETIQKLSSLEVELEDRSHCCEELEGTCLELQLQLESITANPTSADNMDQKGI

### NbL01g03060.1 caffeoyl−CoA_O−methyltransferase_6 [MW = 27.74 kDa]

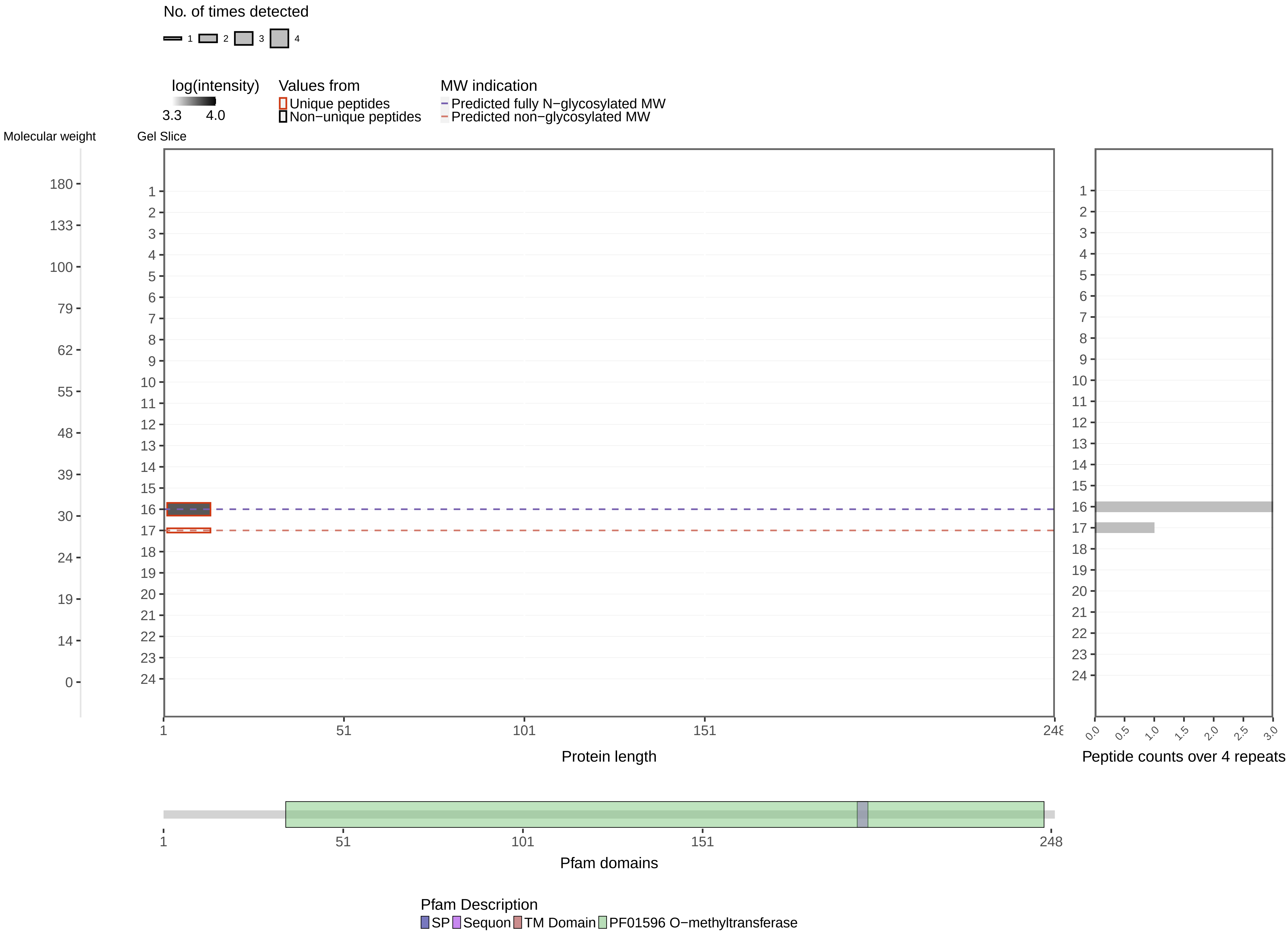


Protein sequence:

MDINRENYEIGLPVIQKAGVAHKIDFREGPALPVLDLMVEDKNNHGTYDFIFVDADKDNYINYHKRIIELVKVGGVIGYDNTLWNGSVAAPPDAPMRKYVRYYRDFVLELNKALAADPRIEICMLPVGDGITLCRRIS*

NbL01g02850.1 probable_quinone_oxidoreductase [MW = 68.33 kDa]

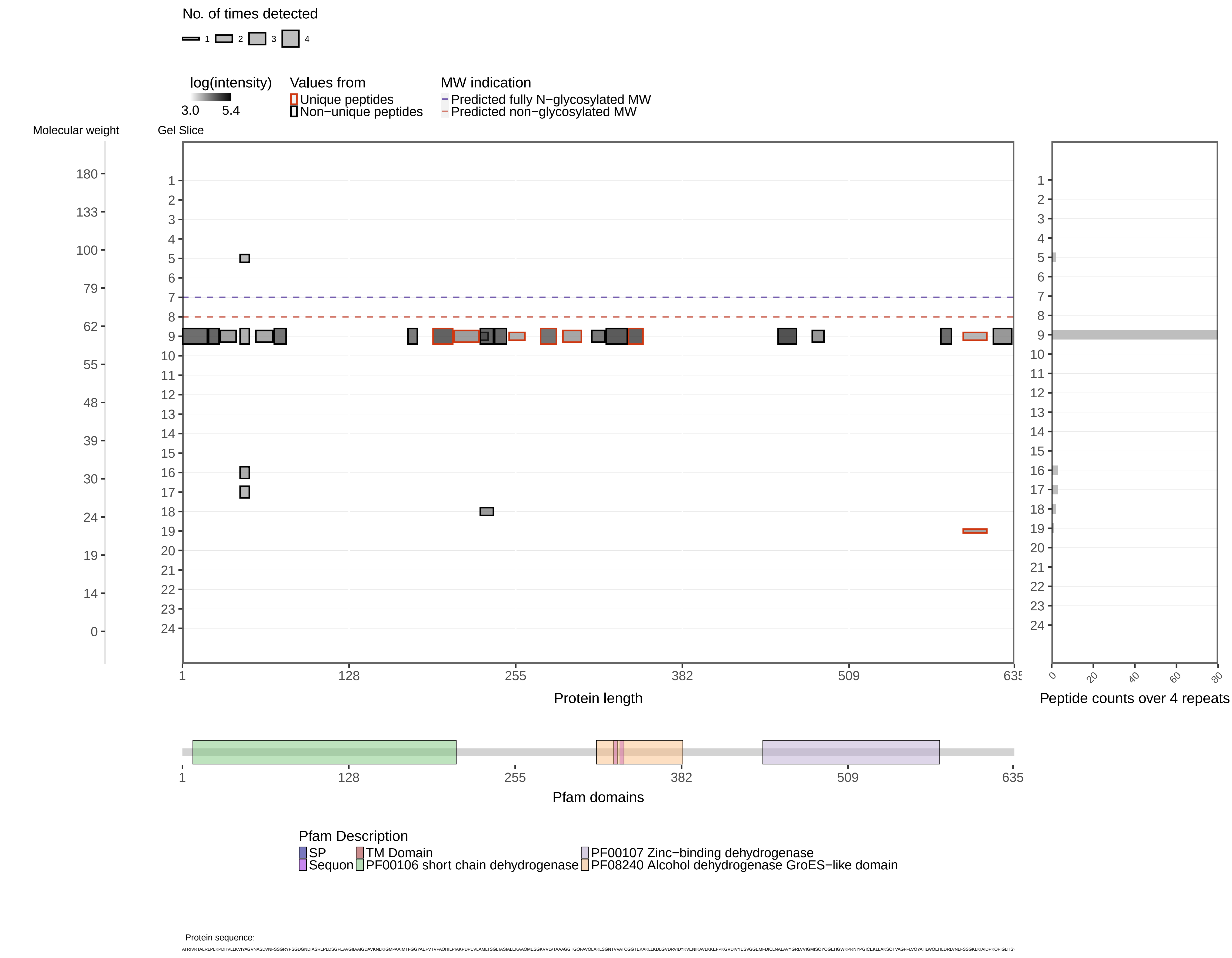

NbL01g02790.1 EKC/KEOPS_complex_subunit_TPRKB [MW = 18.69 kDa]

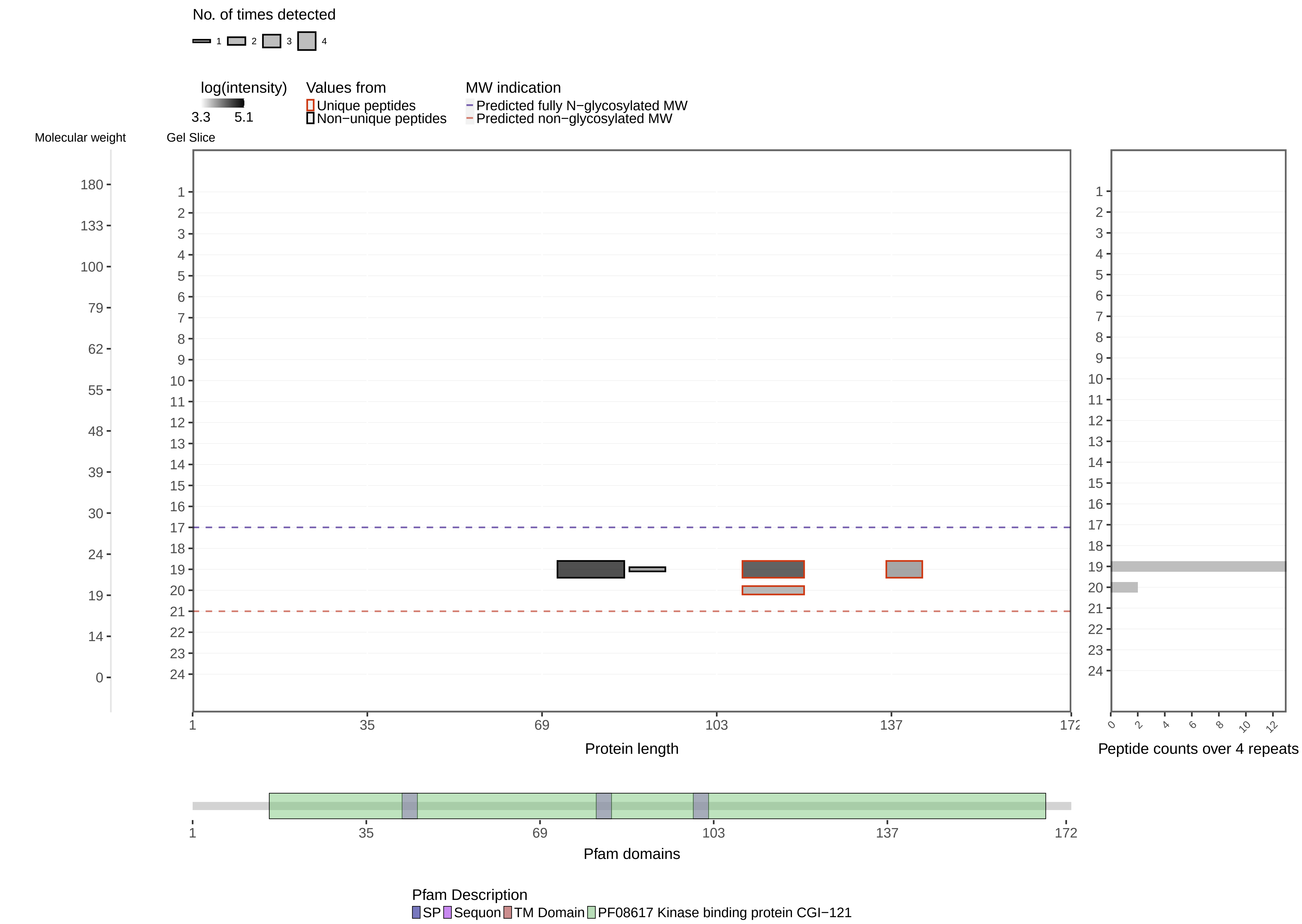


Protein sequence:

TLHSELVYNYSGSKHISESLKRCGIADNTSYILAARFGALADEMVAVEKLINGKEIDLKELEQRINQAQIQKHYKISNLELEISSLADAITCRIAARDAL*

### NbL01g02710.1 ferredoxin−−nitrite_reductase_chloroplastic−like [MW = 68.22 kDa]

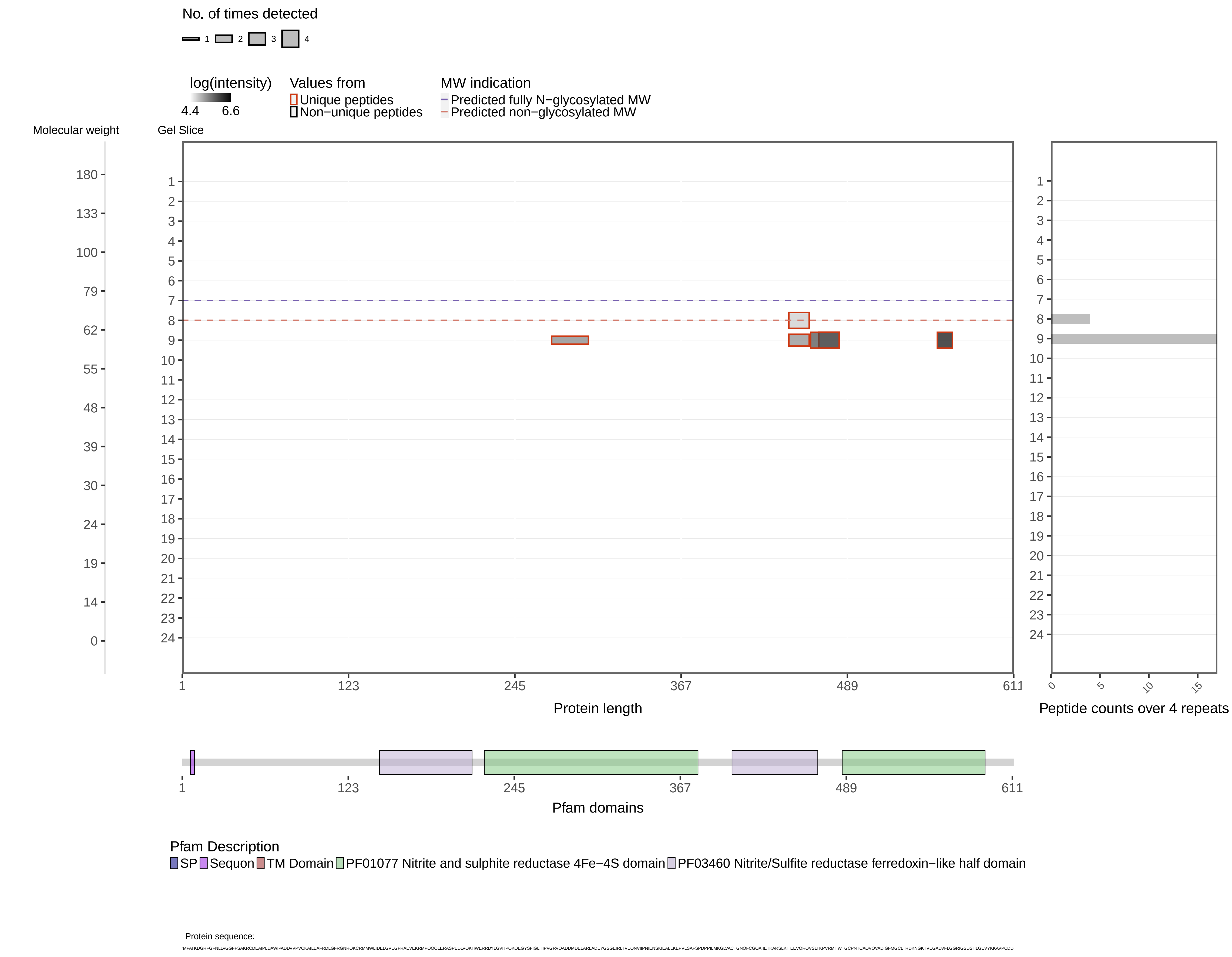

### NbL01g02640.1 presequence_protease_1_chloroplastic/mitochondrial−like [MW = 127.3 kDa]

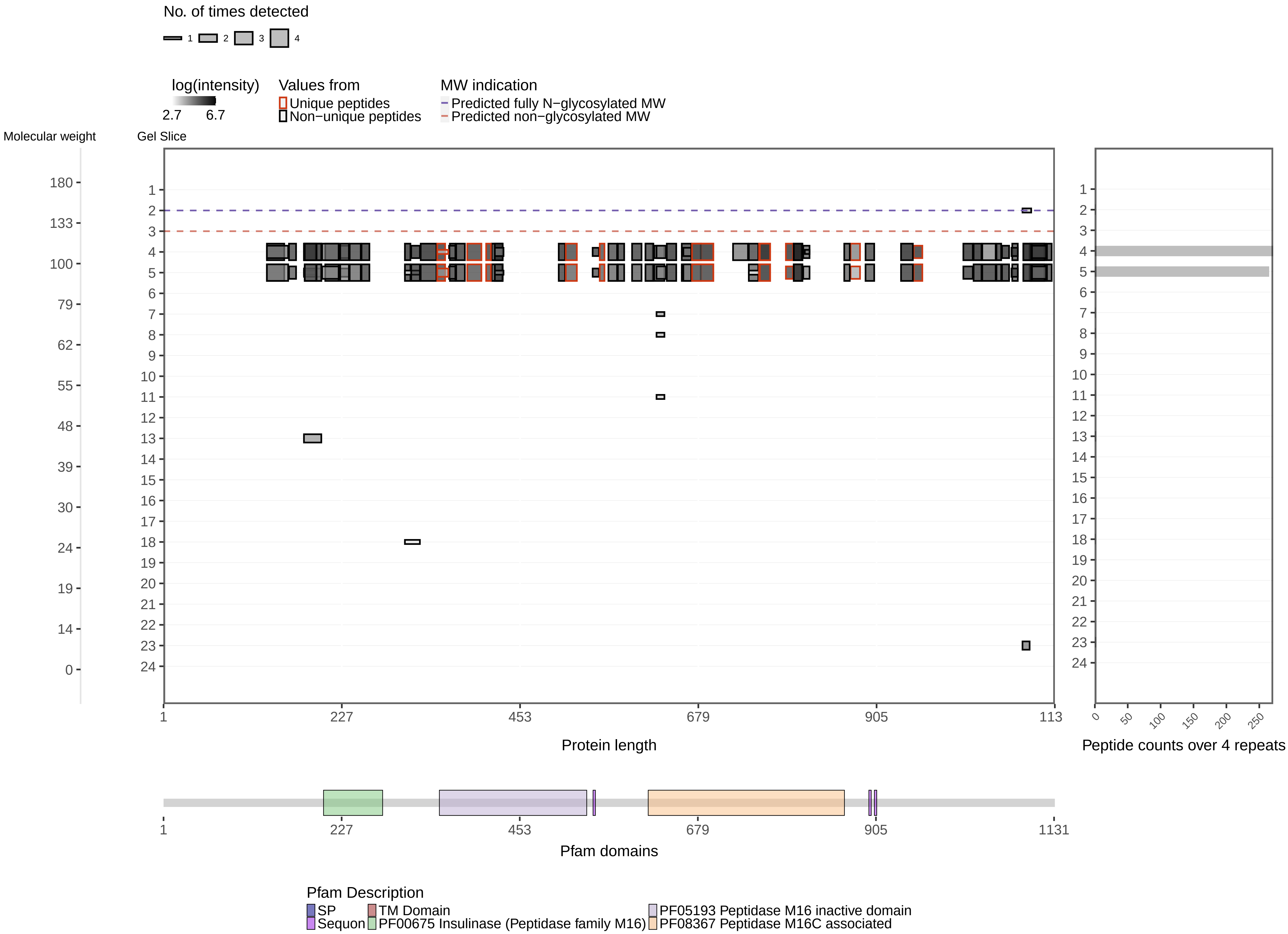


Protein sequence:

GLALMLRSIGKWIYDMDPFEPLKYQKPLEALKARIAKEGSKAVFAPLIDQYILRNPHRVTVEMQPDPKKASREEEIEKETLDKVKASMTQEDLAELARATHELRLKQETPDPPEALKSVPSLSLQDIPREPTHVPTEVGDINGVKVLRHDLFTNDVLYAEVVFDMSSLKQELLPLVPLFCQSLLEMGTKDLDFVQLNQLIGRKTGGISFYPFTSSVRGKVEPCSKIIVRGKAMSQRTDDLFNLINRVLQDVQLNDHKRFKQFVSQSRARMENRLRGSGHSIAASRMGAKLNVAG

NbL01g02630.1 2−hydroxyisoflavanone_dehydratase−like [MW = 35.3 kDa]

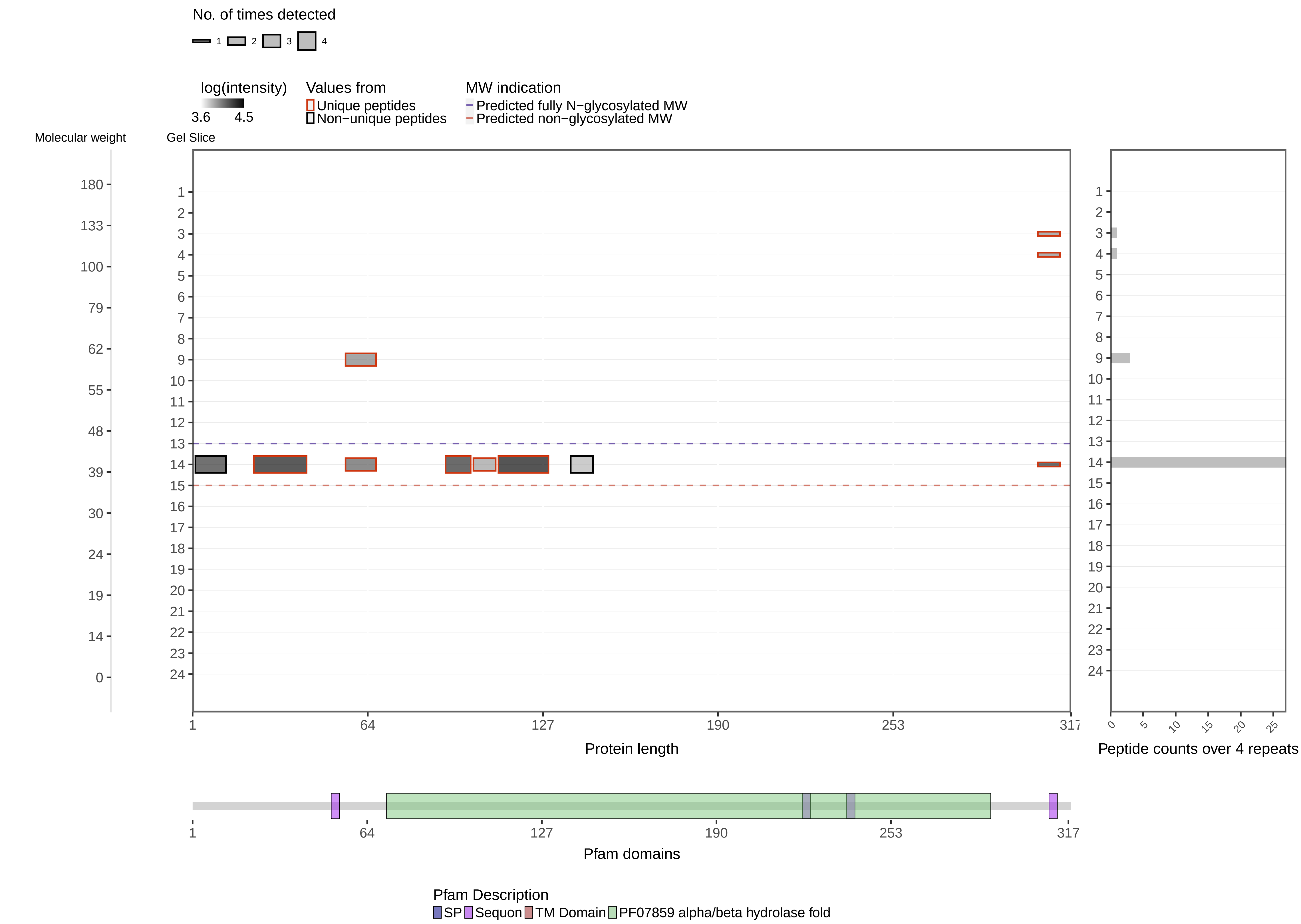


Protein sequence:

HADLNHVYFGGDSAGANIAHNIAIRVGLEKLNGVKLDGIFLACPYFWGIDLIDGEAENLDCKNYIEKLWAFAHPNSSGLDDPVMNPEKDPNLSSLGCDKVLVYYVAGKDPLKYRGLYYKELLEKSGWOGTVEVVEIKDEEHVFHLFAPTSENAMVVLKKLASFLNOSKALR*

### NbL01g02620.1 probable_carboxylesterase_12 [MW = 34.86 kDa]

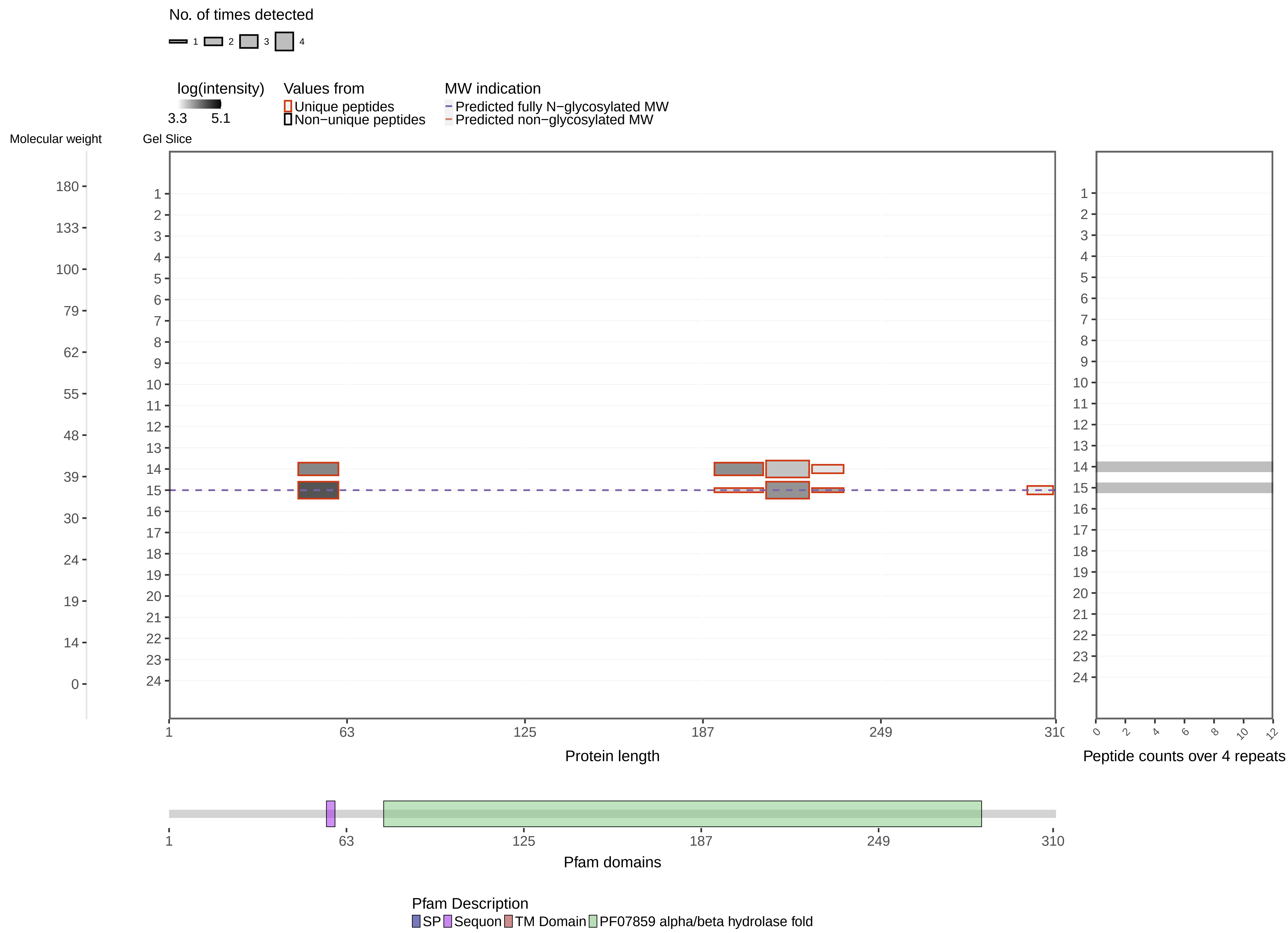


Protein sequence:

ƆGTEPWLGSHADFSRVFLAGDSAGANIAHNMMMQASVDEDKLGDGLKLVGMALVHPYFGNNGPDRIWSYCCPENPNTDDPRFNPAAHPSLLSKLVCSKILICTGEKDFIRDRGWTYYEALKKCGWKGEVEIKETEGEEHVFHLVTPTCDKAKALMKWLADFFQQSS*

NbL01g02610.1 2−hydroxyisoflavanone_dehydratase−like [MW = 43.46 kDa]

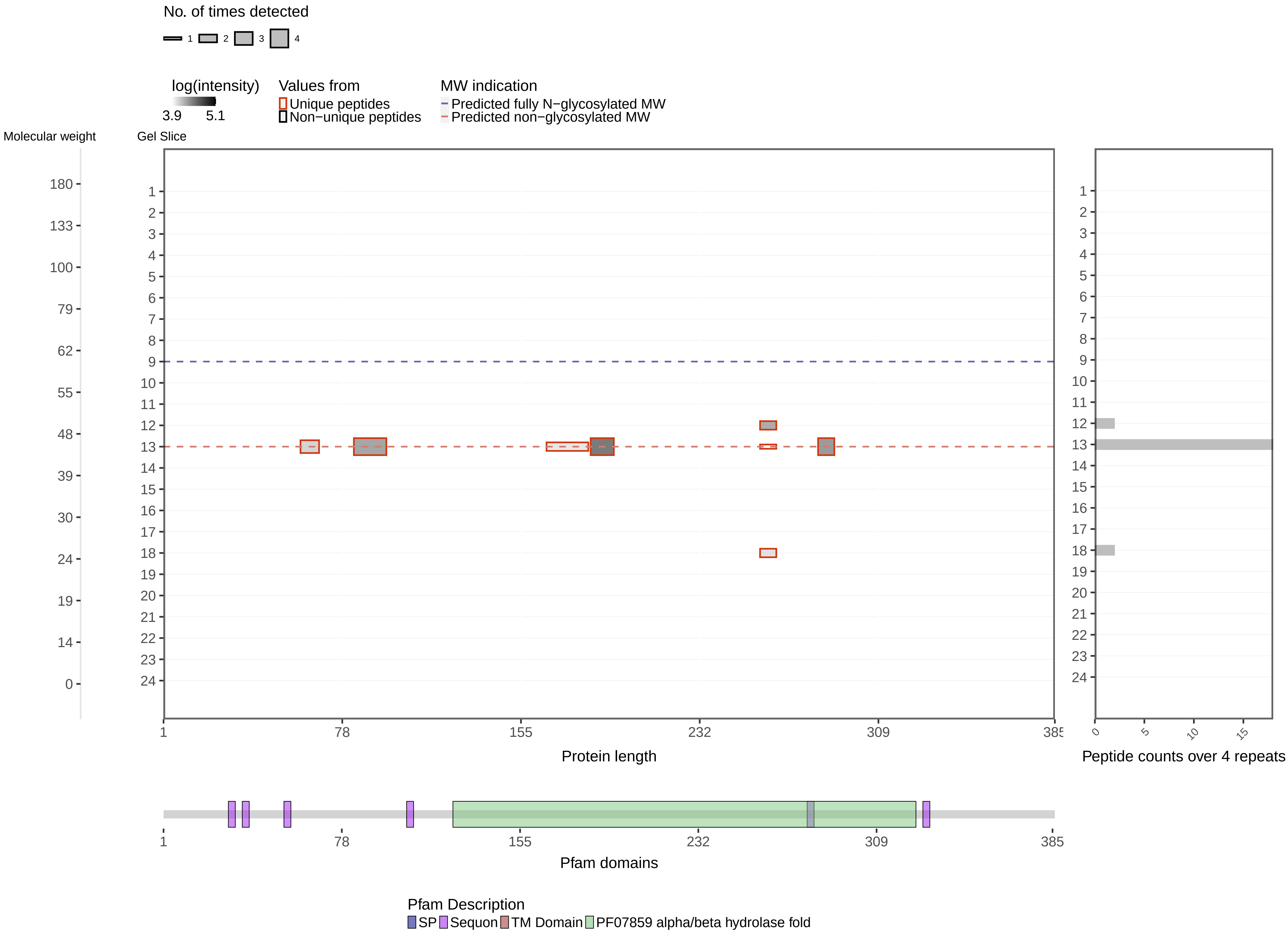


Protein sequence:

IKWVEQGTDPWLKQHADFSRVYLAGDSAGANIAHNMMVRVSEESHFSPSLIGMALIDPYFGNDKPDTLWNYLCPKTTGIYDPRFNPAAHPMLLSKLNCSKVLVCTAGKDFLRDRAWTYYENLKKSGWKGEVDMKEIEGEDHVFHLYNQTSEKANVLMKSVVDFFVYDDSSQHVNLGSKVSEFSMSLIIFAFNLFCICVYKSS*

### NbL01g02600.1 probable_carboxylesterase_2 [MW = 33.84 kDa]

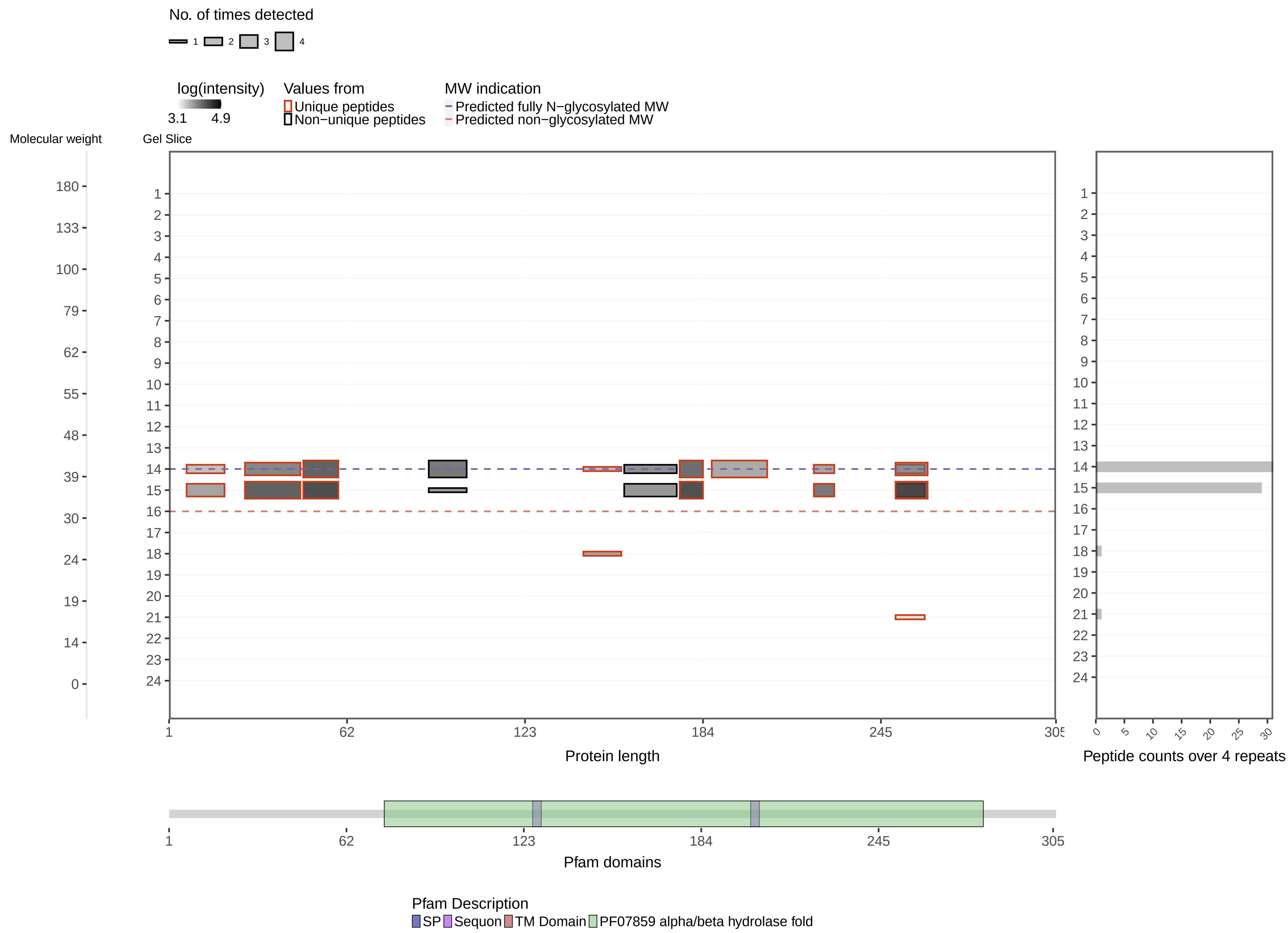


Protein sequence:

SKGPESWLNEYADFGKVFIGGESAGANIANDVAVRAGVSDLDLKVKILGLYLVHPYFGVENDTLYKVLCPTDTGGCFEDPRVNPLLDPRLKSMACKKVLFLVAEKDFLKEGTMNYYEGLKKSEWNGELEIMETKGEGHCFHFINPMSEKAVALMDKLVDFLRKD*

### NbL01g02580.1 polyadenylate−binding_protein_RBP47 [MW = 46.86 kDa]

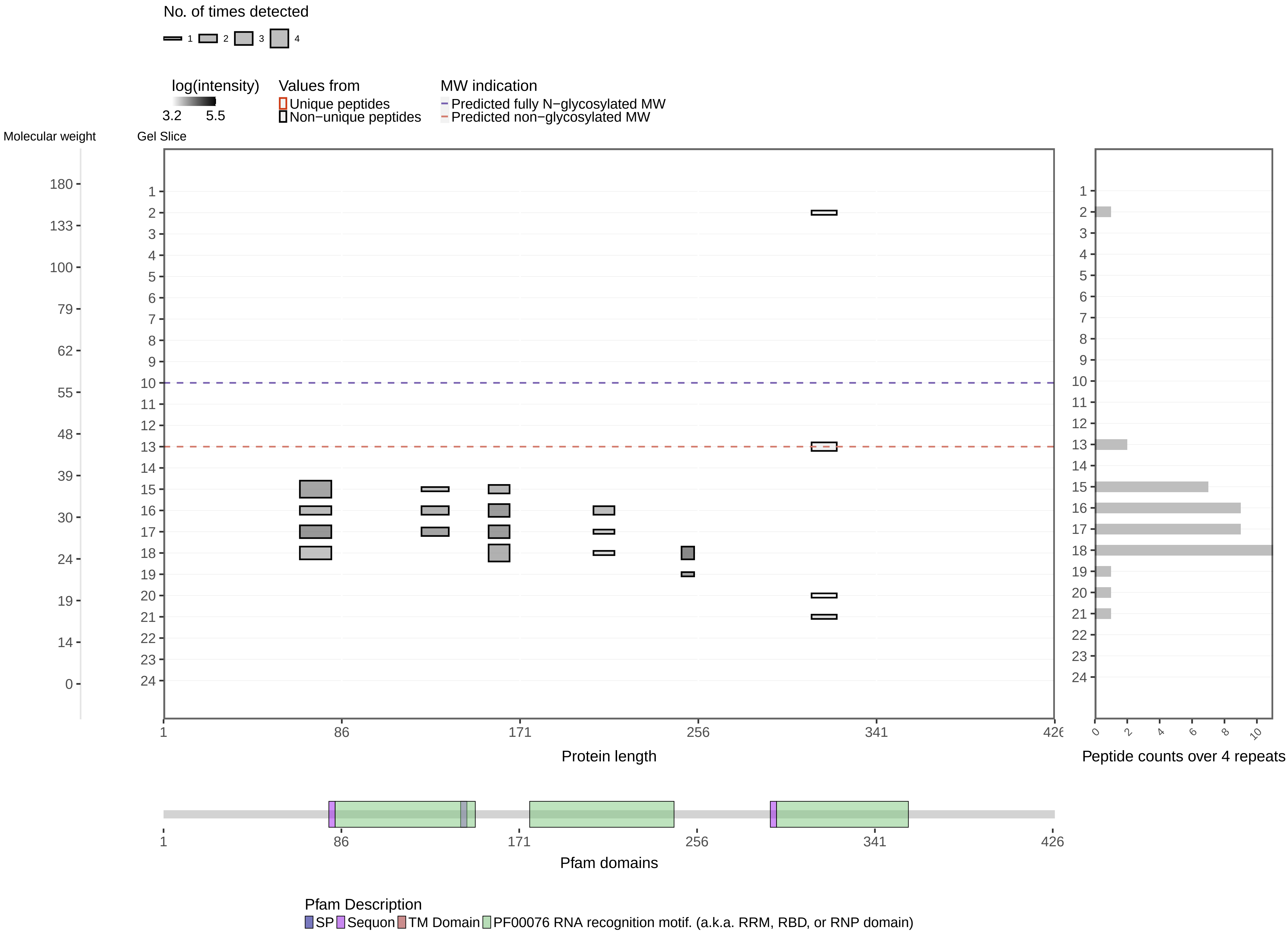


Protein sequence:

YPSLKGAKVVVDANTGHSKGYGFVRFGDESERSRAMTEMNGVYCSSRAMRIGVATPKKPSAQQQYSSQAVILSGGYASNGAAPHGSQSDGDSSNTTIFVGGLDSEVTDEELRQSFNQFGEVVSVKIPAGKGCGFVQFSDRSSAQEAIQKLSGAIIGKQAVRLSWGRSPANKQMRTDSGSQWNGGYNGRQNYGGYGYGTSQNEDSGMYATGAAYGASSNGYGNHQQPVS*

NbL01g02560.1 serine_carboxypeptidase−like_26 [MW = 56.27 kDa]

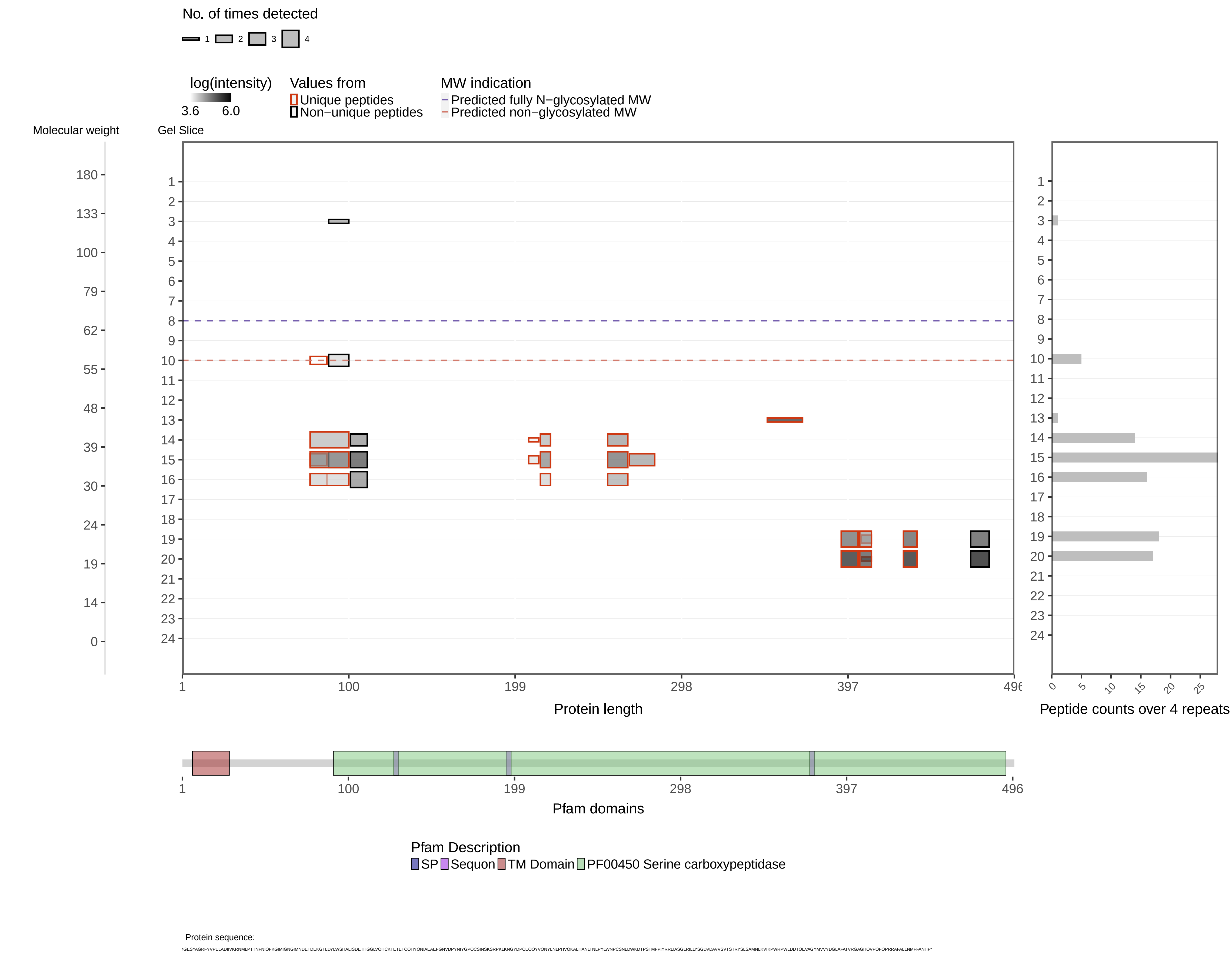

### NbL01g02240.1 CCG−binding_protein_1 [MW = 23.08 kDa]

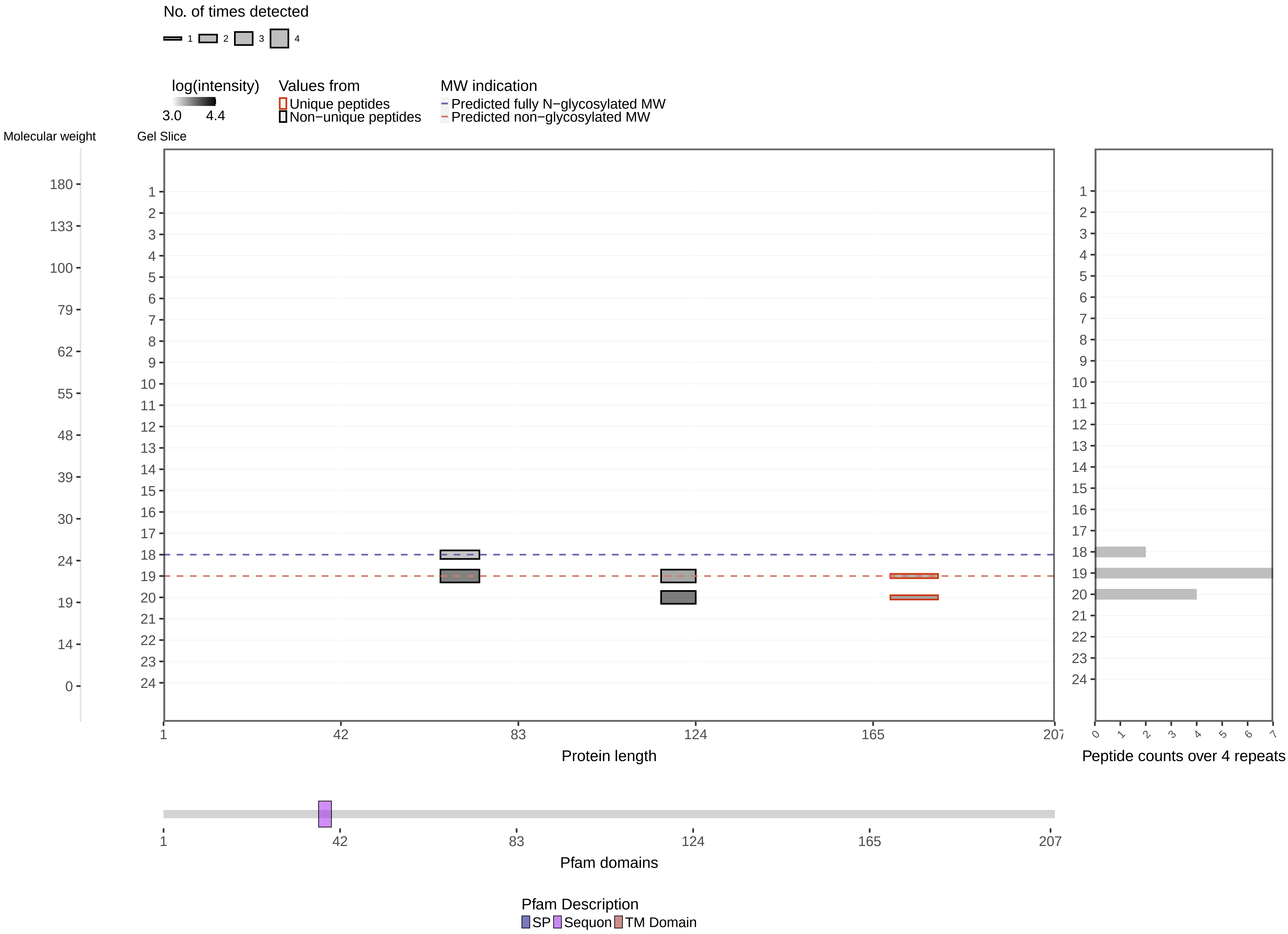


Protein sequence:

YSCWQAYFELKDLEKEAPKEEVERLILQAGGVKTLIGCLHGVSDMHKAKRASKESTKMANLEEKVARPCPVPDGLPKTREELEEEEKGKMPDSPFTRLLRAKGTHPVWYSSALDY*

### NbL01g02140.1 proteasome_assembly_chaperone_2 [MW = 30.54 kDa]

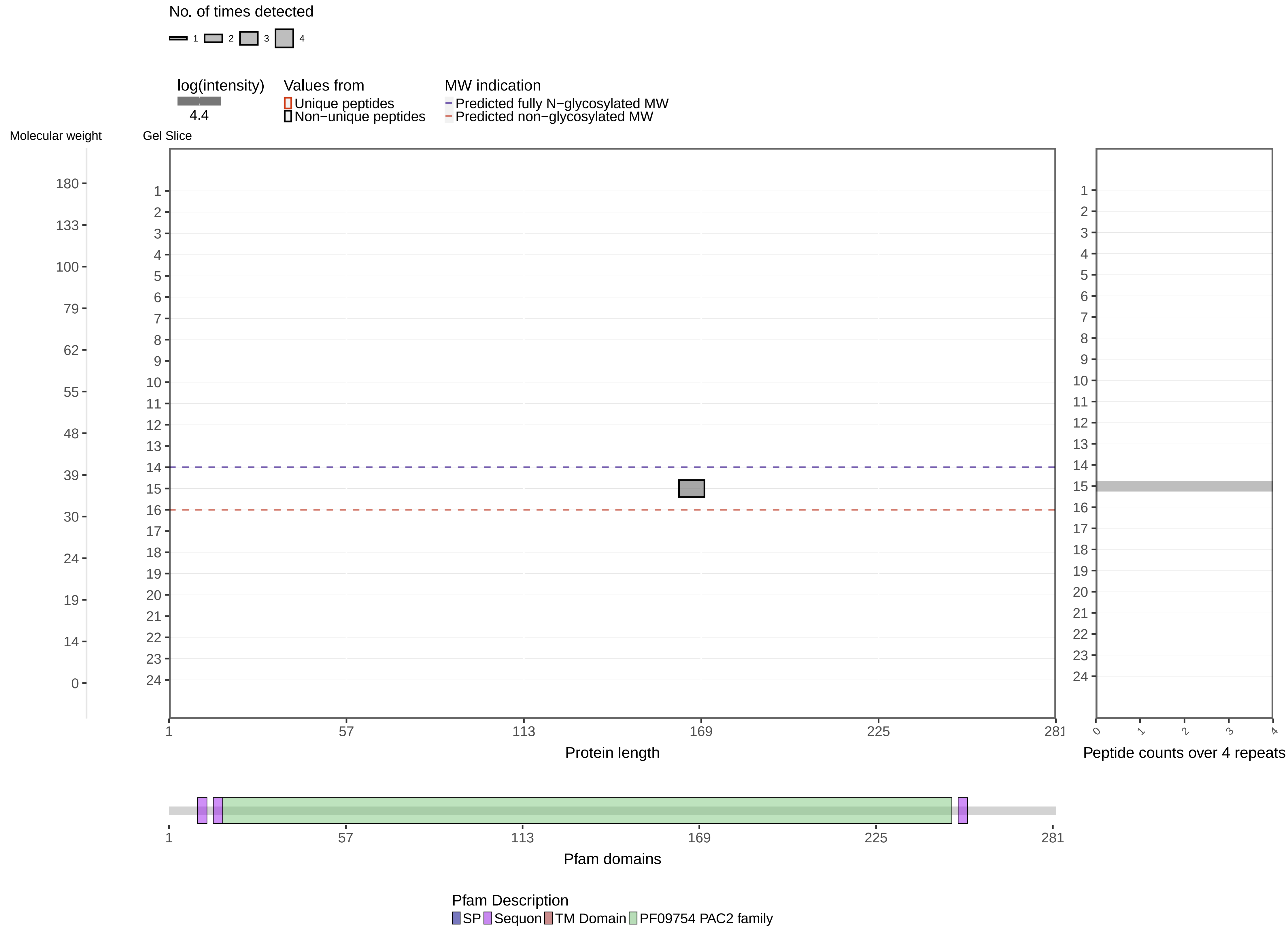


Protein sequence:

RNIDMSSGLQIHYLSSSNSDGTDDQCESQGWKRLPEYDPSQRMWKYLNDLAKNTASEVEDLPYEELGDEDYDASLPYAALFSCLKAKGLKVTCLLCYCSEGDNIPDAFNLADAVSKALGLRPNSSQGNEGGSWTVPFSWKSVYGPPPDMSLF*

### NbL01g02110.1 phosphoribosylamine−−glycine_ligase−like [MW = 54.98 kDa]

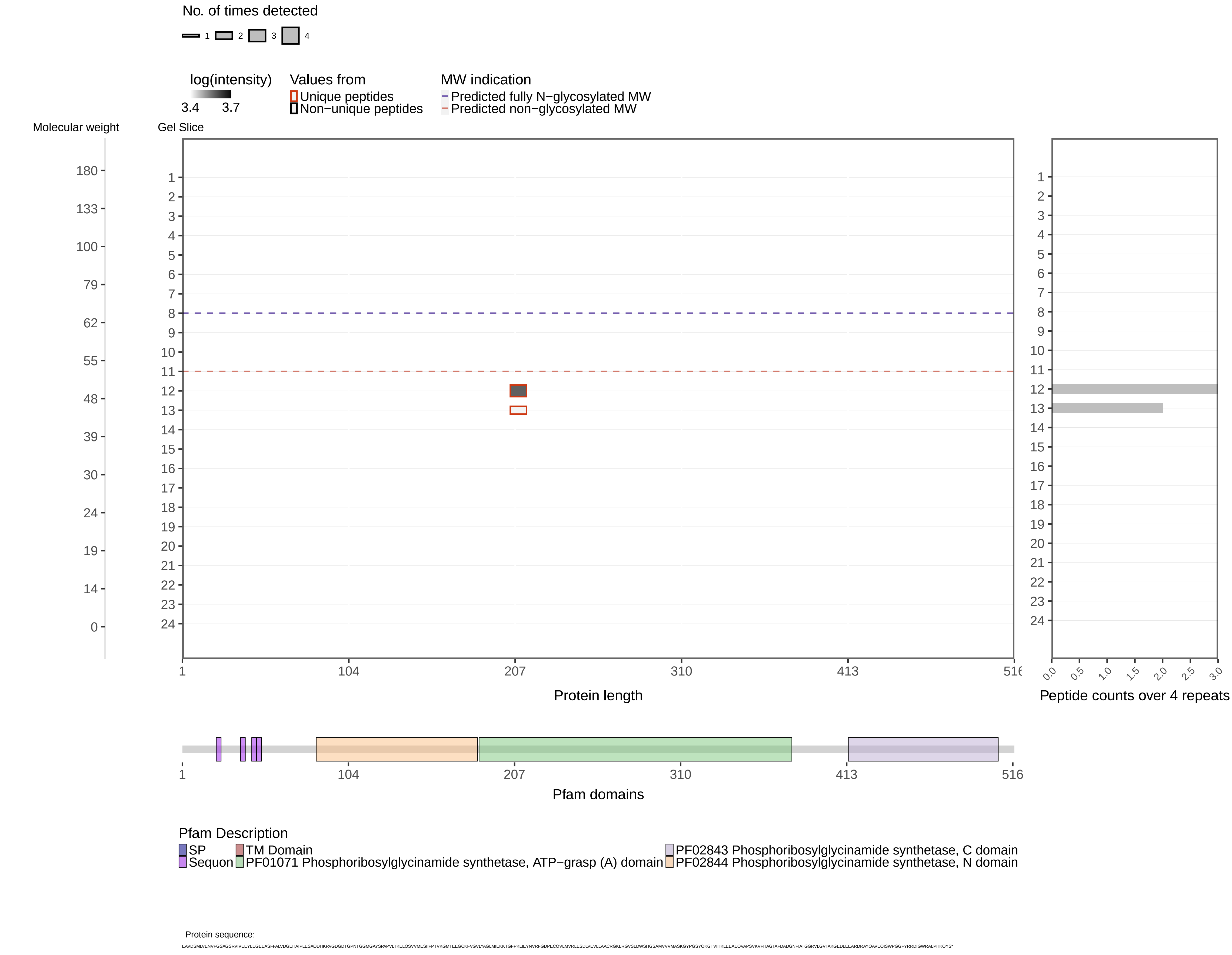

### NbL01g01930.1 tyrosine−−tRNA_ligase_1_cytoplasmic−like [MW = 46.3 kDa]

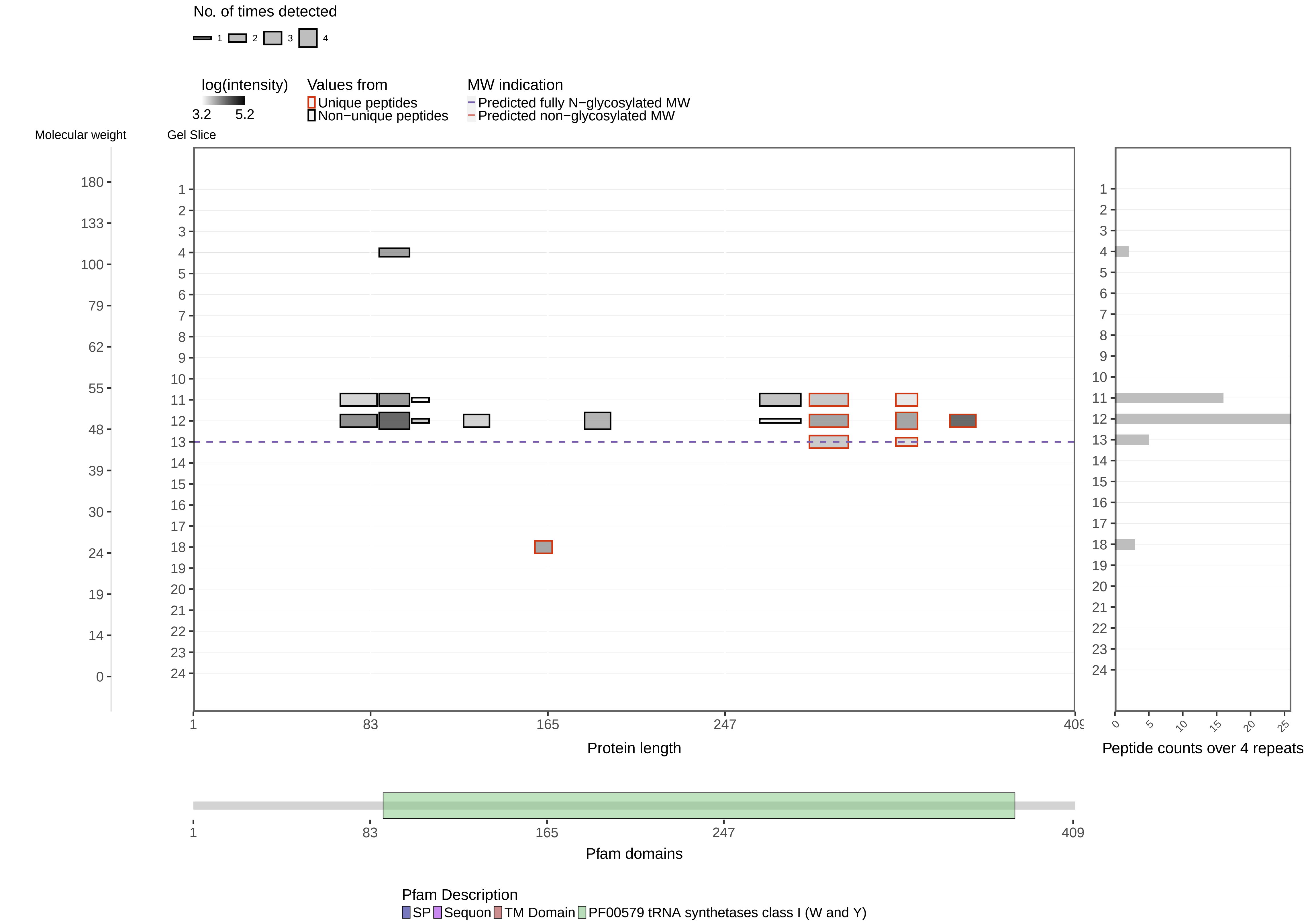


Protein sequence:

DIARRNKLPRILRCCQIMGRSEQDELTAAQIFYPCMQCADIFFLKADICQLGMDQRKVNVLAREYCDDIKRKNKPIILSHHMLPGLQEGQEKMSKSDPSSSIYMEDEEAEVNLKIKKAFCPPKVVEKNPCLEYIKYIVLPWFNEFKIERIAENGGDKTYKNFEELAADYESGKLHPADLKPALSKAINKILQPVRDHFKNDQKAKDLMKRVKSFKVTR*

NbL01g01760.1 ras−related_protein_RABG3f [MW = 23 kDa]

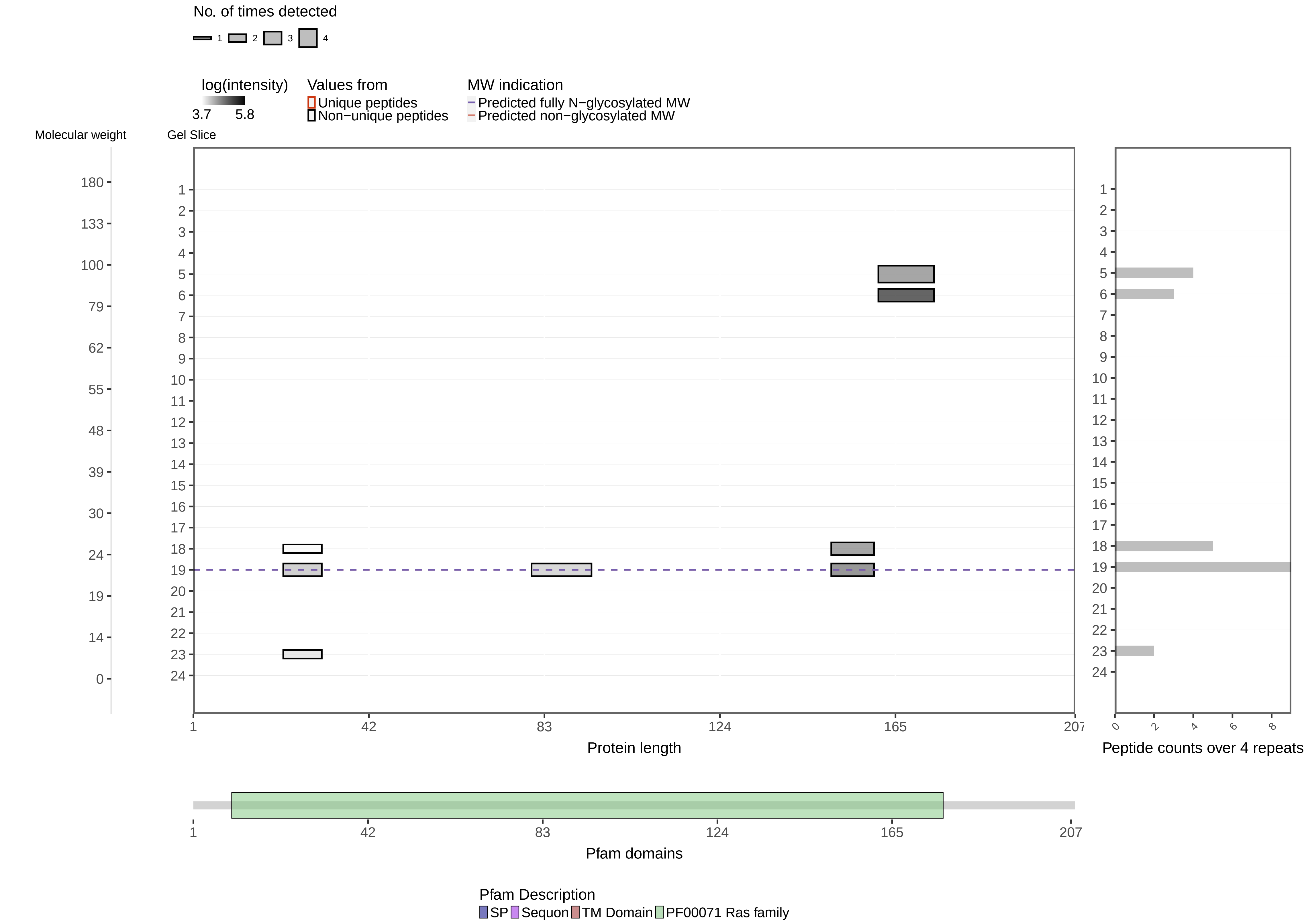


Protein sequence:

ISMKSFENLNNWREEFLIQASPSDPENFPFVVLGNKVDVDGGNSRVVSEKKARAWCASKGNIPYFETSAKEGTNVEEAFQCIAKNALKSGEEEEIYLPDTIDVGTSSQPRTGGCEC*

### NbL01g01740.1 coatomer_subunit_gamma [MW = 98.06 kDa]

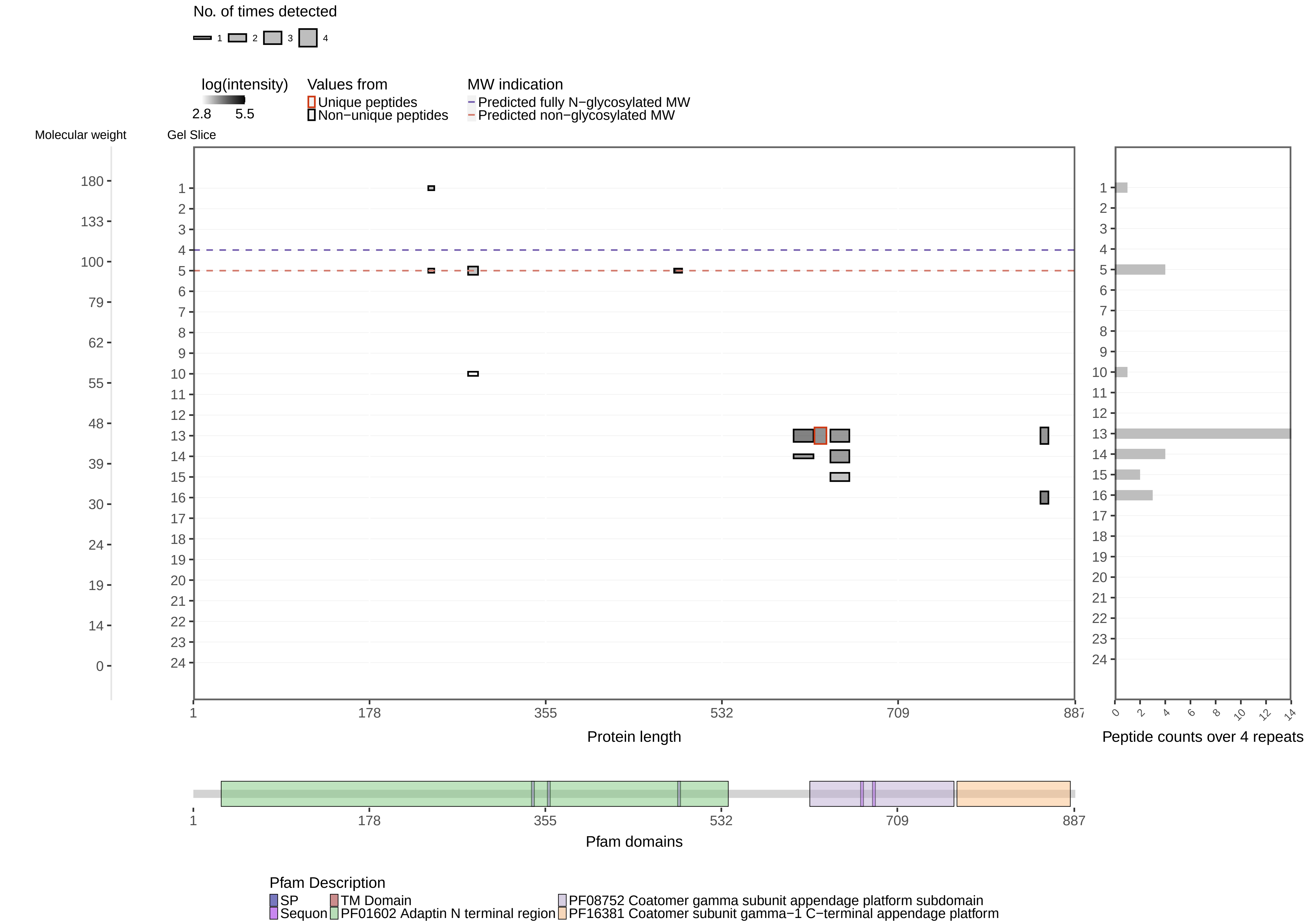


Protein sequence:

AKEGGLLHLCEFIEDCEFTYLSTQILHFLGNEGPKTSDPSKYIRYIYNRVILENATVRASAVSTLAKFGALVDSLKPRIFVLLKRCLFDSDDEVRDRATLYLNTLGGDGAVVETDEEVKEFLFGSLGVPLTNLETSLKNYEPSEEPFDIHSVPKEVKSQPLSEKKAPGKKPTGLAAPPVAPTSTVDAYERLLSSIPEFTSFGKLFKSSAPVELTEAETEYAVNVVKHNFDNHIVFQYNCTNTIPEQLLENVTVIVDASEAEEFSEVASKPLKSLPYDTPGQTFVAFEKPEGVPAVGKFS

NbL01g01540.1 glucose−1−phosphate_adenylyltransferase_large_subunit_1 [MW = 63.07 kDa]

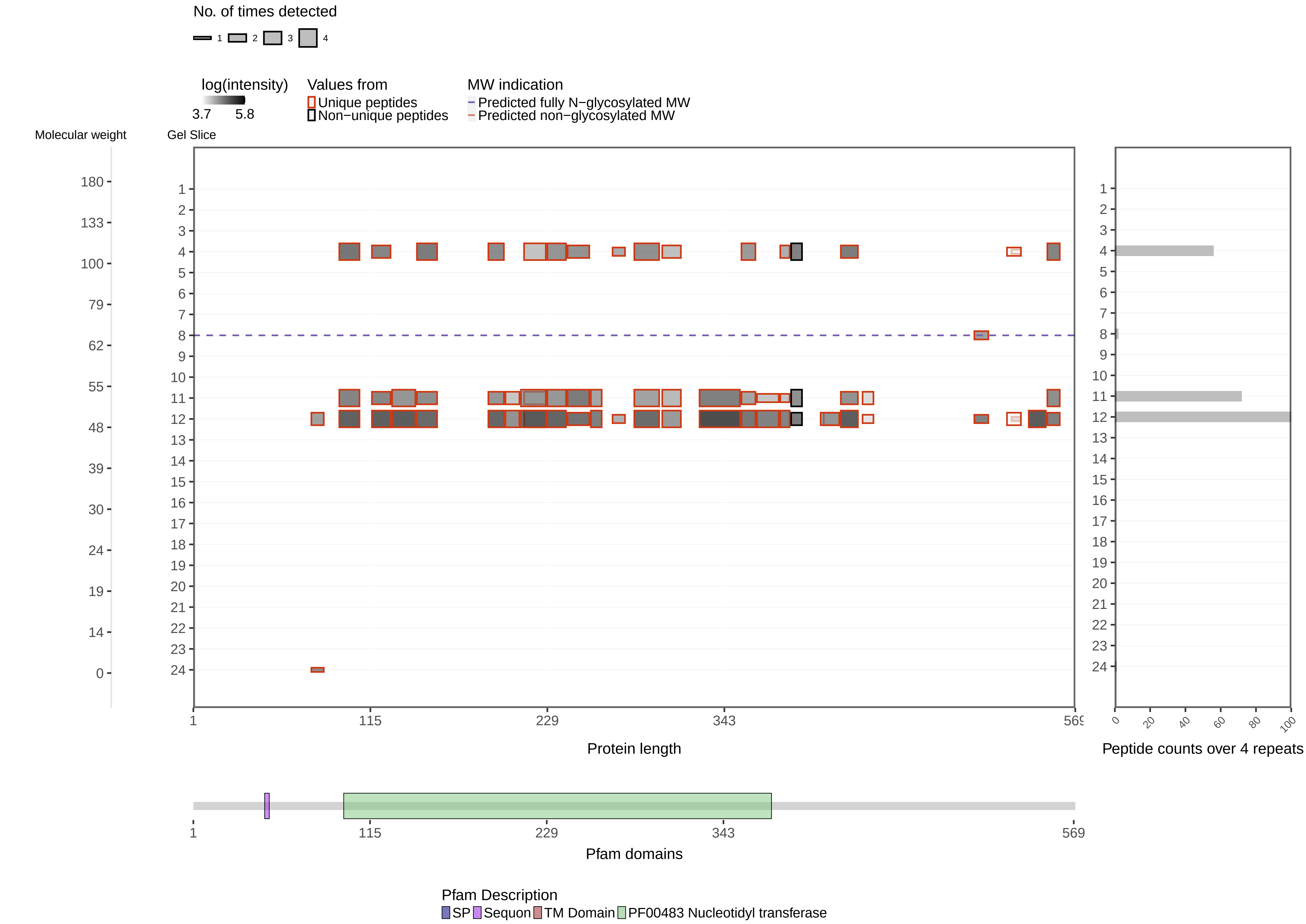


Protein sequence:

GRVVQFAEKPKGFDLKAMQVDTTLIGLSPQEAKRSPYIASMGVYVFKTDVLLKLLKWRYPTANDFGSEIIPAAINEHNVQAYIFRDYWEDIGTIKSFYDANLALTAESPKFEFYDPKTPFYTSPRFLPPTKIDNCKIKDAIISHGCFLRECSVDHSIVGERSRLDCGVELKVCMYLNICIYFLSFMQNAPFFLVHFLVRYGPQTPVLLPFQDTLMMGADYYQTESEIASLLAEGKVPMGIGENTKIRNCIIDKNAKIGKDVVIMNKDGVKEADRPEDGFYIRAGLTIVAEKATIKDGTVI*

NbL01g01530.1 bifunctional_dihydrofolate_reductase−thymidylate_synthase−like [MW = 58.93 kDa]

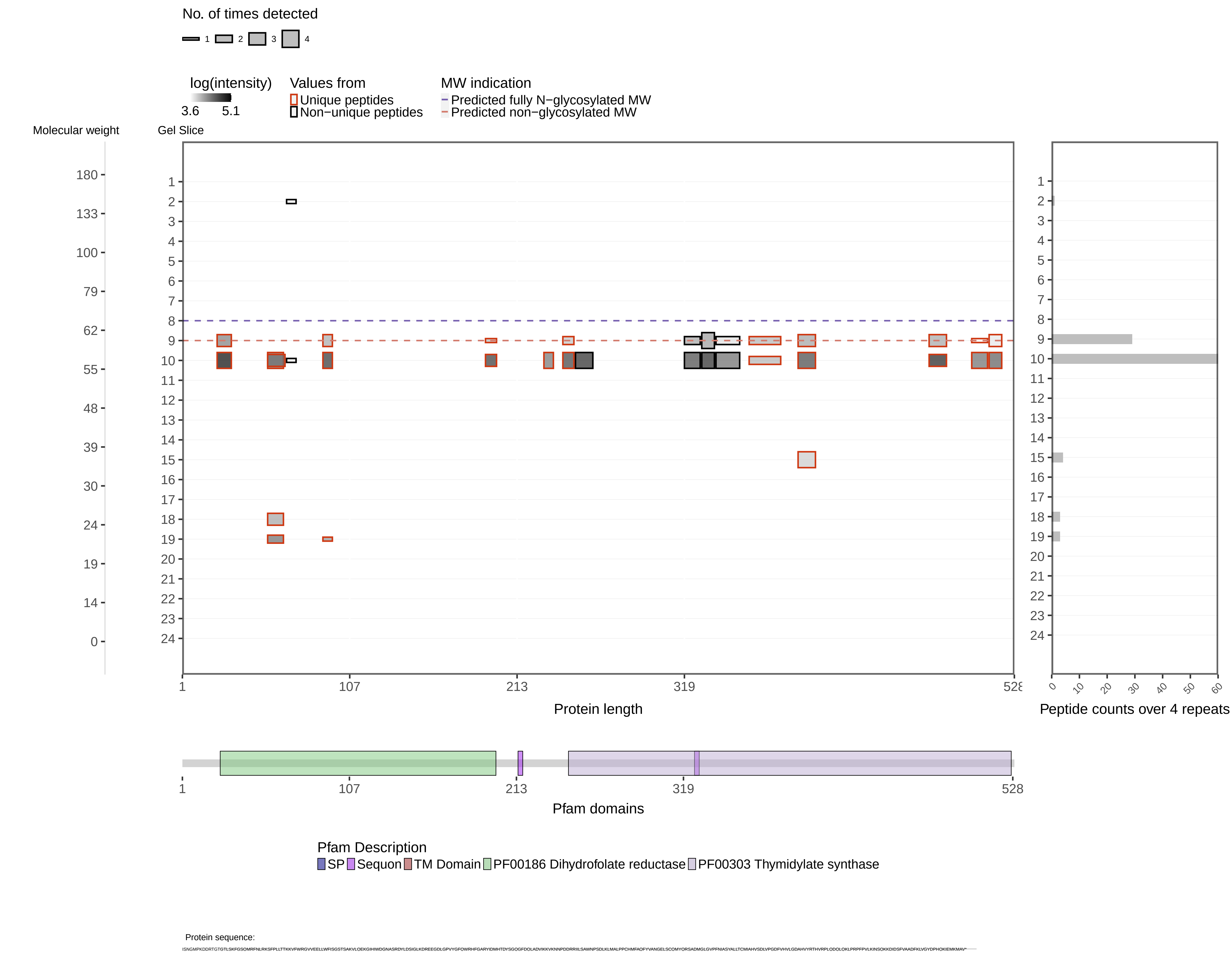

NbL01g01390.1 SUMO−activating_enzyme_subunit_2 [MW = 72.67 kDa]

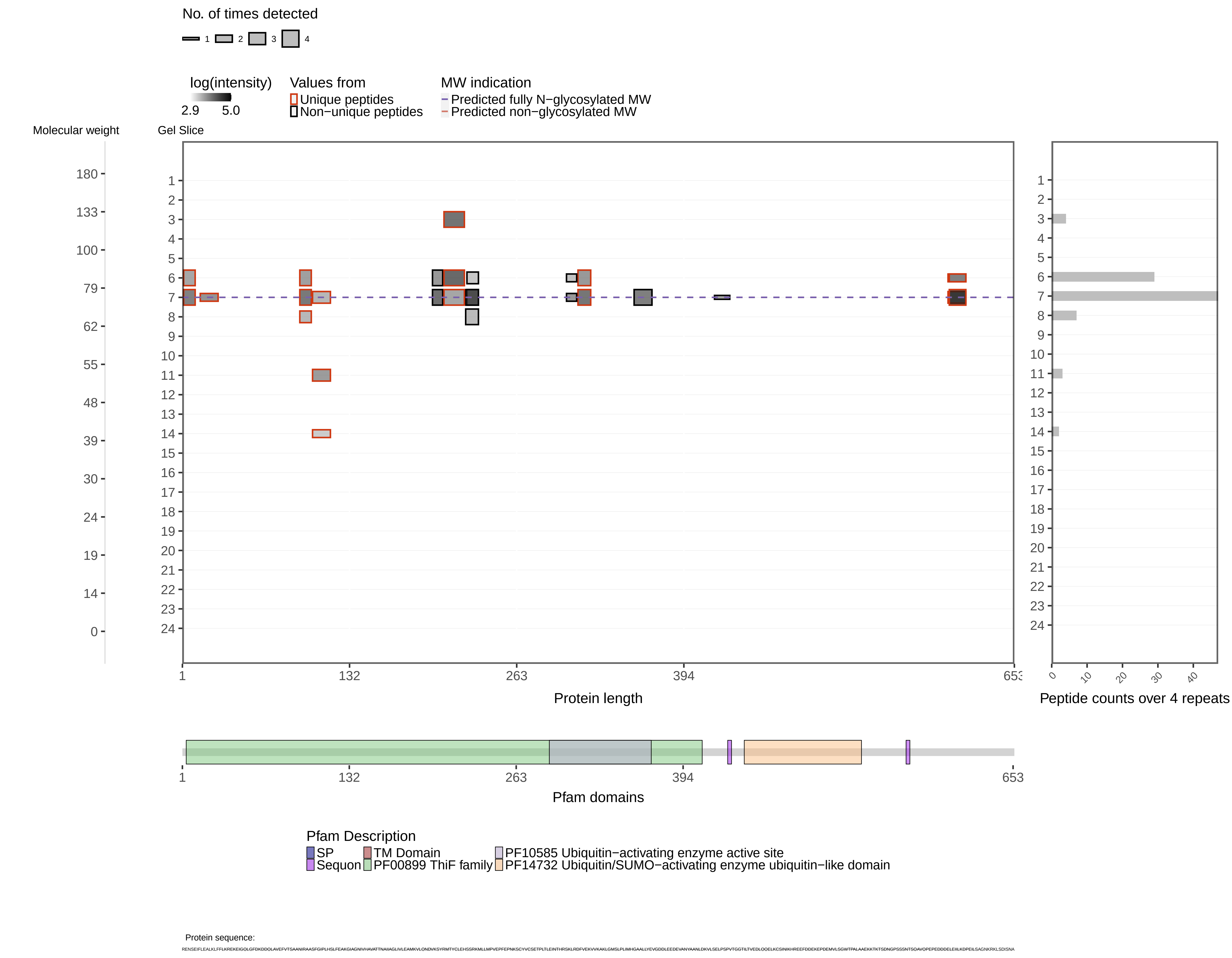

### NbL01g01340.1 probable_receptor_protein_kinase_TMK1 [MW − SP = 98.75 kDa]

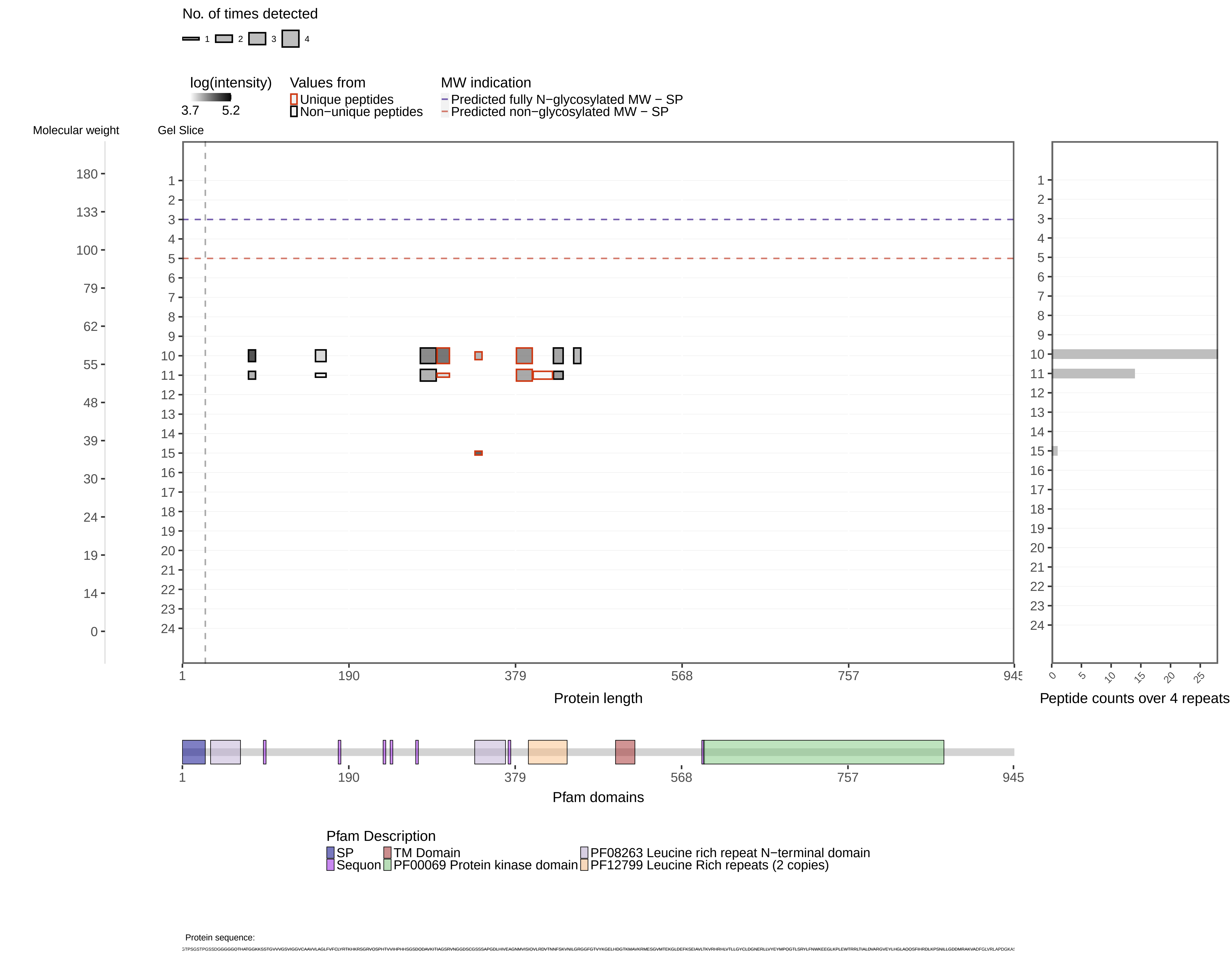

### NbL01g01330.1 beta−galactosidase_3−like [MW − SP = 81.74 kDa]

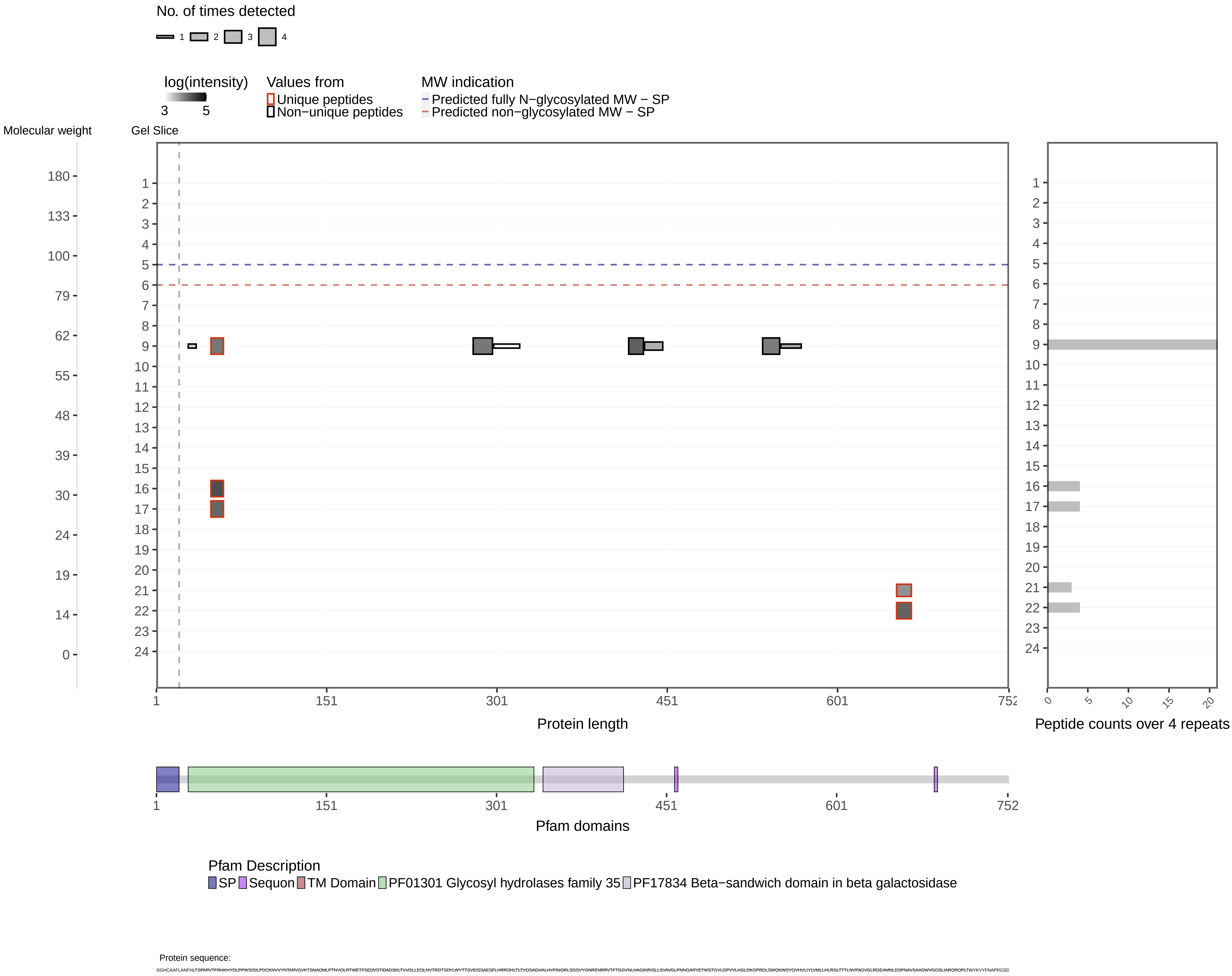

### NbL01g01310.1 purple_acid_phosphatase_2−like [MW − SP = 50.88 kDa]

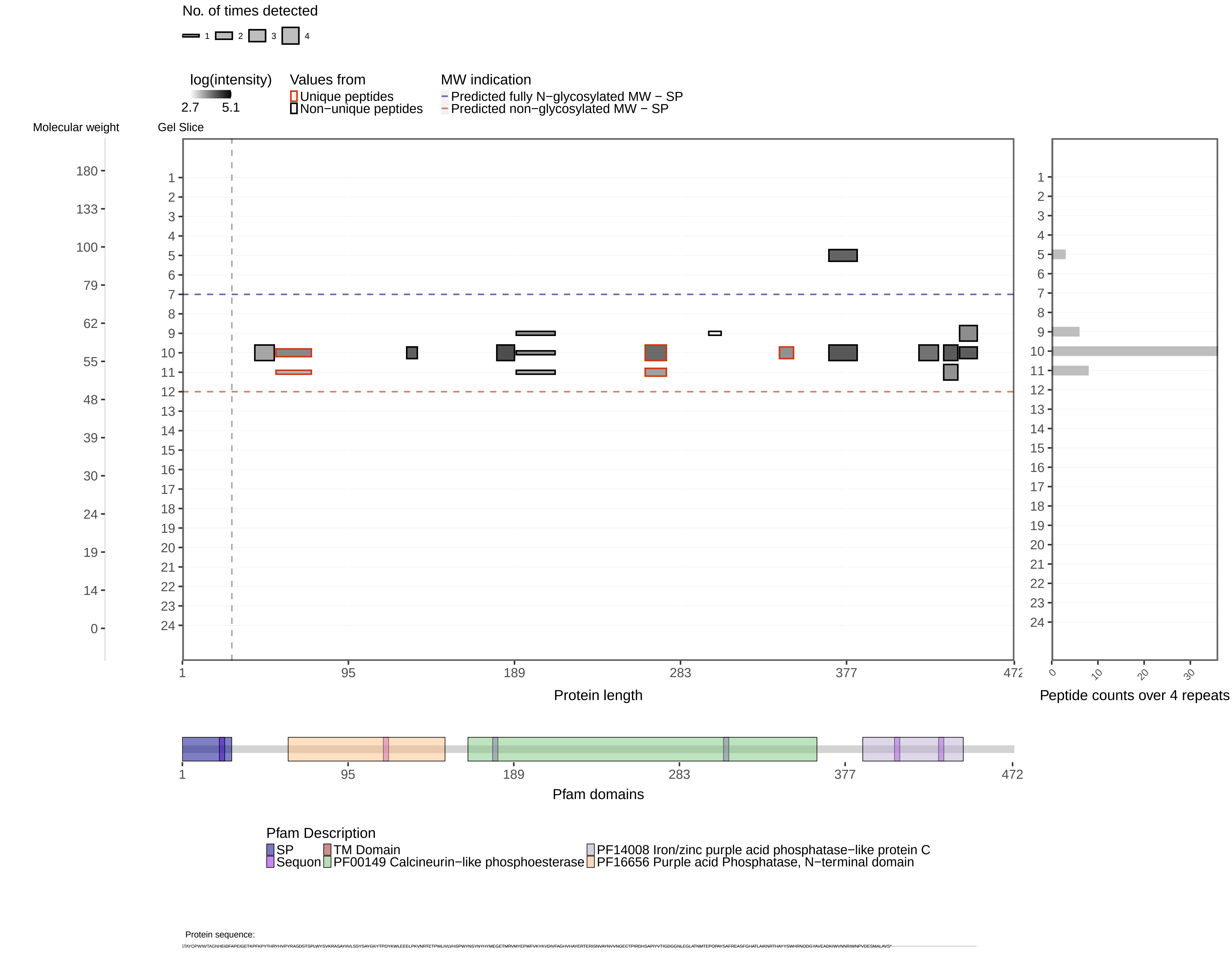

NbL01g01290.1 cysteine_protease_RD19A−like [MW − SP = 38.07 kDa]

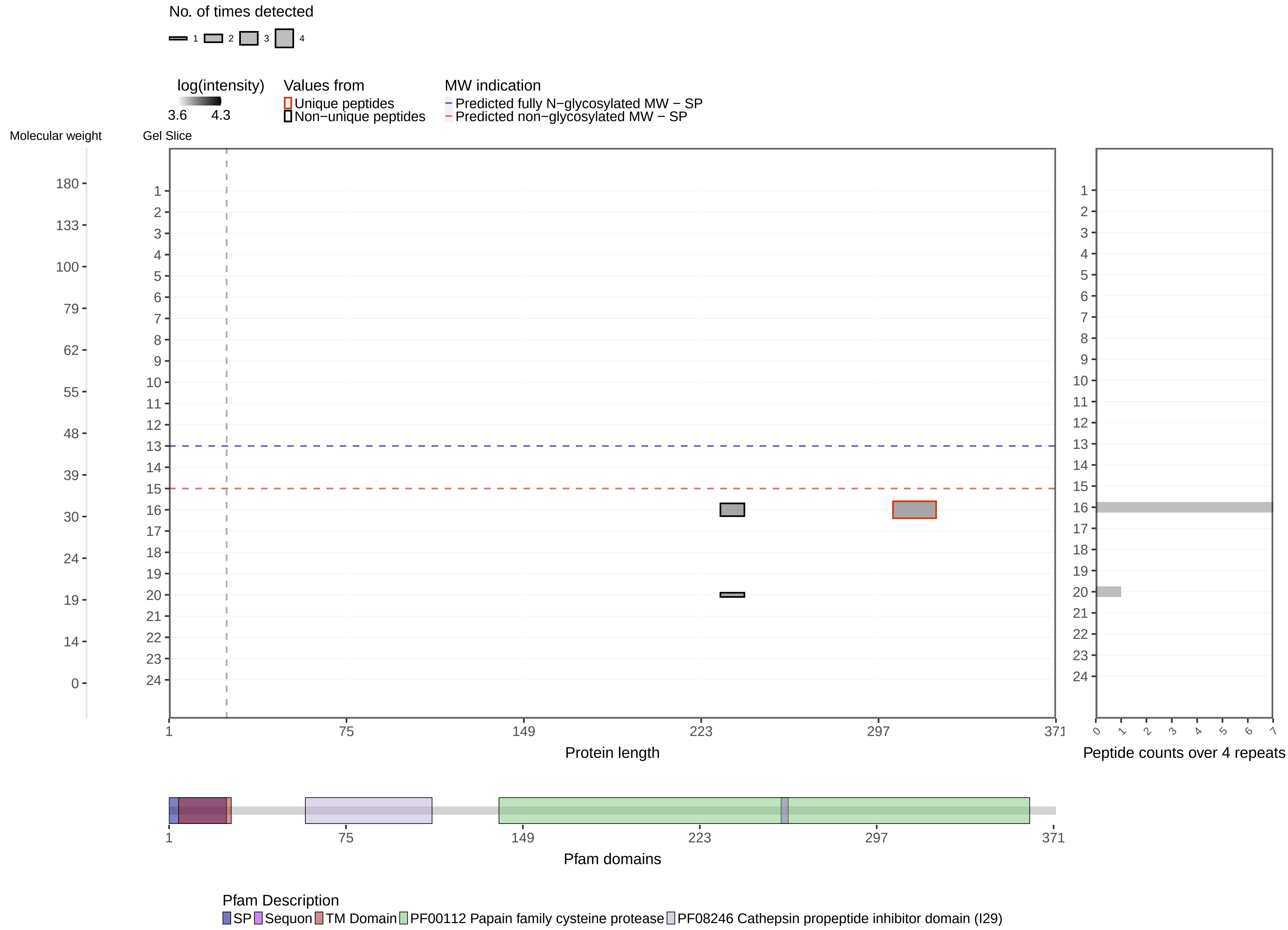


Protein sequence:

GANYLSTGKLVRLSEQQLVDCDHECDPEEKDSCDAGCNGGLMNSAFEYTLKAGGLMREEDYPYTGTDRGTCKFDNTKVAAKVANFSVVSLDEEQIAANLVKNGPLAVAINAVFMQTYVGGVSCPYICSKNLDHGVLLVGYGTGFSPIRMKEKPYWIIKNSWGEKWGENGYYKICRGRNVCGVDSMVSTVAAVSTSSH*

### NbL01g01280.1 V−type_proton_ATPase_subunit_a2−like [MW = 92.56 kDa]

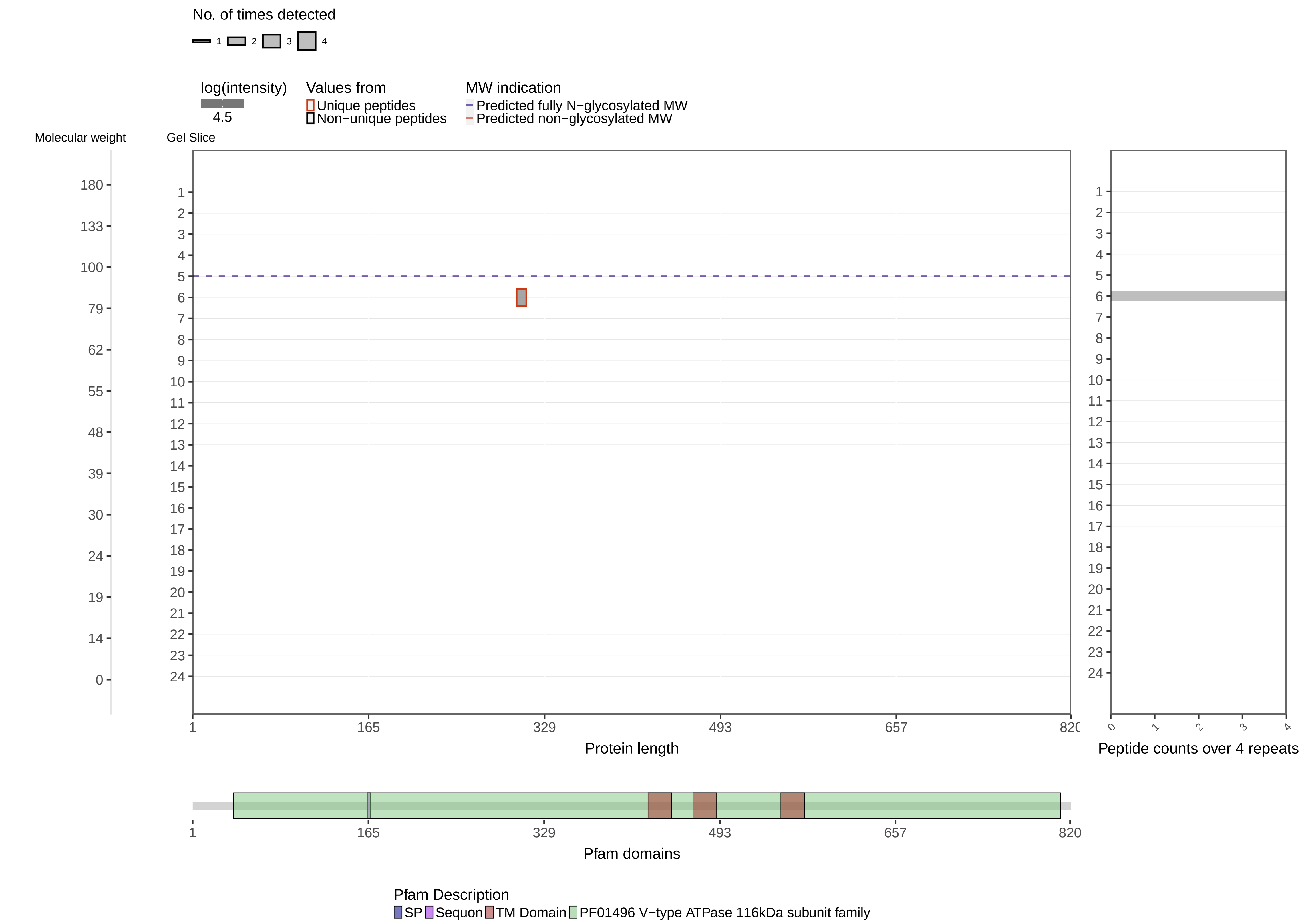


Protein sequence:

[illegible]

NbL01g01250.1 uncharacterized_LOC104113315 [MW − SP = 39.6 kDa]

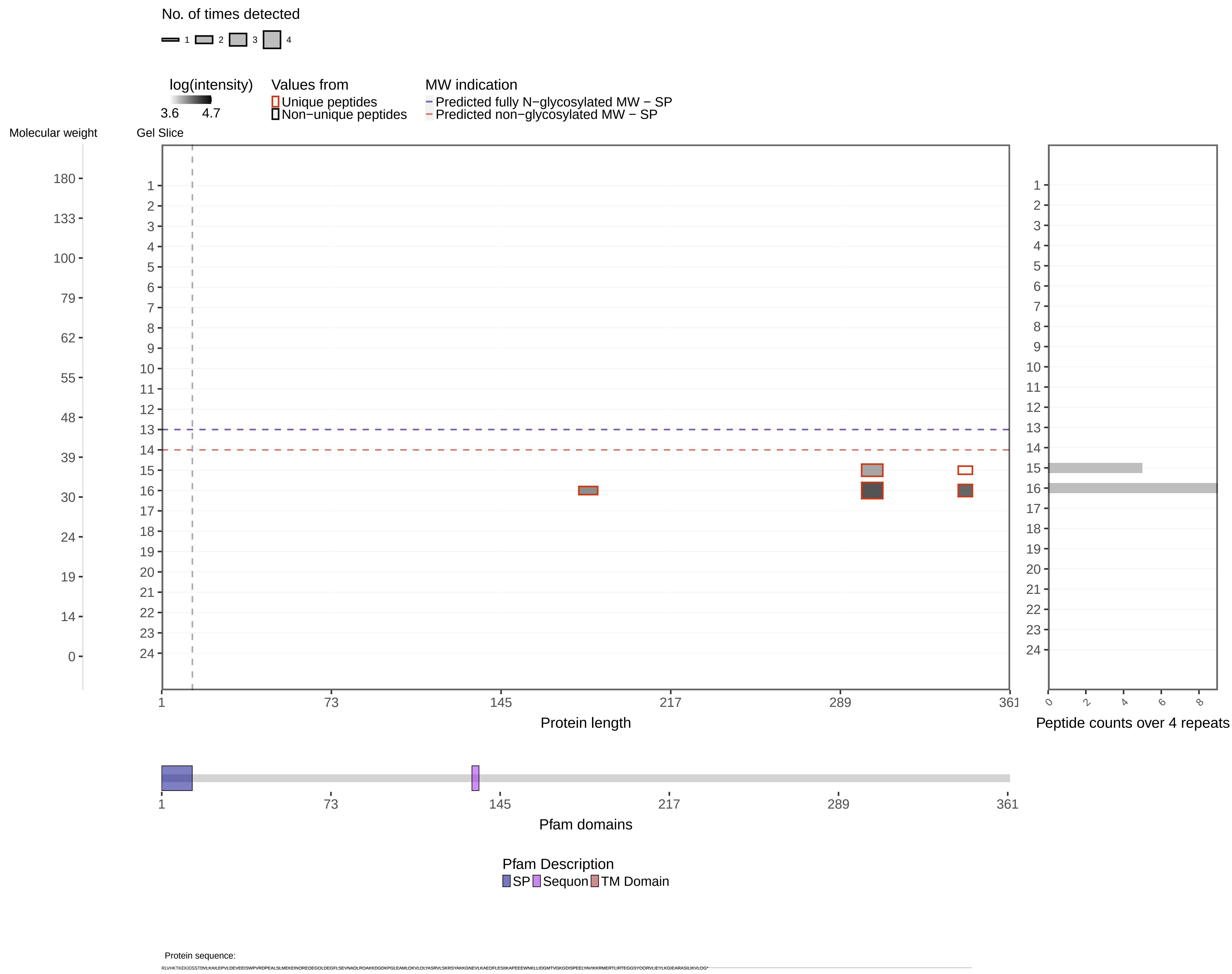

### NbL01g01210.1 uncharacterized_LOC104090505 [MW = 28.12 kDa]

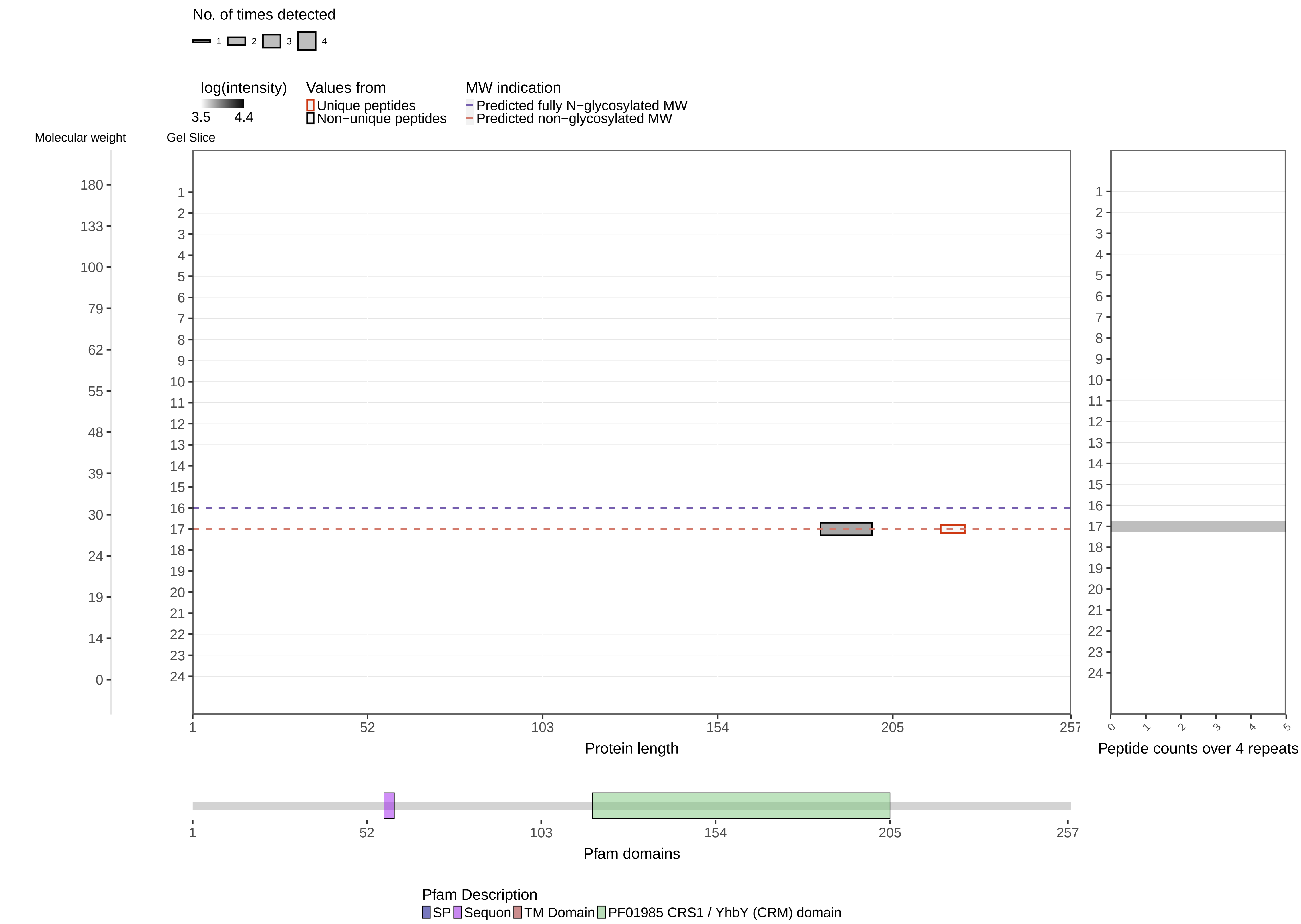


Protein sequence:

SLTVKEKKELASYAHSLGKKLKSQQVGKSGVTDSVIFALDETLEANELLKLKIHGTCPGGELDDVVKHLEEATGSVVVGQIGRTVILYRPSVTKLKAEEKKKQSQNAFMRKKQYAAKQSFESKGQVPKPYARGRRGSSRV*

### NbL01g01190.1 40S_ribosomal_protein_S11 [MW = 17.68 kDa]

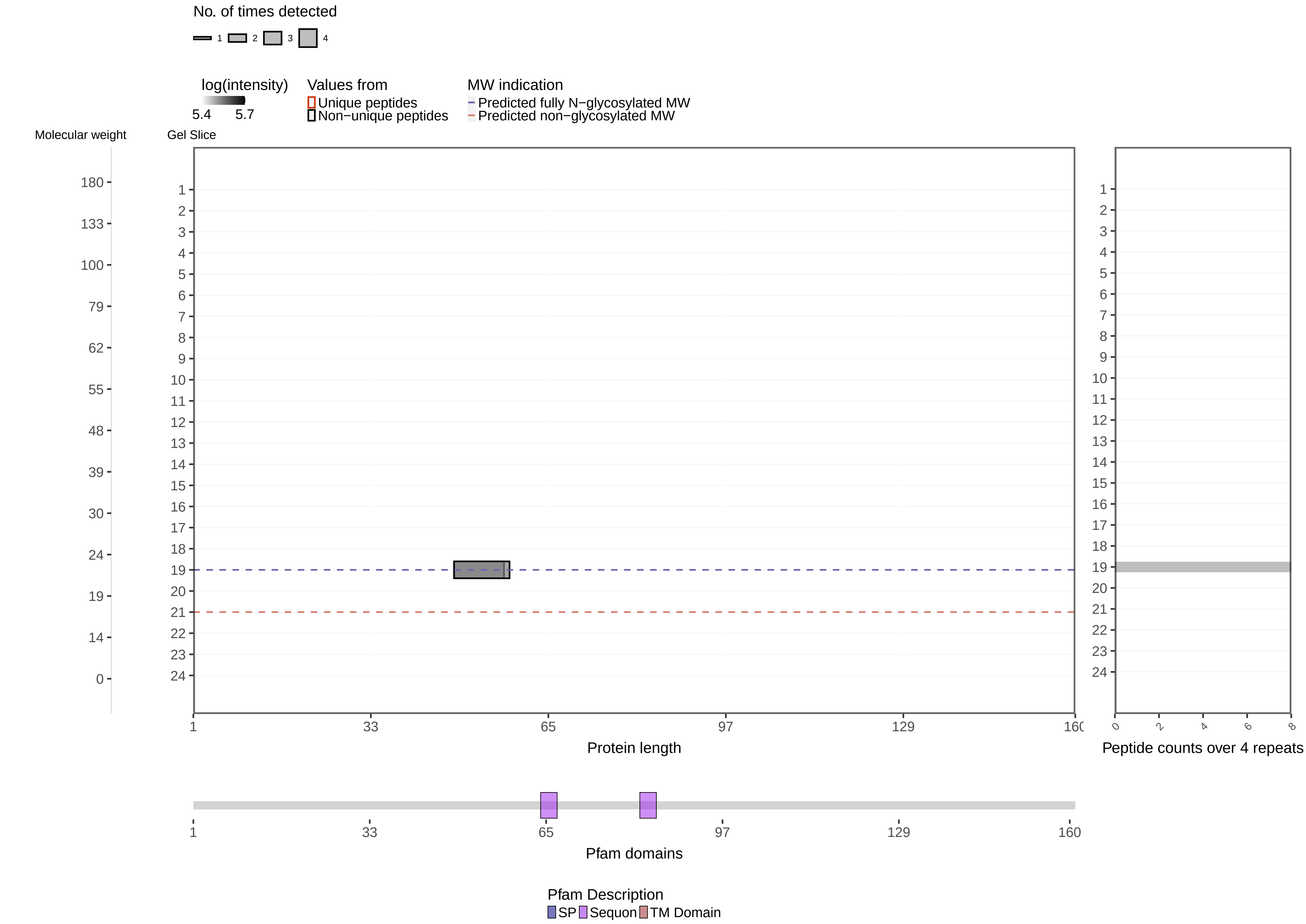


Protein sequence:

SIRGRILAGTCHSAKMNRTIIVRRNYLHYVKKYQRYEKRHSNIPAHISPCFRVKEGDHVIIGQCRPLSKTVRFNVLKVIPAGSAGGGKKAFTGM*

NbL01g01130.1 fructose−bisphosphate_aldolase_1_chloroplastic−like [MW = 42.8 kDa]

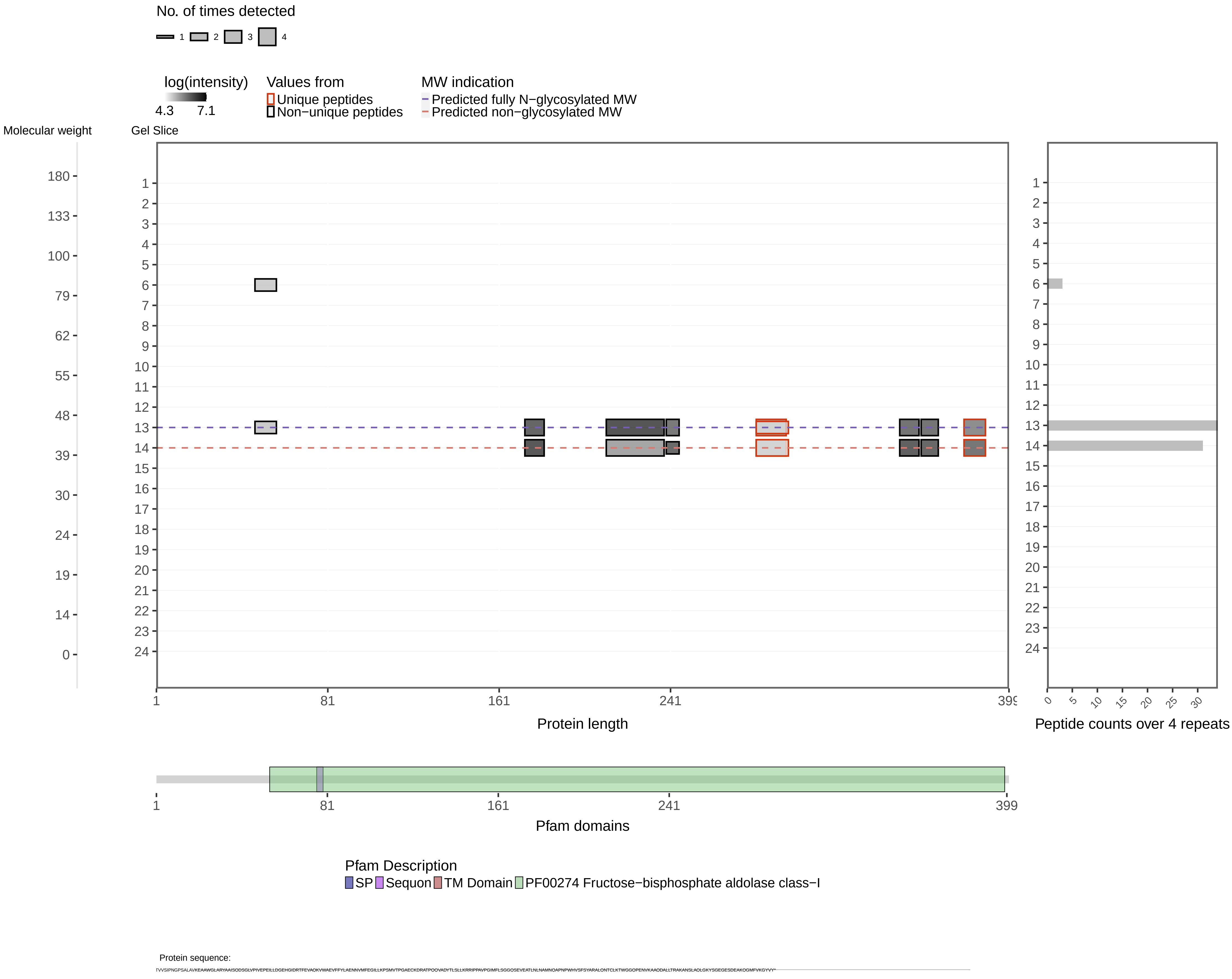

### NbL01g01040.1 NADP−dependent_D−sorbitol−6−phosphate_dehydrogenase−like [MW = 34.64 kDa]

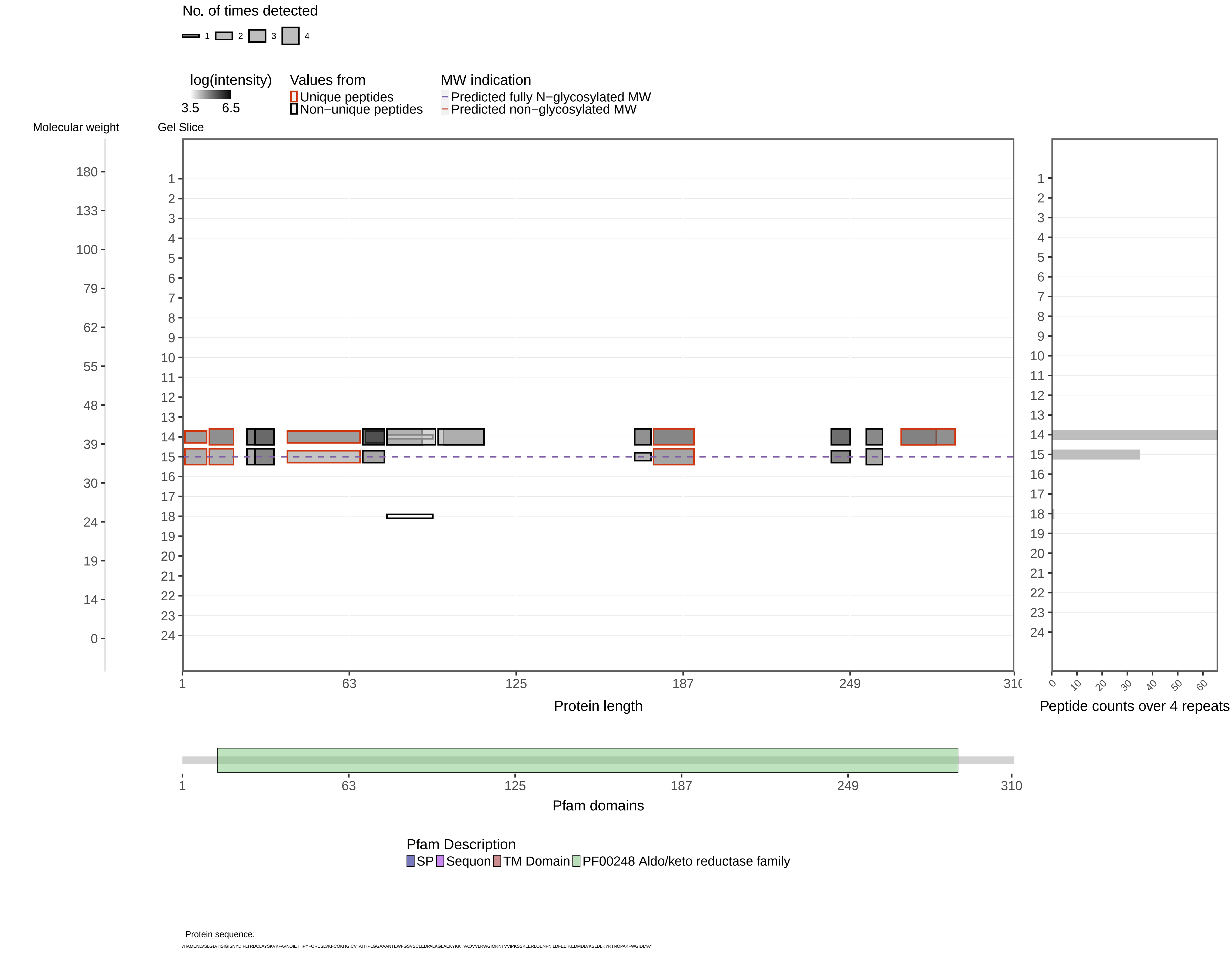

NbL01g01000.1 tRNA−dihydrouridine(47)_synthase_[NAD(P)(+)]−like [MW = 78.35 kDa]

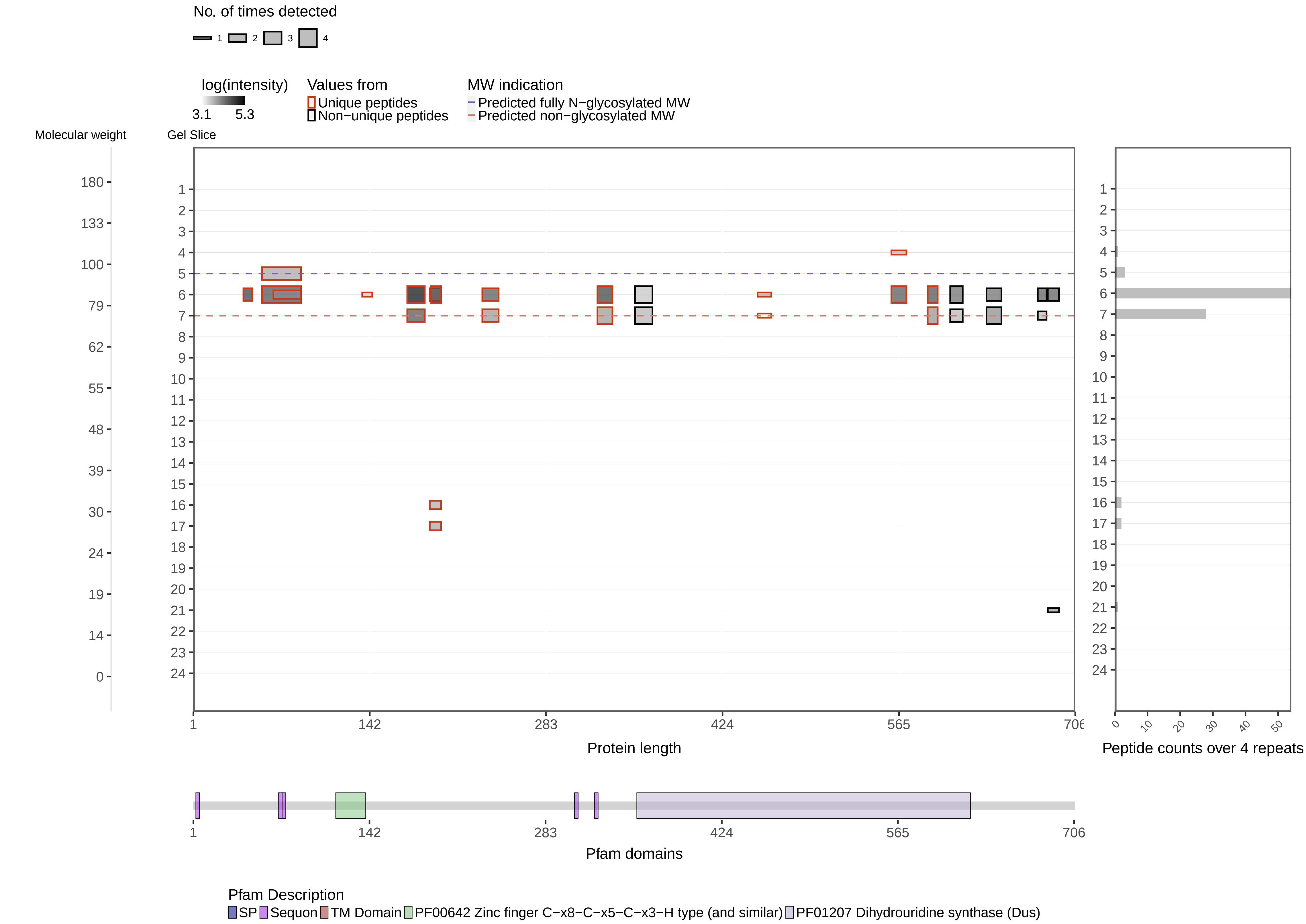


Protein sequence:

[illegible]

NbL01g00990.1 amidophosphoribosyltransferase_chloroplastic−like [MW − SP = 61.8 kDa]

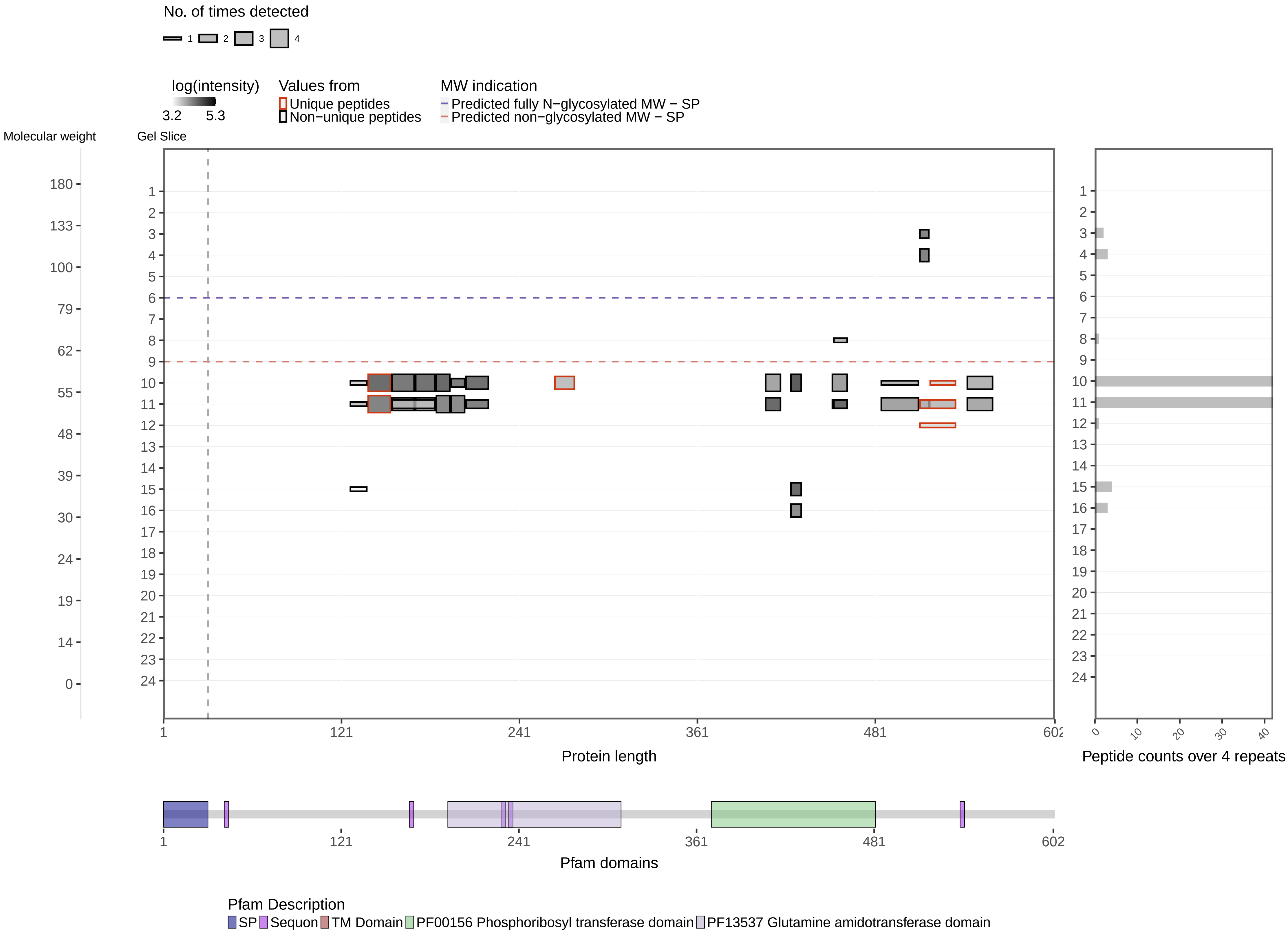


Protein sequence:

RPLVMGRRSNGAVVFASETCALDLIEATYEREVSPGEVVVVDKDGVQSICLMPHPERRSCIFEHIYFALPNSVVFGRSVYESRRAFGEILATEAPVECDVVIAVPDSGVVAALGYAAKAGVPFQQGLIRSHYVGRTFIEPSQKIRDFGVKLKLSPVRAVLEGKRVVVVDDSIVRGTTSSKIVRLLKEAGAKEVHMRIASPPIIASCYYGVDTPSSDELISNRMSVEEIKEFIGSDSLAFLPIDSLNKLLGNDSKSFCYACFSGNYPVEPTGKVKRIGDFMDDGLSGDMDSIDGGWL

### NbL01g00980.1 peptidyl−prolyl_cis−trans_isomerase_Pin1 [MW = 12.95 kDa]

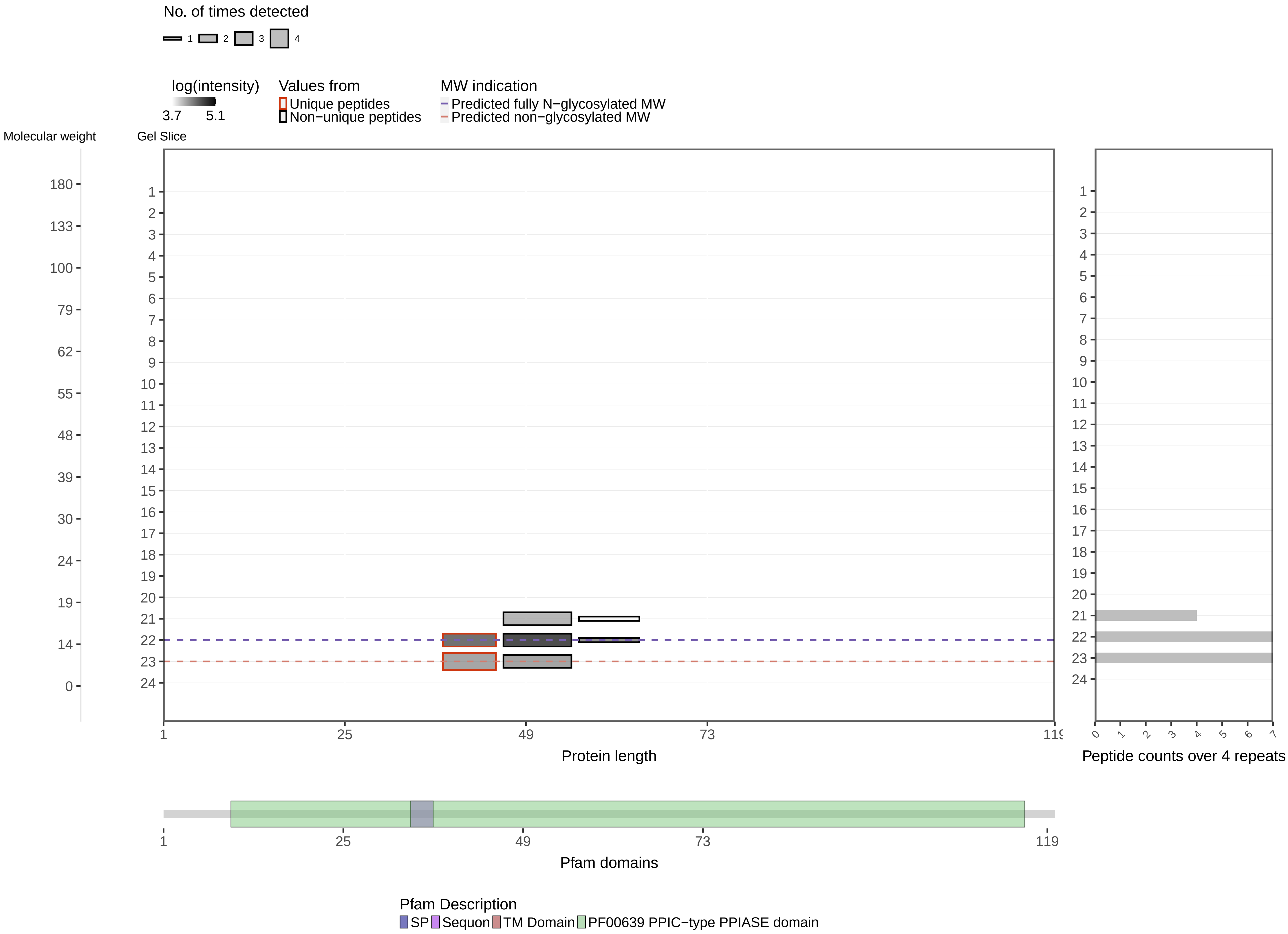


Protein sequence:

AFRDDIISGKSKFEDVASRFSDCSSAKRGGDLGPFGRGQMQKPFEDKTFALKVGEISDIIDTDSGVHIILRTG*

### NbL01g00870.1 pentatricopeptide_repeat−containing_protein_MRL1_chloroplastic [MW = 123.04 kDa]

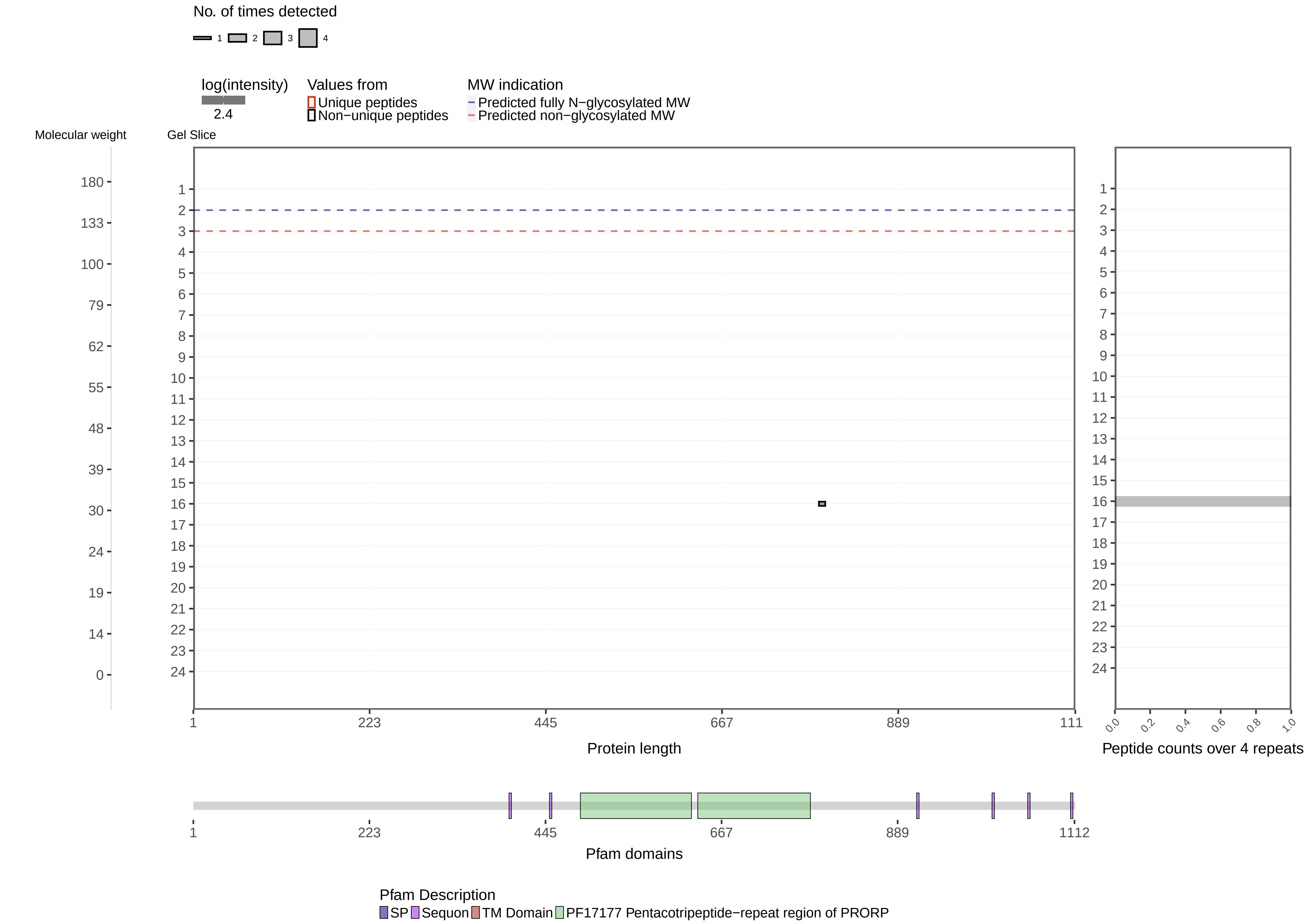


Protein sequence:

EVFHEMVNAGVEPNVNTYGALIDGCAKAGQVAKAFGAYGIMRSKNVKPDRVVFNALITACGQSGAVDRAFDVLSEMKAEARPIEPDQITIGALMKACANAGQVDRALDVYRMIDKYDIKGTAEVYTIAVNCCSQNGNWDFARSIYDDMTRKGVYPDEMFISALVDVAGHAGKLDAAFDVLEEARTKGINVGSMSYSSLMGACSNAKNWQKALELYEDVKGVKLKPTVSMMNALVTALCDADQYQKALEIFSEMKRVDLCPNTITYSTLLVASEKKDELDIGLMLLSHAKKDG

NbL01g00850.1 5'−methylthioadenosine/S−adenosylhomocysteine_nucleosidase_2 [MW = 27.78 kDa]

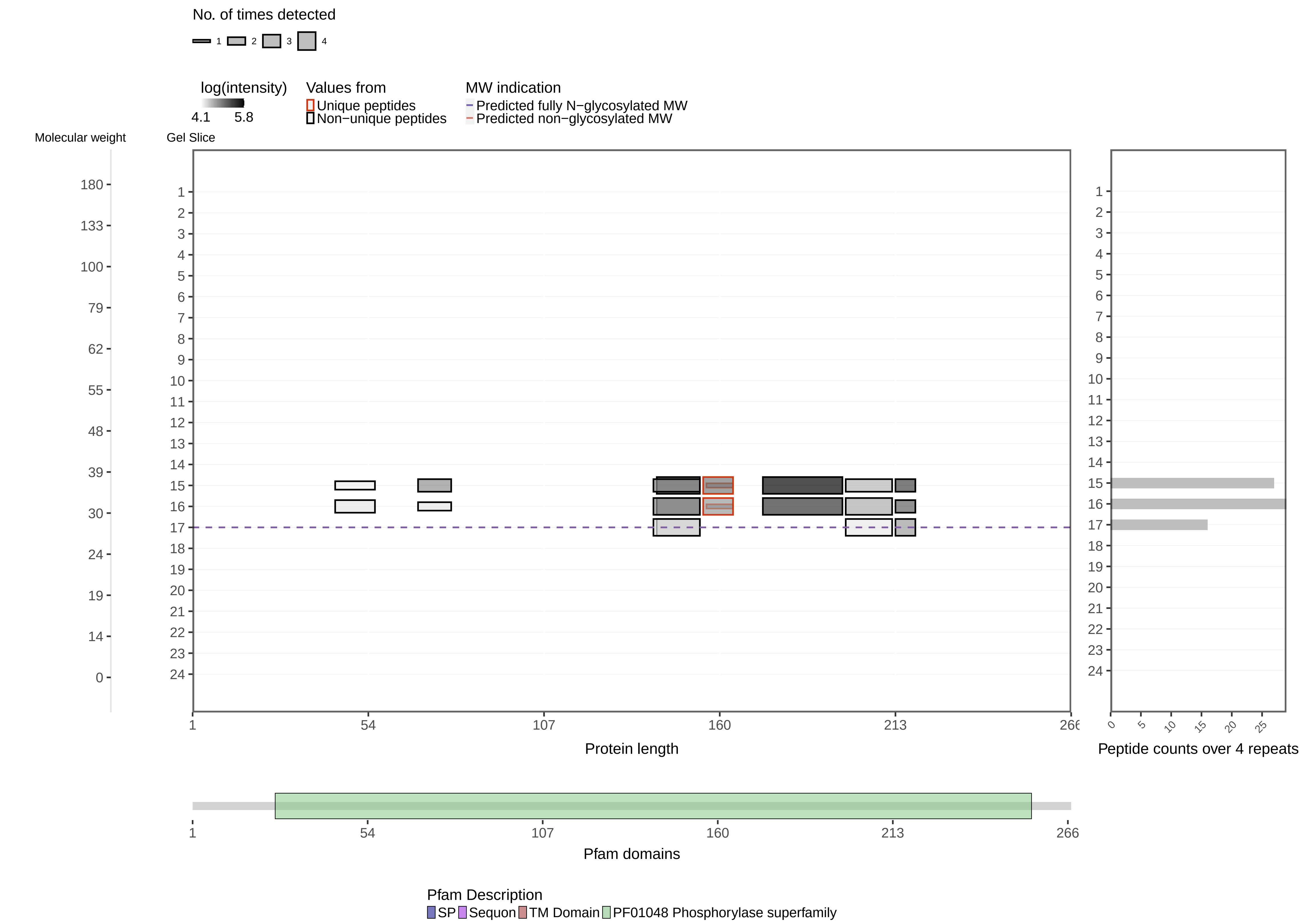


Protein sequence:

AKGASIGDVFLASEVAFHDRRIPIPVFDIYGVGSRKAFATPNLLKELNLKVGKLSTGDSLDMSPMDEASIVANDATVKDMEGAAVAYVADLLKVPVIFLKAVTDIVDGDKPTAEEFLQNLAAVTAALDQTATLVVDCISGKCLSDL*

NbL01g00810.1 triosephosphate_isomerase_chloroplastic [MW = 34.78 kDa]

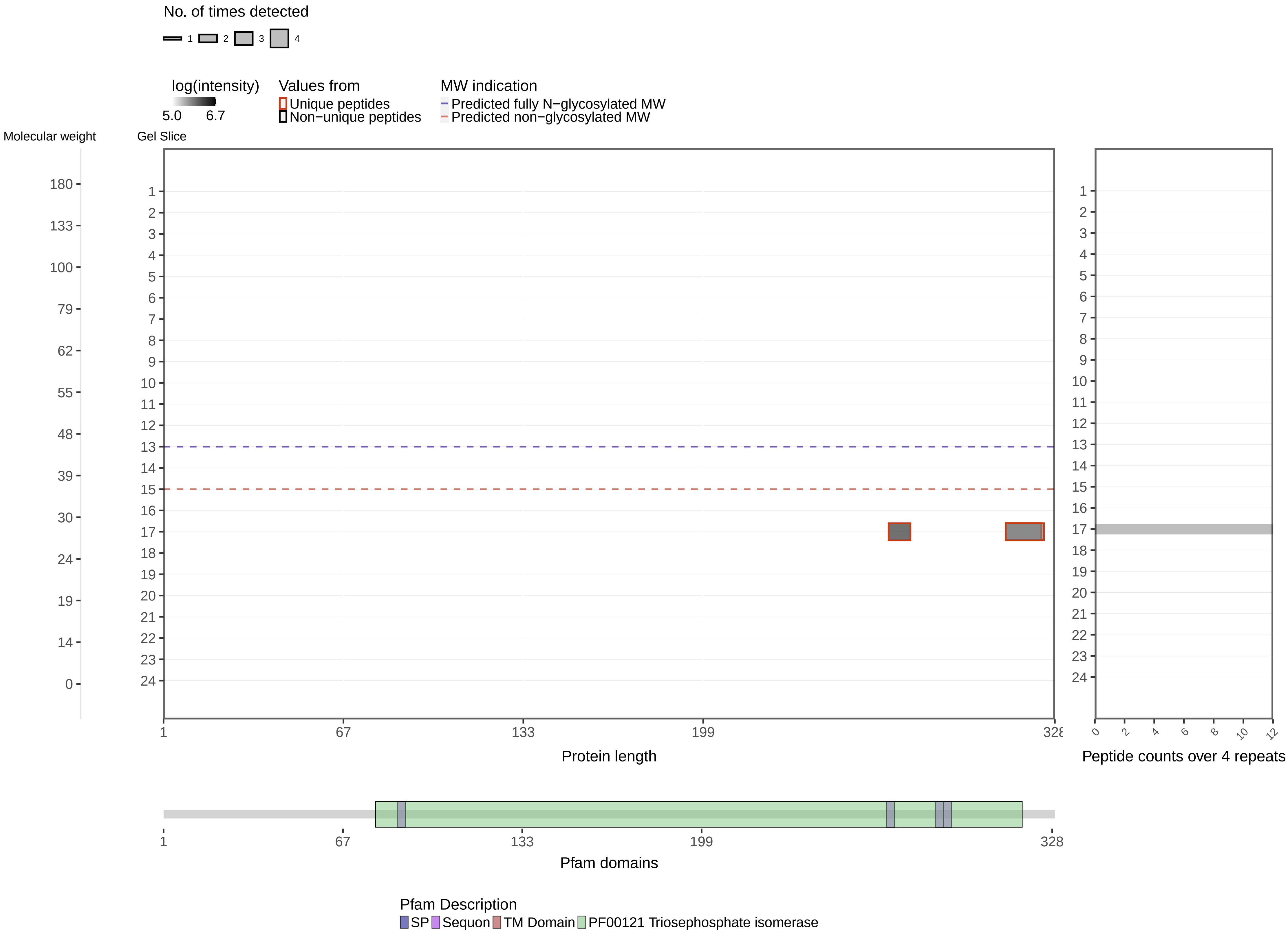


Protein sequence:

EISVEQLKDLGCKWVILGHSERRHVIGENDEFIGKKAAYALSQGVGVIACIGELLQEREAGKTFDVCFQQLKAFADALPSWDNVVIAYEPVWAIGTGKVASPEQAQEVHAAVREWLTKNVSADVASKTRIIYGGSVNGSNSSDLAKKEDIDGFLVGGASLKGTEFATIINSVTSKKVAA*

NbL01g00760.1 peptidyl−prolyl_cis−trans_isomerase−like [MW = 18.1 kDa]

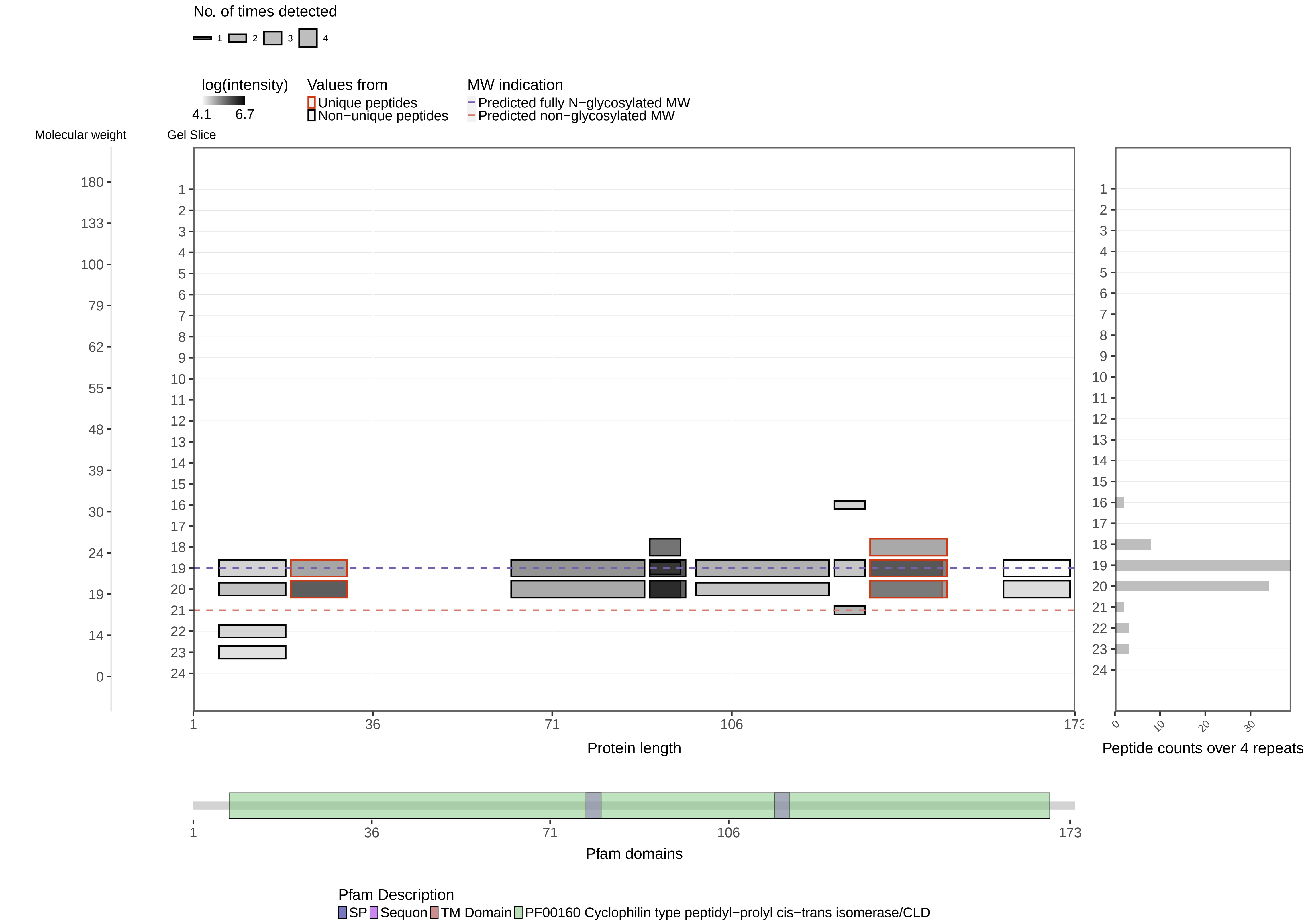


Protein sequence:

FTAGNGTGGESIYGAKFADENFKRKHTGPGILSMANAGPGTNGSQFFICTAKTEWLDGKHVVFGQVVEGYDVIKKAEAVGSGSGRCSKPVVIADCGQLC*

NbL01g00700.1 translocase_of_chloroplast_120_chloroplastic−like [MW = 152 kDa]

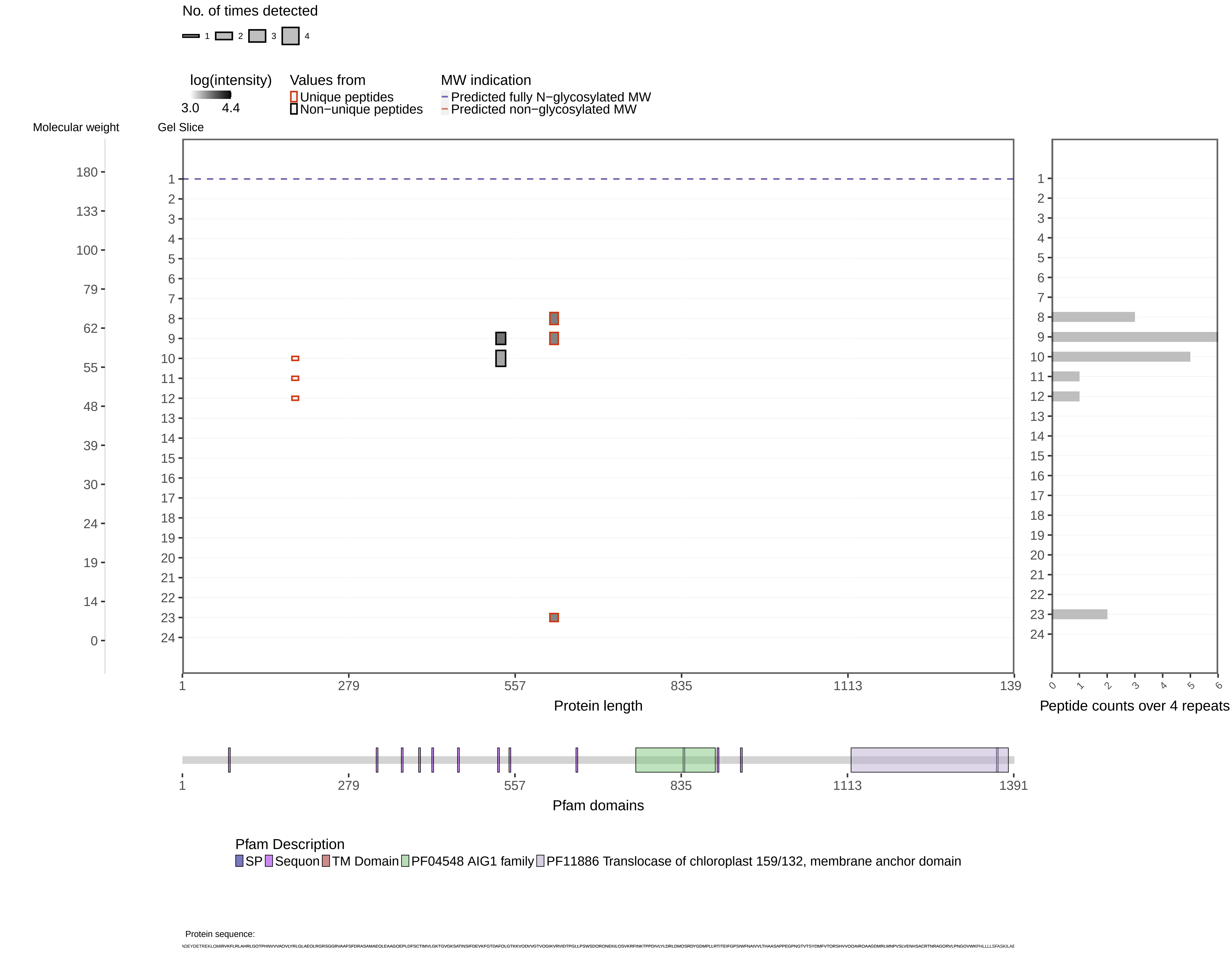

NbL01g00680.1 protein_2_(GRP−2)_mRNA_X60007.1_N.sylvestris_mRNA_for_glycine_rich_protein_2 [MW = 19.67 kDa]

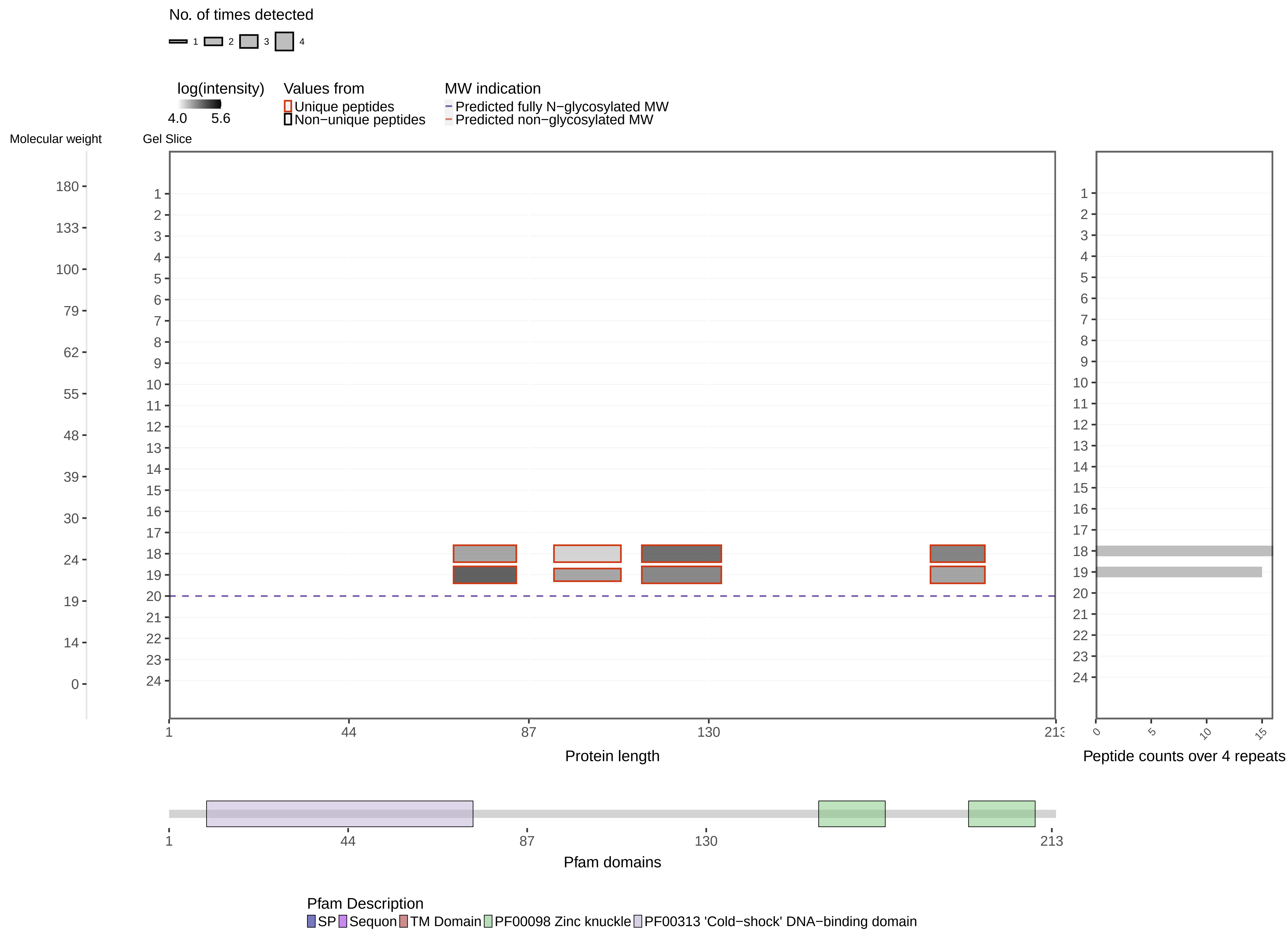


Protein sequence:

GGGGGGYGGGRGGSRGYGGGDGGYGGGGGYGGGSRYGGGGGGYGGGGGYGGGGSGGGNGCFKCGESGHFARDCTQSGGGGGGGRFGGGGGGGGGGCYKCGEDGHFARECTSGGR*

### NbL01g00640.1 thaumatin−like_protein_1b [MW = 41.31 kDa]

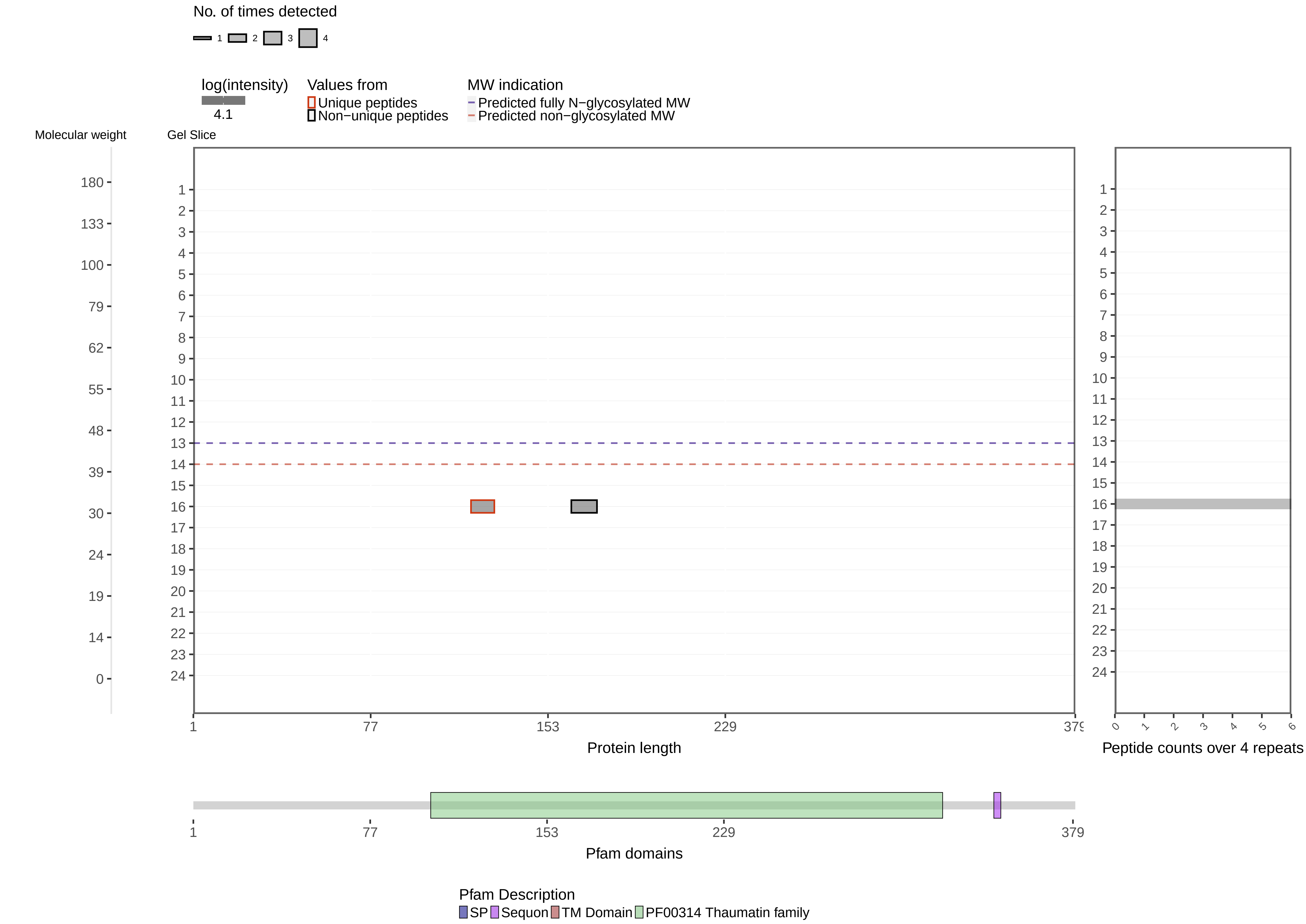


Protein sequence:

CAGGGAIPPATLAEFTLNGDQGLDFYDVSLVDGYNLPMLIVAKGGTRGGCSATGCLVDLNGACPRDLRVASGKGSHSESVGCKSACLAFNDPIYCCSESHNTPDTCQPSVYSLFFKHACPRSYSYAYDDKTSTFTCASADYLIIFCPLPYTSMKVLGLRKDPAELPLVNKTMMYIGRHHRSGESSIRPSLLHPYGVSNNLHL*

NbL01g00540.1
L−ascorbate_peroxidase_3_peroxisomal_(LOC104249656)_transcript_variant_X1_mRNA_XM_009806132.1_PREDICTED:_Nicotiana_sylvestris_L−
[MW = 35.39 kDa]

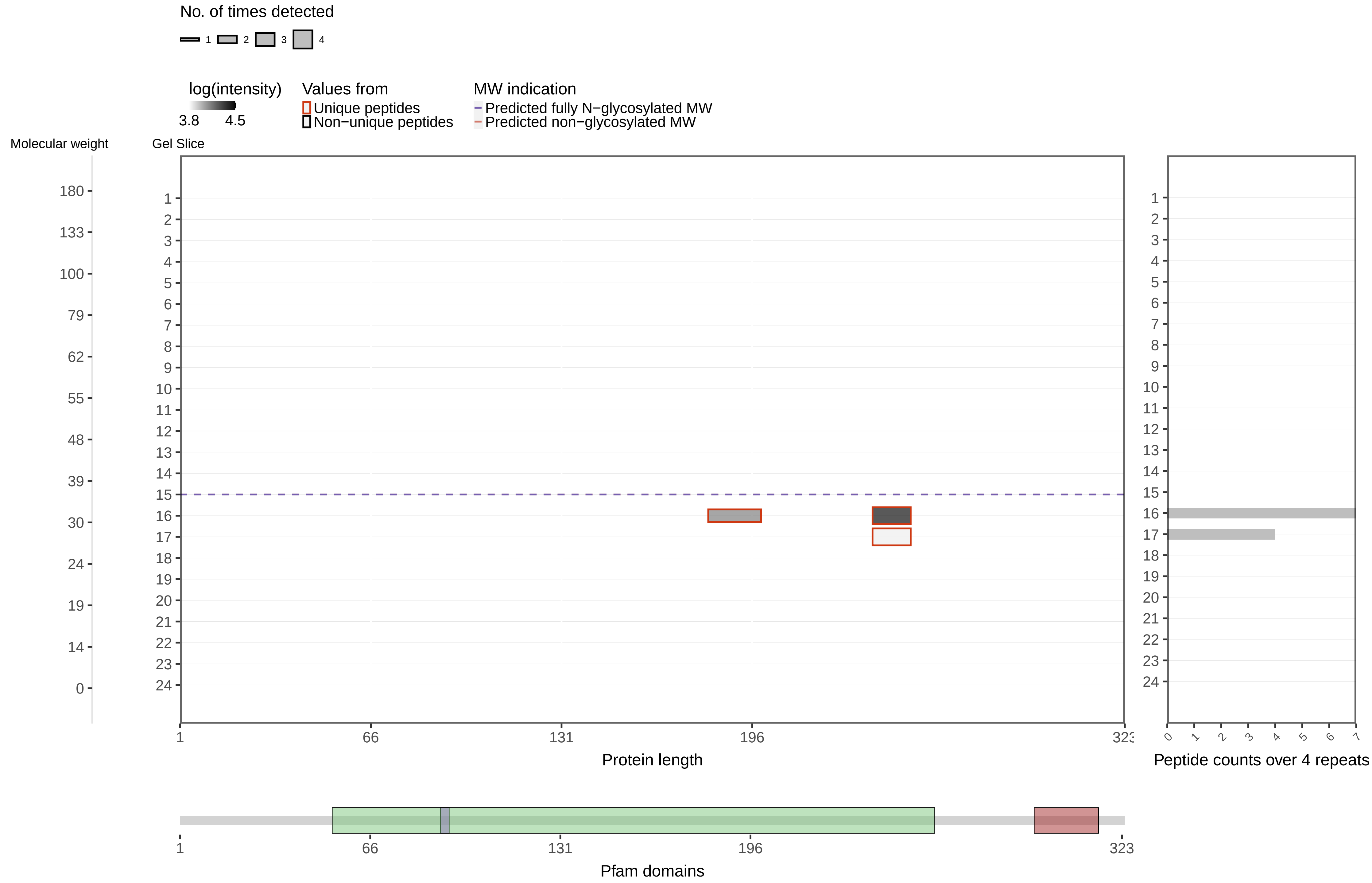


Protein sequence:
GRKDSSVSPKEGRLPDAKQGVPHLKDVFYRMGLSDEDIVALSGGHTLGRAHPERSGFDGPWTKEPLKFDNSYFVELLKGESEGLLKLPTDIALLDDPEFRQYVELYAKDEDAFFRDYAISHKKLSELGFTPSSGSKATVRDGTILAQSAVGVVVAAAVVALSYWYEVRKRMK*

### NbL01g00220.1 L−galactose_dehydrogenase [MW = 34.92 kDa]

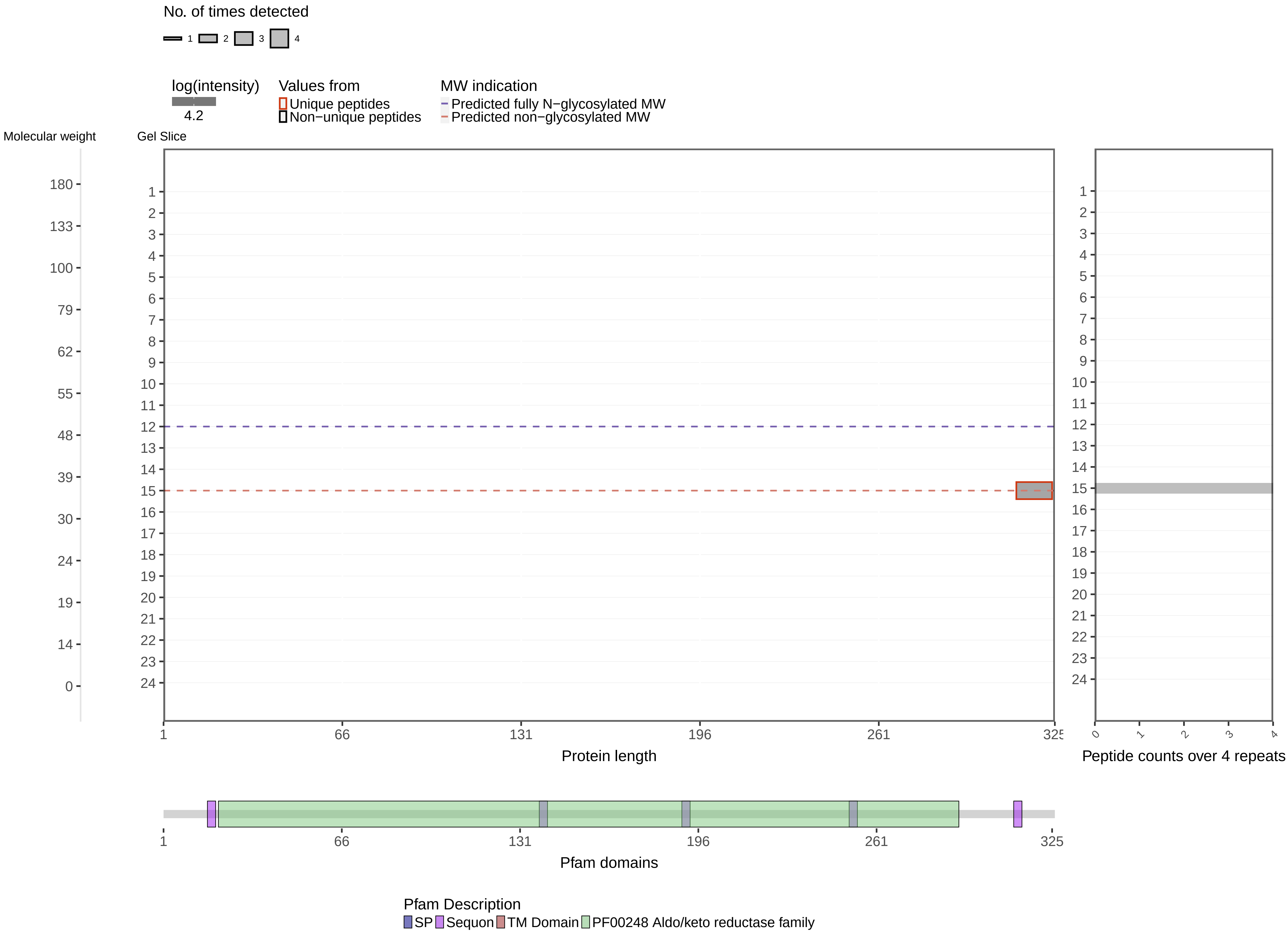


Protein sequence:

KQAGKIRFIGITGLPLGIFTYVLDRVPPGTVDVILSYCHYSINDSTLEDPLPYLKSKGVGVISASPLAMGLLTEAGAPEWHPASSELKAACRAAVDHCKGKGKNISKLALQYSLANTDISSVLVGMKSVKEVEENIGAALELATAGMDEETLSEITDILKPVKNQSWPSGIQQSLFV*

### NbL01g00180.1 peptide_methionine_sulfoxide_reductase_B5−like [MW = 22.45 kDa]

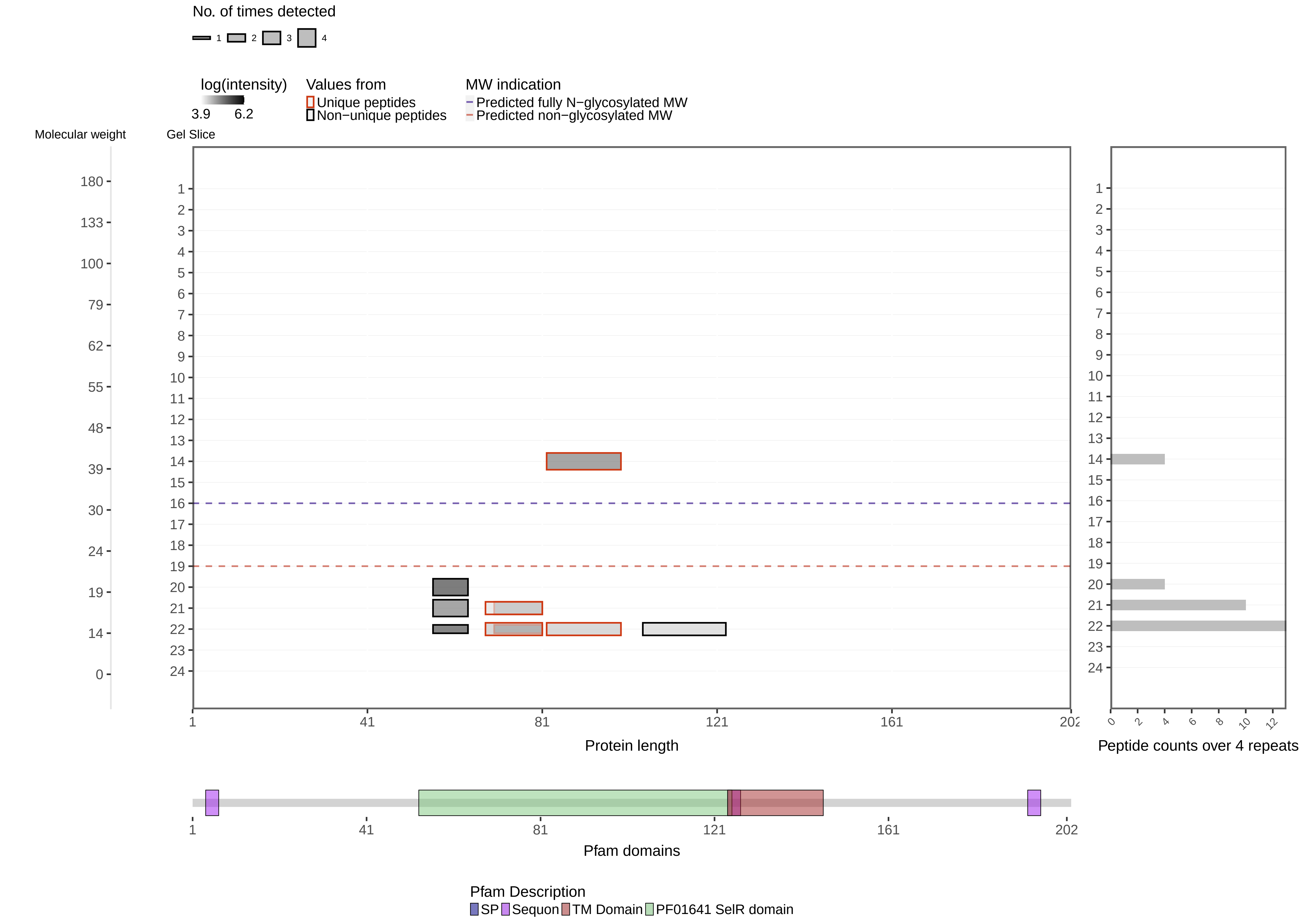


Protein sequence:

TYLCAGCGTPLYRSATKFNSPCGWPSFYEGLPGAINRNVSFSSIIFASSLFFFFNFLNIFRLSYHSSFSFCSFDNSAALTSLLEMSALSTTFLTLELLKCTKQEGKNFSCFPLHN*

### NbL01g00100.1 multiple_organellar_RNA_editing_factor_1_mitochondrial−like [MW = 46.08 kDa]

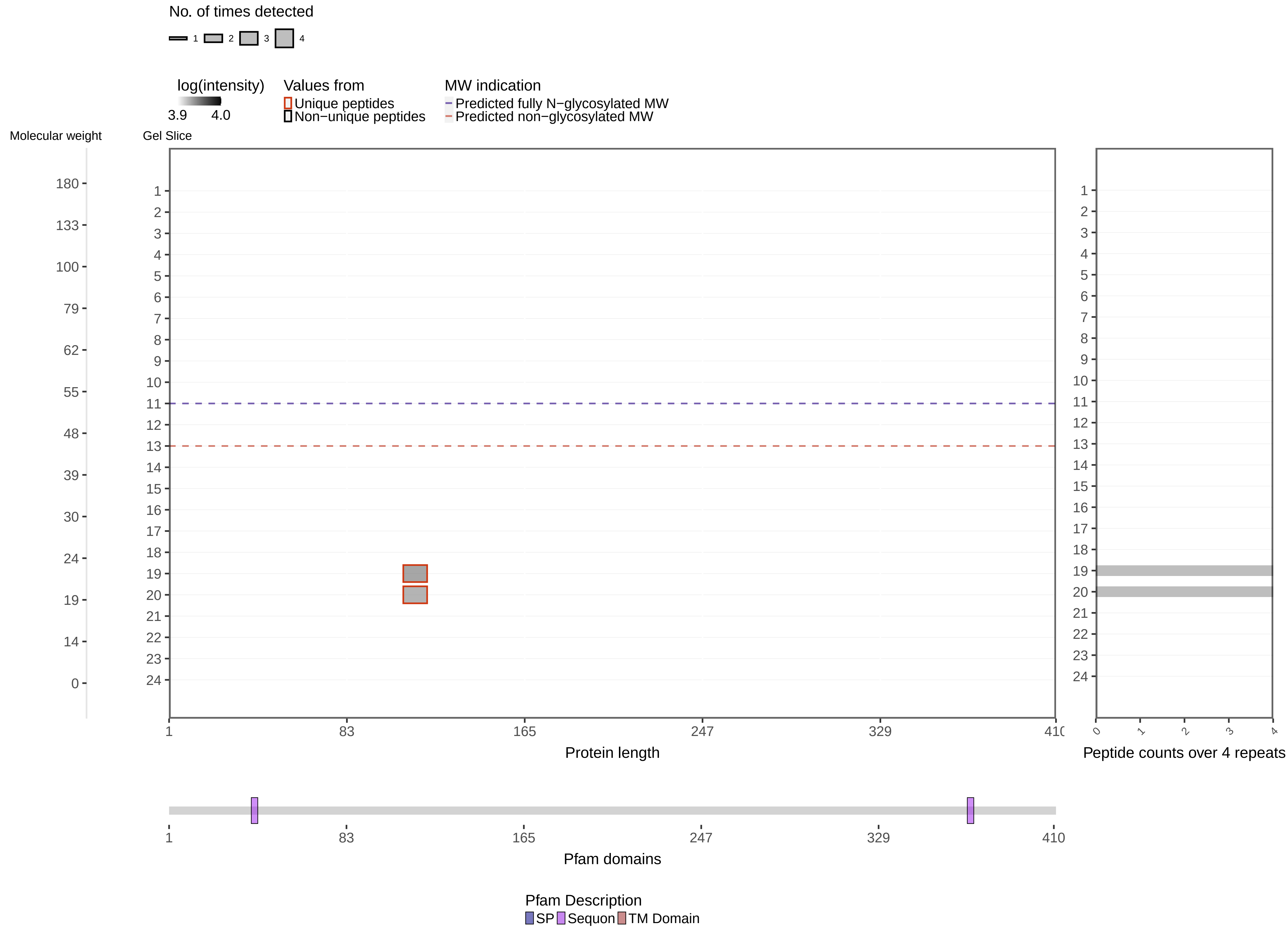


Protein sequence:
SNRPRRGPOFOOGNYGPPHSPPOONFGPPOGAPAOONYGAAOGPPPOONFGOPRYSPAOONYGSOOYPPPPOOIYSAPOSPPOONYGOPONPLPOOSYSAPONVPSOWNHGPOOSFSPOONYGPOGAGDPWHOTPLGSSPGGWDNSOGGRODSRPOYOVNYNOGEOGNYSPORDORGYAPPEERGFRGDNOHYAHPEORGFRGDNOR*

### NbL00g03500.1 stromal_70_kDa_heat_shock−related_protein_chloroplastic [MW = 46.85 kDa]

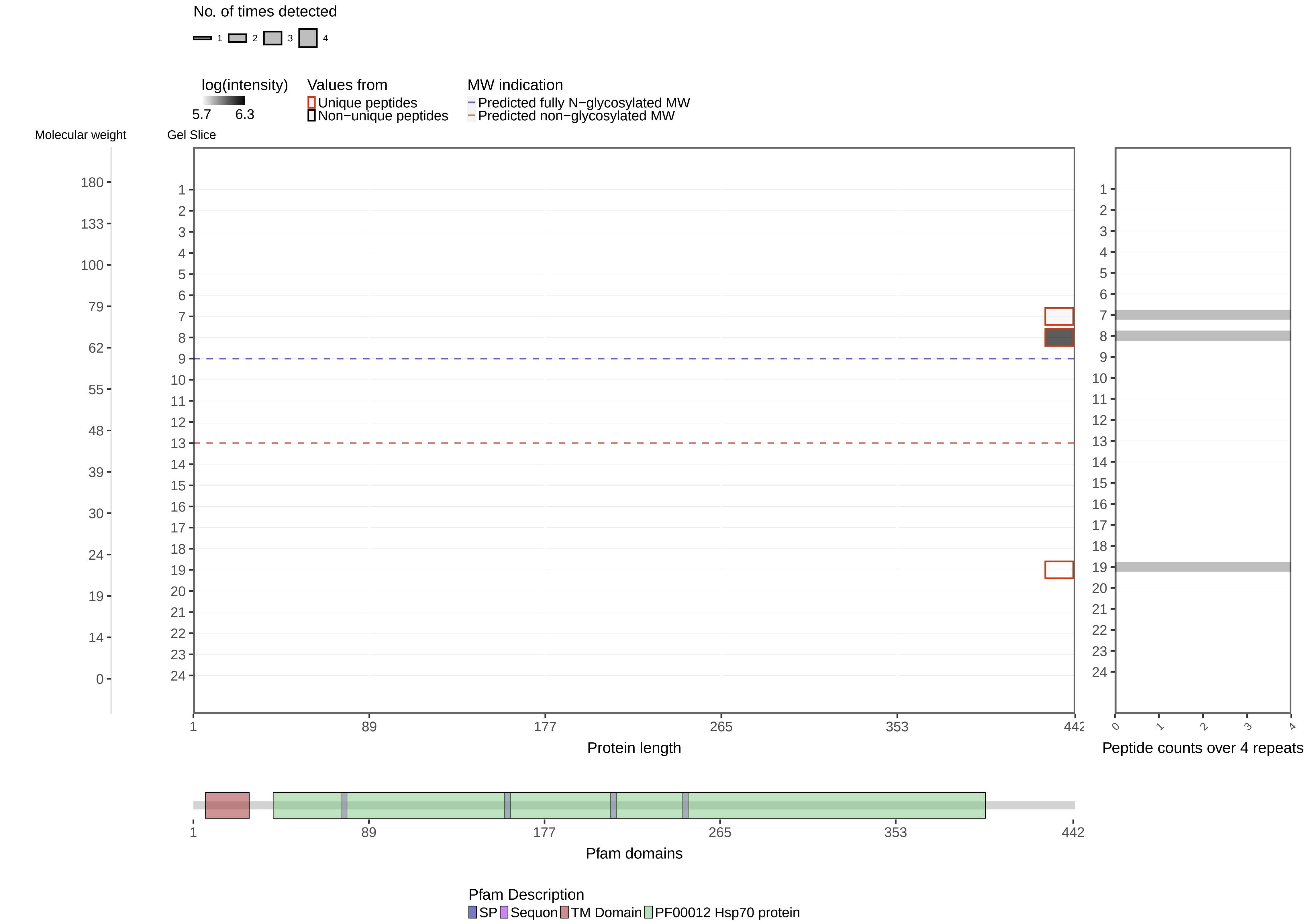


Protein sequence:

TSKSEVFSTAADGOTSVEINVLOGEREFVRDNKSLGSFRLDGIPPAPRGVPOIEVKFDIDANGILSVTAIDKGTGKKODITITGASTLPGDEVERMVKEAERFAOEDKEKRDAIDTKNOADSVVYOTEKOLKELGDKVPGPVKEKVEAKLGELKEAISGGSTOTMKDAMAALNOEVMOLGOSLYNOPGAGAAPGAGPAPGDSAGPSESSSGKGPDGDVIDADFTDSK*

### NbL00g03310.1 DNA−directed_RNA_polymerase_subunit_beta" [MW = 140.46 kDa]

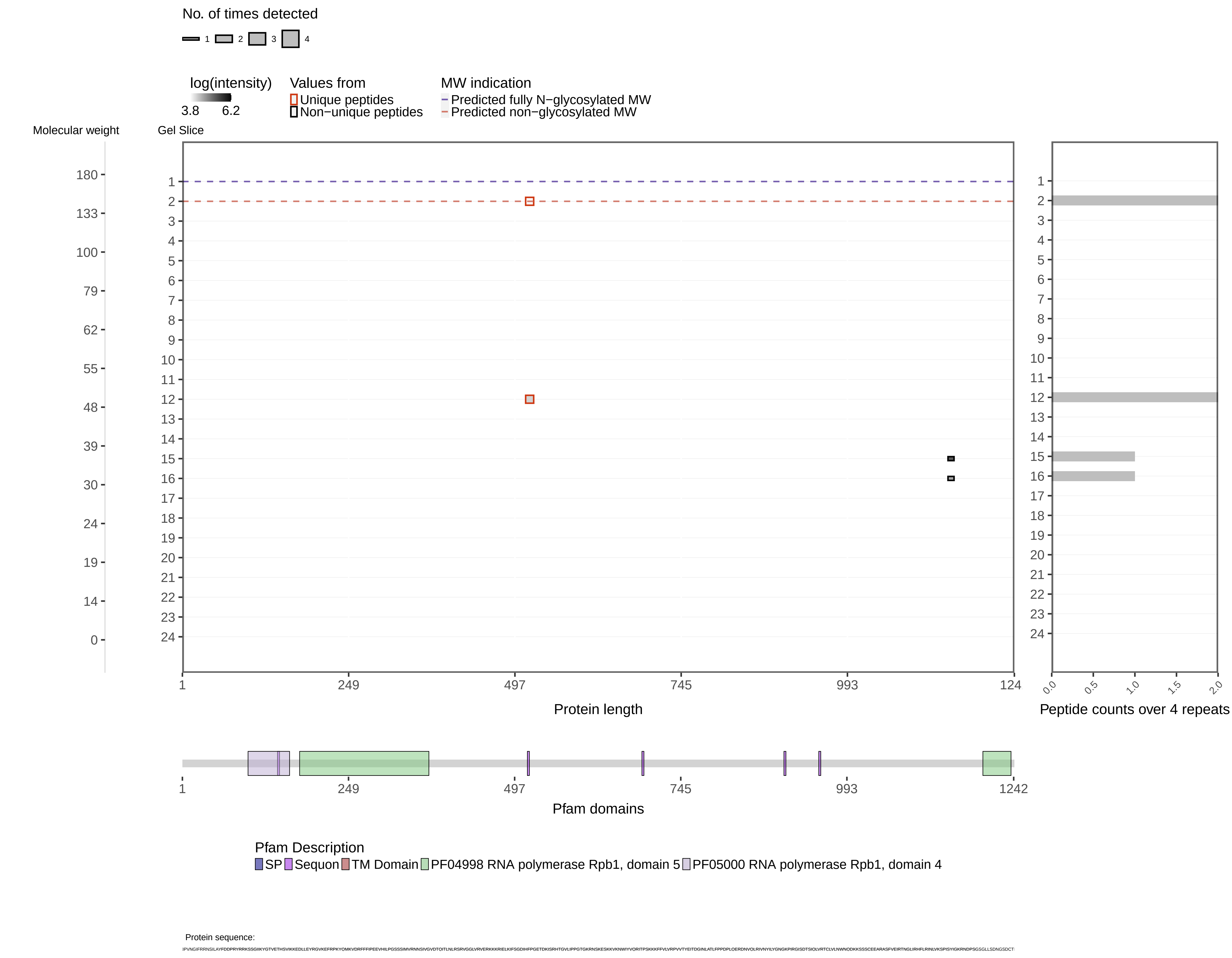

NbL00g03000.1 putative_deoxyribonuclease_TATDN1 [MW = 19.35 kDa]

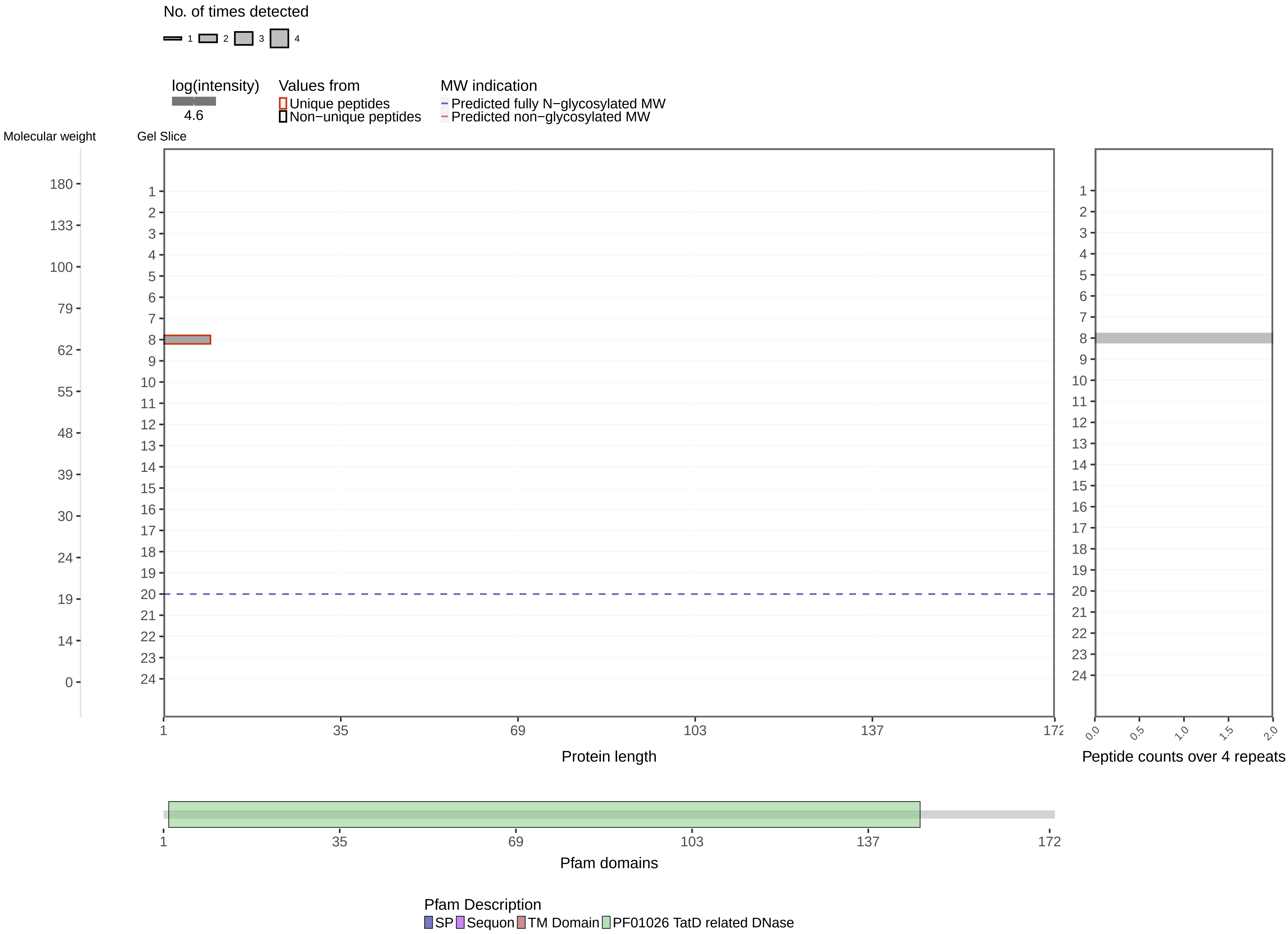


Protein sequence:

DSPYCDIKNSHAGIHFVKSSWPSKKKERHDQECLVKGRNEPCLVRQVLEVVAGTKGITDIDQLGKTLYHNTCRVFFPYDLDSAAEALLVSGREAICKKS*

### NbL00g02860.1 cathepsin_B−like [MW − SP = 39.32 kDa]

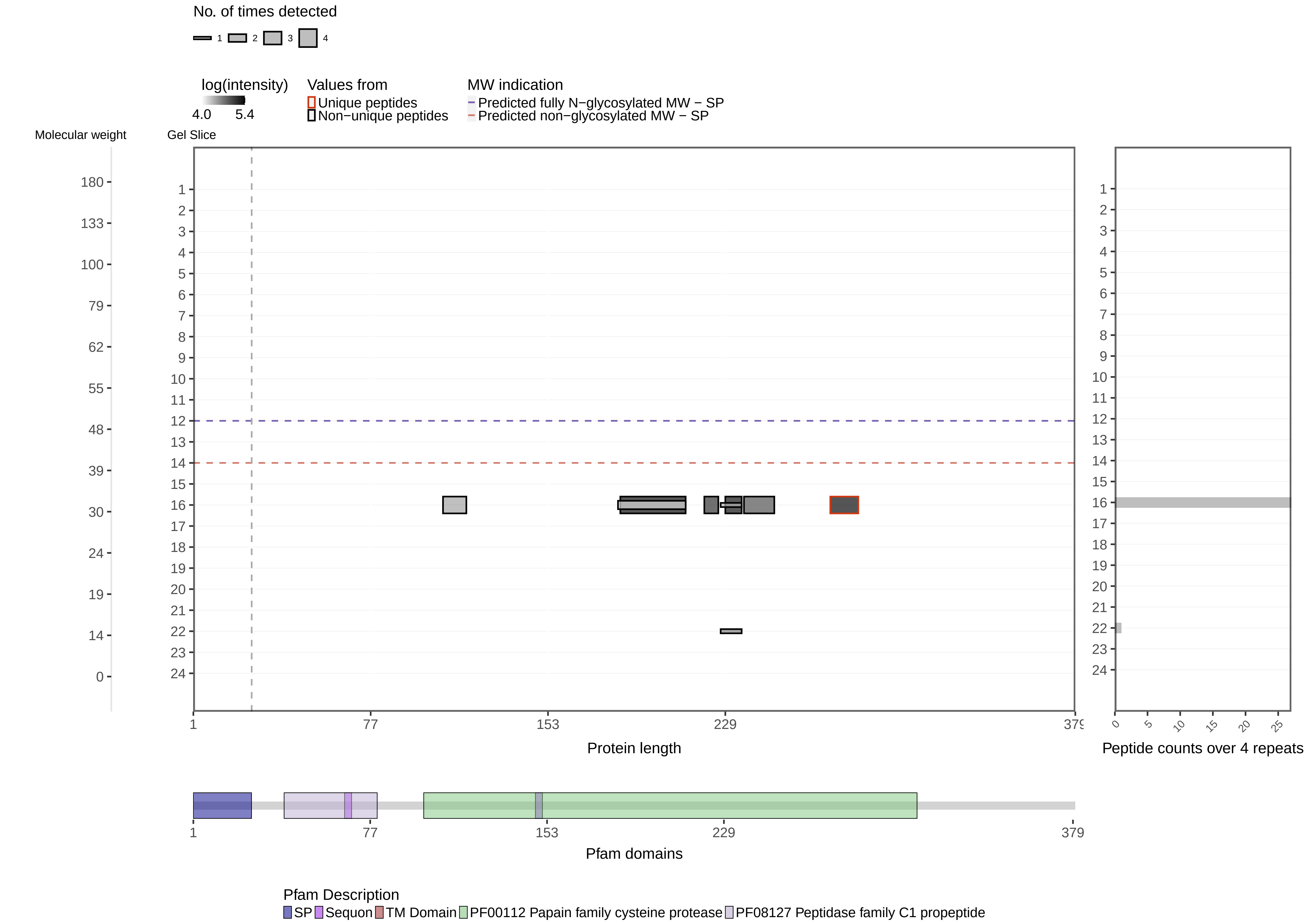


Protein sequence:

IRRGVVTEECDPYFDNEGCSHPGCEPGYPTPKCQRKCVKENLLWGKSKHYGVNAYRIHHDPHSIMTEIYKNGPVEVSFTVYEDFAHYKSGVYKHVIGQSMGGHAVKLIGWGTSEQGEDYWLIANSWNRGWGDVCTLFSPYAFIYEQPHFQPRGCKSESPQEQGGKFLEGSMLGVYLETTSLPKGRGKVCVHTTLPRPH*

### NbL00g02820.1 transaldolase [MW = 28.28 kDa]

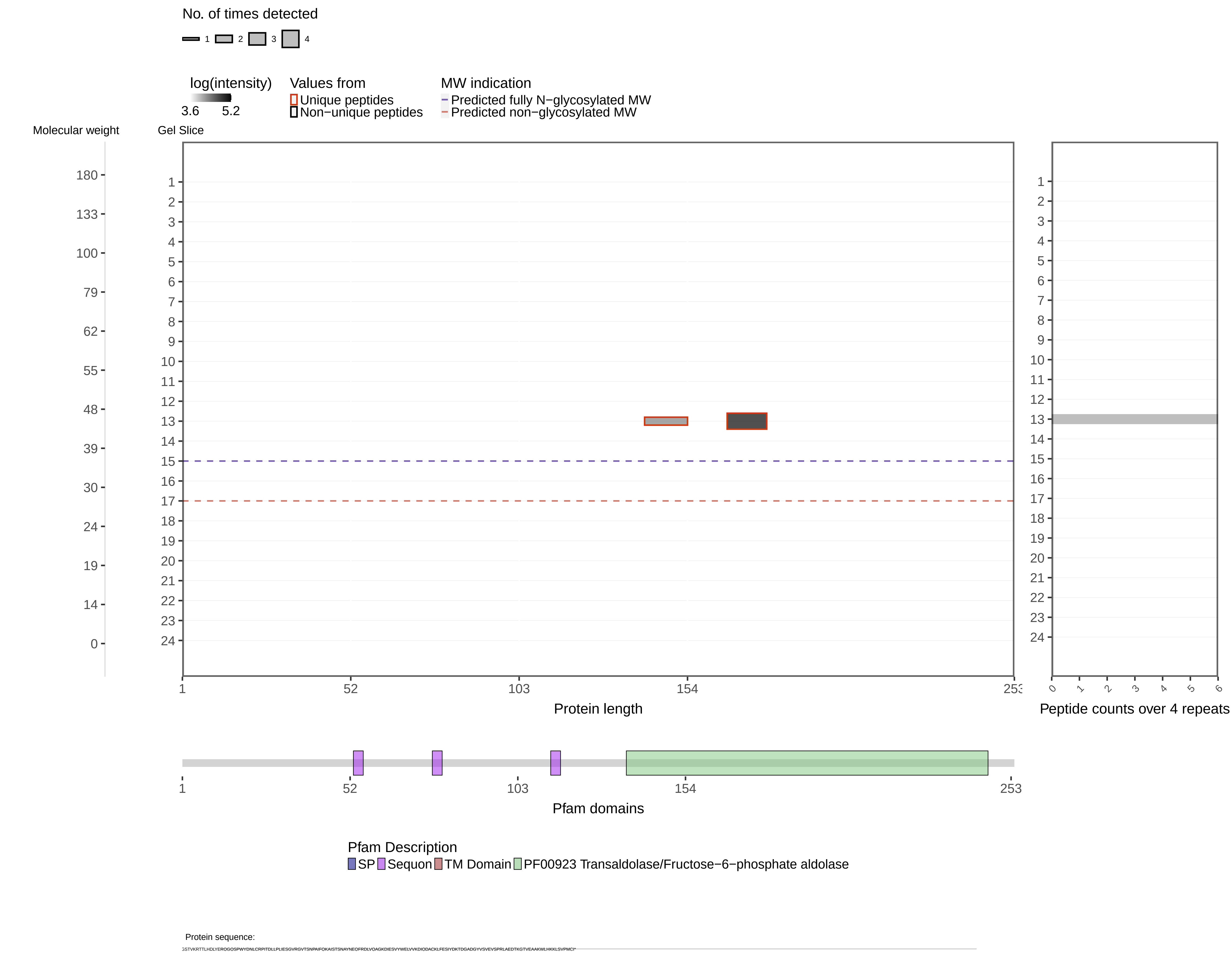

### NbL00g02050.1 ubiquitin−conjugating_enzyme_E2_7−like [MW = 18.95 kDa]

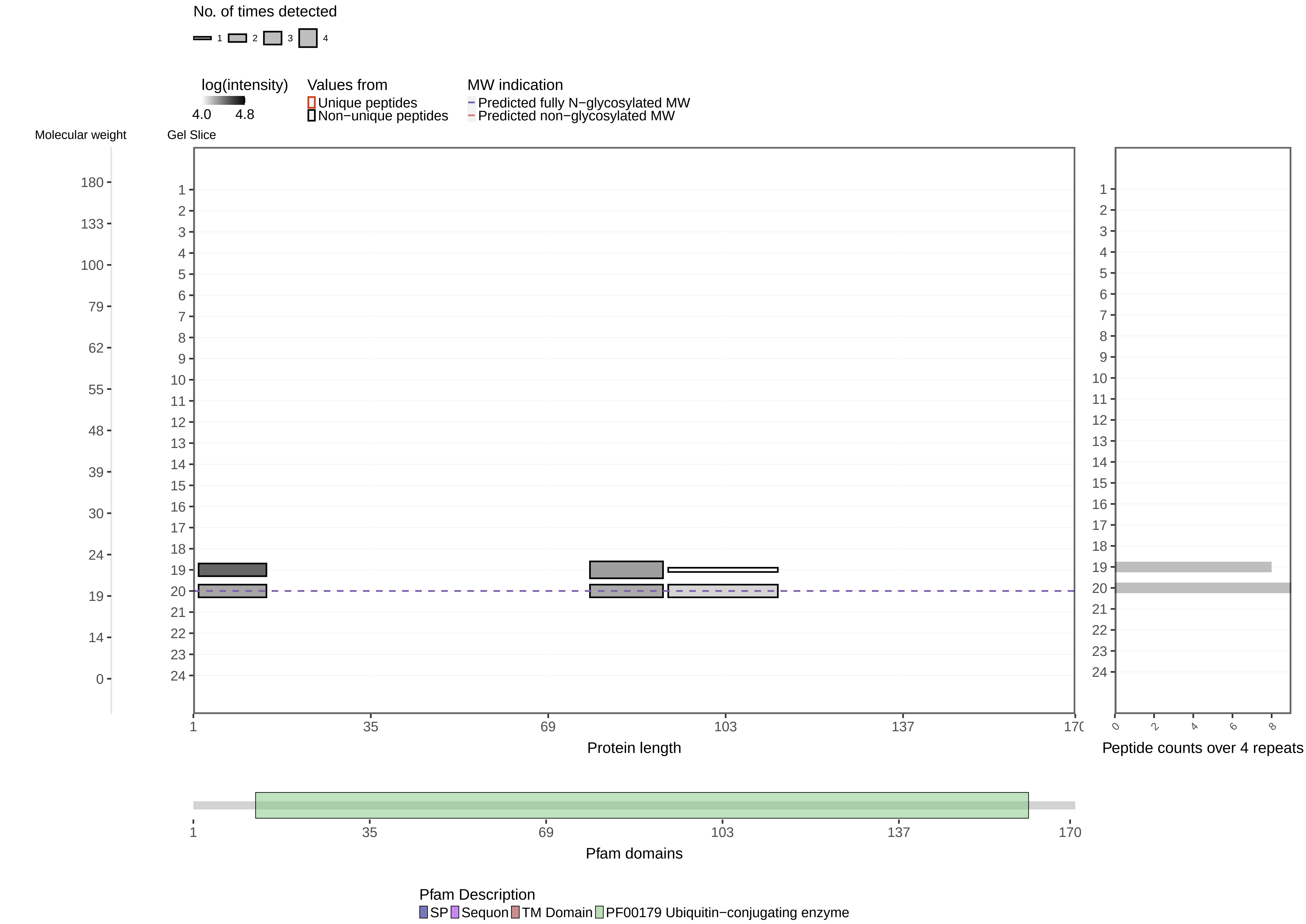


Protein sequence:

PPTVRFTTDIWHPNVYSDGKVCISILHPPGDDPNGYELASERWSPVHTVESIVLSIISMLSSPNDESPANVEAAKEWREKRDEFKKRVSRCVRRSQEI*

NbL00g01620.1 uncharacterized_LOC107788871 [MW = 23.73 kDa]

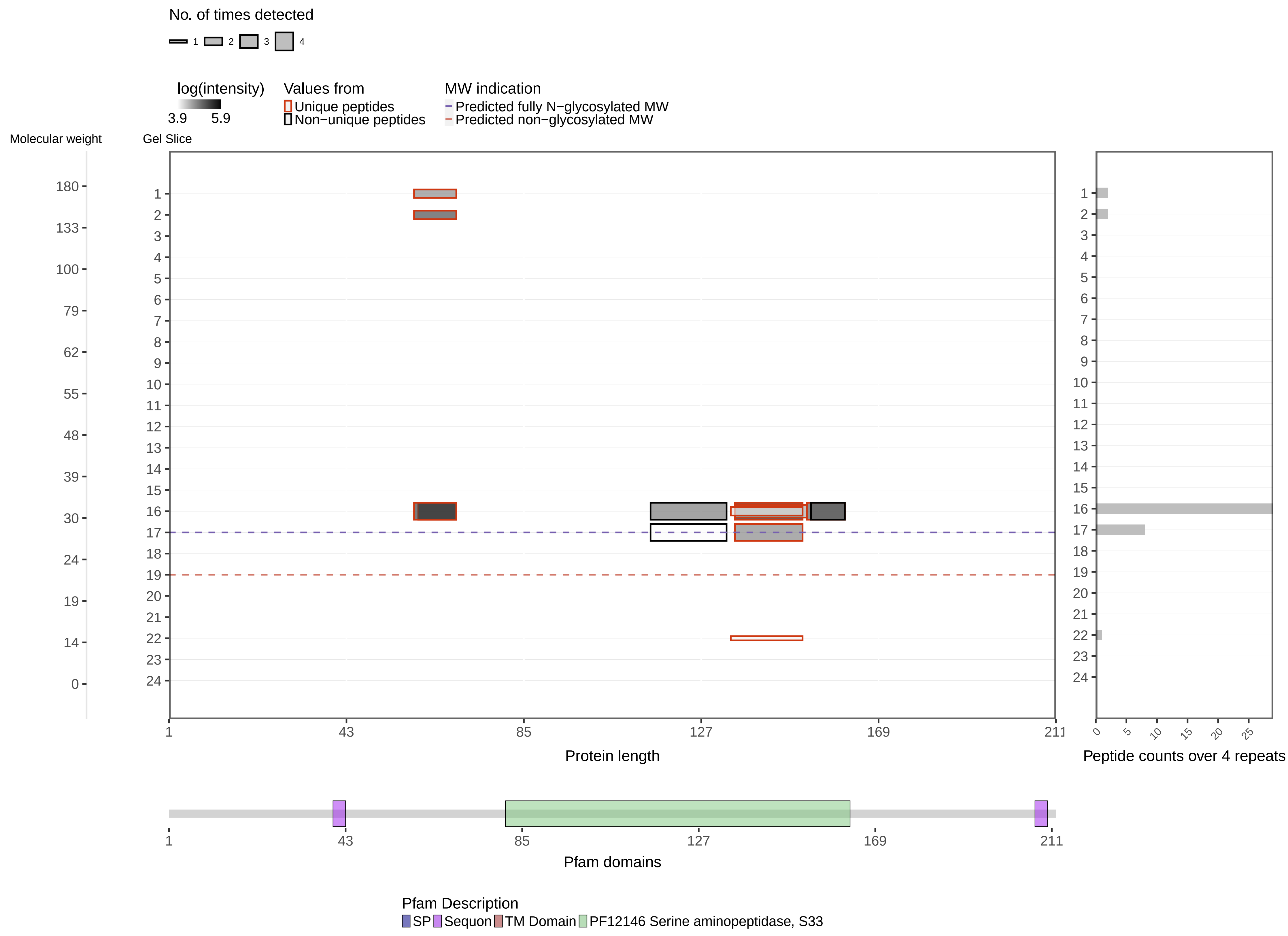


Protein sequence:

DFYTMVNLAVAMEKEGISVFRFDFPGNGESEGSFQYGNYRREADDLHSVVEYFNGANRKVTAVLGHSKGVSTFAKSCVYIESILCTVVKKRFCTRNYCALGSFFFSWVFSGCDNITF*

NbL00g01610.1 DNA−directed_RNA_polymerase_subunit_beta−like [MW = 123.55 kDa]

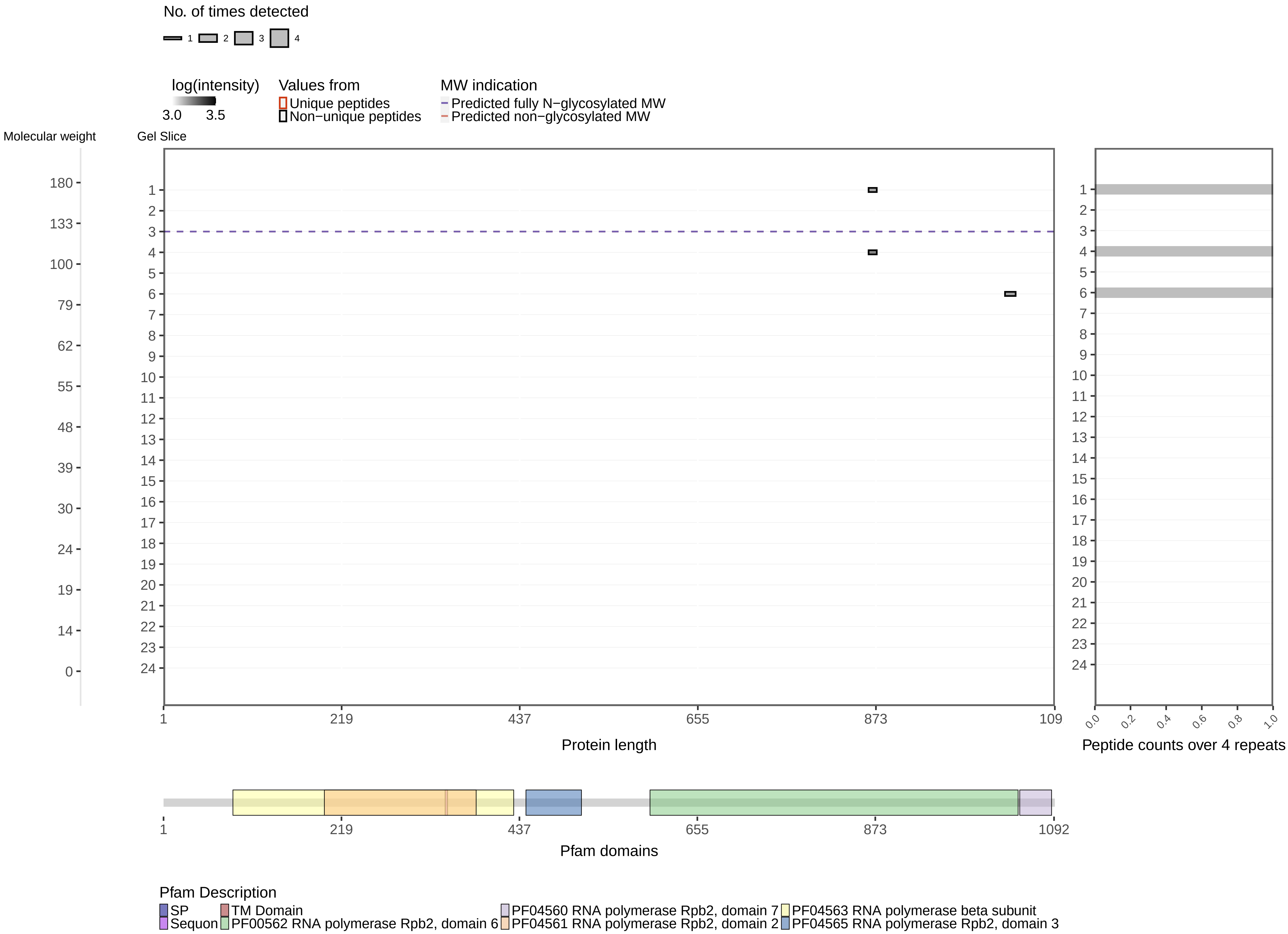


Protein sequence:

### NbL00g01400.1 ras−related_protein_RABF1 [MW = 21.57 kDa]

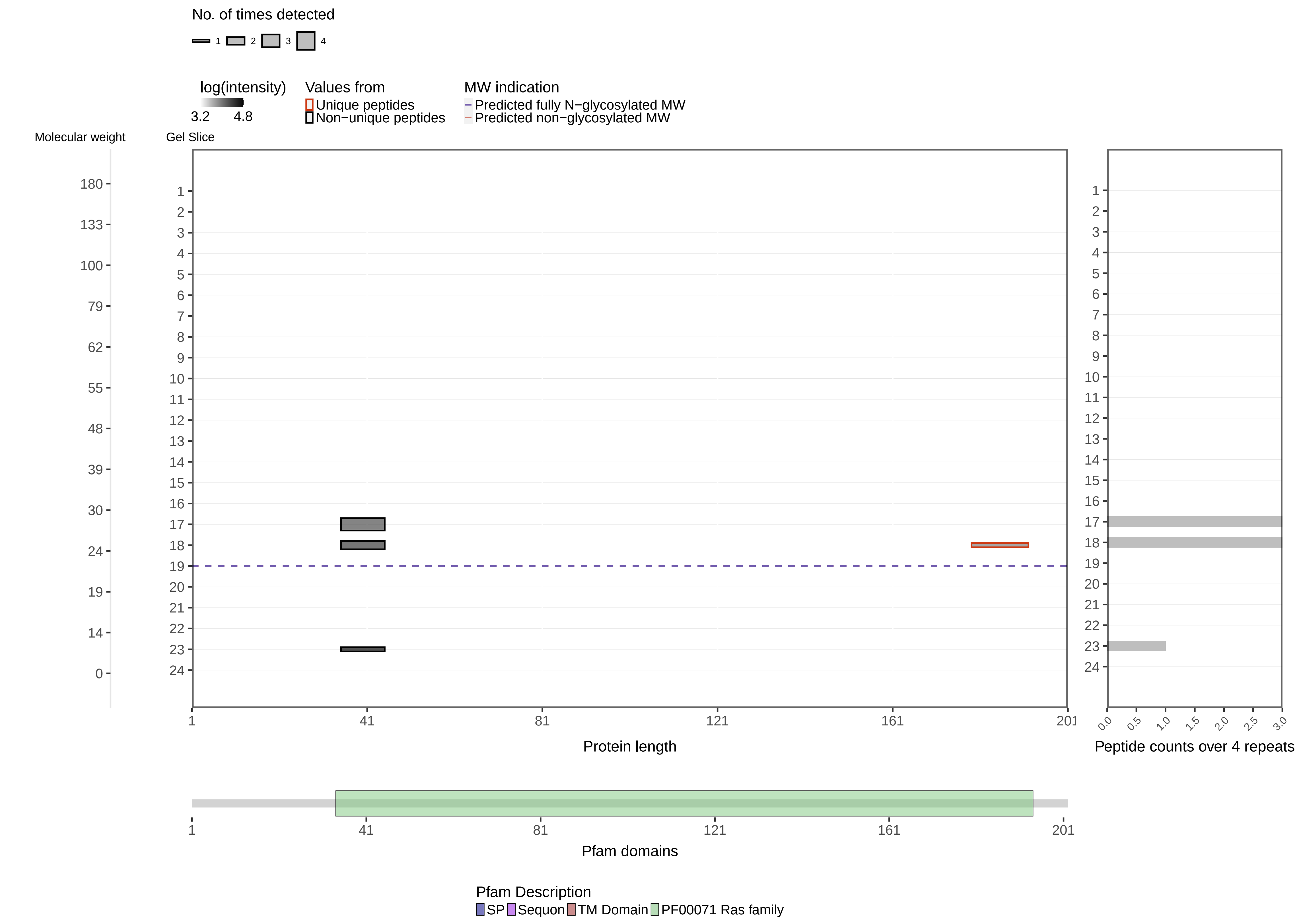


Protein sequence:

WDTAGQERYAALAPLYYRGAEVAVVVYDITSPESFAKAQYWVKELQKHGSPDIVMALVGNKADLHEKREVTTQDGIDCAEKNGMFFIETSAKTADNINQLFEEIAKRLPRPSAA*

### NbL00g01300.1 50S_ribosomal_protein_L2−A_chloroplastic [MW = 30.01 kDa]

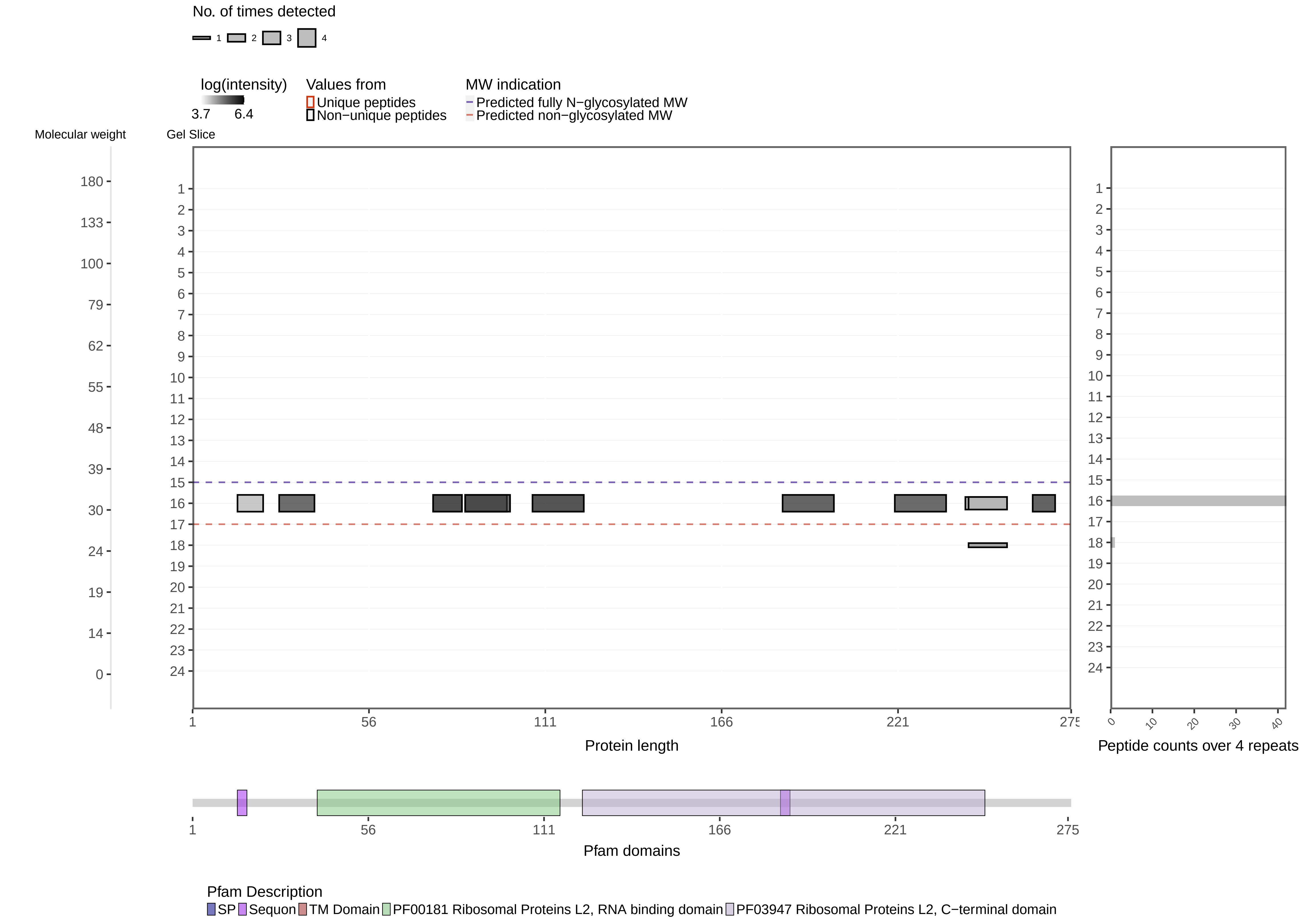


Protein sequence:

ALPLTDMPLGTAIHNIEITLGKGGQLARAAGAVAKLIAKEGKSATLKLPSGEVRLISKNCSATVGQVGNVGVNQKSLGRAGSKRWLGKRPVVRGVVMNPVDHPHGGGEGRAPIGRKKPTTPWGYPALGRRSRKRNKYSDNLILRRRSK*

### NbL00g00910.1 protein_IN2−1_homolog_B−like [MW = 33.24 kDa]

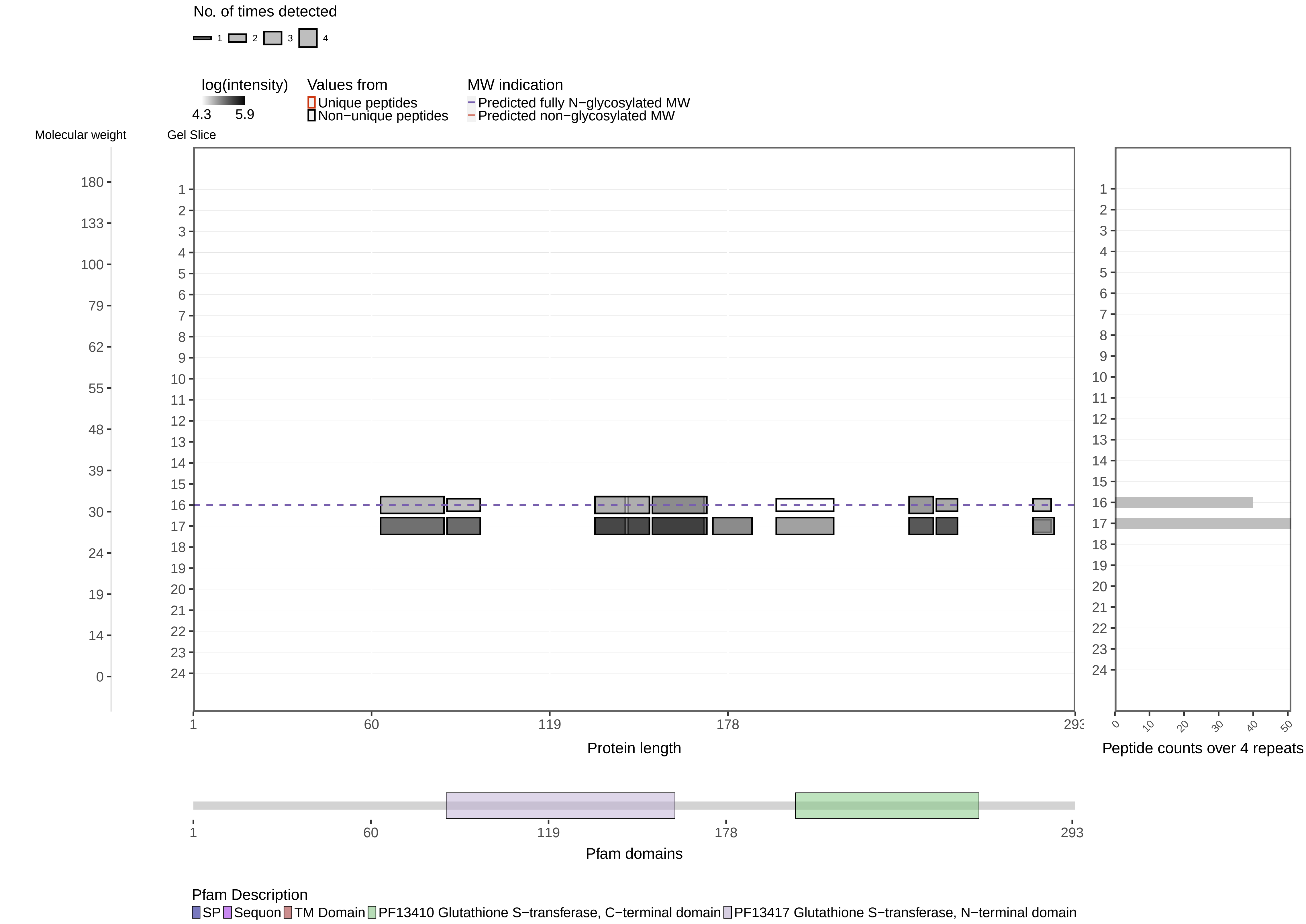


Protein sequence:

VPSLEHNNEVKGESMDLIRYIDSNFEGPSLFPDDPSKRKFAOEMFSHFDSFYKAVISSFKEDTANDANAAFDSIETALSKFVDGPFFLGTFSLVDIAYAPFIERFOPFLLDVKNFDITTGRSKLAAWIEEMNOIEAYTVTKRNPKELLESYKKRFLSOV*

NbL00g00890.1 ras−related_protein_RABE1a [MW = 23.76 kDa]

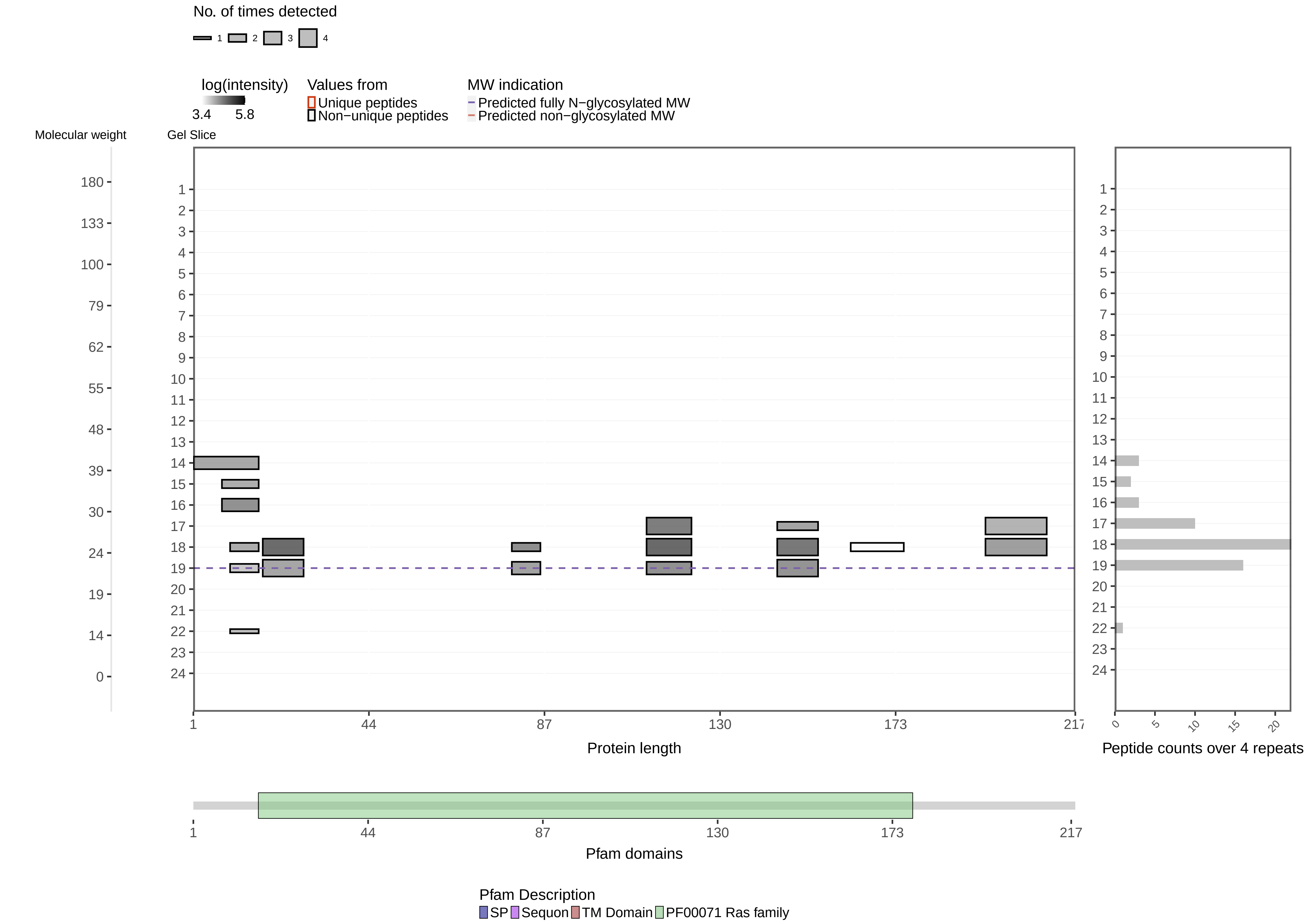


Protein sequence:

DESSFNNIRNWIRNIEQHASDNVNKILVGNKADMDESKRAVPTSKGQALADEYGIKFFETSAKTNLNVEQVFFSIARDIKQRLADTDSKAEPSTLKINQPDAGAGGSQAAQKSACCGS*

### NbL00g00550.1 heat_shock_cognate_70_kDa_protein_2−like [MW = 71.01 kDa]

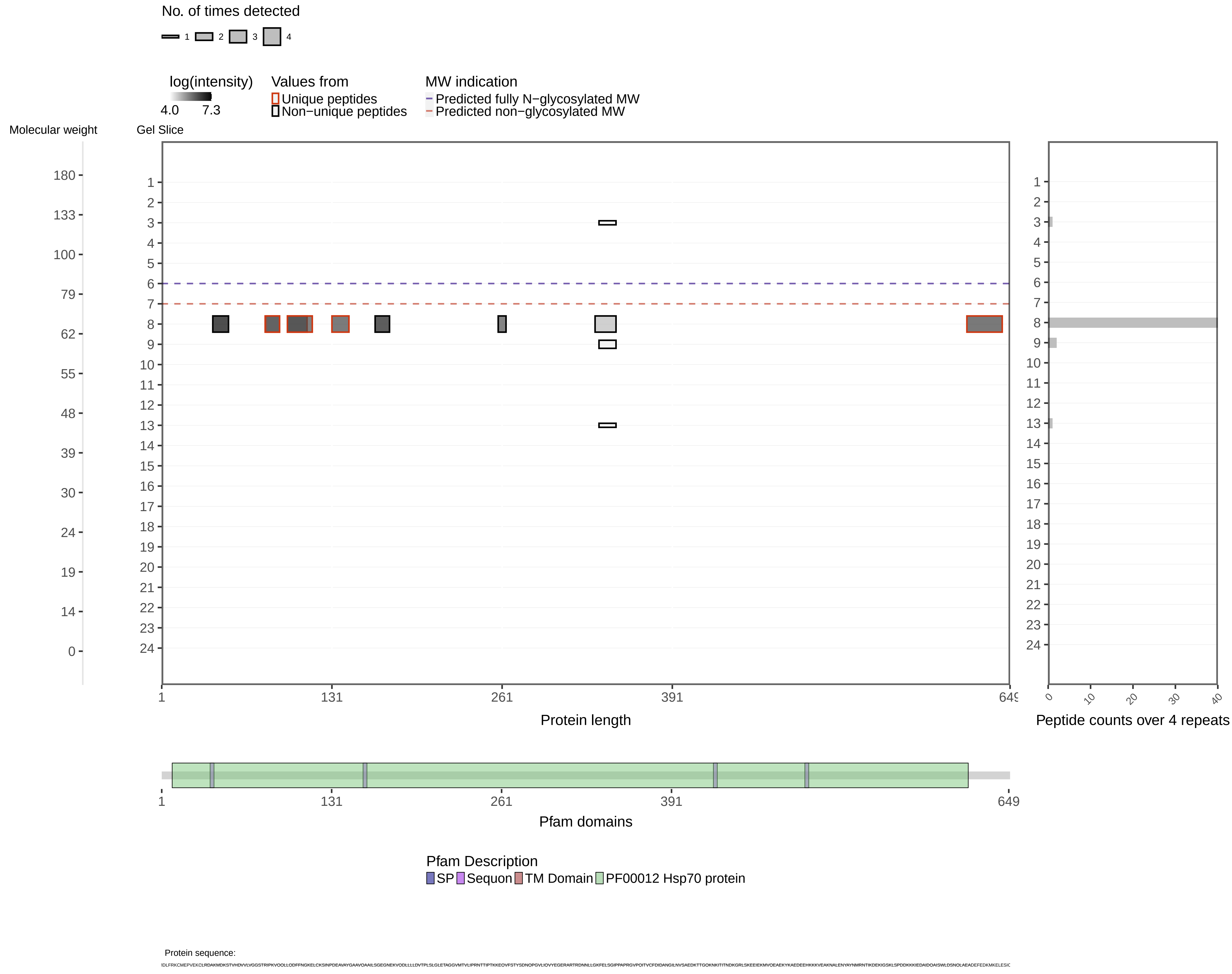

### NbL00g00360.1 glutelin_type−B_2−like [MW = 38.19 kDa]

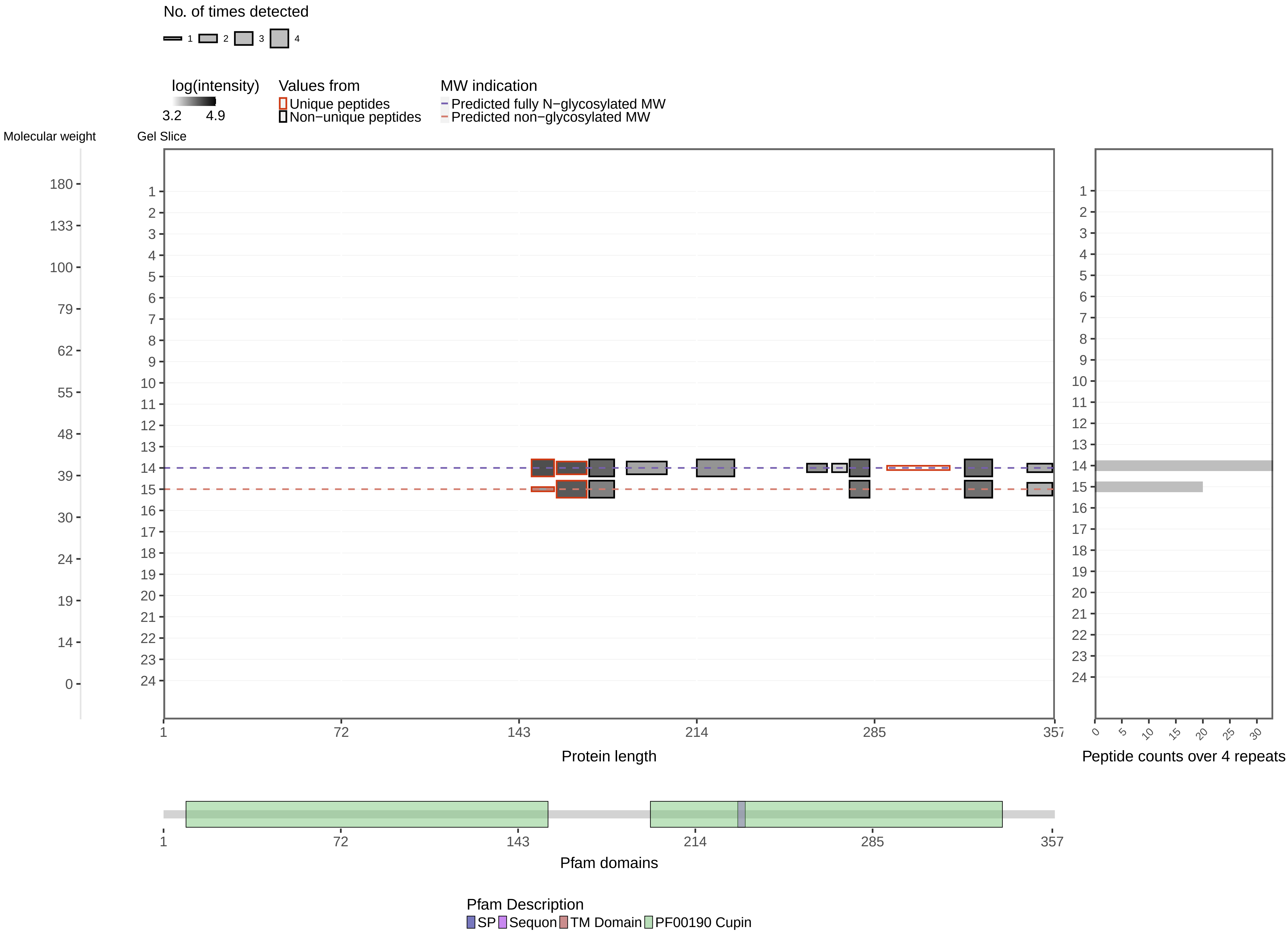


Protein sequence:

QGIVKLDAGFQMPEPKQGHRDGMVLNCLEAPLDVDIKGGGKVVVLNTKNLPLVGEVGLGADLVRLNGSAMCSPGFSCDSALQVTYIVRGSGRVQVVGPDGKRVLETHIKAGNLFIVPRFFVVSKIGDPDGMDWFSIITTPNPIFTHLAGRTSVWKALSPQVLQAAFKVSADVEKQFSSKRTAEEIFFPPPN*

### NbL00g00330.1 photosystem_II_CP43_reaction_center_protein−like [MW = 14.98 kDa]

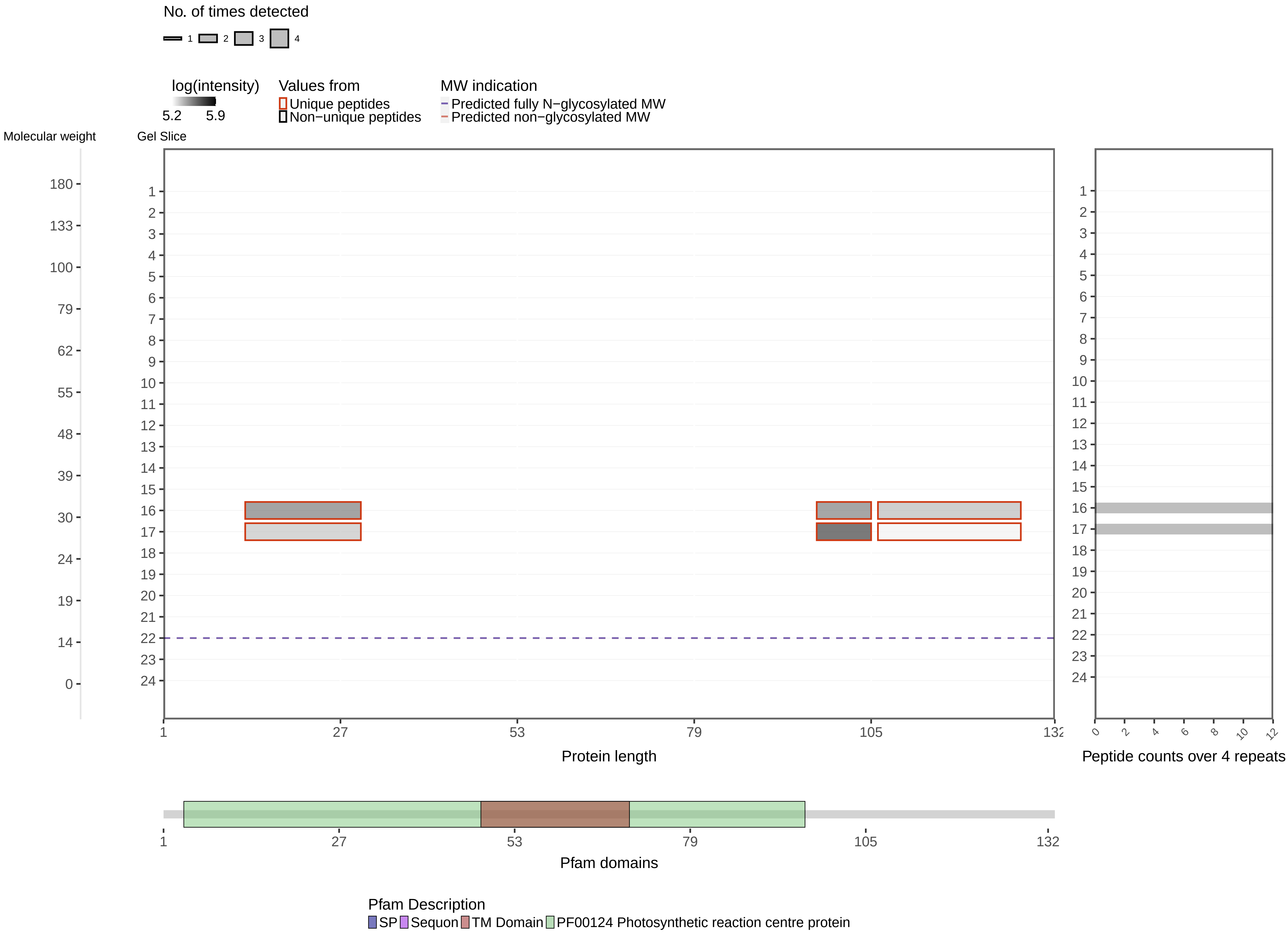


Protein sequence:

VPVTGLWMSALGVVGLALNLRAYDFVSQEIRAAEDPEFETFYTKNILLNEGIRAWMAAQDQPHENLIFPEEVLPRGNAL*

### NbL00g00250.1 GTPase_Der [MW = 72.38 kDa]

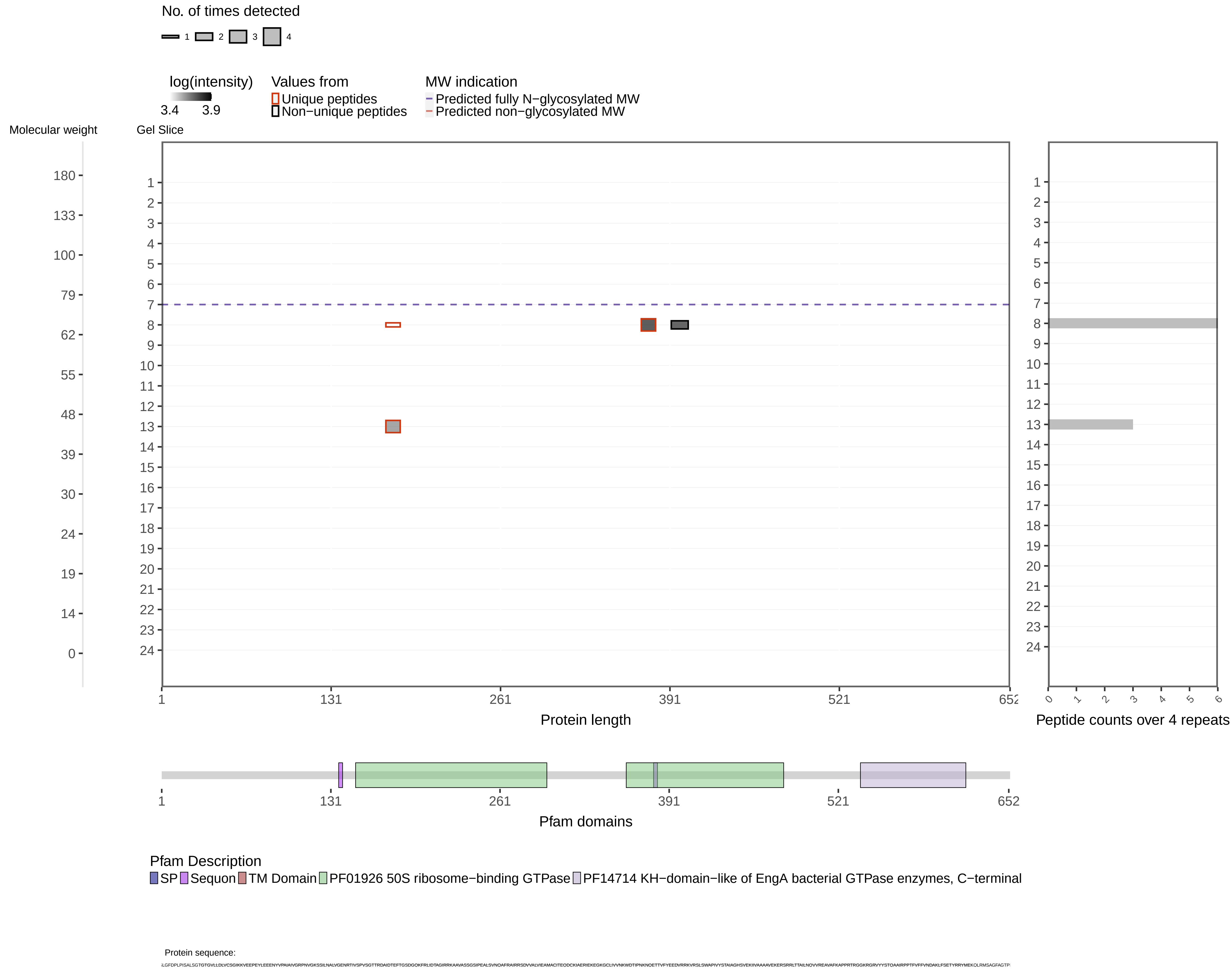

NbL00g00230.1 xanthine_dehydrogenase_1−like [MW = 150.4 kDa]

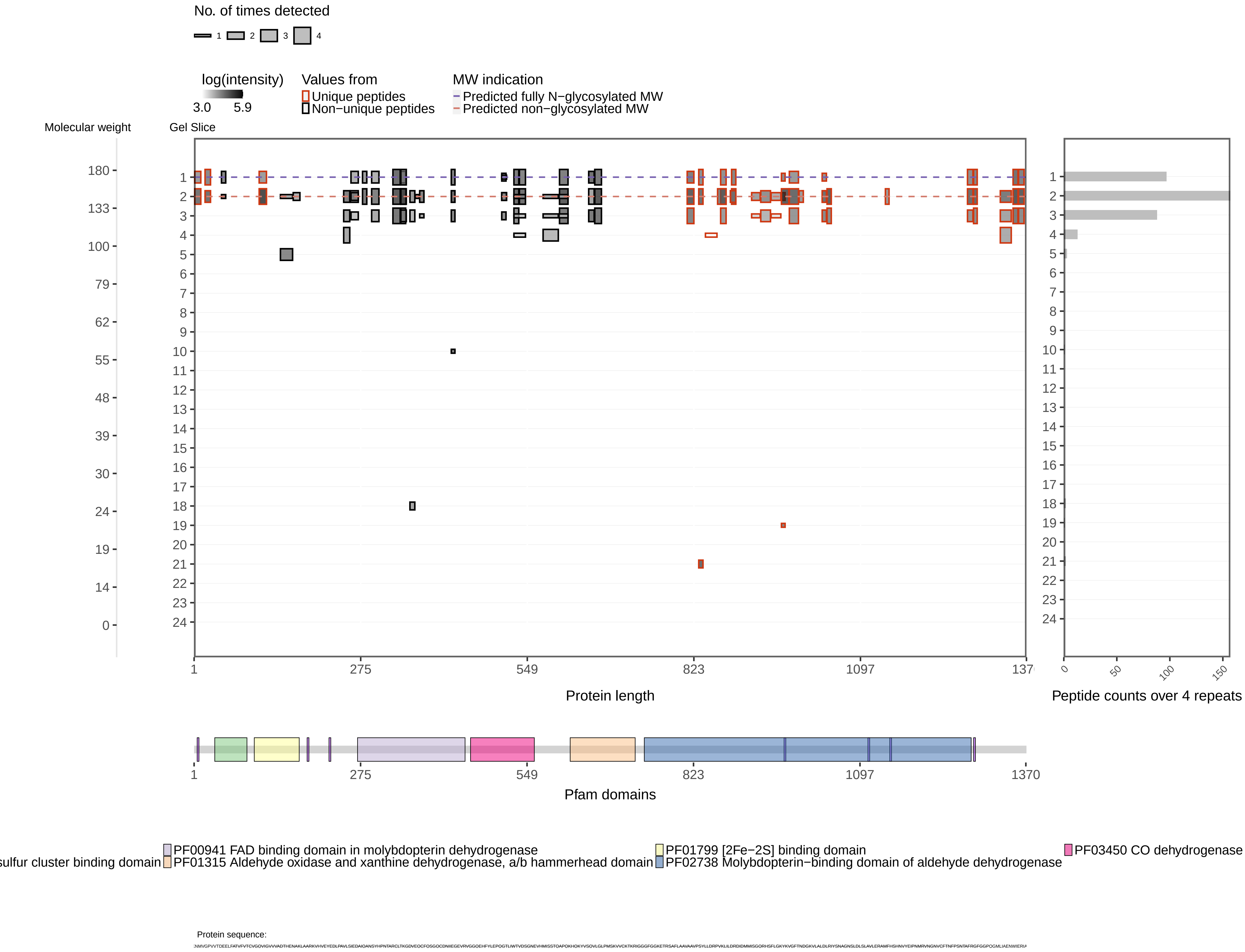
