## Supplementary material for "The proteome of agroinfiltrated *Nicotiana benthamiana* is shaped by extensive protein processing": File S3: functions.R

```
##functions for peptographs 

#for a certain molecular weight, find out the expected gel slice
gel = c(0,11.25,13.75,16.25,18.75,21.5,24.5,27.5,30.5,34.25,38.75,43.25,47.75,
        51.6875,55.0625,58.4375,61.8125,68.6875,79.0625,89.4375,99.8125,114.375,133.125,151.875,800)
exp_gel = function(mw){
  for (i in 1:(length(gel)-1)){
    if(between(mw,gel[i],gel[i+1])){
      return(25-i)}}}

#color vectors for plots
qual_col_pals = brewer.pal.info[brewer.pal.info$category == 'qual',]
col_vector = unlist(mapply(brewer.pal, qual_col_pals$maxcolors, rownames(qual_col_pals)))
col_vector = c("darkblue","purple","brown",col_vector)

##mw scale
scale = 
  ggplot(data.frame(c(gel[1:24],180)),aes(y =factor(round(c.gel.1.24...180.)), xmin = 0,xmax =0.1,x = 0.05))+
  geom_blank()+
  scale_y_discrete(expand = c(0,1.7),breaks = function(x){x[c(TRUE,FALSE)]}) +
  labs(y = "Molecular weight") +
  scale_theme

##peptograph
plot_ppt_gly_f = function(pro){
  
  # subset a dataframe for "pro" that is to be plotted
  test = ppt_1[ppt_1$Leading.razor.protein == pro,]
  m = which(colnames(test) == "Mean 1 ")
  test = test[,!names(test) == "Leading.razor.protein"]
  test.m = melt(test, id = c("Sequence","Start.position","End.position","Unique..Proteins."))
  
  test.m = test.m %>% 
    mutate(gel.slice = gsub("[^0-9]","",variable)) %>%
    mutate(gel.slice = as.numeric(gel.slice)) %>%
    mutate(measure = gsub("[0-9]","",variable)) 
  
  if (any(test.m$Unique..Proteins. == "yes")) {
    test.m[test.m$Unique..Proteins. == "yes",]$Unique..Proteins. <- "Unique peptides"}
  if (!is.null(test.m) && any(test.m$Unique..Proteins. == "no")) {
    test.m[test.m$Unique..Proteins. == "no",]$Unique..Proteins. <- "Non-unique peptides" }
  
  test.m1 = test.m[,!names(test.m) == "variable"]
  test.m1m = test.m1[test.m1$measure == "Mean ",]
  test.m1c = test.m1[test.m1$measure == "Count ",]
  test.m1c = test.m1c[,c(1,5,6)]
  test.m1 = merge(test.m1m, test.m1c, by = c("Sequence","gel.slice"))
  test.m = test.m1[test.m1$value.x>0,]
  test.m$value.y = as.numeric(test.m$value.y)
  test.m$gel.slice = as.numeric(test.m$gel.slice)
  test.m$multiplier = 0.1*test.m$value.y
  names(test.m)[names(test.m) == "value.x"] = "Intensity"
  test.m$Intensity = log10(as.numeric(test.m$Intensity))
  test.m$Unique..Proteins. = factor(test.m$Unique..Proteins., levels = c("Unique peptides","Non-unique peptides"))
  
  # match the expected molecular weight of the protein
  mw = lab360[lab360$Leading.razor.protein == pro,]$exp.mw
  exp_gly_gel = exp_gel(lab360[lab360$Leading.razor.protein == pro,]$gly.mw[1])
  
  if (pro %in% unique(signalp$seqid)){
    sp.end = signalp[signalp$seqid == pro,]$end 
    sp.sequence = substring(lab360[lab360$Leading.razor.protein == pro,]$sequence,1,sp.end)
    sp.mw = round(mw(sp.sequence)/1000,2)
    exp.gel = exp_gel(mw-sp.mw)
    exp.gly.gel = exp_gel(lab360[lab360$Leading.razor.protein == pro,]$gly.mw[1]-sp.mw)}else{
      sp.end = 0
      sp.mw = 0
      exp.gel = exp_gel(mw)
      exp.gly.gel = exp_gly_gel}
  
  if(!sp.end == 0){mw_2="MW - SP"}else{mw_2 = "MW"}
  

  # plot a is for the protomap
  a = ggplot(test.m) + 
    geom_rect(aes(xmin = Start.position, xmax = End.position, 
                  ymin = gel.slice - multiplier,ymax = gel.slice + multiplier,
                  col = Unique..Proteins., fill = Intensity),linewidth = 0.5,alpha = 0.7) + 
    scale_colour_manual(values = c("Non-unique peptides" = "black","Unique peptides" = "#ce3812"), drop = FALSE) +
    scale_fill_gradient(low = "white",high = "black",
                        limits = c(round(min(test.m$Intensity),1),round(max(test.m$Intensity),1)),
                        breaks = c(round(min(test.m$Intensity),1),round(max(test.m$Intensity),1))) +
    geom_hline(aes(yintercept = exp.gel, linetype = paste0("Predicted non-glycosylated ",mw_2)), color = "#d37b6d") +
    geom_hline(aes(yintercept = exp_gly_gel, linetype = paste0("Predicted fully N-glycosylated ", mw_2)), color = "#765faf") +
    scale_linetype_manual(name = "MW indication", values = c(2, 2), 
                          guide = guide_legend(title.position = "top",
                                               override.aes = list(color = c("#765faf","#d37b6d")),
                                               nrow = 2)) +
    geom_vline(aes(xintercept = sp.end), linetype = "dashed", color = "darkgrey") +
    xlab("Protein length") + ylab("Gel Slice") +
    scale_x_continuous(limits = c(1,lab360[lab360$Leading.razor.protein == pro,]$length),
                       breaks = c(seq(1,(lab360[lab360$Leading.razor.protein == pro,]$length)-(round(lab360[lab360$Leading.razor.protein == pro,]$length/5,0))+1,
                                      by =round(lab360[lab360$Leading.razor.protein == pro,]$length/5,0)),
                                  lab360[lab360$Leading.razor.protein == pro,]$length),
                       expand = c(0,0)) +
    scale_y_reverse(limits = c(24.6,0.2),breaks = seq(24,1, by = -1)) +
    guides(fill = guide_colorbar("log(intensity)", title.position = "top", order = 1),
           col = guide_legend("Values from", title.position = "top",override.aes = list(fill = NA), nrow = 2, order = 2)) +
    peptograph_theme
 
  
  # plot b for peptide counts (over 4 repeats) on the right
  b=ggplot(data = test.m %>% mutate(gel.slice = factor(gel.slice, levels = seq(24:1))) %>% mutate(gel.slice = fct_rev(gel.slice)), 
           aes(x = gel.slice, y = value.y)) + 
    geom_col(fill = "grey", width=0.5) +
    #geom_point(aes(col = Unique..Proteins.,),size = 0.8, position = position_jitter()) +
    scale_color_manual(values = c("Non-unique peptides" = "black","Unique peptides" = "#ce3812"),drop = FALSE) +
    coord_flip() + 
    scale_x_discrete(drop = FALSE, expand = expansion(add = 2)) +
    scale_y_continuous(limits = c(0,NA),expand = c(0,0),breaks = pretty_breaks()) +
    ylab("Peptide counts over 4 repeats") +
    guides(col = guide_legend("Values from",title.position = "top", nrow = 2))+
    count_barplot_theme
  
  #plot c for annotation
  #sp
  annot = pfam[pfam$protein.id == pro & pfam$database == "Pfam",]
  if (!is.null(sp.end) && sp.end>0) {
    annot$prediction = paste(annot$number,annot$prediction)
    annot[nrow(annot)+1,] = c(pro,"SignalP","","SP",1,sp.end,
                              "","","","")} else {
                                annot$prediction = paste(annot$number,annot$prediction)
                                annot[nrow(annot)+1,] = c(pro,"SignalP","","SP",NA,NA,"","","","")}
  
  #sequon
  sequon.loc = unlist(gregexpr("N.[ST]",lab360[lab360$Leading.razor.protein == pro,]$sequence))
  npt.loc = unlist(gregexpr("NP[ST]",lab360[lab360$Leading.razor.protein == pro,]$sequence))
  sequon.loc = sequon.loc[!sequon.loc %in% npt.loc]
  annot = annot[,names(annot) %in% c("database","start","stop","prediction")]
  
  if(length(sequon.loc)>0) {
    annot[nrow(annot) + length(sequon.loc),] <- NA
    annot[(nrow(annot)-length(sequon.loc)+1):nrow(annot),"database"] = "Sequon"
    annot[(nrow(annot)-length(sequon.loc)+1):nrow(annot),"start"] <- sequon.loc
    annot[(nrow(annot)-length(sequon.loc)+1):nrow(annot),"stop"] <- sequon.loc+3
    annot[(nrow(annot)-length(sequon.loc)+1):nrow(annot),"prediction"] = "Sequon"}else{
      annot[nrow(annot)+1,] <- NA
      annot[nrow(annot),"database"] = "Sequon"
      annot[nrow(annot),"start"] <- NA
      annot[nrow(annot),"stop"] <- NA
      annot[nrow(annot),"prediction"] = "Sequon" 
    }  
  
  annot = annot[order(nrow(annot):1),]
  
  ##add tm domains
  topo = tm[tm$V1 == pro,]$V6
  topo = sub("Topology=","",topo)
  tm_start = c()
  tm_end = c() 
  len = gregexpr("i",topo)
  len = sum(unlist(len) != -1)
  topo_start = unlist(str_extract_all(topo, "[io](\\d+)"))
  topo_end = unlist(str_extract_all(topo,"(\\d+)[io]"))
  
  if(length(topo_start)>0){
    for (i in 1:len){
      tm_start[i] = as.numeric(str_extract_all(topo_start[i], "\\d+"))
      tm_end[i] = as.numeric(str_extract_all(topo_end[i], "\\d+"))
    }
    annot[nrow(annot)+length(tm_start),] <- NA
    annot[(nrow(annot)-length(tm_start)):nrow(annot),"database"] <- "TMHMM"
    annot[(nrow(annot)-length(tm_start)+1):nrow(annot),"start"] <- tm_start
    annot[(nrow(annot)-length(tm_start)+1):nrow(annot),"stop"] <- tm_end
    annot[(nrow(annot)-length(tm_start)+1):nrow(annot),"prediction"] <- "TM Domain"}else{
    annot[(nrow(annot)+1),"database"] <- "TMHMM"
    annot[nrow(annot),"start"] <- NA
    annot[nrow(annot),"stop"] <- NA
    annot[nrow(annot),"prediction"] <- "TM Domain"
  }
  
  
  c = ggplot(annot, 
             aes(xmin = as.numeric(start), xmax = as.numeric(stop), 
                 ymin = 0.6, ymax = 1.0, y =0.8,
                 fill = fct_relevel(prediction,"SP","Sequon","TM Domain"))) +
    geom_hline(yintercept = 0.8,col = "lightgrey",linewidth = 2.5)+
    geom_rect(linewidth = 0.2, col = "black",alpha = 0.5)+
    scale_fill_manual(values = c(col_vector), drop = FALSE)+
    scale_x_continuous(limits = c(1,lab360[lab360$Leading.razor.protein == pro,]$length+1),
                       breaks = c(seq(1,(lab360[lab360$Leading.razor.protein == pro,]$length)-(round(lab360[lab360$Leading.razor.protein == pro,]$length/5,0))+1,
                                      by =round(lab360[lab360$Leading.razor.protein == pro,]$length/5,0)),
                                  lab360[lab360$Leading.razor.protein == pro,]$length),
                       expand = c(0,0))+
    guides(fill = guide_legend("Pfam Description", title.position = "top"))+ 
    labs(x = "Pfam domains") +
    annot_theme
  
  string = lab360[lab360$Leading.razor.protein == pro,]$sequence
  
  
  d = ggplot(annot, aes(xmin = -1, xmax = lab360[lab360$Leading.razor.protein == pro,"length"], ymin = 0, ymax=0))+ 
    geom_rect(fill = "transparent", col = "transparent") + 
    annotate("text",x = -1, y = 0, label = "Protein sequence:", size = 2,hjust = 0, vjust = 0) +
    annotate("text",x = 1, y = 0,label = str_replace_all(string,"(.{250})",paste0("\\1","\n")), hjust = 0, vjust = 0, size = 1) + 
    theme(axis.text = element_blank(), axis.ticks = element_blank(), 
          axis.title = element_blank(),panel.background = element_blank(),
          panel.grid = element_blank())
  
  
  e = ggplot(data.frame(a = "", b = ""), aes(ymin=4,ymax = 5, xmin=0,xmax=1)) +
    geom_rect(fill = "grey", col = "black") + annotate("text",x = 1.5, y = 4.5, label = "1", size = 2) +
    geom_rect(aes(ymin = 3, ymax = 6, xmin = 2, xmax = 3),fill = "grey", col = "black")+ annotate("text",x = 3.5, y = 4.5,label = "2", size = 2) +
    geom_rect(aes(ymin = 2, ymax = 7, xmin = 4, xmax = 5),fill = "grey", col = "black")+ annotate("text",x = 5.5, y = 4.5, label = "3", size = 2) +
    geom_rect(aes(ymin = 1, ymax = 8, xmin = 6, xmax = 7),fill = "grey", col = "black") + annotate("text",x = 7.5, y = 4.5,label = "4", size = 2) +
    scale_x_continuous(limits = c(0,50), expand = c(0,0))+
    ggtitle("No. of times detected")+
    theme(panel.background = element_blank(), panel.border = element_blank(),
          axis.title = element_blank(),axis.text = element_blank(),axis.ticks = element_blank(),
          title = element_text(size = 8))
  
  if(!sp.end == 0){mw_1="[MW - SP"}else{mw_1 = "[MW"}
  # join the two plots
  plot_spacer()+plot_spacer()+plot_spacer()+
    plot_spacer()+e+plot_spacer()+
    scale + a + b + 
    plot_spacer()+plot_spacer()+plot_spacer() +
    plot_spacer()+c +plot_spacer()+
    plot_spacer()+plot_spacer()+plot_spacer() +
    plot_spacer() + d+
    plot_spacer()+plot_spacer()+plot_spacer() +
    # plot_layout(guides = "collect")  + 
    plot_layout(ncol = 3, nrow = 8, widths = c(0.05,5,1), heights = c(0.05,0.7,20,0.05,1,0.05,2,0.05)) +
    plot_annotation(title = str_wrap(paste(pro,lab360[lab360$Leading.razor.protein == pro,]$protein.annotation," ", mw_1,"=",mw-sp.mw,"kDa]"),120)) 
  
}


##simplified protomap to show gel slices 
protomap_slice = function(pro){
  test = ppt_1[ppt_1$Leading.razor.protein == pro,]
  m = which(colnames(test) == "Mean 1 ")
  test = test[,!names(test) == "Leading.razor.protein"]
  test.m = melt(test, id = c("Sequence","Start.position","End.position","Unique..Proteins."))
  
  test.m = test.m %>% 
    mutate(gel.slice = gsub("[^0-9]","",variable)) %>%
    mutate(gel.slice = as.numeric(gel.slice)) %>%
    mutate(measure = gsub("[0-9]","",variable)) 
  test.c = test.m[test.m$measure == "Count ",]
  test.c$value = as.numeric(test.c$value)
  test.sum = aggregate(value ~ gel.slice,data = test.c, FUN = sum)
  
  # match the expected molecular weight of the protein
  mw = lab360[lab360$Leading.razor.protein == pro,]$exp.mw
  
  #sequon
  sequon.loc = unlist(gregexpr("N.[ST]",lab360[lab360$Leading.razor.protein == pro,]$sequence))
  npt.loc = unlist(gregexpr("NP[ST]",lab360[lab360$Leading.razor.protein == pro,]$sequence))
  sequon.loc = sequon.loc[!sequon.loc %in% npt.loc]
  
  mw = lab360[lab360$Leading.razor.protein == pro,]$exp.mw
  exp_gly_gel = exp_gel(lab360[lab360$Leading.razor.protein == pro,]$gly.mw[1])
  
  if (pro %in% unique(signalp$seqid)){
    sp.end = signalp[signalp$seqid == pro,]$end 
    sp.sequence = substring(lab360[lab360$Leading.razor.protein == pro,]$sequence,1,sp.end)
    sp.mw = round(mw(sp.sequence)/1000,2)
    exp.gel = exp_gel(mw-sp.mw)
    exp.gly.gel = exp_gel(lab360[lab360$Leading.razor.protein == pro,]$gly.mw[1]-sp.mw)}else{
      sp.end = 0
      sp.mw = 0
      exp.gel = exp_gel(mw)
      exp.gly.gel = exp_gly_gel}
  
  ggplot(test.sum, aes(xmin=0,xmax=2,ymin=gel.slice-0.5,ymax=gel.slice+0.5, fill = value))+
    geom_rect()+
    geom_rect(aes(xmin=0,xmax=2,ymin=exp.gel-0.5,ymax=exp.gel+0.5),col="#d37b6d",linewidth = 0.4, fill = "transparent")+
    geom_rect(aes(xmin=0,xmax=2,ymin=exp.gly.gel-0.5,ymax=exp.gly.gel+0.5),col="#765faf",linewidth = 0.4, fill= "transparent")+
    scale_fill_gradient(low="white",high="black")+
    scale_y_reverse(limits = c(24.6,0.5),breaks = seq(24,1, by = -1), expand = c(0,0)) + 
    xlab(pro)+
    protomap_slice_theme
}


##simplifed peptographs to show sequence
protomap_seq = function(pro){
  test = ppt_1[ppt_1$Leading.razor.protein == pro,]
  m = which(colnames(test) == "Mean 1 ")
  test = test[,!names(test) == "Leading.razor.protein"]
  test.m = melt(test, id = c("Sequence","Start.position","End.position","Unique..Proteins."))
  
  test.m = test.m %>% 
    mutate(gel.slice = gsub("[^0-9]","",variable)) %>%
    mutate(gel.slice = as.numeric(gel.slice)) %>%
    mutate(measure = gsub("[0-9]","",variable)) 
  test.c = test.m[test.m$measure == "Count ",]
  test.c$value = as.numeric(test.c$value)
  test.sum = aggregate(value ~ gel.slice,data = test.c, FUN = sum)
  
  # match the expected molecular weight of the protein
  mw = lab360[lab360$Leading.razor.protein == pro,]$exp.mw
  exp_gly_gel = exp_gel(lab360[lab360$Leading.razor.protein == pro,]$gly.mw[1])
  
  if (pro %in% unique(signalp$seqid)){
    sp.end = signalp[signalp$seqid == pro,]$end 
    sp.sequence = substring(lab360[lab360$Leading.razor.protein == pro,]$sequence,1,sp.end)
    sp.mw = round(mw(sp.sequence)/1000,2)
    exp.gel = exp_gel(mw-sp.mw)
    exp.gly.gel = exp_gel(lab360[lab360$Leading.razor.protein == pro,]$gly.mw[1]-sp.mw)}else{
      sp.end = 0
      sp.mw = 0
      exp.gel = exp_gel(mw)
      exp.gly.gel = exp_gly_gel}
  
  #sequon
  sequon.loc = unlist(gregexpr("N.[ST]",lab360[lab360$Leading.razor.protein == pro,]$sequence))
  npt.loc = unlist(gregexpr("NP[ST]",lab360[lab360$Leading.razor.protein == pro,]$sequence))
  sequon.loc = sequon.loc[!sequon.loc %in% npt.loc]
  
  a = ggplot(test.c, aes(xmin=Start.position, xmax = End.position, ymin = -0.1, ymax =0.1, col = Unique..Proteins.),
             linewidth = 1.3,alpha = 0.3)+
    geom_segment(aes(x = 1, xend = lab360[lab360$Leading.razor.protein == pro,]$length,
                     y = 0, yend = 0),col = "lightgrey", linewidth = 2 )+
    geom_rect(fill = "darkgrey", alpha = 0.4,)+
    scale_x_continuous(limits = c(1,max(ppt_1$End.position)),
                       breaks = seq(1,lab360[lab360$Leading.razor.protein == pro,]$length,
                                    by =round(lab360[lab360$Leading.razor.protein == pro,]$length/5,0)),
                       expand = c(0,0)) +
    scale_color_manual(values = c("no" = "black","yes" = "#ce3812"),drop = FALSE) +
    labs(y = paste0(pro)) +
    protomap_seq_a_theme
  
  annot = pfam[pfam$protein.id == pro & pfam$database == "Pfam",]
  if (!is.null(sp.end) && sp.end>0) {
    annot$prediction = paste(annot$number,annot$prediction)
    annot[nrow(annot)+1,] = c(pro,"SignalP","","SP",1,sp.end,
                              "","","","")} else {
                                annot$prediction = paste(annot$number,annot$prediction)
                                annot[nrow(annot)+1,] = c(pro,"SignalP","","SP",NA,NA,"","","","")}
  
  annot = annot[,names(annot) %in% c("database","start","stop","prediction")]
  if(length(sequon.loc)>0) {
    annot[nrow(annot) + length(sequon.loc),] <- NA
    annot[(nrow(annot)-length(sequon.loc)+1):nrow(annot),"database"] = "Sequon"
    annot[(nrow(annot)-length(sequon.loc)+1):nrow(annot),"start"] <- sequon.loc
    annot[(nrow(annot)-length(sequon.loc)+1):nrow(annot),"stop"] <- sequon.loc+3
    annot[(nrow(annot)-length(sequon.loc)+1):nrow(annot),"prediction"] = "Sequon"}else{
      annot[nrow(annot)+1,] <- NA
      annot[nrow(annot),"database"] = "Sequon"
      annot[nrow(annot),"start"] <- NA
      annot[nrow(annot),"stop"] <- NA
      annot[nrow(annot),"prediction"] = "Sequon" 
    }  
  
  annot = annot[order(nrow(annot):1),]
  
  annot = annot[,names(annot) %in% c("database","start","stop","prediction")]
  if(length(sequon.loc)>0) {
    annot[nrow(annot) + length(sequon.loc),] <- NA
    annot[(nrow(annot)-length(sequon.loc)+1):nrow(annot),"database"] = "Sequon"
    annot[(nrow(annot)-length(sequon.loc)+1):nrow(annot),"start"] <- sequon.loc
    annot[(nrow(annot)-length(sequon.loc)+1):nrow(annot),"stop"] <- sequon.loc+3
    annot[(nrow(annot)-length(sequon.loc)+1):nrow(annot),"prediction"] = "Sequon"}else{
      annot[nrow(annot)+1,] <- NA
      annot[nrow(annot),"database"] = "Sequon"
      annot[nrow(annot),"start"] <- NA
      annot[nrow(annot),"stop"] <- NA
      annot[nrow(annot),"prediction"] = "Sequon" 
    }  
  
  annot = annot[order(nrow(annot):1),]
  
  ##add tm domains
  topo = tm[tm$V1 == pro,]$V6
  topo = sub("Topology=","",topo)
  tm_start = c()
  tm_end = c() 
  len = gregexpr("i",topo)
  len = sum(unlist(len) != -1)
  topo_start = unlist(str_extract_all(topo, "[io](\\d+)"))
  topo_end = unlist(str_extract_all(topo,"(\\d+)[io]"))
  
  if(length(topo_start)>0){
    for (i in 1:len){
      tm_start[i] = as.numeric(str_extract_all(topo_start[i], "\\d+"))
      tm_end[i] = as.numeric(str_extract_all(topo_end[i], "\\d+"))
    }
    annot[nrow(annot)+length(tm_start),] <- NA
    annot[(nrow(annot)-length(tm_start)):nrow(annot),"database"] <- "TMHMM"
    annot[(nrow(annot)-length(tm_start)+1):nrow(annot),"start"] <- tm_start
    annot[(nrow(annot)-length(tm_start)+1):nrow(annot),"stop"] <- tm_end
    annot[(nrow(annot)-length(tm_start)+1):nrow(annot),"prediction"] <- "TM Domain"
  }else{
    annot[(nrow(annot)+1),"database"] <- "TMHMM"
    annot[nrow(annot),"start"] <- NA
    annot[nrow(annot),"stop"] <- NA
    annot[nrow(annot),"prediction"] <- "TM Domain"
  }
  
  
  b = ggplot(annot, 
             aes(xmin = as.numeric(start), xmax = as.numeric(stop), 
                 ymin = -0.1, ymax = 0.1,
                 fill = fct_relevel(prediction,"SP","Sequon","TM Domain"))) +
    geom_segment(aes(x = 1, xend = lab360[lab360$Leading.razor.protein == pro,]$length,
                     y = 0, yend = 0),col = "lightgrey", linewidth = 2 )+
    geom_rect(linewidth = 0.2, col = "black",alpha = 0.5)+
    scale_fill_manual(values = c(col_vector), drop = FALSE)+
    scale_x_continuous(limits = c(1,max(ppt_1$End.position)),
                       breaks = seq(1,lab360[lab360$Leading.razor.protein == pro,]$length,
                                    by =round(lab360[lab360$Leading.razor.protein == pro,]$length/5,0)),
                       expand = c(0,0)) +
    guides(fill = guide_legend("Domain description", title.position = "top",nrow = 2))+ 
    labs(y = "Pfam domains") +
    protomap_seq_b_theme
  
  a + b +
    plot_layout(nrow = 2)
}
```
