## Supplementary material for "The proteome of agroinfiltrated *Nicotiana benthamiana* is shaped by extensive protein processing": File S3: lib_requiremens.R

```
##library requirements

##the following libraries are required to process data (prep_files.R)
library(dplyr)
library(Peptides)
library(tidyverse)
library(ape)
library(zeallot)
library(rvest)

##the following libraries are required to generate functions for plots (functions.R)
library(RColorBrewer)
library(gridExtra)
library(patchwork)
library(reshape2)
library(reshape)
library(cowplot)
library(scales)


##plot themes

##scale_theme
scale_theme = 
  theme(axis.ticks.x = element_blank(),
        axis.title.x = element_blank(),
        axis.text.x = element_blank(),
        panel.border = element_blank(), panel.background = element_blank(), 
        panel.grid= element_blank(),
        axis.line.x = element_blank(),
        axis.title.y.left = element_text(hjust = 1, angle = 0,vjust = 1.03,margin = margin(r = -30,l= 20)),
        axis.title.y = element_text(size = 8),       
        axis.line.y = element_line(color = "grey90"))


##theme peptograph
peptograph_theme = 
  theme(panel.background = element_rect(fill = 'transparent'), 
        panel.border = element_rect(colour = alpha("black",0.6),fill = NA,linewidth = 1),
        panel.grid.major.y = element_line(color = "grey90", linewidth = 0.1),
        plot.background = element_rect(fill='transparent', color=NA), 
        legend.position = "top", legend.title.align = 0,
        legend.text = element_text(size = 9),legend.key.size = unit(0.2,"cm"), legend.title = element_text(size = 10),
        legend.spacing.x = unit(0.05,"cm"),
        legend.spacing.y = unit(0.1,"cm"),
        legend.justification = "left",
        panel.grid = element_line(),
        axis.title.x = element_text(size = 10,margin = margin(t = 5)),
        axis.title.y.left = element_text(hjust = 1, angle = 0,vjust = 1.03,margin = margin(r = -30,l= 20)),
        axis.title.y = element_text(size = 8))  


##count_barplot theme
count_barplot_theme = 
  theme(panel.background = element_blank(),  
        panel.border = element_rect(colour = alpha("black",0.6), fill=NA, linewidth = 1), 
        axis.title.y = element_blank(),
        axis.text.x = element_text(angle = 45, size = 7,hjust = 1),
        legend.position = "top",
        legend.spacing.x = unit(0.05,"cm"),legend.spacing.y = unit(0.1,"cm"),
        legend.text = element_text(size = 9),legend.key.size = unit(0.2,"cm"), legend.title = element_text(size = 10),
        axis.title.x = element_text(size = 10,margin = margin(t = 5)),
        panel.grid.major.y = element_line(color = "grey90", linewidth = 0.1),
        panel.grid.major.x = element_blank(),
        legend.key = element_rect(fill = NA))


##annotation plot theme
annot_theme = 
  theme(axis.title.y=element_blank(),
        axis.text.y=element_blank(),
        axis.ticks.y=element_blank(), 
        panel.border = element_blank(), panel.background = element_blank(), 
        panel.grid= element_blank(),
        legend.position = "bottom",
        legend.spacing.x = unit(0.05,"cm"),legend.spacing.y = unit(0.1,"cm"),
        legend.text = element_text(size = 9),legend.key.size = unit(0.2,"cm"), legend.title = element_text(size = 10),
        axis.title.x = element_text(size = 10,margin = margin(t = 5)),
        axis.line.x = element_line(colour = "white"), axis.line.y = element_blank())


##simplified protomap gel slice plot theme
protomap_slice_theme = 
  theme(panel.background = element_blank(),panel.border = element_rect(colour = alpha("black",0.6),fill = NA,linewidth = 0.3),
        axis.ticks.x = element_blank(),
        axis.ticks.y = element_line(size = 0.3),
        axis.ticks.length.y = unit(0.05,"cm"),
        axis.text.x = element_blank(),
        axis.text.y = element_text(size = 3),
        legend.position = "none",
        axis.title.x = element_text( angle = 90, size = 4,vjust = 0.5, margin = margin(t = -2)))


##simplified protomap sequence plot theme
protomap_seq_a_theme = 
  theme(axis.text.y=element_blank(),
        axis.ticks.y=element_blank(),
        axis.text.x = element_text(size = 5),
        panel.border = element_blank(), panel.background = element_blank(),
        panel.grid= element_blank(),
        legend.position = "none",
        axis.title.x = element_blank(), axis.ticks.x = element_line(linewidth = 0.2),
        axis.title.y.left = element_text(hjust = 1, angle = 0, vjust = 0.5, size = 7),
        axis.line.x = element_line(colour = "white"), axis.line.y = element_blank())

protomap_seq_b_theme = 
  theme(axis.title.y=element_blank(),
        axis.text.y=element_blank(),
        axis.ticks.y=element_blank(), 
        panel.border = element_blank(), panel.background = element_blank(), 
        panel.grid= element_blank(),
        legend.position = "bottom", legend.justification = "left",
        legend.spacing.x = unit(0.05,"cm"),legend.spacing.y = unit(0.1,"cm"),
        legend.text = element_text(size = 6),legend.key.size = unit(0.2,"cm"), 
        legend.title = element_text(size = 7),
        axis.title.x = element_blank(), axis.ticks.x = element_line(size = 0.2),
        axis.text.x = element_text(size = 5),
        axis.title.y.left = element_text(hjust = 1, angle = 0, vjust = 0.5, size = 7),
        axis.line.x = element_line(colour = "white"), axis.line.y = element_blank())
```
