## Supplementary material for "The proteome of agroinfiltrated *Nicotiana benthamiana* is shaped by extensive protein processing": File S3: prep_files.R

```
##protomap load files

##directory and file names
dir = "/Users/user/Desktop/Oxford/03_dphil/2_protomap" 
ace0277 = "ACE_0277_Sec._7.7.MQ08_fullyTryp_peptides.txt"     ##raw proteomics data 
pfam = "NbLab360_interproscan.csv"        ##interpro annotation on LAB360 database
signalp = "NbLab360.sl2.gff3"     ##signalP results for LAB360 database
LAB360 = "NbLab360.sl2.fasta"     ##genome 
TMHMM = "tmhmm_annot.txt"

##set directory
setwd(dir)

##load raw proteomics data
ace0277 = read.csv(ace0277,sep = '\t')

##load genome as a dataframe for later use
lab360 <- readLines(LAB360)
lab360.header = lab360[which(str_detect(lab360,">"))]
lab360.sequence = lab360[which(!str_detect(lab360,">"))]
lab360 = data.frame(header = lab360.header, sequence = lab360.sequence) %>%
  mutate(Leading.razor.protein = str_extract(header,"(?<=\\>)Nb.+?(?= )")) %>%
  mutate(protein.annotation = str_extract(header, "\\s(\\S+)")) %>%
  mutate(length = nchar(sequence)) %>%
  mutate(exp.mw = round(as.numeric(mw(sequence)/1000),2)) 
##add sequons
lab360 = lab360 %>% 
  mutate(sequon = str_count(sequence,"N.[ST]")) %>%
  mutate(p_sequon = str_count(sequence,"NP[ST]")) %>%
  mutate(sequon = sequon - p_sequon) %>%
  mutate(gly.mw = exp.mw + 3*sequon) 

##load pfam
pfam <- read.csv(pfam, header = FALSE, sep = ',')
pfam = pfam[,c(1,4:9,12:13)]
colnames(pfam) <- c("protein.id","database","number","prediction","start","stop","evalue","interpro.no","interpro.pre")  
##add length from lab360 
pfam <- merge(pfam, lab360[,c("Leading.razor.protein","length")], by.x = "protein.id", by.y = "Leading.razor.protein")
pfam[,"length"] <- as.numeric(pfam[,"length"])       
pfam[,"start"] = as.numeric(pfam[,"start"])
pfam$stop = as.numeric(pfam$stop)

##load signalP
signalp<- read.gff(signalp, GFF3 = TRUE)

##load TMHMM
tmhmm = read.table(TMHMM)
tmhmm$V6 = sub('\\\\+$','',tmhmm$V6)
tm = tmhmm[!tmhmm$V6 == "Topology=o",]
tm[] <- lapply(tm, function(x) gsub(",", "", x))


##clean up proteomics data and select the data needed for peptographs
first_sample = "LFQ.intensity.097" #intensity for the first sample
slices = 24 #number of slices
reps = 4  #number of repeats
c(s,w,l,p,a,b,u,i) %<-% which(names(ace0277) %in% c("Sequence","Amino.acid.before","Last.amino.acid","Leading.razor.protein","Start.position","End.position","Unique..Proteins.",first_sample))

# There are 96 experiments (97-192, which are i:(i+slices*rep)). 
peptide_groups_1 = ace0277[,c(s,w,l,p,a,b,u,i:(i+slices*reps))]
#select Nb proteins only
peptide_groups_1 = peptide_groups_1[grep("^Nb",peptide_groups_1$Leading.razor.protein), ]
#select proteins with >0 value
q = as.numeric(which(colnames(peptide_groups_1) == first_sample))
ppt_1 = peptide_groups_1[colSums(peptide_groups_1[,q:ncol(peptide_groups_1)])>0.01,]


##make the dataset 
# Count how many times the same peptide is detected among the repeats
for (i in 0:23){
  ppt_1[,ncol(ppt_1)+1] = rowSums(ppt_1[,c(q+i,q+i+24,q+i+48,q+i+72)] != 0) 
  names(ppt_1)[ncol(ppt_1)] = paste("Count",i+1)
}
# Calculate the mean intensities 
for (i in 0:23){
  ppt_1[,ncol(ppt_1)+1] = rowMeans(ppt_1[,c(q+i,q+i+24,q+i+48,q+i+72)]) 
  names(ppt_1)[ncol(ppt_1)] = paste("Mean",i+1)
}
# drop the columns we do not use in later analysis i.e. the individual intensity reads
q = as.numeric(which(colnames(ppt_1) == first_sample)) 
ppt_1 = ppt_1[,-c(q:(q+96))]

#for each peptide, remove the bands that is less than 1% of its total intensity summing up
p = as.numeric(which(colnames(ppt_1) == "Mean 1"))
ppt_1$sum = rowSums(ppt_1[,(p:ncol(ppt_1))])
ppt_1 = ppt_1[ppt_1$sum>0,]
ppt_1[,p:ncol(ppt_1)] = lapply(ppt_1[,p:ncol(ppt_1)], function(x) ifelse(x<ppt_1$sum*0.1,0,x))
ppt_1$sum = rowSums(ppt_1[,(p:ncol(ppt_1))])


##total proteins detected from the dataset
protein = unique(ppt_1$Leading.razor.protein)
protein = sort(protein, decreasing = TRUE)
```
