## Supplementary material for "The proteome of agroinfiltrated *Nicotiana benthamiana* is shaped by extensive protein processing": File S4

### Slide 1
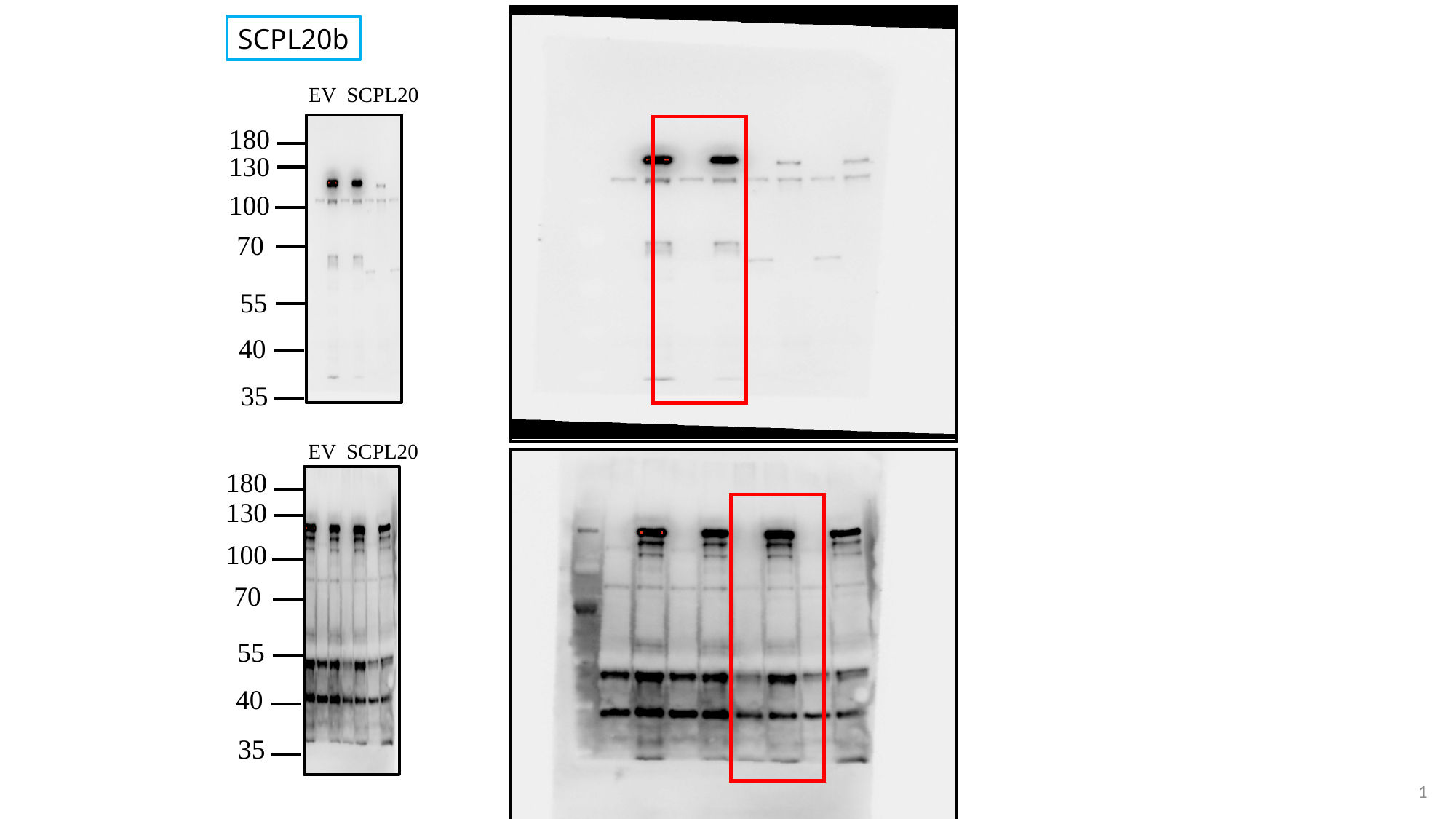

SCPL20b
EV SCPL20
180
130
100
70
55
40
35
EV SCPL20
180
130
100
70
55
40
35
1

### Slide 2
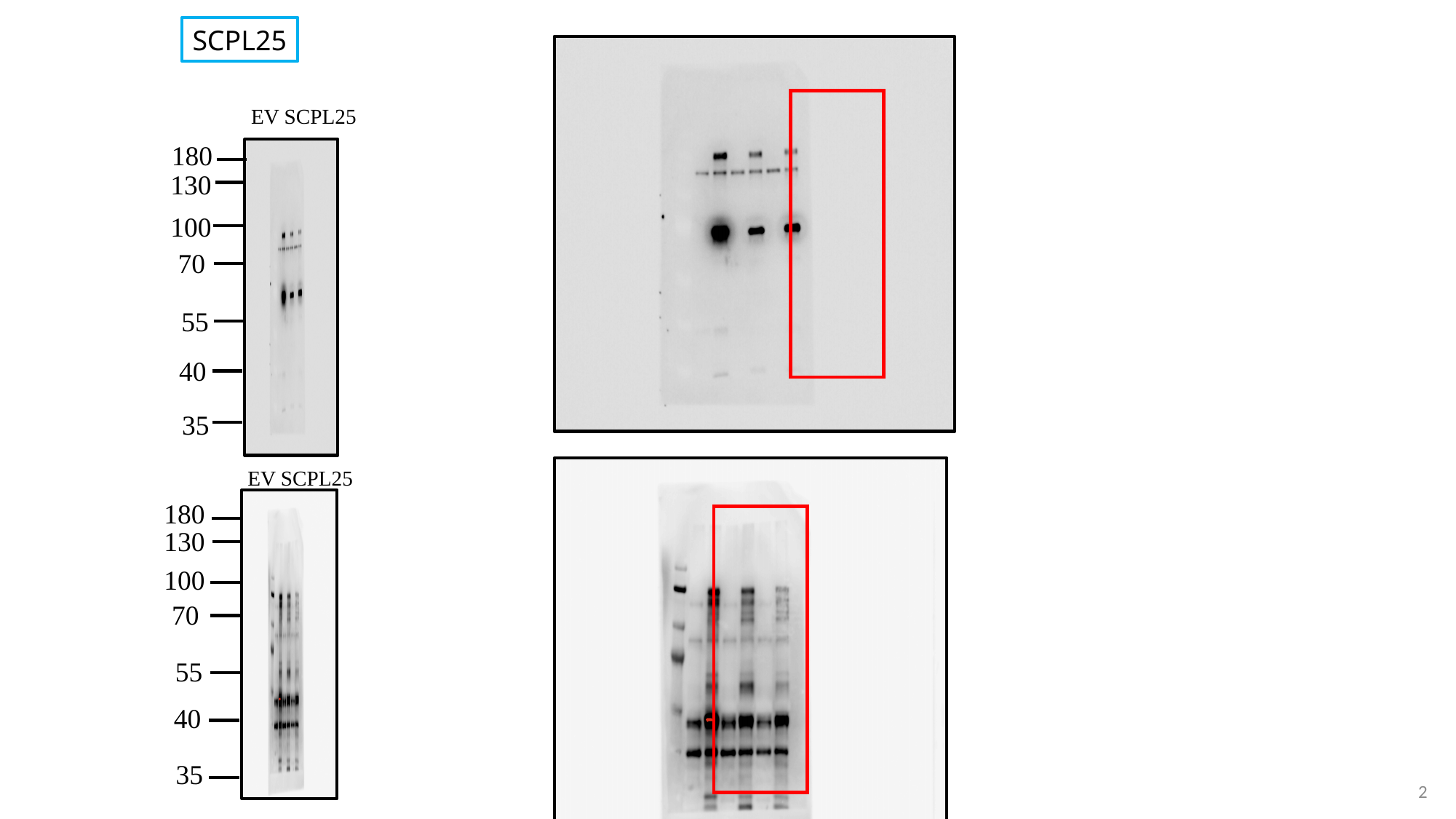

SCPL25
EV SCPL25
180
130
100
70
55
40
35
EV SCPL25
180
130
100
70
55
40
35
2

### Slide 3
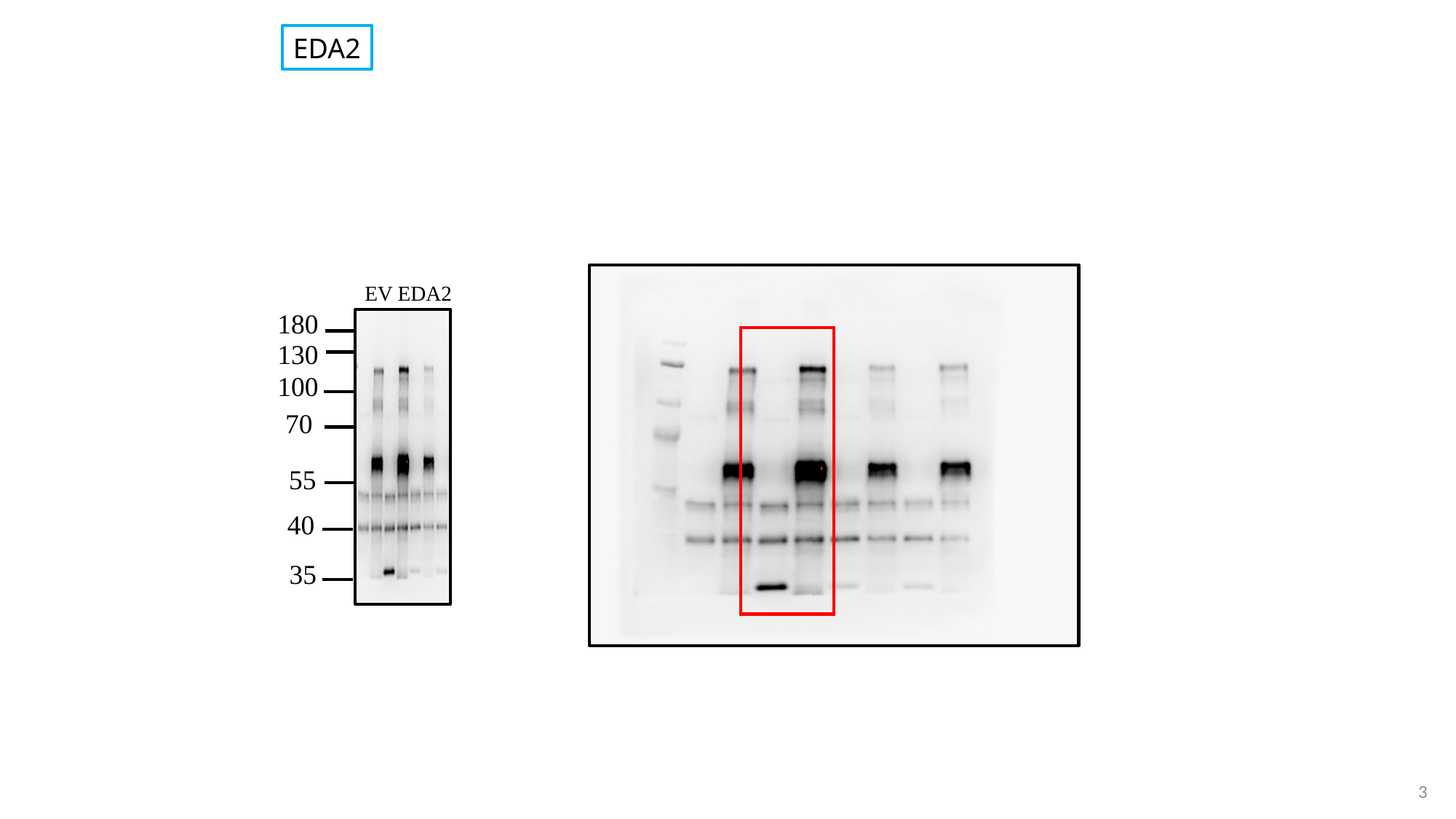

EDA2
EV EDA2
180
130
100
70
55
40
35
3

### Slide 4
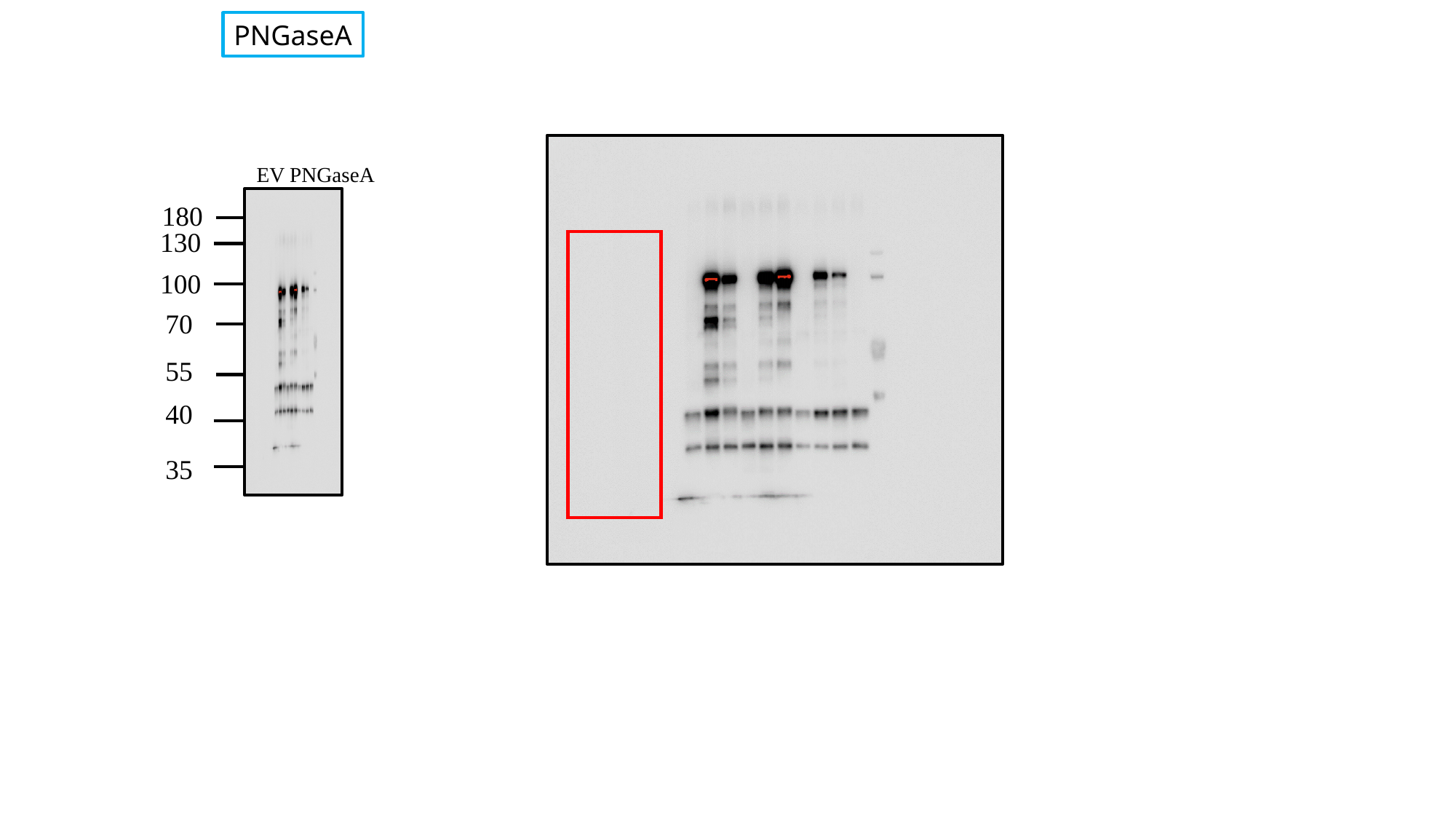

PNGaseA
EV PNGaseA
180
130
100
70
55
40
35
4

### Slide 5
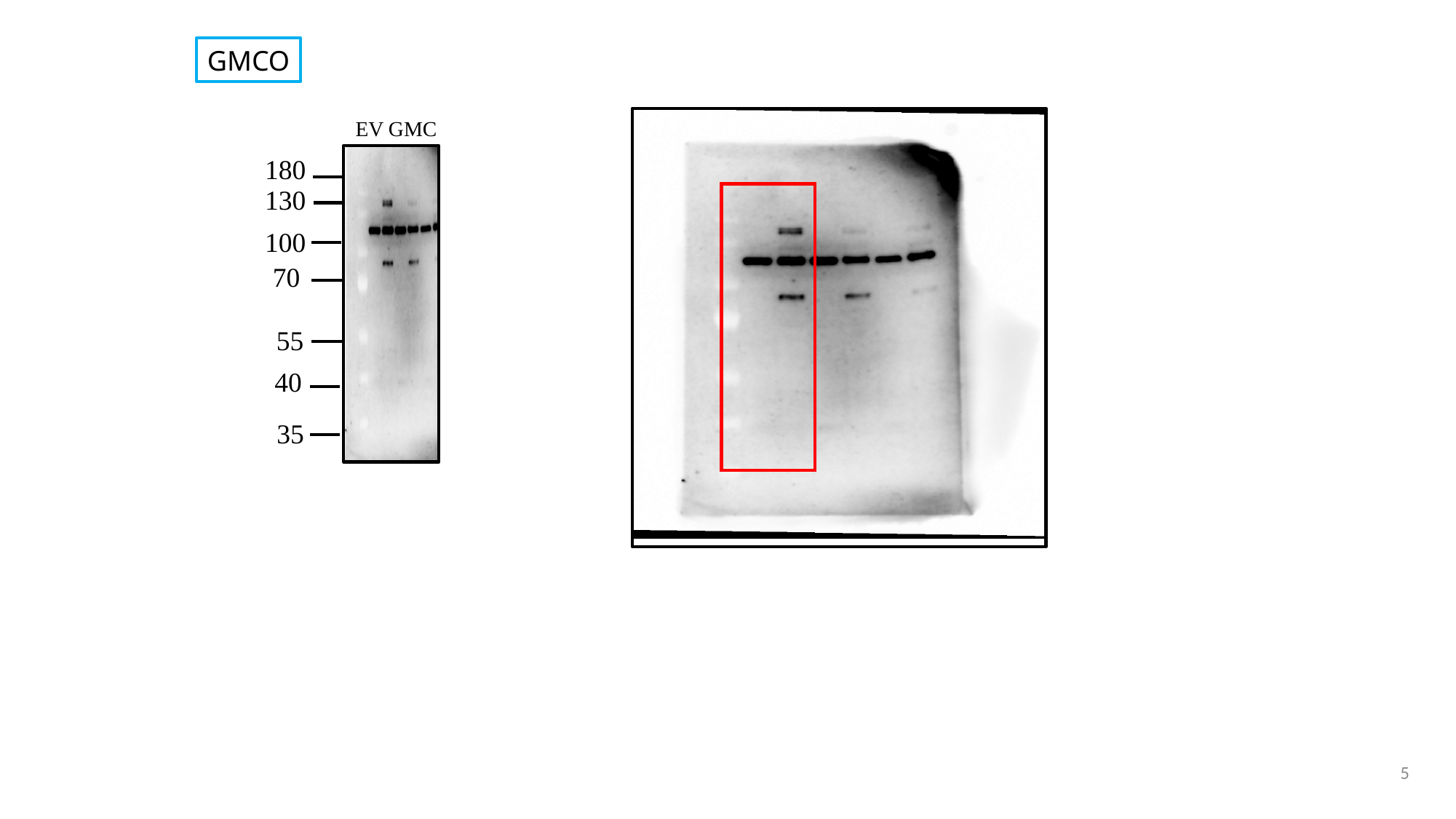

GMCO
EV GMC
180
130
100
70
55
40
35
5
