## Supplementary material for "The proteome of agroinfiltrated *Nicotiana benthamiana* is shaped by extensive protein processing": Figure S1

**Supplemental Figure S1**

Zheng et al.: Protein topography mapping reveals endogenous protease substrates and homomers in *Nicotiana benthamiana.*

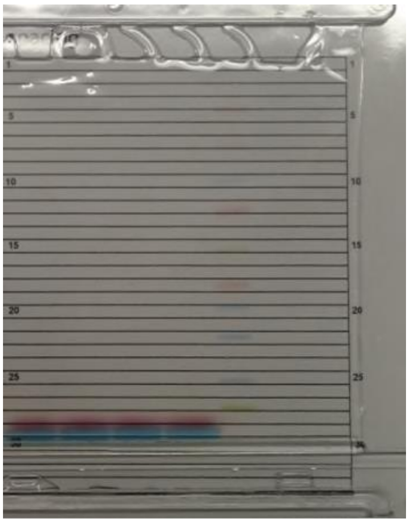

**Figure S1** Image of 4-12% gradient SDS PAGE used for PROTOMAP.

Gel slices were excised in 2.5 mm bands guided by gridlines in one cut. Next, the four biological replicate lanes were separated. The prestained MW marker is just visible.
