## Supplementary material for "The proteome of agroinfiltrated *Nicotiana benthamiana* is shaped by extensive protein processing": Figure S2

**Supplemental Figure S2**

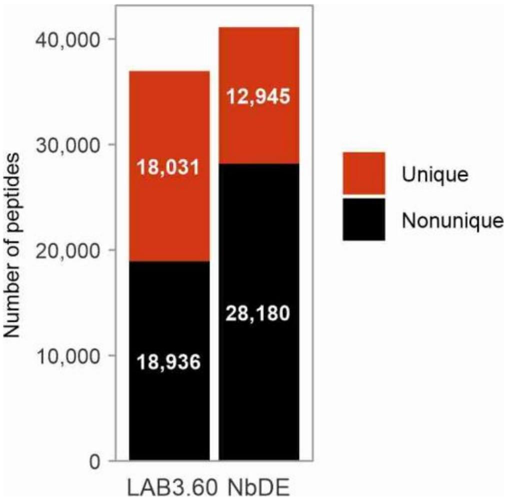


**Figure S2** Number of annotated (unique) tryptic peptides to LAB3.60 vs NbDE proteomes. Spectra were annotated to the tryptic proteomes of the LAB3.60 and NbDE annotations and the number of non-unique and unique peptides were counted.
