## Supplementary material for "The proteome of agroinfiltrated *Nicotiana benthamiana* is shaped by extensive protein processing": Figure S3

**Supplemental Figure S3**

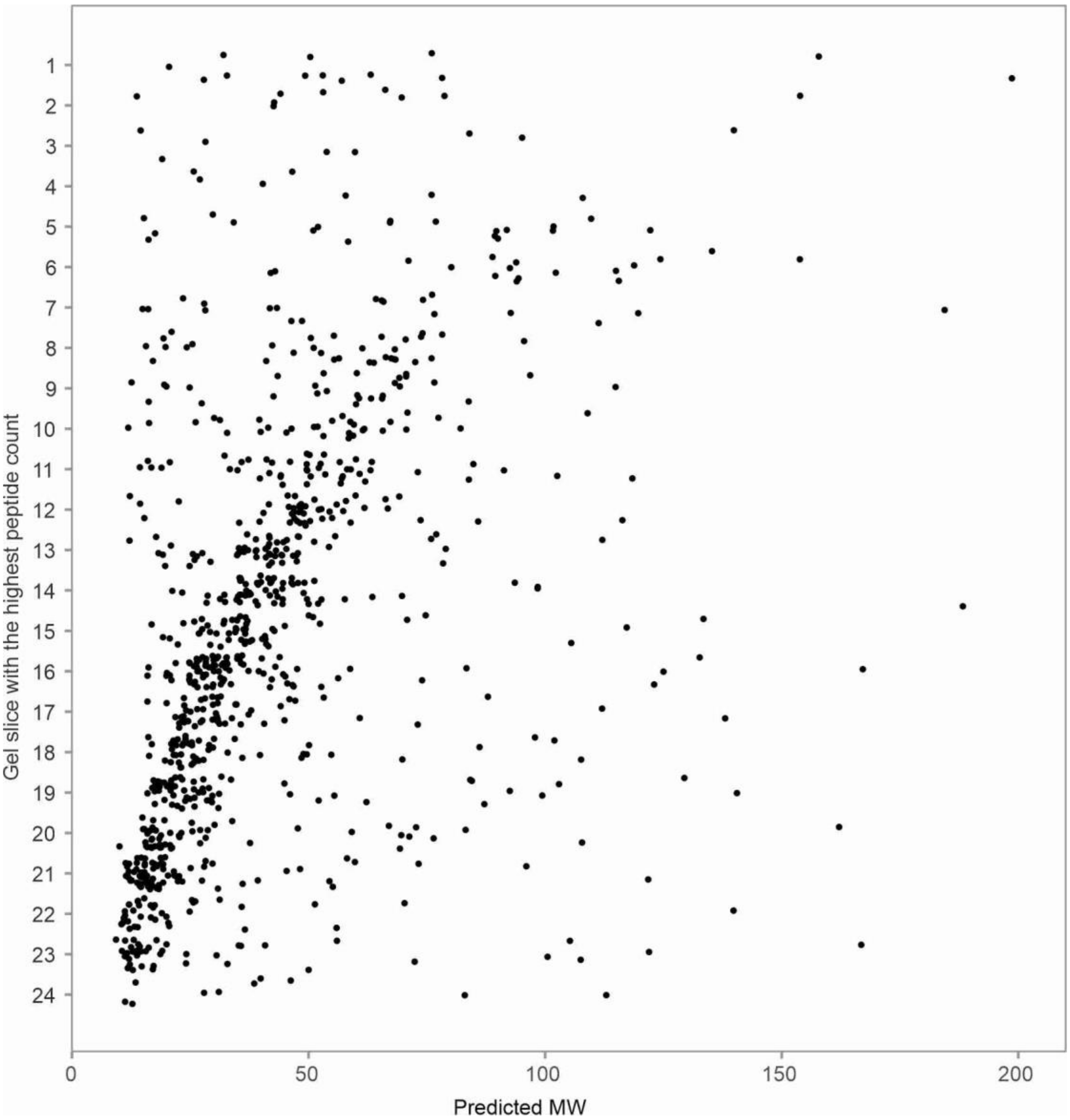


**Figure S3** Correlation between predicted and apparent MW for proteins detected with a single peptide. Slices with the highest count were taken if a single peptide was detected in multiple slices.
