## Supplementary material for "The proteome of agroinfiltrated *Nicotiana benthamiana* is shaped by extensive protein processing": Figure S5

### Supplemental Figure S5

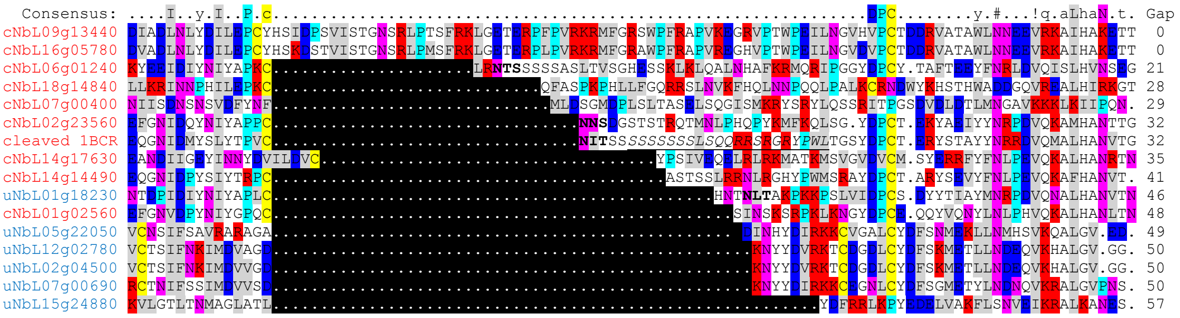


**Figure S5** Alignment of the linker region of SCPLs. Shown are nine processed SCPLs (red annotation) including wheat SCPL1 (1BCR) and six unprocessed SCPLs (blue annotation). Also indicated are the sequons (NxS/T, bold) and the linker peptide that is removed from wheat SCPL1 (1bcr, underlined italics). SCPLs are ranked based on the length of the linker region.
