## Supplementary material for "The proteome of agroinfiltrated *Nicotiana benthamiana* is shaped by extensive protein processing": Figure S7

### Supplemental Figure S6

```
cNbL12g00390 .....YPKWLKKKDRTLLQ.APINQIKFDLVVAKDGSGNFRTINEALSTAPNSS.STRFV
cNbL03g24710 .....WPEWLSAGDRRLLQ.S..STVRPDVVVAADGSGNFKTVSEAVARAPEKS.SKRYV
cNbL13g12920 D....FPKWLSRRDRKLLN.KSVSTIQADIIVAKDGSGTVKTIAEAIKKVPEKS.NRRTI
cNbL11g05910 EDFAPFPKWLNRKERVLLD.TPVSAIHADIIVSKDGNGTFKTIAEAIKKVPQYS.NRRII
cNbL05g08840 .....MPSWVNSRDRKLLE.SPAEDIKANAVVAQDGSGDYQTLTEAVAAAPDKS.KTRYV
cNbL02g21830 .....LPTWV...DRRLLQ.LSANAIKANVIVAKDGSGQYKTVKEAVASAPDNS.KTRYV
cNbL03g24700 .....YPEWVRPGDRRLLQ.A..VNPKPDVTVASDGTRDVLTIQEAVKRVPKKS.KVRFV
cNbL05g15200 .....FPFWLNRKDRRLLQLTPSTGVVADVVVALDGTGNFTRIKDAISAAPQLS.TKRFV
uNbL05g19120 .....WEGSGSDGPCHDIKGGVPSGLKPDVTVCKEGGCDYKIVQEAVNAAPDNGLTRKFV
uNbL12g21760 .....WEGSGS.GESGQAKVGVPSGLKPDVTVCKEGGCNYKTVQEAVNAAPDNEVTRKFV
uNbL05g19090 .....WEPGSVSG..FEFKGGFPSGLDPNVTVCKEGGCDYKMVQEAVNATPDNLGPGKFV
uNbL01g19610 .....FPKWLKKRDRALLQ.ATINETEIDLVVAKDGSGNFSTINEALSAAPNAS.RTRFV
```

**Figure S6** Alignment of the linker region of PMEs. Shown are eight cleaved PMEs (c, red annotation) and four uncleaved PMEs (u, blue annotation). The putative cleavage motif is highlighted in red.
