## Supplementary material for "The proteome of agroinfiltrated *Nicotiana benthamiana* is shaped by extensive protein processing": Figure S8

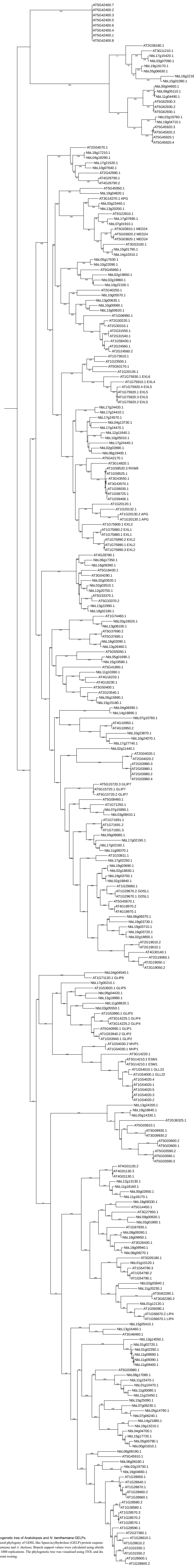

**Figure S7** Phylogenetic tree of Arabidopsis and *N. benthamiana* GELPs.
Maximum likelihood phylogeny of GDSL-like lipase/acylhydrolase (GELP) protein sequences from *N. benthamiana* and *A. thaliana*. Branch support values were calculated using ultrafast bootstrap with 1000 replications. The phylogenetic tree was visualised using iTOL and displayed with midpoint rooting.
