## Supplementary material for "The proteome of agroinfiltrated *Nicotiana benthamiana* is shaped by extensive protein processing": Figure S9

**Supplemental Figure S9**

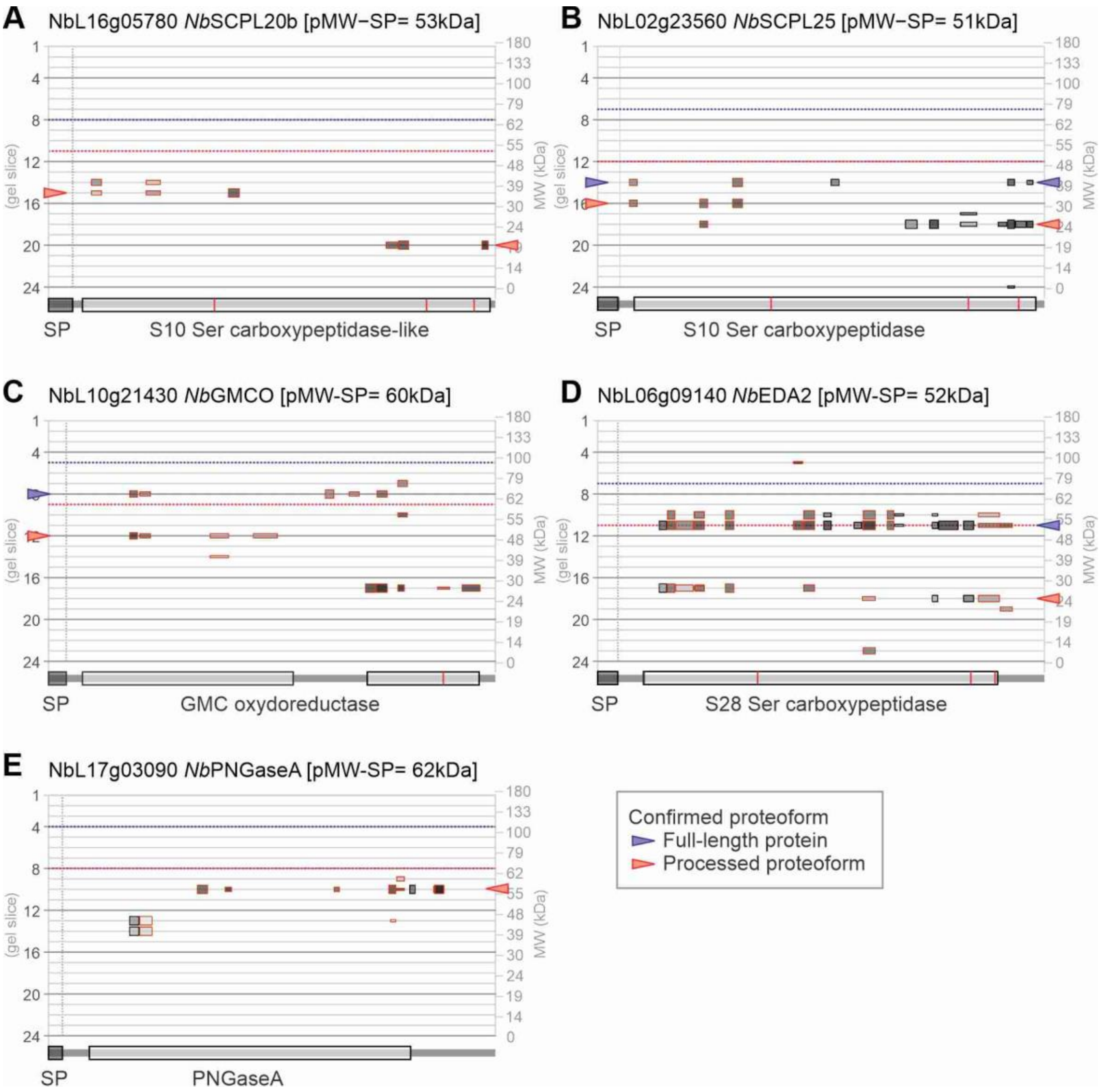


**Figure S9** Peptographs of five processed proteins confirmed by double tagging.

Highlighted with arrowheads are the proteoforms that were detected *in vivo* with double-tagged constructs.
