## Supplementary material for "The proteome of agroinfiltrated *Nicotiana benthamiana* is shaped by extensive protein processing": Table S1

**Supplemental Table S2** Used plasmids

| Plasmid | Description* | Reference |
|---|---|---|
| pJK001c | Backbone for pKZ45 | Paulus et al., 2020 |
| pJK497 | Binary empty vector | Kourelis et al., 2020 |
| pJK497 | Template for 2x35S-SP-intron | Kourelis et al., 2020 |
| pPJ057 | Template for FLAG-RFP | Beritza et al., in prep. |
| pJK227 | Template for GFP-terminator | Kourelis et al., 2020 |
| pKZ45 | Binary 35S: SP-FR-HG | This work |
| pKZ47 | Binary 35S: SP-FR-SCPL25-HG | This work |
| pKZ49 | Binary 35S: SP-FR-EDA2-HG | This work |
| pKZ54 | Binary 35S: SP-FR-GMC-HG | This work |
| pKZ60 | Binary 35s: SP-FR-PNGaseA-HG | This work |
| pKZ67 | Binary 35S: SP-FR-SCPL20b-HG | This work |

*, SP, PR1a Signal Peptide; FR, FLAG-RFP; HG, 2xHis-GFP.

**Kourelis J, Malik S, Mattinson O, Krauter S, Kahlon PS, Paulus JK, van der Hoorn RAL.** (2020) Evolution of a guarded decoy protease and its receptor in solanaceous plants. *Nat Commun.* **11**, 4393.

**Paulus JK, Kourelis J, Ramasubramanian S, Homma F, Godson A, Hörger AC, Hong TN, Krahn D, Ossorio Carballo L, Wang S, Win J, Smoker M, Kamoun S, Dong S, van der Hoorn RAL.** (2020) Extracellular proteolytic cascade in tomato activates immune protease Rcr3. *Proc. Natl. Acad. Sci. USA.* **117**, 17409-17417.
