## Supplementary material for "The proteome of agroinfiltrated *Nicotiana benthamiana* is shaped by extensive protein processing": Table S2

**Supplemental Table S3** Used oligonucleotides

| Name | Sequence (5' – 3') |
|---|---|
| SPintron-EcoRI-F1 | CAAgaattcGGATCCGGAGGTCAACATGGTGGAGCAC |
| 2x35S-SPi-SalI-R1 | CAAgtcgacCTGCACATCAACAAATTTTGGTCATATATTAG |
| FLAGRFP-SalI-F1 | CAAgtcgacGAATGGATTACAAGGATGACGACGATAAGGAGCCGCGGATGGCCT CCTCCGAGGAC |
| FLAGRFP-KpnI-R1 | CAAggtaccACTAGTCTTAAGGGCGCCGGTGGAGTGGCGGCC |
| HisGFPterm-KpnI-F1 | CAAggtaccCCCGGGTCTAGAACGCGTCATCACCATCACCATCACCATCACCATC ACCATCACCTGCAGATGGTGAGCAAGGGCGAGG |
| HisGFPterm-PvuI-R2 | CAAcgatcgACTCTAGCTAGAGAAGCTGATCAATGCATC |
| SCPL25-AflII-F1 | CAActtaagACCAATTACAAGGAAGAAGAAGAAGCTG |
| SCPL25-XbaI-R1 | CAAtctagaTGACTTGGGAAGTGGCTCTC |
| EDA2-KpnI-F1 | CAAggtaccATCTCAACTTCTCATCTTCTTCTTCAG |
| EDA2-XbaI-R1 | CAAtctagaTACATCTGAAACCTGGCACTG |
| GMC-AflII-F2 | CAActtaagGAAAAAGCTCCAAACTACTCATTC |
| GMC-KpnI-R1 | CAAggtaccATTAATATGACTCTTCTCTTTTGCAAG |
| PNGaseA-AflII-F1 | CAActtaagCTTGAACACCATTCCCACTTC |
| PNGaseA-KpnI-R1 | CAAggtaccCAAAGCAGAAAACGAATCAAACTTC |
| SCPL20b-AflII-F1 | CAActtaagGTACCTGAAAATGCATTAATTACTCAAATTCCTG |
| SCPL20b-XbaI-R1 | CAAtctagaTATGTTCTTGCCCTCTAGCCAG |

Restrictions sites for restriction enzymes indicated in the primer name are printed in small case.
